# Selective degradation of BRD9 by a DCAF16-recruiting targeted glue: mode of action elucidation and in vivo proof of concept

**DOI:** 10.1101/2024.12.31.630899

**Authors:** Scott J. Hughes, Wojciech J. Stec, Colin T. R. Davies, David McGarry, Alicia Williams, Ivan Del Barco Barrantes, Rebecca Harris, Dominic D. G. Owens, Alexander Fawcett, John Hellicar, Gregor P. Meier, Andrew C. Runcie, Liliana Greger, Martin O’Rourke, Ian Churcher, Martin Pass, Giles A. Brown, Alessio Ciulli, Louise K. Modis, Andrea Testa

## Abstract

Prospective discovery of molecular glues degraders for a specific therapeutic target protein of interest is an emerging strategy in drug discovery. Modification of pre-existing ligands with fragments that can alter the protein surface can lead to the creation of novel compounds (‘’targeted glues’’) able to induce neo-interactions between the target and an E3 ligase, resulting in targeted protein degradation. By screening a library of potential BRD9 targeted glue compounds, we have discovered a potent and selective, reversibly covalent BRD9 degrader, **AMPTX-1**. Co-immunoprecipitation-mass spectrometry experiments demonstrated that cell treatment with **AMPTX-1** induces selective recruitment of BRD9 to the E3 ligase DCAF16. Degradation is dependent on the engagement of the surface Cys58 of DCAF16 and formation of a covalent adduct to DCAF16 is facilitated by the ternary complex formation with BRD9. BRD9 degradation is achieved *in viv*o after oral dosing, demonstrating that covalent recruitment of DCAF16 is a viable strategy for targeted protein degradation and can be achieved with drug-like, orally bioavailable compounds with promising *in vivo* activity.

## Introduction

Targeted protein degradation with small molecule degraders, designed to remove a specific protein from the cell, is an emerging therapeutic modality in clinical trials for multiple indications, including solid and haematological malignancies and autoimmune diseases^1^. Most small molecule degraders work by recruiting a specific protein of interest (POI) to an E3 ligase, by means of either heterobifunctional compounds (PROTACs) or monovalent compounds (molecular glues). Generally, a PROTAC shows measurable binary binding affinity for both POI and E3 ligase and can be rationally designed for a specific POI. By contrast, a molecular glue degrader can show measurable binary binding affinity only to one of the two proteins, typically an E3 ligase (such as CRBN), with formation of a POI:glue:E3 complex made possible by ternary complex cooperativity. To date, rational design of molecular glues remains a largely unsolved challenge and high throughput screening of compound libraries continues to be the preferred strategy to discover novel, monofunctional compounds able to induce protein degradation by direct recruitment of an E3 ligase^2,3^.

While the advancement of clinical stage degraders has been made possible by the development of high quality E3 ligase binders for VHL^4^ and CRBN^2,3^, challenges remain in the discovery of drug-like ligands for novel E3 ligases^5,6^, limiting the protein degradation toolbox available to medicinal chemists and potentially limiting the range of degradable targets. Recent disclosures of KEAP1^7^, KLHDC2^8^ and DCAF1^9,10^ ligands have demonstrated the feasibility of discovering substrate-competitive small molecules for new PROTAC modalities. However, even when high quality ligands are available, the complex biology of E3 ligases makes it difficult to establish new E3 systems for general use in TPD, as the ubiquitination machinery may be differently regulated in diverse biological systems^6^.

A potential solution to this problem is “ligase agnostic” cellular screening of compound libraries, with degradation of the POI as a phenotypic readout. Such compound libraries can be obtained by chemical modification of POI ligands at their solvent exposed region: upon binding, the ligand could modify the POI surface to potentially enable *de novo* protein-protein interactions (PPI) with an E3 ligase. If the interaction is functional, degradation of the POI would be observed. A proof of concept for this approach comes from serendipitously discovered glue degraders GNE-0011^11,12^, CR8^13^ and UNC8153^14,15^, simple derivatives of POI inhibitors, that present chemical fragments on the POI surface, allowing interactions with components of E3 ligase complexes (Fig 1).

**Fig. 1:**
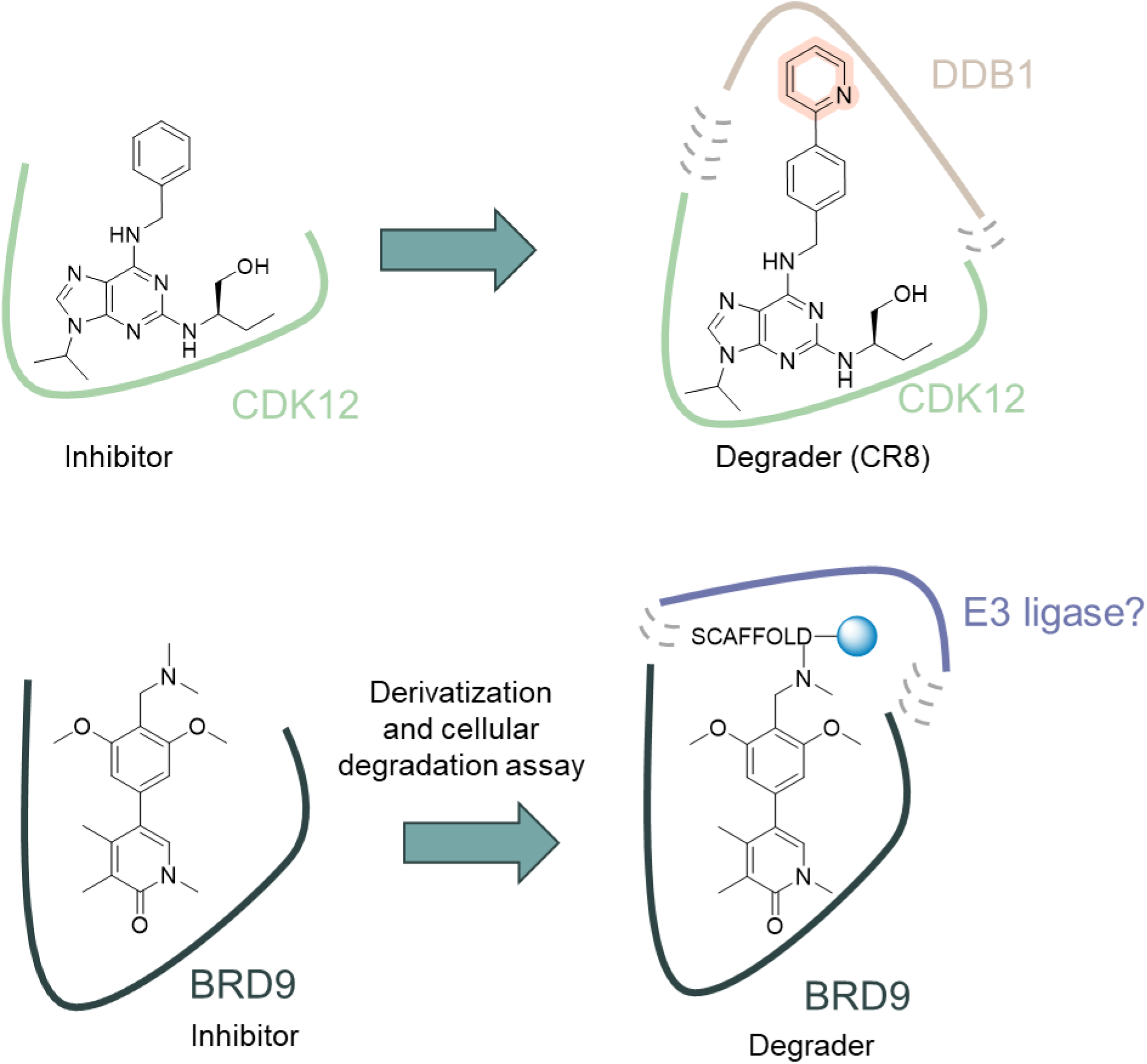
Chemical derivatization of protein inhibitors to facilitate E3 ligase recruitment. **a** Simple derivatization of solvent-exposed regions on CDK12 inhibitors modifies the protein surface to allow recruitment of DDB1. **b** Overview of novel modification approach applied to a BRD9 inhibitor in the present study.

We applied this “ligase agnostic” approach via chemical modification of POI ligands at their solvent exposed region to create new chemical entity degraders of bromodomain containing protein 9 (BRD9), defined hereafter as “targeted glues”. BRD9 is a component of the non-canonical BRG-/BRM-associated factor (ncBAF) chromatin remodelling complex^16^, that has been shown to be degradable by both CRBN and VHL-based PROTACs^17–21^. Two CRBN based PROTAC degraders of BRD9, CFT8634 from C4 Therapeutics and FHD-609 from Foghorn Therapeutics, have shown encouraging efficacy in preclinical models of disease^22^ and have advanced to Phase 1 clinical trials in oncology (NCT05355753 and NCT04965753). However, no molecular glue degrader has been reported for BRD9 to date.

Herein we report the discovery and mode of action elucidation of a potent and selective BRD9 targeted glue degrader that covalently recruits DCAF16. The drug-like properties of the compound allowed a proof-of-concept degradation study *in vivo*, which demonstrates the potential for DCAF16 covalent recruitment by targeted glues to be a viable alternative to traditional CRBN or VHL-based PROTACs.

## Results

### AMPTX-1 is a potent and selective BRD9 degrader

A focussed library of potential targeted glues was prepared by modification of high affinity BRD9 ligands with a proprietary library of molecular scaffolds bearing diverse electrophilic warheads. We reasoned that diverse molecular scaffolds could form non-covalent interactions at the surface of BRD9, facilitating the formation of potential protein-protein interactions with an E3 ligase, potentially aiding covalent E3 ligase engagement and BRD9 degradation. This library was then screened in a HiBiT endpoint degradation assay, in which C-terminally tagged BRD9-HiBit HEK293 cells were treated with compounds for 6 h, and BRD9 protein levels were quantified^19^. Compounds identified as having promising degradation activity were then further optimized.

A novel optimized compound having a warhead with a cyanoacrylamide-bearing tetrahydroisoquinoline scaffold, **AMPTX-1**, was identified as a potent BRD9 degrader (DC_50_ = 0.05 nM, D_max_ = 88%, upon 6 h treatment). We reasoned that the cyanoacrylamide moiety may be important for the degradation activity. To test this hypothesis, a des-cyano analogue, compound **2**, was prepared and tested in the same assay. As expected, **2** did not show protein degradation across the concentration range tested (Fig. 2a, b), suggesting that the cyanoacrylamide moiety was essential to the degradation activity of **AMPTX-1**. Both compounds were subjected to live cell kinetic profiling to better understand their activity over a 24 h period (Fig. 2c, Supplementary Fig. 1a). The results were comparable to those observed in the endpoint assay, with the D_max_ observed at concentrations above 1 nM and reached after 3 h of treatment. The degradation profile was sustained at higher concentrations throughout the duration of the experiment (24 h). As expected, the negative control compound **2** did not produce any degradation of the target (Supplementary Fig. 1a).

**Fig. 2:**
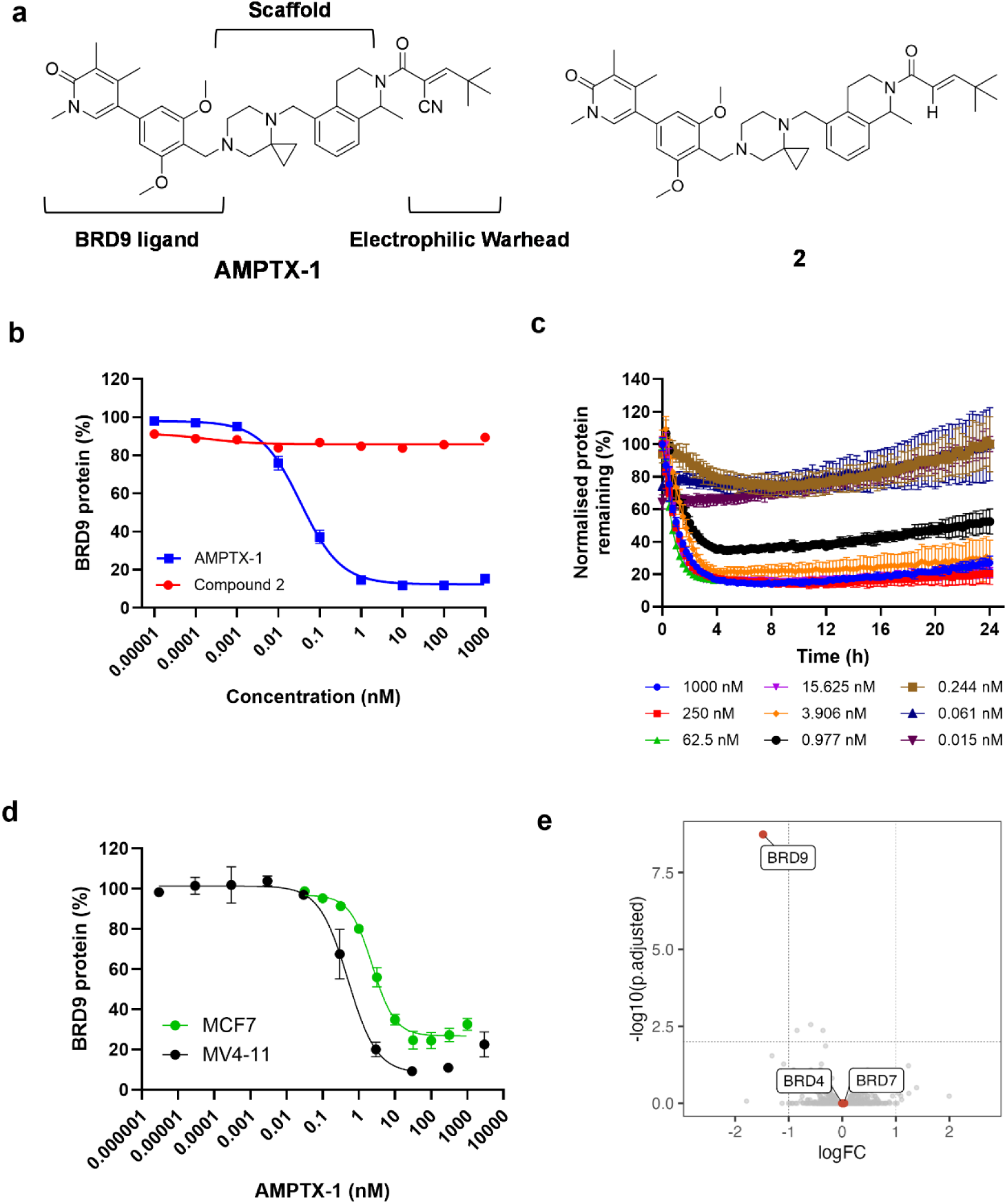
AMPTX-1 is a potent and selective degrader of BRD9. **a** Chemical structures of **AMPTX-1** and **2. b** Degradation of BRD9-HiBiT following 6 h treatment with a range of concentrations of **AMPTX-1** or **2**. **c** Live cell degradation of BRD9- HiBiT by **AMPTX-1** over 24 h. **d** BRD9 degradation in MV4-11 and MCF-7 cells following 6 h treatment with **AMPTX-1**. BRD9 levels were quantified by immunofluorescence. **e** Global expression proteomics following 6 h treatment of MV4-11 cells with 100 nM **AMPTX-1**. 8350 proteins were quantified at 1% FDR. Data are presented as logFC relative to treatment with 100 nM **2**.

**AMPTX-1** induced degradation of endogenous, untagged BRD9 in solid and liquid tumour cancer cell lines (MCF-7 and MV4-11 respectively) within 6 h (Fig. 2d). Potent BRD9 degradation was maintained in both cell lines (MV4-11 - DC_50_ = 0.5 nM, D_max_ = 93%; MCF-7 - DC_50_ = 2 nM, D_max_ = 70%), and a characteristic “hook effect” is noticeable at concentrations above 100 nM in both cell lines. To assess degradation selectivity across the proteome in a disease-relevant cell line (MV4-11), we conducted an unbiased quantitative multiplexed tandem mass tag (TMT) proteomics experiment. MV4-11 cells were treated in triplicate for 6 h with either **AMPTX-1** or **2**, each at a concentration of 100 nM (approximately 100-fold over the DC_50_ of **AMPTX-1**). Out of 8350 quantified proteins, BRD9 was the only protein significantly degraded by **AMPTX-1** (adjusted P-value = 1.84e-9), (Fig. 2e; Table Sup. 2). Conversely, treatment with **2** resulted in no significant changes in any proteins, including all quantified BET family members (Supplementary Fig. 1b). Interestingly, BRD9 ligands containing a pyridone scaffold are known to bind BRD7 with similar affinity (BRD9 IC_50_ = 37 nM; BRD7 IC_50_ = 41 nM for the dimethyl pyridone analogue)^23^, but no degradation of BRD7 was observed in this experiment, suggesting that selectivity of the novel compound **AMPTX- 1** may come from more effective and functional ternary complex formation and target ubiquitination, and not from selective target engagement.

### AMPTX-1 covalently recruits DCAF16:DDB1 complex to BRD9

A close inspection of the degradation data in MV4-11 and MCF-7 cells (Fig. 2d) reveals the presence of a hook effect. This is a characteristic of bifunctional degraders and suggests saturation of two independent binding sites.

We first assessed whether BRD9 degradation is mediated via the proteasome and Cullin RING E3 ligases (CRL), a class of NEDD8-dependent E3 ligases that are the most exploited for TPD approaches. Degradation of BRD9 induced by **AMPTX-1** was blocked by the proteasome inhibitor **Bortezomib** and the NEDD8-activating E1 (NAE1) inhibitor **MLN4924** (Fig. 3a), demonstrating that BRD9 degradation is proteasome and NAE1 dependent, and suggesting the ubiquitination step is CRL-mediated.

**Fig. 3:**
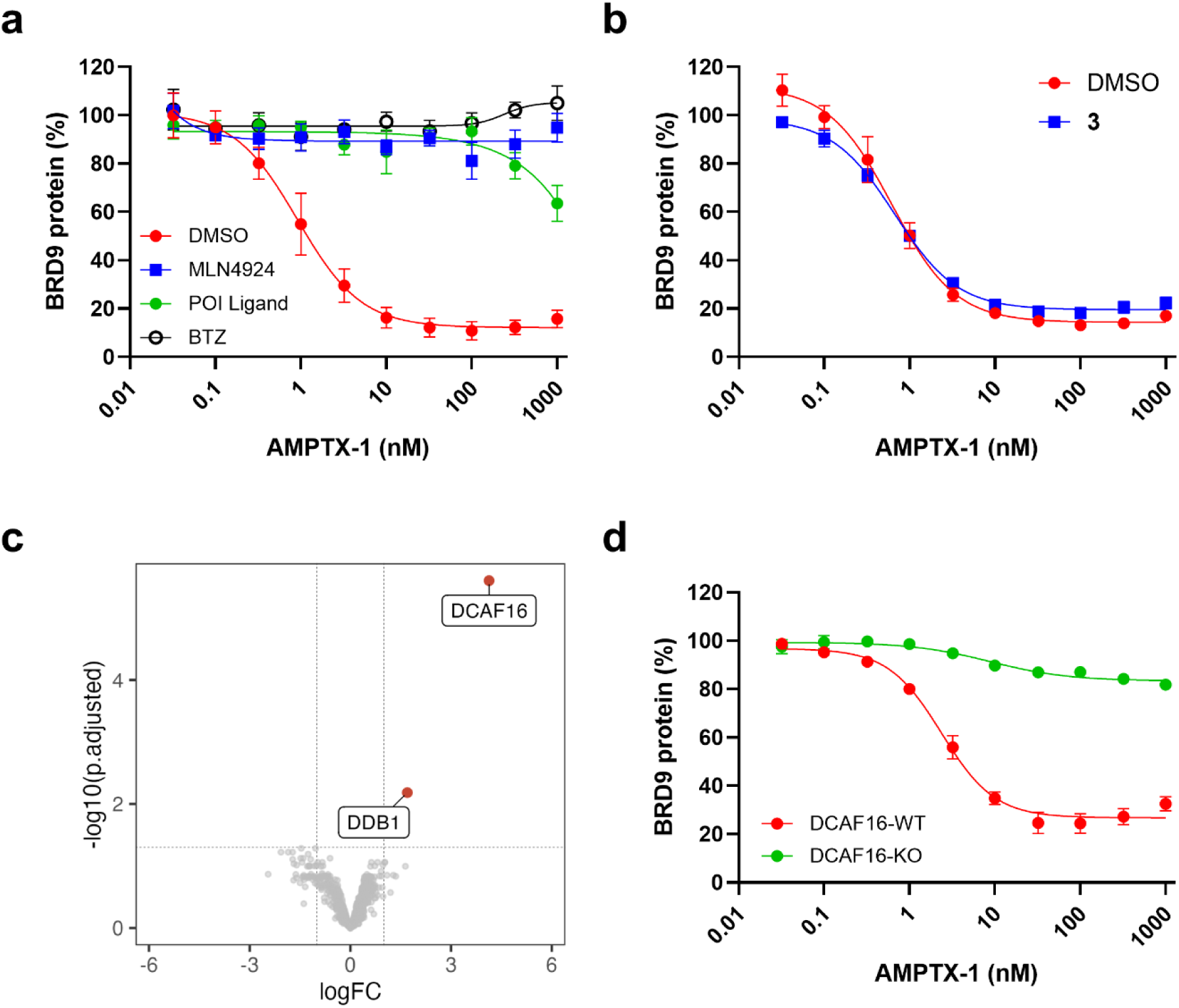
AMPTX-1 induced BRD9 degradation is dependent on the DCAF16 Cullin RING E3 ligase. **a.** Degradation of BRD9-HiBiT after 6 h treatment with **AMPTX-1,** following 1 h pre-treatment with DMSO (red), 3 µM NAE1 inhibitor **MLN4924** (blue), 10 µM proteasome inhibitor **Bortezomib** (BTZ, black), or 10 µM **BI-7273** (BRD9 POI ligand, green). Data are represented as mean ± standard deviation from *n* = 2 biological replicates. **b** Degradation of BRD9-HiBiT after 6 h treatment of **AMPTX-1,** following 1 h pre-treatment of DMSO (red) or 10 µM warhead **3** (blue). Data are represented as mean ± standard deviation from *n* = 2 biological replicates. **c** Volcano plot showing proteins enriched by anti-HiBiT co-immunoprecipitation following 2 h treatment of BRD9-HiBiT HEK293 cells with 100 nM **AMPTX-1**. Data are presented as logFC relative to 100 nM **2**. **d** Degradation of BRD9 following 6 h treatment with a range of concentrations of **AMPTX-1** in parental (red, DCAF16-WT) or knockout (green, DCAF16-KO) MCF-7 cells. Data are represented as mean ± standard deviation from *n* = 2 biological replicates.

We next performed a competition assay to assess whether degradation by **AMPTX-1** could be impaired by an excess of either the BRD9 inhibitor **BI-7273**^23^ or our newly identified warhead **3** (structure in Supplementary Fig. 2). As expected, excess **BI-7273** (10 µM, approximately 300- fold over IC_50_ for BRD9) was able to block BRD9 degradation mediated by **AMPTX-1,** presumably through competition for BRD9 cellular engagement (Fig. 3a). In contrast, a comparable concentration of warhead **3** (10 µM) failed to prevent degradation (Fig. 3b). This suggests that the warhead alone could not efficiently engage with its putative E3 ligase target at a binary level, indicating that the mode of action of **AMPTX-1** shows characteristics of a molecular glue but, with the presence of the hook effect, in a novel, hybrid mechanism.

To identify cellular CRL components engaged by **AMPTX-1**, we performed a ternary complex pulldown that utilizes the HiBiT tag on BRD9 to enrich for proteins that form protein-protein interactions following treatment with **AMPTX-1** or **2**. BRD9-HiBiT HEK293 cells were treated for 1 h with DMSO or increasing concentrations of **AMPTX-1** or **2** (1, 10, 300, and 1000 nM). To ensure maximum retention of the ternary complex, all samples were pre-treated for 1 h with 3 µM of **MLN4924**. Comparisons of protein recruitment to BRD9 by **AMPTX-1** and **2** using proteomics showed significant enrichment of the substrate receptor DCAF16 and its adaptor DDB1, members of the CRL4^DCAF16^ E3 ligase complex (Fig. 3c; Supplementary Table 3). Recruitment of DDB1 and DCAF16 to BRD9 was dose-dependent, with increasing quantification observed across 10, 300, and 1000 nM of **AMPTX-1** (Supplementary Fig. 2b). DDB1 and DCAF16 were the only proteins significantly enriched at all concentrations, indicating excellent specificity of ternary complex formation by the compound.

To functionally validate the dependency on DCAF16, we generated DCAF16-knockout (KO) MCF-7 cells and assessed the impact on BRD9 degradation. **AMPTX-1** showed impaired activity (<10% degradation up to 1 µM at 6 h) in the KO cells compared to WT parental control, consistent with a functional role for DCAF16 in the mode of action of **AMPTX-1** (Fig. 3d).

The importance of stereochemistry of the tetrahydroisoquinoline (THIQ) moiety for degradation potency and ternary complex formation was investigated. Chiral separation of a key intermediate in the synthesis of **AMPTX-1** was performed (see supplementary methods) to obtain the separate enantiomers of **AMPTX-1** (*ent-1*: 97.8% ee and *ent-2*: 91.0% ee). **AMPTX- 1-*ent-1*** and **AMPTX-1-*ent-2*** were tested in MV4-11 cells using an immunofluorescence (IF)- based BRD9 degradation assay (Fig. 4a). Only one of the two enantiomers, **AMPTX-1-*ent-1***, induced potent and deep degradation (DC_50_ 0.2 nM, D_max_ 94%), while **AMPTX-1-*ent-2***, showed significantly weaker activity (DC_50_ 5 nM, D_max_ 43%). Cyanoacrylamides are known to reversibly form covalent bonds, and we confirmed reversible formation of an **AMPTX-1** glutathione adduct in a dilution experiment (Supplementary Fig. 3) ^24,25^. Moreover, multiple cysteines on DCAF16 are known to be engaged by covalent degraders to induce targeted protein degradation^26–29^. To assess whether covalent reaction with DCAF16 occurred, we incubated recombinant DCAF16:DDB1:DDA1 (3.6 µM) complex with an excess of **AMPTX-1-*ent-1*** or **AMPTX-1-*ent-2*** (18 µM, 5 equivalents). Quantification of the modified/unmodified DCAF16 ratio was performed by ESI/MS analysis of the reaction mixtures (Fig. 4b, c). Despite the excess of warhead, the most abundant species was found to be the unmodified E3 ligase, with 71% and 82% unmodified DCAF16 with **AMPTX-1-*ent-1*** and **AMPTX-1-*ent-2***, respectively. The majority of modified DCAF16 showed mass shift consistent with a single adduct, but a small population (∼4%) for both **AMPTX-1-*ent-1*** and **AMPTX-1-*ent-2*** showed mass shifts consistent with >1 modification, suggesting multiple cysteines can react with the warhead.

**Fig. 4:**
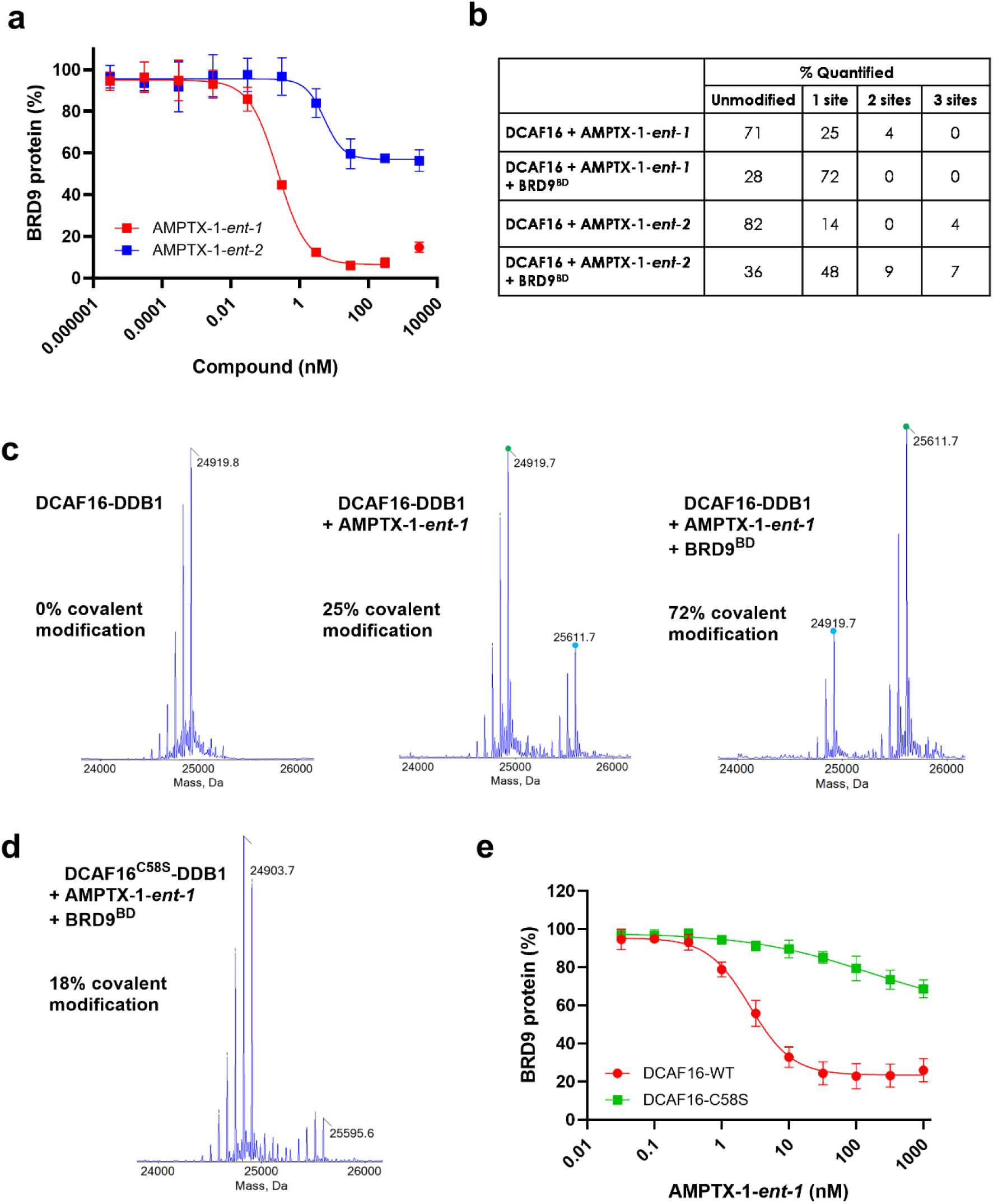
BRD9 and warhead stereochemistry facilitate covalent modification of cysteine 58 on CRL4^DCAF16^. **a** BRD9 degradation in MV4-11 cells following 6 h treatment with **AMPTX-1-*ent-1*** (red) and **AMPTX-1-*ent-2*** (blue). BRD9 levels were quantified by immunofluorescence. Data are represented as mean ± standard deviation from *n* = 3 biological replicates. **b** Table showing relative abundance (%) for each species, calculated using the peak area for unmodified DCAF16 (24,919.7 Da, corresponding to the molecular weight of the most abundant post-translationally modified state of DCAF16) as a reference m/z. Relative abundances for 1, 2 and 3 sites were then calculated using 24,919.7 + 692 Da, 24,919.7 + 1384 Da, and 24,919.7 + 2076 Da, respectively. **c** Intact MS deconvolution spectra of DCAF16 protein from samples prepared using 3.6 µM DCAF16-DDB1 alone; 3.6 µM DCAF16-DDB1 and 18 µM **AMPTX-1-*ent-1***; or 3.6 µM DCAF16-DDB1, 18 µM **AMPTX-*1-ent-1*,** and 3.6 µM BRD9^BD^. Samples were incubated at room temperature for 2 h prior to MS analysis. The different deconvolution peaks of DCAF16 correspond to multiply-phosphorylated protein species. **d** Intact MS spectra corresponding to a sample prepared using 3.6 µM DCAF16^C58S^-DDB1, 18 µM **AMPTX-1-*ent-1***, 3.6 µM BRD9^BD^, 2 h incubation at room temperature prior to MS analysis. **e** BRD9 degradation following 6 h treatment of **AMPTX-1-*ent-1*** in MCF-7 wildtype (DCAF16-WT, red) or MCF-7 cells containing a C58S mutation in DCAF16 (DCAF16-C58S, green). BRD9 levels were quantified by immunofluorescence.

The experiment was repeated in the presence of the bromodomain of BRD9 (residues 134- 250; BRD9^BD^), 3.6 µM, 1 equivalent): under these conditions the main species in solution was modified E3 ligase, with 72% and 64% modified DCAF16 with **AMPTX-1-*ent-1*** and **AMPTX-1- *ent-2***, respectively. These data indicate that the presence of BRD9^BD^ shifts the equilibrium in favour of the covalent adduct and suggests that engagement of BRD9 may be required to achieve sufficient labelling of DCAF16 in a cellular environment. While the molecular weight shift indicates only a single site on DCAF16 is modified by **AMPTX-1-*ent-1***, a significant fraction (16%) of DCAF16 is modified by **AMPTX-1-*ent-2*** at 2 or 3 sites. This suggests that stereochemistry within the tetrahydroisoquinoline scaffold may guide the specificity of adduct formation, which may relate to how the different enantiomers are accommodated within the BRD9-DCAF16 ternary complex. Given the more potent degradation activity observed with **AMPTX-1-*ent-1*** compared to **AMPTX-1-*ent-2***, we postulated that a specific ternary complex, relying on engagement of a single cysteine in DCAF16, can yield much deeper degradation of BRD9 when compared to engaging multiple cysteines sub-optimally.

Modified DCAF16 (**AMPTX-1-*ent-1***) was subjected to trypsin digestion and peptide mapping. This experiment identified DCAF16-Cys58 as the cysteine modified by **AMPTX-1-*ent-1*** (Supplementary Fig. 4 and Supplementary Table 4). Intact MS of C58S mutant DCAF16 (DCAF16^C58S^–DDB1) showed significantly less modification (18%) compared to wildtype DCAF16 (72%) when incubated with **AMPTX-1-*ent-1*** and BRD9^BD^, confirming that Cys58 is the primary cysteine engaged by the compound (Fig. 4d). To validate the cellular dependency on DCAF16-Cys58, we generated a homozygous C58S knock-in mutation of DCAF16 in MCF-7 cells using CRISPR-Cas9 genome editing and assessed the impact on BRD9 degradation. **AMPTX-1-*ent-1*** showed impaired activity compared to WT parental control, which is consistent with a functional role for DCAF16-Cys58 in the degradation of BRD9 (Fig. 4e). The residual degradation (∼25%) observed in the C58S mutant cell line may be the result of a second cysteine that is engaged sub-optimally, consistent with the 18% modification observed in the C58S mutant intact MS experiment.

### Oral administration of AMPTX-1 induces sustained BRD9 protein degradation in a xenograft model

To assess whether the novel BRD9 targeted glue degraders were suitable to demonstrate targeted protein degradation via DCAF16 *in vivo*, the mouse pharmacokinetic (PK) profiles of the single enantiomers were investigated upon intravenous (IV) and oral administration (PO). Oral bioavailability (F) was 20% and 30% for compound **AMPTX-1-*ent-1*** and **AMPTX-1-*ent-2*** respectively, with moderate elimination half-lifes (Supplementary table 1). Free fraction for **AMPTX-1-*ent-1*** and **AMPTX-1-*ent-2*** in plasma was found to be 0.6% and 0.5%. The free drug concentration profile over time, upon oral administration at 10 mg/kg is shown in Fig. 5. At this dose, systemic exposure of free **AMPTX-1*-ent-1*** was higher than the *in vitro* MV4-11 DC_50_ (0.2 nM) for at least 4 h, suggesting that the PK properties of **AMPTX-1-*ent-1*** were adequate for a pharmacodynamic (PD) proof of concept study *in vivo*.

**Fig 5:**
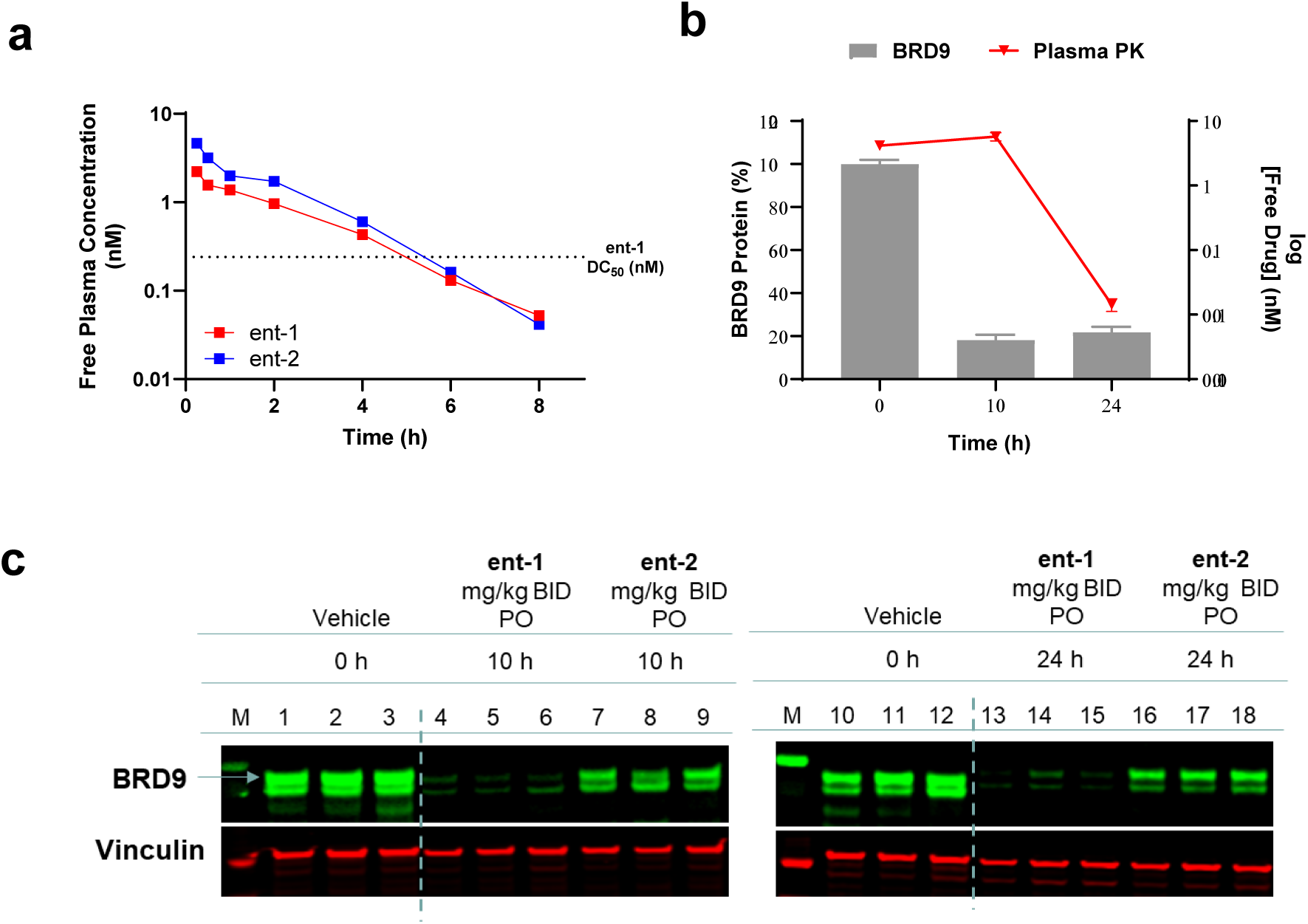
*In vivo* PK in mice and PD effect in a MV4-11 mouse model. **a** Free drug plasma concentration for **AMPTX-1-*ent-1*** and **AMPTX-1-*ent-2*** after oral dosing (10 mg/kg). CD-1 male mice (fasted state) n = 3, PO formulation: 5% DMSO, 5% Solutol, 90% (15%) HPBCD. **b** BRD9 quantification (grey bar, normalised to actin) and free drug concentration in plasma (red line) for tumour samples collected at 10 and 24 h after the first oral dose of **AMPTX-1-*ent-1*** (50 mg/kg). **c** Immunoblot analysis of tumour tissues for detection of BRD9 degradation. Samples were collected at 10 and 24 h after oral dosing (BID, first dose t = 0 h, second dose at t = 8 h) of **AMPTX-1-*ent-1*** or **AMPTX-1-*ent-2*** at 50 mg/kg.

To ensure the free concentration of degrader in plasma was closer to DC_90_ (1.4 nM), a dose of 50 mg/kg PO was chosen for the PD study. Mice subcutaneously implanted with MV4-11 xenografts received two oral doses of AMPTX-1 enantiomers, first dose at t = 0 h, second dose at t = 8 h. Tumour samples were collected at 10 and 24 h after the initial dose (2 and 16 h post last dose), and BRD9 protein levels were quantified by western blot analysis. At 2 h post last dose, 82% BRD9 degradation was observed for the samples derived from the animals treated with **AMPTX-1*-ent-1***, whereas only 31% degradation was achieved by the less active **AMPTX- 1-*ent-2*** at the same time point (Supplementary Fig. 5). Notably, degradation was maintained at 16 hours post last dose, with 78% BRD9 degradation observed **AMPTX-1-*ent-1*** treatment and 28% for **AMPTX-1-*ent-2*** (Supplementary Fig. 5). At this time point the concentration of total drug in plasma being over 3 orders of magnitude lower compared to the levels detected at the 2 h post last dose, highlighting an extended pharmacodynamic effect.

## Discussion

**AMPTX-1** is a potent and selective covalent targeted glue that induces degradation of BRD9 by recruitment of DCAF16, a relatively uncharacterized E3 ligase. While DCAF16’s endogenous functional and biological role remain elusive, it has recently emerged as an E3 ligase hijackable via small molecules. DCAF16-mediated degradation of neo-substrate BRD4 has been demonstrated by means of a variety of electrophilic bifunctional degraders^27–30^.However, these compounds show deep target degradation only at relatively high (micromolar) concentrations and have been shown to react with a wide range of cysteine-containing proteins in the proteome ^29^ limiting the potential for drug development. During the preparation of this manuscript, electrophilic derivatives of JQ1 were reported to recruit DCAF16 (and the off-target Cyclin G-associated kinase, GAK) to BRD4 by a template-assisted covalent modification mechanism^26,30^. Consistently to our findings, DCAF16-Cys58 was shown to be the target of the BRD4-templated modification by the JQ1 derivative MMH2^26^ and cryo-EM data revealed that BRD4^BD2^ acts as a “structural scaffold” facilitating the covalent bond formation^26^. Moreover, inspection of cryo-EM structures available for ternary complexes comprising BRD4 and DCAF16 reveals structural complementarity among the two proteins^26,31^. As degradation of BRD4 by DCAF16 can be induced by diverse degrader chemotypes, it has been suggested DCAF16 may play a role in BRD4 protein quality control^27^. Our work shows for the first time that BRD9 can also be efficiently glued to DCAF16 by a new targeted glue bearing a novel, reversible covalent warhead. The molecular complexity inherent within targeted glues may facilitate additional protein-ligand and protein-protein interactions, expanding the neo-substrate scope of DCAF16 beyond BRD4 as we have shown here. It has been noted that the structural plasticity of DCAF16 may confer high structural flexibility to the DDB1-DCAF16 E3 ligase complex^26^, allowing for recognition of a variety of neo-substrates in addition to the proposed endogenous substrate, SPIN4^32^.

Proteomics experiments indicate that **AMPTX-1** selectively targets BRD9 for degradation by inducing formation of a ternary complex with DCAF16. The absence of BRD7 degradation, which is engaged with similar potency by this class of bromodomain binders^23^ also points to a model where specific protein-protein interactions between ligase and target protein, and/or preferential protein ubiquitinability^33^, add a layer of selectivity beyond the simple target engagement. Selective degradation of BRD9 over BRD7 protein is also observed with CRBN-based PROTACs^17,20^, but not VHL-based PROTACs^18^.

A unique, counterintuitive, feature of the mode of action here described is the presence of a hook effect^34,35^, albeit mild, at high compound concentration in our protein degradation assays. (Fig. 2 and Fig. 3). Interestingly, a similar hook effect was observed by Li et al. in both ternary complex formation and degradation assays of related covalent glue molecules^26^. Hook effects are often observed with hetero-bifunctional PROTACs because they can engage each protein individually. Typically, a hook effect results from saturation of the individual POI and E3 binding sites in 1:1 engagement, out-competing the 1:1:1 ternary complex formation at high degrader concentrations. However, despite the presence of a hook effect, no competition effects of **AMPTX-1**-induced BRD9 degradation were observed in the presence of a large excess of warhead **3**, suggesting that the warhead alone is unable to effectively saturate the E3 ligase. Therefore, the hook effect observed with the targeted glue could be explained by an “E3 ligase covalent loading” process coupled to BRD9 binding site saturation accordingly to the following steps (**Fig. 6**):

1. BRD9 engagement with **AMPTX-1**;
2. ternary complex formation between DCAF16 and **AMPTX-1-**bound BRD9, giving DCAF16 labelling;
3. ubiquitin transfer and BRD9 dissociation from the complex;
4. degradation cycle established: DCAF16 remains labelled and primed for engagement with additional BRD9 proteins; and
5. hook effect: excess of targeted glue leads to saturation of both BRD9 and DCAF16.

**Fig 6:**
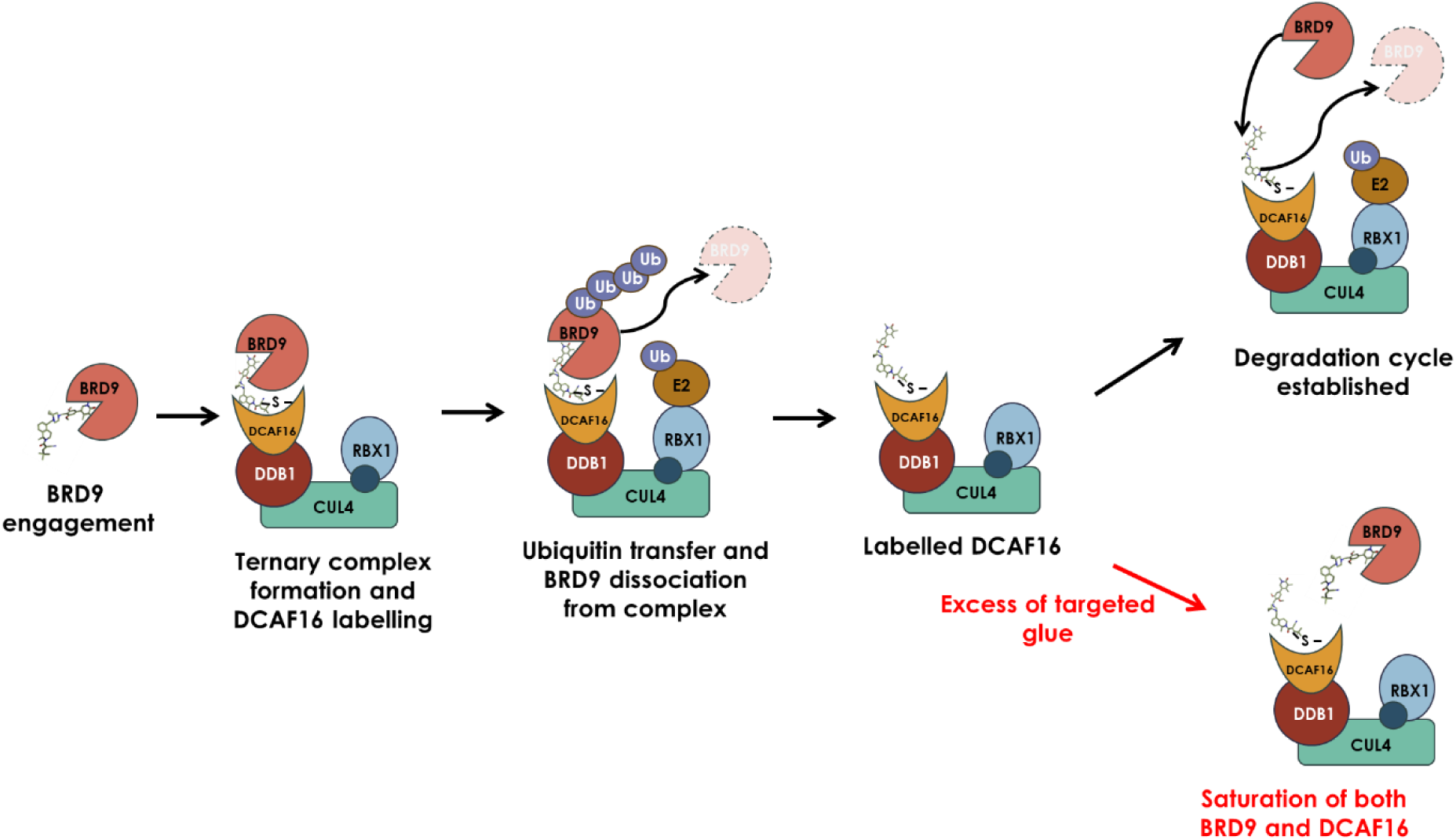
Schematic representation of the “E3 ligase loading” process and hook effect. Labelling of DCAF16 Cys58 is achieved upon formation of the complex with BRD9. Dissociation of BRD9 from the complex leaves DCAF16 covalently modified with the targeted glue, primed to recruit additional copies of BRD9. Excess of targeted glue may block the binding site of BRD9, which cannot be recruited to the labelled DCAF16, causing the hook effect.

The hook effect suggests the persistence of a portion of modified DCAF16, essentially a “neo E3 ligase”^28^ bearing a covalently modified substrate receptor subunit (in this case a DCAF16:**AMPTX-1** covalent adduct). This population of E3 ligase is primed for the target recruitment and is likely to persist inside the cell^36–38^. The half-life of this species will depend on the half-life of DCAF16 (155.4 hours, as reported from proteomics studies^39^) and the reversibility of the covalent bond formed between DCAF16-Cys58 and **AMPTX-1**. Dissociation of the warhead, possible for this class of reversible covalent electrophiles, may be kinetically unfavoured because of protein-compound interactions that could stabilise the adduct and prevent the retro-Michael reaction by steric hindrance^40^. Persistent modification of DCAF16 may therefore underscore the high potency of this compound.

Here we also demonstrate the unprecedented potential of hijacking an E3 ligase via a covalent targeted glue to achieve robust and persistent target protein degradation *in vivo* in an animal model. Deep protein degradation was achieved in the tumour and degradation levels were maintained when plasma concentrations of the drug dropped by over 1000-fold. This apparent disconnect between PK and PD may be a feature of the covalent mode of action as previously demonstrated for reversibly covalent BTK inhibitors^40,41^.

Notably, the degradation potency of **AMPTX-1** and its oral bioavailability suggest the potential for drug development for this class of reversible covalent compounds with further medicinal chemistry efforts.

In summary, we have identified and characterised a potent and selective BRD9 degrader that works via a “targeted glue” mode of action. Covalent modification of the E3 ligase DCAF16 is enabled by a templating effect of the BRD9 protein, which likely positions the covalent warhead in proximity of DCAF16-Cys58. Proteomics studies show that **AMPTX-1** induces selective target degradation through selective recruitment of DCAF16 to BRD9. Importantly, the remarkable bioavailability of **AMPTX-1** *in vivo* supports targeted protein degradation via covalent recruitment of an E3 ligase as a viable strategy for drug development. More generally, this work exemplifies a novel, drug-like small molecule able to selectively engage an undruggable protein surface, the one of DCAF16, to a highly ligandable protein (BRD9) to enable event-driven pharmacology. We envisage this approach could open the way to the recruitment of additional hijackable E3 ligases and expanding the potential for TPD approaches and induced proximity pharmacology in general.

## Acknowledgements

The authors would like to thank Dr Victoria Lovatt (Amphista Therapeutics Ltd) for helpful discussions: Dr Kate Heesom (Director of the Bristol University Proteomic Facility, UK), Dr Joshodeep Boruwa and Dr Kamalkishor Landge (Syngene International Ltd, India), Dr Samuel Bloor, Camila Torres (Sygnature Discovery, UK) and Marieke Lamers, Jason Pembroke (Domainex, UK) for their assistance with this work.

