## Supplementary material for "Selective degradation of BRD9 by a DCAF16-recruiting targeted glue: mode of action elucidation and in vivo proof of concept": Methods

**Experimental Methods**

Cell Culture & Models (incl. Cell Engineering)

HEK293 HiBit-BRD9 KI LgBit cells were obtained from Promega Corp. (CS3023412) and MCF7 cells from ATCC (HTB-22). Briefly, both cell lines were maintained in Dulbecco's Modified Eagle's Medium with glutaMAX (Gibco 31966-021) supplemented with 10% FBS (v/v) (Thermo Fisher A5256701). MV4-11 cells were obtained from ATCC (CRL-9591) and maintained in IMDM (Thermo Scientific 21056023) supplemented with 10% FBS. All lines were grown at 37°C in a humidified 5% $CO_2$ atmosphere.

$BRD9^{BD}$ (residue 134- 250) Protein Production

10 µl of competent cells of *E. coli* BL21 (DE3) were transformed using 1 µl of 50 ng/ul plasmid DNA (pRSET_BRD9_134–250) following standard procedures. One colony was inoculated in 100 mL LB plus 2% glucose and grown overnight at 37 °C, 150 rpm until $OD_{600}$ = 1.5. 50 mL was used to inoculate 1 Litre of Luria Broth (LB) medium plus 0.2% glucose or 1 L Terrific Broth (TB, used readymade EZMix™ powder from Sigma plus 4% glycerol) medium. Two litres of each were set up. Starter $OD_{600}$ = 0.075. Cultures were grown to $OD_{600}$ = 1.1 and induced with 0.5 mM IPTG overnight at 17 °C. Cells were cooled in cold cabinet at 4 °C for 30 mins before induction. Next day the cells were spun down at 4000 rpm (Beckman Avant J-26 XP rotor JLA8.1000) for 30 min, washed once with 25 mL cold PBS and stored at -80 °C. 1 mL of each cell pellets was lysed in 0.5 mL B-per in the presence of lysozyme at 1 mg/mL and DNAse I at 10 µg/mL. Post lysis, whole cell and soluble fractions were analysed on SDS-PAGE followed by Anti-His western blotting.

Cell pellets were thawed in the presence of Lysis Buffer (50 mM Tris pH 8.0, 300 mM NaCl, 10 mM Imidazole, 10% glycerol, 1 mM TCEP, 1 mg/ml lysozyme, (1:1000) DNase I stock for 10 µg/ml final, and 2 protease tablets (SIGMAFAST™ Protease Inhibitor Cocktail Tablets, EDTA-Free Cat# S8830)). Soluble proteins were released from cells pellets by using the cell disruptor (Constant Systems) at 25 PSI. Insoluble proteins were removed by high-speed centrifugation (Beckman Avant J-26 XP, Rotor JA25.50, 20,000 rpm) for 30 min at 4 °C. For each litre of cell culture, 0.8 mL (1.6 mL of slurry) of Ni-NTA agarose beads (Ni-NTA Affinity Resin – Amintra, cat# ab270549) were employed in 50 mL tube. Each 0.8 mL of resin was washed in 25 mL of water followed by 25 mL of Buffer A (50 mM Tris pH 8.0, 300 mM NaCl, 10 mM Imidazole, 10% glycerol, 1 mM TCEP). Binding was performed at 4 °C gently rolling overnight.

The following day, Ni-NTA beads were collected through centrifugation at a low speed (Eppendorf 5810R, Rotor 1200 rpm for 10 min) and collected into one 50 mL Falcon tube and washed twice with 50 mL of buffer A. The beads were transferred to a 10 mL OmniFit column. The column was washed with 20 mL of Buffer A and then 20 mL with 15% (45 mM Imidazole) Buffer B (50 mM Tris pH 8.0, 300 mM NaCl, 300 mM imidazole, 1 mM TCEP, 10% glycerol). Protein elution was then achieved using a step elution at 100% Buffer B. For His-tag cleavage, samples were incubated with TEV protease (1:10) at 20 °C overnight.

For final purification, samples were loaded onto a size exclusion S75 26/60 column pre-equilibrated in 25 mM Hepes pH 7.5, 200 mM NaCl, 10% glycerol, 1 mM TCEP. Fractions were collected and analysed by SDS-PAGE. Highest purity fractions were concentrated, aliquoted and frozen at -80 °C.

DCAF16:DDB1ΔB and DCAF16$^{C58S}$:DDB1ΔB Protein Production

WT and C58S DCAF16(1-216), with N-terminal 6His-TEV affinity tags, were subcloned into individual pOET1 vectors. DDB1(1-1140, Δ396-705, linker) and full length DDA1(1-102) were both subcloned into a single pOET5.1 duet vector for co-expression. SF21 insect cells in suspension were grown in SF-900TM II SFM media (Gibco®) at 28°C and 140rpm in shake flasks with vented caps. Cells were passaged every 2-3 days to a density within the log phase of the growth to maintain optimal growth conditions. The SF21 cell line used was routinely passaged at least 4 times before use in an expression grow and had a viability of above 95%.
On the day of infection, the cell number was adjusted to 2 x 106 cells/mL by the addition of fresh SF900II media prior to co-infection with the two baculovirus constructs at a 1 : 1 ratio and MOI of 2.5. Approximately 72 hours after infection the cell pellet was harvested by centrifugation at 7000 x g for 20 minutes at 4°C before freezing the cell pellet at -80°C prior to purification.
Cells were counted using a Beckman Coulter Vi-Cell BLU Cell Viability Analyzer which automates the Trypan Blue Dye Exclusion method for cell viability analysis.

Harvested cells were resuspended in 50 mM Tris-HCl pH 8.0, 300 mM NaCl, 20 mM Imidazole, 10 % Glycerol, 2 mM TCEP (lysis buffer), supplemented with protease inhibitors (AbCam) and Nuclease A (GenScript). The cells were disrupted by sonication (Q500 QSonica), 5 x 20 s pulse, 40 s rest, and then clarified by centrifugation at 185,000 x g for 1 hour (Ti45 rotor, Optima XPN-80 Ultracentrifuge). The soluble supernatant was filtered using a 0.45 µm membrane prior to nickel affinity chromatography.
The filtered supernatant was applied directly to 10 mL Ni2+ HisTrap Crude FF resin (Cytiva).
The resin was washed using 5 CV lysis buffer and eluted using a 10 CV 0 – 100 % lysis buffer:IMAC elution buffer gradient (50 mM Tris-HCl, pH 8.0, 300 mM NaCl, 600 mM Imidazole, 10 % Glycerol, 2 mM TCEP). Elution fractions containing the DCAF16:DDB1:DDA1 complex were pooled and then treated with TEV protease in a 1:15 ratio. The sample was dialysed overnight at 4 °C into 50 mM Tris-HCl pH 8.0, 300 mM NaCl, 20 mM Imidazole, 10 % Glycerol, 1 mM TCEP.

Following dialysis, the cleaved DCAF16:DDB1:DDA1 sample was passed through 10 mL Ni2+ HisTrap Crude FF resin (Cytiva) and the flow through collected. The subsequent sample was diluted to reduce the NaCl concentration to < 50 mM and applied to a HiTrap Q FF column (Cytiva), which had been equilibrated into 25 mM Tris-HCl, pH 8.2, 50 mM NaCl, 10 2% Glycerol, 1 mM TCEP (IEX buffer). A 40 CV 0 – 0.5 M NaCl IEX buffer gradient, followed by 5 CV of 1 M NaCl IEX buffer, was used to elute the protein.

The protein was then concentrated by ultrafiltration (Amicon) and loaded onto a Superdex 200 column (Cytiva) equilibrated in 25 mM Tris-HCl, pH 8.0, 300 mM NaCl, 10 % Glycerol, 1 mM TCEP. The protein eluted as a single peak representing a 1:1:1 DCAF16:DDB1:DDA1 complex. The protein was snap frozen in liquid N2 and stored at -80 °C prior to use.

BRD9 HiBit Endpoint and Kinetic assays

HiBiT-BRD9 KI HEK293 (LgBiT) Cells (Promega CS3023412) were seeded at 8 x $10^3$ cells per well in 36 µL of media in sterile white bottom 384 well plates (Greiner 781080) and incubated overnight at 37°C with 5% $CO_2$. The following day, compounds were prepared at 1000x final concentration in DMSO and diluted 1:100 in maintenance media. In case of competition or mechanism of action studies (NEDDylation- or proteasome-dependence) plates were pre-treated with DMSO, 3 µM NAE1 inhibitor MLN4924, 10 µM proteasome inhibitor Bortezomib, 10 µM BI-7273 (BRD9 POI ligand) or 10 µM of warhead **3** with a Multidrop Pico8 digital dispenser and incubated for 1 hour. Cells were treated with compound **1** and incubated for 6 h. BRD9 expression was quantified using the NanoGlo Lytic Endpoint assay (Promega N3050). The lytic buffer was equilibrated to room temperature for 10-15 minutes before lytic reagent was prepared by adding substrate (1:50) and LgBit protein (1:100) in lytic buffer. 40 µl lytic reagent mix was added to each well of the cell plate and plates were centrifuged briefly at ~50G. Plates were incubated with shaking (350RPM) for 12-24 minutes before reading using the LUM plus module of a BMG Pherastar FSX. The percentage of BRD9 remaining was calculated by normalisation to average data from high and low control wells (DMSO treated cells and no cells respectively).

For live cell kinetic 8 x $10^3$ Hek293 HiBit-BRD9 KI LgBit cells per well were seeded in 80 µL of phenol red-free OptiMEM media (Gibco 11058021) supplemented with 4% FBS (ATCC 302025) in white opaque 384-welled plates (GBO 781080). Following overnight incubation, 20 µL of assay media supplemented with 1:40 of Nano-Glo Endurazine (Promega N2571) was added to each well and incubated at 37°C for 3 h. Compounds of interest were dispensed into plates using FlexDropIQ non-contact dispenser and normalised for DMSO concentration to 0.1% (vol/vol). Plates were placed immediately in ClarioSTAR microplate reader equipped with ACU (BMG) that was pre-normalised to 5% $CO_2$ and 37°C. Recording was taken every 5 min with 0.8 s integration time.

Resulting measurements were analysed by determination of fractional luminescence and normalisation to first timepoint. Four-parameter variable slope dose response as well as one phase decay fitting was performed in GraphPad Prism.

BRD9 Immunofluorescence assay

A suspension of MV4-11 cells was prepared in phenol red-free assay media supplemented with 10% FBS and cells were seeded at 20,000 cells per well (45 µL) in sterile black poly-D-lysine coated 384 well plates (Greiner 781948). Compounds were prepared at 1000x final concentration in DMSO, diluted 1:100 in assay media, and 5 µL compound was added to each well of the cell plate. Cells were incubated for 24 hours at 37°C with 5% $CO_2$. All the following incubations for immunofluorescence staining were at room temperature. 15 µL of 16% PFA was added to each well (3.7% final concentration) and the cells were fixed for 15 min then washed twice with DPBS. Cells were permeabilised with 0.1% Triton X-100 for 10 min, Triton X-100 was removed, then blocked with 1% BSA in DPBS for 1 hour. Cells were stained with 25 µL anti-BRD9 E4Q3F antibody (CST 48306) diluted 1:25600 in 1% BSA in DPBS for 2-3 hours. Wells were washed twice with DPBS then incubated with 25 µL of 1% BSA containing a 1:1000 dilution of Anti-rabbit Alexa Fluor™ 647 secondary antibody (Thermo Scientific A21244) and 1 µg/mL Hoechst nuclear counter stain (Abcam ab228551) for 1 hour. Wells were washed twice with DPBS prior to imaging on a Perkin Elmer Operetta CLS with 10X air lens. Images were processed using Harmony High-Content Imaging and Analysis Software (Perkin Elmer) and the mean contrast ratio of Alexa Fluor™ 647 in central nuclei was used to quantify BRD9 protein levels. Results were normalised using DMSO and no-primary antibody control and four-parameter variable slope dose response was fitted in GraphPad Prism.

Immunoprecipitation Mass Spectrometry (IP-MS)

HEK293 cells expressing BRD9-HiBiT were seeded in 100 mm plates at 7x$10^6$ cells/dish. The following day, all plates were pre-treated with **MLN4924** at 3 µM for 30 mins before addition of DMSO, **AMPTX-1** or compound **2** at the indicated concentrations for 2h at 37°C. Cell monolayers were washed 1x in ice-cold PBS, scraped into PBS, pelleted at 300 x g for 5 minutes and washed 1x PBS. Cell pellets were resuspended in 500 µL NP-40 lysis buffer (Thermo, J60766.AK) containing protease inhibitors + 250 U benzonase (Thermo Fisher, 88701). Cells were lysed on ice for 30 min and cleared through centrifugation at 16,000 x g for 15 minutes. Supernatants were directly transferred to HiBiT antibody (Promega N7200), bound dynabeads protein G (Thermo Fisher, 10765583) and incubated on an Eppendorf rotator for 30 min. Beads were isolated with a magnetic rack and washed 2x in PBS and 1x in H2O and resuspended in 25 uL of 50 mM TEAB. BRD9-enriched samples were reduced (10mM TCEP, 55°C for 1h), alkylated (18.75mM iodoacetamide, room temperature for 30min.) and then digested from the beads with trypsin (1.25µg trypsin; 37°C, overnight). Beads were placed on a magnetic rack and digested peptides removed and subjected to TMT labelling.

Expression Proteomics

MV4;11 cells were seeded at 8x10^5 for 6 h and 24 h time points and 6.2x10^5 cells for 48 h. After seeding, cells were immediately treated with 100 µM of indicated compounds in triplicate. At indicated time points, cells were washed twice with PBS, pelleted, and resuspended in 100 µL of lysis buffer (50 mM Tris pH 7.6, 150 mM NaCl, 2% SDS, 10% glycerol, protease inhibitors (Pierce A32961), 1 mM PMSF, 250 U benzonase (Thermo Fisher, 88701) and sonicated for 1 minute at 100% power with a probe sonicator. Cell debris was removed by centrifugation at 21,000 × g for 10 min.

FASP-based Tryptic Digestion

246 µg of each sample was used for FASP as follows: Dithiothreitol was added to a final concentration of 83.3 mM, incubated at 99°C for 5 mins, then cooled to room temperature. VIVACON 500 filter units, 30,000 MWCO (Sartorius Stedim, Biotech GmbH, 37079 Goettingen, Germany) were washed with 8 M urea in 100 mM Tris/HCl pH 8.5, centrifuged at 14,000 × g, 20°C for 15 mins. Samples were loaded in 50 µL aliquots plus 200 µL 8 M urea, centrifuged at 14,000 × g, 20°C for 15 mins. The filter units were washed with 8 M urea, centrifuged at 14,000 × g, 20°C for 15 mins. 100 µL 50 mM Iodoacetamide in 8 M urea was added to the filter units and incubated for 30 mins in the dark. Filter units were centrifuged at 14,000 × g, 20°C for 10 mins, then washed with 100 µL 8 M urea, centrifuged at 14,000 × g, 20°C for 15 mins. This wash step was repeated two times, for a total of three washes. Filters were further washed with 100 µL 50 mM Triethylammonium Bicarbonate Buffer pH 8.5 (TEAB, (Thermo Fisher Scientific)), centrifuged at 14,000 × g, 20°C for 10 mins. This wash step was repeated two times, for a total of three washes. Filter units were transferred to new tubes and 1.25% (w/w) trypsin (Pierce MS grade, Thermo Fisher Scientific, Loughborough, LE11 5RG, UK) in 60 µL 50 mM TEAB was added. The Filter units were sealed with Parafilm and digestion performed overnight at 37°C with shaking. Following overnight digestion, the filter units were centrifuged at 14,000 × g, 20°C for 20 min. and the flow through containing the peptides was retained. The filter units were washed with 40 µL 50 mM TEAB, centrifuged at 14,000 × g, 20°C for 10 mins and then 50 µL 0.5M NaCl, centrifuged at 14,000 × g, 20°C for 20 mins and these washes were added to the flow through. Peptide samples were desalted and cleaned up using Sep-Pak cartridges according to the manufacturer's instructions (Waters, Milford, Massachusetts, USA). Eluate from the Sep-Pak cartridge was evaporated to dryness and resuspended in 100 µl 50 mM TEAB and 10µl used for a peptide assay. All chemicals from Merck Life Science UK Limited, Dorset, SP8 4XT unless otherwise stated.

TMT Labelling, High pH reversed-phase chromatography

For expression proteomics samples, approximately 60 µg of each sample was labelled with Tandem Mass Tag (TMTpro) 18-plex reagents according to the manufacturer's protocol (Thermo Fisher Scientific, Loughborough, LE11 5RG, UK). In addition, 5 µg of all 34 samples was combined to make a bridging sample, and two 60 µg aliquots of this bridging sample were labelled with the TMTpro 126C tag. The labelled samples were pooled into experiment one - 3-plex and experiment two - 6-plex experiments and labelled as follows, experiment 1 layout; TMT-127N = DMSO 6h Repeat-1, TMT-127C = compound **2** 6h Repeat 1, TMT-128N = **AMPTX-1** 6h Repeat 1, Experiment 2 layout; TMT-127N = DMSO 6h Repeat 2, TMT-127C = compound **2** 6h Repeat 2, TMT-128N = **AMPTX-1** 6h Repeat 2, TMT-131N = DMSO 6h Repeat 3, TMT-131C = compound **2** 6h Repeat 3, TMT-132N = **AMPTX-1** 6h Repeat 3.

For IP-MS the entire digest was TMT labelled according to the manufacturer's protocol (Thermo Fisher Scientific, Loughborough, LE11 5RG). The labelling was as follows: TMT-126C = DMSO Repeat 1, TMT-127N = 10 nM compound **2** Repeat 1, TMT-127C = DMSO Repeat 2, TMT-128N = 10 nM compound **2** Repeat 2, TMT-128C = DMSO Repeat 3, TMT-129N = 10 nM compound **2** Repeat 3, TMT-129C = 1 nM **AMPTX-1** Repeat 1, TMT-130N = 300 nM compound **2** Repeat 1, TMT-130C = 10 nM AMPTX-1 - Repeat 1, TMT-131N = 300 nM compound **2** Repeat 2, TMT-131C = 10 nM **AMPTX-1** Repeat 2, TMT-132N = 300 nM compound **2** Repeat 3, TMT-132C = 10 nM **AMPTX-1** Repeat 3, TMT-133N = 300 nM **AMPTX-1** Repeat 1, TMT-133C = 300 nM compound **1** Repeat 2, TMT-134N = 300 nM **AMPTX-1** Repeat 3, TMT-135N = 1000 nM AMPH-3340 Repeat 1.

Offline HpRP fractionation

For expression proteomics, samples were combined to a total of 100 µg each TMT plex and desalted using a SepPak cartridge according to the manufacturer's instructions (Waters, Milford, Massachusetts, USA). Eluate from the SepPak cartridge was evaporated to dryness and resuspended in buffer A (20 mM ammonium hydroxide, pH 10) prior to fractionation by high pH reversed-phase chromatography using an Ultimate 3000 liquid chromatography system (Thermo Fisher Scientific). In brief, the sample was loaded onto an XBridge BEH C18 Column (130Å, 3.5 µm, 2.1 mm X 150 mm, Waters, UK) in buffer A, and peptides were eluted with an increasing gradient of buffer B (20 mM Ammonium Hydroxide in acetonitrile, pH 10) from 0-95% over 60 minutes. The resulting fractions (20 in total) were evaporated to dryness and resuspended in 1% formic acid prior to analysis by nano-LC MSMS using an Orbitrap Fusion Lumos mass spectrometer (Thermo Scientific).

For IP-MS, the entire labelled material was combined prior to SepPak desalting as described above.

Nano-LC MS3

High pH RP fractions were further fractionated using an Ultimate 3000 nano-LC system in line with an Orbitrap Fusion Lumos mass spectrometer (Thermo Scientific). In brief, peptides in 1% (vol/vol) formic acid were injected onto an Acclaim PepMap C18 nano-trap column (Thermo Scientific). After washing with 0.5% (vol/vol) acetonitrile 0.1% (vol/vol) formic acid peptides were resolved on a 250 mm × 75 µm Acclaim PepMap C18 reverse phase analytical column (Thermo Scientific) over a 150 min organic gradient, using 7 gradient segments in solvent B (1-6% for 1 min., 6-15% for 58 min., 15-32% for 58 min., 32-40% for 5 min., 40-90% for 1 min, held at 90% for 6 min and then reduced to 1% over 1 min.) with a flow rate of 300 nl min−1. Solvent A was 0.1% formic acid and Solvent B was aqueous 80% acetonitrile in 0.1% formic acid. Peptides were ionized by nano-electrospray ionization at 2.0kV using a stainless-steel emitter with an internal diameter of 30 µm (Thermo Scientific) and a capillary temperature of 300°C. For expression proteomics, 12 samples were collected for SPS-MS3 analysis. For IP-MS six fractions were collected.

All spectra were acquired using an Orbitrap Fusion Lumos mass spectrometer controlled by Xcalibur 3.0 software (Thermo Scientific) and operated in data-dependent acquisition mode using an SPS-MS3 workflow. FTMS1 spectra were collected at a resolution of 120,000, with an automatic gain control (AGC) target of 200,000 and a max injection time of 50 ms. Precursors were filtered with an intensity threshold of 5,000, according to charge state (to include charge states 2-7) and with monoisotopic peak determination set to Peptide. Previously interrogated precursors were excluded using a dynamic window (60s +/-10 ppm). The MS2 precursors were isolated with a quadrupole isolation window of 0.7m/z. ITMS2 spectra were collected with an AGC target of 10,000, max injection time of 70 ms and CID collision energy of 35%.

For FTMS3 analysis, the Orbitrap was operated at 50,000 resolution with an AGC target of 50,000 and a max injection time of 105 ms. Precursors were fragmented by high energy collision dissociation (HCD) at a normalised collision energy of 60% to ensure maximal TMT reporter ion yield. Synchronous Precursor Selection (SPS) was enabled to include up to 10 MS2 fragment ions in the FTMS3 scan.

<u>Data processing</u>

Raw data files were converted to mzML format using msconvert proteowizard (version 3.0.22167). Database searches were performed using MSFragger (version 3.8)[1] within FragPipe (version 20.0). Searches were performed against the reference human proteome from Uniprot (2023-08-17) including only reviewed accessions. For expression proteomics, proteins with less than two unique peptides were excluded. Carbamidomethylation of cysteine (+57.02146 Da) and TMTpro labelling of Lysine (+304.20715 Da) were set as fixed modifications. Oxidation of methionine (+15.9949 Da), N-terminal acetylation (+42.0106 Da), and N-terminal TMTpro labelling (+304.20715 Da) were set as variable modifications. False-discovery rate filtering was set to 1% at the PSM, peptide, and protein level. Each channel intensity was normalized to the median intensity of all channels to account for protein loading differences and log2 transformation was applied. For global proteomics, internal reference scaling normalisation[2] was performed to correct for batch effect between plexes. Statistical analysis was performed in R (v4.3.1) using limma 3.56.2[3]. A single model was constructed to test degrader vs equal concentration of negative control compound. P-values were adjusted using the Benjamini & Hochberg method to account for multiple comparisons. An adjusted p-value threshold of 0.05 and log2 fold-change threshold of 1 was used to identify significantly changed proteins.

<u>Intact MS</u>

DCAF16-DDB1 or DCAF16$^{C58S}$-DDB1 (3.6 µM) was incubated with 18 µM compound in the presence or absence of 3.6 µM BRD9$^{BD}$ (1:1:5) at RT for 2 h. The samples were diluted to approximately 0.1mg/ml with 0.1 % formic acid, 5 % acetonitrile. 10 µl of sample was loaded onto the Sciex Exion LC and a 5 min reverse phase gradient was used.

Buffer A was 0.1 % formic acid and Buffer B was 0.1% formic acid 100% acetonitrile. The flow was set to 500 µl starting at 5% B leading to 45% B over 3 minutes before a 95% B wash and equilibration at 5%B. A Phenomenex bioZen 3.6 µm, XB-C8, 50 x 2.1 mm column was used. The flow from the column was passed into the Sciex X500B mass spectrometer collecting data in positive ion mode. To enable ionisation of the eluate the source was set to 400 °C, 5500 V with gas at 50 psi. A TOF mass window of 500 to 3000 Da was collected scanning at 0.5 seconds. The X500B was calibrated with positive calibration mix, the error for this experiment was estimated at < 1 Da. The resultant TIC was deconvoluted using BioToolKit. The relative abundances were calculated using the peak area for DCAF16 species 1 (24,919.7) in the intact MS spectra as a reference m/z. The relative abundances were then calculated using 24,919.7 + 692 Da, and +1384 Da, respectively for 1:1 and 1:2 ratios.

Peptide Mapping Mass Spectrometry

For sample preparation, 5 µM DCAF16-DDB1 and 5 µM compound, in the presence or absence of 5 µM $BRD9^{BD}$ were incubated at RT for 20 h in 50 mM ammonium bicarbonate. Samples were added to a 2 mL Lo-Bind plate and incubated with 12.5 mM TCEP in 8 M Urea for 45 min at 60 °C while mixing at 500 rpm. Each sample was then incubated with 20 mM Iodoacetamide in 50 mM Ammonium Bicarbonate for 45 min at RT while mixing at 500 rpm. For the digest, samples were incubated overnight at 37 °C in 100 µL of 1 µg/mL trypsin. Digests were quenched using 30% Formic Acid (aq) for 5 min prior to injection on the LC-MS system.

The column used was an Acquity UPLC CSH C18 column, 1.7 µ m, 2.1 mm x 100 mm, which was kept at 60oC during analysis. Buffer A was 0.1% formic acid in water and Buffer B was 0.1% formic acid in Acetonitrile. The flow was set to 500 µl/min starting at 3% A leading to 90% A over 57.5 min, before a 97% B wash and equilibration at 3% A. Data was acquired in data independent acquisition mode (MSe) across the m/z range of 300-1200. A collision energy ramp of 30-60 V was used to generate MS/MS data. Data was processed within the peptide mapping module for Waters UNIFI software. The data within UNIFI was filtered to display only peptides that met the following criteria: i) the peptide had a binder attached; ii) The observed mass of the peptide was within +- 10ppm of its theoretical mass, and iii) The peptide had a MS response of >50000.

Animal Studies and Immunoblotting Procedures

All animal handling, care, and treatments for this study were conducted according to protocols approved by the Institutional Animal Care and Use Committee (IACUC) of WuXi AppTec, in compliance with the standards of the Association for Assessment and Accreditation of Laboratory Animal Care (AAALAC). Female Balb/c nude mice were inoculated subcutaneously in the right flank with MV4-11 tumor cells (10 x 10^6) in 0.2 mL PBS mixed with Matrigel (50:50) to promote tumour growth. Once the tumours reached an average volume of 379 mm³ on day 29 post-inoculation, animals were randomized into treatment groups. The compounds were administered orally twice daily, formulated in a solution containing 5% DMSO, 5% Solutol, and 90% HPβCD (15% v/v), with an 8-hour interval between doses. Plasma and tumor samples were collected at 10 and 24 hours following the first dose.

For immunoblotting, protein lysates were prepared in RIPA buffer and separated using NuPAGE® Novex 4-12% Bis-Tris gels, followed by transfer onto a nitrocellulose membrane with the iBlot®2 Gel Transfer Device. Membranes were blocked for 2 hours at room temperature in Odyssey blocking buffer (LI-COR, 927-60001). Primary antibodies were diluted in Odyssey blocking buffer with 0.1% Tween20 and incubated with the membrane overnight. BRD9 antibody (Cell Signaling, 58906) was used at 1:1000, with Vinculin (Sigma, SAB4200080) at 1:10000 as a loading control. Secondary antibodies, IRDye® 680RD Goat anti-Mouse IgG (LI-COR, 926-68070) and IRDye® 800CW Goat anti-Rabbit IgG (LI-COR, 926-32211), were diluted 1:10000 in Odyssey blocking buffer with 0.1% Tween20 and incubated for 1 hour at room temperature. Band intensities were quantified using Image Studio software.

### GSH Reversibility Study

A solution containing 0.5 mM **AMPTX-1** and 5.0 mM L-glutathione (reduced form) in a 9:1 mixture of PBS buffer and DMSO was allowed to sit at ambient temperature in a sealed well of a 100 uL v-bottom 384-well plate for 2 h (performed in duplicate).

After this time, LC–MS of each well revealed that 38.4% (average of two experiments) of **AMPTX-1** ($[M+H^+]^+$ ion with m/z 692.6) had reacted with glutathione (reduced form) to form the corresponding adduct GSH adduct ($[M+H^+]^+$ *m/z* 999.8 and $[M+H^+]^{2+}$ *m/z* 500.5). This amount of adduct formation was formed in ~2 h and was maintained up to at least 24 h in a sealed well.

The wells were then diluted by 10x by removing 5 uL of the original solution and transferring into a separate well containing 45 uL of 9:1 mixture of PBS buffer and DMSO. This produced a solution containing 0.05 mM **AMPTX-1** and 0.5 mM L-glutathione (reduced form) in a 9:1 mixture of PBS buffer and DMSO, which was allowed to sit at ambient temperature in a sealed well of a 100 uL v-bottom 384-well plate for 4 h (performed in duplicate).

After this time, LC–MS of each well revealed that <2% (average of two experiments) of **AMPTX-1** ($[M+H^+]^+$ ion with *m/z* 692.6) was now an adduct GSH adduct with glutathione (reduced form) ($[M+H^+]^+$ *m/z* 999.8 and $[M+H^+]^{2+}$ *m/z* 500.5).

These data support the dynamic equilibrium and reversible nature of the reaction of **AMPTX-1** with glutathione (reduced form). The amount of adduct formation peaked at ~2 h and was stable up to 24 h, and then after dilution the amount of adduct decreased and reversed to reform **AMPTX-1** (see supplementary figure 3).

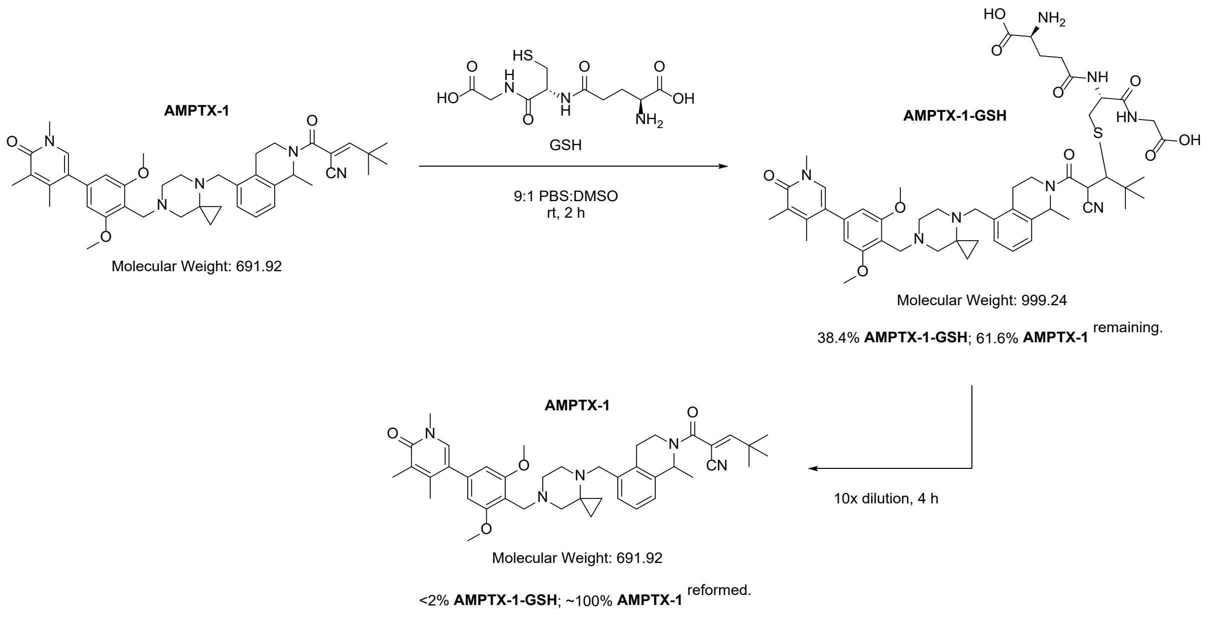


### Synthetic Methods

All chemicals, unless otherwise stated, were commercially available and used without further purification. Commercially available dry solvents were used. Flash silica-gel column chromatography was performed using Biotage Isolera using pre-packed silica (230-400 mesh particle size) with a flow rate suggested by the column manufacturer. Reverse-phase silica gel column chromatography was performed using Grace using pre-packed C18 (particle size = 40 µm) with a flow rate suggested by the column manufacturer. NMR Spectra were recorded on a Bruker Avance Neo 400 MHz. Chemical shifts ($\delta$) are reported in ppm and referenced to the residual solvent signals: $^1$H NMR $\delta$ (ppm) = 2.50 (DMSO-$d_6$). Signal splitting patterns are described as singlet (s), doublet (d), triplet (t), quartet (q), multiplet (m), broad (br) or a combination thereof. Coupling constants ($J$) are measured in Hertz (Hz).

LCMS chromatograms were recorded on a Waters Acquity UPLC using one of the methods below:

A. Acidic method: column = Zorbax Eclipse Plus C18 (2.1 x 50 µm, 1.8 µm particle size). Gradient = 5-100% acetonitrile + formic acid 0.05%/water + 0.1% formic acid. Flow rate = 0.7 mL/min over 5 min.
B. Neutral method: column = BEH C18 (2.1 mm X 50 mm, 1.7 µm particle size). Gradient = 5−98% acetonitrile/water + 10 mM ammonium acetate. Flow rate = 0.6 mL/min over 4 min.

HPLC chromatograms were recorded using the following method: column = X-Bridge C8 (4.6 x 150 mm, 5 µm particle size). Gradient = 10-100% acetonitrile/water + 10 mM ammonium acetate. Flow rate = 1.5 mL/min over 12 min.

SFC conditions are as follows: column: IZ (30 x 250 µm, 5 µm particle size); mobile phase: 60:40 $CO_2$:0.5% isopropylamine in MeOH:MeCN (60:40); total flow: 120 mL/min; back pressure: 100 bar; wavelength: 254 nm; cycle time: 8 min.

**Synthesis of (*E*)-2-cyano-4,4-dimethylpent-2-enoic acid**

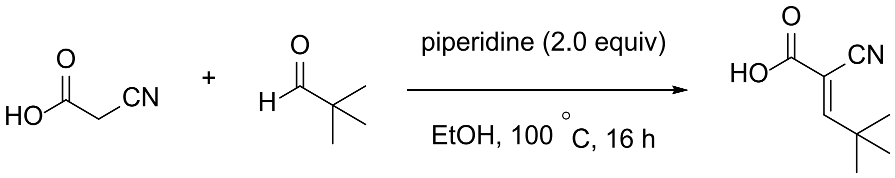


To a stirred solution of 2-cyanoacetic acid (2.00 g, 23.51 mmol, 1.0 equiv) in ethanol (30 mL) was added piperidine (4.00 g, 47.0 mmol, 2.0 equiv) and pivalaldehyde (3.04 g, 35.3 mmol, 1.5 equiv), then the reaction was stirred at 100 °C for 16 h. The reaction was monitored by thin-layer chromatography. Upon completion, the mixture was concentrated under reduced pressure. Water (10 mL) was added to the residue, and the mixture was then acidified with 1.5 N HCl to pH ~2. The mixture was extracted with ethyl acetate (2x 100 mL). The combined organic layers were washed with brine (30 mL), and the organic layer was dried ($Na_2SO_4$), filtered, and concentrated under reduced pressure. The crude material obtained was triturated with hexane and dried under reduced pressure to afford (*E)*-2-cyano-4,4-dimethylpent-2-enoic acid (2.5 g, 16.32 mmol, 69.4 % yield) as a pale brown solid. **LCMS:** $m/z$ = 153.2 ($[M]^+$), purity: 98.8%, $t_R$ = 1.200 min (Method B). **$^1$H NMR (400 MHz, $CDCl_3$):** $\delta$ 7.91 (br s, 1H), 7.72 (s, 1H), 1.33 (s, 9H).

1H NMR (400 MHz, CDCl3)

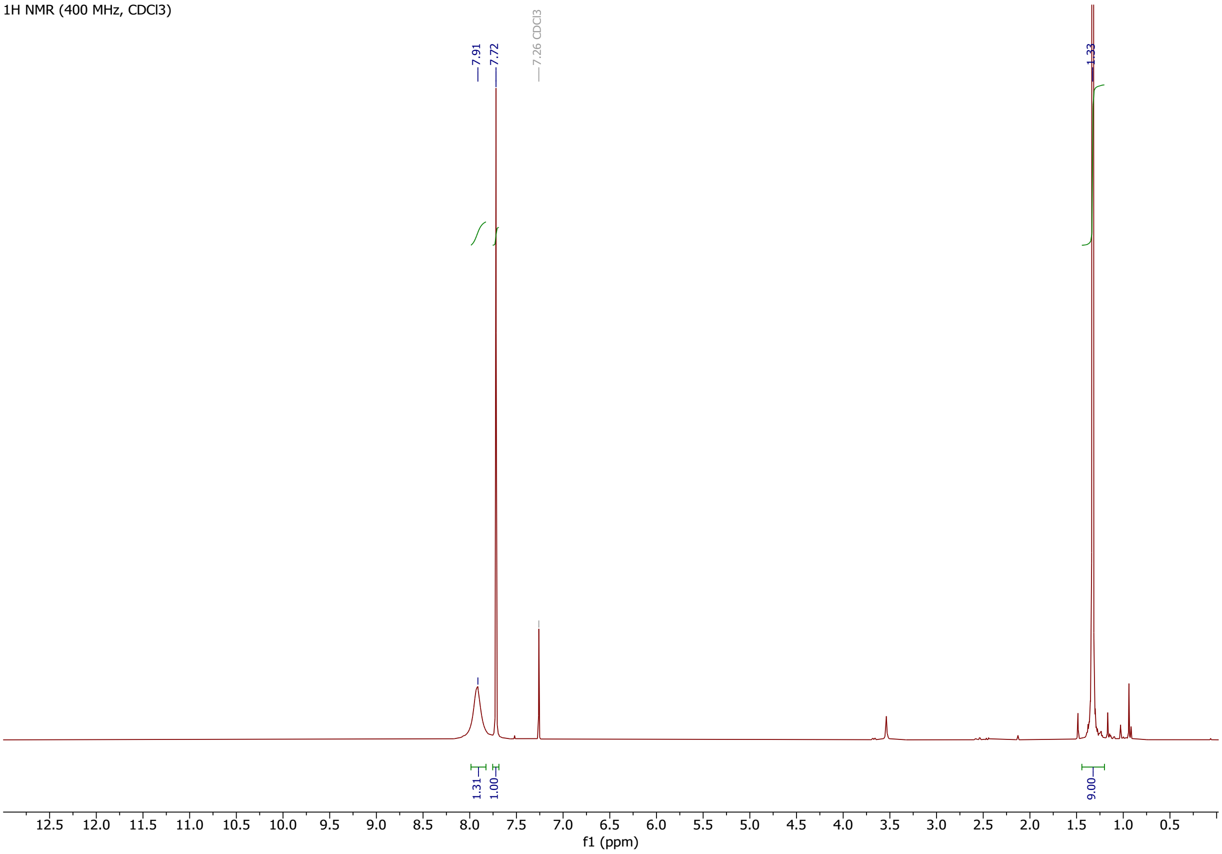

**$^{13}$C NMR (100 MHz, CDCl$_3$):** $\delta$ 174.2 (d, $J$ = 159.7 Hz), 167.1, 113.8 (d, $J$ = 14.5 Hz), 105.5, 35.5, 28.8 (q, $J$ = 128.6 Hz). *Note: This data and spectrum below were measured* *<u>without</u>* *$^1$H decoupling; all peaks except those at 167.1 and 105.5 ppm showed fine splitting which has not been measured.*

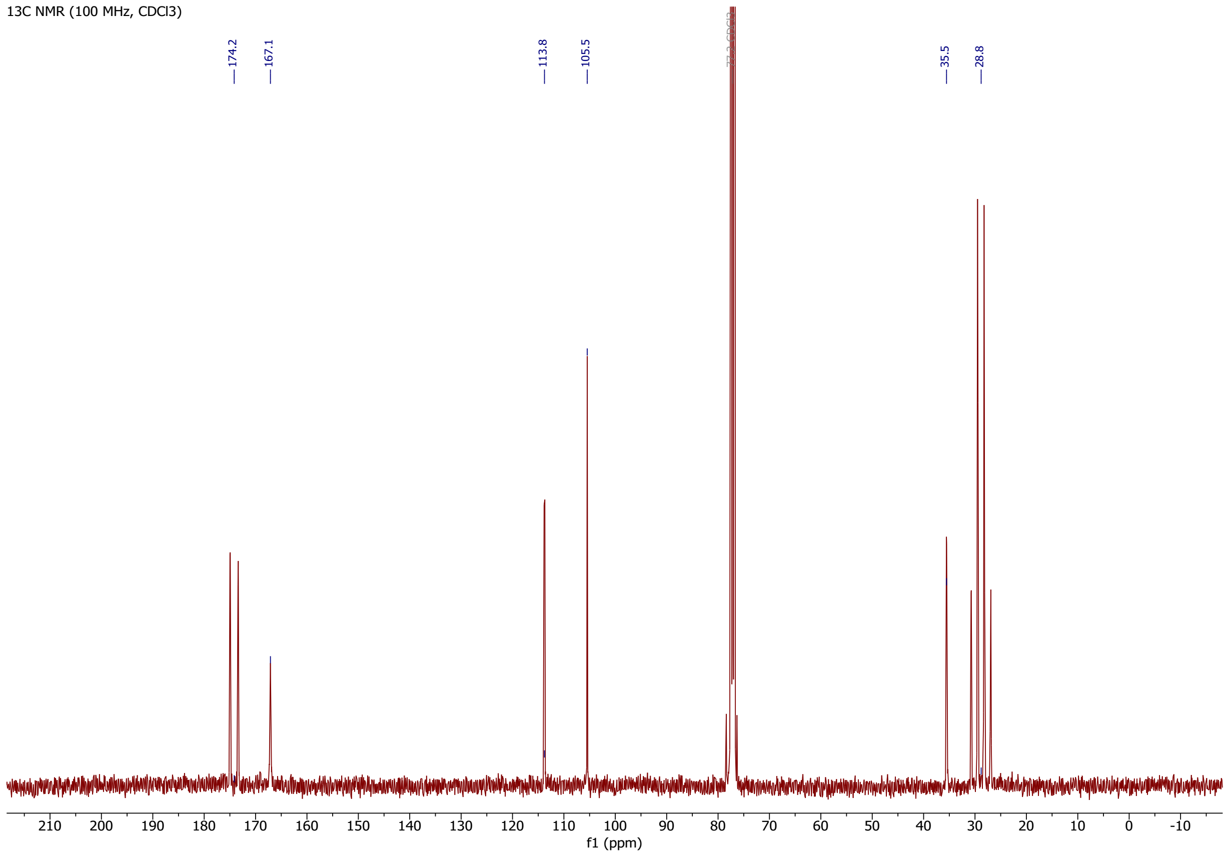


**Confirmation of double bond geometry:** As reported above, the $^{13}$C NMR resonance corresponding to the nitrile C≡N carbon is a doublet with a coupling constant of $J$ = 14.5 Hz. This is consistent with *E* (*trans*) coupling with the only available *E*-proton at 7.72 ppm in the $^1$H NMR spectrum (bound to the carbon at 174.2 ppm in the $^{13}$C spectrum). Correspondingly, the $^{13}$C NMR resonance corresponding to the carboxylic acid COOH carbon is an apparent singlet and thus not subject to any strong coupling with adjacent protons (therefore must be *Z* to the only available *E*-proton).

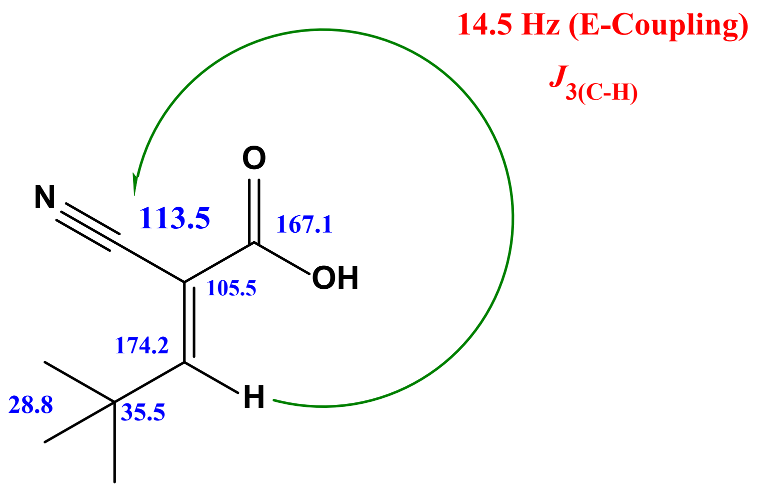

**Synthesis of *tert*-butyl 7-(2,6-dimethoxy-4-(1,4,5-trimethyl-6-oxo-1,6-dihydropyridin-3-yl)benzyl)-4,7-diazaspiro[2.5]octane-4-carboxylate (c)**

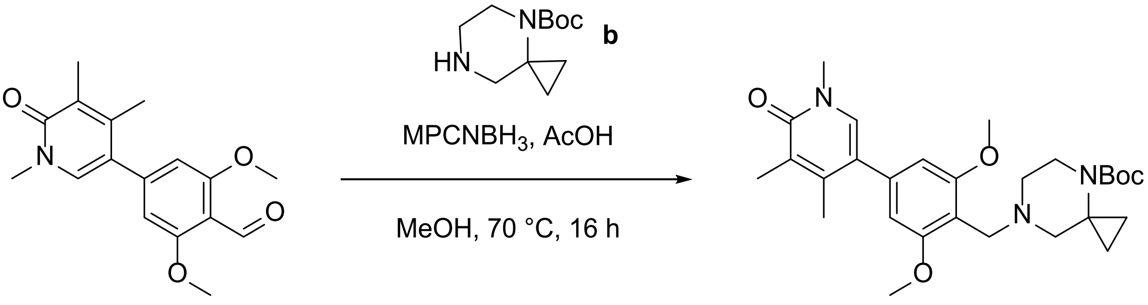


To a stirred solution of 2,6-dimethoxy-4-(1,4,5-trimethyl-6-oxo-1,6-dihydropyridin-3-yl) benzaldehyde[4] (**a**) (2.20 g, 7.30 mmol, 1 equiv) and *tert*-butyl 4,7-diazaspiro[2.5]octane-4-carboxylate (**b**) (2.02 g, 9.49 mmol, 1.3 equiv) in MeOH (15 mL) was added acetic acid (0.438 g, 7.30 mmol, 1 equiv) at ambient temperature and was stirred for 1 h, then polymer supported cyanoborohydride (MPCHN$H_3$, 3.00 g, 7.30 mmol, w/w) was added. The mixture was heated at 70 °C for 16 h, then allowed to cool to ambient temperature and concentrated under reduced pressure. The resulting crude residue was purified by silica gel column chromatography (gradient = 5-10% MeOH in DCM). The pure fractions were combined and concentrated under reduced pressure to afford *tert*-butyl 7-(2,6-dimethoxy-4-(1,4,5-trimethyl-6-oxo-1,6-dihydropyridin-3-yl)benzyl)-4,7-diazaspiro[2.5]octane-4-carboxylate (**c**) (3.4 g, 92% yield) as a colourless gum. **LCMS:** $m/z$ = 498.2 ($[M+H]^+$), $t_R$ = 2.06 min (Method A). **$^1$H NMR (400 MHz, DMSO-$d_6$):** $\delta$ 7.51 (s, 1H), 6.54 (s, 2H), 3.77 (s, 6H), 3.45 (s, 3H), 3.61 – 3.41 (br m, 2H), 2.43 – 2.13 (br m, 4H), 2.08 – 2.00 (br m, 6H), 1.37 (s, 9H), 0.81 (br s, 2H), 0.64 (br s, 2H). *One methylene signal obscured by deuterated solvent.*

**Synthesis of 5-(4-((4,7-diazaspiro[2.5]octan-7-yl)methyl)-3,5-dimethoxyphenyl)-1,3,4-trimethylpyridin-2(1*H*)-one hydrochloride (d)**

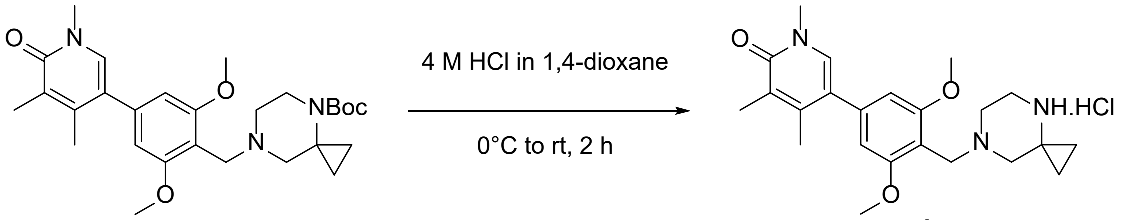


To a stirred solution of *tert*-butyl 7-(2,6-dimethoxy-4-(1,4,5-trimethyl-6-oxo-1,6-dihydropyridin-3-yl)benzyl)-4,7-diazaspiro[2.5]octane-4-carboxylate (**c**) (3.4 g, 6.83 mmol, 1 equiv) in DCM (20 mL) at 0 °C was added HCl (4 M in 1,4-dioxane, 7 mL, 28 mmol, 4.1 equiv). The mixture was then allowed to warm to ambient temperature and then stirred for 2 h before being concentrated under reduced pressure. The resulting residue was triturated with MTBE (2x 20 mL) before being dried under reduced pressure to afford 5-(4-((4,7-diazaspiro[2.5]octan-7-yl)methyl)-3,5-dimethoxyphenyl)-1,3,4-trimethylpyridin-2(1*H*)-one hydrochloride (**d**) (2.8 g, 100% yield) as a yellow solid. **LCMS:** *m/z* = 398.2 ($[M+H]^+$), $t_R$ = 1.45 min (Method B). **$^1$H NMR (400 MHz, DMSO-$d_6$):** $\delta$ 10.31 (br s, 1H), 7.53 (s, 1H), 6.69 (s, 2H), 3.87 (s, 6H), 3.68 – 3.37 (br m, 6H), 3.48 (s, 3H), 3.35 – 3.17 (br m, 2H), 2.09 (s, 3H), 2.06 (s, 3H), 1.31 – 1.18 (m, 2H), 1.12 – 1.01 (m, 2H).

**Synthesis of *tert*-butyl 5-((7-(2,6-dimethoxy-4-(1,4,5-trimethyl-6-oxo-1,6-dihydropyridin-3-yl)benzyl)-4,7-diazaspiro[2.5]octan-4-yl)methyl)-1-methyl-3,4-dihydroisoquinoline-2(1H)-carboxylate (f)**

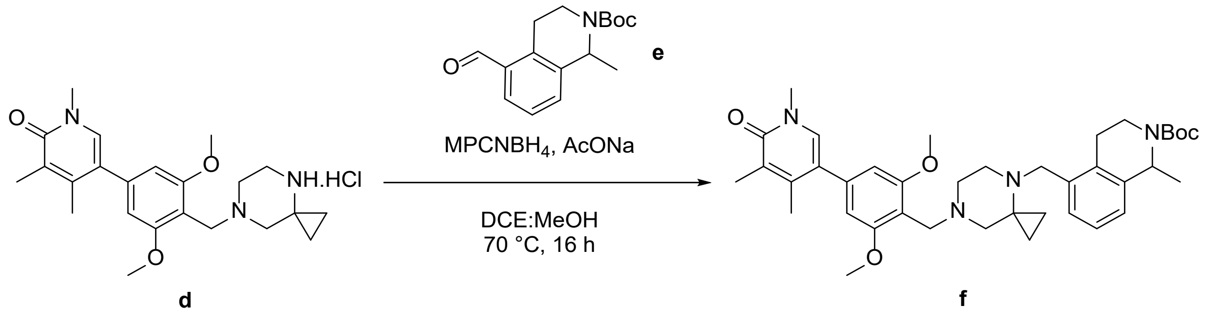


To a stirred solution of 5-(4-((4,7-diazaspiro[2.5]octan-7-yl)methyl)-3,5-dimethoxyphenyl)-1,3,4-trimethylpyridin-2(1*H*)-one hydrochloride (**d**) (2.80 g, 7.04 mmol, 1 equiv) in DCE (40 mL) and MeOH (10 mL) was added *tert*-butyl 5-formyl-1-methyl-3,4-dihydroisoquinoline-2(1*H*)-carboxylate[5] (**e**) (2.91 g, 10.57 mmol, 1.5 equiv) followed by sodium acetate (1.16 g, 14.09 mmol, 2 equiv). The mixture was stirred at ambient temperature for 1 h before polymer supported cyanoborohydride ($MPCHNH_3$, 2.80 g, 6.81 mmol, w/w) was added, then the mixture was stirred at 70 °C for 16 h. The mixture was allowed to cool to ambient temperature and was then concentrated under reduced pressure. The resulting crude residue was purified by silica-gel column chromatography (gradient = 5 – 15% MeOH in DCM) to afford *tert*-butyl 5-((7-(2,6-dimethoxy-4-(1,4,5-trimethyl-6-oxo-1,6-dihydropyridin-3-yl)benzyl)-4,7-diazaspiro[2.5]octan-4-yl)methyl)-1-methyl-3,4-dihydroisoquinoline-2(1*H*)-carboxylate (**f**) (3.2 g, 69% yield) as a brown solid. **LCMS:** *m/z* = 657.4 ($[M+H]^+$), $t_R$ = 1.98 min (Method B). **$^1$H NMR** (400 MHz, DMSO-$d_6$): $\delta$ 7.52 (s, 1H), 7.18-7.08 (m, 3H), 6.68 (s, 2H), 5.13-4.93 (br m, 1H), 4.41-4.30 (m, 1H), 4.27-4.18 (m, 1H), 4.10-3.92 (m, 2H), 3.87 (s, 6H), 3.74-3.61 (br m, 2H), 3.47 (s, 3H), 3.12-2.98 (br m, 4H), 2.84-2.63 (br m, 4H), 2.08 (s, 3H), 2.06 (s, 3H), 1.42 (s, 9H), 1.40-1.32 (br m, 5H), 0.74-0.66 (br m, 2H). *Note: compound isolated with some AcOH, giving the acetate salt of **f**, causing downfield shift in some signals.*

**Synthesis of 5-(3,5-dimethoxy-4-((4-((1-methyl-1,2,3,4-tetrahydroisoquinolin-5-yl)methyl)-4,7-diazaspiro[2.5]octan-7-yl)methyl)phenyl)-1,3,4-trimethylpyridin-2(1H)-one hydrochloride (g)**

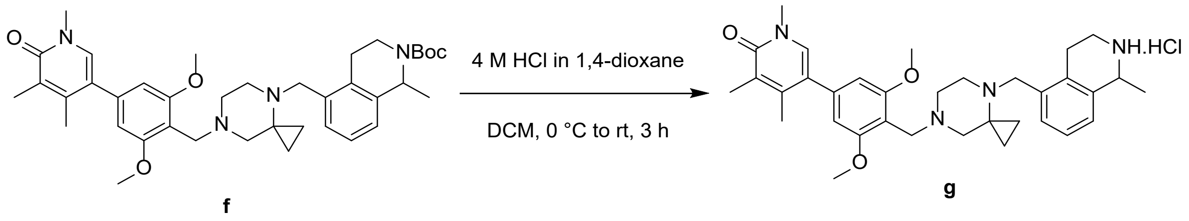

To a stirred solution of *tert*-butyl 5-((7-(2,6-dimethoxy-4-(1,4,5-trimethyl-6-oxo-1,6-dihydropyridin-3-yl)benzyl)-4,7-diazaspiro[2.5]octan-4-yl)methyl)-1-methyl-3,4-dihydroisoquinoline-2(1*H*)-carboxylate (**f**) (3.20 g, 4.87 mmol, 1 equiv) in DCM (25 mL) at 0 °C was added HCl (4 M in 1,4-dioxane, 5 mL). The mixture was allowed to warm to ambient temperature and then stirred for 3 h before being concentrated under reduced pressure. The resulting residue was washed with MTBE (2x 20 mL) and was dried under reduced pressure to afford 5-(3,5-dimethoxy-4-((4-((1-methyl-1,2,3,4-tetrahydroisoquinolin-5-yl)methyl)-4,7-diazaspiro[2.5]octan-7-yl)methyl)phenyl)-1,3,4-trimethylpyridin-2(1*H*)-one hydrochloride (**g**) (3 g, 5.05 mmol, 100% yield) as an off-white solid. **LCMS:** $m/z$ = 557.3 ([M+H]$^+$), $t_R$ = 1.71 min (Method A). **$^1$H NMR** (400 MHz, DMSO-$d_6$): $\delta$ 7.54 (s, 1H), 7.28 (br m, 3H), 6.68 (s, 2H), 4.53 (br s, 1H), 4.37 (br s, 1H), 4.28-4.25 (m, 2H), 3.89 (s, 6H), 3.57 (s, 6H), 3.48 (s, 3H), 3.40-3.39 (m, 2H), 3.27 (br s, 2H), 3.17-2.90 (m, 4H), 2.08 (m, 6H), 1.61-1.60 (m, 3H), 0.83 (br s, 3H).

**Synthesis of (*E*)-2-(5-((7-(2,6-dimethoxy-4-(1,4,5-trimethyl-6-oxo-1,6-dihydropyridin-3-yl)benzyl)-4,7-diazaspiro[2.5]octan-4-yl)methyl)-1-methyl-1,2,3,4-tetrahydroisoquinoline-2-carbonyl)-4,4-dimethylpent-2-enenitrile) (AMPTX-1)**

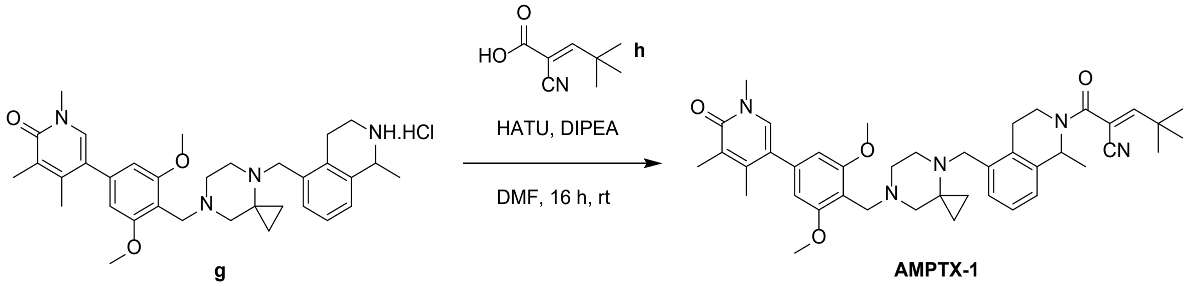

To a stirred solution of 5-(3,5-dimethoxy-4-((4-((1-methyl-1,2,3,4-tetrahydroisoquinolin-5-yl)methyl)-4,7-diazaspiro[2.5]octan-7-yl)methyl)phenyl)-1,3,4-trimethylpyridin-2(1*H*)-one hydrochloride (**g**) (3.0 g, 5.06 mmol, 1.0 equiv), DIPEA (4.4 mL, 25.3 mmol, 5.0 equiv) and (*E)*-2-cyano-4,4-dimethylpent-2-enoic acid (**h**) (1.55 g, 10.1 mmol, 2.0 equiv) in DMF (8 mL) at 0 °C was added HATU (3.85 g, 10.1 mmol, 2.0 equiv). The mixture was stirred for 15 min at 0 °C before being allowed to warm to ambient temperature and then stirred for 16 h. The mixture was quenched with ice water (30 mL) and extracted with ethyl acetate (3x 30 mL). The combined organic layers were dried over sodium sulfate, filtered, and concentrated under reduced pressure. The resulting crude residue was purified by silica-gel column chromatography (gradient = 10-20% MeOH in DCM), and the appropriate fractions were concentrated under reduced pressure. Then, the resulting residue was purified by reverse-phase column chromatography (eluent = 0.1% aqueous $NH_4OAc$/MeCN), and the appropriate fractions were lyophilized to obtain **AMPTX-1** (1.5 g, 42% yield) as a yellow solid. **LCMS:** $m/z$ = 692.4 ([M+H]$^+$), $t_R$ = 2.27 min (Method B). **HPLC:** $t_R$ = 6.94 min. **$^1$H NMR** (400 MHz, DMSO-$d_6$): $\delta$ 7.51 (s, 1H), 7.28 – 7.06 (m, 3H), 6.90 (br s, 1H), 6.65 (br s, 2H), 5.39 – 5.25 (br m, 1H), 4.50-4.11 (br m, 2H), 4.09-3.39 (br m, 5H), 3.85 (s, 6H), 3.47 (s, 3H), 3.20-2.60 (br m, 6H), 2.07 (s, 3H), 2.06 (s, 3H), 1.60-1.34 (br m, 4H), 1.25 (s, 9H), 1.11-0.85 (br m, 2H), 0.82-0.48 (br m, 2H). *Note: compound isolated with some AcOH, giving the acetate salt of AMPTX-1, causing downfield shift in some signals.*

**Synthesis of single enantiomers of (*E)*-2-(5-((7-(2,6-dimethoxy-4-(1,4,5-trimethyl-6-oxo-1,6-dihydropyridin-3-yl)benzyl)-4,7-diazaspiro[2.5]octan-4-yl)methyl)-1-methyl-1,2,3,4-tetrahydroisoquinoline-2-carbonyl)-4,4-dimethylpent-2-enenitrile) (AMPTX-*ent*-1 and AMPTX-*ent*-2)**

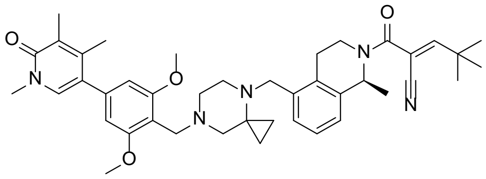

**AMPTX-1-*ent*-1**

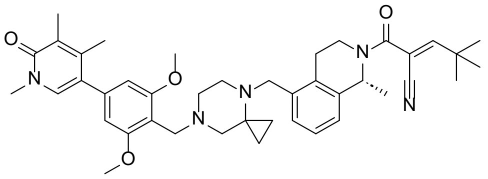

**AMPTX-1-*ent*-2**

*Single enantiomers of unknown absolute stereochemistry (assigned arbitrarily)*

**Synthesis of single enantiomers of *tert*-butyl 5-((7-(2,6-dimethoxy-4-(1,4,5-trimethyl-6-oxo-1,6-dihydropyridin-3-yl)benzyl)-4,7-diazaspiro[2.5]octan-4-yl)methyl)-1-methyl-3,4-dihydroisoquinoline-2(1*H*)-carboxylate (i and j)**

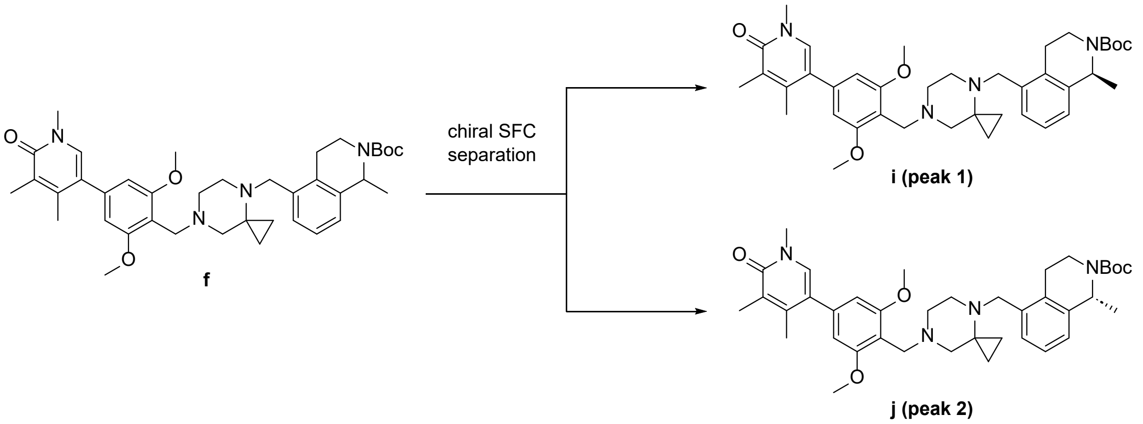


*Single enantiomers of unknown absolute stereochemistry (assigned arbitrarily)*

Racemic *tert*-butyl 5-((7-(2,6-dimethoxy-4-(1,4,5-trimethyl-6-oxo-1,6-dihydropyridin-3-yl)benzyl)-4,7-diazaspiro[2.5]octan-4-yl)methyl)-1-methyl-3,4-dihydroisoquinoline-2(1*H*)-carboxylate (**f**) was separated by chiral SFC to afford single enantiomers of *tert*-butyl 5-((7-(2,6-dimethoxy-4-(1,4,5-

trimethyl-6-oxo-1,6-dihydropyridin-3-yl)benzyl)-4,7-diazaspiro[2.5]octan-4-yl)methyl)-1-methyl-3,4-dihydroisoquinoline-2(1*H*)-carboxylate (**i** and **j**). **SFC:** column: IZ (30 x 250 μm, 5 μm particle size); mobile phase: 60:40 $CO_2$:0.5% isopropylamine in MeOH:MeCN (60:40); total flow: 120 mL/min; back pressure: 100 bar; wavelength: 254 nm; cycle time: 8 min. **Peak 1 of *tert*-butyl 5-((7-(2,6-dimethoxy-4-(1,4,5-trimethyl-6-oxo-1,6-dihydropyridin-3-yl)benzyl)-4,7-diazaspiro[2.5]octan-4-yl)methyl)-1-methyl-3,4-dihydroisoquinoline-2(1H)-carboxylate (i)**: white sold, 170 mg. **LCMS:** *m/z* = 657.4 ($[M+H]^+$), $t_R$ = 2.18 min (Method B). **SFC:** $t_R$ = 2.44 min, 97.8% ee. **$^1$H NMR** (400 MHz, DMSO-$d_6$): $\delta$ 7.52 (s, 1H), 7.13-7.05 (m, 3H), 6.53 (s, 2H), 5.13-4.93 (br m, 1H), 4.09-3.83 (br m, 2H), 3.77 (s, 6H), 3.74-3.61 (br m, 2H), 3.50 (br s, 2H), 3.45 (s, 3H), 2.82-2.54 (br m, 6H), 2.45-2.36 (br m, 2H), 2.05 (br app. s, 6H), 1.42 (s, 9H), 1.39-1.31 (br m, 3H), 0.62-0.52 (br m, 2H), 0.38-0.30 (br m, 2H). **Peak 2 of *tert*-butyl 5-((7-(2,6-dimethoxy-4-(1,4,5-trimethyl-6-oxo-1,6-dihydropyridin-3-yl)benzyl)-4,7-diazaspiro[2.5]octan-4-yl)methyl)-1-methyl-3,4-dihydroisoquinoline-2(1H)-carboxylate (j)**: brown solid, 200 mg. **LCMS:** *m/z* = 657.4 ($[M+H]^+$), $t_R$ = 2.16 min (Method B). **SFC:** $t_R$ = 2.27 min, 91.0% ee **$^1$H NMR** (400 MHz, DMSO-$d_6$): $\delta$ 7.51 (s, 1H), 7.14-7.04 (m, 3H), 6.53 (s, 2H), 5.12-4.93 (br m, 1H), 4.09-3.82 (br m, 2H), 3.77 (s, 6H), 3.73-3.64 (br m, 2H), 3.51 (br s, 2H), 3.45 (s, 3H), 2.82-2.56 (br m, 6H), 2.45-2.36 (br m, 2H), 2.05 (br app. s, 6H), 1.42 (s, 9H), 1.39-1.31 (br m, 3H), 0.64-0.53 (br m, 2H), 0.40-0.30 (br m, 2H).

**Synthesis of single enantiomers of 5-(3,5-dimethoxy-4-((4-((1-methyl-1,2,3,4-tetrahydroisoquinolin-5-yl)methyl)-4,7-diazaspiro[2.5]octan-7-yl)methyl)phenyl)-1,3,4-trimethylpyridin-2(1*H*)-one hydrochloride (k and l)**

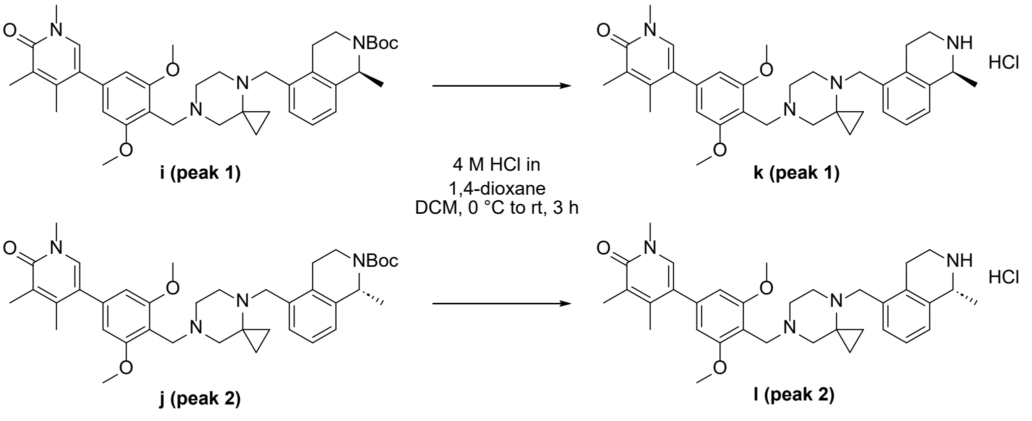


*Single enantiomers of unknown absolute stereochemistry (assigned arbitrarily)*

To a stirred solution of the relevant starting material (**i** or **j**) (1 equiv) in DCM at 0 °C was added HCl (4 M in 1,4-dioxane, 10 equiv). The resultant mixture was allowed to warm to ambient temperature and stir for 3 h and then was concentrated under reduced pressure. The resulting residue was washed with MTBE and dried under high vacuum to afford the corresponding single enantiomers of 5-(3,5-dimethoxy-4-((4-((1-methyl-1,2,3,4-tetrahydroisoquinolin-5-yl)methyl)-4,7-diazaspiro[2.5] octan-7-yl)methyl)phenyl)-1,3,4-trimethylpyridin-2(1*H*)-one hydrochloride (**k** and **l**). **5-(3,5-dimethoxy-4-((4-((1-methyl-1,2,3,4-tetrahydroisoquinolin-5-yl)methyl)-4,7-diazaspiro[2.5] octan-7-yl)methyl)phenyl)-1,3,4-trimethylpyridin-2(1H)-one hydrochloride (k) (from i):** brown solid, 140 mg, 100% yield. **LCMS:** $m/z$ = 557.2 ([M+H]$^+$), $t_R$ = 1.71 min (Method A) **$^1$H NMR** (400 MHz, DMSO-$d_6$): $\delta$ 9.70 (m, 1H), 9.30 (m, 2H), 7.53 (s, 1H), 7.26 (s, 3H), 6.69 (s, 2H), 4.55 - 4.65 (m, 1H), 4.26 - 4.23 (m, 2H), 3.89 (s, 6H), 3.77 (br s, 3H), 3.48 (s, 3H), 3.42 - 3.41 (m, 2H), 3.30 (br s, 1H), 3.10 - 3.00 (m, 4H), 2.81 (br s, 2H), 2.09 - 2.07 (m, 6H), 1.60 - 1.59 (m, 3H), 0.74 (br s, 4H). **5-(3,5-dimethoxy-4-((4-((1-methyl-1,2,3,4-tetrahydroisoquinolin-5-yl)methyl)-4,7-diazaspiro[2.5] octan-7-yl)methyl)phenyl)-1,3,4-trimethylpyridin-2(1H)-one hydrochloride (l) (from j):** brown solid, 120 mg, 72% yield. **LCMS:** $m/z$ = 557.2 ([M+H]$^+$), $t_R$ = 1.71 min (Method A). **$^1$H NMR** (400 MHz, DMSO-$d_6$): $\delta$ 9.41 (m, 3H), 7.52 (s, 1H), 7.26 (s, 3H), 6.69 (s, 2H), 4.55 - 4.65 (m, 1H), 4.35 (br s, 1H), 4.23 (s, 1H), 3.89 (s, 2H), 3.75 (m, 5H), 3.48 (s, 3H), 3.44 - 3.41 (m, 2H), 3.31 - 3.29 (m, 1H), 3.17 - 3.08 (m, 4H), 3.05 - 2.98 (m, 2H), 2.08 (m, 6H), 1.59 (m, 3H), 0.73 (br s, 4H).

**Synthesis of single enantiomers of (*E*)-2-(5-((7-(2,6-dimethoxy-4-(1,4,5-trimethyl-6-oxo-1,6-dihydropyridin-3-yl)benzyl)-4,7-diazaspiro[2.5]octan-4-yl)methyl)-1-methyl-1,2,3,4-tetrahydroisoquinoline-2-carbonyl)-4,4-dimethylpent-2-enenitrile (AMPTX-1-*ent*-1 and AMPTX-*ent*-2)**

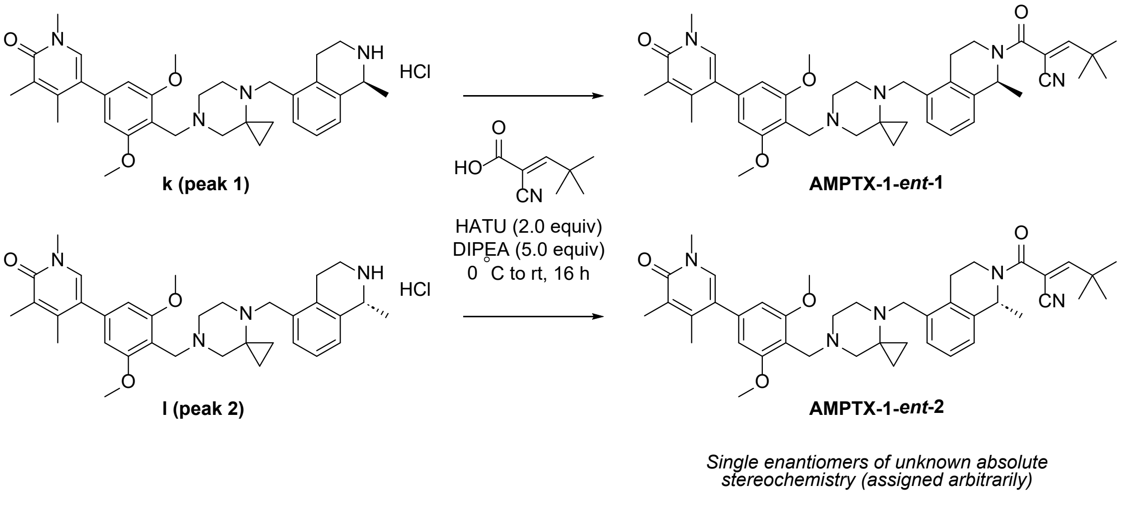


To a stirred solution of the relevant amine hydrochloride salt (**k** or **l**) (1.0 equiv) in DMF at 0 °C was added DIPEA (5.0 equiv) followed by (*E*)-2-cyano-4,4-dimethylpent-2-enoic acid (**h**) (2.0 equiv) and HATU (2.0 equiv). The resultant mixture was stirred for 15 min before being allowed to warm to ambient temperature and then stir for 16 h. Then mixture was quenched with iced water and the resultant solid was collected by vacuum filtration and then purified by reverse-phase column chromatography (eluent = 0.1% aqueous $NH_4OAc$ in MeCN) to give the corresponding single enantiomers of (*E*)-2-(5-((7-(2,6-dimethoxy-4-(1,4,5-trimethyl-6-oxo-1,6-dihydropyridin-3-yl)benzyl)-4,7-diazaspiro[2.5]octan-4-yl)methyl)-1-methyl-1,2,3,4-tetrahydroisoquinoline-2-carbonyl)-4,4-dimethylpent-2-enenitrile (**AMPTX-1-*ent*-1** and **AMPTX-1-*ent*-2**). **AMPTX-1-*ent*-1 (from k)**: off-white solid, 53 mg, 38% yield. **LCMS:** $m/z$ = 692.4 ($[M+H]^+$), $t_R$ = 2.04 min (Method B). **SFC:** $t_R$ = 3.19 min, 98.4% ee. **$^1$H NMR** (400 MHz, DMSO-$d_6$): $\delta$ 7.51 (s, 1H), 7.19-7.06 (m, 3H), 6.87 (br s, 1H), 6.53 (s, 2H), 5.37-5.25 (br m, 1H), 3.96-3.83 (br m, 1H), 3.77 (s, 6H), 3.73-3.64 (br m, 2H), 3.50 (s, 2H), 3.45 (s, 3H), 2.94-2.80 (br m, 2H), 2.64-2.56 (br m, 2H), 2.44-2.37 (br m, 2H), 2.05 (app. s, 6H), 1.59-1.38 (br m, 4H), 1.25 (s, 9H), 1.09-0.82 (m, 2H), 0.63-0.52 (br m, 2H), 0.36 (br s, 2H). **AMPTX-1-*ent*-2 (from l)**: off-white solid, 41 mg, 29% yield. **LCMS:** $m/z$ = 692.4 ($[M+H]^+$), $t_R$ = 2.05 min (Method B). **SFC:** $t_R$ = 1.49 min, 97.1% ee. **$^1$H NMR** (400 MHz, DMSO-$d_6$): $\delta$ 7.51 (s, 1H), 7.19-7.06 (m, 3H), 6.87 (br s, 1H), 6.53 (s, 2H), 5.37-5.26 (br m, 1H), 3.97-3.85 (br m, 1H), 3.77 (s, 6H), 3.70 (br s, 2H), 3.50 (br s, 2H), 3.45 (s, 3H), 2.93-2.80 (br m, 2H), 2.64-2.55 (br m, 2H), 2.44-2.36 (br m, 2H), 2.05 (app. s, 6H), 1.59-1.37 (br m, 4H), 1.25 (9H), 1.13-0.95 (m, 2H), 0.63-0.51 (m, 2H), 0.36 (br s, 2H).

**Synthesis of (*E*)-5-(4-((4-((2-(4,4-dimethylpent-2-enoyl)-1-methyl-1,2,3,4-tetrahydroisoquinolin-5-yl)methyl)-4,7-diazaspiro[2.5]octan-7-yl)methyl)-3,5-dimethoxyphenyl)-1,3,4-trimethylpyridin-2(1H)-one (2)**

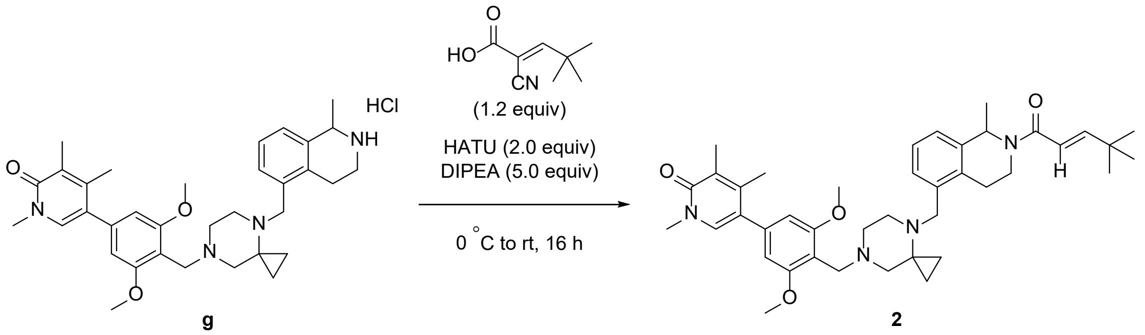

**g** **2**

To a stirred solution of 5-(3,5-dimethoxy-4-((4-((1-methyl-1,2,3,4-tetrahydroisoquinolin-5-yl)methyl)-4,7-diazaspiro[2.5]octan-7-yl)methyl)phenyl)-1,3,4-trimethylpyridin-2(1*H*)-one hydrochloride (**g**) (1.0 equiv) in DMF at 0 °C was added DIPEA (5.0 equiv) followed by (*E*)-4,4-dimethylpent-2-enoic acid (1.2 equiv) (*see below for confirmation of stereochemistry*) and HATU (2.0 equiv). The resultant mixture was stirred for 15 min before being allowed to warm to ambient temperature and then stir for 16 h. Then mixture was quenched with iced water and the resultant solid was collected by vacuum filtration and then purified by reverse-phase column chromatography (eluent = 0.1% aqueous $NH_4OAc$ in MeCN) to give (E)-5-(4-((4-((2-(4,4-dimethylpent-2-enoyl)-1-methyl-1,2,3,4-tetrahydroisoquinolin-5-yl)methyl)-4,7-diazaspiro[2.5]octan-7-yl)methyl)-3,5-dimethoxyphenyl)-1,3,4-trimethylpyridin-2(1*H*)-one (**2**). **LCMS:** $m/z$ = 667.3 ($[M+H]^+$), $t_R$ = 2.38 min (Method A). **$^1$H NMR** (400 MHz, DMSO-$d_6$): $\delta$

7.52 (s, 1H), 7.17-7.11 (m, 3H), 6.75 (dd, $J$ = 8.80, 15.20 Hz, 1H), 6.59 (s, 2H), 6.36 (d, $J$ = 15.20 Hz, 1H), 5.51-5.26 (m, 1H), 4.50-4.10 (m, 1H), 3.82 (s, 9H), 3.47 (s, 3H), 3.41 (s, 2H), 2.85-2.84 (m, 1H), 2.79-2.68 (m, 6H), 2.34-2.33 (m, 1H), 2.07 (d, $J$ = 3.20 Hz, 6H), 1.49-1.37 (m, 3H), 1.08 (s, 9H), 0.64-0.47 (m, 4H).

**Confirmation of double bond geometry of (*E*)-4,4-dimethylpent-2-enoic acid:** The commercially sourced sample of (*E*)-4,4-dimethylpent-2-enoic acid used for this synthesis was assessed by $^1$H NMR. **$^1$H NMR** (400 MHz, DMSO-$d_6$): 12.16 (br s, 1H), 6.84 (d, $J$ = 16.0 Hz, 1H), 5.64 (d, $J$ = 16.0 Hz, 1H), 1.05 (s, 9H). The coupling constant between resonances at 6.84 and 5.64 ppm of 16.0 Hz is only consistent with *E* (*trans*) coupling. No presence of the *Z* (*cis*) geometric isomer was detected (expected $J \approx 10$ Hz).

**Synthesis of *tert*-butyl 1-methyl-5-((4-methylpiperazin-1-yl)methyl)-3,4-dihydroisoquinoline-2(1*H*)-carboxylate (m)**

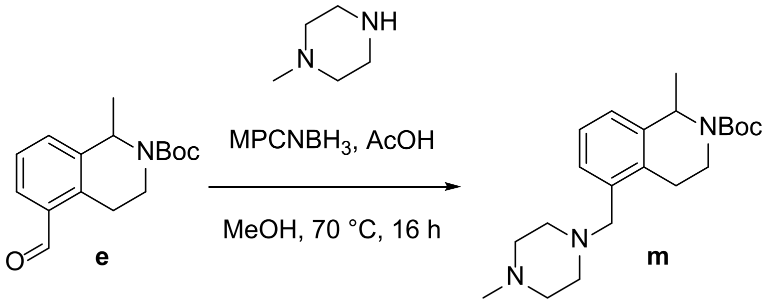


To a stirred solution of *tert*-butyl 5-formyl-1-methyl-3,4-dihydroisoquinoline-2(1*H*)-carboxylate (**e**) (490 mg, 1.78 mmol, 1 equiv) and 1-methylpiperazine (214 mg, 2.14 mmol, 1.2 equiv) in MeOH (5 mL) at ambient temperature was added acetic acid (0.107 g, 1.78 mmol, 1 equiv). The resultant mixture was stirred at ambient temperature for 1 h, then polymer supported cyanoborohydride (MPCNBH$_3$, 0.73 g, 1.78 mmol, w/w) was added. The mixture was heated at 70°C for 16 h before being allowed to cool to ambient temperature and then concentrated under reduced pressure. The resulting residue was purified by silica-gel column chromatography to afford *tert*-butyl 1-methyl-5-((4-methylpiperazin-1-yl)methyl)-3,4-dihydroisoquinoline-2(1*H*)-carboxylate (**m**) (650 mg, 99% yield). **LCMS:** $m/z$ = 360.4 ([M+H]$^+$), $t_R$ = 2.13 min (Method A).

**Synthesis of 1-methyl-5-((4-methylpiperazin-1-yl)methyl)-1,2,3,4-tetrahydroisoquinoline hydrochloride (o)**

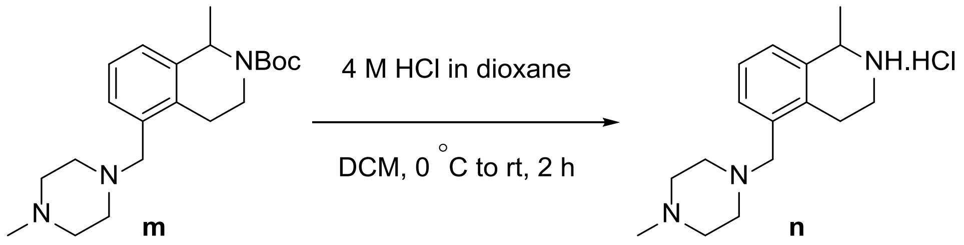

To a stirred solution of *tert*-butyl 1-methyl-5-((4-methylpiperazin-1-yl)methyl)-3,4-dihydroisoquinoline-2(1*H*)-carboxylate (**m**) (650 mg, 1.79 mmol, 1 equiv) in DCM (5 mL) at 0 °C was added HCl (4 M in 1,4-dioxane, 4.5 mL, 17.9 mmol, 10 equiv). The resultant mixture was allowed to warm to ambient temperature and stirred for 2 h before. The mixture was then concentrated under reduced pressure and the residue was washed with MTBE (2x 20 mL) and dried under high vacuum to afford 1-methyl-5-((4-methylpiperazin-1-yl)methyl)-1,2,3,4-tetrahydroisoquinoline hydrochloride (**n**) (530 mg, 99% yield). **LCMS:** $m/z$ = 260.2 ($[M+H]^+$), $t_R$ = 0.26 min (Method A).

**Synthesis of (*E*)-4,4-dimethyl-2-(1-methyl-5-((4-methylpiperazin-1-yl)methyl)-1,2,3,4-tetrahydroisoquinoline-2-carbonyl)pent-2-enenitrile (3)**

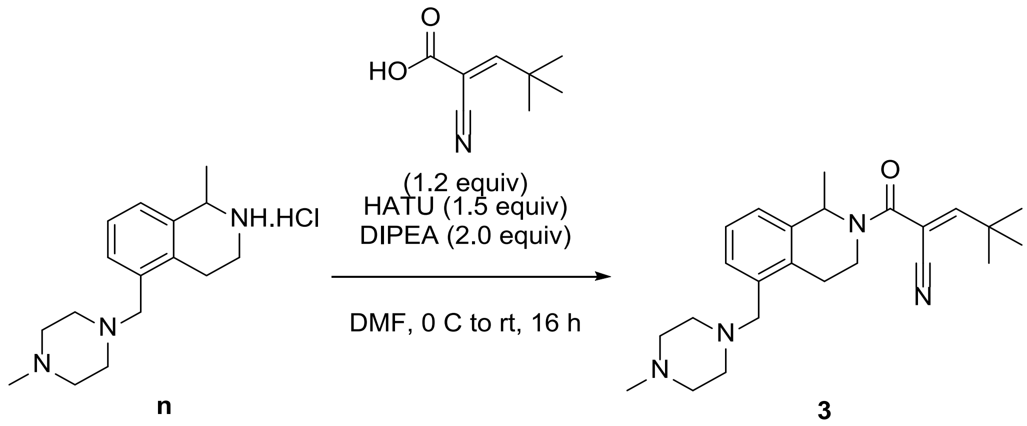

To a stirred solution of 1-methyl-5-((4-methylpiperazin-1-yl)methyl)-1,2,3,4-tetrahydroisoquinoline hydrochloride (**o**) (66 mg, 0.224 mmol, 1.0 equiv) in DMF (1 mL) at 0 °C was added DIPEA (0.08 mL, 0.448 mmol, 2 equiv), (*E*)-2-cyano-4,4-dimethylpent-2-enoic acid (**h**) (*exclusively E geometry, see previous*) (41 mg, 0.269 mmol, 1.2 equiv), and then HATU (128 mg, 0.336 mmol, 1.5 equiv). The resultant mixture was stirred for 15 min before being allowed to warm to ambient temperature and stir for 16 h. The mixture was quenched with iced water (5 mL) and extracted with ethyl acetate (3x 5 mL). The combined organic layers were dried over sodium sulfate, filtered, and concentrated under reduced pressure. The resulting residue was purified by silica-gel column chromatography to afford (*E*)-4,4-dimethyl-2-(1-methyl-5-((4-methylpiperazin-1-yl)methyl)-1,2,3,4-tetrahydroisoquinoline-2-carbonyl)pent-2-enenitrile **3** (37 mg, 42% yield). **LCMS:** $m/z$ = 395.2 ($[M+H]^+$), $t_R$ = 1.94 min (Method

A) **$^1$H NMR** (400 MHz, DMSO-$d_6$): $\delta$ 7.18-7.12 (m, 3H), 6.87 (s, 1H), 5.34 (d, $J$ = 5.60 Hz, 1H), 3.92 (d, $J$ = 12.40 Hz, 1H), 3.51 (s, 2H), 2.97 (s, 2H), 2.36-2.30 (m, 8H), 2.17 (s, 3H), 1.55-1.44 (m, 4H), 1.26 (s, 9H).
