## supplementary figures 1_5 for "Selective degradation of BRD9 by a DCAF16-recruiting targeted glue: mode of action elucidation and in vivo proof of concept"

**Supplementary Figure 1**

**a** Live cell degradation of HiBiT-BRD9 by Compound 2 over 24 h. **b** Global expression proteomics following 6 h treatment of MV4-11 cells with 100 nM **Compound 2.** 8350 proteins were quantified at 1% FDR. Data is presented as logFC relative to DMSO.

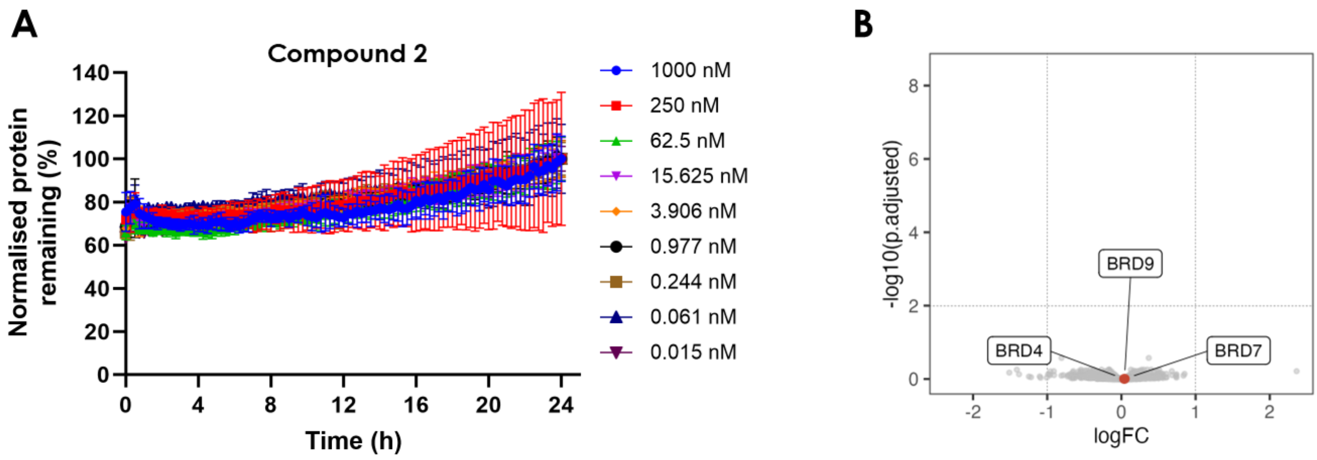


**Supplementary figure 2**

**a.** Chemical structures of BRD9 inhibitor BI-7273 and reversibly covalent warhead **3. B.** Enrichment of DCAF16 and DDB1 to BRD9-HiBiT following co-immunoprecipitation by anti-HiBiT following treatment with AMPTX-1 and compound **2** vs DMSO.

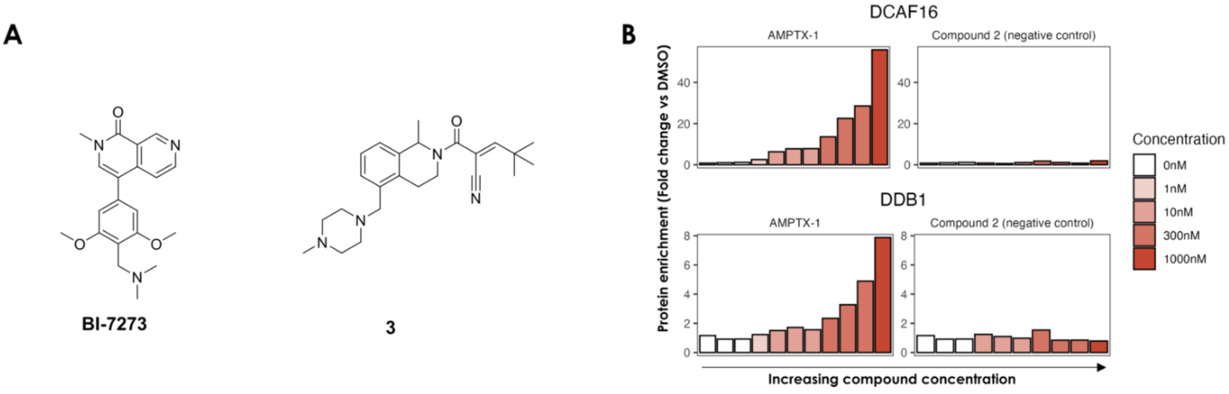


**Supplementary figure 3**

**a.** LC–MS chromatogram of a 0.5 mM **AMPTX-1** and 5.0 mM L-glutathione solution in a 9:1 mixture of PBS buffer and DMSO after 2 h incubation. **b.** LC–MS chromatogram of the solution from **a** following 10-fold dilution and 4 h incubation.

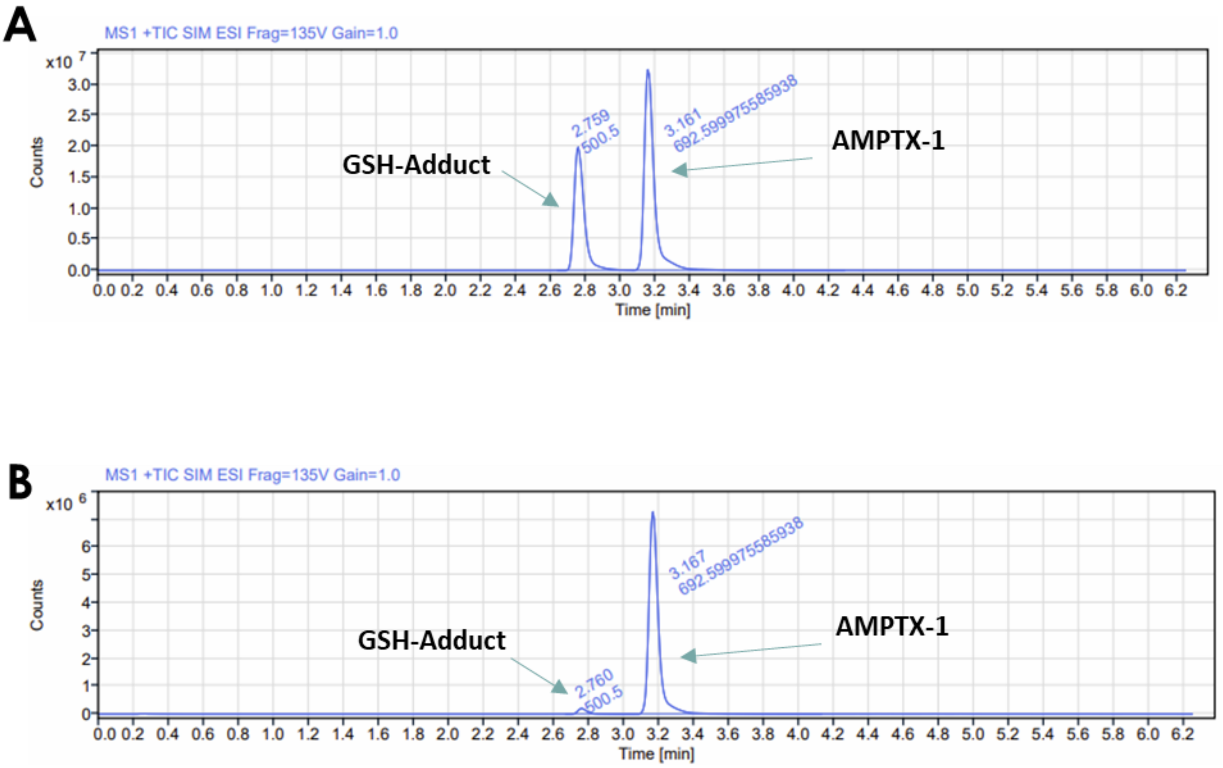


**Supplementary figure 4**

Detection of DCAF16$^{Cys58}$ modification by peptide mapping MS. Recombinant DCAF16-DDB1 was incubated with 5 μM **AMPTX-1** in the presence and absence of BRD9$^{BD}$ for 20 h prior to proteolytic digest.

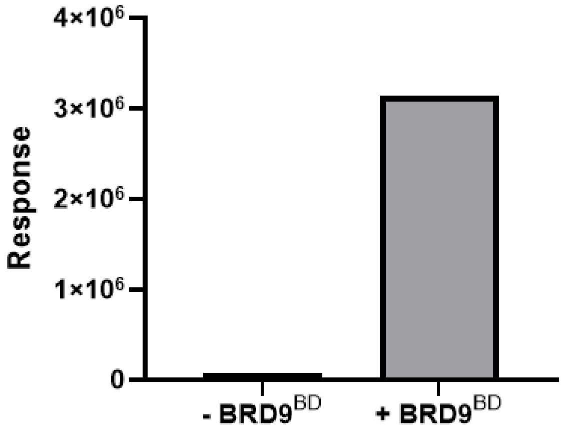


**Supplementary figure 5**

BRD9 quantification (gray bar, normalised to actin) and total drug concentration in plasma (red line) for tumour samples collected at 10 and 24 h after the first oral dose of **AMPTX-1-*ent-2*** (50 mg/kg).

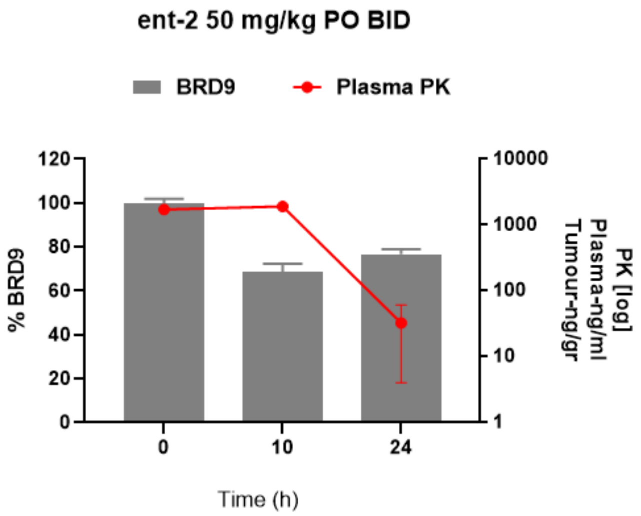


**Supplementary table 1**

In vivo PK data for **AMPTX-1-*ent-1* and AMPTX-1-*ent-2*** dosed intravenously (1 mg/kg) or orally (10 mg/kg) in male CD-1 Mice (n = 3 per time point). IV formulation: 5% DMSO, 95% (15%) HPBCD; PO formulation: 5% DMSO, 5% Solutol, 90% (15%) HPBCD.

| | AMPTX-1-*ent-1* | | AMPTX-1-*ent-2* | |
|---|---|---|---|---|
| | 1 mg/kg (IV) | 10 mg/kg (PO) | 1 mg/kg (IV) | 10 mg/kg (PO) |
| **C_max (ng/mL)** | 878 ± 115 | 508 ± 189 | 1120 ± 103 | 1343 ± 306 |
| **AUC last (h*ng/mL)** | 540 ± 102 | 1056 ± 289 | 744 ± 91 | 2209 ± 345 |
| **Cl (mL/min/Kg)** | 31 ± 6 | - | 23 ± 2.97 | - |
| **t ½ (h)** | 1.9 ± 0.26 | 1.4 ± 0.12 | 0.94 ± 0.10 | 1.1 ± 0.12 |
| **F (%)** | - | 20 | - | 30 |
| **Free fraction (%)** | 0.6 | | 0.5 | |

**Supplementary table 2**

Global proteomics data of **AMPTX-1** treated MV4-11 cells. Intensity values following normalisation are shown. Fold changes and P-values are displayed from Limma analysis.

**Supplementary table 3**

Co-immunoprecipitation mass spectrometry data from **AMPTX-1** and compound **2** treated HEK293 BRD9-HiBiT KI LgBit cells. Intensity values following normalisation are shown. Fold changes and P-values are displayed from Limma analysis.

**Supplementary table 4**

Peptide digest mass spectrometry data of DCAF16-DDB1 treated with **AMPTX-1** in the presence and absence of $BRD9^{BD}$.
