## Supplementary table 2 for "Selective degradation of BRD9 by a DCAF16-recruiting targeted glue: mode of action elucidation and in vivo proof of concept"

| Uniprot Accession | Protein name | DMSO.6h_1_abundance_norm | DMSO.6h_2_abundance_norm | DMSO.6h_3_abundance_norm | AMPH-04647.6h_1_abundance_norm | AMPH-04647.6h_2_abundance_norm |
|---|---|---|---|---|---|---|
| Q9H8M2 | BRD9 | 21.1263972 | 20.9992843 | 20.9339317 | 20.8648324 | 21.1342087 |
| P11926 | ODC1 | 18.47111 | 18.6421207 | 18.5321291 | 18.2967679 | 18.4739786 |
| Q02556 | IRF8 | 22.0637781 | 22.1361385 | 21.9807427 | 21.8529176 | 21.952769 |
| P10242 | MYB | 19.4056325 | 19.5245527 | 19.313945 | 19.2535273 | 19.3829871 |
| Q9HAI6 | TASL | 19.6287353 | 19.6054017 | 19.6854557 | 19.6669704 | 19.5489552 |
| Q8NFL0 | B3GNT7 | 20.9443557 | 20.694379 | 20.8342507 | 20.7905703 | 20.6282371 |
| Q16678 | CYP1B1 | 17.6888019 | 17.8182889 | 17.9750254 | 17.3892411 | 17.7306247 |
| O14880 | MGST3 | 23.1783138 | 23.2209839 | 23.2155075 | 23.039254 | 23.0557379 |
| P61129 | ZC3H6 | 14.7797735 | 15.7143877 | 15.3216867 | 16.2037842 | 15.5193587 |
| Q86WC6 | PPP1R27 | 21.9787793 | 21.9971072 | 21.5104046 | 21.685992 | 21.4978462 |
| Q14409 | GK3 | NA | 16.7959975 | 16.7553309 | NA | 16.9327877 |
| Q6N075 | MFSD5 | 18.3195103 | 18.5992768 | 18.5948511 | 18.2696025 | 18.3026267 |
| Q86UU0 | BCL9L | 15.558826 | 15.7527168 | 15.7796083 | 16.6580893 | 16.0329368 |
| P81605 | DCD | 21.9167856 | 21.9984704 | 19.116115 | 19.87521 | 19.4948703 |
| Q96S97 | MYADM | 23.3899194 | 23.2028925 | 23.4355835 | 23.1211786 | 23.1681738 |
| P20702 | ITGAX | 13.8936724 | NA | NA | 14.4019473 | NA |
| Q7RTV5 | PRXL2C | NA | 14.7159428 | 15.2271803 | NA | 15.2061085 |
| Q8TAA9 | VANGL1 | 15.655241 | NA | NA | 15.8382659 | NA |
| P05089 | ARG1 | 23.2613145 | 23.1684431 | 22.8651556 | 22.8673827 | 22.2997085 |
| Q96QS1 | TSPAN32 | 16.7007555 | 17.0937115 | 17.2614844 | 17.042205 | 16.6479445 |
| Q56A73 | SPIN4 | 17.6597769 | 17.8098233 | 17.616653 | 17.6292386 | 17.8173791 |
| Q9NQG7 | HPS4 | 17.4147439 | 17.4583336 | 17.3190941 | 17.4145577 | 17.3639606 |
| P48651 | PTDSS1 | 21.2797927 | 21.2309074 | 21.4811492 | 21.179711 | 21.2652572 |
| Q5XKP0 | MICOS13 | 22.74593 | 22.6600691 | 22.8177105 | 22.6452897 | 22.7028603 |
| O43677 | NDUFC1 | 20.1720521 | 20.2208888 | 20.1975399 | 19.9630016 | 19.9589656 |
| P36941 | LTBR | 20.5607318 | 20.6589579 | 20.5256712 | 20.6123597 | 20.6029683 |
| O15156 | ZBTB7B | 20.2680794 | 20.4745731 | 20.4279916 | 20.4465607 | 20.3777173 |
| Q7Z5Q1 | CPEB2 | 16.7099881 | 16.8670909 | 17.0077502 | 16.8914361 | 16.9382804 |
| Q8TED4 | SLC37A2 | 18.0558992 | 18.1151395 | 18.1012353 | 17.977145 | 18.2501362 |
| P15104 | GLUL | 21.0357456 | 20.9985685 | 20.8750673 | 20.8355324 | 20.8115504 |
| P03886 | MT-ND1 | 19.7112287 | 19.5270932 | 19.6539674 | 19.5105345 | 19.3737286 |
| P10415 | BCL2 | 21.3632012 | 21.1400069 | 21.0297273 | 20.9921751 | 21.0174403 |
| Q96MD2 | KICS2 | 17.0112519 | 16.8664963 | 17.5828048 | 16.8595895 | 16.4260874 |
| Q8NA72 | POC5 | 19.5284698 | 19.5364104 | 19.6288745 | 19.6274783 | 19.66423 |
| Q5VUA4 | ZNF318 | 19.6504508 | 19.7261352 | 19.6191809 | 19.6250376 | 19.7896706 |
| Q9Y3A4 | RRP7A | 20.5152194 | 20.5967184 | 20.6771901 | 20.4374796 | 20.5201241 |
| Q9H967 | WDR76 | 20.0196058 | 19.9030534 | 19.9365983 | 19.9500088 | 19.9955703 |
| Q8IX21 | SLF2 | 15.6659924 | 14.8521997 | 14.3830629 | 16.0726264 | 14.6275773 |

| | | | | | | |
|---|---|---|---|---|---|---|
| Q6UWP8 | SBSN | 17.2347316 | 16.6636545 | 15.5186989 | 15.524979 | 15.0401312 |
| P53999 | SUB1 | 25.1826055 | 25.1904549 | 25.114056 | 25.0838358 | 25.3118399 |
| Q96G21 | IMP4 | 20.8727083 | 20.9941835 | 20.8591599 | 20.7311318 | 20.9332677 |
| Q9NP58 | ABCB6 | 13.5158528 | 14.2575937 | 14.0665859 | 13.6865075 | 13.4060324 |
| P10620 | MGST1 | 21.426111 | 21.1990414 | 21.5630551 | 21.2119499 | 21.3524777 |
| P17707 | AMD1 | 17.0823273 | 17.1814605 | 16.8928068 | 17.1296439 | 17.0937885 |
| Q9UJJ7 | RPUSD1 | NA | 15.9303064 | 16.070039 | NA | 15.8628267 |
| Q6T4P5 | PLPPR3 | 20.4649348 | 20.3765993 | 20.2782177 | 20.3825524 | 20.2413932 |
| P62244 | RPS15A | 26.8162678 | 26.6757511 | 26.9181727 | 26.5109154 | 26.761941 |
| O43760 | SYNGR2 | 22.7882505 | 22.8122393 | 22.7777446 | 22.840667 | 22.7953088 |
| Q9HBI0 | PARVG | 21.1202379 | 21.1068126 | 21.1657546 | 20.9920693 | 20.9124303 |
| Q8WWK9 | CKAP2 | 20.5388536 | 20.6262142 | 20.5608026 | 20.7037186 | 20.6974993 |
| O43293 | DAPK3 | 18.4960601 | 18.4927004 | 18.5923139 | 18.5543795 | 18.5976786 |
| P14854 | COX6B1 | 24.9129806 | 24.9921193 | 24.876254 | 24.994393 | 25.0392647 |
| Q9Y608 | LRRFIP2 | 19.2015147 | 19.2797672 | 19.2908288 | 19.3466181 | 19.3481916 |
| O95789 | ZMYM6 | 14.4992047 | 14.7614824 | 14.5176256 | 14.6646379 | 14.5689634 |
| O75093 | SLIT1 | 19.0320798 | 18.1455711 | 17.6687545 | 19.0347744 | 18.2545747 |
| Q9BQE4 | SELENOS | 18.8075404 | 18.8661232 | 18.8770096 | 18.7968472 | 18.7156547 |
| Q6NXT4 | SLC30A6 | 18.6335241 | 18.8005525 | 18.7003435 | 18.5160295 | 18.5250418 |
| P62269 | RPS18 | 26.7426081 | 26.6837338 | 26.8172168 | 26.5039499 | 26.5664117 |
| Q96EB6 | SIRT1 | 20.1910232 | 20.337636 | 20.153206 | 20.259442 | 20.3297567 |
| Q9UBB5 | MBD2 | 21.9443682 | 22.1567056 | 21.9566208 | 22.2465976 | 22.2148679 |
| A0AV96 | RBM47 | 16.2318687 | 16.8611198 | 16.2179417 | 16.3254558 | 15.8183306 |
| Q08117 | TLE5 | 18.1804987 | 18.2158542 | 18.2841159 | 18.2625544 | 18.2372215 |
| P55851 | UCP2 | 13.7591678 | NA | NA | 14.2667239 | NA |
| Q96FZ5 | CMTM7 | 14.2236343 | 14.5322244 | 14.5834256 | 14.3591469 | 14.1052537 |
| Q92546 | RGP1 | 17.6311112 | 17.1384796 | 17.4216541 | 17.0369188 | 16.9948199 |
| Q9NR96 | TLR9 | 18.9033803 | 19.3850872 | 19.1311468 | 19.214515 | 19.0381601 |
| P33552 | CKS2 | 20.5291336 | 20.4037525 | 20.3945535 | 20.4791637 | 20.6552781 |
| Q9Y6G3 | MRPL42 | 20.7607752 | 20.9735466 | 20.8252436 | 20.9402244 | 21.1205078 |
| P49207 | RPL34 | 25.4015493 | 25.3113096 | 25.535579 | 25.3093395 | 25.2680449 |
| P16150 | SPN | 24.80037 | 24.7474359 | 24.8544832 | 24.6837541 | 24.8060161 |
| Q8IWB7 | WDFY1 | 22.2712915 | 22.1843026 | 22.2522385 | 22.158975 | 22.1808889 |
| Q86TW2 | ADCK1 | 17.3285802 | 17.158167 | 17.1934027 | 17.3870897 | 17.3215179 |
| Q9NWS9 | ZNF446 | NA | 15.2765586 | 15.2254746 | NA | 15.1537906 |
| Q53EP0 | FNDC3B | 19.7228625 | 19.752678 | 19.7886582 | 19.6707958 | 19.6877062 |
| Q5H9F3 | BCORL1 | 16.33908 | 16.8798057 | 16.5238632 | 16.610116 | 16.7728025 |
| Q16873 | LTC4S | 22.3503694 | 22.2604129 | 22.1379156 | 22.0824966 | 22.0232918 |
| Q6ZT21 | TMPPE | 19.2961969 | 19.4818262 | 19.4790472 | 19.1138455 | 19.1324603 |
| O96015 | DNAL4 | 17.210732 | 17.39649 | 17.2306968 | 17.1830713 | 17.3665854 |
| P35408 | PTGER4 | 16.4630505 | NA | NA | 16.7780168 | NA |
| Q9UKG9 | CROT | 17.0955663 | 17.4506892 | 17.5751878 | 17.1589722 | 16.8853495 |

| | | | | | | |
|---|---|---|---|---|---|---|
| Q96PC5 | MIA2 | 20.2472768 | 20.3199785 | 20.2158755 | 20.276374 | 20.3827397 |
| Q5THK1 | PRR14L | 17.6338202 | 17.7097637 | 17.7155272 | 17.807045 | 17.8905704 |
| Q8IWD4 | CCDC117 | 18.3221601 | 18.642447 | 18.4054531 | 18.5254014 | 18.5368014 |
| Q9P2B4 | CTTNBP2NL | 17.7844222 | 18.0578947 | 17.9809875 | 17.9775361 | 18.1884698 |
| A0A1W2PR95 | IGBP1C | NA | 16.144937 | 15.8174167 | NA | 16.1981206 |
| Q96ES6 | MFSD3 | 19.7735046 | 20.0538584 | 19.6589125 | 19.7453855 | 19.7192432 |
| P62280 | RPS11 | 26.3539093 | 26.2591007 | 26.2906154 | 26.1258627 | 26.1519968 |
| Q00534 | CDK6 | 22.8504411 | 22.7772921 | 22.7142075 | 22.7253473 | 22.6803446 |
| Q9BYN7 | ZNF341 | NA | 11.7267259 | 11.4043533 | NA | 11.7618557 |
| P01040 | CSTA | 23.4711155 | 22.4578237 | 23.7633321 | 22.4877429 | 21.5957528 |
| Q58FF6 | HSP90AB4P | 23.7004444 | 23.4145356 | 23.0236142 | 22.7816065 | 22.8603865 |
| Q13509 | TUBB3 | 22.6998474 | 22.6041926 | 22.6899578 | 22.4574677 | 22.3972164 |
| P62249 | RPS16 | 26.6247139 | 26.4450974 | 26.6998332 | 26.3998363 | 26.3481961 |
| Q8N653 | LZTR1 | 19.246516 | 19.2486311 | 19.1217899 | 19.0964999 | 19.0136754 |
| Q6GTX8 | LAIR1 | 21.4594025 | 21.649153 | 21.3935153 | 21.4368561 | 21.4138632 |
| Q9BRJ7 | NUDT16L1 | 21.0862774 | 21.1115661 | 21.1588405 | 21.1040736 | 21.117022 |
| O00748 | CES2 | 14.8705392 | 14.9841445 | 15.2076936 | 15.1311824 | 15.1831305 |
| Q96MT8 | CEP63 | 17.4702674 | 17.5823814 | 17.2716198 | 17.550176 | 17.7204795 |
| Q8IUD6 | RNF135 | 17.2599634 | 17.1337315 | 17.349572 | 17.4448601 | 17.4312337 |
| C9JLW8 | MCRIP1 | 20.2477952 | 19.986893 | 20.1328393 | 20.0871888 | 20.5401999 |
| Q969W3 | FAM104A | 14.5990229 | 14.656122 | 14.6587491 | 14.7595809 | 14.7802948 |
| Q9Y664 | KPTN | 16.9988536 | 17.1644133 | 16.9154501 | 16.4873876 | 16.823716 |
| P13591 | NCAM1 | 12.9461764 | NA | NA | 13.0300038 | NA |
| Q86WC4 | OSTM1 | 18.6688718 | 18.7436652 | 18.8624396 | 18.8031602 | 18.9690865 |
| P56373 | P2RX3 | NA | 18.0319734 | 17.4081743 | NA | 17.6620343 |
| Q659C4 | LARP1B | 18.6761839 | 18.6814923 | 18.745891 | 18.74301 | 18.8242002 |
| P26651 | ZFP36 | 14.3958659 | NA | NA | 14.5904022 | NA |
| Q9P2K6 | KLHL42 | 16.536382 | 16.5175888 | 16.7042686 | 16.3283495 | 16.4252239 |
| Q8N4H5 | TOMM5 | 23.9459334 | 24.0620714 | 23.9485513 | 23.7812153 | 23.8208283 |
| Q6P589 | TNFAIP8L2 | 21.0995919 | 21.330057 | 21.1847858 | 21.1487753 | 21.0789575 |
| P15529 | CD46 | 19.794544 | 19.8914927 | 19.9976509 | 19.7346693 | 19.8087231 |
| Q00325 | SLC25A3 | 25.9505682 | 25.8767719 | 25.9996537 | 25.8047017 | 25.8987691 |
| Q9ULH1 | ASAP1 | 18.1157586 | 18.0988837 | 18.1578427 | 18.2209991 | 18.2511966 |
| Q6ZSR9 | NA | 18.7002195 | 18.790816 | 18.808656 | 18.8727269 | 18.7489545 |
| Q15428 | SF3A2 | 22.1689509 | 22.279655 | 22.3247594 | 22.3518297 | 22.3728734 |
| Q5T3J3 | LRIF1 | 19.0574807 | 19.1274156 | 19.2222206 | 19.1140116 | 19.3971156 |
| Q9GZX9 | TWSG1 | NA | 15.0067264 | 14.6943385 | NA | 15.4267825 |
| Q9Y5G9 | PCDHGA4 | NA | 17.9241769 | 17.7845979 | NA | 18.0327516 |
| Q5JTW2 | CEP78 | 19.0091457 | 18.7220981 | 19.0229014 | 18.8660633 | 19.1331555 |
| O00232 | PSMD12 | 23.9172721 | 23.8505198 | 23.9003199 | 23.749637 | 23.8187987 |
| O00483 | NDUFA4 | 23.9274821 | 23.8359389 | 23.9325555 | 23.73668 | 23.7620393 |
| P35221 | CTNNA1 | 20.3561794 | 20.3390712 | 20.3413013 | 20.4019957 | 20.4017045 |

| | | | | | | |
|---|---|---|---|---|---|---|
| P83876 | TXNL4A | 21.2127874 | 21.2022704 | 21.245016 | 21.2369675 | 21.3047373 |
| Q9H5K3 | POMK | 19.6029769 | 19.5053097 | 19.699917 | 19.7030195 | 19.853415 |
| Q8IWJ2 | GCC2 | 20.1730314 | 20.3435768 | 20.1834272 | 20.2602866 | 20.3154208 |
| Q9BSK0 | MARVELD1 | 18.6369133 | 18.6118789 | 18.5499355 | 18.843262 | 18.7724636 |
| Q9NPD3 | EXOSC4 | 21.4695113 | 21.5751365 | 21.512679 | 21.5121855 | 21.8673181 |
| Q96Q07 | BTBD9 | 19.9277982 | 19.9888393 | 20.0902366 | 19.9263885 | 19.9285026 |
| Q8N2K1 | UBE2J2 | 18.7856169 | 18.5250476 | 18.6125904 | 18.4363077 | 18.4981622 |
| Q86X95 | CIR1 | 19.1662573 | 19.1251393 | 19.2737856 | 19.1209084 | 19.1004963 |
| P40937 | RFC5 | 21.9128378 | 21.924204 | 22.0483411 | 21.8363158 | 21.8436294 |
| Q16533 | SNAPC1 | 17.2605863 | 17.6202824 | 17.725085 | 17.1967597 | 16.9733145 |
| Q8IVL0 | NAV3 | 19.1917095 | 19.3668127 | 19.2751892 | 19.1972052 | 19.2307435 |
| Q15386 | UBE3C | 22.638496 | 22.5997114 | 22.7431464 | 22.4865095 | 22.583008 |
| Q9NYH9 | UTP6 | 21.5877823 | 21.5674018 | 21.6170883 | 21.4683467 | 21.545018 |
| Q9Y572 | RIPK3 | 19.1028185 | 19.0103949 | 19.0812689 | 19.1311151 | 19.1420623 |
| Q9NZM4 | BICRA | 17.1707578 | 17.2680022 | 17.0418349 | 17.301548 | 17.2806235 |
| P58107 | EPPK1 | 13.318731 | NA | NA | 13.8182474 | NA |
| P32249 | GPR183 | 16.3579151 | 16.6237953 | 16.4470914 | 16.1674413 | 16.3532151 |
| A6NNL5 | C15orf61 | 17.8128826 | 17.6462367 | 17.6907287 | 17.8294572 | 17.5000377 |
| B4DJY2 | TMEM233 | 19.4833785 | 19.4311137 | 19.3661644 | 19.4080731 | 19.3462214 |
| P59998 | ARPC4 | 25.7782672 | 25.5994962 | 25.6708921 | 25.4287669 | 25.5699391 |
| Q5MNZ6 | WDR45B | 19.312491 | 19.3112282 | 19.3734949 | 19.2060337 | 19.2753247 |
| Q9Y320 | TMX2 | 21.6523808 | 21.64361 | 21.8153286 | 21.6826975 | 21.7032082 |
| Q7Z569 | BRAP | 20.4542992 | 20.5079267 | 20.510956 | 20.3864058 | 20.5534727 |
| P82933 | MRPS9 | 22.4058975 | 22.4453795 | 22.4979905 | 22.3884141 | 22.4105092 |
| Q969S9 | GFM2 | 20.3555779 | 20.3880112 | 20.3421037 | 20.3555025 | 20.5012072 |
| O60508 | CDC40 | 20.7472491 | 20.7049433 | 20.7841252 | 20.7556197 | 20.916586 |
| Q9UHY1 | NRBP1 | 20.6309494 | 20.7840213 | 20.6015484 | 20.7334629 | 20.8460124 |
| Q9Y3X0 | CCDC9 | 20.4439461 | 20.5880461 | 20.3417441 | 20.5641073 | 20.6267711 |
| Q9NX08 | COMMD8 | 21.2309115 | 21.2629605 | 21.4106885 | 21.3721657 | 21.4688097 |
| Q6Y288 | B3GLCT | 17.6976349 | 17.9129657 | 17.7679047 | 17.881931 | 18.0592247 |
| O14647 | CHD2 | 18.4794205 | 18.5303992 | 18.5488743 | 18.4861209 | 18.6589477 |
| Q53FT3 | HIKESHI | 20.2662423 | 20.1095331 | 20.2717939 | 20.3137957 | 20.521684 |
| Q8TD26 | CHD6 | 17.3132223 | 17.4507822 | 17.2718406 | 17.4850466 | 17.5590494 |
| Q86UA6 | RPAIN | 17.6200829 | 17.5596213 | 17.5371908 | 17.789793 | 17.7303465 |
| Q63ZY3 | KANK2 | 19.2776244 | 19.4071055 | 19.5060699 | 19.501711 | 19.6466451 |
| Q9BQ69 | MACROD1 | 19.4936761 | 19.4339856 | 19.5112844 | 19.3977837 | 19.6211956 |
| Q9HC77 | CENPJ | 16.2289309 | 16.3957288 | 16.5296462 | 16.4750642 | 16.4732898 |
| Q9UII2 | ATP5IF1 | 23.7809887 | 23.6256093 | 23.5478985 | 23.7988569 | 23.8351294 |
| Q9H254 | SPTBN4 | 24.0603268 | 24.2160884 | 23.8035368 | 24.0749822 | 24.2357553 |
| Q96HU1 | SGSM3 | 16.6795317 | NA | NA | 16.7487846 | NA |
| Q9Y2F5 | ICE1 | 15.0936313 | NA | NA | 15.7723038 | NA |
| Q9ULQ0 | STRIP2 | 15.867096 | NA | NA | 15.8608002 | NA |

| | | | | | | |
|---|---|---|---|---|---|---|
| Q96BV0 | ZNF775 | 15.8188056 | NA | NA | 16.2868168 | NA |
| P54753 | EPHB3 | 13.2491567 | NA | NA | 13.2839927 | NA |
| Q5SVZ6 | ZMYM1 | 13.628091 | NA | NA | 13.9750042 | NA |
| Q8NCF5 | NFATC2IP | 18.5426579 | 18.5637802 | 18.5437269 | 18.5283908 | 18.7420852 |
| P67812 | SEC11A | 23.6446747 | 23.6950951 | 23.5578156 | 23.4539887 | 23.6079956 |
| Q6DKI1 | RPL7L1 | 21.9572586 | 21.9217796 | 21.9387503 | 21.8439241 | 21.8297911 |
| Q16881 | TXNRD1 | 23.1518064 | 23.1308411 | 23.1438112 | 23.0516987 | 23.0573408 |
| Q9BX63 | BRIP1 | 16.9898929 | 17.1700397 | 17.3060902 | 17.1543689 | 17.2978648 |
| Q13277 | STX3 | 18.400437 | 18.5250473 | 18.5488037 | 18.7692209 | 18.8933568 |
| Q14257 | RCN2 | 20.4144613 | 20.6981694 | 20.279986 | 20.6249438 | 20.7249825 |
| Q8NI08 | NCOA7 | 15.2948462 | 15.6140695 | 15.2798351 | 16.0344465 | 15.2675747 |
| Q96JK4 | HHIPL1 | 13.9173228 | 14.6167164 | 14.1913974 | 13.8161546 | 13.9451442 |
| Q92968 | PEX13 | 17.926336 | 17.8626824 | 17.8034277 | 17.7454219 | 17.6133757 |
| O00762 | UBE2C | 21.3958423 | 21.2391537 | 21.3632703 | 21.2842255 | 21.2549949 |
| O15056 | SYNJ2 | 17.100201 | 17.1536363 | 17.1339208 | 17.0952676 | 17.2180175 |
| P52739 | ZNF131 | 15.6667128 | 16.0269972 | 16.2106334 | 16.1303212 | 16.1459914 |
| O95059 | RPP14 | 19.73861 | 19.6635405 | 19.7404681 | 19.5043938 | 19.5672429 |
| Q96A37 | RNF166 | NA | 15.6768399 | 15.6728274 | NA | 16.2016356 |
| A0AVI4 | TMEM129 | 15.2019283 | NA | NA | 15.0205904 | NA |
| P23610 | NA | 19.3124724 | 19.5746726 | 19.4837139 | 19.2756101 | 19.4428756 |
| O14965 | AURKA | 18.6764814 | 18.7050408 | 18.7254485 | 18.6189303 | 18.5425338 |
| O43747 | AP1G1 | 22.9732271 | 22.9744769 | 23.0326672 | 22.8571521 | 22.911844 |
| Q7Z2W9 | MRPL21 | 21.8355484 | 21.8163243 | 21.8916942 | 21.7193924 | 21.734662 |
| Q9NWK9 | ZNHIT6 | 18.8117287 | 18.9182973 | 18.7789428 | 18.8902582 | 19.0340348 |
| Q9UBS8 | RNF14 | 19.6017975 | 19.5355476 | 19.7897588 | 19.6938582 | 19.6943769 |
| P15336 | ATF2 | 19.662437 | 20.017827 | 19.6299746 | 19.8717466 | 20.0146344 |
| Q9P1T7 | MDFIC | 15.2887916 | NA | NA | 15.6321374 | NA |
| O75398 | DEAF1 | 14.6439971 | NA | NA | 15.0191961 | NA |
| Q96DX7 | TRIM44 | 18.1029217 | 18.1156057 | 18.0853356 | 18.2576749 | 18.1909237 |
| Q6ZNK6 | TIFAB | 17.8815825 | 17.8910354 | 18.5439279 | 17.7472518 | 18.2932455 |
| Q6P5R6 | RPL22L1 | 22.5922346 | 22.595987 | 22.5779152 | 22.4676364 | 22.3231566 |
| Q8IZF4 | ADGRG5 | 19.2256715 | 19.6575379 | 19.5074103 | 19.0809582 | 19.064279 |
| P60981 | DSTN | 22.266627 | 22.3957822 | 22.2426255 | 22.3379393 | 22.4791876 |
| P34949 | MPI | 21.3925588 | 21.4045355 | 21.3815913 | 21.3647544 | 21.4674614 |
| P08246 | ELANE | 24.7161573 | 24.2381264 | 24.716481 | 24.3576623 | 24.3976705 |
| Q9HAU8 | RNPEPL1 | 14.9739599 | NA | NA | 15.0927607 | NA |
| A9UHW6 | MIF4GD | 20.6763316 | 20.7493595 | 20.5819146 | 20.6929327 | 20.7588321 |
| Q8N108 | MIER1 | 20.3813485 | 20.3217214 | 20.4043557 | 20.4360278 | 20.4338025 |
| Q8NDA2 | HMCN2 | NA | 15.2214619 | 15.0453222 | NA | 15.062082 |
| O96020 | CCNE2 | 15.2642848 | 15.133091 | 15.3845943 | 15.5962962 | 15.4760075 |
| Q14520 | HABP2 | 11.528158 | NA | NA | 11.5620306 | NA |
| Q13323 | BIK | NA | 14.2926892 | 14.2168503 | NA | 14.257976 |

| | | | | | | |
|---|---|---|---|---|---|---|
| Q14344 | GNA13 | 22.3125264 | 22.2631348 | 22.2880775 | 22.1258583 | 22.1907808 |
| O14508 | SOCS2 | NA | 15.5090524 | 15.5161923 | NA | 15.550052 |
| Q9NQE9 | HINT3 | 19.4816986 | 19.4690186 | 19.3985252 | 19.2538936 | 19.3983069 |
| O95425 | SVIL | 18.5375777 | 18.7834464 | 18.7546301 | 18.6643254 | 18.8064704 |
| Q6ZMG9 | CERS6 | 21.6845163 | 21.5249623 | 21.6436079 | 21.4071428 | 21.5308281 |
| Q15643 | TRIP11 | 22.2852037 | 22.296199 | 22.2119298 | 22.2940802 | 22.4242115 |
| Q6UXZ4 | UNC5D | NA | 14.7008612 | 14.8618339 | NA | 14.2984034 |
| Q9NY74 | ETAA1 | 14.1202446 | NA | NA | 13.8151306 | NA |
| Q9NY12 | GAR1 | 22.8961514 | 23.0556434 | 22.9916349 | 22.7691632 | 22.7855653 |
| P51159 | RAB27A | 23.231102 | 23.3708182 | 23.2329664 | 23.1288099 | 23.078393 |
| P60763 | RAC3 | 19.7164649 | 19.8788794 | 19.9942947 | 19.6036103 | 19.8637393 |
| P27635 | RPL10 | 26.7049078 | 26.7217981 | 26.7661859 | 26.5766007 | 26.5698083 |
| P00749 | PLAU | 18.7184086 | 18.938347 | 18.9020719 | 18.7956327 | 18.8957823 |
| Q02543 | RPL18A | 26.0867633 | 26.0375003 | 26.156024 | 25.8638287 | 25.9026212 |
| P30307 | CDC25C | 18.6944064 | 18.7641383 | 18.795044 | 18.9479702 | 18.9268687 |
| O60346 | PHLPP1 | 19.6109094 | 19.7272314 | 19.8267033 | 19.5681095 | 19.7636211 |
| P30679 | GNA15 | 21.0099841 | 21.0317445 | 21.0809446 | 20.9763026 | 20.9854841 |
| Q14CS0 | UBXN2B | 17.4364026 | 17.6261508 | 17.3262575 | 17.4092763 | 17.5488091 |
| Q9ULD5 | ZNF777 | 15.1468648 | 15.9195801 | 15.5221897 | 15.2104652 | 15.017912 |
| Q9NXB0 | MKS1 | 15.3964603 | 15.4549838 | 15.4188715 | 15.7939547 | 15.4473871 |
| Q9H7X0 | NAA60 | NA | 15.9196979 | 15.7789833 | NA | 15.3982919 |
| Q9NTM9 | CUTC | 20.3055906 | 20.4873371 | 20.3246884 | 20.0997119 | 20.3874819 |
| O00587 | MFNG | 18.0222366 | 18.0716612 | 18.13332 | 18.1095732 | 17.9953564 |
| P35268 | RPL22 | 26.1707059 | 26.105969 | 26.0693241 | 25.8934889 | 25.9615253 |
| Q9Y3B3 | TMED7 | 23.5302551 | 23.5330747 | 23.54906 | 23.4386406 | 23.4320992 |
| Q15025 | TNIP1 | 19.4806802 | 19.447062 | 19.4801014 | 19.592122 | 19.4930361 |
| Q8NEM2 | SHCBP1 | 20.6192795 | 20.6734093 | 20.7224045 | 20.6318996 | 20.7605599 |
| Q9NS00 | C1GALT1 | 17.3342519 | 17.3462592 | 17.4601788 | 17.5226806 | 17.3671787 |
| Q13901 | C1D | 20.365186 | 20.4669527 | 20.6350226 | 20.4922734 | 20.6715878 |
| Q99643 | SDHC | 22.2480489 | 22.2721182 | 22.5159962 | 22.4843396 | 22.655569 |
| P51808 | DYNLT3 | 17.5007341 | 17.5454013 | 17.2706447 | 17.5801783 | 17.5485564 |
| Q9Y3D2 | MSRB2 | 17.2818576 | 17.2523022 | 17.4617193 | 17.6094898 | 17.7411881 |
| Q8WVX9 | FAR1 | 21.1792241 | 21.0559218 | 21.2178952 | 21.050762 | 21.0788744 |
| Q99715 | COL12A1 | NA | 16.0297068 | 16.2148145 | NA | 15.9965216 |
| Q02978 | SLC25A11 | 22.7420147 | 22.8699605 | 22.9613881 | 22.6889867 | 22.7648459 |
| Q9NNW7 | TXNRD2 | 20.5833154 | 20.5775723 | 20.4813096 | 20.5554356 | 20.606105 |
| P18510 | IL1RN | 17.9915633 | 17.9076558 | 18.1192657 | 18.0662667 | 18.2139644 |
| P07996 | THBS1 | 15.9612712 | 15.9008216 | 15.3276143 | 16.0132449 | 15.7767318 |
| Q9Y5Z9 | UBIAD1 | 18.0883228 | 18.4026733 | 18.2068948 | 18.4381848 | 18.2415826 |
| Q9Y6V0 | PCLO | NA | 16.6241636 | 16.4260032 | NA | 15.9294774 |
| P01106 | MYC | 17.8484316 | 18.1488989 | 17.6512348 | 17.5874672 | 17.7265163 |
| Q86XT9 | TMEM219 | 18.0299879 | 17.8086127 | 17.8256064 | 17.5983665 | 17.9569614 |

| | | | | | | |
|---|---|---|---|---|---|---|
| L0R6Q1 | SLC35A4 | 22.0386945 | 21.8938686 | 22.1943309 | 21.8893992 | 22.0027957 |
| P10321 | HLA-C | 23.2218318 | 23.1835509 | 23.1662666 | 23.0536966 | 23.1097579 |
| Q9Y2Z2 | MTO1 | 19.7133289 | 19.6421083 | 19.7838852 | 19.6975327 | 19.8298834 |
| P22033 | MMUT | 20.8349539 | 20.9519734 | 20.8620583 | 20.9057707 | 21.0572314 |
| Q96HR3 | MED30 | 19.6124437 | 19.8027507 | 19.9158349 | 19.9060482 | 20.0922728 |
| Q9Y2V0 | CDIN1 | 16.4109684 | 16.3404809 | 16.6388405 | 16.782559 | 16.6931203 |
| Q9H1N7 | SLC35B3 | 17.0181608 | 17.4622141 | 17.3488781 | 16.9202843 | 17.3941687 |
| O14735 | CDIPT | 22.3315617 | 22.1213692 | 22.0832761 | 21.8879703 | 21.9364201 |
| P23396 | RPS3 | 26.581183 | 26.5834558 | 26.6060018 | 26.3882087 | 26.482136 |
| Q8IZD6 | SLC22A15 | 17.4682236 | 17.4378465 | 17.453927 | 17.3354521 | 17.5454904 |
| Q9NW15 | ANO10 | 18.4420904 | 18.3619018 | 18.3613844 | 18.4576584 | 18.5384252 |
| Q16822 | PCK2 | 19.9939396 | 20.1031473 | 20.0399963 | 19.9548663 | 20.0516934 |
| Q96N96 | SPATA13 | 14.5107642 | 14.2560092 | 14.8959985 | 14.4595121 | 14.9308275 |
| P55061 | TMBIM6 | 16.974292 | 17.0341723 | 17.2942016 | 16.642119 | 17.4070518 |
| Q8NBI2 | CYB561A3 | 17.3695731 | 17.1106457 | 17.1429876 | 17.3055406 | 17.1207386 |
| Q8N726 | CDKN2A | 18.1028083 | 17.7859311 | 17.685493 | 17.5133924 | 17.7412158 |
| Q9NX05 | FAM120C | 19.3671211 | 19.4187543 | 19.5842406 | 19.3450212 | 19.2850974 |
| Q6UWL2 | SUSD1 | 18.4810316 | 18.7848064 | 18.6143802 | 18.5961611 | 18.5338146 |
| O75964 | ATP5MG | 24.3293735 | 24.1814311 | 24.3099919 | 24.1105989 | 24.244636 |
| P50897 | PPT1 | 23.5437007 | 23.5381173 | 23.5415192 | 23.4467672 | 23.596688 |
| Q96GG9 | DCUN1D1 | 21.6597699 | 21.6596972 | 21.7410829 | 21.6358647 | 21.6377097 |
| Q9NUD5 | ZCCHC3 | 21.0155045 | 21.1355211 | 21.1643935 | 20.9728012 | 21.0282267 |
| Q9H490 | PIGU | 20.4651565 | 20.4056887 | 20.4694402 | 20.4083782 | 20.4084798 |
| Q7L945 | ZNF627 | 19.9150063 | 20.0073516 | 20.0621376 | 20.0017011 | 20.025841 |
| Q9HCM1 | RESF1 | 17.6065142 | 17.7301209 | 17.4031889 | 17.6269657 | 17.7554944 |
| Q2KHR2 | RFX7 | 16.4912989 | 16.4812414 | 16.86006 | 16.8745204 | 16.765295 |
| Q99576 | TSC22D3 | 11.073551 | NA | NA | 11.4504639 | NA |
| Q7L0J3 | SV2A | 16.118831 | 16.6385694 | 16.0049367 | 16.3683395 | 16.2419085 |
| P29122 | PCSK6 | 15.2780794 | NA | NA | 15.1928678 | NA |
| Q9Y2C4 | EXOG | 17.2338063 | 17.3873154 | 17.3302744 | 16.9675542 | 17.1713473 |
| P03905 | MT-ND4 | 20.3492027 | 20.4614407 | 20.7648093 | 20.3592297 | 20.5470477 |
| P42574 | CASP3 | 22.6833661 | 22.6841322 | 22.6731089 | 22.5284056 | 22.5101398 |
| Q9H7F4 | TMEM185B | 18.9276352 | 18.8901191 | 18.968566 | 18.8288978 | 18.824286 |
| Q969H6 | POP5 | 20.2596832 | 20.0680523 | 20.225826 | 19.9601967 | 20.0021258 |
| Q9ULT0 | TTC7A | 20.0353379 | 19.9728341 | 19.949435 | 19.8703832 | 19.9497799 |
| P10619 | CTSA | 23.5143483 | 23.3833542 | 23.4633895 | 23.4888047 | 23.4111604 |
| P54920 | NAPA | 24.0728074 | 24.0668157 | 24.0882334 | 23.9653489 | 23.9991301 |
| Q03188 | CENPC | 19.6042326 | 19.667014 | 19.7330778 | 19.6853085 | 19.7811514 |
| Q9H204 | MED28 | 19.9350017 | 20.1879261 | 20.218174 | 20.2273549 | 20.2786018 |
| Q99081 | TCF12 | 18.9392291 | 18.9837129 | 19.0565856 | 18.9933889 | 19.0889097 |
| A0A0U1RRE5 | NBDY | 19.1444632 | 19.1668868 | 19.1912349 | 19.1873723 | 19.4434014 |
| Q8N720 | ZNF655 | 16.5459391 | 16.701809 | 16.7516898 | 16.7210962 | 16.344383 |

| | | | | | | |
|---|---|---|---|---|---|---|
| Q9NYY3 | PLK2 | 10.9123969 | NA | NA | 11.6543051 | NA |
| Q9Y3C0 | WASHC3 | 20.3858483 | 20.3828449 | 20.4095449 | 20.3249989 | 20.3011312 |
| Q8N5C8 | TAB3 | NA | 11.8318433 | 12.0344734 | NA | 11.7090126 |
| Q53RE8 | ANKRD39 | 15.8019601 | 16.3196809 | 16.2929001 | 15.8772143 | 16.1194372 |
| Q9H857 | NT5DC2 | 21.6556885 | 21.6295194 | 21.8689875 | 21.5459202 | 21.7019195 |
| Q92979 | EMG1 | 21.9078289 | 21.8438073 | 21.927909 | 21.7912958 | 21.8649958 |
| Q15058 | KIF14 | 21.3056594 | 21.190679 | 21.379392 | 21.4498887 | 21.347606 |
| Q15119 | PDK2 | 16.7333949 | 16.8892227 | 17.0328603 | 16.9761858 | 17.0678379 |
| Q6UXH1 | CRELD2 | 21.0297175 | 21.2377795 | 20.8567543 | 21.1353971 | 21.2464374 |
| Q14592 | ZNF460 | 18.9244149 | 19.0839947 | 18.8394323 | 19.0519114 | 19.0874251 |
| Q9H0K1 | SIK2 | 16.2157634 | 16.1416806 | 16.5162911 | 16.3638692 | 16.421398 |
| O75629 | CREG1 | 16.8133941 | 16.4421423 | 16.8611746 | 16.8617094 | 16.8993767 |
| Q02413 | DSG1 | 17.5872468 | 16.8219669 | 17.509534 | 17.0046711 | 17.0454841 |
| P61604 | HSPE1 | 26.4742212 | 26.6198298 | 26.3618048 | 26.2571339 | 25.8236418 |
| O95456 | PSMG1 | 21.8765848 | 21.9337563 | 21.9941925 | 21.8307993 | 21.8725232 |
| P63173 | RPL38 | 23.928078 | 23.8779637 | 23.849448 | 23.7775986 | 23.7941428 |
| O14757 | CHEK1 | 19.4849412 | 19.598949 | 19.6564796 | 19.5846956 | 19.6936912 |
| Q02818 | NUCB1 | 21.0097241 | 21.1180655 | 20.9831839 | 21.0832326 | 21.1933548 |
| Q96BW1 | UPRT | 16.780169 | 17.0120889 | 17.1400886 | 16.7731041 | 17.0932847 |
| Q15051 | IQCB1 | 18.2213566 | 18.2678409 | 18.2298778 | 18.0714395 | 18.1391597 |
| Q86T24 | ZBTB33 | 20.8091161 | 20.6367743 | 20.6500744 | 20.7072401 | 20.5677084 |
| Q96AQ6 | PBXIP1 | 20.3432619 | 20.2790691 | 20.2292276 | 20.2907016 | 20.3117058 |
| Q53H96 | PYCR3 | 22.0962344 | 22.248871 | 22.2870231 | 22.2592734 | 22.3885999 |
| Q6IQ49 | SDE2 | 19.4810867 | 19.4867043 | 19.4362933 | 19.5692539 | 19.6801383 |
| Q16864 | ATP6V1F | 21.7517224 | 21.9018057 | 21.6940429 | 21.8021978 | 21.8352663 |
| P22083 | FUT4 | 19.8859574 | 19.6662575 | 19.6977837 | 19.6969761 | 19.8383068 |
| Q96G28 | CFAP36 | 16.2456772 | 16.4091277 | 16.2484939 | 16.0290441 | 16.204826 |
| P47974 | ZFP36L2 | 19.7787805 | 19.9883745 | 19.9084738 | 19.7770291 | 20.0134995 |
| Q12788 | TBL3 | 21.8260075 | 21.833853 | 21.8837366 | 21.7303704 | 21.7944492 |
| Q9BUE0 | MED18 | 20.1637726 | 20.1323814 | 20.2320435 | 20.214355 | 20.2780438 |
| Q96CD2 | PPCDC | 17.966568 | 18.0608412 | 18.0748091 | 18.0146548 | 18.1092845 |
| Q6UB28 | METAP1D | 17.6249488 | 17.6802428 | 17.7564549 | 17.7454359 | 17.9918271 |
| Q7Z5L2 | R3HCC1L | 17.9846518 | 18.2324247 | 18.1595103 | 18.1376543 | 18.3336979 |
| Q9H2D1 | SLC25A32 | 18.9102766 | 19.0533396 | 19.2478674 | 18.7991832 | 19.0257701 |
| P60228 | EIF3E | 24.2131588 | 24.2260761 | 24.2328419 | 24.0591891 | 24.1441314 |
| Q9H910 | JPT2 | 21.2458323 | 21.3231876 | 21.2441712 | 21.3048883 | 21.4350391 |
| Q8TAP9 | MPLKIP | 17.098608 | 17.4165477 | 17.131999 | 17.6819323 | 17.3857232 |
| P52756 | RBM5 | 20.5889233 | 20.6322348 | 20.5590077 | 20.5971315 | 20.6353997 |
| Q7Z2H8 | SLC36A1 | 18.4840945 | 18.123939 | 18.1901602 | 18.1792889 | 17.7941481 |
| Q9Y2D2 | SLC35A3 | 16.6294084 | 16.5105181 | 16.6304009 | 16.3730693 | 16.3741302 |
| Q6TFL4 | KLHL24 | NA | 11.9217069 | 11.8650801 | NA | 11.6538532 |
| P07204 | THBD | 21.068362 | 20.9912789 | 21.205475 | 21.0665895 | 21.0079241 |

| | | | | | | |
|---|---|---|---|---|---|---|
| Q15041 | ARL6IP1 | 23.701844 | 23.4540488 | 23.6377493 | 23.480253 | 23.4023755 |
| P02786 | TFRC | 24.8139106 | 24.5897204 | 24.6444629 | 24.6409376 | 24.5676302 |
| O15347 | HMGB3 | 25.1648808 | 25.1931611 | 25.1189846 | 25.1762431 | 25.2078008 |
| Q96FF9 | CDCA5 | 18.1040472 | 18.3637616 | 18.077048 | 18.2679295 | 18.4066793 |
| Q9NTW7 | ZFP64 | NA | 15.5537031 | 15.5738824 | NA | 15.8964934 |
| O95297 | MPZL1 | 18.5648484 | 18.4853116 | 18.5342973 | 18.3793381 | 18.4562751 |
| P55899 | FCGRT | 20.6970295 | 20.5256744 | 20.6796859 | 20.5867462 | 20.4982545 |
| Q9HC07 | TMEM165 | 20.380071 | 20.2822141 | 20.4324985 | 20.240902 | 20.2864426 |
| P14316 | IRF2 | 19.9815257 | 20.088363 | 20.0016905 | 20.1616476 | 20.2223626 |
| Q9NRW3 | APOBEC3C | 21.2334699 | 21.0891707 | 21.5216244 | 21.1494279 | 21.1707977 |
| Q9UK97 | FBXO9 | 18.1246188 | 18.0059331 | 18.1987455 | 17.8840461 | 18.0923076 |
| P24158 | PRTN3 | 26.1690914 | 26.0628082 | 26.0535902 | 26.0027358 | 25.9776125 |
| Q9P035 | HACD3 | 22.5989989 | 22.6365812 | 22.5253565 | 22.527699 | 22.4752884 |
| O95613 | PCNT | 21.1745564 | 21.314352 | 21.2499407 | 21.2595154 | 21.2866582 |
| O43252 | PAPSS1 | 20.6960207 | 20.8570722 | 20.8658706 | 20.86922 | 20.9768861 |
| Q9Y6G5 | COMMD10 | 20.3604974 | 20.3771735 | 20.4009288 | 20.3755764 | 20.551517 |
| Q02086 | SP2 | 18.6285886 | 18.8112591 | 18.8558388 | 18.8520943 | 18.8057038 |
| Q969G6 | RFK | 18.6860393 | 19.0791223 | 19.0169356 | 18.9969039 | 18.9611263 |
| Q12851 | MAP4K2 | 17.2208125 | 17.1491826 | 17.4309751 | 17.1328959 | 17.3791372 |
| Q9Y2Q5 | LAMTOR2 | 20.5813573 | 20.615405 | 20.6441942 | 20.5046295 | 20.7074671 |
| Q13637 | RAB32 | 22.0523865 | 21.9898976 | 22.0585995 | 21.8748155 | 21.9808613 |
| O95817 | BAG3 | 20.2043633 | 20.4044061 | 20.052777 | 20.2737746 | 20.3676637 |
| Q12968 | NFATC3 | 17.9555921 | 18.0611512 | 17.9267206 | 18.0267571 | 18.1843068 |
| Q96PY5 | FMNL2 | 15.6812314 | NA | NA | 15.4833342 | NA |
| Q9UBX3 | SLC25A10 | 23.2368468 | 23.1662449 | 23.2921251 | 23.1140235 | 23.217374 |
| P24928 | POLR2A | 21.9291018 | 21.9874853 | 21.9123709 | 21.9257166 | 21.9769018 |
| Q15043 | SLC39A14 | 21.3078805 | 21.3446964 | 21.2900637 | 21.1490642 | 21.2764729 |
| Q8ND30 | PPFIBP2 | 19.8077025 | 20.043137 | 19.8963374 | 20.0096386 | 20.1072417 |
| O43505 | B4GAT1 | NA | 11.4333172 | 11.0343333 | NA | 11.7984165 |
| Q14155 | ARHGEF7 | 21.0472243 | 21.1414011 | 21.1061859 | 21.1349493 | 21.2672224 |
| Q96C57 | C12orf43 | 19.6026318 | 19.8936069 | 19.7749175 | 19.806221 | 19.9114139 |
| P62304 | SNRPE | 24.4263852 | 24.2525456 | 24.1658344 | 24.0304592 | 24.1248717 |
| P53609 | PGGT1B | 21.229456 | 21.310327 | 21.3424228 | 21.1341073 | 21.1532853 |
| P62380 | TBPL1 | 17.784561 | 17.4578826 | 17.707207 | 17.3037908 | 17.6259481 |
| P52434 | POLR2H | 21.9071146 | 21.7591057 | 21.8736574 | 21.5089563 | 21.8173152 |
| O75425 | MOSPD3 | 17.4353921 | 17.2795046 | 17.3522141 | 17.3670972 | 17.5408287 |
| Q5TBA9 | FRY | 19.8504212 | 19.8407149 | 19.8966075 | 19.7817179 | 19.862044 |
| Q9BUI4 | POLR3C | 20.255468 | 20.2418215 | 20.3574111 | 20.1802845 | 20.2267281 |
| Q96FT9 | IFT43 | 16.6158135 | 16.950911 | 16.4851642 | 16.7663682 | 16.9510293 |
| Q9H3G5 | CPVL | NA | 13.104657 | 13.9956333 | NA | 13.5881355 |
| Q6UW56 | ATRAID | 19.4457375 | 19.2688059 | 19.6534542 | 19.3953903 | 19.3646106 |
| Q8WTW3 | COG1 | 20.9797757 | 21.0524414 | 21.0830787 | 20.9236943 | 20.9944752 |

| | | | | | | |
|---|---|---|---|---|---|---|
| Q9BTT4 | MED10 | 19.6970848 | 19.7474043 | 19.8226551 | 19.7938767 | 19.9159686 |
| Q96L93 | KIF16B | 16.4991304 | 16.5707821 | 16.8556462 | 16.7038079 | 16.9145887 |
| Q9NQ55 | PPAN | 21.5166543 | 21.5878042 | 21.6910303 | 21.4677601 | 21.5063458 |
| P14621 | ACYP2 | 19.7138212 | 19.6530384 | 19.7509626 | 19.840161 | 19.7311805 |
| Q8TC59 | PIWIL2 | 23.8217981 | NA | NA | 23.0138908 | NA |
| Q8WWB7 | GLMP | 17.382478 | 17.528004 | 17.082073 | 17.1359185 | 16.9780726 |
| Q4G0I0 | UQCC4 | 18.7678965 | 18.8746749 | 18.9219268 | 18.6746542 | 18.5265263 |
| P14324 | FDPS | 23.7610727 | 23.7787429 | 23.6974533 | 23.6317479 | 23.5927931 |
| Q9NV06 | DCAF13 | 21.641281 | 21.7154542 | 21.7534343 | 21.5876363 | 21.7329978 |
| Q9NZM5 | NOP53 | 20.9780278 | 21.0140485 | 21.0093601 | 20.9478387 | 20.9154308 |
| Q9Y4E5 | ZNF451 | 19.6210098 | 19.6637131 | 19.7271232 | 19.7114266 | 19.7701645 |
| Q8NB90 | AFG2A | 20.8379732 | 20.9336321 | 20.8015574 | 20.8240848 | 20.9550136 |
| O75410 | TACC1 | 21.6152498 | 21.7499884 | 21.4255314 | 21.6391372 | 21.730234 |
| Q7Z6M1 | RABEPK | 18.3523443 | 18.4301116 | 18.3257658 | 18.2958162 | 18.4444561 |
| O75391 | SPAG7 | 20.8257818 | 20.8661153 | 20.6615132 | 20.8598632 | 21.0590619 |
| Q8N423 | LILRB2 | 15.3537238 | NA | NA | 15.4387886 | NA |
| Q9NZT1 | CALML5 | 18.2253746 | 17.5509902 | 16.244747 | 16.6901381 | 16.326054 |
| P08138 | NGFR | NA | 14.9146025 | 14.3239637 | NA | 14.4518585 |
| P03928 | MT-ATP8 | 20.1195806 | 20.0185269 | 19.9634419 | 19.4178442 | 19.8464844 |
| Q9BZL1 | UBL5 | 23.6540989 | 23.6599051 | 23.7012725 | 23.5501042 | 23.5812212 |
| P56937 | HSD17B7 | 18.3629942 | 18.649614 | 18.5535241 | 18.4095473 | 18.6167271 |
| Q8TC12 | RDH11 | 22.4016995 | 22.4148395 | 22.4832532 | 22.2706002 | 22.4125171 |
| Q8WWI5 | SLC44A1 | 23.7430232 | 23.7456507 | 23.7884244 | 23.5862287 | 23.6706311 |
| O75153 | CLUH | 23.8431854 | 23.9133826 | 23.9115463 | 23.7765415 | 23.8619837 |
| Q9UL40 | ZNF346 | 19.816531 | 19.7735532 | 19.7783462 | 19.7641256 | 19.8832557 |
| P46013 | MKI67 | 21.8832033 | 21.9704842 | 21.8425878 | 21.8488593 | 21.9556076 |
| Q08378 | GOLGA3 | 20.8009295 | 20.909318 | 20.807636 | 20.8924902 | 21.0280007 |
| P28290 | ITPRID2 | 18.7388575 | 18.8621035 | 18.8481531 | 18.9043407 | 18.942889 |
| Q9NQC7 | CYLD | 19.3163788 | 19.4678141 | 19.4760964 | 19.4145594 | 19.5707337 |
| Q96CS2 | HAUS1 | 19.6917647 | 19.9196178 | 19.5729463 | 19.9160186 | 19.8653923 |
| A6ND36 | FAM83G | NA | 11.3940512 | 11.5822713 | NA | 11.537816 |
| P07738 | BPGM | 17.0835419 | 17.3517135 | 17.320266 | 17.0640411 | 17.4684514 |
| P14649 | MYL6B | 17.796066 | 18.31061 | 17.8826339 | 18.0813328 | 18.2490092 |
| Q7Z6J8 | UBE3D | 13.8698823 | NA | NA | 13.973667 | NA |
| Q9Y330 | ZBTB12 | 14.4553339 | NA | NA | 14.712419 | NA |
| Q09019 | DMWD | 13.1192907 | NA | NA | 13.5412372 | NA |
| O75934 | BCAS2 | 21.9461357 | 21.9704517 | 21.91885 | 21.9729018 | 22.0021305 |
| P34810 | CD68 | 15.6083433 | NA | NA | 16.0033157 | NA |
| P08754 | GNAI3 | 23.0522536 | 22.9532636 | 23.0221882 | 22.9832258 | 23.0837096 |
| Q9UKY7 | CDV3 | 23.0276171 | 23.1822256 | 22.9289163 | 23.1061399 | 23.1082538 |
| O15321 | TM9SF1 | 20.8316765 | 20.8667405 | 20.9554355 | 20.8708135 | 20.7315224 |
| Q9UPQ9 | TNRC6B | 19.9878129 | 20.0020847 | 19.9193529 | 20.0466987 | 20.1473342 |

| | | | | | | |
|---|---|---|---|---|---|---|
| Q92783 | STAM | 20.1421164 | 20.2898541 | 20.1326108 | 20.2348412 | 20.3322585 |
| Q9ULI4 | KIF26A | NA | 16.8833147 | 16.2204071 | NA | 16.0630492 |
| Q8NGE1 | OR6C4 | NA | 14.4283738 | 14.4935012 | NA | 14.5253766 |
| O00470 | MEIS1 | 20.2453405 | 20.3322377 | 20.3117487 | 20.3237012 | 20.5674216 |
| Q9HAW4 | CLSPN | 18.6385169 | 18.6362932 | 18.583943 | 18.6939316 | 18.549617 |
| Q96QZ7 | MAGI1 | NA | 19.2633352 | 18.8658448 | NA | 19.4867043 |
| P24071 | FCAR | NA | 14.249083 | 14.3192228 | NA | 14.8777937 |
| O60486 | PLXNC1 | 13.7691845 | NA | NA | 13.5505724 | NA |
| O96005 | CLPTM1 | 23.1810041 | 23.2476712 | 23.2421502 | 23.1553256 | 23.1231001 |
| P62979 | RPS27A | 28.4465297 | 28.4396511 | 28.4050678 | 28.1263775 | 28.3561772 |
| Q8WVM0 | TFB1M | 19.7540196 | 19.8894327 | 19.7781268 | 19.8235089 | 19.8571447 |
| Q9NUG6 | PDRG1 | 19.8600332 | 19.84197 | 19.7964559 | 19.8294747 | 19.9559664 |
| P48595 | SERPINB10 | NA | 15.872568 | 15.8318904 | NA | 15.7957236 |
| Q9P0M9 | MRPL27 | 21.3751714 | 21.3583288 | 21.4036521 | 21.5491307 | 21.5956499 |
| Q6ZNA5 | FRRS1 | 14.2000997 | NA | NA | 14.3496558 | NA |
| O95373 | IPO7 | 23.0629292 | 23.063243 | 23.1074145 | 23.0025582 | 23.0519781 |
| Q9BW71 | HIRIP3 | 20.7940747 | 20.9650688 | 20.8044387 | 20.8203222 | 20.9860336 |
| P50914 | RPL14 | 26.2077417 | 26.3594505 | 26.4372686 | 26.1232124 | 26.1103679 |
| Q7L211 | ABHD13 | 17.9913329 | 18.0878146 | 18.2280383 | 17.8887883 | 18.0159582 |
| Q8IYI8 | ZNF440 | 17.4792327 | 17.6260067 | 17.6461531 | 17.453023 | 17.4921115 |
| P62081 | RPS7 | 26.3696467 | 26.5342111 | 26.4524825 | 26.2393843 | 26.3381158 |
| O14929 | HAT1 | 24.0147225 | 24.052404 | 24.0206663 | 23.8946961 | 23.847168 |
| P23381 | WARS1 | 23.8950694 | 23.9400999 | 23.930731 | 23.829961 | 23.8981221 |
| Q13813 | SPTAN1 | 18.8490618 | 18.9081419 | 18.8504288 | 18.8219913 | 18.9910007 |
| O00499 | BIN1 | 20.102427 | 20.5101919 | 20.3283998 | 20.470959 | 20.5004349 |
| Q7Z2Z1 | TICRR | 16.4321426 | 16.6352375 | 16.8237783 | 16.6850495 | 16.9079999 |
| P30536 | TSPO | 23.8618521 | 23.7775659 | 23.9989435 | 23.8054375 | 23.7908917 |
| P05141 | SLC25A5 | 27.1211915 | 27.0958226 | 27.0537085 | 26.8773381 | 26.9908868 |
| Q58WW2 | DCAF6 | 18.8939948 | 18.9167807 | 18.8905597 | 18.765759 | 18.8882218 |
| P23921 | RRM1 | 23.3373826 | 23.309084 | 23.3901028 | 23.250814 | 23.3120647 |
| Q8TES7 | FBF1 | 15.8894736 | 16.1228885 | 15.8019398 | 15.8543522 | 15.6409672 |
| Q8N8E2 | ZNF513 | 12.9057283 | NA | NA | 13.5139281 | NA |
| Q4LDG9 | DNAL1 | 15.5500952 | 15.3886587 | 15.6763446 | 15.9007756 | 15.9274027 |
| C9JVW0 | INAFM1 | NA | 15.4026844 | 15.1992135 | NA | 15.0636481 |
| Q9Y240 | CLEC11A | 23.2561923 | 23.2256779 | 23.0184334 | 23.1967573 | 23.0877073 |
| Q96MG8 | PCMTD1 | 16.8338139 | 17.2837343 | 17.4311233 | 17.606931 | 17.3548839 |
| Q96LB3 | IFT74 | 18.4619461 | 18.3981707 | 18.5351731 | 18.6437516 | 18.5252175 |
| P19634 | SLC9A1 | 18.8830283 | 19.0454932 | 19.0131501 | 19.1638615 | 19.1871568 |
| O75995 | SASH3 | 20.5526939 | 20.7136333 | 20.5591333 | 20.6116535 | 20.8564146 |
| Q8N0Y2 | ZNF444 | NA | 16.9707357 | 16.9998989 | NA | 16.8525586 |
| Q92733 | PRCC | 20.7084899 | 20.8980482 | 20.6735203 | 20.8422603 | 20.8861067 |
| Q9BQI7 | PSD2 | 15.7849618 | NA | NA | 16.2839337 | NA |

| | | | | | | |
|---|---|---|---|---|---|---|
| Q9Y2D4 | EXOC6B | 18.6961807 | 18.8559619 | 18.9057241 | 19.0196759 | 18.9419939 |
| Q9Y221 | NIP7 | 21.5359769 | 21.5745179 | 21.6437062 | 21.5247038 | 21.6049374 |
| Q7Z7H8 | MRPL10 | 20.2088955 | 20.3316355 | 20.2287961 | 20.375532 | 20.4307599 |
| P20160 | AZU1 | 24.0242326 | 23.5936556 | 23.777699 | 23.836393 | 23.7452598 |
| P61006 | RAB8A | 23.3975066 | 23.4568614 | 23.4324114 | 23.2442867 | 23.2913866 |
| P50238 | CRIP1 | 21.9780558 | 22.0115401 | 22.1229842 | 22.0669239 | 22.1271178 |
| P31944 | CASP14 | 15.3210112 | 16.1257354 | 14.6758928 | 14.839918 | 14.2653432 |
| Q8IYW5 | RNF168 | 15.0440313 | 15.7860119 | 15.7180429 | 15.4392881 | 15.1335999 |
| O95807 | TMEM50A | 18.0440192 | 18.3139952 | 17.9921153 | 17.9526929 | 17.8769927 |
| Q15836 | VAMP3 | 18.3273583 | 18.330283 | 18.6560391 | 18.2701741 | 18.34236 |
| Q7Z7N9 | TMEM179B | 20.8487394 | 20.8836429 | 20.8041096 | 20.7030559 | 20.7982722 |
| Q15629 | TRAM1 | 20.42181 | 20.4970133 | 20.6806251 | 20.4676575 | 20.4093292 |
| Q9BU23 | LMF2 | 20.0831838 | 20.1923933 | 20.3967697 | 20.0281211 | 19.9642297 |
| P53794 | SLC5A3 | 20.1012729 | 20.1904921 | 20.2042494 | 20.0023079 | 20.0987262 |
| P46779 | RPL28 | 25.7289551 | 25.592001 | 25.8251603 | 25.548722 | 25.5228211 |
| P62424 | RPL7A | 26.4935033 | 26.3600239 | 26.6149313 | 26.3626527 | 26.3208758 |
| Q04837 | SSBP1 | 23.9685108 | 24.002864 | 23.9563099 | 23.8995544 | 24.0008396 |
| Q9NUQ2 | AGPAT5 | 21.7263078 | 21.6720205 | 21.787338 | 21.611832 | 21.7099462 |
| Q8NI36 | WDR36 | 21.1164419 | 21.063166 | 21.2241989 | 21.070879 | 21.1413645 |
| P48047 | ATP5PO | 24.1823696 | 24.2242337 | 24.219222 | 24.1815207 | 24.2165646 |
| Q9UNM6 | PSMD13 | 23.605749 | 23.6117037 | 23.5646127 | 23.486594 | 23.5067661 |
| Q8NBN3 | TMEM87A | 20.6025257 | 20.6200895 | 20.7142127 | 20.5122569 | 20.6101715 |
| Q96GC6 | ZNF274 | 19.8475867 | 19.9198386 | 19.900144 | 19.9631397 | 19.9972645 |
| Q8IXT5 | RBM12B | 21.2170855 | 21.2189243 | 21.2142765 | 21.1581318 | 21.317303 |
| O15037 | KHNYN | 19.5804306 | 19.4559299 | 19.5302249 | 19.5350688 | 19.6503734 |
| P32019 | INPP5B | 19.8868576 | 19.8338887 | 20.0540092 | 19.9611752 | 20.0703901 |
| Q96BD8 | SKA1 | 18.7692221 | 18.8336333 | 18.8591023 | 18.77119 | 18.8965503 |
| Q9HCE5 | METTL14 | 19.4616604 | 19.6841902 | 19.4574104 | 19.569888 | 19.6381472 |
| Q9NYK5 | MRPL39 | 22.0394402 | 22.1912155 | 22.05939 | 22.1348759 | 22.214986 |
| Q9P2K5 | MYEF2 | 19.054506 | 19.2998535 | 19.1830973 | 19.1436543 | 19.2278989 |
| Q01167 | FOXK2 | 19.7988504 | 19.9556562 | 19.9461491 | 20.0325699 | 20.0480967 |
| Q9NUS5 | AP5S1 | 19.1504388 | 18.9686339 | 19.0259022 | 19.1230665 | 19.1146603 |
| Q9H4I2 | ZHX3 | 18.4656317 | 18.5330748 | 18.5334919 | 18.6383331 | 18.6784395 |
| P27816 | MAP4 | 22.4903916 | 22.5632538 | 22.3938615 | 22.4513989 | 22.6328384 |
| Q6ULP2 | AFTPH | 18.8743812 | 18.979248 | 18.8113575 | 18.9365256 | 19.0813657 |
| P42677 | RPS27 | 24.6126408 | 24.7155233 | 24.6797481 | 24.5513211 | 25.0074226 |
| P41236 | PPP1R2 | 20.7225891 | 20.6337352 | 20.37049 | 20.7177257 | 20.7437488 |
| Q86UU1 | PHLDB1 | 17.000705 | 16.8182754 | 17.1934109 | 16.6633658 | 17.0573798 |
| O75570 | MTRF1 | NA | 14.0607485 | 13.5324088 | NA | 14.1401448 |
| Q9UBP9 | GULP1 | 16.771402 | 17.1902127 | 17.1724728 | 16.7150868 | 17.419159 |
| Q71UM5 | RPS27L | 20.1192895 | 20.1554061 | 20.1856311 | 19.8863387 | 20.6486396 |
| O75317 | USP12 | NA | 16.5701758 | 16.7509942 | NA | 16.1739938 |

| | | | | | | |
|---|---|---|---|---|---|---|
| Q9BZE1 | MRPL37 | 23.0037693 | 22.954289 | 23.0201266 | 22.9460149 | 22.9382172 |
| Q9NVC3 | SLC38A7 | 16.0482331 | 16.3289506 | 16.1292783 | 15.8257151 | 16.0907259 |
| Q8TF40 | FNIP1 | 17.9154744 | 17.9481034 | 17.7401759 | 17.7848882 | 17.7737492 |
| P09661 | SNRPA1 | 23.2492709 | 23.3418726 | 23.3856831 | 23.2368268 | 23.4488498 |
| P07602 | PSAP | 24.0200626 | 24.2512821 | 23.8986593 | 24.1348787 | 24.2595352 |
| Q9BVG9 | PTDSS2 | 18.138389 | 17.8792474 | 17.9394828 | 17.9586432 | 17.8914468 |
| P62263 | RPS14 | 26.4527961 | 26.3951174 | 26.3577902 | 26.2664084 | 26.2703769 |
| Q7L7V1 | DHX32 | 18.3033395 | 18.185079 | 18.3599258 | 18.3169692 | 18.5198661 |
| Q5JV73 | FRMPD3 | NA | 19.3835091 | 19.3654436 | NA | 19.8810733 |
| Q9BVK8 | TMEM147 | 19.1845562 | 19.1644748 | 19.2379235 | 19.1161391 | 18.9207564 |
| Q96S99 | PLEKHF1 | 19.4817542 | 19.4728324 | 19.521879 | 19.5152726 | 19.4728185 |
| Q9H4G4 | GLIPR2 | 21.063041 | 21.0336327 | 21.2409627 | 21.0221441 | 21.0843812 |
| Q14019 | COTL1 | 24.5511809 | 24.6050838 | 24.5558655 | 24.3202064 | 24.5278983 |
| O43809 | NUDT21 | 23.7950392 | 23.8319244 | 23.8166312 | 23.6297182 | 23.7204231 |
| Q8TDW0 | LRRC8C | 21.5262794 | 21.6153505 | 21.6268712 | 21.4569742 | 21.4982669 |
| P28715 | ERCC5 | 19.3934146 | 19.3691779 | 19.4230416 | 19.3793793 | 19.4722555 |
| P43353 | ALDH3B1 | 19.1384928 | 19.1334172 | 19.161177 | 19.2310804 | 19.342158 |
| Q9C0B0 | UNK | 19.3793235 | 19.5360923 | 19.5569323 | 19.5144984 | 19.6335848 |
| P82094 | TMF1 | 19.924518 | 19.9946868 | 19.8136182 | 19.9448675 | 20.0985422 |
| Q6UVJ0 | SASS6 | 17.746656 | 17.8700721 | 17.9464607 | 17.8619371 | 18.009043 |
| P37173 | TGFBR2 | 18.1600503 | 18.2766853 | 18.4375898 | 18.58714 | 18.5210278 |
| Q6MZP7 | LIN54 | 17.5699634 | 17.438638 | 17.6034287 | 17.58969 | 17.6796026 |
| Q9Y2G9 | SBNO2 | 18.7772081 | 18.7811423 | 19.0251441 | 18.9739372 | 19.2014301 |
| Q9H5V7 | IKZF5 | 16.4831486 | 16.7649498 | 16.3147071 | 16.4511206 | 16.3700401 |
| Q8IVV7 | GID4 | 17.1359555 | 17.0022715 | 17.2162987 | 16.9688531 | 17.1266764 |
| Q9UKJ3 | GPATCH8 | 19.3812405 | 19.5486231 | 19.3602266 | 19.4129962 | 19.5053243 |
| O75157 | TSC22D2 | 18.1276546 | 18.0732177 | 17.989476 | 18.1887097 | 18.3134975 |
| Q5T6F0 | DCAF12 | 17.1772378 | 17.3839155 | 17.4633779 | 17.3455182 | 16.9733387 |
| Q99808 | SLC29A1 | 21.2441882 | 21.2129339 | 21.1826026 | 21.1261057 | 21.0993839 |
| Q6PCB8 | EMB | 19.0932049 | 19.2281501 | 19.2407016 | 19.239484 | 19.0219709 |
| Q8TEX9 | IPO4 | 22.3863931 | 22.3892154 | 22.4346079 | 22.2860708 | 22.3427849 |
| Q13151 | HNRNPA0 | 24.7805757 | 24.8642822 | 24.8217388 | 24.7510502 | 24.8109134 |
| Q9Y5S1 | TRPV2 | 20.5742898 | 20.6674133 | 20.610041 | 20.5342877 | 20.5092823 |
| Q8N884 | CGAS | 21.9340927 | 21.9295593 | 21.9810817 | 21.9056069 | 21.9470288 |
| P49406 | MRPL19 | 22.190519 | 22.2321913 | 22.2035755 | 22.1262118 | 22.2078889 |
| Q8TBA6 | GOLGA5 | 19.5321949 | 19.6983335 | 19.6744473 | 19.6084532 | 19.6888964 |
| Q9UQL6 | HDAC5 | 19.3488361 | 19.4478782 | 19.337682 | 19.3313828 | 19.4387067 |
| P55145 | MANF | 22.7730804 | 22.9577308 | 22.6942378 | 22.8068566 | 22.8765796 |
| Q13309 | SKP2 | 18.0306938 | 17.8892778 | 18.3623921 | 18.359199 | 18.5028561 |
| Q9NRP4 | SDHAF3 | 18.7280295 | 18.5970896 | 18.7155466 | 18.9628363 | 18.860805 |
| Q8TEY7 | USP33 | 17.5189407 | 17.3889161 | 17.2073434 | 17.5942022 | 17.3783773 |
| Q9UKZ4 | TENM1 | 21.7108164 | 22.05994 | 22.0250425 | 22.0989775 | 22.0386715 |

| | | | | | | |
|---|---|---|---|---|---|---|
| Q9BTX3 | TMEM208 | 18.8230182 | NA | NA | 18.8596092 | NA |
| Q8NHZ8 | CDC26 | 20.0391732 | 20.3538461 | 20.0634162 | 20.1071887 | 20.4033593 |
| O43852 | CALU | 22.9030916 | 23.1096811 | 22.7495911 | 22.9928357 | 23.1677537 |
| A6NLJ0 | C2CD4B | NA | 16.7459787 | 16.0017657 | NA | 15.2783065 |
| Q6ICB0 | DESI1 | 17.5996826 | 17.7183123 | 18.5849139 | 17.6738241 | 17.422841 |
| Q5T3U5 | ABCC10 | 16.9343437 | 17.192496 | 16.7775557 | 16.8218625 | 16.7483872 |
| Q9BU19 | ZNF692 | 16.696497 | 17.239894 | 17.2033791 | 16.640157 | 17.0975463 |
| Q8IX15 | HOMEZ | 15.6499916 | 15.5883286 | 15.5930572 | 15.3546348 | 15.6874701 |
| O00115 | DNASE2 | 19.4677377 | 19.4554504 | 19.5821156 | 19.5128876 | 19.3268001 |
| Q9H0M5 | ZNF700 | 17.0772942 | 17.2037079 | 17.5043397 | 17.0097159 | 16.8066201 |
| Q96AN5 | TMEM143 | 19.0535963 | 19.170886 | 19.0604383 | 19.1474483 | 19.020102 |
| O15127 | SCAMP2 | 23.2169469 | 23.2023124 | 23.3617037 | 23.1611594 | 23.0726036 |
| O43286 | B4GALT5 | 16.5672189 | 16.3622044 | 16.6119607 | 16.5589132 | 16.6026942 |
| Q14558 | PRPSAP1 | 20.1538455 | 19.9492067 | 20.1658127 | 20.0195905 | 20.2624597 |
| Q15057 | ACAP2 | 21.4098512 | 21.545285 | 21.5546037 | 21.4507255 | 21.6073856 |
| Q9BSE5 | AGMAT | 19.1375131 | 19.1517802 | 19.2161161 | 19.2153914 | 19.3040526 |
| Q9UKX7 | NUP50 | 22.5938419 | 22.7150235 | 22.6514211 | 22.662471 | 22.7149874 |
| Q7Z7M0 | MEGF8 | 19.9006366 | 20.0112552 | 19.8910538 | 19.9291935 | 20.0555361 |
| O95551 | TDP2 | 19.6864365 | 19.8169588 | 19.8535112 | 19.7496959 | 19.9888798 |
| Q9BPZ7 | MAPKAP1 | 17.7700796 | 17.7161537 | 17.8902872 | 17.7805189 | 17.9489484 |
| Q9H2G9 | BLZF1 | 16.1432244 | 15.9084227 | 15.9606994 | 15.9305325 | 16.177978 |
| Q8N4S0 | CCDC82 | 14.4520934 | 14.5080532 | 15.0226521 | 15.0114723 | 15.3036863 |
| Q8N187 | CARF | NA | 16.5061447 | 17.081221 | NA | 16.9994047 |
| Q9BT43 | POLR3GL | 17.2074413 | 17.6724474 | 17.2968433 | 17.6372092 | 17.8555731 |
| Q96RD7 | PANX1 | 16.5177938 | 16.2773775 | 16.4849675 | 16.452138 | 16.8107771 |
| Q32M88 | PGGHG | 18.6091584 | 18.4128427 | 18.5422151 | 18.224821 | 18.4643447 |
| Q9Y5Y0 | FLVCR1 | 17.1083431 | 17.1214392 | 17.279132 | 17.0259737 | 16.9400312 |
| A8MQ14 | ZNF850 | 18.5914554 | 18.5839473 | 18.5612095 | 18.4793733 | 18.5152927 |
| Q9HB58 | SP110 | 17.6574793 | 17.8340606 | 17.9873722 | 17.8138045 | 17.9495078 |
| O00154 | ACOT7 | 22.8029384 | 22.8131137 | 22.8997918 | 22.7749268 | 22.7963493 |
| P05388 | RPLP0 | 26.5854139 | 26.6406587 | 26.6666683 | 26.4677388 | 26.6357205 |
| Q96S15 | WDR24 | 17.0215762 | 17.1504412 | 17.0443295 | 17.2817811 | 17.3304378 |
| Q04941 | PLP2 | 21.7233047 | 21.8280042 | 22.3500692 | 21.6136252 | 22.2830937 |
| Q9H649 | NSUN3 | 15.8536623 | 15.9899975 | 15.822891 | 15.7428434 | 15.7152124 |
| Q8NFC6 | BOD1L1 | 22.7454427 | 22.7844869 | 22.6748321 | 22.7154359 | 22.8209256 |
| Q8NE00 | TMEM104 | 19.556542 | 19.5379307 | 19.6995297 | 19.6273919 | 19.2971165 |
| Q6IQ20 | NAPEPLD | NA | 16.811431 | 16.9819938 | NA | 17.1616175 |
| Q9UK23 | NAGPA | 20.2391284 | 20.1531388 | 20.2612678 | 20.3402037 | 20.1747677 |
| Q6PCE3 | PGM2L1 | 17.9183516 | 17.9951458 | 17.9955927 | 17.9358332 | 18.0059164 |
| A0JNW5 | BLTP3B | 19.9486681 | 19.9923591 | 19.9763187 | 19.981561 | 20.0748662 |
| Q6UW78 | UQCC3 | NA | 11.6441968 | 11.6880071 | NA | 11.7932336 |
| Q8N8R3 | SLC25A29 | 21.3653482 | 20.9857676 | 21.2553702 | 20.911601 | 20.911507 |

| | | | | | | |
|---|---|---|---|---|---|---|
| P37108 | SRP14 | 24.7882884 | 24.7148009 | 24.788982 | 24.6363706 | 24.7315565 |
| Q8NBU5 | ATAD1 | 20.9241416 | 20.9606047 | 20.9275119 | 20.9923743 | 21.0595849 |
| Q9NZ72 | STMN3 | 16.5908667 | 16.593071 | 16.6755484 | 16.7049465 | 16.7148471 |
| Q9Y5X1 | SNX9 | 18.3425595 | 18.6362779 | 18.682198 | 18.6590452 | 18.7841785 |
| Q5VST9 | OBSCN | 18.2817184 | 17.7843618 | 18.110372 | 18.3146622 | 18.3528751 |
| Q3T906 | GNPTAB | 19.1220049 | 19.2520742 | 19.2156013 | 19.2451531 | 19.2636198 |
| P83731 | RPL24 | 25.464822 | 25.4590259 | 25.6586724 | 25.3451053 | 25.3412979 |
| P40616 | ARL1 | 21.165917 | 21.1211173 | 21.3215457 | 21.0896155 | 21.1387194 |
| O60890 | OPHN1 | 15.2091105 | NA | NA | 15.2980385 | NA |
| Q8WUH6 | TMEM263 | 17.1872084 | 17.7582436 | 17.6756673 | 17.2235997 | 17.6206169 |
| Q0VG06 | FAAP100 | 17.1567625 | 17.3340875 | 17.2673734 | 17.3231116 | 17.3070705 |
| P54709 | ATP1B3 | 23.3332819 | 23.4174499 | 23.4436846 | 23.2534378 | 23.2878639 |
| Q9UBS3 | DNAJB9 | 17.3022051 | 17.3655623 | 17.3012194 | 17.3062156 | 17.0907787 |
| Q13303 | KCNAB2 | 23.0925237 | 23.0544703 | 23.1829079 | 23.0164813 | 23.1161321 |
| Q96S94 | CCNL2 | 20.6597866 | 20.6361634 | 20.8927983 | 20.6336852 | 20.6953129 |
| P09001 | MRPL3 | 22.4628453 | 22.4244815 | 22.477214 | 22.3860356 | 22.4022795 |
| Q9BY42 | RTF2 | 21.5245046 | 21.5406471 | 21.5767202 | 21.5524949 | 21.51702 |
| Q6P1K2 | PMF1 | 18.8476442 | 19.0333662 | 19.0883667 | 19.0946606 | 19.3123005 |
| Q9P0S3 | ORMDL1 | 20.8504524 | 20.7517564 | 20.6823398 | 20.6129522 | 20.678552 |
| Q96CN5 | LRRC45 | 17.5118932 | 17.4370081 | 17.5924755 | 17.5029133 | 17.5236158 |
| Q96NB1 | CEP20 | 17.2608032 | 17.6289522 | 17.4800936 | 17.8604968 | 17.5882388 |
| Q6P2C8 | MED27 | 20.3298918 | 20.3076492 | 20.2499301 | 20.1948098 | 20.2398352 |
| Q08188 | TGM3 | 16.8185282 | 16.1109271 | 16.8144095 | 16.6278153 | 15.4523368 |
| O94826 | TOMM70 | 23.1163739 | 23.2020347 | 23.2630027 | 23.0693118 | 23.155665 |
| Q8TCG2 | PI4K2B | 17.6600094 | 17.5538876 | 17.6801091 | 17.6841928 | 17.6022105 |
| P0DTL6 | ZFTRAF1 | 19.1507289 | 19.211459 | 19.4265707 | 19.0572794 | 19.3918177 |
| P46778 | RPL21 | 25.4153708 | 25.4130218 | 25.5402041 | 25.2919192 | 25.3932298 |
| Q14159 | SPIDR | 15.3497056 | 15.7242109 | 15.4961522 | 15.1825479 | 15.2544499 |
| Q8NEZ3 | WDR19 | 17.8653071 | 18.130759 | 17.7389767 | 17.9439903 | 17.9738343 |
| Q9NRX1 | PNO1 | 21.7739912 | 21.870219 | 22.0719662 | 21.5935861 | 21.7317107 |
| Q5VST6 | ABHD17B | 20.7517246 | 20.5922648 | 20.7736467 | 20.6701223 | 20.6734233 |
| Q8IWA0 | WDR75 | 21.0334043 | 20.9078779 | 21.0958703 | 20.8840816 | 21.0231464 |
| Q9Y3T9 | NOC2L | 21.7178906 | 21.6431238 | 21.7977714 | 21.6078288 | 21.6863048 |
| P36222 | CHI3L1 | 23.6613212 | 23.6312468 | 23.6769413 | 23.522842 | 23.6462949 |
| Q9NUQ8 | ABCF3 | 21.5791564 | 21.6824488 | 21.5995482 | 21.5367964 | 21.5038981 |
| Q9NX62 | BPNT2 | 22.1186654 | 22.1577236 | 22.2114453 | 22.1645168 | 22.071275 |
| Q9NR30 | DDX21 | 25.1896578 | 25.196848 | 25.1972643 | 25.1124544 | 25.0844247 |
| Q9GZT3 | SLIRP | 23.432356 | 23.4180079 | 23.401257 | 23.3896258 | 23.5037796 |
| Q9H0E9 | BRD8 | 20.0710766 | 20.0235182 | 20.0452743 | 20.144162 | 20.1694123 |
| O43516 | WIPF1 | 22.2413661 | 22.3715373 | 22.300714 | 22.3370215 | 22.3713195 |
| P07311 | ACYP1 | 20.65352 | 20.7212842 | 20.6651722 | 20.6259796 | 20.8029924 |
| P07992 | ERCC1 | 19.3344747 | 19.3969237 | 19.3896788 | 19.4655423 | 19.5912766 |

| | | | | | | |
|---|---|---|---|---|---|---|
| Q96SY0 | INTS14 | 19.8005692 | 19.9349141 | 20.0769112 | 19.8620143 | 20.131707 |
| Q9Y6A5 | TACC3 | 20.3284557 | 20.4726921 | 20.3215477 | 20.3638261 | 20.5646964 |
| Q86W92 | PPFIBP1 | 18.9369285 | 19.1969781 | 19.1216223 | 19.0556657 | 19.3084128 |
| Q96D70 | R3HDM4 | 18.0111882 | 18.1102771 | 18.1655677 | 17.8882544 | 18.0263765 |
| Q6ZSJ8 | C1orf122 | 18.1212135 | 18.3208485 | 18.0946168 | 18.2393059 | 18.4552262 |
| Q9H8Y8 | GORASP2 | 22.7972224 | 22.8435591 | 22.5839874 | 22.7947819 | 22.9367494 |
| Q8TC20 | CAGE1 | NA | 15.8815056 | 16.2117153 | NA | 16.2643397 |
| P80404 | ABAT | 16.5930706 | 16.5564703 | 16.6028643 | 16.3483819 | 16.6889022 |
| Q8TB52 | FBXO30 | 17.624501 | 18.0308065 | 17.9363454 | 17.9589932 | 18.0589925 |
| Q96D15 | RCN3 | 19.2403147 | 19.4086574 | 19.1676125 | 19.263027 | 19.6079967 |
| Q9BQI3 | EIF2AK1 | NA | 13.655509 | 13.9009671 | NA | 14.007628 |
| O15013 | ARHGEF10 | 18.5311278 | 18.7255931 | 18.8744725 | 18.5493748 | 18.79854 |
| Q8TC05 | MDM1 | 16.1735326 | 16.0949438 | 16.0792198 | 16.0923236 | 16.4739386 |
| Q9UHH9 | IP6K2 | NA | 13.0953868 | 13.360064 | NA | 13.284172 |
| Q68DE3 | USF3 | 20.0412174 | 20.2170964 | 19.8060238 | 19.5449394 | 19.894458 |
| Q92626 | PXDN | 15.9987243 | NA | NA | 15.8883567 | NA |
| P08962 | CD63 | 25.088113 | 24.9483509 | 24.9852114 | 24.9333777 | 24.8525686 |
| Q16670 | ZSCAN26 | 15.8383586 | 16.8428823 | 16.1586366 | 15.9402864 | 15.8031496 |
| Q96S16 | JMJD8 | 18.8509974 | 19.1602451 | 18.7533936 | 18.7807357 | 18.6321184 |
| O00562 | PITPNM1 | 16.2308886 | 16.03548 | 16.1824569 | 16.136378 | 16.2819575 |
| Q9UNL2 | SSR3 | 22.0333749 | 22.2020045 | 22.3212068 | 21.9635673 | 21.6202048 |
| Q9Y4U1 | MMACHC | 18.6859491 | 19.0863201 | 19.0363195 | 18.6124027 | 18.8148524 |
| O95478 | NSA2 | 21.0205749 | 20.9819583 | 21.2147078 | 20.9936465 | 20.9219537 |
| P01111 | NRAS | 20.6641342 | 20.8946982 | 20.8111771 | 20.7157228 | 20.6433364 |
| P42575 | CASP2 | 20.7212279 | 20.9183931 | 20.8963042 | 20.7018206 | 20.7918503 |
| Q9HBH1 | PDF | 19.4625302 | 19.5825404 | 19.4695797 | 19.3386453 | 19.4977303 |
| P18621 | RPL17 | 25.5137641 | 25.4412415 | 25.6664598 | 25.4093619 | 25.4944563 |
| Q6P0N0 | MIS18BP1 | 18.9704366 | 19.0953288 | 19.0828981 | 18.9214268 | 18.961921 |
| P14384 | CPM | 21.9656981 | 22.0124136 | 22.0193845 | 21.8365748 | 21.8251979 |
| Q9Y333 | LSM2 | 22.6906187 | 22.7406033 | 22.8578601 | 22.6605357 | 22.6278375 |
| Q9UID3 | VPS51 | 19.9435886 | 19.8521065 | 20.0073881 | 19.8094367 | 19.947202 |
| Q9Y399 | MRPS2 | 22.2709767 | 22.2379514 | 22.2856568 | 22.190615 | 22.1847496 |
| P36404 | ARL2 | 21.7101398 | 21.6444625 | 21.7302096 | 21.5917032 | 21.633279 |
| O00400 | SLC33A1 | 20.3277451 | 20.264105 | 20.3363482 | 20.2167003 | 20.3485561 |
| P08567 | PLEK | 23.9791074 | 24.0214744 | 23.9870067 | 23.8838034 | 23.8850786 |
| Q9UG56 | PISD | 19.0750154 | 19.1077162 | 19.177093 | 19.1758394 | 19.2443636 |
| Q86UW6 | N4BP2 | 19.6383612 | 19.6137314 | 19.5425879 | 19.6103924 | 19.7616682 |
| O60563 | CCNT1 | 20.2939015 | 20.2919217 | 20.3577885 | 20.2906244 | 20.3582766 |
| O15550 | KDM6A | 20.0408487 | 20.1852441 | 20.099903 | 20.1842759 | 20.2523901 |
| O15533 | TAPBP | 21.9947534 | 21.9988202 | 21.8284717 | 21.8731313 | 22.0637844 |
| Q13608 | PEX6 | 19.9043977 | 19.9381713 | 20.0036448 | 19.9508667 | 20.0350582 |
| Q96T23 | RSF1 | 19.9191452 | 20.0734503 | 19.965551 | 19.9131104 | 20.0964167 |

| | | | | | | |
|---|---|---|---|---|---|---|
| Q14596 | NBR1 | 20.2520489 | 20.0076902 | 20.1374229 | 20.2804565 | 20.2831738 |
| Q9HA65 | TBC1D17 | 19.8388104 | 19.9223318 | 20.011147 | 19.9908017 | 20.039822 |
| P10301 | RRAS | 18.5168256 | 18.7018948 | 18.5365665 | 18.650897 | 18.7652741 |
| Q8IXW5 | RPAP2 | 18.6873193 | 18.8742463 | 18.9124926 | 18.8185215 | 18.8941112 |
| O75886 | STAM2 | 20.8351547 | 20.9295002 | 20.8441902 | 20.8701183 | 21.0398659 |
| Q7L3B6 | CDC37L1 | 18.9149702 | 19.0328327 | 19.0000487 | 18.9375558 | 19.1834846 |
| Q4V328 | GRIPAP1 | 20.7336641 | 20.9613832 | 20.6684639 | 20.8024575 | 20.9776133 |
| Q5T3F8 | TMEM63B | NA | 15.3871141 | 15.4599338 | NA | 15.5670315 |
| Q99571 | P2RX4 | 18.0233197 | 18.4049551 | 18.1626333 | 18.2962817 | 18.4088561 |
| Q9BYJ4 | TRIM34 | 16.0177268 | 15.9983884 | 16.1529222 | 16.3028683 | 16.1064776 |
| O00757 | FBP2 | 18.9099951 | 19.1127173 | 18.8296612 | 19.0439484 | 19.0324557 |
| Q16875 | PFKFB3 | 21.8973441 | 21.9480982 | 21.7113628 | 21.7416941 | 21.8879651 |
| P02655 | APOC2 | 22.1865643 | 22.2691321 | 22.1710999 | 22.3785007 | 22.4298624 |
| O43291 | SPINT2 | 19.2885904 | 19.2727444 | 19.2059101 | 19.280324 | 19.4221829 |
| Q8N5D6 | GBGT1 | 17.3261431 | 17.0843497 | 17.2919035 | 17.5302715 | 17.3751639 |
| Q96B45 | BORCS7 | 18.7840914 | 19.0975099 | 19.0525992 | 19.0631636 | 19.271987 |
| Q8WUR7 | C15orf40 | 17.6390012 | 17.708904 | 17.5561322 | 17.8878182 | 17.7710601 |
| Q9NRI5 | DISC1 | 16.2377215 | 16.7470872 | 16.4766051 | 16.5028772 | 16.7846516 |
| O75072 | FKTN | 17.8438619 | 18.0756917 | 17.5718435 | 17.9323977 | 17.9776145 |
| Q8IXQ5 | KLHL7 | 15.873341 | 16.1252633 | 15.9881638 | 16.2305273 | 16.0391439 |
| Q9Y5J5 | PHLDA3 | 17.2734787 | 17.4621008 | 17.3330461 | 18.1286229 | 17.5994189 |
| Q9P2S2 | NRXN2 | NA | 15.0508929 | 15.0161617 | NA | 15.2413109 |
| Q5RL73 | RBM48 | NA | 15.5659144 | 15.1487362 | NA | 15.713742 |
| P58005 | SESN3 | 14.168601 | NA | NA | 14.1642915 | NA |
| Q53T94 | TAF1B | 14.8493817 | 14.6102979 | 14.7222813 | 15.3467473 | 15.4330061 |
| P14927 | UQCRB | 24.5933845 | 24.597244 | 24.5673969 | 24.4546514 | 24.5624062 |
| O00330 | PDHX | 22.1046739 | 22.3073051 | 22.0997594 | 22.1337168 | 22.3253983 |
| Q15080 | NCF4 | 22.8303979 | 22.8286151 | 22.9000442 | 22.7794645 | 22.7812537 |
| Q8WXD5 | GEMIN6 | 18.61163 | 18.7409998 | 18.800891 | 18.6095866 | 19.0613642 |
| Q9NPE3 | NOP10 | 21.5628288 | 21.3302737 | 21.6615827 | 21.4268017 | 21.3474804 |
| P35612 | ADD2 | 22.1592398 | 22.3014679 | 22.10313 | 22.1906333 | 22.2678138 |
| O14545 | TRAFD1 | 20.2600264 | 20.1330764 | 20.2460748 | 20.1836545 | 20.2431844 |
| Q9UL45 | BLOC1S6 | 17.4480002 | 17.2932546 | 17.1691188 | 17.5164375 | 17.4025741 |
| Q99569 | PKP4 | 13.7576191 | NA | NA | 14.2474435 | NA |
| Q9H633 | RPP21 | 16.7504656 | 16.6792193 | 16.6329818 | 16.4852855 | 16.6379375 |
| Q9UGJ0 | PRKAG2 | NA | 16.0174922 | 16.0669484 | NA | 16.2037784 |
| P46977 | STT3A | 24.212487 | 24.223134 | 24.1591653 | 24.0676613 | 24.0358651 |
| P27105 | STOM | 24.115193 | 23.9569782 | 23.8397538 | 23.8896732 | 23.8288312 |
| Q14088 | RAB33A | 20.3811597 | 20.3015157 | 20.4018031 | 20.2082876 | 20.2777825 |
| Q9H6E4 | CCDC134 | 20.2146121 | 20.2613356 | 20.293421 | 20.194917 | 20.305538 |
| Q04206 | RELA | 21.7117454 | 21.7736857 | 21.7417418 | 21.767021 | 21.7795403 |
| Q6PID6 | TTC33 | 17.5270349 | 17.549347 | 17.6100235 | 17.6197088 | 17.6849699 |

| | | | | | | |
|---|---|---|---|---|---|---|
| P12259 | F5 | 20.5190191 | 20.7088481 | 20.7056248 | 20.5120621 | 20.6903277 |
| Q9HBH9 | MKNK2 | 16.2554337 | 16.385409 | 16.1915104 | 16.188578 | 16.3436308 |
| O95235 | KIF20A | 21.7824334 | 21.9027753 | 21.8448493 | 21.7857004 | 21.9044779 |
| P23193 | TCEA1 | 23.4581037 | 23.4733722 | 23.4648456 | 23.4394582 | 23.5071682 |
| Q9H410 | DSN1 | 19.5952233 | 19.6177901 | 19.5730655 | 19.6267296 | 19.6898125 |
| Q9NRA8 | EIF4ENIF1 | 19.3045641 | 19.3707284 | 19.5074186 | 19.550807 | 19.6058124 |
| O75531 | BANF1 | 24.5122764 | 24.6862383 | 24.4929328 | 24.5518727 | 24.6580781 |
| P08865 | RPSA | 26.550242 | 26.5996945 | 26.4710372 | 26.4436668 | 26.5322299 |
| Q8IWP9 | CCDC28A | 16.7484716 | 16.8885668 | 16.8575268 | 16.6785794 | 16.819228 |
| Q15572 | TAF1C | 12.5544517 | NA | NA | 12.8440721 | NA |
| O15519 | CFLAR | 16.1793741 | 16.1823575 | 16.4055883 | 16.1084121 | 16.2465526 |
| P0C1Z6 | TFPT | 18.6152563 | 18.5708202 | 18.5183966 | 18.7849459 | 18.7566152 |
| Q8N128 | FAM177A1 | 17.9271846 | 18.0770253 | 18.1631582 | 18.0275118 | 18.2109929 |
| O14641 | DVL2 | 19.6010393 | 19.7012062 | 19.5148681 | 19.4084303 | 19.6114469 |
| Q96GX2 | ATXN7L3B | 19.6712148 | 19.9566986 | 19.7638165 | 19.7874216 | 19.8093353 |
| Q6ZYL4 | GTF2H5 | 16.726502 | 16.6857148 | 16.627339 | 16.9080326 | 16.9230454 |
| P79483 | HLA-DRB3 | 19.6134977 | 20.1058384 | 19.6395028 | 19.7717988 | 19.8954026 |
| P13056 | NR2C1 | 16.8916099 | 16.8033393 | 16.9914698 | 17.0294591 | 17.1335354 |
| Q9BYX4 | IFIH1 | 15.8070222 | 15.4576573 | 15.4704832 | 15.5046489 | 15.2025728 |
| P80723 | BASP1 | 21.7153419 | 21.7570884 | 21.9678385 | 21.743436 | 21.6873474 |
| Q13564 | NAE1 | 22.4068546 | 22.4750429 | 22.4209368 | 22.3170149 | 22.3448653 |
| Q53GQ0 | HSD17B12 | 23.9994248 | 24.0087592 | 24.0140626 | 23.875062 | 23.9789876 |
| Q9NS87 | KIF15 | 20.5713895 | 20.6954572 | 20.6297952 | 20.6087604 | 20.6647209 |
| Q9H5Z1 | DHX35 | 18.7212545 | 18.8221851 | 18.880708 | 18.8795356 | 18.9262211 |
| Q9H9Q2 | COPS7B | 21.7187306 | 21.6691779 | 21.6232776 | 21.5268217 | 21.6681504 |
| O15083 | ERC2 | 15.6225317 | NA | NA | 15.6482763 | NA |
| Q8IV20 | LACC1 | 19.0996358 | 18.9818949 | 19.1818603 | 18.8944552 | 18.9795644 |
| Q8IUD2 | ERC1 | 19.6081423 | 19.677376 | 19.5812958 | 19.6175486 | 19.7502094 |
| Q01970 | PLCB3 | 20.5373192 | 20.6304938 | 20.5268278 | 20.5490942 | 20.6492091 |
| Q6PCD5 | RFWD3 | 16.550658 | 16.6053778 | 16.5559849 | 16.414233 | 16.1952927 |
| Q9H9Y4 | GPN2 | 18.7573267 | 18.7588589 | 18.871559 | 18.7011381 | 18.7951916 |
| Q8NBP0 | TTC13 | 18.0136243 | 18.0513073 | 18.1429443 | 17.9308366 | 17.9699 |
| P61923 | COPZ1 | 22.3193097 | 22.435291 | 22.3046415 | 22.2228576 | 22.2616142 |
| O95848 | NUDT14 | 19.5118722 | 19.361248 | 19.4999427 | 19.4111266 | 19.5693352 |
| Q5VUG0 | SFMBT2 | 18.6086787 | 18.6991976 | 18.8093296 | 18.5883214 | 18.8101073 |
| P67870 | CSNK2B | 22.6998037 | 22.7875193 | 22.7945258 | 22.6873882 | 22.795926 |
| Q96BZ4 | PLD4 | 24.8553595 | 24.8623344 | 24.7576494 | 24.7311504 | 24.7670284 |
| Q14738 | PPP2R5D | 21.5300104 | 21.5771183 | 21.6225675 | 21.4671906 | 21.5659338 |
| Q4G176 | ACSF3 | 21.7187915 | 21.7423372 | 21.815976 | 21.6733315 | 21.7083682 |
| A8MT69 | CENPX | 18.3438636 | 18.3849366 | 18.3854657 | 18.4158735 | 18.4745192 |
| Q8IUC6 | TICAM1 | 16.6914251 | 16.4893396 | 16.4233203 | 16.4051197 | 16.427415 |
| Q2VPK5 | CTU2 | 20.3055106 | 20.2806022 | 20.3866312 | 20.2709971 | 20.2561836 |

| | | | | | | |
|---|---|---|---|---|---|---|
| P41567 | EIF1 | 22.3571637 | 22.4900643 | 22.3065294 | 22.410091 | 22.5034925 |
| O95258 | SLC25A14 | 15.6697215 | NA | NA | 15.9210367 | NA |
| Q8WU67 | ABHD3 | 15.9091914 | NA | NA | 15.9521602 | NA |
| Q8IWR1 | TRIM59 | 19.2104163 | 19.2477507 | 19.3143864 | 19.1104001 | 19.299336 |
| Q96A49 | SYAP1 | 20.5584497 | 20.6726545 | 20.5339205 | 20.6873145 | 20.6580703 |
| P09429 | HMGB1 | 25.6751488 | 25.7791515 | 25.561054 | 25.6661537 | 25.7096008 |
| Q8NAF0 | ZNF579 | 19.2721589 | 19.5019405 | 19.4325635 | 19.5094102 | 19.500497 |
| Q6P1M3 | LLGL2 | NA | 15.1532285 | 14.7910707 | NA | 14.713376 |
| Q9NW75 | GPATCH2 | 17.6083036 | 17.5549134 | 17.2726198 | 17.5726803 | 17.4558419 |
| P14207 | FOLR2 | 13.808833 | 14.8655285 | 14.2155596 | 14.2326371 | 14.4002161 |
| Q03001 | DST | 19.5965741 | 20.2194842 | 19.8932809 | 20.022608 | 19.8178806 |
| Q9UHB6 | LIMA1 | 21.1948401 | 21.3281937 | 21.2765352 | 21.2913498 | 21.4040217 |
| Q9Y3B6 | EMC9 | NA | 15.2167575 | 15.3468023 | NA | 15.5263805 |
| P61619 | SEC61A1 | 24.0686747 | 24.1729323 | 24.0535716 | 23.9300186 | 24.0046962 |
| Q8N6M0 | OTUD6B | 21.4915453 | 21.5903994 | 21.4461633 | 21.4452757 | 21.5041828 |
| Q8TF74 | WIPF2 | 18.4843011 | NA | NA | 18.4495094 | NA |
| O43257 | ZNHIT1 | 17.394065 | 17.7465396 | 17.5728982 | 17.3545693 | 17.4038604 |
| P56378 | ATP5MJ | 22.7203945 | 22.1847657 | 22.5043101 | 22.2371572 | 22.0429936 |
| Q96E14 | RMI2 | 18.9289248 | 19.3096697 | 19.2205669 | 19.1192995 | 19.0269235 |
| P62851 | RPS25 | 26.7002264 | 26.713569 | 26.8976235 | 26.5892375 | 26.5178861 |
| Q8NG68 | TTL | 17.9332448 | 18.0615755 | 18.0677471 | 17.9574759 | 17.9532794 |
| P19438 | TNFRSF1A | 18.5247097 | 18.7701054 | 18.6596777 | 18.5284361 | 18.4868097 |
| Q8NEM0 | MCPH1 | 15.3511407 | 15.3173868 | 15.4739489 | 15.2523784 | 15.2294595 |
| P62913 | RPL11 | 26.0160886 | 25.9878351 | 25.9826051 | 25.9113032 | 25.9691076 |
| Q9NVR2 | INTS10 | 20.3422769 | 20.4242827 | 20.5114016 | 20.2661997 | 20.369919 |
| O00148 | DDX39A | 25.0333405 | 25.0837341 | 25.0758098 | 24.8995587 | 25.0672975 |
| O14925 | TIMM23 | 21.3272557 | 21.3254524 | 21.4364715 | 21.3052018 | 21.4256646 |
| Q14145 | KEAP1 | 21.0961953 | 21.1003081 | 21.0702664 | 21.0131958 | 21.115994 |
| Q5T1M5 | FKBP15 | 20.8243245 | 20.938371 | 20.8071285 | 20.8361878 | 20.9419716 |
| Q9BRT9 | GINS4 | 20.5580549 | 20.5977471 | 20.4760721 | 20.6113953 | 20.6279387 |
| P48382 | RFX5 | 19.2996864 | 19.4060974 | 19.3185772 | 19.4015429 | 19.2458723 |
| O15118 | NPC1 | 20.6362972 | 20.7231874 | 20.7367153 | 20.632649 | 20.7681847 |
| Q6ZU80 | CEP128 | 18.9426748 | 18.9997492 | 18.8330852 | 19.0402631 | 19.0401734 |
| Q6P2H3 | CEP85 | 19.8232651 | 19.9222227 | 19.7473079 | 19.8081951 | 19.8460593 |
| Q8IXQ4 | GPALPP1 | 19.7577893 | 19.8607964 | 19.8222787 | 19.9853041 | 20.0325163 |
| Q9P2S5 | WRAP73 | 16.6160744 | 16.6576668 | 16.5049317 | 16.7047873 | 16.6861644 |
| O00233 | PSMD9 | 21.9595986 | 21.8805347 | 21.914889 | 22.0109591 | 21.8122075 |
| P51957 | NEK4 | 16.8717988 | 16.866841 | 17.053716 | 17.1353359 | 17.1663757 |
| Q8IYJ3 | SYTL1 | 19.5427498 | 20.124342 | 19.8798298 | 19.9772535 | 20.1137254 |
| Q9NZ09 | UBAP1 | 18.7375765 | 18.8893436 | 18.8409628 | 18.9337505 | 19.0508688 |
| Q15293 | RCN1 | 21.438015 | 21.576814 | 21.324825 | 21.4724883 | 21.5135703 |
| P07225 | PROS1 | 15.8096348 | NA | NA | 15.9440912 | NA |

| | | | | | | |
|---|---|---|---|---|---|---|
| Q15776 | ZKSCAN8 | 18.8098586 | 18.6512626 | 18.7997027 | 18.8932567 | 19.0499686 |
| Q9BU70 | TRMO | 14.3389889 | NA | NA | 14.190602 | NA |
| Q15788 | NCOA1 | 16.0658102 | 16.5679347 | 16.4125063 | 16.4266195 | 16.7651765 |
| Q9NX38 | ABITRAM | 15.8914095 | NA | NA | 16.0059954 | NA |
| Q96E09 | PABIR1 | 15.5735635 | 16.5575571 | 16.308854 | 16.3715041 | 16.3224657 |
| O75503 | CLN5 | 19.7178265 | 19.83626 | 19.8156988 | 19.7975877 | 19.8580691 |
| Q13671 | RIN1 | 19.0920353 | 19.0743322 | 19.1242825 | 19.0627749 | 19.1523854 |
| Q92610 | ZNF592 | 19.0082547 | 19.0899761 | 19.1379207 | 19.0313977 | 19.1829877 |
| P68036 | UBE2L3 | 24.0405046 | 24.1335026 | 23.9179645 | 24.027575 | 24.1763535 |
| O95793 | STAU1 | 20.597626 | 20.6552037 | 20.6386694 | 20.6528248 | 20.7141293 |
| Q8N3Z3 | GTPBP8 | 17.8059601 | 17.9170374 | 18.205819 | 18.0792704 | 18.2078981 |
| P53803 | POLR2K | 21.003325 | 20.861806 | 20.9814914 | 20.7066823 | 20.5344583 |
| Q6P3X8 | PGBD2 | 15.7592873 | 15.9196065 | 15.696868 | 15.6869269 | 15.6951229 |
| P61626 | LYZ | 28.6293904 | 28.4000676 | 28.4219157 | 28.4855701 | 28.3456053 |
| Q5JRA6 | MIA3 | 20.6737514 | 20.8328744 | 20.7324207 | 20.718849 | 20.7795297 |
| Q9UQN3 | CHMP2B | 19.333247 | 19.4211611 | 19.2492972 | 19.3302537 | 19.5072744 |
| Q49AG3 | ZBED5 | 15.2376699 | 15.4350207 | 15.3965978 | 15.4340039 | 15.571524 |
| Q9BQE5 | APOL2 | 18.5413005 | 18.5352448 | 18.3098882 | 18.6881802 | 18.5840307 |
| O76041 | NEBL | NA | 13.3142268 | 13.0662058 | NA | 12.7966154 |
| P62888 | RPL30 | 25.8808464 | 26.0232021 | 25.9529589 | 25.8447544 | 25.8679719 |
| P15924 | DSP | 20.3079871 | 19.7395581 | 19.3444721 | 19.791068 | 19.1060907 |
| O00559 | EBAG9 | 18.0827949 | 18.1005682 | 18.2512069 | 18.1051839 | 18.0205909 |
| Q13362 | PPP2R5C | 21.0942105 | 21.0034611 | 20.9116374 | 20.9178514 | 20.863585 |
| Q9UMZ2 | SYNRG | 19.4614729 | 19.5215424 | 19.4170173 | 19.4771354 | 19.5513805 |
| Q9Y6M5 | SLC30A1 | 20.4371028 | 20.4500587 | 20.5100774 | 20.4520268 | 20.5047173 |
| Q9H1B5 | XYLT2 | 16.5416942 | 16.5534433 | 16.710839 | 16.4467204 | 16.6112225 |
| Q9BVP2 | GNL3 | 22.6372338 | 22.6339591 | 22.7804497 | 22.5903709 | 22.6736338 |
| P08631 | HCK | 22.6824061 | 22.6776538 | 22.7518475 | 22.6211124 | 22.68977 |
| Q7L1Q6 | BZW1 | 23.8869981 | 23.8632285 | 23.8376683 | 23.6922314 | 23.7811374 |
| P98194 | ATP2C1 | 19.8297619 | 19.981093 | 19.9245472 | 19.8551861 | 19.9131917 |
| Q8TDZ2 | MICAL1 | 23.0040909 | 22.9523726 | 22.8921732 | 22.886108 | 22.9088027 |
| P45973 | CBX5 | 23.8443341 | 23.87679 | 23.8542994 | 23.7645726 | 23.9134396 |
| Q16774 | GUK1 | 19.2657451 | 19.4477995 | 19.3513595 | 19.452263 | 19.5019412 |
| Q9BV73 | CEP250 | 19.4907949 | 19.6362032 | 19.506596 | 19.5905573 | 19.6719563 |
| Q96MW5 | COG8 | 20.2052003 | 20.1731336 | 20.1891229 | 20.1995353 | 20.311398 |
| Q9Y2U5 | MAP3K2 | 18.1106121 | 18.3492746 | 18.1713742 | 18.6442178 | 18.240604 |
| Q8IWB1 | ITPRIP | 17.2460818 | 17.7628526 | 17.4598565 | 17.5102772 | 17.7569785 |
| P14210 | HGF | 18.8375582 | 18.9181829 | 18.9736158 | 18.6520655 | 19.6630706 |
| P02765 | AHSG | 25.8350836 | 25.0722103 | 24.9543383 | 24.954596 | 24.9031845 |
| Q13433 | SLC39A6 | 16.6326813 | 17.024565 | 16.7674435 | 16.8586516 | 17.1615628 |
| Q9UPV0 | CEP164 | NA | 11.4418457 | 12.3731105 | NA | 11.7242284 |
| Q8TDN6 | BRIX1 | 22.1646079 | 22.1780541 | 22.3387127 | 22.0289388 | 22.2131202 |

| | | | | | | |
|---|---|---|---|---|---|---|
| Q9BV19 | C1orf50 | 18.8535937 | 18.9735019 | 18.6395322 | 18.9596806 | 19.0889076 |
| P46939 | UTRN | 19.9108895 | 20.0739194 | 19.9981674 | 19.9589951 | 20.1004345 |
| Q969Q6 | PPP2R3C | 15.8082709 | NA | NA | 15.8374243 | NA |
| Q9NS73 | MBIP | 17.4828089 | 17.5729195 | 17.4709074 | 17.4410594 | 17.5978705 |
| Q9UBR2 | CTSZ | 23.5901559 | 23.972689 | 23.651161 | 23.6030729 | 23.6949088 |
| Q15326 | ZMYND11 | 17.876788 | 18.0366372 | 18.1086074 | 17.7753415 | 17.8334837 |
| Q9NZL4 | HSPBP1 | 22.4357267 | 22.4742514 | 22.4967357 | 22.3279938 | 22.5422122 |
| Q9H488 | POFUT1 | 21.2173411 | 21.1364398 | 21.2531597 | 21.2277281 | 21.2556864 |
| Q9Y6V7 | DDX49 | 21.0525992 | 20.9941624 | 21.0895785 | 20.9457323 | 21.069345 |
| Q9BWJ5 | SF3B5 | 23.0564053 | 23.0382709 | 23.0629285 | 23.0673268 | 23.1394051 |
| Q7Z6E9 | RBBP6 | 21.6622371 | 21.7078928 | 21.6561773 | 21.6985404 | 21.6852079 |
| O75528 | TADA3 | 19.3713531 | 19.5623284 | 19.4250739 | 19.3740539 | 19.5838909 |
| P06756 | ITGAV | 14.774497 | NA | NA | 15.2832067 | NA |
| Q9H3T2 | SEMA6C | 18.9227047 | 18.826224 | 18.8904626 | 18.8300745 | 18.593712 |
| P62312 | LSM6 | 23.2270441 | 23.1410595 | 23.1888665 | 23.0000874 | 23.1177313 |
| P13995 | MTHFD2 | 21.1870381 | 21.3537027 | 21.2190357 | 21.2915295 | 21.2968425 |
| P51575 | P2RX1 | 17.6891508 | 17.6711071 | 17.7148622 | 17.8601666 | 17.5656909 |
| Q9BV38 | WDR18 | 21.4351349 | 21.5385479 | 21.6030565 | 21.6302227 | 21.7112967 |
| Q6ZQX7 | LIAT1 | 19.7061422 | 19.9535697 | 19.620753 | 19.8633994 | 20.0263453 |
| Q9NY15 | STAB1 | NA | 12.9834056 | 12.6205534 | NA | 12.7731991 |
| Q5T4B2 | CERCAM | 17.8954015 | 18.2179548 | 18.0209262 | 18.0167651 | 17.768425 |
| Q8ND90 | PNMA1 | NA | 15.3017055 | 15.1185699 | NA | 15.4030826 |
| Q9Y2U9 | KLHDC2 | 17.6411023 | 17.2351197 | 17.4213556 | 17.3526771 | 17.2768363 |
| Q6IN84 | MRM1 | 19.4557869 | 19.601677 | 19.6412761 | 19.35579 | 19.6979852 |
| Q13439 | GOLGA4 | 19.4149153 | 19.5401551 | 19.4067009 | 19.4384453 | 19.5456905 |
| Q9ULM6 | CNOT6 | 18.1488933 | 18.5404859 | 18.3144876 | 18.1504638 | 18.2799648 |
| Q9H3U5 | MFSD1 | 21.7737401 | 21.9002285 | 21.837678 | 21.7265322 | 21.5077431 |
| Q7Z614 | SNX20 | 19.9018045 | 19.9337278 | 20.0504303 | 19.9518699 | 19.9765712 |
| O76021 | RSL1D1 | 24.3597862 | 24.3830757 | 24.4952419 | 24.2782587 | 24.2985364 |
| Q8N766 | EMC1 | 21.6471158 | 21.6539437 | 21.6353176 | 21.5967953 | 21.643239 |
| Q9UKA4 | AKAP11 | 19.2149869 | 19.2030081 | 19.1476117 | 19.2366013 | 19.4217625 |
| P51970 | NDUFA8 | 22.6822571 | 22.8274025 | 23.0052555 | 22.7854855 | 22.8128434 |
| P63208 | SKP1 | 23.8552965 | 23.8612178 | 23.5902222 | 23.7766117 | 23.8539487 |
| Q7Z6B0 | CCDC91 | 18.6738784 | 18.8455083 | 18.5442875 | 18.712706 | 18.8632246 |
| Q9NPB8 | GPCPD1 | 19.7223786 | 19.8030632 | 19.794856 | 19.7706945 | 19.8286819 |
| Q8N9H8 | EXD3 | 14.4519186 | 14.1618551 | 14.2605871 | 13.9988839 | 13.2996779 |
| Q8WV28 | BLNK | 18.6575996 | 18.6622797 | 18.8124277 | 18.3607387 | 18.6222324 |
| Q5JXC2 | MIIP | NA | 13.6580636 | 13.6788723 | NA | 13.3364665 |
| P54259 | ATN1 | 15.9866595 | NA | NA | 15.9481958 | NA |
| Q9HBE1 | PATZ1 | 18.9363722 | 19.0229533 | 19.0974224 | 18.9459792 | 18.8795449 |
| O75064 | DENND4B | 21.9427202 | 22.0877223 | 22.0077296 | 21.9543075 | 21.9553117 |
| Q9Y3B7 | MRPL11 | 22.6565059 | 22.5702003 | 22.6251974 | 22.614637 | 22.6470191 |

| | | | | | | |
|---|---|---|---|---|---|---|
| Q9NR45 | NANS | 23.2673641 | 23.3205399 | 23.2684172 | 23.2740407 | 23.3692866 |
| Q5TGY3 | AHDC1 | 17.3067857 | 17.4593593 | 17.4394735 | 17.5200396 | 17.575 |
| Q13443 | ADAM9 | 17.110621 | 17.5391585 | 17.5717987 | 17.4882312 | 17.4133101 |
| Q9UKT7 | FBXL3 | NA | 15.1820845 | 15.190508 | NA | 15.2623875 |
| P42684 | ABL2 | 14.9998466 | 15.3906714 | 15.4933905 | 15.4321625 | 15.3755389 |
| Q9NXC5 | MIOS | 19.8214478 | 19.9594335 | 19.9706245 | 19.9045712 | 20.0540195 |
| Q96QT6 | PHF12 | 18.0480716 | 18.2440515 | 18.3943507 | 18.3230047 | 18.3626175 |
| O75373 | ZNF737 | 15.4922466 | NA | NA | 15.1988284 | NA |
| Q9BUT9 | MCRIP2 | 19.5804689 | 19.4043814 | 20.03819 | 19.5276196 | 19.7364264 |
| O75477 | ERLIN1 | 22.9577246 | 23.0034343 | 22.9614269 | 22.8788233 | 22.8823152 |
| P05166 | PCCB | 20.7303807 | 20.8482774 | 20.7887453 | 20.7507236 | 20.890464 |
| P51665 | PSMD7 | 23.7167335 | 23.5550024 | 23.7572737 | 23.4822864 | 23.7013252 |
| O43299 | AP5Z1 | 20.2509392 | 20.3150971 | 20.4064665 | 20.2744467 | 20.2233261 |
| Q52LW3 | ARHGAP29 | 17.1373202 | NA | NA | 16.4265138 | NA |
| Q9HBH0 | RHOF | 19.0493127 | 19.225376 | 19.4313671 | 19.1724421 | 19.3860083 |
| P56962 | STX17 | 19.3116208 | 19.4648436 | 19.6000956 | 19.2003124 | 19.3083713 |
| P62273 | RPS29 | 25.9570763 | 25.8813011 | 25.9683405 | 25.9609526 | 25.8925096 |
| O75161 | NPHP4 | 14.7490072 | 15.3681978 | 15.593465 | 15.4215631 | 15.0970328 |
| Q9BS16 | CENPK | 18.1087254 | 18.3857532 | 18.1358597 | 18.1453206 | 18.2991223 |
| A6NKF1 | SAC3D1 | 18.5516265 | 18.4802258 | 18.4873557 | 18.3997082 | 18.5531236 |
| Q7L5D6 | GET4 | 20.8364982 | 20.7270316 | 20.8369291 | 20.7063806 | 20.8641599 |
| Q8WV60 | PTCD2 | 18.7151786 | 18.6280141 | 18.9027983 | 18.7135574 | 18.8412018 |
| Q8IY81 | FTSJ3 | 22.3974738 | 22.5071336 | 22.5001519 | 22.4172298 | 22.3971708 |
| P38398 | BRCA1 | 19.4614537 | 19.6324605 | 19.5129358 | 19.5311868 | 19.6733676 |
| Q9P2X0 | DPM3 | 22.1176404 | 22.0880173 | 21.8733117 | 22.0440515 | 21.9226802 |
| Q8NE86 | MCU | 20.8649577 | 20.99854 | 20.9925135 | 20.8584023 | 21.0213794 |
| Q9Y6B7 | AP4B1 | 19.3037051 | 19.2502259 | 19.3586569 | 19.1690226 | 19.3201145 |
| P43155 | CRAT | 20.5063417 | 20.589169 | 20.6315637 | 20.5246118 | 20.6164577 |
| Q13283 | G3BP1 | 24.1772991 | 24.2719314 | 24.2162828 | 24.2367064 | 24.2909 |
| P15498 | VAV1 | 22.9567118 | 22.9579907 | 23.0536529 | 22.8483248 | 22.9262921 |
| O95239 | KIF4A | 20.9859645 | 21.1101651 | 21.0947969 | 21.0746892 | 21.1073368 |
| Q9BV40 | VAMP8 | 25.6058642 | 25.6160171 | 25.7828402 | 25.5062549 | 25.4610953 |
| Q8WY91 | THAP4 | 17.8680197 | 17.9310231 | 17.9511865 | 17.8127591 | 18.1504697 |
| Q8N7H5 | PAF1 | 20.84459 | 20.8043365 | 20.7645659 | 20.815233 | 20.9251406 |
| Q9UDV7 | ZNF282 | 12.96357 | NA | NA | 13.4088197 | NA |
| P53804 | TTC3 | 18.2078384 | 18.0610838 | 17.5914292 | 17.5227397 | 17.7723016 |
| Q99684 | GFI1 | 20.2039468 | 20.346924 | 20.3293614 | 20.1881633 | 20.3202043 |
| Q9HBH5 | RDH14 | 20.000031 | 20.1594556 | 19.9387497 | 19.9681272 | 19.9822725 |
| Q14197 | MRPL58 | 22.3322379 | 22.3468389 | 22.3489215 | 22.2521324 | 22.3360315 |
| Q9NZ52 | GGA3 | 20.3870096 | 20.508601 | 20.4338767 | 20.4059075 | 20.4897712 |
| Q86UW9 | DTX2 | 17.0347846 | 17.5990905 | 17.7807344 | 17.3601771 | 17.5473769 |
| Q13569 | TDG | 20.3995539 | 20.5965276 | 20.3984923 | 20.4791629 | 20.2683422 |

| | | | | | | |
|---|---|---|---|---|---|---|
| P13716 | ALAD | 21.2664996 | 21.2403075 | 21.3941705 | 21.1955369 | 21.3217403 |
| P51825 | AFF1 | 21.0940456 | 21.1417583 | 21.1360146 | 20.9728683 | 21.0123017 |
| Q8IVH2 | FOXP4 | 18.6899767 | 18.7023118 | 18.6999571 | 18.7132238 | 18.7299017 |
| Q8TEU7 | RAPGEF6 | 19.9240591 | 19.9624792 | 19.920813 | 19.949244 | 20.091165 |
| Q0P6H9 | TMEM62 | 18.7225531 | NA | NA | 19.2849023 | NA |
| Q92564 | DCUN1D4 | 17.6336904 | 17.639779 | 17.7896734 | 17.6080829 | 17.5406052 |
| O15541 | RNF113A | 20.1907449 | 20.3648184 | 20.2210196 | 20.3661702 | 20.4975744 |
| Q96SN8 | CDK5RAP2 | 19.3303153 | 19.4704494 | 19.2100128 | 19.2317527 | 19.4294854 |
| O75665 | OFD1 | 21.3304187 | 21.3198174 | 21.2006028 | 21.2180546 | 21.1933718 |
| Q15291 | RBBP5 | 21.1728193 | 21.2807774 | 21.3901755 | 21.2978202 | 21.4175946 |
| Q3MIT2 | PUS10 | 17.8802445 | 17.7413284 | 18.0507768 | 17.9462282 | 18.0299897 |
| P50053 | KHK | 19.0480075 | 19.065285 | 19.0979355 | 19.1750014 | 19.2489098 |
| P35813 | PPM1A | 20.5321622 | 20.595739 | 20.5171914 | 20.6133962 | 20.5945248 |
| Q8N9N8 | EIF1AD | 19.8174799 | 19.8968988 | 19.9049471 | 20.0296839 | 20.1675567 |
| Q9Y324 | FCF1 | 20.6947547 | 20.7715932 | 20.7833985 | 20.5084257 | 20.7019297 |
| Q96CW1 | AP2M1 | 23.6286152 | 23.5735106 | 23.6867134 | 23.5537947 | 23.587632 |
| O43914 | TYROBP | 19.8083953 | 19.5863158 | 19.6762263 | 19.6903901 | 19.5446354 |
| Q14146 | URB2 | 20.3382157 | 20.3132297 | 20.4533533 | 20.3223161 | 20.3519506 |
| P15559 | NQO1 | 19.8375657 | 19.8682505 | 19.8195063 | 19.7990074 | 19.8606366 |
| Q9Y2G5 | POFUT2 | 19.5043906 | 19.5363904 | 19.5625254 | 19.6071903 | 19.6262863 |
| Q92551 | IP6K1 | 17.4520757 | 17.6726316 | 17.5533427 | 17.5547991 | 17.8980764 |
| Q9BXP2 | SLC12A9 | 20.4174339 | 20.4410137 | 20.477718 | 20.2539983 | 20.4568117 |
| Q9NZQ3 | NCKIPSD | 20.0494849 | 20.0357276 | 20.2156051 | 20.0501088 | 20.1950165 |
| P02750 | LRG1 | 18.3491379 | 18.4155413 | 18.5095659 | 18.1755582 | 18.5051702 |
| Q9Y2R5 | MRPS17 | 20.8978082 | 21.0525689 | 20.9071014 | 20.8289066 | 20.8823549 |
| Q8IWI9 | MGA | 18.8646328 | 19.0429713 | 19.0452984 | 19.0129627 | 19.0986254 |
| Q14005 | IL16 | 20.4302068 | 20.4871469 | 20.5373439 | 20.4548623 | 20.5853191 |
| Q9NV64 | TMEM39A | 16.7387245 | 16.7574562 | 17.206368 | 16.812272 | 16.8896813 |
| Q8TDH9 | BLOC1S5 | 19.4538599 | 19.4669854 | 19.2992708 | 19.3561823 | 19.6021217 |
| Q9NQS1 | AVEN | 20.2032323 | 20.3193389 | 20.1734585 | 20.3044492 | 20.3742007 |
| Q15286 | RAB35 | 22.3010382 | 22.3464754 | 22.3688618 | 22.1660414 | 22.1866821 |
| Q53EZ4 | CEP55 | 19.7415077 | 19.7281067 | 19.734014 | 19.7525864 | 19.8197314 |
| Q9P0B6 | CCDC167 | 20.0304653 | 20.032665 | 20.2041778 | 19.9465831 | 19.9316198 |
| Q6DCA0 | AMMECR1L | 15.1504652 | 15.1098292 | 15.4697959 | 15.1436911 | 15.1340545 |
| Q7L0Q8 | RHOU | 14.5336782 | NA | NA | 14.6872355 | NA |
| O95067 | CCNB2 | 20.148877 | 20.1838884 | 20.1827101 | 19.9827408 | 20.0801913 |
| O43196 | MSH5 | NA | 17.0584256 | 17.3516977 | NA | 17.0746603 |
| Q2QL34 | MPV17L | 18.0536505 | 18.1410827 | 18.3892122 | 17.8769003 | 17.6112 |
| Q6NUK4 | REEP3 | 18.1344466 | 17.6499265 | 17.7556089 | 18.1591913 | 17.4436995 |
| P35613 | BSG | 23.9194323 | 24.0953474 | 24.0457416 | 23.8397688 | 23.9763152 |
| P56182 | RRP1 | 22.301981 | 22.3084878 | 22.3375286 | 22.187799 | 22.2019182 |
| P63218 | GNG5 | 24.6912045 | 24.7329281 | 24.6639644 | 24.5329426 | 24.525403 |

| | | | | | | |
|---|---|---|---|---|---|---|
| P15170 | GSPT1 | 24.247138 | 24.2066957 | 24.1581743 | 24.2248844 | 24.3319381 |
| Q08AF3 | SLFN5 | 15.1164166 | NA | NA | 15.1663873 | NA |
| P52788 | SMS | 22.9078749 | 22.920769 | 22.8327036 | 22.8249855 | 22.8653865 |
| Q9Y4I5 | TESMIN | 13.9484686 | NA | NA | 13.5492284 | NA |
| Q9UKT8 | FBXW2 | 16.3586442 | 16.74297 | 16.7996654 | 16.5430897 | 16.4805494 |
| Q969Z4 | RELT | 14.611388 | 14.693258 | 14.6716988 | 14.8035545 | 14.6629508 |
| Q96IZ5 | RBM41 | 17.1995894 | 16.8739647 | 16.7164286 | 16.7870396 | 16.7638036 |
| Q8IW52 | SLITRK4 | 16.0379533 | 16.1377214 | 16.2057637 | 15.8158372 | 16.1715165 |
| Q9NVH6 | TMLHE | 18.7384069 | 18.8376067 | 18.7304846 | 18.6296212 | 18.4962636 |
| P03915 | MT-ND5 | 19.6995499 | 19.7668597 | 19.9320905 | 19.7078248 | 19.7704072 |
| P62899 | RPL31 | 26.7798109 | 26.6898775 | 26.7918594 | 26.5161744 | 26.6244656 |
| P48509 | CD151 | 22.2839685 | 22.1126923 | 22.0238292 | 22.1115846 | 22.034019 |
| Q6UWP2 | DHRS11 | 17.4393949 | 17.5049961 | 17.5082781 | 17.3546125 | 17.5112258 |
| Q9UBB6 | NCDN | 21.4305253 | 21.5199059 | 21.5759215 | 21.3779114 | 21.4600774 |
| Q9NP79 | VTA1 | 22.4294649 | 22.5800038 | 22.5532311 | 22.4628221 | 22.5276075 |
| Q8IU81 | IRF2BP1 | 21.6197164 | 21.675337 | 21.6645041 | 21.6595884 | 21.7908203 |
| O95825 | CRYZL1 | 20.3338492 | 20.5179327 | 20.4416142 | 20.3852773 | 20.4983401 |
| Q86YR5 | GPSM1 | 18.6660297 | 18.6047827 | 18.7779474 | 18.6545219 | 18.7550029 |
| O95365 | ZBTB7A | 19.3143148 | 19.3623111 | 19.4294925 | 19.5400733 | 19.5113418 |
| Q0VD83 | APOBR | 22.2122258 | 22.4178436 | 22.2741535 | 22.2536662 | 22.4791397 |
| P83881 | RPL36A | 25.0480094 | 25.2461305 | 25.400304 | 25.2387351 | 25.381757 |
| Q9UBJ2 | ABCD2 | 17.0790619 | 16.9867852 | 17.3068499 | 17.2164678 | 17.3815833 |
| Q9H0Y0 | ATG10 | 15.3749577 | 15.2491682 | 15.3165475 | 15.1880699 | 15.4626042 |
| Q9NX63 | CHCHD3 | 23.966432 | 24.0329 | 24.0172513 | 23.91167 | 23.9139944 |
| P78332 | RBM6 | 20.8828188 | 20.932145 | 20.868979 | 20.8693675 | 20.9263032 |
| Q96EA4 | SPDL1 | 19.22381 | 19.5929115 | 19.3245722 | 19.2539443 | 19.4929165 |
| Q96MY1 | NOL4L | 18.1044807 | 18.0905055 | 18.2588059 | 18.3867015 | 18.2271174 |
| Q8N184 | ZNF567 | 17.0980429 | 16.7489937 | 17.0010875 | 17.1963706 | 16.822367 |
| Q9P0K8 | FOXJ2 | 17.0291367 | 17.4294756 | 17.0376195 | 17.0485861 | 17.396443 |
| Q9P1U1 | ACTR3B | 17.5548991 | 17.541866 | 17.8417778 | 17.7294664 | 17.6856332 |
| Q6NZY4 | ZCCHC8 | 19.6832082 | 19.9074098 | 19.7797356 | 19.7472958 | 19.9321464 |
| Q10469 | MGAT2 | 21.7016253 | 21.6094716 | 21.6348282 | 21.6074274 | 21.5896642 |
| Q14676 | MDC1 | 21.2496483 | 21.3836234 | 21.3369828 | 21.2817193 | 21.3976847 |
| Q9BQA9 | CYBC1 | 23.0259696 | 23.2335479 | 22.9322862 | 22.893994 | 23.1632087 |
| Q9BYD3 | MRPL4 | 22.0992728 | 22.0082501 | 22.1213547 | 22.0849412 | 22.0645027 |
| Q969V6 | MRTFA | 19.0088526 | 19.1888015 | 18.9663324 | 18.9765391 | 19.1183113 |
| Q9UDW1 | UQCR10 | 22.0597347 | 21.8265354 | 22.0565335 | 21.670338 | 22.2133156 |
| O76096 | CST7 | 19.6453428 | 19.2245985 | 19.3560054 | 19.4600414 | 19.3033695 |
| Q9Y6A1 | POMT1 | 17.284186 | 17.0517665 | 17.1865682 | 16.6207172 | 17.0184682 |
| Q9NP50 | SINHCAF | 18.1737328 | 18.4133499 | 18.6096706 | 18.3795513 | 18.1435778 |
| P78415 | IRX3 | 16.7856175 | 16.746682 | 16.8098532 | 16.6202845 | 16.8837321 |
| Q99685 | MGLL | 16.9317969 | 16.6710466 | 16.7167619 | 16.8169265 | 16.7602161 |

| | | | | | | |
|---|---|---|---|---|---|---|
| Q4AC94 | C2CD3 | 13.9815834 | 14.3752767 | 14.0881027 | 14.0942364 | 14.2580345 |
| Q8WXK3 | ASB13 | 15.9696532 | 15.8605008 | 15.9373479 | 15.8819668 | 15.8386994 |
| Q8TBF5 | PIGX | 18.735705 | 19.2355564 | 18.8016518 | 18.6798287 | 18.6220786 |
| P19823 | ITIH2 | 19.547466 | 19.4444612 | 19.2817704 | 19.2364374 | 19.4007403 |
| Q96HV5 | TMEM41A | 19.3200902 | 19.3535238 | 19.4669619 | 19.322961 | 19.4086663 |
| P40429 | RPL13A | 26.3918636 | 26.4286598 | 26.5110667 | 26.2394998 | 26.3242592 |
| P62753 | RPS6 | 26.015963 | 26.0360536 | 26.189551 | 25.9221374 | 26.0164652 |
| P60468 | SEC61B | 22.5702695 | 22.6633414 | 22.6467874 | 22.5097457 | 22.6576061 |
| P20138 | CD33 | 21.7483612 | 21.9794593 | 21.9122796 | 21.7054131 | 21.8848651 |
| Q14CZ7 | FASTKD3 | 18.5667694 | 18.7117627 | 18.6739424 | 18.572267 | 18.473418 |
| P83111 | LACTB | 21.0381514 | 21.1093561 | 21.239118 | 20.9750255 | 21.0624904 |
| Q86V21 | AACS | 22.0688759 | 21.9741816 | 22.1342193 | 21.9012494 | 22.0863233 |
| Q8WWY3 | PRPF31 | 22.4586274 | 22.478507 | 22.5180328 | 22.3124209 | 22.3481928 |
| Q9BVV7 | TIMM21 | 20.9019765 | 20.9618265 | 20.9315353 | 20.8241565 | 20.9710807 |
| Q9Y3D6 | FIS1 | 22.352114 | 22.3804963 | 22.4291063 | 22.2364693 | 22.3904313 |
| P52565 | ARHGDIA | 25.830869 | 25.8345881 | 25.7994947 | 25.7303034 | 25.7459254 |
| Q8N2K0 | ABHD12 | 21.5309347 | 21.5340105 | 21.7054602 | 21.555007 | 21.6138268 |
| Q9NVP1 | DDX18 | 22.9942943 | 23.0079554 | 23.0648896 | 22.9535276 | 22.9486991 |
| Q8N5M1 | ATPAF2 | 19.7030122 | 19.6594624 | 19.6221308 | 19.5915738 | 19.6225494 |
| A0FGR8 | ESYT2 | 22.8799629 | 22.9021111 | 22.9198908 | 22.7686813 | 22.8654913 |
| Q5TBB1 | RNASEH2B | 22.2944213 | 22.3455888 | 22.3200693 | 22.2218643 | 22.3013042 |
| O14617 | AP3D1 | 22.4465867 | 22.4788159 | 22.5383724 | 22.3686943 | 22.468624 |
| Q9Y4D7 | PLXND1 | 19.185724 | 19.2907915 | 19.23539 | 19.1997131 | 19.1982446 |
| Q9UNF0 | PACSIN2 | 22.7631113 | 22.8500966 | 22.7609807 | 22.7923567 | 22.8648761 |
| Q53F19 | NCBP3 | 20.7800502 | 20.8398514 | 20.8150822 | 20.8521718 | 20.8852969 |
| Q9UQB8 | BAIAP2 | 19.2923565 | 19.2424049 | 19.3292657 | 19.2815385 | 19.3409065 |
| P61758 | VBP1 | 22.4229125 | 22.4539575 | 22.4174919 | 22.444407 | 22.5106568 |
| Q6BDS2 | BLTP3A | 19.3843866 | 19.2505045 | 19.2524363 | 19.3310574 | 19.3911683 |
| Q96R06 | SPAG5 | 19.0390333 | 19.0977093 | 19.0458213 | 19.0151459 | 19.1401364 |
| Q96BR1 | SGK3 | 19.9188958 | 19.8621129 | 19.9410417 | 19.9747598 | 20.0524736 |
| Q9H4Z3 | PCIF1 | 20.1706218 | 20.2626982 | 20.2740834 | 20.2235777 | 20.2649266 |
| Q5SW79 | CEP170 | 20.8894996 | 20.9886309 | 20.9123287 | 20.8923097 | 20.9621372 |
| Q86YH6 | PDSS2 | 19.0081735 | 19.1330252 | 19.1179517 | 19.1223746 | 19.1074597 |
| Q6ZU65 | UBN2 | 18.8515306 | 19.054575 | 18.9549708 | 18.9010918 | 18.9979193 |
| Q6P2P2 | PRMT9 | 18.9561893 | 18.8797341 | 18.9239046 | 19.0046836 | 19.0067134 |
| Q9Y2Q9 | MRPS28 | 21.7718393 | 21.8823018 | 21.920271 | 21.9323659 | 21.8299771 |
| P30044 | PRDX5 | 23.0051611 | 23.1638798 | 23.0685995 | 23.0613305 | 23.2079247 |
| Q86WA8 | LONP2 | 17.9783322 | 18.0595196 | 18.029167 | 18.1159624 | 18.1048679 |
| Q8WUD4 | CCDC12 | 21.5847148 | 21.7420478 | 21.736952 | 21.7903076 | 21.8334964 |
| Q99676 | ZNF184 | 17.4287852 | 17.5138401 | 17.4741524 | 17.5541102 | 17.4543513 |
| Q86V88 | MDP1 | 19.8356343 | 19.9290028 | 19.7121947 | 19.8308359 | 19.8341736 |
| Q96CN9 | GCC1 | 19.4946806 | 19.762645 | 19.6135062 | 19.6836875 | 19.7654749 |

| | | | | | | |
|---|---|---|---|---|---|---|
| P16333 | NCK1 | 20.1019821 | 20.2228297 | 20.0200762 | 20.1414314 | 20.2581639 |
| Q9NRY5 | FAM114A2 | 21.1168684 | 21.1599491 | 21.0183485 | 21.1118745 | 21.2596995 |
| Q96RF0 | SNX18 | 18.3869379 | 18.3665091 | 18.3206795 | 18.5143844 | 18.5552962 |
| P82932 | MRPS6 | 22.2179663 | 22.0628545 | 22.3392367 | 22.2207049 | 22.1998143 |
| Q8N5Y2 | MSL3 | 17.9446948 | 18.1380686 | 18.1689925 | 18.2076964 | 18.3671645 |
| Q9H0N5 | PCBD2 | 22.402913 | 22.5384712 | 22.3313527 | 22.4715473 | 22.4157234 |
| O95229 | ZWINT | 19.4034259 | 19.4257422 | 19.4605882 | 19.437959 | 19.5336963 |
| Q96C23 | GALM | 21.5897758 | 21.6137287 | 21.6421933 | 21.7114961 | 21.8489192 |
| Q9NRH2 | SNRK | 18.1464382 | 17.9591815 | 18.1138299 | 18.1192065 | 18.0843687 |
| Q15545 | TAF7 | 19.7923134 | 19.9557578 | 19.7456707 | 19.8493279 | 19.9492098 |
| Q8IZ21 | PHACTR4 | 19.0513321 | 19.1321861 | 19.2497578 | 19.3252582 | 19.5593767 |
| Q9UEE9 | CFDP1 | 20.6446261 | 20.8993244 | 20.512524 | 20.6791258 | 20.8145247 |
| Q9H2S1 | KCNN2 | 19.1123071 | 18.9123815 | 18.9511756 | 19.0546411 | 18.9769474 |
| P17096 | HMGA1 | 25.2565893 | 25.3627713 | 25.1163614 | 25.1768917 | 25.3105517 |
| Q9BY12 | SCAPER | 18.4214139 | 18.3368231 | 18.4614593 | 18.5643572 | 18.6170671 |
| Q14CB8 | ARHGAP19 | 17.8304941 | 18.0357396 | 17.9274707 | 17.9445506 | 18.2559366 |
| Q93073 | SECISBP2L | 21.5037798 | 21.7667057 | 21.5977494 | 21.6971197 | 21.8827901 |
| Q96K80 | ZC3H10 | 17.6826681 | 17.9850174 | 17.8528719 | 17.9873255 | 18.0445646 |
| Q3KRA9 | ALKBH6 | 18.5428393 | 18.7143611 | 18.6518703 | 18.5459802 | 18.7144666 |
| Q9H8K7 | C10orf88 | 14.5209321 | NA | NA | 14.7675111 | NA |
| P09466 | PAEP | NA | 14.7133171 | 14.383119 | NA | 14.8461435 |
| P49459 | UBE2A | 18.6916962 | 19.1001598 | 18.8974246 | 19.0050459 | 19.0905979 |
| Q99952 | PTPN18 | 18.9902679 | 19.0947979 | 19.0980832 | 19.0562271 | 19.4598124 |
| Q9NYQ7 | CELSR3 | 17.7074997 | 18.1031154 | 17.7159632 | 17.8144791 | 17.6499952 |
| Q8NI22 | MCFD2 | 19.6389649 | 19.4672733 | 19.3835062 | 19.6544317 | 19.8654455 |
| Q9BY66 | KDM5D | 15.6916892 | NA | NA | 15.6344799 | NA |
| Q9BS18 | ANAPC13 | 11.2253274 | NA | NA | 11.6772643 | NA |
| Q70CQ4 | USP31 | 12.2389385 | NA | NA | 12.4520459 | NA |
| Q9NRM2 | ZNF277 | 16.0515434 | 16.41357 | 16.1054903 | 15.9721467 | 15.8494893 |
| Q9NRX4 | PHPT1 | 23.3154219 | 23.3966643 | 23.172981 | 23.2024398 | 22.8799597 |
| Q9NX07 | TRNAU1AP | 18.3749026 | 18.4844879 | 18.5550777 | 18.3967703 | 18.6001397 |
| P22234 | PAICS | 25.4491726 | 25.4938373 | 25.4610381 | 25.3538301 | 25.452117 |
| Q9Y4C8 | RBM19 | 20.2510747 | 20.2642211 | 20.3032442 | 20.3491617 | 20.3491598 |
| Q9P2N5 | RBM27 | 20.4019695 | 20.5545704 | 20.4283605 | 20.4549951 | 20.5214351 |
| Q9Y520 | PRRC2C | 22.1702082 | 22.327955 | 22.1519542 | 22.1891222 | 22.2730053 |
| Q99590 | SCAF11 | 21.4994576 | 21.6737853 | 21.449291 | 21.4795642 | 21.6459009 |
| O94967 | WDR47 | 18.3565216 | 18.2685592 | 18.2608686 | 18.1441689 | 18.4414275 |
| Q9HD42 | CHMP1A | 22.4404329 | 22.5111821 | 22.2038744 | 22.3850022 | 22.4089387 |
| Q9Y5K6 | CD2AP | 19.6867337 | 19.8639086 | 19.632208 | 19.6855503 | 19.9355842 |
| Q02750 | MAP2K1 | 21.7008076 | 21.8666745 | 21.8908253 | 21.6583433 | 21.8725631 |
| Q9NVT9 | ARMC1 | 20.9686772 | 20.9499588 | 20.906599 | 20.9157734 | 20.8420833 |
| Q9NXH9 | TRMT1 | 21.6711733 | 21.6933405 | 21.6498963 | 21.5918434 | 21.7398281 |

| | | | | | | |
|---|---|---|---|---|---|---|
| Q6ZVM7 | TOM1L2 | 19.123317 | 19.2190003 | 19.1654381 | 19.1927637 | 19.3170896 |
| Q9H477 | RBKS | 21.082371 | 21.1349899 | 21.2316779 | 21.0818985 | 21.1547336 |
| Q9UHD9 | UBQLN2 | 20.9120107 | 21.1127384 | 21.0123632 | 21.0457133 | 21.3051121 |
| Q96ST8 | CEP89 | 18.4994064 | 18.6367061 | 18.4433615 | 18.416348 | 18.7257475 |
| P01024 | C3 | 18.3164799 | 17.9528001 | 18.2737167 | 17.8499719 | 17.3310592 |
| P56192 | MARS1 | 22.7170323 | 22.7309891 | 22.7766557 | 22.6112011 | 22.7319033 |
| O60232 | ZNRD2 | 20.5234239 | 20.6366982 | 20.5213981 | 20.5355655 | 20.6690676 |
| Q86X76 | NIT1 | 20.1915663 | 20.2882478 | 20.2136907 | 20.2007467 | 20.4388619 |
| P49759 | CLK1 | 17.2675989 | 17.2540525 | 17.3497693 | 17.2539931 | 17.5758341 |
| Q5HYA8 | TMEM67 | NA | 22.2496272 | 22.4904432 | NA | 22.4573568 |
| P17544 | ATF7 | 18.3773422 | 18.4874473 | 18.3138358 | 18.379014 | 18.6314775 |
| Q96QA5 | GSDMA | NA | 13.0698407 | 13.1885254 | NA | 12.5287444 |
| Q08AE8 | SPIRE1 | 16.9338053 | 17.1459471 | 17.3789635 | 16.7317807 | 16.8876308 |
| Q13203 | MYBPH | 20.3284306 | 20.7576046 | 20.7408837 | 20.33306 | 20.8334215 |
| P52849 | NDST2 | 17.1898034 | 17.0766903 | 17.2041125 | 16.7827624 | 16.991275 |
| O43741 | PRKAB2 | 16.0390036 | 16.6637408 | 16.1609528 | 16.1003529 | 15.8649851 |
| Q8N5H7 | SH2D3C | 17.9673937 | 18.4072467 | 18.4237676 | 18.1693065 | 18.3297433 |
| Q6UX65 | DRAM2 | 19.4533167 | 19.6023712 | 19.6568889 | 19.4683391 | 19.3290018 |
| P56134 | ATP5MF | 23.0511978 | 23.048119 | 23.1397953 | 22.9563529 | 23.1392432 |
| P01730 | CD4 | 20.6250354 | 20.6285186 | 20.7726028 | 20.5373568 | 20.707589 |
| P20042 | EIF2S2 | 23.3497095 | 23.5388529 | 23.4483279 | 23.2803013 | 23.3092067 |
| P10253 | GAA | 22.0599759 | 22.1207209 | 22.0360745 | 22.0103998 | 22.1329868 |
| P17174 | GOT1 | 22.0130941 | 22.0495251 | 22.0152755 | 21.9416871 | 22.0722848 |
| Q9Y3R5 | DOP1B | 19.8256036 | 19.8422518 | 19.9218159 | 19.8507856 | 19.8233982 |
| Q92530 | PSMF1 | 21.7419817 | 21.8799805 | 21.7628032 | 21.7676049 | 21.8993221 |
| Q9Y4E1 | WASHC2C | 21.4494052 | 21.568253 | 21.423313 | 21.4621107 | 21.5591748 |
| P11182 | DBT | 20.5808179 | 20.7069798 | 20.7339372 | 20.6390541 | 20.7578416 |
| P26583 | HMGB2 | 24.2983093 | 24.3358422 | 24.4039299 | 24.2691115 | 24.3558461 |
| Q8TBX8 | PIP4K2C | 19.9582963 | 19.9908651 | 20.0547355 | 19.8830379 | 20.1090913 |
| O43529 | CHST10 | 17.3532401 | 17.4883637 | 17.7670272 | 17.8108281 | 17.672972 |
| Q96B01 | RAD51AP1 | 17.73482 | 18.1342279 | 17.4789595 | 17.7102033 | 17.8100377 |
| O95872 | GPANK1 | 15.9699522 | 16.3162305 | 15.8408779 | 15.9800647 | 16.284328 |
| Q8N556 | AFAP1 | NA | 14.5326675 | 14.4864691 | NA | 14.6474963 |
| Q9NQT4 | EXOSC5 | 19.8251366 | 20.0026721 | 19.8270385 | 19.8391286 | 20.052528 |
| Q9ULC3 | RAB23 | 16.9363038 | 17.2088732 | 17.0497086 | 17.1649005 | 17.2728417 |
| O75564 | JRK | 15.7902811 | 15.9768011 | 15.7756431 | 15.7304423 | 16.1245788 |
| Q8N5J2 | MINDY1 | 14.6205104 | 15.0948071 | 14.6973492 | 14.7421882 | 14.7333396 |
| Q13404 | UBE2V1 | 24.1569655 | 24.340536 | 24.2121239 | 24.2040994 | 24.2417656 |
| A3KMH1 | VWA8 | 19.3325389 | 19.4133353 | 19.4325949 | 19.3514863 | 19.4245683 |
| Q5UIP0 | RIF1 | 21.346561 | 21.4369994 | 21.3847041 | 21.3635107 | 21.5556779 |
| O95865 | DDAH2 | 20.0759937 | 20.2319209 | 20.115466 | 20.1668123 | 20.2592367 |
| Q9ULW0 | TPX2 | 20.2481818 | 20.3989932 | 20.2479714 | 20.2839199 | 20.4236179 |

| | | | | | | |
|---|---|---|---|---|---|---|
| Q8TF47 | ZFP90 | 17.0827901 | 16.8399513 | 16.9514322 | 17.2377272 | 17.3115572 |
| Q96RI0 | F2RL3 | 12.7961246 | NA | NA | 12.789832 | NA |
| Q68DA7 | FMN1 | NA | 18.0620599 | 18.1551863 | NA | 17.8888179 |
| Q12846 | STX4 | 21.595464 | 21.7237113 | 21.9411542 | 21.569458 | 21.6236748 |
| Q8N442 | GUF1 | 20.6805591 | 20.759312 | 20.7881745 | 20.6633197 | 20.8136793 |
| Q99996 | AKAP9 | 20.4708239 | 20.6328984 | 20.5411401 | 20.5410679 | 20.6826571 |
| P14859 | POU2F1 | 19.3209821 | 19.4657688 | 19.440272 | 19.4509086 | 19.6328832 |
| Q9UH99 | SUN2 | 23.9441694 | 23.958775 | 23.9769264 | 23.8785195 | 23.8993675 |
| Q86YC3 | NRROS | NA | 16.6682601 | 16.1209286 | NA | 16.5132481 |
| Q12982 | BNIP2 | 19.7056323 | 19.7398579 | 19.3948791 | 19.545007 | 19.6875052 |
| Q92844 | TANK | 18.2792358 | 18.4994324 | 18.5272393 | 18.2784706 | 18.5163832 |
| O75071 | EFCAB14 | 15.568201 | 15.79948 | 15.8772419 | 15.7165595 | 15.7194935 |
| Q8NBM4 | UBAC2 | 21.61084 | 21.6288704 | 21.6681669 | 21.6499318 | 21.527646 |
| P39880 | CUX1 | 20.4383945 | 20.6287062 | 20.5298047 | 20.5247126 | 20.6096006 |
| Q15596 | NCOA2 | 18.5643743 | 18.6445044 | 18.7243118 | 18.6971083 | 18.7135203 |
| Q9BW83 | IFT27 | 20.787976 | 20.8033578 | 20.8660237 | 20.8712812 | 20.9430577 |
| P08887 | IL6R | 17.6691277 | 17.5436152 | 17.5976293 | 17.7109626 | 17.677473 |
| Q969N2 | PIGT | 21.4827908 | 21.3863696 | 21.4909447 | 21.3027476 | 21.4421359 |
| P48739 | PITPNB | 22.5406439 | 22.5385673 | 22.6039328 | 22.4905897 | 22.561944 |
| Q9P275 | USP36 | 19.406972 | 19.4366327 | 19.4319253 | 19.3182944 | 19.4356723 |
| Q5VWZ2 | LYPLAL1 | 20.8147389 | 20.87512 | 20.9727543 | 20.9087888 | 20.941171 |
| Q9Y678 | COPG1 | 23.4994367 | 23.4815897 | 23.5036331 | 23.378241 | 23.4651456 |
| Q9Y6D5 | ARFGEF2 | 20.3221879 | 20.2755355 | 20.3192525 | 20.2902695 | 20.4223936 |
| Q9BS26 | ERP44 | 23.8435896 | 23.9192317 | 23.7800498 | 23.8269681 | 23.9339479 |
| Q96C11 | FGGY | 18.5648929 | 18.5068883 | 18.6081724 | 18.6067717 | 18.5154064 |
| Q9NV72 | ZNF701 | 16.0745955 | 16.2260797 | 16.1522367 | 16.3433073 | 16.1527016 |
| Q06481 | APLP2 | 20.6525802 | 20.6283024 | 20.5432712 | 20.6652031 | 20.5916188 |
| A1A4S6 | ARHGAP10 | 18.8470191 | 18.9912557 | 18.9326357 | 18.9530114 | 19.0632351 |
| Q95460 | MR1 | 15.5190394 | 15.4411785 | 15.5787963 | 15.6358396 | 15.653565 |
| P61457 | PCBD1 | 21.6109484 | 21.7805196 | 21.5839161 | 21.6744078 | 21.6555648 |
| Q9NWZ5 | UCKL1 | 18.5311509 | 18.7749495 | 18.7285197 | 18.7237514 | 18.7493954 |
| Q9Y371 | SH3GLB1 | 20.9628811 | 21.1919609 | 21.0646718 | 21.0365614 | 21.2094817 |
| Q9Y6C9 | MTCH2 | 22.7959864 | 22.8734143 | 22.8663124 | 22.7890146 | 22.8910552 |
| Q8TEH3 | DENND1A | 19.6443605 | 19.8209812 | 19.7688399 | 19.6055306 | 19.7958184 |
| Q9Y4G6 | TLN2 | 24.1196415 | 24.2310117 | 23.999749 | 23.955382 | 23.9513081 |
| Q0VF96 | CGNL1 | 17.2655099 | NA | NA | 17.3745907 | NA |
| Q8IZV5 | RDH10 | 15.057385 | 15.7644109 | 15.7682881 | 15.6961137 | 15.7269436 |
| Q7Z698 | SPRED2 | 18.994701 | 19.5874319 | 18.8478757 | 18.7786295 | 18.8300211 |
| Q9BWT1 | CDCA7 | 17.5110701 | 17.5523294 | 17.6668683 | 17.6251472 | 17.7720611 |
| Q8WUY8 | NAT14 | 18.1508429 | 17.871674 | 18.1870401 | 17.9192097 | 18.0880162 |
| P05387 | RPLP2 | 26.6192254 | 26.6635688 | 26.3399732 | 26.4001671 | 26.5234525 |
| Q96EU6 | RRP36 | 19.6227809 | 19.7935585 | 19.8081327 | 19.6511107 | 19.6353405 |

| | | | | | | |
|---|---|---|---|---|---|---|
| Q92902 | HPS1 | 19.7029791 | 19.9274225 | 19.7795607 | 19.7574435 | 19.589185 |
| O75882 | ATRN | 24.3500043 | 24.128875 | 24.1115938 | 24.1966952 | 24.0803478 |
| Q96RR4 | CAMKK2 | 18.5807683 | 18.6405088 | 18.619342 | 18.395652 | 18.5086327 |
| Q9NW13 | RBM28 | 21.627119 | 21.6313415 | 21.710601 | 21.5746765 | 21.5789695 |
| O60832 | DKC1 | 23.2676326 | 23.379277 | 23.3649855 | 23.1890656 | 23.2011448 |
| Q9UJS0 | SLC25A13 | 21.3437137 | 21.412089 | 21.4746132 | 21.3206071 | 21.4454425 |
| Q8WUB8 | PHF10 | 20.2843573 | 20.3699026 | 20.4193909 | 20.3268034 | 20.4059605 |
| P43007 | SLC1A4 | 22.9072598 | 22.9254587 | 23.0592073 | 22.9033299 | 22.8716528 |
| Q9NRK6 | ABCB10 | 22.9029425 | 22.8324953 | 22.8111689 | 22.7879482 | 22.7210167 |
| Q96TA2 | YME1L1 | 22.082365 | 22.0915395 | 22.1083803 | 22.0278008 | 22.0736886 |
| Q96AG4 | LRRC59 | 24.6103148 | 24.6917871 | 24.6217561 | 24.5728674 | 24.6313875 |
| P25098 | GRK2 | 22.929081 | 23.0135997 | 22.9248888 | 22.907031 | 22.9229305 |
| P05161 | ISG15 | 21.3846784 | 21.5019433 | 21.4187485 | 21.548918 | 21.6420339 |
| Q69YQ0 | SPECC1L | 18.4891972 | 18.664419 | 18.5111721 | 18.6998424 | 18.6193075 |
| P84085 | ARF5 | 22.0395593 | 22.2113304 | 22.0920266 | 22.1149649 | 22.2686205 |
| Q6NUQ1 | RINT1 | 20.5497409 | 20.654696 | 20.6420297 | 20.6435071 | 20.5429183 |
| Q9NXN4 | GDAP2 | 19.2345411 | 19.2264286 | 19.2843307 | 19.2764449 | 19.3606403 |
| P42331 | ARHGAP25 | 19.5120747 | 19.7493774 | 19.7504519 | 19.5914492 | 19.7219225 |
| Q92576 | PHF3 | 19.7082463 | 19.8327517 | 19.8889749 | 19.7796413 | 19.8806912 |
| Q9UBU6 | FAM8A1 | 18.3674381 | 18.4607692 | 18.2612259 | 18.3329343 | 18.1966704 |
| Q9UHW9 | SLC12A6 | 19.9348162 | 19.8304727 | 19.9487381 | 20.0306833 | 20.0594293 |
| Q9BQ75 | CMSS1 | 20.2815772 | 20.1805818 | 20.3037019 | 20.3307791 | 20.4236513 |
| Q9NVS9 | PNPO | 21.7056689 | 21.7624353 | 21.72647 | 21.6916902 | 21.8743715 |
| Q9C0C2 | TNKS1BP1 | 19.9553023 | 20.0543651 | 19.8302683 | 19.9731335 | 20.0580018 |
| P51617 | IRAK1 | 19.0162337 | 19.1094027 | 19.1164016 | 19.0922101 | 19.1443641 |
| Q13371 | PDCL | 19.1466557 | 19.3228313 | 19.0672981 | 19.3235012 | 19.2885654 |
| Q09666 | AHNAK | 25.0412422 | 25.1360326 | 24.8384608 | 24.9936979 | 25.1430182 |
| Q9Y508 | RNF114 | 21.6881884 | 21.8740036 | 21.7587978 | 21.8628049 | 21.9451937 |
| Q9H467 | CUEDC2 | 19.2391972 | 19.3036215 | 19.2952669 | 19.2658843 | 19.3521287 |
| Q9NU19 | TBC1D22B | 17.9987015 | 18.2702823 | 17.9267375 | 18.1066849 | 18.1599568 |
| Q9NXG0 | CNTLN | 14.1428091 | 14.6614303 | 14.3333712 | 14.4542788 | 14.4640522 |
| P18846 | ATF1 | 16.860132 | 16.9431619 | 17.0279192 | 17.0150712 | 17.311135 |
| Q8IUG5 | MYO18B | NA | 16.5605474 | 16.7667822 | NA | 16.5571869 |
| C0HLS1 | SCRIB | 16.2411264 | 16.3383348 | 16.372403 | 16.5927203 | 16.6809679 |
| Q9BTX7 | TTPAL | 16.5206381 | 16.9897426 | 16.0662226 | 16.466033 | 16.0532822 |
| Q1L5Z9 | LONRF2 | 17.992383 | 17.9661043 | 17.8050391 | 17.7856118 | 17.9650555 |
| P84095 | RHOG | 24.4437289 | 24.6221273 | 24.642575 | 24.447593 | 24.4790698 |
| Q96P20 | NLRP3 | 18.2174588 | 18.25509 | 18.2424048 | 18.2643168 | 18.1960013 |
| Q9NRX5 | SERINC1 | 19.918582 | 19.819319 | 19.7403354 | 19.8410046 | 19.671611 |
| Q96PE7 | MCEE | 19.299624 | 19.3972174 | 19.3337429 | 19.3728566 | 19.622467 |
| Q6DN14 | MCTP1 | NA | 12.4661358 | 12.5805033 | NA | 11.9197736 |
| Q9UKT4 | FBXO5 | NA | 15.5703719 | 15.4038329 | NA | 14.3342521 |

| | | | | | | |
|---|---|---|---|---|---|---|
| Q6UWY2 | PRSS57 | 21.2872724 | 20.8855404 | 21.0734323 | 21.0306166 | 21.2426388 |
| O43169 | CYB5B | 23.8096275 | 23.9910422 | 24.0467138 | 23.6866084 | 23.7151999 |
| Q8TBZ6 | TRMT10A | 18.4818015 | 18.8503011 | 18.8659324 | 18.6119132 | 18.7714438 |
| Q96S52 | PIGS | 21.7410588 | 21.6974822 | 21.7141421 | 21.6382827 | 21.7019686 |
| Q9NYY8 | FASTKD2 | 19.9562784 | 19.9868029 | 19.9667047 | 19.9270309 | 19.9419012 |
| Q9Y2W1 | THRAP3 | 23.3770138 | 23.4893176 | 23.3532579 | 23.3533219 | 23.4893645 |
| Q9H0R8 | GABARAPL1 | 17.0907732 | 16.6923444 | 16.7889806 | 17.0321617 | 16.9952536 |
| Q9Y592 | CEP83 | 16.5463529 | 16.3633953 | 16.7902426 | 16.7289196 | 16.8234449 |
| Q9H3T3 | SEMA6B | 17.618507 | 17.3520392 | 17.6332153 | 17.7080833 | 17.5876494 |
| Q96D53 | COQ8B | 19.9412916 | 20.0269832 | 19.8962354 | 19.8852699 | 19.8622842 |
| P08311 | CTSG | 25.272514 | 24.9576808 | 24.9707779 | 24.9816231 | 24.7575877 |
| Q13393 | PLD1 | 19.5919845 | 19.4298208 | 19.637911 | 19.4921791 | 19.4783161 |
| P61353 | RPL27 | 25.8288169 | 25.7298778 | 25.9965007 | 25.6496462 | 25.6677172 |
| O15360 | FANCA | 17.9586924 | 17.9723599 | 18.0154138 | 17.8895539 | 17.9555812 |
| P11233 | RALA | 21.027026 | 21.1256303 | 21.2527646 | 20.9902348 | 21.1430788 |
| P16070 | CD44 | 24.5700474 | 24.5589692 | 24.6340768 | 24.4736421 | 24.4788313 |
| Q13951 | CBFB | 22.9585285 | 23.0147681 | 23.0065878 | 22.8672085 | 23.0211874 |
| P27695 | APEX1 | 26.1876112 | 26.1564611 | 26.103317 | 26.07256 | 26.1016282 |
| Q96SB3 | PPP1R9B | 21.3697829 | 21.5400571 | 21.3960025 | 21.3701183 | 21.5136753 |
| Q15276 | RABEP1 | 20.5342907 | 20.7121637 | 20.5535682 | 20.5514732 | 20.7365859 |
| Q5T848 | GPR158 | NA | 15.6914325 | 15.6853122 | NA | 15.6639285 |
| Q8IWZ3 | ANKHD1 | 21.8163924 | 21.9114691 | 21.7958639 | 21.86228 | 21.9615722 |
| Q9HBR0 | SLC38A10 | 21.3729512 | 21.4005757 | 21.2691493 | 21.3182846 | 21.3397936 |
| Q969Z0 | TBRG4 | 21.917271 | 21.8399551 | 21.915932 | 21.7984759 | 21.8519222 |
| Q99623 | PHB2 | 25.2917913 | 25.2580845 | 25.1641621 | 25.1140755 | 25.1658906 |
| A1L188 | NDUFAF8 | 19.1953663 | 19.2106766 | 18.9752799 | 18.8866797 | 18.7534752 |
| Q8N2U9 | SLC66A2 | 19.9972922 | 19.8378605 | 20.0009144 | 19.9210147 | 20.034287 |
| Q86Y82 | STX12 | 20.7531563 | 20.7170956 | 20.9024523 | 20.6964716 | 20.842328 |
| P10153 | RNASE2 | 21.7746329 | 21.6283636 | 21.7147271 | 21.8585503 | 21.6165214 |
| Q567V2 | MPV17L2 | 18.672836 | 18.6820657 | 18.6606472 | 18.7859316 | 18.7499548 |
| O96000 | NDUFB10 | 23.0744036 | 23.2078549 | 23.1052144 | 23.1825674 | 23.1496281 |
| Q9Y6A4 | CFAP20 | 20.7564472 | 20.5933969 | 20.8067397 | 20.6735533 | 20.5921196 |
| O60337 | MARCHF6 | 18.5334637 | 18.4333758 | 18.5141499 | 18.3358848 | 18.4250117 |
| Q8WUW1 | BRK1 | 21.7323116 | 21.9275427 | 21.8366667 | 21.6507358 | 21.8653142 |
| P54802 | NAGLU | 19.5748928 | 19.5069042 | 19.7110367 | 19.5140338 | 19.6214044 |
| Q9BRU9 | UTP23 | 20.4899271 | 20.6626539 | 20.6629802 | 20.5484837 | 20.4219115 |
| Q9Y5Y6 | ST14 | 18.9008388 | 18.881888 | 18.9826947 | 18.8995543 | 18.9996557 |
| P42126 | ECI1 | 22.7858411 | 22.8076347 | 22.7787732 | 22.7152148 | 22.7666258 |
| P61018 | RAB4B | 20.9375816 | 20.9477082 | 21.0318285 | 20.9269632 | 21.0063007 |
| P82914 | MRPS15 | 21.1544034 | 21.0415139 | 21.2075661 | 21.0381833 | 21.0251787 |
| Q9H6Z4 | RANBP3 | 22.5075365 | 22.4721794 | 22.5193613 | 22.3873902 | 22.4589612 |
| P25789 | PSMA4 | 24.7264318 | 24.7518561 | 24.6833434 | 24.6123749 | 24.6822052 |

| | | | | | | |
|---|---|---|---|---|---|---|
| P47897 | QARS1 | 23.7047664 | 23.7517462 | 23.7585124 | 23.6689202 | 23.7548114 |
| Q5W111 | SPRYD7 | 20.0151796 | 20.0736214 | 20.1231116 | 20.1681518 | 20.0930114 |
| Q96EP1 | CHFR | 18.9908992 | 19.0624402 | 19.1794927 | 19.0586618 | 19.0626425 |
| P25490 | YY1 | 22.1661114 | 22.0979575 | 22.2593987 | 22.161248 | 22.3009346 |
| Q96GM5 | SMARCD1 | 19.4522319 | 19.6092891 | 19.5252708 | 19.5753294 | 19.6068074 |
| Q14149 | MORC3 | 21.6921701 | 21.7377664 | 21.6543194 | 21.7036195 | 21.8056702 |
| Q96BK5 | PINX1 | 20.4220477 | 20.4115871 | 20.4833784 | 20.3635395 | 20.5710412 |
| Q9P086 | MED11 | NA | 18.0649165 | 18.1148433 | NA | 18.1646616 |
| Q09028 | RBBP4 | 25.7701893 | 25.8108061 | 25.6729787 | 25.7648799 | 25.8694203 |
| O43303 | CCP110 | 19.1186351 | 19.390102 | 19.3331478 | 19.3199258 | 19.4619732 |
| Q9UHB4 | NDOR1 | 18.9052583 | 19.0852826 | 19.0537779 | 18.9951596 | 19.1313774 |
| O75962 | TRIO | 20.0415829 | 20.1345212 | 20.1996163 | 19.9991421 | 20.1336751 |
| Q96CG3 | TIFA | 18.598961 | 18.7065789 | 18.4006057 | 18.6591125 | 18.527724 |
| Q9H0V1 | TMEM168 | 21.7830451 | 21.7567793 | 21.6967075 | 21.7974537 | 21.800447 |
| Q9NVP2 | ASF1B | 22.139777 | 22.168563 | 21.8997211 | 22.1403292 | 22.2997968 |
| Q9Y6R0 | NUMBL | 16.1428207 | NA | NA | 16.4065886 | NA |
| P00451 | F8 | NA | 19.6323057 | 19.7605857 | NA | 20.2274491 |
| Q96AJ9 | VTI1A | 20.1138706 | 20.3322337 | 20.3519745 | 20.0783733 | 20.0197556 |
| Q6AI08 | HEATR6 | 18.8207615 | 18.9436629 | 18.9486032 | 18.7708374 | 18.9211591 |
| P23497 | SP100 | 20.8113761 | 20.9393731 | 20.8450479 | 20.8183253 | 20.9952836 |
| O75037 | KIF21B | 16.4289118 | 16.5136254 | 16.6747406 | 16.6507574 | 16.7498029 |
| Q96PL5 | ERMAP | 14.9926509 | NA | NA | 15.0295611 | NA |
| Q9H063 | MAF1 | 18.0080456 | 17.9579112 | 18.1996381 | 17.9142811 | 18.0691854 |
| Q9NX47 | MARCHF5 | 20.1411647 | 20.2591376 | 20.2648518 | 20.1276943 | 20.2542265 |
| Q9P258 | RCC2 | 23.7626208 | 23.8023234 | 23.8452553 | 23.640234 | 23.808799 |
| Q9BRR6 | ADPGK | 21.8507728 | 21.7631921 | 21.7894157 | 21.6755589 | 21.8486673 |
| P51571 | SSR4 | 23.5105119 | 23.592675 | 23.4540765 | 23.3460589 | 23.4682571 |
| Q9Y692 | GMEB1 | 19.8459339 | 19.9168092 | 19.7261338 | 19.7325524 | 20.0275287 |
| Q8IYL2 | TRMT44 | 15.6787392 | 15.7018085 | 15.778754 | 15.7157052 | 15.4436998 |
| Q2NL68 | PROSER3 | NA | 13.902245 | 14.258928 | NA | 14.0276901 |
| Q9Y6F1 | PARP3 | 19.62414 | 19.7282108 | 19.5931716 | 19.5019461 | 19.4560734 |
| Q9P0U1 | TOMM7 | 20.1453475 | 19.9750139 | 20.4203417 | 20.1139117 | 20.1860414 |
| Q9BU61 | NDUFAF3 | 20.6688505 | 20.6681962 | 20.7121963 | 20.6773431 | 20.730705 |
| Q8IUH4 | ZDHHC13 | 18.5973286 | 18.4972015 | 18.5954975 | 18.5051592 | 18.5820304 |
| P06737 | PYGL | 24.8132945 | 24.7504486 | 24.740907 | 24.6873298 | 24.682085 |
| P22415 | USF1 | 23.1025915 | 23.1655415 | 23.0575013 | 23.0410251 | 23.0450018 |
| Q8NB46 | ANKRD52 | 19.2058965 | 19.2828041 | 19.2270077 | 19.2600439 | 19.3771626 |
| Q17R31 | TATDN3 | 17.2756387 | 17.4675746 | 17.5515836 | 17.5036653 | 17.5379995 |
| Q9H8G2 | CAAP1 | 19.9911901 | 19.9716517 | 19.7599564 | 20.0261096 | 20.1363732 |
| Q9NUQ6 | SPATS2L | 18.739775 | 18.8895181 | 18.8582197 | 18.7781048 | 18.8522171 |
| Q6XUX3 | DSTYK | 15.2949812 | 15.0444391 | 14.795303 | 14.9479125 | 14.9886439 |
| Q0P6D6 | CCDC15 | 16.9906211 | 17.246981 | 16.4765275 | 16.9840713 | 16.8720908 |

| | | | | | | |
|---|---|---|---|---|---|---|
| Q9P2E5 | CHPF2 | 19.4647373 | 19.4539315 | 19.4252193 | 19.4946783 | 19.3700111 |
| P50750 | CDK9 | 21.3961311 | 21.3455564 | 21.4752279 | 21.2970236 | 21.4147338 |
| Q5JTJ3 | COA6 | 21.1585655 | 21.1248059 | 20.9864069 | 21.0880428 | 21.3524557 |
| Q9P2N6 | KANSL3 | 18.0224316 | 17.982357 | 17.8134796 | 17.8800089 | 18.041424 |
| A2RU54 | HMX2 | 15.8390211 | NA | NA | 15.9362073 | NA |
| P81274 | GPSM2 | 14.4491603 | 14.6531385 | 14.6384277 | 14.6371395 | 14.593172 |
| O75575 | CRCP | 16.4371786 | 16.0732099 | 16.5909659 | 16.4380537 | 16.6375596 |
| P04049 | RAF1 | 19.3522605 | 19.3327411 | 19.5383418 | 19.3631307 | 19.4883715 |
| Q8TAK5 | GABPB2 | 14.7474462 | 14.4283003 | 14.8476939 | 14.4438933 | 14.6755451 |
| Q92618 | ZNF516 | 20.5917152 | 20.58755 | 20.5960878 | 20.5866599 | 20.4850441 |
| Q86SQ7 | SDCCAG8 | 17.8756332 | 18.0973661 | 17.9155353 | 17.9050683 | 18.1712437 |
| P36954 | POLR2I | 18.3420627 | NA | NA | 18.1086113 | NA |
| P98161 | PKD1 | 20.3873945 | NA | NA | 19.9299813 | NA |
| P51687 | SUOX | 17.3044938 | 17.3729922 | 17.2215553 | 17.3884099 | 17.482095 |
| P09132 | SRP19 | 21.667741 | 21.6644983 | 21.9481485 | 21.719723 | 21.763303 |
| Q9UHQ4 | BCAP29 | 19.472193 | 19.7256387 | 19.7903104 | 19.4985989 | 19.5318422 |
| Q04760 | GLO1 | 23.9364655 | 23.9210388 | 23.9024209 | 23.8777585 | 23.9474953 |
| P49768 | PSEN1 | 20.5831305 | 20.6446517 | 20.6477134 | 20.7022782 | 20.6273035 |
| Q8IZ69 | TRMT2A | 21.1026586 | 21.2394869 | 21.1590155 | 21.0598754 | 21.1886329 |
| Q9Y5J1 | UTP18 | 21.2489611 | 21.3139607 | 21.2990006 | 21.208033 | 21.3372809 |
| Q32P28 | P3H1 | 20.9434678 | 20.9441835 | 21.0317297 | 20.9396143 | 20.9822053 |
| P35658 | NUP214 | 21.7702013 | 21.8512223 | 21.8065854 | 21.7716474 | 21.9010869 |
| Q9Y3B8 | REXO2 | 19.7825145 | 19.7549925 | 19.8381764 | 19.8001363 | 19.7442998 |
| Q15185 | PTGES3 | 25.8183193 | 25.9348261 | 25.9491923 | 25.891284 | 25.909274 |
| Q9BWU0 | SLC4A1AP | 20.5335171 | 20.7288576 | 20.5020774 | 20.6534315 | 20.7102998 |
| Q641Q2 | WASHC2A | 19.4254713 | 19.5274746 | 19.2592142 | 19.4632899 | 19.4894484 |
| P27815 | PDE4A | 14.3311257 | 14.4199236 | 14.0757583 | 14.3512949 | 14.458015 |
| Q9Y3C5 | RNF11 | 15.5022731 | 15.6098615 | 15.7752901 | 15.7396774 | 15.9772845 |
| Q96BZ9 | TBC1D20 | 17.5499171 | 16.9971467 | 17.4444917 | 17.182375 | 17.3150273 |
| Q9Y5Z7 | HCFC2 | 17.0177215 | 17.1643887 | 17.1109188 | 17.0429227 | 17.2012849 |
| Q9NVV0 | TMEM38B | 19.1914473 | 19.7758469 | 19.4265293 | 19.2522701 | 19.4526423 |
| P61254 | RPL26 | 26.4436732 | 26.4342137 | 26.5366177 | 26.3204864 | 26.4349289 |
| Q9NVW2 | RLIM | 18.8139237 | 18.8908349 | 18.9058586 | 18.8073116 | 18.8683355 |
| O75420 | GIGYF1 | 19.9642138 | 19.9518779 | 19.9900652 | 19.8585863 | 19.8757727 |
| Q86UX7 | FERMT3 | 23.3355681 | 23.3775209 | 23.4500673 | 23.2397297 | 23.3604649 |
| P42285 | MTREX | 24.6112395 | 24.6034372 | 24.6182226 | 24.4309818 | 24.5004303 |
| O43592 | XPOT | 22.2289465 | 22.2353477 | 22.2052161 | 22.1097871 | 22.2024665 |
| Q9HBL8 | NMRAL1 | 21.1927182 | 21.2425822 | 21.1166769 | 21.1845016 | 21.2744221 |
| Q9BVJ6 | UTP14A | 21.0008362 | 21.1570261 | 20.9968601 | 21.0209774 | 21.1646547 |
| Q8IXI2 | RHOT1 | 19.9445128 | 20.1347481 | 20.0457801 | 20.0368296 | 20.1763681 |
| Q9HCU8 | POLD4 | 18.5000717 | 18.6088586 | 18.4802757 | 18.5903382 | 18.729283 |
| P30622 | CLIP1 | 19.7324994 | 19.8902736 | 19.7668967 | 19.7575383 | 19.9157398 |

| | | | | | | |
|---|---|---|---|---|---|---|
| P30046 | DDT | 24.9102028 | 24.9974335 | 24.8002404 | 24.8618426 | 25.0116201 |
| Q2LD37 | BLTP1 | 17.2776397 | 17.607553 | 17.4128212 | 17.7632253 | 17.7379852 |
| Q6AI39 | BICRAL | 15.1800885 | 15.5859651 | 15.8353959 | 15.0372363 | 15.7883447 |
| P01130 | LDLR | 18.0689249 | 18.754515 | 18.6080907 | 18.5849321 | 18.8258943 |
| Q13751 | LAMB3 | 15.7863153 | NA | NA | 16.0327271 | NA |
| Q14185 | DOCK1 | 19.1759282 | 19.5416546 | 19.1927699 | 19.2937576 | 19.1973282 |
| Q7Z6J9 | TSEN54 | 17.9094696 | 17.9637562 | 17.9796006 | 18.0518228 | 17.8901141 |
| Q7Z5Y7 | KCTD20 | NA | 14.6517985 | 15.0933955 | NA | 15.04183 |
| Q9H3F6 | KCTD10 | 19.6414078 | 19.7290165 | 19.9407059 | 19.5899311 | 19.7291122 |
| Q8N300 | SVBP | 18.7923205 | 18.8376842 | 18.6088122 | 18.5709576 | 18.5682086 |
| O43242 | PSMD3 | 24.0159844 | 24.0322071 | 23.9998189 | 23.8425908 | 23.9131341 |
| O75394 | MRPL33 | 20.4978487 | 20.781558 | 20.9719734 | 20.5474855 | 20.8092795 |
| Q96LJ7 | DHRS1 | 19.0734529 | 19.3148466 | 19.2993768 | 19.2737717 | 19.4020601 |
| Q9P2F9 | ZNF319 | 16.8300853 | 16.3868473 | 16.4753065 | 16.7307208 | 16.5918325 |
| Q8N5A5 | ZGPAT | 19.1374556 | 19.2688215 | 19.2287277 | 19.2145387 | 19.2545652 |
| Q969H8 | MYDGF | 24.9604832 | 24.6509425 | 24.9061148 | 24.6694278 | 24.7406691 |
| Q9H9G7 | AGO3 | 18.2169892 | 18.4263416 | 18.4209092 | 18.3002977 | 18.4067856 |
| Q96CS4 | ZNF689 | 17.7366348 | 17.8279925 | 17.8118014 | 17.7335253 | 17.7852955 |
| P61769 | B2M | 23.8578224 | 23.8053377 | 23.7688706 | 23.6383927 | 23.7392566 |
| Q9NX20 | MRPL16 | 21.6179714 | 21.6259198 | 21.7557603 | 21.5535049 | 21.6153361 |
| Q92544 | TM9SF4 | 22.8241293 | 22.8870683 | 22.8069578 | 22.8276265 | 22.7883163 |
| P41440 | SLC19A1 | 20.9838885 | 20.8940848 | 20.7799688 | 20.9041223 | 20.7871547 |
| O14980 | XPO1 | 23.713772 | 23.6836109 | 23.7812384 | 23.590433 | 23.7012168 |
| Q8NEZ4 | KMT2C | 18.9071445 | 19.0129444 | 18.8884319 | 18.9335164 | 19.1249986 |
| Q99551 | MTERF1 | 16.1768337 | 16.3905852 | 16.4872634 | 16.4053825 | 16.442053 |
| Q8N8U2 | CDYL2 | 16.2394905 | 16.3906731 | 16.2966274 | 16.1093419 | 16.197339 |
| Q8NF64 | ZMIZ2 | 17.1339965 | 17.339053 | 17.2283384 | 17.0293396 | 17.1623633 |
| P82921 | MRPS21 | 20.4752145 | 20.4797456 | 20.5370262 | 20.4750544 | 20.4049827 |
| Q03154 | ACY1 | 21.3124539 | 21.3755428 | 21.3344567 | 21.3486488 | 21.3569305 |
| Q5JSH3 | WDR44 | 19.5694858 | 19.5612706 | 19.6348539 | 19.5968975 | 19.7125227 |
| P09496 | CLTA | 24.3047485 | 24.515999 | 24.1763162 | 24.2453957 | 24.4436248 |
| O14561 | NDUFAB1 | 22.7315368 | 22.9648126 | 22.5831452 | 22.7149472 | 22.8735256 |
| Q9NWB7 | IFT57 | 14.9391841 | 14.8409638 | 15.3047476 | 15.1693314 | 14.9294679 |
| Q8NBW4 | SLC38A9 | 17.4830753 | 17.5348776 | 17.5835654 | 17.5774852 | 17.6025131 |
| Q9Y3U8 | RPL36 | 25.1953685 | 25.2534084 | 25.3468528 | 25.1201226 | 25.0659457 |
| O94887 | FARP2 | 18.5698809 | 18.6095466 | 18.6587826 | 18.581685 | 18.5774061 |
| P49641 | MAN2A2 | 19.3168926 | 19.5132574 | 19.5018873 | 19.5721954 | 19.416427 |
| O94927 | HAUS5 | 19.630097 | 19.8459491 | 19.71196 | 19.6164423 | 19.7063016 |
| P00491 | PNP | 24.884344 | 24.8226472 | 24.7432536 | 24.7893967 | 24.6964335 |
| P53621 | COPA | 23.7152213 | 23.702575 | 23.7342243 | 23.647915 | 23.6968677 |
| P15144 | ANPEP | 20.099886 | 20.1666572 | 20.1373694 | 20.0348182 | 20.1325295 |
| P13612 | ITGA4 | 21.8686281 | 21.852049 | 21.8239511 | 21.8250332 | 21.8181886 |

| | | | | | | |
|---|---|---|---|---|---|---|
| Q9H9P8 | L2HGDH | 20.421967 | 20.3504022 | 20.42516 | 20.4155121 | 20.4884391 |
| P56537 | EIF6 | 23.2720662 | 23.3730577 | 23.2193972 | 23.2819111 | 23.4530891 |
| Q9BSY4 | CHCHD5 | 21.556199 | 21.5622218 | 21.4526335 | 21.587698 | 21.676253 |
| Q96F24 | NRBF2 | 19.3852172 | 19.5724435 | 19.2142036 | 19.506889 | 19.5001148 |
| Q0JRZ9 | FCHO2 | 16.1585926 | 15.4774362 | 15.096338 | 16.5927022 | 15.2922057 |
| Q5T011 | SZT2 | 19.9985809 | 19.7946337 | 19.9874493 | 19.7748035 | 19.7288269 |
| O75044 | SRGAP2 | 20.7379958 | 20.80913 | 20.8008985 | 20.7806912 | 20.824558 |
| Q13503 | MED21 | 20.9190144 | 21.5048907 | 20.576579 | 21.0981666 | 20.8646512 |
| Q9HD40 | SEPSECS | 20.3982568 | 20.5379369 | 20.5056361 | 20.3378528 | 20.455899 |
| Q56NI9 | ESCO2 | 18.7338527 | 18.8493645 | 18.9706227 | 18.8511886 | 18.8529984 |
| Q8N9N5 | BANP | 18.5142218 | 18.6170862 | 18.6759036 | 18.6791513 | 18.744759 |
| Q8NC51 | SERBP1 | 25.7573017 | 25.9925702 | 25.667416 | 25.7978603 | 25.8935564 |
| Q9HAZ1 | CLK4 | NA | 13.6291425 | 14.1379649 | NA | 13.8856919 |
| Q15283 | RASA2 | 19.3310106 | 19.4891653 | 19.4175276 | 19.2240452 | 19.2856786 |
| A5PLL7 | PEDS1 | 19.937438 | 19.9445302 | 20.073994 | 19.8335362 | 20.0279558 |
| P46782 | RPS5 | 26.6491084 | 26.6747734 | 26.6885988 | 26.537896 | 26.605408 |
| Q15853 | USF2 | 20.167952 | 20.0535022 | 20.0483394 | 20.0107806 | 19.9873134 |
| Q99459 | CDC5L | 22.3718592 | 22.4566847 | 22.4192904 | 22.4326069 | 22.4998728 |
| Q8NFD5 | ARID1B | 19.5068631 | 19.5772105 | 19.5472051 | 19.5787048 | 19.5963328 |
| O60307 | MAST3 | 19.748189 | 19.8550515 | 19.8604589 | 19.8650694 | 19.8809951 |
| Q96AT1 | KIAA1143 | 20.3109607 | 20.3864866 | 20.1409181 | 20.3862032 | 20.4340275 |
| Q9Y294 | ASF1A | 20.6997517 | 20.8924158 | 20.7965172 | 20.8109081 | 20.9811231 |
| Q96I36 | COX14 | 19.7793215 | 19.3486305 | 19.701866 | 19.4457872 | 19.5996543 |
| Q9NQ89 | C12orf4 | 19.5041102 | 19.6151424 | 19.7189192 | 19.4155593 | 19.4167368 |
| Q9NS86 | LANCL2 | 21.0767215 | 21.1046928 | 21.0704449 | 21.0085128 | 21.0460255 |
| Q8IY63 | AMOTL1 | 17.7556404 | 17.7440403 | 17.5567117 | 17.8194205 | 17.7223714 |
| Q8N0X4 | CLYBL | 18.50091 | 18.6488454 | 18.6883346 | 18.4971971 | 18.7556754 |
| Q9UK39 | NOCT | 17.7724095 | 17.8923748 | 18.0723595 | 17.8901735 | 18.0386314 |
| P25787 | PSMA2 | 24.2927782 | 24.2672115 | 24.2335609 | 24.233273 | 24.3212211 |
| Q9NWU5 | MRPL22 | 21.9189419 | 21.8335022 | 22.0376866 | 21.8334409 | 21.9229656 |
| P52732 | KIF11 | 20.4014968 | 20.5148427 | 20.446438 | 20.4016317 | 20.5129413 |
| P62993 | GRB2 | 24.8938868 | 24.9557258 | 24.8635909 | 24.8674366 | 24.9915405 |
| P57678 | GEMIN4 | 22.4785384 | 22.6449083 | 22.6233664 | 22.4576052 | 22.5065586 |
| Q9UIJ7 | AK3 | 21.6228467 | 21.7107543 | 21.6629628 | 21.570597 | 21.6497157 |
| O75569 | PRKRA | 21.2624048 | 21.4484423 | 21.2211833 | 21.3359259 | 21.4397496 |
| Q96CN4 | EVI5L | 17.6399828 | 17.5350056 | 17.7527117 | 17.6678853 | 17.7838329 |
| Q9HAB3 | SLC52A2 | 20.7372985 | 20.7468455 | 20.9401554 | 20.5997774 | 20.4110292 |
| P18124 | RPL7 | 26.2536576 | 26.2772775 | 26.3957114 | 26.1860746 | 26.1455558 |
| Q8WUY1 | THEM6 | 22.6389646 | 22.7117663 | 22.6730835 | 22.5058818 | 22.4808804 |
| P31153 | MAT2A | 24.0481582 | 24.1836698 | 24.1296115 | 23.9862001 | 24.0719732 |
| Q9UJ41 | RABGEF1 | 20.1122344 | 20.2436772 | 20.2363921 | 20.1495999 | 20.2334526 |
| Q14151 | SAFB2 | 22.1639082 | 22.2238326 | 22.2265128 | 22.1280881 | 22.1807606 |

| | | | | | | |
|---|---|---|---|---|---|---|
| Q9NYJ8 | TAB2 | 17.7342195 | 17.7336876 | 17.6538653 | 17.7254387 | 17.7700391 |
| Q13451 | FKBP5 | 22.0582608 | 22.172333 | 22.0273398 | 22.1400233 | 22.1935331 |
| Q96KA5 | CLPTM1L | 20.7380483 | 20.7863999 | 20.8085849 | 20.6525573 | 20.7445606 |
| Q6DKJ4 | NXN | 15.3806653 | 15.7297546 | 16.1065245 | 15.8790069 | 15.8309761 |
| P50416 | CPT1A | 23.3642899 | 23.4131914 | 23.3696372 | 23.3085647 | 23.3044374 |
| O95563 | MPC2 | 20.5985314 | 20.4166449 | 20.9070713 | 20.5796265 | 20.6531241 |
| Q7L2H7 | EIF3M | 24.2187589 | 24.146383 | 24.2135586 | 24.0944559 | 24.1278106 |
| O75376 | NCOR1 | 20.6328752 | 20.7806857 | 20.748984 | 20.6849728 | 20.7688658 |
| Q9UKY1 | ZHX1 | 18.916125 | 19.1060674 | 19.0107772 | 18.9976267 | 19.1125317 |
| Q01105 | SET | 25.4658209 | 25.5654027 | 25.2361584 | 25.448995 | 25.4883471 |
| Q9Y6K8 | AK5 | 14.1003242 | NA | NA | 14.0166298 | NA |
| Q9ULR5 | PAIP2B | 16.0106724 | 16.1164565 | 16.0332739 | 16.0919632 | 16.6271139 |
| Q8IW41 | MAPKAPK5 | 16.9835366 | 16.4235412 | 16.8972509 | 16.8106278 | 16.1373556 |
| P39023 | RPL3 | 26.3265982 | 26.3553324 | 26.426649 | 26.1759294 | 26.2547484 |
| P11021 | HSPA5 | 26.8480083 | 26.9292103 | 26.6840994 | 26.8265051 | 26.8915999 |
| P51608 | MECP2 | 20.6271182 | 20.7169153 | 20.6166588 | 20.6203265 | 20.7349871 |
| P55735 | SEC13 | 23.2619411 | 23.4844517 | 23.2807274 | 23.3211485 | 23.4303526 |
| Q5JTV8 | TOR1AIP1 | 22.9272924 | 23.014302 | 23.0280277 | 22.9345504 | 22.9280705 |
| Q2KHM9 | KIAA0753 | NA | 12.8332012 | 13.1306297 | NA | 12.6458076 |
| P57721 | PCBP3 | 22.1985585 | 22.2922392 | 22.2197336 | 22.1801463 | 22.2576917 |
| P49754 | VPS41 | 20.2717801 | 20.2940936 | 20.3203409 | 20.2207873 | 20.3481755 |
| Q5THJ4 | VPS13D | 17.9798871 | 17.9761835 | 18.0407045 | 18.1215845 | 18.076462 |
| Q12888 | TP53BP1 | 20.1493631 | 20.278787 | 20.1597357 | 20.1284119 | 20.2901497 |
| P06748 | NPM1 | 27.4081164 | 27.5051906 | 27.1868584 | 27.3001725 | 27.3913332 |
| Q6VY07 | PACS1 | 16.1502468 | 15.9336045 | 15.9867992 | 15.9652803 | 16.025704 |
| Q86YV9 | HPS6 | 19.7671314 | 19.7423646 | 19.7816156 | 19.886752 | 19.9124431 |
| P31327 | CPS1 | 13.5143958 | NA | NA | 13.2219053 | NA |
| Q8WVX3 | C4orf3 | 20.1717775 | 20.481263 | 19.9475755 | 20.0268524 | 19.8617888 |
| Q9H992 | MARCHF7 | 18.5483653 | 18.7803794 | 18.8494608 | 18.412399 | 18.6663218 |
| Q7Z392 | TRAPPC11 | 21.0283805 | 21.0172801 | 21.0627752 | 20.8963557 | 21.0491052 |
| Q9UBU8 | MORF4L1 | 21.2052406 | 21.303896 | 21.2637104 | 21.2159254 | 21.2237505 |
| Q16658 | FSCN1 | 19.4017551 | 19.6210229 | 19.5868798 | 19.5073735 | 19.6110951 |
| Q7Z591 | AKNA | 19.6538809 | 19.6978219 | 19.6920415 | 19.6723326 | 19.8258693 |
| Q96CV9 | OPTN | 20.0280071 | 20.1582665 | 20.0295099 | 20.0873554 | 20.2170724 |
| Q9BXI6 | TBC1D10A | 18.8452828 | 18.8727116 | 18.9550768 | 18.8705929 | 18.895219 |
| Q8IU68 | TMC8 | 18.8543381 | 18.7744744 | 18.9937983 | 18.8711631 | 18.8717704 |
| Q5T749 | KPRP | 18.3788504 | 17.9792759 | 18.6309036 | 18.2280492 | 17.6070218 |
| P25025 | CXCR2 | 17.6069134 | 17.777331 | 17.8863056 | 17.471381 | 17.8720972 |
| Q8TE04 | PANK1 | 19.3607143 | 19.6398556 | 19.4250607 | 19.3159259 | 19.4422038 |
| Q9UJX4 | ANAPC5 | 20.3934101 | 20.4477295 | 20.5362444 | 20.3038923 | 20.4225304 |
| P16152 | CBR1 | 23.3650786 | 23.431824 | 23.3134478 | 23.2701821 | 23.3389414 |
| Q9Y487 | ATP6V0A2 | 20.977301 | 20.9462678 | 20.9856426 | 20.9479884 | 20.9462955 |

| | | | | | | |
|---|---|---|---|---|---|---|
| P49711 | CTCF | 21.4638475 | 21.5091479 | 21.5338301 | 21.5120995 | 21.552199 |
| P15289 | ARSA | 20.9321826 | 21.0180084 | 21.057886 | 21.0640235 | 21.1653875 |
| Q96IZ7 | RSRC1 | 19.5678651 | 19.7276814 | 19.4945085 | 19.5996669 | 19.6455261 |
| Q14493 | SLBP | 20.3205682 | 20.4680406 | 20.2305126 | 20.4807823 | 20.478851 |
| Q562F6 | SGO2 | 19.1693777 | 19.4242163 | 19.1582525 | 19.2949645 | 19.0918133 |
| Q86X83 | COMMD2 | 20.9371164 | 21.3112818 | 21.3813394 | 21.0103428 | 21.1390874 |
| Q96MW1 | CCDC43 | 20.8516814 | 21.1320863 | 20.8008054 | 20.8814252 | 21.0606245 |
| Q96EX3 | DYNC2I2 | 14.9299494 | 15.3999934 | 15.4402177 | 15.2890616 | 15.307298 |
| Q9Y6I8 | PXMP4 | 18.851237 | 18.7709546 | 18.9618815 | 18.8661734 | 18.8526179 |
| Q7L3T8 | PARS2 | 18.6236816 | 18.7218616 | 18.7409358 | 18.6309166 | 18.6738365 |
| Q9BRX9 | WDR83 | 18.4313164 | 18.7177657 | 18.6423607 | 18.5695618 | 18.7648464 |
| Q8N9Q2 | SREK1IP1 | 16.5250245 | NA | NA | 16.7053819 | NA |
| O95294 | RASAL1 | 14.224903 | NA | NA | 14.4827207 | NA |
| Q9NY35 | CLDND1 | 19.4524765 | 19.3925454 | 19.5028673 | 19.471872 | 19.4000187 |
| P78346 | RPP30 | 21.6018644 | 21.7205671 | 21.6433168 | 21.5453671 | 21.7061079 |
| Q9UGH3 | SLC23A2 | 18.8534104 | 18.743744 | 19.0354469 | 18.9005905 | 18.959493 |
| P61313 | RPL15 | 26.2540867 | 26.3446033 | 26.5109665 | 26.1676587 | 26.3170237 |
| O43290 | SART1 | 22.9170911 | 22.9841063 | 22.8173118 | 22.9307599 | 23.0502366 |
| Q7L4I2 | RSRC2 | 20.4683374 | 20.4868322 | 20.2827466 | 20.4700726 | 20.6583178 |
| Q07890 | SOS2 | 15.7574692 | 15.958278 | 15.6598124 | 16.0397662 | 15.8375021 |
| P52907 | CAPZA1 | 24.9901223 | 25.0580423 | 25.0281834 | 24.8758222 | 24.9910252 |
| Q9H6R4 | NOL6 | 21.4236654 | 21.5102377 | 21.4104825 | 21.4587991 | 21.4817012 |
| Q8IX01 | SUGP2 | 21.4291993 | 21.5088335 | 21.5633317 | 21.442779 | 21.5985635 |
| Q7L7X3 | TAOK1 | 20.2097544 | 20.361818 | 20.2275339 | 20.2461932 | 20.3334014 |
| Q9BUB5 | MKNK1 | 19.525658 | 19.5729621 | 19.65939 | 19.4957667 | 19.6135855 |
| Q8TEK3 | DOT1L | 17.3404433 | 17.5249756 | 17.4264747 | 17.3182189 | 17.4636718 |
| Q6P4Q7 | CNNM4 | 17.6854721 | 17.7056356 | 17.8630244 | 17.826652 | 17.7257539 |
| Q1RMZ1 | BMT2 | NA | 15.4739399 | 15.7287408 | NA | 15.2566717 |
| Q8IZP0 | ABI1 | 20.543468 | 20.6985836 | 20.5384959 | 20.5533375 | 20.4771819 |
| P20810 | CAST | 20.4923129 | 20.6571889 | 20.4558282 | 20.5144941 | 20.5839886 |
| Q99547 | MPHOSPH6 | 20.2362449 | 20.3645919 | 20.0867005 | 20.2535955 | 20.2462877 |
| Q4VC44 | FLYWCH1 | 14.7439135 | NA | NA | 14.0781619 | NA |
| Q9H165 | BCL11A | 16.9013687 | 17.0404969 | 17.2170236 | 16.8896307 | 16.8387683 |
| P48506 | GCLC | 19.0997347 | 19.1618706 | 19.1091547 | 19.1320481 | 19.1874874 |
| Q9UQC2 | GAB2 | 19.4416389 | 19.3827063 | 19.4060377 | 19.4617603 | 19.5645314 |
| Q9BTE7 | DCUN1D5 | 21.0068092 | 21.0874886 | 21.1989767 | 21.0579 | 21.1709648 |
| Q9P0U4 | CXXC1 | 20.2011096 | 20.3092855 | 20.3332396 | 20.2666534 | 20.264785 |
| Q05209 | PTPN12 | 21.9258295 | 22.0621218 | 21.8754947 | 21.9540361 | 21.9198629 |
| Q14789 | GOLGB1 | 23.6803241 | 23.8057222 | 23.5535081 | 23.6439263 | 23.7514824 |
| O60828 | PQBP1 | 21.1856199 | 21.3157812 | 21.1978026 | 21.2949522 | 21.3540696 |
| P26885 | FKBP2 | 22.1345 | 22.1595236 | 22.0748427 | 22.1009256 | 22.2624211 |
| Q13033 | STRN3 | 19.1226578 | 19.2324125 | 19.1731475 | 19.2067757 | 19.3584431 |

| | | | | | | |
|---|---|---|---|---|---|---|
| Q96RL1 | UIMC1 | 20.1436035 | 20.3379534 | 20.2262174 | 20.1989188 | 20.3843884 |
| O15015 | ZNF646 | 17.9316808 | 17.9640124 | 18.1367213 | 17.9908442 | 18.0674644 |
| P35869 | AHR | 17.4250836 | 17.3340582 | 17.600552 | 17.3459335 | 17.6026339 |
| Q1MSJ5 | CSPP1 | 16.7906321 | 16.9865629 | 16.8156013 | 16.97551 | 16.954716 |
| O43716 | GATC | 18.4688475 | 18.2641029 | 18.1542817 | 18.4399041 | 18.6087594 |
| P99999 | CYCS | 23.8218803 | 23.9033723 | 23.9419477 | 23.5300605 | 24.2439375 |
| O76075 | DFFB | 18.6634997 | 18.4956924 | 18.5201244 | 18.5174531 | 18.4010452 |
| Q8TDQ1 | CD300LF | 18.7574062 | 18.6116663 | 18.8808409 | 18.7956932 | 18.8806794 |
| Q9Y6J9 | TAF6L | 20.4732191 | 20.4414174 | 20.3553435 | 20.3084383 | 20.3566447 |
| Q7Z6P3 | RAB44 | 20.0766999 | 20.12039 | 20.0123687 | 20.078224 | 20.1342756 |
| Q8TEP8 | CEP192 | 18.733418 | 19.0250373 | 18.9411774 | 18.9998411 | 18.948836 |
| Q9BV10 | ALG12 | 16.7337051 | 16.6625838 | 17.1457717 | 16.9220948 | 16.8708783 |
| Q9BYD1 | MRPL13 | 22.1241781 | 22.1081828 | 22.0868339 | 22.107672 | 22.1006853 |
| Q92890 | UFD1 | 21.627274 | 21.6499266 | 21.7226816 | 21.5737756 | 21.6661252 |
| Q9Y315 | DERA | 21.7997706 | 21.901149 | 21.9586009 | 21.7288832 | 21.8086785 |
| Q8IZD4 | DCP1B | 19.589931 | 19.763909 | 19.6568296 | 19.6036122 | 19.7133429 |
| Q9BQ04 | RBM4B | 23.4697059 | 23.6015809 | 23.4728109 | 23.4225661 | 23.6231029 |
| Q96TA1 | NIBAN2 | 22.2163634 | 22.3854262 | 22.3904898 | 22.2285378 | 22.3471093 |
| Q8IZR5 | CMTM4 | 14.4717517 | NA | NA | 14.0709923 | NA |
| Q15800 | MSMO1 | 18.6061372 | 18.77408 | 19.1032289 | 18.4970054 | 19.2417161 |
| Q13530 | SERINC3 | 19.2387989 | 19.43755 | 19.4131194 | 19.2946897 | 19.2036652 |
| Q9Y247 | FAM50B | 19.2779924 | 19.0602078 | 19.1266823 | 19.0231307 | 19.1101395 |
| Q12792 | TWF1 | 19.7325735 | 19.7697022 | 19.691673 | 19.5983278 | 19.6466278 |
| Q8WUX9 | CHMP7 | 20.9373337 | 21.0935174 | 20.9697924 | 20.8845616 | 20.988986 |
| P60891 | PRPS1 | 23.3064458 | 23.3555613 | 23.366466 | 23.159794 | 23.3168119 |
| Q15631 | TSN | 24.3254885 | 24.440149 | 24.274778 | 24.2494681 | 24.1830411 |
| Q92734 | TFG | 22.5612817 | 22.5698641 | 22.5545067 | 22.5732463 | 22.594695 |
| P29372 | MPG | 21.8198858 | 21.841256 | 21.8745949 | 21.8127582 | 21.8464572 |
| Q9NPR9 | GPR108 | 20.8394765 | 20.8352528 | 20.824125 | 20.7464343 | 20.7550748 |
| O14907 | TAX1BP3 | 21.374502 | 21.4157376 | 21.3584531 | 21.3460109 | 21.3880663 |
| Q9Y580 | RBM7 | 19.6672215 | 19.6919144 | 19.7104955 | 19.7144871 | 19.7636775 |
| Q01201 | RELB | 17.4442744 | 17.5984577 | 17.6837914 | 17.5223844 | 17.6742414 |
| Q6PK18 | OGFOD3 | 17.9180131 | 17.9555032 | 18.1583891 | 18.0450343 | 18.190067 |
| Q9H444 | CHMP4B | 22.3420163 | 22.503068 | 22.1866799 | 22.3908089 | 22.4674822 |
| Q969F1 | GTF3C6 | 17.1968581 | 17.3551181 | 17.1942783 | 17.3293635 | 17.4519165 |
| Q8TE73 | DNAH5 | 13.5742699 | NA | NA | 13.8268992 | NA |
| Q9BTV4 | TMEM43 | 23.2385879 | 23.1908347 | 23.3651574 | 23.1348596 | 23.1969243 |
| Q9NYV6 | RRN3 | 18.7759728 | 18.9061454 | 18.8378782 | 18.652175 | 18.8386853 |
| Q9UBK8 | MTRR | 19.4489913 | 19.5857358 | 19.533237 | 19.4875256 | 19.5340181 |
| P08133 | ANXA6 | 26.0594797 | 25.961304 | 25.9491975 | 25.8857322 | 25.8932425 |
| Q9GZT4 | SRR | 18.8960348 | 18.9902849 | 19.0299815 | 18.9248194 | 19.1043634 |
| O14562 | UBFD1 | 19.6302668 | 19.8364781 | 19.7089284 | 19.7140538 | 19.9597102 |

| | | | | | | |
|---|---|---|---|---|---|---|
| Q68DQ2 | CRYBG3 | 19.3740498 | 19.4865078 | 19.4179014 | 19.3910799 | 19.6019359 |
| O14607 | UTY | 17.151606 | 17.3601691 | 17.2859206 | 17.332797 | 17.4787786 |
| Q08999 | RBL2 | 19.0307701 | 19.0290031 | 19.1123301 | 19.0870489 | 19.1419234 |
| Q13422 | IKZF1 | 21.9267317 | 21.9374153 | 21.9615912 | 21.8730398 | 21.9104136 |
| Q96N66 | MBOAT7 | 22.1106929 | 22.2437191 | 22.240394 | 22.0053307 | 22.1077109 |
| Q8TCF1 | ZFAND1 | 20.5347386 | 20.4531298 | 20.6411509 | 20.4034011 | 20.6132171 |
| P63146 | UBE2B | 18.3776555 | 18.4759982 | 18.3343521 | 18.4570813 | 18.7036875 |
| Q9H5P4 | PDZD7 | 17.5914408 | 17.8608108 | 17.452198 | 17.4682315 | 17.2485285 |
| Q8WU79 | SMAP2 | 22.2487278 | 22.2757302 | 22.2938639 | 22.0942875 | 22.1917032 |
| Q8N465 | D2HGDH | 19.6254098 | 19.6668549 | 19.6208943 | 19.5574619 | 19.7067305 |
| Q9HCS7 | XAB2 | 21.7638817 | 21.8412441 | 21.8611325 | 21.7796778 | 21.8227708 |
| Q13627 | DYRK1A | 20.6976696 | 20.8838078 | 20.7786605 | 20.6679871 | 20.8866783 |
| Q969P0 | IGSF8 | 15.1439045 | NA | NA | 15.2737189 | NA |
| Q9H6U6 | BCAS3 | 19.17803 | 19.1889682 | 19.4194919 | 19.2023388 | 19.2446128 |
| Q9H7X7 | IFT22 | 13.6936954 | NA | NA | 13.5256721 | NA |
| Q63HM1 | AFMID | NA | 12.8885076 | 12.647146 | NA | 12.5500994 |
| Q9C000 | NLRP1 | 14.3853121 | 14.4455709 | 14.8555058 | 14.4598756 | 14.6121032 |
| Q7Z6K3 | PTAR1 | 17.1160561 | 17.2162989 | 17.2786612 | 16.9784712 | 17.0546485 |
| Q9HCM7 | FBRSL1 | 21.7642736 | 21.948403 | 22.0626353 | 21.6464573 | 21.6949978 |
| P16671 | CD36 | 22.396987 | 22.4700984 | 22.5664466 | 22.3386889 | 22.4684543 |
| Q6P1X6 | C8orf82 | 22.2295838 | 22.0731371 | 22.2221216 | 22.0863945 | 22.0613203 |
| Q5VWN6 | TASOR2 | 18.2108971 | 18.3356025 | 18.2126033 | 18.133902 | 18.2590795 |
| Q13617 | CUL2 | 21.8793593 | 21.9320483 | 21.9387456 | 21.8187519 | 21.8852661 |
| P00966 | ASS1 | 23.4645441 | 23.460763 | 23.4514528 | 23.4359818 | 23.4319669 |
| Q8WW01 | TSEN15 | 17.7927663 | 17.7420976 | 17.7243212 | 17.8297377 | 17.8501031 |
| Q7Z4G1 | COMMD6 | 19.9459898 | 20.1355423 | 20.0451976 | 19.9660642 | 20.2436207 |
| Q15032 | R3HDM1 | 18.0876001 | 18.2973685 | 18.2636141 | 18.2594171 | 18.3561516 |
| Q9BVJ7 | DUSP23 | 21.9758089 | 21.8749904 | 22.1250752 | 22.1408406 | 22.1537802 |
| Q9H765 | ASB8 | 14.3543249 | 14.3044532 | 14.506865 | 14.4440617 | 14.4128891 |
| P19419 | ELK1 | 16.2223244 | 16.7156403 | 16.551605 | 16.3880535 | 16.6708002 |
| Q93096 | PTP4A1 | 13.6645174 | 14.0723921 | 13.9514568 | 14.2048899 | 14.2991161 |
| Q9NRN9 | METTL5 | 19.6728345 | 19.600544 | 19.6828656 | 19.6946332 | 19.7976766 |
| Q86U90 | YRDC | 19.0358031 | 19.1082493 | 19.1183549 | 19.0766323 | 19.1370652 |
| Q96G74 | OTUD5 | 13.9546182 | NA | NA | 13.8796856 | NA |
| Q6PI98 | INO80C | 17.8189758 | 18.3993714 | 18.0597437 | 17.9880467 | 18.0489656 |
| Q9H0E2 | TOLLIP | 21.3182085 | 21.3211571 | 21.2547327 | 21.2464396 | 21.3546768 |
| Q9HCK8 | CHD8 | 20.5700448 | 20.6554593 | 20.6875822 | 20.6054037 | 20.6656453 |
| P48723 | HSPA13 | 19.5574014 | 19.6365263 | 19.5251909 | 19.5538339 | 19.6979543 |
| Q14142 | TRIM14 | 17.6337406 | 17.7485251 | 17.678944 | 17.8345312 | 17.7410391 |
| Q06587 | RING1 | 20.4962492 | 20.534227 | 20.5487919 | 20.561505 | 20.6859297 |
| Q8NDH2 | CCDC168 | 18.7768844 | NA | NA | 18.685129 | NA |
| Q9NUJ7 | PLCXD1 | 16.1182498 | 16.2339188 | 15.7553526 | 16.2115056 | 16.1216957 |

| | | | | | | |
|---|---|---|---|---|---|---|
| P30260 | CDC27 | 19.8928093 | 19.9287092 | 20.0243841 | 19.8556694 | 19.8861445 |
| P17812 | CTPS1 | 23.1896416 | 23.1132941 | 23.1505466 | 23.0359205 | 23.0932493 |
| Q92674 | CENPI | 18.6998925 | 18.9289039 | 19.3523063 | 18.9815979 | 18.7981382 |
| Q6WRX3 | ZYG11A | 14.6962329 | NA | NA | 14.5933531 | NA |
| Q5SQI0 | ATAT1 | 16.2781341 | 16.5345606 | 16.6559043 | 16.3755261 | 16.3706708 |
| Q9Y5W8 | SNX13 | 16.636101 | 16.6733459 | 16.6944564 | 16.5108096 | 16.5402486 |
| Q10589 | BST2 | 22.3792996 | 22.7530942 | 22.6616922 | 22.3372288 | 22.4841617 |
| P26373 | RPL13 | 26.7081018 | 26.6856161 | 26.8159985 | 26.570491 | 26.417018 |
| Q149N8 | SHPRH | 17.8218287 | 17.9509942 | 18.078329 | 17.7771595 | 17.9608942 |
| P01889 | HLA-B | 22.0303666 | 22.0090507 | 22.163155 | 21.8621549 | 22.0179887 |
| Q01081 | U2AF1 | 24.1934785 | 24.2863098 | 24.2515942 | 24.1214677 | 24.1975291 |
| Q6XQN6 | NAPRT | 23.1313766 | 23.222112 | 23.141622 | 23.0736069 | 23.2047804 |
| Q9HD20 | ATP13A1 | 22.8274273 | 22.8102775 | 22.8573961 | 22.7523097 | 22.8114519 |
| Q16531 | DDB1 | 23.6289001 | 23.6859877 | 23.6316204 | 23.5489443 | 23.6487101 |
| Q8NFY9 | KBTBD8 | 19.500923 | 19.5048164 | 19.5260944 | 19.5031915 | 19.4907519 |
| Q86U44 | METTL3 | 21.4100916 | 21.5380658 | 21.4238841 | 21.4038712 | 21.5433533 |
| P61916 | NPC2 | 22.7012841 | 22.646454 | 22.5130384 | 22.6843311 | 22.6639528 |
| Q15648 | MED1 | 20.1916226 | 20.337821 | 20.2105785 | 20.2751365 | 20.4329468 |
| Q8NEZ2 | VPS37A | 19.7166087 | 19.8573588 | 19.7495082 | 19.870124 | 19.8449563 |
| P16220 | CREB1 | 21.5284684 | 21.7503542 | 21.4892003 | 21.6503911 | 21.8121225 |
| O60733 | PLA2G6 | 21.2391265 | 21.5261809 | 21.5839863 | 21.4880555 | 21.4438881 |
| P13929 | ENO3 | 16.0304185 | NA | NA | 16.202256 | NA |
| O15265 | ATXN7 | 14.9879588 | 15.1301942 | 14.9463721 | 15.2730627 | 14.6562884 |
| Q9BT40 | INPP5K | 19.7088474 | 19.666572 | 19.8838569 | 19.812348 | 19.6268486 |
| P57088 | TMEM33 | 22.5369442 | 22.4637447 | 22.5877424 | 22.4029187 | 22.5731478 |
| Q14186 | TFDP1 | 18.957384 | 18.9458864 | 19.1044995 | 18.8684158 | 19.0952594 |
| Q7Z4H3 | HDDC2 | 19.7599447 | 19.7067382 | 19.7309938 | 19.5747106 | 19.7132694 |
| A2A288 | ZC3H12D | 18.6774656 | 18.4657455 | 18.7663356 | 18.6001751 | 18.7301785 |
| P36888 | FLT3 | 21.0568804 | 20.8530248 | 20.9304156 | 20.8731367 | 20.9182775 |
| O00567 | NOP56 | 23.7667933 | 23.8213115 | 23.7857875 | 23.700272 | 23.7316066 |
| Q9Y3S2 | ZNF330 | 20.358813 | 20.4462498 | 20.5255684 | 20.524924 | 20.5313032 |
| P33527 | ABCC1 | 18.0566618 | 18.1979177 | 18.2590813 | 18.2709981 | 18.2878051 |
| Q8N9V3 | WDSUB1 | 18.183785 | 18.3278607 | 18.3356679 | 18.2744619 | 18.5157662 |
| P05204 | HMGN2 | 25.1667143 | 24.7907492 | 24.9643294 | 24.8675379 | 24.7699983 |
| Q15365 | PCBP1 | 25.4390471 | 25.5501146 | 25.4577017 | 25.2834518 | 25.4479552 |
| P23528 | CFL1 | 27.2451219 | 27.4226074 | 27.0185375 | 27.1430711 | 27.303776 |
| O15498 | YKT6 | 21.6054037 | 21.6216445 | 21.7043844 | 21.6038796 | 21.609804 |
| O95154 | AKR7A3 | 16.7827044 | 16.4089846 | 16.6645787 | 16.7394665 | 16.5393225 |
| Q9NTJ4 | MAN2C1 | 19.6941196 | 19.9126703 | 19.9671659 | 19.7616308 | 19.8569607 |
| Q7LDG7 | RASGRP2 | 21.9091551 | 21.7876886 | 21.7895031 | 21.7790736 | 21.8166989 |
| Q8IV53 | DENND1C | 21.4373158 | 21.4823753 | 21.3932635 | 21.4680903 | 21.5066707 |
| Q6NYC8 | PPP1R18 | 21.7578661 | 21.9167113 | 21.9038201 | 21.8784351 | 21.9496513 |

| | | | | | | |
|---|---|---|---|---|---|---|
| Q96C90 | PPP1R14B | 20.6476089 | 20.712927 | 20.6172554 | 20.8974716 | 20.8025621 |
| Q96B54 | ZNF428 | 19.887162 | 19.8670661 | 19.7225742 | 19.9335464 | 20.0550368 |
| Q9H583 | HEATR1 | 21.5327069 | 21.5781527 | 21.6172634 | 21.4978933 | 21.5510632 |
| Q9BPX3 | NCAPG | 22.1239745 | 22.1952245 | 22.1942912 | 22.0701605 | 22.1394308 |
| Q92575 | UBXN4 | 21.8632517 | 21.8945763 | 21.8779624 | 21.723574 | 21.682248 |
| Q86Y91 | KIF18B | 17.7442264 | 17.8638647 | 17.7202899 | 17.7164112 | 18.0006018 |
| Q68DK7 | MSL1 | 19.4947598 | 19.5978138 | 19.565733 | 19.4709045 | 19.6279554 |
| Q14684 | RRP1B | 21.7161872 | 21.6775119 | 21.7739762 | 21.7113359 | 21.6582174 |
| O43464 | HTRA2 | 19.5522923 | 19.6886991 | 19.5124637 | 19.46384 | 19.6577698 |
| Q8ND76 | CCNY | 16.091008 | 16.160576 | 16.3790072 | 16.1863917 | 16.3244183 |
| A0MZ66 | SHTN1 | 21.7329377 | 21.8109467 | 21.7613738 | 21.7436566 | 21.8972397 |
| P55273 | CDKN2D | 17.3741967 | 17.1929301 | 17.4461345 | 17.3694262 | 17.7148346 |
| Q5SSJ5 | HP1BP3 | 24.3787343 | 24.5136762 | 24.5248371 | 24.3539823 | 24.4462536 |
| Q13247 | SRSF6 | 25.3242174 | 25.4577954 | 25.173114 | 25.3025382 | 25.2794069 |
| Q9BYX2 | TBC1D2 | 15.0949869 | 15.3305675 | 15.5206506 | 15.1918033 | 15.3642707 |
| A2RTY3 | HEATR9 | 15.108343 | NA | NA | 14.8607329 | NA |
| O00303 | EIF3F | 24.0770148 | 24.0525391 | 24.1215752 | 23.9570751 | 24.0883865 |
| P49755 | TMED10 | 23.6808371 | 23.7510573 | 23.6436016 | 23.7120343 | 23.7460393 |
| Q7Z3B3 | KANSL1 | 18.0598635 | 17.8471556 | 18.1896135 | 18.2988457 | 18.2733818 |
| Q9H4Z2 | ZNF335 | NA | 16.1180453 | 16.1506613 | NA | 16.0882369 |
| Q9UJT2 | TSKS | 16.4546374 | 16.7703903 | 16.4260013 | 16.492727 | 16.5833752 |
| Q86UV5 | USP48 | 21.0747117 | 21.1474198 | 21.1400653 | 21.0511932 | 21.0971191 |
| Q14582 | MXD4 | 15.7895567 | 15.9647853 | 15.7870889 | 15.9767631 | 15.8010478 |
| Q9Y5P6 | GMPPB | 20.849903 | 20.8885415 | 21.0385032 | 20.8619135 | 20.9294213 |
| Q7Z3T8 | ZFYVE16 | 19.6995946 | 19.8476646 | 19.6927344 | 19.7389659 | 19.8229329 |
| O75365 | PTP4A3 | 16.590278 | 16.8158646 | 16.8423814 | 16.4619789 | 16.6374823 |
| Q9UBU7 | DBF4 | 11.3741712 | NA | NA | 11.2658849 | NA |
| Q96B36 | AKT1S1 | 20.6048727 | 20.2891346 | 20.1064033 | 20.2065794 | 20.1889446 |
| Q96I51 | RCC1L | 20.1622712 | 20.2216727 | 20.1884848 | 20.0912101 | 20.3063251 |
| O95900 | TRUB2 | 17.7185302 | 17.8050196 | 17.7460443 | 18.0070463 | 17.6837844 |
| Q9NQC3 | RTN4 | 22.2962301 | 22.420305 | 22.4080409 | 22.299299 | 22.2576421 |
| Q92989 | CLP1 | 20.7156243 | 20.8174699 | 20.9062042 | 20.7293003 | 20.8895396 |
| Q8NCG7 | DAGLB | 21.3594011 | 21.2966202 | 21.3868752 | 21.3437103 | 21.4097514 |
| Q15532 | SS18 | 20.1265671 | 20.129323 | 20.1313646 | 20.239935 | 20.3958214 |
| Q9NSA3 | CTNNBIP1 | 18.2856302 | 18.0780265 | 18.2682952 | 18.0930867 | 18.4553876 |
| Q9C0K1 | SLC39A8 | 17.9915473 | 17.7724348 | 17.9515065 | 17.8769122 | 17.6513052 |
| O75695 | RP2 | 21.7912581 | 21.8660929 | 21.844956 | 21.7599117 | 21.975518 |
| P09669 | COX6C | 24.1305612 | 23.8496799 | 24.3472337 | 24.104761 | 24.0578389 |
| Q96HR9 | REEP6 | 21.7790331 | 21.9156907 | 21.9430039 | 21.8200072 | 21.6014708 |
| Q9H3K2 | GHITM | 21.4712777 | 21.4657744 | 21.6944485 | 21.3542982 | 21.3925913 |
| Q14331 | FRG1 | 21.3503825 | 21.5481752 | 21.5544129 | 21.3677583 | 21.3120563 |
| Q08426 | EHHADH | 19.0886084 | 19.1126265 | 19.1318642 | 19.0983425 | 19.2011773 |

| | | | | | | |
|---|---|---|---|---|---|---|
| Q14160 | SCRIB | 21.9566602 | 22.0886476 | 21.9548848 | 21.987548 | 22.1487791 |
| Q9BYC5 | FUT8 | 19.2926643 | 19.509316 | 19.4094822 | 19.4394338 | 19.4295481 |
| Q86VU5 | COMTD1 | 17.8318577 | 17.6924286 | 17.7426166 | 17.625593 | 17.7905874 |
| Q92585 | MAML1 | 18.7905472 | 18.9076588 | 18.8226058 | 18.8786354 | 18.7538863 |
| P27449 | ATP6V0C | 20.9003282 | 20.5544853 | 20.9312079 | 20.5859562 | 20.6660329 |
| Q92800 | EZH1 | 13.0623956 | NA | NA | 13.1997249 | NA |
| P42766 | RPL35 | 25.2415181 | 25.4702513 | 25.6858747 | 25.3745909 | 25.2882925 |
| P62995 | TRA2B | 24.4790026 | 24.4767427 | 24.4871657 | 24.4668061 | 24.6326883 |
| Q9NVH2 | INTS7 | 20.2372965 | 20.2379814 | 20.3054083 | 20.0777083 | 20.3343942 |
| O75127 | PTCD1 | 19.263883 | 19.4218392 | 19.4264554 | 19.3241744 | 19.3653915 |
| Q99436 | PSMB7 | 23.9160805 | 23.9792411 | 23.7124784 | 23.8584065 | 23.8392479 |
| P98170 | XIAP | 18.5852181 | 18.7867861 | 18.9064174 | 18.8210663 | 18.8873012 |
| Q8N9M1 | C19orf47 | 19.6084542 | 19.7331522 | 19.7619524 | 19.5718556 | 19.8083707 |
| O43768 | ENSA | 23.2812868 | 23.4486066 | 23.0897485 | 23.2773223 | 23.3947144 |
| Q8WVR3 | TRAPPC14 | 13.9690589 | 14.2918376 | 14.7981773 | 14.5259007 | 14.3671597 |
| Q96T21 | SECISBP2 | 15.811827 | 15.2639303 | 16.1115995 | 15.7046949 | 16.0381524 |
| Q9BTL3 | RAMAC | 20.5912207 | 20.7284894 | 20.7727631 | 20.7752565 | 20.8982167 |
| P26951 | IL3RA | 20.2339676 | 20.1975752 | 20.2801436 | 20.1374257 | 20.2524326 |
| Q15831 | STK11 | 18.9895044 | 19.0584153 | 19.1059806 | 19.0559519 | 19.1715507 |
| Q8WXH0 | SYNE2 | 19.1449532 | 19.5090312 | 18.6022363 | 18.8924132 | 19.4205134 |
| O14782 | KIF3C | 20.8149616 | 20.7751988 | 20.5037342 | 20.2414706 | 19.5246683 |
| Q9Y283 | INVS | 15.8911908 | 16.2074436 | 16.1122761 | 16.0857657 | 16.229453 |
| P01008 | SERPINC1 | 21.3495033 | 21.5199662 | 21.1593584 | 21.0748476 | 21.2562279 |
| Q8IUR0 | TRAPPC5 | 21.5357498 | 21.4876349 | 21.6801021 | 21.5326817 | 21.7247603 |
| Q13395 | TARBP1 | 20.5255261 | 20.6199692 | 20.6635801 | 20.5647604 | 20.6389125 |
| P00374 | DHFR | 22.4323369 | 22.3569873 | 22.5348009 | 22.3186509 | 22.2470548 |
| P06280 | GLA | 22.357171 | 22.3755473 | 22.4040347 | 22.247915 | 22.3991918 |
| Q8IV63 | VRK3 | 20.4590046 | 20.2811914 | 20.5528987 | 20.5342121 | 20.5790117 |
| O94992 | HEXIM1 | 21.525364 | 21.6413615 | 21.3557922 | 21.4885611 | 21.5248778 |
| O95271 | TNKS | 17.0630063 | 16.9727804 | 17.1322701 | 16.8271373 | 16.6168727 |
| Q8N4W9 | ZNF808 | 16.8774604 | 16.8003078 | 16.7445938 | 16.7152416 | 16.6718608 |
| Q8IZA0 | KIAA0319L | 17.726011 | 17.8887908 | 17.961046 | 17.8688023 | 17.7907407 |
| O60499 | STX10 | 19.1878026 | 19.2261718 | 19.168321 | 19.1107361 | 19.0312301 |
| Q9H2C0 | GAN | 20.2001429 | 20.171941 | 20.2569549 | 20.1621843 | 20.1867903 |
| Q68CQ7 | GLT8D1 | 20.2212228 | 20.2175031 | 20.193501 | 20.1819341 | 20.0441569 |
| O00442 | RTCA | 21.2897499 | 21.4202673 | 21.4500887 | 21.2413944 | 21.4476356 |
| Q8WW59 | SPRYD4 | 20.8225435 | 20.7830106 | 20.8872328 | 20.8518632 | 20.7334987 |
| Q86XL3 | ANKLE2 | 19.6582535 | 19.7524189 | 19.6792282 | 19.6948963 | 19.7803228 |
| Q14671 | PUM1 | 21.6870306 | 21.8062491 | 21.74289 | 21.6734827 | 21.8063223 |
| Q9BTZ2 | DHRS4 | 20.2198666 | 20.3274247 | 20.252204 | 20.2161603 | 20.3401463 |
| Q9Y2D5 | PALM2AKAP2 | 21.3501238 | 21.4448855 | 21.400704 | 21.3572957 | 21.5037228 |
| O43823 | AKAP8 | 23.2971021 | 23.3386741 | 23.1530521 | 23.301103 | 23.3037184 |

| | | | | | | |
|---|---|---|---|---|---|---|
| P12270 | TPR | 23.0966736 | 23.1991433 | 23.0202777 | 23.0277687 | 23.1666324 |
| Q9NYG2 | ZDHHC3 | 18.6778305 | 18.7815336 | 18.8892337 | 18.8656779 | 18.819933 |
| P23025 | XPA | 17.3457552 | 17.6177057 | 17.6128149 | 17.6040251 | 17.5866149 |
| O94986 | CEP152 | 17.6866349 | 17.8362753 | 17.5661278 | 17.7609928 | 17.8362095 |
| Q16513 | PKN2 | 17.4629039 | 17.8091739 | 17.6628728 | 17.5518255 | 17.8476251 |
| Q9HC29 | NOD2 | 14.7163594 | 14.3869693 | 14.8569863 | 14.6938905 | 14.6338894 |
| Q96P50 | ACAP3 | 13.452759 | 13.2305734 | 13.4176846 | 13.5762704 | 13.3044374 |
| Q9C073 | FAM117A | 16.0107196 | 16.1289279 | 16.5740387 | 15.8351191 | 15.6724684 |
| Q16854 | DGUOK | 18.3520398 | 18.1302359 | 18.4028809 | 18.2333635 | 18.2533421 |
| Q9Y619 | SLC25A15 | 20.446114 | 20.4862693 | 20.5489285 | 20.3730566 | 20.5415379 |
| Q15075 | EEA1 | 23.1669869 | 23.3155321 | 23.0536899 | 23.165 | 23.242274 |
| O14618 | CCS | 21.6816982 | 21.9213551 | 21.6483078 | 21.6308526 | 21.8715231 |
| Q9H8V3 | ECT2 | 17.1370264 | 16.9903394 | 17.2218231 | 17.131572 | 17.1874991 |
| Q9Y5U2 | TSSC4 | 21.0016282 | 20.9264077 | 21.0280091 | 20.7145722 | 21.2985725 |
| P08048 | ZFY | 14.5805414 | NA | NA | 14.8209638 | NA |
| P49642 | PRIM1 | 20.192392 | 20.1811986 | 20.3123282 | 20.1464838 | 20.1650297 |
| Q9UGV2 | NDRG3 | 19.5194207 | 19.4731075 | 19.595112 | 19.6117594 | 19.8516038 |
| Q96RK4 | BBS4 | NA | 14.2285645 | 14.1811639 | NA | 14.2771115 |
| Q9H0U6 | MRPL18 | 21.5359328 | 21.3613597 | 21.5787279 | 21.3211747 | 21.4723723 |
| O15126 | SCAMP1 | 20.5859485 | 20.6583618 | 20.7297461 | 20.5205761 | 20.5743015 |
| Q9UHA4 | LAMTOR3 | 21.0214539 | 21.2124704 | 21.1297344 | 21.1190429 | 21.2910596 |
| Q9H4K7 | MTG2 | 17.4333216 | 17.6764381 | 17.6113223 | 17.6917746 | 17.6776272 |
| P25054 | APC | 16.745205 | 17.0433128 | 16.568294 | 16.8522409 | 16.845025 |
| Q16626 | MEA1 | 18.0001574 | 18.1378553 | 17.8810634 | 18.0651341 | 18.2147575 |
| Q8TD31 | CCHCR1 | 15.4119759 | 15.376151 | 15.6603765 | 15.5620304 | 15.7666516 |
| O96007 | MOCS2 | 17.0946007 | 17.070931 | 17.5723313 | 17.0793972 | 17.6592697 |
| Q96RQ1 | ERGIC2 | 19.7391043 | 19.7936812 | 19.9832969 | 19.6630661 | 19.6480291 |
| Q92520 | FAM3C | 20.7148645 | 20.8110118 | 20.7859309 | 20.8258361 | 20.7888063 |
| Q8TD10 | MIPOL1 | 18.0998579 | NA | NA | 17.5441769 | NA |
| Q86WR7 | PROSER2 | 18.483859 | 18.5545126 | 18.7111394 | 18.6595442 | 18.5534514 |
| Q7Z388 | DPY19L4 | 16.2367244 | 16.8448633 | 16.6211635 | 16.6550335 | 16.912556 |
| Q99755 | PIP5K1A | 14.8367793 | NA | NA | 15.0346041 | NA |
| Q9NUQ3 | TXLNG | 20.3031484 | 20.2421666 | 20.3624661 | 20.3334928 | 20.4611765 |
| Q9UI12 | ATP6V1H | 21.1796289 | 21.2423635 | 21.2759358 | 21.1102002 | 21.2298207 |
| Q09161 | NCBP1 | 23.1097074 | 23.1092337 | 23.2128085 | 23.0174815 | 23.0635027 |
| Q9P0P8 | MTRES1 | 21.8845019 | 22.1712033 | 21.9207581 | 21.9961326 | 22.192679 |
| P78337 | PITX1 | 21.3524738 | 21.4400336 | 21.3124912 | 21.3400851 | 21.5532587 |
| Q96K49 | TMEM87B | 19.100748 | 19.213428 | 19.2394384 | 19.2101523 | 19.1523771 |
| Q9UHA2 | SS18L2 | 16.6470333 | 16.6068792 | 17.1620748 | 17.2029078 | 17.2945525 |
| O95476 | CTDNEP1 | 18.2388185 | 18.420719 | 18.2951719 | 18.201674 | 18.3686855 |
| Q3SXY8 | ARL13B | 16.7396121 | 16.8784907 | 17.1653811 | 16.8355635 | 16.9767605 |
| Q9NWT1 | PAK1IP1 | 20.960106 | 20.8613845 | 21.0082418 | 20.8862005 | 20.8813804 |

| | | | | | | |
|---|---|---|---|---|---|---|
| P23526 | AHCY | 25.2805955 | 25.2451086 | 25.1718521 | 25.1986024 | 25.2763534 |
| A8MTT3 | CEBPZOS | 20.7021981 | 20.6348877 | 20.8031391 | 20.6912711 | 20.8156601 |
| Q8IXM2 | C17orf49 | 22.017601 | 22.1137656 | 21.8769824 | 21.9985119 | 22.1406736 |
| O94864 | SUPT7L | 19.3141037 | 19.4658842 | 19.4491871 | 19.4850808 | 19.7098395 |
| A0PK00 | TMEM120B | 17.4262531 | 17.5365902 | 17.6330287 | 17.568296 | 17.4481372 |
| Q96AT9 | RPE | 22.1228469 | 22.2654914 | 22.2029691 | 22.1510052 | 22.4925891 |
| Q6PK04 | CCDC137 | 20.6360293 | 20.7364567 | 20.7900984 | 20.743997 | 20.5680023 |
| Q8IXM6 | NRM | 21.1645164 | 21.3046392 | 21.3907254 | 21.1404689 | 21.2432487 |
| Q9BVK2 | ALG8 | 19.7831085 | 19.7200149 | 19.7979666 | 19.581876 | 19.7494137 |
| Q7Z7H5 | TMED4 | 22.4796072 | 22.5506958 | 22.5478597 | 22.4350698 | 22.4941659 |
| Q9BW19 | KIFC1 | 20.7208687 | 20.8133135 | 20.7929191 | 20.8102554 | 20.768994 |
| Q9HAN9 | NMNAT1 | 18.5212849 | 18.6239591 | 18.751169 | 18.6439087 | 18.7439081 |
| P33240 | CSTF2 | 20.6630418 | 20.6736403 | 20.4483478 | 20.6492059 | 20.7699692 |
| Q9H825 | METTL8 | 15.3486244 | 15.6359906 | 15.5284735 | 15.3344342 | 15.4961413 |
| Q8NBK3 | SUMF1 | 19.0371098 | 18.9169019 | 18.8583789 | 18.9455654 | 18.9451662 |
| Q9Y262 | EIF3L | 24.7124817 | 24.7641934 | 24.8099637 | 24.6885143 | 24.6643951 |
| Q9Y2Q3 | GSTK1 | 22.7858321 | 22.7937152 | 22.8026041 | 22.7963656 | 22.8231836 |
| Q9NTZ6 | RBM12 | 22.4714239 | 22.5330587 | 22.4266293 | 22.4684718 | 22.54988 |
| Q8IW92 | GLB1L2 | 18.5242865 | 18.4389869 | 18.4127529 | 18.5319406 | 18.0389944 |
| Q12894 | IFRD2 | 18.1433641 | 18.4680441 | 18.3949789 | 18.4340967 | 18.2432905 |
| Q9NZP8 | C1RL | 15.7952983 | NA | NA | 15.8295117 | NA |
| P35555 | FBN1 | NA | 16.7259989 | 16.5191706 | NA | 16.1911552 |
| Q9NZC7 | WWOX | 18.2657956 | 18.1642331 | 18.0771486 | 18.1766782 | 18.3840609 |
| Q69YL0 | NCBP2AS2 | 19.7376072 | 19.3297115 | 19.7456946 | 19.3821386 | 19.5685379 |
| Q13823 | GNL2 | 21.1104549 | 21.2199872 | 21.2741679 | 21.1276168 | 21.1698599 |
| P35789 | ZNF93 | 20.5722954 | 20.7158575 | 20.7218088 | 20.5771449 | 20.4888924 |
| Q9H7E9 | C8orf33 | 22.1013069 | 22.0812761 | 22.1142593 | 21.9853474 | 21.9540867 |
| Q96D96 | HVCN1 | 21.8801822 | 22.0353889 | 21.8825006 | 21.8433459 | 21.7600149 |
| O14828 | SCAMP3 | 22.5731561 | 22.6105766 | 22.5872444 | 22.5314274 | 22.5993321 |
| Q9Y2W6 | TDRKH | 19.4039844 | 19.3961693 | 19.499573 | 19.443916 | 19.6639673 |
| Q969X5 | ERGIC1 | 18.2013773 | 18.3109377 | 18.3318624 | 18.4663369 | 18.3613764 |
| P22830 | FECH | 20.8840923 | 20.9737175 | 21.0135821 | 20.9279233 | 20.9228133 |
| Q8N5D0 | WDTC1 | 18.5904832 | 18.5264725 | 18.5865644 | 18.5450332 | 18.6387957 |
| Q9NPF2 | CHST11 | 19.8416837 | 19.9494461 | 20.0442297 | 19.8124914 | 20.021115 |
| O75290 | ZNF780A | 18.5003839 | 18.3044912 | 18.5956126 | 18.5945122 | 18.6441874 |
| Q9BQB6 | VKORC1 | 21.3752644 | 21.2397458 | 21.3789749 | 21.2328716 | 21.0640375 |
| Q9Y256 | RCE1 | 18.020084 | 18.1300387 | 18.2120156 | 18.083017 | 18.0768765 |
| O95259 | KCNH1 | 18.1860761 | 18.9234739 | 18.0635682 | 18.4410286 | 18.1978588 |
| Q9NWS6 | FAM118A | 19.505464 | 19.6212547 | 19.6971739 | 19.5932144 | 19.7647611 |
| Q9NZI7 | UBP1 | 19.3332645 | 19.611994 | 19.3643212 | 19.4025396 | 19.6473548 |
| O75386 | TULP3 | 15.6418402 | 15.9299376 | 15.673052 | 15.6458984 | 15.8010002 |
| Q5VW38 | GPR107 | 19.3939674 | 19.3329433 | 19.4967999 | 19.463689 | 19.4990455 |

| | | | | | | |
|---|---|---|---|---|---|---|
| Q9P2R6 | RERE | 17.5125186 | 18.0836277 | 18.3260158 | 17.7763932 | 17.8236945 |
| Q86V87 | FHIP2B | 16.6957569 | 16.7251267 | 16.9312864 | 16.5864547 | 16.892237 |
| P51151 | RAB9A | 20.8937206 | 21.0097333 | 20.9416515 | 20.8313909 | 20.9312545 |
| P53597 | SUCLG1 | 22.2808911 | 22.3600134 | 22.332157 | 22.3719441 | 22.4918604 |
| Q4ZIN3 | TMEM259 | 19.2053983 | 19.0340161 | 19.1639617 | 19.040375 | 19.3478791 |
| P17026 | ZNF22 | 19.7836154 | 20.1362862 | 20.0541339 | 19.9261607 | 19.7425598 |
| Q9BWE0 | REPIN1 | 20.1772482 | 20.2226368 | 20.2131115 | 20.1310981 | 19.712453 |
| Q5JUR7 | TEX30 | 20.9045225 | 20.8253054 | 20.9397816 | 20.7878879 | 20.8208977 |
| Q93074 | MED12 | 20.2258668 | 20.268627 | 20.3885224 | 20.2150467 | 20.283332 |
| O15213 | WDR46 | 21.0926263 | 21.0634845 | 21.0775035 | 20.9782606 | 21.063091 |
| Q9H0G5 | NSRP1 | 20.4436857 | 20.5347927 | 20.4004186 | 20.4690125 | 20.5846366 |
| Q8TBZ8 | ZNF564 | 18.0476475 | 18.1174226 | 17.9694796 | 17.9557196 | 17.8717325 |
| Q9P0W2 | HMG20B | 19.7169902 | 19.7716183 | 19.7425383 | 19.7456417 | 19.8293024 |
| Q9HAV7 | GRPEL1 | 23.819044 | 24.0157084 | 23.7891035 | 23.8361526 | 23.9881297 |
| O95777 | LSM8 | 22.1693166 | 22.2848748 | 22.138161 | 22.1135884 | 22.2613934 |
| Q99747 | NAPG | 21.1902778 | 21.2864189 | 21.3023819 | 21.241762 | 21.376716 |
| Q8N5F7 | NKAP | 19.2661208 | 19.3607948 | 19.4537865 | 19.3477027 | 19.5307091 |
| O75170 | PPP6R2 | 19.5165603 | 19.6552972 | 19.5793113 | 19.6024881 | 19.7015849 |
| Q9NS93 | TM7SF3 | 19.0486953 | 19.1238315 | 19.0235536 | 19.0951196 | 19.1538129 |
| Q9UPU3 | SORCS3 | NA | 16.6896995 | 16.1604122 | NA | 16.3651784 |
| Q7Z5G4 | GOLGA7 | 20.4589619 | 20.591591 | 20.6681713 | 20.6271433 | 20.5069338 |
| Q9H7B2 | RPF2 | 21.7020507 | 21.7916154 | 21.9061716 | 21.8031492 | 21.9448477 |
| Q13158 | FADD | 20.7829742 | 20.8355943 | 20.8854918 | 20.8269594 | 20.9704334 |
| Q7Z7L9 | ZSCAN2 | 15.4735082 | NA | NA | 15.2346035 | NA |
| O15511 | ARPC5 | 24.6392675 | 24.7044099 | 24.6507837 | 24.551273 | 24.567555 |
| Q9BVL4 | SELENOO | 20.1647346 | 20.1129737 | 20.1319231 | 20.1344969 | 20.2419601 |
| Q9P1Z2 | CALCOCO1 | 19.3725436 | 19.4283009 | 19.3386468 | 19.4740553 | 19.4994577 |
| P57060 | RWDD2B | 17.4890206 | 17.5388262 | 17.6530024 | 17.6606211 | 17.7028179 |
| Q8WYH8 | ING5 | 20.4499293 | 20.17952 | 20.3578564 | 20.3001574 | 20.4741028 |
| Q13542 | EIF4EBP2 | 19.4130422 | 19.592878 | 19.2863802 | 19.3329299 | 19.7965328 |
| O75506 | HSBP1 | 21.1446473 | 21.4223272 | 21.0099114 | 21.2219286 | 21.3526375 |
| P08708 | RPS17 | 25.8985002 | 25.9224253 | 26.0529626 | 25.8133537 | 25.9203121 |
| Q8NB16 | MLKL | 22.2360655 | 22.2582038 | 22.4103385 | 22.4361372 | 22.4229781 |
| Q99717 | SMAD5 | 18.5719035 | 18.6107437 | 18.867598 | 18.7733239 | 19.0355932 |
| Q9UIA9 | XPO7 | 22.1846992 | 22.2170465 | 22.248425 | 22.0983263 | 22.2163563 |
| P31350 | RRM2 | 22.0025569 | 22.0493597 | 22.104815 | 22.0192862 | 22.0970443 |
| Q8NG11 | TSPAN14 | 19.3348513 | 19.2720305 | 19.3372688 | 19.2130423 | 19.1788669 |
| Q8N131 | TMEM123 | 21.5414433 | 21.551554 | 21.8527191 | 21.5780102 | 21.6148846 |
| Q68CP4 | HGSNAT | 19.9620823 | 19.9375219 | 19.8848236 | 19.8471042 | 19.7809332 |
| O60879 | DIAPH2 | 20.2526145 | 20.3442839 | 20.3655526 | 20.2933682 | 20.3895493 |
| Q8N5G0 | SMIM20 | 19.0925836 | 19.2576779 | 19.2223137 | 19.060699 | 19.0424744 |
| P49720 | PSMB3 | 24.0217241 | 23.8740251 | 24.0181067 | 23.8892391 | 23.9008316 |

| | | | | | | |
|---|---|---|---|---|---|---|
| Q8N1G4 | LRRC47 | 22.7586777 | 22.8519724 | 22.7737613 | 22.7784588 | 22.8409187 |
| Q71F56 | MED13L | 19.3187001 | 19.3827354 | 19.4626151 | 19.3330733 | 19.2691117 |
| Q96H79 | ZC3HAV1L | 20.4270694 | 20.6466928 | 20.6785113 | 20.4581059 | 20.5287431 |
| Q96N11 | INTS15 | 19.5538385 | 19.3621309 | 19.8442771 | 19.4406148 | 19.5058786 |
| Q8TAP6 | CEP76 | 20.5559269 | 20.6110678 | 20.5370013 | 20.4695145 | 20.5063185 |
| Q9Y6E2 | BZW2 | 23.6953093 | 23.7775438 | 23.6739214 | 23.5969655 | 23.5862021 |
| Q9UHB9 | SRP68 | 22.6900104 | 22.7499269 | 22.7493555 | 22.6537493 | 22.7391003 |
| Q92905 | COPS5 | 22.6569865 | 22.6343971 | 22.6453404 | 22.5102593 | 22.699498 |
| P50395 | GDI2 | 25.8512237 | 25.8262775 | 25.8270233 | 25.7161675 | 25.8095226 |
| Q15154 | PCM1 | 21.799683 | 21.8199506 | 21.7145752 | 21.7702814 | 21.8954125 |
| Q99807 | COQ7 | 20.6910595 | 20.4916103 | 20.566457 | 20.528285 | 20.6061226 |
| Q14004 | CDK13 | 23.3774166 | 23.4015405 | 23.5075558 | 23.3986646 | 23.4466035 |
| Q8IWE5 | PLEKHM2 | 18.669759 | 18.7387435 | 18.5850384 | 18.620617 | 18.7969526 |
| O43399 | TPD52L2 | 22.7958928 | 22.9216687 | 22.6365197 | 22.771824 | 22.9660895 |
| Q86Y22 | COL23A1 | 21.3534413 | 21.0330831 | 21.3380467 | 21.1133733 | 21.2945052 |
| O00716 | E2F3 | 16.3896967 | 16.2486726 | 16.4426219 | 16.4879255 | 16.4520349 |
| Q5T4F4 | ZFYVE27 | 16.930781 | 16.9279342 | 16.9453005 | 16.9976743 | 16.8392272 |
| Q9NXZ1 | SAGE1 | NA | 16.6043363 | 16.2647558 | NA | 16.3765684 |
| Q8TB03 | CXorf38 | 24.1070309 | 24.2202451 | 24.1397328 | 23.9792474 | 24.1315141 |
| P63000 | RAC1 | 23.8310839 | 24.0017705 | 24.0142996 | 23.8641865 | 23.8975357 |
| P29218 | IMPA1 | 21.91297 | 21.9345866 | 21.9286396 | 21.8393166 | 21.908301 |
| Q8NFA0 | USP32 | 19.0279094 | 19.0708199 | 19.2428021 | 19.0879544 | 19.1165399 |
| Q9UBE0 | SAE1 | 24.8968429 | 24.9011 | 24.8282573 | 24.8055487 | 24.8532698 |
| P54652 | HSPA2 | 20.2846024 | 20.3238339 | 20.3808258 | 20.3405587 | 20.5580858 |
| Q7Z7G8 | VPS13B | 18.6653843 | 18.843031 | 18.7788879 | 18.8076072 | 18.7416598 |
| Q9NZV5 | SELENON | 17.8500496 | 17.9635541 | 18.1285222 | 18.0002043 | 18.0182736 |
| Q76L83 | ASXL2 | 18.9253148 | 19.1916345 | 18.9996556 | 19.2013307 | 19.1511472 |
| P20333 | TNFRSF1B | 19.1359587 | 19.2875837 | 19.2834018 | 19.3421818 | 19.3096465 |
| P15923 | TCF3 | 19.8642944 | 20.0389567 | 19.8060263 | 19.8370731 | 20.1195776 |
| Q6NUT3 | MFSD12 | 18.1871619 | 18.7137375 | 18.6245765 | 18.4639641 | 18.6264481 |
| Q9NVJ2 | ARL8B | 24.099367 | 23.9149186 | 24.0570289 | 24.0086594 | 24.1036531 |
| Q9BRT2 | UQCC2 | 20.9817595 | 21.072859 | 21.1032051 | 20.9670352 | 21.0018754 |
| Q15031 | LARS2 | 20.2018525 | 20.2660917 | 20.3663682 | 20.2621715 | 20.2846927 |
| Q9UNL4 | ING4 | 18.49392 | 18.657649 | 18.9554119 | 18.7395059 | 18.7447702 |
| Q9Y3A0 | COQ4 | 17.8099584 | 18.0086212 | 17.8193867 | 17.88884 | 17.8463882 |
| Q16563 | SYPL1 | 21.4535006 | 21.3905835 | 21.4362694 | 21.3713742 | 21.3583625 |
| Q9H9E3 | COG4 | 20.2794789 | 20.4621411 | 20.3504082 | 20.3506264 | 20.4832273 |
| Q8N5I4 | DHRSX | 19.1708657 | 19.0257603 | 18.9713312 | 19.0535224 | 19.0288131 |
| Q8IX90 | SKA3 | 16.3738613 | 16.7055581 | 16.5502803 | 16.7136489 | 16.8245803 |
| Q9HBG7 | LY9 | 15.0860521 | NA | NA | 14.9050131 | NA |
| Q6UW02 | CYP20A1 | 20.396678 | 20.4831495 | 20.5057036 | 20.3665555 | 20.391977 |
| Q9C0E2 | XPO4 | 21.1328585 | 21.0583072 | 21.1900745 | 21.0614613 | 21.1185566 |

| | | | | | | |
|---|---|---|---|---|---|---|
| C9J7I0 | UMAD1 | NA | 15.8274034 | 15.6973158 | NA | 16.0796652 |
| P57086 | SCAND1 | 15.639239 | 15.5066557 | 15.86239 | 15.9140531 | 16.3330302 |
| P28676 | GCA | 22.0849374 | 21.9714069 | 22.046595 | 21.9512595 | 21.9991568 |
| Q6P3S6 | FBXO42 | 16.119098 | 16.4261038 | 16.7015396 | 16.0177462 | 16.4896249 |
| P48729 | CSNK1A1 | 22.3670214 | 22.3695686 | 22.4921815 | 22.3493082 | 22.4815756 |
| Q8N954 | GPATCH11 | 18.8172366 | 18.8715562 | 18.7041573 | 18.7321803 | 19.0305058 |
| Q9NUX5 | POT1 | 18.5083946 | 18.689541 | 18.6564979 | 18.6242632 | 18.7896317 |
| Q8WUJ1 | CYB5D2 | 15.6950182 | NA | NA | 15.4937354 | NA |
| P19447 | ERCC3 | 19.5257255 | 19.5006616 | 19.5841153 | 19.4604433 | 19.5225714 |
| P17098 | ZNF8 | 19.2746219 | 19.1660306 | 19.1954949 | 19.2954188 | 19.5647218 |
| Q9NUE0 | ZDHHC18 | 14.1650027 | NA | NA | 14.3627478 | NA |
| Q9Y672 | ALG6 | 20.4258958 | 20.6299915 | 20.494654 | 20.4746158 | 20.4502165 |
| P40855 | PEX19 | 20.6327142 | 20.802536 | 20.573587 | 20.6776248 | 20.9032408 |
| O60678 | PRMT3 | 20.3093635 | 20.313217 | 20.3023083 | 20.2221389 | 20.2376651 |
| P39656 | DDOST | 24.7783321 | 24.8471733 | 24.7873678 | 24.7637799 | 24.8026077 |
| Q12931 | TRAP1 | 24.2416817 | 24.2208495 | 24.2179585 | 24.1616234 | 24.1931353 |
| O43318 | MAP3K7 | 19.2746559 | 19.3342373 | 19.4905013 | 19.3894246 | 19.5425174 |
| Q9Y448 | KNSTRN | 19.0783532 | 19.2575895 | 19.1857252 | 19.1878993 | 19.2701415 |
| O15121 | DEGS1 | 19.8027918 | 19.9503838 | 20.0220769 | 19.8400724 | 19.874558 |
| P08236 | GUSB | 21.6479053 | 21.6032452 | 21.6640519 | 21.5670733 | 21.6739775 |
| Q8TBE9 | NANP | 15.4805062 | 15.6311965 | 15.6521116 | 15.3342026 | 15.656939 |
| Q8IY95 | TMEM192 | 20.8349651 | 20.9215751 | 20.9320554 | 20.864067 | 20.7706636 |
| Q13616 | CUL1 | 20.9042691 | 20.9880942 | 21.0223449 | 20.8549828 | 20.898705 |
| Q9HCU5 | PREB | 22.1557926 | 22.215288 | 22.2454362 | 22.1453109 | 22.1776572 |
| Q15036 | SNX17 | 21.2068599 | 21.2216235 | 21.238802 | 21.1119913 | 21.181624 |
| Q96FV9 | THOC1 | 20.6783231 | 20.7769979 | 20.7353665 | 20.6928733 | 20.8206099 |
| Q9NW82 | WDR70 | 20.660592 | 20.7822683 | 20.7682545 | 20.729816 | 20.8001289 |
| Q8TF01 | PNISR | 19.960833 | 19.7450431 | 19.8542336 | 19.8142163 | 19.9594152 |
| P38117 | ETFB | 23.8519858 | 24.0009117 | 23.8704527 | 23.8238705 | 24.0158455 |
| Q9BY32 | ITPA | 21.3345169 | 21.5232056 | 21.3880056 | 21.4281685 | 21.5556162 |
| Q9HB09 | BCL2L12 | 16.2096632 | 16.3317605 | 16.5613149 | 16.3030486 | 16.4914373 |
| Q99873 | PRMT1 | 25.0794076 | 24.9712213 | 25.024167 | 24.9242874 | 24.9949407 |
| O96033 | MOCS2 | 19.8600265 | 19.9806724 | 19.9449487 | 19.8924847 | 19.9057669 |
| Q9UHC7 | MKRN1 | 17.5741624 | 17.55549 | 17.6701639 | 17.6236716 | 17.7315781 |
| Q8WVN8 | UBE2Q2 | 18.3702694 | 18.504319 | 18.6662083 | 18.7185389 | 18.9454933 |
| Q9HAV0 | GNB4 | 20.8957683 | 20.7887744 | 20.9750493 | 20.7758932 | 21.0446333 |
| O75843 | AP1G2 | 20.2437767 | 20.1733293 | 20.2696742 | 20.053912 | 20.2343821 |
| Q96SU4 | OSBPL9 | 18.9022283 | 18.7746459 | 19.0845252 | 18.8954814 | 19.1955209 |
| O14949 | UQCRQ | 22.7155884 | 22.5350763 | 22.7996865 | 22.7183269 | 22.5968704 |
| P62805 | NA | 29.7707112 | 29.9146578 | 29.9956332 | 29.6695111 | 29.7487862 |
| Q9NPI1 | BRD7 | 21.4959099 | 21.5697253 | 21.5131966 | 21.5490933 | 21.5697206 |
| Q8IWF2 | FOXRED2 | 17.7793438 | 17.6248829 | 18.0719834 | 17.832873 | 17.6963416 |

| | | | | | | |
|---|---|---|---|---|---|---|
| Q9NX00 | TMEM160 | 21.2703407 | 21.438494 | 21.4685569 | 21.1125393 | 21.407982 |
| Q16526 | CRY1 | 16.7786907 | 16.8190924 | 16.6950619 | 17.06001 | 16.8973284 |
| P61803 | DAD1 | 23.5916032 | 23.7005499 | 23.5336482 | 23.4077568 | 23.4308583 |
| Q96AQ8 | MCUR1 | 19.9858229 | 19.9919795 | 20.1553966 | 19.9579989 | 19.8397973 |
| P35790 | CHKA | 18.7608576 | 18.6683376 | 18.8643699 | 18.7719313 | 18.8271336 |
| Q06330 | RBPJ | 20.122796 | 20.1412042 | 20.1921998 | 20.1575007 | 20.1262881 |
| P11940 | PABPC1 | 25.691543 | 25.7017877 | 25.556133 | 25.5392375 | 25.5991357 |
| P26038 | MSN | 26.7876403 | 26.787923 | 26.7542201 | 26.6323474 | 26.6878453 |
| Q14746 | COG2 | 19.8214212 | 19.9190846 | 19.8986152 | 19.8934467 | 19.9198613 |
| Q6I9Y2 | THOC7 | 21.2918499 | 21.4556872 | 21.3240154 | 21.3326237 | 21.3714086 |
| P07858 | CTSB | 22.5338146 | 22.6917245 | 22.5873777 | 22.6110247 | 22.6995846 |
| Q9NUP7 | TRMT13 | 17.80611 | 18.0292909 | 18.0781892 | 17.9621355 | 18.0320574 |
| O95400 | CD2BP2 | 20.7811196 | 20.9345088 | 20.7770557 | 20.8519346 | 20.9580925 |
| Q15208 | STK38 | 20.7680852 | 20.769961 | 20.9397422 | 20.8342216 | 20.9181279 |
| Q9BVM2 | DPCD | 18.4624717 | 18.6212166 | 18.6076134 | 18.6005014 | 18.6574054 |
| Q9Y6K9 | IKBKG | 21.5384882 | 21.6924886 | 21.4132571 | 21.5011357 | 21.5327024 |
| Q9Y6E0 | STK24 | 19.5547791 | 19.6945782 | 19.672048 | 19.5958642 | 19.6543246 |
| Q9GZV4 | EIF5A2 | 17.0257604 | 17.349083 | 16.9732858 | 17.1220273 | 17.1712651 |
| Q9Y2B0 | CNPY2 | 22.0613413 | 22.2518238 | 21.949264 | 22.0218327 | 22.2160005 |
| Q9Y2M5 | KLHL20 | 12.568812 | NA | NA | 13.4053776 | NA |
| Q16831 | UPP1 | 15.3979734 | 15.1847846 | 15.5744572 | 14.7106062 | 15.0641077 |
| Q96C12 | ARMC5 | 18.2434426 | 18.5040742 | 18.4163791 | 18.2936625 | 18.4149839 |
| P62917 | RPL8 | 26.7435301 | 26.8370373 | 26.9311115 | 26.6854769 | 26.693622 |
| Q9UQ13 | SHOC2 | 20.4164067 | 20.2904016 | 20.6342114 | 20.3877525 | 20.4265309 |
| Q96IU4 | ABHD14B | 23.7400174 | 23.5889182 | 23.7997783 | 23.6026157 | 23.7173001 |
| P04439 | HLA-A | 23.1523387 | 23.2504673 | 23.2741461 | 23.1103021 | 23.1727662 |
| Q92769 | HDAC2 | 22.95251 | 22.9566054 | 22.9732575 | 22.8783655 | 22.9643414 |
| P61247 | RPS3A | 26.5518401 | 26.5265984 | 26.5152295 | 26.4199539 | 26.4143231 |
| Q9BV20 | MRI1 | 22.2749525 | 22.3774769 | 22.331116 | 22.2808642 | 22.3415617 |
| Q9UHD2 | TBK1 | 20.613152 | 20.6990895 | 20.74786 | 20.6282922 | 20.7298061 |
| Q5T200 | ZC3H13 | 20.7722614 | 20.8460272 | 20.8292352 | 20.8123053 | 20.8561629 |
| Q9BZL4 | PPP1R12C | 19.115841 | 19.4429889 | 19.3461228 | 19.3486088 | 19.3681643 |
| Q69YH5 | CDCA2 | 19.3078472 | 19.5717806 | 19.5374746 | 19.4821471 | 19.6131282 |
| P15374 | UCHL3 | 21.9089949 | 21.8403709 | 21.8498583 | 21.8150478 | 22.0217998 |
| B2RTY4 | MYO9A | 16.3731361 | 16.4926564 | 16.4656366 | 16.448976 | 16.6694361 |
| Q96PP9 | GBP4 | 18.620271 | 18.5964706 | 18.3080671 | 18.5425826 | 18.5332311 |
| Q14055 | COL9A2 | 14.8386231 | 15.113048 | 14.6777516 | 14.9890892 | 14.8468258 |
| P30049 | ATP5F1D | 23.2903982 | 23.7092871 | 23.203602 | 23.4433104 | 23.7010378 |
| Q7Z624 | CAMKMT | 16.9244438 | 17.4802232 | 17.552364 | 17.4233447 | 17.1226536 |
| Q9UFN0 | NIPSNAP3A | 23.3124175 | 23.2665846 | 23.4277552 | 23.239394 | 23.3736391 |
| Q00653 | NFKB2 | 20.3083487 | 20.5297691 | 20.4241376 | 20.3650859 | 20.470115 |
| Q9UKD1 | GMEB2 | 19.6816992 | 19.9175797 | 19.6872659 | 19.7631819 | 19.9195226 |

| | | | | | | |
|---|---|---|---|---|---|---|
| Q86TN4 | TRPT1 | 19.3620699 | 19.4348358 | 19.5388325 | 19.3922276 | 19.6245076 |
| Q6ZWJ1 | STXBP4 | 13.2586762 | NA | NA | 13.7477955 | NA |
| P53611 | RABGGTB | 19.9142675 | 20.0648728 | 20.0974649 | 19.9169825 | 19.894947 |
| Q96GN5 | CDCA7L | 18.2531379 | 18.391882 | 18.5483646 | 18.145936 | 18.2735337 |
| Q9Y2R0 | COA3 | 22.088338 | 22.1027702 | 22.0719391 | 21.9863926 | 21.9814549 |
| P50225 | SULT1A1 | 20.9830497 | 21.1458373 | 21.1381907 | 21.0714295 | 21.2985254 |
| Q86YS6 | RAB43 | 19.8687581 | 19.7400889 | 19.9137688 | 19.6514268 | 19.7881149 |
| Q9NZJ0 | DTL | 17.4378693 | 17.3567475 | 17.5562223 | 17.3974307 | 17.1172207 |
| Q9UKR5 | ERG28 | 20.1888521 | 20.3390832 | 20.4466149 | 20.2045463 | 20.2484124 |
| Q9UM00 | TMCO1 | 23.0864464 | 23.0835051 | 22.8802166 | 23.0175207 | 22.9589923 |
| Q96JN8 | NEURL4 | 17.5702393 | 17.6046169 | 17.639348 | 17.5070923 | 17.6166055 |
| Q92973 | TNPO1 | 22.996785 | 23.0353209 | 23.0405732 | 22.9375505 | 23.0761377 |
| O43815 | STRN | 20.5136361 | 20.578243 | 20.5288266 | 20.5725185 | 20.7132033 |
| Q8IVD9 | NUDCD3 | 20.6439168 | 20.8502604 | 20.6659347 | 20.7043226 | 20.7897219 |
| P0C7T5 | ATXN1L | 16.5276503 | 16.271734 | 16.7429759 | 16.6613289 | 16.7998855 |
| Q5SNV9 | C1orf167 | 22.0323685 | 22.4315667 | 21.7525356 | 22.0193584 | 22.1116082 |
| O00401 | WASL | 15.0394654 | 14.9661359 | 15.0171764 | 15.3038609 | 15.0053508 |
| Q16649 | NFIL3 | 16.7638604 | 17.1353663 | 17.0160082 | 17.1047582 | 17.0463324 |
| P78560 | CRADD | 19.619358 | 19.4283201 | 19.4789462 | 19.6371694 | 19.2949305 |
| O15400 | STX7 | 21.0988976 | 21.2262592 | 21.4274493 | 21.139201 | 21.1675158 |
| Q5T653 | MRPL2 | 21.8243688 | 21.7938912 | 22.0653813 | 21.8068773 | 21.8296647 |
| Q9UJQ1 | LAMP5 | 22.4405818 | 22.2030266 | 22.2001068 | 22.337639 | 22.3774856 |
| Q16739 | UGCG | 21.743462 | 21.5181331 | 21.5255523 | 21.6635811 | 21.4160551 |
| Q9Y3L5 | RAP2C | 20.8462461 | 20.8359868 | 20.7733318 | 20.5939412 | 20.8489866 |
| Q9H0K6 | PUS7L | 18.5324046 | 18.5822122 | 18.5615682 | 18.5238385 | 18.472159 |
| O75925 | PIAS1 | 19.3537509 | 19.5492128 | 19.5849783 | 19.437276 | 19.6243808 |
| Q9BSC4 | NOL10 | 21.25347 | 21.3166223 | 21.4261662 | 21.199924 | 21.3235012 |
| Q14914 | PTGR1 | 22.3797244 | 22.336573 | 22.5667656 | 22.3070901 | 22.4599389 |
| P40692 | MLH1 | 21.0167913 | 21.1628029 | 21.1940815 | 20.9704096 | 21.0725676 |
| Q13895 | BYSL | 21.8698843 | 21.9924663 | 21.8387984 | 21.7712697 | 21.9399069 |
| O43865 | AHCYL1 | 24.4609182 | 24.4714288 | 24.5038718 | 24.3745071 | 24.393431 |
| P50990 | CCT8 | 26.1920398 | 26.292864 | 26.2460443 | 26.1286632 | 26.1876355 |
| P60842 | EIF4A1 | 26.906577 | 26.9435471 | 26.8501088 | 26.7513447 | 26.8197119 |
| O95819 | MAP4K4 | 21.1711923 | 21.0449831 | 21.0894275 | 21.009217 | 21.0868499 |
| Q6IC98 | GRAMD4 | 20.8632636 | 20.862382 | 20.8380788 | 20.8670629 | 20.7475119 |
| Q15738 | NSDHL | 22.3161348 | 22.3341228 | 22.3023147 | 22.1865347 | 22.3337186 |
| Q9NNW5 | WDR6 | 20.411592 | 20.5671721 | 20.4701105 | 20.4624296 | 20.5524179 |
| Q9H9J2 | MRPL44 | 22.4275905 | 22.3659961 | 22.4645433 | 22.3939814 | 22.4315654 |
| Q6NUQ4 | TMEM214 | 20.8137719 | 20.8605217 | 20.921019 | 20.846783 | 20.8703009 |
| Q8NE01 | CNNM3 | 19.7447141 | 19.7805494 | 19.751952 | 19.7599542 | 19.7494994 |
| Q8N9T8 | KRI1 | 21.667873 | 21.7189402 | 21.6703046 | 21.6339748 | 21.6876491 |
| Q9ULU4 | ZMYND8 | 20.0275056 | 20.0570589 | 20.0453797 | 20.0327113 | 20.1482737 |

| | | | | | | |
|---|---|---|---|---|---|---|
| Q9NSV4 | DIAPH3 | 18.2941449 | 18.4092861 | 18.405721 | 18.3354435 | 18.446759 |
| Q9UBC2 | EPS15L1 | 21.4449785 | 21.5876767 | 21.4265122 | 21.4526486 | 21.6183941 |
| Q7LBC6 | KDM3B | 21.549656 | 21.603674 | 21.603963 | 21.5197341 | 21.659497 |
| Q8N6N3 | C1orf52 | 19.3446007 | 19.327612 | 19.2355084 | 19.2688928 | 19.2594506 |
| Q99871 | HAUS7 | 18.622684 | 18.7993758 | 18.7321112 | 18.779839 | 18.7864813 |
| Q8WW22 | DNAJA4 | 19.4943648 | 19.5608357 | 19.6732958 | 19.6007275 | 19.7111685 |
| O95376 | ARIH2 | 18.923425 | 19.0441898 | 19.0093623 | 18.9765418 | 19.0253181 |
| P61244 | MAX | 20.6864763 | 20.7698232 | 20.6102415 | 20.6083381 | 20.7716823 |
| O60674 | JAK2 | 17.6350504 | 17.8122572 | 18.007911 | 17.9225331 | 17.9466564 |
| Q04726 | TLE3 | 21.0032037 | 21.1662936 | 21.1860291 | 21.1338605 | 21.2906391 |
| Q8WUH1 | CHURC1 | 19.7401136 | 19.8117657 | 19.8666857 | 19.8454148 | 19.4767463 |
| P00441 | SOD1 | 26.1723218 | 26.3678344 | 26.059529 | 26.1445211 | 26.2484778 |
| P16949 | STMN1 | 25.4192073 | 25.6003211 | 25.309331 | 25.4285638 | 25.580255 |
| P52737 | ZNF136 | 16.4927614 | 16.5742609 | 16.8106933 | 16.8513013 | 16.8655533 |
| O95429 | BAG4 | 20.5160487 | 20.7680877 | 20.4872984 | 20.5462333 | 20.7373274 |
| P08174 | CD55 | 17.3936696 | 16.977187 | 17.3285041 | 17.3303034 | 17.4503377 |
| Q6PJ61 | FBXO46 | 16.8157028 | NA | NA | 16.8077375 | NA |
| A6NHX0 | CASTOR2 | 15.5648872 | 15.8457976 | 15.9831522 | 15.9079524 | 16.0606562 |
| Q8TB05 | UBALD1 | NA | 16.2560582 | 15.7862548 | NA | 16.2154351 |
| P04899 | GNAI2 | 24.9544648 | 24.9637529 | 24.9675415 | 24.8730865 | 24.9316203 |
| P51606 | RENBP | 18.9137755 | 18.8241546 | 18.8971014 | 18.8832188 | 18.7487093 |
| O75896 | TUSC2 | 16.1175712 | 16.2084579 | 16.6164989 | 16.3278233 | 16.4882643 |
| Q9NZ01 | TECR | 24.8601098 | 24.9646354 | 24.8745555 | 24.8160605 | 24.6960527 |
| Q8WVC6 | DCAKD | 20.5965456 | 20.4388809 | 20.6307846 | 20.6379366 | 20.6485597 |
| Q6PML9 | SLC30A9 | 19.7226378 | 19.8461122 | 19.7900711 | 19.7838695 | 19.8533777 |
| Q8N5I2 | ARRDC1 | 20.5145787 | 20.5254783 | 20.6506405 | 20.6378649 | 20.6934336 |
| P55854 | SUMO3 | 21.9636021 | NA | NA | 22.153145 | NA |
| Q5FBB7 | SGO1 | 12.9961604 | NA | NA | 12.7996278 | NA |
| Q96SL1 | SLC49A4 | 16.6201656 | NA | NA | 16.4820751 | NA |
| Q96KF7 | SMIM8 | 18.9821815 | NA | NA | 19.1486506 | NA |
| Q9H8P0 | SRD5A3 | 20.5071354 | 20.4048515 | 20.8082306 | 20.480921 | 20.6148597 |
| Q15018 | ABRAXAS2 | 21.395652 | 21.513098 | 21.4522225 | 21.3989283 | 21.4464859 |
| Q15020 | SART3 | 21.727838 | 21.7907613 | 21.7852921 | 21.7215037 | 21.7990932 |
| A8MW92 | PHF20L1 | 18.3771909 | 18.5799416 | 18.4975808 | 18.6046078 | 18.49983 |
| Q9UPW6 | SATB2 | 19.3775395 | 19.7023345 | 19.5012766 | 19.5834167 | 19.8066774 |
| Q9BPY3 | FAM118B | 18.138052 | 18.3331185 | 18.4488734 | 18.2236978 | 18.4712024 |
| Q8N302 | AGGF1 | 17.7200607 | 17.8578724 | 17.8158113 | 17.7412292 | 17.9232055 |
| Q8NC44 | RETREG2 | 19.7342953 | 19.6942977 | 19.8627649 | 19.7157433 | 19.5942783 |
| Q9NRG7 | SDR39U1 | 21.3456624 | 21.3225826 | 21.3775652 | 21.1679135 | 21.3434026 |
| Q9NSI2 | SLX9 | 20.8475858 | 20.9124298 | 21.0378945 | 20.8755789 | 20.9932429 |
| Q9NS69 | TOMM22 | 22.8905237 | 23.0871417 | 22.9298164 | 22.8445395 | 22.9462396 |
| Q9NVH1 | DNAJC11 | 21.0366687 | 21.0791579 | 21.0431228 | 21.0095181 | 21.0985661 |

| | | | | | | |
|---|---|---|---|---|---|---|
| Q14966 | ZNF638 | 21.2065856 | 21.2506839 | 21.2274961 | 21.1781202 | 21.2688725 |
| P63098 | PPP3R1 | 21.2919969 | 21.3464583 | 21.2848383 | 21.2812431 | 21.4291734 |
| P25398 | RPS12 | 26.0229214 | 26.075208 | 25.8937925 | 25.9653074 | 26.0777736 |
| Q96B97 | SH3KBP1 | 22.2581788 | 22.376786 | 22.2042989 | 22.2674484 | 22.4144146 |
| P30519 | HMOX2 | 21.6453476 | 21.6881929 | 21.7664143 | 21.6375408 | 21.647038 |
| Q8N5M4 | TTC9C | 20.0110802 | 20.1011513 | 19.9839692 | 20.0668205 | 19.8830658 |
| Q8IYQ7 | THNSL1 | 19.308302 | 19.4039391 | 19.3226618 | 19.3842787 | 19.5299088 |
| P00846 | MT-ATP6 | 23.7256388 | 23.5794749 | 23.809575 | 23.5322174 | 23.2918626 |
| P31749 | AKT1 | 21.6537575 | 21.5555618 | 21.6572187 | 21.4787429 | 21.5488106 |
| Q68DI1 | ZNF776 | NA | 14.1439723 | 13.497806 | NA | 13.6589664 |
| Q8NI77 | KIF18A | 14.1895025 | NA | NA | 14.3972393 | NA |
| Q92997 | DVL3 | 15.5555407 | 15.6535769 | 15.9454544 | 15.880067 | 15.7729516 |
| P0DMQ5 | INAFM2 | 15.7691705 | 15.9781649 | 16.3009224 | 16.0170956 | 16.0059333 |
| A1XBS5 | CIBAR1 | 15.4970388 | 15.6358177 | 15.4851827 | 15.8025616 | 15.6063669 |
| Q2TAA5 | ALG11 | 18.7119228 | 18.7901664 | 18.8959591 | 18.9213493 | 18.7734244 |
| Q9Y251 | HPSE | 18.5683678 | 18.7588871 | 18.7425028 | 18.5730986 | 18.6286603 |
| Q96C55 | ZNF524 | 16.4901912 | NA | NA | 16.7921232 | NA |
| Q96HY6 | DDRGK1 | 21.0244482 | 21.2423503 | 21.4645128 | 21.2540374 | 21.1419012 |
| Q86T03 | PIP4P1 | 19.7813593 | 19.7326143 | 19.8287182 | 19.6882824 | 19.7538745 |
| Q8IWT3 | CUL9 | 19.7752379 | 19.7420668 | 19.5286079 | 19.6528106 | 19.559989 |
| P36578 | RPL4 | 26.6194409 | 26.6368818 | 26.9735733 | 26.5542767 | 26.5405801 |
| Q15973 | ZNF124 | 17.8070664 | 17.9247555 | 17.83495 | 17.8565376 | 17.7824314 |
| Q92664 | GTF3A | 18.0046886 | 18.3518994 | 18.3922185 | 18.1467237 | 18.1542605 |
| Q9GZP9 | DERL2 | 19.7996299 | 19.9067291 | 20.0517677 | 19.7716407 | 19.8926554 |
| Q96AA3 | RFT1 | 20.6595016 | 20.7427038 | 20.7958214 | 20.6068791 | 20.5820878 |
| Q6PJ69 | TRIM65 | 20.8395353 | 20.8902035 | 20.8259569 | 20.7903644 | 20.84151 |
| Q9Y6X3 | MAU2 | 19.9832151 | 20.0808641 | 20.0790783 | 19.8989119 | 20.0703885 |
| Q13363 | CTBP1 | 22.4204748 | 22.5222185 | 22.4395031 | 22.3018254 | 22.4534063 |
| O75396 | SEC22B | 23.0543265 | 23.1351995 | 23.1696275 | 23.0049516 | 23.1135287 |
| O75190 | DNAJB6 | 21.0400447 | 21.1554376 | 21.0532915 | 20.9867773 | 20.9901441 |
| Q14667 | BLTP2 | 19.0934605 | 19.0644172 | 19.1691793 | 19.0247201 | 19.2238498 |
| Q96DI7 | SNRNP40 | 22.714625 | 22.7697116 | 22.7362213 | 22.6711073 | 22.6971719 |
| Q9UI26 | IPO11 | 21.3409009 | 21.2817691 | 21.3124512 | 21.2563749 | 21.2959445 |
| O00541 | PES1 | 23.9341909 | 23.9680266 | 23.9793999 | 23.8744095 | 23.9255646 |
| Q9UGP8 | SEC63 | 23.1001402 | 23.1087708 | 23.1310812 | 23.0632951 | 23.0953356 |
| O60749 | SNX2 | 22.6854444 | 22.7877843 | 22.7770894 | 22.6312371 | 22.770208 |
| Q8WVT3 | TRAPPC12 | 20.7671823 | 20.7274079 | 20.7864509 | 20.6907695 | 20.7758266 |
| O00754 | MAN2B1 | 25.0689554 | 25.0970148 | 25.0372529 | 24.9800136 | 25.0448068 |
| Q9HAD4 | WDR41 | 19.9310705 | 19.9010102 | 20.0232912 | 19.9382844 | 19.858212 |
| P49327 | FASN | 25.5394097 | 25.4823507 | 25.4990882 | 25.4342098 | 25.4736938 |
| Q92665 | MRPS31 | 21.7123864 | 21.7710309 | 21.7364222 | 21.7711705 | 21.7683374 |
| Q86Y37 | CACUL1 | 18.6323368 | 18.6482131 | 18.6773896 | 18.5702214 | 18.6545596 |

| | | | | | | |
|---|---|---|---|---|---|---|
| P50579 | METAP2 | 23.0104746 | 23.0763234 | 23.0101859 | 23.0187692 | 23.1326174 |
| Q9NUL3 | STAU2 | 21.2560815 | 21.3762741 | 21.348154 | 21.2958575 | 21.436986 |
| Q8N1G2 | CMTR1 | 19.8582549 | 20.0092859 | 19.8869701 | 19.8852591 | 19.9128151 |
| P20839 | IMPDH1 | 22.0893616 | 22.115894 | 22.1692672 | 22.0476986 | 22.3075795 |
| Q96Q11 | TRNT1 | 21.6553615 | 21.7313156 | 21.7396359 | 21.5670257 | 21.8152889 |
| Q96EV2 | RBM33 | 20.1949102 | 20.1253704 | 20.2782859 | 20.2607873 | 20.294568 |
| Q16584 | MAP3K11 | 16.8826167 | 16.9363716 | 16.9712465 | 16.9776731 | 16.9947601 |
| Q8NEM7 | SUPT20H | 18.6542159 | 18.7915713 | 18.7997615 | 18.7786044 | 18.7912841 |
| Q9H1I8 | ASCC2 | 20.3499701 | 20.3334262 | 20.5226986 | 20.4033784 | 20.4844805 |
| Q9GZS3 | SKIC8 | 21.7582137 | 21.8534638 | 21.6629192 | 21.7072169 | 21.9021906 |
| Q99611 | SEPHS2 | 20.4346531 | 20.4730285 | 20.4036684 | 20.2851576 | 20.4174652 |
| Q13445 | TMED1 | 20.4753526 | 20.591315 | 20.6449015 | 20.5704554 | 20.4763473 |
| P49790 | NUP153 | 21.1982643 | 21.3045655 | 21.2593944 | 21.2639223 | 21.3861536 |
| Q5RI15 | COX20 | 18.4348226 | 18.6850149 | 18.6168414 | 18.5421667 | 18.5479509 |
| Q9UQR0 | SCML2 | 19.1346284 | 19.1887194 | 19.0817205 | 19.1309315 | 19.220253 |
| Q8IW35 | CEP97 | 18.3301944 | 18.3331962 | 18.3441383 | 18.3111565 | 18.3609375 |
| P50542 | PEX5 | 19.8318943 | 19.959448 | 19.8707555 | 19.8432781 | 20.0357207 |
| Q08752 | PPID | 22.2089982 | 22.3887684 | 22.0991418 | 22.1783035 | 22.3392817 |
| Q6JBY9 | RCSD1 | 22.4340817 | 22.7864205 | 22.4727995 | 22.5534192 | 22.5501993 |
| Q9UF56 | FBXL17 | 18.1760855 | 18.2097803 | 18.2712469 | 18.2971691 | 18.512678 |
| Q9H1B7 | IRF2BPL | 17.8261086 | 17.761628 | 17.7461423 | 17.909575 | 18.0497935 |
| Q01826 | SATB1 | 22.9821502 | 23.209412 | 23.0996624 | 22.9632271 | 23.1997723 |
| Q9NW68 | BSDC1 | 18.693113 | 18.9129225 | 18.6310374 | 18.8342706 | 18.9516855 |
| Q9P1Z9 | CCDC180 | 19.9597527 | 19.6514183 | 19.8434319 | 19.7252242 | 19.7604304 |
| P85299 | PRR5 | 15.0829764 | 15.4227861 | 15.9945749 | 16.0107822 | 16.0337291 |
| Q9UPM9 | B9D1 | 15.4120642 | 14.7903649 | 15.7376481 | 15.8898945 | 14.7946593 |
| P61073 | CXCR4 | 18.8155349 | 19.0322581 | 18.8830139 | 18.9580007 | 18.7430243 |
| O95619 | YEATS4 | 19.6822014 | 19.6839131 | 19.9835835 | 19.7809407 | 20.1096774 |
| Q9NZE8 | MRPL35 | 21.4105207 | 21.2343375 | 21.5990327 | 21.284431 | 21.2574925 |
| Q14978 | NOLC1 | 23.7582576 | 23.7253792 | 23.8300057 | 23.6578151 | 23.4810099 |
| Q9UNA4 | POLI | 16.5422166 | 16.9036639 | 16.694559 | 16.5648259 | 16.6481028 |
| O94782 | USP1 | 18.5989169 | 18.4317439 | 18.4653716 | 18.5112835 | 18.4168035 |
| Q7Z7K6 | CENPV | 20.2265336 | 20.1617222 | 20.2506008 | 20.1054849 | 20.1162844 |
| Q9HAB8 | PPCS | 21.4989106 | 21.5054735 | 21.5086104 | 21.3351433 | 21.5978821 |
| P51532 | SMARCA4 | 22.8442868 | 22.8876921 | 22.8575143 | 22.8442997 | 22.8340037 |
| Q9P2J5 | LARS1 | 23.448723 | 23.4297135 | 23.4619203 | 23.3623886 | 23.4134285 |
| P07339 | CTSD | 24.9320752 | 24.8815122 | 24.8155961 | 24.90032 | 24.827069 |
| O14974 | PPP1R12A | 21.012343 | 21.2428271 | 21.0560458 | 21.0711221 | 21.2276132 |
| Q9Y2J2 | EPB41L3 | 22.2125213 | 22.3697259 | 22.3207483 | 22.2041451 | 22.4159494 |
| Q9H999 | PANK3 | 17.0725378 | 17.2402326 | 17.3215137 | 17.2702615 | 17.5159377 |
| O75521 | ECI2 | 16.1816921 | NA | NA | 16.1056325 | NA |
| O60683 | PEX10 | 14.7650097 | NA | NA | 14.8513932 | NA |

| | | | | | | |
|---|---|---|---|---|---|---|
| Q14C87 | TMEM132D | NA | 15.8560794 | 15.5181634 | NA | 15.6359947 |
| Q8TCN5 | ZNF507 | 13.0903374 | NA | NA | 13.5222816 | NA |
| P47989 | XDH | 14.573227 | NA | NA | 14.7451136 | NA |
| Q13114 | TRAF3 | 16.4626095 | 16.4326362 | 16.3346513 | 16.2623759 | 16.5377809 |
| P68402 | PAFAH1B2 | 22.0966367 | 21.8923979 | 21.8894089 | 21.6234153 | 21.8725273 |
| Q2PZI1 | DPY19L1 | 17.8994308 | 17.9828957 | 18.2848612 | 17.9577161 | 18.1006236 |
| O95861 | BPNT1 | 21.2054786 | 21.3311008 | 21.3696155 | 21.2278043 | 21.3536597 |
| P61601 | NCALD | 20.7750159 | 20.7986046 | 20.8039471 | 20.6478238 | 20.7669743 |
| Q9UHR6 | ZNHIT2 | 20.4095073 | 20.2931317 | 20.353152 | 20.330819 | 20.3970041 |
| Q9Y4P3 | TBL2 | 21.4639336 | 21.493345 | 21.590941 | 21.4879832 | 21.5374192 |
| P33897 | ABCD1 | 21.0599926 | 21.1206279 | 21.1095584 | 21.0899313 | 21.0580435 |
| Q5T7W0 | ZNF618 | 19.0046529 | 19.1675728 | 19.0543813 | 19.0633388 | 19.1533732 |
| Q8IY92 | SLX4 | 19.18396 | 19.2824071 | 19.2826087 | 19.2160059 | 19.2588547 |
| P49356 | FNTB | 20.6589532 | 20.7639829 | 20.7470528 | 20.7187283 | 20.8695575 |
| Q14680 | MELK | 18.7211576 | 18.6452097 | 18.4801893 | 18.6917906 | 18.6466726 |
| Q9NPE2 | NGRN | 17.3530302 | 17.2169193 | 17.1235128 | 17.2122568 | 17.2580428 |
| P84101 | SERF2 | 23.1016341 | 23.3235488 | 22.9678412 | 23.2479876 | 23.0283146 |
| Q76MJ5 | ERN2 | NA | 13.9868345 | 13.7383872 | NA | 14.6774035 |
| Q9HCN4 | GPN1 | 19.6863964 | 19.6490185 | 19.8331518 | 19.7207201 | 19.7526183 |
| P11279 | LAMP1 | 24.9476145 | 25.0090924 | 24.8916225 | 24.8100324 | 24.89408 |
| P51398 | DAP3 | 21.5691528 | 21.5537696 | 21.5579549 | 21.5602356 | 21.6155797 |
| P49247 | RPIA | 22.1243138 | 22.1766062 | 22.2047955 | 22.1077249 | 22.175349 |
| P31323 | PRKAR2B | 18.9933779 | 19.1860527 | 19.3392445 | 19.0961444 | 19.3969709 |
| Q02833 | RASSF7 | 15.4417888 | 15.4025677 | 15.3137877 | 15.2127583 | 15.3583585 |
| Q86UX6 | STK32C | 19.6828432 | 19.6809487 | 19.7067247 | 19.6431062 | 19.5915655 |
| P39019 | RPS19 | 25.753725 | 25.8207664 | 25.7769047 | 25.6945328 | 25.7322621 |
| Q9H7P9 | PLEKHG2 | 14.3283125 | 14.7077792 | 14.4218329 | 14.2421759 | 14.705531 |
| Q9P032 | NDUFAF4 | 20.7639719 | 20.7805875 | 20.6693462 | 20.6373616 | 20.7236382 |
| Q9Y316 | MEMO1 | 21.7825297 | 21.8508298 | 21.8952042 | 21.7508665 | 21.8717415 |
| Q16186 | ADRM1 | 22.8897971 | 22.6855259 | 22.776233 | 22.6746198 | 22.6426612 |
| Q8NHQ9 | DDX55 | 20.0231017 | 20.1165142 | 20.2160747 | 20.0401559 | 20.1869495 |
| Q8N4A0 | GALNT4 | 15.3488755 | 15.2473064 | 15.6368988 | 15.4533829 | 15.4324748 |
| Q5SZK8 | FREM2 | NA | 21.0728434 | 20.062443 | NA | 20.6731049 |
| Q6PCB0 | VWA1 | NA | 14.0869526 | 13.6485919 | NA | 13.5695148 |
| Q9HC21 | SLC25A19 | 21.7748928 | 21.7604831 | 21.8199264 | 21.6918581 | 21.7618468 |
| P26641 | EEF1G | 25.7202974 | 25.7394622 | 25.671719 | 25.62042 | 25.6663295 |
| Q9UHR5 | SAP30BP | 20.2713301 | 20.3048954 | 20.3568003 | 20.2224613 | 20.3902084 |
| Q9H4P4 | RNF41 | 18.6563013 | 18.5790508 | 18.7740278 | 18.7727595 | 18.78908 |
| Q9P0T7 | TMEM9 | 17.5387585 | 17.8813244 | 17.6415382 | 17.9427308 | 17.8108997 |
| Q96HA1 | POM121 | 16.1421495 | NA | NA | 16.2214915 | NA |
| P32927 | CSF2RB | 13.3713703 | NA | NA | 13.5102334 | NA |
| Q5VWQ0 | RSBN1 | 18.9383094 | 19.1406969 | 19.1056551 | 18.9663316 | 19.0477586 |

| | | | | | | |
|---|---|---|---|---|---|---|
| O15553 | MEFV | 19.0444738 | 19.0338955 | 19.0642065 | 18.9791585 | 19.1218002 |
| Q9BXV9 | GON7 | 20.5230336 | 20.8025626 | 20.4728012 | 20.641914 | 20.8310597 |
| Q10567 | AP1B1 | 24.1537147 | 24.1382252 | 24.2253969 | 24.0515301 | 24.1670715 |
| Q9NZU5 | LMCD1 | 18.6005331 | 18.9308697 | 18.8947136 | 18.7400937 | 18.7457576 |
| O00264 | PGRMC1 | 21.3815756 | 21.3757802 | 21.5308729 | 21.3195517 | 21.1716067 |
| Q8IVQ6 | ZDHHC21 | NA | 16.0541787 | 15.8936892 | NA | 15.7068247 |
| Q8TAU3 | ZNF417 | 20.0696175 | 20.2753178 | 20.2885591 | 20.0295483 | 19.6385027 |
| Q9C0C4 | SEMA4C | 19.32188 | 19.4826783 | 19.5839107 | 19.3658958 | 19.4881841 |
| Q86UY8 | NT5DC3 | 17.8717822 | 18.0455035 | 18.0704138 | 17.9178517 | 18.0248408 |
| Q86X52 | CHSY1 | 19.8057676 | 19.6795916 | 19.7734838 | 19.7210082 | 19.6958771 |
| O00182 | LGALS9 | 24.2569152 | 24.1759364 | 24.2191497 | 24.381743 | 24.2416207 |
| Q9UBF2 | COPG2 | 21.9270331 | 21.9615053 | 22.0278503 | 21.9171515 | 21.9961771 |
| P18583 | SON | 23.7513392 | 23.9325403 | 23.7774751 | 23.7762592 | 23.8545035 |
| Q5H8A4 | PIGG | 19.128563 | 19.1095272 | 19.1349996 | 19.1171731 | 19.1744775 |
| Q01850 | CDR2 | 15.3831803 | 15.6832063 | 15.9065101 | 16.0194311 | 15.8355957 |
| Q9H6K4 | OPA3 | 21.6160533 | 21.6405288 | 21.7716793 | 21.6034189 | 21.5247605 |
| Q14108 | SCARB2 | 19.35038 | 19.4029871 | 19.553519 | 19.382109 | 19.4051846 |
| P00387 | CYB5R3 | 23.7579763 | 23.7560151 | 23.8410064 | 23.7049712 | 23.8218104 |
| P53350 | PLK1 | 20.8694173 | 20.892911 | 20.9012129 | 20.816652 | 20.8917645 |
| Q6P2Q9 | PRPF8 | 23.8271276 | 23.8418683 | 23.8848949 | 23.7608192 | 23.8061364 |
| Q9BXS5 | AP1M1 | 22.4860439 | 22.4610181 | 22.4665529 | 22.3841254 | 22.4556809 |
| Q9HBF4 | ZFYVE1 | 19.2755243 | 19.2809209 | 19.2463424 | 19.209187 | 19.3424092 |
| Q9H8W4 | PLEKHF2 | 19.9714077 | 19.9502207 | 20.0784536 | 19.9565955 | 20.0313605 |
| Q96K21 | ZFYVE19 | 19.0446205 | 19.2523944 | 19.0648552 | 19.2480344 | 19.151912 |
| O43557 | TNFSF14 | 15.823974 | 15.8596836 | 15.9913733 | 15.9338471 | 15.824553 |
| Q5U5Q3 | MEX3C | 17.4598708 | 17.3384137 | 17.1866475 | 16.9739733 | 17.2384522 |
| P0CG13 | CHTF8 | 18.5327055 | 18.4405835 | 18.5443809 | 18.2469369 | 18.6199641 |
| Q9BQC6 | MRPL57 | 20.4115285 | 20.4444472 | 20.5860117 | 20.3265661 | 20.3497554 |
| O15120 | AGPAT2 | 20.0982956 | 20.1871648 | 20.2130591 | 20.1602292 | 20.0942195 |
| Q9BTX1 | NDC1 | 20.2840012 | 20.3278376 | 20.4691575 | 20.2655552 | 20.3248006 |
| A8CG34 | POM121C | 20.1431591 | 20.2206023 | 20.1057631 | 20.22055 | 20.1390692 |
| Q9BSF4 | TIMM29 | 20.7290797 | 20.8485819 | 20.813194 | 20.7900027 | 20.8581055 |
| Q86UE8 | TLK2 | 22.0858033 | 22.0382535 | 21.9311601 | 21.9438836 | 22.0536347 |
| Q6PI78 | TMEM65 | 14.4806061 | NA | NA | 14.820393 | NA |
| P31949 | S100A11 | 26.2234471 | 26.2719678 | 26.2060709 | 26.0981406 | 26.2124279 |
| Q9BYD6 | MRPL1 | 22.4667428 | 22.5011188 | 22.4349478 | 22.3539677 | 22.4286564 |
| Q08623 | PUDP | 19.9325628 | 20.0548664 | 20.0608692 | 19.9782367 | 20.0657081 |
| O94842 | TOX4 | 20.4165731 | 20.485132 | 20.3923771 | 20.3689829 | 20.5071765 |
| Q15424 | SAFB | 23.9727822 | 24.0217998 | 23.9313176 | 23.8760374 | 23.9480428 |
| Q8IYT2 | CMTR2 | 18.1816108 | 18.1071156 | 18.2528405 | 18.2610053 | 18.2144921 |
| Q9HD36 | BCL2L10 | 16.9354758 | NA | NA | 16.8454298 | NA |
| O95070 | YIF1A | 17.6082356 | 17.4265516 | 17.7491303 | 17.3179832 | 17.5048985 |

| | | | | | | |
|---|---|---|---|---|---|---|
| P55884 | EIF3B | 23.902905 | 23.926022 | 23.9235621 | 23.8394769 | 23.8819574 |
| O00327 | BMAL1 | 15.1873223 | 15.916287 | 15.5926298 | 15.0709274 | 15.1063488 |
| Q99735 | MGST2 | 20.8513321 | 20.7214728 | 20.94028 | 20.9221573 | 20.9536706 |
| Q86WQ0 | NR2C2AP | 20.2703167 | 20.2700376 | 20.4245132 | 20.072339 | 20.289248 |
| P61326 | MAGOH | 24.0974619 | 24.249141 | 24.2746189 | 24.0327847 | 24.0499606 |
| Q96FZ7 | CHMP6 | 21.3722528 | 21.5485985 | 21.4192517 | 21.3283401 | 21.3418555 |
| Q9Y248 | GINS2 | 19.9348049 | 20.1845064 | 20.1408391 | 20.04998 | 20.1199194 |
| Q9BYJ9 | YTHDF1 | 20.4039265 | 20.5579296 | 20.4456958 | 20.3706785 | 20.4717115 |
| Q9NPD8 | UBE2T | 20.7265952 | 20.8490993 | 20.8821497 | 20.7850834 | 20.8525483 |
| Q5GLZ8 | HERC4 | 20.8272568 | 20.9780677 | 20.9783179 | 20.8154117 | 20.9716174 |
| Q96HY7 | DHTKD1 | 21.0851168 | 20.9949653 | 21.1099318 | 20.9715071 | 20.9973253 |
| Q9BZQ6 | EDEM3 | 19.8594101 | 19.9551336 | 19.8326225 | 19.8609736 | 19.9061345 |
| Q96EB1 | ELP4 | 20.5752786 | 20.6783623 | 20.6413847 | 20.6996761 | 20.779945 |
| Q9BZE9 | ASPSCR1 | 20.2670134 | 20.4486137 | 20.4151098 | 20.4073564 | 20.4683221 |
| Q96GC5 | MRPL48 | 21.4887151 | 21.6542002 | 21.593848 | 21.4225701 | 21.620917 |
| Q8N4C8 | MINK1 | 18.0355069 | 18.2410425 | 18.1258102 | 18.049095 | 18.2359917 |
| Q9H8W3 | FAM204A | 17.5879318 | 17.8676135 | 17.8692317 | 17.821846 | 17.9686531 |
| P17900 | GM2A | 22.3328856 | 22.5056145 | 22.4261339 | 22.2769396 | 22.4879727 |
| Q8NC96 | NECAP1 | 17.2522389 | 17.2912102 | 17.2659893 | 17.2011522 | 17.3431508 |
| Q15121 | PEA15 | 21.7546207 | 21.9831805 | 21.7528048 | 21.7711535 | 22.0482937 |
| Q7Z7M8 | B3GNT8 | 15.4988299 | 15.3163766 | 15.709538 | 15.8416481 | 15.6983753 |
| Q9Y6D0 | SELENOK | 18.4947487 | 18.7684483 | 18.494338 | 18.7506824 | 18.9953156 |
| P06703 | S100A6 | 24.3572142 | 24.3820997 | 24.1786853 | 24.3323803 | 24.2220935 |
| Q9ULX9 | MAFF | 13.9299264 | NA | NA | 14.072971 | NA |
| Q7Z4W1 | DCXR | 23.4674991 | 23.5363344 | 23.6222873 | 23.4268863 | 23.4166182 |
| O00453 | LST1 | 20.9678957 | 21.1415349 | 21.2802479 | 21.019863 | 21.0357565 |
| O60281 | ZNF292 | 19.4659583 | 19.3570955 | 19.4987209 | 19.3407125 | 19.3926106 |
| P46781 | RPS9 | 27.2210237 | 27.146013 | 27.2542383 | 27.0016997 | 27.0202212 |
| Q9Y5Q8 | GTF3C5 | 20.3089852 | 20.433692 | 20.4432571 | 20.3027978 | 20.4231516 |
| O15270 | SPTLC2 | 21.8759402 | 22.0151104 | 21.9568776 | 21.7483283 | 21.8738099 |
| Q01804 | OTUD4 | 20.0491052 | 20.128508 | 20.0811174 | 20.0383215 | 20.1047003 |
| Q9HCE6 | ARHGEF10L | 17.3848592 | 17.4890995 | 17.5302653 | 17.4782845 | 17.4477726 |
| Q8N2G6 | ZCCHC24 | 18.0315036 | 18.2480002 | 18.3681691 | 18.3480295 | 18.3784172 |
| O00273 | DFFA | 20.9135766 | 21.0643674 | 20.8740029 | 20.9611795 | 21.1538907 |
| Q8TD57 | DNAH3 | 19.238742 | 19.3702263 | 19.4207978 | 19.2977421 | 19.3015479 |
| P12694 | BCKDHA | 21.4129231 | 21.5814618 | 21.5830238 | 21.411663 | 21.5890713 |
| Q13155 | AIMP2 | 23.0476567 | 23.080957 | 23.0720907 | 22.9677577 | 23.0770821 |
| Q92552 | MRPS27 | 21.7236548 | 21.7695565 | 21.7709705 | 21.6649641 | 21.7855853 |
| Q9Y4D8 | HECTD4 | 17.7636798 | 17.7444879 | 17.9024256 | 17.8552761 | 17.8678634 |
| Q96NY9 | MUS81 | 16.8526588 | 16.7836894 | 16.3074999 | 16.4529051 | 16.5560003 |
| Q93015 | NAA80 | 18.8733723 | 18.5883996 | 18.7754569 | 18.7253245 | 18.680764 |
| P60510 | PPP4C | 21.1333007 | 21.2412547 | 21.1567747 | 21.0946556 | 21.1827786 |

| | | | | | | |
|---|---|---|---|---|---|---|
| Q9H0W8 | SMG9 | 20.4371467 | 20.3744092 | 20.5437848 | 20.4398974 | 20.26011 |
| Q9NWV4 | CZIB | 21.9376667 | 22.0681855 | 21.8476638 | 21.8777783 | 21.9976616 |
| P49756 | RBM25 | 22.9812801 | 23.0337031 | 23.0047863 | 22.8323932 | 22.8642732 |
| P56377 | AP1S2 | 21.770478 | 21.8628434 | 21.7000829 | 21.6193887 | 21.6885237 |
| P61927 | RPL37 | 24.5891803 | 24.561158 | 25.0620107 | 24.538069 | 24.4235643 |
| Q14011 | CIRBP | 23.6013272 | 23.754334 | 23.2379298 | 23.4250125 | 23.4708261 |
| Q96DE0 | NUDT16 | 20.141797 | 20.3407711 | 20.3834476 | 20.3048611 | 20.4096839 |
| Q7Z5K2 | WAPL | 21.5041092 | 21.6246887 | 21.5333799 | 21.4801846 | 21.617509 |
| O00159 | MYO1C | 21.2564484 | 21.3090491 | 21.4246224 | 21.2536328 | 21.3094301 |
| O15381 | NVL | 20.3181277 | 20.3277788 | 20.3641232 | 20.3480464 | 20.525431 |
| Q92562 | FIG4 | 18.256705 | 18.057151 | 18.2882552 | 18.2041856 | 18.2718045 |
| Q9BS92 | NIPSNAP3B | 17.9826892 | 17.9942247 | 18.1190584 | 17.9255151 | 18.1604389 |
| P78411 | IRX5 | 15.8612419 | 15.9645817 | 15.9816075 | 15.7740441 | 16.1491612 |
| O95167 | NDUFA3 | 22.7240186 | 22.7460148 | 22.590982 | 22.5973967 | 22.5432622 |
| Q96ME7 | ZNF512 | 20.1970755 | 20.2764779 | 20.4186976 | 20.2135665 | 20.2524655 |
| Q9UBV8 | PEF1 | 21.5039381 | 21.3810265 | 21.5260127 | 21.3927865 | 21.3856124 |
| Q6ICG6 | KIAA0930 | 21.3347159 | 21.3460171 | 21.4821309 | 21.2937783 | 21.397249 |
| Q7RTS9 | DYM | 20.7611632 | 20.7741379 | 20.7923823 | 20.6660569 | 20.6776864 |
| Q15393 | SF3B3 | 24.1446035 | 24.1359253 | 24.1280594 | 24.0160086 | 24.0869772 |
| O60266 | ADCY3 | 18.2512971 | 18.2967307 | 18.3090262 | 18.2899332 | 18.3571778 |
| Q9Y224 | RTRAF | 23.0656605 | 23.092198 | 23.0008003 | 23.0170585 | 23.1615246 |
| Q6UXD5 | SEZ6L2 | 19.706139 | 19.7331151 | 19.6438384 | 19.6737868 | 19.8158765 |
| P67809 | YBX1 | 25.0329179 | 25.3172392 | 25.102875 | 25.1790525 | 25.2111205 |
| Q07021 | C1QBP | 25.1514235 | 25.2888373 | 25.0206208 | 25.0933259 | 25.2977443 |
| Q7L5Y9 | MAEA | 20.8888008 | 20.9400716 | 20.9778533 | 20.8770767 | 20.9199481 |
| Q8IUH3 | RBM45 | 20.2937169 | 20.357192 | 20.4563353 | 20.3461581 | 20.4959186 |
| Q8N567 | ZCCHC9 | 18.8366636 | 19.0239913 | 18.9712493 | 18.9506677 | 18.8380333 |
| P28799 | GRN | 24.565946 | 24.6814663 | 24.5182407 | 24.5085265 | 24.6745888 |
| Q9H098 | FAM107B | 21.1590245 | 21.3468451 | 21.1137854 | 21.2899566 | 21.3762769 |
| Q5VTT5 | MYOM3 | 16.6813555 | NA | NA | 16.5115048 | NA |
| Q9Y2H1 | STK38L | 19.1682088 | 19.2040164 | 19.5992136 | 19.2000524 | 19.278608 |
| P30038 | ALDH4A1 | 22.0657181 | 22.0775865 | 22.0816054 | 22.1049549 | 22.1415052 |
| Q9NSK0 | KLC4 | 19.5071457 | 19.4698354 | 19.5041414 | 19.5771322 | 19.560629 |
| P26440 | IVD | 22.7846406 | 22.8288033 | 22.8066907 | 22.7770283 | 22.8416456 |
| Q8NF99 | ZNF397 | 15.2177334 | 14.6776456 | 15.9133281 | 14.8112685 | 14.8808683 |
| Q8N336 | ELMOD1 | NA | 12.8426006 | 13.3232996 | NA | 12.6649212 |
| P14923 | JUP | 17.4143108 | 16.9499788 | 17.0345484 | 16.8219313 | 16.5756708 |
| P41231 | P2RY2 | 18.3172672 | 17.8661213 | 17.9348 | 17.0320058 | 18.0044809 |
| O00220 | TNFRSF10A | 17.4697789 | 17.1842977 | 17.6727164 | 17.3504904 | 17.2928658 |
| O75880 | SCO1 | 20.5351456 | 20.760297 | 20.8668709 | 20.5215638 | 20.7929997 |
| Q8TE02 | ELP5 | 19.7738361 | 19.9443946 | 20.0412856 | 19.7976506 | 19.7940671 |
| O43633 | CHMP2A | 23.4656628 | 23.5345958 | 23.4820549 | 23.3961944 | 23.3547124 |

| | | | | | | |
|---|---|---|---|---|---|---|
| O00299 | CLIC1 | 26.4139009 | 26.5530216 | 26.4552899 | 26.3168394 | 26.3912884 |
| Q9UJF2 | RASAL2 | 23.3895916 | 23.3984323 | 23.4523968 | 23.373744 | 23.2400233 |
| Q9H2V7 | SPNS1 | 19.6711053 | 19.7753795 | 19.7805766 | 19.6725485 | 19.7505092 |
| P49841 | GSK3B | 21.8360235 | 22.0126463 | 21.9520585 | 21.8386936 | 22.0345547 |
| Q15819 | UBE2V2 | 24.5075767 | 24.4144616 | 24.3304562 | 24.2770263 | 24.3618435 |
| Q9Y639 | NPTN | 21.8874598 | 21.7766344 | 21.7368803 | 21.7812941 | 21.7799845 |
| P61201 | COPS2 | 22.5036957 | 22.5948774 | 22.6150803 | 22.4622742 | 22.5328978 |
| P13639 | EEF2 | 27.8291292 | 27.8700356 | 27.7539844 | 27.6800827 | 27.7632136 |
| Q99575 | POP1 | 21.6683548 | 21.6610266 | 21.7965078 | 21.6340586 | 21.7255232 |
| P27348 | YWHAQ | 24.8826275 | 24.9632678 | 24.8753369 | 24.7553818 | 24.8635109 |
| Q93091 | RNASE6 | 20.4426482 | 20.4170683 | 20.4292092 | 20.4245944 | 20.4251104 |
| Q6ZRP7 | QSOX2 | 20.4084632 | 20.3976235 | 20.4681284 | 20.4961546 | 20.4855805 |
| Q9NPF5 | DMAP1 | 20.3069053 | 20.4195984 | 20.3612484 | 20.390944 | 20.3785822 |
| P36957 | DLST | 23.6660127 | 23.8004951 | 23.7209012 | 23.7069663 | 23.8292791 |
| Q96MF7 | NSMCE2 | 20.1997169 | 20.4030681 | 20.3500147 | 20.3908539 | 20.4318285 |
| O75151 | PHF2 | 19.9880981 | 20.0559563 | 20.1235508 | 20.0861005 | 20.1981676 |
| Q9Y6R4 | MAP3K4 | 20.8403755 | 20.8337628 | 20.93801 | 20.8161614 | 20.9880162 |
| Q9C0E8 | LNPK | 19.1680249 | 19.2784326 | 19.5378075 | 19.4910756 | 19.6742602 |
| Q9Y535 | POLR3H | 17.7571429 | 18.1159432 | 17.8254009 | 17.9362011 | 18.044127 |
| Q9BXJ8 | TMEM120A | 18.6587635 | 18.4652823 | 18.7254002 | 18.7369919 | 18.7138075 |
| Q9Y5Z4 | HEBP2 | 21.8731052 | 21.982825 | 21.7118347 | 21.9360298 | 22.1229979 |
| Q9BTK6 | PAGR1 | 17.7262608 | NA | NA | 18.1109493 | NA |
| Q9NRL3 | STRN4 | 20.1126568 | 20.2728719 | 20.1154035 | 20.2067762 | 20.3365612 |
| Q9Y6Q2 | STON1 | NA | 12.0029092 | 12.1946291 | NA | 12.5278429 |
| Q5T4D3 | TMTC4 | 15.5601062 | NA | NA | 15.4721068 | NA |
| P51149 | RAB7A | 24.5531256 | 24.5512447 | 24.5811904 | 24.4895977 | 24.5755527 |
| Q9H3P2 | NELFA | 20.3618785 | 20.3944934 | 20.4031941 | 20.3515219 | 20.5073531 |
| O95721 | SNAP29 | 21.3749261 | 21.4932202 | 21.456965 | 21.352509 | 21.4775582 |
| P54198 | HIRA | 19.0892313 | 19.1008783 | 19.340441 | 19.1903192 | 19.3148255 |
| Q9NWW5 | CLN6 | 21.2915087 | 21.1911383 | 21.2602262 | 21.2118286 | 21.2004083 |
| P49006 | MARCKSL1 | 22.7090555 | 22.8595725 | 22.8861011 | 22.7511491 | 22.6625116 |
| O95822 | MLYCD | 21.2719015 | 21.3495177 | 21.3767112 | 21.2254164 | 21.3193307 |
| P26572 | MGAT1 | 20.9937004 | 21.0260652 | 21.1174857 | 21.0503817 | 21.0129713 |
| Q9NSC5 | HOMER3 | 22.6241448 | 22.7265779 | 22.6278204 | 22.6116123 | 22.684478 |
| O43447 | PPIH | 26.9087257 | 27.0521143 | 26.8495252 | 26.9121922 | 27.0625562 |
| P62699 | YPEL5 | 19.8976682 | 19.9486819 | 19.8194987 | 19.8676036 | 20.0785018 |
| Q96ES7 | SGF29 | 18.3358362 | 18.5025455 | 18.407225 | 18.5271429 | 18.5489831 |
| Q9BVC4 | MLST8 | 18.947918 | 19.1666991 | 19.1599515 | 19.0654066 | 19.2428314 |
| Q96SQ9 | CYP2S1 | 16.7074745 | 16.7865853 | 17.1606131 | 16.9285241 | 16.9745201 |
| Q9NRY2 | INIP | 19.9505332 | 20.0087104 | 19.8321831 | 19.757563 | 20.0182945 |
| Q6UW63 | POGLUT2 | NA | 14.8015517 | 14.9380288 | NA | 14.8661198 |
| Q15459 | SF3A1 | 23.6403781 | 23.7312508 | 23.5363078 | 23.5635576 | 23.679958 |

| | | | | | | |
|---|---|---|---|---|---|---|
| Q6ZUT9 | DENND5B | 18.8691352 | 19.0502566 | 19.0985542 | 18.8346677 | 18.9603209 |
| P40938 | RFC3 | 21.4042176 | 21.4757666 | 21.5039149 | 21.3232546 | 21.407422 |
| P49589 | CARS1 | 21.497265 | 21.5399929 | 21.5154296 | 21.4965588 | 21.5543012 |
| Q5QJ74 | TBCEL | 18.0014841 | 18.0887919 | 18.0426912 | 18.1355345 | 18.1245841 |
| Q96FZ2 | HMCES | 18.6842331 | 18.9071336 | 18.9209266 | 18.8432739 | 19.0444479 |
| O43182 | ARHGAP6 | 12.4158038 | 13.0723591 | 13.181112 | 13.2502643 | 13.362351 |
| Q05066 | SRY | 17.181081 | NA | NA | 17.262059 | NA |
| Q96AX1 | VPS33A | 21.5354483 | 21.5738835 | 21.5968772 | 21.5295422 | 21.6466133 |
| O95391 | SLU7 | 20.3956741 | 20.4566471 | 20.4423807 | 20.3755633 | 20.4139396 |
| Q8IZ52 | CHPF | 19.3595717 | 19.5744778 | 19.5644718 | 19.5328997 | 19.2808055 |
| Q9NYP9 | MIS18A | 18.1968066 | 18.380846 | 18.2392637 | 18.1731222 | 18.4426722 |
| P49750 | YLPM1 | 23.6574053 | 23.7522387 | 23.6057176 | 23.6316689 | 23.7562572 |
| Q12974 | PTP4A2 | 20.9031652 | 20.7800639 | 20.7007344 | 20.6620324 | 20.7430154 |
| Q9H707 | ZNF552 | 17.4242361 | 17.7845895 | 17.7209132 | 17.4345716 | 17.5280771 |
| P53778 | MAPK12 | 18.7342067 | 18.822091 | 18.9505549 | 18.9375957 | 18.7837233 |
| P22087 | FBL | 24.0358049 | 24.1155599 | 24.1918355 | 24.026751 | 24.1025459 |
| Q9UHX3 | ADGRE2 | 19.9942209 | 20.0889338 | 20.0370935 | 20.0015339 | 20.0987859 |
| O15234 | CASC3 | 20.2528278 | 20.5050859 | 20.2651057 | 20.427912 | 20.2536309 |
| Q5JPH6 | EARS2 | 20.362094 | 20.4492471 | 20.5695337 | 20.4967505 | 20.5243896 |
| Q06413 | MEF2C | 19.0802024 | 19.3907549 | 19.3575754 | 19.2295077 | 19.375243 |
| Q6ZXV5 | TMTC3 | 19.8681295 | 19.6578393 | 19.6896662 | 19.7421984 | 19.9223002 |
| P84022 | SMAD3 | 19.1596169 | 18.8733409 | 18.9663171 | 19.0669511 | 19.3649036 |
| Q15653 | NFKBIB | 19.69732 | 19.8579093 | 19.8319426 | 19.7191684 | 19.8865628 |
| Q9ULA0 | DNPEP | 22.0324873 | 22.1204652 | 22.1217693 | 21.9592411 | 22.0977436 |
| P43405 | SYK | 23.7479407 | 23.7503985 | 23.8250442 | 23.6821382 | 23.6759719 |
| Q9UNX4 | WDR3 | 21.5809064 | 21.5757642 | 21.6402565 | 21.454793 | 21.5863387 |
| O95352 | ATG7 | 22.0896676 | 22.0892542 | 22.1240181 | 22.0461807 | 22.1537906 |
| P61086 | UBE2K | 22.8188559 | 22.7563569 | 22.8704575 | 22.7634851 | 22.8381501 |
| Q9BRP4 | PAAF1 | 20.8083662 | 20.8256538 | 20.844495 | 20.7474756 | 20.9381036 |
| P20339 | RAB5A | 20.141835 | 20.276305 | 20.2801756 | 20.2064481 | 20.3383347 |
| Q9H0U9 | TSPYL1 | 19.2942648 | 19.0585566 | 19.0137952 | 19.2731975 | 19.1787143 |
| Q8NDV7 | TNRC6A | 17.7978934 | 17.8821939 | 17.9910764 | 18.0474069 | 18.1457011 |
| O95232 | LUC7L3 | 22.5693592 | 22.6633554 | 22.6479775 | 22.4934935 | 22.6164773 |
| Q9UIL8 | PHF11 | 13.2097912 | NA | NA | 13.3379416 | NA |
| P52815 | MRPL12 | 23.6644746 | 23.8999647 | 23.5098879 | 23.7454682 | 23.8189093 |
| Q5VT52 | RPRD2 | 20.3024149 | 20.3629069 | 20.3624891 | 20.26845 | 20.4250623 |
| Q58EX7 | PLEKHG4 | 16.3011113 | 16.3025855 | 16.3955359 | 16.441332 | 16.2890097 |
| P42704 | LRPPRC | 24.8906168 | 24.9228907 | 24.8289612 | 24.8086944 | 24.8385818 |
| Q53ET0 | CRTC2 | 18.0357594 | 17.7998246 | 17.899325 | 17.9405407 | 18.0037053 |
| Q9UDX5 | MTFP1 | 19.1075501 | 19.4780896 | 19.4565073 | 19.3007486 | 19.4468985 |
| Q49A26 | GLYR1 | 20.8050334 | 20.8562724 | 20.9455092 | 20.8486287 | 20.9343223 |
| Q5UE93 | PIK3R6 | 15.3112903 | NA | NA | 15.5026761 | NA |

| | | | | | | |
|---|---|---|---|---|---|---|
| Q9NUU7 | DDX19A | 18.8058007 | 18.4307283 | 19.0408756 | 18.5353219 | 18.9732944 |
| Q96HE9 | PRR11 | NA | 16.5462853 | 16.6855195 | NA | 16.4691377 |
| P22794 | EVI2A | 19.0475232 | 19.1154791 | 19.2795961 | 19.1546744 | 19.099117 |
| Q07020 | RPL18 | 26.2127661 | 26.2022065 | 26.4276157 | 26.114486 | 26.1118172 |
| Q9UKZ1 | CNOT11 | 19.538083 | 19.6590522 | 19.7324469 | 19.6094396 | 19.6701183 |
| P17706 | PTPN2 | 20.8538386 | 20.8804514 | 20.9518661 | 20.7925425 | 20.8462344 |
| Q00577 | PURA | 21.6559749 | 21.6399205 | 21.8464139 | 21.7004089 | 21.7531791 |
| Q6ZN55 | ZNF574 | 19.9752237 | 20.2018462 | 20.2134065 | 20.0027637 | 20.1477102 |
| P51798 | CLCN7 | 21.7349253 | 21.7272293 | 21.704686 | 21.6731848 | 21.6258938 |
| O43566 | RGS14 | 19.4607572 | 19.577506 | 19.5582389 | 19.5166982 | 19.6615298 |
| P19838 | NFKB1 | 20.8403373 | 20.8491305 | 20.8555076 | 20.8381981 | 20.9570351 |
| Q8WZ60 | KLHL6 | 15.7807999 | 16.0463875 | 16.1115498 | 16.0417202 | 16.0902839 |
| Q9H7P6 | MVB12B | 16.9371992 | 16.9720707 | 16.7198167 | 17.0742854 | 17.1280611 |
| Q9HC98 | NEK6 | 20.4745118 | 20.6798762 | 20.6298192 | 20.4982159 | 20.7311362 |
| Q8WTV0 | SCARB1 | 21.5534904 | 21.2385963 | 21.312659 | 21.3286514 | 21.3311073 |
| O43847 | NRDC | 22.5792331 | 22.5505952 | 22.5772151 | 22.530409 | 22.55942 |
| Q86VX9 | MON1A | 14.1221374 | NA | NA | 13.9534353 | NA |
| P09958 | FURIN | 16.954544 | 17.2968738 | 17.046099 | 16.9906927 | 16.8158844 |
| P16401 | H1-5 | 27.6162946 | 27.8159858 | 27.9122034 | 27.5272708 | 27.598392 |
| Q8NBX0 | SCCPDH | 20.9154715 | 20.9479086 | 21.1717417 | 20.9129842 | 21.0551924 |
| Q9H3H1 | TRIT1 | 17.8914539 | 18.1989041 | 18.1769853 | 17.9310232 | 18.117361 |
| Q8N668 | COMMD1 | 20.1899061 | 20.1053001 | 19.8315895 | 19.9589307 | 20.0603683 |
| Q9ULV4 | CORO1C | 22.56281 | 22.5050067 | 22.7271146 | 22.5127821 | 22.6188535 |
| Q9H173 | SIL1 | 19.7508032 | 19.7210097 | 19.8559 | 19.6815545 | 19.8052056 |
| Q6ZRV2 | FAM83H | 18.8600261 | 19.0513125 | 19.0630046 | 18.9287708 | 19.1362976 |
| O15427 | SLC16A3 | 23.1097353 | 23.1964207 | 23.1622509 | 23.0440242 | 23.1467971 |
| P14635 | CCNB1 | 20.4065307 | 20.5079545 | 20.6511408 | 20.4759224 | 20.5042534 |
| O60825 | PFKFB2 | 19.5248997 | 19.5623747 | 19.7120192 | 19.5188003 | 19.6961606 |
| O75683 | SURF6 | 20.819404 | 20.8896729 | 21.1370536 | 20.8617558 | 20.9055809 |
| Q96ME1 | FBXL18 | 17.8654011 | 18.0250416 | 17.9400615 | 17.866783 | 17.9928402 |
| O00186 | STXBP3 | 20.9043932 | 20.9198879 | 20.9437676 | 20.8119586 | 20.8750756 |
| P50281 | MMP14 | 23.3145787 | 23.2652477 | 23.2610139 | 23.3266522 | 23.1623515 |
| Q96GD4 | AURKB | 19.2758549 | 19.4104117 | 19.3537878 | 19.3056445 | 19.3205765 |
| Q2VPB7 | AP5B1 | 18.3595982 | 18.3174354 | 18.47526 | 18.3023701 | 18.431738 |
| O43663 | PRC1 | 19.9281633 | 20.0477919 | 20.0600717 | 19.9464437 | 19.9871551 |
| Q9Y237 | PIN4 | 22.1006101 | 22.0653514 | 22.1851305 | 22.0589467 | 22.1493638 |
| Q6JQN1 | ACAD10 | 19.2591951 | 19.3752638 | 19.3265766 | 19.2864745 | 19.2995588 |
| Q6DKK2 | TTC19 | 19.7734204 | 19.7891848 | 19.7990051 | 19.7748278 | 19.8465742 |
| Q9Y3I0 | RTCB | 24.1305997 | 24.1499927 | 24.1288556 | 24.1155284 | 24.2028708 |
| P78362 | SRPK2 | 20.4156329 | 20.4577697 | 20.51175 | 20.5334373 | 20.4978024 |
| P07954 | FH | 24.889799 | 25.0046279 | 24.8616081 | 24.8470878 | 24.9774587 |
| Q9BW61 | DDA1 | 20.8764775 | 21.0279765 | 21.0051554 | 20.9589213 | 21.0231206 |

| | | | | | | |
|---|---|---|---|---|---|---|
| Q96F15 | GIMAP5 | 18.1418475 | 18.2933883 | 18.2080761 | 18.1623098 | 18.2856377 |
| Q8WUH2 | TGFBRAP1 | 19.4243 | 19.4546755 | 19.3743494 | 19.3709528 | 19.5351809 |
| Q9Y3C4 | TPRKB | 20.1978213 | 20.2792647 | 20.3959725 | 20.2836283 | 20.4289651 |
| Q96MH6 | TMEM68 | 19.0565974 | 19.1502002 | 19.2589495 | 19.0491529 | 19.1459825 |
| Q92993 | KAT5 | 19.396177 | 19.4196904 | 19.3003888 | 19.4208294 | 19.4438911 |
| P42898 | MTHFR | 20.2428699 | 20.3458263 | 20.3893783 | 20.3988145 | 20.331837 |
| Q9BW91 | NUDT9 | 20.4453427 | 20.2049847 | 20.2079197 | 20.2777119 | 20.4363317 |
| P62942 | FKBP1A | 24.012264 | 24.0209009 | 23.7114061 | 23.932719 | 24.0484311 |
| Q7Z6M4 | MTERF4 | 18.6326959 | 18.696885 | 18.8512545 | 18.6893522 | 18.6809441 |
| O60262 | GNG7 | 18.4079835 | 18.5042283 | 18.356747 | 18.5926776 | 18.6182751 |
| Q9P0R6 | GSKIP | 18.2097674 | 18.3849171 | 18.4024225 | 18.4386117 | 18.5949706 |
| Q7Z4H7 | HAUS6 | 19.7211066 | 19.9214571 | 19.8405828 | 19.8398733 | 19.8298559 |
| Q9NQ88 | TIGAR | 21.2280352 | 21.2612295 | 21.3980034 | 21.2225609 | 21.2803015 |
| P13473 | LAMP2 | 24.6013631 | 24.7268054 | 24.5399439 | 24.3717893 | 24.5481848 |
| Q6P2D0 | ZFP1 | 16.7543735 | 16.5352036 | 16.4269736 | 16.8522688 | 16.7241397 |
| Q9UBB4 | ATXN10 | 22.6256898 | 22.6553168 | 22.6235095 | 22.5078963 | 22.6502919 |
| Q15785 | TOMM34 | 21.5640651 | 21.570467 | 21.6740106 | 21.4811645 | 21.6775397 |
| Q96PU5 | NEDD4L | 15.609212 | 15.8837201 | 15.3981357 | 15.7484313 | 16.0549239 |
| Q16656 | NRF1 | 19.7950969 | 19.9380191 | 19.6767594 | 19.7914198 | 19.7541398 |
| P19256 | CD58 | 19.2896616 | 19.5885521 | 19.6808887 | 19.4211758 | 19.685753 |
| Q9NXK8 | FBXL12 | 19.6362317 | 19.7034188 | 19.8587576 | 19.5964396 | 19.7710027 |
| Q56P03 | EAPP | 20.1677435 | 20.3874752 | 20.073374 | 20.0781496 | 20.1274595 |
| Q9Y5J6 | TIMM10B | 17.7322857 | 17.7108369 | 17.8224092 | 17.6750262 | 17.7964111 |
| Q11206 | ST3GAL4 | 17.9866313 | 17.8141206 | 17.9797615 | 17.9982857 | 17.9084699 |
| Q7L2J0 | MEPCE | 21.5135838 | 21.5040229 | 21.4485239 | 21.4636575 | 21.4911364 |
| P09668 | CTSH | 19.6087684 | 19.6311275 | 19.7727475 | 19.6416152 | 19.9006195 |
| Q9NXS2 | QPCTL | 21.3020697 | 21.3198099 | 21.3456469 | 21.3727179 | 21.4286543 |
| P23468 | PTPRD | NA | 18.3214833 | 18.203884 | NA | 18.4137736 |
| Q8WYA6 | CTNNBL1 | 22.0340773 | 22.1063865 | 22.1673262 | 21.9317857 | 22.0325339 |
| O75845 | SC5D | 19.003335 | 19.0549708 | 19.3800287 | 19.1719233 | 19.3048061 |
| Q9BXX0 | EMILIN2 | 21.4194754 | 21.4433848 | 21.3657167 | 21.4463389 | 21.406856 |
| Q7RTP6 | MICAL3 | 19.6105018 | 19.3713398 | 19.5544578 | 19.5190406 | 19.4423956 |
| Q7Z406 | MYH14 | 19.4642455 | 20.5470239 | 19.4840937 | 19.7872742 | 19.8093011 |
| Q8WVD3 | RNF138 | 19.4347729 | 19.3226548 | 19.5467374 | 19.5402273 | 19.5442127 |
| Q8N554 | ZNF276 | 17.3815293 | 17.4210042 | 17.3832308 | 17.2574935 | 17.2616567 |
| Q96HR8 | NAF1 | 19.4810761 | 19.4701992 | 19.5860864 | 19.2986348 | 19.4642162 |
| Q9BZF2 | OSBPL7 | 17.4906342 | 17.5145422 | 17.5350346 | 17.5327188 | 17.5599704 |
| Q969S3 | ZNF622 | 21.4701531 | 21.5089851 | 21.633925 | 21.4315315 | 21.4836925 |
| Q92759 | GTF2H4 | 20.1225579 | 20.1373987 | 20.1968091 | 20.0497281 | 20.1432584 |
| P10768 | ESD | 23.8621911 | 23.7747895 | 23.7981772 | 23.7753765 | 23.7761252 |
| Q9Y617 | PSAT1 | 23.1269069 | 23.1030106 | 23.109125 | 23.0314039 | 23.1611903 |
| Q9P2I0 | CPSF2 | 21.6725378 | 21.7556408 | 21.6819077 | 21.7437649 | 21.7554364 |

| | | | | | | |
|---|---|---|---|---|---|---|
| Q9NV88 | INTS9 | 20.1846997 | 20.1898854 | 20.34166 | 20.2283068 | 20.3495445 |
| Q05932 | FPGS | 19.3034265 | 19.2854528 | 19.2848899 | 19.2666562 | 19.5166057 |
| Q9H814 | PHAX | 22.7201729 | 22.8997754 | 22.7061088 | 22.7295455 | 22.8531809 |
| A6NDU8 | RIMOC1 | 19.4104052 | 19.5925586 | 19.385044 | 19.3477802 | 19.4786636 |
| Q96JG6 | VPS50 | 19.615071 | 19.5617404 | 19.6551175 | 19.5694652 | 19.6410615 |
| P37837 | TALDO1 | 26.3209028 | 26.3764031 | 26.2911676 | 26.1768758 | 26.2696472 |
| Q5VYK3 | ECPAS | 21.7309469 | 21.7176001 | 21.7722357 | 21.6701036 | 21.7626233 |
| Q96T58 | SPEN | 21.3124074 | 21.35241 | 21.2587995 | 21.261034 | 21.3329651 |
| O00418 | EEF2K | 21.1701494 | 21.3865519 | 21.2585164 | 21.2195903 | 21.3759621 |
| Q9UL03 | INTS6 | 19.7882786 | 19.788017 | 19.9360497 | 19.8509254 | 19.992658 |
| Q9UII4 | HERC5 | 15.1498049 | 15.1675602 | 15.3345186 | 15.4581253 | 15.2914134 |
| Q9Y6K1 | DNMT3A | 17.783944 | 18.1578311 | 17.8786013 | 17.8713751 | 18.1434055 |
| Q8N1S5 | SLC39A11 | 19.0855084 | 19.111405 | 19.2191126 | 19.0906141 | 19.1363905 |
| O75438 | NDUFB1 | 21.9869327 | 21.940568 | 22.0347493 | 21.9105099 | 21.7511996 |
| Q9UFG5 | C19orf25 | 19.7272593 | 20.1648177 | 19.8563196 | 19.8881562 | 20.1648895 |
| Q99583 | MNT | 18.9497647 | 19.4440986 | 19.249395 | 19.191144 | 19.0983727 |
| P07686 | HEXB | 22.9946764 | 23.07496 | 23.0892293 | 22.9217251 | 23.0584859 |
| O75909 | CCNK | 21.20849 | 21.2892802 | 21.2999829 | 21.1644652 | 21.1764135 |
| Q96GS4 | BORCS6 | 19.3358685 | 19.5404918 | 19.4413374 | 19.4180481 | 19.4286484 |
| Q9NQ48 | LZTFL1 | 20.9827726 | 21.0587885 | 20.8889232 | 20.9527202 | 20.9308397 |
| Q6UN15 | FIP1L1 | 21.1703716 | 21.252157 | 21.1056619 | 21.1593364 | 21.2929037 |
| Q9BWF3 | RBM4 | 20.1092935 | 20.3696404 | 20.224169 | 20.1382923 | 20.368004 |
| Q03426 | MVK | 20.5698593 | 20.7032255 | 20.677196 | 20.5951912 | 20.6167356 |
| P11474 | ESRRA | 18.2107937 | 18.2458629 | 18.3846814 | 18.2086608 | 18.2264598 |
| P58317 | ZNF121 | 19.1014416 | 19.124858 | 19.1709563 | 19.0412799 | 18.9192618 |
| Q6PCB7 | SLC27A1 | 21.7728371 | 21.9106736 | 21.8429055 | 21.6602353 | 21.9468393 |
| Q16537 | PPP2R5E | 20.2608171 | 20.3450666 | 20.3088048 | 20.2517099 | 20.3455419 |
| P20336 | RAB3A | 16.7629528 | 17.1282942 | 17.0544867 | 17.1254575 | 17.0115861 |
| Q86W50 | METTL16 | 20.2360936 | 20.2667945 | 20.2982416 | 20.1428401 | 20.2889043 |
| P61081 | UBE2M | 24.2457946 | 24.1805005 | 24.2116807 | 24.1794519 | 24.2298209 |
| P56211 | ARPP19 | 21.7481571 | 22.0135011 | 21.9078284 | 21.757802 | 21.9622909 |
| P25325 | MPST | 22.1339856 | 22.3228049 | 22.1912321 | 22.2168083 | 22.3993592 |
| Q13563 | PKD2 | 19.21907 | 19.1188263 | 19.1577529 | 19.1677219 | 19.2726241 |
| P12081 | HARS1 | 22.8884354 | 22.8460525 | 22.8419766 | 22.7919408 | 22.9084609 |
| P16260 | SLC25A16 | 16.1470226 | 16.2341772 | 16.5897584 | 16.2437622 | 16.2204603 |
| Q16181 | SEPTIN7 | 23.7689823 | 23.7667087 | 23.7217952 | 23.6506574 | 23.6684534 |
| P08240 | SRPRA | 23.0652622 | 23.0840506 | 23.0502517 | 23.0375703 | 23.0085007 |
| P19623 | SRM | 23.6754804 | 23.6747418 | 23.7186065 | 23.6272745 | 23.6783661 |
| Q9P2E9 | RRBP1 | 23.4330304 | 23.4477907 | 23.3062772 | 23.3338542 | 23.3826796 |
| Q460N5 | PARP14 | 19.7368569 | 19.7854759 | 19.8351439 | 19.7954254 | 19.8773297 |
| Q6WKZ4 | RAB11FIP1 | 21.1745522 | 21.2563568 | 21.2146216 | 21.1608182 | 21.2768411 |
| Q8WUQ7 | CACTIN | 19.5200735 | 19.6629365 | 19.5245179 | 19.5318757 | 19.6348213 |

| | | | | | | |
|---|---|---|---|---|---|---|
| Q9Y4X0 | AMMECR1 | 17.7506229 | 17.5850345 | 17.7837362 | 17.683144 | 17.6586423 |
| Q9GZY6 | LAT2 | 20.3757943 | 20.319185 | 20.6124066 | 20.4595009 | 20.6435241 |
| Q9Y6I3 | EPN1 | 21.8347918 | 21.930561 | 21.794585 | 21.7717545 | 22.0385782 |
| Q9H2C2 | ARV1 | 20.7355106 | 20.9644117 | 21.0533247 | 20.8805602 | 20.8320118 |
| Q6P6B7 | ANKRD16 | 20.6893416 | 20.5741482 | 20.6812461 | 20.6760882 | 20.5166256 |
| P46736 | BRCC3 | 20.1762326 | 20.1226864 | 19.8990296 | 20.1423079 | 20.0764315 |
| P55327 | TPD52 | 21.2367088 | 21.4017903 | 21.0510233 | 21.3328783 | 21.4001655 |
| Q99674 | CGREF1 | 20.0573252 | 20.0530088 | 19.828778 | 19.8901827 | 20.2005482 |
| Q9NRP2 | CMC2 | 19.5113575 | 19.7056385 | 19.432149 | 19.6583897 | 19.770306 |
| P49450 | CENPA | NA | 19.3294315 | 19.2884149 | NA | 19.1089601 |
| Q69YN2 | CWF19L1 | 21.4489693 | 21.5359285 | 21.4842588 | 21.4002643 | 21.4882293 |
| Q96FV2 | SCRN2 | 18.908354 | 19.1744068 | 19.0014822 | 18.9739706 | 18.8953125 |
| P07948 | LYN | 23.0397803 | 22.8594753 | 23.0354218 | 22.9522401 | 23.0256004 |
| Q9ULF5 | SLC39A10 | 18.5409902 | 18.6312188 | 18.6739373 | 18.5088929 | 18.6938324 |
| O43660 | PLRG1 | 22.1487748 | 22.1947492 | 22.28351 | 22.0552465 | 22.233661 |
| Q96NT1 | NAP1L5 | NA | 16.2100394 | 16.0818175 | NA | 16.3548887 |
| Q02742 | GCNT1 | 16.4472948 | 16.277176 | 16.485589 | 16.470968 | 16.4590369 |
| Q96PE2 | ARHGEF17 | 14.2365775 | NA | NA | 13.5875267 | NA |
| A6NM28 | ZFP92 | 15.8505285 | 15.8103739 | 16.0081131 | 15.8372225 | 15.7797547 |
| P13598 | ICAM2 | 11.5763563 | NA | NA | 11.6357971 | NA |
| P08574 | CYC1 | 24.7065233 | 24.9037307 | 24.9716748 | 24.6794225 | 24.7866533 |
| O15143 | ARPC1B | 25.3107057 | 25.3831975 | 25.4384289 | 25.2632319 | 25.3665913 |
| O14593 | RFXANK | 18.4104499 | 18.6047293 | 18.5138829 | 18.6366698 | 18.5874052 |
| P05771 | PRKCB | 21.7661096 | 21.7761601 | 21.8615113 | 21.62696 | 21.8443131 |
| O94761 | RECQL4 | 19.3396145 | 19.3878989 | 19.5431148 | 19.330741 | 19.317924 |
| Q7Z7E8 | UBE2Q1 | 20.8190758 | 20.9548999 | 20.8989598 | 20.8781644 | 20.8832399 |
| P24864 | CCNE1 | 18.2387598 | 18.5039485 | 18.2327849 | 18.1918901 | 18.2502997 |
| Q9BUF5 | TUBB6 | 22.5883293 | 22.5795233 | 22.6108848 | 22.518296 | 22.6185425 |
| Q9BQ52 | ELAC2 | 21.6758665 | 21.6930934 | 21.728809 | 21.6619455 | 21.7172669 |
| Q9Y618 | NCOR2 | 19.6533298 | 19.654054 | 19.6505307 | 19.6801933 | 19.6982787 |
| Q5JSZ5 | PRRC2B | 20.3507814 | 20.2822668 | 20.4009306 | 20.3075722 | 20.3718276 |
| P38606 | ATP6V1A | 23.5025991 | 23.5307137 | 23.4503696 | 23.4570316 | 23.5755716 |
| Q86YT6 | MIB1 | 19.1957601 | 19.2350976 | 19.2882381 | 19.2905188 | 19.2674689 |
| Q8IVH4 | MMAA | 19.4400465 | 19.5333976 | 19.4019672 | 19.3490082 | 19.5263584 |
| P45984 | MAPK9 | 20.8690472 | 20.962111 | 21.1065806 | 21.0115995 | 21.1109364 |
| Q8NCD3 | HJURP | 18.5644434 | 18.4350942 | 18.4223623 | 18.5302713 | 18.4969077 |
| O94768 | STK17B | 16.988459 | 17.0688807 | 17.2156801 | 17.1314615 | 17.2758538 |
| Q9Y680 | FKBP7 | 18.2543548 | 18.4030526 | 18.1667533 | 18.3666032 | 18.4910507 |
| O75369 | FLNB | 22.0866786 | 22.2205285 | 22.0454554 | 22.0695548 | 22.3468664 |
| Q9BTF0 | THUMPD2 | 17.46392 | 17.8081554 | 17.6711518 | 17.6270345 | 17.6960648 |
| Q7Z4G4 | TRMT11 | 17.852039 | 18.0461996 | 18.1025105 | 17.8953473 | 17.9464677 |
| Q96A08 | H2BC1 | NA | 16.3890508 | 16.0309034 | NA | 17.1139735 |

| | | | | | | |
|---|---|---|---|---|---|---|
| O43924 | PDE6D | 19.6658633 | 19.7328613 | 19.639346 | 19.7376664 | 19.8604795 |
| Q8WUU4 | ZNF296 | 16.3863501 | 16.324714 | 16.3808126 | 16.0038794 | 16.1270208 |
| P14174 | MIF | 27.0589105 | 27.114711 | 27.0494527 | 26.8686941 | 27.0007609 |
| Q8N584 | TTC39C | 18.7715069 | 18.8527321 | 18.8410091 | 18.7254173 | 18.5969413 |
| P30533 | LRPAP1 | 22.1008815 | 22.1480807 | 21.8945584 | 22.1037808 | 22.1099183 |
| P61163 | ACTR1A | 22.518974 | 22.6021338 | 22.6349681 | 22.4330285 | 22.5049125 |
| Q8NI27 | THOC2 | 22.4081562 | 22.4093449 | 22.4471558 | 22.3206904 | 22.3579364 |
| O75152 | ZC3H11A | 20.6117651 | 20.757449 | 20.752589 | 20.6544775 | 20.7536576 |
| Q09328 | MGAT5 | 18.0404433 | 17.8841553 | 18.1251322 | 17.9216256 | 17.9809428 |
| Q15388 | TOMM20 | 22.8270257 | 23.0022705 | 23.0318448 | 22.9641281 | 22.8220303 |
| Q9H0R4 | HDHD2 | 21.7103234 | 21.8358596 | 21.7495542 | 21.7170991 | 21.8482899 |
| Q9H974 | QTRT2 | 19.8503597 | 19.9788255 | 19.8860422 | 19.8509808 | 19.9951035 |
| Q53TQ3 | INO80D | 17.0375584 | 17.0344097 | 17.069003 | 17.1056237 | 17.0526324 |
| Q7Z340 | ZNF551 | 18.1102897 | 18.0825828 | 18.1557904 | 18.0696731 | 18.1407579 |
| Q9H892 | TTC12 | 18.5137996 | 18.7165193 | 19.3469364 | 18.5620389 | 17.7857147 |
| Q14849 | STARD3 | 19.4245451 | 19.6210656 | 19.6655209 | 19.537227 | 19.4831074 |
| Q8NAG6 | ANKLE1 | 17.1621897 | 17.2473781 | 17.1869688 | 17.0143673 | 17.2719557 |
| P00734 | F2 | 20.1399196 | 20.3446824 | 20.2715278 | 20.012051 | 20.1909655 |
| Q15050 | RRS1 | 22.3174945 | 22.398105 | 22.712995 | 22.4217946 | 22.3788372 |
| O14526 | FCHO1 | 20.3247557 | 20.3611897 | 20.3665861 | 20.191576 | 20.2774231 |
| Q9UL46 | PSME2 | 24.6019751 | 24.6779543 | 24.5492568 | 24.530245 | 24.5102649 |
| Q9UGU5 | HMGXB4 | 18.8271497 | 18.8705909 | 18.8892191 | 18.8292378 | 18.9740518 |
| Q6GYQ0 | RALGAPA1 | 18.1439313 | 18.2506604 | 18.4174951 | 18.2464403 | 18.2291233 |
| Q9NPC7 | MYNN | 13.5608331 | NA | NA | 13.9213972 | NA |
| O15554 | KCNN4 | 19.8836392 | 19.9748441 | 19.8691381 | 19.8582041 | 19.8546604 |
| O94822 | LTN1 | 20.6448562 | 20.6328831 | 20.6961808 | 20.5517316 | 20.6772076 |
| Q8NBI5 | SLC43A3 | 22.2957099 | 22.2981874 | 22.23083 | 22.1807558 | 21.9931112 |
| Q5JU69 | TOR2A | 19.4177134 | 19.4664005 | 19.5614868 | 19.4876941 | 19.3934651 |
| Q96P53 | WDFY2 | 18.8206017 | 18.8444343 | 18.8449204 | 18.8097686 | 18.8450254 |
| Q9UMS0 | NFU1 | 19.8606408 | 20.0136964 | 19.8099003 | 19.8481671 | 20.1224251 |
| O43488 | AKR7A2 | 23.9582031 | 24.1051944 | 24.052884 | 23.873271 | 24.0903547 |
| Q5T1R4 | HIVEP3 | NA | 15.2346207 | 15.1327337 | NA | 14.4914249 |
| Q8WYN0 | ATG4A | 15.5881342 | 15.8866709 | 15.8210154 | 15.5914572 | 15.7690355 |
| Q8IWF6 | DENND6A | 18.458859 | 18.5448312 | 18.5212501 | 18.3969553 | 18.7038206 |
| Q9H061 | TMEM126A | 20.3542298 | 20.4608345 | 20.4180615 | 20.3443813 | 20.4209606 |
| O00592 | PODXL | 19.9720113 | 19.8329006 | 19.9994098 | 19.8593938 | 20.0014925 |
| P37268 | FDFT1 | 23.2770408 | 23.197835 | 23.156978 | 23.2169723 | 23.0489487 |
| Q9NV66 | TYW1 | 20.0456088 | 20.0456183 | 20.1831604 | 19.9949126 | 20.0978301 |
| Q14161 | GIT2 | 21.644306 | 21.6903614 | 21.8176746 | 21.6294 | 21.6712787 |
| Q5SW96 | LDLRAP1 | 18.7251414 | 18.72353 | 18.7725307 | 18.7326025 | 18.8257447 |
| Q9Y5M8 | SRPRB | 22.9672621 | 23.0589158 | 23.0586699 | 22.9747145 | 23.023107 |
| P04632 | CAPNS1 | 24.4383844 | 24.4476233 | 24.4194653 | 24.3114411 | 24.396426 |

| | | | | | | |
|---|---|---|---|---|---|---|
| Q16539 | MAPK14 | 22.2524009 | 22.2099113 | 22.3299369 | 22.1379396 | 22.2089811 |
| Q08211 | DHX9 | 25.345176 | 25.3613231 | 25.3992527 | 25.2528942 | 25.3238038 |
| P07384 | CAPN1 | 23.7091507 | 23.8038492 | 23.7798858 | 23.6777157 | 23.7215863 |
| Q86Y56 | DNAAF5 | 21.3655254 | 21.3388555 | 21.3315386 | 21.274514 | 21.3408908 |
| Q9BRZ2 | TRIM56 | 19.9057134 | 20.0016265 | 19.9453119 | 19.9674634 | 19.9964862 |
| Q9UGM6 | WARS2 | 19.8029606 | 19.855002 | 19.8042492 | 19.7712596 | 19.8176835 |
| Q12972 | PPP1R8 | 21.0258432 | 20.9797981 | 21.0625015 | 21.1428862 | 21.0795312 |
| A6NDG6 | PGP | 22.1424053 | 22.2857758 | 22.2720308 | 22.1657583 | 22.3017054 |
| P29084 | GTF2E2 | 20.6570156 | 20.6668596 | 20.8516456 | 20.7729895 | 20.7561917 |
| P61962 | DCAF7 | 20.4967887 | 20.7003837 | 20.5886707 | 20.5763774 | 20.6755764 |
| Q9UHQ9 | CYB5R1 | 21.1793012 | 21.3199262 | 21.3351984 | 21.2113438 | 21.4012643 |
| Q9NRG4 | SMYD2 | 20.0658586 | 20.1300963 | 20.1080964 | 20.1953533 | 20.1525961 |
| Q2KHT3 | CLEC16A | 21.1157777 | 21.299323 | 21.1820482 | 21.1594417 | 21.1871647 |
| O75529 | TAF5L | 18.8300216 | 18.9022015 | 18.9343969 | 18.9321935 | 19.0337408 |
| Q9BSQ5 | CCM2 | 20.7900839 | 20.8067986 | 20.7015893 | 20.7318514 | 20.9425271 |
| Q96PQ6 | ZNF317 | 19.1098919 | 19.1937893 | 19.2484612 | 19.3744217 | 19.3382533 |
| Q969E2 | SCAMP4 | 20.8035571 | 20.834786 | 20.9338083 | 20.8615795 | 20.7166334 |
| Q684P5 | RAP1GAP2 | 18.664643 | 18.7774561 | 18.8127903 | 18.85309 | 18.8806579 |
| Q05519 | SRSF11 | 21.7464349 | 21.8131276 | 21.624754 | 21.6742778 | 21.8820466 |
| Q643R3 | LPCAT4 | 19.4213695 | 19.3337657 | 19.5167022 | 19.4389296 | 19.4772898 |
| Q8WZA0 | LZIC | 21.8151029 | 22.0781092 | 21.6037178 | 21.7970585 | 21.8756554 |
| Q96PY6 | NEK1 | 16.5488788 | 16.4772057 | 16.6233782 | 16.5849054 | 16.3579625 |
| P98172 | EFNB1 | 18.7688101 | 18.7663549 | 19.0046658 | 19.0322819 | 19.0745967 |
| O14686 | KMT2D | 19.3843273 | 19.5069024 | 19.3607079 | 19.3613903 | 19.4505673 |
| Q96H35 | RBM18 | 17.0516271 | 17.2435993 | 17.1286562 | 17.1496216 | 17.2579762 |
| O75822 | EIF3J | 23.4668864 | 23.5421274 | 23.4146233 | 23.4527571 | 23.5104637 |
| Q96LL9 | DNAJC30 | 17.3696454 | 17.6028956 | 17.9007223 | 17.3562407 | 17.7113161 |
| O15392 | BIRC5 | 18.9098809 | 18.8265726 | 18.8815188 | 18.7704171 | 18.939719 |
| P01116 | KRAS | 21.4804277 | 21.5388231 | 21.5840618 | 21.4136855 | 21.4218969 |
| Q5T0D9 | TPRG1L | 19.2428709 | 19.1812901 | 19.2746823 | 19.3886846 | 19.2722605 |
| Q9Y296 | TRAPPC4 | 20.9642283 | 20.9540509 | 20.8968291 | 20.8304179 | 20.8963692 |
| Q9NXX6 | NSMCE4A | 19.9122258 | 20.00325 | 20.1858286 | 19.8915803 | 20.1598828 |
| Q53S33 | BOLA3 | 20.3769771 | 20.3460696 | 20.0975832 | 20.3351232 | 20.3875414 |
| P25208 | NFYB | 19.3188611 | 19.1066944 | 19.1615109 | 19.1563019 | 19.5185822 |
| P63272 | SUPT4H1 | 18.9676195 | 18.9019173 | 18.7900406 | 18.8418607 | 19.210052 |
| Q9P246 | STIM2 | 15.1870115 | NA | NA | 15.3893447 | NA |
| P00492 | HPRT1 | 23.3625305 | 23.4531184 | 23.5170367 | 23.3028065 | 23.4906795 |
| Q9UJY4 | GGA2 | 21.5700405 | 21.6461561 | 21.5677984 | 21.5010526 | 21.5805855 |
| P41223 | BUD31 | 21.7000506 | 21.8628124 | 21.5829986 | 21.6701431 | 21.7013494 |
| Q9C037 | TRIM4 | 20.2904177 | 20.3079652 | 20.3885612 | 20.3301816 | 20.4064081 |
| O75554 | WBP4 | 18.6205995 | 18.8616078 | 18.7964183 | 18.7416915 | 18.9879979 |
| Q96GJ1 | TRMT2B | 15.8320721 | 16.2037738 | 15.5286826 | 15.7734774 | 15.9657602 |

| | | | | | | |
|---|---|---|---|---|---|---|
| Q5TA45 | INTS11 | 20.2961116 | 20.4173927 | 20.5059512 | 20.3441974 | 20.4694412 |
| Q9Y2G2 | CARD8 | 19.8373732 | 19.9640505 | 19.9859263 | 20.0340135 | 19.9980751 |
| Q5W0Z9 | ZDHHC20 | 23.1259388 | 23.0946323 | 23.068193 | 23.0278939 | 23.1123986 |
| Q9BVC3 | DSCC1 | 19.6435876 | 19.8726596 | 19.7425795 | 19.7315079 | 19.8024322 |
| Q9Y2Y9 | KLF13 | 19.6437202 | 19.5442763 | 19.5562429 | 19.6437593 | 19.6594808 |
| P29597 | TYK2 | 20.6104887 | 20.7201762 | 20.8328048 | 20.6978298 | 20.7763501 |
| Q9C0D3 | ZYG11B | 19.1144598 | 19.2467119 | 19.338352 | 19.200736 | 19.3666465 |
| Q9NQX3 | GPHN | 20.3294557 | 20.5043134 | 20.3296335 | 20.3598751 | 20.5966253 |
| Q9NY65 | TUBA8 | 20.7805545 | 20.9037178 | 20.6682109 | 20.670986 | 20.7294804 |
| Q6AZZ1 | TRIM68 | 16.0890717 | 15.7398738 | 15.7531227 | 15.7745262 | 15.7695177 |
| Q6P5X5 | C22orf39 | 16.3371384 | 15.7663715 | 15.8172777 | 16.3357014 | 15.9927754 |
| Q13505 | MTX1 | 20.3658436 | 20.5271883 | 20.4751566 | 20.4349494 | 20.5522373 |
| Q6PJW8 | CNST | 14.3733784 | 14.2196132 | 14.5836075 | 14.8566463 | 14.9592118 |
| Q86U70 | LDB1 | 20.6504811 | 20.7114101 | 20.6323237 | 20.5714273 | 20.5856063 |
| Q8WWQ0 | PHIP | 20.1801974 | 20.283515 | 20.3310364 | 20.2033289 | 20.262164 |
| Q9UMX0 | UBQLN1 | 22.7758829 | 22.8453591 | 22.7041971 | 22.8657602 | 22.8965532 |
| Q96JL9 | ZNF333 | 11.6306877 | NA | NA | 11.8411836 | NA |
| Q8TAV0 | FAM76A | 15.7263119 | 15.9870973 | 15.8763459 | 15.5756235 | 15.505723 |
| P61968 | LMO4 | 16.7395876 | 16.7412598 | 16.9344271 | 16.8934482 | 17.1620695 |
| Q9H668 | STN1 | 15.8403738 | 15.9659122 | 16.4657279 | 15.9229917 | 16.1265185 |
| Q6ZTW0 | TPGS1 | 18.3179194 | 18.2593672 | 18.3251618 | 18.3814806 | 18.3791271 |
| Q96EK4 | THAP11 | 17.8928993 | 17.9001795 | 17.9386124 | 17.9223082 | 18.0561719 |
| O15431 | SLC31A1 | 18.4191494 | 18.3679837 | 18.5275252 | 18.4847544 | 18.4661057 |
| O15389 | SIGLEC5 | 18.4454284 | 18.7251689 | 18.7244267 | 18.453646 | 18.5699116 |
| Q9BPX7 | C7orf25 | 17.07427 | 17.2108764 | 17.3700306 | 17.0537859 | 17.1286193 |
| Q15544 | TAF11 | 17.5521661 | 17.494725 | 17.6286999 | 17.4469176 | 17.6279425 |
| Q9UHW5 | GPN3 | 18.2427344 | 18.4781131 | 18.5054909 | 18.3566702 | 18.3652849 |
| Q53GL7 | PARP10 | 20.0614281 | 20.3094895 | 20.2283774 | 20.0602812 | 20.3206409 |
| Q9Y289 | SLC5A6 | 18.8819429 | 18.8056574 | 18.9696048 | 18.7786675 | 18.9194441 |
| Q96F07 | CYFIP2 | 19.1443596 | 19.3654297 | 19.4016324 | 19.1858 | 19.3031745 |
| O75818 | RPP40 | 20.9328831 | 20.8452695 | 21.1261694 | 20.9512284 | 21.0090326 |
| O95674 | CDS2 | 20.5292362 | 20.5513998 | 20.7337864 | 20.4127822 | 20.5908282 |
| Q8TAF3 | WDR48 | 20.4251914 | 20.5343752 | 20.4500135 | 20.4649128 | 20.4746239 |
| Q8NCE0 | TSEN2 | 19.1243424 | 19.1957781 | 19.2115406 | 19.0782647 | 19.2044374 |
| Q8TCJ2 | STT3B | 23.1289034 | 23.1233932 | 23.22284 | 23.0639806 | 23.1387906 |
| Q8IYS2 | KIAA2013 | 20.7644565 | 20.68333 | 20.9020914 | 20.7653048 | 20.7423703 |
| Q96T60 | PNKP | 21.4090438 | 21.3643244 | 21.3906665 | 21.2705787 | 21.3397473 |
| O95298 | NDUFC2 | 21.8217398 | 21.783509 | 21.7932497 | 21.7291558 | 21.7015568 |
| Q9H0A0 | NAT10 | 22.6749806 | 22.7170057 | 22.7490973 | 22.6149598 | 22.7172364 |
| P63010 | AP2B1 | 22.6395907 | 22.6007294 | 22.5602984 | 22.514893 | 22.6050455 |
| Q8N6G5 | CSGALNACT2 | 18.2196185 | 18.2876193 | 18.3303901 | 18.2594091 | 18.3660802 |
| O43379 | WDR62 | 18.8877865 | 18.8925627 | 18.8184933 | 18.9441991 | 18.8529756 |

| | | | | | | |
|---|---|---|---|---|---|---|
| O00411 | POLRMT | 20.4223335 | 20.5562999 | 20.5104136 | 20.4270288 | 20.5235615 |
| P22307 | SCP2 | 23.7060072 | 23.8140717 | 23.7457869 | 23.771328 | 23.8139213 |
| P13667 | PDIA4 | 24.5986418 | 24.6532193 | 24.4883183 | 24.531757 | 24.5902983 |
| Q9BTU6 | PI4K2A | 19.9861742 | 20.0880083 | 20.0134579 | 20.0492138 | 20.0437668 |
| Q96PE3 | INPP4A | 19.3159435 | 19.3479352 | 19.3536207 | 19.3453706 | 19.3689867 |
| Q12830 | BPTF | 19.9946918 | 20.0450688 | 20.0658298 | 19.9957279 | 20.0582333 |
| Q9H2D6 | TRIOBP | 19.7583248 | 19.843852 | 19.8128183 | 19.7411823 | 19.8736346 |
| Q9P2X3 | IMPACT | 20.5238878 | 20.4425847 | 20.7166349 | 20.5941726 | 20.599599 |
| P55212 | CASP6 | 20.5947782 | 20.62804 | 20.6757052 | 20.5950604 | 20.6069908 |
| P48643 | CCT5 | 25.2091073 | 25.2868054 | 25.127615 | 25.1061336 | 25.2400247 |
| Q9Y383 | LUC7L2 | 24.2860963 | 24.3514662 | 24.1993279 | 24.2301304 | 24.3342969 |
| Q9UGU0 | TCF20 | 19.0230441 | 19.0489031 | 19.096759 | 19.0715601 | 19.1479983 |
| Q96PU8 | QKI | 21.6673588 | 21.8958558 | 21.7391908 | 21.7345073 | 21.8433529 |
| Q8TCD5 | NT5C | 20.6943386 | 20.6092785 | 20.9193484 | 20.8071984 | 20.7914271 |
| Q96C19 | EFHD2 | 23.5158152 | 23.7288436 | 23.4961062 | 23.5480286 | 23.6724082 |
| Q2TBE0 | CWF19L2 | 21.0869221 | 21.1428777 | 21.0311285 | 21.059431 | 21.1914746 |
| Q9BQP7 | MGME1 | 19.2364926 | 19.5716647 | 19.4305168 | 19.4314235 | 19.3894596 |
| Q3L8U1 | CHD9 | 18.3228229 | 18.5326657 | 18.4749509 | 18.3503445 | 18.368741 |
| P21333 | FLNA | 24.8780125 | 24.8372129 | 24.6390495 | 24.7641627 | 24.8864091 |
| Q8WXA9 | SREK1 | 21.4514259 | 21.4402282 | 21.3315613 | 21.2547392 | 21.4836934 |
| Q8WUI4 | HDAC7 | 18.8374199 | 18.682283 | 18.8357613 | 18.7284682 | 18.8764484 |
| Q9NV35 | NUDT15 | 19.5735344 | 19.7600103 | 19.821887 | 19.8339034 | 19.7904101 |
| P25440 | BRD2 | 20.6822521 | 20.8388748 | 20.8759363 | 20.867021 | 20.8522928 |
| Q7RTN6 | STRADA | 19.2147026 | 19.47136 | 19.6196002 | 19.3837511 | 19.5387503 |
| Q86VQ1 | GLCCI1 | 15.9837622 | 15.8126543 | 15.9438313 | 15.9071688 | 15.9766392 |
| Q9BWT6 | MND1 | 17.2748743 | 17.3418118 | 17.1754167 | 17.3653701 | 17.5234606 |
| P0DI83 | RAB34 | 16.2650889 | 16.4038845 | 16.091692 | 16.1081221 | 16.3072288 |
| P51397 | DAP | 22.9096815 | 22.8901153 | 22.732153 | 22.9329552 | 22.9428498 |
| Q66K89 | E4F1 | 16.9531605 | 16.9552784 | 17.323734 | 16.9581733 | 17.4388552 |
| Q8NCA9 | ZNF784 | 14.7805441 | NA | NA | 15.3307555 | NA |
| Q96QE2 | SLC2A13 | 15.5407458 | 16.0492018 | 16.1042682 | 16.4551056 | 16.3526158 |
| Q7Z736 | PLEKHH3 | 17.7074021 | NA | NA | 17.6170486 | NA |
| Q14587 | ZNF268 | 13.3232432 | NA | NA | 13.2680548 | NA |
| Q9H6A0 | DENND2D | 16.6102755 | 16.4302723 | 16.9489479 | 16.5862964 | 16.5671185 |
| Q9BRB3 | PIGQ | 18.2000901 | 17.7204071 | 18.5256363 | 18.1363394 | 18.0134194 |
| Q8N8Q3 | ENDOV | 15.7543318 | NA | NA | 15.7073126 | NA |
| P0DJ93 | SMIM13 | 17.7558659 | 17.9227107 | 17.5541603 | 17.7454492 | 17.4386972 |
| Q14240 | EIF4A2 | 21.745703 | 21.9412198 | 21.8982868 | 21.7215432 | 21.8880296 |
| Q92854 | SEMA4D | 19.5287337 | 19.6879119 | 19.8424243 | 19.6229748 | 19.6701163 |
| Q14314 | FGL2 | 22.2028217 | 22.4172218 | 22.1799427 | 22.1424378 | 22.2747256 |
| Q9NQV6 | PRDM10 | 18.7421879 | 18.783618 | 18.9699715 | 18.7830051 | 18.7663259 |
| O14874 | BCKDK | 19.7392135 | 19.8569147 | 19.9670603 | 19.6924152 | 19.6933426 |

| | | | | | | |
|---|---|---|---|---|---|---|
| P60602 | ROMO1 | 20.8524534 | 20.9724763 | 21.2039744 | 20.9593309 | 21.0571382 |
| Q9BXK1 | KLF16 | 20.2592791 | 20.3144453 | 20.3124715 | 20.2433824 | 20.3474181 |
| Q9BQA1 | WDR77 | 21.5304931 | 21.5901181 | 21.5424181 | 21.4473049 | 21.6465068 |
| Q15649 | ZNHIT3 | 19.0247261 | 19.0083553 | 19.1838952 | 19.025176 | 19.0711449 |
| O95470 | SGPL1 | 20.1665001 | 20.1896877 | 20.2473789 | 20.1124074 | 20.1731262 |
| O75208 | COQ9 | 21.2156007 | 21.1805836 | 21.1534898 | 21.1725378 | 21.2523507 |
| Q14139 | UBE4A | 20.8252183 | 20.7457602 | 20.8480254 | 20.8104281 | 20.8009785 |
| Q8N4Q0 | PTGR3 | 20.6880196 | 20.8422241 | 20.7395814 | 20.6412361 | 20.7292441 |
| B0I1T2 | MYO1G | 24.8827243 | 24.9338142 | 24.9751097 | 24.8444327 | 24.8831972 |
| O95394 | PGM3 | 20.6717453 | 20.6899879 | 20.7059602 | 20.5929142 | 20.6681015 |
| Q9NRP0 | OSTC | 22.7188818 | 22.9090521 | 22.852344 | 22.7751314 | 22.8013165 |
| Q9NRF8 | CTPS2 | 20.3110175 | 20.3808707 | 20.3584698 | 20.2242938 | 20.4281244 |
| Q9HB90 | RRAGC | 20.3480348 | 20.3401923 | 20.4517455 | 20.3461113 | 20.3714974 |
| Q9UIQ6 | LNPEP | 22.8477891 | 22.725362 | 22.7474393 | 22.6998361 | 22.7288598 |
| P14735 | IDE | 21.1360157 | 21.172906 | 21.1882718 | 21.1264492 | 21.2689029 |
| Q6P2E9 | EDC4 | 22.5416418 | 22.5890715 | 22.5685637 | 22.5209242 | 22.6039418 |
| Q9P2R3 | ANKFY1 | 21.4782085 | 21.4205741 | 21.5711282 | 21.5097386 | 21.5477809 |
| Q13573 | SNW1 | 22.239538 | 22.3793455 | 22.2709026 | 22.2542318 | 22.3259132 |
| Q9UJV9 | DDX41 | 21.1643855 | 21.2842287 | 21.1614411 | 21.1553159 | 21.2566622 |
| P82664 | MRPS10 | 20.9701498 | 21.0524637 | 21.0112224 | 20.9729758 | 20.8848266 |
| Q8IYK4 | COLGALT2 | 17.809262 | 17.8352393 | 17.8885616 | 17.8019808 | 17.9904712 |
| P11498 | PC | 20.3174824 | 20.4009901 | 20.4321807 | 20.4055413 | 20.4574857 |
| Q86V48 | LUZP1 | 19.7693522 | 19.8676845 | 19.8710693 | 19.7816274 | 19.8753072 |
| Q9UPN3 | MACF1 | 21.4274867 | 21.576769 | 21.4914455 | 21.4592814 | 21.58602 |
| Q99933 | BAG1 | 20.6088825 | 20.7943945 | 20.4817244 | 20.6939664 | 20.6829526 |
| Q9H1K1 | ISCU | 20.5059694 | 20.8006909 | 20.4233834 | 20.6358065 | 20.7567148 |
| P52655 | GTF2A1 | 21.469027 | 21.5815717 | 21.2898563 | 21.4860151 | 21.6582849 |
| Q9ULR0 | ISY1 | 21.4791966 | 21.6079909 | 21.2894222 | 21.3294594 | 21.5317966 |
| P49116 | NR2C2 | 18.4971766 | 18.6597679 | 18.2873487 | 18.3787078 | 18.5652055 |
| Q9Y2K6 | USP20 | 17.5824866 | 17.5908674 | 17.6215513 | 17.5962389 | 17.7379337 |
| Q8WVD5 | RNF141 | 19.177455 | 19.036403 | 19.0945644 | 19.3154787 | 19.1687694 |
| A6NJ78 | METTL15 | 17.9851274 | 18.0705952 | 17.7562638 | 18.032415 | 18.2774501 |
| Q9BY67 | CADM1 | 12.7302475 | NA | NA | 12.7340038 | NA |
| Q96QD5 | DEPDC7 | 12.2222761 | 13.5215327 | 13.389251 | 13.8432012 | 13.3159712 |
| Q15042 | RAB3GAP1 | 20.1742408 | 20.2866015 | 20.3373769 | 20.2742033 | 20.3389292 |
| Q9HCK5 | AGO4 | NA | 14.3367369 | 14.4186007 | NA | 14.3718456 |
| Q9H4I3 | TRABD | 21.1293172 | 21.1653615 | 21.3326548 | 21.1556782 | 21.3362669 |
| Q13643 | FHL3 | 19.4395049 | 19.5208487 | 19.5040975 | 19.6603549 | 19.7807106 |
| Q9P2P5 | HECW2 | NA | 15.846675 | 14.92979 | NA | 15.4093981 |
| Q14114 | LRP8 | 16.3764025 | 16.6881733 | 16.6665 | 16.7370886 | 16.5755066 |
| P04818 | TYMS | 21.6906216 | 21.7092171 | 21.7329675 | 21.6317924 | 21.8113259 |
| Q8WVM8 | SCFD1 | 22.9134052 | 22.9914586 | 23.0523722 | 22.9100602 | 22.9666159 |

| | | | | | | |
|---|---|---|---|---|---|---|
| Q9Y5X3 | SNX5 | 22.5435657 | 22.5798767 | 22.5063663 | 22.4553117 | 22.5578634 |
| P25786 | PSMA1 | 25.0375219 | 25.0288496 | 25.0476558 | 24.9926245 | 25.0296605 |
| Q9H116 | GZF1 | 17.6757742 | 17.6718977 | 17.9367776 | 17.7981416 | 17.8115666 |
| Q14BN4 | SLMAP | 18.6346569 | 18.8118628 | 18.6315448 | 18.6918852 | 18.7023184 |
| Q92609 | TBC1D5 | 20.9109481 | 21.0008724 | 21.0058836 | 20.9032602 | 21.1201273 |
| Q8N999 | RLIG1 | 20.8855721 | 20.850612 | 20.7951626 | 20.8365973 | 20.8471595 |
| O95834 | EML2 | 21.0465376 | 21.1286529 | 21.0936065 | 21.0467885 | 21.2663175 |
| Q8NCH0 | CHST14 | 19.6276151 | 19.4712259 | 19.649948 | 19.5482643 | 19.5394657 |
| Q8ND04 | SMG8 | 19.3851748 | 19.5613854 | 19.5657909 | 19.5181066 | 19.5811544 |
| Q13761 | RUNX3 | 20.6575024 | 20.5199619 | 20.6005682 | 20.4400455 | 20.5822263 |
| P53680 | AP2S1 | 23.30001 | 23.392333 | 23.3029943 | 23.2279197 | 23.2205458 |
| P43234 | CTSO | 18.6907338 | 18.8753726 | 18.3113369 | 18.7114402 | 18.8411686 |
| Q8TBQ9 | TMEM167A | 21.3189635 | 20.5383703 | 21.4807302 | 20.8083235 | 21.0944071 |
| O43414 | ERI3 | 19.9202199 | 19.8919335 | 19.8909236 | 19.8531571 | 19.9541944 |
| Q8IXJ6 | SIRT2 | 20.6261703 | 20.7505214 | 20.7273836 | 20.6912859 | 20.7832307 |
| Q00535 | CDK5 | 21.4115877 | 21.483669 | 21.476395 | 21.4340684 | 21.5008815 |
| Q9C035 | TRIM5 | 18.8356357 | 18.8849402 | 18.8512489 | 18.8726667 | 18.8461603 |
| Q9GZM5 | YIPF3 | 20.1223875 | 20.1991794 | 20.2619205 | 20.2203565 | 20.3328042 |
| Q15181 | PPA1 | 24.1700057 | 24.2655524 | 24.0814382 | 24.0845233 | 24.26624 |
| Q13112 | CHAF1B | 20.6986527 | 20.8390798 | 20.7491322 | 20.7749668 | 20.6822856 |
| Q8WVJ2 | NUDCD2 | 20.985894 | 21.0563871 | 20.8791928 | 20.9262715 | 21.0926001 |
| Q14693 | LPIN1 | 15.5042058 | 15.8176199 | 15.7003759 | 15.5660937 | 15.7486889 |
| O60831 | PRAF2 | 17.8635557 | 17.8212121 | 17.9760675 | 17.8947741 | 18.098997 |
| Q9Y255 | PRELID1 | 16.3712641 | 16.7426159 | 16.5969795 | 16.6608026 | 16.5499634 |
| O60292 | SIPA1L3 | 17.2422548 | 17.4140763 | 17.2115503 | 17.230914 | 16.6701351 |
| Q8NDH3 | NPEPL1 | 19.8752313 | 19.8211029 | 20.0847888 | 19.842769 | 19.9837589 |
| P13196 | ALAS1 | 17.9418013 | 17.9982543 | 17.9164089 | 17.7966703 | 18.1167153 |
| Q96CX6 | LRRC58 | 20.4260607 | 20.4603634 | 20.5411334 | 20.5679881 | 20.537077 |
| Q8TCZ2 | CD99L2 | 19.8171029 | 20.1561381 | 19.9660079 | 19.8704385 | 19.9174867 |
| Q6UW68 | TMEM205 | 22.3492878 | 22.4105569 | 22.4838021 | 22.2885048 | 22.452767 |
| P09417 | QDPR | 21.5048542 | 21.536802 | 21.5703513 | 21.4991497 | 21.6543509 |
| Q5SY16 | NOL9 | 20.7952133 | 20.9281528 | 20.9494244 | 20.873057 | 20.9660056 |
| Q9Y484 | WDR45 | 18.8188175 | 18.8670382 | 18.8380592 | 18.8231539 | 19.0032123 |
| Q9NP66 | HMG20A | 19.9217357 | 20.1648771 | 19.7882259 | 19.8890928 | 20.0941284 |
| Q9Y4G2 | PLEKHM1 | 15.2223597 | 14.9734507 | 15.5635504 | 15.0677599 | 15.5567501 |
| Q68CR1 | SEL1L3 | NA | 11.8287742 | 11.8421582 | NA | 11.8810514 |
| O94829 | IPO13 | 21.1013662 | 21.06948 | 21.035467 | 20.860366 | 20.9283166 |
| P54578 | USP14 | 23.0152568 | 23.0114122 | 23.0330866 | 22.9258049 | 22.9612901 |
| Q9H6T3 | RPAP3 | 21.3579355 | 21.5110856 | 21.3452819 | 21.3729743 | 21.4253812 |
| P29083 | GTF2E1 | 20.887681 | 20.9506823 | 20.8914787 | 20.9207022 | 20.964906 |
| Q5VWG9 | TAF3 | 18.0322471 | 18.0917534 | 18.1956089 | 18.097702 | 18.3711154 |
| Q96DP5 | MTFMT | 12.2252106 | 12.5838419 | 12.3109168 | 12.6446071 | 12.6835945 |

| | | | | | | |
|---|---|---|---|---|---|---|
| Q9Y6X8 | ZHX2 | 19.2094181 | 19.2622527 | 19.1280963 | 19.053335 | 19.1065721 |
| Q14657 | LAGE3 | 19.8270091 | 19.7755753 | 19.6243061 | 19.7287688 | 19.5009483 |
| P46776 | RPL27A | 26.2699721 | 26.3370204 | 26.5062702 | 26.2661419 | 26.2964673 |
| Q9H9T3 | ELP3 | 21.4599926 | 21.6494128 | 21.5947705 | 21.426413 | 21.5883974 |
| O95985 | TOP3B | 19.8021047 | 19.8040877 | 19.8921868 | 19.8568456 | 19.7455184 |
| Q9UPR3 | SMG5 | 18.5848402 | 18.6480935 | 18.6478595 | 18.5304163 | 18.4507979 |
| P49590 | HARS2 | 19.6881777 | 19.684508 | 19.7576281 | 19.632108 | 19.7487388 |
| Q9H269 | VPS16 | 21.150691 | 21.1643048 | 21.235407 | 21.1020141 | 21.1377489 |
| P62191 | PSMC1 | 23.1446763 | 23.2699923 | 23.1651003 | 23.1741117 | 23.2308832 |
| Q6PIU2 | NCEH1 | 20.1779964 | 20.1832549 | 20.2804685 | 20.1488684 | 20.2429667 |
| Q8WUZ0 | BCL7C | 19.6647649 | 19.7800273 | 19.6601655 | 19.7735139 | 19.8151256 |
| P49441 | INPP1 | 24.4837981 | 24.6309018 | 24.5730878 | 24.4838029 | 24.5174577 |
| Q99653 | CHP1 | 19.8074664 | 20.0352332 | 20.0000903 | 19.9119748 | 19.8542928 |
| Q96A47 | ISL2 | 16.173413 | 16.4147054 | 16.3875941 | 15.8307001 | 16.3825333 |
| P53985 | SLC16A1 | 21.8992529 | 21.8528891 | 21.9391356 | 21.685168 | 21.8063239 |
| Q8N960 | CEP120 | 17.0079354 | 17.1139924 | 17.175774 | 17.1527684 | 17.2364614 |
| P06744 | GPI | 26.6928324 | 26.7234986 | 26.7226459 | 26.5767921 | 26.7290645 |
| Q13772 | NCOA4 | 19.0600381 | 18.9572218 | 19.0203901 | 19.2210397 | 19.1194774 |
| Q96AP0 | ACD | 18.0518465 | 18.0265196 | 18.1038896 | 17.9756431 | 17.9937915 |
| Q9C0B7 | TANGO6 | 17.370224 | 17.3711676 | 17.4515588 | 17.271071 | 17.4777643 |
| Q8N7R7 | CCNYL1 | 19.4079243 | 19.5722528 | 19.4768382 | 19.4156148 | 19.6128025 |
| P57740 | NUP107 | 21.944804 | 22.0038731 | 21.9429795 | 21.8557123 | 21.954551 |
| P50148 | GNAQ | 21.2986831 | 21.2464974 | 21.2232516 | 21.1971106 | 21.2544936 |
| O15305 | PMM2 | 21.5625855 | 21.5979197 | 21.5372911 | 21.4987057 | 21.6427272 |
| Q6P996 | PDXDC1 | 21.3905028 | 21.4274549 | 21.3811269 | 21.386827 | 21.4508075 |
| O75419 | CDC45 | 19.340711 | 19.3349546 | 19.4473699 | 19.2920798 | 19.3614314 |
| Q8TEQ0 | SNX29 | 17.8242446 | 17.9717971 | 18.1359244 | 18.015972 | 18.1920722 |
| Q8WXX5 | DNAJC9 | 22.7885937 | 23.0276229 | 22.7760376 | 22.8446508 | 22.9070055 |
| Q14686 | NCOA6 | 21.2690805 | 21.5586092 | 21.2627176 | 21.2736206 | 21.4335315 |
| Q13127 | REST | 17.1757589 | 17.5824208 | 17.6778836 | 17.5198378 | 17.5438335 |
| Q75T13 | PGAP1 | 20.3285819 | NA | NA | 20.1554919 | NA |
| P62750 | RPL23A | 26.6269324 | 26.6691658 | 26.8569585 | 26.5645186 | 26.5905212 |
| Q9UI10 | EIF2B4 | 21.3194531 | 21.3707971 | 21.3912066 | 21.2449298 | 21.3228429 |
| Q86SQ6 | ADGRA1 | NA | 16.7515812 | 16.6794537 | NA | 16.6731975 |
| Q13057 | COASY | 22.3121852 | 22.2606786 | 22.2721851 | 22.1755806 | 22.2019224 |
| Q9P2W9 | STX18 | 20.0634489 | 19.9941507 | 20.125515 | 20.0946064 | 20.0779712 |
| Q5BKY9 | FAM133B | 20.1567842 | 20.0018837 | 20.3693631 | 20.2664955 | 20.0826225 |
| Q93050 | ATP6V0A1 | 20.5826754 | 20.4818958 | 20.7069442 | 20.6091763 | 20.7673608 |
| Q9HD26 | GOPC | 20.9485906 | 21.1276126 | 20.9459376 | 20.9974232 | 21.0910411 |
| Q6GQQ9 | OTUD7B | 16.8940864 | 17.0615271 | 17.0071146 | 17.1844755 | 17.1558678 |
| Q8IV48 | ERI1 | 20.5468318 | 20.5747225 | 20.6062908 | 20.5291952 | 20.6653218 |
| P13987 | CD59 | 17.7830869 | 18.1744986 | 17.8662285 | 17.6978758 | 17.9879476 |

| | | | | | | |
|---|---|---|---|---|---|---|
| P49773 | HINT1 | 24.431307 | 24.6393683 | 24.4692614 | 24.4884676 | 24.6027794 |
| Q96Q05 | TRAPPC9 | 20.0804442 | 20.2092305 | 20.1703707 | 20.0790972 | 20.1361674 |
| P23219 | PTGS1 | 17.174359 | NA | NA | 17.2776783 | NA |
| O43310 | CTIF | 17.5617887 | 17.704123 | 17.8531369 | 17.6100057 | 17.5156063 |
| Q8TDX7 | NEK7 | 21.2009818 | 21.3413038 | 21.3368824 | 21.1422841 | 21.3037519 |
| O95782 | AP2A1 | 23.5091935 | 23.539918 | 23.5696704 | 23.4596013 | 23.5134476 |
| Q9NYJ1 | COA4 | 21.4201731 | 21.4069978 | 21.2515711 | 21.1667985 | 21.5829066 |
| O14522 | PTPRT | 19.7569272 | NA | NA | 19.609609 | NA |
| Q5TA50 | CPTP | 16.5668426 | 16.5149338 | 17.1755456 | 16.6308303 | 16.8571609 |
| P17036 | ZNF3 | NA | 16.4553281 | 15.9312574 | NA | 15.9801852 |
| P49716 | CEBPD | 19.3005747 | 19.5775523 | 19.3197517 | 19.4695579 | 19.5857374 |
| Q8TBF4 | ZCRB1 | 18.872624 | 18.8865404 | 18.8856036 | 18.9281144 | 18.9090858 |
| Q8TDN4 | CABLES1 | 17.0269611 | 16.8730398 | 17.028264 | 17.0822525 | 17.1038311 |
| P40425 | PBX2 | 18.8709407 | 18.9385706 | 19.0804821 | 18.8526696 | 18.9768736 |
| Q9H0H0 | INTS2 | 19.5198224 | 19.5431937 | 19.5832424 | 19.3992505 | 19.5985566 |
| Q3MIX3 | ADCK5 | 18.0008118 | 18.075767 | 17.9742481 | 17.9868212 | 18.0142323 |
| P31146 | CORO1A | 26.2518325 | 26.2499291 | 26.2546491 | 26.0916532 | 26.0808012 |
| P48059 | LIMS1 | 21.607497 | 21.6215895 | 21.7575027 | 21.5170414 | 21.6528249 |
| Q96ER3 | SAAL1 | 23.808446 | 23.8257778 | 23.74026 | 23.6183307 | 23.695117 |
| Q9Y6M9 | NDUFB9 | 22.4978893 | 22.3982775 | 22.3733809 | 22.3465906 | 22.4969563 |
| O75937 | DNAJC8 | 23.9784024 | 24.0613982 | 23.901221 | 23.8336465 | 23.9136678 |
| O94901 | SUN1 | 19.4392087 | 19.3928497 | 19.4114358 | 19.3232772 | 19.4380961 |
| Q8NI60 | COQ8A | 20.200271 | 20.2825042 | 20.3844715 | 20.2655291 | 20.3229658 |
| Q8WVY7 | UBLCP1 | 22.1327397 | 22.1211706 | 22.1462392 | 22.0618214 | 22.0990972 |
| Q6P3W7 | SCYL2 | 20.7696549 | 20.8841011 | 20.8686638 | 20.8510833 | 20.9014058 |
| Q9UBP0 | SPAST | 19.8790319 | 19.9986663 | 20.0089961 | 19.9791357 | 19.968636 |
| P00813 | ADA | 22.9609372 | 23.0038957 | 22.8454109 | 22.9537539 | 22.9509598 |
| Q68CZ6 | HAUS3 | 19.3634513 | 19.4958564 | 19.3015698 | 19.3800865 | 19.4544026 |
| Q7Z333 | SETX | 21.6297004 | 21.7886594 | 21.6991232 | 21.6309363 | 21.7329176 |
| O94885 | SASH1 | 19.7407119 | 19.9292289 | 19.7328703 | 19.7521633 | 19.890469 |
| P12268 | IMPDH2 | 24.9975424 | 25.1202666 | 25.0159495 | 24.9850999 | 25.1622581 |
| Q86XP1 | DGKH | 17.2424068 | 17.1831889 | 17.2188704 | 17.2256367 | 17.3084413 |
| Q9NPI6 | DCP1A | 18.9365206 | 19.1374661 | 19.0833012 | 19.0642206 | 19.2683791 |
| Q9P0N9 | TBC1D7 | 16.6961902 | 16.7964437 | 16.625611 | 16.764969 | 16.6434832 |
| P02452 | COL1A1 | 17.9526346 | 17.6741592 | 17.9735805 | 17.8827877 | 17.7634939 |
| Q8N490 | PNKD | 18.5242874 | 18.5782443 | 18.6697108 | 18.4230611 | 18.6986068 |
| Q9Y6M7 | SLC4A7 | 18.1392557 | 18.3934537 | 18.5303269 | 18.332665 | 18.5508356 |
| Q7Z2K8 | GPRIN1 | 20.3554925 | 20.3327145 | 20.2871592 | 20.347313 | 20.1644561 |
| Q5VYV7 | SLX4IP | 17.7723483 | 18.0754706 | 18.0195032 | 18.2381505 | 18.1709099 |
| Q66K64 | DCAF15 | 18.1285294 | 18.0337575 | 18.0947383 | 17.7778265 | 17.8877856 |
| Q6UUV7 | CRTC3 | 17.0258181 | 16.8471821 | 17.0927882 | 16.9343213 | 17.0411036 |
| P57775 | FBXW4 | NA | 14.3639722 | 14.1557478 | NA | 14.0177786 |

| | | | | | | |
|---|---|---|---|---|---|---|
| O14640 | DVL1 | 14.9623278 | NA | NA | 15.3268358 | NA |
| Q96EN9 | REX1BD | 17.7492362 | 18.0268181 | 17.9233075 | 18.0778186 | 17.8971817 |
| P51648 | ALDH3A2 | 21.3276389 | 21.5389645 | 21.5651401 | 21.3329352 | 21.4350853 |
| Q14697 | GANAB | 25.5108881 | 25.5609135 | 25.520769 | 25.4186544 | 25.5229029 |
| Q8TDQ7 | GNPDA2 | 18.746226 | 18.7519445 | 18.674816 | 18.7807317 | 18.7904142 |
| Q9Y5U9 | IER3IP1 | 21.9346579 | 21.9549036 | 22.2311118 | 22.0622051 | 22.2696743 |
| Q6UWB1 | IL27RA | NA | 14.8570954 | 13.9413368 | NA | 13.2622157 |
| Q6ZSZ6 | TSHZ1 | 17.2469087 | 17.2978907 | 17.0213674 | 16.9228543 | 16.9618526 |
| Q9H920 | RNF121 | 16.1755221 | 16.3934098 | 16.3637939 | 16.3761403 | 16.2637819 |
| P35626 | GRK3 | 17.9184908 | 18.2797843 | 18.2857089 | 17.818749 | 17.9787454 |
| Q13588 | GRAP | 19.8279402 | 19.6356469 | 19.7018693 | 19.7862488 | 19.6468951 |
| Q96JK2 | DCAF5 | 18.2093599 | 18.3796332 | 18.348043 | 18.3289046 | 18.3236204 |
| Q92611 | EDEM1 | 19.3081801 | 19.2188571 | 19.2557484 | 19.0667861 | 19.2485793 |
| O95210 | STBD1 | 19.6898321 | 19.7247195 | 19.7346186 | 19.6636025 | 19.669998 |
| P78368 | CSNK1G2 | 20.2089164 | 20.386386 | 20.3058612 | 20.3110579 | 20.2863227 |
| Q8NEY8 | PPHLN1 | 21.8427009 | 21.869466 | 21.7915501 | 21.8022442 | 21.9273752 |
| Q2NL82 | TSR1 | 21.7289068 | 21.8085772 | 21.8680179 | 21.7576289 | 21.8219225 |
| P07437 | TUBB | 27.4706533 | 27.5206145 | 27.454291 | 27.3513314 | 27.5014367 |
| Q86UT6 | NLRX1 | 19.2507074 | 19.3049985 | 19.3467589 | 19.200249 | 19.3623159 |
| Q8NEF9 | SRFBP1 | 19.8534192 | 19.8983162 | 19.9228548 | 19.8452538 | 19.8857214 |
| P42679 | MATK | 20.1368279 | 20.2083298 | 20.1674575 | 20.1431448 | 20.1122308 |
| Q99698 | LYST | 19.8542781 | 19.9448585 | 19.9522093 | 19.8303831 | 19.9878797 |
| Q15382 | RHEB | 21.6359116 | 21.7317741 | 21.708954 | 21.5502404 | 21.6062882 |
| Q9NZN5 | ARHGEF12 | 18.4973393 | 18.5394684 | 18.5365533 | 18.5630827 | 18.6546856 |
| P54132 | BLM | 22.8808714 | 23.0096383 | 22.8985647 | 22.9181 | 22.9616012 |
| P04083 | ANXA1 | 23.0397999 | 23.1244755 | 22.9752487 | 22.9419948 | 23.0665591 |
| P12109 | COL6A1 | 18.5820319 | 18.5909571 | 18.5001655 | 18.5684068 | 18.531727 |
| Q96BY7 | ATG2B | 20.3891766 | 20.3440265 | 20.4119611 | 20.3856066 | 20.4160285 |
| Q9Y5J7 | TIMM9 | 21.4674363 | 21.6470729 | 21.5447652 | 21.4385878 | 21.6250452 |
| Q8WZA9 | IRGQ | 21.6039758 | 21.8239168 | 21.6081193 | 21.6443562 | 21.7280016 |
| Q7Z4L5 | TTC21B | 18.1239814 | 18.1663365 | 18.2306794 | 18.1944816 | 18.2242406 |
| Q5T9S5 | CCDC18 | 17.2756748 | 17.3450833 | 17.4408719 | 17.3336856 | 17.4250274 |
| Q9UBC1 | NFKBIL1 | 17.8389845 | 17.7746656 | 18.1828042 | 18.0012246 | 17.8543041 |
| Q8N149 | LILRA2 | 17.9566364 | 18.2408499 | 18.0743433 | 18.1465918 | 18.058481 |
| Q9Y587 | AP4S1 | 17.783542 | 18.0906267 | 18.0867666 | 17.8564924 | 18.1372214 |
| Q86WZ6 | ZNF227 | NA | 15.2469789 | 15.4048603 | NA | 15.3138131 |
| O43504 | LAMTOR5 | 21.1349918 | 21.3871529 | 21.0101127 | 21.172697 | 21.564198 |
| P57772 | EEFSEC | 21.0273622 | 21.0813515 | 21.1666998 | 20.9997118 | 21.0649227 |
| Q96SD1 | DCLRE1C | 19.2046693 | 19.2791116 | 19.161959 | 19.0857607 | 19.1054763 |
| Q9NPA8 | ENY2 | 22.6216073 | 22.75935 | 22.5176542 | 22.3605194 | 22.5227931 |
| Q9NSP4 | CENPM | 18.7818077 | 18.8242902 | 18.8990771 | 19.1142438 | 18.7086839 |
| P00338 | LDHA | 27.5532107 | 27.6679031 | 27.6096982 | 27.4332753 | 27.5841838 |

| | | | | | | |
|---|---|---|---|---|---|---|
| P62873 | GNB1 | 24.7959685 | 24.8630278 | 24.7667558 | 24.6997748 | 24.8301926 |
| P50748 | KNTC1 | 20.3462029 | 20.4376778 | 20.3947762 | 20.2851227 | 20.3711293 |
| Q8IWR0 | ZC3H7A | 20.0027696 | 20.0045461 | 20.0953626 | 19.9792248 | 20.071775 |
| Q75QN2 | INTS8 | 20.8175412 | 20.8390737 | 20.8597507 | 20.8083232 | 20.8959883 |
| Q86TB9 | PATL1 | 20.3801209 | 20.4399712 | 20.438379 | 20.3474578 | 20.5369786 |
| Q5VT06 | CEP350 | 21.6374736 | 21.7586297 | 21.5528211 | 21.5650316 | 21.6884988 |
| Q9UNI6 | DUSP12 | 22.2818015 | 22.2398464 | 22.4136195 | 22.2195612 | 22.3761582 |
| Q9P2L0 | WDR35 | 18.8677118 | 18.8753524 | 18.707188 | 18.8084672 | 18.9011622 |
| Q9UKI2 | CDC42EP3 | 17.9034131 | 18.2736692 | 17.9579173 | 17.9342293 | 18.1729185 |
| Q9H9L4 | KANSL2 | 18.3470624 | 18.5077909 | 18.4167777 | 18.4007974 | 18.4171102 |
| Q9Y4F5 | CEP170B | 14.9338939 | 14.5480049 | 14.9177354 | 14.9173969 | 14.6498553 |
| O94972 | TRIM37 | 14.9481729 | 14.9216747 | 14.8182371 | 14.9195615 | 14.4495415 |
| Q9NS18 | GLRX2 | 16.6090777 | 16.5323033 | 16.3163638 | 16.2539194 | 16.2261211 |
| Q9UHN1 | POLG2 | 19.7602476 | 19.8950354 | 20.0998745 | 19.8773317 | 20.1259512 |
| O15294 | OGT | 21.1999372 | 21.3254524 | 21.3386198 | 21.1724028 | 21.3492159 |
| Q96A26 | FAM162A | 21.0660175 | 21.1447486 | 21.127609 | 20.9458619 | 21.128504 |
| P15153 | RAC2 | 26.2450754 | 26.2974784 | 26.3295224 | 26.1561553 | 26.2231997 |
| Q5C9Z4 | NOM1 | 20.2612667 | 20.2277177 | 20.3629241 | 20.1932315 | 20.293023 |
| Q96KB5 | PBK | 22.1851046 | 22.2099238 | 22.1415678 | 22.0769673 | 22.1565532 |
| Q9BVK6 | TMED9 | 24.4615939 | 24.5697444 | 24.5054608 | 24.4172072 | 24.4277137 |
| Q9UN86 | G3BP2 | 22.8670423 | 22.8343141 | 22.7843443 | 22.7000451 | 22.789022 |
| P42356 | PI4KA | 20.404529 | 20.386158 | 20.4864186 | 20.3781903 | 20.4405313 |
| Q7Z2T5 | TRMT1L | 20.7842549 | 20.83953 | 20.8909471 | 20.8582578 | 20.8795494 |
| Q8NHG8 | ZNRF2 | 19.1482232 | 19.2781126 | 19.2989677 | 19.0959763 | 19.3899078 |
| Q3V6T2 | CCDC88A | 20.7697093 | 20.9330985 | 20.8929857 | 20.8250151 | 20.947982 |
| O95881 | TXNDC12 | 22.7162085 | 22.7460485 | 22.540496 | 22.6932165 | 22.8177413 |
| P53384 | NUBP1 | 21.1638802 | 21.3350899 | 21.1874649 | 21.0963784 | 21.3382424 |
| Q8IZT6 | ASPM | 19.1103106 | 19.14571 | 19.2748161 | 19.1621414 | 19.1547857 |
| Q96GA7 | SDSL | 18.2140838 | 18.311005 | 18.0856776 | 18.1241309 | 18.3776378 |
| P62256 | UBE2H | 20.0223413 | 20.0106261 | 20.2161656 | 20.1144157 | 20.0496625 |
| Q6GMV2 | SMYD5 | 19.6860404 | 19.8332135 | 19.6704958 | 19.7249646 | 19.8695632 |
| Q9Y6Y0 | IVNS1ABP | 18.7798067 | 18.8846652 | 18.9319405 | 18.8984512 | 18.9859542 |
| Q9NX70 | MED29 | 20.5277614 | 20.5627282 | 20.3297561 | 20.4109849 | 20.6480157 |
| Q9Y3Q8 | TSC22D4 | 20.3806838 | 20.6030178 | 20.2266716 | 20.381684 | 20.5899586 |
| Q8N129 | CNPY4 | 18.9263099 | 19.1137949 | 18.8445597 | 19.0002694 | 19.1733189 |
| P13498 | CYBA | 22.993827 | 23.1486602 | 23.0296147 | 22.8697825 | 22.8997581 |
| O15173 | PGRMC2 | 21.9386044 | 22.0514617 | 22.0385117 | 21.9776758 | 22.0909641 |
| Q9Y4E8 | USP15 | 22.2660978 | 22.3419729 | 22.192446 | 22.191456 | 22.3638218 |
| Q96AE4 | FUBP1 | 23.6419766 | 23.7229666 | 23.5190736 | 23.5952527 | 23.7527885 |
| O00214 | LGALS8 | 20.8507122 | 20.862186 | 21.0565024 | 20.80868 | 20.9324818 |
| O00311 | CDC7 | 17.6295097 | 17.6196463 | 17.7779711 | 17.6530602 | 17.541561 |
| Q9Y244 | POMP | 20.3973095 | 20.5217815 | 20.5844335 | 20.5779481 | 20.6114734 |

| | | | | | | |
|---|---|---|---|---|---|---|
| Q8WW33 | GTSF1 | 19.4739954 | 19.2729181 | 19.1702599 | 19.4435821 | 19.5976412 |
| O00241 | SIRPB1 | 18.9664436 | 18.9482586 | 19.0504971 | 18.8340957 | 18.7836219 |
| Q96K12 | FAR2 | 18.0774684 | 18.037417 | 18.1010666 | 18.1024966 | 18.1743721 |
| P23443 | RPS6KB1 | 19.9526172 | 19.7817037 | 19.94627 | 19.8686475 | 19.9134306 |
| P61586 | RHOA | 25.6911714 | 25.5534586 | 25.6280384 | 25.5631899 | 25.570208 |
| O95870 | ABHD16A | 22.6779985 | 22.6383796 | 22.6954953 | 22.6340355 | 22.573826 |
| Q86VS8 | HOOK3 | 20.5403343 | 20.672092 | 20.5898684 | 20.5946179 | 20.6954431 |
| O00584 | RNASET2 | 23.6798598 | 23.8408316 | 23.6008399 | 23.7184027 | 23.7772024 |
| Q6ZPD9 | DPY19L3 | 19.8608756 | 19.9440075 | 20.0541992 | 19.8918255 | 19.7767471 |
| Q15011 | HERPUD1 | 18.8082699 | 18.7327018 | 18.9643356 | 18.6676718 | 19.0288164 |
| Q13287 | NMI | 20.9979845 | 21.1974126 | 21.0426703 | 21.011949 | 21.1483001 |
| Q8IXT1 | DDIAS | 16.5258112 | 17.5258013 | 16.9931822 | 16.752024 | 17.1530791 |
| P41247 | PNPLA4 | 13.5046476 | 13.4146812 | 13.9860061 | 13.5855937 | 14.0934396 |
| Q5JSJ4 | INTS6L | 16.7012021 | 17.0514119 | 16.8929853 | 16.6859911 | 17.008726 |
| P0C7P0 | CISD3 | 20.4553985 | 20.4344716 | 20.469655 | 20.3286151 | 20.4853172 |
| O95602 | POLR1A | 20.7333875 | 20.7736029 | 20.7608214 | 20.6343845 | 20.7951392 |
| P61088 | UBE2N | 25.2536873 | 25.3220358 | 25.1562507 | 25.1442575 | 25.2414173 |
| P27824 | CANX | 25.8309801 | 25.8715844 | 25.6977881 | 25.765222 | 25.8209021 |
| O00221 | NFKBIE | 19.3859489 | 19.4544287 | 19.2979316 | 19.3477546 | 19.3933189 |
| Q5T9A4 | ATAD3B | 20.6053413 | 20.6674264 | 20.7452993 | 20.6488654 | 20.7353318 |
| O00444 | PLK4 | NA | 13.0226138 | 12.9442395 | NA | 12.7624782 |
| P41226 | UBA7 | 21.1037369 | 21.1257977 | 21.1286918 | 21.0185405 | 21.1771926 |
| P30050 | RPL12 | 26.0289172 | 26.0989814 | 26.0349308 | 26.0101281 | 26.042091 |
| P23588 | EIF4B | 23.9642092 | 24.1372028 | 23.8335056 | 23.9111535 | 23.9695519 |
| Q16667 | CDKN3 | 16.0141091 | 16.3433064 | 16.4788627 | 16.6563444 | 16.5677654 |
| Q01658 | DR1 | 21.6050369 | 21.7953111 | 21.5109128 | 21.4936775 | 21.6972664 |
| Q9NW61 | PLEKHJ1 | NA | 15.8093355 | 16.3037591 | NA | 15.8537945 |
| Q9Y2T3 | GDA | NA | 14.3198889 | 14.1529641 | NA | 13.9111358 |
| Q6PIY7 | TENT2 | 18.1794358 | 18.1285228 | 18.1199122 | 17.9219914 | 18.1846092 |
| Q96DH6 | MSI2 | 19.3640747 | 19.5656953 | 19.3976904 | 19.3858048 | 19.5799258 |
| Q86VX2 | COMMD7 | 20.6474688 | 20.7545195 | 20.7207944 | 20.6161366 | 20.5200239 |
| O75448 | MED24 | 20.6225105 | 20.7187939 | 20.7475219 | 20.6245663 | 20.7162568 |
| Q6PCB5 | RSBN1L | 19.5535434 | 19.750241 | 19.6393509 | 19.6290525 | 19.7334429 |
| Q13243 | SRSF5 | 23.6591358 | 23.7337852 | 23.4049553 | 23.6418587 | 23.6150307 |
| O43422 | THAP12 | 18.0523784 | 18.097244 | 18.2167445 | 18.0427524 | 18.285905 |
| Q5XPI4 | RNF123 | 20.299137 | 20.2790818 | 20.4106238 | 20.2448099 | 20.3924071 |
| P40926 | MDH2 | 26.2942161 | 26.3249962 | 26.3439327 | 26.1616849 | 26.3574202 |
| Q14156 | EFR3A | 21.1697299 | 21.1372963 | 21.2389798 | 21.1750307 | 21.2648338 |
| Q3KR37 | GRAMD1B | 11.4561002 | NA | NA | 11.9101602 | NA |
| Q8IVF2 | AHNAK2 | 14.4945509 | 14.429231 | 14.5504968 | 14.7097568 | 14.4439945 |
| Q15012 | LAPTM4A | 17.0654213 | 16.818642 | 16.8149937 | 16.894693 | 16.7561345 |
| Q5VTL7 | FNDC7 | NA | 19.0527122 | 19.079283 | NA | 18.4576092 |

| | | | | | | |
|---|---|---|---|---|---|---|
| Q9NU23 | LYRM2 | 19.2675892 | 19.2715531 | 19.1205083 | 19.176063 | 19.3995446 |
| P02771 | AFP | NA | 17.8381398 | 18.0537898 | NA | 17.9700544 |
| Q9H6P5 | TASP1 | 19.6321305 | 19.2728843 | 19.2215314 | 19.3581341 | 19.2493089 |
| Q5SXM2 | SNAPC4 | 17.2169229 | 17.2465383 | 17.2947398 | 17.1649883 | 17.3477986 |
| Q9HB19 | PLEKHA2 | 19.4713526 | 19.6576577 | 19.5836669 | 19.3382853 | 19.7366747 |
| Q14119 | VEZF1 | 20.1857421 | 20.3203759 | 20.4053114 | 20.3519817 | 20.221827 |
| Q499Z4 | ZNF672 | 18.1078606 | 18.1522736 | 17.870319 | 18.1532496 | 18.0800432 |
| Q9NUN5 | LMBRD1 | 18.7495802 | 18.7692318 | 18.9429589 | 18.7791418 | 18.6889941 |
| Q02878 | RPL6 | 26.4096153 | 26.3723263 | 26.6007075 | 26.277315 | 26.3834489 |
| Q96BY9 | SARAF | 19.176297 | 18.9864892 | 18.9643552 | 18.864066 | 18.9336161 |
| Q9P031 | CCDC59 | 19.9155477 | 20.0068346 | 20.0486364 | 19.847335 | 20.052677 |
| Q9BRG1 | VPS25 | 21.6797818 | 21.6667004 | 21.7905395 | 21.6785292 | 21.6726275 |
| Q13948 | CUX1 | 18.9052804 | 19.0549952 | 18.8913346 | 18.7925761 | 19.0583702 |
| Q12980 | NPRL3 | 18.5057053 | 18.336453 | 18.4801246 | 18.292743 | 18.5404971 |
| Q9UKF7 | PITPNC1 | 18.9739603 | 18.7821347 | 18.8708701 | 18.8701388 | 18.6984929 |
| P26447 | S100A4 | 26.3709553 | 26.6275985 | 26.483655 | 26.3234157 | 26.3897231 |
| Q9Y3D3 | MRPS16 | 21.515778 | 21.6670208 | 21.7612019 | 21.646696 | 21.61294 |
| Q9BXK5 | BCL2L13 | 21.2292808 | 21.3147067 | 21.2886728 | 21.2366829 | 21.1794885 |
| P28288 | ABCD3 | 21.8283219 | 21.9562096 | 21.8937181 | 21.7073322 | 21.9052209 |
| Q9Y570 | PPME1 | 21.4433921 | 21.5557459 | 21.5249317 | 21.3697739 | 21.4606864 |
| Q9Y2Z9 | COQ6 | 21.9814252 | 21.9367224 | 22.0193757 | 21.8748108 | 21.8081308 |
| O60496 | DOK2 | 21.8526617 | 21.9878168 | 22.0539263 | 21.8128015 | 21.9483002 |
| Q9HB40 | SCPEP1 | 21.5961091 | 21.7325951 | 21.7795071 | 21.6022511 | 21.6909355 |
| Q96EE3 | SEH1L | 21.7747042 | 21.7112799 | 21.9006831 | 21.7117448 | 21.8350146 |
| Q9NVA2 | SEPTIN11 | 21.4705631 | 21.6465733 | 21.5286546 | 21.5023871 | 21.5600464 |
| O75083 | WDR1 | 25.1338052 | 25.24756 | 25.2067357 | 25.1121588 | 25.2662783 |
| P20226 | TBP | 19.9599278 | 19.9240311 | 20.0644622 | 19.8527294 | 19.9067232 |
| Q7L1V2 | MON1B | 19.5932903 | 19.6280476 | 19.712899 | 19.5436299 | 19.623598 |
| Q5VW32 | BROX | 22.4217908 | 22.4545837 | 22.5544196 | 22.3399211 | 22.390862 |
| Q9NVU7 | SDAD1 | 24.0050464 | 24.0273302 | 24.0511178 | 23.9548195 | 24.0438631 |
| P22695 | UQCRC2 | 24.2685593 | 24.3180809 | 24.3300219 | 24.1691802 | 24.2590304 |
| Q9Y5L0 | TNPO3 | 21.4019317 | 21.5186157 | 21.5413708 | 21.3959092 | 21.4643496 |
| P30520 | ADSS2 | 23.7183374 | 23.8130664 | 23.7406862 | 23.6609265 | 23.7332916 |
| O15269 | SPTLC1 | 22.0443055 | 22.0631961 | 22.0434935 | 22.0122143 | 22.045794 |
| Q5ST30 | VARS2 | 18.9655007 | 18.8690039 | 18.9470094 | 18.9713855 | 18.8898831 |
| Q6PJG2 | MIDEAS | 20.0561546 | 20.1459378 | 20.0382554 | 20.0103281 | 20.1074235 |
| Q9BVQ7 | AFG2B | 18.9549129 | 19.1221273 | 19.0518163 | 19.0225636 | 19.1910476 |
| P42694 | HELZ | 19.5330127 | 19.6809367 | 19.6493556 | 19.6207151 | 19.7660289 |
| Q9Y6W3 | CAPN7 | 19.9061721 | 19.8880179 | 19.9885352 | 19.8834553 | 20.0333755 |
| Q86UP2 | KTN1 | 23.3731262 | 23.4692381 | 23.3418886 | 23.2782506 | 23.4323006 |
| Q13614 | MTMR2 | 19.2087022 | 19.2835568 | 19.2853805 | 19.2264063 | 19.3848488 |
| P30101 | PDIA3 | 26.1831138 | 26.2143301 | 26.088692 | 26.1092719 | 26.2229784 |

| | | | | | | |
|---|---|---|---|---|---|---|
| Q9NVX0 | HAUS2 | 19.1613096 | 19.2067948 | 19.3468706 | 19.2289782 | 19.2862945 |
| P57081 | WDR4 | 19.9674568 | 20.1644315 | 20.1488915 | 20.1407278 | 20.1591063 |
| P42566 | EPS15 | 20.319728 | 20.4831441 | 20.4117952 | 20.3548148 | 20.5080696 |
| Q96L58 | B3GALT6 | 21.0764609 | 21.1517665 | 21.1915596 | 21.1880507 | 21.2235363 |
| Q9H773 | DCTPP1 | 22.3800642 | 22.5283713 | 22.3612325 | 22.37134 | 22.533088 |
| Q7Z4V5 | HDGFL2 | 23.2680867 | 23.5448405 | 23.3413134 | 23.334122 | 23.3074437 |
| P51795 | CLCN5 | 17.5425968 | 17.4971068 | 17.817107 | 17.6413777 | 17.7518648 |
| Q07955 | SRSF1 | 25.1422236 | 25.2495184 | 25.0739314 | 25.1295452 | 25.1991523 |
| Q86WN1 | FCHSD1 | 19.2060856 | 19.246841 | 19.2073568 | 19.216316 | 19.4832468 |
| Q9NYL9 | TMOD3 | 21.7950422 | 22.0635114 | 21.8627554 | 21.8677137 | 22.0319739 |
| Q32MZ4 | LRRFIP1 | 22.3253644 | 22.4742441 | 22.1621762 | 22.3231121 | 22.4336802 |
| Q03014 | HHEX | 16.8713384 | 16.8443013 | 16.8904388 | 16.8701857 | 17.0228501 |
| Q9BU89 | DOHH | 21.0505807 | 21.0773587 | 20.8443161 | 21.0239259 | 21.0833408 |
| Q8TD22 | SFXN5 | 17.3822202 | 17.4174775 | 17.580993 | 17.3095758 | 17.7105254 |
| Q9BRV8 | SIKE1 | 19.2566214 | 19.5124436 | 19.2192016 | 19.2923444 | 19.4514485 |
| O00628 | PEX7 | 17.0311485 | 16.9500915 | 17.2262443 | 17.0443313 | 17.2989951 |
| P80303 | NUCB2 | 23.7236059 | 23.8710331 | 23.4515759 | 23.6317283 | 23.77196 |
| P04114 | APOB | 19.1238858 | 19.1916927 | 19.1151395 | 19.3065082 | 19.0875314 |
| Q9NP73 | ALG13 | 18.444915 | 18.328754 | 18.4810609 | 18.4194031 | 18.5528142 |
| Q14207 | NPAT | 18.4018827 | 18.4738755 | 18.8934087 | 18.5554733 | 18.6978622 |
| Q96DU7 | ITPKC | 16.4710266 | 17.0344262 | 17.0025008 | 16.7503298 | 17.1278252 |
| P21817 | RYR1 | NA | 14.6852529 | 14.4020917 | NA | 14.9076723 |
| O43715 | TRIAP1 | 20.6457937 | 20.8574505 | 20.6519208 | 20.6013378 | 21.0008006 |
| Q8TF76 | HASPIN | 16.9043418 | 16.7142834 | 17.095942 | 16.9998647 | 17.0692387 |
| Q86WI3 | NLRC5 | 13.0213878 | 13.2594107 | 13.2665065 | 12.9085453 | 13.5435769 |
| P12036 | NEFH | 15.1195225 | NA | NA | 15.1050267 | NA |
| Q9P0I2 | EMC3 | 20.6472161 | 20.5413115 | 20.6545244 | 20.5773733 | 20.7297513 |
| Q9C0J8 | WDR33 | 20.9640342 | 20.9519285 | 21.087114 | 20.9756164 | 21.0690014 |
| Q96EY1 | DNAJA3 | 20.9894233 | 21.0030922 | 20.9412583 | 20.8093146 | 20.9865208 |
| Q96KV7 | WDR90 | 22.3719592 | 22.2315849 | 22.0973712 | 22.2085047 | 22.4167247 |
| O75674 | TOM1L1 | 18.5703618 | NA | NA | 18.6308912 | NA |
| Q86VZ1 | P2RY8 | 18.3377848 | 18.4620534 | 18.6923794 | 18.153942 | 18.5272349 |
| Q9BT73 | PSMG3 | 19.6896824 | 19.496622 | 19.7842387 | 19.5996234 | 19.8058099 |
| Q9NZJ7 | MTCH1 | 20.9646213 | 21.0681426 | 21.2829317 | 21.015625 | 21.1485534 |
| Q9NXH8 | TOR4A | 20.4785792 | 20.5315713 | 20.5275568 | 20.3162471 | 20.446313 |
| Q96DT0 | LGALS12 | 19.5369157 | 19.5104722 | 19.585659 | 19.5164811 | 19.5869289 |
| P20701 | ITGAL | 24.9307006 | 25.0079244 | 24.96135 | 24.8148127 | 24.9685151 |
| O60234 | GMFG | 24.9058151 | 24.984405 | 24.8011874 | 24.7729123 | 24.9408527 |
| Q9NVU0 | POLR3E | 19.8525495 | 19.9146068 | 19.8695492 | 19.8219179 | 19.9967984 |
| Q9UHI6 | DDX20 | 23.3099029 | 23.4049011 | 23.4474755 | 23.3133409 | 23.4083022 |
| Q9UI08 | EVL | 20.7476345 | 20.665196 | 20.7392981 | 20.6106685 | 20.6114808 |
| Q14203 | DCTN1 | 22.3309844 | 22.4048009 | 22.3675661 | 22.2618382 | 22.3457763 |

| | | | | | | |
|---|---|---|---|---|---|---|
| Q9HAU5 | UPF2 | 21.4466955 | 21.4812112 | 21.4980081 | 21.3885152 | 21.4517531 |
| P29590 | PML | 21.346158 | 21.4286877 | 21.4512718 | 21.3977136 | 21.4620818 |
| Q8ND24 | RNF214 | 22.4307418 | 22.4731367 | 22.3413033 | 22.3394759 | 22.4269363 |
| Q8NFH4 | NUP37 | 21.2877694 | 21.3529147 | 21.3532148 | 21.2855879 | 21.31832 |
| P04179 | SOD2 | 22.6352363 | 22.7145426 | 22.728295 | 22.6832584 | 22.7275436 |
| Q86VP1 | TAX1BP1 | 22.2990268 | 22.2065057 | 22.0379715 | 22.2867846 | 22.2073689 |
| Q13444 | ADAM15 | 19.1722975 | 19.2924413 | 19.1295682 | 19.1325793 | 19.2692314 |
| P30043 | BLVRB | 23.6730759 | 23.8316346 | 23.7419958 | 23.6592835 | 23.8780812 |
| Q08AN1 | ZNF616 | 15.9681435 | 16.0434418 | 16.0768353 | 16.1879551 | 15.9902847 |
| O00534 | VWA5A | 18.5872829 | 18.4653618 | 18.544093 | 18.6543301 | 18.5976563 |
| Q9P2W1 | PSMC3IP | 18.1302043 | 18.1985486 | 18.0329053 | 18.2106822 | 18.1346925 |
| Q8IZQ1 | WDFY3 | 17.3429567 | 17.4617914 | 17.422554 | 17.4085179 | 17.6422107 |
| A0JLT2 | MED19 | 18.866923 | 19.012729 | 18.8160311 | 18.7713629 | 18.9435182 |
| Q9NQ92 | COPRS | 17.7868962 | 18.128632 | 17.6656122 | 17.7786318 | 17.6723228 |
| Q9BTV6 | DPH7 | 17.763977 | 17.8909904 | 18.0505607 | 17.8727086 | 18.0957603 |
| P15907 | ST6GAL1 | 14.6549624 | NA | NA | 14.9810381 | NA |
| Q96HA8 | NTAQ1 | 18.7623615 | 18.8256 | 18.773073 | 18.7280159 | 18.6454363 |
| P55055 | NR1H2 | 18.1531881 | 18.5319881 | 18.2862866 | 18.1151672 | 18.3364633 |
| P40337 | VHL | 19.3906184 | 19.5356512 | 19.4807233 | 19.3908406 | 19.3823376 |
| Q06187 | BTK | 21.9803742 | 21.9763105 | 22.060513 | 21.9023689 | 21.9981813 |
| Q15654 | TRIP6 | 21.9948669 | 21.875992 | 21.9792203 | 21.9128075 | 22.0266143 |
| P50213 | IDH3A | 22.4330814 | 22.4517279 | 22.4568613 | 22.4018769 | 22.4750255 |
| Q96MU7 | YTHDC1 | 22.3366935 | 22.2967105 | 22.402429 | 22.1878844 | 22.2671935 |
| P20645 | M6PR | 24.6183446 | 24.6324096 | 24.6059372 | 24.484075 | 24.5303936 |
| P28070 | PSMB4 | 23.236645 | 23.3256469 | 23.4063705 | 23.2746094 | 23.3438402 |
| A4D1P6 | WDR91 | 21.2357038 | 21.2882823 | 21.3389093 | 21.329334 | 21.4247866 |
| P46199 | MTIF2 | 19.8062701 | 19.9278428 | 19.8937393 | 19.8860174 | 19.8518648 |
| Q00613 | HSF1 | 20.1909976 | 20.3208213 | 20.295319 | 20.2348237 | 20.345416 |
| O75717 | WDHD1 | 20.8365679 | 20.8688167 | 20.8871404 | 20.8187295 | 20.9643679 |
| Q01844 | EWSR1 | 23.9884053 | 23.9754918 | 23.8931509 | 23.9030765 | 23.9997571 |
| P49458 | SRP9 | 24.5052727 | 24.5153207 | 24.3022319 | 24.4411467 | 24.5063802 |
| Q96RS0 | TGS1 | 19.9792616 | 19.9039911 | 20.0783868 | 19.9715442 | 20.0642961 |
| Q9H4A3 | WNK1 | 22.3956783 | 22.5355625 | 22.3436985 | 22.364604 | 22.451815 |
| O43181 | NDUFS4 | 21.7662647 | 21.987337 | 21.822922 | 21.8337057 | 21.9536365 |
| Q9Y6Q9 | NCOA3 | 18.6601997 | 18.9017223 | 18.7212789 | 18.7578794 | 18.8398519 |
| O00151 | PDLIM1 | 24.2960031 | 24.4460173 | 24.2359137 | 24.2566899 | 24.4154746 |
| Q9Y2I1 | NISCH | 17.7088979 | 17.7497932 | 17.863206 | 17.7190246 | 17.760818 |
| Q08357 | SLC20A2 | 18.3302223 | 18.4227362 | 18.4742066 | 18.2272582 | 18.4665812 |
| Q96LZ7 | RMDN2 | 15.6447919 | 15.4793186 | 15.810204 | 15.5742632 | 15.8848969 |
| Q9H0R6 | QRSL1 | 19.6076116 | 19.7122313 | 19.6652872 | 19.7069945 | 19.664929 |
| P17029 | ZKSCAN1 | 19.6235361 | 19.7185885 | 19.5938316 | 19.5625748 | 19.5195322 |
| Q9BZ23 | PANK2 | 17.6324088 | 17.6920333 | 17.8335234 | 17.7108206 | 17.8173 |

| | | | | | | |
|---|---|---|---|---|---|---|
| P13984 | GTF2F2 | 21.8627218 | 21.8622427 | 21.9246374 | 21.8003958 | 21.9043324 |
| Q6RI45 | BRWD3 | 23.4783699 | 23.3066589 | 23.1452226 | 23.4252248 | 23.4502238 |
| P49757 | NUMB | 20.5227979 | 20.4059842 | 20.4547837 | 20.4814154 | 20.6281816 |
| Q08462 | ADCY2 | 15.215411 | NA | NA | 15.5762326 | NA |
| Q96QD9 | FYTTD1 | 20.3656079 | 20.3470119 | 20.667595 | 20.1881424 | 20.3453672 |
| P17010 | ZFX | 18.0187404 | 18.0894156 | 18.3030258 | 17.9735072 | 18.0165098 |
| Q9NSU2 | TREX1 | 19.6652829 | 19.5373966 | 19.8524799 | 19.5072262 | 19.6888914 |
| O94973 | AP2A2 | 21.3552456 | 21.3754977 | 21.4804662 | 21.3628344 | 21.4237772 |
| Q32NC0 | C18orf21 | 18.6382496 | 18.5225063 | 18.596486 | 18.5371777 | 18.5400955 |
| Q9ULX3 | NOB1 | 21.6877056 | 21.7722558 | 21.784062 | 21.5998939 | 21.6093445 |
| Q8TAT6 | NPLOC4 | 21.4952605 | 21.5880547 | 21.6431863 | 21.536774 | 21.6225837 |
| O95625 | ZBTB11 | 18.5209334 | 18.5577128 | 18.8705406 | 18.3749016 | 18.6204166 |
| P09543 | CNP | 22.9191889 | 22.9236286 | 22.9686054 | 22.8449555 | 22.9532847 |
| Q13084 | MRPL28 | 20.9948682 | 21.1298682 | 21.0113492 | 21.0965848 | 21.0387337 |
| P49585 | PCYT1A | 21.4724229 | 21.6254726 | 21.4915157 | 21.5338712 | 21.6061907 |
| Q96K17 | BTF3L4 | 23.2976652 | 23.383609 | 23.2107243 | 23.2473048 | 23.334398 |
| Q8N5W9 | RFLNB | 19.4492158 | 19.4577026 | 19.3244816 | 19.384765 | 19.769772 |
| P49914 | MTHFS | 19.4720819 | 19.5270537 | 19.6338523 | 19.4330089 | 19.558117 |
| Q8WY36 | BBX | 17.6600378 | 17.9270058 | 17.75144 | 17.7267839 | 17.7546692 |
| P15848 | ARSB | 21.5356599 | 21.5603611 | 21.6829233 | 21.6105643 | 21.7424878 |
| P23743 | DGKA | 17.5472144 | 17.717347 | 17.7675515 | 17.5257139 | 17.6774726 |
| Q8WV41 | SNX33 | 17.377227 | 17.4268323 | 17.3441765 | 17.4273127 | 17.6023015 |
| O75955 | FLOT1 | 23.2578343 | 23.5727351 | 23.3767469 | 23.2642813 | 23.5508387 |
| Q5VSG8 | MANEAL | 18.0226101 | 17.997364 | 18.1605209 | 18.1805579 | 17.7916154 |
| P62875 | POLR2L | 21.7620556 | 21.8786607 | 21.8432462 | 21.7615651 | 21.8533079 |
| Q7Z417 | NUFIP2 | 21.6792436 | 21.8029617 | 21.5968403 | 21.6821721 | 21.7930915 |
| P07237 | P4HB | 27.7768286 | 27.7777749 | 27.5761621 | 27.6561017 | 27.7237475 |
| Q9H3P7 | ACBD3 | 21.5569618 | 21.5019201 | 21.5234205 | 21.5608341 | 21.5697542 |
| Q9H081 | MIS12 | 19.4146662 | 19.3851695 | 19.4185598 | 19.3991896 | 19.4739745 |
| Q99496 | RNF2 | 19.2848603 | 19.3023962 | 19.3562625 | 19.3141916 | 19.4266545 |
| Q8IX19 | MCEMP1 | 18.929648 | 18.8571476 | 19.0247236 | 19.1263088 | 18.987339 |
| P09497 | CLTB | 23.0956117 | 23.3183039 | 22.9662273 | 22.9775378 | 23.3358511 |
| P08069 | IGF1R | 12.4628846 | 12.9017279 | 12.7774753 | 12.9877883 | 13.1339406 |
| P42892 | ECE1 | 16.7548302 | 16.8115469 | 16.8420752 | 16.9198425 | 16.7859112 |
| Q6IA69 | NADSYN1 | 20.2931147 | 20.3874483 | 20.3652498 | 20.2014445 | 20.3562199 |
| Q9NZJ6 | COQ3 | 20.1707673 | 20.257733 | 20.2006054 | 20.156954 | 20.1399196 |
| Q8WU17 | RNF139 | 16.9769399 | 16.8249994 | 17.113486 | 17.1301796 | 17.1184421 |
| Q8NEL9 | DDHD1 | 17.3103846 | 17.3567915 | 17.4660308 | 17.3264468 | 17.2642422 |
| O00142 | TK2 | 20.3917952 | 20.3926676 | 20.4764308 | 20.3068064 | 20.4711102 |
| Q9P2D0 | IBTK | 17.525484 | 17.6448873 | 17.7463903 | 17.7066851 | 17.8391735 |
| O75618 | DEDD | NA | 14.3896074 | 13.7815564 | NA | 13.8660961 |
| P49411 | TUFM | 24.6942728 | 24.7256874 | 24.6503799 | 24.6308734 | 24.7712675 |

| | | | | | | |
|---|---|---|---|---|---|---|
| O75347 | TBCA | 24.6776837 | 24.9873925 | 24.6056187 | 24.777119 | 24.9696632 |
| Q5VYS8 | TUT7 | 19.9380067 | 20.0855828 | 20.1697674 | 20.0344329 | 20.0817842 |
| Q6UXN9 | WDR82 | 22.9940026 | 23.0534457 | 23.0404692 | 23.0054035 | 23.1304792 |
| O75792 | RNASEH2A | 22.0802443 | 22.0886607 | 22.0191365 | 22.0497317 | 22.2520743 |
| Q96G04 | EEF2KMT | 15.0931668 | 15.075374 | 15.1726 | 14.9943582 | 15.2030024 |
| Q7Z3Z4 | PIWIL4 | 15.443853 | 15.4559834 | 15.7386908 | 15.5975428 | 15.7140998 |
| Q9BT09 | CNPY3 | 22.6386801 | 22.8297504 | 22.5475781 | 22.5730396 | 22.7010311 |
| O15243 | LEPROT | 17.5234093 | 17.3529171 | 17.3620667 | 16.9242382 | 17.2931931 |
| Q96D46 | NMD3 | 20.452175 | 20.5070727 | 20.5804311 | 20.4277968 | 20.5393334 |
| Q99961 | SH3GL1 | 22.76262 | 22.8535575 | 22.6219221 | 22.7543179 | 22.7651282 |
| Q5SXM8 | DNLZ | 12.1107065 | NA | NA | 11.4803126 | NA |
| Q3SYG4 | BBS9 | 17.3847332 | 17.9137054 | 17.9143599 | 17.2258821 | 17.5836655 |
| Q92834 | RPGR | 16.4143727 | 16.4262067 | 16.6114657 | 16.6958596 | 16.6706194 |
| Q5TEA3 | DNAAF9 | 17.0449328 | 17.4196495 | 17.3058654 | 17.1542286 | 17.165224 |
| Q5D0E6 | DALRD3 | 15.334439 | 15.4594066 | 15.4919919 | 15.6135548 | 15.5467399 |
| Q9GZN8 | ADISSP | 22.8754502 | 22.7796319 | 22.8154586 | 22.7938961 | 22.8445941 |
| P51523 | ZNF84 | 17.1100023 | 17.3774501 | 17.4358447 | 17.2416546 | 17.1308062 |
| Q969T9 | WBP2 | 22.3235904 | 22.3750515 | 22.4071774 | 22.3567934 | 22.3798768 |
| Q9Y3Y2 | CHTOP | 22.6041794 | 22.6544235 | 22.8208968 | 22.5719055 | 22.5266659 |
| Q92521 | PIGB | 18.6473625 | 18.4471609 | 18.5841509 | 18.6403962 | 18.6373849 |
| P61019 | RAB2A | 23.4779524 | 23.5429527 | 23.5930596 | 23.4220929 | 23.6124711 |
| P12236 | SLC25A6 | 24.4804026 | 24.4811699 | 24.4635016 | 24.3957279 | 24.4363746 |
| Q9BT78 | COPS4 | 22.0187101 | 22.1189987 | 22.0247911 | 21.9784813 | 22.0782069 |
| P29401 | TKT | 26.3902081 | 26.3920023 | 26.3759873 | 26.3082052 | 26.3293439 |
| O60547 | GMDS | 21.2592543 | 21.2495883 | 21.2891706 | 21.2058055 | 21.316504 |
| Q8NHP8 | PLBD2 | 22.2934722 | 22.2841031 | 22.2826553 | 22.3489315 | 22.2715381 |
| Q9Y2H0 | DLGAP4 | 20.4637929 | 20.6610965 | 20.5079509 | 20.5155525 | 20.5704035 |
| Q7KZ85 | SUPT6H | 21.7175036 | 21.7983207 | 21.8354676 | 21.7682454 | 21.8749032 |
| Q9NP97 | DYNLRB1 | 21.9925118 | 22.068664 | 21.9749054 | 22.056034 | 22.1829782 |
| Q13469 | NFATC2 | 19.6066611 | 19.7697937 | 19.5926002 | 19.6631176 | 19.6745569 |
| Q9Y5Y2 | NUBP2 | 22.0139174 | 22.2163607 | 22.1301106 | 22.2667859 | 22.361888 |
| P19525 | EIF2AK2 | 19.8399496 | 20.1714383 | 20.1165764 | 19.9993761 | 20.1739203 |
| Q8NB49 | ATP11C | 17.1754635 | 17.1301517 | 17.5409194 | 17.3666894 | 17.2974967 |
| O14737 | PDCD5 | 23.7514351 | 23.9422021 | 23.6644694 | 23.7027227 | 23.8922606 |
| Q9Y6A9 | SPCS1 | 21.2274848 | 21.088551 | 21.366276 | 21.1939312 | 21.2955729 |
| Q96AH8 | RAB7B | 13.5644438 | 13.5692635 | 14.0437442 | 14.1344639 | 14.0541314 |
| Q9UK76 | JPT1 | 22.5428937 | 22.6114682 | 22.382973 | 22.5748117 | 22.6694071 |
| Q53GS9 | USP39 | 22.1367048 | 22.044764 | 22.2015621 | 22.0886395 | 22.07976 |
| O14979 | HNRNPDL | 25.3833415 | 25.4461969 | 25.2703182 | 25.2432575 | 25.4100066 |
| Q70HW3 | SLC25A26 | NA | 14.5380555 | 14.6429292 | NA | 14.6600209 |
| P62277 | RPS13 | 26.668664 | 26.7131291 | 26.7522813 | 26.5848028 | 26.6622016 |
| Q15007 | WTAP | 22.2626817 | 22.3333278 | 22.2349207 | 22.149868 | 22.3027864 |

| | | | | | | |
|---|---|---|---|---|---|---|
| O14578 | CIT | 20.487708 | 20.5915552 | 20.4721347 | 20.4167565 | 20.507524 |
| P17568 | NDUFB7 | 21.3077396 | 21.3742371 | 21.5437917 | 21.3753094 | 21.4592573 |
| P54725 | RAD23A | 23.3314601 | 23.5023718 | 23.2163748 | 23.2579171 | 23.3819344 |
| Q9H4H8 | FAM83D | 17.4478872 | 17.4979824 | 17.4902047 | 17.6475372 | 17.5360416 |
| Q9BQ48 | MRPL34 | 20.2166634 | 20.2466646 | 20.4774604 | 20.242168 | 20.2125976 |
| Q9P2D1 | CHD7 | 19.2244567 | 19.2883546 | 19.3592966 | 19.2454616 | 19.3087738 |
| Q9P1U0 | POLR1H | 18.2147462 | 18.3586104 | 18.0187354 | 18.3177901 | 18.3596581 |
| Q9BSA9 | TMEM175 | 17.0207258 | 17.1763884 | 17.3441145 | 17.0305187 | 17.124733 |
| Q15005 | SPCS2 | 23.4050946 | 23.3683166 | 23.4459685 | 23.2618901 | 23.3583278 |
| Q96DE5 | ANAPC16 | NA | 14.8918927 | 14.7974881 | NA | 15.3922281 |
| Q16595 | FXN | 20.3882738 | 20.4720386 | 20.2935479 | 20.2747976 | 20.4357459 |
| A8MVM7 | NA | 20.2195808 | NA | NA | 20.1790151 | NA |
| Q9BY77 | POLDIP3 | 21.9461828 | 21.9643352 | 22.0848139 | 21.9630945 | 22.0301485 |
| O75356 | ENTPD5 | 16.5717321 | NA | NA | 16.2567072 | NA |
| Q9NRA0 | SPHK2 | 15.3571496 | 15.9401111 | 15.9045625 | 15.6106607 | 15.7931016 |
| Q96C92 | ENTR1 | 16.9342443 | 16.8638262 | 16.7541035 | 16.6085619 | 16.6554185 |
| P58557 | YBEY | 18.9473992 | 18.9331897 | 18.8859525 | 18.8628051 | 18.8640787 |
| P05067 | APP | 17.6160035 | 17.4637898 | 17.6527709 | 17.600131 | 17.5851277 |
| P53801 | PTTG1IP | 22.8258451 | 22.8351665 | 22.9543165 | 22.8578417 | 22.7415622 |
| P18077 | RPL35A | 26.0481492 | 26.0563078 | 26.0829015 | 25.9131096 | 25.9345349 |
| Q9UK99 | FBXO3 | 19.162925 | 19.2077562 | 19.2586206 | 19.1951512 | 19.311827 |
| Q13190 | STX5 | 19.911036 | 19.9869751 | 20.1559312 | 20.0463664 | 20.0048667 |
| P33908 | MAN1A1 | 19.9617577 | 19.9061445 | 20.0273592 | 20.0076428 | 20.0390846 |
| O94855 | SEC24D | 21.2364531 | 21.2264397 | 21.2667762 | 21.2403953 | 21.2833964 |
| Q86Y07 | VRK2 | 18.9793331 | 18.9672613 | 19.0680052 | 18.9453384 | 19.0469983 |
| Q969X0 | RILPL2 | 20.5392534 | 20.6881115 | 20.6744323 | 20.604568 | 20.8425654 |
| Q8WVB3 | HEXD | 17.4392908 | 17.6141934 | 17.5450655 | 17.5559918 | 17.4734011 |
| P31943 | HNRNPH1 | 26.4386123 | 26.5191775 | 26.3800091 | 26.3693457 | 26.5319766 |
| Q9NRV9 | HEBP1 | 21.6096238 | 21.5904938 | 21.5948972 | 21.4832668 | 21.7388216 |
| Q9HD34 | LYRM4 | 20.3879256 | 20.7620092 | 20.6169927 | 20.6595069 | 20.7392056 |
| Q9Y597 | KCTD3 | 18.0410773 | 18.135762 | 18.2786256 | 18.1788997 | 18.0967663 |
| Q96A73 | KIAA1191 | 17.0549287 | 17.2778286 | 17.3892444 | 17.2654882 | 17.5820435 |
| Q6ZVC0 | NYAP1 | NA | 14.816305 | 14.6967955 | NA | 14.3412609 |
| Q8WYL5 | SSH1 | 17.6505493 | 17.1071464 | 17.7959115 | 17.563578 | 17.9652237 |
| Q15813 | TBCE | 21.290423 | 21.3778664 | 21.3229937 | 21.3122011 | 21.3058674 |
| O94854 | MACF1 | 15.8673504 | 16.212344 | 15.7104752 | 15.8486195 | 16.0133845 |
| Q8TDD1 | DDX54 | 22.6945862 | 22.6371549 | 22.7936088 | 22.6007863 | 22.6340384 |
| Q92692 | NECTIN2 | 24.7216826 | 24.8773997 | 24.7459491 | 24.6496756 | 24.6124531 |
| Q92985 | IRF7 | 15.5289083 | 15.8793344 | 15.5517895 | 15.5950066 | 15.2276278 |
| P31930 | UQCRC1 | 24.196452 | 24.3139768 | 24.2777397 | 24.1633416 | 24.4938062 |
| Q5TF39 | MFSD4B | 17.8907831 | 18.1197056 | 18.1346439 | 17.7045563 | 18.2741611 |
| O00746 | NME4 | NA | 17.0597723 | 17.4208041 | NA | 17.3035506 |

| | | | | | | |
|---|---|---|---|---|---|---|
| Q5HYI8 | RABL3 | 19.2347631 | 19.3469864 | 19.5017512 | 19.2393863 | 19.3154357 |
| Q16762 | TST | 22.4683473 | 22.5652225 | 22.6544919 | 22.4284791 | 22.538626 |
| Q08379 | GOLGA2 | 20.8558155 | 20.9031027 | 20.858296 | 20.8286015 | 20.9579015 |
| O75379 | VAMP4 | 16.6594541 | 17.0011742 | 16.7761359 | 16.6809571 | 17.5167787 |
| Q9UPV9 | TRAK1 | 19.1954352 | NA | NA | 19.3012201 | NA |
| P56282 | POLE2 | 19.2686166 | 19.3726955 | 19.3415131 | 19.2345877 | 19.2582872 |
| Q9UJX5 | ANAPC4 | 19.4689666 | 19.4819053 | 19.585498 | 19.4244434 | 19.5690364 |
| Q14690 | PDCD11 | 21.846836 | 21.8999086 | 21.9091265 | 21.7897602 | 21.8341515 |
| P52740 | ZNF132 | 17.9561199 | NA | NA | 18.0479975 | NA |
| O95164 | UBL3 | 17.0984985 | 17.1896938 | 17.2750099 | 17.0378172 | 17.1859793 |
| Q9Y4R8 | TELO2 | 21.8059809 | 21.7521954 | 21.7873913 | 21.7070207 | 21.7495 |
| Q96DZ1 | ERLEC1 | 20.1505431 | 20.1877465 | 20.167227 | 20.1143068 | 20.2164992 |
| Q12872 | SFSWAP | 20.2891725 | 20.405484 | 20.4430836 | 20.3262975 | 20.3848545 |
| Q9H3S7 | PTPN23 | 22.7557509 | 22.8939932 | 22.8161997 | 22.8281746 | 22.8873744 |
| Q9UBI9 | HECA | 18.3314789 | 18.2678454 | 18.3783915 | 18.3389792 | 18.2248151 |
| O94813 | SLIT2 | NA | 15.6072731 | 16.2194758 | NA | 15.1661774 |
| P02794 | FTH1 | 22.1462532 | 22.3471285 | 22.3774818 | 22.271229 | 22.3891011 |
| Q9Y295 | DRG1 | 22.5799432 | 22.5903636 | 22.5884567 | 22.5156089 | 22.5791143 |
| Q96RT7 | TUBGCP6 | 19.2043684 | 19.1760182 | 19.1996428 | 19.1311494 | 19.2327271 |
| Q6UB99 | ANKRD11 | 16.7449085 | 16.8354344 | 16.933371 | 17.073013 | 17.1522034 |
| Q2VYF4 | LETM2 | 16.3231827 | 16.0886491 | 16.1793423 | 16.1465461 | 16.2094531 |
| O15182 | CETN3 | NA | 17.5642943 | 17.2276255 | NA | 17.2784981 |
| Q9Y4H4 | GPSM3 | 18.479504 | 18.8466803 | 18.6824588 | 18.4507589 | 18.8655005 |
| Q86Y38 | XYLT1 | 19.5563847 | 19.4696263 | 19.6395829 | 19.4706001 | 19.4906647 |
| Q96ST2 | IWS1 | 22.0439414 | 22.0358383 | 22.1825643 | 21.9724357 | 22.0118293 |
| A1L390 | PLEKHG3 | 18.3770188 | 18.2637513 | 18.2531499 | 18.3187423 | 18.3731384 |
| Q9H7C9 | AAMDC | 19.9557934 | 20.0142291 | 19.8621222 | 19.9364878 | 20.1898335 |
| Q5T1C6 | THEM4 | 18.6016901 | 18.6668969 | 18.6751959 | 18.6090633 | 18.7452021 |
| Q9P2Y5 | UVRAG | 20.1897068 | 20.2811873 | 20.2464445 | 20.110619 | 20.2105244 |
| O94898 | LRIG2 | NA | 15.3136597 | 14.8430568 | NA | 15.0892613 |
| O00337 | SLC28A1 | NA | 10.7757337 | 11.4600892 | NA | 11.6604498 |
| Q6UXV4 | APOOL | 21.7013253 | 21.7147171 | 21.6500476 | 21.550824 | 21.6844663 |
| Q96I23 | PYURF | 14.5327593 | 14.7458241 | 15.2124681 | 14.830214 | 15.3652616 |
| Q7Z3B4 | NUP54 | 21.3643892 | 21.4146016 | 21.4308884 | 21.3145007 | 21.3731996 |
| Q7Z7B1 | PIGW | 17.609324 | 17.4610713 | 17.6409041 | 17.6347454 | 17.5261446 |
| Q15750 | TAB1 | 19.8918639 | 20.170233 | 20.0619795 | 20.0151571 | 20.0618623 |
| Q16540 | MRPL23 | 21.3930142 | 21.468541 | 21.5179346 | 21.4464531 | 21.4709395 |
| P53396 | ACLY | 25.1487029 | 25.2096159 | 25.178103 | 25.0726642 | 25.1030538 |
| P61647 | ST8SIA6 | 18.3618688 | 18.57578 | 18.3918884 | 18.2263965 | 18.2522905 |
| Q969U7 | PSMG2 | 21.4644251 | 21.5404824 | 21.6014316 | 21.4345392 | 21.408433 |
| Q86W42 | THOC6 | 21.1214511 | 21.0949248 | 21.1777715 | 21.0176643 | 21.2328858 |
| Q9NXF1 | TEX10 | 22.090946 | 22.1953433 | 22.2446053 | 22.0989975 | 22.146397 |

| | | | | | | |
|---|---|---|---|---|---|---|
| Q13098 | GPS1 | 22.7873485 | 22.8218801 | 22.8270863 | 22.7203022 | 22.6848471 |
| O60287 | URB1 | 20.4853133 | 20.4888298 | 20.6254447 | 20.4627966 | 20.5787527 |
| Q12981 | BNIP1 | 19.5282732 | 19.6201885 | 19.7520577 | 19.6560617 | 19.7031923 |
| P54819 | AK2 | 24.8263595 | 24.9289816 | 24.8016291 | 24.787893 | 24.8561064 |
| Q6P4I2 | WDR73 | 19.1025841 | 19.3101112 | 19.2458791 | 19.2144994 | 19.3006517 |
| Q9NZD8 | SPG21 | 21.2354304 | 21.2417672 | 21.307521 | 21.2403522 | 21.2824285 |
| Q9NZN8 | CNOT2 | 20.5709901 | 20.4319217 | 20.5176703 | 20.482737 | 20.5666421 |
| P55196 | AFDN | 19.8969975 | 20.0844658 | 19.9513838 | 19.9207543 | 20.058921 |
| P98082 | DAB2 | 11.3866277 | NA | NA | 11.0458529 | NA |
| Q86XH1 | IQCA1 | 13.2652725 | NA | NA | 13.1875251 | NA |
| Q9H8E8 | KAT14 | 17.7227587 | 18.1425638 | 17.9722304 | 17.9019504 | 18.0507949 |
| Q15834 | CCDC85B | 17.7616177 | 18.0748126 | 18.0218032 | 18.024552 | 17.9208861 |
| P27448 | MARK3 | 19.896272 | 19.9204107 | 19.9372836 | 19.9400975 | 19.9475784 |
| Q13724 | MOGS | 22.2529038 | 22.2758533 | 22.3663238 | 22.2231687 | 22.2919121 |
| O60841 | EIF5B | 23.0889856 | 23.2344315 | 23.0926288 | 23.1022534 | 23.174589 |
| Q8TDF6 | RASGRP4 | 15.6463249 | 15.9165834 | 15.8214219 | 16.2126213 | 15.522008 |
| P62306 | SNRPF | 23.356571 | 23.1878803 | 23.4745893 | 23.2366788 | 23.2107471 |
| Q9Y4C4 | MFHAS1 | 15.8442575 | 15.8576442 | 15.9206645 | 15.9898386 | 15.7886439 |
| Q96F44 | TRIM11 | 18.5593663 | 18.6742158 | 18.8678975 | 18.5083224 | 18.6457504 |
| O60502 | OGA | 20.6514506 | 20.7912359 | 20.7210966 | 20.6465977 | 20.7437015 |
| Q14997 | PSME4 | 22.1790582 | 22.1089636 | 22.2521884 | 22.1827913 | 22.1505301 |
| Q96L91 | EP400 | 20.2741183 | 20.3113448 | 20.2570539 | 20.2928525 | 20.3553996 |
| P55072 | VCP | 25.3653714 | 25.4030074 | 25.2683475 | 25.3250473 | 25.4100695 |
| Q9UIM3 | FKBPL | 17.6192034 | 18.0664809 | 17.5928419 | 17.6436056 | 17.7136577 |
| P32926 | DSG3 | NA | 16.6046994 | 16.2670922 | NA | 16.5538773 |
| Q92540 | SMG7 | 19.2330526 | 19.3980547 | 19.3740875 | 19.3461158 | 19.3035272 |
| O43426 | SYNJ1 | 17.8708618 | 18.0484439 | 18.0744433 | 17.8728459 | 17.9574883 |
| P15586 | GNS | 21.921805 | 21.9147489 | 21.8485542 | 21.8778408 | 21.9383395 |
| P15056 | BRAF | 18.1912728 | 18.3152615 | 18.3825449 | 18.385964 | 18.4997439 |
| Q9BZG8 | DPH1 | 19.7256419 | 19.7872832 | 19.7453603 | 19.5976948 | 19.822701 |
| Q9Y3B2 | EXOSC1 | 21.4202269 | 21.326331 | 21.5727848 | 21.4350318 | 21.4792005 |
| O00522 | KRIT1 | 17.9649969 | 17.9903934 | 18.0478848 | 18.0403004 | 18.0509012 |
| P62877 | RBX1 | 22.70263 | 22.88744 | 22.9482892 | 22.760016 | 22.7525957 |
| P62241 | RPS8 | 26.0587576 | 26.116074 | 26.1474941 | 25.9075261 | 26.0604589 |
| Q15052 | ARHGEF6 | 22.1771272 | 22.1760163 | 22.1056577 | 22.0781026 | 22.1776382 |
| P38432 | COIL | 20.6287339 | 20.8317785 | 20.7276129 | 20.6966092 | 20.7752867 |
| P55160 | NCKAP1L | 21.8400106 | 21.9987758 | 22.066919 | 21.8449377 | 21.9225595 |
| Q99805 | TM9SF2 | 23.3769466 | 23.298223 | 23.3398099 | 23.3415592 | 23.3058825 |
| Q8NBJ5 | COLGALT1 | 24.3301138 | 24.3009904 | 24.2944917 | 24.2124032 | 24.2775232 |
| Q8N0V3 | RBFA | 18.8663959 | 19.1006849 | 19.0884195 | 18.9610651 | 19.2210883 |
| Q0VDD7 | BRME1 | NA | 12.6100117 | 12.2500282 | NA | 12.3956013 |
| Q8N680 | ZBTB2 | 15.6876429 | 15.6877513 | 15.9394116 | 15.7494982 | 15.9578896 |

| | | | | | | |
|---|---|---|---|---|---|---|
| Q9BSK2 | SLC25A33 | 16.9872375 | 16.7819014 | 16.8789019 | 17.1308117 | 16.5993785 |
| Q00839 | HNRNPU | 27.3390418 | 27.4065141 | 27.394809 | 27.2311063 | 27.2529917 |
| Q8N6Q8 | METTL25 | 17.2166361 | 17.3792414 | 17.3757793 | 17.1567644 | 17.4293602 |
| P09086 | POU2F2 | 16.5539225 | 16.7601041 | 16.9446924 | 16.584085 | 16.9996182 |
| Q9Y4Y9 | LSM5 | 22.6338585 | 22.7442614 | 22.4501033 | 22.6125274 | 22.8216643 |
| Q9UPG8 | PLAGL2 | 15.1653966 | 15.4233395 | 15.3249667 | 15.326574 | 14.6544291 |
| Q9NTG7 | SIRT3 | 24.933946 | 24.9840679 | 24.8697498 | 24.8147518 | 24.8015338 |
| Q9H0H5 | RACGAP1 | 19.7716059 | 19.7737359 | 19.8013276 | 19.7472977 | 19.8177657 |
| Q15013 | MAD2L1BP | 18.0511254 | 18.4438928 | 18.3391552 | 18.0780943 | 18.3099012 |
| Q12874 | SF3A3 | 23.4713053 | 23.571705 | 23.5539976 | 23.5318313 | 23.5907287 |
| Q96DG6 | CMBL | 18.2467277 | 18.2928491 | 18.0717277 | 18.2040851 | 18.3190588 |
| Q9NYA4 | MTMR4 | 15.1497206 | 15.1872632 | 15.2846929 | 15.3729984 | 15.3551564 |
| F2Z333 | FNDC10 | 16.5458832 | 16.1883461 | 16.5664948 | 16.6442224 | 16.3047428 |
| Q6P3S1 | DENND1B | 16.5433945 | 16.7885749 | 16.578334 | 16.4644193 | 16.3971339 |
| O14727 | APAF1 | 21.3778245 | 21.4593373 | 21.5072597 | 21.3884071 | 21.478433 |
| Q9NUY8 | TBC1D23 | 20.0709449 | 20.2052477 | 20.2367041 | 20.1031949 | 20.2489523 |
| Q9UKK3 | PARP4 | 19.5719415 | 19.6652014 | 19.676767 | 19.6475124 | 19.7060687 |
| P34741 | SDC2 | 20.5741386 | 20.9074648 | 20.9676079 | 20.6686781 | 20.6209013 |
| Q8WZ73 | RFFL | 17.2602444 | 17.4503891 | 17.4363257 | 17.4615693 | 17.2817488 |
| Q96H55 | MYO19 | 17.4080439 | 17.4860882 | 17.8878757 | 17.5348337 | 17.6135076 |
| P46777 | RPL5 | 27.1197351 | 27.027231 | 27.1372445 | 26.9929137 | 27.0113235 |
| P40925 | MDH1 | 25.1289907 | 25.2442886 | 25.1527007 | 25.1050119 | 25.183425 |
| Q9H9B1 | EHMT1 | 22.1589025 | 22.1860504 | 22.1418539 | 22.0612549 | 22.2067403 |
| P14868 | DARS1 | 24.4955001 | 24.591857 | 24.5444111 | 24.411203 | 24.4626106 |
| Q9UJA5 | TRMT6 | 21.8396433 | 21.7743129 | 21.7886708 | 21.7508858 | 21.8591447 |
| P60660 | MYL6 | 25.7763959 | 26.0081665 | 25.4745819 | 25.663834 | 25.8601825 |
| P46926 | GNPDA1 | 21.6514263 | 21.5209578 | 21.6979561 | 21.5967383 | 21.5871015 |
| Q13488 | TCIRG1 | 21.6214992 | 21.5338533 | 21.5571137 | 21.5606321 | 21.5526009 |
| Q7Z3J2 | VPS35L | 20.5811088 | 20.6370279 | 20.7249169 | 20.5834038 | 20.7236101 |
| P09525 | ANXA4 | 22.7754623 | 22.8181674 | 22.7693165 | 22.7226933 | 22.8446937 |
| O94880 | PHF14 | 20.9801354 | 20.9975724 | 20.9951893 | 21.0077615 | 21.0717541 |
| P46108 | CRK | 20.6751251 | 20.8935645 | 20.5913173 | 20.7542896 | 20.9351622 |
| Q8IYA6 | CKAP2L | 17.4968671 | 17.8314478 | 17.7602891 | 17.5420863 | 17.5195695 |
| P07919 | UQCRH | 23.7121907 | 23.9663094 | 23.6100191 | 23.670466 | 23.9006633 |
| A4D1U4 | DENND11 | NA | 16.3754168 | 16.4690996 | NA | 16.5276284 |
| Q70IA6 | MOB2 | 19.9755079 | 19.9772527 | 20.1146392 | 19.9414088 | 19.9674526 |
| P23141 | CES1 | 24.5139879 | 24.5348885 | 24.5458876 | 24.4411279 | 24.5929008 |
| Q3B726 | POLR1F | 20.0905411 | 20.1410229 | 20.0306727 | 20.144615 | 20.1315894 |
| Q08AG7 | MZT1 | 16.8329362 | 17.0227043 | 17.1112531 | 17.3600962 | 16.8553688 |
| O75385 | ULK1 | 16.8172028 | 16.7062704 | 16.5708835 | 16.5582131 | 16.6303055 |
| P12074 | COX6A1 | 19.0094039 | 18.7153331 | 19.2912158 | 19.3425294 | 19.3185743 |
| O95503 | CBX6 | 19.0640417 | 19.0026182 | 18.9429338 | 18.8026982 | 18.7058532 |

| | | | | | | |
|---|---|---|---|---|---|---|
| Q8WV22 | NSMCE1 | 20.1170393 | 20.2810701 | 20.3059658 | 20.0522106 | 20.3368824 |
| Q14807 | KIF22 | 20.9425422 | 20.8992046 | 20.9630173 | 20.7795685 | 20.936279 |
| Q96F46 | IL17RA | 19.7815813 | 19.8279354 | 19.7382794 | 19.7522617 | 19.7108233 |
| Q8N531 | FBXL6 | 20.7864144 | 20.7110386 | 20.7379402 | 20.69463 | 20.6252737 |
| Q9BYG3 | NIFK | 21.9747398 | 22.0303766 | 22.1382541 | 22.0307952 | 22.0161114 |
| O43567 | RNF13 | 20.7823596 | 20.8469579 | 20.9075139 | 20.7423482 | 20.7283562 |
| P21283 | ATP6V1C1 | 22.6996271 | 22.7683379 | 22.6516424 | 22.6182305 | 22.6280516 |
| P82673 | MRPS35 | 21.2559194 | 21.1871368 | 21.2965612 | 21.2165083 | 21.2790914 |
| Q9NWX6 | THG1L | 20.1316184 | 20.166078 | 20.0915854 | 19.9882169 | 20.184052 |
| O43353 | RIPK2 | 18.9213416 | 18.848985 | 18.9217995 | 18.9404668 | 19.0194341 |
| Q6NXT1 | ANKRD54 | 20.7333648 | 20.9786889 | 20.8035396 | 20.7486801 | 20.8803938 |
| Q00059 | TFAM | 23.5656701 | 23.6029432 | 23.413625 | 23.4017906 | 23.5563884 |
| Q969J2 | ZKSCAN4 | 17.689116 | 17.4815117 | 17.7002699 | 17.6167903 | 17.7877592 |
| P35236 | PTPN7 | 21.7543073 | 21.6818604 | 21.7768257 | 21.7440076 | 21.6882176 |
| Q6P1J9 | CDC73 | 21.30054 | 21.3728324 | 21.4535273 | 21.3591018 | 21.4818767 |
| O60237 | PPP1R12B | 17.8572185 | 17.3602826 | 17.2532768 | 17.5213237 | 17.3876528 |
| Q5JTH9 | RRP12 | 22.4395672 | 22.5064294 | 22.5626135 | 22.4161964 | 22.4061717 |
| Q12772 | SREBF2 | 19.9099531 | 20.13734 | 19.7712343 | 20.004835 | 19.9743133 |
| Q96IC2 | REXO5 | 20.8107219 | 21.1231877 | 21.1219034 | 20.3792448 | 20.7487133 |
| Q8N684 | CPSF7 | 22.1955298 | 22.2902434 | 22.2261285 | 22.1217799 | 22.2729403 |
| Q9BZV1 | UBXN6 | 20.1158755 | 20.1706349 | 20.2237078 | 20.0710048 | 20.1352412 |
| P62166 | NCS1 | 20.3340602 | 20.4379451 | 20.4455545 | 20.3921889 | 20.4305071 |
| O15031 | PLXNB2 | 21.6288244 | 21.7286189 | 21.7133522 | 21.5939123 | 21.7410086 |
| Q10570 | CPSF1 | 22.1440433 | 22.2007583 | 22.2207577 | 22.1534259 | 22.2270496 |
| Q8IYB3 | SRRM1 | 22.9017447 | 22.9722614 | 22.9437248 | 22.8474156 | 23.0561943 |
| P20338 | RAB4A | 21.5547517 | 21.6481573 | 21.6864822 | 21.6125461 | 21.7561026 |
| Q9BSI4 | TINF2 | 18.2217658 | 18.3752326 | 18.1823718 | 18.2093091 | 18.2352621 |
| P06746 | POLB | 19.9429703 | 20.0899544 | 20.1005068 | 19.8884569 | 20.2021396 |
| Q03468 | ERCC6 | 18.1221282 | 18.3348303 | 18.3771199 | 18.3448792 | 18.3361609 |
| Q86VI3 | IQGAP3 | 19.5878763 | 19.5763775 | 19.6187614 | 19.4942446 | 19.5960154 |
| Q9UHY8 | FEZ2 | 16.7326388 | 16.9212619 | 16.9491101 | 16.8930127 | 16.9877773 |
| Q9NZ53 | PODXL2 | 14.7719012 | 14.8521498 | 14.7851805 | 14.9441703 | 14.8948302 |
| Q5SYE7 | NHSL1 | 16.0630321 | 16.3533193 | 16.05369 | 16.2596258 | 16.2527248 |
| Q96QP1 | ALPK1 | 15.8447793 | 15.4558299 | 16.2289589 | 15.4857476 | 15.5541381 |
| O75871 | CEACAM4 | 11.1834006 | NA | NA | 11.3105433 | NA |
| Q96MD7 | C9orf85 | NA | 15.8117718 | 15.4875884 | NA | 15.5918008 |
| Q9NPA1 | KCNMB3 | 15.032086 | NA | NA | 15.2562683 | NA |
| O43707 | ACTN4 | 26.1373882 | 26.2614639 | 26.0696062 | 26.0399182 | 26.2168071 |
| Q9P013 | CWC15 | 21.882389 | 21.925538 | 21.8740747 | 21.8838688 | 21.9340684 |
| Q9Y223 | GNE | 20.2065677 | 20.2001825 | 20.2493019 | 20.1869756 | 20.2842362 |
| Q99638 | RAD9A | 17.5089833 | 17.618085 | 17.5552002 | 17.3954437 | 17.2088259 |
| Q96B23 | ARK2N | 19.0605206 | 19.1916915 | 19.168005 | 19.0710751 | 19.0191994 |

| | | | | | | |
|---|---|---|---|---|---|---|
| Q8NDT2 | RBM15B | 19.5726469 | 19.6204791 | 19.6246016 | 19.5891967 | 19.6798213 |
| Q9BUB4 | ADAT1 | 19.0105817 | 19.3839887 | 19.4771787 | 19.3295831 | 19.1579819 |
| Q96I76 | GPATCH3 | 15.0708423 | 15.0534056 | 14.4635514 | 14.3726874 | 14.285634 |
| P10398 | ARAF | 21.481522 | 21.565761 | 21.6321949 | 21.4745073 | 21.5973251 |
| Q6UB35 | MTHFD1L | 23.349287 | 23.3464661 | 23.3117434 | 23.3086761 | 23.3490473 |
| Q5K651 | SAMD9 | 19.8094154 | 19.9974348 | 20.0311471 | 19.8855872 | 20.0019553 |
| P12277 | CKB | 20.9184819 | 21.1763285 | 21.0487308 | 20.983839 | 21.1922506 |
| Q93009 | USP7 | 22.7131085 | 22.7765995 | 22.7377777 | 22.696894 | 22.7598564 |
| P45985 | MAP2K4 | 20.2312974 | 20.2492637 | 20.3056659 | 20.2129626 | 20.4467863 |
| Q9NTJ5 | SACM1L | 23.3984922 | 23.3392325 | 23.4789398 | 23.274484 | 23.3865015 |
| Q9NUL7 | DDX28 | 20.7447739 | 20.6702979 | 20.7576818 | 20.6926785 | 20.7652605 |
| Q5VTR2 | RNF20 | 21.5401194 | 21.6466059 | 21.5256479 | 21.5308151 | 21.6026123 |
| P51689 | ARSD | 17.9694535 | 17.9290794 | 18.0977633 | 18.0437529 | 18.1363096 |
| Q13683 | ITGA7 | 15.7483294 | 15.7147804 | 15.8807166 | 15.3636106 | 15.6639995 |
| Q9NUQ9 | CYRIB | 24.8796903 | 24.8924437 | 24.8428323 | 24.7430978 | 24.8206363 |
| Q8NCA5 | FAM98A | 21.9148709 | 21.8568439 | 22.042665 | 21.8659541 | 21.9236486 |
| O43678 | NDUFA2 | 21.6038131 | 21.8540826 | 21.7698535 | 21.5968204 | 21.8089525 |
| Q99714 | HSD17B10 | 23.1555196 | 23.2427257 | 23.213557 | 23.1099643 | 23.2924101 |
| O43791 | SPOP | 16.1710536 | 16.2887121 | 16.1094706 | 16.2512743 | 16.3258073 |
| P60059 | SEC61G | 18.7266971 | 19.0181975 | 18.4706227 | 18.5866582 | 18.2215774 |
| O00506 | STK25 | 19.0967979 | 19.0304404 | 19.1679267 | 19.1092826 | 19.1078357 |
| P17097 | ZNF7 | 17.3902024 | 17.6779531 | 17.54996 | 17.3557444 | 17.1858566 |
| O60506 | SYNCRIP | 26.2388057 | 26.280473 | 26.1037877 | 26.1423611 | 26.2270455 |
| Q49AR2 | C5orf22 | 19.0183078 | 19.2559272 | 19.1154158 | 19.2090233 | 19.1430949 |
| P20749 | BCL3 | 13.2172374 | NA | NA | 13.3371597 | NA |
| P25445 | FAS | 17.7526403 | 17.7082067 | 17.6015742 | 17.6414647 | 17.7776175 |
| Q7LGA3 | HS2ST1 | 20.6578922 | 20.6348562 | 20.6910635 | 20.6157814 | 20.7077187 |
| Q96PM5 | RCHY1 | 19.2393248 | 19.3628024 | 19.2116738 | 19.1055328 | 19.2285523 |
| Q9Y2P5 | SLC27A5 | 19.4175044 | 19.1272644 | 19.502021 | 19.2353097 | 19.4495257 |
| Q6FI81 | CIAPIN1 | 23.5713012 | 23.6548739 | 23.435774 | 23.4524592 | 23.5896215 |
| O95166 | GABARAP | 23.2735727 | 23.312546 | 23.2001679 | 23.2874448 | 23.2486486 |
| Q9UPN7 | PPP6R1 | 22.1788122 | 22.298191 | 22.1978484 | 22.0457972 | 22.1591345 |
| Q12802 | AKAP13 | 19.6040646 | 19.8261154 | 19.6517005 | 19.6272302 | 19.8384558 |
| Q96LC7 | SIGLEC10 | 14.0091152 | NA | NA | 14.1064247 | NA |
| Q92794 | KAT6A | 14.9025612 | 15.2574892 | 14.9978029 | 15.1540557 | 14.6121558 |
| P01112 | HRAS | 14.4631806 | NA | NA | 14.4898176 | NA |
| Q99942 | RNF5 | 18.7931892 | 18.9713737 | 19.0836932 | 18.8998144 | 18.7547904 |
| Q8N0S6 | CENPL | 20.0290767 | 20.0125775 | 20.2977257 | 19.973242 | 20.170776 |
| P01019 | AGT | 17.9920054 | 17.8021973 | 17.9133766 | 17.9303843 | 17.8385198 |
| Q9H9C1 | VIPAS39 | 21.1848151 | 21.2108809 | 21.2847908 | 21.1008207 | 21.2435585 |
| Q8N3P4 | VPS8 | 19.3194759 | 19.4280438 | 19.3004947 | 19.1889667 | 19.2693818 |
| Q14692 | BMS1 | 20.7193507 | 20.8549235 | 20.8584008 | 20.7447615 | 20.7454048 |

| | | | | | | |
|---|---|---|---|---|---|---|
| Q7Z3D6 | DGLUCY | 20.4645489 | 20.3977428 | 20.5129762 | 20.4398543 | 20.5449474 |
| P29374 | ARID4A | 19.9337664 | 20.0393446 | 19.9909738 | 19.9848895 | 20.0207274 |
| P36543 | ATP6V1E1 | 23.2573821 | 23.2943788 | 23.306106 | 23.2684624 | 23.2687459 |
| P11117 | ACP2 | 20.6103884 | 20.7385004 | 20.7286018 | 20.5759987 | 20.749838 |
| Q96SI1 | KCTD15 | 19.1475765 | 19.2673686 | 19.2729953 | 19.179636 | 19.4294768 |
| Q4ADV7 | RIC1 | 17.177334 | 17.2036109 | 17.2122106 | 17.1470579 | 17.2897369 |
| O95402 | MED26 | 17.852945 | 18.1163434 | 18.0602698 | 17.9947844 | 18.1710681 |
| A5D8V6 | VPS37C | 18.1047172 | 18.1611515 | 18.033434 | 18.0995754 | 18.1883431 |
| Q8N488 | RYBP | 19.5041137 | 19.6082554 | 19.7074896 | 19.7171198 | 19.9047613 |
| Q659A1 | ICE2 | 17.5741692 | 17.6078515 | 17.7851381 | 17.6610364 | 17.6786676 |
| O60927 | PPP1R11 | 19.1085144 | 19.1921895 | 19.2672194 | 19.2126293 | 19.5069588 |
| O14594 | NCAN | 18.0774191 | NA | NA | 17.9118128 | NA |
| P48551 | IFNAR2 | 16.7412029 | 16.6202552 | 16.9530216 | 16.8884933 | 16.4312459 |
| P49795 | RGS19 | 19.8212511 | 19.8978034 | 19.9620628 | 19.8247728 | 20.0064955 |
| Q8NBT0 | POC1A | 17.9624901 | 18.0960891 | 18.3303394 | 18.2120155 | 18.3344693 |
| Q8N3U4 | STAG2 | 22.0695582 | 22.0415196 | 22.1623265 | 22.0262978 | 22.0902324 |
| Q9UGL1 | KDM5B | 18.5971653 | 18.7734124 | 18.8403569 | 18.5306196 | 18.7881929 |
| Q5HY98 | ZNF766 | 15.2999727 | 15.9395838 | 15.372799 | 15.4229351 | 15.1999547 |
| P35914 | HMGCL | 20.5898806 | 20.4564706 | 20.6793178 | 20.4335344 | 20.7077746 |
| Q13496 | MTM1 | 20.3353717 | 20.394853 | 20.4854438 | 20.3461212 | 20.3787351 |
| O15357 | INPPL1 | 20.8255669 | 20.9737352 | 20.9949512 | 20.8562695 | 20.9366128 |
| P38919 | EIF4A3 | 25.0061273 | 25.0526042 | 25.0290458 | 24.9088247 | 24.9080096 |
| Q71RC2 | LARP4 | 20.1851553 | 20.407104 | 20.2398113 | 20.2905584 | 20.3367084 |
| Q9BRJ2 | MRPL45 | 21.3439397 | 21.4067849 | 21.2501576 | 21.3524324 | 21.3421468 |
| Q14444 | CAPRIN1 | 24.0242798 | 24.1336027 | 23.8655943 | 23.9985478 | 24.0258613 |
| Q9BY43 | CHMP4A | 22.5956724 | 22.7936467 | 22.5319804 | 22.5751566 | 22.7682858 |
| Q99720 | SIGMAR1 | 22.0635607 | 22.0423171 | 22.267147 | 22.0679575 | 22.2044033 |
| P63151 | PPP2R2A | 23.0978677 | 23.1725893 | 23.1575184 | 23.0543556 | 23.1943429 |
| Q96N46 | TTC14 | 17.7589879 | 17.6711482 | 17.7047515 | 17.5544475 | 17.773181 |
| Q99741 | CDC6 | 24.0953089 | 24.344376 | 24.263225 | 23.8965696 | 24.2304191 |
| P57076 | CFAP298 | 21.1060964 | 21.2466854 | 21.2043122 | 21.1447917 | 21.1778629 |
| P35606 | COPB2 | 23.4923095 | 23.462413 | 23.4894896 | 23.423733 | 23.5211186 |
| P40306 | PSMB10 | 21.9032753 | 22.0319577 | 21.9089235 | 21.9101029 | 22.0680317 |
| Q1ED39 | KNOP1 | 19.149953 | 19.3666904 | 19.3409275 | 19.2884891 | 19.2255409 |
| O75140 | DEPDC5 | 18.4841913 | 18.7318786 | 18.6015979 | 18.5663424 | 18.6017674 |
| Q96E22 | NUS1 | 17.5906353 | 17.9584872 | 17.7011627 | 17.8480508 | 17.6178183 |
| Q9UPQ8 | DOLK | 11.5737604 | NA | NA | 11.6311833 | NA |
| P51587 | BRCA2 | 17.1563435 | 17.6670448 | 17.202869 | 17.3703904 | 17.4521062 |
| P32942 | ICAM3 | 19.7912198 | 20.2593038 | 20.0186753 | 19.7883761 | 20.2569842 |
| P47755 | CAPZA2 | 22.8873912 | 23.0194337 | 23.0025912 | 22.7984286 | 23.0093709 |
| P11217 | PYGM | 17.5619328 | NA | NA | 17.58451 | NA |
| Q8NB78 | KDM1B | 18.8158025 | 18.7063981 | 18.7454767 | 18.6351197 | 18.6849595 |

| | | | | | | |
|---|---|---|---|---|---|---|
| Q504Q3 | PAN2 | 19.789005 | 19.8342184 | 19.8892937 | 19.8256977 | 19.8490292 |
| P07199 | CENPB | 20.7580855 | 20.9322424 | 20.9474191 | 20.8074142 | 20.8503626 |
| Q6ZVK8 | NUDT18 | 20.1177517 | 20.0514419 | 20.1782215 | 20.1020798 | 20.1735891 |
| Q9BQ90 | KLHDC3 | 19.3781755 | 19.1729218 | 19.4067045 | 19.2534953 | 19.4277635 |
| Q7L5Y1 | ENOSF1 | 19.5913275 | 19.594895 | 19.6087056 | 19.7166705 | 19.8546264 |
| Q92504 | SLC39A7 | 19.2560838 | 19.0537246 | 19.3621796 | 19.2689166 | 18.6878103 |
| Q16829 | DUSP7 | 15.3879544 | 15.7340072 | 15.6607818 | 15.6337539 | 15.907803 |
| Q9NX61 | TMEM161A | 19.6225669 | 19.7614468 | 19.78205 | 19.7338957 | 19.7006379 |
| Q9H2J4 | PDCL3 | 21.932057 | 22.0494492 | 22.0286961 | 21.934252 | 22.0049874 |
| O00461 | GOLIM4 | 21.4450975 | 21.4545056 | 21.4783469 | 21.4003577 | 21.4041043 |
| Q9HC35 | EML4 | 20.9546651 | 21.0530569 | 21.0371644 | 20.9616212 | 21.0472403 |
| Q96JY6 | PDLIM2 | 22.1444201 | 22.2743249 | 22.2922552 | 22.2594392 | 22.3092619 |
| P54727 | RAD23B | 23.6575943 | 23.9097205 | 23.59303 | 23.649881 | 23.8239763 |
| Q9Y2Y0 | ARL2BP | 18.3781564 | 18.2533406 | 18.0272483 | 18.3239596 | 18.311391 |
| Q9HCH0 | NCKAP5L | 18.4630289 | 18.697909 | 18.5030743 | 18.5504259 | 18.7186294 |
| Q96F10 | SAT2 | NA | 18.0222055 | 17.7766997 | NA | 17.748441 |
| P02792 | FTL | 22.3864773 | 22.5430141 | 22.4759876 | 22.3478877 | 22.2683762 |
| Q9Y6N5 | SQOR | 24.162314 | 24.1571974 | 24.2674893 | 24.0841194 | 24.2090307 |
| Q9BYW2 | SETD2 | 19.3934551 | 19.4928841 | 19.5160538 | 19.3752553 | 19.5337292 |
| Q13405 | MRPL49 | 22.2861297 | 22.3904369 | 22.2077805 | 22.3369278 | 22.429886 |
| Q49AM3 | TTC31 | 15.9302887 | 16.2785268 | 15.986731 | 16.0030259 | 15.8542699 |
| O94827 | PLEKHG5 | 13.1694248 | NA | NA | 13.4527244 | NA |
| Q99832 | CCT7 | 25.8237222 | 25.8642774 | 25.7618544 | 25.6903797 | 25.7520549 |
| Q6ZNW5 | GDPGP1 | 16.6474434 | 16.8547211 | 16.7393759 | 16.6289669 | 16.9153249 |
| Q9H840 | GEMIN7 | 18.9104455 | 19.0064443 | 19.0935504 | 19.0112816 | 18.9635916 |
| P25774 | CTSS | 21.7234374 | 21.7570405 | 21.9395994 | 21.7020595 | 21.8794816 |
| Q9Y282 | ERGIC3 | 22.1160056 | 22.1489496 | 22.1073209 | 22.0465188 | 22.1948987 |
| Q8N8J7 | FAM241A | 17.9240577 | 17.7145582 | 17.9691386 | 18.1120569 | 17.9889364 |
| Q8NAV1 | PRPF38A | 20.9200709 | 20.9759728 | 20.8111112 | 20.871468 | 20.9060533 |
| Q32P44 | EML3 | 20.6321396 | 20.7017713 | 20.7446764 | 20.6489356 | 20.6673358 |
| O00203 | AP3B1 | 22.416078 | 22.4200147 | 22.4776854 | 22.302008 | 22.4303596 |
| Q96PE1 | ADGRA2 | 17.6688752 | 17.7888587 | 17.7487547 | 17.7537483 | 17.9192214 |
| Q9H5X1 | CIAO2A | 19.9761848 | 20.0994366 | 20.0941636 | 19.7418698 | 20.1384826 |
| P11388 | TOP2A | 22.4416458 | 22.5001854 | 22.5411826 | 22.357376 | 22.4905074 |
| Q9HAS3 | SLC28A3 | 15.9204732 | NA | NA | 16.2531608 | NA |
| Q9HB96 | FANCE | 16.1681161 | 16.0502988 | 16.2918844 | 16.0964259 | 16.3959962 |
| Q6QNY1 | BLOC1S2 | 19.752809 | 20.0325404 | 19.5686049 | 19.6901529 | 19.4807764 |
| Q96B96 | LDAF1 | 17.7758559 | 17.9880529 | 17.851424 | 17.8189666 | 17.7808878 |
| P11802 | CDK4 | 19.7794723 | 19.685719 | 19.9038384 | 19.8007145 | 19.957977 |
| Q15758 | SLC1A5 | 23.2429232 | 23.3402732 | 23.3890005 | 23.2098932 | 23.3274922 |
| P18433 | PTPRA | 18.9992284 | 19.0496007 | 19.0329501 | 19.0768377 | 19.1932193 |
| P55263 | ADK | 24.1375396 | 24.2225904 | 24.2527185 | 24.1547393 | 24.1529015 |

| | | | | | | |
|---|---|---|---|---|---|---|
| P49848 | TAF6 | 20.3167868 | 20.4936056 | 20.395925 | 20.3487471 | 20.4528737 |
| Q09472 | EP300 | 20.8268677 | 20.8982456 | 20.9784121 | 20.855663 | 20.9248656 |
| O15523 | DDX3Y | 21.3971691 | 21.4816151 | 21.3628537 | 21.3290012 | 21.4840369 |
| P50402 | EMD | 22.1916021 | 22.2985039 | 22.2863554 | 22.1724519 | 22.3509342 |
| Q96T51 | RUFY1 | 21.6581253 | 21.7868533 | 21.6188879 | 21.6661306 | 21.7888051 |
| Q8N3X1 | FNBP4 | 20.1750091 | 20.2571938 | 20.1376328 | 20.1751774 | 20.2843001 |
| Q96Q83 | ALKBH3 | 18.6464912 | 18.6497284 | 18.6612714 | 18.6911426 | 18.7676332 |
| Q9Y3D5 | MRPS18C | 20.4281443 | 20.4413526 | 20.3712501 | 20.4133374 | 20.4618057 |
| Q9NYU1 | UGGT2 | NA | 16.0836329 | 15.5242788 | NA | 15.8784201 |
| Q15742 | NAB2 | 18.289651 | 18.4325948 | 18.275001 | 18.1181809 | 18.4042901 |
| P51812 | RPS6KA3 | 22.9379025 | 23.0725005 | 23.0274575 | 22.8912977 | 23.0540915 |
| Q86TI2 | DPP9 | 22.0572898 | 22.2032368 | 22.1322997 | 22.1069309 | 22.1302183 |
| P19971 | TYMP | 23.4168226 | 23.4670708 | 23.2988846 | 23.340021 | 23.389334 |
| Q9BT22 | ALG1 | 21.9784475 | 22.0375107 | 22.0890285 | 21.948372 | 21.9581786 |
| Q8NFV4 | ABHD11 | 21.2723495 | 21.4043959 | 21.3857785 | 21.3493269 | 21.4358885 |
| Q9BU02 | THTPA | 19.6126193 | 19.693909 | 19.6528708 | 19.6002844 | 19.7336188 |
| Q96J01 | THOC3 | 21.9587949 | 22.1872631 | 22.0166872 | 21.9851196 | 22.1512592 |
| O43298 | ZBTB43 | 17.0955016 | 17.067185 | 16.9550765 | 16.8414039 | 16.889679 |
| Q8NA42 | ZNF383 | 15.1631671 | NA | NA | 15.3510131 | NA |
| Q96S66 | CLCC1 | 18.7717813 | 18.9034698 | 18.987506 | 18.7905657 | 18.7875117 |
| Q8IXI1 | RHOT2 | 21.564751 | 21.6408656 | 21.6562816 | 21.535985 | 21.5233619 |
| P53041 | PPP5C | 22.7790587 | 22.7672458 | 22.806868 | 22.6170858 | 22.7948396 |
| Q9BUL8 | PDCD10 | 22.0007892 | 22.1455608 | 21.9580819 | 21.9641523 | 21.9790247 |
| O43617 | TRAPPC3 | 22.4650891 | 22.5658567 | 22.3668521 | 22.3174328 | 22.5250669 |
| Q9BXP5 | SRRT | 23.6362394 | 23.6973725 | 23.6177708 | 23.5635036 | 23.6467851 |
| Q01469 | FABP5 | 25.551227 | 25.5704812 | 25.50603 | 25.5205933 | 25.6934268 |
| Q5THR3 | EFCAB6 | NA | 15.50255 | 15.7278775 | NA | 15.8930488 |
| P24863 | CCNC | 18.8523484 | 18.733419 | 19.0075124 | 18.6533381 | 18.9095677 |
| Q9Y4P8 | WIPI2 | 20.425473 | 20.5271136 | 20.350589 | 20.4145993 | 20.5406549 |
| Q9H4W6 | EBF3 | 19.1806599 | 19.0345229 | 19.0547262 | 19.1737055 | 19.0981341 |
| Q13426 | XRCC4 | 20.5968648 | 20.7075941 | 20.5941804 | 20.6510717 | 20.6599466 |
| Q99442 | SEC62 | 22.5814419 | 22.7803871 | 22.8161798 | 22.7094966 | 22.7387425 |
| Q76FK4 | NOL8 | 20.220176 | 20.2035248 | 20.1601029 | 20.1432297 | 20.2569437 |
| Q8NHP6 | MOSPD2 | 19.0058647 | 19.0725558 | 19.020339 | 19.0715403 | 19.0287642 |
| Q9BVL2 | NUP58 | 21.6626356 | 21.9300015 | 21.7963943 | 21.7302901 | 21.8987649 |
| A6PVC2 | TTLL8 | 19.4255373 | NA | NA | 19.5902627 | NA |
| P22090 | RPS4Y1 | 24.4209507 | 24.4308571 | 24.4582335 | 24.3613602 | 24.4101597 |
| Q8IV38 | ANKMY2 | 20.2511177 | 20.4723011 | 20.4017994 | 20.2651622 | 20.3948729 |
| Q3KPI0 | CEACAM21 | 15.8555091 | 15.7592764 | 15.9039346 | 15.2839711 | 15.3041903 |
| P61960 | UFM1 | 22.4192272 | 22.5608186 | 22.4456062 | 22.4295435 | 22.7195719 |
| Q9BV68 | RNF126 | 21.0191238 | 21.1220959 | 21.0218188 | 20.9422385 | 21.1238437 |
| Q12769 | NUP160 | 22.5953667 | 22.5502299 | 22.6479049 | 22.4965432 | 22.5934743 |

| | | | | | | |
|---|---|---|---|---|---|---|
| Q9HAT2 | SIAE | 20.237888 | 20.2647533 | 20.3483641 | 20.0785983 | 20.3183716 |
| A1A5D9 | BICDL2 | 17.4930607 | NA | NA | 17.2208322 | NA |
| P02686 | MBP | 20.5489935 | 20.5916456 | 20.8463699 | 20.5705977 | 20.5878857 |
| L0R8F8 | MIEF1 | 19.6473806 | 19.9365352 | 19.7590876 | 19.8110066 | 19.6275506 |
| Q5SNT2 | TMEM201 | 20.0049904 | 20.0041707 | 19.9762111 | 19.8965299 | 19.911229 |
| O43396 | TXNL1 | 23.6560639 | 23.5265589 | 23.5914591 | 23.5100464 | 23.7373916 |
| Q8WUK0 | PTPMT1 | 19.4558621 | 19.7474355 | 19.7396905 | 19.7231245 | 19.561574 |
| P37235 | HPCAL1 | 23.1973056 | 23.2575023 | 23.3195587 | 23.1362746 | 23.2923919 |
| O95639 | CPSF4 | 20.4220933 | 20.5040202 | 20.6359025 | 20.4497751 | 20.5297498 |
| Q13867 | BLMH | 22.2563369 | 22.3122297 | 22.367088 | 22.24604 | 22.3450629 |
| Q8NCE2 | MTMR14 | 20.6829072 | 20.7074497 | 20.7820773 | 20.6882255 | 20.7116882 |
| Q3YEC7 | RABL6 | 21.107853 | 21.1099755 | 21.235056 | 21.0784984 | 21.1204505 |
| Q53H47 | SETMAR | 20.10281 | 20.2567947 | 20.3017963 | 20.2637117 | 20.2213702 |
| Q9Y6B6 | SAR1B | 21.0062775 | 21.0547662 | 21.1693527 | 21.0834382 | 21.2147042 |
| O60711 | LPXN | 21.6662252 | 21.6505747 | 21.8734114 | 21.793446 | 21.883343 |
| Q6XZF7 | DNMBP | 18.5190832 | 18.2710969 | 18.4730583 | 18.7346456 | 18.6417846 |
| Q9UI30 | TRMT112 | 21.6548248 | 21.6601492 | 21.7222367 | 21.7304877 | 21.9035996 |
| Q9NX55 | HYPK | 21.8702686 | 22.0654685 | 21.805885 | 21.9608933 | 21.9808285 |
| Q02487 | DSC2 | 12.8354743 | NA | NA | 12.3332131 | NA |
| O75891 | ALDH1L1 | NA | 11.4548619 | 11.9918415 | NA | 11.9725266 |
| Q8N4B1 | PHETA1 | 17.5475695 | 17.566774 | 17.7358477 | 17.5465808 | 17.5059383 |
| P04920 | SLC4A2 | 20.6410239 | 20.7123854 | 20.7875347 | 20.6309325 | 20.7148253 |
| Q6PD62 | CTR9 | 22.3691121 | 22.4871817 | 22.4776669 | 22.4425004 | 22.5181237 |
| Q8NDB2 | BANK1 | NA | 16.2659257 | 15.400785 | NA | 15.4534121 |
| Q96HD1 | CRELD1 | 18.1645713 | 18.4563904 | 18.4164136 | 18.4084645 | 18.3532163 |
| P02774 | GC | 18.2880159 | 18.0616238 | 18.123648 | 18.1089704 | 18.0150358 |
| Q9Y4J8 | DTNA | 19.5201018 | 19.624551 | 19.5979002 | 19.579161 | 19.5397163 |
| Q9Y2V7 | COG6 | 19.8855421 | 20.0650453 | 19.9882626 | 19.9596867 | 20.0809667 |
| Q9NR56 | MBNL1 | 23.2937394 | 23.3236827 | 23.3853971 | 23.3063941 | 23.406719 |
| Q9UIA0 | CYTH4 | 20.0678185 | 20.2001308 | 20.1834001 | 20.1871005 | 20.1589721 |
| Q9UPW0 | FOXJ3 | 17.8207667 | 17.9449109 | 17.8291037 | 17.9302188 | 18.0847848 |
| P35659 | DEK | 24.022678 | 24.0835973 | 24.0574356 | 23.976788 | 24.018701 |
| P35222 | CTNNB1 | 15.6653933 | 15.8051163 | 16.1531419 | 16.1157376 | 15.8556333 |
| P60604 | UBE2G2 | 20.3147222 | 20.3300805 | 20.5627144 | 20.2633361 | 20.3355556 |
| P51692 | STAT5B | 19.9159671 | 19.9842366 | 20.0814864 | 19.8642116 | 20.0902176 |
| Q9UH65 | SWAP70 | 20.9737536 | 21.0693711 | 21.0681818 | 20.9771675 | 21.0856485 |
| O60447 | EVI5 | 17.14337 | 17.2546598 | 17.2667003 | 17.2237032 | 17.2076217 |
| P17947 | SPI1 | 20.9981503 | 21.3468137 | 21.1277998 | 21.408323 | 21.3498179 |
| Q9H9J4 | USP42 | 15.995064 | 15.9432028 | 15.808831 | 15.5705502 | 15.7220441 |
| Q76N32 | CEP68 | NA | 14.8470558 | 14.6089094 | NA | 14.7414937 |
| Q96DT6 | ATG4C | 18.708161 | 18.7370472 | 18.7705029 | 18.7227797 | 18.7645267 |
| P15509 | CSF2RA | 18.312242 | 18.3656401 | 18.1795679 | 18.2895803 | 18.051127 |

| | | | | | | |
|---|---|---|---|---|---|---|
| Q8WUD1 | RAB2B | 19.26753 | 19.439882 | 19.424467 | 19.2331154 | 19.4050915 |
| P07741 | APRT | 25.0605879 | 25.0567743 | 24.9254341 | 24.9061806 | 25.0000029 |
| P30837 | ALDH1B1 | 22.2342231 | 22.251752 | 22.3377971 | 22.1943432 | 22.3118381 |
| Q99570 | PIK3R4 | 20.8598107 | 20.8882653 | 20.941301 | 20.8611124 | 20.9140061 |
| Q92620 | DHX38 | 21.0587112 | 21.1552778 | 21.1656394 | 21.055409 | 21.2009611 |
| Q8NG08 | HELB | 18.7989704 | 18.8458819 | 18.9747102 | 18.8682003 | 18.9616929 |
| Q13571 | LAPTM5 | 22.0315408 | 21.8761541 | 21.9486522 | 21.9699376 | 21.818822 |
| Q9Y3D7 | PAM16 | 21.6460555 | 21.7074122 | 21.9238727 | 21.7348776 | 21.8293616 |
| O94868 | FCHSD2 | 19.8536457 | 19.8327069 | 19.8187946 | 19.8467832 | 19.8743426 |
| Q13148 | TARDBP | 23.8582423 | 23.9330409 | 23.8395762 | 23.8168117 | 23.9201224 |
| Q8WUM9 | SLC20A1 | 20.4546671 | 20.4417612 | 20.3250762 | 20.5006062 | 20.4973557 |
| Q86SE9 | PCGF5 | 16.4678149 | 16.4095745 | 16.3995085 | 16.3894543 | 16.3283575 |
| P41134 | ID1 | NA | 14.0362095 | 14.0930768 | NA | 14.3407475 |
| Q9BUM1 | G6PC3 | 18.5734119 | 18.1721675 | 18.5339311 | 18.5598851 | 18.4807539 |
| Q92889 | ERCC4 | 19.2473556 | 19.3027953 | 19.4510902 | 19.3368579 | 19.3612913 |
| P19793 | RXRA | 20.4129017 | 20.5822761 | 20.6039605 | 20.511219 | 20.5922499 |
| Q9UK61 | TASOR | 20.6028969 | 20.7715828 | 20.7428043 | 20.6287662 | 20.7836964 |
| Q14050 | COL9A3 | 14.8918514 | 14.8394381 | 15.1576418 | 15.2215006 | 15.2079681 |
| Q9UJJ9 | GNPTG | 18.6982423 | 18.5654136 | 18.4946651 | 18.6387192 | 18.6915456 |
| Q96RP9 | GFM1 | 21.5465438 | 21.6456295 | 21.5862081 | 21.5988187 | 21.6734777 |
| Q8WXG6 | MADD | 19.1757999 | 19.2322863 | 19.2507504 | 19.1790404 | 19.2794298 |
| Q9H582 | ZNF644 | 18.7686758 | 18.8528665 | 18.8962951 | 18.8151543 | 18.8413252 |
| Q2PPJ7 | RALGAPA2 | 18.2831644 | 18.3781862 | 18.3872857 | 18.295227 | 18.4505232 |
| Q17R89 | ARHGAP44 | 18.204475 | 18.1741539 | 18.4346313 | 18.2974006 | 18.4814721 |
| P06753 | TPM3 | 25.2334104 | 25.4050585 | 25.0163761 | 25.1961417 | 25.4029955 |
| O14933 | UBE2L6 | 16.9676093 | 16.9182064 | 17.1814619 | 17.1383218 | 17.2014796 |
| P56545 | CTBP2 | 24.0482673 | 24.1845009 | 24.0932193 | 24.0943404 | 24.2531424 |
| Q9H2Y7 | ZNF106 | 19.986182 | 20.0811839 | 20.0033618 | 19.8526323 | 19.996829 |
| Q6VMQ6 | ATF7IP | 20.2311212 | 20.2883716 | 20.1343481 | 20.2015782 | 20.2915566 |
| Q9Y2E5 | MAN2B2 | 20.4878303 | 20.3792379 | 20.4643902 | 20.410769 | 20.5848888 |
| Q9HCL2 | GPAM | 19.3594097 | 19.7745406 | 20.0173466 | 19.5558868 | 19.7032359 |
| P18054 | ALOX12 | 18.1368628 | 17.7326017 | 17.8301009 | 18.0305492 | 17.9454533 |
| Q53FV1 | ORMDL2 | 20.0306352 | 20.0080848 | 20.1826749 | 20.063747 | 19.9338054 |
| Q15361 | TTF1 | 18.1282428 | 18.5286758 | 18.3688926 | 18.2795292 | 18.2571215 |
| Q9BZC7 | ABCA2 | 18.8219453 | 18.6849668 | 18.8093676 | 18.7895491 | 18.8026579 |
| P46087 | NOP2 | 23.1614038 | 23.1336526 | 23.1798311 | 23.1210959 | 23.1221132 |
| Q9BVG4 | PBDC1 | 21.9594006 | 22.0335177 | 21.8260319 | 21.850573 | 22.0318006 |
| P63220 | RPS21 | 25.5991254 | 25.7044809 | 25.5718278 | 25.5887394 | 25.6427484 |
| Q86WW8 | COA5 | 19.9851168 | 20.1707923 | 20.2771575 | 20.0806262 | 20.3101109 |
| Q9UBN6 | TNFRSF10D | 16.895007 | 17.497493 | 17.7411998 | 17.4084889 | 16.8234383 |
| Q99607 | ELF4 | 16.3190055 | 16.303694 | 16.3305165 | 16.1976877 | 16.8904974 |
| O60861 | GAS7 | 17.3866761 | 17.10697 | 17.5911059 | 17.4815113 | 17.480861 |

| | | | | | | |
|---|---|---|---|---|---|---|
| Q71RG4 | TMUB2 | NA | 16.3459021 | 16.5259199 | NA | 16.569417 |
| O75884 | RBBP9 | 18.4828246 | 18.4083843 | 18.773775 | 18.4517281 | 18.6377964 |
| Q96AX2 | RAB37 | 19.1710027 | 19.1768771 | 19.3246147 | 19.0632529 | 19.1951663 |
| Q86XN6 | ZNF761 | 18.768403 | 18.8906397 | 19.1026522 | 18.8714596 | 18.7801008 |
| P49593 | PPM1F | 20.6498229 | 20.6381964 | 20.6494265 | 20.603201 | 20.6764843 |
| Q96MX3 | ZNF48 | 18.7770789 | 18.7705143 | 18.7846634 | 18.6151254 | 18.6645722 |
| P46783 | RPS10 | 26.3552468 | 26.3402119 | 26.4743425 | 26.2841938 | 26.4176199 |
| Q06265 | EXOSC9 | 21.117031 | 21.0840913 | 21.1061154 | 20.9046952 | 21.1172838 |
| Q9UGJ1 | TUBGCP4 | 19.6730118 | 19.7207829 | 19.7616777 | 19.5778125 | 19.6700526 |
| P29144 | TPP2 | 23.4546855 | 23.4681527 | 23.4513744 | 23.3545505 | 23.500712 |
| A2RUS2 | DENND3 | 22.0739382 | 22.1384727 | 22.1102636 | 21.9944429 | 22.1163796 |
| Q9Y666 | SLC12A7 | 20.1082322 | 20.0237877 | 20.1652131 | 20.101065 | 20.0719368 |
| Q2NKX8 | ERCC6L | 19.4530258 | 19.5282428 | 19.5115016 | 19.4472471 | 19.5944599 |
| Q8TAD4 | SLC30A5 | 19.2802757 | 19.424143 | 19.3617865 | 19.4167314 | 19.4684612 |
| Q86WR0 | CCDC25 | 21.968259 | 21.9895943 | 22.0046219 | 21.9255738 | 21.9832442 |
| Q9Y388 | RBMX2 | 19.1385295 | 19.1852077 | 19.2081924 | 19.1681072 | 19.2198335 |
| O75298 | RTN2 | 19.6015526 | 19.802539 | 19.9238 | 19.6985183 | 19.8698095 |
| Q9Y4H2 | IRS2 | 13.9803766 | NA | NA | 14.1842818 | NA |
| Q5T6F2 | UBAP2 | 20.9135752 | 20.96998 | 20.770901 | 20.9472834 | 21.0759615 |
| Q14774 | HLX | 15.9579758 | 16.2874866 | 16.2622684 | 16.277922 | 16.2239173 |
| Q8N1P7 | CRYBG2 | 17.5049997 | NA | NA | 17.6549403 | NA |
| P07333 | CSF1R | 20.8190838 | 20.7552074 | 20.6952612 | 20.7073606 | 20.6427032 |
| Q9UK59 | DBR1 | 20.8490386 | 20.9564439 | 21.0248774 | 20.9038068 | 20.9363073 |
| Q14191 | WRN | 19.6326135 | 19.8010503 | 19.7460159 | 19.7586919 | 19.7551881 |
| Q8IUI8 | CRLF3 | 18.511025 | 18.7001267 | 18.5593916 | 18.4515997 | 18.5348972 |
| Q5TA31 | RNF187 | 14.8364022 | 15.2589599 | 15.5346353 | 15.581624 | 15.5414572 |
| Q6P161 | MRPL54 | 20.6748396 | 20.7074933 | 20.7222409 | 20.699694 | 20.887236 |
| Q05086 | UBE3A | 22.3901108 | 22.4043702 | 22.4239166 | 22.3566005 | 22.5164679 |
| Q07812 | BAX | 23.2387409 | 23.3798982 | 23.3242139 | 23.2304933 | 23.3135601 |
| Q9UHB7 | AFF4 | 18.0379993 | 17.9675667 | 18.1661131 | 18.0860627 | 18.0348842 |
| Q6AI12 | ANKRD40 | 15.7848031 | 16.1211286 | 16.3113174 | 16.4844605 | 16.2195507 |
| Q9Y2R4 | DDX52 | 20.7202496 | 20.7812598 | 20.8679088 | 20.6887461 | 20.7693781 |
| Q9NZJ4 | SACS | 20.4211786 | 20.5334711 | 20.5409731 | 20.5329681 | 20.5541037 |
| P28065 | PSMB9 | 20.433951 | 20.38867 | 20.4498846 | 20.4849861 | 20.5719619 |
| Q9NSD4 | ZNF275 | 15.5039467 | 15.2123417 | 15.7378854 | 15.5736779 | 15.3504009 |
| Q3KNV8 | PCGF3 | 16.5236492 | 16.692823 | 16.4324411 | 16.5857377 | 16.6636927 |
| P28340 | POLD1 | 23.2099421 | 23.2999136 | 23.2594369 | 23.1372756 | 23.2308311 |
| A0AVF1 | IFT56 | 17.3676652 | 17.1296188 | 17.5252104 | 17.3701612 | 17.3059595 |
| P43250 | GRK6 | 19.5475657 | 19.7065845 | 19.7837787 | 19.560972 | 19.6560549 |
| Q8WU76 | SCFD2 | 19.882337 | 19.8314181 | 19.9149108 | 19.8418149 | 20.0116256 |
| O60476 | MAN1A2 | 19.1859987 | 19.2256612 | 19.4109065 | 19.1895544 | 19.1720852 |
| Q6P158 | DHX57 | 18.6608132 | 18.865317 | 18.8529566 | 18.7712786 | 18.8736641 |

| | | | | | | |
|---|---|---|---|---|---|---|
| Q96JB2 | COG3 | 19.8212553 | 19.8077349 | 19.8877549 | 19.8275307 | 19.8847036 |
| O15084 | ANKRD28 | 19.2417642 | 19.2624686 | 19.2967942 | 19.3098325 | 19.4513281 |
| Q8WUF8 | FAM172A | 18.7549736 | 18.8955011 | 18.8445867 | 18.8036145 | 18.9684606 |
| Q9H0X4 | FAM234A | 19.1854763 | 19.2642924 | 19.0921898 | 19.1093807 | 19.0626702 |
| P59768 | GNG2 | 16.8769568 | 17.0628455 | 17.3006702 | 17.1309496 | 17.0902704 |
| Q9NXR5 | ANKRD10 | 18.2514888 | 18.6309037 | 18.5168957 | 18.2180112 | 18.5787163 |
| P62266 | RPS23 | 26.4490239 | 26.5021337 | 26.5204568 | 26.3261567 | 26.4430929 |
| Q13601 | KRR1 | 21.4111229 | 21.4753598 | 21.504337 | 21.3637677 | 21.4795744 |
| O60762 | DPM1 | 22.6904556 | 22.7812057 | 22.8056465 | 22.7180153 | 22.8136371 |
| Q9NZC3 | GDE1 | 20.1069824 | 20.4613109 | 20.3744841 | 20.2612151 | 20.5172889 |
| Q12979 | ABR | 21.8640501 | 21.9177048 | 21.9197932 | 21.855705 | 21.9564861 |
| Q2TAC6 | KIF19 | 24.3431624 | NA | NA | 23.3803981 | NA |
| Q5VTQ0 | TTC39B | NA | 13.7287272 | 13.5904197 | NA | 13.5092983 |
| Q9HBI1 | PARVB | 18.9850978 | 19.0406579 | 19.0858965 | 18.8525793 | 19.116559 |
| Q6YHU6 | THADA | 19.900148 | 19.9621584 | 20.0451419 | 19.8653873 | 19.9922946 |
| Q96KM6 | ZNF512B | 21.7747771 | 21.9061681 | 21.8209441 | 21.63514 | 21.7546941 |
| Q9Y312 | AAR2 | 21.2197296 | 21.2976788 | 21.2693662 | 21.1870717 | 21.3321095 |
| Q9UH03 | SEPTIN3 | 17.8864645 | 17.847716 | 17.872262 | 17.8503296 | 18.2000265 |
| Q9Y6H1 | CHCHD2 | 22.087595 | 22.3429722 | 22.0625527 | 22.1074863 | 22.3715318 |
| P27482 | CALML3 | NA | 16.5764764 | 16.7009267 | NA | 17.0856924 |
| Q9BSJ6 | PIMREG | 15.4464322 | 15.475293 | 15.5070822 | 15.5116911 | 15.3368714 |
| P47914 | RPL29 | 25.5284011 | 25.441383 | 25.8096659 | 25.4467033 | 25.3074351 |
| Q9NZ71 | RTEL1 | 18.314382 | 18.3649808 | 18.475981 | 18.454727 | 18.4493701 |
| Q9H0U3 | MAGT1 | 22.8473138 | 22.9658824 | 22.8961301 | 22.813705 | 22.7053876 |
| O00471 | EXOC5 | 20.9470546 | 20.9835672 | 21.0139743 | 20.8726045 | 21.0090431 |
| Q99417 | MYCBP | 22.0298022 | 22.2278913 | 22.0441339 | 22.0536345 | 22.1171753 |
| Q9HBL0 | TNS1 | 20.5668552 | 20.7447506 | 20.5945169 | 20.5987398 | 20.7066067 |
| P24534 | EEF1B2 | 25.6919732 | 25.8437399 | 25.5203807 | 25.6665914 | 25.8426049 |
| Q7Z4F1 | LRP10 | 16.6397043 | 16.3425783 | 16.4773345 | 16.5366383 | 16.3792357 |
| Q9UHA3 | RSL24D1 | 21.1193454 | 21.1199491 | 21.2643609 | 21.1320983 | 21.0356591 |
| P05107 | ITGB2 | 24.0927225 | 24.3009161 | 24.2273805 | 24.0109118 | 24.2068309 |
| Q8NFF5 | FLAD1 | 20.0351107 | 19.98058 | 20.070585 | 19.9753912 | 20.0814712 |
| Q8N7Z5 | ANKRD31 | 17.9453513 | NA | NA | 17.9744595 | NA |
| Q96E11 | MRRF | 21.0585397 | 21.2335876 | 21.1433745 | 21.0415427 | 21.2212623 |
| Q9NR09 | BIRC6 | 21.929561 | 21.9787967 | 21.9904025 | 21.9125866 | 21.9706768 |
| Q9BY44 | EIF2A | 22.1920349 | 22.2034303 | 22.2865185 | 22.1805999 | 22.2781691 |
| Q92797 | SYMPK | 22.288522 | 22.2960093 | 22.3095257 | 22.2035132 | 22.2690391 |
| Q8NBJ7 | SUMF2 | 21.1647357 | 21.3832183 | 21.347353 | 21.2442259 | 21.2204613 |
| Q7KZN9 | COX15 | 20.2352224 | 20.4219562 | 20.3897801 | 20.1475717 | 20.418907 |
| P02787 | TF | 22.4853088 | 21.9644546 | 22.0524142 | 22.2814136 | 22.0870119 |
| Q9BPW8 | NIPSNAP1 | 20.9501227 | 20.9690481 | 20.9003244 | 20.8781346 | 20.9518843 |
| Q8N7X0 | ADGB | NA | 15.8188619 | 15.9249556 | NA | 16.093249 |

| | | | | | | |
|---|---|---|---|---|---|---|
| Q9Y3E0 | GOLT1B | 21.0275244 | 21.3875945 | 21.2523397 | 21.045237 | 21.1353902 |
| P30085 | CMPK1 | 23.4130594 | 23.431948 | 23.3583486 | 23.315219 | 23.4311417 |
| Q9Y6I4 | USP3 | 20.1636808 | 20.1324783 | 20.124404 | 20.1260594 | 20.0740079 |
| P29353 | SHC1 | 20.4260771 | 20.593194 | 20.3267408 | 20.4205084 | 20.5387981 |
| O15033 | AREL1 | 14.8768333 | 15.3416671 | 15.5447727 | 14.7307999 | 15.3945924 |
| A4D1E9 | GTPBP10 | 20.6236057 | 20.560087 | 20.4953006 | 20.6299908 | 20.6016511 |
| O75177 | SS18L1 | 15.0928704 | 14.9199489 | 15.2729252 | 15.4188296 | 15.3246117 |
| A0A1W2PQL4 | ZNF722 | 18.1421238 | 17.9462216 | 18.294559 | 17.7495431 | 17.6260131 |
| Q5T6S3 | PHF19 | 18.429329 | 18.3742792 | 18.744747 | 18.5049672 | 18.5915529 |
| Q92882 | OSTF1 | 23.453918 | 23.4974241 | 23.4389245 | 23.3279408 | 23.4767173 |
| Q96NL6 | SCLT1 | 24.281657 | 24.5781996 | 24.4715435 | 24.2513509 | 24.2596082 |
| Q9BTY2 | FUCA2 | 22.0167838 | 21.9675946 | 22.1028217 | 21.9005725 | 22.1266617 |
| P61106 | RAB14 | 24.9724625 | 25.0348091 | 25.0733161 | 24.9068433 | 25.0183612 |
| P15880 | RPS2 | 26.5825318 | 26.5893892 | 26.5449994 | 26.4797409 | 26.5375066 |
| Q9Y265 | RUVBL1 | 23.7631234 | 23.7832396 | 23.776226 | 23.6961235 | 23.8111004 |
| O00468 | AGRN | 19.2515058 | 19.3267555 | 19.2360827 | 19.2244181 | 19.3441406 |
| P21281 | ATP6V1B2 | 23.4915532 | 23.6532109 | 23.4666017 | 23.493574 | 23.5695226 |
| Q9UQ84 | EXO1 | 19.5983827 | 19.5397995 | 19.6131264 | 19.5280318 | 19.4554787 |
| Q9NWS0 | PIH1D1 | 19.9106814 | 20.0913276 | 19.9659301 | 20.0418587 | 20.0973092 |
| Q9H6Y2 | WDR55 | 20.5284585 | 20.6131057 | 20.5053803 | 20.4688257 | 20.6712396 |
| P0DPB5 | POLR1D | 20.1377866 | 20.1045137 | 20.2287597 | 20.1036882 | 20.0551405 |
| Q9NWV8 | BABAM1 | 21.6662647 | 21.7591323 | 21.5652293 | 21.6537978 | 21.8608603 |
| Q5SR56 | MFSD14B | 16.7346634 | 16.404427 | 16.7237728 | 17.0175121 | 16.8723855 |
| O60293 | ZFC3H1 | 18.3621366 | 18.4408978 | 18.5648852 | 18.309965 | 18.5676383 |
| Q96BM9 | ARL8A | 18.5877424 | 18.6454412 | 18.9459052 | 18.5994695 | 19.1656912 |
| Q9UL25 | RAB21 | 22.3758852 | 22.6000204 | 22.5460549 | 22.4682071 | 22.4327306 |
| Q8N1G0 | ZNF687 | 18.7307584 | 18.6732818 | 18.8839178 | 18.6132734 | 18.7064214 |
| Q14181 | POLA2 | 20.841528 | 20.8737 | 20.8222109 | 20.7123542 | 20.8937769 |
| Q5I0G3 | MDH1B | 19.9296096 | 20.7058363 | 20.5714897 | 19.8152109 | 20.3982874 |
| P52790 | HK3 | 19.1496417 | 19.2255066 | 19.3799049 | 19.2007468 | 19.2172091 |
| O43402 | EMC8 | 22.3733531 | 22.659721 | 22.5099986 | 22.537985 | 22.7018334 |
| Q9H3Y8 | PPDPF | 16.8902692 | 17.0389603 | 16.384964 | 16.8255809 | 16.9061656 |
| Q92536 | SLC7A6 | 18.5633522 | 18.5724979 | 18.8757858 | 18.6069743 | 18.7617344 |
| P28074 | PSMB5 | 23.2426065 | 23.1450705 | 23.2054138 | 23.1570988 | 23.136609 |
| Q96BI1 | SLC22A18 | 19.6150309 | 19.8923988 | 20.0234103 | 19.7162182 | 20.028614 |
| Q9Y3C8 | UFC1 | 23.1912598 | 23.218774 | 23.3966963 | 23.1189518 | 23.3118384 |
| Q03169 | TNFAIP2 | 20.6644708 | 20.678771 | 20.7155731 | 20.6205556 | 20.7155549 |
| Q9UPZ3 | HPS5 | 18.3539345 | 18.3339315 | 18.3777899 | 18.3710489 | 18.5817256 |
| Q14689 | DIP2A | 17.4660468 | 17.4596047 | 17.6522836 | 17.5652433 | 17.6348991 |
| O75694 | NUP155 | 22.4581484 | 22.4397911 | 22.4956294 | 22.3086949 | 22.4536639 |
| O60725 | ICMT | 18.4503829 | 18.8381624 | 18.5257653 | 18.0610745 | 18.5580175 |
| O96028 | NSD2 | 19.5700582 | 19.6229856 | 19.6839215 | 19.648463 | 19.7507289 |

| | | | | | | |
|---|---|---|---|---|---|---|
| Q9NWZ3 | IRAK4 | 20.4349041 | 20.5808866 | 20.6298963 | 20.5347636 | 20.7006701 |
| P55197 | MLLT10 | 17.9979739 | 18.0183452 | 17.9762458 | 17.9780325 | 17.9273793 |
| Q9H9L3 | ISG20L2 | 19.2486393 | 19.5121504 | 19.5127618 | 19.4604732 | 19.5719522 |
| Q9H5I1 | SUV39H2 | 17.8125885 | 17.6555921 | 17.9562472 | 17.8906174 | 17.7801524 |
| Q15125 | EBP | 21.3862258 | 21.0626464 | 20.8854414 | 21.1987535 | 21.0535579 |
| Q8WUN7 | UBTD2 | 16.4531131 | 16.6442781 | 16.6605888 | 16.701009 | 16.4799425 |
| Q9NVF7 | FBXO28 | 18.6119192 | 18.6493372 | 18.6510667 | 18.6509133 | 18.5876566 |
| O14548 | COX7A2L | 21.0340426 | 21.2098995 | 21.293593 | 21.1816415 | 21.1410262 |
| Q14165 | MLEC | 23.5013649 | 23.4810926 | 23.5683171 | 23.3614306 | 23.5476333 |
| O15027 | SEC16A | 20.9482893 | 21.0594244 | 21.0260186 | 20.9795078 | 21.1047142 |
| P55957 | BID | 21.1511532 | 21.1485255 | 21.048641 | 21.0677893 | 21.2445604 |
| P0CAP2 | POLR2M | 19.0053734 | 19.1005436 | 19.0157058 | 19.1508311 | 19.1094103 |
| Q9NQ86 | TRIM36 | 13.5315338 | 13.6264743 | 13.6314283 | 13.6123971 | 13.901056 |
| Q49B96 | COX19 | 18.303353 | 18.6181124 | 18.4343702 | 18.5534447 | 18.4984235 |
| Q9H9Y2 | RPF1 | 20.1939265 | 20.2669086 | 20.3188721 | 20.2108193 | 20.1840675 |
| Q9BWW4 | SSBP3 | 14.7077872 | 14.7374279 | 14.308965 | 14.4358813 | 14.7524287 |
| O43813 | LANCL1 | 22.0341549 | 22.083445 | 22.0626626 | 22.0437981 | 22.0702782 |
| O00458 | IFRD1 | 16.4460893 | NA | NA | 16.3825381 | NA |
| Q7Z460 | CLASP1 | 20.5761526 | 20.6539236 | 20.7070096 | 20.6583072 | 20.700514 |
| P43378 | PTPN9 | 20.2395427 | 20.2705171 | 20.3192957 | 20.1947054 | 20.2788157 |
| P30041 | PRDX6 | 24.8092951 | 24.9115794 | 24.8353069 | 24.7686105 | 24.8671128 |
| Q06547 | GABPB1 | 19.257482 | 19.5439078 | 19.352709 | 19.3787228 | 19.6416355 |
| Q9H1Y3 | OPN3 | 17.1165544 | 16.668799 | 16.8368082 | 16.9787706 | 17.01571 |
| Q8N3Y1 | FBXW8 | 21.3938498 | 21.6717176 | 21.529503 | 21.5752387 | 21.5445959 |
| P52888 | THOP1 | 22.9428108 | 22.9787291 | 22.9747821 | 22.8559523 | 22.9133612 |
| P06865 | HEXA | 22.6058474 | 22.6072632 | 22.6073656 | 22.5967905 | 22.6940852 |
| Q9BRN9 | TM2D3 | 16.45388 | 15.8091099 | 16.0460064 | 15.4314677 | 15.8378745 |
| O95359 | TACC2 | 15.5390215 | 15.5246975 | 15.2557773 | 15.6096651 | 15.3399796 |
| O60333 | KIF1B | 12.7346907 | NA | NA | 12.7510948 | NA |
| A8MPS7 | YDJC | 16.7522284 | 16.7827528 | 16.8314194 | 16.7970828 | 16.6675161 |
| Q6UWW8 | CES3 | 15.8859843 | NA | NA | 15.8693262 | NA |
| P30273 | FCER1G | 21.3470158 | 21.4050278 | 21.477241 | 21.1563152 | 21.4012947 |
| Q9NRX6 | TMEM167B | 19.9900092 | 20.1221741 | 20.0560207 | 19.9809666 | 20.1048402 |
| Q9UG22 | GIMAP2 | 18.4892487 | 18.378238 | 18.6998776 | 17.9460626 | 18.4446278 |
| P12931 | SRC | 19.7391263 | 19.7900441 | 20.0690752 | 19.8065767 | 19.937178 |
| Q96IP4 | TENT5A | 15.7156427 | 16.1292043 | 15.75497 | 15.8434587 | 15.8708578 |
| Q13227 | GPS2 | 19.9806084 | 20.0924162 | 20.1447257 | 19.908925 | 20.0540604 |
| Q9Y679 | AUP1 | 21.7823343 | 21.850531 | 21.7110251 | 21.6142777 | 21.7323662 |
| Q9P2M4 | TBC1D14 | 18.0338954 | 18.0797446 | 18.0216883 | 18.131264 | 18.0198625 |
| Q99467 | CD180 | 21.6109011 | 21.6597914 | 21.6699573 | 21.6095551 | 21.6467203 |
| Q9BSB4 | ATG101 | 19.187687 | 19.2740832 | 19.3395521 | 19.0299506 | 19.3156192 |
| P51572 | BCAP31 | 25.0654324 | 25.091216 | 25.0622589 | 24.9501601 | 24.8416666 |

| | | | | | | |
|---|---|---|---|---|---|---|
| Q9H2W6 | MRPL46 | 21.8993318 | 21.9550417 | 21.8563847 | 21.8136056 | 21.8214613 |
| Q9H0S4 | DDX47 | 21.2384102 | 21.2968725 | 21.3623907 | 21.2344396 | 21.3539088 |
| Q6P1L8 | MRPL14 | 21.7631644 | 21.875065 | 21.9057218 | 21.8055484 | 21.5887253 |
| O00459 | PIK3R2 | 19.7543081 | 19.895584 | 19.7973638 | 19.7629511 | 19.8477794 |
| Q9BTE3 | MCMBP | 21.9605628 | 21.9519223 | 22.0019624 | 21.9136374 | 22.0272681 |
| Q15796 | SMAD2 | 21.0134908 | 20.9635456 | 20.9835414 | 20.8258895 | 20.879578 |
| O75935 | DCTN3 | 21.4031021 | 21.4475278 | 21.3825984 | 21.3761737 | 21.302449 |
| O00116 | AGPS | 22.1475909 | 22.2745789 | 22.2530801 | 22.1149423 | 22.2189175 |
| Q92616 | GCN1 | 23.6647856 | 23.7256892 | 23.7138454 | 23.6022885 | 23.6854217 |
| Q96DV4 | MRPL38 | 22.1309583 | 22.0583699 | 22.1663347 | 22.1022549 | 22.2248405 |
| Q9H257 | CARD9 | 21.4412759 | 21.4064999 | 21.3932342 | 21.3250113 | 21.3097572 |
| P26639 | TARS1 | 24.3964512 | 24.3810637 | 24.4092289 | 24.2863484 | 24.3356758 |
| O75874 | IDH1 | 23.9296131 | 23.9076962 | 23.8858351 | 23.853207 | 23.8946082 |
| Q6PJT7 | ZC3H14 | 20.9608644 | 21.0932636 | 21.0399438 | 21.0061499 | 21.0822541 |
| Q13136 | PPFIA1 | 20.889564 | 20.9664828 | 20.9516032 | 20.8992166 | 20.9671797 |
| P85037 | FOXK1 | 21.1763281 | 21.2317646 | 21.1869082 | 21.098307 | 21.2878694 |
| Q8WWM7 | ATXN2L | 22.4653288 | 22.5584772 | 22.391382 | 22.40289 | 22.5506601 |
| Q10713 | PMPCA | 22.1314636 | 22.2893284 | 22.2742968 | 22.1519029 | 22.2894932 |
| P28331 | NDUFS1 | 22.8012479 | 22.9447274 | 22.8623036 | 22.8073297 | 22.9714981 |
| Q00722 | PLCB2 | 20.9938956 | 21.1154342 | 21.0154092 | 20.9750165 | 21.1149219 |
| Q9BRR8 | GPATCH1 | 19.5245975 | 19.6432481 | 19.6293789 | 19.5331206 | 19.5890611 |
| Q9Y5A7 | NUB1 | 19.7325303 | 19.7549661 | 19.7581235 | 19.6681399 | 19.7460169 |
| Q13485 | SMAD4 | 20.7588391 | 20.7903443 | 20.6999504 | 20.7213623 | 20.8593997 |
| Q9UDT6 | CLIP2 | 18.738301 | 18.8502096 | 18.6009526 | 18.7524381 | 18.7576194 |
| Q05D32 | CTDSPL2 | 20.9527843 | 21.0520138 | 21.0912868 | 21.0460905 | 21.1608606 |
| Q13242 | SRSF9 | 23.6231466 | 23.7400091 | 23.5162024 | 23.5576264 | 23.7639425 |
| Q14118 | DAG1 | 22.3913293 | 22.6448187 | 22.5607877 | 22.4508247 | 22.4283728 |
| P02649 | APOE | 19.5533894 | 19.5530033 | 19.3411129 | 19.495154 | 19.6476711 |
| P56181 | NDUFV3 | 19.7016334 | 20.0593124 | 19.9009259 | 19.8870026 | 19.9849171 |
| Q9BSF8 | BTBD10 | 15.9588417 | 16.1899874 | 15.8160612 | 15.9973066 | 15.7854989 |
| Q15306 | IRF4 | 18.9283849 | 19.1747363 | 18.9159822 | 18.8720247 | 19.2601618 |
| P20674 | COX5A | 24.9955274 | 25.1068468 | 25.1281543 | 25.0240393 | 25.3319609 |
| Q9BXX2 | ANKRD30B | NA | 18.3751695 | 18.3935138 | NA | 18.1356694 |
| Q9H015 | SLC22A4 | 13.7914484 | NA | NA | 13.3106565 | NA |
| Q8NB91 | FANCB | 16.1048111 | 16.1093213 | 16.3860288 | 16.0209995 | 15.8399273 |
| Q8N2Z9 | CENPS | 18.3132433 | 18.7344872 | 18.4293359 | 18.2552941 | 18.3470508 |
| Q6P1M0 | SLC27A4 | 20.0650882 | 20.1278443 | 20.0278021 | 20.1341081 | 20.1240077 |
| O14777 | NDC80 | 20.3233539 | 20.4303527 | 20.3000553 | 20.2618706 | 20.3348699 |
| Q04864 | REL | 19.8886556 | 19.8909515 | 19.8850757 | 19.9011505 | 19.9762629 |
| Q9Y3L3 | SH3BP1 | 22.2476986 | 22.3405252 | 22.315784 | 22.2273764 | 22.3084672 |
| Q9NZD2 | GLTP | 18.4831059 | 18.595885 | 18.5685619 | 18.5045774 | 18.6657122 |
| Q53T59 | HS1BP3 | 20.0672553 | 19.9957087 | 20.3024364 | 20.1064252 | 20.2743769 |

| | | | | | | |
|---|---|---|---|---|---|---|
| P58004 | SESN2 | 12.7338347 | NA | NA | 13.1915902 | NA |
| O75781 | PALM | 11.7461704 | NA | NA | 11.4246174 | NA |
| Q9BZ67 | FRMD8 | 18.5692842 | 18.689119 | 18.5467607 | 18.2323316 | 18.596015 |
| Q969V5 | MUL1 | 19.5491589 | 19.6813107 | 19.7961764 | 19.5625762 | 19.6118671 |
| O43683 | BUB1 | 18.6414478 | 18.6698961 | 18.6881706 | 18.6388833 | 18.7576754 |
| Q9Y305 | ACOT9 | 20.2725982 | 20.4306821 | 20.47023 | 20.2342012 | 20.396247 |
| Q9UI14 | RABAC1 | 20.0736986 | 20.1680603 | 20.0700302 | 20.1094408 | 19.9345842 |
| O43861 | ATP9B | 19.20137 | 19.1675081 | 19.193169 | 19.2684751 | 19.1482842 |
| P15121 | AKR1B1 | 23.6911965 | 23.6351385 | 23.7682043 | 23.5972264 | 23.7092958 |
| Q8TCT9 | HM13 | 22.8466717 | 22.9228753 | 22.849744 | 22.7642268 | 22.7710304 |
| P11413 | G6PD | 24.3394829 | 24.38679 | 24.4102869 | 24.293641 | 24.3943745 |
| P27361 | MAPK3 | 20.936061 | 20.9435067 | 20.8762223 | 20.911005 | 20.9780917 |
| Q96A35 | MRPL24 | 22.4412909 | 22.3824317 | 22.4570214 | 22.3753722 | 22.3875302 |
| Q9UBW8 | COPS7A | 20.870843 | 21.0321613 | 20.9258829 | 20.8227887 | 20.9746687 |
| P61764 | STXBP1 | 18.7150238 | 18.7160043 | 18.8063706 | 18.7231764 | 18.921188 |
| P62068 | USP46 | 17.6741093 | 17.5668723 | 17.6919513 | 17.5827832 | 17.8661446 |
| Q9NZ63 | C9orf78 | 21.4563954 | 21.573824 | 21.3255399 | 21.4102017 | 21.4793489 |
| Q96GQ5 | RUSF1 | 18.4719842 | 18.7259658 | 18.8069634 | 18.6116699 | 18.8098695 |
| Q5JTD0 | TJAP1 | 18.5056727 | 18.5649488 | 18.4364411 | 18.7635217 | 18.7260466 |
| Q16514 | TAF12 | 19.7813194 | 19.8681688 | 19.9945608 | 19.8081405 | 20.1881956 |
| O95759 | TBC1D8 | 14.5447456 | 14.6362753 | 14.6147728 | 14.6250329 | 14.6778338 |
| P05109 | S100A8 | 22.9257232 | 22.7215307 | 22.8306247 | 22.8822415 | 22.7117799 |
| Q8IZY2 | ABCA7 | 16.6641393 | 16.5166795 | 16.8755045 | 16.9200376 | 16.6498657 |
| Q9BZX2 | UCK2 | 21.6711397 | 21.6501058 | 21.6832028 | 21.5507047 | 21.6183758 |
| P52948 | NUP98 | 21.9759387 | 21.9723579 | 21.9726253 | 21.9806921 | 22.0340324 |
| Q8IVS2 | MCAT | 21.1119752 | 21.1433741 | 21.1572327 | 21.1042211 | 21.2137071 |
| Q9BSR8 | YIPF4 | 20.287067 | 20.4827156 | 20.3768767 | 20.322725 | 20.5417132 |
| Q9UNZ2 | NSFL1C | 22.6026583 | 22.7824906 | 22.5040796 | 22.5668707 | 22.7670583 |
| Q8TF61 | FBXO41 | 14.3209964 | NA | NA | 14.5228527 | NA |
| Q9BY50 | SEC11C | 21.7979442 | 21.7815033 | 21.7329217 | 21.7739738 | 21.7480421 |
| Q9UGC7 | MTRF1L | 17.359511 | 17.0502098 | 17.716366 | 17.5591165 | 17.1849013 |
| Q96E16 | SMIM19 | 17.3201232 | 17.1595258 | 17.3271222 | 17.3453615 | 17.3496868 |
| A6H8Y1 | BDP1 | NA | 13.6807329 | 13.1299471 | NA | 13.3714921 |
| Q14CW9 | ATXN7L3 | 17.7879663 | 17.1921762 | 17.7555447 | 17.1283403 | 16.8895656 |
| O95382 | MAP3K6 | 16.5466798 | 16.803209 | 16.7428608 | 16.6953253 | 16.9210628 |
| Q8TCY9 | URGCP | 18.0335062 | 18.0286928 | 18.0467842 | 17.9494193 | 18.3184856 |
| P29558 | RBMS1 | 18.4807147 | 18.5068594 | 18.5020126 | 18.4906705 | 18.7210955 |
| Q9H7B4 | SMYD3 | 19.0317004 | 19.0888057 | 19.1748786 | 19.0125709 | 19.0657427 |
| Q9Y616 | IRAK3 | 20.3117999 | 20.3174285 | 20.4424578 | 20.2791208 | 20.4241986 |
| Q9NPH3 | IL1RAP | 21.5558963 | 21.5468846 | 21.5636826 | 21.4546252 | 21.6129785 |
| Q92621 | NUP205 | 22.470865 | 22.3999233 | 22.5007599 | 22.3943683 | 22.4824257 |
| Q8NHM5 | KDM2B | 20.0342896 | 20.0981228 | 20.1623188 | 20.0168962 | 20.0797205 |

| | | | | | | |
|---|---|---|---|---|---|---|
| Q96PZ0 | PUS7 | 20.9087268 | 20.9421662 | 20.9668799 | 20.9341229 | 21.038798 |
| Q9NX01 | TXNL4B | 18.5723306 | 18.4433088 | 18.6317882 | 18.5258391 | 18.6298301 |
| Q8IZM8 | ZNF654 | 18.6266426 | 18.7383239 | 18.7406153 | 18.6671123 | 18.7720704 |
| Q8ND56 | LSM14A | 23.555993 | 23.8342629 | 23.7263866 | 23.6991829 | 23.6304784 |
| O15014 | ZNF609 | 17.1086058 | 17.1263363 | 17.2542814 | 17.0645742 | 17.2394844 |
| Q9H5H4 | ZNF768 | 18.3011039 | 18.4612636 | 18.2727718 | 18.3002174 | 18.5448141 |
| Q8IWU2 | LMTK2 | NA | 13.2341289 | 12.4792214 | NA | 12.9478521 |
| Q9BSJ5 | C17orf80 | 16.3466002 | 16.7778748 | 17.1022755 | 16.6010277 | 16.7723071 |
| Q02880 | TOP2B | 23.5254299 | 23.5161293 | 23.5283326 | 23.445161 | 23.457107 |
| Q5TB80 | CEP162 | 14.566962 | 14.4927531 | 14.57588 | 14.3141316 | 14.7348015 |
| P86452 | ZBED6 | 12.8641618 | 13.2888595 | 13.1804489 | 12.8443658 | 13.3254749 |
| Q96PC3 | AP1S3 | 18.5803801 | 18.681455 | 18.4526077 | 18.6381372 | 17.8990917 |
| P78317 | RNF4 | 16.971668 | NA | NA | 16.9267407 | NA |
| Q8WXA3 | RUFY2 | 17.0276358 | 17.009214 | 17.121945 | 17.233398 | 17.1139831 |
| Q96GY3 | LIN37 | 16.8473194 | 16.8861404 | 17.0249688 | 16.9963458 | 17.009809 |
| Q9NPA3 | MID1IP1 | 18.5191297 | 18.7429414 | 18.7859602 | 18.5831619 | 18.7951851 |
| Q15833 | STXBP2 | 22.799494 | 22.8648549 | 22.9147248 | 22.7361309 | 22.8125769 |
| Q96P48 | ARAP1 | 21.8807011 | 21.9960077 | 22.0107008 | 21.8357556 | 21.9334447 |
| P41240 | CSK | 22.7599537 | 22.8773767 | 22.9076258 | 22.7893592 | 22.9402048 |
| P06730 | EIF4E | 23.190247 | 23.1976529 | 23.3039345 | 23.1456796 | 23.281911 |
| Q13838 | DDX39B | 26.2546348 | 26.3247531 | 26.2262797 | 26.078312 | 26.2064632 |
| P68366 | TUBA4A | 28.1884944 | 28.2538729 | 28.1172876 | 28.0520406 | 28.1059872 |
| Q96PZ2 | FAM111A | 19.6459671 | 19.6788265 | 19.7704836 | 19.590185 | 19.6214089 |
| Q53R41 | FASTKD1 | 19.9139305 | 19.9316522 | 20.0322353 | 19.8303507 | 19.9479169 |
| Q9H4A4 | RNPEP | 24.0872083 | 24.0866175 | 24.1237493 | 23.984147 | 24.0661286 |
| P43304 | GPD2 | 21.3614728 | 21.3903215 | 21.3439661 | 21.3058487 | 21.3410679 |
| Q92896 | GLG1 | 21.9548886 | 21.9814434 | 21.9842272 | 21.9456787 | 21.991544 |
| P08559 | PDHA1 | 23.3442315 | 23.4259533 | 23.3611091 | 23.3090901 | 23.4354454 |
| Q9Y4L1 | HYOU1 | 24.2051293 | 24.2912607 | 24.1100272 | 24.2164482 | 24.1813979 |
| Q9UL15 | BAG5 | 20.4606835 | 20.5498607 | 20.4772395 | 20.4093818 | 20.5153862 |
| P78417 | GSTO1 | 24.7860685 | 24.7354999 | 24.7402992 | 24.7018779 | 24.8609195 |
| Q8N8A2 | ANKRD44 | 20.1991362 | 20.3165737 | 20.298108 | 20.2309499 | 20.4018948 |
| Q6UXT9 | ABHD15 | 20.3899957 | 20.6118257 | 20.5953642 | 20.4547773 | 20.5834749 |
| P78314 | SH3BP2 | 19.0347767 | 18.9486415 | 19.1982409 | 18.9804565 | 19.0970407 |
| O95816 | BAG2 | 22.6245788 | 22.8154922 | 22.6376416 | 22.5734319 | 22.5692031 |
| Q9BT17 | MTG1 | 19.2161659 | 19.3521135 | 19.4752221 | 19.3915487 | 19.5510965 |
| O00764 | PDXK | 22.1740572 | 22.1880017 | 22.1154166 | 22.0739913 | 22.070314 |
| Q8NBZ0 | INO80E | 18.1604001 | 18.0012809 | 18.125492 | 18.2765615 | 18.4073515 |
| Q07666 | KHDRBS1 | 24.2853305 | 24.5169511 | 24.2249511 | 24.2481656 | 24.3710162 |
| Q8IWT0 | ZBTB8OS | 19.9682089 | 20.1622562 | 20.1705287 | 19.9776359 | 19.8145585 |
| O43639 | NCK2 | 20.4340558 | 20.5455622 | 20.3663006 | 20.4841221 | 20.5308273 |
| Q8IVM0 | CCDC50 | 19.9567468 | 20.1561876 | 19.8404802 | 19.9744076 | 20.025742 |

| | | | | | | |
|---|---|---|---|---|---|---|
| Q5T8I3 | EEIG2 | 17.7710654 | 17.9526583 | 18.2069655 | 17.8575944 | 17.810296 |
| Q9H777 | ELAC1 | 16.6171503 | 16.9159902 | 17.0621793 | 16.773858 | 16.8224057 |
| Q9H469 | FBXL15 | 17.2090879 | 17.222893 | 17.1397931 | 17.2145652 | 17.2075514 |
| Q02446 | SP4 | 16.1638782 | 16.5950514 | 16.2423349 | 16.2036205 | 16.143337 |
| O96018 | APBA3 | 14.0736305 | 14.0576109 | 13.604342 | 14.5221514 | 14.9530997 |
| P82912 | MRPS11 | 19.7590678 | 19.8982372 | 20.0330992 | 20.0036517 | 19.8978351 |
| P22059 | OSBP | 21.8510929 | 21.8084607 | 21.8551611 | 21.8100231 | 21.8799795 |
| Q14980 | NUMA1 | 23.1317653 | 23.2636423 | 23.1518883 | 23.0893617 | 23.1746092 |
| Q9UKL3 | CASP8AP2 | 18.0033709 | 18.0530523 | 18.0292724 | 17.9956475 | 17.9378793 |
| Q14166 | TTLL12 | 23.3114958 | 23.3256809 | 23.2958983 | 23.2777221 | 23.3392236 |
| Q9Y385 | UBE2J1 | 20.650648 | 20.7465862 | 20.6230451 | 20.5621595 | 20.6088111 |
| Q9NP87 | POLM | 21.1866975 | 20.7128361 | 21.0847913 | 20.8087971 | 21.0689426 |
| Q9C0I1 | MTMR12 | 19.2653562 | 19.2881623 | 19.3590892 | 19.2336803 | 19.4091801 |
| O75832 | PSMD10 | 23.7631236 | 23.7467053 | 23.834483 | 23.7229954 | 23.8657112 |
| P20936 | RASA1 | 19.6019848 | 19.6818737 | 19.7203743 | 19.5750818 | 19.6862159 |
| A5YKK6 | CNOT1 | 21.8037951 | 21.8263064 | 21.869536 | 21.7456943 | 21.8505229 |
| O15371 | EIF3D | 24.1287861 | 24.113099 | 24.1992002 | 24.107428 | 24.1127549 |
| P30740 | SERPINB1 | 23.7395344 | 23.7446071 | 23.6667023 | 23.658641 | 23.639544 |
| P62195 | PSMC5 | 23.4765737 | 23.5937321 | 23.4633277 | 23.4600478 | 23.5934917 |
| Q9Y276 | BCS1L | 21.7012675 | 21.6305092 | 21.7060274 | 21.6933358 | 21.7695693 |
| Q9NVV4 | MTPAP | 22.1274433 | 22.1922158 | 22.3012135 | 22.2123156 | 22.2175099 |
| P29692 | EEF1D | 25.9788996 | 26.0309447 | 25.8316863 | 25.87738 | 25.942106 |
| P10809 | HSPD1 | 27.1369327 | 27.3015112 | 27.0628907 | 27.0979014 | 27.2557359 |
| Q9UFF9 | CNOT8 | 18.5764902 | 18.7384773 | 18.8130328 | 18.667854 | 18.6277311 |
| Q86TI0 | TBC1D1 | 17.430261 | 17.5489926 | 17.7213227 | 17.5355822 | 17.6976694 |
| Q66GS9 | CEP135 | 18.1964082 | 18.5426108 | 18.4653102 | 18.4862337 | 18.450753 |
| P49321 | NASP | 24.78681 | 24.9891861 | 24.6530406 | 24.7775925 | 24.8599298 |
| O95684 | CEP43 | 19.3133126 | 19.4828957 | 19.3604152 | 19.370049 | 19.4244535 |
| A0A096LP01 | SMIM26 | 18.3032097 | 18.5426441 | 18.2942735 | 18.3208741 | 18.2809773 |
| P11215 | ITGAM | 17.5475974 | 17.141279 | 17.590282 | 17.5291785 | 17.3984261 |
| Q13454 | TUSC3 | 20.1599627 | NA | NA | 20.1485611 | NA |
| O95999 | BCL10 | 19.3096347 | 19.4647283 | 19.2835709 | 19.2114136 | 19.4525162 |
| P84243 | NA | 22.8472682 | 22.7615491 | 22.7339604 | 22.1448919 | 22.4472164 |
| O15085 | ARHGEF11 | 15.559526 | NA | NA | 15.7120905 | NA |
| Q9BST9 | RTKN | 14.7050497 | 14.7842431 | 15.0270645 | 15.0457259 | 15.0507263 |
| O75771 | RAD51D | 14.3808934 | NA | NA | 14.2154034 | NA |
| P15408 | FOSL2 | 22.8655956 | 23.0818587 | 22.6772623 | 22.6714393 | 23.023379 |
| Q5JUQ0 | FAM78A | 18.4767531 | 18.4489952 | 18.6012895 | 18.4689962 | 18.3394599 |
| Q5JXA9 | SIRPB2 | NA | 14.2592908 | 14.2398008 | NA | 14.2660716 |
| Q9H9H4 | VPS37B | 17.0858651 | 17.1639898 | 17.4379995 | 17.3307765 | 17.0509202 |
| Q9BV23 | ABHD6 | NA | 14.8256896 | 14.9521894 | NA | 14.8341062 |
| Q9UIW2 | PLXNA1 | 17.0094992 | 17.1976315 | 17.2726913 | 16.9005711 | 17.0925935 |

| | | | | | | |
|---|---|---|---|---|---|---|
| O95199 | RCBTB2 | NA | 15.0407018 | 15.0026663 | NA | 14.9939555 |
| Q4FZB7 | KMT5B | 17.1039847 | 16.9703009 | 17.0857129 | 16.7880674 | 16.6645325 |
| O14975 | SLC27A2 | 21.8963143 | 21.9842904 | 21.8494086 | 21.7558113 | 21.7479328 |
| O14662 | STX16 | 19.0431785 | 19.2268559 | 19.486336 | 19.195966 | 19.3133744 |
| Q8N0Z6 | TTC5 | 20.7750269 | 20.7261234 | 20.5752608 | 20.6401312 | 20.8137575 |
| P56556 | NDUFA6 | 21.7072893 | 21.7676077 | 21.8600528 | 21.6603609 | 21.6892236 |
| P16104 | H2AX | 29.4466248 | 29.4477964 | 29.4830961 | 29.2547156 | 29.2578239 |
| Q96AY2 | EME1 | 17.1381107 | 16.9552593 | 17.2466853 | 17.0221305 | 16.8972152 |
| Q6WCQ1 | MPRIP | 18.3933048 | 18.7512944 | 18.6986786 | 18.5509725 | 18.6346324 |
| O75352 | MPDU1 | 21.6358419 | 21.5407391 | 21.7722453 | 21.3961868 | 21.7688088 |
| Q9BX59 | TAPBPL | 19.9254727 | 20.1816396 | 20.1296129 | 20.1452419 | 20.0857165 |
| Q02809 | PLOD1 | 21.090618 | 21.2310019 | 21.2874866 | 21.0498744 | 21.2747536 |
| Q9BT30 | ALKBH7 | 18.5965489 | 18.6893455 | 18.6680498 | 18.7124401 | 18.8483304 |
| P01033 | TIMP1 | 23.0871416 | 23.2876625 | 23.2089215 | 23.1481889 | 23.0668954 |
| Q9HCU9 | BRMS1 | 18.1137463 | 18.1826573 | 18.43446 | 18.1952849 | 18.2325417 |
| Q9Y485 | DMXL1 | 18.3574334 | 18.458701 | 18.557161 | 18.4498785 | 18.4096224 |
| Q68CZ2 | TNS3 | 20.1567372 | 20.2729155 | 20.324968 | 20.0947494 | 20.3014035 |
| P82675 | MRPS5 | 21.2432878 | 21.2418289 | 21.4535374 | 21.1991672 | 21.2718824 |
| O75063 | FAM20B | 19.4453118 | 19.4830958 | 19.502017 | 19.4087995 | 19.3734105 |
| Q9H3S1 | SEMA4A | 20.0545217 | 20.0291974 | 20.2424862 | 20.0849602 | 20.1195343 |
| Q9UBM7 | DHCR7 | 21.7653267 | 21.7321097 | 21.7549565 | 21.7065957 | 21.7448521 |
| Q6RW13 | AGTRAP | 21.8556683 | 21.8814078 | 21.8783839 | 21.8744373 | 21.8367857 |
| Q7L5N1 | COPS6 | 21.9195139 | 21.9416034 | 22.0562735 | 21.9280475 | 22.0907334 |
| Q8IUR7 | ARMC8 | 20.783433 | 20.9255736 | 20.8886673 | 20.7743894 | 20.8737732 |
| Q9BWD1 | ACAT2 | 24.0662165 | 23.9435582 | 24.0534417 | 23.905386 | 24.0460003 |
| P07195 | LDHB | 27.1255624 | 27.1192238 | 27.0413827 | 26.9679283 | 27.0658966 |
| Q5J8M3 | EMC4 | 20.4750396 | 20.598428 | 20.5792791 | 20.4383512 | 20.4872235 |
| Q9BQG0 | MYBBP1A | 23.4251383 | 23.425626 | 23.4811102 | 23.3598055 | 23.4256723 |
| Q9NV96 | TMEM30A | 21.5991258 | 21.6332061 | 21.5826383 | 21.5536379 | 21.5663882 |
| Q9BZE4 | GTPBP4 | 22.5598884 | 22.6174487 | 22.663572 | 22.5399077 | 22.5409755 |
| O14920 | IKBKB | 20.7405506 | 20.7817355 | 20.8078004 | 20.6895486 | 20.7311273 |
| P24539 | ATP5PB | 24.8243639 | 24.8678987 | 24.8691777 | 24.721131 | 24.850439 |
| O14818 | PSMA7 | 24.3451549 | 24.4094495 | 24.3375542 | 24.2863985 | 24.3849698 |
| O15228 | GNPAT | 20.8807048 | 20.9821566 | 20.9567065 | 20.8688454 | 20.8938484 |
| Q13630 | GFUS | 22.4679198 | 22.4423178 | 22.4814059 | 22.4145924 | 22.490504 |
| P07332 | FES | 21.6327289 | 21.6616109 | 21.6347077 | 21.5392853 | 21.688526 |
| A3KN83 | SBNO1 | 20.8787041 | 20.7886903 | 20.8843352 | 20.8026703 | 20.8632075 |
| Q13075 | NAIP | 20.3539534 | 20.3214409 | 20.4105377 | 20.3014895 | 20.4171136 |
| P34932 | HSPA4 | 24.5226446 | 24.5398779 | 24.5028317 | 24.4484555 | 24.5105361 |
| O75533 | SF3B1 | 23.7444953 | 23.7820119 | 23.6966924 | 23.682548 | 23.738499 |
| O60701 | UGDH | 20.5294463 | 20.5995395 | 20.5980052 | 20.5308991 | 20.5923452 |
| P23246 | SFPQ | 25.9870164 | 26.0812542 | 25.9096846 | 25.9111199 | 25.9547492 |

| | | | | | | |
|---|---|---|---|---|---|---|
| Q16630 | CPSF6 | 23.4662288 | 23.5001343 | 23.3945474 | 23.3909426 | 23.4707821 |
| O75663 | TIPRL | 21.7229911 | 21.7797595 | 21.759012 | 21.734747 | 21.7382441 |
| P15309 | ACP3 | 21.0829996 | 21.0567289 | 21.1295016 | 21.0155355 | 21.1380735 |
| Q9UGR2 | ZC3H7B | 19.1257151 | 19.1951124 | 19.3876667 | 19.2598371 | 19.2888784 |
| P60900 | PSMA6 | 23.9938222 | 24.0280197 | 24.1225274 | 24.0107196 | 24.0935248 |
| P53701 | HCCS | 21.9134495 | 22.0177924 | 21.9881412 | 22.0038074 | 22.0364679 |
| Q9H871 | RMND5A | 19.8841919 | 20.0260139 | 19.8797034 | 19.9019376 | 20.0728912 |
| P13804 | ETFA | 25.8575784 | 25.8763168 | 25.8791392 | 25.7604302 | 25.9756191 |
| Q9BSD7 | NTPCR | 21.6139021 | 21.6048062 | 21.6765482 | 21.6493193 | 21.6846475 |
| Q96EV8 | DTNBP1 | 19.7368125 | 19.9075282 | 19.9086846 | 19.7704226 | 19.9327836 |
| Q14764 | MVP | 21.5293049 | 21.638056 | 21.5545118 | 21.4753971 | 21.6673189 |
| Q9H334 | FOXP1 | 19.8486559 | 19.9745656 | 19.9843038 | 19.9520051 | 20.093622 |
| Q9H5N1 | RABEP2 | 21.350884 | 21.4185495 | 21.2513563 | 21.2834224 | 21.3877848 |
| P60983 | GMFB | 21.0499747 | 21.0321564 | 20.9390843 | 21.0094861 | 21.0212686 |
| Q9NW64 | RBM22 | 21.967903 | 22.0042452 | 22.0142032 | 21.9202712 | 21.9843756 |
| Q8WZ82 | OVCA2 | 20.98005 | 20.9367714 | 20.8248472 | 20.8132902 | 20.9191433 |
| Q9BSW2 | CRACR2A | 19.6026621 | 19.7943873 | 19.6688496 | 19.5149214 | 19.6992618 |
| Q13188 | STK3 | 19.7010683 | 19.9690798 | 19.7931222 | 19.7729294 | 19.9584387 |
| Q96S19 | METTL26 | 20.7354555 | 20.8062993 | 20.8341647 | 20.7293033 | 20.940778 |
| Q7Z7K0 | CMC1 | 21.0824428 | 21.3437536 | 21.1437773 | 21.1790279 | 21.3403367 |
| Q8N4C6 | NIN | 18.3856243 | 18.5198668 | 18.3202707 | 18.3974084 | 18.359933 |
| Q9NVM4 | PRMT7 | 19.9754016 | 20.1462426 | 20.1845825 | 20.0367494 | 20.1783222 |
| Q6P1R4 | DUS1L | 19.8278839 | 19.8841907 | 19.9938557 | 19.9304318 | 20.0013883 |
| Q15370 | ELOB | 23.2215659 | 23.5007694 | 23.3899839 | 23.3040977 | 23.5464541 |
| Q9UPT8 | ZC3H4 | 21.4399355 | 21.5067 | 21.3685466 | 21.3999907 | 21.5582129 |
| Q9BUR4 | WRAP53 | 18.1467697 | 18.293038 | 18.2318406 | 18.2202077 | 18.4569506 |
| Q16787 | LAMA3 | NA | 18.3692444 | 18.2678876 | NA | 18.4976226 |
| Q96BP2 | CHCHD1 | 19.5791931 | 19.666692 | 19.8951967 | 19.9317806 | 19.7406591 |
| Q5VVQ6 | YOD1 | 18.1050723 | 18.1370164 | 18.1625873 | 18.1108113 | 18.3463933 |
| Q12962 | TAF10 | 20.1467409 | 20.3478488 | 20.1536187 | 20.1721064 | 20.4695275 |
| Q6ICL3 | TANGO2 | 20.2658312 | 20.4571463 | 20.0984782 | 20.1847912 | 20.178031 |
| O43933 | PEX1 | 20.4877452 | 20.8019625 | 20.7117042 | 20.6331619 | 20.6487857 |
| Q8WYR1 | PIK3R5 | 19.4383703 | 19.4300412 | 19.5880652 | 19.5144308 | 19.5300719 |
| Q16637 | NA | 21.4001697 | 21.548 | 21.4253289 | 21.4815551 | 21.5793451 |
| O15116 | LSM1 | 21.8750541 | 21.8417452 | 21.948172 | 21.8347855 | 21.9573204 |
| Q92503 | SEC14L1 | 18.7892126 | 19.0558964 | 19.0683083 | 18.8693888 | 18.8540567 |
| Q5QP82 | DCAF10 | 17.6276495 | 17.515723 | 17.6126209 | 17.6316752 | 17.5419269 |
| Q8WTS6 | SETD7 | 20.343741 | 20.4441388 | 20.3158862 | 20.3044201 | 20.5265207 |
| Q5T6J7 | IDNK | 16.3294701 | 16.5006829 | 16.6458374 | 16.5409662 | 16.4293633 |
| O95786 | RIGI | 18.9245881 | 19.0642068 | 19.0652555 | 19.2562752 | 19.1408138 |
| P51858 | HDGF | 24.831533 | 24.9170418 | 24.601306 | 24.782911 | 24.8765945 |
| Q13322 | GRB10 | 17.3217345 | 17.3783986 | 17.473187 | 17.3565134 | 17.5032985 |

| | | | | | | |
|---|---|---|---|---|---|---|
| Q96C00 | ZBTB9 | 18.6678346 | 18.8996645 | 18.7007582 | 18.6605478 | 18.4653937 |
| O43909 | EXTL3 | NA | 16.2949613 | 16.3511674 | NA | 16.2757992 |
| Q9P021 | CRIPT | 18.0671845 | 18.1742725 | 18.2924243 | 18.2256254 | 18.5011228 |
| Q96QU6 | ACCS | 15.6910429 | 15.0384566 | 15.3822118 | 15.3360474 | 15.519168 |
| O94921 | CDK14 | NA | 15.27346 | 15.2907405 | NA | 15.6586023 |
| Q92567 | FAM168A | 16.7881532 | 16.4084179 | 16.1825982 | 16.7813711 | 16.7074515 |
| P00747 | PLG | 16.9511265 | NA | NA | 16.9921129 | NA |
| Q6P5Z2 | PKN3 | 14.281622 | 14.2298441 | 14.8883326 | 14.4414217 | 14.3079524 |
| P34913 | EPHX2 | NA | 14.4123698 | 14.0875353 | NA | 13.9166084 |
| P41279 | MAP3K8 | 11.9985653 | NA | NA | 12.5965249 | NA |
| Q96T83 | SLC9A7 | 12.8823602 | NA | NA | 13.0085104 | NA |
| B1APH4 | ZNF487 | NA | 13.7160799 | 13.9077673 | NA | 14.0031897 |
| Q9BYX7 | POTEKP | 16.3195791 | 16.4639765 | 16.728432 | 19.5810238 | 19.3310762 |
| Q9UJK0 | TSR3 | 17.4169336 | 17.654087 | 17.7014612 | 17.4559935 | 17.5440032 |
| P32322 | PYCR1 | 21.1119296 | 21.0745519 | 21.1387803 | 21.1828918 | 21.2206041 |
| Q15084 | PDIA6 | 25.8267888 | 25.795786 | 25.7163673 | 25.787023 | 25.8158617 |
| Q99943 | AGPAT1 | 18.5179737 | 18.6088049 | 18.6695724 | 18.51469 | 18.6244314 |
| Q7Z6J0 | SH3RF1 | 13.2525115 | 14.4235898 | 13.7562431 | 13.9985014 | 14.0049537 |
| P40426 | PBX3 | 19.608696 | 19.6936634 | 19.7625153 | 19.6378628 | 19.7704861 |
| Q6ZMZ3 | SYNE3 | 21.0750437 | 21.1682926 | 21.1503488 | 21.0714321 | 21.1101732 |
| P04181 | OAT | 22.2344555 | 22.1424029 | 22.2422121 | 22.147715 | 22.2632188 |
| O00193 | C11orf58 | 24.4022854 | 24.5080911 | 24.3431611 | 24.2754393 | 24.3929894 |
| P31948 | STIP1 | 25.3195877 | 25.4043746 | 25.1564724 | 25.2046647 | 25.3665577 |
| Q9UK41 | VPS28 | 21.1487623 | 21.1339031 | 21.1961833 | 21.1063481 | 21.2190671 |
| Q9UFH2 | DNAH17 | NA | 15.1065204 | 14.8804303 | NA | 15.0544102 |
| Q8WVV9 | HNRNPLL | 21.7946916 | 21.8272756 | 21.8273795 | 21.7470617 | 21.9132906 |
| Q8NAT1 | POMGNT2 | 15.7620579 | 16.0755991 | 16.0650229 | 16.0204078 | 16.1089615 |
| Q8NEN9 | PDZD8 | 17.8136395 | 17.6850119 | 17.7537869 | 17.6965091 | 17.7957332 |
| Q9NTK5 | OLA1 | 24.4975579 | 24.5124427 | 24.4815375 | 24.4239501 | 24.5445648 |
| Q9BTA9 | WAC | 20.4237027 | 20.4429076 | 20.5431125 | 20.4376267 | 20.4382615 |
| P20290 | BTF3 | 23.8299708 | 24.0329056 | 23.7680525 | 23.774023 | 23.9477061 |
| Q9Y2S7 | POLDIP2 | 21.8563301 | 21.8449942 | 21.8543224 | 21.8022203 | 21.8470585 |
| P22670 | RFX1 | 19.3492638 | 19.5019628 | 19.5398007 | 19.4329993 | 19.5390868 |
| P08651 | NFIC | 17.6206734 | 17.7460199 | 17.8555053 | 17.6788492 | 17.8405276 |
| Q9BRR9 | ARHGAP9 | 20.1385304 | 20.1044091 | 20.2346214 | 20.0459733 | 20.1640748 |
| Q9NXV6 | CDKN2AIP | 20.93898 | 21.0104511 | 20.9646486 | 20.970523 | 21.0481662 |
| P36955 | SERPINF1 | 19.3888802 | 19.2550195 | 19.3115506 | 19.3255257 | 19.3332342 |
| O43542 | XRCC3 | 17.1783814 | 17.4810417 | 17.2585696 | 17.3087377 | 17.2688977 |
| Q9P0Z9 | PIPOX | 16.848538 | 17.1615249 | 17.3114266 | 17.2219264 | 17.2312421 |
| Q5RKV6 | EXOSC6 | 21.3819788 | 21.6108751 | 21.5948417 | 21.3801797 | 21.4475558 |
| P42773 | CDKN2C | 17.6599042 | 17.7288296 | 17.7673875 | 17.5319973 | 17.7414388 |
| Q9ULV3 | CIZ1 | 18.6591236 | 18.565167 | 18.7190449 | 18.6306063 | 18.7875179 |

| | | | | | | |
|---|---|---|---|---|---|---|
| Q99986 | VRK1 | 23.4741043 | 23.5079683 | 23.4073772 | 23.4070086 | 23.3848886 |
| Q96AB3 | ISOC2 | 22.2798862 | 22.3380706 | 22.3727731 | 22.3014909 | 22.3081761 |
| Q9P219 | CCDC88C | 17.9335241 | 17.9330716 | 17.9212749 | 18.0024092 | 18.0460882 |
| Q9GZZ1 | NAA50 | 23.1856862 | 23.1044176 | 23.1691748 | 23.0827665 | 23.2419011 |
| P53370 | NUDT6 | 18.5833703 | 18.6318185 | 18.6956341 | 18.6469873 | 18.6959249 |
| O95544 | NADK | 20.3576944 | 20.4552804 | 20.4569414 | 20.3800569 | 20.5616985 |
| Q9BYN0 | SRXN1 | 15.7898275 | 15.7254287 | 15.561485 | 15.9738575 | 15.9043306 |
| Q9GZY4 | COA1 | NA | 16.1311886 | 16.4345638 | NA | 15.3637276 |
| Q53HV7 | SMUG1 | 13.8584925 | 13.6990988 | 13.806369 | 13.6902671 | 13.9879606 |
| Q9UIB8 | CD84 | 15.6003375 | 15.8432736 | 15.6042097 | 15.7072359 | 15.6821621 |
| Q969S8 | HDAC10 | 18.7276615 | 18.9964424 | 18.9929912 | 18.6803105 | 18.9997659 |
| P24557 | TBXAS1 | 18.6343875 | 18.722508 | 18.6717926 | 18.504583 | 18.7887046 |
| Q8NFZ5 | TNIP2 | 18.3527482 | 18.4447935 | 18.3678431 | 18.3461567 | 18.4125478 |
| Q6UWP7 | LCLAT1 | 19.7630723 | 19.7442877 | 19.8020351 | 19.7991662 | 19.7376756 |
| Q9H7F0 | ATP13A3 | 19.1599099 | 18.9302196 | 19.0876101 | 18.9477915 | 19.0785089 |
| P20020 | ATP2B1 | 20.7384551 | 20.7954269 | 20.9184215 | 20.7307592 | 20.7956324 |
| Q9H9Y6 | POLR1B | 20.6546954 | 20.7034402 | 20.7704718 | 20.6391567 | 20.7768394 |
| P50570 | DNM2 | 23.7621925 | 23.7881104 | 23.8467963 | 23.6739775 | 23.8460556 |
| O43237 | DYNC1LI2 | 21.0864198 | 21.1260375 | 21.1363721 | 21.1194037 | 21.1640187 |
| P11387 | TOP1 | 23.701531 | 23.8012307 | 23.7624748 | 23.6816773 | 23.7578227 |
| Q9BQF6 | SENP7 | 17.8598833 | 18.169115 | 18.0214859 | 17.9502184 | 18.0382983 |
| Q92614 | MYO18A | 22.8147286 | 22.9144801 | 22.9162958 | 22.7818225 | 22.9001904 |
| P45880 | VDAC2 | 24.6040088 | 24.5899417 | 24.6085768 | 24.5364929 | 24.8217804 |
| P27918 | CFP | 19.716981 | 19.7384152 | 19.7429852 | 19.7125688 | 19.8412007 |
| Q99614 | TTC1 | 22.7251041 | 22.9384544 | 22.6737141 | 22.7891892 | 22.8327515 |
| O95926 | SYF2 | 20.0842662 | 20.0363262 | 19.9143805 | 20.1144703 | 20.0096967 |
| Q9UKV5 | AMFR | 18.8534026 | 19.2414508 | 19.1485351 | 18.967674 | 19.0315117 |
| Q16621 | NFE2 | 14.8074473 | 15.0177646 | 15.1441466 | 15.1197121 | 15.4083174 |
| P47736 | RAP1GAP | 18.4111038 | 18.1848581 | 18.2590855 | 18.2354345 | 18.322178 |
| O43524 | FOXO3 | 16.5496319 | 16.9423613 | 16.7977227 | 16.6430908 | 16.7502232 |
| Q9NR81 | ARHGEF3 | 15.5352012 | 15.4127459 | 15.2720735 | 15.5870823 | 15.4992322 |
| O15160 | POLR1C | 21.5586418 | 21.6881366 | 21.6901454 | 21.5838974 | 21.7639711 |
| Q8IXB1 | DNAJC10 | 20.1678605 | 20.2129464 | 20.2748154 | 20.346596 | 20.3058309 |
| O43324 | EEF1E1 | 22.4862215 | 22.6337844 | 22.4344798 | 22.4790839 | 22.5430128 |
| Q8N370 | SLC43A2 | 20.2203854 | 20.1030642 | 20.1920711 | 20.2743854 | 20.023818 |
| Q6P9B9 | INTS5 | 19.8183014 | 19.9137885 | 19.9440485 | 19.7464313 | 19.8923521 |
| O60264 | SMARCA5 | 23.0359027 | 23.0827766 | 23.0672933 | 22.970934 | 23.0760387 |
| Q96EY4 | TMA16 | 21.8786374 | 22.0958993 | 21.8820193 | 21.9478721 | 22.0626373 |
| P62906 | RPL10A | 27.1481155 | 27.1970453 | 27.1282319 | 27.053504 | 27.0625145 |
| Q5U3C3 | TMEM164 | 18.422118 | 18.6210034 | 18.4467052 | 18.5864847 | 18.5963413 |
| O14787 | TNPO2 | 21.4092308 | 21.381145 | 21.4153127 | 21.3408694 | 21.3914557 |
| Q0VGE8 | ZNF816 | 17.5448944 | 17.6538609 | 17.7799908 | 17.4725519 | 17.3872442 |

| | | | | | | |
|---|---|---|---|---|---|---|
| Q8N6Y0 | USHBP1 | NA | 14.8246407 | 14.8070122 | NA | 14.764013 |
| Q8N8Z6 | DCBLD1 | 13.7554756 | 13.4770108 | 13.7416176 | 13.7348769 | 13.35032 |
| Q86U06 | RBM23 | 17.9782936 | 17.9397189 | 18.0612271 | 17.9721058 | 18.1245404 |
| Q96P47 | AGAP3 | 19.974796 | 20.130597 | 20.000074 | 19.9816082 | 19.9297524 |
| P53602 | MVD | 21.3742203 | 21.4199028 | 21.4784122 | 21.3935081 | 21.5488023 |
| Q9H7Z7 | PTGES2 | 21.4197046 | 21.4088847 | 21.4692054 | 21.3503876 | 21.4595085 |
| P35269 | GTF2F1 | 21.1689003 | 21.2046546 | 21.1682783 | 21.1107825 | 21.1452412 |
| Q9UJW0 | DCTN4 | 20.5147641 | 20.4377986 | 20.5051431 | 20.4949627 | 20.4283389 |
| Q8WX92 | NELFB | 21.2058274 | 21.2893036 | 21.2262415 | 21.1710026 | 21.1577094 |
| Q96I25 | RBM17 | 22.4113529 | 22.5286793 | 22.4537334 | 22.4328021 | 22.4603038 |
| Q5JTZ9 | AARS2 | 23.0033039 | 23.0966695 | 22.9993854 | 23.0033187 | 23.0320732 |
| Q9H6U8 | ALG9 | 17.84006 | 17.9963655 | 18.0923988 | 17.9766228 | 18.0683276 |
| Q14157 | UBAP2L | 23.0622276 | 23.1032331 | 22.9334003 | 22.9927592 | 23.144375 |
| Q9NWL6 | ASNSD1 | 18.745391 | 18.7984498 | 19.0188419 | 18.8000168 | 18.8160353 |
| Q8IZ81 | ELMOD2 | 19.7562784 | 19.7736147 | 19.8781389 | 19.6795146 | 19.8324986 |
| Q7Z7A4 | PXK | 20.9530084 | 20.8673954 | 20.810942 | 20.8686494 | 20.7033893 |
| Q96B70 | LENG9 | 20.1570427 | 20.2577308 | 20.1819506 | 20.162701 | 20.2029848 |
| Q2TAY7 | SMU1 | 22.9340234 | 22.8436949 | 22.9436661 | 22.780877 | 22.9505637 |
| Q9Y5T5 | USP16 | 19.4442639 | 19.5619712 | 19.5389094 | 19.5289568 | 19.5922655 |
| O60942 | RNGTT | 20.8375767 | 21.0704412 | 20.9767903 | 20.9319666 | 20.9845582 |
| Q9NXR8 | ING3 | 18.1738955 | 18.1773426 | 18.1920747 | 18.1908959 | 18.2348462 |
| Q8NCY6 | MSANTD4 | 12.9054986 | 13.1454426 | 13.2753477 | 12.9011843 | 13.3129457 |
| P16144 | ITGB4 | NA | 19.4431371 | 19.2986232 | NA | 19.5425316 |
| P78559 | MAP1A | 17.9071515 | 18.5134571 | 18.2143979 | 17.5374644 | 18.5077888 |
| Q494V2 | CFAP100 | 22.5973973 | 22.0254221 | 21.8281646 | 22.1581601 | 22.1234645 |
| Q8NCW6 | GALNT11 | 15.2831606 | 15.0134768 | 15.6434177 | 15.1028858 | 15.2824804 |
| O75828 | CBR3 | 15.5533173 | NA | NA | 15.6530169 | NA |
| P26010 | ITGB7 | 20.8695686 | 21.0172759 | 21.1592113 | 20.9614847 | 21.0246059 |
| P61970 | NUTF2 | 24.3850955 | 24.3480015 | 24.4107578 | 24.3233319 | 24.4288934 |
| Q15008 | PSMD6 | 23.4664647 | 23.4988286 | 23.4922265 | 23.3687074 | 23.4191284 |
| Q9BZZ5 | API5 | 23.522203 | 23.6672805 | 23.5867857 | 23.5408792 | 23.5666083 |
| Q14814 | MEF2D | 21.5996936 | 21.7652635 | 21.6306138 | 21.6259121 | 21.7292881 |
| Q96CW6 | SLC7A6OS | 20.0596659 | 20.1165412 | 20.1324428 | 20.095279 | 20.337725 |
| Q02410 | APBA1 | 16.6931387 | 16.8380145 | 16.4474977 | 16.7870686 | 16.8100306 |
| Q6P179 | ERAP2 | 17.7729723 | 17.7935754 | 17.9061001 | 17.6970584 | 17.8144195 |
| Q9NXU5 | ARL15 | NA | 16.0651226 | 16.0970153 | NA | 16.0629426 |
| P54277 | PMS1 | 20.8464755 | 20.8627947 | 20.9715676 | 20.8686975 | 20.9359153 |
| O60518 | RANBP6 | 19.6240983 | 19.6346112 | 19.7792947 | 19.6106639 | 19.7814407 |
| Q96SB8 | SMC6 | 21.2073538 | 21.353379 | 21.2847174 | 21.1574732 | 21.2908444 |
| Q9BUT1 | BDH2 | 15.5664781 | 15.7656728 | 16.031175 | 15.4039515 | 16.0784441 |
| Q9UPQ4 | TRIM35 | 18.6471221 | 18.4275867 | 18.5789329 | 18.6563657 | 18.5036208 |
| Q96IV0 | NGLY1 | 20.1729056 | 20.2923976 | 20.1722757 | 20.1183515 | 20.1610241 |

| | | | | | | |
|---|---|---|---|---|---|---|
| Q6P4F7 | ARHGAP11A | 18.1507556 | 18.2447342 | 18.2430034 | 18.0313203 | 18.2093317 |
| Q14202 | ZMYM3 | 20.8751435 | 20.8943275 | 20.9700991 | 20.7432239 | 20.8746675 |
| O75323 | NIPSNAP2 | 22.1437969 | 22.3193582 | 22.3273259 | 22.16627 | 22.2546084 |
| Q8WVM7 | STAG1 | 20.4636816 | 20.5227266 | 20.5975453 | 20.4317368 | 20.5405513 |
| O15144 | ARPC2 | 25.4981623 | 25.5756817 | 25.5125309 | 25.4257372 | 25.5153313 |
| Q9NPQ8 | RIC8A | 24.1316198 | 24.1761389 | 24.1681905 | 24.0458742 | 24.1609289 |
| Q4G0N4 | NADK2 | 21.9366385 | 21.8055971 | 21.8987107 | 21.897445 | 21.829251 |
| P19387 | POLR2C | 21.4878994 | 21.4694511 | 21.431614 | 21.4632356 | 21.5817464 |
| Q9Y228 | TRAF3IP3 | 21.4120993 | 21.5279229 | 21.6001246 | 21.4053326 | 21.5820361 |
| Q96A54 | ADIPOR1 | 18.5768045 | 18.5994894 | 18.7273108 | 18.5346311 | 18.572827 |
| Q9ULJ3 | ZBTB21 | 19.1414155 | 19.230643 | 19.1952469 | 19.0108658 | 19.2520794 |
| Q13177 | PAK2 | 23.9062971 | 23.9832571 | 23.8694642 | 23.9019907 | 24.0593626 |
| Q9NP77 | SSU72 | 20.7541133 | 20.820025 | 20.7081744 | 20.7658827 | 20.8354545 |
| Q96HP4 | OXNAD1 | 18.3836914 | 18.6144703 | 18.6394511 | 18.5654771 | 18.6604058 |
| P38936 | CDKN1A | 19.8190727 | 19.9902538 | 19.7978345 | 19.7182835 | 19.8182148 |
| P06241 | FYN | 18.1948338 | 18.2822945 | 18.3421751 | 18.0945032 | 18.4528121 |
| O15066 | KIF3B | 17.1570683 | 17.0350718 | 17.2085567 | 17.2799561 | 17.2939893 |
| Q5M7Z0 | RNFT1 | 16.8132803 | 17.1785105 | 17.1905747 | 16.7581195 | 17.0875275 |
| Q9H6S1 | AZI2 | 16.5529056 | 16.7772053 | 16.9636347 | 16.8940396 | 16.9749869 |
| Q9BRQ0 | PYGO2 | 16.5931227 | 16.6578554 | 16.5633043 | 16.6535813 | 16.7512895 |
| P01893 | HLA-H | 21.4376268 | 21.4050196 | 21.6894167 | 21.5280254 | 21.4879598 |
| Q9H497 | TOR3A | 18.7718633 | 18.6561214 | 18.7192827 | 18.6155167 | 18.824427 |
| P82979 | SARNP | 23.483296 | 23.6724877 | 23.3510426 | 23.4704087 | 23.5731048 |
| Q9UHY7 | ENOPH1 | 21.9646604 | 22.1260821 | 22.0097555 | 21.8827829 | 21.9940637 |
| Q9NQW6 | ANLN | 20.7796312 | 20.8175216 | 20.7291544 | 20.7388099 | 20.8484726 |
| Q14116 | IL18 | 18.8448177 | 19.0713865 | 18.8728275 | 18.8122148 | 19.0837179 |
| Q5CZC0 | FSIP2 | 18.745537 | 18.0634236 | 17.9653191 | 17.4607471 | 17.5244234 |
| Q8N6S5 | ARL6IP6 | 19.8645171 | 20.0703069 | 20.0170234 | 19.932675 | 20.099588 |
| Q7Z3E5 | ARMC9 | 16.5899482 | 16.397006 | 16.4731017 | 16.6776409 | 16.61897 |
| Q15120 | PDK3 | 18.4568193 | 18.5569198 | 18.4896901 | 18.478234 | 18.5290944 |
| Q9NUQ7 | UFSP2 | 17.4500595 | 17.5511754 | 17.4815022 | 17.4507765 | 17.6062277 |
| Q15006 | EMC2 | 21.5937456 | 21.8189889 | 21.6495649 | 21.6224976 | 21.7120044 |
| O43819 | SCO2 | 22.1863136 | 22.2204392 | 22.1665362 | 22.1836546 | 22.2208629 |
| P49917 | LIG4 | 18.438156 | 18.4834437 | 18.4815805 | 18.4925039 | 18.5785576 |
| Q9BZ95 | NSD3 | 20.4281369 | 20.542478 | 20.6080612 | 20.4996125 | 20.5418064 |
| P48426 | PIP4K2A | 21.9402015 | 22.0104104 | 22.0539526 | 21.9489411 | 22.0785079 |
| P60484 | PTEN | 22.9105102 | 22.9511437 | 22.9234455 | 22.789265 | 22.7867463 |
| Q9UN70 | PCDHGC3 | 17.2521101 | 17.2168414 | 17.3310583 | 17.2380857 | 17.3261064 |
| Q9UKM9 | RALY | 24.1184727 | 24.1239633 | 24.1818105 | 24.0013821 | 24.063495 |
| P84103 | SRSF3 | 26.2398033 | 26.3332032 | 26.1111037 | 26.1422525 | 26.3098297 |
| Q9UPP1 | PHF8 | 20.4132393 | 20.7048826 | 20.5987933 | 20.457663 | 20.6344773 |
| O14713 | ITGB1BP1 | 17.7880525 | 17.7513816 | 17.5613503 | 17.7390351 | 17.5793465 |

| | | | | | | |
|---|---|---|---|---|---|---|
| Q8N4Q1 | CHCHD4 | 21.0569561 | 21.1555204 | 20.9236038 | 20.9761708 | 21.156337 |
| P49715 | CEBPA | 22.3277614 | 22.420608 | 22.1246003 | 22.1837799 | 22.4642045 |
| Q14147 | DHX34 | 18.1739259 | 18.1615943 | 18.2810712 | 18.1957189 | 18.1909646 |
| O94763 | URI1 | 20.892747 | 20.946666 | 20.7944882 | 20.8969645 | 21.0604434 |
| Q8TEB1 | DCAF11 | 18.6886067 | 18.8782193 | 18.8098329 | 18.8829424 | 18.8887597 |
| P47224 | RABIF | 21.0073889 | 21.1336416 | 21.0409296 | 21.098791 | 21.2061592 |
| P11047 | LAMC1 | 16.4313267 | 16.5833645 | 16.6532924 | 16.5260568 | 16.7119988 |
| P61024 | CKS1B | 20.8347028 | 20.9318587 | 20.5851228 | 20.7326826 | 20.9675258 |
| Q9HCP0 | CSNK1G1 | 18.5632719 | 18.8433061 | 18.8091215 | 18.6466413 | 18.9007238 |
| Q8NBM8 | PCYOX1L | 21.434186 | 21.3872751 | 21.3909934 | 21.4299911 | 21.4053023 |
| Q12912 | IRAG2 | 20.6505957 | 20.8078618 | 20.6709813 | 20.6689 | 20.7991528 |
| Q9UI95 | MAD2L2 | 18.6708346 | 18.6582017 | 18.7882559 | 18.6158998 | 18.8415426 |
| Q9HAP2 | MLXIP | 16.3413638 | 16.4086028 | 16.4380565 | 16.0855111 | 16.2874607 |
| P08238 | HSP90AB1 | 28.0863563 | 28.1403542 | 28.0784183 | 27.9846582 | 28.069032 |
| Q53H82 | LACTB2 | 22.1740528 | 22.2686806 | 22.2365684 | 22.2486974 | 22.2943074 |
| O15397 | IPO8 | 20.484471 | 20.5751561 | 20.6092733 | 20.5139082 | 20.6180243 |
| P11717 | IGF2R | 22.6538028 | 22.6295804 | 22.6444552 | 22.622254 | 22.6698151 |
| Q9UBG0 | MRC2 | 21.6039064 | 21.566312 | 21.5247473 | 21.5383071 | 21.565872 |
| Q99733 | NAP1L4 | 24.6322394 | 24.7589652 | 24.5313112 | 24.5094275 | 24.6354401 |
| Q96QC0 | PPP1R10 | 21.4389885 | 21.5371645 | 21.4380805 | 21.3992915 | 21.4883594 |
| P25685 | DNAJB1 | 23.4137454 | 23.536094 | 23.4808532 | 23.377421 | 23.4962328 |
| O43837 | IDH3B | 21.9758307 | 22.081578 | 22.1642416 | 21.998258 | 22.2156706 |
| Q9UIF8 | BAZ2B | 19.9265036 | 19.8916986 | 19.940852 | 19.7898601 | 19.9369847 |
| Q5TC82 | RC3H1 | 18.3897857 | 18.4903797 | 18.4670549 | 18.4373587 | 18.5438808 |
| O43432 | EIF4G3 | 20.5218477 | 20.5790528 | 20.5261648 | 20.5014815 | 20.560577 |
| O75179 | ANKRD17 | 20.7366047 | 20.7303636 | 20.7881055 | 20.6934902 | 20.8013175 |
| Q9UJ68 | MSRA | 21.6686925 | 21.5019172 | 21.4740939 | 21.5798389 | 21.5671497 |
| Q9UHG3 | PCYOX1 | 22.7219596 | 22.8033297 | 22.7911115 | 22.6622838 | 22.7414261 |
| Q5H9R7 | PPP6R3 | 21.717044 | 21.7691061 | 21.7383237 | 21.6637211 | 21.7919759 |
| P78527 | PRKDC | 24.759864 | 24.7675907 | 24.7750789 | 24.6623055 | 24.7367093 |
| Q53G59 | KLHL12 | 16.7821553 | 16.8689352 | 16.6049463 | 16.7533454 | 16.7796702 |
| Q86V85 | GPR180 | 16.6833078 | 16.8565046 | 16.8479504 | 16.6520085 | 16.9715243 |
| Q8N103 | TAGAP | 14.1503591 | 14.032757 | 13.9101341 | 13.8249558 | 14.4188251 |
| O94851 | MICAL2 | 18.5292435 | 18.5139939 | 18.7265571 | 18.6202023 | 18.3869334 |
| Q9NVS2 | MRPS18A | 22.0148214 | 21.8775692 | 22.1024809 | 21.9376195 | 22.0727207 |
| Q9P243 | ZFAT | 16.8276083 | 16.7198705 | 16.7863451 | 16.6812548 | 16.7927088 |
| Q9UI43 | MRM2 | 19.0871866 | 19.1751158 | 19.145433 | 18.9669256 | 19.2595802 |
| P15813 | CD1D | 16.1566146 | 16.5439238 | 16.8335824 | 16.1486787 | 16.6031473 |
| Q6V1X1 | DPP8 | 17.8873103 | 18.0942735 | 18.3155385 | 18.2241518 | 18.0673846 |
| Q8IWY9 | CDAN1 | 19.3402238 | 19.3496582 | 19.4187762 | 19.3624269 | 19.3801537 |
| Q9P1Y6 | PHRF1 | 20.1003851 | 20.0706586 | 20.1462942 | 20.0570644 | 20.0098403 |
| Q9UBT6 | POLK | 17.5745465 | 17.64699 | 17.5082187 | 17.4565984 | 17.4952419 |

| | | | | | | |
|---|---|---|---|---|---|---|
| O00767 | SCD | 20.4839619 | 20.8721635 | 20.9785558 | 20.335176 | 20.7391757 |
| Q147X3 | NAA30 | 18.5671526 | 18.3481346 | 18.5096506 | 18.3359702 | 18.4810112 |
| Q6NT76 | HMBOX1 | 17.3504646 | 17.491568 | 17.4258385 | 17.2774219 | 17.2620426 |
| Q969R8 | ITFG2 | 18.247187 | 18.3514182 | 18.3991568 | 18.3248486 | 18.3655635 |
| P54284 | CACNB3 | 15.4594769 | NA | NA | 15.3925068 | NA |
| P04040 | CAT | 25.7499663 | 25.7346047 | 25.6816091 | 25.6504192 | 25.7446455 |
| Q969F9 | HPS3 | 22.3180504 | 22.277205 | 22.3726661 | 22.2305919 | 22.3270119 |
| Q9BQ67 | GRWD1 | 21.7334266 | 21.7901817 | 21.7858945 | 21.8027354 | 21.8759705 |
| Q9GZQ3 | COMMD5 | 20.7458545 | 20.8024108 | 20.8780824 | 20.7968228 | 20.9564691 |
| P35232 | PHB1 | 25.0566311 | 25.1292211 | 25.0192214 | 24.9638865 | 25.1028862 |
| P22061 | PCMT1 | 22.9545767 | 23.0092283 | 22.8224005 | 22.8816196 | 22.9524681 |
| Q13029 | PRDM2 | 19.4962547 | 19.6830285 | 19.5823034 | 19.5138607 | 19.5975515 |
| Q7Z6I6 | ARHGAP30 | 21.6212342 | 21.7734776 | 21.6910681 | 21.6992593 | 21.8510379 |
| Q7KZI7 | MARK2 | 19.5958408 | 19.6807018 | 19.8217691 | 19.7380407 | 19.7655629 |
| Q2KHR3 | QSER1 | 18.4356253 | 18.4289417 | 18.4821564 | 18.3539389 | 18.4368377 |
| Q7Z422 | SZRD1 | 20.7446574 | 20.8776125 | 20.8060925 | 20.9088132 | 20.9112757 |
| Q9NUP1 | BLOC1S4 | 17.8945862 | 18.0387455 | 18.0522704 | 17.8007281 | 18.056449 |
| Q9NPI8 | FANCF | 15.2147257 | 15.647376 | 15.6887185 | 15.3433944 | 15.6361759 |
| Q96RY5 | CRAMP1 | 16.1002459 | 16.3140997 | 16.4029933 | 16.1229719 | 16.0229082 |
| P19474 | TRIM21 | 21.3693214 | 21.2937261 | 21.4314659 | 21.2420167 | 21.4090943 |
| P32456 | GBP2 | 18.1570714 | 18.1309618 | 18.212511 | 18.196792 | 18.2979653 |
| Q8TCT8 | SPPL2A | 19.7377845 | 19.5308917 | 19.6081634 | 19.5300931 | 19.4112314 |
| Q9ULC4 | MCTS1 | 22.0717972 | 22.1248101 | 22.1977902 | 22.1044257 | 22.1713656 |
| Q8IWW6 | ARHGAP12 | 17.757801 | 17.7211335 | 17.7579972 | 17.503611 | 17.8180274 |
| Q7Z628 | NET1 | NA | 14.4142257 | 14.8054372 | NA | 14.4268971 |
| Q96QG7 | MTMR9 | 20.3307317 | 20.2354746 | 20.4999502 | 20.3011538 | 20.4571766 |
| O95831 | AIFM1 | 23.7680456 | 23.7823119 | 23.7443675 | 23.7025097 | 23.7818743 |
| Q15274 | QPRT | 19.1268255 | 19.3764386 | 19.3041431 | 19.2544497 | 19.4245744 |
| Q9Y3P4 | RHBDD3 | 18.4250137 | 18.2133728 | 18.3840762 | 18.4464577 | 18.3247021 |
| Q5SWA1 | PPP1R15B | 15.203942 | 15.400176 | 15.7851656 | 14.631495 | 15.0740746 |
| Q8WXD9 | CASKIN1 | NA | 17.3508224 | 17.4016478 | NA | 17.0563254 |
| Q9Y3R0 | GRIP1 | 16.8395418 | 17.2390317 | 16.884757 | 16.8305118 | 16.9669862 |
| Q15560 | TCEA2 | 16.7599783 | 16.7701355 | 16.9952837 | 16.7816132 | 16.9553492 |
| E9PRG8 | C11orf98 | 21.1190524 | 21.2600143 | 21.6867587 | 21.3329976 | 21.2709868 |
| Q8TF62 | ATP8B4 | 17.3994294 | 17.4554042 | 17.4228515 | 17.3901766 | 17.5141563 |
| O15055 | PER2 | 17.9215331 | 17.8976522 | 17.9972212 | 17.8289288 | 17.7082548 |
| Q8N4V1 | MMGT1 | 19.9089188 | 19.7091618 | 19.9587883 | 19.9017708 | 19.878335 |
| Q9H3C7 | GGNBP2 | 17.7272965 | 17.6459865 | 17.780133 | 17.6792922 | 17.6455038 |
| Q9UJ90 | KCNE5 | 19.9466698 | 19.8574342 | 19.6958695 | 19.9715563 | 19.8190217 |
| Q8TBF2 | PRXL2B | 19.2890872 | 19.2955661 | 19.2572266 | 19.2463209 | 19.3170427 |
| Q86SQ9 | DHDDS | 17.9949018 | 18.0899434 | 18.0777881 | 17.9594125 | 17.8523838 |
| Q6ZT12 | UBR3 | 18.577817 | 18.7236249 | 18.8239833 | 18.6055459 | 18.5880684 |

| | | | | | | |
|---|---|---|---|---|---|---|
| Q99490 | AGAP2 | 19.7974006 | 19.9049423 | 19.9172185 | 19.8358184 | 19.8589402 |
| O43493 | TGOLN2 | 21.6532343 | 21.7863896 | 21.7090982 | 21.5605842 | 21.6351134 |
| O00217 | NDUFS8 | 22.4261247 | 22.4625032 | 22.4723153 | 22.3348974 | 22.3288045 |
| P20073 | ANXA7 | 22.7442274 | 22.6513996 | 22.7550561 | 22.586986 | 22.7412941 |
| P41212 | ETV6 | 21.5925726 | 21.4428001 | 21.5303119 | 21.4498846 | 21.545202 |
| P18074 | ERCC2 | 20.4708467 | 20.5870886 | 20.7249127 | 20.4879576 | 20.5848126 |
| P30626 | SRI | 22.2913473 | 22.2198293 | 22.3207179 | 22.3264938 | 22.3337249 |
| Q15392 | DHCR24 | 23.6018289 | 23.5269509 | 23.6366734 | 23.5494936 | 23.5786186 |
| P12956 | XRCC6 | 26.0928503 | 26.1423019 | 26.0542367 | 25.9597563 | 26.0291613 |
| Q9Y673 | ALG5 | 21.4536778 | 21.5068955 | 21.5532777 | 21.4467944 | 21.4511553 |
| P55084 | HADHB | 24.3182813 | 24.353741 | 24.2870795 | 24.2682696 | 24.2843694 |
| Q9BTE6 | AARSD1 | 20.9999224 | 20.9959794 | 20.8995325 | 20.9326754 | 20.9469791 |
| Q70J99 | UNC13D | 22.884355 | 22.9282222 | 22.9479719 | 22.8553157 | 22.9224977 |
| Q9H845 | ACAD9 | 22.233571 | 22.3269779 | 22.3373851 | 22.2161429 | 22.30191 |
| P61421 | ATP6V0D1 | 22.7839538 | 22.623736 | 22.7091394 | 22.6604494 | 22.6645478 |
| P78347 | GTF2I | 23.1213871 | 23.2230274 | 23.0983984 | 23.0692699 | 23.1096105 |
| Q9NSE4 | IARS2 | 22.578388 | 22.618097 | 22.5999115 | 22.5452505 | 22.6314172 |
| O95140 | MFN2 | 20.3773779 | 20.3994149 | 20.3756863 | 20.463855 | 20.4454821 |
| Q9Y263 | PLAA | 22.1340528 | 22.1636123 | 22.1307509 | 22.1018575 | 22.1604213 |
| P55786 | NPEPPS | 23.3624859 | 23.3791881 | 23.3676941 | 23.2976376 | 23.4404901 |
| Q9NWU1 | OXSM | 20.5101256 | 20.5046804 | 20.4852201 | 20.4968661 | 20.5326453 |
| Q8TAG9 | EXOC6 | 22.3400515 | 22.3441754 | 22.4320308 | 22.3467651 | 22.4117867 |
| Q5T8P6 | RBM26 | 21.8543967 | 21.9294418 | 21.8836999 | 21.8025782 | 21.987537 |
| Q5VZE5 | NAA35 | 19.7739224 | 19.9155877 | 19.8490726 | 19.8023331 | 19.8770015 |
| Q9NQH7 | XPNPEP3 | 20.5208814 | 20.6552434 | 20.6374337 | 20.5222221 | 20.6728569 |
| P62140 | PPP1CB | 24.6682005 | 24.6782465 | 24.7606314 | 24.6541247 | 24.7054703 |
| Q13415 | ORC1 | 19.5875612 | 19.6965771 | 19.6951031 | 19.6225402 | 19.6887573 |
| Q9P2R7 | SUCLA2 | 22.4070531 | 22.6440991 | 22.5927967 | 22.4602315 | 22.5680413 |
| Q96HA7 | TONSL | 20.1903243 | 20.1669982 | 20.2643488 | 20.1586292 | 20.3245894 |
| Q9NWT6 | HIF1AN | 20.1494593 | 20.3458191 | 20.3433074 | 20.2232332 | 20.276019 |
| Q9Y2Z0 | SUGT1 | 22.7928707 | 22.8835748 | 22.8124954 | 22.7717501 | 22.9793506 |
| Q13561 | DCTN2 | 22.7806412 | 22.9442747 | 22.6649383 | 22.7999841 | 22.8641499 |
| P50749 | RASSF2 | 21.3730058 | 21.4753792 | 21.5139414 | 21.3899974 | 21.5045156 |
| P54105 | CLNS1A | 21.8975864 | 22.082414 | 21.7829727 | 21.9449046 | 22.0498052 |
| Q9Y3C1 | NOP16 | 21.8291445 | 22.1008304 | 21.9831243 | 21.9175528 | 21.9486436 |
| Q8NC42 | RNF149 | 18.897338 | 18.6839043 | 18.6540881 | 18.8505641 | 18.7756805 |
| Q9Y696 | CLIC4 | 23.0309358 | 23.20881 | 23.0261345 | 22.9832507 | 22.9735993 |
| Q15637 | SF1 | 23.1961887 | 23.2533161 | 23.1427787 | 23.2089074 | 23.2909385 |
| Q52LR7 | EPC2 | 17.18786 | 17.3209975 | 16.979583 | 17.0134463 | 17.276647 |
| P13051 | UNG | 22.4335389 | 22.554635 | 22.7050873 | 22.5485573 | 22.611953 |
| Q9NVH0 | EXD2 | 17.3239147 | 17.6709538 | 17.5857145 | 17.6077874 | 17.5843854 |
| Q9NVK5 | FGFR1OP2 | 19.8976278 | 19.9011602 | 19.9733609 | 19.8526718 | 20.1456626 |

| | | | | | | |
|---|---|---|---|---|---|---|
| Q14332 | FZD2 | 18.0074426 | 18.1340021 | 17.9911288 | 18.0021305 | 18.1106948 |
| Q5T1V6 | DDX59 | 18.1739707 | 18.4125964 | 18.2308056 | 18.2863143 | 18.38543 |
| Q5PSV4 | BRMS1L | 18.9671005 | 18.9008145 | 19.137696 | 18.9259289 | 19.0567421 |
| O14730 | RIOK3 | 18.900429 | 18.8096417 | 18.9617691 | 18.8460541 | 18.7228773 |
| Q9BVW5 | TIPIN | 19.403525 | 19.6095375 | 19.3071086 | 19.3511092 | 19.3287051 |
| Q6P1Q9 | METTL2B | 19.7588779 | 19.9588886 | 19.7991615 | 19.8348642 | 19.9868067 |
| Q96BZ8 | LENG1 | 19.9064624 | 19.8000497 | 19.591774 | 19.7739061 | 19.787219 |
| Q04727 | TLE4 | 19.0108641 | 19.1180179 | 19.2041666 | 19.0401964 | 19.2399374 |
| Q9ULM3 | YEATS2 | 17.7510177 | 18.1747852 | 17.9920266 | 17.9937801 | 18.1933857 |
| O14745 | NHERF1 | 24.1943508 | 24.360479 | 24.0058027 | 24.1500904 | 24.2495935 |
| Q9BY41 | HDAC8 | 15.9874442 | 16.2981913 | 16.3459094 | 16.1020586 | 16.3870728 |
| Q99757 | TXN2 | 19.7051029 | 19.8313428 | 19.6614625 | 19.6639259 | 19.9412352 |
| Q15697 | ZNF174 | 17.1382987 | 17.1085632 | 17.015619 | 16.9382871 | 16.9591508 |
| Q66PJ3 | ARL6IP4 | 20.5989705 | 20.9445015 | 20.6864379 | 20.7328019 | 20.9347228 |
| P82970 | HMGN5 | 19.166672 | 19.3379859 | 18.876108 | 19.0049648 | 19.3415664 |
| Q8IV50 | LYSMD2 | 17.899292 | 17.930824 | 18.3556855 | 18.041995 | 18.5904036 |
| Q5JY77 | GPRASP1 | NA | 17.0024399 | 16.9507803 | NA | 17.3170042 |
| O00194 | RAB27B | 14.5250884 | NA | NA | 14.4462546 | NA |
| Q53F39 | MPPE1 | 14.949759 | 14.7036537 | 14.6155286 | 15.3393076 | 14.9026481 |
| Q70EL1 | USP54 | 17.4043577 | NA | NA | 17.7128368 | NA |
| Q8N8K9 | KIAA1958 | 15.6693207 | 15.1169673 | 16.3686736 | 16.007906 | 15.3043721 |
| Q9NQY0 | BIN3 | 18.3766263 | 18.2537317 | 18.452903 | 18.2441142 | 18.4245963 |
| Q9Y530 | OARD1 | 21.6023175 | 21.5898083 | 21.733059 | 21.6774325 | 21.754533 |
| Q7Z5L9 | IRF2BP2 | 23.4680094 | 23.6473044 | 23.5092616 | 23.410994 | 23.5631889 |
| A5PLN9 | TRAPPC13 | 20.9752055 | 21.0409589 | 20.9691365 | 20.9558786 | 21.0350754 |
| Q9UIG8 | SLCO3A1 | NA | 16.5573002 | 16.2625173 | NA | 16.4702516 |
| P51530 | DNA2 | 16.3912966 | 16.4409857 | 16.5497215 | 16.5640421 | 16.4169975 |
| O60524 | NEMF | 22.4965669 | 22.5255715 | 22.5717117 | 22.4517151 | 22.5026277 |
| O00189 | AP4M1 | 22.533012 | 22.6743871 | 22.7232137 | 22.6475485 | 22.6993921 |
| Q9NP99 | TREM1 | 16.3087368 | 16.3265362 | 16.4330863 | 16.4937485 | 16.3871903 |
| Q5TAX3 | TUT4 | 18.2876904 | 18.2650957 | 18.4711418 | 18.3727555 | 18.433445 |
| Q8NDX6 | ZNF740 | 18.536694 | 18.6658126 | 18.7291263 | 18.7308558 | 18.7387739 |
| Q99608 | NDN | 19.9374086 | 20.035689 | 19.9062705 | 19.9026321 | 20.0056996 |
| Q01518 | CAP1 | 25.8475383 | 25.9827123 | 25.9043979 | 25.7468165 | 25.9160285 |
| Q9H8H0 | NOL11 | 22.9770445 | 23.0172704 | 22.908713 | 22.8502366 | 22.9074031 |
| Q9Y2L5 | TRAPPC8 | 20.4876249 | 20.5610321 | 20.5863116 | 20.5157321 | 20.6213438 |
| P20292 | ALOX5AP | 21.0989945 | 20.8446542 | 21.7280717 | 21.0806286 | 21.5562707 |
| Q6DD88 | ATL3 | 22.7173844 | 22.7727338 | 22.7132778 | 22.6648195 | 22.7069197 |
| P55789 | GFER | 18.8101157 | 18.7401076 | 18.6005577 | 18.8982515 | 18.8367651 |
| Q9BUK6 | MSTO1 | 20.0213441 | 20.0478402 | 19.9918263 | 19.9281217 | 19.9942307 |
| Q12770 | SCAP | 17.877801 | 17.8599553 | 17.9899404 | 17.856068 | 17.9583947 |
| Q16594 | TAF9 | 19.8342209 | 19.8745992 | 19.945895 | 19.8627254 | 19.9229933 |

| | | | | | | |
|---|---|---|---|---|---|---|
| Q5BJF6 | ODF2 | 19.2873213 | 19.402618 | 19.3114812 | 19.251072 | 19.2857349 |
| Q9Y3E5 | PTRH2 | 21.085266 | 20.9493067 | 21.2103505 | 21.0015157 | 21.1296491 |
| Q9NQS7 | INCENP | 19.6515497 | 19.6063533 | 19.7262095 | 19.6825082 | 19.6449301 |
| Q96MP5 | ZSWIM3 | NA | 14.0665594 | 13.9639293 | NA | 14.1243397 |
| Q9H9Q4 | NHEJ1 | 19.8242774 | 19.9352901 | 19.8428094 | 19.6679857 | 19.8442452 |
| Q9Y3C7 | MED31 | 19.3271525 | 19.1825627 | 19.3894639 | 19.367343 | 19.3352462 |
| P36551 | CPOX | 22.5460053 | 22.6023107 | 22.6224703 | 22.5167675 | 22.6214492 |
| Q86YC2 | PALB2 | NA | 15.2699253 | 15.4038781 | NA | 14.7643996 |
| Q9HA92 | RSAD1 | 16.6348013 | 16.8213087 | 16.9581183 | 16.6814484 | 16.5739529 |
| O95568 | METTL18 | 19.1193982 | 19.0657947 | 19.0713543 | 18.9403786 | 18.9181555 |
| O14656 | TOR1A | 20.604809 | 20.640417 | 20.6263963 | 20.5823733 | 20.6142843 |
| P16615 | ATP2A2 | 23.966667 | 24.0080936 | 23.9653601 | 23.9149308 | 23.9193516 |
| Q9H6W3 | RIOX1 | 20.5980801 | 20.4389148 | 20.6486807 | 20.5691147 | 20.6384543 |
| Q99550 | MPHOSPH9 | NA | 13.5894292 | 14.0235175 | NA | 13.4958138 |
| Q53TN4 | CYBRD1 | 19.0816559 | 18.9628472 | 18.5653376 | 18.8739073 | 18.7906356 |
| Q96RN5 | MED15 | 20.0287645 | 20.0382621 | 20.0745961 | 20.0239905 | 20.0689869 |
| Q8N1B4 | VPS52 | 20.2390818 | 20.3091437 | 20.4154963 | 20.2900856 | 20.467315 |
| Q9H0U4 | RAB1B | 25.2850975 | 25.2997261 | 25.3066653 | 25.1474803 | 25.3165192 |
| Q92615 | LARP4B | 21.643827 | 21.7367549 | 21.645928 | 21.5696336 | 21.6930494 |
| Q08945 | SSRP1 | 24.213647 | 24.2797777 | 24.205151 | 24.1719438 | 24.1794579 |
| Q9Y230 | RUVBL2 | 24.2244365 | 24.3028322 | 24.195602 | 24.1456012 | 24.2232889 |
| Q8IYD1 | GSPT2 | 19.4831588 | 19.4679541 | 19.5560556 | 19.6072068 | 19.6377125 |
| Q9Y421 | FAM32A | 21.0792647 | 21.4462123 | 20.9836442 | 21.2246087 | 21.2806427 |
| Q96JM7 | L3MBTL3 | 17.4122462 | 17.4916838 | 17.6832023 | 17.5014453 | 17.6387602 |
| Q96LI5 | CNOT6L | 22.2816361 | 22.3097766 | 22.3858821 | 22.2830192 | 22.3908207 |
| Q9P0P0 | RNF181 | 21.1332002 | 21.2463254 | 21.0099518 | 21.1426658 | 21.1772519 |
| Q9HD45 | TM9SF3 | 22.5289389 | 22.629417 | 22.5729346 | 22.5593628 | 22.5469466 |
| P13497 | BMP1 | 18.6304966 | 18.5767543 | 18.4088447 | 18.5280221 | 18.5320319 |
| Q96QR8 | PURB | 21.1358517 | 21.1462636 | 21.2369335 | 21.1677312 | 21.283652 |
| P63027 | VAMP2 | 21.4220277 | 21.3764509 | 21.6749045 | 21.4265383 | 21.4484912 |
| Q9H1J1 | UPF3A | 18.379884 | 18.3311556 | 18.7312027 | 18.4786354 | 18.6809575 |
| Q9GZT6 | CCDC90B | 19.6830854 | 19.8940347 | 19.837881 | 19.8086205 | 19.9254303 |
| Q9BWL3 | C1orf43 | 16.4312875 | 16.9657013 | 16.8346093 | 16.7784691 | 16.6674296 |
| Q9Y5K8 | ATP6V1D | 21.5158389 | 21.7500731 | 21.552083 | 21.5286665 | 21.5959031 |
| P48553 | TRAPPC10 | 19.6974343 | 19.7677867 | 19.8254365 | 19.7411441 | 19.8140976 |
| O43934 | MFSD11 | 15.7988861 | 16.3474603 | 16.1891836 | 16.0417937 | 16.1483434 |
| O75865 | TRAPPC6A | 21.0634585 | 21.0723432 | 21.2206679 | 20.9342404 | 21.3401361 |
| Q92845 | KIFAP3 | 16.3121506 | 16.4926292 | 16.9870817 | 16.5087815 | 16.9374126 |
| P00414 | MT-CO3 | 17.5887928 | 17.8170096 | 17.5297124 | 17.6151041 | 17.7573463 |
| Q9H5U6 | ZCCHC4 | 17.5461387 | 17.3917667 | 17.715333 | 17.5561978 | 17.2607369 |
| P50151 | GNG10 | 20.6533616 | 20.5542747 | 20.6195944 | 20.6301613 | 20.6755903 |
| Q6IPU0 | CENPP | 17.7827236 | 18.0997216 | 17.8901932 | 17.8672038 | 17.5306932 |

| | | | | | | |
|---|---|---|---|---|---|---|
| P42680 | TEC | 18.4314428 | 18.4219066 | 18.5258778 | 18.396459 | 18.5055989 |
| Q9UBH6 | XPR1 | 17.5940161 | 17.512721 | 17.5773818 | 17.5250434 | 17.4764115 |
| P60709 | ACTB | 25.2484643 | 25.4545228 | 25.1834105 | 25.1998912 | 25.3033466 |
| Q8NDF8 | TENT4B | 19.7644295 | 19.7726142 | 19.8761186 | 19.6712694 | 19.6869089 |
| Q96SZ6 | CDK5RAP1 | 18.5928895 | 18.5989567 | 18.6555091 | 18.5337884 | 18.4867772 |
| Q9NP61 | ARFGAP3 | 21.1407034 | 21.35285 | 21.2194235 | 21.1650881 | 21.2570684 |
| Q9H4M9 | EHD1 | 22.625859 | 22.6914726 | 22.7075053 | 22.5496943 | 22.6993089 |
| O95696 | BRD1 | 20.7928193 | 20.807902 | 20.9501278 | 20.8029705 | 20.8729804 |
| P55769 | SNU13 | 23.8501239 | 23.8342116 | 23.8523892 | 23.8391311 | 23.9915232 |
| Q99543 | DNAJC2 | 21.6348812 | 21.6509012 | 21.6948854 | 21.5653731 | 21.6910822 |
| Q9NY33 | DPP3 | 23.4055744 | 23.4591696 | 23.4108119 | 23.3435938 | 23.405831 |
| O43175 | PHGDH | 22.098903 | 22.1684556 | 22.0413463 | 22.1103823 | 22.0942302 |
| O00566 | MPHOSPH10 | 22.6977418 | 22.7983045 | 22.7218025 | 22.6533281 | 22.7821448 |
| P49662 | CASP4 | 20.2721639 | 20.3396571 | 20.3211441 | 20.2104722 | 20.3359128 |
| Q9UIS9 | MBD1 | 19.4952964 | 19.6173084 | 19.6049272 | 19.5093711 | 19.6794831 |
| P48060 | GLIPR1 | 18.83907 | 18.8611917 | 18.8347827 | 18.8507929 | 18.8143401 |
| Q9NZJ5 | EIF2AK3 | 17.7347524 | 18.0455428 | 18.0336869 | 17.9741955 | 17.831949 |
| Q9H7M9 | VSIR | 21.0551216 | 21.0704615 | 21.2137992 | 21.0460803 | 21.2725648 |
| Q8IYR2 | SMYD4 | 18.0746206 | 18.1234001 | 18.3263752 | 18.3467424 | 18.2701627 |
| Q8WTS1 | ABHD5 | 14.5556113 | 14.633559 | 14.5613956 | 14.4232838 | 14.5624961 |
| P49916 | LIG3 | 20.6983588 | 20.7384034 | 20.7607003 | 20.6437221 | 20.7666966 |
| Q96H22 | CENPN | 16.7905789 | 16.9629612 | 16.8858395 | 16.9642382 | 17.0343321 |
| Q9Y5Y5 | PEX16 | 21.3944998 | 21.3598428 | 21.4092957 | 21.1138354 | 21.1765754 |
| Q05655 | PRKCD | 23.2413024 | 23.2856842 | 23.3451001 | 23.1911079 | 23.339802 |
| Q86XK2 | FBXO11 | 17.8105255 | 17.8154209 | 18.1490069 | 18.0728302 | 17.9761543 |
| Q9UKB1 | FBXW11 | 18.9992358 | 19.0524 | 18.9621362 | 18.9315254 | 18.855497 |
| Q9GZP4 | PITHD1 | 20.5943363 | 20.4854409 | 20.5365137 | 20.5524195 | 20.5336006 |
| Q6VN20 | RANBP10 | 20.195701 | 20.3569988 | 20.3236839 | 20.3543457 | 20.264784 |
| Q96DF8 | ESS2 | 18.549645 | 18.8348473 | 18.8082611 | 18.7712541 | 18.8086535 |
| P22079 | LPO | 19.1866526 | NA | NA | 19.0356267 | NA |
| P17655 | CAPN2 | 23.4506926 | 23.517634 | 23.5192049 | 23.3958906 | 23.5380381 |
| Q8WVK2 | SNRNP27 | 18.9552411 | 19.1487267 | 19.2480898 | 19.0723722 | 19.2529845 |
| Q96HH6 | TMEM19 | 17.9245845 | 17.2327538 | 18.4208456 | 17.9825131 | 17.3963316 |
| Q02539 | H1-1 | 19.195776 | 19.2539703 | 19.7146153 | 19.1459946 | 19.3393923 |
| Q9UGQ3 | SLC2A6 | 17.9933624 | 18.2272763 | 18.1779491 | 18.0050368 | 18.0932981 |
| P16402 | H1-3 | 23.4841164 | 23.7843038 | 23.8784254 | 23.392017 | 23.4894093 |
| Q9NPH2 | ISYNA1 | 19.0345724 | 19.0560388 | 19.2603402 | 18.8487938 | 19.0884415 |
| Q02386 | ZNF45 | 16.7139729 | 16.8104922 | 16.7494027 | 16.8532076 | 16.8731383 |
| Q9HCJ6 | VAT1L | 18.6880407 | 18.8365534 | 18.8200323 | 18.7623045 | 18.9019828 |
| P53671 | LIMK2 | 14.6963729 | 14.9071718 | 14.7585026 | 14.6863662 | 14.6305039 |
| O43765 | SGTA | 22.8395445 | 22.8219197 | 22.7291522 | 22.7762507 | 22.9499212 |
| Q9NXA8 | SIRT5 | 18.6950346 | 18.6354091 | 18.4085394 | 18.5930004 | 18.4897287 |

| | | | | | | |
|---|---|---|---|---|---|---|
| O60637 | TSPAN3 | 20.3163426 | 20.4483812 | 20.1510763 | 20.5021736 | 20.19777 |
| P57105 | SYNJ2BP | 18.3403149 | 18.4156901 | 18.4646342 | 18.6559222 | 18.3575499 |
| Q9BR76 | CORO1B | 23.1105777 | 23.1871794 | 23.0653657 | 23.1233347 | 23.1341422 |
| Q92545 | TMEM131 | 21.9603063 | 22.2250722 | 22.1979576 | 21.9645141 | 22.1315715 |
| Q6P1R3 | MSANTD2 | 19.264315 | 19.3345151 | 19.2804495 | 19.0679132 | 19.2746816 |
| Q9BZD4 | NUF2 | 20.422498 | 20.4985231 | 20.5469694 | 20.3730057 | 20.3151173 |
| Q53HC9 | EIPR1 | 20.3859386 | 20.5142103 | 20.3828858 | 20.5045138 | 20.5946923 |
| Q8N988 | ZNF557 | 17.4825439 | 17.7161393 | 17.8403713 | 17.8200083 | 17.7946075 |
| Q9H0X9 | OSBPL5 | 20.8219547 | 20.8416175 | 20.7806275 | 20.7311085 | 20.7809948 |
| Q9Y2X0 | MED16 | 20.3459764 | 20.3997988 | 20.4996561 | 20.3313343 | 20.339779 |
| Q9H1Z4 | WDR13 | 18.4231612 | 18.6202723 | 18.5927793 | 18.4311431 | 18.5166175 |
| Q12959 | DLG1 | 19.1981958 | 19.3377435 | 19.1370552 | 19.1091779 | 19.3778362 |
| P14317 | HCLS1 | 24.4026584 | 24.5616181 | 24.2259939 | 24.3469237 | 24.3366295 |
| Q03405 | PLAUR | 20.1919339 | 20.4680844 | 20.4751259 | 20.3322003 | 20.3953413 |
| Q9H0C5 | BTBD1 | 15.3288024 | NA | NA | 15.2508876 | NA |
| A2AJT9 | BCLAF3 | 18.0909507 | 18.2962807 | 18.4169831 | 18.3396545 | 18.0432558 |
| Q8NDX5 | PHC3 | 18.5128318 | 18.2529038 | 18.4422141 | 18.5163696 | 18.3261319 |
| Q9UKL6 | PCTP | 20.1535546 | 20.1306167 | 20.3504232 | 20.100314 | 20.1030383 |
| O95185 | UNC5C | 19.5657731 | 19.656364 | 19.7947913 | 19.8597652 | 19.7724098 |
| O95218 | ZRANB2 | 22.4491463 | 22.5465412 | 22.411429 | 22.4202152 | 22.4778246 |
| O00287 | RFXAP | 17.5249877 | 17.5781317 | 17.6308835 | 17.6769031 | 17.7334273 |
| Q8IUZ5 | PHYKPL | 18.1762807 | 18.442164 | 18.4009097 | 18.4190432 | 18.5430988 |
| Q92851 | CASP10 | 18.3784351 | 18.3978014 | 18.8000486 | 18.5664074 | 18.7247809 |
| Q9UM82 | SPATA2 | NA | 16.1696887 | 16.3489479 | NA | 16.2425751 |
| Q96NW4 | ANKRD27 | 19.5260586 | 19.4859473 | 19.5983506 | 19.489519 | 19.5252108 |
| Q9NTI5 | PDS5B | 21.7310958 | 21.7791548 | 21.8512337 | 21.7084411 | 21.7958831 |
| O14497 | ARID1A | 21.5813871 | 21.614542 | 21.5649683 | 21.5902095 | 21.6389859 |
| P35052 | GPC1 | 12.017737 | 12.7001155 | 12.2810629 | 12.185174 | 12.2400947 |
| Q9BQD3 | KXD1 | 19.0873264 | 18.9541012 | 19.0629299 | 18.9408342 | 18.8897501 |
| Q9Y3B1 | PRELID3B | 16.3476718 | 16.3079599 | 16.5200907 | 16.4497458 | 16.3840197 |
| O95025 | SEMA3D | 21.3264226 | NA | NA | 21.0487621 | NA |
| Q92901 | RPL3L | NA | 14.5949079 | 14.7449294 | NA | 14.2622563 |
| P56202 | CTSW | 16.5346515 | 16.6342986 | 16.5471913 | 16.3831507 | 16.6287086 |
| Q3ZAQ7 | VMA21 | 19.6331601 | 19.5935435 | 19.5213924 | 19.3374388 | 19.2480604 |
| Q9NUM4 | TMEM106B | 20.2711676 | 20.3287682 | 20.4398942 | 20.4814578 | 20.3414086 |
| Q8WXE1 | ATRIP | 18.9192413 | 18.9580778 | 18.9414563 | 18.9906043 | 18.9992425 |
| Q5VSL9 | STRIP1 | 19.9963938 | 19.9716202 | 19.9215797 | 19.9093758 | 19.9866149 |
| Q9NV70 | EXOC1 | 21.2736326 | 21.360424 | 21.3902208 | 21.2298948 | 21.3027046 |
| Q14258 | TRIM25 | 23.4396089 | 23.4237854 | 23.5431755 | 23.3685957 | 23.460345 |
| P26374 | CHML | 20.3833515 | 20.4633203 | 20.3990387 | 20.4207416 | 20.3487344 |
| Q96BP3 | PPWD1 | 21.6261372 | 21.6646709 | 21.642952 | 21.6234595 | 21.6198445 |
| O95749 | GGPS1 | 19.8325222 | 19.8600916 | 19.8826028 | 19.7925674 | 19.8562266 |

| | | | | | | |
|---|---|---|---|---|---|---|
| Q16763 | UBE2S | 20.8339768 | 20.9315269 | 20.8548895 | 20.8486076 | 20.9572347 |
| Q5TDH0 | DDI2 | 21.1439715 | 21.3953101 | 21.3421449 | 21.2907877 | 21.3525557 |
| Q9UNQ2 | DIMT1 | 20.6315819 | 20.7167373 | 20.875349 | 20.7299547 | 20.843781 |
| Q05823 | RNASEL | 20.7349003 | 20.7852871 | 20.7976576 | 20.661691 | 20.8233964 |
| Q14164 | IKBKE | 18.0810174 | 18.1296316 | 18.2835568 | 18.2246931 | 18.1958879 |
| Q8WYA0 | IFT81 | 17.3913268 | 17.2895654 | 17.3629127 | 17.481194 | 17.3910256 |
| Q9H3R0 | KDM4C | 16.721882 | 16.9818622 | 17.227108 | 16.9556164 | 17.0512872 |
| Q12766 | HMGXB3 | 15.8850932 | 15.6765463 | 16.2484185 | 16.0107329 | 15.8807241 |
| Q9BRR3 | PGAP4 | 16.6471521 | NA | NA | 16.677893 | NA |
| O75436 | VPS26A | 23.4724118 | 23.498399 | 23.4318612 | 23.333128 | 23.3987862 |
| Q9H981 | ACTR8 | 19.9597027 | 20.0877248 | 20.1525883 | 20.0641946 | 20.1231932 |
| Q9Y4I1 | MYO5A | 19.5869468 | 19.6989184 | 19.7265527 | 19.6155887 | 19.6900345 |
| Q9H944 | MED20 | 20.0155741 | 20.1345934 | 20.1481383 | 20.0321203 | 20.2342489 |
| Q14242 | SELPLG | 20.0410709 | 20.3620081 | 20.2025336 | 20.1526312 | 20.3408832 |
| O00743 | PPP6C | 23.0757667 | 23.0800148 | 23.1499575 | 22.9850257 | 23.043777 |
| P00746 | CFD | 21.609655 | 21.7658176 | 21.6663659 | 21.5869863 | 21.6508252 |
| P13073 | COX4I1 | 25.4176153 | 25.4074721 | 25.3562274 | 25.2773161 | 25.3037187 |
| Q9NSI6 | BRWD1 | 16.6322724 | 16.6756646 | 16.671535 | 16.4941297 | 16.7067119 |
| Q93100 | PHKB | 19.5486487 | 19.6302143 | 19.6672573 | 19.4727113 | 19.5965922 |
| Q6UWE0 | LRSAM1 | 20.697673 | 20.7752108 | 20.7379184 | 20.6455907 | 20.8370879 |
| O95873 | C6orf47 | 18.3247153 | 18.4037371 | 18.684255 | 18.5183203 | 18.3848411 |
| Q13286 | CLN3 | 17.5007458 | NA | NA | 17.6298788 | NA |
| Q9BQG2 | NUDT12 | NA | 15.3636584 | 15.2639716 | NA | 15.1997502 |
| Q9BQ65 | USB1 | 15.8067622 | 15.5924601 | 16.2230555 | 15.9353955 | 15.8844784 |
| Q8WUX2 | CHAC2 | 18.4811955 | 18.7576018 | 18.9327792 | 18.5913177 | 18.4455311 |
| P27487 | DPP4 | 18.4277193 | 18.9006529 | 18.8366634 | 18.5024527 | 18.7231946 |
| Q96GC9 | VMP1 | 18.7326252 | 18.9052706 | 18.9969111 | 18.8379954 | 18.7285278 |
| Q15363 | TMED2 | 24.3953862 | 24.4410467 | 24.3841167 | 24.1706906 | 24.1844647 |
| Q8WUU5 | GATAD1 | 17.2945547 | 17.0279177 | 17.42298 | 17.2313498 | 17.309217 |
| Q86WA6 | BPHL | 19.0095454 | 19.1149252 | 19.1545392 | 18.9330271 | 19.2774764 |
| P29728 | OAS2 | 17.3120484 | 17.6035064 | 17.5134824 | 17.3335328 | 17.5045389 |
| Q8TF50 | ZNF526 | 19.3591621 | 19.2863997 | 19.3205112 | 19.2602725 | 19.2233111 |
| Q9NRC1 | ST7 | 20.8479779 | 20.8903457 | 20.8114458 | 20.7253902 | 20.6597445 |
| O60704 | TPST2 | 17.5090409 | 17.5536804 | 17.5680308 | 17.4186868 | 17.5060354 |
| Q8NBJ9 | SIDT2 | 20.1143951 | 20.1729393 | 20.0967016 | 19.9745585 | 19.9036554 |
| Q9H553 | ALG2 | 19.7702627 | 19.7826649 | 20.0906831 | 19.736686 | 19.9507645 |
| Q5VV42 | CDKAL1 | 19.7378498 | 19.9468216 | 19.9974695 | 19.7180797 | 19.95151 |
| Q9P260 | RELCH | 18.0039609 | 17.986014 | 17.9177378 | 17.9129781 | 17.8868954 |
| Q7L2Z9 | CENPQ | 16.9017496 | 16.7252361 | 17.0160792 | 16.6742196 | 16.8634615 |
| Q96KC8 | DNAJC1 | 18.2506531 | 18.4871275 | 18.2974875 | 18.20256 | 18.292894 |
| Q9H307 | PNN | 23.2031227 | 23.2627104 | 23.259188 | 23.1626553 | 23.1400693 |
| O00160 | MYO1F | 23.656926 | 23.7127724 | 23.7520913 | 23.6445017 | 23.7257614 |

| | | | | | | |
|---|---|---|---|---|---|---|
| Q96I24 | FUBP3 | 22.824268 | 22.8453805 | 22.8094876 | 22.7787518 | 22.7737969 |
| Q9NQG6 | MIEF1 | 19.8972322 | 20.049894 | 19.9384774 | 19.9434621 | 20.0215955 |
| P41091 | EIF2S3 | 23.8541015 | 23.743757 | 23.9276304 | 23.7644312 | 23.8318431 |
| P32321 | DCTD | 20.3267826 | 20.3240578 | 20.3303512 | 20.3174714 | 20.3931178 |
| Q9UNS2 | COPS3 | 22.0948118 | 22.2114518 | 22.1714465 | 22.1321895 | 22.1648956 |
| O95433 | AHSA1 | 23.8790843 | 23.9926665 | 23.9807845 | 23.8622042 | 23.919786 |
| P48735 | IDH2 | 26.1193074 | 26.1350683 | 26.0835593 | 26.0354074 | 26.1323878 |
| Q9NUT2 | ABCB8 | 20.1860283 | 20.2541493 | 20.2321726 | 20.0766443 | 20.2500851 |
| P35579 | MYH9 | 26.1794135 | 26.2539037 | 26.174509 | 26.071529 | 26.1483573 |
| O00231 | PSMD11 | 23.5622606 | 23.6356488 | 23.5066549 | 23.4638937 | 23.543648 |
| P54886 | ALDH18A1 | 22.9363327 | 22.9725653 | 22.964692 | 22.8506512 | 22.9324532 |
| P12955 | PEPD | 23.5270179 | 23.4949815 | 23.468085 | 23.3995689 | 23.4335735 |
| P42696 | RBM34 | 20.6491706 | 20.6950803 | 20.7564441 | 20.6685444 | 20.7377173 |
| O94966 | USP19 | 20.9419344 | 20.9017499 | 20.9549425 | 20.8930781 | 20.8818525 |
| Q8IZQ5 | SELENOH | 22.4339919 | 22.4726264 | 22.3957467 | 22.432913 | 22.4945282 |
| O43913 | ORC5 | 19.2298452 | 19.3140444 | 19.3703414 | 19.2815216 | 19.3736695 |
| Q16644 | MAPKAPK3 | 21.0722617 | 21.0348918 | 21.0556113 | 21.1092707 | 21.1099029 |
| Q01831 | XPC | 20.8838006 | 20.7559029 | 20.8260343 | 20.8711457 | 20.9387056 |
| Q96N64 | PWWP2A | 18.8586834 | 18.8157189 | 18.8347437 | 18.8935683 | 18.7962324 |
| Q13555 | CAMK2G | 19.9318165 | 20.0523319 | 20.0519803 | 19.9971468 | 19.9571007 |
| Q6ZMI0 | PPP1R21 | 20.3622629 | 20.5742869 | 20.4629829 | 20.450942 | 20.5361443 |
| Q13347 | EIF3I | 24.2018606 | 24.2358008 | 24.1593541 | 24.1170444 | 24.1928694 |
| Q9H0L4 | CSTF2T | 22.2415629 | 22.2470911 | 22.091396 | 22.1857381 | 22.2372431 |
| Q16555 | DPYSL2 | 25.1696183 | 25.2317859 | 25.1026386 | 25.0816211 | 25.2594827 |
| P51580 | TPMT | 21.1390071 | 21.1784171 | 21.2451936 | 21.23668 | 21.3204874 |
| P51784 | USP11 | 20.4170936 | 20.5383451 | 20.4525701 | 20.3953718 | 20.5244491 |
| O75475 | PSIP1 | 23.4079307 | 23.5296505 | 23.4808595 | 23.4245767 | 23.4914376 |
| P24941 | CDK2 | 20.1077899 | 20.2632702 | 20.155716 | 20.1193443 | 20.1797625 |
| Q9H993 | ARMT1 | 21.9538074 | 22.0607854 | 22.0706079 | 21.9272623 | 22.0817025 |
| Q96LT9 | RNPC3 | 18.4086915 | 18.3271669 | 18.5679057 | 18.5327561 | 18.5585657 |
| Q9P2K3 | RCOR3 | 18.738108 | 18.8750704 | 18.8475247 | 18.7936209 | 18.9472826 |
| Q96IY1 | NSL1 | 18.9277508 | 19.1244219 | 19.1268863 | 19.0480422 | 19.1447749 |
| P12314 | FCGR1A | 20.9677107 | 20.9154597 | 21.1147665 | 20.857316 | 21.1480993 |
| Q96MN5 | TCEANC2 | 17.3611688 | 17.7345455 | 17.5854202 | 17.4348022 | 17.6069304 |
| O14893 | GEMIN2 | 19.2631806 | 19.3749053 | 19.274875 | 19.3450802 | 19.1430544 |
| Q99595 | TIMM17A | 19.2690095 | 19.3406401 | 19.3907929 | 19.4057167 | 19.4945916 |
| Q9H8G1 | ZNF430 | 18.0449007 | 18.0819141 | 18.2419234 | 18.0669049 | 17.9620618 |
| Q96B77 | TMEM186 | 18.1198173 | 18.0452461 | 18.1850826 | 17.9056784 | 18.2069671 |
| O14867 | BACH1 | 17.6184757 | 17.7716719 | 17.7761489 | 17.5914868 | 17.9413821 |
| Q9NWZ8 | GEMIN8 | 19.7844856 | 19.5915403 | 20.0027118 | 19.689658 | 19.8241189 |
| O00124 | UBXN8 | 18.4781027 | 18.5438352 | 18.4828014 | 18.5284783 | 18.5954533 |
| O15162 | PLSCR1 | 19.5050231 | 19.8000267 | 19.3167925 | 19.5312511 | 19.3728864 |

| | | | | | | |
|---|---|---|---|---|---|---|
| Q96GX5 | MASTL | 17.5920241 | 17.5788198 | 17.8008991 | 17.7014626 | 17.5420958 |
| Q9Y5S9 | RBM8A | 24.3410967 | 24.5245367 | 24.2028395 | 24.2728869 | 24.4131462 |
| Q8TDR0 | TRAF3IP1 | 16.5895328 | 16.8405257 | 16.9989547 | 16.5516137 | 16.9718577 |
| P78549 | NTHL1 | 18.0815717 | 18.2632712 | 18.2437016 | 18.1989266 | 18.4584377 |
| Q13163 | MAP2K5 | 19.2778365 | 19.4057677 | 19.4638434 | 19.3818689 | 19.0111031 |
| P18859 | ATP5PF | 23.6205021 | 23.8780733 | 23.5818472 | 23.5967836 | 23.872386 |
| Q86YD1 | PTOV1 | 16.8024912 | 17.1359512 | 16.947783 | 17.0809139 | 16.6649994 |
| P24311 | COX7B | 21.2823784 | 21.1057429 | 21.624258 | 21.1368013 | 21.357384 |
| O43822 | CFAP410 | 14.8337721 | NA | NA | 14.9041223 | NA |
| Q15506 | SPA17 | 15.8573183 | 15.6043809 | 15.4683827 | 16.013623 | 15.9923514 |
| Q9NRP7 | STK36 | 15.5213193 | NA | NA | 15.6481607 | NA |
| O43151 | TET3 | 15.8665686 | 15.9591799 | 15.9248884 | 15.7546823 | 16.0192522 |
| Q8TCD1 | C18orf32 | 19.1467111 | 19.0422114 | 19.2588358 | 19.0353783 | 18.9825422 |
| Q9BV94 | EDEM2 | 17.7493022 | 17.796389 | 18.0383812 | 17.7225993 | 17.9281615 |
| Q9BW60 | ELOVL1 | 17.7565764 | 17.6683194 | 17.621749 | 17.8523838 | 17.3879967 |
| Q16611 | BAK1 | 20.7893617 | 20.6957503 | 21.0300886 | 20.8780373 | 20.7857634 |
| Q8NCN2 | ZBTB34 | 18.2807688 | 18.3288871 | 18.2040722 | 18.2293012 | 18.2755338 |
| O43296 | ZNF264 | 18.3402731 | 18.6593867 | 18.6438914 | 18.4891518 | 18.3876068 |
| Q9NX40 | OCIAD1 | 22.5643205 | 22.5939374 | 22.6032549 | 22.463262 | 22.4876491 |
| Q96BN2 | TADA1 | 18.7697228 | 19.0054885 | 18.9586877 | 18.812378 | 18.8827402 |
| Q15628 | TRADD | 19.8507272 | 20.013602 | 19.9522791 | 19.9466664 | 19.8911076 |
| P42765 | ACAA2 | 22.9857971 | 23.1199444 | 23.0377742 | 22.8784727 | 22.9979848 |
| O15355 | PPM1G | 23.4170866 | 23.5044533 | 23.3245509 | 23.3433285 | 23.387113 |
| Q9P289 | STK26 | 22.2749875 | 22.3529962 | 22.313035 | 22.229763 | 22.3559737 |
| Q96GA3 | LTV1 | 20.5329708 | 20.5850728 | 20.4620499 | 20.4817011 | 20.6370413 |
| Q9NPL8 | TIMMDC1 | 21.6004132 | 21.6599386 | 21.5633831 | 21.5993332 | 21.6373245 |
| Q9UBS0 | RPS6KB2 | 18.8718469 | 19.0926061 | 19.0260549 | 18.9406868 | 19.0155048 |
| Q92625 | ANKS1A | 19.6129978 | 19.6652304 | 19.768573 | 19.6037141 | 19.8202848 |
| Q16643 | DBN1 | 21.5625959 | 21.6419181 | 21.4929567 | 21.510167 | 21.694583 |
| P06493 | CDK1 | 23.0183921 | 23.1363308 | 23.1606827 | 23.0638803 | 23.192145 |
| Q9H4A5 | GOLPH3L | 19.4668946 | 19.2473443 | 19.493319 | 19.5162325 | 19.4819736 |
| O60613 | SELENOF | 21.4767049 | 21.5827851 | 21.4055589 | 21.4376654 | 21.5938606 |
| Q5M775 | SPECC1 | 20.8769953 | 21.0467178 | 20.9062799 | 20.9060815 | 21.0296366 |
| Q9H2K0 | MTIF3 | 19.849628 | 19.9306694 | 19.9596736 | 19.8843003 | 19.9141941 |
| P55201 | BRPF1 | 19.1887936 | 19.3157127 | 19.3905174 | 19.2367791 | 19.0676013 |
| Q8WXI9 | GATAD2B | 20.4195404 | 20.4298469 | 20.4675718 | 20.3470975 | 20.5725989 |
| Q5VZ89 | DENND4C | 19.2987261 | 19.576899 | 19.3267302 | 19.3836517 | 19.3663948 |
| O00750 | PIK3C2B | 17.6074821 | 17.4411005 | 17.3620432 | 17.5725115 | 17.4484677 |
| Q9H1E3 | NUCKS1 | 24.850841 | 24.9812599 | 24.6505071 | 24.667474 | 24.8401327 |
| Q96Q42 | ALS2 | 15.6163143 | 15.5251752 | 15.8258644 | 15.9301147 | 15.6817427 |
| Q14061 | COX17 | 21.8363114 | 21.7963767 | 21.746005 | 21.7763597 | 22.0737384 |
| Q96KR6 | FAM210B | 16.4113559 | 16.6621125 | 16.3609142 | 16.6165059 | 16.3621863 |

| | | | | | | |
|---|---|---|---|---|---|---|
| P16452 | EPB42 | 19.9027214 | 19.6968006 | 20.2782715 | 20.0424188 | 20.1769451 |
| Q9HCI7 | MSL2 | 17.999626 | 18.366921 | 18.0346411 | 18.2183592 | 18.2724248 |
| Q9NY93 | DDX56 | 20.8406097 | 20.9749998 | 20.9720125 | 20.795152 | 20.9040469 |
| Q86XJ0 | CALHM3 | 15.1638343 | NA | NA | 15.7065745 | NA |
| A0A0U1RRL7 | MMP24OS | 18.8769438 | 18.6376011 | 18.8524362 | 18.8788022 | 18.7656198 |
| Q9NSG2 | FIRRM | 16.6228794 | 16.8837617 | 16.9613377 | 16.7587168 | 16.860188 |
| P30281 | CCND3 | 19.8500034 | 20.0794207 | 19.7309177 | 19.6841057 | 19.8637458 |
| O43709 | BUD23 | 20.7047095 | 20.7192581 | 20.8189639 | 20.6967737 | 20.7458778 |
| Q9BQE3 | TUBA1C | 27.348138 | 27.4830017 | 27.3160842 | 27.1853589 | 27.3138559 |
| P54922 | ADPRH | 19.7294455 | 19.7445463 | 19.6964331 | 19.6988936 | 19.6037377 |
| Q7Z478 | DHX29 | 20.7055792 | 20.7427105 | 20.8149202 | 20.6307595 | 20.703826 |
| Q9BZJ0 | CRNKL1 | 21.2255342 | 21.2716647 | 21.3499027 | 21.2162109 | 21.3393847 |
| O43149 | ZZEF1 | 22.3027963 | 22.3493473 | 22.3466043 | 22.2201671 | 22.3265157 |
| Q96JI7 | SPG11 | 18.8069805 | 18.7656634 | 18.8894523 | 18.7160834 | 18.7940909 |
| Q8NEZ5 | FBXO22 | 21.0600286 | 21.100277 | 21.1962007 | 21.0694365 | 21.1830075 |
| Q00169 | PITPNA | 22.3215958 | 22.2990145 | 22.3507849 | 22.2846157 | 22.3295974 |
| Q330K2 | NDUFAF6 | 18.6219932 | 18.4147176 | 18.5998509 | 18.4014514 | 18.5757462 |
| Q8NHY2 | COP1 | 17.4356972 | 17.6512878 | 17.6550758 | 17.5022064 | 17.7183147 |
| Q5FWF4 | ZRANB3 | 17.2507297 | 17.4841793 | 17.3376966 | 17.2626368 | 17.2168803 |
| Q96CD0 | FBXL8 | 17.5708944 | 17.8390838 | 17.4741422 | 17.7558255 | 17.757149 |
| Q9NP80 | PNPLA8 | 17.4831515 | 17.7272497 | 17.8897617 | 17.6919801 | 17.9134568 |
| Q4VC05 | BCL7A | 19.1298035 | 19.3008579 | 19.2311401 | 19.2758945 | 19.2029402 |
| Q7Z3E2 | CCDC186 | 16.2889747 | 16.3190915 | 16.133716 | 16.4016613 | 16.4178572 |
| Q8NCM8 | DYNC2H1 | 16.1436065 | 16.4790195 | 16.2472183 | 16.1791677 | 16.390517 |
| Q9NSY1 | BMP2K | 18.6716185 | 18.7492088 | 18.7808688 | 18.7010027 | 18.763126 |
| Q9BVG8 | KIFC3 | NA | 14.360805 | 14.3631207 | NA | 14.5830456 |
| O94811 | TPPP | 15.0012741 | NA | NA | 14.7382658 | NA |
| A7MCY6 | TBKBP1 | 15.3808539 | 15.1201745 | 14.8121151 | 14.9451694 | 15.2538309 |
| Q15118 | PDK1 | 18.7301712 | 18.8547234 | 18.7795544 | 18.6250805 | 18.8354907 |
| Q8WUX1 | SLC38A5 | 20.7893249 | 20.842919 | 20.7954671 | 20.6798775 | 20.7814756 |
| Q04446 | GBE1 | 22.283858 | 22.4223014 | 22.4992888 | 22.2496196 | 22.4100121 |
| P07737 | PFN1 | 28.0089079 | 28.0363397 | 28.0387744 | 27.8469792 | 28.0424357 |
| Q3KQV9 | UAP1L1 | 20.264985 | 20.3933212 | 20.3247393 | 20.2553785 | 20.4187526 |
| Q12768 | WASHC5 | 21.5342347 | 21.5694617 | 21.6192774 | 21.4893219 | 21.5127726 |
| Q9UG63 | ABCF2 | 22.975782 | 22.9665417 | 22.9567646 | 22.9009381 | 22.8869147 |
| Q14152 | EIF3A | 24.5520789 | 24.549378 | 24.5869244 | 24.4757815 | 24.5074374 |
| Q13769 | THOC5 | 21.0080217 | 21.049291 | 21.0697734 | 20.9888265 | 21.0995538 |
| Q92556 | ELMO1 | 21.3967 | 21.4462237 | 21.4498219 | 21.2966932 | 21.452519 |
| P35249 | RFC4 | 22.0765685 | 22.2221865 | 22.2691018 | 22.1135459 | 22.1973278 |
| A6NED2 | RCCD1 | 18.0881591 | 18.3375503 | 18.4050553 | 18.4026919 | 18.328512 |
| Q7Z5U6 | WDR53 | 15.836686 | 16.0403325 | 15.9537656 | 15.9699343 | 15.9540042 |
| Q6ZMD2 | SPNS3 | NA | 15.7168232 | 15.5097988 | NA | 15.6181918 |

| | | | | | | |
|---|---|---|---|---|---|---|
| Q8TF32 | ZNF431 | 16.2266864 | 15.9672978 | 16.4705197 | 16.2031845 | 15.8286891 |
| Q4W5G0 | TIGD2 | 14.6143615 | NA | NA | 14.7012769 | NA |
| P04080 | CSTB | 24.6900037 | 24.7413336 | 24.6602353 | 24.4862447 | 24.8975178 |
| Q969X1 | TMBIM1 | 17.2825097 | 17.1094035 | 17.6725576 | 17.3021499 | 17.3250514 |
| Q86WP2 | GPBP1 | 17.214738 | 17.0367934 | 17.2380499 | 17.1373492 | 17.0298526 |
| Q9UBR5 | CKLF | 18.343781 | 18.2761014 | 18.3276658 | 18.2384488 | 18.456907 |
| P54274 | TERF1 | 17.1578162 | 17.249745 | 17.1152191 | 17.1002876 | 17.0912441 |
| Q4G0F5 | VPS26B | 22.1241064 | 22.2164542 | 22.2536213 | 22.0686504 | 22.1970384 |
| Q06787 | FMR1 | 20.5131692 | 20.6210501 | 20.61355 | 20.5223842 | 20.5047751 |
| Q15061 | WDR43 | 21.9756417 | 21.9500934 | 21.9563047 | 21.9758877 | 21.8853625 |
| P78356 | PIP4K2B | 20.9693418 | 21.0648694 | 21.0441015 | 21.060211 | 20.9135067 |
| P61158 | ACTR3 | 24.7811132 | 24.8353046 | 24.8294363 | 24.73115 | 24.8653536 |
| Q9ULK4 | MED23 | 20.9580478 | 21.0527971 | 21.069724 | 20.9813815 | 21.0485836 |
| O94776 | MTA2 | 23.5597555 | 23.5870375 | 23.6478709 | 23.4934537 | 23.5962706 |
| Q9P281 | BAHCC1 | 19.088941 | 19.1973495 | 19.0578528 | 19.0596366 | 19.0914124 |
| Q99615 | DNAJC7 | 23.1201556 | 23.1748694 | 23.1413237 | 23.1562356 | 23.191315 |
| Q66K74 | MAP1S | 21.7687286 | 21.7999857 | 21.8003694 | 21.6926866 | 21.8458204 |
| Q6ZSZ5 | ARHGEF18 | 21.055001 | 21.193994 | 21.1080919 | 21.1186189 | 21.1687179 |
| Q8WYP5 | AHCTF1 | 20.3783852 | 20.4909913 | 20.4317724 | 20.4196955 | 20.5647113 |
| Q15287 | RNPS1 | 22.6919871 | 22.7538981 | 22.7808923 | 22.7004048 | 22.7979234 |
| Q14677 | CLINT1 | 22.7383127 | 22.773125 | 22.7058067 | 22.7494348 | 22.807073 |
| Q15652 | JMJD1C | 19.157416 | 19.1264083 | 19.3085097 | 19.1793233 | 19.2764139 |
| P17050 | NAGA | 21.8451979 | 21.9813083 | 21.9695955 | 21.8821516 | 21.9683174 |
| Q5T5Y3 | CAMSAP1 | 18.5810126 | 18.7327162 | 18.7337004 | 18.6842456 | 18.7075396 |
| P0CJ78 | ZNF865 | 18.552794 | 18.59881 | 18.6116525 | 18.5421255 | 18.7656346 |
| P27707 | DCK | 21.5820052 | 21.6371291 | 21.6433708 | 21.6335483 | 21.648858 |
| P04183 | TK1 | 20.8672426 | 20.9567269 | 20.9847154 | 20.8323512 | 20.9520412 |
| Q9Y5X2 | SNX8 | 20.0043202 | 20.1523212 | 20.2389635 | 20.1068083 | 20.3249409 |
| P21953 | BCKDHB | 19.8344946 | 19.7171603 | 19.8791089 | 19.7867734 | 19.9086709 |
| Q5K4L6 | SLC27A3 | 19.7502086 | 19.923701 | 19.832426 | 19.9019151 | 19.8907603 |
| O15446 | POLR1G | 20.3264496 | 20.4348167 | 20.2967355 | 20.3123786 | 20.4239998 |
| Q96CN7 | ISOC1 | 21.0204108 | 21.1579806 | 21.2100644 | 20.9832172 | 21.1435697 |
| Q17RN3 | FAM98C | 19.4450842 | 19.6815209 | 19.4618716 | 19.5290519 | 19.5934746 |
| Q92581 | SLC9A6 | 19.4691749 | 19.3092004 | 19.3382556 | 19.4492384 | 19.4538222 |
| Q9BZF9 | UACA | 18.7459836 | 18.828754 | 18.8347526 | 18.6960658 | 18.9103631 |
| Q86YQ8 | CPNE8 | 19.1115542 | 19.2102936 | 19.218076 | 19.0987204 | 19.3156896 |
| Q8TBE0 | BAHD1 | 17.9312721 | 18.126321 | 18.0564033 | 18.1605083 | 18.2420746 |
| Q9H305 | CDIP1 | 19.1619292 | 19.2435161 | 19.2259548 | 19.3098224 | 19.0538763 |
| Q96HJ9 | FMC1 | 20.0531457 | 20.0668428 | 20.233806 | 20.0655261 | 20.155603 |
| Q9NXV2 | KCTD5 | 18.5421258 | 18.5441663 | 18.2538348 | 18.6393154 | 18.6999926 |
| P31937 | HIBADH | 20.2420842 | 20.3096507 | 20.4567098 | 20.3795812 | 20.6604254 |
| Q5T7V8 | GORAB | 16.298625 | 16.442193 | 16.6431304 | 16.489479 | 16.6913554 |

| | | | | | | |
|---|---|---|---|---|---|---|
| Q86YA3 | ZGRF1 | 14.1963944 | 14.3580682 | 14.6401389 | 14.3122197 | 14.1815007 |
| P43003 | SLC1A3 | 16.5663048 | 16.2626107 | 16.5114293 | 16.5232978 | 16.278606 |
| P41002 | CCNF | 14.638715 | NA | NA | 14.6503614 | NA |
| Q6UWJ1 | TMCO3 | 16.2058785 | 16.8680314 | 16.9370776 | 16.1100416 | 17.2780967 |
| Q6NTF9 | RHBDD2 | 18.389336 | 18.2966789 | 18.5386386 | 17.8494446 | 18.4449064 |
| P05114 | HMGN1 | 24.8340748 | 25.0190899 | 24.5828781 | 24.8092849 | 24.6248828 |
| Q96EK5 | KIFBP | 19.9951823 | 20.0205688 | 20.1415238 | 20.0038799 | 20.1052845 |
| O60870 | KIN | 20.6257813 | 20.6909712 | 20.7189996 | 20.589386 | 20.6875492 |
| Q96B26 | EXOSC8 | 21.4363539 | 21.6041795 | 21.3194977 | 21.3748148 | 21.4926043 |
| Q9P2G1 | ANKIB1 | 16.3820979 | 16.1663339 | 16.556164 | 16.3729715 | 16.36982 |
| P61026 | RAB10 | 23.3956301 | 23.3487636 | 23.4058327 | 23.3111278 | 23.4281086 |
| P56589 | PEX3 | 20.2939434 | 20.3443535 | 20.4898321 | 20.3366032 | 20.4629002 |
| Q9NQX7 | ITM2C | 21.2042886 | 21.1352961 | 21.3504278 | 21.2543021 | 21.3371938 |
| Q96EY5 | MVB12A | 18.6971381 | 18.697113 | 18.8353321 | 18.7104775 | 18.9241941 |
| Q15573 | TAF1A | 19.9688576 | 20.0957192 | 20.1381321 | 19.9436242 | 20.3061581 |
| Q8TAP8 | PPP1R35 | 16.9759975 | 17.0934129 | 16.9629126 | 16.7722869 | 16.2940869 |
| Q9Y2K2 | SIK3 | 17.9637197 | 17.8207349 | 17.9808324 | 17.7963916 | 17.9207999 |
| O95479 | H6PD | 19.4025367 | 19.3475265 | 19.4067197 | 19.4869086 | 19.3678309 |
| Q6PI47 | KCTD18 | 19.2083754 | 19.2549963 | 19.387161 | 19.2630155 | 19.3032068 |
| Q8WV44 | TRIM41 | 18.6304983 | 18.7765257 | 18.6204866 | 18.6622822 | 18.8498243 |
| Q96L73 | NSD1 | 18.2679028 | 18.3307025 | 18.2562731 | 18.2710653 | 18.2755485 |
| Q13042 | CDC16 | 21.3459916 | 21.3949875 | 21.3731445 | 21.3321294 | 21.366099 |
| O95273 | CCNDBP1 | 16.5467898 | 16.8745876 | 16.8423817 | 16.8806712 | 16.7078264 |
| O75145 | PPFIA3 | 17.5418926 | 17.3203667 | 17.4366196 | 17.3752844 | 17.2755469 |
| A2VDF0 | FUOM | 16.4606555 | 16.7447262 | 16.8655412 | 16.5971219 | 16.7502328 |
| Q9NPA0 | EMC7 | 20.8592611 | 21.0248919 | 20.9446128 | 20.807011 | 21.0386858 |
| Q13884 | SNTB1 | 21.3728598 | 21.570047 | 21.4439512 | 21.3123592 | 21.5220748 |
| Q9BRQ6 | CHCHD6 | 22.6267121 | 22.5905088 | 22.6395213 | 22.4476361 | 22.5042703 |
| Q02156 | PRKCE | 23.691107 | 23.6738865 | 23.5746742 | 23.5876409 | 23.5865067 |
| O95983 | MBD3 | 21.597884 | 21.7914174 | 21.7459093 | 21.5762322 | 21.6735549 |
| Q8NBI6 | XXYLT1 | 19.8167244 | 19.7789905 | 19.8671406 | 19.7723799 | 19.827553 |
| P00367 | GLUD1 | 24.9379843 | 24.9727668 | 24.9945675 | 24.867431 | 24.9137971 |
| Q96MX6 | DNAAF10 | 20.5835537 | 20.6706852 | 20.8118839 | 20.6201296 | 20.7274535 |
| Q96DM3 | RMC1 | 21.6770158 | 21.7563227 | 21.7562555 | 21.6783479 | 21.7705 |
| Q7L1T6 | CYB5R4 | 19.2453179 | 19.2385576 | 19.3190621 | 19.2354847 | 19.2733111 |
| Q13435 | SF3B2 | 23.9171129 | 24.0278939 | 23.8376007 | 23.85966 | 23.9925502 |
| P36897 | TGFBR1 | 19.3866219 | 19.3195707 | 19.3235309 | 19.5071189 | 19.4933255 |
| P09622 | DLD | 24.1879188 | 24.1481225 | 24.0535352 | 24.0322522 | 24.1621901 |
| Q92542 | NCSTN | 23.1589915 | 23.3285777 | 23.1877404 | 23.1605937 | 23.2171541 |
| Q9H7Z3 | NRDE2 | 19.1765187 | 19.2791994 | 19.1894239 | 19.1592776 | 19.222229 |
| Q658Y4 | FAM91A1 | 20.9952611 | 21.0371872 | 21.0805182 | 20.9696036 | 21.0572593 |
| Q96QT4 | TRPM7 | 18.0444625 | 18.0512911 | 18.3495975 | 18.1730543 | 18.1598303 |

| | | | | | | |
|---|---|---|---|---|---|---|
| Q02252 | ALDH6A1 | 24.1368301 | 24.234376 | 24.2090358 | 24.15125 | 24.24577 |
| Q14511 | NEDD9 | 17.9049632 | 17.9386298 | 17.9587742 | 17.8360777 | 18.0811448 |
| Q6UX04 | CWC27 | 19.9596352 | 20.0966229 | 19.9688169 | 19.9734665 | 20.0815402 |
| Q96K19 | RNF170 | 19.594441 | 19.4022983 | 19.7035914 | 19.6753764 | 19.5287484 |
| P46109 | CRKL | 21.3826767 | 21.4626746 | 21.3852791 | 21.3390272 | 21.5533932 |
| Q8NHG7 | SVIP | 17.5910535 | 17.6458786 | 17.8508632 | 17.7917055 | 17.5850268 |
| Q9UKY4 | POMT2 | 18.3087059 | 18.508763 | 18.7779837 | 18.3565343 | 18.4560707 |
| Q68CQ4 | UTP25 | 19.3713196 | 19.4807556 | 19.5575887 | 19.4111442 | 19.525068 |
| O75027 | ABCB7 | 20.5507354 | 20.675369 | 20.6979943 | 20.5497628 | 20.6352199 |
| P21980 | TGM2 | 15.8301116 | 16.6080706 | 16.2282273 | 16.1795998 | 16.4223932 |
| O95243 | MBD4 | 18.8139762 | 18.9020147 | 18.9485122 | 18.8503237 | 18.9283482 |
| O14832 | PHYH | 20.8010705 | 20.7222926 | 20.7698628 | 20.7098272 | 20.854282 |
| Q96JM3 | CHAMP1 | 21.1519549 | 21.1786395 | 21.1807676 | 21.177316 | 21.228327 |
| P06396 | GSN | 26.0227312 | 26.1620821 | 25.9797997 | 26.0197094 | 26.0734584 |
| Q8NBF2 | NHLRC2 | 20.1442752 | 20.2671795 | 20.0836181 | 20.1121931 | 20.2384534 |
| P11441 | UBL4A | 20.9676729 | 21.1267189 | 21.0645021 | 21.00417 | 21.0423868 |
| P16383 | GCFC2 | 17.6571643 | 17.936784 | 17.8528699 | 17.8482416 | 17.9565932 |
| Q7L8J4 | SH3BP5L | 19.2118534 | 19.3421378 | 19.0494107 | 19.1168005 | 19.3617064 |
| Q9BWN1 | PRR14 | 18.710273 | 18.5863369 | 18.3458645 | 18.5207003 | 18.370607 |
| Q4G148 | GXYLT1 | 19.5208635 | 19.4110251 | 19.5990039 | 19.3825579 | 19.5383398 |
| P21796 | VDAC1 | 25.6826647 | 25.7547497 | 25.8265991 | 25.6593059 | 25.7856881 |
| P04062 | GBA1 | 21.4237427 | 21.5180666 | 21.5590139 | 21.4119886 | 21.4797141 |
| Q7Z2K6 | ERMP1 | 22.2370397 | 22.294095 | 22.3283265 | 22.1976842 | 22.2515885 |
| Q01780 | EXOSC10 | 21.8945889 | 22.0228296 | 21.9995816 | 21.8679492 | 21.9746672 |
| Q7Z401 | DENND4A | 19.6198139 | 19.6780177 | 19.7051366 | 19.6467783 | 19.7346141 |
| Q9BV44 | THUMPD3 | 21.0105074 | 21.0681339 | 21.0252884 | 21.021432 | 21.015095 |
| Q13464 | ROCK1 | 21.5731021 | 21.70744 | 21.675348 | 21.6126347 | 21.6704041 |
| O75150 | RNF40 | 21.3867335 | 21.4625722 | 21.4521607 | 21.3584201 | 21.4573778 |
| Q9UBD5 | ORC3 | 19.4710495 | 19.4899165 | 19.5251914 | 19.5228588 | 19.4981722 |
| P12724 | RNASE3 | 21.7114862 | 21.6276943 | 21.5813902 | 21.638647 | 21.6140023 |
| Q6NSJ2 | PHLDB3 | 17.2951251 | 17.5553401 | 17.233493 | 16.8797342 | 17.6444782 |
| O00487 | PSMD14 | 23.0461446 | 23.102662 | 23.105887 | 22.9899415 | 23.210932 |
| Q8N4J0 | CARNMT1 | 20.1370746 | 20.2769884 | 20.2564564 | 20.2245796 | 20.1935972 |
| Q9Y5B8 | NME7 | 21.5102531 | 21.4704541 | 21.4931689 | 21.454109 | 21.5083993 |
| P00390 | GSR | 21.0317333 | 21.1208626 | 21.1812122 | 21.0467902 | 21.2825087 |
| Q9H2L5 | RASSF4 | 18.7426375 | 18.9647225 | 19.0416236 | 18.7667848 | 18.722212 |
| Q9UBL6 | CPNE7 | 17.5760889 | 17.662744 | 17.6812715 | 17.7282295 | 17.6778865 |
| P46060 | RANGAP1 | 22.9163757 | 22.9419332 | 22.9137589 | 22.8430092 | 22.9208222 |
| Q15651 | HMGN3 | 19.1296338 | 19.2481082 | 19.274201 | 19.2201739 | 18.8729799 |
| Q8IU85 | CAMK1D | 19.1500331 | 19.244438 | 19.416309 | 19.3060404 | 19.3055655 |
| Q9NUW8 | TDP1 | 20.5065856 | 20.5039533 | 20.6398161 | 20.4919954 | 20.5424873 |
| P78316 | NOP14 | 21.6063575 | 21.727557 | 21.6745399 | 21.5906059 | 21.6513835 |

| | | | | | | |
|---|---|---|---|---|---|---|
| P46063 | RECQL | 22.4089302 | 22.4879311 | 22.4752758 | 22.3263651 | 22.4871591 |
| Q96EY7 | PTCD3 | 22.0760714 | 22.1083843 | 22.0998246 | 22.0579547 | 22.1846928 |
| Q6Y7W6 | GIGYF2 | 21.8164527 | 21.8806252 | 21.7851065 | 21.7649597 | 21.8455778 |
| Q7Z4H8 | POGLUT3 | 20.7398513 | 20.8512285 | 20.8515679 | 20.8098474 | 20.8805823 |
| Q9P209 | CEP72 | 16.5419583 | 16.7533744 | 16.9298282 | 17.3655419 | 16.7891931 |
| P78383 | SLC35B1 | 17.9031583 | 17.5477655 | 17.6027815 | 17.5643364 | 17.611561 |
| Q13795 | ARFRP1 | 19.2092989 | 19.3633453 | 19.354259 | 19.3132464 | 19.4466542 |
| Q9Y3T6 | R3HCC1 | 19.5301658 | 19.5458738 | 19.6027192 | 19.4262772 | 19.5137607 |
| Q6DHV7 | ADAL | 13.6639436 | 14.203584 | 13.9071825 | 14.0100469 | 13.8736797 |
| Q86UA1 | PRPF39 | 20.3139004 | 20.2447481 | 20.3598208 | 20.2483442 | 20.38259 |
| Q9ULL0 | KIAA1210 | 15.0760187 | NA | NA | 15.2534059 | NA |
| Q5FWF5 | ESCO1 | 14.8693063 | 14.6556718 | 15.0569741 | 14.3855108 | 14.7374252 |
| Q7Z3K3 | POGZ | 21.1931319 | 21.1762634 | 21.2632675 | 21.177159 | 21.2890398 |
| P48634 | PRRC2A | 20.9134131 | 20.9646952 | 20.9410305 | 20.9087476 | 21.0387038 |
| P0C024 | NUDT7 | 18.8791193 | 19.0676282 | 19.0993665 | 18.8344267 | 19.0374381 |
| P51148 | RAB5C | 23.8952801 | 23.8827515 | 23.9676581 | 23.844817 | 23.9399414 |
| Q5SRE5 | NUP188 | 22.1633845 | 22.2188673 | 22.2989071 | 22.1782527 | 22.2423875 |
| Q96SI9 | STRBP | 20.147199 | 20.1282209 | 20.2092322 | 20.0723817 | 20.1137851 |
| Q9H9B4 | SFXN1 | 22.4395695 | 22.3927019 | 22.4667183 | 22.3631999 | 22.5583446 |
| O75191 | XYLB | 18.9576384 | 19.1750866 | 19.197327 | 19.144477 | 19.1688597 |
| Q9Y2H2 | INPP5F | 19.3144572 | 19.3395936 | 19.4392034 | 19.3832843 | 19.4386404 |
| Q9GZR2 | REXO4 | 19.9821884 | 20.0772121 | 20.2312891 | 20.0527571 | 20.1286648 |
| Q99700 | ATXN2 | 20.3657 | 20.5556761 | 20.4584386 | 20.437658 | 20.5319937 |
| Q4KMP7 | TBC1D10B | 20.5214571 | 20.6570772 | 20.6539902 | 20.6087382 | 20.6494819 |
| Q96BR5 | COA7 | 21.1201141 | 21.1955031 | 21.1329091 | 21.1177877 | 21.2648333 |
| Q9NUA8 | ZBTB40 | 18.832767 | 18.9466242 | 19.1455882 | 18.914693 | 18.9624702 |
| P55010 | EIF5 | 23.0305247 | 23.1155108 | 23.0435483 | 23.0320526 | 23.1097303 |
| Q7L2K0 | TEDC2 | 14.253121 | 14.0468012 | 14.7014679 | 14.2855153 | 14.3650461 |
| P49281 | SLC11A2 | 17.3455 | 17.461537 | 17.463184 | 17.5732132 | 17.4016718 |
| P69905 | NA | 23.9544883 | 24.3714555 | 24.2450633 | 23.6632336 | 24.276049 |
| Q9H0F6 | SHARPIN | 20.160288 | 20.3950072 | 20.0070196 | 20.0720274 | 20.1895083 |
| P04843 | RPN1 | 25.2556513 | 25.2575936 | 25.2065963 | 25.1403801 | 25.2107715 |
| P55809 | OXCT1 | 22.5074873 | 22.5857554 | 22.4968709 | 22.3701556 | 22.580867 |
| Q16254 | E2F4 | 19.9949029 | 20.1432006 | 19.9717787 | 19.9682134 | 20.1155126 |
| Q8NCN4 | RNF169 | 18.6472597 | 18.8596809 | 18.691988 | 18.7025173 | 18.7308479 |
| P78330 | PSPH | 19.1489273 | 19.0032156 | 19.2607096 | 19.1285048 | 19.2349003 |
| Q9BYC9 | MRPL20 | 22.3370866 | 22.2803264 | 22.5170456 | 22.3391561 | 22.2583127 |
| Q9Y2L6 | FRMD4B | 17.4927405 | NA | NA | 17.3015962 | NA |
| Q8N5P1 | ZC3H8 | 18.3105876 | 18.2799113 | 18.3896582 | 18.2623835 | 18.1309402 |
| P61020 | RAB5B | 21.671043 | 21.8592524 | 21.7379289 | 21.5686131 | 21.6142843 |
| Q6ZW31 | SYDE1 | NA | 15.3885987 | 15.5287986 | NA | 15.5081517 |
| Q8N4T4 | ARHGEF39 | 16.4523788 | 16.8025635 | 16.7831113 | 16.8882253 | 16.8471997 |

| | | | | | | |
|---|---|---|---|---|---|---|
| Q96S82 | UBL7 | 20.624841 | 20.7354054 | 20.5612243 | 20.5085804 | 20.8698123 |
| Q96S06 | LMF1 | 14.7490365 | 14.9805501 | 15.0172078 | 14.8354912 | 14.7085159 |
| Q8WVL7 | ANKRD49 | NA | 16.2255811 | 16.2565861 | NA | 16.3194205 |
| Q969H0 | FBXW7 | NA | 15.4948508 | 15.7987279 | NA | 15.6938446 |
| Q9BRA0 | NAA38 | 18.0504971 | 17.9769311 | 17.9561449 | 17.8170572 | 18.0358871 |
| Q9NWD8 | TMEM248 | 17.6392054 | 17.7290838 | 17.8901866 | 17.7794498 | 17.8129949 |
| Q9NZM1 | MYOF | 18.1426201 | 18.2125922 | 17.9483611 | 18.0621346 | 18.265079 |
| O94941 | UBOX5 | 16.6007973 | 16.8838477 | 16.8503429 | 16.7432899 | 16.7536034 |
| Q9UNK0 | STX8 | 20.7423388 | 20.8240666 | 21.0599651 | 20.7739878 | 20.8298808 |
| P21912 | SDHB | 22.9855292 | 23.0557494 | 23.0623266 | 22.9056998 | 23.12535 |
| Q13094 | LCP2 | 21.6317245 | 21.6091413 | 21.6434842 | 21.5666858 | 21.6502545 |
| Q9NXD2 | MTMR10 | 17.9972583 | 18.2470978 | 18.2869956 | 18.0089105 | 18.0688997 |
| Q96EK6 | GNPNAT1 | 20.5886158 | 20.5660134 | 20.6686086 | 20.4858951 | 20.7035416 |
| Q8NF50 | DOCK8 | 22.1752081 | 22.2476045 | 22.273898 | 22.1480321 | 22.2806866 |
| P51570 | GALK1 | 23.5324241 | 23.562729 | 23.5399421 | 23.4288315 | 23.4810121 |
| O95299 | NDUFA10 | 21.9506501 | 21.9051165 | 22.0231458 | 21.8668969 | 21.9851977 |
| P42226 | STAT6 | 22.194508 | 22.2412742 | 22.241235 | 22.1599794 | 22.1679708 |
| Q8WVB6 | CHTF18 | 20.0198566 | 20.0699433 | 20.1293237 | 19.9983489 | 20.1057304 |
| P49748 | ACADVL | 23.7690602 | 23.7599074 | 23.8107543 | 23.6494546 | 23.792834 |
| O75116 | ROCK2 | 21.1923251 | 21.2907396 | 21.2808958 | 21.1880065 | 21.2293655 |
| Q9H089 | LSG1 | 20.2024784 | 20.3230694 | 20.3497882 | 20.2794985 | 20.3333357 |
| Q9NXJ5 | PGPEP1 | 20.8818843 | 20.8804885 | 20.9852905 | 20.8948748 | 20.8580304 |
| Q13568 | IRF5 | 20.712211 | 20.621436 | 20.6687044 | 20.7024666 | 20.8401896 |
| O60568 | PLOD3 | 21.1623052 | 21.2312937 | 21.2183655 | 21.1623217 | 21.3121269 |
| O94762 | RECQL5 | 18.5305518 | 18.5813778 | 18.4763573 | 18.4086316 | 18.3547396 |
| P30048 | PRDX3 | 25.7628894 | 25.8735252 | 25.7973013 | 25.6921781 | 25.9119887 |
| Q8IY31 | IFT20 | 17.402599 | 17.5417214 | 17.5515282 | 17.5347876 | 17.6201378 |
| Q08334 | IL10RB | 18.9581351 | 19.0711153 | 19.101036 | 18.9086028 | 19.0543006 |
| Q8NHL6 | LILRB1 | 18.0748147 | 18.2420332 | 18.281164 | 18.3114849 | 18.4348597 |
| Q9NVR5 | DNAAF2 | 17.7326267 | 18.0369869 | 17.7220906 | 17.7448545 | 17.8585018 |
| Q8NF91 | SYNE1 | 19.829735 | 20.4052907 | 19.8918126 | 19.5624036 | 19.9763587 |
| Q07617 | SPAG1 | 17.3775433 | 17.3039127 | 17.1573871 | 17.1370915 | 17.3101842 |
| Q9UDY4 | DNAJB4 | 16.4181068 | 16.8476418 | 16.736537 | 16.6555886 | 16.6471998 |
| P52848 | NDST1 | NA | 15.6737356 | 16.074057 | NA | 15.5174611 |
| P48380 | RFX3 | NA | 17.8402066 | 17.6079524 | NA | 17.7146205 |
| Q6SPF0 | SAMD1 | 19.9903224 | 19.9635621 | 19.9127144 | 19.9499651 | 19.9187511 |
| P13284 | IFI30 | 20.1990693 | 20.1252264 | 20.1406587 | 19.9832107 | 20.2008565 |
| P32969 | RPL9 | 25.6921821 | 25.6526895 | 25.8378673 | 25.5592421 | 25.6126621 |
| Q92900 | UPF1 | 23.4348826 | 23.4959869 | 23.4762422 | 23.3762043 | 23.5103626 |
| Q86WB0 | ZC3HC1 | 20.7735255 | 20.8083177 | 20.8194222 | 20.687862 | 20.8314118 |
| Q9P2K9 | DISP3 | NA | 11.5598778 | 11.5847158 | NA | 11.549052 |
| P13727 | PRG2 | NA | 11.7755967 | 11.7632207 | NA | 11.83963 |

| | | | | | | |
|---|---|---|---|---|---|---|
| O75787 | ATP6AP2 | 20.9111475 | 20.905795 | 20.7358469 | 20.9029593 | 20.9072517 |
| P47756 | CAPZB | 25.1654384 | 25.1931725 | 25.1313955 | 25.0296494 | 25.1204118 |
| P49427 | CDC34 | 21.0441086 | 21.198258 | 21.1539828 | 21.0635118 | 21.0558592 |
| Q9C0C9 | UBE2O | 20.9552527 | 21.0193544 | 21.0154062 | 20.9374554 | 21.0339551 |
| Q8NCR0 | B3GALNT2 | 18.0111085 | 18.3655096 | 18.1796457 | 18.1505441 | 18.2206866 |
| Q9NYT6 | ZNF226 | 16.8348489 | 16.9228434 | 17.0789607 | 16.8552546 | 16.7877899 |
| Q9BTC8 | MTA3 | 16.4598835 | 16.5650244 | 16.5937839 | 16.7337109 | 16.3341797 |
| Q13191 | CBLB | 14.1108867 | 14.4976868 | 14.4006734 | 13.73292 | 14.4312628 |
| Q9H7L9 | SUDS3 | 19.7344434 | 19.9536365 | 19.6334073 | 19.7153113 | 19.7075043 |
| Q9Y547 | IFT25 | 21.1386798 | 21.190874 | 21.0572034 | 21.0439579 | 20.9799805 |
| Q8TF68 | ZNF384 | 19.1760484 | 19.1885398 | 19.5454 | 19.0554831 | 19.1831281 |
| O75586 | MED6 | 19.6596545 | 19.7371856 | 19.6953089 | 19.6194708 | 19.8454159 |
| Q13889 | GTF2H3 | 19.4406482 | 19.3198301 | 19.561488 | 19.448286 | 19.4254747 |
| Q9UBQ5 | EIF3K | 23.8560036 | 23.888856 | 23.8486471 | 23.8201355 | 23.7958669 |
| O00410 | IPO5 | 23.4434988 | 23.4646138 | 23.4646499 | 23.3753571 | 23.443426 |
| Q96P11 | NSUN5 | 20.6018121 | 20.633477 | 20.7032162 | 20.6168474 | 20.6928371 |
| P00488 | F13A1 | 18.9839092 | 18.9343578 | 19.0024025 | 19.0249135 | 18.971153 |
| O14841 | OPLAH | 19.6552756 | 19.6838541 | 19.6875629 | 19.6031707 | 19.7263574 |
| Q8TED0 | UTP15 | 21.6398912 | 21.6861989 | 21.6844524 | 21.6118168 | 21.7783082 |
| Q96C36 | PYCR2 | 21.6036955 | 21.6903687 | 21.5998131 | 21.5590364 | 21.6422844 |
| O15164 | TRIM24 | 22.4590918 | 22.5231166 | 22.4007204 | 22.4148431 | 22.5493539 |
| Q8NEJ9 | NGDN | 21.061895 | 21.1594338 | 21.2220014 | 21.1518614 | 21.1721034 |
| P19404 | NDUFV2 | 22.4814882 | 22.5346161 | 22.3940021 | 22.3913562 | 22.6055018 |
| P53814 | SMTN | 17.9286029 | 17.987527 | 17.9172708 | 17.92772 | 17.8791432 |
| Q8WVE0 | EEF1AKMT1 | 17.9295277 | 18.1149022 | 18.1839505 | 18.1811405 | 18.3367504 |
| Q9C0B9 | ZCCHC2 | 14.872945 | NA | NA | 15.070864 | NA |
| Q86U42 | PABPN1 | 23.4295774 | 23.5334935 | 23.4272538 | 23.3596812 | 23.5114328 |
| Q8IUX1 | TMEM126B | 18.2997549 | 18.1617588 | 18.4382402 | 18.2281246 | 18.3784373 |
| O75460 | ERN1 | 15.168974 | 15.2599732 | 14.8024984 | 14.9518048 | 14.6027402 |
| Q15935 | ZNF77 | 14.4530056 | 15.0643429 | 14.940951 | 14.67232 | 14.5954116 |
| O43502 | RAD51C | 17.0144653 | 17.4094801 | 17.1526965 | 16.9810315 | 17.2666494 |
| Q13952 | NFYC | 20.945636 | 20.8357275 | 20.9986397 | 20.8652153 | 20.9728487 |
| Q6NXR4 | TTI2 | 20.2915041 | 20.2917084 | 20.2774986 | 20.2030934 | 20.3362601 |
| P26640 | VARS1 | 23.8586253 | 23.935291 | 23.9071279 | 23.821627 | 23.8893114 |
| Q9NYV4 | CDK12 | 20.1094318 | 20.1460558 | 20.1254076 | 20.1208429 | 20.2325459 |
| P61978 | HNRNPK | 27.1555711 | 27.2073145 | 27.1106301 | 27.0573288 | 27.1443865 |
| P47985 | UQCRFS1 | 24.0153131 | 24.0819877 | 24.0951025 | 24.0385815 | 24.0793689 |
| Q8NDZ4 | DIPK2A | 19.2036269 | 19.3802753 | 19.4510095 | 19.2078537 | 19.3646066 |
| Q12873 | CHD3 | 18.591162 | 18.6810396 | 18.9307262 | 18.7262651 | 18.8785461 |
| Q9H6R7 | WDCP | 17.6163542 | 17.6965953 | 17.5218539 | 17.5960023 | 17.8515871 |
| Q86U28 | ISCA2 | 19.7281422 | 19.9453467 | 19.6802528 | 19.7781102 | 19.926463 |
| Q6UB98 | ANKRD12 | 17.5754137 | 18.0835411 | 17.8920173 | 17.6238058 | 17.6694566 |

| | | | | | | |
|---|---|---|---|---|---|---|
| Q92879 | CELF1 | 20.3774827 | 20.2497967 | 20.4808306 | 20.3963434 | 20.5439569 |
| Q9UQ16 | DNM3 | 17.6791276 | 17.8647839 | 17.8484933 | 17.7082098 | 17.7166918 |
| O60603 | TLR2 | 20.9432723 | 20.9583153 | 21.1438969 | 21.0288781 | 21.0659216 |
| Q9P0U3 | SENP1 | 18.9011593 | 18.8933048 | 19.1159036 | 19.0409764 | 19.0975379 |
| P80217 | IFI35 | 20.4059523 | 20.6982391 | 20.7003224 | 20.575095 | 20.7295484 |
| Q96D31 | ORAI1 | 18.4167355 | 18.0632646 | 18.4067542 | 18.511911 | 18.2761393 |
| Q86V81 | ALYREF | 23.9089975 | 23.9822525 | 23.8416398 | 23.8927806 | 24.0075768 |
| Q9BRP8 | PYM1 | 22.4717653 | 22.5198773 | 22.1491378 | 22.365668 | 22.5309864 |
| P49902 | NT5C2 | 21.5795951 | 21.6619218 | 21.645956 | 21.5486847 | 21.5543715 |
| Q8IYB8 | SUPV3L1 | 21.9542233 | 21.9623621 | 22.0300275 | 21.9478478 | 21.9994636 |
| Q6P444 | MTFR2 | 18.2052958 | 18.2624311 | 18.2956493 | 18.1099898 | 18.2950023 |
| Q9H6E5 | TUT1 | 19.0600799 | 19.1794374 | 19.1552813 | 19.0496393 | 19.2167663 |
| O95159 | ZFPL1 | 20.600989 | 20.5726116 | 20.7419906 | 20.6461512 | 20.6457478 |
| Q8TB22 | SPATA20 | 18.0595065 | 18.1387479 | 18.1249144 | 18.1438401 | 18.1278885 |
| O43670 | ZNF207 | 23.3510516 | 23.3703615 | 23.4370093 | 23.4159411 | 23.3758481 |
| Q8N9B5 | JMY | 19.9242852 | 20.0569689 | 19.9117951 | 19.9382167 | 19.995181 |
| Q6ZN18 | AEBP2 | 15.4754315 | 15.3364646 | 15.6704663 | 15.8710929 | 15.6730196 |
| Q92995 | USP13 | 19.3052385 | 19.3700173 | 19.4705433 | 19.3000626 | 19.2321988 |
| Q9P2H3 | IFT80 | 17.9816526 | 18.4656596 | 18.0549239 | 18.2189455 | 18.1650793 |
| Q6ZUM4 | ARHGAP27 | 19.8815413 | 19.9399699 | 20.0345759 | 19.8048199 | 19.9408175 |
| Q00536 | CDK16 | 18.0303814 | 18.1521027 | 18.2728754 | 18.0288622 | 18.1973159 |
| Q6GMV3 | PTRHD1 | 21.7028799 | 21.7275983 | 21.7138374 | 21.6296507 | 21.8702392 |
| P04406 | GAPDH | 28.7060462 | 28.8202676 | 28.6745684 | 28.5117756 | 28.7078376 |
| Q9NX14 | NDUFB11 | 22.1716296 | 22.3981829 | 22.2870895 | 22.2585716 | 22.2720462 |
| Q15257 | PTPA | 22.6922525 | 22.8645983 | 22.845859 | 22.7504778 | 22.794214 |
| P34059 | GALNS | 21.1251757 | 21.2361668 | 21.2409968 | 21.1542851 | 21.2614707 |
| Q96CS3 | FAF2 | 21.8387123 | 21.8643135 | 22.0049742 | 21.8468669 | 21.9155497 |
| P63104 | YWHAZ | 26.9626118 | 26.9225749 | 26.8337866 | 26.8037769 | 26.860551 |
| Q96EZ8 | MCRS1 | 19.2799231 | 19.3506449 | 19.4033349 | 19.2728445 | 19.3883634 |
| P36776 | LONP1 | 23.528087 | 23.6143673 | 23.4958746 | 23.4462559 | 23.5335856 |
| Q9UPU5 | USP24 | 20.7394179 | 20.7808739 | 20.7604844 | 20.7135145 | 20.7732134 |
| Q7KZF4 | SND1 | 24.9639671 | 24.9480398 | 24.9015099 | 24.8712592 | 24.9137962 |
| A6NHR9 | SMCHD1 | 21.9925617 | 22.0836815 | 22.0541234 | 21.9773159 | 22.0869857 |
| Q8TDJ6 | DMXL2 | 19.0913198 | 19.1416817 | 19.2668533 | 19.1090132 | 19.221976 |
| P07203 | GPX1 | 24.1430391 | 24.2020153 | 24.1314418 | 24.1053736 | 24.2546097 |
| P10074 | ZBTB48 | 18.3094382 | 18.2376462 | 18.3457979 | 18.3423242 | 18.3577564 |
| O60504 | SORBS3 | 20.4363292 | 20.4592306 | 20.4555841 | 20.43095 | 20.5087923 |
| Q86TJ2 | TADA2B | 18.7531966 | 18.9196612 | 18.7756467 | 18.8597852 | 18.823544 |
| Q9H4E7 | DEF6 | 19.6570312 | 19.6835594 | 19.7587234 | 19.6831748 | 19.8235164 |
| Q5TC84 | OGFRL1 | 18.2274074 | 18.6426785 | 18.5044649 | 18.5609118 | 18.5095265 |
| Q4G0X4 | KCTD21 | 16.0685514 | 16.4096603 | 16.0175193 | 15.9694138 | 16.2370236 |
| Q8IY57 | YAF2 | 16.7594049 | 17.0205761 | 16.9275683 | 17.0704236 | 17.1624355 |

| | | | | | | |
|---|---|---|---|---|---|---|
| O43543 | XRCC2 | 15.4435964 | 15.4669565 | 15.7697839 | 15.9165443 | 15.668994 |
| Q9BQD7 | ANTKMT | 17.9407795 | 18.1495152 | 17.84535 | 17.8402668 | 17.73288 |
| Q99766 | DMAC2L | 20.6525537 | 20.9057599 | 20.8400908 | 20.6666647 | 20.8255449 |
| Q9UJ83 | HACL1 | 21.1685806 | 21.1555457 | 21.2338714 | 21.1788554 | 21.2272753 |
| Q13330 | MTA1 | 21.0528576 | 21.0564646 | 21.1373801 | 21.039149 | 21.1203391 |
| Q16698 | DECR1 | 23.1537898 | 23.2292454 | 23.1923704 | 23.1689841 | 23.3083513 |
| Q9HBM0 | VEZT | 18.6210549 | 18.6006867 | 18.8962936 | 18.7622981 | 18.8110505 |
| Q68D85 | NCR3LG1 | 16.4054424 | 17.3112759 | 16.998023 | 17.1797655 | 17.1397429 |
| O75582 | RPS6KA5 | 19.0066517 | 19.1319921 | 19.2611083 | 19.186958 | 19.150439 |
| P14209 | CD99 | 21.0885225 | 21.1107431 | 20.9263426 | 21.2700403 | 21.0916494 |
| Q04721 | NOTCH2 | 17.7992256 | 17.7660878 | 17.8700069 | 17.7074482 | 17.7203181 |
| Q96MN2 | NLRP4 | 16.868599 | NA | NA | 17.0365314 | NA |
| P61218 | POLR2F | 18.5322988 | 18.2095983 | 18.4351818 | 18.5611872 | 18.8978997 |
| Q96A19 | CCDC102A | 14.2991002 | NA | NA | 14.5871634 | NA |
| O14681 | EI24 | 22.0061312 | 22.0022771 | 21.9713784 | 21.869504 | 21.822847 |
| Q9UQR1 | ZNF148 | 19.5585424 | 19.7071778 | 19.6288686 | 19.6215031 | 19.8439197 |
| O94766 | B3GAT3 | 20.8492666 | 20.9298929 | 20.8048476 | 20.8487254 | 20.7769826 |
| O15145 | ARPC3 | 24.9611808 | 25.015411 | 25.0417784 | 24.9379371 | 24.9193115 |
| P55795 | HNRNPH2 | 22.278744 | 22.3618692 | 22.3963476 | 22.2535775 | 22.3039346 |
| O94923 | GLCE | 19.1084845 | 19.2014322 | 19.2143589 | 19.1755582 | 19.2101346 |
| Q02224 | CENPE | 19.1354239 | 19.2169028 | 19.1743641 | 19.0274095 | 19.0542042 |
| Q9Y620 | RAD54B | 17.6190753 | 17.6752421 | 17.8052604 | 17.7835373 | 17.9437225 |
| Q9BYC8 | MRPL32 | 19.6864277 | 19.9537398 | 19.8748455 | 19.8782242 | 19.9914479 |
| Q9NZR1 | TMOD2 | 22.8095879 | 23.351824 | 22.8172038 | 22.7229859 | 23.1538486 |
| Q9BQ15 | NABP2 | 21.9223221 | 21.7219767 | 21.8779587 | 21.8560122 | 21.6884908 |
| Q9H2H9 | SLC38A1 | 18.9683361 | 18.9350058 | 18.9080525 | 18.7757605 | 18.9366426 |
| Q6UUV9 | CRTC1 | 19.4890839 | 19.6888771 | 19.735704 | 19.4324822 | 19.7542669 |
| P55082 | MFAP3 | 18.6268221 | 18.3615972 | 18.3599183 | 18.4737634 | 18.5299662 |
| O14776 | TCERG1 | 22.5845164 | 22.6475012 | 22.6608891 | 22.5628878 | 22.6420985 |
| Q96S38 | RPS6KC1 | 20.4299896 | 20.4453637 | 20.1993606 | 20.2904835 | 20.4008609 |
| Q9Y314 | NOSIP | 22.5697547 | 22.6531305 | 22.6042304 | 22.4953611 | 22.680141 |
| Q8NEB9 | PIK3C3 | 20.4606451 | 20.5117889 | 20.5244512 | 20.3934789 | 20.5572515 |
| Q8NDZ2 | SIMC1 | 16.1129882 | 16.4913141 | 16.3491178 | 16.708496 | 16.3074284 |
| O43167 | ZBTB24 | 15.9614321 | NA | NA | 15.9567903 | NA |
| Q9Y426 | C2CD2 | 14.7800639 | 14.7306872 | 14.923285 | 14.7473933 | 14.5816309 |
| Q9Y4G8 | RAPGEF2 | 14.70392 | 14.6730355 | 14.4300822 | 14.8138829 | 14.448961 |
| P05556 | ITGB1 | 23.4692382 | 23.5004124 | 23.5167722 | 23.4714894 | 23.5095737 |
| Q6V9R5 | ZNF562 | 17.7652021 | 17.7907756 | 17.7643693 | 17.6804707 | 17.9269571 |
| O95149 | SNUPN | 20.5268155 | 20.860795 | 20.9375348 | 20.7582957 | 20.8888354 |
| Q8NHH9 | ATL2 | 20.6735939 | 20.7821501 | 20.7819847 | 20.7572454 | 20.7316924 |
| Q9NRG9 | AAAS | 20.5348203 | 20.6515401 | 20.6837558 | 20.5661678 | 20.7011854 |
| Q9BW27 | NUP85 | 21.7591772 | 21.8822596 | 21.8700834 | 21.8120892 | 21.7574246 |

| | | | | | | |
|---|---|---|---|---|---|---|
| Q8N8A6 | DDX51 | 20.4042881 | 20.4751169 | 20.4368429 | 20.4117367 | 20.4558015 |
| Q86UE4 | MTDH | 22.7412056 | 22.8592396 | 22.9123706 | 22.7791274 | 22.7970712 |
| P41252 | IARS1 | 23.6351817 | 23.6592926 | 23.6563682 | 23.5557173 | 23.6347604 |
| Q92696 | RABGGTA | 21.0100114 | 21.0819772 | 21.0731271 | 21.0093882 | 21.0599205 |
| Q8N0T1 | RBIS | 19.3180335 | 19.5233999 | 19.5147263 | 19.4607555 | 19.5171627 |
| Q9Y5N5 | N6AMT1 | 17.1698607 | 17.4187378 | 17.2674533 | 17.3640317 | 17.4528829 |
| Q9NPA2 | MMP25 | 20.5023326 | 20.1713131 | 20.548691 | 20.360747 | 20.2304931 |
| P30047 | GCHFR | 18.6062412 | 18.5281211 | 18.3311232 | 18.4302574 | 18.3557993 |
| O14492 | SH2B2 | 17.111255 | 17.2977257 | 17.3402741 | 17.0765173 | 16.859228 |
| P15260 | IFNGR1 | 19.6646198 | 19.5323938 | 19.7347308 | 19.7037299 | 19.8432628 |
| Q96I34 | PPP1R16A | 15.8658932 | 16.2757809 | 15.7910439 | 16.05401 | 16.1255679 |
| Q9HBD1 | RC3H2 | 14.7200069 | 14.3526792 | 14.9788019 | 14.5406719 | 14.6618835 |
| Q8IUN9 | CLEC10A | 18.2141605 | 18.561237 | 18.6741928 | 18.4530145 | 18.6712503 |
| Q9NVZ3 | NECAP2 | 21.4054082 | 21.4594251 | 21.4552044 | 21.3139591 | 21.4684081 |
| Q96FK6 | WDR89 | 19.6612004 | 19.7325531 | 19.8717761 | 19.7524162 | 19.7386239 |
| Q9NS91 | RAD18 | 18.9781347 | 19.1153227 | 19.2470801 | 19.0516406 | 19.274454 |
| Q9NQR1 | KMT5A | 15.6123855 | 15.4546755 | 15.4687691 | 15.4393536 | 15.5063118 |
| Q14232 | EIF2B1 | 22.322603 | 22.3832921 | 22.4730895 | 22.3686163 | 22.4558943 |
| Q9NU53 | GINM1 | 16.694939 | 16.8728032 | 16.3161636 | 16.5174438 | 16.0384214 |
| Q8IZL8 | PELP1 | 21.1543086 | 21.2265769 | 21.1891126 | 21.1158522 | 21.3171259 |
| Q9Y490 | TLN1 | 24.4633782 | 24.6130331 | 24.4707071 | 24.4072008 | 24.5924261 |
| Q99460 | PSMD1 | 23.5379038 | 23.4767423 | 23.5766731 | 23.4650821 | 23.5028791 |
| P22694 | PRKACB | 18.7209129 | 18.8080493 | 18.8051029 | 18.548083 | 18.8580671 |
| Q9UPN6 | SCAF8 | 20.0647588 | 19.996343 | 19.9241474 | 20.0077526 | 19.9767081 |
| Q8IYB7 | DIS3L2 | 22.5513722 | 22.6821426 | 22.5843324 | 22.5215162 | 22.6696228 |
| Q9C0H2 | TTYH3 | 21.7815966 | 21.5081199 | 21.5385853 | 21.7430737 | 21.5438196 |
| Q8N8D1 | PDCD7 | 18.243216 | 18.240601 | 18.4505843 | 18.2717248 | 18.1633882 |
| O43513 | MED7 | 18.1135427 | 18.1335793 | 18.226305 | 18.3679642 | 18.2341488 |
| Q99759 | MAP3K3 | 16.4868882 | 15.9079866 | 16.1066405 | 16.0201025 | 16.3062692 |
| Q9BSM1 | PCGF1 | 18.0595591 | 17.8884595 | 18.1716602 | 18.0016335 | 18.0086026 |
| Q8WV93 | AFG1L | 18.5296673 | 18.4224525 | 18.5721079 | 18.6239418 | 18.6903502 |
| Q96EW2 | HSPBAP1 | 16.2697413 | NA | NA | 16.2694753 | NA |
| O75607 | NPM3 | 22.088354 | 22.1295068 | 22.1277755 | 22.0216775 | 22.0524358 |
| Q8IWV7 | UBR1 | 21.3518438 | 21.4989705 | 21.4050085 | 21.2469321 | 21.2646075 |
| P52657 | GTF2A2 | 21.6583758 | 21.7030388 | 21.7048377 | 21.6147371 | 21.4905976 |
| Q06830 | PRDX1 | 26.5285512 | 26.5607734 | 26.4773734 | 26.4930778 | 26.5580772 |
| Q9HCD5 | NCOA5 | 21.9077841 | 22.0011866 | 21.9627465 | 21.9503847 | 21.8917375 |
| P23469 | PTPRE | 20.0003427 | 20.0888109 | 20.1191354 | 20.0003063 | 20.0718711 |
| O43491 | EPB41L2 | 20.2652329 | 20.4732229 | 20.4504928 | 20.2989214 | 20.5410547 |
| Q96CJ1 | EAF2 | 20.0335923 | 19.9206696 | 20.0275966 | 20.1232967 | 20.1173762 |
| Q9NVA4 | TMEM184C | 17.7329824 | 18.1729896 | 18.2608096 | 18.0654094 | 18.2796545 |
| Q7Z6M3 | MILR1 | 19.185527 | 19.2069374 | 19.3511164 | 19.0877855 | 19.1419526 |

| | | | | | | |
|---|---|---|---|---|---|---|
| P36873 | PPP1CC | 21.0930358 | 21.0447221 | 20.8744117 | 21.0233256 | 21.0500383 |
| O43361 | ZNF749 | 13.4301623 | 13.5172446 | 13.3007857 | 13.70012 | 13.3655566 |
| Q63HQ0 | AP1AR | 15.3104371 | 15.7293289 | 15.4227489 | 15.7083555 | 15.5035564 |
| P10588 | NR2F6 | 18.2763593 | 18.3055061 | 18.2348147 | 18.0247676 | 18.4492982 |
| Q567U6 | CCDC93 | 21.3376344 | 21.2687316 | 21.3900934 | 21.2651507 | 21.3169791 |
| Q14320 | FAM50A | 21.0708048 | 21.0339787 | 21.1272019 | 20.9712684 | 21.1261402 |
| O75821 | EIF3G | 23.7541714 | 23.8932885 | 23.6734758 | 23.7303258 | 23.7416315 |
| Q08170 | SRSF4 | 23.0154037 | 23.0834573 | 22.8767203 | 23.0479784 | 22.8989695 |
| Q9BTM9 | URM1 | 20.0343394 | 20.2597575 | 19.4600341 | 19.9240545 | 19.8103737 |
| O75144 | ICOSLG | 17.9742599 | 18.181688 | 18.276734 | 18.2800382 | 18.5329378 |
| O95183 | VAMP5 | 15.2652795 | 15.1112482 | 15.4576311 | 15.3614559 | 15.590981 |
| Q8N5B7 | CERS5 | 21.7927067 | 21.7951606 | 22.0022061 | 21.6420632 | 21.8465291 |
| Q96EY9 | ADAT3 | 19.0509138 | 18.8732256 | 18.9323352 | 18.8752529 | 19.072586 |
| Q9GZT9 | EGLN1 | 20.4873668 | 20.608947 | 20.6489275 | 20.4579338 | 20.6586341 |
| Q75N03 | CBLL1 | 19.8982898 | 19.9613083 | 19.9380369 | 19.9609652 | 20.0261035 |
| P22392 | NME2 | 27.4842092 | 27.5811108 | 27.4115317 | 27.3435183 | 27.3490128 |
| Q15047 | SETDB1 | 20.5572857 | 20.5928165 | 20.680288 | 20.573564 | 20.6365849 |
| Q86X55 | CARM1 | 22.8999844 | 23.0444038 | 22.9306595 | 22.9121336 | 22.9554886 |
| Q9H2M9 | RAB3GAP2 | 22.3284343 | 22.4125655 | 22.3802461 | 22.2710523 | 22.4500054 |
| P10276 | RARA | 19.8463294 | 19.9579285 | 20.1306611 | 20.0182215 | 20.0299001 |
| P62330 | ARF6 | 22.0165999 | 22.0931716 | 22.1624272 | 21.8985447 | 22.1950677 |
| P46934 | NEDD4 | 17.9880698 | 18.1058343 | 18.2141983 | 17.9531863 | 18.2639363 |
| A6NC98 | CCDC88B | 20.5512012 | 20.7644464 | 20.6867555 | 20.5610301 | 20.7316379 |
| P62714 | PPP2CB | 20.5626043 | 20.5271727 | 20.4410176 | 20.5459262 | 20.6582545 |
| Q92538 | GBF1 | 20.793612 | 20.8648361 | 20.871689 | 20.807135 | 20.9236601 |
| Q9NRY4 | ARHGAP35 | 20.8803656 | 20.9163098 | 20.938593 | 20.8126967 | 20.9860215 |
| Q9H939 | PSTPIP2 | 21.7911837 | 21.8857992 | 21.8981635 | 21.7869047 | 21.8342424 |
| Q8TF46 | DIS3L | 18.8213476 | 19.0671134 | 18.8900163 | 18.8884568 | 18.942516 |
| Q8ND71 | GIMAP8 | 18.8477598 | 18.7780416 | 18.8143444 | 18.7564984 | 18.8429217 |
| P68543 | UBXN2A | NA | 16.9649895 | 16.9047519 | NA | 16.8898542 |
| O95363 | FARS2 | 20.2337797 | 20.2371223 | 20.3250392 | 20.2330723 | 20.2866691 |
| Q06033 | ITIH3 | 19.0619951 | 19.294759 | 19.1357609 | 18.8811755 | 19.4043667 |
| Q12907 | LMAN2 | 24.0546661 | 24.2118235 | 24.1985457 | 24.0751702 | 24.149677 |
| Q9UHF1 | EGFL7 | 19.9222898 | 19.9794788 | 19.9649992 | 20.1461995 | 20.0626044 |
| Q8N2M8 | CLASRP | 20.6032477 | 20.6538242 | 20.608449 | 20.5819177 | 20.6809451 |
| P13611 | VCAN | 14.6935456 | NA | NA | 14.9436477 | NA |
| Q96GX9 | APIP | 20.7050235 | 20.7788516 | 20.6713835 | 20.663889 | 20.7226376 |
| Q9P2C4 | TMEM181 | 18.5689954 | 18.6191227 | 18.8416114 | 18.6662052 | 18.7463858 |
| Q6IQ32 | ADNP2 | 17.9733197 | 18.0077993 | 18.2762516 | 18.0988617 | 18.1879018 |
| Q9BWT3 | PAPOLG | 19.2632795 | 19.3628469 | 19.3767606 | 19.3489659 | 19.4583444 |
| A6NJG2 | SOWAHD | 18.7436748 | 18.969895 | 18.900324 | 18.8738956 | 18.8691603 |
| Q02040 | AKAP17A | 21.0285465 | 21.0630426 | 21.0620332 | 20.9819978 | 21.0212066 |

| | | | | | | |
|---|---|---|---|---|---|---|
| Q2T9J0 | TYSND1 | 18.7815849 | 18.8654549 | 18.7798066 | 18.6895754 | 18.8735945 |
| O75129 | ASTN2 | NA | 14.628746 | 14.5941245 | NA | 14.73809 |
| Q96M94 | KLHL15 | 15.8081753 | 16.4075413 | 16.4173065 | 16.3947115 | 16.4603004 |
| Q96IX5 | ATP5MK | 21.0068446 | 20.8988811 | 21.002323 | 20.9536185 | 20.9192458 |
| Q9UG01 | IFT172 | 17.8572931 | 18.2497308 | 18.5078995 | 18.079497 | 17.8418021 |
| Q5T6C5 | ATXN7L2 | NA | 14.8642845 | 15.0560904 | NA | 15.085705 |
| Q8WWW0 | RASSF5 | 17.0119128 | 17.0955382 | 17.1976306 | 17.0250854 | 17.3889867 |
| Q96K37 | SLC35E1 | 20.100931 | 19.8160049 | 20.1117352 | 19.6632235 | 19.8299338 |
| Q14534 | SQLE | 20.9425041 | 21.0875558 | 21.2019928 | 20.9162636 | 21.19068 |
| Q2M385 | MPEG1 | 21.591478 | 21.4487585 | 21.4275472 | 21.625344 | 21.4536063 |
| Q8N357 | SLC35F6 | 19.7738723 | 19.7160364 | 19.7188257 | 19.5602349 | 19.58989 |
| P17252 | PRKCA | 19.8072558 | 19.7570185 | 20.0026706 | 19.827491 | 19.8334865 |
| Q8N4P3 | HDDC3 | 20.9457714 | 21.1101578 | 20.9459236 | 20.7311381 | 21.1077542 |
| Q9BSH4 | TACO1 | 19.7959833 | 19.8326182 | 19.8503094 | 19.7708326 | 19.7182114 |
| Q9Y478 | PRKAB1 | 18.7132105 | 18.8600034 | 18.9147874 | 18.8152284 | 18.9542202 |
| Q9H6Y7 | RNF167 | 21.6168109 | 21.5525334 | 21.406893 | 21.4399241 | 21.2969831 |
| Q9BUL9 | RPP25 | 19.548004 | 19.5202486 | 19.6042905 | 19.2296882 | 19.4982874 |
| Q13315 | ATM | 22.0598908 | 22.1219922 | 22.1994345 | 22.0243387 | 22.16837 |
| Q9Y483 | MTF2 | 19.5892221 | 19.6830182 | 19.830743 | 19.6679218 | 19.7450802 |
| Q8NDX1 | PSD4 | 19.446789 | 19.4453152 | 19.5419192 | 19.381692 | 19.5027259 |
| Q2TAA2 | IAH1 | 21.2932342 | 21.3787281 | 21.4561416 | 21.3205369 | 21.390705 |
| O95155 | UBE4B | 20.990739 | 21.108098 | 21.0081757 | 20.9136975 | 21.0182591 |
| Q9P215 | POGK | 19.288129 | 19.3864663 | 19.2516274 | 19.2457291 | 19.3103646 |
| Q9Y512 | SAMM50 | 22.6501259 | 22.7858043 | 22.7466882 | 22.6237062 | 22.7595331 |
| P13010 | XRCC5 | 25.669742 | 25.6298485 | 25.5712082 | 25.5471812 | 25.5575651 |
| Q13574 | DGKZ | 21.0148403 | 20.9357318 | 21.1189793 | 21.0303031 | 21.0399492 |
| Q96PU4 | UHRF2 | 20.0818423 | 19.9890708 | 20.0808631 | 20.0016471 | 20.0843846 |
| Q8TAE8 | GADD45GIP1 | 21.6422201 | 21.6322384 | 21.669769 | 21.6193957 | 21.7375309 |
| Q96JC1 | VPS39 | 19.691344 | 19.7082356 | 19.8046894 | 19.7196604 | 19.73555 |
| P51649 | ALDH5A1 | 21.006549 | 21.043146 | 21.0107338 | 20.9797068 | 21.0949892 |
| Q96EP5 | DAZAP1 | 24.2369939 | 24.2057468 | 24.148501 | 24.1664577 | 24.2872748 |
| Q96TC7 | RMDN3 | 19.5722387 | 19.5756318 | 19.6588672 | 19.5580368 | 19.614268 |
| Q6P1N0 | CC2D1A | 21.3190837 | 21.4477966 | 21.3975658 | 21.3405036 | 21.3922174 |
| Q9UPT5 | EXOC7 | 20.6072144 | 20.6613506 | 20.7747856 | 20.7364528 | 20.7732645 |
| P35240 | NF2 | 19.5209813 | 19.507614 | 19.5531736 | 19.4569018 | 19.611742 |
| Q9UBW7 | ZMYM2 | 19.3013408 | 19.4134189 | 19.415359 | 19.3743867 | 19.3440563 |
| Q9BRS2 | RIOK1 | 19.4051318 | 19.3873638 | 19.5204366 | 19.4078745 | 19.4239323 |
| Q86XI2 | NCAPG2 | 20.5632321 | 20.7077309 | 20.7526538 | 20.6181525 | 20.7075829 |
| Q5SWX8 | ODR4 | 19.1356861 | 19.1801872 | 19.3710439 | 19.2628369 | 19.4381881 |
| Q9Y3C6 | PPIL1 | 22.7101221 | 22.6447498 | 22.7218679 | 22.5429382 | 22.7408981 |
| Q9Y3F4 | STRAP | 23.0713538 | 23.1080105 | 22.974634 | 22.9669324 | 23.1710269 |
| Q9NUU6 | OTULINL | 21.8686899 | 21.9424127 | 21.9299046 | 21.7632619 | 21.8187587 |

| | | | | | | |
|---|---|---|---|---|---|---|
| Q96II8 | LRCH3 | 19.7742097 | 19.99112 | 19.9244136 | 19.9110348 | 19.9445063 |
| Q9H832 | UBE2Z | 21.2724431 | 21.4258325 | 21.5010297 | 21.3402805 | 21.4175362 |
| Q13416 | ORC2 | 19.920019 | 20.0095705 | 19.8603822 | 19.8623269 | 20.062084 |
| Q7L273 | KCTD9 | 18.0046386 | 18.2012067 | 18.2137809 | 18.1922674 | 18.248376 |
| Q9BVC5 | C2orf49 | 20.1844422 | 20.2319262 | 20.434031 | 20.2962528 | 20.3362384 |
| Q96AX9 | MIB2 | 18.0907161 | 17.9902477 | 18.1041914 | 18.0262143 | 18.1109854 |
| P04150 | NR3C1 | 21.6377501 | 21.7895208 | 21.540726 | 21.5871801 | 21.7390036 |
| Q9NVM6 | DNAJC17 | 20.3897934 | 20.6430849 | 20.4106852 | 20.4556526 | 20.5854995 |
| Q9NSY0 | NRBP2 | 17.8126178 | 17.6669389 | 17.609306 | 17.4749988 | 17.8149181 |
| Q9HA47 | UCK1 | 17.1314284 | 17.2166645 | 17.0885242 | 17.262848 | 17.5354398 |
| O60925 | PFDN1 | 22.3335572 | 22.4791462 | 22.2170516 | 22.2852169 | 22.5258005 |
| Q9GZR1 | SENP6 | 17.6795915 | 17.9771161 | 17.9637837 | 17.8538815 | 17.7272256 |
| Q9H9A7 | RMI1 | 17.6198694 | 17.5902155 | 17.6483964 | 17.6840165 | 17.6485321 |
| Q96BJ3 | AIDA | 18.5344382 | 18.7769113 | 18.9691534 | 18.7439891 | 18.7700129 |
| Q8NC06 | ACBD4 | 14.2912242 | 14.3433619 | 14.575899 | 14.3753087 | 14.266617 |
| P55008 | AIF1 | 22.9940716 | 23.2610722 | 23.0260539 | 22.9438671 | 23.257663 |
| Q6EEV4 | POLR2M | 19.138606 | 19.2173367 | 19.1358237 | 19.0394339 | 19.3953481 |
| P07205 | PGK2 | 19.7763849 | 19.6148488 | 19.9608643 | 19.6050267 | 20.0623377 |
| P28907 | CD38 | 16.65289 | 16.9018966 | 16.8498922 | 16.7403949 | 16.893323 |
| Q13547 | HDAC1 | 24.0684453 | 24.1608863 | 24.1901476 | 24.0649216 | 24.1304652 |
| Q9UHV7 | MED13 | 17.4921098 | 17.7459145 | 17.8593007 | 17.539594 | 17.9393584 |
| O43805 | SSNA1 | 20.9001372 | 21.0492121 | 21.0917371 | 21.0559603 | 21.2168156 |
| Q92871 | PMM1 | 12.8957729 | 13.9814333 | 13.811681 | 13.2922042 | 13.4504595 |
| O60343 | TBC1D4 | 17.3787305 | 17.2908939 | 17.3291878 | 17.3212544 | 17.6261551 |
| O75164 | KDM4A | 19.5493369 | 19.769922 | 19.8192609 | 19.676236 | 19.863398 |
| Q9GZN7 | ROGDI | 18.8921161 | 18.8983414 | 18.9551384 | 18.8639835 | 18.9099189 |
| Q8TA86 | RP9 | 18.7357335 | 18.8745795 | 18.5888204 | 18.9533547 | 18.7724565 |
| Q7L5N7 | LPCAT2 | 21.7282453 | 21.8290959 | 21.7547315 | 21.6746784 | 21.7812755 |
| Q9Y4K0 | LOXL2 | 14.247856 | NA | NA | 14.4419731 | NA |
| Q9Y3D8 | AK6 | 21.3731717 | 21.303658 | 21.263156 | 21.3084793 | 21.3826707 |
| Q13740 | ALCAM | 18.9294623 | 18.7693117 | 19.0467036 | 18.9612616 | 18.9917068 |
| Q14573 | ITPR3 | 16.4777401 | 16.2858432 | 16.6586519 | 15.8226879 | 16.5666735 |
| P31483 | TIA1 | 21.8956513 | 21.8661373 | 22.0248596 | 21.826461 | 22.1900515 |
| Q14204 | DYNC1H1 | 24.7021104 | 24.732581 | 24.7292236 | 24.6259023 | 24.6964586 |
| P00519 | ABL1 | 18.4409443 | 18.6104929 | 18.7373052 | 18.5308911 | 18.7302223 |
| Q15751 | HERC1 | 19.49225 | 19.5342806 | 19.6092364 | 19.461188 | 19.5610775 |
| P20248 | CCNA2 | 20.9940667 | 21.1399672 | 21.0753873 | 20.9465602 | 21.0542703 |
| Q9BWC9 | CCDC106 | 19.1909208 | 19.4065811 | 19.4056883 | 19.290324 | 19.4391753 |
| Q9H3R5 | CENPH | 20.048234 | 20.0572165 | 20.1105432 | 20.0312967 | 20.0658128 |
| Q9UPY3 | DICER1 | 19.7212567 | 19.7931901 | 19.8022222 | 19.7346043 | 19.8609481 |
| O94817 | ATG12 | 20.1343784 | 20.3380046 | 20.3180996 | 20.0806043 | 19.9894723 |
| Q92925 | SMARCD2 | 22.5383518 | 22.6278082 | 22.5924544 | 22.446487 | 22.5008078 |

| | | | | | | |
|---|---|---|---|---|---|---|
| Q9BYD2 | MRPL9 | 23.3795211 | 23.2877657 | 23.3658506 | 23.3228521 | 23.2919566 |
| Q9Y2R9 | MRPS7 | 21.1987528 | 21.1496407 | 21.2429326 | 21.210054 | 21.3180678 |
| Q9BSA4 | TTYH2 | 19.2068414 | 19.0485518 | 18.9596176 | 19.178923 | 18.9268827 |
| Q14103 | HNRNPD | 26.5236862 | 26.4423542 | 26.460758 | 26.3685512 | 26.4881693 |
| O76095 | JTB | 18.9989355 | 19.0007102 | 19.1874672 | 18.9636599 | 19.1214091 |
| P42695 | NCAPD3 | 24.6215829 | 24.6514341 | 24.6610785 | 24.4948143 | 24.5905932 |
| Q9Y4B4 | RAD54L2 | 17.9913162 | 18.1425678 | 18.0394279 | 18.0516711 | 18.0437716 |
| Q9UJX2 | CDC23 | 21.0055407 | 21.0126808 | 21.1058713 | 20.9955732 | 21.0612269 |
| Q7RTV0 | PHF5A | 22.020391 | 22.0760582 | 22.0375878 | 21.9560211 | 22.1213163 |
| L0R819 | ASDURF | 19.6424158 | 19.8464568 | 19.2225549 | 19.5499604 | 19.7713612 |
| Q96QH2 | PRAM1 | 17.4511857 | 17.5463893 | 17.7272762 | 17.3968238 | 17.7880911 |
| Q5TZA2 | CROCC | 21.3192409 | 21.0310296 | 21.0477411 | 21.0443474 | 20.7545347 |
| Q53GL0 | PLEKHO1 | 15.1010043 | 15.211521 | 15.2842191 | 15.1915275 | 15.3485734 |
| Q8NFI3 | ENGASE | 19.1438763 | 19.2369061 | 19.3414478 | 19.171114 | 19.2780517 |
| Q13011 | ECH1 | 24.5539555 | 24.6652291 | 24.661733 | 24.5511743 | 24.6589963 |
| Q8NDA8 | MROH1 | 20.3133614 | 20.2519227 | 20.3824737 | 20.2478464 | 20.3332131 |
| Q93033 | CD101 | 19.3809885 | 19.5106045 | 19.5457956 | 19.45404 | 19.5007619 |
| O15062 | ZBTB5 | 15.8379906 | 16.3511302 | 16.3789896 | 16.1721905 | 16.1789513 |
| Q96IR7 | HPDL | 19.4915441 | 19.5715639 | 19.4195037 | 19.4729261 | 19.3730715 |
| Q9Y5K5 | UCHL5 | 22.7859837 | 22.8248243 | 22.7664484 | 22.6906006 | 22.807202 |
| Q9HD15 | SRA1 | 21.2107061 | 21.3757085 | 21.2745587 | 21.2217459 | 21.2902408 |
| Q9UNY4 | TTF2 | 19.6828259 | 19.8183605 | 19.8596293 | 19.7352036 | 19.8275358 |
| O95801 | TTC4 | 21.1031512 | 21.1868762 | 21.1892297 | 21.165443 | 21.2663518 |
| Q8WUM4 | PDCD6IP | 23.8566602 | 23.9124407 | 23.8494248 | 23.8084693 | 23.926834 |
| Q9NWX5 | ASB6 | 16.4924386 | 16.6705781 | 16.7089185 | 16.6922866 | 16.5945078 |
| Q14153 | FAM53B | 15.232965 | 15.4332026 | 15.4468384 | 15.4169197 | 15.5905365 |
| Q9H019 | MTFR1L | 18.260615 | 18.2979705 | 18.5964651 | 18.2594476 | 18.3661012 |
| Q96N21 | TEPSIN | 17.3986557 | 17.5706346 | 17.5081491 | 17.4353269 | 17.4909677 |
| Q9Y5U8 | MPC1 | 20.1018801 | 19.9667271 | 19.8888755 | 20.1015234 | 19.8486716 |
| Q9NVE5 | USP40 | 18.4853108 | 18.598685 | 18.4403928 | 18.4980096 | 18.4597935 |
| Q9H1X3 | DNAJC25 | 17.6988877 | 17.8397659 | 18.0086283 | 17.7340933 | 17.846466 |
| Q8NF37 | LPCAT1 | 21.6339186 | 21.7571585 | 21.5790955 | 21.6072768 | 21.7842973 |
| Q99598 | TSNAX | 22.4240555 | 22.4319013 | 22.3896134 | 22.3550141 | 22.3762469 |
| Q7Z4Q2 | HEATR3 | 21.0731818 | 21.0665677 | 21.1567591 | 21.0569395 | 21.2424593 |
| Q8TD19 | NEK9 | 22.3486842 | 22.4325249 | 22.4773129 | 22.3127448 | 22.4540869 |
| Q9H936 | SLC25A22 | 21.2015716 | 21.2933134 | 21.2987401 | 21.2242594 | 21.3276251 |
| Q9HCG8 | CWC22 | 21.315919 | 21.2967858 | 21.4334343 | 21.2207424 | 21.3229193 |
| Q6PIW4 | FIGNL1 | 18.171613 | 18.2585883 | 18.3583848 | 18.0908311 | 18.1305533 |
| Q71F23 | CENPU | 19.0138628 | 19.190676 | 19.113494 | 19.1764651 | 19.1991914 |
| Q13595 | TRA2A | 22.4581156 | 22.4744316 | 22.5218322 | 22.3481849 | 22.5232761 |
| P50336 | PPOX | 19.2434612 | 19.2143612 | 19.2474982 | 19.1943631 | 19.3281223 |
| O43583 | DENR | 21.0572739 | 21.1068381 | 21.0654496 | 21.0518762 | 21.2157267 |

| | | | | | | |
|---|---|---|---|---|---|---|
| Q8TCU4 | ALMS1 | 18.4074554 | 18.5226725 | 18.4279947 | 18.4817267 | 18.5598869 |
| A6NK89 | RASSF10 | NA | 17.8568148 | 18.1935674 | NA | 17.9521862 |
| Q96JC9 | EAF1 | 17.7108717 | 17.4867146 | 17.6359458 | 17.5320822 | 17.5239664 |
| Q674X7 | KAZN | 15.2786666 | 15.0601723 | 15.4967532 | 14.9511547 | 15.1293373 |
| Q9UJ70 | NAGK | 19.065865 | 19.1657772 | 19.1451324 | 19.0881302 | 19.218665 |
| Q12824 | SMARCB1 | 21.8668171 | 21.9491812 | 21.8624105 | 21.8683586 | 21.8774723 |
| Q5SGD2 | PPM1L | 17.9723005 | 17.9852235 | 18.0985883 | 17.767961 | 18.0819316 |
| Q96NJ6 | ZFP3 | 15.4200734 | 15.3351733 | 15.5569299 | 15.2903012 | 15.4858761 |
| Q6YN16 | HSDL2 | 21.5478938 | 21.5477258 | 21.6317958 | 21.6547912 | 21.6159132 |
| O00161 | SNAP23 | 22.2862078 | 22.3792092 | 22.3398413 | 22.2080083 | 22.2024553 |
| P62820 | RAB1A | 22.6498009 | 22.6963082 | 22.8002707 | 22.6479202 | 22.8050587 |
| P19440 | GGT1 | 20.9833299 | 21.0799124 | 21.0052391 | 20.9117496 | 21.0513981 |
| Q9UQ80 | PA2G4 | 25.6853025 | 25.7041949 | 25.6344518 | 25.565545 | 25.6678433 |
| Q9NXW2 | DNAJB12 | 20.7118975 | 20.7617647 | 20.7859002 | 20.8131986 | 20.7847667 |
| P33981 | TTK | 18.8863175 | 18.9893374 | 19.0719207 | 18.8797844 | 19.0151258 |
| Q9BQ24 | ZFYVE21 | 19.0186492 | 18.8513324 | 18.9561283 | 18.9674309 | 18.9201229 |
| O75940 | SMNDC1 | 21.1279258 | 21.2791896 | 21.0011347 | 21.1685285 | 21.1095248 |
| Q96EY8 | MMAB | 21.7021038 | 21.8835463 | 21.7123289 | 21.8424779 | 21.9102032 |
| Q99941 | ATF6B | 17.5432591 | 17.9885791 | 17.7530169 | 17.7151863 | 17.7486987 |
| Q86T82 | USP37 | 16.1731223 | 16.3146735 | 16.6983001 | 16.4312834 | 16.7669893 |
| O14733 | MAP2K7 | 20.5189253 | 20.5729512 | 20.655656 | 20.4016601 | 20.6245479 |
| Q96EC8 | YIPF6 | 18.8490252 | 19.0240916 | 18.9339548 | 19.0095514 | 18.9449961 |
| Q5VTL8 | PRPF38B | 20.9323802 | 21.0334197 | 21.1040455 | 20.8554536 | 21.0459033 |
| P43034 | PAFAH1B1 | 22.7261817 | 22.7156871 | 22.7439844 | 22.6303862 | 22.7675328 |
| Q9NT62 | ATG3 | 22.2795329 | 22.2071017 | 22.2942342 | 22.1234989 | 22.3092819 |
| P35244 | RPA3 | 22.4735298 | 22.4536551 | 22.4945118 | 22.3910936 | 22.5102101 |
| Q6P4R8 | NFRKB | 18.8302013 | 18.9900484 | 18.9063267 | 18.8680549 | 18.9523664 |
| O14763 | TNFRSF10B | 18.2668049 | 18.1236972 | 18.2797344 | 18.3485454 | 18.3040315 |
| Q92688 | ANP32B | 25.4923759 | 25.6280665 | 25.3984215 | 25.4280178 | 25.5630301 |
| Q96NT0 | CCDC115 | 19.260075 | 19.3300691 | 19.3900072 | 19.2550905 | 19.3354721 |
| Q6P4A7 | SFXN4 | 20.006288 | 20.1482855 | 20.1636405 | 20.0260917 | 20.1650739 |
| P17861 | XBP1 | 17.9144038 | 18.4729029 | 18.1105845 | 18.1457134 | 17.6867625 |
| Q9NR31 | SAR1A | 23.5078532 | 23.5453514 | 23.5471388 | 23.370077 | 23.6241577 |
| Q05639 | EEF1A2 | 23.1893702 | 23.1278915 | 23.2039717 | 23.1899005 | 23.1601629 |
| Q96RE7 | NACC1 | 21.3429234 | 21.2418939 | 21.2946297 | 21.201664 | 21.2267342 |
| Q9UP83 | COG5 | 20.8719613 | 20.8355213 | 20.9014161 | 20.8527949 | 20.9539909 |
| Q8IYH5 | ZZZ3 | 17.7008584 | 17.8003216 | 17.757199 | 17.8256257 | 17.8659555 |
| O43820 | HYAL3 | 18.4007414 | 18.6066804 | 18.4687321 | 18.5937062 | 18.4206453 |
| Q8IZD2 | KMT2E | 16.7510041 | 16.8124533 | 16.9873329 | 16.9638174 | 16.8210122 |
| P62987 | UBA52 | 20.6139028 | 19.7361587 | 21.0918229 | 20.5133107 | 19.5565864 |
| O60763 | USO1 | 22.0445861 | 22.0734762 | 22.0377716 | 21.9989806 | 22.0367424 |
| Q969G3 | SMARCE1 | 22.7371666 | 22.9082278 | 22.7261379 | 22.7046278 | 22.8098007 |

| | | | | | | |
|---|---|---|---|---|---|---|
| Q9BSG1 | ZNF2 | NA | 15.3971288 | 15.3499014 | NA | 15.1148279 |
| P12004 | PCNA | 26.350919 | 26.4045596 | 26.3381666 | 26.288999 | 26.3665367 |
| Q92560 | BAP1 | 18.2514816 | 18.2389376 | 18.2686852 | 18.3508397 | 18.274246 |
| O75820 | ZNF189 | 18.3770288 | 18.30111 | 18.3804806 | 18.4843383 | 18.3261916 |
| P17612 | PRKACA | 22.6020893 | 22.6322687 | 22.8817847 | 22.6011048 | 22.7465973 |
| Q6IQ22 | RAB12 | 17.2158202 | 17.1576212 | 17.1915393 | 17.1447207 | 17.2192196 |
| Q86YM7 | HOMER1 | 17.6954841 | 17.6130325 | 17.645434 | 17.8743746 | 17.6363099 |
| Q5T2E6 | ARMH3 | 20.7878224 | 20.7374442 | 20.8390065 | 20.7411855 | 20.9377554 |
| Q8WXI4 | ACOT11 | 20.2180582 | 20.1946462 | 20.1996208 | 20.1209338 | 20.2187422 |
| Q9NQZ2 | UTP3 | 20.7872921 | 20.7721776 | 20.8531203 | 20.7775066 | 20.8145209 |
| O43166 | SIPA1L1 | NA | 15.2970147 | 14.7699236 | NA | 14.8436656 |
| Q15554 | TERF2 | 20.7867797 | 20.8567999 | 20.8129838 | 20.8033006 | 20.8625719 |
| Q9GZY8 | MFF | 19.6478946 | 19.8121779 | 19.733759 | 19.7104353 | 19.7788488 |
| Q9P107 | GMIP | 21.8338512 | 21.9442376 | 21.9351907 | 21.8739614 | 21.9683475 |
| P98179 | RBM3 | 24.9052604 | 24.9431564 | 24.5587349 | 24.7678896 | 24.7923949 |
| Q9BXQ6 | TMEM121B | 16.7466174 | 16.9875138 | 16.8887415 | 16.7626165 | 16.8642687 |
| P31150 | GDI1 | 20.687496 | 20.8739434 | 20.9901585 | 20.7235938 | 20.9019789 |
| Q6PK81 | ZNF773 | 18.6593461 | 18.7726072 | 18.8808595 | 18.6544935 | 18.8353139 |
| Q9NP64 | ZCCHC17 | 19.2740725 | 19.3647442 | 19.3871192 | 19.2918707 | 19.3612575 |
| Q9UKW4 | VAV3 | 19.9331382 | 19.9702414 | 20.1259078 | 20.0556848 | 20.0280925 |
| Q9Y584 | TIMM22 | 18.3949545 | 18.5947575 | 18.6420578 | 18.4928935 | 18.6104727 |
| Q712K3 | UBE2R2 | 19.7227047 | 19.7066483 | 19.6477457 | 19.6803217 | 19.8140582 |
| Q9BRT3 | MIEN1 | 21.7435583 | 21.6719603 | 21.4902922 | 21.5888142 | 21.6797261 |
| O15235 | MRPS12 | 20.9340198 | 20.9556876 | 21.2924192 | 20.9032406 | 21.198907 |
| P12980 | LYL1 | 16.614332 | 16.8022387 | 16.6239957 | 16.4936239 | 16.7531681 |
| P51809 | VAMP7 | 21.9291601 | 21.9272432 | 21.9184107 | 21.8655813 | 21.7938231 |
| Q9NZ32 | ACTR10 | 20.7684068 | 20.7394447 | 20.9202242 | 20.7586135 | 20.8944309 |
| Q9H1A4 | ANAPC1 | 20.5108267 | 20.7336204 | 20.6825529 | 20.6005079 | 20.6966018 |
| P07900 | HSP90AA1 | 27.2563144 | 27.3168494 | 27.2476179 | 27.1903989 | 27.308931 |
| Q96SB4 | SRPK1 | 21.8574838 | 21.9472299 | 21.9430738 | 21.8905564 | 21.8799974 |
| P49815 | TSC2 | 19.7075059 | 19.7188391 | 19.7867856 | 19.6758101 | 19.793348 |
| P16284 | PECAM1 | 21.0593092 | 21.0846721 | 21.0721213 | 21.0825492 | 21.0083736 |
| Q9UL63 | MKLN1 | 19.5847144 | 19.6379767 | 19.6001665 | 19.6118978 | 19.6208224 |
| P78413 | IRX4 | 16.5370603 | 16.6877874 | 16.6407998 | 16.5254209 | 16.7327302 |
| Q9Y3E7 | CHMP3 | 21.2763229 | 21.284989 | 21.5776971 | 21.2937347 | 21.3983971 |
| P10746 | UROS | 20.3641877 | 20.3548113 | 20.4269485 | 20.3655809 | 20.662833 |
| Q8IYD8 | FANCM | 13.0067637 | 12.7873022 | 13.0466486 | 12.612742 | 13.1127126 |
| Q99828 | CIB1 | 19.0109034 | 19.1943153 | 19.2270904 | 19.0303729 | 19.1483582 |
| Q6IQ21 | ZNF770 | 16.9509486 | 17.1257997 | 17.7218719 | 17.4576664 | 16.8748994 |
| Q9BTE1 | DCTN5 | 20.6426438 | 20.6877541 | 20.6197102 | 20.4365689 | 20.9142772 |
| Q9Y2Y1 | POLR3K | 17.255874 | 17.4869486 | 17.4551302 | 17.6314309 | 17.6852284 |
| Q6IE81 | JADE1 | 16.4599163 | 16.3360284 | 16.5608936 | 16.5793398 | 16.6462963 |

| | | | | | | |
|---|---|---|---|---|---|---|
| P30566 | ADSL | 22.8592472 | 22.8143251 | 22.8221088 | 22.7130189 | 22.7935265 |
| E9PAV3 | NACA | 25.3676882 | 25.4036316 | 25.3404646 | 25.3517986 | 25.3554245 |
| Q8WV37 | ZNF480 | 17.3352936 | 17.3698805 | 17.2043198 | 17.3733474 | 17.2668414 |
| P14314 | PRKCSH | 24.9060316 | 25.0745481 | 24.8298374 | 24.8220428 | 24.8951688 |
| Q13418 | ILK | 21.827971 | 21.8866658 | 21.9417482 | 21.7764272 | 21.9040903 |
| O94913 | PCF11 | 20.2325016 | 20.3296564 | 20.2421174 | 20.2629079 | 20.2846733 |
| Q92994 | BRF1 | 19.4770412 | 19.6697422 | 19.6184829 | 19.5837528 | 19.6553434 |
| Q969J3 | BORCS5 | 19.3459857 | 19.4714366 | 19.3736653 | 19.3692717 | 19.4763885 |
| Q8N0U4 | FAM185A | NA | 15.8812352 | 15.4988009 | NA | 15.4962956 |
| Q8N6T7 | SIRT6 | 20.3629973 | 20.3243261 | 20.3777899 | 20.304277 | 20.2780219 |
| Q96ER9 | CCDC51 | 21.0382479 | 21.0516433 | 21.1218707 | 20.9894791 | 21.0879931 |
| Q7Z7F7 | MRPL55 | 20.9398822 | 21.0138429 | 21.0853812 | 20.9742043 | 21.0303459 |
| O75122 | CLASP2 | 21.0271556 | 21.0254065 | 21.0972838 | 21.0360218 | 21.0969697 |
| O15231 | ZNF185 | 21.3284274 | 21.4925644 | 21.3907365 | 21.3306812 | 21.4556792 |
| P42025 | ACTR1B | 21.2652194 | 21.4795519 | 21.388665 | 21.2375133 | 21.4187961 |
| Q9Y253 | POLH | NA | 15.2122868 | 15.1882702 | NA | 15.0430208 |
| A2VDJ0 | TMEM131L | 18.2069537 | 18.1127075 | 18.2947987 | 18.2339728 | 18.3048715 |
| Q96J42 | TXNDC15 | 18.0816241 | 18.1896551 | 18.0888867 | 17.9675444 | 18.0593696 |
| Q86Y97 | KMT5C | 19.3951989 | 19.1752933 | 19.3791939 | 19.6079401 | 19.3772243 |
| P27797 | CALR | 27.5353418 | 27.5607602 | 27.3272012 | 27.4786401 | 27.4841737 |
| Q8TBC3 | SHKBP1 | 20.6277336 | 20.7346285 | 20.7506804 | 20.6702286 | 20.6122339 |
| Q86SZ2 | TRAPPC6B | 19.3495228 | 19.2723532 | 19.490109 | 19.3699633 | 19.5497178 |
| Q8WU90 | ZC3H15 | 22.4652123 | 22.4928022 | 22.4079298 | 22.3776509 | 22.4223275 |
| Q96H40 | ZNF486 | 15.1426744 | 14.8398795 | 15.00118 | 14.9868221 | 14.8271504 |
| Q14696 | MESD | 22.5119003 | 22.7165492 | 22.6097535 | 22.5044211 | 22.5911325 |
| Q9Y5V0 | ZNF706 | 21.2864401 | 21.2819085 | 21.2820776 | 21.1281535 | 21.2021718 |
| O43766 | LIAS | 17.8479586 | 17.9483212 | 17.9935766 | 17.8100406 | 18.0098422 |
| O00255 | MEN1 | 20.2580252 | 20.2131362 | 20.3616798 | 20.2043203 | 20.2010943 |
| P54687 | BCAT1 | 19.0012121 | 19.2217131 | 19.1289274 | 19.1192451 | 19.1842583 |
| Q8N5G2 | MACO1 | 19.0862903 | 19.1557018 | 19.0208881 | 19.1204613 | 18.9657823 |
| Q5T3I0 | GPATCH4 | 20.0253751 | 20.2242995 | 20.2042433 | 20.0911792 | 20.1686039 |
| Q9BQQ7 | RTP3 | 12.9778961 | NA | NA | 13.1022744 | NA |
| Q9H425 | C1orf198 | 13.7767415 | 13.9401207 | 13.9847883 | 13.7236176 | 13.6664239 |
| P09110 | ACAA1 | 21.5116693 | 21.6091013 | 21.5105082 | 21.4892636 | 21.5234747 |
| Q15269 | PWP2 | 22.1743769 | 22.2049252 | 22.2984751 | 22.1520981 | 22.1657835 |
| Q96PK6 | RBM14 | 24.0044713 | 24.050858 | 23.9601429 | 23.9981247 | 23.979836 |
| Q9BTY7 | HGH1 | 22.5210146 | 22.5160217 | 22.3073583 | 22.4515857 | 22.5020511 |
| O95835 | LATS1 | 18.5588915 | 19.0084035 | 19.0753467 | 18.9304445 | 18.96874 |
| Q9UGN4 | CD300A | 18.7503255 | 19.2957758 | 19.2175463 | 19.1064094 | 19.1685252 |
| Q6AHZ1 | ZNF518A | 15.7451559 | 15.8172384 | 15.8978842 | 15.8205218 | 15.5739934 |
| O60443 | GSDME | 18.8547942 | 19.1272654 | 19.0996106 | 18.8823522 | 19.0470997 |
| Q8NEW0 | SLC30A7 | 20.9715042 | 20.9695807 | 21.0173793 | 20.9500997 | 21.0378069 |

| | | | | | | |
|---|---|---|---|---|---|---|
| Q9BQ95 | ECSIT | 20.0414817 | 19.9116262 | 20.0121929 | 19.9712029 | 20.0396706 |
| Q9BZF1 | OSBPL8 | 22.1327056 | 22.0114958 | 22.1426043 | 22.0509055 | 22.1349786 |
| Q2M389 | WASHC4 | 21.3188919 | 21.3269878 | 21.4048955 | 21.2977698 | 21.3860201 |
| Q9BQ70 | TCF25 | 20.562842 | 20.4306117 | 20.5501532 | 20.4916944 | 20.4802665 |
| P27694 | RPA1 | 23.4573915 | 23.409396 | 23.4976221 | 23.3651498 | 23.4446806 |
| P18858 | LIG1 | 22.628695 | 22.7130805 | 22.6656926 | 22.6052173 | 22.6775141 |
| Q9Y2P8 | RCL1 | 20.9821298 | 21.0768376 | 21.0201408 | 21.0105008 | 21.070556 |
| Q9BUQ8 | DDX23 | 22.2391577 | 22.3400071 | 22.2752122 | 22.2319972 | 22.3252147 |
| Q15311 | RALBP1 | 20.9789721 | 20.9955058 | 21.0403195 | 20.9453694 | 21.0039891 |
| Q9Y4B6 | DCAF1 | 20.0448457 | 20.1467383 | 20.1369272 | 20.0478791 | 20.2050761 |
| Q9Y4K4 | MAP4K5 | 18.4593751 | 18.2731698 | 18.3125085 | 18.2679605 | 18.3732896 |
| O00443 | PIK3C2A | 18.6325571 | 18.6979147 | 18.7438985 | 18.6092756 | 18.8004828 |
| Q15468 | STIL | 17.4069496 | 17.6667918 | 17.790163 | 17.712001 | 17.6866553 |
| P17931 | LGALS3 | 18.5470259 | 18.5412501 | 18.9643301 | 18.5879704 | 18.699689 |
| Q5JVS0 | HABP4 | 15.8131367 | 16.1124459 | 16.172869 | 16.0263584 | 16.3679925 |
| Q9BQ89 | FAM110A | 16.909775 | 17.16833 | 16.7807589 | 16.5426092 | 16.8022879 |
| Q96K78 | ADGRG7 | NA | 18.860384 | 18.2104242 | NA | 18.9706055 |
| Q9UKF6 | CPSF3 | 21.7308061 | 21.8497655 | 21.7867695 | 21.7358756 | 21.7286873 |
| Q86UK7 | ZNF598 | 21.592058 | 21.6956069 | 21.6419733 | 21.6011052 | 21.6147165 |
| P84090 | ERH | 25.0005149 | 24.8601462 | 24.8985802 | 24.871632 | 24.9934111 |
| P45983 | MAPK8 | 18.8610725 | 19.0331813 | 19.1006721 | 18.8575032 | 19.0758381 |
| P16219 | ACADS | 20.6672568 | 20.5743825 | 20.8412542 | 20.6064656 | 20.72522 |
| Q969P6 | TOP1MT | 15.1672554 | 15.917939 | 15.67835 | 14.9581208 | 15.7208338 |
| Q6UX71 | PLXDC2 | 17.3211457 | 17.7545044 | 17.9108179 | 17.6622504 | 17.7487845 |
| P17021 | ZNF17 | 16.44309 | 16.5343186 | 16.8091024 | 16.4780946 | 16.4909156 |
| Q9UBZ4 | APEX2 | 16.2054072 | 16.3059247 | 16.5789727 | 16.265523 | 16.0118729 |
| P63172 | DYNLT1 | 19.561088 | 19.7132976 | 19.9773911 | 19.5753038 | 19.7961435 |
| P49366 | DHPS | 20.3574629 | 20.517583 | 20.5414661 | 20.3953903 | 20.5653748 |
| P43897 | TSFM | 22.2184976 | 22.2970619 | 22.298829 | 22.1879204 | 22.3568661 |
| Q8N5N7 | MRPL50 | 21.7266793 | 21.6965867 | 21.5934075 | 21.7157483 | 21.630318 |
| Q6DN90 | IQSEC1 | 19.4441923 | 19.4867368 | 19.3405481 | 19.4514152 | 19.4269998 |
| Q13263 | TRIM28 | 26.0993591 | 26.144213 | 26.0329268 | 25.9776023 | 25.9910062 |
| Q15435 | PPP1R7 | 22.6423857 | 22.5579722 | 22.6334253 | 22.6016403 | 22.6201064 |
| O95785 | WIZ | 21.0039483 | 20.9819675 | 20.9978627 | 20.9402232 | 21.0329313 |
| Q676U5 | ATG16L1 | 19.5485462 | 19.5485329 | 19.6304287 | 19.6007511 | 19.5160103 |
| Q8NC60 | NOA1 | 20.9440665 | 21.0242348 | 20.9425825 | 20.9326291 | 20.9411607 |
| P22681 | CBL | 20.1124621 | 20.1540992 | 20.2136609 | 20.1352077 | 20.2929091 |
| Q15014 | MORF4L2 | 19.2456877 | 19.2043801 | 19.2921643 | 19.2944973 | 19.3482869 |
| Q9NVN8 | GNL3L | 19.984443 | 20.0988253 | 20.0975537 | 20.0308959 | 20.1807428 |
| O60566 | BUB1B | 19.590117 | 19.5960194 | 19.6515369 | 19.5473966 | 19.6925938 |
| Q9H2U1 | DHX36 | 19.7459923 | 19.7795157 | 19.7358643 | 19.7165491 | 19.7936334 |
| Q8WYQ5 | DGCR8 | 19.419202 | 19.5297417 | 19.4080536 | 19.3687321 | 19.4485742 |

| | | | | | | |
|---|---|---|---|---|---|---|
| P12755 | SKI | 17.8055599 | 17.8407793 | 17.878756 | 17.6780356 | 17.9660467 |
| Q9BRD0 | BUD13 | 19.6982378 | 19.921296 | 19.8044084 | 19.785504 | 19.8798233 |
| Q99418 | CYTH2 | 18.3928667 | 18.5137962 | 18.6486518 | 18.4364073 | 18.6559451 |
| Q86XK3 | SFR1 | 15.7573529 | 15.4519001 | 15.5935399 | 15.7654393 | 15.826182 |
| Q702N8 | XIRP1 | 21.3025528 | 21.3668319 | 21.2758215 | 21.0971159 | 21.0501887 |
| Q96FX2 | DPH3 | 16.7356741 | 17.1902048 | 16.681197 | 16.8393883 | 17.1002956 |
| Q9BSV6 | TSEN34 | 19.6851836 | 19.6717237 | 19.6307148 | 19.6470555 | 19.6536477 |
| Q5PRF9 | SAMD4B | 18.8492614 | 18.9140721 | 18.991786 | 18.9264803 | 18.9753102 |
| P49368 | CCT3 | 26.1355439 | 26.1359131 | 26.1135383 | 26.0565032 | 26.1019921 |
| Q03933 | HSF2 | 17.0811109 | 16.4306464 | 16.507961 | 16.9948223 | 15.950107 |
| Q86Y39 | NDUFA11 | 20.9750404 | 21.0538803 | 21.0883877 | 20.8934816 | 20.9063108 |
| Q4U2R6 | MRPL51 | 20.5669147 | 20.6790047 | 20.7283388 | 20.6944736 | 20.5635746 |
| Q8IV08 | PLD3 | 23.7321265 | 23.9640737 | 23.9174046 | 23.6717741 | 23.8308022 |
| Q53QZ3 | ARHGAP15 | 21.7845299 | 21.8977728 | 21.8787705 | 21.7377725 | 21.8883476 |
| P49407 | ARRB1 | 21.5151853 | 21.5767226 | 21.7135965 | 21.5218009 | 21.6057316 |
| Q8NBT2 | SPC24 | 20.4542612 | 20.5123251 | 20.3783169 | 20.3999762 | 20.5178379 |
| O95251 | KAT7 | 20.50688 | 20.3208737 | 20.4932966 | 20.3628535 | 20.3783218 |
| O15111 | CHUK | 19.810269 | 19.9802646 | 19.9274292 | 19.7904593 | 20.0236077 |
| Q01968 | OCRL | 18.2971974 | 18.4387368 | 18.3497265 | 18.3144105 | 18.3937211 |
| P78324 | SIRPA | 21.7646305 | 21.9380738 | 21.8832772 | 21.7503995 | 21.7839355 |
| Q96BW5 | PTER | 20.6940833 | 20.7015433 | 20.7246994 | 20.6999305 | 20.7939035 |
| Q9BQ39 | DDX50 | 22.4299421 | 22.5010744 | 22.499391 | 22.4388253 | 22.4342456 |
| Q16706 | MAN2A1 | 20.6520462 | 20.671653 | 20.7295819 | 20.6159067 | 20.7040526 |
| P17980 | PSMC3 | 23.7472503 | 23.826296 | 23.7627591 | 23.7171663 | 23.8440849 |
| O75592 | MYCBP2 | 21.1043945 | 21.169965 | 21.19615 | 21.108221 | 21.1861431 |
| Q16401 | PSMD5 | 22.0906313 | 22.0785319 | 22.1802993 | 22.0837666 | 22.1286331 |
| Q9H3U1 | UNC45A | 21.4740487 | 21.5744477 | 21.5659115 | 21.4833247 | 21.6055292 |
| Q9UPM8 | AP4E1 | 20.0938581 | 20.1307868 | 20.1370762 | 20.0627428 | 20.1750862 |
| P35251 | RFC1 | 22.1286781 | 22.2198073 | 22.2060246 | 22.1303045 | 22.1845259 |
| Q8TD16 | BICD2 | 20.1797172 | 20.4065291 | 20.1692094 | 20.1913244 | 20.3579016 |
| P31040 | SDHA | 23.5191817 | 23.5801683 | 23.6488689 | 23.494196 | 23.7437334 |
| Q92522 | H1-10 | 22.7641498 | 22.8166351 | 22.9782704 | 22.7374536 | 22.7594577 |
| Q9UKA9 | PTBP2 | 16.5354762 | 16.5618261 | 16.6912731 | 16.4446594 | 16.2430798 |
| O75167 | PHACTR2 | 16.9282816 | 16.873028 | 17.5127407 | 17.0992543 | 17.1699109 |
| Q9H0F7 | ARL6 | 12.3405509 | NA | NA | 12.7738712 | NA |
| Q66K14 | TBC1D9B | 21.0218891 | 21.0502202 | 21.116549 | 20.9992608 | 21.0875808 |
| Q8IWE4 | DCUN1D3 | 14.8826295 | 14.9980387 | 15.3095902 | 15.1180895 | 14.9553992 |
| O15287 | FANCG | 18.4836924 | 18.6902686 | 18.5805411 | 18.6219034 | 18.5386681 |
| Q9ULH7 | MRTFB | 15.6802659 | 15.7167849 | 15.7583557 | 15.5127346 | 15.9545511 |
| Q86UZ6 | ZBTB46 | 15.7500853 | 15.2905359 | 15.593874 | 15.6620516 | 15.4689813 |
| Q9BRQ3 | NUDT22 | 17.6781433 | 17.8869901 | 17.5919379 | 17.5393386 | 18.1035767 |
| Q6NSZ9 | ZSCAN25 | 17.30236 | 17.4412904 | 17.1674586 | 17.2639937 | 17.5953737 |

| | | | | | | |
|---|---|---|---|---|---|---|
| O00213 | APBB1 | 18.022202 | 18.397144 | 18.2926292 | 18.1839991 | 18.2713135 |
| O14972 | VPS26C | 21.8959263 | 21.8649308 | 21.8712372 | 21.7930718 | 21.953972 |
| Q8IWA4 | MFN1 | 18.9749022 | 19.0023415 | 19.0790872 | 19.0239671 | 19.0271791 |
| Q9NVI1 | FANCI | 20.6056307 | 20.6659202 | 20.7841664 | 20.5921513 | 20.7160887 |
| Q9BW85 | YJU2 | 20.7826014 | 20.9141095 | 20.8418167 | 20.7270382 | 20.8439911 |
| P07355 | ANXA2 | 24.8436157 | 24.9162302 | 24.8079363 | 24.6892315 | 24.8384438 |
| P15531 | NME1 | 25.9028422 | 25.9658802 | 25.8537387 | 25.7943729 | 25.8496161 |
| Q9BXW9 | FANCD2 | 20.4887633 | 20.5141679 | 20.430153 | 20.4762556 | 20.5156355 |
| Q9Y232 | CDYL | 19.0103253 | 19.2015514 | 19.0589751 | 18.9385756 | 19.0347746 |
| O14798 | TNFRSF10C | 17.5065058 | 17.39235 | 17.8659741 | 17.6364538 | 17.7520807 |
| P19484 | TFEB | 16.9647729 | 17.3393999 | 17.5308762 | 17.1939825 | 17.6319455 |
| Q2YD98 | UVSSA | 17.5958953 | 17.747261 | 17.855938 | 17.6511305 | 17.6840362 |
| Q0P651 | ABHD18 | NA | 15.999909 | 16.0121355 | NA | 15.9186291 |
| P01137 | TGFB1 | 21.9198013 | 22.0586675 | 22.0173391 | 21.7583242 | 21.9592162 |
| O00257 | CBX4 | 18.6557249 | 18.6603321 | 18.8857762 | 18.5246601 | 18.5942975 |
| Q15390 | MTFR1 | 18.8084665 | 18.8322977 | 18.9583628 | 18.8090747 | 18.8572631 |
| O00422 | SAP18 | 25.1004196 | 25.1228232 | 25.0583048 | 24.9803978 | 25.0890405 |
| Q5TC12 | ATPAF1 | 19.4611285 | 19.4644736 | 19.6504474 | 19.419022 | 19.5483872 |
| O95372 | LYPLA2 | 22.4393215 | 22.3873348 | 22.2701431 | 22.3152891 | 22.4747398 |
| Q13162 | PRDX4 | 23.2254513 | 23.2906166 | 23.3020169 | 23.2884273 | 23.4647794 |
| P19532 | TFE3 | 19.1056578 | 19.1773341 | 19.1377717 | 18.9168214 | 19.1380234 |
| P25686 | DNAJB2 | 18.0500244 | 18.1306887 | 17.8977616 | 18.1410049 | 18.0896882 |
| Q3B7T1 | EDRF1 | 19.9890696 | 20.2175727 | 20.3134534 | 20.1169545 | 20.2082508 |
| P32970 | CD70 | 19.1653723 | 19.154873 | 19.3428545 | 19.3027083 | 19.0567568 |
| P62701 | RPS4X | 26.8599564 | 26.8069483 | 26.9092101 | 26.7378997 | 26.8628188 |
| P62310 | LSM3 | 23.0003751 | 22.9429743 | 22.9890009 | 23.0237436 | 23.0417384 |
| Q8WWN8 | ARAP3 | 18.2295287 | 18.4059963 | 18.1025331 | 17.9236385 | 18.2143116 |
| Q9ULD4 | BRPF3 | 13.7711615 | NA | NA | 13.8933321 | NA |
| Q6PGN9 | PSRC1 | NA | 14.6805136 | 13.7116505 | NA | 14.3454349 |
| O75558 | STX11 | 18.1218092 | 18.1881076 | 18.3367137 | 18.1970279 | 18.055766 |
| P45974 | USP5 | 23.0224936 | 23.1175469 | 23.0336782 | 22.9247485 | 23.0079729 |
| O15379 | HDAC3 | 20.2648755 | 20.3943352 | 20.3338353 | 20.2112908 | 20.455937 |
| Q9Y2K5 | R3HDM2 | 16.4306671 | 16.6139835 | 16.792759 | 16.3135353 | 16.4393727 |
| P61966 | AP1S1 | 19.7043828 | 19.5247935 | 20.2473067 | 19.7396672 | 19.5963861 |
| Q9NQT5 | EXOSC3 | 21.4335658 | 21.573295 | 21.5471684 | 21.606778 | 21.6092537 |
| Q86WJ1 | CHD1L | 20.7194258 | 20.7869412 | 20.8242012 | 20.704085 | 20.7877869 |
| A6NHT5 | HMX3 | 19.9774659 | 19.8762988 | 20.0538733 | 20.0093654 | 20.1148059 |
| Q13686 | ALKBH1 | 17.8544599 | 17.7317853 | 17.9976634 | 17.9990079 | 17.9294995 |
| Q8N5C7 | DTWD1 | 18.2089377 | 18.50316 | 18.3312723 | 18.1492222 | 18.4096988 |
| Q86XR8 | CEP57 | 18.3996464 | 18.7301408 | 18.6228903 | 18.4797305 | 18.550887 |
| Q9UKV8 | AGO2 | 21.3339148 | 21.3740403 | 21.4653308 | 21.3732645 | 21.4461529 |
| Q92804 | TAF15 | 22.3236699 | 22.6192265 | 22.4542487 | 22.2613394 | 22.2528652 |

| | | | | | | |
|---|---|---|---|---|---|---|
| Q14674 | ESPL1 | 16.5540648 | 16.8635769 | 16.8403906 | 16.7979398 | 16.7972732 |
| Q7Z7F0 | KHDC4 | 18.8870544 | 18.9664991 | 18.9060739 | 18.8439531 | 18.8075617 |
| Q9NQR4 | NIT2 | 22.7569745 | 22.8801915 | 22.8385425 | 22.8233848 | 22.9647428 |
| Q00610 | CLTC | 25.4004703 | 25.4177421 | 25.4224163 | 25.3086544 | 25.4036701 |
| Q9NRA2 | SLC17A5 | 17.9596588 | 18.1430942 | 18.0591197 | 18.0796978 | 17.9847417 |
| Q9Y6P5 | SESN1 | 16.0184066 | 15.9519805 | 16.1304717 | 16.1926332 | 16.1073883 |
| O15327 | INPP4B | NA | 16.6237786 | 16.3258 | NA | 16.5269536 |
| P10075 | GLI4 | 14.518164 | NA | NA | 14.8069812 | NA |
| Q96JP2 | MYO15B | 22.8719226 | NA | NA | 22.9772382 | NA |
| Q14376 | GALE | 20.6765246 | 20.7992178 | 20.7324779 | 20.7283337 | 20.6895893 |
| Q96P16 | RPRD1A | 20.1920181 | 20.355881 | 20.2845457 | 20.2059241 | 20.2942897 |
| Q9Y287 | ITM2B | 20.2318761 | 20.406304 | 20.3406116 | 20.3300332 | 20.214169 |
| P48960 | ADGRE5 | 22.9758758 | 23.1634607 | 22.9849097 | 22.9632089 | 23.0839052 |
| O60610 | DIAPH1 | 23.1674817 | 23.2779912 | 23.2294861 | 23.1095743 | 23.2557684 |
| Q96A46 | SLC25A28 | 16.35283 | 16.6542498 | 16.2548029 | 16.3643085 | 16.0108948 |
| O15394 | NCAM2 | 13.6998634 | NA | NA | 13.0112846 | NA |
| Q13370 | PDE3B | 22.1578167 | 22.0748747 | 22.1337812 | 22.0372281 | 22.1085829 |
| Q8N0Z8 | PUSL1 | 16.463252 | 16.8516353 | 16.9093135 | 16.7524015 | 16.7256549 |
| Q9GZU8 | PSME3IP1 | 22.2603675 | 22.3571574 | 22.112885 | 22.197283 | 22.2842005 |
| Q14703 | MBTPS1 | 15.2867567 | 15.400977 | 15.53425 | 15.52623 | 15.5802153 |
| Q9BUA3 | SPINDOC | 16.146692 | 16.0479906 | 16.2132604 | 16.3008292 | 16.3219142 |
| Q9H6R0 | DHX33 | 20.2206087 | 20.29337 | 20.2142605 | 20.1998686 | 20.2824061 |
| Q9NYF8 | BCLAF1 | 23.5115995 | 23.442124 | 23.467794 | 23.4045358 | 23.4730643 |
| P32780 | GTF2H1 | 19.3114407 | 19.4398252 | 19.4333032 | 19.4645559 | 19.5298558 |
| P22626 | HNRNPA2B1 | 27.7411957 | 27.7356692 | 27.6755169 | 27.6116814 | 27.6732611 |
| Q53HC5 | KLHL26 | 18.0265084 | 17.9418955 | 18.1334707 | 17.9921108 | 18.0765599 |
| O60885 | BRD4 | 21.7312956 | 21.7887081 | 21.8039077 | 21.6969112 | 21.8531808 |
| P56277 | CMC4 | 17.7633683 | 18.1481996 | 17.8787168 | 17.9034641 | 18.1426644 |
| Q9BVC6 | TMEM109 | 23.4472481 | 23.4988167 | 23.6213665 | 23.5684144 | 23.447506 |
| O43759 | SYNGR1 | 20.2107777 | 20.1225474 | 19.9124457 | 20.1074423 | 20.0941372 |
| Q86SF2 | GALNT7 | 20.4802572 | 20.6651671 | 20.7153843 | 20.6379169 | 20.6257195 |
| O94964 | SOGA1 | 23.6477091 | 23.8370246 | 23.632475 | 23.5990055 | 23.7221185 |
| Q01543 | FLI1 | 19.9884339 | 20.0834492 | 20.0304883 | 19.9842913 | 19.9792195 |
| Q5MIZ7 | PPP4R3B | 20.1337953 | 20.2112214 | 20.3198766 | 20.2004077 | 20.2243388 |
| Q9H813 | PACC1 | 18.6563785 | 18.7631172 | 18.9638885 | 19.0103796 | 19.0247492 |
| Q96K76 | USP47 | 20.5981044 | 20.6938833 | 20.6588937 | 20.6164269 | 20.7336337 |
| Q9H330 | TMEM245 | 19.8399222 | 20.1391201 | 20.0590964 | 20.2156682 | 20.0531049 |
| Q12913 | PTPRJ | 19.9458107 | 20.0168807 | 20.0384758 | 19.9820917 | 20.0853869 |
| Q9H900 | ZWILCH | 19.8384102 | 19.8777881 | 20.0279461 | 19.8357402 | 19.9652206 |
| Q15811 | ITSN1 | 16.7403606 | 17.0300459 | 16.9961695 | 16.9372117 | 17.0483937 |
| Q96BT7 | ALKBH8 | 16.3984211 | 16.7040823 | 16.60464 | 16.3840313 | 16.6947312 |
| O14678 | ABCD4 | 16.433805 | 16.6280387 | 16.3850028 | 16.6535132 | 16.413199 |

| | | | | | | |
|---|---|---|---|---|---|---|
| P03891 | MT-ND2 | 18.9342109 | 19.1872283 | 19.0265895 | 19.0364738 | 18.8002634 |
| Q6ZS17 | RIPOR1 | 18.6954723 | 18.7978407 | 18.6519408 | 18.6343999 | 18.718406 |
| Q96L50 | LRR1 | 17.897057 | 17.9060504 | 18.0235936 | 18.1003954 | 18.0214904 |
| Q9C0B1 | FTO | 20.7885715 | 20.9319463 | 20.8918657 | 20.8219117 | 20.9313485 |
| P26599 | PTBP1 | 25.9016503 | 25.9931921 | 25.971391 | 25.8458656 | 26.0316486 |
| Q8N587 | ZNF561 | 18.665013 | 18.6917436 | 18.7291341 | 18.7036938 | 18.5678933 |
| Q9UHV9 | PFDN2 | 22.9215658 | 23.0916552 | 22.8445682 | 22.8199217 | 23.0868881 |
| Q05BQ5 | MBTD1 | 16.3940341 | 16.2291178 | 17.5559431 | 16.8592526 | 17.1152114 |
| P55060 | CSE1L | 24.4801868 | 24.5004755 | 24.4918904 | 24.3643844 | 24.4647867 |
| Q4KMQ1 | TPRN | 19.3707015 | 19.3804604 | 19.3151442 | 19.2991849 | 19.3197547 |
| Q9NW81 | DMAC2 | 18.9102744 | 18.7194813 | 18.9588832 | 18.951256 | 18.9446922 |
| Q9Y5W7 | SNX14 | 17.2573688 | 17.4302015 | 17.3617839 | 17.3005307 | 17.210141 |
| Q9C0F1 | CEP44 | 19.3098925 | 19.3392562 | 19.3867854 | 19.2608349 | 19.4314327 |
| Q96KR7 | PHACTR3 | 16.9361914 | 17.3314617 | 16.9072903 | 17.0876158 | 17.2154659 |
| P51797 | CLCN6 | 17.3390444 | 17.7772149 | 17.8704988 | 17.674977 | 17.783399 |
| Q4VC31 | MIX23 | 21.4767722 | 21.5621647 | 21.5756379 | 21.4845723 | 21.5850524 |
| Q6P9A3 | ZNF549 | 16.7486917 | 16.7271186 | 16.720143 | 16.4090907 | 16.7436562 |
| Q9BV79 | MECR | 20.9623919 | 20.9864265 | 21.0465541 | 20.9143596 | 21.1881059 |
| O60318 | MCM3AP | 21.0693138 | 21.0489039 | 21.1102359 | 21.0752687 | 21.1095351 |
| Q9ULJ7 | ANKRD50 | NA | 14.285914 | 14.3824294 | NA | 14.3670324 |
| Q92934 | BAD | 18.5320837 | 18.7746674 | 18.1339068 | 18.4324503 | 18.8000458 |
| P18085 | ARF4 | 24.4509955 | 24.3702328 | 24.3204816 | 24.2877086 | 24.3859707 |
| Q58A45 | PAN3 | 19.8601268 | 19.7284165 | 19.7487645 | 19.8161713 | 19.8973283 |
| O75351 | VPS4B | 23.9556724 | 24.0421257 | 24.0053226 | 23.9216972 | 24.0140278 |
| P16930 | FAH | 22.9016884 | 22.779885 | 22.9038439 | 22.7874788 | 22.9623358 |
| Q7Z589 | EMSY | 18.4674285 | 18.620522 | 18.6731768 | 18.598155 | 18.6719534 |
| Q9H3Z4 | DNAJC5 | 22.2511636 | 22.2412623 | 22.2891253 | 22.1846196 | 22.2783735 |
| P01034 | CST3 | 23.0439868 | 22.90197 | 22.9320745 | 22.978421 | 22.9380585 |
| Q96PV6 | LENG8 | 20.4607831 | 20.4388751 | 20.5393534 | 20.4568043 | 20.5119571 |
| Q9H4L7 | SMARCAD1 | 20.1606072 | 20.2280251 | 20.2471293 | 20.1230127 | 20.3361849 |
| P46459 | NSF | 23.4639824 | 23.5559332 | 23.447571 | 23.4027169 | 23.5805028 |
| Q86SX6 | GLRX5 | 21.754589 | 21.8904038 | 21.7833951 | 21.7023634 | 22.1101606 |
| P62910 | RPL32 | 26.2572005 | 26.1434839 | 26.4117935 | 26.0806794 | 26.148041 |
| Q68CP9 | ARID2 | 20.7321215 | 20.7570286 | 20.7777131 | 20.7117082 | 20.8133984 |
| Q96M27 | PRRC1 | 22.9106857 | 22.8641471 | 22.8236041 | 22.8327423 | 22.9529505 |
| P49069 | CAMLG | 21.3279234 | 21.5866601 | 21.36279 | 21.534806 | 21.4483672 |
| P40818 | USP8 | 21.1345906 | 21.1087134 | 21.1712778 | 21.0358999 | 21.167214 |
| Q8NBL1 | POGLUT1 | 20.7908509 | 20.822659 | 20.896944 | 20.8893967 | 20.8332113 |
| Q15029 | EFTUD2 | 23.3715544 | 23.4335479 | 23.3934271 | 23.3249627 | 23.4379785 |
| Q5NDL2 | EOGT | 19.7942123 | 20.0028799 | 19.9954926 | 19.8561157 | 19.8486953 |
| P61599 | NAA20 | 19.7647437 | 19.9130661 | 19.7072921 | 19.8515219 | 19.9198652 |
| O75367 | MACROH2A1 | 25.318889 | 25.3406103 | 25.4607599 | 25.3850539 | 25.3964736 |

| | | | | | | |
|---|---|---|---|---|---|---|
| P21399 | ACO1 | 22.0168248 | 22.0653431 | 22.0969481 | 21.9632773 | 22.0473641 |
| O60271 | SPAG9 | 24.1728298 | 24.2721487 | 24.1670149 | 24.1395837 | 24.1988658 |
| P34896 | SHMT1 | 20.6802356 | 20.8150357 | 20.7301683 | 20.7173277 | 20.8480208 |
| O60716 | CTNND1 | 17.0786358 | 17.2132425 | 16.7244433 | 17.2943742 | 16.8623258 |
| Q9Y6J0 | CABIN1 | 20.2095223 | 20.3260987 | 20.2971474 | 20.2450129 | 20.3140993 |
| Q96EH3 | MALSU1 | 18.8532216 | 18.6748938 | 18.8006514 | 18.6937797 | 18.9561397 |
| Q5BJH7 | YIF1B | 19.145114 | 19.2443536 | 19.2164459 | 18.9751689 | 19.2040004 |
| O00625 | PIR | 19.4412091 | 19.677813 | 19.6016569 | 19.3345691 | 19.2138061 |
| Q86SE8 | NPM2 | 18.3065929 | 18.2773165 | 18.1751822 | 18.2753575 | 18.2787435 |
| Q03519 | TAP2 | 20.9604316 | 20.8541242 | 20.9395372 | 20.7776706 | 20.9347759 |
| Q9HCJ0 | TNRC6C | NA | 13.0656283 | 12.8771353 | NA | 13.1313122 |
| Q8N344 | MIER2 | 15.8281005 | 15.8804683 | 16.2215981 | 16.3236954 | 16.3615006 |
| Q5VWC8 | HACD4 | 17.6207095 | 17.7649968 | 17.8672045 | 17.685444 | 17.7798438 |
| Q03701 | CEBPZ | 21.6014842 | 21.651689 | 21.7435424 | 21.5997463 | 21.6538334 |
| Q8N883 | ZNF614 | 21.2295235 | 21.033421 | 21.1513272 | 21.249754 | 21.1179831 |
| O95486 | SEC24A | 20.504927 | 20.5473574 | 20.5646105 | 20.5190996 | 20.6188272 |
| Q08722 | CD47 | 21.254715 | 21.5620141 | 21.5000598 | 21.3758263 | 21.489455 |
| Q66LE6 | PPP2R2D | 19.5922293 | 19.8448578 | 19.8316967 | 19.7019505 | 19.8618064 |
| Q9UBW5 | BIN2 | 21.6936822 | 21.899924 | 21.9762338 | 21.7532114 | 21.9506693 |
| Q9P203 | BTBD7 | 13.3039181 | 12.9173283 | 13.1822772 | 13.3277213 | 13.2884339 |
| O43462 | MBTPS2 | 16.2886696 | 17.4383741 | 17.2767655 | 17.1269405 | 17.3547088 |
| Q9NV79 | PCMTD2 | 18.0767562 | 18.0904804 | 18.0779703 | 17.9401644 | 17.9986191 |
| Q969M7 | UBE2F | 18.5862689 | 18.5813828 | 18.84257 | 18.6826248 | 18.8215533 |
| Q01650 | SLC7A5 | 22.8933371 | 23.0040573 | 22.8528852 | 22.8016521 | 22.7771114 |
| Q86YV0 | RASAL3 | 21.4321948 | 21.5886448 | 21.6114518 | 21.3864101 | 21.5464575 |
| Q9NUL5 | SHFL | 20.5062013 | 20.4774662 | 20.5697512 | 20.3998201 | 20.5189439 |
| Q8TB24 | RIN3 | 18.9686589 | 19.1100414 | 19.1275606 | 18.9124524 | 19.041455 |
| P20340 | RAB6A | 22.7995354 | 22.8644111 | 22.8793272 | 22.7524274 | 22.8582085 |
| P23786 | CPT2 | 22.1586276 | 22.2303635 | 22.19271 | 22.088367 | 22.253715 |
| Q15369 | ELOC | 23.3120999 | 23.3828592 | 23.347589 | 23.3168838 | 23.3777071 |
| Q14739 | LBR | 23.5586376 | 23.5738842 | 23.6031307 | 23.5505443 | 23.6127354 |
| Q9Y266 | NUDC | 24.6669141 | 24.6924191 | 24.5617026 | 24.5687585 | 24.6372184 |
| Q15126 | PMVK | 22.3990533 | 22.4395719 | 22.4409299 | 22.4232246 | 22.4467285 |
| Q86VV8 | RTTN | 17.1849153 | 17.3179055 | 17.4110416 | 17.3561177 | 17.3936581 |
| Q16576 | RBBP7 | 21.623849 | 21.7060192 | 21.8733295 | 21.6256246 | 21.8136308 |
| Q9HCE0 | EPG5 | 17.1860696 | 17.2577546 | 17.5519356 | 17.1925992 | 17.3615316 |
| Q9H299 | SH3BGRL3 | 26.1802061 | 26.2643769 | 25.9427061 | 25.9881287 | 26.0540588 |
| Q9ULZ3 | PYCARD | 25.372885 | 25.5244216 | 25.2641075 | 25.2888604 | 25.5091021 |
| Q15223 | NECTIN1 | 20.54663 | 20.8040736 | 20.8562705 | 20.5886975 | 20.78835 |
| Q16610 | ECM1 | 16.506037 | 16.8428126 | 16.3761851 | 16.5673834 | 16.5878745 |
| Q9NQW8 | CNGB3 | 15.483832 | NA | NA | 15.1688703 | NA |
| Q86YP4 | GATAD2A | 21.8430144 | 21.9071982 | 21.822459 | 21.8533608 | 21.819223 |

| | | | | | | |
|---|---|---|---|---|---|---|
| Q15024 | EXOSC7 | 21.5835588 | 21.7083785 | 21.5683458 | 21.5440102 | 21.6996072 |
| Q86X53 | ERICH1 | 18.8040992 | 18.9122818 | 18.7531554 | 18.7370772 | 18.868316 |
| P10412 | H1-4 | 22.7646164 | 22.9359165 | 23.0903309 | 22.7511468 | 22.1236628 |
| O43676 | NDUFB3 | 21.4356363 | 21.2339644 | 21.4121595 | 21.0973792 | 21.6886846 |
| O60512 | B4GALT3 | 19.067652 | 19.1422109 | 19.1594977 | 19.2864364 | 19.0322836 |
| O76031 | CLPX | 21.3041656 | 21.2836239 | 21.2382419 | 21.2097038 | 21.2876839 |
| Q9H2U2 | PPA2 | 23.4143724 | 23.4153877 | 23.4391912 | 23.3774581 | 23.4259023 |
| Q9BTC0 | DIDO1 | 22.3011889 | 22.3406743 | 22.3309425 | 22.2683772 | 22.3645786 |
| O75306 | NDUFS2 | 22.8788474 | 22.9361458 | 23.0220984 | 22.8721443 | 23.0227021 |
| Q15814 | TBCC | 20.9471198 | 21.0107399 | 20.9037871 | 20.9710434 | 21.0738324 |
| Q15542 | TAF5 | 21.8121033 | 21.949374 | 21.8509808 | 21.8568609 | 22.0025378 |
| Q9UNP9 | PPIE | 20.9563101 | 21.0591543 | 21.004813 | 20.9252367 | 21.08348 |
| P29466 | CASP1 | 21.6159655 | 21.7408813 | 21.8254199 | 21.676183 | 21.8536696 |
| Q9Y6H3 | ATP23 | 20.2423914 | 20.3429776 | 20.3860641 | 20.3684622 | 20.2841341 |
| P16035 | TIMP2 | 21.3435401 | 22.0216312 | 21.0781802 | 20.8798067 | 20.3003818 |
| Q2NKJ3 | CTC1 | 16.6007325 | 16.7339038 | 16.742997 | 16.6448801 | 16.8305588 |
| Q9UMY1 | NOL7 | 20.0866477 | 20.1235428 | 20.3113811 | 20.1011207 | 20.071164 |
| Q9BRX5 | GINS3 | 19.9630122 | 19.9925531 | 20.0124581 | 20.0221364 | 20.0839001 |
| Q4LE39 | ARID4B | 18.4287732 | 18.6474375 | 18.6363508 | 18.6261614 | 18.7065077 |
| Q9BUV8 | RAB5IF | 18.1554364 | 18.3405091 | 18.2154154 | 18.2409994 | 18.0852678 |
| Q13356 | PPIL2 | 21.0135036 | 21.1030537 | 21.1014322 | 21.0297118 | 21.1460464 |
| Q9ULR3 | PPM1H | 11.7003729 | NA | NA | 10.5987559 | NA |
| O15151 | MDM4 | NA | 14.9587552 | 15.0385724 | NA | 14.9638917 |
| Q8WV74 | NUDT8 | 20.134478 | 20.4323428 | 20.3991312 | 20.1243917 | 20.3305224 |
| Q53FA7 | TP53I3 | 21.227505 | 21.4813937 | 21.3988341 | 21.2191979 | 21.4095064 |
| Q9ULW3 | ABT1 | 19.1340888 | 19.2922752 | 19.1397563 | 19.2809215 | 19.1915412 |
| Q9UKI8 | TLK1 | 20.2613812 | 20.2793771 | 20.3148658 | 20.1936354 | 20.2441351 |
| P18669 | PGAM1 | 25.1918531 | 25.1797452 | 25.245831 | 25.1197784 | 25.2475096 |
| P31946 | YWHAB | 25.3048596 | 25.247816 | 25.2601758 | 25.1086466 | 25.2337613 |
| Q5XUX1 | FBXW9 | 20.3943102 | 20.307885 | 20.3022221 | 20.310511 | 20.3402648 |
| Q15021 | NCAPD2 | 22.3182979 | 22.3229664 | 22.4242454 | 22.244094 | 22.3601378 |
| O14657 | TOR1B | 20.2385275 | 20.3168458 | 20.3527013 | 20.2943546 | 20.3951846 |
| P13637 | ATP1A3 | 18.729188 | 18.6880416 | 18.7220415 | 18.8061965 | 18.8010631 |
| Q9NVE7 | PANK4 | 19.7651052 | 19.8659721 | 19.8717486 | 19.84228 | 19.8769065 |
| A2RTX5 | TARS3 | 18.8962233 | 19.1364032 | 19.0686148 | 18.9822302 | 19.1066912 |
| P30084 | ECHS1 | 24.0344164 | 24.1372844 | 23.965889 | 24.0059304 | 24.1148759 |
| Q96EL3 | MRPL53 | 21.2748596 | 21.2914781 | 21.2656449 | 21.2702662 | 21.3145196 |
| Q15131 | CDK10 | 15.8567382 | 15.6467067 | 15.8634684 | 15.7980424 | 15.9587913 |
| Q8NFZ0 | FBH1 | 16.5172616 | 16.968596 | 16.8769836 | 16.6573814 | 16.7073779 |
| O43295 | SRGAP3 | 17.4066149 | 17.435993 | 17.5397827 | 17.6391564 | 17.5657828 |
| Q14106 | TOB2 | 14.6854531 | 14.6767937 | 15.0447397 | 14.3370987 | 15.0712523 |
| P0CJ79 | ZNF888 | 17.3816593 | 17.5998731 | 17.5868792 | 17.563385 | 17.5867096 |

| | | | | | | |
|---|---|---|---|---|---|---|
| O14807 | MRAS | 18.2781749 | 18.3828816 | 18.524549 | 18.3355162 | 18.2993565 |
| O95630 | STAMBP | 21.1477415 | 21.1666964 | 21.1329121 | 21.0703457 | 21.2118082 |
| Q15048 | LRRC14 | 18.7729221 | 18.8237849 | 18.9885444 | 18.8087275 | 18.9017621 |
| Q5R3I4 | TTC38 | 19.0573358 | 19.3106597 | 19.2246613 | 19.0887805 | 19.1942286 |
| Q6ZNJ1 | NBEAL2 | 19.6312251 | 19.7217215 | 19.7283795 | 19.6796864 | 19.7612055 |
| Q8IWT6 | LRRC8A | 19.2934593 | 19.3390321 | 19.4620215 | 19.2487941 | 19.3922835 |
| Q6UWD8 | C16orf54 | 19.456623 | 19.4932718 | 19.5532007 | 19.4027542 | 19.5836211 |
| Q9NZ43 | USE1 | 20.0379978 | 19.9717086 | 20.2382312 | 20.0755633 | 20.0506738 |
| Q7Z7L1 | SLFN11 | 17.8320959 | 17.8790446 | 18.0391225 | 17.9429753 | 17.8685958 |
| Q9HAU4 | SMURF2 | 14.4653176 | NA | NA | 14.5989306 | NA |
| P04839 | CYBB | 18.2465715 | 18.480774 | 18.7125351 | 18.5673241 | 18.5324239 |
| Q8NC54 | C5orf15 | 15.5590697 | 15.4036387 | 15.5018849 | 15.4846901 | 15.6746096 |
| P52564 | MAP2K6 | 19.8847467 | 20.1877068 | 19.9964204 | 19.813822 | 19.9934894 |
| Q96HQ2 | CDKN2AIPNL | 19.9606801 | 20.2229526 | 20.0513286 | 20.0727357 | 20.2024713 |
| Q9GZS1 | POLR1E | 22.1750709 | 22.1951345 | 22.2431771 | 22.0582734 | 22.2175664 |
| Q8N201 | INTS1 | 20.4078997 | 20.5222018 | 20.5241271 | 20.4193066 | 20.5176423 |
| P49189 | ALDH9A1 | 22.878785 | 22.9480338 | 22.8723751 | 22.794545 | 22.8922287 |
| Q969V3 | NCLN | 22.3509091 | 22.3960494 | 22.4458502 | 22.304021 | 22.39336 |
| O43708 | GSTZ1 | 21.0343094 | 21.0008575 | 20.9638108 | 20.9244466 | 21.0250329 |
| Q9NRL2 | BAZ1A | 21.3271801 | 21.3094749 | 21.3554197 | 21.2379359 | 21.2925096 |
| Q96RU3 | FNBP1 | 22.2043983 | 22.2856258 | 22.1383408 | 22.1805824 | 22.2743511 |
| O95573 | ACSL3 | 21.5789808 | 21.5649277 | 21.5975138 | 21.5692654 | 21.6575598 |
| P08195 | SLC3A2 | 23.313162 | 23.3241941 | 23.2126505 | 23.2016615 | 23.3152426 |
| O94903 | PLPBP | 22.5687764 | 22.6539338 | 22.6408262 | 22.5499531 | 22.7293207 |
| Q13107 | USP4 | 20.741259 | 20.8016014 | 20.7866884 | 20.6653794 | 20.825849 |
| P51946 | CCNH | 19.6554939 | 19.505409 | 19.6562964 | 19.5903276 | 19.5364499 |
| O15525 | MAFG | 20.2362152 | 20.408334 | 20.1794356 | 20.2616333 | 20.3315342 |
| Q7L8W6 | DPH6 | 18.7836336 | 19.0141795 | 18.9989158 | 18.8942874 | 19.0934052 |
| P51553 | IDH3G | 21.3253313 | 21.3435415 | 21.4855371 | 21.2687411 | 21.4071198 |
| Q9UM22 | EPDR1 | 19.3887035 | 19.6833077 | 19.4966962 | 19.4195268 | 19.4923272 |
| Q9BVT8 | TMUB1 | 19.5232862 | 19.3796386 | 19.3954529 | 19.527125 | 19.5754489 |
| Q7Z6L1 | TECPR1 | 18.1237948 | 18.1960449 | 18.2095856 | 18.3179982 | 18.0717654 |
| Q8IZ07 | ANKRD13A | 19.8585989 | 20.0397923 | 20.1344729 | 19.9402406 | 20.1092894 |
| Q9BV81 | EMC6 | 19.9205769 | 20.2173948 | 20.195329 | 19.959504 | 20.0976898 |
| Q8IVT5 | KSR1 | 16.5386213 | 16.7138214 | 16.720569 | 16.5486253 | 16.7174313 |
| P22732 | SLC2A5 | 19.782315 | 20.025699 | 20.176474 | 19.9120348 | 20.1685406 |
| Q9NTG1 | PKDREJ | NA | 15.9338673 | 16.2416607 | NA | 16.0106281 |
| P32246 | CCR1 | 20.3144513 | 20.5740508 | 20.6309713 | 20.0185921 | 20.5863894 |
| P17014 | ZNF12 | 12.2818221 | NA | NA | 12.4783785 | NA |
| O94979 | SEC31A | 22.5540773 | 22.4867446 | 22.5726337 | 22.4757695 | 22.5714013 |
| Q13586 | STIM1 | 20.3818027 | 20.5579565 | 20.5852968 | 20.37895 | 20.5510968 |
| Q9H2H8 | PPIL3 | 20.9792082 | 21.1206192 | 21.0880714 | 20.9889035 | 21.1726925 |

| | | | | | | |
|---|---|---|---|---|---|---|
| Q8TDB6 | DTX3L | 18.6911604 | 18.7354371 | 18.8027639 | 18.7717973 | 18.7939912 |
| P10073 | ZSCAN22 | 14.720351 | NA | NA | 14.6770998 | NA |
| O75346 | ZNF253 | NA | 15.3629588 | 15.2181593 | NA | 14.9449561 |
| Q9UPW5 | AGTPBP1 | 17.6173505 | 17.7355612 | 17.8448873 | 17.7047283 | 17.8073175 |
| O43164 | PJA2 | 18.5252905 | 18.5116148 | 18.1839069 | 18.4181127 | 18.5263388 |
| Q96AY4 | TTC28 | 21.4153851 | 21.5674128 | 21.5448758 | 21.3902223 | 21.5354472 |
| Q12905 | ILF2 | 24.8736138 | 24.8473868 | 24.8900858 | 24.8045729 | 24.8817843 |
| Q9H0W9 | C11orf54 | 19.0650438 | 19.076397 | 19.3467531 | 19.0770621 | 19.2834956 |
| Q9BVV6 | KIAA0586 | 18.8026862 | NA | NA | 18.8082401 | NA |
| Q2M2I8 | AAK1 | 20.0310569 | 19.9906171 | 20.0640824 | 20.0418507 | 20.0428089 |
| Q58FG1 | HSP90AA4P | 22.4109096 | 22.2185045 | 22.2130459 | 22.350992 | 22.2347921 |
| Q96LT7 | C9orf72 | 19.181199 | 19.1671641 | 19.0216066 | 19.2335638 | 19.2383426 |
| O14569 | CYB561D2 | 20.0529928 | 19.8115329 | 20.1375139 | 19.7158529 | 19.9115953 |
| Q96I99 | SUCLG2 | 22.2100016 | 22.3260856 | 22.1832905 | 22.1505148 | 22.3305483 |
| P46100 | ATRX | 20.3869783 | 20.4669874 | 20.466982 | 20.3966263 | 20.498546 |
| P49354 | FNTA | 21.9223466 | 21.9290213 | 21.9176832 | 21.9049336 | 22.0320777 |
| Q9HC52 | CBX8 | 19.1900962 | 19.4244837 | 19.4365623 | 19.3240019 | 19.4389147 |
| Q8NCP5 | ZBTB44 | 16.9340433 | 17.0163617 | 16.8065602 | 17.0000875 | 17.2776221 |
| Q96BI3 | APH1A | 16.6202678 | 16.9859295 | 16.5565205 | 17.1134437 | 17.1868209 |
| P84157 | MXRA7 | 19.6054915 | 19.6303099 | 19.5029678 | 19.7194369 | 19.437015 |
| Q14CX7 | NAA25 | 21.121656 | 21.1361397 | 21.2210214 | 21.0828413 | 21.254343 |
| P08621 | SNRNP70 | 24.695753 | 24.7540021 | 24.6056843 | 24.5668212 | 24.7110065 |
| Q99574 | SERPINI1 | NA | 15.4780071 | 15.7567827 | NA | 15.8233329 |
| P22692 | IGFBP4 | 17.5219151 | 17.4969787 | 17.6324598 | 17.3505229 | 17.6274565 |
| Q9Y2H6 | FNDC3A | 20.0672977 | 20.1539661 | 20.0700288 | 19.9829749 | 20.1290579 |
| Q9NXE8 | CWC25 | 19.5163622 | 19.6034863 | 19.6146048 | 19.5158661 | 19.5373974 |
| O95361 | TRIM16 | 20.1310909 | 20.0820138 | 20.114112 | 20.1279174 | 20.188117 |
| O14732 | IMPA2 | 19.6263117 | 19.8691468 | 19.8587046 | 19.5987013 | 19.8229378 |
| P15151 | PVR | 18.6787042 | 18.748255 | 18.9903205 | 18.815293 | 18.8814196 |
| Q9NR19 | ACSS2 | 20.3269308 | 20.3749129 | 20.428801 | 20.3053466 | 20.4398963 |
| P30154 | PPP2R1B | 19.4579895 | 19.4873188 | 19.6902003 | 19.5064063 | 19.5734365 |
| Q9NX18 | SDHAF2 | 19.2946644 | 19.3444562 | 19.0748303 | 19.136999 | 19.2581344 |
| Q8NE22 | SETD9 | 15.2070346 | 14.7305664 | 15.7899137 | 15.2066192 | 15.7363373 |
| Q15582 | TGFBI | 19.5339113 | 19.5732237 | 19.5777585 | 19.5514763 | 19.4583448 |
| P42338 | PIK3CB | 19.7538793 | 19.9462683 | 19.9247606 | 19.8301546 | 19.8680663 |
| Q5T8D3 | ACBD5 | 20.1159346 | 20.1464211 | 20.1855939 | 20.0957461 | 20.123296 |
| Q6PI26 | SHQ1 | 16.3672079 | 16.5271726 | 16.3357065 | 15.8425319 | 16.3868154 |
| Q12893 | TMEM115 | 18.4427236 | 18.5690342 | 18.6814649 | 18.5900948 | 18.5873327 |
| O15504 | NUP42 | 19.1953802 | 19.2007496 | 19.3580711 | 19.2208022 | 19.3611241 |
| Q14562 | DHX8 | 21.771576 | 21.7294298 | 21.7540183 | 21.7120332 | 21.8535961 |
| P27708 | CAD | 23.37676 | 23.4923328 | 23.4756096 | 23.3589188 | 23.4606134 |
| Q9Y2I7 | PIKFYVE | 20.0417607 | 20.0155476 | 20.1518045 | 20.087728 | 20.1135605 |

| | | | | | | |
|---|---|---|---|---|---|---|
| Q02241 | KIF23 | 19.1377187 | 19.1797635 | 19.1989907 | 19.1534299 | 19.1890159 |
| P58335 | ANTXR2 | 18.0368132 | 18.0700608 | 18.3583151 | 18.3174151 | 18.2107328 |
| Q9Y496 | KIF3A | 16.2863198 | 16.4208906 | 16.3980136 | 16.3624769 | 16.5867396 |
| Q9H0R3 | TMEM222 | 17.3830614 | 17.6649032 | 17.5250994 | 17.9028626 | 17.6686309 |
| Q93070 | ART4 | NA | 14.5767757 | 14.8117912 | NA | 14.7224649 |
| Q2NL98 | VMAC | 17.6288207 | 17.8222951 | 17.5895869 | 17.6235707 | 17.8889175 |
| Q8IYM0 | FAM186B | 19.38349 | 19.4463502 | 19.4413615 | 19.3657893 | 19.2880188 |
| P57789 | KCNK10 | 19.210513 | 18.8869651 | 19.0878736 | 19.1381137 | 19.0991382 |
| Q15397 | PUM3 | 21.2505318 | 21.4466293 | 21.3400563 | 21.2733013 | 21.3474713 |
| Q86VP6 | CAND1 | 23.3323227 | 23.4170069 | 23.3367783 | 23.2823202 | 23.4102976 |
| Q9NUI1 | DECR2 | 21.2707416 | 21.3452506 | 21.3421809 | 21.2763126 | 21.2813455 |
| O60826 | CCDC22 | 21.7046257 | 21.7565367 | 21.7502564 | 21.7120105 | 21.7106677 |
| Q6N069 | NAA16 | 19.8409949 | 19.9795503 | 19.973993 | 19.8809532 | 20.0126115 |
| P52294 | KPNA1 | 22.2104629 | 22.3983162 | 22.1770581 | 22.208561 | 22.1930598 |
| O60888 | CUTA | 20.6694772 | 20.8661093 | 20.5353981 | 20.6357421 | 20.8332493 |
| Q92966 | SNAPC3 | 17.1977482 | 17.2146451 | 17.325059 | 17.4223358 | 17.1533487 |
| Q9NSD9 | FARSB | 23.514406 | 23.6606894 | 23.5013428 | 23.489105 | 23.59182 |
| Q5F1R6 | DNAJC21 | 20.4050917 | 20.531617 | 20.4034719 | 20.4134452 | 20.5313669 |
| Q9C0D7 | ZC3H12C | NA | 14.4695558 | 14.1678876 | NA | 14.3385685 |
| O95319 | CELF2 | 22.7948737 | 22.7907803 | 22.6592129 | 22.6136224 | 22.6625891 |
| O75676 | RPS6KA4 | 21.2557615 | 21.2868546 | 21.2766637 | 21.2037094 | 21.3062986 |
| Q6NUS6 | TCTN3 | 15.4599252 | 15.341609 | 15.4126825 | 15.484225 | 15.6446194 |
| Q9Y2Z4 | YARS2 | 21.0633425 | 21.0703221 | 21.1024318 | 21.0338984 | 21.1585357 |
| Q14669 | TRIP12 | 22.1066696 | 22.1261751 | 22.18281 | 22.0669033 | 22.1759223 |
| Q13045 | FLII | 21.7964024 | 21.7910783 | 21.8748541 | 21.7583001 | 21.8263423 |
| O00571 | DDX3X | 23.7144052 | 23.8126885 | 23.7544198 | 23.6963521 | 23.7766003 |
| Q7Z3C6 | ATG9A | 20.6240371 | 20.5863542 | 20.773118 | 20.6561454 | 20.6782081 |
| P54577 | YARS1 | 23.9750436 | 24.0053333 | 23.8779469 | 23.9162116 | 23.9588746 |
| Q9UNN5 | FAF1 | 21.6539107 | 21.7708156 | 21.743661 | 21.6625846 | 21.7802067 |
| Q86W56 | PARG | 19.5778762 | 19.6311194 | 19.6086422 | 19.6326577 | 19.7196905 |
| Q9NYL2 | MAP3K20 | 21.7213298 | 21.8278861 | 21.816031 | 21.8016079 | 21.9246563 |
| P32119 | PRDX2 | 26.4670909 | 26.5211596 | 26.3445955 | 26.3214487 | 26.4872898 |
| Q15691 | MAPRE1 | 24.575209 | 24.7108899 | 24.6920752 | 24.6021982 | 24.6989763 |
| Q6P587 | FAHD1 | 20.9191359 | 20.9251613 | 21.068893 | 20.9561255 | 21.0771227 |
| Q9BX70 | BTBD2 | 20.0917073 | 19.9856918 | 19.8554235 | 19.9554729 | 19.952728 |
| Q6NUK1 | SLC25A24 | 21.8033472 | 21.8823089 | 21.9025998 | 21.7616872 | 21.9837115 |
| O94888 | UBXN7 | 20.8092813 | 21.0312566 | 20.9552394 | 20.8891609 | 20.9460719 |
| Q9NWY4 | HPF1 | 19.9426939 | 20.1118459 | 20.157271 | 19.999316 | 20.054573 |
| Q99541 | PLIN2 | 17.3755117 | 17.4538376 | 17.3159366 | 17.659188 | 17.4387894 |
| Q13625 | TP53BP2 | 17.5265723 | 17.7079459 | 17.8249219 | 17.7572036 | 17.6789521 |
| P35520 | CBS | 16.873587 | 17.2692946 | 17.1539849 | 16.9189872 | 17.1089224 |
| Q8N2H3 | PYROXD2 | NA | 18.9901863 | 18.6547549 | NA | 18.6103791 |

| | | | | | | |
|---|---|---|---|---|---|---|
| O94916 | NFAT5 | 14.4965424 | NA | NA | 14.2807824 | NA |
| P84098 | RPL19 | 26.3198458 | 26.3362449 | 26.5881307 | 26.2180421 | 26.3279852 |
| P49760 | CLK2 | 18.9095033 | 19.1114733 | 19.2178962 | 19.0230805 | 19.0402125 |
| O15047 | SETD1A | 19.5206394 | 19.624717 | 19.6110245 | 19.4941269 | 19.575923 |
| Q96EK9 | KTI12 | 20.4505119 | 20.5154314 | 20.4633568 | 20.4023028 | 20.4408515 |
| Q9P015 | MRPL15 | 22.0760115 | 22.0776099 | 22.1620003 | 22.017292 | 22.0559575 |
| P49137 | MAPKAPK2 | 22.3149062 | 22.3391398 | 22.3201858 | 22.2780452 | 22.2597022 |
| O60493 | SNX3 | 23.9342789 | 24.0516955 | 23.9188858 | 23.940106 | 23.8757891 |
| Q7Z6M2 | FBXO33 | NA | 16.1994495 | 16.179488 | NA | 16.3177543 |
| Q5T0F9 | CC2D1B | 19.6404818 | 19.6316063 | 19.6740106 | 19.5672392 | 19.6386464 |
| O75312 | ZPR1 | 21.8050533 | 21.9557171 | 21.7002087 | 21.7719772 | 21.9370839 |
| Q6PL24 | TMED8 | 21.1371728 | 21.2751822 | 21.2557526 | 21.1886921 | 21.3836076 |
| Q13541 | EIF4EBP1 | 22.2519303 | 22.2033755 | 21.8564733 | 22.0422193 | 22.2509699 |
| Q9H6H4 | REEP4 | 21.1279157 | 21.2624343 | 21.2869199 | 21.1489322 | 21.1497561 |
| P52298 | NCBP2 | 22.2174533 | 22.3088186 | 22.1848782 | 22.156477 | 22.1872907 |
| Q9Y3A3 | MOB4 | 20.858557 | 21.1076012 | 21.1248661 | 20.9435771 | 21.081487 |
| Q9UL54 | TAOK2 | 19.297535 | 19.3757695 | 19.3956431 | 19.3363615 | 19.6018851 |
| P07108 | DBI | 24.7658631 | 24.9042518 | 24.6819237 | 24.6282029 | 24.9725546 |
| Q53EQ6 | TIGD5 | 12.5035367 | 13.5611005 | 13.923127 | 12.4231728 | 14.1727096 |
| Q9UK58 | CCNL1 | 19.2261734 | 19.2242018 | 19.2739726 | 19.2794638 | 19.2997814 |
| Q8ND82 | ZNF280C | 18.54827 | 18.6779646 | 18.5065876 | 18.6133869 | 18.5962723 |
| Q8NHJ6 | LILRB4 | 21.3281271 | 21.2493297 | 21.361283 | 21.3502605 | 21.380962 |
| Q9NY27 | PPP4R2 | 21.4647728 | 21.4772944 | 21.4780676 | 21.4556775 | 21.5961565 |
| Q9C005 | DPY30 | 21.8515802 | 21.8761963 | 21.7243349 | 21.6023504 | 21.6761564 |
| O14717 | TRDMT1 | NA | 17.0034907 | 17.311152 | NA | 17.2275256 |
| Q9Y2X9 | ZNF281 | 18.912128 | 18.8338464 | 19.0644061 | 18.9101939 | 18.9580921 |
| Q99447 | PCYT2 | 22.3277441 | 22.3350792 | 22.3075487 | 22.295241 | 22.3518851 |
| Q9BXW6 | OSBPL1A | 21.2493887 | 21.3408904 | 21.2601179 | 21.2472326 | 21.3452088 |
| Q9BX69 | CARD6 | 17.8794669 | 17.8633783 | 17.9506363 | 17.9338469 | 18.0260927 |
| P31689 | DNAJA1 | 23.6513312 | 23.6879705 | 23.6858242 | 23.6114497 | 23.6917425 |
| P28749 | RBL1 | 19.0527459 | 19.0091592 | 19.1720119 | 19.0245308 | 19.1279996 |
| Q8WVK7 | SKA2 | 18.4864598 | 18.4747345 | 18.7087545 | 18.6168327 | 18.608391 |
| Q9UID6 | ZNF639 | 16.1257498 | 16.6690124 | 16.4380711 | 16.3518806 | 16.0895359 |
| Q8N8Q9 | NIPA2 | 18.247338 | 17.9914497 | 18.4944502 | 18.1072933 | 17.9102906 |
| P00395 | MT-CO1 | 19.1325661 | 18.7010326 | 19.0090812 | 18.9529071 | 18.8700705 |
| P61011 | SRP54 | 22.4725702 | 22.5574766 | 22.4112894 | 22.4270385 | 22.5162256 |
| Q15345 | LRRC41 | 19.5976817 | 19.6349782 | 19.8037416 | 19.6515961 | 19.6753402 |
| Q96RK0 | CIC | 18.8671601 | 18.9863951 | 18.7902864 | 18.8325445 | 18.9564988 |
| Q14588 | ZNF234 | 14.666233 | 14.6307659 | 15.064016 | 14.6269748 | 14.6771015 |
| O75380 | NDUFS6 | 21.7786731 | 21.7518973 | 21.5742721 | 21.5122733 | 21.8621924 |
| Q96T68 | SETDB2 | 17.2519015 | 17.3773178 | 17.5611607 | 17.4359725 | 17.325903 |
| Q9HBG6 | IFT122 | 18.358097 | 18.4620663 | 18.4689086 | 18.5174227 | 18.5264866 |

| | | | | | | |
|---|---|---|---|---|---|---|
| Q15404 | RSU1 | 22.4300642 | 22.4056101 | 22.5196795 | 22.3527773 | 22.4078459 |
| Q8IYI6 | EXOC8 | 20.7592136 | 20.8425109 | 20.8110541 | 20.7510113 | 20.8787258 |
| Q96GZ6 | SLC41A3 | 17.5759738 | 17.8563821 | 17.5196018 | 17.6750028 | 17.5684952 |
| P41227 | NAA10 | 22.8064989 | 22.8612845 | 22.7562508 | 22.7857915 | 22.8579279 |
| Q9UKD2 | MRTO4 | 22.6144395 | 22.6396364 | 22.6495109 | 22.5813646 | 22.6435436 |
| Q8N6R0 | METTL13 | 21.2319822 | 21.2972166 | 21.2297297 | 21.1181122 | 21.3163831 |
| Q13043 | STK4 | 21.8459707 | 21.9611721 | 21.9580165 | 21.8152303 | 21.9821145 |
| Q9NY61 | AATF | 20.88742 | 20.8290492 | 20.9121005 | 20.8592752 | 20.920921 |
| P08758 | ANXA5 | 25.1773026 | 25.2665266 | 25.1246455 | 25.0319386 | 25.2545323 |
| Q9NR12 | PDLIM7 | 21.6422155 | 21.6480215 | 21.7086147 | 21.5829596 | 21.6393124 |
| P52701 | MSH6 | 22.301384 | 22.4033323 | 22.355681 | 22.2854255 | 22.3380792 |
| Q9H211 | CDT1 | 18.2636471 | 18.55 | 18.5139696 | 18.5351402 | 18.4329264 |
| Q07864 | POLE | 21.5186862 | 21.680835 | 21.7701134 | 21.5154683 | 21.7138606 |
| Q08J23 | NSUN2 | 22.2769422 | 22.275763 | 22.3246062 | 22.2547214 | 22.3102615 |
| P34897 | SHMT2 | 24.3544225 | 24.4633883 | 24.3901258 | 24.270727 | 24.4532449 |
| Q7Z6I8 | C5orf24 | 18.8875914 | 18.5636176 | 19.0136482 | 18.7187085 | 19.0358102 |
| Q53FZ2 | ACSM3 | 16.9003063 | NA | NA | 15.8939208 | NA |
| P10599 | TXN | 25.4055268 | 25.548525 | 25.1799699 | 25.2724625 | 25.4438867 |
| P04035 | HMGCR | 18.2392441 | 18.4266852 | 18.4848748 | 18.3977745 | 18.549415 |
| Q99683 | MAP3K5 | 14.206268 | NA | NA | 14.6007933 | NA |
| Q8TF42 | UBASH3B | 16.5060471 | 16.7241993 | 17.0950936 | 16.8042669 | 16.9645023 |
| Q9BRT6 | LLPH | 21.1743966 | 21.2519624 | 21.5263065 | 21.4265252 | 21.3996373 |
| P23434 | GCSH | 20.4405706 | 20.5634746 | 20.1909554 | 20.2500022 | 20.5863771 |
| Q19AV6 | ZSWIM7 | 13.5859617 | 13.4124236 | 13.9242485 | 13.3511659 | 13.5334637 |
| Q6ZW76 | ANKS3 | NA | 14.1370712 | 13.9326713 | NA | 13.9542887 |
| Q12955 | ANK3 | 11.1068 | NA | NA | 11.1658957 | NA |
| Q86UL3 | GPAT4 | 19.6229613 | 19.5699771 | 19.7008196 | 19.4195796 | 19.6475495 |
| Q13620 | CUL4B | 23.4169872 | 23.4249832 | 23.3531002 | 23.2920463 | 23.3431211 |
| Q96G25 | MED8 | 20.5179852 | 20.5592136 | 20.4325474 | 20.4792757 | 20.5231711 |
| Q9UHP3 | USP25 | 19.2217662 | 19.3886765 | 19.2686405 | 19.2749122 | 19.3266863 |
| Q9BX40 | LSM14B | 22.3175684 | 22.4525112 | 22.2565834 | 22.17344 | 22.4765741 |
| Q9ULQ1 | TPCN1 | 16.259954 | 16.2141625 | 16.3378688 | 16.1520116 | 16.3746026 |
| O14523 | C2CD2L | 18.4879546 | 18.5938204 | 18.4804161 | 18.360744 | 18.4089441 |
| P55347 | PKNOX1 | 19.1338703 | 19.0545355 | 19.1925934 | 19.1341674 | 19.2567956 |
| Q8WWI1 | LMO7 | 17.2743605 | 17.3380268 | 17.5914882 | 17.2775874 | 17.3409874 |
| P13861 | PRKAR2A | 22.6797519 | 22.7386579 | 22.6833849 | 22.5835681 | 22.7167365 |
| Q969M1 | TOMM40L | 16.9489653 | NA | NA | 16.9904289 | NA |
| P08243 | ASNS | 20.8430495 | 20.8030259 | 20.8428396 | 20.8564151 | 20.8240859 |
| O95671 | ASMTL | 22.7568577 | 22.8213302 | 22.8178912 | 22.6987533 | 22.7354777 |
| Q9NZL6 | RGL1 | 19.2796366 | 18.916193 | 19.3367261 | 19.0466838 | 18.9798357 |
| Q9HB20 | PLEKHA3 | 15.2698898 | 15.9810505 | 15.7660062 | 15.7113499 | 15.4384455 |
| Q9H0Z9 | RBM38 | 18.0937019 | 18.3328044 | 18.250534 | 18.258028 | 18.2308726 |

| | | | | | | |
|---|---|---|---|---|---|---|
| Q5MCW4 | ZNF569 | 17.4319169 | 17.2817545 | 17.4843739 | 17.2975095 | 17.1918764 |
| P78536 | ADAM17 | 19.6619942 | 19.8160938 | 19.7057529 | 19.5892255 | 19.6987149 |
| O60315 | ZEB2 | 22.0025663 | 22.1120327 | 21.9245087 | 21.9830415 | 22.1008285 |
| Q53HL2 | CDCA8 | 22.1480012 | 22.3309105 | 22.2679034 | 22.1470313 | 22.2168235 |
| Q9NXF7 | DCAF16 | 18.9904141 | 18.9579612 | 19.0593944 | 19.0380536 | 19.0358733 |
| O00267 | SUPT5H | 22.4258612 | 22.5004486 | 22.532912 | 22.4204882 | 22.5242813 |
| Q06124 | PTPN11 | 21.9291553 | 21.9650278 | 21.922875 | 21.8833467 | 21.9669752 |
| Q14008 | CKAP5 | 22.5538141 | 22.6401674 | 22.6213902 | 22.5502942 | 22.6279182 |
| O43719 | HTATSF1 | 22.2709219 | 22.3080317 | 22.2001607 | 22.2287177 | 22.2816701 |
| O14734 | ACOT8 | 20.6434686 | 20.558987 | 20.6910151 | 20.6325124 | 20.6722737 |
| P35754 | GLRX | 22.6743258 | 22.6548043 | 22.4377041 | 22.5318144 | 22.6301549 |
| Q8N8Q8 | COX18 | 19.4384116 | 19.2483386 | 19.5001059 | 19.4777755 | 19.190375 |
| O75414 | NME6 | 19.0616147 | 19.0388669 | 19.1085896 | 19.3714136 | 19.1002448 |
| P04733 | MT1F | 22.4129846 | 22.1382992 | 22.1841461 | 22.1314223 | 22.5184415 |
| Q99487 | PAFAH2 | 18.1028052 | 18.4346249 | 18.5277125 | 18.2558155 | 18.3066051 |
| O75390 | CS | 25.1678601 | 25.1005791 | 25.1354976 | 25.0744011 | 25.0800108 |
| P29275 | ADORA2B | NA | 14.8864295 | 15.0269986 | NA | 14.6166216 |
| Q9UKU7 | ACAD8 | 19.5211273 | 19.7182477 | 19.5297628 | 19.486034 | 19.5972461 |
| Q9Y697 | NFS1 | 20.6420017 | 20.6904501 | 20.7494992 | 20.5869995 | 20.7415937 |
| Q8WWW8 | GAB3 | 16.2688905 | 16.1430742 | 16.9864666 | 16.5131341 | 16.4712646 |
| P14652 | HOXB2 | NA | 15.2867707 | 15.3845626 | NA | 15.5709633 |
| Q4J6C6 | PREPL | 19.5249471 | 19.6567848 | 19.6135882 | 19.6691054 | 19.7668757 |
| Q9NSI8 | SAMSN1 | 18.9663626 | 18.9949804 | 18.8904777 | 18.9931091 | 19.0067613 |
| O60331 | PIP5K1C | 19.8801636 | 19.899426 | 19.9940952 | 19.8719684 | 19.9752069 |
| Q8NFW8 | CMAS | 20.9578502 | 21.112947 | 21.0854794 | 20.9573629 | 21.0342422 |
| P08047 | SP1 | 19.5268242 | 19.6384444 | 19.7656478 | 19.6255812 | 19.7239483 |
| O96019 | ACTL6A | 22.4461606 | 22.5941397 | 22.3313963 | 22.3303477 | 22.5615777 |
| Q9H3L0 | MMADHC | 15.6994495 | 15.8365459 | 15.8356874 | 15.747907 | 15.9972609 |
| Q8N9A8 | CNEP1R1 | 17.7703984 | 17.3529475 | 17.953072 | 17.8176272 | 17.343955 |
| Q9H930 | SP140L | 16.4739206 | 16.2559607 | 16.4637832 | 16.5728723 | 16.258675 |
| Q9Y257 | KCNK6 | 21.5671334 | 21.4495721 | 21.3549569 | 21.3179775 | 21.3797171 |
| P63165 | SUMO1 | 23.2514779 | 23.4211531 | 23.284158 | 23.1593079 | 23.2358125 |
| Q9UJY5 | GGA1 | 20.5712456 | 20.6325143 | 20.5627334 | 20.5985185 | 20.6555869 |
| P00167 | CYB5A | 21.1192773 | 21.1245779 | 21.15806 | 21.1956936 | 21.1318852 |
| Q9UNW1 | MINPP1 | 18.6780099 | 18.5264373 | 18.6058576 | 18.5003496 | 18.5993115 |
| Q6EMB2 | TTLL5 | 15.6800863 | 15.8399377 | 15.9105427 | 15.942619 | 15.4974482 |
| P63279 | UBE2I | 23.1080916 | 23.1410461 | 23.2204174 | 23.154035 | 23.1192101 |
| Q9HA64 | FN3KRP | 21.2981351 | 21.359103 | 21.3397902 | 21.3094374 | 21.2917587 |
| Q9NQ66 | PLCB1 | 15.3807396 | NA | NA | 15.3570884 | NA |
| O00623 | PEX12 | 15.5859937 | 16.108065 | 16.1613368 | 16.0503601 | 15.8127678 |
| Q6ZRS2 | SRCAP | 19.4607598 | 19.6365727 | 19.6134512 | 19.4826492 | 19.5996957 |
| Q14527 | HLTF | 18.5719887 | 18.6900655 | 18.8102792 | 18.7136899 | 18.73476 |

| | | | | | | |
|---|---|---|---|---|---|---|
| Q9Y375 | NDUFAF1 | 21.0323935 | 20.9516015 | 21.2352071 | 21.0107005 | 21.0916631 |
| P43686 | PSMC4 | 23.6009523 | 23.7130909 | 23.5419928 | 23.4871347 | 23.6503171 |
| Q96SL8 | FIZ1 | 17.565578 | 17.5573312 | 17.946481 | 17.5872511 | 17.5741328 |
| Q96Q89 | KIF20B | 20.2411637 | 20.3057473 | 20.2878745 | 20.1912532 | 20.3463671 |
| Q9UHJ3 | SFMBT1 | 18.6858764 | 18.8534485 | 18.7533697 | 18.6527921 | 18.7613177 |
| Q9NYM9 | BET1L | 19.4058275 | 19.5030266 | 19.5836972 | 19.5988696 | 19.3113379 |
| Q38SD2 | LRRK1 | 19.0816595 | 18.6984519 | 19.1079897 | 18.8389811 | 18.891635 |
| Q92508 | PIEZO1 | 19.3737414 | 19.3712479 | 19.5129191 | 19.3892667 | 19.382259 |
| O60921 | HUS1 | 18.8484501 | 18.8235881 | 18.9953671 | 18.7815808 | 18.886202 |
| Q9NPJ3 | ACOT13 | 21.4066603 | 21.4696599 | 21.3079428 | 21.2718094 | 21.2264581 |
| A8MXV4 | NUDT19 | 20.7103974 | 20.7638561 | 20.7551448 | 20.7966633 | 20.727309 |
| P02788 | LTF | 22.7829563 | 22.4838602 | 22.382077 | 22.6047889 | 21.8708594 |
| Q9UNE7 | STUB1 | 22.3368903 | 22.4784631 | 22.4353238 | 22.3209598 | 22.4538855 |
| Q5EBL2 | ZNF628 | 11.5969817 | NA | NA | 12.3391634 | NA |
| Q96L34 | MARK4 | 13.8366571 | NA | NA | 13.9497902 | NA |
| Q9H8X2 | IPPK | 16.5630483 | 17.0122831 | 16.6313185 | 16.9698714 | 16.5889581 |
| P09769 | FGR | 17.5753511 | 17.6108481 | 18.0627116 | 17.5663995 | 18.0167445 |
| Q5BKX6 | SLC45A4 | 18.8137693 | 18.3533366 | 18.8602484 | 18.4812829 | 18.2221561 |
| O75716 | STK16 | 17.2700719 | 17.2898724 | 17.2634015 | 17.4838983 | 17.1367504 |
| Q07954 | LRP1 | NA | 20.1902671 | 20.2888223 | NA | 20.2964923 |
| Q9Y394 | DHRS7 | 20.8522873 | 20.778199 | 20.8800031 | 20.6416384 | 20.6806614 |
| Q9P0T4 | ZNF581 | 18.7038381 | 18.7636296 | 18.7447488 | 18.8409379 | 18.7490988 |
| Q9BWU1 | CDK19 | 19.868603 | 19.9605476 | 20.0314424 | 19.8463692 | 20.039887 |
| Q5BJF2 | TMEM97 | 22.5762935 | 22.663159 | 22.7549003 | 22.561279 | 22.6724775 |
| Q68D91 | MBLAC2 | 20.6875995 | 20.8134056 | 20.759272 | 20.6460936 | 20.9061203 |
| Q7L775 | EPM2AIP1 | 19.998415 | 19.9640415 | 20.0890386 | 19.9629493 | 20.0321643 |
| Q9H0P0 | NT5C3A | 21.01023 | 21.0141144 | 21.1459439 | 20.9810963 | 21.0607716 |
| Q07889 | SOS1 | 18.8245704 | 18.9335604 | 18.8375792 | 18.7231367 | 18.8812667 |
| Q7Z7A3 | CTU1 | 19.873387 | 19.9814923 | 19.9494103 | 19.8794138 | 19.9104674 |
| Q9BYI3 | HYCC1 | 18.1385384 | 18.2818884 | 18.2255589 | 18.1678602 | 18.2108582 |
| P07910 | HNRNPC | 27.1594638 | 27.223779 | 27.1255695 | 27.0019623 | 27.0233264 |
| Q9NXE4 | SMPD4 | 20.4916017 | 20.554474 | 20.6086414 | 20.5345145 | 20.6214736 |
| P11310 | ACADM | 22.7031032 | 22.754226 | 22.6882716 | 22.6721061 | 22.7193004 |
| Q96GM8 | TOE1 | 20.8976902 | 20.8501116 | 20.908615 | 20.8349024 | 20.9090929 |
| Q13144 | EIF2B5 | 21.0488736 | 21.0269717 | 21.0735438 | 20.8612672 | 20.9082225 |
| Q13217 | DNAJC3 | 22.2143809 | 22.2313487 | 22.1702389 | 22.1183428 | 22.0827616 |
| Q06210 | GFPT1 | 21.5574465 | 21.5788848 | 21.6274682 | 21.5279622 | 21.5675022 |
| Q13200 | PSMD2 | 23.8964119 | 23.942456 | 23.9080342 | 23.861963 | 23.9561427 |
| P16455 | MGMT | 19.8669388 | 19.8342701 | 19.9186597 | 19.815058 | 19.9113915 |
| Q9HDC9 | APMAP | 23.2446283 | 23.3586829 | 23.2014302 | 23.2027564 | 23.2124583 |
| P41218 | MNDA | 22.1265198 | 22.1780679 | 22.2406595 | 22.1316537 | 22.1792843 |
| Q9UIV1 | CNOT7 | 20.5494699 | 20.650506 | 20.7688612 | 20.6112772 | 20.7787101 |

| | | | | | | |
|---|---|---|---|---|---|---|
| Q9H223 | EHD4 | 20.4499256 | 20.5144054 | 20.5818615 | 20.4160074 | 20.5104866 |
| P12814 | ACTN1 | 25.0782889 | 25.187938 | 25.0525052 | 25.0226923 | 25.164824 |
| O96006 | ZBED1 | 19.0573038 | 19.0726421 | 19.1654143 | 19.0632888 | 19.1956138 |
| P16298 | PPP3CB | 20.1112763 | 20.1910772 | 20.0706802 | 20.0934942 | 20.2054196 |
| Q9Y3I1 | FBXO7 | 20.3082167 | 20.3394379 | 20.4485451 | 20.3549963 | 20.3667549 |
| P62136 | PPP1CA | 23.6418189 | 23.7306471 | 23.7047413 | 23.6316936 | 23.6690242 |
| Q9Y4E6 | WDR7 | 22.4988488 | 22.5111611 | 22.5265125 | 22.4795071 | 22.5665198 |
| Q9H4L4 | SENP3 | 21.0217255 | 20.9982776 | 21.1210557 | 21.133262 | 21.1130861 |
| Q5VWJ9 | SNX30 | 18.6875265 | 18.9421422 | 18.9341314 | 18.7570242 | 18.8553728 |
| Q01196 | RUNX1 | 21.3639163 | 21.4260696 | 21.427548 | 21.3905297 | 21.5040688 |
| P52758 | RIDA | 20.4776135 | 20.6258159 | 20.6553683 | 20.5164381 | 20.5431752 |
| P43487 | RANBP1 | 24.9318784 | 25.0593109 | 24.9918639 | 24.976307 | 25.0156149 |
| Q14254 | FLOT2 | 20.0212039 | 20.2439924 | 20.1084227 | 20.0106159 | 20.2765823 |
| P17024 | ZNF20 | 17.1269316 | 17.3305405 | 17.4544548 | 17.278463 | 17.1127918 |
| O95772 | STARD3NL | 19.7408015 | 19.867805 | 19.9611085 | 19.9550563 | 19.972568 |
| Q14872 | MTF1 | 16.5803604 | 16.9152001 | 17.0122195 | 16.7057148 | 17.0528353 |
| Q6ZMW3 | EML6 | 16.0699649 | 17.1692825 | 16.8075086 | 16.2740709 | 16.7613681 |
| P50993 | ATP1A2 | 14.7676655 | NA | NA | 14.3441047 | NA |
| Q5SQN1 | SNAP47 | 16.1430632 | 15.9792216 | 16.1706577 | 15.7657272 | 16.1877584 |
| P56385 | ATP5ME | 23.2167339 | 23.2620813 | 23.3069576 | 23.1935056 | 23.3086402 |
| Q96FJ2 | DYNLL2 | 21.2578899 | 21.3623972 | 21.3794565 | 21.2156665 | 21.4295387 |
| Q6PIJ6 | FBXO38 | 19.3905801 | 19.5726413 | 19.4933947 | 19.3865899 | 19.5444527 |
| Q9BR61 | ACBD6 | 20.0620118 | 20.0716742 | 19.9902536 | 19.993961 | 20.0655482 |
| Q92945 | KHSRP | 25.6236444 | 25.7732607 | 25.4600977 | 25.5605865 | 25.6280989 |
| P57737 | CORO7 | 22.5781152 | 22.6611214 | 22.5390653 | 22.5504232 | 22.7046354 |
| Q0VAK6 | LMOD3 | 18.0674682 | 18.0818543 | 18.0749805 | 18.0642677 | 18.2394096 |
| Q9NYB9 | ABI2 | NA | 11.5763999 | 11.4366554 | NA | 11.6163274 |
| Q8N5H3 | FAM89B | 15.6508833 | 15.1746588 | 15.5117949 | 15.5021162 | 15.6715048 |
| Q9UKB3 | DNAJC12 | 16.1194122 | 15.947044 | 16.3075347 | 16.2207126 | 16.3163642 |
| P16591 | FER | 18.0481101 | 18.0760641 | 18.0008662 | 18.176112 | 18.0964925 |
| Q13105 | ZBTB17 | 15.667522 | 15.8529996 | 15.7824917 | 15.7872903 | 15.8824691 |
| Q15427 | SF3B4 | 21.3425524 | 21.5544176 | 21.3678679 | 21.4081622 | 21.6934651 |
| Q8TB36 | GDAP1 | 19.9634185 | 19.8548138 | 20.0202395 | 19.9251272 | 19.9367915 |
| P60520 | GABARAPL2 | 23.0861603 | 23.1598469 | 23.0128095 | 22.9768402 | 23.0981855 |
| Q13905 | RAPGEF1 | 18.478811 | 18.8069322 | 18.6406933 | 18.5842758 | 18.6410324 |
| Q9H5Q4 | TFB2M | 19.7578015 | 19.6615621 | 19.9223422 | 19.8191704 | 19.9034981 |
| Q9BV57 | ADI1 | 22.692163 | 22.8406089 | 22.6999039 | 22.7044496 | 22.8738333 |
| Q86XA9 | HEATR5A | 17.1959558 | 17.3683366 | 17.6173633 | 17.3637735 | 17.5070756 |
| P49005 | POLD2 | 21.1994193 | 21.2236612 | 21.3578627 | 21.209123 | 21.3263064 |
| Q7L8L6 | FASTKD5 | 20.5773012 | 20.5612211 | 20.6170244 | 20.577475 | 20.6376276 |
| P27701 | CD82 | 19.5703612 | 19.2548821 | 19.4116988 | 19.5811611 | 19.337951 |
| Q8NI37 | PPTC7 | 16.9774875 | 17.0874698 | 17.0735628 | 16.9723718 | 17.2924049 |

| | | | | | | |
|---|---|---|---|---|---|---|
| Q8IVB5 | LIX1L | 18.4178234 | 18.9909854 | 18.4715149 | 18.645989 | 18.4757207 |
| P24666 | ACP1 | 22.6859152 | 22.761867 | 22.7927079 | 22.6233016 | 22.7731844 |
| O60306 | AQR | 23.004713 | 23.0411761 | 23.118554 | 23.0079341 | 23.1134853 |
| O15042 | U2SURP | 22.3226677 | 22.345067 | 22.4068236 | 22.3121107 | 22.4085243 |
| Q9UK53 | ING1 | 18.677461 | 18.7493512 | 18.8437405 | 18.6388948 | 18.7193954 |
| Q92466 | DDB2 | 22.0530243 | 22.0951559 | 22.1854491 | 22.1525015 | 22.1510654 |
| Q92833 | JARID2 | 18.5608448 | 18.7469914 | 18.7461996 | 18.7265485 | 18.6517785 |
| Q70CQ2 | USP34 | 20.239092 | 20.3089272 | 20.3658034 | 20.24225 | 20.3321173 |
| P28838 | LAP3 | 23.5844588 | 23.5840487 | 23.5490049 | 23.5009372 | 23.5284088 |
| Q9NQC1 | JADE2 | 16.6574202 | 16.8735869 | 16.5311448 | 16.5809636 | 16.6114034 |
| Q9UEW8 | STK39 | 18.632381 | 18.7584281 | 18.7997506 | 18.6956906 | 18.5883208 |
| O43264 | ZW10 | 21.2809843 | 21.368198 | 21.3948079 | 21.2922193 | 21.3913598 |
| Q9UL33 | TRAPPC2L | 20.4379223 | 20.467639 | 20.4421707 | 20.4671725 | 20.4337919 |
| Q9H0V9 | LMAN2L | 19.7264501 | 19.8894397 | 19.984834 | 19.9470269 | 20.0420478 |
| P23142 | FBLN1 | 17.6181707 | 17.269469 | 17.2035019 | 17.3236169 | 17.259138 |
| Q15438 | CYTH1 | 18.1908175 | 18.5795689 | 18.4419671 | 18.2041157 | 18.4311922 |
| P20592 | MX2 | 16.1508577 | 16.0116719 | 16.189376 | 15.9793883 | 16.3878593 |
| Q14964 | RAB39A | 18.3884019 | 18.5333113 | 18.4565202 | 18.315248 | 18.4411612 |
| Q8N137 | CNTROB | 16.6496136 | 16.9284563 | 16.804846 | 16.6250728 | 16.4860299 |
| Q6GPH4 | XAF1 | 17.5774479 | 17.217606 | 17.8593702 | 17.1778831 | 17.3418589 |
| Q9H6F5 | CCDC86 | 20.4590354 | 20.7952708 | 20.8198625 | 20.6388422 | 20.6963656 |
| O60830 | TIMM17B | 21.5742531 | 21.8391166 | 21.7080472 | 21.5990821 | 21.6363897 |
| Q96GS6 | ABHD17A | 18.7857627 | 18.5785946 | 18.7719498 | 18.6426695 | 18.6628631 |
| Q9BT25 | HAUS8 | 18.873684 | 18.8740968 | 19.0544164 | 18.9835331 | 19.0954125 |
| Q9NVR0 | KLHL11 | 18.0027839 | 18.0512574 | 18.12479 | 18.0083795 | 17.97996 |
| O75223 | GGCT | 22.1148398 | 22.2821726 | 22.1484419 | 22.0661825 | 22.2312559 |
| Q9HB07 | MYG1 | 21.6385122 | 21.6251629 | 21.6512891 | 21.6251707 | 21.6754556 |
| Q6NUM9 | RETSAT | 20.3230694 | 20.2228055 | 20.473716 | 20.3005319 | 20.3736477 |
| O15530 | PDPK1 | 19.8952186 | 19.9615696 | 20.0315983 | 19.8225001 | 20.0187075 |
| O95396 | MOCS3 | 19.8113931 | 19.8804855 | 19.8362914 | 19.8588725 | 19.9792172 |
| P50583 | NUDT2 | 20.209388 | 20.3632194 | 20.4441036 | 20.1820571 | 20.3651206 |
| O60869 | EDF1 | 23.9231189 | 23.8718032 | 24.0302466 | 23.8583554 | 23.5387594 |
| P06702 | S100A9 | 22.2776688 | 22.1057875 | 21.9227023 | 22.2350196 | 22.075536 |
| P49842 | STK19 | 16.2482259 | 16.3584331 | 16.3725151 | 16.1540464 | 16.3008372 |
| Q9BQ49 | SMIM7 | 19.9195518 | 20.0093943 | 19.6814169 | 20.0257352 | 19.4552853 |
| Q99519 | NEU1 | 20.8520446 | 20.7125973 | 20.7617078 | 20.6956525 | 20.8106315 |
| Q53H12 | AGK | 21.3781172 | 21.406826 | 21.395565 | 21.3609844 | 21.3528363 |
| Q9NPF4 | OSGEP | 21.8683958 | 21.8061172 | 21.8738116 | 21.7916315 | 21.9573748 |
| Q9BRX8 | PRXL2A | 18.9402792 | 18.7354452 | 18.9167332 | 18.8712955 | 18.9978393 |
| P28072 | PSMB6 | 23.9269889 | 23.8930667 | 23.8030647 | 23.896215 | 23.9727484 |
| P43243 | MATR3 | 24.6230583 | 24.7029981 | 24.6182383 | 24.588263 | 24.6229428 |
| P30086 | PEBP1 | 25.4552 | 25.5410149 | 25.4776691 | 25.4740635 | 25.5484629 |

| | | | | | | |
|---|---|---|---|---|---|---|
| P63167 | DYNLL1 | 22.9014943 | 22.910175 | 23.0596617 | 22.8694779 | 23.165169 |
| Q6N043 | ZNF280D | 17.6521046 | 17.6799705 | 17.6180667 | 17.5571061 | 17.7522296 |
| P52429 | DGKE | 17.837511 | 18.0750652 | 17.5557829 | 17.9429654 | 17.7892288 |
| Q96E39 | RBMXL1 | 20.1985972 | 19.8920258 | 20.1039174 | 19.9714211 | 19.7264908 |
| Q9UN36 | NDRG2 | 18.0074938 | 18.3747624 | 18.4319026 | 18.2042537 | 18.5195142 |
| Q86SX3 | TEDC1 | 16.2673793 | NA | NA | 16.3394811 | NA |
| Q8IY67 | RAVER1 | 21.6540708 | 21.7865869 | 21.7109624 | 21.6991197 | 21.6473958 |
| Q96QE3 | ATAD5 | 18.044095 | 18.2884841 | 18.4258027 | 18.1639788 | 18.1050557 |
| Q6IAN0 | DHRS7B | 20.1020151 | 20.1574838 | 20.2655407 | 20.0156727 | 20.260434 |
| O75431 | MTX2 | 21.0170312 | 21.0303895 | 20.9068435 | 21.0935938 | 20.9659464 |
| Q96C86 | DCPS | 22.7505649 | 22.8850542 | 22.7755364 | 22.7531486 | 22.8682871 |
| P51003 | PAPOLA | 21.2217918 | 21.2608583 | 21.2666345 | 21.226572 | 21.3166157 |
| Q9BU76 | C1orf35 | 19.9202199 | 20.0598691 | 19.80871 | 19.9293723 | 19.9261561 |
| Q00978 | IRF9 | 17.7114285 | 17.7865267 | 17.624089 | 17.6909749 | 17.8110682 |
| Q9Y4C2 | TCAF1 | 16.4427527 | 16.6732024 | 16.7586386 | 16.6777508 | 16.7231576 |
| Q04724 | TLE1 | 17.452778 | 17.6336802 | 17.7409494 | 17.6509903 | 17.7795288 |
| Q8IZC7 | ZNF101 | 15.7503885 | 15.6990292 | 16.0586469 | 15.5970733 | 16.0383596 |
| Q9Y2G3 | ATP11B | 20.0031353 | 19.9874501 | 20.1169793 | 19.9874424 | 20.1062505 |
| Q9HD67 | MYO10 | 15.0134516 | 15.2476763 | 15.6597505 | 15.5615155 | 15.2505038 |
| Q9C026 | TRIM9 | NA | 17.4895327 | 17.0815328 | NA | 16.9297349 |
| O00409 | FOXN3 | 16.1844959 | 16.5246629 | 16.6579798 | 16.3475352 | 16.5715369 |
| Q9NV92 | NDFIP2 | 19.1806683 | 19.1095502 | 19.2184913 | 18.9206693 | 19.1662365 |
| Q86UD0 | SAPCD2 | 17.009858 | 16.9681126 | 16.8772046 | 16.702551 | 17.135935 |
| Q96LD8 | SENP8 | 17.3081327 | 17.4088381 | 17.4714522 | 17.3597428 | 17.5400346 |
| O95685 | PPP1R3D | 19.3176612 | 19.5511766 | 19.502704 | 19.4124838 | 19.5427399 |
| P61225 | RAP2B | 20.3933922 | 20.522405 | 20.5176702 | 20.4344374 | 20.5220813 |
| Q6IBS0 | TWF2 | 21.3487909 | 21.3244875 | 21.4313445 | 21.3691708 | 21.445121 |
| Q9BTT0 | ANP32E | 23.5450134 | 23.6157757 | 23.6444498 | 23.4982977 | 23.6340083 |
| Q13428 | TCOF1 | 22.0191104 | 22.0905645 | 22.1093317 | 21.9859971 | 21.9976947 |
| Q9H2F5 | EPC1 | 18.3920248 | 18.4189432 | 18.4855957 | 18.3547418 | 18.4879932 |
| Q3KP31 | ZNF791 | 20.4608326 | 20.5708628 | 20.6426766 | 20.5569184 | 20.58781 |
| Q9Y2X3 | NOP58 | 23.644199 | 23.7335929 | 23.6163988 | 23.6018428 | 23.6017389 |
| Q9H0D6 | XRN2 | 22.4310012 | 22.4529901 | 22.5099082 | 22.4323054 | 22.4586333 |
| Q99536 | VAT1 | 23.1911377 | 23.2618954 | 23.3525729 | 23.2411512 | 23.3468079 |
| Q9UJZ1 | STOML2 | 22.7804537 | 22.9105604 | 22.7898433 | 22.7600458 | 22.8309967 |
| Q92917 | GPKOW | 21.210802 | 21.2796869 | 21.2505789 | 21.2012648 | 21.2620925 |
| P08582 | MELTF | 18.8532745 | 18.8408984 | 18.8893747 | 18.9129014 | 18.8420901 |
| Q52LJ0 | FAM98B | 21.5173277 | 21.4248841 | 21.6184565 | 21.4931139 | 21.4986878 |
| P10155 | RO60 | 22.1595715 | 22.1666655 | 22.1486876 | 22.1095664 | 22.2505283 |
| O95905 | ECD | 20.3599036 | 20.3430995 | 20.5474041 | 20.3376979 | 20.4722953 |
| Q9UBV7 | B4GALT7 | 18.7274261 | 19.1167892 | 18.8308511 | 19.0072585 | 19.0536758 |
| Q96RU2 | USP28 | 16.989831 | 17.2181441 | 16.928447 | 16.8333918 | 17.122994 |

| | | | | | | |
|---|---|---|---|---|---|---|
| Q8N697 | SLC15A4 | 19.1977949 | 19.3383124 | 19.0385903 | 19.1350329 | 19.1229449 |
| Q6P4F2 | FDX2 | 18.4256226 | 18.6611361 | 18.5266402 | 18.4194998 | 18.5955197 |
| P35247 | SFTPD | 22.4141481 | 22.4090994 | 22.4918055 | 21.9272835 | 21.6507475 |
| P0DPB6 | POLR1D | 20.0348945 | 20.1793383 | 19.9852116 | 19.7877329 | 20.194484 |
| P79522 | PRR3 | 17.1359966 | 17.4801303 | 17.5653791 | 17.1642611 | 17.4272644 |
| Q8N612 | FHIP1B | 16.8810034 | 16.8204872 | 16.9003863 | 16.699157 | 16.9352719 |
| Q9HC44 | GPBP1L1 | NA | 15.5336467 | 15.3505817 | NA | 15.562278 |
| Q9H8H3 | TMT1A | 19.4023786 | 19.6264243 | 19.7216434 | 19.4568822 | 19.5905676 |
| Q8N1G1 | REXO1 | 18.8443324 | 18.5311147 | 18.6873876 | 18.6240503 | 18.5250708 |
| Q96PN7 | TRERF1 | 16.7791852 | 16.991703 | 17.0149039 | 16.941995 | 16.844454 |
| O95219 | SNX4 | 19.5083984 | 19.4872474 | 19.4583058 | 19.5258142 | 19.4454002 |
| Q92539 | LPIN2 | 17.6559052 | 17.8670032 | 17.8395513 | 17.8436855 | 17.8509801 |
| P14678 | SNRPB | 24.9307966 | 24.883796 | 24.9168431 | 24.7753778 | 24.8934015 |
| Q6ZW49 | PAXIP1 | 20.4689089 | 20.5833744 | 20.6180481 | 20.5052123 | 20.595236 |
| Q9UL18 | AGO1 | 19.2191077 | 19.2198851 | 19.2942697 | 19.171755 | 19.3009654 |
| Q16543 | CDC37 | 24.3698136 | 24.4227864 | 24.3859315 | 24.3301667 | 24.35818 |
| Q96G03 | PGM2 | 23.8195733 | 23.8834206 | 23.8684094 | 23.741847 | 23.7965085 |
| P61221 | ABCE1 | 23.9388208 | 23.963504 | 23.9725599 | 23.9010134 | 23.9778728 |
| Q96G46 | DUS3L | 23.1872959 | 23.2375717 | 23.2400043 | 23.1314573 | 23.1951924 |
| O14908 | GIPC1 | 20.6097752 | 20.6825416 | 20.5505296 | 20.6032528 | 20.5942261 |
| P38159 | RBMX | 25.7707267 | 25.7768732 | 25.7408888 | 25.6320377 | 25.6862794 |
| Q8N1F8 | STK11IP | 20.7649181 | 20.7900627 | 20.8835948 | 20.722011 | 20.768107 |
| Q7Z2W4 | ZC3HAV1 | 22.4292222 | 22.4500099 | 22.4132867 | 22.3499802 | 22.4505669 |
| O95248 | SBF1 | 19.746634 | 19.7299374 | 19.787999 | 19.7789472 | 19.7443482 |
| Q8NHU6 | TDRD7 | 19.83369 | 19.7959956 | 19.7732485 | 19.8611039 | 19.8060926 |
| Q07866 | KLC1 | 21.9316594 | 22.032566 | 21.9228471 | 21.9087742 | 21.9689651 |
| O43156 | TTI1 | 20.510194 | 20.5055957 | 20.5416503 | 20.4755766 | 20.5519225 |
| P59780 | AP3S2 | 20.8944207 | 20.9987903 | 20.9556717 | 21.0135078 | 20.981716 |
| Q9NV31 | IMP3 | 20.9371841 | 20.92851 | 21.0217277 | 21.0048507 | 21.0352741 |
| Q13546 | RIPK1 | 20.0928154 | 19.9593061 | 20.1251168 | 20.0303412 | 20.087786 |
| Q14318 | FKBP8 | 23.0368368 | 23.0990484 | 23.0718769 | 23.0335776 | 23.077409 |
| Q96T66 | NMNAT3 | 18.3955561 | 18.2796726 | 18.3583207 | 18.4462888 | 18.4313413 |
| Q8N4T8 | CBR4 | 20.5642715 | 20.4862927 | 20.6612428 | 20.6304018 | 20.6034408 |
| Q9UQ35 | SRRM2 | 22.3938049 | 22.5604619 | 22.4016968 | 22.4243599 | 22.5093913 |
| Q14847 | LASP1 | 23.7741895 | 23.9109428 | 23.8005602 | 23.7906707 | 23.9265769 |
| P78406 | RAE1 | 22.3277532 | 22.3284248 | 22.360373 | 22.2753559 | 22.506972 |
| Q96ED9 | HOOK2 | 18.1419419 | 18.2893374 | 18.3794559 | 18.3850516 | 18.3295332 |
| Q9UNZ5 | C19orf53 | 22.4744671 | 22.7261431 | 22.6851004 | 22.5974461 | 22.555867 |
| P39687 | ANP32A | 25.3597515 | 25.5721638 | 25.2781114 | 25.372776 | 25.4023195 |
| P47712 | PLA2G4A | 20.4262724 | 20.5027846 | 20.6130206 | 20.4377978 | 20.5119047 |
| O95628 | CNOT4 | 18.9410089 | 19.0435455 | 19.2046342 | 19.0821502 | 19.2580907 |
| Q7Z403 | TMC6 | 17.5051485 | 17.5190536 | 17.5336749 | 17.3142201 | 17.4325942 |

| | | | | | | |
|---|---|---|---|---|---|---|
| Q9Y4C1 | KDM3A | 21.1683061 | 21.4037481 | 21.3910906 | 21.1412435 | 21.3871921 |
| Q9UHQ1 | NARF | 18.8302235 | 19.052878 | 19.1975228 | 18.8404019 | 19.2071266 |
| Q9NYP7 | ELOVL5 | 19.1145559 | 19.5765475 | 19.5004225 | 19.2464682 | 19.3568026 |
| Q96AJ1 | CLUAP1 | NA | 11.4776045 | 11.3648223 | NA | 11.3628795 |
| O15440 | ABCC5 | 15.0853789 | 15.4380328 | 15.7207588 | 15.7265869 | 15.357207 |
| Q12983 | BNIP3 | 15.7079814 | 15.908468 | 15.6987344 | 15.6831029 | 16.1312766 |
| Q9BT67 | NDFIP1 | 19.9152213 | 19.7257898 | 19.7682599 | 19.9155168 | 19.7049119 |
| P62487 | POLR2G | 20.6311026 | 20.6865254 | 20.6999067 | 20.6499992 | 20.7866875 |
| Q8WTW4 | NPRL2 | 16.6831995 | 16.8726485 | 16.933626 | 16.8386224 | 16.6758784 |
| Q9BSJ8 | ESYT1 | 23.843898 | 23.9497584 | 23.8890274 | 23.8078929 | 23.8824588 |
| Q9NPJ6 | MED4 | 19.8683523 | 19.8523068 | 19.7291867 | 19.7385122 | 19.8946965 |
| Q9BWH6 | RPAP1 | 20.1692326 | 20.2205957 | 20.216806 | 20.1096969 | 20.2514446 |
| Q8N806 | UBR7 | 20.9605126 | 21.1865014 | 21.0455508 | 20.9974625 | 21.1188718 |
| Q96EM0 | L3HYPDH | 19.1864252 | 19.3316685 | 19.4266242 | 19.2161718 | 19.3742987 |
| O94810 | RGS11 | NA | 15.5017386 | 15.6845168 | NA | 15.6157879 |
| O75319 | DUSP11 | 17.4344115 | 17.8696494 | 17.7414287 | 17.7601751 | 17.7415727 |
| Q12996 | CSTF3 | 21.7965241 | 21.8137889 | 21.8794193 | 21.7594438 | 21.8181142 |
| Q8WVQ1 | CANT1 | 20.5762097 | 20.5555017 | 20.607188 | 20.6420071 | 20.5687266 |
| P21964 | COMT | 23.6325836 | 23.7526634 | 23.6161482 | 23.559014 | 23.7335361 |
| O15021 | MAST4 | NA | 13.8343836 | 14.4782843 | NA | 14.0628734 |
| P11142 | HSPA8 | 27.5574746 | 27.6965732 | 27.540535 | 27.550883 | 27.6325735 |
| P08910 | ABHD2 | 16.1919462 | 15.7880223 | 16.0328663 | 16.3206226 | 15.9011856 |
| Q3SXZ3 | ZNF718 | NA | 15.6100848 | 15.2723343 | NA | 15.3748191 |
| O95125 | ZNF202 | 17.197003 | 17.3381162 | 16.9703906 | 17.1925625 | 17.1147755 |
| Q9NWH2 | TMEM242 | 15.4632647 | NA | NA | 15.4468836 | NA |
| Q6ZTN6 | ANKRD13D | 16.9426023 | 17.0119798 | 17.0880983 | 16.9254896 | 17.1003843 |
| Q70Z53 | FRA10AC1 | 16.7506153 | 17.0029694 | 17.0426645 | 17.0697123 | 16.9924793 |
| Q99728 | BARD1 | 17.1976029 | 17.4288417 | 17.2960959 | 17.0448637 | 17.0579435 |
| Q7L4E1 | MIGA2 | 14.9514803 | 15.134838 | 15.3095886 | 15.1988402 | 15.3330921 |
| Q96AD5 | PNPLA2 | 17.2669613 | 17.3351675 | 17.4395444 | 17.1989562 | 17.4365752 |
| Q86W33 | TPRA1 | 19.1484604 | 19.2852853 | 19.2643235 | 19.1110416 | 19.0895208 |
| Q9BSY9 | DESI2 | 20.1278323 | 19.9999866 | 20.3120702 | 20.1681626 | 20.2046478 |
| Q9Y4K1 | CRYBG1 | 17.6601203 | 17.7113754 | 17.8627541 | 17.6984317 | 17.91002 |
| Q15942 | ZYX | 23.7588339 | 23.8312013 | 23.7114979 | 23.6799503 | 23.8772412 |
| Q8N0Z3 | SPICE1 | 17.9763082 | 18.1745673 | 18.1399843 | 17.9193703 | 18.0472603 |
| Q9Y5T4 | DNAJC15 | 19.0206772 | 19.1872806 | 19.3260791 | 19.1695633 | 19.1804637 |
| Q92624 | APPBP2 | 18.4450124 | 18.6108457 | 18.6472726 | 18.4547174 | 18.5157147 |
| P42167 | TMPO | 24.3407908 | 24.3523481 | 24.3991349 | 24.3321593 | 24.3537542 |
| P54098 | POLG | 19.0839269 | 19.2249239 | 19.1481076 | 19.1017992 | 19.1651834 |
| P37802 | TAGLN2 | 26.1538904 | 26.1517854 | 26.1020942 | 26.0223712 | 26.1982566 |
| P46734 | MAP2K3 | 21.0471924 | 21.1475151 | 21.2255637 | 21.0180006 | 21.1754562 |
| Q9H3M7 | TXNIP | 21.561807 | 21.5738631 | 21.5616288 | 21.5607762 | 21.5205939 |

| | | | | | | |
|---|---|---|---|---|---|---|
| P53618 | COPB1 | 23.902462 | 23.9153217 | 23.9216055 | 23.8591361 | 23.9262189 |
| O60341 | KDM1A | 22.7598474 | 22.8295205 | 22.7544805 | 22.7129489 | 22.7506623 |
| P49736 | MCM2 | 23.7198808 | 23.7604779 | 23.6535361 | 23.62076 | 23.6722362 |
| Q5VZL5 | ZMYM4 | 20.2531877 | 20.3544374 | 20.2909102 | 20.2392315 | 20.3626418 |
| Q7L592 | NDUFAF7 | 18.6491053 | 18.6783998 | 18.8174142 | 18.7277147 | 18.8091304 |
| O00139 | KIF2A | 21.6184349 | 21.7387821 | 21.6804586 | 21.6352554 | 21.6500082 |
| O43929 | ORC4 | 19.7049343 | 19.8035772 | 19.7860757 | 19.7172717 | 19.8280019 |
| Q6PD74 | AAGAB | 20.6968205 | 20.8253323 | 20.8638164 | 20.7368611 | 20.896371 |
| Q96JP5 | ZFP91 | 20.6540255 | 20.9278736 | 20.7728933 | 20.7651759 | 20.7650412 |
| P26368 | U2AF2 | 24.6957462 | 24.8209783 | 24.6156599 | 24.6667952 | 24.7175418 |
| Q6PL18 | ATAD2 | 21.4156904 | 21.5661156 | 21.545332 | 21.4432215 | 21.5746243 |
| P0C2W1 | FBXO45 | 20.4044654 | 20.3627538 | 20.3550836 | 20.4298895 | 20.343999 |
| Q9H7N4 | SCAF1 | 20.961372 | 21.1049991 | 20.9534137 | 21.0078972 | 21.117248 |
| Q5JSP0 | FGD3 | 20.5609885 | 20.6016777 | 20.6851925 | 20.5087459 | 20.6835317 |
| O43586 | PSTPIP1 | 18.884951 | 19.0825144 | 19.0269445 | 19.0453902 | 19.061636 |
| O96013 | PAK4 | 19.010915 | 19.0174284 | 19.1575587 | 18.9164773 | 19.1312437 |
| Q6P1M9 | ARMCX5 | 16.4620001 | 16.4495347 | 16.5332241 | 16.4924031 | 16.6297865 |
| Q8N8R7 | ARL14EP | 19.0806854 | 19.0807215 | 18.9205288 | 19.0210194 | 18.8805241 |
| Q14353 | GAMT | 20.577506 | 20.614536 | 20.6957451 | 20.6263838 | 20.8817545 |
| P50135 | HNMT | 22.672336 | 22.8179898 | 22.8318664 | 22.6419563 | 22.8315946 |
| Q13685 | AAMP | 21.7371173 | 21.864584 | 21.6507926 | 21.6314012 | 21.8918756 |
| Q9UL26 | RAB22A | 19.5782871 | 19.3227873 | 19.7039535 | 19.5038863 | 19.665003 |
| Q13153 | PAK1 | 20.3473425 | 20.457411 | 20.2662253 | 20.3043551 | 20.5630732 |
| Q6NXT6 | TAPT1 | 16.5398635 | 16.6943754 | 16.9255932 | 16.4658982 | 16.6916306 |
| Q6P0Q8 | MAST2 | 13.7034723 | 13.2653472 | 13.9785346 | 13.752343 | 13.4926761 |
| Q96MH2 | HEXIM2 | 15.3524374 | NA | NA | 15.400246 | NA |
| O43439 | CBFA2T2 | NA | 14.4348433 | 14.5354789 | NA | 14.6653788 |
| Q8TDM6 | DLG5 | NA | 18.013155 | 19.2313497 | NA | 18.8271207 |
| Q9NVN3 | RIC8B | 18.3946706 | 18.2370811 | 18.3868152 | 18.2920607 | 18.2215256 |
| Q96SZ5 | ADO | 20.2498645 | 20.2144342 | 20.1854556 | 20.128645 | 20.2656629 |
| Q8TEA7 | TBCK | 18.9454639 | 19.1041671 | 19.0490361 | 18.9238789 | 19.1240542 |
| Q14289 | PTK2B | 22.0134609 | 22.1130485 | 22.0971371 | 21.9961367 | 22.0969926 |
| Q9H0B6 | KLC2 | 18.2228467 | 18.4139006 | 18.3840038 | 18.3154044 | 18.3173885 |
| P50452 | SERPINB8 | 23.7346399 | 23.5460538 | 23.6092929 | 23.6580838 | 23.6154032 |
| Q96KG9 | SCYL1 | 20.8169942 | 20.9008793 | 20.9041045 | 20.8353011 | 20.8773648 |
| Q00341 | HDLBP | 22.819728 | 22.9063409 | 22.7933768 | 22.7784414 | 22.8538875 |
| P18615 | NELFE | 20.7500542 | 20.8100303 | 20.6653177 | 20.7418691 | 20.7692143 |
| Q9NQG5 | RPRD1B | 21.9096754 | 22.1459966 | 21.873536 | 21.9323091 | 22.0966854 |
| P35716 | SOX11 | 16.6024462 | 16.7675264 | 16.6344907 | 16.8994685 | 16.5680574 |
| Q01995 | TAGLN | 15.0029221 | NA | NA | 15.0298736 | NA |
| Q9BR77 | CCDC77 | 18.6882933 | 18.8359251 | 18.6071336 | 18.9438291 | 18.7321248 |
| Q15111 | PLCL1 | 19.4543088 | NA | NA | 19.5864667 | NA |

| | | | | | | |
|---|---|---|---|---|---|---|
| Q9NZ45 | CISD1 | 21.3070112 | 21.5163826 | 21.5898538 | 21.2139623 | 21.5514721 |
| P18065 | IGFBP2 | 20.1745644 | 19.9984065 | 19.7401535 | 20.0797589 | 20.151754 |
| Q5T0B9 | ZNF362 | 16.8591968 | 16.8829481 | 17.070443 | 16.8661426 | 16.965631 |
| P05091 | ALDH2 | 18.5570088 | 18.7794965 | 18.7995767 | 18.4501309 | 18.7323652 |
| Q9UJ98 | STAG3 | 17.485325 | 17.4296268 | 17.7672135 | 17.6551599 | 17.6630517 |
| P67775 | PPP2CA | 23.4656459 | 23.5132883 | 23.5424771 | 23.4630928 | 23.6208121 |
| Q9Y6X9 | MORC2 | 21.4980491 | 21.4622863 | 21.6066433 | 21.4486075 | 21.4921745 |
| Q5R372 | RABGAP1L | 21.6641297 | 21.6318272 | 21.6328887 | 21.6002147 | 21.6593181 |
| P29350 | PTPN6 | 25.4868869 | 25.5647475 | 25.4285308 | 25.4457991 | 25.5563788 |
| Q9NQW7 | XPNPEP1 | 22.5516249 | 22.5721558 | 22.5723 | 22.4818242 | 22.6945194 |
| P40222 | TXLNA | 25.353053 | 25.4749373 | 25.2651477 | 25.2998128 | 25.3285114 |
| P49959 | MRE11 | 21.9041692 | 22.0077627 | 22.0802132 | 21.9370014 | 22.0529416 |
| Q6RFH5 | WDR74 | 21.0803727 | 21.1354596 | 21.1890959 | 21.0853638 | 21.2615021 |
| Q9NXR1 | NDE1 | 19.5197282 | 19.629408 | 19.5455214 | 19.5535152 | 19.6184311 |
| Q9Y646 | CPQ | 20.6243742 | 20.6418437 | 20.6446164 | 20.519986 | 20.8240829 |
| P56270 | MAZ | 20.2593567 | 20.4387224 | 20.302041 | 20.3910417 | 20.2392082 |
| Q96BT3 | CENPT | 16.6598634 | 16.6217818 | 16.97235 | 16.6704 | 16.7468924 |
| Q6DHV5 | CC2D2B | 15.9150076 | NA | NA | 15.7775971 | NA |
| Q8IWZ6 | BBS7 | 15.7314568 | NA | NA | 16.1192322 | NA |
| Q15555 | MAPRE2 | 20.728971 | 20.881665 | 20.9090849 | 20.7600045 | 20.8759789 |
| P07947 | YES1 | 20.937144 | 20.8497178 | 20.9874903 | 20.9047703 | 21.0476151 |
| O14936 | CASK | NA | 13.9031102 | 13.3348468 | NA | 13.6525058 |
| P50995 | ANXA11 | 23.6591272 | 23.7076232 | 23.5947007 | 23.5512216 | 23.6263964 |
| Q96N67 | DOCK7 | 22.5741921 | 22.6344348 | 22.7059088 | 22.5563316 | 22.7165825 |
| Q6FIF0 | ZFAND6 | 19.4706725 | 19.5431074 | 19.5289846 | 19.4741174 | 19.5717962 |
| Q9HCL3 | ZFP14 | NA | 14.6976653 | 14.7810152 | NA | 14.3930721 |
| Q9BS40 | LXN | 19.5615233 | 19.4944614 | 19.6739566 | 19.4721137 | 19.5546066 |
| Q8TDY2 | RB1CC1 | 19.6561198 | 19.8373422 | 19.7841454 | 19.7627341 | 19.6957476 |
| P21580 | TNFAIP3 | 18.2928563 | 18.4306853 | 18.5092055 | 18.3202392 | 18.1914851 |
| Q9H300 | PARL | 19.861213 | 19.9590077 | 20.1164756 | 19.968105 | 19.9679346 |
| Q16772 | GSTA3 | 16.7251138 | NA | NA | 16.9177143 | NA |
| O60427 | FADS1 | 21.4419037 | 21.5093991 | 21.601512 | 21.4642355 | 21.6149199 |
| P32189 | GK | 21.5727569 | 21.6343316 | 21.6561961 | 21.5487843 | 21.7243217 |
| Q96S59 | RANBP9 | 21.0287532 | 21.0616278 | 21.1051972 | 20.9847294 | 21.0627424 |
| Q68EM7 | ARHGAP17 | 22.3622551 | 22.4136752 | 22.3669613 | 22.2366427 | 22.3772792 |
| P28066 | PSMA5 | 23.7749715 | 23.8633738 | 23.7415001 | 23.7290789 | 23.8889347 |
| Q9NXW9 | ALKBH4 | 19.5314031 | 19.3903729 | 19.7235902 | 19.4982524 | 19.5171103 |
| Q06136 | KDSR | 19.0602691 | 19.333649 | 19.1972569 | 19.2506362 | 19.2248627 |
| Q13615 | MTMR3 | 18.299593 | 18.4394605 | 18.3853672 | 18.3662801 | 18.3195558 |
| O75113 | N4BP1 | 19.1289805 | 19.1156938 | 19.1778685 | 19.0731431 | 19.3125732 |
| Q02083 | NAAA | 20.4124177 | 20.2619879 | 20.5227887 | 20.4323662 | 20.5071738 |
| O75943 | RAD17 | 18.7239603 | 18.9028685 | 18.8268051 | 18.8454554 | 18.7454468 |

| | | | | | | |
|---|---|---|---|---|---|---|
| Q8TF05 | PPP4R1 | 20.6989906 | 20.6756872 | 20.6123775 | 20.5377119 | 20.6489976 |
| Q7Z739 | YTHDF3 | 22.4602029 | 22.557349 | 22.5451296 | 22.4528668 | 22.4962718 |
| O60934 | NBN | 22.0119739 | 22.0082568 | 22.0530692 | 21.9152051 | 22.0635908 |
| P49454 | CENPF | 20.7252999 | 20.8017453 | 20.5894503 | 20.6494334 | 20.7557476 |
| Q03111 | MLLT1 | 20.7258897 | 20.8332666 | 20.7621164 | 20.7478501 | 20.7627879 |
| Q9BPX5 | ARPC5L | 22.4771436 | 22.426559 | 22.4767711 | 22.3996291 | 22.4151645 |
| Q9NUJ1 | ABHD10 | 22.9394225 | 22.976658 | 23.0357312 | 22.8627005 | 23.0241926 |
| Q8N3Z6 | ZCCHC7 | 18.468759 | 18.4891375 | 18.6499026 | 18.6073681 | 18.4164093 |
| O75368 | SH3BGRL | 24.3271845 | 24.2757537 | 24.2020518 | 24.1991046 | 24.3239331 |
| Q16877 | PFKFB4 | 18.4661165 | 18.6542234 | 18.6674888 | 18.4447233 | 18.7022963 |
| B7ZAP0 | RABGAP1L | 16.2381779 | 16.4267759 | 16.4192436 | 16.1649637 | 16.1172213 |
| Q15233 | NONO | 25.6725548 | 25.763481 | 25.649924 | 25.5911995 | 25.6535845 |
| O95466 | FMNL1 | 22.0486025 | 22.1529539 | 22.1487152 | 22.0650193 | 22.1537975 |
| Q9BRP1 | PDCD2L | 19.7348947 | 19.9479436 | 19.6405264 | 19.8113132 | 19.6843093 |
| P39210 | MPV17 | 20.5159411 | 20.5887819 | 20.7702962 | 20.4922784 | 20.7167478 |
| Q9Y5W9 | SNX11 | 18.8938808 | 18.9887557 | 19.0542817 | 19.0009096 | 18.9610345 |
| Q5JS37 | NHLRC3 | 18.4865593 | 18.529519 | 18.5529316 | 18.4112053 | 18.5491847 |
| Q96RT8 | TUBGCP5 | 18.8631748 | 18.8771177 | 18.7847301 | 18.8039985 | 18.8754215 |
| Q9P270 | SLAIN2 | 19.5936259 | 19.6332517 | 19.6060639 | 19.5681881 | 19.651397 |
| Q99816 | TSG101 | 21.0129423 | 21.0557814 | 21.1054265 | 20.9235703 | 20.978429 |
| O14646 | CHD1 | 20.7096932 | 20.7591861 | 20.7508569 | 20.662062 | 20.6983591 |
| Q15650 | TRIP4 | 20.2879579 | 20.3532802 | 20.4187826 | 20.3696759 | 20.4477374 |
| Q06323 | PSME1 | 25.5116913 | 25.5954447 | 25.3790365 | 25.3952192 | 25.5422323 |
| Q13619 | CUL4A | 22.0556064 | 22.1480294 | 22.0779778 | 21.9931553 | 22.051804 |
| P52747 | ZNF143 | 18.924014 | 19.1613241 | 19.1277981 | 19.0417724 | 19.1244213 |
| Q6PII5 | HAGHL | 19.7026767 | 19.6112617 | 19.6591978 | 19.5602376 | 19.7076171 |
| Q9P1F3 | ABRACL | 24.0273604 | 24.1251066 | 23.8988233 | 23.9065649 | 24.2140006 |
| Q6ZN44 | UNC5A | NA | 14.5432323 | 14.8929574 | NA | 14.7680839 |
| O43159 | RRP8 | 20.2123964 | 20.2921128 | 20.5143414 | 20.3244237 | 20.4006865 |
| Q16560 | SNRNP35 | 15.3586625 | 15.5727172 | 15.2993457 | 15.6549846 | 15.9300362 |
| Q9NRD5 | PICK1 | 15.2738773 | NA | NA | 14.0547385 | NA |
| P63313 | TMSB10 | 25.3336948 | 25.4518479 | 25.1797813 | 25.1317721 | 25.4328081 |
| Q00537 | CDK17 | 14.1885155 | NA | NA | 14.3127019 | NA |
| Q14624 | ITIH4 | 19.3589295 | 19.0996939 | 19.5147935 | 18.837554 | 18.8271493 |
| Q96JE9 | MAP6 | NA | 15.4773915 | 15.5432492 | NA | 15.1579774 |
| O75326 | SEMA7A | 13.6269914 | 13.9967332 | 13.6896041 | 13.7791405 | 13.5640999 |
| Q6P9B6 | MEAK7 | 18.5790935 | 18.7817902 | 18.9976994 | 18.5247741 | 18.7788396 |
| Q9UGN5 | PARP2 | 18.2187969 | 18.3303424 | 18.337389 | 18.1752616 | 18.5696464 |
| Q9H6Q4 | CIAO3 | 20.3490223 | 20.7340842 | 20.6984007 | 20.5338831 | 20.6640959 |
| P19544 | WT1 | 18.2246711 | 17.9103923 | 18.2153704 | 18.0485038 | 18.1716789 |
| P17535 | JUND | 20.965652 | 21.1601619 | 20.9972347 | 20.9213983 | 21.1870197 |
| Q6NTE8 | MRNIP | 14.9019868 | 15.1343239 | 15.2719886 | 15.0388192 | 15.3364913 |

| | | | | | | |
|---|---|---|---|---|---|---|
| Q93063 | EXT2 | 16.7577902 | 16.6910444 | 16.8356146 | 16.6195553 | 16.7024217 |
| Q13131 | PRKAA1 | 20.536852 | 20.7138299 | 20.8095408 | 20.6520223 | 20.816149 |
| Q9P0J7 | KCMF1 | 20.163641 | 20.2371248 | 20.0460302 | 20.1891649 | 20.1984756 |
| Q5TKA1 | LIN9 | 17.6130143 | 17.6749513 | 17.8648117 | 17.6173334 | 17.6311881 |
| P36871 | PGM1 | 21.8995644 | 22.0886775 | 22.0655524 | 21.8984266 | 22.0493391 |
| Q96J02 | ITCH | 19.1829298 | 19.3568939 | 19.4408419 | 19.2909009 | 19.3618362 |
| O95716 | RAB3D | 23.0064721 | 23.0724066 | 23.0952849 | 22.9163604 | 23.0370096 |
| P28062 | PSMB8 | 22.589961 | 22.7012338 | 22.7390091 | 22.5473976 | 22.6936124 |
| Q9UET6 | FTSJ1 | 20.3175896 | 20.3547349 | 20.2508128 | 20.2881582 | 20.3135063 |
| Q9Y2I8 | WDR37 | 19.5717647 | 19.5996362 | 19.7402896 | 19.5484673 | 19.7029917 |
| Q8TC07 | TBC1D15 | 21.8176721 | 21.8930614 | 21.9274962 | 21.7242666 | 21.8785829 |
| O00329 | PIK3CD | 21.288676 | 21.3751845 | 21.431173 | 21.2411448 | 21.3489812 |
| Q12904 | AIMP1 | 23.123118 | 23.3273422 | 23.1997728 | 23.1346915 | 23.2532793 |
| P42345 | MTOR | 20.5209538 | 20.5497645 | 20.6663488 | 20.4630754 | 20.5940209 |
| Q5TAQ9 | DCAF8 | 20.3914931 | 20.3827264 | 20.3570988 | 20.3663679 | 20.3611347 |
| P11586 | MTHFD1 | 24.6400599 | 24.6649913 | 24.613727 | 24.5312305 | 24.6979369 |
| O14744 | PRMT5 | 22.4305129 | 22.4617729 | 22.4864455 | 22.4247915 | 22.5054394 |
| Q04637 | EIF4G1 | 23.7622881 | 23.8397294 | 23.798248 | 23.7242005 | 23.7766354 |
| Q8TBC4 | UBA3 | 22.3018813 | 22.4339617 | 22.3731271 | 22.3136512 | 22.4361044 |
| Q14498 | RBM39 | 23.7172041 | 23.7006344 | 23.6982877 | 23.6050453 | 23.7362066 |
| Q9P2E3 | ZNFX1 | 19.2916646 | 19.3936974 | 19.3830909 | 19.2934654 | 19.3571394 |
| O14672 | ADAM10 | 21.7842792 | 21.7960431 | 21.8360641 | 21.7383918 | 21.8056288 |
| P26196 | DDX6 | 23.433929 | 23.4603249 | 23.4605049 | 23.4012363 | 23.4829848 |
| P51610 | HCFC1 | 22.3347835 | 22.4521028 | 22.3655177 | 22.3160646 | 22.4656 |
| P50851 | LRBA | 19.5893847 | 19.6423032 | 19.6902108 | 19.5372909 | 19.6671978 |
| Q6P3X3 | TTC27 | 20.6253519 | 20.6428178 | 20.6317876 | 20.5798448 | 20.7189877 |
| Q9BYN8 | MRPS26 | 21.9834739 | 22.0731722 | 22.0816468 | 22.0504414 | 22.0351781 |
| Q8N392 | ARHGAP18 | 20.3524364 | 20.4719708 | 20.4385318 | 20.3465556 | 20.3762785 |
| O75947 | ATP5PD | 24.2138039 | 24.3283978 | 24.2984748 | 24.2378243 | 24.3176109 |
| Q8IYS1 | PM20D2 | 21.7457495 | 21.8311295 | 21.7671229 | 21.7463338 | 21.8463636 |
| Q9UPN4 | CEP131 | 20.6819011 | 20.757801 | 20.74603 | 20.7160119 | 20.709984 |
| Q9NQ50 | MRPL40 | 21.3476258 | 21.3691075 | 21.2473605 | 21.2413704 | 21.2918582 |
| Q9UIC8 | LCMT1 | 19.5223755 | 19.6411615 | 19.6723071 | 19.5272308 | 19.6319877 |
| P51828 | ADCY7 | 20.0311304 | 20.1397621 | 20.2980394 | 20.1616057 | 20.1823527 |
| Q9UEU0 | VTI1B | 20.6022987 | 20.6137256 | 20.7070148 | 20.5925951 | 20.6782285 |
| P25705 | ATP5F1A | 26.9390348 | 27.0253964 | 26.8098027 | 26.8032755 | 26.9471398 |
| Q14694 | USP10 | 21.4727159 | 21.5269385 | 21.5594424 | 21.4031234 | 21.6026809 |
| Q99798 | ACO2 | 23.5313363 | 23.6598593 | 23.6725329 | 23.5261323 | 23.7156902 |
| P49336 | CDK8 | 17.9854937 | 17.9644319 | 18.053826 | 18.0285351 | 18.0738907 |
| Q969Y2 | GTPBP3 | 19.0755319 | 19.2656414 | 19.3098478 | 19.2276554 | 19.3827089 |
| O95379 | TNFAIP8 | 19.2706682 | 19.2845706 | 19.1438944 | 19.156406 | 19.2015841 |
| Q9UJU6 | DBNL | 23.3419643 | 23.5506644 | 23.3845846 | 23.3209815 | 23.4025852 |

| | | | | | | |
|---|---|---|---|---|---|---|
| Q9BQC3 | DPH2 | 19.7366329 | 19.8562046 | 19.7284201 | 19.7116115 | 19.8562811 |
| O15212 | PFDN6 | 23.2341445 | 23.3332616 | 23.0845107 | 23.1220475 | 23.1954139 |
| P53365 | ARFIP2 | 22.371176 | 22.3370446 | 22.4299341 | 22.4570774 | 22.3557984 |
| Q13501 | SQSTM1 | 22.4861732 | 22.3653966 | 22.4205013 | 22.5951311 | 22.3683789 |
| P61513 | RPL37A | 25.3287597 | 25.1763606 | 25.5688044 | 25.3456704 | 25.3934117 |
| Q9H082 | RAB33B | 18.2025584 | 18.4021629 | 18.4207412 | 18.4254261 | 18.4221888 |
| Q96RQ9 | IL4I1 | 20.3738916 | 20.6096361 | 20.6700975 | 20.3931451 | 20.6267695 |
| Q8IYN0 | ZNF100 | 16.9298764 | 17.3052933 | 17.1371259 | 17.0363337 | 17.340572 |
| P08514 | ITGA2B | 17.3252816 | 17.192239 | 17.1715097 | 17.4023096 | 17.1880312 |
| P42336 | PIK3CA | 15.9769344 | 16.1150998 | 16.3096016 | 15.799276 | 16.0970609 |
| O43482 | OIP5 | 18.0529255 | 18.0050993 | 18.1320437 | 18.1005848 | 18.0335166 |
| Q92613 | JADE3 | 16.3403241 | 15.8451737 | 16.2182851 | 16.212044 | 15.8381322 |
| Q8NB50 | ZFP62 | 16.5557112 | 16.9832307 | 16.9679794 | 16.6660551 | 16.5024421 |
| Q14012 | CAMK1 | 19.3038048 | 19.5686277 | 19.4806247 | 19.3300863 | 19.613731 |
| Q8WVZ9 | KBTBD7 | 16.3839308 | 16.5326534 | 16.6959254 | 16.5174276 | 16.5768693 |
| Q9HAR2 | ADGRL3 | 18.0972356 | 17.9253027 | 17.920586 | 17.999002 | 17.7027917 |
| Q9BQQ3 | GORASP1 | 16.1100453 | 16.5293053 | 16.0179924 | 16.472894 | 16.3943826 |
| O95670 | ATP6V1G2 | 15.9195676 | 16.3222538 | 15.5204537 | 15.9105651 | 15.6550786 |
| Q8NBS3 | SLC4A11 | 17.9105751 | 18.2322355 | 18.2494102 | 18.1768216 | 18.6385923 |
| O60347 | TBC1D12 | 15.0959845 | 14.4127303 | 15.1290473 | 14.7039384 | 14.7932871 |
| P27144 | AK4 | 17.582319 | 18.3123966 | 18.1433044 | 17.7205118 | 18.1672286 |
| Q9UJ14 | GGT7 | 14.9570997 | 14.8339534 | 14.6790817 | 14.9400505 | 15.44101 |
| P33241 | LSP1 | 21.0466297 | 21.866434 | 21.5105608 | 21.0833576 | 21.8243664 |
| O43674 | NDUFB5 | 21.4901112 | 21.6424873 | 21.6887502 | 21.4906323 | 21.627421 |
| Q8IZE3 | SCYL3 | 17.6291918 | 17.6748523 | 17.7694167 | 17.709214 | 17.7061047 |
| Q63HN8 | RNF213 | 19.7678957 | 19.8454177 | 19.9161144 | 19.7357436 | 19.8382192 |
| Q9Y4Z0 | LSM4 | 23.3099062 | 23.3125524 | 23.4073109 | 23.2133779 | 23.2986062 |
| O43390 | HNRNPR | 24.1345616 | 24.1873954 | 24.0934198 | 24.0604943 | 24.1443099 |
| Q9BPZ3 | PAIP2 | 22.1390645 | 22.1497334 | 21.9330263 | 22.1236614 | 22.0908416 |
| P35475 | IDUA | 20.2649089 | 20.4745462 | 20.3633617 | 20.3356639 | 20.4305359 |
| P13994 | YJU2B | 17.8312082 | 18.1124085 | 18.0958776 | 17.9781467 | 18.2594161 |
| Q86VP3 | PACS2 | 18.5542532 | 18.4577537 | 18.7141551 | 18.5645135 | 18.6755727 |
| Q9NVX2 | NLE1 | 19.6298889 | 19.5900918 | 19.7056422 | 19.5428001 | 19.728073 |
| O95644 | NFATC1 | 20.9810948 | 21.1160304 | 21.0361961 | 21.0377853 | 21.0705861 |
| Q14687 | GSE1 | 19.9553103 | 20.1181565 | 20.1233956 | 19.9672949 | 20.0873705 |
| Q0VDF9 | HSPA14 | 22.0221778 | 22.2136487 | 21.9912594 | 22.0095764 | 22.0868912 |
| Q6R327 | RICTOR | 19.6754609 | 19.7883609 | 19.746946 | 19.7135003 | 19.5639684 |
| Q5EBL8 | PDZD11 | 16.642986 | 16.701386 | 16.5385555 | 16.5891116 | 16.5605269 |
| Q70CQ3 | USP30 | 18.2095223 | 18.2487465 | 18.2631055 | 18.3293808 | 18.2093374 |
| P53567 | CEBPG | 21.5498363 | 21.5439806 | 21.6373769 | 21.602409 | 21.619165 |
| Q8TEA8 | DTD1 | 20.3420074 | 20.5985083 | 20.422428 | 20.3658235 | 20.4292766 |
| Q8TEA1 | NSUN6 | 18.4030768 | 18.1581598 | 18.4740124 | 18.3528741 | 18.2851174 |

| | | | | | | |
|---|---|---|---|---|---|---|
| Q8N6M3 | FITM2 | 16.1586954 | 16.2646811 | 16.4458468 | 16.3316124 | 16.3833113 |
| Q9NX57 | RAB20 | 18.2523968 | 18.3731896 | 18.6423644 | 18.4268515 | 18.4369465 |
| Q658P3 | STEAP3 | 19.3186368 | 19.2954944 | 19.5224014 | 19.3088895 | 19.3762833 |
| P11274 | BCR | 18.8255629 | 18.9013635 | 18.9413356 | 18.6709974 | 18.9270752 |
| Q8IY47 | KBTBD2 | 17.7752829 | 17.8381152 | 17.9312089 | 17.84047 | 17.9075712 |
| Q9NRX2 | MRPL17 | 22.5113834 | 22.5165455 | 22.4325991 | 22.4155614 | 22.5203033 |
| Q92830 | KAT2A | 19.1850807 | 19.4145306 | 19.5023569 | 19.3661868 | 19.3282198 |
| P16435 | POR | 21.9859609 | 22.062863 | 22.0753594 | 21.9261448 | 22.1223296 |
| Q8NC56 | LEMD2 | 20.569222 | 20.6796053 | 20.7071397 | 20.5080257 | 20.6042907 |
| Q9P016 | THYN1 | 21.7029905 | 21.78187 | 21.7767618 | 21.6768296 | 21.7870062 |
| P82930 | MRPS34 | 22.0945054 | 22.0928583 | 22.1760487 | 22.0951259 | 22.0623918 |
| Q8N543 | OGFOD1 | 20.9474484 | 21.0049274 | 20.921551 | 20.850096 | 20.8969735 |
| Q92841 | DDX17 | 25.2401832 | 25.3664318 | 25.2445931 | 25.180633 | 25.2620288 |
| O75746 | SLC25A12 | 19.9423226 | 19.9884982 | 19.9956616 | 19.966311 | 20.095326 |
| Q8IXH7 | NELFCD | 20.6692583 | 20.7201949 | 20.8408422 | 20.7414711 | 20.8372658 |
| Q14141 | SEPTIN6 | 23.2600326 | 23.2622733 | 23.2955964 | 23.1890005 | 23.3480446 |
| O95347 | SMC2 | 23.278457 | 23.3169118 | 23.2982467 | 23.1666539 | 23.2411585 |
| O43776 | NARS1 | 24.0774265 | 24.1720042 | 24.1609084 | 24.0632546 | 24.122664 |
| P14373 | TRIM27 | 20.3040212 | 20.2395822 | 20.2884998 | 20.2722801 | 20.3196411 |
| Q92785 | DPF2 | 22.0196793 | 22.0876148 | 22.0261778 | 22.0453016 | 22.042412 |
| P55265 | ADAR | 23.0631627 | 23.050657 | 23.0340476 | 23.0194486 | 23.0661939 |
| Q15907 | RAB11B | 24.8563213 | 24.9038436 | 24.7968769 | 24.8136808 | 24.8894257 |
| Q96GQ7 | DDX27 | 21.7179032 | 21.8072475 | 21.726576 | 21.7483096 | 21.8380944 |
| Q71SY5 | MED25 | 20.3271669 | 20.2577982 | 20.2794495 | 20.2399687 | 20.3381554 |
| P48736 | PIK3CG | 23.7066585 | 23.6834361 | 23.6883104 | 23.5941042 | 23.5860339 |
| Q5VV52 | ZNF691 | 18.75251 | 18.8105684 | 18.7236404 | 18.9409858 | 18.8194285 |
| Q86YV5 | PRAG1 | 21.2301559 | 21.3449586 | 21.3856401 | 21.2840461 | 21.4806865 |
| Q96BD5 | PHF21A | 18.0300331 | 18.1759311 | 18.1575199 | 18.0379267 | 18.1603545 |
| O75600 | GCAT | 19.4857509 | 19.6620322 | 19.5380698 | 19.5535099 | 19.5642475 |
| P51688 | SGSH | 17.4153121 | 17.5121024 | 17.4936083 | 17.6261506 | 17.6104245 |
| Q9GZN1 | ACTR6 | 19.0812171 | 19.1330248 | 18.7871366 | 18.9909914 | 19.0223886 |
| P17152 | TMEM11 | 20.944536 | 20.8403217 | 21.0942923 | 20.8288986 | 21.0127649 |
| Q96CS7 | PLEKHB2 | 19.6863714 | 19.7339925 | 19.4882725 | 19.8503894 | 19.4537243 |
| Q15527 | SURF2 | 16.3126027 | 16.4766961 | 16.5989291 | 16.6152861 | 16.7198721 |
| Q8IXS8 | HYCC2 | 17.8336342 | 17.721913 | 17.8421244 | 17.8267061 | 17.7421066 |
| O75496 | GMNN | 19.3348124 | 19.42079 | 19.315977 | 19.4307498 | 19.4732015 |
| Q8IWE2 | FAM114A1 | 17.8001739 | 17.8481888 | 17.9448401 | 17.7897172 | 18.1357762 |
| O15211 | RGL2 | 17.2446233 | 17.6114996 | 17.4470167 | 17.3037106 | 17.488195 |
| Q502W6 | VWA3B | 18.7286973 | 19.2562487 | 18.7530099 | 18.9064696 | 18.9278452 |
| Q9P2D8 | UNC79 | 14.2849685 | 13.4122859 | 13.9622911 | 14.4838224 | 13.4279972 |
| Q6ZQQ6 | WDR87 | 19.3636333 | 19.1983056 | 18.887662 | 18.8860847 | 17.8746543 |
| P04004 | VTN | 21.8778826 | 22.2512302 | 21.699876 | 21.3631693 | 22.0321877 |

| | | | | | | |
|---|---|---|---|---|---|---|
| Q06418 | TYRO3 | 11.4639268 | NA | NA | 11.7221935 | NA |
| Q562R1 | ACTBL2 | 22.3166101 | 22.770931 | 22.6186797 | 22.3688566 | 22.7923482 |
| Q13099 | IFT88 | 16.7793041 | 17.1022732 | 16.8750752 | 16.8332657 | 16.94694 |
| Q3KQV3 | ZNF792 | NA | 15.1522206 | 15.0147171 | NA | 14.6525421 |
| Q8NCL4 | GALNT6 | 17.8082381 | 18.0672413 | 18.1584258 | 18.1050084 | 18.1574385 |
| P52789 | HK2 | 22.8034664 | 23.0173544 | 22.9613381 | 22.7461563 | 22.9356799 |
| Q13907 | IDI1 | 23.1206637 | 23.1234686 | 23.1641439 | 23.0130673 | 23.1259976 |
| Q13557 | CAMK2D | 21.4867215 | 21.6496208 | 21.614029 | 21.5170388 | 21.6297368 |
| P11908 | PRPS2 | 21.817348 | 21.8651546 | 21.8169844 | 21.6923824 | 21.8046258 |
| O14524 | NEMP1 | 19.6979177 | 19.8938996 | 19.9032999 | 19.7679515 | 19.8605317 |
| Q8WWV3 | RTN4IP1 | 18.6687346 | 18.6422998 | 18.7684274 | 18.7485443 | 18.8629934 |
| Q9NVI7 | ATAD3A | 24.2223874 | 24.2270149 | 24.3023188 | 24.1046976 | 24.1914201 |
| Q9H2G2 | SLK | 21.4471323 | 21.5794161 | 21.5781981 | 21.4280873 | 21.4713501 |
| O14981 | BTAF1 | 20.8472149 | 20.9232687 | 20.9611322 | 20.8234128 | 20.9393682 |
| Q86W74 | ANKRD46 | 18.3247491 | 18.404555 | 18.3774564 | 18.3270285 | 18.3845134 |
| O96008 | TOMM40 | 24.2675628 | 24.3791764 | 24.2988697 | 24.2753131 | 24.4354314 |
| P13798 | APEH | 23.5909488 | 23.5619018 | 23.5690134 | 23.4858669 | 23.5175004 |
| Q9H0C8 | ILKAP | 22.4578992 | 22.5152985 | 22.5677379 | 22.4267479 | 22.5241082 |
| P48556 | PSMD8 | 24.0876509 | 24.0952586 | 24.0278859 | 24.0120861 | 24.059341 |
| Q5T4S7 | UBR4 | 24.0062224 | 24.0214982 | 24.0354569 | 23.9248277 | 24.0108769 |
| Q96KP4 | CNDP2 | 23.9238973 | 23.9330709 | 23.8790865 | 23.8460295 | 23.9404218 |
| Q14642 | INPP5A | 19.454399 | 19.4304721 | 19.4766397 | 19.4078268 | 19.3950221 |
| Q9Y4W6 | AFG3L2 | 22.7252309 | 22.7800223 | 22.6811806 | 22.6466204 | 22.788496 |
| Q9ULE6 | PALD1 | 19.4943857 | 19.5059333 | 19.581264 | 19.5601719 | 19.5153463 |
| Q12789 | GTF3C1 | 21.0288818 | 21.0840634 | 21.0949916 | 21.0163216 | 21.0937508 |
| Q96EK7 | FAM120B | 18.8811535 | 18.9215911 | 18.8815524 | 18.9193076 | 18.9486596 |
| P78345 | RPP38 | 20.3238749 | 20.3473118 | 20.2980686 | 20.3221024 | 20.29289 |
| Q9NTX5 | ECHDC1 | 22.1936837 | 22.2000636 | 22.2826991 | 22.1792408 | 22.2816766 |
| Q93034 | CUL5 | 21.1956634 | 21.1523331 | 21.2114098 | 21.1160734 | 21.2010487 |
| Q0VDG4 | SCRN3 | 19.6967221 | 19.7317474 | 19.7116641 | 19.7067798 | 19.7946226 |
| Q9GZL7 | WDR12 | 22.0850657 | 22.116439 | 22.1581697 | 22.1106744 | 22.1026462 |
| Q9H7Z6 | KAT8 | 19.8735574 | 19.9818366 | 19.973041 | 19.8265377 | 19.9487547 |
| P15927 | RPA2 | 23.0000687 | 23.1994321 | 23.023031 | 23.011147 | 23.1201414 |
| O00170 | AIP | 23.1576495 | 23.2033989 | 23.212575 | 23.0860145 | 23.1537298 |
| Q15717 | ELAVL1 | 24.2896407 | 24.3451626 | 24.150429 | 24.1676843 | 24.2997121 |
| Q13472 | TOP3A | 19.0001194 | 18.9263297 | 19.0874692 | 18.8967738 | 18.9528708 |
| Q8TEQ8 | PIGO | 18.1535445 | 18.3879453 | 18.3816572 | 18.2812312 | 18.396453 |
| Q92820 | GGH | 22.7917075 | 22.8978355 | 22.8374314 | 22.7517938 | 22.9304026 |
| Q9HD23 | MRS2 | 18.61104 | 18.5934553 | 18.7871156 | 18.6759692 | 18.8060577 |
| Q9NRD1 | FBXO6 | 20.1872169 | 20.222101 | 20.3225828 | 20.2153773 | 20.3972788 |
| P63241 | EIF5A | 26.5940642 | 26.659978 | 26.4441028 | 26.5108417 | 26.5802146 |
| Q9NQZ5 | STARD7 | 19.4222547 | 19.5562953 | 19.7295163 | 19.4570037 | 19.4787485 |

| | | | | | | |
|---|---|---|---|---|---|---|
| P67936 | TPM4 | 26.5504458 | 26.6912308 | 26.3455498 | 26.4682984 | 26.5618681 |
| Q8N0W3 | FCSK | 17.97473 | 17.8006001 | 17.9176622 | 17.8365952 | 17.9476721 |
| Q9NWQ9 | C14orf119 | 18.4194388 | 18.4662832 | 18.4487525 | 18.4289151 | 18.5670784 |
| Q15569 | TESK1 | NA | 16.6896056 | 16.5241321 | NA | 16.6560323 |
| Q9NW38 | FANCL | 15.9220811 | 16.1077688 | 16.2233225 | 15.9327345 | 16.2271348 |
| P51826 | AFF3 | 17.631547 | 17.3876644 | 17.5930809 | 17.5737439 | 17.5420368 |
| Q9NQT8 | KIF13B | 17.8545896 | 17.9989206 | 18.0025416 | 17.8931444 | 17.9707217 |
| Q9Y3E2 | BOLA1 | 19.6747551 | 19.6936051 | 19.3473958 | 19.5496029 | 19.6145133 |
| Q06609 | RAD51 | 20.2364828 | 20.274239 | 20.4300971 | 20.363026 | 20.5238759 |
| P31641 | SLC6A6 | 18.1226038 | 17.9925776 | 18.2458542 | 18.3110033 | 17.9658858 |
| P63096 | GNAI1 | 16.0645232 | 15.8585036 | 16.0156975 | 15.9623667 | 16.0762362 |
| Q49AN0 | CRY2 | 16.6814827 | 16.8242096 | 17.027623 | 16.9189032 | 16.9207071 |
| Q9Y6B2 | EID1 | 17.4176495 | 17.1408364 | 17.4414967 | 17.4245612 | 17.3584333 |
| Q9NR48 | ASH1L | NA | 16.3656284 | 16.0991443 | NA | 16.0739087 |
| P42081 | CD86 | 16.4900861 | 16.9286497 | 17.1995419 | 16.5474947 | 17.2216895 |
| O43189 | PHF1 | 15.9080904 | 16.024934 | 16.2288194 | 16.0565021 | 16.4482207 |
| Q9NYF3 | FAM53C | 13.9254856 | NA | NA | 13.7376077 | NA |
| Q9UQE7 | SMC3 | 22.9449495 | 23.0684133 | 23.0271971 | 22.9245809 | 23.0148633 |
| Q92685 | ALG3 | 20.1080591 | 20.2233154 | 20.0451802 | 20.1890886 | 20.0540623 |
| Q9Y376 | CAB39 | 23.1428268 | 23.1302522 | 23.1786465 | 23.0685289 | 23.2139939 |
| Q7L014 | DDX46 | 22.9545574 | 23.0552049 | 22.9371119 | 22.8797616 | 22.9321134 |
| Q92805 | GOLGA1 | 14.3527574 | 14.3005524 | 14.1543179 | 14.0198236 | 14.2424939 |
| P62847 | RPS24 | 26.0837719 | 25.9626044 | 26.338138 | 25.9961377 | 25.9949206 |
| Q9H3S4 | TPK1 | 19.7669961 | 19.8996138 | 19.9603372 | 19.8482948 | 19.9574152 |
| Q96CB9 | NSUN4 | 20.345103 | 20.4006969 | 20.3888811 | 20.3557668 | 20.3141626 |
| Q9Y277 | VDAC3 | 24.348869 | 24.4281834 | 24.4400935 | 24.3157279 | 24.5071874 |
| Q96RS6 | NUDCD1 | 22.4198893 | 22.3906726 | 22.4461771 | 22.3474725 | 22.3364288 |
| Q14566 | MCM6 | 24.0083157 | 24.0475975 | 23.9526145 | 23.9030221 | 23.9372533 |
| Q86VN1 | VPS36 | 21.0786366 | 21.209542 | 21.1914151 | 21.1508167 | 21.0739733 |
| Q9Y4W2 | LAS1L | 20.7732142 | 20.9219327 | 20.8457173 | 20.7735038 | 20.8721668 |
| P01009 | SERPINA1 | 19.0742857 | 19.3818696 | 19.1507487 | 19.0068161 | 19.220926 |
| P00156 | MT-CYB | 22.3391325 | 22.4356365 | 22.1012133 | 22.3925647 | 22.15132 |
| Q14568 | HSP90AA2P | 11.7051437 | 11.4945237 | 11.5413963 | 11.7635824 | 11.6463323 |
| Q96BW9 | TAMM41 | 19.010049 | 18.941754 | 19.1237691 | 19.1562565 | 19.0844443 |
| O15297 | PPM1D | 20.774586 | 20.98839 | 20.6465346 | 20.6661596 | 20.6283416 |
| Q9UL12 | SARDH | 20.0299091 | 19.9570939 | 20.0622967 | 19.9862886 | 20.0105371 |
| Q9NZC9 | SMARCAL1 | 19.0918464 | 19.3346508 | 19.3360717 | 19.2432086 | 19.2163177 |
| Q9H792 | PEAK1 | NA | 12.2589612 | 12.1457665 | NA | 12.2698589 |
| Q8IXK0 | PHC2 | 19.3288691 | 19.2670904 | 19.4049216 | 19.2933708 | 19.4119654 |
| A2RUC4 | TYW5 | NA | 15.8011307 | 15.5054946 | NA | 15.8031925 |
| O00244 | ATOX1 | 21.8791394 | 21.9434252 | 21.8456367 | 21.8605174 | 22.1090668 |
| Q9BUN8 | DERL1 | 21.9601145 | 22.2722622 | 21.953441 | 22.0793955 | 21.9783238 |

| | | | | | | |
|---|---|---|---|---|---|---|
| Q8NCW5 | NAXE | 21.3799498 | 21.3074384 | 21.4536322 | 21.3941421 | 21.3156849 |
| Q68DH5 | LMBRD2 | 20.6360547 | 20.5380488 | 20.716538 | 20.5223888 | 20.4451508 |
| Q96PM9 | ZNF385A | 19.6969398 | 19.917504 | 19.9692357 | 19.9609081 | 19.7856313 |
| Q13523 | PRPF4B | 21.9737476 | 21.9908333 | 22.0587534 | 21.9402031 | 21.9630377 |
| O75446 | SAP30 | 20.4579576 | 20.2539435 | 20.457988 | 20.3809187 | 20.5206126 |
| O43325 | LYRM1 | 19.165094 | 19.3248255 | 19.5006713 | 19.2101894 | 19.3984083 |
| O60675 | MAFK | 14.2983945 | NA | NA | 14.3082447 | NA |
| Q99549 | MPHOSPH8 | 20.0405299 | 20.1572559 | 20.1356353 | 20.0054811 | 20.1137833 |
| O15527 | OGG1 | 15.1378889 | 14.8412699 | 15.3278831 | 14.9341645 | 14.9287375 |
| Q16527 | CSRP2 | NA | 14.0878613 | 15.9387973 | NA | 14.3511077 |
| Q9H4A6 | GOLPH3 | 21.515094 | 21.5277522 | 21.5678073 | 21.5522186 | 21.5856623 |
| Q9NX46 | ADPRS | 21.0812184 | 21.1639063 | 21.1150469 | 21.0209646 | 21.2462877 |
| Q9NUP9 | LIN7C | 21.0980533 | 21.1885923 | 21.0878337 | 21.1038274 | 21.1160319 |
| P13797 | PLS3 | 16.0323194 | NA | NA | 15.943165 | NA |
| Q92918 | MAP4K1 | 17.7260869 | 17.9459138 | 18.0107749 | 17.7745258 | 17.9660727 |
| O95249 | GOSR1 | 21.4879246 | 21.543212 | 21.4492483 | 21.4259434 | 21.482202 |
| P23284 | PPIB | 26.4221791 | 26.4146035 | 26.3675356 | 26.2900244 | 26.357318 |
| P33992 | MCM5 | 23.8350669 | 23.811453 | 23.8323638 | 23.7388434 | 23.8104055 |
| P04075 | ALDOA | 27.7226381 | 27.909504 | 27.8024275 | 27.6717077 | 27.7698168 |
| Q9NX94 | WBP1L | NA | 15.9864838 | 16.2918243 | NA | 16.068851 |
| Q9NQP4 | PFDN4 | 23.3383481 | 23.4199133 | 23.2424174 | 23.2762207 | 23.493221 |
| O60884 | DNAJA2 | 23.7992542 | 23.9195626 | 23.7594639 | 23.6978218 | 23.8705024 |
| Q8N2A8 | PLD6 | 16.4444151 | 16.9245668 | 17.0319348 | 16.8790639 | 16.7745503 |
| O14966 | RAB29 | 18.7155708 | 18.5607365 | 18.6666711 | 18.5789686 | 18.7581977 |
| O75362 | ZNF217 | 21.0082227 | 21.143338 | 21.2027086 | 21.0646262 | 21.1477061 |
| Q5U4P2 | ASPHD1 | 15.1296341 | 15.1539182 | 14.9241522 | 14.6872082 | 15.0997654 |
| Q6NSJ0 | MYORG | 11.5399499 | NA | NA | 11.9698127 | NA |
| Q8N983 | MRPL43 | 22.5948791 | 22.5607245 | 22.4576912 | 22.5746601 | 22.7907267 |
| Q10471 | GALNT2 | 21.2012013 | 21.1400652 | 21.3194767 | 21.3018109 | 21.214994 |
| Q13137 | CALCOCO2 | 22.5819806 | 22.5269905 | 22.4314408 | 22.5776296 | 22.4291785 |
| P53582 | METAP1 | 22.4291638 | 22.5057845 | 22.5601776 | 22.4363337 | 22.527984 |
| P62328 | TMSB4X | 28.4131398 | 28.5593287 | 28.1615892 | 28.2087229 | 28.5747141 |
| Q99622 | C12orf57 | 18.9631475 | 19.1278288 | 18.8743459 | 19.0383109 | 19.1729431 |
| P06733 | ENO1 | 28.3004185 | 28.3650487 | 28.2625366 | 28.1865577 | 28.3304232 |
| Q96RR1 | TWNK | 20.0690763 | 20.1377014 | 20.0564798 | 19.9950007 | 20.124051 |
| Q93008 | USP9X | 21.5271148 | 21.6364976 | 21.6364037 | 21.4858364 | 21.6392951 |
| P33993 | MCM7 | 24.2015021 | 24.2776262 | 24.2165909 | 24.1077281 | 24.1581227 |
| Q02218 | OGDH | 22.530493 | 22.5506789 | 22.5763012 | 22.5064196 | 22.5720271 |
| Q2TAZ0 | ATG2A | 18.9534079 | 19.0151916 | 19.0349933 | 18.9791125 | 18.9803377 |
| Q5VW36 | FOCAD | 22.4254455 | 22.5163905 | 22.566569 | 22.3963139 | 22.4441516 |
| O75449 | KATNA1 | 20.0133108 | 20.0519234 | 19.9718473 | 19.9789223 | 20.0164704 |
| P61289 | PSME3 | 23.6723454 | 23.7835372 | 23.6679047 | 23.6288734 | 23.5853496 |

| | | | | | | |
|---|---|---|---|---|---|---|
| Q14137 | BOP1 | 21.9482841 | 21.9042299 | 21.931898 | 21.8497431 | 21.9732894 |
| P52735 | VAV2 | 21.0916375 | 21.1710666 | 21.2982615 | 21.1343182 | 21.2127546 |
| Q16204 | CCDC6 | 21.0621686 | 21.2669121 | 20.9677213 | 21.0910741 | 21.1974242 |
| P05023 | ATP1A1 | 24.2987372 | 24.3635895 | 24.3482491 | 24.2219984 | 24.2424989 |
| P56557 | TMEM50B | 18.4156961 | 18.3397671 | 18.4455286 | 18.2314158 | 18.3986738 |
| Q9Y225 | RNF24 | 18.0804071 | 17.8412531 | 17.888312 | 18.2521744 | 17.7810367 |
| O95976 | IGSF6 | 16.1398818 | 15.8799709 | 15.8746041 | 15.8037169 | 15.7960411 |
| Q9Y6N1 | COX11 | 16.8664203 | 16.8092313 | 16.8054886 | 16.9544943 | 16.9580207 |
| Q8NB37 | GATD1 | 19.4956297 | 19.63601 | 19.4089661 | 19.3069617 | 19.620873 |
| Q96DN5 | TBC1D31 | 17.5623792 | 17.6191291 | 17.8157241 | 17.5411354 | 17.452298 |
| P49761 | CLK3 | 19.3329644 | 19.2653465 | 19.5232134 | 19.3860573 | 19.3826775 |
| O95707 | POP4 | 20.6248376 | 20.7208698 | 20.8269014 | 20.7820937 | 20.6608686 |
| Q9BUR5 | APOO | 19.9279117 | 19.9510744 | 20.2690464 | 19.9692357 | 20.0447973 |
| P45954 | ACADSB | 22.8608484 | 22.9039092 | 22.9591568 | 22.8559451 | 23.0525901 |
| Q99973 | TEP1 | 18.7425809 | 18.8816085 | 18.8171814 | 18.7021143 | 18.9430293 |
| Q9GZZ9 | UBA5 | 21.6914748 | 21.8516622 | 21.793859 | 21.5304046 | 21.7919194 |
| Q96I15 | SCLY | 20.4455691 | 20.4199727 | 20.5186479 | 20.4104719 | 20.5083707 |
| P35611 | ADD1 | 21.8754874 | 21.9262687 | 21.9654863 | 21.8931004 | 21.9917707 |
| Q9Y3D9 | MRPS23 | 21.6883211 | 21.7130343 | 21.7073747 | 21.668227 | 21.8053604 |
| P11172 | UMPS | 22.8898063 | 22.9042487 | 22.926102 | 22.8050742 | 22.8994668 |
| Q8IWZ8 | SUGP1 | 20.9504177 | 20.8887148 | 20.9593435 | 20.9039854 | 21.0023174 |
| Q13123 | IK | 22.0741584 | 22.0650496 | 22.0806407 | 22.0018449 | 22.1364858 |
| Q9NWS8 | RMND1 | 20.2753103 | 20.2929639 | 20.2553304 | 20.2332571 | 20.3373825 |
| Q9UN37 | VPS4A | 22.0826319 | 22.0739121 | 22.153911 | 22.138004 | 22.2025083 |
| Q13576 | IQGAP2 | 22.9324105 | 23.0218413 | 23.0295813 | 22.9305193 | 23.0481394 |
| P51452 | DUSP3 | 22.6459222 | 22.8151806 | 22.7406377 | 22.7362803 | 22.868781 |
| O95292 | VAPB | 21.4705278 | 21.5067289 | 21.6345626 | 21.4753271 | 21.546262 |
| Q6PFW1 | PPIP5K1 | 18.1531069 | 18.3333295 | 18.3285453 | 18.3285101 | 18.4268873 |
| P30414 | NKTR | 18.8917409 | 19.068833 | 18.9333534 | 18.9388762 | 19.0013452 |
| Q9Y605 | MRFAP1 | 19.9185831 | 20.071056 | 19.9934283 | 19.9632426 | 19.9997767 |
| Q96LW4 | PRIMPOL | 17.3489262 | 17.5051637 | 17.5994798 | 17.4969573 | 17.3566299 |
| Q6UWZ7 | ABRAXAS1 | 19.9161148 | 19.9529449 | 19.8699078 | 19.8762012 | 19.9233369 |
| Q8N8N7 | PTGR2 | 19.4720332 | 19.5234132 | 19.5037391 | 19.4450197 | 19.7395639 |
| O76080 | ZFAND5 | 19.2191022 | 19.5093316 | 19.3671416 | 19.3794508 | 19.4898958 |
| O60239 | SH3BP5 | 14.7796006 | 15.1984415 | 14.7906991 | 14.77405 | 15.3045594 |
| Q9NY59 | SMPD3 | 15.0155146 | NA | NA | 15.3483979 | NA |
| Q5JPI3 | C3orf38 | 19.0996039 | 19.1527746 | 19.2747462 | 19.1567829 | 19.3585678 |
| Q4KMQ2 | ANO6 | 19.6867995 | 19.661632 | 19.8327401 | 19.7468221 | 19.5889092 |
| Q9HAJ7 | SAP30L | 19.1442897 | 18.975641 | 19.2065126 | 18.9275576 | 19.1393886 |
| Q8WWH5 | TRUB1 | 19.1019242 | 19.1307227 | 19.1479393 | 19.0426839 | 19.1816349 |
| Q92506 | HSD17B8 | 20.9719395 | 20.938408 | 20.8478993 | 20.8557058 | 21.0035509 |
| Q8N5C6 | SRBD1 | 21.0146625 | 21.018353 | 21.1840014 | 20.9866221 | 21.0778678 |

| | | | | | | |
|---|---|---|---|---|---|---|
| Q9NX74 | DUS2 | 19.380486 | 19.5212608 | 19.4641921 | 19.4169027 | 19.5033048 |
| Q9Y5B6 | PAXBP1 | 20.1681245 | 20.2624706 | 20.2884331 | 20.1744024 | 20.3420754 |
| Q13492 | PICALM | 21.9153464 | 22.009618 | 21.9875544 | 21.9598026 | 21.985282 |
| Q99661 | KIF2C | 21.4995577 | 21.518928 | 21.6404444 | 21.5452849 | 21.577588 |
| O95104 | SCAF4 | 21.2875852 | 21.248701 | 21.227053 | 21.1893902 | 21.3171212 |
| Q9HC16 | APOBEC3G | 20.5775061 | 20.6262391 | 20.7459817 | 20.613153 | 20.7301539 |
| Q9HCG7 | GBA2 | 16.7476021 | 16.7876564 | 16.8609619 | 16.580707 | 16.8782285 |
| Q8IXL6 | FAM20C | 16.8810825 | 16.878823 | 17.1363046 | 16.8150979 | 16.9803413 |
| Q15700 | DLG2 | 14.7182795 | NA | NA | 14.5234117 | NA |
| Q16891 | IMMT | 24.3085186 | 24.3361582 | 24.3228964 | 24.2370406 | 24.2939891 |
| Q9Y5P4 | CERT1 | 20.432097 | 20.5245814 | 20.5124083 | 20.5332843 | 20.5428251 |
| Q8N386 | LRRC25 | 19.3844916 | 19.3779453 | 19.6478565 | 19.3790945 | 19.5025783 |
| P19397 | CD53 | 21.1104822 | 21.639335 | 21.6402993 | 21.4261776 | 21.2234415 |
| Q53H80 | AKIRIN2 | 17.2521242 | 17.4085433 | 17.4835651 | 17.4185061 | 17.4869765 |
| P17676 | CEBPB | 19.5991394 | 19.9154781 | 19.6827981 | 19.5978113 | 19.9139608 |
| Q01459 | CTBS | 19.8118583 | 19.8287861 | 19.9425862 | 19.804199 | 19.8466382 |
| Q9BWH2 | FUNDC2 | 21.0185504 | 21.2679409 | 21.4440891 | 21.0232688 | 21.1596002 |
| Q00796 | SORD | 22.9935693 | 23.0076422 | 22.9927418 | 22.8958171 | 23.0647782 |
| Q12834 | CDC20 | 20.1076709 | 20.1498229 | 20.1428316 | 20.0364665 | 20.0245625 |
| Q9UJX3 | ANAPC7 | 20.5180441 | 20.5175206 | 20.616567 | 20.5450484 | 20.5779905 |
| Q9NWF9 | RNF216 | 17.6492314 | 17.7140012 | 17.5899822 | 17.6029814 | 17.6539093 |
| Q86WV6 | STING1 | 21.5059932 | 21.7179097 | 21.6011838 | 21.501043 | 21.5721648 |
| P04424 | ASL | 21.3676027 | 21.5406315 | 21.51572 | 21.3015726 | 21.4769044 |
| Q8NFH5 | NUP35 | 21.3369249 | 21.3676102 | 21.3880871 | 21.2554154 | 21.5458722 |
| Q6ZRI6 | C15orf39 | 18.3882971 | 18.4734143 | 18.3903202 | 18.3890699 | 18.554273 |
| P49840 | GSK3A | 20.5203579 | 20.5474977 | 20.5778308 | 20.4541751 | 20.7392351 |
| Q02447 | SP3 | 20.0646641 | 20.3192541 | 20.3547601 | 20.1777749 | 20.2612367 |
| Q14571 | ITPR2 | 20.1216222 | 20.313675 | 20.3505262 | 20.1775936 | 20.3506634 |
| Q5BKU9 | OXLD1 | 17.0522834 | 16.855528 | 17.0520471 | 17.0389962 | 17.0499618 |
| Q9Y519 | TMEM184B | NA | 16.1612794 | 15.8252626 | NA | 15.9390615 |
| Q6PJI9 | WDR59 | 18.3970881 | 18.6052259 | 18.4889894 | 18.4925913 | 18.579754 |
| Q9NUB1 | ACSS1 | 20.4705833 | 20.6112669 | 20.6502195 | 20.5049419 | 20.5466331 |
| Q9H1P3 | OSBPL2 | 18.7632283 | 18.6904025 | 18.8755497 | 18.6690357 | 18.6497558 |
| Q06546 | GABPA | 20.7213279 | 20.8002391 | 20.7109797 | 20.7161428 | 20.7618996 |
| Q96AH0 | NABP1 | 12.8272256 | NA | NA | 12.1473768 | NA |
| Q01581 | HMGCS1 | 22.4993427 | 22.6968833 | 22.658969 | 22.5285722 | 22.7788252 |
| P07305 | H1-0 | 21.088026 | 21.2977933 | 21.4680502 | 21.1629205 | 21.2515472 |
| Q7Z7L7 | ZER1 | 18.9955317 | 18.9045972 | 19.1399043 | 18.9715642 | 19.0553072 |
| Q5VVJ2 | MYSM1 | 18.694414 | 18.8435267 | 18.6642298 | 18.6973112 | 18.6805705 |
| P78537 | BLOC1S1 | 19.8957366 | 20.0147369 | 20.0285674 | 19.9018352 | 19.8899428 |
| Q3SXM5 | HSDL1 | 19.8816197 | 20.233759 | 19.9132657 | 19.9606692 | 20.039509 |
| Q9Y657 | SPIN1 | 20.2048418 | 20.2291787 | 20.3143733 | 20.0630652 | 20.3144865 |

| | | | | | | |
|---|---|---|---|---|---|---|
| Q15019 | SEPTIN2 | 23.2125122 | 23.2063452 | 23.2733835 | 23.2032474 | 23.2446256 |
| Q15070 | OXA1L | 22.2866607 | 22.3961567 | 22.3212212 | 22.2622913 | 22.2709827 |
| Q6PKC3 | TXNDC11 | 18.4027085 | 18.6443548 | 18.4234957 | 18.3208553 | 18.271928 |
| Q86X10 | RALGAPB | 21.2786136 | 21.2844925 | 21.3375287 | 21.223492 | 21.3003095 |
| Q96K62 | ZBTB45 | 16.7002979 | 16.7917814 | 16.5705063 | 16.6733442 | 16.4296359 |
| P40763 | STAT3 | 21.0465115 | 21.2095561 | 21.3362453 | 20.9594396 | 21.152186 |
| O94915 | FRYL | 20.6693967 | 20.765607 | 20.8079569 | 20.6361851 | 20.7720172 |
| Q13535 | ATR | 19.408927 | 19.3862348 | 19.4483925 | 19.3782911 | 19.4182993 |
| B1ANS9 | WDR64 | 17.0216868 | NA | NA | 17.8721945 | NA |
| Q3KQZ1 | SLC25A35 | 18.2219223 | 18.4406228 | 18.3254106 | 18.3699635 | 18.4271147 |
| Q9UK45 | LSM7 | 23.1562914 | 23.2889596 | 23.1255035 | 23.1491456 | 23.1851228 |
| Q8N2F6 | ARMC10 | 20.127784 | 20.3392153 | 20.1829078 | 20.107748 | 20.0899593 |
| O95336 | PGLS | 24.1255898 | 24.2376024 | 24.1687328 | 24.1747159 | 24.2033442 |
| Q9H7D0 | DOCK5 | 21.1724388 | 21.2406379 | 21.2992057 | 21.165686 | 21.2352698 |
| Q9BTD8 | RBM42 | 22.3299243 | 22.326789 | 22.2427666 | 22.3085212 | 22.3775785 |
| P42785 | PRCP | 21.2087414 | 21.3688357 | 21.3636644 | 21.3602181 | 21.3683278 |
| Q14249 | ENDOG | 19.6463882 | 19.529189 | 19.4747534 | 19.4565985 | 19.7612245 |
| P38935 | IGHMBP2 | 18.527236 | 18.7055383 | 18.7852656 | 18.6688432 | 18.7231526 |
| Q86TM6 | SYVN1 | 20.9257747 | 21.0983153 | 20.9886291 | 20.8171089 | 20.9019421 |
| Q96B02 | UBE2W | 18.96675 | 19.0920409 | 19.3313596 | 19.0865261 | 19.2563157 |
| Q8N511 | TMEM199 | 18.5838918 | 18.6896716 | 18.6997648 | 18.6528595 | 18.6799913 |
| O94953 | KDM4B | 19.6072384 | 19.7955242 | 19.8169149 | 19.5061106 | 19.8790701 |
| Q9UN19 | DAPP1 | 16.8879688 | 17.2966121 | 17.1229288 | 17.2451259 | 17.4590475 |
| O95562 | SFT2D2 | 21.2117949 | 21.2859497 | 21.2049938 | 21.2833802 | 21.1936441 |
| P50443 | SLC26A2 | 18.1962411 | 18.1327384 | 18.2799665 | 18.3602445 | 18.1599897 |
| P16278 | GLB1 | 21.5106469 | 21.6029619 | 21.6978617 | 21.5316635 | 21.6696539 |
| Q9P2K8 | EIF2AK4 | 20.5971745 | 20.6896643 | 20.7769984 | 20.6350766 | 20.6526719 |
| P27986 | PIK3R1 | 21.3007681 | 21.365401 | 21.4084006 | 21.2866047 | 21.3240211 |
| Q8IY18 | SMC5 | 21.0258471 | 21.0656297 | 21.1071648 | 20.9391837 | 21.0262718 |
| Q969K4 | ABTB1 | 16.2443816 | 16.4256404 | 16.4066283 | 16.4967016 | 16.3565467 |
| Q96DC7 | TMCO6 | 18.6862614 | 18.1144927 | 18.9227522 | 18.6663733 | 18.2908957 |
| A6NCS6 | C2orf72 | 13.4189788 | 13.723183 | 13.5684607 | 13.6553196 | 13.9267081 |
| Q9H9D4 | ZNF408 | 17.4592446 | 17.5716043 | 17.6932449 | 17.2629163 | 17.4083292 |
| Q13825 | AUH | 18.3997708 | 18.4050983 | 18.6443471 | 18.5959901 | 18.5969865 |
| Q8IZ40 | RCOR2 | 17.2518914 | 17.4165459 | 17.4991724 | 17.3305534 | 17.3578452 |
| Q96LX8 | ZNF597 | NA | 14.0899488 | 14.0501689 | NA | 14.1232962 |
| Q9UK73 | FEM1B | 18.234689 | 17.8562442 | 18.0051205 | 17.9603477 | 18.0848904 |
| Q96E52 | OMA1 | 17.7757154 | 17.7420052 | 17.9581304 | 17.9120438 | 17.8076084 |
| O75131 | CPNE3 | 22.3085771 | 22.4408119 | 22.4987339 | 22.364684 | 22.5126743 |
| P15735 | PHKG2 | 19.5065567 | 19.4638237 | 19.5777 | 19.3582682 | 19.5867212 |
| O60858 | TRIM13 | 18.8583335 | 19.0735122 | 19.0776371 | 18.9406073 | 19.1750315 |
| P13807 | GYS1 | 22.036064 | 22.0756812 | 22.121446 | 22.0105721 | 22.1458861 |

| | | | | | | |
|---|---|---|---|---|---|---|
| Q96FS4 | SIPA1 | 20.6559736 | 20.8406039 | 20.7736152 | 20.6422536 | 20.7380424 |
| Q99836 | MYD88 | 20.7436701 | 20.8679121 | 20.9974041 | 20.7844041 | 20.8696552 |
| Q9BYE7 | PCGF6 | 18.7307149 | 19.0366813 | 18.8179366 | 18.7874018 | 18.9616828 |
| Q96T76 | MMS19 | 20.8209784 | 20.7965291 | 20.9546905 | 20.7351292 | 20.7999395 |
| P15311 | EZR | 25.0118293 | 25.0119399 | 24.9555915 | 24.8847146 | 25.0152915 |
| Q96KR1 | ZFR | 21.6473772 | 21.762655 | 21.7017692 | 21.6555623 | 21.7570645 |
| Q96A33 | CCDC47 | 22.3410365 | 22.2997735 | 22.3119824 | 22.3067887 | 22.3525972 |
| Q6YP21 | KYAT3 | 21.360616 | 21.2658224 | 21.3332725 | 21.3087716 | 21.4091222 |
| Q9UBT2 | UBA2 | 24.1073525 | 24.1426633 | 24.1164663 | 24.0176873 | 24.1431744 |
| Q9BWS9 | CHID1 | 22.811033 | 22.7540886 | 22.8400332 | 22.8099194 | 22.8389584 |
| Q13438 | OS9 | 20.8640763 | 20.87012 | 20.7888778 | 20.8240891 | 20.8353877 |
| Q9Y6D6 | ARFGEF1 | 22.1171187 | 22.1082408 | 22.1450013 | 22.0431588 | 22.1772227 |
| P34910 | EVI2B | 20.7667121 | 20.8362988 | 20.9346683 | 20.7702369 | 20.8656478 |
| Q96DA6 | DNAJC19 | 21.2716762 | 21.3454458 | 21.4851439 | 21.3199265 | 21.3170468 |
| P49903 | SEPHS1 | 21.3223812 | 21.3984958 | 21.3780123 | 21.4048761 | 21.5293113 |
| Q17RS7 | GEN1 | 17.8257362 | 17.9748757 | 18.0455754 | 17.9520918 | 17.9088769 |
| Q16134 | ETFDH | 20.6647339 | 20.7241495 | 20.8480715 | 20.7453987 | 20.8530263 |
| Q9ULL5 | PRR12 | 19.3854731 | 19.4793293 | 19.4859955 | 19.3846249 | 19.413322 |
| P78318 | IGBP1 | 20.7384431 | 20.7717106 | 20.9252154 | 20.7723723 | 20.8941989 |
| Q96HN2 | AHCYL2 | 18.3891826 | 18.3779625 | 18.4081014 | 18.3825387 | 18.540396 |
| Q9Y3Z3 | SAMHD1 | 24.2247656 | 24.4089104 | 24.3314772 | 24.1718701 | 24.4106044 |
| P55211 | CASP9 | 18.6572266 | 18.9580746 | 18.7950135 | 18.7405185 | 18.7699092 |
| Q14699 | RFTN1 | 20.7422166 | 20.9114837 | 20.8981029 | 20.7495504 | 20.854795 |
| Q93052 | LPP | 17.6775963 | 17.8368429 | 17.8392433 | 17.845817 | 18.036811 |
| P17275 | JUNB | 19.215318 | 19.3522748 | 19.2097518 | 19.2105705 | 19.3764397 |
| O00488 | ZNF593 | 19.4300451 | 19.4306781 | 19.4996714 | 19.4921425 | 19.5988499 |
| Q5BJD5 | TMEM41B | 20.2683632 | 20.5906414 | 20.4573295 | 20.6206215 | 20.5458263 |
| Q6Q0C0 | TRAF7 | 18.5028319 | 18.5500459 | 18.4602741 | 18.5116367 | 18.7023954 |
| Q16270 | IGFBP7 | 16.6460048 | 16.4486439 | 16.4100271 | 16.6990023 | 16.6523024 |
| P05164 | MPO | 17.8788753 | 17.7261073 | 17.613683 | 17.9178617 | 17.9092019 |
| Q2QGD7 | ZXDC | 17.8472522 | 18.2634177 | 18.0115122 | 18.3395564 | 17.5945273 |
| Q8N4N3 | KLHL36 | 16.9041973 | 16.8248619 | 16.7234319 | 17.3079454 | 16.9539217 |
| Q6ZTR5 | CFAP47 | 16.8009266 | 17.2578266 | 17.1730926 | 17.1560326 | 17.0809632 |
| Q9HC62 | SENP2 | 14.5270474 | 14.8331548 | 14.7804187 | 14.7380418 | 14.9225902 |
| O15258 | RER1 | 22.3432481 | 22.4640624 | 22.3336786 | 22.2930185 | 22.1872452 |
| O94919 | ENDOD1 | 19.9310526 | 20.1787043 | 20.1714025 | 20.0839277 | 20.0812539 |
| Q9Y3A2 | UTP11 | 20.5390167 | 20.7052712 | 20.7542049 | 20.6311894 | 20.6937008 |
| Q9H5V9 | STEEP1 | 19.8491608 | 19.8728232 | 19.9686614 | 19.8943906 | 19.9687535 |
| Q13618 | CUL3 | 21.7697681 | 21.7722002 | 21.867019 | 21.7351541 | 21.8329154 |
| Q9NX58 | LYAR | 22.1981837 | 22.1920578 | 22.2691095 | 22.1471149 | 22.0950992 |
| Q7Z3V4 | UBE3B | 18.2482132 | 18.4231297 | 18.3004866 | 18.3080158 | 18.3306348 |
| Q96NC0 | ZMAT2 | 21.8263681 | 22.0320311 | 22.0694838 | 21.9686744 | 22.0836937 |

| | | | | | | |
|---|---|---|---|---|---|---|
| Q9Y286 | SIGLEC7 | 19.1581021 | 19.2652639 | 19.4912827 | 19.2563961 | 19.4921963 |
| Q6IA17 | SIGIRR | 17.7854045 | 17.9516744 | 17.9559642 | 17.6834506 | 17.9404399 |
| Q8WUA4 | GTF3C2 | 20.0580759 | 19.9902778 | 20.1996376 | 19.9900219 | 20.1451488 |
| P20700 | LMNB1 | 26.0636703 | 26.1745156 | 26.0774121 | 25.9854266 | 26.0816345 |
| Q9H000 | MKRN2 | 20.4013425 | 20.4739064 | 20.6466721 | 20.5364355 | 20.5865094 |
| Q7Z407 | CSMD3 | NA | 11.8107475 | 10.9333368 | NA | 11.6819909 |
| Q9GZM8 | NDEL1 | 21.7098582 | 22.136154 | 21.940011 | 21.5373154 | 21.8502488 |
| Q9NY26 | SLC39A1 | 19.126732 | 19.0620331 | 18.5297776 | 18.8210514 | 18.3743606 |
| Q8NEP9 | ZNF555 | 15.9054329 | 15.8620473 | 16.1676217 | 16.2247184 | 16.3363651 |
| Q5JS54 | PSMG4 | 19.6118854 | 19.4693626 | 19.6369734 | 19.3246148 | 19.7524773 |
| Q9P2J8 | ZNF624 | 15.2301515 | 15.6213983 | 15.2676701 | 15.4167933 | 15.0780859 |
| P49674 | CSNK1E | 17.3113911 | 17.5716123 | 17.52619 | 17.7048183 | 17.5705422 |
| O15078 | CEP290 | NA | 22.9548127 | 22.4543633 | NA | 22.4070859 |
| Q6IEG0 | SNRNP48 | 17.6239627 | 17.647734 | 17.837003 | 17.6350672 | 17.6429742 |
| Q9NPF0 | CD320 | 19.5849262 | 19.6272294 | 19.9264627 | 19.7470336 | 19.6835578 |
| Q9NX76 | CMTM6 | 20.850361 | 20.8653133 | 20.8790786 | 20.811033 | 20.7103083 |
| O76062 | TM7SF2 | 19.1598607 | 18.9467611 | 18.833959 | 19.0407653 | 18.8771635 |
| Q96G23 | CERS2 | 21.0376029 | 21.2181053 | 21.5248192 | 21.2824577 | 21.2497642 |
| O75915 | ARL6IP5 | 23.054928 | 23.28316 | 23.2695805 | 23.1518723 | 23.1087573 |
| Q5ZPR3 | CD276 | 19.3892926 | 19.7153128 | 19.71103 | 19.5518316 | 19.7529585 |
| P36956 | SREBF1 | 17.8586424 | 17.7835711 | 18.0995886 | 17.7973152 | 17.9007028 |
| P51790 | CLCN3 | 18.3257188 | 18.5996033 | 18.6815441 | 18.3762377 | 18.529631 |
| Q6NUJ5 | PWWP2B | 16.4630729 | 16.5655845 | 16.5177544 | 16.3418598 | 16.50923 |
| P62314 | SNRPD1 | 24.4187561 | 24.4500703 | 24.5782652 | 24.3027435 | 24.2738872 |
| Q969M3 | YIPF5 | 21.2021821 | 21.1979526 | 21.3822002 | 21.2358546 | 21.3617993 |
| Q8N9L9 | ACOT4 | 17.598811 | 17.8470197 | 17.7778937 | 17.6464408 | 18.0056975 |
| Q68DK2 | ZFYVE26 | 18.9297026 | 18.9338866 | 18.9389133 | 18.9622666 | 18.9886761 |
| Q9NRS6 | SNX15 | 18.4071951 | 18.6558744 | 18.6250352 | 18.6104522 | 18.6788301 |
| O75808 | CAPN15 | 19.0838582 | 19.0880879 | 19.266906 | 19.2258575 | 19.1153095 |
| P62834 | RAP1A | 21.1006307 | 21.153277 | 21.3100789 | 21.0310719 | 21.0447561 |
| Q92802 | N4BP2L2 | 19.1796779 | 19.1136625 | 19.1511039 | 19.045288 | 19.1045643 |
| O60671 | RAD1 | 19.4914831 | 19.4877863 | 19.7209437 | 19.5802314 | 19.4852786 |
| Q9NRN7 | AASDHPPT | 23.1185546 | 22.966906 | 23.1313409 | 22.9781011 | 23.184863 |
| P14618 | PKM | 27.1351548 | 27.2924433 | 27.1538659 | 27.0267597 | 27.1781127 |
| Q7Z7A1 | CNTRL | 19.358581 | 19.5593256 | 19.3532091 | 19.2725558 | 19.387669 |
| Q9BX68 | HINT2 | 22.6324609 | 22.734567 | 22.7360697 | 22.6873915 | 22.8144334 |
| P60953 | CDC42 | 24.6665202 | 24.6764821 | 24.8190181 | 24.6454543 | 24.7776629 |
| Q8N5M9 | JAGN1 | 21.3912667 | 21.399904 | 21.4670754 | 21.3769921 | 21.3643801 |
| P49643 | PRIM2 | 20.4674324 | 20.4573559 | 20.6184996 | 20.3903424 | 20.5646751 |
| O75489 | NDUFS3 | 23.210168 | 23.273821 | 23.3163987 | 23.1887449 | 23.3244482 |
| Q92990 | GLMN | 20.918186 | 21.0340066 | 21.0752034 | 20.8894951 | 21.0328449 |
| Q12899 | TRIM26 | 19.6354835 | 19.6904409 | 19.7709668 | 19.6899744 | 19.7666917 |

| | | | | | | |
|---|---|---|---|---|---|---|
| Q8IYU8 | MICU2 | 19.6796992 | 19.5275715 | 19.6941392 | 19.6102427 | 19.5556447 |
| Q3KQU3 | MAP7D1 | 20.6774383 | 20.7371331 | 20.8928648 | 20.6943144 | 20.823269 |
| Q9UNF1 | MAGED2 | 20.5857702 | 20.5783629 | 20.6100115 | 20.5063647 | 20.5828429 |
| Q8NBS9 | TXNDC5 | 23.7361762 | 23.7452499 | 23.6268121 | 23.628534 | 23.7935506 |
| Q9BRX2 | PELO | 20.3481749 | 20.4065501 | 20.408491 | 20.2463514 | 20.4542961 |
| Q9UBL3 | ASH2L | 21.7905694 | 21.8983893 | 21.9624047 | 21.8218773 | 21.8610055 |
| Q27J81 | INF2 | 20.8908534 | 21.0078961 | 20.9788109 | 20.8629235 | 20.9750994 |
| Q9NZI8 | IGF2BP1 | 22.2360318 | 22.3447873 | 22.3944428 | 22.2030311 | 22.3291325 |
| Q96IG2 | FBXL20 | 20.4067475 | 20.464435 | 20.5359455 | 20.4076424 | 20.5109916 |
| O15439 | ABCC4 | 20.5707405 | 20.603808 | 20.6353641 | 20.5217649 | 20.6075518 |
| Q9BZG1 | RAB34 | 21.1184083 | 21.3115919 | 21.1812985 | 21.1314216 | 21.2245328 |
| Q8WXW3 | PIBF1 | 19.4512473 | 19.5121961 | 19.4116113 | 19.476436 | 19.4804471 |
| Q8TCU6 | PREX1 | 21.8966999 | 21.9250567 | 22.0024549 | 21.8999856 | 21.93733 |
| Q8WWC4 | MAIP1 | 21.2220568 | 21.348146 | 21.3119082 | 21.2615235 | 21.372541 |
| P18887 | XRCC1 | 21.4093097 | 21.4471398 | 21.4455886 | 21.4451166 | 21.5002616 |
| Q14643 | ITPR1 | 21.5314802 | 21.5546628 | 21.5952096 | 21.5015127 | 21.5828819 |
| Q9UKM7 | MAN1B1 | 19.4965581 | 19.5395331 | 19.4503872 | 19.453443 | 19.5297868 |
| Q9P287 | BCCIP | 21.9093074 | 21.9088292 | 21.9528539 | 21.8825469 | 21.9721293 |
| Q969X6 | UTP4 | 21.0945054 | 21.0724222 | 21.1971129 | 21.0965217 | 21.1183803 |
| P24386 | CHM | 19.3761546 | 19.398939 | 19.4282097 | 19.469872 | 19.4299454 |
| P61160 | ACTR2 | 25.3953189 | 25.4688526 | 25.4436042 | 25.3317808 | 25.4047506 |
| Q8N163 | CCAR2 | 23.8784047 | 23.9195796 | 23.8570894 | 23.8288558 | 23.843225 |
| Q9H9A6 | LRRC40 | 21.5106743 | 21.581588 | 21.6449168 | 21.544444 | 21.6135282 |
| Q6KC79 | NIPBL | 21.5348318 | 21.6520827 | 21.6331068 | 21.5101676 | 21.5827459 |
| O94905 | ERLIN2 | 24.0256306 | 24.0522004 | 23.9758868 | 23.9402458 | 24.0355863 |
| Q92974 | ARHGEF2 | 23.1769274 | 23.2611188 | 23.2744016 | 23.1380021 | 23.303534 |
| Q96HS1 | PGAM5 | 23.9023657 | 23.8696743 | 23.8327566 | 23.8049379 | 23.8524737 |
| Q9H7D7 | WDR26 | 21.2541263 | 21.2857153 | 21.2921024 | 21.2463653 | 21.3285946 |
| P09467 | FBP1 | 21.5402212 | 21.4694654 | 21.4697139 | 21.5014372 | 21.590086 |
| P21359 | NF1 | 19.4313247 | 19.4325347 | 19.4646474 | 19.4015562 | 19.5121094 |
| Q9Y2G8 | DNAJC16 | 19.2551049 | 19.3314626 | 19.4476317 | 19.2703594 | 19.2903973 |
| Q96BX8 | MOB3A | 22.9074781 | 22.8836267 | 22.9793863 | 22.8490513 | 22.8560006 |
| Q9NZ08 | ERAP1 | 20.6920886 | 20.5594986 | 20.6774675 | 20.5759413 | 20.6676566 |
| Q5JSL3 | DOCK11 | 20.6859385 | 20.7695879 | 20.7817532 | 20.7027608 | 20.7989025 |
| Q96FW1 | OTUB1 | 23.0774118 | 23.1169559 | 23.1551974 | 23.0906371 | 23.1483829 |
| Q96JH7 | VCPIP1 | 20.5555055 | 20.6326166 | 20.585796 | 20.5529567 | 20.6342747 |
| Q6PKG0 | LARP1 | 22.8164191 | 22.8777122 | 22.7912251 | 22.7473344 | 22.8404177 |
| O94906 | PRPF6 | 23.1766966 | 23.2557785 | 23.2022224 | 23.151394 | 23.2417434 |
| P39748 | FEN1 | 23.7946677 | 23.8338893 | 23.8072917 | 23.742885 | 23.7978438 |
| O60216 | RAD21 | 21.7154442 | 21.6373932 | 21.7188674 | 21.6148788 | 21.7030809 |
| Q14C86 | GAPVD1 | 21.5628457 | 21.6757972 | 21.6350719 | 21.5726352 | 21.6786656 |
| Q8WXF1 | PSPC1 | 23.1916558 | 23.2932424 | 23.1202068 | 23.1338149 | 23.2113008 |

| | | | | | | |
|---|---|---|---|---|---|---|
| Q96QU8 | XPO6 | 20.6266864 | 20.6920002 | 20.6737548 | 20.6864072 | 20.717982 |
| P17987 | TCP1 | 25.4793458 | 25.5375724 | 25.5159459 | 25.3852653 | 25.4768741 |
| Q9UIF9 | BAZ2A | 20.7914087 | 20.8401314 | 20.8581871 | 20.7760146 | 20.9001158 |
| Q5VIR6 | VPS53 | 20.178533 | 20.3021315 | 20.3153875 | 20.2965187 | 20.272558 |
| Q7Z2Z2 | EFL1 | 20.705993 | 20.7708815 | 20.8589628 | 20.7472106 | 20.8111947 |
| Q9H2P0 | ADNP | 22.4227352 | 22.4134684 | 22.4473036 | 22.3781067 | 22.4582313 |
| O43172 | PRPF4 | 22.1930442 | 22.2015013 | 22.27563 | 22.1441682 | 22.2498746 |
| Q9Y2T2 | AP3M1 | 21.6001646 | 21.7093756 | 21.7970663 | 21.7110555 | 21.7425848 |
| Q5T447 | HECTD3 | 20.6914967 | 20.7871258 | 20.740876 | 20.677483 | 20.790698 |
| Q9Y4X5 | ARIH1 | 20.915731 | 21.0545607 | 21.0699549 | 20.9268794 | 20.9783481 |
| Q86YS7 | C2CD5 | 21.715946 | 21.7917365 | 21.7441874 | 21.6556627 | 21.7498717 |
| Q8NCJ5 | SPRYD3 | 18.6245995 | 18.7010051 | 18.6472466 | 18.6132808 | 18.7272965 |
| Q9UBB9 | TFIP11 | 20.2727137 | 20.3156446 | 20.3280279 | 20.2496038 | 20.2941297 |
| P55199 | ELL | 21.269326 | 21.3519106 | 21.3743402 | 21.3341903 | 21.2811923 |
| Q9P2T1 | GMPR2 | 22.6197588 | 22.6740678 | 22.777659 | 22.6630548 | 22.7362321 |
| Q6NVY1 | HIBCH | 21.8660612 | 21.835407 | 21.8815639 | 21.7173051 | 21.8617807 |
| Q6IA86 | ELP2 | 20.3647685 | 20.403941 | 20.6452307 | 20.4551848 | 20.5734015 |
| P10606 | COX5B | 24.7581874 | 24.8263188 | 24.6334345 | 24.6811596 | 24.8181654 |
| Q92598 | HSPH1 | 24.3667839 | 24.5019526 | 24.3942443 | 24.3609543 | 24.4455709 |
| P20585 | MSH3 | 19.8170084 | 19.967526 | 20.0026204 | 19.9050998 | 19.9634331 |
| Q9NPG3 | UBN1 | 19.1202573 | 19.2499768 | 19.2560442 | 19.1184797 | 19.1472402 |
| P56524 | HDAC4 | 20.8340113 | 21.0515025 | 20.8942651 | 20.8717277 | 21.0506285 |
| Q7Z2E3 | APTX | 19.2591932 | 19.2930634 | 19.3465025 | 19.3208734 | 19.3258162 |
| Q9BUB7 | TMEM70 | 21.088869 | 21.0210159 | 21.0487131 | 20.8676407 | 21.0394047 |
| Q8TE77 | SSH3 | 20.3288165 | 20.4381839 | 20.5031266 | 20.3527464 | 20.5890305 |
| Q8NDI1 | EHBP1 | 19.4568736 | 19.4544834 | 19.450664 | 19.3355599 | 19.5714201 |
| Q9UFC0 | LRWD1 | 19.8172907 | 19.9386988 | 19.9841177 | 19.923734 | 20.0463532 |
| Q6ZRQ5 | MMS22L | 18.3506639 | 18.5117548 | 18.5517989 | 18.3726446 | 18.4750459 |
| Q96ND0 | FAM210A | 20.746964 | 20.9899372 | 20.9905153 | 20.8474631 | 20.9466999 |
| Q08257 | CRYZ | 20.4299298 | 20.434486 | 20.4647982 | 20.4508593 | 20.5965105 |
| P21675 | TAF1 | 18.9689692 | 19.305086 | 19.1804256 | 19.0796747 | 19.1090158 |
| P54803 | GALC | 19.9924063 | 19.7869238 | 19.9675137 | 19.8277579 | 19.9137848 |
| Q15059 | BRD3 | 21.4267213 | 21.588009 | 21.6025963 | 21.4872353 | 21.5968871 |
| O00629 | KPNA4 | 23.5539012 | 23.648327 | 23.5081595 | 23.4518879 | 23.6453439 |
| Q9Y3B9 | RRP15 | 21.1456586 | 21.2286042 | 21.0053493 | 21.0773285 | 21.1376939 |
| P25791 | LMO2 | 18.4131019 | 18.7020186 | 18.7693407 | 18.5762293 | 18.5357115 |
| Q5VU43 | PDE4DIP | 22.078872 | 22.3936117 | 22.1211765 | 22.137012 | 22.1555661 |
| Q8IYB5 | SMAP1 | 20.7090999 | 20.5428014 | 20.800414 | 20.4924954 | 20.7201113 |
| Q8WV99 | ZFAND2B | 18.6963474 | 18.8991453 | 18.9689389 | 18.7434545 | 18.8811807 |
| Q5BJH2 | TMEM128 | 18.5072538 | 18.5929577 | 18.5432417 | 18.6954831 | 18.5250918 |
| O60930 | RNASEH1 | 19.5782098 | 19.5112898 | 19.3231827 | 19.5283319 | 19.3923297 |
| P13747 | HLA-E | 18.1834518 | 18.2663407 | 18.3092913 | 18.3649224 | 18.2339577 |

| | | | | | | |
|---|---|---|---|---|---|---|
| Q96F63 | CCDC97 | 20.5766315 | 20.7877034 | 20.4916936 | 20.5607769 | 20.7510974 |
| P31260 | HOXA10 | 19.0547247 | 19.3333274 | 19.3921682 | 19.1537223 | 19.3799958 |
| P53805 | RCAN1 | 18.9040697 | 19.1083497 | 18.8709088 | 19.0094879 | 19.0658504 |
| P06132 | UROD | 21.8618585 | 21.886035 | 21.8529676 | 21.8004616 | 21.9708823 |
| Q9H1K0 | RBSN | 17.0911999 | 17.0571601 | 17.2018483 | 17.0356534 | 17.3874199 |
| Q9NZZ3 | CHMP5 | 20.9836483 | 21.1899895 | 20.8934812 | 20.9645157 | 21.0868889 |
| Q9H160 | ING2 | 18.3617157 | 18.277776 | 18.365514 | 18.282406 | 18.4293225 |
| Q6ZS30 | NBEAL1 | 18.5512231 | 18.8428869 | 18.8562055 | 18.6861429 | 18.4325288 |
| Q9H6R6 | ZDHHC6 | 18.1711455 | 18.4173232 | 18.3471657 | 18.4074731 | 18.3262974 |
| Q96IK1 | BOD1 | 19.6591614 | 19.5076825 | 19.6103527 | 19.5068308 | 19.6950215 |
| P62072 | TIMM10 | 21.178171 | 21.3748461 | 21.2680404 | 21.2112623 | 21.3530643 |
| Q96NA8 | TSNARE1 | 15.2310496 | 15.5774805 | 15.7318936 | 15.5957345 | 15.8908261 |
| Q8TET4 | GANC | 16.7305181 | 16.268771 | 16.4889723 | 16.3869561 | 16.3595996 |
| Q96GE4 | CEP95 | 16.7923405 | 16.9739683 | 16.4894843 | 16.7758124 | 16.5447234 |
| O60783 | MRPS14 | 20.0049641 | 20.0358028 | 20.1720772 | 20.0872284 | 19.9927814 |
| Q9Y250 | LZTS1 | 20.7324571 | 20.7785614 | 20.704681 | 20.7082659 | 20.5345449 |
| Q14296 | FASTK | 15.3962583 | 15.6419967 | 15.5334904 | 15.8450011 | 15.6261461 |
| Q14781 | CBX2 | 15.94212 | 16.116655 | 16.0216459 | 16.1962806 | 15.8761579 |
| Q9H6A9 | PCNX3 | 16.4613761 | 16.6664956 | 16.5920844 | 16.4744521 | 16.746494 |
| Q5TEC6 | H3-7 | 25.8382686 | 25.9896921 | 25.9562936 | 25.8504414 | 25.6184501 |
| P10109 | FDX1 | 19.3437659 | 19.2709881 | 19.068643 | 19.3164958 | 19.2136387 |
| Q8IYL3 | C1orf174 | 18.9986037 | 18.6815226 | 18.4130759 | 18.6285423 | 18.7154073 |
| Q9NYG5 | ANAPC11 | 18.1036976 | 17.5981759 | 17.8559309 | 18.3408336 | 18.2643396 |
| Q9H6K1 | ILRUN | NA | 14.1784392 | 14.4319004 | NA | 14.6940534 |
| O75417 | POLQ | 14.6723493 | 15.0042886 | 15.5422594 | 14.9578797 | 15.2146307 |
| Q9Y219 | JAG2 | NA | 13.8201547 | 13.8392672 | NA | 13.039038 |
| Q9Y2R2 | PTPN22 | 14.4046595 | 15.1399897 | 15.5846656 | 14.7824308 | 15.6384637 |
| Q9P1Z0 | ZBTB4 | 15.2768951 | 16.0756203 | 15.6779318 | 14.980967 | 15.6990546 |
| Q5T5X7 | BEND3 | 15.4141956 | 15.4100476 | 15.5696277 | 15.6244205 | 15.5277501 |
| Q86UY6 | NAA40 | 18.9049075 | 18.855684 | 19.0394161 | 18.8059322 | 19.0312183 |
| O75182 | SIN3B | 17.6016339 | 17.7605471 | 17.7424588 | 17.5372725 | 17.7192641 |
| P09211 | GSTP1 | 25.9861774 | 26.0404568 | 25.8954728 | 25.87368 | 25.9780866 |
| Q8IX12 | CCAR1 | 22.7104266 | 22.7347856 | 22.7022857 | 22.660287 | 22.7708688 |
| Q9NXG2 | THUMPD1 | 21.1603946 | 21.2973492 | 21.1928808 | 21.1808354 | 21.2577628 |
| Q92541 | RTF1 | 21.479289 | 21.5357129 | 21.5654412 | 21.4707374 | 21.5422493 |
| O75400 | PRPF40A | 22.7050691 | 22.7639166 | 22.6768804 | 22.6345346 | 22.7550138 |
| P33121 | ACSL1 | 22.0905968 | 22.1473152 | 22.1765917 | 22.1030097 | 22.2359765 |
| P30876 | POLR2B | 22.4493713 | 22.3956463 | 22.4712468 | 22.4068851 | 22.4470059 |
| P20618 | PSMB1 | 24.7085835 | 24.7918759 | 24.7549256 | 24.68427 | 24.8422257 |
| P30419 | NMT1 | 22.2471811 | 22.2095194 | 22.2121294 | 22.1983831 | 22.2572593 |
| O60488 | ACSL4 | 22.6511249 | 22.7956083 | 22.8058081 | 22.6646598 | 22.824951 |
| Q9Y5A9 | YTHDF2 | 21.9435884 | 22.0095583 | 22.155447 | 21.9942605 | 22.0502046 |

| | | | | | | |
|---|---|---|---|---|---|---|
| Q9NVM9 | INTS13 | 21.1431468 | 21.2572059 | 21.1990749 | 21.1184809 | 21.2523824 |
| O94925 | GLS | 20.7968097 | 20.9736694 | 20.8231553 | 20.7854669 | 20.9635806 |
| Q8TCG1 | CIP2A | 21.5675044 | 21.7383352 | 21.7082171 | 21.6313926 | 21.6861029 |
| Q99497 | PARK7 | 26.2396947 | 26.4054166 | 26.1528729 | 26.1305719 | 26.4131581 |
| Q9H446 | RWDD1 | 19.8599664 | 20.086572 | 19.7138575 | 19.8135564 | 20.1311188 |
| Q12882 | DPYD | 19.4565747 | 19.8643079 | 20.0194919 | 19.6227482 | 20.0092903 |
| P04271 | S100B | 14.3028687 | 14.717514 | 13.8779001 | 14.0769603 | 14.2081947 |
| Q96EX1 | SMIM12 | 20.3653395 | 20.3055999 | 20.3834086 | 20.2544722 | 20.2299972 |
| Q9NVP4 | DZANK1 | 16.9334389 | NA | NA | 17.2868556 | NA |
| P10244 | MYBL2 | 19.0346106 | 19.0574951 | 18.7890283 | 18.9410307 | 19.0865145 |
| Q9Y5R8 | TRAPPC1 | 20.7591533 | 20.7919111 | 20.7201829 | 20.8051031 | 21.0086155 |
| Q5VTE6 | ANGEL2 | 17.7254323 | 17.9516436 | 18.2716985 | 18.0466754 | 18.0251336 |
| Q8IUH5 | ZDHHC17 | 18.1619579 | 18.2119011 | 18.0968513 | 18.1104085 | 18.3427765 |
| Q9UGY1 | NOL12 | 19.6426525 | 20.0578269 | 20.1043606 | 19.9164462 | 19.8317065 |
| Q9Y281 | CFL2 | 17.8335381 | 17.8668771 | 17.9487866 | 17.782134 | 17.9287792 |
| Q9BZQ8 | NIBAN1 | 17.5831158 | 17.5170381 | 17.5377042 | 17.5520342 | 17.5521125 |
| Q765P7 | MTSS2 | 19.9932475 | 20.114148 | 20.035082 | 19.9533704 | 20.0203065 |
| Q92793 | CREBBP | 19.3976153 | 19.4790717 | 19.4124795 | 19.3936245 | 19.4636027 |
| O00165 | HAX1 | 19.5075172 | 19.7070091 | 19.7021342 | 19.5555757 | 19.5455723 |
| O75143 | ATG13 | 20.2769029 | 20.3383398 | 20.2504867 | 20.2367689 | 20.278551 |
| O75132 | ZBED4 | 18.758147 | 18.8303594 | 18.6451366 | 18.708154 | 18.6456051 |
| Q8N8R5 | C2orf69 | 18.9658558 | 18.9337466 | 18.9348583 | 18.865815 | 18.9898393 |
| O60664 | PLIN3 | 24.7967481 | 24.8881708 | 24.7299186 | 24.7192478 | 24.8034377 |
| A0AVT1 | UBA6 | 21.8340994 | 21.9155824 | 21.9109518 | 21.8080623 | 21.9930203 |
| Q9NWB6 | ARGLU1 | 22.7964925 | 22.877139 | 22.7008547 | 22.6347935 | 22.7109326 |
| Q9H147 | DNTTIP1 | 20.5617134 | 20.6143506 | 20.6249092 | 20.6118243 | 20.6326075 |
| P43246 | MSH2 | 22.4961036 | 22.5910114 | 22.5442806 | 22.4819771 | 22.5944338 |
| P46379 | BAG6 | 22.3114187 | 22.390537 | 22.2898774 | 22.2640766 | 22.3645999 |
| Q16851 | UGP2 | 23.4148626 | 23.4419819 | 23.4231754 | 23.4037603 | 23.4780453 |
| Q08AM6 | VAC14 | 20.4362784 | 20.4877835 | 20.4022454 | 20.4236314 | 20.4347685 |
| Q6IN85 | PPP4R3A | 21.562107 | 21.6452478 | 21.5950401 | 21.5409418 | 21.6164939 |
| Q6L8Q7 | PDE12 | 21.5470823 | 21.6381523 | 21.5399032 | 21.5574752 | 21.6404729 |
| Q01433 | AMPD2 | 20.8375113 | 20.9506485 | 20.9190587 | 20.8402021 | 21.0340853 |
| Q8TEV9 | SMCR8 | 19.184151 | 19.3568751 | 19.4799604 | 19.2897707 | 19.4509573 |
| O75340 | PDCD6 | 23.1268628 | 23.1618541 | 23.136124 | 23.1127652 | 23.0990112 |
| Q9NR33 | POLE4 | 20.1973083 | 20.3560597 | 20.2206618 | 20.2677205 | 20.4237655 |
| P62861 | FAU | 24.5507509 | 24.828972 | 24.8078218 | 24.5943656 | 24.7856501 |
| Q99471 | PFDN5 | 23.4923378 | 23.6353899 | 23.3915589 | 23.4649177 | 23.6450353 |
| Q9P299 | COPZ2 | 16.8652297 | 17.0030356 | 17.1299511 | 16.7412822 | 17.1790623 |
| Q13064 | MKRN3 | 15.0283584 | NA | NA | 14.8971247 | NA |
| Q9UHN6 | CEMIP2 | NA | 18.8775443 | 18.9490762 | NA | 18.7303197 |
| Q53S58 | TMEM177 | 16.3273136 | NA | NA | 16.3123752 | NA |

| | | | | | | |
|---|---|---|---|---|---|---|
| Q9C0A6 | SETD5 | 18.077347 | 18.45844 | 18.222269 | 18.1550608 | 18.0638077 |
| Q9BSJ2 | TUBGCP2 | 21.0629635 | 21.0953232 | 21.154264 | 21.0670612 | 21.1421094 |
| P23511 | NFYA | 19.1630651 | 19.3370958 | 19.1457832 | 19.1912432 | 19.2670181 |
| Q8N3C0 | ASCC3 | 22.6068791 | 22.6733131 | 22.6659837 | 22.5745482 | 22.6644372 |
| Q5TBK1 | N4BP2L1 | 17.9969864 | 18.0140469 | 17.8868328 | 18.0852918 | 18.1088176 |
| O75354 | ENTPD6 | 16.0064166 | 16.06627 | 16.0667817 | 16.222342 | 16.0280847 |
| Q8N594 | MPND | 15.1901651 | 15.2370265 | 15.4501397 | 15.3976918 | 15.062025 |
| Q96CP6 | GRAMD1A | 16.5333798 | 16.4567857 | 16.900061 | 16.4075159 | 16.6489195 |
| Q9BWG4 | SSBP4 | 19.0239246 | 18.9956549 | 19.0953433 | 19.0360015 | 19.2049844 |
| P15291 | B4GALT1 | 17.9099042 | 18.0686137 | 17.946997 | 17.7116718 | 17.9598633 |
| Q68D10 | SPTY2D1 | 18.7419502 | 18.9831191 | 19.0231727 | 18.817507 | 18.7037313 |
| Q96EL2 | MRPS24 | 20.5439275 | 20.5644185 | 20.5783885 | 20.5071958 | 20.5831819 |
| Q9UDY2 | TJP2 | 17.6378534 | 17.7514983 | 17.758992 | 17.7386978 | 17.8015278 |
| O14773 | TPP1 | 24.1362212 | 24.2422886 | 24.1625464 | 24.0738811 | 24.1684679 |
| Q5HYJ3 | FAM76B | 19.5950826 | 19.7949578 | 19.7230271 | 19.6574742 | 19.8714153 |
| O43752 | STX6 | 20.0688203 | 20.2127215 | 20.1320672 | 19.9969611 | 20.0716254 |
| P46976 | GYG1 | 22.8684049 | 23.0150427 | 23.1016733 | 22.905529 | 22.8538495 |
| Q96HW7 | INTS4 | 20.5208291 | 20.5688301 | 20.6519323 | 20.476871 | 20.5719986 |
| Q6P087 | RPUSD3 | 19.8964151 | 19.8368374 | 19.9225011 | 19.8378268 | 19.8938036 |
| Q6ZUJ8 | PIK3AP1 | 22.4409881 | 22.5093908 | 22.5515994 | 22.438864 | 22.5381416 |
| Q9HA77 | CARS2 | 21.5630312 | 21.5270563 | 21.5290183 | 21.4920328 | 21.5927818 |
| P17858 | PFKL | 23.8865332 | 23.9158861 | 23.9415622 | 23.8486494 | 23.977706 |
| Q92947 | GCDH | 22.4003504 | 22.452583 | 22.389089 | 22.3434956 | 22.4367257 |
| O00291 | HIP1 | 21.4979104 | 21.5279473 | 21.501112 | 21.4538532 | 21.574051 |
| Q7L523 | RRAGA | 22.079885 | 22.0783053 | 22.0727541 | 22.0084144 | 22.027984 |
| Q86U38 | NOP9 | 20.3891837 | 20.2932429 | 20.4680764 | 20.3515946 | 20.4566941 |
| P26358 | DNMT1 | 22.7158687 | 22.7657921 | 22.7670085 | 22.6732454 | 22.7508745 |
| O60313 | OPA1 | 22.033828 | 22.1056176 | 22.0986753 | 22.0464185 | 22.105365 |
| P14550 | AKR1A1 | 22.2772301 | 22.2579663 | 22.2002273 | 22.2176965 | 22.2141541 |
| P17813 | ENG | 19.9971849 | 19.9338456 | 19.8825571 | 20.0152207 | 19.9314521 |
| Q8NBZ7 | UXS1 | 20.075017 | 20.0947144 | 20.2058922 | 20.1491678 | 20.0891219 |
| Q9UMR2 | DDX19B | 22.9730248 | 22.9979963 | 22.9756238 | 22.8822518 | 22.9393843 |
| Q04917 | YWHAH | 23.9485336 | 23.9614204 | 23.9443544 | 23.8040825 | 23.9401196 |
| Q8NCN5 | PDPR | 20.5491749 | 20.6776912 | 20.6802226 | 20.6032013 | 20.6775609 |
| P49591 | SARS1 | 23.2196873 | 23.2956893 | 23.3389927 | 23.2175228 | 23.3194075 |
| Q15843 | NEDD8 | 23.8086922 | 23.895786 | 23.7959729 | 23.667195 | 23.9522007 |
| Q9UBN7 | HDAC6 | 20.4399734 | 20.6135559 | 20.6610365 | 20.5252217 | 20.6264239 |
| Q9BQS8 | FYCO1 | 19.2830879 | 19.3094556 | 19.4227686 | 19.25216 | 19.4320464 |
| Q6P1Q0 | LETMD1 | 19.1507125 | 19.1749901 | 19.1831507 | 19.1589489 | 19.1834495 |
| Q14790 | CASP8 | 19.4918398 | 19.6089711 | 19.6492087 | 19.505334 | 19.7523576 |
| O96011 | PEX11B | 21.1575871 | 21.0882406 | 21.1875764 | 21.0511887 | 21.1581772 |
| Q9NRR4 | DROSHA | 18.4613282 | 18.5245594 | 18.7167667 | 18.3289755 | 18.4439274 |

| | | | | | | |
|---|---|---|---|---|---|---|
| Q9Y4F3 | MARF1 | 18.8331094 | 18.7002128 | 18.9064505 | 18.8543486 | 18.9011444 |
| Q9Y5S2 | CDC42BPB | 20.3007475 | 20.8368404 | 20.6958767 | 20.4431624 | 20.5879616 |
| Q96DY7 | MTBP | 15.4975325 | 15.5985659 | 15.4332432 | 15.4047928 | 15.5383148 |
| Q8IY33 | MICALL2 | 17.6375714 | 17.6341062 | 17.5320822 | 17.8462187 | 17.6261404 |
| P10914 | IRF1 | 15.8040295 | NA | NA | 15.6812287 | NA |
| P62341 | SELENOT | 19.5896673 | 19.5983906 | 19.7095539 | 19.627577 | 19.6211834 |
| Q92628 | KIAA0232 | 17.0652586 | 17.492375 | 17.0947795 | 17.1179204 | 17.3259179 |
| Q9NUJ3 | TCP11L1 | 17.3801553 | 17.8501779 | 17.6779745 | 17.550232 | 17.7169706 |
| Q9NRZ9 | HELLS | 21.1940872 | 21.2373634 | 21.279377 | 21.2091648 | 21.334635 |
| Q9HCE3 | ZNF532 | 12.8483767 | 13.2179906 | 13.0497729 | 12.7023919 | 12.066609 |
| Q86UQ4 | ABCA13 | NA | 17.0110822 | 17.4525521 | NA | 17.3207935 |
| Q12906 | ILF3 | 25.5737186 | 25.6452494 | 25.5270379 | 25.4641161 | 25.5324604 |
| Q9P265 | DIP2B | 20.7298866 | 20.8188856 | 20.9195812 | 20.7838938 | 20.9574459 |
| Q9Y3P9 | RABGAP1 | 19.6153303 | 19.6909403 | 19.7596016 | 19.596072 | 19.6770712 |
| Q5VTB9 | RNF220 | 17.9300794 | 18.1564922 | 18.2454671 | 18.2001574 | 18.1720785 |
| Q9Y3Q3 | TMED3 | 22.2665725 | 22.4521009 | 22.185039 | 22.2385865 | 22.1532115 |
| Q86XN8 | MEX3D | 15.3175802 | 15.8359254 | 15.8660222 | 16.8641768 | 15.9004172 |
| Q5T2R2 | PDSS1 | 17.9921325 | 17.9956117 | 17.978672 | 17.965214 | 18.0418026 |
| P62495 | ETF1 | 23.402776 | 23.407449 | 23.4031654 | 23.3609523 | 23.4035606 |
| P23368 | ME2 | 23.086896 | 23.071251 | 23.0928986 | 23.0120694 | 23.2032327 |
| P48200 | IREB2 | 20.8757568 | 20.9901496 | 21.0376241 | 20.8621128 | 21.0644123 |
| P16403 | H1-2 | 26.9932883 | 27.1044762 | 27.3939851 | 26.9573852 | 26.9707972 |
| P55081 | MFAP1 | 22.2836237 | 22.3019217 | 22.2361341 | 22.1326186 | 22.2678548 |
| Q8N5K1 | CISD2 | 22.0931583 | 22.0918646 | 22.2459042 | 22.0524795 | 22.1915898 |
| Q8IV32 | CCDC71 | NA | 14.7973599 | 15.1907573 | NA | 14.5682677 |
| O75204 | TMEM127 | 17.9630512 | 17.8496986 | 18.322763 | 18.0340384 | 18.2683675 |
| Q9H0Q0 | CYRIA | 19.6889518 | 20.0205627 | 20.0457069 | 19.8820318 | 20.0259801 |
| Q6IAA8 | LAMTOR1 | 21.8922822 | 21.897177 | 22.0010466 | 21.9030672 | 21.9588136 |
| P46527 | CDKN1B | 18.6085275 | 18.7332106 | 18.6607926 | 18.5976291 | 18.7859063 |
| Q8IUW5 | RELL1 | 19.2596052 | 19.3478136 | 19.4196306 | 19.5253962 | 19.4131841 |
| Q9UH17 | APOBEC3B | 20.0932835 | 20.1932084 | 20.2689503 | 20.1355863 | 20.2788383 |
| Q9BZD2 | SLC29A3 | 17.6723463 | 17.8953647 | 17.8583635 | 17.9909813 | 17.9276404 |
| Q7LG56 | RRM2B | 21.1829874 | 21.154217 | 21.2861288 | 21.1347111 | 21.1690729 |
| P46019 | PHKA2 | 19.6766163 | 19.6815699 | 19.8460898 | 19.6521443 | 19.768584 |
| Q8IX18 | DHX40 | 17.955747 | 18.0461397 | 18.1373908 | 17.9824148 | 18.1786311 |
| Q76I76 | SSH2 | 17.8140844 | 17.8297907 | 17.7662377 | 17.7777226 | 17.8434465 |
| Q9NVX7 | KBTBD4 | 18.0351016 | 18.1383117 | 18.0844536 | 18.135348 | 18.042749 |
| Q12800 | TFCP2 | 23.8223227 | 23.8848277 | 23.856678 | 23.791398 | 23.8097947 |
| Q6PJG6 | BRAT1 | 21.0656294 | 21.0378669 | 21.0973255 | 20.9912067 | 21.1710762 |
| Q15155 | NOMO1 | 24.9497109 | 24.95286 | 24.8680511 | 24.8869155 | 24.9762088 |
| Q8IZ83 | ALDH16A1 | 22.5692909 | 22.6292035 | 22.605542 | 22.5410151 | 22.6100812 |
| P53992 | SEC24C | 22.3615415 | 22.3901446 | 22.4003457 | 22.3074714 | 22.4842039 |

| | | | | | | |
|---|---|---|---|---|---|---|
| Q8N6T3 | ARFGAP1 | 21.0212204 | 21.134695 | 21.0140534 | 21.0160436 | 21.0920443 |
| Q99848 | EBNA1BP2 | 22.2882563 | 22.4061352 | 22.4774128 | 22.2904143 | 22.3221986 |
| O75616 | ERAL1 | 19.1831907 | 19.0453296 | 19.1649212 | 19.1315769 | 19.1261254 |
| Q9BUH6 | PAXX | 20.8703466 | 21.0860888 | 20.9157674 | 20.8361694 | 21.0846954 |
| O43684 | BUB3 | 23.7969876 | 23.6473967 | 23.7808752 | 23.6746239 | 23.7135726 |
| Q6N063 | OGFOD2 | 18.9823481 | 18.9805992 | 19.041031 | 18.8848126 | 19.0166364 |
| P23258 | TUBG1 | 22.1466074 | 22.3754584 | 22.2851771 | 22.2419802 | 22.3720875 |
| Q587I9 | SFT2D3 | 18.8334091 | 18.8948011 | 19.0360131 | 19.0437317 | 18.7795874 |
| Q9ULG6 | CCPG1 | 21.7056718 | 21.4741004 | 21.379746 | 21.606108 | 21.3807374 |
| Q9UH92 | MLX | 16.7588205 | 16.8072536 | 16.7667136 | 16.9176002 | 16.3866966 |
| O75146 | HIP1R | 13.8714262 | 13.7606966 | 13.9823566 | 13.8988832 | 13.9558824 |
| Q6NUN9 | ZNF746 | 16.0131519 | 16.4726509 | 15.7705916 | 16.1413954 | 16.1706158 |
| Q9BXS6 | NUSAP1 | 21.0047298 | 21.1085125 | 21.1987664 | 20.9675717 | 21.0021093 |
| Q5JPB2 | ZNF831 | 17.9518545 | 18.0987665 | 18.2329647 | 18.1214525 | 17.4828008 |
| Q7Z412 | PEX26 | 16.8413771 | NA | NA | 16.9377082 | NA |
| Q7L5Y6 | DET1 | NA | 15.3236358 | 15.2291265 | NA | 15.165811 |
| Q96EU7 | C1GALT1C1 | 15.1504001 | NA | NA | 15.034264 | NA |
| Q9BT23 | LIMD2 | 20.9078902 | 20.9598887 | 21.2578757 | 20.711922 | 21.0577266 |
| P83369 | LSM11 | 16.6765973 | 16.5200624 | 16.3315806 | 16.4186955 | 16.6083633 |
| Q92599 | SEPTIN8 | 19.4837081 | 19.6938614 | 19.6347014 | 19.6084845 | 19.7069407 |
| Q96IR2 | ZNF845 | 18.9719629 | 19.1284412 | 19.1690576 | 19.3305479 | 18.9932942 |
| Q9BVM4 | GGACT | 19.2555228 | 19.191227 | 19.2697261 | 19.2699032 | 19.4270733 |
| Q7Z7C8 | TAF8 | 19.2155522 | 19.2007594 | 19.1242508 | 19.1656552 | 19.1652703 |
| Q9UER7 | DAXX | 19.7953247 | 19.9434879 | 19.896477 | 19.8290897 | 19.847454 |
| Q15528 | MED22 | 19.7423666 | 19.9000919 | 19.8268333 | 19.757177 | 19.819291 |
| P60866 | RPS20 | 26.2773837 | 26.1792928 | 26.1843692 | 26.05462 | 26.1724415 |
| Q8NBQ5 | HSD17B11 | 22.8063814 | 22.9047321 | 22.9485029 | 22.6881793 | 22.8309103 |
| O75381 | PEX14 | 21.0198699 | 21.0627405 | 21.0539912 | 20.9898453 | 21.0982338 |
| Q8TB61 | SLC35B2 | 20.4701949 | 20.4688615 | 20.5101446 | 20.4422878 | 20.5003609 |
| Q16795 | NDUFA9 | 24.5642166 | 24.6201987 | 24.6809858 | 24.5009017 | 24.5573993 |
| Q9ULH0 | KIDINS220 | 18.4693947 | 18.575976 | 18.4718536 | 18.6182361 | 18.4880851 |
| Q7Z3U7 | MON2 | 19.9871996 | 19.9712117 | 20.1097733 | 19.9840546 | 20.0903936 |
| P33176 | KIF5B | 22.8898973 | 22.8972107 | 22.8622486 | 22.7557344 | 22.8548171 |
| P23634 | ATP2B4 | 21.5947158 | 21.6508846 | 21.6588186 | 21.5623111 | 21.6326795 |
| Q96RQ3 | MCCC1 | 21.4693376 | 21.5922625 | 21.5131486 | 21.4969296 | 21.5877734 |
| Q96D71 | REPS1 | 21.4611454 | 21.6327449 | 21.5052202 | 21.4949211 | 21.5538404 |
| O60684 | KPNA6 | 22.088647 | 22.2111154 | 22.1815618 | 22.1025637 | 22.166774 |
| P24752 | ACAT1 | 24.2484846 | 24.3359774 | 24.2474416 | 24.2239356 | 24.3228245 |
| Q96KQ7 | EHMT2 | 23.1477232 | 23.3031687 | 23.2392897 | 23.1871162 | 23.1895624 |
| P62829 | RPL23 | 25.4801998 | 25.5969408 | 25.5836137 | 25.4599094 | 25.6309878 |
| P46531 | NOTCH1 | 20.0089965 | 20.219877 | 20.038679 | 19.9152133 | 20.0707676 |
| Q0VGL1 | LAMTOR4 | 20.6859222 | 20.6699771 | 20.5828383 | 20.7319798 | 20.8487116 |

| | | | | | | |
|---|---|---|---|---|---|---|
| Q8WUJ0 | STYX | 17.675366 | 17.6112211 | 17.6722052 | 17.6454843 | 17.7130937 |
| Q13432 | UNC119 | 18.7669262 | 19.0602097 | 19.077181 | 18.8143676 | 18.9379813 |
| Q15049 | MLC1 | 23.3841972 | 23.2402524 | 23.0955889 | 23.2302466 | 23.1058349 |
| P58546 | MTPN | 23.4786839 | 23.4203332 | 23.2878646 | 23.2175382 | 23.5496751 |
| Q96BD0 | SLCO4A1 | 15.3187607 | 15.326841 | 15.4280967 | 15.4281253 | 15.4488596 |
| Q9H4L5 | OSBPL3 | 16.5647555 | 16.6260428 | 16.1347172 | 16.235521 | 16.5956169 |
| P20591 | MX1 | 13.5101713 | 13.1721725 | 13.2046636 | 13.5488866 | 13.3282407 |
| P09104 | ENO2 | 18.2288888 | NA | NA | 18.3540035 | NA |
| P0C7X2 | ZNF688 | 12.7750519 | NA | NA | 12.9865779 | NA |
| Q8N9N7 | LRRC57 | 20.4728791 | 20.5893052 | 20.6459823 | 20.5517853 | 20.6474196 |
| Q9HB71 | CACYBP | 24.8587181 | 24.8710805 | 24.8780921 | 24.7700372 | 24.9162241 |
| Q07960 | ARHGAP1 | 23.9417187 | 24.1000849 | 24.0714307 | 23.9140501 | 24.1057762 |
| Q96JJ7 | TMX3 | 22.5354388 | 22.7156012 | 22.5486589 | 22.517054 | 22.6164358 |
| O75864 | PPP1R37 | 18.0348189 | 17.9659134 | 18.0410789 | 17.9508026 | 17.9786106 |
| Q16222 | UAP1 | 21.4432772 | 21.4767196 | 21.4124416 | 21.3877882 | 21.3926485 |
| Q9BUP3 | HTATIP2 | 21.4668294 | 21.5222852 | 21.5076092 | 21.4557868 | 21.4690342 |
| Q96T37 | RBM15 | 21.7099925 | 21.8195331 | 21.7659209 | 21.6593642 | 21.7280806 |
| P55209 | NAP1L1 | 24.059523 | 24.0733615 | 24.0748962 | 24.0540109 | 24.2143165 |
| P07099 | EPHX1 | 21.7485475 | 21.8125558 | 21.8941216 | 21.7976337 | 21.8526052 |
| Q5T5C0 | STXBP5 | 14.0902222 | NA | NA | 14.1282276 | NA |
| Q9BXC9 | BBS2 | 16.3645594 | 16.1063237 | 15.9182255 | 15.71915 | 16.2314241 |
| O95980 | RECK | 16.301861 | 16.7018116 | 16.1433243 | 16.1409317 | 16.5386556 |
| P62253 | UBE2G1 | 21.187808 | 21.2535158 | 21.0890282 | 21.0942845 | 21.2767964 |
| Q9Y2V2 | CARHSP1 | 21.3501023 | 21.4412003 | 21.2379191 | 21.2877551 | 21.4616744 |
| Q5UCC4 | EMC10 | 19.644078 | 19.5869748 | 19.6322523 | 19.6187749 | 19.6248704 |
| Q9BSK4 | FEM1A | 17.4677845 | 17.292167 | 17.5241182 | 17.4596539 | 17.6825184 |
| P31321 | PRKAR1B | 19.0002748 | 19.1471842 | 19.3198408 | 19.1190365 | 19.1323451 |
| O43278 | SPINT1 | 17.6935833 | 18.04163 | 17.8942682 | 17.703306 | 17.8109916 |
| Q9P0S2 | COX16 | 18.0678321 | 17.9521198 | 18.1361645 | 18.0457095 | 18.0892788 |
| P00558 | PGK1 | 27.0058065 | 27.1653053 | 27.0316697 | 26.8968727 | 27.0555293 |
| O43598 | DNPH1 | 23.7089548 | 23.7127481 | 23.7509188 | 23.6698466 | 23.7677209 |
| Q9NWU2 | GID8 | 22.9266622 | 23.0023349 | 22.9679324 | 22.8128284 | 23.0195874 |
| Q5T280 | SPOUT1 | 20.8008783 | 20.8341554 | 20.8594229 | 20.6708058 | 20.8521243 |
| Q9H0R1 | AP5M1 | 20.4279677 | 20.3606386 | 20.4489321 | 20.3724044 | 20.4041802 |
| Q9NZM3 | ITSN2 | 20.5377542 | 20.7203139 | 20.674272 | 20.5839618 | 20.6646993 |
| Q9H2K8 | TAOK3 | 22.4874027 | 22.5732804 | 22.587452 | 22.4496023 | 22.4795259 |
| Q9Y6M1 | IGF2BP2 | 23.0934263 | 23.162529 | 23.1320551 | 23.0336013 | 23.1847554 |
| O15091 | PRORP | 20.1782384 | 20.2208524 | 20.2825553 | 20.2043932 | 20.2425528 |
| Q7L0Y3 | TRMT10C | 22.0319936 | 22.0279878 | 21.9926427 | 21.9971146 | 21.9954172 |
| Q5W0V3 | FHIP2A | 19.4960252 | 19.542333 | 19.5900862 | 19.500675 | 19.6181333 |
| Q969R5 | L3MBTL2 | 19.2923656 | 19.2868498 | 19.4420606 | 19.334376 | 19.3150786 |
| Q9ULI0 | ATAD2B | 19.4246264 | 19.5727992 | 19.5272495 | 19.4208413 | 19.4937335 |

| | | | | | | |
|---|---|---|---|---|---|---|
| P28482 | MAPK1 | 23.5512451 | 23.7096125 | 23.6174435 | 23.5318763 | 23.59532 |
| Q13427 | PPIG | 21.1405076 | 21.1602308 | 21.2115992 | 21.0605313 | 21.2199486 |
| Q8WW12 | PCNP | 23.0235183 | 22.9935265 | 22.9852317 | 22.903401 | 23.0388233 |
| Q3LXA3 | TKFC | 22.0236893 | 22.0939815 | 22.1501764 | 21.95487 | 22.0892981 |
| Q96AE7 | TTC17 | 18.356595 | 18.3561297 | 18.3922994 | 18.3078225 | 18.4687128 |
| Q01813 | PFKP | 24.2652071 | 24.3974218 | 24.3879766 | 24.2464855 | 24.4426668 |
| O95295 | SNAPIN | 20.4841321 | 20.6143212 | 20.5805304 | 20.4867087 | 20.586363 |
| Q9H875 | PRKRIP1 | 20.3264041 | 20.5154893 | 20.3644238 | 20.4941038 | 20.4649427 |
| Q9NRG0 | CHRAC1 | 21.6528293 | 21.4248815 | 21.6723791 | 21.607375 | 21.6446944 |
| Q86UT8 | CENATAC | 17.5163555 | 17.5848513 | 17.29716 | 17.4483328 | 17.8660235 |
| Q9UMX5 | NENF | 21.8062787 | 21.9667657 | 21.698054 | 21.7550723 | 22.0917413 |
| A6NK58 | LIPT2 | 16.9092315 | 17.0319261 | 16.6882101 | 16.8128076 | 16.7770601 |
| O00763 | ACACB | NA | 15.2322135 | 15.6369618 | NA | 15.2731497 |
| P54278 | PMS2 | 18.9649271 | 19.1917154 | 19.2290567 | 19.1127691 | 19.2100625 |
| Q14919 | DRAP1 | 22.4800427 | 22.6239688 | 22.5500065 | 22.4175243 | 22.5129939 |
| P09651 | HNRNPA1 | 27.5075231 | 27.5466595 | 27.4708608 | 27.4147353 | 27.4858238 |
| Q96KP1 | EXOC2 | 21.1541426 | 21.2150833 | 21.1974198 | 21.1386697 | 21.1949565 |
| Q5JRX3 | PITRM1 | 21.9807864 | 21.9746607 | 22.0258401 | 21.9715197 | 22.0433533 |
| P41214 | EIF2D | 19.7099495 | 19.7778862 | 19.8157949 | 19.7281149 | 19.8237183 |
| Q9NRR8 | CDC42SE1 | 18.748904 | 18.1803324 | 18.1833059 | 18.4652467 | 17.9633172 |
| H7BZ55 | CROCC2 | 15.2707735 | NA | NA | 14.6212126 | NA |
| Q86VR2 | RETREG3 | 19.9894394 | 20.0969993 | 20.1020059 | 20.0149354 | 20.1425412 |
| P11166 | SLC2A1 | 23.0908959 | 23.193699 | 23.2097496 | 22.9367894 | 23.1405514 |
| P32519 | ELF1 | 21.3734182 | 21.4575296 | 21.3657487 | 21.3395046 | 21.446511 |
| Q8NFH3 | NUP43 | 21.1150013 | 21.1337778 | 21.1650366 | 21.1587688 | 21.2479969 |
| Q0PNE2 | ELP6 | 20.0972082 | 20.0568767 | 20.0742201 | 19.967305 | 20.0463135 |
| Q8IY22 | CMIP | 19.5486872 | 19.6574059 | 19.6075424 | 19.5203494 | 19.6619634 |
| Q8N122 | RPTOR | 20.2912948 | 20.3716675 | 20.3287846 | 20.2756367 | 20.3446297 |
| O95747 | OXSR1 | 22.5456706 | 22.6979419 | 22.6255042 | 22.5011245 | 22.7233002 |
| Q15022 | SUZ12 | 20.3654259 | 20.4932778 | 20.5046526 | 20.4579711 | 20.4344124 |
| Q16836 | HADH | 23.3597512 | 23.4961422 | 23.477001 | 23.3463607 | 23.5160294 |
| P00403 | MT-CO2 | 25.2510868 | 25.3399046 | 25.3772048 | 25.2126158 | 25.4268446 |
| Q56VL3 | OCIAD2 | 17.9075032 | 17.6678331 | 17.9265454 | 17.7620781 | 17.7581406 |
| Q01085 | TIAL1 | 21.1071513 | 21.0501833 | 21.021912 | 20.9826006 | 21.1355487 |
| Q99594 | TEAD3 | 19.2756653 | 19.2262889 | 19.3562887 | 19.2441738 | 19.1812165 |
| P30040 | ERP29 | 26.1128066 | 26.0918645 | 25.816755 | 26.0004259 | 26.0683872 |
| Q8TCT7 | SPPL2B | 20.5127854 | 20.381344 | 20.3403477 | 20.4800876 | 20.3767603 |
| Q99470 | SDF2 | 17.9169816 | 17.6140712 | 18.0985151 | 17.8934439 | 18.2965736 |
| P06454 | PTMA | 25.8508117 | 26.0702877 | 25.6366027 | 25.7665486 | 25.8352286 |
| P07814 | EPRS1 | 24.5922231 | 24.6279063 | 24.5836383 | 24.5345548 | 24.6130321 |
| Q86Y79 | PTRH1 | 19.6775772 | 19.7934005 | 19.641729 | 19.7291235 | 19.6495515 |
| Q4KWH8 | PLCH1 | 20.6934988 | 20.892847 | 20.6423186 | 20.5476993 | 21.0588096 |

| | | | | | | |
|---|---|---|---|---|---|---|
| Q8NHV4 | NEDD1 | 18.3663515 | 18.4088507 | 18.3652088 | 18.2968249 | 18.264478 |
| Q9Y336 | SIGLEC9 | 15.4688811 | NA | NA | 15.793267 | NA |
| Q9Y259 | CHKB | 20.0086984 | 19.9661661 | 19.9592231 | 20.0699055 | 19.763624 |
| Q6NS38 | ALKBH2 | 16.64796 | 17.0751326 | 17.0492409 | 16.9957667 | 17.0021891 |
| Q9Y6R9 | CCDC61 | 18.6352291 | 18.6787838 | 18.5771513 | 18.5388004 | 18.4944102 |
| Q9UPS6 | SETD1B | 18.4794312 | 18.5012944 | 18.443954 | 18.4628212 | 18.5057708 |
| Q96QE5 | TEFM | 18.6549352 | 18.4817868 | 18.7580342 | 18.7094467 | 18.7423364 |
| Q9BX10 | GTPBP2 | 19.1504582 | 19.1997243 | 19.1876209 | 19.1613895 | 19.1356535 |
| Q9Y546 | LRRC42 | NA | 14.8131845 | 14.9927012 | NA | 15.0704193 |
| P78382 | SLC35A1 | 19.3091224 | 19.237849 | 19.1625009 | 19.209449 | 18.9704782 |
| Q9H1D9 | POLR3F | 23.2317752 | 23.3483416 | 23.3114467 | 23.2098615 | 23.2604823 |
| Q5TEJ8 | THEMIS2 | 18.9755373 | 18.9456894 | 19.0127366 | 18.9463697 | 18.9495076 |
| Q969Q5 | RAB24 | 21.0754053 | 21.2163726 | 21.2137555 | 21.1288807 | 21.2609124 |
| Q8IUF8 | RIOX2 | 21.2227605 | 21.222632 | 21.4045233 | 21.3022903 | 21.2896638 |
| Q9UHD1 | CHORDC1 | 22.3687627 | 22.4070384 | 22.4115219 | 22.3481202 | 22.3902374 |
| P09884 | POLA1 | 20.2339971 | 20.3054855 | 20.3583873 | 20.2339664 | 20.358162 |
| O75691 | UTP20 | 21.6582437 | 21.6409431 | 21.7260455 | 21.6008443 | 21.7203561 |
| Q9ULT8 | HECTD1 | 20.6743279 | 20.7497073 | 20.7832433 | 20.6970561 | 20.8233891 |
| P62826 | RAN | 25.7831695 | 25.6758763 | 25.7588477 | 25.7389984 | 25.7980663 |
| Q9Y5B9 | SUPT16H | 24.1392405 | 24.1634233 | 24.2107174 | 24.0723548 | 24.1290519 |
| P48449 | LSS | 22.0340284 | 22.0513935 | 22.0988591 | 22.0500075 | 22.1066265 |
| P83436 | COG7 | 20.0104625 | 20.1288746 | 20.1252544 | 20.0314048 | 20.1041926 |
| P48147 | PREP | 23.1458962 | 23.2047117 | 23.1780105 | 23.1012457 | 23.1909987 |
| P51659 | HSD17B4 | 23.6642838 | 23.6436957 | 23.6468096 | 23.5972601 | 23.6810464 |
| Q9BYT8 | NLN | 21.4958701 | 21.5200819 | 21.5531134 | 21.5324687 | 21.5568127 |
| Q8WUM0 | NUP133 | 22.0665939 | 22.082999 | 22.1366099 | 22.0555042 | 22.1531866 |
| Q96GD0 | PDXP | 21.9121825 | 22.0656272 | 21.9725044 | 21.9454181 | 21.9917746 |
| P18031 | PTPN1 | 22.0497944 | 22.1161394 | 22.1594994 | 22.0041525 | 22.0499067 |
| O43818 | RRP9 | 21.5928171 | 21.707563 | 21.5496533 | 21.5439606 | 21.6851269 |
| O76071 | CIAO1 | 20.8604443 | 20.9101559 | 21.0498979 | 20.9059001 | 20.9877953 |
| Q6IBW4 | NCAPH2 | 20.0916554 | 20.2516156 | 20.2077236 | 20.1275427 | 20.2562996 |
| Q9H0E3 | SAP130 | 18.9911014 | 19.1930971 | 19.2759976 | 19.0707302 | 19.1795814 |
| P35226 | BMI1 | 19.1618395 | 19.241647 | 19.1371145 | 19.0143642 | 19.1576247 |
| Q9H9V9 | JMJD4 | 15.5331659 | 15.813167 | 15.47464 | 15.5036708 | 15.7897284 |
| Q15696 | ZRSR2 | 17.701479 | 18.0027638 | 17.843615 | 18.0006587 | 17.8960705 |
| Q96IW7 | SEC22A | 16.7061035 | 16.6858736 | 16.932368 | 16.8745433 | 16.9471439 |
| Q96IQ9 | ZNF414 | 17.7644714 | 17.2138191 | 17.4234472 | 17.5681087 | 17.6899315 |
| P57764 | GSDMD | 21.454751 | 21.4813114 | 21.5811427 | 21.4833618 | 21.5561768 |
| Q9NZB2 | FAM120A | 22.0854176 | 22.1313728 | 22.1781038 | 22.0281908 | 22.1288282 |
| Q99627 | COPS8 | 21.2345004 | 21.442107 | 21.3212772 | 21.3039161 | 21.4634 |
| O14924 | RGS12 | 16.3593635 | 15.734816 | 16.3556315 | 15.6754204 | 16.3223849 |
| P09234 | SNRPC | 23.1955918 | 23.1160251 | 23.1555518 | 23.1017922 | 23.1512616 |

| | | | | | | |
|---|---|---|---|---|---|---|
| Q9Y291 | MRPS33 | 20.9052326 | 21.0785057 | 21.2076997 | 20.9079986 | 20.8886794 |
| Q6ZNB6 | NFXL1 | 18.8035832 | 18.9165062 | 19.1162835 | 18.8938654 | 18.907232 |
| Q15418 | RPS6KA1 | 22.1509751 | 22.3442171 | 22.372524 | 22.2140646 | 22.3499856 |
| Q9C0C7 | AMBRA1 | 18.2777101 | 18.3837969 | 18.3656155 | 18.2290054 | 18.5716348 |
| Q96IJ6 | GMPPA | 21.4659679 | 21.4636091 | 21.5103026 | 21.47486 | 21.5625399 |
| P25788 | PSMA3 | 25.049438 | 24.9861872 | 24.9447732 | 24.9267442 | 24.9937551 |
| Q9BXS4 | TMEM59 | 19.9653415 | 20.1025861 | 20.00923 | 19.7170761 | 20.0474512 |
| Q9HAV4 | XPO5 | 21.84531 | 21.9526895 | 21.9417239 | 21.8193623 | 21.971004 |
| Q9NX78 | TMEM260 | 16.5103266 | 16.6283276 | 16.6143818 | 16.5399003 | 16.6611745 |
| Q5T8I9 | HENMT1 | 18.1896806 | 18.3001484 | 18.2088607 | 18.11775 | 18.3331205 |
| Q96CW5 | TUBGCP3 | 22.4660976 | 22.5911246 | 22.6355124 | 22.4936619 | 22.5541219 |
| Q9BYK8 | HELZ2 | 19.2003921 | 19.3317384 | 19.366426 | 19.1974086 | 19.3223503 |
| Q9UKT5 | FBXO4 | 20.1139485 | 20.1264903 | 20.1577598 | 20.0671339 | 20.2244831 |
| Q99797 | MIPEP | 20.6336648 | 20.7083614 | 20.5939711 | 20.6100289 | 20.69723 |
| Q99618 | CDCA3 | 18.543518 | 18.582371 | 18.6142576 | 18.6792535 | 18.6019792 |
| Q15046 | KARS1 | 24.1227211 | 24.1597077 | 24.1036143 | 24.0903698 | 24.0804368 |
| P29120 | PCSK1 | 15.3033809 | NA | NA | 15.4089237 | NA |
| P81877 | SSBP2 | 19.6725066 | 19.4659229 | 19.4592547 | 19.8573848 | 19.5239967 |
| P35270 | SPR | 21.3096206 | 21.3748768 | 21.4818599 | 21.3201949 | 21.4859079 |
| Q86W34 | AMZ2 | 16.6345293 | 16.760066 | 16.8329547 | 16.7289835 | 16.2754643 |
| Q9Y5V3 | MAGED1 | 15.9975142 | 16.0014498 | 15.8303278 | 15.8786702 | 15.8411485 |
| Q9UBC3 | DNMT3B | 16.9075516 | 16.8681541 | 16.8856687 | 16.9441368 | 16.8334378 |
| O14786 | NRP1 | 17.2948954 | 17.1640999 | 17.3788987 | 17.3781569 | 17.3849617 |
| P07902 | GALT | 20.2103682 | 20.0869196 | 20.2856484 | 20.1576394 | 20.2847025 |
| Q99729 | HNRNPAB | 24.2004891 | 24.2276875 | 24.1487853 | 24.0489103 | 24.1417519 |
| O60573 | EIF4E2 | 20.4668435 | 20.4769665 | 20.6229781 | 20.4819102 | 20.4843239 |
| Q99829 | CPNE1 | 22.9293163 | 23.0368221 | 22.9358792 | 22.8128967 | 22.9534086 |
| Q9Y227 | ENTPD4 | 18.2533541 | 18.2633908 | 18.3295887 | 18.3724559 | 18.2805412 |
| Q92766 | RREB1 | 20.5998223 | 20.7250569 | 20.5957232 | 20.6314703 | 20.7123574 |
| Q9UMS4 | PRPF19 | 24.6051328 | 24.6087285 | 24.5767629 | 24.4718591 | 24.627286 |
| P51965 | UBE2E1 | 21.5852586 | 21.532626 | 21.6106611 | 21.5308513 | 21.5504302 |
| Q9P253 | VPS18 | 20.7646889 | 20.7565362 | 20.9069843 | 20.7752788 | 20.8306479 |
| Q6P1X5 | TAF2 | 19.5268676 | 19.6072606 | 19.6047979 | 19.5591257 | 19.5976983 |
| Q05397 | PTK2 | 19.5321733 | 19.5842322 | 19.5727451 | 19.5105156 | 19.674174 |
| Q8NBF6 | AVL9 | 19.7433292 | 19.7821715 | 19.8766265 | 19.754973 | 19.89551 |
| Q9NWH9 | SLTM | 22.4767689 | 22.5755198 | 22.614495 | 22.4228956 | 22.4947796 |
| O00178 | GTPBP1 | 20.5118349 | 20.6588401 | 20.6602609 | 20.5010394 | 20.6572575 |
| P49821 | NDUFV1 | 22.8123903 | 22.8532794 | 22.9020726 | 22.7520572 | 22.8625557 |
| P17028 | ZNF24 | 20.2589388 | 20.2838048 | 20.3258354 | 20.3102041 | 20.3747411 |
| O95427 | PIGN | 18.3571746 | 18.4479823 | 18.6632184 | 18.5027666 | 18.5781981 |
| Q13636 | RAB31 | 20.4295242 | 20.645351 | 20.6682601 | 20.5196373 | 20.7599089 |
| P48378 | RFX2 | 16.0444529 | 15.9406492 | 16.1117671 | 16.2663253 | 16.0595674 |

| | | | | | | |
|---|---|---|---|---|---|---|
| P10124 | SRGN | 22.6778234 | 22.7620944 | 22.5131681 | 22.6567863 | 22.8096122 |
| Q3KRA6 | C2orf76 | 19.011409 | 19.3507665 | 19.2278749 | 19.0014305 | 19.3751582 |
| O95081 | AGFG2 | 18.1108649 | 18.2154675 | 18.787484 | 18.1185637 | 18.2724291 |
| Q8NHS3 | MFSD8 | 16.4960205 | 16.0390242 | 16.2323567 | 16.2641549 | 16.0628467 |
| Q8N142 | ADSS1 | 19.2196685 | 19.6550245 | 19.7963835 | 19.232361 | 19.642768 |
| Q9H2W1 | MS4A6A | 19.3484069 | 19.0553407 | 19.2253465 | 19.2883934 | 19.1620904 |
| P62879 | GNB2 | 23.3420309 | 23.5745672 | 23.3462117 | 23.2916836 | 23.5042117 |
| Q8IXZ2 | ZC3H3 | 18.4155952 | 18.7585102 | 18.4541496 | 18.3349556 | 18.4932772 |
| Q01664 | TFAP4 | 20.1519884 | 20.2075783 | 20.2102552 | 20.1880531 | 20.3562066 |
| Q13442 | PDAP1 | 23.2223781 | 23.2913059 | 23.1761273 | 23.1679058 | 23.082275 |
| Q8WVP7 | LMBR1 | 18.4249407 | 18.7318447 | 18.6545866 | 18.4673249 | 18.6021168 |
| Q8NDZ6 | TMEM161B | 15.598205 | NA | NA | 15.5218608 | NA |
| Q96S21 | RAB40C | 17.701181 | 17.8913547 | 17.9773888 | 17.8473134 | 17.8208339 |
| P32121 | ARRB2 | 20.572965 | 20.4497281 | 20.836512 | 20.6133106 | 20.7246558 |
| Q14728 | MFSD10 | 22.4121575 | 22.4611712 | 22.4009234 | 22.3334775 | 22.3181306 |
| Q13946 | PDE7A | 18.0473664 | 18.1403003 | 17.9864499 | 18.164505 | 17.9623171 |
| Q9Y4P1 | ATG4B | 20.2591752 | 20.1935943 | 20.0502414 | 20.1674142 | 20.285543 |
| Q99426 | TBCB | 23.4161204 | 23.6079774 | 23.4478034 | 23.3118699 | 23.4577725 |
| Q96HE7 | ERO1A | 21.4874993 | 21.5986625 | 21.583773 | 21.471123 | 21.6057991 |
| Q03518 | TAP1 | 19.9805419 | 20.2225069 | 20.1173994 | 20.0808703 | 20.1691462 |
| P08648 | ITGA5 | 21.2935818 | 21.2846385 | 21.2492781 | 21.2533527 | 21.3210318 |
| Q8NDD1 | C1orf131 | 20.3165969 | 20.3404776 | 20.336463 | 20.2428187 | 20.2405683 |
| Q9NZW5 | PALS2 | 19.0562167 | 19.1051224 | 19.1281122 | 18.8952701 | 19.100485 |
| O75879 | GATB | 19.0289338 | 19.179189 | 19.0475555 | 19.1007957 | 19.0730637 |
| Q9HC38 | GLOD4 | 23.6369231 | 23.6793534 | 23.5874861 | 23.5738936 | 23.7153432 |
| Q13950 | RUNX2 | 21.7432859 | 21.8372076 | 21.723485 | 21.7674827 | 21.8135414 |
| P52597 | HNRNPF | 25.7789653 | 25.8696978 | 25.6952816 | 25.702837 | 25.7887465 |
| Q9UI09 | NDUFA12 | 22.3236427 | 22.3585189 | 22.458721 | 22.3321475 | 22.4740984 |
| Q96L92 | SNX27 | 22.0655806 | 22.1261379 | 22.1412124 | 22.0115695 | 22.1182436 |
| Q9NZL9 | MAT2B | 23.282185 | 23.3131899 | 23.2584818 | 23.1144352 | 23.2513052 |
| Q13409 | DYNC1I2 | 21.9247966 | 21.9907023 | 21.8940617 | 21.8903209 | 22.0128208 |
| Q9BXS9 | SLC26A6 | 19.8316496 | 19.93469 | 19.9738727 | 19.88425 | 19.8885262 |
| O43681 | GET3 | 22.8015628 | 22.8938689 | 22.824636 | 22.7304856 | 22.8947598 |
| Q96QK1 | VPS35 | 24.0274963 | 23.9966175 | 24.0164543 | 23.9078926 | 24.0150101 |
| Q9BPX6 | MICU1 | 20.4639839 | 20.5165237 | 20.6293966 | 20.4941652 | 20.5413341 |
| Q9H270 | VPS11 | 21.6252854 | 21.6249547 | 21.7053374 | 21.6485515 | 21.679073 |
| P40939 | HADHA | 24.355645 | 24.4460887 | 24.403558 | 24.3196893 | 24.477166 |
| Q96P70 | IPO9 | 21.974566 | 22.0726361 | 22.0796271 | 21.9689282 | 22.0618707 |
| Q8IWX8 | CHERP | 22.3073481 | 22.3202826 | 22.2755975 | 22.2469333 | 22.3680165 |
| Q9Y285 | FARSA | 22.8791275 | 22.9149702 | 22.945092 | 22.8378209 | 22.8837437 |
| Q9Y2K1 | ZBTB1 | 18.9232486 | 19.0402296 | 18.9936586 | 18.9569862 | 19.0013156 |
| Q15054 | POLD3 | 20.8049065 | 20.8714749 | 20.8319082 | 20.8258739 | 20.8943132 |

| | | | | | | |
|---|---|---|---|---|---|---|
| Q04323 | UBXN1 | 22.6063921 | 22.7021771 | 22.4916669 | 22.5889772 | 22.5003331 |
| Q15723 | ELF2 | 20.9018505 | 20.9605533 | 20.8489762 | 20.8869212 | 21.0616897 |
| Q9NR46 | SH3GLB2 | 20.5950147 | 20.6567735 | 20.6769226 | 20.6028265 | 20.7390512 |
| Q9NWQ8 | PAG1 | 18.5940462 | 18.7376932 | 18.7056478 | 18.6654221 | 18.9139031 |
| Q8NG31 | KNL1 | 17.6214846 | 17.9345569 | 17.8042827 | 17.7855205 | 17.8199236 |
| Q8IUA7 | ABCA9 | NA | 20.4504342 | 20.2841648 | NA | 20.1745324 |
| Q9UGK8 | SERGEF | 19.104305 | 19.01093 | 19.2500272 | 19.2222841 | 19.1364267 |
| Q6ZS11 | RINL | 19.0803252 | 19.2012226 | 19.3202719 | 19.0525127 | 19.3601568 |
| O95302 | FKBP9 | 18.5004367 | 18.3540378 | 18.3942364 | 18.4939764 | 18.6015352 |
| Q969S2 | NEIL2 | 18.1116953 | 17.9352829 | 18.022407 | 17.9990813 | 18.1149884 |
| Q86X19 | TMEM17 | NA | 15.3290376 | 15.4297103 | NA | 15.5275451 |
| P54253 | ATXN1 | 16.5007372 | 16.7747779 | 16.6892846 | 16.6494166 | 16.7375627 |
| Q96GR4 | ZDHHC12 | 18.2726763 | 17.9010931 | 18.1800316 | 18.1102674 | 18.2386035 |
| Q969W8 | ZNF566 | 15.8959124 | 15.8325191 | 15.7993472 | 16.1483172 | 15.8864585 |
| Q04759 | PRKCQ | 18.5113736 | 18.7298536 | 18.6106492 | 18.4313068 | 18.6158194 |
| Q9BXY0 | MAK16 | 21.1769824 | 21.1696109 | 21.2502159 | 21.1668047 | 21.2508189 |
| Q9ULX6 | AKAP8L | 20.9443661 | 21.0138145 | 20.9779028 | 20.9413169 | 20.9227419 |
| O76003 | GLRX3 | 23.0707289 | 23.254755 | 23.1173424 | 23.111829 | 23.2558998 |
| O96017 | CHEK2 | 21.1379418 | 21.1907541 | 21.1966094 | 21.1838467 | 21.1961215 |
| Q9NXG6 | P4HTM | 22.371097 | 22.3273137 | 22.0966122 | 22.275024 | 22.1169935 |
| P21854 | CD72 | 18.0552555 | 17.8532823 | 17.803806 | 17.7872954 | 17.9240561 |
| Q5T7W7 | TSTD2 | 16.7541038 | 16.6402397 | 16.9145122 | 16.6919067 | 16.7982775 |
| Q9H2I8 | LRMDA | 21.0761389 | 21.159174 | 21.2529998 | 21.1374606 | 21.2601994 |
| Q5T6V5 | QNG1 | 21.0772984 | 21.1023619 | 21.2074227 | 21.0089417 | 21.1934043 |
| Q9P2B2 | PTGFRN | 21.0943812 | 21.20691 | 21.1901705 | 21.0356022 | 21.1716712 |
| Q9UNA1 | ARHGAP26 | 20.3423524 | 20.3234081 | 20.3029832 | 20.2914798 | 20.3301619 |
| Q9UPN9 | TRIM33 | 20.2787599 | 20.2905285 | 20.349582 | 20.2942996 | 20.3496443 |
| Q9Y6G9 | DYNC1LI1 | 21.8998392 | 21.9989292 | 21.9466721 | 21.8845579 | 21.984635 |
| Q9H8Y5 | ANKZF1 | 20.5369663 | 20.5667585 | 20.5327685 | 20.5172 | 20.6270399 |
| Q13185 | CBX3 | 24.7095112 | 24.8472052 | 24.5951714 | 24.6112881 | 24.7862839 |
| Q96FX7 | TRMT61A | 18.9436758 | 19.0733556 | 19.0420467 | 19.1106685 | 19.2549724 |
| Q8TE68 | EPS8L1 | 15.4161624 | 15.5152078 | 15.7843945 | 15.8606098 | 15.659888 |
| Q9NS56 | TOPORS | 17.8662251 | 18.1014662 | 18.1248854 | 18.0493302 | 18.1134737 |
| Q9H9A5 | CNOT10 | 20.7976724 | 20.8301955 | 20.8405057 | 20.7895891 | 20.8726026 |
| Q9UBF6 | RNF7 | 18.7598113 | 19.1920119 | 19.083367 | 19.1060958 | 19.209916 |
| Q9P2B7 | CFAP97 | 16.5328619 | 16.7815565 | 16.9575526 | 16.3021443 | 16.6222191 |
| Q15398 | DLGAP5 | 19.6097532 | 19.7504469 | 19.5940931 | 19.6734307 | 19.716039 |
| Q9NVA1 | UQCC1 | 21.4781634 | 21.5530321 | 21.5550149 | 21.5071246 | 21.668703 |
| Q15436 | SEC23A | 21.064159 | 21.1573321 | 21.267478 | 21.0353498 | 21.2128352 |
| Q9BUL5 | PHF23 | 19.8197371 | 19.8990062 | 19.7161295 | 19.7356566 | 19.8603445 |
| O75689 | ADAP1 | 21.051908 | 21.0824677 | 21.1996326 | 21.0520355 | 21.152862 |
| Q58DX5 | NAALADL2 | 16.3976028 | NA | NA | 16.2430334 | NA |

| | | | | | | |
|---|---|---|---|---|---|---|
| Q69YN4 | VIRMA | 20.2880157 | 20.3267676 | 20.3146964 | 20.2689338 | 20.3583081 |
| Q9Y624 | F11R | 18.6547563 | 18.8415963 | 18.9294637 | 18.7104988 | 18.9159578 |
| Q2TAM9 | TUSC1 | 17.5732232 | 17.7486557 | 17.6144276 | 17.6998335 | 17.8752231 |
| Q9P2Q2 | FRMD4A | 15.2347825 | 15.4116297 | 15.1770795 | 15.5663645 | 15.0812265 |
| Q9NQ34 | TMEM9B | 20.0974559 | 19.9171913 | 20.1520387 | 20.0424015 | 20.3753577 |
| Q8NFG4 | FLCN | 19.0886044 | 19.1365133 | 19.2471834 | 19.1241057 | 19.191443 |
| P49770 | EIF2B2 | 21.1301938 | 21.3266207 | 21.1535966 | 21.0752832 | 21.1917445 |
| Q9P0L0 | VAPA | 23.2996577 | 23.4588538 | 23.350195 | 23.2617101 | 23.2659379 |
| Q9H267 | VPS33B | 20.6664346 | 20.7199942 | 20.6907837 | 20.6285859 | 20.6643118 |
| Q96Q15 | SMG1 | 20.3960251 | 20.4660314 | 20.4658138 | 20.3289731 | 20.4293734 |
| Q13085 | ACACA | 21.3473554 | 21.3846422 | 21.3943903 | 21.3517663 | 21.4191567 |
| Q96C03 | MIEF2 | 13.7106814 | NA | NA | 13.2659806 | NA |
| Q5JPI9 | EEF1AKMT2 | 17.3561607 | 17.3080764 | 17.491854 | 17.3615783 | 17.5516635 |
| Q01064 | PDE1B | 17.2734498 | 17.5680058 | 17.6690838 | 17.3456269 | 17.642808 |
| P00568 | AK1 | 18.3614669 | 18.5320522 | 18.3543499 | 18.1692875 | 18.5471457 |
| Q13510 | ASAH1 | 22.0321471 | 21.9949021 | 22.0033745 | 22.0197864 | 21.9500669 |
| Q92499 | DDX1 | 23.3205991 | 23.3849064 | 23.3250868 | 23.278348 | 23.3772223 |
| P98196 | ATP11A | 20.3359818 | 20.3707459 | 20.4862539 | 20.3960828 | 20.4575614 |
| Q6P1L5 | FAM117B | 17.954415 | 17.9610054 | 17.8675476 | 17.8612373 | 17.8434658 |
| Q6P1A2 | LPCAT3 | 20.1805506 | 20.1337685 | 20.2676302 | 20.1654479 | 20.1748536 |
| Q9BW66 | CINP | 18.3498299 | 18.6235217 | 18.4750425 | 18.4169021 | 18.5369754 |
| Q9H3H5 | DPAGT1 | 20.0134091 | 20.0537448 | 20.1503391 | 19.9115861 | 20.1847234 |
| Q8IX05 | CD302 | 19.5138119 | 19.4430128 | 19.7172306 | 19.5466054 | 19.7054867 |
| P38435 | GGCX | 18.8001583 | 18.7147138 | 18.6807704 | 18.8477246 | 18.5735936 |
| P78312 | FAM193A | 18.3333485 | 18.4016558 | 18.4274333 | 18.4905928 | 18.3561218 |
| Q9C0D5 | TANC1 | 17.8222828 | 18.0430078 | 17.7485351 | 17.7916144 | 17.7705927 |
| Q8N4L2 | PIP4P2 | 19.0536733 | 18.9314329 | 19.2565461 | 19.1029117 | 18.904594 |
| P36639 | NUDT1 | 21.7310673 | 21.6903297 | 21.7372655 | 21.636599 | 21.7518132 |
| Q9BW92 | TARS2 | 21.2789595 | 21.3045181 | 21.2555169 | 21.2210749 | 21.2397792 |
| Q8N0X7 | SPART | 22.5032755 | 22.4579789 | 22.3453605 | 22.3985997 | 22.4279153 |
| P46940 | IQGAP1 | 23.7649896 | 23.8084397 | 23.8225689 | 23.6946874 | 23.80846 |
| P32320 | CDA | 17.4921242 | 17.552372 | 17.3706572 | 17.3381462 | 17.4804358 |
| P82663 | MRPS25 | 21.3385391 | 21.2459134 | 21.4121357 | 21.2626611 | 21.4101723 |
| Q9UJX6 | ANAPC2 | 21.1263434 | 21.2672629 | 21.1506774 | 21.2139193 | 21.2185148 |
| Q02338 | BDH1 | 22.5257502 | 22.5286041 | 22.6321612 | 22.493179 | 22.6472065 |
| P31942 | HNRNPH3 | 23.9891142 | 24.1098444 | 24.0731383 | 23.9996223 | 24.0735031 |
| Q8N6H7 | ARFGAP2 | 22.1182514 | 22.2774405 | 22.3427838 | 22.136686 | 22.2419606 |
| O14579 | COPE | 23.7703938 | 23.897705 | 23.7304891 | 23.7075974 | 23.8042136 |
| P42858 | HTT | 21.1587661 | 21.1606667 | 21.1957358 | 21.0804148 | 21.1666579 |
| Q9NYU2 | UGGT1 | 22.8650995 | 22.8770229 | 22.9190462 | 22.8327152 | 22.8874032 |
| P31939 | ATIC | 24.2157439 | 24.2847303 | 24.2339727 | 24.1956126 | 24.2850826 |
| Q8NFQ8 | TOR1AIP2 | 20.2116878 | 20.4358644 | 20.3707049 | 20.3503731 | 20.3139943 |

| | | | | | | |
|---|---|---|---|---|---|---|
| O95057 | DIRAS1 | 17.4382472 | 17.1822741 | 17.4041679 | 17.4075532 | 17.2130175 |
| Q96GE5 | ZNF799 | 16.9700035 | 17.2267272 | 17.2935845 | 17.091995 | 17.0569046 |
| O75817 | POP7 | 19.8025819 | 19.7406539 | 20.1280144 | 19.9580006 | 19.9590386 |
| Q3T8J9 | GON4L | 16.8709552 | 16.8950814 | 16.7623242 | 16.9137522 | 16.807166 |
| P51114 | FXR1 | 23.1580053 | 23.2733895 | 23.2491977 | 23.1655659 | 23.2308194 |
| Q92636 | NSMAF | 21.2901525 | 21.2487942 | 21.2953519 | 21.2552656 | 21.2302814 |
| O94788 | ALDH1A2 | 18.0470159 | 18.5402514 | 18.2541295 | 17.8613106 | 18.2837989 |
| Q9HAH7 | FBRS | 18.7366148 | 18.7855841 | 19.0939212 | 19.0196075 | 18.4855103 |
| Q8N5U6 | RNF10 | 17.9796064 | 17.996115 | 18.1672059 | 17.9145281 | 17.8867187 |
| Q969Q4 | ARL11 | 17.0924051 | 17.1960818 | 17.0009626 | 17.0319303 | 17.2284133 |
| Q96HC4 | PDLIM5 | 19.1539855 | 19.4290487 | 19.3193174 | 19.3003527 | 19.3837109 |
| O95139 | NDUFB6 | 22.5317982 | 22.6554729 | 22.6909173 | 22.529721 | 22.556604 |
| Q99439 | CNN2 | 23.6906518 | 23.8430261 | 23.7318252 | 23.6681916 | 23.8147154 |
| Q9BQ61 | TRIR | 23.6154822 | 23.7060269 | 23.570798 | 23.5626137 | 23.6811986 |
| Q9BYM8 | RBCK1 | 19.0690458 | 19.3716912 | 19.285423 | 19.1671405 | 19.2968498 |
| Q6PCT2 | FBXL19 | 18.7698784 | 18.6828221 | 18.8085006 | 18.6152115 | 18.7422578 |
| O43808 | SLC25A17 | 19.3246339 | 19.500537 | 19.4572931 | 19.3343014 | 19.3582986 |
| O43920 | NDUFS5 | 22.3198419 | 22.4574104 | 22.4984129 | 22.3749231 | 22.5049233 |
| Q9H8H2 | DDX31 | 20.0159858 | 19.9832704 | 20.2127476 | 19.9862038 | 20.0987692 |
| P36915 | GNL1 | 20.7500943 | 20.7450436 | 20.8341151 | 20.7489535 | 20.842808 |
| Q00688 | FKBP3 | 24.1309713 | 24.2207543 | 24.1567549 | 24.0352384 | 24.1925597 |
| Q96F86 | EDC3 | 20.4457954 | 20.6244752 | 20.5216751 | 20.5159384 | 20.6265914 |
| Q9UPT9 | USP22 | 19.1573128 | 19.2052124 | 19.2542221 | 19.23139 | 19.2702517 |
| Q03164 | KMT2A | 19.6088071 | 19.7441632 | 19.7448411 | 19.5747879 | 19.7052951 |
| Q9UFW8 | CGGBP1 | 20.8453438 | 20.9404069 | 20.9854177 | 20.816261 | 20.9397451 |
| Q3ZCQ8 | TIMM50 | 22.7124115 | 22.796242 | 22.7366398 | 22.8157055 | 22.7717772 |
| O43427 | FIBP | 22.2680202 | 22.3106093 | 22.3254435 | 22.1994725 | 22.3396385 |
| P31751 | AKT2 | 21.7476211 | 21.8330989 | 21.8584591 | 21.7590971 | 21.843066 |
| P06576 | ATP5F1B | 26.3775244 | 26.4749257 | 26.3276147 | 26.276195 | 26.3716702 |
| P50747 | HLCS | 18.9345541 | 18.9760971 | 18.9277717 | 18.9214834 | 19.0197032 |
| P35998 | PSMC2 | 23.9706649 | 24.050997 | 23.9715558 | 23.9163565 | 24.0298457 |
| P49588 | AARS1 | 23.8869924 | 23.926618 | 23.8634116 | 23.8144344 | 23.9734973 |
| Q9UBQ7 | GRHPR | 22.9372239 | 22.9365945 | 22.993964 | 22.9170588 | 23.0529697 |
| Q00403 | GTF2B | 24.1513608 | 24.2600097 | 24.0903521 | 24.09544 | 24.1826309 |
| Q9BXW7 | HDHD5 | 22.6488566 | 22.6826986 | 22.6712443 | 22.6879394 | 22.7042713 |
| O43314 | PPIP5K2 | 21.5369691 | 21.5614996 | 21.5755054 | 21.4647932 | 21.6018501 |
| O95989 | NUDT3 | 21.6237142 | 21.5986039 | 21.6502259 | 21.672569 | 21.6737608 |
| Q6UX53 | TMT1B | 20.81825 | 20.7293696 | 20.7531775 | 20.8179535 | 20.6504598 |
| P53677 | AP3M2 | 18.3207847 | 18.4167467 | 18.3830926 | 18.2700183 | 18.3544178 |
| Q8TBM8 | DNAJB14 | 19.2436666 | 19.526627 | 19.3561238 | 19.3944651 | 19.4664406 |
| Q9BRJ6 | C7orf50 | 20.1503683 | 20.2902319 | 20.1836473 | 20.1498278 | 20.2021553 |
| Q9H869 | YY1AP1 | 17.487097 | 17.6357336 | 17.5802525 | 17.4905505 | 17.7301532 |

| | | | | | | |
|---|---|---|---|---|---|---|
| P98160 | HSPG2 | 19.3536486 | 19.5471421 | 19.2650396 | 19.3346707 | 19.3475674 |
| O75251 | NDUFS7 | 22.2218296 | 22.24927 | 22.2045652 | 22.129943 | 22.2731579 |
| O14772 | FPGT | 19.336172 | 19.2988866 | 19.57796 | 19.3940169 | 19.5399392 |
| Q8TEL6 | TRPC4AP | 16.866754 | 16.8123018 | 16.8696012 | 16.9018968 | 16.9861718 |
| P13489 | RNH1 | 25.9501993 | 26.0386405 | 25.8264147 | 25.8384296 | 26.0408169 |
| P23919 | DTYMK | 22.9633412 | 22.8185515 | 22.9331593 | 22.850054 | 23.055515 |
| A6NFI3 | ZNF316 | 18.5885383 | 18.5949897 | 18.7063114 | 18.6845515 | 18.728521 |
| Q9BU64 | CENPO | 19.0875882 | 19.1128592 | 19.2614189 | 19.0114912 | 19.2993328 |
| P09382 | LGALS1 | 26.5849676 | 26.7226046 | 26.5074971 | 26.5181686 | 26.8098172 |
| P78552 | IL13RA1 | 16.8304414 | 17.0180085 | 16.913864 | 17.1356533 | 17.0266072 |
| Q9UHC1 | MLH3 | 13.7228387 | 14.1663623 | 14.6117159 | 14.3658668 | 14.2632113 |
| Q16635 | TAFAZZIN | 14.0387433 | 14.3281144 | 14.4094646 | 13.9051843 | 14.5196529 |
| P41229 | KDM5C | 21.0182055 | 21.3200305 | 21.321778 | 21.0852739 | 21.3485401 |
| Q9GZU1 | MCOLN1 | 18.0922511 | 18.0222818 | 17.923981 | 18.0753343 | 18.0555837 |
| Q16718 | NDUFA5 | 22.3591127 | 22.5165037 | 22.405947 | 22.3364294 | 22.4730105 |
| P11234 | RALB | 22.5621549 | 22.6489931 | 22.65198 | 22.4201689 | 22.5564532 |
| Q29RF7 | PDS5A | 22.950739 | 22.96399 | 23.0332788 | 22.9183076 | 22.989709 |
| P62318 | SNRPD3 | 24.715409 | 24.7624504 | 24.7927511 | 24.6502016 | 24.7188334 |
| O60503 | ADCY9 | 19.5190939 | 19.5300186 | 19.5276575 | 19.5612575 | 19.5366582 |
| P98171 | ARHGAP4 | 22.6244629 | 22.634543 | 22.6234871 | 22.5392597 | 22.6269266 |
| P22314 | UBA1 | 25.3673693 | 25.4536521 | 25.3586511 | 25.3189668 | 25.3875388 |
| Q53EL6 | PDCD4 | 24.0515392 | 24.0689837 | 23.933098 | 24.0046401 | 24.0460219 |
| Q15067 | ACOX1 | 22.145654 | 22.1537111 | 22.276759 | 22.1266803 | 22.2357408 |
| O14964 | HGS | 21.7130982 | 21.8740406 | 21.767195 | 21.6904257 | 21.8671099 |
| Q9Y2L9 | LRCH1 | 18.5194339 | 18.4821411 | 18.6328437 | 18.6106313 | 18.7254525 |
| P49721 | PSMB2 | 23.875267 | 23.9290265 | 23.8385695 | 23.7816375 | 23.9386044 |
| Q5U5X0 | LYRM7 | 20.3115783 | 20.5314978 | 20.2225085 | 20.3116957 | 20.4083766 |
| Q9BSU1 | PHAF1 | 16.9881635 | 17.1276424 | 16.7526016 | 17.033167 | 17.0817813 |
| Q07002 | CDK18 | NA | 13.3784158 | 14.4666439 | NA | 13.3965863 |
| Q9UBK9 | UXT | 19.9966915 | 20.2265243 | 20.08704 | 20.0735172 | 20.1064472 |
| Q86TU7 | SETD3 | 21.2469696 | 21.3011595 | 21.3422787 | 21.2716844 | 21.3077186 |
| Q6DD87 | ZNF787 | 21.8185592 | 21.8496782 | 21.612749 | 21.7086159 | 21.5765697 |
| O75976 | CPD | 20.8519078 | 20.879781 | 20.8073268 | 20.8014619 | 20.8619081 |
| O75718 | CRTAP | 21.4105309 | 21.331154 | 21.3087534 | 21.2539727 | 21.2414141 |
| O15260 | SURF4 | 23.315174 | 23.4296691 | 23.1675755 | 23.18575 | 23.2177539 |
| Q9NYZ3 | GTSE1 | 18.811092 | 18.8469721 | 18.7589393 | 18.6368054 | 18.8109972 |
| O94804 | STK10 | 22.3397308 | 22.3500017 | 22.3172216 | 22.2616541 | 22.3124319 |
| Q15102 | PAFAH1B3 | 23.7382376 | 23.8407338 | 23.7465092 | 23.6990696 | 23.7236442 |
| P39060 | COL18A1 | 16.0368562 | 15.8550344 | 15.6334066 | 15.7804756 | 15.7261234 |
| Q96K58 | ZNF668 | 20.8531396 | 20.9799135 | 20.9269627 | 20.9557599 | 21.0416505 |
| Q15642 | TRIP10 | 19.8989886 | 20.1075485 | 20.0020792 | 19.8105566 | 20.045618 |
| P52594 | AGFG1 | 20.9222027 | 20.9464795 | 20.8361828 | 20.8601714 | 20.9967889 |

| | | | | | | |
|---|---|---|---|---|---|---|
| A1L0T0 | ILVBL | 22.0844901 | 22.1376212 | 22.242832 | 22.137318 | 22.2350412 |
| Q10472 | GALNT1 | 21.1905047 | 21.2229557 | 21.3425847 | 21.2519138 | 21.3133982 |
| P51991 | HNRNPA3 | 25.8926951 | 26.024808 | 26.0180442 | 25.9430523 | 26.0646252 |
| Q8N9N2 | ASCC1 | 21.3119332 | 21.4893742 | 21.4954047 | 21.2681572 | 21.4246689 |
| Q9NTX7 | RNF146 | 17.3355971 | 17.5208169 | 17.608106 | 17.6384343 | 17.4341962 |
| Q96H78 | SLC25A44 | 16.0705824 | NA | NA | 16.1508658 | NA |
| Q96CT7 | CCDC124 | 23.3576169 | 23.4190948 | 23.1672214 | 23.2834256 | 23.3465715 |
| Q9UKK9 | NUDT5 | 24.8856199 | 25.0055635 | 24.9427848 | 24.8737511 | 24.9067538 |
| Q9NR11 | ZNF302 | 13.637777 | NA | NA | 12.5476867 | NA |
| Q6NW34 | NEPRO | 19.1001401 | 19.0571706 | 19.1247614 | 19.015163 | 19.0235825 |
| Q9NP92 | MRPS30 | 21.6187879 | 21.5169177 | 21.7551667 | 21.575502 | 21.5765924 |
| Q16512 | PKN1 | 22.8617343 | 22.9223047 | 22.8173257 | 22.7271571 | 22.8764619 |
| P53367 | ARFIP1 | 21.1205863 | 21.1837817 | 21.1442049 | 21.0849164 | 21.2114864 |
| Q9HAF1 | MEAF6 | 20.2717544 | 20.4389977 | 20.2752433 | 20.1979475 | 20.3515893 |
| Q13129 | RLF | 19.4407634 | 19.6276871 | 19.8005732 | 19.4567434 | 19.7438395 |
| Q9H501 | ESF1 | 20.5817972 | 20.7422226 | 20.625056 | 20.6089853 | 20.6386628 |
| Q8N3F0 | MTURN | 16.1305011 | 15.9408111 | 16.2198666 | 16.038717 | 16.1523263 |
| Q8IVW6 | ARID3B | 19.8037692 | 19.832728 | 19.8726468 | 19.7699494 | 19.8533281 |
| Q9NPP4 | NLRC4 | 13.5298363 | NA | NA | 13.7131 | NA |
| P01903 | HLA-DRA | 20.8815361 | 20.9528796 | 21.1099356 | 20.8167492 | 21.0895916 |
| P10644 | PRKAR1A | 23.1477711 | 23.23827 | 23.2151183 | 23.1471797 | 23.2087309 |
| P21127 | CDK11B | 21.6409523 | 21.7127772 | 21.6906655 | 21.6089246 | 21.7053707 |
| Q9BVI4 | NOC4L | 21.7887271 | 21.9011209 | 21.9147673 | 21.7920666 | 21.8274314 |
| O14976 | GAK | 21.534703 | 21.5876594 | 21.6335563 | 21.480462 | 21.629835 |
| P62258 | YWHAE | 28.1180812 | 28.1644862 | 28.0155066 | 27.9651479 | 28.0844341 |
| Q9ULJ6 | ZMIZ1 | 20.9327279 | 20.8817022 | 20.9069841 | 20.9741539 | 20.9120403 |
| Q5D1E8 | ZC3H12A | 17.3125154 | 17.4450415 | 17.4167975 | 17.3941452 | 17.6707764 |
| Q9H7H0 | METTL17 | 21.0525738 | 21.1041227 | 21.2588733 | 20.7430137 | 21.0874533 |
| Q9Y4B5 | MTCL1 | NA | 16.1196388 | 15.7866784 | NA | 16.3077554 |
| Q13126 | MTAP | 22.2312411 | 22.4496516 | 22.3312568 | 22.3027511 | 22.4018995 |
| P15954 | COX7C | 24.4488836 | 24.2411807 | 24.4008102 | 24.1876299 | 24.3198866 |
| P19878 | NCF2 | 21.1830472 | 21.3178055 | 21.3494183 | 21.1730294 | 21.1573218 |
| Q8IZ73 | RPUSD2 | 21.0156645 | 21.0605676 | 21.0419361 | 21.0235821 | 21.123869 |
| Q8NBR6 | MINDY2 | 15.5318069 | 15.485371 | 15.6873279 | 15.6209555 | 15.7504401 |
| O75844 | ZMPSTE24 | 21.7756976 | 21.6508717 | 21.8076807 | 21.7608528 | 21.8011842 |
| Q9UBI1 | COMMD3 | 20.5732886 | 20.6040829 | 20.6652496 | 20.5660029 | 20.7321307 |
| Q12797 | ASPH | 22.3618052 | 22.4592866 | 22.4293196 | 22.327004 | 22.4381035 |
| P63244 | RACK1 | 26.2287535 | 26.1991884 | 26.2413826 | 26.1282824 | 26.2948458 |
| Q9H2P9 | DPH5 | 18.8282681 | 18.9131121 | 18.9867507 | 18.9419919 | 18.9903317 |
| P52292 | KPNA2 | 22.5923728 | 22.6261322 | 22.5985181 | 22.6334424 | 22.5906573 |
| P37198 | NUP62 | 21.8585092 | 22.1500323 | 22.015201 | 21.8431127 | 22.0812197 |
| Q8NB14 | USP38 | 15.4387711 | 15.4912622 | 15.4742327 | 15.3274895 | 15.8892015 |

| | | | | | | |
|---|---|---|---|---|---|---|
| P52824 | DGKQ | 18.1306418 | 18.0918231 | 18.0560629 | 18.2674384 | 18.0950163 |
| Q00013 | MPP1 | 20.3521547 | 20.3848886 | 20.3921952 | 20.3064539 | 20.4738854 |
| Q86VE0 | MYPOP | 14.5903411 | NA | NA | 14.4953962 | NA |
| O60645 | EXOC3 | 21.5998978 | 21.7192524 | 21.7030357 | 21.6499422 | 21.6682184 |
| Q15773 | MLF2 | 19.9807443 | 20.0808021 | 20.1522792 | 19.9529578 | 19.9648297 |
| Q70UQ0 | IKBIP | 20.8337241 | 20.8463158 | 20.8293097 | 20.798669 | 20.8606238 |
| A0PJW6 | TMEM223 | 18.1185513 | 18.1493842 | 18.3150492 | 18.2052014 | 18.2115543 |
| Q9NRR5 | UBQLN4 | 21.2220192 | 21.2791911 | 21.1528947 | 21.2050035 | 21.3635103 |
| P08575 | PTPRC | 22.2723259 | 22.2982688 | 22.3664712 | 22.2039264 | 22.3580679 |
| Q5SRI9 | MANEA | 18.1468734 | 17.9592048 | 18.0763264 | 18.1646354 | 17.9791994 |
| O75643 | SNRNP200 | 23.5651076 | 23.6201812 | 23.56891 | 23.5006154 | 23.5669902 |
| Q9UKG1 | APPL1 | 21.0439141 | 21.1390043 | 21.2525181 | 21.1569082 | 21.2420607 |
| Q14393 | GAS6 | 18.6288587 | 19.0142546 | 19.1984933 | 18.5903349 | 19.1815464 |
| Q8IUW3 | SPATA2L | 17.2482469 | 17.3828535 | 17.2926335 | 17.356589 | 17.3481627 |
| Q96AG3 | SLC25A46 | 19.598082 | 19.6566929 | 19.6852504 | 19.6134347 | 19.685992 |
| O15067 | PFAS | 23.127851 | 23.1858756 | 23.0712737 | 23.0588903 | 23.1764629 |
| Q9Y5L4 | TIMM13 | 22.7721848 | 22.8144887 | 22.7854464 | 22.7787862 | 22.7778101 |
| O75534 | CSDE1 | 23.7855794 | 23.9044419 | 23.7339492 | 23.7543811 | 23.8717138 |
| Q9UKN8 | GTF3C4 | 20.8792612 | 20.90125 | 20.9529275 | 20.8348928 | 20.9878152 |
| P09960 | LTA4H | 21.2144741 | 21.1877059 | 21.2675152 | 21.2085886 | 21.3607437 |
| O14530 | TXNDC9 | 20.9182078 | 21.1109334 | 21.0333151 | 20.9564627 | 21.0816433 |
| O60906 | SMPD2 | 15.2066215 | NA | NA | 15.3972322 | NA |
| P15884 | TCF4 | 17.042207 | 17.3864375 | 17.2183964 | 17.368505 | 17.415349 |
| P05362 | ICAM1 | 18.7173078 | 18.9876496 | 19.0220181 | 18.6934955 | 18.9508204 |
| Q05193 | DNM1 | 19.0809656 | 19.3207344 | 19.3558211 | 19.0972396 | 19.2482003 |
| P43490 | NAMPT | 22.0615653 | 22.0355021 | 22.0236086 | 21.9930428 | 22.0619619 |
| Q9UNH7 | SNX6 | 22.3286353 | 22.4463571 | 22.3832855 | 22.3179964 | 22.3786511 |
| O94886 | TMEM63A | 13.6020511 | NA | NA | 13.5018134 | NA |
| C9JLR9 | ZFTA | 12.690837 | NA | NA | 12.1676903 | NA |
| A7KAX9 | ARHGAP32 | 15.0929653 | NA | NA | 15.1394471 | NA |
| Q9NVG8 | TBC1D13 | 20.742359 | 20.8517297 | 20.8788569 | 20.8140712 | 20.8856614 |
| Q4G0J3 | LARP7 | 21.3805913 | 21.4040873 | 21.4949895 | 21.3838114 | 21.3876499 |
| Q96JN0 | LCOR | 13.876376 | NA | NA | 13.9292574 | NA |
| P55210 | CASP7 | 16.354576 | 16.9718011 | 16.9399005 | 17.1311082 | 16.9296982 |
| Q969Q0 | RPL36AL | 24.0175967 | 24.037967 | 23.9303146 | 23.7883622 | 23.7493812 |
| Q05469 | LIPE | 16.6699523 | 16.7980794 | 16.8798945 | 16.7202522 | 16.9118307 |
| Q9BUE6 | ISCA1 | 18.7622686 | 18.7772483 | 18.8234553 | 18.8061647 | 18.9801601 |
| P11171 | EPB41 | 20.393715 | 20.3582169 | 20.3949814 | 20.3415839 | 20.3623301 |
| P49792 | RANBP2 | 26.744365 | 26.812233 | 26.705814 | 26.6404711 | 26.7216889 |
| Q96BN8 | OTULIN | 22.2219131 | 22.2846061 | 22.3584239 | 22.2402434 | 22.3740412 |
| Q96NB2 | SFXN2 | 19.5448431 | 19.5781621 | 19.6038476 | 19.4433293 | 19.6874409 |
| P60903 | S100A10 | 22.4431132 | 22.7379586 | 22.575333 | 22.5512747 | 22.5942776 |

| | | | | | | |
|---|---|---|---|---|---|---|
| O00192 | ARVCF | NA | 13.2245826 | 13.3362149 | NA | 12.7662017 |
| O15054 | KDM6B | NA | 14.5595308 | 14.4766302 | NA | 14.6472698 |
| Q12965 | MYO1E | 19.786272 | 20.0644637 | 20.1248787 | 19.851298 | 20.1279113 |
| O60583 | CCNT2 | 17.538579 | 17.7089593 | 17.728806 | 17.7243691 | 17.9595817 |
| Q9Y2U8 | LEMD3 | 20.7571776 | 20.8401719 | 20.8415889 | 20.742685 | 20.8130897 |
| Q96CM8 | ACSF2 | 19.7321878 | 19.7084284 | 19.7447155 | 19.69222 | 19.8424067 |
| Q14839 | CHD4 | 23.061088 | 23.0861417 | 23.0781494 | 22.9817048 | 23.0347615 |
| O75330 | HMMR | 19.6830067 | 19.7056349 | 19.6702197 | 19.6537242 | 19.749048 |
| Q9ULG1 | INO80 | 20.0156799 | 20.1562965 | 20.1492003 | 20.0523706 | 20.1453997 |
| Q9H078 | CLPB | 21.6599001 | 21.7942022 | 21.6908566 | 21.661493 | 21.7018032 |
| Q13868 | EXOSC2 | 21.0980153 | 21.1869129 | 21.2655325 | 21.0847848 | 21.2195763 |
| Q9UHK0 | NUFIP1 | 18.9680102 | 19.0410731 | 19.0819766 | 18.9990389 | 19.1082545 |
| Q96E29 | MTERF3 | 19.1845586 | 19.4237012 | 19.4458531 | 19.3638061 | 19.3790479 |
| Q5TEU4 | NDUFAF5 | 16.5774415 | 16.6701297 | 16.7022886 | 16.7182603 | 16.7567565 |
| Q9BSL1 | UBAC1 | 20.9270053 | 21.031272 | 20.8531001 | 20.935252 | 21.0895299 |
| P08253 | MMP2 | 17.4517828 | 17.453158 | 17.4397093 | 17.4967056 | 17.3015338 |
| O00258 | GET1 | 17.9846062 | 17.981693 | 18.0416521 | 17.8768505 | 17.9871313 |
| P41208 | CETN2 | 21.0752112 | 21.0162335 | 20.853144 | 20.8966515 | 21.1229588 |
| P08123 | COL1A2 | 11.7780413 | NA | NA | 11.8811151 | NA |
| P30305 | CDC25B | 12.6531537 | NA | NA | 12.6111852 | NA |
| Q92622 | RUBCN | 15.1756725 | 16.481373 | 17.0645524 | 16.0199573 | 16.4339809 |
| Q8NHW5 | RPLP0P6 | 19.1271994 | 19.1922049 | 19.3047501 | 19.0524794 | 19.1340616 |
| Q9UP95 | SLC12A4 | 18.7107828 | 18.6239555 | 18.8350431 | 18.6281729 | 18.8474821 |
| Q9BW72 | HIGD2A | 21.3038285 | 21.2453471 | 21.5487946 | 21.2997107 | 21.4992548 |
| Q9P0M6 | MACROH2A2 | 23.0725339 | 23.2663764 | 23.1125386 | 23.0396026 | 22.9753007 |
| Q96S55 | WRNIP1 | 20.1459695 | 20.3278832 | 20.2803058 | 20.1564094 | 20.3162019 |
| Q5JVF3 | PCID2 | 21.2775584 | 21.3456507 | 21.3499333 | 21.203049 | 21.3406947 |
| Q7Z6Z7 | HUWE1 | 22.712104 | 22.7760576 | 22.7268327 | 22.6407591 | 22.7935684 |
| P82909 | MRPS36 | 21.1589934 | 21.4096902 | 21.4424279 | 21.2565871 | 21.3462778 |
| Q9H4D5 | NXF3 | 19.1035847 | 19.0207469 | 19.1700871 | 19.0470737 | 19.1036119 |
| Q86XN7 | PROSER1 | NA | 16.0796926 | 16.1498399 | NA | 16.067364 |
| O15409 | FOXP2 | 17.8928437 | 18.1197514 | 17.9803061 | 17.7039488 | 17.8142765 |
| O60312 | ATP10A | 14.3176478 | 14.6555334 | 14.9226895 | 14.6035287 | 14.6623354 |
| Q08380 | LGALS3BP | 18.4162376 | 18.6092583 | 18.5859403 | 18.4910376 | 18.7595869 |
| Q9HBM1 | SPC25 | 21.0285016 | 21.0265944 | 21.0241941 | 20.8943485 | 20.9108537 |
| Q15437 | SEC23B | 22.6863361 | 22.6303015 | 22.7434397 | 22.6468936 | 22.7441521 |
| Q5T5P2 | KIAA1217 | 16.1665519 | NA | NA | 16.1965808 | NA |
| P28845 | HSD11B1 | 13.891386 | 13.7151239 | 13.608386 | 13.9181651 | 14.2211636 |
| O15315 | RAD51B | 15.691364 | NA | NA | 15.6411914 | NA |
| Q15034 | HERC3 | 17.7262914 | 17.3632519 | 17.4603216 | 17.1061055 | 17.4165311 |
| Q8IWY8 | ZSCAN29 | 16.4762805 | 16.8043278 | 16.7714878 | 16.4867491 | 16.7036894 |
| Q9NRY6 | PLSCR3 | 18.6906657 | 18.5025637 | 18.5468091 | 18.3796567 | 18.6667653 |

| | | | | | | |
|---|---|---|---|---|---|---|
| Q6N021 | TET2 | 16.3584049 | 16.306772 | 16.2151091 | 16.1416713 | 16.4724115 |
| Q8IYU2 | HACE1 | 17.170126 | 17.271157 | 17.2457465 | 17.1169024 | 16.9333648 |
| Q7Z3D4 | LYSMD3 | 15.7490849 | 15.6022133 | 15.6328222 | 15.7472087 | 15.7035798 |
| Q9UEY8 | ADD3 | 18.7466009 | 18.8812299 | 19.070649 | 18.6456092 | 18.9978683 |
| O43572 | AKAP10 | 18.655416 | 18.5346528 | 18.6045465 | 18.6095342 | 18.6917746 |
| Q03113 | GNA12 | 18.6419246 | 18.5549317 | 18.7993918 | 18.7006834 | 18.6268165 |
| P21291 | CSRP1 | 22.1715521 | 22.0520165 | 22.3385638 | 22.0870907 | 22.2158596 |
| Q2TB10 | ZNF800 | 24.0183616 | 23.9612715 | 24.1678965 | 23.993961 | 23.9500288 |
| Q8WTT2 | NOC3L | 21.6736106 | 21.7358598 | 21.8671396 | 21.6888945 | 21.7811739 |
| P51116 | FXR2 | 23.1294809 | 23.292431 | 23.2838329 | 23.1853044 | 23.1651532 |
| Q96CB8 | INTS12 | 20.3661475 | 20.3251752 | 20.4232028 | 20.3690654 | 20.324938 |
| Q9UNS1 | TIMELESS | 19.8064022 | 19.8424885 | 19.9307926 | 19.7490379 | 19.8130467 |
| Q12986 | NFX1 | 19.2047463 | 19.3991783 | 19.3600011 | 19.283934 | 19.2897607 |
| Q96T88 | UHRF1 | 22.6426864 | 22.6992127 | 22.7271312 | 22.5764097 | 22.7229916 |
| P08397 | HMBS | 21.0342088 | 21.0260279 | 21.0656109 | 21.0036032 | 21.1339664 |
| Q9Y4A5 | TRRAP | 24.35732 | 24.3798086 | 24.3993445 | 24.2915498 | 24.3530596 |
| O14653 | GOSR2 | 20.6691579 | 20.6510785 | 20.7127036 | 20.6081974 | 20.7096671 |
| Q5BKZ1 | ZNF326 | 22.0529078 | 22.1287039 | 22.1068737 | 22.0107743 | 22.1656328 |
| P08579 | SNRPB2 | 23.1156443 | 23.199887 | 23.1735614 | 23.1089017 | 23.1839875 |
| Q9Y3A6 | TMED5 | 22.2803169 | 22.3704251 | 22.440744 | 22.312976 | 22.3580039 |
| Q9BVA0 | KATNB1 | 20.4791923 | 20.5117922 | 20.4874798 | 20.5258102 | 20.6321204 |
| Q9Y217 | MTMR6 | 20.0182964 | 20.0047164 | 20.1305121 | 20.0674931 | 20.1136002 |
| O00505 | KPNA3 | 21.6620775 | 21.6086723 | 21.7121749 | 21.637984 | 21.7186189 |
| Q99856 | ARID3A | 21.4472609 | 21.6198733 | 21.4747586 | 21.424473 | 21.4072831 |
| Q13596 | SNX1 | 22.2970215 | 22.327333 | 22.3212033 | 22.2725396 | 22.371996 |
| P05455 | SSB | 25.2364922 | 25.2754268 | 25.1310787 | 25.1060274 | 25.1325972 |
| P36542 | ATP5F1C | 24.6484237 | 24.6568823 | 24.6683872 | 24.55952 | 24.6251778 |
| O43395 | PRPF3 | 22.0073101 | 22.0606921 | 22.0601405 | 21.9861641 | 21.9721124 |
| Q6PGP7 | SKIC3 | 20.8282738 | 20.8366205 | 20.8767925 | 20.8116429 | 20.8638451 |
| P49915 | GMPS | 23.7829408 | 23.7748737 | 23.7654418 | 23.6987793 | 23.8121365 |
| Q9NU22 | MDN1 | 21.5034998 | 21.553631 | 21.5414173 | 21.4697237 | 21.5383494 |
| Q9UHD8 | SEPTIN9 | 23.4034648 | 23.5073371 | 23.3791229 | 23.349908 | 23.3554929 |
| P38571 | LIPA | 22.5157937 | 22.4585986 | 22.4938405 | 22.3872097 | 22.5787068 |
| P41743 | PRKCI | 19.6615952 | 19.8299635 | 19.9133168 | 19.715972 | 19.8421672 |
| Q92835 | INPP5D | 21.3212543 | 21.5552944 | 21.5279525 | 21.3555487 | 21.5067577 |
| Q86VM9 | ZC3H18 | 21.7588306 | 21.8513304 | 21.8045479 | 21.7357715 | 21.6637568 |
| Q14653 | IRF3 | 21.7758876 | 21.9319954 | 21.9679539 | 21.8586687 | 21.8685843 |
| Q969K3 | RNF34 | NA | 17.4384176 | 17.3906502 | NA | 17.5134987 |
| Q9C086 | INO80B | 17.9644603 | 18.0724292 | 17.9394024 | 17.9242629 | 18.1010792 |
| P48454 | PPP3CC | 18.980816 | 19.1347579 | 19.2821138 | 18.9907563 | 19.2065607 |
| P33316 | DUT | 24.6754852 | 24.8577932 | 24.6162827 | 24.6088246 | 24.861863 |
| Q96CP2 | FLYWCH2 | 20.7739362 | 20.9614202 | 20.7525779 | 20.8178081 | 20.8908632 |

| | | | | | | |
|---|---|---|---|---|---|---|
| Q69YI7 | NAIF1 | 16.6106485 | 16.8885121 | 16.7824677 | 16.8223339 | 16.7376886 |
| Q9H8N7 | ZNF395 | 17.6579029 | 17.688687 | 17.7198241 | 17.3785995 | 18.0598481 |
| Q9Y6F7 | NA | 16.0498403 | NA | NA | 15.4244518 | NA |
| Q96GF1 | RNF185 | 18.7499845 | 18.908762 | 18.6660282 | 18.5452251 | 18.7307376 |
| Q9UNK9 | ANGEL1 | 14.6305386 | 14.3466154 | 15.1452895 | 14.9808239 | 14.5661254 |
| P78563 | ADARB1 | 18.2789136 | 18.1840834 | 18.239436 | 18.3363206 | 18.0404659 |
| P51948 | MNAT1 | 20.1585145 | 20.1228467 | 19.9861628 | 20.1210547 | 20.2387072 |
| O75928 | PIAS2 | 13.541871 | NA | NA | 13.5241276 | NA |
| Q96HP0 | DOCK6 | 20.5301565 | 20.5331224 | 20.5842724 | 20.4507196 | 20.6469367 |
| Q9NP72 | RAB18 | 23.1113649 | 23.1679313 | 23.1956807 | 23.1271247 | 23.2184751 |
| Q8WUA2 | PPIL4 | 20.7971185 | 20.9483021 | 20.9421996 | 20.7091178 | 20.9904981 |
| P23458 | JAK1 | 25.0739114 | 25.1832556 | 25.1980737 | 25.0348929 | 25.1482714 |
| Q16666 | IFI16 | 20.3431956 | 20.5194357 | 20.6104472 | 20.4577114 | 20.5232095 |
| P11177 | PDHB | 23.046928 | 23.1397149 | 23.0635593 | 23.03123 | 23.1391099 |
| P53007 | SLC25A1 | 24.8988795 | 24.977232 | 24.8473534 | 24.8024125 | 24.8614965 |
| Q8IWV8 | UBR2 | 19.8632712 | 19.9107719 | 20.0488754 | 19.9073696 | 19.98448 |
| Q03252 | LMNB2 | 24.216214 | 24.3924142 | 24.1614106 | 24.1329334 | 24.2009698 |
| Q96A65 | EXOC4 | 21.2342152 | 21.2892757 | 21.2985875 | 21.219363 | 21.2555981 |
| P55011 | SLC12A2 | 18.2942299 | 18.4428123 | 18.4758731 | 18.3555702 | 18.4618268 |
| Q13423 | NNT | 23.6369976 | 23.7239508 | 23.5785998 | 23.5307335 | 23.6904587 |
| O95714 | HERC2 | 21.1798578 | 21.19023 | 21.2668733 | 21.1816398 | 21.2479592 |
| P09874 | PARP1 | 24.0207914 | 24.019274 | 24.0509894 | 23.9832277 | 24.0075901 |
| Q9H074 | PAIP1 | 21.9226349 | 21.9277656 | 21.8825756 | 21.892759 | 21.8331798 |
| Q96BY6 | DOCK10 | 20.8057701 | 20.8215676 | 20.9359741 | 20.7822819 | 20.932026 |
| O75794 | CDC123 | 21.1407865 | 21.0686439 | 21.2516446 | 21.1325701 | 21.2661361 |
| Q6NW29 | RWDD4 | 20.3052238 | 20.3228696 | 20.2046888 | 20.1655859 | 20.4432686 |
| Q9H3H3 | C11orf68 | 20.4518009 | 20.5044725 | 20.5608189 | 20.4503684 | 20.4868574 |
| P35580 | MYH10 | NA | 12.2120315 | 13.7181155 | NA | 13.1104223 |
| Q8IX07 | ZFPM1 | 15.8860963 | 16.2325943 | 15.8554492 | 16.0423171 | 15.4663257 |
| Q1ZZU3 | SWI5 | 17.6996302 | NA | NA | 17.6459396 | NA |
| Q9C010 | PKIB | 18.0067113 | 18.0437366 | 18.3753486 | 17.9508658 | 18.5170047 |
| Q8WZA1 | POMGNT1 | 15.9933481 | 15.9969927 | 16.1847096 | 15.9504266 | 16.0206161 |
| Q01082 | SPTBN1 | 18.7642381 | 18.8880105 | 18.786049 | 18.7178095 | 18.9155495 |
| Q9NX24 | NHP2 | 21.8077857 | 21.8690631 | 21.9616775 | 21.9677397 | 22.129419 |
| Q15027 | ACAP1 | 19.6988439 | 19.9739177 | 20.0220344 | 19.8876278 | 19.9577388 |
| P19338 | NCL | 27.4941339 | 27.598648 | 27.3482829 | 27.3750715 | 27.4386692 |
| Q9H665 | IGFLR1 | 20.1842722 | 20.1979645 | 20.1420933 | 20.0595978 | 20.2864189 |
| Q96FN4 | CPNE2 | 19.5903851 | 19.6312126 | 19.7365597 | 19.54601 | 19.6565318 |
| P18440 | NAT1 | 19.8334963 | 19.8083636 | 20.0802585 | 19.8768068 | 19.805679 |
| Q9NXR7 | BABAM2 | 21.5458461 | 21.4649762 | 21.5931655 | 21.5199301 | 21.6576842 |
| O75427 | LRCH4 | 21.0831476 | 21.0807082 | 21.1526253 | 21.062303 | 21.0350951 |
| Q08209 | PPP3CA | 22.8101455 | 22.980492 | 22.8617691 | 22.7995559 | 22.9689995 |

| | | | | | | |
|---|---|---|---|---|---|---|
| P50613 | CDK7 | 20.0437188 | 20.1166035 | 20.1204559 | 20.050832 | 20.0552718 |
| Q6PI48 | DARS2 | 21.8269917 | 21.820041 | 21.8764832 | 21.7045105 | 21.8419423 |
| O60244 | MED14 | 19.9017473 | 19.9076942 | 20.0033566 | 19.8652708 | 19.9130045 |
| Q9UBU9 | NXF1 | 21.3342513 | 21.3415223 | 21.369605 | 21.2648283 | 21.3889198 |
| Q9NY97 | B3GNT2 | 20.1767515 | 20.2400883 | 20.2555956 | 20.2212563 | 20.1843051 |
| O00560 | SDCBP | 23.1789574 | 23.2524046 | 23.1386167 | 23.1518359 | 23.2078841 |
| Q68E01 | INTS3 | 21.3651136 | 21.4515325 | 21.4358048 | 21.3462063 | 21.4097979 |
| O43143 | DHX15 | 24.1172298 | 24.2215543 | 24.1912233 | 24.0725329 | 24.1630905 |
| Q92747 | ARPC1A | 21.8470097 | 21.8272578 | 21.8773401 | 21.8022859 | 21.7533257 |
| P35573 | AGL | 19.9769987 | 20.0311435 | 20.0920727 | 19.9561369 | 20.0718408 |
| P18206 | VCL | 23.196778 | 23.2794446 | 23.1859679 | 23.1303636 | 23.2368434 |
| Q86TS9 | MRPL52 | 19.0136677 | 19.0221927 | 19.0819285 | 19.161563 | 19.1324195 |
| P00505 | GOT2 | 23.1930733 | 23.2728033 | 23.3938425 | 23.2098247 | 23.3266963 |
| Q8TB72 | PUM2 | 21.4906465 | 21.6580952 | 21.5246967 | 21.4969603 | 21.5995105 |
| Q9Y3A5 | SBDS | 21.2055085 | 21.3675135 | 21.4287837 | 21.2766786 | 21.4366497 |
| Q9BZE2 | PUS3 | 19.8407045 | 19.8663214 | 19.9163786 | 19.7587586 | 19.8749222 |
| Q04756 | HGFAC | 18.1200561 | 18.121813 | 18.2095697 | 18.1655299 | 18.2441961 |
| Q8TBB5 | KLHDC4 | 20.3315098 | 20.2911496 | 20.5471359 | 20.3962623 | 20.5538213 |
| Q11201 | ST3GAL1 | 17.9749686 | 18.1433918 | 17.9827641 | 18.1416595 | 18.0742042 |
| Q96C01 | FAM136A | 20.8077704 | 20.9003373 | 20.7690736 | 20.7130482 | 20.9835128 |
| O00479 | HMGN4 | 19.4163711 | 19.5048671 | 19.714121 | 19.5935675 | 19.5727925 |
| Q86XE0 | SNX32 | 15.6652575 | 15.9670911 | 15.7958649 | 16.0058591 | 15.9161618 |
| Q05048 | CSTF1 | 21.7168606 | 21.761036 | 21.917577 | 21.6975844 | 21.8583186 |
| Q9Y463 | DYRK1B | 16.9225384 | 17.0790031 | 17.1864204 | 17.1471259 | 17.5235923 |
| Q13487 | SNAPC2 | NA | 17.5513801 | 17.6528828 | NA | 17.6730047 |
| P19367 | HK1 | 21.8730048 | 22.0112448 | 22.0197037 | 21.8066173 | 21.9756267 |
| O95571 | ETHE1 | 23.5446026 | 23.5332781 | 23.5348061 | 23.4698781 | 23.686027 |
| Q9H6V9 | LDAH | 20.0101241 | 20.1656973 | 20.2001991 | 19.9240352 | 20.197142 |
| Q96EF0 | MTMR8 | 15.9138883 | 15.9192673 | 15.7014224 | 15.4865352 | 15.6298334 |
| O15304 | SIVA1 | 17.6289193 | 18.0359741 | 18.0875646 | 17.7400929 | 17.9369947 |
| Q9UDW3 | ZMAT5 | 15.9091532 | 15.8790039 | 15.987808 | 15.9856495 | 15.9174732 |
| P98164 | LRP2 | 18.7767922 | NA | NA | 18.8506265 | NA |
| Q15678 | PTPN14 | 16.5260144 | 16.4273125 | 16.611143 | 16.6158682 | 16.7663321 |
| Q6IPR3 | TYW3 | 18.8961357 | 18.9945928 | 18.9950396 | 18.904225 | 18.7704752 |
| P05386 | RPLP1 | 25.1739688 | 25.2546489 | 25.0841517 | 25.1025107 | 25.1762432 |
| Q7LBR1 | CHMP1B | 22.2922755 | 22.2565259 | 21.9278596 | 22.0350446 | 22.1770771 |
| Q96LW7 | CARD19 | 20.036903 | 19.9936142 | 20.1047587 | 19.959137 | 20.1507093 |
| Q8N2W9 | PIAS4 | 20.9189156 | 21.0394787 | 21.0611948 | 20.9415032 | 21.0689946 |
| Q8TDP1 | RNASEH2C | 21.4599183 | 21.4629809 | 21.4165233 | 21.5927294 | 21.6378516 |
| P42166 | TMPO | 23.6273022 | 23.7298203 | 23.5624445 | 23.6125128 | 23.6966392 |
| Q8TAD8 | SNIP1 | 20.1449995 | 20.1737103 | 20.256867 | 20.1143329 | 20.0549991 |
| Q7L1W4 | LRRC8D | 19.456027 | 19.5922165 | 19.6385202 | 19.4360534 | 19.6306553 |

| | | | | | | |
|---|---|---|---|---|---|---|
| Q92922 | SMARCC1 | 23.3790443 | 23.3906055 | 23.3190343 | 23.3120136 | 23.2728584 |
| P41250 | GARS1 | 23.8063165 | 23.7175522 | 23.7651428 | 23.7367711 | 23.8028498 |
| P54136 | RARS1 | 23.4976468 | 23.6091678 | 23.5754528 | 23.5110677 | 23.6231238 |
| P48444 | ARCN1 | 23.1154059 | 23.0965348 | 23.0548465 | 23.0452874 | 23.0316081 |
| Q86XP3 | DDX42 | 22.5543771 | 22.6448623 | 22.6139328 | 22.562406 | 22.5947629 |
| Q9GZT8 | NIF3L1 | 21.5689795 | 21.5346317 | 21.5099504 | 21.4865826 | 21.5275426 |
| Q9BXB4 | OSBPL11 | 19.5095447 | 19.4630173 | 19.5024956 | 19.4540272 | 19.464177 |
| O75165 | DNAJC13 | 21.2475662 | 21.2587301 | 21.3151763 | 21.2412819 | 21.3053322 |
| Q92619 | ARHGAP45 | 23.2858699 | 23.4561547 | 23.3885865 | 23.3064478 | 23.4071993 |
| Q9UBF8 | PI4KB | 19.0942245 | 19.2007 | 19.2381509 | 19.1742327 | 19.2622451 |
| Q8TEQ6 | GEMIN5 | 23.4458141 | 23.5492333 | 23.5563204 | 23.4496363 | 23.5060414 |
| Q8IWC1 | MAP7D3 | 20.4330876 | 20.4075546 | 20.4088374 | 20.3666565 | 20.4948418 |
| Q96RG2 | PASK | 21.0453 | 21.2038679 | 21.1063942 | 21.0508983 | 21.210506 |
| Q9BYB4 | GNB1L | 19.9361639 | 20.1114698 | 20.0626433 | 19.9289354 | 20.0896666 |
| Q9Y613 | FHOD1 | 20.4726771 | 20.5353798 | 20.580348 | 20.423744 | 20.5808277 |
| P51531 | SMARCA2 | 21.2109101 | 21.3639284 | 21.4159475 | 21.2315242 | 21.3082374 |
| P38646 | HSPA9 | 25.7475564 | 25.8459638 | 25.604394 | 25.6319346 | 25.7164467 |
| Q15910 | EZH2 | 20.3254104 | 20.4147607 | 20.5075307 | 20.3829948 | 20.4396529 |
| Q96NB3 | ZNF830 | 20.0454952 | 20.1067407 | 20.0767908 | 20.0395781 | 20.1843455 |
| Q7Z6J4 | FGD2 | 17.6450426 | 17.8601394 | 18.0782387 | 17.6504575 | 17.8560928 |
| P18754 | RCC1 | 23.4360794 | 23.610947 | 23.4520454 | 23.4412309 | 23.584769 |
| Q8N9R8 | SCAI | 18.5870304 | 18.7513033 | 18.8132038 | 18.7551564 | 18.7252706 |
| Q9UM11 | FZR1 | 17.9072308 | 18.0649709 | 18.0878933 | 17.8962188 | 18.2983742 |
| O15417 | TNRC18 | 20.5920572 | 20.7952716 | 20.5920351 | 20.5669547 | 20.5477879 |
| Q6PID8 | KLHDC10 | 15.6941352 | 16.0457522 | 15.9611981 | 15.917166 | 15.9006891 |
| Q96PV7 | FAM193B | 17.2043282 | 17.153315 | 17.05139 | 16.9279148 | 17.1982139 |
| Q9BRQ8 | AIFM2 | 16.8768508 | NA | NA | 16.909251 | NA |
| Q96JH8 | RADIL | 15.2961686 | 15.3943541 | 15.2527565 | 15.5846258 | 15.5343781 |
| P37287 | PIGA | 17.5037924 | 17.3100579 | 17.4688297 | 17.371875 | 17.3943001 |
| Q8TB37 | NUBPL | 18.9378102 | 19.0958567 | 19.0889037 | 19.077798 | 19.0307338 |
| Q96JJ3 | ELMO2 | 22.1909633 | 22.2426844 | 22.2253439 | 22.1283668 | 22.2156263 |
| Q9NP81 | SARS2 | 23.7154108 | 23.7593828 | 23.7665729 | 23.6832051 | 23.8334381 |
| Q5T160 | RARS2 | 19.8817185 | 19.8776127 | 19.864146 | 19.8503089 | 19.9217398 |
| Q96JB5 | CDK5RAP3 | 20.8485484 | 21.0338947 | 20.9447449 | 20.8954899 | 21.0844827 |
| P62316 | SNRPD2 | 25.4251771 | 25.4457084 | 25.4165609 | 25.3875493 | 25.396893 |
| Q15904 | ATP6AP1 | 22.2282851 | 22.2859049 | 22.0709211 | 22.2821396 | 22.3193884 |
| Q8IY26 | PLPP6 | 19.4586559 | 19.3536428 | 19.5482874 | 19.3186522 | 19.3726434 |
| Q8N5I9 | NOPCHAP1 | 17.4295101 | 17.2583957 | 17.2632094 | 17.2636264 | 17.4502525 |
| Q9BUZ4 | TRAF4 | 17.7266752 | 17.8981742 | 18.1821038 | 17.6460204 | 17.8703196 |
| Q86YN1 | DOLPP1 | 19.5770446 | 19.3280088 | 19.1219101 | 19.4281914 | 19.2275668 |
| P32929 | CTH | 15.4455745 | 15.1354769 | 15.369102 | 15.156103 | 15.0277361 |
| Q9Y3M9 | ZNF337 | NA | 12.2693259 | 13.0982918 | NA | 11.5348449 |

| | | | | | | |
|---|---|---|---|---|---|---|
| P22570 | FDXR | 22.8107752 | 22.9384289 | 22.755339 | 22.7710461 | 22.8568987 |
| Q9C0H5 | ARHGAP39 | NA | 18.6261136 | 18.7528447 | NA | 18.7248435 |
| Q92878 | RAD50 | 22.4475259 | 22.5623552 | 22.5238088 | 22.3972618 | 22.5050717 |
| O95178 | NDUFB2 | 13.3860194 | NA | NA | 13.5892244 | NA |
| Q8WV92 | MITD1 | 18.7655927 | 19.0931241 | 19.0344757 | 18.9159047 | 18.982192 |
| O15514 | POLR2D | 20.6750536 | 20.912804 | 20.8028039 | 20.6758637 | 20.6596582 |
| Q8TER5 | ARHGEF40 | 18.821022 | 19.035444 | 18.9651622 | 18.8847555 | 19.0576348 |
| P17661 | DES | 14.8752979 | NA | NA | 14.9543153 | NA |
| Q7L591 | DOK3 | 22.0024227 | 21.8865989 | 21.9054192 | 21.820226 | 21.9682797 |
| Q9H8U3 | ZFAND3 | 16.5247793 | 16.5181169 | 16.622338 | 16.5064249 | 16.4694241 |
| Q96NG3 | ODAD4 | 21.5989303 | 21.8266733 | 21.9049844 | 21.7378661 | 21.8294335 |
| Q9UBP6 | METTL1 | 20.2949821 | 20.2524468 | 20.3590009 | 20.2682186 | 20.3719589 |
| Q9H1E5 | TMX4 | 19.1783372 | 19.262235 | 19.3928985 | 19.2586683 | 19.3554974 |
| Q7L9B9 | EEPD1 | 19.5473025 | 19.6651683 | 19.6595436 | 19.5459251 | 19.6156804 |
| Q9NV56 | MRGBP | 19.1514583 | 19.452838 | 19.2908329 | 19.3056731 | 19.3424747 |
| O43824 | GTPBP6 | 19.6264685 | 19.6692088 | 19.6779724 | 19.6985424 | 19.6732133 |
| Q6QEF8 | CORO6 | NA | 16.1837247 | 16.2609193 | NA | 16.3035329 |
| Q7L576 | CYFIP1 | 21.5816896 | 21.6876884 | 21.74313 | 21.5702916 | 21.7022468 |
| Q96SW2 | CRBN | 19.2667848 | 19.1289855 | 19.4178297 | 19.2810433 | 19.194299 |
| P48552 | NRIP1 | 17.8013136 | 17.7839835 | 17.9371435 | 17.6446161 | 17.8012421 |
| Q14554 | PDIA5 | 21.9927976 | 22.050753 | 21.9989882 | 21.9686741 | 21.9570701 |
| P13796 | LCP1 | 26.5239234 | 26.5936339 | 26.4787271 | 26.4270514 | 26.563699 |
| Q9C0I4 | THSD7B | 18.9857339 | NA | NA | 18.8036793 | NA |
| Q13526 | PIN1 | 22.9388033 | 23.1124587 | 23.0515053 | 23.0394831 | 23.0102543 |
| Q96S44 | TP53RK | 21.1014886 | 21.0433881 | 21.0953983 | 21.0984492 | 21.1180263 |
| P62937 | PPIA | 28.1907572 | 28.2301603 | 28.0604827 | 28.0529003 | 28.1771966 |
| Q16775 | HAGH | 21.8135875 | 21.8573346 | 21.842616 | 21.741942 | 21.91688 |
| Q9Y450 | HBS1L | 21.1748177 | 21.2425123 | 21.2608505 | 21.0830432 | 21.137761 |
| Q9Y6W5 | WASF2 | 21.478445 | 21.6260337 | 21.5087838 | 21.515235 | 21.5140111 |
| Q86VK4 | ZNF410 | 15.5362608 | 15.932774 | 15.5919137 | 15.8931297 | 15.612206 |
| Q9NRW7 | VPS45 | 19.9054367 | 20.0002819 | 20.0525079 | 19.9132563 | 20.0098737 |
| Q6NYC1 | JMJD6 | 19.6304787 | 19.7669497 | 19.8911834 | 19.688847 | 19.8030078 |
| Q5JVG8 | ZNF506 | 16.9592039 | 16.8739418 | 17.0420472 | 16.5849617 | 17.1381954 |
| Q6Y1H2 | HACD2 | 19.6272257 | 19.9962156 | 19.9855973 | 19.6019356 | 19.0140216 |
| Q8IYD2 | KLHDC8A | NA | 15.7895907 | 15.9451036 | NA | 15.9182444 |
| Q9Y252 | RNF6 | 16.966009 | 16.8980255 | 17.0128929 | 17.0889009 | 16.9733927 |
| Q10588 | BST1 | 19.6926303 | 19.8642563 | 19.8657045 | 19.7844771 | 19.7622787 |
| Q9NX04 | AIRIM | 18.3072143 | 18.4511204 | 18.467448 | 18.3340082 | 18.3403132 |
| Q9C0G0 | ZNF407 | 18.1354679 | 18.2900424 | 18.2067472 | 18.1412286 | 18.1159935 |
| Q9BV86 | NTMT1 | 20.3671448 | 20.5000504 | 20.3623975 | 20.3681303 | 20.3882474 |
| P78371 | CCT2 | 25.7158672 | 25.7466255 | 25.6600796 | 25.6182012 | 25.7530565 |
| Q9NTJ3 | SMC4 | 23.0157468 | 23.1065916 | 23.0192625 | 22.9192407 | 23.03485 |

| | | | | | | |
|---|---|---|---|---|---|---|
| Q96ST3 | SIN3A | 21.4434658 | 21.523471 | 21.5630638 | 21.4468699 | 21.5145805 |
| P98175 | RBM10 | 21.5424836 | 21.6223126 | 21.5967691 | 21.5090898 | 21.5693035 |
| P62333 | PSMC6 | 23.9719394 | 24.0291243 | 23.9376159 | 23.872696 | 23.9401914 |
| P54252 | ATXN3 | 20.9174609 | 21.136563 | 20.9597692 | 20.9889054 | 21.0411258 |
| Q92574 | TSC1 | 18.5337125 | 18.6197615 | 18.5932546 | 18.4814258 | 18.5113373 |
| Q9UEG4 | ZNF629 | 19.5105969 | 19.5625544 | 19.5845188 | 19.6027603 | 19.5253113 |
| Q9UM13 | ANAPC10 | 17.1744395 | 17.388487 | 17.2938245 | 17.2055449 | 17.016069 |
| O43826 | SLC37A4 | 17.4258108 | 17.3127777 | 17.400486 | 17.3961495 | 17.5370197 |
| A6NIH7 | UNC119B | 17.9581879 | 17.9946999 | 18.2213122 | 17.9303558 | 18.1540208 |
| O60220 | TIMM8A | 20.7450886 | 20.9199589 | 20.755452 | 20.6916116 | 21.0664638 |
| Q9UQ88 | CDK11A | 17.4466362 | 17.6725489 | 17.5988318 | 17.4699509 | 17.4846151 |
| O43292 | GPAA1 | 21.7812513 | 21.8318114 | 22.0474432 | 21.7817476 | 21.8920933 |
| Q15003 | NCAPH | 21.7030045 | 21.8709215 | 21.6915418 | 21.6185683 | 21.708936 |
| Q00765 | REEP5 | 20.8558637 | 20.9382145 | 20.9717104 | 20.8427042 | 20.8537968 |
| Q15172 | PPP2R5A | 20.047303 | 20.1669288 | 20.1041757 | 20.1385432 | 20.1670629 |
| Q8TB45 | DEPTOR | 19.8707152 | 19.8698755 | 19.8432704 | 19.8662008 | 19.8209709 |
| Q9H3K6 | NA | 23.3004813 | 23.2116544 | 23.2382324 | 23.2365772 | 23.3908016 |
| A7E2V4 | ZSWIM8 | 19.5723206 | 19.6149472 | 19.4603808 | 19.4692235 | 19.4859383 |
| Q99567 | NUP88 | 21.5172517 | 21.5556477 | 21.5088182 | 21.514653 | 21.550202 |
| P48507 | GCLM | 20.4782025 | 20.5209076 | 20.5470323 | 20.4197549 | 20.6113637 |
| Q8IY17 | PNPLA6 | 22.4719952 | 22.6083702 | 22.6516029 | 22.5165315 | 22.5877896 |
| P17480 | UBTF | 22.590145 | 22.5709998 | 22.6821064 | 22.5472377 | 22.6655385 |
| O75348 | ATP6V1G1 | 22.6102731 | 22.8109701 | 22.5938241 | 22.7076023 | 22.5122675 |
| Q9Y606 | PUS1 | 22.0656425 | 22.0421873 | 22.1027733 | 22.0020136 | 21.9964081 |
| Q15149 | PLEC | 24.2457115 | 24.3058372 | 24.2041448 | 24.1698118 | 24.2387668 |
| Q6PJF5 | RHBDF2 | 18.8923403 | 19.1728818 | 19.1761108 | 19.0986638 | 18.9779179 |
| O75147 | OBSL1 | 19.117966 | 19.2545737 | 19.1012614 | 19.0735262 | 19.0256659 |
| Q8IXM3 | MRPL41 | 21.3449322 | 21.2797648 | 21.3708922 | 21.210124 | 21.1937936 |
| Q53GT1 | KLHL22 | 19.0736441 | 18.953585 | 19.0684137 | 18.9349807 | 19.1266536 |
| Q9H0A8 | COMMD4 | 20.4403119 | 20.5905392 | 20.592251 | 20.4853668 | 20.681287 |
| Q12891 | HYAL2 | 14.7849156 | 15.2322471 | 15.5112321 | 15.257673 | 15.3039352 |
| Q13232 | NME3 | 23.0544979 | 23.0593772 | 23.0626592 | 22.7850637 | 22.9868361 |
| Q9UHX1 | PUF60 | 24.040637 | 24.0698565 | 23.9985371 | 23.9815691 | 24.0563968 |
| Q8TB96 | ITFG1 | 18.4610056 | 18.5606795 | 18.5311775 | 18.619014 | 18.6068456 |
| O43463 | SUV39H1 | 17.8776019 | 18.1982461 | 18.477491 | 18.1692189 | 18.1222823 |
| Q9NR50 | EIF2B3 | 21.4020462 | 21.3099997 | 21.3694462 | 21.331136 | 21.429352 |
| Q5T440 | IBA57 | 20.3885234 | 20.3626698 | 20.5851431 | 20.3641649 | 20.4276606 |
| O94759 | TRPM2 | 20.1293179 | 20.0920959 | 20.066518 | 20.0844205 | 20.0317367 |
| P20933 | AGA | 20.7743757 | 20.9680469 | 20.9086905 | 20.7891445 | 20.9157789 |
| Q9UBS4 | DNAJB11 | 22.9102361 | 23.0643563 | 22.9429112 | 22.9997371 | 22.9279168 |
| O75494 | SRSF10 | 23.5882179 | 23.7132231 | 23.4589874 | 23.5277877 | 23.5905573 |
| Q8IWS0 | PHF6 | 21.6403278 | 21.502833 | 21.6179019 | 21.4629269 | 21.5812119 |

| | | | | | | |
|---|---|---|---|---|---|---|
| Q9UGP4 | LIMD1 | 19.6003398 | 19.6283337 | 19.6030803 | 19.5957654 | 19.8057659 |
| Q9UNX3 | RPL26L1 | 21.8492248 | 21.6198371 | 21.8396838 | 21.8726994 | 21.3693153 |
| Q9P2K2 | TXNDC16 | 16.3022446 | 16.3953055 | 16.4902288 | 16.6424653 | 16.2945391 |
| Q15477 | SKIC2 | 21.5985611 | 21.6378862 | 21.6375675 | 21.576051 | 21.6019832 |
| Q14974 | KPNB1 | 24.0713893 | 24.0973304 | 24.0684732 | 23.945638 | 24.122038 |
| P16885 | PLCG2 | 21.9858006 | 22.1193157 | 22.102734 | 21.9827813 | 22.1067458 |
| Q86TP1 | PRUNE1 | 18.593965 | 18.607446 | 18.7011005 | 18.489132 | 18.7126526 |
| Q13164 | MAPK7 | 18.60879 | 18.7870412 | 18.7853146 | 18.704654 | 18.6810457 |
| Q13572 | ITPK1 | 19.296896 | 19.5839699 | 19.4573222 | 19.4814003 | 19.481879 |
| Q6UX07 | DHRS13 | 18.9448553 | 19.1890373 | 19.1556223 | 18.8806131 | 19.08502 |
| Q92673 | SORL1 | 17.4487154 | 17.4801573 | 17.4234297 | 17.8350228 | 17.4724742 |
| P51684 | CCR6 | 17.6176055 | 17.5069737 | 17.2298856 | 17.3947265 | 17.3088139 |
| Q8NCC3 | PLA2G15 | 20.7129426 | 20.7662913 | 20.7715293 | 20.7274975 | 20.7443538 |
| P17405 | SMPD1 | 17.057088 | 17.3877514 | 17.3144539 | 17.1617358 | 17.4306549 |
| A6NFY7 | SDHAF1 | 17.2925899 | 17.0089064 | 17.5748762 | 17.2205198 | 17.2717308 |
| P30405 | PPIF | 24.4224634 | 24.5092072 | 24.4198759 | 24.4625836 | 24.5244494 |
| P36969 | GPX4 | 24.2513609 | 24.204369 | 24.1758739 | 24.1975154 | 24.250929 |
| Q5HYI7 | MTX3 | 19.0439348 | 19.14165 | 19.3417628 | 19.1250465 | 19.2536351 |
| O95169 | NDUFB8 | 21.539368 | 21.6039032 | 21.674033 | 21.6128545 | 21.599615 |
| P40121 | CAPG | 23.6203662 | 23.763677 | 23.6464941 | 23.5670861 | 23.6931558 |
| P43251 | BTD | 19.0246163 | 19.2461556 | 19.2633148 | 19.1678859 | 19.1113536 |
| P50991 | CCT4 | 25.650096 | 25.7504139 | 25.6292326 | 25.583229 | 25.7002601 |
| Q9Y2X7 | GIT1 | 21.6781983 | 21.7051862 | 21.7883116 | 21.6519176 | 21.8284932 |
| Q16236 | NFE2L2 | 16.491708 | NA | NA | 16.1346963 | NA |
| Q8IWL3 | HSCB | 19.5342288 | 19.6840434 | 19.5708047 | 19.6245527 | 19.6105547 |
| P50454 | SERPINH1 | 23.4058736 | 23.44511 | 23.4237972 | 23.365416 | 23.4489541 |
| Q9UKV3 | ACIN1 | 23.2923846 | 23.39835 | 23.2961886 | 23.2543323 | 23.3195721 |
| Q9H3N1 | TMX1 | 22.8985232 | 22.9737762 | 23.0634168 | 22.9173612 | 23.0693687 |
| Q9Y5B0 | CTDP1 | 21.2189989 | 21.3066438 | 21.2329016 | 21.2022915 | 21.2267161 |
| Q86WV5 | TEN1 | 12.5967366 | NA | NA | 12.5979905 | NA |
| P06400 | RB1 | 22.5000042 | 22.5642531 | 22.5601509 | 22.4061024 | 22.5269863 |
| Q9NW08 | POLR3B | 19.9432997 | 19.934958 | 20.0738845 | 19.9133814 | 20.0018225 |
| P55036 | PSMD4 | 23.9426944 | 23.9208662 | 23.8692801 | 23.8649265 | 23.9627748 |
| Q9BW62 | KATNAL1 | 16.0464261 | 16.2231216 | 16.333802 | 16.137859 | 16.4211498 |
| Q6P4E1 | GOLM2 | 19.9208012 | 20.0158489 | 20.015727 | 20.0274684 | 19.8848031 |
| Q6NXE6 | ARMC6 | 22.5719252 | 22.6027846 | 22.6566025 | 22.5524353 | 22.6429117 |
| P35241 | RDX | 21.4255502 | 21.4644096 | 21.4598885 | 21.3901247 | 21.4762777 |
| Q99633 | PRPF18 | 19.8547944 | 19.9190976 | 19.8104046 | 19.8296351 | 19.8952084 |
| Q9NZT2 | OGFR | 21.5079708 | 21.6736373 | 21.6178464 | 21.5311391 | 21.66496 |
| Q9NPR2 | SEMA4B | 15.1119238 | 15.6232308 | 15.5694614 | 14.662504 | 14.3760544 |
| Q8TAT5 | NEIL3 | 14.9781198 | NA | NA | 14.9667511 | NA |
| Q9BRA2 | TXNDC17 | 23.1831867 | 23.1333247 | 22.9788873 | 23.0608967 | 23.3072091 |

| | | | | | | |
|---|---|---|---|---|---|---|
| Q9Y5J9 | TIMM8B | 22.0078993 | 22.0259046 | 21.8717248 | 21.8451298 | 21.9231533 |
| P11766 | ADH5 | 23.6389614 | 23.7474302 | 23.6765962 | 23.6224292 | 23.6629661 |
| Q92572 | AP3S1 | 21.2556251 | 21.2501572 | 21.2724022 | 21.2519543 | 21.3769487 |
| Q8N573 | OXR1 | 19.7718223 | 19.9141671 | 19.8991046 | 19.7591965 | 19.8997795 |
| Q9NRW4 | DUSP22 | 17.0458346 | 16.919725 | 17.3056788 | 17.2440131 | 17.2622204 |
| Q9UJA3 | MCM8 | 15.0734632 | 15.6678402 | 15.86003 | 15.3559985 | 15.7023255 |
| P0CG12 | DERPC | 19.807512 | 19.9771849 | 20.2953073 | 19.9525564 | 19.7003375 |
| Q8N5S9 | CAMKK1 | 18.0504281 | 18.2850893 | 18.2101503 | 18.0494993 | 18.2563499 |
| Q8TD30 | GPT2 | 20.37024 | 20.3056437 | 20.2681129 | 20.3377896 | 20.2384823 |
| P14625 | HSP90B1 | 26.4613228 | 26.488382 | 26.3888299 | 26.3779495 | 26.4354335 |
| O15254 | ACOX3 | 20.9807731 | 20.8712205 | 21.0191815 | 20.9283437 | 21.0340988 |
| Q9Y2W2 | WBP11 | 21.8937632 | 22.0442067 | 21.9017961 | 21.9584439 | 21.9823901 |
| Q13111 | CHAF1A | 20.7532596 | 20.7803037 | 20.8145415 | 20.7558911 | 20.786696 |
| Q5EBM0 | CMPK2 | 17.8187878 | 17.8381303 | 17.8825802 | 17.7491982 | 17.8477774 |
| Q6ZS72 | PEAK3 | 19.217637 | 19.3728917 | 19.4018147 | 19.2093533 | 19.3879171 |
| P62633 | CNBP | 23.7083308 | 23.9283708 | 23.6737734 | 23.6155571 | 23.8199618 |
| O95445 | APOM | 16.9942048 | 17.5686761 | 17.9316242 | 16.7909862 | 17.6287512 |
| Q9BS34 | ZNF670 | 21.5833481 | 21.6336756 | 21.5811554 | 21.6471354 | 21.6539014 |
| Q96T49 | PPP1R16B | 16.6489915 | 16.6653591 | 16.4071891 | 16.5638259 | 16.5603859 |
| Q9BQE9 | BCL7B | 21.0428058 | 21.1616064 | 20.9554459 | 20.9817391 | 21.1224632 |
| P60174 | TPI1 | 26.9423552 | 27.1057421 | 26.9012841 | 26.8436256 | 27.0656887 |
| Q9NPJ8 | NXT2 | 21.0046874 | 20.9202019 | 20.9838683 | 21.0493716 | 20.9129472 |
| P49257 | LMAN1 | 23.4004974 | 23.5146607 | 23.446603 | 23.3872205 | 23.407336 |
| P48730 | CSNK1D | 20.966292 | 20.9706953 | 20.9566569 | 20.8661588 | 20.9597541 |
| Q9ULP9 | TBC1D24 | 20.7230111 | 20.6959718 | 20.9009137 | 20.6890829 | 20.8114683 |
| O75387 | SLC43A1 | 19.6318132 | 19.6375274 | 19.7218632 | 19.4665363 | 19.6480498 |
| P05423 | POLR3D | 18.6943518 | 18.7680328 | 18.7709161 | 18.8307972 | 18.6935998 |
| P62841 | RPS15 | 23.9754202 | 24.036829 | 24.0698376 | 23.9524294 | 23.8264731 |
| O75648 | TRMU | 20.21545 | 20.2025787 | 20.3008577 | 20.1785113 | 20.3029269 |
| P50440 | GATM | 21.9684609 | 22.0998707 | 22.0988022 | 22.0263231 | 22.131477 |
| Q9P227 | ARHGAP23 | NA | 12.7451275 | 12.4594327 | NA | 10.8251004 |
| P53667 | LIMK1 | 18.8265876 | 19.0738533 | 18.9183332 | 18.7991817 | 19.1530257 |
| Q8WU10 | PYROXD1 | 19.5716466 | 19.6499511 | 19.587235 | 19.679213 | 19.6104224 |
| O95487 | SEC24B | 20.2245438 | 20.2118158 | 20.306574 | 20.2814613 | 20.3280825 |
| P61981 | YWHAG | 27.9531249 | 27.929084 | 27.8145176 | 27.7535928 | 27.8004946 |
| Q96FH0 | BORCS8 | 18.8852301 | 18.8187586 | 19.1537082 | 19.1271494 | 19.0250589 |
| O75123 | ZNF623 | 16.3774034 | 16.3684868 | 16.4308855 | 16.4243614 | 16.3993899 |
| Q8TCT0 | CERK | 17.1120283 | 17.1086916 | 17.6148477 | 16.9751424 | 17.3375283 |
| Q9NWW6 | NMRK1 | 15.4640233 | 15.3931326 | 15.6123571 | 15.7853871 | 15.7279024 |
| Q99640 | PKMYT1 | 18.4624966 | 18.5127299 | 18.6761883 | 18.4402726 | 18.6218138 |
| O00268 | TAF4 | 19.258731 | 19.3011166 | 19.2575807 | 19.3141792 | 19.3548233 |
| Q02127 | DHODH | 20.6959102 | 20.8328329 | 20.7981593 | 20.6721585 | 20.7300525 |

| | | | | | | |
|---|---|---|---|---|---|---|
| O95182 | NDUFA7 | 21.9559997 | 21.803173 | 22.0865342 | 21.878413 | 21.9239519 |
| Q92930 | RAB8B | 20.1643518 | 20.2625817 | 20.55514 | 20.4973578 | 20.6274515 |
| Q9H8M5 | CNNM2 | NA | 13.4791786 | 14.5496909 | NA | 13.5174743 |
| Q8WVC0 | LEO1 | 21.0033465 | 21.0968221 | 21.2031841 | 20.9792301 | 21.0400299 |
| Q9Y2K7 | KDM2A | 20.1279182 | 20.1620456 | 20.1942437 | 20.0287634 | 20.2054226 |
| P46952 | HAAO | 18.5505516 | 18.7799889 | 18.9040612 | 18.5763113 | 18.8428002 |
| Q9NYB0 | TERF2IP | 21.0033421 | 21.239924 | 20.9332611 | 21.0284031 | 21.1211066 |
| P19237 | TNNI1 | NA | 12.8241866 | 13.2495913 | NA | 13.1015541 |
| Q9BRK4 | LZTS2 | 16.2246877 | 16.1286515 | 16.1728973 | 16.6308485 | 16.136959 |
| Q96A57 | TMEM230 | 18.0191524 | 17.947585 | 18.3743203 | 18.072432 | 18.0074733 |
| Q9H1C4 | UNC93B1 | 21.0668642 | 21.1324757 | 21.264954 | 21.0255366 | 21.2255633 |
| Q9HCN8 | SDF2L1 | 21.9708392 | 22.0688841 | 22.0850124 | 22.0221757 | 22.0462319 |
| Q9UH62 | ARMCX3 | 20.4547108 | 20.5298185 | 20.4758902 | 20.4875259 | 20.4705622 |
| O95197 | RTN3 | 23.2214006 | 23.2485733 | 23.1596522 | 23.1463385 | 23.0188344 |
| Q14691 | GINS1 | 19.4752637 | 19.6002073 | 19.4255611 | 19.543804 | 19.5570321 |
| Q9UBQ0 | VPS29 | 23.959515 | 23.9859723 | 24.0708677 | 23.9183121 | 23.9970255 |
| O43772 | SLC25A20 | 21.0252152 | 21.1863986 | 21.1273708 | 20.9520043 | 21.0025438 |
| P06213 | INSR | 21.2502968 | 21.3195417 | 21.2783997 | 21.2197056 | 21.2974905 |
| O95758 | PTBP3 | 22.637101 | 22.6206429 | 22.7360231 | 22.6077649 | 22.7294656 |
| O60231 | DHX16 | 21.1104071 | 21.3022227 | 21.2058833 | 21.1212133 | 21.2213527 |
| P61009 | SPCS3 | 23.1839588 | 23.0786835 | 23.1330127 | 23.0774 | 23.0971062 |
| Q14457 | BECN1 | 20.188763 | 20.316248 | 20.2601445 | 20.2068281 | 20.2437497 |
| O43615 | TIMM44 | 22.6392551 | 22.7062716 | 22.6075481 | 22.5858752 | 22.5716375 |
| Q709C8 | VPS13C | 20.7216198 | 20.779918 | 20.7968177 | 20.6999857 | 20.7923445 |
| Q92608 | DOCK2 | 22.2046907 | 22.2458999 | 22.2714026 | 22.1713355 | 22.2560785 |
| Q9BXR0 | QTRT1 | 21.2959712 | 21.2790255 | 21.2856236 | 21.1632472 | 21.2778704 |
| O75439 | PMPCB | 22.3786486 | 22.291387 | 22.350522 | 22.3191934 | 22.4229381 |
| Q7Z5J4 | RAI1 | 20.6227234 | 20.7457201 | 20.7849803 | 20.657276 | 20.7036048 |
| Q13490 | BIRC2 | 18.5010409 | 18.6800811 | 18.4491476 | 18.5230388 | 18.4687863 |
| Q96EP0 | RNF31 | 19.8938947 | 20.1908895 | 20.0987297 | 20.0564762 | 20.178036 |
| Q96GK7 | FAHD2A | 21.0993292 | 21.2820724 | 21.193236 | 21.1526813 | 21.2616266 |
| P04233 | CD74 | 21.8836506 | 22.0027209 | 21.9945414 | 21.8810794 | 21.9934405 |
| Q13257 | MAD2L1 | 22.2320853 | 22.2773442 | 22.2688076 | 22.1346473 | 22.193401 |
| Q9NRF9 | POLE3 | 23.0336323 | 23.1526493 | 22.8818509 | 22.920793 | 22.9239831 |
| Q8IU60 | DCP2 | 20.0526301 | 19.9941234 | 20.1880533 | 20.0109747 | 20.0675953 |
| Q13233 | MAP3K1 | 18.8752368 | 19.1723775 | 19.1304162 | 18.9869233 | 19.1399702 |
| O95864 | FADS2 | 23.3442799 | 23.308957 | 23.3880173 | 23.3313312 | 23.4230959 |
| Q5W0B1 | OBI1 | 18.7315809 | 18.9132275 | 18.8626317 | 18.6516885 | 18.8077211 |
| Q9H008 | LHPP | 18.6533524 | 18.6453176 | 18.7888822 | 18.5216825 | 18.6332772 |
| Q6QNY0 | BLOC1S3 | 20.3662431 | 20.42093 | 20.2203778 | 20.3554461 | 20.5011024 |
| Q93075 | TATDN2 | 18.4129025 | 18.7692975 | 18.61003 | 18.5320363 | 18.5962823 |
| P17181 | IFNAR1 | 16.6079173 | 16.8284053 | 16.8841319 | 16.762801 | 16.8866712 |

| | | | | | | |
|---|---|---|---|---|---|---|
| P0C870 | JMJD7 | 16.1917767 | 16.2995429 | 16.5038191 | 16.1753653 | 16.4628291 |
| Q16850 | CYP51A1 | 20.8993408 | 21.0263705 | 21.1361414 | 20.7786894 | 21.2051575 |
| Q3LIE5 | ADPRM | 12.8014655 | 12.765152 | 13.1426467 | 12.8841161 | 13.2013598 |
| Q92597 | NDRG1 | 20.9123291 | 21.3474029 | 21.1640944 | 20.8546995 | 21.4369748 |
| Q9BV29 | CCDC32 | 18.8274618 | 19.0878988 | 19.0719497 | 18.8162283 | 18.8436773 |
| O75478 | TADA2A | 16.6050928 | 16.7301843 | 16.9140213 | 16.4955229 | 16.7658827 |
| Q8TBZ3 | WDR20 | 18.0895855 | 18.3577175 | 18.2585573 | 18.3182913 | 18.3425491 |
| P29375 | KDM5A | 18.7555182 | 18.8457615 | 18.9663248 | 18.8167876 | 18.8743146 |
| O14628 | ZNF195 | 20.05711 | 20.1428823 | 20.1641125 | 20.1041938 | 20.1833467 |
| P53990 | IST1 | 22.5047203 | 22.6463543 | 22.6385329 | 22.4841517 | 22.5433701 |
| P61964 | WDR5 | 21.7281137 | 21.7402124 | 21.7303071 | 21.6873585 | 21.7525201 |
| Q9Y689 | ARL5A | 16.9769028 | 17.2089315 | 17.1402602 | 17.1386749 | 17.1975658 |
| Q5TFE4 | NT5DC1 | 19.1550297 | 19.2105433 | 19.4345582 | 19.1257836 | 19.2903302 |
| Q01151 | CD83 | 17.1561989 | 17.1927628 | 17.0306161 | 17.10421 | 17.3698938 |
| Q15072 | ZNF146 | 19.0322332 | 19.2890384 | 19.2815794 | 19.1067621 | 19.0373002 |
| Q9Y4D1 | DAAM1 | 17.5618817 | 17.6659407 | 17.7006115 | 17.5551569 | 17.526223 |
| Q96RL7 | VPS13A | 18.6856655 | 18.7391137 | 18.7795995 | 18.6837639 | 18.8245135 |
| Q9BVS4 | RIOK2 | 20.1796691 | 20.3099296 | 20.2981223 | 20.2972437 | 20.375433 |
| P22102 | GART | 24.0366566 | 24.0887699 | 23.9968386 | 23.9746602 | 24.0390951 |
| Q12933 | TRAF2 | 20.532793 | 20.5177028 | 20.5696558 | 20.4491716 | 20.5042317 |
| P08670 | VIM | 27.0855348 | 27.1786756 | 27.0043171 | 26.9564319 | 27.093976 |
| Q86VW0 | SESTD1 | 16.7294554 | 16.7111692 | 16.6601933 | 16.6008763 | 16.5948823 |
| Q92888 | ARHGEF1 | 22.2927063 | 22.4016498 | 22.3834783 | 22.2567709 | 22.4034425 |
| O00462 | MANBA | 20.6374085 | 20.5660811 | 20.642286 | 20.6438661 | 20.7547332 |
| Q96SK2 | TMEM209 | 19.7718435 | 19.7259685 | 19.8124462 | 19.7978168 | 19.8408197 |
| Q9UHJ6 | SHPK | 19.9296249 | 19.735226 | 19.8454442 | 19.8173011 | 19.9003051 |
| Q9NZK5 | ADA2 | 20.6022687 | 20.7763468 | 20.9456457 | 20.6187809 | 20.9426483 |
| Q7L2E3 | DHX30 | 21.5990796 | 21.6006251 | 21.6474136 | 21.5595137 | 21.6218859 |
| P05165 | PCCA | 21.0769102 | 21.1806036 | 21.1053289 | 21.0380351 | 21.1345866 |
| Q8N335 | GPD1L | 21.0854207 | 21.3038136 | 21.1707456 | 21.0578282 | 21.2434011 |
| Q9GZR7 | DDX24 | 22.304123 | 22.4182653 | 22.4194202 | 22.3163276 | 22.4292209 |
| P49619 | DGKG | 15.6313869 | NA | NA | 15.7222231 | NA |
| P04792 | HSPB1 | 24.0298864 | 24.1475364 | 24.0491036 | 24.0017731 | 24.1171892 |
| Q14644 | RASA3 | 20.0774883 | 20.1968069 | 20.1935441 | 20.1706746 | 20.1795523 |
| O00391 | QSOX1 | 21.0151233 | 20.9721498 | 20.8126824 | 20.9851186 | 20.7067715 |
| O43829 | ZBTB14 | 17.8015212 | 18.0826173 | 17.8956133 | 17.8670928 | 17.8291076 |
| O95168 | NDUFB4 | 22.2508747 | 22.3069865 | 22.2827291 | 22.1691186 | 22.277049 |
| Q9H6S0 | YTHDC2 | 21.9927207 | 21.997667 | 22.0036822 | 21.9512439 | 22.0418637 |
| Q5QJE6 | DNTTIP2 | 21.0864564 | 21.1628667 | 21.1599868 | 21.157051 | 21.2223452 |
| P55039 | DRG2 | 20.8277713 | 20.8357362 | 20.964611 | 20.7585716 | 20.9292978 |
| P98173 | FAM3A | 15.5182415 | 15.4962917 | 15.7138048 | 15.605223 | 15.5862657 |
| Q86XZ4 | SPATS2 | 15.1330168 | 15.3365435 | 15.4593718 | 15.4951778 | 15.3444893 |

| | | | | | | |
|---|---|---|---|---|---|---|
| Q8N5Y8 | PARP16 | 15.099086 | 15.4485969 | 15.7380095 | 15.2149852 | 15.6059908 |
| Q9Y5N6 | ORC6 | 18.5410501 | 18.7357998 | 18.7055689 | 18.6177574 | 18.4429337 |
| Q9UIG0 | BAZ1B | 21.8637115 | 21.9281388 | 21.9921608 | 21.822951 | 21.8997479 |
| Q8IW45 | NAXD | 19.5708524 | 19.6253676 | 19.6483986 | 19.6261261 | 19.6621428 |
| Q6AW86 | ZNF324B | 13.5503312 | NA | NA | 13.8810803 | NA |
| P18850 | ATF6 | 16.4776928 | NA | NA | 16.4877591 | NA |
| Q9NYR9 | NKIRAS2 | 19.2013332 | 19.4092181 | 19.2906356 | 19.21686 | 19.3944369 |
| Q9HBL7 | PLGRKT | 19.3217317 | 19.6801563 | 19.4939077 | 19.5275554 | 19.5362399 |
| P04066 | FUCA1 | 21.8735774 | 21.8936275 | 22.0169188 | 21.8011821 | 21.9474375 |
| Q9Y3D0 | CIAO2B | 19.7082093 | 19.7100118 | 19.7794131 | 19.6651838 | 19.7872783 |
| O75608 | LYPLA1 | 22.990727 | 23.0692895 | 23.0595893 | 22.9781898 | 23.1311239 |
| Q9P2D3 | HEATR5B | 21.7544514 | 21.7323396 | 21.7750259 | 21.6822937 | 21.7626138 |
| P52306 | RAP1GDS1 | 23.2074904 | 23.2800962 | 23.2217397 | 23.1393943 | 23.2515222 |
| Q9Y5Q9 | GTF3C3 | 20.666572 | 20.7376747 | 20.8007128 | 20.6966174 | 20.7191482 |
| Q9BVS5 | TRMT61B | 18.453173 | 18.4576358 | 18.5181365 | 18.4787124 | 18.4605776 |
| Q9BRK5 | SDF4 | 21.7070095 | 21.8196013 | 21.691758 | 21.6849507 | 21.7771606 |
| Q52LA3 | LIN52 | 16.9380937 | 17.2000067 | 16.6265162 | 16.800815 | 17.2776112 |
| Q5TGZ0 | MICOS10 | 20.6705191 | 21.0226658 | 21.1626712 | 20.7647018 | 20.8710836 |
| Q96RY7 | IFT140 | 17.2814227 | 17.3222839 | 17.1645568 | 17.1864314 | 17.4588271 |
| P52566 | ARHGDIB | 27.254429 | 27.2971031 | 27.2198627 | 27.1212507 | 27.2353009 |
| P42229 | STAT5A | 21.5787184 | 21.5472409 | 21.5986473 | 21.5146929 | 21.5520603 |
| P53634 | CTSC | 23.0157062 | 22.9411947 | 23.1491474 | 22.9950751 | 23.126194 |
| O15382 | BCAT2 | 22.1310548 | 22.1731731 | 22.2068815 | 22.0689055 | 22.1700967 |
| Q9HCE1 | MOV10 | 16.7278104 | 16.7595434 | 16.7808235 | 16.5785173 | 16.5924944 |
| P11049 | CD37 | 20.0656553 | 20.2993052 | 20.1695158 | 20.2401117 | 20.2065709 |
| Q9Y6Y8 | SEC23IP | 22.1725866 | 22.1451622 | 22.218219 | 22.1178908 | 22.2511054 |
| P36405 | ARL3 | 20.6068549 | 20.7148328 | 20.7197605 | 20.5083094 | 20.8336656 |
| Q9Y2S2 | CRYL1 | 18.6055702 | 18.5504276 | 18.8728049 | 18.6253911 | 18.6921383 |
| Q8N183 | NDUFAF2 | 20.400327 | 20.3193709 | 20.3063073 | 20.3160795 | 20.4181236 |
| P57071 | PRDM15 | 16.6576541 | 16.7506084 | 16.8302193 | 16.8234334 | 16.7857437 |
| P18084 | ITGB5 | 17.5365025 | 17.5669723 | 17.3860731 | 17.5055829 | 17.5791579 |
| Q15165 | PON2 | 18.3304657 | 18.4708922 | 18.4291587 | 18.2918265 | 18.4830714 |
| P54619 | PRKAG1 | 20.9803641 | 20.9300139 | 20.913496 | 20.8745304 | 21.0213024 |
| Q6ZS81 | WDFY4 | 20.9130486 | 20.9674199 | 20.9519952 | 20.8636269 | 20.9743057 |
| Q7Z434 | MAVS | 20.9356349 | 20.9550507 | 21.0029951 | 20.9330237 | 20.9971849 |
| Q9Y676 | MRPS18B | 20.4562764 | 20.5581455 | 20.5357224 | 20.4130431 | 20.4176673 |
| Q8TCC3 | MRPL30 | 21.5716356 | 21.5477189 | 21.6195722 | 21.6269057 | 21.6187879 |
| Q8IYM9 | TRIM22 | 22.1181962 | 22.1149744 | 22.1152765 | 22.0362014 | 22.1479316 |
| Q2M296 | MTHFSD | 20.3473798 | 20.4191091 | 20.3207185 | 20.4755324 | 20.46308 |
| P36894 | BMPR1A | NA | 13.6739466 | 13.7809548 | NA | 13.9929012 |
| O75175 | CNOT3 | 21.4724661 | 21.599724 | 21.4775283 | 21.4378382 | 21.5153214 |
| Q9H1Y0 | ATG5 | 19.844773 | 20.0439686 | 19.8996614 | 19.8276551 | 20.0167136 |

| | | | | | | |
|---|---|---|---|---|---|---|
| P25942 | CD40 | NA | 14.8123314 | 14.5092454 | NA | 14.4723014 |
| Q92686 | NRGN | 21.7665192 | 22.0436749 | 22.0443596 | 21.8955097 | 21.9380493 |
| P52209 | PGD | 25.9563477 | 25.8992552 | 25.8649579 | 25.8270933 | 25.889315 |
| Q8N0U8 | VKORC1L1 | 20.8428573 | 20.8201447 | 21.0029761 | 20.8473586 | 20.8302879 |
| O75907 | DGAT1 | 19.9792536 | 20.1802983 | 20.0588318 | 20.0934051 | 20.0391788 |
| O75081 | CBFA2T3 | 19.2905304 | 19.3994895 | 19.4381564 | 19.257519 | 19.3459309 |
| Q96GW9 | MARS2 | 19.9608225 | 19.9249845 | 19.9658425 | 19.9431651 | 20.119387 |
| Q9Y6K0 | CEPT1 | 20.0593418 | 20.125706 | 20.151381 | 20.0610148 | 20.1588034 |
| Q8N3F8 | MICALL1 | 23.9964615 | 23.9700092 | 23.9820622 | 23.8681565 | 23.905552 |
| P0DJ07 | PET100 | 20.6012574 | 20.4965888 | 20.5759589 | 20.3232862 | 20.4688274 |
| Q96CM3 | RPUSD4 | 19.9797605 | 19.7719925 | 19.9375683 | 19.8942639 | 19.8124986 |
| P05198 | EIF2S1 | 23.9305151 | 23.9484698 | 23.8944454 | 23.9051111 | 23.9562208 |
| Q12849 | GRSF1 | 21.4170699 | 21.5576721 | 21.4367212 | 21.4425051 | 21.5695332 |
| O94874 | UFL1 | 22.2850431 | 22.3912852 | 22.4020038 | 22.323363 | 22.4222415 |
| Q15334 | LLGL1 | 19.3952203 | 19.4738654 | 19.5620819 | 19.4525875 | 19.5037854 |
| Q6ZNE5 | ATG14 | 17.3614354 | 17.2451237 | 17.2769343 | 17.4890084 | 17.2166694 |
| Q96CX2 | KCTD12 | 18.8853212 | 19.2727307 | 19.4757679 | 18.9174531 | 19.3502435 |
| Q8IX04 | UEVLD | 17.5835085 | 17.8941811 | 17.8832111 | 17.7764619 | 17.9195936 |
| P07307 | ASGR2 | 16.80806 | NA | NA | 16.8226255 | NA |
| O60449 | LY75 | 20.4995147 | 20.5612837 | 20.6459334 | 20.4637374 | 20.6376597 |
| Q562E7 | WDR81 | 19.9252088 | 19.9893134 | 20.0589919 | 20.0655589 | 20.0791621 |
| O43148 | RNMT | 21.5807483 | 21.5961864 | 21.5814586 | 21.5719356 | 21.6333014 |
| Q9UPU7 | TBC1D2B | 20.2546554 | 20.3451086 | 20.4040196 | 20.3097971 | 20.2430045 |
| Q07065 | CKAP4 | 21.0684173 | 21.2341462 | 21.0856811 | 21.0411746 | 21.1848007 |
| Q96JB3 | HIC2 | 15.3844953 | 15.8856184 | 15.857247 | 15.9951398 | 15.6980582 |
| Q5VUM1 | SDHAF4 | 17.1906579 | 17.077997 | 16.8923543 | 17.2382626 | 17.0141221 |
| Q9HBU6 | ETNK1 | 20.1194137 | 20.2125604 | 20.2783505 | 20.1681295 | 20.3178408 |
| Q9NRC8 | SIRT7 | 18.1270914 | 18.1490106 | 18.4170472 | 18.3116353 | 18.2403411 |
| O75563 | SKAP2 | 20.7062719 | 20.6677168 | 20.6199651 | 20.6198529 | 20.8158559 |
| O60907 | TBL1X | 19.4051026 | 19.4189193 | 19.4666645 | 19.3683979 | 19.521163 |
| Q9HD33 | MRPL47 | 21.9132414 | 21.8169895 | 21.9714399 | 21.8587465 | 21.8268278 |
| Q9UQ90 | SPG7 | 21.7892457 | 21.7716694 | 21.8732273 | 21.7737748 | 21.8333882 |
| Q15366 | PCBP2 | 26.0990989 | 26.1687072 | 26.0949221 | 26.0008788 | 26.0637084 |
| P17844 | DDX5 | 25.1272033 | 25.169939 | 25.0819685 | 25.0645219 | 25.1100186 |
| Q5HYK3 | COQ5 | 20.881289 | 20.7858933 | 20.8955993 | 20.7515608 | 20.8542663 |
| O95071 | UBR5 | 20.7185333 | 20.7488863 | 20.7533044 | 20.7180596 | 20.8008561 |
| Q9UBV2 | SEL1L | 21.1907363 | 21.2065462 | 21.1260307 | 21.1985735 | 21.2259432 |
| O95453 | PARN | 20.7607371 | 20.673946 | 20.7543804 | 20.6575977 | 20.7305975 |
| P52272 | HNRNPM | 26.4325288 | 26.5116854 | 26.4131851 | 26.3615177 | 26.4112421 |
| P40227 | CCT6A | 25.7495474 | 25.7791315 | 25.7391675 | 25.6903521 | 25.8252587 |
| P35637 | FUS | 25.5210484 | 25.6261575 | 25.4937123 | 25.4918464 | 25.5591943 |
| Q14683 | SMC1A | 23.081027 | 23.1224911 | 23.0904286 | 22.993548 | 23.0756431 |

| | | | | | | |
|---|---|---|---|---|---|---|
| P02545 | LMNA | 25.6051635 | 25.663906 | 25.5402687 | 25.4711723 | 25.5882073 |
| Q02790 | FKBP4 | 25.3502644 | 25.4137308 | 25.3342617 | 25.2776168 | 25.3328293 |
| Q99704 | DOK1 | 21.0765281 | 21.2927314 | 21.2044934 | 21.0994782 | 21.1247702 |
| Q7L590 | MCM10 | 18.9458363 | 18.8921408 | 18.8515521 | 18.8391146 | 18.9143031 |
| Q9BVI0 | PHF20 | 16.1362823 | 16.1409358 | 16.2492438 | 16.3198194 | 16.0912923 |
| Q13613 | MTMR1 | 16.2840078 | 16.505037 | 16.3797505 | 16.6033442 | 16.5180175 |
| P36269 | GGT5 | 18.2618568 | 18.6251824 | 18.4369216 | 18.3396208 | 18.4658486 |
| Q96MX0 | CMTM3 | 20.7258373 | 21.0092413 | 20.8964931 | 20.7224703 | 20.8775646 |
| Q8IY37 | DHX37 | 20.3996414 | 20.5001489 | 20.534929 | 20.4141394 | 20.5644688 |
| Q13206 | DDX10 | 20.6984171 | 20.8336852 | 20.9293472 | 20.7911768 | 20.9074557 |
| O95757 | HSPA4L | 23.1484923 | 23.2294368 | 23.1895932 | 23.0893564 | 23.1737922 |
| Q92643 | PIGK | 20.6757624 | 20.7500572 | 20.8905175 | 20.633627 | 20.8273374 |
| O60294 | LCMT2 | 17.7804625 | 17.6077617 | 17.6351001 | 17.671081 | 17.7910858 |
| P35610 | SOAT1 | 19.5903304 | 19.6374835 | 19.7259911 | 19.6540268 | 19.6764839 |
| Q9Y4K3 | TRAF6 | 19.9236289 | 20.0723211 | 20.0649669 | 19.9491712 | 20.1422143 |
| Q9BZK7 | TBL1XR1 | 22.260487 | 22.2827051 | 22.3882409 | 22.2247954 | 22.385042 |
| O95202 | LETM1 | 23.7862572 | 23.8634288 | 23.8126404 | 23.7643242 | 23.7117749 |
| Q13459 | MYO9B | 21.0253594 | 21.1859175 | 21.1860459 | 21.0554848 | 21.1693694 |
| Q8N3A8 | PARP8 | 18.3581748 | 18.4948135 | 18.5101751 | 18.32695 | 18.6370472 |
| O95260 | ATE1 | 19.6817639 | 19.9361148 | 20.0730311 | 19.8800274 | 19.9213496 |
| Q6P1N9 | TATDN1 | 21.6072121 | 21.7502009 | 21.6031817 | 21.5295034 | 21.5750785 |
| Q9HCE9 | ANO8 | 14.8348029 | 15.3153261 | 15.0511379 | 14.9265931 | 15.0029526 |
| Q16719 | KYNU | 21.4277076 | 21.545567 | 21.5179888 | 21.3377896 | 21.5152794 |
| Q8NEB7 | ACRBP | 17.321134 | 17.2356334 | 17.4903255 | 17.2954435 | 17.5281031 |
| Q9BUK0 | CHCHD7 | 19.2251508 | 19.4063972 | 19.0864412 | 19.2910604 | 19.3165232 |
| Q2TAL8 | QRICH1 | 20.9214843 | 20.9630028 | 20.8673832 | 20.8965331 | 20.9762382 |
| Q96QD8 | SLC38A2 | 19.0223055 | 19.3128659 | 18.9985872 | 19.1235294 | 19.139523 |
| Q96LA8 | PRMT6 | 19.0217622 | 19.0100535 | 19.1705024 | 19.0486605 | 19.2137022 |
| P42768 | WAS | 21.3410018 | 21.5196458 | 21.3913201 | 21.3725705 | 21.4781942 |
| P04844 | RPN2 | 24.6458508 | 24.6599548 | 24.6445779 | 24.6077107 | 24.6161075 |
| Q9UHL4 | DPP7 | 23.1293123 | 23.1396932 | 23.1251952 | 23.0315471 | 23.0746623 |
| O94830 | DDHD2 | 18.6912391 | 18.8120788 | 18.8803746 | 18.7188905 | 18.915375 |
| O15155 | BET1 | 19.340341 | 19.5190117 | 19.6605764 | 19.5596148 | 19.4104238 |
| Q8TBP6 | SLC25A40 | 18.0258367 | 18.3550682 | 18.1666533 | 18.108553 | 18.2905525 |
| Q86V24 | ADIPOR2 | 15.6290375 | 16.4615655 | 16.5066669 | 15.8335336 | 16.4420425 |
| Q9NRM0 | SLC2A9 | 16.2352696 | 16.0283471 | 15.5945069 | 16.1589995 | 16.2793363 |
| P09972 | ALDOC | 24.5999635 | 24.9984423 | 24.936273 | 24.5511794 | 25.0143178 |
| Q2M2Z5 | KIZ | 16.533908 | 16.3663901 | 16.4977859 | 16.6760021 | 16.6004511 |
| B2RBV5 | MRFAP1L2 | 15.5865973 | 14.9531866 | 16.1603554 | 15.8386319 | 15.7248985 |
| P41162 | ETV3 | 17.5383264 | 17.2147189 | 17.6726834 | 17.5915474 | 17.3084657 |
| P83916 | CBX1 | 22.3966219 | 22.8633452 | 22.5742006 | 22.4886759 | 22.6657055 |
| P15407 | FOSL1 | 16.6366628 | 16.7296377 | 16.723548 | 16.5947127 | 16.888866 |

| | | | | | | |
|---|---|---|---|---|---|---|
| Q96B49 | TOMM6 | 20.5976834 | 20.6677642 | 20.7093843 | 20.5932002 | 20.7578625 |
| P08842 | STS | 17.0240898 | 17.2623508 | 17.2847421 | 16.8980003 | 17.248569 |
| Q13216 | ERCC8 | 16.8194305 | 16.5637767 | 16.8266526 | 16.7156195 | 16.4259716 |
| P51157 | RAB28 | 17.4707119 | 17.8628849 | 17.6425994 | 17.7147583 | 17.7821084 |
| Q86UB9 | TMEM135 | 19.5273932 | 19.7248574 | 19.8091647 | 19.7190032 | 19.7247997 |
| O00635 | TRIM38 | 19.6571417 | 19.8099842 | 19.8371424 | 19.7751861 | 19.7742926 |
| Q15633 | TARBP2 | 18.6757536 | 18.7455671 | 18.6457696 | 18.7864212 | 18.7346631 |
| Q96F25 | ALG14 | NA | 14.4628546 | 14.9454257 | NA | 14.7833136 |
| Q969F2 | NKD2 | 19.3410523 | 19.4309536 | 19.4894533 | 19.4683983 | 19.386219 |
| Q6ZUT1 | NKAPD1 | 21.727762 | 21.7772732 | 21.6669727 | 21.5400646 | 21.7402207 |
| Q8N755 | SLC66A3 | 17.1743016 | 17.2308723 | 17.2517087 | 17.2237548 | 17.325822 |
| Q8NBN7 | RDH13 | 20.8874972 | 21.0410892 | 20.8547471 | 20.871339 | 20.9600783 |
| P27540 | ARNT | 18.0672608 | 18.3897588 | 18.3685405 | 18.1795229 | 18.4089331 |
| Q9HAS0 | C17orf75 | 19.3528403 | 19.4648327 | 19.5233879 | 19.3858672 | 19.561884 |
| Q92547 | TOPBP1 | 20.3201042 | 20.4520696 | 20.4661509 | 20.2970109 | 20.4600669 |
| O95163 | ELP1 | 20.6374868 | 20.7788764 | 20.8777665 | 20.6571093 | 20.9399545 |
| Q8NE71 | ABCF1 | 22.9245637 | 22.9321762 | 22.9909144 | 22.8282885 | 22.8609871 |
| O15040 | TECPR2 | 16.6916294 | 16.8362312 | 16.7685331 | 16.8107578 | 16.6526443 |
| Q6P6C2 | ALKBH5 | 20.4429842 | 20.5594108 | 20.6634973 | 20.6269804 | 20.7033107 |
| Q96H20 | SNF8 | 20.6266144 | 20.6173736 | 20.7097744 | 20.6566801 | 20.5842519 |
| Q9H6D7 | HAUS4 | 21.3447045 | 21.3092092 | 21.4047946 | 21.3114673 | 21.3992985 |
| P35237 | SERPINB6 | 22.8922048 | 22.7344095 | 22.9273307 | 22.8406274 | 22.8784155 |
| O14802 | POLR3A | 20.041565 | 20.0573095 | 20.1049368 | 20.0184351 | 20.0290327 |
| Q9P000 | COMMD9 | 20.5231024 | 20.573395 | 20.5102792 | 20.4648955 | 20.5705853 |
| P14866 | HNRNPL | 25.9431107 | 25.9999498 | 25.9634929 | 25.9032359 | 25.947662 |
| Q15056 | EIF4H | 24.6489026 | 24.7655759 | 24.666566 | 24.686133 | 24.7843019 |
| O60784 | TOM1 | 21.3767163 | 21.3438675 | 21.4745597 | 21.2431504 | 21.3522178 |
| Q6UWU4 | C6orf89 | 17.7116884 | 17.6557683 | 17.6784107 | 17.6685148 | 17.7369005 |
| Q9NR28 | DIABLO | 22.6318782 | 22.819272 | 22.6544586 | 22.6257237 | 22.7841068 |
| P98155 | VLDLR | 20.8603913 | 20.7109514 | 20.836847 | 20.8531905 | 20.8885337 |
| Q8TCS8 | PNPT1 | 22.4087113 | 22.4211382 | 22.4206302 | 22.3632352 | 22.4468165 |
| P19174 | PLCG1 | 19.1201631 | 19.3486976 | 19.2371468 | 19.2077901 | 19.2190728 |
| Q9BTW9 | TBCD | 21.5423875 | 21.6754591 | 21.6543988 | 21.5950352 | 21.602578 |
| Q96DX4 | RSPRY1 | 18.7485011 | 18.72177 | 18.7626751 | 18.7941419 | 18.7484017 |
| O60256 | PRPSAP2 | 21.0580393 | 21.244918 | 21.3208226 | 21.1709809 | 21.2321637 |
| P19388 | POLR2E | 22.901761 | 22.9899666 | 22.9789543 | 22.9691139 | 22.8969638 |
| Q5BKX5 | ACTMAP | 19.6529561 | 19.6074513 | 19.6637178 | 19.5971664 | 19.7598065 |
| Q8N3D4 | EHBP1L1 | 22.4736776 | 22.5164159 | 22.6273986 | 22.4672412 | 22.5559925 |
| O15226 | NKRF | 22.4191826 | 22.4990756 | 22.4729958 | 22.4050749 | 22.4227113 |
| P48637 | GSS | 22.8647872 | 22.9518468 | 22.8292888 | 22.7953241 | 22.8888605 |
| Q32P41 | TRMT5 | 20.1291457 | 20.2633284 | 20.2675843 | 20.1434387 | 20.2271294 |
| Q9H7E2 | TDRD3 | 19.7252854 | 19.7642436 | 19.7692534 | 19.7041743 | 19.7544372 |

| | | | | | | |
|---|---|---|---|---|---|---|
| O94819 | KBTBD11 | 20.9471797 | 20.9158429 | 20.8743053 | 20.9875722 | 20.9770044 |
| A2RRP1 | NBAS | 20.3857609 | 20.4160366 | 20.471011 | 20.3904902 | 20.5057095 |
| Q96DB5 | RMDN1 | 22.0468964 | 22.1064216 | 22.0203385 | 21.9504876 | 22.0664606 |
| Q7Z5R6 | APBB1IP | 21.7778645 | 21.8538914 | 21.7743019 | 21.7764765 | 21.8311124 |
| P16083 | NQO2 | 21.1388879 | 21.2885255 | 21.3158363 | 21.1557243 | 21.2988125 |
| O00429 | DNM1L | 22.264447 | 22.383647 | 22.3520772 | 22.255094 | 22.3193455 |
| P78344 | EIF4G2 | 22.7847614 | 22.7883384 | 22.8362322 | 22.7630485 | 22.7876733 |
| Q9UMN6 | KMT2B | 19.2219141 | 19.361296 | 19.3510701 | 19.268715 | 19.371489 |
| Q8TEM1 | NUP210 | 23.6393375 | 23.7508995 | 23.6770848 | 23.5759688 | 23.7572812 |
| P43307 | SSR1 | 23.3011901 | 23.4942386 | 23.2587685 | 23.255807 | 23.2885636 |
| Q99707 | MTR | 19.3444591 | 19.351433 | 19.4480759 | 19.2864827 | 19.4360911 |
| Q9Y3B4 | SF3B6 | 23.1186021 | 23.2448682 | 23.2103275 | 23.103094 | 23.1908549 |
| P14406 | COX7A2 | 24.3107099 | 24.2867908 | 24.2762935 | 24.0934086 | 24.2939354 |
| P30825 | SLC7A1 | 21.0736968 | 21.0479565 | 21.0100669 | 20.9180068 | 20.8557096 |
| P10515 | DLAT | 22.7296105 | 22.8275583 | 22.8109397 | 22.7012649 | 22.8631766 |
| Q16342 | PDCD2 | 19.688319 | 19.6267908 | 19.5160797 | 19.6193963 | 19.5492596 |
| Q53GS7 | GLE1 | 19.8901625 | 19.9462241 | 19.9502323 | 19.9235061 | 20.0073385 |
| Q86US8 | SMG6 | 19.4612905 | 19.5688258 | 19.5821536 | 19.4440214 | 19.5338104 |
| Q8NEC7 | GSTCD | 18.3626718 | 18.399798 | 18.4705772 | 18.320989 | 18.3354256 |
| Q96DT7 | ZBTB10 | 17.5458506 | 17.7710115 | 17.8529132 | 17.6013447 | 17.6490212 |
| Q9NQ29 | LUC7L | 20.4813833 | 20.6060248 | 20.584482 | 20.4063348 | 20.5818281 |
| Q16740 | CLPP | 23.0491856 | 23.2099313 | 23.0875564 | 23.0597131 | 23.0431117 |
| Q9P0J1 | PDP1 | 20.7504613 | 20.8323732 | 20.7955519 | 20.7322735 | 20.8271585 |
| Q8IZH2 | XRN1 | 20.3610844 | 20.230172 | 20.3470282 | 20.215743 | 20.2962489 |
| Q9HC36 | MRM3 | 21.9076679 | 21.8762927 | 21.9669223 | 21.8751981 | 21.9585896 |
| Q9UMY4 | SNX12 | 21.9137345 | 21.9860954 | 22.0618441 | 21.8441045 | 21.9968801 |
| P62857 | RPS28 | 25.2476817 | 25.1958406 | 25.1806871 | 25.1084306 | 25.2083936 |
| Q8IXQ3 | C9orf40 | 19.2228502 | 19.3677497 | 19.3334175 | 19.1876017 | 19.2643437 |
| Q6PHR2 | ULK3 | 19.0882221 | 19.1503676 | 19.2032863 | 19.2001144 | 19.2427889 |
| P50548 | ERF | 18.3744723 | 18.3347814 | 18.2801813 | 18.3243994 | 18.3838516 |
| Q99504 | EYA3 | 22.3263007 | 22.4080358 | 22.3428588 | 22.2976109 | 22.3446748 |
| Q01432 | AMPD3 | 17.4619572 | 17.5245168 | 17.7427403 | 17.6096612 | 17.7291223 |
| Q96BH1 | RNF25 | 18.7051121 | 18.8650578 | 18.8132952 | 18.8125661 | 18.9747835 |
| Q92526 | CCT6B | 16.3932478 | 16.4665571 | 16.8311331 | 16.4744279 | 16.6150775 |
| Q15526 | SURF1 | 20.7667077 | 20.9622101 | 20.9881871 | 20.88965 | 20.8449244 |
| Q9H1A3 | METTL9 | 18.3430965 | 18.375285 | 18.3629951 | 18.3754734 | 18.3113636 |
| P52630 | STAT2 | 24.5705411 | 24.6094913 | 24.6066274 | 24.4603604 | 24.584259 |
| Q969E8 | TSR2 | 21.2497651 | 21.3244063 | 20.9124012 | 21.1185735 | 21.167137 |
| Q969U6 | FBXW5 | 18.0245324 | 18.2269048 | 18.0158659 | 18.1874487 | 18.2394938 |
| Q9UDY8 | MALT1 | 17.1320272 | 17.2922315 | 17.1535318 | 17.0645441 | 17.2514875 |
| Q8N6I1 | EID2 | 18.1753613 | 18.4610648 | 18.5752186 | 18.4351336 | 18.5544506 |
| P10114 | RAP2A | 15.7016526 | 15.486438 | 15.9728568 | 15.7874922 | 15.936538 |

| | | | | | | |
|---|---|---|---|---|---|---|
| P10721 | KIT | 19.8167658 | 20.1113186 | 19.9925517 | 19.8038217 | 19.9770404 |
| Q8N371 | KDM8 | 16.4189108 | 16.4704393 | 16.4394962 | 16.3647224 | 16.4201235 |
| P30039 | PBLD | 15.6083341 | 15.7178655 | 15.5968739 | 15.5946398 | 15.6750026 |
| Q8WXB1 | METTL21A | 16.254795 | 16.3778827 | 16.5943335 | 16.6366934 | 16.4232976 |
| Q9NS23 | RASSF1 | 17.6422752 | 17.9908935 | 17.8295595 | 17.9509203 | 17.9332958 |
| Q969Z3 | MTARC2 | 17.8148339 | 18.0518892 | 17.9537543 | 17.7697085 | 18.0496018 |
| P55198 | MLLT6 | 15.2476058 | 15.3983482 | 15.4735764 | 14.4154383 | 14.9584868 |
| P02458 | COL2A1 | 18.1589997 | 17.9637243 | 17.9774183 | 18.3709903 | 17.8111855 |
| A6NGG8 | PCARE | 16.3583846 | NA | NA | 15.9780675 | NA |
| P20908 | COL5A1 | NA | 11.4752133 | 11.1515909 | NA | 12.3211334 |
| Q7Z4V0 | ZNF438 | 16.3160997 | 16.2771946 | 16.042135 | 16.334846 | 16.0294474 |
| Q8WWP7 | GIMAP1 | 19.783045 | 20.1895097 | 19.9264668 | 19.8891444 | 19.9927917 |
| Q16629 | SRSF7 | 24.4701986 | 24.6664467 | 24.4311848 | 24.4264295 | 24.5546061 |
| Q86XS8 | RNF130 | 18.8366616 | 18.6574439 | 18.8304706 | 18.922882 | 18.7412876 |
| Q13310 | PABPC4 | 22.9317012 | 22.980663 | 22.9016042 | 22.8886074 | 22.970734 |
| P82650 | MRPS22 | 21.8768534 | 22.0356212 | 21.9810227 | 21.8830518 | 21.9879771 |
| Q9C0B5 | ZDHHC5 | 19.6806795 | 19.6000576 | 19.7831772 | 19.6134607 | 19.7487482 |
| Q1KMD3 | HNRNPUL2 | 24.3884138 | 24.4984027 | 24.4155083 | 24.326891 | 24.4182803 |
| P35250 | RFC2 | 21.8814731 | 21.982739 | 21.9941297 | 21.8761705 | 21.9098218 |
| Q8IWB9 | TEX2 | 19.1517358 | 19.2077132 | 19.2281824 | 19.1668643 | 19.2359673 |
| Q6Y2X3 | DNAJC14 | 17.7682393 | 17.7515282 | 17.9689785 | 17.7594115 | 17.7890418 |
| Q9BZL6 | PRKD2 | 20.3210622 | 20.268955 | 20.3684354 | 20.3158817 | 20.4315454 |
| Q8N5L8 | RPP25L | 19.8257621 | 20.0326312 | 19.7117723 | 20.1076648 | 19.9697342 |
| Q9H9F9 | ACTR5 | 19.7555018 | 19.966963 | 19.9414705 | 19.8261625 | 19.9299002 |
| Q96MG7 | NSMCE3 | 19.8136188 | 19.9177333 | 19.9175197 | 19.8725753 | 19.8535348 |
| P50552 | VASP | 23.6377009 | 23.8131882 | 23.7313218 | 23.6642058 | 23.7442506 |
| Q07820 | MCL1 | 17.563965 | 17.8083136 | 17.5568591 | 17.7318895 | 17.8911993 |
| Q9H9S5 | FKRP | 16.6045107 | 16.8159936 | 16.4534972 | 16.8062478 | 16.6448702 |
| Q9BXJ9 | NAA15 | 23.2654846 | 23.3462194 | 23.26427 | 23.218658 | 23.2434145 |
| Q96I59 | NARS2 | 20.3777348 | 20.5116338 | 20.5673001 | 20.4516863 | 20.5309072 |
| Q9NVC6 | MED17 | 20.1343273 | 20.2187896 | 20.2059081 | 20.1828868 | 20.2886335 |
| Q3MHD2 | LSM12 | 21.5947858 | 21.6818855 | 21.5811993 | 21.3906206 | 21.6947453 |
| P04637 | TP53 | 17.0245495 | 17.7373277 | 17.3104069 | 17.0885469 | 17.026391 |
| P11831 | SRF | 18.2145632 | 18.0515424 | 18.1764217 | 17.9346389 | 18.2494249 |
| Q96GE9 | DMAC1 | 19.5202836 | 19.5328632 | 19.4713226 | 19.4214674 | 19.4700206 |
| Q8NB15 | ZNF511 | 16.7485718 | 17.3778999 | 16.8063129 | 16.9578181 | 16.8649691 |
| Q9UK33 | ZNF580 | 18.7575174 | 18.9754098 | 18.628331 | 18.9124555 | 19.1046983 |
| P13693 | TPT1 | 24.9907459 | 25.1079139 | 25.0068317 | 25.0084764 | 25.071059 |
| Q93084 | ATP2A3 | 21.655837 | 21.6785605 | 21.7121066 | 21.644081 | 21.6795769 |
| Q92600 | CNOT9 | 21.2346293 | 21.1572687 | 21.3146728 | 21.1274477 | 21.2874992 |
| Q9UKK6 | NXT1 | 17.5313227 | 17.6162182 | 17.7842613 | 17.7556738 | 17.8306355 |
| O15117 | FYB1 | 18.2210994 | 18.2937451 | 18.3393177 | 18.3583953 | 18.3438617 |

| | | | | | | |
|---|---|---|---|---|---|---|
| Q15906 | VPS72 | 19.9881037 | 19.9575288 | 19.8824071 | 19.9647194 | 19.7860213 |
| P41273 | TNFSF9 | 17.8087951 | 18.0368962 | 18.2691426 | 18.0974205 | 18.0096382 |
| Q9NRZ7 | AGPAT3 | 20.238335 | 20.3854923 | 20.4042867 | 20.1928148 | 20.290217 |
| Q96HI0 | SENP5 | 16.5377692 | 16.1719398 | 16.3859489 | 16.3940685 | 16.3977259 |
| P42224 | STAT1 | 22.2526876 | 22.344227 | 22.2898545 | 22.2642777 | 22.3120785 |
| Q96BS2 | TESC | 22.230285 | 22.3023564 | 22.1918911 | 22.2534472 | 22.3154308 |
| P19784 | CSNK2A2 | 21.5527641 | 21.5314572 | 21.6896845 | 21.5242914 | 21.6187856 |
| Q6P2H8 | TMEM53 | 16.3317877 | 17.1594398 | 17.0447178 | 16.9238113 | 16.8441028 |
| Q14435 | GALNT3 | 15.8299295 | 15.9297311 | 15.8637975 | 16.1132404 | 15.7645059 |
| Q96AB6 | NTAN1 | 18.0296785 | 18.2554658 | 18.008419 | 18.1666635 | 18.116331 |
| O43896 | KIF1C | 18.7906583 | 18.8916757 | 18.8893876 | 18.7964865 | 18.8829299 |
| Q9HCC0 | MCCC2 | 22.0854753 | 22.1808945 | 22.1463745 | 22.1079137 | 22.1611536 |
| O76094 | SRP72 | 23.106485 | 23.2203849 | 23.1555129 | 23.0984124 | 23.1646959 |
| Q14241 | ELOA | 20.4964294 | 20.5050711 | 20.6096901 | 20.4878074 | 20.5333686 |
| Q9UBQ6 | EXTL2 | 17.3173734 | 17.2755204 | 17.3265622 | 17.3302518 | 17.5229545 |
| Q9NPG8 | ZDHHC4 | 16.6909996 | 17.1176592 | 16.4566207 | 16.6918957 | 16.8220241 |
| Q7LFL8 | CXXC5 | 18.4056333 | 17.8257048 | 18.1583266 | 18.0716655 | 18.3312114 |
| P25205 | MCM3 | 23.972764 | 24.0212808 | 23.961711 | 23.876704 | 24.0088295 |
| P13674 | P4HA1 | 22.2139281 | 22.5769799 | 22.4725571 | 22.1821959 | 22.5011252 |
| P30153 | PPP2R1A | 24.5049266 | 24.5966168 | 24.4803965 | 24.4256947 | 24.61283 |
| P33991 | MCM4 | 23.7806767 | 23.870864 | 23.7904121 | 23.7683139 | 23.8777832 |
| Q9BWM7 | SFXN3 | 22.9828514 | 23.1023853 | 23.0514302 | 23.0017792 | 23.1740018 |
| Q01130 | SRSF2 | 26.1734677 | 26.3186778 | 25.9646015 | 26.0348753 | 26.2533151 |
| P30291 | WEE1 | 18.3853417 | 18.3786345 | 18.4968304 | 18.2940608 | 18.2382882 |
| Q9H977 | WDR54 | 17.6865465 | 18.0412165 | 18.3641172 | 17.7801378 | 18.1816521 |
| Q9Y3M8 | STARD13 | NA | 18.0035704 | 18.2824303 | NA | 18.5911298 |
| O00399 | DCTN6 | 19.5661236 | 19.8867897 | 19.4942599 | 19.5599347 | 19.6603417 |
| Q9P0J0 | NDUFA13 | 22.9923395 | 23.169989 | 23.1256886 | 23.0688022 | 23.0215839 |
| Q96RT1 | ERBIN | 22.7303187 | 22.8867472 | 22.9038453 | 22.7624795 | 22.9967991 |
| O15372 | EIF3H | 23.7532585 | 23.7534045 | 23.7599127 | 23.6944494 | 23.7458084 |
| Q8N1F7 | NUP93 | 22.9607432 | 22.9933732 | 22.9910572 | 22.8969499 | 22.9685828 |
| Q9BUJ2 | HNRNPUL1 | 24.2012348 | 24.2029019 | 24.2769035 | 24.1238475 | 24.1846236 |
| O43665 | RGS10 | 22.5407925 | 22.4730111 | 22.4092076 | 22.4040946 | 22.4536893 |
| O60291 | MGRN1 | 20.1808265 | 20.0648708 | 20.1734157 | 20.1507673 | 20.1621929 |
| Q6PII3 | CCDC174 | 18.0381737 | 17.8642387 | 17.7325175 | 17.8425335 | 17.9016546 |
| Q9H4B0 | OSGEPL1 | 19.1784735 | 19.3202619 | 19.4268383 | 19.337154 | 19.410742 |
| Q13610 | PWP1 | 20.8597638 | 20.8536781 | 20.9410066 | 20.8853122 | 21.0557055 |
| Q9NR83 | SLC2A4RG | 15.5209191 | 15.4334113 | 15.6801632 | 15.3124407 | 15.6932675 |
| P28702 | RXRB | 19.3600591 | 19.5367478 | 19.6541861 | 19.3549991 | 19.4788547 |
| Q15004 | PCLAF | 20.8070769 | 20.9620255 | 20.9572426 | 21.00024 | 20.9660606 |
| Q8WY22 | BRI3BP | 23.9662965 | 23.8233081 | 24.0171838 | 23.9125896 | 23.9414516 |
| Q8TAQ2 | SMARCC2 | 22.0157177 | 22.0924528 | 22.045614 | 21.9430096 | 21.9974378 |

| | | | | | | |
|---|---|---|---|---|---|---|
| Q15645 | TRIP13 | 21.4575102 | 21.5438444 | 21.5526623 | 21.5246924 | 21.5677555 |
| Q14999 | CUL7 | 18.8228124 | 18.8887177 | 19.0012235 | 18.8138967 | 18.8235194 |
| Q86WX3 | RPS19BP1 | 19.7878991 | 19.8337141 | 19.8636763 | 19.7786911 | 19.8134173 |
| Q9BYV8 | CEP41 | 19.2465408 | 19.3721848 | 19.3481646 | 19.3355757 | 19.4666937 |
| Q9Y6I9 | TEX264 | 21.3090068 | 21.4075513 | 21.4945204 | 21.3032246 | 21.4148023 |
| Q8TD55 | PLEKHO2 | 20.7933869 | 20.9119737 | 21.0010361 | 20.7762364 | 20.893312 |
| Q96CU9 | FOXRED1 | 20.4543178 | 20.4274082 | 20.4868843 | 20.4644641 | 20.5306074 |
| Q7Z309 | PABIR2 | 12.8256864 | NA | NA | 13.3262827 | NA |
| P16989 | YBX3 | 22.8266456 | 23.0067505 | 22.6279572 | 22.8321176 | 22.8198353 |
| O75888 | TNFSF13 | 17.5179165 | 17.9618709 | 17.6146032 | 17.4743849 | 17.2900946 |
| Q53SF7 | COBLL1 | 22.7421043 | NA | NA | 22.6983984 | NA |
| Q8IWB6 | TEX14 | 18.2727475 | NA | NA | 18.3481935 | NA |
| Q14586 | ZNF267 | 17.3538205 | 17.5143882 | 17.7061863 | 17.4963601 | 17.5240511 |
| P60033 | CD81 | 21.479379 | 21.6397897 | 21.6753745 | 21.4922179 | 21.5656074 |
| Q92667 | AKAP1 | 19.8454525 | 19.9765941 | 19.9388899 | 19.8386844 | 19.9302143 |
| Q3SY69 | ALDH1L2 | 20.269753 | 20.2079469 | 20.1997823 | 20.3152677 | 20.136863 |
| Q86U86 | PBRM1 | 22.5405271 | 22.5478288 | 22.5902257 | 22.5542603 | 22.5824324 |
| Q9UKL0 | RCOR1 | 21.4082097 | 21.4569353 | 21.5122909 | 21.3932802 | 21.4873491 |
| Q9BZH6 | WDR11 | 20.8250659 | 20.8702809 | 20.8810424 | 20.8447428 | 20.8935678 |
| P09012 | SNRPA | 24.4881411 | 24.5346936 | 24.4322881 | 24.4152122 | 24.5740738 |
| Q9Y6D9 | MAD1L1 | 21.3169689 | 21.4532542 | 21.4285249 | 21.360967 | 21.3632348 |
| Q06203 | PPAT | 20.9521676 | 21.1850248 | 21.2921575 | 21.0117236 | 21.252521 |
| Q9UFB7 | ZBTB47 | 16.7804624 | 17.3207127 | 16.9978241 | 17.0476239 | 17.0202009 |
| Q99501 | GAS2L1 | 16.7980179 | 16.7111103 | 16.5662637 | 16.3786701 | 16.241403 |
| P11473 | VDR | NA | 15.5445115 | 15.5639711 | NA | 15.1750055 |
| P68400 | CSNK2A1 | 22.9045619 | 22.9978432 | 22.9976922 | 22.844923 | 22.9995053 |
| Q9Y2L1 | DIS3 | 23.4659442 | 23.4674658 | 23.4201919 | 23.4080859 | 23.455572 |
| Q8N2G8 | GHDC | 21.5326744 | 21.6171643 | 21.5230784 | 21.5297447 | 21.5224021 |
| Q9BZI7 | UPF3B | 20.158861 | 20.1270131 | 20.3975968 | 20.2196433 | 20.2452413 |
| O75688 | PPM1B | 20.6353211 | 20.6869693 | 20.785432 | 20.6786623 | 20.7734967 |
| Q8WUA7 | TBC1D22A | 20.4642713 | 20.3689289 | 20.3825287 | 20.3503845 | 20.3615871 |
| P11216 | PYGB | 22.5461349 | 22.5865083 | 22.6190875 | 22.5335029 | 22.5400844 |
| O75530 | EED | 20.4402813 | 20.4347545 | 20.5029578 | 20.3896275 | 20.4428 |
| Q01415 | GALK2 | 23.7235421 | 23.7977684 | 23.8026945 | 23.6909065 | 23.8096424 |
| Q2TAK8 | PWWP3A | 18.5376273 | 18.4520421 | 18.6844921 | 18.5779935 | 18.2338494 |
| P36507 | MAP2K2 | 22.621603 | 22.7586922 | 22.6469893 | 22.5978323 | 22.7406578 |
| O00308 | WWP2 | 21.9193311 | 21.8428334 | 21.9230701 | 21.886659 | 21.8899156 |
| Q6W2J9 | BCOR | 18.9430501 | 18.981784 | 18.9151605 | 18.8466878 | 19.042834 |

| AMPH-04647.6h_3_abundance_norm | AMPH-03340.6h_1_abundance_norm | AMPH-03340.6h_2_abundance_norm | AMPH-03340.6h_3_abundance_norm | AMPH-03340.6h - AMPH-04647.6h__logFC | AMPH-03340.6h - AMPH-04647.6h__P.Value |
|---|---|---|---|---|---|
| 21.1838154 | 19.478449 | 19.7089473 | 19.5602672 | -1.478397684 | 2.20E-13 |
| 18.3891049 | 18.0263718 | 18.0493036 | 18.0028164 | -0.360453216 | 1.74E-06 |
| 21.9541546 | 21.6618712 | 21.755614 | 21.7676985 | -0.191552508 | 2.93E-04 |
| 19.4826864 | 18.701092 | 18.8401293 | 18.8145897 | -0.587796638 | 6.62E-07 |
| 19.4431338 | 19.2849657 | 19.2058455 | 19.2302412 | -0.312669023 | 8.27E-06 |
| 20.6240176 | 20.3526614 | 20.2369363 | 20.2820704 | -0.390385609 | 5.09E-05 |
| 17.7471588 | 17.2090252 | 16.9173584 | 17.1840079 | -0.51887771 | 2.80E-04 |
| 23.0409371 | 22.8246954 | 22.9489616 | 22.9370216 | -0.141750152 | 0.021990522 |
| 15.7719844 | 16.015584 | 16.3765172 | 16.5168165 | 0.471263487 | 0.025691936 |
| 21.5195252 | 21.2858869 | 21.2985353 | 21.2607308 | -0.286070148 | 0.012678257 |
| 17.0475528 | NA | 16.1117752 | 16.1771647 | -0.845700339 | 2.06E-06 |
| 18.2391698 | 18.0373184 | 18.0205052 | 18.1190808 | -0.211498205 | 0.024749186 |
| 16.0090513 | 16.2006504 | 16.8251636 | 16.7679114 | 0.364549335 | 0.057687763 |
| 19.1266937 | 18.6956379 | 18.771978 | 18.5021466 | -0.842337171 | 0.104805347 |
| 23.2052008 | 23.0181739 | 23.0436031 | 22.9337128 | -0.166354427 | 0.036368875 |
| NA | 14.5359554 | NA | NA | 0.134008112 | 0.193488741 |
| 15.3173904 | NA | 13.8977374 | 13.999632 | -1.313064775 | 2.06E-05 |
| NA | 14.7432504 | NA | NA | -1.095015571 | 4.55E-05 |
| 22.4170479 | 22.251342 | 22.5631912 | 21.7529732 | -0.338877597 | 0.118235297 |
| 16.6026769 | 16.5114217 | 16.6702446 | 16.1302216 | -0.326979492 | 0.021813445 |
| 17.5975108 | 17.3092325 | 17.3873852 | 17.3489624 | -0.332849455 | 3.22E-04 |
| 17.2877937 | 17.1936985 | 17.1692658 | 17.0600743 | -0.214424499 | 0.001131125 |
| 21.2385487 | 20.9447579 | 21.1184841 | 21.0605634 | -0.186570498 | 0.010690022 |
| 22.7055149 | 22.3175762 | 22.5709078 | 22.4828494 | -0.227443832 | 0.002212638 |
| 19.9617471 | 19.6701012 | 19.8651304 | 19.6949668 | -0.217838629 | 0.052957388 |
| 20.7056302 | 20.7260658 | 20.8542612 | 20.8276632 | 0.16234398 | 0.004904257 |
| 20.5271347 | 20.5847923 | 20.673139 | 20.7284909 | 0.211669816 | 0.003170266 |
| 16.9931584 | 17.2063038 | 17.0775833 | 17.3517716 | 0.270927918 | 0.003422253 |
| 18.1977358 | 17.6203663 | 17.6323229 | 17.4132732 | -0.586351511 | 1.25E-04 |
| 20.7450473 | 20.7648217 | 20.8144406 | 20.7559869 | -0.018960259 | 0.685489622 |
| 19.6499723 | 19.3508674 | 19.2165008 | 19.2688759 | -0.23266375 | 0.012003956 |
| 20.9150645 | 20.8353521 | 20.8587433 | 20.8764912 | -0.118031112 | 0.143643647 |
| 16.9201092 | 16.6285523 | 16.1290185 | 16.3350886 | -0.371042243 | 0.066388187 |
| 19.599139 | 19.6670487 | 19.8106432 | 19.8484617 | 0.145102148 | 0.010200278 |
| 19.7230868 | 19.8406624 | 19.8434743 | 19.8668585 | 0.137733375 | 0.006209056 |
| 20.5348787 | 20.4113056 | 20.3734325 | 20.3575478 | -0.116732163 | 0.041555828 |
| 19.9710566 | 20.113079 | 20.1947982 | 20.1171501 | 0.169463828 | 0.001494349 |
| 14.8350272 | 15.976474 | 16.5678568 | 16.7169389 | 1.242012953 | 0.002639716 |

| | | | | | |
|---|---|---|---|---|---|
| 14.7109914 | 14.7446773 | 13.6993084 | NA | -0.87004101 | 0.143929066 |
| 25.275462 | 25.0321375 | 24.7520504 | 24.6434642 | -0.414495174 | 1.20E-04 |
| 20.8696434 | 20.6594867 | 20.6149911 | 20.7075631 | -0.184000592 | 0.008333205 |
| 13.9762338 | 13.5233266 | 13.2278118 | 12.7588037 | -0.519610547 | 0.018881601 |
| 21.3129903 | 20.76223 | 21.0620081 | 21.0079602 | -0.348406541 | 0.008285976 |
| 16.962492 | 16.6997146 | 16.8348156 | 16.5830051 | -0.35612972 | 7.34E-04 |
| 15.7387659 | NA | 15.6474982 | 15.660835 | -0.146629696 | 0.085382065 |
| 20.2795993 | 20.1976063 | 20.0156899 | 20.0385952 | -0.217217843 | 0.008955806 |
| 26.7045261 | 26.4641696 | 26.5529311 | 26.5548168 | -0.135155016 | 0.079165554 |
| 22.6435176 | 22.4866533 | 22.6241934 | 22.6215494 | -0.182365757 | 0.004130178 |
| 20.9954996 | 20.9600809 | 21.0578358 | 20.9112783 | 0.009731945 | 0.814121786 |
| 20.7111352 | 20.6714221 | 20.736293 | 20.8121214 | 0.035827795 | 0.421388363 |
| 18.5606434 | 18.7134777 | 18.7157727 | 18.6707519 | 0.129100281 | 0.010211012 |
| 25.009294 | 25.1213934 | 25.2256043 | 25.1272764 | 0.143774134 | 0.027194739 |
| 19.4213694 | 19.4507382 | 19.5503896 | 19.5724007 | 0.152449848 | 0.042523362 |
| 14.5166392 | 14.8964517 | 14.723799 | 15.1360194 | 0.335343195 | 7.54E-04 |
| 18.1506396 | 19.0322112 | 19.4610784 | 19.7628889 | 0.938729924 | 0.004822083 |
| 18.7554563 | 18.6933527 | 18.5758936 | 18.6034894 | -0.131740848 | 0.041073145 |
| 18.4708209 | 18.5722506 | 18.4197644 | 18.4589338 | -0.020314493 | 0.748412436 |
| 26.6692296 | 26.5248783 | 26.5946361 | 26.5414016 | -0.026225006 | 0.62770041 |
| 20.3691231 | 20.4412031 | 20.4457128 | 20.4179696 | 0.115521237 | 0.053764563 |
| 22.2739489 | 22.2039907 | 22.3202433 | 22.2488215 | 0.012547024 | 0.849587781 |
| 16.2788612 | 17.4577719 | 17.4502518 | 17.219514 | 1.234963373 | 6.51E-05 |
| 18.4220239 | 17.9369399 | 17.909157 | 17.8877325 | -0.395990123 | 1.27E-04 |
| NA | 14.4333801 | NA | NA | 0.166656214 | 0.272133624 |
| 14.4265664 | 13.6448899 | 13.9604191 | 14.2373384 | -0.34943987 | 0.018568493 |
| 17.2772269 | 16.8183983 | 16.9200761 | 17.0875218 | -0.160989803 | 0.217798327 |
| 19.1335853 | 19.7748143 | 19.8385801 | 19.9050985 | 0.71074414 | 0.001294553 |
| 20.5740306 | 20.6428053 | 20.8366676 | 20.7885483 | 0.186516261 | 0.045145411 |
| 21.1163602 | 21.1599607 | 21.2047562 | 21.0574899 | 0.081704797 | 0.325305377 |
| 25.3585152 | 25.1869793 | 25.1982539 | 25.1750911 | -0.125191776 | 0.067466165 |
| 24.7635802 | 24.5753629 | 24.5924616 | 24.5960289 | -0.163165671 | 0.012501373 |
| 22.1658904 | 22.1064237 | 22.1078802 | 22.078818 | -0.070877426 | 0.085130189 |
| 17.5455968 | 17.5192584 | 17.3427925 | 17.5100706 | 0.03930568 | 0.554858975 |
| 15.3433952 | NA | 15.7958924 | 15.7467616 | 0.522734068 | 0.00171442 |
| 19.7445105 | 19.5586982 | 19.6198004 | 19.6370824 | -0.095810495 | 0.03647804 |
| 16.6430174 | 16.9391018 | 17.1641939 | 16.9954622 | 0.357607345 | 0.011380703 |
| 21.9734287 | 21.8745168 | 22.0387745 | 21.9960126 | -0.056637734 | 0.492634171 |
| 19.2804317 | 19.0191223 | 19.2394868 | 18.9951988 | -0.090976516 | 0.359532609 |
| 17.2419466 | 17.004952 | 16.8936642 | 17.0155585 | -0.292476181 | 0.003263095 |
| NA | 15.8705383 | NA | NA | -0.907478437 | 1.34E-04 |
| 16.8634972 | 17.2069195 | 16.7633935 | 16.7619732 | -0.058510907 | 0.671222389 |

| | | | | | |
|---|---|---|---|---|---|
| 20.3770313 | 20.3835697 | 20.4172682 | 20.4191876 | 0.061293528 | 0.165943281 |
| 17.7843822 | 17.8584969 | 17.8860804 | 17.9094816 | 0.057353741 | 0.336387731 |
| 18.7905906 | 18.7451615 | 18.9679035 | 18.7234559 | 0.194575834 | 0.074843817 |
| 18.1094028 | 18.2728743 | 18.1099874 | 18.153017 | 0.086823359 | 0.228379344 |
| 16.2223352 | NA | 16.3317507 | 16.5068709 | 0.209082908 | 0.08837599 |
| 19.7212808 | 19.6568772 | 19.4186917 | 19.5630147 | -0.182441978 | 0.03961803 |
| 26.2383435 | 26.1025655 | 26.1787778 | 26.1325463 | -0.034104487 | 0.488489776 |
| 22.678231 | 22.6471004 | 22.6384703 | 22.6349743 | -0.054459332 | 0.205453452 |
| 12.0246358 | NA | 12.1050364 | 12.1627346 | 0.240639747 | 0.128666631 |
| 21.8679652 | 21.473257 | 21.1884265 | 21.3008988 | -0.662959562 | 0.260015759 |
| 23.2266548 | 21.799597 | 22.9917343 | 22.8047012 | -0.424205099 | 0.108813484 |
| 22.5695452 | 22.1718847 | 22.3904263 | 22.5636141 | -0.099434752 | 0.263420589 |
| 26.4189814 | 26.3625031 | 26.3385478 | 26.3812106 | -0.028250811 | 0.687723295 |
| 19.117599 | 18.8976517 | 19.123182 | 18.9183109 | -0.096209925 | 0.172018485 |
| 21.4982664 | 21.1688806 | 21.4008427 | 21.266588 | -0.170891476 | 0.016627838 |
| 21.1007868 | 20.9674707 | 20.9792732 | 20.9761882 | -0.132983421 | 0.005452163 |
| 15.0725109 | 15.1787052 | 15.3480935 | 15.3604783 | 0.16681773 | 0.056559304 |
| 17.5981784 | 17.7214003 | 17.8273307 | 17.684286 | 0.121394395 | 0.197900072 |
| 17.6387552 | 17.624286 | 17.7170655 | 17.4155725 | 0.080691673 | 0.43688501 |
| 20.3401421 | 20.5210266 | 20.5507006 | 20.4098045 | 0.171333623 | 0.140027198 |
| 14.5636967 | 14.7269759 | 15.0687707 | 15.5278654 | 0.406679889 | 0.008521657 |
| 16.5081989 | 16.5968578 | 16.6941972 | 16.7295675 | 0.067106677 | 0.537217298 |
| NA | 13.6489736 | NA | NA | 0.618969826 | 0.006147908 |
| 18.9041948 | 18.8970289 | 19.0555396 | 18.9670922 | 0.081073061 | 0.22853709 |
| 17.550504 | NA | 18.6436197 | 18.5406602 | 0.985870779 | 0.001324258 |
| 18.801325 | 18.8310873 | 18.9854004 | 18.8928635 | 0.113605324 | 0.079139895 |
| NA | 15.2002686 | NA | NA | 0.609866383 | 0.014092527 |
| 16.4599734 | 16.3410754 | 16.0969889 | 16.2231582 | -0.18410809 | 0.117212725 |
| 23.8332863 | 23.5682101 | 23.8944745 | 23.6806958 | -0.097316531 | 0.256702192 |
| 21.1590276 | 20.9698764 | 20.9323803 | 20.9956854 | -0.162939449 | 0.036864049 |
| 19.7706046 | 19.5774066 | 19.680447 | 19.7538907 | -0.100750839 | 0.159240698 |
| 25.8142317 | 25.7556255 | 25.7350084 | 25.8029085 | -0.074720073 | 0.187591692 |
| 18.2255205 | 18.2028239 | 18.2930237 | 18.3916885 | 0.063273322 | 0.246507677 |
| 19.0012574 | 18.9886002 | 18.908261 | 18.9882871 | 0.087403175 | 0.160461715 |
| 22.4499638 | 22.4704942 | 22.4256351 | 22.4013851 | 0.040949177 | 0.458567288 |
| 19.3135628 | 19.310948 | 19.3126817 | 19.3970503 | 0.065329975 | 0.315666105 |
| 15.0749795 | NA | 15.4987248 | 15.3755438 | 0.186253302 | 0.268739358 |
| 18.2971706 | NA | 18.5012945 | 18.562612 | 0.366992153 | 0.072312557 |
| 19.0976971 | 19.0753181 | 19.2359987 | 19.2796485 | 0.164683088 | 0.070461266 |
| 23.7816685 | 23.7317171 | 23.7828145 | 23.7098247 | -0.041915953 | 0.373690082 |
| 23.7960659 | 23.6395875 | 23.7614726 | 23.7739378 | -0.039929111 | 0.469770852 |
| 20.4102752 | 20.4708504 | 20.4477947 | 20.4279196 | 0.044196459 | 0.185773608 |

| | | | | | |
|---|---|---|---|---|---|
| 21.319813 | 21.4389521 | 21.4379996 | 21.3131561 | 0.109530004 | 0.059596595 |
| 19.7472852 | 19.7396093 | 19.7762343 | 19.8700127 | 0.027378876 | 0.655561059 |
| 20.3321592 | 20.3484651 | 20.4161325 | 20.4072569 | 0.087995929 | 0.088435504 |
| 18.8217933 | 18.7186445 | 18.8367802 | 18.8962962 | 0.004734007 | 0.945702894 |
| 21.6846692 | 21.6933187 | 21.7791139 | 21.7692711 | 0.059176984 | 0.419245539 |
| 19.855423 | 19.8578405 | 19.578935 | 19.734883 | -0.179551845 | 0.052424633 |
| 18.485753 | 18.3648895 | 18.4441485 | 18.2872971 | -0.107962587 | 0.228680325 |
| 19.2664294 | 19.4324931 | 19.3053925 | 19.4575435 | 0.235864994 | 0.001654306 |
| 21.8308124 | 21.8585667 | 21.8255223 | 21.7872487 | -0.013140003 | 0.768027892 |
| 16.952587 | 17.1322269 | 16.8771887 | 17.3112342 | 0.065996208 | 0.635844898 |
| 19.3162565 | 19.0587342 | 19.1166259 | 19.1468497 | -0.140665095 | 0.016631185 |
| 22.6248467 | 22.3899844 | 22.5106598 | 22.5873313 | -0.068796221 | 0.204780276 |
| 21.5070626 | 21.4831822 | 21.4713205 | 21.464147 | -0.033925833 | 0.378876525 |
| 19.2330928 | 19.2310047 | 19.2003743 | 19.238833 | 0.054647232 | 0.291268574 |
| 17.1575795 | 17.4046573 | 17.4910569 | 17.3525864 | 0.16951655 | 0.048902773 |
| NA | 14.626859 | NA | NA | 0.808611642 | 0.043078774 |
| 16.1142753 | 16.1833173 | 15.9535271 | 16.1311399 | -0.122315822 | 0.341508758 |
| 17.3231909 | 17.3733085 | 17.3642577 | 17.4179178 | -0.165733925 | 0.142115223 |
| 19.3381166 | 19.3360676 | 19.1647196 | 19.1642377 | -0.142462068 | 0.044573614 |
| 25.5882037 | 25.5048883 | 25.5506819 | 25.4556707 | -0.02522291 | 0.67130209 |
| 19.2641613 | 19.1828581 | 19.2499693 | 19.0985261 | -0.071388714 | 0.171916561 |
| 21.7415058 | 21.5847798 | 21.4976658 | 21.6263754 | -0.139530165 | 0.004033876 |
| 20.4724069 | 20.3347133 | 20.3907354 | 20.3731191 | -0.104572556 | 0.016221164 |
| 22.4466653 | 22.2925053 | 22.3455114 | 22.359074 | -0.082832598 | 0.040399859 |
| 20.4254788 | 20.4003863 | 20.5450798 | 20.5502159 | 0.071164551 | 0.127461123 |
| 20.9051298 | 20.8435908 | 20.9115095 | 20.9273012 | 0.035022017 | 0.477276813 |
| 20.8634386 | 20.8303082 | 20.9095209 | 20.8067777 | 0.034564311 | 0.550303733 |
| 20.6730716 | 20.6013664 | 20.6232317 | 20.6907118 | 0.017119959 | 0.775111973 |
| 21.3119857 | 21.4272702 | 21.5556387 | 21.4788003 | 0.102915966 | 0.104181515 |
| 17.9426999 | 17.9791432 | 18.0332706 | 17.9462454 | 0.024934543 | 0.698272561 |
| 18.6696678 | 18.7375629 | 18.7399684 | 18.7541903 | 0.138995073 | 0.066780929 |
| 20.5275762 | 20.3126832 | 20.4543364 | 20.5615238 | -0.011504173 | 0.878969634 |
| 17.5261212 | 17.4114315 | 17.6417978 | 17.6830955 | 0.055369183 | 0.47774963 |
| 17.742265 | 17.7727583 | 17.8501045 | 17.7996091 | 0.053355801 | 0.495831009 |
| 19.4671708 | 19.4994527 | 19.6850171 | 19.75662 | 0.108520954 | 0.193094541 |
| 19.5244526 | 19.7693882 | 19.6687985 | 19.7752123 | 0.223322375 | 0.014592565 |
| 16.6856219 | 16.559007 | 16.6336156 | 16.872499 | 0.143715271 | 0.163144956 |
| 23.7361899 | 23.8625838 | 24.0821959 | 24.0294689 | 0.201357513 | 0.084991404 |
| 24.2624065 | 24.3722834 | 24.3053519 | 24.4553395 | 0.186610282 | 0.120221318 |
| NA | 17.089779 | NA | NA | 0.340994398 | 0.01373257 |
| NA | 15.6666817 | NA | NA | -0.105622101 | 0.49947896 |
| NA | 16.467767 | NA | NA | 0.606966868 | 0.004660013 |

| | | | | | |
|---|---|---|---|---|---|
| NA | 16.8294141 | NA | NA | 0.54259733 | 0.077096584 |
| NA | 15.2788302 | NA | NA | 1.9948375 | 0.005824284 |
| NA | 14.0983124 | NA | NA | 0.123308133 | 0.35847634 |
| 18.7869565 | 18.6613333 | 18.7937875 | 18.7703505 | 0.056012938 | 0.386887791 |
| 23.5189188 | 23.3837356 | 23.4829105 | 23.5147534 | -0.066501213 | 0.257373832 |
| 21.8630502 | 21.7739832 | 21.8408806 | 21.7533466 | -0.056184993 | 0.271837077 |
| 23.0951897 | 23.0573191 | 23.0616357 | 23.0335551 | -0.017239797 | 0.577076809 |
| 17.2629922 | 17.3718568 | 17.3931768 | 17.3431811 | 0.130996244 | 0.079149895 |
| 18.7257883 | 18.6111127 | 18.8804339 | 18.6594748 | -0.079114864 | 0.304419634 |
| 20.7843015 | 20.7405107 | 20.8451129 | 20.8266991 | 0.092698327 | 0.421836936 |
| 15.317471 | 15.9281001 | 16.2863058 | 16.0642451 | 0.553052934 | 0.025477983 |
| 14.3453892 | 13.8740216 | 13.5473261 | 12.8932879 | -0.597350768 | 0.036118694 |
| 17.9707483 | 17.3669325 | 17.4795391 | 17.7033693 | -0.259901688 | 0.035234368 |
| 21.2469397 | 21.1518989 | 21.1511952 | 21.2124001 | -0.090221969 | 0.107182699 |
| 17.3415602 | 17.2304456 | 17.4840133 | 17.3745687 | 0.144727416 | 0.077339263 |
| 16.3511015 | 16.3363642 | 16.2291103 | 16.4531719 | 0.130410798 | 0.305610707 |
| 19.5614099 | 19.4425886 | 19.4946369 | 19.537347 | -0.052824681 | 0.489034296 |
| 15.9520276 | NA | 16.0156466 | 15.9388737 | -0.099571434 | 0.296665906 |
| NA | 14.7985358 | NA | NA | -0.222054604 | 0.084159789 |
| 19.3222754 | 19.1595833 | 19.3717999 | 19.160458 | -0.116306625 | 0.142315826 |
| 18.5222356 | 18.549926 | 18.5548804 | 18.4919637 | -0.028976531 | 0.620903129 |
| 22.9070668 | 22.8466139 | 22.9288104 | 22.8410086 | -0.019876635 | 0.634059756 |
| 21.7496403 | 21.7757468 | 21.7172535 | 21.6915079 | -0.006395491 | 0.876708892 |
| 18.9875556 | 18.9844057 | 18.9760378 | 19.0175139 | 0.022036277 | 0.682819506 |
| 19.7831096 | 19.738749 | 19.7435809 | 19.9400089 | 0.083664711 | 0.14720842 |
| 19.9376909 | 19.894512 | 20.1376921 | 20.0867718 | 0.098301313 | 0.293569222 |
| NA | 15.6424136 | NA | NA | 0.010276135 | 0.919306243 |
| NA | 15.0528313 | NA | NA | 0.033635193 | 0.774907753 |
| 17.8271164 | 18.2871713 | 18.4128143 | 18.5200878 | 0.314786134 | 0.005804574 |
| 18.1405376 | 17.9712101 | 17.4899587 | 17.512513 | -0.402451008 | 0.014811212 |
| 22.3007272 | 22.413626 | 22.3896479 | 22.3618195 | 0.02452441 | 0.724641623 |
| 19.1201691 | 19.0826582 | 19.2708797 | 19.2248263 | 0.104319303 | 0.27372004 |
| 22.4290324 | 22.4085878 | 22.5371085 | 22.4783245 | 0.059287194 | 0.328921041 |
| 21.512191 | 21.476062 | 21.6364565 | 21.5576039 | 0.108571873 | 0.066050253 |
| 24.3625869 | 24.0216899 | 24.1605371 | 24.1327855 | -0.267635734 | 0.099975013 |
| NA | 14.5934359 | NA | NA | -0.499324867 | 0.001704462 |
| 20.7169652 | 20.7112356 | 20.8638752 | 20.9158585 | 0.107413118 | 0.065453698 |
| 20.4436967 | 20.4854443 | 20.5496131 | 20.6143456 | 0.111958679 | 0.085839687 |
| 14.7835497 | NA | 15.3636403 | 15.6383022 | 0.578155399 | 3.38E-04 |
| 15.5223676 | 15.3816914 | 15.473095 | 15.640006 | -0.033292972 | 0.689837455 |
| NA | 11.1426056 | NA | NA | -0.419424978 | 0.005154326 |
| 14.0379339 | NA | 13.8971746 | 14.0473589 | -0.175688185 | 0.070531567 |

| | | | | | |
|---|---|---|---|---|---|
| 22.2405807 | 22.1389767 | 22.214374 | 22.1651421 | -0.012909034 | 0.750114327 |
| 15.5390818 | NA | 15.355049 | 15.317631 | -0.208226913 | 0.002982643 |
| 19.2493922 | 19.3196085 | 19.2923966 | 19.2213983 | -0.022729733 | 0.70806893 |
| 18.8496946 | 18.8198593 | 18.8821617 | 18.9455712 | 0.109033929 | 0.113505365 |
| 21.3544841 | 21.2660057 | 21.422548 | 21.5231141 | -0.026929063 | 0.721678265 |
| 22.4309998 | 22.3456112 | 22.4338259 | 22.5165257 | 0.048890426 | 0.412403576 |
| 14.4480689 | NA | 14.2383535 | 14.2512468 | -0.128436006 | 0.461756394 |
| NA | 13.2715569 | NA | NA | -0.5435737 | 0.058522694 |
| 22.7682974 | 22.8076716 | 22.8487698 | 22.7322987 | 0.021904726 | 0.738824293 |
| 23.0849948 | 23.2359979 | 23.1175138 | 23.0170687 | 0.026127555 | 0.634819644 |
| 19.8016667 | 19.5608734 | 19.6514778 | 19.5428971 | -0.171255975 | 0.092206079 |
| 26.6509693 | 26.4807756 | 26.605149 | 26.5654086 | -0.048681688 | 0.451716331 |
| 18.7611378 | 18.6136881 | 18.7175198 | 18.6919453 | -0.143133194 | 0.031724469 |
| 25.944253 | 25.9525589 | 25.9050373 | 25.9519749 | 0.032956107 | 0.55622404 |
| 19.0071391 | 18.8887273 | 18.969779 | 19.0213095 | -7.21E-04 | 0.99226886 |
| 19.7160625 | 19.5300999 | 19.561151 | 19.5080281 | -0.149504708 | 0.03372533 |
| 20.9639392 | 20.9368445 | 20.9521391 | 20.8392593 | -0.065827669 | 0.169085342 |
| 17.3137193 | 17.715595 | 17.8909162 | 17.7318371 | 0.355514528 | 0.003920496 |
| 15.202114 | 15.0926617 | 15.2922198 | 15.0167372 | -0.009624142 | 0.945823086 |
| 15.5457871 | 15.6659334 | 16.023498 | 16.1583934 | 0.353565311 | 0.069238263 |
| 15.5501911 | NA | 15.3760044 | 15.6414316 | 0.034476461 | 0.758313299 |
| 20.2498835 | 20.0474083 | 20.209165 | 20.1370556 | -0.114482812 | 0.194311679 |
| 18.0573669 | 17.9535188 | 17.857292 | 17.8271394 | -0.174782126 | 0.019689686 |
| 25.9352228 | 25.982679 | 25.9376223 | 25.9217937 | 0.017285965 | 0.774665836 |
| 23.5215177 | 23.2682034 | 23.5325538 | 23.3787283 | -0.070924035 | 0.180186529 |
| 19.5693751 | 19.5607452 | 19.6004458 | 19.6342322 | 0.046963323 | 0.316377726 |
| 20.7194548 | 20.8571315 | 20.813931 | 20.7605004 | 0.106549558 | 0.041160754 |
| 17.3743033 | 17.6600629 | 17.5321344 | 17.5098026 | 0.145945771 | 0.037773796 |
| 20.6481403 | 20.6506648 | 20.6984561 | 20.7192853 | 0.085468206 | 0.243097085 |
| 22.6625982 | 22.4883416 | 22.6294198 | 22.6027325 | -0.027337593 | 0.740116856 |
| 17.4897192 | 17.5039843 | 17.6783139 | 17.8674159 | 0.143753392 | 0.111418729 |
| 17.6891632 | 17.4970761 | 17.6218926 | 17.6217138 | -0.099719535 | 0.26999915 |
| 21.0788241 | 21.0797578 | 21.0351358 | 20.9886758 | -0.034963687 | 0.409183554 |
| 15.8741184 | NA | 15.6933589 | 15.6774409 | -0.249920097 | 0.10350241 |
| 22.8633691 | 22.6271628 | 22.7206279 | 22.6283115 | -0.113699811 | 0.123895248 |
| 20.581578 | 20.6793712 | 20.6970158 | 20.6995707 | 0.110946343 | 0.042179973 |
| 18.0402366 | 18.0921531 | 18.2593214 | 18.3163032 | 0.115769998 | 0.14854822 |
| 15.829172 | 16.414152 | 16.0594769 | 15.9597033 | 0.271394483 | 0.079464208 |
| 18.2743594 | 18.5451039 | 18.5805627 | 18.3039044 | 0.158481417 | 0.083484357 |
| 16.0311591 | NA | 16.3109639 | 16.0392544 | 0.194790891 | 0.115186131 |
| 17.5992492 | 17.3691834 | 17.4683997 | 17.836223 | -0.079808851 | 0.504031236 |
| 17.8829601 | 17.5392685 | 17.6701081 | 17.5954753 | -0.211145345 | 0.052417255 |

| | | | | | |
|---|---|---|---|---|---|
| 22.0539109 | 21.6102385 | 21.7969132 | 21.9916347 | -0.182439798 | 0.048742336 |
| 23.0697174 | 22.9114066 | 23.1283374 | 23.0463058 | -0.049040771 | 0.410494831 |
| 19.7943366 | 19.4916958 | 19.6065229 | 19.5655398 | -0.219331356 | 7.95E-04 |
| 21.0052107 | 20.9769002 | 21.0586941 | 21.0370993 | 0.034826929 | 0.502863277 |
| 20.0144736 | 19.7980496 | 20.0839405 | 20.1703141 | 0.013169914 | 0.881106037 |
| 16.5827886 | 16.7595475 | 16.6732451 | 16.7267092 | 0.033677983 | 0.719342206 |
| 17.3174036 | 16.7798587 | 17.1623659 | 16.8844395 | -0.268397528 | 0.036453629 |
| 21.905625 | 21.8635071 | 22.0430888 | 22.0506301 | 0.075736874 | 0.290745142 |
| 26.492369 | 26.5268995 | 26.4115166 | 26.4455331 | 0.007078475 | 0.881410283 |
| 17.3952664 | 17.3496763 | 17.193797 | 17.3258185 | -0.135639039 | 0.032162575 |
| 18.48783 | 18.4836517 | 18.5091652 | 18.5772624 | 0.028721891 | 0.565947398 |
| 20.0143236 | 19.8650709 | 19.9849351 | 19.9687932 | -0.067361399 | 0.095576097 |
| 15.0347666 | 14.9162191 | 14.87094 | 15.1634123 | 0.175155077 | 0.276154792 |
| 17.279866 | 16.128524 | 16.7013192 | 16.5464948 | -0.650899595 | 0.010893658 |
| 17.2090877 | 16.6097371 | 16.9042844 | 17.0364709 | -0.361624804 | 0.01120926 |
| 17.7500604 | 17.5749337 | 17.5155856 | 17.6571944 | -0.08565167 | 0.409030614 |
| 19.454541 | 19.1536216 | 19.1431027 | 19.3223073 | -0.155209337 | 0.105887095 |
| 18.4833949 | 18.2662049 | 18.2885853 | 18.5920946 | -0.155495249 | 0.097789622 |
| 24.1085619 | 24.1005548 | 24.067725 | 24.0782085 | -0.072436158 | 0.316688734 |
| 23.651664 | 23.3211988 | 23.4074815 | 23.4042737 | -0.187388401 | 0.004594677 |
| 21.6844035 | 21.4547923 | 21.5374286 | 21.6463473 | -0.106469917 | 0.051353846 |
| 21.0757568 | 20.9237452 | 20.9942937 | 21.0363214 | -0.040808135 | 0.366048336 |
| 20.3982006 | 20.3972764 | 20.3065708 | 20.3066556 | -0.068185233 | 0.105458236 |
| 19.9972589 | 20.0472671 | 20.0956459 | 20.2200316 | 0.112714527 | 0.023363777 |
| 17.8099299 | 17.7804677 | 17.8169964 | 17.7155996 | 0.040224612 | 0.57432446 |
| 16.9593374 | 17.0053081 | 16.8441668 | 16.8385693 | 0.029630476 | 0.780873048 |
| NA | 11.5526405 | NA | NA | 0.10217662 | 0.5107934 |
| 16.4459137 | 16.607183 | 16.9137614 | 16.6811 | 0.381960879 | 0.040969593 |
| NA | 14.8461693 | NA | NA | -0.346698454 | 0.033321652 |
| 17.2449098 | 16.8822886 | 17.1964167 | 17.0565865 | -0.08283986 | 0.421045987 |
| 20.5311086 | 20.285202 | 20.360664 | 20.166252 | -0.208422636 | 0.038012291 |
| 22.6357753 | 22.3977601 | 22.5701072 | 22.609762 | -0.032230504 | 0.580746827 |
| 18.9120049 | 18.773069 | 18.8179761 | 18.734076 | -0.080022558 | 0.175012407 |
| 20.0174586 | 20.0375364 | 20.0033006 | 20.0743979 | 0.045151227 | 0.416268993 |
| 19.8675947 | 19.8438631 | 19.8446083 | 19.8831532 | -0.038711069 | 0.427109362 |
| 23.4904386 | 23.3934897 | 23.3368936 | 23.270446 | -0.1298581 | 0.007580102 |
| 24.0145155 | 23.9557342 | 23.9753847 | 23.9825318 | -0.02178132 | 0.582878148 |
| 19.8280013 | 19.7501536 | 19.8432731 | 19.8439741 | 0.047646548 | 0.385369808 |
| 20.2817091 | 20.2168684 | 20.2971878 | 20.3263478 | 0.017579378 | 0.779533089 |
| 19.1181507 | 19.1327989 | 19.3023571 | 19.2814057 | 0.17203748 | 0.07228317 |
| 19.2783174 | 19.5047719 | 19.5707015 | 19.252487 | 0.139623084 | 0.184704384 |
| 16.7523142 | 16.9019659 | 17.1064687 | 16.8249521 | 0.338531054 | 0.003088014 |

| | | | | | |
|---|---|---|---|---|---|
| NA | 11.6485317 | NA | NA | -0.005773311 | 0.980792932 |
| 20.3267935 | 20.2597246 | 20.199848 | 20.1657405 | -0.109203492 | 0.126216987 |
| 11.0960632 | NA | 11.3555642 | 11.3434875 | -0.053012021 | 0.7944963 |
| 16.1490427 | 15.9133615 | 15.7827078 | 15.5764822 | -0.291047554 | 0.052596323 |
| 21.6914637 | 21.4469517 | 21.5311436 | 21.6408604 | -0.106782549 | 0.123294022 |
| 21.877651 | 21.8047109 | 21.7046852 | 21.7707494 | -0.084599038 | 0.103300748 |
| 21.3883599 | 21.4560176 | 21.3954001 | 21.5629577 | 0.076173609 | 0.270276997 |
| 17.0257274 | 17.0198724 | 17.0790352 | 17.1398145 | 0.056323664 | 0.448747826 |
| 21.1485613 | 21.3188184 | 21.2189185 | 21.200226 | 0.069189 | 0.376825499 |
| 19.0926245 | 19.1381534 | 19.2154064 | 19.2686876 | 0.130095466 | 0.191419042 |
| 16.645475 | 16.5379132 | 16.4143867 | 16.7631002 | 0.094885955 | 0.377533107 |
| 17.189419 | 15.5991497 | 15.7835135 | 16.2103542 | -1.119162531 | 0.001621127 |
| 17.0533303 | 16.961725 | 16.5108426 | 16.2371219 | -0.464598639 | 0.107337661 |
| 26.0876967 | 26.1480816 | 25.6276617 | 25.6566275 | -0.245367159 | 0.344920433 |
| 21.9273092 | 21.7516284 | 21.873934 | 21.809522 | -0.065182453 | 0.173438408 |
| 23.7908603 | 23.4555415 | 23.8109025 | 23.776499 | -0.106552866 | 0.179709245 |
| 19.6263047 | 19.6535745 | 19.7918174 | 19.813831 | 0.118177131 | 0.083344723 |
| 21.2337965 | 21.2099487 | 21.2635038 | 21.4123636 | 0.125144034 | 0.212575726 |
| 16.7043674 | 16.7357368 | 16.5344695 | 16.5384614 | -0.254029518 | 0.08644429 |
| 18.1184602 | 18.0189769 | 18.0656283 | 17.9571235 | -0.095776932 | 0.275012515 |
| 20.4868606 | 20.6407491 | 20.5248382 | 20.4454636 | -0.050252724 | 0.42025186 |
| 20.3533888 | 20.3936319 | 20.3739906 | 20.3922178 | 0.068014684 | 0.094058452 |
| 22.3164458 | 22.3408851 | 22.3492237 | 22.323139 | 0.016309553 | 0.737948769 |
| 19.7075031 | 19.5595207 | 19.6202399 | 19.7337904 | -0.014448063 | 0.824543462 |
| 21.9977739 | 21.8338325 | 22.0094972 | 22.036364 | 0.081485273 | 0.2383624 |
| 19.6460936 | 19.571463 | 19.511087 | 19.4221049 | -0.225573835 | 0.025349243 |
| 16.1632034 | 16.0948438 | 15.970161 | 16.1385801 | -0.064496202 | 0.47456649 |
| 19.9154237 | 19.624221 | 19.8092048 | 19.698229 | -0.191432459 | 0.010543228 |
| 21.8472401 | 21.6537753 | 21.7850329 | 21.7697535 | -0.054499303 | 0.212749691 |
| 20.3415453 | 20.2292296 | 20.2853178 | 20.3770653 | 0.019222857 | 0.681323699 |
| 18.0918504 | 18.0623871 | 18.2483451 | 18.2048501 | 0.099930859 | 0.06931204 |
| 17.8805716 | 17.8569045 | 17.9655053 | 17.8255435 | 0.010039584 | 0.893940132 |
| 18.3632677 | 18.2048885 | 18.4446536 | 18.376793 | 0.063905087 | 0.447289345 |
| 19.0776242 | 18.3726465 | 18.9311629 | 18.9504633 | -0.216101602 | 0.08887158 |
| 24.165642 | 24.1375168 | 24.1226874 | 24.0683051 | -0.013484386 | 0.76100692 |
| 21.4636487 | 21.4035282 | 21.5346648 | 21.4728941 | 0.069170332 | 0.373667883 |
| 17.3069997 | 17.6236267 | 17.783804 | 17.4775881 | 0.170121205 | 0.292767199 |
| 20.7157452 | 20.6270638 | 20.7611124 | 20.7574184 | 0.065772728 | 0.172094999 |
| 18.0329708 | 17.2576083 | 17.9691822 | 17.9326114 | -0.282335317 | 0.192250275 |
| 16.4499653 | 16.2190811 | 16.2819762 | 16.3250908 | -0.123672209 | 0.317519408 |
| 11.6570896 | NA | 11.471478 | 11.7320336 | -0.053715567 | 0.608885987 |
| 20.8416571 | 20.8628971 | 20.9105614 | 20.7579782 | -0.128244666 | 0.18475154 |

| | | | | | |
|---|---|---|---|---|---|
| 23.4716061 | 23.2654549 | 23.4555528 | 23.4891413 | -0.048028561 | 0.528002721 |
| 24.5454087 | 24.5830512 | 24.5481199 | 24.4914584 | -0.043782338 | 0.431292593 |
| 25.1351407 | 25.3718396 | 25.2749541 | 25.2708739 | 0.132827651 | 0.02694433 |
| 18.3467444 | 18.2928259 | 18.4225114 | 18.4444777 | 0.046153969 | 0.564584568 |
| 15.688645 | NA | 15.7315025 | 15.8796368 | 0.01300045 | 0.880787156 |
| 18.5157252 | 18.1307125 | 18.4658368 | 18.4127508 | -0.114012755 | 0.13653585 |
| 20.6409531 | 20.4750375 | 20.4657297 | 20.4041026 | -0.127028003 | 0.089142948 |
| 20.3591302 | 20.3611898 | 20.1504514 | 20.0812715 | -0.097854038 | 0.142820259 |
| 20.2770966 | 20.1230373 | 20.1937549 | 20.2289342 | -0.038460148 | 0.534940245 |
| 21.310693 | 21.0126619 | 21.0801377 | 21.0033389 | -0.17826005 | 0.077541868 |
| 18.1421667 | 17.9916586 | 17.7933171 | 17.9221798 | -0.137121621 | 0.100178035 |
| 26.0288685 | 25.9285079 | 25.9441608 | 25.9567944 | -0.059917894 | 0.318886092 |
| 22.4522844 | 22.465536 | 22.4448355 | 22.4147524 | -0.043382666 | 0.44895007 |
| 21.2892826 | 21.3074008 | 21.40472 | 21.3922088 | 0.089624441 | 0.069859418 |
| 20.9911742 | 20.8852663 | 20.9436537 | 20.9898806 | -0.006159878 | 0.906309284 |
| 20.4300052 | 20.4869696 | 20.571917 | 20.5569834 | 0.086257166 | 0.174623664 |
| 18.9365961 | 18.8177633 | 19.0538498 | 18.9925479 | 0.089922265 | 0.233604003 |
| 19.087699 | 19.2418152 | 19.0450166 | 19.1136166 | 0.118239727 | 0.151994836 |
| 17.1593418 | 17.6010864 | 17.4929103 | 17.4163263 | 0.279649372 | 0.00528584 |
| 20.5270584 | 20.519971 | 20.4968161 | 20.3897257 | -0.110880761 | 0.060488953 |
| 21.9255487 | 21.9793547 | 21.8567842 | 21.9010975 | -0.014663021 | 0.758608773 |
| 20.3580164 | 20.4402247 | 20.4486877 | 20.4281485 | 0.105868726 | 0.224641524 |
| 18.1524589 | 18.1338765 | 18.2721078 | 18.2539166 | 0.098792672 | 0.298860282 |
| NA | 16.0414086 | NA | NA | 0.55807442 | 0.001567078 |
| 23.2641343 | 23.036886 | 23.1070297 | 23.1419111 | -0.103235056 | 0.063976053 |
| 21.9904845 | 22.0304732 | 22.0169155 | 22.0482725 | 0.067519434 | 0.062987674 |
| 21.2040068 | 21.1565129 | 21.1441182 | 21.2464931 | -0.027473224 | 0.598496784 |
| 20.1820263 | 20.0715392 | 20.0459889 | 20.2686713 | 0.029097598 | 0.729587833 |
| 11.4523241 | NA | 11.5827418 | 11.92023 | 0.126115592 | 0.508988915 |
| 21.2347717 | 21.1655765 | 21.2294449 | 21.2488304 | 0.002302794 | 0.960101135 |
| 19.8768149 | 19.9708684 | 19.9411546 | 19.8304641 | 0.049345744 | 0.430427671 |
| 24.1350573 | 23.873554 | 24.1662238 | 24.2035523 | -0.015686003 | 0.843660142 |
| 21.2312769 | 21.0956393 | 21.1630919 | 21.2039889 | -0.018649853 | 0.736968444 |
| 17.534014 | 17.358632 | 17.355625 | 17.4858127 | -0.087894426 | 0.379829974 |
| 21.7454094 | 21.6639257 | 21.6838001 | 21.7519194 | 0.009321426 | 0.873092582 |
| 17.4586216 | 17.5247072 | 17.549226 | 17.5585095 | 0.088631709 | 0.243214731 |
| 19.7900388 | 19.7626946 | 19.7781957 | 19.7849498 | -0.035986874 | 0.305551138 |
| 20.1790941 | 20.1647417 | 20.1449044 | 20.195633 | -0.026942514 | 0.562932707 |
| 16.724716 | 16.9710452 | 17.0255582 | 16.8987917 | 0.151093883 | 0.18547227 |
| 13.6159872 | NA | 12.8586785 | 13.0652572 | -0.640093477 | 0.01189227 |
| 19.3517596 | 19.1068923 | 19.3391015 | 19.1731303 | -0.164212077 | 0.109532416 |
| 20.9886342 | 20.9603226 | 20.9436458 | 20.9318713 | -0.023654694 | 0.527839012 |

| | | | | | |
|---|---|---|---|---|---|
| 19.8423027 | 19.7873904 | 20.0613397 | 19.9069914 | 0.067857833 | 0.302948755 |
| 16.7566256 | 16.8696195 | 16.7685282 | 17.0843568 | 0.115827415 | 0.282842286 |
| 21.5640435 | 21.475435 | 21.4466025 | 21.3942615 | -0.07395011 | 0.255058205 |
| 19.5726423 | 19.86404 | 19.9561902 | 20.1633788 | 0.279875059 | 0.022349661 |
| NA | 23.5208029 | NA | NA | 0.50691205 | 0.001148653 |
| 17.1022608 | 17.1316711 | 17.1021947 | 16.9189454 | -0.021146862 | 0.851999029 |
| 18.6707113 | 18.6168796 | 18.5167154 | 18.7775326 | 0.013078571 | 0.882004097 |
| 23.7108169 | 23.5542489 | 23.6713796 | 23.6158523 | -0.031292363 | 0.557791881 |
| 21.6765877 | 21.5346592 | 21.5898949 | 21.5929903 | -0.093225796 | 0.087467339 |
| 20.8359357 | 20.8668195 | 20.8765855 | 20.8866352 | -0.023055007 | 0.646211457 |
| 19.7257035 | 19.7457686 | 19.8200766 | 19.7846893 | 0.047746659 | 0.299622235 |
| 20.825019 | 20.9944576 | 21.018576 | 20.9625837 | 0.123833301 | 0.029879466 |
| 21.7832149 | 21.679452 | 21.8004022 | 21.8151599 | 0.047476017 | 0.485989634 |
| 18.4531679 | 18.4962469 | 18.5946493 | 18.5404325 | 0.145962859 | 0.046644745 |
| 20.9827423 | 20.9465521 | 20.9943523 | 21.0739553 | 0.037730757 | 0.672175577 |
| NA | 15.65839 | NA | NA | 0.219601336 | 0.068634116 |
| 16.0374285 | 16.260182 | 16.4631993 | 16.513548 | 0.061102892 | 0.871575245 |
| 14.2672456 | NA | 14.2720017 | 14.0025984 | -0.222252024 | 0.236708051 |
| 19.7163125 | 19.6197452 | 19.7614356 | 19.8059771 | 0.068838924 | 0.580777495 |
| 23.4561656 | 23.3650987 | 23.5772405 | 23.5756079 | -0.023181339 | 0.731761623 |
| 18.5518838 | 18.3305315 | 18.3956083 | 18.3731554 | -0.159620978 | 0.017359126 |
| 22.3995313 | 22.257102 | 22.3105663 | 22.2984912 | -0.072163032 | 0.226517338 |
| 23.7206018 | 23.6567366 | 23.6573724 | 23.6022339 | -0.020372919 | 0.677130999 |
| 23.8745808 | 23.7180388 | 23.820943 | 23.787727 | -0.062132373 | 0.188169967 |
| 19.7058814 | 19.6240374 | 19.690726 | 19.7161137 | -0.107461893 | 0.026106985 |
| 21.9718724 | 21.9953499 | 22.0134017 | 22.0403168 | 0.090909703 | 0.067479422 |
| 21.0088513 | 20.9317212 | 20.9744047 | 21.0150089 | -0.002735783 | 0.960034912 |
| 19.0628154 | 18.8612731 | 18.9758518 | 19.0574068 | -0.005171134 | 0.931548489 |
| 19.4568998 | 19.5182596 | 19.7216464 | 19.5414836 | 0.113065595 | 0.120187157 |
| 19.8298277 | 19.8044666 | 20.0547374 | 19.9924456 | 0.080137001 | 0.378624306 |
| 11.6529809 | NA | 11.7403332 | 11.6935244 | 0.12153034 | 0.177584282 |
| 17.2578719 | 17.6443601 | 17.3417493 | 17.4587185 | 0.218154519 | 0.027148263 |
| 18.0726223 | 18.3555766 | 18.2270285 | 18.2221473 | 0.133929372 | 0.231247525 |
| NA | 14.2043266 | NA | NA | 0.230659567 | 0.082843317 |
| NA | 14.9120813 | NA | NA | 0.19966233 | 0.244364478 |
| NA | 13.8759524 | NA | NA | 0.334715285 | 0.238338253 |
| 22.0441723 | 22.0045496 | 22.0550165 | 22.1018511 | 0.047404207 | 0.293399304 |
| NA | 16.4737169 | NA | NA | 0.470401134 | 0.159174259 |
| 22.9154713 | 22.8835151 | 22.7947077 | 22.8786739 | -0.141836665 | 0.035626937 |
| 23.2020096 | 23.2555995 | 23.2692632 | 23.2534179 | 0.120625745 | 0.176725441 |
| 20.7873585 | 20.8219032 | 20.7752258 | 20.7369265 | -0.018546272 | 0.673506299 |
| 20.1149044 | 20.0954093 | 20.1536011 | 20.1413499 | 0.027140946 | 0.68215376 |

| | | | | | |
|---|---|---|---|---|---|
| 20.4476961 | 20.2800522 | 20.4166453 | 20.3548272 | 0.012242993 | 0.854918961 |
| 16.4916743 | NA | 17.246895 | 17.4950074 | 1.093589481 | 0.004368112 |
| 14.4938017 | NA | 14.5424173 | 14.7272939 | 0.12526644 | 0.080838969 |
| 20.4424972 | 20.3524112 | 20.4635108 | 20.439584 | -0.026038031 | 0.606675467 |
| 18.738496 | 18.7503713 | 18.8294945 | 18.8658316 | 0.15455092 | 0.065087509 |
| 19.3361045 | NA | 19.5128513 | 19.2941649 | -0.007896294 | 0.951659833 |
| 14.2527295 | NA | 14.8710519 | 14.8316731 | 0.2861009 | 0.205916664 |
| NA | 13.450308 | NA | NA | -0.100264352 | 0.400440067 |
| 23.2206593 | 23.1485363 | 23.123314 | 23.0992279 | -0.042668918 | 0.307643926 |
| 28.3872938 | 28.2935367 | 28.1826469 | 28.1803841 | -0.071093574 | 0.42201257 |
| 19.6818307 | 19.6231299 | 19.6506958 | 19.7338176 | -0.118280328 | 0.047509394 |
| 20.0485877 | 19.9822095 | 19.9912041 | 20.1520199 | 0.097134887 | 0.269198594 |
| 15.6853997 | NA | 16.055247 | 16.0777208 | 0.325922278 | 0.001955496 |
| 21.5447214 | 21.5370589 | 21.5966552 | 21.4142914 | -0.047165493 | 0.411210199 |
| NA | 14.5630809 | NA | NA | 0.213425162 | 0.137456204 |
| 23.0528935 | 22.9413242 | 23.0204599 | 22.9780116 | -0.055878018 | 0.178576248 |
| 21.0259673 | 20.8886725 | 21.0912349 | 21.0642002 | 0.070594805 | 0.295521852 |
| 26.1769493 | 26.0498799 | 26.1844507 | 26.1336857 | -0.01417112 | 0.873278159 |
| 18.0567043 | 17.9983536 | 17.951383 | 17.7521784 | -0.086511965 | 0.31026215 |
| 17.6865277 | 17.4968025 | 17.2687298 | 17.4853456 | -0.126928089 | 0.078223177 |
| 26.3537984 | 26.3011054 | 26.3308914 | 26.3399961 | 0.013564801 | 0.800446918 |
| 23.9455932 | 23.8595479 | 23.9933792 | 23.94081 | 0.035426576 | 0.390713645 |
| 23.8991057 | 23.7877407 | 23.8521622 | 23.8525068 | -0.044926395 | 0.245570682 |
| 19.0346614 | 18.9297034 | 19.0510656 | 19.0923051 | 0.075140224 | 0.252599845 |
| 20.5533146 | 20.4722732 | 20.5095729 | 20.6382881 | 0.031808619 | 0.736891221 |
| 16.7025383 | 16.8323601 | 16.761148 | 17.0522625 | 0.116727663 | 0.272044238 |
| 23.7949608 | 23.7515431 | 23.7086482 | 23.5897588 | -0.113779976 | 0.174806163 |
| 27.025902 | 26.8183874 | 26.9808107 | 27.0211592 | -0.024589879 | 0.696774244 |
| 18.7838809 | 18.6920133 | 18.7721196 | 18.8647074 | -0.036340451 | 0.486900683 |
| 23.3295177 | 23.2704955 | 23.1946941 | 23.2864506 | -0.04691875 | 0.245257909 |
| 15.6590794 | 16.063881 | 16.2603246 | 16.249957 | 0.47325458 | 1.36E-04 |
| NA | 13.6777148 | NA | NA | 0.16378665 | 0.572331298 |
| 15.8924455 | 15.6629147 | 15.797253 | 15.7097188 | -0.183579041 | 0.024931085 |
| 14.8626309 | NA | 14.9343156 | 14.8908758 | -0.0505438 | 0.740493759 |
| 23.1390453 | 23.0735278 | 22.9905283 | 22.9773836 | -0.127356719 | 0.056625826 |
| 17.4859527 | 17.4494476 | 17.4258403 | 17.4580245 | -0.038151778 | 0.729588752 |
| 18.5660151 | 18.579831 | 18.6913626 | 18.6510952 | 0.062434872 | 0.402665851 |
| 19.1678419 | 19.1226313 | 19.0094526 | 19.2612309 | -0.041848471 | 0.512168105 |
| 20.8222758 | 20.7843503 | 20.7174415 | 20.9036901 | 0.038379349 | 0.638997168 |
| 16.7697504 | NA | 16.6782846 | 16.8749023 | -0.034561097 | 0.675462225 |
| 21.0061208 | 20.8775696 | 20.9971136 | 20.9574515 | 0.032548993 | 0.676504709 |
| NA | 16.3898865 | NA | NA | 0.10595278 | 0.643218987 |

| | | | | | |
|---|---|---|---|---|---|
| 18.9563932 | 18.9646164 | 19.1080419 | 19.1153378 | 0.089977703 | 0.386155635 |
| 21.6181293 | 21.5673717 | 21.3428042 | 21.4285363 | -0.13635283 | 0.027106227 |
| 20.3988027 | 20.3934507 | 20.427931 | 20.5323344 | 0.049540507 | 0.550610563 |
| 23.7795779 | 23.5418766 | 23.3841762 | 23.3242055 | -0.370324156 | 0.029780359 |
| 23.2941559 | 23.2285808 | 23.3261845 | 23.3368017 | 0.020579251 | 0.713667459 |
| 21.9959059 | 22.4961365 | 22.2573404 | 22.0474446 | 0.203657949 | 0.045403197 |
| 14.2922474 | 14.706262 | 14.6149338 | 14.3057739 | 0.076487052 | 0.83017853 |
| 15.2420897 | 14.9580113 | 15.3807357 | 14.9186793 | -0.185850499 | 0.319616752 |
| 18.014691 | 17.8160694 | 17.8459677 | 17.9188604 | -0.087826355 | 0.42900693 |
| 18.4126184 | 18.2684858 | 18.2928913 | 17.9913411 | -0.15747809 | 0.158209989 |
| 20.8481934 | 20.7739811 | 20.7260207 | 20.4126852 | -0.145611476 | 0.112998074 |
| 20.4871861 | 20.3286068 | 20.3331104 | 20.3308561 | -0.123866504 | 0.163682934 |
| 20.1644096 | 20.0353426 | 20.0854008 | 20.0027454 | -0.011090551 | 0.88729088 |
| 20.0876834 | 19.961207 | 20.0021958 | 20.013585 | -0.070576527 | 0.346746451 |
| 25.6077355 | 25.5230211 | 25.5669392 | 25.567487 | -0.007277073 | 0.91729447 |
| 26.3851815 | 26.3513422 | 26.3158188 | 26.3182767 | -0.027757436 | 0.686973598 |
| 23.9879801 | 23.9369025 | 23.8029103 | 23.7786483 | -0.123304342 | 0.043164557 |
| 21.6870032 | 21.474923 | 21.6078045 | 21.6999103 | -0.075381222 | 0.197957175 |
| 21.1436953 | 20.9372818 | 21.0710429 | 21.0500701 | -0.099181293 | 0.054440029 |
| 24.2279863 | 24.0724562 | 24.1281889 | 24.1208867 | -0.101513262 | 0.027486755 |
| 23.5105858 | 23.4513912 | 23.5478439 | 23.5143597 | 0.003216303 | 0.933056741 |
| 20.5279085 | 20.5691553 | 20.5047456 | 20.5959608 | 0.006508243 | 0.865296173 |
| 19.9781127 | 19.983852 | 20.0781925 | 19.9333624 | 0.018963356 | 0.687773003 |
| 21.2915684 | 21.3151402 | 21.3089539 | 21.3630521 | 0.073380968 | 0.13694131 |
| 19.626909 | 19.5876241 | 19.6056059 | 19.7229476 | 0.034608814 | 0.493203594 |
| 20.0694431 | 19.9969081 | 20.1251381 | 20.0277504 | 0.016262736 | 0.763237378 |
| 18.8434329 | 18.9282739 | 18.966306 | 18.9489335 | 0.110780094 | 0.049822117 |
| 19.6158335 | 19.6066471 | 19.6944063 | 19.6853177 | 0.054167473 | 0.32440649 |
| 22.2583031 | 22.172969 | 22.2754039 | 22.2303758 | 0.023527878 | 0.67472791 |
| 19.1638806 | 19.2709153 | 19.2493644 | 19.4075385 | 0.130794781 | 0.026498177 |
| 20.0356423 | 19.8995731 | 20.0719447 | 20.1461742 | 4.61E-04 | 0.993814903 |
| 19.0420949 | 19.1753135 | 19.2179066 | 19.2173456 | 0.11024797 | 0.105949902 |
| 18.7523858 | 18.6857427 | 18.7111318 | 18.7089174 | 0.012211192 | 0.88203036 |
| 22.6296892 | 22.626198 | 22.7139288 | 22.6847359 | 0.103645407 | 0.219450486 |
| 19.1081939 | 18.9649873 | 19.1681657 | 19.1359979 | 0.047688569 | 0.583892041 |
| 24.7303394 | 24.9010767 | 24.9197532 | 24.8569771 | 0.129574643 | 0.183954424 |
| 20.8165683 | 20.7708067 | 20.8858436 | 20.8763792 | 0.084995566 | 0.461463419 |
| 16.950612 | 17.7498129 | 17.2577828 | 17.0864111 | 0.474216406 | 0.004894732 |
| 14.1720568 | NA | 14.2301592 | 14.2205683 | 0.069262968 | 0.686797118 |
| 17.1564979 | 17.5730238 | 17.4036576 | 17.4740565 | 0.386664702 | 0.049317235 |
| 20.3239491 | 20.8442152 | 20.4634641 | 20.4721245 | 0.306958767 | 0.114966857 |
| 16.0793412 | NA | 16.2529689 | 16.3482329 | 0.173933415 | 0.246591939 |

| | | | | | |
|---|---|---|---|---|---|
| 22.9220707 | 22.8904418 | 22.8790149 | 22.8982123 | -0.046211313 | 0.307364828 |
| 15.7222528 | 16.0143356 | 15.9548873 | 15.5610304 | -0.036146831 | 0.797845076 |
| 17.7528139 | 18.0172078 | 18.0192136 | 17.9653668 | 0.230112294 | 4.42E-04 |
| 23.3301026 | 23.1912209 | 23.1973136 | 23.2132193 | -0.13800849 | 0.01651862 |
| 24.2410211 | 24.1862913 | 24.3429842 | 24.328137 | 0.073992503 | 0.459235579 |
| 18.0891794 | 17.6807103 | 17.7714116 | 17.9489424 | -0.179401683 | 0.033533955 |
| 26.222866 | 26.2148174 | 26.276379 | 26.2454673 | -0.007662557 | 0.910234152 |
| 18.4235735 | 18.3212997 | 18.4681632 | 18.490754 | 0.006602736 | 0.915925563 |
| 19.5487389 | NA | 19.8748607 | 19.6599956 | 0.052522073 | 0.743080221 |
| 18.8939309 | 19.078006 | 18.989149 | 18.7354753 | -0.04273204 | 0.708730785 |
| 19.2932224 | 19.4040082 | 19.3867542 | 19.1806183 | -0.103310908 | 0.168390763 |
| 21.077373 | 20.8804332 | 20.9827795 | 20.9817274 | -0.112986049 | 0.123864552 |
| 24.5227881 | 24.3070241 | 24.4932503 | 24.4214603 | -0.049719402 | 0.488504534 |
| 23.7747859 | 23.6475253 | 23.7213686 | 23.7194005 | -0.012210976 | 0.813536739 |
| 21.5565952 | 21.470284 | 21.4903447 | 21.4939929 | -0.019071529 | 0.675952957 |
| 19.4442427 | 19.4671589 | 19.4903696 | 19.4868391 | 0.049496728 | 0.196402164 |
| 19.2160274 | 19.1503712 | 19.3312325 | 19.3290093 | 0.007115708 | 0.896814549 |
| 19.642523 | 19.4957144 | 19.6583901 | 19.6988188 | 0.02077238 | 0.708557824 |
| 20.0609825 | 19.9555458 | 20.0892355 | 20.0946251 | 0.011671441 | 0.843894706 |
| 17.8863835 | 17.947203 | 18.0282969 | 18.0850951 | 0.101077108 | 0.169969671 |
| 18.5865499 | 18.3805654 | 18.5687614 | 18.4338142 | -0.103858892 | 0.168177013 |
| 17.8400436 | 17.5914684 | 17.779407 | 17.8446861 | 0.035408438 | 0.687290849 |
| 18.9509346 | 19.0643706 | 19.0493306 | 19.1572065 | 0.048201933 | 0.630174883 |
| 16.6685736 | 16.7986039 | 16.7754412 | 16.9844391 | 0.356249968 | 0.020274583 |
| 17.0629262 | 16.8984335 | 16.8846398 | 17.0362734 | -0.11303633 | 0.156242288 |
| 19.5353939 | 19.4845426 | 19.602879 | 19.6160983 | 0.083268517 | 0.175599953 |
| 18.1506069 | 18.2733911 | 18.201746 | 18.3618799 | 0.061400983 | 0.517426513 |
| 16.8383188 | 17.2427372 | 17.1883032 | 16.7976236 | 0.023829385 | 0.837995666 |
| 21.0900329 | 20.7713263 | 21.0841708 | 21.1991033 | -0.086974021 | 0.315418508 |
| 19.1228709 | 18.9746642 | 19.075965 | 19.0753048 | -0.0861306 | 0.18582826 |
| 22.3541042 | 22.2047576 | 22.335578 | 22.3488624 | -0.031253944 | 0.509103891 |
| 24.8825546 | 24.6625267 | 24.7426672 | 24.7604771 | -0.09294905 | 0.043369427 |
| 20.5996493 | 20.5002367 | 20.5027202 | 20.5520269 | -0.029411817 | 0.501768745 |
| 21.8565255 | 21.8701642 | 21.8554945 | 21.8436163 | -0.046628756 | 0.25458699 |
| 22.2515024 | 22.1008497 | 22.1069081 | 22.1455674 | -0.077426018 | 0.062381978 |
| 19.6254223 | 19.6756964 | 19.7470765 | 19.7932507 | 0.097750542 | 0.040335686 |
| 19.4902294 | 19.4641898 | 19.524904 | 19.4848506 | 0.071208501 | 0.161428047 |
| 22.9083741 | 22.9117179 | 22.9309835 | 22.9706046 | 0.073831906 | 0.201059668 |
| 18.2764023 | 18.1304074 | 18.375312 | 18.4971629 | -0.045191668 | 0.669173042 |
| 18.7851356 | 18.9315262 | 19.0057131 | 18.8718024 | 0.066754909 | 0.554683865 |
| 17.3254899 | 17.447257 | 17.9685455 | 17.6637967 | 0.26050991 | 0.074683442 |
| 22.054651 | 22.3381618 | 22.3928471 | 22.1993767 | 0.246028525 | 0.147637993 |

| | | | | | |
|---|---|---|---|---|---|
| NA | 18.3075749 | NA | NA | -0.552034307 | 0.023020138 |
| 20.3229945 | 20.4007335 | 20.2746592 | 20.4519165 | 0.097922258 | 0.325997971 |
| 23.185007 | 23.1648317 | 23.217256 | 23.2349272 | 0.090472787 | 0.474430076 |
| 15.553766 | NA | 15.6925348 | 15.7808198 | 0.320641082 | 0.242963017 |
| 17.6686449 | 17.6238901 | 16.9535475 | 17.5797767 | -0.202698546 | 0.435247649 |
| 16.3237297 | 16.6826527 | 16.5240224 | 16.6535915 | -0.011237568 | 0.942156531 |
| 16.7640023 | 16.5290514 | 16.9339397 | 16.7074243 | -0.110430081 | 0.444713144 |
| 15.5124865 | 15.1273745 | 15.4000636 | 15.4361279 | -0.197008525 | 0.134640994 |
| 19.5301539 | 19.491527 | 19.1103465 | 19.1549253 | -0.204347607 | 0.073097217 |
| 16.9708059 | 16.954987 | 17.1562005 | 16.9671005 | 0.097048756 | 0.357126086 |
| 19.2046002 | 18.6778283 | 18.9343776 | 19.0000567 | -0.253295953 | 0.016678135 |
| 23.3120972 | 22.9060426 | 22.9412236 | 23.2653383 | -0.144418566 | 0.152581602 |
| 16.7126245 | 16.2745513 | 16.1993088 | 16.4011333 | -0.333079464 | 0.002169805 |
| 20.098134 | 19.8474122 | 19.991705 | 19.8578596 | -0.227735815 | 0.011533431 |
| 21.6102956 | 21.5408794 | 21.6790573 | 21.6347491 | 0.062093004 | 0.230752232 |
| 19.1679569 | 19.1829979 | 19.3919893 | 19.2888014 | 0.058795904 | 0.273511391 |
| 22.8057232 | 22.685058 | 22.8085315 | 22.829749 | 0.046718932 | 0.387768757 |
| 19.8241853 | 19.96059 | 20.1488766 | 20.1183007 | 0.139617468 | 0.033325497 |
| 19.9362826 | 19.9126164 | 19.9763857 | 19.9560923 | 0.056745418 | 0.43515753 |
| 18.0395849 | 17.7334863 | 18.0963776 | 18.2223813 | 0.094397648 | 0.349910025 |
| 15.840374 | 16.2606107 | 16.3365063 | 16.2392352 | 0.295822584 | 0.02177475 |
| 14.8407542 | 15.1719947 | 15.4351 | 15.1826879 | 0.211289918 | 0.433202311 |
| 16.9993867 | NA | 16.0670088 | 16.433845 | -0.748968803 | 0.006122991 |
| 17.7464627 | 17.9386543 | 17.8134273 | 17.844841 | 0.11922584 | 0.573142588 |
| 16.7574509 | 16.6376378 | 16.6547962 | 16.7406133 | 0.004227131 | 0.969866398 |
| 18.308903 | 18.0267634 | 18.3469754 | 18.3811813 | -0.081049587 | 0.503532023 |
| 16.9772724 | 16.8418745 | 17.0021292 | 17.0337262 | -0.021849124 | 0.816162783 |
| 18.4520384 | 18.3282313 | 18.5006923 | 18.4315751 | -0.06206856 | 0.385173813 |
| 17.9038557 | 17.5903292 | 17.6829987 | 17.7643403 | -0.209833235 | 0.003516283 |
| 22.8422734 | 22.7891545 | 22.7292157 | 22.6612204 | -0.077986268 | 0.128655073 |
| 26.635584 | 26.5617923 | 26.5196729 | 26.5219662 | -0.045203926 | 0.299193663 |
| 17.2380983 | 17.3057886 | 17.3159527 | 17.3833427 | 0.051588954 | 0.661524623 |
| 21.9454434 | 21.7032072 | 21.42337 | 21.179634 | -0.511983671 | 0.039530407 |
| 15.7380283 | 15.7050204 | 15.781725 | 15.519469 | -0.063289869 | 0.522891941 |
| 22.8826487 | 22.8628733 | 22.8851671 | 22.9280039 | 0.085678025 | 0.230332516 |
| 19.4664151 | 19.7818527 | 19.7640407 | 19.9869214 | 0.380630425 | 0.00185093 |
| 16.8237324 | NA | 17.1122377 | 17.211284 | 0.169085917 | 0.149884243 |
| 20.28115 | 20.3603041 | 20.2769974 | 20.4164991 | 0.085893097 | 0.160357501 |
| 17.9799214 | 17.7790078 | 17.7873874 | 17.8655553 | -0.163240139 | 0.029668269 |
| 20.0170263 | 20.0129695 | 20.0670591 | 20.0983407 | 0.03497194 | 0.375712537 |
| 11.6349095 | NA | 11.327777 | 11.3459441 | -0.377210993 | 0.018266012 |
| 21.2512352 | 20.9106748 | 20.8564598 | 20.9638209 | -0.114462588 | 0.388298786 |

| | | | | | |
|---|---|---|---|---|---|
| 24.7878904 | 24.6519719 | 24.6490746 | 24.6296808 | -0.075030099 | 0.175482821 |
| 21.0539799 | 21.0089977 | 21.0556843 | 21.0717962 | 0.010179744 | 0.834532349 |
| 16.5698973 | 16.8866837 | 16.6390161 | 16.8341834 | 0.123397445 | 0.109742764 |
| 18.7332341 | 18.7842327 | 18.6711519 | 18.7269158 | 0.001947547 | 0.980164946 |
| 18.2109494 | 18.1980373 | 18.5147117 | 18.3501914 | 0.061484564 | 0.644895591 |
| 19.3630243 | 19.2733839 | 19.2667798 | 19.3916526 | 0.020006379 | 0.698132483 |
| 25.3622426 | 25.2488902 | 25.3743604 | 25.3535462 | -0.023949674 | 0.793197863 |
| 21.149525 | 20.9733439 | 21.1388895 | 21.1174701 | -0.049385483 | 0.39077847 |
| NA | 15.6573184 | NA | NA | 0.359279927 | 0.076300945 |
| 17.818518 | 17.3139061 | 17.2764322 | 17.1745857 | -0.299270193 | 0.027507885 |
| 17.3945733 | 17.2819329 | 17.3950213 | 17.5142629 | 0.055487216 | 0.399149169 |
| 23.2924456 | 23.2645977 | 23.3269044 | 23.246444 | 0.001399572 | 0.979255469 |
| 17.0914665 | 17.2878418 | 16.9157262 | 17.1207744 | -0.054706146 | 0.575734457 |
| 23.0609102 | 22.8732357 | 23.0154278 | 23.0235603 | -0.09376657 | 0.146072538 |
| 20.6653187 | 20.5385362 | 20.6803578 | 20.5803091 | -0.065037905 | 0.274873276 |
| 22.4795272 | 22.3491423 | 22.3843992 | 22.3777277 | -0.052191029 | 0.181138214 |
| 21.6152108 | 21.6687682 | 21.6483573 | 21.6954849 | 0.109294924 | 0.060263419 |
| 19.1874218 | 19.073411 | 19.3048984 | 19.2246287 | 0.002851729 | 0.97626242 |
| 20.5834858 | 20.6758023 | 20.7309586 | 20.4422888 | -0.008646755 | 0.8958068 |
| 17.5877303 | 17.5740046 | 17.6718833 | 17.6705477 | 0.100725442 | 0.085871083 |
| 17.6974084 | 17.680625 | 17.7349495 | 17.5871437 | -0.04780861 | 0.619351797 |
| 20.182619 | 20.0662645 | 20.3031073 | 20.1764925 | -0.023799925 | 0.646908283 |
| 15.6284967 | 15.6086 | 16.0454267 | 15.9723958 | -0.027408762 | 0.932117409 |
| 23.2127742 | 23.0329225 | 23.1060375 | 23.118113 | -0.060226014 | 0.228772236 |
| 17.4993568 | 17.8217763 | 17.679901 | 17.7244613 | 0.146792836 | 0.007044338 |
| 19.3565609 | 19.0235219 | 19.1344607 | 19.0525904 | -0.198361679 | 0.031623339 |
| 25.380759 | 25.2978469 | 25.3448486 | 25.3504057 | -0.024268902 | 0.671857342 |
| 15.4729898 | 15.0340753 | 15.2773215 | 15.4093883 | -0.063067501 | 0.6295123 |
| 17.8216916 | 17.6514176 | 17.7945485 | 17.5982445 | -0.231768533 | 0.036897188 |
| 21.9499296 | 21.6962549 | 21.7388533 | 21.7748465 | -0.021757234 | 0.778612152 |
| 20.6978538 | 20.6428565 | 20.4586588 | 20.6374678 | -0.100805427 | 0.092366514 |
| 20.9397748 | 20.8512631 | 20.9288055 | 20.8914733 | -0.058486943 | 0.299401078 |
| 21.6916693 | 21.5566531 | 21.6337951 | 21.6462446 | -0.049703347 | 0.318221762 |
| 23.6827835 | 23.5262317 | 23.5710695 | 23.5695534 | -0.061688565 | 0.19042768 |
| 21.5795303 | 21.4723282 | 21.5341209 | 21.5614133 | -0.017454167 | 0.69849684 |
| 22.1410388 | 22.0075615 | 22.0927076 | 22.1233044 | -0.05108573 | 0.21518121 |
| 25.1452116 | 25.0809362 | 25.1245797 | 25.1171819 | -0.006464314 | 0.872088462 |
| 23.4446068 | 23.5347 | 23.4809651 | 23.4960788 | 0.057910571 | 0.156363773 |
| 20.1393035 | 20.122037 | 20.1466875 | 20.1408615 | -0.014430608 | 0.72695574 |
| 22.2745939 | 22.3820352 | 22.3492382 | 22.5266909 | 0.091676492 | 0.090785124 |
| 20.7887268 | 20.8087391 | 20.8347231 | 20.8052577 | 0.077007031 | 0.225818641 |
| 19.4942413 | 19.4917322 | 19.5587711 | 19.5000908 | -1.55E-04 | 0.99812209 |

| | | | | | |
|---|---|---|---|---|---|
| 20.0615981 | 20.0833876 | 20.1527671 | 20.0416348 | 0.074156703 | 0.304932167 |
| 20.4901784 | 20.4949907 | 20.5837198 | 20.5152469 | 0.058418835 | 0.421368956 |
| 19.1410333 | 19.2043275 | 19.2862102 | 19.2605039 | 0.081976571 | 0.284979994 |
| 17.9898966 | 18.4507968 | 18.1666444 | 18.1682692 | 0.293727664 | 6.37E-04 |
| 18.2169827 | 18.3281689 | 18.4301419 | 18.3528536 | 0.066549884 | 0.450047345 |
| 22.9712255 | 22.8376777 | 22.9763226 | 22.9906953 | 0.033979559 | 0.701463642 |
| 16.284775 | NA | 16.2564894 | 16.2358212 | -0.028402062 | 0.743771038 |
| 16.5551776 | 16.6385954 | 16.8518578 | 16.9020248 | 0.266672096 | 0.011062921 |
| 18.0133288 | 17.9773936 | 18.1097338 | 18.1483693 | 0.068060736 | 0.493762161 |
| 19.4414524 | 19.379476 | 19.5175196 | 19.575497 | 0.053338842 | 0.597972188 |
| 13.7139434 | NA | 13.957347 | 14.0988368 | 0.167306192 | 0.143598855 |
| 18.9170602 | 19.2629563 | 18.8532363 | 18.8206494 | 0.223955665 | 0.079911415 |
| 16.0958139 | 16.1631309 | 16.4727747 | 16.6494358 | 0.207755039 | 0.158104391 |
| 13.328195 | NA | 13.8573943 | 13.4243984 | 0.334712882 | 0.081295971 |
| 20.388657 | 20.4087561 | 20.3542974 | 20.8364813 | 0.590493488 | 0.01783571 |
| NA | 16.5733591 | NA | NA | 0.68500247 | 0.018036854 |
| 24.8254369 | 24.918591 | 24.817114 | 24.7650174 | -0.036886942 | 0.646386241 |
| 15.8306011 | 16.1203557 | 15.9605581 | 15.750006 | 0.08562758 | 0.585112029 |
| 18.4114498 | 18.7156713 | 18.7062916 | 18.4721927 | 0.023283942 | 0.863777585 |
| 16.1021515 | 15.3844103 | 16.178582 | 16.0200252 | -0.312489856 | 0.026273609 |
| 21.8984223 | 21.8948062 | 21.9923702 | 21.9983646 | 0.134448909 | 0.201668804 |
| 18.7115168 | 18.7392378 | 18.6939745 | 18.7616469 | 0.018695754 | 0.844538076 |
| 20.916114 | 20.8824515 | 20.9418321 | 20.8435066 | -0.054641376 | 0.521116535 |
| 20.6961009 | 20.4460402 | 20.7083264 | 20.7200159 | -0.060259193 | 0.439633288 |
| 20.8466539 | 20.5962472 | 20.7117976 | 20.7350725 | -0.099069137 | 0.2023165 |
| 19.3385301 | 19.3368528 | 19.3756111 | 19.3794379 | -0.02766798 | 0.672594718 |
| 25.4790162 | 25.3180828 | 25.4251633 | 25.4694666 | -0.056707194 | 0.377305747 |
| 19.0079205 | 18.8291313 | 18.990864 | 18.9253261 | -0.048648979 | 0.442513954 |
| 21.9385499 | 21.7765563 | 21.8674995 | 21.9508556 | -0.001803736 | 0.977134681 |
| 22.6455675 | 22.6666832 | 22.628394 | 22.6124992 | -0.008788145 | 0.882282697 |
| 19.9981632 | 19.7471989 | 19.7997903 | 19.8925137 | -0.105099643 | 0.069706357 |
| 22.1066449 | 22.1549249 | 22.2181334 | 22.1102892 | 4.46E-04 | 0.992623908 |
| 21.6997983 | 21.6081263 | 21.5912005 | 21.5910959 | -0.044785919 | 0.335735041 |
| 20.3046484 | 20.1737622 | 20.1986578 | 20.2680303 | -0.076484803 | 0.096619252 |
| 23.9287487 | 23.848963 | 23.9278839 | 23.9375026 | 0.005572949 | 0.896349566 |
| 19.235011 | 19.2683654 | 19.2324874 | 19.1274869 | -0.008958124 | 0.829637942 |
| 19.7060652 | 19.6609677 | 19.6890797 | 19.7870578 | 0.019659822 | 0.712727866 |
| 20.377894 | 20.3602567 | 20.5334787 | 20.3963469 | 0.087762404 | 0.110161844 |
| 20.2130252 | 20.2326137 | 20.260878 | 20.1838724 | 0.009224303 | 0.866891237 |
| 21.8976238 | 22.1118434 | 22.0978309 | 21.9788357 | 0.117990175 | 0.045935714 |
| 20.0841992 | 20.0236811 | 20.0936603 | 20.0972313 | 0.048149535 | 0.405844504 |
| 20.047391 | 20.0242823 | 20.1044761 | 20.2292777 | 0.100372672 | 0.116500733 |

| | | | | | |
|---|---|---|---|---|---|
| 20.2272706 | 20.3594747 | 20.2181031 | 20.2462723 | 0.010983065 | 0.869147536 |
| 20.0509754 | 19.9972844 | 20.0679173 | 20.1357412 | 0.039781235 | 0.55410606 |
| 18.6820233 | 18.7566979 | 18.748008 | 18.6796905 | 0.028733959 | 0.66722433 |
| 18.768476 | 18.8995641 | 18.9297993 | 19.0768332 | 0.141695953 | 0.043135941 |
| 21.0751485 | 20.9427543 | 21.0448584 | 21.0784941 | 0.0269913 | 0.70320391 |
| 19.1206712 | 19.0775094 | 19.2157709 | 19.1289501 | 0.06017296 | 0.415408327 |
| 20.9640606 | 20.9010741 | 21.0002164 | 20.957584 | 0.038247704 | 0.621485861 |
| 15.6176985 | NA | 15.5756577 | 15.6061661 | -0.001453095 | 0.984244519 |
| 18.2876999 | 18.3795553 | 18.4064319 | 18.3575993 | 0.050249612 | 0.559426293 |
| 16.2754656 | 16.1117644 | 16.3082646 | 16.3032185 | 0.012811979 | 0.882659713 |
| 19.1827458 | 19.236936 | 19.0512274 | 19.1307469 | 0.053253502 | 0.546160503 |
| 21.7992642 | 21.9436977 | 22.0523532 | 22.1339012 | 0.233676215 | 0.014009369 |
| 22.3135647 | 22.3911967 | 22.3857581 | 22.4451715 | 0.03339949 | 0.718255121 |
| 19.0639834 | 19.4844393 | 19.3767202 | 19.520214 | 0.20496104 | 0.040439647 |
| 17.6327012 | 17.4180318 | 17.4426053 | 17.4578377 | -0.073220601 | 0.449193254 |
| 19.1697543 | 19.1418022 | 19.3157311 | 19.1196746 | 0.024100968 | 0.808988724 |
| 17.7006015 | 17.8474697 | 17.9245237 | 17.8046242 | 0.072379259 | 0.490590736 |
| 16.7987796 | 16.7896217 | 16.8072713 | 16.6563952 | 0.055659939 | 0.653813396 |
| 17.7919998 | 18.0373638 | 18.2605914 | 18.059358 | 0.21843373 | 0.11127875 |
| 16.1924966 | 16.4416252 | 16.2466381 | 16.1729692 | 0.133021622 | 0.333366856 |
| 17.6404174 | 17.6727294 | 17.6564187 | 17.6146294 | -0.141560574 | 0.302695963 |
| 15.3326317 | NA | 15.4117994 | 15.3569707 | 0.097413789 | 0.533175027 |
| 15.4629938 | NA | 15.6212162 | 15.8994248 | 0.171952545 | 0.343898492 |
| NA | 14.6250878 | NA | NA | 0.460796293 | 0.039830649 |
| 15.4495158 | 14.6729249 | 15.5069429 | 15.7064295 | -0.114323986 | 0.668927148 |
| 24.6027374 | 24.3543419 | 24.4701874 | 24.5368604 | -0.086135081 | 0.173739408 |
| 22.3293884 | 22.2490191 | 22.3375551 | 22.3755744 | 0.057881674 | 0.415488393 |
| 22.7492178 | 22.7758944 | 22.7114071 | 22.7946635 | -0.009323689 | 0.83030629 |
| 18.9232557 | 18.7646634 | 19.1494227 | 19.0936405 | 0.137840047 | 0.308976992 |
| 21.428569 | 21.11412 | 21.3692411 | 21.3296063 | -0.129961227 | 0.270918861 |
| 22.3230481 | 22.3175847 | 22.3187555 | 22.2814241 | 0.045423081 | 0.41735649 |
| 20.3000573 | 20.3156567 | 20.3538524 | 20.3706191 | 0.104410644 | 0.106605275 |
| 17.3992216 | 17.4273591 | 17.4595103 | 17.4749538 | 0.014530036 | 0.837714515 |
| NA | 14.0365562 | NA | NA | -0.210887348 | 0.105072374 |
| 16.7244352 | 16.2958931 | 16.4475782 | 16.632976 | -0.157070298 | 0.154554956 |
| 15.9182386 | NA | 15.8148535 | 15.8497432 | -0.228710175 | 0.029488142 |
| 24.0791846 | 24.0304544 | 24.1224881 | 24.0643857 | 0.011539066 | 0.846152549 |
| 23.7933583 | 23.9319368 | 23.849785 | 23.7784381 | 0.016099096 | 0.771554303 |
| 20.2943533 | 20.229988 | 20.2287848 | 20.2938977 | -0.009250958 | 0.859616122 |
| 20.2582991 | 20.3473005 | 20.3304846 | 20.3464041 | 0.088478378 | 0.035111949 |
| 21.841954 | 21.8414427 | 21.8368436 | 21.8361018 | 0.041957584 | 0.358127112 |
| 17.9008619 | 17.6998079 | 17.6592999 | 17.8820127 | 0.011859953 | 0.892073833 |

| | | | | | |
|---|---|---|---|---|---|
| 20.5571568 | 20.2055923 | 20.467487 | 20.5644182 | -0.174016358 | 0.121881126 |
| 16.30027 | 16.4162274 | 16.4370083 | 16.4318227 | 0.150859866 | 0.042773021 |
| 21.9072584 | 21.9254877 | 21.9662218 | 21.9352099 | 0.076494226 | 0.111587008 |
| 23.564093 | 23.510921 | 23.5795131 | 23.5978293 | 0.059181333 | 0.206812217 |
| 19.693721 | 19.5944034 | 19.8121442 | 19.7771634 | 0.057815997 | 0.361183403 |
| 19.6255021 | 19.4544222 | 19.5278062 | 19.6456309 | -0.051420688 | 0.467016014 |
| 24.6536654 | 24.6920451 | 24.698825 | 24.7546156 | 0.093956518 | 0.197871702 |
| 26.5777107 | 26.3743203 | 26.4673154 | 26.4381214 | -0.091283444 | 0.09989613 |
| 16.7141359 | 17.0081704 | 17.14589 | 16.9799619 | 0.307359694 | 0.00508498 |
| NA | 12.7919992 | NA | NA | -0.052072813 | 0.6142226 |
| 16.0669881 | 16.2131847 | 16.0102054 | 15.9062379 | -0.097441564 | 0.338956706 |
| 18.5327743 | 18.7279372 | 18.8260238 | 18.8547687 | 0.111464796 | 0.322253631 |
| 18.1694508 | 18.1933475 | 18.1993971 | 18.1927871 | 0.059192059 | 0.375459802 |
| 19.6450372 | 19.7756203 | 19.7350529 | 19.7482225 | 0.197993783 | 0.008488603 |
| 19.8303043 | 19.9641794 | 19.9524239 | 19.9180359 | 0.135859294 | 0.061947653 |
| 16.6558815 | 16.7210701 | 17.0509289 | 17.0950854 | 0.126708279 | 0.338829791 |
| 19.8868953 | 19.9490128 | 20.0087009 | 20.23219 | 0.211935684 | 0.116733574 |
| 16.7584968 | 17.022978 | 16.9807832 | 17.2400665 | 0.107445452 | 0.231331061 |
| 15.4586215 | 15.1349796 | 15.347965 | 15.2259132 | -0.15232844 | 0.354376895 |
| 21.6587215 | 21.7418636 | 21.6591715 | 21.595658 | -0.030937274 | 0.661527202 |
| 22.4054537 | 22.3197929 | 22.3638937 | 22.3125576 | -0.023696576 | 0.627651918 |
| 23.9614526 | 23.9134644 | 23.9544134 | 23.859711 | -0.029304487 | 0.533366681 |
| 20.6816922 | 20.6791613 | 20.781146 | 20.7504059 | 0.08517988 | 0.097219978 |
| 18.9291067 | 18.8397816 | 18.9416045 | 18.946461 | -0.002338773 | 0.96140074 |
| 21.5703926 | 21.609992 | 21.5904263 | 21.5048714 | -0.020024982 | 0.68184695 |
| NA | 15.9355832 | NA | NA | 0.28730688 | 0.060918386 |
| 18.9839065 | 18.8400093 | 19.0062185 | 18.9108792 | -0.033606355 | 0.67800065 |
| 19.7370387 | 19.6515366 | 19.8046415 | 19.8266162 | 0.059332552 | 0.375187431 |
| 20.6552223 | 20.6207151 | 20.6911203 | 20.686117 | 0.048142249 | 0.325133763 |
| 16.5151691 | 16.1020454 | 16.2805888 | 16.4534535 | -0.096202354 | 0.494032788 |
| 18.8616139 | 18.576218 | 18.6481355 | 18.7613988 | -0.124063765 | 0.062404616 |
| 18.0246644 | 17.9485 | 17.9796542 | 17.9026313 | -0.031538531 | 0.601917063 |
| 22.2836547 | 22.2292091 | 22.265586 | 22.2932321 | 0.006633571 | 0.878535332 |
| 19.485405 | 18.9849254 | 19.4296472 | 19.1974302 | -0.284621336 | 0.026616198 |
| 18.7366192 | 18.4063703 | 18.5295799 | 18.6942991 | -0.168266174 | 0.038838744 |
| 22.7013644 | 22.5577369 | 22.705484 | 22.6827723 | -0.079561829 | 0.148066297 |
| 24.8325603 | 24.6344071 | 24.781459 | 24.7267085 | -0.062721497 | 0.246123218 |
| 21.5629122 | 21.3950721 | 21.549413 | 21.4699532 | -0.060532768 | 0.237358137 |
| 21.7821189 | 21.7126135 | 21.5990814 | 21.6667028 | -0.06180696 | 0.204494333 |
| 18.5131051 | 18.4921012 | 18.6886208 | 18.4867017 | 0.087975293 | 0.325446451 |
| 16.2878733 | 16.4720887 | 17.0456123 | 16.7921632 | 0.396485421 | 0.001596593 |
| 20.3803922 | 20.1655161 | 20.1646779 | 20.3082168 | -0.089720656 | 0.103661009 |

| | | | | | |
|---|---|---|---|---|---|
| 22.5750457 | 22.5187032 | 22.4631707 | 22.6060647 | 0.033103114 | 0.635697576 |
| NA | 16.23796 | NA | NA | 0.316923263 | 0.226087752 |
| NA | 16.2505858 | NA | NA | 0.298425634 | 0.074015895 |
| 19.2649264 | 19.1555473 | 19.116182 | 19.1576273 | -0.081768635 | 0.147505138 |
| 20.7359181 | 20.6458208 | 20.7590734 | 20.8065717 | 0.043387668 | 0.547533199 |
| 25.7584295 | 25.7615794 | 25.8331467 | 25.8322251 | 0.097589103 | 0.150163948 |
| 19.4061048 | 19.6490201 | 19.4483353 | 19.533211 | 0.071518128 | 0.300883213 |
| 14.72344 | NA | 14.7732462 | 14.535926 | -0.063821897 | 0.662993412 |
| 17.7371581 | 17.5705469 | 17.797671 | 17.8162819 | 0.139606499 | 0.254445302 |
| 14.5927503 | 14.4853848 | 14.9169702 | 14.5018337 | 0.226195044 | 0.175310637 |
| 19.9916736 | 20.1364915 | 20.0950523 | 20.1950552 | 0.198145611 | 0.096538156 |
| 21.3482559 | 21.3422409 | 21.4130751 | 21.4238807 | 0.045189771 | 0.464118928 |
| 15.4924591 | NA | 15.4907912 | 15.3888686 | -0.069589876 | 0.348334243 |
| 23.971735 | 24.0265602 | 23.9951137 | 23.9290382 | 0.014754067 | 0.791432613 |
| 21.5972102 | 21.5412345 | 21.6837472 | 21.7074444 | 0.128585769 | 0.058349434 |
| NA | 17.9774003 | NA | NA | -0.472109091 | 0.063679072 |
| 17.1763956 | 17.1521822 | 17.4687564 | 17.2566719 | -0.019071601 | 0.889043699 |
| 22.0843079 | 22.3039126 | 22.0125436 | 22.2795148 | 0.077170764 | 0.560946398 |
| 19.1013491 | 19.0034171 | 19.0554836 | 18.7738678 | -0.138267878 | 0.182702224 |
| 26.5794707 | 26.5780692 | 26.6167572 | 26.5001142 | 0.002782121 | 0.977935529 |
| 17.9767417 | 17.9228361 | 17.8286536 | 17.8087083 | -0.109099682 | 0.191486657 |
| 18.5426857 | 18.4566175 | 18.4341541 | 18.6047848 | -0.020791721 | 0.781231214 |
| 15.25286 | 15.2162824 | 15.2780922 | 15.204976 | -0.011782422 | 0.870680639 |
| 26.0632672 | 25.726054 | 25.9022164 | 25.9398364 | -0.125190416 | 0.075216011 |
| 20.3698439 | 20.2862897 | 20.2619325 | 20.3750577 | -0.027560851 | 0.635029468 |
| 25.079229 | 24.9813855 | 24.9731207 | 24.9375409 | -0.051346068 | 0.298253945 |
| 21.3315828 | 21.28419 | 21.2352139 | 21.2691164 | -0.091309656 | 0.07203156 |
| 21.0800987 | 20.990431 | 21.0446913 | 20.9510627 | -0.07436787 | 0.112429336 |
| 20.8874166 | 20.9173496 | 20.9973744 | 20.9329287 | 0.060692253 | 0.187868287 |
| 20.6430833 | 20.5709303 | 20.6873675 | 20.6515239 | 0.00913477 | 0.840260337 |
| 19.4244093 | 19.3867409 | 19.4767594 | 19.4459265 | 0.079200741 | 0.097475537 |
| 20.7780649 | 20.7450327 | 20.8076738 | 20.8320374 | 0.068615072 | 0.153373163 |
| 18.984809 | 18.9922748 | 19.0462646 | 19.090712 | 0.021335312 | 0.711140419 |
| 19.9440698 | 19.9579427 | 19.9425135 | 19.9488962 | 0.083676077 | 0.157830184 |
| 20.0257511 | 19.8597156 | 19.944004 | 20.0069449 | -0.07763565 | 0.20648346 |
| 16.6530372 | 16.749322 | 16.6809425 | 16.7395179 | 0.04193115 | 0.513797383 |
| 22.1507661 | 21.960494 | 22.0453143 | 22.2027352 | 0.078203627 | 0.29506337 |
| 17.0518455 | 17.0272121 | 17.1162552 | 17.1087805 | -0.033769806 | 0.652622601 |
| 20.1467407 | 20.0545858 | 19.9899063 | 20.0452918 | -0.049311865 | 0.579743798 |
| 19.0413707 | 18.9580266 | 19.0639685 | 19.0047554 | 2.54E-04 | 0.997785153 |
| 21.6181249 | 21.6211961 | 21.6787472 | 21.6259264 | 0.10722874 | 0.269150911 |
| NA | 16.0186452 | NA | NA | 0.074553947 | 0.432536529 |

| | | | | | |
|---|---|---|---|---|---|
| 18.6202082 | 18.8273374 | 19.186876 | 18.9516068 | 0.134128905 | 0.24965106 |
| NA | 14.6717763 | NA | NA | 0.481174295 | 0.009710714 |
| 16.485984 | 16.5410499 | 16.8634772 | 16.6600335 | 0.128926872 | 0.438108116 |
| NA | 16.2419872 | NA | NA | 0.235991842 | 0.158081508 |
| 16.5499879 | 16.5094286 | 16.5027779 | 16.6681326 | 0.145460473 | 0.475491584 |
| 19.9256083 | 19.8066808 | 19.9361834 | 19.988809 | 0.050136021 | 0.400687695 |
| 18.9983562 | 18.9541673 | 19.0177444 | 19.0416656 | -0.066646414 | 0.150597845 |
| 19.0989204 | 19.1385396 | 19.1845225 | 19.2649566 | 0.091570997 | 0.121131194 |
| 24.2177782 | 24.1509439 | 24.2211999 | 24.208564 | 0.053000376 | 0.509789589 |
| 20.8278376 | 20.6875005 | 20.7874544 | 20.7867286 | 0.022297254 | 0.713760005 |
| 18.1024246 | 18.0906408 | 18.4562801 | 18.1067222 | 0.088016645 | 0.461524257 |
| 20.6516323 | 20.0573037 | 20.8007205 | 20.6551287 | -0.126540027 | 0.564889359 |
| 15.3705261 | 15.7201563 | 15.6161739 | 15.0706819 | -0.115187952 | 0.472270145 |
| 28.3419502 | 28.4307217 | 28.2836853 | 28.2956354 | -0.054361029 | 0.456641773 |
| 20.800455 | 20.8311973 | 20.861023 | 20.8654345 | 0.086273709 | 0.10884049 |
| 19.2825965 | 19.3576354 | 19.6551891 | 19.5285302 | 0.140410053 | 0.1213615 |
| 15.4810292 | 15.4089845 | 15.5832439 | 15.6558353 | 0.053835546 | 0.57273177 |
| 18.5176454 | 18.6535798 | 18.7078683 | 18.6643335 | 0.078641812 | 0.457034882 |
| 12.6812705 | NA | 13.8855617 | 13.6956831 | 1.051679442 | 0.002743017 |
| 25.8863243 | 25.7791146 | 25.9258821 | 25.8266443 | -0.022469905 | 0.675374147 |
| 19.2297671 | 19.1079715 | 19.3739432 | 19.2864356 | -0.11952519 | 0.655678794 |
| 18.0886067 | 17.9403679 | 17.8874163 | 17.9630881 | -0.141169732 | 0.190857008 |
| 20.9209947 | 20.8127935 | 20.8883674 | 20.9582117 | -0.014352832 | 0.803589785 |
| 19.5587149 | 19.5242867 | 19.5948729 | 19.6325999 | 0.054842916 | 0.348175508 |
| 20.5155397 | 20.5208902 | 20.5554418 | 20.6053918 | 0.069813329 | 0.14548102 |
| 16.5174276 | 16.5178853 | 16.4136341 | 16.4957387 | -0.049370786 | 0.434294782 |
| 22.6475125 | 22.5493413 | 22.6324282 | 22.5723856 | -0.052454048 | 0.291985981 |
| 22.7243737 | 22.5978633 | 22.6077023 | 22.6269535 | -0.06757897 | 0.152436504 |
| 23.7887422 | 23.7775318 | 23.7820064 | 23.7608755 | 0.019434209 | 0.661550754 |
| 19.9531094 | 19.8252574 | 19.7993879 | 19.8524808 | -0.081453707 | 0.065615624 |
| 22.9280609 | 22.8591796 | 22.8668377 | 22.8828373 | -0.038039028 | 0.342427766 |
| 23.9139231 | 23.8850428 | 24.011772 | 24.0314402 | 0.112106581 | 0.063902175 |
| 19.5091758 | 19.4422499 | 19.5149445 | 19.4839736 | -0.007404004 | 0.905276601 |
| 19.6570399 | 19.5948349 | 19.6477543 | 19.7974998 | 0.040178503 | 0.552060976 |
| 20.2899666 | 20.2908728 | 20.3163939 | 20.3793711 | 0.061912581 | 0.376118893 |
| 18.3061327 | 18.2913763 | 18.5547672 | 18.3887268 | 0.014638649 | 0.883563193 |
| 17.8401291 | 17.566302 | 17.924572 | 17.6675011 | 0.016996787 | 0.881423309 |
| 19.2877837 | 19.0859857 | 19.544911 | 19.5568317 | 0.19493617 | 0.422930532 |
| 24.8334668 | 24.9773323 | 25.067487 | 24.7145113 | 0.022694411 | 0.901214704 |
| 16.8311783 | 17.2239602 | 17.2061511 | 17.1804835 | 0.253067393 | 0.205406661 |
| 11.5199316 | NA | 11.2459031 | 11.4191567 | -0.289550063 | 0.299592606 |
| 22.2305454 | 22.0357582 | 22.1067742 | 22.1100582 | -0.073337896 | 0.308233889 |

| | | | | | |
|---|---|---|---|---|---|
| 19.0561398 | 18.9811108 | 19.0988227 | 19.2153549 | 0.063520104 | 0.645196598 |
| 20.093245 | 20.0480179 | 20.1346978 | 20.1330332 | 0.054358126 | 0.330273597 |
| NA | 15.3982643 | NA | NA | -0.439160046 | 0.041956217 |
| 17.5574872 | 17.0851383 | 17.337699 | 17.4261572 | -0.249140854 | 0.0350986 |
| 23.6526301 | 23.4350345 | 23.6516289 | 23.5284028 | -0.111848586 | 0.26820102 |
| 17.8498193 | 17.9001754 | 17.8040414 | 17.9206717 | 0.055414635 | 0.405938993 |
| 22.5191411 | 22.3358495 | 22.3635319 | 22.3433509 | -0.115538266 | 0.065559195 |
| 21.2398754 | 21.0147566 | 21.1399544 | 21.1243182 | -0.148086875 | 0.011205914 |
| 21.0127436 | 20.9005699 | 20.936309 | 20.9967972 | -0.064714945 | 0.206541586 |
| 23.1755173 | 23.1507043 | 23.1679414 | 23.1470758 | 0.02782407 | 0.588314046 |
| 21.8300795 | 21.7195307 | 21.7963621 | 21.8881205 | 0.063395144 | 0.318306422 |
| 19.578082 | 19.5106725 | 19.7300335 | 19.5810389 | 0.095239396 | 0.224015608 |
| NA | 15.0989529 | NA | NA | -0.184253745 | 0.236536341 |
| 18.9095163 | 18.6228604 | 18.668631 | 18.6979287 | -0.114627564 | 0.295708151 |
| 23.0863608 | 23.0281442 | 23.0792713 | 23.0977833 | 3.40E-04 | 0.995387795 |
| 21.4324748 | 21.3359822 | 21.3641093 | 21.3596593 | 0.012968001 | 0.796116732 |
| 17.6091332 | 17.8970029 | 17.8859303 | 17.8340942 | 0.194012246 | 0.040241921 |
| 21.656438 | 21.6284522 | 21.6507404 | 21.6083498 | -0.036805032 | 0.481570143 |
| 19.9349641 | 19.7905675 | 20.0437995 | 20.005927 | 0.005195062 | 0.955809775 |
| 13.0729199 | NA | 13.2607708 | 13.2118988 | 0.313275317 | 0.150296969 |
| 18.1621906 | 17.7952775 | 17.8732816 | 17.782051 | -0.165590224 | 0.156732834 |
| 15.4375533 | NA | 15.5361735 | 15.3664785 | 0.031008026 | 0.788515012 |
| 17.3414798 | 16.9537579 | 17.2694978 | 17.2989203 | -0.149605702 | 0.256671447 |
| 19.4086938 | 19.4198457 | 19.3770549 | 19.398988 | -0.088860156 | 0.297955075 |
| 19.5274679 | 19.4699499 | 19.5929165 | 19.6664471 | 0.072569984 | 0.246190503 |
| 18.1089874 | 18.1760662 | 18.155417 | 18.2145532 | 0.002206816 | 0.977183642 |
| 21.6509846 | 21.7895751 | 21.712665 | 21.6188103 | 0.078596805 | 0.238403433 |
| 20.0558937 | 19.8435211 | 19.8375197 | 19.8538675 | -0.149808851 | 0.016985102 |
| 24.3714004 | 24.2532647 | 24.3044005 | 24.3607843 | -0.009915335 | 0.853913621 |
| 21.6666233 | 21.5427738 | 21.6046764 | 21.5882661 | -0.056980408 | 0.100612718 |
| 19.3133709 | 19.2491947 | 19.3045464 | 19.3622863 | -0.018569094 | 0.753402124 |
| 22.9800218 | 22.9391666 | 22.988588 | 22.9647955 | 0.104733169 | 0.108760126 |
| 23.9511795 | 23.8765117 | 23.9151618 | 23.9286182 | 0.046183942 | 0.508844215 |
| 18.8464292 | 18.7684409 | 18.8591965 | 18.8762506 | 0.027176075 | 0.714480141 |
| 19.7592713 | 19.8563635 | 19.8265935 | 19.8710362 | 0.065115203 | 0.107982192 |
| 14.6071046 | 15.1849945 | 15.0543284 | 14.989428 | 1.107694822 | 0.009363475 |
| 18.6224628 | 18.0642604 | 18.5213378 | 18.5218866 | -0.165983014 | 0.342667388 |
| 13.6363615 | NA | 13.2015764 | 13.4770972 | -0.147077171 | 0.364029258 |
| NA | 15.743645 | NA | NA | -0.204550838 | 0.099782126 |
| 18.9243474 | 18.9737144 | 18.8523986 | 18.8519194 | -0.023946391 | 0.709335101 |
| 21.9166445 | 21.826808 | 21.9706784 | 21.9301482 | -0.032876362 | 0.53277402 |
| 22.60758 | 22.5319709 | 22.5584805 | 22.5156992 | -0.08769519 | 0.042955046 |

| | | | | | |
|---|---|---|---|---|---|
| 23.3992639 | 23.3269039 | 23.3485341 | 23.4550754 | 0.029307403 | 0.529011221 |
| 17.5194577 | 17.5315664 | 17.5288725 | 17.5624682 | 0.002803264 | 0.968334362 |
| 17.3109164 | 17.4615075 | 17.6288172 | 17.6794527 | 0.185773238 | 0.054022372 |
| 15.2727865 | NA | 15.3282758 | 15.4790376 | 0.136069659 | 0.211830609 |
| 15.6136283 | 15.4474211 | 15.5149824 | 15.6808135 | 0.073962432 | 0.566301206 |
| 19.9819091 | 19.9735238 | 19.9922631 | 20.0734971 | 0.032928009 | 0.501685063 |
| 18.1448085 | 18.2452782 | 18.5812011 | 18.4240453 | 0.140031333 | 0.151830606 |
| NA | 15.1498522 | NA | NA | -0.04897617 | 0.761125483 |
| 19.6069924 | 19.4289707 | 19.3369533 | 19.4932829 | -0.203943824 | 0.125047697 |
| 22.9637895 | 22.8713349 | 22.9182583 | 22.92614 | -0.003064976 | 0.930535636 |
| 20.8754372 | 20.8016523 | 20.9348994 | 20.9260813 | 0.048669404 | 0.336486541 |
| 23.6833624 | 23.4353068 | 23.5220755 | 23.6741155 | -0.078492062 | 0.251813227 |
| 20.1947294 | 20.1921982 | 20.2295333 | 20.2580474 | -0.004241104 | 0.932167347 |
| NA | 16.5932946 | NA | NA | 0.166780833 | 0.52025157 |
| 19.2459045 | 19.080681 | 19.0364702 | 19.0001646 | -0.229013031 | 0.029605441 |
| 19.5420113 | 18.8763868 | 19.2743724 | 19.4559521 | -0.147994554 | 0.265560114 |
| 25.8597241 | 25.7994084 | 25.8876667 | 25.8289805 | -0.065710264 | 0.192470637 |
| 15.5194196 | 16.0000555 | 15.5232434 | 15.4824706 | 0.322584689 | 0.152639352 |
| 18.2226828 | 18.2632559 | 18.430875 | 18.3497998 | 0.125601672 | 0.084253503 |
| 18.4967777 | 18.4358087 | 18.3329621 | 18.3449116 | -0.111975693 | 0.114818728 |
| 20.7973795 | 20.6835697 | 20.69492 | 20.7213529 | -0.089359167 | 0.090897853 |
| 18.796841 | 18.8732014 | 18.8427931 | 18.9189078 | 0.09443401 | 0.163478622 |
| 22.4507945 | 22.3489644 | 22.3572283 | 22.3766261 | -0.06079212 | 0.275113515 |
| 19.7022693 | 19.5693643 | 19.657679 | 19.7597227 | 0.026647461 | 0.682225728 |
| 22.0884811 | 21.8199615 | 21.7407446 | 21.9296603 | -0.188282132 | 0.070763232 |
| 20.8471142 | 20.7481593 | 20.8406448 | 20.905645 | -0.077482285 | 0.217823801 |
| 19.3144225 | 19.1688027 | 19.2063236 | 19.2626697 | -0.05525456 | 0.246871378 |
| 20.5905327 | 20.6299784 | 20.6132769 | 20.7480653 | 0.086572819 | 0.064764568 |
| 24.3505456 | 24.2885071 | 24.3090855 | 24.3774631 | 0.032301249 | 0.54377825 |
| 22.9555314 | 22.8591767 | 22.9514674 | 22.9106818 | -0.002940812 | 0.944674483 |
| 21.1748495 | 21.0824234 | 21.2068182 | 21.2174886 | 0.049951523 | 0.360488864 |
| 25.5040918 | 25.5533244 | 25.4509657 | 25.514544 | 0.015797327 | 0.849893969 |
| 18.0899492 | 17.7232031 | 17.7995383 | 17.7829442 | -0.249164158 | 0.002917232 |
| 20.9897842 | 20.8483122 | 20.9386351 | 20.9706708 | 0.009153451 | 0.876913334 |
| NA | 13.4591243 | NA | NA | 0.05030454 | 0.831102466 |
| 17.8950521 | 17.615715 | 17.762883 | 17.6484909 | -0.054334797 | 0.705227341 |
| 20.3612859 | 19.9996541 | 20.2282552 | 20.1773908 | -0.154784489 | 0.067266075 |
| 20.0448951 | 19.8607745 | 19.9887322 | 19.8956104 | -0.083392559 | 0.178032703 |
| 22.3174999 | 22.1954463 | 22.325556 | 22.2152292 | -0.05647746 | 0.266755742 |
| 20.5362172 | 20.4762253 | 20.5309097 | 20.7027901 | 0.092676393 | 0.165056074 |
| 17.7165962 | 18.3306382 | 17.9058759 | 17.5093749 | 0.373912923 | 0.111873562 |
| 20.243846 | 20.221931 | 20.4744211 | 20.2454272 | -0.016523924 | 0.832481547 |

| | | | | | |
|---|---|---|---|---|---|
| 21.3685772 | 21.0743611 | 21.2153282 | 21.212324 | -0.127947058 | 0.071868921 |
| 21.0338441 | 20.8962404 | 21.0775853 | 21.0008546 | -0.014777947 | 0.829187902 |
| 18.8675792 | 18.7609933 | 18.8309621 | 18.8266996 | 0.03598347 | 0.524344359 |
| 20.038274 | 19.9171005 | 20.1392533 | 20.1329321 | 0.036867636 | 0.576622997 |
| NA | 19.6485604 | NA | NA | 0.363658098 | 0.420596928 |
| 17.571308 | 17.2371501 | 17.6274197 | 17.6536424 | -0.067261308 | 0.476905882 |
| 20.4457555 | 20.313519 | 20.5011854 | 20.4763936 | -0.006134059 | 0.944959343 |
| 19.5023595 | 19.4725481 | 19.4649465 | 19.5478216 | 0.107239517 | 0.198342411 |
| 21.1346504 | 21.1462714 | 21.1501488 | 21.2401364 | -0.00317342 | 0.953416039 |
| 21.418922 | 21.3245161 | 21.3950799 | 21.4365215 | 0.007260218 | 0.893098206 |
| 18.0836447 | 18.1447474 | 17.99619 | 18.0102807 | 0.030451805 | 0.713605264 |
| 19.3945347 | 19.2917499 | 19.0865374 | 19.3985337 | -0.013874964 | 0.887272136 |
| 20.6007329 | 20.6113179 | 20.6857937 | 20.6586075 | 0.049021713 | 0.366086203 |
| 20.1637062 | 19.9593044 | 20.0756474 | 20.1305512 | -0.065147924 | 0.493050091 |
| 20.5542973 | 20.6301721 | 20.630914 | 20.5835965 | 0.026676641 | 0.704627479 |
| 23.5907602 | 23.533586 | 23.6090862 | 23.5090948 | -0.026806633 | 0.516287082 |
| 19.7430841 | 19.5655289 | 19.3874284 | 19.530053 | -0.16503308 | 0.115102622 |
| 20.3736518 | 20.2191722 | 20.3214906 | 20.308615 | -0.066213536 | 0.144231733 |
| 19.8276214 | 19.7487295 | 19.8074326 | 19.7305222 | -0.066860342 | 0.115760555 |
| 19.6371881 | 19.5887588 | 19.6730134 | 19.6516665 | 0.01425798 | 0.791408427 |
| 17.8602683 | 17.6894655 | 17.7549423 | 17.7632548 | -0.035160403 | 0.702777628 |
| 20.4392195 | 20.3075629 | 20.3235544 | 20.3684903 | -0.050140608 | 0.394944921 |
| 20.1996196 | 20.1229408 | 20.2383063 | 20.2864879 | 0.067663371 | 0.268013377 |
| 18.5935042 | 18.0473211 | 18.2572245 | 18.3338616 | -0.211941788 | 0.064487184 |
| 20.9150173 | 20.825187 | 20.8057359 | 20.9365025 | -0.019617791 | 0.698178638 |
| 19.1333995 | 19.0664255 | 19.0573078 | 19.1438049 | 0.007516889 | 0.890852915 |
| 20.6732707 | 20.5120227 | 20.6570578 | 20.6438075 | 0.033145269 | 0.596198031 |
| 17.1705419 | 16.7816764 | 16.5844208 | 16.6070541 | -0.299781337 | 0.025054367 |
| 19.4062741 | 19.5513145 | 19.6566064 | 19.482312 | 0.108551567 | 0.192929062 |
| 20.4454032 | 20.3257887 | 20.4613504 | 20.3882533 | 0.01711309 | 0.837790302 |
| 22.2371084 | 22.1957147 | 22.2291932 | 22.1626258 | -7.66E-04 | 0.991830075 |
| 19.8213889 | 19.7744847 | 19.8775193 | 19.8718976 | 0.043398279 | 0.440125244 |
| 19.7706498 | 19.9183576 | 19.9471515 | 19.8503996 | 0.022351999 | 0.816472619 |
| 15.0148523 | 14.9630433 | 15.1044489 | 14.8851264 | -0.113326401 | 0.407524323 |
| NA | 14.2942493 | NA | NA | -0.392986191 | 0.007945826 |
| 20.1359685 | 20.073005 | 20.0612651 | 20.0337558 | -0.010291558 | 0.865431852 |
| 17.2543849 | NA | 17.1233759 | 16.7385107 | -0.233579345 | 0.109481756 |
| 17.8368926 | 17.880858 | 18.0796527 | 17.8326353 | 0.15605105 | 0.265401493 |
| 17.324808 | 17.0740734 | 17.3606947 | 17.2893555 | -0.401191752 | 0.214357479 |
| 23.9723113 | 23.802364 | 23.9282811 | 23.9503322 | -0.035805981 | 0.591463724 |
| 22.1701601 | 22.2158973 | 22.1383547 | 22.2264838 | 0.006952808 | 0.914029733 |
| 24.5492732 | 24.6211733 | 24.5617094 | 24.5458504 | 0.04037144 | 0.522903727 |

| | | | | | |
|---|---|---|---|---|---|
| 24.3294346 | 24.221139 | 24.3225391 | 24.3545732 | 0.003998078 | 0.936493479 |
| NA | 15.4725995 | NA | NA | 0.306212235 | 0.105242372 |
| 22.9056348 | 22.9247581 | 23.0473214 | 22.9810578 | 0.119043472 | 0.027140333 |
| NA | 13.4146896 | NA | NA | -0.134538799 | 0.609998257 |
| 16.6937603 | 16.3946313 | 16.0917255 | 16.5079077 | -0.241044974 | 0.139970243 |
| 14.4725147 | 14.1837302 | 14.4136528 | 14.4953967 | -0.282080062 | 0.078909417 |
| 16.8044257 | 16.2881356 | 16.8168503 | 16.8314211 | -0.139620604 | 0.356800957 |
| 16.0345379 | 15.814538 | 16.1326063 | 15.7828893 | -0.097286 | 0.400076582 |
| 18.6654398 | 18.4869656 | 18.5182938 | 18.7218052 | -0.021420029 | 0.83352714 |
| 19.7061246 | 19.6534469 | 19.5270663 | 19.6999014 | -0.101314011 | 0.273892067 |
| 26.7597625 | 26.4637883 | 26.7044477 | 26.6886216 | -0.014514952 | 0.83912286 |
| 22.003846 | 22.1066874 | 22.0004067 | 21.9292542 | -0.037700408 | 0.578506545 |
| 17.4817649 | 17.4113761 | 17.4563638 | 17.231305 | -0.082852762 | 0.190577626 |
| 21.4739437 | 21.3803576 | 21.4766623 | 21.3818801 | -0.02434418 | 0.631057569 |
| 22.5151701 | 22.4036283 | 22.4206823 | 22.4632572 | -0.072677241 | 0.142318366 |
| 21.7536461 | 21.681035 | 21.7239395 | 21.7711756 | -0.009301526 | 0.807680836 |
| 20.5143093 | 20.4373662 | 20.6111962 | 20.5353279 | 0.061987888 | 0.233511074 |
| 18.7563638 | 18.8084877 | 18.8573689 | 18.6782921 | 0.059420027 | 0.259563012 |
| 19.562147 | 19.5057767 | 19.4536613 | 19.49135 | -0.054258055 | 0.376386017 |
| 22.4722214 | 22.3295436 | 22.4833478 | 22.4548481 | 0.020904064 | 0.74497991 |
| 25.2833485 | 25.4110475 | 25.2915409 | 25.4147662 | 0.071171331 | 0.34262941 |
| 17.1739432 | 17.1763122 | 17.3456445 | 17.335834 | 0.028598817 | 0.738346447 |
| 15.5652626 | 15.3315357 | 15.5263237 | 15.5896823 | 0.077201695 | 0.391313062 |
| 23.8573527 | 23.8033413 | 23.9842554 | 23.8946074 | -2.71E-04 | 0.996345424 |
| 20.9565088 | 20.9147435 | 21.0444712 | 20.9885269 | 0.065187404 | 0.170687352 |
| 19.5197123 | 19.4937975 | 19.5503305 | 19.5036481 | 0.093734327 | 0.200098365 |
| 18.3023906 | 18.1663445 | 18.3213134 | 18.3745521 | -0.017999858 | 0.803474934 |
| 16.7880703 | 17.0161589 | 17.2463995 | 17.2101037 | 0.221951398 | 0.053222265 |
| 17.1516771 | 17.443886 | 17.4534791 | 17.3845924 | 0.228417115 | 0.109265354 |
| 17.5931376 | 17.8918641 | 17.7672083 | 17.7440073 | 0.131614115 | 0.118527996 |
| 19.9069078 | 19.8997648 | 19.8588386 | 19.9970451 | 0.056432822 | 0.411525337 |
| 21.5793595 | 21.542492 | 21.5678003 | 21.5462541 | -0.039968254 | 0.438664076 |
| 21.395761 | 21.3810622 | 21.4622713 | 21.4634687 | 0.077212412 | 0.203025103 |
| 22.9431048 | 22.8232824 | 22.9110492 | 23.0046591 | -0.087105632 | 0.283931416 |
| 22.167826 | 22.1955664 | 22.1435357 | 22.201669 | 0.074500392 | 0.185911911 |
| 19.1404826 | 19.1075457 | 19.2222014 | 19.2406999 | 0.111704682 | 0.130304784 |
| 22.0704698 | 21.7136613 | 21.6402588 | 21.7335913 | -0.288870717 | 0.070667099 |
| 19.3394739 | 19.5057729 | 19.0015646 | 18.9615542 | -0.211331023 | 0.131820326 |
| 16.7930186 | 16.7449461 | 16.9607244 | 17.0941648 | 0.122543758 | 0.34593956 |
| 18.2695701 | 17.9973515 | 18.0541857 | 18.460424 | -0.093579274 | 0.44640341 |
| 16.5652065 | 16.443656 | 16.6129909 | 16.6178655 | -0.131570192 | 0.273165095 |
| 16.6382183 | 16.6985611 | 16.4932419 | 16.5005244 | -0.174344484 | 0.127396653 |

| | | | | | |
|---|---|---|---|---|---|
| 14.1659852 | 13.9609018 | 13.8576302 | 14.0290907 | -0.223544458 | 0.044159572 |
| 15.8168035 | 15.7492599 | 15.7011502 | 15.7614939 | -0.108521935 | 0.283776122 |
| 18.8971909 | 18.7201566 | 18.6349655 | 18.8666965 | 0.007573469 | 0.938907671 |
| 19.3133399 | 19.2428579 | 19.1892151 | 19.3085666 | -0.069959327 | 0.468085945 |
| 19.3064079 | 19.1819946 | 19.2734324 | 19.2208877 | -0.120573466 | 0.154965224 |
| 26.3625564 | 26.2571425 | 26.3127755 | 26.3567524 | 1.18E-04 | 0.998699549 |
| 26.0181333 | 25.9042114 | 25.962838 | 25.9884493 | -0.033745745 | 0.628280381 |
| 22.4325977 | 22.5803637 | 22.4645201 | 22.4535335 | -0.033844047 | 0.62394114 |
| 21.7157068 | 21.745756 | 21.6607296 | 21.870296 | -0.009734467 | 0.881401386 |
| 18.5592437 | 18.635674 | 18.4481876 | 18.5060944 | -0.004990945 | 0.938674432 |
| 21.1761298 | 21.001424 | 20.9555021 | 21.083357 | -0.057787553 | 0.353920428 |
| 22.0593558 | 21.8417995 | 22.0190031 | 21.9755458 | -0.07019342 | 0.259768545 |
| 22.436759 | 22.2782783 | 22.445249 | 22.4063118 | 0.010822143 | 0.853344151 |
| 20.9372585 | 20.8142973 | 20.8621139 | 20.8113566 | -0.081575959 | 0.149229701 |
| 22.3810157 | 22.3299709 | 22.2909138 | 22.2504164 | -0.045538422 | 0.387941866 |
| 25.767555 | 25.7434235 | 25.7196361 | 25.7174537 | -0.021090222 | 0.680377388 |
| 21.5543984 | 21.4459565 | 21.533842 | 21.5121734 | -0.077086727 | 0.1314683 |
| 22.9734469 | 22.9165633 | 22.9648758 | 22.9083083 | -0.02864206 | 0.569229079 |
| 19.5838127 | 19.6354029 | 19.5533929 | 19.5284528 | -0.026895795 | 0.578383283 |
| 22.883596 | 22.8130776 | 22.8346387 | 22.8017426 | -0.022769871 | 0.618141987 |
| 22.2800645 | 22.1868841 | 22.2561456 | 22.2759509 | -0.02808413 | 0.519186793 |
| 22.4750627 | 22.3604217 | 22.4476307 | 22.4266462 | -0.025894132 | 0.531525247 |
| 19.1692156 | 19.1030863 | 19.2175816 | 19.1809299 | -0.021858481 | 0.561676724 |
| 22.8293747 | 22.8366282 | 22.8796485 | 22.8918299 | 0.040499688 | 0.335506685 |
| 20.8484698 | 20.825264 | 20.8990304 | 20.9554282 | 0.031261358 | 0.475814437 |
| 19.3899551 | 19.4557462 | 19.3051802 | 19.3488559 | 0.032460723 | 0.463919749 |
| 22.4970209 | 22.4679336 | 22.5565425 | 22.5237861 | 0.032059165 | 0.48645886 |
| 19.3371884 | 19.3530082 | 19.4323108 | 19.3577148 | 0.027873212 | 0.548005392 |
| 19.0888081 | 19.0873311 | 19.1875014 | 19.1798288 | 0.070190351 | 0.16215504 |
| 19.9335058 | 19.9770337 | 19.9946359 | 20.0251799 | 0.012036794 | 0.808152196 |
| 20.3334562 | 20.2743319 | 20.3783803 | 20.338968 | 0.056573241 | 0.276150058 |
| 20.9962563 | 20.9543134 | 21.067631 | 21.0578234 | 0.076354859 | 0.150661996 |
| 19.1604782 | 19.1258662 | 19.2402085 | 19.1928409 | 0.056201052 | 0.305485184 |
| 19.0959499 | 18.9836552 | 19.1360514 | 19.052641 | 0.059128916 | 0.292500933 |
| 18.9824027 | 19.0178698 | 19.058791 | 19.000202 | 0.027687708 | 0.628568585 |
| 21.9587695 | 21.8612378 | 22.0309256 | 22.0040131 | 0.058354668 | 0.31827285 |
| 23.2029436 | 23.2156142 | 23.2083248 | 23.1373813 | 0.029707174 | 0.608263034 |
| 18.1140716 | 18.0624332 | 18.032204 | 18.3024256 | 0.020720342 | 0.727387789 |
| 21.8505393 | 21.7854804 | 21.7594099 | 21.8546054 | -0.024949215 | 0.68022869 |
| 17.5472502 | 17.5118273 | 17.6545019 | 17.591372 | 0.067329839 | 0.273146137 |
| 19.809868 | 19.9220596 | 19.9017164 | 19.9951718 | 0.114690098 | 0.069057202 |
| 19.6960436 | 19.6579081 | 19.7509578 | 19.823289 | 0.028983004 | 0.657538421 |

| | | | | | |
|---|---|---|---|---|---|
| 20.248475 | 20.2186621 | 20.2381912 | 20.2566094 | 0.02179745 | 0.73996049 |
| 21.2016126 | 21.1530955 | 21.2543901 | 21.2589568 | 0.031085344 | 0.641431767 |
| 18.5537337 | 18.3660726 | 18.4924367 | 18.5973091 | -0.05586527 | 0.419874602 |
| 22.3080935 | 22.2868154 | 22.4086516 | 22.3068091 | 0.091221165 | 0.187993656 |
| 18.2699351 | 18.1388714 | 18.1777888 | 18.3191777 | -0.069652717 | 0.321069462 |
| 22.4191274 | 22.5047361 | 22.6351991 | 22.5206193 | 0.118052138 | 0.101633347 |
| 19.4602242 | 19.5689653 | 19.5918051 | 19.5202313 | 0.083040742 | 0.243386246 |
| 21.7526407 | 21.6690519 | 21.8563633 | 21.7324184 | -0.018407464 | 0.801785026 |
| 18.0562302 | 18.241109 | 18.2811514 | 18.1126114 | 0.125022113 | 0.101455739 |
| 19.993666 | 19.9325406 | 19.9591279 | 20.0182473 | 0.039237373 | 0.600423932 |
| 19.3360534 | 19.2259144 | 19.3233178 | 19.3253155 | -0.115380193 | 0.150000333 |
| 20.8916215 | 20.775729 | 20.8681796 | 20.8608338 | 0.039823498 | 0.620644157 |
| 18.8506812 | 19.1078742 | 19.1755522 | 19.1503195 | 0.183825432 | 0.032046177 |
| 25.3550957 | 25.4021269 | 25.4458837 | 25.3550358 | 0.120169107 | 0.158553949 |
| 18.4945156 | 18.4498453 | 18.6441132 | 18.6193075 | 0.012442076 | 0.889107976 |
| 17.9882833 | 18.0341845 | 18.2239673 | 18.0354008 | 0.034927367 | 0.695900035 |
| 21.8041041 | 21.6818157 | 21.8505988 | 21.8882472 | 0.012215918 | 0.901474134 |
| 17.9120632 | 18.0077945 | 17.9560331 | 18.1316946 | 0.050522963 | 0.628008424 |
| 19.0068708 | 18.6861032 | 18.9212052 | 18.8881298 | 0.076040205 | 0.469588974 |
| NA | 14.7197054 | NA | NA | -0.047805671 | 0.632886889 |
| 14.7066651 | NA | 14.8608166 | 14.6989622 | 0.003485101 | 0.976897941 |
| 19.2338633 | 19.0160959 | 19.2932059 | 19.091257 | 0.023683899 | 0.851952619 |
| 19.4727772 | 19.4360325 | 19.3854142 | 19.1991176 | 0.010582541 | 0.943841142 |
| 17.8857585 | 18.0332427 | 18.2473818 | 18.0984588 | 0.342950156 | 0.031548834 |
| 19.8696914 | 19.7477538 | 19.9371293 | 19.7317192 | 0.009011233 | 0.956960621 |
| NA | 16.0060824 | NA | NA | 0.371602429 | 0.040473379 |
| NA | 11.5546013 | NA | NA | -0.122662952 | 0.466167097 |
| NA | 12.7060647 | NA | NA | 0.254018821 | 0.285789948 |
| 16.0469764 | 16.0375969 | 16.1608123 | 15.7344294 | 0.021408765 | 0.853310899 |
| 23.1199234 | 22.9646396 | 23.3193736 | 23.0797728 | 0.053821054 | 0.570733986 |
| 18.4285621 | 18.3690509 | 18.389506 | 18.25625 | -0.13688841 | 0.067863954 |
| 25.5231922 | 25.3179232 | 25.4045809 | 25.4098858 | -0.065583139 | 0.191120921 |
| 20.3531183 | 20.2955382 | 20.3391332 | 20.4437933 | 0.00900827 | 0.848359486 |
| 20.5694562 | 20.499537 | 20.5998056 | 20.5761116 | 0.043189279 | 0.415973489 |
| 22.322599 | 22.2672436 | 22.3378134 | 22.3391973 | 0.053175954 | 0.323187821 |
| 21.6208846 | 21.6164745 | 21.6816265 | 21.6702694 | 0.074006906 | 0.24423517 |
| 18.3082256 | 18.4380941 | 18.3888497 | 18.4561704 | 0.129764081 | 0.080834206 |
| 22.4581245 | 22.4415492 | 22.5438834 | 22.5782053 | 0.103857515 | 0.16902436 |
| 19.9116683 | 19.8459551 | 19.9075978 | 19.9174938 | 0.046081274 | 0.604086656 |
| 21.91067 | 21.6189979 | 21.7525185 | 21.7159576 | -0.118034112 | 0.088896064 |
| 20.8351191 | 20.8266393 | 20.8891181 | 20.8229016 | -0.018105568 | 0.728577603 |
| 21.6539093 | 21.563229 | 21.6018565 | 21.642254 | -0.059413795 | 0.123758243 |

| | | | | | |
|---|---|---|---|---|---|
| 19.1879788 | 19.2025768 | 19.2757732 | 19.3275454 | 0.036021113 | 0.508342648 |
| 21.2185147 | 21.1507173 | 21.3786671 | 21.2257782 | 0.10000523 | 0.08169051 |
| 21.2773852 | 21.0853623 | 21.2318183 | 21.2112721 | -0.033252692 | 0.710086787 |
| 18.5455783 | 18.6320085 | 19.0222708 | 18.7456748 | 0.237426735 | 0.120689896 |
| 17.9211071 | 18.0799902 | 17.0815044 | 18.0491685 | 0.036174976 | 0.881714573 |
| 22.7337713 | 22.6034852 | 22.6830557 | 22.6784524 | -0.037294126 | 0.433946932 |
| 20.6272576 | 20.701057 | 20.659078 | 20.5940794 | 0.040774521 | 0.415241702 |
| 20.371534 | 20.2824976 | 20.4172631 | 20.425693 | 0.038103695 | 0.629441743 |
| 17.4641456 | 17.5481143 | 17.5924701 | 17.4332921 | 0.093301231 | 0.469007193 |
| 22.368673 | NA | 23.0928079 | 22.4160673 | 0.341422711 | 0.110787206 |
| 18.5448433 | 18.6608496 | 18.5690553 | 18.5284831 | 0.06768441 | 0.524200067 |
| 12.6455168 | NA | 12.382133 | 12.2552758 | -0.26842618 | 0.532611859 |
| 16.7950125 | 17.1793561 | 15.8316432 | 17.0424198 | -0.120334953 | 0.639850609 |
| 20.7927706 | 19.9794068 | 20.4445273 | 20.4254001 | -0.369972638 | 0.047129097 |
| 17.0101036 | 16.3833958 | 17.1116373 | 17.1892718 | -0.033278702 | 0.817140108 |
| 16.0396765 | 15.9232842 | 16.2477785 | 16.0024631 | 0.056170447 | 0.65664151 |
| 18.2296953 | 18.2607872 | 17.8438081 | 18.0813286 | -0.180940383 | 0.116547282 |
| 19.4959317 | 19.436641 | 19.3483596 | 19.4014976 | -0.035591466 | 0.712803448 |
| 23.0615374 | 22.7057307 | 23.0355063 | 23.0524475 | -0.121149657 | 0.1460446 |
| 20.7165466 | 20.4491633 | 20.5008743 | 20.639239 | -0.124071912 | 0.130550334 |
| 23.3990921 | 23.3786095 | 23.3270973 | 23.3311446 | 0.01608376 | 0.77050945 |
| 22.0935405 | 21.9870396 | 22.0110061 | 22.001205 | -0.079225479 | 0.055687916 |
| 22.0738286 | 22.0866099 | 22.1180042 | 22.1293138 | 0.082042441 | 0.090035954 |
| 19.9369079 | 19.9139484 | 19.9508988 | 19.9982738 | 0.084009744 | 0.103232029 |
| 21.8773507 | 21.9092084 | 21.9104101 | 21.8805684 | 0.051969689 | 0.372107064 |
| 21.6441323 | 21.5482982 | 21.6044861 | 21.6049882 | 0.030784903 | 0.597418479 |
| 20.8305712 | 20.7063485 | 20.8249712 | 20.8279462 | 0.043932975 | 0.480364648 |
| 24.4957335 | 24.3536789 | 24.5338561 | 24.5275155 | 0.098119779 | 0.16447698 |
| 20.0483111 | 20.0852997 | 20.1663464 | 20.1337092 | 0.114971654 | 0.108269855 |
| 17.7041483 | 17.6762247 | 17.8457637 | 17.6377298 | -0.009410073 | 0.925663595 |
| 17.7265359 | 18.0965747 | 18.0556513 | 17.9387455 | 0.281398218 | 0.046491739 |
| 16.5262597 | 16.3700803 | 16.0610649 | 16.4707787 | 0.037090505 | 0.794152402 |
| 14.4536316 | NA | 14.7369081 | 14.841277 | 0.238528591 | 0.125229915 |
| 20.0048479 | 19.8256554 | 20.2378069 | 20.1439065 | 0.103621416 | 0.312387696 |
| 17.1997528 | 17.0784566 | 17.195369 | 17.4434689 | 0.026599851 | 0.782030258 |
| 16.1170427 | 15.9642981 | 15.897725 | 16.3525358 | 0.080831695 | 0.514823289 |
| NA | 14.6941133 | 14.1945924 | 13.689802 | -0.544928015 | 0.156717207 |
| 24.0951252 | 24.0676104 | 24.1951962 | 24.1262979 | -0.050628563 | 0.394201529 |
| 19.449621 | 19.4409104 | 19.4860389 | 19.5199025 | 0.073725377 | 0.144569855 |
| 21.5542598 | 21.4012023 | 21.5414306 | 21.569362 | 0.012848853 | 0.839371422 |
| 20.3745265 | 20.2671935 | 20.247904 | 20.282583 | -9.65E-04 | 0.988842329 |
| 20.4336959 | 20.3584373 | 20.4822439 | 20.461744 | 0.053730512 | 0.476372188 |

| | | | | | |
|---|---|---|---|---|---|
| 17.0957327 | 17.144113 | 17.2457828 | 17.2139767 | -0.013714848 | 0.918759634 |
| NA | 12.529761 | NA | NA | -0.260071026 | 0.086252549 |
| 17.8764866 | NA | 17.9213602 | 17.798146 | -0.022899157 | 0.862612711 |
| 21.7175165 | 21.5876282 | 21.6448814 | 21.5732287 | -0.034970289 | 0.677187736 |
| 20.7827657 | 20.5918027 | 20.6412939 | 20.6639146 | -0.120917805 | 0.056577239 |
| 20.6234059 | 20.5721969 | 20.6601026 | 20.7003079 | 0.028492185 | 0.592875394 |
| 19.5897499 | 19.419747 | 19.5775005 | 19.5624428 | -0.037950489 | 0.537997268 |
| 23.9754452 | 23.8059385 | 23.9122125 | 23.9334191 | -0.033920676 | 0.424238242 |
| 16.3771174 | NA | 16.0779428 | 15.8896996 | -0.461361545 | 0.053425356 |
| 19.750349 | 19.7223383 | 19.8173798 | 19.8237299 | 0.126862284 | 0.197067206 |
| 18.4218003 | 18.4779528 | 18.6919754 | 18.6911306 | 0.214801553 | 0.044686042 |
| 15.7141557 | 15.400327 | 15.6571729 | 15.6983872 | -0.131440528 | 0.155129785 |
| 21.5712374 | 21.5104053 | 21.5938641 | 21.5383494 | -0.035398803 | 0.473127834 |
| 20.5877737 | 20.5787224 | 20.6429299 | 20.6291667 | 0.042910705 | 0.365556491 |
| 18.6784293 | 18.7028343 | 18.7589633 | 18.7371979 | 0.036645849 | 0.458923137 |
| 21.0562137 | 21.0167023 | 21.0351618 | 20.8683143 | 0.016541936 | 0.846992804 |
| 17.7003454 | 17.485558 | 17.5452043 | 17.3777581 | -0.226753589 | 0.005101852 |
| 21.4561604 | 21.3335578 | 21.4070032 | 21.3109118 | -0.049856977 | 0.387424676 |
| 22.5532233 | 22.3735873 | 22.5280602 | 22.5000269 | -0.068027573 | 0.200124234 |
| 19.3420356 | 19.3608369 | 19.2331347 | 19.4199079 | -0.027374223 | 0.57456178 |
| 20.8785105 | 20.7563047 | 20.8280525 | 20.831219 | -0.104298 | 0.030224124 |
| 23.4573567 | 23.393521 | 23.4738531 | 23.3947883 | -0.012860307 | 0.755978131 |
| 20.3623386 | 20.3955547 | 20.3816618 | 20.3878649 | 0.030026541 | 0.515699941 |
| 23.9507164 | 23.8957277 | 23.9361039 | 23.9768318 | 0.032343703 | 0.513987426 |
| 18.5935071 | 18.6977231 | 18.7352671 | 18.6512705 | 0.122858537 | 0.110919593 |
| 16.2911369 | 16.370626 | 16.328212 | 16.3225856 | 0.078092606 | 0.462182122 |
| 20.5058004 | 20.5381851 | 20.5793463 | 20.4832135 | -0.053959091 | 0.201526078 |
| 18.936834 | 18.9621668 | 19.1326884 | 18.9930138 | 0.044929467 | 0.448549036 |
| 15.6461721 | 15.6300604 | 15.7386326 | 15.6803668 | 0.037827703 | 0.690590136 |
| 21.6475098 | 21.4459371 | 21.5054617 | 21.4182014 | -0.202627449 | 0.081620775 |
| 18.6310904 | 18.5409748 | 18.5474379 | 18.5485842 | -0.155746766 | 0.043621227 |
| 21.1164347 | 21.1031523 | 21.2191134 | 21.2720053 | 0.077264361 | 0.274002639 |
| 22.9285884 | 22.6345988 | 22.8095831 | 22.8066168 | -0.119286517 | 0.032099956 |
| 19.6232542 | 19.577991 | 19.599161 | 19.6865235 | -0.053642554 | 0.440185187 |
| 24.0842765 | 24.0023834 | 23.963021 | 24.0582429 | 0.010893555 | 0.858291859 |
| NA | 16.3705813 | NA | NA | -1.004009455 | 0.057248455 |
| 14.9492909 | 15.4514638 | 14.9654647 | 15.1940101 | -0.253803231 | 0.174494755 |
| 18.472986 | 18.8528318 | 18.8793203 | 18.8635103 | 0.171341905 | 0.279843463 |
| 17.8305459 | 17.2824033 | 17.1790534 | 17.5457713 | -0.40684207 | 0.005533626 |
| 18.0718085 | 17.6518458 | 17.9454396 | 18.0588786 | -0.140956785 | 0.184987606 |
| 26.5708171 | 26.288125 | 26.4675081 | 26.372901 | -0.121967528 | 0.197914256 |
| 19.7979897 | 19.5629144 | 19.5298995 | 19.6671584 | -0.108156201 | 0.223752491 |

| | | | | | |
|---|---|---|---|---|---|
| 19.6941898 | 19.7005526 | 19.7242639 | 19.5263035 | -0.029899455 | 0.729639743 |
| 24.0975033 | 24.142248 | 24.0717034 | 24.0210917 | -0.046501028 | 0.487775852 |
| 18.5782725 | 18.4125063 | 18.5361792 | 18.5634394 | 0.009855883 | 0.873272248 |
| 21.5807733 | 21.4564639 | 21.6160237 | 21.5802998 | -0.027210653 | 0.649252108 |
| 23.2650574 | 23.167014 | 23.2748462 | 23.2738534 | 0.020148631 | 0.718781776 |
| 21.3616228 | 21.2848495 | 21.3547486 | 21.3293809 | -0.052897795 | 0.285398049 |
| 20.3671234 | 20.2169052 | 20.3439452 | 20.2547181 | -0.094772925 | 0.060243245 |
| 22.9808028 | 22.8655242 | 22.870384 | 22.9063669 | -0.03783681 | 0.4250637 |
| 22.7274983 | 22.7969163 | 22.753398 | 22.7821787 | 0.032009919 | 0.430718028 |
| 22.0477343 | 22.0990624 | 22.024944 | 21.9518793 | -0.024446022 | 0.528316035 |
| 24.5862195 | 24.5643478 | 24.5535345 | 24.6033522 | -0.023079979 | 0.546172501 |
| 22.9864222 | 22.8814632 | 22.8480674 | 22.9356154 | -0.050412594 | 0.19348819 |
| 21.6014315 | 21.5415536 | 21.5130602 | 21.4909792 | -0.082263465 | 0.078441978 |
| 18.6433925 | 18.6304034 | 18.6927155 | 18.6105125 | -0.009636976 | 0.84943424 |
| 22.2436045 | 22.0978607 | 22.2923924 | 22.2519598 | 0.005007649 | 0.929276391 |
| 20.6233007 | 20.6322085 | 20.7170746 | 20.8087688 | 0.116108604 | 0.056622802 |
| 19.4772421 | 19.4009702 | 19.2672529 | 19.3966143 | -0.016496627 | 0.784058945 |
| 19.7181815 | 19.7091556 | 19.8347389 | 19.7952848 | 0.102542077 | 0.104387658 |
| 19.9356779 | 19.8522753 | 19.923732 | 19.9969765 | 0.058991125 | 0.36365565 |
| 18.3168767 | 18.375004 | 18.4590105 | 18.6062444 | 0.197925833 | 0.005549757 |
| 20.0067096 | 19.9170223 | 20.056548 | 20.0954095 | -0.009280846 | 0.889877965 |
| 20.5210175 | 20.2663505 | 20.3435784 | 20.5113607 | -0.051386066 | 0.446130519 |
| 21.7362608 | 21.8585116 | 21.8942151 | 21.8157672 | 0.088723822 | 0.213834164 |
| 20.016597 | 20.0492873 | 20.0811394 | 20.0840714 | 0.055588602 | 0.431980138 |
| 19.1391135 | 19.0796059 | 19.2641232 | 19.2746567 | 0.080899423 | 0.257077205 |
| 19.3393062 | 19.343056 | 19.2286412 | 19.3494796 | -0.010065324 | 0.889004721 |
| 25.1445574 | 25.1055115 | 25.155332 | 25.1493231 | 0.042964363 | 0.563040349 |
| 21.9866272 | 21.8065236 | 21.9777025 | 21.9528586 | -0.019180322 | 0.806122113 |
| 19.6172728 | 19.3636822 | 19.4588873 | 19.4862524 | 0.024512027 | 0.782211007 |
| 18.0660708 | 18.1563889 | 18.5481686 | 17.9984411 | 0.123428679 | 0.201383123 |
| 14.4025664 | 14.493666 | 14.6476461 | 14.5558286 | 0.125414443 | 0.240101904 |
| 17.3349441 | 16.9973272 | 17.2860777 | 17.1847979 | -0.06431582 | 0.591653505 |
| 16.9408311 | NA | 16.8969133 | 16.9118973 | 0.155396334 | 0.247107411 |
| 16.4626633 | 16.5351634 | 16.7021614 | 16.5443086 | 0.015093959 | 0.922745307 |
| 16.8693179 | 16.6268832 | 16.9962082 | 16.9486212 | 0.394359811 | 0.042418648 |
| 17.8728211 | 18.1794542 | 18.1170035 | 18.4649362 | 0.37930185 | 0.050740827 |
| 24.5430651 | 24.3737296 | 24.4996712 | 24.464329 | -0.043999367 | 0.531603982 |
| 18.110586 | 18.0691945 | 18.2095889 | 18.0299256 | -0.087398366 | 0.261393301 |
| 19.7741059 | 19.7668109 | 19.5930386 | 19.7293553 | -0.065838899 | 0.374572131 |
| 19.3853616 | 19.431464 | 19.5877815 | 19.4378009 | 0.025453718 | 0.752696293 |
| 11.9549374 | NA | 12.1867601 | 12.0052411 | 0.158645108 | 0.501952405 |
| 14.8156299 | NA | 15.2779155 | 14.8504255 | 0.489229513 | 0.053294454 |

| | | | | | |
|---|---|---|---|---|---|
| 20.9479409 | 21.0406629 | 20.9222117 | 20.7015704 | -0.185583779 | 0.101654232 |
| 23.8958227 | 23.7598092 | 23.7971997 | 23.8567097 | 0.03869587 | 0.639016883 |
| 18.5672516 | 18.5325643 | 18.6820124 | 18.5686637 | -0.055789392 | 0.480603425 |
| 21.7410322 | 21.5697481 | 21.6473207 | 21.6681536 | -0.065353702 | 0.204727304 |
| 19.9546775 | 19.9126385 | 19.8766745 | 19.881883 | -0.05080457 | 0.267848596 |
| 23.4999538 | 23.5031939 | 23.4949044 | 23.4977534 | 0.051070529 | 0.333704826 |
| 16.9696358 | 17.1450357 | 16.8884953 | 17.1080963 | 0.048192078 | 0.656161069 |
| 16.6721006 | 16.8132399 | 16.6586487 | 16.8367552 | 0.028059554 | 0.808030048 |
| 17.6383977 | 17.8655033 | 17.8086965 | 17.5761486 | 0.105405996 | 0.393063564 |
| 19.8803013 | 19.8539327 | 19.8518108 | 19.9183788 | -0.001244332 | 0.978253031 |
| 24.843945 | 25.0418219 | 24.8609449 | 24.8669591 | 0.062190066 | 0.450343378 |
| 19.558071 | 19.34636 | 19.5200176 | 19.4973193 | -0.054956452 | 0.333242588 |
| 25.8193005 | 25.6226794 | 25.6687277 | 25.7636812 | -0.027191917 | 0.775415127 |
| 17.9975486 | 17.8098642 | 17.8006621 | 17.9205611 | -0.103865429 | 0.196179593 |
| 21.067966 | 20.980141 | 21.0791614 | 20.9557145 | -0.062087568 | 0.407045511 |
| 24.5359989 | 24.5069215 | 24.4004213 | 24.5570596 | -0.008023296 | 0.887724696 |
| 22.995864 | 22.8486111 | 22.9318586 | 22.9041141 | -0.066558713 | 0.243021478 |
| 26.2014559 | 26.0478385 | 26.0731981 | 26.0976243 | -0.052327704 | 0.235617344 |
| 21.5364413 | 21.4422785 | 21.5764719 | 21.6307179 | 0.076411098 | 0.249096945 |
| 20.7336119 | 20.6444613 | 20.762238 | 20.763048 | 0.049358774 | 0.48537906 |
| 15.6571754 | NA | 15.8986988 | 16.0561593 | 0.316877076 | 0.065733994 |
| 21.9357052 | 21.9456011 | 21.9417855 | 21.9305007 | 0.019443258 | 0.729114431 |
| 21.3602887 | 21.4467731 | 21.4471767 | 21.3643148 | 0.079965897 | 0.060601642 |
| 21.7902331 | 21.8124685 | 21.8205262 | 21.7929473 | -0.004896403 | 0.917337346 |
| 25.1658868 | 25.1184661 | 25.2157272 | 25.1478677 | 0.012069377 | 0.785227965 |
| 19.0844572 | 19.003031 | 18.9757527 | 18.7665974 | 0.006922984 | 0.954632489 |
| 20.0058312 | 19.6894222 | 19.8847227 | 19.814456 | -0.190844006 | 0.033698078 |
| 20.7174652 | 20.56261 | 20.7735445 | 20.6073944 | -0.104238657 | 0.21173224 |
| 21.6853876 | 21.5749812 | 21.5554332 | 21.5714294 | -0.15287184 | 0.064769651 |
| 18.8222501 | 18.5228141 | 18.6482796 | 18.4569259 | -0.243372301 | 0.002908342 |
| 23.1024269 | 22.8974374 | 23.0904898 | 23.0153228 | -0.143790779 | 0.059773168 |
| 20.5446271 | 20.6660328 | 20.5894508 | 20.5216657 | -0.011050179 | 0.878877912 |
| 18.4756267 | 18.3443036 | 18.2647706 | 18.5135183 | -0.037976931 | 0.581650984 |
| 21.7256236 | 21.6165734 | 21.7971989 | 21.7384478 | -0.029817854 | 0.651064308 |
| 19.5977352 | 19.5069961 | 19.4720346 | 19.4944814 | -0.086553748 | 0.166287152 |
| 20.5493435 | 20.4603052 | 20.5278525 | 20.5202991 | -0.00376064 | 0.948882006 |
| 18.9321618 | 18.8362394 | 18.800259 | 18.8242493 | -0.123541369 | 0.042917739 |
| 22.7963917 | 22.6148054 | 22.7188668 | 22.7434548 | -0.067035124 | 0.242243321 |
| 20.9242327 | 20.9156028 | 20.8785907 | 20.8432425 | -0.073353576 | 0.180642558 |
| 21.043085 | 21.1006487 | 21.0296134 | 21.0256747 | 0.016496583 | 0.728656819 |
| 22.5035624 | 22.4202044 | 22.3943937 | 22.4598257 | -0.025163336 | 0.560665735 |
| 24.6796668 | 24.5799106 | 24.7186721 | 24.6507414 | -0.008307604 | 0.838637237 |

| | | | | | |
|---|---|---|---|---|---|
| 23.7528228 | 23.6804826 | 23.6676415 | 23.6674984 | -0.053643965 | 0.168747065 |
| 20.238497 | 20.0675415 | 20.24448 | 20.1719312 | -0.00523584 | 0.91978201 |
| 19.1065977 | 19.192495 | 19.1229666 | 19.2042086 | 0.097256073 | 0.086590484 |
| 22.2868428 | 22.2367887 | 22.2734401 | 22.3033599 | 0.021521092 | 0.698769624 |
| 19.5627528 | 19.650882 | 19.5983681 | 19.6285669 | 0.044309134 | 0.431093195 |
| 21.832631 | 21.7499901 | 21.7561738 | 21.8753579 | 0.013200394 | 0.817180513 |
| 20.5906036 | 20.536926 | 20.594084 | 20.5104358 | 0.038753821 | 0.536092322 |
| 18.234319 | NA | 18.2755706 | 18.1456934 | 0.011141712 | 0.868087071 |
| 25.9495758 | 25.7850619 | 25.8970772 | 25.9371069 | 0.011789976 | 0.866713821 |
| 19.4709552 | 19.2677532 | 19.4549174 | 19.4905755 | -0.013202717 | 0.852530032 |
| 19.1959451 | 19.0101563 | 19.2017619 | 19.2361487 | 0.041861598 | 0.589424547 |
| 20.1346491 | 20.1377783 | 20.2680513 | 20.3744901 | 0.170951119 | 0.035701458 |
| 18.5369865 | 18.67396 | 18.716472 | 18.7279708 | 0.131526574 | 0.104484116 |
| 21.738269 | 21.9947778 | 21.835474 | 21.832408 | 0.108830049 | 0.18924388 |
| 22.0371558 | 22.1973611 | 22.2459937 | 22.2085339 | 0.058202248 | 0.494940311 |
| NA | 16.3548449 | NA | NA | -0.051743634 | 0.651943454 |
| 20.1113902 | NA | 19.9328534 | 20.0849235 | -0.160531258 | 0.360999004 |
| 20.1334874 | 20.1494059 | 20.0838605 | 20.1188037 | 0.040151252 | 0.640771866 |
| 18.8401839 | 18.7791488 | 18.8435945 | 18.8350608 | -0.024792143 | 0.614806682 |
| 21.0020453 | 20.8934103 | 21.0057417 | 21.0041331 | 0.029210269 | 0.622612368 |
| 16.9227427 | 16.6176055 | 16.7211833 | 16.7751612 | -0.069784352 | 0.467949548 |
| NA | 14.7080992 | NA | NA | -0.321461884 | 0.063238647 |
| 18.2899517 | 17.9157567 | 17.8543789 | 17.8052077 | -0.232691643 | 0.049333922 |
| 20.341755 | 20.1575783 | 20.1275081 | 20.0665975 | -0.123997311 | 0.048752709 |
| 23.8214904 | 23.6765918 | 23.7651246 | 23.672078 | -0.052243019 | 0.364371825 |
| 21.8184387 | 21.5336695 | 21.8191772 | 21.754297 | -0.078507049 | 0.177305025 |
| 23.5211727 | 23.3880194 | 23.43974 | 23.4446209 | -0.021036138 | 0.70209697 |
| 19.8322879 | 19.9398403 | 20.0044073 | 19.9877431 | 0.113207237 | 0.192950806 |
| 15.0717361 | 15.1263363 | 15.5885115 | 15.5461716 | 0.009959424 | 0.95418218 |
| 13.9731242 | NA | 13.7582988 | 13.8560954 | -0.193210044 | 0.222250188 |
| 19.2732912 | 19.5244073 | 19.4818169 | 19.210879 | -0.004735861 | 0.973224983 |
| 20.3120624 | 19.8874603 | 19.9621456 | 20.1006174 | -0.220597415 | 0.062604376 |
| 20.4637655 | 20.4535759 | 20.5909433 | 20.621909 | -0.0684618 | 0.359594661 |
| 18.5620859 | 18.3922034 | 18.4927123 | 18.4519796 | -0.104126713 | 0.136592775 |
| 24.695128 | 24.724691 | 24.6999487 | 24.6894255 | 0.016507493 | 0.655076441 |
| 23.1147733 | 23.1482024 | 23.2285496 | 23.185386 | 0.120445955 | 0.013463986 |
| 19.3320353 | 19.2311048 | 19.4212696 | 19.3398676 | 0.007666725 | 0.886187202 |
| 17.5475076 | 17.6027506 | 17.4589129 | 17.5551844 | 0.009225182 | 0.882411542 |
| 20.0726542 | 19.9517025 | 20.0271879 | 20.1338099 | -0.040812233 | 0.589970004 |
| 19.0142595 | 18.8505056 | 19.0636615 | 19.0282501 | 0.099278607 | 0.265009594 |
| 15.2234482 | 15.1059323 | 15.326645 | 15.3132574 | 0.195276719 | 0.107373308 |
| 16.859553 | 17.1392903 | 17.4128538 | 17.3427036 | 0.393044209 | 0.094230046 |

| | | | | | |
|---|---|---|---|---|---|
| 19.3752746 | 19.3142041 | 19.4304186 | 19.3232565 | -0.057361554 | 0.288977765 |
| 21.4357462 | 21.3308277 | 21.3118542 | 21.2831044 | -0.073905766 | 0.197793441 |
| 21.2735602 | 21.3369299 | 21.4339634 | 21.1759133 | 0.077582626 | 0.556215113 |
| 17.8746222 | 18.1169032 | 18.0046691 | 18.0663897 | 0.130635641 | 0.077948779 |
| NA | 15.6364269 | NA | NA | -0.299780396 | 0.023148962 |
| 14.5341755 | 14.7906242 | 14.6358081 | 14.7638501 | 0.141931794 | 0.113477542 |
| 16.6955436 | 16.5708453 | 16.5417589 | 16.6447083 | -0.0046148 | 0.971132554 |
| 19.4466647 | 19.4331004 | 19.4790163 | 19.614026 | 0.075991939 | 0.204397288 |
| 14.7313994 | 14.5960945 | 14.7269941 | 13.9781508 | -0.183199484 | 0.19988393 |
| 20.5685549 | 20.5106032 | 20.5736232 | 20.5160725 | -0.013319983 | 0.696346439 |
| 17.9382677 | 18.0043135 | 18.2220626 | 18.1287107 | 0.113502362 | 0.218384677 |
| NA | 17.7207513 | NA | NA | -0.38786001 | 0.274292515 |
| NA | 19.9858236 | NA | NA | 0.055842305 | 0.800997996 |
| 16.6973627 | 17.11752 | 17.1092707 | 17.0748067 | -0.088756774 | 0.449218655 |
| 21.867744 | 21.5808531 | 21.6494875 | 21.6447323 | -0.158565676 | 0.053879023 |
| 19.5692641 | 19.537799 | 19.5169887 | 19.5731066 | 0.009396392 | 0.89354028 |
| 24.0412857 | 23.8270089 | 23.7947989 | 23.8153909 | -0.143113627 | 0.030345542 |
| 20.5394565 | 20.583446 | 20.4551705 | 20.5364616 | -0.097986679 | 0.103865848 |
| 21.1784704 | 21.0260108 | 21.1121731 | 21.1126417 | -0.058717696 | 0.236792009 |
| 21.2709209 | 21.250154 | 21.1980296 | 21.1847159 | -0.061111711 | 0.179700684 |
| 20.9297702 | 20.906113 | 20.9311977 | 20.8705277 | -0.047917095 | 0.251753358 |
| 21.9216566 | 21.8364359 | 21.9246802 | 21.8940893 | 0.020271471 | 0.648786159 |
| 19.7465266 | 19.7990038 | 19.9396757 | 19.8852024 | 0.110973053 | 0.029323278 |
| 25.9769118 | 25.9531859 | 25.9863024 | 26.0711815 | 0.077733313 | 0.20401748 |
| 20.7538266 | 20.626661 | 20.7492305 | 20.7352883 | -0.002126026 | 0.974964149 |
| 19.5913515 | 19.5370325 | 19.4820564 | 19.6322314 | 0.035743476 | 0.67743506 |
| 14.4097141 | 14.4105084 | 14.6381567 | 14.317172 | 0.048937665 | 0.642264626 |
| 15.8251347 | 16.0438452 | 15.8008042 | 15.8575805 | 0.053377759 | 0.737576366 |
| 17.2801369 | 16.6843272 | 17.1682383 | 17.2239896 | -0.23366139 | 0.201026039 |
| 16.9896742 | 16.8750629 | 17.0484157 | 16.8673748 | -0.14767611 | 0.142032211 |
| 19.539454 | 19.291002 | 19.3732064 | 19.2451974 | -0.111653562 | 0.246760215 |
| 26.42448 | 26.3428616 | 26.3640704 | 26.3862007 | -0.028920897 | 0.646889773 |
| 19.0023138 | 18.7603646 | 18.7313249 | 18.7984858 | -0.129261891 | 0.047639787 |
| 19.9917378 | 19.8668416 | 19.9200639 | 19.8371109 | -0.034026837 | 0.539762827 |
| 23.3549189 | 23.2421131 | 23.3365697 | 23.3123428 | -0.021362667 | 0.689643306 |
| 24.5724689 | 24.4763603 | 24.5380476 | 24.5511849 | 0.020570606 | 0.695699556 |
| 22.2316217 | 22.1296236 | 22.170058 | 22.1591004 | -0.028364469 | 0.495236538 |
| 21.2478674 | 21.2149331 | 21.3184928 | 21.237957 | 0.021530605 | 0.618114984 |
| 21.1183015 | 21.1098841 | 21.1661779 | 21.1621843 | 0.044770835 | 0.423697815 |
| 20.009161 | 20.1314349 | 20.1490521 | 20.1411912 | 0.066439861 | 0.259774267 |
| 18.6769489 | 18.64894 | 18.6303278 | 18.6439998 | -0.024434129 | 0.709520672 |
| 19.9181472 | 19.8667011 | 19.9020143 | 19.9560273 | 0.044439126 | 0.501299676 |

| | | | | | |
|---|---|---|---|---|---|
| 25.0542436 | 24.9667607 | 25.0373647 | 25.048162 | 0.041527035 | 0.539706349 |
| 17.453389 | 17.4453548 | 17.5766432 | 17.7933929 | -0.046402868 | 0.648321428 |
| 15.7124686 | 15.8548667 | 15.7835206 | 15.8886228 | 0.329653534 | 0.078025382 |
| 18.7621681 | 18.505951 | 18.9412698 | 19.0159807 | 0.096735664 | 0.632688816 |
| NA | 16.7927731 | NA | NA | 0.760046041 | 0.195073736 |
| 19.1222292 | 19.3275731 | 19.739396 | 19.7411296 | 0.398261235 | 0.031362366 |
| 18.0484433 | 18.066583 | 18.0665623 | 18.1081176 | 0.083627603 | 0.279144105 |
| 14.8647023 | NA | 14.3879676 | 14.7718798 | -0.373342445 | 0.042216479 |
| 19.7476995 | 19.6494707 | 19.5978378 | 19.7118171 | -0.035872407 | 0.605261484 |
| 19.1914985 | 19.1036264 | 19.1497741 | 18.7020057 | 0.208247174 | 0.148146923 |
| 23.927863 | 23.922874 | 23.9006562 | 23.9522984 | 0.030746909 | 0.567377336 |
| 20.8112847 | 20.3957206 | 20.577494 | 20.7198212 | -0.15833796 | 0.158356798 |
| 19.3433051 | 19.0182242 | 19.117439 | 19.126074 | -0.252466594 | 0.005475533 |
| 16.7162433 | 17.0607557 | 16.5151536 | 16.785126 | 0.107412893 | 0.418010584 |
| 19.2711637 | 19.2076864 | 19.3447014 | 19.297723 | 0.036614376 | 0.391759635 |
| 24.8146217 | 24.5902813 | 24.7357087 | 24.7564493 | -0.047426443 | 0.582511261 |
| 18.4270065 | 18.1419603 | 18.4116337 | 18.0990628 | -0.160477617 | 0.057057759 |
| 17.4615063 | 17.6398097 | 17.7807523 | 17.5456476 | -0.004705846 | 0.953753674 |
| 23.5650301 | 23.7229461 | 23.7527008 | 23.558812 | 0.030593153 | 0.697492075 |
| 21.6562432 | 21.5865981 | 21.5228511 | 21.6080721 | -0.035854306 | 0.522304019 |
| 22.7893937 | 22.7703757 | 22.7277775 | 22.7396633 | -0.055840014 | 0.317906774 |
| 20.7639522 | 20.8510525 | 20.7483791 | 20.7847404 | -0.023685777 | 0.662087373 |
| 23.7114483 | 23.5648765 | 23.6827827 | 23.6754925 | -0.02664882 | 0.598413697 |
| 18.8859019 | 19.0364955 | 19.011067 | 19.0950814 | 0.066075698 | 0.321287739 |
| 16.3513 | 16.5736112 | 16.4254256 | 16.524791 | 0.108364112 | 0.249287199 |
| 16.3470639 | 16.5229677 | 16.5137052 | 16.4382224 | 0.273716827 | 0.017182746 |
| 17.1760845 | 17.0384 | 17.1844905 | 17.0225911 | -0.0407686 | 0.652437445 |
| 20.3595304 | 20.443127 | 20.4056079 | 20.4042685 | 0.004478623 | 0.924630231 |
| 21.3194866 | 21.3261436 | 21.2875657 | 21.2291576 | -0.060732968 | 0.0969066 |
| 19.6200984 | 19.6068024 | 19.674912 | 19.7435628 | 0.031919527 | 0.536527632 |
| 24.4684162 | 24.4525113 | 24.5410905 | 24.4527402 | 0.096301773 | 0.284659911 |
| 22.921657 | 22.9236649 | 23.0046422 | 23.0808509 | 0.16634273 | 0.255629555 |
| 14.7114375 | 15.1111991 | 15.4362394 | 15.327975 | 0.355058916 | 0.030587743 |
| 17.5884023 | 17.4077147 | 17.4396343 | 17.2690481 | -0.217334484 | 0.031360879 |
| 25.1888405 | 25.1347797 | 25.1148486 | 25.2053184 | 0.026679379 | 0.693351983 |
| 18.6916987 | 18.4511907 | 18.5311359 | 18.5563235 | -0.104046545 | 0.088888674 |
| 19.4516358 | 19.3297691 | 19.3267538 | 19.3766551 | -0.13569337 | 0.029239299 |
| 19.596468 | 19.6078122 | 19.6623748 | 19.6445441 | -0.001493609 | 0.97804801 |
| 24.711326 | 24.6941829 | 24.811215 | 24.7238069 | 0.010682885 | 0.807614705 |
| 23.7246023 | 23.6154688 | 23.6497602 | 23.67899 | -0.041721974 | 0.317211064 |
| 20.1130598 | 20.0948352 | 20.060968 | 20.0566544 | -0.022649948 | 0.553071948 |
| 21.7960174 | 21.7581841 | 21.8265232 | 21.7934502 | -0.020360547 | 0.539446753 |

| | | | | | |
|---|---|---|---|---|---|
| 20.5708185 | 20.3728304 | 20.5412577 | 20.532484 | -0.009399205 | 0.849389462 |
| 23.3533324 | 23.2536427 | 23.5409373 | 23.4310544 | 0.045767268 | 0.525078818 |
| 21.7039084 | 21.5850742 | 21.7320455 | 21.6183086 | -0.010810386 | 0.881144117 |
| 19.3590986 | 19.4936993 | 19.5614289 | 19.5072157 | 0.065413844 | 0.402057093 |
| 15.4998756 | 15.9301586 | 16.1389322 | 16.1959914 | 0.293432893 | 0.339954709 |
| 19.8784958 | 19.5866157 | 19.7694886 | 19.837736 | -0.062761941 | 0.592182557 |
| 20.9115093 | 20.8311976 | 20.8408546 | 20.8997609 | 0.018351549 | 0.680877904 |
| 20.7986761 | 21.2462086 | 21.173316 | 21.3624377 | 0.34015611 | 0.036617352 |
| 20.347945 | 20.3558708 | 20.377934 | 20.4104838 | 8.64E-04 | 0.988377187 |
| 18.885817 | 18.9544133 | 18.902399 | 18.9661426 | 0.077650301 | 0.156215719 |
| 18.6725729 | 18.60269 | 18.7711566 | 18.7283645 | 0.00190932 | 0.974107009 |
| 25.950738 | 25.8682967 | 26.0060529 | 25.9557311 | 0.062641999 | 0.448971575 |
| 13.7189302 | NA | 13.6851338 | 13.5734219 | -0.173033215 | 0.253013836 |
| 19.3183826 | 19.2092148 | 19.327171 | 19.3964655 | 0.034914957 | 0.567666592 |
| 19.9169971 | 19.7352161 | 19.9272187 | 19.8941751 | -0.073959724 | 0.358846354 |
| 26.6559632 | 26.5410639 | 26.5687186 | 26.6103958 | -0.026362958 | 0.652836534 |
| 19.9854134 | 19.9728014 | 20.1006011 | 19.9412975 | 0.010397544 | 0.838684316 |
| 22.4788993 | 22.4254262 | 22.5064748 | 22.530309 | 0.016943697 | 0.692999836 |
| 19.5560211 | 19.603844 | 19.671395 | 19.5803443 | 0.041508178 | 0.358945025 |
| 19.9503213 | 19.9061556 | 19.8836132 | 19.9286138 | 0.007332262 | 0.885633721 |
| 20.3265073 | 20.455312 | 20.3920994 | 20.4305473 | 0.043740251 | 0.619408521 |
| 20.8473592 | 21.0957448 | 20.9446065 | 20.935717 | 0.112225974 | 0.344056219 |
| 19.7805634 | 19.3333206 | 19.1415159 | 19.5678539 | -0.261104837 | 0.106875763 |
| 19.2852214 | 19.3911912 | 19.3445943 | 19.5721384 | 0.063468749 | 0.551432751 |
| 21.1984598 | 20.9172525 | 21.0584924 | 21.0223776 | -0.084958527 | 0.103842327 |
| 17.7126214 | 17.7975018 | 17.754608 | 17.8501948 | 0.049297095 | 0.479023271 |
| 18.6153204 | 18.6663322 | 18.7600942 | 18.7159348 | 0.091389412 | 0.1427043 |
| 17.9910469 | 17.8384621 | 17.6231679 | 17.8745428 | -0.194559667 | 0.021918423 |
| 24.3285657 | 24.1759654 | 24.2285759 | 24.1734367 | -0.101693951 | 0.025505016 |
| 21.9030055 | 21.8447159 | 21.8587862 | 21.8608105 | -0.031699773 | 0.48660695 |
| 20.4862133 | 20.5095165 | 20.5772 | 20.5253322 | 0.070420824 | 0.16750143 |
| 25.0303848 | 24.9939235 | 24.9882957 | 25.0189074 | 0.03725493 | 0.521130412 |
| 22.5983237 | 22.4463124 | 22.5166919 | 22.4868281 | -0.037551669 | 0.531195384 |
| 21.7148183 | 21.5443039 | 21.6337876 | 21.6049732 | -0.05068876 | 0.244296608 |
| 21.3893077 | 21.3556171 | 21.4269105 | 21.434743 | 0.017429112 | 0.761489313 |
| 17.6871358 | 17.6819992 | 17.8555769 | 17.7348958 | 0.044539283 | 0.522208557 |
| 20.7227987 | 20.67057 | 20.6905714 | 20.576126 | 0.067887356 | 0.491056865 |
| 26.2031394 | 26.0889549 | 26.2474564 | 26.2267368 | 0.009459445 | 0.897187577 |
| 22.6359513 | 22.4909651 | 22.6755775 | 22.5845939 | 0.042807651 | 0.438570705 |
| 24.0740411 | 24.0405308 | 24.025025 | 24.0546707 | -0.003995946 | 0.934392708 |
| 20.2430386 | 20.234646 | 20.2775528 | 20.3090568 | 0.065054819 | 0.1664665 |
| 22.2486076 | 22.2306511 | 22.3470562 | 22.2791445 | 0.099798513 | 0.049665319 |

| | | | | | |
|---|---|---|---|---|---|
| 17.8043154 | 17.7706021 | 17.9156755 | 17.8465554 | 0.077679931 | 0.352177021 |
| 22.1914923 | 22.1298974 | 22.168169 | 22.1636636 | -0.021106224 | 0.608356056 |
| 20.79342 | 20.6430608 | 20.7294123 | 20.7236993 | -0.031455155 | 0.512149782 |
| 15.8742788 | 15.5912255 | 15.3553122 | 15.4140104 | -0.407904608 | 0.024999765 |
| 23.3228159 | 23.285978 | 23.3323117 | 23.3177804 | 8.40E-05 | 0.998427543 |
| 20.6244725 | 20.5043229 | 20.4730475 | 20.5126637 | -0.122396328 | 0.169855103 |
| 24.2015931 | 24.0653694 | 24.1700409 | 24.1085728 | -0.026625489 | 0.576320038 |
| 20.816937 | 20.7210276 | 20.8470339 | 20.8616453 | 0.052977026 | 0.331513193 |
| 19.1212663 | 19.051617 | 19.2400985 | 19.0830402 | 0.047776998 | 0.491434389 |
| 25.4990688 | 25.499246 | 25.6064134 | 25.5415492 | 0.070265884 | 0.364690769 |
| NA | 14.3310802 | NA | NA | 0.314450451 | 0.038933496 |
| 16.3050467 | 16.3756162 | 16.4709814 | 16.259695 | 0.027389613 | 0.886186529 |
| 16.8448309 | 16.3274745 | 16.4777736 | 16.5717721 | -0.138598066 | 0.463305299 |
| 26.293375 | 26.2745812 | 26.2542096 | 26.261969 | 0.022235641 | 0.730297971 |
| 26.9531738 | 26.8719957 | 26.9549367 | 26.9423866 | 0.032680028 | 0.60167064 |
| 20.835585 | 20.7051252 | 20.7902509 | 20.8654641 | 0.056647222 | 0.486821213 |
| 23.4442496 | 23.3140038 | 23.4732546 | 23.5172649 | 0.036257509 | 0.52117058 |
| 22.9827542 | 22.830813 | 22.9521901 | 22.9351328 | -0.04241311 | 0.409203882 |
| 12.860706 | NA | 12.5610514 | 12.6047111 | -0.170375569 | 0.473823129 |
| 22.3421019 | 22.1656158 | 22.2203511 | 22.0410471 | -0.117642 | 0.049691153 |
| 20.3128264 | 20.4221786 | 20.4023395 | 20.311316 | 0.084681677 | 0.104140284 |
| 18.1867865 | 17.9794824 | 18.1023695 | 18.1653253 | -0.045885308 | 0.370781615 |
| 20.2368469 | 20.2226632 | 20.3337256 | 20.3408864 | 0.080622212 | 0.207181606 |
| 27.4263309 | 27.4144516 | 27.4912489 | 27.5091756 | 0.099013162 | 0.130135355 |
| 16.0689965 | 16.3175038 | 16.0522265 | 16.1560193 | 0.155256299 | 0.102361418 |
| 19.8759598 | 19.800064 | 19.8166187 | 19.8448782 | -0.071197959 | 0.048401823 |
| NA | 13.0977049 | NA | NA | -0.124200446 | 0.607753574 |
| 19.9514124 | 19.9169762 | 20.0674932 | 20.0648725 | 0.069762768 | 0.536017568 |
| 18.6303513 | 18.6246368 | 18.4254773 | 18.6956973 | 0.012246483 | 0.889497832 |
| 20.9997582 | 20.8995237 | 21.0157943 | 20.9414155 | -0.029495185 | 0.566677152 |
| 21.1798571 | 21.1948066 | 21.2197506 | 21.1272042 | -0.025923851 | 0.582769577 |
| 19.5131431 | 19.5716327 | 19.6459141 | 19.6835729 | 0.089836017 | 0.138890135 |
| 19.7695581 | 19.7538026 | 19.7701451 | 19.8126154 | 0.02293437 | 0.700906002 |
| 20.1934108 | 20.0608592 | 20.2617434 | 20.2235484 | 0.016104135 | 0.811047184 |
| 18.9933059 | 18.8937436 | 19.0555798 | 19.1191837 | 0.103129762 | 0.207769879 |
| 19.0347256 | 19.032416 | 18.9864643 | 19.1764683 | 0.139229848 | 0.239568644 |
| 17.7013966 | 18.2700439 | 17.7849719 | 17.8618195 | 0.12678923 | 0.563550082 |
| 17.7408267 | 17.6026715 | 17.6365643 | 17.5351307 | -0.103312791 | 0.312876785 |
| 19.333697 | 19.291876 | 19.4760366 | 19.2388026 | -0.028370562 | 0.740553078 |
| 20.4219453 | 20.3368303 | 20.3802846 | 20.4163902 | -0.004954334 | 0.920800428 |
| 23.3662089 | 23.2610933 | 23.3287893 | 23.3041536 | -0.02709873 | 0.541329871 |
| 20.9185789 | 20.9245756 | 20.8902531 | 20.8891678 | -0.036288755 | 0.39012863 |

| | | | | | |
|---|---|---|---|---|---|
| 21.6280384 | 21.5443805 | 21.6195138 | 21.5899264 | 0.020494621 | 0.685404369 |
| 21.1199668 | 21.025657 | 21.1384734 | 21.1459948 | -0.013084182 | 0.832291133 |
| 19.7589286 | 19.6591832 | 19.7765778 | 19.7496833 | 0.060440904 | 0.456831809 |
| 20.5492275 | 20.375902 | 20.622194 | 20.4564869 | -0.01809259 | 0.839011636 |
| 19.5438067 | 19.4435525 | 19.5065961 | 19.3090285 | 0.109530875 | 0.296270244 |
| 21.3857544 | 21.4732014 | 21.2238 | 21.4617776 | 0.20786482 | 0.063395425 |
| 21.1635826 | 20.9825127 | 21.1479667 | 21.1904745 | 0.071773872 | 0.514571067 |
| 15.3773434 | 15.3788707 | 15.424559 | 15.5650141 | 0.13158028 | 0.287662365 |
| 18.7940884 | 18.6433493 | 18.7862493 | 18.8028677 | -0.093471143 | 0.201792015 |
| 18.5940519 | 18.5795463 | 18.6322596 | 18.6582636 | -0.009578453 | 0.828913876 |
| 18.5201585 | 18.5788424 | 18.9038035 | 18.7329848 | 0.120354672 | 0.173810547 |
| NA | 16.7296697 | NA | NA | 0.024287737 | 0.838030363 |
| NA | 14.7091164 | NA | NA | 0.226395733 | 0.429524545 |
| 19.4414834 | 19.3670219 | 19.3049269 | 19.2698352 | -0.123863358 | 0.145395193 |
| 21.8031696 | 21.6608946 | 21.8312952 | 21.7604621 | 0.066002443 | 0.267701589 |
| 18.9481947 | 18.5159054 | 18.7616739 | 18.8230239 | -0.235891682 | 0.038673034 |
| 26.2725569 | 26.1960516 | 26.2630379 | 26.2374762 | -0.020224494 | 0.811553262 |
| 23.0033742 | 22.9433592 | 23.0243812 | 23.0454186 | 0.009596055 | 0.87398033 |
| 20.6610008 | 20.4398524 | 20.5883832 | 20.6409363 | -0.040073088 | 0.651612036 |
| 15.959211 | 16.0532374 | 16.3011914 | 16.3850043 | 0.300984622 | 0.28802662 |
| 24.9846403 | 24.8631388 | 24.9392133 | 25.0013569 | -0.015926178 | 0.776496857 |
| 21.4662822 | 21.3441512 | 21.4273124 | 21.3806003 | -0.084906221 | 0.039838771 |
| 21.5442776 | 21.5439187 | 21.5675938 | 21.5993178 | 0.041736739 | 0.336736736 |
| 20.4103877 | 20.2851868 | 20.3517585 | 20.3896107 | 0.012191228 | 0.794521513 |
| 19.6892285 | 19.5873949 | 19.6797433 | 19.7814085 | 0.08332198 | 0.171448277 |
| 17.5706909 | 17.502902 | 17.6147232 | 17.5521223 | 0.105722001 | 0.181446401 |
| 17.9492362 | 17.9070544 | 17.7666318 | 18.0837496 | 0.085264538 | 0.413428737 |
| 15.5268957 | NA | 15.4520767 | 15.2401088 | -0.045690923 | 0.764829855 |
| 20.6431609 | 20.6287588 | 20.7607416 | 20.7260101 | 0.147276732 | 0.040771907 |
| 20.7076947 | 20.5968424 | 20.6900285 | 20.7321923 | 0.070961922 | 0.408081243 |
| 20.3816614 | 20.3117014 | 20.3177414 | 20.4223125 | 0.056736857 | 0.452088985 |
| NA | 14.2698849 | NA | NA | 0.191723011 | 0.496600938 |
| 17.1317415 | 16.5128786 | 16.6967847 | 17.2255476 | -0.141643194 | 0.34733906 |
| 19.1325775 | 18.9256247 | 19.1007363 | 19.0625258 | -0.121075426 | 0.045864862 |
| 19.4844717 | 19.4770239 | 19.5588731 | 19.450377 | -0.008163109 | 0.877144763 |
| 21.1438727 | 21.2401342 | 21.1364626 | 21.1730003 | 0.058953174 | 0.271302233 |
| 20.3737151 | 20.3081246 | 20.3647078 | 20.4611739 | 0.076284257 | 0.209959241 |
| 22.0256373 | 21.9663045 | 22.0372135 | 22.1680029 | 0.090661543 | 0.162216395 |
| 23.729758 | 23.7693941 | 23.7596682 | 23.8221681 | 0.075354575 | 0.248756251 |
| 21.3831352 | 21.2794642 | 21.3295381 | 21.4040237 | -0.006377015 | 0.921683475 |
| 22.3166264 | 22.2686472 | 22.2726634 | 22.1635613 | 0.008299579 | 0.904811742 |
| 19.4174816 | 19.2092963 | 19.276219 | 19.3837712 | -0.037804648 | 0.593009231 |

| | | | | | |
|---|---|---|---|---|---|
| 20.3610628 | 20.2504545 | 20.4725786 | 20.3278091 | 0.035490751 | 0.617355102 |
| 18.1801524 | 18.0438325 | 18.1831976 | 18.1789403 | 0.05583646 | 0.471027589 |
| 17.5223464 | 17.5568693 | 17.5756709 | 17.6373015 | 0.099642604 | 0.246549619 |
| 17.0505047 | 16.9096999 | 17.0055955 | 17.0983191 | 0.010961295 | 0.899750023 |
| 18.4936326 | 18.4265474 | 18.5499079 | 18.3763736 | -0.063155703 | 0.512706838 |
| 24.0772117 | 23.9202016 | 24.1788318 | 24.2465707 | 0.164798166 | 0.246693146 |
| 18.360424 | 18.2781854 | 18.4912767 | 18.4646712 | -0.014929643 | 0.871307647 |
| 18.9883563 | 18.9505639 | 18.9055741 | 18.7466335 | -0.020652412 | 0.77765731 |
| 20.3979294 | 20.2104179 | 20.3621119 | 20.3649792 | -0.041834511 | 0.545142674 |
| 20.1302338 | 20.0828809 | 20.1948983 | 20.166298 | 0.033781249 | 0.489245527 |
| 18.9782281 | 18.9527492 | 19.00148 | 19.0294531 | 0.018925666 | 0.748147751 |
| 16.9939138 | 16.6251142 | 16.6962113 | 16.638771 | -0.275596734 | 0.029700958 |
| 22.1241829 | 21.9619794 | 22.0496432 | 22.0788575 | -0.080686757 | 0.098240358 |
| 21.7581039 | 21.545774 | 21.6571683 | 21.5543688 | -0.080231203 | 0.120011026 |
| 21.8452847 | 21.7390602 | 21.8310526 | 21.8352983 | 0.007521577 | 0.886667141 |
| 19.5636009 | 19.848161 | 19.7435712 | 19.7173313 | 0.142835832 | 0.028250173 |
| 23.5808514 | 23.5744367 | 23.637269 | 23.6322678 | 0.072484316 | 0.250899536 |
| 22.4209831 | 22.3588369 | 22.4868593 | 22.4531221 | 0.100729356 | 0.122732787 |
| NA | 13.9093906 | NA | NA | -0.161601732 | 0.630053936 |
| 19.1559889 | 18.4600764 | 18.6131061 | 18.4026431 | -0.472961575 | 0.031853828 |
| 19.0158089 | 19.362286 | 19.1304039 | 19.1809242 | 0.053150072 | 0.540746387 |
| 19.1547045 | 19.1206356 | 18.9640434 | 19.010459 | -0.064278891 | 0.408002173 |
| 19.6139173 | 19.5928375 | 19.5981234 | 19.6382503 | -0.009887273 | 0.896648296 |
| 21.0325429 | 20.7935862 | 20.9983086 | 20.9286323 | -0.061854474 | 0.294422908 |
| 23.3108678 | 23.2010247 | 23.2802008 | 23.2688101 | -0.012479391 | 0.829925557 |
| 24.1672026 | 24.3014282 | 24.2590118 | 24.2345072 | 0.065078455 | 0.210774689 |
| 22.5699246 | 22.4918642 | 22.5340578 | 22.4236521 | -0.09609728 | 0.059712394 |
| 21.8237332 | 21.7655892 | 21.7377425 | 21.8096034 | -0.056671138 | 0.22803366 |
| 20.7114496 | 20.7818001 | 20.803785 | 20.6943342 | 0.02232023 | 0.625673695 |
| 21.417968 | 21.4416804 | 21.525732 | 21.4292916 | 0.081552914 | 0.12371999 |
| 19.713291 | 19.68319 | 19.837264 | 19.8380302 | 0.0556762 | 0.358369676 |
| 17.5707664 | 17.6095534 | 17.7651685 | 17.6570924 | 0.088140732 | 0.173547946 |
| 18.054981 | 18.0883097 | 18.1498269 | 18.2293253 | 0.059126554 | 0.516203384 |
| 22.4190186 | 22.5268613 | 22.4882268 | 22.4723374 | 0.070038619 | 0.463004803 |
| 17.5830986 | 17.4159904 | 17.4324197 | 17.4988841 | -0.005694795 | 0.963692876 |
| NA | 14.4960206 | NA | NA | 0.669121393 | 0.240824657 |
| 23.216659 | 23.1777432 | 23.1480381 | 23.1741976 | -0.016154654 | 0.792965465 |
| 18.9127633 | 18.6873937 | 18.751458 | 18.7029187 | -0.087284434 | 0.275229873 |
| 19.4418323 | 19.3923841 | 19.5270372 | 19.374984 | -0.056323583 | 0.329310428 |
| 25.938517 | 25.9409675 | 25.9038878 | 25.9235709 | 0.016978161 | 0.687476227 |
| 19.1224382 | 19.0648636 | 19.0624996 | 19.0572385 | 0.010993527 | 0.844611018 |
| 19.8891153 | 19.7601499 | 19.8554475 | 19.927659 | -0.006540933 | 0.932171296 |

| | | | | | |
|---|---|---|---|---|---|
| 19.6188142 | 19.4621494 | 19.5906467 | 19.5746161 | 0.005194054 | 0.943275887 |
| 17.4209391 | 17.3213321 | 17.5017276 | 17.4609524 | 0.017165817 | 0.866026914 |
| 19.1605555 | 19.1971271 | 19.1330792 | 19.0926142 | 0.011097562 | 0.832538988 |
| 21.9704258 | 21.9654206 | 22.0404 | 22.0305494 | 0.094163604 | 0.040727495 |
| 22.1210503 | 22.0200622 | 22.0974986 | 22.1412069 | 0.008225258 | 0.906979355 |
| 20.5962614 | 20.3691748 | 20.4475239 | 20.4951691 | -0.100337271 | 0.139732586 |
| 18.63794 | 18.4104953 | 18.808353 | 18.5754209 | -0.001479897 | 0.990711031 |
| 17.3679872 | 17.0951347 | 17.3513061 | 17.7780677 | 0.046587082 | 0.744597378 |
| 22.262928 | 22.0811835 | 22.2158177 | 22.2036967 | -0.016073604 | 0.809699791 |
| 19.595562 | 19.5321264 | 19.5654716 | 19.5934608 | -0.05623184 | 0.234748316 |
| 21.8132513 | 21.7168691 | 21.772975 | 21.7616178 | -0.054745993 | 0.231447276 |
| 20.8157762 | 20.8205313 | 20.9262011 | 20.9290568 | 0.101782555 | 0.133610044 |
| NA | 15.3986652 | NA | NA | 0.124946285 | 0.420366539 |
| 19.3431699 | 19.0689502 | 19.1275826 | 19.2553012 | -0.112762521 | 0.117959635 |
| NA | 13.3167573 | NA | NA | -0.2089148 | 0.367041576 |
| 12.6905458 | NA | 12.3847744 | 12.4324033 | -0.211733745 | 0.346207041 |
| 14.3428304 | 13.5866706 | 14.6023982 | 14.5118275 | -0.237970988 | 0.257048793 |
| 17.2628356 | 17.1936177 | 16.983996 | 17.0095621 | -0.036259785 | 0.685502505 |
| 21.7834654 | 21.8216077 | 21.7996025 | 21.7664004 | 0.087563376 | 0.28968491 |
| 22.4970094 | 22.2382214 | 22.4183144 | 22.4114895 | -0.078709105 | 0.311347524 |
| 22.0884388 | 21.9200826 | 22.1347615 | 22.1587218 | -0.00752923 | 0.908744992 |
| 18.2229068 | 18.1857203 | 18.1531018 | 18.1781384 | -0.032975937 | 0.520001041 |
| 21.8877367 | 21.8738744 | 21.8171674 | 21.8520517 | -0.016220444 | 0.710333003 |
| 23.3933511 | 23.422192 | 23.3927944 | 23.3561817 | -0.030043933 | 0.489296048 |
| 17.9289784 | 17.6721774 | 18.0016655 | 17.9320886 | -9.63E-04 | 0.989494075 |
| 20.2061176 | 20.0670947 | 20.1938486 | 20.2416174 | 0.028919383 | 0.715719573 |
| 18.4037858 | 18.3519285 | 18.3120374 | 18.3684993 | 0.004370256 | 0.956861184 |
| 22.3397512 | 22.085499 | 22.1061397 | 22.1790913 | -0.087880647 | 0.295770611 |
| 14.4455594 | 14.5918255 | 14.4758526 | 14.5897244 | 0.118297421 | 0.259756825 |
| 16.5950013 | 16.5379706 | 16.6365647 | 16.838007 | 0.119562403 | 0.28418451 |
| 14.153538 | 14.0663666 | 14.3086658 | 14.2076054 | -0.024968742 | 0.894610681 |
| 19.6935576 | 19.7871274 | 19.8169522 | 19.6753217 | 0.031177992 | 0.648385958 |
| 19.0165344 | 18.9787963 | 19.0533515 | 18.9538219 | -0.081420726 | 0.171186282 |
| NA | 13.4749914 | NA | NA | -0.404694189 | 0.185541754 |
| 18.2253721 | 17.8689681 | 18.0101355 | 17.7648117 | -0.206156381 | 0.133359096 |
| 21.3128307 | 21.3401968 | 21.4022592 | 21.3918744 | 0.07346111 | 0.15651141 |
| 20.6771269 | 20.6887416 | 20.706746 | 20.771366 | 0.072892591 | 0.182567956 |
| 19.7002924 | 19.5054182 | 19.7502651 | 19.7172193 | 0.006940722 | 0.897187836 |
| 17.7592888 | 17.8200762 | 17.8062053 | 17.7344988 | 0.008640369 | 0.891613753 |
| 20.7493442 | 20.583549 | 20.6416051 | 20.7123404 | -0.019761469 | 0.794416739 |
| NA | 18.9557731 | NA | NA | 0.270644029 | 0.031526702 |
| 16.6345415 | 16.2917639 | 16.3511881 | 16.7346896 | 0.136632951 | 0.612140802 |

| | | | | | |
|---|---|---|---|---|---|
| 19.870463 | 19.8681544 | 19.880322 | 19.8847149 | 0.006971452 | 0.877064773 |
| 23.1472278 | 23.0372206 | 23.0823649 | 23.1271685 | -0.009881189 | 0.821605584 |
| 18.9319702 | 18.7253078 | 18.8287081 | 18.858345 | -0.099781812 | 0.412729235 |
| NA | 14.5337471 | NA | NA | -0.059606033 | 0.548825787 |
| 16.1389699 | 16.3868913 | 16.3240011 | 16.2715549 | 0.032426863 | 0.753917307 |
| 16.2547917 | 16.5285924 | 16.5391639 | 16.4658962 | 0.075934157 | 0.449373413 |
| 22.305975 | 22.3707732 | 22.4848806 | 22.4934445 | 0.073910923 | 0.436923539 |
| 26.5638678 | 26.6248268 | 26.581765 | 26.5811838 | 0.078799579 | 0.383191812 |
| 17.8338861 | 17.7967487 | 17.8494308 | 17.8794795 | -0.015426951 | 0.823843625 |
| 22.0084876 | 22.0101532 | 21.9906518 | 21.9080727 | 0.006748806 | 0.913807645 |
| 24.2102867 | 24.1062172 | 24.237762 | 24.1423672 | -0.014312352 | 0.783078319 |
| 23.1681532 | 23.0116291 | 23.1445935 | 23.1000661 | -0.063417275 | 0.21801386 |
| 22.8000572 | 22.7532181 | 22.7882082 | 22.7528556 | -0.023178937 | 0.587851979 |
| 23.6461002 | 23.5318191 | 23.6468634 | 23.5772343 | -0.029279271 | 0.472500495 |
| 19.5334426 | 19.5264787 | 19.5400152 | 19.67377 | 0.070959264 | 0.117728792 |
| 21.490688 | 21.4890247 | 21.5557608 | 21.5641159 | 0.056996274 | 0.26427714 |
| 22.781472 | 22.7110226 | 22.7065693 | 22.7001045 | -0.004019827 | 0.941178662 |
| 20.3849114 | 20.2516444 | 20.4213569 | 20.3487267 | -0.023755555 | 0.691621981 |
| 19.9358959 | 19.8189146 | 19.991094 | 19.8419719 | 3.35E-04 | 0.996165521 |
| 21.8306039 | 21.7173558 | 21.7634125 | 21.7960728 | -0.005425526 | 0.959967207 |
| 21.1933775 | 21.6404614 | 21.5949567 | 21.7075104 | 0.272535777 | 0.038934661 |
| NA | 16.2288566 | NA | NA | 0.026600528 | 0.824828485 |
| 14.5104806 | 15.310473 | 15.6925 | 15.221912 | 0.595017778 | 0.022214235 |
| 19.7083036 | 19.8845157 | 20.0363391 | 19.7762154 | 0.183190027 | 0.05820712 |
| 22.4953828 | 22.328475 | 22.4165801 | 22.4982852 | -0.076036321 | 0.306073977 |
| 19.125406 | 18.8772915 | 18.9112522 | 18.8375648 | -0.154324253 | 0.066702433 |
| 19.7048405 | 19.5727553 | 19.5873601 | 19.6804757 | -0.050743097 | 0.506925018 |
| 18.737782 | 18.6182737 | 18.5106843 | 18.4446757 | -0.164833996 | 0.028459285 |
| 20.8804277 | 20.9223622 | 20.8394394 | 20.8431068 | -0.02231117 | 0.657333726 |
| 23.7884883 | 23.6620534 | 23.7421362 | 23.7533054 | -0.02095729 | 0.650497418 |
| 20.6252042 | 20.472994 | 20.6349649 | 20.5197216 | -0.017916982 | 0.777534633 |
| 18.3291204 | 17.9857291 | 18.0608572 | 18.1418258 | -0.23317054 | 0.002313016 |
| 18.4989197 | 18.2635682 | 18.4278658 | 18.5106849 | -0.029009682 | 0.702556367 |
| 25.0794843 | 25.1803551 | 25.1811517 | 25.1583386 | 0.267608326 | 0.044606447 |
| 25.5034703 | 25.303932 | 25.4607657 | 25.4371511 | -0.011009504 | 0.833964455 |
| 27.3393468 | 27.3783649 | 27.4063521 | 27.390587 | 0.129703358 | 0.222851099 |
| 21.5726814 | 21.5153975 | 21.5941161 | 21.5690136 | -0.035945964 | 0.509204827 |
| 16.8241655 | 16.7082139 | 16.7995788 | 16.8500393 | 0.08495919 | 0.432621399 |
| 19.7301688 | 19.7828888 | 19.751472 | 19.7449621 | -0.023145797 | 0.715132708 |
| 21.8700149 | 21.7207314 | 21.783809 | 21.7852318 | -0.05867176 | 0.172785552 |
| 21.5721056 | 21.4689485 | 21.4967495 | 21.5470714 | -0.011365696 | 0.791407275 |
| 21.9069325 | 21.8524128 | 21.9789382 | 21.9837865 | 0.026706162 | 0.600381087 |

| | | | | | |
|---|---|---|---|---|---|
| 20.7962308 | 20.7256752 | 20.8087408 | 20.9134683 | -0.016126747 | 0.873230631 |
| 20.049787 | 19.8285901 | 20.0605692 | 20.0993333 | -0.016625864 | 0.879830227 |
| 21.5912041 | 21.4251004 | 21.5376175 | 21.5426012 | -0.044947145 | 0.352431927 |
| 22.1153709 | 22.1347407 | 22.0727044 | 22.1091094 | -0.002802568 | 0.947261599 |
| 21.8744394 | 21.7167009 | 21.783707 | 21.8728218 | 0.030989451 | 0.584634377 |
| 17.8547663 | 17.8600904 | 18.0988064 | 17.9214312 | 0.102849534 | 0.390293041 |
| 19.5919281 | 19.4670622 | 19.3544366 | 19.582803 | -0.095495415 | 0.090032233 |
| 21.6821574 | 21.6702994 | 21.6529496 | 21.6577746 | -0.02356232 | 0.559470032 |
| 19.893601 | 19.6055815 | 19.7743947 | 19.721226 | 0.028663793 | 0.702567439 |
| 16.3925654 | 16.366608 | 16.2866869 | 16.3733313 | 0.041083592 | 0.631123041 |
| 21.868883 | 21.7937326 | 21.8448073 | 21.9325584 | 0.020439645 | 0.721772118 |
| 17.3206252 | 17.4989059 | 17.5635292 | 17.5076587 | 0.055069301 | 0.647353505 |
| 24.3980359 | 24.2587789 | 24.4108175 | 24.432603 | -0.03202415 | 0.638463326 |
| 25.3576549 | 25.3090703 | 25.5022979 | 25.4193734 | 0.097047236 | 0.112008272 |
| 15.4380663 | 15.5498539 | 15.5636298 | 15.4704662 | 0.196603177 | 0.161721284 |
| NA | 14.1739731 | NA | NA | -0.686759718 | 0.252786536 |
| 24.0969978 | 23.9014014 | 24.0614103 | 24.0654505 | -0.038065718 | 0.432530055 |
| 23.7113211 | 23.7133929 | 23.7582919 | 23.8065461 | 0.036278739 | 0.411230921 |
| 18.2751749 | 18.0960391 | 18.1417216 | 18.1847703 | -0.141623816 | 0.053077069 |
| 16.0474662 | NA | 16.1799422 | 16.3089943 | 0.17661674 | 0.024093441 |
| 16.6379618 | 16.6087705 | 16.7672239 | 16.7430651 | 0.13499852 | 0.189857712 |
| 21.1441827 | 21.035256 | 21.0640016 | 21.0874392 | -0.035266093 | 0.357114331 |
| 16.0689004 | 15.57423 | 15.8735326 | 15.3284194 | -0.356843124 | 0.039202782 |
| 20.9372049 | 20.6363931 | 20.849694 | 20.9376983 | -0.101584766 | 0.191697966 |
| 19.7931241 | 19.825238 | 19.7878196 | 19.9280592 | 0.062031336 | 0.345872496 |
| 16.6831022 | 16.4885353 | 16.5118478 | 16.7582786 | -0.007967245 | 0.940196479 |
| NA | 11.1670878 | NA | NA | -0.098797098 | 0.448624097 |
| 20.1751081 | 20.1530755 | 20.1760748 | 20.1456198 | -0.031953996 | 0.779671426 |
| 20.3306109 | 20.0704264 | 20.0276804 | 20.1370716 | -0.164322555 | 0.032377511 |
| 17.8248162 | 17.588765 | 17.7522733 | 17.6193769 | -0.185077218 | 0.010070503 |
| 22.3835028 | 22.1914663 | 22.275929 | 22.3680254 | -0.035007717 | 0.578573453 |
| 20.9406048 | 20.7878935 | 20.6737957 | 20.7055695 | -0.130728643 | 0.034619024 |
| 21.2942983 | 21.3439094 | 21.2598433 | 21.2583753 | -0.061877324 | 0.123479928 |
| 20.3318453 | 20.216995 | 20.2758449 | 20.2779563 | -0.065601856 | 0.43347493 |
| 18.3287819 | 18.3882033 | 18.3832796 | 18.2547716 | 0.049666076 | 0.563429332 |
| 17.9882612 | 18.1173643 | 17.9074861 | 18.1080493 | 0.205473667 | 0.030733898 |
| 21.8800412 | 21.8441077 | 22.0083752 | 21.9225078 | 0.053173214 | 0.373553079 |
| 24.1110426 | 23.6846393 | 23.8959919 | 23.9225097 | -0.256833838 | 0.160670512 |
| 21.7344664 | 21.7245318 | 21.7830208 | 21.7102123 | 0.02060684 | 0.821828065 |
| 21.366097 | 21.3954711 | 21.4548299 | 21.4637713 | 0.067028628 | 0.3362058 |
| 21.3697026 | 21.4256649 | 21.4129706 | 21.4034106 | 0.064176274 | 0.170218172 |
| 19.2561393 | 19.1652017 | 19.1189643 | 19.3119907 | 0.013499192 | 0.813595866 |

| | | | | | |
|---|---|---|---|---|---|
| 22.1086268 | 22.0070086 | 22.1234546 | 22.1383046 | 0.007937954 | 0.891960686 |
| 19.4615568 | 19.4854481 | 19.4618562 | 19.5430455 | 0.053270352 | 0.384055067 |
| 17.6086825 | 18.203911 | 17.8354841 | 17.797606 | 0.270712745 | 0.03699797 |
| 18.6920356 | 18.9049886 | 18.9508709 | 18.9564056 | 0.162569254 | 0.015907283 |
| 20.6502073 | 20.4027966 | 20.5598286 | 20.6084573 | -0.110371286 | 0.536094217 |
| NA | 12.6876448 | NA | NA | -0.512080122 | 0.052934763 |
| 25.3072705 | 25.1463018 | 25.4134222 | 25.3213763 | -0.029684507 | 0.792519905 |
| 24.5160598 | 24.258853 | 24.457974 | 24.4160008 | -0.160908779 | 0.024138193 |
| 20.1901417 | 20.082711 | 20.241387 | 20.1798317 | -0.032771534 | 0.589133534 |
| 19.4471507 | 19.2529521 | 19.309979 | 19.3185306 | -0.085084948 | 0.100558385 |
| 23.8386552 | 23.9376215 | 23.986144 | 24.0067563 | 0.131404067 | 0.071152248 |
| 18.6721612 | 18.7938695 | 18.8998064 | 18.9170809 | 0.076742729 | 0.296013622 |
| 19.5228235 | 19.7126543 | 19.8229913 | 19.950461 | 0.19435226 | 0.027076333 |
| 23.3557922 | 23.3540415 | 23.4519907 | 23.4763651 | 0.084856122 | 0.40474286 |
| 14.3944313 | 14.6593003 | 14.6804876 | 14.5822772 | 0.21152448 | 0.268135122 |
| 15.8255216 | 15.936094 | 16.0619779 | 16.2350409 | 0.221581315 | 0.336572641 |
| 20.803065 | 20.7791872 | 20.8478246 | 20.8198383 | -0.009896033 | 0.898457807 |
| 20.2224602 | 20.0427192 | 20.2240008 | 20.0728817 | -0.09090558 | 0.270184602 |
| 19.1202332 | 19.1364657 | 19.1942398 | 19.0635978 | 0.015522472 | 0.768328759 |
| 19.3087164 | 19.1878929 | 19.4789473 | 19.5708847 | 0.205360615 | 0.343959105 |
| 20.3570522 | 20.4096123 | 20.6076381 | 20.1002694 | 0.331442861 | 0.130632916 |
| 16.4447738 | 15.7308837 | 16.1398265 | 15.6479107 | -0.413790549 | 0.010845577 |
| 21.0755768 | 21.0475936 | 21.2025791 | 21.2539389 | 0.03248643 | 0.778019076 |
| 21.5698102 | 21.4584522 | 21.5676318 | 21.393 | -0.136056057 | 0.036956886 |
| 20.6326705 | 20.4712645 | 20.4778964 | 20.6143346 | -0.090949344 | 0.100818077 |
| 22.3650783 | 22.3215543 | 22.4017525 | 22.3818686 | 0.058130487 | 0.23328801 |
| 22.4084243 | 22.3148303 | 22.3450095 | 22.2871079 | -0.036194442 | 0.388562366 |
| 20.5718382 | 20.4680283 | 20.4767116 | 20.6036133 | -0.045569574 | 0.41923471 |
| 21.5337349 | 21.4870015 | 21.7150218 | 21.6991795 | 0.118009689 | 0.163759691 |
| 16.5266883 | 16.7553232 | 17.0743306 | 16.7783375 | 0.212430964 | 0.094450461 |
| 16.6718118 | 16.614818 | 16.7542745 | 16.5669608 | -0.040953668 | 0.70239292 |
| 17.7848576 | 17.7495765 | 17.7348035 | 17.6951057 | -0.088304999 | 0.316816862 |
| 19.0133258 | 19.1378442 | 18.9051719 | 19.152574 | 0.013432749 | 0.874504174 |
| 20.1542024 | 20.2029 | 19.9558342 | 20.1561285 | -0.062771393 | 0.368408972 |
| 20.1727983 | 20.1273688 | 20.1003493 | 20.1413818 | -0.009929816 | 0.864144421 |
| 21.4777134 | 21.2731204 | 21.3454188 | 21.3004602 | -0.082581291 | 0.127951908 |
| 20.8184209 | 20.7775489 | 20.7500419 | 20.7726694 | -0.034507535 | 0.419274815 |
| 19.7852681 | 19.7445373 | 19.7384821 | 19.8047047 | 0.009078962 | 0.834679951 |
| 21.856688 | 21.7593785 | 21.8609017 | 21.8458098 | 0.043199061 | 0.396639388 |
| 20.340971 | 20.2774669 | 20.3691413 | 20.3886273 | 0.045986029 | 0.380087778 |
| 21.5054377 | 21.431415 | 21.4937559 | 21.5139395 | 0.024218021 | 0.651357483 |
| 23.4136613 | 23.3104955 | 23.3467196 | 23.3827587 | 0.00716369 | 0.897034119 |

| | | | | | |
|---|---|---|---|---|---|
| 23.1975181 | 23.1179545 | 23.2290006 | 23.2425867 | 0.065874173 | 0.279689773 |
| 18.939531 | 18.7858354 | 18.8610719 | 18.9863094 | 0.002691553 | 0.96577848 |
| 17.5674415 | 17.6165678 | 17.6201695 | 17.7009785 | 0.059878125 | 0.453991249 |
| 17.6914892 | 17.6855703 | 17.8849193 | 17.9027472 | 0.061515097 | 0.469263119 |
| 17.7213807 | 17.6938875 | 17.8206466 | 17.8168649 | 0.070189244 | 0.424323294 |
| 14.6985336 | 14.8897683 | 14.7013559 | 14.9425502 | 0.169120278 | 0.188357395 |
| 13.465839 | 13.4862817 | 13.854443 | 13.441019 | 0.145065657 | 0.338492834 |
| 16.3491204 | 15.7245085 | 16.1651095 | 15.8696786 | -0.032470458 | 0.877633892 |
| 18.2372959 | 18.2159237 | 18.1270273 | 18.1146435 | -0.088802354 | 0.351430165 |
| 20.4797088 | 20.2449917 | 20.3749731 | 20.529303 | -0.081678523 | 0.271540153 |
| 23.285387 | 23.2150553 | 23.275817 | 23.2930475 | 0.030419568 | 0.579543872 |
| 21.9021903 | 21.693029 | 22.0439493 | 21.90881 | 0.08040744 | 0.359979895 |
| 17.2074516 | 17.2655112 | 17.2080575 | 17.2729771 | 0.073341024 | 0.406865184 |
| 21.1947225 | 21.1766469 | 21.1657998 | 21.1755958 | 0.103391748 | 0.40810473 |
| NA | 14.8225267 | NA | NA | 0.001562981 | 0.991826789 |
| 20.2524892 | 20.0741625 | 20.1821352 | 20.1631084 | -0.048198946 | 0.416831919 |
| 19.7372551 | 19.528563 | 19.7339114 | 19.6672657 | -0.090292737 | 0.239578411 |
| 14.154424 | NA | 14.0152745 | 14.0473571 | -0.184451929 | 0.120042236 |
| 21.3351308 | 21.3852741 | 21.2650741 | 21.433136 | -0.015064516 | 0.862587058 |
| 20.5868895 | 20.5592155 | 20.538024 | 20.6255517 | 0.013674687 | 0.806075371 |
| 21.1470597 | 21.2779369 | 21.2985704 | 21.1293376 | 0.049560887 | 0.51531616 |
| 17.5768975 | 17.6946924 | 17.8036434 | 17.5750011 | 0.042345878 | 0.588529454 |
| 16.868314 | 16.8646169 | 17.059703 | 16.9009255 | 0.086555203 | 0.407860501 |
| 18.1809503 | 18.0691704 | 18.1532643 | 18.3216089 | 0.027733908 | 0.811577185 |
| 15.9009217 | 15.4635628 | 15.8110773 | 15.8105218 | -0.048147301 | 0.733271107 |
| 17.5851525 | 17.3287135 | 16.70828 | 16.7960743 | -0.496917228 | 0.01926744 |
| 19.5840137 | 19.7862973 | 19.6363258 | 19.7499135 | 0.092475903 | 0.231613342 |
| 20.836098 | 20.8516729 | 20.8512334 | 20.8239888 | 0.025384889 | 0.595845151 |
| NA | 17.6415139 | NA | NA | 0.097337049 | 0.738779906 |
| 18.6233011 | 18.5316356 | 18.7345139 | 18.9462077 | 0.125353491 | 0.229511806 |
| 16.7199147 | 17.0063549 | 16.601436 | 16.7566008 | 0.025629174 | 0.861628728 |
| NA | 15.1431075 | NA | NA | 0.108503425 | 0.580425194 |
| 20.4192557 | 20.3956712 | 20.4157768 | 20.3749947 | -0.009160777 | 0.882427044 |
| 21.2005518 | 21.0614631 | 21.1653295 | 21.2213485 | -0.030810552 | 0.580983844 |
| 23.0890071 | 22.9878257 | 23.1205933 | 23.1038078 | 0.014078515 | 0.773683053 |
| 22.1131296 | 22.0311508 | 22.0250123 | 22.251989 | 0.002070303 | 0.977653374 |
| 21.4492645 | 21.3511244 | 21.5853175 | 21.5023809 | 0.032071527 | 0.666821651 |
| 19.1078563 | 19.2675069 | 19.3704315 | 19.291696 | 0.153082893 | 0.076782825 |
| 17.1404781 | 17.0909458 | 16.8087889 | 17.2237419 | -0.171487265 | 0.282321015 |
| 18.2060625 | 18.1580819 | 18.1718578 | 18.1479287 | -0.099517848 | 0.354314685 |
| 16.9724422 | 16.7709685 | 16.9366076 | 16.6609675 | -0.138740859 | 0.142988595 |
| 20.8413943 | 20.8639368 | 20.8905092 | 20.8776302 | 0.007700322 | 0.861484566 |

| | | | | | |
|---|---|---|---|---|---|
| 25.2582509 | 25.2602584 | 25.2883395 | 25.3424973 | 0.0526295 | 0.229809073 |
| 20.6380866 | 20.8081092 | 20.7885439 | 20.8433362 | 0.098323808 | 0.150466287 |
| 22.1618564 | 22.0479711 | 22.1521677 | 22.1263782 | 0.008491732 | 0.904909383 |
| 19.4374002 | 19.4982547 | 19.5614274 | 19.5272974 | -0.015113672 | 0.849913207 |
| 17.5094365 | 17.5495767 | 17.638089 | 17.8105703 | 0.157455451 | 0.08876146 |
| 22.4748437 | 22.2519585 | 22.3628214 | 22.4574942 | -0.015387989 | 0.886106624 |
| 20.7015426 | 20.5531002 | 20.5843156 | 20.6625735 | -0.071184187 | 0.383913125 |
| 21.1317131 | 21.1419918 | 21.2152475 | 21.1520231 | -0.002056105 | 0.979099764 |
| 19.7984595 | 19.6089019 | 19.7308877 | 19.6533811 | -0.045526194 | 0.510706445 |
| 22.5575315 | 22.3876994 | 22.492481 | 22.4660377 | -0.0468497 | 0.370571094 |
| 20.90972 | 20.7928837 | 20.8326834 | 20.9521476 | 0.029581714 | 0.598960963 |
| 18.7894175 | 18.6225322 | 18.7972862 | 18.7347927 | -0.007541073 | 0.896157898 |
| 20.7661134 | 20.6337765 | 20.6520325 | 20.8569709 | -0.014169501 | 0.859632287 |
| 15.6307725 | 15.3870632 | 15.2391376 | 15.4306835 | -0.134821223 | 0.194881359 |
| 18.9290813 | 18.7101494 | 18.828249 | 18.8778342 | -0.13452676 | 0.138762452 |
| 24.7465725 | 24.6823105 | 24.7134126 | 24.677242 | -0.008838945 | 0.853791243 |
| 22.8471284 | 22.7952283 | 22.8433085 | 22.9045163 | 0.025458521 | 0.482799697 |
| 22.5552708 | 22.4607749 | 22.5806007 | 22.5924491 | 0.020067401 | 0.659699264 |
| 18.380523 | 18.5994171 | 18.5714543 | 18.6580846 | 0.292499308 | 0.007517799 |
| 18.3067107 | 18.1081415 | 18.1965595 | 18.3826681 | -0.098909558 | 0.175200999 |
| NA | 15.5385268 | NA | NA | -0.290984899 | 0.105533165 |
| 16.2862431 | NA | 16.3433949 | 16.1714273 | 0.018711939 | 0.937654895 |
| 18.2961486 | 18.2257595 | 18.3007178 | 18.3953852 | 0.021658254 | 0.816245144 |
| 19.3989312 | 19.3872854 | 19.4865894 | 19.4305047 | -0.015076065 | 0.895067765 |
| 21.2021486 | 21.045259 | 21.1889507 | 21.1264325 | -0.046327688 | 0.401685025 |
| 20.595918 | 20.4908918 | 20.5931409 | 20.635168 | 0.019081793 | 0.770732279 |
| 22.1021105 | 22.0053618 | 21.9453716 | 22.0624844 | -0.009442251 | 0.882442309 |
| 21.8182606 | 21.7538644 | 21.8629813 | 21.9493809 | 0.048201709 | 0.359780915 |
| 22.6127511 | 22.5819986 | 22.6881833 | 22.7070652 | 0.077912153 | 0.10256215 |
| 19.583553 | 19.4767249 | 19.5913685 | 19.547223 | -0.025373256 | 0.721289262 |
| 18.3705712 | 18.3169877 | 18.3745159 | 18.4683145 | -0.012822113 | 0.856876841 |
| 21.0942322 | 20.9150059 | 21.126078 | 21.1894588 | 0.095191325 | 0.245478685 |
| 18.6233578 | 18.5578195 | 18.6982862 | 18.6797645 | 0.042894458 | 0.415616714 |
| 19.6477578 | 19.8076824 | 19.8589716 | 19.8089177 | -0.001930818 | 0.980999763 |
| 18.6158735 | 18.5025654 | 18.5836976 | 18.7173598 | -0.016983499 | 0.851825859 |
| 21.0123002 | 20.949853 | 21.2984747 | 21.2252634 | 0.054793928 | 0.641323341 |
| 18.152537 | 18.0673288 | 18.0304839 | 17.9908804 | -0.074579167 | 0.233600251 |
| 18.3713303 | 18.798841 | 18.6706984 | 18.717898 | 0.392406581 | 0.095654098 |
| 19.5981245 | 19.3452854 | 19.6574418 | 19.5241736 | -0.143066409 | 0.040921102 |
| 19.4577513 | 19.4239743 | 19.5830501 | 19.6382858 | 0.045888163 | 0.546517055 |
| 15.7160737 | 15.9178401 | 15.869637 | 15.8746887 | 0.166397851 | 0.086607202 |
| 19.4836483 | 19.5356044 | 19.4959665 | 19.5075842 | 0.03092418 | 0.665016153 |

| | | | | | |
|---|---|---|---|---|---|
| 17.980422 | 17.7920969 | 17.50622 | 17.8896917 | -0.130833705 | 0.433435587 |
| 16.742798 | 16.7480654 | 16.6869335 | 16.5504802 | -0.078670228 | 0.347272542 |
| 20.7896169 | 20.9294935 | 20.8985458 | 20.817638 | 0.031138325 | 0.492218657 |
| 22.4202118 | 22.3169755 | 22.458756 | 22.4905245 | -0.005920082 | 0.928886591 |
| 19.219344 | 19.0965507 | 19.3011591 | 19.3757687 | 0.055293423 | 0.510670508 |
| 19.7913976 | 19.7516721 | 19.8572518 | 19.9167154 | 0.021840388 | 0.830148503 |
| 20.0170244 | 20.0252022 | 20.0260016 | 20.1450221 | 0.111883462 | 0.24282729 |
| 20.8382643 | 20.8670945 | 20.7669536 | 20.8190335 | 0.002010562 | 0.96735297 |
| 20.3168316 | 20.1504051 | 20.2341318 | 20.29673 | -0.0446478 | 0.333294724 |
| 21.0656276 | 21.022936 | 20.9443701 | 21.0827539 | -0.018973063 | 0.649097048 |
| 20.5672995 | 20.4232753 | 20.6007834 | 20.5972199 | 1.10E-04 | 0.998403401 |
| 18.0053505 | 18.0519988 | 18.162294 | 18.1855046 | 0.18899822 | 0.003941601 |
| 19.8664842 | 19.7242067 | 19.8963021 | 19.9028244 | 0.027301692 | 0.680942774 |
| 23.9677421 | 23.9155747 | 24.0413245 | 23.988787 | 0.051220585 | 0.484717475 |
| 22.3781154 | 22.2799992 | 22.3363728 | 22.3192583 | 0.06084437 | 0.437634555 |
| 21.3812141 | 21.3342748 | 21.321245 | 21.328856 | -0.005105435 | 0.912841886 |
| 19.4301258 | 19.3227039 | 19.520039 | 19.4946033 | 0.00960286 | 0.869245649 |
| 19.7131418 | 19.6093581 | 19.7238101 | 19.6830741 | -3.24E-04 | 0.995704011 |
| 19.0605724 | 19.1371681 | 19.106739 | 19.2216605 | 0.052020892 | 0.398695961 |
| 16.3716088 | NA | 16.7766056 | 16.6863879 | 0.363103115 | 0.094430359 |
| 20.4795483 | 20.5252088 | 20.4516057 | 20.4556887 | -0.060374039 | 0.357522422 |
| 21.8819602 | 21.8434095 | 21.8910309 | 21.8917934 | -0.001241083 | 0.980763778 |
| 20.9861415 | 20.8243267 | 21.0121772 | 20.9619177 | 0.004962419 | 0.940902987 |
| NA | 14.8878753 | NA | NA | -0.346728211 | 0.375515792 |
| 24.6286259 | 24.4768562 | 24.6044809 | 24.6364599 | -0.009885649 | 0.875212327 |
| 20.1069237 | 20.1662937 | 20.2098272 | 20.1847217 | 0.025820658 | 0.456052889 |
| 19.5060975 | 19.4090556 | 19.4359256 | 19.5172996 | -0.039109899 | 0.443460181 |
| 17.7785823 | 17.5950719 | 17.7466684 | 17.6689478 | -0.04377774 | 0.56130143 |
| 20.3304622 | 20.3179492 | 20.5637449 | 20.4570901 | 0.078020587 | 0.334428748 |
| 19.7037948 | 19.4762134 | 19.7209048 | 19.6847838 | 0.016214829 | 0.90388706 |
| 21.3251979 | 21.3400765 | 21.5115775 | 21.3937555 | 0.115215161 | 0.451530812 |
| 26.0388505 | 25.7210635 | 25.8661682 | 25.9474793 | -0.079268464 | 0.310462073 |
| 22.4814803 | 22.3006112 | 22.4292993 | 22.4797753 | -0.043636582 | 0.532164767 |
| 18.8415981 | 18.729039 | 18.8170927 | 18.8131101 | -0.09709112 | 0.176162726 |
| 22.1996344 | 22.1002809 | 22.1856906 | 22.1698899 | -0.019485259 | 0.660890827 |
| 22.0318258 | 21.9422773 | 22.0051522 | 21.9742075 | -0.075506397 | 0.167472693 |
| 19.3034358 | 19.187218 | 19.1789069 | 19.2629976 | -0.022074149 | 0.759097843 |
| 21.6184014 | 21.6695544 | 21.4860825 | 21.3325879 | -0.10769048 | 0.308062262 |
| 19.7732804 | 19.758305 | 19.866748 | 19.9253121 | 0.049682429 | 0.35706284 |
| 20.3985304 | 20.3755556 | 20.3890531 | 20.4266301 | 0.036596998 | 0.485997457 |
| 19.1603251 | 19.1002202 | 18.9642179 | 19.1478227 | -0.017079263 | 0.835899545 |
| 23.9106036 | 23.8459805 | 23.919577 | 23.8832993 | -0.017272549 | 0.775789244 |

| | | | | | |
|---|---|---|---|---|---|
| 22.7783986 | 22.8471569 | 22.8760221 | 22.8492489 | 0.058217292 | 0.183591302 |
| 19.4612896 | 19.4906209 | 19.4185407 | 19.5318873 | 0.125858103 | 0.055735424 |
| 20.5048192 | 20.3679637 | 20.5274817 | 20.5055912 | -0.030210572 | 0.707829259 |
| 19.5954436 | 19.3631382 | 19.5575367 | 19.4992612 | -0.040666963 | 0.603081354 |
| 20.6380529 | 20.3889223 | 20.4968666 | 20.5322807 | -0.065272089 | 0.324427537 |
| 23.5538276 | 23.6609579 | 23.6500371 | 23.5885766 | 0.054192127 | 0.343615351 |
| 22.6856587 | 22.5725648 | 22.7135536 | 22.6632158 | -0.043058038 | 0.437003575 |
| 22.682764 | 22.60257 | 22.6157806 | 22.5040054 | -0.056721735 | 0.254740444 |
| 25.8070182 | 25.7621201 | 25.7626342 | 25.7909605 | -0.005664473 | 0.895818055 |
| 21.7931474 | 21.7915309 | 21.8691108 | 21.8691589 | 0.023653063 | 0.59861082 |
| 20.6436476 | 20.6839464 | 20.5674683 | 20.7163569 | 0.063238779 | 0.214316758 |
| 23.5452546 | 23.4565979 | 23.5161442 | 23.5525732 | 0.044930884 | 0.415438941 |
| 18.8972577 | 18.5955666 | 18.9022069 | 18.8084194 | -0.002878135 | 0.967973941 |
| 22.9396903 | 22.8079684 | 22.915684 | 22.9543473 | 1.32E-04 | 0.998582925 |
| 21.3140078 | 21.2803137 | 21.4554917 | 21.5039678 | 0.172629 | 0.15268296 |
| 16.4372206 | 16.325524 | 16.7288162 | 16.6441968 | 0.107118696 | 0.451815051 |
| 16.9681343 | 16.6704083 | 17.0496417 | 16.4767952 | -0.202730179 | 0.15487517 |
| 16.3182528 | NA | 16.3454643 | 16.25759 | -0.045883452 | 0.609942394 |
| 24.1974662 | 24.0366575 | 24.0048323 | 24.1218619 | -0.048291978 | 0.491148631 |
| 23.9015499 | 23.8265525 | 23.8638333 | 23.9119706 | -0.020305242 | 0.719004847 |
| 21.8672484 | 21.7917746 | 21.8801116 | 21.8993452 | -0.014544845 | 0.758112528 |
| 19.0995846 | 18.9945469 | 19.0829471 | 19.0653668 | -0.053739331 | 0.246206192 |
| 24.8659986 | 24.7446214 | 24.8812248 | 24.8316151 | -0.022451946 | 0.564046798 |
| 20.5013001 | 20.3987323 | 20.4442723 | 20.3720137 | -0.061642108 | 0.241706506 |
| 18.8680867 | 18.8284637 | 18.7996457 | 18.8985843 | 0.036446627 | 0.510262277 |
| 18.0247561 | 18.0255402 | 18.0196449 | 18.1534231 | 0.051791367 | 0.383409998 |
| 19.1266478 | 19.14519 | 19.120778 | 19.1418499 | -0.023769307 | 0.722983671 |
| 19.1932629 | 19.331857 | 19.4527532 | 19.264207 | 0.067908666 | 0.391778641 |
| 20.2077297 | 19.9427107 | 20.1099106 | 20.0207499 | -0.030336391 | 0.717423161 |
| 18.7911611 | 18.8483405 | 18.6009246 | 18.6079698 | 0.058553854 | 0.633153763 |
| 24.0852603 | 23.9591406 | 23.9887445 | 23.8032592 | -0.148809473 | 0.052218414 |
| 20.9932947 | 20.8621483 | 21.0458793 | 21.017292 | -0.012295258 | 0.818606244 |
| 20.3588245 | 20.1919948 | 20.2467245 | 20.184721 | -0.094082791 | 0.061789177 |
| 18.7930894 | 18.6970686 | 18.3969108 | 18.5133037 | -0.223360794 | 0.061652156 |
| 17.865361 | 17.6330601 | 17.8755176 | 17.6974155 | -0.131532001 | 0.194885551 |
| 21.3693848 | 21.3676369 | 21.4082222 | 21.2748482 | -0.0161381 | 0.760987075 |
| 20.4262675 | 20.3934758 | 20.4816989 | 20.4309714 | 0.015341656 | 0.756467036 |
| 19.0925708 | 19.2215212 | 19.2273173 | 19.1827009 | 0.152211065 | 0.163309334 |
| 16.7552702 | 16.4715805 | 16.8502294 | 16.8114933 | -0.053398734 | 0.646712506 |
| NA | 15.4530721 | NA | NA | 0.548058973 | 0.047889144 |
| 20.418052 | 20.3820472 | 20.4439408 | 20.3637874 | 0.004396974 | 0.922606932 |
| 21.0686841 | 21.0404627 | 21.058156 | 21.0878597 | -0.020741183 | 0.64556278 |

| | | | | | |
|---|---|---|---|---|---|
| 16.0504806 | NA | 16.1299227 | 15.9656445 | -0.017289322 | 0.92852134 |
| 16.422318 | 16.0268305 | 15.7023307 | 16.3565813 | -0.194552932 | 0.437488431 |
| 22.0926875 | 21.8858732 | 21.9875876 | 21.9719277 | -0.065905098 | 0.274376808 |
| 16.5639238 | 15.6756003 | 16.5938927 | 16.1448883 | -0.21897118 | 0.261865265 |
| 22.4598567 | 22.3376955 | 22.3354199 | 22.3347797 | -0.094281796 | 0.074210814 |
| 18.9602696 | 18.8148192 | 18.967751 | 18.9885827 | 0.016065713 | 0.854451945 |
| 18.7359665 | 18.7238719 | 18.6864347 | 18.6972821 | -0.014090888 | 0.810137014 |
| NA | 15.5118135 | NA | NA | 0.018078062 | 0.882017619 |
| 19.4458811 | 19.4339157 | 19.5192683 | 19.4504218 | -0.008430015 | 0.860503758 |
| 19.3852576 | 19.2742741 | 19.3211577 | 19.3596721 | -0.096764765 | 0.198159402 |
| NA | 14.4489247 | NA | NA | 0.08617694 | 0.650252809 |
| 20.3627816 | 20.4213013 | 20.4147391 | 20.4526295 | 3.52E-04 | 0.995378145 |
| 20.829003 | 20.6892897 | 20.9459374 | 20.9020943 | 0.042484255 | 0.729453091 |
| 20.2322574 | 20.1900837 | 20.2884905 | 20.2474124 | 0.011308393 | 0.806761051 |
| 24.8408085 | 24.7430682 | 24.7616958 | 24.7206682 | -0.060587962 | 0.172642879 |
| 24.2592958 | 24.1389649 | 24.1947171 | 24.210953 | -0.023139804 | 0.466384578 |
| 19.5525298 | 19.4216969 | 19.4577712 | 19.4862275 | -0.039592111 | 0.524370077 |
| 19.2477986 | 19.1743243 | 19.2479732 | 19.4589351 | 0.058464378 | 0.486539939 |
| 19.8022073 | 19.7846813 | 19.7803189 | 19.8086169 | -0.047740223 | 0.610544769 |
| 21.6620701 | 21.4625021 | 21.6067696 | 21.6120019 | -0.073949075 | 0.182261619 |
| 15.3664008 | 15.7929572 | 15.7074344 | 15.7117279 | 0.284859034 | 0.010686161 |
| 20.7601815 | 20.7849487 | 20.8755342 | 20.7627333 | 0.009434701 | 0.878893667 |
| 20.9255519 | 20.7751273 | 20.9133542 | 20.9803036 | -0.003484854 | 0.951567059 |
| 22.1647987 | 22.1184597 | 22.1765239 | 22.0968049 | -0.031992784 | 0.542860992 |
| 21.2418079 | 21.1635207 | 21.184609 | 21.1140503 | -0.024414401 | 0.610336953 |
| 20.7861031 | 20.8142748 | 20.8045497 | 20.7438327 | 0.021023654 | 0.601114344 |
| 20.8058358 | 20.7092634 | 20.8531803 | 20.8590125 | 0.028558452 | 0.562035208 |
| 19.9598877 | 19.8367227 | 19.9625339 | 20.020176 | 0.028637789 | 0.636718813 |
| 24.0269207 | 23.9294736 | 24.0598912 | 24.0168066 | 0.046511543 | 0.48317044 |
| 21.5346167 | 21.4755231 | 21.5837614 | 21.5473108 | 0.029397948 | 0.727105592 |
| 16.2975863 | 16.4554659 | 16.5500956 | 16.5073609 | 0.140283399 | 0.150949993 |
| 24.995967 | 24.8981172 | 25.0226553 | 24.9512867 | -0.014378606 | 0.76164325 |
| 20.0378727 | 19.9241589 | 20.1150867 | 20.0557964 | 0.086305905 | 0.239137816 |
| 17.8671994 | 17.5850117 | 17.7409202 | 17.8209495 | -0.025189227 | 0.75620891 |
| 18.6833197 | 18.7552231 | 18.5721435 | 18.6201526 | -0.133277568 | 0.168846073 |
| 21.0110603 | 20.7438438 | 20.8443793 | 20.7426372 | -0.166908803 | 0.037798605 |
| 20.1785017 | 20.0179261 | 20.1923905 | 20.1614162 | -0.03168766 | 0.667602893 |
| 19.0417849 | 18.9533082 | 19.059762 | 19.0822874 | -0.01247651 | 0.873111965 |
| 22.8240857 | 22.416523 | 22.552813 | 22.6176094 | -0.184112506 | 0.098823935 |
| 29.8448937 | 29.7762167 | 29.779579 | 29.7748668 | 0.022490436 | 0.784089465 |
| 21.6001609 | 21.5154649 | 21.6417036 | 21.6620207 | 0.033404819 | 0.554046517 |
| 17.7870885 | 17.5925308 | 17.6033607 | 17.5092293 | -0.203727428 | 0.267051046 |

| | | | | | |
|---|---|---|---|---|---|
| 21.3572161 | 21.2290556 | 21.2329791 | 21.3688737 | -0.015609651 | 0.847648389 |
| 16.7580706 | 16.6457494 | 16.6112294 | 16.7038125 | -0.251539234 | 0.003190846 |
| 23.4768355 | 23.4653003 | 23.5354181 | 23.504617 | 0.063294914 | 0.403203266 |
| 20.0084357 | 19.8610525 | 20.0060862 | 19.9750372 | 0.011981327 | 0.860533033 |
| 18.8489867 | 18.7099837 | 18.6408508 | 18.6643483 | -0.144289631 | 0.03462149 |
| 20.1036759 | 20.0614394 | 20.0872854 | 20.0494567 | -0.063094372 | 0.303056398 |
| 25.6531232 | 25.4996914 | 25.5999208 | 25.5919156 | -0.033322887 | 0.583891348 |
| 26.7425858 | 26.6477338 | 26.7526899 | 26.7114504 | 0.016365187 | 0.749540895 |
| 19.9012704 | 19.8824066 | 19.9672159 | 19.9648616 | 0.033301868 | 0.422168524 |
| 21.4491116 | 21.3599389 | 21.4626854 | 21.4636625 | 0.044380974 | 0.381332404 |
| 22.6336011 | 22.5856134 | 22.7817212 | 22.7211429 | 0.048089053 | 0.459545772 |
| 18.0382068 | 18.0069509 | 17.9909204 | 18.1966153 | 0.054028916 | 0.415791525 |
| 20.9008024 | 20.8182214 | 21.0110884 | 20.9494837 | 0.022654694 | 0.736561239 |
| 20.9899809 | 20.8455543 | 20.9103984 | 21.0121651 | 0.00859574 | 0.899623076 |
| 18.5917799 | 18.6838188 | 18.6601538 | 18.6566894 | 0.05032511 | 0.490681121 |
| 21.5789086 | 21.5339423 | 21.7708634 | 21.6919089 | 0.127989292 | 0.131372548 |
| 19.6736784 | 19.7210101 | 19.7697708 | 19.7995573 | 0.122157008 | 0.16689453 |
| 17.3711732 | 17.1066567 | 17.3547313 | 17.3208779 | 0.039266761 | 0.700328976 |
| 22.1451865 | 22.1364759 | 22.3097385 | 22.2794179 | 0.114204212 | 0.298788011 |
| NA | 13.0384775 | NA | NA | -0.366900075 | 0.265513999 |
| 15.3878273 | 14.9560324 | 15.6042674 | 14.8080793 | 0.068612615 | 0.712596504 |
| 18.2867837 | 18.2347349 | 18.4200865 | 18.112179 | -0.076143241 | 0.418082119 |
| 26.7580111 | 26.7214552 | 26.7329511 | 26.7532996 | 0.023531988 | 0.743025418 |
| 20.4059974 | 20.2558827 | 20.4282333 | 20.3629015 | -0.057754449 | 0.407042073 |
| 23.7698623 | 23.5960161 | 23.6007304 | 23.651144 | -0.080629215 | 0.230848091 |
| 23.1646181 | 23.1168366 | 23.1536205 | 23.1733533 | -0.001291989 | 0.981277391 |
| 22.8198281 | 22.8866658 | 22.9181323 | 22.8904613 | 0.01090817 | 0.804752513 |
| 26.5296506 | 26.4826393 | 26.4578134 | 26.474754 | 0.01709305 | 0.68534714 |
| 22.3068947 | 22.2142291 | 22.3388149 | 22.2666538 | -0.036540901 | 0.347900414 |
| 20.7683555 | 20.6968467 | 20.7970854 | 20.7793965 | 0.048958271 | 0.334810221 |
| 20.9124702 | 20.7729593 | 20.9385673 | 20.9548175 | 0.028468597 | 0.582500279 |
| 19.4728512 | 19.342634 | 19.4398706 | 19.407676 | 1.85E-04 | 0.997802828 |
| 19.5912154 | 19.4384654 | 19.6827075 | 19.5959816 | 0.010221303 | 0.885304684 |
| 22.0592643 | 21.9045518 | 21.9860415 | 22.0181514 | 0.004210903 | 0.954015073 |
| 16.4026339 | 16.5739973 | 16.5385418 | 16.6105441 | 0.067345766 | 0.467724396 |
| 18.7012424 | 18.5069176 | 18.6459809 | 18.7970546 | 0.057632361 | 0.566595725 |
| 14.7422024 | 14.9447383 | 15.0150701 | 15.2844466 | 0.222045902 | 0.133811178 |
| 23.5946062 | 23.449797 | 23.7417352 | 23.6718168 | 0.041464832 | 0.790125411 |
| 17.5243069 | 17.382753 | 17.5377369 | 18.0336618 | 0.294615526 | 0.217320864 |
| 23.3770565 | 23.1535657 | 23.3280158 | 23.272666 | -0.078614045 | 0.195761437 |
| 20.4378476 | 20.5570781 | 20.4194791 | 20.5136991 | 0.072402667 | 0.187350643 |
| 19.8279191 | 19.7738947 | 19.9047993 | 19.9201119 | 0.029394139 | 0.691097084 |

| | | | | | |
|---|---|---|---|---|---|
| 19.5698734 | 19.4965515 | 19.6593636 | 19.4961902 | 0.02183224 | 0.770740761 |
| NA | 13.5404172 | NA | NA | -0.207378315 | 0.295754979 |
| 19.9399301 | 19.9478877 | 19.9292282 | 19.9668643 | 0.0307069 | 0.578243162 |
| 18.2656622 | 18.3170027 | 18.2760742 | 18.220321 | 0.042755378 | 0.635863602 |
| 22.0611086 | 21.9768248 | 21.9866664 | 22.0717721 | 0.002102407 | 0.968944659 |
| 21.2290459 | 21.1179063 | 21.2472184 | 21.2441079 | 0.003410582 | 0.966468526 |
| 19.8255829 | 19.7470538 | 19.7372904 | 19.8107571 | 0.009992261 | 0.853191653 |
| 17.4873938 | 17.7442249 | 17.4737044 | 17.6987965 | 0.304893524 | 0.030486227 |
| 20.0739402 | 20.1289053 | 20.3038255 | 20.1774678 | 0.027766585 | 0.748766091 |
| 22.898875 | 22.7727124 | 23.0358671 | 22.88311 | -0.061232852 | 0.474594447 |
| 17.6787096 | 17.4092556 | 17.5869592 | 17.5363117 | -0.089960313 | 0.186982358 |
| 23.0486656 | 22.84027 | 23.0153656 | 23.0020017 | -0.068238791 | 0.189517614 |
| 20.6405517 | 20.5556616 | 20.6665932 | 20.6575715 | -0.015482364 | 0.801532658 |
| 20.8005519 | 20.6990972 | 20.8298753 | 20.919522 | 0.051299353 | 0.456685789 |
| 16.8153702 | 16.6249497 | 16.7397984 | 16.6333245 | -0.092837299 | 0.39500845 |
| 22.308264 | 22.4040688 | 22.4412203 | 22.4078278 | 0.271295434 | 0.276824782 |
| 14.9545515 | 14.5150102 | 14.7832337 | 14.857032 | -0.369495793 | 0.082847362 |
| 17.0685393 | 16.6639254 | 16.7339103 | 16.9219999 | -0.299931436 | 0.043724068 |
| 19.2663458 | 19.2814397 | 19.4735728 | 19.3145943 | -0.042946276 | 0.69555722 |
| 21.273104 | 21.1570237 | 21.1289285 | 21.0771727 | -0.072231958 | 0.441619469 |
| 21.8436127 | 21.8109881 | 21.6886615 | 21.8303065 | -0.050066175 | 0.556550872 |
| 22.136685 | 22.1988737 | 22.2442752 | 22.063634 | -0.115008924 | 0.16136086 |
| 21.4377602 | 21.6288614 | 21.4615716 | 21.3726848 | -0.018092858 | 0.816171214 |
| 20.8397862 | 20.6708768 | 20.8005196 | 20.6924161 | -0.039633825 | 0.571674385 |
| 18.4390883 | 18.4226392 | 18.3139797 | 18.668537 | -0.009976649 | 0.877350431 |
| 19.5201945 | 19.3924243 | 19.3774949 | 19.4501845 | -0.120582537 | 0.069321236 |
| 21.2753138 | 21.2332867 | 21.2531624 | 21.2659929 | -0.015432321 | 0.792022146 |
| 22.4320269 | 22.3243485 | 22.356038 | 22.3727679 | -0.048633814 | 0.381335889 |
| 21.0706287 | 21.0166036 | 21.0528598 | 21.0767532 | 0.010870187 | 0.841104068 |
| 21.9431757 | 21.8143718 | 21.8437698 | 21.8204933 | -0.058572454 | 0.27491884 |
| 24.4896235 | 24.3639626 | 24.4721061 | 24.3837494 | -0.012581189 | 0.807105104 |
| 26.2232711 | 26.0992088 | 26.2223067 | 26.2020179 | -0.005345462 | 0.914067862 |
| 26.8986764 | 26.7781425 | 26.8565025 | 26.8608034 | 0.00857182 | 0.861172965 |
| 21.137834 | 21.0042893 | 21.0300987 | 21.0714547 | -0.042686032 | 0.375710846 |
| 20.7181678 | 20.8500858 | 20.8228931 | 20.6930744 | 0.011103583 | 0.814861627 |
| 22.3693705 | 22.2089149 | 22.2937413 | 22.2527103 | -0.044752483 | 0.347852572 |
| 20.510074 | 20.4046847 | 20.4402923 | 20.4156841 | -0.088086776 | 0.058758675 |
| 22.3992132 | 22.31401 | 22.3480244 | 22.4132597 | -0.049821977 | 0.2593456 |
| 20.8598954 | 20.8142502 | 20.8214165 | 20.7834954 | -0.052605752 | 0.220082823 |
| 19.8483776 | 19.763711 | 19.8124752 | 19.8780662 | 0.0321404 | 0.450747984 |
| 21.6959902 | 21.7021464 | 21.7480195 | 21.7841436 | 0.072231848 | 0.097097546 |
| 20.1116354 | 20.0475518 | 20.1451363 | 20.1341585 | 0.011408703 | 0.808139905 |

| | | | | | |
|---|---|---|---|---|---|
| 18.3649236 | 18.4407293 | 18.4632116 | 18.4271672 | 0.061327338 | 0.252650689 |
| 21.5819814 | 21.5156453 | 21.5813766 | 21.5904427 | 0.011480147 | 0.834037843 |
| 21.6825223 | 21.5561115 | 21.7081966 | 21.7263959 | 0.042983542 | 0.445093416 |
| 19.2845559 | 19.4417844 | 19.2791187 | 19.4205896 | 0.109531169 | 0.059424258 |
| 18.8801485 | 18.8133199 | 18.806878 | 18.7688128 | -0.019152728 | 0.734256279 |
| 19.6821756 | 19.6418983 | 19.6843909 | 19.6634303 | -0.001450709 | 0.981542117 |
| 19.028775 | 19.1129083 | 19.1576112 | 18.9690447 | 0.069643072 | 0.274697657 |
| 20.7556004 | 20.7324497 | 20.8361215 | 20.7633182 | 0.065422874 | 0.30897112 |
| 17.8359886 | 17.9621562 | 17.8830873 | 17.9056716 | 0.015245674 | 0.829287051 |
| 21.3354697 | 21.2020054 | 21.2640262 | 21.2056822 | -0.029418481 | 0.6963048 |
| 19.805032 | 19.8777278 | 19.9809461 | 19.8832405 | 0.204907072 | 0.012613384 |
| 26.2313674 | 26.3144961 | 26.3802128 | 26.3852161 | 0.151852942 | 0.191886834 |
| 25.5835919 | 25.5534396 | 25.6806191 | 25.5857872 | 0.075811738 | 0.521027974 |
| 16.8087574 | 16.5254492 | 16.7772221 | 17.0794314 | -0.047836411 | 0.69155639 |
| 20.6991334 | 20.6668987 | 20.7879264 | 20.8254955 | 0.099208791 | 0.417262972 |
| 17.2857138 | 17.4827986 | 17.5328004 | 17.223731 | 0.057658327 | 0.656806624 |
| NA | 17.0518448 | NA | NA | 0.244107323 | 0.158503675 |
| 15.9849089 | 16.2539736 | 15.9630036 | 15.8915511 | 0.051670272 | 0.762288625 |
| 16.3637663 | NA | 16.290185 | 16.231714 | -0.028651173 | 0.86472005 |
| 25.0603924 | 24.7964201 | 24.937945 | 24.9278801 | -0.067617982 | 0.21489636 |
| 18.8776155 | 19.1758127 | 18.881781 | 18.8927103 | 0.146920159 | 0.061324802 |
| 16.4874539 | 16.4653282 | 16.3715371 | 16.5743981 | 0.035907245 | 0.750101506 |
| 24.8304289 | 24.656993 | 24.8866646 | 24.8649814 | 0.022032241 | 0.752859021 |
| 20.6152468 | 20.6040384 | 20.658539 | 20.6060895 | -0.011025419 | 0.820965146 |
| 19.9203484 | 19.8203546 | 19.8717748 | 19.9075664 | 0.01403341 | 0.808613934 |
| 20.650946 | 20.6849876 | 20.6515381 | 20.622465 | -0.007751272 | 0.90439356 |
| NA | 21.0302983 | NA | NA | -1.122846663 | 0.110440544 |
| NA | 12.6878466 | NA | NA | -0.111781196 | 0.60433317 |
| NA | 16.4168818 | NA | NA | -0.065193377 | 0.645367374 |
| NA | 18.839144 | NA | NA | -0.309506601 | 0.01200602 |
| 20.5138858 | 20.5008315 | 20.3711193 | 20.4386154 | -0.099700137 | 0.31712527 |
| 21.4091207 | 21.377239 | 21.4065873 | 21.3801892 | -0.030173162 | 0.526088766 |
| 21.8292865 | 21.7881956 | 21.8499378 | 21.8622339 | 0.050161305 | 0.294993824 |
| 18.425102 | 18.4533521 | 18.7354 | 18.5760041 | 0.078405497 | 0.29909455 |
| 19.474947 | 19.6260401 | 19.690227 | 19.6381611 | 0.029795685 | 0.740856932 |
| 18.3393174 | 18.364402 | 18.5195958 | 18.4100599 | 0.086613361 | 0.34079496 |
| 17.8169166 | 18.0387523 | 17.8861382 | 17.84312 | 0.095553058 | 0.294892736 |
| 19.6853803 | 19.5964735 | 19.6859716 | 19.6711452 | -0.013937186 | 0.86428159 |
| 21.3116351 | 21.213382 | 21.2712495 | 21.287476 | -0.016947868 | 0.79760887 |
| 20.9434247 | 20.7084138 | 20.9435712 | 20.8766718 | -0.094529873 | 0.154198015 |
| 22.9173697 | 22.7623764 | 22.9660708 | 22.9099758 | -0.023241929 | 0.720591731 |
| 21.2184529 | 21.0816174 | 21.1527979 | 21.12602 | 0.011299438 | 0.816431954 |

| | | | | | |
|---|---|---|---|---|---|
| 21.3012819 | 21.2034344 | 21.3407259 | 21.368423 | 0.054769567 | 0.324081231 |
| 21.4879544 | 21.433403 | 21.5104836 | 21.3934702 | 0.0463286 | 0.643600648 |
| 26.1419257 | 26.0474457 | 26.1243679 | 26.1267309 | 0.03784594 | 0.610362839 |
| 22.3442452 | 22.2893296 | 22.391151 | 22.3628977 | 0.00575669 | 0.907146359 |
| 21.6683011 | 21.6446875 | 21.6039217 | 21.6445522 | -0.019906172 | 0.691073952 |
| 20.0709726 | 19.5989345 | 20.0090766 | 19.9870274 | -0.141940158 | 0.24809198 |
| 19.4089304 | 19.3607298 | 19.4006294 | 19.466229 | -0.031843244 | 0.496802063 |
| 23.3747108 | 23.5614512 | 23.4352339 | 23.6756528 | 0.157849043 | 0.149048752 |
| 21.4918859 | 21.5394298 | 21.5765715 | 21.5418794 | 0.046147119 | 0.36559357 |
| 13.6337714 | NA | 13.4715468 | 13.3964696 | -0.212360721 | 0.451074285 |
| NA | 13.8544617 | NA | NA | -0.54277769 | 0.044875404 |
| 15.6118272 | 15.5409743 | 15.6044321 | 15.2871959 | -0.277414482 | 0.124754514 |
| 16.1224099 | 15.7472363 | 15.8648918 | 15.778151 | -0.251719865 | 0.126561482 |
| 15.684561 | 15.2932805 | 15.3528382 | 15.4697917 | -0.325859698 | 0.01252484 |
| 18.7415213 | 18.3176889 | 18.8081091 | 18.7888914 | -0.173868587 | 0.148820689 |
| 18.7780909 | 18.1639085 | 18.8638692 | 18.56408 | -0.129330716 | 0.275224366 |
| NA | 16.3368354 | NA | NA | -0.455287803 | 0.002409316 |
| 21.1809405 | 21.1593779 | 21.1437135 | 20.9700454 | -0.101247457 | 0.367259925 |
| 19.5691762 | 19.7386744 | 19.660358 | 19.5488255 | -0.021158401 | 0.826704836 |
| 19.4478264 | 19.4596211 | 19.6757412 | 19.5204987 | -0.001588325 | 0.986752204 |
| 26.6983191 | 26.6006544 | 26.6086968 | 26.6465329 | 0.020902752 | 0.818406415 |
| 17.6835728 | 17.7537793 | 17.7838018 | 17.6815703 | -0.034463484 | 0.686069467 |
| 17.9527908 | 18.1235888 | 18.0578572 | 18.2295145 | 0.052395173 | 0.525496863 |
| 19.8712342 | 19.7633998 | 19.7484396 | 19.9461328 | -0.025852701 | 0.725711731 |
| 20.6350841 | 20.5072595 | 20.7217399 | 20.6825543 | 0.029167589 | 0.675795111 |
| 20.8056607 | 20.7312329 | 20.8926015 | 20.6709636 | -0.047579011 | 0.454728222 |
| 19.9634369 | 19.9008558 | 20.0072153 | 19.9889571 | -0.011903057 | 0.843724804 |
| 22.4540787 | 22.2843925 | 22.4097599 | 22.4434307 | -0.023909071 | 0.688756341 |
| 23.0825315 | 23.0189733 | 23.0633044 | 23.0534092 | -0.02177499 | 0.689605062 |
| 20.9429207 | 20.9175535 | 21.1443215 | 20.9709486 | 0.037660526 | 0.477632112 |
| 19.2215511 | 18.9606288 | 19.0960022 | 19.0588629 | -0.118209035 | 0.029728311 |
| 22.7728919 | 22.6165878 | 22.7043739 | 22.6979897 | -0.040739924 | 0.411381444 |
| 21.311243 | 21.2332357 | 21.1732265 | 21.3302508 | -0.042283147 | 0.388193378 |
| 23.9151517 | 23.9087811 | 23.9230913 | 23.8591248 | -0.008042866 | 0.862050023 |
| 23.0786124 | 23.1093818 | 23.0109456 | 23.0361929 | -0.026907615 | 0.550841531 |
| 22.7575401 | 22.6690119 | 22.7106597 | 22.6893697 | -0.029981318 | 0.500622045 |
| 20.733547 | 20.6615357 | 20.7233609 | 20.7298299 | -0.028472205 | 0.486482352 |
| 25.070685 | 24.9385891 | 25.0653717 | 25.0387215 | -0.017607683 | 0.653802412 |
| 19.8860034 | 19.9262641 | 19.8654076 | 19.9035793 | 0.004250378 | 0.91331436 |
| 25.4945375 | 25.4530007 | 25.4696095 | 25.4548538 | -0.008325712 | 0.81190276 |
| 21.8444246 | 21.7459829 | 21.8223945 | 21.8303792 | 0.004941381 | 0.909426175 |
| 18.7115607 | 18.7014729 | 18.719245 | 18.7274263 | 0.070600808 | 0.137673487 |

| | | | | | |
|---|---|---|---|---|---|
| 23.1486124 | 23.0505128 | 23.1134934 | 23.1326312 | -0.001120529 | 0.98157042 |
| 21.4279756 | 21.3343794 | 21.4072493 | 21.4543802 | 0.011729967 | 0.823475063 |
| 20.0411768 | 19.951609 | 20.037666 | 19.9922386 | 0.047420863 | 0.397285792 |
| 22.1939432 | 22.1610388 | 22.2334941 | 22.2079206 | 0.017744094 | 0.750495397 |
| 21.8025145 | 21.7720133 | 21.7986604 | 21.7876887 | 0.057844431 | 0.309752218 |
| 20.3873318 | 20.1957643 | 20.3028256 | 20.3352119 | -0.036295115 | 0.527140385 |
| 16.8689033 | 16.9509881 | 17.0689773 | 17.0082846 | 0.06230449 | 0.290539261 |
| 18.8741452 | 18.7987452 | 18.8212536 | 18.8685362 | 0.014833745 | 0.802520342 |
| 20.4848574 | 20.31079 | 20.5338178 | 20.6074818 | 0.026457765 | 0.659747715 |
| 21.8499628 | 21.7667887 | 21.8857842 | 21.8720961 | 0.021766216 | 0.720177442 |
| 20.5086966 | 20.500507 | 20.5179506 | 20.5487247 | 0.118620946 | 0.067549683 |
| 20.5485311 | 20.6760807 | 20.6298851 | 20.6633491 | 0.124660376 | 0.055765144 |
| 21.3945444 | 21.2792694 | 21.3678793 | 21.3885194 | -0.002984079 | 0.964345902 |
| 18.6256377 | 18.6426451 | 18.6292657 | 18.739102 | 0.098419136 | 0.152170212 |
| 19.3021722 | 19.0996083 | 19.2598366 | 19.3341807 | 0.013422974 | 0.849547747 |
| 18.413946 | 18.2439741 | 18.4984255 | 18.5581486 | 0.071502695 | 0.320946813 |
| 20.0448762 | 19.8715398 | 20.036261 | 20.0475908 | 0.010505494 | 0.883080587 |
| 22.3302129 | 22.2886738 | 22.3461701 | 22.3632695 | 0.050105136 | 0.499723652 |
| 22.6335675 | 22.6777802 | 22.6766493 | 22.6666029 | 0.094615464 | 0.240610067 |
| 18.1382779 | 18.2603434 | 18.353975 | 18.3775649 | 0.014586097 | 0.858679428 |
| 18.020158 | 17.8202624 | 17.9624522 | 17.9192191 | -0.092530936 | 0.309873984 |
| 23.2172365 | 23.0860244 | 23.2851799 | 23.2945623 | 0.095176887 | 0.301822014 |
| 18.7153343 | 18.926422 | 18.7806778 | 18.9329044 | 0.04623792 | 0.637118588 |
| 19.5124151 | 19.7455265 | 20.1366224 | 20.0695531 | 0.317877416 | 0.014150689 |
| 15.6693684 | 15.8405601 | 15.8462848 | 15.535575 | -0.163819906 | 0.358202099 |
| 14.8845996 | 15.5070563 | 15.6659635 | 15.9661423 | 0.523336254 | 0.08374305 |
| 18.7124749 | 18.6069409 | 18.9256403 | 18.6882202 | -0.064232828 | 0.609471178 |
| 19.8832911 | 19.6768287 | 19.6931107 | 19.4795028 | -0.308155644 | 0.018135653 |
| 21.4286796 | 21.1964315 | 21.3481005 | 21.3212288 | -0.034947446 | 0.707822879 |
| 23.7382208 | 23.6691202 | 23.6121338 | 23.6851501 | 0.029786074 | 0.727729731 |
| 16.7846164 | 16.6203128 | 16.6067162 | 16.5691647 | -0.067117081 | 0.430780742 |
| 18.492518 | 18.2609354 | 18.5469015 | 18.3825937 | -0.076724842 | 0.313069662 |
| 20.2574222 | 20.0691235 | 20.1419592 | 20.1958859 | -0.024074309 | 0.673804231 |
| 21.4889882 | 21.3791345 | 21.4637672 | 21.4582641 | -0.040282604 | 0.44297661 |
| 22.8644914 | 22.7456033 | 22.8191704 | 22.8381944 | -0.046608906 | 0.31544141 |
| 23.4173841 | 23.3732386 | 23.4086609 | 23.3919223 | -0.006459795 | 0.874955178 |
| 24.8176151 | 24.8620135 | 24.7886986 | 24.8263529 | -0.022646399 | 0.546555529 |
| 21.1946473 | 21.140238 | 21.2189515 | 21.2303888 | 0.032065258 | 0.640369386 |
| 22.4029763 | 22.262723 | 22.4786567 | 22.4633705 | 0.06055982 | 0.416980161 |
| 17.4154165 | 17.4005274 | 17.3554735 | 17.2480372 | -0.065859223 | 0.470932041 |
| NA | 16.3600886 | NA | NA | 0.254456086 | 0.071182349 |
| NA | 14.9666739 | NA | NA | 0.115280644 | 0.429441775 |

| | | | | | |
|---|---|---|---|---|---|
| 15.6317687 | NA | 15.7775626 | 16.0630188 | 0.286408969 | 0.10907557 |
| NA | 13.5534002 | NA | NA | 0.031118601 | 0.924552323 |
| NA | 15.1320396 | NA | NA | 0.386925948 | 0.343752635 |
| 16.600952 | 16.0771378 | 16.1998551 | 16.488799 | -0.211772303 | 0.073676959 |
| 21.8764849 | 21.6724553 | 21.8453427 | 21.999676 | 0.048348809 | 0.588905049 |
| 18.0631011 | 17.8680207 | 17.8509169 | 18.0993918 | -0.101037121 | 0.247718921 |
| 21.426723 | 21.222552 | 21.2473797 | 21.1967704 | -0.113828319 | 0.063301493 |
| 20.736543 | 20.5746292 | 20.7705433 | 20.8093093 | 0.001046919 | 0.984843695 |
| 20.239133 | 20.2360565 | 20.3527547 | 20.2520087 | -0.042045418 | 0.431161268 |
| 21.505256 | 21.3988893 | 21.4914651 | 21.4775246 | -0.05425979 | 0.229203536 |
| 21.0436722 | 20.9882025 | 21.0561036 | 21.0721732 | -0.025055919 | 0.560955058 |
| 19.020388 | 19.0706874 | 19.1705757 | 19.2039776 | 0.069380255 | 0.205858095 |
| 19.3026167 | 19.2954894 | 19.300744 | 19.3854581 | 0.068071407 | 0.243073007 |
| 20.7003308 | 20.6683089 | 20.8213453 | 20.9434017 | 0.048146443 | 0.460622217 |
| 18.5314294 | 18.6665761 | 18.8015449 | 18.6860547 | 0.094761017 | 0.21957095 |
| 17.3269298 | 17.4003465 | 17.3791852 | 17.3184349 | 0.100245722 | 0.321005225 |
| 23.1981833 | 23.3128626 | 23.2561916 | 23.2891193 | 0.127895998 | 0.271043526 |
| 13.6645943 | NA | 14.3305602 | 14.0757321 | 0.032147252 | 0.896272779 |
| 19.7421266 | 19.4359936 | 19.7862331 | 19.5612529 | -0.143995146 | 0.139218941 |
| 24.8765265 | 24.8294135 | 24.9244934 | 24.8546276 | 0.009298539 | 0.87550489 |
| 21.5748162 | 21.6024492 | 21.5903013 | 21.6494931 | 0.030537339 | 0.44707641 |
| 22.1644743 | 21.978511 | 22.1421512 | 22.1558886 | -0.056999155 | 0.319785 |
| 19.3033856 | 19.0568075 | 19.4643422 | 19.4887924 | 0.071147081 | 0.559902586 |
| 15.2529371 | 14.9811578 | 15.3409081 | 15.2751853 | -0.075600896 | 0.587331971 |
| 19.6297768 | 19.5118307 | 19.5488281 | 19.7795253 | -0.00808815 | 0.887326561 |
| 25.8307133 | 25.6864668 | 25.6889833 | 25.7600355 | -0.040674183 | 0.44968761 |
| 14.2670163 | 14.5557888 | 14.9239732 | 14.6177834 | 0.29427402 | 0.072345811 |
| 20.7315334 | 20.5828809 | 20.7000342 | 20.6999302 | -0.036562635 | 0.524860409 |
| 21.874424 | 21.7398652 | 21.8067709 | 21.7391276 | -0.070422754 | 0.249109866 |
| 22.7577293 | 22.5655787 | 22.7672826 | 22.781937 | 0.013262675 | 0.821624201 |
| 20.1570535 | 19.9691662 | 20.077709 | 20.0878781 | -0.083135179 | 0.138406868 |
| 15.2173326 | 15.4832402 | 15.5235256 | 15.750002 | 0.217859186 | 0.103028659 |
| 20.6294905 | NA | 21.0499297 | 20.7878307 | 0.267582492 | 0.307042528 |
| 12.5980762 | NA | 13.3231523 | 13.6335755 | 0.394568409 | 0.182518722 |
| 21.8040484 | 21.7279934 | 21.7079012 | 21.7287172 | -0.031047213 | 0.514041554 |
| 25.721502 | 25.624605 | 25.6571873 | 25.6745595 | -0.017299876 | 0.691518557 |
| 20.4993574 | 20.3659284 | 20.4094996 | 20.4359187 | 0.033106563 | 0.633259818 |
| 18.7804701 | 18.7434084 | 18.7310801 | 18.8857124 | 0.005963743 | 0.945468454 |
| 17.8880986 | 17.6494251 | 17.9574565 | 18.0311729 | -0.001224865 | 0.993182123 |
| NA | 16.5389298 | NA | NA | 0.317438305 | 0.282635239 |
| NA | 12.9786588 | NA | NA | -0.531574618 | 0.087232712 |
| 19.0969611 | 19.0329652 | 19.2283917 | 19.1846895 | 0.11166504 | 0.095201747 |

| | | | | | |
|---|---|---|---|---|---|
| 19.1134097 | 19.0770701 | 19.1916245 | 19.2188426 | 0.091056276 | 0.294373587 |
| 20.7620254 | 20.7373979 | 20.95165 | 20.7944742 | 0.082840995 | 0.62798005 |
| 24.1162319 | 24.0741233 | 24.1385206 | 24.142492 | 0.006767461 | 0.86703744 |
| 18.9543568 | 18.7764585 | 18.9719954 | 19.0084943 | 0.105580046 | 0.207590344 |
| 21.2681111 | 21.2443063 | 21.433934 | 21.1306158 | 0.016528863 | 0.890136955 |
| 15.5555165 | NA | 15.7327759 | 15.938436 | 0.20443538 | 0.063091472 |
| 19.9534047 | 19.8950521 | 20.2337498 | 20.0912676 | 0.199537956 | 0.061773766 |
| 19.4659178 | 19.2672928 | 19.4158658 | 19.4001271 | -0.078903966 | 0.305558558 |
| 18.062539 | 17.7552474 | 18.0461513 | 17.8974793 | -0.102117811 | 0.164699283 |
| 19.7963538 | 19.607246 | 19.6658593 | 19.7697785 | -0.056785109 | 0.296912286 |
| 24.2578789 | 24.2189916 | 24.1685474 | 24.0594043 | -0.144766448 | 0.00850939 |
| 21.9816589 | 21.8911873 | 21.9046938 | 21.9587781 | -0.04677612 | 0.252697448 |
| 23.910031 | 23.8175446 | 23.9163341 | 23.945285 | 0.046123317 | 0.399819128 |
| 19.1604674 | 19.1569071 | 19.2164501 | 19.2754099 | 0.06554969 | 0.344848203 |
| 15.7024982 | 15.8143435 | 15.8849784 | 16.0393923 | 0.060396422 | 0.752283994 |
| 21.6792664 | 21.5310211 | 21.5799795 | 21.6425506 | -0.017964858 | 0.793655579 |
| 19.4314564 | 19.1325101 | 19.4483559 | 19.4249671 | -0.070972334 | 0.350075534 |
| 23.8039306 | 23.6553307 | 23.7451987 | 23.7472645 | -0.060972735 | 0.246193782 |
| 20.8073444 | 20.7995567 | 20.7930706 | 20.8757747 | -0.015786341 | 0.746724218 |
| 23.8421149 | 23.7400449 | 23.8201839 | 23.820004 | -0.00961258 | 0.824944523 |
| 22.4610505 | 22.3857072 | 22.4189227 | 22.4361354 | -0.020030478 | 0.643940056 |
| 19.3192498 | 19.2606047 | 19.3520315 | 19.3613628 | 0.034384316 | 0.42471012 |
| 20.0961529 | 19.9652386 | 20.1437237 | 20.1555839 | 0.060145765 | 0.367615768 |
| 19.266032 | 19.133976 | 19.3201134 | 19.3353699 | 0.041160298 | 0.700875597 |
| 16.0148784 | 16.0241036 | 16.0123923 | 16.2658374 | 0.176351636 | 0.267334809 |
| 17.400583 | 16.6172506 | 17.2492868 | 17.3880625 | -0.119469502 | 0.51620147 |
| 18.4986357 | 18.2775274 | 18.4540395 | 18.3050627 | -0.109635687 | 0.367247765 |
| 20.4268568 | 20.378751 | 20.3191249 | 20.3915248 | -0.004592524 | 0.958597742 |
| 20.1801435 | 20.1488971 | 20.0190967 | 20.0951401 | -0.057152771 | 0.33753488 |
| 20.3124586 | 20.2714677 | 20.3496868 | 20.2410602 | -0.013533254 | 0.805253442 |
| 20.2282488 | 20.2202227 | 20.2269094 | 20.2735024 | 0.04425549 | 0.484393545 |
| 20.8463156 | 20.8617456 | 20.9082575 | 20.955123 | 0.07690077 | 0.362742046 |
| 22.1418599 | 22.1045486 | 22.1910903 | 22.0945305 | 0.083597066 | 0.324549171 |
| NA | 14.6571459 | NA | NA | -0.163247056 | 0.223165219 |
| 26.1404328 | 25.9583009 | 26.2609458 | 26.189941 | -0.01393784 | 0.849457723 |
| 22.3984588 | 22.35192 | 22.4591442 | 22.4300465 | 0.020009258 | 0.62287304 |
| 20.0495545 | 20.0284254 | 20.1162485 | 20.0975683 | 0.049580985 | 0.313591116 |
| 20.5526799 | 20.4085854 | 20.4898643 | 20.6215677 | 0.030392674 | 0.593205267 |
| 24.0240559 | 23.980747 | 24.0602659 | 24.0561448 | 0.083007193 | 0.062617033 |
| 18.3366275 | 17.9509075 | 18.130656 | 18.1925797 | -0.179327233 | 0.012374879 |
| NA | 16.637863 | NA | NA | -0.207566733 | 0.350342435 |
| 17.5202034 | 17.1787087 | 17.5502238 | 17.4696036 | -0.048182979 | 0.743724452 |

| | | | | | |
|---|---|---|---|---|---|
| 23.8738649 | 23.8301279 | 23.856571 | 23.903756 | -0.001614761 | 0.968388282 |
| 15.1353193 | 15.3588812 | 15.6985546 | 14.9021222 | 0.215654162 | 0.253108395 |
| 21.0444467 | 20.7823098 | 20.7499827 | 20.5991698 | -0.26293746 | 0.010625111 |
| 20.208841 | 20.1296055 | 20.3107468 | 20.2217327 | 0.030552367 | 0.690282138 |
| 24.1637272 | 24.0530359 | 24.1293231 | 24.1516365 | 0.029174289 | 0.687734316 |
| 21.3030206 | 21.3441455 | 21.3649731 | 21.3609546 | 0.032285682 | 0.635938382 |
| 20.1263956 | 19.9029649 | 20.0560255 | 20.0477771 | -0.096509124 | 0.139885404 |
| 20.4387209 | 20.3547164 | 20.4183246 | 20.3896694 | -0.039466816 | 0.524129208 |
| 20.9612503 | 20.6891354 | 20.7505201 | 20.7791152 | -0.126703783 | 0.043820927 |
| 20.9000476 | 20.7625335 | 20.8704778 | 20.9278111 | -0.042084771 | 0.457409936 |
| 21.0143173 | 20.9561337 | 20.9898879 | 21.0591616 | 0.007344513 | 0.874673734 |
| 19.8983494 | 19.8944882 | 19.9999913 | 19.9429897 | 0.057337254 | 0.239734735 |
| 20.7573675 | 20.6636874 | 20.6944677 | 20.7452994 | -0.044511347 | 0.401492007 |
| 20.4874792 | 20.4546472 | 20.394519 | 20.5012422 | -0.004249734 | 0.939004876 |
| 21.601884 | 21.6644511 | 21.6530597 | 21.6812128 | 0.117784171 | 0.084343718 |
| 18.1984339 | 18.1804383 | 18.2226709 | 18.2760289 | 0.065205855 | 0.355383672 |
| 17.9840401 | 17.8786399 | 17.8751707 | 17.8695451 | -0.05039446 | 0.505420317 |
| 22.5750906 | 22.4659727 | 22.465875 | 22.63375 | 0.075198275 | 0.326886249 |
| 17.4007695 | 17.296713 | 17.397306 | 17.4708423 | 0.073262884 | 0.417532308 |
| 21.9840556 | 21.8453802 | 22.0584128 | 22.0391528 | 0.046480996 | 0.68468739 |
| 15.5335204 | 15.7733817 | 15.8993768 | 15.5113706 | 0.036861752 | 0.824966319 |
| 18.6129404 | 18.9795346 | 18.6999051 | 18.9740596 | 0.09818696 | 0.664674981 |
| 24.3318785 | 24.3773597 | 24.3844946 | 24.4740625 | 0.116521481 | 0.155653151 |
| NA | 14.0732409 | NA | NA | 2.70E-04 | 0.997944191 |
| 23.5423286 | 23.3640892 | 23.4738989 | 23.5216083 | -0.008745617 | 0.89701299 |
| 21.0595831 | 20.954311 | 21.142591 | 20.9164033 | -0.033965717 | 0.722544136 |
| 19.5011099 | 19.2395107 | 19.3375274 | 19.4648116 | -0.064194418 | 0.370304958 |
| 27.1564893 | 27.1150386 | 27.1106602 | 27.144736 | 0.064008172 | 0.319460356 |
| 20.4422206 | 20.2061877 | 20.3677288 | 20.3653437 | -0.076303238 | 0.229767251 |
| 21.8731275 | 21.8766159 | 21.9076041 | 21.8719591 | 0.053637831 | 0.276306283 |
| 20.1235777 | 20.0816035 | 20.1319049 | 20.2334912 | 0.060133326 | 0.215588771 |
| 17.4365207 | 17.4910185 | 17.6494108 | 17.5474986 | 0.108450035 | 0.140393298 |
| 18.2850475 | 18.2184292 | 18.4608895 | 18.2938543 | -0.012773739 | 0.877041 |
| 21.1117875 | 20.9746461 | 21.1841542 | 21.1289721 | 0.020304898 | 0.854473748 |
| 19.770306 | 19.4111462 | 19.4731733 | 19.8333799 | 0.116034437 | 0.5085672 |
| 21.5307045 | 21.4206782 | 21.4106653 | 21.4955818 | -0.068171168 | 0.289912584 |
| 23.0443941 | 22.9324082 | 23.0183609 | 23.0287252 | -0.036579859 | 0.517242681 |
| 21.7697708 | 21.6846245 | 21.6965953 | 21.7165592 | -0.040847035 | 0.338604704 |
| 17.854419 | 17.7517009 | 17.994918 | 17.9186092 | 0.029223218 | 0.652361123 |
| 16.4480296 | 16.2552865 | 16.5024403 | 16.4996728 | -0.066511765 | 0.703771811 |
| 18.5709813 | 18.6170767 | 18.6483826 | 18.7012472 | -0.003454433 | 0.959981215 |
| 21.1244497 | 21.1286457 | 21.1385339 | 21.0898502 | -0.014951375 | 0.736306604 |

| | | | | | |
|---|---|---|---|---|---|
| 20.4334679 | 20.3633608 | 20.3201952 | 20.4267207 | -0.007732838 | 0.901393681 |
| 22.0347121 | 21.9953319 | 22.0644266 | 22.0159892 | 0.05519856 | 0.334087783 |
| 22.9156723 | 22.9255257 | 22.9265541 | 22.9763398 | 0.072027004 | 0.148401252 |
| 21.6785454 | 21.613849 | 21.7218964 | 21.7083249 | 0.01920416 | 0.794832636 |
| 24.5933779 | 24.5035426 | 24.520444 | 24.6199644 | 0.029646569 | 0.838112052 |
| 23.6125164 | 23.7016431 | 23.6713644 | 23.6449977 | 0.169883407 | 0.125661972 |
| 20.3610139 | 20.0544179 | 20.2158029 | 20.3040849 | -0.167084434 | 0.032322815 |
| 21.6096216 | 21.5823985 | 21.6090516 | 21.6620526 | 0.048729173 | 0.323337011 |
| 21.3214802 | 21.1709509 | 21.2873318 | 21.2828432 | -0.047805719 | 0.454907666 |
| 20.368084 | 20.3057344 | 20.4140956 | 20.4751918 | -0.015513215 | 0.743054874 |
| 18.1949292 | 18.3203543 | 18.1732594 | 18.3395155 | 0.054069981 | 0.363853976 |
| 17.951703 | 17.6722402 | 17.9613716 | 18.0506319 | -0.117804465 | 0.270157075 |
| 16.0455019 | 15.8932025 | 15.6937162 | 15.816499 | -0.188429807 | 0.077880453 |
| 22.6552528 | 22.6362089 | 22.4928384 | 22.6331734 | -0.01123035 | 0.883376916 |
| 20.2848166 | 20.2623896 | 20.2303407 | 20.1051164 | -0.051000676 | 0.500977483 |
| 21.4117256 | 21.1895681 | 21.4966767 | 21.4337662 | -0.023371159 | 0.753756882 |
| 21.3464442 | 21.2351223 | 21.3362031 | 21.3502746 | -0.03862381 | 0.533205866 |
| 20.7704097 | 20.6829575 | 20.7596157 | 20.6949586 | 0.007792906 | 0.872764538 |
| 24.0772007 | 24.0513264 | 24.0889146 | 24.1234661 | 0.027840195 | 0.45536694 |
| 18.1986045 | 18.3007546 | 18.3881303 | 18.3474996 | 0.063556295 | 0.174341472 |
| 23.152706 | 23.0467974 | 23.1800343 | 23.1120758 | 0.002539415 | 0.956149941 |
| 19.7349163 | 19.6864412 | 19.8071588 | 19.8395984 | 0.036206259 | 0.573083938 |
| 25.221869 | 25.3033528 | 25.2098566 | 25.265387 | 0.055518142 | 0.507153672 |
| 25.3203427 | 25.2306546 | 25.2501866 | 25.3697013 | 0.046376574 | 0.642749367 |
| 20.8910381 | 20.8005493 | 20.9171337 | 20.9167621 | -0.017872566 | 0.686310382 |
| 20.4999969 | 20.4504669 | 20.3900477 | 20.4913395 | -0.003406507 | 0.952838829 |
| 18.9558966 | 18.9434136 | 19.0456272 | 19.1277778 | 0.124073656 | 0.098660638 |
| 24.679418 | 24.687405 | 24.7221293 | 24.6528836 | 0.066628233 | 0.385187613 |
| 21.4269653 | 21.2444007 | 21.3638896 | 21.3882648 | -0.032214577 | 0.739413806 |
| NA | 17.0667911 | NA | NA | 0.555286326 | 0.079691697 |
| 19.3594376 | 19.0919807 | 19.3105394 | 19.2098511 | -0.075242279 | 0.417722735 |
| 22.1278315 | 22.0384297 | 22.2184899 | 22.1225123 | 0.001713399 | 0.965571456 |
| 19.5188937 | 19.4955999 | 19.5553868 | 19.6008378 | -0.001610161 | 0.970712216 |
| 22.8506704 | 22.7811534 | 22.9193777 | 22.9132444 | 0.048143709 | 0.337550717 |
| 15.2559985 | 14.9590331 | 14.8611112 | 15.0304127 | -0.032526137 | 0.895229834 |
| 12.8585083 | NA | 12.6125724 | 12.9834891 | 0.036315992 | 0.866190833 |
| 16.6354288 | 16.8354282 | 16.9713441 | 16.7774976 | 0.183746314 | 0.386440013 |
| 17.8913218 | 17.5768049 | 17.8190228 | 17.9488747 | 0.13896461 | 0.486915113 |
| 17.6397847 | 17.2473409 | 17.1226238 | 17.4128979 | -0.166759422 | 0.241676598 |
| 20.4641521 | 20.3606502 | 20.7270067 | 20.682862 | -0.002732242 | 0.978401002 |
| 19.9234622 | 19.7192498 | 19.8094506 | 19.9501136 | -0.012121971 | 0.867021027 |
| 23.4646392 | 23.2657704 | 23.5014584 | 23.4390796 | -0.003079223 | 0.965501117 |

| | | | | | |
|---|---|---|---|---|---|
| 26.513102 | 26.3285254 | 26.4002776 | 26.4314896 | -0.020312412 | 0.76439254 |
| 23.4220335 | 23.3258159 | 23.3045777 | 23.3862863 | -0.006373669 | 0.911758034 |
| 19.7814145 | 19.7027534 | 19.6959684 | 19.6131079 | -0.064214189 | 0.253581078 |
| 21.8910915 | 21.7961214 | 21.9109689 | 21.8785845 | -0.059554965 | 0.287611517 |
| 24.4515975 | 24.2902791 | 24.3686675 | 24.382312 | -0.01640288 | 0.763676438 |
| 21.856048 | 21.6766802 | 21.7803454 | 21.7364701 | -0.074610321 | 0.172202148 |
| 22.536737 | 22.4989594 | 22.5303452 | 22.48653 | -0.005358138 | 0.916288598 |
| 27.8097651 | 27.7054747 | 27.7886899 | 27.7799901 | 0.007031146 | 0.879085593 |
| 21.6958685 | 21.575189 | 21.6711143 | 21.7010971 | -0.03601663 | 0.435617624 |
| 24.8633606 | 24.8537821 | 24.7818231 | 24.9072851 | 0.020212289 | 0.661619941 |
| 20.395372 | 20.3823093 | 20.3930896 | 20.3725426 | -0.032378449 | 0.376223837 |
| 20.4800953 | 20.4657821 | 20.50282 | 20.4816499 | -0.003859513 | 0.932406643 |
| 20.3744908 | 20.4078532 | 20.388764 | 20.4942097 | 0.048936647 | 0.354878851 |
| 23.7865656 | 23.7615803 | 23.8249511 | 23.8093568 | 0.024359046 | 0.651591111 |
| 20.4359881 | 20.3405418 | 20.3502061 | 20.4967667 | -0.023718621 | 0.695210826 |
| 20.1712414 | 20.0303351 | 20.1603116 | 20.2121309 | -0.017577315 | 0.772420059 |
| 20.9771288 | 20.8601133 | 21.0099303 | 20.9883879 | 0.025708371 | 0.686790327 |
| 19.5050958 | 19.4167426 | 19.4361673 | 19.4256367 | -0.130628331 | 0.094105162 |
| 18.0437122 | 18.0247532 | 18.0017049 | 17.9750751 | -0.007502379 | 0.923560412 |
| 18.8780669 | 18.7717449 | 18.6181346 | 18.8849951 | -0.017997189 | 0.869436652 |
| 22.016164 | 21.877708 | 22.0493427 | 22.1245352 | -0.007868661 | 0.949698495 |
| NA | 17.9590309 | NA | NA | -0.151918414 | 0.391298585 |
| 20.3608943 | 20.1966644 | 20.329823 | 20.2194454 | -0.052766325 | 0.408395408 |
| 12.4435404 | NA | 12.4104859 | 12.4609141 | -0.049991629 | 0.845315858 |
| NA | 15.43259 | NA | NA | -0.039516742 | 0.680744781 |
| 24.5737256 | 24.4529863 | 24.5653102 | 24.5081534 | -0.037475344 | 0.367003838 |
| 20.511087 | 20.3782357 | 20.4929925 | 20.5096744 | 0.003646891 | 0.949255835 |
| 21.4656465 | 21.2740229 | 21.3478551 | 21.437336 | -0.078833258 | 0.258801073 |
| 19.2752084 | 19.2566114 | 19.2709071 | 19.2681482 | 0.005104524 | 0.940820663 |
| 21.2212576 | 21.0770077 | 21.246864 | 21.0773916 | -0.077410433 | 0.387008724 |
| 22.6633973 | 22.7168671 | 22.7477813 | 22.6917626 | 0.026451041 | 0.73328758 |
| 21.2965007 | 21.1711294 | 21.3311618 | 21.3053584 | -0.011199417 | 0.820956682 |
| 21.0531998 | 20.997688 | 20.9925952 | 20.9787191 | -0.049183481 | 0.266994963 |
| 22.5704376 | 22.7603237 | 22.6741085 | 22.7171529 | 0.095019046 | 0.042880193 |
| 27.0367076 | 26.9997515 | 27.0512042 | 26.9850423 | 0.008180615 | 0.889091204 |
| 19.8861183 | 19.8942684 | 20.0399126 | 19.9857659 | 0.029241103 | 0.658536441 |
| 18.5150343 | 18.4424641 | 18.5540525 | 18.5074987 | -0.029048366 | 0.666177983 |
| 19.3131327 | 19.0764582 | 19.1973248 | 19.2754543 | -0.024044477 | 0.736230101 |
| 17.1458232 | 16.9637278 | 17.0473943 | 17.075156 | 0.012470199 | 0.911548185 |
| 19.8502463 | 20.081031 | 20.1003199 | 20.0449798 | 0.200075599 | 0.086072409 |
| 15.0197237 | NA | 15.0604807 | 15.0604743 | 0.117555766 | 0.426079167 |
| 23.6723408 | 23.6480579 | 23.7301821 | 23.7271208 | 0.063168098 | 0.225650591 |

| | | | | | |
|---|---|---|---|---|---|
| 18.9459672 | 18.8450254 | 18.9131061 | 18.7927105 | -0.063371244 | 0.60356365 |
| 21.4060268 | 21.383305 | 21.3671518 | 21.4140543 | 0.009269215 | 0.871280318 |
| 21.5893501 | 21.5458732 | 21.5889578 | 21.5629149 | 0.019178534 | 0.612632223 |
| 18.1419045 | 18.0303121 | 18.1876068 | 18.2170819 | 0.010992559 | 0.888980932 |
| 19.0653243 | 18.8982241 | 19.0095514 | 18.9235172 | -0.040584511 | 0.626125577 |
| 12.8015141 | 13.3103209 | 13.2057629 | 12.9175714 | 0.006508624 | 0.973942211 |
| NA | 16.9193839 | NA | NA | -0.342675121 | 0.110895557 |
| 21.6018411 | 21.6109044 | 21.6043621 | 21.6365882 | 0.024619386 | 0.518681538 |
| 20.4858253 | 20.4544333 | 20.4881033 | 20.5230365 | 0.063414956 | 0.163001763 |
| 19.426956 | 19.3977175 | 19.3831582 | 19.4519883 | -0.002599049 | 0.970114822 |
| 18.4114425 | 18.2884981 | 18.5489508 | 18.34498 | 0.051730688 | 0.588941141 |
| 23.7462542 | 23.6707869 | 23.7587735 | 23.7704814 | 0.021953836 | 0.649726611 |
| 20.6704679 | 20.6805887 | 20.8204388 | 20.6490419 | 0.024851249 | 0.684773954 |
| 17.376744 | 17.4372544 | 17.4892341 | 17.5793477 | 0.055481156 | 0.617739805 |
| 18.8673642 | 18.6590918 | 18.7852828 | 18.7945537 | -0.116584921 | 0.105386767 |
| 24.1734091 | 23.9556612 | 24.0505494 | 24.1113615 | -0.061711295 | 0.301095703 |
| 20.0337204 | 20.1590733 | 20.0574937 | 20.1545418 | 0.0790229 | 0.235191933 |
| 20.4521011 | 20.4129797 | 20.4565224 | 20.4214518 | 0.052436641 | 0.456720067 |
| 20.5220391 | 20.4988867 | 20.50076 | 20.5484976 | 0.001655071 | 0.969840776 |
| 19.1180943 | 19.3057576 | 19.4184334 | 19.4269343 | 0.142760065 | 0.10013062 |
| 19.9941144 | 19.7200376 | 19.4582796 | 19.7166654 | -0.254543461 | 0.005195813 |
| 18.9182703 | 18.8147995 | 18.8732015 | 18.8850499 | -0.259024677 | 0.027550178 |
| 19.7810975 | 19.7728428 | 19.655296 | 19.6968972 | -0.087264252 | 0.212099859 |
| 22.0714051 | 21.9323839 | 22.060895 | 22.0783834 | -0.01890919 | 0.723208482 |
| 23.732867 | 23.7016604 | 23.6667941 | 23.7795763 | 0.019017934 | 0.680030964 |
| 21.5896711 | 21.5098451 | 21.5502968 | 21.5758018 | 0.00171366 | 0.967606811 |
| 22.1354228 | 22.0092777 | 22.0953098 | 22.0390786 | -0.063909325 | 0.135133334 |
| 22.7547268 | 22.7043841 | 22.7695129 | 22.8127183 | -0.023248891 | 0.578468096 |
| 20.9887481 | 20.8337874 | 20.974248 | 20.9119163 | 0.015208189 | 0.81044602 |
| 20.3741852 | 20.2495756 | 20.3795142 | 20.3344732 | 0.014865002 | 0.831026063 |
| 19.360227 | 19.2085387 | 19.1009888 | 19.3661535 | -0.045485963 | 0.576218103 |
| 18.0088035 | 17.8290655 | 18.1311382 | 18.071918 | -0.056596605 | 0.551948829 |
| 22.6166223 | 22.5234393 | 22.6200045 | 22.5695579 | -0.004530447 | 0.918114371 |
| NA | 13.5238256 | NA | NA | 0.185884017 | 0.445202242 |
| 23.8702785 | 23.7371238 | 23.9585705 | 23.8929886 | 0.051342308 | 0.704495083 |
| 20.4142751 | 20.335597 | 20.4661082 | 20.4272269 | 0.040381604 | 0.447662972 |
| 16.2756789 | 15.9397524 | 16.4390492 | 16.1350484 | -0.164056884 | 0.206369095 |
| 24.8478494 | 24.8393625 | 24.8633167 | 24.826062 | 0.011205153 | 0.708493402 |
| 18.222763 | 18.00076 | 17.8968104 | 18.198906 | -0.023510847 | 0.804929735 |
| 19.4078158 | 19.1562479 | 19.3280014 | 19.243989 | -0.142408188 | 0.094044008 |
| 20.9384945 | 20.8867984 | 20.93486 | 21.000085 | 0.03343263 | 0.556114237 |
| NA | 15.5675997 | NA | NA | 0.0649236 | 0.740685038 |

| | | | | | |
|---|---|---|---|---|---|
| 18.7217588 | 18.1123647 | 18.7030226 | 18.6194649 | -0.265174289 | 0.239367237 |
| 16.4551507 | NA | 16.3830715 | 16.5736129 | 0.016198057 | 0.879364322 |
| 19.1421588 | 18.8427273 | 19.140933 | 19.0937495 | -0.106180128 | 0.276398181 |
| 26.1582701 | 26.1196634 | 26.1810185 | 26.186061 | 0.034056551 | 0.71729358 |
| 19.7210163 | 19.514849 | 19.6308496 | 19.4955148 | -0.119786957 | 0.124678869 |
| 20.8442902 | 20.7289717 | 20.8133296 | 20.8645368 | -0.025409656 | 0.730840939 |
| 21.867637 | 21.6512271 | 21.5554495 | 21.6714129 | -0.147711812 | 0.042292393 |
| 20.0830543 | 19.9787125 | 20.1255099 | 20.0564681 | -0.024279225 | 0.688989582 |
| 21.6394256 | 21.6678707 | 21.6229235 | 21.6680267 | 0.006772262 | 0.901922156 |
| 19.5802408 | 19.4926459 | 19.703445 | 19.6239572 | 0.020526434 | 0.728415054 |
| 20.9820711 | 20.8478527 | 20.9379346 | 21.0036366 | 0.004039868 | 0.950100331 |
| 16.038745 | 16.1998644 | 16.0836793 | 16.0744844 | 0.062426315 | 0.574215761 |
| 16.7014718 | 17.0941777 | 17.0974401 | 16.9973972 | 0.09506556 | 0.521821713 |
| 20.788974 | 20.4393142 | 20.5039895 | 20.4909815 | -0.19468033 | 0.043505114 |
| 21.4005616 | 21.1366666 | 21.3171146 | 21.3642542 | -0.080761661 | 0.292032702 |
| 22.567193 | 22.4670829 | 22.5194441 | 22.5827152 | -0.029259935 | 0.424909984 |
| NA | 13.8048386 | NA | NA | -0.148596696 | 0.54885091 |
| 17.18614 | 16.8029838 | 16.8314917 | 17.0116149 | -0.115542211 | 0.507253009 |
| 27.6287004 | 27.6739718 | 27.6928257 | 27.6156507 | 0.076028361 | 0.431479265 |
| 21.0587815 | 20.7705276 | 20.9524708 | 20.9927829 | -0.103725598 | 0.228471681 |
| 18.0807147 | 17.787327 | 18.0553424 | 18.1078073 | -0.059540771 | 0.482608151 |
| 19.8700049 | 19.8711067 | 20.0645373 | 19.8836455 | -0.023338078 | 0.775514407 |
| 22.6943979 | 22.452916 | 22.4673789 | 22.5823534 | -0.107795058 | 0.172723134 |
| 19.7164621 | 19.6485335 | 19.7590477 | 19.6322743 | -0.054455596 | 0.477003593 |
| 19.017934 | 18.8728774 | 18.9772727 | 18.8418459 | -0.130335429 | 0.091975883 |
| 23.0677839 | 22.9427705 | 23.0882718 | 23.1649187 | -0.020881374 | 0.772598675 |
| 20.5443883 | 20.3532384 | 20.5133389 | 20.4337859 | -0.074733621 | 0.29281817 |
| 19.6229438 | 19.5346056 | 19.475762 | 19.5286663 | -0.099623581 | 0.157685765 |
| 20.9024057 | 20.885593 | 20.9189533 | 20.8082504 | -0.018981942 | 0.759302273 |
| 18.008461 | 17.7900999 | 17.9152324 | 17.9053923 | -0.085786528 | 0.150023892 |
| 20.9167341 | 20.6939067 | 20.9602105 | 20.8960305 | -0.017873527 | 0.758271674 |
| 23.1955613 | 23.2911118 | 23.1708487 | 23.1683379 | -0.018088881 | 0.745771551 |
| 19.2239404 | 19.2588202 | 19.2980901 | 19.2760523 | -0.005732911 | 0.916744701 |
| 18.3853694 | 18.254816 | 18.3525887 | 18.3401138 | -0.057319627 | 0.297416895 |
| 20.040284 | 19.9394607 | 19.9127076 | 19.9856032 | -0.045370481 | 0.392115553 |
| 22.1689306 | 21.9710223 | 22.0661821 | 22.115799 | -0.07474588 | 0.163630711 |
| 19.3296786 | 19.2427013 | 19.3097929 | 19.2500021 | -0.037738538 | 0.372131767 |
| 19.8140333 | 19.8241483 | 19.8712761 | 19.8291439 | 0.029711018 | 0.495362789 |
| 24.1614467 | 24.1459173 | 24.2146188 | 24.2172774 | 0.032655876 | 0.466688039 |
| 20.579457 | 20.4831395 | 20.4768285 | 20.6117218 | -0.013002265 | 0.793698503 |
| 24.9966658 | 24.8981952 | 25.0745723 | 24.9856502 | 0.045735119 | 0.397949317 |
| 21.0468905 | 21.0343862 | 21.056588 | 21.0254087 | 0.029150168 | 0.597080419 |

| | | | | | |
|---|---|---|---|---|---|
| 18.3273361 | 18.2982728 | 18.2630818 | 18.3160512 | 0.03404075 | 0.585382552 |
| 19.453396 | 19.4111051 | 19.5098226 | 19.5668019 | 0.042733324 | 0.49273578 |
| 20.4665012 | 20.3458166 | 20.3988306 | 20.3794759 | -0.018323844 | 0.784068576 |
| 19.2915289 | 19.1463403 | 19.2953235 | 19.2793377 | 0.078112398 | 0.256323679 |
| 19.3762444 | 19.5147879 | 19.4389379 | 19.4277868 | 0.046849226 | 0.507687912 |
| 20.4004046 | 20.4314322 | 20.4219944 | 20.393645 | 0.038671862 | 0.589252198 |
| 20.4566388 | 20.3105999 | 20.3880723 | 20.4605752 | -0.003811665 | 0.962061414 |
| 24.0129682 | 24.0693948 | 23.9843927 | 23.9999405 | 0.019869916 | 0.808463456 |
| 18.6924483 | 18.6488031 | 18.8644234 | 18.9923255 | 0.147602472 | 0.096376318 |
| 18.6287773 | 18.6574536 | 18.4235821 | 18.5655512 | -0.064380977 | 0.523212748 |
| 18.527335 | 18.3899503 | 18.5169543 | 18.4719884 | -0.060674745 | 0.551428373 |
| 19.943619 | 19.832783 | 19.9260045 | 19.9522383 | 0.032559199 | 0.592864887 |
| 21.3153014 | 21.2562758 | 21.2638205 | 21.1834123 | -0.038218346 | 0.437944703 |
| 24.5424793 | 24.7553323 | 24.7454876 | 24.6156855 | 0.218017354 | 0.002716692 |
| 16.9547388 | 16.7674688 | 16.6214475 | 16.7417081 | -0.133507621 | 0.233450951 |
| 22.6625658 | 22.4858658 | 22.5992414 | 22.6147455 | -0.040300432 | 0.463146102 |
| 21.6275419 | 21.5473763 | 21.5528337 | 21.5290416 | -0.052331489 | 0.279892752 |
| 15.9277455 | 15.5884682 | 15.8369361 | 15.9485502 | -0.119048717 | 0.35982739 |
| 20.0564152 | 19.8169913 | 20.134921 | 19.9264526 | 0.092130077 | 0.463992404 |
| 19.6532477 | 19.4140688 | 19.469978 | 19.295986 | -0.193381257 | 0.065219469 |
| 19.6776756 | 19.7212103 | 19.4761419 | 19.7016804 | -0.048695115 | 0.544199357 |
| 20.3197237 | 20.2692183 | 20.3840827 | 20.3281597 | 0.152042626 | 0.115968592 |
| 17.7260363 | 17.4885202 | 17.6185991 | 17.7282212 | -0.120711029 | 0.299542304 |
| 17.9306385 | 17.9160575 | 17.8020552 | 17.7790382 | -0.113414423 | 0.143170088 |
| 21.5867583 | 21.4655373 | 21.6434306 | 21.6000837 | 0.055833145 | 0.393543138 |
| 19.8458285 | 19.6273138 | 19.9150079 | 19.7828143 | -0.020975743 | 0.801884152 |
| 21.4437327 | 21.4201863 | 21.4543633 | 21.408424 | 0.012622862 | 0.881140111 |
| 18.3723225 | NA | 18.3817887 | 18.3773283 | -0.013489573 | 0.883458438 |
| 22.0308936 | 21.9829955 | 22.0713117 | 22.0686214 | 0.042571783 | 0.393347532 |
| 19.2812432 | 19.262818 | 19.2610643 | 19.3148885 | 0.026932709 | 0.801818349 |
| 21.4844422 | 21.4453441 | 21.4641653 | 21.4598778 | 0.010583408 | 0.779307654 |
| 19.449894 | 19.2894353 | 19.4137987 | 19.5198738 | -0.062740794 | 0.457409741 |
| 19.8755754 | 20.4946191 | 20.1841491 | 20.2987015 | 0.501772997 | 0.213928854 |
| 19.3081626 | 19.3073488 | 19.2255733 | 19.3311081 | -0.17619083 | 0.14424278 |
| 17.1016052 | 17.180571 | 17.687533 | 16.806435 | 0.01792787 | 0.896286916 |
| 19.5388902 | 19.2812314 | 19.4031029 | 19.4662514 | -0.050385157 | 0.629027427 |
| 17.4769804 | 17.4786863 | 17.2137798 | 17.5300199 | -0.11572788 | 0.183607678 |
| 21.5646211 | 21.446551 | 21.4500563 | 21.5039284 | -0.026436502 | 0.64413006 |
| 20.1694441 | 20.0860914 | 20.1096112 | 20.0699291 | -0.032266291 | 0.531674825 |
| 23.7907638 | 23.7098051 | 23.772135 | 23.7769971 | -0.027776119 | 0.559056603 |
| 23.1425793 | 22.9773173 | 23.1166114 | 23.096855 | -0.048129907 | 0.234032969 |
| 21.8092387 | 21.6890045 | 21.7866686 | 21.8072399 | -0.008508994 | 0.854707687 |

| | | | | | |
|---|---|---|---|---|---|
| 20.3791297 | 20.3205637 | 20.2978743 | 20.309719 | -0.009607986 | 0.866221064 |
| 19.3526669 | 19.3923266 | 19.3586972 | 19.3405766 | -0.014776119 | 0.80071037 |
| 22.9509634 | 22.7914119 | 22.8292386 | 22.9838626 | 0.023607772 | 0.752426265 |
| 19.4880449 | 19.528003 | 19.5619908 | 19.6212191 | 0.132241366 | 0.136432387 |
| 19.6146189 | 19.5977638 | 19.5135664 | 19.530216 | -0.06119979 | 0.239788857 |
| 26.2981156 | 26.2378637 | 26.318628 | 26.2607863 | 0.024213119 | 0.600858764 |
| 21.7488663 | 21.6354663 | 21.7137685 | 21.7222498 | -0.036702923 | 0.366118721 |
| 21.3503333 | 21.3242999 | 21.3468028 | 21.4355548 | 0.054108365 | 0.278273568 |
| 21.3770088 | 21.2514428 | 21.3888228 | 21.3861667 | 0.017957026 | 0.752809236 |
| 19.9740755 | 19.8234872 | 19.9372187 | 19.9641126 | -0.03094681 | 0.590067732 |
| 15.2169398 | 15.2526287 | 15.3361238 | 15.3890425 | 0.00377219 | 0.96577393 |
| 18.0113956 | 18.1023409 | 18.0764126 | 18.0246917 | 0.059089647 | 0.568270807 |
| 19.3749567 | 19.2404126 | 19.177292 | 19.2667101 | 0.027484454 | 0.704817633 |
| 21.7724939 | 21.7816384 | 21.961615 | 21.8548542 | 0.054634733 | 0.579601188 |
| 20.1503079 | 20.048952 | 20.0267089 | 20.0662564 | -0.020478736 | 0.84724725 |
| 19.1377717 | 19.0643574 | 19.2010392 | 18.9746062 | -0.062428573 | 0.568422391 |
| 23.0214201 | 22.8973813 | 23.0343784 | 23.0004863 | -0.023128375 | 0.706333854 |
| 21.2683092 | 21.1931297 | 21.2009037 | 21.2220025 | 0.002282621 | 0.96296241 |
| 19.4763926 | 19.4163521 | 19.5483725 | 19.535509 | 0.059048109 | 0.238337692 |
| 20.9401228 | 21.0681042 | 21.0162255 | 21.0327114 | 0.097786119 | 0.061564177 |
| 21.3013735 | 21.2183415 | 21.3127395 | 21.2512915 | 0.009586316 | 0.889175059 |
| 20.4769849 | 20.2365807 | 20.3378224 | 20.4995244 | 0.030215448 | 0.763438774 |
| 20.6665681 | 20.5307125 | 20.6542746 | 20.5832779 | -0.036743291 | 0.45851298 |
| 18.3290216 | 18.0799548 | 18.1234728 | 18.2930119 | -0.089234242 | 0.344106432 |
| 18.9615375 | 18.9127873 | 19.0789229 | 19.1101506 | 0.059927173 | 0.456725971 |
| 21.7698698 | 21.6327842 | 21.7980692 | 21.8390054 | -0.035695202 | 0.608501295 |
| 20.3540015 | 20.1916727 | 20.2692232 | 20.2635136 | -0.075614596 | 0.151090706 |
| 16.9469693 | 16.8829184 | 16.891068 | 16.8977799 | -0.137415574 | 0.073495193 |
| 20.2717381 | 20.1041281 | 20.1107498 | 20.3280789 | -0.053508553 | 0.448157619 |
| 24.2847445 | 24.1229775 | 24.2602011 | 24.0843417 | -0.075498997 | 0.112158672 |
| 22.1472618 | 21.9181484 | 22.1091179 | 22.0706032 | 0.076838256 | 0.511040605 |
| 22.2869153 | 22.1515617 | 22.3845246 | 22.3650368 | -6.53E-04 | 0.992427581 |
| 19.2133039 | 19.2027772 | 19.2141587 | 19.230682 | -0.002010653 | 0.961211027 |
| 22.900163 | 22.8396528 | 22.9477096 | 22.9325036 | 0.039767093 | 0.313119887 |
| 16.4523014 | 16.0606558 | 16.3361962 | 16.1583664 | -0.12043514 | 0.294252464 |
| 23.7165416 | 23.6583228 | 23.7309251 | 23.7132723 | 0.022289228 | 0.599078244 |
| 23.0125524 | 23.0399136 | 23.0538004 | 22.9899499 | 0.008346829 | 0.792088685 |
| 23.7467918 | 23.7603 | 23.7451123 | 23.7109895 | 0.054656468 | 0.182401522 |
| 23.3999998 | 23.4196783 | 23.4847349 | 23.4327837 | 0.073554402 | 0.081895577 |
| 19.9126966 | 19.8027712 | 19.8574277 | 19.8663452 | -0.019635845 | 0.670440476 |
| 21.3684148 | 21.2219559 | 21.2975475 | 21.3011307 | 0.004853326 | 0.918957016 |
| 19.6468061 | 19.5722205 | 19.6443076 | 19.689253 | 0.03075936 | 0.571027457 |

| | | | | | |
|---|---|---|---|---|---|
| 17.6269227 | 17.7915525 | 17.8507055 | 17.7056709 | 0.126406595 | 0.050923196 |
| 20.473461 | 20.4179728 | 20.5591486 | 20.583463 | -0.005300552 | 0.938891894 |
| 21.9489273 | 21.8147477 | 21.9679104 | 22.0342123 | 0.019203488 | 0.784525817 |
| 20.9830836 | 21.1088316 | 21.0252384 | 20.9290083 | 0.122474222 | 0.155627947 |
| 20.5865635 | 20.7905636 | 20.7571754 | 20.7073271 | 0.15859628 | 0.069997605 |
| 20.1026349 | 20.127429 | 20.0974323 | 20.2888183 | 0.064101775 | 0.459322773 |
| 21.4528331 | 21.3165504 | 21.3974264 | 21.4024517 | -0.023149453 | 0.842242103 |
| 20.1842565 | 20.008099 | 20.1865628 | 20.1727379 | 0.030804112 | 0.791926031 |
| 20.0143943 | 19.6054481 | 19.9805525 | 19.5921047 | -0.088328286 | 0.541661395 |
| 19.2370892 | NA | 19.4530136 | 19.533231 | 0.320097629 | 0.048171111 |
| 21.4644093 | 21.3957313 | 21.4882207 | 21.4157999 | -0.017717043 | 0.702305668 |
| 18.8733877 | 18.8800371 | 19.0379358 | 18.8930291 | 0.022777102 | 0.760750129 |
| 22.9461344 | 22.8487862 | 22.9781121 | 22.8745224 | -0.07418475 | 0.251192573 |
| 18.6559406 | 18.5909256 | 18.4875327 | 18.501466 | -0.0929139 | 0.209522137 |
| 22.3714431 | 22.0380023 | 22.1427662 | 22.2180369 | -0.087181751 | 0.170877567 |
| 16.4243013 | NA | 16.4345167 | 16.3905971 | 0.022961955 | 0.914850067 |
| 16.3923273 | 16.4106507 | 16.2786726 | 16.2306108 | -0.134132695 | 0.100479884 |
| NA | 14.0465238 | NA | NA | 0.45899716 | 0.013734724 |
| 15.7054313 | 15.7193734 | 15.4909655 | 16.0306239 | -0.027148546 | 0.817815738 |
| NA | 11.4347177 | NA | NA | -0.201079463 | 0.103739932 |
| 24.9094398 | 24.7649973 | 24.719664 | 24.7853298 | -0.035174856 | 0.681916967 |
| 25.394973 | 25.1861648 | 25.3579413 | 25.2874213 | -0.064422941 | 0.438729382 |
| 18.4430623 | 18.3748498 | 18.4023513 | 18.4628084 | -0.142375937 | 0.081533251 |
| 21.8844125 | 21.6167149 | 21.7413184 | 21.784892 | -0.070920094 | 0.32789703 |
| 19.3677573 | 19.2239493 | 19.3613882 | 19.4271654 | -0.001306464 | 0.985260179 |
| 20.9645382 | 20.6574488 | 20.8629692 | 20.8984016 | -0.102374303 | 0.151806446 |
| 18.2139849 | 18.19074 | 18.2341531 | 18.3202791 | 0.02966582 | 0.639546054 |
| 22.5810525 | 22.4027254 | 22.5959969 | 22.563534 | -0.051878206 | 0.386687818 |
| 21.6609457 | 21.6281775 | 21.6700034 | 21.6507309 | -0.030415438 | 0.460011474 |
| 19.6802067 | 19.7011315 | 19.7239616 | 19.7142577 | 0.026890713 | 0.590662565 |
| 20.3994648 | 20.4116035 | 20.3415432 | 20.4669484 | 0.047076848 | 0.361649259 |
| 23.550217 | 23.4531254 | 23.5988815 | 23.6226045 | 0.030597087 | 0.560295376 |
| 19.2894501 | 19.232135 | 19.3763348 | 19.3028551 | 0.021295677 | 0.687828188 |
| 19.455679 | 19.4729373 | 19.5532413 | 19.5592111 | 0.084781383 | 0.150041127 |
| 21.0571939 | 21.0206465 | 21.0681691 | 21.0773414 | -0.004524265 | 0.942601958 |
| 18.5923592 | 18.513244 | 18.5208427 | 18.6440769 | 0.019541761 | 0.781875116 |
| 17.1628469 | 17.1830624 | 17.1512211 | 17.2072113 | -0.009555831 | 0.896865165 |
| 18.3342512 | 18.4819686 | 18.3954706 | 18.2395861 | -0.024959889 | 0.756734246 |
| 22.2781398 | 22.1332189 | 22.2827989 | 22.2462343 | -0.010769626 | 0.899381214 |
| 17.6725639 | 17.7499746 | 17.7973906 | 17.714365 | 0.088688998 | 0.314882414 |
| 17.9724645 | 17.9967685 | 18.0187218 | 18.3759433 | 0.192384667 | 0.08182959 |
| 16.1428789 | NA | 16.5670258 | 16.595137 | -0.047344875 | 0.874508683 |

| | | | | | |
|---|---|---|---|---|---|
| 19.7915413 | 19.6595036 | 19.8449451 | 19.7994475 | -0.028596956 | 0.696515974 |
| 16.2533256 | 15.7591571 | 16.4678929 | 16.2609189 | 0.034581068 | 0.835490181 |
| 26.816874 | 26.9310771 | 27.0852212 | 26.9820735 | 0.104014241 | 0.102130649 |
| 18.6463099 | 18.7540158 | 18.7074634 | 18.8397136 | 0.110841462 | 0.02035974 |
| 22.1418462 | 22.1444768 | 22.1214685 | 22.1402472 | 0.016882421 | 0.815739712 |
| 22.5952343 | 22.4287904 | 22.5559333 | 22.5925836 | 0.014710717 | 0.765676666 |
| 22.3386903 | 22.375064 | 22.3781851 | 22.3629262 | 0.032952712 | 0.424306277 |
| 20.7990846 | 20.7257488 | 20.7718775 | 20.7981497 | 0.029518764 | 0.540267575 |
| 18.1210251 | 17.9413877 | 18.319918 | 18.1170042 | 0.118238822 | 0.199857789 |
| 22.9109025 | 22.8986431 | 22.8451548 | 22.8847986 | -0.022821448 | 0.72272601 |
| 21.8357035 | 21.6088657 | 21.7539192 | 21.7036055 | -0.111567334 | 0.087418654 |
| 19.8950464 | 19.8127667 | 19.8435321 | 19.8544101 | -0.076807252 | 0.181867269 |
| 17.1051881 | 17.0910775 | 17.0638009 | 17.2436584 | 0.045030844 | 0.528515611 |
| 18.1997148 | 18.1342817 | 18.15922 | 18.292581 | 0.058645597 | 0.375325605 |
| 18.6579861 | 18.9056346 | 18.1869737 | 18.4953626 | 0.194077032 | 0.480871056 |
| 19.6019774 | 19.4232493 | 19.5498963 | 19.4127238 | -0.078814158 | 0.385303093 |
| 17.2757699 | 17.0171578 | 17.0856947 | 17.1849044 | -0.091445329 | 0.290534906 |
| 20.2204093 | 20.0583156 | 20.2111654 | 20.1778844 | 0.007979861 | 0.925669734 |
| 22.42813 | 22.3221472 | 22.3907672 | 22.4119589 | -0.034629516 | 0.680973761 |
| 20.3267838 | 20.2497477 | 20.3453893 | 20.254378 | 0.017910717 | 0.750035569 |
| 24.6245181 | 24.5461982 | 24.4960243 | 24.5994474 | -0.00778602 | 0.880842559 |
| 18.9803054 | 18.846569 | 19.0320507 | 18.9840257 | 0.026350166 | 0.730331966 |
| 18.3383916 | 18.2527105 | 18.5866869 | 18.3297093 | 0.118383781 | 0.237568061 |
| NA | 13.7994709 | NA | NA | -0.121926306 | 0.530372748 |
| 19.895697 | 19.7694619 | 19.9094571 | 19.8565671 | -0.024358526 | 0.648026246 |
| 20.7045721 | 20.5211672 | 20.6078932 | 20.6536805 | -0.050256784 | 0.34724718 |
| 22.1480968 | 22.164653 | 22.0834589 | 22.2968298 | 0.07432598 | 0.342453469 |
| 19.4639288 | 19.3536268 | 19.483447 | 19.372898 | -0.045038711 | 0.492852755 |
| 18.7896153 | 18.8942832 | 19.0069591 | 18.8833102 | 0.113381079 | 0.145279879 |
| 20.0920804 | 19.9240744 | 20.0877844 | 20.0355469 | -0.005088971 | 0.959736663 |
| 24.0658197 | 23.8959098 | 24.0015853 | 24.0106022 | -0.040449338 | 0.486226235 |
| 14.905795 | NA | 14.9950355 | 14.9386768 | 0.268246155 | 0.150912483 |
| 15.5493142 | 15.5518692 | 15.6924608 | 15.68648 | 0.007000998 | 0.945199483 |
| 18.4924456 | 18.4303352 | 18.3631064 | 18.4030012 | -0.132259557 | 0.157703732 |
| 20.3829021 | 20.2655703 | 20.2830716 | 20.4159555 | -0.061215559 | 0.417875551 |
| 19.9460949 | 19.8499279 | 19.8949947 | 19.8338017 | -0.076085627 | 0.232675345 |
| 23.1023529 | 23.2084574 | 23.1378662 | 23.0688445 | 0.015631405 | 0.795836604 |
| 20.0681912 | 19.9348693 | 20.0794501 | 20.0557903 | -0.030274696 | 0.59681924 |
| 21.6773481 | 21.5976208 | 21.6743529 | 21.6781524 | -0.009300206 | 0.868856234 |
| 18.6766908 | 18.7097597 | 18.6660859 | 18.6481345 | -0.070352623 | 0.206895822 |
| 23.0484454 | 22.9460362 | 23.0087059 | 22.950781 | -0.046914586 | 0.350644632 |
| 24.3933945 | 24.2954202 | 24.4351213 | 24.3974445 | 0.00890814 | 0.857236248 |

| | | | | | |
|---|---|---|---|---|---|
| 22.2552371 | 22.143535 | 22.2683208 | 22.206856 | 0.00551799 | 0.909344083 |
| 25.3759346 | 25.2600267 | 25.3309315 | 25.3505238 | -0.003716897 | 0.935145278 |
| 23.7437468 | 23.68276 | 23.7376789 | 23.7088161 | -0.004597945 | 0.919625186 |
| 21.2990269 | 21.267127 | 21.2782889 | 21.3273544 | -0.013887185 | 0.760900704 |
| 19.9472764 | 19.850942 | 19.9467513 | 19.9056496 | -0.06929438 | 0.105913793 |
| 19.8557183 | 19.9185455 | 19.9038599 | 19.7946978 | 0.057480589 | 0.190778502 |
| 21.1664874 | 21.0610984 | 21.0904723 | 21.0804786 | -0.052285156 | 0.257964404 |
| 22.2688209 | 22.2076228 | 22.3691506 | 22.308932 | 0.049806952 | 0.340406595 |
| 20.85122 | 20.7270434 | 20.7659661 | 20.8766732 | -0.003572834 | 0.947359656 |
| 20.592046 | 20.5822499 | 20.6607002 | 20.7475438 | 0.048831364 | 0.395802571 |
| 21.3084962 | 21.3422183 | 21.3272497 | 21.3705146 | 0.039626126 | 0.491677522 |
| 20.2246937 | 20.0962405 | 20.1433397 | 20.2745157 | -0.019515747 | 0.738905326 |
| 21.2595781 | 21.1359405 | 21.389945 | 21.2821703 | 0.067290459 | 0.259467053 |
| 19.0507858 | 18.9353302 | 18.9489898 | 18.9985875 | -0.044604231 | 0.461635486 |
| 20.9532519 | 20.7922175 | 20.8958039 | 20.8333703 | -0.035412919 | 0.568828836 |
| 19.3783346 | 19.2198046 | 19.1978737 | 19.3829749 | -0.096785467 | 0.171181965 |
| 20.7394173 | 20.8663735 | 21.0058678 | 20.9596903 | 0.171433781 | 0.024157574 |
| 18.6923218 | 18.7121396 | 18.8947863 | 18.9255791 | 0.035478448 | 0.647415557 |
| 21.8697858 | 21.6921156 | 21.8652611 | 21.9318647 | 0.021043753 | 0.804354009 |
| 19.6510879 | 19.372491 | 19.6709906 | 19.5949596 | 0.023711343 | 0.816874819 |
| 21.9939529 | 21.9187849 | 21.9931262 | 22.0561707 | 0.100471685 | 0.44809809 |
| 16.6594317 | 16.2781094 | 16.8914242 | 16.9777775 | 0.181670515 | 0.197483594 |
| 19.036332 | 18.8769238 | 18.9496753 | 18.9522306 | -0.12146027 | 0.078255493 |
| 19.5181405 | 19.4098891 | 19.4871188 | 19.5522864 | 0.039732081 | 0.473931411 |
| 17.2150415 | 16.7107333 | 17.290431 | 17.0567119 | -0.188254365 | 0.075131212 |
| 23.5842773 | 23.5351327 | 23.5364468 | 23.5536559 | 0.025912425 | 0.646889585 |
| 17.5230662 | 17.5770092 | 17.2460715 | 17.4838499 | -0.094564117 | 0.553313747 |
| 18.855614 | 18.8203158 | 18.6951415 | 18.8040704 | -0.082074126 | 0.33169328 |
| 21.5389728 | 21.4774013 | 21.4450467 | 21.4501357 | -6.57E-04 | 0.991894725 |
| 19.3914785 | 19.2082656 | 19.1579116 | 19.1113132 | -0.191644384 | 0.004516097 |
| 20.9051608 | 20.823946 | 20.9426757 | 20.8842853 | 0.006319689 | 0.890866011 |
| 19.9631491 | 20.032602 | 20.182252 | 20.1228799 | 0.107707234 | 0.113529612 |
| 20.4713141 | 20.4622746 | 20.3673413 | 20.365081 | 2.39E-04 | 0.998180152 |
| 19.40157 | 19.2917363 | 19.3742559 | 19.3043936 | -0.035356139 | 0.742726628 |
| 18.9433115 | 18.8462557 | 19.1583815 | 19.043213 | 0.017541993 | 0.872066468 |
| NA | 15.3336041 | NA | NA | -0.05574055 | 0.643402301 |
| 23.4844456 | 23.2589684 | 23.3732757 | 23.4673292 | -0.05945279 | 0.367695379 |
| 21.5636205 | 21.482447 | 21.5735291 | 21.5809608 | -0.002773861 | 0.946451404 |
| 21.7283778 | 21.4721539 | 21.7468483 | 21.6365665 | -0.081433863 | 0.322875462 |
| 20.4668072 | 20.340394 | 20.3842573 | 20.4353533 | -0.014464039 | 0.766357792 |
| 18.7708054 | 18.7244428 | 18.9749851 | 18.9043434 | 0.034425511 | 0.706744221 |
| 15.2462098 | 15.4620691 | 15.9829222 | 15.2103946 | -0.110020495 | 0.667732043 |

| | | | | | |
|---|---|---|---|---|---|
| 20.4669293 | 20.3161773 | 20.3772106 | 20.3510602 | -0.078706599 | 0.118820348 |
| 20.0196407 | 19.9697651 | 20.0395637 | 19.994553 | -0.015949148 | 0.7936481 |
| 23.154852 | 23.123798 | 23.1570529 | 23.2443911 | 0.076699181 | 0.255289165 |
| 19.9155662 | 19.7373813 | 19.8390593 | 19.953902 | 0.026945406 | 0.724651031 |
| 19.6389614 | 19.6233259 | 19.6534027 | 19.6743147 | 0.002947266 | 0.959675717 |
| 20.7320919 | 20.7298695 | 20.8098559 | 20.8379476 | 0.057133749 | 0.348183532 |
| 19.3447177 | 19.2119952 | 19.3101598 | 19.3979621 | 0.00267234 | 0.965707837 |
| 20.5276503 | 20.4164104 | 20.4443175 | 20.5440327 | -0.026463351 | 0.69772254 |
| 20.7872653 | 20.7443171 | 21.0920231 | 20.8459114 | 0.164840011 | 0.085071704 |
| 15.7155465 | 16.0945192 | 16.1156947 | 15.8352106 | 0.261944716 | 0.053467919 |
| 15.9520257 | 16.3072735 | 15.9434412 | 16.3575021 | 0.109238078 | 0.574116258 |
| 20.4177547 | 20.2959969 | 20.4720722 | 20.2778592 | -0.119670979 | 0.19539784 |
| 14.3861936 | 13.9441858 | 14.3589243 | 14.0937133 | -0.601742779 | 0.010564508 |
| 20.6547918 | 20.597885 | 20.5917072 | 20.6049008 | -0.0057775 | 0.918356893 |
| 20.3390458 | 20.3005925 | 20.3621676 | 20.3033251 | 0.05384886 | 0.271296833 |
| 22.9514671 | 22.8103712 | 22.9302469 | 22.8701508 | -0.034337148 | 0.670177405 |
| NA | 10.9015811 | NA | NA | -0.93960244 | 0.145599757 |
| 14.9799696 | 15.6818475 | 15.4147149 | 15.6703393 | 0.235195227 | 0.319814058 |
| 16.8489644 | 16.6721063 | 16.7373327 | 16.5244088 | -0.323544785 | 0.024347111 |
| 15.8837591 | 15.8683807 | 16.0330131 | 16.0000624 | -0.010604345 | 0.920074043 |
| 18.1875911 | 18.1212022 | 18.0665024 | 18.3498901 | -0.136868037 | 0.191955832 |
| 18.0231104 | 17.611349 | 17.8762353 | 17.8885317 | -0.208491482 | 0.048310424 |
| 18.3762014 | 18.4424076 | 18.3291851 | 18.1920521 | -0.121138921 | 0.232167752 |
| 18.6684592 | 18.5530588 | 18.534384 | 18.4899981 | -0.038191951 | 0.672950644 |
| 17.0637328 | 17.0410267 | 17.2604229 | 17.0540957 | 0.036469033 | 0.668109822 |
| 17.4554236 | 17.2829174 | 17.5363105 | 17.5642979 | -0.048919284 | 0.555282522 |
| 18.5750892 | 18.1971427 | 18.3278543 | 18.4218521 | -0.116731741 | 0.150931941 |
| 20.2493077 | 19.9768226 | 20.2027761 | 20.1602606 | -0.096790146 | 0.198677668 |
| 18.8550041 | 18.8335721 | 18.7988904 | 18.775482 | -0.048390408 | 0.496379137 |
| 19.3717134 | 19.269687 | 19.1476916 | 19.2490193 | -0.064763359 | 0.354281316 |
| 21.0181134 | 20.8007591 | 20.9330346 | 20.9373892 | -0.102397188 | 0.131082546 |
| 20.5810896 | 20.507024 | 20.4713368 | 20.6037234 | -8.72E-04 | 0.989494485 |
| 20.4639969 | 20.2880947 | 20.4208285 | 20.4735885 | -0.073673973 | 0.259061874 |
| 19.20024 | 19.1279098 | 19.06671 | 19.1111288 | -0.059064523 | 0.360111239 |
| 23.0828796 | 23.0864569 | 23.0921508 | 23.0971923 | -0.003283625 | 0.953651678 |
| 20.8013687 | 20.7465961 | 20.6808609 | 20.7297555 | -0.050610454 | 0.357561303 |
| 21.45706 | 21.2471292 | 21.3734257 | 21.3530238 | -0.031269102 | 0.564222077 |
| 21.6782059 | 21.734155 | 21.7378512 | 21.7396996 | 0.034262481 | 0.520233375 |
| 22.7575168 | 22.6049398 | 22.6654167 | 22.6922786 | -0.042359326 | 0.407958048 |
| 22.591279 | 22.4868162 | 22.5594382 | 22.5971781 | -0.022595031 | 0.614730593 |
| 18.3129127 | 18.2792871 | 18.2080796 | 18.1973725 | -0.084554239 | 0.05921525 |
| 18.8622778 | 18.754574 | 18.8287383 | 18.8681806 | -0.069319891 | 0.10654498 |

| | | | | | |
|---|---|---|---|---|---|
| 20.5453587 | 20.4595967 | 20.4676617 | 20.4205304 | -0.04938675 | 0.224800749 |
| 23.7724067 | 23.8321419 | 23.8142341 | 23.7713378 | 0.020019268 | 0.641643595 |
| 24.6559984 | 24.6224651 | 24.6364838 | 24.6357189 | 0.038871364 | 0.380638787 |
| 20.0834505 | 20.0702404 | 20.082487 | 20.0920531 | 0.022783125 | 0.610961902 |
| 19.4211717 | 19.2995358 | 19.4606639 | 19.4287831 | 0.017817939 | 0.715019367 |
| 20.103052 | 20.0483874 | 20.0968194 | 20.1335961 | 0.040596571 | 0.413456896 |
| 19.8858557 | 19.7563382 | 19.9224071 | 19.912684 | 0.030252181 | 0.545911915 |
| 20.5331601 | 20.6269124 | 20.6060442 | 20.628746 | 0.044923585 | 0.37967007 |
| 20.6788323 | 20.6692213 | 20.704341 | 20.7072295 | 0.066636092 | 0.203351346 |
| 25.2586699 | 25.2164398 | 25.3175119 | 25.2739852 | 0.067702898 | 0.205568883 |
| 24.3282061 | 24.2494759 | 24.4503628 | 24.3263522 | 0.044519142 | 0.413124394 |
| 19.0951765 | 19.0248743 | 19.0927055 | 19.2581857 | 0.020343539 | 0.73090314 |
| 21.7833833 | 21.7721321 | 21.9055733 | 21.844787 | 0.053749662 | 0.394042429 |
| 20.8095129 | 20.8188778 | 20.8086678 | 20.8164794 | 0.011962202 | 0.849485221 |
| 23.613552 | 23.6245231 | 23.7259111 | 23.615413 | 0.043952803 | 0.495686472 |
| 20.892379 | 21.1293068 | 21.1949443 | 21.1696436 | 0.116870062 | 0.086151546 |
| 19.4683511 | 19.4207223 | 19.5083204 | 19.5481854 | 0.062664604 | 0.360771611 |
| 18.3760991 | 18.5859816 | 18.4480046 | 18.5525983 | 0.16379994 | 0.031081841 |
| 24.8546707 | 24.886231 | 24.8793093 | 24.8543763 | 0.038224698 | 0.614119871 |
| 21.5418413 | 21.3706475 | 21.5883318 | 21.5317875 | 0.070164284 | 0.362374535 |
| 18.691186 | 18.6896126 | 18.9907974 | 18.9612513 | 0.115186243 | 0.162981195 |
| 19.9351669 | 19.7353183 | 19.8316604 | 19.874754 | -0.039249266 | 0.630365313 |
| 20.9475714 | 20.8531637 | 20.8948674 | 20.9464148 | 0.009186891 | 0.91324902 |
| 19.6055581 | 19.4892933 | 19.5054247 | 19.6303384 | 0.03233228 | 0.721600858 |
| 16.025444 | 16.0439778 | 16.0593968 | 15.9596908 | 0.051271114 | 0.577057279 |
| 17.4234228 | 17.4523404 | 17.3486715 | 17.4317337 | -0.026502606 | 0.832438285 |
| 16.404508 | 16.1386654 | 16.4178684 | 16.7009843 | 0.145886418 | 0.306565704 |
| 22.9371846 | 22.9700092 | 23.0706369 | 23.0080016 | 0.07855269 | 0.592840048 |
| 17.3641045 | 17.1195469 | 17.3142853 | 17.3256633 | -5.46E-04 | 0.997075871 |
| NA | 15.0089245 | NA | NA | -0.321830926 | 0.117933568 |
| 15.4775138 | 16.2229358 | 16.1415634 | 16.202833 | 0.094032346 | 0.704271634 |
| NA | 18.0421964 | NA | NA | 0.425147773 | 0.151211925 |
| NA | 13.0911991 | NA | NA | -0.176855664 | 0.374682236 |
| 16.6158001 | 16.4710463 | 16.3545534 | 16.6636994 | -0.093305304 | 0.519032542 |
| 18.2582386 | 17.8608597 | 17.8914144 | 18.1958457 | -0.153292521 | 0.287080381 |
| NA | 15.6009755 | NA | NA | -0.106337086 | 0.418691415 |
| 17.6459324 | 17.6347544 | 17.5247873 | 17.66483 | -0.001902392 | 0.987101359 |
| 21.8739021 | 21.544597 | 21.7978693 | 21.9049079 | -0.078700227 | 0.420665408 |
| 19.7031676 | 19.4640273 | 19.6724752 | 19.6312282 | -0.076175995 | 0.36587524 |
| 22.2393471 | 22.1397093 | 22.2600261 | 22.1094475 | -0.049109233 | 0.558022423 |
| 18.8083546 | 18.5903146 | 18.8147467 | 18.8078958 | -0.048242815 | 0.552803982 |
| 19.9652847 | 19.6795811 | 19.798865 | 19.8068086 | -0.021929245 | 0.784076935 |

| | | | | | |
|---|---|---|---|---|---|
| 21.073753 | 20.9329922 | 20.9685367 | 20.8776179 | -0.103691768 | 0.155531771 |
| 20.2751375 | 20.18475 | 20.1994309 | 20.2666995 | -0.071685911 | 0.294005484 |
| 21.6150774 | 21.3806669 | 21.5538481 | 21.5000277 | -0.091448751 | 0.171141648 |
| 19.1355906 | 19.0275103 | 18.9299449 | 19.0346807 | -0.079925201 | 0.223594774 |
| 20.1460383 | 20.1292777 | 20.0757986 | 20.1825976 | -0.014632615 | 0.813017627 |
| 21.1786875 | 21.0330227 | 21.2021131 | 21.1118026 | -0.085545887 | 0.150511305 |
| 20.885611 | 20.5803537 | 20.8272989 | 20.8097919 | -0.093190998 | 0.115329626 |
| 20.6606569 | 20.6877892 | 20.720247 | 20.6794158 | 0.018771661 | 0.721156719 |
| 24.9175384 | 24.8065105 | 24.8979786 | 24.9055631 | -0.011705349 | 0.822807144 |
| 20.6734388 | 20.5244679 | 20.6883059 | 20.6797443 | -0.013978828 | 0.780717495 |
| 22.719251 | 22.7944976 | 22.726201 | 22.7844817 | 0.003160496 | 0.949753682 |
| 20.3059429 | 20.3145537 | 20.2941219 | 20.2777933 | -0.023964072 | 0.612795808 |
| 20.4277125 | 20.353902 | 20.3621036 | 20.2670202 | -0.054098477 | 0.237811453 |
| 22.7873696 | 22.7207765 | 22.7808691 | 22.6627213 | -0.017232874 | 0.700809326 |
| 21.1812307 | 21.1860882 | 21.2026196 | 21.2467939 | 0.019639651 | 0.620997359 |
| 22.6190592 | 22.5957494 | 22.6121075 | 22.6344429 | 0.032791553 | 0.425909872 |
| 21.6271795 | 21.496511 | 21.5470556 | 21.5933733 | -0.015919697 | 0.74097228 |
| 22.3768941 | 22.3129217 | 22.3832787 | 22.3646293 | 0.03459683 | 0.48151532 |
| 21.283719 | 21.2037585 | 21.3475069 | 21.2486812 | 0.034749841 | 0.527089193 |
| 21.0199045 | 21.0506391 | 21.0738746 | 21.0999683 | 0.115591697 | 0.041974841 |
| 17.870102 | 17.8365509 | 17.9708055 | 17.9232006 | 0.022667669 | 0.690868553 |
| 20.4330632 | 20.3555293 | 20.4842416 | 20.5096757 | 0.017785461 | 0.756040317 |
| 19.9128491 | 19.8704397 | 19.8819044 | 19.9612432 | 0.047934531 | 0.418109826 |
| 21.5815851 | 21.4675865 | 21.605237 | 21.632232 | 0.026056348 | 0.665422371 |
| 20.7092744 | 20.65548 | 20.7366887 | 20.7634346 | 0.023136636 | 0.766033855 |
| 20.6968232 | 20.674885 | 20.6405165 | 20.6899123 | -0.028010253 | 0.723526068 |
| 21.4795345 | 21.4525555 | 21.6335349 | 21.542035 | 0.001430321 | 0.986214477 |
| 21.5658538 | 21.509679 | 21.5311696 | 21.6251139 | 0.079617538 | 0.343953706 |
| 18.4999517 | 18.5767555 | 18.5684182 | 18.591678 | 0.097662255 | 0.251451284 |
| 17.5517973 | 17.6283645 | 17.6266507 | 17.8340442 | 0.067696506 | 0.426280522 |
| 19.1895695 | 19.2646849 | 19.2142731 | 19.1630939 | -0.010588574 | 0.911940983 |
| 18.2259558 | 17.6486165 | 18.290985 | 18.3203843 | -0.091945049 | 0.475656506 |
| NA | 12.9587425 | NA | NA | 0.2247387 | 0.260244769 |
| 13.4976749 | 13.4756477 | 13.1393206 | 13.3499506 | -0.230642758 | 0.338540789 |
| 20.3617174 | 20.2494246 | 20.3928715 | 20.3642292 | 0.010558466 | 0.860482876 |
| 14.3637251 | NA | 14.6460068 | 14.4145901 | 0.162513081 | 0.226180154 |
| 21.2065227 | 21.0721796 | 21.2257625 | 21.1334173 | -0.089036126 | 0.124075054 |
| 19.6771763 | 19.456077 | 19.6680316 | 19.65127 | -0.11428774 | 0.210021271 |
| 15.3671846 | NA | 15.8382001 | 15.5605252 | 0.311071296 | 0.254465991 |
| 16.51651 | 16.3645631 | 16.4938897 | 16.5215168 | -0.149711855 | 0.148729529 |
| 21.7247474 | 21.5067939 | 21.682738 | 21.6877802 | -0.096851174 | 0.197270887 |
| 23.0644901 | 22.8817144 | 22.9326984 | 22.9360847 | -0.063556281 | 0.292535623 |

| | | | | | |
|---|---|---|---|---|---|
| 22.4940449 | 22.4535176 | 22.5318937 | 22.5055005 | -0.005436051 | 0.892318242 |
| 25.0451439 | 25.0393927 | 25.1199232 | 25.0793111 | 0.057066025 | 0.121874325 |
| 17.9290801 | 17.696036 | 17.9066502 | 17.9552784 | 0.006392101 | 0.935543383 |
| 18.7495998 | 18.7388054 | 18.7607973 | 18.7590108 | 0.038270034 | 0.466516709 |
| 21.150166 | 20.9517892 | 21.0608582 | 21.1224393 | -0.012822249 | 0.838588691 |
| 21.0625046 | 20.9000538 | 20.9365598 | 20.938472 | 0.009608058 | 0.891715889 |
| 21.1380232 | 21.0498324 | 21.2131378 | 21.2635894 | 0.025143485 | 0.73633426 |
| 19.5557787 | 19.4860902 | 19.5040619 | 19.5569255 | -0.032143711 | 0.583563037 |
| 19.5100024 | 19.5341623 | 19.5805678 | 19.5785646 | 0.028010406 | 0.594228989 |
| 20.5943344 | 20.5117313 | 20.4437004 | 20.625518 | -0.011885445 | 0.835323313 |
| 23.26812 | 23.3156788 | 23.2563566 | 23.2477403 | 0.03439673 | 0.501285877 |
| 18.4289095 | 18.611324 | 18.8140047 | 18.8221629 | 0.088657753 | 0.411830678 |
| 21.3260608 | 20.3000861 | 20.9557491 | 21.1181997 | -0.284918854 | 0.314752669 |
| 19.9479687 | 19.7278887 | 19.8981274 | 19.8553054 | -0.091332893 | 0.164843146 |
| 20.7514637 | 20.5694853 | 20.6670014 | 20.6793987 | -0.103364979 | 0.067991565 |
| 21.5914552 | 21.4729652 | 21.5489723 | 21.5101513 | 0.001894559 | 0.967602596 |
| 18.8145466 | 18.8996899 | 18.9243572 | 18.9121705 | 0.067614663 | 0.165176458 |
| 20.2377741 | 20.2298203 | 20.2967762 | 20.2283018 | -0.012012204 | 0.809770046 |
| 24.2654334 | 24.1412052 | 24.3060047 | 24.2649805 | 0.031997884 | 0.574408609 |
| 20.8456214 | 20.7108647 | 20.8877927 | 20.8931474 | 0.062976954 | 0.295300002 |
| 20.9579838 | 21.0041605 | 21.0522178 | 21.1290922 | 0.069538384 | 0.368531198 |
| 15.6880183 | 15.872261 | 15.8223663 | 15.6267261 | 0.106184169 | 0.228638525 |
| 18.0156402 | 17.8292686 | 18.0420791 | 18.0894131 | -0.016216855 | 0.852368235 |
| 16.6350063 | 16.6955048 | 16.6558981 | 16.7239309 | 0.076520529 | 0.47277762 |
| 17.1256687 | 17.4037246 | 17.4795748 | 17.5535461 | 0.470042576 | 0.008142667 |
| 19.9396596 | 19.8106752 | 19.8859358 | 19.8745409 | -0.065011869 | 0.292960499 |
| 18.1757043 | 17.9885744 | 18.046459 | 18.1062434 | 0.0173956 | 0.833885711 |
| 20.4993812 | 20.4236428 | 20.3814079 | 20.4697499 | -0.109881866 | 0.019554511 |
| 19.8990919 | 20.1575492 | 20.0613773 | 20.1342495 | 0.22205296 | 0.074606497 |
| 22.4109735 | 22.3723671 | 22.3167344 | 22.3205713 | -0.047524178 | 0.488162693 |
| 21.6232878 | 21.5603398 | 21.6022081 | 21.6069671 | -0.002424427 | 0.957896539 |
| 20.9961924 | 20.8726366 | 20.9932257 | 20.9691426 | -8.34E-05 | 0.998593459 |
| 18.9642017 | 18.8467643 | 18.8900776 | 18.9815282 | -0.024065949 | 0.671793266 |
| 20.0463897 | 19.9818821 | 20.1482993 | 20.0580363 | 0.052868892 | 0.563866237 |
| 15.3111774 | 15.1136612 | 14.8910687 | 15.0898576 | -0.280366634 | 0.15711555 |
| 11.4748166 | NA | 11.8123061 | 11.4915883 | -0.025986789 | 0.869399147 |
| 20.9946818 | 20.9474768 | 20.9566064 | 21.1289089 | 0.083209275 | 0.109074987 |
| 23.0446761 | 22.9532799 | 22.9551644 | 22.9885421 | -0.011594909 | 0.807462831 |
| 21.4838778 | 21.4335125 | 21.4599158 | 21.4827157 | 0.031303574 | 0.509496786 |
| 21.0025335 | 20.9217447 | 20.9808385 | 21.0157099 | 0.010050479 | 0.855140504 |
| 18.3286716 | 18.2327685 | 18.1384316 | 18.2378399 | -0.062816339 | 0.459797897 |
| 12.3325141 | 12.8404958 | 12.3318353 | 12.4476783 | -0.013568779 | 0.926003028 |

| | | | | | |
|---|---|---|---|---|---|
| 19.3191357 | 18.9542494 | 19.0265628 | 19.2872549 | -0.070325252 | 0.471577269 |
| 19.636469 | 19.6373634 | 19.8288247 | 19.461702 | 0.020567987 | 0.814160119 |
| 26.2460798 | 26.25443 | 26.3108738 | 26.2553426 | 0.00398583 | 0.96292 |
| 21.5552153 | 21.4372029 | 21.5631078 | 21.4566458 | -0.037689741 | 0.603871476 |
| 19.8178767 | 19.7054674 | 19.7846685 | 19.8080486 | -0.040685415 | 0.489148688 |
| 18.4993328 | 18.4645599 | 18.6087196 | 18.6119537 | 0.06822871 | 0.23971589 |
| 19.7468268 | 19.5876214 | 19.7458663 | 19.6425065 | -0.050559768 | 0.26929808 |
| 21.1546793 | 21.097112 | 21.1579093 | 21.1629685 | 0.007849116 | 0.839704185 |
| 23.2542883 | 23.2312116 | 23.2670698 | 23.2234118 | 0.020803397 | 0.617766985 |
| 20.3275294 | 20.1903135 | 20.3035041 | 20.3283722 | 0.034275135 | 0.518206714 |
| 19.8508812 | 19.820415 | 19.6997693 | 19.7735524 | -0.048594637 | 0.383281421 |
| 24.6001398 | 24.514205 | 24.681224 | 24.708042 | 0.100690173 | 0.119745107 |
| 19.8199866 | 19.8674644 | 20.0803499 | 20.1669496 | 0.176169921 | 0.034303425 |
| 16.4035246 | 16.4194595 | 16.5963135 | 16.4979299 | 0.298981636 | 0.066430888 |
| 21.9224907 | 21.8161623 | 21.7269376 | 21.885877 | 0.004998109 | 0.948185201 |
| 17.2897969 | 17.044528 | 17.2322146 | 17.3221919 | -0.026697387 | 0.762652542 |
| 26.734324 | 26.5928277 | 26.6891909 | 26.6797167 | -0.026148413 | 0.615910298 |
| 19.0818177 | 19.1147331 | 19.0825069 | 19.0619557 | -0.054379693 | 0.405971107 |
| 17.806631 | 17.8443581 | 17.8713955 | 18.110543 | 0.016743638 | 0.872760703 |
| 17.4373345 | 17.490805 | 17.3746206 | 17.563441 | 0.080898912 | 0.249185678 |
| 19.5268627 | 19.3678952 | 19.4557597 | 19.4230295 | -0.102865164 | 0.105662469 |
| 21.9750517 | 21.8823022 | 21.9569148 | 21.9090865 | -0.012337163 | 0.770128724 |
| 21.2280211 | 21.2280359 | 21.2063002 | 21.2111335 | -0.011385259 | 0.752960252 |
| 21.6229606 | 21.5566921 | 21.6625458 | 21.6193787 | 0.024741049 | 0.551610864 |
| 21.5142536 | 21.3702513 | 21.4508932 | 21.54872 | 0.005992161 | 0.90507048 |
| 19.2846814 | 19.3764971 | 19.4503897 | 19.5145197 | 0.134404615 | 0.044601472 |
| 18.1382437 | 17.9708886 | 18.0552522 | 18.1248678 | -0.065093099 | 0.317073738 |
| 22.9459536 | 22.8949682 | 22.957502 | 22.9662019 | 0.040354105 | 0.546227673 |
| 21.4595242 | 21.4069827 | 21.4846451 | 21.4899368 | 0.071629412 | 0.405931526 |
| 17.7104965 | 17.660797 | 17.4310423 | 17.6965966 | 0.004756033 | 0.963420518 |
| NA | 20.6023584 | NA | NA | 0.446866496 | 0.084582795 |
| 26.6240994 | 26.6122903 | 26.6289127 | 26.6896201 | 0.05056126 | 0.4424227 |
| 21.3503012 | 21.2575369 | 21.3392582 | 21.3429671 | 0.007229463 | 0.862453139 |
| 16.7638026 | NA | 16.885626 | 16.8150509 | 0.131838423 | 0.274078779 |
| 22.2811496 | 22.0749421 | 22.2658909 | 22.2866328 | -0.010395583 | 0.871283964 |
| 20.0092197 | 20.0845049 | 20.1522968 | 20.1248756 | 0.059959985 | 0.260950388 |
| 20.2298884 | 19.9795386 | 20.1022773 | 20.1473106 | -0.116626665 | 0.193852307 |
| 20.7793616 | 20.6147847 | 20.6315594 | 20.7205173 | -0.06301239 | 0.280108068 |
| 21.2469314 | 21.0200233 | 21.141151 | 21.1508488 | -0.007790884 | 0.927440027 |
| 16.9338064 | 16.916939 | 17.2306422 | 17.1287557 | 7.29E-04 | 0.993716074 |
| 20.5880867 | 20.553151 | 20.541265 | 20.4730305 | -0.071719015 | 0.138639979 |
| 17.9395948 | 17.5582338 | 17.9600781 | 17.7654006 | -0.11390188 | 0.477818626 |

| | | | | | |
|---|---|---|---|---|---|
| 24.6163939 | 24.3710076 | 24.4852354 | 24.4521808 | -0.133072422 | 0.06067361 |
| 20.1060214 | 20.0672887 | 20.1114323 | 20.0914846 | -0.017026816 | 0.761757022 |
| NA | 17.2996956 | NA | NA | 0.022017246 | 0.838765637 |
| 17.7403693 | 17.561248 | 17.7912043 | 17.357445 | -0.052028016 | 0.667834811 |
| 21.3126157 | 21.0985524 | 21.2678528 | 21.2580886 | -0.044719317 | 0.554126868 |
| 23.5136123 | 23.4451465 | 23.5269155 | 23.5052566 | -0.003114201 | 0.940695897 |
| 21.4247602 | 20.8814058 | 21.1846447 | 21.3302503 | -0.259388195 | 0.2075512 |
| NA | 19.5336213 | NA | NA | -0.075987706 | 0.69628559 |
| 16.7679922 | 16.6231506 | 16.5177479 | 16.4747001 | -0.213461574 | 0.26968941 |
| 15.8200612 | NA | 16.1175663 | 15.908609 | 0.112964463 | 0.479468474 |
| 19.5727925 | 19.1325451 | 19.2828403 | 19.3670977 | -0.28186823 | 0.030048538 |
| 18.8594561 | 18.7423601 | 18.9118186 | 18.6716512 | -0.123608795 | 0.199698263 |
| 16.9558053 | 16.7116137 | 16.9644039 | 16.979889 | -0.161994104 | 0.053384758 |
| 19.0043548 | 18.9286393 | 18.8898831 | 18.811331 | -0.068014878 | 0.383262828 |
| 19.6599615 | 19.2978312 | 19.5359652 | 19.5748427 | -0.083043138 | 0.247428757 |
| 17.9269128 | 17.8900872 | 18.0587315 | 17.869064 | -0.036694555 | 0.596550098 |
| 26.2459281 | 26.0752109 | 26.2385179 | 26.2125286 | 0.035958301 | 0.601225496 |
| 21.6977426 | 21.5460166 | 21.6698744 | 21.5539506 | -0.032589093 | 0.613612291 |
| 23.7929379 | 23.6378484 | 23.7350315 | 23.796379 | 0.020957793 | 0.730687404 |
| 22.4807537 | 22.2440702 | 22.4340956 | 22.3922362 | -0.08463283 | 0.160201664 |
| 23.9813832 | 23.8598222 | 23.9148791 | 23.9816515 | 0.009218457 | 0.866961441 |
| 19.492938 | 19.3379269 | 19.328906 | 19.396097 | -0.063793823 | 0.241801405 |
| 20.2803607 | 20.2163946 | 20.2250653 | 20.2504431 | -0.05898418 | 0.26192109 |
| 22.0803077 | 22.0810139 | 22.0645115 | 22.1044201 | 0.00290638 | 0.948227895 |
| 20.937114 | 20.8065277 | 20.9407396 | 20.9401065 | -7.43E-04 | 0.987924526 |
| 20.033876 | 19.9417933 | 20.0582466 | 20.0585714 | 0.025654559 | 0.61746421 |
| 22.9597995 | 22.9557282 | 23.0312589 | 22.9966693 | 0.039714422 | 0.44369435 |
| 19.4558219 | 19.3455971 | 19.4880753 | 19.5117078 | 0.018356392 | 0.738034613 |
| 21.7277669 | 21.6960455 | 21.7669929 | 21.856709 | 0.076042231 | 0.213893055 |
| 19.9125659 | 19.8033871 | 19.8887239 | 19.9265044 | 0.021139041 | 0.742725496 |
| 25.1498286 | 25.0795905 | 25.1255872 | 25.1546165 | 0.020869161 | 0.756193106 |
| 17.0921499 | 17.333708 | 17.2143916 | 17.3428166 | 0.088229421 | 0.235417607 |
| 19.1584619 | 19.0258055 | 19.2171851 | 19.1637036 | -0.028122484 | 0.704374729 |
| 16.5450597 | 16.6956573 | 16.8042695 | 16.8688723 | 0.138429071 | 0.072815861 |
| 17.9459884 | 17.9867161 | 17.902862 | 17.9648606 | 0.087389568 | 0.251783136 |
| 18.6228852 | 18.6047257 | 18.641097 | 18.7858014 | 0.09569034 | 0.22036155 |
| 18.413682 | 18.3190197 | 18.4677029 | 18.5567969 | 0.015445645 | 0.853313717 |
| 20.4439356 | 20.3904702 | 20.4004542 | 20.5009611 | 0.112060298 | 0.239232563 |
| 18.1542647 | 17.8050207 | 18.2728916 | 18.1449974 | -0.11347184 | 0.288535094 |
| 18.2061349 | 18.3370057 | 18.0628296 | 18.2674555 | 0.265181253 | 0.036930052 |
| 17.3997052 | 17.2142884 | 17.2347889 | 16.9634908 | 0.012479346 | 0.925265454 |
| 13.8671765 | NA | 14.6542243 | 14.3827355 | 0.576002402 | 0.024899149 |

| | | | | | |
|---|---|---|---|---|---|
| NA | 15.3447442 | NA | NA | 0.0179084 | 0.957032174 |
| 17.7572581 | 17.8612766 | 17.7329619 | 17.7515364 | -0.128827831 | 0.228703027 |
| 21.4495964 | 21.3205855 | 21.4664971 | 21.4070986 | -0.007811891 | 0.912208523 |
| 25.5013411 | 25.464965 | 25.5071874 | 25.4593113 | -0.00381156 | 0.936819276 |
| 18.8496177 | 18.7466176 | 18.7755712 | 18.8196079 | -0.026322302 | 0.60229884 |
| 21.9561814 | 22.0202758 | 22.2545642 | 22.1439362 | 0.04357177 | 0.625016697 |
| 13.591548 | NA | 13.9327521 | NA | 0.505870266 | 0.235273765 |
| 17.3105753 | 16.7936876 | 17.1097648 | 17.1558112 | -0.04533952 | 0.765449963 |
| 16.1904429 | 16.1333617 | 16.097052 | 16.2960952 | -0.10128536 | 0.40791276 |
| 18.0479416 | 17.7837672 | 18.1169748 | 18.199577 | 0.084961024 | 0.462511163 |
| 19.6288454 | 19.4083892 | 19.6387231 | 19.7938491 | -0.073675963 | 0.45196697 |
| 18.3014806 | 18.0576266 | 18.3157286 | 18.2864607 | -0.098063223 | 0.245207359 |
| 19.0933415 | 19.0998298 | 19.0909045 | 19.3366714 | 0.039566295 | 0.605590705 |
| 19.686857 | 19.6286454 | 19.5606206 | 19.7181603 | -0.03767708 | 0.603867087 |
| 20.1930007 | 20.3101346 | 20.1737902 | 20.1869023 | -0.039851371 | 0.564230738 |
| 21.9003264 | 21.6825526 | 21.7976758 | 21.7976551 | -0.117354099 | 0.09190863 |
| 21.8273668 | 21.6346174 | 21.7530087 | 21.7996806 | -0.073203791 | 0.268363164 |
| 27.5134133 | 27.3316654 | 27.4306742 | 27.4741963 | -0.043215167 | 0.492943979 |
| 19.3382357 | 19.2050594 | 19.2297397 | 19.2724582 | -0.064514411 | 0.276849301 |
| 19.7678216 | 19.7507664 | 19.8108986 | 19.9370552 | -2.55E-05 | 0.999614837 |
| 20.138486 | 20.0326146 | 20.1107438 | 20.1962712 | -0.018077336 | 0.728161651 |
| 19.9348937 | 19.7872324 | 19.8912544 | 19.9007943 | -0.057958423 | 0.26677746 |
| 21.6404429 | 21.5752047 | 21.6459687 | 21.6972201 | 0.04047405 | 0.397395235 |
| 18.5031209 | 18.5721982 | 18.594916 | 18.5667641 | 0.004329731 | 0.928180769 |
| 22.9497769 | 22.9390844 | 23.0020431 | 23.0119725 | 0.041207292 | 0.404123476 |
| 23.1622749 | 23.0917007 | 23.1032455 | 23.1183921 | 0.047503168 | 0.366787412 |
| 18.6318682 | 18.5451576 | 18.5790144 | 18.7247609 | 0.038976974 | 0.462163011 |
| 20.4481107 | 20.4945544 | 20.403937 | 20.4365054 | 0.028417012 | 0.617538386 |
| 21.6710216 | 21.6476565 | 21.6431137 | 21.5588374 | 0.038317651 | 0.503147666 |
| 21.7643983 | 21.6590754 | 21.810947 | 21.7760191 | 0.036428456 | 0.563683878 |
| 18.1067239 | 18.3142337 | 18.1158116 | 18.3138481 | 0.072815781 | 0.281208556 |
| 17.3375365 | 17.4176409 | 17.3396668 | 17.5461735 | 0.069077245 | 0.343826145 |
| 17.8679599 | 17.8977177 | 17.9762958 | 18.2322332 | 0.127586004 | 0.176286175 |
| 18.2658677 | 18.0718802 | 18.2270142 | 18.304906 | 0.044286611 | 0.656217017 |
| 17.9764341 | 18.1936689 | 17.9029738 | 18.2163328 | 0.114275897 | 0.28387841 |
| 15.5302417 | NA | 15.3694269 | 15.5531859 | 0.039279001 | 0.743589018 |
| 21.352551 | 21.3313153 | 21.446979 | 21.4127785 | 0.033875617 | 0.864070047 |
| 21.1182083 | 21.0051311 | 21.0405272 | 21.0503181 | -0.028955469 | 0.592271441 |
| 19.1992371 | 19.3022929 | 19.2163105 | 19.382816 | 0.17031504 | 0.034227445 |
| 22.5491892 | 22.4631041 | 22.639015 | 22.4966953 | 0.055437585 | 0.539912522 |
| 18.8537001 | 18.5805891 | 18.796911 | 18.7923024 | -0.168941778 | 0.103042823 |
| 27.6138597 | 27.5000682 | 27.5602134 | 27.543101 | -0.009312069 | 0.891875134 |

| | | | | | |
|---|---|---|---|---|---|
| 24.7871322 | 24.6652071 | 24.8601263 | 24.7223298 | -0.023145473 | 0.66675754 |
| 20.4231154 | 20.2897872 | 20.3701075 | 20.357188 | -0.020761571 | 0.67035555 |
| 19.9200493 | 19.9210768 | 20.0268034 | 20.0086411 | -0.004842627 | 0.912407294 |
| 20.8390157 | 20.8404597 | 20.889308 | 20.9380207 | 0.041487091 | 0.367170673 |
| 20.529175 | 20.4442303 | 20.4784161 | 20.5105822 | 0.006539051 | 0.901128242 |
| 21.7053195 | 21.6551454 | 21.7349292 | 21.7471162 | 0.059446922 | 0.29936163 |
| 22.5157353 | 22.3427836 | 22.3779813 | 22.4357721 | 0.01502745 | 0.821581369 |
| 18.9174387 | 18.8724439 | 18.9866544 | 18.91664 | 0.049556705 | 0.614143882 |
| 18.1241854 | 17.8524622 | 18.0717306 | 17.9093127 | -0.132609261 | 0.152925664 |
| 18.4478856 | 18.5579222 | 18.4552694 | 18.4757533 | 0.07438391 | 0.262308275 |
| 14.5583059 | 14.0516493 | 14.6447245 | 15.0472682 | -0.127305369 | 0.523390224 |
| 14.738804 | 14.5235652 | 14.5582199 | 14.9727772 | -0.01778159 | 0.926072695 |
| 16.3405371 | 16.0542008 | 16.5949088 | 16.3934037 | 0.07397855 | 0.556057576 |
| 20.1002676 | 19.7363629 | 19.811476 | 19.8784214 | -0.225763402 | 0.030375796 |
| 21.3605821 | 21.0725735 | 21.2682758 | 21.2498914 | -0.097153347 | 0.245940624 |
| 21.0403337 | 20.9168184 | 21.03117 | 21.1414347 | -0.008425487 | 0.911073076 |
| 26.260538 | 26.1844882 | 26.2250329 | 26.2300064 | -1.22E-04 | 0.998620149 |
| 20.2945462 | 20.1079732 | 20.2541509 | 20.2942344 | -0.041480735 | 0.485842577 |
| 22.0996602 | 22.015845 | 22.1831178 | 22.1443827 | 0.0033883 | 0.953775296 |
| 24.5727795 | 24.387154 | 24.4945091 | 24.4664699 | -0.023189114 | 0.684489946 |
| 22.8956496 | 22.7314318 | 22.7804897 | 22.8097571 | -0.021012704 | 0.672260482 |
| 20.4246079 | 20.3245155 | 20.4144159 | 20.3884218 | -0.038658769 | 0.396370385 |
| 20.9572969 | 20.8105697 | 20.9425824 | 20.9099632 | -0.010662976 | 0.812408121 |
| 19.351892 | 19.3226772 | 19.3749662 | 19.2295556 | 0.029807597 | 0.626387969 |
| 20.9744482 | 20.8526969 | 20.9555203 | 21.0087315 | 0.0231678 | 0.729591002 |
| 22.8174246 | 22.6747989 | 22.7999469 | 22.7501464 | -0.034496694 | 0.607737253 |
| 21.3696588 | 21.2454185 | 21.265699 | 21.3991463 | 0.035328068 | 0.602233423 |
| 19.1701641 | 19.1940267 | 19.2319804 | 19.3351077 | 0.091341219 | 0.197900361 |
| 18.2685381 | 18.3731247 | 18.2061572 | 18.2646855 | 0.024553526 | 0.72839806 |
| 20.3475209 | 20.1037946 | 20.0569546 | 20.3261509 | -0.008233017 | 0.908975548 |
| 19.7544561 | 19.6943214 | 19.8947269 | 19.8408065 | 0.026956949 | 0.711301289 |
| 18.9671854 | 18.8897799 | 18.9638692 | 18.9858071 | -0.004044855 | 0.956059635 |
| 20.5053785 | 20.5496537 | 20.6418032 | 20.544722 | 0.057266612 | 0.550869552 |
| 20.5053704 | 20.4751657 | 20.5511074 | 20.5045509 | 0.017937051 | 0.853290008 |
| 19.1124086 | 18.9751758 | 19.0272322 | 19.2390493 | -0.014846503 | 0.89074276 |
| 22.9275872 | 22.9098761 | 22.9545438 | 22.9872688 | 0.051520292 | 0.596815543 |
| 21.9905776 | 21.9443108 | 21.9684337 | 21.8918916 | -0.084860444 | 0.218688564 |
| 22.3144165 | 22.2806759 | 22.3412056 | 22.3521345 | 0.034773887 | 0.511094733 |
| 23.7293866 | 23.625825 | 23.7370035 | 23.7495113 | 0.011637341 | 0.866736086 |
| 21.0731505 | 20.9220021 | 21.0048103 | 21.0942181 | 0.068906046 | 0.370940373 |
| 17.6629703 | 17.6672538 | 17.9031919 | 17.7118858 | 0.141580026 | 0.076791525 |
| 20.695622 | 20.6181564 | 20.5801132 | 20.567211 | -0.039854286 | 0.617021302 |

| | | | | | |
|---|---|---|---|---|---|
| 19.482503 | 19.2887735 | 19.5384799 | 19.5185137 | -0.059319707 | 0.64906994 |
| 18.9741141 | 18.8069526 | 18.9587471 | 18.9425491 | 0.038805672 | 0.620498008 |
| 18.0748515 | 17.9596538 | 18.0138111 | 17.9937515 | -0.128167922 | 0.099026433 |
| 19.8838662 | 19.6896156 | 19.827151 | 19.9468981 | -0.067426575 | 0.311494748 |
| 25.6257148 | 25.469173 | 25.6309824 | 25.5823688 | -0.025529466 | 0.659631749 |
| 22.7163478 | 22.6183564 | 22.5684933 | 22.6587647 | -0.026198329 | 0.605557781 |
| 20.7142107 | 20.5758387 | 20.6897237 | 20.7008625 | -0.012615578 | 0.800553801 |
| 23.8199694 | 23.7039097 | 23.8414702 | 23.7485283 | -0.007222099 | 0.890413901 |
| 19.9326047 | 20.0600746 | 20.0006578 | 20.063052 | 0.174202352 | 0.038500382 |
| 18.858088 | 18.8185506 | 18.91883 | 19.0331923 | 0.071998895 | 0.375813296 |
| 21.1824624 | 21.0682233 | 21.3002394 | 21.2588312 | 0.094860843 | 0.42580116 |
| 16.8453288 | 17.2651102 | 17.3290273 | 16.9779322 | 0.273879262 | 0.096927493 |
| 13.271317 | 12.9848534 | 13.7772916 | 13.4909416 | -0.232421234 | 0.248513508 |
| 16.7749782 | 16.7968351 | 16.739141 | 16.7936952 | -0.046674716 | 0.628426422 |
| 20.4604432 | 20.3728894 | 20.3585775 | 20.3495516 | -0.064452306 | 0.450118536 |
| 20.7607945 | 20.6556548 | 20.7221308 | 20.7219048 | -0.03020929 | 0.556691095 |
| 25.246843 | 25.2420783 | 25.3474397 | 25.3075894 | 0.088196503 | 0.088795872 |
| 25.8642937 | 25.8102775 | 25.897365 | 25.8763543 | 0.044526354 | 0.429459408 |
| 19.4066801 | 19.4312758 | 19.4314421 | 19.4622007 | 0.059054979 | 0.304527898 |
| 20.7949244 | 20.6426402 | 20.8708266 | 20.7000046 | 0.011449928 | 0.847601175 |
| 12.3680942 | NA | 13.0574239 | 13.3651811 | 0.646016275 | 0.008046927 |
| 21.2395951 | 20.9804455 | 21.0471498 | 21.1427811 | -0.088317299 | 0.132411216 |
| 26.1344223 | 25.9408791 | 26.0404582 | 26.0031836 | -0.06737353 | 0.222843525 |
| 24.0052851 | 24.0219158 | 24.1136226 | 24.0589475 | 0.10283182 | 0.201948212 |
| 16.4127037 | 16.3411863 | 16.3294073 | 16.4364687 | -0.176583725 | 0.040689925 |
| 21.6939495 | 21.6030834 | 21.7412052 | 21.8622133 | 0.107202833 | 0.240679592 |
| 16.0506507 | NA | 15.4131794 | 16.213853 | -0.138706389 | 0.530937189 |
| 13.9335541 | NA | 14.0905391 | 14.0921478 | 0.168998486 | 0.21330996 |
| 17.8111381 | 17.9675392 | 17.910222 | 18.2210996 | 0.060374064 | 0.550215776 |
| 19.5340574 | 19.4532599 | 19.5818922 | 19.5078159 | 0.014393348 | 0.827349844 |
| 20.6767216 | 20.5900989 | 20.7529499 | 20.5512543 | 0.027140298 | 0.698504949 |
| 20.6544551 | 20.5547957 | 20.719177 | 20.6497367 | -0.023856301 | 0.638156715 |
| 19.7534496 | 19.6487425 | 19.7153123 | 19.7667852 | 0.004964993 | 0.931232418 |
| 23.7211754 | 23.6136981 | 23.7035907 | 23.7180398 | 0.019087903 | 0.793295815 |
| 18.1741592 | 18.1396031 | 18.2496499 | 18.1822518 | 0.02289608 | 0.716319696 |
| 20.3419962 | 20.20163 | 20.2740547 | 20.3373534 | -0.055391669 | 0.309527103 |
| 26.3629198 | 26.1678608 | 26.3344019 | 26.2831753 | -0.032195647 | 0.556498361 |
| 21.1925518 | 21.1205923 | 21.1544792 | 21.138302 | -0.073014256 | 0.082054202 |
| NA | 10.9358011 | NA | NA | -0.974359084 | 0.066300002 |
| 14.4105888 | 13.4852165 | 14.2673451 | 14.4908261 | -0.440317469 | 0.253663772 |
| 16.5164271 | 16.5022678 | 16.7966259 | 16.7394848 | -0.042958692 | 0.83247419 |
| 18.564566 | NA | 19.1696205 | 18.6251372 | 0.386291269 | 0.026084513 |

| | | | | | |
|---|---|---|---|---|---|
| 19.310336 | 18.8173763 | 19.2730241 | 19.1732553 | -0.207429293 | 0.099993341 |
| 17.7342218 | NA | 17.8395037 | 17.8002994 | -0.032236602 | 0.77893289 |
| 19.0942161 | 19.0219992 | 19.3266206 | 19.4240003 | 0.023653654 | 0.829474952 |
| 17.252021 | 16.9621728 | 17.2593642 | 17.2082483 | -0.111674233 | 0.274613503 |
| 19.6876488 | 19.4004474 | 19.450308 | 19.5799216 | -0.110643978 | 0.213385729 |
| 20.2406215 | 20.0651015 | 20.3291236 | 20.2427243 | -0.059160295 | 0.490338323 |
| 17.977721 | 17.8385473 | 18.0435677 | 17.9830535 | -0.115281804 | 0.170760488 |
| 18.8254603 | 18.8492969 | 18.565096 | 18.7828081 | -0.032131748 | 0.695384798 |
| 26.4234194 | 26.3077503 | 26.3992543 | 26.4251809 | 0.01600076 | 0.836628354 |
| 19.0605969 | 19.0073319 | 19.0143252 | 18.8553085 | 0.006228813 | 0.936244482 |
| 19.8639738 | 19.9057314 | 19.9018353 | 19.9234059 | -0.011004433 | 0.882233395 |
| 21.7014377 | 21.5414317 | 21.6595422 | 21.7010274 | -0.050197678 | 0.492713017 |
| 18.8009119 | 18.7901901 | 18.9292427 | 18.902205 | -0.010073409 | 0.887000757 |
| 18.4264969 | 18.3584398 | 18.411145 | 18.3271312 | -0.054340351 | 0.44081732 |
| 18.7246176 | 18.951974 | 18.7389121 | 18.7131332 | 0.036923336 | 0.590972766 |
| 26.4701684 | 26.3885051 | 26.4159064 | 26.4568391 | 0.025981151 | 0.704620013 |
| 21.5853104 | 21.6459945 | 21.5857272 | 21.4960732 | -0.039050526 | 0.559514959 |
| 21.2440205 | 21.2451958 | 21.1451351 | 21.2334649 | -0.012132064 | 0.852145421 |
| 21.8298258 | 21.7280582 | 21.9007849 | 21.8464419 | 0.010968677 | 0.861954783 |
| 21.536158 | 21.3408148 | 21.472564 | 21.5085298 | -0.014903266 | 0.812328271 |
| 21.8542386 | 21.8695745 | 21.938362 | 21.9325439 | 0.067766723 | 0.27452247 |
| 22.0692932 | 21.8706136 | 21.943052 | 21.8845647 | -0.044054859 | 0.467977443 |
| 21.6461348 | 21.6226717 | 21.6633271 | 21.6268756 | -0.00881568 | 0.884290181 |
| 21.7737888 | 21.6942431 | 21.7611584 | 21.7408905 | -0.041418758 | 0.48272299 |
| 21.5437295 | 21.4299928 | 21.4896282 | 21.5383473 | -0.049398216 | 0.399051665 |
| 25.1850877 | 25.0715838 | 25.1804494 | 25.1509423 | -0.053516434 | 0.351586993 |
| 19.8733142 | 19.9099899 | 19.9423983 | 19.9135567 | 0.044392662 | 0.434902674 |
| 19.6003407 | 19.6107901 | 19.612196 | 19.5308963 | -0.004562038 | 0.934942237 |
| 22.439213 | 22.4135706 | 22.4414366 | 22.4086808 | 0.031230631 | 0.546472051 |
| 23.9752623 | 23.9222218 | 23.9810538 | 24.0268503 | -0.014606322 | 0.759889714 |
| 24.2788085 | 24.2394906 | 24.2717802 | 24.2552008 | 0.019817484 | 0.671693934 |
| 21.4033691 | 21.4461117 | 21.4500702 | 21.4175179 | 0.016690664 | 0.716625504 |
| 23.7786916 | 23.7108045 | 23.7228698 | 23.6996752 | -0.013186733 | 0.758712541 |
| 22.1016916 | 22.0784473 | 22.1042911 | 22.1051169 | 0.042718448 | 0.314852904 |
| 18.9662539 | 18.9576276 | 19.0175064 | 18.9473997 | 0.031670388 | 0.470382379 |
| 20.1373023 | 20.0786619 | 20.1623783 | 20.1542644 | 0.046750197 | 0.336522849 |
| 19.1066494 | 19.147022 | 19.0821967 | 19.059649 | -0.010464279 | 0.833075803 |
| 19.6921588 | 19.6113438 | 19.7059823 | 19.7117408 | -0.016611985 | 0.746516298 |
| 20.0205737 | 19.9337511 | 20.0068367 | 20.0111757 | 0.0047863 | 0.927357145 |
| 23.4567142 | 23.4017586 | 23.4720511 | 23.4809939 | 0.062512703 | 0.243781445 |
| 19.2753374 | 19.2496539 | 19.3771301 | 19.3235236 | 0.021238343 | 0.692580991 |
| 26.2628096 | 26.1888478 | 26.2114915 | 26.2603486 | 0.021876027 | 0.68651866 |

| | | | | | |
|---|---|---|---|---|---|
| 19.2947269 | 19.3306086 | 19.2997359 | 19.2607738 | 0.027039541 | 0.618854844 |
| 20.2153351 | 20.1739556 | 20.1624114 | 20.1205951 | -0.019402301 | 0.723999278 |
| 20.5476654 | 20.3948567 | 20.5044148 | 20.5185198 | 0.002413767 | 0.969531716 |
| 21.1223053 | 21.1617321 | 21.1850081 | 21.276239 | 0.029695596 | 0.635821927 |
| 22.5276293 | 22.460198 | 22.489134 | 22.5355811 | 0.017618575 | 0.792704662 |
| 23.4762359 | 23.4665057 | 23.4546236 | 23.4546486 | 0.085992081 | 0.217317304 |
| 17.6844616 | 17.5882783 | 17.6986277 | 17.8013266 | 0.003509483 | 0.960988159 |
| 25.2486449 | 25.1482643 | 25.2993765 | 25.2694081 | 0.04656884 | 0.5528311 |
| 19.3491398 | 19.2303086 | 19.308147 | 19.3764918 | -0.044585061 | 0.571156819 |
| 22.0439036 | 21.8904813 | 22.0319895 | 22.0558792 | 0.011586295 | 0.883696737 |
| 22.4341749 | 22.3779332 | 22.3868938 | 22.4609909 | 0.011616858 | 0.886248807 |
| 16.9311212 | 16.7958641 | 17.0853647 | 16.9959704 | 0.017680703 | 0.832849087 |
| 21.0178526 | 21.0989429 | 21.0893571 | 21.0641303 | 0.042437008 | 0.624201201 |
| 17.5332347 | 17.5337923 | 17.6072134 | 17.5278204 | 0.038496701 | 0.66649966 |
| 19.2723365 | 19.3223343 | 19.5237019 | 19.4314249 | 0.087110512 | 0.333781028 |
| 17.2093198 | 17.2028571 | 17.1651808 | 17.1366084 | -0.015999971 | 0.861195197 |
| 23.8142785 | 23.7171571 | 23.8184567 | 23.8363203 | 0.051322439 | 0.612290945 |
| 19.1052607 | 19.1659533 | 19.2923814 | 19.3006101 | 0.086548142 | 0.398999163 |
| 18.679197 | 18.3333245 | 18.5333506 | 18.7241169 | -0.020207435 | 0.846928788 |
| 18.7636909 | 18.6483457 | 18.7286939 | 18.7306025 | 0.030205221 | 0.77279308 |
| 16.8705198 | 16.6780954 | 17.1319813 | 17.1018067 | 0.054402864 | 0.662825902 |
| 14.678139 | NA | 14.5946239 | 14.779287 | -0.105950174 | 0.427509247 |
| 20.8114522 | 20.8603833 | 20.9335941 | 20.8025931 | 0.060993266 | 0.656828415 |
| 16.9936735 | 17.1623949 | 16.9929459 | 17.0026768 | 0.031746888 | 0.81633006 |
| 12.9921541 | 13.2371609 | 13.3982623 | 13.3870211 | 0.192722689 | 0.20039635 |
| NA | 15.3208722 | NA | NA | 0.215845495 | 0.258209924 |
| 20.6419367 | 20.5762733 | 20.5504228 | 20.5345276 | -0.095945828 | 0.099726718 |
| 21.1003567 | 21.0299544 | 21.0554566 | 21.0855758 | 0.008670786 | 0.868471932 |
| 20.9699496 | 20.6897351 | 21.0043293 | 20.9945801 | -0.025713533 | 0.736959773 |
| 21.9393528 | 22.3758891 | 22.4479392 | 22.2572442 | 0.172163459 | 0.155190971 |
| NA | 18.29407 | NA | NA | -0.336821214 | 0.205707469 |
| 18.5063163 | 18.1581598 | 18.3164802 | 18.4915383 | -0.07377165 | 0.654425213 |
| 19.7783221 | 19.5535419 | 19.6069571 | 19.5538717 | -0.156461546 | 0.059985628 |
| 21.2573159 | 20.9838542 | 21.0087736 | 21.084736 | -0.114710176 | 0.132739823 |
| 20.4455851 | 20.3336357 | 20.4451763 | 20.5321526 | 0.034273145 | 0.630138438 |
| 19.5565723 | 19.3025504 | 19.5831961 | 19.5264413 | -0.08259817 | 0.238521236 |
| 24.9387173 | 24.8179063 | 24.9300244 | 24.960344 | -0.004590124 | 0.938934516 |
| 24.9221904 | 24.786496 | 24.8884181 | 24.8314592 | -0.043194059 | 0.458737794 |
| 19.9347018 | 19.7497476 | 19.8885415 | 19.8324235 | -0.094235169 | 0.079633033 |
| 23.3750752 | 23.2879432 | 23.3401157 | 23.3717769 | -0.032294179 | 0.528197735 |
| 20.6327252 | 20.6920354 | 20.6411552 | 20.6629213 | 0.047079114 | 0.34059823 |
| 22.3893958 | 22.3040103 | 22.3220643 | 22.3457401 | -0.008398527 | 0.838699821 |

| | | | | | |
|---|---|---|---|---|---|
| 21.486394 | 21.38165 | 21.4239861 | 21.4898371 | -0.010396381 | 0.798833092 |
| 21.5004477 | 21.4311665 | 21.486522 | 21.4379255 | -0.001543053 | 0.969666953 |
| 22.4692781 | 22.4173523 | 22.5044518 | 22.4920495 | 0.059387731 | 0.267202725 |
| 21.4167358 | 21.2866611 | 21.4872136 | 21.4192371 | 0.057489391 | 0.360295894 |
| 22.7752606 | 22.6545468 | 22.8469012 | 22.7925479 | 0.035977735 | 0.594849285 |
| 22.2236023 | 22.3510217 | 22.2366625 | 22.1850444 | 0.018324276 | 0.798930306 |
| 19.2416746 | 19.2172409 | 19.2247 | 19.3886108 | 0.062355481 | 0.402791189 |
| 23.8280074 | 23.6977886 | 23.9627704 | 23.8774392 | 0.057542037 | 0.530020945 |
| 15.9032705 | 16.1830252 | 16.0824243 | 16.1167402 | 0.100226472 | 0.282681001 |
| 18.6101681 | 18.5085476 | 18.547196 | 18.8467569 | 0.013448697 | 0.888585772 |
| 18.2302781 | 18.1193591 | 18.3421952 | 18.2154387 | 0.033780058 | 0.733010615 |
| 17.5216229 | 17.4315795 | 17.4377853 | 17.6868272 | -0.005386448 | 0.958346716 |
| 19.1296358 | 18.9950514 | 19.1330916 | 18.9006538 | 0.061426608 | 0.557917575 |
| 17.7699553 | 18.0405136 | 18.1431697 | 17.7550051 | 0.239259476 | 0.040253675 |
| 18.0595728 | 18.0219133 | 18.0257851 | 18.0450132 | 0.021556673 | 0.858709868 |
| NA | 14.7924993 | NA | NA | -0.188538867 | 0.161168879 |
| 18.73777 | 18.5637062 | 18.7100621 | 18.7206649 | -0.038929693 | 0.73586817 |
| 18.3492961 | 18.3969389 | 18.0304668 | 18.2076985 | -0.055274089 | 0.603380657 |
| 19.4564268 | 19.3295637 | 19.3235201 | 19.4687009 | -0.035940088 | 0.689877481 |
| 22.0906599 | 21.827438 | 21.964614 | 22.0109893 | -0.062722926 | 0.357102648 |
| 22.0316008 | 21.920949 | 21.8908446 | 21.8471614 | -0.104022554 | 0.092952545 |
| 22.4695314 | 22.3366217 | 22.426194 | 22.4007102 | -0.060969295 | 0.282481322 |
| 22.2665862 | 22.2085136 | 22.33739 | 22.3193803 | 0.047873246 | 0.377308605 |
| 24.5718151 | 24.5158349 | 24.5844401 | 24.5897243 | 0.034571889 | 0.511577817 |
| 23.2767347 | 23.2994228 | 23.2508245 | 23.2743256 | -0.023537153 | 0.605047509 |
| 21.4321286 | 21.2223387 | 21.3897416 | 21.4179233 | -0.052081891 | 0.327697432 |
| 19.9651235 | 19.8173382 | 19.9128501 | 20.0680699 | 0.031750827 | 0.555776327 |
| 20.3058387 | 20.2761308 | 20.3472957 | 20.3543216 | 0.030556588 | 0.5714087 |
| 21.014045 | 20.8429863 | 20.9520952 | 20.9765373 | -0.008507902 | 0.879953413 |
| 24.1175371 | 23.9767107 | 24.0635114 | 23.9964468 | 0.005432757 | 0.923369124 |
| 24.4829861 | 24.5015363 | 24.5361548 | 24.46616 | 0.024446046 | 0.668052593 |
| 20.1269176 | 20.0528566 | 20.0673736 | 20.0374183 | -0.001703174 | 0.977997275 |
| 22.4805437 | 22.3844058 | 22.5465823 | 22.5452717 | 0.059765712 | 0.350616464 |
| 21.9152374 | 21.895727 | 21.9878444 | 21.9134006 | 0.031464122 | 0.650374944 |
| 18.9589274 | 18.7268621 | 18.8914484 | 18.8897408 | -0.016202487 | 0.819126124 |
| 24.4493347 | 24.3378229 | 24.4361626 | 24.4413475 | 0.031277944 | 0.676156647 |
| 17.8220994 | 17.7471991 | 17.8629719 | 17.949763 | 0.085997283 | 0.257032164 |
| 18.471541 | 18.5832996 | 18.4629429 | 18.5035265 | 0.128129579 | 0.214044691 |
| 15.719474 | 15.5702511 | 15.7992593 | 15.9670474 | 0.052641258 | 0.677441543 |
| 19.816104 | 19.6816675 | 19.7706612 | 19.7081823 | -0.009172157 | 0.868435664 |
| 19.6065161 | 19.6625734 | 19.688137 | 19.7705372 | 0.144208125 | 0.019715448 |
| 17.9637097 | 17.7506645 | 17.7329401 | 17.9154929 | -0.030910947 | 0.685038622 |

| | | | | | |
|---|---|---|---|---|---|
| 21.9301674 | 21.7807373 | 21.8410393 | 21.8693449 | -0.04792468 | 0.342726871 |
| 23.4582148 | 23.6785272 | 23.372291 | 23.3330683 | 0.016741003 | 0.906979441 |
| 20.5503533 | 20.4715763 | 20.4922245 | 20.6073551 | -0.029598154 | 0.618422363 |
| NA | 15.4018376 | NA | NA | -0.174395019 | 0.32712614 |
| 20.3405998 | 20.3749126 | 20.2998624 | 20.4018657 | 0.06751046 | 0.48431306 |
| 18.0392514 | 17.8952784 | 18.097631 | 18.1224597 | 0.028700229 | 0.759281582 |
| 19.7214933 | 19.548637 | 19.5965467 | 19.6799452 | -0.030827368 | 0.672457013 |
| 21.4447815 | 21.4006854 | 21.4674581 | 21.4988812 | 0.045210501 | 0.36290676 |
| 18.4835799 | 18.5668419 | 18.6805712 | 18.7159807 | 0.134180259 | 0.048005861 |
| 21.7363095 | 21.619379 | 21.7375376 | 21.7115401 | 0.040969573 | 0.463431821 |
| 21.6363971 | 21.5402247 | 21.6544324 | 21.6793696 | 0.026090643 | 0.577511217 |
| 18.7005572 | 18.4088684 | 18.4685806 | 18.7418104 | -0.02553869 | 0.807206811 |
| 22.9794337 | 22.8460485 | 22.9132368 | 22.9046276 | -0.03792035 | 0.420447827 |
| 21.1194811 | 21.0371983 | 21.1587254 | 21.1008667 | 0.01399694 | 0.783509619 |
| 21.6228609 | 21.515662 | 21.6284278 | 21.6183476 | -1.62E-04 | 0.997645467 |
| 23.3484392 | 23.4217241 | 23.36449 | 23.4277653 | 0.094612464 | 0.357906107 |
| 19.5011311 | 19.4990808 | 19.5604288 | 19.6102241 | 0.004688548 | 0.973066925 |
| 19.6220869 | 19.3507797 | 19.5788248 | 19.4300315 | -0.084525588 | 0.334249251 |
| 17.9048351 | 17.8524064 | 17.8941752 | 17.8799669 | 0.080086772 | 0.38459104 |
| 21.7132701 | 21.56175 | 21.6790642 | 21.6975274 | -0.0426602 | 0.402482267 |
| 17.5442036 | 17.4602373 | 17.7017551 | 17.5983338 | 0.00431204 | 0.960169708 |
| 17.4182041 | 17.3633616 | 17.5733809 | 17.4247017 | -0.028791375 | 0.671327518 |
| 23.5642904 | 23.4183594 | 23.5741247 | 23.5392226 | 0.050765439 | 0.623008307 |
| 17.98077 | 18.1544655 | 18.248698 | 18.2173307 | 0.222516994 | 0.120232936 |
| 21.8201238 | 21.8374853 | 21.7818202 | 21.7099125 | -0.035259622 | 0.475826823 |
| 21.7694024 | 21.6840975 | 21.8056624 | 21.7631563 | 0.002750104 | 0.960330904 |
| 27.779107 | 27.7105363 | 27.8071503 | 27.7961045 | 0.051611618 | 0.379353021 |
| 21.6775357 | 21.4617909 | 21.618942 | 21.6851238 | -0.014089092 | 0.809242531 |
| 19.6494348 | 19.3896122 | 19.5267579 | 19.4934067 | -0.037607387 | 0.538259027 |
| 19.4426175 | 19.244205 | 19.4599917 | 19.4451734 | -0.011364511 | 0.862334919 |
| 19.1200664 | 18.9331877 | 19.0482761 | 19.0531747 | -0.066358561 | 0.353648872 |
| 23.2681489 | 23.1170347 | 23.349532 | 23.2473588 | 0.044129205 | 0.678331533 |
| 13.1741363 | 13.2212067 | 12.7132166 | 12.8275891 | -0.177950927 | 0.370822486 |
| 16.8876447 | 16.6403091 | 16.7267414 | 16.7882296 | -0.146039459 | 0.079245705 |
| 20.2449672 | 20.2576417 | 20.3165076 | 20.3327536 | 0.034757106 | 0.435058727 |
| 20.2675339 | 20.0553096 | 20.2064077 | 20.2013204 | -0.033789927 | 0.524869721 |
| 17.0909712 | 16.655052 | 16.8342759 | 16.9507613 | -0.299834593 | 0.056738362 |
| 17.4608221 | 17.3800737 | 17.2097266 | 17.2332567 | -0.076151383 | 0.444000052 |
| 20.5645537 | 20.2734959 | 20.429397 | 20.3190428 | -0.106844849 | 0.169065105 |
| 17.7046454 | 17.6717701 | 17.7974083 | 17.6318705 | -0.049818322 | 0.400532285 |
| 13.5081457 | NA | 13.8508744 | 13.7786058 | 0.127619183 | 0.619896736 |
| 24.8096175 | 24.7058622 | 24.745182 | 24.7701808 | 0.003155582 | 0.947823108 |

| | | | | | |
|---|---|---|---|---|---|
| 24.960188 | 24.8123528 | 24.9071118 | 24.9511619 | -0.012114588 | 0.924494996 |
| 20.0758205 | 20.0949796 | 20.127751 | 20.1226824 | 0.051125161 | 0.29863101 |
| 23.1509335 | 23.029368 | 23.0912797 | 23.1551693 | -0.003666414 | 0.951366308 |
| 22.2367516 | 22.0761449 | 22.1629937 | 22.1526416 | -0.048925775 | 0.45536902 |
| 15.1275904 | 15.2132748 | 15.3902552 | 15.0481506 | 0.108909908 | 0.278747773 |
| 15.7990959 | 15.3516974 | 16.0223077 | 15.6755683 | -0.020388366 | 0.876813617 |
| 22.7647358 | 22.7056088 | 22.7241603 | 22.8192426 | 0.070068409 | 0.351730644 |
| 17.0959176 | 17.3801764 | 17.1148859 | 17.3302397 | 0.170651075 | 0.204371454 |
| 20.54586 | 20.4457038 | 20.4654596 | 20.475268 | -0.042186229 | 0.393582555 |
| 22.752062 | 22.7688752 | 22.8081861 | 22.8399664 | 0.048506492 | 0.400788105 |
| NA | 11.6669167 | NA | NA | 0.186604066 | 0.655162832 |
| 17.6060976 | 17.6357596 | 17.5691268 | 17.5166917 | 0.101977634 | 0.520571053 |
| 16.6725634 | 16.6784796 | 15.9325554 | 16.4155902 | -0.337472379 | 0.01965304 |
| 17.1952284 | 17.2289694 | 16.9674598 | 17.1502323 | -0.056006533 | 0.681164642 |
| 15.2117927 | 15.047365 | 15.3822311 | 15.4865228 | -0.151989493 | 0.208942843 |
| 22.7976638 | 22.6887877 | 22.7274719 | 22.7359831 | -0.094637108 | 0.359211169 |
| 17.2241643 | 17.2586811 | 17.2713744 | 17.0765763 | 0.003335553 | 0.973877519 |
| 22.4284365 | 22.2128699 | 22.3462401 | 22.2606705 | -0.115108736 | 0.218608658 |
| 22.6089979 | 22.58195 | 22.6528553 | 22.6276663 | 0.051634084 | 0.46248146 |
| 18.5528339 | 18.4361091 | 18.5072509 | 18.539144 | -0.116037009 | 0.074648573 |
| 23.5833301 | 23.4865865 | 23.5197749 | 23.4306581 | -0.060291496 | 0.294072597 |
| 24.5272218 | 24.3919241 | 24.4758147 | 24.3995837 | -0.030667299 | 0.546422256 |
| 22.0328582 | 21.9786194 | 22.0437177 | 22.0127217 | -0.018162541 | 0.658718234 |
| 26.3882864 | 26.3399202 | 26.3408671 | 26.3516106 | 0.002187469 | 0.956869414 |
| 21.3298639 | 21.2783317 | 21.3234564 | 21.3252971 | 0.024970596 | 0.549353351 |
| 22.29066 | 22.3097349 | 22.3183364 | 22.3719281 | 0.029623222 | 0.512342171 |
| 20.640066 | 20.5593382 | 20.5572811 | 20.6800625 | 0.023553304 | 0.656115963 |
| 21.8665596 | 21.7673337 | 21.8761945 | 21.8757178 | 0.003179227 | 0.953106353 |
| 22.1763994 | 21.9524354 | 22.118744 | 22.1616807 | -0.060850505 | 0.341196777 |
| 19.7603464 | 19.5931877 | 19.7404971 | 19.8562996 | 0.030654483 | 0.666686356 |
| 22.3485733 | 22.0463613 | 22.3084947 | 22.2428623 | -0.126509606 | 0.106413607 |
| 20.2448384 | 20.0822122 | 20.1247963 | 20.1704887 | -0.013545839 | 0.866223914 |
| 17.3688558 | 17.3029068 | 17.5078634 | 17.289089 | 0.022272429 | 0.784950935 |
| 23.8820133 | 23.8393748 | 23.9195892 | 23.91643 | 0.066132439 | 0.517709508 |
| 21.3711331 | 21.4595775 | 21.2617394 | 21.34372 | 0.068133204 | 0.580610218 |
| 14.3259489 | 13.8854909 | 13.8667383 | 13.8278819 | -0.311477678 | 0.022518455 |
| 22.7326209 | 22.5583889 | 22.678484 | 22.589009 | -0.050319296 | 0.589625129 |
| 22.1295892 | 21.9986257 | 22.1063677 | 22.1113679 | -0.027209123 | 0.613623703 |
| 25.4304557 | 25.4106838 | 25.4824161 | 25.424849 | 0.07807639 | 0.272795001 |
| 14.3318154 | NA | 14.4641652 | 14.3386629 | -0.09450411 | 0.602796677 |
| 26.6627932 | 26.6161277 | 26.6811982 | 26.6682306 | 0.0185863 | 0.733028352 |
| 22.3654124 | 22.3198871 | 22.3639033 | 22.2991526 | 0.05495871 | 0.268625997 |

| | | | | | |
|---|---|---|---|---|---|
| 20.4959945 | 20.500021 | 20.5765914 | 20.6395728 | 0.098636754 | 0.072969571 |
| 21.5280021 | 21.4598209 | 21.4440944 | 21.548398 | 0.029914835 | 0.683210959 |
| 23.3665614 | 23.3545208 | 23.4826661 | 23.4658721 | 0.09888203 | 0.232620947 |
| 17.3318257 | 17.6139061 | 17.5123874 | 17.613569 | 0.074819345 | 0.44830175 |
| 20.2360115 | 20.2072366 | 20.2702477 | 20.2334156 | 0.006707559 | 0.928080514 |
| 19.2899615 | 19.3404226 | 19.3367035 | 19.3883378 | 0.073755658 | 0.244345272 |
| 18.344967 | 18.0002878 | 18.1704787 | 17.9279588 | -0.307896654 | 0.062581332 |
| 17.1857867 | 17.0732892 | 17.1023213 | 17.1452263 | -0.006733898 | 0.924763369 |
| 23.3258441 | 23.3506668 | 23.3401903 | 23.3704679 | 0.038421054 | 0.454866179 |
| 15.0536203 | NA | 15.120755 | 15.1418597 | -0.091616787 | 0.739047733 |
| 20.5143383 | 20.3938658 | 20.5395617 | 20.5074416 | 0.071995793 | 0.441079198 |
| NA | 20.3228954 | NA | NA | 0.143880265 | 0.167883555 |
| 22.026672 | 21.9303652 | 21.9130719 | 22.0129384 | -0.054513148 | 0.232372128 |
| NA | 16.4033123 | NA | NA | 0.146605155 | 0.371733653 |
| 15.3096895 | 15.7313132 | 15.5478082 | 15.4831027 | 0.016257439 | 0.909272196 |
| 16.9880056 | 16.6258656 | 16.6934728 | 16.7968273 | -0.045273446 | 0.749580635 |
| 18.8559273 | 18.7387503 | 18.8833765 | 18.8778613 | -0.027607693 | 0.749871118 |
| 17.5492443 | 17.3497228 | 17.5648399 | 17.5569241 | -0.087672039 | 0.30714714 |
| 22.7594565 | 22.7694187 | 22.7763844 | 22.8558429 | 0.014261867 | 0.837514257 |
| 26.001228 | 25.9607771 | 26.0140849 | 26.0315469 | 0.052512138 | 0.37585895 |
| 19.1694311 | 19.0842986 | 19.1492428 | 19.2277324 | -0.071711864 | 0.197105215 |
| 20.0677805 | 20.0155973 | 19.9453447 | 19.9418049 | -0.0720889 | 0.150320479 |
| 20.0866806 | 19.9230082 | 19.8638976 | 19.9678248 | -0.12622581 | 0.009896282 |
| 21.2707753 | 21.1285073 | 21.2695149 | 21.2148111 | -0.06057787 | 0.119963936 |
| 19.0560305 | 19.0434353 | 19.0084226 | 19.1217232 | 0.041737935 | 0.421553941 |
| 20.6730834 | 20.6729886 | 20.7180701 | 20.7182396 | -0.003639534 | 0.956924898 |
| 17.5263211 | 17.4876635 | 17.611826 | 17.7074647 | 0.083746748 | 0.22297437 |
| 26.5653216 | 26.4243277 | 26.5434673 | 26.5789574 | 0.026702859 | 0.694569906 |
| 21.7021881 | 21.5493471 | 21.7697695 | 21.6876716 | 0.027503935 | 0.689715695 |
| 20.6495221 | 20.7188441 | 20.6434306 | 20.6472306 | -0.012909741 | 0.870118967 |
| 18.3352427 | 18.0537465 | 18.2718896 | 18.4055921 | 0.040106531 | 0.654646872 |
| 17.3264686 | 17.2435636 | 17.4611884 | 17.3145616 | -0.051562237 | 0.594695472 |
| 14.2838981 | NA | 14.6663595 | 15.1247297 | 0.58296506 | 9.33E-04 |
| 17.5589025 | 17.6412343 | 17.5459212 | 17.8204981 | -0.0266835 | 0.856206184 |
| 21.3947596 | 21.2481216 | 21.448898 | 21.4601084 | 0.048099912 | 0.377006547 |
| 15.2665436 | 16.0456127 | 16.0339917 | 16.2014143 | 0.384157052 | 0.022747905 |
| 22.6677557 | 22.5898127 | 22.6988333 | 22.6636323 | 0.016565952 | 0.76918181 |
| 24.7017062 | 24.6776119 | 24.7318789 | 24.7385855 | 0.061413804 | 0.343186776 |
| 15.6100285 | 15.6034573 | 15.9212602 | 15.7961273 | 0.296060633 | 0.017605301 |
| 24.422792 | 24.2727145 | 24.3512398 | 24.3792902 | -0.025565123 | 0.715999841 |
| 18.1362671 | 18.0420756 | 18.1657761 | 18.2801704 | 0.124345873 | 0.272458782 |
| 17.2185872 | NA | 17.0349075 | 17.1757684 | -0.155730959 | 0.247374014 |

| | | | | | |
|---|---|---|---|---|---|
| 19.3447815 | 19.2823347 | 19.2739708 | 19.3398095 | -0.001162848 | 0.984842782 |
| 22.5949859 | 22.4852311 | 22.477728 | 22.5147113 | -0.028140225 | 0.683216591 |
| 20.9859174 | 20.8787466 | 20.9289404 | 20.9632436 | -4.97E-04 | 0.992121461 |
| 17.0384657 | 17.1031327 | 17.0290506 | 16.9657384 | -0.046093251 | 0.831284006 |
| NA | 19.5282901 | NA | NA | 0.227070048 | 0.483943166 |
| 19.2640976 | 19.0768611 | 19.392792 | 19.307639 | 0.006773195 | 0.919858348 |
| 19.4722466 | 19.4692148 | 19.463639 | 19.4474511 | -0.02847383 | 0.578478255 |
| 21.8606339 | 21.7762698 | 21.84181 | 21.9075602 | 0.013698168 | 0.748282085 |
| NA | 18.0847158 | NA | NA | 0.036718278 | 0.766875707 |
| 16.9613351 | 16.8203883 | 17.2508497 | 17.0496666 | -0.021408986 | 0.882718509 |
| 21.8312923 | 21.6880174 | 21.7510834 | 21.7623802 | -0.028777286 | 0.543941295 |
| 20.1861407 | 20.1077062 | 20.1390707 | 20.1396827 | -0.043495689 | 0.271256249 |
| 20.4056718 | 20.3547704 | 20.4250654 | 20.5011852 | 0.05473242 | 0.250446843 |
| 22.8962635 | 22.8598205 | 22.8436044 | 22.9191502 | 0.003587512 | 0.944285187 |
| 18.419734 | 18.3590305 | 18.4304887 | 18.4480486 | 0.084679828 | 0.325207697 |
| 16.145227 | NA | 15.9459676 | 16.649647 | 0.642105107 | 0.105863984 |
| 22.5426797 | 22.2969578 | 22.4239794 | 22.4467304 | -0.011780732 | 0.903758519 |
| 22.5838945 | 22.5325713 | 22.5441429 | 22.5692698 | -0.010877854 | 0.769230959 |
| 19.1437408 | 19.1767741 | 19.2805015 | 19.3226361 | 0.090764773 | 0.174911729 |
| 17.1589347 | 16.9301626 | 16.8640618 | 17.0153031 | -0.191541163 | 0.057569092 |
| 16.2568356 | 16.2649389 | 16.3884402 | 16.2891153 | 0.10988656 | 0.34524274 |
| 17.4153986 | NA | 17.5266242 | 17.6514124 | 0.24206998 | 0.213891275 |
| 18.7230133 | 18.5429255 | 18.4759089 | 18.5680207 | -0.150805842 | 0.282450868 |
| 19.5541513 | 19.4237939 | 19.3989412 | 19.6361937 | -0.018829089 | 0.781988738 |
| 21.9531334 | 21.9558847 | 22.0583598 | 22.0559049 | 0.044250323 | 0.486157166 |
| 18.3589451 | 18.416828 | 18.295281 | 18.3885953 | 0.016626192 | 0.80700254 |
| 20.0483128 | 20.0557014 | 20.0678039 | 19.9521097 | -0.033006398 | 0.680742523 |
| 18.7556891 | 18.6373514 | 18.7360914 | 18.7200652 | -0.005482168 | 0.911431634 |
| 20.1907924 | 20.2080397 | 20.1677098 | 20.2000419 | 0.021285205 | 0.648529514 |
| 14.9256025 | NA | 15.2632225 | 15.2084333 | 0.228395936 | 0.155924866 |
| 12.0475085 | NA | 11.7223618 | 11.0616796 | -0.461958476 | 0.104503988 |
| 21.7041567 | 21.5748148 | 21.6769767 | 21.6586868 | -0.009656232 | 0.85067769 |
| 14.7900165 | 14.430414 | 14.3543262 | 14.394835 | -0.601972318 | 0.171979288 |
| 21.4600244 | 21.3448587 | 21.3857955 | 21.3368895 | -0.026727024 | 0.570358548 |
| 17.5610154 | 17.6488919 | 17.6305652 | 17.6370192 | 0.064856962 | 0.342227052 |
| 20.0356707 | 20.0673775 | 20.119369 | 20.1601755 | 0.078077324 | 0.293288358 |
| 21.381324 | 21.374033 | 21.4189074 | 21.4556818 | -0.016698104 | 0.69949451 |
| 25.138299 | 25.1082045 | 25.1510539 | 25.1476498 | 0.03096371 | 0.471408688 |
| 17.9681748 | 18.3347651 | 18.2874076 | 18.2296564 | 0.134989094 | 0.396405945 |
| 21.4253484 | 21.461958 | 21.4814462 | 21.4530583 | 0.042713979 | 0.539705458 |
| 21.1875951 | 21.0439398 | 21.1277573 | 21.0340451 | -0.077467659 | 0.22081627 |
| 22.1637052 | 22.0513277 | 22.1872572 | 22.1219794 | -0.016178462 | 0.773972437 |

| | | | | | |
|---|---|---|---|---|---|
| 22.728777 | 22.7515117 | 22.828785 | 22.6952654 | 0.04721197 | 0.378238732 |
| 20.5834636 | 20.4453448 | 20.4863578 | 20.5264833 | -0.055608991 | 0.240530332 |
| 19.7290791 | 19.711097 | 19.7439356 | 19.6539145 | 0.006871357 | 0.920585311 |
| 24.852022 | 24.8305581 | 24.9690261 | 24.8824215 | 0.061994776 | 0.142670475 |
| 19.4734245 | 19.3199526 | 19.2292812 | 19.3427542 | -0.032195884 | 0.677359549 |
| 21.2655792 | 21.2333447 | 21.1727508 | 21.2465677 | -0.045232283 | 0.305913719 |
| 20.4228304 | 20.4804669 | 20.5031279 | 20.3281027 | -0.053504011 | 0.442291281 |
| 20.0767231 | 19.9550837 | 20.0816498 | 20.0811743 | 0.020503115 | 0.738399381 |
| NA | 11.5166525 | NA | NA | 0.470799606 | 0.004789876 |
| NA | 13.6083412 | NA | NA | 0.420816071 | 0.229183617 |
| 17.8621636 | 17.6043904 | 17.9529284 | 17.9580807 | -0.099836454 | 0.354706386 |
| 17.7206229 | 17.6421995 | 17.9850046 | 17.9454926 | -0.031121421 | 0.742856555 |
| 19.9227655 | 19.7864472 | 19.9264783 | 19.8880024 | -0.069837835 | 0.177130677 |
| 22.2974016 | 22.2250769 | 22.2568465 | 22.2615291 | -0.02300997 | 0.648031284 |
| 23.1729169 | 23.1493214 | 23.1778696 | 23.2246472 | 0.034026307 | 0.452698849 |
| 16.1517159 | 15.8585245 | 15.9213918 | 16.0868969 | -0.006510683 | 0.967584913 |
| 23.2758906 | 23.415563 | 23.1796549 | 23.0559548 | -0.024047925 | 0.844120136 |
| 15.9579572 | 15.6509028 | 15.9518958 | 15.6942903 | -0.146450269 | 0.184124839 |
| 18.5542443 | 18.5346062 | 18.5089311 | 18.8259495 | 0.053723184 | 0.488289655 |
| 20.745489 | 20.6354066 | 20.64551 | 20.7335169 | -0.040451608 | 0.418648679 |
| 22.1292578 | 22.1142031 | 22.1502933 | 22.1604744 | -0.012536118 | 0.744066758 |
| 20.3320602 | 20.263115 | 20.3530298 | 20.3622737 | -6.31E-04 | 0.988867098 |
| 25.4348072 | 25.3625301 | 25.3952051 | 25.430357 | 0.006056021 | 0.904284416 |
| 18.1853682 | 17.8498526 | 17.6871032 | 18.1230879 | 0.039137429 | 0.758013779 |
| 16.2812885 | NA | 16.8090824 | 16.6473653 | 0.310640998 | 0.293201616 |
| 19.4264212 | 19.3829399 | 19.3806301 | 19.4114397 | 0.032981825 | 0.561053625 |
| 18.0658953 | 17.8531616 | 17.9053502 | 18.0278745 | -0.036614399 | 0.597117349 |
| 21.9385187 | 21.7482036 | 21.8739064 | 21.8939649 | -0.079541395 | 0.16608578 |
| 18.2862424 | 18.3198049 | 18.4815987 | 18.3697813 | -2.55E-04 | 0.997834819 |
| 19.7867676 | 19.5929624 | 19.7854384 | 19.6689865 | -0.053258693 | 0.451656736 |
| 21.6232403 | 21.3476866 | 21.3770923 | 21.4095705 | -0.134374367 | 0.037275802 |
| 17.9857143 | 17.9408315 | 17.8800732 | 17.9506341 | -0.101792343 | 0.195751834 |
| 22.8468465 | 22.8770517 | 22.7220906 | 22.7308229 | -0.009831024 | 0.887662068 |
| 26.0441073 | 26.0803474 | 26.0640174 | 25.9852284 | 0.039166986 | 0.544197912 |
| 22.210441 | 22.0502085 | 22.1371221 | 22.1469835 | -0.043955889 | 0.295386831 |
| 20.7737002 | 20.7202769 | 20.7993989 | 20.8111788 | 0.028419496 | 0.552555421 |
| 21.9599224 | 21.8358844 | 21.914344 | 21.9499592 | -0.009077337 | 0.894905921 |
| 23.359411 | 23.2789163 | 23.3328564 | 23.2774085 | -0.039223841 | 0.354985808 |
| 24.2587491 | 24.2431336 | 24.2904947 | 24.2696865 | 0.018213084 | 0.656595045 |
| 19.142187 | 18.9935486 | 19.1405096 | 19.1226368 | -0.02254847 | 0.737651705 |
| 11.3705416 | NA | 12.8765771 | 12.6411505 | 0.875792303 | 0.01868421 |
| 16.0935296 | 15.894833 | 15.8156882 | 16.0410741 | -0.016440682 | 0.910416851 |

| | | | | | |
|---|---|---|---|---|---|
| 16.3554481 | 16.9363743 | 17.2636407 | 17.0269949 | 0.380457197 | 0.058336895 |
| 27.3903528 | 27.2515585 | 27.3256835 | 27.3752605 | 0.026017252 | 0.679613366 |
| 17.4883928 | 17.4108716 | 17.4459408 | 17.3693735 | 0.050556114 | 0.55442081 |
| 17.0693125 | 16.8009372 | 16.8409133 | 16.9934649 | -0.005900089 | 0.962712898 |
| 22.8156908 | 22.571701 | 22.8815961 | 22.7974758 | 2.97E-04 | 0.998327845 |
| 15.0804185 | 15.3035812 | 16.0549027 | 15.2492272 | 0.515429831 | 0.034345774 |
| 24.9524462 | 24.8566239 | 24.8712281 | 24.8936684 | 0.017596195 | 0.752562806 |
| 19.7530737 | 19.7742928 | 19.8387408 | 19.9070519 | 0.067316142 | 0.253400796 |
| 18.2929021 | 18.1106748 | 18.29268 | 18.1276422 | -0.049966854 | 0.624128934 |
| 23.5327975 | 23.5040164 | 23.6206587 | 23.6027939 | 0.024037156 | 0.584111473 |
| 18.3041082 | 18.30709 | 18.3348818 | 18.2451191 | 0.019946245 | 0.82945879 |
| 15.5041287 | 15.3384649 | 15.3704245 | 15.3044978 | -0.072965481 | 0.580198682 |
| 16.1822236 | 16.2443655 | 16.2580914 | 16.4108144 | -0.072639186 | 0.578068196 |
| 16.4032113 | 16.435694 | 16.6549034 | 16.5324376 | 0.119423523 | 0.222888088 |
| 21.4978019 | 21.3100763 | 21.4181688 | 21.4597853 | -0.058870494 | 0.268405409 |
| 20.2568961 | 20.1785633 | 20.2528791 | 20.2380843 | 0.020161147 | 0.702114597 |
| 19.6828114 | 19.6771871 | 19.6830362 | 19.680197 | 0.001342612 | 0.974812914 |
| 20.8623843 | 20.5653279 | 20.7087671 | 20.7441855 | -0.044561089 | 0.758868187 |
| 17.4925738 | 17.2144378 | 17.3236242 | 17.3253615 | -0.124156132 | 0.200472028 |
| 17.5126131 | 17.551962 | 17.5180421 | 17.4597453 | -0.043734991 | 0.607406642 |
| 27.0269585 | 27.0223721 | 27.0273481 | 27.0705 | 0.029674862 | 0.591733571 |
| 25.1957773 | 25.199765 | 25.254849 | 25.2297169 | 0.066705503 | 0.21759187 |
| 22.215488 | 22.1346326 | 22.237592 | 22.2736651 | 0.054135512 | 0.316799639 |
| 24.5307892 | 24.4396733 | 24.5183089 | 24.5260021 | 0.026460475 | 0.596133617 |
| 21.767527 | 21.7152089 | 21.8152933 | 21.7385976 | -0.036152575 | 0.424453425 |
| 25.8840511 | 25.7672491 | 25.9706335 | 25.886837 | 0.072217331 | 0.559026587 |
| 21.6652388 | 21.6569056 | 21.5364435 | 21.5276333 | -0.042698749 | 0.400054434 |
| 21.5787287 | 21.5519223 | 21.4646697 | 21.5500236 | -0.041782003 | 0.399461716 |
| 20.6901486 | 20.5844617 | 20.6434038 | 20.5798439 | -0.063151042 | 0.174860989 |
| 22.7986238 | 22.7659224 | 22.7585557 | 22.7174141 | -0.041372894 | 0.316264175 |
| 21.0315615 | 20.9493042 | 21.0382651 | 21.1489769 | 0.008489708 | 0.877895188 |
| 20.9143097 | 20.7009962 | 20.8616654 | 20.8549336 | -0.06205541 | 0.47775502 |
| 17.692009 | 17.5999768 | 17.9248052 | 17.8388835 | 0.203333578 | 0.036176826 |
| 23.9091947 | 23.9219118 | 23.9686267 | 23.8950448 | 0.101753086 | 0.545677343 |
| 16.4635933 | NA | 16.3658302 | 16.3116344 | -0.156878567 | 0.080884786 |
| 19.9712454 | 19.9220542 | 20.0067729 | 19.9041306 | -0.015716367 | 0.842795727 |
| 24.5645024 | 24.4348226 | 24.5121471 | 24.5049453 | -0.048872029 | 0.316288416 |
| 20.1887845 | 20.0082395 | 20.1939196 | 20.2369132 | -0.008638813 | 0.885066819 |
| 16.9879928 | 16.997865 | 17.0834719 | 17.2500119 | 0.042630329 | 0.729625091 |
| 16.6842021 | 16.8915836 | 16.7136696 | 16.8128318 | 0.181788075 | 0.105858235 |
| 19.3287475 | 18.9415768 | 19.2679181 | 19.2787593 | -0.167198967 | 0.300467104 |
| 19.0165005 | 18.9151307 | 18.8827264 | 18.9329398 | 0.06858167 | 0.46981988 |

| | | | | | |
|---|---|---|---|---|---|
| 20.2740761 | 20.0598515 | 20.2253093 | 20.2099464 | -0.056020613 | 0.431528471 |
| 20.9599212 | 20.7596453 | 20.9388376 | 20.9158347 | -0.020483675 | 0.751390375 |
| 19.6397841 | 19.7221751 | 19.7898793 | 19.666681 | 0.025288736 | 0.659716816 |
| 20.6746903 | 20.6198187 | 20.6645883 | 20.7825244 | 0.024112469 | 0.67300785 |
| 22.0757959 | 21.9620149 | 21.9894631 | 22.0249163 | -0.048769398 | 0.392456218 |
| 20.8076014 | 20.7451715 | 20.7932903 | 20.8332221 | 0.03112601 | 0.579127277 |
| 22.6628686 | 22.6553906 | 22.6805439 | 22.6584305 | 0.028404764 | 0.505641688 |
| 21.3043285 | 21.2971372 | 21.2738479 | 21.3004702 | 0.023842388 | 0.594680055 |
| 20.1727166 | 20.1491693 | 20.2224147 | 20.1836052 | 0.07006787 | 0.219196385 |
| 19.0315175 | 18.8503974 | 19.0208998 | 19.0310626 | -0.029686183 | 0.677525108 |
| 20.9379611 | 20.7567592 | 20.9593714 | 21.0103536 | 0.053149714 | 0.459567437 |
| 23.6143803 | 23.4600815 | 23.7417165 | 23.6428281 | 0.090688896 | 0.313147479 |
| 17.9252207 | 17.5546467 | 17.9935876 | 17.7115799 | -0.023318643 | 0.859612985 |
| 21.6799242 | 21.6802195 | 21.6470582 | 21.6949407 | -0.029976962 | 0.644051417 |
| 21.4601602 | 21.3712389 | 21.4768663 | 21.4443149 | -0.002906232 | 0.958764282 |
| 17.3279162 | 17.3883544 | 17.167318 | 17.4017024 | -0.093172678 | 0.594302507 |
| 22.4413993 | 22.4424864 | 22.4261488 | 22.4804246 | 0.02843085 | 0.600999994 |
| 20.1264531 | 19.977152 | 20.1057026 | 20.0113092 | -0.003812527 | 0.967530521 |
| 20.7528538 | 21.0615735 | 20.6452858 | 20.7767317 | 0.200926388 | 0.307888529 |
| 22.3130989 | 22.1180796 | 22.2021313 | 22.2124977 | -0.058370156 | 0.34236007 |
| 20.255482 | 20.0545436 | 20.1220357 | 20.1628713 | -0.040759119 | 0.48627773 |
| 20.4372125 | 20.3304807 | 20.3701245 | 20.3561626 | -0.067713586 | 0.22323706 |
| 21.6900663 | 21.5718064 | 21.6646812 | 21.6949028 | -0.031198939 | 0.513047296 |
| 22.2233589 | 22.0968866 | 22.2095184 | 22.14368 | -0.051249802 | 0.199381752 |
| 23.0074693 | 22.9542009 | 23.0060529 | 23.0116955 | 0.020290037 | 0.700168628 |
| 21.6522716 | 21.705428 | 21.6679418 | 21.6745415 | 0.008996965 | 0.867636284 |
| 18.3278393 | 18.2134793 | 18.277654 | 18.4615936 | 0.060105489 | 0.314437751 |
| 20.1148899 | 19.9963854 | 20.1487152 | 20.1780604 | 0.039224823 | 0.54523852 |
| 18.2845946 | 18.2636781 | 18.3687849 | 18.3997763 | 0.022201512 | 0.742406363 |
| 19.6503981 | 19.5711897 | 19.6436848 | 19.771241 | 0.081819102 | 0.244009167 |
| 17.0129036 | 16.9629731 | 17.0603422 | 16.9513341 | 0.026985266 | 0.831403805 |
| 14.9405659 | 14.6930198 | 15.0332796 | 15.055771 | 8.35E-04 | 0.994759678 |
| 16.0599679 | 16.4871714 | 16.1842929 | 16.1784285 | 0.092524724 | 0.475993681 |
| 16.029543 | 15.9298729 | 15.9168315 | 16.2294964 | 0.335590686 | 0.080593258 |
| NA | 11.529137 | NA | NA | 0.218593753 | 0.532354811 |
| 15.9830094 | NA | 15.9921563 | 16.0491973 | 0.233271739 | 0.536472675 |
| NA | 15.478211 | NA | NA | 0.221942694 | 0.621619503 |
| 26.2021124 | 26.1340158 | 26.2557617 | 26.2543762 | 0.061771983 | 0.308048229 |
| 22.0468001 | 21.9266913 | 21.966439 | 22.0504507 | 0.026281259 | 0.768787513 |
| 20.3444076 | 20.2456062 | 20.2604144 | 20.3128456 | 0.001082265 | 0.984473925 |
| 17.3690062 | 17.3295363 | 17.3122839 | 17.6883776 | 0.11897398 | 0.327450515 |
| 19.0154131 | 19.1161976 | 19.038589 | 19.0726239 | 0.040574278 | 0.539637302 |

| | | | | | |
|---|---|---|---|---|---|
| 19.7101967 | 19.5610052 | 19.7019655 | 19.6884715 | -0.009257521 | 0.839580618 |
| 19.2892391 | 19.2746726 | 19.3741108 | 19.5508096 | 0.140929654 | 0.215703039 |
| 14.7682282 | 14.2812856 | 14.6847918 | 14.957828 | 0.165785287 | 0.467780823 |
| 21.5838013 | 21.5437702 | 21.6675192 | 21.6277367 | 0.061130798 | 0.268582831 |
| 23.3896266 | 23.3224207 | 23.3986388 | 23.4111342 | 0.028281215 | 0.5094124 |
| 20.0309637 | 19.9910527 | 20.0003444 | 20.0029729 | 0.025287927 | 0.637290148 |
| 21.1644339 | 20.9753544 | 21.168752 | 21.2543256 | 0.019302823 | 0.825007142 |
| 22.7220914 | 22.7307251 | 22.817624 | 22.7894997 | 0.05300234 | 0.16888265 |
| 20.2091837 | 20.1571013 | 20.274931 | 20.1696077 | -0.089097528 | 0.167167217 |
| 23.4399004 | 23.2470913 | 23.4064026 | 23.4018982 | -0.015164633 | 0.783905431 |
| 20.727012 | 20.7935511 | 20.7918032 | 20.7098129 | 0.03673874 | 0.384812961 |
| 21.5447548 | 21.5532122 | 21.641594 | 21.6472172 | 0.054613755 | 0.225991755 |
| 18.0478679 | 18.0286944 | 18.1306412 | 18.073533 | 0.001646027 | 0.983798878 |
| 16.0457007 | 15.6811675 | 15.5577128 | 15.6420843 | -0.064115411 | 0.686813763 |
| 24.8689151 | 24.7439436 | 24.8060964 | 24.8859703 | 0.001120347 | 0.985434404 |
| 22.0008604 | 21.8992576 | 21.854206 | 21.9098743 | -0.042375052 | 0.416554376 |
| 21.8257206 | 21.430447 | 21.7296405 | 21.7411737 | -0.110077416 | 0.330770356 |
| 23.2364468 | 23.0795168 | 23.1916041 | 23.1840497 | -0.061216878 | 0.261216519 |
| 16.2812633 | 16.1788042 | 16.2826123 | 16.3326769 | -0.021417182 | 0.781686949 |
| 18.3484608 | 18.5316281 | 18.702575 | 18.2072991 | 0.094935297 | 0.72151599 |
| 19.0266482 | 18.9492205 | 19.1015868 | 19.0328219 | -0.05337916 | 0.465128531 |
| 17.1992298 | 17.5524325 | 17.4979982 | 17.2767232 | 0.195440998 | 0.059613554 |
| 26.2572099 | 26.1892655 | 26.2821768 | 26.3036061 | 0.049477335 | 0.348208995 |
| 19.2761322 | 19.0973425 | 19.1647311 | 19.2923077 | -0.024623021 | 0.66437422 |
| NA | 12.6825072 | NA | NA | -0.654652499 | 0.247073066 |
| 17.6379904 | 17.3357295 | 17.8113982 | 17.678152 | -0.077264269 | 0.348286703 |
| 20.6861383 | 20.6342571 | 20.5047138 | 20.6857776 | -0.061629979 | 0.266608317 |
| 19.3640932 | 19.321433 | 19.3542554 | 19.3435615 | 0.107023834 | 0.139135811 |
| 19.5149848 | 19.251604 | 19.5527254 | 19.5535901 | 0.052699762 | 0.623727924 |
| 23.673598 | 23.523658 | 23.6774172 | 23.6442649 | 0.043220459 | 0.496068802 |
| 23.2794825 | 23.550763 | 23.2597837 | 23.3369857 | 0.110652141 | 0.377901978 |
| 22.2051293 | 22.0995048 | 22.1628525 | 22.2184682 | 0.023588141 | 0.724756543 |
| 19.6592494 | 19.7437646 | 19.8411076 | 19.7092158 | 0.056384168 | 0.443679279 |
| NA | 13.8083929 | NA | NA | -0.298031878 | 0.169677286 |
| 14.8771944 | 14.9531781 | 14.6153518 | 15.1176471 | 0.014257074 | 0.930400679 |
| NA | 14.3143586 | NA | NA | -0.175458956 | 0.266036634 |
| 18.5709051 | 18.7301958 | 18.9832912 | 18.7602069 | 0.082727971 | 0.524931157 |
| 20.3423267 | 19.9846697 | 19.8964502 | 20.095443 | -0.169927296 | 0.184052841 |
| 17.9502937 | 17.7860473 | 17.8402342 | 17.781149 | -0.103922454 | 0.322357505 |
| 21.3186034 | 20.986676 | 21.1957736 | 21.2985083 | -0.06067492 | 0.383470339 |
| 19.2962114 | 19.2494524 | 19.3377053 | 19.307311 | 0.046636254 | 0.382554946 |
| 20.834908 | 20.7591716 | 20.7406651 | 20.8036016 | -0.007211993 | 0.87182995 |

| | | | | | |
|---|---|---|---|---|---|
| 20.5409671 | 20.4287406 | 20.5095295 | 20.5716431 | -0.005285212 | 0.90951835 |
| 20.0623119 | 20.0367723 | 19.9940296 | 20.0681979 | 0.010356973 | 0.824219035 |
| 23.3363247 | 23.2426875 | 23.3855673 | 23.3783544 | 0.044358739 | 0.392416748 |
| 20.7143453 | 20.6224114 | 20.8415788 | 20.7970872 | 0.073631811 | 0.253085879 |
| 19.2733306 | 19.2359243 | 19.3653301 | 19.280378 | -2.70E-04 | 0.996785715 |
| 17.2894148 | 17.1490617 | 17.3002134 | 17.3641281 | 0.029064541 | 0.702595344 |
| 18.1186316 | 18.0479026 | 18.0579991 | 18.1578655 | -0.00690563 | 0.932163708 |
| 17.7334441 | 18.1616131 | 18.2449834 | 18.1662841 | 0.183839349 | 0.061109592 |
| 19.5930872 | 19.6445965 | 19.7603298 | 19.7113218 | -0.032906735 | 0.747863596 |
| 17.7761587 | 17.7406533 | 17.7859048 | 17.7716322 | 0.060775874 | 0.596648354 |
| 19.43953 | 19.2453303 | 19.3415531 | 19.3186068 | -0.084542638 | 0.470670135 |
| NA | 18.2055564 | NA | NA | 0.293743599 | 0.048234751 |
| 16.733696 | 16.9289088 | 17.0213242 | 16.7887179 | 0.228505281 | 0.128606289 |
| 19.9508855 | 19.9752321 | 19.9770911 | 19.9264538 | 0.032207741 | 0.638945803 |
| 18.4580439 | 18.0854915 | 18.3239593 | 18.2175533 | -0.125841541 | 0.136003591 |
| 22.0580311 | 22.0681802 | 22.0479496 | 22.0260257 | -0.010801915 | 0.812451025 |
| 18.8204404 | 18.4318588 | 18.7141931 | 18.7493288 | -0.081290739 | 0.460426814 |
| 15.3384759 | 15.3326159 | 15.5581913 | 15.279019 | 0.069486841 | 0.652079273 |
| 20.738035 | 20.3872467 | 20.477366 | 20.5431488 | -0.157194169 | 0.163051168 |
| 20.4121662 | 20.3036134 | 20.3682037 | 20.3828455 | -0.027453256 | 0.624392521 |
| 21.0193784 | 20.7975994 | 20.922881 | 20.914698 | -0.059027461 | 0.290943523 |
| 24.9579458 | 24.9424717 | 25.010537 | 25.0213233 | 0.066517262 | 0.100606472 |
| 20.3163591 | 20.2992271 | 20.3448228 | 20.3647812 | 0.021735041 | 0.723907651 |
| 21.3632975 | 21.3835238 | 21.404695 | 21.3313495 | 0.02056385 | 0.619310428 |
| 24.0522292 | 23.9525472 | 24.1323645 | 24.1455512 | 0.051274858 | 0.4785609 |
| 22.6982335 | 22.623411 | 22.7643246 | 22.7650821 | 0.037047305 | 0.64626999 |
| 22.057661 | 22.0243519 | 22.0833685 | 22.0710819 | -0.0504065 | 0.458798564 |
| 23.1903331 | 23.0536657 | 23.1060039 | 23.1224275 | -0.052311522 | 0.308364174 |
| 17.8217475 | 17.725361 | 17.8982425 | 17.7083788 | 0.060868782 | 0.379259343 |
| 24.1876555 | 24.26307 | 24.508075 | 24.3465248 | 0.267675212 | 0.073193545 |
| 21.1912204 | 21.0604011 | 21.0965256 | 21.2027841 | -0.051388056 | 0.458477674 |
| 23.5092199 | 23.500903 | 23.5038769 | 23.5390597 | 0.029922706 | 0.393141429 |
| 22.0152298 | 21.9236897 | 22.0786184 | 22.0419871 | 0.016976926 | 0.808017078 |
| 19.341692 | 19.2790898 | 19.3486589 | 19.4386553 | 0.070227326 | 0.339566803 |
| 18.6277403 | 18.5545516 | 18.6964954 | 18.7986735 | 0.084623443 | 0.301386122 |
| 17.9500202 | 17.7307626 | 17.9510517 | 18.1284611 | 0.131462036 | 0.503300698 |
| NA | 11.8483772 | NA | NA | 0.217193879 | 0.449385503 |
| 17.4664925 | 17.378197 | 17.6952445 | 17.2966693 | 0.027040532 | 0.82200501 |
| 19.9733252 | 19.8011775 | 20.103727 | 19.8079477 | -0.101944421 | 0.416482248 |
| 22.9480388 | 22.9085875 | 22.9080156 | 22.9333229 | -0.001970787 | 0.971823326 |
| NA | 17.3120486 | NA | NA | -0.272461397 | 0.306465804 |
| 18.7994255 | 18.6402343 | 18.7113692 | 18.7495904 | -0.00610362 | 0.916678752 |

| | | | | | |
|---|---|---|---|---|---|
| 19.8785488 | 19.7372246 | 19.8133361 | 19.8442288 | -0.052828744 | 0.207496703 |
| 20.8812806 | 20.8743633 | 20.9526657 | 20.9520011 | 0.079990885 | 0.115001379 |
| 20.2517065 | 20.1389577 | 20.2134086 | 20.1620446 | -0.004321512 | 0.941139238 |
| 19.2518238 | 19.3292744 | 19.4267111 | 19.3914158 | 0.07143954 | 0.287812992 |
| 19.6174488 | 19.5745384 | 19.7710736 | 19.6705057 | -0.057542705 | 0.459768421 |
| 19.4470855 | 19.3420619 | 19.1852312 | 19.6626526 | 0.262044396 | 0.156916765 |
| 15.6104963 | 15.5683261 | 15.5074371 | 15.3463187 | -0.243323777 | 0.063489534 |
| 19.7492241 | 19.62136 | 19.7192272 | 19.6193205 | -0.074616655 | 0.307149902 |
| 22.1262274 | 21.9287791 | 21.9386502 | 21.9629986 | -0.078346301 | 0.221583137 |
| 21.4499714 | 21.3742209 | 21.4270609 | 21.4398151 | -0.004445466 | 0.926280573 |
| 21.0625812 | 21.0078564 | 21.1032353 | 21.0673705 | 0.03567315 | 0.449287288 |
| 22.4198604 | 22.1846975 | 22.3934857 | 22.3398174 | -0.023520296 | 0.746269494 |
| 23.7801804 | 23.7131725 | 23.8616391 | 23.8457803 | 0.055518099 | 0.54479429 |
| 18.3269045 | 18.2349784 | 18.3516047 | 18.4078747 | 0.010734221 | 0.927448359 |
| 18.5932188 | 18.5964706 | 18.6227482 | 18.7921718 | 0.049705523 | 0.68373008 |
| 17.7452473 | NA | 17.7438058 | 17.6536176 | -0.048132442 | 0.818281086 |
| 22.5825884 | 22.2849992 | 22.378102 | 22.456993 | -0.02625268 | 0.793799168 |
| 24.2331124 | 24.1524718 | 24.1392867 | 24.1430125 | -0.030497183 | 0.57040038 |
| 19.5347495 | 19.4319702 | 19.5253115 | 19.6163256 | 0.043291062 | 0.474524715 |
| 22.4171829 | 22.346414 | 22.4158524 | 22.3257674 | -0.0319876 | 0.656044694 |
| 16.1474612 | 15.9863447 | 16.2901285 | 16.4645452 | 0.245420461 | 0.208160332 |
| NA | 13.3799439 | NA | NA | -0.072780452 | 0.739274579 |
| 25.7991363 | 25.7159956 | 25.8040766 | 25.7975552 | 0.025352139 | 0.587126538 |
| 16.932349 | 16.7254901 | 16.5410633 | 16.6507648 | -0.186440842 | 0.111654134 |
| 19.1483927 | 18.8874034 | 19.0698365 | 18.7486658 | -0.139120049 | 0.202167331 |
| 21.8781876 | 21.6059818 | 21.8018408 | 21.7893986 | -0.087502472 | 0.271129443 |
| 22.2509501 | 22.1954923 | 22.1560515 | 22.155154 | 0.004776738 | 0.919709612 |
| 17.8762307 | 17.9677881 | 17.6361965 | 17.7276029 | -0.21521217 | 0.034920792 |
| 20.9452735 | 20.9072103 | 20.9972362 | 20.9761943 | 0.052615358 | 0.393665988 |
| 20.7379191 | 20.674172 | 20.7265978 | 20.8755813 | 0.074053513 | 0.294306523 |
| 22.4014974 | 22.3638977 | 22.408519 | 22.4151504 | 0.017900731 | 0.688468098 |
| 17.7480119 | 17.8610392 | 17.8557954 | 17.7809732 | 0.025608695 | 0.803609371 |
| 20.015645 | 19.8986871 | 20.0163821 | 20.047217 | 0.022096245 | 0.763409876 |
| 22.5016812 | 22.3694595 | 22.4737755 | 22.4876822 | -0.006215812 | 0.908029581 |
| NA | 15.6254822 | NA | NA | -0.627678561 | 0.068021527 |
| 16.3386897 | 15.6880317 | 16.1691101 | 16.158502 | -0.27182271 | 0.130343269 |
| 19.6989533 | 19.7741329 | 19.3874443 | 19.7714114 | 0.021035315 | 0.88795161 |
| 17.5499882 | 17.7581794 | 17.7985366 | 17.7632797 | 0.056717682 | 0.589067462 |
| 19.7466199 | 19.7385308 | 19.7770085 | 19.6680305 | -0.107247227 | 0.112306262 |
| 23.2726324 | 23.2429945 | 23.2968973 | 23.2475784 | -0.007515873 | 0.908695594 |
| 19.147794 | 18.9479235 | 18.8870712 | 19.0709812 | -0.170625029 | 0.010533566 |
| 24.2403247 | 24.0962862 | 24.1946967 | 24.1795009 | -0.025827204 | 0.609808472 |

| | | | | | |
|---|---|---|---|---|---|
| 20.4543143 | 20.401589 | 20.4302262 | 20.5097506 | 0.028543556 | 0.552824578 |
| 20.9107742 | 20.8806369 | 20.9692256 | 20.9893444 | 0.049301372 | 0.311054634 |
| 21.4931436 | 21.4715929 | 21.4200819 | 21.494887 | 0.026793396 | 0.602492552 |
| 22.3325192 | 22.282268 | 22.3266883 | 22.3169516 | 0.023334199 | 0.659623346 |
| 21.8008779 | 21.6632671 | 21.7802368 | 21.7745829 | -0.012575591 | 0.818024175 |
| 20.2418436 | 20.1881739 | 20.305157 | 20.2387829 | 0.010264267 | 0.858364484 |
| 18.7046711 | 18.5657169 | 18.7146626 | 18.8565507 | -0.008838922 | 0.88953575 |
| 20.5708815 | 20.3865558 | 20.506666 | 20.5464443 | -0.002119468 | 0.976039843 |
| 15.8947167 | NA | 15.9701065 | 16.1269621 | 0.161965921 | 0.532945575 |
| 18.1935632 | 18.2466318 | 18.2737191 | 18.2495807 | 0.01796579 | 0.823946167 |
| 23.0038666 | 22.8596406 | 23.048101 | 22.9594421 | -0.027357372 | 0.652925969 |
| 22.1473409 | 22.0280315 | 22.1081505 | 22.1183608 | -0.043315791 | 0.382332705 |
| 23.3946447 | 23.3305058 | 23.3997766 | 23.3164362 | -0.025760375 | 0.595682052 |
| 21.9991593 | 21.9537475 | 21.9696785 | 22.0479323 | 0.021882796 | 0.645902369 |
| 21.4516265 | 21.3978837 | 21.3985693 | 21.4281574 | -0.004077183 | 0.943587541 |
| 19.7822136 | 19.6842368 | 19.7070356 | 19.7543798 | 0.009845174 | 0.881837215 |
| 22.2203888 | 22.1106746 | 22.1206568 | 22.1296294 | 0.001397733 | 0.984188174 |
| 17.0511413 | 17.1269559 | 17.0616789 | 17.1703465 | 0.19225236 | 0.032348944 |
| NA | 15.2554548 | NA | NA | -0.095558298 | 0.337132374 |
| 18.8095301 | 18.744525 | 18.8502038 | 18.8626869 | 0.023269392 | 0.75047795 |
| 21.6490242 | 21.6419468 | 21.7084881 | 21.6515294 | 0.097864436 | 0.058470236 |
| 22.8793598 | 22.665029 | 22.7308248 | 22.7761361 | -0.03976505 | 0.539585023 |
| 22.0875283 | 21.9892216 | 21.952549 | 21.9850842 | -0.034616861 | 0.58530981 |
| 22.375483 | 22.4506941 | 22.4214118 | 22.3720927 | 0.008738588 | 0.873115069 |
| 23.6793976 | 23.5595823 | 23.6124367 | 23.6488932 | -0.022924704 | 0.622923404 |
| 25.6070635 | 25.5825404 | 25.5893037 | 25.6192458 | -0.009997925 | 0.863682908 |
| 15.6957169 | NA | 15.6960316 | 15.7531269 | -0.069803579 | 0.549263831 |
| 18.7602334 | 18.9410018 | 18.6843314 | 18.7390036 | 0.013732562 | 0.866512188 |
| 20.4761228 | 20.4341582 | 20.3205457 | 20.3708083 | -0.101954899 | 0.116805868 |
| 19.1169011 | 19.0923639 | 19.0627348 | 18.9663812 | -0.089086959 | 0.101776603 |
| 20.7527999 | 20.5980274 | 20.6580804 | 20.7949708 | -0.004246573 | 0.937765969 |
| 22.8240745 | 22.8322782 | 22.7754966 | 22.7853599 | 0.04027364 | 0.600965879 |
| 20.2576062 | 20.12837 | 20.2753541 | 20.3361632 | 0.027369258 | 0.624373371 |
| 19.0802879 | 19.0988014 | 19.1205782 | 19.0487936 | 0.029193624 | 0.630367027 |
| 21.8269834 | 21.7473701 | 21.8805649 | 21.9368864 | 0.036261013 | 0.564869828 |
| NA | 19.5306912 | NA | NA | -0.059571569 | 0.593311645 |
| 24.429472 | 24.3824487 | 24.3863646 | 24.4160436 | -0.005378351 | 0.90434918 |
| 20.4729872 | 20.3698625 | 20.4470448 | 20.4781221 | 0.054002384 | 0.377526632 |
| 15.6816785 | 15.6454505 | 15.7407109 | 15.6287925 | 0.248371295 | 0.17644636 |
| 22.5810141 | 22.4895496 | 22.4146317 | 22.2823146 | -0.181211165 | 0.042502581 |
| 21.0853914 | 20.8864744 | 20.9700666 | 21.0858686 | -0.069687969 | 0.381281272 |
| 22.6339287 | 22.4529407 | 22.5984546 | 22.6004986 | -0.024017437 | 0.636461619 |

| | | | | | |
|---|---|---|---|---|---|
| 20.3537092 | 20.0907566 | 20.2693896 | 20.2511633 | -0.04645655 | 0.589533357 |
| NA | 17.2848998 | NA | NA | 0.064067657 | 0.771578333 |
| 20.7961144 | 20.3974194 | 20.5460769 | 20.6909903 | -0.106703754 | 0.403120374 |
| 19.7465965 | 19.4403525 | 19.8921456 | 19.7170663 | -0.04519641 | 0.668841331 |
| 19.9676045 | 19.7914524 | 19.9050103 | 20.0348629 | -0.01467929 | 0.872121784 |
| 23.6769864 | 23.4756914 | 23.5645159 | 23.5217082 | -0.12083628 | 0.121104668 |
| 19.6862802 | 19.5954003 | 19.5740196 | 19.563607 | -0.079317277 | 0.297586145 |
| 23.2305645 | 23.1222459 | 23.2040975 | 23.2696823 | -0.021068414 | 0.741927592 |
| 20.5158992 | 20.4534606 | 20.541652 | 20.3985894 | -0.03390736 | 0.576107291 |
| 22.2977202 | 22.2077237 | 22.2665584 | 22.3026628 | -0.037292747 | 0.514127413 |
| 20.7366907 | 20.6475578 | 20.6901119 | 20.6941976 | -0.034912404 | 0.491167233 |
| 21.1646365 | 21.1420918 | 21.1059461 | 21.0857914 | -0.00991866 | 0.816460313 |
| 20.1706419 | 20.2423696 | 20.3397111 | 20.2416342 | 0.055997047 | 0.341518611 |
| 21.2347502 | 21.1613286 | 21.0977955 | 21.1403526 | -0.044471979 | 0.465954071 |
| 21.8781779 | 21.7791273 | 21.797417 | 21.8236401 | -0.051594177 | 0.49633588 |
| 18.6175462 | 18.4854599 | 18.5446586 | 18.4546851 | -0.169724281 | 0.040949597 |
| 21.8760424 | 21.7105018 | 21.7615444 | 21.8045987 | -0.077828294 | 0.368710747 |
| 22.084326 | 22.0218293 | 22.0051905 | 22.0335707 | 0.011514257 | 0.92001478 |
| NA | 12.621286 | NA | NA | 0.288072928 | 0.226044288 |
| 11.8884279 | NA | 11.4956664 | 11.5485992 | -0.408344407 | 0.077560581 |
| 17.7370954 | 17.2487442 | 17.5463808 | 17.6600263 | -0.11148772 | 0.436124082 |
| 20.6340569 | 20.6268063 | 20.6413219 | 20.7215262 | 0.003279932 | 0.952004434 |
| 22.554271 | 22.4630776 | 22.5096527 | 22.5289982 | -0.004388837 | 0.942105514 |
| 15.5515237 | NA | 16.5019785 | 16.079197 | 0.788119879 | 0.126686024 |
| 18.5522358 | 18.3659849 | 18.444825 | 18.4338853 | -0.023073817 | 0.757607362 |
| 18.0879484 | 17.9221683 | 17.993263 | 18.272478 | -0.008015094 | 0.937917251 |
| 19.6640892 | 19.5060507 | 19.7002535 | 19.6823399 | 0.035225897 | 0.505569538 |
| 20.0372285 | 19.9795471 | 20.0604879 | 20.0400181 | 0.000723745 | 0.988661523 |
| 23.4435382 | 23.4003081 | 23.4146009 | 23.3198052 | -0.007312346 | 0.877910657 |
| 20.242439 | 20.0999621 | 20.240359 | 20.2764074 | 0.00940561 | 0.874826871 |
| 17.9964642 | 17.9084811 | 17.9442253 | 17.9884455 | -0.056771934 | 0.525375162 |
| 24.0502776 | 23.8710758 | 24.0371755 | 24.0841554 | -0.017786675 | 0.774095006 |
| 15.8706017 | 15.4401495 | 15.8114149 | 15.8726231 | -0.239261706 | 0.192897308 |
| 20.4439847 | 20.2612789 | 20.2822623 | 20.3935643 | -0.035256958 | 0.719053927 |
| 20.046423 | 19.9483474 | 19.9564279 | 19.8391388 | -0.085646031 | 0.324222597 |
| 21.0442045 | 21.0201985 | 21.0920643 | 21.1111671 | 0.038803095 | 0.343224103 |
| 17.2407463 | 17.3247382 | 17.3305951 | 17.2657012 | 0.08298773 | 0.374704446 |
| 21.2102273 | 21.2751407 | 21.3951981 | 21.0678791 | -0.076716761 | 0.427650344 |
| 15.9562385 | 16.0745239 | 16.1862005 | 15.8253732 | 0.279088258 | 0.030182229 |
| 14.8184028 | NA | 14.839788 | 14.8967216 | 0.088306565 | 0.561024577 |
| 18.8261493 | 18.7385694 | 18.8424155 | 18.7861597 | 0.017896254 | 0.744562766 |
| 18.2844051 | 18.3327833 | 18.0040702 | 18.2239262 | -0.021444233 | 0.842178399 |

| | | | | | |
|---|---|---|---|---|---|
| 19.440077 | 19.2847356 | 19.1948385 | 19.3849011 | -0.071269563 | 0.464998118 |
| 25.0299547 | 24.8921316 | 24.9789326 | 25.0091181 | -0.018651959 | 0.752117379 |
| 22.2897595 | 22.1600249 | 22.2758336 | 22.2441055 | -0.038658923 | 0.46092132 |
| 20.9207274 | 20.8271086 | 20.8850338 | 20.881793 | -0.033970155 | 0.33115839 |
| 21.2071675 | 21.1221021 | 21.1766887 | 21.2350296 | 0.023427615 | 0.675668532 |
| 18.9543066 | 18.8416432 | 18.8967287 | 19.0356702 | -0.003385893 | 0.95180769 |
| 21.6927446 | 22.1067706 | 21.8509701 | 21.6136248 | 0.02995382 | 0.772576919 |
| 21.8739327 | 21.7300051 | 21.7343967 | 21.600944 | -0.124275405 | 0.115859032 |
| 19.8681274 | 19.8618678 | 19.8608277 | 19.8951705 | 0.009537587 | 0.816028262 |
| 23.8824978 | 23.8871309 | 23.9582401 | 23.909561 | 0.045166694 | 0.321434899 |
| 20.3942447 | 20.4643427 | 20.5360679 | 20.3735948 | -0.006067064 | 0.912788615 |
| 16.3373359 | 16.3698979 | 16.5764672 | 16.5295632 | 0.140260208 | 0.061194202 |
| 14.2773451 | NA | 14.0833032 | 14.4100975 | -0.062345903 | 0.751444478 |
| 18.6179215 | 18.5096733 | 18.1543565 | 18.2771231 | -0.239135836 | 0.060905891 |
| 19.4432047 | 19.2330645 | 19.3201975 | 19.3066391 | -0.093817592 | 0.07707962 |
| 20.6291799 | 20.5433642 | 20.5721225 | 20.6157238 | -4.79E-04 | 0.992038361 |
| 20.7840814 | 20.6597118 | 20.8084243 | 20.8348636 | 0.03548522 | 0.600587803 |
| 14.4993108 | 14.5591854 | 15.0980251 | 14.6310225 | -0.213515487 | 0.334386244 |
| 18.7727964 | 18.5341082 | 18.5027542 | 18.5470775 | -0.17304039 | 0.010908641 |
| 21.6663605 | 21.591756 | 21.6772 | 21.6331781 | -0.012174263 | 0.787375977 |
| 19.309231 | 19.1845519 | 19.3188893 | 19.295001 | 0.010246979 | 0.840511412 |
| 18.8897248 | 18.9115874 | 18.8505654 | 18.9116715 | 0.042539986 | 0.457101167 |
| 18.4007621 | 18.3246764 | 18.457543 | 18.4494032 | 0.028370089 | 0.671147194 |
| 18.3747631 | 18.1087752 | 18.4826919 | 18.4920156 | -0.023384364 | 0.812355535 |
| 25.3357174 | 25.2166068 | 25.392203 | 25.3674669 | 0.013807369 | 0.906277497 |
| 17.1671093 | 17.0134887 | 17.1359846 | 17.2983376 | -0.019699942 | 0.887115223 |
| 24.2461442 | 24.0947587 | 24.229462 | 24.1872365 | -0.027389948 | 0.686225342 |
| 19.9613074 | 19.9243341 | 20.0171177 | 19.9964232 | 0.042368816 | 0.386057059 |
| 20.3198321 | 20.2606644 | 20.2894097 | 20.2587993 | -0.001364486 | 0.980775992 |
| 20.4812798 | 20.4617053 | 20.5051707 | 20.4852811 | -0.008260189 | 0.85150416 |
| 19.5695079 | 19.8515454 | 19.8481592 | 19.8147977 | 0.228623909 | 0.094093132 |
| 18.0328391 | 17.8804775 | 17.7597472 | 17.8238499 | -0.181589029 | 0.043310362 |
| 20.10009 | 20.1754555 | 20.0318086 | 19.7815555 | -0.036274328 | 0.670315319 |
| 18.2136578 | 18.3682544 | 18.1309125 | 18.3189664 | 0.022608237 | 0.766371814 |
| 18.7879513 | 18.7538657 | 18.6530339 | 18.7627218 | -0.070178976 | 0.198150519 |
| 23.2303957 | 23.086211 | 23.1394761 | 23.1314303 | -0.038829123 | 0.371383992 |
| 22.0573822 | 21.9798899 | 21.9863076 | 22.0130583 | 0.013166653 | 0.822571901 |
| 25.6270461 | 25.8162129 | 25.6602512 | 25.5699534 | 0.062627882 | 0.321882124 |
| 20.2987462 | 20.1592085 | 20.2971471 | 20.2075596 | -0.008522681 | 0.919666306 |
| 17.1852899 | 16.9106228 | 17.4415708 | 17.2489889 | 0.061321816 | 0.753555777 |
| 16.3793855 | 16.0099071 | 16.3398535 | 16.1735551 | -0.314751623 | 0.054505808 |
| 17.1419827 | 17.3999231 | 16.8442333 | 17.4118161 | -0.149460839 | 0.347088479 |

| | | | | | |
|---|---|---|---|---|---|
| 16.3989608 | NA | 16.2961223 | 16.4096459 | -0.131304783 | 0.164576666 |
| 18.6602897 | 18.4237192 | 18.3989329 | 18.6032056 | -0.107985515 | 0.224729973 |
| 19.3871021 | 19.0761139 | 19.1081277 | 19.2625663 | -0.066237743 | 0.427092324 |
| 18.8797504 | 18.8041087 | 18.835718 | 18.9025492 | 0.003688377 | 0.963426078 |
| 20.6816642 | 20.4875897 | 20.548608 | 20.6996713 | -0.075160145 | 0.315814485 |
| 18.9535133 | 18.6043688 | 18.7858589 | 18.7549397 | -0.029347814 | 0.66995793 |
| 26.4623122 | 26.2515989 | 26.3846091 | 26.3564152 | -0.057167568 | 0.382539031 |
| 21.0448298 | 21.0817932 | 21.0708502 | 21.0185271 | 0.034787203 | 0.488552559 |
| 19.6788064 | 19.6470539 | 19.6683026 | 19.7125487 | 0.033744586 | 0.474236773 |
| 23.4891884 | 23.3463608 | 23.4538047 | 23.4526022 | -0.03056107 | 0.495653671 |
| 22.1192971 | 22.0592032 | 22.0524404 | 22.0935351 | -0.008313643 | 0.847434212 |
| 20.1457398 | 20.1564157 | 20.1733992 | 20.0863398 | 0.03247099 | 0.459598502 |
| 19.5020591 | 19.5106587 | 19.5723369 | 19.5328635 | 0.02403106 | 0.595577257 |
| 19.4465133 | 19.3740386 | 19.3982453 | 19.4507644 | -0.036219242 | 0.530722124 |
| 22.0752386 | 22.0157038 | 22.0892792 | 22.0201093 | 0.047011866 | 0.434918251 |
| 19.0978896 | 19.2417326 | 19.2950709 | 19.1696393 | 0.07353751 | 0.256732603 |
| 19.7585028 | 19.7872861 | 19.8467719 | 19.9553078 | 0.087511709 | 0.365872637 |
| NA | 14.0703216 | NA | NA | -0.113960263 | 0.260996487 |
| 21.0931525 | 20.977373 | 21.0148458 | 20.9451186 | -0.059686688 | 0.567646036 |
| 16.1090061 | 16.2717501 | 16.1161035 | 16.4149638 | 0.063990673 | 0.556829868 |
| NA | 17.613539 | NA | NA | -0.041401328 | 0.725631996 |
| 20.6584076 | 20.8269908 | 20.6459806 | 20.646283 | 0.036927661 | 0.50644353 |
| 21.0367863 | 20.9640082 | 21.0334466 | 20.9989199 | 0.039824785 | 0.516624796 |
| 19.7138586 | 19.7571006 | 19.7799252 | 19.7996016 | 0.036296276 | 0.5319367 |
| 18.606363 | 18.4414202 | 18.6291258 | 18.4801619 | -0.014050626 | 0.86241292 |
| 15.4944321 | 15.3224755 | 15.4353075 | 15.2665708 | -0.19771984 | 0.181737326 |
| 20.7884337 | 20.797683 | 20.8002768 | 20.6720605 | -0.035114484 | 0.566439562 |
| 22.4729647 | 22.411483 | 22.4756592 | 22.4536611 | -0.001743269 | 0.969229009 |
| 23.2760375 | 23.2078797 | 23.2812589 | 23.2957486 | -0.011734569 | 0.8406299 |
| 17.9898755 | 18.0844707 | 18.1572377 | 18.0981125 | 0.076332864 | 0.226007904 |
| 15.8037613 | 15.9498705 | 16.2903824 | 16.4139554 | 0.048811933 | 0.762557677 |
| 20.6847826 | 20.6913667 | 20.7036741 | 20.830327 | 0.027487026 | 0.606691236 |
| 20.5192603 | 20.4616894 | 20.5982953 | 20.5601873 | 0.004613336 | 0.920202705 |
| 20.5344664 | 20.4409459 | 20.5406709 | 20.457541 | -0.050752185 | 0.413186123 |
| 15.6275378 | 15.658364 | 15.5055725 | 15.6533728 | 0.088564218 | 0.511565716 |
| 16.5586875 | 16.4743709 | 16.3917258 | 16.48454 | -0.152493717 | 0.175243763 |
| 23.2970955 | 23.1587273 | 23.2287276 | 23.2157481 | -0.020666391 | 0.737091947 |
| 17.2408137 | 17.4103961 | 17.3513519 | 17.4949058 | 0.11323977 | 0.19885124 |
| 19.7344737 | 19.558112 | 19.6234979 | 19.6715313 | -0.032786516 | 0.633061889 |
| 19.8597893 | 19.7588232 | 19.8158459 | 19.8955793 | -0.080993785 | 0.176255612 |
| 19.2542653 | 19.1334635 | 19.2721175 | 19.2637792 | 0.017818425 | 0.754070314 |
| 18.7968939 | 18.7380343 | 18.715535 | 18.7754463 | -0.070940348 | 0.209907475 |

| | | | | | |
|---|---|---|---|---|---|
| 19.9043918 | 19.8040235 | 19.9246749 | 19.9103718 | 0.007481357 | 0.869091483 |
| 19.4499921 | 19.2493111 | 19.3269338 | 19.3682911 | -0.088872274 | 0.103949725 |
| 19.0144062 | 18.8886239 | 18.8040575 | 18.9733434 | -0.040152203 | 0.52820947 |
| 19.3225145 | 19.2870939 | 19.3613102 | 19.1336164 | 0.095818385 | 0.287441243 |
| 17.039627 | 17.1579656 | 17.3940715 | 17.0810637 | 0.124084571 | 0.39771007 |
| 18.3398719 | 18.4399013 | 18.2581016 | 18.4413231 | 9.09E-04 | 0.992487785 |
| 26.5716394 | 26.3528069 | 26.4754582 | 26.4633977 | -0.016408715 | 0.806143064 |
| 21.426222 | 21.380311 | 21.4363657 | 21.4319121 | -0.006991756 | 0.894701926 |
| 22.8189793 | 22.684 | 22.7171143 | 22.7369236 | -0.070864617 | 0.178649501 |
| 20.5737269 | 20.2813125 | 20.4477776 | 20.4537507 | -0.056463387 | 0.528429879 |
| 21.9212099 | 21.8771187 | 21.9819674 | 21.9736114 | 0.033098832 | 0.499119367 |
| NA | 24.0370378 | NA | NA | 0.656639705 | 0.077710649 |
| 13.3299208 | NA | 13.4845589 | 13.6138782 | 0.129609007 | 0.299637377 |
| 19.1346349 | 18.9090248 | 18.9567017 | 19.0312508 | -0.068931914 | 0.391562213 |
| 19.9936166 | 19.8100688 | 19.9741474 | 19.9626845 | -0.034799311 | 0.561462611 |
| 21.7497422 | 21.6950104 | 21.8398739 | 21.7804271 | 0.058578393 | 0.402746624 |
| 21.2770894 | 21.1678649 | 21.2505997 | 21.2264159 | -0.050463382 | 0.344025774 |
| 17.9495568 | 17.9494811 | 17.8865372 | 17.9884525 | -0.058480687 | 0.473600492 |
| 22.292258 | 22.0477414 | 22.3316201 | 22.4129031 | 0.00699617 | 0.949927562 |
| 17.1650006 | NA | 17.0070151 | 16.8111519 | -0.216262968 | 0.474798405 |
| 15.5280809 | 15.2960891 | 15.6077627 | 15.178847 | -0.097981538 | 0.450263538 |
| 25.5312395 | 25.4629421 | 25.4844052 | 25.5318219 | 0.064597097 | 0.565138649 |
| 18.5550274 | 18.4013991 | 18.4955188 | 18.4794164 | -0.027596732 | 0.737979185 |
| 22.8393 | 22.7693964 | 22.8506853 | 22.8799803 | 0.047223146 | 0.546774317 |
| 20.9279659 | 20.9158477 | 20.9163412 | 20.9697906 | -0.002544706 | 0.961870682 |
| 22.2740069 | 22.0543848 | 22.1878349 | 22.2167403 | 0.004714426 | 0.935921002 |
| 20.8597723 | 20.6096275 | 20.7913033 | 20.7344243 | -0.009921304 | 0.907664665 |
| 25.8123329 | 25.7076975 | 25.8742464 | 25.8055229 | 0.021979195 | 0.859116654 |
| 16.3188415 | 16.2082637 | 16.4676299 | 16.3823007 | -0.058840421 | 0.695538622 |
| 21.1368834 | 20.8908227 | 21.1446437 | 21.1656426 | -0.034510615 | 0.760591282 |
| 24.2697317 | 23.99436 | 24.1987989 | 24.1956623 | -0.032884397 | 0.705450402 |
| 20.1310039 | 19.9542937 | 19.9679814 | 20.0009967 | -0.088198159 | 0.156426754 |
| NA | 17.8551888 | NA | NA | -0.119270743 | 0.251293557 |
| 21.246511 | 21.0125072 | 21.1578531 | 21.0150142 | -0.107980507 | 0.255085722 |
| 21.9810741 | 21.8812324 | 21.9185013 | 21.9576584 | -0.035648474 | 0.502614967 |
| 22.2297409 | 22.1914665 | 22.186369 | 22.182663 | -0.042670447 | 0.352625672 |
| 22.3114309 | 22.2144359 | 22.3169454 | 22.2579654 | 0.001787823 | 0.963653966 |
| 21.3487419 | 21.3308525 | 21.3761018 | 21.3351943 | 0.07623987 | 0.173552122 |
| 20.3703454 | 20.1653927 | 20.3387593 | 20.344788 | -0.029294699 | 0.693498456 |
| 21.9082203 | 22.2399615 | 21.9972645 | 21.9375443 | -0.033958518 | 0.782286833 |
| 20.822721 | 20.7485892 | 20.9439383 | 20.9156895 | -0.014840957 | 0.851489427 |
| 15.9821666 | NA | 15.4081545 | 16.0110356 | -0.328112746 | 0.090332352 |

| | | | | | |
|---|---|---|---|---|---|
| 21.0037505 | 21.2361914 | 21.1780154 | 20.9054541 | 0.045094361 | 0.730048877 |
| 23.4300003 | 23.3370198 | 23.4681579 | 23.5162932 | 0.048369965 | 0.28097105 |
| 20.1481456 | 20.1293961 | 20.193833 | 20.235546 | 0.070187417 | 0.183726072 |
| 20.55515 | 20.5059046 | 20.4592329 | 20.5819411 | 0.010874036 | 0.885494031 |
| 15.4654623 | 15.6469321 | 15.3474836 | 15.3052272 | 0.236262765 | 0.247249176 |
| 20.6141087 | 20.6120654 | 20.626472 | 20.5513052 | -0.018636002 | 0.655234011 |
| 15.1999098 | 15.4238084 | 15.0802124 | 15.2782625 | -0.053689253 | 0.773769947 |
| 18.0275851 | 17.8255382 | 18.1476775 | 17.8098045 | 0.126626298 | 0.575762489 |
| 18.5046909 | 18.1694688 | 18.6481101 | 18.4370475 | -0.11552824 | 0.302558101 |
| 23.4402718 | 23.2845824 | 23.4952281 | 23.4377666 | -0.009117604 | 0.888479396 |
| 24.373941 | 24.3860296 | 24.4274401 | 24.3469203 | 0.091829995 | 0.162526566 |
| 22.1290185 | 21.9753482 | 21.9961357 | 21.9690573 | -0.071903783 | 0.200393556 |
| 25.0554408 | 24.9579834 | 25.0136272 | 24.9631214 | -0.015304464 | 0.781131576 |
| 26.5750104 | 26.451149 | 26.5441161 | 26.5809275 | -0.005355076 | 0.919678084 |
| 23.8516684 | 23.7929804 | 23.8314215 | 23.8126872 | 0.026065604 | 0.547100214 |
| 19.2667958 | 19.2764029 | 19.3450189 | 19.3196693 | 0.035245526 | 0.462533154 |
| 23.6040526 | 23.5359477 | 23.5802345 | 23.6415426 | 0.030191907 | 0.585121344 |
| 19.5942453 | 19.5998974 | 19.6463453 | 19.6676751 | 0.112054042 | 0.076614411 |
| 20.1413228 | 19.9328079 | 20.0921583 | 20.1168609 | -0.046221177 | 0.482965279 |
| 20.6129339 | 20.5366951 | 20.6527821 | 20.6467388 | 0.027738936 | 0.697950938 |
| 20.2760536 | 20.1720194 | 20.2172128 | 20.2878001 | 0.080716674 | 0.304027458 |
| 21.8504187 | 21.7700469 | 21.7518174 | 21.7005415 | -0.04755699 | 0.587574518 |
| 16.8058726 | 16.9873555 | 16.564777 | 16.6284517 | -0.171728703 | 0.160387491 |
| 18.4869014 | 18.393117 | 18.3853546 | 18.4273396 | -0.052897844 | 0.391173656 |
| 18.8037754 | 18.5674517 | 18.656673 | 18.5976089 | -0.249067503 | 0.074266574 |
| 22.4902421 | 22.3779884 | 22.4865568 | 22.4865036 | -0.013377003 | 0.83595646 |
| 18.75565 | 18.6673961 | 18.7785388 | 18.6792183 | 0.01660281 | 0.787417506 |
| 20.97158 | 20.8620856 | 20.8720626 | 20.9540829 | 0.036839994 | 0.519918336 |
| 20.5004942 | 20.1092735 | 20.4956127 | 20.0755106 | -0.011198558 | 0.955182194 |
| 19.2281256 | 19.2521802 | 19.391715 | 19.3424231 | 0.113412286 | 0.203097934 |
| 22.5769763 | 22.443542 | 22.7231162 | 22.6461326 | -0.001334649 | 0.989583439 |
| 16.7577519 | 16.8471054 | 16.7439557 | 17.1571776 | 0.086246772 | 0.600911759 |
| 18.5984322 | 18.2373945 | 18.5366553 | 18.8988189 | -0.098090682 | 0.448003014 |
| 23.2094758 | 23.141459 | 23.1709588 | 23.1466914 | -0.014691452 | 0.772769869 |
| 19.8572965 | 19.8115505 | 19.6745017 | 19.8145487 | -0.100509275 | 0.256479612 |
| 23.3785047 | 23.0543137 | 23.2646767 | 23.2577148 | -0.077529935 | 0.378641226 |
| 20.7364191 | 20.7006353 | 20.7603104 | 20.7279574 | 0.038791213 | 0.43522384 |
| 18.4285374 | 18.3471186 | 18.3885965 | 18.4817946 | -0.054600751 | 0.347739339 |
| 17.6106242 | 17.5939117 | 17.5353164 | 17.6144436 | -0.022364993 | 0.722634938 |
| 22.4300039 | 22.368664 | 22.4352685 | 22.467673 | 0.026414292 | 0.569946803 |
| 18.4948748 | 18.4867303 | 18.5822395 | 18.3435927 | 0.099531915 | 0.516652763 |
| 19.6939601 | 19.5777099 | 19.7439771 | 19.6910675 | -0.026799157 | 0.604777785 |

| | | | | | |
|---|---|---|---|---|---|
| 20.6984798 | 20.5066844 | 20.6903491 | 20.6221893 | -0.038230219 | 0.563767414 |
| 17.9938769 | 18.0233682 | 18.0835074 | 18.0207985 | 0.07612846 | 0.147406912 |
| 19.461652 | 19.4328395 | 19.510912 | 19.5204154 | -0.009970175 | 0.890752545 |
| 17.8378646 | 17.8185469 | 17.8427127 | 17.9782005 | 0.043608552 | 0.595216187 |
| 21.0657318 | 20.8286198 | 20.9531164 | 21.1210406 | -0.138422136 | 0.403928237 |
| 16.6019371 | 16.1530079 | 16.4681182 | 16.7341532 | -0.14253642 | 0.357318021 |
| 18.6135795 | 18.5147575 | 18.5639463 | 18.6445328 | -0.042970916 | 0.552670862 |
| 21.3460576 | 21.0922514 | 21.1556319 | 21.1025818 | -0.106086727 | 0.145783836 |
| 23.4876444 | 23.4865799 | 23.5026075 | 23.4317537 | 0.008077609 | 0.870492873 |
| 21.1063321 | 20.9718458 | 21.0694857 | 21.1081024 | -0.013706719 | 0.756158803 |
| 21.2285713 | 21.1229618 | 21.1852777 | 21.2168121 | -0.005289802 | 0.937391211 |
| 19.1712646 | 18.920184 | 19.1711014 | 19.2271601 | -0.037686849 | 0.616420155 |
| 13.9536596 | 13.6315023 | 13.771361 | 13.6556548 | -0.13619817 | 0.191568843 |
| 18.5113533 | 18.7655398 | 18.6307786 | 18.2526456 | 0.028580844 | 0.798397208 |
| 20.1449787 | 20.2005987 | 20.1776759 | 20.209644 | 0.016017687 | 0.826956831 |
| 14.7861031 | 14.670222 | 14.8936296 | 14.9697035 | 0.18638067 | 0.532701387 |
| 22.0316381 | 21.9936506 | 22.0336627 | 22.0348479 | -0.027851054 | 0.538624699 |
| NA | 16.5947231 | NA | NA | 0.212185063 | 0.227253653 |
| 20.6787012 | 20.6544118 | 20.6752451 | 20.7347546 | 0.008963023 | 0.85395211 |
| 20.2850317 | 20.2117475 | 20.2716186 | 20.2359219 | -0.013088238 | 0.756189204 |
| 24.8817883 | 24.8885999 | 24.8864804 | 24.9262879 | 0.061285559 | 0.275581051 |
| 19.6021058 | 19.3368886 | 19.537942 | 19.5481117 | -0.066507312 | 0.519720731 |
| 16.7762766 | 17.0054173 | 16.8222561 | 17.0696077 | 0.042174651 | 0.689060299 |
| 21.6542013 | 21.6203153 | 21.6220309 | 21.6680787 | 0.045462976 | 0.706878604 |
| 22.9250481 | 22.8982676 | 22.9723694 | 22.9210158 | 0.032430417 | 0.421833629 |
| 22.625754 | 22.585485 | 22.6613588 | 22.680066 | 0.003426703 | 0.933043922 |
| 16.0229755 | 16.0690118 | 15.8234217 | 15.9156148 | 0.171910211 | 0.377629407 |
| 15.2754051 | 15.2156059 | 15.0049054 | 15.6234778 | -0.127020241 | 0.488512026 |
| NA | 12.5977313 | NA | NA | -0.153363503 | 0.339269929 |
| 16.8704788 | 16.6867845 | 16.5751773 | 16.6975941 | -0.125173961 | 0.426557475 |
| NA | 15.7761036 | NA | NA | -0.093222556 | 0.462581961 |
| 21.3529155 | 21.3147253 | 21.3158566 | 21.2953335 | 0.005129975 | 0.964825245 |
| 19.9999708 | 20.0014153 | 19.9542266 | 19.9153252 | -0.071603504 | 0.533756068 |
| 18.2793303 | 18.1696586 | 18.616965 | 18.5064592 | 0.207687362 | 0.057922738 |
| 19.9549114 | 19.7656948 | 19.7797275 | 19.82995 | -0.107764612 | 0.218746043 |
| 15.7810684 | 15.6757985 | 15.8110869 | 15.8907231 | -0.039258807 | 0.648542041 |
| 20.0419223 | 19.968052 | 19.913456 | 20.117452 | -0.001982547 | 0.98125672 |
| 21.5746795 | 21.7218357 | 21.7747288 | 21.6335984 | 0.069613185 | 0.399579719 |
| 17.9787742 | 17.980412 | 17.9765475 | 17.9804796 | -0.064153869 | 0.404467199 |
| 21.7180446 | 21.5437431 | 21.5826419 | 21.6244734 | -0.074487192 | 0.315148095 |
| 19.2333702 | 19.2987679 | 19.0778874 | 19.2444666 | 0.014060649 | 0.840248404 |
| 24.9482476 | 24.949473 | 25.0363609 | 25.0540337 | 0.099931088 | 0.157696683 |

| | | | | | |
|---|---|---|---|---|---|
| 21.7695623 | 21.7407976 | 21.9255923 | 21.8903805 | 0.050713718 | 0.394987732 |
| 21.3305376 | 21.1994689 | 21.2902649 | 21.2577645 | -0.057129268 | 0.329018496 |
| 21.8329262 | 21.7992648 | 21.7939385 | 21.8048242 | 0.056942536 | 0.31829507 |
| 19.8033608 | 19.741339 | 19.820536 | 19.7456144 | -0.035533971 | 0.509540862 |
| 22.0149317 | 21.8576056 | 21.9363675 | 21.9856245 | -0.058746499 | 0.264399289 |
| 20.9561649 | 20.8817288 | 20.975076 | 20.9689335 | 0.054701923 | 0.298183298 |
| 21.3617505 | 21.2918684 | 21.3249362 | 21.4842267 | 0.020219336 | 0.691740299 |
| 22.3046732 | 22.1369233 | 22.2336619 | 22.1843993 | -0.027849494 | 0.550015601 |
| 23.7430782 | 23.6200583 | 23.6999805 | 23.6650692 | -0.015226752 | 0.739917064 |
| 22.1683101 | 22.0616278 | 22.0813735 | 22.1083553 | -0.081349625 | 0.05233346 |
| 21.4131176 | 21.343394 | 21.3927493 | 21.4022152 | 0.030157505 | 0.447174915 |
| 24.3228308 | 24.3069093 | 24.38447 | 24.3938596 | 0.046794621 | 0.240509773 |
| 23.9509442 | 23.9292998 | 23.9778516 | 23.9241916 | 0.044194593 | 0.293423815 |
| 21.1097692 | 20.9939987 | 21.1031654 | 21.1208292 | 0.006606671 | 0.889687451 |
| 20.9976047 | 20.9010946 | 21.0370531 | 21.0037222 | 0.025956256 | 0.616763897 |
| 21.2726548 | 21.2239127 | 21.2641152 | 21.2492969 | 0.026164554 | 0.635213363 |
| 22.5393985 | 22.4544195 | 22.527718 | 22.5763019 | 0.021830256 | 0.693370601 |
| 22.2875798 | 22.2640442 | 22.2528569 | 22.3262017 | 0.038042262 | 0.506352881 |
| 22.9572644 | 22.8323813 | 22.9357022 | 22.9888268 | 0.006939352 | 0.903448427 |
| 21.0810796 | 20.9973973 | 21.1604655 | 21.1158939 | 0.034246182 | 0.551148834 |
| 19.6468083 | 19.5081942 | 19.6943281 | 19.7449065 | 0.059479601 | 0.310398726 |
| 19.7636505 | 19.6666492 | 19.8723657 | 19.8677425 | 0.076316685 | 0.2279649 |
| 20.8807988 | 20.7729235 | 20.8257588 | 20.8218621 | -0.013672135 | 0.836899471 |
| 18.883308 | 18.7803186 | 18.792722 | 18.7982343 | -0.007363532 | 0.916039171 |
| 21.0601841 | 21.0170202 | 21.112602 | 21.1581183 | 0.006868458 | 0.926266557 |
| 23.672412 | 23.668816 | 23.7437746 | 23.661576 | 0.026728594 | 0.721381414 |
| 22.573117 | 22.5550301 | 22.5551534 | 22.7032275 | 0.120365457 | 0.156577437 |
| 19.4877225 | 19.4723398 | 19.561856 | 19.6399417 | 0.014529984 | 0.868106086 |
| 20.0747275 | 19.8884067 | 20.0560332 | 19.9679825 | -0.011408256 | 0.906387583 |
| 15.9483988 | 16.0998466 | 16.0457325 | 16.1256206 | 0.179998475 | 0.137028911 |
| 19.1109449 | 18.8962193 | 19.1902532 | 19.2402048 | 0.02784863 | 0.814757359 |
| 25.2606163 | 25.105666 | 25.2643226 | 25.1727331 | -0.024631601 | 0.837278051 |
| 18.3317294 | NA | 18.3921935 | 18.5929281 | 0.258861369 | 0.056664725 |
| NA | 13.5947888 | NA | NA | 0.28413226 | 0.229293633 |
| 16.2319147 | 16.030558 | 16.2068966 | 16.0534359 | 0.066016327 | 0.583441592 |
| 18.4281247 | 18.4088958 | 18.4594629 | 18.3501825 | 0.062690531 | 0.533463549 |
| 20.1397519 | 20.1360901 | 20.0695421 | 20.1161955 | -0.025346698 | 0.519938501 |
| 20.3958719 | 20.3379276 | 20.4056404 | 20.4301092 | 0.06035491 | 0.202049371 |
| 19.9755515 | 19.8401008 | 19.9588737 | 19.9945999 | -0.019796811 | 0.692664744 |
| 22.3330659 | 22.2746119 | 22.3742953 | 22.3874966 | 0.055831409 | 0.283091363 |
| 18.5844721 | 18.5420578 | 18.634103 | 18.6165625 | 0.012653915 | 0.822433963 |
| 20.0918943 | 20.2932572 | 20.0721914 | 20.217776 | 0.036842702 | 0.663195453 |

| | | | | | |
|---|---|---|---|---|---|
| NA | 12.9796901 | NA | NA | -0.211900129 | 0.459442994 |
| NA | 11.2693453 | NA | NA | -0.155272088 | 0.777804338 |
| 18.5512582 | 18.4846463 | 18.5351519 | 18.4783609 | 0.03951802 | 0.741740657 |
| 19.6880304 | 19.6582392 | 19.5116316 | 19.651633 | -0.013656657 | 0.864598214 |
| 18.8059027 | 18.62596 | 18.5327985 | 18.6477023 | -0.13200018 | 0.088896309 |
| 20.3985778 | 20.2089001 | 20.3757528 | 20.4124257 | -0.010649099 | 0.877072012 |
| 20.0000427 | 20.0860893 | 19.9898681 | 20.0632154 | 0.031701694 | 0.637678468 |
| 19.1456256 | 19.1181126 | 19.0621282 | 19.2241732 | -0.052656944 | 0.395498287 |
| 23.7507899 | 23.5936113 | 23.6713328 | 23.6909543 | -0.033804545 | 0.532437425 |
| 22.8202365 | 22.7773818 | 22.836557 | 22.8785234 | 0.045656196 | 0.359369508 |
| 24.4198659 | 24.2994751 | 24.3802341 | 24.3359727 | -0.030733163 | 0.515556955 |
| 20.9247935 | 21.021588 | 20.9392921 | 20.9237016 | 0.023563822 | 0.639063797 |
| 22.4622103 | 22.4435886 | 22.4754612 | 22.492133 | 0.062023357 | 0.22990254 |
| 20.9885255 | 20.9842973 | 21.0116901 | 20.9696613 | 0.059888596 | 0.26649748 |
| 18.8702961 | 18.7496207 | 18.7917609 | 18.8445569 | -0.042907327 | 0.46134669 |
| 17.6116416 | 17.6598334 | 17.7548921 | 17.7044931 | 0.019549719 | 0.787889534 |
| 21.5304355 | 21.5182845 | 21.4538473 | 21.570741 | 0.040962233 | 0.575840872 |
| 18.6757292 | 18.7554955 | 18.7225649 | 18.7166795 | 0.032490484 | 0.661529335 |
| 18.6430439 | 18.4208292 | 18.7101652 | 18.6067189 | -0.131632983 | 0.151059275 |
| 19.9018573 | 19.8170034 | 20.077021 | 20.0046276 | 1.53E-04 | 0.998771851 |
| 14.8141777 | 14.7324152 | 14.6297366 | 14.7765971 | 0.007234844 | 0.956874121 |
| 22.8799969 | 22.7792536 | 22.7761459 | 22.7443422 | -0.058092205 | 0.40681129 |
| 16.6328271 | 16.8564591 | 16.5866194 | 16.278214 | -0.160479284 | 0.227101595 |
| 21.7096061 | 21.5893918 | 21.6182558 | 21.6501808 | -0.006952756 | 0.903407314 |
| 22.0389035 | 21.9463631 | 22.0600197 | 22.0350778 | -0.004055797 | 0.931379234 |
| 21.2079109 | 21.094834 | 21.2351432 | 21.2057524 | 0.003296805 | 0.945407193 |
| 20.3848183 | 20.4085056 | 20.3328301 | 20.587113 | 0.026397372 | 0.710954697 |
| 22.6989543 | 22.6209702 | 22.7500511 | 22.7041965 | 0.01411153 | 0.846311669 |
| NA | 14.6489316 | NA | NA | 0.12607885 | 0.740268746 |
| 21.7164253 | 21.8453425 | 21.6878583 | 21.650719 | -0.018173788 | 0.71823713 |
| 17.7328057 | 17.5056328 | 17.1161518 | 18.0327839 | 0.059248354 | 0.775147985 |
| 17.2136973 | 17.0522514 | 17.3059322 | 17.2460486 | -0.10150446 | 0.208733488 |
| 12.5283483 | NA | 13.5853851 | 12.6563462 | 0.17094546 | 0.609725253 |
| 17.3157589 | 17.3551249 | 17.262999 | 17.3548064 | 0.213088497 | 0.480301376 |
| 16.3863542 | 16.5167388 | 16.8009627 | 16.3527697 | -0.110757026 | 0.506196791 |
| 18.0601871 | 17.8394613 | 18.0301226 | 17.9934532 | -0.155018321 | 0.118019979 |
| 18.5423223 | 18.2591898 | 18.4908782 | 18.5490634 | -0.151652328 | 0.051424384 |
| 19.1435358 | 19.0114875 | 19.0025939 | 19.128917 | -0.026283653 | 0.661471799 |
| 20.4232992 | 20.2080052 | 20.3654217 | 20.3477842 | -0.068469143 | 0.253297933 |
| 21.516745 | 21.3957857 | 21.5442841 | 21.589409 | -0.018289984 | 0.734403285 |
| 22.4562519 | 22.3260283 | 22.4711395 | 22.44647 | -0.029802703 | 0.554891194 |
| 20.1169368 | 20.036943 | 20.046331 | 20.0997796 | -0.010166603 | 0.817143195 |

| | | | | | |
|---|---|---|---|---|---|
| 20.9990664 | 20.9463825 | 20.9711397 | 21.0321733 | -0.007430554 | 0.886310984 |
| 18.6498347 | 18.5819402 | 18.6408277 | 18.5662971 | -0.005479639 | 0.921760072 |
| 18.7267301 | 18.6606509 | 18.7794171 | 18.8320962 | 0.035417126 | 0.589231847 |
| 23.7779444 | 23.7571981 | 23.7361025 | 23.8127807 | 0.066158509 | 0.378776474 |
| 17.2770748 | 17.13029 | 17.330461 | 17.2634204 | 0.047679375 | 0.606302339 |
| 18.4265543 | 18.4515173 | 18.5052109 | 18.390865 | 0.025335813 | 0.836913507 |
| 12.880006 | NA | 12.9164506 | 13.2049059 | 0.146749178 | 0.541891036 |
| 16.5224621 | 16.5151562 | 16.7800979 | 16.5639995 | -0.012181127 | 0.932968247 |
| 23.5364153 | 23.4689508 | 23.4840813 | 23.523537 | 0.012628597 | 0.731640965 |
| 14.3289521 | 14.439756 | 14.5683387 | 14.1543833 | -0.071802384 | 0.701578021 |
| 13.2540551 | 13.2045424 | 12.8304311 | 12.8421124 | -0.182269993 | 0.317647368 |
| 18.5234424 | 18.5368476 | 18.2509529 | 18.5250627 | 0.084064025 | 0.598123319 |
| NA | 16.8584909 | NA | NA | -0.068249786 | 0.609971725 |
| 17.2057876 | 16.8776566 | 16.9439621 | 17.0086254 | -0.240974883 | 0.072326912 |
| 17.0578672 | 16.8086262 | 16.9264039 | 16.8136019 | -0.171796652 | 0.046284384 |
| 18.7587807 | 18.611832 | 18.5999961 | 18.6298978 | -0.098467268 | 0.234397744 |
| 22.8878621 | 22.6849792 | 22.8684287 | 22.8708606 | -0.004100505 | 0.946800516 |
| 21.9750513 | 21.8406834 | 21.9913335 | 21.9077335 | -0.001500379 | 0.979583356 |
| 22.9036652 | 22.6877073 | 22.8510775 | 22.8632046 | -0.077079919 | 0.18056463 |
| 23.2042785 | 23.1656804 | 23.2248534 | 23.1620685 | -0.026422271 | 0.632672474 |
| 26.2987648 | 26.1468943 | 26.2160183 | 26.3059011 | 0.02842455 | 0.600009421 |
| 28.1819523 | 28.0753077 | 28.1935134 | 28.1552091 | 0.028016739 | 0.603491289 |
| 19.5817161 | 19.615376 | 19.685116 | 19.6596095 | 0.055597203 | 0.304632373 |
| 19.9635643 | 19.9414245 | 19.9343844 | 19.8782513 | 0.004076078 | 0.933955135 |
| 24.1370405 | 24.0509974 | 24.0404571 | 24.0880531 | -0.002602796 | 0.955720172 |
| 21.3530859 | 21.2922222 | 21.3655245 | 21.3252734 | -0.005660798 | 0.899431909 |
| 21.9999411 | 21.9943886 | 22.0045577 | 22.0059359 | 0.022572803 | 0.500836093 |
| 23.4160455 | 23.3858597 | 23.4104157 | 23.4440303 | 0.026574891 | 0.540361345 |
| 24.2051601 | 24.2135996 | 24.2897793 | 24.2197456 | 0.040039407 | 0.389102497 |
| 20.5031001 | 20.4717726 | 20.5469013 | 20.5919014 | 0.06090243 | 0.219103228 |
| 24.8515702 | 24.7417953 | 24.8376784 | 24.808799 | -0.008698278 | 0.862092908 |
| 20.4201679 | 20.2227423 | 20.3966309 | 20.3392558 | -0.031461183 | 0.585429689 |
| 20.4538353 | 20.5455477 | 20.5764433 | 20.6284672 | 0.086123556 | 0.163895194 |
| 19.1387335 | 19.0739886 | 19.1235986 | 19.1510749 | 0.044143807 | 0.50718618 |
| 22.7169305 | 22.6887415 | 22.8912203 | 22.6655555 | 0.128650626 | 0.061190094 |
| 19.4313873 | 19.3804469 | 19.5447688 | 19.2932059 | -0.051870318 | 0.455698092 |
| 22.202271 | 22.0443035 | 22.2709882 | 22.3391577 | 0.102624326 | 0.152673285 |
| 18.3298415 | 18.1509997 | 18.3719086 | 17.9601451 | -0.176900369 | 0.029203773 |
| 24.4447899 | 24.4348078 | 24.424484 | 24.3642046 | 0.053174879 | 0.494371045 |
| 20.1144955 | 20.0227157 | 20.2743002 | 20.2076571 | 0.199327722 | 0.019671602 |
| 20.5001477 | 20.4371427 | 20.5538335 | 20.5757551 | 0.017211448 | 0.844229988 |
| 20.080058 | 20.0422545 | 20.1163529 | 20.0159851 | 0.031461631 | 0.720746696 |

| | | | | | |
|---|---|---|---|---|---|
| 17.8777586 | 18.0893776 | 17.976119 | 18.0943266 | 0.204724742 | 0.031245706 |
| 17.0772631 | 16.9587792 | 16.9344043 | 16.946479 | 0.055378573 | 0.567277435 |
| 17.2361933 | 17.2084089 | 17.3487727 | 17.2706976 | 0.056523109 | 0.577636232 |
| 16.4143726 | 16.505668 | 16.4590456 | 16.3133804 | 0.17225461 | 0.123555088 |
| 14.078051 | 14.2040244 | 13.8390611 | 14.2401047 | -0.423370622 | 0.059578546 |
| 20.1338171 | 19.8610068 | 19.7975753 | 19.8464258 | -0.176765364 | 0.023021019 |
| 21.8909183 | 21.8306885 | 21.9111892 | 21.8840252 | 0.014994014 | 0.734871294 |
| 23.2042243 | 23.1561162 | 23.2552594 | 23.2563965 | 0.066525581 | 0.173379619 |
| 18.0208312 | 17.9363485 | 18.0788521 | 18.2329689 | 0.097937141 | 0.13875681 |
| 23.3647039 | 23.1524723 | 23.324251 | 23.2979474 | -0.068993014 | 0.279864672 |
| 20.579048 | 20.5897527 | 20.6298239 | 20.6377114 | 0.035756486 | 0.583216943 |
| 21.174303 | 20.4729609 | 20.9983997 | 21.0904487 | -0.163411144 | 0.338554456 |
| 19.3210699 | 19.1522009 | 19.3035827 | 19.2846521 | -0.074498206 | 0.284263201 |
| 23.9442073 | 23.6476548 | 23.7903164 | 23.7390672 | -0.118625158 | 0.085649959 |
| 19.6659492 | 19.6024638 | 19.6252997 | 19.6307121 | -0.022923773 | 0.694810279 |
| 21.8786241 | 21.7232083 | 21.8407704 | 21.8141188 | -0.032247909 | 0.508870762 |
| 24.1718279 | 24.0785155 | 24.1278601 | 24.1158895 | -0.023248595 | 0.625836036 |
| 23.6957488 | 23.6940787 | 23.689982 | 23.6778366 | 0.022654488 | 0.52501141 |
| 23.585915 | 23.529465 | 23.58017 | 23.536268 | 0.002149478 | 0.961808815 |
| 21.7038971 | 21.6866641 | 21.7940671 | 21.6917143 | 0.001881122 | 0.972088173 |
| 22.2003154 | 22.189297 | 22.3183566 | 22.2558519 | 0.044454864 | 0.439139939 |
| 26.0403249 | 25.9210649 | 26.0376022 | 26.03171 | 0.043522059 | 0.466166639 |
| 27.2713826 | 27.1594995 | 27.2596143 | 27.2638293 | 0.019307723 | 0.790408898 |
| 18.806468 | 18.7177926 | 18.8190463 | 18.7772134 | 0.0706664 | 0.346027767 |
| 17.7103138 | 17.5688912 | 17.6282222 | 17.6965865 | -0.016621797 | 0.829254931 |
| 18.3798956 | 18.4359851 | 18.4642386 | 18.5121648 | 0.03183539 | 0.701706351 |
| 24.9031438 | 24.8562457 | 24.8908897 | 24.8965818 | 0.034350381 | 0.689097343 |
| 19.3805576 | 19.3678924 | 19.4882925 | 19.5191246 | 0.066749824 | 0.448379036 |
| 18.3291115 | 18.5003896 | 18.4120049 | 18.495007 | 0.15881285 | 0.145818939 |
| 17.5238054 | 17.2808135 | 17.7354852 | 17.6198505 | 0.061579706 | 0.666948871 |
| NA | 20.4573163 | NA | NA | 0.30875527 | 0.391297839 |
| 19.4569468 | 19.3465828 | 19.453109 | 19.4551857 | 0.044666962 | 0.572472562 |
| 22.2746166 | 22.4704564 | 22.6867867 | 22.4305975 | 0.240371862 | 0.438148423 |
| NA | 15.4234267 | NA | NA | -0.288663812 | 0.105852838 |
| 14.9402645 | 14.7273533 | 14.7052555 | 14.7003746 | -0.301244437 | 0.059559912 |
| NA | 14.2601772 | NA | NA | 0.044773764 | 0.755142457 |
| 23.0992232 | 22.5812695 | 22.8744653 | 22.8092044 | -0.176367424 | 0.234910531 |
| 18.5650037 | 18.2895711 | 18.4148071 | 18.5086319 | -0.053483255 | 0.672963699 |
| 14.2158796 | NA | 14.1320465 | 14.1648546 | -0.092524991 | 0.455015409 |
| 17.1320805 | 16.7774308 | 17.3214599 | 17.2936668 | -0.04040657 | 0.738886581 |
| 14.8135255 | NA | 14.7183518 | 14.866992 | -0.031143942 | 0.791056511 |
| 17.2062742 | 17.2057914 | 16.9334558 | 17.0686808 | 0.002829724 | 0.97973546 |

| | | | | | |
|---|---|---|---|---|---|
| 14.9191402 | NA | 14.9383689 | 14.9364059 | -0.019160397 | 0.85083778 |
| 16.6994994 | 16.812638 | 17.1155785 | 16.9797698 | 0.251962372 | 0.019167093 |
| 22.0935705 | 21.8828632 | 21.7713718 | 21.8406888 | -0.034130222 | 0.719490429 |
| 19.2234496 | 19.0824546 | 19.2434631 | 19.2055701 | -0.067100752 | 0.463958648 |
| 20.6974061 | 20.3844008 | 20.7335036 | 20.7352721 | -0.099372746 | 0.278252245 |
| 21.6656843 | 21.759919 | 21.681995 | 21.6734911 | 0.033378754 | 0.708275351 |
| 29.3762685 | 29.34075 | 29.4195509 | 29.3988178 | 0.09010359 | 0.315207082 |
| 16.8331247 | 16.9860185 | 17.1039404 | 17.0380321 | 0.125173524 | 0.154690908 |
| 18.6884573 | 18.4644162 | 18.5631292 | 18.6081967 | -0.07944005 | 0.347067854 |
| 21.4655495 | 21.3859272 | 21.6680256 | 21.6886306 | 0.037346111 | 0.654529961 |
| 20.159539 | 20.0692516 | 20.0698837 | 19.9082719 | -0.114363412 | 0.146436632 |
| 21.2686143 | 21.0299285 | 21.1751919 | 21.216339 | -0.057260951 | 0.456239848 |
| 18.5852691 | 18.5803415 | 18.5399739 | 18.6471315 | -0.126197548 | 0.104021258 |
| 23.2620466 | 23.1441795 | 23.0790208 | 23.1759856 | -0.025981684 | 0.731447183 |
| 18.1521323 | 18.0941442 | 18.2200817 | 18.2329052 | -0.010942612 | 0.884294452 |
| 18.4646252 | 18.2337868 | 18.4632169 | 18.4941028 | -0.044339887 | 0.545645978 |
| 20.2018559 | 20.1303385 | 20.283151 | 20.1634909 | -0.007009481 | 0.922727096 |
| 21.2565764 | 21.308903 | 21.2488531 | 21.2208887 | 0.017006306 | 0.795093654 |
| 19.4090839 | 19.3365066 | 19.4002153 | 19.5365928 | 0.027340258 | 0.669911462 |
| 20.0888383 | 20.059359 | 20.0974443 | 20.0268101 | -0.036573115 | 0.53244546 |
| 21.6960646 | 21.574096 | 21.7739174 | 21.7641482 | -0.011783598 | 0.834801912 |
| 21.7295585 | 21.7586966 | 21.8032744 | 21.9160249 | 0.012404797 | 0.824887059 |
| 21.9742789 | 21.9029332 | 21.8843456 | 21.9931307 | -0.070883442 | 0.214143326 |
| 20.8919067 | 20.7496087 | 20.8549663 | 20.8590462 | -0.025482668 | 0.637333991 |
| 24.106734 | 23.9147526 | 24.023203 | 23.9932117 | -0.04231763 | 0.435098145 |
| 27.0845631 | 26.9995537 | 27.0896263 | 27.0669317 | 0.01257462 | 0.810370619 |
| 20.4448976 | 20.5090338 | 20.5718982 | 20.4448078 | 0.05175582 | 0.319171112 |
| 23.4074637 | 23.3315123 | 23.4283832 | 23.4529013 | 0.006618457 | 0.890975409 |
| 21.6210305 | 21.5008635 | 21.6653976 | 21.5300819 | -0.014904573 | 0.759285758 |
| 22.6386747 | 22.5635019 | 22.5572249 | 22.6018055 | 9.91E-04 | 0.983442634 |
| 20.7886697 | 20.6761007 | 20.7401312 | 20.7957628 | 8.83E-04 | 0.985253189 |
| 24.766173 | 24.7332029 | 24.8726055 | 24.8437798 | 0.03728179 | 0.411267339 |
| 24.335491 | 24.3860366 | 24.2953239 | 24.3065003 | -0.006332802 | 0.880519446 |
| 20.9687515 | 20.903386 | 20.9034959 | 20.9109354 | -0.004542695 | 0.911322615 |
| 22.4517866 | 22.4229237 | 22.4432337 | 22.4249337 | -0.021930628 | 0.591884905 |
| 21.6566963 | 21.59044 | 21.6175049 | 21.6227856 | -0.017925732 | 0.651308289 |
| 20.8517536 | 20.8026601 | 20.8147019 | 20.8369958 | -0.021091186 | 0.594165345 |
| 20.4506941 | 20.3412731 | 20.3162786 | 20.3329979 | -0.059582538 | 0.130424472 |
| 24.522713 | 24.4765529 | 24.4990257 | 24.5094773 | 0.001117106 | 0.972912186 |
| 23.7876511 | 23.7426732 | 23.7593782 | 23.8194036 | 0.037585589 | 0.353304699 |
| 20.6581371 | 20.6097336 | 20.6151305 | 20.6009965 | 0.014826401 | 0.712078806 |
| 25.9641139 | 25.981492 | 26.0550111 | 26.0446439 | 0.083721358 | 0.055505143 |

| | | | | | |
|---|---|---|---|---|---|
| 23.4663955 | 23.4584407 | 23.5064962 | 23.5005298 | 0.045782156 | 0.286954247 |
| 21.7983556 | 21.717474 | 21.7773851 | 21.8806097 | 0.034707398 | 0.45509818 |
| 21.1882446 | 21.1257138 | 21.1223796 | 21.1448445 | 0.017028106 | 0.732703208 |
| 19.3058916 | 19.2391483 | 19.2750017 | 19.3241195 | -0.005445885 | 0.917436357 |
| 24.0907225 | 24.0781068 | 24.1477827 | 24.0483949 | 0.026439176 | 0.615255023 |
| 22.091133 | 21.9735731 | 22.0350545 | 22.0422949 | -0.026828615 | 0.618603697 |
| 19.989074 | 19.9741909 | 19.9858776 | 19.9624175 | -0.013805595 | 0.797320338 |
| 25.9960445 | 25.8395572 | 25.9567423 | 25.9521607 | 0.005455468 | 0.921294477 |
| 21.6464232 | 21.5899481 | 21.7397864 | 21.7026083 | 0.017317578 | 0.756559442 |
| 19.9283315 | 19.7861041 | 19.926414 | 19.979034 | 0.020004845 | 0.720848286 |
| 21.6392597 | 21.5650227 | 21.642105 | 21.6543065 | 0.026486149 | 0.643553289 |
| 20.0007138 | 19.9149468 | 20.0627182 | 19.9699311 | -0.032914957 | 0.561761169 |
| 21.3650157 | 21.3428106 | 21.447136 | 21.3714505 | 0.0417247 | 0.469128312 |
| 21.051068 | 21.0082359 | 21.030665 | 21.1236547 | 0.026910914 | 0.636671895 |
| 22.0082429 | 21.9126829 | 22.1131538 | 22.1047736 | 0.072573533 | 0.224537339 |
| 20.9855989 | 20.8858659 | 21.058582 | 20.9451799 | 0.05719843 | 0.34996776 |
| 19.6725725 | 19.6915403 | 19.7656473 | 19.7579354 | 0.109455751 | 0.078093243 |
| 19.9082665 | 19.826724 | 19.8714859 | 19.9192784 | -0.00738211 | 0.906725733 |
| 20.8896929 | 20.8331336 | 20.9217639 | 20.7761998 | -0.009558955 | 0.878700528 |
| 21.3198047 | 21.2555166 | 21.3079944 | 21.162658 | -0.03766679 | 0.552668592 |
| 18.4956014 | 18.401425 | 18.499451 | 18.4858015 | 0.044578207 | 0.49467014 |
| 20.1842258 | 20.140385 | 20.1492612 | 20.1815497 | 0.023966198 | 0.72200974 |
| 20.0700281 | 19.9219974 | 19.9288816 | 20.0248419 | -0.042042465 | 0.544638171 |
| 23.5219973 | 23.4096099 | 23.5076809 | 23.3649354 | -0.030107669 | 0.662091535 |
| 21.6124487 | 21.4660662 | 21.5167034 | 21.5043779 | -0.027834938 | 0.690785956 |
| 18.2086236 | 18.2651298 | 18.286238 | 18.2927525 | -0.013887177 | 0.842584699 |
| 18.3240588 | NA | 18.4031097 | 18.3498713 | -0.034350171 | 0.620916059 |
| 19.7722133 | 19.7904266 | 19.7316616 | 19.8000077 | -0.040852363 | 0.583268697 |
| 18.2956325 | 18.0993769 | 18.244308 | 18.2472877 | -0.053954871 | 0.479899188 |
| 20.2793993 | 20.2958284 | 20.2831191 | 20.2555854 | -0.028833458 | 0.704718678 |
| 20.2295801 | 20.1929066 | 20.4386452 | 20.382457 | 0.140535512 | 0.07768686 |
| 20.6130557 | 20.6514303 | 20.8005825 | 20.7447094 | 0.100572959 | 0.212450881 |
| 19.5165982 | 19.5808999 | 19.5102273 | 19.5635257 | 0.031184009 | 0.700320085 |
| 21.5341145 | 21.4049198 | 21.5828066 | 21.5850699 | -0.007406129 | 0.926608869 |
| 21.9191819 | 21.8392439 | 21.9588575 | 22.0754058 | 0.05407318 | 0.524363646 |
| 18.9389531 | 19.0685621 | 18.9274276 | 19.1294988 | 0.154363291 | 0.080605829 |
| 17.4421479 | 17.7045029 | 17.6389712 | 17.6458318 | 0.124518682 | 0.191319628 |
| 20.5286789 | 20.4362824 | 20.4785802 | 20.4232898 | -0.007155746 | 0.940660357 |
| 16.6743245 | 16.5339216 | 16.6985011 | 16.4846554 | 0.024141399 | 0.805294267 |
| 19.1551601 | 18.8103919 | 19.1704112 | 19.3149015 | -0.085514812 | 0.390590361 |
| 24.8959393 | 24.8707785 | 24.8548598 | 24.8871234 | 0.019105617 | 0.858532276 |
| 17.569335 | 17.5224676 | 17.4744587 | 17.4510523 | 0.006277223 | 0.955020404 |

| | | | | | |
|---|---|---|---|---|---|
| 18.8766832 | 18.5369341 | 19.2031183 | 18.8528166 | 0.196748091 | 0.142647343 |
| 16.1785275 | NA | 16.4917868 | 16.3743017 | 0.205880909 | 0.137218447 |
| 18.3735601 | 18.244183 | 18.3492368 | 18.2755499 | -0.077112874 | 0.573445781 |
| 15.6486902 | 15.5916098 | 15.5900501 | 15.2950986 | -0.009048992 | 0.951711046 |
| 15.3236334 | NA | 15.3552093 | 15.4693907 | -0.078817833 | 0.620708433 |
| 16.9165813 | 16.4757385 | 16.724245 | 16.5760201 | -0.209800067 | 0.200255525 |
| NA | 17.1104232 | NA | NA | 0.118310346 | 0.544305168 |
| 14.2355524 | 14.4415526 | 14.6803417 | 14.7768781 | 0.304615301 | 0.138295638 |
| 14.3126307 | NA | 14.3363012 | 14.5474495 | 0.32725582 | 0.174960597 |
| NA | 12.2684663 | NA | NA | -0.32805868 | 0.326257499 |
| NA | 13.1987758 | NA | NA | 0.190265349 | 0.620237911 |
| 13.1993421 | NA | 14.2765225 | 13.989272 | 0.531631287 | 0.182946947 |
| 17.6851902 | 17.4396486 | 17.451951 | 16.3434474 | -1.787414397 | 0.01523996 |
| 17.5174237 | 17.442544 | 17.6649148 | 17.3578827 | -0.017359654 | 0.890484906 |
| 21.1587206 | 21.1444496 | 21.1387983 | 21.1496951 | -0.043091201 | 0.334310285 |
| 25.8879455 | 25.8481577 | 25.776158 | 25.8377819 | -0.009577553 | 0.849801444 |
| 18.6885971 | 18.5572426 | 18.8913533 | 18.575199 | 0.065358786 | 0.485915859 |
| 14.1685603 | 13.9812107 | 13.6643418 | 14.3906684 | -0.045264855 | 0.855189097 |
| 19.7258853 | 19.5756539 | 19.7038016 | 19.6512056 | -0.067857721 | 0.225607889 |
| 21.1412302 | 21.0097361 | 21.1494383 | 21.103649 | -0.020004028 | 0.710446389 |
| 22.2451775 | 22.1652943 | 22.1871775 | 22.1541117 | -0.049842596 | 0.28683648 |
| 24.5372906 | 24.3642537 | 24.5106672 | 24.5383342 | 0.06917859 | 0.297535949 |
| 25.3490989 | 25.2535631 | 25.4149643 | 25.384753 | 0.044319686 | 0.535104216 |
| 21.1539301 | 21.0679168 | 21.1103651 | 21.156883 | -0.048060177 | 0.420260198 |
| 15.132993 | NA | 15.2069452 | 15.0277038 | 0.023622911 | 0.876518456 |
| 21.8817561 | 21.8107977 | 21.882264 | 21.9118124 | 0.020921925 | 0.744799646 |
| 16.0621456 | 16.1034415 | 16.1019197 | 16.0056102 | 0.006485503 | 0.959364259 |
| 17.7062003 | 17.6657746 | 17.748448 | 17.6541954 | -0.043341553 | 0.569935177 |
| 24.5534246 | 24.5070578 | 24.5122999 | 24.5745445 | 0.023987539 | 0.57184058 |
| 20.621185 | 20.4494564 | 20.5108376 | 20.5843834 | 0.015868075 | 0.776062922 |
| 23.9716917 | 23.8045627 | 23.7871459 | 23.8323263 | -0.089795305 | 0.299182456 |
| 21.8547996 | 21.8437657 | 21.8140088 | 21.8162611 | -0.010014234 | 0.766875878 |
| 19.5384611 | 19.4911038 | 19.4972512 | 19.5474814 | 0.008429718 | 0.888029118 |
| 17.848192 | 17.6167547 | 17.7020011 | 17.7058427 | -0.114323381 | 0.170145726 |
| 20.2065432 | 19.9477969 | 20.171829 | 20.1853781 | -0.037195803 | 0.603540801 |
| 20.9950556 | 20.9884929 | 21.0204305 | 21.0010636 | -0.001252572 | 0.974813256 |
| 19.2148421 | 19.3255339 | 19.3328662 | 19.4005287 | 0.061775583 | 0.15775636 |
| 17.3290394 | 17.2931635 | 17.3326834 | 17.5383249 | 0.085832344 | 0.403157134 |
| 17.2059849 | 17.15197 | 17.2178528 | 17.2134218 | -0.025302957 | 0.81548112 |
| 21.5318159 | 21.4862406 | 21.6588826 | 21.61351 | 0.133027257 | 0.069404529 |
| 17.8804494 | 17.8212922 | 17.6900154 | 17.9025155 | 0.086645866 | 0.420352854 |
| 18.7135508 | 18.5020086 | 18.7005857 | 18.5525361 | -0.125514896 | 0.117011149 |

| | | | | | |
|---|---|---|---|---|---|
| 23.4576043 | 23.4283522 | 23.3875903 | 23.4578419 | 0.008094309 | 0.866148752 |
| 22.4387124 | 22.3697138 | 22.3241171 | 22.4257193 | 0.023723615 | 0.657971297 |
| 18.0994925 | 17.9744756 | 17.9666715 | 18.0185931 | -0.062749916 | 0.382877778 |
| 23.4000006 | 23.0637927 | 23.1706832 | 23.0761173 | -0.138025016 | 0.033009754 |
| 18.6177984 | 18.4838441 | 18.6363852 | 18.6567879 | -0.061231094 | 0.276642492 |
| 20.4755859 | 20.433741 | 20.4097817 | 20.5460835 | -0.009245007 | 0.852630733 |
| 15.8348702 | 15.966488 | 15.7788701 | 15.6648813 | -0.100939622 | 0.468718196 |
| 15.9051799 | NA | 16.0921419 | 16.1076461 | 0.46544023 | 0.059960032 |
| 13.5867409 | 13.4943078 | 13.4782887 | 14.0292977 | -0.087691436 | 0.563108308 |
| 15.8308196 | 15.6588618 | 15.5756448 | 15.5526405 | -0.14435684 | 0.193019745 |
| 19.0381012 | 18.6653129 | 18.8034224 | 18.9967949 | -0.0842158 | 0.425367452 |
| 18.6851749 | 18.5350492 | 18.772427 | 18.5224029 | -0.049527798 | 0.55247024 |
| 18.4781945 | 18.3590612 | 18.2823371 | 18.3479729 | -0.082509277 | 0.267599414 |
| 19.7023792 | 19.6737331 | 19.7745344 | 19.6925139 | -0.032813204 | 0.640891165 |
| 19.1358217 | 18.9478787 | 19.0002708 | 19.0928551 | -0.040372518 | 0.481679276 |
| 20.8494697 | 20.776953 | 20.7282344 | 20.8121701 | -0.019501263 | 0.729367009 |
| 20.8187682 | 20.6337441 | 20.7634235 | 20.6065374 | -0.077019739 | 0.147515178 |
| 23.8195353 | 23.7193973 | 23.7705187 | 23.787135 | -0.020839115 | 0.678055393 |
| 21.1483864 | 21.0075209 | 21.0802629 | 21.1474593 | -0.065521908 | 0.175412637 |
| 23.7778218 | 23.7312271 | 23.8100296 | 23.8496318 | 0.057855565 | 0.27649582 |
| 17.9790435 | 18.0682941 | 18.0840135 | 18.0396697 | 0.074805674 | 0.212771375 |
| 22.9138229 | 22.8678834 | 22.9569964 | 22.9626013 | 0.063881758 | 0.286169182 |
| 24.7702019 | 24.6417007 | 24.7220133 | 24.582934 | -0.060609064 | 0.318541754 |
| 19.8934581 | 19.797862 | 19.7730162 | 19.791774 | -0.028191837 | 0.681871069 |
| 22.9289694 | 22.7791711 | 22.8745171 | 22.8620233 | -0.011732853 | 0.875101032 |
| 20.1400473 | 20.0460731 | 20.0528091 | 20.1255661 | -0.013255303 | 0.866656433 |
| 19.1519285 | 19.1493394 | 19.1492417 | 19.1390207 | 0.095495836 | 0.243823318 |
| 15.3188673 | 15.0140703 | 15.1899198 | 15.0680114 | -0.191631762 | 0.13867746 |
| 18.1461676 | 18.5657989 | 18.3558408 | 18.2594541 | 0.159104581 | 0.248612383 |
| 16.9839697 | 16.5551976 | 16.9102495 | 17.177198 | 0.088453797 | 0.550414891 |
| 15.0239789 | 15.6155299 | 15.6142837 | 15.4317825 | 0.183767526 | 0.322521927 |
| 21.6646794 | 21.6329512 | 21.7483708 | 21.6964284 | 0.021734126 | 0.712578845 |
| 20.3293544 | 20.254517 | 20.2877624 | 20.2388946 | -0.066869106 | 0.21046787 |
| 22.4931834 | 22.5369443 | 22.5868813 | 22.568157 | 0.0589009 | 0.311057322 |
| 20.1204743 | 20.1167926 | 20.0362754 | 20.1586434 | -0.03565543 | 0.676722023 |
| 19.9242349 | 19.7323465 | 19.8499308 | 19.9457219 | -0.01167303 | 0.850900783 |
| 23.0855221 | 22.972359 | 23.061152 | 23.0431511 | -0.018610894 | 0.685051744 |
| 21.9254847 | 21.8866472 | 21.9097683 | 21.9025864 | -0.078997344 | 0.238518992 |
| 27.0668154 | 27.1204197 | 27.131676 | 27.1116985 | 0.060320096 | 0.196914345 |
| 18.4105429 | 18.31666 | 18.3507814 | 18.5001094 | -0.141939384 | 0.299394084 |
| 21.3337764 | 21.2608208 | 21.3788172 | 21.4191258 | -0.002445899 | 0.96837444 |
| 17.484415 | 17.3038353 | 17.7231781 | 17.5593308 | 0.080711003 | 0.625181401 |

| | | | | | |
|---|---|---|---|---|---|
| 15.3002171 | NA | 14.316582 | 14.8125196 | -0.467564249 | 0.156584916 |
| 13.7986336 | 13.4559784 | 13.49369 | 13.6198168 | -0.104781748 | 0.540336441 |
| 17.94458 | 17.7628227 | 18.0206117 | 17.9398061 | -0.105995208 | 0.329655734 |
| 19.8800521 | 19.9571691 | 20.0444858 | 19.9668171 | 0.059019742 | 0.311530201 |
| 21.5111736 | 21.3148084 | 21.3643035 | 21.4668223 | -0.102516598 | 0.063451398 |
| 21.489949 | 21.3376697 | 21.3994334 | 21.4400085 | -0.040911155 | 0.422899318 |
| 21.2564142 | 21.0693231 | 21.1115825 | 21.2470411 | -0.028163742 | 0.557766474 |
| 20.5200316 | 20.4063783 | 20.492388 | 20.4552812 | -0.029761893 | 0.496895957 |
| 21.1963976 | 21.2084051 | 21.2040453 | 21.2215413 | 0.036294057 | 0.328677861 |
| 22.4755377 | 22.4806742 | 22.534155 | 22.478149 | 0.041444837 | 0.325131458 |
| 23.1432364 | 22.9957123 | 23.1100362 | 23.1233311 | 0.016817099 | 0.758384448 |
| 17.8652542 | 18.0166718 | 18.109149 | 17.9733966 | 0.063004317 | 0.383660317 |
| 23.1347741 | 23.0805675 | 23.0994389 | 23.1034863 | 0.003861445 | 0.960390359 |
| 18.8064661 | 18.8332545 | 18.6946812 | 18.8305415 | -0.021347002 | 0.804249841 |
| 19.7832442 | 19.6883635 | 19.7480882 | 19.8253101 | -0.011165185 | 0.856493079 |
| 20.7667907 | 20.8522442 | 20.7796113 | 20.8758709 | 0.056298976 | 0.286161717 |
| 20.2588183 | 20.0951359 | 20.0986115 | 20.2839336 | -0.048941025 | 0.334629753 |
| 22.9473151 | 22.8270475 | 22.8863886 | 22.8964981 | -0.022940534 | 0.626197581 |
| 19.6108242 | 19.5269079 | 19.5996113 | 19.5515512 | -0.01799205 | 0.748542052 |
| 21.0590857 | 20.9814957 | 21.0464263 | 20.9942157 | 0.015509085 | 0.789094468 |
| 18.2885251 | 18.1478866 | 18.1933336 | 18.3771387 | 0.001363889 | 0.985267079 |
| 13.3559115 | 13.211915 | 13.2645045 | 13.2009719 | 0.035783315 | 0.809420075 |
| 19.1990466 | NA | 19.1475812 | 19.2873708 | -0.153313103 | 0.433252264 |
| 18.3036159 | 17.5684446 | 17.9729274 | 18.5177691 | -0.096575988 | 0.692337448 |
| 21.6708213 | 22.0614223 | 21.9065177 | 22.0590281 | 0.02484074 | 0.889974696 |
| 15.1171675 | 15.1885867 | 15.1511942 | 15.2097731 | 0.015673394 | 0.924481805 |
| NA | 15.4460589 | NA | NA | -0.206957929 | 0.150477844 |
| 21.0622508 | 20.8702124 | 20.9456206 | 21.0791589 | -0.051116494 | 0.426844224 |
| 24.4048811 | 24.2811355 | 24.3692555 | 24.3665503 | -0.046721689 | 0.388287232 |
| 23.4578053 | 23.4368037 | 23.4385751 | 23.4641211 | 0.031286257 | 0.532822388 |
| 23.5641946 | 23.5487154 | 23.5775083 | 23.5489585 | 0.001166714 | 0.978249168 |
| 21.7579075 | 21.6476979 | 21.7600576 | 21.7467833 | 0.013810369 | 0.83745437 |
| 20.3501045 | 20.2220162 | 20.135283 | 20.1878741 | -0.079311759 | 0.431526968 |
| 16.4429013 | 16.702384 | 16.7686371 | 16.8399474 | 0.090322696 | 0.523060325 |
| 17.6300474 | 17.6943715 | 17.6748657 | 17.8357407 | 0.021150908 | 0.852368412 |
| 15.9580637 | NA | 15.9856127 | 16.0333429 | -0.001025401 | 0.99094849 |
| 20.9133673 | 20.775172 | 20.8282546 | 20.9303723 | -0.06139374 | 0.329785565 |
| 19.6933649 | 19.6815332 | 19.7484569 | 19.7272853 | 0.023935307 | 0.637137385 |
| 21.3818486 | 21.2951738 | 21.343493 | 21.3443769 | 0.050959186 | 0.386216145 |
| 15.826849 | 15.4169508 | 15.9412292 | 15.6173961 | -0.111222848 | 0.501951503 |
| 18.4758493 | 18.1928177 | 18.6180698 | 18.5375577 | -0.095796846 | 0.462393385 |
| 20.1508825 | 20.0289341 | 20.17657 | 20.260647 | 0.011964296 | 0.8701128 |

| | | | | | |
|---|---|---|---|---|---|
| 18.1410158 | 18.1212866 | 18.1295838 | 18.2176988 | 0.028967159 | 0.688749356 |
| 20.8769055 | 20.7874766 | 20.9126207 | 20.9027736 | 0.036024699 | 0.536946207 |
| 22.2783885 | 22.2327431 | 22.2189108 | 22.2034026 | -0.014736779 | 0.798445152 |
| 20.49135 | 20.4549489 | 20.5020156 | 20.504739 | -6.45E-04 | 0.990102085 |
| 25.5399933 | 25.4444112 | 25.4993435 | 25.5218489 | -0.005152738 | 0.920238197 |
| 24.0923418 | 24.0634083 | 24.1582493 | 24.1427239 | 0.021745554 | 0.648452866 |
| 21.8301365 | 21.8640924 | 21.8079675 | 21.8684267 | -0.005448581 | 0.898730116 |
| 21.5214216 | 21.4550744 | 21.5034351 | 21.5382934 | -0.023200208 | 0.614016968 |
| 21.5927733 | 21.5160969 | 21.5740726 | 21.5839447 | 0.03132409 | 0.584286056 |
| 18.6605101 | 18.7250709 | 18.6426515 | 18.6750825 | 0.09161223 | 0.130999977 |
| 19.2531183 | 19.2305232 | 19.3402471 | 19.144585 | 0.066430577 | 0.297050384 |
| 24.0144266 | 23.8876623 | 24.0278221 | 23.9960673 | -0.021409404 | 0.742391902 |
| 20.823814 | 20.7442989 | 20.8484185 | 20.8575354 | 0.008367174 | 0.907021432 |
| 18.5241329 | 18.5302476 | 18.6794644 | 18.6117102 | 0.023802155 | 0.76128568 |
| 19.7951356 | 19.8249501 | 19.9857274 | 19.9810069 | 0.15335013 | 0.060182202 |
| 18.1528452 | 18.4367934 | 18.2140035 | 18.3651507 | 0.105262375 | 0.216636466 |
| 17.4707025 | 17.1202103 | 17.125055 | 17.3909743 | -0.136136061 | 0.183504391 |
| 17.2782948 | 16.9909335 | 17.2645935 | 17.2409968 | 0.124193976 | 0.357686038 |
| 17.0631995 | 16.636564 | 16.7755342 | 17.2263214 | -0.097935467 | 0.505765655 |
| 16.699294 | 16.5474107 | 16.4259085 | 16.58408 | -0.182255175 | 0.105908859 |
| 21.3666489 | 21.485771 | 21.5477679 | 21.2932049 | -0.018630125 | 0.831815974 |
| 18.6585109 | 18.7638916 | 18.7129453 | 18.7910294 | 0.056470544 | 0.27880762 |
| 23.5797138 | 23.5047195 | 23.5813266 | 23.5813438 | 0.014720856 | 0.830153897 |
| 21.9830644 | 21.9730493 | 22.0230817 | 21.9955577 | 0.04392591 | 0.34928684 |
| 20.8029259 | 20.759336 | 20.8762625 | 20.8092138 | 0.018201305 | 0.719688988 |
| 19.0756324 | 18.8824244 | 19.0595028 | 19.0626737 | 0.011011888 | 0.904932913 |
| 17.3630786 | 18.1538975 | 18.0969563 | 17.7083502 | 0.536984981 | 0.133050945 |
| 19.7947193 | 19.9730286 | 19.8799177 | 19.8435925 | -0.043481178 | 0.691446129 |
| 16.4290544 | 16.4168828 | 16.4650851 | 16.346408 | -0.165763105 | 0.104325912 |
| 18.4746971 | 18.3319579 | 18.4760549 | 18.4729361 | -0.067025508 | 0.483821852 |
| 17.7507969 | 17.4452863 | 17.3322757 | 17.5514697 | -0.159589833 | 0.021847729 |
| 21.568512 | 21.5923447 | 21.7197811 | 21.5995176 | 0.002876481 | 0.964478658 |
| 22.2350164 | 22.1340934 | 22.1949977 | 22.125797 | -0.061548624 | 0.232348116 |
| 18.5619723 | 18.4046434 | 18.4467279 | 18.4584436 | -0.107739641 | 0.011576054 |
| 20.5631308 | 20.5024102 | 20.5781544 | 20.6247575 | 0.033590799 | 0.537391367 |
| 22.0240473 | 21.9922728 | 22.0783586 | 22.0859081 | 0.035014382 | 0.591509668 |
| 22.800814 | 22.8638588 | 22.9964667 | 23.0898463 | 0.191115487 | 0.01142913 |
| 17.381277 | 17.4927138 | 17.2466295 | 17.2901205 | 0.027998239 | 0.776030931 |
| 24.1311962 | 24.0685757 | 24.1297631 | 24.1222284 | 0.041497981 | 0.355218869 |
| 26.2967871 | 26.2312933 | 26.3899911 | 26.2755249 | 0.049313296 | 0.590151068 |
| 20.5196493 | 20.4548415 | 20.5657124 | 20.541484 | -0.016583906 | 0.803250805 |
| 17.8917253 | 17.6303047 | 17.762781 | 17.9206102 | 0.034529668 | 0.706247932 |

| | | | | | |
|---|---|---|---|---|---|
| 21.055159 | 20.9991265 | 21.2571597 | 21.2537312 | 0.107450196 | 0.505570268 |
| 22.2622993 | 22.1592865 | 22.1974996 | 22.3180069 | -0.078496896 | 0.361072196 |
| 18.2485929 | 18.2028578 | 18.3128519 | 18.2524749 | 0.044302722 | 0.498731835 |
| 20.979082 | 20.8472977 | 20.989868 | 20.9671341 | -0.044063366 | 0.549084153 |
| 18.8681991 | 18.7867715 | 18.8793825 | 18.8430013 | -0.043581983 | 0.447585285 |
| 21.2882719 | 21.0261829 | 21.1350755 | 21.1653204 | -0.088881131 | 0.161993462 |
| 16.5802462 | 16.5426825 | 16.6016702 | 16.7046608 | 0.01023721 | 0.895569993 |
| 21.017633 | 20.8705308 | 20.8860093 | 20.8805177 | -0.026927863 | 0.826850982 |
| 18.6847734 | 18.759796 | 18.5811879 | 18.6398154 | -0.083779741 | 0.411951763 |
| 21.4162094 | 21.2088715 | 21.5161798 | 21.3548699 | -0.05719385 | 0.322087022 |
| 20.7936337 | 20.6998712 | 20.6976706 | 20.8706452 | 0.002166867 | 0.971154168 |
| 18.8524663 | 18.8553426 | 18.764583 | 18.6365838 | -0.017799743 | 0.767254904 |
| 16.6643614 | 16.3298316 | 16.580423 | 16.5556891 | 0.142870182 | 0.240408972 |
| 28.1235735 | 27.9848978 | 28.0853399 | 28.1006868 | -0.002113047 | 0.970932219 |
| 22.229479 | 22.2651725 | 22.2744908 | 22.2433917 | 0.003523723 | 0.937351602 |
| 20.5966822 | 20.466069 | 20.5076625 | 20.5811409 | -0.057914058 | 0.246703563 |
| 22.6689013 | 22.6449969 | 22.6902153 | 22.6672266 | 0.013822821 | 0.668861744 |
| 21.4949079 | 21.5189118 | 21.5343876 | 21.547229 | 4.80E-04 | 0.99061754 |
| 24.6347279 | 24.5097168 | 24.6117285 | 24.6484139 | -0.003245428 | 0.960860509 |
| 21.5177765 | 21.464226 | 21.527065 | 21.5394656 | 0.041776397 | 0.411037222 |
| 23.5196235 | 23.5143022 | 23.5407849 | 23.5247843 | 0.062198071 | 0.340352105 |
| 22.2023366 | 22.077731 | 22.1456504 | 22.1260934 | -0.022263504 | 0.688252874 |
| 19.880995 | 19.800555 | 19.8826569 | 19.9525239 | 0.009298652 | 0.86197267 |
| 18.5809252 | 18.4011579 | 18.4655638 | 18.3504571 | -0.114995323 | 0.050067891 |
| 20.6341801 | 20.4944075 | 20.6044845 | 20.6467431 | 0.01646546 | 0.749189502 |
| 20.8284642 | 20.7167768 | 20.8212708 | 20.8486923 | 0.021156027 | 0.71153729 |
| 21.6188524 | 21.5060248 | 21.5302461 | 21.4485657 | -0.093668113 | 0.184896376 |
| 22.7527635 | 22.7495446 | 22.7822739 | 22.6941897 | 0.023178285 | 0.556225226 |
| 21.8276564 | 21.7358404 | 21.8058772 | 21.7996113 | 0.019325166 | 0.703692492 |
| 24.7733647 | 24.6555133 | 24.7705678 | 24.7716453 | 0.00844895 | 0.852822033 |
| 17.0608739 | 16.8165893 | 16.7577666 | 16.9306508 | -0.029627566 | 0.784315961 |
| 16.8430098 | 16.5737946 | 16.9298288 | 16.6032644 | -0.119884944 | 0.330715051 |
| 13.8715068 | 13.9542963 | 14.6830661 | 14.2216725 | 0.247915721 | 0.458941605 |
| 18.7011131 | 18.6889714 | 18.6263053 | 18.6234505 | 0.076826174 | 0.300855964 |
| 21.8932693 | 21.9495223 | 21.9553236 | 21.9560797 | -0.014227918 | 0.807244352 |
| 16.8058569 | 16.7334579 | 16.9335109 | 16.822803 | 0.069983754 | 0.307601105 |
| 19.3368781 | 19.2484049 | 19.30631 | 19.0888624 | 0.02673112 | 0.794644308 |
| 16.5415746 | 16.4679331 | 16.6575141 | 16.7558175 | 0.195954737 | 0.201788185 |
| 18.3624138 | 18.0383393 | 18.1582765 | 18.2902187 | -0.055705216 | 0.501990575 |
| 19.3208309 | 19.2762771 | 19.3832018 | 19.3133775 | -0.030185007 | 0.610874725 |
| 20.2139392 | 19.9133757 | 20.0550315 | 20.1974403 | -0.03833213 | 0.562916557 |
| 17.5777346 | 17.4684426 | 17.7229187 | 17.7468927 | 0.13622635 | 0.14247815 |

| | | | | | |
|---|---|---|---|---|---|
| 20.7506984 | 20.4603041 | 20.7759019 | 20.672734 | 0.027963293 | 0.880755518 |
| 18.6358582 | 18.4180644 | 18.5015594 | 18.2259025 | -0.102437794 | 0.405679908 |
| 17.3500387 | 17.209598 | 17.4080993 | 17.4246394 | 0.050944493 | 0.606930804 |
| 18.4386595 | 18.2671735 | 18.3415344 | 18.183557 | -0.112268887 | 0.219206392 |
| NA | 15.3987799 | NA | NA | 0.006273194 | 0.936332728 |
| 25.6961832 | 25.749629 | 25.7625695 | 25.7466033 | 0.055851362 | 0.175819818 |
| 22.3950768 | 22.3483927 | 22.3655834 | 22.3515325 | 0.03760935 | 0.381061374 |
| 21.8331758 | 21.780038 | 21.8131861 | 21.8312256 | -0.029144017 | 0.563479761 |
| 20.926339 | 20.7794184 | 20.900117 | 20.8638995 | -0.045398691 | 0.379073213 |
| 25.1455298 | 25.0534057 | 25.1429234 | 25.1260651 | 0.036697226 | 0.476338722 |
| 22.9748261 | 22.9999267 | 22.9673633 | 22.9366641 | 0.03168008 | 0.539920389 |
| 19.6340295 | 19.526695 | 19.6267604 | 19.7388544 | 0.048956075 | 0.395639263 |
| 21.7596163 | 21.6270411 | 21.8429515 | 21.7782264 | -0.020564817 | 0.772536193 |
| 19.8306126 | 19.5949281 | 19.7798591 | 19.8877104 | -0.023906215 | 0.738958752 |
| 18.5839881 | 18.3552576 | 18.578004 | 18.5878952 | 0.048797369 | 0.523674292 |
| 20.9904124 | 20.8836474 | 20.8203269 | 20.909371 | -0.065718683 | 0.419181785 |
| 17.7664314 | 18.0078758 | 18.1661782 | 18.0413655 | 0.197270343 | 0.058479494 |
| 15.6813106 | 15.6287087 | 15.4921465 | 15.6899664 | 0.049980233 | 0.661029434 |
| 16.4217944 | 16.3370299 | 16.0337649 | 16.7703932 | 0.191171186 | 0.185119961 |
| 21.4092423 | 21.274054 | 21.353661 | 21.3495726 | -0.027688551 | 0.590963049 |
| 18.3212642 | 18.2264132 | 18.2110778 | 18.1994301 | -0.059700131 | 0.322567276 |
| 19.5377261 | 19.6360888 | 19.5589618 | 19.4745807 | 0.063526925 | 0.486750795 |
| 22.1291815 | 22.0869148 | 22.2013017 | 22.2325336 | 0.038592465 | 0.488558329 |
| 17.7786311 | 17.9187411 | 17.797633 | 17.7518685 | 0.122657656 | 0.233644012 |
| 14.6265848 | NA | 14.5979501 | 14.9026411 | 0.223554622 | 0.238484726 |
| 20.349767 | 20.2702196 | 20.324576 | 20.3422704 | -0.057010449 | 0.319533865 |
| 23.7391956 | 23.6958458 | 23.7650084 | 23.751148 | -0.003859136 | 0.915405056 |
| 19.4437918 | 19.1514956 | 19.4187939 | 19.4275244 | -0.041667281 | 0.619356852 |
| 18.5458918 | 18.3460167 | 18.6253298 | 18.3683065 | 0.007533774 | 0.956908014 |
| 15.5316853 | 15.2532102 | 14.9899388 | 15.5886075 | 0.198167165 | 0.425176658 |
| 17.2470081 | NA | 17.1740217 | 17.3303753 | 0.100531768 | 0.546071361 |
| 16.7564518 | 16.871766 | 16.9246052 | 16.8466874 | 0.029702972 | 0.833349234 |
| 17.0187576 | 16.6792801 | 16.7615131 | 16.8114673 | -0.167819857 | 0.174229582 |
| 21.2782945 | 21.2928043 | 21.2455592 | 21.2715379 | -0.024125873 | 0.831803606 |
| 17.3916043 | 17.4091419 | 17.5104705 | 17.1342257 | -0.080699695 | 0.422510527 |
| 17.927736 | 17.9510924 | 17.957062 | 17.691469 | 0.044901267 | 0.641486303 |
| 19.8321319 | 19.5957517 | 19.9092055 | 19.8951811 | -0.070699826 | 0.37410126 |
| 17.7675558 | 17.4869435 | 17.819242 | 17.6862137 | -0.033317554 | 0.638943157 |
| 19.7985392 | 19.9315741 | 19.6543336 | 19.7563294 | -0.082293387 | 0.245979916 |
| 19.2557106 | 19.1750295 | 19.3314687 | 19.1790754 | -0.044500219 | 0.52351851 |
| 17.935791 | 17.9507884 | 18.05157 | 18.0053132 | 0.086694775 | 0.215000811 |
| 18.6445722 | 18.5681443 | 18.7081062 | 18.6976827 | 0.045248884 | 0.499576926 |

| | | | | | |
|---|---|---|---|---|---|
| 19.777338 | 19.7472729 | 19.8657134 | 19.8671975 | 0.002695736 | 0.965519456 |
| 21.5928059 | 21.6454765 | 21.6702343 | 21.6969893 | 0.074732231 | 0.219396001 |
| 22.416788 | 22.3312859 | 22.4593831 | 22.4370535 | 0.049077567 | 0.41160849 |
| 22.7502678 | 22.5873434 | 22.6985281 | 22.7321661 | -0.020170084 | 0.730063431 |
| 21.518144 | 21.3529116 | 21.572622 | 21.5081919 | -0.026501737 | 0.65032693 |
| 20.5435002 | 20.5333793 | 20.5881086 | 20.5314021 | 0.012206572 | 0.8338003 |
| 22.1961633 | 22.0817129 | 22.3136559 | 22.3073344 | -0.051226312 | 0.377469378 |
| 23.6071511 | 23.56093 | 23.5287375 | 23.5546722 | -0.030307904 | 0.572158301 |
| 26.0729423 | 26.0188652 | 26.0672559 | 26.0831026 | 0.035787887 | 0.502040767 |
| 21.4954197 | 21.4805146 | 21.4582105 | 21.4563059 | 5.54E-04 | 0.991688148 |
| 24.3355898 | 24.1842329 | 24.3485338 | 24.3201791 | -0.011761013 | 0.803475977 |
| 20.9488109 | 20.9016753 | 20.9709816 | 20.9225839 | -0.011074893 | 0.804509479 |
| 22.9349922 | 22.8434608 | 22.905911 | 22.9159198 | -0.015837992 | 0.707515207 |
| 22.2881279 | 22.2507252 | 22.2656826 | 22.2894231 | -1.17E-04 | 0.997725661 |
| 22.7177751 | 22.6789105 | 22.7057844 | 22.642453 | -0.005208118 | 0.896752395 |
| 23.1268016 | 23.0720579 | 23.1730877 | 23.1101528 | 0.016538792 | 0.668237209 |
| 22.6028549 | 22.5828765 | 22.678922 | 22.6270473 | 0.036441036 | 0.380934421 |
| 20.4181895 | 20.359 | 20.4195207 | 20.4668412 | -0.027388217 | 0.508704448 |
| 22.186436 | 22.1578174 | 22.2036966 | 22.1657381 | 0.026179123 | 0.552723891 |
| 23.4154331 | 23.3419126 | 23.468359 | 23.4086984 | 0.021803072 | 0.653687556 |
| 20.6048086 | 20.4958538 | 20.6228175 | 20.4942625 | -0.007128759 | 0.887595253 |
| 22.4354653 | 22.3390018 | 22.4758037 | 22.4159194 | 0.012235937 | 0.810978319 |
| 21.9931458 | 21.8476548 | 21.9998498 | 21.9414785 | 0.001907349 | 0.971631085 |
| 19.8963081 | 19.8031792 | 19.887249 | 19.9702922 | 0.028359247 | 0.599770555 |
| 20.6763244 | 20.5912842 | 20.6921459 | 20.653136 | 0.021720916 | 0.690829139 |
| 24.8036507 | 24.6981729 | 24.7535062 | 24.7787838 | 0.022405747 | 0.682441047 |
| 19.733481 | 19.6262068 | 19.7169617 | 19.7602433 | 0.0195444 | 0.724373575 |
| 22.6180031 | 22.5304762 | 22.5817701 | 22.6580046 | 0.041324995 | 0.46239428 |
| 20.2356362 | 20.1604325 | 20.298325 | 20.2919925 | 0.010631736 | 0.851803951 |
| 20.2881583 | 20.2586195 | 20.4018057 | 20.3102236 | 0.061079437 | 0.301896772 |
| 22.9047593 | 22.7606625 | 22.9313291 | 22.9347545 | -0.009704654 | 0.874361282 |
| 22.8022541 | 22.784388 | 22.8966634 | 22.8479039 | 0.020855765 | 0.735512304 |
| 21.5190174 | 21.4172586 | 21.5066328 | 21.5805448 | 0.030301965 | 0.634019839 |
| 21.9554923 | 21.8846362 | 22.0881045 | 21.9338067 | -0.014551578 | 0.820218544 |
| 22.0931638 | 21.9185854 | 22.0255651 | 22.1277416 | 0.037510639 | 0.593407718 |
| 18.7579821 | 18.7429744 | 18.8267828 | 18.8248244 | 0.003451624 | 0.961365148 |
| 23.115025 | 23.1715078 | 23.1142595 | 23.1425977 | 0.118830027 | 0.109333528 |
| 23.3729361 | 23.1883819 | 23.303465 | 23.2653412 | -0.038531297 | 0.598030635 |
| 17.1391423 | 17.2297122 | 17.2886062 | 17.1386454 | 0.075909452 | 0.316083898 |
| 22.648583 | 22.6426404 | 22.5135481 | 22.7105788 | 0.019224664 | 0.804563423 |
| 17.5820663 | 17.5060687 | 17.5912915 | 17.6591423 | -0.005912197 | 0.939277317 |
| 20.0546175 | 19.9040033 | 20.0337706 | 20.0134381 | -0.033913332 | 0.669718351 |

| | | | | | |
|---|---|---|---|---|---|
| 17.9448973 | 18.13059 | 18.1885876 | 17.9984105 | 0.086621849 | 0.296857903 |
| 18.3723407 | 18.2416157 | 18.3675612 | 18.3951697 | -0.013246149 | 0.873828223 |
| 18.9151668 | 19.1164015 | 18.9308092 | 19.1461643 | 0.098512392 | 0.243393964 |
| 19.0662471 | 18.9928895 | 18.8520129 | 19.0220609 | 0.077261567 | 0.3795019 |
| 19.3295688 | 19.480306 | 19.6452141 | 19.3948248 | 0.170320579 | 0.064316166 |
| 20.0129725 | 19.7893366 | 20.0050831 | 19.9369293 | -0.03443142 | 0.719052777 |
| 19.6510693 | 19.8474549 | 19.7651584 | 19.9047209 | 0.101713267 | 0.300861297 |
| 19.2060539 | 19.1110831 | 19.2738408 | 19.168512 | 0.022416099 | 0.819067899 |
| 18.1037177 | 17.8319304 | 18.1012481 | 18.2184019 | -0.046434389 | 0.657024566 |
| 24.2853833 | 24.2617238 | 24.2957082 | 24.2383503 | 0.036905039 | 0.722776438 |
| 16.3204311 | 16.2221866 | 16.1622585 | 16.4830932 | 0.01932526 | 0.854118633 |
| 19.8431048 | 19.6854157 | 19.9464673 | 19.8421774 | 0.00859816 | 0.944373182 |
| 17.1065074 | 16.9307253 | 17.4448359 | 17.1638196 | 0.178478517 | 0.152565573 |
| 20.9326985 | 20.6968503 | 20.893905 | 20.9215473 | -0.029306889 | 0.81411221 |
| 19.165722 | 19.3464689 | 19.1557193 | 19.1952481 | 0.061727682 | 0.660944979 |
| 18.4619663 | 18.1357283 | 18.1236091 | 18.253015 | -0.194004186 | 0.190049562 |
| 16.6374519 | NA | 17.0670182 | 17.1364105 | 0.12448631 | 0.454729724 |
| NA | 14.6518941 | NA | NA | 0.205639476 | 0.236338409 |
| 14.8239851 | 14.9621279 | 14.7519227 | 14.9787132 | -0.124392317 | 0.510688707 |
| NA | 17.6140581 | NA | NA | -0.098778789 | 0.722076348 |
| 15.9581569 | 15.6274515 | 16.2424475 | 15.9711129 | 0.190192299 | 0.530242542 |
| 18.3060688 | 18.0723027 | 18.5073216 | 18.3009893 | -0.031388589 | 0.72991244 |
| 21.8055738 | 21.466847 | 21.6432809 | 21.6640109 | -0.154466838 | 0.029614354 |
| 23.6064691 | 23.5397528 | 23.6223939 | 23.5911322 | 0.057542294 | 0.322993716 |
| 21.1034778 | 21.0903929 | 20.989371 | 21.0987176 | 0.028016537 | 0.746458034 |
| 16.3579072 | NA | 16.3795583 | 16.2775304 | -0.085535041 | 0.437808551 |
| 16.326138 | 16.3578307 | 16.2694371 | 16.5133106 | -0.055533106 | 0.609108177 |
| 22.461436 | 22.4361139 | 22.5147497 | 22.5367674 | 0.023950679 | 0.616121845 |
| 22.662317 | 22.7630204 | 22.8222194 | 22.5223707 | 0.032784309 | 0.68014862 |
| 16.5477291 | 16.4684143 | 16.5543374 | 16.3474237 | -0.019497486 | 0.885579309 |
| 18.2455712 | 18.580592 | 18.7553225 | 18.1484287 | 0.144190498 | 0.487481672 |
| 18.7793941 | 18.6286981 | 18.7639755 | 18.6857369 | -0.056871076 | 0.391806567 |
| 19.9190533 | 20.0445885 | 19.8758965 | 19.7841094 | -0.040930193 | 0.603459739 |
| 25.922088 | 25.7446575 | 25.9379057 | 25.9060139 | 0.001214712 | 0.985250912 |
| 22.9459221 | 22.8440191 | 22.991274 | 22.9551372 | 0.028956167 | 0.568588393 |
| 20.5610943 | 20.4451777 | 20.5915284 | 20.4872276 | -0.058078864 | 0.251119172 |
| 21.4681992 | 21.0334796 | 21.4614171 | 21.5865721 | -0.007876566 | 0.965911331 |
| 22.8102158 | 22.7403755 | 22.780327 | 22.8000735 | 0.046273703 | 0.384873508 |
| 18.8405456 | 18.5741694 | 18.6450047 | 18.7544352 | -0.200650931 | 0.017793366 |
| 20.0704044 | 19.9698394 | 19.9832355 | 19.9836535 | -0.018676133 | 0.739269607 |
| 17.8781628 | 17.8722864 | 17.8244023 | 17.908214 | -0.029240922 | 0.598514306 |
| 19.9540508 | 19.7958663 | 19.8452607 | 19.9012986 | -0.065781321 | 0.201134701 |

| | | | | | |
|---|---|---|---|---|---|
| 19.3760635 | 19.2732533 | 19.4964265 | 19.3605385 | 0.072449308 | 0.219057507 |
| 21.1068564 | 20.9424796 | 21.0279255 | 21.1394085 | -0.042735844 | 0.485549409 |
| 19.6207369 | 19.7181173 | 19.7237534 | 19.6798491 | 0.057848223 | 0.356094617 |
| 13.8181148 | NA | 14.0303896 | 13.7580594 | -0.077002713 | 0.638155454 |
| 19.8021669 | 19.6869365 | 19.8154002 | 19.9173832 | 0.035107345 | 0.671085531 |
| 19.3228056 | 19.3624787 | 19.2403234 | 19.163258 | -0.086444883 | 0.159096067 |
| 22.6249545 | 22.5243666 | 22.6100061 | 22.5308556 | -0.032647624 | 0.4960072 |
| 15.3863026 | NA | 15.4240624 | 15.5128466 | 0.393103423 | 0.04411753 |
| 16.3892615 | 16.6778224 | 17.1813733 | 16.9299294 | 0.381487457 | 0.031960078 |
| 18.9571346 | 19.107001 | 19.0038217 | 18.9876564 | 0.094270128 | 0.195435408 |
| 20.677618 | 20.5671857 | 20.6010119 | 20.6243193 | -0.027252898 | 0.449440619 |
| 23.9296155 | 23.9111091 | 23.9885731 | 23.9479279 | 0.027903989 | 0.507676202 |
| 20.6502326 | 20.5155316 | 20.6500189 | 20.6817522 | -0.003499623 | 0.961938427 |
| 12.8021446 | NA | 13.2493889 | 13.428958 | 0.190194297 | 0.764291864 |
| 18.8078099 | 18.7197765 | 19.0680074 | 18.5059983 | -0.059523559 | 0.679043842 |
| 20.1105046 | 19.8702603 | 20.0506628 | 20.0544865 | -0.076024138 | 0.319477815 |
| 20.3962119 | 20.2003109 | 20.3190213 | 20.3117999 | -0.107493493 | 0.083631783 |
| 25.2915141 | 25.2033102 | 25.3267087 | 25.2391473 | 0.004550865 | 0.934801683 |
| 21.6477336 | 21.5437509 | 21.6650176 | 21.7040777 | 8.10E-04 | 0.987486485 |
| 24.2264837 | 24.084862 | 24.2508182 | 24.2582114 | 0.005335415 | 0.910628099 |
| 24.1841082 | 24.1831262 | 24.2543983 | 24.195662 | 0.0267294 | 0.514072974 |
| 19.7988379 | 19.5583944 | 19.5366997 | 19.5824496 | -0.122071204 | 0.123963469 |
| 21.2799156 | 21.12471 | 21.3191957 | 21.290926 | -0.016778407 | 0.870076412 |
| 17.6104531 | 17.4193614 | 17.4506582 | 17.532343 | -0.116098693 | 0.175450017 |
| 22.4301771 | 22.4006968 | 22.341146 | 22.3338628 | -0.009437082 | 0.833230126 |
| 21.0308889 | 21.1306 | 21.2728776 | 21.223127 | 0.091932681 | 0.397632795 |
| 22.5843702 | 22.5650775 | 22.5049466 | 22.5383371 | -0.027439481 | 0.624841961 |
| 18.3693009 | 18.4528425 | 18.53863 | 18.4458545 | 0.002657337 | 0.973971441 |
| 21.3289907 | 21.2288051 | 21.1738897 | 21.2320608 | -0.048539415 | 0.361356891 |
| 21.3026427 | 21.42455 | 21.4026073 | 21.3802313 | 0.009905512 | 0.934939229 |
| 18.71024 | 18.4930199 | 18.3765248 | 18.3609658 | -0.213107447 | 0.035014315 |
| 19.9317874 | 19.7737043 | 19.9600125 | 19.8270907 | -0.035010252 | 0.599562653 |
| 16.7446945 | 16.9912425 | 16.8737432 | 16.6766862 | 0.11702625 | 0.411154055 |
| 21.5769901 | 21.5610684 | 21.5435919 | 21.5655564 | -0.010447708 | 0.876953781 |
| 19.851 | 19.706704 | 19.8405872 | 19.8770458 | 0.00603176 | 0.921264725 |
| 16.2108237 | 16.1817762 | 15.9711618 | 15.8753703 | -0.124217463 | 0.379873229 |
| 21.1309175 | 21.1948122 | 21.0968178 | 20.8468125 | -0.088950508 | 0.375869929 |
| 16.6738254 | 16.3510349 | 16.6054629 | 16.4445113 | -0.239670131 | 0.18828986 |
| 17.9084861 | 17.6225348 | 17.4621976 | 17.5105231 | -0.228560341 | 0.151476954 |
| 17.5148344 | 17.4112293 | 17.6806888 | 17.2699719 | 0.010040302 | 0.940152935 |
| 20.6727523 | 20.5662024 | 20.4828817 | 20.5686114 | -0.120269425 | 0.217605467 |
| 17.7778791 | 17.8849958 | 17.8912039 | 17.8152581 | 0.138560581 | 0.104570905 |

| | | | | | |
|---|---|---|---|---|---|
| 18.4668531 | 18.4339859 | 18.311073 | 18.465653 | -0.052733012 | 0.498337724 |
| 17.4582266 | 17.4904864 | 17.426011 | 17.6092514 | 0.022022464 | 0.761838597 |
| 25.3436869 | 25.2098891 | 25.2566467 | 25.2669585 | -0.037810143 | 0.590945154 |
| 19.8943792 | 19.6290471 | 19.7494241 | 19.8931348 | 0.006349461 | 0.922324787 |
| 18.5119698 | 18.5135383 | 18.5117915 | 18.6831803 | 0.058658223 | 0.360923876 |
| 21.2045165 | 21.1974995 | 21.2092757 | 21.179599 | -0.01343297 | 0.817647186 |
| 22.7084917 | 22.536648 | 22.6631735 | 22.7010935 | -0.01885995 | 0.740712238 |
| 20.9330665 | 20.7772337 | 20.7989271 | 20.8536411 | -0.059738522 | 0.288408932 |
| 23.9068905 | 23.7819122 | 23.8381381 | 23.8030549 | -0.1048132 | 0.053115106 |
| 21.664564 | 21.6005511 | 21.6361619 | 21.6521755 | -0.010710246 | 0.799214775 |
| 23.413689 | 23.4015488 | 23.4192421 | 23.3774326 | 0.011703211 | 0.741731856 |
| 22.097448 | 22.0976138 | 22.1455083 | 22.1524595 | 0.031173691 | 0.437581221 |
| 22.8029247 | 22.746061 | 22.7868991 | 22.7984403 | 0.031000916 | 0.552823567 |
| 20.3600297 | 20.2552329 | 20.3833171 | 20.4188637 | 0.050333017 | 0.381900057 |
| 19.6431372 | 19.5571419 | 19.7236717 | 19.5933925 | 0.014071538 | 0.844715353 |
| 18.8499479 | 18.959183 | 18.8450999 | 18.8944141 | 0.061205367 | 0.41889198 |
| 18.0512971 | 17.8588379 | 18.0436333 | 18.087212 | 0.044080527 | 0.586306861 |
| 21.2769705 | 21.0669078 | 21.2629812 | 21.1940418 | -0.023894926 | 0.777704689 |
| 18.3341041 | 18.232957 | 18.2524944 | 18.2388746 | -0.075561024 | 0.413054984 |
| 14.4648563 | 14.7210841 | 14.5198572 | 14.8246534 | 0.204986174 | 0.165410419 |
| 20.7201526 | 20.7330442 | 20.7861858 | 20.7761032 | 0.05492067 | 0.229762778 |
| 16.9497751 | 16.9184762 | 16.9685896 | 16.9076844 | -0.051198411 | 0.476612513 |
| 21.2384412 | 21.4442047 | 21.4443995 | 21.4436854 | 0.267812532 | 0.001832897 |
| 23.3041664 | 23.1663997 | 23.2866587 | 23.2906484 | -0.030456549 | 0.607622052 |
| 18.0737758 | 17.9155208 | 18.0422621 | 18.0128718 | -0.050701842 | 0.575282575 |
| 18.9021285 | 18.9829438 | 18.9590231 | 18.9238299 | 0.058881977 | 0.391976925 |
| 20.6169645 | 20.4613838 | 20.4595673 | 20.5488709 | -0.077720875 | 0.256369783 |
| 20.3850764 | 20.3135321 | 20.3259614 | 20.3553727 | -0.003113319 | 0.954580948 |
| 18.6505113 | 18.7901951 | 18.7438568 | 18.812674 | 0.038768988 | 0.586274091 |
| NA | 19.3419763 | NA | NA | 0.306349565 | 0.175270957 |
| 23.5119789 | 23.3863172 | 23.4614094 | 23.5133627 | -0.028272789 | 0.629267918 |
| 19.2495263 | 19.0471896 | 18.9682204 | 19.0502419 | -0.169743705 | 0.207134383 |
| 18.4874664 | 18.2548864 | 17.7645303 | 18.3725016 | 0.175202376 | 0.642213878 |
| 19.4178623 | 18.9099103 | 19.3909861 | 19.5162691 | -0.028694551 | 0.858306181 |
| 18.0919572 | 18.1247263 | 18.1631631 | 18.3288155 | 0.142137588 | 0.167319478 |
| 23.7919637 | 23.3793341 | 23.6914082 | 23.7267226 | 0.041358239 | 0.798516257 |
| 18.9602934 | 18.9324418 | 19.1361085 | 19.0846184 | 0.085213329 | 0.3563515 |
| 16.727425 | 16.7537181 | 16.5893582 | 16.7635285 | -0.115722039 | 0.144026671 |
| 18.7811868 | 18.8361108 | 18.8040848 | 18.8567613 | 0.017160917 | 0.807921558 |
| 14.6536096 | 14.9226364 | 14.8373045 | 14.8133536 | 0.200938285 | 0.048523923 |
| 22.945934 | 22.7775187 | 22.9322937 | 22.8488829 | -0.0378035 | 0.628452258 |
| 18.4138356 | 18.5727503 | 18.6884085 | 18.6659469 | 0.143513674 | 0.10905838 |

| | | | | | |
|---|---|---|---|---|---|
| 20.206704 | 20.7568157 | 20.207759 | 20.2046658 | 0.087530959 | 0.458758297 |
| 18.370057 | 18.3766526 | 18.2757403 | 18.3553391 | -0.125265714 | 0.212034405 |
| 23.1676099 | 23.1325763 | 23.1487925 | 23.1851112 | 0.013797701 | 0.773905483 |
| 22.1601786 | 22.2020798 | 22.212502 | 22.1344215 | 0.097579703 | 0.211750279 |
| 19.1232873 | 19.3622759 | 19.2826389 | 19.3700652 | 0.183032679 | 0.007100216 |
| 20.4761082 | 20.4464169 | 20.5531174 | 20.6210169 | 0.152106645 | 0.039943732 |
| 20.4874787 | 20.3151036 | 20.594597 | 20.5345876 | -0.047465578 | 0.528077499 |
| 17.698079 | 17.8013959 | 17.630744 | 17.8362926 | -0.014754097 | 0.88994824 |
| 20.8173737 | 20.7498326 | 20.7979084 | 20.7799835 | -5.84E-04 | 0.991394073 |
| 20.4132494 | 20.3940021 | 20.4675478 | 20.4941251 | 0.090437456 | 0.08716324 |
| 18.4774686 | 18.6335931 | 18.5488097 | 18.5972288 | 0.118134171 | 0.086104904 |
| 19.3504981 | 19.3693969 | 19.126663 | 19.3539207 | 0.004156143 | 0.959785364 |
| 24.4713432 | 24.4162389 | 24.4681311 | 24.4900303 | 0.073167961 | 0.397091067 |
| 20.531518 | 20.3072712 | 20.5584073 | 20.4571173 | 0.021245406 | 0.808061658 |
| NA | 15.4106509 | NA | NA | 0.159763212 | 0.182589802 |
| 18.3851867 | 17.7917837 | 18.5046486 | 18.1844236 | -0.095747057 | 0.527460099 |
| 18.4194868 | 18.2321631 | 18.4279669 | 18.2989162 | -0.100980705 | 0.388196806 |
| 20.2959954 | 19.9086141 | 20.0225382 | 20.4574897 | -0.036901888 | 0.748084913 |
| 19.7380911 | 19.6099505 | 19.6167022 | 19.6394317 | -0.168060539 | 0.023532939 |
| 22.4805311 | 22.4789738 | 22.5428086 | 22.5096577 | 0.050956414 | 0.383651329 |
| 17.7645238 | 17.5960369 | 17.6089794 | 17.708192 | -0.087215308 | 0.30240969 |
| 18.4092802 | 18.4191832 | 18.3949855 | 18.4056742 | -0.050526393 | 0.589856645 |
| 18.6951932 | 18.5746289 | 18.488019 | 18.7236821 | -0.066683827 | 0.497815282 |
| 16.1974964 | NA | 16.3113865 | 16.3900539 | 0.130684434 | 0.315172751 |
| 19.565916 | 19.4409272 | 19.5740692 | 19.5070834 | -0.019521994 | 0.636599827 |
| 21.8125651 | 21.6455087 | 21.792858 | 21.795366 | -0.027718832 | 0.643186867 |
| 21.6593095 | 21.5802053 | 21.6033415 | 21.6735743 | -0.010461239 | 0.816133942 |
| 12.1972272 | 12.0651397 | 12.4622604 | 12.0584658 | -0.01220997 | 0.949583669 |
| 19.0304457 | 18.8971458 | 19.1475065 | 18.9059893 | 0.029870572 | 0.678412473 |
| 16.1840381 | 16.0117299 | 16.2442007 | 16.4526694 | -0.103067854 | 0.639014959 |
| NA | 21.1991534 | NA | NA | 0.150391285 | 0.407390436 |
| 14.1278749 | NA | 14.5859454 | 14.537966 | 0.366890065 | 0.030101206 |
| 16.5903033 | 16.4050586 | 16.5860006 | 16.512319 | -0.032928115 | 0.741791567 |
| 19.4162786 | 19.4051575 | 19.5345578 | 19.6004513 | 0.179462918 | 0.074359192 |
| 20.3458418 | 20.3099023 | 20.3128175 | 20.2839586 | -0.087343213 | 0.171118 |
| 18.8999382 | 18.8631779 | 18.8648536 | 18.9771789 | -0.061524881 | 0.25393434 |
| 20.0324551 | 19.7787523 | 19.9599444 | 20.0387597 | -0.050329799 | 0.343380078 |
| 21.3682959 | 21.2854904 | 21.2770802 | 21.3499965 | 0.003890629 | 0.940962799 |
| 23.4785984 | 23.415345 | 23.4209197 | 23.4690643 | -7.37E-04 | 0.987646024 |
| 20.4101931 | 20.3835582 | 20.3271352 | 20.4438911 | -0.00836157 | 0.845041038 |
| 21.6308944 | 21.638051 | 21.695306 | 21.6992849 | 0.052814486 | 0.261146002 |
| 19.8479159 | 19.8311756 | 19.9387099 | 19.914677 | 0.062617534 | 0.230127585 |

| | | | | | |
|---|---|---|---|---|---|
| 20.9323075 | 20.8725022 | 20.9308609 | 20.9282978 | -0.002162944 | 0.967027538 |
| 21.4055636 | 21.2871605 | 21.335018 | 21.3790133 | -0.015905064 | 0.777418445 |
| 20.8608953 | 20.8245966 | 20.7583071 | 20.7715422 | -0.026728393 | 0.664378282 |
| 20.8440767 | 20.7136798 | 20.8699488 | 20.871113 | 0.041859174 | 0.516920504 |
| 18.1704184 | 18.2016369 | 18.1891778 | 18.2580118 | 0.019275677 | 0.790521864 |
| 17.4954899 | 17.3667465 | 17.3291591 | 17.5261331 | -0.048556958 | 0.56376973 |
| 16.8865401 | 17.0118384 | 17.0842174 | 17.050219 | 0.084277031 | 0.406852983 |
| 16.218696 | 15.8925973 | 15.9939348 | 16.2113104 | -0.004103541 | 0.975768093 |
| NA | 16.499581 | NA | NA | -0.178312005 | 0.398168542 |
| 23.4309251 | 23.3916078 | 23.4210325 | 23.4648762 | 0.038225759 | 0.518404715 |
| 20.1371022 | 19.9579768 | 20.1141574 | 20.0138539 | -0.079500657 | 0.147084637 |
| 19.7674094 | 19.6512016 | 19.6952382 | 19.7806299 | 0.018012399 | 0.738884613 |
| 20.0698339 | 20.1169581 | 20.1623097 | 20.1751709 | 0.039411899 | 0.593109053 |
| 20.405539 | 20.3146183 | 20.2018615 | 20.3439283 | -0.012881744 | 0.914630075 |
| 23.0846466 | 23.0561401 | 23.1286405 | 23.0223627 | 0.031231307 | 0.504046555 |
| 21.6536436 | 21.6366955 | 21.6632826 | 21.5897875 | -5.63E-04 | 0.993742472 |
| 25.3620704 | 25.3262127 | 25.3462265 | 25.3835115 | 0.037615172 | 0.526927507 |
| 16.6702095 | 16.6973392 | 16.4093472 | 16.6462241 | -0.03938018 | 0.713386571 |
| 19.5545614 | 19.4330199 | 19.6678544 | 19.596408 | 0.024472457 | 0.72834555 |
| 20.7185466 | 20.7393325 | 20.8065343 | 20.7783148 | 0.040985487 | 0.447189778 |
| 18.5745895 | 18.6518643 | 18.473843 | 18.5851017 | 0.077686012 | 0.582753748 |
| NA | 17.3317015 | NA | NA | -0.298177324 | 0.236491357 |
| 15.333045 | NA | 15.2972893 | 15.1287802 | -0.053362846 | 0.709766842 |
| 15.670159 | 15.8229108 | 15.7031918 | 15.8171047 | -0.048941843 | 0.716275135 |
| 18.7992682 | 18.4764956 | 18.570527 | 18.8491673 | 0.020024287 | 0.879486833 |
| 18.6477782 | 18.5165195 | 18.6709501 | 18.7331007 | 0.015714901 | 0.893061827 |
| 18.6743149 | 18.84325 | 18.8581957 | 18.6905785 | 0.050395355 | 0.663327033 |
| 24.2394325 | 24.2907532 | 24.3071626 | 24.4433141 | 0.14888072 | 0.090303974 |
| 17.420265 | 17.1874388 | 17.1622208 | 17.2169835 | -0.131396212 | 0.133985812 |
| 19.0248857 | 18.9372698 | 19.107177 | 19.0623268 | -0.042871895 | 0.604153996 |
| 17.2385592 | 17.3893023 | 17.3767147 | 17.4916822 | 0.060356093 | 0.464846741 |
| 19.1840845 | 19.2025226 | 19.2521406 | 19.3452988 | 0.044098001 | 0.579037412 |
| 20.8642813 | 20.9316784 | 20.6726427 | 20.7816729 | 0.045525949 | 0.564804668 |
| 17.5064757 | 17.3336767 | 17.5815087 | 17.5534111 | 0.012466169 | 0.871793123 |
| 20.0946662 | 20.0423893 | 20.1316671 | 20.0492236 | 0.083466589 | 0.283858483 |
| 19.9555987 | 19.8514718 | 19.7488919 | 19.8825404 | -0.053381748 | 0.488514324 |
| 19.9693664 | 19.780617 | 19.8717528 | 19.8699826 | -0.038867898 | 0.608913811 |
| 17.9989257 | 17.9424821 | 18.0248192 | 17.7985718 | -0.010975408 | 0.87207504 |
| 16.8946479 | 16.8834702 | 16.8422053 | 16.7818773 | 0.02507464 | 0.698269802 |
| 18.2371333 | 18.1860723 | 18.3448389 | 18.3763971 | 0.058240341 | 0.344068787 |
| 23.199094 | 23.1430686 | 23.1975846 | 23.2578948 | 0.032243151 | 0.595757169 |
| 23.7662408 | 23.619399 | 23.7087277 | 23.6753761 | -0.044333678 | 0.433916549 |

| | | | | | |
|---|---|---|---|---|---|
| 22.8445948 | 22.7463709 | 22.7932219 | 22.8224256 | -0.011708395 | 0.835483724 |
| 20.0673305 | 19.8781886 | 19.956552 | 19.9365985 | -0.087016349 | 0.120177616 |
| 23.8476724 | 23.7861434 | 23.8299612 | 23.7995437 | -0.009432764 | 0.856473753 |
| 20.4099095 | 20.2295143 | 20.345091 | 20.2971723 | -0.082907043 | 0.123031855 |
| 22.1398431 | 22.0814252 | 22.1792609 | 22.1115798 | -0.021554093 | 0.667036524 |
| 23.9425291 | 23.8692089 | 23.9410317 | 23.9382126 | 0.007977988 | 0.872856504 |
| 26.1571028 | 25.9837997 | 26.1413473 | 26.1103248 | -0.029808714 | 0.547250066 |
| 20.1693481 | 20.1699728 | 20.1581217 | 20.2431083 | 0.025041776 | 0.603907384 |
| 26.1924631 | 26.1309129 | 26.1777442 | 26.2009705 | 0.032426067 | 0.491760323 |
| 23.5303246 | 23.5145094 | 23.5554872 | 23.5503842 | 0.027504799 | 0.494817893 |
| 22.9776361 | 22.9014387 | 22.9579209 | 22.9332077 | 0.010608923 | 0.785678685 |
| 23.4582125 | 23.471527 | 23.4719927 | 23.4698805 | 0.040681742 | 0.274339127 |
| 20.6894409 | 20.7367096 | 20.7316785 | 20.7211936 | 0.031293057 | 0.467465536 |
| 20.8732322 | 20.9444893 | 20.9749737 | 20.9707574 | 0.080685817 | 0.07438081 |
| 22.4751695 | 22.3577848 | 22.4893317 | 22.5527675 | -9.09E-04 | 0.984531024 |
| 19.3697759 | 19.3131719 | 19.3486154 | 19.3505712 | -0.004202826 | 0.929109913 |
| 21.14886 | 21.0419315 | 21.1081868 | 21.1144492 | -0.034488718 | 0.480706925 |
| 20.9342868 | 20.8324739 | 20.8912555 | 20.8441831 | -0.058741872 | 0.234530968 |
| 18.8876397 | 18.8633336 | 18.9371694 | 18.8168222 | 0.013294944 | 0.796756562 |
| 20.000145 | 19.9515227 | 20.1823691 | 20.0124782 | 0.063992463 | 0.233224567 |
| 20.5149409 | 20.4301608 | 20.5205686 | 20.5629525 | 0.003884917 | 0.94276758 |
| 24.219359 | 24.1485404 | 24.3007771 | 24.2637091 | 0.061251252 | 0.271042693 |
| 22.2336568 | 22.1805341 | 22.2329556 | 22.2874552 | 0.014768995 | 0.796957813 |
| 25.2259908 | 25.131031 | 25.2725083 | 25.2223643 | 0.019603009 | 0.73881579 |
| 21.3152013 | 21.1261421 | 21.2910219 | 21.2678826 | -0.062440722 | 0.294212475 |
| 20.5085791 | 20.4010304 | 20.5977615 | 20.5316604 | 0.034017409 | 0.563048745 |
| 23.4716673 | 23.4302781 | 23.5975121 | 23.5151775 | 0.051762035 | 0.389553809 |
| 20.2525948 | 20.1126648 | 20.2463778 | 20.2941319 | 0.033824343 | 0.579545765 |
| 22.0671647 | 21.9795564 | 22.13087 | 22.1022725 | 0.045523147 | 0.457376835 |
| 18.4966284 | 18.3950045 | 18.4994474 | 18.5373854 | -0.052037625 | 0.401476164 |
| 18.898237 | 18.7823451 | 19.016074 | 18.7989485 | -0.013924314 | 0.831486016 |
| 19.1344868 | 18.9749974 | 19.1796842 | 19.1632467 | -0.00312521 | 0.962554531 |
| 21.0952439 | 21.0203921 | 20.9641562 | 21.1691686 | 0.017685927 | 0.811187514 |
| 17.5518174 | 17.6731328 | 17.5581816 | 17.6255102 | 0.087758223 | 0.299783692 |
| 19.2953813 | 19.3150087 | 19.3420661 | 19.4360092 | 0.103189412 | 0.239548016 |
| 19.3738311 | 19.2649389 | 19.4223201 | 19.4978419 | -0.029679479 | 0.737932631 |
| 18.2332631 | 18.1004571 | 18.2551819 | 18.1985937 | 0.097334292 | 0.279017593 |
| 18.2512454 | 18.2091939 | 18.0987324 | 18.2358182 | 0.059951223 | 0.519408175 |
| 17.8467552 | 17.6542339 | 17.7560882 | 17.9576494 | -0.003884163 | 0.967730834 |
| 19.8868773 | 19.7417438 | 19.9143014 | 19.9314675 | 0.062286198 | 0.536251023 |
| 18.5672929 | 18.6695615 | 18.6431411 | 18.403219 | 0.008232365 | 0.935219673 |
| 19.5182643 | 19.6382018 | 19.5005175 | 19.6965758 | 0.137631095 | 0.188540657 |

| | | | | | |
|---|---|---|---|---|---|
| 17.7530352 | 17.6633113 | 17.6667856 | 17.8674057 | 0.066969669 | 0.534148077 |
| 24.5167053 | 24.4340519 | 24.3795347 | 24.481194 | 0.030680753 | 0.778197339 |
| 16.8801828 | 16.9094667 | 16.8521418 | 16.8964555 | 0.084803283 | 0.438714779 |
| 18.2024822 | 18.2223686 | 18.0524799 | 18.5531669 | -0.010610388 | 0.926064778 |
| 19.2707278 | 19.3761568 | 19.455702 | 19.5859468 | 0.251368602 | 0.062293975 |
| 23.8637626 | 23.6707749 | 23.8940882 | 23.7875527 | 0.006494533 | 0.960065462 |
| 17.1157909 | 17.1145136 | 16.7806496 | 17.2985569 | 0.110671979 | 0.456458805 |
| 21.5090639 | 21.5347051 | 21.3267883 | 21.4691189 | 0.109121031 | 0.473253232 |
| NA | 14.9581515 | NA | NA | 0.05402911 | 0.759489491 |
| 15.7861135 | 16.3797708 | 15.5150319 | 15.501679 | -0.131868783 | 0.557268769 |
| NA | 15.6875878 | NA | NA | 0.039427094 | 0.866404605 |
| 16.0166299 | 15.8039609 | 16.0044907 | 15.6583539 | -0.107919607 | 0.434729689 |
| 18.8913365 | 19.1044197 | 19.1634983 | 18.9670204 | 0.108560456 | 0.296011422 |
| 17.5071783 | 17.8359539 | 17.7581595 | 17.7915883 | 0.07592083 | 0.433811481 |
| 17.6240925 | 17.6124151 | 17.7326899 | 17.5095541 | -0.003271311 | 0.97200943 |
| 20.781391 | 20.7789327 | 20.6534797 | 20.9005376 | -0.037413897 | 0.673298604 |
| 18.2952598 | 18.0698675 | 18.2596099 | 18.3123451 | -0.052757442 | 0.529275006 |
| 18.6257808 | 18.4551203 | 18.3710563 | 18.650615 | -0.008582593 | 0.915628743 |
| 22.5697481 | 22.3532086 | 22.6127766 | 22.6360219 | 0.027116024 | 0.726231804 |
| 18.8430859 | 18.8208117 | 18.8298262 | 18.9243413 | 0.012258318 | 0.873849559 |
| 19.9302087 | 19.8183179 | 19.9128333 | 19.9423832 | -0.031482744 | 0.649763888 |
| 23.0787376 | 22.9690431 | 23.0125456 | 23.0508725 | 0.025755333 | 0.632402579 |
| 23.4323102 | 23.3642006 | 23.3907019 | 23.3919878 | -0.005287157 | 0.912510761 |
| 22.3351863 | 22.2388786 | 22.4299253 | 22.369298 | 0.039059649 | 0.410158487 |
| 20.5226358 | 20.4859311 | 20.5972386 | 20.6015805 | 0.01445737 | 0.774932218 |
| 21.6015976 | 21.6744251 | 21.5759343 | 21.6888051 | 0.033636384 | 0.550067648 |
| 19.01962 | 19.003766 | 19.0413108 | 19.069086 | 0.046117076 | 0.4431061 |
| 19.8515716 | 19.730106 | 19.6687664 | 19.7726353 | -0.034687618 | 0.567466733 |
| 21.6054787 | 21.5855332 | 21.6714364 | 21.5654512 | 0.004064035 | 0.94659583 |
| 23.2020064 | 23.0868092 | 23.1803008 | 23.1746932 | -0.005409505 | 0.929739553 |
| 19.5765378 | 19.4981789 | 19.3467051 | 19.499104 | -0.076918609 | 0.250105976 |
| 21.545269 | 21.4353803 | 21.6442452 | 21.5231393 | 0.008656615 | 0.89644041 |
| 21.0259681 | 20.9143212 | 21.0113935 | 21.0420045 | 0.002011013 | 0.975983137 |
| 20.0327075 | 19.807066 | 20.033961 | 20.0375659 | 0.015796954 | 0.814561628 |
| 19.274995 | 19.3157264 | 19.3387498 | 19.3811199 | 0.152073559 | 0.032835195 |
| 20.5070356 | 20.4623604 | 20.4447522 | 20.5583551 | 0.012911906 | 0.857442243 |
| 19.5975991 | 19.4031351 | 19.3808551 | 19.5722541 | 0.002866257 | 0.969372292 |
| 17.5237033 | 17.4427044 | 17.5347909 | 17.6208977 | 0.017903491 | 0.844530917 |
| 25.0389071 | 24.8120105 | 24.8527067 | 25.0157363 | 0.044646594 | 0.640769227 |
| 15.6669766 | 15.7031511 | 15.7061045 | 15.7657114 | -0.034622332 | 0.731313913 |
| 22.1102972 | 21.7876109 | 21.8738616 | 22.040088 | -0.086278266 | 0.583151228 |
| 16.0946102 | 16.5099355 | 16.4462164 | 16.9324623 | 0.271770544 | 0.225103587 |

| | | | | | |
|---|---|---|---|---|---|
| 20.0131303 | 19.7431345 | 19.8737025 | 19.897887 | -0.239256696 | 0.183382204 |
| 18.0169862 | 18.1707139 | 18.070179 | 17.9543266 | -0.104183591 | 0.303663122 |
| 20.9548023 | 20.8447622 | 20.9396843 | 20.8905295 | 0.006991659 | 0.898491521 |
| NA | 14.9660246 | NA | NA | -0.740549893 | 0.028342879 |
| 18.8437726 | 18.7650497 | 18.7093085 | 18.6566021 | -0.119078084 | 0.305879352 |
| 16.9004592 | 16.6547973 | 16.6898839 | 16.9314338 | -0.081082952 | 0.389884663 |
| 19.9763206 | 19.8885685 | 19.8279523 | 19.7896026 | -0.006016229 | 0.936413749 |
| 20.6695206 | 20.734776 | 20.6563276 | 20.7017003 | -0.006456073 | 0.929691867 |
| 27.3695516 | 27.2754496 | 27.3721086 | 27.3615314 | 0.046774379 | 0.489680766 |
| 19.7694085 | 19.6252877 | 19.7417823 | 19.6984551 | -0.002171573 | 0.966210357 |
| 20.7320551 | 20.6818167 | 20.7553625 | 20.722133 | 0.030890523 | 0.544244371 |
| 21.3053665 | 21.1673285 | 21.2957299 | 21.2881336 | -0.036589994 | 0.438032165 |
| 22.3648264 | 22.2634591 | 22.3258021 | 22.3225375 | 9.65E-05 | 0.998188812 |
| 18.8570184 | 18.7948306 | 18.9344633 | 18.8380775 | 0.066726226 | 0.202312953 |
| 21.1743184 | 21.0891404 | 21.1780019 | 21.1963233 | 0.012234422 | 0.814813534 |
| 22.3863354 | 22.3300772 | 22.3532738 | 22.3965618 | 0.026454783 | 0.618792169 |
| 18.5102032 | 18.7255615 | 18.5664797 | 18.49014 | 0.098260122 | 0.173978206 |
| 17.5941424 | 17.4909923 | 17.7289709 | 17.6953367 | 0.033545424 | 0.692020301 |
| 17.2707002 | 17.4808469 | 17.4809035 | 17.3065356 | 0.172689534 | 0.079267229 |
| 17.7211009 | 17.4800444 | 17.737873 | 17.8681335 | -0.04934153 | 0.617794829 |
| 17.8487779 | 17.8108207 | 17.7269756 | 17.7886281 | -0.042596804 | 0.699950917 |
| 19.0547901 | 19.3819073 | 19.4612836 | 19.0478836 | 0.119149918 | 0.291830658 |
| 16.5955348 | 16.4180593 | 16.4081766 | 16.1467442 | -0.14735777 | 0.198080609 |
| 15.9316676 | 16.2073188 | 16.2637203 | 16.0749316 | 0.01487279 | 0.925231049 |
| 18.7758201 | 18.5874884 | 18.6926009 | 18.7905649 | -0.056431545 | 0.382644816 |
| 14.3319633 | NA | 14.4538226 | 14.5202942 | 0.029553994 | 0.871469798 |
| NA | 15.258897 | NA | NA | 0.520631288 | 0.189299925 |
| 14.7512912 | 14.6682641 | 15.0372864 | 15.1353842 | -0.036452266 | 0.875237491 |
| 18.7783731 | 18.605093 | 18.813122 | 18.7489487 | -0.023926861 | 0.804915833 |
| 20.8547796 | 20.718333 | 20.7129202 | 20.8283774 | -0.018834028 | 0.819289604 |
| 22.4092593 | 22.2862121 | 22.3911431 | 22.3746645 | -0.005623701 | 0.940524537 |
| 28.0713949 | 27.8786148 | 28.0588329 | 28.0125798 | -0.003594113 | 0.956457075 |
| 20.359808 | 20.1882391 | 20.3930483 | 20.2931295 | -0.053174062 | 0.324032675 |
| 21.5722819 | 21.5096594 | 21.5520678 | 21.5744733 | 0.020608036 | 0.628893944 |
| 22.8896133 | 22.9228751 | 22.9462369 | 22.9433812 | 0.045009032 | 0.295718283 |
| 24.5437549 | 24.500797 | 24.5460974 | 24.5561131 | 0.025344578 | 0.547156354 |
| 21.0291262 | 21.0020609 | 21.1224246 | 21.0823926 | 0.02979051 | 0.450070427 |
| 21.463144 | 21.4093951 | 21.4933088 | 21.4896982 | 0.060015299 | 0.226544842 |
| 22.2840945 | 22.1755583 | 22.253264 | 22.2593892 | 0.031081075 | 0.599500731 |
| 18.3105343 | 18.2762615 | 18.3700712 | 18.3509641 | -0.014813793 | 0.856467592 |
| 16.1229583 | 15.9568138 | 15.9655911 | 16.0942663 | -0.010075174 | 0.912040804 |
| 15.2312579 | NA | 15.4435814 | 15.5294039 | 0.061767731 | 0.741153414 |

| | | | | | |
|---|---|---|---|---|---|
| 15.8650537 | 16.3039885 | 16.0316295 | 16.0078095 | 0.148833398 | 0.355331812 |
| NA | 14.5145938 | NA | NA | -0.186683174 | 0.225610325 |
| 24.8431783 | 24.4644298 | 24.6636319 | 24.6759767 | -0.140967485 | 0.324484894 |
| 17.3719406 | 17.3677281 | 17.3664352 | 17.0761193 | -0.062953056 | 0.6189717 |
| 17.1160198 | 16.9810293 | 17.0436552 | 17.2399743 | -0.006187607 | 0.955785728 |
| 18.3765575 | 18.1899751 | 18.4374649 | 18.1590813 | -0.095130653 | 0.238314902 |
| 16.8508531 | 17.0659742 | 17.098047 | 17.2070403 | 0.109558914 | 0.151484601 |
| 22.1961071 | 22.0067581 | 22.2204071 | 22.2299746 | -0.001552028 | 0.981771832 |
| 20.5550944 | 20.487863 | 20.5323297 | 20.5995583 | 0.012499084 | 0.844462527 |
| 22.0081708 | 21.8686054 | 21.9695407 | 21.931054 | -0.033406928 | 0.55209064 |
| 21.0603443 | 20.9406427 | 20.991444 | 21.0348882 | -0.022362379 | 0.685535333 |
| 24.852724 | 24.7190091 | 24.789068 | 24.8306627 | -0.036829303 | 0.490487292 |
| 21.1024841 | 20.9216548 | 21.0404592 | 21.0202992 | -0.050011998 | 0.306637215 |
| 23.5850399 | 23.5162465 | 23.6217924 | 23.5610555 | 0.008110087 | 0.863672185 |
| 19.1494043 | 19.0903686 | 19.0588264 | 19.1048391 | -0.01547309 | 0.730335285 |
| 23.2072133 | 23.1429456 | 23.1762935 | 23.2073186 | -0.009402048 | 0.832804976 |
| 21.8230338 | 21.7529351 | 21.8728706 | 21.843612 | 0.035958931 | 0.471464552 |
| 21.1943983 | 21.0689639 | 21.1932807 | 21.1957084 | -0.00792736 | 0.873637235 |
| 20.4773848 | 20.4220824 | 20.4967054 | 20.4890943 | -0.017969856 | 0.73538984 |
| 22.7959253 | 22.7034521 | 22.7811138 | 22.8489973 | 0.013103197 | 0.805261993 |
| 22.8400611 | 22.6877074 | 22.8547726 | 22.7837038 | -0.023461694 | 0.664413829 |
| 19.3397412 | 19.1811144 | 19.2755939 | 19.2477872 | -0.030327658 | 0.585579709 |
| 21.952173 | 21.8789006 | 22.0138315 | 22.0159771 | 0.035355734 | 0.528356279 |
| 18.619506 | 18.6751036 | 18.6712226 | 18.8160408 | 0.050358592 | 0.382375541 |
| 18.670991 | 18.5482498 | 18.6527849 | 18.6781672 | -0.033183053 | 0.56511725 |
| 21.7082588 | 21.5962064 | 21.690231 | 21.6937298 | -0.00349931 | 0.952108684 |
| 20.9690488 | 20.9583072 | 20.956498 | 21.0120634 | 0.057809131 | 0.33031802 |
| 20.285608 | 20.0937397 | 20.2096517 | 20.2166881 | -0.065759238 | 0.290752272 |
| 19.9101931 | 19.7038667 | 19.9599094 | 19.898468 | -0.01446445 | 0.825019229 |
| 19.8730075 | 19.8810746 | 19.8099898 | 19.9563999 | -0.006072861 | 0.931009272 |
| 20.4588836 | 20.2861215 | 20.4221352 | 20.4915955 | 0.001530051 | 0.98252648 |
| 21.1845772 | 21.126146 | 21.2589354 | 21.1456626 | 0.073126606 | 0.305570165 |
| 19.5997738 | 19.4581378 | 19.6661117 | 19.610123 | 0.004024104 | 0.955384016 |
| 19.4902935 | 19.3805058 | 19.4677587 | 19.4184429 | -0.042215548 | 0.572733412 |
| 18.9063043 | 18.6912788 | 18.9312172 | 18.9410884 | 0.016950439 | 0.824515974 |
| 19.2470446 | 19.1017232 | 19.2890736 | 19.3045483 | 0.011296804 | 0.882923858 |
| 18.0296882 | 17.9871779 | 18.1390497 | 18.1528858 | -0.051052507 | 0.533322979 |
| 19.2587499 | 19.3758478 | 19.0910349 | 19.3490588 | 0.064497618 | 0.484664619 |
| 20.100413 | 20.0594287 | 20.2147206 | 20.2730891 | 0.075232126 | 0.433433715 |
| 18.6542307 | 18.445431 | 18.3773339 | 18.7417897 | -0.14299467 | 0.2028281 |
| 20.6924786 | 20.255206 | 20.4481969 | 20.5353502 | -0.164577394 | 0.159128595 |
| 16.6545847 | 16.6169514 | 16.3793348 | 16.6562355 | -0.060965782 | 0.645962125 |

| | | | | | |
|---|---|---|---|---|---|
| 14.4751061 | 14.2832779 | 14.5458224 | 14.6491117 | 0.169795208 | 0.235433409 |
| 16.2461192 | 16.6254647 | 16.6366274 | 16.3969154 | 0.203661527 | 0.201891425 |
| NA | 14.8316068 | NA | NA | 0.18124545 | 0.529322467 |
| 17.0501955 | 16.3479534 | 17.2097818 | 17.1153267 | 0.078242762 | 0.811669201 |
| 18.5750003 | 18.3640627 | 18.3210175 | 18.2253634 | 0.013697427 | 0.930279555 |
| 24.6747524 | 24.5272131 | 24.8318085 | 24.7780737 | 0.009391719 | 0.949749657 |
| 20.04948 | 20.0253604 | 19.9640998 | 20.0673726 | -0.033937197 | 0.500229428 |
| 20.7196412 | 20.5964081 | 20.6214505 | 20.7259828 | -0.017578347 | 0.701885673 |
| 21.4959653 | 21.4473616 | 21.5333166 | 21.4985641 | 0.038619285 | 0.517872438 |
| 16.4928482 | 16.5908721 | 16.3476542 | 16.3973375 | 0.033408098 | 0.772335268 |
| 23.4307891 | 23.4020815 | 23.3161433 | 23.2954643 | -0.052112123 | 0.447630982 |
| 20.4157689 | 20.3694777 | 20.4010743 | 20.4771336 | 0.01080442 | 0.85651914 |
| 21.1984529 | 21.3146489 | 21.320479 | 21.1740923 | 0.006423761 | 0.91416331 |
| 18.8722414 | 18.7044434 | 18.9246463 | 18.7496464 | -0.042725687 | 0.568055658 |
| 19.8722801 | 19.7780714 | 20.0076706 | 20.1258502 | -0.070156716 | 0.631222353 |
| 16.8791971 | 16.8626551 | 17.1340769 | 16.8000385 | 0.283733167 | 0.022573178 |
| 17.6860916 | 17.8840375 | 17.8348224 | 17.9077942 | 0.074457015 | 0.28899026 |
| 19.455713 | 19.3356687 | 19.2808083 | 19.4435165 | -0.083486351 | 0.094181302 |
| 19.42594 | 19.2332217 | 19.3228878 | 19.4184879 | -0.005854933 | 0.924925248 |
| 18.6512827 | 18.6569162 | 18.7655046 | 18.7309439 | -0.003341491 | 0.957677168 |
| 18.2846491 | 18.1607234 | 18.3982616 | 18.423891 | 0.050537707 | 0.433745191 |
| 21.4428153 | 21.32729 | 21.4499276 | 21.4515318 | 0.029235238 | 0.611952437 |
| 16.8955091 | 16.6466449 | 16.7289226 | 16.6303115 | -0.159375881 | 0.227224519 |
| 17.3785137 | 17.3116969 | 17.3815015 | 17.3716145 | 0.011822628 | 0.919947344 |
| 16.7476238 | 16.3513105 | 16.68137 | 16.8049713 | -0.085775515 | 0.466741598 |
| 20.9455094 | 20.7951613 | 20.894656 | 20.976889 | -0.041499962 | 0.612753005 |
| 21.4957184 | 21.2732455 | 21.4851302 | 21.478481 | -0.031098547 | 0.681475129 |
| 22.4092559 | 22.5349107 | 22.6067056 | 22.5916735 | 0.124042462 | 0.054943406 |
| 23.6644126 | 23.5887864 | 23.6059402 | 23.6239837 | -0.006616649 | 0.913760033 |
| 21.7712387 | 21.6197493 | 21.7328763 | 21.6641733 | -0.001408936 | 0.981107386 |
| 19.7774498 | 19.8025636 | 19.7724853 | 19.7782578 | -0.008025332 | 0.884301645 |
| 24.9550382 | 24.9124074 | 24.9439549 | 24.959558 | 0.02655135 | 0.55665591 |
| 20.6979573 | 20.6457412 | 20.6831579 | 20.648092 | -0.022849756 | 0.611433099 |
| 21.8141354 | 21.6730187 | 21.7861963 | 21.8140907 | 0.003440792 | 0.934957993 |
| 19.3419232 | 19.2748085 | 19.352003 | 19.2704136 | 0.015502029 | 0.744277845 |
| 23.9644244 | 23.9187342 | 23.9592479 | 24.0069517 | 0.02276636 | 0.660079529 |
| 19.4032509 | 19.4399401 | 19.3757636 | 19.3171745 | -0.090272426 | 0.092353426 |
| 24.1161525 | 24.1133399 | 24.2041692 | 24.175261 | 0.060725081 | 0.247665015 |
| 23.2533819 | 23.248696 | 23.2466681 | 23.2966083 | 0.05361425 | 0.366843854 |
| 19.1474763 | 19.2575707 | 19.2416144 | 19.2627616 | 0.077654601 | 0.194061804 |
| 21.1434524 | 21.0244448 | 21.1004332 | 21.1054877 | 0.020016782 | 0.735921586 |
| 18.1102724 | 18.1678605 | 18.1208947 | 18.2862095 | 0.043935902 | 0.501904438 |

| | | | | | |
|---|---|---|---|---|---|
| 24.3007593 | 24.1393419 | 24.2844071 | 24.2871217 | 0.004363821 | 0.947161993 |
| 17.9801234 | 17.8770307 | 17.9692984 | 18.0894645 | 0.012815893 | 0.848693863 |
| 19.9780009 | 19.9827417 | 20.1188733 | 20.0642106 | 0.044272637 | 0.534396137 |
| 19.6364338 | 19.5127197 | 19.67924 | 19.6719331 | 0.007778115 | 0.924730508 |
| 21.6056655 | 21.3956482 | 21.4756211 | 21.5261231 | -0.033564522 | 0.68983288 |
| 17.3381432 | 17.6422362 | 17.8301737 | 17.8441182 | 0.200550874 | 0.091447017 |
| 18.5301463 | 18.5543215 | 18.5315844 | 18.774434 | 0.172529567 | 0.202221848 |
| 19.4821779 | 19.4991531 | 19.4941288 | 19.5252499 | 0.033380548 | 0.545449613 |
| 20.6092064 | 20.5773169 | 20.6469063 | 20.5919862 | 0.007340062 | 0.8929813 |
| 16.3548681 | 16.3476489 | 15.9538306 | 16.0295293 | -0.208617476 | 0.226668365 |
| 18.9322387 | 18.802236 | 18.9740162 | 18.7763934 | -0.052754968 | 0.355921151 |
| 20.7572267 | 20.7783798 | 20.7904547 | 20.6237108 | -0.042930222 | 0.402696905 |
| 21.2418333 | 21.1617029 | 21.2133509 | 21.2103698 | -0.020684274 | 0.582564318 |
| 26.1180315 | 26.0384498 | 26.0873384 | 26.1398995 | 0.018162799 | 0.723254837 |
| 20.3310648 | 20.0757793 | 20.2732187 | 20.2717781 | -0.020311729 | 0.750010511 |
| 21.0797472 | 20.9601274 | 21.1253333 | 21.2001724 | 0.053109718 | 0.410705559 |
| 17.7210881 | 17.8358676 | 17.8681837 | 17.8775688 | 0.018565705 | 0.785699708 |
| 19.2740508 | 19.0918603 | 19.3986653 | 19.3069363 | 0.014968056 | 0.879030545 |
| 18.5384578 | 18.6638337 | 18.6178714 | 18.5918204 | 0.147920147 | 0.213382389 |
| 19.4671594 | 19.4266643 | 19.4507734 | 19.5230065 | 0.004129002 | 0.950250447 |
| 25.7818926 | 25.7133811 | 25.7530633 | 25.6878226 | -0.024206562 | 0.664323563 |
| 21.4852167 | 21.5387983 | 21.4216579 | 21.4635986 | 0.0157118 | 0.687143688 |
| 22.326777 | 22.1893935 | 22.3022236 | 22.2466411 | -0.01259718 | 0.838142427 |
| 21.9875549 | 21.9194929 | 21.9781468 | 21.9345861 | 6.85E-04 | 0.987321175 |
| 19.6312475 | 19.612489 | 19.6620678 | 19.6500996 | -0.029327824 | 0.464076011 |
| 21.0639574 | 20.9930155 | 21.0655076 | 21.1305786 | 0.029539117 | 0.497479277 |
| 21.7005008 | 21.6264936 | 21.6943126 | 21.7310863 | 0.022784273 | 0.641351701 |
| 21.4949875 | 21.4023409 | 21.4912851 | 21.5080094 | 0.030283304 | 0.55343586 |
| 19.6050936 | 19.4136421 | 19.6008969 | 19.5861782 | -0.008469095 | 0.88428691 |
| 21.5488006 | 21.7994525 | 21.6533242 | 21.5892911 | 0.080205986 | 0.201301559 |
| 17.4852516 | 17.3326098 | 17.6205914 | 17.4749725 | 0.139569924 | 0.428136048 |
| 23.1171224 | 22.9343192 | 23.1538954 | 23.0434991 | -0.062094098 | 0.326187347 |
| 20.2233055 | 20.124228 | 20.2196711 | 20.2163995 | -0.027061231 | 0.630522335 |
| 21.4986706 | 21.4551893 | 21.5844141 | 21.5704438 | 0.049622738 | 0.476404188 |
| 21.2160884 | 21.0690467 | 21.267315 | 21.1540543 | -0.018323803 | 0.818452502 |
| 18.8944796 | 18.7235798 | 18.8760333 | 18.9824506 | 0.066195746 | 0.439535652 |
| 17.5168553 | 17.521461 | 17.5273572 | 17.7338852 | -0.046755969 | 0.506393986 |
| 22.9572297 | 22.8174494 | 22.9405575 | 22.9183718 | -0.014894101 | 0.760399019 |
| 19.1824059 | 19.2009503 | 19.1393819 | 19.0809315 | 0.048567996 | 0.680639852 |
| 19.243118 | 19.0968943 | 19.2202479 | 19.3494167 | -0.062721643 | 0.398338994 |
| 20.4808874 | 20.436373 | 20.5293687 | 20.5562567 | 0.002209437 | 0.973108236 |
| 21.6558695 | 21.619218 | 21.6570995 | 21.6406289 | 0.006362516 | 0.891715232 |

| | | | | | |
|---|---|---|---|---|---|
| 22.4099308 | 22.4055668 | 22.4577726 | 22.422919 | 0.020934458 | 0.634088152 |
| 22.1285981 | 22.0368034 | 22.0811325 | 22.0867386 | -0.055523712 | 0.180085328 |
| 21.8613991 | 21.7987953 | 21.8995348 | 21.8813411 | 0.035911538 | 0.473374116 |
| 20.8749436 | 20.7499434 | 20.9352482 | 20.8833382 | 0.001052207 | 0.986943249 |
| 16.9298056 | 16.580378 | 16.8803758 | 17.0723439 | -0.183814303 | 0.248563358 |
| 17.521833 | 17.5159971 | 17.6852847 | 17.6287751 | 0.044108844 | 0.700452155 |
| 19.3828301 | 19.2922128 | 19.3615359 | 19.3714426 | -0.039179807 | 0.438841933 |
| 19.6616695 | 19.5450321 | 19.6052466 | 19.7153568 | 0.087976038 | 0.36227558 |
| 14.1655327 | 13.9225186 | 14.4038887 | 13.7550392 | 0.010729034 | 0.945582459 |
| 20.2687352 | 20.2179022 | 20.292783 | 20.3116342 | -0.02578333 | 0.601864274 |
| NA | 15.3991187 | NA | NA | 0.145712794 | 0.767926418 |
| 14.7973242 | 14.7921152 | 14.8410608 | 14.5854534 | 0.099456392 | 0.594928647 |
| 21.2546931 | 21.1791646 | 21.2572752 | 21.2953077 | 0.003618575 | 0.94331518 |
| 20.9806128 | 20.9145766 | 20.9474455 | 21.0524974 | -0.00451487 | 0.926603785 |
| 19.0573935 | 18.8433674 | 18.9376473 | 19.0840751 | -0.021389486 | 0.818302588 |
| 23.923707 | 23.8326167 | 23.9584236 | 23.8585514 | -0.019624614 | 0.69169734 |
| 22.229147 | 22.1133151 | 22.1756067 | 22.2765839 | -0.02809386 | 0.637673113 |
| 20.0525798 | 20.1537753 | 20.1173225 | 20.1092638 | 0.047205021 | 0.382754932 |
| 22.4733595 | 22.3879481 | 22.5009795 | 22.5240545 | 0.006026068 | 0.918154462 |
| 19.1329597 | 19.107436 | 19.1534494 | 19.2332325 | 0.01594051 | 0.850326422 |
| 19.4180227 | 19.3276083 | 19.4134037 | 19.4538085 | -0.015042316 | 0.773848107 |
| 20.1812594 | 19.9575768 | 20.1685272 | 20.0143023 | -0.074091672 | 0.342976409 |
| 20.5623293 | 20.4233362 | 20.4949787 | 20.5738227 | -0.013281112 | 0.81880327 |
| 20.6426681 | 20.5878022 | 20.721774 | 20.6436822 | 0.017456771 | 0.779375288 |
| 21.2567791 | 21.1636462 | 21.2413567 | 21.1680487 | -0.022116158 | 0.730905674 |
| 18.9463076 | 18.9641326 | 19.0124651 | 19.0816998 | 0.07827556 | 0.261032057 |
| 23.1388733 | 23.0490179 | 23.1012611 | 23.1373947 | 0.002339124 | 0.963148107 |
| 14.1281574 | 14.3204295 | 14.4794361 | 14.4971567 | 0.172767834 | 0.263883773 |
| 17.311026 | 17.6147961 | 17.5512818 | 17.4065774 | 0.095581411 | 0.542093489 |
| 24.1274896 | 23.7159278 | 24.2736122 | 24.1822881 | 0.035018612 | 0.865372045 |
| 20.1918153 | 20.1085843 | 20.1539866 | 20.0899145 | -0.033621888 | 0.756700216 |
| 25.2298238 | 25.2094248 | 25.230388 | 25.2102724 | 0.023036588 | 0.52472818 |
| 22.6139023 | 22.4804405 | 22.5925388 | 22.6258415 | 0.044631954 | 0.43063327 |
| 20.1092952 | 20.0447342 | 20.0565296 | 20.1339416 | 0.014061427 | 0.828514968 |
| 18.7794147 | 18.7039379 | 18.8416564 | 18.7825159 | 0.038443431 | 0.566455006 |
| 19.0815637 | 19.0440647 | 19.2501978 | 19.26221 | 0.037167881 | 0.617747868 |
| 22.4558367 | 22.4202281 | 22.3682293 | 22.5099557 | 0.081702528 | 0.339816522 |
| NA | 17.6644788 | NA | NA | 0.362882583 | 0.193871695 |
| 18.3929976 | 18.1986932 | 18.2976108 | 18.3120791 | 0.007353905 | 0.934155412 |
| 21.7260061 | 21.6749595 | 21.7950961 | 21.6545344 | 0.071895461 | 0.338409241 |
| 15.4834751 | NA | 15.4697327 | 15.6983849 | 0.088245409 | 0.650616761 |
| 16.714915 | 16.5862798 | 16.7771178 | 16.4601898 | -0.208917521 | 0.069368405 |

| | | | | | |
|---|---|---|---|---|---|
| 20.4090467 | 20.5327909 | 20.7519938 | 20.3288201 | -0.057944838 | 0.717829639 |
| 14.5976706 | 14.6662931 | 14.8206665 | 15.0040005 | 0.116427476 | 0.38747118 |
| 16.1283469 | NA | 16.1295862 | 16.2001597 | -0.059010763 | 0.621719764 |
| 15.7099438 | NA | 15.5493549 | 15.5922026 | -0.131115453 | 0.280521345 |
| 17.9319703 | 17.916083 | 17.953831 | 17.9132773 | -5.74E-04 | 0.995620849 |
| 17.7743537 | 17.7811815 | 17.665331 | 17.6177293 | -0.100852174 | 0.323744486 |
| 18.3987597 | 17.9287683 | 18.012231 | 18.1687325 | -0.205413829 | 0.050928108 |
| 16.9295837 | 16.7288906 | 16.6358827 | 16.8012973 | -0.086802123 | 0.329291428 |
| 20.9176872 | 20.7894698 | 20.8084696 | 20.8681539 | -0.018487485 | 0.825468423 |
| 23.0770381 | 22.8371892 | 23.0885113 | 23.0391199 | -0.047755813 | 0.511535131 |
| 21.6573126 | 21.5146782 | 21.5340954 | 21.7069283 | -0.039517021 | 0.557486903 |
| 18.0695889 | 18.1279648 | 18.0894161 | 18.1890644 | 0.086348764 | 0.191749746 |
| 20.6390729 | 20.5388279 | 20.565144 | 20.6126783 | -0.037286483 | 0.503588328 |
| 22.2771971 | 22.1217543 | 22.2486863 | 22.2252604 | -0.036738304 | 0.489100557 |
| 23.5846501 | 23.4438335 | 23.5210583 | 23.5724981 | 0.014298708 | 0.779784974 |
| 22.0574344 | 21.857356 | 21.9827251 | 21.9413639 | -0.042694651 | 0.402816706 |
| 22.2301745 | 22.1299857 | 22.2452977 | 22.2073017 | 0.008153414 | 0.869036453 |
| 20.0891647 | 19.9909978 | 20.0691234 | 20.0708183 | -0.020768192 | 0.652665191 |
| 23.7968024 | 23.7300223 | 23.7607753 | 23.7657842 | 0.005830304 | 0.893459213 |
| 21.2758512 | 21.2336651 | 21.3065508 | 21.3106783 | 0.052556992 | 0.251234836 |
| 20.3424298 | 20.2456007 | 20.3870479 | 20.3317556 | 0.003046762 | 0.947863604 |
| 20.9663039 | 20.9520526 | 20.944887 | 20.9405228 | 0.039417773 | 0.401714941 |
| 20.7908173 | 20.6502024 | 20.6772344 | 20.7670307 | -0.079668658 | 0.10749099 |
| 21.2181375 | 21.1608389 | 21.2722827 | 21.2712282 | 0.003921184 | 0.935191956 |
| 18.5482966 | 18.5403484 | 18.5121975 | 18.6489397 | 0.129939231 | 0.035001389 |
| 25.9071112 | 25.7702577 | 25.9077886 | 25.872607 | 0.013125072 | 0.829057792 |
| 17.5561926 | 17.5415877 | 17.4490709 | 17.6522261 | -0.022744422 | 0.767701959 |
| 19.0781033 | 18.9382448 | 19.2552237 | 19.0890116 | 0.08049113 | 0.314011944 |
| 18.2128293 | 18.2270857 | 18.3128182 | 18.2349032 | -0.061455579 | 0.505968571 |
| 17.9871758 | 17.7007932 | 17.967747 | 18.0425289 | 0.04017904 | 0.725835459 |
| 19.9008776 | 19.9943733 | 20.2110128 | 20.1504521 | 0.305399464 | 0.016132827 |
| 17.1282399 | 17.270331 | 17.500446 | 17.3077546 | 0.167671971 | 0.186270836 |
| 16.2878367 | 16.6830014 | 16.8066111 | 16.8283861 | 0.242457829 | 0.14840945 |
| 15.83302 | NA | 15.9534302 | 16.0070447 | 0.304996942 | 0.086246482 |
| 17.70325 | NA | 17.9651896 | 17.819814 | 0.183566545 | 0.488725155 |
| 19.9344874 | 19.9034154 | 20.0109511 | 20.0901105 | 0.067091109 | 0.357111279 |
| 20.1521285 | 19.9758006 | 20.1752092 | 20.1785301 | -0.002218586 | 0.975069287 |
| 25.678475 | 25.6414331 | 25.6685345 | 25.7676575 | 0.075748652 | 0.178140458 |
| 23.5285497 | 23.352098 | 23.4723771 | 23.4878704 | -0.034256996 | 0.493406314 |
| 20.8997925 | 20.720871 | 20.9041302 | 20.8938549 | 0.033263238 | 0.591370651 |
| 11.1676116 | NA | 11.6491866 | 11.7233536 | 0.327938323 | 0.086852736 |
| 11.6126754 | NA | 11.615345 | 11.7064713 | -0.065244508 | 0.70261426 |

| | | | | | |
|---|---|---|---|---|---|
| 20.9954947 | 20.7917426 | 20.82049 | 20.8283217 | -0.121717122 | 0.04793894 |
| 25.1886039 | 25.0552846 | 25.1380716 | 25.1860242 | 0.013571787 | 0.815972191 |
| 21.0856534 | 21.0466182 | 21.0870377 | 21.1645693 | 0.031066922 | 0.549030691 |
| 21.0305751 | 20.9742722 | 21.0296502 | 21.0730363 | 0.024991011 | 0.585961125 |
| 18.1874747 | 18.2459935 | 18.1937772 | 18.3270493 | 0.06937155 | 0.533247459 |
| 17.0116975 | 17.1259996 | 17.0522385 | 16.9181421 | 0.147212742 | 0.286938962 |
| 16.4007655 | 16.2460348 | 16.8828804 | 16.7682847 | 0.14284787 | 0.334762438 |
| 13.9634064 | 13.4696701 | 14.4129801 | 14.5136931 | 0.089584702 | 0.78244345 |
| 19.8142056 | 19.6941054 | 19.6812114 | 19.7905461 | -0.023719432 | 0.772876849 |
| 21.1616706 | 21.201635 | 20.9578148 | 21.0723045 | 0.015381816 | 0.851217448 |
| 19.2435683 | 19.1505484 | 19.3309904 | 19.2842758 | 0.094545012 | 0.222374831 |
| 19.79705 | 19.6080753 | 19.5847767 | 19.766345 | -0.100913258 | 0.159815043 |
| 19.4261963 | 19.2544332 | 19.4651021 | 19.4905462 | -0.029958506 | 0.613950629 |
| 23.8232924 | 23.8395053 | 23.863685 | 23.8025982 | 0.022164563 | 0.633135884 |
| 23.4770693 | 23.3505104 | 23.469994 | 23.4698872 | -0.001820241 | 0.966671078 |
| 20.6997402 | 20.640089 | 20.6978323 | 20.6703398 | -3.88E-04 | 0.991614996 |
| 18.9658568 | 19.0927383 | 18.9382306 | 18.9670264 | 0.01202399 | 0.768805249 |
| 19.7279101 | 19.6637087 | 19.6921085 | 19.751275 | 0.016551319 | 0.697151276 |
| 21.7259263 | 21.6757184 | 21.7329745 | 21.6928261 | -0.004844099 | 0.91969009 |
| 21.5882335 | 21.6340355 | 21.6979081 | 21.6600671 | 0.067485482 | 0.20044209 |
| 22.5173821 | 22.4057649 | 22.5520983 | 22.5255363 | 6.07E-04 | 0.990896563 |
| 21.2473151 | 21.0985563 | 21.2051204 | 21.2826979 | 0.005031571 | 0.94681751 |
| 22.6310173 | 22.4204421 | 22.5927525 | 22.5721529 | -0.014175934 | 0.878608278 |
| 17.9612217 | 17.9634634 | 17.9260501 | 18.1294986 | 0.083642351 | 0.39649777 |
| 18.0074803 | 18.0827728 | 18.2002946 | 18.1398134 | -0.034163428 | 0.739549304 |
| NA | 15.0984806 | NA | NA | 0.027616614 | 0.937986697 |
| 23.4889596 | 23.352068 | 23.4325358 | 23.441214 | -0.044751931 | 0.60849634 |
| 18.3449463 | 18.2972258 | 18.4357389 | 18.3082017 | 0.029886105 | 0.690652275 |
| 15.4178254 | 15.1644248 | 14.6699096 | 14.8826015 | -0.08514483 | 0.755530092 |
| 14.4624552 | 14.7006129 | 15.1510435 | 14.255677 | 0.125715545 | 0.502477338 |
| 17.3688918 | 16.9657156 | 17.240219 | 17.1784186 | -0.077406503 | 0.451052618 |
| 20.9976766 | 20.8157773 | 20.9068419 | 20.938446 | -0.058225148 | 0.361319502 |
| 20.3945282 | 20.1749413 | 20.3195328 | 20.2619175 | -0.059163351 | 0.291623221 |
| 23.8979212 | 23.8132839 | 23.9001113 | 23.911999 | 0.005511573 | 0.891054579 |
| 20.2061309 | 20.1003919 | 20.1338456 | 20.2272192 | -0.032687679 | 0.448339835 |
| 27.1970578 | 27.117535 | 27.2067712 | 27.2493832 | 0.05830543 | 0.279723717 |
| 24.0269841 | 24.1354412 | 24.0503239 | 24.1069054 | 0.049245321 | 0.358876378 |
| 19.2066209 | 19.3278654 | 19.3770141 | 19.4484463 | 0.124748215 | 0.056369045 |
| 18.8541306 | 18.7142437 | 18.7346955 | 18.8816793 | -0.042774408 | 0.530138022 |
| 17.8007979 | 17.6393711 | 17.7851183 | 17.5710097 | -0.084296066 | 0.327943254 |
| 19.8132072 | 19.8092016 | 19.8538952 | 19.9275554 | 0.024290569 | 0.846910668 |
| 17.5193934 | 17.6728524 | 17.8760956 | 17.7188591 | 0.151717083 | 0.319231274 |

| | | | | | |
|---|---|---|---|---|---|
| 20.4789378 | 20.4052324 | 20.3869177 | 20.4241448 | -0.067647783 | 0.246814583 |
| 17.8451944 | 17.7366236 | 17.9648318 | 17.8506806 | 0.094013326 | 0.274860677 |
| 21.0908267 | 20.8778447 | 20.9986556 | 21.0557404 | -0.084461899 | 0.169669498 |
| 19.022269 | 18.994239 | 19.0221618 | 19.0067986 | -0.045861267 | 0.448627752 |
| 20.7382609 | 20.6198197 | 20.7189953 | 20.6200643 | -0.028008363 | 0.734025121 |
| 18.3622641 | 18.0487635 | 18.3920899 | 18.6487994 | -0.020220551 | 0.851826858 |
| 24.0443256 | 23.9709926 | 23.9989373 | 23.9094311 | -0.02177399 | 0.780781684 |
| 22.4160821 | 22.4786525 | 22.4593824 | 22.4146935 | 0.013330641 | 0.906217294 |
| 21.5639946 | 21.4995799 | 21.5915588 | 21.6783198 | 0.034135868 | 0.589229758 |
| 22.0047398 | 21.9783092 | 21.9985169 | 22.0603604 | 0.028378435 | 0.558904879 |
| 18.3908594 | 18.2258322 | 18.3741959 | 18.3378372 | 0.047337937 | 0.612499067 |
| 19.1957516 | 18.9651497 | 19.1925155 | 19.1058023 | -0.066229909 | 0.349921113 |
| 20.659213 | 20.6042435 | 20.6269136 | 20.5947466 | -0.041736129 | 0.387738161 |
| 17.9671838 | 18.0314315 | 18.0631492 | 18.1085961 | -0.011911857 | 0.852817637 |
| 23.4953462 | 23.2089539 | 23.4878065 | 23.332198 | -0.086059014 | 0.221562803 |
| 19.9457971 | 19.9753537 | 20.0728381 | 19.9564965 | 0.041831177 | 0.486524282 |
| 15.6235501 | 15.5921956 | 15.5426922 | 15.5441017 | -0.162891064 | 0.131064964 |
| 19.3001216 | 19.2995963 | 19.3582057 | 19.3644516 | 0.063290209 | 0.343826765 |
| 18.4130007 | 17.9973627 | 18.1951276 | 18.1196299 | -0.161635096 | 0.122103959 |
| 19.8904088 | 19.9656828 | 19.8687776 | 19.872988 | 0.023800784 | 0.766290306 |
| 18.267006 | 18.0842275 | 18.0416137 | 18.1890847 | -0.059419422 | 0.434944007 |
| 21.7782439 | 21.5967912 | 21.7313515 | 21.6851897 | -0.088267118 | 0.217650298 |
| 28.7268547 | 28.6040475 | 28.7615376 | 28.7144787 | 0.044531998 | 0.496506804 |
| 22.2828214 | 22.25974 | 22.2999274 | 22.1806257 | -0.024382014 | 0.698902265 |
| 22.7947086 | 22.7674343 | 22.7681452 | 22.7510719 | -0.017583015 | 0.778856964 |
| 21.2386088 | 21.0939742 | 21.2076532 | 21.1975425 | -0.051731558 | 0.356950188 |
| 21.8938191 | 21.8156486 | 21.8614487 | 21.933022 | -0.015372147 | 0.771022394 |
| 26.8910223 | 26.8811082 | 26.8561122 | 26.8912355 | 0.02436857 | 0.61746564 |
| 19.4100612 | 19.2433432 | 19.374768 | 19.3262964 | -0.042287177 | 0.38277323 |
| 23.6128512 | 23.4603379 | 23.5321701 | 23.5628144 | -0.01245677 | 0.78061008 |
| 20.7715688 | 20.7050387 | 20.744324 | 20.7485383 | -0.02013194 | 0.652259475 |
| 24.9032015 | 24.9201879 | 24.9114591 | 24.9193559 | 0.020915304 | 0.537063849 |
| 22.1013057 | 22.0080268 | 22.1163033 | 22.0962632 | 0.01832865 | 0.706997814 |
| 19.1964194 | 19.1442649 | 19.1964699 | 19.250522 | 0.021282715 | 0.665716288 |
| 24.2118402 | 24.0851854 | 24.2429106 | 24.2492701 | 0.001847501 | 0.972831041 |
| 18.3723532 | 18.1943207 | 18.4149789 | 18.3971295 | -0.022001537 | 0.718577251 |
| 20.5259034 | 20.3613652 | 20.4469151 | 20.65747 | 3.49E-05 | 0.999550555 |
| 18.7935281 | 18.8430446 | 18.8562116 | 18.885708 | 0.036035648 | 0.623501569 |
| 19.8231019 | 19.686661 | 19.7261112 | 19.8292231 | -0.029265913 | 0.704010907 |
| 18.3950363 | 18.3558122 | 18.6098082 | 18.572984 | 0.02437657 | 0.782649397 |
| 16.1499943 | 16.1271872 | 16.3226578 | 16.255462 | 0.116291748 | 0.308349512 |
| 16.9117414 | 16.7303147 | 17.1076038 | 17.0975686 | -0.06970447 | 0.570572759 |

| | | | | | |
|---|---|---|---|---|---|
| 15.7185331 | 15.5220462 | 15.5878406 | 15.8089635 | -0.128407013 | 0.322508062 |
| 17.9907147 | 17.9524748 | 17.9890101 | 17.7226161 | 0.033413168 | 0.820021451 |
| 20.9492133 | 20.6551064 | 20.8070449 | 20.8057676 | -0.057834668 | 0.415362832 |
| 21.2604757 | 21.0759041 | 21.2269058 | 21.1424995 | -0.07376565 | 0.233031837 |
| 21.0874861 | 21.0446112 | 21.0430345 | 21.0723089 | -0.029006545 | 0.537805076 |
| 23.1971902 | 23.1782329 | 23.3158537 | 23.1720634 | -0.002791863 | 0.954642108 |
| 18.8925338 | 18.8228013 | 18.8185592 | 18.7094653 | -0.03835222 | 0.76091798 |
| 16.8785876 | 16.9345394 | 16.9596379 | 17.0669524 | -0.078988731 | 0.554296432 |
| 19.1753961 | 19.1524474 | 19.1397864 | 19.2194688 | -3.63E-04 | 0.995215329 |
| 21.2353043 | 21.068246 | 20.8013302 | 21.0839124 | -0.214501775 | 0.028567584 |
| 17.9557323 | 17.7583023 | 17.8366828 | 18.024241 | 0.078575808 | 0.433271254 |
| NA | 16.9842325 | NA | NA | -0.052298879 | 0.77955117 |
| 18.6175849 | 18.2457331 | 18.1744836 | 18.3311012 | -0.441784622 | 0.06506117 |
| NA | 14.2303436 | NA | NA | -0.356819809 | 0.010450274 |
| 21.7895084 | 21.9478121 | 21.9085366 | 21.9779451 | 0.117478114 | 0.145413427 |
| 19.7931832 | 19.5005672 | 19.5544069 | 19.6949178 | -0.169571352 | 0.03893221 |
| 20.7759718 | 20.8577602 | 20.8299022 | 20.7765116 | 0.02083138 | 0.748821915 |
| 24.9990908 | 24.9616694 | 24.9652868 | 24.9949756 | 0.021864143 | 0.676747587 |
| 22.3914577 | 22.3360719 | 22.3979212 | 22.3939887 | 0.059670723 | 0.233596091 |
| 19.2018346 | 19.1827345 | 19.2981654 | 19.140238 | 0.011203537 | 0.831190948 |
| 18.8857842 | 19.1900465 | 19.2872401 | 19.1835842 | 0.231157612 | 0.003641994 |
| 17.7670827 | 17.7836528 | 17.5860197 | 17.9160882 | -0.06952728 | 0.493876736 |
| 20.0254523 | 19.9426574 | 19.8111179 | 19.957314 | -0.061345045 | 0.566167284 |
| 22.8560935 | 22.6059253 | 23.0011352 | 23.0247126 | -0.033718261 | 0.858182262 |
| 22.0252895 | 21.7654533 | 21.4776236 | 22.0142582 | -0.104152492 | 0.47208292 |
| 19.1213648 | 18.73551 | 18.8242966 | 19.0782987 | -0.065220877 | 0.491332312 |
| 19.8012898 | 19.6799923 | 19.6210683 | 19.7567195 | 0.023247043 | 0.767083351 |
| 18.7596815 | 18.4700347 | 18.5118805 | 18.5226287 | -0.086289086 | 0.315137724 |
| 22.6634837 | 22.592262 | 22.693577 | 22.6909959 | 0.036121662 | 0.432208736 |
| 20.4713355 | 20.3789763 | 20.5404147 | 20.3756614 | 0.04412418 | 0.713565712 |
| 22.6646446 | 22.5912917 | 22.6687773 | 22.6611704 | 0.027030885 | 0.599979276 |
| 20.5096314 | 20.4860856 | 20.5443581 | 20.5610386 | 0.043706827 | 0.400488519 |
| 16.2676694 | 16.3293129 | 16.4650796 | 16.3850272 | -0.034724702 | 0.778423371 |
| NA | 16.0622895 | NA | NA | 0.105499189 | 0.525127054 |
| 14.8768253 | 14.4475971 | 15.2548213 | 15.0508127 | 0.182460542 | 0.300207607 |
| 14.2275102 | 15.0864512 | 14.7075569 | 14.396874 | 0.233509328 | 0.270927573 |
| 23.5097951 | 23.4070563 | 23.5083086 | 23.4909542 | -0.028179691 | 0.521401389 |
| 17.6314952 | 17.78195 | 17.5749855 | 17.7563618 | -0.041875253 | 0.712604303 |
| 20.7659416 | 20.8118197 | 20.7238553 | 20.6368755 | -0.080174062 | 0.342495719 |
| 20.761979 | 20.6180676 | 20.7506368 | 20.7635996 | -0.039537568 | 0.495911004 |
| 20.6902357 | 20.530738 | 20.632946 | 20.6059559 | -0.06264966 | 0.260498207 |
| 21.8330207 | 21.793241 | 21.7750925 | 21.8441337 | 0.003310884 | 0.951400456 |

| | | | | | |
|---|---|---|---|---|---|
| 20.5488744 | 20.3293866 | 20.4305471 | 20.4585907 | -0.065962732 | 0.225692922 |
| 22.8446905 | 22.7677619 | 22.8542037 | 22.8108297 | 0.003968737 | 0.927951473 |
| 23.663241 | 23.5829562 | 23.6253832 | 23.6672014 | 0.007274061 | 0.860264363 |
| 21.0457174 | 20.9968865 | 21.1789418 | 21.0804765 | 0.047092896 | 0.349799158 |
| 19.5686587 | 19.3639646 | 19.5466488 | 19.549624 | -0.028779807 | 0.614709241 |
| 17.2848336 | 17.2690087 | 17.2328955 | 17.546013 | -0.017943693 | 0.864612481 |
| 20.6034267 | 20.4601937 | 20.5673546 | 20.393019 | 0.075300181 | 0.490901035 |
| 18.3499014 | 18.6476712 | 18.4513391 | 18.5749057 | 0.179319271 | 0.125864063 |
| 16.8384724 | 17.3193849 | 17.2962347 | 17.3853838 | 0.408928597 | 0.006123105 |
| 19.8469864 | 19.7829832 | 19.6518007 | 19.7519937 | -0.069067169 | 0.621347383 |
| 16.1738915 | 15.9579451 | 16.0480159 | 16.1961586 | -0.050449962 | 0.733292662 |
| 15.0522211 | 14.7206 | 14.8715191 | 14.7461567 | 0.027833078 | 0.85953338 |
| 18.5619141 | 18.4155098 | 18.4826066 | 18.3414814 | -0.148860391 | 0.204152795 |
| 21.4631774 | 21.3650106 | 21.4574777 | 21.4034488 | -0.006535858 | 0.899411559 |
| 19.9024289 | 19.6797175 | 19.8539499 | 19.8551154 | -0.001562065 | 0.981609301 |
| 19.1160456 | 19.1485034 | 19.1449807 | 19.1918164 | 0.014386766 | 0.856595336 |
| 15.403219 | 15.6221404 | 15.5571095 | 15.5568305 | 0.129065347 | 0.247393307 |
| 22.4238751 | 22.4777009 | 22.3586133 | 22.4209068 | 0.002945052 | 0.94547672 |
| 16.3590097 | 16.769368 | 16.4194144 | 16.3578867 | 0.210598061 | 0.263216286 |
| 21.2973591 | 21.1327325 | 21.2659615 | 21.2787075 | -0.017645288 | 0.765764451 |
| 24.5965173 | 24.4393733 | 24.6285885 | 24.6083604 | 0.026726014 | 0.70776917 |
| 23.5779455 | 23.4238864 | 23.5194717 | 23.5625976 | -0.013316981 | 0.777443395 |
| 18.8140669 | 18.7673918 | 18.6193232 | 18.7729472 | -0.020184946 | 0.834086558 |
| 19.9987858 | 19.9387218 | 20.1052321 | 20.0441712 | 0.034959524 | 0.539849901 |
| 22.6805935 | 22.555981 | 22.6632962 | 22.7054304 | 0.017658354 | 0.765063742 |
| 21.5472367 | 21.6889625 | 21.5767094 | 21.746666 | 0.059402608 | 0.560872248 |
| 18.160344 | 18.1673762 | 18.4527717 | 18.1301956 | 0.051628878 | 0.613316182 |
| 18.1686575 | 18.1015583 | 18.2955015 | 18.2603698 | -0.037780291 | 0.711144911 |
| 16.1840903 | 15.8576293 | 16.193434 | 16.2142066 | -0.081730691 | 0.533916164 |
| 18.0561928 | 18.0016888 | 18.1128399 | 18.1252215 | 0.057773713 | 0.388663542 |
| 18.6744369 | 18.3887744 | 18.6273597 | 18.6480904 | -0.108168127 | 0.171384025 |
| NA | 16.1455914 | NA | NA | -0.123883945 | 0.55079106 |
| 22.1666804 | 22.0126193 | 22.0665558 | 22.115534 | -0.015361513 | 0.854866241 |
| 21.3384465 | 21.3376754 | 21.3933479 | 21.4088478 | 0.09662828 | 0.141597052 |
| 21.5393968 | 21.6973674 | 21.658054 | 21.6038722 | 0.10485403 | 0.086473244 |
| 26.5744464 | 26.4181567 | 26.5295804 | 26.5413732 | -0.045497037 | 0.297078366 |
| 22.1104831 | 21.9493391 | 21.9662485 | 22.0523472 | 0.005109862 | 0.923873491 |
| 20.028839 | 20.0674849 | 20.0801098 | 20.1715936 | 0.072723989 | 0.244732107 |
| 20.4518589 | 20.3686265 | 20.4764283 | 20.4921222 | 0.015113977 | 0.854293765 |
| 19.8856137 | 20.0177054 | 20.073971 | 20.0412345 | 0.002208124 | 0.979025025 |
| 18.1824709 | 18.1060073 | 18.1084161 | 18.1079245 | -0.068395631 | 0.431839506 |
| 19.2386795 | 19.203177 | 19.3596552 | 19.3650288 | 0.153147781 | 0.143049488 |

| | | | | | |
|---|---|---|---|---|---|
| 21.1385085 | 21.0076792 | 21.1884214 | 21.0238506 | 0.002692908 | 0.981351835 |
| 13.2963018 | 13.3334472 | 13.6279905 | 13.5555933 | 0.051684163 | 0.729809353 |
| 15.4256891 | 15.3884692 | 15.3132084 | 15.4949259 | -0.146999174 | 0.325401657 |
| 18.2407713 | 18.3998514 | 18.389865 | 18.2048499 | 0.093243026 | 0.350663645 |
| 21.3688468 | 21.35738 | 21.3214081 | 21.4125227 | 0.046778038 | 0.379974455 |
| 21.2313475 | 21.1120143 | 21.0744733 | 21.164409 | 0.007380155 | 0.91131857 |
| 23.8133615 | 23.7896295 | 23.7975632 | 23.8659429 | 0.055938971 | 0.450288801 |
| 23.0520798 | 22.8851004 | 23.259527 | 22.9946173 | 0.046739019 | 0.609585928 |
| 19.6515192 | 19.6567839 | 20.2814378 | 20.1897149 | 0.247329749 | 0.242008567 |
| 18.2906455 | 18.019209 | 18.3289629 | 18.3099946 | -0.148485005 | 0.244396671 |
| 15.3090097 | 14.8869894 | 15.3666351 | 15.3540545 | -0.21792251 | 0.094516147 |
| 22.0544101 | 21.5887046 | 21.7723768 | 22.0358846 | -0.048678749 | 0.652683852 |
| 19.05133 | 18.7842076 | 18.9477947 | 18.9728612 | -0.098101781 | 0.253989014 |
| 20.7114815 | 20.4366102 | 20.5708657 | 20.6041114 | -0.072154076 | 0.339302057 |
| 19.8695873 | 19.9503849 | 19.8819239 | 19.8458616 | -0.059495174 | 0.377597241 |
| 27.4799516 | 27.3603892 | 27.5038802 | 27.5041054 | 0.065297356 | 0.288037007 |
| 20.6238658 | 20.6015088 | 20.6466147 | 20.6652093 | 0.026439347 | 0.570072869 |
| 23.017423 | 22.9485497 | 23.0609773 | 22.9509296 | 0.025137152 | 0.60020097 |
| 22.4638633 | 22.2700882 | 22.4384721 | 22.5241536 | 0.015931002 | 0.798512555 |
| 19.9249413 | 20.011249 | 20.0097505 | 20.0373467 | 0.028427837 | 0.681263402 |
| 22.1683313 | 22.0516476 | 22.2023374 | 22.1526088 | 0.048216708 | 0.522220755 |
| 17.9719767 | 18.134497 | 18.1035547 | 18.206423 | 0.085125135 | 0.271119054 |
| 20.7300806 | 20.6193655 | 20.7368013 | 20.7932677 | 0.042228664 | 0.608232837 |
| 20.5656147 | 20.3747052 | 20.6311487 | 20.6780102 | -0.028643769 | 0.738641395 |
| 20.8807107 | 20.788771 | 20.8874456 | 20.9312534 | -0.001345258 | 0.975263149 |
| 20.9111665 | 20.8685372 | 20.9656482 | 20.990691 | 0.038330535 | 0.448333634 |
| 21.9559315 | 21.7989846 | 21.9552983 | 21.9482524 | 0.041818938 | 0.559675022 |
| 18.852031 | 18.8424754 | 18.9527976 | 18.8860215 | -5.70E-04 | 0.991666764 |
| 18.82527 | 18.9384444 | 18.7862755 | 18.8141325 | 0.038054058 | 0.494502772 |
| 16.7355096 | NA | 16.771814 | 16.7794183 | -0.037065802 | 0.889736951 |
| 20.374095 | 20.2517235 | 20.2831484 | 20.3478456 | -0.003706284 | 0.939424 |
| 19.1883968 | 19.096312 | 18.9008487 | 18.9408616 | -0.178638869 | 0.568623007 |
| 24.1922652 | 24.0566085 | 24.1478612 | 24.1483923 | -0.021416773 | 0.734869602 |
| 19.9728305 | 20.0242246 | 19.839458 | 19.8989038 | -0.139682644 | 0.023948668 |
| 20.7163018 | 20.6082831 | 20.6705248 | 20.6887061 | -0.003883519 | 0.946131692 |
| NA | 14.9330562 | NA | NA | -0.010591516 | 0.978959332 |
| 20.7602649 | 20.6284789 | 20.8087103 | 20.8145752 | 0.034990966 | 0.52182211 |
| 18.6764267 | 18.6134133 | 18.7801287 | 18.4948122 | -0.06688781 | 0.404754624 |
| 18.2196498 | 17.9311759 | 17.9869827 | 18.2064228 | -0.127277305 | 0.09863891 |
| 19.3199586 | 19.2088395 | 19.3309908 | 19.3352331 | -0.084068514 | 0.249977788 |
| 18.8426845 | 18.8797482 | 18.8653262 | 18.7610428 | -0.026541069 | 0.663445178 |
| 21.0461569 | 21.0500301 | 21.1171057 | 21.0647712 | 0.060848559 | 0.176617035 |

| | | | | | |
|---|---|---|---|---|---|
| 18.8292146 | 18.8271928 | 18.8999951 | 18.8452802 | 0.060027842 | 0.467799039 |
| 14.8629623 | NA | 14.7186566 | 14.6562337 | -0.113081007 | 0.386133493 |
| 15.8889459 | 15.9773623 | 16.2506472 | 16.0704553 | -0.148497626 | 0.4393052 |
| 20.82274 | 20.8783629 | 20.8550462 | 20.8820916 | -0.026701186 | 0.872031714 |
| 18.0726652 | 18.0831041 | 17.9072085 | 18.3749006 | 0.123749619 | 0.389907415 |
| 15.2288312 | NA | 14.9269144 | 14.8429472 | -0.272337279 | 0.052888963 |
| 17.0434045 | 17.0758628 | 17.2251372 | 16.8000755 | -0.118800346 | 0.311332842 |
| 19.9591291 | 19.9050684 | 19.9029909 | 20.0342209 | 0.129997892 | 0.227701781 |
| 21.1543004 | 20.8794133 | 21.0457473 | 21.1275475 | -0.069511967 | 0.499645737 |
| 21.4191704 | 21.6405105 | 21.3569896 | 21.3061449 | -0.064825244 | 0.491646358 |
| 19.6099275 | 19.6680369 | 19.7754113 | 19.6100565 | 0.097817434 | 0.271393994 |
| 19.9595938 | 19.7603688 | 19.8028988 | 19.8635828 | -0.064573649 | 0.418974948 |
| 20.9707441 | 20.8598362 | 21.0147627 | 20.988541 | 0.017834528 | 0.821733375 |
| 19.8604337 | 19.7964983 | 19.7839135 | 19.7692057 | 4.66E-05 | 0.999498539 |
| 18.866936 | 18.7397059 | 18.8581693 | 18.7653658 | -0.091047841 | 0.210229329 |
| 21.5111116 | 21.5954826 | 21.4324633 | 21.4271573 | 0.069028145 | 0.325002486 |
| 19.5874706 | 19.4987747 | 19.5376966 | 19.5263848 | 0.082469985 | 0.197606692 |
| 22.1899662 | 22.0003584 | 22.1255641 | 22.1458156 | -0.036978941 | 0.554854129 |
| 19.6763787 | 19.6363691 | 19.7247655 | 19.6344452 | -0.031266993 | 0.61065689 |
| 19.4838547 | 19.3648132 | 19.4971879 | 19.466196 | -0.013358508 | 0.825892505 |
| 21.5021679 | 21.3511191 | 21.3059106 | 21.3682236 | -0.062718794 | 0.289156948 |
| 21.0011939 | 20.9358018 | 21.1198604 | 20.9563898 | 0.026300531 | 0.630311367 |
| 19.2735058 | 19.2229286 | 19.3221641 | 19.2880607 | 0.001184624 | 0.982295334 |
| 22.7386924 | 22.6047615 | 22.7114306 | 22.7737646 | -0.01065831 | 0.838926553 |
| 25.6029898 | 25.5850298 | 25.589689 | 25.6168187 | 0.027933805 | 0.536929325 |
| 21.06837 | 20.9574086 | 20.9912863 | 21.0416719 | -0.049418549 | 0.277991625 |
| 20.070537 | 19.9535151 | 20.0220576 | 20.1026189 | -0.02612569 | 0.537836754 |
| 21.6237158 | 21.6157976 | 21.6495007 | 21.6136256 | -0.033906161 | 0.366238397 |
| 19.7507608 | 19.7345049 | 19.7513874 | 19.7810788 | 0.020333282 | 0.570327676 |
| 21.0721083 | 21.0007741 | 21.1047842 | 21.0279518 | -0.004431427 | 0.914943696 |
| 24.264143 | 24.1386127 | 24.2414055 | 24.2954697 | -0.014129183 | 0.76774962 |
| 19.5686435 | 19.5790344 | 19.7098151 | 19.6025341 | 0.05014509 | 0.302653765 |
| 21.4464675 | 21.3486189 | 21.4535876 | 21.452249 | 0.025088995 | 0.627474087 |
| 20.7394169 | 20.6305028 | 20.7061769 | 20.799599 | -0.037618506 | 0.476588487 |
| 19.5954336 | 19.4800567 | 19.5637742 | 19.6317469 | 0.003833461 | 0.943222933 |
| 19.4536502 | 19.3348089 | 19.4205498 | 19.4706923 | 0.01798592 | 0.742850464 |
| 19.3802055 | 19.3401706 | 19.6147447 | 19.4595331 | 0.067478738 | 0.24887509 |
| 20.6590459 | 20.6521152 | 20.7363199 | 20.7390597 | 0.047571134 | 0.423792369 |
| 19.3485821 | 19.2035146 | 19.2671189 | 19.3247188 | -0.084751615 | 0.17768443 |
| 22.7272511 | 22.7928211 | 22.6633991 | 22.7290384 | 0.058057046 | 0.354459653 |
| 23.1107086 | 23.0039491 | 23.1447489 | 23.1147573 | 0.004929107 | 0.936688463 |
| 21.8783617 | 21.8889046 | 21.9573222 | 22.0068252 | 0.130889911 | 0.047690561 |

| | | | | | |
|---|---|---|---|---|---|
| 20.0444008 | 19.8710843 | 19.978785 | 19.9525134 | -0.032519741 | 0.611062467 |
| 21.4014631 | 21.3932147 | 21.4108482 | 21.5103773 | 0.051720148 | 0.43050833 |
| 19.95696 | 19.9138015 | 19.943582 | 20.0572393 | 0.011084023 | 0.875179334 |
| 18.2973206 | 18.1358283 | 18.1619559 | 18.2471334 | -0.064348812 | 0.372520796 |
| 20.2371255 | 20.2357223 | 20.3744532 | 20.3683018 | 0.036286845 | 0.618107242 |
| 18.1783613 | 17.9649246 | 18.1456256 | 18.2075856 | 8.58E-04 | 0.990964259 |
| 21.6914557 | 21.6760928 | 21.6664656 | 21.7678738 | 0.030930965 | 0.702191754 |
| 20.5717215 | 20.3299542 | 20.6287218 | 20.6287375 | -0.008486698 | 0.917562036 |
| 17.6876786 | 17.5877759 | 17.8203336 | 17.8248181 | 0.085110687 | 0.307898328 |
| 17.2938783 | 17.180015 | 17.1689914 | 17.238533 | -0.168208883 | 0.060629816 |
| 22.41264 | 22.3522035 | 22.4290838 | 22.399739 | -0.014210375 | 0.869789031 |
| 17.5416219 | 17.8792897 | 17.8670754 | 18.0261866 | 0.216607544 | 0.018800702 |
| 17.6099864 | 17.4335238 | 17.6915929 | 17.8987857 | 0.027122446 | 0.772049554 |
| 18.9483326 | 18.914454 | 18.696603 | 18.8469447 | -0.00144432 | 0.988634404 |
| 14.4515994 | 14.543539 | 14.429367 | 14.4370144 | 0.105465085 | 0.358118863 |
| 23.2816523 | 22.9723328 | 23.274852 | 23.2425388 | 0.00218039 | 0.985455196 |
| 19.3300859 | 19.2388908 | 19.2362654 | 19.2619949 | -0.00923893 | 0.947280294 |
| 19.9389424 | 19.6501229 | 19.9180171 | 20.0332901 | -0.001625551 | 0.990914358 |
| 16.6645496 | 16.6475706 | 16.9405353 | 17.0773067 | 0.122381704 | 0.413642437 |
| 24.1904396 | 24.057483 | 24.1517618 | 24.1189025 | -0.019226396 | 0.714322348 |
| 17.9331592 | 17.4606311 | 17.7088174 | 17.7732648 | -0.156466093 | 0.087395131 |
| 21.3053902 | 20.8840239 | 21.1175701 | 21.2401998 | -0.112124044 | 0.335418636 |
| 12.6062693 | 13.1315459 | 13.3618066 | 13.6030575 | 0.249158989 | 0.466938094 |
| 17.4948982 | 17.2417658 | 17.32747 | 17.6148338 | -0.086079407 | 0.422111852 |
| 19.7539808 | 19.5338239 | 19.7000901 | 19.7477847 | -0.103972033 | 0.255408483 |
| 18.9054867 | 18.8633536 | 18.9444387 | 18.8550514 | -0.005515135 | 0.907804575 |
| 18.8179885 | 18.8653759 | 18.8588226 | 18.7769586 | -0.014214189 | 0.934834427 |
| 21.786851 | 21.6007652 | 21.8283062 | 21.7832514 | -0.010160702 | 0.859627159 |
| NA | 14.3915263 | NA | NA | -0.050446774 | 0.837988285 |
| 21.3044125 | 21.1694234 | 21.4097488 | 21.4752437 | 0.019617775 | 0.766293286 |
| 18.957597 | 18.7711794 | 19.0596876 | 19.0567136 | -0.007661636 | 0.92547306 |
| 16.5531215 | 16.3751584 | 16.3942555 | 16.4102855 | 0.079072211 | 0.573407306 |
| 22.0852646 | 21.8567498 | 21.9103951 | 21.8704166 | -0.154738516 | 0.081257518 |
| 24.7109985 | 24.6407147 | 24.7157498 | 24.7336694 | 0.018924793 | 0.657139143 |
| 18.7126814 | 18.5462431 | 18.6827451 | 18.6777139 | -0.02236422 | 0.7427928 |
| 19.5701841 | 19.4191936 | 19.5168168 | 19.6076821 | -0.01625239 | 0.760073572 |
| 21.1229772 | 21.0939183 | 21.1061085 | 21.0952717 | 0.057163599 | 0.255318665 |
| 19.3846709 | 19.1979957 | 19.3356392 | 19.3712966 | -0.069746189 | 0.226093918 |
| 20.0485788 | 19.9825574 | 20.0587109 | 20.0939999 | -0.003473344 | 0.940654568 |
| 19.8833232 | 19.7161768 | 19.8526079 | 19.8337078 | -0.025461013 | 0.608750652 |
| 20.2286528 | 20.0393128 | 20.0327702 | 20.4881214 | 0.08715834 | 0.514914375 |
| 22.5795703 | 22.533184 | 22.5305597 | 22.6074966 | 0.048125118 | 0.345266613 |

| | | | | | |
|---|---|---|---|---|---|
| 23.3247266 | 23.2932082 | 23.2979313 | 23.3680696 | 0.006557946 | 0.878196521 |
| 21.2721111 | 21.2203276 | 21.2009118 | 21.2584098 | -0.040194582 | 0.434553511 |
| 19.1045754 | 18.9997178 | 19.0015517 | 19.0701611 | -0.046316831 | 0.578767673 |
| 26.5095789 | 26.362857 | 26.4879321 | 26.469205 | -0.015435122 | 0.803045218 |
| 19.1922796 | 18.9664254 | 19.109747 | 18.9567917 | -0.081461507 | 0.36633718 |
| 24.6244554 | 24.5072866 | 24.6856079 | 24.6343465 | 0.039126014 | 0.529421216 |
| 18.0521938 | 17.9647379 | 18.0695274 | 18.0594651 | -0.017968696 | 0.697521651 |
| 21.0697069 | 21.0220001 | 21.0313805 | 20.9990194 | -0.024702328 | 0.553911866 |
| 22.0255501 | 22.1404342 | 22.0363644 | 22.0566841 | 0.043531682 | 0.453326606 |
| 19.672542 | 19.6202324 | 19.5802768 | 19.7354373 | -0.019305717 | 0.882398159 |
| 17.7463635 | 17.4955988 | 17.5421771 | 17.433779 | -0.153241163 | 0.303336974 |
| 20.8905912 | 20.9470461 | 21.0737865 | 21.1455598 | 0.15897305 | 0.244245635 |
| 15.4141145 | 15.1694099 | 15.2893492 | 14.9252756 | -0.190060264 | 0.133850593 |
| 19.2165225 | 19.1041786 | 19.2577134 | 19.2599435 | -0.014617559 | 0.802806645 |
| 24.6651349 | 24.5556284 | 24.6136437 | 24.6162654 | -0.029922651 | 0.591319818 |
| 20.3060177 | 20.2165166 | 20.3296325 | 20.323445 | -0.005827664 | 0.898065049 |
| 19.5022041 | 19.3399173 | 19.621509 | 19.6361536 | 0.046857979 | 0.61626951 |
| 16.2799851 | 16.270229 | 16.512284 | 16.0680927 | 0.073159623 | 0.657291103 |
| 19.4620829 | 19.2138488 | 19.5735812 | 19.5544844 | 0.011277923 | 0.890716196 |
| 22.7625986 | 22.7809448 | 22.7790278 | 22.746797 | 0.015456136 | 0.707336407 |
| 21.3138951 | 21.2439489 | 21.3667893 | 21.334089 | 0.03964843 | 0.420759062 |
| 19.8184596 | 19.7154174 | 19.8608393 | 19.8801621 | 0.025073258 | 0.653701126 |
| 21.2512139 | 21.0872133 | 21.0888046 | 21.2152719 | -0.097239615 | 0.067345668 |
| 23.9147184 | 23.8327333 | 23.9448771 | 23.9156927 | 0.014427124 | 0.741357822 |
| 16.7243467 | 16.6946431 | 16.751527 | 16.5983507 | 0.011126556 | 0.912117389 |
| 15.631633 | 15.2870674 | 15.6205694 | 15.4294326 | -0.100673213 | 0.44456195 |
| 18.3894534 | 18.2489489 | 18.3677615 | 18.3889734 | -0.003106155 | 0.971615962 |
| 17.5474799 | 17.1721204 | 17.6186036 | 17.4990742 | -0.061325405 | 0.579077327 |
| 19.8707007 | 19.8328533 | 19.9780548 | 19.9737589 | -0.01207624 | 0.905047492 |
| 18.7083915 | 18.4267134 | 18.3798406 | 18.5927205 | -0.088973376 | 0.232363977 |
| 17.7715297 | 17.7676922 | 17.7880455 | 17.8685836 | 0.024077432 | 0.738281424 |
| 21.7559489 | 21.5410072 | 21.7148043 | 21.6114312 | -0.09342675 | 0.129982645 |
| 22.3716547 | 22.3718402 | 22.4471768 | 22.353578 | 0.023226416 | 0.589265957 |
| 21.1816734 | 21.0966221 | 21.1632966 | 21.1138541 | -0.035766479 | 0.432777697 |
| 22.4679831 | 22.3800111 | 22.4920397 | 22.479159 | 0.038798385 | 0.477919175 |
| 21.3250802 | 21.3527612 | 21.2477126 | 21.2882227 | 0.00391058 | 0.944203645 |
| 21.3075237 | 21.3614849 | 21.3327781 | 21.449589 | 0.097555545 | 0.097739561 |
| 18.1344593 | 18.2310042 | 18.2713274 | 18.3876256 | 0.178037832 | 0.005638769 |
| 19.2308197 | 19.0799193 | 19.2935924 | 19.0528794 | -0.060028383 | 0.349296452 |
| 22.52069 | 22.4066726 | 22.6188446 | 22.5386182 | 0.057328133 | 0.377823165 |
| 19.27113 | 19.2502483 | 19.1870844 | 19.3796606 | 0.007792616 | 0.905042597 |
| 21.19275 | 21.0900802 | 21.1646309 | 21.1369878 | -0.022884659 | 0.810344766 |

| | | | | | |
|---|---|---|---|---|---|
| 18.7780729 | 18.3865474 | 18.564527 | 18.5964371 | -0.090725035 | 0.416989873 |
| 18.2141967 | NA | 18.0395641 | 18.1530326 | 0.013106853 | 0.916382846 |
| 17.5658397 | 17.6497842 | 17.795941 | 17.6623067 | 0.162047869 | 0.318055337 |
| 15.2101764 | 15.3080697 | 15.7702173 | 15.1823878 | 0.323335494 | 0.202320914 |
| 19.2197475 | 19.0912475 | 19.1822188 | 19.2298001 | -0.0077588 | 0.916947236 |
| 21.9921566 | 21.9578064 | 21.9003983 | 21.9186142 | 0.012943825 | 0.82320751 |
| 18.0943064 | 18.0353337 | 18.0729764 | 18.0779114 | 0.080674182 | 0.297478836 |
| 15.4082013 | 15.3380304 | 15.4819267 | 15.7038901 | 0.113156187 | 0.367923979 |
| 21.6747495 | 21.4355573 | 21.5365939 | 21.6373127 | -0.111996658 | 0.116404906 |
| 22.3356338 | 22.1671094 | 22.3851942 | 22.3513323 | 0.052512811 | 0.385689027 |
| 22.6840112 | 22.6193725 | 22.7548117 | 22.6748708 | -0.0293117 | 0.610591232 |
| 21.0033978 | 21.0097032 | 21.0283228 | 20.9538504 | 0.008443674 | 0.851863901 |
| 25.662087 | 25.594144 | 25.6729804 | 25.6864199 | 0.019356332 | 0.642498404 |
| 20.7775761 | 20.7651108 | 20.7913133 | 20.7803968 | -0.01290684 | 0.777110449 |
| 18.9986107 | 19.0245642 | 19.007873 | 19.0114562 | 0.050124194 | 0.382427484 |
| 19.0590327 | 18.9127582 | 19.0414579 | 19.0203097 | 0.009313097 | 0.915347669 |
| 21.2122612 | 21.1772272 | 21.1716269 | 21.2107293 | 0.023089649 | 0.796463943 |
| 21.9151102 | 21.8327785 | 21.9020304 | 21.7257625 | -0.069073314 | 0.474254323 |
| 17.766189 | 17.8474293 | 17.9684099 | 17.6383803 | 0.074715169 | 0.457786324 |
| 16.8412681 | 16.4274897 | 16.4855158 | 16.4965356 | -0.209999905 | 0.120282346 |
| 20.5810576 | 20.5515006 | 20.5475431 | 20.5351806 | 0.008986244 | 0.893343464 |
| 18.9061506 | 18.8340231 | 18.9823112 | 18.853808 | -0.063518637 | 0.436476542 |
| 21.0386349 | 20.863325 | 20.9798373 | 21.1050245 | 0.002731615 | 0.969818885 |
| 22.8278994 | 22.6373857 | 22.7575702 | 22.703017 | -0.042615119 | 0.416480145 |
| 22.2560856 | 22.180316 | 22.2566341 | 22.2628661 | 0.003649952 | 0.939460157 |
| 22.4972934 | 22.3841169 | 22.5453081 | 22.5911888 | 0.040672207 | 0.49052443 |
| 18.9662248 | 18.9250932 | 18.9581982 | 18.9422563 | 0.012967195 | 0.824987937 |
| 18.201205 | 18.3769067 | 18.1616006 | 18.2374363 | -0.025946093 | 0.679488501 |
| 25.6162271 | 25.4517529 | 25.589031 | 25.5883155 | 0.007274804 | 0.911472284 |
| 19.294614 | 19.3246992 | 19.3869472 | 19.3927193 | 0.073063012 | 0.326222019 |
| 20.0699486 | 20.0576581 | 20.1471814 | 20.239099 | 0.060941439 | 0.41622227 |
| 17.6452775 | 18.0922742 | 18.5852426 | 18.2874656 | 0.495742999 | 0.082091361 |
| 23.5851565 | 23.4145497 | 23.6188599 | 23.6814932 | 0.045170513 | 0.508529854 |
| 23.2848788 | 23.1752578 | 23.1215727 | 23.1384603 | -0.066550457 | 0.20004874 |
| 21.2156658 | 21.2500992 | 21.2758269 | 21.269401 | 0.05042106 | 0.318685723 |
| 20.9292851 | 20.8378205 | 20.9222165 | 20.9300762 | -0.015319213 | 0.751355144 |
| 17.8149612 | 17.6334201 | 17.805158 | 17.9442849 | -0.041226455 | 0.578894758 |
| 18.2838598 | 18.5105264 | 18.48353 | 18.6419985 | 0.112614508 | 0.242503037 |
| 16.8660222 | 16.8299494 | 16.8773009 | 17.0200879 | 0.025495457 | 0.808165594 |
| 21.0563471 | 20.996609 | 19.9871281 | 21.0598657 | 0.305786182 | 0.396539167 |
| 22.0636994 | 21.9945366 | 22.0925058 | 22.1337922 | 0.040470745 | 0.300840138 |
| 22.777662 | 22.7591396 | 22.8454951 | 22.8568698 | 0.056471324 | 0.297286253 |

| | | | | | |
|---|---|---|---|---|---|
| 15.2599496 | NA | 15.3300085 | 15.6024555 | 0.278843178 | 0.111639549 |
| 26.3985183 | 26.277877 | 26.366342 | 26.3689582 | -0.013625611 | 0.776588232 |
| 18.2951156 | 18.3337941 | 18.2445706 | 18.2768741 | -0.02165413 | 0.70606754 |
| 18.4585534 | 18.4034933 | 18.2746586 | 18.4856033 | -0.035109341 | 0.576668911 |
| 22.7543574 | 22.6530448 | 22.5996012 | 22.7561347 | -0.031092939 | 0.628271904 |
| 16.9930879 | 17.1119944 | 17.2490777 | 17.3776748 | 0.127239558 | 0.226654347 |
| 17.9103391 | 17.5503609 | 17.7926106 | 17.7317628 | -0.115429707 | 0.118376829 |
| 20.8859847 | 20.7703211 | 20.9093307 | 20.781437 | -0.034612255 | 0.550930075 |
| 20.254813 | 20.2129266 | 20.2663463 | 20.2281774 | 0.037653775 | 0.509757093 |
| 20.809064 | 20.7203807 | 20.7709515 | 20.8246211 | -0.028379384 | 0.624244929 |
| 14.2912784 | NA | 14.8012418 | 15.0177198 | 0.342008779 | 0.142570886 |
| 20.8226183 | 20.8385623 | 20.8467639 | 20.8348753 | 0.010570274 | 0.781680022 |
| 19.7957876 | 19.7144457 | 19.7640262 | 19.8124903 | 0.001963458 | 0.973072719 |
| 21.9527208 | 21.9074699 | 21.9398596 | 21.9454059 | -7.65E-04 | 0.987199772 |
| 24.7880067 | 24.8998224 | 24.8975291 | 24.8212807 | 0.090113647 | 0.479702655 |
| 16.5890277 | 17.0809939 | 16.6305259 | 16.7045792 | 0.066728686 | 0.592461452 |
| 20.9223443 | 20.6608664 | 20.8533116 | 20.9032811 | -0.04348598 | 0.590673055 |
| 18.8222649 | 18.5742733 | 18.7848696 | 18.8367081 | -0.03874039 | 0.582279558 |
| 19.3843642 | 19.4422207 | 19.1726907 | 19.3133563 | -0.036408227 | 0.536363922 |
| 20.0848898 | 20.0304954 | 19.9949904 | 20.0859513 | -0.019076651 | 0.699510235 |
| 18.5966301 | 18.4312975 | 18.6690947 | 18.6472556 | 0.015883834 | 0.81972812 |
| 19.6321573 | 19.6419235 | 19.6471802 | 19.6796114 | -0.052607375 | 0.422736115 |
| 21.5772397 | 21.3254308 | 21.6803827 | 21.6802575 | -0.053236331 | 0.688343437 |
| 21.1655653 | 21.0621872 | 20.8626725 | 21.0818979 | -0.086985097 | 0.414373453 |
| 16.9402116 | 16.6156645 | 16.6455079 | 16.6079795 | -0.105950553 | 0.311316417 |
| 21.8593078 | 21.8897638 | 21.9138566 | 21.8634353 | 0.049447839 | 0.450212291 |
| 20.9789662 | 20.7165288 | 20.7815375 | 20.830365 | -0.101193062 | 0.101102512 |
| 20.6866133 | 20.6026326 | 20.6188376 | 20.6109018 | -0.050450325 | 0.379739981 |
| 27.3142592 | 27.1826873 | 27.2555475 | 27.2953823 | -0.026657337 | 0.613737263 |
| 21.947759 | 21.842171 | 21.9037806 | 21.9152479 | -0.019037779 | 0.716435665 |
| 19.7084676 | 19.6632639 | 19.7058705 | 19.7702859 | -0.012735148 | 0.775018858 |
| 21.0395388 | 21.0920021 | 21.0477845 | 21.0058046 | 0.005043206 | 0.905580805 |
| 19.7205674 | 19.6153224 | 19.6413191 | 19.6654932 | -0.010384263 | 0.862698185 |
| 16.6363144 | 16.5926546 | 16.6754025 | 16.7142671 | 0.029286204 | 0.67748781 |
| 21.452193 | 21.3049321 | 21.4563064 | 21.510983 | 0.042632231 | 0.596995603 |
| 20.6379943 | 20.4630165 | 20.4650783 | 20.3884685 | -0.116614961 | 0.263262422 |
| 13.0350311 | 13.145755 | 12.9072542 | 13.0051124 | 0.099211954 | 0.452548278 |
| 19.1717342 | 19.117872 | 19.2097517 | 19.208122 | 0.061760134 | 0.328496589 |
| 17.5724424 | 17.3400773 | 17.175975 | 17.705951 | 0.105665051 | 0.680491327 |
| 20.8180434 | 20.4720802 | 20.6131742 | 20.6395549 | -0.148026716 | 0.284201664 |
| 17.6949805 | 17.1982437 | 17.6142624 | 17.1928702 | -0.335421161 | 0.007626495 |
| 16.437374 | 16.3453394 | 16.4193257 | 16.4520122 | -0.148777621 | 0.089423965 |

| | | | | | |
|---|---|---|---|---|---|
| 22.8243458 | 22.7333921 | 22.8253237 | 22.8508491 | 0.026224559 | 0.61668876 |
| 25.3612859 | 25.3108298 | 25.4760148 | 25.4071717 | 0.041835778 | 0.404611357 |
| 17.4861634 | 17.2187103 | 17.1804761 | 17.6777951 | -0.016456909 | 0.87135117 |
| 24.9276893 | 24.9316968 | 24.9417644 | 25.043673 | 0.09074445 | 0.169396738 |
| 21.9513128 | 21.8476639 | 21.9386065 | 21.9592856 | 0.037908552 | 0.486087204 |
| 20.2893173 | 20.2543305 | 20.2757814 | 20.3467599 | 0.013324422 | 0.76313171 |
| 19.7171955 | 19.5028446 | 19.6871134 | 19.6771415 | -0.029730705 | 0.632340343 |
| 19.427332 | 19.4114071 | 19.374751 | 19.4868205 | -4.53E-06 | 0.999927563 |
| 15.5445367 | NA | 15.7769117 | 15.4631269 | 0.099603152 | 0.441115352 |
| 20.355533 | 20.2746752 | 20.3664724 | 20.3314861 | 0.011600565 | 0.837202006 |
| 20.992133 | 20.9345198 | 21.0401253 | 21.1506782 | 0.01857267 | 0.725063251 |
| 21.0068903 | 20.9866884 | 20.9938051 | 20.9799487 | -0.016999441 | 0.723676385 |
| 21.1016014 | 20.9893382 | 21.1202162 | 21.1179264 | -0.002370694 | 0.960116251 |
| 21.4879092 | 21.3438857 | 21.4925018 | 21.5065066 | 0.022874841 | 0.775288699 |
| 21.4571794 | 21.3368465 | 21.4394302 | 21.4928073 | 0.051865086 | 0.532834541 |
| 15.5857362 | NA | 15.093373 | 15.5267282 | -0.004327872 | 0.982754815 |
| 18.3510909 | 18.1999772 | 18.2139658 | 18.287718 | -0.062758007 | 0.245401851 |
| 17.9255328 | 18.200491 | 18.2278636 | 18.0806729 | 0.185526889 | 0.049919862 |
| 19.5154473 | 19.4823864 | 19.4617146 | 19.2216457 | -0.111621661 | 0.401331197 |
| 27.590403 | 27.5000511 | 27.5107688 | 27.5192921 | -0.007701602 | 0.906283431 |
| 20.7015507 | 20.6517844 | 20.794251 | 20.7401161 | 0.067379414 | 0.141456564 |
| 19.5003131 | 19.3807356 | 19.4843994 | 19.3529735 | -0.067295238 | 0.306340212 |
| 22.4363874 | 22.4212028 | 22.5218639 | 22.4888362 | 0.065179059 | 0.116270025 |
| 14.7276209 | 14.8049062 | 14.8826577 | 15.0465263 | 0.06416563 | 0.676092501 |
| 22.6833685 | 22.5436571 | 22.5599301 | 22.6278527 | -0.01582739 | 0.809354224 |
| 21.2493489 | 21.1920008 | 21.3511859 | 21.214274 | 0.0592622 | 0.304748202 |
| 17.9781887 | 18.0038515 | 17.9362741 | 17.9497407 | 0.030598292 | 0.619375968 |
| 20.2267095 | 20.204489 | 20.2588012 | 20.2858098 | 0.038991963 | 0.451851827 |
| 19.1554499 | 19.1827672 | 19.1793697 | 19.0721152 | -0.008233767 | 0.871149627 |
| 19.0054759 | 19.0738011 | 19.1702209 | 19.1059243 | 0.086075559 | 0.117589396 |
| 20.2634567 | 20.2082722 | 20.1992615 | 20.151785 | 0.012026283 | 0.853197155 |
| NA | 13.0780889 | NA | NA | -0.024185562 | 0.895515532 |
| 13.8151709 | 13.9066466 | 13.6093963 | 13.9722955 | 0.094375342 | 0.475917153 |
| 21.5707453 | 21.4444808 | 21.5112311 | 21.5690349 | -0.019578907 | 0.76621773 |
| 22.20512 | 22.1242208 | 22.2457608 | 22.2229872 | 0.023322391 | 0.656585881 |
| 23.9579331 | 23.9586402 | 24.0142133 | 23.9589282 | -0.001370728 | 0.978828854 |
| 22.516853 | 22.4156551 | 22.5329848 | 22.4981758 | -0.007891337 | 0.900728537 |
| 18.9576574 | 19.0008433 | 18.9112129 | 18.9189575 | -0.008609376 | 0.940967038 |
| 19.2373445 | 18.9510278 | 19.0587489 | 19.0523624 | -0.150046665 | 0.237478184 |
| 15.7157513 | 15.8941458 | 15.7594756 | 15.6127674 | 0.052040772 | 0.666435086 |
| 19.1358557 | 18.9243306 | 18.9566337 | 19.0566599 | -0.042561152 | 0.635277899 |
| 20.9519671 | 20.9069406 | 20.9696101 | 20.9644514 | -0.032957218 | 0.651175291 |

| | | | | | |
|---|---|---|---|---|---|
| 19.9989271 | 19.9074648 | 19.9567792 | 20.0119669 | -0.044529908 | 0.423847422 |
| 22.1189313 | 21.9556527 | 22.1517989 | 22.0911894 | -0.035391495 | 0.519960564 |
| 21.3435269 | 21.2651209 | 21.3099663 | 21.3932057 | -0.01967463 | 0.701162927 |
| 20.5106111 | 20.4516056 | 20.5081218 | 20.5071785 | -0.005222061 | 0.912312105 |
| 23.4448316 | 23.3789039 | 23.4438653 | 23.4700153 | 0.012707489 | 0.775136223 |
| 22.7196998 | 22.6708824 | 22.6888237 | 22.7142758 | 0.023850233 | 0.565047284 |
| 21.114993 | 21.0139761 | 21.0829015 | 21.0540714 | -0.015033584 | 0.736628999 |
| 22.3653956 | 22.2612533 | 22.3240038 | 22.3449086 | 0.002519347 | 0.957139323 |
| 21.1074878 | 20.9681171 | 21.0825454 | 21.045305 | 0.013040402 | 0.795222748 |
| 20.1847272 | 20.0892833 | 20.1364159 | 20.1918813 | -0.006700665 | 0.903676426 |
| 18.3681943 | 18.27334 | 18.353257 | 18.5356164 | 0.050923001 | 0.485474082 |
| 18.7724486 | 18.6197176 | 18.773946 | 18.8000763 | 0.003844278 | 0.958634117 |
| 17.616951 | 17.6633231 | 17.7103299 | 17.6215898 | -0.006788184 | 0.933455267 |
| 18.7696292 | 18.7916807 | 18.4992718 | 18.9567667 | 0.063476879 | 0.599556663 |
| 15.630939 | 16.1521575 | 16.155871 | 16.1468083 | 0.143182292 | 0.518173456 |
| 16.988087 | 17.1848467 | 16.9646483 | 17.0802357 | 0.298915513 | 0.200634142 |
| 18.7173339 | NA | 18.8409586 | 18.4839087 | -0.181536083 | 0.446448164 |
| 21.8710456 | 21.7303634 | 21.775032 | 21.7857141 | -0.014832946 | 0.754731819 |
| 21.6391613 | 21.574239 | 21.6803734 | 21.6046773 | 0.001435579 | 0.973841945 |
| 25.0522361 | 24.8867666 | 24.9798439 | 24.9866948 | -0.021324567 | 0.71588217 |
| 19.0997368 | 18.9441166 | 19.0557892 | 19.1294918 | 0.032106471 | 0.701764349 |
| 20.6055703 | 20.62636 | 20.6701338 | 20.6756737 | 0.011637207 | 0.866051829 |
| 15.4595165 | 15.1145098 | 15.6831025 | 15.666509 | 0.10855006 | 0.563983097 |
| 17.7130366 | 17.6296085 | 17.6704265 | 17.4477837 | -0.125417629 | 0.405813584 |
| 16.4060481 | 16.2792933 | 16.5957108 | 16.7324786 | 0.077474811 | 0.492115927 |
| 16.11678 | 16.3182974 | 16.3088347 | 16.3140675 | 0.182341232 | 0.058519695 |
| 19.8034566 | 19.6884633 | 19.7535757 | 19.6761696 | -0.018898436 | 0.821385505 |
| 20.5977559 | 20.3158946 | 20.4477315 | 20.5193279 | -0.091855707 | 0.278189656 |
| 22.3760847 | 22.126486 | 22.3363173 | 22.2419655 | -0.072034138 | 0.300267634 |
| 21.7498023 | 21.6099275 | 21.6361209 | 21.6700195 | -0.059933574 | 0.345561626 |
| 19.4328798 | 19.3113979 | 19.4400734 | 19.4333954 | -0.042142695 | 0.439190865 |
| 26.0865637 | 26.0527374 | 26.0366483 | 26.1132731 | 0.049162187 | 0.291343872 |
| 22.6169516 | 22.5993029 | 22.6381403 | 22.6644082 | 0.021051056 | 0.622727516 |
| 21.029299 | 21.0050699 | 21.0561402 | 20.99629 | 0.018348837 | 0.690763048 |
| 19.5770955 | 19.6174497 | 19.5879245 | 19.6007058 | 0.037407679 | 0.450637522 |
| 20.9175523 | 21.0583097 | 20.8992998 | 21.0438295 | 0.0700323 | 0.224076786 |
| 20.2791727 | 20.1708035 | 20.207518 | 20.1927508 | -0.045405768 | 0.42824342 |
| 19.293696 | 19.2358166 | 19.2493262 | 19.3534007 | -0.032645564 | 0.59017991 |
| 20.1689766 | 20.0017907 | 20.1483639 | 20.1316041 | -0.032952192 | 0.602738938 |
| 19.7060687 | 19.5137662 | 19.6846008 | 19.7535707 | 0.00195954 | 0.978120567 |
| 19.7929012 | 19.6665926 | 19.8610468 | 19.8498984 | 0.024818045 | 0.733541694 |
| 19.3696703 | 19.4885945 | 19.5071657 | 19.492832 | 0.100538534 | 0.227833777 |

| | | | | | |
|---|---|---|---|---|---|
| 17.7351648 | 17.8739537 | 17.9808681 | 17.8037205 | 0.09309843 | 0.27025298 |
| 20.0321286 | 19.7583991 | 19.8691003 | 19.9357674 | -0.04472969 | 0.608260742 |
| 18.4797092 | 18.3976583 | 18.6863565 | 18.6133661 | 0.041773106 | 0.638848895 |
| 15.5592935 | 15.6560619 | 15.7937548 | 15.5065137 | -0.064861472 | 0.501699451 |
| 21.1627334 | 21.5676084 | 21.2765498 | 21.3230962 | 0.28573882 | 0.048852884 |
| 17.0903717 | 16.8350682 | 17.015404 | 17.0653964 | -0.038062334 | 0.843643046 |
| 19.4999339 | 19.5554096 | 19.6416233 | 19.695393 | 0.030596239 | 0.609877077 |
| 18.9805895 | 18.8228199 | 19.0015002 | 19.0187112 | -0.013116282 | 0.812533908 |
| 26.1683275 | 26.0469784 | 26.1133531 | 26.1449149 | -0.007192143 | 0.886108321 |
| 16.6086625 | 15.9173978 | 16.5723364 | 16.9594025 | -0.034818379 | 0.923049788 |
| 20.9128593 | 21.2288924 | 21.0146912 | 20.7382445 | 0.089725479 | 0.29954517 |
| 20.577166 | 20.6187631 | 20.6300854 | 20.617134 | 0.010256154 | 0.881323124 |
| 23.9055462 | 23.8085551 | 23.8207657 | 23.8815003 | 0.034232905 | 0.598913775 |
| 21.8461763 | 21.7673445 | 21.8569724 | 21.8384339 | -0.003181895 | 0.959304599 |
| 21.6218321 | 21.5928975 | 21.5265565 | 21.5922787 | -0.012543993 | 0.832095494 |
| 20.4435887 | 20.3946145 | 20.4323487 | 20.4242794 | -0.036720059 | 0.538041823 |
| 20.3711399 | 20.3620756 | 20.4548503 | 20.4158906 | 0.040167054 | 0.472649456 |
| 19.8916236 | 19.9287299 | 19.8137052 | 19.8875812 | -0.025224766 | 0.651877882 |
| 18.3664364 | 18.2678425 | 18.339623 | 18.3933034 | -0.024599723 | 0.646574703 |
| 21.8551771 | 21.7697379 | 21.8555198 | 21.8773213 | 0.037688996 | 0.478569842 |
| 20.6841403 | 20.6034341 | 20.712997 | 20.7288826 | -0.044220189 | 0.356608672 |
| 22.4568285 | 22.4058376 | 22.4566897 | 22.4956785 | 0.009435454 | 0.83592536 |
| 20.723837 | 20.6168017 | 20.6965696 | 20.6869952 | -0.01447662 | 0.664837618 |
| 23.8077777 | 23.7061944 | 23.8602778 | 23.8437603 | 0.013734518 | 0.768837379 |
| 21.2060171 | 21.1279123 | 21.2048817 | 21.2137622 | 0.015391686 | 0.749081487 |
| 22.0970713 | 22.065222 | 22.1711585 | 22.1894059 | 0.038771783 | 0.425400935 |
| 21.6426491 | 21.5156079 | 21.637923 | 21.5378379 | -0.013378057 | 0.784246471 |
| 20.2530261 | 20.1062064 | 20.1421282 | 20.1973626 | -0.015052642 | 0.777489961 |
| 22.2370429 | 22.1452234 | 22.2347583 | 22.2603355 | 0.029481336 | 0.586975723 |
| 20.2988344 | 20.2001839 | 20.3196883 | 20.3487126 | 0.006841485 | 0.924033579 |
| 23.6833856 | 23.5077297 | 23.7023858 | 23.6540561 | -0.01904778 | 0.794877308 |
| 22.8571785 | 22.8041942 | 22.9594296 | 22.918766 | 0.109433344 | 0.170050233 |
| 16.4227381 | 16.5124604 | 16.7450989 | 16.7185965 | 0.288559542 | 0.021557035 |
| 17.3145693 | 16.9983923 | 17.249912 | 17.348494 | 0.004354586 | 0.980607324 |
| NA | 12.5361168 | NA | NA | -0.237754393 | 0.52491113 |
| 21.083288 | 20.9961161 | 21.1407285 | 21.1330986 | 0.033271234 | 0.520780939 |
| 14.9288282 | 14.838669 | 15.1680538 | 14.9570597 | -0.012844821 | 0.928859837 |
| 18.5928817 | 18.4768327 | 18.6780253 | 18.7333996 | 0.044934821 | 0.598018523 |
| 15.7210657 | 15.9504542 | 15.7217779 | 15.1598353 | -0.118761309 | 0.565968352 |
| 15.3011482 | 15.1504196 | 15.5475704 | 15.6462717 | -0.029306484 | 0.87429535 |
| 17.9393806 | 17.6019718 | 17.7016444 | 17.6230559 | -0.218541284 | 0.146022807 |
| 17.682056 | 17.1195887 | 17.2680316 | 17.3435402 | -0.270087658 | 0.025156401 |

| | | | | | |
|---|---|---|---|---|---|
| 18.2951081 | 18.088561 | 18.2204922 | 18.2497607 | -0.063868916 | 0.514406569 |
| 21.938782 | 21.7746312 | 21.9272117 | 21.8228321 | -0.05371692 | 0.435560981 |
| 18.9764202 | 18.933415 | 19.0243601 | 19.0047824 | -0.02166963 | 0.717855333 |
| 20.6924489 | 20.6041235 | 20.690387 | 20.6698723 | -0.012102026 | 0.835427968 |
| 20.8983382 | 20.7413069 | 20.825977 | 20.8822882 | -0.006598454 | 0.907340532 |
| 24.8680251 | 24.7460063 | 24.8850696 | 24.8601648 | 0.031846785 | 0.516422175 |
| 25.9200193 | 25.8484445 | 25.9997341 | 25.9459078 | 0.076692702 | 0.102248895 |
| 20.5000756 | 20.4169343 | 20.5787406 | 20.521463 | 0.00839043 | 0.875382808 |
| 19.143828 | 19.1137289 | 19.0913254 | 19.1915482 | 0.093141402 | 0.251945259 |
| 17.7528355 | 17.6964407 | 17.5473635 | 17.6679069 | -0.076552984 | 0.417288734 |
| 17.1363059 | 16.9138128 | 17.3863085 | 17.3128047 | -0.116435986 | 0.41726824 |
| 17.6321915 | 17.5752167 | 17.8531471 | 17.6080696 | 0.023025073 | 0.825344625 |
| 15.9335777 | NA | 15.8769641 | 16.0419605 | 0.033358887 | 0.709569604 |
| 22.0022704 | 21.8206402 | 22.0446353 | 22.0013133 | 0.048926011 | 0.555464074 |
| 18.7531633 | 18.6842574 | 18.6383795 | 18.7517049 | 0.067406961 | 0.413128736 |
| 18.9514633 | 18.8926678 | 18.7875173 | 18.8110247 | -0.042197099 | 0.544054039 |
| 25.0990552 | 25.016353 | 25.1108542 | 25.0479095 | 0.002207705 | 0.974166513 |
| 19.4812103 | 19.3966129 | 19.525974 | 19.5588432 | 0.010936854 | 0.856974291 |
| 22.3725161 | 22.367288 | 22.4085873 | 22.4141984 | 0.009176263 | 0.878112602 |
| 23.435062 | 23.2361251 | 23.2926574 | 23.3965441 | -0.087647375 | 0.210101505 |
| 19.0921176 | 19.2072935 | 19.2133873 | 19.113562 | 0.129093495 | 0.084661578 |
| 18.1249169 | 18.1016897 | 18.1209128 | 17.9779709 | -0.051678862 | 0.511588576 |
| 20.2169916 | 20.1302742 | 20.2396939 | 20.2806783 | 0.036149792 | 0.666100097 |
| 19.1969295 | 19.1164378 | 19.4804231 | 19.2030968 | 0.081187706 | 0.359662248 |
| 26.8833626 | 26.7626747 | 26.8526539 | 26.871162 | 8.03E-04 | 0.988879269 |
| 22.9077977 | 22.628162 | 23.1817306 | 22.9288732 | -0.078171299 | 0.531785965 |
| 18.4056778 | 17.8618361 | 18.3407681 | 18.2870701 | -0.017984519 | 0.910513126 |
| NA | 13.8825387 | NA | NA | -0.010793423 | 0.959804714 |
| 14.534457 | NA | 14.9647152 | 13.9095282 | -0.002824277 | 0.995148058 |
| 18.1366401 | 18.1184503 | 18.1642559 | 18.2423447 | 0.04520564 | 0.566156149 |
| 23.0733027 | 22.9658301 | 23.0411271 | 23.0833956 | 0.028109589 | 0.602699573 |
| 20.4018004 | 20.273048 | 20.3967871 | 20.43204 | 0.010948924 | 0.876302104 |
| 16.7033051 | 16.3669061 | 16.6746767 | 16.6008733 | 0.062080958 | 0.623168434 |
| 20.0820727 | 19.7487811 | 19.432855 | 20.018656 | -0.07261127 | 0.685162734 |
| 21.5679809 | 21.4970234 | 21.5209887 | 21.4464842 | -0.10650542 | 0.07529607 |
| 20.8316596 | 20.7489141 | 20.824948 | 20.8254366 | 0.025255742 | 0.571328522 |
| 20.1052586 | 19.9312178 | 20.0182603 | 20.0673788 | -0.070857634 | 0.321193654 |
| 17.8745923 | 17.8230431 | 17.7822402 | 17.8419941 | -0.118607417 | 0.188320131 |
| 18.2774482 | 18.2370418 | 18.4258294 | 18.248522 | 0.025008022 | 0.770279766 |
| 18.5192942 | 18.3725318 | 18.5614195 | 18.6877311 | 0.023923496 | 0.778238516 |
| 21.4272262 | 21.3479937 | 21.3686665 | 21.3892755 | -0.04690264 | 0.288154681 |
| 22.3799027 | 22.3063752 | 22.4733465 | 22.4397274 | 0.108447258 | 0.351094722 |

| | | | | | |
|---|---|---|---|---|---|
| 16.7418615 | 16.6239989 | 16.7641126 | 16.7421271 | -0.068945284 | 0.408942645 |
| 18.8806945 | 18.8047349 | 18.932537 | 18.9124232 | 0.039161937 | 0.584669065 |
| 22.8915855 | 22.7542117 | 22.8066217 | 22.8362873 | -0.094197499 | 0.074040056 |
| 25.420593 | 25.3489889 | 25.4208944 | 25.4067681 | 0.014577962 | 0.726235358 |
| 17.8243555 | 18.3189074 | 18.0727005 | 17.93783 | 0.146880949 | 0.185203788 |
| 16.0501062 | 16.0621419 | 16.2167387 | 16.0543914 | -0.005618559 | 0.970300224 |
| 16.3449676 | NA | 16.5123029 | 16.6206529 | 0.130517317 | 0.468998339 |
| NA | 14.6462778 | NA | NA | -0.160703379 | 0.52354192 |
| NA | 23.0500884 | NA | NA | 0.072850224 | 0.833386793 |
| 20.6217141 | 20.6188671 | 20.7055852 | 20.7998351 | 0.028216756 | 0.606932403 |
| 20.3282145 | 20.2086594 | 20.3727921 | 20.3494987 | 0.034173949 | 0.595687669 |
| 20.1717843 | 20.2249279 | 20.5027404 | 20.0139638 | 0.008548577 | 0.955973153 |
| 23.134032 | 22.9511581 | 23.0381701 | 23.0523399 | -0.046492634 | 0.392444465 |
| 23.2364504 | 23.1431629 | 23.2244137 | 23.2271403 | -0.002358763 | 0.964106527 |
| 16.5340891 | 16.6077176 | 16.2959439 | 16.586819 | 0.193729354 | 0.202012324 |
| NA | 13.5897799 | NA | NA | 0.578495291 | 0.023599588 |
| 22.0881153 | 22.079838 | 22.0765691 | 22.1446952 | 0.022392005 | 0.601543824 |
| 16.6949753 | 16.6563342 | 16.7431225 | 16.6029077 | -0.056889095 | 0.69577938 |
| 22.3228358 | 22.212665 | 22.3020926 | 22.3251753 | 0.011871256 | 0.868518363 |
| 15.5987803 | 15.4533831 | 15.3634625 | 15.6164125 | -0.090655824 | 0.514025249 |
| 16.175548 | 15.6636501 | 16.1285399 | 16.24604 | -0.253353805 | 0.30172975 |
| 20.1843746 | 20.0941783 | 20.2503839 | 20.2890255 | -0.011020508 | 0.859274007 |
| 23.5461554 | 23.4467613 | 23.4895184 | 23.5598576 | 0.024127256 | 0.622933634 |
| 19.5213792 | 19.4114714 | 19.4393796 | 19.4117725 | -0.08438917 | 0.108392399 |
| 27.7600567 | 27.7040205 | 27.7594105 | 27.7687716 | 0.062401125 | 0.23964923 |
| 18.167843 | 18.0334764 | 18.1421131 | 18.0190873 | -0.013945617 | 0.818985566 |
| 21.8771443 | 21.716501 | 21.8516691 | 21.851862 | -0.002401415 | 0.969611215 |
| 18.1001166 | 18.017945 | 18.0538166 | 17.9378147 | -0.04555622 | 0.751421309 |
| 23.482397 | 23.3728685 | 23.6229108 | 23.3869435 | -0.038531557 | 0.751253654 |
| 20.0993124 | 19.9789277 | 20.0968931 | 20.0802046 | -0.048288857 | 0.416078473 |
| 20.6317794 | 20.717473 | 20.6022317 | 20.6300545 | 0.018114477 | 0.756818768 |
| 23.7156484 | 23.603505 | 23.8159479 | 23.7961466 | 0.059609041 | 0.360862354 |
| 20.1732449 | 19.9285576 | 20.0961058 | 20.1917143 | 0.026540684 | 0.723660749 |
| 20.2267846 | 20.1874426 | 20.1899924 | 20.2155847 | -0.01950384 | 0.68068997 |
| 18.7988568 | 18.7625523 | 18.7531941 | 18.7543726 | -0.187955541 | 0.01754578 |
| 20.754691 | 20.5912082 | 20.6999003 | 20.7425693 | -0.023691283 | 0.664829405 |
| 20.053538 | 20.1580961 | 20.0050051 | 20.0048645 | -0.051448469 | 0.548968886 |
| 20.0395664 | 19.9175794 | 20.0111628 | 19.9869146 | -0.063796071 | 0.264364553 |
| 19.9917658 | 19.9137957 | 19.9280984 | 19.9907171 | 0.013294867 | 0.820008576 |
| 17.0193064 | 16.9005788 | 17.1042054 | 16.8887556 | -0.037124004 | 0.65853376 |
| 16.3542579 | 16.615511 | 16.5971318 | 16.6724744 | 0.150698937 | 0.206990555 |
| 16.3843829 | 16.6857738 | 16.4964213 | 16.4475855 | 0.059561841 | 0.622573544 |

| | | | | | |
|---|---|---|---|---|---|
| 18.995934 | 19.2405235 | 18.8428778 | 18.891673 | 0.047467697 | 0.678396871 |
| 18.7053527 | 18.616048 | 18.6375571 | 18.7850085 | -0.006514989 | 0.926349988 |
| 18.0068722 | 17.9623141 | 17.9232861 | 18.0299074 | -0.071083459 | 0.233161628 |
| 20.987694 | 20.7741878 | 20.9767319 | 20.9577788 | -0.01075187 | 0.866115226 |
| 26.0502948 | 25.8759544 | 25.9780038 | 25.9280094 | -0.048613794 | 0.387272253 |
| 18.7090511 | 18.6969025 | 18.6554325 | 18.6505113 | 0.007402709 | 0.893059619 |
| 23.0765133 | 22.9609957 | 22.9638424 | 23.0388492 | -0.006545316 | 0.925729245 |
| 17.4088394 | 16.682478 | 17.0455486 | 16.9127983 | -0.247492824 | 0.422060376 |
| 24.4678119 | 24.4131646 | 24.5129628 | 24.4705761 | 0.033240184 | 0.511562465 |
| 19.3884975 | 19.3591616 | 19.4231858 | 19.3506794 | 0.04186325 | 0.349922238 |
| 18.9197125 | 18.9494353 | 18.8992669 | 18.8786655 | -0.029430973 | 0.750079975 |
| 17.2294793 | 17.236613 | 17.439644 | 17.2621018 | 0.066069256 | 0.375285714 |
| 19.3565207 | 19.371963 | 19.3838242 | 19.3840238 | 0.030340906 | 0.661493348 |
| 17.3736157 | 17.0290249 | 16.8625991 | 17.5028782 | -0.094065036 | 0.521887764 |
| 17.9436363 | 17.721759 | 17.3935732 | 17.6982262 | -0.19615131 | 0.098394001 |
| 21.6249584 | 21.5380862 | 21.6006739 | 21.5616646 | 0.001947227 | 0.972858513 |
| 16.4347533 | 16.5044347 | 16.7402676 | 16.7644242 | 0.140542113 | 0.265481249 |
| 21.0574009 | 20.9549019 | 20.9465828 | 20.9772572 | -0.093708135 | 0.235373329 |
| 21.1487053 | 21.0835089 | 21.0951714 | 21.1147651 | -0.013354565 | 0.758445484 |
| 14.3333127 | NA | 14.2394656 | 14.5664631 | 0.052791783 | 0.701734745 |
| 18.419846 | 18.7058368 | 18.3235167 | 18.6093558 | -0.004544241 | 0.972603585 |
| 24.3331322 | 24.3154122 | 24.3858588 | 24.3602817 | 0.018247144 | 0.733786781 |
| 19.8752517 | 19.6991749 | 19.85947 | 19.8460635 | -0.061347643 | 0.181405749 |
| 24.0356852 | 23.9286172 | 24.0753194 | 24.0822553 | 0.038260526 | 0.493035297 |
| 22.9021581 | 22.7562034 | 22.8506061 | 22.8741726 | -0.056996908 | 0.418173569 |
| 18.6423734 | 18.5313959 | 18.6497841 | 18.6740428 | -0.019086317 | 0.762068426 |
| 22.3156667 | 22.3224726 | 22.2536987 | 22.3286627 | 0.04205809 | 0.611398174 |
| 22.8725367 | 22.9957712 | 23.0225142 | 22.9319311 | 0.053733462 | 0.27370067 |
| 20.5601529 | 20.360377 | 20.5334994 | 20.4719074 | -0.054376829 | 0.275639993 |
| 20.255193 | 20.2173335 | 20.2306434 | 20.2557663 | -0.003549183 | 0.938155453 |
| 23.5480971 | 23.4338901 | 23.5700962 | 23.5362683 | 0.002979234 | 0.951472302 |
| 22.1105222 | 21.8675326 | 22.0024531 | 21.7870245 | -0.088678664 | 0.5654604 |
| 26.1638091 | 26.1774459 | 26.2538739 | 26.2508992 | 0.096563197 | 0.276678238 |
| 20.8169829 | 20.6406308 | 20.7624174 | 20.7917777 | -0.049087878 | 0.316810955 |
| 22.9387405 | 22.83815 | 22.9571664 | 22.8723893 | -0.018909157 | 0.68612984 |
| 21.4999408 | 21.3112575 | 21.5632053 | 21.5261961 | -0.027485084 | 0.741216949 |
| 21.1639537 | 21.0597594 | 21.1507877 | 21.1185163 | -0.012668113 | 0.826328889 |
| 20.8727786 | 20.823908 | 20.8754271 | 20.888558 | -0.00249784 | 0.96189778 |
| 23.4741517 | 23.371955 | 23.4500999 | 23.4434296 | 0.009463873 | 0.83431013 |
| 19.840363 | 19.9477609 | 19.8760864 | 19.8682704 | 0.048981207 | 0.472980211 |
| 19.7990559 | 19.7790915 | 19.6559025 | 19.8153379 | -0.10670368 | 0.248553584 |
| 25.4061053 | 25.3006229 | 25.3747122 | 25.3750338 | -0.045754617 | 0.338050322 |

| | | | | | |
|---|---|---|---|---|---|
| 22.0508841 | 21.9514944 | 22.0882603 | 22.0758097 | 0.018012997 | 0.676001939 |
| 24.2257255 | 24.195754 | 24.2342516 | 24.2505807 | 0.038803786 | 0.406337148 |
| 20.7147964 | 20.7458999 | 20.73975 | 20.8285639 | 0.01135629 | 0.850212873 |
| 16.9081261 | 16.9032322 | 17.1202378 | 17.3064618 | 0.08836853 | 0.677832427 |
| 20.3322199 | 20.3177572 | 20.2985237 | 20.2815284 | 0.002159093 | 0.960956819 |
| 18.9485398 | 18.8576767 | 18.8282328 | 18.7679697 | -0.048193292 | 0.571399145 |
| 19.2335323 | 19.2311609 | 19.195642 | 19.3428036 | 0.118968263 | 0.289628345 |
| 19.2064858 | 19.5684414 | 19.5035398 | 19.4505683 | 0.255896178 | 0.066323066 |
| 18.3705129 | 18.2284337 | 18.2218518 | 18.1250173 | -0.116436998 | 0.355594078 |
| 20.9600662 | 20.8006341 | 21.0156761 | 21.0493403 | 0.064379242 | 0.399409054 |
| 13.0420085 | NA | 12.962333 | 13.1313295 | -0.039829108 | 0.795490374 |
| 16.3745066 | 15.7124311 | 16.2458223 | 16.2402302 | -0.287072995 | 0.12410572 |
| 17.8499731 | 17.600344 | 17.7332124 | 17.7639345 | -0.072589989 | 0.49452486 |
| 21.6849027 | 21.5288024 | 21.6595061 | 21.7164295 | -0.011248144 | 0.857538331 |
| 21.0796411 | 21.0444311 | 21.1333987 | 21.3624521 | 0.030967913 | 0.71868012 |
| 20.6659214 | 20.4721373 | 20.6347142 | 20.5833476 | -0.037883045 | 0.451129747 |
| 21.4673672 | 21.4615626 | 21.4953544 | 21.4555775 | 0.026615339 | 0.684456533 |
| 19.9700266 | 19.7318761 | 19.8225114 | 19.824176 | -0.051739986 | 0.492368691 |
| 21.9523458 | 21.7339775 | 21.9684025 | 21.9908983 | 0.012350573 | 0.883595019 |
| 12.8301281 | 13.0681415 | 13.0254192 | 13.5151141 | 0.054130524 | 0.699776163 |
| 17.0336042 | 16.9055815 | 17.3021221 | 17.125434 | -0.060705332 | 0.787458685 |
| 17.9605361 | 18.0510386 | 18.1392217 | 18.1486236 | 0.146521478 | 0.029345303 |
| 18.9772957 | 18.5397205 | 18.6326195 | 18.7047619 | -0.201457282 | 0.035379023 |
| 22.834188 | 22.8900587 | 22.8551862 | 22.8833605 | 0.071884657 | 0.392658535 |
| 21.6248108 | 21.4371569 | 21.5223596 | 21.5631437 | -0.011672777 | 0.877016073 |
| 20.5358264 | 20.3934399 | 20.5161336 | 20.5529297 | 0.002637607 | 0.966174377 |
| 19.0849248 | 18.9065302 | 19.0789241 | 19.1306616 | 0.025761218 | 0.677722119 |
| 22.8178029 | 22.7103751 | 22.8545064 | 22.8949934 | 0.010478675 | 0.854968106 |
| 22.2847117 | 22.0896461 | 22.1788095 | 22.2368945 | -0.040481198 | 0.442428168 |
| 23.3402656 | 23.2354832 | 23.3672925 | 23.3729267 | -0.019718047 | 0.667528158 |
| 23.6370603 | 23.5067549 | 23.5940785 | 23.5680281 | -0.043826131 | 0.34237289 |
| 24.6285079 | 24.6282855 | 24.6693963 | 24.6868351 | 0.050010714 | 0.258372386 |
| 22.542829 | 22.4123514 | 22.4787089 | 22.4720588 | -0.016554339 | 0.773554073 |
| 17.4113831 | 17.2938488 | 17.3928289 | 17.3377177 | -0.045587817 | 0.549750251 |
| 21.8836959 | 21.7655152 | 21.7939224 | 21.7563332 | -0.002393489 | 0.975342953 |
| 17.4045689 | 17.3362511 | 17.450465 | 17.3258542 | 0.051290197 | 0.524605144 |
| 26.1669071 | 26.0781309 | 26.2286011 | 26.2002391 | 0.099292197 | 0.233486123 |
| 25.4941589 | 25.3561538 | 25.4833957 | 25.4510062 | -5.22E-04 | 0.995294836 |
| 20.7431002 | 20.6872986 | 20.7792744 | 20.8998093 | 0.082078231 | 0.455511088 |
| 16.7619848 | 16.7478301 | 16.7204973 | 16.4644706 | 0.005185073 | 0.971027471 |
| NA | 15.7578682 | NA | NA | 0.588997841 | 0.309069168 |
| 21.8805135 | 21.7750982 | 21.8789269 | 21.8559506 | -0.014373823 | 0.739486333 |

| | | | | | |
|---|---|---|---|---|---|
| 21.6612376 | 21.6268651 | 21.6349759 | 21.6778104 | 0.011598847 | 0.832021737 |
| 18.8624884 | 18.8557337 | 18.821544 | 18.8809279 | 0.030108003 | 0.622910907 |
| 22.8305408 | 22.7302097 | 22.9672275 | 22.8466466 | 0.279577821 | 0.110301165 |
| 21.3835571 | 21.4476799 | 21.422632 | 21.0337525 | -0.088518843 | 0.473516142 |
| 19.0413192 | 19.1941693 | 19.0057518 | 19.0551268 | -0.034997081 | 0.658642553 |
| 21.2095952 | 21.2564936 | 21.2575106 | 21.2636309 | 0.023550761 | 0.482959351 |
| 23.4295068 | 23.3622123 | 23.4788574 | 23.4959742 | 0.034725572 | 0.461991213 |
| 22.3999799 | 22.2817834 | 22.3808231 | 22.3794756 | 0.003048841 | 0.949171909 |
| 23.0050164 | 22.8995877 | 23.0462118 | 22.9640432 | 0.003326598 | 0.947148395 |
| 21.0977211 | 20.9611124 | 21.0044697 | 20.9718291 | -0.068395237 | 0.198309291 |
| 21.9004847 | 21.8363551 | 21.9549112 | 21.9008687 | -0.022582796 | 0.682496135 |
| 21.099833 | 20.9747616 | 21.1073559 | 21.0289427 | 8.37E-04 | 0.989353094 |
| 21.7410278 | 21.7188862 | 21.7668357 | 21.7886109 | 0.001150807 | 0.985580997 |
| 20.3694652 | 20.3363191 | 20.3465646 | 20.4512428 | 0.037355003 | 0.739902133 |
| 21.2860843 | 21.0343074 | 21.3047603 | 21.7132284 | 0.528674383 | 0.059887285 |
| 16.6786779 | 16.6005857 | 16.7066655 | 16.6165237 | -0.076780662 | 0.474217079 |
| 20.1249255 | 20.0699418 | 20.3125931 | 20.0189205 | 0.034748391 | 0.677114778 |
| 20.0961944 | 19.9972518 | 20.0235535 | 20.0323031 | -0.049707484 | 0.402603787 |
| 18.7750024 | 18.5016449 | 18.6479137 | 18.6804531 | -0.092553242 | 0.262022599 |
| 18.1644333 | 18.2507942 | 18.2352388 | 18.3741271 | 0.123153215 | 0.238805454 |
| 21.1554971 | 21.0176658 | 21.15753 | 21.1231646 | -0.010964955 | 0.844254559 |
| NA | 11.484118 | NA | NA | 0.885362087 | 0.075735786 |
| 14.8980524 | NA | 14.7225369 | 15.1597209 | 0.010156846 | 0.932452252 |
| 20.5367813 | 20.4655421 | 20.1820334 | 20.1765054 | -0.055871537 | 0.572292439 |
| 21.4473633 | 21.1439777 | 21.3785896 | 21.4541321 | -0.033122712 | 0.717779808 |
| 19.2782142 | 19.1333387 | 19.218217 | 19.1144311 | -0.094896723 | 0.181179615 |
| 20.2550197 | 20.239779 | 20.2556488 | 20.2719748 | 0.024870809 | 0.687108044 |
| 25.255875 | 25.110161 | 25.2021845 | 25.2196755 | -0.030380676 | 0.611528277 |
| 25.2937875 | 25.182457 | 25.2786027 | 25.2664905 | 0.030451599 | 0.608734445 |
| 20.2145338 | 20.2475806 | 20.3679587 | 20.3153122 | 0.021847278 | 0.671085423 |
| 22.4027853 | 22.2752477 | 22.3497217 | 22.3672956 | -0.004917376 | 0.923150006 |
| 20.3319761 | 20.3289829 | 20.3088614 | 20.3395773 | -0.014697901 | 0.761679993 |
| 18.8320634 | 18.6364655 | 18.7893628 | 18.783605 | -0.076629889 | 0.12633844 |
| 19.8546143 | 19.7280352 | 19.983243 | 19.8715159 | 0.002997748 | 0.957161793 |
| 19.0774847 | 19.0627531 | 19.0916772 | 19.0323855 | 0.006803208 | 0.909256767 |
| 24.0857166 | 24.0443102 | 24.1099794 | 24.0718733 | 0.006546653 | 0.915609677 |
| 21.4479681 | 21.4238315 | 21.2772816 | 21.2565437 | -0.02503235 | 0.775526952 |
| 15.7925646 | 15.9467217 | 15.6819729 | 15.8662575 | -0.018148751 | 0.839165791 |
| 16.7452991 | 16.8269409 | 16.7545987 | 16.9200345 | 0.130505198 | 0.184987744 |
| 17.3442774 | 17.4585262 | 17.4840334 | 17.5705476 | -0.012036449 | 0.895216929 |
| 14.9224939 | 14.6999827 | 14.574431 | 14.8769149 | -0.059838735 | 0.738339425 |
| 17.5785513 | 17.4298557 | 17.3951914 | 17.6323119 | -0.09042897 | 0.251677077 |

| | | | | | |
|---|---|---|---|---|---|
| 18.5135304 | 18.3242176 | 18.3085336 | 18.4519053 | -0.021248864 | 0.76412647 |
| 21.1730932 | 21.0933844 | 21.1797073 | 21.110008 | -0.024049101 | 0.594045274 |
| 18.9973008 | 18.8042798 | 18.9524272 | 18.9296616 | -0.007140588 | 0.919726154 |
| 19.2101665 | 18.9678864 | 19.1907346 | 19.2657407 | -0.02293796 | 0.846033312 |
| 19.7324699 | 19.68844 | 19.7302963 | 19.7377626 | -0.00562098 | 0.915188835 |
| 19.3411327 | 19.3825456 | 19.3021464 | 19.3247186 | 0.009066746 | 0.879525308 |
| 19.5800323 | 19.4564673 | 19.4867296 | 19.4829616 | -0.046749693 | 0.391020405 |
| 20.0273161 | 20.0646178 | 20.0959156 | 20.005269 | 0.004083039 | 0.943710333 |
| 18.0670636 | 17.9069847 | 17.9409484 | 18.0285366 | -7.22E-04 | 0.993521063 |
| NA | 14.3139776 | NA | NA | -0.284953053 | 0.385527074 |
| 18.4799635 | 18.3643872 | 18.4823766 | 18.3950565 | -0.112630451 | 0.423565762 |
| 15.4566055 | 15.2460297 | 15.5489863 | 15.5206083 | -0.100093621 | 0.347774238 |
| 19.9022849 | 19.9629892 | 19.9133722 | 20.0773611 | 0.081375369 | 0.323770307 |
| 20.0609013 | 19.9775683 | 20.0410427 | 20.1068266 | -0.070223553 | 0.368303007 |
| 22.1856612 | 22.0826228 | 22.1959658 | 22.2531572 | 0.02341493 | 0.685769465 |
| 20.465833 | 20.4185881 | 20.5215114 | 20.4419887 | -0.006897956 | 0.893057974 |
| 22.8754597 | 22.8053013 | 22.9316207 | 22.8937977 | 0.02282882 | 0.638117957 |
| 22.4260312 | 22.2767772 | 22.4403933 | 22.4094563 | 0.001071539 | 0.981840022 |
| 20.9457625 | 21.0055208 | 20.9959502 | 20.9346822 | 0.013637099 | 0.759745282 |
| 21.3291606 | 21.2788927 | 21.3332249 | 21.3171399 | 0.023217117 | 0.603829207 |
| 22.2983797 | 22.1855819 | 22.2097412 | 22.1702564 | -0.062577947 | 0.166775765 |
| 21.5954036 | 21.5648976 | 21.6026111 | 21.6220589 | -0.01088702 | 0.748568309 |
| 23.3256455 | 23.3004373 | 23.2600401 | 23.3563425 | 0.024756725 | 0.600866424 |
| 22.6904743 | 22.5881059 | 22.7217436 | 22.6261224 | -0.01125871 | 0.82584369 |
| 20.8149527 | 20.7178805 | 20.8834045 | 20.8083259 | 0.034476599 | 0.544236286 |
| 19.6205128 | 19.5603515 | 19.6635598 | 19.6741745 | 0.050265146 | 0.381988081 |
| 20.3194795 | 20.3362924 | 20.2920465 | 20.2785251 | -0.001927622 | 0.973901248 |
| 18.9802194 | 18.9405224 | 18.9613359 | 18.981269 | -0.028261609 | 0.644597495 |
| 21.4753459 | 21.3992729 | 21.425231 | 21.4223667 | 0.031887916 | 0.627012277 |
| 19.5863122 | 19.5441309 | 19.60691 | 19.5227388 | 0.058537846 | 0.434633562 |
| 19.5123912 | 19.3911282 | 19.5698265 | 19.4453125 | -0.069565929 | 0.366069857 |
| 18.2280311 | 18.2321896 | 18.2119504 | 18.1970473 | 0.007797558 | 0.921779531 |
| 20.0938595 | 19.8508481 | 20.1257404 | 20.1774686 | 0.003555876 | 0.966926537 |
| 20.0210612 | 20.2812147 | 20.1808124 | 20.0125624 | 0.132111554 | 0.19590435 |
| 16.6548093 | 16.5242805 | 16.6764427 | 16.9486433 | 0.076166859 | 0.544745316 |
| 20.1038787 | 19.9740054 | 20.0646355 | 20.1294019 | -0.005470408 | 0.966385698 |
| 16.0842076 | NA | 16.1448564 | 16.2100513 | 0.130036065 | 0.499443032 |
| 20.5225574 | 20.2731905 | 20.6857473 | 20.8297468 | 0.220381891 | 0.255614708 |
| NA | 12.4050321 | NA | NA | -0.073346396 | 0.779019046 |
| 22.5982509 | 22.4453953 | 22.5296356 | 22.5745876 | -0.031934422 | 0.484398391 |
| 20.5910935 | 20.5043372 | 20.533176 | 20.5828189 | 0.03306388 | 0.626713814 |
| 21.1822613 | 20.9934509 | 21.0820054 | 21.0309278 | -0.079157742 | 0.180792757 |

| | | | | | |
|---|---|---|---|---|---|
| 18.7897015 | 18.7449114 | 18.7364323 | 18.8143425 | -0.019934606 | 0.673823105 |
| NA | 14.5407426 | NA | NA | -0.136357178 | 0.722810122 |
| 15.2901497 | NA | 15.1735539 | 15.3001925 | 0.119320268 | 0.309411661 |
| 17.6876308 | 17.7795549 | 17.7500894 | 17.7543931 | 0.028120269 | 0.648549087 |
| 18.504708 | 18.4072306 | 18.4630127 | 18.4593038 | -0.039870819 | 0.60852564 |
| 21.6149339 | 21.4349608 | 21.5930956 | 21.6106201 | 0.032691001 | 0.680550382 |
| 24.9146777 | 24.8373991 | 24.8823917 | 24.8318919 | -0.016450732 | 0.698976934 |
| 19.3253475 | 19.1392858 | 19.1479347 | 19.3318049 | -0.022293255 | 0.811764706 |
| NA | 18.7522998 | NA | NA | -0.055940266 | 0.606613576 |
| 20.0660795 | 19.9788106 | 20.0560122 | 19.9939218 | -0.040664854 | 0.323892539 |
| 22.2250929 | 22.1369095 | 22.326881 | 22.2761175 | -0.023656329 | 0.747696523 |
| 19.2374266 | 19.1033044 | 19.1642529 | 19.1951181 | -0.082219185 | 0.223246981 |
| 20.0213506 | 20.0178602 | 20.064547 | 20.0669569 | 0.166855068 | 0.123391391 |
| 22.289092 | 22.2689021 | 22.2539633 | 22.2689347 | 0.007214981 | 0.889674203 |
| 20.5284829 | 20.3793264 | 20.4651114 | 20.5552736 | -0.007981298 | 0.887808024 |
| 21.9432497 | 21.8374943 | 21.9807586 | 21.8833257 | -0.05956082 | 0.226226434 |
| 19.3720349 | 19.3146672 | 19.4082322 | 19.4085868 | -0.001155101 | 0.984039954 |
| 17.2392418 | 16.9283247 | 16.953272 | 16.9900883 | -0.215088821 | 0.014266469 |
| 16.9735664 | 16.6283098 | 17.2031736 | 16.6058551 | -0.278830818 | 0.166075725 |
| 19.6114339 | 19.4762601 | 19.5064364 | 19.6181087 | -0.055693545 | 0.576087908 |
| 21.1740682 | 21.1022875 | 21.1282517 | 21.1803082 | -0.033468331 | 0.496222737 |
| 24.7774499 | 24.6568181 | 24.7504318 | 24.723027 | 0.024999768 | 0.643480961 |
| 15.7811339 | NA | 15.9519341 | 15.4708661 | -0.090833284 | 0.655218162 |
| 17.621704 | 17.3744033 | 17.552617 | 17.5525961 | -0.040022357 | 0.746868457 |
| 20.1142738 | 20.0057972 | 20.1144378 | 20.1070379 | 3.22E-04 | 0.99443242 |
| 19.6594807 | 19.5296194 | 19.5982652 | 19.5418789 | -0.014326926 | 0.759225373 |
| 20.1158914 | 20.2045165 | 20.0240475 | 20.0264022 | -0.058986534 | 0.264363548 |
| 19.5770581 | 19.6321895 | 19.8463295 | 19.7374208 | 0.072414179 | 0.470943342 |
| 18.9126148 | 18.7422054 | 18.7324809 | 18.818997 | -0.105214718 | 0.245860585 |
| 20.375925 | 20.3363772 | 20.3731649 | 20.3535153 | -0.019370144 | 0.692561759 |
| 19.5725598 | 19.5502397 | 19.5685401 | 19.599291 | 0.021889434 | 0.714525338 |
| 19.4446364 | 19.2120996 | 19.289751 | 19.3630148 | 0.0083652 | 0.938902337 |
| 15.3109273 | 15.226593 | 15.4422511 | 15.3007623 | -0.094759149 | 0.589276413 |
| 19.456569 | 19.5383489 | 19.5788237 | 19.4762639 | 0.042348826 | 0.524211123 |
| 19.8836535 | 19.7570263 | 19.8887095 | 19.9077004 | -0.009479421 | 0.854860456 |
| 20.1338888 | 20.1503817 | 20.1729509 | 20.2019272 | 0.057442929 | 0.310319913 |
| 16.3236463 | 16.2087507 | 16.3580258 | 16.4041783 | 0.139320421 | 0.46208027 |
| 18.6761598 | 18.6442458 | 18.4202766 | 18.5366358 | -0.084143029 | 0.215701606 |
| 19.2386401 | 19.1926429 | 19.184538 | 19.2864445 | -0.052313684 | 0.429878849 |
| 21.8550503 | 21.6695815 | 21.7807053 | 21.7376407 | -0.077584013 | 0.120944137 |
| 23.4932066 | 23.3754733 | 23.4567508 | 23.4454879 | -0.011675613 | 0.810416384 |
| 20.125541 | 19.9786836 | 20.1098706 | 20.0562946 | -0.060660173 | 0.201954881 |

| | | | | | |
|---|---|---|---|---|---|
| 19.171075 | 19.1031774 | 19.2050137 | 19.2812412 | 0.025303817 | 0.633823462 |
| 18.092419 | 18.170661 | 18.2458244 | 18.1459046 | -0.019392317 | 0.784068099 |
| 16.5599698 | 16.2194383 | 16.3059718 | 16.7297754 | -0.084666956 | 0.440538538 |
| 17.6525358 | 17.3950618 | 17.6985977 | 17.6644753 | -0.155298155 | 0.254221316 |
| 14.225068 | NA | 14.8495357 | 14.7651067 | 0.333554801 | 0.192095146 |
| 17.7607218 | 17.1684248 | 17.8341409 | 17.8150517 | -0.1518642 | 0.355185253 |
| 19.3239602 | 19.3120945 | 19.4338346 | 19.3914279 | 0.053196219 | 0.587127173 |
| 19.1444235 | 18.9257515 | 19.1240704 | 19.0197192 | -0.104044766 | 0.225709289 |
| 21.3686686 | 21.2967729 | 21.3465447 | 21.3143798 | -0.010581236 | 0.85525342 |
| 23.371418 | 23.2854639 | 23.3892868 | 23.3495351 | -0.013249982 | 0.7689512 |
| 21.3071009 | 21.2944998 | 21.337454 | 21.2657562 | 0.010983665 | 0.803705677 |
| 21.731022 | 21.697517 | 21.7928976 | 21.7763877 | 0.037700683 | 0.355568647 |
| 19.9334156 | 19.9464154 | 19.9488531 | 19.9611632 | 0.009817172 | 0.827766148 |
| 22.126216 | 22.1990084 | 22.3482596 | 22.3133585 | 0.110929898 | 0.050817474 |
| 20.8201088 | 20.7237588 | 20.805485 | 20.6898569 | -0.023333202 | 0.829200786 |
| 17.4690984 | 17.2865875 | 17.1954241 | 17.4397222 | -0.041016381 | 0.760434686 |
| 23.6127728 | 23.5424288 | 23.6348407 | 23.5666311 | 0.016734266 | 0.734785551 |
| 20.5746956 | 20.4071873 | 20.5044714 | 20.5068135 | -0.033678479 | 0.557674953 |
| 14.2024743 | NA | 14.4014888 | 14.4158968 | 0.138171377 | 0.487726368 |
| 22.6986436 | 22.6247297 | 22.8156099 | 22.7194238 | 0.061636116 | 0.327156932 |
| 21.2328118 | 21.2655129 | 21.2956658 | 21.3246134 | 0.047657461 | 0.332142318 |
| 15.5896602 | 15.0647366 | 15.3608952 | 15.6216425 | -0.223743455 | 0.077778555 |
| 21.1266894 | 21.0075461 | 21.068362 | 21.082515 | -0.053566825 | 0.352334854 |
| 22.0878688 | 22.0416282 | 22.1676739 | 22.1332453 | 0.003951004 | 0.941588752 |
| 21.8574887 | 21.73351 | 21.8360687 | 21.8298406 | -0.014237249 | 0.759086589 |
| 23.7516922 | 23.6953781 | 23.7553513 | 23.7684182 | -0.001832343 | 0.968174771 |
| 20.7348312 | 20.6184375 | 20.6243762 | 20.6848197 | -0.047183754 | 0.257264126 |
| 23.9756662 | 23.9548958 | 23.9871216 | 23.965205 | 0.01882328 | 0.602821151 |
| 21.8347273 | 21.6937172 | 21.7746338 | 21.7691471 | -0.013340164 | 0.793634155 |
| 19.6651547 | 19.5267799 | 19.725646 | 19.6346269 | -0.043483328 | 0.398923568 |
| 21.8868765 | 21.7874757 | 21.832256 | 21.8153883 | -0.059340226 | 0.255624161 |
| 26.4900757 | 26.4328722 | 26.4904987 | 26.483762 | 0.036106208 | 0.510433352 |
| 24.770396 | 24.6188961 | 24.7107781 | 24.7240404 | -0.005951961 | 0.914907519 |
| 20.9427295 | 21.0067019 | 21.0332513 | 20.9527414 | 0.005572315 | 0.923857042 |
| 19.9418233 | 19.9720872 | 20.0286761 | 20.0176451 | 0.056128073 | 0.377418244 |
| 21.8775406 | 21.767974 | 21.9506493 | 21.956091 | 0.017258345 | 0.786612244 |
| 20.9751643 | 20.8811596 | 20.9658326 | 21.0370656 | 0.024553549 | 0.704760222 |
| 20.2315741 | 20.0624044 | 20.1567016 | 20.0946787 | 0.009440576 | 0.899787685 |
| 17.4178927 | 17.5557164 | 17.3669204 | 17.3260658 | -0.089055794 | 0.246656075 |
| 17.7107208 | 17.6277266 | 17.7677044 | 17.7931834 | 0.013912592 | 0.883501584 |
| 17.3212806 | 16.9561211 | 17.146596 | 17.3566383 | 0.036721718 | 0.758124572 |
| 18.6026139 | NA | 19.1120646 | 18.6954437 | 0.297257609 | 0.116395689 |

| | | | | | |
|---|---|---|---|---|---|
| NA | 14.6315034 | NA | NA | 0.35072099 | 0.256464478 |
| 26.3326893 | 26.3292923 | 26.3611585 | 26.4136108 | 0.075114997 | 0.470630025 |
| 19.0221402 | 18.9107499 | 19.0923615 | 19.1199718 | 0.012550025 | 0.883480226 |
| 19.5444184 | 19.4466301 | 19.5946612 | 19.6436198 | 0.023480906 | 0.657660199 |
| 20.4809519 | 20.4687176 | 20.4173147 | 20.4790398 | 0.013655329 | 0.77362138 |
| 22.1002873 | 22.1178825 | 22.1833294 | 22.0807837 | 0.069486248 | 0.165355075 |
| 22.3584483 | 22.2849035 | 22.3849809 | 22.3713322 | 0.04834031 | 0.333120059 |
| 23.9886905 | 23.9441239 | 24.0483757 | 23.9842655 | 0.057393199 | 0.286276159 |
| 16.0597816 | NA | 16.107215 | 16.3990992 | 0.064389195 | 0.6495712 |
| 19.6783262 | 19.6351806 | 19.6940451 | 19.6854857 | 0.043499882 | 0.395433128 |
| 21.9142878 | 21.7586696 | 21.8845185 | 21.9188673 | -0.020431162 | 0.785246423 |
| 21.3011865 | 21.1326847 | 21.2811642 | 21.3718917 | -0.029248542 | 0.737411926 |
| 22.1779508 | 21.9588391 | 22.3114041 | 22.2689437 | 0.022682286 | 0.892968859 |
| 21.2944012 | 21.1873906 | 21.2563642 | 21.3254083 | 0.058691197 | 0.392768633 |
| 22.1470396 | 22.0831929 | 22.2438208 | 22.3055359 | 0.047247424 | 0.421012237 |
| 20.9929235 | 21.0376204 | 21.0066812 | 20.9665577 | -0.002376114 | 0.968174547 |
| 19.3621456 | 19.3235158 | 19.4727626 | 19.3670578 | -0.045685323 | 0.516542437 |
| 24.9956519 | 24.9436932 | 24.7879803 | 24.8666013 | 6.22E-04 | 0.997287293 |
| 13.4997939 | 12.2539084 | 13.6794867 | 13.2669237 | -0.29845251 | 0.611304986 |
| 19.1536416 | 19.1728027 | 19.24987 | 19.2162823 | -0.031310541 | 0.622913378 |
| 18.6888106 | 18.4748877 | 18.6040687 | 18.5695112 | -0.083334052 | 0.19193043 |
| 21.3080366 | 21.3513946 | 21.3109331 | 21.2219062 | -0.051675056 | 0.210032479 |
| 21.5654355 | 21.4622223 | 21.5413707 | 21.4833568 | -0.043439869 | 0.385987433 |
| 21.8209622 | 21.5849628 | 22.0353087 | 21.9549228 | 0.158575085 | 0.092770994 |
| 17.2536074 | NA | 17.2841717 | 16.8935384 | -0.15171149 | 0.332223457 |
| 18.9739568 | 18.7878598 | 18.900153 | 19.0271923 | -0.042345872 | 0.552937292 |
| 22.3656229 | 22.2656755 | 22.3194248 | 22.3205327 | -0.03570534 | 0.462607884 |
| 21.285853 | 21.2975519 | 21.3444015 | 21.2719252 | 0.011861388 | 0.802101112 |
| 18.0014773 | 17.9224687 | 17.8575387 | 17.9839925 | -0.065805643 | 0.218794814 |
| 23.8033807 | 23.6178606 | 23.750274 | 23.7387595 | 1.07E-04 | 0.998599958 |
| 19.0944326 | 19.0730542 | 19.177719 | 19.0725876 | 0.02546594 | 0.702876321 |
| 18.8223143 | 18.6763754 | 18.4401269 | 18.6768049 | -0.084743599 | 0.361837514 |
| 16.2342586 | 16.4977466 | 16.6841731 | 16.2768431 | 0.261029208 | 0.130714444 |
| 18.2900556 | 18.4445918 | 18.2852983 | 18.2458442 | 0.22269821 | 0.225868299 |
| 19.1145974 | 18.6930213 | 18.765469 | 19.179975 | -0.099703277 | 0.515859949 |
| 22.4895492 | 22.3950993 | 22.489357 | 22.4961959 | -0.017387028 | 0.70233394 |
| 19.7094683 | 19.749535 | 19.6479374 | 19.7124365 | 0.024501384 | 0.656877319 |
| 18.9937078 | 18.8495604 | 18.9107339 | 18.9779364 | -0.014840137 | 0.833585075 |
| 15.0359051 | 14.8109646 | 14.949587 | 14.8291396 | 0.083236578 | 0.627642955 |
| 21.8516386 | 21.5298898 | 21.744215 | 21.5963469 | -0.118550884 | 0.501836866 |
| 17.4464028 | 17.2629075 | 17.4359248 | 17.3645562 | -0.048296565 | 0.612780251 |
| 18.4872187 | 18.4082365 | 18.4530568 | 18.5154929 | -0.05144729 | 0.436485463 |

| | | | | | |
|---|---|---|---|---|---|
| 22.4655554 | 22.3496699 | 22.4941119 | 22.4506342 | 0.022745781 | 0.619732622 |
| 20.9055283 | 20.7629972 | 20.8262292 | 20.8926366 | -0.017800815 | 0.731516264 |
| 17.4696386 | 17.6255972 | 17.8301564 | 17.6187427 | 0.120453189 | 0.1982651 |
| 22.8493572 | 22.6902203 | 22.8209478 | 22.8485339 | -0.044458184 | 0.362343549 |
| 22.6122151 | 22.6177024 | 22.6220686 | 22.6037488 | 0.002132183 | 0.962378332 |
| 21.3015674 | 21.2910625 | 21.2483222 | 21.2815878 | 0.028303245 | 0.54587127 |
| 21.9310477 | 21.8600913 | 21.9708421 | 22.0106802 | 0.037740329 | 0.513605528 |
| 20.9199249 | 20.8456816 | 20.8869262 | 20.9643394 | -0.001058016 | 0.983603509 |
| 25.2091724 | 25.0673941 | 25.2095734 | 25.2044028 | -0.004757667 | 0.942253158 |
| 21.6116801 | 21.5460665 | 21.6504375 | 21.7352632 | 0.032605015 | 0.520892567 |
| 22.3882848 | 22.3039476 | 22.3475919 | 22.3489446 | -0.003768456 | 0.933520382 |
| 18.5710208 | 18.4624788 | 18.4741372 | 18.4780263 | -0.041481731 | 0.528784591 |
| 21.6819869 | 21.5720323 | 21.6481517 | 21.6559024 | -0.011743134 | 0.86812804 |
| 22.3422487 | 22.1965505 | 22.3023002 | 22.3269089 | -0.027157359 | 0.48808921 |
| 24.3980527 | 24.2774622 | 24.4447943 | 24.4051716 | 0.001801125 | 0.97641765 |
| 19.2215293 | 18.7522262 | 18.750967 | 19.1363599 | -0.112164986 | 0.399507066 |
| NA | 16.6286019 | NA | NA | 0.734681158 | 0.252216241 |
| 25.4558217 | 25.3077064 | 25.5042902 | 25.481855 | 0.040560226 | 0.738073292 |
| 18.4017298 | 18.2301325 | 18.4492439 | 18.3544568 | -0.105028694 | 0.24289409 |
| NA | 14.083319 | NA | NA | -0.517474214 | 0.091960141 |
| 16.8732076 | 16.7242968 | 16.9281131 | 16.8042146 | -0.061784071 | 0.537771012 |
| 21.5605782 | 21.405781 | 21.3704517 | 21.354796 | -0.085237384 | 0.531295281 |
| 20.64726 | 20.4856075 | 20.403134 | 20.5541132 | -0.013594868 | 0.942503054 |
| 13.6474668 | 13.5763032 | 13.8804479 | 13.6485715 | 0.191075413 | 0.177437533 |
| 13.7953679 | NA | 13.9913967 | 13.8331243 | 0.037432222 | 0.893510361 |
| NA | 10.991971 | NA | NA | -0.173924693 | 0.512312098 |
| 19.602535 | 19.4944683 | 19.6100219 | 19.6972699 | 0.044032035 | 0.531869253 |
| 23.3596979 | 23.3347342 | 23.4275502 | 23.3662252 | 0.044548118 | 0.383995573 |
| 20.4916243 | 20.4702486 | 20.6300204 | 20.4863345 | 0.030844152 | 0.599645202 |
| 19.2762069 | 19.2820283 | 19.3309843 | 19.3476367 | 0.027614642 | 0.657536082 |
| 22.3775448 | 22.3558041 | 22.4124219 | 22.3402567 | 0.026974626 | 0.666667211 |
| 16.3104279 | 16.2332128 | 16.326243 | 16.1078815 | -0.056568283 | 0.60980537 |
| 18.4686139 | 18.3004712 | 18.657311 | 18.4874665 | 0.068982249 | 0.443594847 |
| 19.1885764 | 19.0237945 | 19.1504069 | 19.0952239 | -0.103371378 | 0.233437225 |
| 17.4055925 | 17.289634 | 17.3881235 | 17.4146919 | 0.0227607 | 0.790012827 |
| 22.719903 | 22.6453397 | 22.7013559 | 22.6905176 | 0.005668508 | 0.908951235 |
| NA | 17.0108266 | NA | NA | 0.020397696 | 0.885789872 |
| 20.8977836 | 20.8185884 | 20.7878294 | 20.8372186 | -0.044882776 | 0.204705508 |
| 22.7248684 | 22.7232952 | 22.8041876 | 22.8007204 | 0.056367914 | 0.285972693 |
| 19.0711977 | 19.2220999 | 18.993409 | 19.1081792 | 0.075323635 | 0.640445016 |
| 15.6963964 | 15.5518952 | 15.6142242 | 15.6471146 | -0.010985915 | 0.944235873 |
| 18.2820892 | 17.975402 | 18.3082197 | 18.2410009 | -0.082122387 | 0.486297603 |

| | | | | | |
|---|---|---|---|---|---|
| 17.3298635 | 17.3224194 | 17.3568409 | 17.3919966 | 0.084002479 | 0.395519096 |
| 19.7302541 | 19.6014352 | 19.7588115 | 19.7361345 | 0.026062193 | 0.699614782 |
| 22.0940139 | 21.8870888 | 22.0537281 | 22.0150386 | -0.074009494 | 0.256888 |
| 22.2290477 | 22.2120776 | 22.2376022 | 22.2191669 | 0.02531473 | 0.675355583 |
| 19.0130233 | 18.9495291 | 19.0499603 | 18.9434837 | -0.04799236 | 0.34309113 |
| 22.5266489 | 22.4747153 | 22.5211098 | 22.5179173 | 0.014108015 | 0.737777905 |
| 22.0198843 | 21.9622664 | 21.9945763 | 21.9155927 | 7.43E-04 | 0.98616371 |
| 22.6536371 | 22.5777468 | 22.6195132 | 22.6757554 | 0.013721945 | 0.758400872 |
| 22.2628974 | 22.3113116 | 22.2639674 | 22.2684139 | 0.023469251 | 0.637651458 |
| 20.7499203 | 20.6106756 | 20.6839888 | 20.6659458 | -0.031365386 | 0.547468352 |
| 22.6417528 | 22.4800305 | 22.6646668 | 22.7178161 | 0.019597082 | 0.791228106 |
| 19.3205957 | 19.4095783 | 19.3216003 | 19.5559846 | 0.099472337 | 0.20750966 |
| 19.1293963 | 19.0469412 | 19.2141152 | 19.0664613 | -0.091178995 | 0.324372655 |
| 22.1139508 | 22.5719213 | 22.2437308 | 22.1832201 | 0.078352547 | 0.699970256 |
| 18.2454952 | 18.3536689 | 18.2426117 | 18.3599707 | 0.04944518 | 0.559776399 |
| 25.1453962 | 25.03508 | 25.1535914 | 25.1537063 | 0.014189854 | 0.76659245 |
| 14.8971214 | NA | 14.9837637 | 14.8240216 | 0.147021192 | 0.246144217 |
| 19.5984859 | 19.5330306 | 19.5562545 | 19.5916633 | -2.73E-04 | 0.996828259 |
| 20.8026551 | 20.6424319 | 20.7415201 | 20.7767034 | 0.009802376 | 0.872604922 |
| 16.4973602 | 16.4595736 | 16.401112 | 16.7386398 | 0.039188801 | 0.801915181 |
| 15.4505991 | NA | 15.3916919 | 15.4257826 | -0.102043952 | 0.551664137 |
| 19.6981394 | 19.6291243 | 19.6322919 | 19.6036314 | -0.089690968 | 0.107244423 |
| 18.9811172 | 18.8284179 | 18.9986302 | 18.9482632 | -0.068558781 | 0.256048318 |
| 20.0089123 | 19.8779649 | 19.9010748 | 19.9201915 | -0.052285439 | 0.370750041 |
| 21.0146577 | 20.9939268 | 21.0502346 | 21.0543783 | 0.030758943 | 0.495974417 |
| 19.7002808 | 19.5509756 | 19.6401932 | 19.8653443 | 0.002234254 | 0.981782029 |
| 22.4984293 | 22.4220283 | 22.5764924 | 22.4842244 | 0.030796818 | 0.722446762 |
| 15.6976193 | 15.892229 | 15.8944069 | 15.7501622 | 0.031336967 | 0.807800706 |
| 18.0817242 | 17.8829479 | 17.393733 | 18.0543792 | 0.029251258 | 0.882964241 |
| 16.373594 | 16.2893828 | 16.2093945 | 16.4945337 | -0.070610084 | 0.652276821 |
| 21.2776556 | 21.3359699 | 21.55793 | 21.3618315 | 0.093460439 | 0.30790053 |
| 23.3274024 | 23.3306135 | 23.2595314 | 23.2879034 | 0.051841842 | 0.403266983 |
| 20.5765151 | 20.5427809 | 20.6147426 | 20.6693487 | -0.001249405 | 0.97888844 |
| 21.143363 | 21.0768333 | 21.1701805 | 21.2266216 | 8.98E-04 | 0.987220004 |
| 18.8386317 | 18.5408482 | 18.677536 | 18.7015952 | -0.006104467 | 0.943205249 |
| 15.6226631 | 15.7212489 | 15.9807619 | 15.8845329 | 0.174604472 | 0.160529098 |
| 23.1964176 | 23.0702849 | 23.2027488 | 23.1229896 | -0.024546462 | 0.670814951 |
| 21.2716991 | 21.2166455 | 21.385113 | 21.3310354 | 0.019966243 | 0.69267839 |
| NA | 15.4654033 | NA | NA | 0.108314962 | 0.589272137 |
| 15.8988047 | 16.1357653 | 15.6107202 | 16.3567857 | 0.113779523 | 0.56052902 |
| 19.6178201 | 19.5216791 | 19.6453783 | 19.6122173 | 0.026369924 | 0.625244149 |
| 18.7447838 | 18.5309167 | 18.6399767 | 18.8066451 | -0.071898384 | 0.339747318 |

| | | | | | |
|---|---|---|---|---|---|
| 21.0462774 | 21.0099938 | 21.0413693 | 21.0935051 | -0.001257602 | 0.982877751 |
| 23.6896828 | 23.5792539 | 23.6650392 | 23.6888981 | 0.035352147 | 0.562585372 |
| 17.5218146 | 17.7270415 | 17.7465205 | 17.7449648 | 0.178442767 | 0.137999109 |
| 20.3205006 | 20.2427001 | 20.3219758 | 20.3361031 | 0.014219389 | 0.785090758 |
| 18.7407145 | 18.8656832 | 18.6210623 | 18.929025 | 0.086982078 | 0.374581502 |
| 19.4089111 | 19.465465 | 19.393485 | 19.4768125 | 0.005547958 | 0.964354427 |
| 18.8249021 | 18.7728015 | 19.0328286 | 18.9550148 | 0.068375605 | 0.499598685 |
| 19.4252586 | 19.3175206 | 19.4087426 | 19.4671088 | -0.001137423 | 0.982241084 |
| 18.9753715 | 18.7878648 | 18.9557244 | 19.0196005 | 0.040011822 | 0.598653582 |
| 21.2563104 | 21.465591 | 21.3972899 | 21.4058623 | 0.17138844 | 0.015916378 |
| 20.6978533 | 20.7378725 | 20.7353785 | 20.8441544 | 0.031859998 | 0.648999096 |
| 22.0928736 | 22.5523653 | 22.5623494 | 22.3476153 | 0.297935967 | 0.052946444 |
| 22.3895 | 22.3373614 | 22.4587558 | 22.3955114 | 0.009094405 | 0.846393967 |
| NA | 11.414627 | NA | NA | -0.924536335 | 0.061775248 |
| NA | 13.7607461 | NA | NA | -0.189044074 | 0.312653741 |
| 16.5700847 | 16.5982242 | 16.7939 | 16.6224153 | -0.03812489 | 0.80531261 |
| 17.8244959 | 17.5344693 | 17.8183212 | 17.7192524 | -0.111865668 | 0.432500994 |
| 18.6680602 | 18.4382138 | 18.5168233 | 18.898284 | 0.160607319 | 0.253350012 |
| 17.1406228 | 17.1367323 | 17.2287508 | 17.2838382 | -0.037316697 | 0.789729066 |
| 20.2246591 | NA | 20.2340135 | 20.1635235 | -0.061807167 | 0.531287405 |
| 20.7616366 | 20.5305297 | 20.9338915 | 20.9320897 | 0.104191489 | 0.263783056 |
| 18.4808938 | 18.6507491 | 18.7884844 | 18.6624274 | 0.010243486 | 0.908734009 |
| 20.020753 | 19.8364705 | 19.9065827 | 20.0308442 | -0.044370622 | 0.526875911 |
| 22.6770791 | 22.6382708 | 22.7047181 | 22.5719661 | 0.001373144 | 0.982908316 |
| 20.8345285 | 20.6155422 | 20.7402654 | 20.8265005 | -0.068144768 | 0.284939244 |
| 19.9406196 | 19.9949788 | 19.9531332 | 20.0254768 | 0.012618528 | 0.839787826 |
| 21.1069517 | 20.9624211 | 21.0810432 | 21.0532243 | -0.017376989 | 0.768630913 |
| 18.8552633 | 18.8057806 | 18.8789 | 18.8374953 | 0.020836363 | 0.726284275 |
| 20.0944338 | 19.8588101 | 19.9042521 | 19.9690388 | -0.050738012 | 0.388682205 |
| 18.259465 | 18.1794341 | 18.171194 | 18.2244868 | -0.021022811 | 0.71166209 |
| 27.1329363 | 27.1042711 | 27.1478367 | 27.1867176 | 0.093533457 | 0.103109511 |
| 20.5274284 | 20.497086 | 20.5878191 | 20.501818 | -0.032231112 | 0.558173823 |
| 22.7319246 | 22.6058474 | 22.7316615 | 22.7403839 | -0.015146069 | 0.779217505 |
| 20.959622 | 20.8299315 | 20.8605779 | 20.9051342 | -0.03599122 | 0.465681148 |
| 20.9656455 | 20.9863858 | 21.079292 | 21.0251529 | 0.118565189 | 0.016777669 |
| 22.1638961 | 22.1932105 | 22.1778411 | 22.1955523 | 0.067201157 | 0.099593742 |
| 21.6142584 | 21.5628637 | 21.5698622 | 21.5825717 | 0.001858268 | 0.961862869 |
| 23.9640524 | 23.8661632 | 23.9589834 | 23.9720625 | 0.005017019 | 0.901461184 |
| 19.9247484 | 19.8706607 | 19.8424227 | 19.9618539 | 0.007913092 | 0.85753319 |
| 23.2869542 | 23.2321025 | 23.326362 | 23.3061236 | 0.054139697 | 0.262353855 |
| 22.1771271 | 22.1491196 | 22.2091891 | 22.2476099 | 0.03928449 | 0.42545732 |
| 20.6659598 | 20.6389068 | 20.676862 | 20.7202213 | -0.006652356 | 0.901591715 |

| | | | | | |
|---|---|---|---|---|---|
| 20.5722272 | 20.4376235 | 20.5263205 | 20.6503549 | 0.038525871 | 0.481481684 |
| 25.1227045 | 25.0484767 | 25.1483416 | 25.1903521 | 0.025649857 | 0.64089328 |
| 19.1331769 | 18.9930734 | 19.1955885 | 19.1766295 | -0.008929349 | 0.873469775 |
| 20.2544243 | 20.1254447 | 20.096592 | 20.221636 | -0.036555145 | 0.524076717 |
| 20.4266267 | 20.3490697 | 20.4162166 | 20.4038725 | 0.006926902 | 0.905811408 |
| 23.7742202 | 23.6765946 | 23.7750804 | 23.7010818 | 0.025939614 | 0.670012814 |
| 22.6515949 | 22.4184529 | 22.5703177 | 22.6253569 | -0.027831438 | 0.653515684 |
| 21.1205549 | 20.9207563 | 21.0527851 | 21.2489855 | -0.048125379 | 0.466651155 |
| 18.8559359 | 18.8994989 | 18.7950038 | 18.954519 | 0.060229591 | 0.382379568 |
| 21.4759783 | 21.3069302 | 21.4558685 | 21.5406599 | -0.022372712 | 0.745376762 |
| 20.6654788 | 20.5899257 | 20.6712361 | 20.5893978 | 0.041822486 | 0.572805323 |
| 25.0722951 | 25.0110934 | 24.9885093 | 25.0812363 | 0.005540669 | 0.943980804 |
| 20.1769807 | 20.0295863 | 20.2663844 | 20.1768591 | 0.002883592 | 0.971170208 |
| 17.2591719 | 17.4213031 | 17.3598865 | 17.2333141 | 0.121358988 | 0.151514874 |
| 19.7813499 | 19.890215 | 19.8554995 | 19.9391299 | -0.008043281 | 0.930833076 |
| 16.8341701 | 16.6699641 | 17.0438054 | 17.0016843 | 0.040911223 | 0.807680985 |
| 16.3472654 | 16.7077109 | 16.9328607 | 16.6921721 | 0.316679745 | 0.177880312 |
| NA | 14.9467506 | NA | NA | 0.60264593 | 0.184405719 |
| 15.8975531 | 16.1488474 | 15.8819262 | 16.0787904 | 0.086175115 | 0.563923609 |
| 23.4148556 | 23.2528212 | 23.2485734 | 23.1629627 | -0.08421473 | 0.396520028 |
| 21.3007181 | 21.1902275 | 21.3049949 | 21.4039134 | -0.015595864 | 0.848802626 |
| 19.4151506 | 19.4447235 | 19.4731714 | 19.4528778 | 0.008193136 | 0.906447052 |
| 20.0899999 | 19.9196322 | 20.016036 | 20.1360365 | -0.025934749 | 0.542568815 |
| 25.6666806 | 25.5815488 | 25.7223938 | 25.6347001 | 0.027758923 | 0.675560228 |
| 22.661138 | 22.5228594 | 22.7055492 | 22.6327591 | -0.018342951 | 0.785079618 |
| 18.2316451 | 18.0031935 | 18.2142648 | 18.1292569 | -0.062869102 | 0.528713236 |
| 11.5990794 | NA | 11.5233521 | 11.5785549 | -0.056749896 | 0.601057066 |
| 15.3134743 | 15.1728163 | 15.6574035 | 15.6871731 | 0.010099207 | 0.944816241 |
| 15.9995528 | 16.3171214 | 15.9938088 | 16.2809893 | 0.018429968 | 0.917144716 |
| 18.0798122 | 18.1292189 | 17.9583307 | 18.1297774 | -0.045029873 | 0.549920958 |
| 15.8092722 | 15.8912411 | 15.7790045 | 15.7248143 | -0.027990557 | 0.708948796 |
| 21.5859354 | 21.292096 | 21.4814268 | 21.5977463 | -0.105431236 | 0.230550278 |
| 19.9776305 | 19.8154137 | 19.9598728 | 19.9831562 | -0.027035512 | 0.679410514 |
| 22.996316 | 23.1721161 | 23.1082733 | 23.0489063 | 0.085984626 | 0.14323149 |
| 18.5079828 | 18.6091035 | 18.7338743 | 18.6621534 | 0.090613406 | 0.166489794 |
| 19.8170132 | 19.7123755 | 19.7856402 | 19.9308646 | -0.036933792 | 0.60457348 |
| 22.8835465 | 22.6541175 | 22.8350625 | 22.8500387 | -0.040870252 | 0.638652568 |
| 17.4073394 | 17.3106576 | 17.3516988 | 17.4144313 | -0.06713363 | 0.436261784 |
| 21.2722794 | 21.0110278 | 21.3897445 | 21.2758937 | -0.043680972 | 0.609143446 |
| 20.6575445 | 20.6515554 | 20.5745758 | 20.5745567 | -0.023986428 | 0.517490622 |
| 19.3907048 | 19.6285449 | 19.4382915 | 19.3297675 | 0.028928956 | 0.824655857 |
| 16.8198737 | 17.0478063 | 16.9063043 | 17.0206701 | -0.036623226 | 0.784741963 |

| | | | | | |
|---|---|---|---|---|---|
| 18.3682597 | 18.4425659 | 18.9133487 | 18.3592607 | 0.075068644 | 0.580315478 |
| 22.7988853 | 22.6329908 | 22.7918089 | 22.7473445 | -0.007742356 | 0.890049438 |
| 23.1244906 | 23.0294671 | 23.0468677 | 23.033347 | -0.045409416 | 0.31640324 |
| 22.385307 | 22.2893407 | 22.3279334 | 22.4052605 | -0.027802402 | 0.516404876 |
| 18.7275262 | 18.7348197 | 18.7957353 | 18.8227408 | 0.089159815 | 0.196365012 |
| 22.1752412 | 22.0614375 | 22.0957214 | 22.1201239 | -0.067175092 | 0.154804657 |
| 18.7108498 | 18.5274505 | 18.6920622 | 18.7284915 | -0.047057581 | 0.590046342 |
| 20.3158009 | 20.2046454 | 20.2719044 | 20.3711605 | -0.014152647 | 0.794566759 |
| 23.602302 | 23.48034 | 23.6088705 | 23.5729891 | 0.010183879 | 0.822936515 |
| 16.8110924 | 16.634122 | 16.6913702 | 16.5907977 | -0.029056481 | 0.809203745 |
| 18.6439008 | 18.5942488 | 18.7971358 | 18.7190699 | 0.060847404 | 0.360731856 |
| 21.4046066 | 21.331773 | 21.3649268 | 21.4000844 | 0.00286615 | 0.947502368 |
| 20.5097596 | 20.4101932 | 20.5440107 | 20.4646642 | 0.002714724 | 0.96311398 |
| 19.9863882 | 19.8953201 | 19.8629228 | 19.9269301 | -0.096763277 | 0.172907121 |
| 17.4250233 | 17.2881654 | 17.3108853 | 17.3119025 | -0.032274985 | 0.828359438 |
| 18.3827936 | 18.2995287 | 18.3176498 | 18.4884194 | 0.029165519 | 0.74129002 |
| 16.3252938 | 15.9776116 | 16.1218586 | 16.4125027 | -0.060189538 | 0.649815858 |
| 18.415092 | 18.3683643 | 18.576568 | 18.5368876 | 0.103439606 | 0.234144489 |
| 16.7163495 | 16.7996764 | 16.5231946 | 16.9108177 | 0.135412129 | 0.279511297 |
| 17.5936481 | 17.531069 | 17.3976274 | 17.5768652 | 0.130723821 | 0.294386621 |
| 20.6552657 | 20.6572378 | 20.6346359 | 20.6586667 | -0.013311037 | 0.896680651 |
| 21.6006984 | 21.6920612 | 21.7411364 | 21.5898116 | 0.062279641 | 0.448134453 |
| 18.5663976 | 18.6697067 | 18.6396892 | 18.7627342 | 0.066733299 | 0.216612009 |
| 19.1059183 | 18.7806094 | 19.0521683 | 19.0665716 | -0.095171546 | 0.243965446 |
| 18.2175964 | 17.8504237 | 18.0137486 | 18.2033146 | -0.046149661 | 0.619590767 |
| 22.269915 | 22.040591 | 22.2313379 | 22.1994679 | -0.031985549 | 0.604545215 |
| 21.7365448 | 21.5514159 | 21.6431681 | 21.6598673 | -0.060906608 | 0.23399579 |
| 20.3649174 | 20.3632817 | 20.3411599 | 20.3788923 | 0.014745624 | 0.781570995 |
| 19.9791068 | 19.9684134 | 20.0192292 | 19.9669187 | 0.044748969 | 0.419761584 |
| 19.9455776 | 19.8276684 | 19.8592093 | 19.9181984 | -0.059530381 | 0.357043884 |
| 20.2907133 | 20.2811563 | 20.3320808 | 20.4893525 | 0.088232869 | 0.223777832 |
| 23.9258713 | 23.9367785 | 23.9578212 | 24.0293901 | 0.200334593 | 0.021063955 |
| 22.1212331 | 22.252958 | 22.1141642 | 22.0470504 | -0.005872025 | 0.947915416 |
| 16.1635071 | 16.2376475 | 16.3004187 | 16.5809736 | 0.166883018 | 0.160262314 |
| 20.1908233 | 20.0991933 | 20.0800329 | 19.6898975 | 0.065759997 | 0.760037074 |
| 20.6861468 | 20.7773816 | 20.7640771 | 20.7316734 | 0.026900476 | 0.546506276 |
| 21.3856785 | 21.3069986 | 21.4105347 | 21.4110257 | 0.009686637 | 0.823374239 |
| 21.9230381 | 21.8342716 | 21.8507153 | 21.9217227 | -0.021778242 | 0.655942998 |
| 18.9891392 | 18.9061751 | 18.7972712 | 18.8163506 | -0.112825695 | 0.073181171 |
| 23.8739472 | 23.801462 | 23.8931788 | 23.8697602 | -0.059503242 | 0.234672907 |
| 24.6661112 | 24.611213 | 24.6241565 | 24.6562078 | 0.004753443 | 0.914429478 |
| 25.5247757 | 25.4422213 | 25.4971622 | 25.4900138 | -0.039301574 | 0.298830392 |

| | | | | | |
|---|---|---|---|---|---|
| 23.1238244 | 22.979545 | 22.9027095 | 23.0959781 | -0.060079587 | 0.505060556 |
| 17.6509279 | 17.5142264 | 17.8020069 | 17.7475103 | 0.034493306 | 0.71806465 |
| 17.8877001 | 17.6777235 | 17.9588014 | 17.9603352 | -0.00767808 | 0.943247067 |
| 20.1911576 | 19.9796291 | 20.2561282 | 20.1020828 | 0.14959019 | 0.221567174 |
| 18.4555641 | 18.1762594 | 18.3425241 | 18.4652427 | -0.065101927 | 0.649110956 |
| NA | 16.3447786 | NA | NA | 0.005297544 | 0.978249851 |
| 21.8402618 | 21.6676842 | 21.7429352 | 21.8080289 | 0.010623684 | 0.850701918 |
| 18.2970055 | 18.2504492 | 18.2816064 | 18.32702 | 0.097678537 | 0.255398799 |
| 20.1745875 | 20.1203263 | 20.1591808 | 20.1627014 | -0.002828544 | 0.967692871 |
| 20.942766 | 21.0174971 | 20.9143099 | 20.9424004 | -0.042699596 | 0.52944603 |
| 22.9086138 | 22.7546559 | 22.8195337 | 22.777079 | -0.059593603 | 0.245079767 |
| 21.306029 | 21.2330176 | 21.2750422 | 21.2803181 | -0.020279599 | 0.539711213 |
| 20.0217701 | 19.9680993 | 20.0508928 | 19.885971 | 0.009221493 | 0.925068917 |
| 17.7191372 | 17.566215 | 17.8167995 | 17.861901 | 0.007911723 | 0.939224067 |
| 16.7362649 | 16.6847697 | 16.7072144 | 16.607997 | -0.045730725 | 0.666447508 |
| 18.0082565 | 17.4574851 | 17.7617898 | 17.7562793 | -0.154407107 | 0.223937612 |
| 15.8204679 | 15.6731632 | 16.0415702 | 15.953323 | 0.070718545 | 0.601322361 |
| 20.0254395 | 19.9741715 | 20.0391322 | 20.0353653 | -0.023487835 | 0.637877637 |
| 15.2058146 | 14.7586728 | 15.402603 | 15.5377415 | -0.10627219 | 0.576158647 |
| 17.1351505 | NA | 17.0433001 | 17.3870528 | 0.182733705 | 0.320866018 |
| 16.3886608 | 16.4850287 | 16.2327205 | 16.4557197 | -0.044754594 | 0.785558317 |
| 19.0267574 | 19.2265426 | 19.1479527 | 18.9753284 | 0.078720164 | 0.559533155 |
| 16.8160682 | 16.7454444 | 16.9947843 | 16.961586 | 0.015753491 | 0.903679965 |
| 17.3336689 | 17.3029357 | 17.4594701 | 17.3037079 | -0.055777555 | 0.59400119 |
| 19.4261566 | 19.441803 | 19.3319327 | 19.510758 | -0.03229559 | 0.662630492 |
| 20.5461835 | 20.3186603 | 20.5467043 | 20.4917986 | -0.048512994 | 0.45891295 |
| 21.4066289 | 21.2569165 | 21.4240685 | 21.3534096 | -0.062175369 | 0.301079146 |
| 23.569992 | 23.5062077 | 23.6095988 | 23.6272272 | 0.013578586 | 0.797641017 |
| 22.0392336 | 22.0253737 | 22.054422 | 22.0798619 | 0.045577418 | 0.369617574 |
| 18.4928461 | 18.4108336 | 18.3911366 | 18.4367335 | -0.032292481 | 0.514751127 |
| 20.5846835 | 20.5627192 | 20.5835451 | 20.4737109 | -0.036478884 | 0.433858381 |
| 23.6254669 | 23.596362 | 23.6714644 | 23.6736833 | 0.037487066 | 0.406521848 |
| 22.5192047 | 22.4374161 | 22.500272 | 22.5016392 | 0.00972792 | 0.801297329 |
| 23.3370889 | 23.2894211 | 23.2698954 | 23.3011914 | -0.021513349 | 0.646025105 |
| 22.8232023 | 22.7957208 | 22.858432 | 22.8835377 | 0.041148592 | 0.399134867 |
| 21.2725221 | 21.1705071 | 21.3159963 | 21.3119858 | 0.020869925 | 0.670179045 |
| 18.8157123 | 18.8599358 | 18.8877297 | 18.8986543 | 0.025205355 | 0.638119235 |
| 21.566642 | 21.5303632 | 21.5031073 | 21.5907841 | 0.021936969 | 0.685886299 |
| 22.1902947 | 22.0551028 | 22.2305629 | 22.2578439 | -0.002293232 | 0.968711992 |
| 20.4103929 | 20.4426377 | 20.4449667 | 20.4351103 | 0.03410957 | 0.581867634 |
| 19.070948 | 18.8894887 | 18.9696459 | 18.9161349 | -0.118870924 | 0.172963926 |
| 16.9805529 | 17.0317589 | 17.0380616 | 17.1710519 | 0.101311249 | 0.26014228 |

| | | | | | |
|---|---|---|---|---|---|
| 19.2344593 | 19.3322894 | 19.1806031 | 19.1832756 | 0.067910388 | 0.512489535 |
| 18.7237385 | 18.5410466 | 18.5106328 | 18.7095523 | 0.007491212 | 0.952384271 |
| 22.4561911 | 22.4033618 | 22.48574 | 22.2585359 | 0.371138544 | 0.015949112 |
| 20.2573376 | 19.8276333 | 20.1543493 | 20.0735073 | -0.061354861 | 0.620827148 |
| 17.4315994 | 17.3541737 | 17.4266983 | 17.2606015 | 0.006116207 | 0.959946177 |
| 16.915973 | 16.6719288 | 16.8364269 | 16.9598068 | -0.027413124 | 0.812538475 |
| 15.4142217 | NA | 15.4506174 | 15.3628934 | -0.081494492 | 0.382024724 |
| 19.5890997 | 19.4243796 | 19.5994776 | 19.6215609 | 0.002956223 | 0.973963738 |
| 18.7621484 | 18.5371411 | 18.660315 | 18.7769669 | 0.021051176 | 0.781209049 |
| 16.7742635 | 16.7847164 | 16.9689577 | 16.9485669 | 0.047176202 | 0.513855622 |
| 19.4428454 | 19.3557422 | 19.4860763 | 19.5353342 | -0.012302364 | 0.852032776 |
| 17.8104335 | 17.8405013 | 17.8203185 | 17.6296305 | -0.071549647 | 0.257768213 |
| 24.9030146 | 24.8156232 | 24.9491658 | 24.8979781 | 0.030324407 | 0.608603704 |
| 20.638351 | 20.4672731 | 20.5361567 | 20.601982 | -0.044462463 | 0.433941975 |
| 19.2447853 | 19.1454442 | 19.3325144 | 19.1932801 | -0.015422384 | 0.771799045 |
| 24.3810604 | 24.3287074 | 24.4168713 | 24.372396 | 0.016189217 | 0.758887438 |
| 23.8555234 | 23.7529485 | 23.8722646 | 23.8866522 | 0.039328762 | 0.449740603 |
| 23.9984006 | 23.9237937 | 23.9615226 | 23.933175 | -0.019598513 | 0.685727127 |
| 23.246957 | 23.0889373 | 23.2485728 | 23.2710413 | 0.011648219 | 0.810995616 |
| 20.6156601 | 20.5369465 | 20.6258188 | 20.6267217 | -0.007883983 | 0.865010511 |
| 25.7459758 | 25.6967669 | 25.7420376 | 25.7974818 | 0.057331129 | 0.210575031 |
| 20.8055863 | 20.7596324 | 20.8077686 | 20.8200699 | 0.030588862 | 0.489892838 |
| 22.4192549 | 22.3937932 | 22.4162033 | 22.4359683 | 0.008720918 | 0.828525763 |
| 19.7860656 | 19.7689022 | 19.7775422 | 19.7570097 | -0.001968975 | 0.953281289 |
| 19.7138502 | 19.8858328 | 19.7654079 | 19.8069985 | 0.025730813 | 0.590111564 |
| 21.9752528 | 21.9115804 | 22.0229146 | 22.0092047 | 0.030235851 | 0.536422098 |
| 20.5776452 | 20.5472203 | 20.574868 | 20.495156 | 0.004033314 | 0.937842725 |
| 21.0099877 | 20.9153598 | 21.010425 | 20.9837808 | -0.03188197 | 0.544678439 |
| 21.0189158 | 20.8943591 | 21.0493377 | 21.0063234 | -0.03634013 | 0.503704827 |
| 20.1039806 | 20.103444 | 20.0588441 | 20.0780989 | 0.006093087 | 0.910471092 |
| 23.1138533 | 23.1145247 | 23.1107425 | 23.053267 | 0.017898139 | 0.769949676 |
| 18.4054571 | 18.4547768 | 18.4156339 | 18.2340511 | -0.059541813 | 0.331800655 |
| 20.6986119 | 20.5081811 | 20.6148492 | 20.6610686 | -0.04945188 | 0.428517904 |
| 22.5039885 | 22.4113263 | 22.5053306 | 22.5161449 | -0.001645988 | 0.980113302 |
| 23.8584019 | 23.7544489 | 23.8661716 | 23.9424573 | -0.004190575 | 0.950333883 |
| 22.4598308 | 22.359249 | 22.3862004 | 22.3498748 | -0.048944873 | 0.472969889 |
| 18.2316265 | 18.2837517 | 18.2574282 | 18.351985 | -0.017682102 | 0.803652737 |
| 22.6887504 | 22.6670761 | 22.6886894 | 22.6202685 | 0.044656852 | 0.570483756 |
| 25.3892309 | 25.3635017 | 25.5386125 | 25.4026735 | 0.046820415 | 0.571067101 |
| 20.5296111 | 20.4277676 | 20.5578865 | 20.6540149 | 0.053451807 | 0.528682611 |
| 19.1316425 | 19.0669804 | 19.1488925 | 19.0718341 | -0.061392148 | 0.475791562 |
| 17.342325 | 17.4192443 | 17.6264359 | 17.6239829 | 0.193507964 | 0.054593764 |

| | | | | | |
|---|---|---|---|---|---|
| 21.3999672 | 21.2056457 | 21.4814331 | 21.4010139 | 0.053229959 | 0.62529248 |
| 19.0492377 | 18.9083334 | 19.1745641 | 19.1277366 | 0.037955946 | 0.736792344 |
| 19.2546382 | 19.2948187 | 19.5698541 | 19.4613415 | 0.156035057 | 0.189944851 |
| 11.6033195 | NA | 11.6138069 | 11.3246421 | -0.013874973 | 0.911299759 |
| 15.1423086 | 15.4933127 | 15.4381042 | 15.5611922 | 0.088835514 | 0.68077073 |
| 15.720418 | 15.635586 | 15.9454529 | 16.0379549 | 0.028065464 | 0.914038681 |
| 19.8420804 | 20.0676356 | 19.5281383 | 19.6653955 | -0.067113176 | 0.603792098 |
| 20.8057383 | 20.6381809 | 20.6340406 | 20.6785999 | -0.097201146 | 0.103608065 |
| 16.8973035 | 16.8020487 | 16.6744881 | 16.8967634 | -0.012834706 | 0.899329992 |
| 23.9491931 | 23.7897759 | 23.8827968 | 23.9435804 | -0.007797245 | 0.893290837 |
| 19.8479308 | 19.7611625 | 19.8297953 | 19.7974598 | -0.030907344 | 0.566380769 |
| 20.2461097 | 20.1465224 | 20.221909 | 20.1874869 | -0.017110957 | 0.699566778 |
| 21.0996925 | 21.0418609 | 21.1029669 | 21.1101038 | 0.012968298 | 0.81206844 |
| 19.3787772 | 19.3432553 | 19.3310558 | 19.3392191 | 0.014760847 | 0.806372215 |
| 15.6148795 | NA | 15.6156414 | 15.6739969 | 0.029485418 | 0.827618375 |
| 17.7257294 | 17.6401559 | 17.611122 | 17.7048149 | -0.090461455 | 0.254102837 |
| 21.8626454 | 21.7551181 | 21.8583185 | 21.8305043 | 0.001245864 | 0.975222868 |
| 20.5402018 | 20.6432364 | 20.5665369 | 20.4812013 | -0.019986986 | 0.635054397 |
| 23.7581627 | 23.5104095 | 23.7717364 | 23.8003847 | 0.010605962 | 0.881503785 |
| 13.7352894 | NA | 14.0100626 | 14.1507924 | 0.181346091 | 0.372145176 |
| 27.6787407 | 27.5664869 | 27.6461125 | 27.6444768 | -0.001706997 | 0.975181402 |
| 16.0075662 | 16.3284127 | 15.7631228 | 15.6996751 | -0.14605457 | 0.462334351 |
| 15.6751145 | NA | 15.4097674 | 15.3531384 | -0.143513922 | 0.375782242 |
| 16.7224196 | 17.0898923 | 17.130928 | 17.1458411 | 0.11230127 | 0.365110669 |
| NA | 15.4183961 | NA | NA | -0.028487535 | 0.811374923 |
| 17.0581848 | 17.1137066 | 16.9203997 | 16.8747922 | -0.058386742 | 0.62204336 |
| 16.9961607 | 16.7261683 | 16.9416844 | 17.0025398 | -0.129319982 | 0.256066712 |
| 17.1940446 | 17.2817732 | 17.2194541 | 17.2998423 | 0.168072608 | 0.124642893 |
| 15.3786171 | 14.8854145 | 15.112642 | 15.2799811 | -0.210837207 | 0.051630105 |
| 17.3355768 | 17.1814327 | 17.3340619 | 17.4176874 | -0.012642097 | 0.894995551 |
| 19.1344444 | 19.1080478 | 19.2054387 | 19.2902054 | 0.089561681 | 0.291744709 |
| 20.4014962 | 20.2190465 | 19.9422689 | 20.1858314 | -0.142386609 | 0.091244751 |
| 17.7136554 | 17.6058395 | 17.6926204 | 17.8447745 | -0.05962425 | 0.462153737 |
| 23.8787995 | 23.6579229 | 23.7770602 | 23.7814595 | -0.073182783 | 0.339317369 |
| 18.0897025 | 18.1083422 | 18.0529741 | 18.050755 | 0.051912737 | 0.457331614 |
| 19.0627227 | 19.1282916 | 19.1830314 | 19.1464182 | 0.014997165 | 0.825901755 |
| 18.5117356 | 18.613375 | 18.574573 | 18.4427738 | 0.049518045 | 0.441171067 |
| 24.4091202 | 24.2964221 | 24.3910296 | 24.3358844 | -0.023899206 | 0.696685484 |
| 19.1668239 | 19.1477491 | 19.1802751 | 19.0654036 | -0.013459581 | 0.810954541 |
| 26.1720393 | 26.0876598 | 26.1347976 | 26.1223343 | -0.015958491 | 0.773347795 |
| 21.203762 | 21.1132783 | 21.1111 | 21.1364984 | -0.01211401 | 0.818423515 |
| 21.6320949 | 21.5865293 | 21.5387409 | 21.5212051 | -0.022329903 | 0.62449424 |

| | | | | | |
|---|---|---|---|---|---|
| 23.8941632 | 23.8486014 | 23.9468477 | 23.8962333 | 0.004054719 | 0.924025365 |
| 22.7803816 | 22.7351025 | 22.8193587 | 22.7462955 | 0.01892133 | 0.620160087 |
| 23.718046 | 23.6997601 | 23.7505014 | 23.725952 | 0.055057107 | 0.154339118 |
| 20.357983 | 20.2999662 | 20.316488 | 20.3395541 | -0.001282699 | 0.980081035 |
| 18.7565228 | 18.680337 | 18.7056654 | 18.816451 | -0.030304818 | 0.555716913 |
| 21.6945942 | 21.6414174 | 21.7282505 | 21.7261988 | 0.038669624 | 0.459572658 |
| 19.7629427 | 19.7653995 | 19.7854559 | 19.8142523 | 0.018963846 | 0.760013208 |
| 20.8267077 | 20.8535941 | 20.7884032 | 20.8158381 | -7.02E-04 | 0.991272761 |
| 20.7685578 | 20.73472 | 20.7991939 | 20.8934141 | 0.042851034 | 0.505852952 |
| 24.8040556 | 24.657441 | 24.7788454 | 24.7695429 | 0.00581221 | 0.929324514 |
| 21.5538649 | 21.4199144 | 21.5822602 | 21.5986561 | 0.009706647 | 0.88257255 |
| 20.3481708 | 20.366493 | 20.3749435 | 20.455603 | 0.024993412 | 0.706233915 |
| 21.0721323 | 20.9077642 | 21.1198204 | 21.0690865 | -0.033535472 | 0.625284597 |
| 20.6947009 | 20.4989893 | 20.712674 | 20.7152298 | 0.013304851 | 0.850532591 |
| 18.9986935 | 18.9447829 | 19.0625175 | 19.0668807 | -0.010512917 | 0.882492679 |
| 19.2455639 | 19.0976372 | 19.0721371 | 19.1061527 | -0.005785964 | 0.941984654 |
| 16.3622477 | 16.5933304 | 16.5177907 | 16.4248326 | 0.017172122 | 0.83333906 |
| 19.0400582 | 19.0237588 | 18.9871271 | 19.1634434 | 0.07757583 | 0.346002811 |
| 20.8670267 | 20.606365 | 20.6416423 | 20.7402095 | -0.128982748 | 0.158535001 |
| 22.7868769 | 22.7273376 | 22.8549179 | 22.843403 | 0.055076907 | 0.550690109 |
| 21.8870918 | 21.6374529 | 21.8804556 | 21.8393167 | -0.017714454 | 0.848548256 |
| 19.8745716 | 19.5685047 | 19.5483357 | 19.5989368 | -0.109227868 | 0.276281701 |
| 20.5024846 | 20.3265051 | 20.4635719 | 20.3953806 | -0.06148514 | 0.547947441 |
| 16.8158605 | 16.8255927 | 16.7842691 | 16.7059935 | 0.11415531 | 0.408679061 |
| 13.9466395 | 13.8343812 | 13.6497128 | 13.6437078 | -0.02128556 | 0.895213951 |
| NA | 15.4180527 | NA | NA | 0.01780673 | 0.919144285 |
| 14.1170226 | NA | 14.46228 | 14.7310551 | 0.205466827 | 0.492057938 |
| 18.530056 | NA | 18.3024715 | 18.6038145 | -0.225445373 | 0.622268649 |
| 18.1495823 | 18.0572209 | 18.3256133 | 18.5172755 | 0.078980366 | 0.459882002 |
| 20.1666821 | 20.1506922 | 20.1227995 | 20.2963098 | 0.002937168 | 0.96744487 |
| 19.0817355 | 18.9023296 | 19.0325179 | 19.090006 | -0.034938398 | 0.598384165 |
| 22.1080206 | 22.0253858 | 22.0662011 | 22.0757521 | -0.011270331 | 0.824137627 |
| 18.3527633 | 18.3001052 | 18.3392463 | 18.3253505 | -0.006951443 | 0.890147547 |
| 23.6516164 | 23.6177355 | 23.5914168 | 23.6291423 | -0.028936268 | 0.535075351 |
| 20.8549026 | 20.8395504 | 20.9220026 | 20.9044775 | 0.032820651 | 0.411132753 |
| 22.8588796 | 22.8209523 | 22.8699102 | 22.8751606 | 0.024938234 | 0.551887376 |
| 20.7989197 | 20.7004789 | 20.7618334 | 20.8308541 | -0.005612281 | 0.926682825 |
| 22.0668247 | 21.907469 | 22.0903741 | 22.0125457 | -0.02847682 | 0.696505154 |
| 16.5393634 | 16.8497161 | 16.5473977 | 16.7586592 | 0.049627931 | 0.715875188 |
| NA | 15.1054985 | NA | NA | 0.07562485 | 0.784219105 |
| 18.5265784 | 18.3506238 | 18.6918535 | 18.7968974 | -0.121052524 | 0.648051647 |
| NA | 19.3963397 | NA | NA | -0.190126989 | 0.244897353 |

| | | | | | |
|---|---|---|---|---|---|
| 21.4557904 | 21.1520294 | 21.6090497 | 21.5280454 | 0.022633233 | 0.841021475 |
| 20.1600383 | 19.962635 | 19.9325809 | 19.9069016 | -0.196477844 | 0.060319957 |
| 16.9215701 | 16.8167783 | 16.9490655 | 16.9502144 | -0.012428479 | 0.886814892 |
| 18.7192989 | 18.7326688 | 18.616375 | 18.6932619 | 0.046836848 | 0.58163382 |
| 17.6420623 | 17.4846667 | 17.5233732 | 17.5850924 | -0.122380519 | 0.139050186 |
| 23.5748182 | 23.3902311 | 23.565412 | 23.5105874 | -0.064164204 | 0.207908048 |
| 21.5105828 | 21.5046118 | 21.4531785 | 21.5585349 | 0.021653435 | 0.63792015 |
| 21.6887501 | 21.6819357 | 21.6469487 | 21.6412995 | 0.00730039 | 0.845516869 |
| 25.5412852 | 25.4649236 | 25.5223129 | 25.5431285 | -0.004366024 | 0.923609277 |
| 22.6575817 | 22.5525083 | 22.6256254 | 22.5767347 | -0.026352294 | 0.622156483 |
| 25.3947407 | 25.3689072 | 25.3624671 | 25.425802 | 0.044703827 | 0.443682111 |
| 22.0640449 | 22.0063627 | 22.0291734 | 22.0216801 | 0.001076114 | 0.985479107 |
| 21.2430883 | 21.1647142 | 21.1189949 | 21.1879351 | -0.039436685 | 0.515299295 |
| 19.5327856 | 19.5832345 | 19.5127011 | 19.6741923 | 0.021798656 | 0.749752472 |
| 20.7806039 | 20.4973101 | 20.7962442 | 20.7133698 | -0.039249545 | 0.653050437 |
| 20.324402 | 20.3468643 | 20.4340863 | 20.3290909 | 0.051796533 | 0.602445822 |
| 17.0564917 | 16.6981827 | 16.6907659 | 16.996585 | -0.029416867 | 0.805479493 |
| NA | 15.9837372 | NA | NA | 0.206140082 | 0.287082633 |
| NA | 15.8787207 | NA | NA | -0.24051147 | 0.550851373 |
| 20.9205424 | 20.7355131 | 20.8132979 | 20.8977165 | -0.036666065 | 0.582857011 |
| 20.9734307 | 20.8553918 | 20.9832428 | 20.9890545 | -0.032709006 | 0.502372611 |
| 13.7514241 | NA | 13.7847073 | 13.5834056 | -0.017908482 | 0.919596699 |
| 23.6595355 | 23.6180366 | 23.7056481 | 23.6895659 | 0.058699033 | 0.220674451 |
| 22.7433482 | 22.548988 | 22.6868169 | 22.7427757 | -0.012560564 | 0.829721863 |
| 19.5877116 | 19.5291921 | 19.4336224 | 19.6866804 | 0.005289911 | 0.956624215 |
| 14.351403 | NA | 14.6938792 | 14.889798 | 0.419601042 | 0.012095323 |
| 19.4094687 | 19.5328779 | 19.5890913 | 19.5180582 | 0.067946105 | 0.411178598 |
| 19.7778541 | 19.6905952 | 19.7849313 | 19.8598768 | 0.033022505 | 0.533152615 |
| 18.371873 | 18.4378689 | 18.5161684 | 18.3744274 | 0.148289121 | 0.099050871 |
| 19.9959981 | 19.849225 | 20.0059811 | 20.01866 | -0.019390532 | 0.736107536 |
| NA | 16.6339325 | NA | NA | -0.283781816 | 0.275016964 |
| 21.6496368 | 21.3293036 | 21.5424306 | 21.5647956 | -0.097420806 | 0.365524746 |
| 21.6860714 | 21.5785133 | 21.577417 | 21.639117 | -0.054710012 | 0.385875124 |
| 21.1111415 | 20.9465904 | 21.1000291 | 21.0911343 | -0.006953209 | 0.895720985 |
| 22.4138987 | 22.2978465 | 22.3655067 | 22.427155 | 0.020895881 | 0.664067136 |
| 23.9180906 | 23.7192028 | 23.866408 | 23.8572565 | -0.031079019 | 0.592190618 |
| 19.5942995 | 19.6002956 | 19.5317009 | 19.6064853 | 0.042939877 | 0.616335112 |
| 19.3485541 | 19.129382 | 19.2616527 | 19.0989296 | -0.1113629 | 0.237905685 |
| 18.3631948 | 18.3130691 | 18.3869933 | 18.3389572 | -0.00333704 | 0.966119423 |
| 19.2480789 | 19.0454644 | 19.1755345 | 19.2674432 | -0.048451026 | 0.427745751 |
| 20.5701256 | 20.323794 | 20.4287154 | 20.5302343 | -0.075640605 | 0.341493741 |
| 18.808087 | 18.7503857 | 18.7022498 | 18.9121248 | -0.01140969 | 0.889235007 |

| | | | | | |
|---|---|---|---|---|---|
| 20.6642883 | 20.5681839 | 20.6304972 | 20.7244056 | 0.024029685 | 0.684357722 |
| 22.5678635 | 22.4470859 | 22.5729085 | 22.5878065 | 0.030266237 | 0.469476029 |
| 22.0410764 | 21.9618828 | 22.062 | 22.1070501 | 0.03702026 | 0.488795014 |
| 20.755887 | 20.6760919 | 20.7496647 | 20.7572827 | 0.007323769 | 0.905132806 |
| 20.7691843 | 20.73987 | 20.7401764 | 20.7522659 | -0.015836701 | 0.847348919 |
| 22.4764354 | 22.4090337 | 22.4675836 | 22.4409796 | 0.008789312 | 0.880085389 |
| 23.0079444 | 22.9157267 | 22.9936971 | 22.9908519 | 0.001812769 | 0.969685758 |
| 18.5104538 | 18.4893685 | 18.6227505 | 18.5699588 | 0.049282205 | 0.476149374 |
| 24.4107844 | 24.2216792 | 24.3651739 | 24.3036694 | -0.014433194 | 0.855312039 |
| 18.6645332 | 18.4602131 | 18.7495357 | 18.7004824 | 0.032892826 | 0.771585083 |
| 16.1582671 | 16.6416383 | 16.455876 | 16.1156119 | 0.257558074 | 0.039018152 |
| 25.7035058 | 25.6980996 | 25.6951176 | 25.7388224 | 0.06124992 | 0.160168067 |
| 22.1203172 | 22.1175963 | 22.1662296 | 22.1191002 | 0.021264046 | 0.663882835 |
| 19.7714137 | 19.6933242 | 19.9174103 | 19.7961415 | 0.046613269 | 0.548847263 |
| 20.7114993 | 20.4036812 | 20.6095642 | 20.7443823 | -0.054299242 | 0.619972023 |
| 18.8955762 | 18.9087405 | 18.8378433 | 19.1160565 | 0.001706647 | 0.980318907 |
| 18.4898135 | 18.4688363 | 18.6336999 | 18.3952677 | 0.015866803 | 0.810817312 |
| 18.8477784 | 18.6506375 | 18.8512308 | 18.9597416 | -0.02186285 | 0.712313224 |
| 19.7459457 | 19.5255182 | 19.6295316 | 19.6148479 | -0.065210997 | 0.273031097 |
| 21.0259401 | 20.9886494 | 21.0313972 | 21.0970019 | 0.063036399 | 0.242987338 |
| 20.7707526 | 20.6387509 | 20.7522824 | 20.7717662 | 0.010541916 | 0.842220778 |
| 20.4146851 | 20.3813383 | 20.3454822 | 20.3779646 | -0.042437738 | 0.314786254 |
| 25.5488092 | 25.4658489 | 25.5710533 | 25.512746 | 0.021129167 | 0.720529929 |
| 22.120173 | 22.1258503 | 22.0996219 | 22.1251079 | 0.061815921 | 0.339904098 |
| 19.1440796 | 19.0119115 | 19.1306289 | 19.1594264 | -0.002768812 | 0.973277424 |
| 19.6399929 | 19.7196851 | 19.6759721 | 19.6692077 | 0.052339108 | 0.542080487 |
| 24.1852795 | 23.8703514 | 24.2306434 | 24.1167353 | -0.029371645 | 0.849861809 |
| 14.6805187 | NA | 14.5254685 | 15.0965744 | 0.086720139 | 0.739152808 |
| 20.3720335 | 20.220879 | 20.3295932 | 20.3922181 | -0.051484472 | 0.471916904 |
| 15.4856283 | 15.1055475 | 15.6455892 | 15.638195 | -0.227105794 | 0.134289459 |
| NA | 15.077344 | NA | NA | 1.022605542 | 0.099209758 |
| 25.4194264 | 25.0496575 | 25.349183 | 25.3674167 | -0.07258318 | 0.703878068 |
| NA | 14.1252897 | NA | NA | -0.187412225 | 0.310143396 |
| 19.3283175 | 19.1486463 | 19.2201067 | 19.4170053 | 0.264245849 | 0.144686538 |
| 15.4374306 | NA | 15.5650067 | 15.3527543 | 0.161176524 | 0.285546568 |
| 13.6195273 | 13.5881852 | 13.9648193 | 13.6067808 | 0.065672531 | 0.649380067 |
| 18.7945823 | 18.6857819 | 18.6731668 | 18.8810753 | 0.047275998 | 0.678451043 |
| 18.4725543 | 18.1490869 | 18.4525079 | 18.1731043 | -0.147587743 | 0.172142457 |
| 20.6793151 | 20.4855048 | 20.582422 | 20.6046076 | -0.068253235 | 0.508913851 |
| 18.1522828 | 17.8913705 | 18.1021881 | 18.2551556 | -0.041250413 | 0.675285891 |
| 21.1525796 | 20.8417324 | 21.0340997 | 21.145725 | -0.079813545 | 0.410124287 |
| 15.1505516 | 15.0312044 | 15.1372669 | 15.043026 | -0.104788267 | 0.266057056 |

| | | | | | |
|---|---|---|---|---|---|
| 16.8189625 | 16.6414409 | 16.7159536 | 16.8427791 | 0.019744712 | 0.806149579 |
| 20.8403552 | 20.5009195 | 20.7326554 | 20.7446578 | -0.110097943 | 0.172117614 |
| 20.0510724 | 20.0520169 | 20.1740533 | 20.138913 | -0.024576569 | 0.752589506 |
| 17.4575807 | 17.7328689 | 17.6836561 | 17.65512 | 0.121847635 | 0.12784784 |
| 22.0570196 | 21.8399192 | 22.0755433 | 22.0574496 | -0.010624397 | 0.891178091 |
| 19.3340639 | 19.2622062 | 19.3405562 | 19.3105722 | -0.02448884 | 0.704726485 |
| 23.0086686 | 22.9575994 | 23.1056221 | 23.0455371 | 0.048906633 | 0.436680449 |
| 22.7091322 | 22.5917127 | 22.7284447 | 22.6464027 | 0.005472669 | 0.928149211 |
| 20.3326157 | 20.2031493 | 20.2911875 | 20.3673596 | -0.024194607 | 0.684336461 |
| 19.6324026 | 19.5364458 | 19.6662601 | 19.6477833 | -0.011124113 | 0.85007924 |
| 21.9272976 | 21.7976004 | 21.9135213 | 21.8676947 | 0.016223104 | 0.773023107 |
| 21.3649152 | 21.2828372 | 21.3903348 | 21.3665875 | 0.028239383 | 0.595205138 |
| 23.2518971 | 23.1481408 | 23.2258966 | 23.2225316 | -0.014432932 | 0.777205014 |
| 20.58415 | 20.4964376 | 20.5843402 | 20.6036455 | 0.014392316 | 0.775093388 |
| 20.3384284 | 20.2858016 | 20.3789259 | 20.4141197 | 0.004305387 | 0.93205988 |
| 24.6769151 | 24.5517636 | 24.6799599 | 24.6349403 | -0.013139507 | 0.791310566 |
| 22.547593 | 22.4297396 | 22.4186184 | 22.4810637 | -0.049467374 | 0.298084224 |
| 23.8113511 | 23.7104955 | 23.8044301 | 23.8375816 | 0.013440068 | 0.767793558 |
| 22.4593025 | 22.3128924 | 22.3706041 | 22.3797983 | -0.048587743 | 0.267384837 |
| 23.7038345 | 23.6379132 | 23.7332859 | 23.6995748 | 0.008562517 | 0.8431233 |
| 19.3628077 | 19.3259348 | 19.3247887 | 19.3724735 | 0.00326149 | 0.938820978 |
| 21.7890186 | 21.7741393 | 21.7735006 | 21.8262342 | 0.013611614 | 0.740737967 |
| 23.4943117 | 23.3952748 | 23.4450136 | 23.4768157 | -0.02047624 | 0.564501868 |
| 22.439218 | 22.3737597 | 22.4044594 | 22.4243248 | -0.006112875 | 0.899779972 |
| 19.5842873 | 19.6108206 | 19.6547322 | 19.707989 | 0.061588601 | 0.21932935 |
| 20.6846872 | 20.593622 | 20.6629464 | 20.695343 | -0.01053608 | 0.831487473 |
| 22.0275017 | 22.0921656 | 22.024091 | 22.0756424 | 0.026259268 | 0.612151565 |
| 20.3768188 | 20.3930235 | 20.4450752 | 20.4790557 | 0.072500474 | 0.168729466 |
| 24.3683378 | 24.2953469 | 24.2783606 | 24.3239605 | -0.008701676 | 0.872163393 |
| 21.8347765 | 21.7523338 | 21.8150951 | 21.8336163 | -0.00880952 | 0.870469168 |
| 20.8147338 | 20.7004321 | 20.7651243 | 20.7778423 | 8.90E-04 | 0.987224003 |
| 21.3270935 | 21.3851269 | 21.2573982 | 21.3818633 | 0.054688746 | 0.338052665 |
| 19.6042267 | 19.6628254 | 19.6917383 | 19.5428751 | 0.044664515 | 0.456190332 |
| 20.1973014 | 20.1871864 | 20.1559602 | 20.1878404 | -0.003424314 | 0.953751275 |
| 20.7007921 | 20.659125 | 20.6866997 | 20.6431803 | 0.0057964 | 0.926920426 |
| 26.9694398 | 26.8811595 | 27.0288427 | 26.9312992 | 0.040482112 | 0.531917136 |
| 21.5742239 | 21.4042383 | 21.6054294 | 21.6176949 | 0.015778127 | 0.808980507 |
| 23.7552756 | 23.5429529 | 23.7032774 | 23.6873374 | -0.021176782 | 0.750189842 |
| 17.8922886 | 18.0603139 | 17.9642017 | 18.0500008 | 0.026600671 | 0.695317931 |
| 19.3777144 | 19.0756822 | 19.3032493 | 19.3443656 | -0.088260529 | 0.208073782 |
| 19.2930994 | 19.1076672 | 19.2698951 | 19.3942228 | 0.040231869 | 0.556815096 |
| 23.4950859 | 23.329564 | 23.5300256 | 23.4925219 | 0.044486297 | 0.532881437 |

| | | | | | |
|---|---|---|---|---|---|
| 19.7839053 | 19.7198143 | 19.8863103 | 19.7917326 | 0.015353118 | 0.8333113 |
| 23.2069591 | 23.2113353 | 23.2606289 | 23.2607162 | 0.069419982 | 0.367844437 |
| 22.5210861 | 22.3454023 | 22.4456248 | 22.4290208 | -0.037971322 | 0.630666353 |
| 22.4921581 | 22.5279803 | 22.3632045 | 22.4632855 | -0.03373262 | 0.671709093 |
| 25.3900133 | 25.2438251 | 25.4536084 | 25.4666858 | 0.011674631 | 0.892864999 |
| 18.4213041 | 18.3476875 | 18.4639829 | 18.3155685 | -0.047226653 | 0.626734849 |
| 20.7005603 | 20.4353849 | 20.6721855 | 20.6488207 | 0.011972081 | 0.902487491 |
| 16.9702999 | 17.0440177 | 17.3043277 | 17.1307701 | 0.043969957 | 0.668991771 |
| 16.8993014 | 17.3354423 | 17.2046481 | 17.2644453 | 0.104964464 | 0.341938706 |
| 16.2783577 | 16.2387778 | 16.1649577 | 16.1147192 | 0.11458666 | 0.306319821 |
| 18.2600026 | 18.207404 | 17.9841667 | 18.1180605 | -0.028157641 | 0.804956469 |
| 16.2510696 | 16.0582678 | 16.0617559 | 16.4060772 | 0.074951711 | 0.517278028 |
| 16.2478429 | 16.8683958 | 17.0305486 | 16.7351021 | 0.405902136 | 0.002415573 |
| 19.6247943 | 19.2519611 | 19.6167234 | 19.61198 | -0.029315673 | 0.812058554 |
| 16.7294106 | 16.681143 | 16.5553046 | 16.5306879 | -0.018857303 | 0.896921306 |
| 17.9042409 | 17.9286532 | 18.0159803 | 18.1555164 | 0.164705138 | 0.283979776 |
| 16.321402 | 16.2892759 | 16.2425213 | 16.298007 | -0.119624801 | 0.473673088 |
| 15.9000376 | 15.7120499 | 16.0477034 | 16.1858967 | 0.159989571 | 0.369809018 |
| 18.2440019 | 18.2831521 | 18.2034728 | 18.0948286 | -0.159320747 | 0.381864717 |
| 15.3761899 | 14.8552977 | 14.8686982 | 15.1411968 | -0.002740846 | 0.98976328 |
| 18.1670322 | 17.5893896 | 18.1823533 | 18.4994791 | 0.072149792 | 0.741447326 |
| 15.1429855 | 14.9331066 | 14.3668472 | 15.4391537 | -0.261646144 | 0.311030216 |
| 21.821409 | 21.1395945 | 21.6752872 | 21.8881306 | -0.008706889 | 0.974262264 |
| 21.5620294 | 21.6083505 | 21.6171848 | 21.6679705 | 0.071141003 | 0.313835183 |
| 17.7517473 | 17.6241618 | 17.8219888 | 17.6900201 | -0.010298446 | 0.865688196 |
| 19.8943242 | 19.7321703 | 19.8707249 | 19.8630952 | -7.66E-04 | 0.99008605 |
| 23.2784099 | 23.2439935 | 23.3533921 | 23.3739877 | 0.060326409 | 0.293694398 |
| 24.1920264 | 24.0869535 | 24.1907634 | 24.1867448 | 0.022543686 | 0.63744117 |
| 22.1595615 | 22.0398355 | 22.1774461 | 22.1349009 | -0.007293978 | 0.954120042 |
| 20.3972148 | 20.3089645 | 20.3277862 | 20.5234672 | -0.001065545 | 0.984786042 |
| 18.292838 | 17.9971786 | 17.7980583 | 18.152498 | -0.194221993 | 0.038147287 |
| 18.6723983 | 18.5293565 | 18.5114255 | 18.6028809 | -0.089607179 | 0.272424282 |
| 19.7122211 | 19.5343725 | 19.6899695 | 19.6379822 | -0.040256699 | 0.516583749 |
| 21.0656458 | 20.9768905 | 21.0251952 | 21.0735527 | -0.032792888 | 0.56219107 |
| 20.1277337 | 19.9611722 | 20.1403516 | 20.0387043 | -0.014056992 | 0.799216164 |
| 22.1015115 | 22.0291411 | 22.1229006 | 22.1341634 | 0.029408688 | 0.611705711 |
| 19.7038867 | 19.7058094 | 19.761803 | 19.8071807 | 0.097812588 | 0.127053936 |
| 16.4552704 | 16.5980511 | 16.6375612 | 16.7535973 | 0.128100231 | 0.224282437 |
| 18.2567736 | 18.2267183 | 18.4964337 | 17.832937 | -0.079800919 | 0.628954862 |
| 21.8604238 | 21.4333121 | 21.4764277 | 21.7248298 | -0.149142774 | 0.130280282 |
| 20.2767355 | 20.3914437 | 20.4944541 | 20.3901963 | 0.068086207 | 0.434592499 |
| 18.4944986 | 18.2854824 | 18.2752913 | 18.388262 | -0.061151498 | 0.478631036 |

| | | | | | |
|---|---|---|---|---|---|
| 16.1922549 | 16.2651784 | 16.2675716 | 16.2533218 | -0.040368932 | 0.626296767 |
| 18.4927279 | 18.3210952 | 18.4355354 | 18.4347901 | -0.055035117 | 0.46920724 |
| 19.3930295 | 19.4232336 | 19.3526359 | 19.2843069 | -0.006008612 | 0.936808071 |
| 18.9581447 | 18.901619 | 18.7675282 | 18.9281119 | 0.013680635 | 0.845481825 |
| 17.6888001 | 17.8440094 | 17.7796599 | 17.8514776 | 0.012768585 | 0.853546047 |
| 22.4453659 | 22.4181468 | 22.4796248 | 22.4940359 | 0.003525652 | 0.958766654 |
| 19.4315608 | 19.2681484 | 19.3636184 | 19.401652 | -0.030849567 | 0.652181024 |
| 22.0698199 | 21.9518928 | 22.0734314 | 22.0316382 | -0.020443974 | 0.759726145 |
| 20.6607624 | 20.5737278 | 20.6441182 | 20.6746496 | 0.039805662 | 0.527576887 |
| 21.7900257 | 21.6812021 | 21.8030413 | 21.714232 | -0.018462074 | 0.768965642 |
| 22.129134 | 22.0257003 | 22.1145327 | 22.1625309 | 0.00537076 | 0.928391186 |
| 20.9032289 | 20.8656881 | 20.9858643 | 20.9673437 | 0.056199238 | 0.306561869 |
| 25.3524429 | 25.1714847 | 25.3163174 | 25.3090977 | 5.98E-04 | 0.991171216 |
| 19.9724116 | 19.8804888 | 20.0073488 | 19.9843762 | -0.053944939 | 0.320346764 |
| 20.800934 | 20.6841143 | 20.7551107 | 20.7374984 | -0.067649119 | 0.207901336 |
| 23.3001685 | 23.1999431 | 23.2938104 | 23.2721371 | -0.023774338 | 0.646936592 |
| 23.3197228 | 23.2004319 | 23.3316774 | 23.3104007 | 0.038324905 | 0.447394938 |
| 24.1455911 | 24.0464389 | 24.1584727 | 24.1619013 | 0.011767721 | 0.78273109 |
| 20.2663206 | 20.2717888 | 20.3274349 | 20.2707507 | 0.003910878 | 0.916076557 |
| 22.0776931 | 22.0093723 | 22.0998491 | 22.0663977 | 0.00340415 | 0.935372639 |
| 23.0946838 | 23.0082564 | 23.0923952 | 23.0907979 | 0.00370774 | 0.931334778 |
| 24.8497249 | 24.8154938 | 24.874936 | 24.9193742 | 0.018990849 | 0.715355557 |
| 21.8221391 | 21.7214671 | 21.7783117 | 21.8057199 | -0.034348117 | 0.519230709 |
| 20.4051584 | 20.301347 | 20.303417 | 20.3198163 | -0.019567432 | 0.744597396 |
| 23.7123868 | 23.7249875 | 23.6839503 | 23.7302886 | 0.082233863 | 0.178483567 |
| 18.7838687 | 18.7579232 | 18.8236957 | 18.772537 | -0.063375737 | 0.34894101 |
| 21.3524535 | 21.2413377 | 21.3924438 | 21.3945901 | -0.029604828 | 0.659257516 |
| 18.2140331 | 18.1193389 | 18.156336 | 18.1588531 | 0.007404532 | 0.916309883 |
| 19.6177128 | 19.5658753 | 19.6155913 | 19.5765412 | 0.007512552 | 0.915867178 |
| 17.4826422 | 17.4183498 | 17.4124756 | 17.663447 | -0.074981627 | 0.309730584 |
| 19.0723385 | 19.0033586 | 19.0504295 | 19.0286231 | -0.001102477 | 0.989054485 |
| 20.8014783 | 20.8529595 | 21.0473188 | 21.0614491 | 0.106195215 | 0.202434304 |
| 19.6486033 | 19.8132364 | 19.6835461 | 19.4997364 | 0.014600635 | 0.866302749 |
| 16.4430148 | 16.3339578 | 16.5165825 | 16.629444 | -0.099396265 | 0.28043426 |
| 17.6214385 | 17.7041636 | 17.8487017 | 17.9458747 | 0.102829608 | 0.308843666 |
| 19.5071238 | 19.3211912 | 19.4028484 | 19.4498057 | -0.079076654 | 0.436516504 |
| 18.1506928 | 17.6138125 | 18.0601164 | 18.0291601 | -0.124365737 | 0.259104752 |
| 17.4225071 | 17.3763069 | 17.5464085 | 17.5156786 | 0.074660437 | 0.576687304 |
| 18.9353625 | 18.6105387 | 19.2905462 | 19.0857336 | 0.072380413 | 0.767956614 |
| 13.5004257 | 14.4786368 | 13.0297573 | 13.5766181 | -0.109077732 | 0.852165139 |
| 18.5276067 | 18.7527732 | 19.1673429 | 19.2543433 | 0.628704562 | 0.033493485 |
| 21.7384599 | 21.5469315 | 21.9121622 | 22.1746847 | 0.166653815 | 0.404771396 |

| | | | | | |
|---|---|---|---|---|---|
| NA | 11.4062907 | NA | NA | -0.315902718 | 0.098944917 |
| 22.7895863 | 22.1671235 | 22.6756694 | 22.701822 | -0.13539204 | 0.414198769 |
| 16.6628824 | 16.7913899 | 17.1623019 | 16.6809424 | 0.063848716 | 0.611056158 |
| 14.5825677 | NA | 15.0336669 | 15.060554 | 0.429555517 | 0.00208963 |
| 17.9876101 | 17.8749167 | 18.0169195 | 18.0492553 | -0.102988511 | 0.283450285 |
| 22.9741881 | 22.7868483 | 22.9532214 | 22.9551515 | 0.013065626 | 0.882240958 |
| 23.1873335 | 23.020747 | 23.1446907 | 23.1750345 | 0.004691271 | 0.945612979 |
| 21.6440763 | 21.4364539 | 21.6201868 | 21.6276856 | -0.03550858 | 0.598983475 |
| 21.8379492 | 21.726073 | 21.8818083 | 21.8271953 | 0.033373094 | 0.610192755 |
| 19.8730092 | 19.792516 | 19.7956804 | 19.84618 | -0.022371986 | 0.717645964 |
| 18.7417389 | 18.6434292 | 18.7097708 | 18.6667164 | -0.111120066 | 0.075796687 |
| 24.2540339 | 24.1616331 | 24.2758165 | 24.2582225 | 0.048506816 | 0.401384112 |
| 21.5428481 | 21.4754551 | 21.5279286 | 21.5506173 | 0.037238501 | 0.474505598 |
| 20.9762843 | 20.8059798 | 20.9544578 | 20.9219806 | -0.018882348 | 0.702499036 |
| 18.3295542 | 18.3927074 | 18.3745266 | 18.2909257 | 0.00568788 | 0.909243873 |
| 24.3037117 | 24.2489951 | 24.2798484 | 24.3684167 | -0.039065371 | 0.426109749 |
| 23.5814229 | 23.5143379 | 23.5954772 | 23.5640809 | 0.029701892 | 0.546513108 |
| 22.553984 | 22.4181747 | 22.5467901 | 22.529599 | -0.003425415 | 0.942204748 |
| 24.0636921 | 24.0491763 | 24.1034118 | 24.0135296 | 0.010332827 | 0.82036711 |
| 24.0381311 | 23.9594534 | 24.0241694 | 24.0359767 | 0.015254626 | 0.728134069 |
| 23.9599837 | 23.8501879 | 23.9314092 | 23.9120047 | -0.017611042 | 0.68233796 |
| 19.4792242 | 19.467158 | 19.4577451 | 19.3951857 | 0.012671884 | 0.761493956 |
| 22.7511527 | 22.7084924 | 22.7671739 | 22.7513259 | 0.013574371 | 0.744074429 |
| 19.5250006 | 19.5269939 | 19.5119293 | 19.5842486 | 0.007551022 | 0.859290265 |
| 21.092136 | 21.0450055 | 21.087866 | 21.1213104 | 0.017324505 | 0.709922665 |
| 18.9536444 | 18.857475 | 18.9180584 | 18.9579742 | -0.029367968 | 0.554015675 |
| 20.3209452 | 20.2976129 | 20.4188775 | 20.3036438 | 0.028065532 | 0.59096229 |
| 22.266274 | 22.1758288 | 22.284958 | 22.2669585 | 1.85E-04 | 0.997154564 |
| 21.1962749 | 21.1666322 | 21.2179484 | 21.2262329 | 0.032472183 | 0.539710851 |
| 19.8254052 | 19.6732962 | 19.7558138 | 19.7643696 | -0.044442658 | 0.412384502 |
| 22.1761059 | 22.1197027 | 22.1174775 | 22.1770033 | 0.008252341 | 0.881100577 |
| 19.8825265 | 20.0483019 | 19.8703125 | 19.9654623 | 0.07541929 | 0.191854711 |
| 23.0951229 | 23.0513917 | 23.1443657 | 23.083146 | 0.017497352 | 0.761565934 |
| 23.1709551 | 23.0668442 | 23.2800494 | 23.2841839 | 0.073459367 | 0.209042009 |
| 24.2819489 | 24.2186136 | 24.3307649 | 24.2944448 | 0.031492602 | 0.597972208 |
| 19.0233128 | 18.9822647 | 19.0154871 | 19.0782109 | 0.067668401 | 0.290975679 |
| 18.3340771 | 18.2281167 | 18.33801 | 18.4205865 | -0.008349397 | 0.89556779 |
| 22.9202172 | 22.7355389 | 22.9683927 | 22.890196 | -0.002761997 | 0.967217892 |
| 18.7616556 | 18.6301143 | 18.7464339 | 18.6825135 | -0.061540254 | 0.36766853 |
| 20.4151393 | 20.163792 | 20.3668352 | 20.2717145 | -0.075151276 | 0.29178476 |
| 26.6270102 | 26.5604886 | 26.6247947 | 26.5874057 | 0.018207522 | 0.808472829 |
| 19.6401584 | 19.5400733 | 19.6113835 | 19.631767 | 0.069104433 | 0.372505102 |

| | | | | | |
|---|---|---|---|---|---|
| 26.6005133 | 26.5598736 | 26.5596962 | 26.545356 | 0.011415364 | 0.883582483 |
| 17.8294189 | 17.8344131 | 18.0527682 | 17.8874659 | 0.05365366 | 0.518755804 |
| 18.5375858 | 18.5048042 | 18.5816551 | 18.3316 | -0.038506677 | 0.651926312 |
| 16.5583197 | NA | 16.6518043 | 16.6250863 | 0.031269278 | 0.743730996 |
| 16.091037 | 16.0952685 | 16.2369629 | 16.0174338 | 0.032919594 | 0.737814485 |
| 17.4397333 | 17.6232809 | 17.6245032 | 17.4628859 | 0.051718657 | 0.60258354 |
| 17.8866366 | 17.7557925 | 18.1065247 | 18.0923226 | 0.068045692 | 0.49337589 |
| 19.4883047 | 19.4481653 | 19.6589799 | 19.7169824 | 0.057235542 | 0.601512085 |
| 20.5769519 | 20.2859088 | 20.4191152 | 20.348696 | -0.136711292 | 0.236653745 |
| 18.0443622 | 18.1299551 | 18.1568576 | 18.1934268 | 0.052996077 | 0.663471014 |
| 16.1257854 | 16.0116069 | 16.1326458 | 15.9176169 | -0.034172899 | 0.782254687 |
| 16.8809117 | 16.9895036 | 16.7313694 | 16.9363509 | -0.021099349 | 0.86482172 |
| 17.6895074 | 17.3980465 | 17.2984505 | 17.4352515 | -0.113584495 | 0.404242831 |
| 15.9970522 | NA | 16.3259326 | 16.2500792 | 0.252525404 | 0.152861072 |
| 17.0301245 | 16.5554348 | 17.1284229 | 17.1494272 | 0.011325395 | 0.95829627 |
| 16.2341603 | 16.5065704 | 15.2656141 | 16.606726 | -0.119990845 | 0.587402317 |
| NA | 14.012727 | NA | NA | 0.275119312 | 0.319277008 |
| 23.0120633 | 22.9086726 | 23.0309366 | 23.0531322 | 0.013744608 | 0.779760506 |
| 20.1828528 | 20.0360241 | 20.0631508 | 20.1743176 | -0.050837103 | 0.632239214 |
| 23.1393699 | 23.0944869 | 23.2020197 | 23.1947833 | 0.023132427 | 0.571320845 |
| 23.0130818 | 22.9545464 | 23.0101569 | 23.0269643 | 0.055570264 | 0.235083255 |
| 14.3593004 | 14.2369581 | 14.3327462 | 14.1196581 | 0.022581503 | 0.853808481 |
| 26.038782 | 26.1275116 | 26.0852386 | 26.0964359 | 0.093115281 | 0.238357182 |
| 19.9446153 | 19.7854935 | 19.7882759 | 19.9918288 | -0.061575728 | 0.336361728 |
| 20.224435 | 20.2820955 | 20.4163401 | 20.3817807 | 0.061950603 | 0.27680464 |
| 24.4760367 | 24.3814087 | 24.4318702 | 24.3552022 | -0.043490338 | 0.389959278 |
| 22.3825286 | 22.3387611 | 22.4509041 | 22.4198618 | 0.047698995 | 0.333234416 |
| 23.9762328 | 23.9692379 | 23.997361 | 24.0054901 | 0.051860291 | 0.177515473 |
| 21.1292016 | 21.1129729 | 21.2042627 | 21.2111211 | 0.058121723 | 0.254390349 |
| 20.9239083 | 20.8126908 | 20.870149 | 20.9086378 | 0.007299571 | 0.889241837 |
| 19.195365 | 19.219826 | 19.2221436 | 19.2366265 | 0.085163032 | 0.257174785 |
| 22.3536858 | 22.1226905 | 22.411634 | 22.4670521 | 0.034602009 | 0.789608815 |
| 11.6318719 | 11.4731692 | 11.4759449 | 11.6751306 | -0.139180645 | 0.257348906 |
| 19.0831703 | 19.0853056 | 18.8285833 | 19.0900932 | -0.106629664 | 0.160444322 |
| 20.795078 | 20.7916784 | 20.8956235 | 20.8239777 | 0.140566832 | 0.191719949 |
| 20.078923 | 19.9828928 | 20.0827769 | 20.0423169 | 0.010745974 | 0.860376876 |
| 19.3210167 | 19.2060438 | 19.301609 | 19.3170084 | 0.014706062 | 0.820357288 |
| 12.0109065 | NA | 12.2576061 | 12.064107 | 0.020473875 | 0.874848362 |
| 19.2285657 | 19.3293847 | 19.3685917 | 19.352052 | 0.038708838 | 0.452912323 |
| 15.5008155 | NA | 15.7270519 | 15.651766 | 0.037404929 | 0.741145915 |
| 22.0264867 | 21.8337087 | 22.0077503 | 22.0049224 | -0.049896513 | 0.788704391 |
| 22.189392 | 21.9123651 | 22.242687 | 22.1258965 | 0.011279064 | 0.909701455 |

| | | | | | |
|---|---|---|---|---|---|
| 21.4367597 | 21.3077474 | 21.3901628 | 21.3964013 | -0.017425068 | 0.721663963 |
| 20.6732633 | 20.5424695 | 20.6358447 | 20.6286192 | 0.055376858 | 0.529943889 |
| 19.8471676 | 19.8101286 | 19.833357 | 19.8753939 | -0.024942497 | 0.714059701 |
| 21.9901492 | 21.9939876 | 21.9931352 | 21.9902791 | 0.028004 | 0.562560937 |
| 20.544107 | 20.2889121 | 20.4430845 | 20.519761 | -0.064626902 | 0.454411031 |
| 19.3410423 | 19.2773577 | 19.3129988 | 19.4878773 | 0.042864635 | 0.641452743 |
| NA | 14.3659883 | NA | NA | 0.057743618 | 0.786242059 |
| 20.1988518 | 20.1993073 | 20.1181555 | 20.0777589 | 0.025701845 | 0.692652117 |
| 15.0957004 | 14.8475636 | 15.2312385 | 15.0949309 | 0.071710179 | 0.609803957 |
| 15.81452 | NA | 14.2329494 | 15.3350202 | -0.298829042 | 0.680769384 |
| 21.5787261 | 21.5625987 | 21.5459658 | 21.5416198 | -0.022140885 | 0.595754747 |
| 21.2032999 | 21.030358 | 21.2230891 | 21.1628821 | -0.018074311 | 0.760128492 |
| 21.1587877 | 21.0144923 | 21.2248765 | 21.1900544 | 0.016925381 | 0.77036745 |
| NA | 16.0839007 | NA | NA | 0.140735679 | 0.398261255 |
| 18.1096177 | 17.6389294 | 17.9836669 | 17.9367202 | -0.096966552 | 0.459539056 |
| 21.4413332 | 21.3527022 | 21.5832995 | 21.4885738 | 0.025032255 | 0.67169476 |
| 26.450726 | 26.346256 | 26.4052273 | 26.4046945 | 0.019369806 | 0.703703422 |
| 23.8077826 | 23.7348804 | 23.8540607 | 23.849547 | 0.027152239 | 0.526752845 |
| 27.8236246 | 27.7401965 | 27.8138405 | 27.8213128 | 0.03673355 | 0.559118287 |
| 15.9507152 | NA | 16.0885042 | 16.092725 | 0.080831501 | 0.602723296 |
| 23.4344089 | 23.2017888 | 23.4511797 | 23.4510439 | -0.033279366 | 0.761753262 |
| 23.895019 | 23.8048117 | 23.8807458 | 23.8469075 | 0.02304056 | 0.689495823 |
| 16.7962081 | 17.0236456 | 16.6884816 | 16.814945 | 0.025749984 | 0.847671449 |
| 18.7453838 | 18.530241 | 18.681715 | 18.6550522 | -0.071847295 | 0.378371039 |
| 21.1855517 | 21.0247032 | 21.1274788 | 21.1541135 | -0.03052945 | 0.551750503 |
| 14.8081167 | 15.3177774 | 14.6116868 | 15.4974632 | 0.277279045 | 0.241896705 |
| NA | 11.6599036 | NA | NA | -0.309909089 | 0.427200194 |
| 22.5701236 | 22.3927404 | 22.6401935 | 22.5226049 | -0.126657189 | 0.048536465 |
| 21.2002092 | 21.2064183 | 21.2178247 | 21.1909396 | -0.033943782 | 0.487801034 |
| 22.4988042 | 22.5535881 | 22.5385874 | 22.4975697 | 0.028044287 | 0.595860781 |
| 22.5469126 | 22.4137238 | 22.5228608 | 22.6132155 | 0.012856553 | 0.825825659 |
| 28.4518744 | 28.2889985 | 28.5570441 | 28.4871971 | 0.032642764 | 0.878025073 |
| 19.1317031 | 18.9631201 | 19.1096804 | 19.0094898 | -0.086888928 | 0.489645586 |
| 28.3897278 | 28.1956865 | 28.3426706 | 28.3233895 | -0.014987395 | 0.833136221 |
| 20.0614813 | 20.0207734 | 20.0344435 | 20.1427215 | 0.005801812 | 0.933972841 |
| 21.6455747 | 21.5073594 | 21.6147434 | 21.6321818 | -0.005473886 | 0.911215734 |
| 24.1855344 | 24.1674668 | 24.2649362 | 24.2214034 | 0.067473752 | 0.142162918 |
| 22.554772 | 22.5603765 | 22.5632753 | 22.5704636 | 0.020298881 | 0.606858665 |
| 19.0657153 | 18.9624022 | 19.0633495 | 19.0297869 | 0.010124374 | 0.855870401 |
| 22.5648265 | 22.4131919 | 22.5487482 | 22.4953681 | 0.017338737 | 0.752680519 |
| 20.0517632 | 19.9703988 | 20.0337069 | 19.9951463 | -0.015967927 | 0.695352816 |
| 23.6139512 | 23.6755383 | 23.7582812 | 23.6565406 | 0.087395303 | 0.021544265 |

| | | | | | |
|---|---|---|---|---|---|
| 21.9837105 | 21.8730159 | 21.9741269 | 21.9885143 | 0.00963803 | 0.861411456 |
| 21.275819 | 21.1553929 | 21.1907587 | 21.2663752 | -0.003455033 | 0.950324181 |
| 21.1497851 | 21.0053339 | 21.1911157 | 21.1731368 | -0.02289904 | 0.770312211 |
| 24.3544765 | 24.2696828 | 24.3536184 | 24.3358465 | 0.046724626 | 0.402975064 |
| 18.5139892 | 18.3249315 | 18.3931732 | 18.3390318 | -0.028980778 | 0.853052435 |
| 17.8204705 | 18.1117603 | 17.7964341 | 17.7778665 | -0.05587358 | 0.680988047 |
| 15.7864924 | 16.0539913 | 15.8564287 | 15.8779945 | 0.134054671 | 0.255223872 |
| 16.9860572 | 16.585033 | 17.0173796 | 16.7782095 | -0.172650024 | 0.125299329 |
| 19.6191833 | 19.4050498 | 19.4620107 | 19.5784554 | -0.033834032 | 0.743902093 |
| 17.2882781 | 17.6710819 | 17.6332237 | 17.6085987 | 0.210397578 | 0.0280592 |
| 19.3555175 | 19.2756389 | 19.3918909 | 19.377372 | -0.026450226 | 0.750605187 |
| 20.7866561 | 20.8057282 | 20.7015047 | 20.5922966 | -0.043362917 | 0.588367868 |
| 20.123475 | 19.9515333 | 20.0113613 | 20.1135632 | -0.020350067 | 0.793392607 |
| 23.0387076 | 22.829185 | 22.8780217 | 22.9499213 | -0.096704915 | 0.191986329 |
| 18.8467942 | 18.7372616 | 18.7416303 | 18.8983971 | -0.038216289 | 0.584747756 |
| 21.7862612 | 21.772886 | 21.8134991 | 21.690026 | 0.055941945 | 0.40015011 |
| 20.5514913 | 20.4113237 | 20.4355487 | 20.4812184 | -0.047414333 | 0.443037398 |
| 22.0089577 | 21.8491886 | 21.9736463 | 21.8957004 | -0.05843116 | 0.277974655 |
| 21.7335401 | 21.7009298 | 21.6972115 | 21.6651982 | -0.047929338 | 0.340053764 |
| 22.9108386 | 22.8647273 | 22.9113755 | 22.899366 | 0.020029725 | 0.681629333 |
| 21.0416196 | 20.9131121 | 20.9566637 | 20.9658812 | -0.037421774 | 0.360095076 |
| 22.1235357 | 22.0562597 | 22.0666244 | 22.1364646 | -8.39E-04 | 0.984406361 |
| 20.3623313 | 20.3345667 | 20.2937678 | 20.2412198 | -0.021138842 | 0.672531719 |
| 22.1981122 | 22.1053331 | 22.1260288 | 22.1255048 | -0.06058598 | 0.240267521 |
| 23.0663391 | 22.9500004 | 23.047384 | 23.0362147 | -0.003799577 | 0.94431503 |
| 22.776439 | 22.7127512 | 22.8078114 | 22.7358645 | -0.041691068 | 0.484309486 |
| 21.7207758 | 21.5247772 | 21.5658403 | 21.5791369 | -0.024203526 | 0.701014257 |
| 18.3680154 | 18.30574 | 18.2742141 | 18.2994822 | -0.081325478 | 0.250387619 |
| 19.0328622 | 18.9538095 | 19.0038275 | 19.0016384 | -0.004602761 | 0.948068518 |
| 20.0589955 | 19.8773505 | 19.9712166 | 20.2081264 | 0.011559592 | 0.886170221 |
| 17.4725166 | 17.4908605 | 17.4906452 | 17.5468637 | 0.067421804 | 0.408851277 |
| 19.9692274 | 19.9708912 | 19.8972217 | 19.9529098 | 0.017419069 | 0.845015001 |
| 19.7547558 | 19.337216 | 19.6332708 | 19.6175361 | -0.117105481 | 0.232071953 |
| 19.5855113 | 19.3732973 | 19.4096474 | 19.408428 | -0.087828405 | 0.403026118 |
| 15.0609578 | 14.5984778 | 15.235145 | 15.0842561 | -0.073896101 | 0.647958452 |
| NA | 15.0783483 | NA | NA | -0.27004961 | 0.212283376 |
| 19.2065451 | 19.0946825 | 19.2557199 | 19.0779647 | -0.097842897 | 0.374006663 |
| 19.9440033 | 19.6494081 | 19.7088231 | 19.7390718 | -0.060810499 | 0.513602483 |
| 19.0454959 | 18.9482185 | 19.0912142 | 19.2109703 | 0.045986987 | 0.584215192 |
| 19.1508812 | 18.9530649 | 19.2490536 | 19.1146822 | -0.019466447 | 0.782528182 |
| 21.0068632 | 20.7831934 | 21.0394046 | 20.8776387 | -0.055294403 | 0.39090564 |
| 21.1620663 | 21.0155107 | 21.0345431 | 21.1163565 | -0.020048604 | 0.719072268 |

| | | | | | |
|---|---|---|---|---|---|
| 19.5190429 | 19.4277846 | 19.3746355 | 19.5183186 | -0.039503894 | 0.429068667 |
| 20.2882876 | 20.2626433 | 20.2120957 | 20.2014115 | -0.042871657 | 0.368974362 |
| 21.9659871 | 21.8938096 | 22.0230648 | 21.9586174 | -0.011859952 | 0.771487831 |
| 21.6099065 | 21.5049345 | 21.5823154 | 21.6103838 | -0.011715237 | 0.784350708 |
| 21.3204903 | 21.2045837 | 21.314415 | 21.2959324 | -0.004023539 | 0.943601953 |
| 20.8541376 | 20.6090311 | 20.7138201 | 20.6905805 | -0.061337601 | 0.387313391 |
| 16.7974884 | 16.7563545 | 16.9023802 | 16.8258521 | 0.076054278 | 0.437004947 |
| 17.0088623 | 16.8572331 | 17.1764599 | 16.9529687 | 0.06078671 | 0.543715462 |
| NA | 14.7567512 | NA | NA | 0.233339499 | 0.094306404 |
| 24.3238356 | 24.2717668 | 24.3295122 | 24.3289817 | 0.025131813 | 0.544388764 |
| 20.4552582 | 20.4207447 | 20.5227002 | 20.5648689 | -0.007684635 | 0.859773922 |
| 19.6084543 | 19.2675758 | 19.5672242 | 19.4579527 | -0.065791445 | 0.61555463 |
| 21.3478907 | 21.2690123 | 21.5186687 | 21.5003498 | 0.096840342 | 0.397613628 |
| 17.2550101 | 17.2733148 | 17.2216089 | 17.5581667 | -0.035800788 | 0.724414329 |
| 19.912547 | 19.629711 | 19.720413 | 19.7647284 | -0.103155571 | 0.267844925 |
| 19.9659265 | 19.6278319 | 19.9082768 | 19.9742316 | -0.035474469 | 0.663457194 |
| 21.1592625 | 21.2984991 | 21.1095823 | 21.2543605 | 0.106770077 | 0.169913509 |
| 23.0923417 | 22.8995913 | 23.0604691 | 22.9805374 | -0.037446422 | 0.532530015 |
| 20.1587744 | 20.1443675 | 20.1401558 | 20.0659844 | 0.043568085 | 0.436068021 |
| 20.5799503 | 20.4732498 | 20.6167637 | 20.6059102 | -0.002355212 | 0.961608605 |
| 17.5990568 | 17.6370574 | 17.6720549 | 17.6926204 | 0.048595008 | 0.374298236 |
| 21.6746329 | 21.4754138 | 21.6855081 | 21.7183248 | 0.043802007 | 0.469148959 |
| 21.5085762 | 21.4860345 | 21.4679645 | 21.5242976 | 0.06374784 | 0.296495072 |
| 21.5825012 | 21.4054219 | 21.3640688 | 21.3811528 | -0.077715079 | 0.238192794 |
| 18.4298438 | 18.3537964 | 18.4481537 | 18.5083482 | -0.020962807 | 0.747829195 |
| 20.6712317 | 20.5171661 | 20.5862101 | 20.6042998 | -0.052321962 | 0.452218207 |
| 20.3648385 | 20.2817559 | 20.1736964 | 20.3469914 | -4.69E-04 | 0.994738671 |
| 20.3939952 | 20.2044096 | 20.2587885 | 20.3936245 | -0.021809864 | 0.783511928 |
| 16.8589502 | 17.0600939 | 16.9193377 | 17.0739546 | 0.035159284 | 0.736976759 |
| 16.2628369 | NA | 16.002387 | 16.0545558 | -0.072477784 | 0.541927142 |
| 18.5220698 | 18.4018395 | 18.5983229 | 18.5531169 | -0.013711957 | 0.843632691 |
| 20.5774465 | 20.4920731 | 20.6107235 | 20.6877936 | 0.053856272 | 0.415188402 |
| 18.7707053 | 18.6715761 | 18.8212025 | 18.7839527 | 0.062411523 | 0.295053212 |
| 20.8203319 | 20.6715348 | 20.7491015 | 20.8581984 | -0.006513202 | 0.900384904 |
| NA | 12.9642634 | NA | NA | 0.816886653 | 0.104634017 |
| 22.7578873 | 22.4755559 | 22.7825573 | 22.697499 | -0.036557493 | 0.746980822 |
| 21.2000528 | 21.2783718 | 21.2511653 | 21.2552871 | 0.056767902 | 0.468587968 |
| 19.0162923 | 19.047079 | 18.9207308 | 19.0140341 | -0.020439915 | 0.756409429 |
| 18.8184627 | 18.7583859 | 18.7666155 | 18.7318983 | 0.020185132 | 0.744660656 |
| 20.0260455 | 19.9070749 | 20.1014851 | 19.9868551 | 0.059197236 | 0.357041787 |
| 20.0283781 | 20.0672706 | 20.0649285 | 19.9642858 | 0.02264288 | 0.76787812 |
| 20.3143096 | 20.1280785 | 20.2162079 | 20.3308933 | -0.005560529 | 0.946543285 |

| | | | | | |
|---|---|---|---|---|---|
| 23.3101999 | 23.199822 | 23.2218182 | 23.2333244 | -0.034369413 | 0.419371819 |
| 22.2845079 | 22.3842087 | 22.3666753 | 22.3118685 | 0.081656863 | 0.227015011 |
| 18.4487757 | 18.3584872 | 18.730749 | 18.4816205 | 0.176432536 | 0.12961359 |
| 21.3790642 | 21.2476048 | 21.3094135 | 21.3007052 | -0.015047355 | 0.757611359 |
| 16.7407099 | 16.4497494 | 16.8146388 | 16.6753485 | 0.032015531 | 0.818998347 |
| 21.1971807 | 21.1227755 | 21.2196772 | 21.1849108 | 0.072852422 | 0.32897444 |
| 20.753757 | 20.6306206 | 20.7638102 | 20.7978151 | 0.010095508 | 0.861086031 |
| 19.4104687 | 19.4074097 | 19.4090301 | 19.4705477 | 0.026642809 | 0.590770585 |
| NA | 17.0977967 | NA | NA | -0.774397837 | 0.0147466 |
| 18.5711267 | 18.1249744 | 18.2549213 | 18.511643 | -0.158888748 | 0.15846236 |
| 23.3481663 | 23.0981885 | 23.2182829 | 23.1951388 | -0.056941486 | 0.402306708 |
| 20.1822346 | 20.1527805 | 20.2014823 | 20.2464433 | 0.073588101 | 0.198865947 |
| 24.2516054 | 24.0604526 | 24.1789439 | 24.2549212 | -0.045115941 | 0.30078334 |
| 21.2397127 | 21.1947828 | 21.2853114 | 21.2665343 | 0.035319982 | 0.371079774 |
| 22.3712801 | 22.2507085 | 22.3440216 | 22.3436743 | -0.039658467 | 0.377920569 |
| 21.3157809 | 21.2801828 | 21.3647998 | 21.3451297 | -0.01807154 | 0.746816222 |
| 19.5395147 | 19.5887565 | 19.5330691 | 19.583605 | -0.017302363 | 0.783861998 |
| 18.7021403 | 18.6000197 | 18.7179171 | 18.7586986 | -0.005833573 | 0.930686176 |
| 20.9671589 | 21.0748521 | 21.020756 | 20.981273 | 0.130223745 | 0.084370475 |
| 19.3554883 | 19.06501 | 19.1723646 | 19.2198543 | -0.080367032 | 0.301017809 |
| 18.7185606 | 18.6933128 | 18.5975015 | 18.7706133 | 0.003338749 | 0.97348153 |
| 19.7933456 | 19.6635855 | 19.8356931 | 19.8242076 | 0.048319973 | 0.685404784 |
| 17.2385233 | 16.9709073 | 17.329818 | 17.1235098 | -0.172820534 | 0.203803855 |
| 21.0509015 | 21.3099052 | 21.2923056 | 21.0255731 | 0.033285993 | 0.699701943 |
| 18.1651918 | 18.2394563 | 18.167674 | 18.1395507 | -0.046248355 | 0.520492691 |
| 21.722717 | 21.5864015 | 21.5868687 | 21.5876655 | -0.054366246 | 0.354328803 |
| 20.6576002 | 20.6557038 | 20.6447285 | 20.7161194 | 0.023734332 | 0.663325844 |
| 21.3811801 | 21.3110265 | 21.3944757 | 21.4053989 | 0.039698394 | 0.346389663 |
| 21.0447146 | 21.032879 | 21.0680363 | 21.0575778 | 0.049441013 | 0.290694147 |
| 16.2128804 | 16.3379734 | 16.2183539 | 16.4402741 | -0.023175806 | 0.80188595 |
| 18.877443 | 18.7830452 | 18.067434 | 18.6698939 | -0.104779632 | 0.66187508 |
| 13.49771 | 13.3224421 | 13.8404966 | 13.3956371 | -0.173720633 | 0.333478714 |
| 17.3322138 | 17.5576112 | 17.3962132 | 17.6686275 | 0.206330863 | 0.090829951 |
| 18.6150537 | 18.3153706 | 18.3455651 | 18.6932714 | -0.151274386 | 0.181859526 |
| 17.3540271 | 17.3179672 | 17.258766 | 17.4963312 | 0.010212921 | 0.926491951 |
| 14.0379633 | NA | 14.0136115 | 14.066535 | -0.040556466 | 0.698653633 |
| 18.0629916 | 18.0251136 | 17.9357066 | 18.0609656 | -0.028814655 | 0.742288168 |
| 17.9669447 | 17.5986895 | 17.9331935 | 17.8703583 | -0.094785215 | 0.280787537 |
| 22.4872511 | 22.2827431 | 22.4506906 | 22.4505625 | -0.060204422 | 0.423887735 |
| 19.5215026 | 19.4608094 | 19.5213782 | 19.5040995 | 0.006598338 | 0.926860229 |
| 18.9585438 | 19.0434998 | 19.0763184 | 18.8285058 | -0.041952853 | 0.558650266 |
| 22.0694987 | 21.9459227 | 22.1038944 | 22.1286044 | -0.015845142 | 0.805678877 |

| | | | | | |
|---|---|---|---|---|---|
| 20.7514708 | 20.6591478 | 20.7763236 | 20.7799632 | 0.027889208 | 0.663607364 |
| 20.8832321 | 20.7859433 | 20.8574717 | 20.911577 | 0.005900199 | 0.924787024 |
| 18.8330148 | 18.7726349 | 18.8385393 | 18.9224487 | -0.016158803 | 0.788166759 |
| 20.9513373 | 20.7772902 | 20.8689517 | 20.8797668 | 0.013200929 | 0.806199875 |
| 24.9852605 | 24.8760578 | 25.0558311 | 25.0062313 | 0.017617825 | 0.716946282 |
| 21.775833 | 21.6492653 | 21.736928 | 21.6858383 | -0.038809389 | 0.409464428 |
| 22.4334731 | 22.2362243 | 22.366688 | 22.3104318 | -0.059838259 | 0.199127442 |
| 21.368389 | 21.2711922 | 21.3555485 | 21.2944041 | -0.055046002 | 0.231079577 |
| 24.1552596 | 24.1125818 | 24.0852655 | 24.1338194 | 0.005181775 | 0.897745879 |
| 22.8554309 | 22.808752 | 22.7741805 | 22.789144 | -0.044077423 | 0.259359472 |
| 20.8168882 | 20.8604536 | 20.8129802 | 20.8788065 | 0.025291751 | 0.464564478 |
| 22.1694478 | 22.0467169 | 22.1590715 | 22.2076793 | 0.007879476 | 0.876890722 |
| 20.869459 | 20.7432026 | 20.8537081 | 20.9873031 | 0.026290031 | 0.627080404 |
| 21.3170156 | 21.3481511 | 21.3620357 | 21.439142 | 0.065113277 | 0.246647836 |
| 21.4235693 | 21.3405008 | 21.4224154 | 21.3870783 | -0.069254034 | 0.24850766 |
| 17.9969879 | 17.9545303 | 17.9015259 | 18.0419333 | 0.013344311 | 0.824669212 |
| 20.7963547 | 20.7031929 | 20.777103 | 20.8088076 | -0.035225396 | 0.565548639 |
| 19.5141543 | 19.3800095 | 19.5468615 | 19.4795943 | 0.031454699 | 0.632867855 |
| 20.8688963 | 20.7904359 | 20.9052419 | 20.7963841 | -0.014468526 | 0.827615403 |
| 18.2347611 | 18.481489 | 18.3778993 | 18.3741557 | 0.025282706 | 0.712228046 |
| 24.4069861 | 24.2181843 | 24.3843993 | 24.4228743 | 0.011999106 | 0.864155314 |
| 18.8691962 | 18.681338 | 18.8600509 | 18.9303748 | 0.03071326 | 0.666242673 |
| 20.9315774 | 20.8501753 | 20.80455 | 20.963026 | 0.02727621 | 0.721722229 |
| 17.9777832 | 17.7138163 | 17.8094245 | 17.8985608 | -0.146203192 | 0.076314911 |
| 19.4109669 | 19.0840531 | 19.2883537 | 19.4761745 | -0.049798617 | 0.551018358 |
| 19.6628335 | 19.4991233 | 19.4439683 | 19.4935582 | -0.105725353 | 0.240030344 |
| 20.2134141 | 20.4276715 | 20.3479023 | 20.6184606 | 0.00472419 | 0.95878852 |
| 18.5232538 | 18.3545788 | 18.5980702 | 18.6465098 | -0.046042433 | 0.648865307 |
| 16.5377716 | 16.6324385 | 16.4405001 | 16.5224249 | -0.097904243 | 0.360167834 |
| 17.7099842 | 17.8786818 | 17.8011694 | 17.6338319 | -0.074454911 | 0.507893504 |
| 17.805195 | 17.8852812 | 18.2719748 | 18.1458137 | 0.18793034 | 0.377319463 |
| 16.7877882 | 16.6663416 | 16.886277 | 16.774576 | -0.240820235 | 0.114691013 |
| 16.9535233 | 17.1055212 | 16.9424807 | 17.0678785 | -0.024879586 | 0.856786562 |
| 14.5361776 | 14.4843225 | 14.6615958 | 14.8919633 | -0.05297601 | 0.664602159 |
| 22.2898477 | 22.3525566 | 22.344896 | 22.3586714 | 0.095337534 | 0.348597226 |
| 20.1944827 | 19.9968686 | 20.0697256 | 20.1383025 | -0.051589204 | 0.568965665 |
| 20.6409594 | 20.5315072 | 20.6447094 | 20.7629103 | -0.008907567 | 0.899128106 |
| 19.9452545 | 19.8370814 | 19.8918735 | 19.9127369 | -0.055568924 | 0.340957678 |
| 21.76658 | 21.7452998 | 21.8113082 | 21.8163979 | 0.012785481 | 0.764740702 |
| 22.1423383 | 22.2229388 | 22.3211951 | 22.1561912 | 0.105257576 | 0.038569851 |
| 18.3919214 | 18.3866516 | 18.3616927 | 18.2778053 | -0.001474119 | 0.981742312 |
| 22.1685175 | 21.8914959 | 21.9969101 | 22.1029675 | -0.076504021 | 0.313820669 |

| | | | | | |
|---|---|---|---|---|---|
| 19.463545 | 19.1379579 | 19.4256792 | 19.4438352 | -0.06822173 | 0.53611426 |
| 17.9132969 | 17.9768587 | 17.9403852 | 17.8844558 | 0.08817078 | 0.494306557 |
| 20.1910617 | 19.9626341 | 20.0700916 | 20.1597537 | -0.044584356 | 0.501981853 |
| 26.075772 | 26.0236393 | 26.1183489 | 26.1343073 | 0.044487436 | 0.346293534 |
| 20.531348 | 20.4519004 | 20.5877311 | 20.5331466 | -0.027171615 | 0.654589573 |
| 11.6532288 | NA | 11.2934562 | 11.3056406 | -0.368061524 | 0.176754541 |
| 21.7382208 | 21.6990423 | 22.4064769 | 21.4909667 | 0.156900289 | 0.50985842 |
| 18.9384731 | 19.0902563 | 19.0376275 | 18.4167699 | 0.136922854 | 0.530798993 |
| 15.8614067 | 15.8382852 | 15.8763645 | 16.0802231 | -0.209205807 | 0.246796817 |
| 19.7423657 | 19.4514517 | 19.5521412 | 19.5764982 | -0.079788889 | 0.63460392 |
| 15.5045972 | 15.1773159 | 15.4173761 | 15.4073686 | 8.61E-04 | 0.995340067 |
| 17.5517261 | 17.1386496 | 17.3955122 | 17.7643621 | -0.17618759 | 0.197266877 |
| 22.5636583 | NA | 22.674959 | 22.6606185 | 0.182416681 | 0.197260837 |
| 17.7252797 | 17.4936499 | 17.8970157 | 17.620093 | 0.002479131 | 0.983254465 |
| 19.7064311 | 19.5976888 | 19.6406751 | 19.80326 | -0.031799544 | 0.790614453 |
| 20.9204331 | 20.9255141 | 20.8325294 | 20.7398758 | 0.018714959 | 0.872640347 |
| 19.0564815 | 18.9389105 | 19.0533853 | 18.8581995 | -0.041305008 | 0.709265054 |
| 21.3766541 | 21.2480178 | 21.1695439 | 21.2764475 | -0.071622301 | 0.524877907 |
| 23.428386 | 23.1454458 | 23.1903896 | 23.1893171 | -0.054621061 | 0.604298688 |
| 19.6323857 | 19.6094468 | 19.7020643 | 19.4232859 | -0.067459585 | 0.509893367 |
| 17.9057206 | 17.8132417 | 17.9042915 | 17.9439526 | 0.019249043 | 0.84461611 |
| 18.5740809 | 18.5850557 | 18.4452856 | 18.4985423 | 0.016311306 | 0.86698333 |
| 16.4735474 | 16.5123066 | 16.4378084 | 16.5190231 | 0.048166941 | 0.614832913 |
| 24.4386703 | 24.5616196 | 24.4282713 | 24.3849666 | 0.119852169 | 0.199839614 |
| 21.2912928 | 21.0376892 | 21.355878 | 21.3178308 | -0.059182951 | 0.515613975 |
| 17.7733788 | 17.5582176 | 17.7697944 | 17.8268252 | -0.090226647 | 0.312801553 |
| 18.837044 | 18.7477258 | 18.9995712 | 18.9916328 | -0.016352298 | 0.836383596 |
| 18.5410759 | 18.5085512 | 18.6121641 | 18.5039608 | -0.06856068 | 0.371981413 |
| 18.9877934 | 19.0755251 | 19.1152308 | 19.1851722 | 0.01565588 | 0.841736138 |
| 21.0941606 | 21.0639789 | 21.2258451 | 21.2115752 | 0.110470203 | 0.160699374 |
| 19.0872484 | 19.0673556 | 19.1231025 | 19.1934398 | 0.04893243 | 0.514746378 |
| 19.6493738 | 19.5391464 | 19.496774 | 19.6049958 | -0.024655822 | 0.74392504 |
| 23.147253 | 22.9352215 | 23.1990943 | 23.0233099 | -0.050863803 | 0.50348367 |
| 27.2369986 | 27.0901213 | 27.2487782 | 27.1873783 | 0.028135593 | 0.675452665 |
| 19.3863131 | 19.2780257 | 19.4572599 | 19.4809132 | 0.056553619 | 0.402078085 |
| 22.7256496 | 22.689919 | 22.7286048 | 22.6303008 | -0.059549952 | 0.36452892 |
| 24.7835196 | 24.615916 | 24.7817362 | 24.7115597 | -0.032474976 | 0.635583759 |
| 21.4213646 | 21.3158768 | 21.4991242 | 21.3910157 | 0.014426603 | 0.826838741 |
| 20.6091971 | 20.4884202 | 20.4922963 | 20.5111487 | -0.024116478 | 0.699731879 |
| 23.3227948 | 23.2292676 | 23.2443124 | 23.2772559 | -0.028383984 | 0.636095373 |
| 20.9309594 | 20.8989685 | 21.0367814 | 21.0420964 | 0.041515628 | 0.509461667 |
| 19.6788781 | 19.6840803 | 19.8256325 | 19.5379137 | -0.029305882 | 0.639021816 |

| | | | | | |
|---|---|---|---|---|---|
| 19.5281308 | 19.622553 | 19.505287 | 19.7245484 | 0.052790048 | 0.40232406 |
| 20.8879272 | 20.7153236 | 20.7198557 | 20.8242717 | -0.04868651 | 0.434725705 |
| 20.6466792 | 20.5183405 | 20.6134893 | 20.5944932 | -0.003187965 | 0.958861686 |
| 23.7571499 | 23.5529558 | 23.7384926 | 23.769863 | -0.03930771 | 0.512192772 |
| 20.4301603 | 20.3046464 | 20.3978064 | 20.4142257 | -0.00470977 | 0.937157481 |
| 21.8609808 | 21.7986774 | 21.8918227 | 21.9146513 | 0.020429323 | 0.712464577 |
| 20.940746 | 20.8728017 | 20.9855939 | 20.9744623 | 0.018029715 | 0.744193006 |
| 22.4125415 | 22.2397039 | 22.3379566 | 22.3532176 | -0.004609038 | 0.931194019 |
| 20.4814699 | 20.4170611 | 20.4965249 | 20.4494681 | -0.012349949 | 0.822078342 |
| 20.5794335 | 20.5263818 | 20.5802252 | 20.6594014 | 0.019086035 | 0.724313611 |
| 21.142784 | 21.1343175 | 21.2459963 | 21.1871064 | 0.022893925 | 0.678744279 |
| 19.5641227 | 19.391748 | 19.4603829 | 19.4793828 | -0.063164067 | 0.252586277 |
| 22.001393 | 21.8529516 | 21.9543656 | 21.9734221 | -0.019323075 | 0.715493776 |
| 21.3076749 | 21.2439792 | 21.2718301 | 21.3228487 | -0.034360436 | 0.51345543 |
| 21.5667353 | 21.3667868 | 21.4422256 | 21.4498426 | -0.084419506 | 0.118755447 |
| 21.5993188 | 21.448984 | 21.6041285 | 21.5851952 | -0.015135238 | 0.777676137 |
| 19.4779577 | 19.3930115 | 19.476999 | 19.5740953 | -0.005693884 | 0.916256573 |
| 21.9432111 | 21.8986403 | 21.8998012 | 21.9315876 | -0.022619389 | 0.645406578 |
| 21.0846332 | 21.0797582 | 21.1750781 | 21.0686247 | 0.007975285 | 0.874587327 |
| 19.4122443 | 19.3501324 | 19.3416972 | 19.4709408 | -0.049763808 | 0.344037108 |
| 25.4543628 | 25.3398874 | 25.4699695 | 25.457794 | 0.025585535 | 0.601263149 |
| 23.8907615 | 23.8409265 | 23.8908341 | 23.8835994 | 0.017505956 | 0.718929966 |
| 21.6590751 | 21.5368261 | 21.5998501 | 21.5608961 | -0.039825009 | 0.409882078 |
| 21.5858823 | 21.5368197 | 21.6176383 | 21.6262255 | 0.033962552 | 0.505667488 |
| 24.0330749 | 23.9376318 | 24.0107564 | 24.0661547 | 0.001878631 | 0.968377644 |
| 23.3232655 | 23.17753 | 23.2542453 | 23.2418993 | -0.030375699 | 0.546538969 |
| 23.8924562 | 23.8255856 | 23.8730057 | 23.8685825 | 0.005768648 | 0.904921834 |
| 21.3314265 | 21.2224655 | 21.2583472 | 21.3149252 | -0.03688284 | 0.428199883 |
| 21.5429312 | 21.4449486 | 21.5142729 | 21.4841934 | -0.063679837 | 0.172053014 |
| 19.4440641 | 19.3879716 | 19.4405628 | 19.46443 | -0.021588399 | 0.627678255 |
| 19.2707103 | 19.350006 | 19.3118638 | 19.3383741 | 0.056258957 | 0.202787759 |
| 22.8838227 | 22.8894413 | 22.8943948 | 22.9531388 | 0.049366729 | 0.243215498 |
| 20.6886079 | 20.6590957 | 20.6025716 | 20.6343766 | -0.01205399 | 0.773722345 |
| 20.793773 | 20.7120109 | 20.7256358 | 20.7674703 | -0.030106433 | 0.463814812 |
| 23.1189081 | 23.0694905 | 23.1201489 | 23.1280822 | -0.013402172 | 0.740607209 |
| 20.6122727 | 20.5333273 | 20.6530737 | 20.6185647 | 0.001820523 | 0.962624455 |
| 22.8609662 | 22.8088769 | 22.8642884 | 22.8432733 | 0.022573429 | 0.572998037 |
| 23.1462408 | 23.1732066 | 23.2563071 | 23.2395618 | 0.043232418 | 0.30985134 |
| 23.8730661 | 23.7816636 | 23.81186 | 23.8771657 | 0.018964818 | 0.653228873 |
| 21.7789675 | 21.6557911 | 21.6984676 | 21.7526088 | 0.003313462 | 0.939252758 |
| 21.6724546 | 21.5960824 | 21.6417629 | 21.672022 | -0.00462938 | 0.915945194 |
| 23.2204726 | 23.1701618 | 23.2048851 | 23.266324 | 0.025260871 | 0.577113187 |

| | | | | | |
|---|---|---|---|---|---|
| 20.6632096 | 20.6620586 | 20.6373668 | 20.7298537 | -0.012773216 | 0.776270725 |
| 25.4889546 | 25.4781974 | 25.5485826 | 25.5434279 | 0.073037984 | 0.121860649 |
| 20.9188067 | 20.8074713 | 20.8606317 | 20.8595023 | -0.022443948 | 0.641165627 |
| 20.2752692 | 20.2773495 | 20.2982309 | 20.2592083 | -0.003185734 | 0.949283418 |
| 20.8451314 | 20.7801633 | 20.8057069 | 20.7894001 | -0.009422118 | 0.852986748 |
| 22.4835182 | 22.3728896 | 22.4677425 | 22.4832774 | 0.001351096 | 0.979157994 |
| 22.2906247 | 22.1887484 | 22.2498557 | 22.2725136 | 0.008816746 | 0.858606741 |
| 21.7262508 | 21.6378539 | 21.7368563 | 21.7730288 | -0.010717382 | 0.834245445 |
| 20.8114277 | 20.6768763 | 20.7614675 | 20.8225109 | -0.006251316 | 0.90372497 |
| 21.004192 | 20.9533194 | 21.0728978 | 21.0555606 | 0.057452798 | 0.274701369 |
| 21.8157903 | 21.7437505 | 21.7564874 | 21.7934183 | 0.024110484 | 0.656024987 |
| 18.6442634 | 18.5947425 | 18.7146312 | 18.7054929 | 0.010008623 | 0.852214537 |
| 20.3905638 | 20.2010493 | 20.3808869 | 20.376531 | 0.008056647 | 0.875931871 |
| 21.393605 | 21.2840668 | 21.3118761 | 21.4419792 | 0.009644787 | 0.856159639 |
| 22.6402981 | 22.702375 | 22.7297895 | 22.6842324 | 0.025604021 | 0.637503648 |
| 21.9284353 | 21.7722662 | 21.9693189 | 21.8864512 | 0.04017174 | 0.480504158 |
| 20.5839666 | 20.3937126 | 20.5202047 | 20.5456277 | -0.051002611 | 0.390555038 |
| 24.7893229 | 24.6744538 | 24.8074154 | 24.7817087 | -0.008356679 | 0.879639206 |
| 24.4940465 | 24.3755369 | 24.4731484 | 24.460549 | 0.00288753 | 0.959332128 |
| 19.9151612 | 19.8756361 | 20.0385305 | 19.9193801 | 0.016617523 | 0.768197577 |
| 19.3558427 | 19.1683657 | 19.2579473 | 19.2466583 | 0.017136231 | 0.764766193 |
| 21.0395697 | 20.9123371 | 20.9809704 | 20.9333282 | -0.045096732 | 0.436173969 |
| 19.2593024 | 19.2634781 | 19.4154244 | 19.2670715 | 0.013327336 | 0.825389488 |
| 20.9306903 | 21.1503872 | 21.0420038 | 21.0138549 | 0.122836716 | 0.049865015 |
| 20.4674057 | 20.3928241 | 20.478763 | 20.44633 | -0.030421837 | 0.614959087 |
| 19.5069674 | 19.4474783 | 19.4803916 | 19.4829169 | -0.001053521 | 0.985676339 |
| 20.0219387 | 19.8558369 | 19.9720974 | 19.962762 | -0.06710988 | 0.278240154 |
| 18.5170733 | 18.4190553 | 18.5523339 | 18.4943152 | 0.033646867 | 0.592363946 |
| 21.045527 | 20.8768369 | 20.954635 | 20.9475585 | -0.020219858 | 0.758550702 |
| 20.5179346 | 20.4028974 | 20.5054204 | 20.4725785 | -0.061469359 | 0.358492595 |
| 19.1590506 | 19.1399698 | 19.1173579 | 19.2503795 | 0.053322034 | 0.419427641 |
| 19.8814049 | 19.8526637 | 19.9362245 | 20.0116136 | 0.059184743 | 0.382600571 |
| 21.6835343 | 21.4828623 | 21.5976444 | 21.5905783 | -0.032190584 | 0.62569011 |
| 23.7496254 | 23.4609686 | 23.6280887 | 23.6751208 | -0.027559679 | 0.671514966 |
| 21.1561172 | 21.0703623 | 21.1587088 | 21.2068352 | 0.021588952 | 0.761269029 |
| 18.4185711 | 18.5262179 | 18.7433713 | 18.6757038 | 0.138260394 | 0.075779725 |
| 22.2444641 | 22.202212 | 22.2366895 | 22.215995 | 0.039284785 | 0.613289508 |
| 20.5808007 | 20.7137544 | 20.7372643 | 20.6668763 | 0.108162537 | 0.200004845 |
| 18.8633335 | 18.8674413 | 18.8747027 | 18.8918778 | 0.048684418 | 0.567255731 |
| 18.608603 | 18.5547944 | 18.6708221 | 18.4897803 | -0.037927027 | 0.664425816 |
| 19.4408132 | 19.4485793 | 19.4568442 | 19.5807782 | 0.041575633 | 0.655795032 |
| 18.2619842 | 18.0848531 | 18.2353539 | 18.5128315 | -0.009275256 | 0.922945788 |

| | | | | | |
|---|---|---|---|---|---|
| 20.6325584 | 20.6793439 | 20.6237154 | 20.6269478 | -0.004808526 | 0.957923795 |
| 19.4261698 | 19.1571816 | 19.3098112 | 19.3884643 | -0.034810243 | 0.704388767 |
| 18.8923438 | 19.0205822 | 19.0559083 | 18.8821438 | -0.003015883 | 0.97482959 |
| 22.0263565 | 21.8646402 | 21.8985632 | 21.913182 | -0.040438295 | 0.679995283 |
| 17.2172176 | 16.9765039 | 17.3942074 | 17.057382 | -0.070732557 | 0.478974533 |
| 21.0702676 | 20.9780416 | 21.1164125 | 21.0518937 | 0.008225215 | 0.935250552 |
| 18.3649491 | 18.3522261 | 18.5460163 | 18.187202 | 0.002922313 | 0.975944139 |
| 18.6882244 | 18.8201761 | 18.7050987 | 18.8085668 | 0.175648515 | 0.08759725 |
| 18.2707459 | 18.2602999 | 18.3740534 | 18.3862137 | 0.005350218 | 0.960543771 |
| 19.6978265 | 19.4342389 | 19.8397488 | 19.5979094 | -0.009260611 | 0.939166554 |
| 21.4352659 | 21.0359731 | 21.456563 | 21.4233948 | -0.027887216 | 0.8076345 |
| 15.7974982 | 15.4979723 | 15.5198763 | 15.6182365 | -0.215991238 | 0.082342421 |
| 16.591576 | 16.4641763 | 16.6517899 | 16.4697124 | 0.082515602 | 0.487641427 |
| 17.0402306 | 16.8741881 | 16.8851258 | 16.5973832 | -0.00135641 | 0.991570091 |
| 20.002023 | 20.1133079 | 20.1289434 | 20.0728124 | 0.077677006 | 0.557895455 |
| 20.3627635 | 20.813671 | 20.6304973 | 20.8780884 | 0.238894151 | 0.070615107 |
| 15.4152002 | 15.8386341 | 15.585958 | 15.2581658 | -0.067863187 | 0.626334349 |
| 15.7618622 | 15.9864812 | 15.9494007 | 16.2564646 | 0.119348649 | 0.378566202 |
| 16.6338643 | 16.6969355 | 16.7925542 | 16.3463191 | -0.006333892 | 0.966367184 |
| 25.8926558 | 25.9838149 | 25.8294393 | 26.0880213 | 0.179909417 | 0.239307779 |
| 19.3981669 | 19.070724 | 19.4519376 | 19.2884088 | -0.039077013 | 0.807773324 |
| 18.775637 | 18.6928281 | 18.7585449 | 18.7763219 | 0.036036107 | 0.83238256 |
| 18.0304051 | 17.9081972 | 17.9063931 | 17.8819556 | -0.313010835 | 0.075005079 |
| 14.4921165 | NA | 14.5597065 | 14.1521115 | -0.237175954 | 0.219767216 |
| 14.9208283 | 15.0299255 | 15.104187 | 15.2605962 | 0.100456667 | 0.659374653 |
| 13.4146996 | NA | 14.0427533 | 13.737519 | 0.663267326 | 0.012433811 |
| 15.5280491 | 14.6302879 | 15.3176621 | 15.3902358 | -0.203585945 | 0.417747819 |
| 15.6494317 | 15.1593347 | 15.8543021 | 16.2263044 | 0.303495994 | 0.236794718 |
| 15.5975095 | 15.4812216 | 15.6952995 | 15.4329244 | -0.04674492 | 0.862381785 |
| 18.8828612 | 18.8366035 | 18.954408 | 18.9495477 | 0.006849179 | 0.930080544 |
| 17.7532917 | 17.6356501 | 17.7625652 | 17.658022 | 0.015469628 | 0.807276855 |
| 25.9558761 | 25.9470129 | 25.9684467 | 25.9703485 | 0.026055145 | 0.583850671 |
| 22.7277087 | 22.6786177 | 22.7025817 | 22.733 | -0.014888359 | 0.733048664 |
| 21.2361481 | 21.1820758 | 21.2050999 | 21.2337315 | -0.017946336 | 0.645797922 |
| 21.5166231 | 21.4799348 | 21.5845444 | 21.5473449 | 0.027404737 | 0.508033748 |
| 22.7689288 | 22.6688174 | 22.7201911 | 22.791598 | 0.00737643 | 0.871207108 |
| 22.2184645 | 22.0871014 | 22.2071267 | 22.1552641 | -0.035986198 | 0.43357155 |
| 22.4760556 | 22.3836302 | 22.4697899 | 22.4994074 | 0.007626974 | 0.873127189 |
| 24.8052602 | 24.7412687 | 24.7931664 | 24.7582099 | -0.013037016 | 0.788867312 |
| 22.2151289 | 22.1833605 | 22.2480651 | 22.2757692 | 0.012141123 | 0.807779505 |
| 22.8183565 | 22.6924907 | 22.7877804 | 22.8156381 | -0.00401936 | 0.943741894 |
| 22.1313426 | 22.0071481 | 22.0973653 | 22.0478521 | -0.007814046 | 0.891099415 |

| | | | | | |
|---|---|---|---|---|---|
| 21.2212008 | 21.1446578 | 21.2352416 | 21.2684297 | 0.018754988 | 0.769926865 |
| 20.9519158 | 20.7847469 | 20.984825 | 20.8736163 | -0.019258391 | 0.765614873 |
| 21.7303683 | 21.6234498 | 21.7724602 | 21.6689882 | 0.005678083 | 0.932199888 |
| 26.4004189 | 26.2766765 | 26.3460522 | 26.2479032 | -0.024505632 | 0.797483736 |
| 19.892844 | 20.0008675 | 19.7430726 | 19.9904629 | -0.034372072 | 0.722781563 |
| 19.9580774 | 19.5465702 | 19.944112 | 19.983732 | -0.038567234 | 0.825252617 |
| 13.9115825 | 14.5867797 | 14.2143527 | 14.3011758 | 0.30185691 | 0.263250157 |
| 20.3517473 | 20.3489879 | 20.3298804 | 20.3307958 | 0.057815825 | 0.330056806 |
| NA | 16.8840603 | NA | NA | -0.402795253 | 0.06509262 |
| 19.0419481 | 18.9656806 | 18.8186377 | 19.1588874 | -0.042095881 | 0.607584429 |
| 20.758147 | 20.7668013 | 20.7349471 | 20.7165987 | -0.117839512 | 0.10052456 |
| 18.1587969 | 17.5873457 | 18.1064638 | 18.1445268 | -0.130756537 | 0.376284759 |
| 17.9643422 | 18.0048286 | 18.165009 | 18.2292763 | -0.006137786 | 0.94850665 |
| 19.8970954 | 19.8474275 | 19.9209594 | 19.9709862 | 0.03137499 | 0.718287853 |
| 17.9555674 | 17.749866 | 17.869295 | 17.9719888 | -0.025110291 | 0.747437917 |
| 17.5843039 | 17.4981049 | 17.4260793 | 17.6561307 | -0.036045234 | 0.640116422 |
| 20.0475831 | 20.0161407 | 19.9885763 | 20.0841361 | 0.022531038 | 0.75111587 |
| 19.5216613 | 19.4319483 | 19.402109 | 19.4039078 | -0.046974479 | 0.492929152 |
| 19.7062719 | 19.5273847 | 19.7079817 | 19.6307351 | 0.019560536 | 0.771401743 |
| 20.3524708 | 20.1766884 | 20.3011542 | 20.3422395 | -0.015902862 | 0.792239621 |
| 18.7048429 | 18.7795347 | 18.6950549 | 18.7139081 | 0.043298591 | 0.473746794 |
| 18.9979496 | 18.8687607 | 18.9125185 | 19.0099194 | -0.020801787 | 0.71821712 |
| 24.8292588 | 24.7358543 | 24.829875 | 24.8068836 | 0.00688954 | 0.902908306 |
| 21.9963588 | 21.8061737 | 21.903005 | 21.9099197 | -0.059447652 | 0.291571413 |
| 22.6654017 | 22.7690774 | 22.8477175 | 22.7163712 | 0.107346095 | 0.062015518 |
| 20.6851916 | 20.5123479 | 20.6406726 | 20.6135663 | -0.054345512 | 0.243233716 |
| 22.5938524 | 22.47797 | 22.5409167 | 22.5803247 | -0.023684015 | 0.582592724 |
| 22.3575354 | 22.2818933 | 22.3263216 | 22.3523694 | -0.008542487 | 0.837340667 |
| 23.4786098 | 23.4467643 | 23.436105 | 23.4260429 | -0.017167732 | 0.65834076 |
| 20.427374 | 20.3907314 | 20.4934484 | 20.4752943 | 0.024566754 | 0.579604612 |
| 21.6765767 | 21.5758782 | 21.6060427 | 21.655709 | 0.001205864 | 0.979677457 |
| 21.5840915 | 21.5377608 | 21.6568112 | 21.5669078 | -0.006853241 | 0.887109184 |
| 21.0463044 | 20.9189446 | 20.9327893 | 20.8944815 | -0.058125525 | 0.266537136 |
| 19.3594263 | 19.3482074 | 19.3198803 | 19.4018439 | -0.010074272 | 0.878307926 |
| 23.1060216 | 23.0916632 | 23.1461603 | 23.2365642 | 0.052196574 | 0.436160948 |
| 20.3527298 | 20.2867102 | 20.2665115 | 20.2779738 | -0.071006803 | 0.358638777 |
| 24.7502117 | 24.6938927 | 24.8404051 | 24.7117291 | 0.038599838 | 0.619701162 |
| 23.4990057 | 23.3820766 | 23.6513322 | 23.5608611 | -0.00489626 | 0.961258638 |
| 16.7532292 | 17.0049968 | 17.2498349 | 16.8444132 | 0.141890384 | 0.300390134 |
| NA | 15.0737146 | NA | NA | 0.176589916 | 0.346437943 |
| 18.8625295 | NA | 18.9190237 | 18.9983527 | 0.162263563 | 0.38199494 |
| NA | 16.4212435 | NA | NA | 0.108868266 | 0.77271275 |

| | | | | | |
|---|---|---|---|---|---|
| 18.2062557 | 18.3695952 | 18.1721555 | 18.1281057 | 0.081577417 | 0.49408878 |
| 21.1704181 | 21.0507711 | 21.1448747 | 21.1502727 | -0.011223394 | 0.803412997 |
| 19.3235169 | 19.272586 | 19.182445 | 19.2637569 | -0.020996783 | 0.830991617 |
| 22.6680535 | 22.5915935 | 22.6721007 | 22.6498131 | 0.002156108 | 0.961063589 |
| 17.4176502 | 18.1660663 | 17.7698494 | 17.8478062 | 0.057320781 | 0.717152359 |
| 15.8067634 | 16.0685709 | 15.9746898 | 15.9932141 | -0.006905065 | 0.96084683 |
| 15.5115939 | 15.3366106 | 15.1164645 | 15.3222717 | -0.065321283 | 0.636976136 |
| 16.5298186 | 16.6198448 | 16.395696 | 16.7811979 | 0.070161596 | 0.58949301 |
| 18.9885008 | 18.8328786 | 19.0800525 | 19.1154529 | -0.067034217 | 0.568901862 |
| 17.9173862 | 18.0297753 | 17.958534 | 17.8709733 | 0.090120417 | 0.326728909 |
| 18.8511626 | 18.8913904 | 19.0236648 | 18.7772468 | 0.106633685 | 0.175414979 |
| 20.4956655 | 20.5140543 | 20.5256382 | 20.5957741 | 0.016474447 | 0.817048643 |
| 17.7591697 | 17.72874 | 17.731972 | 17.6365714 | -0.067370619 | 0.345229984 |
| 24.1922266 | 24.0789449 | 24.1768169 | 24.2376297 | 0.019605308 | 0.764057092 |
| 19.8063482 | 19.7065355 | 19.6723653 | 19.6896535 | -0.088894514 | 0.150784592 |
| 19.9242626 | 20.1215324 | 20.1603685 | 20.0879122 | 0.125654656 | 0.04368495 |
| 22.9710773 | 22.9928927 | 22.9201013 | 23.0284284 | 0.070322179 | 0.248226039 |
| 20.5252776 | 20.4705606 | 20.5950768 | 20.633574 | 0.041688099 | 0.472775389 |
| 19.9169259 | 19.8975825 | 19.8162965 | 19.9007539 | -0.011307827 | 0.841297127 |
| 22.5395288 | 22.4722841 | 22.5364004 | 22.4532779 | -0.018190675 | 0.737973643 |
| 21.5516163 | 21.4632706 | 21.5773606 | 21.538551 | -0.019082909 | 0.729340486 |
| 23.9401985 | 23.821122 | 23.9567146 | 23.9278128 | -0.020301515 | 0.698892459 |
| 22.350837 | 22.3628177 | 22.4810668 | 22.3598882 | 0.024238115 | 0.647497013 |
| 21.5887799 | 21.4208875 | 21.5671023 | 21.5011234 | -0.042523613 | 0.421328153 |
| 22.1080857 | 22.0624589 | 22.0101396 | 22.121866 | 0.016660125 | 0.739673607 |
| 20.3959074 | 20.3339325 | 20.3848196 | 20.3954417 | -0.030000777 | 0.546137293 |
| 22.7829626 | 22.724739 | 22.7401643 | 22.7500734 | 0.002631403 | 0.954844712 |
| 22.1279534 | 22.0042899 | 22.0716449 | 22.1315653 | -0.024078911 | 0.572037823 |
| 22.2316318 | 22.2491634 | 22.2378597 | 22.2784971 | 0.034012606 | 0.412933948 |
| 19.923943 | 19.9209716 | 19.9822039 | 19.9407511 | -0.008896431 | 0.829852052 |
| 20.0398443 | 20.1315881 | 20.151765 | 20.1232548 | 0.042824594 | 0.314562092 |
| 23.0242521 | 22.9083691 | 23.066292 | 23.0048992 | 0.044557322 | 0.328537779 |
| 23.9599789 | 23.9409955 | 23.9672335 | 23.9817116 | 0.061919865 | 0.211735691 |
| 20.6246369 | 20.5786401 | 20.6849082 | 20.6814846 | 0.013211243 | 0.79723512 |
| 23.3624037 | 23.2494642 | 23.3390885 | 23.3055311 | -0.001750046 | 0.974640528 |
| 23.9090204 | 23.817 | 23.8075188 | 23.9158828 | 0.003995169 | 0.941361725 |
| 20.5430432 | 20.468243 | 20.6609831 | 20.627402 | 0.020646395 | 0.718782566 |
| 19.5224643 | 19.3219016 | 19.3310017 | 19.4055945 | -0.049390985 | 0.405567678 |
| 19.1869445 | 19.1565549 | 19.1354416 | 19.2605273 | 0.00772693 | 0.896501309 |
| 19.6354594 | 19.5998112 | 19.5878957 | 19.6067109 | -0.032911051 | 0.591307779 |
| 21.2228449 | 21.1720442 | 21.190167 | 21.1170061 | 0.015668809 | 0.802978056 |
| 18.5490609 | 18.6040185 | 18.5093139 | 18.6399973 | 0.143788609 | 0.048643188 |

| | | | | | |
|---|---|---|---|---|---|
| 18.8889438 | 18.7789094 | 18.7951312 | 18.9249217 | -0.048491464 | 0.549820536 |
| 20.5389433 | 20.6106429 | 20.7310325 | 20.5662358 | 0.112614612 | 0.277287688 |
| 15.591602 | 15.5345142 | 15.4457253 | 15.62444 | 0.023323297 | 0.822315838 |
| 17.6023706 | 17.3309204 | 17.6061274 | 17.9460981 | -0.063861268 | 0.560926686 |
| NA | 15.8495804 | NA | NA | 0.168351711 | 0.385123152 |
| 19.6061661 | 19.8121165 | 19.6108395 | 19.5264468 | 0.031492092 | 0.662243869 |
| 17.5002218 | 17.0948618 | 17.3853979 | 17.0892172 | -0.124861046 | 0.284643445 |
| 17.7364401 | 17.4982446 | 17.6826951 | 17.6590043 | -0.05456622 | 0.567437347 |
| 21.3269114 | 21.2001936 | 21.2575222 | 21.2886842 | -0.041437085 | 0.4060834 |
| 12.7459053 | 13.2002584 | 12.9246167 | 12.834443 | 0.481470649 | 0.035854769 |
| 17.3556875 | NA | 17.3092939 | 17.0719198 | -0.147633689 | 0.400732101 |
| 25.5909627 | 25.5584437 | 25.5781564 | 25.5772363 | 0.042099056 | 0.352648512 |
| 20.8910643 | 20.7854327 | 20.8506833 | 20.8691291 | -0.042386276 | 0.41302389 |
| 19.7722563 | 19.6296381 | 19.7276217 | 19.7461688 | 0.019342994 | 0.712405623 |
| 18.1706384 | 18.1360738 | 18.132846 | 18.1109968 | -0.054319274 | 0.420625382 |
| 22.2660634 | 22.4175388 | 22.302821 | 22.2347451 | 0.099081157 | 0.176464226 |
| 15.8901568 | 15.6229261 | 15.8365483 | 15.7656078 | -0.476556205 | 0.106552384 |
| 18.0966339 | 17.9791893 | 17.771873 | 18.2917088 | -0.020293094 | 0.849600447 |
| 23.3937059 | 23.3655426 | 23.4368537 | 23.3867933 | 0.010323613 | 0.761733492 |
| 23.133888 | 23.0438276 | 23.1262835 | 23.1116149 | -0.022488019 | 0.603043565 |
| 21.0235248 | 20.8277662 | 21.0660365 | 21.0648395 | 0.002864135 | 0.970527349 |
| 27.1181394 | 27.0519522 | 27.1702247 | 27.1852715 | 0.120375551 | 0.315488765 |
| 22.2986518 | 22.1638858 | 22.2546258 | 22.453665 | 0.057683802 | 0.421168718 |
| 22.1942317 | 22.0012455 | 22.0825427 | 22.2680575 | -0.028818427 | 0.796469457 |
| 14.9754593 | NA | 14.933196 | 14.9695713 | 0.179520141 | 0.335629513 |
| 18.1633489 | 17.9360986 | 18.1346804 | 17.975789 | -0.139728884 | 0.275938253 |
| 19.9607355 | 19.8338359 | 19.9094781 | 19.9417092 | -0.061241361 | 0.541975367 |
| 22.1207309 | 21.9167045 | 21.9443193 | 21.8669555 | -0.084877492 | 0.34669069 |
| 18.7436892 | 18.7510728 | 18.5682078 | 18.622347 | -0.061865674 | 0.479259485 |
| 19.3908721 | 19.2491403 | 19.411681 | 19.3055449 | -0.121028782 | 0.169560794 |
| 20.1451395 | 20.207566 | 20.1718221 | 20.1168777 | -0.021099432 | 0.804077765 |
| 17.9857294 | 17.6965561 | 17.8716292 | 17.8002886 | -0.178625756 | 0.038783483 |
| 21.123472 | 21.1816707 | 21.2556693 | 21.1388086 | 0.049630835 | 0.464604247 |
| 19.7635651 | 19.6395513 | 19.7493316 | 19.7714244 | -0.007995369 | 0.898151902 |
| 18.0256916 | 17.9856087 | 18.1160558 | 17.9947084 | -0.030121508 | 0.622799234 |
| 17.9390742 | 17.7973546 | 17.6715837 | 17.8988581 | -0.064148957 | 0.297578586 |
| 18.1347631 | 18.0354004 | 18.0237779 | 18.1579329 | -0.031916274 | 0.581899483 |
| 23.948467 | 23.8030382 | 23.8302791 | 23.893219 | -0.007707824 | 0.884723902 |
| 21.1264199 | 21.0289431 | 21.0642551 | 21.0767237 | -0.03959361 | 0.374190837 |
| 24.9544178 | 24.8506825 | 24.9335753 | 24.9600027 | -0.024427195 | 0.521001672 |
| 22.582533 | 22.5564547 | 22.6316375 | 22.645457 | 0.033306662 | 0.435913748 |
| 22.4831603 | 22.3624771 | 22.4246956 | 22.3960576 | -0.030535079 | 0.496653047 |

| | | | | | |
|---|---|---|---|---|---|
| 21.0948685 | 21.0083337 | 21.0314342 | 21.1634467 | 8.60E-05 | 0.998561407 |
| 22.3135066 | 22.3655727 | 22.4190083 | 22.4269627 | 0.095141419 | 0.102616259 |
| 19.16291 | 19.106492 | 19.141605 | 19.1850974 | 0.004194049 | 0.941403478 |
| 21.0131087 | 20.9406263 | 21.0128607 | 20.9607102 | -0.006592091 | 0.911980242 |
| 23.7751369 | 23.6552963 | 23.8112517 | 23.8042421 | 0.035818892 | 0.585967697 |
| 19.0599048 | 19.0074564 | 18.8824631 | 19.1653705 | 0.031312077 | 0.667488819 |
| 22.3705228 | 22.1699248 | 22.29265 | 22.3997787 | -0.040745688 | 0.604098094 |
| 18.9300465 | 19.0652459 | 18.7860165 | 18.9735049 | 0.023800596 | 0.781361408 |
| 21.3873469 | 21.6967975 | 21.4733143 | 21.4527392 | 0.082886237 | 0.366392884 |
| 16.9886119 | 16.6396733 | 16.8309546 | 16.9948572 | 0.05752547 | 0.761884218 |
| 13.5487509 | 13.702819 | 14.3589453 | 13.6868157 | 0.115021144 | 0.548004037 |
| 15.7567803 | 16.2682273 | 16.0019293 | 16.1260821 | 0.109149088 | 0.587026439 |
| 21.0634558 | 21.0410647 | 21.1005031 | 21.1194648 | 0.075965271 | 0.306197293 |
| 17.5094253 | 17.9812403 | 18.4615664 | 17.6893466 | 0.339491588 | 0.132631512 |
| NA | 16.7953509 | NA | NA | -0.142357317 | 0.491166637 |
| 14.8455425 | NA | 15.5161894 | 14.9484729 | 0.226654422 | 0.254759357 |
| NA | 15.1153938 | NA | NA | 0.081129842 | 0.602862757 |
| 21.0145612 | 21.1660264 | 21.0442663 | 20.8333008 | 0.086461241 | 0.474833434 |
| 16.4222845 | 16.5832755 | 16.354637 | 16.5125847 | 3.85E-04 | 0.997307972 |
| 19.8186722 | 19.4124242 | 19.7057499 | 19.6405521 | -0.12512375 | 0.121899896 |
| 19.0540458 | 18.9577759 | 19.1145096 | 19.1437002 | -0.053967374 | 0.494897605 |
| 19.3737279 | 19.1235706 | 19.3662197 | 19.1751422 | -0.135257269 | 0.080157786 |
| 19.2841942 | 19.1704581 | 19.1187336 | 19.2017984 | -0.041376506 | 0.569440759 |
| 19.8751863 | 19.7816359 | 20.0061946 | 19.8003444 | 0.012148292 | 0.858528678 |
| 19.8892449 | 19.713643 | 19.9179337 | 19.7926521 | -0.013828067 | 0.833824823 |
| 26.1830751 | 26.1980736 | 26.2514263 | 26.1510267 | 0.063463343 | 0.29447511 |
| 22.8632642 | 22.8131853 | 22.9144886 | 22.891886 | 0.079068714 | 0.187035247 |
| 21.1761172 | 21.0126274 | 21.1529485 | 20.9318409 | -0.05559318 | 0.333157265 |
| 20.4417325 | 20.4151797 | 20.5037245 | 20.4914047 | 0.008642548 | 0.87779148 |
| 24.6317521 | 24.6417615 | 24.626025 | 24.5587653 | 0.045499579 | 0.429520952 |
| 18.6242232 | 18.4158637 | 18.5715228 | 18.4913976 | -0.083920117 | 0.146490462 |
| 20.0512361 | 19.8815895 | 20.0819735 | 20.0664736 | -0.031882564 | 0.572866884 |
| 22.8815092 | 22.7914349 | 22.9146859 | 22.9078238 | 0.040627968 | 0.436276981 |
| 21.6946327 | 21.5524337 | 21.6974212 | 21.6860181 | 0.015416596 | 0.739432029 |
| 21.5855948 | 21.4982791 | 21.5600395 | 21.5478948 | -0.021361467 | 0.646417246 |
| 21.5656152 | 21.4645801 | 21.5699756 | 21.5967516 | 0.005643524 | 0.905103122 |
| 22.182268 | 22.1339924 | 22.1730114 | 22.2069978 | 0.020798587 | 0.664817611 |
| 24.3166643 | 24.2279425 | 24.3122155 | 24.3266119 | 0.001115162 | 0.982452868 |
| 23.2918201 | 23.1925025 | 23.2636918 | 23.2704711 | 0.019388871 | 0.718491096 |
| 25.7105609 | 25.435481 | 25.6433152 | 25.6295564 | -0.031035188 | 0.653610152 |
| 20.210024 | 20.0128163 | 20.1057796 | 20.1971133 | 0.039901474 | 0.567144745 |
| 20.7288786 | 20.6958697 | 20.6894978 | 20.6059628 | -0.106079907 | 0.175990974 |

| | | | | | |
|---|---|---|---|---|---|
| 17.6729107 | 17.6358609 | 17.662592 | 17.7303881 | -8.83E-04 | 0.993055341 |
| 18.600794 | 18.8347422 | 18.9872131 | 19.1524696 | 0.207094019 | 0.051370852 |
| 23.2933862 | 23.2744407 | 23.2833104 | 23.2367802 | 0.055021214 | 0.617575201 |
| 23.5074701 | 23.2828532 | 23.5772415 | 23.4104383 | -0.001383437 | 0.990885227 |
| 15.2900993 | 15.466835 | 15.3573207 | 15.3365481 | -0.002126804 | 0.986664161 |
| 16.770642 | 16.6748455 | 16.0775284 | 16.6944085 | -0.051665844 | 0.770914895 |
| 13.1440883 | 13.2717878 | 13.3000687 | 13.4563261 | 0.00232232 | 0.990914202 |
| NA | 18.2760079 | NA | NA | -0.077995578 | 0.705179665 |
| NA | 12.8464174 | NA | NA | -0.140160516 | 0.65633531 |
| 20.6947915 | 20.527153 | 20.5310976 | 20.6062835 | -0.076487443 | 0.243287755 |
| 24.8821923 | 24.7991248 | 24.8879549 | 24.8821413 | 2.56E-04 | 0.99644083 |
| 24.1069444 | 24.0123045 | 24.055407 | 24.0890029 | 0.009981245 | 0.876878288 |
| 22.6810017 | 22.5561602 | 22.6490052 | 22.6396345 | 0.010102784 | 0.879982997 |
| 17.9601197 | 17.9979413 | 18.0865604 | 18.0043664 | 0.066445093 | 0.344676862 |
| 21.5322534 | 21.3771858 | 21.4984588 | 21.4879893 | 0.016981282 | 0.714271438 |
| 21.6236993 | 21.4222228 | 21.5028244 | 21.5315779 | -0.030631719 | 0.608873149 |
| 21.7665415 | 21.6913482 | 21.7682683 | 21.8028577 | 0.036162626 | 0.463753188 |
| 24.2023018 | 24.0176345 | 24.1566432 | 24.0860432 | -0.070102754 | 0.373593522 |
| 21.9163169 | 21.7502292 | 21.814226 | 21.8525779 | -0.049840881 | 0.385164295 |
| NA | 14.0259524 | NA | NA | -0.102275171 | 0.724388902 |
| 15.8717164 | 15.5041975 | 16.5257098 | 16.2199741 | 0.142530298 | 0.502037125 |
| 16.5291637 | 16.490482 | 16.3798623 | 16.1775019 | -0.053634946 | 0.723098931 |
| 21.368176 | 21.0038863 | 21.190375 | 21.2450691 | -0.099975482 | 0.468499735 |
| 21.3744162 | 21.330676 | 21.5114383 | 21.1031398 | -0.059530528 | 0.643479193 |
| 19.4719659 | 19.5321996 | 19.6951687 | 19.5659185 | 0.025891859 | 0.808155761 |
| 17.4412366 | 17.6540596 | 17.2468059 | 17.3205183 | -0.120675004 | 0.214406411 |
| 19.3080504 | 19.1534729 | 19.0715582 | 19.1856258 | -0.049591711 | 0.567381577 |
| 17.6997154 | 17.7859958 | 17.7710971 | 18.0196415 | 0.120907092 | 0.142175963 |
| 18.152757 | 18.1136686 | 17.9676053 | 18.0279607 | -0.05950356 | 0.40922246 |
| 27.0947795 | 26.9569892 | 27.1041578 | 27.0954252 | 0.036463544 | 0.601261831 |
| 23.7817746 | 23.6360618 | 23.7148768 | 23.7808541 | -0.029183127 | 0.633693653 |
| 22.9940568 | 22.8800928 | 22.9700559 | 23.0063439 | 0.010006688 | 0.869891491 |
| 20.7728309 | 20.6878451 | 20.9138691 | 20.8527249 | 0.052892728 | 0.390809229 |
| 20.5767649 | 20.3159407 | 20.3858026 | 20.5000362 | -0.050523321 | 0.35139661 |
| 20.5933999 | 20.5918166 | 20.6559926 | 20.6497057 | 0.018484624 | 0.727381242 |
| 22.5448311 | 22.4501913 | 22.5708218 | 22.5929266 | 0.046660143 | 0.374176896 |
| 23.1880956 | 23.0976994 | 23.1746322 | 23.084592 | -0.016509546 | 0.728145911 |
| 20.2004246 | 20.1891387 | 20.1853498 | 20.2762771 | 0.001131644 | 0.980569835 |
| 21.9796671 | 21.9781336 | 22.0576027 | 22.0385016 | 0.03401298 | 0.307221114 |
| 19.5919673 | 19.4868155 | 19.5984235 | 19.5734879 | -0.017349555 | 0.704640436 |
| 19.4220502 | 19.2972452 | 19.3517029 | 19.4069719 | -0.005194911 | 0.920324934 |
| 19.5102398 | 19.4715448 | 19.5164888 | 19.5719774 | 0.04506549 | 0.398427454 |

| | | | | | |
|---|---|---|---|---|---|
| 23.6389866 | 23.5704627 | 23.6298195 | 23.7142977 | 0.049465646 | 0.371029843 |
| 21.2163006 | 21.0654823 | 21.3001123 | 21.1832729 | 0.017362298 | 0.755815188 |
| 23.040129 | 22.8889796 | 23.1182024 | 23.0350291 | 0.019952574 | 0.741315348 |
| 22.1484701 | 21.9946897 | 22.1655721 | 22.1484952 | 0.03870626 | 0.535021739 |
| 18.3803495 | 18.311431 | 18.3572076 | 18.4782448 | -0.003333814 | 0.958190098 |
| 24.4354426 | 24.2885247 | 24.4138836 | 24.39194 | -0.010082182 | 0.877375942 |
| 20.589005 | 20.5073411 | 20.6160398 | 20.5998977 | 0.020400629 | 0.761639865 |
| 20.5632435 | 20.4032908 | 20.3618937 | 20.4926808 | -0.088141572 | 0.265756977 |
| 21.8507575 | 21.6109488 | 21.6966189 | 21.502268 | -0.097663728 | 0.28243308 |
| 17.456673 | 17.6403217 | 17.490465 | 17.3515569 | -0.096228527 | 0.44847451 |
| 21.9771875 | 21.8867375 | 21.7166454 | 21.9746511 | -0.081989098 | 0.609420156 |
| 17.0036244 | 16.922682 | 16.8413691 | 16.9754993 | 0.04868612 | 0.771026574 |
| 15.5292417 | NA | 15.0754111 | 15.8963276 | 0.084673636 | 0.717693392 |
| 19.1977779 | 19.0521322 | 19.163909 | 19.125657 | -0.059637122 | 0.380802496 |
| 22.5607926 | 22.5244036 | 22.5156718 | 22.5700748 | 0.039613088 | 0.55670415 |
| 27.4789358 | 27.4179964 | 27.5230776 | 27.5511191 | 0.037566067 | 0.458887872 |
| 21.2142054 | 21.1321547 | 21.2037448 | 21.2014468 | -0.003495102 | 0.938008783 |
| 22.0234492 | 21.9342871 | 22.0295577 | 22.0461804 | -0.009432338 | 0.83031841 |
| 19.7764783 | 19.7829295 | 19.7034122 | 19.8553071 | 0.00444583 | 0.939322155 |
| 18.1955847 | 18.2304102 | 18.5649492 | 18.447292 | 0.206167635 | 0.307093044 |
| NA | 15.1976874 | NA | NA | 0.576474707 | 0.11779843 |
| 20.0776518 | 20.1843852 | 19.888677 | 20.06287 | -0.033065399 | 0.686549983 |
| 23.1918646 | 23.1159506 | 23.1820146 | 23.1489603 | 0.059240053 | 0.424735576 |
| 21.4782928 | 21.2854364 | 21.4445122 | 21.4286041 | -0.035251901 | 0.551615708 |
| 21.2893198 | 20.9782359 | 21.255923 | 21.1419164 | -0.106670022 | 0.077522902 |
| 20.0477968 | 19.9826592 | 20.0642241 | 20.1446791 | 0.043382378 | 0.448783521 |
| 19.6687665 | 19.5826299 | 19.5609506 | 19.6398876 | -0.022537081 | 0.631109672 |
| 20.3265551 | 20.240814 | 20.3619802 | 20.3595993 | 0.00519068 | 0.909196493 |
| 22.7296771 | 22.5655786 | 22.6446113 | 22.6965903 | -0.015773877 | 0.786486174 |
| 20.6305973 | 20.3727811 | 20.5204254 | 20.5094793 | -0.04009833 | 0.509963793 |
| 23.4990939 | 23.3903532 | 23.4824938 | 23.5005168 | 0.003959916 | 0.949550313 |
| 25.3277944 | 25.2405683 | 25.4032687 | 25.3667747 | 0.014452305 | 0.825164256 |
| 17.8767157 | 17.8011887 | 17.8377228 | 17.9058463 | 0.049274462 | 0.460248538 |
| 21.0846996 | 20.8781799 | 21.254096 | 21.0915571 | 0.006994685 | 0.919552517 |
| 19.2944487 | 19.3596308 | 19.251079 | 19.2965965 | 0.062489114 | 0.411830723 |
| 26.0872018 | 26.0388149 | 26.0146829 | 26.0198787 | -0.027546154 | 0.733299553 |
| 20.2780557 | 20.5417466 | 20.3578342 | 20.3879963 | 0.050891111 | 0.539744651 |
| 17.7327779 | 17.9833185 | 17.9833808 | 17.7440255 | -0.070690207 | 0.577111215 |
| 25.9711218 | 25.8710794 | 25.8440624 | 25.9247014 | 0.022314725 | 0.860827013 |
| 24.6286617 | 24.5543534 | 24.6326651 | 24.6419854 | 0.017585131 | 0.655868104 |
| 19.635836 | 19.6937456 | 19.7078401 | 19.7385907 | 0.041888464 | 0.333408892 |
| 20.7992803 | 20.5332962 | 20.8912172 | 20.9271133 | -0.018054176 | 0.92508695 |

| | | | | | |
|---|---|---|---|---|---|
| 18.2257957 | 18.330222 | 18.3125206 | 18.4527299 | 0.102791339 | 0.152389137 |
| NA | 15.5532806 | NA | NA | -0.239986408 | 0.548834425 |
| 19.9078276 | 19.8897385 | 19.9783279 | 20.0203271 | 0.049012119 | 0.496055841 |
| 17.1449971 | 17.051873 | 16.9994354 | 16.8112727 | -0.093457246 | 0.511026104 |
| 18.6679885 | 18.4564715 | 18.8707289 | 18.5095527 | 0.045184689 | 0.59847834 |
| 18.4382587 | 18.4944183 | 18.439071 | 18.4595949 | -0.00458883 | 0.926369139 |
| 18.6329881 | 18.6594701 | 18.6164937 | 18.6564803 | -0.050775703 | 0.39530657 |
| 19.2251908 | 19.1588199 | 19.1246652 | 19.2107115 | -0.009345678 | 0.89184142 |
| 14.9342778 | NA | 15.0586562 | 14.680414 | -0.132813443 | 0.411725968 |
| 19.1713121 | 19.1040867 | 19.2960726 | 19.24979 | 0.099569972 | 0.304756712 |
| 23.4413954 | 23.2395721 | 23.4018931 | 23.2017378 | -0.022845404 | 0.769990274 |
| 18.9825294 | 18.9141385 | 18.9797706 | 18.9981406 | 0.004547648 | 0.946233716 |
| 21.2150046 | 21.0472995 | 21.191901 | 21.2280015 | -0.04586522 | 0.459936811 |
| 21.3076922 | 21.3176532 | 21.2702874 | 21.2258193 | -0.028628779 | 0.624647373 |
| 22.3658818 | 22.3319636 | 22.4186296 | 22.4011328 | 0.015828863 | 0.781357357 |
| 20.4145779 | 20.2155145 | 20.3019846 | 20.3457846 | -0.047807553 | 0.389371287 |
| 21.6887953 | 21.6254695 | 21.6981312 | 21.668161 | -0.006077997 | 0.909590509 |
| 20.7509717 | 20.6368685 | 20.7205157 | 20.8165104 | -0.032507468 | 0.545923314 |
| 25.8401269 | 25.6792179 | 25.7706231 | 25.7347161 | -0.064211488 | 0.237381518 |
| 24.1258732 | 24.130949 | 24.1695425 | 24.1837103 | 0.052307349 | 0.271373829 |
| 22.1082313 | 22.0206914 | 22.1239098 | 22.0109195 | -0.036448181 | 0.431783539 |
| 20.0813838 | 20.062884 | 20.1136992 | 20.0606967 | 0.006766229 | 0.877543891 |
| 23.2214604 | 23.1233967 | 23.1577591 | 23.2207809 | -0.003922708 | 0.926714134 |
| 23.6623849 | 23.5680069 | 23.67645 | 23.6845881 | -0.003882131 | 0.924967816 |
| 21.5756099 | 21.4983501 | 21.5265393 | 21.5657885 | -0.024737816 | 0.477642847 |
| 22.1416604 | 22.0636832 | 22.1104769 | 22.1390121 | -0.012392984 | 0.773831284 |
| 22.0019008 | 21.9491868 | 22.0688698 | 21.9655675 | 0.014843576 | 0.782415328 |
| 22.1241267 | 22.1265419 | 22.0688508 | 22.164053 | 0.060419931 | 0.271272416 |
| 21.593895 | 21.6211915 | 21.6480615 | 21.6167819 | 0.021017445 | 0.717572835 |
| 20.9765532 | 20.9206344 | 20.8778972 | 21.0641213 | -0.002531899 | 0.970114922 |
| 20.3509361 | 20.1552378 | 20.1721718 | 20.2658477 | -0.047173672 | 0.489825708 |
| 19.2421325 | 19.0562771 | 19.2380677 | 19.2081659 | 0.003355533 | 0.960088792 |
| 19.0150124 | 19.1854063 | 19.1569326 | 19.2430828 | 0.132806783 | 0.075465213 |
| 15.6918974 | 15.6242956 | 15.5469371 | 15.7286448 | -0.02847304 | 0.821407148 |
| 17.9749739 | 17.9728279 | 17.9472386 | 17.7087459 | -0.080963599 | 0.535156039 |
| 16.5755134 | 16.998524 | 16.6870238 | 16.7232602 | 0.003869167 | 0.977330432 |
| 17.6133459 | 17.0197211 | 17.7975769 | 17.6958015 | -0.11942887 | 0.504869139 |
| 21.5825422 | 21.4095104 | 21.6005811 | 21.4644846 | -0.049168211 | 0.478751883 |
| 22.1544456 | 22.0747318 | 22.1560787 | 22.1311822 | 0.016842684 | 0.752225519 |
| 21.4289696 | 21.1657172 | 21.4766838 | 21.4000724 | -0.051270729 | 0.479670744 |
| 16.3664907 | 16.0034506 | 16.1872113 | 16.3735689 | 0.06664492 | 0.719169688 |
| 23.1539773 | 23.23402 | 23.16683 | 23.0225988 | 0.005472592 | 0.938497281 |

| | | | | | |
|---|---|---|---|---|---|
| 21.0170889 | 20.9783718 | 21.0219904 | 21.1011974 | 0.09593089 | 0.513928324 |
| 18.8668766 | 18.955365 | 18.9589162 | 18.8797501 | 0.04201912 | 0.543744596 |
| 22.3931412 | 22.1624458 | 22.3300895 | 22.3355679 | -0.043029394 | 0.506301756 |
| 18.2818992 | 18.3021668 | 18.4673078 | 18.2114737 | -0.033863683 | 0.654207944 |
| 21.5378778 | 21.4610175 | 21.5292964 | 21.4795413 | -0.035140813 | 0.475896224 |
| 24.95792 | 24.9493708 | 25.0692345 | 24.9923745 | 0.044186828 | 0.380950345 |
| 19.848535 | 19.995289 | 20.0595528 | 20.0721823 | 0.171320592 | 0.043885594 |
| 21.9334925 | 21.8351084 | 21.9455568 | 21.9271772 | -0.005338768 | 0.918616128 |
| 16.6209461 | 16.4837467 | 16.6510465 | 16.5654356 | -0.040597396 | 0.640073927 |
| 18.1742439 | 18.2606614 | 18.2423883 | 18.1485178 | 0.00881768 | 0.909591349 |
| 22.6011243 | 22.5170947 | 22.5605711 | 22.5810925 | 0.003283423 | 0.953151039 |
| 19.2862011 | 19.3345282 | 19.2390242 | 19.2937106 | 0.020434361 | 0.692430555 |
| 20.1969801 | 20.1020465 | 20.168418 | 20.1569951 | -0.020379182 | 0.673047764 |
| 20.7064646 | 20.6060102 | 20.662518 | 20.6990009 | -0.015398138 | 0.766400991 |
| 18.5525078 | 18.506962 | 18.6166078 | 18.6547784 | -0.018464114 | 0.768896909 |
| 24.1023268 | 24.0693819 | 24.1630566 | 24.1292409 | 0.029515333 | 0.465622659 |
| NA | 15.2813384 | NA | NA | -0.127585286 | 0.262579528 |
| 19.9082621 | 19.5702092 | 19.3826438 | 19.727042 | -0.203249504 | 0.143653771 |
| 21.4894331 | 21.3895688 | 21.3930593 | 21.3413806 | -0.057175724 | 0.418474337 |
| 16.2021425 | 16.6628244 | 16.6782183 | 16.7923416 | 0.308931342 | 0.060419736 |
| 16.2040112 | 15.9506874 | 15.8489833 | 15.9687561 | -0.051801028 | 0.616553523 |
| 17.0759293 | 16.8809597 | 16.7648119 | 16.9551083 | -0.08420798 | 0.414267047 |
| 17.3686241 | 17.4139665 | 17.1985991 | 17.1666981 | -0.117493018 | 0.23602489 |
| 20.2754094 | 20.1670345 | 20.1090924 | 20.265162 | -0.058820785 | 0.406724397 |
| 24.3155095 | 24.0643783 | 24.2438242 | 24.231275 | 0.011101928 | 0.860107726 |
| 20.5186972 | 20.4069161 | 20.575974 | 20.5473057 | 0.015088166 | 0.805377957 |
| 22.9951931 | 22.8611574 | 23.0118242 | 22.9927992 | 0.034760772 | 0.569951927 |
| 18.1039367 | 18.2495405 | 18.3706225 | 18.1925302 | 0.018586466 | 0.743515989 |
| 20.7277488 | 20.5699641 | 20.6323415 | 20.6859256 | -0.061115091 | 0.267269516 |
| 24.6287754 | 24.5319476 | 24.6058228 | 24.6220163 | 0.010622098 | 0.838389863 |
| 21.6495698 | 21.4502889 | 21.6174433 | 21.6321577 | -0.010320447 | 0.83157408 |
| 20.7941207 | 20.7700799 | 20.7918286 | 20.8412694 | 0.00104349 | 0.980198355 |
| 19.6180027 | 19.5489607 | 19.6237904 | 19.5429605 | -0.019705066 | 0.611628311 |
| 19.6497772 | 19.4948782 | 19.6093572 | 19.6152523 | -0.038326359 | 0.455775559 |
| 19.7367162 | 19.7468987 | 19.8808983 | 19.8061015 | 0.015566444 | 0.769702592 |
| 22.5552357 | 22.5720198 | 22.5529263 | 22.5752416 | 0.075758958 | 0.18730484 |
| 20.6913178 | 20.5928494 | 20.6502724 | 20.6219819 | 0.005163027 | 0.928973349 |
| 22.90931 | 22.7780922 | 22.8882913 | 22.936724 | 0.02639488 | 0.657332652 |
| 20.403316 | 20.218508 | 20.3090846 | 20.3826344 | -0.059344726 | 0.39736678 |
| 18.539541 | 18.3711316 | 18.5725748 | 18.5708324 | -0.035322302 | 0.651713713 |
| 20.6500577 | 20.6028066 | 20.6046575 | 20.5822025 | -0.046645764 | 0.552162686 |
| 15.985057 | 16.0994906 | 15.9682882 | 16.0787019 | -0.054823006 | 0.509451835 |

| | | | | | |
|---|---|---|---|---|---|
| 22.6617539 | 22.6152882 | 22.7432138 | 22.6511094 | -0.039513647 | 0.679827326 |
| 19.3927313 | 19.0650094 | 19.3372226 | 19.2545641 | -0.037507978 | 0.738949063 |
| 18.7736353 | 18.3611791 | 18.4509931 | 18.389025 | 0.012189691 | 0.933246024 |
| 16.1144307 | 16.0426487 | 16.1164018 | 16.5154561 | 0.077691465 | 0.631734239 |
| 19.6940621 | 19.3889581 | 19.5953782 | 19.6053002 | 0.006815191 | 0.961683502 |
| 19.1918456 | 19.3978717 | 19.0752135 | 19.0753899 | -0.031284761 | 0.823376929 |
| 23.3397123 | 23.2327397 | 23.5196439 | 23.4451951 | 0.020657039 | 0.857550422 |
| 18.2234042 | 18.593158 | 18.5404061 | 18.4350685 | 0.172331845 | 0.102475436 |
| 20.418528 | 20.0149816 | 20.252305 | 20.2440576 | -0.150481148 | 0.154521703 |
| 23.3883683 | 23.1209258 | 23.1436647 | 23.3682547 | -0.001901332 | 0.984488578 |
| 18.6471212 | 18.5053974 | 18.6138154 | 18.6353971 | 0.012682292 | 0.896254875 |
| NA | 15.5802884 | NA | NA | 0.058427611 | 0.531312102 |
| 17.7637563 | 17.8199661 | 17.8424655 | 17.8615579 | 0.030695288 | 0.704963473 |
| 20.6827349 | 20.4830554 | 20.6313783 | 20.7000695 | -0.068732718 | 0.377603393 |
| 22.440544 | 22.2881276 | 22.4978412 | 22.4436309 | 0.045815855 | 0.559030079 |
| 18.0203028 | 18.026171 | 18.0315313 | 18.0737097 | -0.005237644 | 0.942875965 |
| 20.4161973 | 20.102733 | 20.2140805 | 20.1436058 | -0.136245084 | 0.080166897 |
| 23.4932164 | 23.3871706 | 23.5327007 | 23.5099086 | 0.055640341 | 0.458358396 |
| 21.6408992 | 21.4019047 | 21.6170205 | 21.6118209 | -0.02902507 | 0.676393887 |
| 20.1832561 | 20.101634 | 20.0951383 | 20.0870324 | -0.049822654 | 0.436620501 |
| 21.2181715 | 21.2480582 | 21.2703184 | 21.2728689 | -4.37E-04 | 0.994436757 |
| 20.4844822 | 20.2832324 | 20.329857 | 20.3444711 | -0.003436252 | 0.956802582 |
| 19.0399007 | 18.9860917 | 19.1582452 | 19.1099547 | 0.072878616 | 0.230810587 |
| 19.0936694 | 19.0207043 | 19.0846954 | 19.1159879 | -0.015380417 | 0.797903476 |
| 23.7062512 | 23.5125291 | 23.7419467 | 23.6150347 | -0.04199251 | 0.483213206 |
| 21.8837762 | 21.7147633 | 21.766631 | 21.7904943 | -0.06430392 | 0.26080646 |
| 25.7926705 | 25.6702058 | 25.8177541 | 25.8242328 | 0.009312876 | 0.866909963 |
| 22.4830136 | 22.3579151 | 22.4064423 | 22.3453748 | -0.059842421 | 0.268884528 |
| 22.1639218 | 22.0664723 | 22.0862041 | 22.15042 | 0.003120527 | 0.952555776 |
| 23.2625309 | 23.3001538 | 23.2312423 | 23.2926873 | 0.06527071 | 0.217500184 |
| 22.0159347 | 21.9443 | 21.9667338 | 21.8703397 | -0.045900983 | 0.356774993 |
| 19.9038897 | 19.8151309 | 19.9561043 | 19.9409195 | 0.011829615 | 0.806328693 |
| 22.8935807 | 22.8075606 | 22.8534014 | 22.8310801 | -0.008928003 | 0.856513652 |
| 24.0457989 | 23.974874 | 24.0354257 | 24.003223 | 0.014940389 | 0.754755008 |
| 20.5460075 | 20.4751797 | 20.5418828 | 20.5677568 | 0.001104143 | 0.979886326 |
| 21.6540305 | 21.5884517 | 21.6919454 | 21.701195 | -2.10E-05 | 0.99962332 |
| 24.4458666 | 24.3239952 | 24.4648993 | 24.4436225 | -0.003401636 | 0.942651082 |
| 22.0848857 | 21.9621108 | 22.0952872 | 22.0976781 | 0.013130495 | 0.78893071 |
| 22.389512 | 22.2437214 | 22.3424959 | 22.3454921 | -0.0242508 | 0.622895974 |
| 22.9382359 | 22.9022329 | 22.8886345 | 22.9785018 | 0.036522876 | 0.485791694 |
| 18.9963559 | 18.9761493 | 19.0234684 | 18.9879726 | 0.010977525 | 0.837137692 |
| 20.892589 | 20.8070623 | 20.894832 | 20.8368736 | -0.024669378 | 0.64035945 |

| | | | | | |
|---|---|---|---|---|---|
| 22.570686 | 22.6271226 | 22.6337794 | 22.5699192 | 0.056941624 | 0.286328056 |
| 21.0337021 | 20.8439807 | 20.9199161 | 20.98317 | -0.078415383 | 0.209629872 |
| 20.7077158 | 20.574165 | 20.7659798 | 20.6277078 | -0.027246941 | 0.689428986 |
| 18.6845405 | 18.7113603 | 18.6507089 | 18.7149991 | -0.062265807 | 0.366054796 |
| 17.7619728 | 17.8524713 | 17.7385459 | 17.8108508 | 0.011483702 | 0.873201873 |
| 20.2625989 | NA | 20.3604958 | 20.4061253 | 0.164744856 | 0.069770781 |
| 19.2606287 | 19.0733929 | 19.1111445 | 19.2287596 | -0.068680868 | 0.413229367 |
| 19.2875157 | 19.0844178 | 19.3840916 | 19.1876374 | -0.014679488 | 0.878694615 |
| 18.4409526 | 18.4175127 | 18.254916 | 18.6325704 | -0.077155069 | 0.441842936 |
| 18.2243179 | 17.9441521 | 17.9292156 | 18.2556803 | -0.069779882 | 0.507708795 |
| 15.3878773 | NA | 15.4789599 | 15.3283008 | -0.054080838 | 0.667597977 |
| 16.7631256 | 16.6305521 | 16.5120565 | 16.8992031 | -0.036097787 | 0.783943136 |
| 17.8793856 | 17.8399285 | 18.2941171 | 18.3160004 | 0.073929811 | 0.656979216 |
| 15.664916 | 15.7606842 | 15.9750602 | 15.9199688 | -0.014659515 | 0.946694473 |
| 18.6493913 | 18.5689857 | 18.6127174 | 18.7189748 | 0.068053476 | 0.434519234 |
| 21.1614081 | 21.129148 | 21.201078 | 21.229243 | -0.006520903 | 0.921702385 |
| 20.9944046 | 20.8592672 | 21.0044783 | 21.0443353 | 0.01653913 | 0.740609141 |
| 23.3145697 | 23.0677684 | 23.2092281 | 23.2026756 | -0.067542123 | 0.308490158 |
| 21.1810936 | 21.1689377 | 21.1643943 | 21.2344222 | 0.00223079 | 0.976476017 |
| 22.2890819 | 22.1913308 | 22.2232028 | 22.3366254 | 0.023353227 | 0.765959516 |
| 17.859678 | 17.7900986 | 17.8917903 | 17.9653105 | 0.025389982 | 0.828624507 |
| 16.7868617 | 16.673629 | 16.8539931 | 16.7188342 | -0.010196546 | 0.928059721 |
| 21.0434029 | 21.1081347 | 21.1019488 | 21.2380125 | 0.002344355 | 0.974159443 |
| 21.1483934 | 21.0687864 | 21.1590179 | 21.1264173 | 0.001160704 | 0.984433142 |
| 21.2022399 | 21.0571571 | 21.1801653 | 21.2263786 | 0.018062577 | 0.719107924 |
| 20.3394666 | 20.3127267 | 20.3826362 | 20.250706 | -0.005013102 | 0.902443659 |
| 20.355817 | 20.2465466 | 20.3846036 | 20.3123599 | -0.018750264 | 0.672792803 |
| 21.9856204 | 21.8662228 | 21.9782679 | 22.0289197 | 0.006199065 | 0.901882425 |
| 20.6894526 | 20.4298506 | 20.6088488 | 20.6350938 | -0.053299767 | 0.432098896 |
| 24.8065346 | 24.6590762 | 24.7672554 | 24.7653882 | -0.004128928 | 0.954356309 |
| 19.1478226 | 19.0280121 | 19.133936 | 18.9477452 | -0.134590077 | 0.149089089 |
| 15.714081 | 15.5343805 | 15.5421137 | 15.6960443 | -0.154013463 | 0.139642105 |
| 17.7689693 | 17.9601318 | 18.1373827 | 18.0581932 | 0.074644795 | 0.515241066 |
| 20.8682064 | 20.8142565 | 20.7684276 | 20.8638256 | -0.027962805 | 0.482825079 |
| 19.245477 | 19.0254809 | 19.0751331 | 18.9793158 | -0.16051967 | 0.056829401 |
| 16.7324752 | 16.7634347 | 16.5352597 | 16.8990866 | 0.180314095 | 0.188910656 |
| 19.6491445 | 19.6572087 | 19.6824987 | 19.5822825 | -0.038874757 | 0.511439686 |
| 21.584871 | 21.4879621 | 21.5167114 | 21.5522169 | -0.067936053 | 0.21125784 |
| 21.169491 | 21.1408388 | 21.1398449 | 21.1811533 | 0.014720313 | 0.766154706 |
| 19.9101793 | 19.744983 | 19.8785326 | 19.8413797 | -0.013761695 | 0.801409051 |
| 21.1562458 | 21.0341037 | 21.1741058 | 21.1561757 | 0.001080623 | 0.98443987 |
| NA | 16.4213631 | NA | NA | 0.178329676 | 0.194668599 |

| | | | | | |
|---|---|---|---|---|---|
| 20.3829944 | 20.2507784 | 20.3761697 | 20.3264286 | -0.018953216 | 0.66551598 |
| 18.8149763 | 18.6618699 | 18.9200397 | 18.8827794 | 0.00775204 | 0.913338974 |
| 17.5790409 | 17.5064491 | 17.827721 | 17.6629724 | -0.052318344 | 0.639392488 |
| 15.2954186 | 15.3011796 | 15.2293149 | 15.2283465 | -0.06138955 | 0.606071137 |
| 19.9369447 | 19.9219353 | 20.056436 | 20.1312064 | -0.081708724 | 0.438425156 |
| 19.1952599 | 19.0683112 | 19.1916831 | 19.1778486 | -0.024321901 | 0.702556199 |
| 21.2403103 | 21.1893407 | 21.2181368 | 21.1714657 | 0.023868389 | 0.681759701 |
| 23.309647 | 23.3003358 | 23.4125601 | 23.3668606 | 0.08082052 | 0.138550204 |
| 20.7234118 | 20.6549725 | 20.6887925 | 20.7065806 | 0.011345379 | 0.818508744 |
| 20.3957406 | 20.3963115 | 20.4781105 | 20.4275397 | 0.049291572 | 0.307601563 |
| 21.4023532 | 21.3707873 | 21.3878829 | 21.3860262 | -0.009526572 | 0.778197878 |
| NA | 13.7594599 | NA | NA | 0.493479275 | 0.094487745 |
| 17.4181919 | 17.18897 | 17.4598616 | 17.5744136 | -0.036062851 | 0.771643661 |
| 17.8744848 | 17.1117729 | 17.7612909 | 17.5312553 | -0.152866862 | 0.441903203 |
| 18.2484248 | 18.4210473 | 18.3337657 | 18.4481034 | 0.079352761 | 0.347261345 |
| 22.0581711 | 21.9307053 | 22.0310424 | 22.0447159 | -0.007186977 | 0.871596608 |
| 23.3610696 | 23.3158922 | 23.41905 | 23.3180328 | 0.01211171 | 0.771789299 |
| 20.4422783 | 20.3735044 | 20.3311644 | 20.5193462 | -0.023969211 | 0.678448293 |
| 17.8549501 | 18.0016065 | 17.9168189 | 17.8961392 | 0.084970455 | 0.157339328 |
| 20.1803463 | 20.2252499 | 20.1725296 | 20.2218118 | 0.032981152 | 0.63733215 |
| 18.5470579 | 18.5494052 | 18.4456055 | 18.5032985 | -8.75E-04 | 0.992480556 |
| 20.2496717 | 19.9765834 | 20.1951688 | 20.0967833 | -0.025815216 | 0.78639119 |
| 19.7242322 | 19.4602894 | 19.5525928 | 19.6142075 | -0.116411521 | 0.19289945 |
| 18.8033402 | 18.7376391 | 18.6660458 | 18.8488603 | 0.009295649 | 0.930517464 |
| 18.3668742 | 18.4379623 | 18.3531991 | 18.4030525 | -0.006458305 | 0.913865355 |
| 17.4774924 | 17.9912271 | 17.9172006 | 17.6319054 | 0.166877874 | 0.234935718 |
| 19.1410116 | 19.2936348 | 19.0138055 | 18.8775834 | 0.012168775 | 0.909368405 |
| 21.8893367 | 21.6921563 | 21.6693371 | 21.7585924 | -0.052554338 | 0.471939974 |
| 21.2733236 | 21.2476535 | 21.26414 | 21.3020908 | 0.026568818 | 0.576136456 |
| 22.4593788 | 22.4195144 | 22.4155385 | 22.4510467 | 6.86E-05 | 0.998585637 |
| 23.8156899 | 23.7470422 | 23.8222345 | 23.8020838 | 0.017507723 | 0.707328244 |
| 17.5033838 | 17.5148055 | 17.5404332 | 17.3092937 | 0.014188835 | 0.883496395 |
| 21.3369534 | 21.2925474 | 21.3732943 | 21.2979903 | -0.015318236 | 0.805609516 |
| 21.2166979 | 21.1084443 | 21.1900087 | 21.2136347 | -0.045681432 | 0.459315545 |
| 22.6901818 | 22.4960223 | 22.5561325 | 22.6041511 | -0.058087137 | 0.317101752 |
| 24.1194258 | 23.9893101 | 24.0620838 | 24.0912797 | -0.01662585 | 0.766216804 |
| 22.2857342 | 22.2342914 | 22.2379924 | 22.2387128 | 0.015538603 | 0.767480087 |
| 23.7951406 | 23.745354 | 23.7803796 | 23.8472815 | 0.022021184 | 0.650327704 |
| 21.1699982 | 21.1135043 | 21.1930946 | 21.1841131 | 0.024547011 | 0.597780696 |
| 22.9262155 | 22.8537335 | 22.8976141 | 22.8931503 | -6.12E-04 | 0.984704622 |
| 24.3172274 | 24.2295921 | 24.2711335 | 24.257846 | -0.013117002 | 0.776135261 |
| 20.4337389 | 20.3782631 | 20.303039 | 20.378135 | -0.012889749 | 0.869826018 |

| | | | | | |
|---|---|---|---|---|---|
| 17.5289537 | 17.4063526 | 17.3594754 | 17.3151041 | -0.022864134 | 0.830788717 |
| 17.3405346 | 17.1068522 | 17.1071966 | 17.3384466 | 0.021020387 | 0.859376016 |
| 20.0049002 | 20.0131024 | 19.6737136 | 20.0475133 | -0.062536681 | 0.604902878 |
| 16.7633553 | 16.7500873 | 16.9839111 | 16.8754681 | 0.041731008 | 0.788259712 |
| 23.2611495 | 23.2115245 | 23.2796278 | 23.2195693 | 0.017728934 | 0.75912888 |
| 21.2016474 | 21.3033417 | 21.282431 | 21.228433 | 0.042337104 | 0.279257437 |
| 18.3223937 | 18.1294346 | 18.2194198 | 18.4120505 | 0.097800601 | 0.536101995 |
| 18.8335435 | 18.8123156 | 18.667418 | 19.0673609 | 0.069477717 | 0.607172515 |
| 18.095129 | 17.996115 | 17.9902794 | 18.1139943 | 0.068004265 | 0.425921689 |
| 17.0076968 | 16.8962423 | 17.3110025 | 17.0398231 | -0.006990841 | 0.933756663 |
| 19.4425327 | 19.121488 | 19.4034402 | 19.3356656 | -0.088667508 | 0.282066035 |
| 22.5845195 | 22.5290531 | 22.6806712 | 22.6302462 | 0.056375324 | 0.461651999 |
| 23.787114 | 23.5955815 | 23.827293 | 23.805963 | -0.013727846 | 0.849716756 |
| 23.7472147 | 23.5284578 | 23.6682985 | 23.6599624 | -0.044769444 | 0.530980582 |
| 19.3191108 | 19.1206545 | 19.2653092 | 19.305155 | -0.030660851 | 0.652248606 |
| 18.7259668 | 18.6563509 | 18.7716217 | 18.7991749 | 0.047903784 | 0.477970985 |
| 19.3127779 | 19.3740142 | 19.4074802 | 19.4672676 | 0.081128056 | 0.237803096 |
| 22.4302551 | 22.442379 | 22.4347779 | 22.3652114 | -0.022577747 | 0.727548209 |
| 20.0459992 | 19.9935536 | 20.0697859 | 20.1159927 | 0.016120002 | 0.803354779 |
| 20.7822349 | 20.6701391 | 20.793322 | 20.8333368 | -0.025732871 | 0.692656511 |
| 24.199909 | 24.1022654 | 24.20385 | 24.1700747 | 0.016160994 | 0.799608247 |
| 20.5775825 | 20.4076779 | 20.6015409 | 20.551346 | -0.053182504 | 0.399705107 |
| 19.2505047 | 19.1120831 | 19.229185 | 19.2442698 | -0.055536166 | 0.37590123 |
| 19.7665501 | 19.5847133 | 19.74073 | 19.7416669 | 0.006825676 | 0.911508284 |
| 20.9184099 | 20.8261694 | 20.9443015 | 20.9702553 | 0.022103435 | 0.713709155 |
| 22.8034961 | 22.6482913 | 22.7616524 | 22.8065847 | -0.058150154 | 0.304746534 |
| 22.3879206 | 22.2423366 | 22.2762275 | 22.3577852 | -0.016894118 | 0.755687708 |
| 21.8416669 | 21.808813 | 21.7782451 | 21.8253634 | -0.010469508 | 0.844401346 |
| 26.3947281 | 26.3019988 | 26.4276169 | 26.4255788 | 0.037533734 | 0.445929398 |
| 18.8972663 | 18.8818749 | 18.9487241 | 18.983177 | -0.008225656 | 0.867915815 |
| 24.0229899 | 23.9600648 | 24.0354316 | 24.0187346 | 0.015012939 | 0.713647314 |
| 23.9387609 | 23.8378035 | 23.929359 | 23.9317854 | -0.009248197 | 0.833239506 |
| 23.0062839 | 22.9526495 | 22.9201028 | 23.018455 | -0.028368354 | 0.541236253 |
| 24.2074985 | 24.1500668 | 24.2004424 | 24.175274 | 0.013404621 | 0.778815219 |
| 22.7248152 | 22.6286568 | 22.6721186 | 22.7283583 | -0.02929739 | 0.578881385 |
| 21.597973 | 21.4844062 | 21.6254928 | 21.5909259 | 0.012069487 | 0.822859557 |
| 21.6327138 | 21.5876423 | 21.638209 | 21.6740057 | -0.026395488 | 0.627065707 |
| 20.7392899 | 20.7850027 | 20.7513193 | 20.7920772 | 0.040232006 | 0.456266194 |
| 18.3178074 | 18.234887 | 18.4484647 | 18.4656951 | 0.068934397 | 0.220782038 |
| 19.4142051 | 19.3972438 | 19.3717821 | 19.3880267 | -0.03935272 | 0.522442727 |
| 20.3585012 | 20.2542446 | 20.1137419 | 20.2879719 | -0.018175304 | 0.773616856 |
| 17.7127492 | 17.5504026 | 17.5664437 | 17.6192598 | -0.065782295 | 0.311159908 |

| | | | | | |
|---|---|---|---|---|---|
| 19.2943522 | 19.4051312 | 19.3127022 | 19.4821315 | 0.074458216 | 0.277450873 |
| 22.3022653 | 22.1943901 | 22.2590464 | 22.2565257 | 0.001532027 | 0.982172682 |
| 19.5106494 | 19.3180248 | 19.5006547 | 19.4289828 | -0.065647727 | 0.348644995 |
| 16.7776142 | 16.9741139 | 16.8832435 | 16.7260226 | -0.027434219 | 0.6831582 |
| 26.0261701 | 25.8724924 | 25.9915655 | 25.9870013 | -0.018119145 | 0.795369492 |
| 22.9440934 | 22.8512865 | 22.9529278 | 22.9474476 | -0.032666812 | 0.645967813 |
| 18.5546026 | 18.5215279 | 18.6703399 | 18.7386594 | -0.012382607 | 0.879671024 |
| 19.1238043 | 18.909221 | 19.2725943 | 19.3329785 | 0.026721867 | 0.798183632 |
| 26.7894811 | 26.5072494 | 26.678479 | 26.683164 | -0.082858152 | 0.436614054 |
| 16.725565 | 17.4162549 | 16.774878 | 16.640387 | -0.018768527 | 0.891727271 |
| 14.6884744 | 14.4128515 | 13.9205359 | 14.281368 | -0.234265694 | 0.310615038 |
| 14.4806505 | 14.1681196 | 14.4965597 | 14.3076913 | 0.022294284 | 0.953858304 |
| 21.3112401 | 21.0259598 | 21.2467338 | 21.3387977 | -0.044520968 | 0.650437738 |
| 17.908317 | 18.0771404 | 18.0181804 | 17.8992991 | -0.014871676 | 0.867892368 |
| 22.4405493 | 22.3107785 | 22.4753879 | 22.4600291 | -0.001264557 | 0.986055245 |
| 22.5891155 | 22.5609818 | 22.6110069 | 22.6578163 | 0.088022463 | 0.201962518 |
| 23.0033071 | 22.9006223 | 23.0098789 | 23.0150851 | 0.004754211 | 0.917425852 |
| 24.7394297 | 24.7380315 | 24.7797985 | 24.7304068 | 0.046590702 | 0.31312617 |
| 19.5931178 | 19.534632 | 19.4899578 | 19.5332071 | -0.044412211 | 0.255537649 |
| 22.5865883 | 22.5894033 | 22.6646203 | 22.649367 | 0.050205323 | 0.243450263 |
| 25.3937526 | 25.3250577 | 25.4468969 | 25.4295026 | 0.033733002 | 0.450717377 |
| 24.0479185 | 23.985538 | 24.0621636 | 24.0305494 | -0.006776516 | 0.891622735 |
| 22.2304016 | 22.1718541 | 22.1687675 | 22.2619391 | 0.003246032 | 0.951703039 |
| 21.8418796 | 21.7532207 | 21.82409 | 21.8035406 | -0.006187988 | 0.909006295 |
| 18.6496527 | 18.531247 | 18.4951192 | 18.6365344 | -0.107611982 | 0.072052111 |
| 23.8434741 | 23.8049407 | 23.9289784 | 23.9384867 | 0.036229963 | 0.544706959 |
| 20.5031189 | 20.1908658 | 20.4462943 | 20.4894692 | -0.032187307 | 0.795037444 |
| 17.0666591 | 16.9254575 | 17.1264284 | 16.8815178 | -0.082734582 | 0.533847886 |
| 14.5993882 | NA | 13.4528674 | 14.579745 | 0.018318968 | 0.974559998 |
| 20.0080058 | 20.0693168 | 20.1250835 | 20.0867896 | 0.031073253 | 0.603096502 |
| 21.2839031 | 21.2753103 | 21.2815187 | 21.3122903 | 0.001937727 | 0.964587806 |
| 21.7557354 | 21.7118772 | 21.8075683 | 21.7168724 | 0.06513231 | 0.481060342 |
| 20.8451648 | 20.7704274 | 20.8637574 | 20.8864745 | 0.004041504 | 0.914937579 |
| 21.3376233 | 21.3362966 | 21.3625559 | 21.3776048 | 0.081149046 | 0.138997783 |
| 23.3455231 | 23.2492453 | 23.3080674 | 23.3873925 | 0.065226093 | 0.330708927 |
| 18.8054186 | 18.6550392 | 18.8429657 | 18.8787251 | 0.041169597 | 0.621826912 |
| 22.303067 | 22.3329233 | 22.3170323 | 22.3428549 | 0.038552475 | 0.194366944 |
| 23.768059 | 23.6748961 | 23.7954781 | 23.8855555 | 0.055052312 | 0.385190056 |
| 15.8667598 | 15.8979731 | 15.8638467 | 15.709749 | 0.032736674 | 0.768856107 |
| 21.0696606 | 20.7699379 | 20.95323 | 21.07921 | -0.088231021 | 0.319379626 |
| 20.0144795 | 19.8583811 | 20.1229386 | 19.9832028 | 0.031289453 | 0.732826653 |
| 20.973578 | 20.7966233 | 20.8946586 | 20.9813043 | -0.052650715 | 0.437806514 |

| | | | | | |
|---|---|---|---|---|---|
| 22.2020295 | 22.0212747 | 22.163942 | 22.2515703 | -0.045867208 | 0.437946579 |
| 21.3339835 | 21.1874105 | 21.2597208 | 21.3305071 | -0.04055237 | 0.370629409 |
| 26.0214125 | 25.9735262 | 26.0104276 | 25.9773362 | -0.022599945 | 0.674897659 |
| 21.3894463 | 21.4598034 | 21.4263966 | 21.4364624 | 0.080129996 | 0.146636692 |
| 17.5702709 | 17.4315106 | 17.4387269 | 17.6439931 | -0.042890277 | 0.678973198 |
| NA | 16.0873053 | NA | NA | -0.063560559 | 0.548069069 |
| 23.3594298 | 23.2094753 | 23.3937365 | 23.3053714 | -0.026947916 | 0.716200444 |
| 25.0441252 | 24.8804319 | 24.9707757 | 24.9604091 | -0.004337726 | 0.926166597 |
| NA | 13.5291458 | NA | NA | 0.981459068 | 0.177823028 |
| 19.0734481 | 19.0451507 | 19.080316 | 19.131538 | 0.048270369 | 0.365849276 |
| 21.6984738 | 21.6242527 | 21.62168 | 21.6205578 | 0.00530744 | 0.917810182 |
| 22.8160315 | 22.8416565 | 22.8915773 | 22.8926172 | 0.068733485 | 0.191442595 |
| 21.2038291 | 21.0711115 | 21.203654 | 21.1990363 | -0.008810069 | 0.868584745 |
| 20.3335279 | 20.2606291 | 20.3774725 | 20.3849202 | 0.04665237 | 0.55182684 |
| 19.822961 | 19.4827699 | 19.7109901 | 19.7278738 | -0.03397008 | 0.760350555 |
| 20.722311 | 20.5788212 | 20.6572638 | 20.7425346 | 0.002886833 | 0.963225139 |
| 16.2376207 | 16.1985142 | 15.9322665 | 16.2160522 | -0.027277009 | 0.817181332 |
| 19.8270736 | 19.7570804 | 19.8568745 | 19.9173236 | 0.026975773 | 0.56696138 |
| NA | 13.5022151 | NA | NA | -0.210884909 | 0.254459469 |
| 21.0566138 | 20.8008726 | 21.0545838 | 21.0430744 | -0.021474567 | 0.826342485 |
| 23.2762179 | 23.1752903 | 23.1455418 | 23.2585891 | -0.017569108 | 0.705989576 |
| 21.7300876 | 21.6080559 | 21.7392782 | 21.7176301 | 0.006860457 | 0.876536509 |
| 21.8848529 | 21.8409624 | 21.8387935 | 21.9492343 | 0.04154644 | 0.427604293 |
| 21.6413262 | 21.5326995 | 21.5966695 | 21.6511391 | 0.009628282 | 0.853924476 |
| 28.1265667 | 28.0914138 | 28.0920982 | 28.1403542 | 0.049239163 | 0.376528749 |
| 20.9830247 | 20.8226449 | 20.9493847 | 20.9788241 | -0.039455087 | 0.532713006 |
| 17.5762956 | 17.345408 | 17.3580985 | 17.5136782 | -0.141344188 | 0.132387703 |
| 21.0486863 | 21.178809 | 21.1499654 | 21.1310187 | 0.193546621 | 0.049943636 |
| 16.2321166 | NA | 15.8434906 | 16.1339505 | -0.281215399 | 0.233735759 |
| 22.3678381 | 22.2756481 | 22.3919773 | 22.3198364 | -0.028342337 | 0.594205145 |
| 24.2909542 | 24.3958748 | 24.4032596 | 24.2532409 | 0.084634869 | 0.312077869 |
| 21.2556775 | 21.2429399 | 21.3629234 | 21.2730758 | 0.097636814 | 0.122994528 |
| 21.14022 | 21.0684651 | 21.0002998 | 21.0282305 | -0.063558612 | 0.172322288 |
| 15.5255162 | 15.3908828 | 15.5897315 | 15.6350554 | -0.093747351 | 0.627420472 |
| 21.7848352 | 21.6945131 | 21.7718197 | 21.7466997 | -0.044613201 | 0.337004294 |
| 20.6588519 | 20.5462986 | 20.6385403 | 20.684208 | -0.029312866 | 0.609146574 |
| 22.527703 | 22.3239521 | 22.4653094 | 22.4920095 | -0.003846466 | 0.954108491 |
| 26.2885398 | 26.1830639 | 26.271348 | 26.1911438 | -0.022037417 | 0.671022159 |
| 18.9744584 | 18.8283083 | 18.9428905 | 18.9836042 | -0.050659651 | 0.387720766 |
| 22.6420735 | 22.5418671 | 22.6157211 | 22.6823761 | -0.008736313 | 0.86177165 |
| 22.0840379 | 21.9973517 | 22.0041599 | 22.061667 | 0.018269422 | 0.832726726 |
| 15.4413942 | 15.5419612 | 15.2250088 | 15.5656588 | -0.108485489 | 0.492470792 |

| | | | | | |
|---|---|---|---|---|---|
| 18.1396563 | 17.9453243 | 18.0670529 | 18.2278488 | -0.087295031 | 0.304724132 |
| 20.3609242 | 20.3415828 | 20.3640416 | 20.4442348 | 0.002865222 | 0.949645115 |
| NA | 14.424901 | NA | NA | -0.070495215 | 0.948653029 |
| 21.794801 | 21.6113742 | 21.7004625 | 21.7356708 | -0.021818036 | 0.696462464 |
| 20.0501266 | 20.1268296 | 20.1155233 | 19.9990922 | 0.09117696 | 0.143947113 |
| 20.8533054 | 20.7852008 | 20.8418306 | 20.8621641 | -0.007800872 | 0.861371115 |
| 18.1530716 | 18.2210256 | 18.2343181 | 18.1592708 | 0.014929037 | 0.831592547 |
| 21.3768959 | 21.2026461 | 21.2825739 | 21.2025487 | -0.0858803 | 0.256250721 |
| 22.3892558 | 22.2441657 | 22.3498676 | 22.3757844 | 0.006189241 | 0.932274592 |
| 18.0431994 | 18.0315141 | 18.14158 | 18.0483169 | 0.011458895 | 0.894908768 |
| 23.5820152 | 23.5327693 | 23.5905715 | 23.6128932 | 0.0288711 | 0.473528725 |
| 21.2830579 | 21.1453445 | 21.1580787 | 21.1574465 | -0.073719026 | 0.20169513 |
| 18.9417995 | 18.5509572 | 19.1198407 | 19.0771593 | 0.011425476 | 0.956546917 |
| 17.3056628 | 16.9776725 | 17.3626608 | 17.5256097 | -0.048157192 | 0.713085706 |
| 19.6634001 | 19.5862652 | 19.6386708 | 19.6891932 | -0.016232507 | 0.781874183 |
| 23.1957419 | 23.0646674 | 23.148137 | 23.1478036 | -0.023495725 | 0.669341206 |
| 22.7603516 | 22.7731223 | 22.8264167 | 22.7535947 | 0.012061918 | 0.778123422 |
| 23.9050447 | 23.7671317 | 23.8297641 | 23.8506973 | -0.027848842 | 0.601939371 |
| 21.0800517 | 20.8713187 | 20.9059272 | 20.9800237 | -0.048496692 | 0.368624358 |
| 21.334226 | 21.1483507 | 21.2792705 | 21.2689114 | -0.069008544 | 0.258122897 |
| 21.0213088 | 20.9733258 | 21.0979578 | 21.0192526 | 0.010373805 | 0.869843595 |
| NA | 15.2245976 | NA | NA | -0.172634554 | 0.181236622 |
| 17.4417499 | 16.9629826 | 17.3817877 | 17.3590134 | -0.173940059 | 0.181994941 |
| 19.0349002 | 18.8189707 | 18.9881199 | 18.980077 | 0.035983841 | 0.791553418 |
| 19.3870801 | 19.1104709 | 19.361835 | 19.2464626 | -0.004583826 | 0.95856401 |
| 22.0640639 | 22.0072553 | 22.0691565 | 22.0265835 | -0.005357742 | 0.89409748 |
| 22.4637795 | 22.3391735 | 22.4723002 | 22.3685312 | 0.006525966 | 0.895110172 |
| NA | 13.6300692 | NA | NA | 0.128255868 | 0.508062153 |
| NA | 12.7186726 | NA | NA | 0.550982278 | 0.017250676 |
| NA | 15.0524978 | NA | NA | -0.086949229 | 0.754991898 |
| 20.8060572 | 20.7471908 | 20.7946056 | 20.9069109 | -0.019027507 | 0.731865485 |
| 21.4190263 | 21.3928349 | 21.3933196 | 21.4737086 | 0.023125139 | 0.612165254 |
| NA | 13.8373032 | NA | NA | -0.091954178 | 0.735631523 |
| 16.831189 | 16.5000665 | 16.8707637 | 16.8182086 | -0.234318865 | 0.196282841 |
| 23.9453186 | 23.9904772 | 24.0471969 | 23.9007 | 0.151770711 | 0.175794175 |
| 16.8946217 | 16.663011 | 16.8646176 | 16.7901145 | -0.069653822 | 0.323813668 |
| 18.7876909 | 18.7398281 | 18.8687694 | 18.7247407 | -0.080225833 | 0.249868913 |
| 20.3545458 | 20.3703656 | 20.3679129 | 20.3924467 | 0.024088479 | 0.520052488 |
| 26.7812924 | 26.7227863 | 26.7769919 | 26.7837487 | 0.046691518 | 0.341932719 |
| 22.4561229 | 22.2890477 | 22.2788397 | 22.324549 | -0.059323716 | 0.357405965 |
| 19.8050436 | 19.5430848 | 19.6080132 | 19.6040719 | -0.060214638 | 0.364164654 |
| 22.7028524 | 22.5165272 | 22.6802748 | 22.597934 | -0.017889588 | 0.840796772 |

| | | | | | |
|---|---|---|---|---|---|
| 13.2469554 | NA | 13.4105404 | 13.0879336 | 0.242658478 | 0.2836031 |
| 14.4878512 | NA | 14.4919893 | 14.4952581 | -0.073936809 | 0.672023332 |
| 20.1119552 | 19.8841871 | 20.0350255 | 20.0163796 | -0.051857447 | 0.58548416 |
| 17.6276727 | 17.5484858 | 17.6697011 | 17.7202922 | -0.124381465 | 0.167045383 |
| 20.8038081 | 20.7557563 | 20.8029661 | 20.8579496 | 0.019029736 | 0.718186141 |
| 19.7874227 | 19.6730469 | 19.7235707 | 19.7679749 | -0.052485633 | 0.287526675 |
| 23.0616847 | 23.000226 | 23.0967544 | 23.1089804 | 0.042603258 | 0.352988561 |
| 19.7173259 | 19.6029376 | 19.7380855 | 19.7388246 | -0.013416834 | 0.787537328 |
| 20.0194018 | 20.0675942 | 20.1409411 | 20.1341365 | 0.041833191 | 0.41278187 |
| 21.7275619 | 21.6440468 | 21.7292418 | 21.7933378 | 0.025256079 | 0.619104179 |
| 21.2400706 | 21.1764253 | 21.2243245 | 21.1727401 | 0.009686113 | 0.857886762 |
| 19.1452803 | 18.9407822 | 19.0151887 | 19.1618628 | -0.044913337 | 0.474521198 |
| 19.434498 | 19.3759624 | 19.3079768 | 19.4012565 | -0.030718739 | 0.67746457 |
| 16.68283 | 16.5537588 | 16.6715112 | 16.7575039 | -0.058357607 | 0.45793494 |
| 21.087983 | 20.754265 | 21.0684714 | 21.0241082 | -0.08864007 | 0.297645689 |
| 17.4810093 | 17.3597932 | 17.5009652 | 17.5265567 | 0.036022147 | 0.722752746 |
| 18.1043293 | 17.9038412 | 17.8933215 | 18.2538879 | 0.027579858 | 0.785604236 |
| 21.0789484 | 20.868821 | 21.1078541 | 21.0167034 | -0.035060094 | 0.76111496 |
| NA | 11.7979601 | NA | NA | -0.083155012 | 0.559330645 |
| NA | 12.6805329 | NA | NA | 0.069347722 | 0.719468297 |
| 16.7214728 | 15.3974403 | 16.6861308 | 16.8421477 | -0.083230745 | 0.862646227 |
| 19.2726898 | 19.3367035 | 19.0615873 | 19.1741899 | 0.037749933 | 0.761230683 |
| 18.546798 | 18.511132 | 18.9503671 | 18.657935 | 0.032327012 | 0.788724562 |
| 21.1664083 | 21.3110302 | 21.3674776 | 21.3758507 | 0.029661554 | 0.775419115 |
| 23.06445 | 23.0933088 | 23.0884782 | 23.2393492 | 0.113927585 | 0.127028249 |
| 20.2590538 | 20.1574768 | 20.3176003 | 20.2553253 | -4.21E-04 | 0.994104198 |
| 21.2925064 | 21.3012639 | 21.348823 | 21.3017273 | 0.038521349 | 0.453511915 |
| 22.7837559 | 22.7001894 | 22.7613299 | 22.7728464 | 0.005427421 | 0.907065086 |
| 21.3538708 | 21.3591372 | 21.3450092 | 21.3341748 | 0.027195162 | 0.676833511 |
| 19.0840956 | 19.0669047 | 19.0392592 | 19.2159326 | 0.029105097 | 0.662956591 |
| 16.0938773 | NA | 16.2307816 | 16.0230388 | 0.046289556 | 0.600001281 |
| 17.9639479 | 17.9667688 | 18.1218527 | 17.9512019 | 0.185883377 | 0.108761007 |
| 14.7969031 | 14.4747299 | 14.798474 | 14.7101012 | -0.026487345 | 0.899048953 |
| 18.758023 | 18.3473003 | 18.6210785 | 18.6058357 | -0.144810988 | 0.116164764 |
| 20.9368366 | 21.0270239 | 21.041814 | 20.9856743 | 0.104157825 | 0.091154891 |
| 22.7525398 | 22.6840914 | 22.6775906 | 22.7182621 | -0.021213797 | 0.659877765 |
| NA | 16.1195365 | NA | NA | -0.077044287 | 0.826249739 |
| 13.9627664 | 13.9978588 | 14.1427555 | 12.974834 | -0.328882292 | 0.204244706 |
| NA | 15.6682258 | NA | NA | 0.027034448 | 0.879380552 |
| 17.5226144 | 17.6952037 | 17.4724089 | 17.3170632 | 0.146474882 | 0.383251122 |
| 16.7961252 | 16.5790036 | 16.564005 | 16.8563643 | 0.00426972 | 0.97370117 |
| 18.6113695 | 18.5988276 | 18.6104134 | 18.4813176 | 0.010922364 | 0.928235191 |

| | | | | | |
|---|---|---|---|---|---|
| 16.3073013 | 16.0148807 | 16.3845327 | 16.4382479 | -0.027907573 | 0.792977865 |
| 17.2147526 | 17.0191925 | 17.2066925 | 17.4189995 | 0.126621573 | 0.2482255 |
| 15.6149251 | 15.6361371 | 15.6813286 | 15.6251349 | -0.04103769 | 0.697114882 |
| 19.0332878 | 18.7275479 | 18.898822 | 19.0306246 | -0.006590266 | 0.950442365 |
| 18.7217016 | 18.5232096 | 18.5957849 | 18.6396539 | -0.088120697 | 0.332274576 |
| 18.6688407 | 18.6656316 | 18.7206734 | 18.5778369 | -0.010732877 | 0.895251114 |
| 22.1981259 | 22.1498389 | 22.1839758 | 22.2014159 | 0.011384779 | 0.864226469 |
| 24.1060609 | 23.9338719 | 24.0689826 | 24.1181988 | 0.023667502 | 0.72440215 |
| 21.8448881 | 21.6599117 | 21.7637583 | 21.8274662 | -0.021273414 | 0.737066728 |
| 23.3068077 | 23.2153769 | 23.2248214 | 23.240688 | 0.007873641 | 0.897970234 |
| 20.4331124 | 20.2166205 | 20.4335873 | 20.4402635 | -0.012214832 | 0.843002605 |
| 19.8732781 | 19.7469553 | 19.8706669 | 19.9388035 | 0.040354349 | 0.494316834 |
| 19.3186518 | 19.264051 | 19.3790127 | 19.2981742 | 0.016297138 | 0.775189041 |
| 22.7843074 | 22.5862502 | 22.7232015 | 22.7368935 | -0.01245451 | 0.826554185 |
| 21.0860814 | 20.9922927 | 21.0611298 | 21.0501438 | -0.040028248 | 0.470854421 |
| 24.3972148 | 24.2864225 | 24.4189016 | 24.4094012 | 0.024300349 | 0.65353631 |
| 20.6507041 | 20.6373444 | 20.7369425 | 20.6370526 | 0.014256913 | 0.78607448 |
| 22.119172 | 22.0488326 | 22.1573886 | 22.0609148 | -0.009481054 | 0.859547109 |
| 23.1862122 | 23.0805304 | 23.2190219 | 23.168797 | -0.003584068 | 0.94465159 |
| 22.3860516 | 22.3326369 | 22.334592 | 22.403938 | 0.004711792 | 0.927345367 |
| 20.6551284 | 20.4032014 | 20.5384844 | 20.5170432 | -0.11811001 | 0.019908619 |
| 20.1239568 | 20.064453 | 20.0144991 | 20.0553808 | -0.056905741 | 0.251621795 |
| 21.6895899 | 21.5988892 | 21.6879732 | 21.6774936 | -0.027278878 | 0.548769103 |
| 21.5202228 | 21.5111437 | 21.4819687 | 21.5303129 | 0.057148769 | 0.223858878 |
| 22.345303 | 22.2373723 | 22.346745 | 22.3429935 | -0.020909249 | 0.646317613 |
| 25.1720617 | 25.1969077 | 25.1896707 | 25.2382574 | 0.071383156 | 0.124500609 |
| 24.6467372 | 24.6413754 | 24.7136991 | 24.6004808 | 0.041373465 | 0.363076077 |
| 21.9962061 | 22.0074199 | 22.053239 | 22.0517349 | 0.052637066 | 0.187578217 |
| 20.8639088 | 20.7760864 | 20.8712853 | 20.9097283 | 0.005901054 | 0.877079142 |
| 23.8433459 | 23.7321514 | 23.8021702 | 23.8057947 | -0.004715119 | 0.910308478 |
| 21.5537318 | 21.4909486 | 21.5490169 | 21.576697 | 0.018285861 | 0.68210272 |
| 23.4197856 | 23.4227941 | 23.4261039 | 23.4595959 | 0.061102509 | 0.20081855 |
| 22.465336 | 22.5492133 | 22.4767346 | 22.4616289 | 0.018774748 | 0.695918713 |
| 19.8130168 | 19.8080041 | 19.8097784 | 19.807601 | 0.018075818 | 0.72588823 |
| 21.4948124 | 21.3695034 | 21.5221312 | 21.540828 | 0.025114603 | 0.719437204 |
| 21.8840375 | 21.8075989 | 21.7833789 | 21.8521079 | 0.053173271 | 0.465589447 |
| 21.8739838 | 21.8469478 | 21.8735127 | 21.9847092 | 0.034644313 | 0.640727269 |
| 17.4541828 | NA | 17.399261 | 17.4499835 | -0.059218487 | 0.430410896 |
| 18.0612245 | 17.9110479 | 18.0197025 | 18.0773276 | -0.026162873 | 0.744940474 |
| 19.0513335 | 19.1484434 | 19.0968551 | 19.1845614 | 0.060403128 | 0.448676191 |
| 24.9004333 | 24.6336074 | 24.8199051 | 24.73403 | -0.06119278 | 0.530175768 |
| 20.9506693 | 20.7661067 | 20.8739884 | 20.8871576 | -0.044029321 | 0.662727759 |

| | | | | | |
|---|---|---|---|---|---|
| 16.6154569 | 16.7657885 | 16.7987447 | 16.7682865 | 0.052446774 | 0.683561058 |
| 17.6532721 | 17.3987372 | 17.9189558 | 17.8177947 | 0.014589352 | 0.933337073 |
| NA | 16.1012266 | NA | NA | 0.676774837 | 0.108864917 |
| 18.7104141 | 18.6083339 | 18.8175403 | 18.8631813 | 0.10089291 | 0.277883838 |
| 14.8988416 | 14.7462554 | 14.3930004 | 14.9163151 | -0.130073325 | 0.454420866 |
| 18.1473906 | 18.3249122 | 18.2017261 | 18.1420613 | 0.048174172 | 0.58208494 |
| 20.2700432 | 19.9590034 | 20.1405248 | 20.1372529 | -0.131007989 | 0.10827365 |
| NA | 13.5875916 | NA | NA | 0.063464079 | 0.858221256 |
| 20.4747129 | 20.549257 | 20.4970064 | 20.5823766 | 0.01875694 | 0.703409136 |
| 23.1859707 | 23.1498391 | 23.216157 | 23.124609 | -0.013655124 | 0.737323526 |
| 20.9294535 | 20.8463462 | 20.9495152 | 20.8659308 | 0.010907605 | 0.873498001 |
| 25.178317 | 25.1231462 | 25.1250241 | 25.1820318 | 0.022906913 | 0.72732628 |
| 20.5875228 | 20.4726936 | 20.4459838 | 20.5315388 | -0.039409173 | 0.517207281 |
| 23.1966473 | 22.9955963 | 23.1348601 | 23.0977486 | -0.046260764 | 0.425307047 |
| 24.8460623 | 24.8757205 | 24.9382303 | 24.8879275 | 0.063969017 | 0.266059831 |
| 19.9232109 | 19.8420801 | 19.9671064 | 19.9926936 | -0.004393436 | 0.937163175 |
| 24.2621522 | 24.1935711 | 24.2617781 | 24.2944299 | 0.051241248 | 0.342518908 |
| 21.3924479 | 21.2344603 | 21.2984651 | 21.2695562 | -0.021642457 | 0.677719234 |
| 18.4010362 | 18.4415419 | 18.381324 | 18.3707824 | -0.008261627 | 0.871484796 |
| 23.6725527 | 23.5830073 | 23.696694 | 23.6413246 | 0.009093668 | 0.852786382 |
| 21.2277392 | 21.1798508 | 21.1978788 | 21.2435661 | -0.012014189 | 0.769446741 |
| 24.0252314 | 23.9641534 | 24.04056 | 24.0708424 | 0.019835525 | 0.626218939 |
| 21.8402895 | 21.8539261 | 21.9183622 | 21.977132 | 0.061063977 | 0.166177024 |
| 20.9165671 | 20.7935789 | 20.8955737 | 20.8979576 | -0.014588278 | 0.81537448 |
| 21.2848221 | 21.1475453 | 21.1254274 | 21.2139117 | -0.06554795 | 0.339693905 |
| 20.2741189 | 20.1464565 | 20.359239 | 20.3558019 | -0.007158699 | 0.924435519 |
| 20.4370914 | 20.4702995 | 20.5200936 | 20.5422288 | 0.052768224 | 0.208462615 |
| 12.229987 | NA | 13.212799 | 12.8564796 | 0.364434687 | 0.515909492 |
| 15.98886 | 16.0497274 | 15.6948016 | 16.2857223 | 0.177582832 | 0.241662506 |
| NA | 17.6669363 | NA | NA | 0.020996727 | 0.938020524 |
| 18.4416011 | 17.7500622 | 18.3588511 | 18.234931 | -0.188542459 | 0.408251821 |
| 16.254868 | 16.0635745 | 16.0010482 | 16.0683542 | -0.030977926 | 0.783625136 |
| 18.7669817 | 18.7807614 | 18.7012289 | 18.9205985 | 7.49E-04 | 0.993856441 |
| 22.0098571 | 21.6758138 | 21.9843487 | 21.9448097 | -0.167347849 | 0.081480032 |
| 19.8828753 | 19.7029586 | 19.9317976 | 20.0283886 | -0.021698967 | 0.805714931 |
| 27.5469428 | 27.429904 | 27.4945243 | 27.4894653 | 0.017736735 | 0.81192957 |
| 20.2266815 | 20.117741 | 20.1943807 | 20.1862285 | -0.024782675 | 0.725662946 |
| 19.7508029 | 19.6424808 | 19.6615105 | 19.6302629 | -0.006363478 | 0.922132902 |
| 19.9193184 | 19.9560958 | 19.8795569 | 19.8627826 | 0.032210349 | 0.615116144 |
| 21.649232 | 21.405696 | 21.6115038 | 21.5642162 | -0.081810149 | 0.194312968 |
| 21.1572892 | 21.0356885 | 21.1337541 | 21.12525 | 0.01333507 | 0.824833084 |
| 22.9100185 | 22.8246919 | 22.9433889 | 22.8629667 | -0.015842155 | 0.788544844 |

| | | | | | |
|---|---|---|---|---|---|
| 20.122834 | 20.0262638 | 20.1644871 | 20.0690617 | 0.010291608 | 0.856972792 |
| 21.8448113 | 21.7990142 | 21.8419141 | 21.8620654 | 0.03724317 | 0.514872222 |
| 19.9156212 | 19.8636229 | 19.9469755 | 19.9831208 | 0.033274267 | 0.533605969 |
| 21.4199117 | 21.2041474 | 21.3904104 | 21.431935 | -0.01572231 | 0.760652861 |
| 20.1975902 | 20.1049075 | 20.298592 | 20.2501869 | 0.016844893 | 0.745948259 |
| 23.1443103 | 23.1872616 | 23.1866304 | 23.1791628 | 0.016341537 | 0.724438624 |
| 21.3989201 | 21.3582703 | 21.4423745 | 21.4352637 | 0.026994738 | 0.555591305 |
| 24.1356428 | 24.1030368 | 24.1951729 | 24.215727 | 0.047556857 | 0.293103713 |
| 21.7582867 | 21.8932049 | 21.8289394 | 21.814105 | 0.07411697 | 0.088332205 |
| 20.0408164 | 19.9875221 | 20.0749258 | 20.0555095 | 0.01638779 | 0.734945506 |
| 23.2623475 | 23.1380534 | 23.2747805 | 23.2680226 | 0.017100676 | 0.741277466 |
| 19.0601383 | 18.9925987 | 19.0599279 | 19.0853627 | -0.072077175 | 0.206315817 |
| 23.4218265 | 23.2498638 | 23.3050484 | 23.3257807 | -0.025884819 | 0.657738516 |
| 21.6297547 | 21.4906814 | 21.5808244 | 21.6249076 | -0.009937346 | 0.87656401 |
| 21.4784005 | 21.3323173 | 21.3065102 | 21.3859754 | -0.055641963 | 0.372420887 |
| 19.8568672 | 19.7954365 | 19.902851 | 19.9491761 | 0.052305208 | 0.423015116 |
| 18.1753021 | 18.2452081 | 18.0914731 | 18.1403523 | -0.035998188 | 0.614109861 |
| 20.5020619 | 20.282713 | 20.3863793 | 20.5275596 | -0.085164555 | 0.255691449 |
| 18.0202391 | 18.0706976 | 18.0125711 | 18.044757 | -0.036025714 | 0.629128878 |
| 21.00253 | 20.7689401 | 20.9277225 | 20.8248401 | -0.059196088 | 0.628189566 |
| 19.2272772 | 19.6122368 | 19.8536652 | 19.2427666 | 0.105010455 | 0.60031767 |
| 15.5662922 | 15.9292655 | 15.8536577 | 15.7197678 | 0.004792649 | 0.981521845 |
| 21.8528776 | 21.748359 | 21.8987945 | 21.771741 | 0.003371248 | 0.958932335 |
| 17.0941299 | 17.0241746 | 17.0571403 | 17.1558754 | -0.175885983 | 0.21011369 |
| 17.6114649 | NA | 17.6386581 | 17.5865591 | -0.029626203 | 0.73800435 |
| 21.9947337 | 21.8995638 | 22.0354702 | 21.9442764 | 0.034110944 | 0.626121744 |
| 23.6556179 | 23.4214432 | 23.6383347 | 23.5293839 | -0.074120455 | 0.273752453 |
| 20.2645699 | 20.0614777 | 20.1748121 | 20.1651872 | 0.005243284 | 0.942275785 |
| 15.8508054 | 15.6838998 | 15.8355059 | 15.9599288 | 0.170720167 | 0.305240107 |
| 17.5450995 | 17.9533244 | 17.8217787 | 17.922713 | 0.158542998 | 0.322555418 |
| 16.08672 | 15.9870586 | 15.8874218 | 15.8503235 | -0.088346247 | 0.547278184 |
| NA | 18.7602062 | NA | NA | -0.090420352 | 0.54009396 |
| 16.7862666 | 16.5859814 | 16.5000781 | 16.4392931 | -0.214371429 | 0.077044218 |
| 18.9401491 | 18.7446516 | 18.972669 | 19.1367478 | 0.079739763 | 0.390406719 |
| 25.2956397 | 25.0706276 | 25.1707449 | 25.2401393 | -0.030960612 | 0.729927261 |
| 21.974103 | 22.1079481 | 22.1859179 | 22.1561185 | 0.087919937 | 0.259229749 |
| 20.1997304 | 19.899487 | 20.0974846 | 20.1136784 | -0.066308911 | 0.373756587 |
| 20.8631068 | 21.0486569 | 21.0056078 | 20.9415331 | 0.040731068 | 0.562930556 |
| 21.614532 | 21.430866 | 21.5161374 | 21.370764 | -0.175781855 | 0.009940467 |
| 23.7312712 | 23.5930263 | 23.6390437 | 23.6668266 | -0.047175516 | 0.442260825 |
| 20.2180027 | 20.1541816 | 20.1994873 | 20.2014743 | 0.0559362 | 0.362591612 |
| 19.5612853 | 19.4798883 | 19.5974605 | 19.5915439 | 0.01363289 | 0.795672028 |

| | | | | | |
|---|---|---|---|---|---|
| 23.3691507 | 23.3498252 | 23.3597299 | 23.3616155 | 0.039049308 | 0.460508509 |
| 23.8111064 | 23.7053356 | 23.7910989 | 23.7763404 | -0.025984133 | 0.577411758 |
| 23.6079288 | 23.4669984 | 23.5919434 | 23.6077517 | -0.025142319 | 0.578729544 |
| 23.0843318 | 23.0468414 | 23.0938025 | 23.1118709 | 0.030429147 | 0.468853557 |
| 22.6261311 | 22.5487316 | 22.6033656 | 22.6476195 | 0.005472247 | 0.891188624 |
| 21.5654389 | 21.4630862 | 21.5886565 | 21.5782284 | 0.016802331 | 0.724617569 |
| 19.4770192 | 19.4640861 | 19.4916378 | 19.5357624 | 0.032087674 | 0.513521748 |
| 21.3127175 | 21.2399048 | 21.2869779 | 21.3110536 | -0.00713176 | 0.879707144 |
| 23.3848477 | 23.3321095 | 23.4404425 | 23.3759004 | 0.016652541 | 0.753972408 |
| 19.2032861 | 19.1094545 | 19.1796852 | 19.2630009 | -0.029207745 | 0.611350093 |
| 23.5790476 | 23.4651507 | 23.5657781 | 23.5403022 | 0.012168612 | 0.834780627 |
| 20.4532136 | 20.3416898 | 20.4731458 | 20.4554212 | -0.014818381 | 0.811249001 |
| 21.177379 | 21.0657489 | 21.1668418 | 21.1438573 | -0.020778446 | 0.739221487 |
| 20.0514995 | 19.994473 | 20.0884109 | 20.0482827 | 0.020355053 | 0.736297062 |
| 20.5504449 | 20.4603827 | 20.5850544 | 20.5642557 | 0.01822543 | 0.769992328 |
| 21.363375 | 21.2853805 | 21.3627836 | 21.3643436 | 0.036457005 | 0.558082269 |
| 25.7950944 | 25.677348 | 25.7621707 | 25.7808534 | 0.025632141 | 0.693344177 |
| 20.5383659 | 20.3725563 | 20.4601395 | 20.4379762 | -0.030113841 | 0.651420776 |
| 20.1614235 | 19.9231054 | 20.1339645 | 20.1960585 | -0.044072945 | 0.533886671 |
| 17.8403671 | 17.7920473 | 17.7915767 | 18.0279906 | 0.088232441 | 0.297261518 |
| 23.574713 | 23.3983992 | 23.5201885 | 23.6102968 | -0.023942823 | 0.783279531 |
| 18.9548713 | 18.6693847 | 18.7304107 | 18.7822926 | -0.084403463 | 0.350451769 |
| 18.0909228 | 17.8715576 | 18.1001682 | 18.1202848 | -0.064501725 | 0.489224844 |
| 20.7853016 | 20.5876717 | 20.8086017 | 20.6160684 | 0.037432522 | 0.705626912 |
| 15.7790186 | 16.0604574 | 15.9257136 | 15.7484738 | 0.045923719 | 0.641753961 |
| 17.0884728 | 17.2193489 | 17.1731092 | 17.0581219 | 0.078659503 | 0.529551666 |
| NA | 16.8932928 | NA | NA | -0.015958181 | 0.911311079 |
| 15.0747731 | 15.2583183 | 15.5701535 | 15.1776762 | -0.06254299 | 0.740063438 |
| 17.3270039 | 17.2997381 | 17.4621008 | 17.4890212 | 0.052560362 | 0.590061561 |
| 19.1404007 | 19.05465 | 19.1626058 | 18.8839405 | -0.04924538 | 0.455031302 |
| 22.2486558 | 22.1253118 | 22.260035 | 22.2556128 | 0.016103558 | 0.768986502 |
| 23.8366295 | 23.6539719 | 23.7649014 | 23.8052898 | -0.043036519 | 0.417128892 |
| 19.8809703 | 19.9131695 | 19.7976518 | 19.8961252 | -0.015357485 | 0.761406236 |
| 20.9521427 | 20.873297 | 20.9527116 | 20.9857924 | -0.040104785 | 0.396291617 |
| 25.3843032 | 25.3635445 | 25.4478376 | 25.4627321 | 0.035122867 | 0.390538331 |
| 22.2179367 | 22.2153621 | 22.2218176 | 22.1683547 | -0.071310085 | 0.260437116 |
| 19.5656984 | 19.4437559 | 19.3902363 | 19.5572962 | 0.044764795 | 0.634650049 |
| 17.4625551 | 17.2006402 | 17.3705167 | 17.4129544 | -0.064107583 | 0.525777662 |
| 18.175971 | 17.799121 | 18.0003012 | 18.0509726 | 0.05269464 | 0.691217507 |
| 19.4144715 | 19.0657408 | 19.4788747 | 19.527178 | 5.21E-04 | 0.996948626 |
| 15.0703111 | 15.0544922 | 15.4328791 | 15.5174841 | 0.250235091 | 0.138328798 |
| 12.3798253 | NA | 12.6441933 | 12.6555175 | 0.692520257 | 0.045091352 |

| | | | | | |
|---|---|---|---|---|---|
| 22.8753963 | 22.7605415 | 22.853627 | 22.9056243 | 0.005483897 | 0.907142573 |
| 18.6471451 | NA | 18.6544781 | 18.7432033 | 0.012846421 | 0.882049991 |
| 22.517975 | 22.4524099 | 22.5540628 | 22.5463288 | 0.044164328 | 0.456647585 |
| NA | 13.4084611 | NA | NA | -0.180763252 | 0.396237243 |
| 18.9621774 | 18.8303886 | 19.0700419 | 19.0174022 | 0.019186144 | 0.801770631 |
| 20.8598674 | 20.6457682 | 20.8980307 | 20.8153452 | 0.054584943 | 0.579213421 |
| 19.0237576 | 18.8643816 | 18.9267645 | 19.0516348 | -0.041122291 | 0.534246235 |
| NA | 14.9024849 | NA | NA | -0.051830377 | 0.838192096 |
| 21.9573259 | 21.9092691 | 21.9000947 | 21.9678481 | 0.010460119 | 0.847603243 |
| 16.5278812 | 16.3820933 | 16.6497313 | 16.6631338 | 0.06374274 | 0.499858875 |
| 21.7872498 | 21.7794644 | 21.7485712 | 21.773066 | -0.017815924 | 0.849741773 |
| 20.5178257 | 20.2138812 | 20.2735901 | 20.3956773 | -0.091618198 | 0.224151934 |
| 19.4135181 | 19.1066368 | 19.3613519 | 19.3430583 | -0.07221227 | 0.315720183 |
| 19.6117266 | 19.6084731 | 19.6265489 | 19.6520036 | 0.0378978 | 0.430938151 |
| 19.3678429 | 19.2567929 | 19.2623735 | 19.3550295 | -0.047264931 | 0.484835792 |
| 19.7067569 | 19.607582 | 19.7286262 | 19.6551513 | -0.029051017 | 0.610876659 |
| 16.3759375 | NA | 16.2449015 | 16.2134502 | -0.110559369 | 0.114630598 |
| 21.7429483 | 21.5927492 | 21.74641 | 21.6942335 | 0.005968694 | 0.929105704 |
| 19.2798818 | 19.1689113 | 19.2422551 | 19.428799 | 0.028247103 | 0.739525166 |
| 17.8138847 | 17.6562628 | 17.8780396 | 18.0241446 | 0.099568041 | 0.392241541 |
| 22.0690855 | 22.0145652 | 22.0163085 | 22.0004598 | 0.012167948 | 0.736580745 |
| 26.5956626 | 26.4475258 | 26.5709657 | 26.5606209 | -0.002433531 | 0.965104413 |
| NA | 19.0074081 | NA | NA | 0.203728736 | 0.346915312 |
| 23.1292276 | 22.9112977 | 23.1060342 | 23.0621528 | -0.033160117 | 0.6624764 |
| 21.1078464 | 21.0023683 | 21.1007684 | 21.1186636 | -0.034173914 | 0.573561651 |
| 28.2050812 | 28.1180973 | 28.1971887 | 28.1491149 | 0.009740931 | 0.860739481 |
| 21.8408749 | 21.7818392 | 21.8638052 | 21.8812163 | 0.0090546 | 0.83556585 |
| 21.1739041 | 21.2141439 | 21.2485947 | 21.2335615 | 0.100530614 | 0.099725576 |
| 21.5663905 | 21.4735589 | 21.5359407 | 21.6246699 | 0.01284432 | 0.85115575 |
| 15.6162841 | 15.44839 | 15.7950648 | 15.7656317 | -0.037511103 | 0.826937178 |
| 20.0036002 | 19.9582113 | 19.974342 | 20.0107544 | 0.005525833 | 0.910751683 |
| 19.8562227 | 19.6313368 | 19.8407389 | 19.8403981 | -0.011867937 | 0.880126857 |
| 17.0497286 | 17.166163 | 16.7767239 | 16.8531138 | 0.007704989 | 0.976862396 |
| 19.2534328 | 19.6839293 | 20.2957984 | 19.5742494 | 0.561529008 | 0.004924159 |
| 16.2712717 | NA | 15.6622218 | 16.0381607 | -0.244566803 | 0.172782789 |
| 16.8256616 | 16.9097009 | 17.0961724 | 16.8455387 | -0.012181029 | 0.885434826 |
| 19.7650855 | 19.7277661 | 19.7664417 | 19.904361 | 0.028909139 | 0.723650743 |
| 18.4746378 | 18.201681 | 18.5756107 | 18.4245918 | 0.017641455 | 0.827732196 |
| 18.1494709 | 18.1084043 | 18.0796757 | 18.4221934 | 0.06786013 | 0.367744909 |
| 20.6166919 | 20.3578809 | 20.3754525 | 20.4758675 | -0.054622893 | 0.431513908 |
| 25.7745131 | 25.6532087 | 25.7185798 | 25.7351856 | -0.012932222 | 0.806418426 |
| 23.0455641 | 22.9917648 | 23.0753421 | 23.0601392 | 0.042530444 | 0.386899618 |

| | | | | | |
|---|---|---|---|---|---|
| 21.5808562 | 21.4307649 | 21.5370465 | 21.548731 | -0.008588049 | 0.84968429 |
| 21.5631161 | 21.5447907 | 21.6077802 | 21.6205198 | 0.043860446 | 0.267662564 |
| 23.9439084 | 23.9248134 | 24.0044524 | 24.0225755 | 0.065015153 | 0.153880391 |
| 21.036572 | 20.9856867 | 20.9999175 | 21.0440708 | -0.012309375 | 0.816385288 |
| 18.5090576 | 18.613295 | 18.6016601 | 18.5487425 | 0.087292319 | 0.139400665 |
| 19.5127942 | 19.4465735 | 19.5748013 | 19.6537794 | 0.01142946 | 0.845254922 |
| 17.1671655 | 17.2631907 | 17.3580796 | 17.2558635 | 0.162784827 | 0.023633413 |
| 17.288352 | 17.5405381 | 17.3504456 | 17.2735339 | -0.019001178 | 0.824117205 |
| 18.282079 | 17.8392763 | 18.1627329 | 18.2108332 | -0.051204398 | 0.692205343 |
| 20.9535722 | 20.6848532 | 20.8006363 | 20.9794663 | -0.082230605 | 0.588343039 |
| 17.4733983 | 17.6294654 | 17.6427977 | 17.4196888 | 0.087995882 | 0.331803336 |
| 22.0100106 | 21.9212095 | 21.8903701 | 21.8285972 | -0.014558238 | 0.835007153 |
| 21.7640247 | 21.6886636 | 21.7973384 | 21.7964732 | 0.063648707 | 0.281624394 |
| 20.9391205 | 20.8479997 | 20.9430467 | 20.9922869 | 0.049237228 | 0.41748005 |
| 20.1723345 | 19.9799954 | 20.1575432 | 20.1997212 | -0.046893586 | 0.472224163 |
| 20.0063564 | 19.7549742 | 19.839621 | 19.9699142 | -0.043006274 | 0.526269045 |
| 23.2964493 | 23.1918343 | 23.2151146 | 23.3240918 | -0.064262446 | 0.34171968 |
| 19.5986871 | 19.5630192 | 19.4999364 | 19.5676897 | 0.025598774 | 0.666505298 |
| 21.6073281 | 21.5032303 | 21.5188751 | 21.5444456 | -0.035210698 | 0.505531869 |
| 20.578879 | 20.4866109 | 20.5057241 | 20.5680474 | -0.016538381 | 0.73988129 |
| 22.5991923 | 22.5058181 | 22.6079476 | 22.6351275 | 0.015126579 | 0.797413528 |
| 22.6598795 | 22.5343635 | 22.6543336 | 22.671831 | -0.004042585 | 0.946116833 |
| 22.6743596 | 22.6677176 | 22.7575734 | 22.6134408 | 0.048167451 | 0.559578964 |
| 22.0195101 | 22.0295243 | 22.0892752 | 22.102913 | 0.067926895 | 0.090785071 |
| 24.2805451 | 24.2045477 | 24.2880819 | 24.2751258 | 0.026210545 | 0.539582836 |
| 19.2047139 | 19.0719165 | 19.0229841 | 19.1705548 | -0.005280075 | 0.950536957 |
| 19.249158 | 18.9889573 | 19.3308436 | 19.1770328 | 0.049494503 | 0.545037417 |
| 21.269154 | 21.3318668 | 21.3653427 | 21.2813335 | 0.101823764 | 0.102008147 |
| 19.1126375 | 19.0559589 | 19.0617726 | 18.9952682 | -0.020424068 | 0.741726068 |
| 20.6475788 | 20.4360014 | 20.5831046 | 20.6221373 | -0.057663154 | 0.37666886 |
| 14.9663031 | 15.1053551 | 14.8465414 | 15.4821846 | -0.031276734 | 0.92622601 |
| 23.0448918 | 22.8824946 | 23.0790972 | 23.1915832 | 0.11212778 | 0.188447242 |
| 24.0927718 | 23.9988885 | 24.0461077 | 24.0516589 | -0.011360847 | 0.79855445 |
| 18.5839978 | 18.5655071 | 18.5097576 | 18.4908632 | -0.081243171 | 0.097591638 |
| 18.1808524 | 18.1091282 | 18.2205394 | 18.1921489 | 0.016487613 | 0.88551994 |
| 21.3907985 | 21.3130923 | 21.3526386 | 21.4027134 | -0.027614078 | 0.562844264 |
| 20.5156598 | 20.4214668 | 20.4346654 | 20.4961927 | 0.014946504 | 0.79812974 |
| 20.0834005 | 20.1407984 | 20.1195543 | 20.0405824 | 0.033792463 | 0.4816705 |
| 20.9895589 | 20.80268 | 20.8331264 | 21.0326269 | -0.00868297 | 0.89156138 |
| 22.9598939 | 22.958663 | 22.9297316 | 23.0175064 | 0.006117739 | 0.886431287 |
| 23.4602566 | 23.6036949 | 23.694227 | 23.4350935 | 0.051471285 | 0.614352059 |
| 21.6046459 | 21.5212463 | 21.5855531 | 21.6400088 | 0.032674463 | 0.538817149 |

| | | | | | |
|---|---|---|---|---|---|
| 19.8525099 | 19.5367349 | 19.679277 | 19.5841505 | -0.151292937 | 0.205786458 |
| 21.9447166 | 21.6297805 | 21.5934213 | 22.0289543 | 0.021808301 | 0.918903791 |
| 16.4768361 | 16.5025596 | 16.1611786 | 16.4978739 | -0.084076107 | 0.399846043 |
| 21.6448833 | 21.5600512 | 21.6482017 | 21.654236 | 0.01319046 | 0.761097023 |
| 24.1002149 | 23.9915863 | 24.1381355 | 24.1210645 | 0.027631825 | 0.587524956 |
| 22.101657 | 22.0086901 | 22.112893 | 22.1002566 | 0.010218506 | 0.847062683 |
| 18.6290184 | 18.5864032 | 18.6822196 | 18.65002 | 0.02927995 | 0.628915224 |
| 18.770187 | 18.6568579 | 18.7634911 | 18.7769639 | 0.013808732 | 0.821499324 |
| 19.4994292 | 19.3670088 | 19.4828042 | 19.5061606 | -0.035578305 | 0.595925348 |
| 18.9854667 | 19.0543737 | 19.039557 | 19.2210256 | 0.121285518 | 0.21591403 |
| 17.5854903 | 17.3296963 | 17.4011654 | 17.6610878 | -0.16701257 | 0.272774702 |
| 17.5804827 | 17.4089667 | 17.5317866 | 17.4575826 | 0.038104265 | 0.817060439 |
| 20.7407552 | 20.6687939 | 20.8191564 | 20.7465027 | 0.007282177 | 0.906747655 |
| 17.199759 | 17.2031787 | 16.7873677 | 17.7212047 | -0.026799519 | 0.883388433 |
| 17.5165029 | 17.123771 | 17.366207 | 17.3445279 | -0.058082542 | 0.717312948 |
| 24.5284112 | 24.3994471 | 24.4704441 | 24.4702337 | -0.058439758 | 0.190194622 |
| 24.1763651 | 24.1912039 | 24.2411326 | 24.2077891 | 0.005105406 | 0.876007862 |
| 19.2044341 | 19.1930612 | 19.1418698 | 19.1745986 | -0.024528694 | 0.722016755 |
| 21.634997 | 21.5370664 | 21.550162 | 21.7135364 | -0.015567266 | 0.808836044 |
| 23.6639501 | 23.6526751 | 23.7028876 | 23.6887241 | 0.040031592 | 0.457968749 |
| 19.0829483 | 19.1567951 | 19.2984278 | 19.0975787 | 0.063537942 | 0.388476941 |
| 25.7016621 | 25.6360265 | 25.7000167 | 25.6811108 | 0.01066758 | 0.829170142 |
| 21.7929539 | 21.6659097 | 21.7387912 | 21.7527256 | -0.038646057 | 0.493910048 |
| NA | 16.5035399 | NA | NA | 0.368843629 | 0.025705922 |
| 19.5779771 | 19.5421639 | 19.5842 | 19.648253 | -0.012822516 | 0.823156102 |
| 23.4330799 | 23.3984888 | 23.4384764 | 23.449871 | 0.013128746 | 0.783977975 |
| 23.3311395 | 23.2840369 | 23.3339052 | 23.3814667 | 0.031454975 | 0.526181172 |
| 23.0092901 | 22.9083693 | 23.0424723 | 23.0079758 | -0.012400854 | 0.891960648 |
| 21.1854246 | 21.3055489 | 21.2185655 | 21.2444348 | 0.051372356 | 0.205466065 |
| NA | 12.5713196 | NA | NA | -0.026670958 | 0.930686291 |
| 22.5001474 | 22.4570344 | 22.5736733 | 22.5782296 | 0.058567082 | 0.350899954 |
| 19.9920148 | 19.9453997 | 20.0186626 | 19.9757166 | 0.010853401 | 0.826330811 |
| 23.8988115 | 23.9097522 | 23.9198592 | 23.9147885 | 0.005962378 | 0.897575585 |
| 16.4023951 | 16.3797199 | 16.0438249 | 16.2085917 | -0.109755842 | 0.34796902 |
| 19.9044046 | 19.9809804 | 20.0003281 | 19.9572026 | 0.040611703 | 0.470294431 |
| 22.6308124 | 22.5768329 | 22.6471828 | 22.6177068 | 0.005187681 | 0.902159238 |
| 21.4339683 | 21.3416627 | 21.5613299 | 21.4608749 | 0.021165608 | 0.709965312 |
| 19.9363486 | 19.8002182 | 19.8627668 | 19.9348975 | -0.021103199 | 0.703507368 |
| 21.6835662 | 21.5302531 | 21.6167558 | 21.6682718 | -0.021461549 | 0.740592455 |
| 15.4780717 | 15.3213269 | 15.5923628 | NA | 0.617968127 | 0.029181562 |
| NA | 15.0024599 | NA | NA | 0.035708754 | 0.905035859 |
| 23.2651598 | 22.925785 | 23.1834479 | 23.1601554 | -0.12129244 | 0.264991386 |

| | | | | | |
|---|---|---|---|---|---|
| 21.9838501 | 21.9766407 | 21.8838361 | 22.0285431 | 0.045628875 | 0.504905659 |
| 23.7435902 | 23.7044165 | 23.6234724 | 23.7233718 | 0.007425083 | 0.878407458 |
| 21.3485106 | 21.1711513 | 21.3056858 | 21.3131265 | -0.062483343 | 0.206497027 |
| 19.8280719 | 19.8071898 | 19.8895995 | 19.9017007 | 0.037147394 | 0.507139801 |
| 17.1139462 | 16.9972506 | 17.2058205 | 17.0946284 | -0.10749339 | 0.333333204 |
| 16.0515987 | 15.4389877 | 15.7301145 | 15.4785186 | -0.154100632 | 0.425833735 |
| 20.0520362 | 20.0187896 | 19.9157513 | 20.1212837 | 0.116964825 | 0.252331949 |
| 18.2895022 | 18.2334953 | 18.2279418 | 18.0642337 | -0.023226902 | 0.783660999 |
| 20.2720489 | 20.3640175 | 20.2538383 | 20.3106711 | 0.026735382 | 0.685246984 |
| 26.4606715 | 26.4240972 | 26.4728551 | 26.4303797 | 0.01775915 | 0.708180497 |
| 21.0521675 | 20.9654102 | 20.9891674 | 20.9266893 | -0.044447709 | 0.299061073 |
| 22.0772837 | 21.9300328 | 21.9722688 | 21.9482865 | -0.055843155 | 0.232403056 |
| 20.8044012 | 20.7046003 | 20.8478697 | 20.8074843 | 0.004321986 | 0.930879174 |
| 17.7789565 | 17.7823147 | 17.8382066 | 17.9322123 | 0.058933823 | 0.301547593 |
| 19.4166827 | 19.2450651 | 19.3489704 | 19.415716 | -0.001400528 | 0.984932782 |
| 23.8242157 | 23.6700049 | 23.8410759 | 23.8334482 | 0.028264825 | 0.845672451 |
| 17.6431339 | 17.081735 | 17.5893792 | 17.8872726 | 0.16517183 | 0.542396717 |
| 21.4112386 | 21.6143707 | 21.550058 | 21.6049626 | 0.019038596 | 0.878342482 |
| 16.6794422 | 16.4052013 | 16.6824447 | 16.6132738 | -0.034244709 | 0.704071346 |
| 21.1962363 | 20.9447165 | 21.0335861 | 21.1629387 | -0.053065771 | 0.514965685 |
| 27.0755805 | 26.875359 | 27.0441613 | 27.0113025 | -0.018023982 | 0.823555913 |
| 20.9260199 | 20.8606351 | 21.0652465 | 20.9654294 | 9.91E-04 | 0.989543984 |
| 23.4644194 | 23.3929617 | 23.4636172 | 23.4908647 | 0.02948925 | 0.634966446 |
| 20.9990706 | 20.9973205 | 20.8853064 | 20.9969099 | 0.018184373 | 0.766619455 |
| 20.7763489 | 20.7552677 | 20.784622 | 20.7676807 | 0.010223486 | 0.849192158 |
| 19.7323848 | 19.571686 | 19.6615138 | 19.7727957 | 0.05300818 | 0.414189802 |
| 18.9298149 | 18.7374502 | 18.7928212 | 18.7185476 | -0.068464301 | 0.312835343 |
| 23.7799469 | 23.9612879 | 24.0596289 | 24.0398373 | 0.16730156 | 0.083074748 |
| 20.2662136 | 20.1841974 | 20.2364232 | 20.286641 | -0.013463414 | 0.792517058 |
| 22.1135977 | 22.0890555 | 22.0227334 | 22.0431178 | -0.038830359 | 0.475050176 |
| 11.5592887 | NA | 13.0118325 | 12.1457741 | 1.386608764 | 9.06E-04 |
| 19.0484051 | 18.8004873 | 19.0085181 | 18.9916545 | -0.06665091 | 0.416666907 |
| 19.6281009 | 19.5852549 | 19.6425674 | 19.5913047 | -0.03286977 | 0.4784756 |
| 20.3361931 | 20.1740336 | 20.299028 | 20.2832958 | -0.063126462 | 0.298767149 |
| 27.8746312 | 27.8689981 | 27.8972349 | 27.9443848 | 0.093966394 | 0.140594205 |
| 19.1418551 | 19.0127899 | 18.920435 | 18.9429084 | -0.139310042 | 0.100641932 |
| 16.3906674 | 16.4996811 | 16.120375 | 16.5783139 | -0.005349578 | 0.955642624 |
| 17.4338248 | 17.2781225 | 17.2345006 | 17.3521388 | 0.039422116 | 0.763187173 |
| 15.5431761 | 15.3044346 | 15.5106526 | 15.623767 | -0.205870499 | 0.14858178 |
| 18.5613211 | 18.5601574 | 18.5220187 | 18.5573483 | 0.005372281 | 0.920893151 |
| 19.3628642 | 19.1760147 | 19.3164425 | 19.334482 | -0.068309135 | 0.123018117 |
| 20.8011521 | 20.770056 | 20.8304024 | 20.7394102 | 0.045501841 | 0.442673597 |

| | | | | | |
|---|---|---|---|---|---|
| 21.9339067 | 21.8772065 | 21.9644646 | 22.0217177 | 0.042372384 | 0.599549094 |
| 20.4862043 | 20.2018165 | 20.4157632 | 20.3897662 | -0.201222556 | 0.089774158 |
| 14.2277928 | NA | 13.3499642 | 14.6098091 | 0.107253089 | 0.821302466 |
| 21.0500299 | 21.0257248 | 21.1685736 | 21.0945034 | 0.073170617 | 0.280446606 |
| 20.1547903 | 20.1101811 | 20.2137776 | 20.1494223 | 0.028134887 | 0.570762289 |
| 18.8173756 | 18.6601308 | 18.7820388 | 18.8111441 | 0.005608828 | 0.947988685 |
| 21.1258825 | 21.0805143 | 20.9801771 | 21.1347045 | -0.026665468 | 0.75830338 |
| 13.2862808 | NA | 12.8134392 | 13.2273964 | -0.173499659 | 0.481109144 |
| 16.19461 | 15.7715332 | 16.3058866 | 16.4071605 | -0.159279052 | 0.424464098 |
| 17.9151034 | 18.0512869 | 18.1211575 | 18.1446923 | 0.107376012 | 0.345821584 |
| 21.0211043 | 21.0787046 | 21.2111325 | 21.1565627 | 0.058065198 | 0.509173204 |
| 22.0804384 | 21.9836712 | 22.0597291 | 22.0650608 | -0.013461656 | 0.869033155 |
| 20.3170155 | 20.5084297 | 20.4904394 | 20.4484064 | 0.057390657 | 0.357583786 |
| 23.1311439 | 23.1812684 | 23.2685902 | 23.1668134 | 0.10678508 | 0.107690779 |
| 19.5592093 | 19.4682414 | 19.5148597 | 19.5051763 | -0.057255958 | 0.347949721 |
| 24.0902471 | 23.94429 | 24.0270207 | 24.0330334 | -4.14E-04 | 0.994276518 |
| 21.0530165 | 21.01151 | 21.1776055 | 21.1386211 | 0.106724023 | 0.066617547 |
| 21.2530244 | 21.2320719 | 21.3069772 | 21.2981215 | 0.022316678 | 0.685238529 |
| 22.6675916 | 22.6368022 | 22.6203504 | 22.7257511 | -0.007306142 | 0.886012369 |
| 21.2453899 | 21.122017 | 21.243482 | 21.2421705 | 0.006571172 | 0.899872909 |
| 23.1065119 | 23.1445873 | 23.1807676 | 23.0601345 | 0.034823741 | 0.471366697 |
| 20.228603 | 20.2806381 | 20.3068819 | 20.1675178 | 0.025285686 | 0.607096535 |
| 22.5930467 | 22.5750199 | 22.6892414 | 22.6793597 | 0.064353865 | 0.17030415 |
| 20.7966718 | 20.7365023 | 20.794626 | 20.7749323 | 0.005686198 | 0.880209489 |
| 22.2764134 | 22.1791063 | 22.2794243 | 22.2723684 | 0.009023909 | 0.83105353 |
| 21.2732222 | 21.1940097 | 21.3377693 | 21.3393435 | 0.05226091 | 0.325763054 |
| 22.415145 | 22.2805774 | 22.4078752 | 22.3428338 | -0.041996668 | 0.42649644 |
| 20.729763 | 20.7171939 | 20.6415092 | 20.8065747 | 0.024877985 | 0.667805769 |
| 18.3259445 | 18.4802004 | 18.5803705 | 18.5815896 | 0.108130313 | 0.064156961 |
| 20.1101005 | 19.9835755 | 20.0761425 | 20.1357364 | -0.049719446 | 0.404409951 |
| 21.2221374 | 21.2300721 | 21.1546781 | 21.2024856 | -0.016403154 | 0.791109508 |
| 21.9066684 | 21.9287339 | 21.9868669 | 21.9785616 | 0.037658025 | 0.556248416 |
| 22.2840084 | 22.1539108 | 22.2800561 | 22.3576275 | 0.059845917 | 0.349866989 |
| 23.0062442 | 22.9947248 | 23.0241685 | 23.062657 | 0.076843354 | 0.243543592 |
| 20.0665048 | 20.0079637 | 20.0341592 | 20.2069762 | 0.034674811 | 0.625355287 |
| 19.0225374 | 19.0587192 | 18.9893761 | 19.144265 | 0.014309821 | 0.832814154 |
| 23.4322666 | 23.2393129 | 23.4180759 | 23.3985164 | -0.043596156 | 0.531658051 |
| 18.9897219 | 18.8389999 | 18.8870427 | 18.7963563 | 0.024422474 | 0.741053473 |
| 18.6284844 | 18.5429631 | 18.7801098 | 18.7807255 | 0.106784768 | 0.165980887 |
| 20.5134216 | 20.2171846 | 20.4046773 | 20.402619 | -0.115163057 | 0.16667421 |
| 18.6787049 | 18.5196565 | 18.6245774 | 18.6653316 | 8.47E-04 | 0.992142951 |
| 16.9203747 | 16.7303103 | 16.7973272 | 16.8113411 | -0.076956118 | 0.373171603 |

| | | | | | |
|---|---|---|---|---|---|
| 16.4698968 | 16.2603423 | 16.3582695 | 16.4015659 | -0.029304487 | 0.810513268 |
| 21.232687 | 20.8245383 | 21.0974751 | 21.1701189 | -0.041467211 | 0.773474034 |
| 12.9361213 | 13.3672019 | 12.5943897 | 12.7815718 | -0.092811293 | 0.579327579 |
| 21.2858006 | 20.9674829 | 21.2859008 | 21.2099653 | -0.038041944 | 0.847307673 |
| 18.8929837 | 18.957393 | 19.1167249 | 18.8901059 | 0.137111511 | 0.246702609 |
| 16.7225397 | 16.7927379 | 16.6800697 | 16.7543539 | 0.081072066 | 0.477293606 |
| 18.2565817 | 18.110162 | 18.3039038 | 18.2778352 | -0.075173691 | 0.296110564 |
| 18.8425205 | 18.8346209 | 18.8559146 | 18.8652154 | 0.007376077 | 0.902035271 |
| 20.1436755 | 20.0563269 | 20.0792349 | 20.2169599 | -0.026231411 | 0.655908618 |
| 22.5858019 | 22.6294974 | 22.5789319 | 22.5720695 | 0.055725054 | 0.234537383 |
| 21.7400106 | 21.7125707 | 21.7683548 | 21.7266949 | 0.009243728 | 0.83990441 |
| 17.2135164 | 17.0741595 | 16.9624887 | 17.304382 | -0.069575605 | 0.365828993 |
| 19.319129 | 19.166216 | 19.3078518 | 19.3451119 | 0.027978987 | 0.774253341 |
| 17.065199 | 17.0372549 | 17.1540702 | 17.1588021 | -0.063058563 | 0.683257048 |
| 19.1906894 | 19.057526 | 19.2296578 | 19.3011558 | 0.084529302 | 0.270873563 |
| 17.7451707 | 17.6710347 | 17.5057064 | 17.7384876 | 0.029559368 | 0.670316757 |
| 18.7344491 | 18.6816548 | 18.7576803 | 18.7553795 | -0.016003996 | 0.748515928 |
| 20.2676112 | 20.2116899 | 20.2213358 | 20.3451481 | -0.05403803 | 0.28431351 |
| 24.048519 | 23.9798472 | 24.0752793 | 24.0752069 | 0.022686336 | 0.593548325 |
| 20.5946021 | 20.5215281 | 20.5512376 | 20.5563909 | 0.027050414 | 0.568382111 |
| 27.1190648 | 27.0288045 | 27.1513817 | 27.1008628 | 0.037192087 | 0.567105327 |
| 16.815524 | 16.7708131 | 16.4242403 | 16.9409746 | 0.04158182 | 0.821254374 |
| 22.3830941 | 22.2874739 | 22.378159 | 22.4002212 | 0.007515525 | 0.90569868 |
| 20.6954874 | 20.6031341 | 20.6408612 | 20.609799 | -0.080097502 | 0.06875124 |
| 19.848016 | 19.7404663 | 19.7675424 | 19.8105981 | -0.056015198 | 0.216702478 |
| 19.7626361 | 19.7115338 | 19.8800894 | 19.9058166 | 0.005732517 | 0.933708968 |
| 20.9302809 | 20.6944554 | 20.8130557 | 20.7985464 | -0.061884241 | 0.529600183 |
| 21.6266265 | 21.5734903 | 21.6271604 | 21.6387775 | 0.01046737 | 0.800755647 |
| 21.1649914 | 21.1001292 | 21.1233235 | 21.1490865 | 0.011642011 | 0.824640195 |
| 21.2672635 | 21.0992949 | 21.1877335 | 21.2622468 | -0.006405868 | 0.91229491 |
| 22.4290492 | 22.2864416 | 22.44656 | 22.418049 | -0.007848993 | 0.876162048 |
| NA | 15.62571 | NA | NA | -0.096513055 | 0.316040819 |
| 24.1796085 | 24.0662665 | 24.0905049 | 24.059612 | -0.027395835 | 0.624057306 |
| 20.1790373 | 20.0742856 | 20.2009187 | 20.1839556 | -0.023368086 | 0.624891811 |
| 20.7601099 | 21.1452856 | 20.8523833 | 20.785936 | 0.110534925 | 0.230292588 |
| 17.8011968 | 17.8834342 | 17.8820107 | 17.9980571 | 0.088701581 | 0.33502253 |
| 22.1494418 | 22.1890341 | 22.2644739 | 22.3758501 | 0.077916232 | 0.224533944 |
| 22.0795565 | 21.9357586 | 22.0381316 | 22.0127035 | -0.028690136 | 0.495211572 |
| 21.1824191 | 21.0943153 | 21.1892638 | 21.1337774 | -0.048152907 | 0.289152726 |
| 20.8842073 | 20.8716636 | 20.8821182 | 20.882787 | 0.021497392 | 0.651082589 |
| 15.3820921 | 15.6798362 | 15.6104045 | 15.4568856 | 0.05784852 | 0.583526933 |
| 15.3354256 | 15.0145552 | 15.5470125 | 15.3890507 | -0.074824768 | 0.540888367 |

| | | | | | |
|---|---|---|---|---|---|
| 15.5711152 | 15.1288009 | 15.4796356 | 15.7028094 | -0.026948435 | 0.849292619 |
| 18.5441276 | 18.6823734 | 18.6292448 | 18.657246 | 0.121348495 | 0.123928922 |
| 21.9501835 | 21.8521021 | 21.9486219 | 21.9725594 | 0.033467011 | 0.583067111 |
| 19.6787891 | 19.5630842 | 19.6563827 | 19.616274 | -0.04377236 | 0.387016603 |
| NA | 13.5379314 | NA | NA | -0.343148964 | 0.144838606 |
| NA | 16.4715127 | NA | NA | -0.016246426 | 0.879358659 |
| 19.2053344 | 19.2201452 | 19.3827935 | 19.2827131 | 0.023006797 | 0.799844945 |
| 19.6419622 | 19.5220415 | 19.4397577 | 19.5191583 | -0.074933343 | 0.393635191 |
| 21.9594861 | 21.7831255 | 22.0229079 | 21.9652536 | 0.021060412 | 0.781937372 |
| 19.7710769 | 19.6914046 | 19.7596991 | 19.7372742 | -0.011720382 | 0.830511273 |
| 23.087969 | 22.9873771 | 23.0965711 | 23.0269956 | -0.028779591 | 0.576195584 |
| 21.8139965 | 21.7155209 | 21.7626447 | 21.7755052 | -0.001744383 | 0.970655821 |
| 23.2336761 | 23.1647487 | 23.2521156 | 23.2845721 | 0.025614605 | 0.577659243 |
| 20.7682652 | 20.6888631 | 20.7478786 | 20.7758166 | 0.009509188 | 0.829813334 |
| 18.5236315 | 18.443738 | 18.4639339 | 18.5297108 | -0.008512949 | 0.863672722 |
| 21.8556749 | 21.6999022 | 21.8089227 | 21.7207179 | -0.029414471 | 0.658450851 |
| 16.6534574 | 16.7139345 | 16.9158099 | 17.1595607 | 0.019140428 | 0.895571017 |
| 20.8965352 | 21.195173 | 20.9293979 | 20.7601169 | 0.11745573 | 0.495026851 |
| 17.234831 | 17.2760794 | 17.3628778 | 17.1146295 | -0.04216761 | 0.634308602 |
| 27.2841167 | 27.1747556 | 27.2611973 | 27.3282297 | 0.04117143 | 0.347897437 |
| 21.5435002 | 21.5330469 | 21.5672331 | 21.6313539 | 0.04046017 | 0.342933158 |
| 23.1443236 | 23.0195843 | 23.1050755 | 22.993778 | -0.049051628 | 0.510220693 |
| 22.1700411 | 22.1083929 | 22.2055703 | 22.1885522 | 0.031157409 | 0.55042685 |
| 16.6887128 | 16.6832762 | 16.6791667 | 16.886512 | 0.129743483 | 0.272705796 |
| 20.1633934 | 20.1365379 | 20.2880478 | 20.0981871 | -0.029101078 | 0.685440535 |
| 22.2495702 | 22.1085498 | 22.2486735 | 22.1868588 | -0.024828108 | 0.61918112 |
| 20.7260676 | 20.5784595 | 20.7258972 | 20.7490245 | -0.004887184 | 0.946651807 |
| 18.8443436 | 18.6818942 | 18.6557786 | 18.70473 | -0.039823401 | 0.633831056 |
| 20.5518494 | 20.1935173 | 20.4691699 | 20.3770836 | -0.082093891 | 0.339534449 |
| 16.8417247 | 16.5223554 | 16.9015877 | 16.8359398 | -0.063672981 | 0.632247211 |
| 17.3807842 | 17.3559028 | 17.3839301 | 17.7260424 | 1.17E-04 | 0.999369935 |
| 18.4973747 | 18.4021379 | 18.2849236 | 18.5318255 | -0.017795199 | 0.806764244 |
| 20.9666769 | 20.8928716 | 20.9887179 | 20.9505559 | -0.010121427 | 0.844976138 |
| 20.976745 | 20.8505406 | 20.989082 | 21.0010622 | 0.008669051 | 0.867102873 |
| 21.0537884 | 20.8692225 | 20.9707529 | 21.0447404 | -0.033093695 | 0.56466 |
| 20.5496334 | 20.5200045 | 20.4543368 | 20.5668812 | 0.053626268 | 0.357836078 |
| 21.5725043 | 21.5233913 | 21.6331484 | 21.5765578 | -0.028366783 | 0.455923772 |
| 22.0093782 | 22.0870807 | 22.1466021 | 22.1212942 | 0.053821919 | 0.207775142 |
| 20.457989 | 20.3861982 | 20.3239827 | 20.386591 | -0.099943192 | 0.117130757 |
| 13.7136661 | NA | 13.58836 | 13.8847053 | -0.11675105 | 0.513062655 |
| 21.4847053 | 21.4836191 | 21.5195977 | 21.5544504 | 0.039934089 | 0.445687589 |
| 19.9268238 | 19.7826818 | 20.0342739 | 19.9837128 | 0.009825379 | 0.904722249 |

| | | | | | |
|---|---|---|---|---|---|
| 14.4828045 | NA | 14.6623761 | 14.676062 | 0.19166606 | 0.271974929 |
| 22.2526046 | 21.9054645 | 21.9155099 | 22.0590013 | -0.06872931 | 0.687940865 |
| 25.9149486 | 25.873345 | 25.9125729 | 25.9410478 | 0.031869626 | 0.46394006 |
| 20.8330453 | 20.8243595 | 20.9277856 | 20.901198 | 0.047550448 | 0.580132067 |
| 20.1250024 | 20.0593063 | 20.0306016 | 20.1146584 | -0.017673322 | 0.851495187 |
| 19.3839838 | 19.3594407 | 19.3736904 | 19.386218 | 0.043971782 | 0.466971206 |
| 19.9925651 | 19.8825539 | 19.9677484 | 20.0101348 | -0.064893399 | 0.286412951 |
| 20.0245967 | 20.0282379 | 20.1407421 | 20.1582743 | 0.027613156 | 0.663375069 |
| 24.0157501 | 23.8997965 | 23.9764741 | 24.081595 | 0.056135665 | 0.386987397 |
| 20.4357682 | 20.4494006 | 20.5572309 | 20.6493931 | 0.142714266 | 0.253171896 |
| 19.8894088 | 19.9153194 | 19.8672459 | 19.8991399 | 0.028511326 | 0.593448813 |
| 23.9071212 | 23.8220327 | 23.9949195 | 23.9502812 | -4.07E-04 | 0.99253213 |
| 21.527666 | 21.4713746 | 21.4556121 | 21.4767606 | -0.045318985 | 0.414424594 |
| 22.3660006 | 22.361208 | 22.401046 | 22.3224636 | -0.008962464 | 0.843963765 |
| 19.4801608 | 19.3825983 | 19.5438601 | 19.5146155 | 0.001513438 | 0.982818592 |
| 17.3214719 | 17.4311361 | 17.1539664 | 17.3138225 | -0.042741526 | 0.696783351 |
| 19.303921 | 18.8831482 | 19.4629315 | 19.3136968 | 0.029386331 | 0.873657331 |
| 17.8680772 | 17.798296 | 17.7985849 | 17.7740837 | -0.064389358 | 0.380701832 |
| NA | 16.8170495 | NA | NA | -0.005575981 | 0.977264362 |
| 20.6566265 | 20.4736869 | 20.6044255 | 20.618744 | -0.020389043 | 0.777945605 |
| 20.084036 | 19.9423258 | 20.0192742 | 20.0060697 | -0.087029101 | 0.049412047 |
| 21.6026821 | 21.5738142 | 21.6195993 | 21.5700531 | -0.01481752 | 0.691757388 |
| 20.3512521 | 20.256362 | 20.3771053 | 20.3770093 | 0.035474307 | 0.472654455 |
| 21.1531517 | 21.0565433 | 21.1984198 | 21.1428865 | 0.006240862 | 0.9300245 |
| 15.4626689 | 15.9433359 | 15.2543215 | 15.9027646 | -0.018481673 | 0.93218895 |
| 17.291506 | 17.0052247 | 17.0222815 | 17.112759 | -0.134541824 | 0.413246961 |
| 20.2905667 | 20.0792399 | 20.2882016 | 20.2303073 | -0.05959605 | 0.546805246 |
| 18.3749412 | 18.1569038 | 18.2003373 | 18.3256191 | -0.081352466 | 0.328500802 |
| 20.7977194 | 20.6510332 | 20.6877959 | 20.6460951 | -0.08283474 | 0.254628494 |
| 19.4724618 | 19.2483046 | 19.563326 | 19.4705064 | -0.026628561 | 0.68873862 |
| 21.8690333 | 21.8676291 | 21.9450439 | 21.881577 | 0.046547435 | 0.423459523 |
| 21.9345477 | 21.8339057 | 21.7740308 | 21.8188177 | -0.038318821 | 0.498559588 |
| 26.1054104 | 26.0609042 | 26.1569516 | 26.1382614 | 0.062039903 | 0.217045336 |
| 25.1515979 | 25.070333 | 25.1260658 | 25.1766487 | 0.015636358 | 0.736095185 |
| 20.8903964 | 20.8421297 | 20.8498649 | 20.8649175 | 0.020229568 | 0.648398078 |
| 20.7890277 | 20.6931546 | 20.7692442 | 20.7531593 | -0.030795065 | 0.463153622 |
| 21.218263 | 21.1072921 | 21.2286436 | 21.1929091 | -0.037978281 | 0.369708664 |
| 20.7640084 | 20.6915322 | 20.7905692 | 20.712785 | 0.0142276 | 0.755747927 |
| 26.4721051 | 26.4049777 | 26.4712719 | 26.487191 | 0.039525222 | 0.389319766 |
| 25.8545214 | 25.6928981 | 25.7978639 | 25.7831284 | -0.032080626 | 0.498035256 |
| 25.5927495 | 25.497647 | 25.5967997 | 25.5528067 | 0.00115442 | 0.981640889 |
| 23.0774238 | 23.0431709 | 23.1281984 | 23.1289644 | 0.051239622 | 0.309371812 |

| | | | | | |
|---|---|---|---|---|---|
| 25.5995349 | 25.5394452 | 25.6341804 | 25.6424058 | 0.052372314 | 0.314188266 |
| 25.4007769 | 25.3069487 | 25.4022978 | 25.3959772 | 0.031333556 | 0.576966298 |
| 21.2561003 | 21.1137855 | 21.2188061 | 21.2491563 | 0.033799714 | 0.573605916 |
| 18.9507099 | 18.8869252 | 18.8674197 | 18.945218 | -0.00152157 | 0.984747768 |
| 16.1847297 | 16.240777 | 16.0920858 | 16.204023 | -0.019651915 | 0.813601395 |
| 16.4572145 | 16.4098935 | 16.4142791 | 16.3552374 | -0.13305541 | 0.111652714 |
| 18.2813183 | 18.2426951 | 18.4614477 | 18.6336912 | 0.083682061 | 0.454048511 |
| 20.7563349 | 21.0498974 | 20.8070961 | 20.7626831 | 0.087768934 | 0.377890347 |
| 20.5764095 | 20.4503868 | 20.4820396 | 20.4958395 | -0.042250633 | 0.434053609 |
| 20.8544919 | 20.7593259 | 20.7969708 | 20.8991058 | -0.03257401 | 0.51929212 |
| 23.2510282 | 23.1592184 | 23.2032858 | 23.2102687 | 0.019532033 | 0.653353431 |
| 20.8028859 | 20.6906401 | 20.7727401 | 20.8614233 | 0.020317724 | 0.771648923 |
| 17.6059656 | 17.6284188 | 17.7374799 | 17.6674543 | -0.011593148 | 0.890779569 |
| 19.6910835 | 19.5199218 | 19.6875476 | 19.7351897 | -0.026311667 | 0.787723644 |
| 20.0967641 | 19.9120086 | 20.0099286 | 20.13144 | -0.044924132 | 0.493604012 |
| 22.3329296 | 22.2507286 | 22.3527491 | 22.3224649 | -0.005608151 | 0.906957158 |
| 23.7907129 | 23.8252652 | 23.8242884 | 23.8182453 | 0.066995625 | 0.164693215 |
| 21.181102 | 21.0480312 | 21.1928521 | 21.163806 | -4.22E-04 | 0.99466282 |
| 18.3451667 | 18.2735281 | 18.5999293 | 18.4990777 | 0.021123726 | 0.795573097 |
| 19.9621631 | 19.893312 | 19.776929 | 20.0303887 | -0.020970181 | 0.80347816 |
| 21.6624995 | 21.5252942 | 21.7393332 | 21.7027282 | 0.066758047 | 0.27052906 |
| 15.265381 | 14.9095263 | 15.1171142 | 15.1967294 | 0.009481048 | 0.961356278 |
| 21.5381277 | 21.4548069 | 21.5254495 | 21.5177589 | 0.035606139 | 0.557597106 |
| 17.3936792 | 17.3610495 | 17.2556756 | 17.4403708 | -0.05337662 | 0.553156326 |
| 18.9220094 | 19.1184303 | 19.2976647 | 19.2892558 | 0.058585931 | 0.610196256 |
| 20.9606147 | 20.9236272 | 20.9128014 | 20.9218545 | -0.025034296 | 0.667135482 |
| 19.1194755 | 19.2989066 | 18.9565657 | 19.063376 | -0.021226509 | 0.876612738 |
| 19.1062533 | 18.9047412 | 19.1293635 | 19.1583902 | -0.05870702 | 0.51733922 |
| 21.5173634 | 21.39093 | 21.454798 | 21.4015416 | -0.040286192 | 0.364299678 |
| 24.6517449 | 24.665727 | 24.623243 | 24.6569187 | 0.023441854 | 0.57656591 |
| 23.1008878 | 23.0504552 | 23.1437501 | 23.2051347 | 0.06408099 | 0.184855876 |
| 18.8798279 | 18.7825579 | 18.7744043 | 18.8339394 | -0.04106395 | 0.546441564 |
| 19.3630512 | 19.6320755 | 19.5256552 | 19.3715464 | 0.065395809 | 0.457055517 |
| 18.2792103 | 18.1327084 | 18.3257258 | 18.1002028 | -0.039892955 | 0.703028386 |
| 16.2959061 | 15.6735016 | 16.4607162 | 16.4356114 | -5.51E-04 | 0.998532676 |
| 16.5523643 | 15.9493879 | 15.5047094 | 16.3837827 | -0.384273334 | 0.110244421 |
| 25.0117638 | 24.5815673 | 24.9860479 | 24.9510739 | -0.019523951 | 0.90620012 |
| 16.4015237 | 16.3653679 | 16.4708427 | 16.546763 | -0.098334422 | 0.537356493 |
| 15.461809 | 15.5803737 | 15.6907312 | 15.4143407 | -0.113297961 | 0.551315765 |
| 17.8165662 | 17.4342605 | 17.2201221 | 17.7566717 | -0.101841648 | 0.488389793 |
| 22.7894811 | 22.5597234 | 22.6201132 | 22.6438946 | -0.04004376 | 0.703076422 |
| 16.8213636 | 16.577771 | 16.8378345 | 16.6647468 | -0.074863359 | 0.48294928 |

| | | | | | |
|---|---|---|---|---|---|
| 20.4413309 | 20.7422011 | 20.6512453 | 20.572662 | 0.057904899 | 0.523310947 |
| 17.247349 | 17.4122065 | 17.1132521 | 17.037039 | 0.056193085 | 0.627108268 |
| 16.8024786 | 16.7986997 | 16.5497799 | 16.8527393 | 0.085716415 | 0.543552283 |
| 17.6161263 | 17.7093107 | 17.6022752 | 17.65653 | -0.048292413 | 0.559595163 |
| 19.8434242 | 19.5643905 | 19.7723973 | 19.7168409 | -0.077866143 | 0.382062227 |
| 19.7015175 | 19.6405886 | 19.8508625 | 19.8052727 | 0.015242557 | 0.86736297 |
| 18.7621987 | 18.7292961 | 18.522985 | 18.8076391 | -0.074454279 | 0.42048638 |
| 14.7741546 | NA | 14.6968133 | 14.7067929 | -0.07693098 | 0.508935015 |
| 19.3895109 | 19.5999028 | 19.3004386 | 19.3541447 | 0.003452634 | 0.967467877 |
| 21.8413977 | 21.6806462 | 21.7637093 | 21.7209929 | 0.014555155 | 0.865844788 |
| 17.2406708 | 17.1940699 | 17.1956067 | 17.2607774 | -0.046597897 | 0.58133189 |
| 20.9602667 | 20.8092226 | 21.0149908 | 20.9527404 | -0.004910063 | 0.936072246 |
| 18.3356383 | 18.184981 | 18.3491297 | 18.2852157 | -0.034922623 | 0.639004573 |
| 19.5098963 | 19.3057045 | 19.5117371 | 19.5176325 | -0.040857802 | 0.546505667 |
| 20.4490698 | 20.3727447 | 20.3783932 | 20.4812038 | 0.008731353 | 0.879512056 |
| 20.866127 | 20.5840725 | 20.8349755 | 20.8691869 | -0.05831865 | 0.494368091 |
| 22.9800324 | 22.875256 | 22.9694319 | 22.9971568 | 0.057512208 | 0.31036681 |
| 16.6563931 | 16.8402192 | 16.7838245 | 16.6667214 | 0.056989969 | 0.435521474 |
| 20.6426118 | 20.507123 | 20.5802384 | 20.57308 | -0.104153813 | 0.067623533 |
| 20.7048602 | 20.5615764 | 20.6947012 | 20.6925506 | 0.00101199 | 0.985325615 |
| 21.3599603 | 21.2858952 | 21.3584046 | 21.410083 | -0.005447778 | 0.90532093 |
| 22.8443914 | 22.85407 | 22.9011818 | 22.7943763 | -0.004602102 | 0.924677537 |
| 20.0223462 | 20.0326007 | 20.0768025 | 20.0902313 | 0.04327349 | 0.347609741 |
| 20.6070554 | 20.4889063 | 20.5477883 | 20.5660128 | -0.013276243 | 0.811317218 |
| 26.0191245 | 25.9150157 | 26.0074915 | 25.9800169 | 0.010833879 | 0.805143574 |
| 24.7429755 | 24.5965177 | 24.691926 | 24.7887241 | -0.045414211 | 0.450507903 |
| 21.4210205 | 21.4225248 | 21.3874285 | 21.38132 | 0.058294846 | 0.34648207 |
| 17.6036869 | 17.6159595 | 17.716182 | 17.7099562 | 0.010998497 | 0.849524666 |
| 22.6234979 | 22.6912777 | 22.6895251 | 22.7210898 | 0.022854703 | 0.693613543 |
| 20.8853526 | 20.7309003 | 20.839533 | 20.8342423 | -0.074133736 | 0.144444524 |
| 22.4408627 | 22.3316652 | 22.4294641 | 22.485863 | -0.001307392 | 0.976068755 |
| 19.1947079 | 19.2691283 | 19.2115203 | 19.2218888 | 0.026988847 | 0.557930598 |
| 21.6814112 | 21.5417746 | 21.6098343 | 21.7172298 | -0.003395229 | 0.948136158 |
| 18.6386135 | 18.6665895 | 18.7671346 | 18.7958207 | 0.01612929 | 0.75997731 |
| 21.2350006 | 21.1933813 | 21.2351841 | 21.1918764 | -0.00590114 | 0.915693393 |
| 23.0127023 | 22.9256912 | 22.9593243 | 22.9824072 | -0.003785821 | 0.940998925 |
| 19.6369183 | 19.5701219 | 19.732093 | 19.6186543 | -0.02434062 | 0.643552459 |
| 22.5689681 | 22.5143268 | 22.5467306 | 22.553421 | 0.007425513 | 0.879984801 |
| 22.4371777 | 22.3886498 | 22.5249943 | 22.4800911 | 0.042923811 | 0.292819821 |
| 22.8977075 | 22.8259514 | 22.9314864 | 22.8912579 | 0.022267877 | 0.629446884 |
| 20.2540675 | 20.2159154 | 20.2157977 | 20.231151 | 0.012742799 | 0.751015374 |
| 19.8575651 | 19.7055754 | 19.7635671 | 19.7928151 | -0.018073021 | 0.709679855 |

| | | | | | |
|---|---|---|---|---|---|
| 20.9086445 | 20.8953324 | 20.8986625 | 20.9465566 | -0.044223181 | 0.282792078 |
| 20.4354472 | 20.3913426 | 20.415186 | 20.4695146 | -0.018534541 | 0.65075032 |
| 22.0396892 | 21.9516807 | 22.094687 | 22.1309409 | 0.040223738 | 0.438460851 |
| 21.8186205 | 21.7675099 | 21.8268941 | 21.8153109 | -0.005498204 | 0.912160286 |
| 21.2704576 | 21.1891082 | 21.3152362 | 21.2427057 | 0.007351923 | 0.884993973 |
| 22.3736025 | 22.3083004 | 22.3503968 | 22.3455657 | 0.018740296 | 0.710773722 |
| 22.7934231 | 22.7344231 | 22.8681634 | 22.8108848 | 0.023108804 | 0.600419671 |
| 19.2912025 | 19.259277 | 19.2878263 | 19.3914557 | 0.002384162 | 0.96372307 |
| 23.7401479 | 23.6342075 | 23.7254806 | 23.7119237 | -5.95E-04 | 0.99050177 |
| 23.3943627 | 23.3355743 | 23.3052952 | 23.4176522 | 0.039929468 | 0.488027144 |
| 19.4935252 | 19.2973744 | 19.427921 | 19.4230964 | -0.022569071 | 0.698467285 |
| 23.2564412 | 23.1240953 | 23.2290165 | 23.2251553 | 0.009292344 | 0.870417271 |
| 24.2741529 | 24.251064 | 24.3151486 | 24.3121318 | 0.072282508 | 0.320224426 |
| 20.9622335 | 21.0725502 | 21.0086979 | 21.0552545 | 0.133517552 | 0.058978383 |
| 22.8047852 | 22.733651 | 22.7971048 | 22.8421446 | 0.001224538 | 0.979602617 |
| 19.4191369 | 19.5448571 | 19.5920539 | 19.6990872 | 0.082735135 | 0.303331317 |
| 20.0798307 | 19.8335439 | 19.9797672 | 19.9781447 | -0.073073152 | 0.170103818 |
| 19.4668474 | 19.4641882 | 19.5312835 | 19.6216416 | 0.057478079 | 0.323387495 |
| 18.3632822 | 18.4471766 | 18.3692276 | 18.4215383 | 0.072748575 | 0.289849552 |
| 17.6560882 | 17.702813 | 17.6696153 | 17.8023196 | 0.089431253 | 0.218706738 |
| 20.5704959 | 20.3512718 | 20.6595727 | 20.6660531 | 0.039412926 | 0.580854408 |
| 23.1466622 | 23.048643 | 23.1603609 | 23.1427619 | 0.034092917 | 0.501562133 |
| 20.7940937 | 20.7471791 | 20.8093021 | 20.8270476 | 0.010001037 | 0.857066805 |
| 20.3952987 | 20.2217585 | 20.3101448 | 20.4116412 | 0.012084637 | 0.851095233 |
| 21.9835254 | 21.9577855 | 21.8769129 | 21.9214793 | -0.02037845 | 0.761393254 |
| 21.9957579 | 21.9744156 | 22.0439061 | 21.9486793 | 0.043419499 | 0.485511727 |
| 25.1345906 | 25.2823754 | 25.2228217 | 25.124478 | 0.05942014 | 0.423902402 |
| 19.421857 | 19.1494 | 19.3703466 | 19.4097856 | 0.018576596 | 0.837829825 |
| 19.2751658 | 19.0905797 | 19.1842379 | 19.1726307 | -0.090206913 | 0.120258773 |
| 18.3137961 | 18.4281909 | 18.2290312 | 18.3379486 | -0.008958797 | 0.898643859 |
| 22.4844205 | 22.3553856 | 22.2945233 | 22.4331685 | -0.014542951 | 0.806803873 |
| 17.4264344 | 17.4932418 | 17.7172646 | 17.5246038 | -0.010035909 | 0.907119585 |
| 18.9515425 | 18.722715 | 18.9084814 | 18.7583046 | -0.116463698 | 0.079681099 |
| 16.5357466 | 16.4953509 | 16.7604059 | 16.4412223 | 0.02390908 | 0.809095611 |
| 21.1266539 | 20.8845422 | 20.8032034 | 21.0354254 | -0.04601908 | 0.573823907 |
| 18.3508435 | 18.3278217 | 18.4006089 | 18.359122 | 0.016624036 | 0.815658889 |
| 24.6501989 | 24.5418175 | 24.6151332 | 24.6361361 | 0.032756176 | 0.620703005 |
| 21.0594535 | 21.0953855 | 21.1939753 | 21.2038004 | 0.04933243 | 0.652460295 |
| 18.2227704 | 18.2155 | 17.9336256 | 18.1250344 | -0.125184274 | 0.094208256 |
| 17.1751014 | 17.0673035 | 17.2404639 | 17.2772918 | 0.031308766 | 0.655043705 |
| 18.3261741 | 18.4161757 | 18.3584436 | 18.4444755 | -0.032221164 | 0.715251327 |
| 15.7598286 | 15.6818472 | 15.9211536 | 15.5654407 | -0.105139056 | 0.297754085 |

| | | | | | |
|---|---|---|---|---|---|
| 20.0307233 | 19.9745624 | 20.013931 | 19.9397877 | 0.038898552 | 0.696125498 |
| 16.5976513 | 16.4262279 | 16.4719104 | 16.4394109 | -0.014982673 | 0.870292099 |
| 15.6510286 | 15.6414418 | 15.6760104 | 15.6154942 | 0.004091857 | 0.971130362 |
| 16.3310471 | 16.4513071 | 16.5493247 | 16.2370836 | -0.051107576 | 0.697709886 |
| 17.9232685 | 17.7729191 | 17.8255359 | 17.8753067 | -0.111240933 | 0.406546675 |
| 17.738216 | 17.8544316 | 17.9067489 | 18.071854 | 0.091836069 | 0.466670998 |
| 15.1615888 | 15.3467333 | 15.2441607 | 15.5426613 | 0.532680509 | 0.025112573 |
| 17.9433494 | 18.3903493 | 17.8619856 | 17.8660215 | -0.002389628 | 0.989785536 |
| NA | 16.3681379 | NA | NA | 0.390070412 | 0.21949259 |
| 11.779769 | NA | 11.2414263 | 11.4073792 | -0.726048445 | 0.042124896 |
| 15.7789747 | 16.0807617 | 16.1333391 | 16.4121853 | 0.161005978 | 0.32679907 |
| 19.8510138 | 19.9236598 | 19.9233461 | 20.0453492 | 0.053135039 | 0.641340847 |
| 24.5818533 | 24.4980034 | 24.5950895 | 24.4702189 | 1.41E-04 | 0.998631394 |
| 18.694139 | 18.9010886 | 18.7175374 | 18.7015232 | -0.012719811 | 0.867714814 |
| 22.9666879 | 22.8880448 | 22.9857295 | 22.9371486 | -0.005035469 | 0.923189596 |
| 21.9259327 | 21.819513 | 22.0757519 | 21.9952541 | 0.031185792 | 0.563711756 |
| 19.6506314 | 19.6702459 | 19.6508789 | 19.7398671 | 0.016050543 | 0.750268105 |
| 24.4378192 | 24.3515774 | 24.4927084 | 24.4553531 | 0.038882761 | 0.419783483 |
| 21.9671832 | 21.9123399 | 21.9687043 | 21.9748167 | 0.034228492 | 0.437464128 |
| 19.2260145 | 19.158254 | 19.2731669 | 19.159532 | -0.01263107 | 0.83478344 |
| 17.9661632 | 17.7335032 | 17.7727379 | 17.9894791 | -0.006298773 | 0.959462898 |
| 20.3438167 | 20.3087693 | 20.3162883 | 20.3303405 | -0.045281944 | 0.42489935 |
| 19.8258664 | 19.7872129 | 19.9380269 | 19.838669 | -0.113118862 | 0.346685821 |
| 19.9094369 | 19.7609161 | 19.9360417 | 19.9639509 | -0.001530304 | 0.979013571 |
| 19.9301732 | 19.8169728 | 19.9229512 | 19.9116102 | -0.00158303 | 0.975644235 |
| 23.7959192 | 23.7498552 | 23.7932435 | 23.642398 | -0.006292984 | 0.921402699 |
| 17.6249051 | 17.4494784 | 17.884644 | 17.6014768 | -0.104131538 | 0.410513275 |
| 16.2699899 | 16.4425734 | 16.6346682 | 16.8040204 | 0.053384667 | 0.696153989 |
| 23.2129323 | 23.2642395 | 23.2890636 | 23.3206712 | 0.066323155 | 0.108570196 |
| 20.4757802 | 20.4490505 | 20.4788149 | 20.5310925 | 1.95E-04 | 0.996615957 |
| 20.2987385 | 20.1105902 | 20.1838346 | 20.2675047 | -0.069443105 | 0.238660876 |
| 21.5558363 | 21.7087071 | 21.5928261 | 21.552894 | 0.071074996 | 0.316578837 |
| 17.1722449 | 16.9858978 | 17.6048361 | 17.4713725 | 0.258307815 | 0.255973278 |
| 18.1296425 | 18.0605918 | 18.2452881 | 18.1290699 | 0.040414486 | 0.809689385 |
| 19.5119362 | 19.4084203 | 19.4788014 | 19.6307409 | 0.038179478 | 0.77656518 |
| 16.8949184 | 16.7894515 | 17.0551379 | 17.082136 | 0.069673279 | 0.61736133 |
| 18.7108493 | 18.7441345 | 18.8160139 | 18.7956018 | -0.124084297 | 0.32914637 |
| 25.110828 | 24.9206635 | 25.0554145 | 25.1263349 | -0.029316808 | 0.661473927 |
| 21.6591061 | 21.6356468 | 21.7408667 | 21.6674291 | 0.020392871 | 0.715781317 |
| 21.2998355 | 21.1626106 | 21.2951639 | 21.2513485 | -0.001886462 | 0.972886625 |
| 17.8433107 | 17.6295341 | 17.6419021 | 17.6631356 | -0.16501608 | 0.013713116 |
| 18.3613725 | 18.2673797 | 18.3401265 | 18.2496052 | -0.068839403 | 0.310539789 |

| | | | | | |
|---|---|---|---|---|---|
| 19.9306077 | 19.8516408 | 19.9542488 | 20.0252908 | 0.049944042 | 0.46579003 |
| 17.9502882 | 17.8261196 | 18.1082147 | 18.186369 | 0.02111877 | 0.868543068 |
| 20.3239278 | 20.3073786 | 20.4126575 | 20.3049798 | 0.07268542 | 0.323734745 |
| 16.3986371 | 16.3186337 | 16.3984431 | 16.3714569 | -0.033965909 | 0.843241801 |
| 22.3267573 | 22.2654809 | 22.320879 | 22.3019266 | -0.004942344 | 0.893227478 |
| 22.2536026 | 22.2238804 | 22.3340707 | 22.1638184 | -0.033570372 | 0.626893548 |
| 21.5977254 | 21.5142924 | 21.5873443 | 21.6748821 | 0.011905472 | 0.855926589 |
| 16.6847295 | 17.0334683 | 16.8068674 | 16.6893882 | 0.025693429 | 0.881588831 |
| 15.6065874 | 15.8231208 | 15.6587252 | 16.1354963 | 0.044336156 | 0.796549178 |
| 18.1000868 | 18.0218423 | 18.0800614 | 18.1886408 | -0.030845596 | 0.690857638 |
| 18.8704049 | 18.8367067 | 18.907968 | 18.8248563 | 0.006569881 | 0.914280565 |
| 22.1978193 | 22.0719681 | 22.1922725 | 22.146874 | -0.018590653 | 0.665591695 |
| 23.2024647 | 23.1505258 | 23.1388351 | 23.1945766 | 0.006121474 | 0.88445752 |
| 20.585246 | 20.5032671 | 20.49941 | 20.6106899 | 0.002314972 | 0.967363403 |
| 17.5011029 | 17.1387428 | 17.3608055 | 17.4236267 | -0.143711387 | 0.172836462 |
| 17.0072021 | 16.6487767 | 16.7751379 | 16.847472 | -0.083245078 | 0.62450583 |
| 18.1953268 | 18.0995268 | 18.071465 | 18.2148169 | -0.070798303 | 0.521232442 |
| 23.9898186 | 23.9358478 | 24.0224139 | 23.9990234 | 0.027310989 | 0.539795154 |
| 22.5054278 | 22.1896818 | 22.4926156 | 22.5771117 | 0.023553354 | 0.854488726 |
| 24.5956101 | 24.4097553 | 24.6186229 | 24.5514459 | -0.018103548 | 0.778757892 |
| 23.9248458 | 23.7763639 | 23.8101866 | 23.8539163 | -0.043492037 | 0.338978472 |
| 23.1544996 | 23.0893649 | 23.0395191 | 23.0094805 | -0.063971984 | 0.23973681 |
| 26.1069289 | 26.0956687 | 26.2104937 | 26.1534533 | 0.021498829 | 0.810446552 |
| 18.5348414 | 18.3089059 | 18.3690345 | 18.5803884 | 0.063712781 | 0.428292886 |
| 18.1342594 | 17.8699515 | 18.04428 | 18.1734792 | -0.002779554 | 0.984099691 |
| 18.3940824 | NA | 18.2238644 | 18.0571703 | -0.352088728 | 0.209572287 |
| 19.8615473 | 19.5979226 | 19.7459273 | 19.6002927 | -0.045893713 | 0.705080005 |
| 23.0639234 | 23.095569 | 23.1311849 | 23.0594806 | 0.043974979 | 0.544000139 |
| 22.8851145 | 22.738957 | 22.8961631 | 22.8840513 | -0.041740573 | 0.542298993 |
| 23.782045 | 23.7121306 | 23.8215349 | 23.7317357 | 0.014366157 | 0.755661588 |
| 22.9383036 | 22.9089444 | 23.0032625 | 23.0319193 | 0.046763255 | 0.262616433 |
| 24.2536525 | 24.1715677 | 24.2536452 | 24.257075 | 0.040054763 | 0.419154007 |
| 22.4193472 | 22.4002889 | 22.4981937 | 22.525897 | 0.049082846 | 0.383553221 |
| 20.3437159 | 20.1596044 | 20.1385303 | 20.1223541 | -0.078729168 | 0.1374318 |
| 17.8441424 | 17.8355288 | 17.8849183 | 17.9160356 | 0.016050743 | 0.802483608 |
| 19.4181688 | 19.3365364 | 19.3517166 | 19.2389602 | -0.079617184 | 0.179057418 |
| 21.0682967 | 20.7739307 | 21.0209536 | 20.8618749 | -0.117518436 | 0.150340084 |
| 15.7884543 | 15.5048079 | 15.8582869 | 15.2761183 | -0.051649814 | 0.806175734 |
| 19.5064235 | 19.482951 | 19.491592 | 19.5746554 | 0.069640359 | 0.397713225 |
| 20.9894291 | 20.9133847 | 20.8633279 | 20.9483569 | -0.076886739 | 0.231096758 |
| 23.898334 | 23.8726067 | 23.964466 | 23.9684638 | 0.017720454 | 0.785576218 |
| 22.1258325 | 21.9996248 | 22.0737897 | 22.0792185 | 0.028784344 | 0.634962553 |

| | | | | | |
|---|---|---|---|---|---|
| 21.6492882 | 21.5013301 | 21.539583 | 21.5120854 | -0.062912554 | 0.211620632 |
| 18.904808 | 18.9006097 | 18.8941981 | 18.9189279 | 0.057170531 | 0.269532298 |
| 19.7681746 | 19.7279799 | 19.821127 | 19.9377513 | 0.042191767 | 0.624681065 |
| 19.466736 | 19.2052073 | 19.3482433 | 19.4124416 | -0.101037773 | 0.08814336 |
| 21.3854091 | 21.3801017 | 21.390882 | 21.4392848 | 0.035610806 | 0.463931693 |
| 20.9026719 | 20.8653008 | 20.9157357 | 20.9263323 | 0.045049522 | 0.444789417 |
| 20.4669634 | 20.5089839 | 20.4682842 | 20.3922175 | -0.030849754 | 0.590314014 |
| NA | 12.8280029 | NA | NA | -0.498279776 | 0.287741403 |
| 22.9158708 | 22.721587 | 22.850883 | 22.8874393 | -0.035971474 | 0.727082569 |
| 17.5474223 | 17.7314227 | 17.750123 | 17.6116326 | 0.260425492 | 0.014667644 |
| NA | 22.741728 | NA | NA | 0.043329543 | 0.787290686 |
| NA | 18.2724473 | NA | NA | -0.075746181 | 0.518414193 |
| 17.7486504 | 17.4515803 | 17.6737304 | 17.4482049 | -0.065181973 | 0.523305628 |
| 21.6508363 | 21.4444252 | 21.6389358 | 21.7106254 | 0.028441591 | 0.717974841 |
| 19.9453005 | 19.8651043 | 19.936142 | 19.9591479 | 0.01539835 | 0.776742285 |
| 20.2046647 | 20.2299049 | 20.2283929 | 20.2187047 | 0.006735691 | 0.887633063 |
| 22.6239196 | 22.5174782 | 22.5840893 | 22.5765558 | -0.027496337 | 0.566401525 |
| 21.5100445 | 21.4759572 | 21.457064 | 21.4439995 | -0.00455102 | 0.903461722 |
| 20.915653 | 20.8056545 | 20.8731133 | 20.897252 | -0.025981289 | 0.509148164 |
| 24.4636864 | 24.4518371 | 24.5097742 | 24.4939685 | 8.69E-04 | 0.989038328 |
| 21.3988521 | 21.3537361 | 21.4149863 | 21.4305341 | 0.025400876 | 0.559488242 |
| 21.2050153 | 21.0360381 | 21.2754812 | 21.1188378 | -0.01296759 | 0.894396209 |
| 16.8789425 | 17.028145 | 16.9477851 | 17.1242843 | 0.051149051 | 0.698065024 |
| 16.3338794 | 16.6521187 | 16.668871 | 16.7556693 | 0.374235457 | 0.012721417 |
| 15.3388036 | NA | 15.5992916 | 15.5109098 | 0.298196136 | 0.190315661 |
| 22.9419616 | 22.9170157 | 23.046623 | 22.9360944 | 0.037781047 | 0.528883076 |
| 23.4619591 | 23.4367136 | 23.4470872 | 23.4700579 | 0.00941388 | 0.837494219 |
| 21.513425 | 21.4195252 | 21.6387644 | 21.6145733 | 0.035763713 | 0.527898658 |
| 20.2831499 | 20.1814548 | 20.2119128 | 20.2901765 | -0.021496767 | 0.759569271 |
| 20.7822065 | 20.6314271 | 20.7671367 | 20.7092523 | -0.042183163 | 0.486567338 |
| 20.4300732 | 20.3038334 | 20.4240556 | 20.487953 | 0.024599058 | 0.676479477 |
| 22.6296007 | 22.528332 | 22.6041993 | 22.6193265 | 0.016223214 | 0.713689549 |
| 20.5120795 | 20.4037209 | 20.4406015 | 20.533802 | 0.01120583 | 0.786462721 |
| 23.7760395 | 23.7926955 | 23.7653989 | 23.7660619 | 0.015855997 | 0.719255943 |
| 18.6764142 | 18.6122048 | 18.5367 | 18.525447 | 0.062031572 | 0.490180365 |
| 22.7504251 | 22.5966836 | 22.7118407 | 22.7189735 | -0.020472506 | 0.699220347 |
| 21.7955354 | 21.9215852 | 21.9003947 | 21.8632779 | 0.03771595 | 0.415018866 |
| 19.0122252 | 18.9070482 | 18.976555 | 18.9563915 | -0.020584099 | 0.688064525 |

| AMPH-03340.6h - AMPH-04647.6h__adj.P.Val | AMPH-03340.6h - DMSO.6h__logFC | AMPH-03340.6h - DMSO.6h__P.Value | AMPH-03340.6h - DMSO.6h__adj.P.Val | AMPH-04647.6h - DMSO.6h__logFC | AMPH-04647.6h - DMSO.6h__P.Value |
|---|---|---|---|---|---|
| 1.84E-09 | -1.4373166 | 4.13E-13 | 3.45E-09 | 0.04108113 | 0.70025644 |
| 0.004303476 | -0.5222893 | 2.61E-09 | 1.09E-05 | -0.1618361 | 0.01015322 |
| 0.152848295 | -0.3318252 | 1.22E-07 | 3.41E-04 | -0.1402727 | 0.00500729 |
| 0.002763355 | -0.6294398 | 2.12E-07 | 3.54E-04 | -0.0416431 | 0.64509131 |
| 0.013813838 | -0.3995134 | 1.75E-07 | 3.54E-04 | -0.0868444 | 0.13383476 |
| 0.053114259 | -0.5337724 | 5.38E-07 | 7.48E-04 | -0.1433868 | 0.0854502 |
| 0.152848295 | -0.7239082 | 3.77E-06 | 0.00449866 | -0.2050305 | 0.10794277 |
| 0.956358655 | -0.3013755 | 2.24E-05 | 0.02093488 | -0.1596254 | 0.01089335 |
| 0.971241857 | 1.03102327 | 2.26E-05 | 0.02093488 | 0.55975978 | 0.00930861 |
| 0.781057588 | -0.547046 | 2.60E-05 | 0.02171931 | -0.2609759 | 0.02164434 |
| 0.004303476 | -0.6311942 | 3.32E-05 | 0.02354006 | 0.2145061 | 0.04436081 |
| 0.971241857 | -0.4455779 | 3.38E-05 | 0.02354006 | -0.2340797 | 0.0139334 |
| 0.999490048 | 0.9008581 | 4.58E-05 | 0.02940416 | 0.53630876 | 0.00718106 |
| 0.999490048 | -2.3538695 | 7.96E-05 | 0.0474658 | -1.5115323 | 0.00573869 |
| 0.987721587 | -0.3443019 | 1.11E-04 | 0.06157795 | -0.1779474 | 0.02606058 |
| 0.999490048 | 0.64228301 | 1.28E-04 | 0.06686435 | 0.5082749 | 6.28E-04 |
| 0.028599359 | -1.0228769 | 1.86E-04 | 0.07763131 | 0.29018789 | 0.14842875 |
| 0.053114259 | -0.9119906 | 1.68E-04 | 0.07763131 | 0.18302498 | 0.22532077 |
| 0.999490048 | -0.9091356 | 2.04E-04 | 0.07763131 | -0.570258 | 0.01161614 |
| 0.956358655 | -0.5813545 | 2.00E-04 | 0.07763131 | -0.254375 | 0.06864059 |
| 0.156991024 | -0.346891 | 2.05E-04 | 0.07763131 | -0.0140416 | 0.86198621 |
| 0.355231519 | -0.2563777 | 1.78E-04 | 0.07763131 | -0.0419532 | 0.4788704 |
| 0.771925038 | -0.2893479 | 2.40E-04 | 0.08725481 | -0.1027774 | 0.14120486 |
| 0.439893572 | -0.2841254 | 2.53E-04 | 0.08806958 | -0.0566816 | 0.40377846 |
| 0.999490048 | -0.4534275 | 2.73E-04 | 0.09121806 | -0.2355888 | 0.03749431 |
| 0.578467169 | 0.22087643 | 2.96E-04 | 0.09252184 | 0.05853245 | 0.27660896 |
| 0.493399408 | 0.27192603 | 3.00E-04 | 0.09252184 | 0.06025621 | 0.36182274 |
| 0.510282354 | 0.35027649 | 3.10E-04 | 0.09252184 | 0.07934857 | 0.35290687 |
| 0.081295872 | -0.5354372 | 3.47E-04 | 0.09653315 | 0.05091433 | 0.69748425 |
| 0.999490048 | -0.1913774 | 3.47E-04 | 0.09653315 | -0.1724172 | 9.92E-04 |
| 0.781057588 | -0.3520151 | 3.83E-04 | 0.099862 | -0.1193513 | 0.17699016 |
| 0.999490048 | -0.3207829 | 3.77E-04 | 0.099862 | -0.2027518 | 0.01563785 |
| 0.999490048 | -0.7892979 | 3.97E-04 | 0.10052966 | -0.4182556 | 0.04021987 |
| 0.771925038 | 0.21079966 | 4.49E-04 | 0.11016686 | 0.06569751 | 0.22007166 |
| 0.595926661 | 0.18507608 | 4.75E-04 | 0.11321152 | 0.0473427 | 0.31435333 |
| 0.987721587 | -0.215614 | 5.35E-04 | 0.1209638 | -0.0988819 | 0.08078462 |
| 0.397650117 | 0.18858992 | 5.36E-04 | 0.1209638 | 0.01912609 | 0.69090836 |
| 0.477170731 | 1.45333823 | 6.28E-04 | 0.1380249 | 0.21132528 | 0.57525202 |

| | | | | | |
|---|---|---|---|---|---|
| 0.999490048 | -2.2503688 | 6.93E-04 | 0.14350408 | -1.3803278 | 0.01324256 |
| 0.081295872 | -0.3531548 | 6.81E-04 | 0.14350408 | 0.0613404 | 0.50725209 |
| 0.711558326 | -0.2480036 | 7.05E-04 | 0.14350408 | -0.064003 | 0.32871255 |
| 0.90609983 | -0.7766968 | 9.29E-04 | 0.15351299 | -0.2570862 | 0.22578532 |
| 0.711558326 | -0.4520031 | 0.00101116 | 0.15351299 | -0.1035965 | 0.4021643 |
| 0.286147223 | -0.3463531 | 9.61E-04 | 0.15351299 | 0.00977666 | 0.91680716 |
| 0.999490048 | -0.3460061 | 9.42E-04 | 0.15351299 | -0.1993764 | 0.02597497 |
| 0.740405708 | -0.2892868 | 8.79E-04 | 0.15351299 | -0.072069 | 0.35618232 |
| 0.999490048 | -0.2794247 | 8.57E-04 | 0.15351299 | -0.1442697 | 0.06201365 |
| 0.556241665 | -0.2152795 | 9.95E-04 | 0.15351299 | -0.0329137 | 0.57411678 |
| 0.999490048 | -0.1545367 | 8.80E-04 | 0.15351299 | -0.1642686 | 4.78E-04 |
| 0.999490048 | 0.16465537 | 9.17E-04 | 0.15351299 | 0.12882757 | 0.00696215 |
| 0.771925038 | 0.17297597 | 0.00100339 | 0.15351299 | 0.04387569 | 0.35421599 |
| 0.972489992 | 0.23097341 | 8.91E-04 | 0.15351299 | 0.08719927 | 0.167168 |
| 0.987721587 | 0.26713929 | 9.46E-04 | 0.15351299 | 0.11468944 | 0.12041685 |
| 0.286147223 | 0.3259858 | 9.91E-04 | 0.15351299 | -0.0093574 | 0.91566889 |
| 0.578467169 | 1.13659105 | 9.49E-04 | 0.15351299 | 0.19786113 | 0.52056389 |
| 0.987721587 | -0.2259792 | 0.00107582 | 0.16041259 | -0.0942383 | 0.13596396 |
| 0.999490048 | -0.2278238 | 0.00124339 | 0.17142031 | -0.2075093 | 0.0028037 |
| 0.999490048 | -0.1942142 | 0.00124547 | 0.17142031 | -0.1679892 | 0.0042324 |
| 0.999490048 | 0.20767343 | 0.00123977 | 0.17142031 | 0.0921522 | 0.11895857 |
| 0.999490048 | 0.23845363 | 0.00122838 | 0.17142031 | 0.2259066 | 0.00199066 |
| 0.060355187 | 0.93886922 | 0.00125229 | 0.17142031 | -0.2960942 | 0.26257628 |
| 0.081295872 | -0.3155465 | 0.00135046 | 0.1791543 | 0.08044365 | 0.36693335 |
| 0.999490048 | 0.6742123 | 0.0013517 | 0.1791543 | 0.50755608 | 0.00691381 |
| 0.90609983 | -0.498879 | 0.00138624 | 0.18086136 | -0.1494391 | 0.29192839 |
| 0.999490048 | -0.4550829 | 0.0014597 | 0.18751564 | -0.2940931 | 0.02943091 |
| 0.381294976 | 0.6996262 | 0.00149375 | 0.18898177 | -0.0111179 | 0.95528093 |
| 0.992011001 | 0.3135272 | 0.00157202 | 0.19591544 | 0.12701094 | 0.16343328 |
| 0.999490048 | 0.28754716 | 0.00162488 | 0.19952609 | 0.20584237 | 0.0181426 |
| 0.999490048 | -0.2293712 | 0.00174961 | 0.2030142 | -0.1041794 | 0.12426786 |
| 0.781057588 | -0.2128119 | 0.00171639 | 0.2030142 | -0.0496462 | 0.42070995 |
| 0.999490048 | -0.1382369 | 0.00177485 | 0.2030142 | -0.0673594 | 0.10083905 |
| 0.999490048 | 0.23065723 | 0.00170251 | 0.2030142 | 0.19135155 | 0.00739746 |
| 0.397650117 | 0.52031041 | 0.00177113 | 0.2030142 | -0.0024237 | 0.98516927 |
| 0.987721587 | -0.1495392 | 0.00199805 | 0.22244928 | -0.0537287 | 0.226792 |
| 0.773280431 | 0.45200302 | 0.00198173 | 0.22244928 | 0.09439568 | 0.47765675 |
| 0.999490048 | -0.279798 | 0.00204164 | 0.22431198 | -0.2231603 | 0.01105011 |
| 0.999490048 | -0.3344208 | 0.00208801 | 0.2264275 | -0.2434443 | 0.01941194 |
| 0.495397118 | -0.3079147 | 0.002131 | 0.22812579 | -0.0154385 | 0.86511958 |
| 0.081295872 | -0.5925122 | 0.00217099 | 0.22946565 | 0.31496623 | 0.04666664 |
| 0.999490048 | -0.4630524 | 0.00226064 | 0.23324154 | -0.4045415 | 0.00647353 |

| | | | | | |
|---|---|---|---|---|---|
| 0.999490048 | 0.14563157 | 0.00231186 | 0.23324154 | 0.08433804 | 0.06077442 |
| 0.999490048 | 0.19831592 | 0.00231845 | 0.23324154 | 0.14096218 | 0.02365642 |
| 0.999490048 | 0.35548687 | 0.0022806 | 0.23324154 | 0.16091104 | 0.13679576 |
| 0.999490048 | 0.23752477 | 0.00238421 | 0.23421323 | 0.15070141 | 0.04197836 |
| 0.999490048 | 0.43813398 | 0.0023659 | 0.23421323 | 0.22905107 | 0.06514677 |
| 0.987721587 | -0.282564 | 0.00248132 | 0.23807223 | -0.100122 | 0.24472703 |
| 0.999490048 | -0.1632453 | 0.00245928 | 0.23807223 | -0.1291408 | 0.0133432 |
| 0.999490048 | -0.1404652 | 0.00253754 | 0.23807223 | -0.0860059 | 0.05064135 |
| 0.999490048 | 0.56834589 | 0.00252201 | 0.23807223 | 0.32770614 | 0.04682787 |
| 0.999490048 | -1.9098964 | 0.00275299 | 0.24228775 | -1.2469368 | 0.03987422 |
| 0.999490048 | -0.8475205 | 0.00273987 | 0.24228775 | -0.4233154 | 0.10951996 |
| 0.999490048 | -0.2893576 | 0.00269126 | 0.24228775 | -0.1899228 | 0.03842137 |
| 0.999490048 | -0.2291277 | 0.00290608 | 0.24228775 | -0.2008769 | 0.00780204 |
| 0.999490048 | -0.2259308 | 0.00288203 | 0.24228775 | -0.1297209 | 0.06958308 |
| 0.881091405 | -0.2219198 | 0.00266943 | 0.24228775 | -0.0510284 | 0.45068723 |
| 0.57925586 | -0.144584 | 0.00284805 | 0.24228775 | -0.0116006 | 0.79297181 |
| 0.999490048 | 0.2749666 | 0.00293067 | 0.24228775 | 0.10814887 | 0.2068182 |
| 0.999490048 | 0.30291615 | 0.00289055 | 0.24228775 | 0.18152175 | 0.05903845 |
| 0.999490048 | 0.33788572 | 0.0028328 | 0.24228775 | 0.25719405 | 0.01852749 |
| 0.999490048 | 0.37133471 | 0.00287215 | 0.24228775 | 0.20000109 | 0.08736508 |
| 0.711558326 | 0.46990599 | 0.00290689 | 0.24228775 | 0.0632261 | 0.66103768 |
| 0.999490048 | -0.3526982 | 0.00297838 | 0.24381874 | -0.4198048 | 6.14E-04 |
| 0.595926661 | 0.70279724 | 0.00303116 | 0.24573024 | 0.08382742 | 0.63254502 |
| 0.999490048 | 0.21489469 | 0.00310284 | 0.24674981 | 0.13382162 | 0.05228488 |
| 0.381294976 | 0.87206609 | 0.00308215 | 0.24674981 | -0.1138047 | 0.63308198 |
| 0.999490048 | 0.20192806 | 0.00325004 | 0.25601755 | 0.08832273 | 0.16711301 |
| 0.826281516 | 0.80440274 | 0.00329068 | 0.25679619 | 0.19453636 | 0.34940796 |
| 0.999490048 | -0.3656723 | 0.00350319 | 0.25832207 | -0.1815642 | 0.12211569 |
| 0.999490048 | -0.2710586 | 0.00341396 | 0.25832207 | -0.1737421 | 0.04866446 |
| 0.987721587 | -0.2388309 | 0.00342637 | 0.25832207 | -0.0758914 | 0.31431258 |
| 0.999490048 | -0.2239811 | 0.00347943 | 0.25832207 | -0.1232302 | 0.08812373 |
| 0.999490048 | -0.1778172 | 0.00349056 | 0.25832207 | -0.1030971 | 0.07330814 |
| 0.999490048 | 0.17168368 | 0.00352679 | 0.25832207 | 0.10841036 | 0.05274541 |
| 0.999490048 | 0.19515225 | 0.00344382 | 0.25832207 | 0.10774907 | 0.0866827 |
| 0.999490048 | 0.17471637 | 0.00359304 | 0.25847947 | 0.13376719 | 0.02115333 |
| 0.999490048 | 0.20452106 | 0.00365276 | 0.25847947 | 0.13919108 | 0.03871632 |
| 0.999490048 | 0.58660187 | 0.00364032 | 0.25847947 | 0.40034856 | 0.02916009 |
| 0.999490048 | 0.6775659 | 0.00364733 | 0.25847947 | 0.31057374 | 0.12077295 |
| 0.999490048 | 0.27894003 | 0.00370122 | 0.25970761 | 0.11425694 | 0.20181265 |
| 0.999490048 | -0.1479185 | 0.00373453 | 0.25986122 | -0.1060025 | 0.03065026 |
| 0.999490048 | -0.1736595 | 0.00377222 | 0.26031455 | -0.1337304 | 0.02119963 |
| 0.999490048 | 0.1033376 | 0.00387765 | 0.26323904 | 0.05914114 | 0.08062058 |

| | | | | | |
|---|---|---|---|---|---|
| 0.999490048 | 0.17667798 | 0.00386255 | 0.26323904 | 0.06714798 | 0.23771873 |
| 0.999490048 | 0.19255094 | 0.00393622 | 0.26505992 | 0.16517207 | 0.01158168 |
| 0.999490048 | 0.15727302 | 0.0040024 | 0.26647528 | 0.06927709 | 0.17509171 |
| 0.999490048 | 0.21766438 | 0.00405298 | 0.26647528 | 0.21293038 | 0.00479306 |
| 0.999490048 | 0.22812565 | 0.00402361 | 0.26647528 | 0.16894866 | 0.0272485 |
| 0.999490048 | -0.2784052 | 0.00411245 | 0.26827312 | -0.0988533 | 0.27292758 |
| 0.999490048 | -0.2756399 | 0.00418077 | 0.27061543 | -0.1676773 | 0.06676973 |
| 0.397650117 | 0.2100823 | 0.00430131 | 0.27627655 | -0.0257827 | 0.70335096 |
| 0.999490048 | -0.1380151 | 0.00437207 | 0.27867776 | -0.1248751 | 0.00894485 |
| 0.999490048 | -0.4284346 | 0.00459054 | 0.27884255 | -0.4944308 | 0.00139551 |
| 0.881091405 | -0.1705006 | 0.00460842 | 0.27884255 | -0.0298355 | 0.59107397 |
| 0.999490048 | -0.1644594 | 0.00458711 | 0.27884255 | -0.0956632 | 0.08223188 |
| 0.999490048 | -0.1178742 | 0.00458426 | 0.27884255 | -0.0839484 | 0.03592765 |
| 0.999490048 | 0.15857652 | 0.00441042 | 0.27884255 | 0.10392929 | 0.05091377 |
| 0.999490048 | 0.25590192 | 0.00445072 | 0.27884255 | 0.08638537 | 0.30158364 |
| 0.987721587 | 1.30812807 | 0.00456988 | 0.27884255 | 0.49951642 | 0.17680706 |
| 0.999490048 | -0.3869392 | 0.00511616 | 0.28460799 | -0.2646233 | 0.0461602 |
| 0.999490048 | -0.3314547 | 0.00558418 | 0.28460799 | -0.1657208 | 0.14214558 |
| 0.989730632 | -0.2052105 | 0.00538917 | 0.28460799 | -0.0627485 | 0.36056616 |
| 0.999490048 | -0.1791382 | 0.00533813 | 0.28460799 | -0.1539153 | 0.01469431 |
| 0.999490048 | -0.1552869 | 0.00522103 | 0.28460799 | -0.0838982 | 0.11085629 |
| 0.552178084 | -0.1341661 | 0.00542295 | 0.28460799 | 0.00536403 | 0.9041336 |
| 0.879524171 | -0.1248714 | 0.00499379 | 0.28460799 | -0.0202988 | 0.62128116 |
| 0.987721587 | -0.1173922 | 0.00516826 | 0.28460799 | -0.0345596 | 0.37577033 |
| 0.999490048 | 0.13666309 | 0.00563615 | 0.28460799 | 0.06549854 | 0.15920314 |
| 0.999490048 | 0.14869462 | 0.00516503 | 0.28460799 | 0.11367261 | 0.02741734 |
| 0.999490048 | 0.17669593 | 0.00477963 | 0.28460799 | 0.14213162 | 0.01975434 |
| 0.999490048 | 0.18052453 | 0.00539515 | 0.28460799 | 0.16340457 | 0.01072238 |
| 0.999490048 | 0.18571618 | 0.00543205 | 0.28460799 | 0.08280022 | 0.18702653 |
| 0.999490048 | 0.19338464 | 0.00544748 | 0.28460799 | 0.1684501 | 0.01375181 |
| 0.999490048 | 0.22434254 | 0.00480734 | 0.28460799 | 0.08534746 | 0.2503797 |
| 0.999490048 | 0.22699137 | 0.00552524 | 0.28460799 | 0.23849554 | 0.0037995 |
| 0.999490048 | 0.23349327 | 0.00547408 | 0.28460799 | 0.17812408 | 0.02884275 |
| 0.999490048 | 0.23519232 | 0.00537916 | 0.28460799 | 0.18183652 | 0.02661444 |
| 0.999490048 | 0.25009666 | 0.00493502 | 0.28460799 | 0.14157571 | 0.09326112 |
| 0.835632585 | 0.25815093 | 0.00556529 | 0.28460799 | 0.03482856 | 0.68592555 |
| 0.999490048 | 0.30360524 | 0.00552932 | 0.28460799 | 0.15988997 | 0.12246018 |
| 0.999490048 | 0.33991737 | 0.00563649 | 0.28460799 | 0.13855986 | 0.22864732 |
| 0.999490048 | 0.35100761 | 0.00565807 | 0.28460799 | 0.16439733 | 0.16875626 |
| 0.819178176 | 0.41024728 | 0.00533719 | 0.28460799 | 0.06925288 | 0.54253007 |
| 0.999490048 | 0.57305045 | 0.00482978 | 0.28460799 | 0.67867255 | 0.00181573 |
| 0.578467169 | 0.60067107 | 0.00493551 | 0.28460799 | -0.0062958 | 0.96903413 |

| | | | | | |
|---|---|---|---|---|---|
| 0.999490048 | 1.01060858 | 0.00531402 | 0.28460799 | 0.46801125 | 0.11851353 |
| 0.58593703 | 2.02967348 | 0.00530305 | 0.28460799 | 0.03483598 | 0.94997849 |
| 0.999490048 | 0.47022131 | 0.00576818 | 0.28840914 | 0.34691318 | 0.02512191 |
| 0.999490048 | 0.19176876 | 0.00580571 | 0.28855766 | 0.13575582 | 0.04275976 |
| 0.999490048 | -0.172062 | 0.00603697 | 0.29451651 | -0.1055607 | 0.07766077 |
| 0.999490048 | -0.1498594 | 0.00606454 | 0.29451651 | -0.0936744 | 0.07271844 |
| 0.999490048 | -0.0913163 | 0.00612209 | 0.29451651 | -0.0740765 | 0.0226837 |
| 0.999490048 | 0.214064 | 0.00615863 | 0.29451651 | 0.08306775 | 0.25679664 |
| 0.999490048 | 0.22557782 | 0.0061725 | 0.29451651 | 0.30469268 | 4.46E-04 |
| 0.999490048 | 0.33990204 | 0.00610051 | 0.29451651 | 0.24720372 | 0.03902335 |
| 0.971241857 | 0.69663345 | 0.00613606 | 0.29451651 | 0.14358052 | 0.54302412 |
| 0.987721587 | -0.8036003 | 0.00633517 | 0.29656577 | -0.2062495 | 0.45171327 |
| 0.987721587 | -0.3475351 | 0.00637164 | 0.29656577 | -0.0876334 | 0.45995191 |
| 0.999490048 | -0.1609241 | 0.0063129 | 0.29656577 | -0.0707021 | 0.20229 |
| 0.999490048 | 0.23375647 | 0.00639303 | 0.29656577 | 0.08902906 | 0.26792222 |
| 0.999490048 | 0.37143435 | 0.00633372 | 0.29656577 | 0.24102355 | 0.06459977 |
| 0.999490048 | -0.222682 | 0.00662389 | 0.30533056 | -0.1698573 | 0.03289109 |
| 0.999490048 | 0.30242647 | 0.00665511 | 0.30533056 | 0.4019979 | 9.97E-04 |
| 0.999490048 | -0.4033925 | 0.00706264 | 0.30742622 | -0.1813379 | 0.1462786 |
| 0.999490048 | -0.2263392 | 0.00681992 | 0.30742622 | -0.1100326 | 0.16417404 |
| 0.999490048 | -0.1700669 | 0.00696449 | 0.30742622 | -0.1410903 | 0.02216181 |
| 0.999490048 | -0.1213128 | 0.00692982 | 0.30742622 | -0.1014361 | 0.02115067 |
| 0.999490048 | -0.1196862 | 0.00706298 | 0.30742622 | -0.1132908 | 0.01023304 |
| 0.999490048 | 0.15632955 | 0.00706896 | 0.30742622 | 0.13429328 | 0.01848009 |
| 0.999490048 | 0.1650783 | 0.0067677 | 0.30742622 | 0.08141359 | 0.15794626 |
| 0.999490048 | 0.26957909 | 0.00691561 | 0.30742622 | 0.17127777 | 0.07329562 |
| 0.999490048 | 0.35362198 | 0.00682738 | 0.30742622 | 0.34334585 | 0.00796543 |
| 0.999490048 | 0.40883411 | 0.00685123 | 0.30742622 | 0.37519891 | 0.01063672 |
| 0.58593703 | 0.3054035 | 0.00720048 | 0.31152331 | -0.0093826 | 0.92910867 |
| 0.835632585 | -0.4476213 | 0.00745534 | 0.31944805 | -0.0451703 | 0.7714015 |
| 0.999490048 | -0.2003478 | 0.00746016 | 0.31944805 | -0.2248722 | 0.00314193 |
| 0.999490048 | -0.2707518 | 0.00756542 | 0.32066644 | -0.3750711 | 4.63E-04 |
| 0.999490048 | 0.17299538 | 0.00754947 | 0.32066644 | 0.11370819 | 0.06767011 |
| 0.999490048 | 0.16381224 | 0.00765282 | 0.32273277 | 0.05524037 | 0.33751881 |
| 0.999490048 | -0.4519174 | 0.0079406 | 0.32662078 | -0.1842816 | 0.2506162 |
| 0.397650117 | -0.3805241 | 0.00790155 | 0.32662078 | 0.11880081 | 0.30622149 |
| 0.999490048 | 0.16112125 | 0.00783777 | 0.32662078 | 0.05370813 | 0.34464693 |
| 0.999490048 | 0.18065912 | 0.00791729 | 0.32662078 | 0.06870045 | 0.28297411 |
| 0.156991024 | 0.36757921 | 0.00784756 | 0.32662078 | -0.2105762 | 0.09065234 |
| 0.999490048 | 0.23760746 | 0.00799083 | 0.32707582 | 0.27090043 | 0.00300093 |
| 0.578467169 | -0.3855524 | 0.00806462 | 0.32848585 | 0.03387254 | 0.7672865 |
| 0.999490048 | -0.282503 | 0.00810438 | 0.32850286 | -0.1068148 | 0.24923082 |

| | | | | | |
|---|---|---|---|---|---|
| 0.999490048 | -0.1150819 | 0.00819542 | 0.33058839 | -0.1021729 | 0.01729778 |
| 0.488334682 | -0.1762824 | 0.00833037 | 0.33123154 | 0.03194455 | 0.57354268 |
| 0.999490048 | -0.1719463 | 0.00830432 | 0.33123154 | -0.1492166 | 0.01990567 |
| 0.999490048 | 0.19064602 | 0.00826206 | 0.33123154 | 0.0816121 | 0.23104883 |
| 0.999490048 | -0.2138062 | 0.0084006 | 0.33244072 | -0.1868772 | 0.01930606 |
| 0.999490048 | 0.16754344 | 0.00847563 | 0.33382796 | 0.11865301 | 0.05384448 |
| 0.999490048 | -0.5365474 | 0.00856786 | 0.3358761 | -0.4081114 | 0.03363192 |
| 0.999490048 | -0.8486877 | 0.00868241 | 0.33610034 | -0.305114 | 0.25152826 |
| 0.999490048 | -0.1848966 | 0.00869433 | 0.33610034 | -0.2068013 | 0.00385966 |
| 0.999490048 | -0.1547687 | 0.00864482 | 0.33610034 | -0.1808963 | 0.00269189 |
| 0.999490048 | -0.2781302 | 0.00874552 | 0.33652126 | -0.1068742 | 0.28497243 |
| 0.999490048 | -0.1805195 | 0.00892142 | 0.33729993 | -0.1318378 | 0.04884877 |
| 0.972489992 | -0.1785581 | 0.00886328 | 0.33729993 | -0.0354249 | 0.57846406 |
| 0.999490048 | -0.1569055 | 0.00881861 | 0.33729993 | -0.1898616 | 0.00206084 |
| 0.999490048 | 0.20874238 | 0.00892734 | 0.33729993 | 0.20946312 | 0.00872341 |
| 0.984638128 | -0.1885217 | 0.00897235 | 0.33747338 | -0.039017 | 0.56298255 |
| 0.999490048 | -0.1314768 | 0.00906572 | 0.33794607 | -0.0656491 | 0.1702014 |
| 0.548539439 | 0.31651247 | 0.00906586 | 0.33794607 | -0.0390021 | 0.73039234 |
| 0.999490048 | -0.3956719 | 0.00919787 | 0.33983298 | -0.3860478 | 0.01080441 |
| 0.999490048 | 0.52583639 | 0.00917614 | 0.33983298 | 0.17227108 | 0.36387592 |
| 0.999490048 | -0.3406226 | 0.00968743 | 0.3408255 | -0.3750991 | 0.00551451 |
| 0.999490048 | -0.2413291 | 0.00943152 | 0.3408255 | -0.1268463 | 0.15194617 |
| 0.909524814 | -0.1964225 | 0.0096903 | 0.3408255 | -0.0216404 | 0.76029009 |
| 0.999490048 | -0.167968 | 0.00941418 | 0.3408255 | -0.185254 | 0.00470909 |
| 0.999490048 | -0.1443015 | 0.00957378 | 0.3408255 | -0.0733774 | 0.16610895 |
| 0.999490048 | 0.12919316 | 0.00940548 | 0.3408255 | 0.08222983 | 0.08551118 |
| 0.987721587 | 0.13882319 | 0.0095835 | 0.3408255 | 0.03227363 | 0.52035328 |
| 0.987721587 | 0.18710336 | 0.00942435 | 0.3408255 | 0.04115759 | 0.54185129 |
| 0.999490048 | 0.20041495 | 0.00961215 | 0.3408255 | 0.11494675 | 0.12039803 |
| 0.999490048 | 0.22811022 | 0.00971455 | 0.3408255 | 0.25544781 | 0.00434912 |
| 0.999490048 | 0.24431137 | 0.00959221 | 0.3408255 | 0.10055798 | 0.25931909 |
| 0.999490048 | 0.24826778 | 0.00949684 | 0.3408255 | 0.34798732 | 5.79E-04 |
| 0.999490048 | -0.1164905 | 0.00976049 | 0.34100469 | -0.0815269 | 0.06152015 |
| 0.999490048 | -0.4368608 | 0.00997274 | 0.34205756 | -0.1869407 | 0.21126899 |
| 0.999490048 | -0.1990871 | 0.00996975 | 0.34205756 | -0.0853872 | 0.24303547 |
| 0.987721587 | 0.14458681 | 0.00991516 | 0.34205756 | 0.03364047 | 0.52216268 |
| 0.999490048 | 0.21643099 | 0.00999545 | 0.34205756 | 0.10066099 | 0.20677192 |
| 0.999490048 | 0.41454171 | 0.00986537 | 0.34205756 | 0.14314723 | 0.34417664 |
| 0.999490048 | 0.2438934 | 0.01029439 | 0.35084956 | 0.08541198 | 0.34060771 |
| 0.999490048 | -0.3499743 | 0.01051734 | 0.35339926 | -0.5447651 | 5.53E-04 |
| 0.999490048 | -0.3249197 | 0.01063574 | 0.35339926 | -0.2451109 | 0.047679 |
| 0.999490048 | -0.2864517 | 0.01057891 | 0.35339926 | -0.0753064 | 0.47445278 |

| | | | | | |
|---|---|---|---|---|---|
| 0.999490048 | -0.2427025 | 0.0107448 | 0.35339926 | -0.0602627 | 0.5001176 |
| 0.999490048 | -0.1618665 | 0.01056895 | 0.35339926 | -0.1128258 | 0.06554002 |
| 0.288776683 | -0.1585213 | 0.01075011 | 0.35339926 | 0.06081008 | 0.30058715 |
| 0.999490048 | 0.14123599 | 0.01071423 | 0.35339926 | 0.10640906 | 0.0481774 |
| 0.999490048 | 0.24042496 | 0.01067331 | 0.35339926 | 0.22725504 | 0.01513564 |
| 0.999490048 | 0.25640403 | 0.01046618 | 0.35339926 | 0.22272604 | 0.02394425 |
| 0.987721587 | -0.3341963 | 0.01084683 | 0.3549624 | -0.0657988 | 0.59277913 |
| 0.999490048 | -0.1929936 | 0.01088268 | 0.3549624 | -0.2687305 | 7.62E-04 |
| 0.999490048 | -0.1288972 | 0.01102863 | 0.35832312 | -0.1359756 | 0.0077335 |
| 0.972489992 | -0.1635684 | 0.01126524 | 0.36459197 | -0.0279294 | 0.64449488 |
| 0.999490048 | 0.13490092 | 0.0113461 | 0.36579131 | 0.10617903 | 0.04132762 |
| 0.999490048 | -0.1060947 | 0.0114692 | 0.36692661 | -0.0387333 | 0.32885417 |
| 0.999490048 | 0.42926649 | 0.01145049 | 0.36692661 | 0.25411142 | 0.11879262 |
| 0.771925038 | -0.6421092 | 0.01187716 | 0.36988254 | 0.00879035 | 0.97067219 |
| 0.773280431 | -0.3575713 | 0.01203807 | 0.36988254 | 0.00405348 | 0.9757645 |
| 0.999490048 | -0.2755063 | 0.01220456 | 0.36988254 | -0.1898546 | 0.07446276 |
| 0.999490048 | -0.2503615 | 0.01207084 | 0.36988254 | -0.0951521 | 0.31362614 |
| 0.999490048 | -0.2444444 | 0.01213149 | 0.36988254 | -0.0889492 | 0.334638 |
| 0.999490048 | -0.1914361 | 0.01222606 | 0.36988254 | -0.1189999 | 0.10566825 |
| 0.578467169 | -0.1634611 | 0.01183255 | 0.36988254 | 0.02392734 | 0.69450955 |
| 0.999490048 | -0.1406606 | 0.0121406 | 0.36988254 | -0.0341907 | 0.51713455 |
| 0.999490048 | -0.1203529 | 0.0118116 | 0.36988254 | -0.0795448 | 0.08479833 |
| 0.999490048 | -0.1099275 | 0.01201351 | 0.36988254 | -0.0417423 | 0.31368281 |
| 0.967348048 | 0.12614969 | 0.01216314 | 0.36988254 | 0.01343516 | 0.77586672 |
| 0.999490048 | 0.19107994 | 0.01212657 | 0.36988254 | 0.15085532 | 0.04274416 |
| 0.999490048 | 0.285148 | 0.01205846 | 0.36988254 | 0.25551752 | 0.02283449 |
| 0.999490048 | 0.47908948 | 0.01188721 | 0.36988254 | 0.37691286 | 0.0344299 |
| 0.987721587 | 0.47990242 | 0.01202759 | 0.36988254 | 0.09794154 | 0.58550541 |
| 0.984638128 | -0.4319101 | 0.01259669 | 0.37305198 | -0.0852117 | 0.54685811 |
| 0.999490048 | -0.2720348 | 0.01262297 | 0.37305198 | -0.1891949 | 0.0734344 |
| 0.987721587 | -0.2544449 | 0.01299531 | 0.37305198 | -0.0460223 | 0.63286696 |
| 0.999490048 | -0.154326 | 0.01282924 | 0.37305198 | -0.1220955 | 0.0440842 |
| 0.999490048 | -0.1537331 | 0.01276624 | 0.37305198 | -0.0737105 | 0.2102976 |
| 0.999490048 | -0.1461089 | 0.01296239 | 0.37305198 | -0.1912601 | 0.00174997 |
| 0.999490048 | -0.1286608 | 0.01271911 | 0.37305198 | -0.0899497 | 0.07235932 |
| 0.692187333 | -0.1200876 | 0.01265785 | 0.37305198 | 0.00977053 | 0.82883798 |
| 0.999490048 | -0.1047353 | 0.0129448 | 0.37305198 | -0.082954 | 0.04414991 |
| 0.999490048 | 0.14435881 | 0.01290483 | 0.37305198 | 0.09671226 | 0.08498339 |
| 0.999490048 | 0.16643407 | 0.01283925 | 0.37305198 | 0.1488547 | 0.02434104 |
| 0.999490048 | 0.24567805 | 0.01282472 | 0.37305198 | 0.07364057 | 0.42943801 |
| 0.999490048 | 0.27512513 | 0.01258642 | 0.37305198 | 0.13550205 | 0.19755025 |
| 0.493399408 | 0.27798288 | 0.01257238 | 0.37305198 | -0.0605482 | 0.56345565 |

| | | | | | |
|---|---|---|---|---|---|
| 0.999490048 | 0.73613486 | 0.01300097 | 0.37305198 | 0.74190817 | 0.01251954 |
| 0.999490048 | -0.1843083 | 0.013118 | 0.37512092 | -0.0751048 | 0.28699291 |
| 0.999490048 | -0.5836324 | 0.01334892 | 0.37531014 | -0.5306204 | 0.0215652 |
| 0.999490048 | -0.3806632 | 0.01338741 | 0.37531014 | -0.0896156 | 0.53670869 |
| 0.999490048 | -0.1784133 | 0.01327718 | 0.37531014 | -0.0716307 | 0.2949245 |
| 0.999490048 | -0.1331333 | 0.01340031 | 0.37531014 | -0.0485342 | 0.34137396 |
| 0.999490048 | 0.17954833 | 0.01350073 | 0.37531014 | 0.10337472 | 0.13855379 |
| 0.999490048 | 0.19441474 | 0.01352666 | 0.37531014 | 0.13809108 | 0.07080304 |
| 0.999490048 | 0.20457048 | 0.01340681 | 0.37531014 | 0.13538148 | 0.09050254 |
| 0.999490048 | 0.25813517 | 0.01329059 | 0.37531014 | 0.1280397 | 0.19829313 |
| 0.999490048 | 0.28055498 | 0.01352914 | 0.37531014 | 0.18566903 | 0.090951 |
| 0.397650117 | -0.8412312 | 0.01357711 | 0.37539361 | 0.27793137 | 0.38876547 |
| 0.999490048 | -0.7363527 | 0.01382561 | 0.37906063 | -0.2717541 | 0.33798136 |
| 0.999490048 | -0.674495 | 0.01384593 | 0.37906063 | -0.4291278 | 0.1046583 |
| 0.999490048 | -0.1231498 | 0.01381953 | 0.37906063 | -0.0579673 | 0.22428034 |
| 0.999490048 | -0.2041822 | 0.0138933 | 0.37911469 | -0.0976293 | 0.21764411 |
| 0.999490048 | 0.17295105 | 0.01407094 | 0.38146862 | 0.05477392 | 0.41114082 |
| 0.999490048 | 0.25828085 | 0.01405082 | 0.38146862 | 0.13313681 | 0.18568584 |
| 0.999490048 | -0.3745596 | 0.01433562 | 0.38162555 | -0.1205301 | 0.40519402 |
| 0.999490048 | -0.2257822 | 0.01434656 | 0.38162555 | -0.1300053 | 0.14231814 |
| 0.999490048 | -0.1616379 | 0.01418241 | 0.38162555 | -0.1113852 | 0.08128012 |
| 0.999490048 | 0.10276054 | 0.01439504 | 0.38162555 | 0.03474586 | 0.38244018 |
| 0.999490048 | 0.12703979 | 0.01420045 | 0.38162555 | 0.11073024 | 0.03023341 |
| 0.999490048 | 0.16982227 | 0.0142601 | 0.38162555 | 0.18427033 | 0.00845753 |
| 0.999490048 | 0.17737423 | 0.01439665 | 0.38162555 | 0.09588896 | 0.16751604 |
| 0.971241857 | -0.2484478 | 0.01474908 | 0.38329043 | -0.022874 | 0.81144504 |
| 0.999490048 | -0.233238 | 0.01452284 | 0.38329043 | -0.1687418 | 0.06905248 |
| 0.771925038 | -0.1813247 | 0.01477897 | 0.38329043 | 0.01010781 | 0.88514808 |
| 0.999490048 | -0.1116785 | 0.01470462 | 0.38329043 | -0.0571792 | 0.19179324 |
| 0.999490048 | 0.1211384 | 0.0147613 | 0.38329043 | 0.10191555 | 0.0370002 |
| 0.999490048 | 0.13778805 | 0.01482668 | 0.38329043 | 0.03785719 | 0.47888197 |
| 0.999490048 | 0.19543561 | 0.01464397 | 0.38329043 | 0.18539603 | 0.01987369 |
| 0.999490048 | 0.21658278 | 0.01481743 | 0.38329043 | 0.15267769 | 0.07692603 |
| 0.999490048 | -0.3190703 | 0.01492956 | 0.38475868 | -0.1029687 | 0.4069953 |
| 0.999490048 | -0.1145225 | 0.01499057 | 0.38514224 | -0.1010381 | 0.02976592 |
| 0.999490048 | 0.1992987 | 0.01505066 | 0.38550006 | 0.13012837 | 0.10072004 |
| 0.999490048 | 0.41262138 | 0.01516667 | 0.38728346 | 0.24250018 | 0.13806766 |
| 0.999490048 | 0.12180959 | 0.01528139 | 0.38902317 | 0.05603686 | 0.24228076 |
| 0.999490048 | -0.5462639 | 0.01565307 | 0.38929204 | -0.2639286 | 0.22188325 |
| 0.999490048 | -0.3147264 | 0.0155534 | 0.38929204 | -0.1910542 | 0.12764526 |
| 0.999490048 | -0.2916377 | 0.01538671 | 0.38929204 | -0.2379221 | 0.03946797 |
| 0.999490048 | -0.2445597 | 0.01536976 | 0.38929204 | -0.116315 | 0.2275768 |

| | | | | | |
|---|---|---|---|---|---|
| 0.999490048 | -0.1944977 | 0.01568969 | 0.38929204 | -0.1464692 | 0.06221586 |
| 0.999490048 | -0.1418215 | 0.01571155 | 0.38929204 | -0.0980391 | 0.08533486 |
| 0.972489992 | 0.14688034 | 0.01545 | 0.38929204 | 0.01405269 | 0.80570018 |
| 0.999490048 | 0.20498609 | 0.01564796 | 0.38929204 | 0.15883212 | 0.05540004 |
| 0.999490048 | 0.24177689 | 0.01551272 | 0.38929204 | 0.22877644 | 0.02044053 |
| 0.999490048 | -0.1917191 | 0.01587794 | 0.38961537 | -0.0777063 | 0.30449622 |
| 0.999490048 | -0.18584 | 0.01585089 | 0.38961537 | -0.058812 | 0.42064827 |
| 0.999490048 | -0.1672903 | 0.01588278 | 0.38961537 | -0.0694363 | 0.29328056 |
| 0.999490048 | 0.15804904 | 0.01591124 | 0.38961537 | 0.19650919 | 0.00357033 |
| 0.999490048 | -0.2493755 | 0.01630426 | 0.39027457 | -0.0711155 | 0.46960743 |
| 0.999490048 | -0.2073806 | 0.01605578 | 0.39027457 | -0.070259 | 0.39009509 |
| 0.999490048 | -0.1520089 | 0.01611242 | 0.39027457 | -0.092091 | 0.13057007 |
| 0.999490048 | -0.1452709 | 0.01626538 | 0.39027457 | -0.1018882 | 0.08282612 |
| 0.999490048 | 0.12182683 | 0.01633505 | 0.39027457 | 0.03220239 | 0.50251139 |
| 0.999490048 | 0.13327905 | 0.01639022 | 0.39027457 | 0.13943893 | 0.01247685 |
| 0.999490048 | 0.15909011 | 0.0162321 | 0.39027457 | 0.07283295 | 0.24920753 |
| 0.999490048 | 0.18949146 | 0.01640555 | 0.39027457 | 0.0995692 | 0.18859597 |
| 0.999490048 | 0.20611707 | 0.01627687 | 0.39027457 | 0.08787734 | 0.28260821 |
| 0.57925586 | 0.23645093 | 0.01603604 | 0.39027457 | -0.0431984 | 0.64120516 |
| 0.999490048 | -0.1448146 | 0.01658643 | 0.39123361 | -0.0339338 | 0.55286914 |
| 0.999490048 | -0.1212157 | 0.01654181 | 0.39123361 | -0.1065527 | 0.03290269 |
| 0.999490048 | 0.2185048 | 0.01649895 | 0.39123361 | 0.11263608 | 0.19725318 |
| 0.999490048 | 0.23881231 | 0.01669148 | 0.39149953 | 0.14001964 | 0.14517029 |
| 0.397650117 | 0.36017726 | 0.01667826 | 0.39149953 | -0.1978972 | 0.13697438 |
| 0.999490048 | -0.1364633 | 0.01680473 | 0.39195379 | -0.0332283 | 0.53845919 |
| 0.999490048 | 0.08890107 | 0.01679863 | 0.39195379 | 0.02138163 | 0.54338904 |
| 0.999490048 | -0.1318388 | 0.01688272 | 0.3919716 | -0.1043656 | 0.05350394 |
| 0.999490048 | 0.21300751 | 0.01689937 | 0.3919716 | 0.18390991 | 0.03648531 |
| 0.999490048 | 0.51766061 | 0.0170707 | 0.39484857 | 0.39154501 | 0.05743524 |
| 0.999490048 | 0.11634686 | 0.01713036 | 0.39513408 | 0.11404406 | 0.01920051 |
| 0.999490048 | 0.15711029 | 0.01718572 | 0.39531885 | 0.10776454 | 0.09230227 |
| 0.999490048 | -0.2004784 | 0.01737101 | 0.39779013 | -0.1847924 | 0.02707502 |
| 0.999490048 | -0.1398286 | 0.01738843 | 0.39779013 | -0.1211787 | 0.03669163 |
| 0.999490048 | -0.2498603 | 0.01758756 | 0.39984838 | -0.1619659 | 0.11194119 |
| 0.999490048 | -0.1467442 | 0.01762206 | 0.39984838 | -0.1560656 | 0.01217607 |
| 0.999490048 | 0.18844396 | 0.0176021 | 0.39984838 | 0.09981225 | 0.19037439 |
| 0.999490048 | -0.0873012 | 0.01774468 | 0.40153954 | -0.0513143 | 0.14830076 |
| 0.999490048 | -0.1164739 | 0.01786192 | 0.40294878 | -0.0895314 | 0.06264479 |
| 0.999490048 | 0.28116883 | 0.01790347 | 0.40294878 | 0.13007495 | 0.25218672 |
| 0.781057588 | -0.5881773 | 0.01842475 | 0.4102578 | 0.05191616 | 0.81208104 |
| 0.999490048 | -0.2496245 | 0.01837015 | 0.4102578 | -0.0854124 | 0.39633458 |
| 0.999490048 | -0.093152 | 0.01840746 | 0.4102578 | -0.0694973 | 0.07162025 |

| | | | | | |
|---|---|---|---|---|---|
| 0.999490048 | 0.16285915 | 0.01828447 | 0.4102578 | 0.09500132 | 0.15333272 |
| 0.999490048 | 0.26564858 | 0.01857728 | 0.41255406 | 0.14982116 | 0.16802179 |
| 0.999490048 | -0.1597299 | 0.01862773 | 0.41257707 | -0.0857797 | 0.18871118 |
| 0.961663458 | 0.28859559 | 0.01886127 | 0.41664448 | 0.00872053 | 0.94013631 |
| 0.355231519 | -0.3009952 | 0.0194021 | 0.41882304 | -0.8079073 | 4.79E-05 |
| 0.999490048 | -0.2799146 | 0.01951576 | 0.41882304 | -0.2587677 | 0.02959679 |
| 0.999490048 | -0.2177902 | 0.01943443 | 0.41882304 | -0.2308688 | 0.01382001 |
| 0.999490048 | -0.1319294 | 0.0191079 | 0.41882304 | -0.100637 | 0.06754509 |
| 0.999490048 | -0.1308751 | 0.01946107 | 0.41882304 | -0.0376493 | 0.47929634 |
| 0.999490048 | -0.1237988 | 0.01952863 | 0.41882304 | -0.1007438 | 0.0530435 |
| 0.999490048 | 0.11289614 | 0.0191191 | 0.41882304 | 0.06514948 | 0.16077201 |
| 0.972489992 | 0.13415152 | 0.01956179 | 0.41882304 | 0.01031822 | 0.84941704 |
| 0.999490048 | 0.16808148 | 0.01915157 | 0.41882304 | 0.12060547 | 0.08446669 |
| 0.999490048 | 0.17436901 | 0.01931223 | 0.41882304 | 0.02840615 | 0.68728378 |
| 0.999490048 | 0.22048317 | 0.01921413 | 0.41882304 | 0.18275241 | 0.04848492 |
| 0.999490048 | 0.30466612 | 0.01924002 | 0.41882304 | 0.08506478 | 0.43965637 |
| 0.999490048 | -0.9280608 | 0.02016429 | 0.41939083 | -0.9891637 | 0.01391767 |
| 0.999490048 | -0.481983 | 0.02001334 | 0.41939083 | -0.259731 | 0.17150618 |
| 0.999490048 | -0.3047972 | 0.02031275 | 0.41939083 | -0.3736361 | 0.00550617 |
| 0.999490048 | -0.1657765 | 0.02024349 | 0.41939083 | -0.1425951 | 0.04294426 |
| 0.889255854 | -0.1556123 | 0.0200503 | 0.41939083 | 0.00400864 | 0.94950428 |
| 0.999490048 | -0.1445442 | 0.02004192 | 0.41939083 | -0.0723812 | 0.22515714 |
| 0.999490048 | -0.1202518 | 0.01989089 | 0.41939083 | -0.0998789 | 0.04934684 |
| 0.999490048 | -0.1138018 | 0.02029022 | 0.41939083 | -0.0516694 | 0.27122773 |
| 0.972489992 | -0.1125178 | 0.02040192 | 0.41939083 | -0.0050559 | 0.91232092 |
| 0.999490048 | 0.1175977 | 0.02055602 | 0.41939083 | 0.026688 | 0.57977188 |
| 0.999490048 | 0.13441711 | 0.01988706 | 0.41939083 | 0.1371529 | 0.01774469 |
| 0.999490048 | 0.14847255 | 0.01969289 | 0.41939083 | 0.15364368 | 0.01618605 |
| 0.999490048 | 0.17370008 | 0.02058563 | 0.41939083 | 0.06063449 | 0.3964757 |
| 0.999490048 | 0.22244028 | 0.01984032 | 0.41939083 | 0.14230328 | 0.12399367 |
| 0.999490048 | 0.2287675 | 0.02011774 | 0.41939083 | 0.10723716 | 0.22998088 |
| 0.972489992 | 0.22976885 | 0.02059284 | 0.41939083 | 0.01161433 | 0.90158786 |
| 0.999490048 | 0.27181415 | 0.01974396 | 0.41939083 | 0.13788478 | 0.21815105 |
| 0.999490048 | 0.33444427 | 0.02057965 | 0.41939083 | 0.10378471 | 0.39927306 |
| 0.999490048 | 0.45674734 | 0.0204433 | 0.41939083 | 0.25708501 | 0.14416726 |
| 0.999490048 | 0.75666179 | 0.02025243 | 0.41939083 | 0.4219465 | 0.14670573 |
| 0.999490048 | 0.10865991 | 0.02111793 | 0.42799686 | 0.06125571 | 0.17770256 |
| 0.999490048 | 0.86537352 | 0.02107893 | 0.42799686 | 0.39497238 | 0.22882055 |
| 0.987721587 | -0.1569362 | 0.02125989 | 0.42923914 | -0.0150995 | 0.81504854 |
| 0.999490048 | 0.21317385 | 0.02128204 | 0.42923914 | 0.09254811 | 0.2963647 |
| 0.999490048 | -0.106599 | 0.02157118 | 0.4330451 | -0.0880527 | 0.05371738 |
| 0.999490048 | 0.16036989 | 0.02167819 | 0.4330451 | 0.13322894 | 0.05265326 |

| | | | | | |
|---|---|---|---|---|---|
| 0.999490048 | 0.16231454 | 0.02162766 | 0.4330451 | 0.15007155 | 0.03242232 |
| 0.578467169 | 0.8190903 | 0.02164787 | 0.4330451 | -0.2744992 | 0.39009939 |
| 0.999490048 | 0.17391813 | 0.02175959 | 0.43363386 | 0.04865169 | 0.47131855 |
| 0.999490048 | 0.12205969 | 0.02187697 | 0.43379404 | 0.14809772 | 0.00653761 |
| 0.999490048 | 0.19564809 | 0.02197543 | 0.43379404 | 0.04109717 | 0.61245748 |
| 0.999490048 | 0.3389181 | 0.02192986 | 0.43379404 | 0.3468144 | 0.01961925 |
| 0.999490048 | 0.56720957 | 0.02185591 | 0.43379404 | 0.28110867 | 0.21337783 |
| 0.999490048 | -0.3188765 | 0.02206076 | 0.43445137 | -0.2186121 | 0.08849988 |
| 0.999490048 | -0.0999158 | 0.02215352 | 0.43525151 | -0.0572468 | 0.17460264 |
| 0.999490048 | -0.2115603 | 0.02261722 | 0.44021853 | -0.1404667 | 0.11927509 |
| 0.999490048 | -0.1379786 | 0.02253521 | 0.44021853 | -0.0196983 | 0.73146869 |
| 0.999490048 | 0.20899148 | 0.02252139 | 0.44021853 | 0.11185659 | 0.20502137 |
| 0.4186768 | 0.21425468 | 0.02258069 | 0.44021853 | -0.1116676 | 0.19510692 |
| 0.999490048 | 0.1369511 | 0.02274666 | 0.44075744 | 0.18411659 | 0.00317474 |
| 0.999490048 | 0.36298127 | 0.02275047 | 0.44075744 | 0.14955611 | 0.28111576 |
| 0.999490048 | -0.0979303 | 0.022809 | 0.44086835 | -0.0420523 | 0.30758821 |
| 0.999490048 | 0.16017511 | 0.0228737 | 0.44109791 | 0.0895803 | 0.18726619 |
| 0.999490048 | -0.2121482 | 0.02347859 | 0.44354344 | -0.1979771 | 0.0333677 |
| 0.999490048 | -0.2017569 | 0.02332947 | 0.44354344 | -0.115245 | 0.17988601 |
| 0.999490048 | -0.1668382 | 0.02342632 | 0.44354344 | -0.0399101 | 0.56888829 |
| 0.999490048 | -0.1281158 | 0.02345066 | 0.44354344 | -0.1416806 | 0.01315841 |
| 0.999490048 | -0.0980185 | 0.02325517 | 0.44354344 | -0.1334451 | 0.00296433 |
| 0.999490048 | -0.0911635 | 0.02345906 | 0.44354344 | -0.0462371 | 0.23243605 |
| 0.999490048 | 0.15514717 | 0.02319265 | 0.44354344 | 0.08000695 | 0.22396731 |
| 0.999490048 | 0.22637184 | 0.02319653 | 0.44354344 | 0.19456322 | 0.04807757 |
| 0.999490048 | 0.25153742 | 0.02307741 | 0.44354344 | 0.13480976 | 0.20641838 |
| 0.999490048 | -0.1961371 | 0.02374347 | 0.44372834 | -0.0823572 | 0.32176803 |
| 0.999490048 | -0.1501218 | 0.02377903 | 0.44372834 | -0.1255319 | 0.05502148 |
| 0.999490048 | -0.1241649 | 0.0235431 | 0.44372834 | -0.0878245 | 0.10046231 |
| 0.999490048 | -0.0949764 | 0.02367752 | 0.44372834 | -0.0480577 | 0.23429513 |
| 0.081295872 | 0.25328689 | 0.02380722 | 0.44372834 | -0.2199677 | 0.04694635 |
| 0.999490048 | 0.77198643 | 0.02380634 | 0.44372834 | 0.60819978 | 0.05988753 |
| 0.971241857 | 0.18492938 | 0.02398433 | 0.44603369 | 0.36850842 | 6.38E-05 |
| 0.999490048 | -0.3883533 | 0.02407694 | 0.44676092 | -0.3378095 | 0.04406353 |
| 0.999490048 | -0.1529546 | 0.0241327 | 0.44680282 | -0.0255979 | 0.6913588 |
| 0.999490048 | 0.26154695 | 0.02431854 | 0.44924745 | 0.29969872 | 0.01098916 |
| 0.999490048 | 0.17566627 | 0.0243848 | 0.44947701 | 0.1132314 | 0.13512766 |
| 0.999490048 | 0.15054774 | 0.02455373 | 0.45060143 | 0.19239621 | 0.00524491 |
| 0.999490048 | 0.19334047 | 0.02454621 | 0.45060143 | 0.15496113 | 0.06663919 |
| 0.999490048 | -0.2087239 | 0.02463521 | 0.45110525 | -0.1741628 | 0.05281078 |
| 0.999490048 | 0.18402542 | 0.02483691 | 0.45281257 | 0.15147643 | 0.06060622 |
| 0.999490048 | 0.60492464 | 0.02481656 | 0.45281257 | 0.49897186 | 0.05276546 |

| | | | | | |
|---|---|---|---|---|---|
| 0.999490048 | 0.24337642 | 0.0249017 | 0.4530047 | 0.15339872 | 0.14514311 |
| 0.972489992 | -0.1384963 | 0.02499724 | 0.45375427 | -0.0021435 | 0.97086091 |
| 0.999490048 | 0.19479632 | 0.02527293 | 0.45776356 | 0.14525581 | 0.08820222 |
| 0.972489992 | -0.3817763 | 0.0254877 | 0.45965943 | -0.0114521 | 0.94376825 |
| 0.999490048 | -0.1317375 | 0.02545249 | 0.45965943 | -0.1523167 | 0.01096671 |
| 0.995056951 | 0.22944712 | 0.02564916 | 0.46157429 | 0.02578917 | 0.79193671 |
| 0.999490048 | -0.8318899 | 0.02651194 | 0.46216069 | -0.908377 | 0.0163366 |
| 0.999490048 | -0.4302199 | 0.02690228 | 0.46216069 | -0.2443694 | 0.19382771 |
| 0.999490048 | -0.2564108 | 0.02710677 | 0.46216069 | -0.1685844 | 0.13531486 |
| 0.999490048 | -0.253654 | 0.02736237 | 0.46216069 | -0.096176 | 0.38283388 |
| 0.999490048 | -0.2079349 | 0.02721493 | 0.46216069 | -0.0623235 | 0.48858447 |
| 0.999490048 | -0.2022917 | 0.02733113 | 0.46216069 | -0.0784252 | 0.37229235 |
| 0.999490048 | -0.1829527 | 0.02623126 | 0.46216069 | -0.1718622 | 0.03576023 |
| 0.999490048 | -0.1730089 | 0.02689006 | 0.46216069 | -0.1024323 | 0.17620628 |
| 0.999490048 | -0.1628897 | 0.02704986 | 0.46216069 | -0.1556126 | 0.0339516 |
| 0.999490048 | -0.1610069 | 0.02607465 | 0.46216069 | -0.1332495 | 0.06154359 |
| 0.987721587 | -0.1364079 | 0.02657093 | 0.46216069 | -0.0131035 | 0.82277937 |
| 0.999490048 | -0.1343429 | 0.0265345 | 0.46216069 | -0.0589616 | 0.3108369 |
| 0.999490048 | -0.1151373 | 0.02739665 | 0.46216069 | -0.015956 | 0.74819326 |
| 0.972489992 | -0.1014312 | 0.02760063 | 0.46216069 | 8.21E-05 | 0.9985052 |
| 0.999490048 | -0.0894902 | 0.02629967 | 0.46216069 | -0.0927065 | 0.02180222 |
| 0.999490048 | -0.0889887 | 0.02732036 | 0.46216069 | -0.095497 | 0.01867346 |
| 0.999490048 | 0.10927921 | 0.02734908 | 0.46216069 | 0.09031586 | 0.06415131 |
| 0.999490048 | 0.11228661 | 0.02687035 | 0.46216069 | 0.03890564 | 0.42297372 |
| 0.999490048 | 0.11653077 | 0.02743853 | 0.46216069 | 0.08192196 | 0.11221251 |
| 0.999490048 | 0.12501371 | 0.02746644 | 0.46216069 | 0.10875097 | 0.05237693 |
| 0.999490048 | 0.12718525 | 0.02600491 | 0.46216069 | 0.01640516 | 0.76275439 |
| 0.999490048 | 0.1277034 | 0.025787 | 0.46216069 | 0.07353593 | 0.1844872 |
| 0.999490048 | 0.12956767 | 0.02761894 | 0.46216069 | 0.10603979 | 0.06710486 |
| 0.972489992 | 0.13012045 | 0.02721282 | 0.46216069 | -6.74E-04 | 0.99039919 |
| 0.999490048 | 0.13901214 | 0.02631174 | 0.46216069 | 0.13855106 | 0.02676718 |
| 0.999490048 | 0.15519697 | 0.02630508 | 0.46216069 | 0.044949 | 0.50042014 |
| 0.999490048 | 0.19119783 | 0.02708676 | 0.46216069 | 0.17898664 | 0.03742578 |
| 0.999490048 | 0.19245192 | 0.02760981 | 0.46216069 | 0.08880651 | 0.29078814 |
| 0.999490048 | 0.20138805 | 0.0273297 | 0.46216069 | 0.15369948 | 0.08578579 |
| 0.999490048 | 0.22329827 | 0.02666039 | 0.46216069 | 0.09372363 | 0.33245099 |
| 0.999490048 | 0.26873838 | 0.02607168 | 0.46216069 | 0.18374282 | 0.11840451 |
| 0.578467169 | 0.36053848 | 0.02717716 | 0.46216069 | -0.1136779 | 0.46638671 |
| 0.999490048 | 0.42878514 | 0.02617859 | 0.46216069 | 0.35952217 | 0.05449199 |
| 0.999490048 | 0.43888348 | 0.02724535 | 0.46216069 | 0.05221878 | 0.78255625 |
| 0.999490048 | 0.43982568 | 0.02754323 | 0.46216069 | 0.13286691 | 0.48620664 |
| 0.999490048 | -0.3599841 | 0.02768553 | 0.46234835 | -0.5339175 | 0.00312006 |

| | | | | | |
|---|---|---|---|---|---|
| 0.999490048 | -0.1035053 | 0.02789989 | 0.46499814 | -0.057294 | 0.20817637 |
| 0.999490048 | -0.3254029 | 0.0281034 | 0.4667526 | -0.2892561 | 0.04872013 |
| 0.194125657 | 0.13267818 | 0.02811695 | 0.4667526 | -0.0974341 | 0.09935873 |
| 0.881091405 | -0.1250243 | 0.0282993 | 0.46791908 | 0.01298419 | 0.81098223 |
| 0.999490048 | 0.22913616 | 0.02826892 | 0.46791908 | 0.15514366 | 0.12751883 |
| 0.984638128 | -0.1853516 | 0.02853878 | 0.46817059 | -0.0059499 | 0.94113828 |
| 0.999490048 | -0.1563467 | 0.02851689 | 0.46817059 | -0.1486841 | 0.03644675 |
| 0.999490048 | 0.14395755 | 0.02844108 | 0.46817059 | 0.13735481 | 0.03579441 |
| 0.999490048 | 0.39295181 | 0.02853539 | 0.46817059 | 0.34042973 | 0.05173876 |
| 0.999490048 | -0.2614414 | 0.02927706 | 0.46952258 | -0.2187094 | 0.0644754 |
| 0.999490048 | -0.1683616 | 0.02929658 | 0.46952258 | -0.0650507 | 0.38031219 |
| 0.999490048 | -0.1642321 | 0.02911307 | 0.46952258 | -0.0512461 | 0.47689338 |
| 0.999490048 | -0.1634652 | 0.02929314 | 0.46952258 | -0.1137458 | 0.1202509 |
| 0.999490048 | -0.1184335 | 0.02926416 | 0.46952258 | -0.1062225 | 0.0485268 |
| 0.999490048 | -0.1046265 | 0.02874005 | 0.46952258 | -0.0855549 | 0.06934643 |
| 0.999490048 | 0.08624454 | 0.02922724 | 0.46952258 | 0.03674781 | 0.33387453 |
| 0.999490048 | 0.125842 | 0.02894642 | 0.46952258 | 0.11872629 | 0.03835238 |
| 0.999490048 | 0.12685839 | 0.02945249 | 0.46952258 | 0.10608601 | 0.06486143 |
| 0.999490048 | 0.13552781 | 0.02935991 | 0.46952258 | 0.12385636 | 0.04485413 |
| 0.999490048 | 0.16580206 | 0.02890221 | 0.46952258 | 0.06472495 | 0.37421086 |
| 0.999490048 | 0.16960524 | 0.02891706 | 0.46952258 | 0.27346413 | 9.65E-04 |
| 0.999490048 | 0.20117711 | 0.02914484 | 0.46952258 | 0.16576867 | 0.06808343 |
| 0.999490048 | 0.22913774 | 0.02892827 | 0.46952258 | 0.1809358 | 0.07918269 |
| 0.923933853 | 0.33189287 | 0.02946465 | 0.46952258 | -0.0243571 | 0.86681503 |
| 0.999490048 | -0.178393 | 0.02959252 | 0.46981081 | -0.0653567 | 0.40603807 |
| 0.999490048 | 0.13780991 | 0.02959527 | 0.46981081 | 0.05454139 | 0.36996777 |
| 0.999490048 | 0.21555624 | 0.02970278 | 0.47062279 | 0.15415525 | 0.11171588 |
| 0.999490048 | -0.2652891 | 0.03002945 | 0.47081712 | -0.2891185 | 0.01902193 |
| 0.999490048 | -0.1950414 | 0.03020869 | 0.47081712 | -0.1080674 | 0.2147342 |
| 0.999490048 | -0.1453742 | 0.03029473 | 0.47081712 | -0.0592436 | 0.3583709 |
| 0.999490048 | -0.1070062 | 0.03047167 | 0.47081712 | -0.0757522 | 0.11698138 |
| 0.987721587 | -0.1003086 | 0.03029185 | 0.47081712 | -0.0073595 | 0.86757906 |
| 0.999490048 | -0.0989201 | 0.03053801 | 0.47081712 | -0.0695083 | 0.11974542 |
| 0.999490048 | -0.0918196 | 0.03029196 | 0.47081712 | -0.0451908 | 0.26915449 |
| 0.999490048 | -0.0909868 | 0.03056082 | 0.47081712 | -0.0135608 | 0.73550844 |
| 0.987721587 | 0.10368262 | 0.03047144 | 0.47081712 | 0.00593208 | 0.89666053 |
| 0.999490048 | 0.11318271 | 0.03046747 | 0.47081712 | 0.04197421 | 0.40310079 |
| 0.999490048 | 0.12941899 | 0.02992945 | 0.47081712 | 0.05558709 | 0.33232592 |
| 0.999490048 | 0.24017292 | 0.03017122 | 0.47081712 | 0.28536459 | 0.01140027 |
| 0.999490048 | 0.25612532 | 0.03019614 | 0.47081712 | 0.18937041 | 0.10174163 |
| 0.999490048 | 0.3214663 | 0.03036867 | 0.47081712 | 0.06095639 | 0.66718321 |
| 0.999490048 | 0.37819552 | 0.0302533 | 0.47081712 | 0.132167 | 0.42975834 |

| | | | | | |
|---|---|---|---|---|---|
| 0.965957333 | -0.5154433 | 0.03073875 | 0.47268611 | 0.03659098 | 0.85759859 |
| 0.999490048 | 0.22362454 | 0.0308313 | 0.47323786 | 0.12570228 | 0.21017165 |
| 0.999490048 | 0.28488368 | 0.03090469 | 0.47349392 | 0.19441089 | 0.13115331 |
| 0.999490048 | -0.6371949 | 0.03198147 | 0.47452066 | -0.957836 | 0.00353196 |
| 0.999490048 | -0.5818982 | 0.03160606 | 0.47452066 | -0.3791996 | 0.15041168 |
| 0.999490048 | -0.3480429 | 0.03204667 | 0.47452066 | -0.3368053 | 0.03748861 |
| 0.999490048 | -0.3231183 | 0.03190055 | 0.47452066 | -0.2126882 | 0.14723573 |
| 0.999490048 | -0.2892705 | 0.03205146 | 0.47452066 | -0.092262 | 0.47578681 |
| 0.999490048 | -0.2495016 | 0.03109216 | 0.47452066 | -0.045154 | 0.68287343 |
| 0.999490048 | -0.2356846 | 0.03147703 | 0.47452066 | -0.3327333 | 0.00355493 |
| 0.881091405 | -0.2242193 | 0.03201806 | 0.47452066 | 0.02907662 | 0.77086504 |
| 0.999490048 | -0.2227861 | 0.03164699 | 0.47452066 | -0.0783676 | 0.43092684 |
| 0.439893572 | -0.2221302 | 0.03150547 | 0.47452066 | 0.11094928 | 0.26598052 |
| 0.773280431 | -0.1906293 | 0.03127885 | 0.47452066 | 0.03710647 | 0.66082787 |
| 0.999490048 | 0.11498194 | 0.03173903 | 0.47452066 | 0.05288893 | 0.30545516 |
| 0.999490048 | 0.11945971 | 0.0317227 | 0.47452066 | 0.06066381 | 0.2589268 |
| 0.999490048 | 0.1210173 | 0.03157701 | 0.47452066 | 0.07429837 | 0.17439578 |
| 0.984638128 | 0.1416072 | 0.0310989 | 0.47452066 | 0.00198973 | 0.97465435 |
| 0.999490048 | 0.1627293 | 0.0317404 | 0.47452066 | 0.10598389 | 0.15097264 |
| 0.999490048 | 0.22524154 | 0.03184552 | 0.47452066 | 0.1308439 | 0.19864356 |
| 0.956358655 | 0.27466854 | 0.03195261 | 0.47452066 | -0.021154 | 0.86254296 |
| 0.999490048 | 0.60232795 | 0.03199583 | 0.47452066 | 0.39103804 | 0.15287163 |
| 0.595926661 | -0.5432559 | 0.03222334 | 0.47591856 | 0.20571286 | 0.37395335 |
| 0.999490048 | 0.47339681 | 0.03225987 | 0.47591856 | 0.35417097 | 0.10226058 |
| 0.999490048 | 0.2509695 | 0.03236663 | 0.47665145 | 0.24674237 | 0.03512789 |
| 0.999490048 | -0.2697654 | 0.03280638 | 0.47871946 | -0.1887158 | 0.12654636 |
| 0.999490048 | -0.2103948 | 0.03257292 | 0.47871946 | -0.1885457 | 0.05335378 |
| 0.999490048 | -0.1587045 | 0.03274539 | 0.47871946 | -0.0966359 | 0.18095058 |
| 0.515104687 | -0.1470813 | 0.03290838 | 0.47871946 | 0.06275194 | 0.34463745 |
| 0.999490048 | -0.1120844 | 0.032869 | 0.47871946 | -0.0340981 | 0.4984033 |
| 0.999490048 | -0.0964365 | 0.03262741 | 0.47871946 | -0.0512326 | 0.24077675 |
| 0.999490048 | 0.26291242 | 0.03288382 | 0.47871946 | 0.21132346 | 0.08150123 |
| 0.987721587 | -0.5317223 | 0.03307004 | 0.48023456 | -0.0197386 | 0.93394281 |
| 0.999490048 | -0.2201121 | 0.0331967 | 0.4804028 | -0.1568223 | 0.12087072 |
| 0.999490048 | 0.15709419 | 0.03315728 | 0.4804028 | 0.07141617 | 0.3152677 |
| 0.406717515 | 0.24627077 | 0.03338135 | 0.48223921 | -0.1343597 | 0.23080994 |
| 0.999490048 | 0.26504845 | 0.03349308 | 0.48301758 | 0.09596254 | 0.39861426 |
| 0.999490048 | 0.13342188 | 0.03365304 | 0.48448777 | 0.04752878 | 0.43092982 |
| 0.972489992 | -0.1590465 | 0.03371405 | 0.48453073 | 0.00419361 | 0.95323904 |
| 0.999490048 | 0.08700782 | 0.03390256 | 0.48640272 | 0.05203588 | 0.19164152 |
| 0.897183549 | -0.3292414 | 0.03426187 | 0.48647314 | 0.04796959 | 0.7318715 |
| 0.999490048 | -0.2918435 | 0.03428519 | 0.48647314 | -0.1773809 | 0.18574497 |

| | | | | | |
|---|---|---|---|---|---|
| 0.999490048 | -0.120448 | 0.03431529 | 0.48647314 | -0.0454179 | 0.40668413 |
| 0.999490048 | 0.1080734 | 0.03410748 | 0.48647314 | 0.09789366 | 0.05308903 |
| 0.999490048 | 0.16679904 | 0.03408402 | 0.48647314 | 0.04340159 | 0.56498071 |
| 0.999490048 | 0.17375497 | 0.03414741 | 0.48647314 | 0.17180743 | 0.03603057 |
| 0.999490048 | 0.29549604 | 0.03403343 | 0.48647314 | 0.23401148 | 0.08794204 |
| 0.999490048 | 0.11404532 | 0.034437 | 0.48737103 | 0.09403894 | 0.07699349 |
| 0.999490048 | -0.2019079 | 0.03462836 | 0.48833432 | -0.1779582 | 0.06009284 |
| 0.999490048 | -0.1262922 | 0.03467089 | 0.48833432 | -0.0769067 | 0.18592491 |
| 0.999490048 | 0.44820788 | 0.03468051 | 0.48833432 | 0.08892795 | 0.62871729 |
| 0.972489992 | -0.2853984 | 0.03478041 | 0.48877897 | 0.01387176 | 0.91448159 |
| 0.999490048 | 0.1443312 | 0.03482916 | 0.48877897 | 0.08884399 | 0.18176833 |
| 0.999490048 | -0.1188235 | 0.0349343 | 0.48943196 | -0.120223 | 0.03302139 |
| 0.999490048 | -0.2148815 | 0.03512063 | 0.48957808 | -0.1601753 | 0.10930555 |
| 0.999490048 | -0.139226 | 0.03511845 | 0.48957808 | -0.0454595 | 0.47381093 |
| 0.999490048 | -0.1298484 | 0.0350149 | 0.48957808 | -0.0648105 | 0.27651502 |
| 0.999490048 | -0.0844239 | 0.03529559 | 0.48988759 | -0.0322329 | 0.40365121 |
| 0.999490048 | 0.12357952 | 0.03531884 | 0.48988759 | 0.01428459 | 0.79916684 |
| 0.999490048 | 0.21118701 | 0.03524175 | 0.48988759 | 0.20833528 | 0.03757169 |
| 0.999490048 | -0.1451663 | 0.03558869 | 0.49281194 | -0.1365195 | 0.04701862 |
| 0.999490048 | 0.12501963 | 0.03573945 | 0.49326347 | 0.02429419 | 0.66999599 |
| 0.999490048 | 0.21095641 | 0.03570595 | 0.49326347 | 0.25876502 | 0.01160537 |
| 0.999490048 | -0.1138689 | 0.03580354 | 0.49333265 | -0.090069 | 0.09152165 |
| 0.999490048 | -0.7058141 | 0.03599874 | 0.49520512 | -0.6784053 | 0.04317737 |
| 0.999490048 | -0.1081128 | 0.03609135 | 0.49566251 | -0.0478868 | 0.3360199 |
| 0.666143899 | 0.11071081 | 0.03620275 | 0.49637598 | -0.036082 | 0.47742164 |
| 0.972489992 | -0.1927285 | 0.03632527 | 0.49723937 | 0.00563314 | 0.94897589 |
| 0.999490048 | -0.1251651 | 0.03639114 | 0.49732577 | -0.1008962 | 0.08684821 |
| 0.999490048 | -0.2830946 | 0.03783487 | 0.49747228 | -0.2200271 | 0.10074254 |
| 0.987721587 | -0.2302774 | 0.03802556 | 0.49747228 | 0.00149117 | 0.9887979 |
| 0.999490048 | -0.1687405 | 0.03691088 | 0.49747228 | -0.1469833 | 0.06631334 |
| 0.999490048 | -0.1262177 | 0.03794893 | 0.49747228 | -0.0254122 | 0.66291801 |
| 0.999490048 | -0.1218702 | 0.03659449 | 0.49747228 | -0.0633832 | 0.26165168 |
| 0.999490048 | -0.1073643 | 0.03723932 | 0.49747228 | -0.0576609 | 0.24851278 |
| 0.999490048 | -0.1008849 | 0.03716298 | 0.49747228 | -0.0391963 | 0.40061923 |
| 0.999490048 | -0.0977637 | 0.03764328 | 0.49747228 | -0.0803095 | 0.08342095 |
| 0.999490048 | -0.088087 | 0.03781429 | 0.49747228 | -0.0370012 | 0.36582519 |
| 0.999490048 | -0.0870241 | 0.03803557 | 0.49747228 | -0.0805598 | 0.05338381 |
| 0.999490048 | 0.08670768 | 0.03819797 | 0.49747228 | 0.02879711 | 0.47419436 |
| 0.999490048 | 0.08990562 | 0.03722362 | 0.49747228 | 0.10433623 | 0.01712564 |
| 0.999490048 | 0.11478229 | 0.03705368 | 0.49747228 | 0.0231058 | 0.66135835 |
| 0.999490048 | 0.13624783 | 0.03746318 | 0.49747228 | 0.0592408 | 0.34856936 |
| 0.999490048 | 0.1431723 | 0.03799037 | 0.49747228 | 0.14332768 | 0.03779923 |

| | | | | | |
|---|---|---|---|---|---|
| 0.999490048 | 0.15513163 | 0.03794419 | 0.49747228 | 0.08097493 | 0.26357152 |
| 0.999490048 | 0.15708728 | 0.03737185 | 0.49747228 | 0.09866844 | 0.17955283 |
| 0.999490048 | 0.1651709 | 0.0371209 | 0.49747228 | 0.08319433 | 0.27804898 |
| 0.266102584 | 0.16622583 | 0.03671398 | 0.49747228 | -0.1275018 | 0.10293369 |
| 0.999490048 | 0.19149518 | 0.0365956 | 0.49747228 | 0.1249453 | 0.1620404 |
| 0.999490048 | 0.19330891 | 0.03675058 | 0.49747228 | 0.15932935 | 0.08097651 |
| 0.999490048 | 0.19954484 | 0.03792741 | 0.49747228 | 0.22794691 | 0.02084468 |
| 0.773280431 | 0.21335762 | 0.03764524 | 0.49747228 | -0.0533145 | 0.58826002 |
| 0.999490048 | 0.21461459 | 0.03802795 | 0.49747228 | 0.14655385 | 0.1472675 |
| 0.999490048 | 0.21863603 | 0.03807631 | 0.49747228 | 0.16529719 | 0.11032899 |
| 0.999490048 | 0.24985384 | 0.03824877 | 0.49747228 | 0.08254765 | 0.45375789 |
| 0.999490048 | 0.26854956 | 0.03814654 | 0.49747228 | 0.0445939 | 0.71935352 |
| 0.999490048 | 0.31254839 | 0.03812869 | 0.49747228 | 0.10479335 | 0.46985816 |
| 0.999490048 | 0.41317096 | 0.03716418 | 0.49747228 | 0.07845808 | 0.6618652 |
| 0.891785516 | 0.51173243 | 0.03745955 | 0.49747228 | -0.0787611 | 0.73807293 |
| 0.89605147 | 0.57463481 | 0.03786455 | 0.49747228 | -0.1103677 | 0.64688687 |
| 0.999490048 | -0.173651 | 0.03833411 | 0.49780692 | -0.136764 | 0.09741154 |
| 0.999490048 | -0.3363192 | 0.03947194 | 0.49844986 | -0.4219468 | 0.01152867 |
| 0.999490048 | -0.2901601 | 0.04049971 | 0.49844986 | -0.3134441 | 0.0279209 |
| 0.972489992 | -0.2886027 | 0.03877235 | 0.49844986 | 0.02388718 | 0.85821396 |
| 0.999490048 | -0.2236817 | 0.03872177 | 0.49844986 | -0.3581306 | 0.00178926 |
| 0.999490048 | -0.2045765 | 0.03982582 | 0.49844986 | -0.2232722 | 0.02598931 |
| 0.999490048 | -0.1831502 | 0.03872985 | 0.49844986 | -0.1285089 | 0.13839198 |
| 0.999490048 | -0.165209 | 0.04111133 | 0.49844986 | -0.1049498 | 0.18352241 |
| 0.999490048 | -0.1642693 | 0.03959292 | 0.49844986 | -0.0652002 | 0.39704989 |
| 0.999490048 | -0.1409162 | 0.039015 | 0.49844986 | -0.1132482 | 0.09229091 |
| 0.999490048 | -0.1362509 | 0.04054872 | 0.49844986 | -0.0795437 | 0.21899048 |
| 0.999490048 | -0.1344473 | 0.04081326 | 0.49844986 | -0.0857984 | 0.18092364 |
| 0.999490048 | -0.1341949 | 0.04104653 | 0.49844986 | -0.1323912 | 0.04361715 |
| 0.999490048 | -0.1271686 | 0.04011096 | 0.49844986 | -0.1183805 | 0.05471384 |
| 0.999490048 | -0.1211935 | 0.03844352 | 0.49844986 | -0.0160938 | 0.774107 |
| 0.999490048 | -0.1037458 | 0.03949548 | 0.49844986 | -0.1041918 | 0.03872309 |
| 0.999490048 | -0.0981297 | 0.04133216 | 0.49844986 | -0.0533438 | 0.25338548 |
| 0.999490048 | -0.095916 | 0.04012091 | 0.49844986 | -0.0194312 | 0.66477144 |
| 0.999490048 | -0.0910797 | 0.04130178 | 0.49844986 | -0.0966526 | 0.03119455 |
| 0.999490048 | 0.08950501 | 0.03946367 | 0.49844986 | 0.09846313 | 0.02467335 |
| 0.999490048 | 0.11414156 | 0.04033343 | 0.49844986 | 0.09448174 | 0.08557135 |
| 0.999490048 | 0.11549018 | 0.03889855 | 0.49844986 | 0.02772777 | 0.60542623 |
| 0.999490048 | 0.11712277 | 0.04136273 | 0.49844986 | 0.10789847 | 0.05866671 |
| 0.999490048 | 0.12215495 | 0.03932204 | 0.49844986 | 0.00416478 | 0.94150021 |
| 0.999490048 | 0.1227863 | 0.04085157 | 0.49844986 | 0.07463677 | 0.20187162 |
| 0.999490048 | 0.13329655 | 0.0406006 | 0.49844986 | 0.03292388 | 0.59855471 |

| | | | | | |
|---|---|---|---|---|---|
| 0.999490048 | 0.14222933 | 0.04090783 | 0.49844986 | 0.13124626 | 0.05770583 |
| 0.999490048 | 0.14288455 | 0.04103623 | 0.49844986 | 0.10310331 | 0.13264961 |
| 0.999490048 | 0.14303647 | 0.04001482 | 0.49844986 | 0.11430251 | 0.09576659 |
| 0.987721587 | 0.1440461 | 0.04004836 | 0.49844986 | 0.00235015 | 0.97209694 |
| 0.999490048 | 0.1524205 | 0.03914163 | 0.49844986 | 0.1254292 | 0.08537127 |
| 0.999490048 | 0.15812628 | 0.03914785 | 0.49844986 | 0.09795332 | 0.18967184 |
| 0.999490048 | 0.1651211 | 0.04065858 | 0.49844986 | 0.1268734 | 0.10968919 |
| 0.999490048 | 0.16738797 | 0.0398517 | 0.49844986 | 0.16884106 | 0.03845165 |
| 0.999490048 | 0.1842261 | 0.03975886 | 0.49844986 | 0.13397649 | 0.12724384 |
| 0.999490048 | 0.1847367 | 0.04136835 | 0.49844986 | 0.17192472 | 0.05631593 |
| 0.999490048 | 0.1888456 | 0.03969479 | 0.49844986 | 0.1355921 | 0.1317927 |
| 0.826281516 | 0.19104897 | 0.04057052 | 0.49844986 | -0.0426272 | 0.63413753 |
| 0.999490048 | 0.19844329 | 0.03983281 | 0.49844986 | 0.1650438 | 0.08338288 |
| 0.987721587 | 0.20470953 | 0.04066626 | 0.49844986 | -2.52E-04 | 0.99790539 |
| 0.999490048 | 0.20535947 | 0.04083438 | 0.49844986 | 0.27858007 | 0.00720219 |
| 0.999490048 | 0.21433579 | 0.03944622 | 0.49844986 | 0.19023483 | 0.06519241 |
| 0.999490048 | 0.22419337 | 0.03990602 | 0.49844986 | 0.15181411 | 0.15465952 |
| 0.999490048 | 0.26395812 | 0.04115508 | 0.49844986 | 0.20829818 | 0.10174952 |
| 0.999490048 | 0.28863869 | 0.03874705 | 0.49844986 | 0.07020496 | 0.60043631 |
| 0.999490048 | 0.29148816 | 0.04038917 | 0.49844986 | 0.15846653 | 0.25100193 |
| 0.999490048 | 0.29171731 | 0.03979292 | 0.49844986 | 0.43327788 | 0.00351674 |
| 0.999490048 | 0.35085777 | 0.04057395 | 0.49844986 | 0.25344398 | 0.12211622 |
| 0.999490048 | 0.40299513 | 0.04054495 | 0.49844986 | 0.23104259 | 0.2106916 |
| 0.987721587 | 0.45648687 | 0.04128746 | 0.49844986 | -0.0043094 | 0.98231631 |
| 0.999490048 | 0.56811216 | 0.04134907 | 0.49844986 | 0.68243614 | 0.01600123 |
| 0.999490048 | -0.1322119 | 0.04146262 | 0.49886581 | -0.0460768 | 0.46086638 |
| 0.999490048 | 0.15013671 | 0.04157567 | 0.49950622 | 0.09225503 | 0.19884866 |
| 0.999490048 | -0.0923641 | 0.04178208 | 0.50081286 | -0.0830404 | 0.06513516 |
| 0.999490048 | 0.28473524 | 0.04180438 | 0.50081286 | 0.1468952 | 0.27894534 |
| 0.999490048 | -0.2472393 | 0.04210489 | 0.50082032 | -0.1172781 | 0.31937659 |
| 0.999490048 | 0.11797559 | 0.04209404 | 0.50082032 | 0.07255251 | 0.19968674 |
| 0.999490048 | 0.13365018 | 0.04203979 | 0.50082032 | 0.02923953 | 0.64333248 |
| 0.999490048 | 0.15048319 | 0.04194876 | 0.50082032 | 0.13595316 | 0.06413949 |
| 0.999490048 | 0.27893704 | 0.04189199 | 0.50082032 | 0.48982439 | 0.002748 |
| 0.999490048 | -0.2287398 | 0.04245986 | 0.50101105 | -0.0716695 | 0.50912253 |
| 0.972489992 | -0.209922 | 0.04244357 | 0.50101105 | 0.01878818 | 0.8412503 |
| 0.999490048 | -0.1258194 | 0.04245626 | 0.50101105 | -0.1373585 | 0.02790428 |
| 0.999490048 | -0.117255 | 0.04245628 | 0.50101105 | -0.1333541 | 0.02251492 |
| 0.999490048 | -0.1106027 | 0.0425694 | 0.50101105 | -0.1013517 | 0.0614595 |
| 0.987721587 | 0.08494018 | 0.04240136 | 0.50101105 | -0.0035382 | 0.92973348 |
| 0.999490048 | 0.0957384 | 0.04260094 | 0.50101105 | 0.05378082 | 0.24137013 |
| 0.999490048 | 0.18490506 | 0.04252051 | 0.50101105 | 0.17304511 | 0.05643454 |

| | | | | | |
|---|---|---|---|---|---|
| 0.999490048 | -0.2319981 | 0.04274439 | 0.50128598 | -0.0579818 | 0.59841914 |
| 0.987721587 | 0.1509018 | 0.04271987 | 0.50128598 | 4.19E-05 | 0.99953143 |
| 0.999490048 | 0.09895382 | 0.04285184 | 0.50184135 | 0.0224596 | 0.63245632 |
| 0.999490048 | 0.09731399 | 0.04307106 | 0.50188525 | 0.03813265 | 0.41161728 |
| 0.999490048 | 0.13254407 | 0.04294161 | 0.50188525 | 0.07472807 | 0.24051777 |
| 0.999490048 | 0.14838276 | 0.04302826 | 0.50188525 | 0.19980345 | 0.00821106 |
| 0.999490048 | 0.15134605 | 0.04309602 | 0.50188525 | 0.05738953 | 0.42674802 |
| 0.999490048 | -0.1137389 | 0.04326774 | 0.5022784 | -0.0224554 | 0.67787748 |
| 0.578467169 | 0.21315239 | 0.04324985 | 0.5022784 | -0.0942073 | 0.35573184 |
| 0.999490048 | 0.2375475 | 0.04331023 | 0.5022784 | 0.28962031 | 0.01909833 |
| 0.999490048 | -0.2125639 | 0.04346205 | 0.50264278 | -0.1151224 | 0.26029877 |
| 0.999490048 | 0.23475224 | 0.04345416 | 0.50264278 | 0.12328744 | 0.27463709 |
| 0.999490048 | 0.13938783 | 0.04361254 | 0.5031469 | 0.08019577 | 0.23286527 |
| 0.711558326 | 0.1472607 | 0.04368387 | 0.5031469 | -0.0507331 | 0.47109187 |
| 0.999490048 | 0.14763643 | 0.04376795 | 0.5031469 | 0.01177713 | 0.86686835 |
| 0.999490048 | 0.27584283 | 0.04380692 | 0.5031469 | 0.14913455 | 0.26193228 |
| 0.999490048 | 0.2770216 | 0.04373461 | 0.5031469 | 0.06508591 | 0.62218665 |
| 0.999490048 | 0.18580289 | 0.0439438 | 0.50402578 | 0.07835744 | 0.37951581 |
| 0.999490048 | -0.3421016 | 0.04413788 | 0.50513563 | -0.1897732 | 0.25080906 |
| 0.999490048 | -0.1478586 | 0.04428944 | 0.50513563 | -0.1169213 | 0.10645538 |
| 0.999490048 | -0.1021967 | 0.04425799 | 0.50513563 | -0.0785001 | 0.11627131 |
| 0.999490048 | -0.0982193 | 0.04434304 | 0.50513563 | -0.0689148 | 0.1499083 |
| 0.999490048 | 0.10469045 | 0.04429798 | 0.50513563 | 0.01951057 | 0.69647096 |
| 0.999490048 | 0.10123318 | 0.04444045 | 0.50555558 | 0.10357196 | 0.04010727 |
| 0.999490048 | -0.1019655 | 0.04480738 | 0.50892219 | -0.0819405 | 0.10202709 |
| 0.999490048 | 0.31305146 | 0.04485829 | 0.50892219 | 0.02574458 | 0.85035746 |
| 0.999490048 | -0.1687613 | 0.04502817 | 0.50946512 | -0.135155 | 0.10350722 |
| 0.999490048 | 0.13866008 | 0.04498873 | 0.50946512 | 0.07932753 | 0.2386375 |
| 0.999490048 | 0.10110384 | 0.04519892 | 0.51070502 | 0.05296159 | 0.28001985 |
| 0.999490048 | -0.2919777 | 0.04532056 | 0.5108587 | -0.1957753 | 0.17010502 |
| 0.999490048 | -0.1339974 | 0.04533734 | 0.5108587 | -0.0099337 | 0.87717018 |
| 0.999490048 | -0.1256968 | 0.04539607 | 0.5108587 | -0.0941583 | 0.12723564 |
| 0.999490048 | -0.090405 | 0.04553778 | 0.51176379 | -0.0970386 | 0.03284118 |
| 0.972489992 | -0.2536867 | 0.04599572 | 0.5121633 | 0.03093462 | 0.80004449 |
| 0.987721587 | -0.1623188 | 0.04565739 | 0.5121633 | 0.00594736 | 0.93919572 |
| 0.999490048 | -0.1119519 | 0.04594409 | 0.5121633 | -0.03239 | 0.54895403 |
| 0.999490048 | -0.1109229 | 0.04591471 | 0.5121633 | -0.0482014 | 0.37011935 |
| 0.999490048 | -0.105086 | 0.04579359 | 0.5121633 | -0.0445532 | 0.38141688 |
| 0.999490048 | -0.099569 | 0.04606403 | 0.5121633 | -0.037762 | 0.43354741 |
| 0.999490048 | 0.18438593 | 0.04573146 | 0.5121633 | 0.09641064 | 0.28213717 |
| 0.397650117 | 0.23525971 | 0.04600795 | 0.5121633 | -0.1612257 | 0.1626081 |
| 0.999490048 | -0.1114444 | 0.04616805 | 0.51263729 | -0.0217237 | 0.68610526 |

| | | | | | |
|---|---|---|---|---|---|
| 0.999490048 | 0.14472708 | 0.04625546 | 0.5127334 | 0.11162396 | 0.11835422 |
| 0.999490048 | 0.56823842 | 0.04629952 | 0.5127334 | 0.25131516 | 0.32890507 |
| 0.999490048 | 0.34139442 | 0.04658936 | 0.51525983 | 0.04296879 | 0.77535025 |
| 0.999490048 | -0.1143989 | 0.04681084 | 0.51634147 | -0.0326303 | 0.55625999 |
| 0.999490048 | 0.14881374 | 0.04677594 | 0.51634147 | 0.10542607 | 0.15090463 |
| 0.999490048 | 0.13719896 | 0.04720933 | 0.51936486 | 0.03960986 | 0.55227108 |
| 0.999490048 | 0.14130119 | 0.04718895 | 0.51936486 | 0.06978306 | 0.31251478 |
| 0.999490048 | -0.3175635 | 0.04745988 | 0.52006563 | -0.2537416 | 0.10238768 |
| 0.999490048 | 0.24955431 | 0.04739977 | 0.52006563 | 0.10994781 | 0.36707965 |
| 0.999490048 | 0.33808917 | 0.04741676 | 0.52006563 | 0.11189412 | 0.49655969 |
| 0.999490048 | 0.2390866 | 0.04755994 | 0.52047906 | 0.04094099 | 0.72427798 |
| 0.999490048 | 0.12654256 | 0.04769895 | 0.52131711 | 0.08135279 | 0.19276518 |
| 0.999490048 | 0.15805001 | 0.04777291 | 0.52144284 | 0.22763989 | 0.00828986 |
| 0.999490048 | -0.1148222 | 0.04786366 | 0.52175138 | -0.1295763 | 0.02707965 |
| 0.999490048 | 0.13477269 | 0.04799136 | 0.52246134 | 0.00618693 | 0.92472939 |
| 0.999490048 | -0.5069008 | 0.04967946 | 0.52275926 | -0.0347917 | 0.87823862 |
| 0.999490048 | -0.2786307 | 0.04999251 | 0.52275926 | -0.2595591 | 0.0665251 |
| 0.999490048 | -0.2711664 | 0.04881692 | 0.52275926 | -0.3483371 | 0.01341506 |
| 0.999490048 | -0.2087976 | 0.04894397 | 0.52275926 | -0.0705298 | 0.49097427 |
| 0.999490048 | -0.2054927 | 0.04956073 | 0.52275926 | -0.2082749 | 0.04677655 |
| 0.999490048 | -0.1674565 | 0.04985887 | 0.52275926 | -0.0583568 | 0.47937455 |
| 0.999490048 | -0.1529788 | 0.04934429 | 0.52275926 | -0.1321871 | 0.08639202 |
| 0.999490048 | -0.1477086 | 0.04970985 | 0.52275926 | -0.1359262 | 0.0693295 |
| 0.999490048 | -0.139474 | 0.04910255 | 0.52275926 | -0.0142836 | 0.83400281 |
| 0.999490048 | -0.118227 | 0.04977278 | 0.52275926 | -0.0906662 | 0.1264538 |
| 0.999490048 | -0.1002791 | 0.04842807 | 0.52275926 | -0.048933 | 0.32105897 |
| 0.999490048 | -0.1002198 | 0.04976602 | 0.52275926 | -0.0089101 | 0.85606699 |
| 0.999490048 | -0.0935283 | 0.04899968 | 0.52275926 | -0.0191604 | 0.67530869 |
| 0.999490048 | 0.09260957 | 0.04931558 | 0.52275926 | 0.03191731 | 0.48307429 |
| 0.999490048 | 0.09264918 | 0.04932058 | 0.52275926 | 0.08351441 | 0.07434221 |
| 0.999490048 | 0.09502192 | 0.04936153 | 0.52275926 | 0.01582118 | 0.73381448 |
| 0.999490048 | 0.09618137 | 0.04951597 | 0.52275926 | 0.0275663 | 0.55948964 |
| 0.999490048 | 0.11791407 | 0.04887433 | 0.52275926 | 0.09657876 | 0.10234249 |
| 0.999490048 | 0.11885219 | 0.04910034 | 0.52275926 | 0.03517611 | 0.54604268 |
| 0.999490048 | 0.12326673 | 0.04999857 | 0.52275926 | 0.20090238 | 0.0025177 |
| 0.999490048 | 0.13036981 | 0.04997328 | 0.52275926 | 0.08843867 | 0.17465132 |
| 0.999490048 | 0.15117378 | 0.04915964 | 0.52275926 | 0.07297015 | 0.32788077 |
| 0.999490048 | 0.15329733 | 0.04907952 | 0.52275926 | 0.18706714 | 0.01830633 |
| 0.999490048 | 0.18095414 | 0.05002211 | 0.52275926 | 0.230266 | 0.01470788 |
| 0.999490048 | 0.18628919 | 0.0498707 | 0.52275926 | 0.18603568 | 0.05015972 |
| 0.999490048 | 0.19540524 | 0.04997676 | 0.52275926 | 0.0881765 | 0.36161854 |
| 0.999490048 | 0.20901044 | 0.04879676 | 0.52275926 | 0.13445649 | 0.17416877 |

| | | | | | |
|---|---|---|---|---|---|
| 0.999490048 | 0.23499879 | 0.04941264 | 0.52275926 | 0.10086988 | 0.38388793 |
| 0.771925038 | 0.33278743 | 0.0481818 | 0.52275926 | -0.1483869 | 0.33013397 |
| 0.999490048 | 0.33943646 | 0.04842823 | 0.52275926 | 0.21050959 | 0.20998794 |
| 0.999490048 | 0.35057773 | 0.04923667 | 0.52275926 | 0.11458589 | 0.4717065 |
| 0.999490048 | 0.41345484 | 0.04998545 | 0.52275926 | 0.26799437 | 0.19394555 |
| 0.999490048 | 0.12062928 | 0.05021377 | 0.52410624 | 0.07049326 | 0.24056229 |
| 0.999490048 | -0.0923576 | 0.05044886 | 0.52459279 | -0.0257111 | 0.57241459 |
| 0.999490048 | 0.11728909 | 0.05041817 | 0.52459279 | 0.02571809 | 0.65615122 |
| 0.999490048 | 0.16291206 | 0.05036817 | 0.52459279 | 0.10991168 | 0.17770361 |
| 0.999490048 | 0.1233948 | 0.05061423 | 0.52535152 | 0.10109754 | 0.10494091 |
| 0.999490048 | 0.24160884 | 0.05064766 | 0.52535152 | 0.1535922 | 0.20373833 |
| 0.999490048 | -0.4444898 | 0.05104491 | 0.52756873 | -0.3179498 | 0.15514908 |
| 0.999490048 | -0.3229166 | 0.05136687 | 0.52756873 | -0.2077286 | 0.20002302 |
| 0.999490048 | -0.1471104 | 0.05134969 | 0.52756873 | -0.0927494 | 0.20880385 |
| 0.999490048 | 0.10620278 | 0.05130029 | 0.52756873 | 0.01992907 | 0.70418805 |
| 0.999490048 | 0.17921647 | 0.05130847 | 0.52756873 | 0.03880642 | 0.66147067 |
| 0.999490048 | 0.19292509 | 0.0511363 | 0.52756873 | 0.13908955 | 0.15221663 |
| 0.999490048 | 0.21311606 | 0.05123867 | 0.52756873 | 0.13447425 | 0.20821636 |
| 0.477170731 | 0.60040611 | 0.05109133 | 0.52756873 | -0.4512733 | 0.12838573 |
| 0.999490048 | -0.1084555 | 0.05144007 | 0.52767149 | -0.0859856 | 0.11739894 |
| 0.999490048 | -0.5412223 | 0.05163749 | 0.52777021 | -0.4216971 | 0.12387098 |
| 0.999490048 | -0.2145659 | 0.05164568 | 0.52777021 | -0.0733961 | 0.49101715 |
| 0.999490048 | -0.1166454 | 0.05170252 | 0.52777021 | -0.1022926 | 0.08528688 |
| 0.999490048 | 0.11724231 | 0.05159272 | 0.52777021 | 0.06239939 | 0.28703668 |
| 0.999490048 | 0.0948283 | 0.05192699 | 0.52941434 | 0.02501498 | 0.5951121 |
| 0.999490048 | -0.1262395 | 0.05287899 | 0.53092376 | -0.0768687 | 0.22746878 |
| 0.999490048 | -0.0991626 | 0.05254286 | 0.53092376 | -0.0467085 | 0.3469562 |
| 0.999490048 | -0.0931294 | 0.05266894 | 0.53092376 | -0.0255504 | 0.58181838 |
| 0.999490048 | -0.0891604 | 0.05280271 | 0.53092376 | -0.1085946 | 0.02039591 |
| 0.999490048 | -0.086092 | 0.05256549 | 0.53092376 | -0.0046383 | 0.91358361 |
| 0.999490048 | -0.0799274 | 0.05271201 | 0.53092376 | -0.0418884 | 0.29673594 |
| 0.999490048 | 0.11761051 | 0.05269737 | 0.53092376 | 0.00550393 | 0.92494314 |
| 0.999490048 | 0.12542129 | 0.05239282 | 0.53092376 | 0.1328253 | 0.04080955 |
| 0.999490048 | 0.1354983 | 0.05276332 | 0.53092376 | 0.09531979 | 0.16502431 |
| 0.999490048 | 0.13972698 | 0.05268573 | 0.53092376 | 0.0778144 | 0.26812531 |
| 0.999490048 | 0.20120317 | 0.05268524 | 0.53092376 | 0.18656452 | 0.07096207 |
| 0.999490048 | 0.22986142 | 0.05221355 | 0.53092376 | 0.21286463 | 0.07073155 |
| 0.999490048 | 0.48612381 | 0.05290162 | 0.53092376 | 0.29118764 | 0.23495448 |
| 0.999490048 | -0.3674339 | 0.05305724 | 0.53120858 | -0.3901283 | 0.04089286 |
| 0.999490048 | 0.39530167 | 0.05300023 | 0.53120858 | 0.14223428 | 0.47175777 |
| 0.999490048 | -0.5749482 | 0.05337634 | 0.53306389 | -0.2853981 | 0.30624178 |
| 0.999490048 | -0.142928 | 0.05340549 | 0.53306389 | -0.0695901 | 0.33311254 |

| | | | | | |
|---|---|---|---|---|---|
| 0.999490048 | 0.27622019 | 0.05343407 | 0.53306389 | 0.21270008 | 0.13112689 |
| 0.999490048 | 0.11092421 | 0.05351217 | 0.53320597 | 0.05656608 | 0.31138762 |
| 0.987721587 | -0.4100067 | 0.0539587 | 0.53348809 | 0.02915338 | 0.87675275 |
| 0.987721587 | -0.2258804 | 0.05421596 | 0.53348809 | 0.02326047 | 0.83693352 |
| 0.999490048 | -0.1996466 | 0.05404108 | 0.53348809 | -0.087798 | 0.38253163 |
| 0.999490048 | -0.1323814 | 0.05424328 | 0.53348809 | -0.187796 | 0.00831401 |
| 0.999490048 | -0.1213271 | 0.05395127 | 0.53348809 | -0.0057889 | 0.92391133 |
| 0.773280431 | -0.1093038 | 0.05410115 | 0.53348809 | 0.03878309 | 0.48002628 |
| 0.999490048 | -0.100888 | 0.05401096 | 0.53348809 | -0.0361731 | 0.47529525 |
| 0.999490048 | 0.10270554 | 0.05372551 | 0.53348809 | 0.07488147 | 0.15245594 |
| 0.999490048 | 0.12590204 | 0.05393727 | 0.53348809 | 0.06250689 | 0.32500055 |
| 0.999490048 | 0.15432984 | 0.05414723 | 0.53348809 | 0.05909044 | 0.44645357 |
| 0.999490048 | 0.32445593 | 0.05400045 | 0.53348809 | 0.50870967 | 0.00754395 |
| 0.999490048 | -0.2166571 | 0.0543326 | 0.53373786 | -0.1020295 | 0.35088554 |
| 0.999490048 | -0.1172571 | 0.05472299 | 0.53694124 | -0.1175968 | 0.05407459 |
| 0.999490048 | 0.09999142 | 0.05488446 | 0.53726287 | 0.08702342 | 0.09190085 |
| 0.987721587 | 0.18063576 | 0.05483598 | 0.53726287 | -0.0133765 | 0.88263574 |
| 0.999490048 | 0.10360103 | 0.05517723 | 0.53877737 | 0.14040606 | 0.0115286 |
| 0.999490048 | 0.18660968 | 0.05523275 | 0.53877737 | 0.18141461 | 0.06185883 |
| 0.999490048 | 0.43435529 | 0.05511426 | 0.53877737 | 0.12107998 | 0.56238605 |
| 0.999490048 | -0.2278908 | 0.05538953 | 0.53967626 | -0.0623006 | 0.58769913 |
| 0.999490048 | 0.24118831 | 0.05556478 | 0.54075278 | 0.21018028 | 0.08909733 |
| 0.999490048 | -0.2584672 | 0.05581254 | 0.54212802 | -0.1088615 | 0.40630947 |
| 0.999490048 | -0.1676171 | 0.05586071 | 0.54212802 | -0.078757 | 0.3551557 |
| 0.999490048 | 0.1225141 | 0.05590087 | 0.54212802 | 0.04994411 | 0.42151755 |
| 0.999490048 | -0.1526101 | 0.05669296 | 0.54473332 | -0.1548169 | 0.05345062 |
| 0.999490048 | -0.1301988 | 0.0563459 | 0.54473332 | -0.2087956 | 0.00362504 |
| 0.886410033 | -0.1170181 | 0.05675665 | 0.54473332 | 0.03279074 | 0.5807009 |
| 0.999490048 | -0.1065515 | 0.05651688 | 0.54473332 | -0.0966361 | 0.08177822 |
| 0.999490048 | -0.0668869 | 0.05631402 | 0.54473332 | -0.0099065 | 0.76939943 |
| 0.999490048 | 0.11680693 | 0.05665522 | 0.54473332 | 0.13537602 | 0.02902883 |
| 0.999490048 | 0.12587833 | 0.05657706 | 0.54473332 | 0.02114516 | 0.73986448 |
| 0.999490048 | 0.13785173 | 0.05636616 | 0.54473332 | 0.09166779 | 0.19538562 |
| 0.999490048 | 0.14673797 | 0.05666356 | 0.54473332 | 0.11956189 | 0.11603077 |
| 0.999490048 | 0.07789848 | 0.05711434 | 0.54702031 | 0.01278328 | 0.74621969 |
| 0.766519803 | 0.78479668 | 0.05712595 | 0.54702031 | -0.3228981 | 0.4197298 |
| 0.999490048 | -0.3416074 | 0.05750886 | 0.54735518 | -0.1756244 | 0.31588507 |
| 0.999490048 | -0.3291311 | 0.05748196 | 0.54735518 | -0.1820539 | 0.26586694 |
| 0.999490048 | -0.2430146 | 0.05788199 | 0.54735518 | -0.0384638 | 0.73582647 |
| 0.999490048 | -0.1262385 | 0.05787014 | 0.54735518 | -0.1022921 | 0.11978682 |
| 0.999490048 | -0.1035125 | 0.05740429 | 0.54735518 | -0.0706361 | 0.18623365 |
| 0.987721587 | -0.0819177 | 0.05738802 | 0.54735518 | 0.00577746 | 0.88941506 |

| | | | | | |
|---|---|---|---|---|---|
| 0.999490048 | 0.0913974 | 0.05748083 | 0.54735518 | 0.06209 | 0.1882827 |
| 0.999490048 | 0.1390962 | 0.05763348 | 0.54735518 | 0.13629293 | 0.06249048 |
| 0.999490048 | 0.18273307 | 0.05778442 | 0.54735518 | -0.0030402 | 0.97387376 |
| 0.999490048 | 0.21736042 | 0.05766823 | 0.54735518 | 0.08129076 | 0.44526028 |
| 0.999490048 | 0.2531028 | 0.05771371 | 0.54735518 | 0.17914037 | 0.17149008 |
| 0.999490048 | 0.09592603 | 0.05806176 | 0.54843406 | 0.06299802 | 0.203995 |
| 0.999490048 | 0.18801699 | 0.05822103 | 0.54931706 | 0.04798566 | 0.61693622 |
| 0.999490048 | -0.3423944 | 0.05861831 | 0.55094685 | -0.2934182 | 0.09574628 |
| 0.999490048 | -0.2546111 | 0.05862208 | 0.55094685 | -0.0506673 | 0.69673427 |
| 0.999490048 | -0.0689509 | 0.05865769 | 0.55094685 | -0.0658859 | 0.06996314 |
| 0.999490048 | 0.09840986 | 0.05859509 | 0.55094685 | 0.04974045 | 0.32615949 |
| 0.999490048 | -0.132504 | 0.05874183 | 0.55111716 | -0.0540119 | 0.42716528 |
| 0.999490048 | -0.0975746 | 0.05898589 | 0.55278581 | -0.0933335 | 0.070058 |
| 0.999490048 | -0.5440256 | 0.05919646 | 0.55351672 | -0.7108065 | 0.02066537 |
| 0.972489992 | -0.1962467 | 0.05917121 | 0.55351672 | 0.03276632 | 0.74413356 |
| 0.999490048 | -0.2566162 | 0.05941025 | 0.55427436 | -0.1086217 | 0.41116144 |
| 0.999490048 | -0.0968874 | 0.05938433 | 0.55427436 | -0.0311772 | 0.53088989 |
| 0.999490048 | 0.43169984 | 0.05948317 | 0.55433538 | 0.10911515 | 0.62218174 |
| 0.999490048 | 0.13786413 | 0.05956695 | 0.55449725 | 0.01226246 | 0.86208855 |
| 0.999490048 | -0.1351752 | 0.05970948 | 0.55510644 | -0.0231995 | 0.73783973 |
| 0.999490048 | -0.1002054 | 0.05976535 | 0.55510644 | -0.0108463 | 0.83273758 |
| 0.999490048 | 0.1296371 | 0.05987787 | 0.55553357 | 0.03520309 | 0.5972719 |
| 0.999490048 | -0.1073135 | 0.06001624 | 0.55619931 | -0.0465214 | 0.4012933 |
| 0.999490048 | 0.12663869 | 0.06012735 | 0.55661132 | 0.09999123 | 0.1326112 |
| 0.999490048 | -0.196201 | 0.06039311 | 0.5572596 | -0.0079189 | 0.93737191 |
| 0.999490048 | -0.1205207 | 0.06042158 | 0.5572596 | -0.0430384 | 0.48906356 |
| 0.999490048 | -0.0915973 | 0.06050966 | 0.5572596 | -0.0363427 | 0.44278897 |
| 0.999490048 | 0.08808208 | 0.06053107 | 0.5572596 | 0.00150926 | 0.97340361 |
| 0.999490048 | 0.10318084 | 0.06043424 | 0.5572596 | 0.07087959 | 0.18887665 |
| 0.999490048 | -0.0823432 | 0.06084973 | 0.5589607 | -0.0794024 | 0.06999288 |
| 0.999490048 | 0.10526789 | 0.06081326 | 0.5589607 | 0.05531637 | 0.31214229 |
| 0.999490048 | -0.1619625 | 0.06111769 | 0.55959459 | -0.1777598 | 0.0412232 |
| 0.487177805 | -0.1481812 | 0.06111979 | 0.55959459 | 0.10098293 | 0.1935238 |
| 0.999490048 | 0.11470858 | 0.06106859 | 0.55959459 | 0.10555513 | 0.08317119 |
| 0.999490048 | 0.49555429 | 0.06128092 | 0.56045525 | 0.44524975 | 0.08645255 |
| 0.999490048 | -0.2777542 | 0.06168337 | 0.56095018 | -0.2234194 | 0.12816595 |
| 0.999490048 | -0.1583107 | 0.06165106 | 0.56095018 | -0.0035262 | 0.96559108 |
| 0.999490048 | -0.1177064 | 0.06169241 | 0.56095018 | -0.0343139 | 0.57363762 |
| 0.999490048 | -0.0972556 | 0.06168107 | 0.56095018 | -0.0407782 | 0.41988308 |
| 0.999490048 | 0.12681259 | 0.06159093 | 0.56095018 | 0.0341362 | 0.60302727 |
| 0.999490048 | 0.44375982 | 0.06173811 | 0.56095018 | 0.0698469 | 0.7607876 |
| 0.999490048 | -0.1509315 | 0.06213284 | 0.56148186 | -0.1344076 | 0.09435727 |

| | | | | | |
|---|---|---|---|---|---|
| 0.999490048 | -0.1329881 | 0.06201682 | 0.56148186 | -0.0050411 | 0.94156697 |
| 0.999490048 | -0.1323794 | 0.06205803 | 0.56148186 | -0.1176015 | 0.09501703 |
| 0.999490048 | 0.10880317 | 0.06212394 | 0.56148186 | 0.0728197 | 0.20316369 |
| 0.999490048 | 0.12731155 | 0.06198442 | 0.56148186 | 0.09044391 | 0.17739488 |
| 0.999490048 | 0.92600728 | 0.06224735 | 0.56190852 | 0.56234918 | 0.22546011 |
| 0.999490048 | -0.1816435 | 0.06240782 | 0.56258246 | -0.1143822 | 0.2308387 |
| 0.999490048 | 0.17150505 | 0.06245676 | 0.56258246 | 0.17763911 | 0.05423051 |
| 0.999490048 | 0.15817954 | 0.06256919 | 0.56298786 | 0.05094003 | 0.5359421 |
| 0.999490048 | -0.1047608 | 0.0626872 | 0.56344251 | -0.1015874 | 0.07050774 |
| 0.999490048 | 0.10411507 | 0.06282032 | 0.56399872 | 0.09685486 | 0.08210781 |
| 0.999490048 | 0.15962278 | 0.06295171 | 0.56399872 | 0.12917097 | 0.12786911 |
| 0.999490048 | 0.18853097 | 0.06293959 | 0.56399872 | 0.20240593 | 0.04698711 |
| 0.999490048 | 0.10354214 | 0.0631499 | 0.56480462 | 0.05452043 | 0.31572914 |
| 0.999490048 | 0.18205907 | 0.06317695 | 0.56480462 | 0.24720699 | 0.01405569 |
| 0.999490048 | -0.1350212 | 0.0635913 | 0.56729418 | -0.1616979 | 0.02849171 |
| 0.999490048 | -0.0790241 | 0.0635726 | 0.56729418 | -0.0522174 | 0.21138391 |
| 0.999490048 | -0.1959757 | 0.06390358 | 0.56748614 | -0.0309426 | 0.76210416 |
| 0.999490048 | -0.0851736 | 0.06383737 | 0.56748614 | -0.0189601 | 0.66972965 |
| 0.999490048 | -0.0795461 | 0.0639072 | 0.56748614 | -0.0126858 | 0.75979438 |
| 0.999490048 | 0.10337742 | 0.06391259 | 0.56748614 | 0.08911944 | 0.10715801 |
| 0.999490048 | 0.17653751 | 0.06395263 | 0.56748614 | 0.21169792 | 0.02851036 |
| 0.999490048 | -0.112186 | 0.06412344 | 0.56839782 | -0.0620454 | 0.29434301 |
| 0.999490048 | 0.11563915 | 0.06420351 | 0.56850402 | 0.04797578 | 0.42941589 |
| 0.999490048 | -0.211946 | 0.06448228 | 0.56918978 | -4.16E-06 | 0.99996999 |
| 0.999490048 | -0.0966843 | 0.06455362 | 0.56918978 | -0.0770666 | 0.13583134 |
| 0.999490048 | 0.10487856 | 0.0644317 | 0.56918978 | 0.09736167 | 0.08460715 |
| 0.999490048 | 0.11939677 | 0.06453826 | 0.56918978 | 0.0862515 | 0.17474467 |
| 0.971241857 | -0.2431325 | 0.06463762 | 0.56932924 | 0.05664883 | 0.65637569 |
| 0.999490048 | 0.15670561 | 0.06479544 | 0.56951782 | 0.04815404 | 0.55816976 |
| 0.999490048 | 0.1597876 | 0.06477905 | 0.56951782 | 0.14267451 | 0.09688148 |
| 0.999490048 | -0.1429472 | 0.06499854 | 0.57022427 | -0.1421811 | 0.06635402 |
| 0.999490048 | 0.10675773 | 0.0650124 | 0.57022427 | 0.06335945 | 0.26290778 |
| 0.999490048 | -0.1837998 | 0.0651539 | 0.57086572 | -0.2061518 | 0.04024232 |
| 0.999490048 | -0.2591572 | 0.06543968 | 0.57157044 | -0.1458308 | 0.28866139 |
| 0.711558326 | -0.2394289 | 0.06540448 | 0.57157044 | 0.15355724 | 0.20882734 |
| 0.999490048 | -0.1158165 | 0.06539602 | 0.57157044 | -0.105525 | 0.09135843 |
| 0.999490048 | -0.2741184 | 0.06563857 | 0.57270849 | -0.040539 | 0.76829858 |
| 0.999490048 | -0.2635998 | 0.06572743 | 0.57288525 | -0.4196508 | 0.00516425 |
| 0.999490048 | -0.6052862 | 0.06601899 | 0.57407037 | -0.2040944 | 0.52282829 |
| 0.999490048 | -0.1265147 | 0.06614969 | 0.57407037 | -0.0907087 | 0.18056691 |
| 0.999490048 | -0.1224205 | 0.06626618 | 0.57407037 | -0.1293734 | 0.05316258 |
| 0.999490048 | -0.1197879 | 0.06594344 | 0.57407037 | -0.1601594 | 0.0164609 |

| | | | | | |
|---|---|---|---|---|---|
| 0.999490048 | 0.09541446 | 0.06618915 | 0.57407037 | 0.09141638 | 0.07758101 |
| 0.999490048 | 0.35618296 | 0.06627591 | 0.57407037 | 0.04997072 | 0.77362314 |
| 0.972489992 | 0.09726322 | 0.06666234 | 0.57681918 | -0.0217803 | 0.67136856 |
| 0.999490048 | -0.533779 | 0.06788922 | 0.57733003 | -0.3992402 | 0.15350705 |
| 0.999490048 | -0.3023384 | 0.06738726 | 0.57733003 | -0.0612934 | 0.70161285 |
| 0.999490048 | -0.2945217 | 0.06730645 | 0.57733003 | -0.0124416 | 0.93625702 |
| 0.999490048 | -0.2845252 | 0.06720538 | 0.57733003 | -0.1449046 | 0.33921033 |
| 0.999490048 | -0.217135 | 0.06753168 | 0.57733003 | -0.119849 | 0.30166479 |
| 0.999490048 | -0.1931446 | 0.06693897 | 0.57733003 | -0.1717245 | 0.10097648 |
| 0.999490048 | -0.1726952 | 0.06805251 | 0.57733003 | -0.0713812 | 0.43800444 |
| 0.999490048 | -0.1348967 | 0.06809191 | 0.57733003 | -0.1203818 | 0.1012313 |
| 0.999490048 | -0.1280472 | 0.06736905 | 0.57733003 | -0.0903468 | 0.18939729 |
| 0.999490048 | -0.1178747 | 0.06709884 | 0.57733003 | -0.035022 | 0.57466654 |
| 0.999490048 | -0.0958176 | 0.06713145 | 0.57733003 | -0.0714734 | 0.16577703 |
| 0.999490048 | -0.0917106 | 0.06741819 | 0.57733003 | -0.0190334 | 0.69494535 |
| 0.999490048 | 0.07219756 | 0.06768481 | 0.57733003 | 0.08149909 | 0.04089682 |
| 0.999490048 | 0.0968314 | 0.06802126 | 0.57733003 | 0.03484352 | 0.49887942 |
| 0.999490048 | 0.09846298 | 0.06741925 | 0.57733003 | 0.03904295 | 0.45554928 |
| 0.999490048 | 0.11488985 | 0.06803222 | 0.57733003 | 0.16914791 | 0.00951471 |
| 0.999490048 | 0.12117223 | 0.0681042 | 0.57733003 | 0.10026816 | 0.12719642 |
| 0.999490048 | 0.14097026 | 0.06682214 | 0.57733003 | 0.06979893 | 0.3518518 |
| 0.999490048 | 0.16169793 | 0.06764517 | 0.57733003 | 0.13309911 | 0.12846526 |
| 0.999490048 | 0.1689561 | 0.06774381 | 0.57733003 | 0.09175441 | 0.30974443 |
| 0.999490048 | -0.1114597 | 0.06860189 | 0.57867406 | -0.1111887 | 0.06923426 |
| 0.999490048 | 0.08793297 | 0.06862475 | 0.57867406 | 0.02274557 | 0.62691051 |
| 0.999490048 | 0.13549415 | 0.06867856 | 0.57867406 | 0.04175982 | 0.56294948 |
| 0.999490048 | 0.13613928 | 0.06866812 | 0.57867406 | 0.15413914 | 0.04104897 |
| 0.999490048 | 0.20817934 | 0.06859585 | 0.57867406 | -0.0137721 | 0.90083473 |
| 0.999490048 | 0.26190858 | 0.06844228 | 0.57867406 | 0.03349146 | 0.80963274 |
| 0.999490048 | 0.15484559 | 0.06875653 | 0.57874699 | 0.02323148 | 0.77776461 |
| 0.999490048 | 0.12843164 | 0.06888598 | 0.5792527 | 0.07199882 | 0.29679744 |
| 0.999490048 | -0.0964596 | 0.06906586 | 0.57984619 | -0.0564913 | 0.27653869 |
| 0.999490048 | 0.11218256 | 0.06909544 | 0.57984619 | 0.03497015 | 0.55915417 |
| 0.999490048 | -0.1509377 | 0.06933359 | 0.58036399 | -0.0638321 | 0.42985863 |
| 0.999490048 | 0.10396447 | 0.06926049 | 0.58036399 | 0.02946408 | 0.59540829 |
| 0.999490048 | 0.13548685 | 0.06936566 | 0.58036399 | 0.02378217 | 0.74190071 |
| 0.999490048 | -0.2850974 | 0.07418576 | 0.580377 | 0.00377328 | 0.9805232 |
| 0.999490048 | -0.2523516 | 0.07462769 | 0.580377 | -0.0410206 | 0.76487503 |
| 0.999490048 | -0.2408952 | 0.07059439 | 0.580377 | -0.3634389 | 0.00863095 |
| 0.999490048 | -0.228264 | 0.07077191 | 0.580377 | -0.1346847 | 0.27608847 |
| 0.999490048 | -0.2225468 | 0.06965022 | 0.580377 | -0.0909766 | 0.4457613 |
| 0.999490048 | -0.2090927 | 0.07023149 | 0.580377 | -0.0347482 | 0.75594463 |

| | | | | | |
|---|---|---|---|---|---|
| 0.988558788 | -0.1991133 | 0.07072491 | 0.580377 | 0.02443111 | 0.81875024 |
| 0.999490048 | -0.1851993 | 0.0732935 | 0.580377 | -0.0766774 | 0.44621676 |
| 0.999490048 | -0.1836982 | 0.07207232 | 0.580377 | -0.1912717 | 0.06177499 |
| 0.999490048 | -0.177686 | 0.07299054 | 0.580377 | -0.1077267 | 0.26728518 |
| 0.999490048 | -0.1547537 | 0.07146748 | 0.580377 | -0.0341802 | 0.68118007 |
| 0.999490048 | -0.1349732 | 0.07202594 | 0.580377 | -0.1350915 | 0.07179245 |
| 0.999490048 | -0.1286897 | 0.07329141 | 0.580377 | -0.0949439 | 0.18003177 |
| 0.999490048 | -0.127327 | 0.07354093 | 0.580377 | -0.0934829 | 0.18246835 |
| 0.999490048 | -0.1211062 | 0.07244615 | 0.580377 | -0.1113717 | 0.09692165 |
| 0.999490048 | -0.1208395 | 0.07153613 | 0.580377 | -0.1158486 | 0.08326518 |
| 0.999490048 | -0.1154475 | 0.07078157 | 0.580377 | -0.0576599 | 0.35496627 |
| 0.999490048 | -0.1136429 | 0.07365662 | 0.580377 | -0.0434494 | 0.48199677 |
| 0.999490048 | -0.1084427 | 0.07296248 | 0.580377 | -0.1192648 | 0.05007562 |
| 0.999490048 | -0.1025235 | 0.07318755 | 0.580377 | -0.0209475 | 0.70562966 |
| 0.999490048 | -0.0968052 | 0.07350379 | 0.580377 | -0.0512668 | 0.33203246 |
| 0.999490048 | -0.0948129 | 0.07264711 | 0.580377 | -0.0737226 | 0.15752635 |
| 0.999490048 | -0.0928112 | 0.07214808 | 0.580377 | -0.0157244 | 0.75308424 |
| 0.999490048 | -0.0924639 | 0.07434231 | 0.580377 | -0.0638219 | 0.21044605 |
| 0.999490048 | -0.089119 | 0.07379665 | 0.580377 | -0.0622232 | 0.20450903 |
| 0.999490048 | -0.0841686 | 0.0737887 | 0.580377 | -0.0613987 | 0.18558803 |
| 0.999490048 | -0.0803663 | 0.07304573 | 0.580377 | -0.0522821 | 0.2348684 |
| 0.999490048 | -0.0763588 | 0.07304509 | 0.580377 | -0.0504647 | 0.22771918 |
| 0.999490048 | -0.0701026 | 0.07086244 | 0.580377 | -0.0482441 | 0.20603087 |
| 0.999490048 | 0.07797267 | 0.07028449 | 0.580377 | 0.03747298 | 0.37223363 |
| 0.999490048 | 0.08157958 | 0.07047148 | 0.580377 | 0.05031822 | 0.25488498 |
| 0.999490048 | 0.08191839 | 0.07217792 | 0.580377 | 0.04945767 | 0.26782355 |
| 0.999490048 | 0.08463349 | 0.07393482 | 0.580377 | 0.05257432 | 0.25759823 |
| 0.999490048 | 0.08523543 | 0.07430017 | 0.580377 | 0.05736222 | 0.2216458 |
| 0.999490048 | 0.09069916 | 0.07448142 | 0.580377 | 0.02050881 | 0.67747363 |
| 0.999490048 | 0.09159972 | 0.07355469 | 0.580377 | 0.07956293 | 0.1172854 |
| 0.999490048 | 0.09475894 | 0.07395423 | 0.580377 | 0.0381857 | 0.45939367 |
| 0.999490048 | 0.0964362 | 0.0728465 | 0.580377 | 0.02008134 | 0.69991491 |
| 0.999490048 | 0.09992176 | 0.07464969 | 0.580377 | 0.04372071 | 0.42341798 |
| 0.999490048 | 0.10375708 | 0.07080212 | 0.580377 | 0.04462817 | 0.42468941 |
| 0.999490048 | 0.10567828 | 0.07328654 | 0.580377 | 0.07799057 | 0.17989573 |
| 0.999490048 | 0.10725477 | 0.07296172 | 0.580377 | 0.0489001 | 0.40157781 |
| 0.999490048 | 0.1078933 | 0.07101996 | 0.580377 | 0.07818613 | 0.18400687 |
| 0.999490048 | 0.11001467 | 0.07292921 | 0.580377 | 0.08929433 | 0.14119561 |
| 0.999490048 | 0.11192705 | 0.07307596 | 0.580377 | 0.13687627 | 0.03083715 |
| 0.999490048 | 0.11364115 | 0.07020423 | 0.580377 | 0.04631131 | 0.44811401 |
| 0.999490048 | 0.11403869 | 0.07054824 | 0.580377 | -6.51E-04 | 0.99147814 |
| 0.999490048 | 0.12044103 | 0.07399297 | 0.580377 | 0.09145803 | 0.16913409 |

| | | | | | |
|---|---|---|---|---|---|
| 0.999490048 | 0.12285823 | 0.07012836 | 0.580377 | 0.10106078 | 0.13222316 |
| 0.999490048 | 0.12375883 | 0.07221358 | 0.580377 | 0.09267349 | 0.17216508 |
| 0.999490048 | 0.12723069 | 0.07350335 | 0.580377 | 0.18309596 | 0.0125759 |
| 0.999490048 | 0.12740625 | 0.07031842 | 0.580377 | 0.03618509 | 0.59610235 |
| 0.999490048 | 0.12802735 | 0.07454197 | 0.580377 | 0.19768007 | 0.00815238 |
| 0.999490048 | 0.12927251 | 0.07452948 | 0.580377 | 0.01122037 | 0.87299208 |
| 0.999490048 | 0.13041513 | 0.07229338 | 0.580377 | 0.04737439 | 0.50177051 |
| 0.999490048 | 0.13737862 | 0.06986049 | 0.580377 | 0.15578609 | 0.04161177 |
| 0.999490048 | 0.13847403 | 0.07131989 | 0.580377 | 0.01345192 | 0.85632527 |
| 0.999490048 | 0.13872466 | 0.07237378 | 0.580377 | 0.09948728 | 0.19060818 |
| 0.999490048 | 0.14709053 | 0.06992367 | 0.580377 | 0.26247072 | 0.00238257 |
| 0.999490048 | 0.14942267 | 0.07170633 | 0.580377 | 0.10959917 | 0.17999519 |
| 0.972489992 | 0.15262728 | 0.07109366 | 0.580377 | -0.0311982 | 0.70330658 |
| 0.999490048 | 0.15577473 | 0.0712227 | 0.580377 | 0.03560563 | 0.67045677 |
| 0.999490048 | 0.16452326 | 0.07425593 | 0.580377 | 0.15208118 | 0.09741351 |
| 0.999490048 | 0.16661608 | 0.07088799 | 0.580377 | 0.13168872 | 0.14847915 |
| 0.999490048 | 0.18414226 | 0.07102887 | 0.580377 | 0.17192634 | 0.09056785 |
| 0.999490048 | 0.19165491 | 0.07450656 | 0.580377 | 0.14113195 | 0.18271443 |
| 0.999490048 | 0.19545583 | 0.07071275 | 0.580377 | 0.11941563 | 0.2597203 |
| 0.999490048 | 0.19877325 | 0.072414 | 0.580377 | 0.24657892 | 0.03322718 |
| 0.999490048 | 0.23167135 | 0.07437799 | 0.580377 | 0.22818625 | 0.07824121 |
| 0.999490048 | 0.23709271 | 0.07067564 | 0.580377 | 0.21340881 | 0.10165449 |
| 0.999490048 | 0.27913844 | 0.07220683 | 0.580377 | 0.2685559 | 0.08297126 |
| 0.972489992 | 0.28416835 | 0.07078745 | 0.580377 | -0.0587818 | 0.69960483 |
| 0.999490048 | 0.30895264 | 0.07333877 | 0.580377 | 0.29994141 | 0.08157808 |
| 0.987721587 | 0.31439316 | 0.07288152 | 0.580377 | -0.0572093 | 0.71748222 |
| 0.999490048 | 0.32927392 | 0.07366371 | 0.580377 | 0.45193687 | 0.02222974 |
| 0.999490048 | 0.4671262 | 0.07079652 | 0.580377 | 0.21310738 | 0.36474725 |
| 0.999490048 | -0.2125884 | 0.07536215 | 0.58054944 | -0.2339971 | 0.05179684 |
| 0.999490048 | -0.1737604 | 0.07539522 | 0.58054944 | -0.2275814 | 0.0226067 |
| 0.999490048 | -0.1332204 | 0.07507222 | 0.58054944 | 0.00366797 | 0.9596278 |
| 0.999490048 | -0.0905527 | 0.07536428 | 0.58054944 | -0.0249696 | 0.61346211 |
| 0.999490048 | 0.08664152 | 0.07489874 | 0.58054944 | 0.07763325 | 0.10834009 |
| 0.999490048 | 0.09685125 | 0.07540335 | 0.58054944 | 0.05366197 | 0.31390895 |
| 0.999490048 | 0.09804563 | 0.07491297 | 0.58054944 | 0.04486967 | 0.40317599 |
| 0.999490048 | 0.11527885 | 0.07500901 | 0.58054944 | 0.04127194 | 0.51211865 |
| 0.999490048 | 0.13238829 | 0.07523556 | 0.58054944 | 0.00262421 | 0.97094556 |
| 0.999490048 | 0.13604951 | 0.0753595 | 0.58054944 | 0.03219199 | 0.66446012 |
| 0.999490048 | 0.16273211 | 0.07543666 | 0.58054944 | 0.11665084 | 0.19567172 |
| 0.999490048 | -0.1236112 | 0.07559944 | 0.58126643 | -0.005577 | 0.93402048 |
| 0.999490048 | -0.0955253 | 0.07589137 | 0.5817663 | -0.0774198 | 0.14593886 |
| 0.999490048 | -0.0690235 | 0.07608249 | 0.5817663 | -0.0096097 | 0.79882497 |

| | | | | | |
|---|---|---|---|---|---|
| 0.999490048 | 0.09937998 | 0.07591957 | 0.5817663 | 0.06335886 | 0.24895366 |
| 0.999490048 | 0.10204126 | 0.07602001 | 0.5817663 | 0.00203603 | 0.97083707 |
| 0.999490048 | 0.16378012 | 0.07583277 | 0.5817663 | 0.19703281 | 0.03507441 |
| 0.999490048 | 0.27349333 | 0.07605093 | 0.5817663 | 0.0360666 | 0.80920155 |
| 0.999490048 | -0.4441112 | 0.07679143 | 0.5842681 | -0.4802862 | 0.05690825 |
| 0.999490048 | -0.0865612 | 0.07675045 | 0.5842681 | -0.0492671 | 0.30349188 |
| 0.999490048 | 0.09089806 | 0.07659095 | 0.5842681 | 0.05012354 | 0.31821906 |
| 0.999490048 | 0.14398295 | 0.07672522 | 0.5842681 | 0.10587925 | 0.18673052 |
| 0.999490048 | 0.23415194 | 0.07682951 | 0.5842681 | 0.14085071 | 0.27753854 |
| 0.999490048 | 0.38440238 | 0.07674576 | 0.5842681 | 0.04297967 | 0.83132097 |
| 0.999490048 | 0.19325426 | 0.07697497 | 0.5848417 | 0.12556985 | 0.24198313 |
| 0.999490048 | -0.8104786 | 0.07751916 | 0.58529955 | -0.5420524 | 0.21970144 |
| 0.999490048 | -0.4684322 | 0.07739474 | 0.58529955 | -0.3480973 | 0.18305564 |
| 0.999490048 | -0.3258616 | 0.07780821 | 0.58529955 | 0.04411107 | 0.80546397 |
| 0.999490048 | -0.2621005 | 0.07756009 | 0.58529955 | -0.2288218 | 0.12062354 |
| 0.999490048 | -0.2300571 | 0.07720717 | 0.58529955 | -0.2862276 | 0.03051392 |
| 0.999490048 | -0.2041614 | 0.07851735 | 0.58529955 | -0.023221 | 0.83644342 |
| 0.999490048 | -0.1753596 | 0.0784834 | 0.58529955 | -0.1397681 | 0.15614368 |
| 0.999490048 | -0.1484759 | 0.07783953 | 0.58529955 | -0.0273262 | 0.73790262 |
| 0.999490048 | -0.1456267 | 0.07843335 | 0.58529955 | -0.0215548 | 0.7881925 |
| 0.999490048 | -0.100013 | 0.07857734 | 0.58529955 | -0.1160967 | 0.04327481 |
| 0.999490048 | -0.0725068 | 0.07817928 | 0.58529955 | 0.00671864 | 0.86625526 |
| 0.999490048 | 0.0853444 | 0.07850839 | 0.58529955 | 0.00330196 | 0.94399118 |
| 0.999490048 | 0.09114988 | 0.07843637 | 0.58529955 | 0.00714013 | 0.88687765 |
| 0.999490048 | 0.10514047 | 0.07782947 | 0.58529955 | 0.05317078 | 0.36133655 |
| 0.999490048 | 0.10560041 | 0.07838889 | 0.58529955 | 0.07481551 | 0.20541147 |
| 0.999490048 | 0.11251032 | 0.07842831 | 0.58529955 | 0.06857735 | 0.27404434 |
| 0.999490048 | 0.12565635 | 0.07854843 | 0.58529955 | 0.02753657 | 0.69117997 |
| 0.999490048 | 0.12715277 | 0.07734374 | 0.58529955 | 0.01218112 | 0.86136507 |
| 0.999490048 | 0.18369572 | 0.07764694 | 0.58529955 | 0.19310579 | 0.06445059 |
| 0.999490048 | 0.24765472 | 0.07715148 | 0.58529955 | -0.0337435 | 0.80376771 |
| 0.999490048 | 0.25828781 | 0.07815141 | 0.58529955 | 0.2211973 | 0.12829185 |
| 0.999490048 | 0.27952423 | 0.07775457 | 0.58529955 | 0.04099564 | 0.78093257 |
| 0.999490048 | 0.18417388 | 0.07877073 | 0.58621712 | 0.08055247 | 0.43039767 |
| 0.999490048 | 0.17413634 | 0.07901556 | 0.58737434 | 0.14753649 | 0.13337469 |
| 0.999490048 | 0.22394458 | 0.07906692 | 0.58737434 | 0.14311288 | 0.25307427 |
| 0.999490048 | -0.6113863 | 0.07922579 | 0.58753475 | -0.0664583 | 0.85987187 |
| 0.999490048 | -0.1068403 | 0.07922924 | 0.58753475 | -0.0562118 | 0.3449716 |
| 0.999490048 | 0.08946089 | 0.07955895 | 0.58868266 | 0.01573552 | 0.75056493 |
| 0.999490048 | 0.11457678 | 0.079701 | 0.58868266 | 0.10172793 | 0.11738306 |
| 0.999490048 | 0.12476659 | 0.07973654 | 0.58868266 | 0.12573162 | 0.07757156 |
| 0.999490048 | 0.13575962 | 0.07964217 | 0.58868266 | 0.08202911 | 0.28015988 |

| | | | | | |
|---|---|---|---|---|---|
| 0.999490048 | 0.24323297 | 0.07957051 | 0.58868266 | 0.25694782 | 0.06493849 |
| 0.999490048 | -0.2663636 | 0.08014319 | 0.58893708 | -0.0062926 | 0.96348842 |
| 0.999490048 | -0.2488699 | 0.08018056 | 0.58893708 | -0.2259708 | 0.10801067 |
| 0.999490048 | -0.1515304 | 0.07988952 | 0.58893708 | -0.1165601 | 0.17257876 |
| 0.999490048 | -0.1103448 | 0.08008944 | 0.58893708 | 0.01057304 | 0.8626711 |
| 0.999490048 | 0.09591499 | 0.08019419 | 0.58893708 | 0.06742281 | 0.21168681 |
| 0.999490048 | 0.11088911 | 0.07999558 | 0.58893708 | 0.1488396 | 0.0216446 |
| 0.999490048 | -0.0761002 | 0.08031148 | 0.58928019 | -0.0421796 | 0.32207549 |
| 0.999490048 | -0.4107732 | 0.0804143 | 0.5895166 | 0.05058839 | 0.81709341 |
| 0.999490048 | 0.17435958 | 0.08050364 | 0.58965385 | 0.04749729 | 0.62412942 |
| 0.989730632 | 0.18505043 | 0.08061512 | 0.5899529 | -0.0297511 | 0.77216803 |
| 0.999490048 | -0.1630119 | 0.08104179 | 0.59070342 | -0.0315714 | 0.72773335 |
| 0.999490048 | -0.0884195 | 0.08079894 | 0.59070342 | -0.0530207 | 0.28564776 |
| 0.999490048 | 0.08463786 | 0.08107139 | 0.59070342 | 0.04172716 | 0.37871649 |
| 0.999490048 | 0.08860165 | 0.08090273 | 0.59070342 | 0.0519558 | 0.29636063 |
| 0.999490048 | 0.15427365 | 0.08096661 | 0.59070342 | 0.13773171 | 0.11700849 |
| 0.578467169 | -0.1339506 | 0.08186662 | 0.59208149 | 0.09280295 | 0.22075626 |
| 0.999490048 | -0.1028774 | 0.08148321 | 0.59208149 | -0.0530204 | 0.35849363 |
| 0.999490048 | -0.0938232 | 0.0815208 | 0.59208149 | -0.0257956 | 0.62216466 |
| 0.999490048 | -0.0872168 | 0.0819272 | 0.59208149 | -0.0598426 | 0.22520119 |
| 0.972489992 | -0.0823457 | 0.08171435 | 0.59208149 | 0.02195235 | 0.63295895 |
| 0.999490048 | -0.0741657 | 0.08196961 | 0.59208149 | -0.0613054 | 0.14666499 |
| 0.999490048 | 0.08270182 | 0.08133301 | 0.59208149 | 0.05267528 | 0.25830994 |
| 0.999490048 | 0.08859743 | 0.08172533 | 0.59208149 | 0.05625373 | 0.26040073 |
| 0.999490048 | 0.13476901 | 0.08188026 | 0.59208149 | 0.01191047 | 0.87402135 |
| 0.999490048 | 0.1895039 | 0.08195886 | 0.59208149 | 0.11141129 | 0.296884 |
| 0.999490048 | -0.0744696 | 0.08225469 | 0.59290931 | -0.0205105 | 0.62244139 |
| 0.999490048 | 0.10565283 | 0.08220718 | 0.59290931 | 0.06072337 | 0.30800962 |
| 0.999490048 | 0.17001519 | 0.08229723 | 0.59290931 | 0.13218749 | 0.17163094 |
| 0.999490048 | -0.201928 | 0.08261697 | 0.59367619 | 6.99E-04 | 0.99505306 |
| 0.987721587 | -0.1325411 | 0.08256809 | 0.59367619 | 0.02320567 | 0.75418356 |
| 0.999490048 | 0.12491904 | 0.08257482 | 0.59367619 | 0.04765468 | 0.49666827 |
| 0.972489992 | -0.0949715 | 0.08281832 | 0.59461134 | 0.02431505 | 0.64768513 |
| 0.999490048 | -0.123502 | 0.08294975 | 0.59504328 | -0.0698595 | 0.31678445 |
| 0.999490048 | -0.1089184 | 0.08328858 | 0.59696108 | -0.1198119 | 0.05827425 |
| 0.999490048 | -0.8949287 | 0.08339201 | 0.5971898 | 0.1090808 | 0.81593737 |
| 0.999490048 | -0.3263818 | 0.08435273 | 0.59891412 | -0.0725786 | 0.69281388 |
| 0.999490048 | -0.2781154 | 0.0850553 | 0.59891412 | -0.4494573 | 0.00769293 |
| 0.57925586 | -0.2410133 | 0.08423979 | 0.59891412 | 0.1658288 | 0.22754142 |
| 0.999490048 | -0.1844643 | 0.08656459 | 0.59891412 | -0.0435075 | 0.67756207 |
| 0.999490048 | -0.1647444 | 0.08611795 | 0.59891412 | -0.0427769 | 0.64676411 |
| 0.999490048 | -0.1548333 | 0.08621524 | 0.59891412 | -0.0466771 | 0.59508065 |

| | | | | | |
|---|---|---|---|---|---|
| 0.999490048 | -0.1529475 | 0.08588289 | 0.59891412 | -0.123048 | 0.16269797 |
| 0.999490048 | -0.1184767 | 0.08481809 | 0.59891412 | -0.0719756 | 0.28605433 |
| 0.999490048 | -0.1094981 | 0.08546886 | 0.59891412 | -0.1193539 | 0.0622404 |
| 0.999490048 | -0.1054247 | 0.08659866 | 0.59891412 | -0.0782141 | 0.19770984 |
| 0.999490048 | -0.0987271 | 0.08643848 | 0.59891412 | -0.1188758 | 0.04149295 |
| 0.999490048 | -0.0871456 | 0.08416668 | 0.59891412 | -0.0342478 | 0.48626229 |
| 0.999490048 | -0.0860274 | 0.08616692 | 0.59891412 | 0.00874549 | 0.85738046 |
| 0.999490048 | -0.083217 | 0.08660915 | 0.59891412 | -0.0453802 | 0.34006599 |
| 0.999490048 | -0.0713712 | 0.08622826 | 0.59891412 | -0.1033811 | 0.01586991 |
| 0.999490048 | -0.0687997 | 0.08393383 | 0.59891412 | -0.0443537 | 0.25687405 |
| 0.999490048 | -0.0675412 | 0.08545308 | 0.59891412 | -0.0444612 | 0.24961491 |
| 0.999490048 | -0.0674745 | 0.08578991 | 0.59891412 | -0.0170619 | 0.65500641 |
| 0.999490048 | 0.08007429 | 0.08622094 | 0.59891412 | 0.16233775 | 0.00129645 |
| 0.999490048 | 0.08961435 | 0.08656487 | 0.59891412 | 0.09925133 | 0.05929633 |
| 0.999490048 | 0.09976549 | 0.08616086 | 0.59891412 | 0.09475784 | 0.10216117 |
| 0.999490048 | 0.10386176 | 0.08593285 | 0.59891412 | -0.0122468 | 0.83477056 |
| 0.999490048 | 0.10651234 | 0.08577765 | 0.59891412 | 0.12300897 | 0.04934615 |
| 0.999490048 | 0.10909177 | 0.08509628 | 0.59891412 | 0.00654969 | 0.91515224 |
| 0.999490048 | 0.11433694 | 0.08497556 | 0.59891412 | 0.05534581 | 0.39358711 |
| 0.57925586 | 0.11694186 | 0.08513105 | 0.59891412 | -0.080984 | 0.22599389 |
| 0.999490048 | 0.11831759 | 0.0866453 | 0.59891412 | 0.12759843 | 0.0658782 |
| 0.999490048 | 0.11847627 | 0.08639832 | 0.59891412 | 0.16986233 | 0.01690754 |
| 0.999490048 | 0.12463989 | 0.08523334 | 0.59891412 | 0.03591607 | 0.61016732 |
| 0.999490048 | 0.12485412 | 0.08488787 | 0.59891412 | 0.06926552 | 0.32915317 |
| 0.999490048 | 0.12544928 | 0.08415189 | 0.59891412 | 0.04454986 | 0.52885066 |
| 0.999490048 | 0.12813056 | 0.08473068 | 0.59891412 | 0.13819589 | 0.06422486 |
| 0.999490048 | 0.13147701 | 0.08492966 | 0.59891412 | 0.08851265 | 0.23850491 |
| 0.999490048 | 0.13869832 | 0.0849147 | 0.59891412 | 0.15787864 | 0.05181326 |
| 0.999490048 | 0.15691209 | 0.08572747 | 0.59891412 | 0.13240006 | 0.14370209 |
| 0.999490048 | 0.16909246 | 0.08430348 | 0.59891412 | 0.04566378 | 0.63161021 |
| 0.999490048 | 0.18651003 | 0.08561274 | 0.59891412 | 0.06109558 | 0.56300349 |
| 0.999490048 | 0.21232991 | 0.08486752 | 0.59891412 | 0.27664573 | 0.02768411 |
| 0.999490048 | 0.2407405 | 0.08470321 | 0.59891412 | 0.08534416 | 0.51597262 |
| 0.999490048 | 0.2765897 | 0.08469271 | 0.59891412 | 0.26149574 | 0.10204319 |
| 0.987721587 | 0.33170307 | 0.08414401 | 0.59891412 | -0.0626567 | 0.73690783 |
| 0.999490048 | 0.3326225 | 0.08398581 | 0.59891412 | -0.0466794 | 0.80269963 |
| 0.999490048 | -0.1235671 | 0.08693312 | 0.60040655 | -0.0795677 | 0.26214523 |
| 0.999490048 | -0.1354149 | 0.08713928 | 0.60083647 | -0.0480165 | 0.53358885 |
| 0.999490048 | -0.1296772 | 0.08707633 | 0.60083647 | -0.0638383 | 0.38903571 |
| 0.999490048 | 0.142154 | 0.08732581 | 0.60162581 | 0.11670028 | 0.1565372 |
| 0.999490048 | -0.427319 | 0.08811482 | 0.6023758 | -0.5859641 | 0.02615006 |
| 0.999490048 | -0.4229319 | 0.08822822 | 0.6023758 | -0.9121614 | 0.00193496 |

| | | | | | |
|---|---|---|---|---|---|
| 0.999490048 | -0.1939334 | 0.08792618 | 0.6023758 | -0.0083496 | 0.93967255 |
| 0.999490048 | -0.144555 | 0.08821913 | 0.6023758 | -0.1832509 | 0.03354736 |
| 0.999490048 | -0.1382649 | 0.08810472 | 0.6023758 | -0.0824755 | 0.29991265 |
| 0.999490048 | -0.0891536 | 0.08781971 | 0.6023758 | -0.0237999 | 0.6394908 |
| 0.999490048 | -0.07953 | 0.08820772 | 0.6023758 | -0.0287255 | 0.52752306 |
| 0.999490048 | 0.09208749 | 0.08763176 | 0.6023758 | 0.04101696 | 0.43599679 |
| 0.999490048 | 0.18984303 | 0.08802046 | 0.6023758 | 0.14165096 | 0.19729807 |
| 0.999490048 | 0.20288432 | 0.08794309 | 0.6023758 | 0.17482476 | 0.13854164 |
| 0.999490048 | 0.215529 | 0.08768769 | 0.6023758 | 0.110123 | 0.37255378 |
| 0.999490048 | -0.0801293 | 0.0883615 | 0.60279291 | -0.0788849 | 0.09313172 |
| 0.999490048 | -0.1437489 | 0.08844726 | 0.60288537 | -0.205939 | 0.01766891 |
| 0.999490048 | -0.0986732 | 0.08861695 | 0.60354937 | -0.0437167 | 0.43989554 |
| 0.999490048 | -0.1667024 | 0.0892011 | 0.60374505 | -0.1395105 | 0.15140555 |
| 0.999490048 | -0.1384596 | 0.08885154 | 0.60374505 | -0.0345941 | 0.66209534 |
| 0.999490048 | -0.1301346 | 0.08929642 | 0.60374505 | -0.0680471 | 0.36417901 |
| 0.999490048 | -0.0995637 | 0.08888094 | 0.60374505 | -0.0915404 | 0.1161584 |
| 0.999490048 | -0.0984335 | 0.08917655 | 0.60374505 | -0.0318748 | 0.57203158 |
| 0.999490048 | -0.0762428 | 0.08878313 | 0.60374505 | -0.0239151 | 0.58357626 |
| 0.999490048 | 0.11454191 | 0.08908647 | 0.60374505 | 0.03813081 | 0.56126178 |
| 0.999490048 | 0.1232416 | 0.08918119 | 0.60374505 | 0.07388282 | 0.2992383 |
| 0.999490048 | 0.28905667 | 0.0892768 | 0.60374505 | -0.0278204 | 0.86104059 |
| 0.999490048 | 0.09805392 | 0.08953492 | 0.60486781 | 0.07861067 | 0.16909367 |
| 0.999490048 | 0.07186281 | 0.08961618 | 0.60492736 | -0.0081031 | 0.8438017 |
| 0.999490048 | -0.0824054 | 0.08981793 | 0.60531049 | -0.077509 | 0.10943241 |
| 0.999490048 | -0.0773256 | 0.08976411 | 0.60531049 | -0.089395 | 0.05197436 |
| 0.999490048 | -0.2119806 | 0.09067024 | 0.60540554 | -0.2189035 | 0.08117482 |
| 0.984638128 | -0.1491554 | 0.09130975 | 0.60540554 | 0.04168863 | 0.62787373 |
| 0.999490048 | -0.1430518 | 0.09081738 | 0.60540554 | -0.0388131 | 0.63738166 |
| 0.999490048 | -0.1386266 | 0.0920511 | 0.60540554 | 0.01424523 | 0.85860722 |
| 0.487177805 | -0.1291764 | 0.09226372 | 0.60540554 | 0.11419589 | 0.1342838 |
| 0.999490048 | -0.1280743 | 0.09127139 | 0.60540554 | 0.01571646 | 0.83113016 |
| 0.999490048 | -0.1264782 | 0.09021214 | 0.60540554 | -0.115428 | 0.12029372 |
| 0.999490048 | -0.1194657 | 0.09127949 | 0.60540554 | -0.0814887 | 0.24219116 |
| 0.999490048 | -0.114767 | 0.0902536 | 0.60540554 | -0.0849491 | 0.20401529 |
| 0.999490048 | -0.1064405 | 0.09176739 | 0.60540554 | -0.0198868 | 0.74596697 |
| 0.999490048 | -0.1023682 | 0.09011251 | 0.60540554 | -0.0986075 | 0.10187146 |
| 0.987721587 | -0.101558 | 0.09179141 | 0.60540554 | 0.02198341 | 0.70753535 |
| 0.999490048 | -0.0983741 | 0.09100992 | 0.60540554 | -0.0313389 | 0.58051758 |
| 0.999490048 | -0.0932275 | 0.09229716 | 0.60540554 | -0.0198739 | 0.71221428 |
| 0.999490048 | -0.0825155 | 0.09148986 | 0.60540554 | -0.0990121 | 0.0454011 |
| 0.999490048 | -0.0748845 | 0.09146564 | 0.60540554 | -0.0497211 | 0.25488485 |
| 0.999490048 | -0.0707691 | 0.09183173 | 0.60540554 | -0.0624615 | 0.13435919 |

| | | | | | |
|---|---|---|---|---|---|
| 0.999490048 | -0.0664675 | 0.09130571 | 0.60540554 | -0.0128235 | 0.73761754 |
| 0.999490048 | 0.09068 | 0.09026454 | 0.60540554 | 0.09591584 | 0.07411317 |
| 0.999490048 | 0.09561273 | 0.09173369 | 0.60540554 | -0.0016433 | 0.97619718 |
| 0.999490048 | 0.09670699 | 0.09075018 | 0.60540554 | 0.0751859 | 0.18356093 |
| 0.999490048 | 0.0970084 | 0.0919803 | 0.60540554 | 0.05269926 | 0.35030077 |
| 0.999490048 | 0.09908864 | 0.09168889 | 0.60540554 | 0.08588825 | 0.14099646 |
| 0.999490048 | 0.10814419 | 0.09220752 | 0.60540554 | 0.06939037 | 0.27193115 |
| 0.999490048 | 0.12075207 | 0.0922321 | 0.60540554 | 0.10961036 | 0.12233701 |
| 0.999490048 | 0.12175733 | 0.09212893 | 0.60540554 | 0.10996735 | 0.12626268 |
| 0.999490048 | 0.12378704 | 0.09041328 | 0.60540554 | 0.13698976 | 0.06257103 |
| 0.999490048 | 0.13458269 | 0.09100323 | 0.60540554 | 0.09272109 | 0.23721562 |
| 0.987721587 | 0.13486643 | 0.0921408 | 0.60540554 | -0.0360847 | 0.64346675 |
| 0.999490048 | 0.13741904 | 0.09059841 | 0.60540554 | 0.00589246 | 0.94044475 |
| 0.999490048 | 0.14204266 | 0.09036886 | 0.60540554 | 0.03321261 | 0.68397501 |
| 0.999490048 | 0.14794251 | 0.09050854 | 0.60540554 | 0.08974026 | 0.29574855 |
| 0.999490048 | 0.21202423 | 0.0907225 | 0.60540554 | 0.26376787 | 0.04358371 |
| 0.999490048 | 0.3124427 | 0.0903766 | 0.60540554 | 0.47297395 | 0.01688466 |
| 0.999490048 | -0.1486696 | 0.09251367 | 0.60542919 | -0.1888208 | 0.03557176 |
| 0.999490048 | -0.0850745 | 0.09257283 | 0.60542919 | -0.0602824 | 0.22679282 |
| 0.999490048 | 0.10249599 | 0.09259079 | 0.60542919 | 0.07328572 | 0.22275511 |
| 0.999490048 | 0.16555736 | 0.09258948 | 0.60542919 | 0.23534171 | 0.01994324 |
| 0.999490048 | -0.2845517 | 0.09293863 | 0.6056734 | 0.0369102 | 0.81115926 |
| 0.999490048 | -0.1967505 | 0.09294537 | 0.6056734 | 0.03594113 | 0.75234104 |
| 0.999490048 | -0.1044901 | 0.09313589 | 0.6056734 | 0.01950725 | 0.74724103 |
| 0.999490048 | -0.0988017 | 0.09282989 | 0.6056734 | -0.0465587 | 0.41808637 |
| 0.999490048 | -0.0987457 | 0.09304627 | 0.6056734 | -0.0202386 | 0.7234593 |
| 0.999490048 | -0.094961 | 0.09293935 | 0.6056734 | -0.0739249 | 0.18603208 |
| 0.999490048 | 0.14770459 | 0.09312596 | 0.6056734 | 0.03449736 | 0.68697899 |
| 0.999490048 | -0.2994274 | 0.09325161 | 0.60595404 | -0.3093869 | 0.08343136 |
| 0.999490048 | -0.2733894 | 0.094228 | 0.60755636 | -0.0801794 | 0.6021655 |
| 0.999490048 | -0.2428064 | 0.09444409 | 0.60755636 | -0.2380705 | 0.10069028 |
| 0.999490048 | -0.1968266 | 0.09435631 | 0.60755636 | 0.02377082 | 0.83539078 |
| 0.999490048 | -0.1276049 | 0.09420163 | 0.60755636 | -0.0591431 | 0.42770206 |
| 0.999490048 | -0.1177107 | 0.09437508 | 0.60755636 | -0.013584 | 0.84261376 |
| 0.999490048 | -0.0635283 | 0.09411286 | 0.60755636 | -0.0800358 | 0.03785575 |
| 0.819178176 | 0.07883457 | 0.09411122 | 0.60755636 | -0.0416114 | 0.36715616 |
| 0.999490048 | 0.0921779 | 0.09437133 | 0.60755636 | 0.08451118 | 0.12346991 |
| 0.999490048 | 0.10735035 | 0.09430555 | 0.60755636 | 0.09812517 | 0.12447593 |
| 0.999490048 | 0.12996735 | 0.09439062 | 0.60755636 | 0.17077958 | 0.03106985 |
| 0.999490048 | 0.15163479 | 0.09386114 | 0.60755636 | 0.05235618 | 0.5530725 |
| 0.999490048 | 0.2037038 | 0.09376457 | 0.60755636 | 0.00842708 | 0.94312603 |
| 0.999490048 | 0.39357272 | 0.09381234 | 0.60755636 | 5.29E-04 | 0.99815247 |

| | | | | | |
|---|---|---|---|---|---|
| 0.999490048 | -0.0920029 | 0.09454806 | 0.60775699 | -0.0346414 | 0.51892994 |
| 0.999490048 | -0.097043 | 0.09462968 | 0.6078137 | -0.0231373 | 0.68228593 |
| 0.999490048 | 0.22567611 | 0.09499575 | 0.60969603 | 0.14809349 | 0.26564064 |
| 0.999490048 | 0.12323128 | 0.09524384 | 0.61081877 | -0.0074044 | 0.91786729 |
| 0.966469153 | -0.2025942 | 0.09540548 | 0.61138586 | 0.09718621 | 0.39162906 |
| 0.999490048 | 0.14985196 | 0.09563189 | 0.61189751 | 0.00792017 | 0.92781119 |
| 0.999490048 | 0.21865267 | 0.0955718 | 0.61189751 | 0.22326747 | 0.08914428 |
| 0.999490048 | 0.10093309 | 0.09578194 | 0.61238836 | 0.02494115 | 0.67255265 |
| 0.999490048 | -0.2407337 | 0.09590085 | 0.61242892 | -0.0575342 | 0.68278301 |
| 0.999490048 | -0.0583513 | 0.09600832 | 0.61242892 | -0.0450314 | 0.19397249 |
| 0.999490048 | 0.1555174 | 0.09594144 | 0.61242892 | 0.04201504 | 0.64430508 |
| 0.999490048 | -0.6213114 | 0.09662005 | 0.61305532 | -0.2334514 | 0.50017199 |
| 0.999490048 | -0.4015709 | 0.09735465 | 0.61305532 | -0.4574132 | 0.06488385 |
| 0.999490048 | -0.199148 | 0.09683503 | 0.61305532 | -0.1103912 | 0.34812576 |
| 0.999490048 | -0.135105 | 0.09711283 | 0.61305532 | 0.02346071 | 0.76720965 |
| 0.999490048 | -0.1200826 | 0.09631956 | 0.61305532 | -0.129479 | 0.07421138 |
| 0.972489992 | -0.1075755 | 0.09677266 | 0.61305532 | 0.03553808 | 0.57379244 |
| 0.999490048 | -0.1001391 | 0.09687019 | 0.61305532 | -0.0021525 | 0.97072168 |
| 0.999490048 | -0.0834451 | 0.0972874 | 0.61305532 | -0.0247274 | 0.61423103 |
| 0.999490048 | -0.0763409 | 0.09697708 | 0.61305532 | -0.0152292 | 0.73375873 |
| 0.999490048 | -0.0705142 | 0.09656319 | 0.61305532 | -0.0225971 | 0.58500168 |
| 0.999490048 | 0.07573211 | 0.09733146 | 0.61305532 | 0.05546064 | 0.21883148 |
| 0.972489992 | 0.0827328 | 0.09720517 | 0.61305532 | -0.0282403 | 0.56173262 |
| 0.999490048 | 0.10277737 | 0.09694197 | 0.61305532 | 0.02504406 | 0.67794446 |
| 0.999490048 | 0.11557593 | 0.09714882 | 0.61305532 | 0.11770196 | 0.09147412 |
| 0.999490048 | 0.14638671 | 0.09705706 | 0.61305532 | 0.11064324 | 0.20446588 |
| 0.999490048 | 0.17967648 | 0.09655398 | 0.61305532 | 0.13073881 | 0.22060652 |
| 0.999490048 | 0.27160173 | 0.0970052 | 0.61305532 | 0.21822397 | 0.17819101 |
| 0.999490048 | -0.3050001 | 0.0988469 | 0.61384274 | -0.0713387 | 0.69189163 |
| 0.999490048 | -0.1673919 | 0.09805033 | 0.61384274 | -0.0197158 | 0.84125147 |
| 0.999490048 | -0.1614726 | 0.09864835 | 0.61384274 | -0.049819 | 0.6013578 |
| 0.999490048 | -0.107124 | 0.09820942 | 0.61384274 | -0.0782031 | 0.2214786 |
| 0.999490048 | -0.106814 | 0.09758706 | 0.61384274 | 0.02244789 | 0.72065251 |
| 0.999490048 | -0.0940468 | 0.09805386 | 0.61384274 | -0.06002 | 0.28323329 |
| 0.999490048 | -0.0907103 | 0.09857748 | 0.61384274 | -0.0693476 | 0.20157365 |
| 0.999490048 | -0.0891022 | 0.09887647 | 0.61384274 | -0.1096728 | 0.04503345 |
| 0.999490048 | -0.0702428 | 0.09885977 | 0.61384274 | -0.0418783 | 0.31660995 |
| 0.999490048 | 0.07313521 | 0.09872101 | 0.61384274 | 0.0516046 | 0.23759479 |
| 0.999490048 | 0.09450791 | 0.09832218 | 0.61384274 | 0.04973708 | 0.37484483 |
| 0.999490048 | 0.0988791 | 0.09848424 | 0.61384274 | 0.03243924 | 0.57846896 |
| 0.999490048 | 0.11135392 | 0.09828341 | 0.61384274 | 0.13578805 | 0.04653033 |
| 0.999490048 | 0.11169101 | 0.09868388 | 0.61384274 | 0.06725188 | 0.3116557 |

| | | | | | |
|---|---|---|---|---|---|
| 0.999490048 | 0.11480354 | 0.09793474 | 0.61384274 | 0.0732765 | 0.28299456 |
| 0.999490048 | 0.17245901 | 0.09852818 | 0.61384274 | 0.21886187 | 0.03905303 |
| 0.999490048 | 0.3085202 | 0.09782315 | 0.61384274 | -0.0211333 | 0.90717724 |
| 0.999490048 | 0.34389031 | 0.09771873 | 0.61384274 | 0.24715464 | 0.22781745 |
| 0.999490048 | 1.00645777 | 0.0978478 | 0.61384274 | 0.24641173 | 0.65912462 |
| 0.972489992 | 0.29924868 | 0.09901203 | 0.61422768 | -0.0990126 | 0.57587774 |
| 0.999490048 | 0.12947884 | 0.0990969 | 0.61429777 | 0.04585124 | 0.54964262 |
| 0.987721587 | -0.2926733 | 0.09927735 | 0.6143931 | 0.08066915 | 0.62978835 |
| 0.999490048 | -0.1173349 | 0.09919477 | 0.6143931 | -0.0814625 | 0.24568686 |
| 0.999490048 | 0.23886306 | 0.09933302 | 0.6143931 | 0.03061588 | 0.82815071 |
| 0.999490048 | -0.0907273 | 0.09956073 | 0.61534571 | -0.1214742 | 0.03070369 |
| 0.999490048 | -0.1861148 | 0.0997878 | 0.61538265 | -0.0277768 | 0.80076169 |
| 0.57925586 | -0.1419797 | 0.09975661 | 0.61538265 | 0.11048689 | 0.19546645 |
| 0.999490048 | 0.22293207 | 0.09977234 | 0.61538265 | 0.11551918 | 0.38424413 |
| 0.999490048 | 0.07170202 | 0.10023216 | 0.6176668 | 0.03508764 | 0.4115254 |
| 0.999490048 | -0.1450337 | 0.10088682 | 0.61800803 | -0.0976073 | 0.26251808 |
| 0.999490048 | -0.1371944 | 0.10051353 | 0.61800803 | 0.02328325 | 0.77467511 |
| 0.999490048 | -0.1367397 | 0.10108475 | 0.61800803 | -0.1320339 | 0.11273731 |
| 0.999490048 | -0.1325239 | 0.10085775 | 0.61800803 | -0.1631171 | 0.04626881 |
| 0.999490048 | -0.0940434 | 0.10110167 | 0.61800803 | -0.0581891 | 0.30232257 |
| 0.999490048 | -0.0934463 | 0.10047255 | 0.61800803 | -0.0376063 | 0.49882687 |
| 0.999490048 | -0.0912567 | 0.10072903 | 0.61800803 | -0.0675709 | 0.21865156 |
| 0.999490048 | -0.0851565 | 0.10059707 | 0.61800803 | -0.0585077 | 0.25251096 |
| 0.999490048 | 0.11137436 | 0.10044921 | 0.61800803 | 0.04529866 | 0.4942764 |
| 0.999490048 | 0.15638184 | 0.10109528 | 0.61800803 | 0.04801773 | 0.60582665 |
| 0.889155227 | 0.18270141 | 0.10088315 | 0.61800803 | -0.0910154 | 0.40420718 |
| 0.999490048 | -0.1519688 | 0.10164731 | 0.61817557 | -0.1112002 | 0.22523803 |
| 0.999490048 | -0.079661 | 0.10153734 | 0.61817557 | -0.0841396 | 0.08465182 |
| 0.999490048 | -0.0598622 | 0.10152821 | 0.61817557 | 8.71E-04 | 0.98046905 |
| 0.999490048 | 0.08655562 | 0.10160247 | 0.61817557 | 0.05463609 | 0.29361591 |
| 0.999490048 | 0.14975946 | 0.10145705 | 0.61817557 | 0.05345769 | 0.54938803 |
| 0.999490048 | 0.24322112 | 0.10129262 | 0.61817557 | 0.07687839 | 0.59555152 |
| 0.972489992 | 0.263506 | 0.10136516 | 0.61817557 | -0.0915529 | 0.55989792 |
| 0.972489992 | -0.1617071 | 0.1021741 | 0.61820532 | 0.05562741 | 0.56472928 |
| 0.999490048 | -0.1135609 | 0.10188872 | 0.61820532 | -0.1402403 | 0.04624295 |
| 0.999490048 | -0.0998533 | 0.10176168 | 0.61820532 | 0.0041932 | 0.9437031 |
| 0.972489992 | -0.0996198 | 0.10194879 | 0.61820532 | 0.03607358 | 0.54428419 |
| 0.999490048 | -0.0910917 | 0.10244309 | 0.61820532 | -0.0895981 | 0.10789938 |
| 0.999490048 | -0.07368 | 0.10199792 | 0.61820532 | -0.0843629 | 0.06325279 |
| 0.999490048 | -0.0692672 | 0.10257398 | 0.61820532 | -0.0275452 | 0.50662161 |
| 0.999490048 | -0.0638183 | 0.10261913 | 0.61820532 | -0.0411684 | 0.28484987 |
| 0.999490048 | -0.0554902 | 0.10231612 | 0.61820532 | -0.0351297 | 0.29326929 |

| | | | | | |
|---|---|---|---|---|---|
| 0.999490048 | 0.08301432 | 0.10251379 | 0.61820532 | 0.09241353 | 0.07084084 |
| 0.999490048 | 0.12037111 | 0.10240588 | 0.61820532 | 0.07460384 | 0.30343465 |
| 0.999490048 | 0.12145801 | 0.10203444 | 0.61820532 | 0.13226839 | 0.0763854 |
| 0.999490048 | 0.13015989 | 0.10223051 | 0.61820532 | 0.06474605 | 0.40681191 |
| 0.999490048 | 0.51090514 | 0.10268872 | 0.61820532 | 0.21747224 | 0.47760946 |
| 0.999490048 | -0.1956078 | 0.1031795 | 0.62026555 | -0.1328459 | 0.26157176 |
| 0.999490048 | 0.07459626 | 0.10313703 | 0.62026555 | 0.05624471 | 0.21388408 |
| 0.987721587 | 0.26049271 | 0.10327742 | 0.62040753 | -0.0796634 | 0.6096641 |
| 0.999490048 | -0.0991804 | 0.10357333 | 0.6212398 | -0.1000444 | 0.10075409 |
| 0.999490048 | 0.08970499 | 0.10366376 | 0.6212398 | 0.01205469 | 0.82232321 |
| 0.999490048 | 0.09833317 | 0.10371357 | 0.6212398 | 0.09642385 | 0.11024213 |
| 0.999490048 | 0.1375976 | 0.10349217 | 0.6212398 | 0.0749556 | 0.36614229 |
| 0.999490048 | -0.2542758 | 0.1038116 | 0.62138127 | -0.0812426 | 0.58262744 |
| 0.999490048 | -0.1016174 | 0.10427461 | 0.62370556 | -0.1365323 | 0.03241264 |
| 0.999490048 | -0.1331174 | 0.10489437 | 0.62395686 | -0.0591577 | 0.46162898 |
| 0.999490048 | -0.0974341 | 0.10484454 | 0.62395686 | -0.0710711 | 0.23108067 |
| 0.999490048 | -0.0850312 | 0.10491442 | 0.62395686 | -0.0954287 | 0.07063642 |
| 0.999490048 | 0.07145856 | 0.10450762 | 0.62395686 | 0.05451486 | 0.21053762 |
| 0.999490048 | 0.07476822 | 0.10470242 | 0.62395686 | 0.03326004 | 0.46093257 |
| 0.999490048 | 0.08489438 | 0.10489352 | 0.62395686 | 0.07756212 | 0.13678621 |
| 0.999490048 | 0.1465311 | 0.10441444 | 0.62395686 | 0.10279085 | 0.24832781 |
| 0.999490048 | 0.19579456 | 0.10486379 | 0.62395686 | 0.08356859 | 0.47931725 |
| 0.999490048 | -0.2623758 | 0.10526584 | 0.62456675 | -0.001271 | 0.99356939 |
| 0.999490048 | -0.1767494 | 0.1050927 | 0.62456675 | -0.2402181 | 0.03101439 |
| 0.999490048 | -0.0845789 | 0.10531617 | 0.62456675 | 3.80E-04 | 0.99404241 |
| 0.999490048 | 0.11530406 | 0.10519861 | 0.62456675 | 0.06600697 | 0.34513468 |
| 0.999490048 | 0.10142369 | 0.10547644 | 0.62507328 | 0.01003428 | 0.86937324 |
| 0.956358655 | -0.133657 | 0.10566064 | 0.62572079 | 0.06090265 | 0.45152072 |
| 0.971241857 | -0.0718575 | 0.10574454 | 0.62577383 | 0.02983642 | 0.49237077 |
| 0.999490048 | -0.0752727 | 0.10599201 | 0.6263266 | -0.0435729 | 0.34099623 |
| 0.999490048 | 0.0830904 | 0.10592127 | 0.6263266 | 0.01266957 | 0.80024849 |
| 0.999490048 | 0.0959744 | 0.10606297 | 0.6263266 | 0.05871947 | 0.3148319 |
| 0.999490048 | -0.0989936 | 0.10655682 | 0.62746786 | -0.0614419 | 0.30873652 |
| 0.999490048 | -0.0711664 | 0.10654907 | 0.62746786 | -0.0204776 | 0.63421868 |
| 0.999490048 | 0.09508003 | 0.10654795 | 0.62746786 | 0.07765092 | 0.18370628 |
| 0.999490048 | 0.11492393 | 0.10639784 | 0.62746786 | 0.07038465 | 0.31482925 |
| 0.999490048 | -0.162344 | 0.10708122 | 0.62772105 | -0.2302314 | 0.02576441 |
| 0.999490048 | -0.1211661 | 0.10694055 | 0.62772105 | -0.1306256 | 0.08347614 |
| 0.999490048 | -0.0908926 | 0.10707223 | 0.62772105 | -0.1337003 | 0.02118754 |
| 0.999490048 | -0.0804043 | 0.10671805 | 0.62772105 | -0.0764084 | 0.12435518 |
| 0.999490048 | 0.07631731 | 0.10699629 | 0.62772105 | 0.01126249 | 0.80714867 |
| 0.999490048 | 0.08086606 | 0.10712605 | 0.62772105 | -0.0189325 | 0.6989328 |

| | | | | | |
|---|---|---|---|---|---|
| 0.999490048 | 0.13702021 | 0.10690684 | 0.62772105 | 0.05934028 | 0.47565979 |
| 0.999490048 | 0.06793213 | 0.1072901 | 0.62824145 | 0.08903835 | 0.03808677 |
| 0.999490048 | -0.0789535 | 0.10752687 | 0.62918665 | -0.0474984 | 0.32491371 |
| 0.971241857 | -0.2854654 | 0.10767649 | 0.62962094 | 0.12243918 | 0.4808752 |
| 0.999490048 | -0.0703495 | 0.10798231 | 0.6309673 | -0.0704335 | 0.1075833 |
| 0.999490048 | -0.1440712 | 0.10863522 | 0.63174435 | -0.0216748 | 0.80441792 |
| 0.999490048 | -0.0782391 | 0.10861172 | 0.63174435 | -0.0516137 | 0.28284175 |
| 0.999490048 | 0.08905395 | 0.10842672 | 0.63174435 | 0.03607693 | 0.50626406 |
| 0.999490048 | 0.11392869 | 0.108324 | 0.63174435 | 0.06615169 | 0.34286259 |
| 0.999490048 | 0.12660885 | 0.1086449 | 0.63174435 | 0.05634297 | 0.46600372 |
| 0.987721587 | 0.23075607 | 0.10824607 | 0.63174435 | -0.0836944 | 0.53096205 |
| 0.999490048 | 0.31529659 | 0.10843607 | 0.63174435 | 0.28790698 | 0.14102072 |
| 0.999490048 | -0.3091029 | 0.10912908 | 0.63302832 | -0.1705048 | 0.36824327 |
| 0.999490048 | -0.10594 | 0.10916896 | 0.63302832 | -0.1281756 | 0.05534195 |
| 0.999490048 | 0.10266697 | 0.10916457 | 0.63302832 | 0.06998694 | 0.26823819 |
| 0.999490048 | 0.13338264 | 0.10897132 | 0.63302832 | 0.07673542 | 0.34815219 |
| 0.999490048 | 0.0924677 | 0.10946627 | 0.63431183 | 0.05621019 | 0.32272129 |
| 0.999490048 | -0.0838287 | 0.10956495 | 0.63444337 | -0.0414156 | 0.42016116 |
| 0.999490048 | -0.3990342 | 0.11005177 | 0.63456215 | -0.2286587 | 0.34088242 |
| 0.999490048 | -0.0945058 | 0.1100619 | 0.63456215 | 0.02313622 | 0.68852606 |
| 0.999490048 | 0.08320656 | 0.10999407 | 0.63456215 | -0.0014751 | 0.97679981 |
| 0.999490048 | 0.08346735 | 0.10979957 | 0.63456215 | 0.12935266 | 0.01651252 |
| 0.999490048 | 0.10312975 | 0.11011743 | 0.63456215 | 0.02250754 | 0.72078171 |
| 0.999490048 | 0.10490356 | 0.10977052 | 0.63456215 | 0.00589039 | 0.92656934 |
| 0.999490048 | 0.15169973 | 0.11004788 | 0.63456215 | -0.0035566 | 0.96932737 |
| 0.999490048 | 0.05681647 | 0.11023071 | 0.63477685 | 0.12801443 | 9.80E-04 |
| 0.999490048 | -0.4166909 | 0.11045704 | 0.63519259 | -0.2924904 | 0.24360443 |
| 0.999490048 | -0.183758 | 0.11081884 | 0.63519259 | -0.2535208 | 0.03133814 |
| 0.999490048 | -0.1441313 | 0.11099942 | 0.63519259 | -0.1563778 | 0.08514198 |
| 0.999490048 | -0.0839007 | 0.11106362 | 0.63519259 | -0.0544055 | 0.2943647 |
| 0.999490048 | -0.0770285 | 0.11063672 | 0.63519259 | -0.0511047 | 0.28299268 |
| 0.999490048 | 0.09715396 | 0.110779 | 0.63519259 | 0.00731794 | 0.90190573 |
| 0.999490048 | 0.09760627 | 0.11066932 | 0.63519259 | 0.0746719 | 0.2174854 |
| 0.999490048 | 0.11012254 | 0.11094744 | 0.63519259 | 0.0940184 | 0.17064147 |
| 0.999490048 | 0.13181195 | 0.11084948 | 0.63519259 | 0.02868219 | 0.72214612 |
| 0.999490048 | 0.19091263 | 0.11101741 | 0.63519259 | 0.05168278 | 0.65858777 |
| 0.999490048 | -0.3573982 | 0.11140002 | 0.63522295 | -0.4841874 | 0.03451996 |
| 0.999490048 | -0.1653945 | 0.11164497 | 0.63522295 | -0.0620817 | 0.54157539 |
| 0.999490048 | -0.1396385 | 0.11182339 | 0.63522295 | -0.111268 | 0.20101942 |
| 0.999490048 | -0.081293 | 0.11182063 | 0.63522295 | -0.0763387 | 0.13425147 |
| 0.999490048 | -0.0721047 | 0.11186205 | 0.63522295 | -0.045006 | 0.31353522 |
| 0.999490048 | -0.0684049 | 0.1117074 | 0.63522295 | -0.0321162 | 0.44618357 |

| | | | | | |
|---|---|---|---|---|---|
| 0.999490048 | 0.08233173 | 0.11213397 | 0.63522295 | 0.06183711 | 0.22768499 |
| 0.999490048 | 0.10068274 | 0.11209663 | 0.63522295 | 0.11376692 | 0.07454833 |
| 0.999490048 | 0.13179643 | 0.11180884 | 0.63522295 | 0.07135552 | 0.38074283 |
| 0.999490048 | 0.14515384 | 0.11202977 | 0.63522295 | 0.16324643 | 0.07578359 |
| 0.999490048 | 0.16911022 | 0.11203603 | 0.63522295 | 0.05957935 | 0.56694392 |
| 0.999490048 | 0.17634714 | 0.11185418 | 0.63522295 | -0.0315177 | 0.7708019 |
| 0.999490048 | 0.1787936 | 0.11204329 | 0.63522295 | 0.10701973 | 0.33366178 |
| 0.999490048 | 0.19942773 | 0.11212732 | 0.63522295 | 0.06784745 | 0.58035249 |
| 0.999490048 | -0.1172023 | 0.11272403 | 0.63640678 | -0.0237312 | 0.74205777 |
| 0.999490048 | -0.0721365 | 0.11252795 | 0.63640678 | -0.062558 | 0.16613622 |
| 0.999490048 | 0.141396 | 0.11253122 | 0.63640678 | 0.02104132 | 0.80866449 |
| 0.999490048 | 0.20464514 | 0.11259707 | 0.63640678 | 0.18035741 | 0.155042 |
| 0.999490048 | 0.48421344 | 0.11266671 | 0.63640678 | 0.25781771 | 0.37106671 |
| 0.999490048 | -0.1353684 | 0.11300208 | 0.63711503 | -0.0115051 | 0.89011088 |
| 0.999490048 | 0.09563453 | 0.11296339 | 0.63711503 | 0.02963208 | 0.61526901 |
| 0.987721587 | -0.1773327 | 0.11340895 | 0.6383873 | 0.05855902 | 0.59283206 |
| 0.999490048 | -0.1376969 | 0.11336764 | 0.6383873 | -0.1174724 | 0.17388215 |
| 0.999490048 | 0.09821658 | 0.11346666 | 0.6383873 | 0.08862052 | 0.15136629 |
| 0.999490048 | 0.14375188 | 0.11361 | 0.6383873 | 0.18382497 | 0.0462848 |
| 0.999490048 | 0.45462449 | 0.11356309 | 0.6383873 | 0.15363987 | 0.58442585 |
| 0.999490048 | -0.0908797 | 0.11399173 | 0.63896405 | -0.0749535 | 0.18888577 |
| 0.987721587 | -0.0641073 | 0.1141315 | 0.63896405 | 0.02079897 | 0.60004894 |
| 0.999490048 | 0.06982193 | 0.11380141 | 0.63896405 | 0.02808519 | 0.51587469 |
| 0.999490048 | 0.07581658 | 0.11409181 | 0.63896405 | 0.06362536 | 0.18164216 |
| 0.999490048 | 0.09684554 | 0.1142483 | 0.63896405 | 0.01352356 | 0.82111657 |
| 0.999490048 | 0.12595133 | 0.11400846 | 0.63896405 | 0.02022933 | 0.79470588 |
| 0.999490048 | 0.16776786 | 0.11421686 | 0.63896405 | 0.08250332 | 0.42851999 |
| 0.999490048 | -0.2552476 | 0.11445224 | 0.6389912 | -0.2095567 | 0.18699572 |
| 0.987721587 | 0.11165431 | 0.1144736 | 0.6389912 | -0.0356224 | 0.60645226 |
| 0.999490048 | 0.13791111 | 0.11448274 | 0.6389912 | 0.06694919 | 0.43475344 |
| 0.999490048 | 0.12140599 | 0.11463526 | 0.63941509 | 0.06466913 | 0.39221099 |
| 0.999490048 | -0.4740286 | 0.11591378 | 0.64034277 | -0.6657516 | 0.03816636 |
| 0.999490048 | -0.2412261 | 0.11535717 | 0.64034277 | -0.0995829 | 0.50689423 |
| 0.999490048 | -0.0939578 | 0.11553604 | 0.64034277 | 0.02711767 | 0.64206231 |
| 0.999490048 | 0.08529703 | 0.11517578 | 0.64034277 | 0.09346014 | 0.08583321 |
| 0.999490048 | 0.08544088 | 0.1155348 | 0.64034277 | 0.0264877 | 0.61770112 |
| 0.999490048 | 0.09679054 | 0.11505063 | 0.64034277 | 0.02050628 | 0.73233005 |
| 0.999490048 | 0.10269166 | 0.11534977 | 0.64034277 | 0.01203011 | 0.85000371 |
| 0.999490048 | 0.10389202 | 0.11601621 | 0.64034277 | 0.02853744 | 0.65859006 |
| 0.999490048 | 0.10460739 | 0.11578493 | 0.64034277 | 0.1109844 | 0.09620213 |
| 0.999490048 | 0.11200183 | 0.11553801 | 0.64034277 | 0.10370226 | 0.14367673 |
| 0.999490048 | 0.11368955 | 0.11597439 | 0.64034277 | 0.15149419 | 0.03970152 |

| | | | | | |
|---|---|---|---|---|---|
| 0.999490048 | 0.11435595 | 0.11557341 | 0.64034277 | 0.0788652 | 0.27158579 |
| 0.999490048 | 0.12451865 | 0.11512993 | 0.64034277 | 0.06868219 | 0.37654615 |
| 0.999490048 | 0.13671598 | 0.11602923 | 0.64034277 | 0.03707337 | 0.66268061 |
| 0.999490048 | 0.14027277 | 0.11588629 | 0.64034277 | 0.12931148 | 0.14574706 |
| 0.999490048 | 0.15519894 | 0.1151824 | 0.64034277 | 0.21835465 | 0.03029403 |
| 0.999490048 | 0.22613459 | 0.11610526 | 0.64034277 | 0.06133642 | 0.66267082 |
| 0.999490048 | -0.1483944 | 0.11627056 | 0.64083112 | -0.1334647 | 0.15584571 |
| 0.999490048 | 0.11761941 | 0.11651883 | 0.64177586 | 0.13827183 | 0.06747578 |
| 0.999490048 | -0.1108237 | 0.11666306 | 0.64187982 | -0.0689892 | 0.32141212 |
| 0.999490048 | 0.0782062 | 0.11669145 | 0.64187982 | 0.04442495 | 0.36481636 |
| 0.999490048 | 0.09468322 | 0.11683173 | 0.6422284 | 0.07575756 | 0.2055781 |
| 0.972489992 | -0.193988 | 0.11718513 | 0.64332404 | 0.08160873 | 0.50113671 |
| 0.999490048 | -0.0762383 | 0.11716697 | 0.64332404 | 0.0044485 | 0.92532409 |
| 0.999490048 | -0.080857 | 0.11728427 | 0.64344522 | -6.26E-04 | 0.99008243 |
| 0.999490048 | -0.0847031 | 0.11741597 | 0.64361305 | -0.0922247 | 0.08965219 |
| 0.972489992 | 0.09946466 | 0.11746902 | 0.64361305 | -0.0433712 | 0.48614005 |
| 0.999490048 | 0.09995856 | 0.11755177 | 0.6436441 | 0.02747425 | 0.65979274 |
| 0.999490048 | 0.10217961 | 0.1176843 | 0.64394753 | 0.00145026 | 0.98183599 |
| 0.999490048 | -0.5623611 | 0.11916858 | 0.64446739 | -0.4007593 | 0.2492158 |
| 0.972489992 | -0.3358735 | 0.11902918 | 0.64446739 | 0.13708807 | 0.51605959 |
| 0.999490048 | -0.1386181 | 0.11828197 | 0.64446739 | -0.1917681 | 0.03435359 |
| 0.999490048 | -0.1232482 | 0.11912282 | 0.64446739 | -0.0589693 | 0.44731774 |
| 0.999490048 | -0.1215792 | 0.11911778 | 0.64446739 | -0.1116919 | 0.15069023 |
| 0.999490048 | -0.0933721 | 0.11847778 | 0.64446739 | -0.0315177 | 0.5901278 |
| 0.999490048 | -0.0928125 | 0.11896435 | 0.64446739 | -0.0803331 | 0.17457357 |
| 0.999490048 | -0.0818228 | 0.11887154 | 0.64446739 | -0.1469012 | 0.00766323 |
| 0.999490048 | -0.0786928 | 0.1188344 | 0.64446739 | 0.01740447 | 0.72396183 |
| 0.999490048 | -0.0742672 | 0.1178764 | 0.64446739 | -0.017596 | 0.70446839 |
| 0.999490048 | -0.0729783 | 0.11883389 | 0.64446739 | -0.0952986 | 0.04510173 |
| 0.999490048 | 0.08267045 | 0.11889189 | 0.64446739 | 0.00111754 | 0.98275947 |
| 0.999490048 | 0.09628427 | 0.1181371 | 0.64446739 | 0.04060807 | 0.50122241 |
| 0.999490048 | 0.10176362 | 0.11833597 | 0.64446739 | 0.01362289 | 0.83036134 |
| 0.999490048 | 0.14518547 | 0.11837098 | 0.64446739 | 0.08605891 | 0.34696465 |
| 0.999490048 | 0.15188707 | 0.11851962 | 0.64446739 | 0.08184845 | 0.39201409 |
| 0.999490048 | 0.20034657 | 0.11826094 | 0.64446739 | 0.20604137 | 0.10864517 |
| 0.999490048 | 0.9217507 | 0.1188823 | 0.64446739 | 0.2526293 | 0.64544581 |
| 0.999490048 | -0.0982004 | 0.11933675 | 0.64495915 | -0.0820457 | 0.18992252 |
| 0.999490048 | -0.1260754 | 0.11964404 | 0.64532803 | -0.0387909 | 0.62440885 |
| 0.999490048 | -0.0911863 | 0.1197619 | 0.64532803 | -0.0348627 | 0.54359034 |
| 0.999490048 | -0.067185 | 0.11979143 | 0.64532803 | -0.0841631 | 0.05445056 |
| 0.999490048 | 0.08943344 | 0.11966017 | 0.64532803 | 0.07843991 | 0.16992032 |
| 0.999490048 | 0.1225277 | 0.11979108 | 0.64532803 | 0.12906863 | 0.102161 |

| | | | | | |
|---|---|---|---|---|---|
| 0.999490048 | 0.11631774 | 0.12000894 | 0.64566664 | 0.11112368 | 0.13664739 |
| 0.999490048 | 0.16210548 | 0.11997952 | 0.64566664 | 0.14493966 | 0.16245068 |
| 0.999490048 | 0.0835724 | 0.12010963 | 0.64579229 | 0.07247484 | 0.17512462 |
| 0.987721587 | 0.0702106 | 0.12019278 | 0.64582348 | -0.023953 | 0.58799244 |
| 0.999490048 | -0.1120128 | 0.12045706 | 0.64682732 | -0.1202381 | 0.096725 |
| 0.999490048 | -0.1057172 | 0.12058564 | 0.64710158 | -0.0053799 | 0.93547736 |
| 0.999490048 | 0.20208777 | 0.1208304 | 0.6479986 | 0.20356766 | 0.11826042 |
| 0.999490048 | -0.2266471 | 0.12155125 | 0.64936479 | -0.2732342 | 0.06472841 |
| 0.999490048 | -0.1058747 | 0.12142832 | 0.64936479 | -0.0898011 | 0.186018 |
| 0.999490048 | -0.0740334 | 0.1214651 | 0.64936479 | -0.0178015 | 0.70317991 |
| 0.999490048 | -0.0715988 | 0.12124855 | 0.64936479 | -0.0168528 | 0.7089696 |
| 0.999490048 | 0.10521715 | 0.12155176 | 0.64936479 | 0.0034346 | 0.95869655 |
| 0.999490048 | 0.25476065 | 0.12119623 | 0.64936479 | 0.12981436 | 0.40325133 |
| 0.999490048 | -0.1115521 | 0.12176596 | 0.65009318 | 0.00121046 | 0.98627255 |
| 0.999490048 | -0.3769381 | 0.1225015 | 0.65009629 | -0.1680233 | 0.46410691 |
| 0.999490048 | -0.3592379 | 0.12299666 | 0.65009629 | -0.1475041 | 0.50722248 |
| 0.999490048 | -0.3284975 | 0.12188677 | 0.65009629 | -0.0905265 | 0.66285445 |
| 0.999490048 | -0.1412801 | 0.12291954 | 0.65009629 | -0.1050203 | 0.24647831 |
| 0.999490048 | -0.1292338 | 0.12290398 | 0.65009629 | -0.2167971 | 0.01286438 |
| 0.999490048 | -0.1218355 | 0.12224502 | 0.65009629 | -0.0431264 | 0.57633576 |
| 0.999490048 | -0.1037589 | 0.12308044 | 0.65009629 | -0.0962296 | 0.15135298 |
| 0.999490048 | -0.0807141 | 0.12273827 | 0.65009629 | -0.0477382 | 0.35385247 |
| 0.999490048 | -0.0690199 | 0.12244721 | 0.65009629 | -0.0527995 | 0.23275899 |
| 0.999490048 | -0.0685306 | 0.12194878 | 0.65009629 | -0.0384867 | 0.37722488 |
| 0.999490048 | 0.1155821 | 0.12278395 | 0.65009629 | 0.11654468 | 0.11984165 |
| 0.999490048 | 0.12527699 | 0.12309009 | 0.65009629 | 0.09635761 | 0.23111175 |
| 0.999490048 | 0.12796085 | 0.12215757 | 0.65009629 | 0.12359059 | 0.13482564 |
| 0.999490048 | 0.13161853 | 0.12227116 | 0.65009629 | 0.21949918 | 0.01319589 |
| 0.999490048 | 0.16391982 | 0.12261349 | 0.65009629 | 0.0456224 | 0.66036763 |
| 0.999490048 | 0.17432416 | 0.12308445 | 0.65009629 | 0.05476176 | 0.62057439 |
| 0.999490048 | 0.29809052 | 0.12262455 | 0.65009629 | 0.32305926 | 0.09563164 |
| 0.999490048 | 0.10771908 | 0.12330334 | 0.65081094 | 0.07654108 | 0.2678855 |
| 0.999490048 | -0.0921458 | 0.12338615 | 0.65083663 | -0.0107251 | 0.85428307 |
| 0.999490048 | -0.4796268 | 0.12426629 | 0.65183852 | -0.0749326 | 0.79554931 |
| 0.999490048 | -0.2113919 | 0.12420518 | 0.65183852 | -0.0052355 | 0.96888793 |
| 0.999490048 | 0.08007737 | 0.1238489 | 0.65183852 | 0.00661626 | 0.89638827 |
| 0.999490048 | 0.08458911 | 0.12417431 | 0.65183852 | 0.01169652 | 0.82768315 |
| 0.999490048 | 0.08459473 | 0.12413197 | 0.65183852 | 0.07765401 | 0.15657553 |
| 0.999490048 | 0.09985687 | 0.12414847 | 0.65183852 | 0.0912165 | 0.15854646 |
| 0.999490048 | 0.11940881 | 0.12402281 | 0.65183852 | 0.13917028 | 0.07539924 |
| 0.972489992 | 0.17888871 | 0.12427867 | 0.65183852 | -0.0917553 | 0.40459352 |
| 0.999490048 | 0.42337347 | 0.12413245 | 0.65183852 | 0.28674051 | 0.29150066 |

| | | | | | |
|---|---|---|---|---|---|
| 0.999490048 | -0.0709038 | 0.12446813 | 0.6524224 | -0.0778752 | 0.09307946 |
| 0.999490048 | -0.0689094 | 0.12456364 | 0.65251344 | -0.0590282 | 0.18554941 |
| 0.999490048 | -0.1895806 | 0.12603172 | 0.65255562 | -0.0897988 | 0.46045671 |
| 0.999490048 | -0.1624858 | 0.12596866 | 0.65255562 | -0.1028798 | 0.31132542 |
| 0.999490048 | -0.1620506 | 0.12573848 | 0.65255562 | -0.1944774 | 0.06886216 |
| 0.999490048 | -0.1567503 | 0.12516575 | 0.65255562 | -0.2326844 | 0.02664408 |
| 0.999490048 | -0.1483292 | 0.12537614 | 0.65255562 | -0.2222402 | 0.02546585 |
| 0.999490048 | -0.140647 | 0.12565474 | 0.65255562 | -0.2194465 | 0.02056523 |
| 0.999490048 | -0.1084976 | 0.12614921 | 0.65255562 | -0.0930707 | 0.18682074 |
| 0.999490048 | -0.0978982 | 0.12524821 | 0.65255562 | -0.104647 | 0.10237303 |
| 0.999490048 | -0.0816787 | 0.12477009 | 0.65255562 | -0.0673663 | 0.20210873 |
| 0.999490048 | -0.0796073 | 0.12523482 | 0.65255562 | -0.01619 | 0.74969965 |
| 0.999490048 | -0.0669396 | 0.12535946 | 0.65255562 | -0.0437607 | 0.30984743 |
| 0.999490048 | -0.0635305 | 0.1260016 | 0.65255562 | -0.0342512 | 0.40157402 |
| 0.999490048 | 0.06947668 | 0.12522354 | 0.65255562 | -0.0014826 | 0.97326745 |
| 0.999490048 | 0.07895329 | 0.12621285 | 0.65255562 | 0.02195702 | 0.66378182 |
| 0.999490048 | 0.08563997 | 0.12482821 | 0.65255562 | 0.08965979 | 0.10886483 |
| 0.999490048 | 0.09390195 | 0.12521646 | 0.65255562 | 0.11765751 | 0.05788111 |
| 0.999490048 | 0.1095016 | 0.12482949 | 0.65255562 | 0.10916686 | 0.12593311 |
| 0.999490048 | 0.16960602 | 0.12550263 | 0.65255562 | 0.17503155 | 0.11439699 |
| 0.987721587 | 0.19787829 | 0.12617858 | 0.65255562 | -0.0746575 | 0.55596016 |
| 0.999490048 | 0.19843809 | 0.12590153 | 0.65255562 | 0.17183756 | 0.17738186 |
| 0.96108219 | 0.38678667 | 0.12563731 | 0.65255562 | -0.2082311 | 0.40176797 |
| 0.999490048 | 0.14593135 | 0.12639233 | 0.65307917 | -0.0372587 | 0.68988948 |
| 0.999490048 | -0.1150303 | 0.12648116 | 0.65313401 | -0.038994 | 0.59681211 |
| 0.999490048 | -0.1272205 | 0.1265801 | 0.65324092 | 0.02710378 | 0.73918318 |
| 0.999490048 | -0.1190285 | 0.1267761 | 0.65342346 | -0.0682854 | 0.37353257 |
| 0.972489992 | -0.111971 | 0.12681389 | 0.65342346 | 0.05286295 | 0.46287414 |
| 0.999490048 | -0.0784708 | 0.12685912 | 0.65342346 | -0.0561596 | 0.26914886 |
| 0.999490048 | -0.0721324 | 0.12700548 | 0.65342346 | -0.0511751 | 0.27341686 |
| 0.999490048 | 0.0990164 | 0.12700674 | 0.65342346 | 0.11693338 | 0.07406319 |
| 0.448222939 | -0.1084163 | 0.1274947 | 0.65553 | 0.12475427 | 0.08168521 |
| 0.999490048 | 0.11826843 | 0.12782495 | 0.65641963 | 0.14727812 | 0.06105046 |
| 0.989730632 | 0.19935085 | 0.12780739 | 0.65641963 | -0.0682575 | 0.59448826 |
| 0.999490048 | -0.0816715 | 0.12862706 | 0.66013275 | -0.070662 | 0.18607666 |
| 0.999490048 | 0.16301244 | 0.12871494 | 0.66017796 | 0.03330908 | 0.75087592 |
| 0.999490048 | -0.0843018 | 0.1288214 | 0.6603184 | -0.0483559 | 0.3762457 |
| 0.999490048 | 0.1671881 | 0.12905795 | 0.66112508 | 0.08222891 | 0.4473785 |
| 0.999490048 | -0.098211 | 0.12980259 | 0.66250102 | -0.0750652 | 0.24243723 |
| 0.999490048 | -0.0655249 | 0.12959132 | 0.66250102 | -0.0068531 | 0.87111962 |
| 0.999490048 | 0.06660493 | 0.12973136 | 0.66250102 | 0.07797063 | 0.07854547 |
| 0.999490048 | 0.07891332 | 0.12944671 | 0.66250102 | 0.05220716 | 0.30952914 |

| | | | | | |
|---|---|---|---|---|---|
| 0.999490048 | 0.15669766 | 0.12979458 | 0.66250102 | 0.17282441 | 0.09635836 |
| 0.999490048 | 0.17056342 | 0.12965312 | 0.66250102 | 0.18718928 | 0.09780026 |
| 0.999490048 | -0.0742679 | 0.12996884 | 0.6629443 | -0.0293208 | 0.54209865 |
| 0.999490048 | -0.0656452 | 0.13010843 | 0.66325117 | -0.0628427 | 0.14656209 |
| 0.999490048 | -0.0875202 | 0.13033676 | 0.66377885 | -0.1185096 | 0.04426378 |
| 0.999490048 | 0.18398232 | 0.13037094 | 0.66377885 | 0.08113279 | 0.49676281 |
| 0.999490048 | -0.0846683 | 0.13050074 | 0.66403482 | 0.01082711 | 0.84316618 |
| 0.999490048 | -0.0622172 | 0.1308213 | 0.66445124 | -0.0386549 | 0.34110533 |
| 0.999490048 | 0.11591566 | 0.13080184 | 0.66445124 | 0.08725186 | 0.25072597 |
| 0.999490048 | 0.13201169 | 0.13081769 | 0.66445124 | 0.0909281 | 0.29223044 |
| 0.999490048 | 0.08861334 | 0.13097592 | 0.66483219 | 0.06817369 | 0.24090673 |
| 0.999490048 | 0.18561086 | 0.13114129 | 0.66526718 | 0.13054155 | 0.28281618 |
| 0.999490048 | -0.1050161 | 0.13134746 | 0.66590848 | -0.0729919 | 0.28863751 |
| 0.999490048 | 0.09187166 | 0.1314642 | 0.66597985 | -0.0051756 | 0.93070446 |
| 0.999490048 | 0.21258163 | 0.13152105 | 0.66597985 | 0.01597845 | 0.90765393 |
| 0.999490048 | -0.9343699 | 0.1319141 | 0.66633499 | -0.2476102 | 0.66877921 |
| 0.999490048 | -0.074289 | 0.13203366 | 0.66633499 | -0.0362233 | 0.45487256 |
| 0.999490048 | 0.06757831 | 0.13206999 | 0.66633499 | 0.03129957 | 0.4775927 |
| 0.999490048 | 0.10863284 | 0.13196407 | 0.66633499 | 0.25025666 | 0.00141022 |
| 0.969787208 | 0.11011497 | 0.13178462 | 0.66633499 | -0.0665018 | 0.34682128 |
| 0.999490048 | 0.15601016 | 0.13193309 | 0.66633499 | 0.02101164 | 0.83557602 |
| 0.999490048 | -0.0585 | 0.13217269 | 0.66645044 | -0.0232339 | 0.5421056 |
| 0.987721587 | -0.255083 | 0.13242537 | 0.66726219 | 0.10176017 | 0.54059667 |
| 0.999490048 | -0.1177208 | 0.13249838 | 0.66726219 | -0.016136 | 0.83295667 |
| 0.999490048 | 0.10037443 | 0.13257341 | 0.66726219 | 0.0383431 | 0.55794652 |
| 0.999490048 | -0.1632875 | 0.13291522 | 0.66857956 | -0.1553202 | 0.15202619 |
| 0.999490048 | -0.2070834 | 0.13313629 | 0.66886311 | -0.1082863 | 0.40796153 |
| 0.999490048 | -0.1752135 | 0.13352279 | 0.66886311 | -0.1432595 | 0.21651272 |
| 0.972489992 | -0.1124168 | 0.13373829 | 0.66886311 | 0.05190576 | 0.48090112 |
| 0.771925038 | -0.1030596 | 0.13368635 | 0.66886311 | 0.08201758 | 0.22884292 |
| 0.999490048 | -0.0963851 | 0.1337702 | 0.66886311 | -0.0613774 | 0.33319896 |
| 0.987721587 | -0.0906799 | 0.13374922 | 0.66886311 | 0.04004875 | 0.50001061 |
| 0.999490048 | -0.0602562 | 0.13315373 | 0.66886311 | 0.00162117 | 0.96702241 |
| 0.999490048 | 0.12784714 | 0.1333281 | 0.66886311 | 0.193449 | 0.027083 |
| 0.999490048 | 0.13143419 | 0.13377262 | 0.66886311 | 0.08176812 | 0.34425434 |
| 0.972489992 | 0.13913698 | 0.133606 | 0.66886311 | -0.0663367 | 0.4667119 |
| 0.999490048 | 0.09089455 | 0.13395934 | 0.66939588 | 0.03772133 | 0.52623742 |
| 0.999490048 | -0.274778 | 0.13448428 | 0.66954518 | -0.0179441 | 0.92035792 |
| 0.999490048 | -0.1399876 | 0.1346307 | 0.66954518 | -0.1605944 | 0.08827244 |
| 0.999490048 | -0.1058094 | 0.13419021 | 0.66954518 | -0.172838 | 0.01811558 |
| 0.999490048 | -0.0703082 | 0.13438963 | 0.66954518 | -0.1344845 | 0.00664087 |
| 0.999490048 | 0.08768583 | 0.13417432 | 0.66954518 | 0.07418664 | 0.20220853 |

| | | | | | |
|---|---|---|---|---|---|
| 0.999490048 | 0.08952507 | 0.1342584 | 0.66954518 | 0.08158711 | 0.17072852 |
| 0.999490048 | 0.09296241 | 0.13459464 | 0.66954518 | 0.03969205 | 0.5151883 |
| 0.987721587 | 0.19003273 | 0.1345363 | 0.66954518 | -0.08068 | 0.51747792 |
| 0.876151891 | 0.09715109 | 0.13482626 | 0.67011863 | -0.0654182 | 0.30805895 |
| 0.999490048 | -0.2716463 | 0.13508279 | 0.67099422 | -0.161275 | 0.3680263 |
| 0.999490048 | -0.3747507 | 0.13566187 | 0.67134509 | 0.13732938 | 0.56048342 |
| 0.999490048 | -0.1721813 | 0.13555643 | 0.67134509 | -0.1424968 | 0.21355065 |
| 0.969787208 | -0.1033611 | 0.13570191 | 0.67134509 | 0.05754771 | 0.39887724 |
| 0.999490048 | -0.0922522 | 0.13603162 | 0.67134509 | -0.0594807 | 0.33017535 |
| 0.999490048 | -0.0769053 | 0.1358299 | 0.67134509 | 0.00817968 | 0.87112553 |
| 0.999490048 | 0.10757394 | 0.1354467 | 0.67134509 | -0.0238301 | 0.73541604 |
| 0.999490048 | 0.11077839 | 0.13592361 | 0.67134509 | 0.03403567 | 0.64007141 |
| 0.972489992 | 0.1275159 | 0.13601361 | 0.67134509 | -0.0668364 | 0.42708546 |
| 0.999490048 | 0.1542518 | 0.13593544 | 0.67134509 | 0.06939568 | 0.49463502 |
| 0.999490048 | 0.28766376 | 0.13603783 | 0.67134509 | 0.07613928 | 0.68699742 |
| 0.999490048 | 0.34858534 | 0.13575725 | 0.67134509 | 0.12700402 | 0.57940724 |
| 0.999490048 | 0.11812566 | 0.13636923 | 0.67258304 | 0.12802169 | 0.10780382 |
| 0.999490048 | -0.1240282 | 0.13646907 | 0.67267814 | -0.0331226 | 0.68468983 |
| 0.999490048 | 0.08013432 | 0.13673503 | 0.67359143 | 0.06461185 | 0.22662859 |
| 0.999490048 | 0.32716806 | 0.13686925 | 0.67385509 | 0.12180744 | 0.57232777 |
| 0.999490048 | -0.3254583 | 0.13740906 | 0.67444044 | -0.6569011 | 0.00475671 |
| 0.771925038 | -0.2307632 | 0.13740859 | 0.67444044 | 0.18302737 | 0.2348927 |
| 0.999490048 | -0.1749054 | 0.13748559 | 0.67444044 | -0.2073919 | 0.08083043 |
| 0.987721587 | -0.0948009 | 0.13722685 | 0.67444044 | 0.04125513 | 0.51016929 |
| 0.999490048 | -0.08186 | 0.13771508 | 0.67444044 | 0.00908938 | 0.86618356 |
| 0.999490048 | -0.0729832 | 0.13766736 | 0.67444044 | -0.1311137 | 0.01076362 |
| 0.999490048 | -0.0632684 | 0.13757938 | 0.67444044 | -0.027074 | 0.517575 |
| 0.999490048 | 0.08508616 | 0.13767257 | 0.67444044 | 0.13065574 | 0.02665426 |
| 0.999490048 | 0.12622838 | 0.13742648 | 0.67444044 | 0.00821869 | 0.92121363 |
| 0.999490048 | -0.1866885 | 0.13915311 | 0.67476149 | -0.3991195 | 0.0031498 |
| 0.999490048 | -0.1621029 | 0.13856182 | 0.67476149 | -0.1211492 | 0.26369073 |
| 0.999490048 | -0.1321207 | 0.13896585 | 0.67476149 | -0.0438157 | 0.61672511 |
| 0.999490048 | -0.1289018 | 0.138225 | 0.67476149 | -0.1423345 | 0.10325972 |
| 0.999490048 | -0.1047254 | 0.13897504 | 0.67476149 | -0.041954 | 0.54592342 |
| 0.999490048 | -0.087709 | 0.13927881 | 0.67476149 | -0.0777792 | 0.18778219 |
| 0.999490048 | -0.0803688 | 0.13801378 | 0.67476149 | 0.00221249 | 0.96669277 |
| 0.999490048 | -0.0641755 | 0.1392077 | 0.67476149 | -0.029668 | 0.48668252 |
| 0.999490048 | 0.06594114 | 0.13809875 | 0.67476149 | 0.05686218 | 0.19847797 |
| 0.999490048 | 0.07664016 | 0.13852057 | 0.67476149 | 0.0334411 | 0.51049038 |
| 0.999490048 | 0.07858008 | 0.13931603 | 0.67476149 | 0.03259405 | 0.53228056 |
| 0.999490048 | 0.08113237 | 0.13798377 | 0.67476149 | 0.05691435 | 0.29268257 |
| 0.999490048 | 0.08371514 | 0.13908638 | 0.67476149 | 0.07655145 | 0.17464802 |

| | | | | | |
|---|---|---|---|---|---|
| 0.999490048 | 0.09114902 | 0.13881174 | 0.67476149 | 0.02527485 | 0.67523616 |
| 0.999490048 | 0.09487295 | 0.13914293 | 0.67476149 | 0.09218139 | 0.15021805 |
| 0.999490048 | 0.12048 | 0.1384241 | 0.67476149 | 0.06060188 | 0.44861591 |
| 0.999490048 | 0.12806627 | 0.13856646 | 0.67476149 | 0.06655117 | 0.43408299 |
| 0.999490048 | 0.13214945 | 0.13872548 | 0.67476149 | 0.0619602 | 0.48001797 |
| 0.999490048 | 0.19111976 | 0.13897965 | 0.67476149 | 0.02199948 | 0.86178613 |
| 0.999490048 | 0.22690892 | 0.13943854 | 0.67496336 | 0.08184326 | 0.58684192 |
| 0.999490048 | -0.3181299 | 0.14000253 | 0.67521382 | -0.2856594 | 0.18329033 |
| 0.999490048 | -0.1425207 | 0.14004259 | 0.67521382 | -0.0537184 | 0.57083999 |
| 0.999490048 | -0.1106813 | 0.14013719 | 0.67521382 | -0.0290028 | 0.69312165 |
| 0.999490048 | 0.08257027 | 0.14007651 | 0.67521382 | 0.05215071 | 0.34504991 |
| 0.999490048 | 0.13147575 | 0.13983036 | 0.67521382 | 0.05106831 | 0.55899983 |
| 0.999490048 | 0.1324523 | 0.1401234 | 0.67521382 | 0.05911127 | 0.50278167 |
| 0.999490048 | 0.18733251 | 0.13990425 | 0.67521382 | 0.08394076 | 0.50083014 |
| 0.999490048 | 0.24198536 | 0.13997726 | 0.67521382 | 0.24042237 | 0.14222504 |
| 0.999490048 | -0.0888376 | 0.14059333 | 0.67695283 | -0.0406387 | 0.49278457 |
| 0.999490048 | 0.11403329 | 0.14066026 | 0.67695283 | 0.20432603 | 0.01151168 |
| 0.999490048 | -0.1735484 | 0.14113461 | 0.67720189 | 0.01090357 | 0.92249085 |
| 0.999490048 | -0.1308454 | 0.14128752 | 0.67720189 | -0.1157809 | 0.19097002 |
| 0.999490048 | -0.0837551 | 0.14110368 | 0.67720189 | -0.0974298 | 0.08925886 |
| 0.999490048 | 0.11406204 | 0.14130591 | 0.67720189 | 0.06450116 | 0.39854725 |
| 0.999490048 | 0.11741831 | 0.14111538 | 0.67720189 | 0.07507243 | 0.34055235 |
| 0.999490048 | 0.15614457 | 0.14136083 | 0.67720189 | 0.06958937 | 0.50473653 |
| 0.999490048 | 0.17498915 | 0.14097661 | 0.67720189 | 0.14725525 | 0.21252914 |
| 0.999490048 | 0.21221945 | 0.14128064 | 0.67720189 | 0.26036675 | 0.07412521 |
| 0.909524814 | -0.3015984 | 0.14153315 | 0.67725032 | 0.19531882 | 0.33498982 |
| 0.999490048 | -0.1145153 | 0.14148934 | 0.67725032 | -0.2069912 | 0.01104033 |
| 0.999490048 | 0.071696 | 0.14167305 | 0.67753146 | 0.04631111 | 0.33639228 |
| 0.999490048 | -0.458344 | 0.14218607 | 0.67843069 | -0.555681 | 0.08372756 |
| 0.999490048 | 0.15428207 | 0.14205214 | 0.67843069 | 0.02892858 | 0.77857454 |
| 0.999490048 | 0.22054679 | 0.14215381 | 0.67843069 | 0.19491762 | 0.19246696 |
| 0.999490048 | 0.30632826 | 0.14214412 | 0.67843069 | 0.19782483 | 0.32410903 |
| 0.999490048 | 0.09288724 | 0.14226991 | 0.67844304 | 0.10204802 | 0.10846793 |
| 0.999490048 | -0.0832624 | 0.14322258 | 0.68104168 | -0.0524518 | 0.35018385 |
| 0.999490048 | -0.0731743 | 0.14330422 | 0.68104168 | -0.0872528 | 0.0836325 |
| 0.999490048 | 0.11056295 | 0.14329221 | 0.68104168 | 0.10849264 | 0.15061665 |
| 0.999490048 | 0.11127475 | 0.14314137 | 0.68104168 | 0.07920323 | 0.29222146 |
| 0.999490048 | 0.12533998 | 0.14325633 | 0.68104168 | -0.0277429 | 0.74081577 |
| 0.999490048 | 0.23582981 | 0.14328265 | 0.68104168 | 0.40731708 | 0.01505536 |
| 0.999490048 | -0.158947 | 0.14421942 | 0.68201334 | -0.0594291 | 0.57806101 |
| 0.999490048 | -0.1383135 | 0.14417214 | 0.68201334 | 4.27E-04 | 0.99632037 |
| 0.999490048 | -0.0658854 | 0.14394309 | 0.68201334 | -0.0735857 | 0.10444593 |

| | | | | | |
|---|---|---|---|---|---|
| 0.999490048 | 0.064513 | 0.1438562 | 0.68201334 | 0.0118835 | 0.78336398 |
| 0.999490048 | 0.09992144 | 0.14421361 | 0.68201334 | 0.00159763 | 0.98096388 |
| 0.999490048 | 0.10605599 | 0.14424378 | 0.68201334 | 0.09756426 | 0.17778366 |
| 0.999490048 | 0.1192682 | 0.1438291 | 0.68201334 | 0.13438187 | 0.10148434 |
| 0.999490048 | 0.13412136 | 0.14397661 | 0.68201334 | -0.0233341 | 0.79516064 |
| 0.999490048 | 0.16032218 | 0.14414467 | 0.68201334 | 0.17571017 | 0.11093034 |
| 0.999490048 | -0.120865 | 0.1448673 | 0.68345746 | -0.0496808 | 0.5418157 |
| 0.999490048 | -0.1168729 | 0.14504031 | 0.68345746 | -0.1148168 | 0.15194882 |
| 0.999490048 | -0.1026398 | 0.14497169 | 0.68345746 | -0.0571136 | 0.41045189 |
| 0.999490048 | -0.0773149 | 0.14488687 | 0.68345746 | -0.0304652 | 0.55856034 |
| 0.999490048 | 0.08353776 | 0.14500312 | 0.68345746 | 0.05395605 | 0.34054779 |
| 0.999490048 | 0.08606602 | 0.14485752 | 0.68345746 | 0.0936071 | 0.11418146 |
| 0.999490048 | 0.11925006 | 0.14513341 | 0.68351041 | 0.13341956 | 0.10489104 |
| 0.999490048 | -0.1520681 | 0.14551675 | 0.68492948 | -0.0172468 | 0.86608536 |
| 0.999490048 | -0.1320527 | 0.14585478 | 0.68522175 | 0.00247409 | 0.97778922 |
| 0.999490048 | -0.0712246 | 0.14598916 | 0.68522175 | -0.0623856 | 0.20066967 |
| 0.999490048 | 0.05363389 | 0.14587758 | 0.68522175 | 0.02817537 | 0.43784383 |
| 0.999490048 | 0.06757095 | 0.1459597 | 0.68522175 | 0.04750355 | 0.30153732 |
| 0.692187333 | 0.15097657 | 0.14595815 | 0.68522175 | -0.1415227 | 0.17182378 |
| 0.999490048 | -0.1063393 | 0.14610703 | 0.68538971 | -0.0074297 | 0.91737225 |
| 0.999490048 | -0.2567715 | 0.1461962 | 0.68542295 | 0.03421341 | 0.83577097 |
| 0.999490048 | -0.3651737 | 0.14640305 | 0.6856228 | -0.3838856 | 0.12864548 |
| 0.999490048 | 0.13822838 | 0.14639886 | 0.6856228 | 0.11657013 | 0.21786144 |
| 0.999490048 | -0.1695446 | 0.14649276 | 0.68565836 | -0.1544685 | 0.18430716 |
| 0.999490048 | -0.0813226 | 0.14673349 | 0.68640037 | -0.0349949 | 0.52516758 |
| 0.999490048 | -0.0969203 | 0.14704115 | 0.68644151 | -0.1160021 | 0.0853723 |
| 0.999490048 | -0.0945415 | 0.14717455 | 0.68644151 | -0.0850993 | 0.1902075 |
| 0.999490048 | -0.0772817 | 0.1472059 | 0.68644151 | -0.1254834 | 0.02264866 |
| 0.999490048 | 0.06875668 | 0.14730835 | 0.68644151 | -0.0091555 | 0.84378707 |
| 0.999490048 | 0.10519658 | 0.14717705 | 0.68644151 | 0.13056984 | 0.07514258 |
| 0.999490048 | 0.10521358 | 0.14731775 | 0.68644151 | 0.11803569 | 0.10585923 |
| 0.999490048 | 0.11971694 | 0.14721437 | 0.68644151 | 0.02452562 | 0.76183234 |
| 0.999490048 | 0.07745002 | 0.14748475 | 0.68683639 | 0.03455556 | 0.51096853 |
| 0.999490048 | -0.1199293 | 0.14761066 | 0.6870396 | -0.1179985 | 0.15396959 |
| 0.999490048 | 0.1343783 | 0.1478246 | 0.68765204 | 0.1513618 | 0.10496244 |
| 0.999490048 | -0.1734647 | 0.1479404 | 0.68780754 | -0.2282586 | 0.06064041 |
| 0.999490048 | -0.0911484 | 0.14823177 | 0.68839559 | -0.0165693 | 0.78848289 |
| 0.999490048 | 0.33810641 | 0.14821236 | 0.68839559 | -0.0543002 | 0.81260322 |
| 0.987721587 | -0.0989972 | 0.14833066 | 0.68839796 | 0.04406917 | 0.51282785 |
| 0.999490048 | 0.11191018 | 0.14847961 | 0.68839796 | 0.06602202 | 0.38748548 |
| 0.999490048 | 0.13911202 | 0.1484232 | 0.68839796 | -0.0272858 | 0.77235429 |
| 0.999490048 | 0.1051482 | 0.14871472 | 0.68910541 | 0.07422402 | 0.30296204 |

| | | | | | |
|---|---|---|---|---|---|
| 0.999490048 | -0.2447179 | 0.14891653 | 0.68947465 | -0.1138841 | 0.49469164 |
| 0.999490048 | -0.1222303 | 0.14914205 | 0.68947465 | -0.0435601 | 0.60052131 |
| 0.999490048 | -0.066476 | 0.14920727 | 0.68947465 | -0.0976144 | 0.03842765 |
| 0.999490048 | 0.09773152 | 0.14914891 | 0.68947465 | 0.10365161 | 0.12699968 |
| 0.999490048 | 0.12336744 | 0.14897829 | 0.68947465 | 0.06807401 | 0.41907349 |
| 0.999490048 | -0.1494654 | 0.15040595 | 0.69121819 | -0.1713058 | 0.10144885 |
| 0.999490048 | -0.1389235 | 0.14993902 | 0.69121819 | -0.250807 | 0.01276876 |
| 0.999490048 | -0.0721759 | 0.15024833 | 0.69121819 | -0.0741865 | 0.13966628 |
| 0.999490048 | -0.0672497 | 0.14974712 | 0.69121819 | -0.0226019 | 0.6218496 |
| 0.999490048 | -0.0611848 | 0.14994562 | 0.69121819 | -0.0422117 | 0.31531488 |
| 0.999490048 | 0.08079387 | 0.15009309 | 0.69121819 | 0.08068388 | 0.15062621 |
| 0.548539439 | 0.08841587 | 0.149999 | 0.69121819 | -0.1005823 | 0.1035392 |
| 0.999490048 | 0.09739551 | 0.15023596 | 0.69121819 | 0.07009382 | 0.29566073 |
| 0.999490048 | 0.10727677 | 0.14987567 | 0.69121819 | 0.05605619 | 0.4448904 |
| 0.999490048 | 0.11442596 | 0.15041239 | 0.69121819 | 0.0535816 | 0.49367706 |
| 0.999490048 | 0.06843241 | 0.15078135 | 0.6914818 | 0.07353784 | 0.12376761 |
| 0.999490048 | 0.08554802 | 0.15088381 | 0.6914818 | 0.07594516 | 0.20029522 |
| 0.999490048 | 0.08835783 | 0.15068471 | 0.6914818 | 0.08868199 | 0.14925326 |
| 0.999490048 | 0.0898291 | 0.15062243 | 0.6914818 | 0.03780821 | 0.53825473 |
| 0.999490048 | 0.30644087 | 0.15080631 | 0.6914818 | -0.0566622 | 0.78067641 |
| 0.999490048 | -0.095407 | 0.15098437 | 0.6915393 | -0.0350329 | 0.59131374 |
| 0.999490048 | 0.07546538 | 0.15109224 | 0.6915393 | 0.07670646 | 0.14475359 |
| 0.999490048 | 0.09812042 | 0.15114482 | 0.6915393 | 0.09315801 | 0.17205386 |
| 0.999490048 | -0.5856329 | 0.15135848 | 0.69175879 | -0.2389047 | 0.53611607 |
| 0.999490048 | -0.0922214 | 0.1512937 | 0.69175879 | -0.0823357 | 0.19828062 |
| 0.999490048 | 0.05040377 | 0.1518737 | 0.69185575 | 0.02458311 | 0.47769886 |
| 0.999490048 | 0.07426315 | 0.15169428 | 0.69185575 | 0.11337305 | 0.03293521 |
| 0.999490048 | 0.10994628 | 0.15168177 | 0.69185575 | 0.15372402 | 0.04915005 |
| 0.999490048 | 0.1171595 | 0.15187684 | 0.69185575 | 0.03913891 | 0.62581243 |
| 0.999490048 | 0.19653382 | 0.15171247 | 0.69185575 | 0.18031899 | 0.18701111 |
| 0.999490048 | 0.22284117 | 0.15152124 | 0.69185575 | 0.10762601 | 0.48156671 |
| 0.999490048 | -0.1130591 | 0.15225693 | 0.69267636 | -0.0337906 | 0.66279865 |
| 0.999490048 | 0.10169268 | 0.15230584 | 0.69267636 | 0.14532926 | 0.04501261 |
| 0.999490048 | 0.1029989 | 0.15221835 | 0.69267636 | 0.20009002 | 0.00823838 |
| 0.999490048 | -0.0647698 | 0.15245673 | 0.69298512 | -0.0452845 | 0.31203165 |
| 0.999490048 | -0.0783648 | 0.15255687 | 0.69306305 | -0.0028585 | 0.95751024 |
| 0.999490048 | -0.1050093 | 0.15267515 | 0.69322324 | -0.0829352 | 0.25503522 |
| 0.999490048 | -0.1524972 | 0.15309198 | 0.69398373 | -0.0448067 | 0.66879711 |
| 0.999490048 | -0.0780209 | 0.15306614 | 0.69398373 | -0.1277033 | 0.0235115 |
| 0.999490048 | 0.07626258 | 0.15302227 | 0.69398373 | 0.03966559 | 0.45057127 |
| 0.999490048 | -0.1201049 | 0.15347028 | 0.69447241 | -0.1030256 | 0.21833767 |
| 0.999490048 | -0.088333 | 0.15334967 | 0.69447241 | -0.0710605 | 0.24732874 |

| | | | | | |
|---|---|---|---|---|---|
| 0.999490048 | 0.06267215 | 0.15341071 | 0.69447241 | 0.00445486 | 0.91749357 |
| 0.999490048 | 0.09233277 | 0.15353246 | 0.69447241 | -0.0335253 | 0.59780451 |
| 0.999490048 | -0.117079 | 0.15423557 | 0.69485044 | -0.0868684 | 0.28607389 |
| 0.999490048 | -0.1134368 | 0.15425626 | 0.69485044 | -0.0727699 | 0.35499783 |
| 0.999490048 | -0.0953088 | 0.15460903 | 0.69485044 | -0.0300367 | 0.64768546 |
| 0.999490048 | -0.082401 | 0.1545893 | 0.69485044 | -0.1365931 | 0.02242325 |
| 0.999490048 | -0.0799862 | 0.15470803 | 0.69485044 | -0.0369282 | 0.50433709 |
| 0.999490048 | -0.071456 | 0.15442355 | 0.69485044 | -0.0147343 | 0.76455957 |
| 0.999490048 | -0.0629366 | 0.15404827 | 0.69485044 | -0.0572721 | 0.19306043 |
| 0.999490048 | 0.06519726 | 0.15408103 | 0.69485044 | 0.04154419 | 0.35796312 |
| 0.999490048 | 0.07288159 | 0.15443574 | 0.69485044 | 0.00964281 | 0.84753152 |
| 0.999490048 | 0.07960075 | 0.15478106 | 0.69485044 | 0.03466986 | 0.52864165 |
| 0.999490048 | 0.10421732 | 0.1544081 | 0.69485044 | 0.10709546 | 0.14378747 |
| 0.999490048 | 0.10797285 | 0.15459707 | 0.69485044 | 0.10784088 | 0.15508158 |
| 0.999490048 | 0.17173405 | 0.15473595 | 0.69485044 | -8.95E-04 | 0.99396004 |
| 0.999490048 | 0.20584857 | 0.15431566 | 0.69485044 | 0.09872987 | 0.48764207 |
| 0.999490048 | -0.2023902 | 0.15554086 | 0.69540895 | 3.40E-04 | 0.99805676 |
| 0.999490048 | -0.1330189 | 0.15595171 | 0.69540895 | -0.0871355 | 0.34001769 |
| 0.999490048 | -0.101219 | 0.15545836 | 0.69540895 | -0.052927 | 0.45094403 |
| 0.999490048 | -0.0815992 | 0.15607142 | 0.69540895 | -0.061294 | 0.28245684 |
| 0.999490048 | -0.0683216 | 0.15601124 | 0.69540895 | -0.0537768 | 0.26052084 |
| 0.999490048 | -0.0662235 | 0.15582463 | 0.69540895 | -0.0124842 | 0.78493863 |
| 0.999490048 | -0.0562463 | 0.15546374 | 0.69540895 | -0.0337944 | 0.38725446 |
| 0.999490048 | 0.07525206 | 0.15566021 | 0.69540895 | 0.13689417 | 0.01332529 |
| 0.999490048 | 0.07979679 | 0.15602656 | 0.69540895 | 0.04335016 | 0.43436976 |
| 0.999490048 | 0.08549408 | 0.15546703 | 0.69540895 | 0.03370271 | 0.56887113 |
| 0.999490048 | 0.097071 | 0.15560057 | 0.69540895 | 0.12084031 | 0.08030842 |
| 0.999490048 | 0.11395767 | 0.15604693 | 0.69540895 | 0.046049 | 0.5598616 |
| 0.999490048 | 0.12136458 | 0.15551977 | 0.69540895 | 0.15170097 | 0.07909406 |
| 0.999490048 | 0.177253 | 0.15595308 | 0.69540895 | 0.11869915 | 0.33667564 |
| 0.999490048 | -0.1067234 | 0.15626449 | 0.69589786 | 0.04208609 | 0.56950866 |
| 0.999490048 | -0.0775014 | 0.1564522 | 0.69624899 | -0.0652061 | 0.23042793 |
| 0.999490048 | -0.0702907 | 0.1565101 | 0.69624899 | 0.0237921 | 0.62532477 |
| 0.999490048 | -0.1665659 | 0.15702658 | 0.69691335 | 0.05679487 | 0.62324 |
| 0.999490048 | -0.143991 | 0.15724368 | 0.69691335 | -0.012459 | 0.90061793 |
| 0.999490048 | -0.0765487 | 0.15704739 | 0.69691335 | -0.0604106 | 0.26051222 |
| 0.999490048 | 0.07137267 | 0.15705911 | 0.69691335 | 0.05603101 | 0.26296627 |
| 0.999490048 | 0.15452741 | 0.15724095 | 0.69691335 | 0.00231635 | 0.9827368 |
| 0.999490048 | 0.16786781 | 0.15712815 | 0.69691335 | 0.22126654 | 0.06597836 |
| 0.999490048 | 0.36702001 | 0.15719019 | 0.69691335 | -0.181039 | 0.46387479 |
| 0.999490048 | -0.0652519 | 0.15769411 | 0.69742892 | -0.0696489 | 0.13258865 |
| 0.999490048 | -0.0649206 | 0.15746465 | 0.69742892 | -0.0441794 | 0.33084868 |

| | | | | | |
|---|---|---|---|---|---|
| 0.999490048 | 0.28542399 | 0.1576809 | 0.69742892 | 0.30271331 | 0.13611387 |
| 0.999490048 | 0.35915259 | 0.15762341 | 0.69742892 | 0.55370552 | 0.03376684 |
| 0.999490048 | -0.0858503 | 0.15787828 | 0.69787382 | -0.0199452 | 0.73804075 |
| 0.999490048 | -0.2774533 | 0.15822883 | 0.69831436 | -0.0584822 | 0.76171385 |
| 0.999490048 | -0.0736255 | 0.15812405 | 0.69831436 | 0.02065632 | 0.68663222 |
| 0.999490048 | 0.12606759 | 0.15822534 | 0.69831436 | 0.11000187 | 0.21606483 |
| 0.999490048 | 0.08438506 | 0.15832204 | 0.6983566 | 0.09847595 | 0.10208842 |
| 0.999490048 | -0.1832047 | 0.15864293 | 0.69937512 | -0.2012828 | 0.12593735 |
| 0.999490048 | -0.0689655 | 0.15873667 | 0.69937512 | -0.0605355 | 0.2139899 |
| 0.999490048 | 0.10631882 | 0.15880422 | 0.69937512 | 0.20308358 | 0.01029037 |
| 0.999490048 | 0.28392204 | 0.15895782 | 0.69968256 | 0.1977451 | 0.31112553 |
| 0.999490048 | -0.0872905 | 0.15918089 | 0.70029527 | -0.0876425 | 0.15757165 |
| 0.999490048 | 0.17616141 | 0.15937559 | 0.70078259 | 0.13367716 | 0.28156921 |
| 0.999490048 | -0.0663007 | 0.15960796 | 0.701435 | -0.0776091 | 0.10213678 |
| 0.999490048 | -0.0624804 | 0.16010334 | 0.70225729 | -0.0018924 | 0.96537612 |
| 0.999490048 | -0.0452849 | 0.16013148 | 0.70225729 | -0.0221451 | 0.48556748 |
| 0.999490048 | 0.08876699 | 0.1601077 | 0.70225729 | 0.12835911 | 0.04658519 |
| 0.999490048 | 0.11985489 | 0.16002931 | 0.70225729 | 0.06139051 | 0.46521647 |
| 0.999490048 | -0.1338785 | 0.16045097 | 0.70255144 | -0.0861383 | 0.36093723 |
| 0.999490048 | -0.0779762 | 0.16040511 | 0.70255144 | -0.0040272 | 0.94104178 |
| 0.771925038 | 0.14943505 | 0.16032491 | 0.70255144 | -0.135424 | 0.20167688 |
| 0.999490048 | -0.0884598 | 0.16129604 | 0.70331115 | -0.0978945 | 0.12272063 |
| 0.999490048 | -0.0819744 | 0.1613398 | 0.70331115 | -0.0784895 | 0.1792066 |
| 0.999490048 | -0.0749095 | 0.16097801 | 0.70331115 | -0.0429167 | 0.41573322 |
| 0.999490048 | -0.0683684 | 0.16078679 | 0.70331115 | -0.043954 | 0.36170358 |
| 0.999490048 | 0.05732328 | 0.16117177 | 0.70331115 | 0.03629962 | 0.36929084 |
| 0.999490048 | 0.07011376 | 0.16146676 | 0.70331115 | 0.0415553 | 0.40061014 |
| 0.999490048 | 0.08644096 | 0.16134632 | 0.70331115 | 0.05780318 | 0.34374477 |
| 0.999490048 | 0.0942737 | 0.16146539 | 0.70331115 | 0.04776216 | 0.47160307 |
| 0.999490048 | 0.12028907 | 0.16109383 | 0.70331115 | 0.09089113 | 0.2855864 |
| 0.999490048 | 0.13672797 | 0.16115176 | 0.70331115 | -0.0035554 | 0.97034832 |
| 0.999490048 | -0.0675789 | 0.16187145 | 0.70405627 | -0.0532003 | 0.26726516 |
| 0.999490048 | 0.1031315 | 0.1619751 | 0.70405627 | 0.0168256 | 0.81606168 |
| 0.999490048 | 0.11568836 | 0.16177543 | 0.70405627 | 0.14087759 | 0.09137033 |
| 0.999490048 | 0.13557415 | 0.1618958 | 0.70405627 | 0.26885172 | 0.00844739 |
| 0.987721587 | -0.1095772 | 0.16234175 | 0.70454968 | 0.05733155 | 0.45834722 |
| 0.999490048 | -0.1050158 | 0.16217547 | 0.70454968 | -0.0733281 | 0.32420532 |
| 0.999490048 | 0.11131941 | 0.16232794 | 0.70454968 | 0.12379592 | 0.12189062 |
| 0.999490048 | -0.1544686 | 0.1627451 | 0.70529942 | 0.02964392 | 0.78481918 |
| 0.999490048 | -0.1167799 | 0.1627679 | 0.70529942 | -0.1392704 | 0.0986601 |
| 0.999490048 | 0.08011917 | 0.16264422 | 0.70529942 | 0.04671435 | 0.40954384 |
| 0.999490048 | -0.2570298 | 0.16444691 | 0.70539007 | -0.0533023 | 0.7689249 |

| | | | | | |
|---|---|---|---|---|---|
| 0.999490048 | -0.1154944 | 0.16326971 | 0.70539007 | -0.0998848 | 0.22566175 |
| 0.493399408 | -0.1106846 | 0.16374081 | 0.70539007 | 0.14085465 | 0.07981867 |
| 0.999490048 | -0.106822 | 0.16361767 | 0.70539007 | -0.1701169 | 0.03099855 |
| 0.999490048 | -0.0970077 | 0.16299387 | 0.70539007 | -0.108989 | 0.11885878 |
| 0.987721587 | -0.0927941 | 0.16315429 | 0.70539007 | 0.05149551 | 0.43277413 |
| 0.999490048 | -0.0860061 | 0.16410352 | 0.70539007 | -0.0229118 | 0.70580575 |
| 0.999490048 | -0.0859786 | 0.16456286 | 0.70539007 | -0.0526557 | 0.38890932 |
| 0.999490048 | -0.0726364 | 0.16429334 | 0.70539007 | -0.0890016 | 0.09137866 |
| 0.999490048 | 0.05845436 | 0.16437511 | 0.70539007 | 0.02515249 | 0.5432309 |
| 0.999490048 | 0.07157813 | 0.16300532 | 0.70539007 | 0.02719715 | 0.58983313 |
| 0.999490048 | 0.09185358 | 0.16369106 | 0.70539007 | 0.04376453 | 0.50048861 |
| 0.999490048 | 0.09363216 | 0.16392382 | 0.70539007 | 0.03960324 | 0.54962725 |
| 0.999490048 | 0.09536981 | 0.16446669 | 0.70539007 | 0.07271511 | 0.28526613 |
| 0.999490048 | 0.09677645 | 0.16373728 | 0.70539007 | 0.08818071 | 0.20299583 |
| 0.999490048 | 0.10312009 | 0.16410256 | 0.70539007 | 0.05279498 | 0.469861 |
| 0.999490048 | 0.11749357 | 0.16456261 | 0.70539007 | -0.0104957 | 0.8992544 |
| 0.999490048 | 0.12297763 | 0.16416962 | 0.70539007 | 8.21E-04 | 0.9924458 |
| 0.999490048 | 0.14471221 | 0.16368911 | 0.70539007 | 0.10544545 | 0.30577576 |
| 0.999490048 | 0.15440108 | 0.16377862 | 0.70539007 | 0.04019686 | 0.71195491 |
| 0.999490048 | 0.46966553 | 0.16378373 | 0.70539007 | 0.8365656 | 0.02559287 |
| 0.999490048 | -0.2629454 | 0.1656614 | 0.70590285 | -0.331558 | 0.08381891 |
| 0.999490048 | -0.1322985 | 0.16489506 | 0.70590285 | -0.0561552 | 0.54917951 |
| 0.999490048 | -0.1013243 | 0.16579401 | 0.70590285 | -0.1248563 | 0.09078071 |
| 0.999490048 | -0.0980008 | 0.16478271 | 0.70590285 | -0.0402463 | 0.56202956 |
| 0.999490048 | -0.0936078 | 0.16628873 | 0.70590285 | -0.0129786 | 0.84490438 |
| 0.999490048 | -0.0777139 | 0.16625698 | 0.70590285 | -0.0764219 | 0.17314559 |
| 0.999490048 | -0.0623711 | 0.16546692 | 0.70590285 | -0.0732793 | 0.10568181 |
| 0.999490048 | -0.059487 | 0.16601216 | 0.70590285 | -0.0765801 | 0.07814411 |
| 0.999490048 | -0.0546158 | 0.16510957 | 0.70590285 | -0.0180749 | 0.64018161 |
| 0.999490048 | 0.07107569 | 0.16571983 | 0.70590285 | 0.02211742 | 0.66067393 |
| 0.999490048 | 0.07294016 | 0.16590515 | 0.70590285 | 0.04447156 | 0.39253484 |
| 0.999490048 | 0.09507596 | 0.16615192 | 0.70590285 | 0.09489053 | 0.16694836 |
| 0.999490048 | 0.10001737 | 0.16619896 | 0.70590285 | 0.08979607 | 0.21217062 |
| 0.999490048 | 0.1031735 | 0.16586731 | 0.70590285 | 0.0989626 | 0.18316575 |
| 0.999490048 | 0.13055136 | 0.16519699 | 0.70590285 | 0.0632056 | 0.49525546 |
| 0.999490048 | 0.14171479 | 0.16560653 | 0.70590285 | 0.08408243 | 0.40485218 |
| 0.999490048 | 0.20494414 | 0.16503922 | 0.70590285 | -0.0171018 | 0.90596651 |
| 0.999490048 | 0.22002057 | 0.16577949 | 0.70590285 | 0.17855574 | 0.25760156 |
| 0.999490048 | 0.33237357 | 0.1657101 | 0.70590285 | 0.03775804 | 0.87246216 |
| 0.999490048 | -0.0841699 | 0.16702792 | 0.70757698 | -0.0055559 | 0.9259099 |
| 0.999490048 | 0.07600034 | 0.1671068 | 0.70757698 | 0.00359767 | 0.94683713 |
| 0.999490048 | 0.10408703 | 0.16690272 | 0.70757698 | 0.07469289 | 0.31676653 |

| | | | | | |
|---|---|---|---|---|---|
| 0.999490048 | 0.10545571 | 0.16706971 | 0.70757698 | 0.08362348 | 0.26985245 |
| 0.999490048 | 0.28174105 | 0.16686506 | 0.70757698 | 0.48911936 | 0.0294969 |
| 0.999490048 | -0.0775416 | 0.16720891 | 0.70765049 | -0.1082485 | 0.05808316 |
| 0.999490048 | -0.1266622 | 0.16790873 | 0.70965624 | -0.1694176 | 0.06906545 |
| 0.999490048 | -0.075928 | 0.16786555 | 0.70965624 | -0.0780304 | 0.1568361 |
| 0.999490048 | 0.11405166 | 0.16793781 | 0.70965624 | 0.11064108 | 0.18053579 |
| 0.999490048 | -0.0758381 | 0.16818631 | 0.71034683 | -0.0858304 | 0.12077765 |
| 0.972489992 | 0.18862888 | 0.1683005 | 0.71046977 | -0.1162646 | 0.39019535 |
| 0.999490048 | -0.1214506 | 0.16895846 | 0.71093283 | -0.1492172 | 0.09407038 |
| 0.999490048 | -0.1194929 | 0.16887629 | 0.71093283 | -0.05826 | 0.49609896 |
| 0.999490048 | -0.0938926 | 0.16909133 | 0.71093283 | -0.0039323 | 0.95318498 |
| 0.999490048 | -0.0716806 | 0.1688937 | 0.71093283 | -0.0034418 | 0.9463113 |
| 0.999490048 | 0.08637355 | 0.16868975 | 0.71093283 | 0.10185591 | 0.10707238 |
| 0.999490048 | 0.09612753 | 0.16887315 | 0.71093283 | 0.04482818 | 0.51486541 |
| 0.999490048 | 0.15190413 | 0.16905128 | 0.71093283 | 0.24474143 | 0.03125904 |
| 0.999490048 | 0.3455487 | 0.16908918 | 0.71093283 | 0.07425327 | 0.76344452 |
| 0.999490048 | -0.2891673 | 0.16972463 | 0.71131125 | 0.08032852 | 0.69784679 |
| 0.987721587 | -0.1984664 | 0.17220789 | 0.71131125 | 0.10146504 | 0.47914211 |
| 0.999490048 | -0.1523391 | 0.1725087 | 0.71131125 | -0.1093929 | 0.32291615 |
| 0.999490048 | -0.1298271 | 0.17219315 | 0.71131125 | -0.0575951 | 0.53863824 |
| 0.999490048 | -0.117895 | 0.17276417 | 0.71131125 | -0.0678289 | 0.42703536 |
| 0.999490048 | -0.1123108 | 0.17103673 | 0.71131125 | 0.00269814 | 0.97326123 |
| 0.999490048 | -0.1080099 | 0.17301475 | 0.71131125 | -0.089917 | 0.25395898 |
| 0.999490048 | -0.0972507 | 0.17190297 | 0.71131125 | -0.0576169 | 0.41262292 |
| 0.999490048 | -0.0903431 | 0.17026179 | 0.71131125 | -0.0803664 | 0.22069535 |
| 0.999490048 | -0.0892795 | 0.17231045 | 0.71131125 | 0.03130305 | 0.62659972 |
| 0.999490048 | -0.0812722 | 0.17269305 | 0.71131125 | -0.0658398 | 0.26625337 |
| 0.999490048 | -0.0766362 | 0.17251014 | 0.71131125 | -0.0280024 | 0.61238459 |
| 0.999490048 | -0.0758198 | 0.16994529 | 0.71131125 | -0.0866899 | 0.11869072 |
| 0.999490048 | -0.0741714 | 0.16980631 | 0.71131125 | -0.0155989 | 0.76871964 |
| 0.999490048 | -0.0721336 | 0.16941513 | 0.71131125 | -0.0595524 | 0.25377275 |
| 0.999490048 | -0.0691382 | 0.17084953 | 0.71131125 | -0.0637928 | 0.20512287 |
| 0.999490048 | -0.0682615 | 0.17166769 | 0.71131125 | -0.0768333 | 0.12576407 |
| 0.999490048 | -0.0665867 | 0.17173892 | 0.71131125 | -0.0239007 | 0.61799414 |
| 0.999490048 | -0.0658904 | 0.17249503 | 0.71131125 | -0.076994 | 0.11328775 |
| 0.999490048 | -0.0657353 | 0.17220195 | 0.71131125 | -0.0209828 | 0.65760588 |
| 0.999490048 | -0.0627378 | 0.17074687 | 0.71131125 | 0.02534897 | 0.5738912 |
| 0.999490048 | -0.0609452 | 0.17030673 | 0.71131125 | -0.0111232 | 0.79877716 |
| 0.999490048 | -0.0587169 | 0.17263312 | 0.71131125 | -0.0061111 | 0.88501028 |
| 0.999490048 | 0.05901233 | 0.17177991 | 0.71131125 | 0.02687193 | 0.5276274 |
| 0.999490048 | 0.05906391 | 0.17102158 | 0.71131125 | -0.0131679 | 0.75597935 |
| 0.999490048 | 0.06563414 | 0.17027459 | 0.71131125 | 0.05422543 | 0.25440815 |

| | | | | | |
|---|---|---|---|---|---|
| 0.999490048 | 0.07398539 | 0.17007026 | 0.71131125 | 0.01265806 | 0.81100483 |
| 0.999490048 | 0.07609908 | 0.17273146 | 0.71131125 | 0.06461893 | 0.24451548 |
| 0.999490048 | 0.07780368 | 0.17246596 | 0.71131125 | 0.03482014 | 0.53536078 |
| 0.999490048 | 0.07792389 | 0.17237098 | 0.71131125 | -0.0316073 | 0.57391233 |
| 0.999490048 | 0.07827991 | 0.17286088 | 0.71131125 | 0.09743264 | 0.09298441 |
| 0.999490048 | 0.0870744 | 0.17298763 | 0.71131125 | 0.08852511 | 0.16619984 |
| 0.999490048 | 0.08752902 | 0.17271526 | 0.71131125 | 0.01788595 | 0.77662717 |
| 0.999490048 | 0.08844949 | 0.17258977 | 0.71131125 | 0.02302662 | 0.7178105 |
| 0.999490048 | 0.09856551 | 0.17120851 | 0.71131125 | 0.08331984 | 0.24489128 |
| 0.999490048 | 0.1053958 | 0.16948185 | 0.71131125 | 0.13481428 | 0.0823755 |
| 0.781057588 | 0.10778311 | 0.16987994 | 0.71131125 | -0.097124 | 0.21453214 |
| 0.999490048 | 0.16007998 | 0.16970435 | 0.71131125 | 0.00822704 | 0.94265442 |
| 0.999490048 | 0.16366222 | 0.17222259 | 0.71131125 | 0.08785048 | 0.45771207 |
| 0.999490048 | 0.16812903 | 0.17061817 | 0.71131125 | 0.21596545 | 0.08195814 |
| 0.999490048 | 0.16962861 | 0.17073638 | 0.71131125 | 0.07041981 | 0.56347781 |
| 0.999490048 | 0.17998975 | 0.17264271 | 0.71131125 | 0.12233142 | 0.3491515 |
| 0.999490048 | 0.236142 | 0.17090057 | 0.71131125 | -0.0079653 | 0.96081843 |
| 0.999490048 | 0.23823044 | 0.17084142 | 0.71131125 | 0.18656017 | 0.28002657 |
| 0.999490048 | 0.23979304 | 0.17213127 | 0.71131125 | 0.26844422 | 0.13027758 |
| 0.999490048 | -0.0745047 | 0.17316603 | 0.71158286 | -0.0068867 | 0.8979271 |
| 0.999490048 | 0.10509084 | 0.17341642 | 0.71191105 | -0.0418293 | 0.58200225 |
| 0.999490048 | 0.15624508 | 0.17337517 | 0.71191105 | 0.12033783 | 0.29074652 |
| 0.999490048 | -0.0968872 | 0.17376619 | 0.71212839 | -0.1189195 | 0.09804957 |
| 0.999490048 | 0.06748528 | 0.1738105 | 0.71212839 | 0.0785107 | 0.11590423 |
| 0.999490048 | 0.08029162 | 0.17358695 | 0.71212839 | 0.06625821 | 0.25866291 |
| 0.999490048 | 0.08943109 | 0.17377369 | 0.71212839 | 0.09718236 | 0.14069015 |
| 0.999490048 | -0.9333038 | 0.17427492 | 0.7123509 | 0.18954282 | 0.76998767 |
| 0.999490048 | -0.3083138 | 0.17480323 | 0.7123509 | -0.1965326 | 0.37061062 |
| 0.999490048 | -0.2032838 | 0.17396548 | 0.7123509 | -0.1380905 | 0.34065667 |
| 0.781057588 | -0.1430375 | 0.17468287 | 0.7123509 | 0.16646914 | 0.12112065 |
| 0.999490048 | -0.1365505 | 0.17434365 | 0.7123509 | -0.0368503 | 0.709148 |
| 0.999490048 | -0.0656524 | 0.17405208 | 0.7123509 | -0.0354792 | 0.45667844 |
| 0.999490048 | 0.06549196 | 0.17480131 | 0.7123509 | 0.01533066 | 0.74646696 |
| 0.999490048 | 0.10334769 | 0.17445056 | 0.7123509 | 0.02494219 | 0.73869316 |
| 0.999490048 | 0.12442584 | 0.17479897 | 0.7123509 | 0.09463015 | 0.29832273 |
| 0.999490048 | 0.12467126 | 0.17442586 | 0.7123509 | 0.03805789 | 0.67325806 |
| 0.999490048 | 0.12475536 | 0.17471673 | 0.7123509 | 0.0292023 | 0.74642558 |
| 0.999490048 | -0.1125892 | 0.1752174 | 0.71243446 | -0.098652 | 0.23292252 |
| 0.999490048 | -0.0912342 | 0.17515731 | 0.71243446 | -0.0742863 | 0.26665728 |
| 0.999490048 | -0.0897511 | 0.17525772 | 0.71243446 | 0.00477878 | 0.94138349 |
| 0.999490048 | -0.0896863 | 0.17499082 | 0.71243446 | -0.0664444 | 0.31095387 |
| 0.999490048 | 0.06716197 | 0.17542099 | 0.71243446 | 0.05586253 | 0.25705048 |

| | | | | | |
|---|---|---|---|---|---|
| 0.999490048 | 0.0759392 | 0.17533768 | 0.71243446 | 0.02116963 | 0.70071569 |
| 0.999490048 | 0.13802108 | 0.17517025 | 0.71243446 | 0.09169248 | 0.36281237 |
| 0.999490048 | 0.10220752 | 0.17566113 | 0.71306293 | 0.06436158 | 0.38851316 |
| 0.999490048 | 0.06803821 | 0.17596189 | 0.71393671 | 0.06228152 | 0.21407371 |
| 0.999490048 | -0.0689311 | 0.17609172 | 0.71411651 | -0.049025 | 0.33157358 |
| 0.999490048 | -0.1670541 | 0.17621769 | 0.71428046 | -0.0251139 | 0.83596721 |
| 0.999490048 | 0.06422846 | 0.17647336 | 0.71496969 | 0.0960717 | 0.0478392 |
| 0.999490048 | -0.1474502 | 0.1765741 | 0.71503091 | -0.3052993 | 0.00803339 |
| 0.999490048 | -0.0695524 | 0.17709326 | 0.71678563 | -0.1156996 | 0.02936923 |
| 0.999490048 | -0.386881 | 0.18217536 | 0.71685402 | -0.1745203 | 0.53395906 |
| 0.991295302 | -0.3350409 | 0.18133783 | 0.71685402 | 0.2077368 | 0.39082292 |
| 0.999490048 | -0.2406565 | 0.18043584 | 0.71685402 | 0.03675794 | 0.83502726 |
| 0.999490048 | -0.2193262 | 0.18069744 | 0.71685402 | 0.03239363 | 0.84049423 |
| 0.781057588 | -0.1673763 | 0.17928811 | 0.71685402 | 0.15848341 | 0.20268188 |
| 0.999490048 | -0.1611196 | 0.17967834 | 0.71685402 | 0.01274896 | 0.91390215 |
| 0.999490048 | -0.1593 | 0.1816069 | 0.71685402 | -0.0299693 | 0.79814909 |
| 0.448222939 | -0.1533558 | 0.18209547 | 0.71685402 | 0.30193205 | 0.0203924 |
| 0.999490048 | -0.1527248 | 0.17822637 | 0.71685402 | -0.0514774 | 0.64464763 |
| 0.999490048 | -0.1316113 | 0.18093353 | 0.71685402 | -0.1104529 | 0.2590102 |
| 0.999490048 | -0.1300172 | 0.18194143 | 0.71685402 | -0.1284289 | 0.18714957 |
| 0.999490048 | -0.1246706 | 0.1785853 | 0.71685402 | -0.1455734 | 0.11861782 |
| 0.999490048 | -0.1158735 | 0.18131347 | 0.71685402 | -0.08141 | 0.34328125 |
| 0.999490048 | -0.1126154 | 0.1785604 | 0.71685402 | -0.1650105 | 0.05331134 |
| 0.999490048 | -0.1000515 | 0.18186808 | 0.71685402 | -0.0741988 | 0.31826126 |
| 0.999490048 | -0.095491 | 0.17813429 | 0.71685402 | -0.1246586 | 0.08251911 |
| 0.999490048 | -0.0869659 | 0.17735177 | 0.71685402 | -0.0393869 | 0.53529834 |
| 0.999490048 | -0.0820431 | 0.18191866 | 0.71685402 | -0.07014 | 0.25152174 |
| 0.999490048 | -0.0815378 | 0.17916827 | 0.71685402 | -0.0576287 | 0.33803767 |
| 0.999490048 | -0.0744889 | 0.17908919 | 0.71685402 | -0.0527139 | 0.33734375 |
| 0.999490048 | -0.0719834 | 0.18038369 | 0.71685402 | -0.1096439 | 0.04604498 |
| 0.972489992 | -0.070521 | 0.18130869 | 0.71685402 | 0.04768801 | 0.36136443 |
| 0.999490048 | -0.0672022 | 0.1803558 | 0.71685402 | -0.0264623 | 0.59218038 |
| 0.999490048 | -0.0661361 | 0.18172598 | 0.71685402 | -0.0238529 | 0.62464204 |
| 0.999490048 | -0.06354 | 0.177644 | 0.71685402 | -0.0554971 | 0.23702247 |
| 0.999490048 | -0.0611573 | 0.18149378 | 0.71685402 | -0.0342497 | 0.44869989 |
| 0.999490048 | -0.0604256 | 0.18056104 | 0.71685402 | -0.0304443 | 0.49409213 |
| 0.999490048 | -0.0554382 | 0.18117752 | 0.71685402 | -0.026966 | 0.5096197 |
| 0.999490048 | -0.0535136 | 0.1798986 | 0.71685402 | -0.0359059 | 0.36346469 |
| 0.999490048 | -0.0533737 | 0.17949169 | 0.71685402 | -0.0576241 | 0.14846455 |
| 0.999490048 | -0.0477949 | 0.17956984 | 0.71685402 | -0.0394692 | 0.26500588 |
| 0.999490048 | 0.05963907 | 0.17763804 | 0.71685402 | 0.05469769 | 0.21500772 |
| 0.999490048 | 0.06340156 | 0.18064714 | 0.71685402 | -0.0071992 | 0.87699811 |

| | | | | | |
|---|---|---|---|---|---|
| 0.999490048 | 0.06655115 | 0.17801282 | 0.71685402 | 0.06767168 | 0.17107888 |
| 0.999490048 | 0.07183309 | 0.17960123 | 0.71685402 | 0.06010312 | 0.25894026 |
| 0.999490048 | 0.07566756 | 0.18153022 | 0.71685402 | 0.02824669 | 0.61245603 |
| 0.999490048 | 0.07597691 | 0.18078356 | 0.71685402 | 0.05823282 | 0.30142526 |
| 0.999490048 | 0.0773498 | 0.17784125 | 0.71685402 | 0.01950537 | 0.72956961 |
| 0.999490048 | 0.0784117 | 0.17809138 | 0.71685402 | 0.11470681 | 0.05342301 |
| 0.999490048 | 0.07933842 | 0.18130934 | 0.71685402 | 0.01703393 | 0.77028059 |
| 0.999490048 | 0.08099542 | 0.17972446 | 0.71685402 | 0.06616168 | 0.27026211 |
| 0.999490048 | 0.08199821 | 0.17949377 | 0.71685402 | 0.05554044 | 0.35854798 |
| 0.999490048 | 0.08335744 | 0.17750431 | 0.71685402 | 0.06159123 | 0.31507643 |
| 0.999490048 | 0.08527743 | 0.18174918 | 0.71685402 | -0.0333435 | 0.59598472 |
| 0.999490048 | 0.08591528 | 0.17903499 | 0.71685402 | -0.0387451 | 0.53865025 |
| 0.999490048 | 0.09114795 | 0.18002949 | 0.71685402 | 0.09413203 | 0.1666876 |
| 0.999490048 | 0.09144458 | 0.1821632 | 0.71685402 | -0.0069746 | 0.91748499 |
| 0.999490048 | 0.09618578 | 0.18175608 | 0.71685402 | 0.08276281 | 0.24837042 |
| 0.999490048 | 0.09767309 | 0.17880327 | 0.71685402 | 0.02617039 | 0.71405474 |
| 0.999490048 | 0.09776457 | 0.17895666 | 0.71685402 | 0.08725908 | 0.22861392 |
| 0.999490048 | 0.10040168 | 0.18210551 | 0.71685402 | 0.05029655 | 0.49810195 |
| 0.999490048 | 0.10924356 | 0.1774078 | 0.71685402 | 0.01462809 | 0.854079 |
| 0.999490048 | 0.11159019 | 0.18087931 | 0.71685402 | 0.09700409 | 0.24271682 |
| 0.999490048 | 0.12268493 | 0.18151863 | 0.71685402 | 0.21521586 | 0.02356289 |
| 0.999490048 | 0.12484733 | 0.17887927 | 0.71685402 | 0.02967044 | 0.74515423 |
| 0.999490048 | 0.13431043 | 0.17759378 | 0.71685402 | 0.08807251 | 0.3716921 |
| 0.826281516 | 0.16569969 | 0.18145113 | 0.71685402 | -0.1521777 | 0.21845091 |
| 0.999490048 | 0.2406942 | 0.18121465 | 0.71685402 | 0.4045141 | 0.02930827 |
| 0.999490048 | 0.39969496 | 0.18115384 | 0.71685402 | -0.1236413 | 0.67412717 |
| 0.999490048 | -0.1700018 | 0.18307638 | 0.7176442 | -0.105769 | 0.40236349 |
| 0.89605147 | -0.1667519 | 0.18348019 | 0.7176442 | 0.14140375 | 0.25699743 |
| 0.999490048 | -0.1260434 | 0.18367851 | 0.7176442 | -0.0910959 | 0.33252412 |
| 0.999490048 | -0.1157461 | 0.18339753 | 0.7176442 | -0.1455322 | 0.09770021 |
| 0.999490048 | -0.1147486 | 0.18310172 | 0.7176442 | -0.0476315 | 0.57489118 |
| 0.999490048 | -0.1018673 | 0.18378718 | 0.7176442 | -0.0251425 | 0.73865458 |
| 0.999490048 | -0.077296 | 0.18358662 | 0.7176442 | -0.0532217 | 0.35538355 |
| 0.999490048 | -0.0706096 | 0.18394089 | 0.7176442 | -0.030327 | 0.56255013 |
| 0.999490048 | -0.0621751 | 0.18390787 | 0.7176442 | -0.0155662 | 0.7351225 |
| 0.999490048 | -0.0555117 | 0.183981 | 0.7176442 | -0.0490519 | 0.23859248 |
| 0.999490048 | -0.0507062 | 0.18326308 | 0.7176442 | -0.0280598 | 0.45591052 |
| 0.999490048 | 0.09278746 | 0.18328887 | 0.7176442 | 0.0607222 | 0.37900355 |
| 0.999490048 | 0.10058492 | 0.18258108 | 0.7176442 | 0.0400251 | 0.59025991 |
| 0.999490048 | 0.12325132 | 0.1828273 | 0.7176442 | 0.18911055 | 0.04574336 |
| 0.999490048 | 0.17839651 | 0.18336765 | 0.7176442 | -0.0760596 | 0.55224149 |
| 0.999490048 | 0.20166416 | 0.18327141 | 0.7176442 | 0.08638352 | 0.55032305 |

| | | | | | |
|---|---|---|---|---|---|
| 0.999490048 | 0.23316924 | 0.1835784 | 0.7176442 | -0.0532397 | 0.75212085 |
| 0.999490048 | 0.46306285 | 0.18353891 | 0.7176442 | 0.43194425 | 0.21158366 |
| 0.999490048 | 0.55881258 | 0.18400913 | 0.7176442 | 0.17188664 | 0.66684851 |
| 0.999490048 | -0.1547017 | 0.18481036 | 0.71799857 | 0.0570706 | 0.61933784 |
| 0.999490048 | -0.1203232 | 0.18515973 | 0.71799857 | -0.168672 | 0.06763039 |
| 0.999490048 | -0.1162861 | 0.18527557 | 0.71799857 | -0.015249 | 0.85967961 |
| 0.999490048 | -0.0798309 | 0.18507786 | 0.71799857 | 0.03399739 | 0.56687385 |
| 0.999490048 | -0.0743619 | 0.18502593 | 0.71799857 | -0.0754089 | 0.17913019 |
| 0.999490048 | -0.071657 | 0.18479485 | 0.71799857 | -0.0296116 | 0.57810251 |
| 0.999490048 | -0.0601135 | 0.18423733 | 0.71799857 | -0.0058537 | 0.89528776 |
| 0.999490048 | -0.0578999 | 0.18538981 | 0.71799857 | -0.032844 | 0.44707978 |
| 0.999490048 | 0.07287789 | 0.18462667 | 0.71799857 | 0.00349764 | 0.94831982 |
| 0.999490048 | 0.07757193 | 0.18519245 | 0.71799857 | 0.00950053 | 0.86882813 |
| 0.999490048 | 0.08768898 | 0.18443146 | 0.71799857 | 0.03954254 | 0.5438155 |
| 0.999490048 | 0.10253968 | 0.18514655 | 0.71799857 | 0.00777867 | 0.91850239 |
| 0.999490048 | 0.13483475 | 0.18537721 | 0.71799857 | 0.03458903 | 0.72976448 |
| 0.999490048 | 0.15504978 | 0.18454677 | 0.71799857 | 0.02715378 | 0.81309042 |
| 0.999490048 | 0.34053532 | 0.18501922 | 0.71799857 | 0.30838807 | 0.22672988 |
| 0.999490048 | -0.1283624 | 0.18549076 | 0.7180565 | 0.01563276 | 0.86964839 |
| 0.999490048 | -0.0799317 | 0.18572998 | 0.7183568 | -0.0892302 | 0.14128353 |
| 0.999490048 | 0.05378875 | 0.1857404 | 0.7183568 | 0.02325141 | 0.56170134 |
| 0.999490048 | -0.0763882 | 0.18582769 | 0.71836167 | -0.0193891 | 0.73277098 |
| 0.999490048 | 0.16375568 | 0.18593174 | 0.71843131 | 0.0926086 | 0.44901923 |
| 0.999490048 | -0.1869643 | 0.18603 | 0.71847848 | -0.1113634 | 0.42563873 |
| 0.999490048 | -0.0767775 | 0.1863096 | 0.71856128 | -0.0686893 | 0.23544297 |
| 0.999490048 | -0.0719701 | 0.18628326 | 0.71856128 | -0.031296 | 0.55988829 |
| 0.999490048 | 0.2132069 | 0.18620507 | 0.71856128 | -0.0810671 | 0.60985297 |
| 0.999490048 | -0.0770201 | 0.18639955 | 0.71857627 | -0.0404575 | 0.48207635 |
| 0.999490048 | -0.0809333 | 0.18717834 | 0.72025681 | -0.0105106 | 0.86163402 |
| 0.999490048 | -0.0789192 | 0.18725808 | 0.72025681 | -0.0921819 | 0.12580869 |
| 0.999490048 | -0.0736458 | 0.18724677 | 0.72025681 | 0.0094894 | 0.86272584 |
| 0.999490048 | 0.17456239 | 0.18726677 | 0.72025681 | -0.0432968 | 0.73947922 |
| 0.999490048 | 0.35123698 | 0.18724222 | 0.72025681 | 0.08365449 | 0.74409026 |
| 0.999490048 | -0.3894084 | 0.18785585 | 0.72045619 | -0.7839768 | 0.01634669 |
| 0.999490048 | -0.0635635 | 0.18741563 | 0.72045619 | -0.0325163 | 0.49451033 |
| 0.999490048 | -0.0583756 | 0.18764175 | 0.72045619 | -0.0410757 | 0.34963709 |
| 0.999490048 | 0.09277366 | 0.18792258 | 0.72045619 | 0.0596671 | 0.39224332 |
| 0.999490048 | 0.11694033 | 0.18757708 | 0.72045619 | 0.11097658 | 0.21030908 |
| 0.999490048 | 0.19214447 | 0.18783011 | 0.72045619 | 0.19336934 | 0.18512778 |
| 0.999490048 | 0.3967803 | 0.18775036 | 0.72045619 | 0.07934199 | 0.78087689 |
| 0.999490048 | -0.3927115 | 0.18832761 | 0.7206854 | 0.13886316 | 0.6251072 |
| 0.999490048 | 0.08712831 | 0.18815282 | 0.7206854 | -0.0245367 | 0.7064196 |

| | | | | | |
|---|---|---|---|---|---|
| 0.999490048 | 0.11498715 | 0.18829144 | 0.7206854 | 0.02393088 | 0.78066391 |
| 0.999490048 | 0.22837487 | 0.18828314 | 0.7206854 | 0.14553388 | 0.39692244 |
| 0.999490048 | -0.054067 | 0.18862638 | 0.72148086 | -0.0608344 | 0.14087133 |
| 0.999490048 | 0.11027727 | 0.18870829 | 0.72148086 | 0.00469723 | 0.9545633 |
| 0.999490048 | -0.1597908 | 0.18937434 | 0.72171583 | -0.1763197 | 0.14905654 |
| 0.999490048 | -0.1383279 | 0.18970652 | 0.72171583 | -0.3427633 | 0.00523041 |
| 0.999490048 | -0.1378083 | 0.18895923 | 0.72171583 | -0.3373463 | 0.00285291 |
| 0.999490048 | -0.1017278 | 0.1895405 | 0.72171583 | -0.0228238 | 0.76473054 |
| 0.999490048 | -0.0962739 | 0.18928167 | 0.72171583 | 0.00584396 | 0.93537332 |
| 0.999490048 | -0.0719864 | 0.18893738 | 0.72171583 | -0.0152013 | 0.77786399 |
| 0.711558326 | -0.0683527 | 0.18972051 | 0.72171583 | 0.07641379 | 0.14431718 |
| 0.999490048 | -0.0539098 | 0.18928724 | 0.72171583 | -0.0071337 | 0.85972948 |
| 0.999490048 | 0.07260302 | 0.18961659 | 0.72171583 | 0.0264797 | 0.62718693 |
| 0.999490048 | 0.0918924 | 0.18916116 | 0.72171583 | 0.02634271 | 0.70208614 |
| 0.999490048 | 0.25527251 | 0.1894491 | 0.72171583 | 0.19487609 | 0.31299055 |
| 0.999490048 | -0.0915701 | 0.18983811 | 0.72183434 | -0.0736052 | 0.28902561 |
| 0.999490048 | -0.100351 | 0.19021444 | 0.72224217 | -0.0293787 | 0.69679557 |
| 0.999490048 | -0.069068 | 0.19064219 | 0.72224217 | -0.0080952 | 0.87599941 |
| 0.999490048 | -0.0650464 | 0.19049014 | 0.72224217 | -0.0492601 | 0.31794692 |
| 0.999490048 | -0.0578861 | 0.19032921 | 0.72224217 | -0.0482735 | 0.27227817 |
| 0.999490048 | -0.0576165 | 0.19042051 | 0.72224217 | -0.0375861 | 0.38833953 |
| 0.999490048 | 0.05707046 | 0.19003939 | 0.72224217 | 0.02268614 | 0.59708097 |
| 0.999490048 | 0.08815473 | 0.19072383 | 0.72224217 | 0.02800896 | 0.67283363 |
| 0.999490048 | 0.14252975 | 0.190478 | 0.72224217 | 0.10136945 | 0.34771279 |
| 0.999490048 | 0.2091008 | 0.19062056 | 0.72224217 | 0.03274916 | 0.83485117 |
| 0.999490048 | -0.243444 | 0.19164472 | 0.7227793 | -0.1239745 | 0.50064618 |
| 0.999490048 | -0.1603467 | 0.19130949 | 0.7227793 | -0.0507111 | 0.67466897 |
| 0.999490048 | -0.1175289 | 0.19164243 | 0.7227793 | -0.1129364 | 0.20899368 |
| 0.999490048 | -0.0784619 | 0.19151541 | 0.7227793 | -0.0213091 | 0.71850841 |
| 0.999490048 | -0.0729272 | 0.19136377 | 0.7227793 | -0.0593939 | 0.28452334 |
| 0.999490048 | 0.08370332 | 0.19146785 | 0.7227793 | 0.03944783 | 0.53267946 |
| 0.999490048 | 0.11142352 | 0.19122382 | 0.7227793 | 0.03452275 | 0.68082884 |
| 0.999490048 | 0.11165082 | 0.19157764 | 0.7227793 | 0.02805375 | 0.73876023 |
| 0.999490048 | 0.17653979 | 0.19105411 | 0.7227793 | 0.33978685 | 0.02485931 |
| 0.999490048 | -0.0974327 | 0.19203451 | 0.72294328 | -0.0834949 | 0.2614354 |
| 0.999490048 | -0.0538996 | 0.19183378 | 0.72294328 | -0.0739089 | 0.07773307 |
| 0.999490048 | 0.06464793 | 0.19197129 | 0.72294328 | 0.01506695 | 0.75724364 |
| 0.999490048 | 0.0753117 | 0.19197346 | 0.72294328 | 0.04491903 | 0.43136866 |
| 0.999490048 | 0.05708603 | 0.19225157 | 0.72343426 | -0.0259212 | 0.5483611 |
| 0.781057588 | -0.0891412 | 0.19238329 | 0.7236038 | 0.090186 | 0.18739735 |
| 0.999490048 | -0.2976127 | 0.19260123 | 0.72371661 | -0.090046 | 0.67848888 |
| 0.999490048 | -0.1951271 | 0.1926733 | 0.72371661 | -0.1469442 | 0.32302205 |

| | | | | | |
|---|---|---|---|---|---|
| 0.999490048 | -0.0540114 | 0.19261026 | 0.72371661 | -0.0523966 | 0.20580136 |
| 0.999490048 | -0.2455604 | 0.19485497 | 0.72536669 | -0.4612145 | 0.01923609 |
| 0.771925038 | -0.1272075 | 0.19370926 | 0.72536669 | 0.13572994 | 0.16649073 |
| 0.999490048 | -0.1009275 | 0.19493687 | 0.72536669 | -0.1314798 | 0.09503881 |
| 0.999490048 | -0.0957421 | 0.19403852 | 0.72536669 | -0.1249164 | 0.09391125 |
| 0.999490048 | -0.0900099 | 0.1935274 | 0.72536669 | -0.1222956 | 0.08144687 |
| 0.999490048 | -0.084461 | 0.19412632 | 0.72536669 | 0.01204817 | 0.85059551 |
| 0.999490048 | -0.0816138 | 0.19355643 | 0.72536669 | -0.042147 | 0.49659157 |
| 0.987721587 | -0.0796912 | 0.19385337 | 0.72536669 | 0.04701261 | 0.43832811 |
| 0.999490048 | -0.0742734 | 0.19480244 | 0.72536669 | -0.0321886 | 0.56888884 |
| 0.999490048 | -0.0616102 | 0.19331033 | 0.72536669 | -0.0689548 | 0.14710697 |
| 0.999490048 | 0.0634344 | 0.19473543 | 0.72536669 | 0.00609714 | 0.89912769 |
| 0.999490048 | 0.0694763 | 0.19479309 | 0.72536669 | 0.11398765 | 0.0383815 |
| 0.999490048 | 0.07322385 | 0.19490194 | 0.72536669 | 0.07747359 | 0.17108249 |
| 0.999490048 | 0.08732008 | 0.1947145 | 0.72536669 | -0.0304641 | 0.64606815 |
| 0.999490048 | 0.09225954 | 0.19474774 | 0.72536669 | 0.02705368 | 0.69934984 |
| 0.999490048 | 0.09952628 | 0.194033 | 0.72536669 | 0.14992074 | 0.05526982 |
| 0.999490048 | 0.10032122 | 0.19417766 | 0.72536669 | 0.02512294 | 0.74110769 |
| 0.999490048 | 0.11847428 | 0.19450203 | 0.72536669 | 0.0452114 | 0.61541632 |
| 0.999490048 | 0.15077992 | 0.19460022 | 0.72536669 | 0.10429892 | 0.36537789 |
| 0.999490048 | 0.2197949 | 0.19463204 | 0.72536669 | 0.18293315 | 0.27789723 |
| 0.999490048 | 0.29865476 | 0.19410469 | 0.72536669 | 0.2004678 | 0.37898748 |
| 0.999490048 | 0.10597252 | 0.19505718 | 0.72549106 | -0.010549 | 0.89562994 |
| 0.999490048 | 0.14331446 | 0.19538897 | 0.72640155 | 0.14304456 | 0.19614687 |
| 0.999490048 | -0.0888415 | 0.19603859 | 0.7284923 | -0.0800959 | 0.24229316 |
| 0.999490048 | -0.1254577 | 0.19667585 | 0.72854123 | -0.091492 | 0.34265749 |
| 0.999490048 | -0.0933083 | 0.19677102 | 0.72854123 | -0.0291139 | 0.68258141 |
| 0.999490048 | -0.0836134 | 0.1965119 | 0.72854123 | -0.1476216 | 0.02737356 |
| 0.999490048 | -0.0822247 | 0.19663808 | 0.72854123 | -0.0059215 | 0.92465878 |
| 0.999490048 | -0.0639164 | 0.19673326 | 0.72854123 | -0.1175542 | 0.02212661 |
| 0.999490048 | 0.06275635 | 0.19683701 | 0.72854123 | 0.00262302 | 0.95624729 |
| 0.999490048 | 0.09456798 | 0.19626848 | 0.72854123 | -0.0138821 | 0.8470596 |
| 0.999490048 | 0.10850001 | 0.19622338 | 0.72854123 | 0.12127375 | 0.15025109 |
| 0.999490048 | 0.14527516 | 0.19681931 | 0.72854123 | 0.12497026 | 0.26482031 |
| 0.999490048 | 0.22931108 | 0.19698042 | 0.72874902 | 0.11327664 | 0.51860848 |
| 0.999490048 | -0.0834945 | 0.19737951 | 0.72898388 | -0.0153233 | 0.80997157 |
| 0.999490048 | -0.0737367 | 0.19739312 | 0.72898388 | -0.0371569 | 0.51071247 |
| 0.999490048 | -0.0554676 | 0.19719733 | 0.72898388 | -0.0146206 | 0.72987231 |
| 0.999490048 | 0.08487826 | 0.19734708 | 0.72898388 | 0.05565504 | 0.39344854 |
| 0.999490048 | -0.2288161 | 0.19771224 | 0.72938667 | -0.1623044 | 0.35691657 |
| 0.999490048 | -0.0901741 | 0.19776424 | 0.72938667 | -0.0867197 | 0.21492407 |
| 0.999490048 | -0.0581001 | 0.19768674 | 0.72938667 | -0.0431488 | 0.33513249 |

| | | | | | |
|---|---|---|---|---|---|
| 0.999490048 | -0.0816879 | 0.19802478 | 0.73002512 | -0.0739551 | 0.24247841 |
| 0.999490048 | 0.07407724 | 0.19818141 | 0.73028013 | 0.01887868 | 0.7390419 |
| 0.999490048 | -0.0637833 | 0.19861089 | 0.73153991 | -0.1358103 | 0.00942555 |
| 0.999490048 | -0.0964447 | 0.19876482 | 0.73178407 | -0.1156489 | 0.12598873 |
| 0.999490048 | -0.189466 | 0.19895809 | 0.73197236 | -0.2191126 | 0.13955812 |
| 0.999490048 | 0.14147137 | 0.19899129 | 0.73197236 | -0.028412 | 0.79321672 |
| 0.972489992 | -0.0972367 | 0.19917528 | 0.73222894 | 0.06984774 | 0.35254415 |
| 0.999490048 | 0.06377496 | 0.19923643 | 0.73222894 | 0.01504579 | 0.75832631 |
| 0.999490048 | -0.082998 | 0.19947127 | 0.73276952 | -0.0351923 | 0.58128046 |
| 0.999490048 | 0.06166402 | 0.19956124 | 0.73277766 | 0.07717723 | 0.11160643 |
| 0.999490048 | 0.077006 | 0.19967982 | 0.73289079 | 0.02293602 | 0.69814324 |
| 0.999490048 | -0.1372429 | 0.20084273 | 0.73297062 | -0.0194384 | 0.85389663 |
| 0.999490048 | -0.1346711 | 0.2007921 | 0.73297062 | 0.05375868 | 0.60457229 |
| 0.999490048 | -0.0995982 | 0.20063612 | 0.73297062 | -0.0883679 | 0.25454265 |
| 0.999490048 | -0.0981348 | 0.2007661 | 0.73297062 | -0.0471341 | 0.53371386 |
| 0.999490048 | -0.0969888 | 0.20001368 | 0.73297062 | -0.0736176 | 0.32732972 |
| 0.999490048 | -0.0804213 | 0.20019304 | 0.73297062 | -0.0417975 | 0.50039365 |
| 0.999490048 | -0.0633839 | 0.20007388 | 0.73297062 | -0.0711768 | 0.15190454 |
| 0.999490048 | -0.0482937 | 0.20028972 | 0.73297062 | -0.0761339 | 0.04852207 |
| 0.999490048 | 0.05977683 | 0.20043891 | 0.73297062 | -0.0037795 | 0.93440563 |
| 0.999490048 | 0.06008288 | 0.20070973 | 0.73297062 | 0.05754346 | 0.21978539 |
| 0.999490048 | 0.08336866 | 0.20046735 | 0.73297062 | 0.0471624 | 0.46391831 |
| 0.999490048 | 0.10852141 | 0.2002782 | 0.73297062 | 0.05300326 | 0.52642086 |
| 0.999490048 | 0.12988699 | 0.20039031 | 0.73297062 | 0.08351042 | 0.40583243 |
| 0.999490048 | -0.0574269 | 0.20110684 | 0.73311763 | -0.0395543 | 0.37445675 |
| 0.999490048 | 0.07487 | 0.20107862 | 0.73311763 | 0.0782765 | 0.18199098 |
| 0.999490048 | 0.09497149 | 0.20114641 | 0.73311763 | -0.0291022 | 0.69090136 |
| 0.999490048 | 0.09892165 | 0.20133007 | 0.73346688 | 0.03229342 | 0.67204221 |
| 0.999490048 | 0.12563339 | 0.20149218 | 0.73353862 | 0.15784797 | 0.11180146 |
| 0.999490048 | 0.38543561 | 0.20152546 | 0.73353862 | -0.1698507 | 0.55705845 |
| 0.999490048 | -0.1196892 | 0.20182275 | 0.73360905 | -0.0444469 | 0.63067666 |
| 0.999490048 | 0.05150729 | 0.20186183 | 0.73360905 | 0.04979389 | 0.21678834 |
| 0.999490048 | 0.05690062 | 0.20189624 | 0.73360905 | 0.05851078 | 0.18983426 |
| 0.999490048 | 0.06454694 | 0.20178058 | 0.73360905 | 0.01640323 | 0.74181556 |
| 0.999490048 | -0.3193834 | 0.20329496 | 0.73392707 | -0.2868573 | 0.25170539 |
| 0.999490048 | -0.2849193 | 0.20293317 | 0.73392707 | -0.3212353 | 0.15507382 |
| 0.999490048 | -0.2715227 | 0.20455941 | 0.73392707 | -0.455269 | 0.03850884 |
| 0.999490048 | -0.2578287 | 0.20231869 | 0.73392707 | -0.3967933 | 0.05474166 |
| 0.999490048 | -0.1813102 | 0.20414566 | 0.73392707 | -0.0145507 | 0.91750209 |
| 0.999490048 | -0.1305982 | 0.20303263 | 0.73392707 | -0.127866 | 0.21234042 |
| 0.999490048 | -0.0935674 | 0.20346044 | 0.73392707 | -0.0814455 | 0.26642963 |
| 0.999490048 | -0.0920017 | 0.20364156 | 0.73392707 | -0.0889225 | 0.21869969 |

| | | | | | |
|---|---|---|---|---|---|
| 0.999490048 | -0.0873066 | 0.20462062 | 0.73392707 | -0.0669942 | 0.32716696 |
| 0.999490048 | -0.0745803 | 0.20208285 | 0.73392707 | -0.0682066 | 0.24211654 |
| 0.999490048 | -0.0717439 | 0.20355187 | 0.73392707 | -0.0075298 | 0.89211095 |
| 0.999490048 | -0.0716845 | 0.20282148 | 0.73392707 | -0.0121295 | 0.82667339 |
| 0.999490048 | -0.070412 | 0.2038121 | 0.73392707 | -0.0540091 | 0.32647655 |
| 0.999490048 | -0.0691596 | 0.20454076 | 0.73392707 | 0.00545072 | 0.91904923 |
| 0.999490048 | -0.0659396 | 0.20317684 | 0.73392707 | -0.0605815 | 0.2411637 |
| 0.999490048 | -0.0596648 | 0.20400335 | 0.73392707 | -0.066696 | 0.15727128 |
| 0.999490048 | -0.0594962 | 0.20245615 | 0.73392707 | -0.0234796 | 0.61002791 |
| 0.999490048 | -0.0594474 | 0.20449888 | 0.73392707 | -0.0796597 | 0.0930993 |
| 0.999490048 | -0.0469947 | 0.20287706 | 0.73392707 | -0.0146163 | 0.68770143 |
| 0.999490048 | 0.05867898 | 0.20454642 | 0.73392707 | 0.06253849 | 0.17726003 |
| 0.999490048 | 0.06769162 | 0.20411871 | 0.73392707 | 0.01875497 | 0.72094234 |
| 0.999490048 | 0.06949305 | 0.20412313 | 0.73392707 | 0.045134 | 0.40511213 |
| 0.999490048 | 0.0782383 | 0.20293835 | 0.73392707 | 0.10195692 | 0.10078243 |
| 0.999490048 | 0.07839079 | 0.2041143 | 0.73392707 | 0.09596811 | 0.12295122 |
| 0.999490048 | 0.08209436 | 0.20452292 | 0.73392707 | 0.05638599 | 0.37944771 |
| 0.999490048 | 0.09809388 | 0.20318685 | 0.73392707 | 0.22872221 | 0.00537807 |
| 0.999490048 | 0.1010154 | 0.20373237 | 0.73392707 | 0.10851778 | 0.17318468 |
| 0.999490048 | 0.14180956 | 0.20254454 | 0.73392707 | 0.15980675 | 0.15274753 |
| 0.999490048 | 0.16127362 | 0.20339417 | 0.73392707 | 0.16914228 | 0.18290195 |
| 0.999490048 | 0.23277008 | 0.20219667 | 0.73392707 | 0.3846885 | 0.05052906 |
| 0.999490048 | 0.08166689 | 0.20496049 | 0.73483044 | 0.13443322 | 0.0420605 |
| 0.999490048 | 0.33693082 | 0.20508574 | 0.73496392 | 0.38692245 | 0.15012251 |
| 0.999490048 | -0.1275161 | 0.20533947 | 0.73496396 | -0.0879994 | 0.36949247 |
| 0.999490048 | -0.0530369 | 0.20534981 | 0.73496396 | -0.0155616 | 0.70605756 |
| 0.999490048 | 0.07377888 | 0.205253 | 0.73496396 | 0.07013199 | 0.22783226 |
| 0.999490048 | -0.0886324 | 0.20573601 | 0.73603073 | -0.0097992 | 0.88694878 |
| 0.999490048 | 0.08837201 | 0.20601477 | 0.73671236 | 0.08326749 | 0.23260403 |
| 0.999490048 | -0.11387 | 0.20709274 | 0.73745263 | -0.0364596 | 0.68192515 |
| 0.999490048 | -0.0994394 | 0.20696997 | 0.73745263 | -0.1258904 | 0.113479 |
| 0.999490048 | -0.0634936 | 0.20656146 | 0.73745263 | -0.0522942 | 0.29573264 |
| 0.999490048 | -0.056083 | 0.20727151 | 0.73745263 | -0.0068995 | 0.87474535 |
| 0.987721587 | 0.05768065 | 0.20717492 | 0.73745263 | -0.0373384 | 0.40985812 |
| 0.999490048 | 0.07521091 | 0.20702733 | 0.73745263 | 0.06703029 | 0.25923159 |
| 0.999490048 | 0.0846994 | 0.20693533 | 0.73745263 | 0.0554583 | 0.40432174 |
| 0.999490048 | 0.08613616 | 0.2072816 | 0.73745263 | 0.11518452 | 0.09571707 |
| 0.999490048 | 0.09155623 | 0.20642468 | 0.73745263 | 0.11560071 | 0.11397191 |
| 0.999490048 | 0.14386837 | 0.20723671 | 0.73745263 | 0.13139817 | 0.24812888 |
| 0.999490048 | 0.14496798 | 0.20725922 | 0.73745263 | -0.0551076 | 0.62692895 |
| 0.999490048 | 0.19068724 | 0.20692684 | 0.73745263 | 0.07313147 | 0.61734417 |
| 0.999490048 | 0.06580802 | 0.20741237 | 0.7376036 | 0.00263992 | 0.95900962 |

| | | | | | |
|---|---|---|---|---|---|
| 0.999490048 | -0.1557013 | 0.20806433 | 0.73883848 | -0.09233 | 0.45067677 |
| 0.999490048 | -0.0731293 | 0.20829051 | 0.73883848 | -0.0823985 | 0.15799895 |
| 0.999490048 | 0.04835277 | 0.20789339 | 0.73883848 | 0.02917424 | 0.4427162 |
| 0.999490048 | 0.10067785 | 0.20826926 | 0.73883848 | 0.08968529 | 0.26075922 |
| 0.999490048 | 0.10633311 | 0.20800516 | 0.73883848 | 0.14691762 | 0.08624458 |
| 0.999490048 | 0.25479346 | 0.20827774 | 0.73883848 | 0.24828484 | 0.21976892 |
| 0.999490048 | -0.261697 | 0.20858471 | 0.73894031 | 0.08097807 | 0.6835378 |
| 0.999490048 | 0.04854857 | 0.20851417 | 0.73894031 | 0.02392918 | 0.53037665 |
| 0.999490048 | 0.05695707 | 0.20851431 | 0.73894031 | -0.0064579 | 0.88480882 |
| 0.999490048 | -0.0885524 | 0.20905842 | 0.7399949 | -0.0859534 | 0.22231573 |
| 0.999490048 | 0.12183751 | 0.20905964 | 0.7399949 | 0.07010683 | 0.46506423 |
| 0.999490048 | 0.06156005 | 0.2091834 | 0.74011922 | 0.03960621 | 0.41478592 |
| 0.999490048 | -0.0779647 | 0.20931782 | 0.74028114 | -0.1028159 | 0.10163311 |
| 0.999490048 | -0.1413008 | 0.20983716 | 0.74078986 | -0.196782 | 0.08513587 |
| 0.999490048 | -0.0893081 | 0.20989695 | 0.74078986 | 0.02727682 | 0.69762409 |
| 0.999490048 | -0.0752094 | 0.20990525 | 0.74078986 | -0.0134981 | 0.81923278 |
| 0.999490048 | 0.08362021 | 0.20982136 | 0.74078986 | 0.00459731 | 0.94415105 |
| 0.999490048 | 0.08931151 | 0.20964407 | 0.74078986 | 0.03687487 | 0.59967884 |
| 0.999490048 | 0.05575655 | 0.21001414 | 0.74086102 | 0.05410148 | 0.2234423 |
| 0.999490048 | 0.10753084 | 0.21022305 | 0.74128482 | -0.0352292 | 0.67714437 |
| 0.578467169 | -0.1068841 | 0.21043172 | 0.74170741 | 0.14765932 | 0.08792245 |
| 0.972489992 | -0.1420747 | 0.21112175 | 0.74240831 | 0.11695003 | 0.30088245 |
| 0.999490048 | -0.0873787 | 0.21151968 | 0.74240831 | -1.14E-04 | 0.99867385 |
| 0.999490048 | -0.0676865 | 0.21151315 | 0.74240831 | -0.0487773 | 0.36431631 |
| 0.999490048 | -0.0584508 | 0.21139239 | 0.74240831 | -0.0774688 | 0.10153262 |
| 0.999490048 | -0.0536611 | 0.21072035 | 0.74240831 | -0.0553748 | 0.19696053 |
| 0.999490048 | -0.0530913 | 0.21137531 | 0.74240831 | 0.01081805 | 0.79597333 |
| 0.999490048 | -0.0530183 | 0.21093795 | 0.74240831 | -0.0297694 | 0.47767355 |
| 0.999490048 | 0.0804789 | 0.21130805 | 0.74240831 | 0.06527071 | 0.30811008 |
| 0.999490048 | 0.08841578 | 0.21137723 | 0.74240831 | 0.07355077 | 0.29617635 |
| 0.999490048 | 0.10302146 | 0.21133566 | 0.74240831 | 0.14850743 | 0.07626967 |
| 0.999490048 | 0.12031934 | 0.21160937 | 0.74241101 | 0.17691595 | 0.07107703 |
| 0.999490048 | -0.0558968 | 0.21179106 | 0.7424246 | -0.0513663 | 0.25005294 |
| 0.999490048 | 0.31403436 | 0.21172861 | 0.7424246 | 0.12815034 | 0.59507235 |
| 0.999490048 | 0.17145193 | 0.21188157 | 0.74243018 | 0.12010962 | 0.37801717 |
| 0.999490048 | 0.06704044 | 0.21200661 | 0.7425567 | 0.02665883 | 0.61501508 |
| 0.999490048 | -0.1617942 | 0.21249834 | 0.74334359 | 0.00226264 | 0.98586816 |
| 0.999490048 | -0.0379092 | 0.21245525 | 0.74334359 | -0.0491143 | 0.10984674 |
| 0.999490048 | 0.1205225 | 0.21241164 | 0.74334359 | 0.14403335 | 0.13873306 |
| 0.999490048 | -0.1046362 | 0.21269034 | 0.74336867 | 0.03777195 | 0.64835126 |
| 0.999490048 | 0.07164278 | 0.21277259 | 0.74336867 | 0.03821015 | 0.50157063 |
| 0.999490048 | 0.25630947 | 0.21275647 | 0.74336867 | 0.19138587 | 0.34179875 |

| | | | | | |
|---|---|---|---|---|---|
| 0.999490048 | -0.2808508 | 0.2134813 | 0.74405172 | -0.0156765 | 0.94376594 |
| 0.999490048 | -0.1375602 | 0.21368915 | 0.74405172 | -0.1537582 | 0.16813349 |
| 0.999490048 | -0.1217295 | 0.21377378 | 0.74405172 | -0.0155494 | 0.87187132 |
| 0.999490048 | -0.1186151 | 0.21383962 | 0.74405172 | -0.1526717 | 0.11313688 |
| 0.999490048 | -0.0961229 | 0.21408308 | 0.74405172 | 0.02366402 | 0.75625877 |
| 0.999490048 | -0.093106 | 0.21409679 | 0.74405172 | -0.0676964 | 0.36285908 |
| 0.987721587 | -0.0880733 | 0.2137414 | 0.74405172 | 0.05963856 | 0.39586343 |
| 0.999490048 | -0.0765953 | 0.2131659 | 0.74405172 | -0.0523161 | 0.39123375 |
| 0.999490048 | -0.0693399 | 0.2141187 | 0.74405172 | -0.0761121 | 0.17401576 |
| 0.999490048 | 0.07451536 | 0.21407765 | 0.74405172 | 0.05398893 | 0.36449964 |
| 0.999490048 | 0.08148282 | 0.21400395 | 0.74405172 | 0.07744295 | 0.23689888 |
| 0.999490048 | 0.13976363 | 0.2141265 | 0.74405172 | 0.07733731 | 0.48713177 |
| 0.999490048 | 0.18664281 | 0.21381873 | 0.74405172 | 0.09157725 | 0.53708994 |
| 0.987721587 | -0.1166407 | 0.21440446 | 0.74457549 | 0.07803967 | 0.40214156 |
| 0.999490048 | -0.0955701 | 0.21445558 | 0.74457549 | -0.0148084 | 0.84514644 |
| 0.999490048 | -0.0459337 | 0.21454723 | 0.7445841 | -0.0166738 | 0.64790059 |
| 0.999490048 | -0.3172988 | 0.21789434 | 0.74474734 | -0.1687021 | 0.49754191 |
| 0.999490048 | -0.2171421 | 0.2177221 | 0.74474734 | -0.1015999 | 0.55942962 |
| 0.999490048 | -0.1206785 | 0.21604282 | 0.74474734 | -0.1967069 | 0.04901722 |
| 0.999490048 | -0.1064468 | 0.21684674 | 0.74474734 | -0.0027212 | 0.97442177 |
| 0.999490048 | -0.1056222 | 0.2177359 | 0.74474734 | -0.0460814 | 0.58610928 |
| 0.999490048 | -0.102502 | 0.21710837 | 0.74474734 | -0.079164 | 0.33750067 |
| 0.999490048 | -0.0974277 | 0.21625982 | 0.74474734 | 0.01036739 | 0.89370508 |
| 0.999490048 | -0.0959525 | 0.21499643 | 0.74474734 | -0.0414969 | 0.58708631 |
| 0.999490048 | -0.0941157 | 0.21744256 | 0.74474734 | 0.03621972 | 0.63057669 |
| 0.999490048 | -0.0908153 | 0.21556657 | 0.74474734 | -0.0699339 | 0.33724184 |
| 0.999490048 | -0.0884209 | 0.21527517 | 0.74474734 | -0.0136873 | 0.84557681 |
| 0.999490048 | -0.0867532 | 0.21635856 | 0.74474734 | 0.01287034 | 0.85227775 |
| 0.999490048 | -0.077778 | 0.21599441 | 0.74474734 | -0.058796 | 0.34646993 |
| 0.999490048 | -0.0732599 | 0.21641457 | 0.74474734 | 0.01252666 | 0.83010198 |
| 0.999490048 | -0.0726337 | 0.21770807 | 0.74474734 | -0.0547602 | 0.34956078 |
| 0.999490048 | -0.0701806 | 0.21507864 | 0.74474734 | -0.0520917 | 0.3541501 |
| 0.999490048 | -0.0690306 | 0.21521205 | 0.74474734 | -0.0632977 | 0.25461997 |
| 0.999490048 | -0.0682583 | 0.21672592 | 0.74474734 | -0.0109387 | 0.84072143 |
| 0.999490048 | -0.0660852 | 0.21629499 | 0.74474734 | -0.0207147 | 0.69430388 |
| 0.999490048 | -0.0660295 | 0.21655611 | 0.74474734 | 0.00871639 | 0.8684468 |
| 0.999490048 | -0.0528464 | 0.21484076 | 0.74474734 | -0.0151079 | 0.71906038 |
| 0.999490048 | 0.05431935 | 0.21750521 | 0.74474734 | 0.02460833 | 0.57170707 |
| 0.999490048 | 0.05612183 | 0.21563911 | 0.74474734 | 0.02346595 | 0.60000463 |
| 0.999490048 | 0.06217907 | 0.21783968 | 0.74474734 | 0.07518134 | 0.13893277 |
| 0.999490048 | 0.06746089 | 0.21626171 | 0.74474734 | 0.02172577 | 0.68636464 |
| 0.999490048 | 0.06892449 | 0.21714604 | 0.74474734 | 0.03977432 | 0.47182068 |

| | | | | | |
|---|---|---|---|---|---|
| 0.999490048 | 0.0780313 | 0.21682443 | 0.74474734 | 0.04399055 | 0.48169051 |
| 0.999490048 | 0.07813493 | 0.21472806 | 0.74474734 | 0.03540161 | 0.56927519 |
| 0.999490048 | 0.08368823 | 0.21748442 | 0.74474734 | 0.10201207 | 0.13558482 |
| 0.999490048 | 0.08508479 | 0.21738607 | 0.74474734 | 0.0069724 | 0.91824417 |
| 0.999490048 | 0.08841878 | 0.21629531 | 0.74474734 | 0.04156955 | 0.55630365 |
| 0.999490048 | 0.08966571 | 0.216375 | 0.74474734 | 0.05099385 | 0.4774453 |
| 0.999490048 | 0.10033346 | 0.21760299 | 0.74474734 | 0.10414512 | 0.20115152 |
| 0.999490048 | 0.10305235 | 0.21552795 | 0.74474734 | 0.08318244 | 0.3148684 |
| 0.999490048 | 0.10823885 | 0.21688112 | 0.74474734 | -0.0393636 | 0.64900446 |
| 0.999490048 | 0.12587607 | 0.21716621 | 0.74474734 | 0.19025704 | 0.06716211 |
| 0.999490048 | 0.12726197 | 0.21701293 | 0.74474734 | 0.18793672 | 0.07316742 |
| 0.999490048 | 0.07595977 | 0.21799146 | 0.74477442 | 0.04340057 | 0.47699732 |
| 0.999490048 | -0.0612531 | 0.21808193 | 0.74477879 | -0.0230348 | 0.63884189 |
| 0.477170731 | 0.08279768 | 0.21824777 | 0.74504043 | -0.1352197 | 0.04965428 |
| 0.999490048 | 0.13802459 | 0.21845835 | 0.74545453 | 0.27153221 | 0.02004788 |
| 0.999490048 | -0.0682211 | 0.21880003 | 0.74584558 | -0.0279207 | 0.61023492 |
| 0.999490048 | -0.0597637 | 0.21881173 | 0.74584558 | -0.0074322 | 0.87662056 |
| 0.999490048 | 0.16096226 | 0.21884092 | 0.74584558 | 0.28001098 | 0.03770338 |
| 0.999490048 | 0.15616317 | 0.21907091 | 0.7463248 | 0.06403309 | 0.60978057 |
| 0.999490048 | -0.1263565 | 0.21931537 | 0.74654282 | 0.06702471 | 0.51006676 |
| 0.999490048 | -0.0997919 | 0.21925477 | 0.74654282 | -0.0510968 | 0.52467964 |
| 0.999490048 | 0.11762268 | 0.21940312 | 0.74654282 | -0.0344199 | 0.7155262 |
| 0.999490048 | -0.1433971 | 0.21982218 | 0.74655953 | -0.0226861 | 0.84380225 |
| 0.999490048 | -0.0944542 | 0.21970849 | 0.74655953 | 0.01896025 | 0.80258397 |
| 0.999490048 | 0.08097367 | 0.21959261 | 0.74655953 | 0.02514052 | 0.69916821 |
| 0.999490048 | 0.1041642 | 0.21950714 | 0.74655953 | 0.12513995 | 0.14279465 |
| 0.999490048 | 0.10514904 | 0.21985508 | 0.74655953 | 0.09252618 | 0.27868142 |
| 0.999490048 | 0.11687479 | 0.22003274 | 0.74685911 | 0.13036436 | 0.17477454 |
| 0.999490048 | -0.0616205 | 0.22027899 | 0.74708755 | -0.1041923 | 0.04354159 |
| 0.999490048 | 0.13347877 | 0.22025052 | 0.74708755 | 0.10654606 | 0.32518334 |
| 0.999490048 | 0.0469368 | 0.22059998 | 0.74787244 | 0.03635339 | 0.33985566 |
| 0.999490048 | -0.1043972 | 0.22071109 | 0.74794545 | -0.0416564 | 0.62065027 |
| 0.999490048 | 0.49403554 | 0.22086058 | 0.74814843 | -0.0077375 | 0.98446783 |
| 0.999490048 | -0.1467117 | 0.22103361 | 0.7484309 | 0.02947915 | 0.80296513 |
| 0.999490048 | -0.1704084 | 0.22222578 | 0.74929297 | -0.1883363 | 0.17873267 |
| 0.999490048 | -0.1289253 | 0.22229215 | 0.74929297 | -0.0785402 | 0.45293323 |
| 0.999490048 | -0.1059083 | 0.22232328 | 0.74929297 | 0.00981958 | 0.90855597 |
| 0.999490048 | -0.0708425 | 0.22176441 | 0.74929297 | -0.044406 | 0.43957954 |
| 0.999490048 | -0.0637113 | 0.22199649 | 0.74929297 | -0.031445 | 0.54209214 |
| 0.999490048 | -0.0587402 | 0.2220198 | 0.74929297 | -0.0309641 | 0.51520294 |
| 0.999490048 | -0.0494196 | 0.22208905 | 0.74929297 | -0.0012897 | 0.97419275 |
| 0.999490048 | 0.05760888 | 0.22189143 | 0.74929297 | 0.06611788 | 0.16288751 |

| | | | | | |
|---|---|---|---|---|---|
| 0.999490048 | 0.0706373 | 0.22236503 | 0.74929297 | 0.08024528 | 0.16747783 |
| 0.999490048 | 0.07261041 | 0.22150632 | 0.74929297 | 0.08738653 | 0.14382799 |
| 0.999490048 | 0.09281867 | 0.22142275 | 0.74929297 | 0.0692109 | 0.35870334 |
| 0.999490048 | 0.10773505 | 0.22161018 | 0.74929297 | -0.0245063 | 0.77790492 |
| 0.999490048 | -0.0634609 | 0.22334177 | 0.7508364 | -0.0022611 | 0.96486974 |
| 0.999490048 | -0.0570652 | 0.22325575 | 0.7508364 | -0.0812783 | 0.08739931 |
| 0.999490048 | -0.0497661 | 0.22354243 | 0.7508364 | -0.0130632 | 0.74594134 |
| 0.999490048 | 0.0610135 | 0.22295348 | 0.7508364 | 0.00690514 | 0.8886632 |
| 0.999490048 | 0.0704049 | 0.22339489 | 0.7508364 | 0.05244787 | 0.36122929 |
| 0.999490048 | 0.07082442 | 0.2231962 | 0.7508364 | 0.10177123 | 0.08474464 |
| 0.999490048 | 0.10863712 | 0.22348312 | 0.7508364 | 0.10486493 | 0.23950309 |
| 0.999490048 | 0.1276896 | 0.22300303 | 0.7508364 | 0.06859996 | 0.50817425 |
| 0.999490048 | 0.08946293 | 0.22381186 | 0.75143908 | 0.06197847 | 0.39572814 |
| 0.999490048 | -0.1213808 | 0.22392681 | 0.75149797 | -0.1760156 | 0.08256378 |
| 0.999490048 | 0.13117362 | 0.22400939 | 0.75149797 | 0.15165235 | 0.16184119 |
| 0.999490048 | -0.1344185 | 0.22480417 | 0.75242725 | -0.0719899 | 0.511165 |
| 0.999490048 | -0.0755399 | 0.22472073 | 0.75242725 | -0.0524115 | 0.39596932 |
| 0.999490048 | -0.0605724 | 0.22476056 | 0.75242725 | -0.062855 | 0.20828176 |
| 0.999490048 | 0.06084529 | 0.22474528 | 0.75242725 | 0.00179718 | 0.97096466 |
| 0.999490048 | 0.06218559 | 0.22482916 | 0.75242725 | -0.0356005 | 0.48277358 |
| 0.999490048 | 0.08472734 | 0.22491718 | 0.75242725 | 0.07514102 | 0.28030816 |
| 0.999490048 | 0.12360821 | 0.2247322 | 0.75242725 | 0.09339277 | 0.35595325 |
| 0.999490048 | -0.0606719 | 0.22526801 | 0.75329912 | -0.0239287 | 0.62816906 |
| 0.999490048 | -0.1149661 | 0.22558992 | 0.7535025 | -0.0257319 | 0.78324508 |
| 0.999490048 | -0.0984651 | 0.22568979 | 0.7535025 | -0.1583922 | 0.05665134 |
| 0.999490048 | -0.0855191 | 0.22543595 | 0.7535025 | -0.0498239 | 0.47573813 |
| 0.999490048 | -0.0634264 | 0.22566471 | 0.7535025 | 0.01218824 | 0.81328442 |
| 0.999490048 | -0.0913225 | 0.2259658 | 0.75412248 | 0.04609311 | 0.5366332 |
| 0.999490048 | -0.0860576 | 0.22669257 | 0.75605602 | -0.0325491 | 0.64330863 |
| 0.999490048 | -0.0568185 | 0.22681681 | 0.75605602 | 0.01868047 | 0.68721584 |
| 0.999490048 | 0.14279428 | 0.22680138 | 0.75605602 | 0.06595603 | 0.57221945 |
| 0.999490048 | 0.0843668 | 0.22717847 | 0.75695939 | 0.08502007 | 0.2237044 |
| 0.999490048 | 0.05065629 | 0.22731699 | 0.75711881 | 0.05266694 | 0.20993123 |
| 0.999490048 | 0.04780051 | 0.2274144 | 0.75714126 | 0.00803342 | 0.83695133 |
| 0.999490048 | -0.1385799 | 0.22912969 | 0.75747669 | -0.0181447 | 0.87307774 |
| 0.999490048 | -0.0516554 | 0.22861805 | 0.75747669 | -0.0739446 | 0.08946109 |
| 0.999490048 | -0.0386335 | 0.22893301 | 0.75747669 | -0.0469804 | 0.14618505 |
| 0.999490048 | 0.04919106 | 0.22855341 | 0.75747669 | -0.0054654 | 0.89202557 |
| 0.999490048 | 0.05003286 | 0.22906202 | 0.75747669 | -0.0235215 | 0.56737107 |
| 0.999490048 | 0.05635582 | 0.22795864 | 0.75747669 | 0.07599166 | 0.10805174 |
| 0.999490048 | 0.05836787 | 0.22788037 | 0.75747669 | 0.05351454 | 0.26780841 |
| 0.999490048 | 0.06608441 | 0.22886015 | 0.75747669 | 0.03532506 | 0.51571614 |

| | | | | | |
|---|---|---|---|---|---|
| 0.999490048 | 0.07617841 | 0.22807704 | 0.75747669 | -0.0502282 | 0.4229474 |
| 0.999490048 | 0.08439947 | 0.22899962 | 0.75747669 | 0.08970002 | 0.20191263 |
| 0.999490048 | 0.08564419 | 0.22914804 | 0.75747669 | 0.06644071 | 0.34808659 |
| 0.999490048 | 0.10327709 | 0.22846848 | 0.75747669 | -0.0191971 | 0.82037544 |
| 0.999490048 | 0.10344337 | 0.22848461 | 0.75747669 | -0.0551529 | 0.51644156 |
| 0.999490048 | 0.10524365 | 0.22867097 | 0.75747669 | 0.04114187 | 0.63374027 |
| 0.999490048 | 0.14230205 | 0.22782735 | 0.75747669 | 0.16545151 | 0.16297835 |
| 0.999490048 | 0.14276256 | 0.22798526 | 0.75747669 | 0.11195845 | 0.34173306 |
| 0.999490048 | 0.17632008 | 0.22821256 | 0.75747669 | 0.26464836 | 0.07553424 |
| 0.999490048 | 0.18419909 | 0.22763067 | 0.75747669 | -0.1358985 | 0.36615635 |
| 0.999490048 | -0.0564683 | 0.22928212 | 0.75761997 | -0.0387513 | 0.40576254 |
| 0.999490048 | -0.0910803 | 0.22974297 | 0.75854243 | -0.1138574 | 0.13635796 |
| 0.999490048 | -0.0777522 | 0.22971505 | 0.75854243 | -0.0035675 | 0.95540448 |
| 0.999490048 | -0.0887407 | 0.23007613 | 0.75874236 | 0.00417323 | 0.95432979 |
| 0.999490048 | -0.0760762 | 0.23003372 | 0.75874236 | 0.01110551 | 0.85892679 |
| 0.999490048 | 0.26662847 | 0.22995925 | 0.75874236 | 0.24366652 | 0.27004885 |
| 0.999490048 | -0.0967086 | 0.23028302 | 0.75912483 | 0.03742412 | 0.63834371 |
| 0.819178176 | -0.1900537 | 0.23061883 | 0.75952861 | -0.6490508 | 0.00212133 |
| 0.999490048 | -0.1426842 | 0.23251286 | 0.75952861 | -0.1155357 | 0.33129869 |
| 0.999490048 | -0.1416387 | 0.23243024 | 0.75952861 | 0.05944078 | 0.60269881 |
| 0.999490048 | -0.1039792 | 0.23166116 | 0.75952861 | -0.0688044 | 0.42493333 |
| 0.999490048 | -0.1002683 | 0.23206094 | 0.75952861 | -0.0358453 | 0.66526705 |
| 0.999490048 | -0.0963509 | 0.23081295 | 0.75952861 | 0.04602508 | 0.56270059 |
| 0.999490048 | -0.0869519 | 0.23255082 | 0.75952861 | -0.0160318 | 0.82337398 |
| 0.999490048 | -0.0860418 | 0.23049842 | 0.75952861 | -0.0847353 | 0.23745301 |
| 0.999490048 | -0.0847053 | 0.23267954 | 0.75952861 | 0.01766898 | 0.80071722 |
| 0.999490048 | -0.0767737 | 0.23124447 | 0.75952861 | -0.1064395 | 0.10129837 |
| 0.999490048 | -0.0721604 | 0.23179324 | 0.75952861 | -0.0202822 | 0.73339858 |
| 0.999490048 | -0.0496191 | 0.23226368 | 0.75952861 | -0.0192036 | 0.63976525 |
| 0.999490048 | 0.06047875 | 0.231788 | 0.75952861 | 0.03358804 | 0.50238271 |
| 0.999490048 | 0.06203878 | 0.23213423 | 0.75952861 | 0.01496194 | 0.77018909 |
| 0.999490048 | 0.06364296 | 0.23092116 | 0.75952861 | 0.03304587 | 0.52957338 |
| 0.999490048 | 0.06407634 | 0.23264932 | 0.75952861 | 0.04278066 | 0.42184098 |
| 0.999490048 | 0.06999283 | 0.23164602 | 0.75952861 | -0.0147886 | 0.79775314 |
| 0.999490048 | 0.0761394 | 0.23235008 | 0.75952861 | 0.08066367 | 0.20654871 |
| 0.999490048 | 0.08542122 | 0.23255045 | 0.75952861 | 0.06587946 | 0.3544221 |
| 0.999490048 | 0.08949166 | 0.23146049 | 0.75952861 | 0.09904749 | 0.18677631 |
| 0.999490048 | 0.09762157 | 0.23198369 | 0.75952861 | 0.12258146 | 0.13653548 |
| 0.999490048 | 0.10319655 | 0.23233592 | 0.75952861 | 0.11396618 | 0.18853124 |
| 0.999490048 | 0.10616769 | 0.23095564 | 0.75952861 | 0.01747869 | 0.84145802 |
| 0.999490048 | 0.13022815 | 0.23116003 | 0.75952861 | -0.0621565 | 0.56331657 |
| 0.999490048 | 0.37110426 | 0.2310977 | 0.75952861 | 0.41844914 | 0.18062885 |

| | | | | | |
|---|---|---|---|---|---|
| 0.999490048 | 0.08860858 | 0.2328689 | 0.75984967 | 0.11720554 | 0.11835973 |
| 0.999490048 | -0.2013026 | 0.23324028 | 0.76008038 | -0.2358837 | 0.16465963 |
| 0.999490048 | -0.0749008 | 0.23315667 | 0.76008038 | -0.178915 | 0.00731747 |
| 0.923933853 | -0.0546851 | 0.23316301 | 0.76008038 | -0.1655266 | 0.00106453 |
| 0.999490048 | 0.08755729 | 0.23330372 | 0.76008038 | 0.07067487 | 0.33361877 |
| 0.999490048 | -0.0595895 | 0.23362932 | 0.76084431 | -0.0743002 | 0.14056323 |
| 0.999490048 | -0.0494939 | 0.23382999 | 0.76099444 | -0.0824466 | 0.05284019 |
| 0.999490048 | 0.05799093 | 0.23388136 | 0.76099444 | 0.02847217 | 0.55462051 |
| 0.999490048 | 0.10952635 | 0.23394883 | 0.76099444 | -0.0087125 | 0.92348568 |
| 0.999490048 | -0.0775148 | 0.23424961 | 0.76101769 | -0.0546934 | 0.39794949 |
| 0.999490048 | -0.0764489 | 0.23427863 | 0.76101769 | 0.03511841 | 0.58055233 |
| 0.999490048 | -0.0681728 | 0.23427923 | 0.76101769 | 0.00863441 | 0.87856516 |
| 0.999490048 | 0.08585523 | 0.23432054 | 0.76101769 | 0.04082439 | 0.56745399 |
| 0.999490048 | 0.07913991 | 0.23454345 | 0.7614455 | 0.02049431 | 0.7550541 |
| 0.999490048 | -0.3297615 | 0.23535997 | 0.76180381 | -0.5238385 | 0.06479228 |
| 0.999490048 | -0.1084208 | 0.23547623 | 0.76180381 | -0.0296066 | 0.74270699 |
| 0.999490048 | -0.1029265 | 0.23556616 | 0.76180381 | -0.0114812 | 0.89325116 |
| 0.999490048 | -0.1029214 | 0.23550259 | 0.76180381 | -0.1109013 | 0.20216036 |
| 0.999490048 | -0.1012404 | 0.23526142 | 0.76180381 | -0.0666109 | 0.43115037 |
| 0.999490048 | -0.0676722 | 0.23490075 | 0.76180381 | -0.0855829 | 0.13626851 |
| 0.999490048 | -0.0625054 | 0.2353958 | 0.76180381 | -0.0547194 | 0.29732776 |
| 0.999490048 | 0.09189527 | 0.23545152 | 0.76180381 | 0.0655451 | 0.39414417 |
| 0.999490048 | 0.11900656 | 0.2351774 | 0.76180381 | 6.23E-04 | 0.99497175 |
| 0.999490048 | 0.23863783 | 0.23498 | 0.76180381 | 0.36056414 | 0.08802926 |
| 0.999490048 | -0.0640451 | 0.23571942 | 0.76198259 | -0.0396866 | 0.45856917 |
| 0.999490048 | -0.0637264 | 0.23580395 | 0.76198259 | -0.0134696 | 0.79947966 |
| 0.999490048 | -0.0932619 | 0.23597944 | 0.76225466 | -0.1675879 | 0.0386845 |
| 0.999490048 | -0.078543 | 0.23616384 | 0.76252611 | -0.0335042 | 0.60918915 |
| 0.999490048 | 0.09153204 | 0.23624611 | 0.76252611 | -0.021849 | 0.77446038 |
| 0.999490048 | 0.12105609 | 0.23642761 | 0.76281707 | 0.12614506 | 0.21784982 |
| 0.999490048 | -0.0693947 | 0.23659916 | 0.7630757 | -0.0289454 | 0.61744384 |
| 0.999490048 | -0.2168211 | 0.23810623 | 0.76323692 | -0.4850672 | 0.01743164 |
| 0.999490048 | -0.1216702 | 0.23883362 | 0.76323692 | -0.1286712 | 0.21362416 |
| 0.999490048 | -0.1094992 | 0.23918355 | 0.76323692 | 0.02276037 | 0.80415848 |
| 0.999490048 | -0.0895095 | 0.23966553 | 0.76323692 | -0.028294 | 0.70662398 |
| 0.999490048 | -0.0751991 | 0.23800974 | 0.76323692 | 8.87E-04 | 0.98874341 |
| 0.999490048 | -0.0722285 | 0.23816576 | 0.76323692 | -0.0878599 | 0.15401388 |
| 0.999490048 | -0.0680926 | 0.23940398 | 0.76323692 | -0.0378179 | 0.50942078 |
| 0.999490048 | -0.0674053 | 0.23792225 | 0.76323692 | -0.0581051 | 0.30725476 |
| 0.999490048 | -0.0657407 | 0.23729477 | 0.76323692 | 0.00461194 | 0.93298959 |
| 0.999490048 | -0.0597749 | 0.23688412 | 0.76323692 | -0.0128603 | 0.79644893 |
| 0.999490048 | -0.0591624 | 0.23863978 | 0.76323692 | -0.0680705 | 0.17706933 |

| | | | | | |
|---|---|---|---|---|---|
| 0.999490048 | -0.0578457 | 0.23913898 | 0.76323692 | -0.0633637 | 0.19847519 |
| 0.999490048 | -0.0547566 | 0.23722141 | 0.76323692 | -0.0510397 | 0.26970628 |
| 0.999490048 | -0.0545436 | 0.2378776 | 0.76323692 | -0.0499456 | 0.27870441 |
| 0.999490048 | -0.0543831 | 0.23954439 | 0.76323692 | -0.0404959 | 0.37819868 |
| 0.999490048 | -0.0497696 | 0.23959195 | 0.76323692 | 0.01952475 | 0.64060987 |
| 0.999490048 | 0.05163047 | 0.23840012 | 0.76323692 | -0.0058501 | 0.89224565 |
| 0.999490048 | 0.05463549 | 0.23772675 | 0.76323692 | 0.10692064 | 0.02594252 |
| 0.999490048 | 0.06183117 | 0.23888943 | 0.76323692 | 0.01202421 | 0.81641229 |
| 0.999490048 | 0.06472061 | 0.23829585 | 0.76323692 | 0.06829344 | 0.21408342 |
| 0.999490048 | 0.06821694 | 0.23871205 | 0.76323692 | 0.01938557 | 0.73447539 |
| 0.999490048 | 0.06851896 | 0.23879118 | 0.76323692 | 0.02889283 | 0.61531147 |
| 0.999490048 | 0.07001488 | 0.23788619 | 0.76323692 | 0.08953062 | 0.13458297 |
| 0.999490048 | 0.07030228 | 0.23928951 | 0.76323692 | 0.00301182 | 0.95922468 |
| 0.999490048 | 0.07209581 | 0.23814346 | 0.76323692 | 0.11670004 | 0.06166563 |
| 0.999490048 | 0.07430664 | 0.2369775 | 0.76323692 | 0.10971956 | 0.08569857 |
| 0.999490048 | 0.08283694 | 0.23917647 | 0.76323692 | 0.17962241 | 0.01489272 |
| 0.969787208 | 0.08659341 | 0.23680972 | 0.76323692 | -0.0848404 | 0.24617291 |
| 0.999490048 | 0.09253852 | 0.23851871 | 0.76323692 | 0.05706007 | 0.46348377 |
| 0.999490048 | 0.10164165 | 0.23784454 | 0.76323692 | 0.0805979 | 0.34677548 |
| 0.999490048 | 0.12220129 | 0.23904476 | 0.76323692 | 0.09848995 | 0.34031283 |
| 0.999490048 | 0.15705061 | 0.2395668 | 0.76323692 | 0.05657893 | 0.66800653 |
| 0.999490048 | 0.16594948 | 0.23782121 | 0.76323692 | -0.015721 | 0.90970717 |
| 0.999490048 | 0.07966629 | 0.23978163 | 0.76331551 | 0.20112656 | 0.00546587 |
| 0.999490048 | 0.06578556 | 0.2399013 | 0.76340544 | 0.02605348 | 0.63767457 |
| 0.999490048 | -0.1220021 | 0.24011547 | 0.76379589 | 0.06625227 | 0.51944457 |
| 0.999490048 | 0.0671994 | 0.24041601 | 0.76446066 | 0.04128697 | 0.46690503 |
| 0.999490048 | -0.1887775 | 0.24152182 | 0.76517263 | -0.0942134 | 0.55477577 |
| 0.999490048 | -0.0994815 | 0.24141937 | 0.76517263 | -0.0174074 | 0.83541065 |
| 0.999490048 | -0.0769097 | 0.2410626 | 0.76517263 | -0.0762524 | 0.24499708 |
| 0.578467169 | -0.0737843 | 0.24108834 | 0.76517263 | 0.11786009 | 0.06656092 |
| 0.999490048 | -0.0547338 | 0.24115819 | 0.76517263 | -0.0610535 | 0.19253834 |
| 0.999490048 | 0.07880983 | 0.24142056 | 0.76517263 | -0.0288974 | 0.66375633 |
| 0.999490048 | 0.12468904 | 0.24142529 | 0.76517263 | 0.12444962 | 0.24230552 |
| 0.999490048 | 0.12777311 | 0.2415563 | 0.76517263 | 0.16312925 | 0.13818784 |
| 0.999490048 | 0.1294243 | 0.24121557 | 0.76517263 | 0.11188231 | 0.30933189 |
| 0.999490048 | 0.14659268 | 0.24118416 | 0.76517263 | 0.20233323 | 0.11852962 |
| 0.999490048 | -0.0777041 | 0.24173926 | 0.76528877 | -0.0182513 | 0.78055152 |
| 0.999490048 | -0.0490193 | 0.24177626 | 0.76528877 | -0.0462455 | 0.26873647 |
| 0.999490048 | -0.0967643 | 0.24200402 | 0.76540998 | -0.0153305 | 0.85096195 |
| 0.999490048 | 0.05768689 | 0.24208955 | 0.76540998 | 0.07215093 | 0.14648747 |
| 0.999490048 | 0.10838192 | 0.24207442 | 0.76540998 | 0.07395641 | 0.42132017 |
| 0.999490048 | -0.3030475 | 0.24269099 | 0.76588807 | -0.193027 | 0.45310915 |

| | | | | | |
|---|---|---|---|---|---|
| 0.999490048 | -0.0583358 | 0.24250489 | 0.76588807 | 0.02037077 | 0.67952412 |
| 0.999490048 | 0.07217728 | 0.24269938 | 0.76588807 | 0.08812643 | 0.15646238 |
| 0.999490048 | 0.07882599 | 0.24268706 | 0.76588807 | 0.0021268 | 0.97450275 |
| 0.999490048 | 0.09050529 | 0.24267524 | 0.76588807 | 0.06355988 | 0.40868564 |
| 0.999490048 | 0.06893462 | 0.24346064 | 0.76628705 | 0.06598736 | 0.26366213 |
| 0.999490048 | 0.07140111 | 0.24335011 | 0.76628705 | 0.01426736 | 0.81325247 |
| 0.999490048 | 0.07353114 | 0.24331767 | 0.76628705 | 0.0708588 | 0.26041186 |
| 0.999490048 | 0.08045266 | 0.24346821 | 0.76628705 | 0.10691601 | 0.12494899 |
| 0.999490048 | 0.10992283 | 0.24307738 | 0.76628705 | -0.0549172 | 0.55569516 |
| 0.999490048 | 0.15445211 | 0.24332246 | 0.76628705 | -0.1074926 | 0.41350727 |
| 0.999490048 | 0.22914305 | 0.24344362 | 0.76628705 | 0.11990498 | 0.53757996 |
| 0.999490048 | -0.10742 | 0.24361684 | 0.76646595 | 0.01225093 | 0.89276825 |
| 0.771925038 | -0.2599252 | 0.24386442 | 0.76657748 | 0.34181756 | 0.12906728 |
| 0.999490048 | -0.066574 | 0.24397893 | 0.76657748 | -0.0607965 | 0.28625185 |
| 0.999490048 | 0.05711214 | 0.24401951 | 0.76657748 | 0.00326328 | 0.94619424 |
| 0.999490048 | 0.09510993 | 0.24374583 | 0.76657748 | 0.12944708 | 0.1167149 |
| 0.999490048 | -0.7291066 | 0.24650153 | 0.76662298 | 0.21049585 | 0.72764996 |
| 0.999490048 | -0.2742845 | 0.2476447 | 0.76662298 | -0.5094797 | 0.03733308 |
| 0.971241857 | -0.1604756 | 0.24582653 | 0.76662298 | 0.16306922 | 0.2384819 |
| 0.999490048 | -0.1235192 | 0.24882717 | 0.76662298 | -0.1129149 | 0.29074126 |
| 0.999490048 | -0.1216179 | 0.24460017 | 0.76662298 | 0.01525016 | 0.88242833 |
| 0.999490048 | -0.118525 | 0.24906183 | 0.76662298 | 0.08996645 | 0.37893664 |
| 0.999490048 | -0.1170045 | 0.24804373 | 0.76662298 | 0.00413442 | 0.96699873 |
| 0.999490048 | -0.105861 | 0.24756177 | 0.76662298 | -0.0676691 | 0.4562506 |
| 0.999490048 | -0.0998773 | 0.24588496 | 0.76662298 | -0.1363463 | 0.11728696 |
| 0.999490048 | -0.0973551 | 0.24523803 | 0.76662298 | -0.0484358 | 0.55917105 |
| 0.999490048 | -0.0931631 | 0.24815557 | 0.76662298 | 0.02356861 | 0.76732488 |
| 0.999490048 | -0.0864786 | 0.24923628 | 0.76662298 | 0.01031157 | 0.8892725 |
| 0.999490048 | -0.0830869 | 0.24705443 | 0.76662298 | -0.0346965 | 0.62497151 |
| 0.999490048 | -0.0816746 | 0.24509367 | 0.76662298 | -0.0169112 | 0.80734698 |
| 0.999490048 | -0.077713 | 0.24728622 | 0.76662298 | 0.02468414 | 0.70989443 |
| 0.999490048 | -0.0774461 | 0.24857393 | 0.76662298 | -0.0765741 | 0.25380014 |
| 0.999490048 | -0.0756894 | 0.24658934 | 0.76662298 | -0.0020155 | 0.97504898 |
| 0.999490048 | -0.0753042 | 0.24575158 | 0.76662298 | -0.0162396 | 0.79980487 |
| 0.999490048 | -0.0664456 | 0.24597177 | 0.76662298 | -0.0631619 | 0.26947197 |
| 0.999490048 | -0.0642218 | 0.24544053 | 0.76662298 | -0.0136114 | 0.80304371 |
| 0.999490048 | -0.0634853 | 0.24659185 | 0.76662298 | -0.0322162 | 0.5525635 |
| 0.999490048 | -0.0622642 | 0.24705924 | 0.76662298 | -0.0965267 | 0.078025 |
| 0.999490048 | -0.0594828 | 0.24835065 | 0.76662298 | -0.0171235 | 0.73652945 |
| 0.999490048 | -0.0523954 | 0.24832085 | 0.76662298 | -0.0298003 | 0.50759204 |
| 0.999490048 | -0.0509629 | 0.24476157 | 0.76662298 | 0.03359136 | 0.43975666 |
| 0.999490048 | -0.0491165 | 0.2466894 | 0.76662298 | 0.02020336 | 0.62987201 |

| | | | | | |
|---|---|---|---|---|---|
| 0.999490048 | -0.0470861 | 0.2465147 | 0.76662298 | 0.00230066 | 0.95422807 |
| 0.999490048 | 0.05061601 | 0.24475094 | 0.76662298 | 0.03059675 | 0.47817413 |
| 0.999490048 | 0.05149614 | 0.24817119 | 0.76662298 | 0.01262477 | 0.77431699 |
| 0.999490048 | 0.05238003 | 0.24736271 | 0.76662298 | 0.0295969 | 0.50947294 |
| 0.999490048 | 0.05716112 | 0.24723948 | 0.76662298 | 0.03934318 | 0.42251864 |
| 0.999490048 | 0.0577375 | 0.24797699 | 0.76662298 | 0.01714093 | 0.72841728 |
| 0.999490048 | 0.05881133 | 0.24512003 | 0.76662298 | 0.02855915 | 0.5684548 |
| 0.999490048 | 0.05953168 | 0.24708409 | 0.76662298 | 0.0146081 | 0.77359806 |
| 0.999490048 | 0.0607561 | 0.24484396 | 0.76662298 | -0.00588 | 0.90916359 |
| 0.999490048 | 0.06146976 | 0.24910629 | 0.76662298 | -0.0062331 | 0.90572046 |
| 0.999490048 | 0.06310015 | 0.24921972 | 0.76662298 | 0.01858101 | 0.73122544 |
| 0.999490048 | 0.06901976 | 0.24908323 | 0.76662298 | 0.04867623 | 0.41317144 |
| 0.999490048 | 0.07336234 | 0.2476383 | 0.76662298 | 0.01961268 | 0.75429094 |
| 0.999490048 | 0.0736865 | 0.24861107 | 0.76662298 | 0.0617243 | 0.33190802 |
| 0.999490048 | 0.07502742 | 0.24899974 | 0.76662298 | 0.03107462 | 0.62923584 |
| 0.999490048 | 0.07765548 | 0.24640348 | 0.76662298 | -0.0392146 | 0.55432986 |
| 0.999490048 | 0.07951798 | 0.24856589 | 0.76662298 | 0.01685337 | 0.80434748 |
| 0.972489992 | 0.08538168 | 0.24495098 | 0.76662298 | -0.0784183 | 0.28456183 |
| 0.999490048 | 0.08854721 | 0.24799874 | 0.76662298 | 0.05032251 | 0.50762798 |
| 0.999490048 | 0.08918381 | 0.24935905 | 0.76662298 | 0.01901952 | 0.80348849 |
| 0.999490048 | 0.09539906 | 0.24503086 | 0.76662298 | -0.0197872 | 0.80699444 |
| 0.999490048 | 0.09543366 | 0.24728582 | 0.76662298 | 0.13468293 | 0.10699102 |
| 0.999490048 | 0.09912756 | 0.24617818 | 0.76662298 | 0.08994067 | 0.29156159 |
| 0.999490048 | 0.10646453 | 0.24650001 | 0.76662298 | 0.07413225 | 0.41646196 |
| 0.999490048 | 0.10760589 | 0.24675924 | 0.76662298 | 0.05633477 | 0.54029316 |
| 0.999490048 | 0.14688096 | 0.24716505 | 0.76662298 | 0.17338357 | 0.17414431 |
| 0.999490048 | 0.16561759 | 0.24716157 | 0.76662298 | 0.01973117 | 0.88887002 |
| 0.999490048 | 0.17223262 | 0.24610317 | 0.76662298 | 0.09367993 | 0.52417307 |
| 0.999490048 | 0.17577419 | 0.24441923 | 0.76662298 | 0.17632005 | 0.24299684 |
| 0.999490048 | 0.22838041 | 0.24935657 | 0.76662298 | 0.55021134 | 0.01703727 |
| 0.999490048 | 0.29103882 | 0.24584268 | 0.76662298 | 0.19700647 | 0.42874701 |
| 0.999490048 | 0.33479433 | 0.24696137 | 0.76662298 | -0.0903534 | 0.74490762 |
| 0.999490048 | -0.2320441 | 0.25227526 | 0.76743033 | -0.0551884 | 0.77680408 |
| 0.999490048 | -0.1667322 | 0.25348134 | 0.76743033 | -0.0734269 | 0.61126588 |
| 0.999490048 | -0.1660046 | 0.24990079 | 0.76743033 | -0.0127121 | 0.92884864 |
| 0.999490048 | -0.1533562 | 0.25362715 | 0.76743033 | -0.0470192 | 0.71607569 |
| 0.999490048 | -0.1361218 | 0.25363072 | 0.76743033 | -0.1342194 | 0.26015472 |
| 0.999490048 | -0.1126118 | 0.25240674 | 0.76743033 | -0.0339116 | 0.72720665 |
| 0.999490048 | -0.0971131 | 0.25141211 | 0.76743033 | -0.0209371 | 0.8022489 |
| 0.999490048 | -0.0969344 | 0.2522392 | 0.76743033 | -0.0478252 | 0.56829315 |
| 0.999490048 | -0.0942734 | 0.25073612 | 0.76743033 | -0.0460306 | 0.57103653 |
| 0.999490048 | -0.0926446 | 0.25294814 | 0.76743033 | -0.0707154 | 0.3802697 |

| | | | | | |
|---|---|---|---|---|---|
| 0.999490048 | -0.0832524 | 0.25077416 | 0.76743033 | 0.02043938 | 0.77523898 |
| 0.999490048 | -0.0784385 | 0.25189838 | 0.76743033 | -0.0067526 | 0.92038475 |
| 0.999490048 | -0.0761622 | 0.25166856 | 0.76743033 | 0.01528658 | 0.8157313 |
| 0.999490048 | -0.0749469 | 0.252935 | 0.76743033 | 0.0049783 | 0.93866358 |
| 0.999490048 | -0.0719643 | 0.25077543 | 0.76743033 | -0.0573316 | 0.35790076 |
| 0.999490048 | -0.0675786 | 0.25238126 | 0.76743033 | 0.01796729 | 0.75801158 |
| 0.999490048 | -0.0671864 | 0.25062391 | 0.76743033 | 0.02600455 | 0.65290093 |
| 0.999490048 | -0.060791 | 0.25340981 | 0.76743033 | -0.0795627 | 0.13854329 |
| 0.999490048 | -0.060532 | 0.25289263 | 0.76743033 | -0.0488267 | 0.35419557 |
| 0.999490048 | -0.0583918 | 0.25084963 | 0.76743033 | -0.044413 | 0.37979693 |
| 0.999490048 | -0.0583659 | 0.25084267 | 0.76743033 | -0.0615264 | 0.22676751 |
| 0.999490048 | -0.0546297 | 0.25366559 | 0.76743033 | -0.0306656 | 0.51792168 |
| 0.999490048 | -0.0523156 | 0.25320621 | 0.76743033 | 0.00178286 | 0.96852564 |
| 0.999490048 | -0.0520745 | 0.2512332 | 0.76743033 | -0.0348416 | 0.43934555 |
| 0.999490048 | 0.04610276 | 0.25094237 | 0.76743033 | 0.02646311 | 0.50612196 |
| 0.999490048 | 0.04767426 | 0.25034686 | 0.76743033 | 0.01488271 | 0.71644802 |
| 0.999490048 | 0.05567634 | 0.25339364 | 0.76743033 | 0.07159604 | 0.14525322 |
| 0.999490048 | 0.05701456 | 0.25004943 | 0.76743033 | 0.02241773 | 0.64737626 |
| 0.999490048 | 0.06329712 | 0.25366435 | 0.76743033 | 0.02854728 | 0.60289012 |
| 0.987721587 | 0.06354872 | 0.24975849 | 0.76743033 | -0.052043 | 0.34380338 |
| 0.999490048 | 0.06583139 | 0.25363122 | 0.76743033 | 0.04316372 | 0.45078277 |
| 0.999490048 | 0.06626443 | 0.25287731 | 0.76743033 | 0.04847897 | 0.40000223 |
| 0.999490048 | 0.06849377 | 0.25047951 | 0.76743033 | 0.02055923 | 0.72696302 |
| 0.999490048 | 0.06978481 | 0.25220211 | 0.76743033 | 0.04372846 | 0.46951839 |
| 0.999490048 | 0.09020062 | 0.25189495 | 0.76743033 | 0.06706399 | 0.39149985 |
| 0.999490048 | 0.0917567 | 0.25223042 | 0.76743033 | 0.11976696 | 0.13859766 |
| 0.999490048 | 0.09589014 | 0.25299723 | 0.76743033 | 0.09445982 | 0.25996156 |
| 0.999490048 | 0.09645091 | 0.25356998 | 0.76743033 | 0.01683337 | 0.84004648 |
| 0.999490048 | 0.09751947 | 0.25212694 | 0.76743033 | -1.43E-04 | 0.99864422 |
| 0.999490048 | 0.09805137 | 0.25245914 | 0.76743033 | 0.03035486 | 0.71992416 |
| 0.999490048 | 0.1112098 | 0.25168766 | 0.76743033 | 0.12179837 | 0.21051591 |
| 0.999490048 | 0.14933315 | 0.25055685 | 0.76743033 | 0.24127819 | 0.06892194 |
| 0.999490048 | 0.22849503 | 0.25298106 | 0.76743033 | 0.00375633 | 0.98435143 |
| 0.999490048 | 0.27728638 | 0.25181419 | 0.76743033 | 0.50792914 | 0.04148956 |
| 0.999490048 | 0.06943536 | 0.25386195 | 0.76746823 | 0.05887689 | 0.33151946 |
| 0.999490048 | 0.1526296 | 0.25384522 | 0.76746823 | -0.0098835 | 0.93927884 |
| 0.999490048 | -0.0653247 | 0.25400115 | 0.76761115 | 0.02371143 | 0.67535762 |
| 0.999490048 | 0.10364247 | 0.25432431 | 0.76820422 | 0.21793021 | 0.02144391 |
| 0.999490048 | 0.31113012 | 0.2543814 | 0.76820422 | 5.88E-05 | 0.99982273 |
| 0.999490048 | -0.1170354 | 0.25511554 | 0.76875306 | 0.03267648 | 0.7477526 |
| 0.999490048 | -0.0851647 | 0.25510167 | 0.76875306 | 0.01168651 | 0.87430493 |
| 0.999490048 | -0.0689129 | 0.25466778 | 0.76875306 | -0.0053566 | 0.92851907 |

| | | | | | |
|---|---|---|---|---|---|
| 0.999490048 | -0.046299 | 0.25507568 | 0.76875306 | -0.0408629 | 0.31376426 |
| 0.999490048 | 0.04153325 | 0.25487116 | 0.76875306 | -0.0155328 | 0.66672225 |
| 0.999490048 | 0.09117171 | 0.25491518 | 0.76875306 | 0.0847796 | 0.2889067 |
| 0.999490048 | 0.06018301 | 0.25584572 | 0.76984207 | 0.02191297 | 0.67562686 |
| 0.999490048 | 0.07246093 | 0.2556987 | 0.76984207 | 0.08528318 | 0.18312313 |
| 0.999490048 | 0.08124629 | 0.25580232 | 0.76984207 | 0.07163823 | 0.31496328 |
| 0.999490048 | 0.08592086 | 0.25557239 | 0.76984207 | 0.06077738 | 0.41825699 |
| 0.999490048 | -0.0672371 | 0.25628274 | 0.77060169 | -0.0350934 | 0.54974259 |
| 0.999490048 | 0.06031449 | 0.25619387 | 0.77060169 | 0.03230408 | 0.53934825 |
| 0.999490048 | -0.0656943 | 0.25655019 | 0.77077092 | -0.0538088 | 0.35068509 |
| 0.999490048 | -0.0585205 | 0.25661595 | 0.77077092 | -0.0929172 | 0.07713178 |
| 0.999490048 | 0.12334941 | 0.25650379 | 0.77077092 | 0.03469165 | 0.7467046 |
| 0.999490048 | -0.321343 | 0.25816996 | 0.77182928 | -0.0364242 | 0.89670859 |
| 0.999490048 | -0.0739185 | 0.25779912 | 0.77182928 | 0.0174144 | 0.78724885 |
| 0.999490048 | -0.06273 | 0.25798707 | 0.77182928 | 0.04063501 | 0.46037142 |
| 0.999490048 | 0.05347903 | 0.25776767 | 0.77182928 | 0.05158447 | 0.27458676 |
| 0.999490048 | 0.05479758 | 0.25754442 | 0.77182928 | -0.0128171 | 0.78862386 |
| 0.999490048 | 0.05713692 | 0.25809516 | 0.77182928 | 0.06914913 | 0.17361492 |
| 0.999490048 | 0.06506468 | 0.25814974 | 0.77182928 | 0.0330668 | 0.56177624 |
| 0.999490048 | 0.06831333 | 0.25718588 | 0.77182928 | 0.00533638 | 0.92855346 |
| 0.999490048 | 0.08799885 | 0.25743657 | 0.77182928 | 0.01846046 | 0.80990886 |
| 0.999490048 | 0.09971725 | 0.25743286 | 0.77182928 | -0.0064669 | 0.94068996 |
| 0.999490048 | 0.0999751 | 0.25729169 | 0.77182928 | 0.11619195 | 0.1899468 |
| 0.999490048 | 0.12149145 | 0.25801627 | 0.77182928 | 0.04497092 | 0.67201258 |
| 0.711558326 | 0.18965469 | 0.25721353 | 0.77182928 | -0.2803879 | 0.09883961 |
| 0.999490048 | -0.0699904 | 0.25838909 | 0.77193164 | -0.0049785 | 0.93509455 |
| 0.999490048 | 0.09493744 | 0.25834438 | 0.77193164 | 0.07754184 | 0.3538533 |
| 0.909524814 | -0.050919 | 0.25862439 | 0.77208212 | 0.0589629 | 0.19274493 |
| 0.999490048 | 0.13797569 | 0.25862287 | 0.77208212 | -0.0840773 | 0.48771183 |
| 0.999490048 | -0.0779913 | 0.25910027 | 0.77212249 | -0.0304672 | 0.65581748 |
| 0.999490048 | 0.0525025 | 0.25906027 | 0.77212249 | 0.05492692 | 0.23826964 |
| 0.999490048 | 0.05407147 | 0.25907694 | 0.77212249 | 0.05415485 | 0.25836182 |
| 0.999490048 | 0.06481838 | 0.2590949 | 0.77212249 | 0.08888433 | 0.12584451 |
| 0.999490048 | 0.10445966 | 0.25873186 | 0.77212249 | 0.05159077 | 0.57326686 |
| 0.999490048 | -0.2215911 | 0.25989567 | 0.77261407 | 0.05877554 | 0.76232481 |
| 0.999490048 | -0.183519 | 0.25946651 | 0.77261407 | -0.1575322 | 0.32938951 |
| 0.999490048 | -0.0577737 | 0.2595657 | 0.77261407 | -0.1409829 | 0.00935028 |
| 0.999490048 | -0.0542564 | 0.25998172 | 0.77261407 | -0.0426615 | 0.37345648 |
| 0.999490048 | 0.053947 | 0.25968167 | 0.77261407 | 0.02264342 | 0.6325533 |
| 0.999490048 | 0.06281707 | 0.25982078 | 0.77261407 | 0.05276659 | 0.34207475 |
| 0.999490048 | 0.09647688 | 0.25976427 | 0.77261407 | 0.15929322 | 0.06845946 |
| 0.999490048 | 0.16668007 | 0.26000545 | 0.77261407 | 0.18024885 | 0.22418238 |

| | | | | | |
|---|---|---|---|---|---|
| 0.999490048 | -0.1105666 | 0.26129485 | 0.77297053 | -0.0402414 | 0.67931269 |
| 0.999490048 | -0.0996668 | 0.26053353 | 0.77297053 | -0.1202348 | 0.1771314 |
| 0.999490048 | -0.0975388 | 0.261348 | 0.77297053 | -0.1015246 | 0.24285122 |
| 0.999490048 | -0.0824064 | 0.26142141 | 0.77297053 | -0.0447167 | 0.53862327 |
| 0.999490048 | -0.0667316 | 0.26041945 | 0.77297053 | -0.0260462 | 0.65700193 |
| 0.999490048 | -0.0651867 | 0.26074798 | 0.77297053 | -0.1334154 | 0.02664469 |
| 0.999490048 | -0.0514399 | 0.26125447 | 0.77297053 | -8.80E-04 | 0.98446715 |
| 0.999490048 | -0.0441377 | 0.26126231 | 0.77297053 | -0.0519868 | 0.18789562 |
| 0.999490048 | 0.04730813 | 0.26136125 | 0.77297053 | 0.02650473 | 0.52556769 |
| 0.999490048 | 0.06015669 | 0.26089398 | 0.77297053 | 0.02588155 | 0.62500433 |
| 0.999490048 | 0.06292633 | 0.26133091 | 0.77297053 | 0.11152097 | 0.05236455 |
| 0.999490048 | 0.07189442 | 0.26095768 | 0.77297053 | -0.0287957 | 0.64903357 |
| 0.987721587 | 0.09065801 | 0.26024797 | 0.77297053 | -0.0855119 | 0.28760954 |
| 0.999490048 | 0.17933013 | 0.26059955 | 0.77297053 | -0.1196515 | 0.44969856 |
| 0.999490048 | -0.0874336 | 0.26168385 | 0.77319893 | -0.0924317 | 0.23608031 |
| 0.999490048 | 0.10041094 | 0.26167476 | 0.77319893 | 0.12710833 | 0.15844254 |
| 0.999490048 | -0.0590805 | 0.26189848 | 0.77328583 | -0.0329321 | 0.52812977 |
| 0.999490048 | 0.07384856 | 0.26185796 | 0.77328583 | 0.12822826 | 0.05722262 |
| 0.999490048 | -0.1186531 | 0.26236298 | 0.77410986 | -0.1353967 | 0.20259218 |
| 0.999490048 | 0.07863875 | 0.26230883 | 0.77410986 | -0.0022602 | 0.97396917 |
| 0.999490048 | -0.0701102 | 0.26342329 | 0.77422895 | 0.03275492 | 0.5977318 |
| 0.999490048 | -0.0477844 | 0.26337139 | 0.77422895 | -0.0354472 | 0.404063 |
| 0.999490048 | -0.0409875 | 0.26277285 | 0.77422895 | -0.0296023 | 0.41582831 |
| 0.999490048 | 0.04694012 | 0.26302863 | 0.77422895 | 0.02219907 | 0.59295972 |
| 0.999490048 | 0.05692664 | 0.26321183 | 0.77422895 | 0.05093448 | 0.31559057 |
| 0.989730632 | 0.07279034 | 0.26302636 | 0.77422895 | -0.0616143 | 0.34171222 |
| 0.999490048 | 0.07301416 | 0.2629865 | 0.77422895 | 0.13810726 | 0.04000987 |
| 0.999490048 | 0.07547261 | 0.26338372 | 0.77422895 | 0.03511851 | 0.59916526 |
| 0.999490048 | 0.09705244 | 0.26287904 | 0.77422895 | 0.02542302 | 0.76663758 |
| 0.999490048 | 0.11745754 | 0.26341477 | 0.77422895 | 0.1127015 | 0.28276289 |
| 0.999490048 | 0.27377649 | 0.26282003 | 0.77422895 | -0.17309 | 0.46838852 |
| 0.999490048 | -0.0740778 | 0.26363584 | 0.77430856 | -0.1246391 | 0.06576609 |
| 0.999490048 | -0.0472315 | 0.26358459 | 0.77430856 | -0.054461 | 0.19919574 |
| 0.999490048 | 0.13482099 | 0.2639555 | 0.77497485 | 0.00298256 | 0.97969753 |
| 0.999490048 | -0.0725277 | 0.26420291 | 0.77542857 | -0.0621321 | 0.33722789 |
| 0.999490048 | 0.05952092 | 0.26437055 | 0.77564797 | -4.39E-04 | 0.99334728 |
| 0.999490048 | -0.0996349 | 0.26484031 | 0.77593566 | 0.01699181 | 0.84735036 |
| 0.999490048 | 0.06511536 | 0.26474539 | 0.77593566 | 0.12812775 | 0.03383738 |
| 0.999490048 | 0.09662739 | 0.26468887 | 0.77593566 | 0.10441828 | 0.22904673 |
| 0.999490048 | 0.10453631 | 0.26481365 | 0.77593566 | 0.10380723 | 0.26807819 |
| 0.999490048 | -0.0534662 | 0.26495118 | 0.7759882 | 0.01825285 | 0.70036683 |
| 0.999490048 | -0.1800338 | 0.26546395 | 0.77640069 | -0.066132 | 0.67922978 |

| | | | | | |
|---|---|---|---|---|---|
| 0.999490048 | -0.077171 | 0.26544771 | 0.77640069 | 0.05590143 | 0.41709747 |
| 0.999490048 | -0.0632799 | 0.26541657 | 0.77640069 | -0.0462531 | 0.41292641 |
| 0.999490048 | 0.12533659 | 0.26539954 | 0.77640069 | 0.10331935 | 0.35260281 |
| 0.999490048 | -0.1363837 | 0.26575378 | 0.77670426 | -0.0843557 | 0.48789163 |
| 0.999490048 | -0.0848914 | 0.26574158 | 0.77670426 | -0.0401721 | 0.59486838 |
| 0.999490048 | -0.0471544 | 0.26593562 | 0.77696377 | -0.0440402 | 0.29804303 |
| 0.999490048 | -0.2274805 | 0.26728523 | 0.77709668 | 0.03190774 | 0.87483616 |
| 0.999490048 | -0.2233059 | 0.2681091 | 0.77709668 | -0.1473182 | 0.45516222 |
| 0.999490048 | -0.2139078 | 0.2687139 | 0.77709668 | -4.46E-04 | 0.99813626 |
| 0.999490048 | -0.1802051 | 0.26750137 | 0.77709668 | -0.2931695 | 0.08377315 |
| 0.972489992 | -0.1384652 | 0.26928034 | 0.77709668 | 0.14340304 | 0.2529925 |
| 0.999490048 | -0.1063127 | 0.26799981 | 0.77709668 | 0.01729609 | 0.8552636 |
| 0.999490048 | -0.0907861 | 0.26665998 | 0.77709668 | 0.07120798 | 0.38135624 |
| 0.999490048 | -0.0867133 | 0.26856828 | 0.77709668 | -0.0186985 | 0.80923558 |
| 0.999490048 | -0.0792065 | 0.26937338 | 0.77709668 | 0.00383667 | 0.95680286 |
| 0.999490048 | -0.0776481 | 0.26725347 | 0.77709668 | -0.0409535 | 0.55492234 |
| 0.999490048 | -0.0767178 | 0.26942454 | 0.77709668 | -0.1126761 | 0.1096153 |
| 0.999490048 | -0.0722492 | 0.26769536 | 0.77709668 | -0.0396601 | 0.53939857 |
| 0.999490048 | -0.0684083 | 0.26661919 | 0.77709668 | -0.0893661 | 0.15018926 |
| 0.999490048 | -0.0663818 | 0.26704069 | 0.77709668 | 0.01825098 | 0.75755073 |
| 0.999490048 | -0.0615562 | 0.26909871 | 0.77709668 | -0.0707747 | 0.20561495 |
| 0.999490048 | -0.0601881 | 0.26872958 | 0.77709668 | 0.00360571 | 0.94651642 |
| 0.999490048 | -0.0584479 | 0.26617526 | 0.77709668 | 5.36E-04 | 0.99175687 |
| 0.999490048 | -0.050068 | 0.26922239 | 0.77709668 | -0.0529744 | 0.24308911 |
| 0.999490048 | 0.05498473 | 0.26865814 | 0.77709668 | 0.05572777 | 0.26240059 |
| 0.999490048 | 0.05730568 | 0.26929552 | 0.77709668 | 0.03165112 | 0.53830169 |
| 0.999490048 | 0.05780422 | 0.26802119 | 0.77709668 | 0.0180898 | 0.72593294 |
| 0.999490048 | 0.06150089 | 0.26789027 | 0.77709668 | 0.04314449 | 0.43413878 |
| 0.999490048 | 0.06742143 | 0.26887654 | 0.77709668 | -0.0086208 | 0.88620375 |
| 0.999490048 | 0.07193478 | 0.269395 | 0.77709668 | 0.05079574 | 0.43262729 |
| 0.999490048 | 0.07534523 | 0.26779655 | 0.77709668 | 0.05447607 | 0.42027522 |
| 0.999490048 | 0.08215003 | 0.26837025 | 0.77709668 | -0.0060794 | 0.93390877 |
| 0.999490048 | 0.08313544 | 0.26728709 | 0.77709668 | 0.11125792 | 0.14142308 |
| 0.999490048 | 0.08351808 | 0.26925762 | 0.77709668 | -0.054911 | 0.46450929 |
| 0.999490048 | 0.08468812 | 0.26635939 | 0.77709668 | -0.0027014 | 0.97136047 |
| 0.999490048 | 0.08646056 | 0.26682702 | 0.77709668 | -0.0092298 | 0.90447354 |
| 0.999490048 | 0.09349438 | 0.26883442 | 0.77709668 | 0.07804873 | 0.35418316 |
| 0.999490048 | 0.10550645 | 0.26708049 | 0.77709668 | -0.0065538 | 0.94435032 |
| 0.999490048 | 0.11852914 | 0.26807234 | 0.77709668 | 0.23200098 | 0.03590283 |
| 0.987721587 | 0.13675521 | 0.26649965 | 0.77709668 | -0.128426 | 0.29602233 |
| 0.999490048 | 0.14892655 | 0.26884624 | 0.77709668 | 0.13644721 | 0.3100795 |
| 0.971241857 | 0.25861991 | 0.26911832 | 0.77709668 | -0.3173825 | 0.18091618 |

| | | | | | |
|---|---|---|---|---|---|
| 0.999490048 | 0.38241633 | 0.26893155 | 0.77709668 | 0.36450793 | 0.2903331 |
| 0.999490048 | -0.1178623 | 0.26966195 | 0.77724448 | 0.01096553 | 0.91719739 |
| 0.999490048 | -0.0791874 | 0.26959065 | 0.77724448 | -0.0713755 | 0.31858567 |
| 0.999490048 | -0.0537023 | 0.26994358 | 0.77725133 | -0.0498907 | 0.30461018 |
| 0.999490048 | 0.05627009 | 0.26989055 | 0.77725133 | 0.08259239 | 0.11016445 |
| 0.999490048 | 0.09936758 | 0.26976958 | 0.77725133 | 0.05579581 | 0.53202922 |
| 0.999490048 | -0.466464 | 0.27127176 | 0.77743413 | -0.9723342 | 0.01383789 |
| 0.999490048 | -0.1689677 | 0.27189062 | 0.77743413 | -0.1236282 | 0.41872924 |
| 0.999490048 | -0.1354056 | 0.27113507 | 0.77743413 | -0.0341203 | 0.77909032 |
| 0.999490048 | -0.1278883 | 0.2720006 | 0.77743413 | -0.2128494 | 0.07328245 |
| 0.999490048 | -0.108165 | 0.27260226 | 0.77743413 | -0.034489 | 0.72358943 |
| 0.999490048 | -0.0924067 | 0.27277744 | 0.77743413 | 0.00565648 | 0.94582309 |
| 0.999490048 | -0.0851266 | 0.27116843 | 0.77743413 | -0.1246929 | 0.11178428 |
| 0.999490048 | -0.0805813 | 0.27170591 | 0.77743413 | -0.0429043 | 0.55495211 |
| 0.999490048 | -0.0767788 | 0.2709191 | 0.77743413 | -0.0369274 | 0.59298285 |
| 0.999490048 | -0.0752779 | 0.27161285 | 0.77743413 | 0.04207624 | 0.53548679 |
| 0.999490048 | -0.0727317 | 0.27138708 | 0.77743413 | 4.72E-04 | 0.99423322 |
| 0.999490048 | -0.0696743 | 0.27255048 | 0.77743413 | -0.0264592 | 0.67372786 |
| 0.999490048 | -0.0650692 | 0.2728372 | 0.77743413 | -5.55E-04 | 0.99244935 |
| 0.99974305 | -0.0586234 | 0.2727816 | 0.77743413 | -0.0585979 | 0.2729852 |
| 0.999490048 | -0.0576618 | 0.27283787 | 0.77743413 | -0.0395845 | 0.44868912 |
| 0.999490048 | -0.0573549 | 0.27166782 | 0.77743413 | 6.04E-04 | 0.99065741 |
| 0.999490048 | -0.0527487 | 0.27243561 | 0.77743413 | -0.0932228 | 0.05839014 |
| 0.999490048 | 0.05350577 | 0.27118489 | 0.77743413 | 0.04917604 | 0.31096624 |
| 0.999490048 | 0.0546752 | 0.27083234 | 0.77743413 | 0.01346791 | 0.7837839 |
| 0.999490048 | 0.05793803 | 0.27289335 | 0.77743413 | 0.01043486 | 0.8416261 |
| 0.999490048 | 0.05859282 | 0.27225216 | 0.77743413 | 0.01961584 | 0.71022033 |
| 0.999490048 | 0.06327753 | 0.27086052 | 0.77743413 | 0.03486051 | 0.54067539 |
| 0.999490048 | 0.06344439 | 0.27131212 | 0.77743413 | 0.02512673 | 0.65980171 |
| 0.999490048 | 0.07000987 | 0.27142113 | 0.77743413 | 0.03358142 | 0.59436429 |
| 0.999490048 | 0.0742987 | 0.27175414 | 0.77743413 | 0.00148292 | 0.98228365 |
| 0.999490048 | 0.08061706 | 0.27080156 | 0.77743413 | 0.01153981 | 0.87324048 |
| 0.999490048 | 0.10326412 | 0.27072496 | 0.77743413 | -0.0243219 | 0.79297933 |
| 0.999490048 | 0.1106569 | 0.27095698 | 0.77743413 | 0.06637029 | 0.50575498 |
| 0.999490048 | 0.11734677 | 0.27143631 | 0.77743413 | 0.00307087 | 0.97675682 |
| 0.999490048 | 0.13538683 | 0.27191488 | 0.77743413 | 0.09610783 | 0.42911694 |
| 0.999490048 | 0.21960515 | 0.27278745 | 0.77743413 | 0.18572953 | 0.35201893 |
| 0.999490048 | -0.0598124 | 0.27306509 | 0.77745157 | -0.0308569 | 0.5683442 |
| 0.987721587 | 0.08522644 | 0.27308568 | 0.77745157 | -0.0850886 | 0.27384363 |
| 0.999490048 | -0.0999324 | 0.27321978 | 0.77756823 | -0.1553699 | 0.0938004 |
| 0.999490048 | -0.1117909 | 0.27351205 | 0.77813479 | 0.05715091 | 0.57218862 |
| 0.999490048 | -0.0758098 | 0.27418328 | 0.77839665 | -0.0664978 | 0.33614555 |

| | | | | | |
|---|---|---|---|---|---|
| 0.999490048 | -0.059363 | 0.27433552 | 0.77839665 | -0.0362175 | 0.50159506 |
| 0.999490048 | -0.0538581 | 0.27443355 | 0.77839665 | -0.0330965 | 0.4986245 |
| 0.999490048 | -0.048719 | 0.27419199 | 0.77839665 | -0.0438764 | 0.3236453 |
| 0.999490048 | 0.05047428 | 0.27444308 | 0.77839665 | 0.00898718 | 0.84391341 |
| 0.999490048 | 0.05825252 | 0.27414638 | 0.77839665 | 0.05171347 | 0.33042003 |
| 0.999490048 | 0.06275542 | 0.27385839 | 0.77839665 | 0.00330849 | 0.95343706 |
| 0.999490048 | 0.07375653 | 0.27394011 | 0.77839665 | 0.05872908 | 0.38159383 |
| 0.999490048 | 0.10849537 | 0.27427103 | 0.77839665 | 0.05893867 | 0.54918626 |
| 0.999490048 | -0.100498 | 0.27456287 | 0.77847197 | 0.03211121 | 0.72414503 |
| 0.999490048 | 0.07243795 | 0.27468725 | 0.77856026 | -0.001946 | 0.97630516 |
| 0.999490048 | -0.2186641 | 0.27683048 | 0.77857808 | -0.0913587 | 0.64633747 |
| 0.999490048 | -0.2111741 | 0.27621884 | 0.77857808 | -0.1933925 | 0.31775934 |
| 0.999490048 | -0.1384105 | 0.27482557 | 0.77857808 | -0.2123891 | 0.09901431 |
| 0.972489992 | -0.1096324 | 0.27569668 | 0.77857808 | 0.11613101 | 0.24891455 |
| 0.999490048 | -0.0910895 | 0.27588391 | 0.77857808 | 0.00606381 | 0.94147928 |
| 0.999490048 | -0.082984 | 0.27702461 | 0.77857808 | -0.0745585 | 0.32763709 |
| 0.999490048 | -0.0775162 | 0.27697646 | 0.77857808 | -0.0773944 | 0.27771447 |
| 0.999490048 | -0.0651833 | 0.27684083 | 0.77857808 | -0.0237026 | 0.68948354 |
| 0.999490048 | -0.0644168 | 0.27620364 | 0.77857808 | -0.0678051 | 0.25233083 |
| 0.999490048 | -0.0628887 | 0.27545011 | 0.77857808 | -0.0396996 | 0.48803607 |
| 0.999490048 | -0.054674 | 0.27591758 | 0.77857808 | -0.0336613 | 0.49916957 |
| 0.999490048 | -0.0499175 | 0.27575115 | 0.77857808 | -0.0112587 | 0.80362921 |
| 0.999490048 | 0.04946106 | 0.27646491 | 0.77857808 | 0.06012403 | 0.18832313 |
| 0.999490048 | 0.0672985 | 0.27633751 | 0.77857808 | 0.0374909 | 0.54090031 |
| 0.999490048 | 0.07371841 | 0.27656249 | 0.77857808 | 0.05055061 | 0.45274867 |
| 0.999490048 | 0.0740464 | 0.27506899 | 0.77857808 | 0.1085431 | 0.11438481 |
| 0.999490048 | 0.07460962 | 0.27549507 | 0.77857808 | 0.03928155 | 0.56249305 |
| 0.999490048 | 0.07675939 | 0.27691547 | 0.77857808 | -0.0145818 | 0.83448847 |
| 0.999490048 | 0.07773365 | 0.27677966 | 0.77857808 | 0.05318012 | 0.45399415 |
| 0.999490048 | 0.07925571 | 0.27676229 | 0.77857808 | 0.08748872 | 0.23109956 |
| 0.999490048 | 0.08003504 | 0.27692589 | 0.77857808 | 0.05307809 | 0.46791897 |
| 0.999490048 | 0.08101457 | 0.27554816 | 0.77857808 | 0.08505943 | 0.25285931 |
| 0.999490048 | 0.10531105 | 0.27673665 | 0.77857808 | 0.04804444 | 0.61641601 |
| 0.999490048 | 0.10681696 | 0.27639918 | 0.77857808 | 0.08887991 | 0.36336107 |
| 0.999490048 | 0.11893094 | 0.27683197 | 0.77857808 | 0.13377744 | 0.22268383 |
| 0.999490048 | -0.1068044 | 0.27720275 | 0.77881661 | -0.1583247 | 0.1121073 |
| 0.999490048 | -0.0746472 | 0.27762019 | 0.77943528 | 0.01021324 | 0.88053064 |
| 0.999490048 | 0.05783308 | 0.27807637 | 0.77943528 | 0.0230592 | 0.66223982 |
| 0.999490048 | 0.07610767 | 0.27804473 | 0.77943528 | 0.06447033 | 0.35656938 |
| 0.999490048 | 0.0838766 | 0.27795124 | 0.77943528 | 0.01497055 | 0.84469348 |
| 0.999490048 | 0.08506817 | 0.27801033 | 0.77943528 | -0.0565119 | 0.46818909 |
| 0.999490048 | 0.0873187 | 0.27776126 | 0.77943528 | 0.12717298 | 0.11863192 |

| | | | | | |
|---|---|---|---|---|---|
| 0.999490048 | 0.14286456 | 0.27783539 | 0.77943528 | 0.20218427 | 0.12896866 |
| 0.999490048 | -0.0856502 | 0.27897717 | 0.77945962 | -0.1244558 | 0.12035547 |
| 0.999490048 | -0.0829119 | 0.27829858 | 0.77945962 | 0.04525606 | 0.55069939 |
| 0.999490048 | -0.0723087 | 0.27849125 | 0.77945962 | -0.0048821 | 0.9409756 |
| 0.999490048 | -0.0633814 | 0.27897889 | 0.77945962 | -0.0378519 | 0.5147369 |
| 0.999490048 | -0.0554197 | 0.27905602 | 0.77945962 | -0.0292214 | 0.56486001 |
| 0.999490048 | 0.05471009 | 0.27867048 | 0.77945962 | 0.06732566 | 0.185103 |
| 0.999490048 | 0.057459 | 0.27862779 | 0.77945962 | 0.0646811 | 0.22406063 |
| 0.987721587 | 0.08823404 | 0.27911189 | 0.77945962 | -0.0859683 | 0.29135759 |
| 0.999490048 | 0.08842189 | 0.27865278 | 0.77945962 | 0.01642299 | 0.83871175 |
| 0.999490048 | 0.12974218 | 0.27868284 | 0.77945962 | 0.03488134 | 0.7683963 |
| 0.999490048 | 0.17575832 | 0.27896111 | 0.77945962 | -0.0981209 | 0.54228879 |
| 0.999490048 | -0.2174161 | 0.27958274 | 0.77965764 | 0.01500515 | 0.93980871 |
| 0.999490048 | -0.1053094 | 0.27944642 | 0.77965764 | -0.0586346 | 0.5437705 |
| 0.999490048 | -0.0928355 | 0.27961037 | 0.77965764 | -0.0283832 | 0.73828229 |
| 0.999490048 | -0.0560405 | 0.2795827 | 0.77965764 | -0.0258312 | 0.6149277 |
| 0.999490048 | 0.05504451 | 0.27966552 | 0.77965764 | -0.033152 | 0.5118189 |
| 0.999490048 | 0.06121471 | 0.28002128 | 0.77965764 | 0.01668835 | 0.76589736 |
| 0.999490048 | 0.06220315 | 0.28002315 | 0.77965764 | 0.00314817 | 0.95588091 |
| 0.999490048 | 0.0651348 | 0.27980078 | 0.77965764 | 0.05368487 | 0.37125312 |
| 0.711558326 | 0.22787583 | 0.27997891 | 0.77965764 | -0.4181404 | 0.06099347 |
| 0.999490048 | -0.0626167 | 0.28064667 | 0.7802439 | 0.02570062 | 0.65475143 |
| 0.999490048 | -0.0594362 | 0.28062079 | 0.7802439 | 0.00793737 | 0.88410634 |
| 0.999490048 | 0.08652276 | 0.28070092 | 0.7802439 | -0.0163091 | 0.83705223 |
| 0.987721587 | 0.09026139 | 0.2804837 | 0.7802439 | 0.26684512 | 0.00318545 |
| 0.999490048 | 0.09841369 | 0.28040003 | 0.7802439 | -0.0087891 | 0.92229877 |
| 0.999490048 | -0.2430311 | 0.2809857 | 0.78051583 | -0.1043247 | 0.63612128 |
| 0.999490048 | -0.145083 | 0.28092149 | 0.78051583 | -0.3140815 | 0.03181291 |
| 0.999490048 | -0.10967 | 0.28172931 | 0.78221404 | -0.170044 | 0.10040141 |
| 0.999490048 | 0.07183586 | 0.28178441 | 0.78221404 | 0.05744252 | 0.38743721 |
| 0.999490048 | -0.0761599 | 0.28198478 | 0.7825101 | -0.1033002 | 0.14833478 |
| 0.999490048 | -0.055039 | 0.28249274 | 0.7832551 | -0.0311827 | 0.53939111 |
| 0.999490048 | 0.06256822 | 0.28243097 | 0.7832551 | 0.05760323 | 0.32152902 |
| 0.999490048 | 0.07915071 | 0.28253466 | 0.7832551 | 0.0600628 | 0.41250022 |
| 0.999490048 | 0.06837931 | 0.28280293 | 0.78373862 | 0.04548322 | 0.47211002 |
| 0.999490048 | -0.0586015 | 0.28291866 | 0.7837992 | -0.0032098 | 0.9525493 |
| 0.999490048 | -0.0592356 | 0.28322118 | 0.78411698 | -0.02704 | 0.62103068 |
| 0.999490048 | -0.0442108 | 0.28313179 | 0.78411698 | 0.02880346 | 0.48146581 |
| 0.999490048 | -0.5202991 | 0.28997196 | 0.78449206 | 0.45406 | 0.35175528 |
| 0.999490048 | -0.410297 | 0.28665447 | 0.78449206 | 0.03002046 | 0.93714575 |
| 0.999490048 | -0.2202262 | 0.28358449 | 0.78449206 | -0.1772675 | 0.38613876 |
| 0.972489992 | -0.1686187 | 0.28653621 | 0.78449206 | -0.55491 | 0.00352895 |

| | | | | | |
|---|---|---|---|---|---|
| 0.999490048 | -0.1319983 | 0.28744815 | 0.78449206 | 0.075431 | 0.54016324 |
| 0.999490048 | -0.1260633 | 0.28437731 | 0.78449206 | -0.0938267 | 0.42014532 |
| 0.999490048 | -0.1179754 | 0.28802213 | 0.78449206 | -0.141629 | 0.20435771 |
| 0.999490048 | -0.1094719 | 0.28396624 | 0.78449206 | 0.0022023 | 0.98260161 |
| 0.999490048 | -0.0940001 | 0.28838003 | 0.78449206 | 0.01664389 | 0.8492486 |
| 0.999490048 | -0.0914933 | 0.2892549 | 0.78449206 | -0.032333 | 0.70524109 |
| 0.999490048 | -0.0884283 | 0.28967848 | 0.78449206 | 0.02685352 | 0.74527779 |
| 0.999490048 | -0.08819 | 0.28730342 | 0.78449206 | -0.0560582 | 0.49588313 |
| 0.999490048 | -0.0834879 | 0.28728113 | 0.78449206 | -0.0994886 | 0.20691268 |
| 0.999490048 | -0.083392 | 0.28965511 | 0.78449206 | -0.0896208 | 0.25595328 |
| 0.999490048 | -0.0800154 | 0.28685254 | 0.78449206 | -0.0690109 | 0.35691976 |
| 0.999490048 | -0.0783402 | 0.28758549 | 0.78449206 | -0.0281425 | 0.6996078 |
| 0.999490048 | -0.0766574 | 0.28501888 | 0.78449206 | -0.066584 | 0.35173302 |
| 0.999490048 | -0.0751889 | 0.28877874 | 0.78449206 | -0.0208486 | 0.76627166 |
| 0.999490048 | -0.0743153 | 0.2836054 | 0.78449206 | -0.1112386 | 0.11344704 |
| 0.999490048 | -0.0736527 | 0.28735644 | 0.78449206 | -0.0996339 | 0.15385453 |
| 0.999490048 | -0.0720686 | 0.2854069 | 0.78449206 | -0.0330181 | 0.62137817 |
| 0.999490048 | -0.0696215 | 0.29011457 | 0.78449206 | -0.0574894 | 0.38067691 |
| 0.999490048 | -0.0676549 | 0.28883491 | 0.78449206 | -0.0786236 | 0.21950486 |
| 0.999490048 | -0.0673871 | 0.28826713 | 0.78449206 | -0.0524838 | 0.40610693 |
| 0.999490048 | -0.065681 | 0.28921781 | 0.78449206 | -0.1334477 | 0.03723267 |
| 0.999490048 | -0.0653915 | 0.28438575 | 0.78449206 | -0.0213366 | 0.72413218 |
| 0.999490048 | -0.0651123 | 0.28783854 | 0.78449206 | -0.0562966 | 0.35673917 |
| 0.999490048 | -0.0634584 | 0.28535355 | 0.78449206 | -0.0220396 | 0.70777632 |
| 0.999490048 | -0.0626075 | 0.28721531 | 0.78449206 | -0.0132093 | 0.82040862 |
| 0.999490048 | -0.0617085 | 0.28414397 | 0.78449206 | -0.008192 | 0.88564115 |
| 0.999490048 | -0.0608254 | 0.28724035 | 0.78449206 | -0.1052181 | 0.07163652 |
| 0.999490048 | -0.0601182 | 0.28756098 | 0.78449206 | -0.0555561 | 0.32491548 |
| 0.999490048 | -0.055702 | 0.28590293 | 0.78449206 | -0.0869326 | 0.10115503 |
| 0.999490048 | -0.0511228 | 0.28977848 | 0.78449206 | -0.0365165 | 0.44705421 |
| 0.999490048 | -0.0500635 | 0.28893824 | 0.78449206 | -0.069881 | 0.14295883 |
| 0.999490048 | -0.0494061 | 0.28751506 | 0.78449206 | -0.0660968 | 0.15839908 |
| 0.999490048 | -0.0462468 | 0.28645067 | 0.78449206 | -0.0330601 | 0.44351781 |
| 0.999490048 | 0.04562008 | 0.28378867 | 0.78449206 | 0.00290163 | 0.94501333 |
| 0.999490048 | 0.04700653 | 0.28698514 | 0.78449206 | 0.01533614 | 0.72561395 |
| 0.999490048 | 0.05165225 | 0.28929045 | 0.78449206 | 0.00490205 | 0.91898628 |
| 0.999490048 | 0.0533371 | 0.28801359 | 0.78449206 | 0.06380138 | 0.2059256 |
| 0.999490048 | 0.05525396 | 0.28733991 | 0.78449206 | 0.07186595 | 0.16967589 |
| 0.999490048 | 0.05634606 | 0.2886059 | 0.78449206 | 0.05155976 | 0.3306422 |
| 0.999490048 | 0.05685025 | 0.28801229 | 0.78449206 | -0.0056625 | 0.9147617 |
| 0.999490048 | 0.05755599 | 0.28872589 | 0.78449206 | 0.03631765 | 0.50029835 |
| 0.999490048 | 0.05818399 | 0.28781337 | 0.78449206 | 0.03630796 | 0.50418785 |

| | | | | | |
|---|---|---|---|---|---|
| 0.999490048 | 0.0587144 | 0.28443168 | 0.78449206 | 0.03167486 | 0.56042205 |
| 0.999490048 | 0.05872746 | 0.29001444 | 0.78449206 | 0.07812977 | 0.16277587 |
| 0.999490048 | 0.06770797 | 0.28946332 | 0.78449206 | 0.06529421 | 0.30662156 |
| 0.999490048 | 0.0677307 | 0.28460493 | 0.78449206 | 0.03803511 | 0.54473976 |
| 0.999490048 | 0.07174833 | 0.28967449 | 0.78449206 | 0.05412976 | 0.42214224 |
| 0.999490048 | 0.07384577 | 0.28748082 | 0.78449206 | -0.0121463 | 0.85957264 |
| 0.999490048 | 0.07714065 | 0.28781816 | 0.78449206 | 0.07363117 | 0.30982938 |
| 0.999490048 | 0.08379181 | 0.28932427 | 0.78449206 | 0.03722297 | 0.63477389 |
| 0.999490048 | 0.08488804 | 0.28492193 | 0.78449206 | 0.1294731 | 0.10803021 |
| 0.999490048 | 0.08568035 | 0.28483708 | 0.78449206 | 0.07409405 | 0.35363515 |
| 0.999490048 | 0.08801102 | 0.28396564 | 0.78449206 | 0.07639416 | 0.35099272 |
| 0.999490048 | 0.09037357 | 0.28603871 | 0.78449206 | 0.07269287 | 0.38891016 |
| 0.999490048 | 0.0933916 | 0.28537862 | 0.78449206 | 0.05095459 | 0.55679588 |
| 0.999490048 | 0.09604514 | 0.28692682 | 0.78449206 | 0.05754843 | 0.52038507 |
| 0.999490048 | 0.09639815 | 0.28581577 | 0.78449206 | 0.00928764 | 0.91715377 |
| 0.999490048 | 0.09905399 | 0.28447858 | 0.78449206 | 0.11505396 | 0.21560858 |
| 0.999490048 | 0.10857309 | 0.28792094 | 0.78449206 | 0.05725065 | 0.57208078 |
| 0.999490048 | 0.10940889 | 0.28837775 | 0.78449206 | 0.02286075 | 0.82254311 |
| 0.999490048 | 0.11202068 | 0.2898647 | 0.78449206 | 0.13222812 | 0.21353042 |
| 0.999490048 | 0.11282506 | 0.28598348 | 0.78449206 | 0.08261984 | 0.43217156 |
| 0.999490048 | 0.13464329 | 0.2851943 | 0.78449206 | 0.08024042 | 0.52110086 |
| 0.999490048 | 0.14328311 | 0.28888105 | 0.78449206 | 0.24923328 | 0.07849501 |
| 0.999490048 | 0.14713517 | 0.28836677 | 0.78449206 | 0.0861419 | 0.53115052 |
| 0.999490048 | 0.14781679 | 0.28481616 | 0.78449206 | 0.11606991 | 0.3989192 |
| 0.999490048 | 0.15837978 | 0.29012113 | 0.78449206 | -0.0343429 | 0.81663271 |
| 0.999490048 | 0.20134964 | 0.28909495 | 0.78449206 | -0.0144959 | 0.93690518 |
| 0.999490048 | -0.0606094 | 0.29051351 | 0.7850446 | 0.03533645 | 0.53469924 |
| 0.999490048 | 0.05597006 | 0.29042637 | 0.7850446 | 0.04729927 | 0.3701079 |
| 0.999490048 | -0.0817098 | 0.29073344 | 0.78538474 | -0.0559962 | 0.46639328 |
| 0.999490048 | 0.1267191 | 0.29098969 | 0.78582274 | -0.0454444 | 0.70214101 |
| 0.999490048 | -0.2762918 | 0.29141801 | 0.78600696 | 0.06052943 | 0.81091228 |
| 0.999490048 | -0.1753464 | 0.2918137 | 0.78600696 | -0.1015748 | 0.53841433 |
| 0.999490048 | -0.0853908 | 0.2925901 | 0.78600696 | 0.07107074 | 0.37942791 |
| 0.999490048 | -0.0794439 | 0.29210986 | 0.78600696 | 0.03526625 | 0.63699987 |
| 0.999490048 | -0.0755809 | 0.29257242 | 0.78600696 | -0.109854 | 0.13066949 |
| 0.999490048 | -0.0736197 | 0.29208326 | 0.78600696 | 0.00897846 | 0.8966274 |
| 0.999490048 | -0.0639001 | 0.29161464 | 0.78600696 | -0.05931 | 0.32691881 |
| 0.999490048 | -0.061678 | 0.29279568 | 0.78600696 | -0.018484 | 0.75010222 |
| 0.999490048 | -0.055331 | 0.29363915 | 0.78600696 | 0.03890421 | 0.45775638 |
| 0.999490048 | -0.0541479 | 0.29369362 | 0.78600696 | -0.0218537 | 0.66881846 |
| 0.999490048 | -0.0520056 | 0.29343679 | 0.78600696 | -0.0990847 | 0.05151344 |
| 0.999490048 | -0.0438456 | 0.29312832 | 0.78600696 | -0.0354471 | 0.39355913 |

| | | | | | |
|---|---|---|---|---|---|
| 0.999490048 | -0.0434805 | 0.29160154 | 0.78600696 | -0.0330842 | 0.4201087 |
| 0.999490048 | 0.04316548 | 0.29285421 | 0.78600696 | 0.04470853 | 0.27632564 |
| 0.999490048 | 0.05622392 | 0.29291156 | 0.78600696 | -0.0031638 | 0.95227518 |
| 0.999490048 | 0.06640431 | 0.29200933 | 0.78600696 | 0.00891492 | 0.8862601 |
| 0.999490048 | 0.07197395 | 0.29143279 | 0.78600696 | 0.03599621 | 0.59466116 |
| 0.999490048 | 0.07640824 | 0.29324703 | 0.78600696 | 0.05808396 | 0.42213874 |
| 0.999490048 | 0.07874825 | 0.29269519 | 0.78600696 | 0.01639277 | 0.82477794 |
| 0.999490048 | 0.0970973 | 0.29279482 | 0.78600696 | 0.03955526 | 0.66531565 |
| 0.999490048 | 0.09792302 | 0.29363803 | 0.78600696 | -0.0023035 | 0.98006991 |
| 0.999490048 | 0.10192092 | 0.29367396 | 0.78600696 | 0.08847223 | 0.36070846 |
| 0.999490048 | 0.10511162 | 0.29332126 | 0.78600696 | 0.07133156 | 0.47310815 |
| 0.999490048 | 0.10962997 | 0.29317746 | 0.78600696 | 0.11501642 | 0.27063344 |
| 0.999490048 | 0.1110379 | 0.2932538 | 0.78600696 | 0.04961129 | 0.63564024 |
| 0.987721587 | 0.11918268 | 0.29152057 | 0.78600696 | -0.1200768 | 0.2879962 |
| 0.999490048 | 0.12906123 | 0.29179887 | 0.78600696 | 0.10750455 | 0.37826136 |
| 0.999490048 | 0.13753685 | 0.292919 | 0.78600696 | 0.32607571 | 0.02812623 |
| 0.999490048 | -0.1222005 | 0.29448078 | 0.78634904 | -0.0832708 | 0.47237955 |
| 0.999490048 | -0.1121195 | 0.29597888 | 0.78634904 | -0.0568454 | 0.5931816 |
| 0.999490048 | -0.0950694 | 0.29577438 | 0.78634904 | -0.0591293 | 0.51266845 |
| 0.999490048 | -0.0713855 | 0.2958036 | 0.78634904 | -0.0086626 | 0.89794243 |
| 0.999490048 | -0.0637081 | 0.29497497 | 0.78634904 | 0.04031448 | 0.50469771 |
| 0.999490048 | -0.0593815 | 0.29491871 | 0.78634904 | 0.00158778 | 0.97740769 |
| 0.999490048 | -0.0568497 | 0.29598743 | 0.78634904 | -0.1047229 | 0.06042984 |
| 0.999490048 | -0.055564 | 0.29472475 | 0.78634904 | -0.0901359 | 0.09481105 |
| 0.999490048 | -0.0480299 | 0.29534282 | 0.78634904 | -0.0244927 | 0.59055586 |
| 0.999490048 | 0.05570276 | 0.29585071 | 0.78634904 | 0.10778466 | 0.04930146 |
| 0.999490048 | 0.05680195 | 0.29557272 | 0.78634904 | 0.02505113 | 0.6416233 |
| 0.999490048 | 0.05687012 | 0.29595963 | 0.78634904 | 0.02631353 | 0.62569587 |
| 0.999490048 | 0.0596979 | 0.29453693 | 0.78634904 | 0.0682058 | 0.23262076 |
| 0.999490048 | 0.05987363 | 0.29443465 | 0.78634904 | 0.05444088 | 0.33951397 |
| 0.999490048 | 0.06034194 | 0.29431845 | 0.78634904 | 0.0358959 | 0.52978985 |
| 0.999490048 | 0.0653363 | 0.29541198 | 0.78634904 | 0.06703947 | 0.28327983 |
| 0.999490048 | 0.06710686 | 0.29570993 | 0.78634904 | 0.00734114 | 0.90792804 |
| 0.999490048 | 0.07348279 | 0.29420288 | 0.78634904 | 0.04201867 | 0.54564348 |
| 0.999490048 | 0.07495011 | 0.29526262 | 0.78634904 | 0.0911526 | 0.2053962 |
| 0.999490048 | 0.07913298 | 0.29511397 | 0.78634904 | 0.04785504 | 0.52373531 |
| 0.999490048 | 0.07934563 | 0.29474961 | 0.78634904 | -0.0066517 | 0.92923136 |
| 0.999490048 | 0.10753468 | 0.29483992 | 0.78634904 | -0.0205949 | 0.83929902 |
| 0.999490048 | 0.13408106 | 0.29393381 | 0.78634904 | 0.0814398 | 0.52090019 |
| 0.999490048 | 0.0584603 | 0.29628198 | 0.78638097 | 0.06763246 | 0.22862868 |
| 0.909524814 | 0.06176379 | 0.29624481 | 0.78638097 | -0.0824443 | 0.16677991 |
| 0.999490048 | 0.08037731 | 0.29612081 | 0.78638097 | 0.11128825 | 0.15201154 |

| | | | | | |
|---|---|---|---|---|---|
| 0.999490048 | -0.0528268 | 0.29656534 | 0.78663297 | -0.0049021 | 0.92198762 |
| 0.999490048 | 0.15121168 | 0.29650016 | 0.78663297 | 0.13447068 | 0.35210875 |
| 0.999490048 | 0.06253 | 0.29679435 | 0.78699041 | 0.09212816 | 0.12896707 |
| 0.999490048 | 0.18642654 | 0.29689509 | 0.78700763 | 0.36082156 | 0.06250611 |
| 0.999490048 | -0.1011914 | 0.29742699 | 0.78720403 | -0.1687018 | 0.08819274 |
| 0.999490048 | -0.0986042 | 0.29731008 | 0.78720403 | -0.1273044 | 0.18154581 |
| 0.999490048 | -0.0766769 | 0.29744056 | 0.78720403 | -0.0458495 | 0.53037739 |
| 0.999490048 | 0.05193838 | 0.29718682 | 0.78720403 | 0.00672788 | 0.89141753 |
| 0.999490048 | 0.06871733 | 0.29720274 | 0.78720403 | -0.0654629 | 0.32018506 |
| 0.999490048 | -0.0585222 | 0.29763102 | 0.78720905 | -0.0994918 | 0.08256234 |
| 0.999490048 | 0.04917504 | 0.29758038 | 0.78720905 | 0.0230844 | 0.62188785 |
| 0.999490048 | -0.1099758 | 0.29833388 | 0.78757127 | -0.0844371 | 0.42251418 |
| 0.999490048 | -0.04917 | 0.29830967 | 0.78757127 | -0.0112496 | 0.80998687 |
| 0.999490048 | 0.05356827 | 0.29796823 | 0.78757127 | 0.03957133 | 0.43972458 |
| 0.999490048 | 0.05767541 | 0.29815342 | 0.78757127 | 0.05783721 | 0.29682928 |
| 0.999490048 | 0.107327 | 0.29815546 | 0.78757127 | 0.01271454 | 0.90084604 |
| 0.999490048 | 0.14611125 | 0.29803614 | 0.78757127 | 0.1414227 | 0.31347946 |
| 0.999490048 | -0.0911173 | 0.29865278 | 0.78816395 | -0.0065917 | 0.93940911 |
| 0.999490048 | 0.09602165 | 0.29878595 | 0.78826626 | 0.01593488 | 0.86164901 |
| 0.999490048 | 0.05313246 | 0.29893931 | 0.78842174 | 0.09579266 | 0.0673078 |
| 0.999490048 | -0.0905955 | 0.29933147 | 0.78845987 | -0.0949076 | 0.27742045 |
| 0.999490048 | 0.07106946 | 0.29930614 | 0.78845987 | 0.09986084 | 0.14888434 |
| 0.999490048 | 0.10813013 | 0.2991618 | 0.78845987 | 0.05736469 | 0.57881149 |
| 0.999490048 | 0.1466664 | 0.2991586 | 0.78845987 | -0.0758506 | 0.58834337 |
| 0.999490048 | -0.0515815 | 0.29968869 | 0.78857258 | -0.0163219 | 0.74034098 |
| 0.999490048 | 0.05795691 | 0.2998005 | 0.78857258 | 0.0552068 | 0.32283747 |
| 0.999490048 | 0.06100846 | 0.30021508 | 0.78857258 | 0.00939684 | 0.87188029 |
| 0.999490048 | 0.06118478 | 0.29947228 | 0.78857258 | 0.07527387 | 0.20424943 |
| 0.999490048 | 0.06379377 | 0.29990562 | 0.78857258 | 0.10140116 | 0.10486225 |
| 0.999490048 | 0.06861704 | 0.30022422 | 0.78857258 | 0.07998155 | 0.22900119 |
| 0.999490048 | 0.07437307 | 0.29960528 | 0.78857258 | 0.14073163 | 0.05594004 |
| 0.999490048 | 0.11126086 | 0.3001372 | 0.78857258 | 0.06713166 | 0.52902564 |
| 0.999490048 | 0.20664157 | 0.30004414 | 0.78857258 | 0.38459249 | 0.0600212 |
| 0.999490048 | -0.0843908 | 0.30053319 | 0.78913589 | 0.06164869 | 0.4471929 |
| 0.999490048 | -0.0463033 | 0.30067237 | 0.78925316 | -0.0810604 | 0.07610263 |
| 0.999490048 | -0.055356 | 0.30082845 | 0.7894147 | -0.021566 | 0.6841464 |
| 0.999490048 | -0.1584454 | 0.3012815 | 0.78985887 | 0.14138923 | 0.35526449 |
| 0.999490048 | -0.1033833 | 0.3010938 | 0.78985887 | -0.0272319 | 0.78320253 |
| 0.999490048 | -0.0796526 | 0.30119996 | 0.78985887 | 0.02719221 | 0.72158885 |
| 0.999490048 | 0.0614291 | 0.30169203 | 0.7906869 | 0.11124742 | 0.06765266 |
| 0.999490048 | -0.2708418 | 0.30182574 | 0.79078911 | -0.398461 | 0.13915423 |
| 0.999490048 | 0.05029498 | 0.30213828 | 0.79120095 | 0.0471394 | 0.33286 |

| | | | | | |
|---|---|---|---|---|---|
| 0.999490048 | 0.13331052 | 0.30217243 | 0.79120095 | 0.14542511 | 0.26131426 |
| 0.999490048 | 0.0506854 | 0.30271752 | 0.79138738 | -4.40E-04 | 0.99278806 |
| 0.999490048 | 0.06263318 | 0.30264647 | 0.79138738 | 0.06629959 | 0.27583401 |
| 0.999490048 | 0.06791294 | 0.30259528 | 0.79138738 | 0.11683872 | 0.08215743 |
| 0.999490048 | 0.10351331 | 0.30271411 | 0.79138738 | -0.0053966 | 0.95668672 |
| 0.999490048 | 0.13701542 | 0.30263993 | 0.79138738 | 0.15740379 | 0.23792343 |
| 0.999490048 | 0.07766774 | 0.30291658 | 0.79165993 | 0.00759933 | 0.91885237 |
| 0.999490048 | -0.137697 | 0.30307216 | 0.79181869 | -0.3083481 | 0.02666367 |
| 0.999490048 | -0.0510824 | 0.3032148 | 0.79189286 | -0.0088962 | 0.85622527 |
| 0.999490048 | 0.05964272 | 0.30329022 | 0.79189286 | 0.01113623 | 0.84601204 |
| 0.999490048 | -0.4437899 | 0.30377501 | 0.79255765 | -0.6303939 | 0.15867126 |
| 0.999490048 | -0.1637401 | 0.30540497 | 0.79255765 | -0.2657178 | 0.10188431 |
| 0.909524814 | -0.1418066 | 0.30500133 | 0.79255765 | 0.19566576 | 0.16085207 |
| 0.999490048 | -0.1412621 | 0.30439879 | 0.79255765 | -0.0852556 | 0.53259161 |
| 0.999490048 | -0.1232395 | 0.30568327 | 0.79255765 | 0.02874999 | 0.80926479 |
| 0.999490048 | -0.1060993 | 0.30502872 | 0.79255765 | -0.0114622 | 0.9108353 |
| 0.999490048 | -0.1055551 | 0.30523787 | 0.79255765 | -0.1088907 | 0.29053271 |
| 0.999490048 | -0.0953462 | 0.30589787 | 0.79255765 | 0.0197625 | 0.83024498 |
| 0.999490048 | -0.0723427 | 0.30576726 | 0.79255765 | -0.1239768 | 0.08524123 |
| 0.999490048 | -0.0653901 | 0.30457741 | 0.79255765 | 0.05064695 | 0.42454362 |
| 0.999490048 | -0.0589817 | 0.30445841 | 0.79255765 | 0.00130981 | 0.98161034 |
| 0.999490048 | -0.0525839 | 0.30445422 | 0.79255765 | -0.0219166 | 0.66588319 |
| 0.999490048 | -0.0424803 | 0.30577038 | 0.79255765 | -0.0243178 | 0.5548887 |
| 0.999490048 | -0.0419333 | 0.30499918 | 0.79255765 | -0.0441208 | 0.2810013 |
| 0.999490048 | 0.04302401 | 0.30565178 | 0.79255765 | 0.01805342 | 0.66456326 |
| 0.999490048 | 0.04658956 | 0.30584145 | 0.79255765 | 0.01696634 | 0.70665675 |
| 0.999490048 | 0.05461388 | 0.30601268 | 0.79255765 | 0.03106058 | 0.55764418 |
| 0.999490048 | 0.05598469 | 0.30554423 | 0.79255765 | 0.05280546 | 0.33326922 |
| 0.999490048 | 0.06559297 | 0.30560642 | 0.79255765 | 0.12644347 | 0.05460768 |
| 0.999490048 | 0.07364313 | 0.30506318 | 0.79255765 | 0.04298865 | 0.54656912 |
| 0.999490048 | 0.07910991 | 0.30498823 | 0.79255765 | 0.20561952 | 0.01162366 |
| 0.999490048 | 0.08317762 | 0.30595131 | 0.79255765 | 0.09672346 | 0.23557061 |
| 0.999490048 | 0.08444149 | 0.30566573 | 0.79255765 | 0.06216907 | 0.44855709 |
| 0.999490048 | 0.10576248 | 0.30402485 | 0.79255765 | 0.03963004 | 0.69750109 |
| 0.999490048 | 0.12757504 | 0.30457536 | 0.79255765 | 0.05944184 | 0.62954346 |
| 0.961663458 | 0.13421989 | 0.30480742 | 0.79255765 | 0.44569756 | 0.00185241 |
| 0.999490048 | 0.09618231 | 0.30625682 | 0.79294402 | 0.14650161 | 0.12416728 |
| 0.999490048 | -0.0555566 | 0.30645352 | 0.79296154 | -0.0283475 | 0.59894663 |
| 0.999490048 | 0.07269744 | 0.30643664 | 0.79296154 | -0.0053789 | 0.93904099 |
| 0.999490048 | -0.1890783 | 0.30661138 | 0.79312423 | -0.0945742 | 0.602531 |
| 0.999490048 | -0.0561726 | 0.3071392 | 0.79332009 | -0.0747589 | 0.17751788 |
| 0.999490048 | 0.05067093 | 0.30687812 | 0.79332009 | -0.0042878 | 0.93037076 |

| | | | | | |
|---|---|---|---|---|---|
| 0.999490048 | 0.0549291 | 0.30725715 | 0.79332009 | -0.0437077 | 0.41476404 |
| 0.999490048 | 0.07551498 | 0.30724319 | 0.79332009 | 0.04560014 | 0.53480943 |
| 0.999490048 | 0.08428414 | 0.30707269 | 0.79332009 | -0.0145979 | 0.85815226 |
| 0.999490048 | 0.10126273 | 0.30708186 | 0.79332009 | 0.02644339 | 0.78765608 |
| 0.999490048 | -0.0766295 | 0.30753337 | 0.79354255 | -0.0833371 | 0.26801986 |
| 0.999490048 | 0.06445198 | 0.3074447 | 0.79354255 | -0.0093037 | 0.88166747 |
| 0.999490048 | -0.1644556 | 0.30786311 | 0.7936355 | 0.14344108 | 0.37259011 |
| 0.999490048 | -0.073464 | 0.30794958 | 0.7936355 | -0.0667301 | 0.35352893 |
| 0.999490048 | -0.0526849 | 0.30783749 | 0.7936355 | -0.0911059 | 0.08388082 |
| 0.999490048 | 0.28661699 | 0.30790103 | 0.7936355 | 0.37823378 | 0.18583129 |
| 0.999490048 | 0.09566962 | 0.3081711 | 0.79371643 | 0.02367383 | 0.79897334 |
| 0.999490048 | 0.10331456 | 0.30815895 | 0.79371643 | -0.0405657 | 0.6805758 |
| 0.999490048 | -0.0463188 | 0.30831019 | 0.79382981 | 0.00819437 | 0.85551945 |
| 0.999490048 | -0.1684198 | 0.30862522 | 0.79439599 | -0.3150249 | 0.07611351 |
| 0.999490048 | -0.146533 | 0.30936963 | 0.79481838 | -0.1627905 | 0.25990416 |
| 0.999490048 | -0.1453361 | 0.31007877 | 0.79481838 | -0.1000627 | 0.48225591 |
| 0.999490048 | -0.0888512 | 0.30944547 | 0.79481838 | -0.0612435 | 0.48117829 |
| 0.999490048 | -0.0870258 | 0.31065258 | 0.79481838 | 6.46E-04 | 0.99392996 |
| 0.999490048 | -0.0712274 | 0.31058513 | 0.79481838 | -0.0854892 | 0.22567178 |
| 0.999490048 | -0.0603166 | 0.31028233 | 0.79481838 | -0.1128287 | 0.06405254 |
| 0.999490048 | -0.0560093 | 0.31060052 | 0.79481838 | 0.01570252 | 0.77410367 |
| 0.999490048 | -0.0503985 | 0.30948297 | 0.79481838 | 0.02169043 | 0.65911499 |
| 0.771925038 | -0.0468436 | 0.31023568 | 0.79481838 | 0.07938225 | 0.09144114 |
| 0.999490048 | -0.0389452 | 0.31060105 | 0.79481838 | 0.02163264 | 0.57052864 |
| 0.999490048 | 0.05299382 | 0.30940655 | 0.79481838 | 0.01125589 | 0.8273762 |
| 0.999490048 | 0.06916703 | 0.30971491 | 0.79481838 | 0.07280656 | 0.28548126 |
| 0.999490048 | 0.06946819 | 0.30983557 | 0.79481838 | -0.0142786 | 0.83301105 |
| 0.999490048 | 0.06965118 | 0.31003967 | 0.79481838 | 0.04294832 | 0.52869653 |
| 0.999490048 | 0.07059113 | 0.30979442 | 0.79481838 | 0.04308719 | 0.53263496 |
| 0.999490048 | 0.08085927 | 0.31069308 | 0.79481838 | 0.09376901 | 0.24139723 |
| 0.999490048 | 0.09192112 | 0.30935442 | 0.79481838 | 0.05181458 | 0.56385808 |
| 0.999490048 | 0.09910399 | 0.31016485 | 0.79481838 | 0.15066622 | 0.12783195 |
| 0.31159876 | 0.13899437 | 0.31006367 | 0.79481838 | -0.4439707 | 0.00585938 |
| 0.999490048 | 0.15134881 | 0.3089636 | 0.79481838 | 0.17803231 | 0.23323276 |
| 0.999490048 | 0.05528161 | 0.31115697 | 0.79570567 | 0.0071817 | 0.89425171 |
| 0.964187865 | 0.16361636 | 0.31123051 | 0.79570567 | -0.2205407 | 0.17581414 |
| 0.999490048 | -0.0576905 | 0.31142802 | 0.79596694 | -0.0742565 | 0.19557003 |
| 0.999490048 | -0.0656517 | 0.31153701 | 0.79600186 | -0.1270655 | 0.05653991 |
| 0.890934946 | 0.12027087 | 0.31182166 | 0.79648543 | -0.1757898 | 0.1438591 |
| 0.999490048 | 0.07169199 | 0.31198974 | 0.79667105 | 0.09725712 | 0.17381146 |
| 0.999490048 | 0.11429649 | 0.31220379 | 0.79697391 | -0.0100494 | 0.92846625 |
| 0.999490048 | -0.1349502 | 0.31250511 | 0.79727624 | 0.02078071 | 0.8735121 |

| | | | | | |
|---|---|---|---|---|---|
| 0.999490048 | -0.0624619 | 0.31251319 | 0.79727624 | -0.061299 | 0.32142566 |
| 0.999490048 | -0.0701304 | 0.31332731 | 0.79837748 | -0.0419902 | 0.54340277 |
| 0.999490048 | 0.0512388 | 0.31328238 | 0.79837748 | 0.05173541 | 0.30871092 |
| 0.999490048 | 0.22038582 | 0.31312646 | 0.79837748 | 0.26647907 | 0.22475264 |
| 0.999490048 | 0.33285493 | 0.31322413 | 0.79837748 | 0.10578489 | 0.74127911 |
| 0.999490048 | -0.068511 | 0.31375512 | 0.79856583 | -0.0752842 | 0.26931054 |
| 0.999490048 | -0.0520217 | 0.31353371 | 0.79856583 | -0.0235478 | 0.64555885 |
| 0.999490048 | -0.0434104 | 0.31364751 | 0.79856583 | -0.0571085 | 0.18820971 |
| 0.999490048 | 0.12859585 | 0.31378377 | 0.79856583 | 0.09187757 | 0.46470249 |
| 0.999490048 | -0.1474325 | 0.31452645 | 0.79875178 | -0.1260235 | 0.38864928 |
| 0.999490048 | -0.0480289 | 0.31429971 | 0.79875178 | -0.0192516 | 0.68414193 |
| 0.999490048 | -0.0396857 | 0.31449404 | 0.79875178 | 0.00381004 | 0.9222876 |
| 0.999490048 | 0.04776031 | 0.31440368 | 0.79875178 | -0.0069721 | 0.8820718 |
| 0.999490048 | 0.05221041 | 0.31407442 | 0.79875178 | 0.0486229 | 0.34783499 |
| 0.999490048 | 0.08661736 | 0.31447646 | 0.79875178 | 0.00193753 | 0.98186382 |
| 0.999490048 | 0.38443284 | 0.3144163 | 0.79875178 | -0.2576723 | 0.49460337 |
| 0.999490048 | 0.09893468 | 0.31478326 | 0.79916091 | 0.11071541 | 0.2618241 |
| 0.999490048 | -0.0375931 | 0.31515939 | 0.79946287 | -0.0267153 | 0.47312568 |
| 0.999490048 | 0.06662741 | 0.31522624 | 0.79946287 | -0.0241374 | 0.71356031 |
| 0.999490048 | 0.09860458 | 0.31537758 | 0.79946287 | 0.29014574 | 0.00581005 |
| 0.999490048 | 0.11710679 | 0.31505945 | 0.79946287 | 0.00722022 | 0.95010292 |
| 0.999490048 | 0.19305839 | 0.31538092 | 0.79946287 | -0.0490116 | 0.79442237 |
| 0.999490048 | -0.140596 | 0.31558379 | 0.79953903 | 0.01020982 | 0.94131109 |
| 0.999490048 | -0.0688884 | 0.31588973 | 0.79953903 | -0.0500593 | 0.46398814 |
| 0.999490048 | -0.0640649 | 0.31587886 | 0.79953903 | -0.1083152 | 0.09584988 |
| 0.999490048 | 0.06892813 | 0.31580961 | 0.79953903 | 0.05230194 | 0.44461862 |
| 0.999490048 | 0.08115677 | 0.31580133 | 0.79953903 | 0.11416317 | 0.16225787 |
| 0.999490048 | 0.04990834 | 0.31610601 | 0.799844 | 0.05539051 | 0.26698234 |
| 0.999490048 | -0.0471824 | 0.31622997 | 0.79986101 | -0.0684676 | 0.15025468 |
| 0.999490048 | 0.15746964 | 0.31630432 | 0.79986101 | -0.0709263 | 0.64563212 |
| 0.999490048 | 0.27410925 | 0.31652244 | 0.80017026 | 0.73606773 | 0.01659928 |
| 0.999490048 | -0.0518706 | 0.31664748 | 0.80024408 | -0.0422143 | 0.41351241 |
| 0.999490048 | -0.4371588 | 0.31697242 | 0.80030145 | 0.16481351 | 0.70352372 |
| 0.999490048 | -0.0474451 | 0.31705356 | 0.80030145 | -0.0207181 | 0.65957432 |
| 0.999490048 | 0.06839229 | 0.31705045 | 0.80030145 | 0.00353533 | 0.95833259 |
| 0.999490048 | 0.07428185 | 0.31687846 | 0.80030145 | -0.0037955 | 0.95879786 |
| 0.999490048 | -0.0436225 | 0.31743784 | 0.80102929 | -0.0269244 | 0.53465795 |
| 0.999490048 | -0.0431712 | 0.31767769 | 0.80139235 | -0.0741349 | 0.09219137 |
| 0.999490048 | -0.159236 | 0.3184813 | 0.8017241 | -0.2942251 | 0.07166476 |
| 0.999490048 | -0.0699589 | 0.31822339 | 0.8017241 | -0.1126728 | 0.11344483 |
| 0.999490048 | -0.0628017 | 0.31843552 | 0.8017241 | 0.01466595 | 0.81403932 |
| 0.999490048 | -0.0567768 | 0.3180221 | 0.8017241 | -0.0405983 | 0.47306063 |

| | | | | | |
|---|---|---|---|---|---|
| 0.999490048 | -0.0535842 | 0.31835302 | 0.8017241 | -0.1007962 | 0.06693305 |
| 0.999490048 | -0.047134 | 0.31796292 | 0.8017241 | 0.00847504 | 0.85610799 |
| 0.999490048 | 0.06947589 | 0.31828315 | 0.8017241 | 0.06260453 | 0.36760175 |
| 0.999490048 | 0.04167848 | 0.31866014 | 0.80179936 | -0.0203163 | 0.62425206 |
| 0.999490048 | 0.07780451 | 0.31870324 | 0.80179936 | 0.1100004 | 0.16269421 |
| 0.999490048 | -0.0440185 | 0.31880723 | 0.80181939 | 0.00121378 | 0.97784768 |
| 0.999490048 | -0.0696282 | 0.31934401 | 0.80248852 | -0.0161242 | 0.81591039 |
| 0.999490048 | 0.06168688 | 0.31930953 | 0.80248852 | 0.04118377 | 0.50377373 |
| 0.578467169 | 0.13002473 | 0.3193616 | 0.80248852 | -0.3407749 | 0.023552 |
| 0.999490048 | 0.34306864 | 0.31963694 | 0.80293875 | -0.0777474 | 0.81611896 |
| 0.999490048 | -0.1073845 | 0.32015625 | 0.80303536 | -0.007548 | 0.94373366 |
| 0.999490048 | -0.0951789 | 0.32002022 | 0.80303536 | -0.0640575 | 0.50099909 |
| 0.999490048 | -0.0510128 | 0.32009718 | 0.80303536 | 0.018825 | 0.71131503 |
| 0.999490048 | -0.0505428 | 0.3198302 | 0.80303536 | -0.0275328 | 0.58524883 |
| 0.999490048 | 0.04526409 | 0.31992126 | 0.80303536 | 0.01123778 | 0.80315233 |
| 0.999490048 | 0.16082766 | 0.32030036 | 0.80315556 | 0.16733835 | 0.30149457 |
| 0.999490048 | -0.1226226 | 0.32069813 | 0.80391155 | -0.0985747 | 0.42308628 |
| 0.999490048 | -0.1084925 | 0.32130187 | 0.80396901 | 0.03795781 | 0.72631048 |
| 0.999490048 | -0.077331 | 0.32096782 | 0.80396901 | -0.1310541 | 0.09849611 |
| 0.999490048 | -0.0497832 | 0.32119471 | 0.80396901 | -0.0093316 | 0.85107678 |
| 0.999490048 | -0.0384131 | 0.32143249 | 0.80396901 | -0.025877 | 0.50186576 |
| 0.999490048 | 0.0453005 | 0.32149132 | 0.80396901 | 0.04593176 | 0.31492742 |
| 0.999490048 | 0.05045527 | 0.32117107 | 0.80396901 | 0.04439925 | 0.38161529 |
| 0.999490048 | 0.12717251 | 0.3211442 | 0.80396901 | 0.08803508 | 0.48998123 |
| 0.999490048 | 0.29232807 | 0.32120268 | 0.80396901 | -0.0183129 | 0.9492976 |
| 0.999490048 | 0.05660497 | 0.32163259 | 0.80408148 | 0.02362315 | 0.67665627 |
| 0.999490048 | -0.0691209 | 0.32182567 | 0.80428065 | -0.0325065 | 0.63869044 |
| 0.999490048 | -0.0563444 | 0.32196022 | 0.80428065 | 0.02319696 | 0.68097242 |
| 0.999490048 | 0.09403528 | 0.32200122 | 0.80428065 | 0.09429039 | 0.3207159 |
| 0.999490048 | -0.0702993 | 0.32250562 | 0.80505888 | -0.0170406 | 0.80873203 |
| 0.987721587 | -0.0616644 | 0.32248023 | 0.80505888 | 0.07270996 | 0.24516113 |
| 0.999490048 | -0.0772455 | 0.32277959 | 0.80509859 | 0.02454689 | 0.75123177 |
| 0.999490048 | -0.0694647 | 0.32290721 | 0.80509859 | -0.0596336 | 0.39488524 |
| 0.999490048 | -0.0642442 | 0.32283789 | 0.80509859 | -0.1034112 | 0.11700239 |
| 0.999490048 | -0.0414957 | 0.32266919 | 0.80509859 | 0.00246018 | 0.95277904 |
| 0.999490048 | 0.04757644 | 0.32314347 | 0.80544717 | 0.01915694 | 0.68831921 |
| 0.999490048 | -0.068506 | 0.32349214 | 0.8054496 | -0.0594286 | 0.39058511 |
| 0.999490048 | -0.0419328 | 0.32333025 | 0.8054496 | -0.0027089 | 0.94862421 |
| 0.999490048 | -0.0407604 | 0.32353029 | 0.8054496 | -0.0589735 | 0.15750627 |
| 0.999490048 | 0.0670649 | 0.32335685 | 0.8054496 | 0.08961337 | 0.19029632 |
| 0.90609983 | 0.32884383 | 0.32364009 | 0.80548279 | -0.5469485 | 0.11349815 |
| 0.999490048 | 0.1455965 | 0.3237738 | 0.80557544 | 0.16203718 | 0.27324465 |

| | | | | | |
|---|---|---|---|---|---|
| 0.999490048 | 0.1929897 | 0.32393614 | 0.80573929 | -0.1874675 | 0.33770888 |
| 0.999490048 | -0.0626208 | 0.32414685 | 0.80597793 | -0.0886381 | 0.16687682 |
| 0.999490048 | 0.08484301 | 0.32422513 | 0.80597793 | 0.0342869 | 0.68792064 |
| 0.999490048 | 0.12553214 | 0.32465406 | 0.80671215 | 0.13143223 | 0.30288178 |
| 0.999490048 | 0.14084995 | 0.32473973 | 0.80671215 | 0.14055313 | 0.32574064 |
| 0.987721587 | 0.23133612 | 0.32481033 | 0.80671215 | -0.2840937 | 0.22886487 |
| 0.999490048 | -0.0554144 | 0.32505875 | 0.80684916 | -0.0730106 | 0.19791063 |
| 0.999490048 | 0.05780537 | 0.32505426 | 0.80684916 | -0.0095108 | 0.87014588 |
| 0.999490048 | -0.1010588 | 0.32523676 | 0.8068914 | -0.0510919 | 0.61638188 |
| 0.999490048 | 0.04348701 | 0.32530207 | 0.8068914 | 0.01944986 | 0.6574861 |
| 0.999490048 | 0.09192881 | 0.32536567 | 0.8068914 | 0.07198257 | 0.43952381 |
| 0.999490048 | 0.13057016 | 0.32554143 | 0.80708757 | 0.20353565 | 0.13053333 |
| 0.999490048 | -0.1291509 | 0.32591973 | 0.80719735 | -0.0565118 | 0.66483159 |
| 0.999490048 | -0.0957561 | 0.32584262 | 0.80719735 | -0.2151796 | 0.03330929 |
| 0.999490048 | -0.0521304 | 0.32579343 | 0.80719735 | 0.00674013 | 0.89793067 |
| 0.999490048 | 0.05221004 | 0.32597239 | 0.80719735 | 0.03204889 | 0.54410266 |
| 0.999490048 | 0.04217012 | 0.32608982 | 0.80724874 | 0.0408275 | 0.34143745 |
| 0.999490048 | -0.1436436 | 0.32667086 | 0.80726526 | -0.0990825 | 0.49649951 |
| 0.999490048 | -0.0945119 | 0.3264548 | 0.80726526 | 0.02964424 | 0.75617903 |
| 0.999490048 | -0.0840861 | 0.32667657 | 0.80726526 | -0.0403511 | 0.63534715 |
| 0.999490048 | -0.0546635 | 0.32649114 | 0.80726526 | -0.0843383 | 0.13508538 |
| 0.999490048 | 0.05278359 | 0.32649914 | 0.80726526 | -0.0139219 | 0.79397479 |
| 0.999490048 | 0.05302766 | 0.32660874 | 0.80726526 | -0.0011079 | 0.98348874 |
| 0.999490048 | -0.0492614 | 0.32715519 | 0.8078697 | -0.0757218 | 0.13700037 |
| 0.999490048 | -0.0445091 | 0.32702723 | 0.8078697 | -0.0083565 | 0.85263872 |
| 0.999490048 | 0.12185846 | 0.32721141 | 0.8078697 | 0.04964112 | 0.68741788 |
| 0.999490048 | -0.0497859 | 0.32772185 | 0.80836306 | -0.0070872 | 0.88814278 |
| 0.999490048 | -0.0486168 | 0.32809316 | 0.80836306 | -0.0068348 | 0.88960765 |
| 0.999490048 | -0.0451147 | 0.32805519 | 0.80836306 | 0.0180363 | 0.69343383 |
| 0.999490048 | -0.0403514 | 0.3281246 | 0.80836306 | 0.00102153 | 0.98005706 |
| 0.999490048 | 0.05454968 | 0.32812941 | 0.80836306 | 0.04605997 | 0.4076859 |
| 0.999490048 | 0.08586278 | 0.32818572 | 0.80836306 | 0.14791819 | 0.09812043 |
| 0.987721587 | 0.09168722 | 0.32777962 | 0.80836306 | -0.1116464 | 0.23555128 |
| 0.999490048 | 0.16568808 | 0.32805955 | 0.80836306 | 0.06393499 | 0.70355987 |
| 0.999490048 | -0.0835259 | 0.32845846 | 0.80852096 | 0.07335263 | 0.38842146 |
| 0.999490048 | -0.0781474 | 0.32856975 | 0.80852096 | -0.062431 | 0.43349612 |
| 0.999490048 | -0.0476163 | 0.32863714 | 0.80852096 | 0.00125571 | 0.97924802 |
| 0.999490048 | 0.05894523 | 0.32856605 | 0.80852096 | 0.06758404 | 0.26405538 |
| 0.999490048 | 0.12148503 | 0.32875058 | 0.8085618 | 0.0788547 | 0.52382689 |
| 0.999490048 | 0.10790944 | 0.32888388 | 0.80865148 | -0.0738786 | 0.50166633 |
| 0.999490048 | 0.15743381 | 0.32899375 | 0.80868349 | 0.32463278 | 0.0506507 |
| 0.999490048 | -0.0929322 | 0.32948859 | 0.80879744 | -0.1615139 | 0.09620095 |

| | | | | | |
|---|---|---|---|---|---|
| 0.999490048 | -0.069656 | 0.32968793 | 0.80879744 | -0.0136354 | 0.84728462 |
| 0.999490048 | -0.0634821 | 0.33033936 | 0.80879744 | -0.0429985 | 0.50749165 |
| 0.999490048 | -0.0563536 | 0.33020806 | 0.80879744 | -0.0816423 | 0.16262625 |
| 0.999490048 | -0.0561539 | 0.32948857 | 0.80879744 | -0.0802664 | 0.1674764 |
| 0.999490048 | -0.0556588 | 0.3301432 | 0.80879744 | -0.0068894 | 0.90314806 |
| 0.999490048 | -0.0550492 | 0.32976297 | 0.80879744 | -0.0861752 | 0.13225525 |
| 0.999490048 | -0.0417475 | 0.33039618 | 0.80879744 | -0.0701523 | 0.10775755 |
| 0.999490048 | 0.04394599 | 0.33001084 | 0.80879744 | 0.0201036 | 0.65343018 |
| 0.999490048 | 0.05530244 | 0.32941843 | 0.80879744 | -0.0147654 | 0.79275365 |
| 0.999490048 | 0.0700779 | 0.33008069 | 0.80879744 | 0.09976409 | 0.16964939 |
| 0.999490048 | 0.07029698 | 0.32997236 | 0.80879744 | 0.01714726 | 0.81049763 |
| 0.999490048 | 0.08746257 | 0.33033105 | 0.80879744 | -0.0032263 | 0.97107909 |
| 0.999490048 | 0.12963889 | 0.32997115 | 0.80879744 | 0.15295754 | 0.25214088 |
| 0.999490048 | -0.0635916 | 0.33060957 | 0.80908263 | -0.0336147 | 0.60458142 |
| 0.999490048 | 0.05517341 | 0.33092989 | 0.80962924 | 0.05807965 | 0.30658541 |
| 0.999490048 | -0.1711344 | 0.33118354 | 0.80966271 | -0.0779617 | 0.65556678 |
| 0.999490048 | -0.0531834 | 0.33123447 | 0.80966271 | -0.0816143 | 0.14088184 |
| 0.999490048 | 0.09187879 | 0.33122861 | 0.80966271 | 0.09569132 | 0.31194267 |
| 0.999490048 | -0.1907406 | 0.33256237 | 0.80974557 | -0.391667 | 0.05319785 |
| 0.999490048 | -0.059731 | 0.33143018 | 0.80974557 | -0.0013609 | 0.98217645 |
| 0.999490048 | -0.0569225 | 0.33310048 | 0.80974557 | -0.0161634 | 0.7815925 |
| 0.999490048 | -0.0535974 | 0.3323387 | 0.80974557 | 0.0141162 | 0.79672222 |
| 0.999490048 | -0.0464684 | 0.33245373 | 0.80974557 | -0.0152694 | 0.74806988 |
| 0.999490048 | -0.0384914 | 0.33165304 | 0.80974557 | 0.01275836 | 0.74554277 |
| 0.999490048 | 0.05140616 | 0.33310262 | 0.80974557 | 0.03111612 | 0.55560897 |
| 0.999490048 | 0.05284002 | 0.33214644 | 0.80974557 | 0.04384306 | 0.41964206 |
| 0.999490048 | 0.05778557 | 0.3331109 | 0.80974557 | -0.0023199 | 0.96870919 |
| 0.999490048 | 0.06324314 | 0.33222557 | 0.80974557 | 0.02401831 | 0.7104454 |
| 0.999490048 | 0.06605361 | 0.33223186 | 0.80974557 | 0.0438521 | 0.51754084 |
| 0.999490048 | 0.06770008 | 0.33298593 | 0.80974557 | -0.014119 | 0.83855197 |
| 0.999490048 | 0.12387955 | 0.3327734 | 0.80974557 | 0.09689428 | 0.44705345 |
| 0.999490048 | 0.12427961 | 0.33268272 | 0.80974557 | 0.12344499 | 0.33587911 |
| 0.999490048 | 0.12661713 | 0.33150361 | 0.80974557 | 0.0340924 | 0.79193508 |
| 0.999490048 | 0.18221088 | 0.33232452 | 0.80974557 | -0.1533798 | 0.41319003 |
| 0.999490048 | 0.34573637 | 0.33207591 | 0.80974557 | 0.12714262 | 0.71420514 |
| 0.999490048 | 0.37099671 | 0.33191265 | 0.80974557 | 0.13772497 | 0.71355951 |
| 0.999490048 | 0.44612506 | 0.332175 | 0.80974557 | 0.22418236 | 0.61816109 |
| 0.999490048 | 0.05856511 | 0.33333275 | 0.81004903 | -0.0032069 | 0.9573517 |
| 0.999490048 | 0.08719314 | 0.33358704 | 0.81043112 | 0.06091188 | 0.49730646 |
| 0.999490048 | 0.05427137 | 0.33371564 | 0.81050773 | 0.0531891 | 0.34326732 |
| 0.999490048 | -0.1173569 | 0.33394321 | 0.81054784 | -0.2363309 | 0.05829374 |
| 0.999490048 | -0.0642689 | 0.33400762 | 0.81054784 | -0.1048432 | 0.12057494 |

| | | | | | |
|---|---|---|---|---|---|
| 0.999490048 | 0.04457152 | 0.33412044 | 0.81054784 | 0.05382904 | 0.24543107 |
| 0.999490048 | 0.10928131 | 0.33409482 | 0.81054784 | -0.0316483 | 0.77782043 |
| 0.999490048 | -0.2212979 | 0.33441957 | 0.81080238 | -0.3870832 | 0.09745956 |
| 0.999490048 | 0.05318274 | 0.33436629 | 0.81080238 | -0.0079481 | 0.88422968 |
| 0.999490048 | 0.04156573 | 0.33464993 | 0.81088999 | 0.01328451 | 0.75582359 |
| 0.999490048 | 0.05212422 | 0.33460197 | 0.81088999 | 0.02683629 | 0.61692108 |
| 0.999490048 | 0.08496358 | 0.33475753 | 0.8109154 | 0.06566076 | 0.45431188 |
| 0.999490048 | 0.03678768 | 0.33487434 | 0.81096308 | -0.0162147 | 0.66844807 |
| 0.999490048 | -0.061529 | 0.33528194 | 0.81099227 | 0.02756852 | 0.66358642 |
| 0.999490048 | -0.0537575 | 0.33516938 | 0.81099227 | -0.0385928 | 0.48703963 |
| 0.999490048 | 0.04080452 | 0.33527573 | 0.81099227 | 0.00406578 | 0.92280038 |
| 0.999490048 | 0.04321669 | 0.33537201 | 0.81099227 | -0.0113971 | 0.79773016 |
| 0.999490048 | 0.07885743 | 0.33518541 | 0.81099227 | 0.0772114 | 0.34518631 |
| 0.999490048 | -0.1542873 | 0.33568619 | 0.81114287 | -0.0901719 | 0.57130549 |
| 0.999490048 | -0.059652 | 0.33557425 | 0.81114287 | -0.0607724 | 0.32674657 |
| 0.999490048 | -0.0503473 | 0.33572572 | 0.81114287 | -0.0079723 | 0.87774149 |
| 0.999490048 | -0.1088293 | 0.33617977 | 0.8118037 | 0.0012481 | 0.99111689 |
| 0.999490048 | -0.0522105 | 0.33628688 | 0.8118037 | 0.00900634 | 0.86707178 |
| 0.999490048 | 0.07495236 | 0.3362909 | 0.8118037 | 0.09636954 | 0.21910551 |
| 0.999490048 | -0.258005 | 0.33661119 | 0.81210732 | -0.3529403 | 0.19221 |
| 0.999490048 | -0.0705119 | 0.33652562 | 0.81210732 | -0.0171328 | 0.81375928 |
| 0.999490048 | -0.0969872 | 0.33691368 | 0.81235724 | -0.2924282 | 0.00675444 |
| 0.999490048 | 0.05066069 | 0.33700665 | 0.81235724 | 0.00118336 | 0.98193986 |
| 0.999490048 | 0.05491016 | 0.33685898 | 0.81235724 | 0.07953318 | 0.16842406 |
| 0.999490048 | -0.5347302 | 0.33771268 | 0.81288581 | 0.11992233 | 0.82493997 |
| 0.999490048 | -0.0790471 | 0.33747482 | 0.81288581 | -0.0017829 | 0.98257873 |
| 0.999490048 | -0.0530211 | 0.33766053 | 0.81288581 | 0.00860883 | 0.87516986 |
| 0.999490048 | 0.06848295 | 0.33768887 | 0.81288581 | -0.0385409 | 0.58713538 |
| 0.999490048 | 0.10370994 | 0.33766312 | 0.81288581 | 0.05101017 | 0.63484516 |
| 0.999490048 | 0.06113035 | 0.33790363 | 0.81301925 | 0.0179099 | 0.77705147 |
| 0.999490048 | 0.12041525 | 0.33796285 | 0.81301925 | 0.00976311 | 0.93750084 |
| 0.999490048 | -0.0646754 | 0.33821636 | 0.81316055 | -0.0882635 | 0.19468932 |
| 0.999490048 | 0.07073582 | 0.3381917 | 0.81316055 | 0.01435165 | 0.84456522 |
| 0.999490048 | -0.2007223 | 0.33943856 | 0.81327743 | 0.09730957 | 0.63574923 |
| 0.999490048 | -0.1572254 | 0.33988077 | 0.81327743 | -0.1714824 | 0.29873187 |
| 0.999490048 | -0.148822 | 0.34026301 | 0.81327743 | 0.026637 | 0.86056756 |
| 0.999490048 | -0.1248541 | 0.33979353 | 0.81327743 | -0.207582 | 0.11818437 |
| 0.999490048 | -0.120939 | 0.34023396 | 0.81327743 | 0.04898828 | 0.69705782 |
| 0.999490048 | -0.1000496 | 0.3403738 | 0.81327743 | 0.00387284 | 0.97028756 |
| 0.999490048 | -0.0665097 | 0.3401495 | 0.81327743 | -0.0058347 | 0.93269826 |
| 0.999490048 | -0.0511819 | 0.33865832 | 0.81327743 | -0.0978182 | 0.07401862 |
| 0.999490048 | -0.0430789 | 0.33954386 | 0.81327743 | -0.0358669 | 0.42521199 |

| | | | | | |
|---|---|---|---|---|---|
| 0.999490048 | 0.04488177 | 0.33892414 | 0.81327743 | 0.05016698 | 0.28620563 |
| 0.999490048 | 0.04497162 | 0.33903892 | 0.81327743 | 0.03461465 | 0.46012339 |
| 0.999490048 | 0.04958079 | 0.33998924 | 0.81327743 | 0.00522205 | 0.91921667 |
| 0.999490048 | 0.06119561 | 0.34023505 | 0.81327743 | -0.0124362 | 0.84497658 |
| 0.999490048 | 0.064564 | 0.34040774 | 0.81327743 | 0.06483432 | 0.33842666 |
| 0.999490048 | 0.0734159 | 0.33858972 | 0.81327743 | 0.04435135 | 0.56089843 |
| 0.999490048 | 0.07806966 | 0.34030158 | 0.81327743 | 0.08497529 | 0.30011707 |
| 0.999490048 | 0.09119261 | 0.34007103 | 0.81327743 | -0.0926467 | 0.33256789 |
| 0.999490048 | 0.09879648 | 0.33847038 | 0.81327743 | 0.13170321 | 0.20512873 |
| 0.999490048 | 0.11034381 | 0.33972058 | 0.81327743 | 0.04956794 | 0.66571374 |
| 0.999490048 | 0.11252233 | 0.33892115 | 0.81327743 | 0.19706496 | 0.1000901 |
| 0.999490048 | 0.12813735 | 0.34025541 | 0.81327743 | -0.1656062 | 0.22624256 |
| 0.999490048 | 0.14149042 | 0.33983591 | 0.81327743 | -0.0870149 | 0.55494846 |
| 0.999490048 | 0.06588654 | 0.34054189 | 0.81335892 | 0.0336788 | 0.62377965 |
| 0.999490048 | 0.07936186 | 0.34063666 | 0.81335892 | 0.2052034 | 0.01880736 |
| 0.999490048 | -0.0437496 | 0.34074076 | 0.81337488 | -0.0329477 | 0.47133006 |
| 0.999490048 | -0.1051846 | 0.34129259 | 0.81445931 | -0.0238939 | 0.82738266 |
| 0.999490048 | -0.1475098 | 0.34192699 | 0.8148089 | -0.2169966 | 0.16645241 |
| 0.999490048 | -0.1059692 | 0.34191453 | 0.8148089 | 0.05122501 | 0.64362364 |
| 0.999490048 | -0.0536686 | 0.34172598 | 0.8148089 | -0.0262154 | 0.64003044 |
| 0.999490048 | -0.053025 | 0.34175257 | 0.8148089 | 0.00600246 | 0.91352074 |
| 0.999490048 | -0.0378151 | 0.3417539 | 0.8148089 | -0.1043324 | 0.01299518 |
| 0.999490048 | 0.05892015 | 0.3420855 | 0.81495405 | 0.03718511 | 0.54658917 |
| 0.999490048 | 0.03956201 | 0.34236725 | 0.81509986 | 0.01899816 | 0.64611361 |
| 0.999490048 | 0.06899537 | 0.34228845 | 0.81509986 | 0.01772051 | 0.80568559 |
| 0.999490048 | 0.07717276 | 0.34243956 | 0.81509986 | 0.04012546 | 0.61925932 |
| 0.999490048 | -0.0647408 | 0.34306806 | 0.81613058 | -0.0143343 | 0.8323016 |
| 0.999490048 | -0.0486261 | 0.34301253 | 0.81613058 | 0.00368544 | 0.94218637 |
| 0.999490048 | 0.06569823 | 0.34322296 | 0.81626651 | 0.00482944 | 0.94394883 |
| 0.999490048 | 0.13825333 | 0.34335602 | 0.81635045 | -0.1294219 | 0.37453393 |
| 0.999490048 | -0.0657944 | 0.34421531 | 0.81676551 | -0.0144063 | 0.83452407 |
| 0.999490048 | 0.03320917 | 0.34408432 | 0.81676551 | 0.00328646 | 0.92473079 |
| 0.999490048 | 0.06671287 | 0.34375918 | 0.81676551 | 0.04973595 | 0.47853225 |
| 0.999490048 | 0.06961105 | 0.343745 | 0.81676551 | -6.16E-04 | 0.99325113 |
| 0.999490048 | 0.07735092 | 0.34399057 | 0.81676551 | -0.0072725 | 0.92846504 |
| 0.999490048 | 0.18666339 | 0.34412599 | 0.81676551 | 0.05520136 | 0.77787128 |
| 0.999490048 | 0.27461676 | 0.34397417 | 0.81676551 | 0.05742288 | 0.83876537 |
| 0.999490048 | 0.11461784 | 0.34442618 | 0.8170337 | 0.08757731 | 0.46838385 |
| 0.999490048 | -0.1187822 | 0.34490655 | 0.81794083 | -0.0168378 | 0.89254858 |
| 0.999490048 | -0.0531634 | 0.34503663 | 0.818017 | -0.0511926 | 0.36290353 |
| 0.999490048 | -0.2498843 | 0.34578889 | 0.81813874 | 0.02257712 | 0.9301436 |
| 0.999490048 | -0.0554945 | 0.34580108 | 0.81813874 | -0.0493909 | 0.40055584 |

| | | | | | |
|---|---|---|---|---|---|
| 0.999490048 | -0.0392425 | 0.34567272 | 0.81813874 | 0.01358622 | 0.74209526 |
| 0.999490048 | 0.04709433 | 0.34559005 | 0.81813874 | -0.0328966 | 0.50806042 |
| 0.999490048 | 0.05566527 | 0.34587182 | 0.81813874 | 0.05998678 | 0.31041255 |
| 0.999490048 | 0.06319984 | 0.34586155 | 0.81813874 | -0.0082397 | 0.90132245 |
| 0.999490048 | 0.07372985 | 0.34523627 | 0.81813874 | 0.13127256 | 0.09907573 |
| 0.999490048 | 0.17265258 | 0.34552567 | 0.81813874 | -0.0893918 | 0.62298544 |
| 0.999490048 | -0.1202205 | 0.34657578 | 0.81833194 | 0.12310326 | 0.33534261 |
| 0.999490048 | -0.0687186 | 0.3461983 | 0.81833194 | 0.00589801 | 0.93498498 |
| 0.999490048 | -0.0599248 | 0.34683554 | 0.81833194 | 0.0184215 | 0.77063002 |
| 0.999490048 | -0.0456177 | 0.346677 | 0.81833194 | -0.0411722 | 0.39491031 |
| 0.999490048 | 0.04452527 | 0.34652725 | 0.81833194 | 0.00885212 | 0.85027195 |
| 0.999490048 | 0.06900016 | 0.34633857 | 0.81833194 | 0.09252046 | 0.20986824 |
| 0.999490048 | 0.0867491 | 0.34656922 | 0.81833194 | 0.031231 | 0.73270861 |
| 0.999490048 | 0.11190415 | 0.34678137 | 0.81833194 | 0.10116993 | 0.39422528 |
| 0.999490048 | 0.11579283 | 0.34614312 | 0.81833194 | 0.06608731 | 0.58854438 |
| 0.999490048 | -0.2007409 | 0.34735129 | 0.81910892 | -0.1526085 | 0.47118043 |
| 0.999490048 | -0.0951283 | 0.34755579 | 0.81910892 | -0.0688756 | 0.49461428 |
| 0.999490048 | -0.0507432 | 0.34781587 | 0.81910892 | -0.020246 | 0.70587326 |
| 0.999490048 | 0.05707141 | 0.34761053 | 0.81910892 | 0.01378035 | 0.81910023 |
| 0.999490048 | 0.06789559 | 0.3478242 | 0.81910892 | 0.09988319 | 0.17154084 |
| 0.999490048 | 0.18182396 | 0.3476369 | 0.81910892 | -0.0635965 | 0.74058276 |
| 0.999490048 | 0.21051911 | 0.34785152 | 0.81910892 | 0.28329956 | 0.21626858 |
| 0.999490048 | -0.0440755 | 0.34802068 | 0.8192762 | -0.0694277 | 0.14443285 |
| 0.999490048 | -0.108074 | 0.34832691 | 0.8196597 | 0.07836681 | 0.49466548 |
| 0.999490048 | -0.1015115 | 0.34837991 | 0.8196597 | 0.0376086 | 0.7262668 |
| 0.999490048 | -0.0742853 | 0.34854921 | 0.81979697 | 0.01321715 | 0.86640076 |
| 0.999490048 | 0.04480722 | 0.34863462 | 0.81979697 | 0.04003048 | 0.40157365 |
| 0.987721587 | -0.0920557 | 0.34926508 | 0.82081717 | 0.12315648 | 0.21360428 |
| 0.999490048 | 0.05782865 | 0.34922738 | 0.82081717 | 0.00521329 | 0.93215327 |
| 0.999490048 | 0.06592125 | 0.34943379 | 0.82098259 | -0.0081323 | 0.90730152 |
| 0.999490048 | -0.0420703 | 0.34954436 | 0.82101137 | -0.0599711 | 0.18626505 |
| 0.999490048 | 0.09710642 | 0.34966932 | 0.82107392 | 0.07149772 | 0.48933092 |
| 0.999490048 | -0.0691663 | 0.34991077 | 0.82140987 | -0.0912625 | 0.22038029 |
| 0.999490048 | -0.0506989 | 0.35025618 | 0.82198963 | -0.0444831 | 0.41151974 |
| 0.999490048 | -0.294991 | 0.35142982 | 0.82204238 | 0.33268757 | 0.29681074 |
| 0.999490048 | -0.1648852 | 0.35183062 | 0.82204238 | 0.10693754 | 0.54391747 |
| 0.999490048 | -0.1403219 | 0.35142016 | 0.82204238 | -0.1613572 | 0.28527278 |
| 0.999490048 | -0.0984457 | 0.35126597 | 0.82204238 | -0.1551634 | 0.14684996 |
| 0.999490048 | -0.06182 | 0.35188917 | 0.82204238 | 0.0454272 | 0.49219297 |
| 0.999490048 | -0.0615756 | 0.35161047 | 0.82204238 | -0.0540597 | 0.41255388 |
| 0.771925038 | -0.0586011 | 0.35151057 | 0.82204238 | 0.11202393 | 0.08152218 |
| 0.999490048 | -0.0474549 | 0.35136635 | 0.82204238 | -0.0216277 | 0.66886975 |

| | | | | | |
|---|---|---|---|---|---|
| 0.999490048 | 0.04508281 | 0.35105821 | 0.82204238 | 0.01653926 | 0.7302972 |
| 0.999490048 | 0.04522715 | 0.35195228 | 0.82204238 | -0.0040742 | 0.93259187 |
| 0.999490048 | 0.04830796 | 0.3506507 | 0.82204238 | 0.02151457 | 0.67549791 |
| 0.999490048 | 0.0498155 | 0.35039592 | 0.82204238 | 0.0264813 | 0.61738937 |
| 0.999490048 | 0.05140678 | 0.35092724 | 0.82204238 | 0.06398237 | 0.24788805 |
| 0.999490048 | 0.05409273 | 0.35107261 | 0.82204238 | 0.04382846 | 0.44855055 |
| 0.999490048 | 0.05981304 | 0.35142754 | 0.82204238 | 0.06865196 | 0.28615379 |
| 0.999490048 | 0.06630637 | 0.35165658 | 0.82204238 | 0.06842584 | 0.33674741 |
| 0.999490048 | 0.24457846 | 0.3518411 | 0.82204238 | 0.08261254 | 0.74884191 |
| 0.999490048 | -0.0757717 | 0.35208476 | 0.82212184 | -0.0937375 | 0.25181201 |
| 0.999490048 | -0.0568923 | 0.35291575 | 0.82231674 | -0.0295349 | 0.62744078 |
| 0.999490048 | -0.0460945 | 0.35303913 | 0.82231674 | -0.0027787 | 0.95496287 |
| 0.999490048 | -0.0453531 | 0.35304143 | 0.82231674 | -0.0195928 | 0.68616758 |
| 0.999490048 | -0.0445428 | 0.35285025 | 0.82231674 | -0.0664256 | 0.17031109 |
| 0.999490048 | 0.05402884 | 0.35259646 | 0.82231674 | 0.05810603 | 0.31810261 |
| 0.999490048 | 0.06208438 | 0.35269582 | 0.82231674 | 0.05223921 | 0.43305342 |
| 0.999490048 | 0.06607184 | 0.35305455 | 0.82231674 | 0.06467411 | 0.36315855 |
| 0.972489992 | 0.08040605 | 0.35240386 | 0.82231674 | -0.1118463 | 0.1993916 |
| 0.999490048 | 0.09228779 | 0.35299972 | 0.82231674 | 0.18784609 | 0.07930174 |
| 0.999490048 | -0.0684471 | 0.35328355 | 0.82239754 | -0.0917165 | 0.21667828 |
| 0.999490048 | 0.04668872 | 0.35328623 | 0.82239754 | -0.0511757 | 0.30978019 |
| 0.999490048 | -0.0603942 | 0.35386482 | 0.8225217 | -0.0206291 | 0.74963288 |
| 0.999490048 | -0.0591924 | 0.35356347 | 0.8225217 | -0.0245755 | 0.698055 |
| 0.999490048 | -0.0511998 | 0.35350377 | 0.8225217 | -0.0599384 | 0.27892725 |
| 0.999490048 | -0.0434902 | 0.35393059 | 0.8225217 | -0.0205655 | 0.65898276 |
| 0.999490048 | 0.0544505 | 0.35382048 | 0.8225217 | 0.06444843 | 0.27408644 |
| 0.999490048 | 0.10936549 | 0.3537777 | 0.8225217 | 0.17916907 | 0.14088136 |
| 0.999490048 | -0.0763143 | 0.35429939 | 0.82288497 | -0.0900469 | 0.27601531 |
| 0.999490048 | -0.0592212 | 0.35443967 | 0.82288497 | 0.04273373 | 0.50223606 |
| 0.999490048 | -0.0494764 | 0.35453126 | 0.82288497 | 0.03961055 | 0.45717036 |
| 0.999490048 | 0.0508131 | 0.3543306 | 0.82288497 | 0.05505967 | 0.31627832 |
| 0.999490048 | 0.07170858 | 0.35457966 | 0.82288497 | 0.03143494 | 0.68278248 |
| 0.999490048 | 0.05202786 | 0.35495148 | 0.8229109 | 0.0246586 | 0.65894779 |
| 0.999490048 | 0.05647125 | 0.35498504 | 0.8229109 | 0.02727762 | 0.65288251 |
| 0.999490048 | 0.0585967 | 0.35483671 | 0.8229109 | 0.02233568 | 0.72236216 |
| 0.999490048 | 0.10515387 | 0.35483679 | 0.8229109 | 0.16472544 | 0.16225796 |
| 0.999490048 | -0.0417281 | 0.35526343 | 0.82329896 | -0.0363498 | 0.41969846 |
| 0.999490048 | 0.05660377 | 0.35534964 | 0.82329896 | 0.00260138 | 0.96582449 |
| 0.999490048 | -0.1679221 | 0.35597339 | 0.8240914 | -0.4162934 | 0.02804944 |
| 0.987721587 | -0.0797187 | 0.35608644 | 0.8240914 | 0.10149248 | 0.24249676 |
| 0.999490048 | -0.073543 | 0.35592039 | 0.8240914 | -0.003855 | 0.96107108 |
| 0.999490048 | -0.0472025 | 0.3559981 | 0.8240914 | -0.0231851 | 0.64814457 |

| | | | | | |
|---|---|---|---|---|---|
| 0.999490048 | -0.0798986 | 0.35620307 | 0.82413291 | -0.0334421 | 0.69733377 |
| 0.999490048 | -0.2081608 | 0.35753102 | 0.82432267 | -0.2722285 | 0.23736482 |
| 0.999490048 | -0.1175075 | 0.35791483 | 0.82432267 | -0.0108037 | 0.93206531 |
| 0.999490048 | -0.097813 | 0.35783355 | 0.82432267 | -0.0526166 | 0.61873965 |
| 0.999490048 | -0.0846822 | 0.35712121 | 0.82432267 | -0.0700029 | 0.4452895 |
| 0.999490048 | -0.0707221 | 0.35664407 | 0.82432267 | 0.05011419 | 0.51184575 |
| 0.999490048 | -0.0699871 | 0.35688992 | 0.82432267 | 0.0093302 | 0.90142227 |
| 0.999490048 | -0.059447 | 0.35642599 | 0.82432267 | -0.0383786 | 0.54960816 |
| 0.999490048 | -0.0561047 | 0.35753145 | 0.82432267 | -0.0221973 | 0.71387265 |
| 0.999490048 | -0.0529033 | 0.35676177 | 0.82432267 | -0.0156105 | 0.78402932 |
| 0.999490048 | -0.0468556 | 0.35732649 | 0.82432267 | -0.0119432 | 0.81303457 |
| 0.999490048 | -0.0396851 | 0.35695117 | 0.82432267 | -0.0297664 | 0.48797246 |
| 0.999490048 | 0.05410462 | 0.35779405 | 0.82432267 | -0.0018924 | 0.97412053 |
| 0.999490048 | 0.05636008 | 0.35704771 | 0.82432267 | 0.10083206 | 0.10562409 |
| 0.999490048 | 0.06999105 | 0.35796335 | 0.82432267 | 0.12158523 | 0.11626898 |
| 0.987721587 | 0.07385507 | 0.3573427 | 0.82432267 | 0.24357935 | 0.00481092 |
| 0.999490048 | 0.07981139 | 0.35683589 | 0.82432267 | 0.15763968 | 0.07550873 |
| 0.999490048 | 0.10632282 | 0.35787843 | 0.82432267 | 0.09480856 | 0.4114917 |
| 0.999490048 | -0.2141883 | 0.35819683 | 0.82463291 | -0.5022612 | 0.0512443 |
| 0.999490048 | -0.2012189 | 0.35838517 | 0.82483908 | 0.20712551 | 0.34500018 |
| 0.999490048 | -0.13168 | 0.35882735 | 0.82500198 | -0.0201922 | 0.8871704 |
| 0.999490048 | -0.0504298 | 0.35882722 | 0.82500198 | -0.0537098 | 0.32897505 |
| 0.999490048 | 0.05592261 | 0.35885116 | 0.82500198 | 0.06031144 | 0.32304869 |
| 0.999490048 | 0.45723244 | 0.35855768 | 0.82500198 | -0.3308874 | 0.5024811 |
| 0.999490048 | 0.06910664 | 0.35895028 | 0.82500271 | 0.09218046 | 0.22407844 |
| 0.999490048 | -0.0951261 | 0.35934308 | 0.82545108 | -0.087111 | 0.40061302 |
| 0.999490048 | 0.04869702 | 0.35932755 | 0.82545108 | 0.01347113 | 0.79827984 |
| 0.999490048 | 0.0470677 | 0.35947381 | 0.82552429 | 0.04634395 | 0.36678668 |
| 0.999490048 | 0.04396505 | 0.35965864 | 0.82572165 | 0.05127739 | 0.28680295 |
| 0.999490048 | 0.05512639 | 0.3598237 | 0.82587353 | 0.04572078 | 0.44638036 |
| 0.999490048 | 0.08212355 | 0.3603771 | 0.82691641 | 0.13889549 | 0.12759624 |
| 0.999490048 | -0.0571014 | 0.36049335 | 0.8269559 | -0.0393147 | 0.52713651 |
| 0.999490048 | -0.166488 | 0.36071575 | 0.82708862 | 0.07277367 | 0.68749333 |
| 0.999490048 | -0.0901372 | 0.36118224 | 0.82708862 | -0.0548803 | 0.57622874 |
| 0.999490048 | -0.0792587 | 0.36094214 | 0.82708862 | 0.00638733 | 0.94081566 |
| 0.999490048 | 0.03737448 | 0.36097376 | 0.82708862 | -0.0014286 | 0.97190851 |
| 0.999490048 | 0.08543477 | 0.36101097 | 0.82708862 | 0.00244704 | 0.9789502 |
| 0.999490048 | 0.08848464 | 0.36124457 | 0.82708862 | 0.1652014 | 0.09476893 |
| 0.972489992 | 0.11299989 | 0.36108243 | 0.82708862 | -0.1660884 | 0.18363019 |
| 0.999490048 | 0.14027222 | 0.36138543 | 0.82718431 | 0.05196565 | 0.73095918 |
| 0.999490048 | 0.05047782 | 0.36163659 | 0.82753234 | 0.03258157 | 0.55404303 |
| 0.999490048 | -0.0988901 | 0.36236927 | 0.82762127 | -0.0774458 | 0.47421089 |

| | | | | | |
|---|---|---|---|---|---|
| 0.999490048 | -0.0891346 | 0.36227118 | 0.82762127 | -0.017865 | 0.85395167 |
| 0.999490048 | -0.0542047 | 0.36223316 | 0.82762127 | -0.0355528 | 0.54820259 |
| 0.999490048 | -0.0479361 | 0.3619758 | 0.82762127 | -0.0092772 | 0.85883228 |
| 0.999490048 | -0.0318139 | 0.36221362 | 0.82762127 | 0.0021563 | 0.95034261 |
| 0.999490048 | 0.05139732 | 0.36188131 | 0.82762127 | 0.02796971 | 0.6176939 |
| 0.999490048 | 0.05149322 | 0.36205371 | 0.82762127 | 0.05487911 | 0.33189811 |
| 0.999490048 | -0.0949938 | 0.36300832 | 0.82796957 | -0.1249476 | 0.23430284 |
| 0.999490048 | -0.0706649 | 0.36321587 | 0.82796957 | 0.05361052 | 0.488779 |
| 0.999490048 | 0.03757294 | 0.36314665 | 0.82796957 | 0.02803535 | 0.49583858 |
| 0.999490048 | 0.04135752 | 0.36317232 | 0.82796957 | -0.0038092 | 0.93269691 |
| 0.999490048 | 0.05083362 | 0.36278093 | 0.82796957 | 0.05690069 | 0.30935616 |
| 0.999490048 | 0.06634346 | 0.36274537 | 0.82796957 | -0.0739167 | 0.31152666 |
| 0.999490048 | 0.18205723 | 0.36320336 | 0.82796957 | 0.24440314 | 0.22902962 |
| 0.999490048 | -0.1127859 | 0.36354326 | 0.8280377 | 0.12634997 | 0.30973539 |
| 0.999490048 | -0.0471133 | 0.36337367 | 0.8280377 | 0.04670427 | 0.36748615 |
| 0.999490048 | 0.04402406 | 0.36352922 | 0.8280377 | 0.04450349 | 0.35841462 |
| 0.999490048 | 0.06190525 | 0.36373551 | 0.82824966 | 0.02642003 | 0.69632446 |
| 0.999490048 | -0.2002328 | 0.36478642 | 0.82874756 | 0.01328272 | 0.95166187 |
| 0.771925038 | -0.058127 | 0.3645593 | 0.82874756 | 0.11491336 | 0.07981835 |
| 0.999490048 | 0.0412509 | 0.36441399 | 0.82874756 | 0.05342516 | 0.24272766 |
| 0.999490048 | 0.04653518 | 0.36457303 | 0.82874756 | 0.0362882 | 0.47812067 |
| 0.999490048 | 0.0519956 | 0.36470158 | 0.82874756 | 0.00945562 | 0.86802095 |
| 0.999490048 | 0.06099546 | 0.36445795 | 0.82874756 | 0.03262537 | 0.62557053 |
| 0.999490048 | 0.09007414 | 0.36424158 | 0.82874756 | 0.11345851 | 0.25537082 |
| 0.999490048 | 0.10714392 | 0.36484743 | 0.82874756 | 0.09333655 | 0.42896668 |
| 0.999490048 | 0.12684441 | 0.36460505 | 0.82874756 | 0.14654436 | 0.29622327 |
| 0.999490048 | 0.06182325 | 0.36504634 | 0.82897388 | 0.0892132 | 0.19507832 |
| 0.999490048 | -0.0442842 | 0.36528165 | 0.8292827 | -0.086653 | 0.08320176 |
| 0.999490048 | 0.05167751 | 0.36543695 | 0.82940978 | 0.053042 | 0.35312551 |
| 0.999490048 | 0.0402329 | 0.36568754 | 0.82952756 | 0.04849309 | 0.27734957 |
| 0.999490048 | 0.12106848 | 0.36559375 | 0.82952756 | -0.1075554 | 0.42072444 |
| 0.987721587 | -0.078497 | 0.36635149 | 0.8296141 | 0.10309207 | 0.23822432 |
| 0.999490048 | -0.0775251 | 0.36600397 | 0.8296141 | -0.0412508 | 0.62848524 |
| 0.999490048 | -0.069226 | 0.36647154 | 0.8296141 | -0.0918342 | 0.23376474 |
| 0.999490048 | -0.0488861 | 0.36594406 | 0.8296141 | 0.02129286 | 0.69178422 |
| 0.999490048 | -0.0392567 | 0.36622527 | 0.8296141 | -4.28E-04 | 0.99208309 |
| 0.999490048 | 0.05343522 | 0.36652053 | 0.8296141 | 0.04026857 | 0.49465686 |
| 0.999490048 | 0.05699445 | 0.36651828 | 0.8296141 | -0.0056334 | 0.92829584 |
| 0.999490048 | 0.07694956 | 0.36642805 | 0.8296141 | 0.08547224 | 0.31667557 |
| 0.999490048 | -0.1775058 | 0.36693951 | 0.82967322 | -0.2388276 | 0.22781302 |
| 0.999490048 | -0.1433001 | 0.36721304 | 0.82967322 | 0.17145153 | 0.28233416 |
| 0.999490048 | -0.1429265 | 0.36821344 | 0.82967322 | 0.00653432 | 0.96691599 |

| | | | | | |
|---|---|---|---|---|---|
| 0.999490048 | -0.0830269 | 0.36680168 | 0.82967322 | 0.0482779 | 0.59516462 |
| 0.999490048 | -0.0797087 | 0.36695861 | 0.82967322 | 0.02827679 | 0.74717667 |
| 0.999490048 | -0.0752289 | 0.36794357 | 0.82967322 | -0.0089911 | 0.91361505 |
| 0.999490048 | -0.0731063 | 0.36735675 | 0.82967322 | -0.0767947 | 0.34408712 |
| 0.999490048 | -0.0671922 | 0.36888148 | 0.82967322 | 0.0079679 | 0.91445658 |
| 0.999490048 | -0.0623631 | 0.36816336 | 0.82967322 | -0.0330152 | 0.63177275 |
| 0.999490048 | -0.0590593 | 0.36723669 | 0.82967322 | -0.0018918 | 0.97676849 |
| 0.999490048 | -0.0453558 | 0.36811542 | 0.82967322 | -0.080143 | 0.11783252 |
| 0.999490048 | -0.0425224 | 0.36863146 | 0.82967322 | -0.0762669 | 0.11304194 |
| 0.999490048 | -0.0404816 | 0.36845782 | 0.82967322 | -0.0099206 | 0.82423183 |
| 0.999490048 | -0.0391653 | 0.36849656 | 0.82967322 | -0.0308516 | 0.47733952 |
| 0.999490048 | 0.03964054 | 0.36792072 | 0.82967322 | 0.00716955 | 0.86961729 |
| 0.999490048 | 0.04102965 | 0.36740219 | 0.82967322 | 0.01699859 | 0.70693188 |
| 0.999490048 | 0.052281 | 0.36755077 | 0.82967322 | 0.08850025 | 0.13285481 |
| 0.999490048 | 0.05420566 | 0.36893134 | 0.82967322 | 0.00719379 | 0.90432333 |
| 0.999490048 | 0.05817107 | 0.36732686 | 0.82967322 | -0.0153664 | 0.81035511 |
| 0.999490048 | 0.08715809 | 0.36778131 | 0.82967322 | -3.54E-04 | 0.99706039 |
| 0.999490048 | 0.089945 | 0.3678471 | 0.82967322 | 0.20390526 | 0.06208847 |
| 0.999490048 | 0.09429375 | 0.36891734 | 0.82967322 | 0.15398044 | 0.14760149 |
| 0.999490048 | 0.0983622 | 0.36869502 | 0.82967322 | 0.03437153 | 0.75171364 |
| 0.999490048 | 0.1085393 | 0.36829253 | 0.82967322 | 0.14994063 | 0.22410754 |
| 0.999490048 | -0.0500993 | 0.36916943 | 0.82998513 | -0.087027 | 0.12467777 |
| 0.999490048 | 0.05533829 | 0.3693556 | 0.83018014 | 0.0155135 | 0.79983959 |
| 0.999490048 | 0.05231589 | 0.36964214 | 0.8306006 | 0.01601962 | 0.78197201 |
| 0.999490048 | -0.0732784 | 0.36984417 | 0.83062082 | -0.0592278 | 0.46731769 |
| 0.999490048 | 0.13145214 | 0.36985009 | 0.83062082 | 0.32917198 | 0.03090921 |
| 0.999490048 | 0.05514885 | 0.37021305 | 0.83121241 | 0.09026334 | 0.14778262 |
| 0.999490048 | 0.04080189 | 0.3704815 | 0.83159153 | 0.04254516 | 0.35072574 |
| 0.999490048 | -0.0526553 | 0.3705826 | 0.83159492 | -0.0409207 | 0.48513598 |
| 0.999490048 | 0.05604733 | 0.3707868 | 0.83160617 | -0.0202855 | 0.74425603 |
| 0.999490048 | 0.14565308 | 0.37075469 | 0.83160617 | 0.09684114 | 0.54997992 |
| 0.999490048 | -0.0480168 | 0.37108944 | 0.83173127 | -0.0755038 | 0.16444157 |
| 0.999490048 | 0.04151641 | 0.3711414 | 0.83173127 | 0.03690307 | 0.42585909 |
| 0.999490048 | 0.05555074 | 0.37106397 | 0.83173127 | 0.10630292 | 0.0935765 |
| 0.999490048 | 0.12104513 | 0.37143342 | 0.83216235 | 0.03248092 | 0.80905525 |
| 0.999490048 | -0.0994255 | 0.37173217 | 0.83232229 | 0.05306821 | 0.63157096 |
| 0.999490048 | -0.0553632 | 0.37180385 | 0.83232229 | -0.0346968 | 0.57380488 |
| 0.999490048 | 0.07805311 | 0.37168014 | 0.83232229 | -0.0351867 | 0.68524666 |
| 0.999490048 | -0.0615959 | 0.37250925 | 0.8327498 | -0.0288094 | 0.67468356 |
| 0.999490048 | -0.0528059 | 0.3728924 | 0.8327498 | 0.02818793 | 0.63237252 |
| 0.999490048 | -0.0510687 | 0.37263409 | 0.8327498 | -0.0688872 | 0.23212846 |
| 0.999490048 | -0.0500238 | 0.37283795 | 0.8327498 | 0.0209166 | 0.70758186 |

| | | | | | |
|---|---|---|---|---|---|
| 0.999490048 | 0.04077501 | 0.37276605 | 0.8327498 | 0.03329365 | 0.46556927 |
| 0.999490048 | 0.04783631 | 0.37244157 | 0.8327498 | 0.13670858 | 0.01555644 |
| 0.999490048 | 0.05698777 | 0.37260403 | 0.8327498 | 0.09713997 | 0.1342612 |
| 0.999490048 | 0.08002068 | 0.3727037 | 0.8327498 | -0.0157977 | 0.85923238 |
| 0.999490048 | 0.13087608 | 0.37277392 | 0.8327498 | 0.00679151 | 0.96281264 |
| 0.999490048 | -0.0866541 | 0.37320432 | 0.83280616 | -0.087563 | 0.36829232 |
| 0.999490048 | -0.0599838 | 0.37321684 | 0.83280616 | -0.0435751 | 0.5161326 |
| 0.999490048 | -0.0474103 | 0.37321375 | 0.83280616 | -0.0404185 | 0.44677548 |
| 0.999490048 | -0.0464233 | 0.37333841 | 0.83285485 | 0.0244413 | 0.63734201 |
| 0.999490048 | 0.08002113 | 0.37354467 | 0.83309241 | 0.13648452 | 0.13478113 |
| 0.999490048 | 0.04371647 | 0.37364881 | 0.83310215 | 0.01061764 | 0.82765328 |
| 0.999490048 | -0.3061246 | 0.37391793 | 0.83314021 | -0.9627643 | 0.01784268 |
| 0.999490048 | -0.1103549 | 0.37394969 | 0.83314021 | -0.2399639 | 0.06879316 |
| 0.999490048 | -0.0715583 | 0.37404329 | 0.83314021 | -0.0026264 | 0.97376405 |
| 0.999490048 | -0.0535159 | 0.37406499 | 0.83314021 | -0.0187166 | 0.75422986 |
| 0.999490048 | -0.0621926 | 0.37477729 | 0.83361491 | -0.120771 | 0.09149045 |
| 0.999490048 | -0.047298 | 0.37461982 | 0.83361491 | 0.00316537 | 0.95224075 |
| 0.999490048 | 0.07267611 | 0.37448613 | 0.83361491 | 0.13115679 | 0.11519953 |
| 0.999490048 | 0.0997149 | 0.37456611 | 0.83361491 | 0.09271873 | 0.40849992 |
| 0.999490048 | 0.27038199 | 0.37468344 | 0.83361491 | 0.48664496 | 0.12387167 |
| 0.999490048 | -0.1153695 | 0.37507876 | 0.83377651 | -0.017388 | 0.89282486 |
| 0.999490048 | -0.1000936 | 0.37497201 | 0.83377651 | -0.1646907 | 0.1496708 |
| 0.999490048 | 0.07366349 | 0.3751495 | 0.83377651 | 0.10126022 | 0.22600568 |
| 0.999490048 | -0.0697548 | 0.37544924 | 0.83388983 | -0.1169779 | 0.1427039 |
| 0.999490048 | -0.0475389 | 0.37559996 | 0.83388983 | -0.0449942 | 0.40130567 |
| 0.999490048 | 0.05237753 | 0.37551114 | 0.83388983 | 0.0476631 | 0.41936439 |
| 0.999490048 | 0.07641078 | 0.37544865 | 0.83388983 | 0.08633208 | 0.31768432 |
| 0.999490048 | 0.11045767 | 0.37601985 | 0.83460014 | 0.08847848 | 0.47701749 |
| 0.999490048 | -0.1338076 | 0.37653323 | 0.83534887 | -0.0749672 | 0.618392 |
| 0.999490048 | -0.1008488 | 0.37657495 | 0.83534887 | -0.0663382 | 0.55905373 |
| 0.999490048 | -0.0773993 | 0.37675735 | 0.83534887 | -0.0445149 | 0.60932158 |
| 0.999490048 | -0.0543346 | 0.3766626 | 0.83534887 | 0.03386355 | 0.57982118 |
| 0.999490048 | -0.0901625 | 0.37737801 | 0.83549414 | 0.02910822 | 0.77066354 |
| 0.999490048 | -0.0833758 | 0.37703386 | 0.83549414 | 0.02460473 | 0.7928768 |
| 0.999490048 | -0.0471227 | 0.37713458 | 0.83549414 | -0.0114742 | 0.82847506 |
| 0.999490048 | -0.0404951 | 0.37730119 | 0.83549414 | 0.00217536 | 0.9618736 |
| 0.999490048 | -0.0349035 | 0.37742322 | 0.83549414 | -0.0366913 | 0.35387109 |
| 0.999490048 | 0.04894722 | 0.37706961 | 0.83549414 | -0.0272926 | 0.62045367 |
| 0.999490048 | -0.0660062 | 0.37755595 | 0.83556645 | -0.0367115 | 0.62169332 |
| 0.999490048 | -0.1091358 | 0.37787991 | 0.83589739 | -0.0751773 | 0.54190657 |
| 0.999490048 | -0.0704261 | 0.37790571 | 0.83589739 | -0.0555851 | 0.4851763 |
| 0.999490048 | -0.1623137 | 0.37839997 | 0.83590847 | 0.16579901 | 0.36861296 |

| | | | | | |
|---|---|---|---|---|---|
| 0.999490048 | -0.1159326 | 0.37820021 | 0.83590847 | -0.1610269 | 0.22425784 |
| 0.999490048 | 0.03937162 | 0.37830295 | 0.83590847 | -0.0089983 | 0.8392412 |
| 0.999490048 | 0.04607066 | 0.37807624 | 0.83590847 | -0.0241168 | 0.64261215 |
| 0.999490048 | 0.06702224 | 0.37841126 | 0.83590847 | 0.0561482 | 0.45952218 |
| 0.999490048 | 0.17878995 | 0.37851846 | 0.83592414 | -0.0574728 | 0.77558145 |
| 0.999490048 | 0.03694977 | 0.37878633 | 0.83629452 | 0.05558578 | 0.18974094 |
| 0.999490048 | 0.16551295 | 0.37888764 | 0.83629706 | 0.2192022 | 0.24659335 |
| 0.999490048 | -0.1999614 | 0.37910459 | 0.8365548 | -0.3265877 | 0.15604959 |
| 0.999490048 | -0.0979097 | 0.38083358 | 0.83705197 | 0.01761857 | 0.87375506 |
| 0.999490048 | -0.0575632 | 0.37962082 | 0.83705197 | -0.0484456 | 0.45862922 |
| 0.999490048 | -0.0570033 | 0.38023806 | 0.83705197 | -0.1488333 | 0.02800356 |
| 0.999490048 | -0.0488863 | 0.37979084 | 0.83705197 | 0.02301751 | 0.67737695 |
| 0.999490048 | -0.0486186 | 0.38076766 | 0.83705197 | -0.0333141 | 0.54647629 |
| 0.999490048 | -0.0469093 | 0.3807222 | 0.83705197 | -0.0415542 | 0.43669338 |
| 0.999490048 | 0.0381667 | 0.37994466 | 0.83705197 | 0.0121011 | 0.77921811 |
| 0.999490048 | 0.04224904 | 0.37962124 | 0.83705197 | 0.00700352 | 0.88332581 |
| 0.999490048 | 0.04878634 | 0.37998906 | 0.83705197 | 0.01859443 | 0.73623243 |
| 0.999490048 | 0.0542031 | 0.38020485 | 0.83705197 | -0.0578509 | 0.34949271 |
| 0.999490048 | 0.05796267 | 0.38033572 | 0.83705197 | 0.10418384 | 0.12096849 |
| 0.999490048 | 0.0630905 | 0.38039963 | 0.83705197 | 0.03535156 | 0.62121722 |
| 0.999490048 | 0.06865744 | 0.38057587 | 0.83705197 | -0.0120592 | 0.87668123 |
| 0.999490048 | 0.07725984 | 0.38056234 | 0.83705197 | 0.12481683 | 0.16165782 |
| 0.999490048 | 0.10590699 | 0.38080903 | 0.83705197 | 0.2776357 | 0.0276047 |
| 0.999490048 | -0.0540361 | 0.38121334 | 0.83766617 | -0.0011383 | 0.98516801 |
| 0.999490048 | -0.1191184 | 0.38151796 | 0.83767419 | 0.1299491 | 0.34044614 |
| 0.999490048 | -0.0569706 | 0.38134127 | 0.83767419 | -0.0435936 | 0.50157328 |
| 0.999490048 | -0.0542683 | 0.38143238 | 0.83767419 | -0.0708711 | 0.25556896 |
| 0.999490048 | 0.05026404 | 0.38167196 | 0.83779201 | 0.01342404 | 0.81395553 |
| 0.999490048 | -0.1755129 | 0.38209038 | 0.83849006 | -0.1643143 | 0.41275393 |
| 0.999490048 | 0.07708838 | 0.38281123 | 0.83943986 | -0.0363239 | 0.67909525 |
| 0.999490048 | 0.08990603 | 0.38282479 | 0.83943986 | 0.09124068 | 0.37591032 |
| 0.999490048 | 0.14468178 | 0.38267429 | 0.83943986 | 0.05843501 | 0.72261739 |
| 0.999490048 | -0.1129224 | 0.3832956 | 0.84010199 | -0.0148317 | 0.90814288 |
| 0.999490048 | -0.0446605 | 0.38332798 | 0.84010199 | -0.0299691 | 0.55688856 |
| 0.999490048 | -0.0767464 | 0.38367531 | 0.8401224 | 0.02376286 | 0.7859211 |
| 0.999490048 | -0.076675 | 0.38384035 | 0.8401224 | 8.55E-04 | 0.99219265 |
| 0.999490048 | 0.04336273 | 0.38380058 | 0.8401224 | 0.00457152 | 0.92628741 |
| 0.999490048 | 0.05061791 | 0.38344302 | 0.8401224 | 0.10521866 | 0.0770068 |
| 0.999490048 | 0.05524552 | 0.38366968 | 0.8401224 | 0.07761052 | 0.22439777 |
| 0.999490048 | -0.0406544 | 0.38399399 | 0.84023841 | -0.0670687 | 0.1562061 |
| 0.999490048 | -0.133916 | 0.38452953 | 0.84074929 | -0.2334479 | 0.13537165 |
| 0.999490048 | 0.04526306 | 0.3844141 | 0.84074929 | 0.07206222 | 0.17097525 |

| | | | | | |
|---|---|---|---|---|---|
| 0.999490048 | 0.0578453 | 0.38447031 | 0.84074929 | 0.09607552 | 0.15393253 |
| 0.999490048 | 0.04503641 | 0.38490412 | 0.84100001 | -0.031092 | 0.54700293 |
| 0.999490048 | 0.06353847 | 0.38494635 | 0.84100001 | 0.07350864 | 0.31605682 |
| 0.999490048 | 0.07167743 | 0.38480294 | 0.84100001 | 0.02806888 | 0.731944 |
| 0.999490048 | -0.1438456 | 0.38602733 | 0.8411608 | -0.0054234 | 0.97372854 |
| 0.999490048 | -0.1342335 | 0.38555836 | 0.8411608 | 0.00830288 | 0.95687119 |
| 0.999490048 | -0.0630288 | 0.38572712 | 0.8411608 | -0.0200579 | 0.78106643 |
| 0.999490048 | -0.0623566 | 0.38595396 | 0.8411608 | 0.04373009 | 0.54163194 |
| 0.999490048 | -0.0432778 | 0.38589288 | 0.8411608 | -0.0513555 | 0.30499106 |
| 0.999490048 | 0.03856721 | 0.3854224 | 0.8411608 | 0.05227392 | 0.24236682 |
| 0.999490048 | 0.0589106 | 0.38530107 | 0.8411608 | 0.0642004 | 0.34476534 |
| 0.999490048 | 0.06560756 | 0.38560134 | 0.8411608 | 0.10329441 | 0.1766486 |
| 0.999490048 | 0.08969389 | 0.38516387 | 0.8411608 | 0.22589206 | 0.03521668 |
| 0.999490048 | 0.09770946 | 0.38585757 | 0.8411608 | 0.06912862 | 0.53800728 |
| 0.999490048 | -0.0639295 | 0.38632329 | 0.84136658 | -0.0799472 | 0.28067154 |
| 0.999490048 | 0.25979162 | 0.38625012 | 0.84136658 | 0.07341095 | 0.80525401 |
| 0.999490048 | -0.0393671 | 0.38656017 | 0.84147481 | -0.011516 | 0.79867882 |
| 0.999490048 | 0.14863385 | 0.38657454 | 0.84147481 | -0.0635512 | 0.70572028 |
| 0.999490048 | 0.04244187 | 0.38684338 | 0.84184055 | 0.03347885 | 0.49362682 |
| 0.999490048 | -0.0366891 | 0.38723669 | 0.84195957 | -0.0236009 | 0.5763959 |
| 0.999490048 | 0.0483956 | 0.38712019 | 0.84195957 | -0.01289 | 0.81655653 |
| 0.999490048 | 0.08961447 | 0.3873014 | 0.84195957 | 0.15612178 | 0.13782929 |
| 0.999490048 | 0.09170651 | 0.38709892 | 0.84195957 | 0.04953186 | 0.63860914 |
| 0.999490048 | 0.10511815 | 0.38749236 | 0.84215544 | 0.05965518 | 0.6220574 |
| 0.999490048 | -0.0348897 | 0.387984 | 0.84278523 | -0.0673201 | 0.10240906 |
| 0.999490048 | 0.03547785 | 0.38794011 | 0.84278523 | 0.03205115 | 0.43477741 |
| 0.999490048 | -0.1669827 | 0.39124532 | 0.84283684 | -0.3388929 | 0.08876323 |
| 0.999490048 | -0.1585024 | 0.38865938 | 0.84283684 | -0.0314822 | 0.86307151 |
| 0.999490048 | -0.1369595 | 0.39069582 | 0.84283684 | 0.01640403 | 0.916165 |
| 0.999490048 | -0.1356149 | 0.38952987 | 0.84283684 | -0.010441 | 0.94678592 |
| 0.999490048 | -0.1098807 | 0.38967323 | 0.84283684 | -0.0166581 | 0.89378839 |
| 0.999490048 | -0.1011231 | 0.38825983 | 0.84283684 | -0.1062531 | 0.36501808 |
| 0.999490048 | -0.099079 | 0.3908821 | 0.84283684 | -0.0274754 | 0.81065671 |
| 0.999490048 | -0.0914272 | 0.39001255 | 0.84283684 | -0.2991145 | 0.00836953 |
| 0.999490048 | -0.0742911 | 0.3927543 | 0.84283684 | 0.03347351 | 0.69849949 |
| 0.999490048 | -0.0740695 | 0.39234186 | 0.84283684 | -0.0348107 | 0.68597152 |
| 0.999490048 | -0.0729301 | 0.39099582 | 0.84283684 | -0.0709476 | 0.40381691 |
| 0.999490048 | -0.0712425 | 0.38879858 | 0.84283684 | -0.1408557 | 0.0951828 |
| 0.999490048 | -0.0659631 | 0.39153397 | 0.84283684 | -0.0018092 | 0.98111084 |
| 0.999490048 | -0.0632638 | 0.39225508 | 0.84283684 | 0.01122339 | 0.87849982 |
| 0.999490048 | -0.0600668 | 0.39248464 | 0.84283684 | -0.0741274 | 0.29312864 |
| 0.999490048 | -0.0596799 | 0.39269472 | 0.84283684 | -0.159611 | 0.02837632 |

| | | | | | |
|---|---|---|---|---|---|
| 0.999490048 | -0.0513293 | 0.38935974 | 0.84283684 | -0.102043 | 0.09384681 |
| 0.999490048 | -0.0500584 | 0.39134575 | 0.84283684 | 0.0070709 | 0.902926 |
| 0.999490048 | -0.0486412 | 0.3926717 | 0.84283684 | -0.1055837 | 0.07063287 |
| 0.999490048 | -0.0465888 | 0.38866318 | 0.84283684 | -0.0110549 | 0.83678395 |
| 0.999490048 | -0.04495 | 0.39068126 | 0.84283684 | 0.01379654 | 0.79081654 |
| 0.999490048 | -0.0449465 | 0.39098969 | 0.84283684 | -0.0996485 | 0.0642901 |
| 0.999490048 | -0.0440657 | 0.39031836 | 0.84283684 | -0.0642851 | 0.21388757 |
| 0.999490048 | -0.0400885 | 0.39137067 | 0.84283684 | -0.012239 | 0.79220146 |
| 0.999490048 | -0.0397374 | 0.38934612 | 0.84283684 | -0.0245106 | 0.59374198 |
| 0.999490048 | -0.0347687 | 0.39225546 | 0.84283684 | 0.0465809 | 0.25445429 |
| 0.999490048 | -0.0342171 | 0.38922317 | 0.84283684 | -0.0643747 | 0.11170926 |
| 0.999490048 | -0.033835 | 0.39295375 | 0.84283684 | -0.0806296 | 0.04873708 |
| 0.999490048 | 0.03606623 | 0.38947021 | 0.84283684 | -0.0081284 | 0.845129 |
| 0.999490048 | 0.04130717 | 0.38930636 | 0.84283684 | 0.0347005 | 0.46859084 |
| 0.999490048 | 0.04473992 | 0.39069914 | 0.84283684 | 0.01878367 | 0.71691163 |
| 0.999490048 | 0.0474413 | 0.3921271 | 0.84283684 | 0.02127675 | 0.69944061 |
| 0.999490048 | 0.04775047 | 0.39123164 | 0.84283684 | 0.02592021 | 0.6398884 |
| 0.999490048 | 0.04933798 | 0.39020221 | 0.84283684 | 0.01129572 | 0.84295062 |
| 0.999490048 | 0.04954384 | 0.38993454 | 0.84283684 | 0.04260448 | 0.45884612 |
| 0.999490048 | 0.04967254 | 0.38915763 | 0.84283684 | 0.01542636 | 0.78773068 |
| 0.999490048 | 0.05006809 | 0.39174984 | 0.84283684 | -0.0094115 | 0.87118907 |
| 0.999490048 | 0.05371246 | 0.39266444 | 0.84283684 | -0.0226042 | 0.71740874 |
| 0.999490048 | 0.05713685 | 0.39292014 | 0.84283684 | 0.07080898 | 0.2915834 |
| 0.999490048 | 0.06060389 | 0.38909022 | 0.84283684 | 0.06796742 | 0.33500895 |
| 0.999490048 | 0.06388523 | 0.39283141 | 0.84283684 | 0.05701677 | 0.44500848 |
| 0.999490048 | 0.06493619 | 0.38923755 | 0.84283684 | 0.0382076 | 0.61069234 |
| 0.999490048 | 0.07215843 | 0.38960265 | 0.84283684 | -0.048207 | 0.56384029 |
| 0.999490048 | 0.07554399 | 0.39131656 | 0.84283684 | 0.061014 | 0.48759788 |
| 0.999490048 | 0.08351692 | 0.39273845 | 0.84283684 | 0.09492518 | 0.33237395 |
| 0.999490048 | 0.10210314 | 0.39184051 | 0.84283684 | -0.0778953 | 0.51228488 |
| 0.999490048 | 0.10252464 | 0.39164736 | 0.84283684 | 0.074676 | 0.53123921 |
| 0.999490048 | 0.10406438 | 0.38893792 | 0.84283684 | 0.12869598 | 0.28854742 |
| 0.999490048 | 0.10821912 | 0.39289124 | 0.84283684 | -0.1506423 | 0.24144531 |
| 0.999490048 | -0.1966597 | 0.39416896 | 0.84327718 | -0.4807919 | 0.05849335 |
| 0.999490048 | -0.1030902 | 0.39386798 | 0.84327718 | -0.1691066 | 0.16701635 |
| 0.999490048 | -0.086175 | 0.39362606 | 0.84327718 | -0.1488656 | 0.14622862 |
| 0.999490048 | 0.03369767 | 0.39381447 | 0.84327718 | 0.05904437 | 0.14093751 |
| 0.999490048 | 0.03997177 | 0.39383709 | 0.84327718 | -0.0203831 | 0.66197816 |
| 0.999490048 | 0.04296391 | 0.39376054 | 0.84327718 | 0.06276073 | 0.21658077 |
| 0.999490048 | 0.04413201 | 0.39412731 | 0.84327718 | -0.0116994 | 0.82006884 |
| 0.999490048 | 0.04839018 | 0.39402979 | 0.84327718 | 0.03573626 | 0.52768386 |
| 0.999490048 | 0.07260804 | 0.39332796 | 0.84327718 | 0.03576533 | 0.67242876 |

| | | | | | |
|---|---|---|---|---|---|
| 0.999490048 | 0.2458554 | 0.39352517 | 0.84327718 | 0.45775553 | 0.13135236 |
| 0.999490048 | -0.4768251 | 0.39607254 | 0.8439655 | -0.321553 | 0.56223057 |
| 0.999490048 | -0.102335 | 0.39636811 | 0.8439655 | -0.141853 | 0.24282753 |
| 0.999490048 | -0.0683808 | 0.39650089 | 0.8439655 | -0.0547241 | 0.49630352 |
| 0.999490048 | -0.0643512 | 0.39641696 | 0.8439655 | 0.06764899 | 0.37302397 |
| 0.999490048 | -0.0588105 | 0.39630282 | 0.8439655 | -0.0481614 | 0.48623838 |
| 0.999490048 | -0.0575388 | 0.39508251 | 0.8439655 | -0.0892405 | 0.19160064 |
| 0.999490048 | -0.0525443 | 0.39649497 | 0.8439655 | 1.13E-04 | 0.99853957 |
| 0.999490048 | -0.0462136 | 0.3950083 | 0.8439655 | -0.0124091 | 0.81813154 |
| 0.999490048 | -0.0422763 | 0.39547387 | 0.8439655 | -0.0879325 | 0.08417127 |
| 0.999490048 | -0.0402926 | 0.39539861 | 0.8439655 | -0.0095595 | 0.83911277 |
| 0.999490048 | 0.04293056 | 0.39525135 | 0.8439655 | 0.01936673 | 0.69968315 |
| 0.999490048 | 0.04347958 | 0.39651217 | 0.8439655 | -0.0185438 | 0.716032 |
| 0.999490048 | 0.04558714 | 0.39519501 | 0.8439655 | -0.0143015 | 0.7883157 |
| 0.999490048 | 0.04951325 | 0.39615285 | 0.8439655 | 0.09242057 | 0.119588 |
| 0.999490048 | 0.06209519 | 0.39589026 | 0.8439655 | 0.04254548 | 0.55926057 |
| 0.999490048 | 0.06237116 | 0.39623217 | 0.8439655 | 0.02140892 | 0.76946073 |
| 0.999490048 | 0.06327556 | 0.39639568 | 0.8439655 | 0.03078508 | 0.67821168 |
| 0.999490048 | 0.07688356 | 0.39524172 | 0.8439655 | 0.20851654 | 0.02717003 |
| 0.999490048 | 0.0848677 | 0.39626173 | 0.8439655 | 0.08471483 | 0.39710081 |
| 0.999490048 | 0.1143184 | 0.39627891 | 0.8439655 | 0.10708355 | 0.42643702 |
| 0.999490048 | -0.0593789 | 0.39664392 | 0.84403078 | -0.0012867 | 0.98523821 |
| 0.999490048 | -0.1116769 | 0.39704167 | 0.84423172 | 0.0488024 | 0.70968636 |
| 0.999490048 | -0.0488733 | 0.39698713 | 0.84423172 | -0.0419205 | 0.46667337 |
| 0.999490048 | 0.04017959 | 0.39702437 | 0.84423172 | 0.04423539 | 0.35184102 |
| 0.999490048 | 0.04104918 | 0.39728928 | 0.84454315 | 0.03775237 | 0.43577218 |
| 0.999490048 | 0.06059645 | 0.39771937 | 0.84502716 | 0.03419908 | 0.63147884 |
| 0.999490048 | 0.06199646 | 0.39771276 | 0.84502716 | 0.04788493 | 0.5123953 |
| 0.999490048 | 0.32793514 | 0.39788199 | 0.84515762 | 0.20185629 | 0.59760949 |
| 0.999490048 | -0.0428164 | 0.39808606 | 0.84537604 | -0.0246426 | 0.62504812 |
| 0.999490048 | 0.17616058 | 0.39872952 | 0.84652721 | 0.11691223 | 0.57389863 |
| 0.999490048 | -0.067513 | 0.39893252 | 0.84674288 | 0.03399144 | 0.66939008 |
| 0.999490048 | -0.2844743 | 0.40055311 | 0.84696341 | -0.4554198 | 0.18896411 |
| 0.999490048 | -0.2542523 | 0.40069038 | 0.84696341 | -0.4673408 | 0.12860238 |
| 0.999490048 | -0.1407594 | 0.39947849 | 0.84696341 | -0.0300024 | 0.85650563 |
| 0.999490048 | -0.081982 | 0.40009121 | 0.84696341 | 0.07303632 | 0.4528088 |
| 0.999490048 | -0.0634851 | 0.39996707 | 0.84696341 | 0.08816725 | 0.24551152 |
| 0.999490048 | -0.0507954 | 0.39988857 | 0.84696341 | -0.0245118 | 0.68293193 |
| 0.999490048 | -0.0501583 | 0.39980951 | 0.84696341 | 0.0183108 | 0.75710606 |
| 0.999490048 | -0.0456616 | 0.39982944 | 0.84696341 | -0.0273716 | 0.61215067 |
| 0.999490048 | -0.0426368 | 0.39999134 | 0.84696341 | -0.0128341 | 0.79872425 |
| 0.999490048 | -0.0372259 | 0.40032452 | 0.84696341 | -0.0270593 | 0.53960818 |

| | | | | | |
|---|---|---|---|---|---|
| 0.999490048 | 0.04397423 | 0.40076077 | 0.84696341 | 0.05140479 | 0.32722644 |
| 0.999490048 | 0.04721246 | 0.40074974 | 0.84696341 | 0.0526921 | 0.34917712 |
| 0.999490048 | 0.05552747 | 0.39925651 | 0.84696341 | 0.02011035 | 0.75868844 |
| 0.999490048 | 0.06314621 | 0.40053927 | 0.84696341 | -0.0030123 | 0.96778224 |
| 0.999490048 | 0.07831602 | 0.39939794 | 0.84696341 | 0.03063664 | 0.74013951 |
| 0.999490048 | 0.10415131 | 0.40062249 | 0.84696341 | 0.0788155 | 0.52348444 |
| 0.999490048 | 0.20400311 | 0.40019572 | 0.84696341 | 0.05725393 | 0.81054041 |
| 0.999490048 | -0.122499 | 0.40096872 | 0.84715042 | -0.1103178 | 0.44881779 |
| 0.999490048 | -0.0311076 | 0.40105217 | 0.84715042 | -0.0437362 | 0.24094579 |
| 0.999490048 | -0.1577057 | 0.40263493 | 0.8472391 | -0.0859033 | 0.64684341 |
| 0.999490048 | -0.1521281 | 0.40282466 | 0.8472391 | 0.03014185 | 0.86745685 |
| 0.999490048 | -0.1338598 | 0.40329061 | 0.8472391 | -0.2179238 | 0.17853239 |
| 0.999490048 | -0.1131771 | 0.40429172 | 0.8472391 | -0.0449273 | 0.73581973 |
| 0.999490048 | -0.1095169 | 0.40195786 | 0.8472391 | 0.13145801 | 0.31588722 |
| 0.999490048 | -0.0699322 | 0.40154238 | 0.8472391 | 0.10186443 | 0.22535391 |
| 0.999490048 | -0.0687685 | 0.40286227 | 0.8472391 | 0.02969877 | 0.71634337 |
| 0.999490048 | -0.0516017 | 0.40437817 | 0.8472391 | -0.0475012 | 0.44225819 |
| 0.999490048 | -0.0492197 | 0.40448857 | 0.8472391 | -0.0477193 | 0.41876949 |
| 0.999490048 | -0.0476556 | 0.40251773 | 0.8472391 | 0.02942434 | 0.60366074 |
| 0.999490048 | -0.0464107 | 0.40340131 | 0.8472391 | -0.0199884 | 0.71739684 |
| 0.999490048 | -0.045618 | 0.40206472 | 0.8472391 | -0.0740425 | 0.1786882 |
| 0.999490048 | -0.0452082 | 0.40401566 | 0.8472391 | -0.0732249 | 0.18132879 |
| 0.999490048 | -0.0450586 | 0.4037253 | 0.8472391 | -0.1006558 | 0.06933792 |
| 0.999490048 | -0.0412526 | 0.40491276 | 0.8472391 | -0.0453287 | 0.36079054 |
| 0.999490048 | -0.0393558 | 0.40444738 | 0.8472391 | -0.036753 | 0.43582525 |
| 0.999490048 | -0.0375801 | 0.40470941 | 0.8472391 | -0.0319193 | 0.47826024 |
| 0.999490048 | 0.02810769 | 0.40307148 | 0.8472391 | 0.00553489 | 0.86833273 |
| 0.999490048 | 0.03633726 | 0.40402258 | 0.8472391 | 0.00976237 | 0.82146818 |
| 0.999490048 | 0.03890244 | 0.40251491 | 0.8472391 | -0.001137 | 0.98034171 |
| 0.999490048 | 0.04093059 | 0.40494985 | 0.8472391 | -0.0199718 | 0.68287557 |
| 0.999490048 | 0.04213502 | 0.40327607 | 0.8472391 | 0.0508333 | 0.31483362 |
| 0.999490048 | 0.04827038 | 0.40453944 | 0.8472391 | 0.07973156 | 0.17361584 |
| 0.999490048 | 0.05109085 | 0.40297305 | 0.8472391 | -0.0350327 | 0.56488035 |
| 0.999490048 | 0.05566766 | 0.40417278 | 0.8472391 | 0.01152386 | 0.86197828 |
| 0.999490048 | 0.05593493 | 0.402203 | 0.8472391 | -0.0727157 | 0.27846265 |
| 0.999490048 | 0.0583067 | 0.40243462 | 0.8472391 | 0.11017702 | 0.12003051 |
| 0.999490048 | 0.05899132 | 0.40456391 | 0.8472391 | -0.043633 | 0.53627597 |
| 0.972489992 | 0.06529345 | 0.40136998 | 0.8472391 | 0.24219382 | 0.00401734 |
| 0.999490048 | 0.06542123 | 0.40161484 | 0.8472391 | 0.01224635 | 0.87438652 |
| 0.909524814 | 0.06789308 | 0.40408454 | 0.8472391 | -0.1314346 | 0.1128534 |
| 0.999490048 | 0.07360426 | 0.40389797 | 0.8472391 | 0.05639282 | 0.52129219 |
| 0.999490048 | 0.073726 | 0.40491877 | 0.8472391 | 0.04226437 | 0.63143896 |

| | | | | | |
|---|---|---|---|---|---|
| 0.972489992 | 0.07637802 | 0.40269288 | 0.8472391 | -0.1283467 | 0.16492336 |
| 0.999490048 | 0.08144756 | 0.40193872 | 0.8472391 | 0.02606899 | 0.78716838 |
| 0.999490048 | 0.08536839 | 0.40221207 | 0.8472391 | 0.02884528 | 0.77577408 |
| 0.999490048 | 0.09227644 | 0.40138097 | 0.8472391 | -0.0799782 | 0.46624729 |
| 0.999490048 | 0.18253559 | 0.40297343 | 0.8472391 | 0.60590621 | 0.00915456 |
| 0.965957333 | -0.0617988 | 0.40529627 | 0.84753916 | 0.11496659 | 0.12783918 |
| 0.999490048 | 0.03706273 | 0.40529365 | 0.84753916 | 0.02206872 | 0.61864614 |
| 0.999490048 | 0.04015865 | 0.40561923 | 0.84800216 | -0.0263669 | 0.58356053 |
| 0.999490048 | 0.05415794 | 0.40580521 | 0.8481786 | -0.0437792 | 0.50055936 |
| 0.999490048 | -0.0528014 | 0.4059499 | 0.84826869 | 0.01619157 | 0.79759698 |
| 0.999490048 | -0.0543304 | 0.40629358 | 0.84877442 | -0.0900869 | 0.17359289 |
| 0.999490048 | -0.1408386 | 0.40836283 | 0.84894332 | 0.02257257 | 0.89387152 |
| 0.999490048 | -0.0573907 | 0.40723728 | 0.84894332 | 0.01710752 | 0.80365808 |
| 0.999490048 | -0.0557578 | 0.40830157 | 0.84894332 | 0.06286735 | 0.35206301 |
| 0.999490048 | -0.0485857 | 0.40819344 | 0.84894332 | -0.0256619 | 0.66064255 |
| 0.999490048 | -0.0405133 | 0.40777229 | 0.84894332 | -0.0082654 | 0.86499683 |
| 0.999490048 | -0.0396067 | 0.40827765 | 0.84894332 | -0.0163581 | 0.73122398 |
| 0.999490048 | -0.0296489 | 0.40682515 | 0.84894332 | -0.0523034 | 0.14911922 |
| 0.999490048 | 0.03742316 | 0.40771423 | 0.84894332 | 0.03527368 | 0.43482124 |
| 0.999490048 | 0.04488047 | 0.40710074 | 0.84894332 | 0.04299934 | 0.42678294 |
| 0.999490048 | 0.0475443 | 0.40840782 | 0.84894332 | 0.00308944 | 0.95686286 |
| 0.999490048 | 0.04961548 | 0.40685784 | 0.84894332 | 0.00609342 | 0.91830655 |
| 0.999490048 | 0.06053618 | 0.40759768 | 0.84894332 | 0.04122845 | 0.57132916 |
| 0.999490048 | 0.06201733 | 0.40728449 | 0.84894332 | -0.0086491 | 0.90736656 |
| 0.999490048 | 0.06437454 | 0.40665117 | 0.84894332 | 0.08099634 | 0.29838774 |
| 0.999490048 | 0.06935311 | 0.40663511 | 0.84894332 | 0.03751772 | 0.65190026 |
| 0.999490048 | 0.07156017 | 0.4070925 | 0.84894332 | 0.03720979 | 0.66482392 |
| 0.999490048 | 0.07289534 | 0.40823624 | 0.84894332 | 0.00614552 | 0.94403241 |
| 0.999490048 | 0.08909141 | 0.40771479 | 0.84894332 | -0.0697214 | 0.51591709 |
| 0.999490048 | 0.11899694 | 0.40801629 | 0.84894332 | 0.05741723 | 0.6881636 |
| 0.999490048 | 0.29735367 | 0.40824246 | 0.84894332 | -0.0114016 | 0.97413539 |
| 0.999490048 | 0.06564788 | 0.40853856 | 0.84900372 | 0.02098092 | 0.79040703 |
| 0.999490048 | -0.2516457 | 0.41724015 | 0.84902284 | -0.4920176 | 0.11930373 |
| 0.999490048 | -0.1360993 | 0.41574914 | 0.84902284 | 0.15256447 | 0.36416247 |
| 0.999490048 | -0.1277913 | 0.41044299 | 0.84902284 | 0.1734531 | 0.26660451 |
| 0.999490048 | -0.1207162 | 0.40936643 | 0.84902284 | -0.16549 | 0.26674263 |
| 0.999490048 | -0.1199258 | 0.41572648 | 0.84902284 | 0.05644162 | 0.70021238 |
| 0.999490048 | -0.1046759 | 0.41111375 | 0.84902284 | -0.0511927 | 0.68616273 |
| 0.999490048 | -0.1010953 | 0.41549097 | 0.84902284 | -0.0085703 | 0.94411356 |
| 0.999490048 | -0.0984323 | 0.41934292 | 0.84902284 | -0.0580257 | 0.632575 |
| 0.999490048 | -0.0962676 | 0.41917894 | 0.84902284 | -0.0651237 | 0.58164078 |
| 0.999490048 | -0.0906314 | 0.41898866 | 0.84902284 | -0.0934611 | 0.4048083 |

| | | | | | |
|---|---|---|---|---|---|
| 0.999490048 | -0.0842966 | 0.41494144 | 0.84902284 | -0.0651362 | 0.52619247 |
| 0.909524814 | -0.084004 | 0.41183789 | 0.84902284 | -0.3359664 | 0.00263351 |
| 0.999490048 | -0.0783631 | 0.41218422 | 0.84902284 | -0.0442329 | 0.64190579 |
| 0.999490048 | -0.0749609 | 0.41390496 | 0.84902284 | -0.0078601 | 0.93127008 |
| 0.999490048 | -0.0744115 | 0.41443039 | 0.84902284 | 0.02496121 | 0.78300189 |
| 0.999490048 | -0.0731816 | 0.41450095 | 0.84902284 | -0.1065603 | 0.23822182 |
| 0.999490048 | -0.0727995 | 0.41545621 | 0.84902284 | -0.1629031 | 0.07568153 |
| 0.999490048 | -0.0706881 | 0.41511581 | 0.84902284 | -0.1958616 | 0.03028941 |
| 0.999490048 | -0.0691786 | 0.41188086 | 0.84902284 | 0.01026145 | 0.90247543 |
| 0.999490048 | -0.0687476 | 0.41233184 | 0.84902284 | -0.1060938 | 0.20998693 |
| 0.999490048 | -0.063106 | 0.41606321 | 0.84902284 | 0.05125741 | 0.5079264 |
| 0.999490048 | -0.062549 | 0.41622674 | 0.84902284 | -0.0052881 | 0.94483781 |
| 0.999490048 | -0.0621658 | 0.41384569 | 0.84902284 | 0.06403175 | 0.40016356 |
| 0.999490048 | -0.0615133 | 0.4190026 | 0.84902284 | -0.0355316 | 0.63916658 |
| 0.999490048 | -0.0612441 | 0.41835362 | 0.84902284 | -0.0503015 | 0.50533999 |
| 0.999490048 | -0.0607296 | 0.40935798 | 0.84902284 | -0.0163897 | 0.82269111 |
| 0.999490048 | -0.0592135 | 0.41564127 | 0.84902284 | -0.052204 | 0.47231842 |
| 0.999490048 | -0.0533364 | 0.41813786 | 0.84902284 | -0.0703427 | 0.28792611 |
| 0.999490048 | -0.05237 | 0.41646867 | 0.84902284 | -0.0797103 | 0.22012566 |
| 0.999490048 | -0.0475306 | 0.41843703 | 0.84902284 | -0.0109575 | 0.85110465 |
| 0.999490048 | -0.0467437 | 0.4110821 | 0.84902284 | -0.0349601 | 0.53748449 |
| 0.999490048 | -0.0458214 | 0.4166367 | 0.84902284 | -0.0582262 | 0.30394518 |
| 0.999490048 | -0.0456604 | 0.41937406 | 0.84902284 | 0.025223 | 0.65406947 |
| 0.999490048 | -0.0446842 | 0.41059121 | 0.84902284 | -0.0192015 | 0.72216044 |
| 0.999490048 | -0.0440163 | 0.41712319 | 0.84902284 | -0.0016987 | 0.97485148 |
| 0.999490048 | -0.0433524 | 0.41103186 | 0.84902284 | -0.055927 | 0.29105749 |
| 0.999490048 | -0.0423356 | 0.41362138 | 0.84902284 | -0.0940915 | 0.07645951 |
| 0.999490048 | -0.0396925 | 0.41412166 | 0.84902284 | -0.046311 | 0.3418348 |
| 0.999490048 | -0.0395424 | 0.41889254 | 0.84902284 | -0.0246378 | 0.61307189 |
| 0.999490048 | -0.0394589 | 0.41203661 | 0.84902284 | -0.0404504 | 0.40055034 |
| 0.999490048 | -0.0393639 | 0.41314094 | 0.84902284 | -0.040247 | 0.40288341 |
| 0.999490048 | -0.037284 | 0.4112396 | 0.84902284 | -0.0745658 | 0.10708805 |
| 0.999490048 | -0.0347659 | 0.41235015 | 0.84902284 | -0.0284331 | 0.50162058 |
| 0.999490048 | -0.0339169 | 0.40886333 | 0.84902284 | -0.0293742 | 0.47367988 |
| 0.999490048 | -0.0335175 | 0.41437936 | 0.84902284 | -0.0115868 | 0.77653288 |
| 0.999490048 | -0.0327724 | 0.41089728 | 0.84902284 | -0.0148466 | 0.70798971 |
| 0.999490048 | -0.0324573 | 0.41406448 | 0.84902284 | -0.0113661 | 0.77356251 |
| 0.999490048 | -0.0317941 | 0.41194986 | 0.84902284 | 0.02778841 | 0.47260751 |
| 0.999490048 | -0.0267661 | 0.41895425 | 0.84902284 | -0.0278832 | 0.40006779 |
| 0.999490048 | 0.03275179 | 0.41764802 | 0.84902284 | -0.0048338 | 0.90416375 |
| 0.999490048 | 0.0329565 | 0.41451397 | 0.84902284 | 0.0181301 | 0.65199242 |
| 0.999490048 | 0.0343973 | 0.41713679 | 0.84902284 | -0.0493241 | 0.24787805 |

| | | | | | |
|---|---|---|---|---|---|
| 0.999490048 | 0.03485206 | 0.41536049 | 0.84902284 | -0.0109301 | 0.79718742 |
| 0.999490048 | 0.03790211 | 0.41517582 | 0.84902284 | 0.00319471 | 0.94487805 |
| 0.999490048 | 0.04123596 | 0.41085586 | 0.84902284 | 0.02420785 | 0.62770936 |
| 0.999490048 | 0.04325842 | 0.41338102 | 0.84902284 | 0.04870431 | 0.35794561 |
| 0.999490048 | 0.04330506 | 0.41238743 | 0.84902284 | 0.01686588 | 0.74814088 |
| 0.999490048 | 0.04384644 | 0.41779816 | 0.84902284 | 0.07067505 | 0.19615249 |
| 0.999490048 | 0.04419228 | 0.4138811 | 0.84902284 | 0.05799787 | 0.28587709 |
| 0.999490048 | 0.04514192 | 0.41668077 | 0.84902284 | 0.03968646 | 0.47453869 |
| 0.999490048 | 0.04569542 | 0.41603486 | 0.84902284 | 0.02837785 | 0.61203968 |
| 0.999490048 | 0.04617563 | 0.41207869 | 0.84902284 | 0.02617079 | 0.64047607 |
| 0.999490048 | 0.04652049 | 0.41843959 | 0.84902284 | 0.02003434 | 0.72608242 |
| 0.999490048 | 0.04669025 | 0.4120693 | 0.84902284 | 0.0796052 | 0.16727382 |
| 0.999490048 | 0.0468691 | 0.41677224 | 0.84902284 | 0.0051444 | 0.92850978 |
| 0.999490048 | 0.04711339 | 0.41041635 | 0.84902284 | 0.02020248 | 0.72262308 |
| 0.999490048 | 0.04808626 | 0.41709856 | 0.84902284 | -0.0244873 | 0.6779452 |
| 0.999490048 | 0.04931972 | 0.41925014 | 0.84902284 | -0.0078787 | 0.89667285 |
| 0.999490048 | 0.04974135 | 0.41174061 | 0.84902284 | -0.0597144 | 0.32585639 |
| 0.999490048 | 0.05140597 | 0.41758502 | 0.84902284 | 0.05878808 | 0.35488511 |
| 0.999490048 | 0.05172596 | 0.41197218 | 0.84902284 | 0.06128492 | 0.33233974 |
| 0.999490048 | 0.05206513 | 0.41329815 | 0.84902284 | 0.08973192 | 0.16396778 |
| 0.999490048 | 0.05363853 | 0.41222752 | 0.84902284 | 0.00906032 | 0.88911357 |
| 0.999490048 | 0.05498978 | 0.41683858 | 0.84902284 | 0.03102358 | 0.64541364 |
| 0.999490048 | 0.05659685 | 0.41615681 | 0.84902284 | 0.09863932 | 0.16195489 |
| 0.999490048 | 0.05663566 | 0.41329067 | 0.84902284 | 0.08674333 | 0.21425999 |
| 0.999490048 | 0.05732182 | 0.41507583 | 0.84902284 | 0.08515675 | 0.2296787 |
| 0.999490048 | 0.05749067 | 0.41398035 | 0.84902284 | 0.07137784 | 0.31221058 |
| 0.999490048 | 0.0579245 | 0.40910991 | 0.84902284 | 0.09227467 | 0.19876223 |
| 0.999490048 | 0.06033801 | 0.41942745 | 0.84902284 | 0.10119037 | 0.18075113 |
| 0.999490048 | 0.06209886 | 0.41691753 | 0.84902284 | 0.11605373 | 0.13544225 |
| 0.999490048 | 0.06210818 | 0.41677353 | 0.84902284 | 0.09094164 | 0.23793262 |
| 0.999490048 | 0.06418436 | 0.40876808 | 0.84902284 | -0.0763511 | 0.32714484 |
| 0.999490048 | 0.06510345 | 0.41536952 | 0.84902284 | -0.0354695 | 0.65571365 |
| 0.999490048 | 0.06605877 | 0.41729265 | 0.84902284 | 0.03487476 | 0.66699173 |
| 0.999490048 | 0.06643255 | 0.41182179 | 0.84902284 | 0.07383868 | 0.36243162 |
| 0.999490048 | 0.06951197 | 0.41436356 | 0.84902284 | 0.01543879 | 0.85522539 |
| 0.999490048 | 0.07069038 | 0.41229962 | 0.84902284 | -0.0836729 | 0.3331358 |
| 0.999490048 | 0.07777086 | 0.40953771 | 0.84902284 | -0.0467478 | 0.61854966 |
| 0.999490048 | 0.07812878 | 0.41940553 | 0.84902284 | 0.08528453 | 0.3786932 |
| 0.999490048 | 0.08036258 | 0.41475775 | 0.84902284 | 0.05622118 | 0.56697685 |
| 0.999490048 | 0.08055142 | 0.41829096 | 0.84902284 | 0.16606623 | 0.10213234 |
| 0.999490048 | 0.08762694 | 0.41661096 | 0.84902284 | 0.06852133 | 0.5242265 |
| 0.999490048 | 0.09155281 | 0.41386017 | 0.84902284 | 0.08527559 | 0.44619725 |

| | | | | | |
|---|---|---|---|---|---|
| 0.999490048 | 0.10820393 | 0.4130666 | 0.84902284 | -0.0885442 | 0.50203889 |
| 0.999490048 | 0.1099799 | 0.41028725 | 0.84902284 | -0.095901 | 0.47111571 |
| 0.999490048 | 0.11169611 | 0.4164286 | 0.84902284 | 0.18880898 | 0.17471427 |
| 0.999490048 | 0.12168241 | 0.41852759 | 0.84902284 | 0.1307314 | 0.38525136 |
| 0.999490048 | 0.13019976 | 0.41816547 | 0.84902284 | 0.20901759 | 0.20389719 |
| 0.999490048 | 0.13227815 | 0.41471833 | 0.84902284 | 0.34207821 | 0.04182769 |
| 0.999490048 | 0.15929674 | 0.4186518 | 0.84902284 | 0.04098639 | 0.83190693 |
| 0.999490048 | 0.16632454 | 0.41110539 | 0.84902284 | -0.1382908 | 0.49344128 |
| 0.999490048 | 0.19192282 | 0.41340782 | 0.84902284 | -0.135333 | 0.56107622 |
| 0.999490048 | 0.26990098 | 0.41472468 | 0.84902284 | 0.59795966 | 0.09277593 |
| 0.999490048 | 0.3164156 | 0.41623932 | 0.84902284 | 0.12615025 | 0.74140347 |
| 0.999490048 | 0.32097364 | 0.40928591 | 0.84902284 | -0.2106576 | 0.58476853 |
| 0.854051452 | 0.57435313 | 0.41057427 | 0.84902284 | 2.36176753 | 0.00203502 |
| 0.999490048 | -0.1023801 | 0.41974802 | 0.84904604 | -0.0850204 | 0.50195958 |
| 0.999490048 | 0.03589372 | 0.41994732 | 0.84904604 | 0.07898492 | 0.08317464 |
| 0.999490048 | 0.0410518 | 0.41986605 | 0.84904604 | 0.05062936 | 0.3214446 |
| 0.999490048 | 0.07581461 | 0.41970312 | 0.84904604 | 0.01045583 | 0.91081583 |
| 0.999490048 | 0.20129221 | 0.41994174 | 0.84904604 | 0.24655706 | 0.32477801 |
| 0.999490048 | -0.0447379 | 0.42048122 | 0.84973524 | 0.02311986 | 0.67571275 |
| 0.999490048 | -0.0436206 | 0.42060187 | 0.84973524 | -0.0236166 | 0.66134127 |
| 0.999490048 | -0.0374956 | 0.42069527 | 0.84973524 | 0.01234695 | 0.7896961 |
| 0.999490048 | 0.05323914 | 0.42066116 | 0.84973524 | -0.0159395 | 0.80836255 |
| 0.999490048 | 0.05761523 | 0.42130568 | 0.85076238 | 0.01329555 | 0.85187613 |
| 0.999490048 | -0.0478946 | 0.42184188 | 0.85163919 | 1.66E-04 | 0.99777046 |
| 0.999490048 | 0.12384917 | 0.42203586 | 0.85182486 | 0.10022625 | 0.51370035 |
| 0.999490048 | 0.05184247 | 0.42244633 | 0.85233004 | 0.03092054 | 0.63090377 |
| 0.999490048 | 0.10276385 | 0.4224903 | 0.85233004 | 0.09627835 | 0.45200293 |
| 0.999490048 | -0.0613401 | 0.42285347 | 0.85244483 | -0.0179986 | 0.8129873 |
| 0.999490048 | 0.03412136 | 0.42281361 | 0.85244483 | 0.01013382 | 0.81074055 |
| 0.999490048 | 0.04498491 | 0.42265364 | 0.85244483 | 0.02911684 | 0.60240348 |
| 0.999490048 | -0.0689647 | 0.42319175 | 0.85257561 | 0.02083064 | 0.8077415 |
| 0.999490048 | -0.0272037 | 0.42322466 | 0.85257561 | -0.0171894 | 0.61145025 |
| 0.999490048 | 0.04826973 | 0.42303556 | 0.85257561 | 0.03984001 | 0.5075776 |
| 0.999490048 | -0.0658666 | 0.42349091 | 0.85263315 | 0.04845674 | 0.5548096 |
| 0.999490048 | -0.057519 | 0.42368199 | 0.85263315 | -0.0203232 | 0.77618964 |
| 0.999490048 | 0.03196908 | 0.42336335 | 0.85263315 | 0.03322165 | 0.40566788 |
| 0.999490048 | 0.03449283 | 0.42385469 | 0.85263315 | -0.0272828 | 0.52601232 |
| 0.999490048 | 0.08205968 | 0.42386589 | 0.85263315 | -0.0037727 | 0.97048024 |
| 0.999490048 | 0.08725165 | 0.42374919 | 0.85263315 | 0.1125546 | 0.30415729 |
| 0.999490048 | 0.05697918 | 0.42416613 | 0.8530316 | -0.0760481 | 0.28850639 |
| 0.999490048 | 0.08590056 | 0.42430956 | 0.85311457 | -7.45E-04 | 0.99443314 |
| 0.999490048 | -0.0627351 | 0.4247885 | 0.85378069 | 0.06277985 | 0.42446212 |

| | | | | | |
|---|---|---|---|---|---|
| 0.999490048 | -0.0385552 | 0.42491529 | 0.85378069 | -0.0466495 | 0.33574877 |
| 0.999490048 | 0.0429401 | 0.42501428 | 0.85378069 | 0.01921648 | 0.71970692 |
| 0.999490048 | 0.05728983 | 0.42504986 | 0.85378069 | 0.12003975 | 0.10165888 |
| 0.984398018 | -0.0495618 | 0.42547706 | 0.85400378 | 0.08846322 | 0.16053638 |
| 0.999490048 | -0.0446019 | 0.42552317 | 0.85400378 | 0.01662917 | 0.76512988 |
| 0.999490048 | 0.03989671 | 0.42557003 | 0.85400378 | 0.04914172 | 0.32796349 |
| 0.999490048 | 0.11116606 | 0.42540422 | 0.85400378 | 0.21210568 | 0.13461394 |
| 0.999490048 | -0.1829822 | 0.42774372 | 0.85469049 | -0.6484225 | 0.01385564 |
| 0.999490048 | -0.1206887 | 0.42752563 | 0.85469049 | -0.0329972 | 0.82726258 |
| 0.999490048 | -0.0868912 | 0.42831284 | 0.85469049 | 0.0574656 | 0.59908821 |
| 0.999490048 | -0.083855 | 0.42732836 | 0.85469049 | 3.61E-04 | 0.99725725 |
| 0.999490048 | -0.0662697 | 0.42797826 | 0.85469049 | -0.0167419 | 0.84034929 |
| 0.999490048 | -0.0586712 | 0.42769515 | 0.85469049 | 0.02383805 | 0.74596156 |
| 0.999490048 | -0.0562046 | 0.42625697 | 0.85469049 | -0.0233914 | 0.73922666 |
| 0.999490048 | -0.0455784 | 0.42763244 | 0.85469049 | -0.0052058 | 0.92735384 |
| 0.999490048 | -0.044982 | 0.42725568 | 0.85469049 | -0.0254807 | 0.65151222 |
| 0.999490048 | -0.0416341 | 0.42676877 | 0.85469049 | 0.03538563 | 0.49862759 |
| 0.999490048 | -0.0400161 | 0.4275218 | 0.85469049 | -0.0191769 | 0.7023803 |
| 0.999490048 | -0.0378621 | 0.42795321 | 0.85469049 | 0.02765985 | 0.56136695 |
| 0.999490048 | 0.04188398 | 0.42819622 | 0.85469049 | -0.0159716 | 0.7613075 |
| 0.999490048 | 0.04716437 | 0.42777504 | 0.85469049 | -0.0276413 | 0.64072023 |
| 0.999490048 | 0.04732553 | 0.42706464 | 0.85469049 | -0.0165562 | 0.77993011 |
| 0.999490048 | 0.04804025 | 0.42745677 | 0.85469049 | 0.10864931 | 0.08002325 |
| 0.999490048 | 0.05475697 | 0.42808889 | 0.85469049 | 0.0829488 | 0.23371094 |
| 0.999490048 | 0.05947965 | 0.42836883 | 0.85469049 | 0.0712125 | 0.34433087 |
| 0.999490048 | 0.0631585 | 0.42652241 | 0.85469049 | 0.0764138 | 0.33744346 |
| 0.999490048 | 0.06473775 | 0.42606403 | 0.85469049 | -0.0307581 | 0.70391016 |
| 0.999490048 | 0.10088097 | 0.4283279 | 0.85469049 | 0.29251273 | 0.02786065 |
| 0.999490048 | 0.10868212 | 0.42734855 | 0.85469049 | -0.0504225 | 0.71131183 |
| 0.999490048 | 0.11764308 | 0.42835296 | 0.85469049 | 0.02918928 | 0.84329002 |
| 0.999490048 | 0.14719181 | 0.42650717 | 0.85469049 | -0.0365757 | 0.84243047 |
| 0.999490048 | 0.04694213 | 0.42852326 | 0.85479437 | 0.02520801 | 0.66932434 |
| 0.999490048 | 0.04185058 | 0.42876594 | 0.85487001 | 0.10871969 | 0.04694844 |
| 0.999490048 | 0.04583235 | 0.42867745 | 0.85487001 | -0.0130685 | 0.82042922 |
| 0.999490048 | -0.0679364 | 0.42906449 | 0.85526104 | -0.032281 | 0.7057274 |
| 0.999490048 | -0.0493797 | 0.42928257 | 0.85536092 | -0.0377067 | 0.54507597 |
| 0.999490048 | -0.0364368 | 0.42931947 | 0.85536092 | -0.0178259 | 0.69762675 |
| 0.999490048 | -0.052518 | 0.42968165 | 0.85567416 | 0.02647933 | 0.68911442 |
| 0.999490048 | -0.0365329 | 0.42960944 | 0.85567416 | -0.0968529 | 0.04328848 |
| 0.999490048 | -0.1074253 | 0.43009214 | 0.85625271 | 0.03451408 | 0.79876884 |
| 0.999490048 | -0.0489749 | 0.43017726 | 0.85625271 | -0.046529 | 0.45328734 |
| 0.999490048 | -0.1308006 | 0.43032984 | 0.85635229 | -0.2115116 | 0.20670254 |

| | | | | | |
|---|---|---|---|---|---|
| 0.999490048 | -0.2512756 | 0.43137505 | 0.85652116 | 0.21628866 | 0.49664071 |
| 0.999490048 | -0.1348729 | 0.43174821 | 0.85652116 | -0.0300912 | 0.85993702 |
| 0.999490048 | -0.0853331 | 0.43103482 | 0.85652116 | 0.02066213 | 0.84790466 |
| 0.999490048 | -0.045665 | 0.43171525 | 0.85652116 | -0.1046847 | 0.07883989 |
| 0.999490048 | -0.0422004 | 0.43159287 | 0.85652116 | 0.06031618 | 0.26403103 |
| 0.999490048 | -0.0402277 | 0.43059065 | 0.85652116 | 6.83E-04 | 0.98924836 |
| 0.999490048 | -0.0379622 | 0.4307694 | 0.85652116 | -0.0097984 | 0.83791952 |
| 0.999490048 | -0.0345528 | 0.43100332 | 0.85652116 | -0.0047909 | 0.91251963 |
| 0.999490048 | -0.029127 | 0.43153389 | 0.85652116 | -0.065421 | 0.08459671 |
| 0.999490048 | 0.03307083 | 0.43073646 | 0.85652116 | -0.008374 | 0.84093887 |
| 0.999490048 | 0.04324026 | 0.43144593 | 0.85652116 | 0.02642316 | 0.629357 |
| 0.999490048 | 0.05679771 | 0.43159054 | 0.85652116 | -0.0062066 | 0.93108806 |
| 0.999490048 | 0.06154386 | 0.43150696 | 0.85652116 | 0.05768241 | 0.46062518 |
| 0.999490048 | -0.0680685 | 0.43194238 | 0.85654166 | -0.0467215 | 0.58837427 |
| 0.999490048 | -0.0487568 | 0.43242607 | 0.85654166 | -0.0375916 | 0.5439849 |
| 0.999490048 | -0.0412065 | 0.43256729 | 0.85654166 | -0.0975055 | 0.07071094 |
| 0.999490048 | -0.039681 | 0.43257919 | 0.85654166 | 0.00926 | 0.8538398 |
| 0.999490048 | -0.03715 | 0.43201116 | 0.85654166 | -0.0142095 | 0.76253799 |
| 0.999490048 | 0.04430862 | 0.43225419 | 0.85654166 | 0.06230067 | 0.27235676 |
| 0.999490048 | 0.04577651 | 0.43235738 | 0.85654166 | 0.03026742 | 0.60238754 |
| 0.999490048 | 0.05834871 | 0.43240513 | 0.85654166 | 0.05698482 | 0.44311093 |
| 0.999490048 | 0.11703417 | 0.43283297 | 0.856841 | 0.08125085 | 0.58486766 |
| 0.999490048 | -0.1534042 | 0.43298682 | 0.8569424 | -9.11E-05 | 0.99962329 |
| 0.999490048 | -0.1919551 | 0.43372264 | 0.85737369 | -0.0953791 | 0.69594766 |
| 0.999490048 | -0.1413386 | 0.43398566 | 0.85737369 | -0.1661794 | 0.3587503 |
| 0.999490048 | -0.1301671 | 0.43396199 | 0.85737369 | -0.1458404 | 0.38142548 |
| 0.999490048 | -0.1072583 | 0.43416902 | 0.85737369 | 0.09969961 | 0.46607769 |
| 0.999490048 | -0.0503546 | 0.43368203 | 0.85737369 | 7.62E-04 | 0.99048977 |
| 0.999490048 | -0.0423045 | 0.43408872 | 0.85737369 | 0.00441721 | 0.93450602 |
| 0.999490048 | -0.03934 | 0.43393069 | 0.85737369 | -0.0706262 | 0.16570113 |
| 0.999490048 | -0.0336956 | 0.43389566 | 0.85737369 | -0.0348624 | 0.41834924 |
| 0.999490048 | 0.05298931 | 0.4338555 | 0.85737369 | 0.03917894 | 0.561737 |
| 0.999490048 | 0.07884112 | 0.43423154 | 0.85737369 | 0.15815287 | 0.12340452 |
| 0.999490048 | 0.11077254 | 0.43444039 | 0.85758328 | 0.02044984 | 0.88458463 |
| 0.999490048 | -0.0892233 | 0.43510891 | 0.85792607 | -0.1103742 | 0.33585342 |
| 0.999490048 | -0.0715912 | 0.43489661 | 0.85792607 | -0.0705658 | 0.44132288 |
| 0.999490048 | -0.049013 | 0.43497189 | 0.85792607 | 0.01238077 | 0.84276968 |
| 0.999490048 | 0.03975705 | 0.43512777 | 0.85792607 | 0.01582175 | 0.75488712 |
| 0.999490048 | 0.04586454 | 0.43480953 | 0.85792607 | -0.0050946 | 0.93044623 |
| 0.999490048 | -0.1292499 | 0.43597241 | 0.85834115 | -0.0180271 | 0.91295731 |
| 0.999490048 | -0.1017322 | 0.43544684 | 0.85834115 | -0.0059353 | 0.9634841 |
| 0.999490048 | -0.0571426 | 0.43750137 | 0.85834115 | -0.0691069 | 0.34908121 |

| | | | | | |
|---|---|---|---|---|---|
| 0.999490048 | -0.0566414 | 0.43557776 | 0.85834115 | -0.0856085 | 0.24226275 |
| 0.999490048 | -0.0455664 | 0.43586934 | 0.85834115 | -0.0815911 | 0.16851305 |
| 0.999490048 | -0.0451415 | 0.43660075 | 0.85834115 | -0.0304047 | 0.59907604 |
| 0.999490048 | -0.04075 | 0.4359039 | 0.85834115 | -0.0401051 | 0.44311371 |
| 0.999490048 | -0.0402571 | 0.43679215 | 0.85834115 | -0.0351044 | 0.49708864 |
| 0.999490048 | -0.0371892 | 0.43740488 | 0.85834115 | -0.0589348 | 0.22260106 |
| 0.999490048 | -0.0334866 | 0.43685406 | 0.85834115 | -0.028038 | 0.51427752 |
| 0.999490048 | 0.03594614 | 0.43618739 | 0.85834115 | 0.05914635 | 0.20473005 |
| 0.999490048 | 0.04465581 | 0.43681204 | 0.85834115 | 0.01333172 | 0.81541831 |
| 0.999490048 | 0.04640005 | 0.4365292 | 0.85834115 | -0.0452122 | 0.44822277 |
| 0.999490048 | 0.04934994 | 0.43629742 | 0.85834115 | -0.0170806 | 0.78647309 |
| 0.999490048 | 0.05084446 | 0.43734698 | 0.85834115 | 0.07225386 | 0.27263885 |
| 0.999490048 | 0.05598003 | 0.43729824 | 0.85834115 | 0.04761286 | 0.50811691 |
| 0.999490048 | 0.06126983 | 0.43659964 | 0.85834115 | 0.03746767 | 0.63296863 |
| 0.999490048 | 0.06150778 | 0.43727006 | 0.85834115 | -0.0918424 | 0.2495566 |
| 0.999490048 | 0.06554809 | 0.43737067 | 0.85834115 | -0.0397143 | 0.63666433 |
| 0.999490048 | 0.0785143 | 0.43759979 | 0.85834115 | 0.21465036 | 0.04080121 |
| 0.999490048 | 0.10471946 | 0.43688735 | 0.85834115 | -0.0194745 | 0.88436345 |
| 0.999490048 | 0.11489134 | 0.43577144 | 0.85834115 | 0.2128268 | 0.15474535 |
| 0.999490048 | -0.0856277 | 0.43808885 | 0.85849375 | 0.09662748 | 0.38233695 |
| 0.999490048 | -0.0684398 | 0.43787481 | 0.85849375 | -0.0498097 | 0.57124347 |
| 0.999490048 | 0.04019961 | 0.43807406 | 0.85849375 | -0.0162709 | 0.75240772 |
| 0.999490048 | 0.05352116 | 0.43804169 | 0.85849375 | 0.0388003 | 0.57287483 |
| 0.999490048 | -0.0362697 | 0.43833897 | 0.85858091 | -0.0801956 | 0.093861 |
| 0.999490048 | 0.03950171 | 0.43825898 | 0.85858091 | 0.02130041 | 0.67464776 |
| 0.999490048 | 0.07185637 | 0.43850274 | 0.85870026 | 0.06084448 | 0.51110357 |
| 0.999490048 | -0.2716919 | 0.43951155 | 0.85904297 | -0.8086769 | 0.02764381 |
| 0.999490048 | -0.0851029 | 0.43931388 | 0.85904297 | -0.0416217 | 0.70396724 |
| 0.999490048 | -0.0772266 | 0.43972551 | 0.85904297 | 0.08853647 | 0.37661418 |
| 0.999490048 | -0.0741601 | 0.43902207 | 0.85904297 | -0.0071346 | 0.94029689 |
| 0.956358655 | -0.0512351 | 0.43988751 | 0.85904297 | 0.10835473 | 0.10938024 |
| 0.999490048 | -0.0502187 | 0.43962969 | 0.85904297 | -0.0530952 | 0.41420896 |
| 0.999490048 | -0.039467 | 0.4399123 | 0.85904297 | 0.02208164 | 0.66435072 |
| 0.773280431 | -0.0311217 | 0.43880197 | 0.85904297 | 0.0766179 | 0.0640866 |
| 0.999490048 | 0.04221532 | 0.43928117 | 0.85904297 | 0.00862452 | 0.8737274 |
| 0.999490048 | 0.05065834 | 0.43894704 | 0.85904297 | 0.01564396 | 0.8100765 |
| 0.773280431 | 0.05502413 | 0.43938112 | 0.85904297 | -0.1360914 | 0.0632577 |
| 0.999490048 | 0.07648465 | 0.43945417 | 0.85904297 | 0.04848641 | 0.62280631 |
| 0.999490048 | -0.0345598 | 0.44013355 | 0.85907321 | -0.0760578 | 0.09656909 |
| 0.999490048 | 0.07089968 | 0.44011852 | 0.85907321 | 0.02158638 | 0.81317367 |
| 0.999490048 | -0.0516258 | 0.44046778 | 0.85946523 | -0.0350419 | 0.59937446 |
| 0.999490048 | 0.07097049 | 0.44063843 | 0.85946523 | 0.03644082 | 0.69085437 |

| | | | | | |
|---|---|---|---|---|---|
| 0.999490048 | 0.12464571 | 0.44064319 | 0.85946523 | 0.01719551 | 0.91478218 |
| 0.999490048 | -0.0660589 | 0.44101144 | 0.85959815 | 0.01243797 | 0.88398815 |
| 0.999490048 | 0.05053107 | 0.44098766 | 0.85959815 | 0.00622834 | 0.92388794 |
| 0.999490048 | 0.05679952 | 0.44102018 | 0.85959815 | 0.10086289 | 0.17666011 |
| 0.999490048 | 0.04416543 | 0.44160017 | 0.86052775 | 0.08774742 | 0.13283095 |
| 0.999490048 | 0.04820623 | 0.44185302 | 0.86061878 | 0.13708736 | 0.0354979 |
| 0.999490048 | 0.06034328 | 0.44176268 | 0.86061878 | 0.05010607 | 0.52223502 |
| 0.999490048 | 0.09512451 | 0.44219978 | 0.86109331 | 0.12205237 | 0.32596063 |
| 0.999490048 | -0.0783001 | 0.44281006 | 0.86153527 | 0.00547964 | 0.95691252 |
| 0.999490048 | -0.0441778 | 0.44255921 | 0.86153527 | 0.0130161 | 0.82005318 |
| 0.999490048 | 0.04624942 | 0.44294263 | 0.86153527 | 0.04408255 | 0.46436211 |
| 0.999490048 | 0.04640578 | 0.44266274 | 0.86153527 | 0.06420552 | 0.29074139 |
| 0.999490048 | 0.09264017 | 0.44286495 | 0.86153527 | -0.05023 | 0.67607874 |
| 0.999490048 | -0.0447348 | 0.44315781 | 0.86155244 | -0.0426217 | 0.46475074 |
| 0.999490048 | 0.0345844 | 0.44314492 | 0.86155244 | 0.03106067 | 0.49048093 |
| 0.999490048 | -0.0380093 | 0.44360364 | 0.86201777 | 0.01990476 | 0.68699666 |
| 0.999490048 | 0.02486681 | 0.44350293 | 0.86201777 | 0.01104399 | 0.73238432 |
| 0.999490048 | -0.0314791 | 0.44376586 | 0.86213236 | -0.0319596 | 0.43692361 |
| 0.999490048 | -0.0508855 | 0.44432777 | 0.86274651 | -0.0476401 | 0.47359599 |
| 0.999490048 | 0.03884103 | 0.44426496 | 0.86274651 | -0.0029354 | 0.95362387 |
| 0.999490048 | 0.04972629 | 0.44439194 | 0.86274651 | -0.0124718 | 0.84704886 |
| 0.999490048 | 0.04260817 | 0.44451942 | 0.86279338 | 0.06487168 | 0.24799118 |
| 0.999490048 | -0.0411062 | 0.44469378 | 0.86293122 | -0.0504049 | 0.35008012 |
| 0.999490048 | -0.0433472 | 0.44506272 | 0.86304546 | 0.07164816 | 0.21139077 |
| 0.999490048 | 0.03952327 | 0.44505317 | 0.86304546 | 0.02305781 | 0.65468141 |
| 0.999490048 | 0.0438887 | 0.44505805 | 0.86304546 | 0.02273267 | 0.6911712 |
| 0.999490048 | -0.053289 | 0.44524481 | 0.86305768 | 0.04037912 | 0.56199568 |
| 0.999490048 | -0.0301308 | 0.44537911 | 0.86305768 | -0.0533091 | 0.18226061 |
| 0.999490048 | 0.03895168 | 0.44533338 | 0.86305768 | 0.01962651 | 0.69930866 |
| 0.999490048 | -0.0349357 | 0.44554536 | 0.86312931 | -0.0433847 | 0.34497693 |
| 0.999490048 | 0.08298996 | 0.44562281 | 0.86312931 | 0.11261753 | 0.30300977 |
| 0.999490048 | -0.093625 | 0.44579851 | 0.86324902 | 0.02625994 | 0.82975864 |
| 0.999490048 | 0.25526157 | 0.44589138 | 0.86324902 | 0.00734585 | 0.98239517 |
| 0.999490048 | 0.05631091 | 0.44599486 | 0.86324922 | -0.0205153 | 0.78019026 |
| 0.999490048 | -0.0446486 | 0.44629097 | 0.86327061 | -0.0304206 | 0.60268538 |
| 0.999490048 | 0.05198263 | 0.44641946 | 0.86327061 | -0.0180011 | 0.79098643 |
| 0.999490048 | 0.07861393 | 0.44635103 | 0.86327061 | 0.05188281 | 0.61409472 |
| 0.999490048 | 0.11571464 | 0.44613436 | 0.86327061 | -0.0802401 | 0.59617054 |
| 0.999490048 | 0.0632374 | 0.44657031 | 0.86336237 | 0.11894261 | 0.1581959 |
| 0.999490048 | -0.0452673 | 0.44689806 | 0.86379602 | -0.0150823 | 0.79892417 |
| 0.999490048 | -0.0504968 | 0.44711786 | 0.86382095 | -0.0121647 | 0.85389459 |
| 0.999490048 | 0.06949961 | 0.4470499 | 0.86382095 | -0.0667267 | 0.46518354 |

| | | | | | |
|---|---|---|---|---|---|
| 0.999490048 | -0.1419137 | 0.44900591 | 0.86397111 | -0.169877 | 0.36599194 |
| 0.999490048 | -0.0931372 | 0.44908945 | 0.86397111 | 0.00930062 | 0.93940121 |
| 0.999490048 | -0.0751782 | 0.44916151 | 0.86397111 | -0.1261226 | 0.20885462 |
| 0.999490048 | -0.0684991 | 0.44915826 | 0.86397111 | 0.04376983 | 0.62749759 |
| 0.999490048 | -0.0606969 | 0.44802141 | 0.86397111 | -0.0669701 | 0.40435947 |
| 0.999490048 | 0.03087396 | 0.44845896 | 0.86397111 | -0.0249774 | 0.53892004 |
| 0.999490048 | 0.03252904 | 0.44779839 | 0.86397111 | -0.0050803 | 0.9050932 |
| 0.999490048 | 0.03831564 | 0.44867345 | 0.86397111 | 0.06745966 | 0.18744239 |
| 0.999490048 | 0.03902906 | 0.44861444 | 0.86397111 | 0.08442775 | 0.10829872 |
| 0.999490048 | 0.03910684 | 0.4481848 | 0.86397111 | 0.00240961 | 0.96250967 |
| 0.999490048 | 0.03924948 | 0.44853806 | 0.86397111 | 0.0075694 | 0.88315128 |
| 0.999490048 | 0.04357444 | 0.4489318 | 0.86397111 | -0.0053816 | 0.92506672 |
| 0.999490048 | 0.0541464 | 0.44886264 | 0.86397111 | 0.07471121 | 0.29854148 |
| 0.999490048 | 0.05472862 | 0.44768443 | 0.86397111 | 0.07863483 | 0.27822398 |
| 0.999490048 | 0.05814442 | 0.44812705 | 0.86397111 | 0.00934705 | 0.9023969 |
| 0.999490048 | 0.06166098 | 0.44811772 | 0.86397111 | 0.12737966 | 0.12389613 |
| 0.999490048 | 0.07660581 | 0.44860546 | 0.86397111 | -0.1206645 | 0.23659204 |
| 0.999490048 | 0.0866671 | 0.44879598 | 0.86397111 | 0.03668687 | 0.74733614 |
| 0.999490048 | 0.10794969 | 0.44883908 | 0.86397111 | -0.0832215 | 0.55839035 |
| 0.999490048 | -0.0390753 | 0.44947816 | 0.86418205 | -0.0113867 | 0.82465989 |
| 0.999490048 | 0.04545899 | 0.44944154 | 0.86418205 | 0.10515912 | 0.08760676 |
| 0.999490048 | -0.0690694 | 0.44995086 | 0.86471089 | -0.1325964 | 0.15306789 |
| 0.999490048 | 0.04211753 | 0.45014567 | 0.86471089 | 0.00352506 | 0.9493139 |
| 0.999490048 | 0.07710361 | 0.45014157 | 0.86471089 | -0.045554 | 0.65426356 |
| 0.999490048 | 0.14046412 | 0.45016746 | 0.86471089 | -0.0830905 | 0.65231063 |
| 0.999490048 | -0.0430302 | 0.45052349 | 0.86499681 | 0.01398028 | 0.8053621 |
| 0.999490048 | -0.0275742 | 0.45042577 | 0.86499681 | -0.0237151 | 0.51566012 |
| 0.999490048 | 0.06346893 | 0.45071886 | 0.86517299 | 0.10513621 | 0.21601811 |
| 0.999490048 | 0.10573012 | 0.4508551 | 0.86523561 | 0.09819634 | 0.48341868 |
| 0.999490048 | -0.1858424 | 0.45418869 | 0.86545623 | -0.3840095 | 0.12871419 |
| 0.999490048 | -0.1240366 | 0.45835249 | 0.86545623 | -0.2245684 | 0.19144427 |
| 0.999490048 | -0.1067573 | 0.4518979 | 0.86545623 | -0.1364603 | 0.33800229 |
| 0.999490048 | -0.0910457 | 0.45513284 | 0.86545623 | 0.07677419 | 0.52814996 |
| 0.999490048 | -0.085308 | 0.45500064 | 0.86545623 | -0.0611821 | 0.59107255 |
| 0.999490048 | -0.0746157 | 0.45790598 | 0.86545623 | 0.00608404 | 0.95146881 |
| 0.999490048 | -0.0722611 | 0.45521114 | 0.86545623 | -0.1171623 | 0.23015938 |
| 0.999490048 | -0.0589102 | 0.45784474 | 0.86545623 | 0.0117896 | 0.88125899 |
| 0.999490048 | -0.0536723 | 0.45135566 | 0.86545623 | -0.0203547 | 0.77408585 |
| 0.999490048 | -0.0525789 | 0.45493131 | 0.86545623 | 0.02971453 | 0.67163236 |
| 0.999490048 | -0.0521021 | 0.45580139 | 0.86545623 | -0.0076019 | 0.91287456 |
| 0.999490048 | -0.0516539 | 0.45555856 | 0.86545623 | -0.1383487 | 0.05307213 |
| 0.999490048 | -0.0504973 | 0.45167675 | 0.86545623 | -0.0957462 | 0.15953963 |

| | | | | | |
|---|---|---|---|---|---|
| 0.999490048 | -0.0464592 | 0.45875651 | 0.86545623 | -0.0491549 | 0.43340952 |
| 0.999490048 | -0.0453407 | 0.45186905 | 0.86545623 | -0.1200729 | 0.05374805 |
| 0.999490048 | -0.0444069 | 0.45697478 | 0.86545623 | -0.0934844 | 0.12423304 |
| 0.999490048 | -0.0442152 | 0.45148278 | 0.86545623 | -0.0240451 | 0.68098362 |
| 0.999490048 | -0.0439864 | 0.45345588 | 0.86545623 | -0.0174846 | 0.76461752 |
| 0.999490048 | -0.0433193 | 0.45877987 | 0.86545623 | -0.0555259 | 0.34400602 |
| 0.999490048 | -0.0430638 | 0.45709293 | 0.86545623 | 0.00816252 | 0.88730388 |
| 0.999490048 | -0.0403712 | 0.45290983 | 0.86545623 | -0.0100633 | 0.85079992 |
| 0.999490048 | -0.0400551 | 0.4529743 | 0.86545623 | -0.075843 | 0.1612575 |
| 0.999490048 | -0.0396066 | 0.45880079 | 0.86545623 | -0.0401605 | 0.45261276 |
| 0.999490048 | -0.0353853 | 0.4562795 | 0.86545623 | -0.0236243 | 0.61783448 |
| 0.999490048 | -0.0333978 | 0.45765329 | 0.86545623 | -0.0223229 | 0.61851333 |
| 0.999490048 | -0.0317525 | 0.45385351 | 0.86545623 | -0.0159145 | 0.70617194 |
| 0.999490048 | -0.030701 | 0.45559375 | 0.86545623 | -0.0305844 | 0.45728857 |
| 0.999490048 | -0.0298938 | 0.45879792 | 0.86545623 | -0.0246857 | 0.53998478 |
| 0.999490048 | -0.0291715 | 0.45145749 | 0.86545623 | -0.0457103 | 0.24188139 |
| 0.999490048 | 0.03081644 | 0.45774257 | 0.86545623 | -0.0056246 | 0.89161047 |
| 0.999490048 | 0.03096099 | 0.45557344 | 0.86545623 | 0.0583492 | 0.16551573 |
| 0.999490048 | 0.03294537 | 0.45591711 | 0.86545623 | 0.00676624 | 0.8776455 |
| 0.999490048 | 0.03653399 | 0.45383956 | 0.86545623 | 0.01473091 | 0.76154272 |
| 0.999490048 | 0.03763592 | 0.45794473 | 0.86545623 | 0.04476468 | 0.3784202 |
| 0.999490048 | 0.03815575 | 0.4580421 | 0.86545623 | 0.02591981 | 0.61312821 |
| 0.999490048 | 0.04048155 | 0.45295543 | 0.86545623 | 0.0385742 | 0.47429489 |
| 0.999490048 | 0.04071258 | 0.4525917 | 0.86545623 | 0.01235333 | 0.81879932 |
| 0.999490048 | 0.04100254 | 0.45461096 | 0.86545623 | 0.01928163 | 0.72395049 |
| 0.999490048 | 0.04112818 | 0.45444109 | 0.86545623 | 0.01872243 | 0.73231281 |
| 0.999490048 | 0.0413901 | 0.45716444 | 0.86545623 | 0.0218457 | 0.69356885 |
| 0.999490048 | 0.04210066 | 0.45412813 | 0.86545623 | 7.76E-04 | 0.98893378 |
| 0.999490048 | 0.04302623 | 0.45205754 | 0.86545623 | 0.0323945 | 0.5703254 |
| 0.999490048 | 0.04402101 | 0.45452625 | 0.86545623 | -0.0170584 | 0.77089124 |
| 0.999490048 | 0.0459351 | 0.45661341 | 0.86545623 | 0.05563976 | 0.36844965 |
| 0.999490048 | 0.04636705 | 0.45468198 | 0.86545623 | 0.02551128 | 0.67962719 |
| 0.999490048 | 0.04736995 | 0.45821117 | 0.86545623 | 0.01706798 | 0.78825593 |
| 0.999490048 | 0.04785809 | 0.45708695 | 0.86545623 | 0.06240967 | 0.33403128 |
| 0.999490048 | 0.05293095 | 0.45249785 | 0.86545623 | 0.01542031 | 0.82586338 |
| 0.999490048 | 0.05308378 | 0.4587711 | 0.86545623 | 0.04963216 | 0.48819685 |
| 0.999490048 | 0.05416158 | 0.45630728 | 0.86545623 | -0.0646684 | 0.37488742 |
| 0.999490048 | 0.0549682 | 0.45326732 | 0.86545623 | 0.09349949 | 0.20681542 |
| 0.999490048 | 0.05617446 | 0.4560933 | 0.86545623 | -0.019735 | 0.79245252 |
| 0.999490048 | 0.05783536 | 0.45884727 | 0.86545623 | 0.03861069 | 0.61986667 |
| 0.999490048 | 0.05863984 | 0.45246843 | 0.86545623 | 0.06455203 | 0.40876726 |
| 0.999490048 | 0.05968765 | 0.45455803 | 0.86545623 | 0.09360098 | 0.24470466 |

| | | | | | |
|---|---|---|---|---|---|
| 0.999490048 | 0.06167153 | 0.45522448 | 0.86545623 | -0.0249503 | 0.76148399 |
| 0.999490048 | 0.06232464 | 0.45737111 | 0.86545623 | 0.07557079 | 0.36875961 |
| 0.999490048 | 0.06258802 | 0.45493109 | 0.86545623 | -0.0359244 | 0.66682124 |
| 0.999490048 | 0.06504112 | 0.45839875 | 0.86545623 | -0.0122204 | 0.88860739 |
| 0.999490048 | 0.0667246 | 0.45546428 | 0.86545623 | -0.103596 | 0.25023793 |
| 0.999490048 | 0.07147369 | 0.45716095 | 0.86545623 | 0.10590511 | 0.2737311 |
| 0.999490048 | 0.07301605 | 0.45532065 | 0.86545623 | -0.0286972 | 0.76812743 |
| 0.999490048 | 0.07346242 | 0.45577239 | 0.86545623 | 0.05104632 | 0.60323606 |
| 0.999490048 | 0.07791693 | 0.45784172 | 0.86545623 | 0.12435132 | 0.24004066 |
| 0.999490048 | 0.07838332 | 0.45317433 | 0.86545623 | 0.04147828 | 0.69020432 |
| 0.999490048 | 0.07866449 | 0.45657395 | 0.86545623 | 0.05933923 | 0.57346593 |
| 0.999490048 | 0.09205072 | 0.45756133 | 0.86545623 | 0.08345256 | 0.50021903 |
| 0.999490048 | 0.09229999 | 0.45261384 | 0.86545623 | -0.0861785 | 0.48282717 |
| 0.999490048 | 0.09413092 | 0.45246544 | 0.86545623 | 0.12343781 | 0.32647178 |
| 0.999490048 | 0.10555681 | 0.45490586 | 0.86545623 | 0.04382912 | 0.7552457 |
| 0.999490048 | 0.10885032 | 0.45686765 | 0.86545623 | 0.3028545 | 0.04568706 |
| 0.999490048 | 0.12510425 | 0.45255134 | 0.86545623 | 6.18E-04 | 0.99700027 |
| 0.999490048 | 0.1268057 | 0.45283964 | 0.86545623 | -0.0788338 | 0.63699434 |
| 0.999490048 | 0.14127418 | 0.45560999 | 0.86545623 | 0.26566649 | 0.16643672 |
| 0.999490048 | 0.20970034 | 0.45648172 | 0.86545623 | 0.30847913 | 0.2827979 |
| 0.999490048 | 0.22868343 | 0.45125204 | 0.86545623 | 0.03849113 | 0.89853867 |
| 0.999490048 | -0.0675491 | 0.45944875 | 0.8658084 | -0.0361605 | 0.69094572 |
| 0.972489992 | -0.0503487 | 0.45918138 | 0.8658084 | 0.10411816 | 0.13254724 |
| 0.999490048 | 0.04290115 | 0.45925525 | 0.8658084 | -0.0146411 | 0.79964299 |
| 0.999490048 | 0.06439352 | 0.45943314 | 0.8658084 | 0.03637699 | 0.67485797 |
| 0.999490048 | -0.0813644 | 0.4598854 | 0.86602905 | 0.00417065 | 0.96939439 |
| 0.999490048 | -0.0804752 | 0.4599807 | 0.86602905 | -0.0249421 | 0.8179746 |
| 0.999490048 | -0.0354064 | 0.45994149 | 0.86602905 | -0.0593571 | 0.21990962 |
| 0.999490048 | 0.05899918 | 0.45985449 | 0.86602905 | 0.02621487 | 0.74151018 |
| 0.999490048 | 0.1006054 | 0.46016438 | 0.86602921 | 0.12010289 | 0.37903732 |
| 0.999490048 | 0.15347176 | 0.46018822 | 0.86602921 | 0.00928126 | 0.9641792 |
| 0.999490048 | 0.04892586 | 0.46044113 | 0.86630993 | 0.10579693 | 0.1175006 |
| 0.999490048 | -0.0582579 | 0.46095067 | 0.86668203 | -0.0173277 | 0.82556528 |
| 0.999490048 | -0.0486905 | 0.46115788 | 0.86668203 | -0.0499052 | 0.45017778 |
| 0.999490048 | -0.0375325 | 0.46090385 | 0.86668203 | -0.0664887 | 0.19657837 |
| 0.999490048 | -0.0370116 | 0.46104517 | 0.86668203 | 0.02106724 | 0.67365561 |
| 0.999490048 | 0.13658277 | 0.46110971 | 0.86668203 | 0.14445934 | 0.43596775 |
| 0.999490048 | 0.03912671 | 0.46152878 | 0.86718391 | -0.007147 | 0.89244104 |
| 0.891785516 | -0.0590572 | 0.4623217 | 0.86769751 | 0.14159372 | 0.08556419 |
| 0.999490048 | -0.0414274 | 0.46230496 | 0.86769751 | -0.0227513 | 0.68531367 |
| 0.999490048 | -0.0409313 | 0.46228896 | 0.86769751 | -0.0116904 | 0.83288375 |
| 0.999490048 | -0.0374298 | 0.46202616 | 0.86769751 | 0.02835149 | 0.5765599 |

| | | | | | |
|---|---|---|---|---|---|
| 0.999490048 | 0.04293263 | 0.46201233 | 0.86769751 | -0.0295167 | 0.61205131 |
| 0.999490048 | -0.0450365 | 0.46261189 | 0.86804703 | -0.0023007 | 0.96990359 |
| 0.999490048 | 0.04586913 | 0.46291071 | 0.86841258 | -0.0119791 | 0.84720718 |
| 0.999490048 | -0.1210198 | 0.46317039 | 0.86844092 | -0.0440171 | 0.78734579 |
| 0.999490048 | -0.0608857 | 0.46306724 | 0.86844092 | -0.095993 | 0.25105357 |
| 0.999490048 | -0.044373 | 0.46323783 | 0.86844092 | 0.0420719 | 0.48651236 |
| 0.999490048 | -0.035186 | 0.46345691 | 0.86846167 | -0.0025384 | 0.95758893 |
| 0.988558788 | 0.13155284 | 0.46345421 | 0.86846167 | -0.2615506 | 0.15898617 |
| 0.972489992 | 0.12496562 | 0.46380104 | 0.86891154 | -0.2565218 | 0.13921001 |
| 0.999490048 | -0.0526894 | 0.46412808 | 0.86913422 | -0.1469595 | 0.0485236 |
| 0.999490048 | -0.0263685 | 0.46412521 | 0.86913422 | 8.84E-04 | 0.98030401 |
| 0.999490048 | -0.0308369 | 0.46454879 | 0.869727 | -0.0587409 | 0.16940787 |
| 0.999490048 | 0.05387574 | 0.4649565 | 0.87029517 | 0.05737536 | 0.43679538 |
| 0.999490048 | -0.4672999 | 0.4659145 | 0.87039865 | -0.6574942 | 0.31059442 |
| 0.999490048 | -0.1053529 | 0.46559363 | 0.87039865 | -0.0458293 | 0.74987877 |
| 0.999490048 | -0.0554043 | 0.46605417 | 0.87039865 | 0.02061979 | 0.78520809 |
| 0.999490048 | -0.0441966 | 0.46573999 | 0.87039865 | 0.06329691 | 0.29886587 |
| 0.999490048 | -0.0407742 | 0.46596475 | 0.87039865 | -0.0453251 | 0.41827623 |
| 0.999490048 | -0.0378879 | 0.46550641 | 0.87039865 | -0.0386977 | 0.4561282 |
| 0.999490048 | -0.0348947 | 0.46523606 | 0.87039865 | -0.0402301 | 0.4006672 |
| 0.999490048 | -0.0298947 | 0.46600318 | 0.87039865 | -0.0566241 | 0.17310443 |
| 0.999490048 | 0.05679174 | 0.46586515 | 0.87039865 | 0.17886294 | 0.02799726 |
| 0.999490048 | 0.07523685 | 0.46558193 | 0.87039865 | 0.09201526 | 0.37343309 |
| 0.999490048 | -0.0615899 | 0.46637414 | 0.8704923 | 0.05450882 | 0.51867233 |
| 0.999490048 | 0.03280364 | 0.46641707 | 0.8704923 | 0.04224072 | 0.35000193 |
| 0.999490048 | 0.07904244 | 0.46623428 | 0.8704923 | -0.0128902 | 0.90492142 |
| 0.999490048 | -0.0409765 | 0.46655747 | 0.87055976 | -0.013537 | 0.80902469 |
| 0.999490048 | -0.0595896 | 0.46676297 | 0.87074862 | -0.0622469 | 0.44734939 |
| 0.999490048 | 0.03856895 | 0.46690125 | 0.87081202 | 0.08710837 | 0.1076319 |
| 0.999490048 | -0.0886648 | 0.46732154 | 0.87081786 | -0.0985703 | 0.41964475 |
| 0.987721587 | -0.0705773 | 0.46730593 | 0.87081786 | 0.14253013 | 0.14852771 |
| 0.999490048 | 0.04860213 | 0.46722993 | 0.87081786 | 0.08361238 | 0.21573048 |
| 0.999490048 | 0.10335793 | 0.4672523 | 0.87081786 | -0.0136683 | 0.92301109 |
| 0.999490048 | -0.0492594 | 0.46765687 | 0.87105394 | -0.0388117 | 0.56636942 |
| 0.999490048 | 0.04455984 | 0.46761228 | 0.87105394 | 0.03852808 | 0.52942247 |
| 0.999490048 | -0.1024072 | 0.46799481 | 0.871489 | 0.02181028 | 0.87654128 |
| 0.999490048 | -0.0726757 | 0.46820149 | 0.87167948 | 0.01627479 | 0.87030594 |
| 0.999490048 | -0.1302841 | 0.46904589 | 0.87207202 | 0.10938602 | 0.54262931 |
| 0.999490048 | -0.1134198 | 0.46968393 | 0.87207202 | 0.11514056 | 0.46304336 |
| 0.999490048 | -0.0971161 | 0.47004183 | 0.87207202 | -0.1071564 | 0.42592091 |
| 0.999490048 | -0.0698451 | 0.46949128 | 0.87207202 | 0.05042436 | 0.60062017 |
| 0.999490048 | -0.0603935 | 0.46975706 | 0.87207202 | -0.1989541 | 0.02318339 |

| | | | | | |
|---|---|---|---|---|---|
| 0.999490048 | -0.0561717 | 0.47102333 | 0.87207202 | -0.0034387 | 0.9646183 |
| 0.999490048 | -0.05279 | 0.46948687 | 0.87207202 | -0.0748125 | 0.30787436 |
| 0.999490048 | -0.0509678 | 0.46981039 | 0.87207202 | -0.0131576 | 0.85125689 |
| 0.999490048 | -0.0471855 | 0.47098896 | 0.87207202 | -0.0535349 | 0.41412707 |
| 0.999490048 | -0.0462817 | 0.46960167 | 0.87207202 | -0.10494 | 0.10836788 |
| 0.999490048 | -0.0422009 | 0.47094156 | 0.87207202 | -0.0287679 | 0.62212271 |
| 0.999490048 | -0.0413073 | 0.47044869 | 0.87207202 | -0.0224474 | 0.69382286 |
| 0.999490048 | -0.040349 | 0.47061359 | 0.87207202 | 0.01938948 | 0.72775846 |
| 0.999490048 | -0.0378732 | 0.47012654 | 0.87207202 | 0.06694002 | 0.20668488 |
| 0.999490048 | -0.0305931 | 0.46955286 | 0.87207202 | -0.0198829 | 0.63731475 |
| 0.999490048 | -0.0257775 | 0.46977048 | 0.87207202 | -0.0374807 | 0.2960377 |
| 0.999490048 | 0.02895892 | 0.47046013 | 0.87207202 | -0.0022148 | 0.95574978 |
| 0.999490048 | 0.03785056 | 0.46942198 | 0.87207202 | 0.00684964 | 0.89531119 |
| 0.999490048 | 0.0414829 | 0.47002917 | 0.87207202 | -0.0088501 | 0.87689575 |
| 0.999490048 | 0.0522247 | 0.46945371 | 0.87207202 | 0.03815316 | 0.59627584 |
| 0.999490048 | 0.05455084 | 0.47064439 | 0.87207202 | -0.0066545 | 0.92950806 |
| 0.999490048 | 0.05856704 | 0.4706655 | 0.87207202 | 0.01448651 | 0.85768522 |
| 0.999490048 | 0.06151618 | 0.46935122 | 0.87207202 | 0.08541111 | 0.3174397 |
| 0.999490048 | 0.06664341 | 0.46967534 | 0.87207202 | 0.14220443 | 0.12974685 |
| 0.999490048 | 0.10500962 | 0.47101687 | 0.87207202 | -0.0999766 | 0.49232726 |
| 0.999490048 | 0.03262357 | 0.47144541 | 0.87253141 | -0.0222971 | 0.62161874 |
| 0.999490048 | 0.05179015 | 0.47158494 | 0.87253141 | 0.10298856 | 0.15845794 |
| 0.406717515 | 0.05621707 | 0.47148711 | 0.87253141 | -0.2115955 | 0.01083953 |
| 0.999490048 | -0.0427933 | 0.47177017 | 0.87268076 | -0.0123368 | 0.83487639 |
| 0.999490048 | 0.06523382 | 0.4719006 | 0.87272868 | 0.11593567 | 0.20604262 |
| 0.999490048 | -0.049325 | 0.47234208 | 0.87283765 | -0.108207 | 0.12193767 |
| 0.999490048 | -0.048823 | 0.47237765 | 0.87283765 | 0.02889792 | 0.66953507 |
| 0.999490048 | 0.0394942 | 0.47228827 | 0.87283765 | 0.04260752 | 0.43847987 |
| 0.999490048 | 0.05132418 | 0.47221179 | 0.87283765 | 0.0125552 | 0.85972477 |
| 0.999490048 | 0.15532367 | 0.4727797 | 0.87338728 | -0.1510259 | 0.48472748 |
| 0.999490048 | -0.0421474 | 0.47299603 | 0.87359365 | -0.0138746 | 0.81239858 |
| 0.999490048 | -0.0954686 | 0.47317983 | 0.87373985 | 0.07427515 | 0.57601257 |
| 0.999490048 | 0.27124481 | 0.47329939 | 0.87376738 | 0.09604244 | 0.79866403 |
| 0.999490048 | -0.115732 | 0.47364948 | 0.87415685 | -0.0870375 | 0.58912499 |
| 0.999490048 | 0.0727057 | 0.47371973 | 0.87415685 | -0.0694319 | 0.49366543 |
| 0.999490048 | -0.1164603 | 0.47426329 | 0.87419392 | -0.1578185 | 0.33427264 |
| 0.999490048 | -0.0659275 | 0.47396284 | 0.87419392 | -0.1511409 | 0.10800918 |
| 0.999490048 | -0.0557543 | 0.47422201 | 0.87419392 | 0.05996773 | 0.44184137 |
| 0.999490048 | 0.05077679 | 0.47397517 | 0.87419392 | 0.03361587 | 0.63449351 |
| 0.999490048 | 0.07041575 | 0.47417175 | 0.87419392 | -0.1305225 | 0.19004314 |
| 0.999490048 | 0.05602633 | 0.47451241 | 0.8742058 | 0.09382983 | 0.23529788 |
| 0.999490048 | 0.06270756 | 0.47458382 | 0.8742058 | -0.0808061 | 0.35845499 |

| | | | | | |
|---|---|---|---|---|---|
| 0.999490048 | 0.08448012 | 0.47439918 | 0.8742058 | -0.0030508 | 0.9792835 |
| 0.999490048 | -0.070969 | 0.47484291 | 0.87449014 | 0.05429668 | 0.5837555 |
| 0.999490048 | 0.0344524 | 0.47510112 | 0.87457989 | 0.0206547 | 0.66749377 |
| 0.999490048 | 0.0552224 | 0.47504654 | 0.87457989 | -0.0423573 | 0.58297441 |
| 0.666143899 | 0.04523351 | 0.47555702 | 0.87464783 | -0.1377992 | 0.03669939 |
| 0.987721587 | 0.05085353 | 0.47553552 | 0.87464783 | -0.1012531 | 0.16157745 |
| 0.999490048 | 0.05375114 | 0.47541919 | 0.87464783 | 0.10121672 | 0.18454195 |
| 0.999490048 | 0.07645932 | 0.47552658 | 0.87464783 | 0.09121342 | 0.3956766 |
| 0.999490048 | -0.038825 | 0.47569859 | 0.87471553 | -0.0382409 | 0.48229614 |
| 0.999490048 | 0.03674792 | 0.47617866 | 0.87510862 | -0.0536895 | 0.30067934 |
| 0.999490048 | 0.04780628 | 0.47640733 | 0.87510862 | -0.0703279 | 0.29767612 |
| 0.999490048 | 0.0589954 | 0.47640725 | 0.87510862 | 0.05483926 | 0.50772401 |
| 0.999490048 | 0.06137664 | 0.47654119 | 0.87510862 | -0.0117913 | 0.89068384 |
| 0.999490048 | 0.06255052 | 0.47647302 | 0.87510862 | 0.04130511 | 0.63727152 |
| 0.999490048 | 0.08184844 | 0.47640889 | 0.87510862 | -0.0779148 | 0.49720811 |
| 0.999490048 | -0.1077863 | 0.47734474 | 0.87512251 | -0.0120392 | 0.93642013 |
| 0.999490048 | -0.0829678 | 0.47727263 | 0.87512251 | 0.01801286 | 0.87678165 |
| 0.999490048 | -0.0819842 | 0.47731478 | 0.87512251 | -0.0450823 | 0.69493975 |
| 0.967348048 | -0.0502813 | 0.47732866 | 0.87512251 | 0.1177792 | 0.10343691 |
| 0.999490048 | 0.04144124 | 0.477492 | 0.87512251 | -0.0095152 | 0.86981755 |
| 0.999490048 | 0.05973515 | 0.47747871 | 0.87512251 | 0.14695046 | 0.08817449 |
| 0.999490048 | 0.06682952 | 0.47682499 | 0.87512251 | 0.11735591 | 0.21632802 |
| 0.999490048 | 0.07001494 | 0.47679904 | 0.87512251 | 0.13669877 | 0.17074614 |
| 0.999490048 | 0.09140189 | 0.47715383 | 0.87512251 | -0.0392825 | 0.75772438 |
| 0.999490048 | -0.0294256 | 0.47763463 | 0.87519182 | -0.0099036 | 0.81026711 |
| 0.999490048 | -0.0425838 | 0.47795454 | 0.87539381 | -0.014865 | 0.80350492 |
| 0.999490048 | 0.03207456 | 0.47790452 | 0.87539381 | 0.0425358 | 0.34847095 |
| 0.999490048 | -0.1376832 | 0.47807667 | 0.87542549 | -0.1254732 | 0.51760382 |
| 0.999490048 | -0.0512386 | 0.47841159 | 0.87584669 | -0.0811092 | 0.26540136 |
| 0.999490048 | -0.1557075 | 0.47978786 | 0.87607339 | -0.0526396 | 0.81035767 |
| 0.999490048 | -0.1272692 | 0.48052564 | 0.87607339 | -0.2776605 | 0.14480512 |
| 0.972489992 | -0.107963 | 0.47985391 | 0.87607339 | -0.474853 | 0.00810459 |
| 0.999490048 | -0.0709211 | 0.47966779 | 0.87607339 | -0.037993 | 0.7039282 |
| 0.999490048 | -0.0693098 | 0.47874227 | 0.87607339 | -0.2487727 | 0.01607473 |
| 0.999490048 | -0.0443838 | 0.48063078 | 0.87607339 | 0.04295938 | 0.49468992 |
| 0.999490048 | -0.037855 | 0.47914475 | 0.87607339 | 0.02366988 | 0.6571438 |
| 0.999490048 | -0.0373791 | 0.47984449 | 0.87607339 | 0.0129507 | 0.80576812 |
| 0.999490048 | -0.0372368 | 0.48062655 | 0.87607339 | -0.0411274 | 0.43649044 |
| 0.999490048 | -0.033747 | 0.48035191 | 0.87607339 | -0.0330103 | 0.48989258 |
| 0.999490048 | -0.0303754 | 0.47974859 | 0.87607339 | -0.0220138 | 0.60767553 |
| 0.999490048 | 0.03296058 | 0.47973 | 0.87607339 | -0.0198539 | 0.66932457 |
| 0.999490048 | 0.03644863 | 0.48065439 | 0.87607339 | -0.0261689 | 0.61174903 |

| | | | | | |
|---|---|---|---|---|---|
| 0.999490048 | 0.03708923 | 0.4806402 | 0.87607339 | 0.03925218 | 0.45569404 |
| 0.999490048 | 0.03992173 | 0.47985727 | 0.87607339 | 0.0558268 | 0.32544277 |
| 0.999490048 | 0.04359258 | 0.4805526 | 0.87607339 | 0.07032097 | 0.258908 |
| 0.999490048 | 0.04563222 | 0.4802154 | 0.87607339 | 0.00377305 | 0.95321932 |
| 0.999490048 | 0.05154024 | 0.47945545 | 0.87607339 | 0.03226456 | 0.6569992 |
| 0.999490048 | 0.05941122 | 0.48073872 | 0.87607339 | 0.10796817 | 0.20514418 |
| 0.999490048 | 0.07180752 | 0.47890736 | 0.87607339 | -0.0124695 | 0.90166134 |
| 0.999490048 | 0.09592815 | 0.47987916 | 0.87607339 | 0.10003169 | 0.46148942 |
| 0.999490048 | -0.1475711 | 0.48128418 | 0.87626725 | 0.0307409 | 0.88157213 |
| 0.999490048 | -0.0417185 | 0.48127679 | 0.87626725 | -0.0799442 | 0.18283434 |
| 0.999490048 | -0.0380093 | 0.48103143 | 0.87626725 | 0.04149139 | 0.44221066 |
| 0.999490048 | 0.03821729 | 0.48113358 | 0.87626725 | 0.02020489 | 0.70855614 |
| 0.999490048 | 0.05204434 | 0.48136981 | 0.87626725 | 0.01263244 | 0.86366425 |
| 0.999490048 | 0.08493183 | 0.48184263 | 0.87693678 | 0.09781358 | 0.41868181 |
| 0.999490048 | -0.0328652 | 0.48221035 | 0.87741479 | -0.0640965 | 0.17635553 |
| 0.999490048 | -0.050691 | 0.48244596 | 0.87762335 | -0.0501278 | 0.48727693 |
| 0.999490048 | -0.041788 | 0.48253519 | 0.87762335 | -0.0794032 | 0.18765287 |
| 0.999490048 | -0.0755205 | 0.48276897 | 0.87785734 | -0.0361403 | 0.7359926 |
| 0.999490048 | -0.0496127 | 0.48299939 | 0.87787424 | -0.0740851 | 0.29777124 |
| 0.999490048 | 0.03779311 | 0.48298315 | 0.87787424 | -0.0031924 | 0.95251363 |
| 0.999490048 | 0.09936719 | 0.48309367 | 0.87787424 | 0.02168118 | 0.87780406 |
| 0.999490048 | -0.1690443 | 0.48904044 | 0.87852036 | 0.12913298 | 0.59481413 |
| 0.999490048 | -0.1007803 | 0.48569026 | 0.87852036 | -0.0474174 | 0.74069909 |
| 0.999490048 | -0.0930235 | 0.49119884 | 0.87852036 | -0.0440817 | 0.743341 |
| 0.999490048 | -0.0917955 | 0.48902736 | 0.87852036 | -0.1118198 | 0.40044464 |
| 0.999490048 | -0.0814884 | 0.48779606 | 0.87852036 | -0.0972033 | 0.40884084 |
| 0.999490048 | -0.0809276 | 0.48582042 | 0.87852036 | -0.1313229 | 0.26180675 |
| 0.999490048 | -0.0597733 | 0.48592441 | 0.87852036 | -0.208654 | 0.02070094 |
| 0.999490048 | -0.0596031 | 0.48885341 | 0.87852036 | 0.0717931 | 0.40547209 |
| 0.999490048 | -0.0574121 | 0.48844676 | 0.87852036 | -0.0145402 | 0.86009509 |
| 0.999490048 | -0.0571127 | 0.48893662 | 0.87852036 | -0.1174688 | 0.16094538 |
| 0.999490048 | -0.0553703 | 0.4868256 | 0.87852036 | -0.0994683 | 0.21647116 |
| 0.999490048 | -0.0545918 | 0.49059607 | 0.87852036 | -0.1001178 | 0.21118362 |
| 0.999490048 | -0.0540519 | 0.48612515 | 0.87852036 | -0.066518 | 0.39256176 |
| 0.999490048 | -0.0535853 | 0.48848784 | 0.87852036 | -0.1370519 | 0.08417675 |
| 0.999490048 | -0.0535689 | 0.48700595 | 0.87852036 | -1.87E-04 | 0.99805286 |
| 0.999490048 | -0.0532628 | 0.48424433 | 0.87852036 | -0.0143949 | 0.84937477 |
| 0.999490048 | -0.0472799 | 0.48988208 | 0.87852036 | -0.0363045 | 0.59524331 |
| 0.999490048 | -0.0451707 | 0.48643878 | 0.87852036 | -0.0702453 | 0.28238741 |
| 0.999490048 | -0.0426533 | 0.48653415 | 0.87852036 | -0.1008936 | 0.10721393 |
| 0.999490048 | -0.0421577 | 0.48877964 | 0.87852036 | -0.0744008 | 0.22645865 |
| 0.999490048 | -0.039429 | 0.48590751 | 0.87852036 | 0.00490471 | 0.9305903 |

| | | | | | |
|---|---|---|---|---|---|
| 0.999490048 | -0.0390393 | 0.49055796 | 0.87852036 | -0.0273309 | 0.62850212 |
| 0.999490048 | -0.0380882 | 0.48782124 | 0.87852036 | 0.04892817 | 0.37430059 |
| 0.999490048 | -0.0366135 | 0.48467785 | 0.87852036 | -0.0271807 | 0.60312337 |
| 0.999490048 | -0.0364713 | 0.48912913 | 0.87852036 | 0.0464357 | 0.37991052 |
| 0.999490048 | -0.0351481 | 0.48430467 | 0.87852036 | -0.013594 | 0.78589795 |
| 0.999490048 | -0.034694 | 0.48847513 | 0.87852036 | -0.042672 | 0.39535959 |
| 0.999490048 | -0.0341544 | 0.49091936 | 0.87852036 | -0.0043457 | 0.92982805 |
| 0.999490048 | -0.0337158 | 0.48582964 | 0.87852036 | -0.0587576 | 0.22905747 |
| 0.999490048 | -0.0327329 | 0.48771112 | 0.87852036 | -0.0651589 | 0.17313515 |
| 0.999490048 | -0.0280612 | 0.48621475 | 0.87852036 | -0.055566 | 0.17387968 |
| 0.999490048 | -0.0270076 | 0.49054324 | 0.87852036 | -0.0376165 | 0.33907244 |
| 0.999490048 | -0.0255614 | 0.48902642 | 0.87852036 | -0.0662431 | 0.08073675 |
| 0.999490048 | 0.0296289 | 0.49123492 | 0.87852036 | -0.0016642 | 0.96901199 |
| 0.999490048 | 0.03053122 | 0.48762749 | 0.87852036 | -0.0501546 | 0.25805904 |
| 0.999490048 | 0.03250634 | 0.49013196 | 0.87852036 | 0.03341523 | 0.47819905 |
| 0.999490048 | 0.03270917 | 0.49075022 | 0.87852036 | 0.03691199 | 0.43736565 |
| 0.999490048 | 0.03393423 | 0.48772026 | 0.87852036 | 0.06842295 | 0.16786073 |
| 0.999490048 | 0.03405823 | 0.48656204 | 0.87852036 | 0.0928001 | 0.06572114 |
| 0.999490048 | 0.03605975 | 0.48668291 | 0.87852036 | 0.02276481 | 0.65962855 |
| 0.999490048 | 0.03674709 | 0.4894706 | 0.87852036 | -0.0272454 | 0.6075651 |
| 0.999490048 | 0.03804973 | 0.48411233 | 0.87852036 | 0.03416481 | 0.52943866 |
| 0.999490048 | 0.03867036 | 0.48392138 | 0.87852036 | -0.0225809 | 0.68173716 |
| 0.999490048 | 0.04029829 | 0.4845352 | 0.87852036 | 0.0255293 | 0.65693506 |
| 0.999490048 | 0.04062026 | 0.4912232 | 0.87852036 | 0.02101725 | 0.72078011 |
| 0.999490048 | 0.04080959 | 0.49024883 | 0.87852036 | 0.10325031 | 0.08860629 |
| 0.999490048 | 0.04081449 | 0.48840333 | 0.87852036 | 0.00679708 | 0.90770722 |
| 0.999490048 | 0.04150898 | 0.48904498 | 0.87852036 | -0.0102531 | 0.86369988 |
| 0.999490048 | 0.04213281 | 0.49084206 | 0.87852036 | 0.00830847 | 0.89143248 |
| 0.999490048 | 0.04249935 | 0.48749165 | 0.87852036 | -0.0030238 | 0.96040956 |
| 0.999490048 | 0.04269107 | 0.49026312 | 0.87852036 | 0.0947287 | 0.1327835 |
| 0.999490048 | 0.0455548 | 0.48823275 | 0.87852036 | 0.05947911 | 0.36709544 |
| 0.999490048 | 0.04628977 | 0.48891539 | 0.87852036 | 0.04941498 | 0.46035547 |
| 0.999490048 | 0.05192668 | 0.48495713 | 0.87852036 | 0.03424076 | 0.64423917 |
| 0.999490048 | 0.05856338 | 0.48635706 | 0.87852036 | -0.0291948 | 0.72761158 |
| 0.999490048 | 0.06004107 | 0.48977583 | 0.87852036 | -0.0431483 | 0.61883605 |
| 0.999490048 | 0.06155282 | 0.48934618 | 0.87852036 | 0.0912323 | 0.30825663 |
| 0.999490048 | 0.06183149 | 0.48860339 | 0.87852036 | -0.0355028 | 0.68994076 |
| 0.999490048 | 0.06453284 | 0.48828753 | 0.87852036 | 0.00458161 | 0.96056719 |
| 0.999490048 | 0.06722505 | 0.48596425 | 0.87852036 | 0.07110922 | 0.46138266 |
| 0.999490048 | 0.06959168 | 0.48995813 | 0.87852036 | 0.00730548 | 0.9419495 |
| 0.999490048 | 0.07039409 | 0.48912603 | 0.87852036 | 0.06216173 | 0.54089811 |
| 0.999490048 | 0.07115095 | 0.49113157 | 0.87852036 | -0.0664801 | 0.519769 |

| | | | | | |
|---|---|---|---|---|---|
| 0.999490048 | 0.07525317 | 0.48525769 | 0.87852036 | 0.0082835 | 0.93846968 |
| 0.999490048 | 0.07543593 | 0.49033586 | 0.87852036 | 0.04475518 | 0.6814314 |
| 0.999490048 | 0.07635026 | 0.48519327 | 0.87852036 | -0.008453 | 0.93810428 |
| 0.999490048 | 0.07982364 | 0.48719115 | 0.87852036 | 0.09043402 | 0.43183411 |
| 0.999490048 | 0.09011935 | 0.49069698 | 0.87852036 | -0.1612493 | 0.22227194 |
| 0.999490048 | 0.09066441 | 0.48667701 | 0.87852036 | 0.08416988 | 0.51814338 |
| 0.999490048 | 0.10249824 | 0.49004456 | 0.87852036 | -0.0081737 | 0.95589337 |
| 0.999490048 | 0.10607768 | 0.48555769 | 0.87852036 | -0.0030433 | 0.98395823 |
| 0.999490048 | 0.12437935 | 0.48649425 | 0.87852036 | 0.07035024 | 0.69077667 |
| 0.999490048 | 0.15546656 | 0.48956785 | 0.87852036 | 0.28733535 | 0.2067077 |
| 0.999490048 | 0.16626848 | 0.48460954 | 0.87852036 | 0.12684139 | 0.59149649 |
| 0.999490048 | -0.0946105 | 0.4928615 | 0.87917481 | 0.01330914 | 0.92279004 |
| 0.999490048 | -0.0709399 | 0.49195818 | 0.87917481 | -0.1795004 | 0.08977676 |
| 0.999490048 | -0.0661236 | 0.49479747 | 0.87917481 | -0.1420444 | 0.14913395 |
| 0.999490048 | -0.0639952 | 0.4944889 | 0.87917481 | -0.0607239 | 0.51661852 |
| 0.999490048 | -0.0607502 | 0.49485271 | 0.87917481 | -0.0233363 | 0.79234199 |
| 0.999490048 | -0.0573019 | 0.4947231 | 0.87917481 | -0.0045444 | 0.95661105 |
| 0.999490048 | -0.0555865 | 0.49461238 | 0.87917481 | -0.0470039 | 0.56305449 |
| 0.999490048 | -0.0531685 | 0.49387842 | 0.87917481 | -0.0802845 | 0.30445177 |
| 0.999490048 | -0.0529733 | 0.4945725 | 0.87917481 | -0.0652316 | 0.40141785 |
| 0.999490048 | -0.0476913 | 0.49269239 | 0.87917481 | -0.0162086 | 0.81487487 |
| 0.999490048 | -0.0370182 | 0.49280809 | 0.87917481 | -0.0627735 | 0.24894614 |
| 0.999490048 | -0.0330668 | 0.49397756 | 0.87917481 | -0.0277797 | 0.5650025 |
| 0.999490048 | 0.0323611 | 0.49408286 | 0.87917481 | -0.0066986 | 0.88692887 |
| 0.999490048 | 0.03488558 | 0.4919424 | 0.87917481 | 0.02042821 | 0.68644418 |
| 0.999490048 | 0.03847652 | 0.494668 | 0.87917481 | 0.00484014 | 0.93122344 |
| 0.999490048 | 0.04121828 | 0.49252142 | 0.87917481 | -0.0048988 | 0.93468512 |
| 0.999490048 | 0.04156882 | 0.49382688 | 0.87917481 | 0.07625644 | 0.21439685 |
| 0.999490048 | 0.04165001 | 0.49444452 | 0.87917481 | 0.03758597 | 0.53715204 |
| 0.999490048 | 0.04213257 | 0.49425666 | 0.87917481 | 0.04754207 | 0.44110369 |
| 0.999490048 | 0.04547673 | 0.49278204 | 0.87917481 | 0.12239534 | 0.0726933 |
| 0.999490048 | 0.04590534 | 0.49205082 | 0.87917481 | 0.03724873 | 0.57657395 |
| 0.999490048 | 0.04590875 | 0.49396257 | 0.87917481 | 0.04389774 | 0.51290538 |
| 0.999490048 | 0.04620724 | 0.49447598 | 0.87917481 | 0.03041029 | 0.65210481 |
| 0.982702077 | 0.04685744 | 0.49286621 | 0.87917481 | -0.1052161 | 0.13063885 |
| 0.999490048 | 0.04950284 | 0.49294092 | 0.87917481 | 0.03659093 | 0.61152815 |
| 0.999490048 | 0.05129633 | 0.49402927 | 0.87917481 | 0.04843007 | 0.5182602 |
| 0.999490048 | 0.06258908 | 0.49486486 | 0.87917481 | 0.04468558 | 0.62522287 |
| 0.999490048 | 0.06594853 | 0.49177871 | 0.87917481 | 0.02130194 | 0.82352315 |
| 0.999490048 | 0.06920438 | 0.49402011 | 0.87917481 | 0.10382671 | 0.30766387 |
| 0.999490048 | 0.10762247 | 0.49435344 | 0.87917481 | 0.19390074 | 0.22300408 |
| 0.999490048 | 0.15141051 | 0.49474482 | 0.87917481 | -0.12036 | 0.58664819 |

| | | | | | |
|---|---|---|---|---|---|
| 0.999490048 | -0.1210232 | 0.4952599 | 0.87934217 | 0.11823354 | 0.50516232 |
| 0.999490048 | -0.0686562 | 0.495275 | 0.87934217 | 0.03552738 | 0.723264 |
| 0.999490048 | -0.0375487 | 0.4952442 | 0.87934217 | -0.0445403 | 0.41940855 |
| 0.972489992 | -0.1978097 | 0.49711029 | 0.87944266 | 0.54274022 | 0.08636726 |
| 0.999490048 | -0.0786736 | 0.49617579 | 0.87944266 | 0.04040449 | 0.72581247 |
| 0.999490048 | -0.0639546 | 0.49644094 | 0.87944266 | 0.01712838 | 0.85484634 |
| 0.999490048 | -0.0514061 | 0.4974276 | 0.87944266 | -0.0453899 | 0.54867765 |
| 0.999490048 | -0.0500425 | 0.4960221 | 0.87944266 | -0.0435864 | 0.55277721 |
| 0.999490048 | -0.0460448 | 0.49641448 | 0.87944266 | -0.0928192 | 0.17633735 |
| 0.999490048 | -0.0349666 | 0.49717416 | 0.87944266 | -0.032795 | 0.52403828 |
| 0.999490048 | -0.0346326 | 0.49699728 | 0.87944266 | -0.0655231 | 0.20405808 |
| 0.999490048 | -0.0319698 | 0.49741233 | 0.87944266 | 0.00462015 | 0.92152548 |
| 0.999490048 | -0.0289831 | 0.49743036 | 0.87944266 | -0.0290796 | 0.49601269 |
| 0.999490048 | 0.03509174 | 0.49743805 | 0.87944266 | -0.0316345 | 0.5403996 |
| 0.999490048 | 0.0356531 | 0.4966636 | 0.87944266 | 0.02341868 | 0.65438625 |
| 0.999490048 | 0.03617254 | 0.49722279 | 0.87944266 | 0.00971776 | 0.85466592 |
| 0.999490048 | 0.04853982 | 0.49578373 | 0.87944266 | -0.0497203 | 0.48546244 |
| 0.999490048 | 0.05774634 | 0.49664271 | 0.87944266 | 0.02420092 | 0.77488634 |
| 0.999490048 | 0.06522678 | 0.49596894 | 0.87944266 | -0.1074628 | 0.26574859 |
| 0.999490048 | 0.06731016 | 0.4969942 | 0.87944266 | 0.11665169 | 0.24344674 |
| 0.999490048 | 0.0754205 | 0.49638735 | 0.87944266 | 0.11801731 | 0.29038242 |
| 0.999490048 | 0.07642434 | 0.49613093 | 0.87944266 | -0.0427256 | 0.70267304 |
| 0.999490048 | 0.07706592 | 0.49566701 | 0.87944266 | 0.22442369 | 0.05490217 |
| 0.999490048 | -0.1079579 | 0.49770539 | 0.87964382 | -0.1228307 | 0.44101712 |
| 0.999490048 | -0.0436806 | 0.49786787 | 0.87964382 | 0.01275091 | 0.84248528 |
| 0.999490048 | 0.12509559 | 0.49781865 | 0.87964382 | 0.09554159 | 0.60301942 |
| 0.999490048 | 0.25762293 | 0.49825283 | 0.88013774 | -0.2630084 | 0.48958128 |
| 0.999490048 | -0.1574029 | 0.49966148 | 0.88018368 | -0.1209507 | 0.60329114 |
| 0.999490048 | -0.0657617 | 0.49896349 | 0.88018368 | -0.0418349 | 0.66624773 |
| 0.999490048 | -0.0560268 | 0.49851827 | 0.88018368 | -0.0371928 | 0.65237544 |
| 0.999490048 | -0.0511428 | 0.49940205 | 0.88018368 | -0.0455191 | 0.54735129 |
| 0.999490048 | -0.0446648 | 0.49940862 | 0.88018368 | -0.0410707 | 0.53426355 |
| 0.999490048 | -0.0362096 | 0.49960969 | 0.88018368 | 0.0169645 | 0.75091617 |
| 0.999490048 | -0.0289244 | 0.4985502 | 0.88018368 | -0.0495325 | 0.25061989 |
| 0.999490048 | -0.0288651 | 0.49970092 | 0.88018368 | -0.0738741 | 0.09183523 |
| 0.999490048 | -0.0284579 | 0.49946036 | 0.88018368 | -0.0538025 | 0.20691122 |
| 0.999490048 | 0.02659733 | 0.49959699 | 0.88018368 | -0.0031932 | 0.93510606 |
| 0.999490048 | 0.03321882 | 0.49888869 | 0.88018368 | -0.0267965 | 0.58480636 |
| 0.999490048 | 0.04011822 | 0.49868938 | 0.88018368 | 0.00903715 | 0.87833985 |
| 0.999490048 | 0.0555107 | 0.49975459 | 0.88018368 | 0.0703245 | 0.39386847 |
| 0.999490048 | 0.0619624 | 0.49884804 | 0.88018368 | 0.07203757 | 0.43245062 |
| 0.999490048 | -0.1268184 | 0.50108619 | 0.8805958 | -0.1885861 | 0.3230543 |

| | | | | | |
|---|---|---|---|---|---|
| 0.999490048 | -0.1070255 | 0.50450183 | 0.8805958 | -0.2558589 | 0.11797161 |
| 0.999490048 | -0.0997678 | 0.503042 | 0.8805958 | 0.08691542 | 0.55824489 |
| 0.999490048 | -0.0958447 | 0.50067364 | 0.8805958 | 0.04512274 | 0.75035034 |
| 0.999490048 | -0.0847294 | 0.5041422 | 0.8805958 | -0.0217764 | 0.863119 |
| 0.999490048 | -0.0749742 | 0.5036386 | 0.8805958 | -0.0687866 | 0.5392029 |
| 0.999490048 | -0.0536757 | 0.50175288 | 0.8805958 | 0.04145499 | 0.60323835 |
| 0.999490048 | -0.0505729 | 0.50094676 | 0.8805958 | -0.1601318 | 0.04032095 |
| 0.999490048 | -0.0456807 | 0.50322094 | 0.8805958 | -0.0441287 | 0.51770909 |
| 0.999490048 | -0.0426728 | 0.50473962 | 0.8805958 | -0.0551719 | 0.38986606 |
| 0.999490048 | -0.0376132 | 0.50358209 | 0.8805958 | -0.0042063 | 0.94010778 |
| 0.999490048 | -0.0371126 | 0.5028134 | 0.8805958 | -0.0147502 | 0.78921038 |
| 0.999490048 | -0.0357048 | 0.50370419 | 0.8805958 | 0.00112449 | 0.98312219 |
| 0.999490048 | -0.0327185 | 0.5010474 | 0.8805958 | 0.01729347 | 0.72123617 |
| 0.999490048 | -0.0318565 | 0.50185636 | 0.8805958 | -0.0399666 | 0.40070519 |
| 0.999490048 | -0.0300364 | 0.50484371 | 0.8805958 | -0.0145633 | 0.74559116 |
| 0.999490048 | 0.03006972 | 0.50134265 | 0.8805958 | 0.03947176 | 0.37900251 |
| 0.999490048 | 0.03344466 | 0.50271768 | 0.8805958 | -0.0025143 | 0.95963356 |
| 0.999490048 | 0.03362205 | 0.50181953 | 0.8805958 | 0.04154941 | 0.40766346 |
| 0.999490048 | 0.03557772 | 0.50484576 | 0.8805958 | 0.05354757 | 0.318225 |
| 0.999490048 | 0.03559523 | 0.50468285 | 0.8805958 | 0.02249204 | 0.67251862 |
| 0.999490048 | 0.03631316 | 0.50305301 | 0.8805958 | 0.05977485 | 0.2739447 |
| 0.999490048 | 0.03738717 | 0.50213681 | 0.8805958 | 0.06771483 | 0.22887359 |
| 0.999490048 | 0.03753587 | 0.50345927 | 0.8805958 | 0.00218013 | 0.96886198 |
| 0.999490048 | 0.03831264 | 0.50494523 | 0.8805958 | -0.012046 | 0.83329386 |
| 0.999490048 | 0.03864846 | 0.50336061 | 0.8805958 | 0.07183152 | 0.21855929 |
| 0.999490048 | 0.03922068 | 0.50278226 | 0.8805958 | 0.04271999 | 0.46582166 |
| 0.999490048 | 0.03939457 | 0.50491214 | 0.8805958 | -0.0184146 | 0.75445071 |
| 0.999490048 | 0.04149155 | 0.50214462 | 0.8805958 | 0.10725079 | 0.09055447 |
| 0.999490048 | 0.04382677 | 0.50462923 | 0.8805958 | 0.05829122 | 0.37647949 |
| 0.999490048 | 0.04704289 | 0.50434006 | 0.8805958 | 0.05311575 | 0.45148446 |
| 0.999490048 | 0.04728346 | 0.50048439 | 0.8805958 | 0.04575341 | 0.51432635 |
| 0.999490048 | 0.04742939 | 0.50374739 | 0.8805958 | -0.0256972 | 0.71630053 |
| 0.999490048 | 0.04863194 | 0.50091191 | 0.8805958 | 0.04460784 | 0.53670098 |
| 0.999490048 | 0.05002552 | 0.50444355 | 0.8805958 | 0.09224107 | 0.22326604 |
| 0.999490048 | 0.05136474 | 0.5033488 | 0.8805958 | 0.0344143 | 0.6530701 |
| 0.999490048 | 0.05180709 | 0.50132054 | 0.8805958 | 0.04051029 | 0.59838664 |
| 0.999490048 | 0.05503901 | 0.50211863 | 0.8805958 | 0.10609152 | 0.20117359 |
| 0.999490048 | 0.06151379 | 0.50490858 | 0.8805958 | -0.0029838 | 0.97408197 |
| 0.999490048 | 0.06448133 | 0.50127155 | 0.8805958 | -0.0107508 | 0.9103297 |
| 0.999490048 | 0.07480924 | 0.50016576 | 0.8805958 | 0.21780391 | 0.05736804 |
| 0.999490048 | 0.07676947 | 0.50467127 | 0.8805958 | 0.24134686 | 0.04332648 |
| 0.999490048 | 0.08952445 | 0.50099913 | 0.8805958 | 0.15049023 | 0.26188733 |

| | | | | | |
|---|---|---|---|---|---|
| 0.999490048 | 0.09453687 | 0.50471029 | 0.8805958 | -0.0752583 | 0.59477348 |
| 0.999490048 | 0.10622093 | 0.50055973 | 0.8805958 | -0.0974406 | 0.53632801 |
| 0.999490048 | 0.1928918 | 0.5039106 | 0.8805958 | 0.01164635 | 0.96734337 |
| 0.999490048 | 0.22069149 | 0.50323085 | 0.8805958 | 0.14244873 | 0.66485073 |
| 0.999490048 | -0.1047366 | 0.50536987 | 0.88115231 | -0.1184341 | 0.45190736 |
| 0.999490048 | -0.0996492 | 0.50559794 | 0.88126692 | -0.1090409 | 0.46672985 |
| 0.999490048 | -0.0334807 | 0.50595947 | 0.88126692 | 4.56E-04 | 0.9927318 |
| 0.999490048 | -0.0306369 | 0.50596331 | 0.88126692 | -0.0130586 | 0.77599242 |
| 0.999490048 | 0.03973709 | 0.50593504 | 0.88126692 | 0.00111781 | 0.9850042 |
| 0.999490048 | 0.07708937 | 0.50592912 | 0.88126692 | 0.04368128 | 0.70538377 |
| 0.999490048 | -0.0455124 | 0.50661748 | 0.88203836 | 0.00659968 | 0.92294572 |
| 0.999490048 | 0.03985219 | 0.50659098 | 0.88203836 | 0.02904777 | 0.62760187 |
| 0.999490048 | 0.03973589 | 0.5067767 | 0.88213164 | 0.03331213 | 0.57733727 |
| 0.999490048 | 0.04971761 | 0.50696149 | 0.88226937 | 0.0924433 | 0.22219129 |
| 0.999490048 | -0.0970389 | 0.50763074 | 0.8823307 | -0.0268822 | 0.85378457 |
| 0.961663458 | -0.0785175 | 0.50743944 | 0.8823307 | -0.3622507 | 0.00469633 |
| 0.999490048 | -0.046211 | 0.50753113 | 0.8823307 | -0.120668 | 0.09135182 |
| 0.999490048 | -0.0322631 | 0.50757435 | 0.8823307 | 0.05122322 | 0.29599403 |
| 0.999490048 | 0.04135493 | 0.50753014 | 0.8823307 | 0.04720986 | 0.44994856 |
| 0.999490048 | 0.04195136 | 0.50712692 | 0.8823307 | 0.04529285 | 0.47422075 |
| 0.999490048 | 0.04266582 | 0.50794624 | 0.88269534 | -0.0078719 | 0.90236721 |
| 0.999490048 | 0.03820865 | 0.50811079 | 0.88279757 | 0.00897341 | 0.87597491 |
| 0.999490048 | -0.08596 | 0.51046183 | 0.88314033 | 0.07341587 | 0.57360969 |
| 0.999490048 | -0.078022 | 0.50900799 | 0.88314033 | -0.0898446 | 0.4476329 |
| 0.999490048 | -0.077757 | 0.50900906 | 0.88314033 | 0.00801853 | 0.94546829 |
| 0.999490048 | -0.0540198 | 0.51075159 | 0.88314033 | -0.0125199 | 0.87835365 |
| 0.999490048 | -0.0500004 | 0.51043656 | 0.88314033 | -0.0189019 | 0.80278663 |
| 0.999490048 | -0.0411508 | 0.51034514 | 0.88314033 | -0.1651933 | 0.0127817 |
| 0.999490048 | -0.0403191 | 0.5111057 | 0.88314033 | -0.0337025 | 0.58231947 |
| 0.999490048 | -0.0394706 | 0.50894627 | 0.88314033 | -0.0380617 | 0.52406042 |
| 0.999490048 | -0.0365163 | 0.50966495 | 0.88314033 | -0.0284909 | 0.60632098 |
| 0.999490048 | -0.0297994 | 0.50981199 | 0.88314033 | -0.0563508 | 0.21767711 |
| 0.999490048 | -0.0297106 | 0.50964775 | 0.88314033 | -0.0068608 | 0.87847561 |
| 0.999490048 | 0.02790391 | 0.50991838 | 0.88314033 | 0.02446312 | 0.56306595 |
| 0.999490048 | 0.03142911 | 0.50980753 | 0.88314033 | 0.01592708 | 0.73753639 |
| 0.999490048 | 0.03410876 | 0.51097128 | 0.88314033 | 0.0113424 | 0.82631019 |
| 0.999490048 | 0.03438487 | 0.51116374 | 0.88314033 | 0.12465729 | 0.02326706 |
| 0.999490048 | 0.03439784 | 0.50870203 | 0.88314033 | -0.0263272 | 0.61234507 |
| 0.999490048 | 0.03888756 | 0.51113083 | 0.88314033 | -0.0147267 | 0.80276714 |
| 0.999490048 | 0.03893486 | 0.50958902 | 0.88314033 | -0.0387197 | 0.51191048 |
| 0.999490048 | 0.03913305 | 0.51103693 | 0.88314033 | 0.01911627 | 0.74736253 |
| 0.999490048 | 0.04320456 | 0.50898593 | 0.88314033 | -7.31E-04 | 0.99104099 |

| | | | | | |
|---|---|---|---|---|---|
| 0.999490048 | 0.04354295 | 0.51028271 | 0.88314033 | 0.03917913 | 0.55324401 |
| 0.999490048 | 0.04447547 | 0.50962415 | 0.88314033 | 0.03165957 | 0.63805298 |
| 0.999490048 | 0.04691686 | 0.51047169 | 0.88314033 | 0.00264422 | 0.97028325 |
| 0.999490048 | 0.0545207 | 0.50964863 | 0.88314033 | 0.04674259 | 0.57141349 |
| 0.999490048 | 0.05558732 | 0.50983596 | 0.88314033 | 0.08915184 | 0.29375778 |
| 0.999490048 | 0.07624426 | 0.510727 | 0.88314033 | -0.1243066 | 0.28701432 |
| 0.999490048 | 0.08829579 | 0.50887598 | 0.88314033 | -0.0842338 | 0.52840192 |
| 0.999490048 | 0.03628927 | 0.51128578 | 0.88316844 | 0.00290872 | 0.95782997 |
| 0.999490048 | -0.0359631 | 0.51156488 | 0.88346779 | -0.0433032 | 0.43020481 |
| 0.999490048 | -0.1118003 | 0.51262695 | 0.88347472 | 0.09681722 | 0.57029451 |
| 0.999490048 | -0.0372858 | 0.51229019 | 0.88347472 | 0.01546917 | 0.7849759 |
| 0.999490048 | -0.0335602 | 0.51184684 | 0.88347472 | 0.00937002 | 0.8541058 |
| 0.999490048 | 0.02468717 | 0.51236019 | 0.88347472 | 0.04537144 | 0.23325478 |
| 0.999490048 | 0.03369158 | 0.51256162 | 0.88347472 | 0.01552878 | 0.76199574 |
| 0.999490048 | 0.04190109 | 0.51239352 | 0.88347472 | 0.06221282 | 0.3332007 |
| 0.999490048 | 0.04224644 | 0.51180993 | 0.88347472 | -0.0108633 | 0.86549967 |
| 0.999490048 | 0.04493397 | 0.51201733 | 0.88347472 | 0.02636826 | 0.69958824 |
| 0.999490048 | 0.06468669 | 0.51243582 | 0.88347472 | 0.04971863 | 0.61400299 |
| 0.999490048 | 0.07701706 | 0.51227813 | 0.88347472 | -0.0709031 | 0.54606191 |
| 0.999490048 | -0.0434828 | 0.51295623 | 0.88358513 | -0.0476118 | 0.47414302 |
| 0.999490048 | -0.0365822 | 0.51297043 | 0.88358513 | -0.0123756 | 0.82418111 |
| 0.999490048 | -0.0255894 | 0.51300847 | 0.88358513 | -0.0413012 | 0.29434848 |
| 0.999490048 | -0.040401 | 0.51402643 | 0.88399416 | -0.0278038 | 0.65262325 |
| 0.999490048 | -0.0282581 | 0.51383461 | 0.88399416 | -0.0289429 | 0.50377567 |
| 0.999490048 | -0.0261039 | 0.51414381 | 0.88399416 | 0.00322391 | 0.93550898 |
| 0.999490048 | 0.02839068 | 0.51419876 | 0.88399416 | -0.0011484 | 0.97885803 |
| 0.999490048 | 0.03200078 | 0.51385316 | 0.88399416 | 0.00921651 | 0.85027184 |
| 0.999490048 | 0.03338967 | 0.51381347 | 0.88399416 | 0.00310637 | 0.95135664 |
| 0.999490048 | 0.03818657 | 0.51341803 | 0.88399416 | 0.04665567 | 0.42557685 |
| 0.999490048 | 0.040499 | 0.51362278 | 0.88399416 | -0.039707 | 0.52182222 |
| 0.999490048 | 0.11473853 | 0.5139212 | 0.88399416 | -0.0248314 | 0.88720087 |
| 0.999490048 | -0.0409933 | 0.51456906 | 0.8840847 | 0.02110078 | 0.73646759 |
| 0.999490048 | -0.0367403 | 0.51456198 | 0.8840847 | -0.0096791 | 0.86313175 |
| 0.999490048 | 0.04539037 | 0.51446248 | 0.8840847 | -0.0042324 | 0.95132266 |
| 0.999490048 | 0.05220258 | 0.5147455 | 0.88420591 | 0.07052638 | 0.38045167 |
| 0.999490048 | -0.05564 | 0.51513479 | 0.88432884 | -0.1218357 | 0.16063275 |
| 0.999490048 | -0.0458003 | 0.51504726 | 0.88432884 | 9.56E-04 | 0.98911562 |
| 0.999490048 | -0.0318964 | 0.51511404 | 0.88432884 | -0.0170023 | 0.72782295 |
| 0.999490048 | -0.0768931 | 0.51571604 | 0.88446248 | -0.1254611 | 0.29228337 |
| 0.999490048 | -0.0480737 | 0.51616595 | 0.88446248 | 0.01464791 | 0.84257853 |
| 0.999490048 | -0.0427855 | 0.51568207 | 0.88446248 | -0.044995 | 0.49443303 |
| 0.999490048 | -0.0305027 | 0.51570535 | 0.88446248 | -0.0368652 | 0.43302387 |

| | | | | | |
|---|---|---|---|---|---|
| 0.999490048 | -0.0286262 | 0.51596413 | 0.88446248 | -0.0495607 | 0.26473863 |
| 0.999490048 | -0.0265352 | 0.51590779 | 0.88446248 | 0.02898847 | 0.47823297 |
| 0.999490048 | 0.03249557 | 0.51607612 | 0.88446248 | -0.003416 | 0.94534386 |
| 0.999490048 | 0.04196073 | 0.51580129 | 0.88446248 | 0.04090852 | 0.5262967 |
| 0.999490048 | 0.10264561 | 0.51549132 | 0.88446248 | 0.28645992 | 0.07747454 |
| 0.999490048 | -0.0745495 | 0.51671564 | 0.88504114 | -0.1186583 | 0.30514169 |
| 0.999490048 | 0.03276269 | 0.51664233 | 0.88504114 | 0.0719425 | 0.16096157 |
| 0.999490048 | 0.06229225 | 0.51713893 | 0.88543694 | -0.0256838 | 0.78868265 |
| 0.999490048 | 0.10224542 | 0.5171588 | 0.88543694 | 0.09151639 | 0.56174652 |
| 0.999490048 | -0.03205 | 0.51726949 | 0.88544491 | -0.0062667 | 0.8988344 |
| 0.999490048 | 0.32310003 | 0.51743389 | 0.88554478 | 0.17738724 | 0.71978247 |
| 0.999490048 | -0.1211076 | 0.5179101 | 0.88617814 | -0.220564 | 0.24348569 |
| 0.999490048 | 0.03302824 | 0.51809305 | 0.88630957 | 0.02940966 | 0.56462746 |
| 0.999490048 | 0.03179354 | 0.5182689 | 0.88642878 | 0.03630841 | 0.46123469 |
| 0.999490048 | -0.0603414 | 0.51847562 | 0.88654678 | -0.0389519 | 0.67606531 |
| 0.999490048 | -0.0320327 | 0.51855024 | 0.88654678 | -0.0124081 | 0.80183904 |
| 0.999490048 | -0.0385511 | 0.51893693 | 0.88656545 | -0.0104572 | 0.86057699 |
| 0.999490048 | -0.0347635 | 0.51889829 | 0.88656545 | -0.0819685 | 0.13541741 |
| 0.999490048 | 0.03799747 | 0.51873939 | 0.88656545 | 0.0319714 | 0.58670437 |
| 0.999490048 | 0.05468862 | 0.51898586 | 0.88656545 | 0.03874811 | 0.6470325 |
| 0.999490048 | 0.0338554 | 0.51922237 | 0.88678805 | 0.04889772 | 0.35406106 |
| 0.999490048 | -0.0500945 | 0.51936815 | 0.88685563 | 0.02399721 | 0.75682404 |
| 0.999490048 | 0.03744094 | 0.51992913 | 0.88708791 | 0.05072206 | 0.38501469 |
| 0.999490048 | 0.04024466 | 0.51981936 | 0.88708791 | 0.02278788 | 0.71475375 |
| 0.999490048 | 0.04150843 | 0.5198657 | 0.88708791 | 0.06362459 | 0.32660863 |
| 0.999490048 | 0.04443932 | 0.51981282 | 0.88708791 | -0.0338362 | 0.62347879 |
| 0.999490048 | 0.03269664 | 0.52012446 | 0.88723988 | 0.03035752 | 0.55018343 |
| 0.999490048 | 0.09854405 | 0.5204183 | 0.88737856 | -0.0742238 | 0.62770607 |
| 0.999490048 | 0.10081139 | 0.52039362 | 0.88737856 | 0.00522997 | 0.97328871 |
| 0.999490048 | -0.1330597 | 0.52106526 | 0.887478 | -0.1680783 | 0.41874556 |
| 0.999490048 | -0.0699431 | 0.52058527 | 0.887478 | -0.0363213 | 0.7378992 |
| 0.999490048 | -0.023252 | 0.52089051 | 0.887478 | -0.0462886 | 0.20669267 |
| 0.999490048 | 0.0362357 | 0.52138473 | 0.887478 | -0.0083963 | 0.88142513 |
| 0.999490048 | 0.04177443 | 0.52143318 | 0.887478 | 0.027713 | 0.66987643 |
| 0.999490048 | 0.04306054 | 0.52113815 | 0.887478 | 0.00461711 | 0.94492771 |
| 0.999490048 | 0.04787334 | 0.52105155 | 0.887478 | 0.01070546 | 0.8854434 |
| 0.999490048 | 0.05465151 | 0.52104374 | 0.887478 | -0.027051 | 0.74999772 |
| 0.999490048 | 0.1717383 | 0.52130945 | 0.887478 | -0.1911443 | 0.47672039 |
| 0.999490048 | -0.057258 | 0.52175339 | 0.88766113 | -0.0646119 | 0.4702254 |
| 0.999490048 | -0.0478781 | 0.52165473 | 0.88766113 | -0.1197736 | 0.11646451 |
| 0.999490048 | 0.12536018 | 0.52196838 | 0.887846 | 0.03711477 | 0.84834045 |
| 0.999490048 | -0.0714888 | 0.52213411 | 0.887947 | 0.13742875 | 0.22356773 |

| | | | | | |
|---|---|---|---|---|---|
| 0.999490048 | -0.102622 | 0.52335038 | 0.88817808 | -0.0446771 | 0.78042236 |
| 0.999490048 | -0.0852781 | 0.52529266 | 0.88817808 | -0.2017056 | 0.14007587 |
| 0.999490048 | -0.0762107 | 0.52552316 | 0.88817808 | -0.0171999 | 0.88505447 |
| 0.999490048 | -0.0760106 | 0.52419218 | 0.88817808 | 0.05510488 | 0.6427982 |
| 0.999490048 | -0.0667939 | 0.52507355 | 0.88817808 | -0.0662195 | 0.5286148 |
| 0.999490048 | -0.0647447 | 0.52402021 | 0.88817808 | 0.03610752 | 0.72158271 |
| 0.999490048 | -0.0646139 | 0.52486794 | 0.88817808 | 0.14079996 | 0.17220446 |
| 0.999490048 | -0.0563058 | 0.52456337 | 0.88817808 | 0.03049635 | 0.72960929 |
| 0.999490048 | -0.0534257 | 0.52544557 | 0.88817808 | -0.0349382 | 0.6772525 |
| 0.999490048 | -0.0462616 | 0.52472449 | 0.88817808 | 0.00149421 | 0.98354158 |
| 0.999490048 | -0.0428827 | 0.52468069 | 0.88817808 | -0.0033657 | 0.96001685 |
| 0.999490048 | -0.0416354 | 0.52359505 | 0.88817808 | -0.1279842 | 0.05776457 |
| 0.999490048 | -0.0355292 | 0.52373666 | 0.88817808 | 0.00175727 | 0.9747391 |
| 0.999490048 | -0.0336699 | 0.52578494 | 0.88817808 | 0.00306844 | 0.9537057 |
| 0.999490048 | -0.0325684 | 0.52558659 | 0.88817808 | -0.0468671 | 0.36307955 |
| 0.999490048 | -0.0324891 | 0.52313859 | 0.88817808 | 0.01020551 | 0.84045002 |
| 0.999490048 | -0.0314774 | 0.5259893 | 0.88817808 | -0.0396308 | 0.42573072 |
| 0.999490048 | -0.0293947 | 0.52496741 | 0.88817808 | -0.0086265 | 0.85145615 |
| 0.999490048 | -0.0277134 | 0.52599289 | 0.88817808 | -0.0335437 | 0.44361336 |
| 0.999490048 | 0.02897786 | 0.52316168 | 0.88817808 | -0.0235791 | 0.60288993 |
| 0.999490048 | 0.02968941 | 0.52566906 | 0.88817808 | 0.02664265 | 0.56870866 |
| 0.999490048 | 0.02993303 | 0.52301976 | 0.88817808 | -0.0094847 | 0.83903035 |
| 0.999490048 | 0.03070536 | 0.52584016 | 0.88817808 | 0.11037402 | 0.0292201 |
| 0.999490048 | 0.03079509 | 0.52475281 | 0.88817808 | 0.02687391 | 0.57849233 |
| 0.987721587 | 0.0377329 | 0.52336264 | 0.88817808 | -0.0922063 | 0.1262516 |
| 0.999490048 | 0.03897917 | 0.52290603 | 0.88817808 | 0.0258541 | 0.67103355 |
| 0.999490048 | 0.04901206 | 0.52576038 | 0.88817808 | 0.07175649 | 0.35513541 |
| 0.999490048 | 0.05073126 | 0.52314772 | 0.88817808 | -0.0297599 | 0.70723727 |
| 0.999490048 | 0.05893175 | 0.52344682 | 0.88817808 | 0.12038733 | 0.19811572 |
| 0.999490048 | 0.07312168 | 0.52453918 | 0.88817808 | 0.03294264 | 0.77362956 |
| 0.879524171 | 0.07633332 | 0.5249733 | 0.88817808 | -0.2290661 | 0.06436872 |
| 0.999490048 | 0.07989614 | 0.52318909 | 0.88817808 | -0.0877758 | 0.48342236 |
| 0.999490048 | 0.10523766 | 0.52339924 | 0.88817808 | -0.1372202 | 0.40672451 |
| 0.999490048 | 0.10634112 | 0.5250825 | 0.88817808 | -0.1986558 | 0.24561196 |
| 0.999490048 | 0.16842231 | 0.52456177 | 0.88817808 | -0.0151442 | 0.95394224 |
| 0.999490048 | 0.04595934 | 0.5262783 | 0.88848034 | -0.0211318 | 0.77005436 |
| 0.999490048 | -0.0451382 | 0.52657137 | 0.88879543 | -0.0429196 | 0.54693671 |
| 0.999490048 | -0.0350379 | 0.52747502 | 0.88960137 | -0.1107866 | 0.05349804 |
| 0.999490048 | -0.0315887 | 0.52738652 | 0.88960137 | 0.00266827 | 0.95725441 |
| 0.999490048 | 0.03919687 | 0.52744915 | 0.88960137 | 0.00593363 | 0.92349164 |
| 0.999490048 | 0.11397333 | 0.52728012 | 0.88960137 | -0.213965 | 0.2458276 |
| 0.999490048 | -0.1085005 | 0.52792484 | 0.8900596 | -0.043256 | 0.79980114 |

| | | | | | |
|---|---|---|---|---|---|
| 0.999490048 | -0.0374117 | 0.52852702 | 0.8900596 | 0.0843054 | 0.16214403 |
| 0.999490048 | -0.0368753 | 0.52859947 | 0.8900596 | -0.0504471 | 0.39029116 |
| 0.999490048 | -0.032708 | 0.52828896 | 0.8900596 | -0.063775 | 0.22392927 |
| 0.999490048 | 0.02898179 | 0.52802038 | 0.8900596 | 0.00399078 | 0.93048318 |
| 0.999490048 | 0.07018542 | 0.52849307 | 0.8900596 | 8.14E-04 | 0.99414478 |
| 0.999490048 | 0.08657576 | 0.52807679 | 0.8900596 | -0.060637 | 0.65790037 |
| 0.999490048 | 0.09283599 | 0.52851883 | 0.8900596 | -0.0500119 | 0.73347842 |
| 0.999490048 | -0.2043012 | 0.53019024 | 0.89011075 | -0.2938859 | 0.36856024 |
| 0.999490048 | -0.0518748 | 0.52921261 | 0.89011075 | -0.0281554 | 0.73199591 |
| 0.999490048 | -0.0516676 | 0.53016921 | 0.89011075 | -0.0670494 | 0.4165408 |
| 0.999490048 | -0.0480579 | 0.53052345 | 0.89011075 | -0.1426029 | 0.07072995 |
| 0.999490048 | -0.0443174 | 0.5304208 | 0.89011075 | 0.05659588 | 0.42418001 |
| 0.999490048 | -0.0372949 | 0.53059983 | 0.89011075 | -0.0073364 | 0.90144982 |
| 0.999490048 | -0.0292394 | 0.52960467 | 0.89011075 | -0.051404 | 0.27303535 |
| 0.999490048 | -0.027457 | 0.53014936 | 0.89011075 | -0.0256367 | 0.55756774 |
| 0.999490048 | 0.02325189 | 0.53031671 | 0.89011075 | 0.02363975 | 0.52353546 |
| 0.999490048 | 0.02577523 | 0.52991412 | 0.89011075 | 0.01375124 | 0.73681655 |
| 0.999490048 | 0.02679985 | 0.52961899 | 0.89011075 | 0.01024853 | 0.80940028 |
| 0.999490048 | 0.03032549 | 0.52952433 | 0.89011075 | 0.03516959 | 0.46651138 |
| 0.999490048 | 0.03271114 | 0.52968655 | 0.89011075 | -0.0347743 | 0.50427379 |
| 0.999490048 | 0.03349022 | 0.53051512 | 0.89011075 | 0.03288342 | 0.53793213 |
| 0.999490048 | 0.04768144 | 0.52894849 | 0.89011075 | 0.04264987 | 0.57301146 |
| 0.999490048 | 0.05841368 | 0.53065525 | 0.89011075 | 0.07258962 | 0.43686375 |
| 0.999490048 | 0.06187043 | 0.52918735 | 0.89011075 | -0.0217719 | 0.82415233 |
| 0.999490048 | 0.06483348 | 0.52929274 | 0.89011075 | 0.09899691 | 0.33930238 |
| 0.999490048 | 0.22553555 | 0.53037656 | 0.89011075 | 0.19791894 | 0.58081451 |
| 0.999490048 | -0.0548356 | 0.53078548 | 0.89015038 | -0.0100837 | 0.90788471 |
| 0.999490048 | 0.04713755 | 0.53117741 | 0.89062879 | 0.01725145 | 0.81811552 |
| 0.999490048 | -0.1715032 | 0.53178415 | 0.89076791 | -0.0863584 | 0.75216591 |
| 0.999490048 | -0.1169887 | 0.53234899 | 0.89076791 | -0.2427042 | 0.2008619 |
| 0.999490048 | -0.0640962 | 0.53184678 | 0.89076791 | 0.01331027 | 0.89631354 |
| 0.999490048 | -0.039646 | 0.53238172 | 0.89076791 | 0.01857914 | 0.76914055 |
| 0.999490048 | -0.0347731 | 0.53237452 | 0.89076791 | 0.02439021 | 0.66078769 |
| 0.999490048 | -0.0252166 | 0.53241902 | 0.89076791 | -0.0307282 | 0.4476529 |
| 0.999490048 | 0.02685381 | 0.53259964 | 0.89076791 | 0.05954149 | 0.17284767 |
| 0.999490048 | 0.03339123 | 0.53264721 | 0.89076791 | -0.0249142 | 0.64094789 |
| 0.999490048 | 0.0334224 | 0.53160369 | 0.89076791 | -0.0158229 | 0.76641726 |
| 0.999490048 | 0.03947137 | 0.53248708 | 0.89076791 | -0.0852768 | 0.18358533 |
| 0.999490048 | 0.04256356 | 0.53215153 | 0.89076791 | 0.08533796 | 0.21569091 |
| 0.999490048 | 0.05356523 | 0.53183023 | 0.89076791 | 0.1378613 | 0.11522103 |
| 0.999490048 | 0.07897014 | 0.53170156 | 0.89076791 | 0.05467957 | 0.66433913 |
| 0.999490048 | -0.0943883 | 0.53297 | 0.89088126 | -0.2461054 | 0.1117231 |

| | | | | | |
|---|---|---|---|---|---|
| 0.999490048 | 0.03606161 | 0.53303506 | 0.89088126 | 0.10370939 | 0.08102905 |
| 0.999490048 | 0.05324371 | 0.53296808 | 0.89088126 | -0.0407696 | 0.63251826 |
| 0.999490048 | -0.0377479 | 0.53316718 | 0.89092375 | 0.04671395 | 0.44153477 |
| 0.999490048 | 0.0376106 | 0.53356933 | 0.89113677 | 0.08347187 | 0.1734391 |
| 0.999490048 | 0.05145517 | 0.53361483 | 0.89113677 | 0.07946353 | 0.33896633 |
| 0.999490048 | 0.06763283 | 0.53359098 | 0.89113677 | 0.08785338 | 0.41995346 |
| 0.999490048 | 0.04882377 | 0.5338894 | 0.89130549 | 0.07059776 | 0.37039959 |
| 0.999490048 | 0.07064939 | 0.53392935 | 0.89130549 | 0.05731875 | 0.61333053 |
| 0.999490048 | -0.0393381 | 0.53417684 | 0.89131981 | -0.073474 | 0.25011266 |
| 0.999490048 | 0.03019121 | 0.53425601 | 0.89131981 | 0.00181277 | 0.97011524 |
| 0.999490048 | 0.05816304 | 0.53425816 | 0.89131981 | 0.0108251 | 0.90756347 |
| 0.999490048 | -0.043777 | 0.53465123 | 0.89161929 | 0.02245287 | 0.7494375 |
| 0.999490048 | -0.0298958 | 0.53462616 | 0.89161929 | 0.01184029 | 0.80510797 |
| 0.999490048 | -0.0399973 | 0.53477077 | 0.89164056 | -0.0280854 | 0.66230477 |
| 0.999490048 | -0.0431547 | 0.53530388 | 0.89235125 | 0.04290435 | 0.53765579 |
| 0.999490048 | 0.03721306 | 0.53548411 | 0.89247352 | -0.0046181 | 0.93846944 |
| 0.999490048 | 0.06554235 | 0.53564732 | 0.89256737 | 0.22843341 | 0.03810563 |
| 0.999490048 | -0.0411818 | 0.53580369 | 0.89264981 | -0.104472 | 0.12374638 |
| 0.999490048 | -0.063372 | 0.5361263 | 0.89278625 | 0.09826314 | 0.33998897 |
| 0.999490048 | -0.0495462 | 0.53728843 | 0.89278625 | -0.073347 | 0.36327302 |
| 0.999490048 | -0.0468112 | 0.53754924 | 0.89278625 | 0.01260823 | 0.8676461 |
| 0.999490048 | -0.0436611 | 0.53729401 | 0.89278625 | 0.04460606 | 0.52859193 |
| 0.999490048 | -0.0402728 | 0.53823747 | 0.89278625 | -0.0848048 | 0.20070059 |
| 0.999490048 | -0.0388696 | 0.53841606 | 0.89278625 | -0.0144876 | 0.8180452 |
| 0.999490048 | -0.0386862 | 0.53792188 | 0.89278625 | -0.0211031 | 0.73618555 |
| 0.999490048 | -0.0343898 | 0.53834789 | 0.89278625 | 0.01734177 | 0.75567055 |
| 0.999490048 | -0.0326269 | 0.53797501 | 0.89278625 | -0.0172547 | 0.74396466 |
| 0.999490048 | -0.0301725 | 0.53685692 | 0.89278625 | -0.054541 | 0.26837222 |
| 0.999490048 | -0.0298317 | 0.53651071 | 0.89278625 | 0.01245543 | 0.79571663 |
| 0.999490048 | -0.0276689 | 0.53739237 | 0.89278625 | -0.0152121 | 0.73385638 |
| 0.999490048 | -0.0276251 | 0.53707963 | 0.89278625 | -0.0074931 | 0.8665675 |
| 0.999490048 | -0.020838 | 0.5385586 | 0.89278625 | -0.0417533 | 0.22304249 |
| 0.999490048 | 0.0300756 | 0.53834247 | 0.89278625 | 0.01174695 | 0.8094669 |
| 0.999490048 | 0.03046735 | 0.53706051 | 0.89278625 | 0.00918463 | 0.8518703 |
| 0.999490048 | 0.03362326 | 0.53693873 | 0.89278625 | 0.03177575 | 0.55938103 |
| 0.999490048 | 0.03784896 | 0.53635055 | 0.89278625 | 0.0598505 | 0.33094667 |
| 0.99974305 | 0.03820211 | 0.53850155 | 0.89278625 | 0.03816726 | 0.53886936 |
| 0.999490048 | 0.04548653 | 0.53608246 | 0.89278625 | 0.00945088 | 0.89732862 |
| 0.999490048 | 0.04756042 | 0.53793963 | 0.89278625 | 0.07682633 | 0.32270715 |
| 0.999490048 | 0.0546845 | 0.53726406 | 0.89278625 | 0.03030793 | 0.73171201 |
| 0.999490048 | 0.06985863 | 0.53786002 | 0.89278625 | -0.0464331 | 0.68154943 |
| 0.999490048 | 0.07597929 | 0.53664637 | 0.89278625 | 0.14568377 | 0.2408591 |

| | | | | | |
|---|---|---|---|---|---|
| 0.999490048 | 0.07950446 | 0.53770792 | 0.89278625 | 0.20791147 | 0.11472011 |
| 0.999490048 | -0.0905146 | 0.53901404 | 0.89287792 | -0.1239278 | 0.40186172 |
| 0.999490048 | -0.0434952 | 0.53899876 | 0.89287792 | 0.01433949 | 0.83896157 |
| 0.999490048 | -0.0375627 | 0.53936242 | 0.89287792 | 0.03620291 | 0.55404006 |
| 0.999490048 | -0.0289159 | 0.53906652 | 0.89287792 | 9.06E-05 | 0.9984582 |
| 0.999490048 | 0.03024815 | 0.53927094 | 0.89287792 | 0.03304001 | 0.50280089 |
| 0.999490048 | 0.07759685 | 0.53930667 | 0.89287792 | 0.11594907 | 0.36123233 |
| 0.999490048 | 0.08212951 | 0.53877147 | 0.89287792 | 0.16111824 | 0.23283753 |
| 0.999490048 | 0.03731687 | 0.53966645 | 0.89320413 | 0.03768035 | 0.53575461 |
| 0.972489992 | -0.0573732 | 0.53999266 | 0.89321655 | 0.15712857 | 0.10117362 |
| 0.999490048 | 0.06130189 | 0.53999487 | 0.89321655 | -0.0172739 | 0.86242468 |
| 0.999490048 | 0.11563348 | 0.53992454 | 0.89321655 | 0.16793236 | 0.37957362 |
| 0.999490048 | -0.1419203 | 0.54099358 | 0.89329745 | 0.29986433 | 0.20220988 |
| 0.771925038 | -0.0687566 | 0.54116929 | 0.89329745 | 0.2880632 | 0.02785707 |
| 0.999490048 | -0.0484976 | 0.54065111 | 0.89329745 | -0.1659757 | 0.04379 |
| 0.987721587 | -0.0482323 | 0.54102396 | 0.89329745 | 0.12133906 | 0.13149254 |
| 0.999490048 | -0.0399444 | 0.54035106 | 0.89329745 | -0.0607757 | 0.35391475 |
| 0.999490048 | -0.0321462 | 0.54072296 | 0.89329745 | -0.0540103 | 0.30733463 |
| 0.999490048 | 0.03034036 | 0.54045826 | 0.89329745 | -0.0293304 | 0.55392462 |
| 0.999490048 | 0.03228747 | 0.54033846 | 0.89329745 | 0.02108393 | 0.68863676 |
| 0.524321571 | 0.04472665 | 0.5405749 | 0.89329745 | -0.186431 | 0.01588824 |
| 0.999490048 | 0.06206096 | 0.54101994 | 0.89329745 | 0.13158824 | 0.20071355 |
| 0.999490048 | 0.06535873 | 0.54122058 | 0.89329745 | 0.12670378 | 0.24102926 |
| 0.999490048 | -0.1156142 | 0.54147644 | 0.89336659 | -0.0818959 | 0.66478205 |
| 0.999490048 | -0.0883075 | 0.54147162 | 0.89336659 | 0.01584499 | 0.91244217 |
| 0.999490048 | -0.057763 | 0.54177895 | 0.89339507 | 0.00745784 | 0.93697996 |
| 0.999490048 | 0.04803836 | 0.54166452 | 0.89339507 | 0.02479132 | 0.75213195 |
| 0.999490048 | 0.05206877 | 0.54181469 | 0.89339507 | 0.13835786 | 0.11273711 |
| 0.999490048 | 0.02797604 | 0.54198391 | 0.89349766 | -0.0081456 | 0.85859038 |
| 0.999490048 | 0.07344617 | 0.54216713 | 0.89362328 | 0.02932199 | 0.8071404 |
| 0.999490048 | 0.03137457 | 0.54310888 | 0.89411656 | 0.00434368 | 0.93265792 |
| 0.999490048 | 0.03153239 | 0.54276678 | 0.89411656 | -0.0121744 | 0.81363454 |
| 0.999490048 | 0.0753332 | 0.54277148 | 0.89411656 | 0.1100579 | 0.37591486 |
| 0.999490048 | 0.10085742 | 0.5430716 | 0.89411656 | -0.0046418 | 0.97740125 |
| 0.999490048 | 0.10639835 | 0.5428911 | 0.89411656 | -0.0760622 | 0.66300903 |
| 0.999490048 | 0.12794813 | 0.54290752 | 0.89411656 | -0.1055612 | 0.61526961 |
| 0.999490048 | -0.0267012 | 0.54331789 | 0.89428432 | 0.00147846 | 0.97305252 |
| 0.999490048 | -0.0690166 | 0.54471867 | 0.89435358 | -0.0271413 | 0.81115065 |
| 0.999490048 | -0.050865 | 0.54484137 | 0.89435358 | 0.02930909 | 0.72651406 |
| 0.999490048 | -0.0351415 | 0.54463413 | 0.89435358 | 0.00439602 | 0.93936932 |
| 0.999490048 | -0.0334921 | 0.54381995 | 0.89435358 | 0.02915753 | 0.59683531 |
| 0.999490048 | -0.0330177 | 0.54479749 | 0.89435358 | -0.0363285 | 0.5055504 |

| | | | | | |
|---|---|---|---|---|---|
| 0.999490048 | -0.0325745 | 0.54515656 | 0.89435358 | 0.03338825 | 0.53523592 |
| 0.999490048 | -0.0266735 | 0.54483398 | 0.89435358 | -0.0306422 | 0.48718313 |
| 0.999490048 | -0.0251006 | 0.54491825 | 0.89435358 | -0.0323746 | 0.4360193 |
| 0.999490048 | 0.03039639 | 0.54411823 | 0.89435358 | -0.0166965 | 0.73833024 |
| 0.999490048 | 0.03469256 | 0.54446332 | 0.89435358 | 0.06347236 | 0.271586 |
| 0.999490048 | 0.06395512 | 0.54475121 | 0.89435358 | 0.08189882 | 0.43908119 |
| 0.999490048 | 0.06607685 | 0.54507137 | 0.89435358 | -0.0092233 | 0.93243706 |
| 0.999490048 | 0.0694768 | 0.54510362 | 0.89435358 | -0.1098425 | 0.34137147 |
| 0.595926661 | 0.08391619 | 0.54464946 | 0.89435358 | -0.3250124 | 0.02528921 |
| 0.999490048 | 0.08501103 | 0.54371929 | 0.89435358 | 0.1540782 | 0.27511588 |
| 0.999490048 | 0.08980055 | 0.5451808 | 0.89435358 | 0.14025051 | 0.34725143 |
| 0.999490048 | 0.09559591 | 0.54467691 | 0.89435358 | 0.06776283 | 0.66706491 |
| 0.999490048 | -0.0699975 | 0.54536724 | 0.89448368 | 0.07886289 | 0.49610424 |
| 0.999490048 | -0.0313669 | 0.54551179 | 0.89450773 | -0.024831 | 0.63179665 |
| 0.999490048 | 0.0410844 | 0.54581041 | 0.89450773 | 0.04264647 | 0.53076527 |
| 0.999490048 | 0.04825432 | 0.54579092 | 0.89450773 | 0.03386755 | 0.67100488 |
| 0.999490048 | 0.0667501 | 0.54572907 | 0.89450773 | -0.0623152 | 0.57255159 |
| 0.999490048 | 0.02607881 | 0.54625205 | 0.89505586 | 0.02313376 | 0.59219037 |
| 0.999490048 | -0.1124122 | 0.54676218 | 0.89536462 | -0.3230103 | 0.09138312 |
| 0.999490048 | 0.03580112 | 0.54665443 | 0.89536462 | 0.0534464 | 0.37032187 |
| 0.999490048 | 0.04306795 | 0.54670727 | 0.89536462 | 0.01634193 | 0.81854822 |
| 0.999490048 | -0.0284545 | 0.54702884 | 0.89562564 | -0.0151375 | 0.7480459 |
| 0.999490048 | -0.0581343 | 0.54761724 | 0.89623754 | -0.0379494 | 0.69404835 |
| 0.999490048 | 0.03429196 | 0.5475604 | 0.89623754 | -6.68E-04 | 0.99062445 |
| 0.999490048 | 0.03562011 | 0.54773884 | 0.89626089 | 0.01796175 | 0.76115613 |
| 0.999490048 | 0.06134535 | 0.54821979 | 0.89687211 | 0.00194274 | 0.98477375 |
| 0.999490048 | -0.0613526 | 0.5485821 | 0.89716577 | -0.1129815 | 0.27338158 |
| 0.999490048 | 0.06133421 | 0.54861418 | 0.89716577 | 0.0991145 | 0.33517691 |
| 0.999490048 | -0.0787485 | 0.54883342 | 0.89717288 | 0.00298222 | 0.98182088 |
| 0.999490048 | 0.04002377 | 0.54882228 | 0.89717288 | -0.0177499 | 0.78973253 |
| 0.999490048 | 0.04666563 | 0.54901545 | 0.89729478 | 0.15483376 | 0.05467193 |
| 0.999490048 | -0.1241499 | 0.54995547 | 0.89735688 | -2.66E-04 | 0.99896622 |
| 0.999490048 | -0.050309 | 0.55045053 | 0.89735688 | -0.0349475 | 0.67774494 |
| 0.999490048 | -0.0386506 | 0.54928102 | 0.89735688 | -0.1352789 | 0.04362919 |
| 0.999490048 | -0.0356529 | 0.54951656 | 0.89735688 | -0.140507 | 0.02461444 |
| 0.999490048 | -0.0258626 | 0.55041065 | 0.89735688 | 0.01963447 | 0.64980395 |
| 0.999490048 | 0.03207258 | 0.55006986 | 0.89735688 | 0.02696272 | 0.61498773 |
| 0.999490048 | 0.03696649 | 0.55030059 | 0.89735688 | -0.0357575 | 0.56332601 |
| 0.999490048 | 0.04940948 | 0.54958447 | 0.89735688 | 0.0342955 | 0.67732162 |
| 0.999490048 | 0.05035083 | 0.5502737 | 0.89735688 | 0.0481427 | 0.56777636 |
| 0.999490048 | 0.05185545 | 0.55014972 | 0.89735688 | 0.12025108 | 0.17238292 |
| 0.999490048 | 0.06142673 | 0.5496973 | 0.89735688 | -0.0917211 | 0.37382965 |

| | | | | | |
|---|---|---|---|---|---|
| 0.999490048 | 0.06926053 | 0.54917462 | 0.89735688 | 0.06656762 | 0.56470546 |
| 0.999490048 | 0.08961278 | 0.55024178 | 0.89735688 | 0.03792862 | 0.79980552 |
| 0.999490048 | -0.0886372 | 0.55072771 | 0.89751246 | 0.05836202 | 0.69382423 |
| 0.999490048 | 0.05929537 | 0.55076094 | 0.89751246 | -0.0339477 | 0.73204808 |
| 0.999490048 | 0.03161709 | 0.55123741 | 0.89776329 | -0.0151609 | 0.77446549 |
| 0.999490048 | 0.03963705 | 0.55110941 | 0.89776329 | 0.0322569 | 0.62718254 |
| 0.999490048 | 0.04406662 | 0.55117209 | 0.89776329 | -0.0118724 | 0.87200622 |
| 0.999490048 | 0.05455446 | 0.55151682 | 0.89786809 | 0.00781544 | 0.93177663 |
| 0.999490048 | 0.1246019 | 0.55148714 | 0.89786809 | -0.1227278 | 0.55744642 |
| 0.999490048 | 0.07516152 | 0.55166655 | 0.89793678 | 0.22364652 | 0.08462842 |
| 0.999490048 | -0.0754933 | 0.55259798 | 0.89814063 | 0.14242925 | 0.2668069 |
| 0.999490048 | -0.0643691 | 0.55238612 | 0.89814063 | -0.0156904 | 0.88445454 |
| 0.999490048 | -0.050537 | 0.55295669 | 0.89814063 | 0.04756475 | 0.57639036 |
| 0.999490048 | -0.0445513 | 0.55293446 | 0.89814063 | 0.02760275 | 0.71254455 |
| 0.999490048 | -0.0398215 | 0.55309508 | 0.89814063 | 0.01967365 | 0.76889439 |
| 0.999490048 | -0.036159 | 0.55318242 | 0.89814063 | -0.1014563 | 0.10407777 |
| 0.999490048 | 0.02764749 | 0.55270194 | 0.89814063 | 0.00120814 | 0.97922794 |
| 0.999490048 | 0.02846963 | 0.55309387 | 0.89814063 | 0.00333248 | 0.94445437 |
| 0.999490048 | 0.03715599 | 0.55274947 | 0.89814063 | 0.02122499 | 0.73391057 |
| 0.999490048 | 0.04114243 | 0.55295112 | 0.89814063 | 0.0127146 | 0.85404662 |
| 0.999490048 | 0.04479842 | 0.55191433 | 0.89814063 | -0.0034183 | 0.96366521 |
| 0.999490048 | 0.04545745 | 0.55329765 | 0.89814063 | -0.0396677 | 0.60462791 |
| 0.999490048 | 0.04901048 | 0.55226194 | 0.89814063 | 0.00678181 | 0.93422101 |
| 0.999490048 | 0.0510232 | 0.55325279 | 0.89814063 | 0.07966697 | 0.35700961 |
| 0.999490048 | 0.02577763 | 0.5538023 | 0.89843583 | 0.02712289 | 0.53345839 |
| 0.999490048 | 0.0298693 | 0.5537343 | 0.89843583 | -0.0084612 | 0.86635002 |
| 0.999490048 | 0.04246296 | 0.55368674 | 0.89843583 | 6.44E-04 | 0.99280769 |
| 0.999490048 | -0.0323942 | 0.55403629 | 0.89864083 | -0.0318245 | 0.56097984 |
| 0.999490048 | 0.03290217 | 0.55426817 | 0.89884234 | -0.0051519 | 0.92596777 |
| 0.999490048 | -0.1592546 | 0.55446443 | 0.89887045 | -0.1221888 | 0.64909217 |
| 0.999490048 | 0.02892541 | 0.5545008 | 0.89887045 | 0.0326317 | 0.50533341 |
| 0.999490048 | -0.1848308 | 0.55538861 | 0.89972244 | -0.006192 | 0.98418285 |
| 0.999490048 | -0.0373911 | 0.55528732 | 0.89972244 | -0.0159743 | 0.80048039 |
| 0.969787208 | -0.0347271 | 0.55553061 | 0.89972244 | 0.10495555 | 0.08294493 |
| 0.999490048 | 0.03399773 | 0.55556514 | 0.89972244 | 0.03788125 | 0.51168313 |
| 0.999490048 | 0.2395106 | 0.55537556 | 0.89972244 | 0.25010211 | 0.53843056 |
| 0.999490048 | 0.0321686 | 0.55569362 | 0.89975601 | -0.0028224 | 0.9586247 |
| 0.999490048 | -0.0471251 | 0.55581761 | 0.89978229 | 0.0197627 | 0.80432808 |
| 0.999490048 | -0.044263 | 0.55621356 | 0.89979031 | 0.08301426 | 0.27388143 |
| 0.999490048 | -0.0426079 | 0.55594897 | 0.89979031 | 0.04146064 | 0.56656346 |
| 0.999490048 | -0.0359255 | 0.55646912 | 0.89979031 | -0.0093845 | 0.87751238 |
| 0.999490048 | 0.02609491 | 0.55634435 | 0.89979031 | -0.0347536 | 0.43457757 |

| | | | | | |
|---|---|---|---|---|---|
| 0.999490048 | 0.04854057 | 0.5563977 | 0.89979031 | -0.0114873 | 0.88892843 |
| 0.999490048 | 0.07600991 | 0.55645962 | 0.89979031 | 0.18909092 | 0.15929708 |
| 0.999490048 | -0.1115194 | 0.56035505 | 0.89979762 | 0.03697822 | 0.84642851 |
| 0.999490048 | -0.097516 | 0.55761493 | 0.89979762 | -0.0708148 | 0.66972463 |
| 0.999490048 | -0.0832367 | 0.5614597 | 0.89979762 | -0.2069864 | 0.15575378 |
| 0.999490048 | -0.0752567 | 0.56152712 | 0.89979762 | 0.19708063 | 0.14503247 |
| 0.999490048 | -0.068002 | 0.55951411 | 0.89979762 | 0.05079831 | 0.66240795 |
| 0.999490048 | -0.0621303 | 0.55982106 | 0.89979762 | -0.1921282 | 0.07952543 |
| 0.999490048 | -0.0597815 | 0.5610744 | 0.89979762 | 0.00973046 | 0.92437585 |
| 0.999490048 | -0.0547129 | 0.56109723 | 0.89979762 | 0.01011233 | 0.9141718 |
| 0.999490048 | -0.0517432 | 0.55724505 | 0.89979762 | -0.1495607 | 0.09782471 |
| 0.999490048 | -0.0466982 | 0.55766743 | 0.89979762 | 0.01787548 | 0.82190231 |
| 0.999490048 | -0.0462376 | 0.56026487 | 0.89979762 | -0.0640721 | 0.42105716 |
| 0.99974305 | -0.0430978 | 0.56260536 | 0.89979762 | -0.0431444 | 0.56218565 |
| 0.999490048 | -0.0415868 | 0.56223992 | 0.89979762 | 0.04946109 | 0.49127339 |
| 0.999490048 | -0.0403781 | 0.56229354 | 0.89979762 | -0.1094062 | 0.12408576 |
| 0.999490048 | -0.0365623 | 0.56261996 | 0.89979762 | -0.1190323 | 0.06758678 |
| 0.999490048 | -0.0365265 | 0.55967369 | 0.89979762 | 4.52E-04 | 0.99421451 |
| 0.999490048 | -0.0358011 | 0.5602332 | 0.89979762 | -0.0045342 | 0.9409931 |
| 0.999490048 | -0.0352754 | 0.56246146 | 0.89979762 | -0.0219169 | 0.71839522 |
| 0.999490048 | -0.0342835 | 0.55920056 | 0.89979762 | 0.02843529 | 0.62773064 |
| 0.999490048 | -0.0316535 | 0.56283149 | 0.89979762 | -0.057954 | 0.29318031 |
| 0.999490048 | -0.0310231 | 0.56246588 | 0.89979762 | -0.0322077 | 0.54774604 |
| 0.999490048 | -0.0308872 | 0.55703648 | 0.89979762 | -0.0202289 | 0.69996222 |
| 0.999490048 | -0.0264204 | 0.55905503 | 0.89979762 | -0.0543542 | 0.23446956 |
| 0.999490048 | -0.0263949 | 0.55897627 | 0.89979762 | 0.02302369 | 0.60994827 |
| 0.999490048 | -0.0245282 | 0.56280579 | 0.89979762 | 0.00159745 | 0.96983513 |
| 0.999490048 | -0.0217679 | 0.5599671 | 0.89979762 | 0.01213827 | 0.74457367 |
| 0.999490048 | 0.02090067 | 0.55969158 | 0.89979762 | 5.67E-04 | 0.98732263 |
| 0.999490048 | 0.02436039 | 0.55843806 | 0.89979762 | 0.02879181 | 0.48977442 |
| 0.999490048 | 0.02808207 | 0.55825325 | 0.89979762 | 0.04221125 | 0.38091719 |
| 0.999490048 | 0.02821529 | 0.55905592 | 0.89979762 | -0.0219298 | 0.64931631 |
| 0.999490048 | 0.03000312 | 0.56208613 | 0.89979762 | 0.00491412 | 0.92410063 |
| 0.999490048 | 0.03097604 | 0.5571513 | 0.89979762 | 0.06859455 | 0.19951155 |
| 0.999490048 | 0.03126964 | 0.56258761 | 0.89979762 | 0.02743618 | 0.61116202 |
| 0.999490048 | 0.03197738 | 0.56060967 | 0.89979762 | 0.01399146 | 0.79847004 |
| 0.999490048 | 0.03383873 | 0.55916578 | 0.89979762 | -0.03364 | 0.56145991 |
| 0.999490048 | 0.03462598 | 0.55924261 | 0.89979762 | -0.0129452 | 0.82667447 |
| 0.999490048 | 0.03614503 | 0.55950926 | 0.89979762 | 0.12089665 | 0.05896237 |
| 0.999490048 | 0.03617295 | 0.56191771 | 0.89979762 | -0.0218841 | 0.72510433 |
| 0.999490048 | 0.03648566 | 0.55789912 | 0.89979762 | 0.03155655 | 0.6119831 |
| 0.999490048 | 0.03734825 | 0.55779633 | 0.89979762 | -0.0935417 | 0.14923841 |

| | | | | | |
|---|---|---|---|---|---|
| 0.999490048 | 0.03754648 | 0.55743267 | 0.89979762 | 0.07006622 | 0.27768486 |
| 0.999490048 | 0.03837829 | 0.55746409 | 0.89979762 | -0.0133419 | 0.83791949 |
| 0.999490048 | 0.04155039 | 0.55723726 | 0.89979762 | 0.03046637 | 0.66641968 |
| 0.999490048 | 0.04176381 | 0.5609275 | 0.89979762 | 0.10611262 | 0.14678029 |
| 0.999490048 | 0.04269262 | 0.55788987 | 0.89979762 | 0.00640578 | 0.92970034 |
| 0.999490048 | 0.0443269 | 0.55998798 | 0.89979762 | 0.04346859 | 0.56756025 |
| 0.999490048 | 0.04747846 | 0.55804231 | 0.89979762 | 0.0165475 | 0.83774919 |
| 0.999490048 | 0.04794998 | 0.55998647 | 0.89979762 | 0.05643668 | 0.49326727 |
| 0.999490048 | 0.04802157 | 0.5622955 | 0.89979762 | -0.0370891 | 0.65405597 |
| 0.999490048 | 0.05030745 | 0.56210277 | 0.89979762 | 0.21851634 | 0.01716787 |
| 0.999490048 | 0.05042373 | 0.56201322 | 0.89979762 | 0.0646341 | 0.45829466 |
| 0.90609983 | 0.05068676 | 0.56194658 | 0.89979762 | -0.1659208 | 0.06575298 |
| 0.999490048 | 0.05514037 | 0.55693535 | 0.89979762 | 0.02801792 | 0.76475497 |
| 0.999490048 | 0.05916621 | 0.5608659 | 0.89979762 | 0.06061053 | 0.55140763 |
| 0.999490048 | 0.06647839 | 0.56041178 | 0.89979762 | -0.0389867 | 0.73218487 |
| 0.999490048 | 0.06950862 | 0.56245912 | 0.89979762 | 0.06732823 | 0.57470122 |
| 0.999490048 | 0.08179492 | 0.55961457 | 0.89979762 | 0.09103385 | 0.51647917 |
| 0.999490048 | 0.08311069 | 0.56175794 | 0.89979762 | 0.08473624 | 0.55418419 |
| 0.999490048 | 0.08691128 | 0.56022882 | 0.89979762 | -0.0354704 | 0.81156107 |
| 0.999490048 | -0.030444 | 0.5629561 | 0.89982454 | -0.0112176 | 0.83072937 |
| 0.999490048 | -0.0515372 | 0.56319559 | 0.89986283 | 0.10492885 | 0.24408804 |
| 0.999490048 | 0.06690247 | 0.56310713 | 0.89986283 | 0.17902652 | 0.12936793 |
| 0.999490048 | -0.1974924 | 0.56342439 | 0.89993961 | -0.4466514 | 0.19735195 |
| 0.999490048 | 0.06175244 | 0.56345919 | 0.89993961 | 0.14783185 | 0.17327516 |
| 0.999490048 | -0.0522737 | 0.56366262 | 0.89997694 | 0.05169833 | 0.56793432 |
| 0.999490048 | -0.0275841 | 0.5637069 | 0.89997694 | -0.0220689 | 0.64374451 |
| 0.999490048 | 0.10067461 | 0.56380591 | 0.89997694 | 0.1148888 | 0.51052166 |
| 0.999490048 | -0.03325 | 0.56396836 | 0.90001225 | -0.0230893 | 0.688166 |
| 0.999490048 | 0.14367037 | 0.56404361 | 0.90001225 | 0.19411715 | 0.43969859 |
| 0.999490048 | 0.03814338 | 0.56429034 | 0.90012028 | 0.01852561 | 0.77893538 |
| 0.999490048 | 0.04736761 | 0.56432691 | 0.90012028 | 0.05502924 | 0.50357105 |
| 0.999490048 | -0.0808453 | 0.56489744 | 0.90034237 | -0.1599175 | 0.25944237 |
| 0.999490048 | -0.0496956 | 0.56485946 | 0.90034237 | 0.10504295 | 0.22897109 |
| 0.999490048 | -0.0245937 | 0.56458829 | 0.90034237 | -0.0435185 | 0.31146369 |
| 0.999490048 | 0.03931992 | 0.56485297 | 0.90034237 | 0.06168414 | 0.36880808 |
| 0.999490048 | -0.0306915 | 0.56508182 | 0.90036644 | -0.0144391 | 0.78610969 |
| 0.999490048 | 0.02862574 | 0.56512821 | 0.90036644 | -0.0285379 | 0.5663152 |
| 0.999490048 | -0.0327529 | 0.56526338 | 0.90041001 | 0.03699331 | 0.51640575 |
| 0.999490048 | -0.0269085 | 0.56538631 | 0.90041023 | -0.0234352 | 0.61632915 |
| 0.999490048 | 0.02860783 | 0.56547919 | 0.90041023 | 0.05406885 | 0.28143316 |
| 0.999490048 | -0.0767594 | 0.56592823 | 0.90050371 | -0.1639177 | 0.22560054 |
| 0.999490048 | -0.0291247 | 0.56566844 | 0.90050371 | -0.0772498 | 0.13508178 |

| | | | | | |
|---|---|---|---|---|---|
| 0.999490048 | -0.0246428 | 0.56589493 | 0.90050371 | -0.0312007 | 0.46818129 |
| 0.999490048 | 0.02944103 | 0.56596928 | 0.90050371 | 0.06963561 | 0.1810193 |
| 0.999490048 | -0.04786 | 0.56626355 | 0.90062869 | -0.0015432 | 0.9851956 |
| 0.999490048 | -0.0356014 | 0.56616261 | 0.90062869 | -0.0201663 | 0.74464223 |
| 0.999490048 | -0.0513829 | 0.56689275 | 0.90094299 | 0.03007856 | 0.73691679 |
| 0.999490048 | -0.0356182 | 0.56671434 | 0.90094299 | -0.0747442 | 0.23443272 |
| 0.999490048 | -0.0265271 | 0.56680561 | 0.90094299 | -0.0085584 | 0.85296203 |
| 0.999490048 | -0.0238976 | 0.56681027 | 0.90094299 | 8.05E-04 | 0.98456118 |
| 0.999490048 | 0.03314857 | 0.56706394 | 0.90104356 | -0.0103831 | 0.85729136 |
| 0.999490048 | 0.07483969 | 0.56753147 | 0.90161487 | 0.09414541 | 0.47288535 |
| 0.999490048 | -0.0844321 | 0.56775605 | 0.90162857 | 0.06880902 | 0.64110707 |
| 0.999490048 | -0.0772064 | 0.56768227 | 0.90162857 | -0.2361795 | 0.08860657 |
| 0.999490048 | -0.0709033 | 0.56846483 | 0.90187702 | 0.11915698 | 0.34069485 |
| 0.999490048 | -0.0334649 | 0.56856056 | 0.90187702 | -0.0188473 | 0.74757343 |
| 0.999490048 | -0.0317934 | 0.56848874 | 0.90187702 | -0.0018707 | 0.97314406 |
| 0.999490048 | -0.0260546 | 0.56803347 | 0.90187702 | -0.0202269 | 0.65716958 |
| 0.999490048 | 0.05339711 | 0.56823734 | 0.90187702 | 0.00653913 | 0.94410591 |
| 0.999490048 | 0.09416514 | 0.56847678 | 0.90187702 | 0.02100551 | 0.89845363 |
| 0.999490048 | -0.0468991 | 0.56894684 | 0.9019696 | -0.058177 | 0.48059969 |
| 0.999490048 | -0.0234956 | 0.56891026 | 0.9019696 | -0.0389517 | 0.34769592 |
| 0.999490048 | 0.02795129 | 0.569051 | 0.9019696 | -0.0116971 | 0.81114407 |
| 0.999490048 | 0.03186768 | 0.56902609 | 0.9019696 | 0.00679442 | 0.90305771 |
| 0.999490048 | -0.0293224 | 0.56937492 | 0.90210127 | 0.06791717 | 0.19371227 |
| 0.999490048 | 0.0249258 | 0.56935636 | 0.90210127 | 0.01049867 | 0.81009204 |
| 0.999490048 | 0.05752853 | 0.56945818 | 0.90210127 | 0.04640197 | 0.64598119 |
| 0.999490048 | 0.07468783 | 0.56957125 | 0.90210925 | 0.17536104 | 0.18810432 |
| 0.999490048 | -0.0497889 | 0.56970447 | 0.90214912 | -0.0466828 | 0.59387389 |
| 0.999490048 | -0.0625471 | 0.57158876 | 0.90237645 | -0.0012217 | 0.99115545 |
| 0.999490048 | -0.0576052 | 0.57054496 | 0.90237645 | -0.045529 | 0.65351916 |
| 0.999490048 | -0.0417047 | 0.57130265 | 0.90237645 | 0.04726865 | 0.52147373 |
| 0.999490048 | -0.0409869 | 0.57032261 | 0.90237645 | -0.0650643 | 0.36993191 |
| 0.999490048 | -0.03431 | 0.57047479 | 0.90237645 | 0.05911677 | 0.33134519 |
| 0.999490048 | -0.0243251 | 0.57188514 | 0.90237645 | -0.0475515 | 0.27343433 |
| 0.999490048 | 0.02575471 | 0.5710176 | 0.90237645 | 0.06152119 | 0.18239439 |
| 0.999490048 | 0.03089596 | 0.57128408 | 0.90237645 | -0.0079024 | 0.88451364 |
| 0.999490048 | 0.0316905 | 0.5718294 | 0.90237645 | 0.02777992 | 0.61991959 |
| 0.999490048 | 0.03257098 | 0.57091689 | 0.90237645 | -0.0649846 | 0.26271754 |
| 0.581280463 | 0.03379039 | 0.57074637 | 0.90237645 | -0.1442474 | 0.0214724 |
| 0.999490048 | 0.03611941 | 0.57115426 | 0.90237645 | 0.0961478 | 0.1390947 |
| 0.999490048 | 0.03658537 | 0.57177622 | 0.90237645 | -0.0207428 | 0.74799142 |
| 0.999490048 | 0.03722421 | 0.56998553 | 0.90237645 | 0.02943159 | 0.65292728 |
| 0.999490048 | 0.05404579 | 0.57190134 | 0.90237645 | 0.07693045 | 0.42257463 |

| | | | | | |
|---|---|---|---|---|---|
| 0.999490048 | 0.06312958 | 0.57090818 | 0.90237645 | 0.15385462 | 0.17388404 |
| 0.999490048 | 0.07110721 | 0.57173011 | 0.90237645 | 0.05800036 | 0.64378524 |
| 0.999490048 | 0.09149995 | 0.57026911 | 0.90237645 | -0.0705479 | 0.66120606 |
| 0.999490048 | 0.14169424 | 0.57126227 | 0.90237645 | -0.1816413 | 0.4688712 |
| 0.999490048 | 0.04216394 | 0.57212366 | 0.90255669 | 0.04992274 | 0.50412055 |
| 0.999490048 | 0.03280336 | 0.57229512 | 0.90265665 | 0.01985954 | 0.73193042 |
| 0.999490048 | 0.04336978 | 0.5724326 | 0.90270297 | -0.0373044 | 0.6270129 |
| 0.999490048 | 0.07055686 | 0.57260196 | 0.90279954 | -0.0425993 | 0.73283873 |
| 0.999490048 | -0.0393171 | 0.5731721 | 0.90313901 | 0.07267951 | 0.30135466 |
| 0.999490048 | -0.0338741 | 0.57408012 | 0.90313901 | -0.0863869 | 0.15878003 |
| 0.999490048 | -0.0324416 | 0.57324808 | 0.90313901 | -0.0031299 | 0.95652349 |
| 0.999490048 | -0.025535 | 0.57335161 | 0.90313901 | -0.0339787 | 0.45476391 |
| 0.999490048 | -0.0234683 | 0.57394629 | 0.90313901 | -0.0428246 | 0.30844174 |
| 0.999490048 | 0.02575285 | 0.57310586 | 0.90313901 | 0.03865969 | 0.39949128 |
| 0.999490048 | 0.03210594 | 0.57411519 | 0.90313901 | -0.0180182 | 0.75193248 |
| 0.999490048 | 0.04947193 | 0.57349737 | 0.90313901 | 0.04015883 | 0.64735554 |
| 0.999490048 | 0.05044443 | 0.57406454 | 0.90313901 | 0.02735479 | 0.75999484 |
| 0.999490048 | 0.05419744 | 0.57372103 | 0.90313901 | 0.12327075 | 0.2065772 |
| 0.999490048 | 0.0564548 | 0.57386427 | 0.90313901 | -0.0182604 | 0.85525249 |
| 0.999490048 | 0.07448177 | 0.57347987 | 0.90313901 | 0.28448168 | 0.03894557 |
| 0.999490048 | -0.0377694 | 0.57423578 | 0.90315855 | -0.0467557 | 0.48747957 |
| 0.999490048 | -0.0456431 | 0.57492064 | 0.90338679 | 0.01787551 | 0.82569234 |
| 0.999490048 | -0.0405529 | 0.57553095 | 0.90338679 | -0.0432845 | 0.55025493 |
| 0.999490048 | -0.0292934 | 0.57513734 | 0.90338679 | 0.01332172 | 0.79830487 |
| 0.999490048 | -0.0270175 | 0.57518613 | 0.90338679 | -0.0306675 | 0.52506144 |
| 0.999490048 | 0.03297238 | 0.57557099 | 0.90338679 | -0.0076998 | 0.89566894 |
| 0.999490048 | 0.03299045 | 0.57473921 | 0.90338679 | 0.02002325 | 0.73293006 |
| 0.999490048 | 0.03523572 | 0.57529214 | 0.90338679 | 0.06118182 | 0.33368084 |
| 0.999490048 | 0.03674515 | 0.57556707 | 0.90338679 | 0.02947034 | 0.6530458 |
| 0.999490048 | 0.04140475 | 0.57544998 | 0.90338679 | -0.0316583 | 0.66808716 |
| 0.999490048 | 0.04190817 | 0.57476695 | 0.90338679 | -0.0190333 | 0.79837399 |
| 0.999490048 | 0.15569708 | 0.57453143 | 0.90338679 | -0.3400459 | 0.22559201 |
| 0.999490048 | 0.03818647 | 0.57574504 | 0.90349014 | -0.006984 | 0.91823055 |
| 0.999490048 | -0.0286475 | 0.57608386 | 0.90360404 | 0.03790293 | 0.46050637 |
| 0.999490048 | -0.0280399 | 0.57657513 | 0.90360404 | -0.078461 | 0.12590185 |
| 0.999490048 | 0.0270715 | 0.57634557 | 0.90360404 | 0.04239071 | 0.38378072 |
| 0.999490048 | 0.04149465 | 0.57644312 | 0.90360404 | 0.0827211 | 0.26986796 |
| 0.999490048 | 0.05330034 | 0.57607583 | 0.90360404 | -0.0593142 | 0.53410933 |
| 0.999490048 | 0.05884927 | 0.57617187 | 0.90360404 | 0.03335381 | 0.7508689 |
| 0.999490048 | 0.20057277 | 0.5765672 | 0.90360404 | -0.1052134 | 0.76907126 |
| 0.999490048 | 0.0216669 | 0.5767081 | 0.90364283 | -0.0188038 | 0.62780743 |
| 0.999490048 | 0.02999072 | 0.57691084 | 0.90379091 | -0.0264806 | 0.62205035 |

| | | | | | |
|---|---|---|---|---|---|
| 0.999490048 | 0.09271684 | 0.57707913 | 0.90388496 | -0.1861263 | 0.27297661 |
| 0.999490048 | -0.0268227 | 0.57731449 | 0.90390267 | -0.0131971 | 0.78342596 |
| 0.999490048 | 0.03204482 | 0.57741519 | 0.90390267 | 0.05369895 | 0.35317284 |
| 0.999490048 | 0.03504528 | 0.57736022 | 0.90390267 | 0.07015462 | 0.26906047 |
| 0.999490048 | -0.0357874 | 0.57764192 | 0.90391867 | -0.0046944 | 0.94159262 |
| 0.999490048 | 0.05792209 | 0.57762584 | 0.90391867 | -0.0693175 | 0.50572225 |
| 0.999490048 | 0.04026127 | 0.57776942 | 0.90394878 | 0.15569098 | 0.0387548 |
| 0.999490048 | 0.0322719 | 0.57800366 | 0.90414586 | 0.06688416 | 0.25371699 |
| 0.999490048 | 0.03170836 | 0.57832361 | 0.90447689 | -0.0059454 | 0.91674435 |
| 0.999490048 | -0.0322122 | 0.57847579 | 0.90454548 | -0.0038328 | 0.94712755 |
| 0.999490048 | -0.1239884 | 0.57868339 | 0.90470067 | -0.4659972 | 0.05436691 |
| 0.999490048 | 0.02121274 | 0.57899459 | 0.90484837 | 0.01064246 | 0.78022948 |
| 0.999490048 | 0.03237687 | 0.57897596 | 0.90484837 | 0.03041342 | 0.60207338 |
| 0.999490048 | 0.02648533 | 0.57965006 | 0.90553377 | 0.02725009 | 0.56882248 |
| 0.999490048 | 0.0704935 | 0.57959479 | 0.90553377 | -0.0196201 | 0.8771145 |
| 0.999490048 | -0.0689246 | 0.58041373 | 0.90572234 | -0.1356533 | 0.28080567 |
| 0.999490048 | -0.0447129 | 0.5803014 | 0.90572234 | -0.001227 | 0.98785649 |
| 0.999490048 | -0.0389873 | 0.57989236 | 0.90572234 | -2.47E-04 | 0.99719447 |
| 0.999490048 | -0.0325561 | 0.58003979 | 0.90572234 | 0.00385215 | 0.94763911 |
| 0.999490048 | 0.02738322 | 0.58014699 | 0.90572234 | 0.04645987 | 0.35073047 |
| 0.999490048 | 0.038626 | 0.58042159 | 0.90572234 | 0.02274217 | 0.74433682 |
| 0.999490048 | -0.0361279 | 0.58064671 | 0.90590434 | 0.0164795 | 0.80057074 |
| 0.999490048 | -0.0732466 | 0.5815843 | 0.9063934 | -0.0200102 | 0.87999462 |
| 0.999490048 | -0.0584563 | 0.58194426 | 0.9063934 | 0.02852875 | 0.78768792 |
| 0.999490048 | -0.0571381 | 0.58229658 | 0.9063934 | 0.04881242 | 0.63815913 |
| 0.999490048 | -0.0359195 | 0.58233628 | 0.9063934 | -0.0853673 | 0.19735258 |
| 0.999490048 | -0.0332148 | 0.58135437 | 0.9063934 | 0.06797828 | 0.26367959 |
| 0.999490048 | -0.0315427 | 0.58109323 | 0.9063934 | 0.01890767 | 0.74033325 |
| 0.999490048 | -0.0290549 | 0.58240371 | 0.9063934 | -0.0023975 | 0.96369884 |
| 0.999490048 | -0.0288627 | 0.58250434 | 0.9063934 | -0.0098249 | 0.85116211 |
| 0.999490048 | -0.0245701 | 0.58218462 | 0.9063934 | -0.0118349 | 0.79051449 |
| 0.999490048 | -0.0235038 | 0.58150838 | 0.9063934 | -0.028547 | 0.50382731 |
| 0.999490048 | 0.03309239 | 0.58258843 | 0.9063934 | 0.04347665 | 0.47122611 |
| 0.999490048 | 0.03889223 | 0.58123941 | 0.9063934 | 0.00960602 | 0.89132456 |
| 0.999490048 | 0.04440417 | 0.58192847 | 0.9063934 | 0.00177194 | 0.98241783 |
| 0.999490048 | 0.05687196 | 0.5817245 | 0.9063934 | 0.17348692 | 0.10101243 |
| 0.999490048 | 0.07246904 | 0.58221686 | 0.9063934 | -0.0267429 | 0.83869998 |
| 0.999490048 | 0.03447883 | 0.58290819 | 0.90655306 | -0.0272813 | 0.66356117 |
| 0.999490048 | 0.14112772 | 0.58282643 | 0.90655306 | 0.03546267 | 0.88991332 |
| 0.999490048 | -0.0750996 | 0.5836812 | 0.90715614 | 0.07292713 | 0.59454965 |
| 0.692187333 | -0.0641922 | 0.58368885 | 0.90715614 | 0.271229 | 0.02717824 |
| 0.999490048 | -0.0467203 | 0.58394425 | 0.90715614 | 0.10205731 | 0.23682237 |

| | | | | | |
|---|---|---|---|---|---|
| 0.999490048 | -0.0287054 | 0.58394782 | 0.90715614 | -0.05493 | 0.2985042 |
| 0.999490048 | 0.02741067 | 0.58352924 | 0.90715614 | -0.0144251 | 0.77246733 |
| 0.999490048 | 0.05582923 | 0.58378771 | 0.90715614 | 0.07228614 | 0.47900069 |
| 0.999490048 | 0.03557236 | 0.58406385 | 0.90716762 | -0.0551721 | 0.3978155 |
| 0.999490048 | 0.02972368 | 0.58425293 | 0.90729254 | -0.0081849 | 0.87989859 |
| 0.999490048 | 0.02419882 | 0.58494297 | 0.9081952 | 0.0108744 | 0.80565756 |
| 0.999490048 | 0.0339444 | 0.58516113 | 0.90836501 | 0.06367511 | 0.30946375 |
| 0.999927563 | 0.02729702 | 0.58528817 | 0.90839335 | 0.02730154 | 0.58522632 |
| 0.999490048 | -0.0699988 | 0.58566416 | 0.90857669 | -0.1696019 | 0.20067411 |
| 0.999490048 | -0.0308266 | 0.58602298 | 0.90857669 | -0.0424271 | 0.45472369 |
| 0.999490048 | -0.0288129 | 0.58598831 | 0.90857669 | -0.0473856 | 0.37280497 |
| 0.999490048 | -0.0262214 | 0.58616641 | 0.90857669 | -0.0092219 | 0.84776838 |
| 0.999490048 | 0.02587829 | 0.58626297 | 0.90857669 | 0.02824899 | 0.55265958 |
| 0.999490048 | 0.04372194 | 0.58614066 | 0.90857669 | 0.0208471 | 0.79471311 |
| 0.999490048 | 0.04521593 | 0.58627679 | 0.90857669 | -0.0066492 | 0.93602466 |
| 0.999490048 | 0.10977209 | 0.58611424 | 0.90857669 | 0.11409997 | 0.57164454 |
| 0.999490048 | 0.02906708 | 0.58658121 | 0.90859746 | 0.09182508 | 0.09404338 |
| 0.999490048 | 0.0496205 | 0.58657018 | 0.90859746 | -0.1359064 | 0.14380426 |
| 0.999490048 | 0.0720202 | 0.58661664 | 0.90859746 | 0.18364186 | 0.17239783 |
| 0.999490048 | 0.03560294 | 0.587345 | 0.90955688 | 0.04330454 | 0.5098023 |
| 0.999490048 | 0.02436968 | 0.58779641 | 0.91008715 | -0.0430097 | 0.3417467 |
| 0.999490048 | 0.03537451 | 0.58792722 | 0.91012093 | 0.10266974 | 0.12364363 |
| 0.999490048 | 0.0219862 | 0.58803686 | 0.91012193 | -0.0431929 | 0.29127436 |
| 0.999490048 | -0.0832146 | 0.58837977 | 0.91031956 | -0.1473802 | 0.34082984 |
| 0.999490048 | -0.0355877 | 0.5883826 | 0.91031956 | -0.0197603 | 0.76333038 |
| 0.999490048 | -0.0309885 | 0.58862979 | 0.91036465 | -0.0902507 | 0.12308779 |
| 0.999490048 | 0.03333664 | 0.58855019 | 0.91036465 | 0.00273835 | 0.96445417 |
| 0.999490048 | -0.0279137 | 0.58914018 | 0.91098529 | -0.0669057 | 0.20160523 |
| 0.999490048 | 0.02746647 | 0.58946046 | 0.91127631 | 0.03570023 | 0.48394579 |
| 0.999490048 | 0.02902203 | 0.58960542 | 0.91127631 | -0.0570535 | 0.29295411 |
| 0.999490048 | 0.03513361 | 0.58976493 | 0.91127631 | 0.02310733 | 0.72243451 |
| 0.999490048 | 0.1001928 | 0.58971564 | 0.91127631 | 0.12437837 | 0.50535107 |
| 0.999490048 | -0.071104 | 0.59037668 | 0.91134005 | -0.1654793 | 0.21606963 |
| 0.999490048 | -0.0355106 | 0.59046104 | 0.91134005 | -0.0159317 | 0.80877042 |
| 0.999490048 | -0.0282695 | 0.59027145 | 0.91134005 | -0.0515919 | 0.32903004 |
| 0.999490048 | -0.0278969 | 0.59022977 | 0.91134005 | -0.0265261 | 0.60852022 |
| 0.999490048 | 0.03414036 | 0.59045724 | 0.91134005 | 0.0420317 | 0.50824321 |
| 0.999490048 | 0.06279066 | 0.59019912 | 0.91134005 | 0.07140004 | 0.54062651 |
| 0.999490048 | -0.0671695 | 0.59277251 | 0.91145177 | 0.08287717 | 0.50995889 |
| 0.999490048 | -0.06463 | 0.59280392 | 0.91145177 | -0.1166707 | 0.3374619 |
| 0.999490048 | -0.0480153 | 0.59281339 | 0.91145177 | -0.0054542 | 0.95142382 |
| 0.999490048 | -0.039154 | 0.59148339 | 0.91145177 | -0.0061968 | 0.93211673 |

| | | | | | |
|---|---|---|---|---|---|
| 0.999490048 | -0.0296967 | 0.592463 | 0.91145177 | 0.01483324 | 0.78874174 |
| 0.999490048 | -0.0293883 | 0.59267636 | 0.91145177 | 0.00600323 | 0.91274094 |
| 0.999490048 | -0.0274941 | 0.59229615 | 0.91145177 | -0.0078195 | 0.87862794 |
| 0.999490048 | -0.025567 | 0.59082025 | 0.91145177 | -0.020345 | 0.66843911 |
| 0.999490048 | -0.023875 | 0.59223882 | 0.91145177 | -0.0365825 | 0.41364516 |
| 0.999490048 | 0.02217132 | 0.59257993 | 0.91145177 | -0.0016789 | 0.96758506 |
| 0.999490048 | 0.02394696 | 0.59279675 | 0.91145177 | 0.03898054 | 0.38625671 |
| 0.999490048 | 0.0252629 | 0.59101779 | 0.91145177 | 0.02274355 | 0.62835739 |
| 0.999490048 | 0.02705673 | 0.59106912 | 0.91145177 | 0.01401632 | 0.78029985 |
| 0.999490048 | 0.02968972 | 0.5928257 | 0.91145177 | 0.03639039 | 0.51279802 |
| 0.999490048 | 0.03905335 | 0.59194443 | 0.91145177 | -0.0118696 | 0.8702466 |
| 0.999490048 | 0.03978987 | 0.59243415 | 0.91145177 | 0.03594559 | 0.62849232 |
| 0.999490048 | 0.04377944 | 0.59128159 | 0.91145177 | 0.05056762 | 0.53549475 |
| 0.999490048 | 0.06503772 | 0.59066398 | 0.91145177 | 0.00156084 | 0.98967063 |
| 0.999490048 | 0.11879508 | 0.59139858 | 0.91145177 | -0.0243872 | 0.91200573 |
| 0.999490048 | 0.12362228 | 0.59155532 | 0.91145177 | -0.1752932 | 0.4480643 |
| 0.999490048 | 0.12702954 | 0.59167416 | 0.91145177 | 0.30856562 | 0.20646758 |
| 0.999490048 | -0.0254105 | 0.59322342 | 0.91166387 | -0.0105776 | 0.82362212 |
| 0.999490048 | -0.0234495 | 0.59329119 | 0.91166387 | -0.0248851 | 0.57101082 |
| 0.999490048 | 0.03135469 | 0.59311103 | 0.91166387 | 0.05267926 | 0.37181182 |
| 0.999490048 | 0.04482389 | 0.59342845 | 0.911707 | 0.01271742 | 0.87928727 |
| 0.999490048 | -0.0369086 | 0.59362741 | 0.91184489 | -0.0485459 | 0.48370063 |
| 0.999490048 | -0.0998077 | 0.5955951 | 0.91189021 | -0.2083578 | 0.27236319 |
| 0.999490048 | -0.0795498 | 0.59648162 | 0.91189021 | 0.04586782 | 0.75970789 |
| 0.999490048 | -0.0596761 | 0.59600401 | 0.91189021 | -0.1371509 | 0.22857438 |
| 0.999490048 | -0.0497016 | 0.59379435 | 0.91189021 | -0.2320429 | 0.01836228 |
| 0.999490048 | -0.0445227 | 0.59567206 | 0.91189021 | -0.0256243 | 0.75962051 |
| 0.999490048 | -0.0445193 | 0.59585676 | 0.91189021 | 0.04733637 | 0.57294505 |
| 0.999490048 | -0.0365399 | 0.59632421 | 0.91189021 | 0.03549425 | 0.60682 |
| 0.999490048 | -0.0335352 | 0.59538969 | 0.91189021 | 0.02639835 | 0.67559646 |
| 0.999490048 | -0.0288701 | 0.59498063 | 0.91189021 | 0.01327255 | 0.80648734 |
| 0.999490048 | -0.0246134 | 0.59420497 | 0.91189021 | -0.0737756 | 0.11826135 |
| 0.999490048 | 0.02268942 | 0.59606348 | 0.91189021 | 0.00163836 | 0.96938171 |
| 0.999490048 | 0.02457382 | 0.59464993 | 0.91189021 | 0.00622498 | 0.89248965 |
| 0.999490048 | 0.0261907 | 0.59631916 | 0.91189021 | -0.011217 | 0.82012271 |
| 0.999490048 | 0.03018508 | 0.59581494 | 0.91189021 | -0.0398472 | 0.48470833 |
| 0.999490048 | 0.03028334 | 0.59596114 | 0.91189021 | 0.07568911 | 0.19150882 |
| 0.999490048 | 0.0321038 | 0.59632798 | 0.91189021 | 0.06474936 | 0.28949773 |
| 0.999490048 | 0.03364557 | 0.59517228 | 0.91189021 | 0.06659776 | 0.29686148 |
| 0.999490048 | 0.03808814 | 0.5950347 | 0.91189021 | 0.0361286 | 0.61401466 |
| 0.999490048 | 0.0387218 | 0.59594431 | 0.91189021 | 0.01390375 | 0.84863139 |
| 0.999490048 | 0.04386499 | 0.59440637 | 0.91189021 | -0.0566735 | 0.4923154 |

| | | | | | |
|---|---|---|---|---|---|
| 0.999490048 | 0.04448238 | 0.59487822 | 0.91189021 | -0.048616 | 0.56132406 |
| 0.999490048 | 0.04644154 | 0.59468164 | 0.91189021 | 0.09117123 | 0.30012596 |
| 0.999490048 | 0.04735543 | 0.59494188 | 0.91189021 | 0.00558232 | 0.94988339 |
| 0.999490048 | 0.05117918 | 0.59541093 | 0.91189021 | 0.11604065 | 0.23397766 |
| 0.999490048 | 0.07401607 | 0.59649633 | 0.91189021 | -0.2117227 | 0.13753132 |
| 0.999490048 | 0.10293088 | 0.59468276 | 0.91189021 | 0.14099322 | 0.46722587 |
| 0.999490048 | -0.0317321 | 0.5967458 | 0.9121046 | -0.0623283 | 0.30253973 |
| 0.999490048 | 0.02930391 | 0.59704507 | 0.912395 | 0.04242019 | 0.44550312 |
| 0.999490048 | -0.026583 | 0.59749026 | 0.91290826 | -0.0193908 | 0.69974535 |
| 0.999490048 | -0.1901939 | 0.59874435 | 0.9129313 | -0.1553755 | 0.66699556 |
| 0.999490048 | -0.0451601 | 0.5986119 | 0.9129313 | -0.1348856 | 0.12378625 |
| 0.999490048 | -0.0360919 | 0.60026005 | 0.9129313 | -0.0463481 | 0.50175276 |
| 0.999490048 | -0.0342612 | 0.59861305 | 0.9129313 | -0.0684941 | 0.29661904 |
| 0.999490048 | -0.0327742 | 0.60018431 | 0.9129313 | -0.0295923 | 0.63586238 |
| 0.999490048 | -0.0312573 | 0.59815369 | 0.9129313 | -0.0187133 | 0.75191873 |
| 0.999490048 | -0.0312202 | 0.60020836 | 0.9129313 | 0.0054999 | 0.92623941 |
| 0.999490048 | -0.0294113 | 0.59810033 | 0.9129313 | -0.0695783 | 0.21816108 |
| 0.999490048 | -0.0293155 | 0.6003185 | 0.9129313 | -0.0040907 | 0.94155851 |
| 0.999490048 | -0.0282973 | 0.59813428 | 0.9129313 | -0.0036975 | 0.94493883 |
| 0.999490048 | -0.0278008 | 0.600348 | 0.9129313 | -0.0654898 | 0.22283316 |
| 0.999490048 | -0.0250041 | 0.60006764 | 0.9129313 | 0.01921612 | 0.68665755 |
| 0.999490048 | -0.0240672 | 0.59817872 | 0.9129313 | -0.0335027 | 0.46433706 |
| 0.999490048 | -0.0176382 | 0.59793116 | 0.9129313 | -0.0031616 | 0.92448287 |
| 0.999490048 | 0.02464236 | 0.59868722 | 0.9129313 | 0.01090784 | 0.81538485 |
| 0.999490048 | 0.02534893 | 0.59899835 | 0.9129313 | 0.00995724 | 0.83597601 |
| 0.999490048 | 0.02544133 | 0.59962132 | 0.9129313 | -0.0133305 | 0.7828469 |
| 0.999490048 | 0.02565362 | 0.60030901 | 0.9129313 | 0.03903168 | 0.42701094 |
| 0.999490048 | 0.02799205 | 0.59994974 | 0.9129313 | 0.04304469 | 0.42169637 |
| 0.999490048 | 0.02860241 | 0.59810978 | 0.9129313 | -8.79E-04 | 0.9870401 |
| 0.999490048 | 0.03770975 | 0.60015798 | 0.9129313 | 0.03086826 | 0.66758279 |
| 0.999490048 | 0.03865092 | 0.59859138 | 0.9129313 | 0.0576987 | 0.43351913 |
| 0.999490048 | 0.04111154 | 0.6003109 | 0.9129313 | -0.0683218 | 0.38619313 |
| 0.956358655 | 0.06252683 | 0.60006577 | 0.9129313 | -0.2260327 | 0.0663238 |
| 0.999490048 | 0.09424932 | 0.59983072 | 0.9129313 | 0.08989474 | 0.61670247 |
| 0.999490048 | 0.19556592 | 0.59946886 | 0.9129313 | 0.43332032 | 0.2600324 |
| 0.999490048 | 0.02709502 | 0.60050661 | 0.91300623 | -0.0061762 | 0.9047337 |
| 0.999490048 | -0.0754919 | 0.60076154 | 0.91320248 | -0.0626471 | 0.66382388 |
| 0.999490048 | 0.04458518 | 0.60085442 | 0.91320248 | -3.50E-04 | 0.99671737 |
| 0.999490048 | -0.1077797 | 0.6022081 | 0.91374081 | 0.01098161 | 0.95753091 |
| 0.999490048 | -0.0967445 | 0.6023986 | 0.91374081 | -0.067438 | 0.71611541 |
| 0.999490048 | -0.0767997 | 0.60268084 | 0.91374081 | 0.14174158 | 0.33982857 |
| 0.971241857 | -0.0599828 | 0.60159925 | 0.91374081 | 0.21010484 | 0.07578054 |

| | | | | | |
|---|---|---|---|---|---|
| 0.999490048 | -0.0510538 | 0.60168559 | 0.91374081 | 0.01281512 | 0.89548922 |
| 0.999490048 | -0.0358064 | 0.6020113 | 0.91374081 | 0.01791051 | 0.7937842 |
| 0.999490048 | -0.0312578 | 0.60274064 | 0.91374081 | -0.0095882 | 0.87284441 |
| 0.999490048 | -0.0304448 | 0.60207989 | 0.91374081 | -0.0183428 | 0.75299159 |
| 0.999490048 | -0.0296518 | 0.60189486 | 0.91374081 | -0.0230534 | 0.68472136 |
| 0.999490048 | -0.0255138 | 0.60262604 | 0.91374081 | -0.0573606 | 0.24693366 |
| 0.999490048 | 0.02387508 | 0.6021563 | 0.91374081 | -0.0528176 | 0.25377312 |
| 0.999490048 | 0.02801792 | 0.60141496 | 0.91374081 | 0.01962749 | 0.71402992 |
| 0.999490048 | 0.0419169 | 0.6023244 | 0.91374081 | -0.0512245 | 0.52483822 |
| 0.999490048 | 0.04896035 | 0.60250007 | 0.91374081 | 0.12551333 | 0.18835924 |
| 0.999490048 | -0.074041 | 0.60455474 | 0.91434266 | 0.04239496 | 0.7664194 |
| 0.999490048 | -0.0542203 | 0.6041098 | 0.91434266 | -0.0772453 | 0.46134312 |
| 0.999490048 | -0.0465599 | 0.60428936 | 0.91434266 | -0.0799188 | 0.37935706 |
| 0.999490048 | -0.043073 | 0.6034338 | 0.91434266 | -0.091999 | 0.27176698 |
| 0.999490048 | -0.0424971 | 0.60440948 | 0.91434266 | -0.1099041 | 0.18688082 |
| 0.999490048 | -0.0359724 | 0.60465228 | 0.91434266 | 0.00622471 | 0.92842767 |
| 0.999490048 | -0.035477 | 0.60377886 | 0.91434266 | -0.0376847 | 0.58158051 |
| 0.999490048 | -0.0315398 | 0.60410998 | 0.91434266 | -0.0424767 | 0.48595044 |
| 0.999490048 | 0.03109144 | 0.6042201 | 0.91434266 | 0.02191517 | 0.71447088 |
| 0.999490048 | 0.03574724 | 0.60447658 | 0.91434266 | 0.12339461 | 0.08214829 |
| 0.999490048 | 0.0378264 | 0.60349147 | 0.91434266 | -0.0912671 | 0.21601279 |
| 0.999490048 | 0.0406996 | 0.60461655 | 0.91434266 | 0.09237846 | 0.24509082 |
| 0.999490048 | 0.04351693 | 0.60375514 | 0.91434266 | 0.00736714 | 0.92979905 |
| 0.999490048 | 0.04561927 | 0.60467068 | 0.91434266 | -0.0355684 | 0.68616476 |
| 0.999490048 | -0.0298748 | 0.60510182 | 0.91482893 | -0.0306779 | 0.59548771 |
| 0.999490048 | -0.0645282 | 0.60529529 | 0.91495576 | 0.01364314 | 0.91276689 |
| 0.999490048 | -0.0827946 | 0.60579536 | 0.91554593 | -0.0648101 | 0.68591704 |
| 0.999490048 | 0.1113772 | 0.60610253 | 0.91567869 | 0.12217062 | 0.57231662 |
| 0.999490048 | 0.24103965 | 0.60600902 | 0.91567869 | 0.24386393 | 0.60184684 |
| 0.999490048 | -0.0405265 | 0.60678201 | 0.91641072 | -0.0857322 | 0.28064614 |
| 0.999490048 | -0.0277886 | 0.606825 | 0.91641072 | -0.0558982 | 0.30446879 |
| 0.999490048 | 0.03627636 | 0.60691632 | 0.91641072 | 0.02532744 | 0.71907728 |
| 0.999490048 | -0.0649845 | 0.60709372 | 0.91651284 | -0.1270655 | 0.31826214 |
| 0.999490048 | -0.0920636 | 0.60758438 | 0.91671617 | -0.0194524 | 0.91337631 |
| 0.999490048 | -0.0298443 | 0.60766755 | 0.91671617 | 0.07666112 | 0.19367322 |
| 0.999490048 | 0.02291016 | 0.60738161 | 0.91671617 | -0.0023456 | 0.95793548 |
| 0.999490048 | 0.03640631 | 0.60765014 | 0.91671617 | 0.10726394 | 0.13803211 |
| 0.999490048 | -0.0455437 | 0.60811732 | 0.9167376 | 0.07306371 | 0.4126701 |
| 0.999490048 | -0.0439923 | 0.60812091 | 0.9167376 | -0.0690003 | 0.42304054 |
| 0.999490048 | -0.0436651 | 0.60791834 | 0.9167376 | -0.0675886 | 0.42881426 |
| 0.999490048 | -0.02245 | 0.60802028 | 0.9167376 | 0.02445259 | 0.57658728 |
| 0.999490048 | -0.059232 | 0.60834017 | 0.9167521 | -0.1676793 | 0.15421892 |

| | | | | | |
|---|---|---|---|---|---|
| 0.999490048 | -0.0425979 | 0.60839354 | 0.9167521 | 0.02634743 | 0.7509198 |
| 0.999490048 | -0.0366441 | 0.60891154 | 0.9167521 | -0.075806 | 0.29398308 |
| 0.999490048 | -0.0261959 | 0.60873017 | 0.9167521 | 0.06800156 | 0.19043168 |
| 0.999490048 | -0.0213258 | 0.60900885 | 0.9167521 | -0.0359038 | 0.39143747 |
| 0.999490048 | 0.05585503 | 0.60897008 | 0.9167521 | -0.0910259 | 0.40651114 |
| 0.999490048 | 0.07747107 | 0.60864104 | 0.9167521 | 0.08308963 | 0.58305345 |
| 0.999490048 | 0.09168864 | 0.6087735 | 0.9167521 | -0.0388287 | 0.8275554 |
| 0.999490048 | 0.1281138 | 0.60938427 | 0.91707935 | 0.28881718 | 0.26470958 |
| 0.999490048 | 0.17816578 | 0.60944591 | 0.91707935 | 0.10531556 | 0.76140646 |
| 0.999490048 | -0.0279776 | 0.60996702 | 0.91769813 | -0.0561944 | 0.30934953 |
| 0.999490048 | 0.03283515 | 0.6100964 | 0.91772742 | -0.0013388 | 0.98336812 |
| 0.999490048 | -0.0790532 | 0.61060464 | 0.91831529 | -0.0876017 | 0.57279532 |
| 0.999490048 | -0.027526 | 0.61092712 | 0.91831529 | 0.01896662 | 0.72556379 |
| 0.999490048 | -0.0267474 | 0.61077568 | 0.91831529 | -0.0243886 | 0.64244225 |
| 0.999490048 | 0.07619931 | 0.61084922 | 0.91831529 | -0.11753 | 0.43417468 |
| 0.967348048 | -0.1100835 | 0.61114577 | 0.91847861 | -0.6885788 | 0.01049354 |
| 0.999490048 | -0.0217901 | 0.61129954 | 0.91854439 | -0.0441821 | 0.30664963 |
| 0.999490048 | -0.0739454 | 0.61164924 | 0.91857395 | -0.0170564 | 0.90654226 |
| 0.999490048 | 0.03650767 | 0.61152277 | 0.91857395 | 0.02463641 | 0.73142342 |
| 0.999490048 | 0.07042478 | 0.61158372 | 0.91857395 | 0.16108061 | 0.25054175 |
| 0.999490048 | -0.1232376 | 0.61250893 | 0.918611 | 0.13011616 | 0.59292166 |
| 0.999490048 | -0.0315505 | 0.61255402 | 0.918611 | -0.02053 | 0.74137832 |
| 0.999490048 | 0.02487325 | 0.61227957 | 0.918611 | 7.46E-04 | 0.98784068 |
| 0.999490048 | 0.02601815 | 0.61226358 | 0.918611 | 0.11040732 | 0.03899898 |
| 0.999490048 | 0.02660691 | 0.61205653 | 0.918611 | -0.0357942 | 0.49597967 |
| 0.999490048 | 0.03093405 | 0.61244629 | 0.918611 | 0.04487967 | 0.46363453 |
| 0.999490048 | 0.03204021 | 0.61217496 | 0.918611 | 0.03444163 | 0.58592579 |
| 0.999490048 | 0.07309723 | 0.61183312 | 0.918611 | 0.11865345 | 0.41206871 |
| 0.999490048 | -0.0615695 | 0.61305518 | 0.91867309 | -0.023038 | 0.84958368 |
| 0.999490048 | -0.0299152 | 0.61293825 | 0.91867309 | 0.01837369 | 0.7556401 |
| 0.999490048 | 0.02965018 | 0.61282938 | 0.91867309 | 0.0115357 | 0.84358286 |
| 0.999490048 | 0.03279693 | 0.61307132 | 0.91867309 | -0.0268121 | 0.67904368 |
| 0.999490048 | 0.03800209 | 0.61314553 | 0.91867309 | 0.01146141 | 0.8785213 |
| 0.999490048 | -0.0239579 | 0.61351617 | 0.91906351 | -0.004454 | 0.92500567 |
| 0.890934946 | -0.0377551 | 0.61403225 | 0.91934181 | 0.15020046 | 0.05288948 |
| 0.999490048 | 0.0275988 | 0.61399908 | 0.91934181 | 0.05129008 | 0.35158469 |
| 0.999490048 | 0.04327569 | 0.61381308 | 0.91934181 | 0.09472416 | 0.27393081 |
| 0.999490048 | -0.0285035 | 0.61445079 | 0.91937438 | 0.03529258 | 0.5333741 |
| 0.999490048 | 0.0294889 | 0.61450372 | 0.91937438 | 0.01619403 | 0.78171766 |
| 0.999490048 | 0.04232127 | 0.61460453 | 0.91937438 | 0.07944527 | 0.34763845 |
| 0.999490048 | 0.05932462 | 0.61456452 | 0.91937438 | -0.0913743 | 0.43957038 |
| 0.999490048 | 0.060978 | 0.61436684 | 0.91937438 | 0.00141616 | 0.99063978 |

| | | | | | |
|---|---|---|---|---|---|
| 0.999490048 | -0.0576515 | 0.61482564 | 0.91946838 | -0.1051191 | 0.36169754 |
| 0.999490048 | -0.0355467 | 0.61488759 | 0.91946838 | -0.0290317 | 0.68088992 |
| 0.999490048 | 0.02960219 | 0.61535673 | 0.92000514 | 0.10068565 | 0.09581157 |
| 0.999490048 | 0.03210502 | 0.61546744 | 0.92000592 | 0.04285689 | 0.5033865 |
| 0.999490048 | -0.0280886 | 0.6156254 | 0.92007733 | 0.02052521 | 0.71339065 |
| 0.999490048 | -0.0276815 | 0.61601157 | 0.92033128 | -0.0350842 | 0.52565413 |
| 0.999490048 | 0.03529934 | 0.61601575 | 0.92033128 | 0.04184466 | 0.55258802 |
| 0.999490048 | 0.15390999 | 0.61618112 | 0.92041366 | 0.40140282 | 0.19749719 |
| 0.999490048 | -0.0252831 | 0.6169414 | 0.92138449 | -0.0585233 | 0.25205425 |
| 0.999490048 | 0.02224023 | 0.61725551 | 0.92152733 | -0.019623 | 0.65908236 |
| 0.999490048 | 0.04624293 | 0.61725777 | 0.92152733 | 0.07567391 | 0.41543023 |
| 0.999490048 | -0.0369985 | 0.61762994 | 0.92170671 | -0.1030677 | 0.17134542 |
| 0.999490048 | 0.03462562 | 0.61750209 | 0.92170671 | 0.00428471 | 0.95060207 |
| 0.999490048 | 0.07318628 | 0.61770907 | 0.92170671 | 0.16725131 | 0.25899676 |
| 0.999490048 | -0.0577333 | 0.61783283 | 0.92172666 | 0.13841805 | 0.23708557 |
| 0.999490048 | 0.02861661 | 0.61794906 | 0.92173538 | 0.02666939 | 0.64194993 |
| 0.999490048 | -0.0622756 | 0.61819388 | 0.92193586 | -0.2028177 | 0.11271768 |
| 0.999490048 | -0.0388768 | 0.61838602 | 0.92194166 | 0.05483129 | 0.48340349 |
| 0.999490048 | 0.02166395 | 0.61841859 | 0.92194166 | 0.03501852 | 0.42259936 |
| 0.999490048 | 0.06879263 | 0.61856869 | 0.92200081 | 0.01600084 | 0.90730418 |
| 0.999490048 | 0.06601713 | 0.61870891 | 0.92204523 | 0.07056138 | 0.59489945 |
| 0.999490048 | -0.026719 | 0.61895684 | 0.92209803 | -0.0449662 | 0.40474081 |
| 0.999490048 | 0.02246688 | 0.61907564 | 0.92209803 | 0.08381452 | 0.07204028 |
| 0.999490048 | 0.0276904 | 0.61902556 | 0.92209803 | -0.0105701 | 0.84914323 |
| 0.999490048 | -0.0348118 | 0.61952263 | 0.9223053 | 0.02218514 | 0.75131997 |
| 0.999490048 | 0.03136519 | 0.61935511 | 0.9223053 | 0.0504515 | 0.42607391 |
| 0.999490048 | 0.04109425 | 0.61954616 | 0.9223053 | -9.64E-04 | 0.99068573 |
| 0.999490048 | 0.02406174 | 0.62065655 | 0.92379361 | -0.0296717 | 0.54217434 |
| 0.999490048 | -0.0244092 | 0.62124805 | 0.92401518 | 0.02996759 | 0.54462272 |
| 0.999490048 | 0.02266052 | 0.62117384 | 0.92401518 | 0.02620971 | 0.56792188 |
| 0.999490048 | 0.02425594 | 0.62110785 | 0.92401518 | 0.02127671 | 0.66442115 |
| 0.999490048 | 0.07620743 | 0.62105842 | 0.92401518 | 0.16488609 | 0.28910336 |
| 0.999490048 | -0.0434196 | 0.62143011 | 0.92412136 | -0.1399828 | 0.1194591 |
| 0.999490048 | -0.0240124 | 0.62168529 | 0.92433621 | 0.02507544 | 0.60641743 |
| 0.999490048 | 0.0230896 | 0.62197329 | 0.9244352 | 0.04199876 | 0.37248345 |
| 0.999490048 | 0.04109508 | 0.62194121 | 0.9244352 | 0.06858016 | 0.41259814 |
| 0.999490048 | -0.0285061 | 0.62218514 | 0.92458551 | -0.015838 | 0.78390008 |
| 0.999490048 | 0.02581307 | 0.62237524 | 0.92470342 | 0.02831091 | 0.58928292 |
| 0.999490048 | 0.02231838 | 0.62250338 | 0.92472926 | 0.01285451 | 0.77640037 |
| 0.999490048 | -0.0334891 | 0.62270688 | 0.92486703 | -0.0824703 | 0.23128212 |
| 0.999490048 | -0.0449233 | 0.623283 | 0.92488423 | 0.06178037 | 0.50026994 |
| 0.999490048 | -0.0232968 | 0.62331575 | 0.92488423 | 0.02245784 | 0.63582022 |

| | | | | | |
|---|---|---|---|---|---|
| 0.999490048 | -0.0211839 | 0.62331187 | 0.92488423 | -0.0391969 | 0.36625108 |
| 0.999490048 | 0.02286432 | 0.62307432 | 0.92488423 | -0.0159395 | 0.73154629 |
| 0.999490048 | 0.02959141 | 0.62338305 | 0.92488423 | 0.01823512 | 0.76184076 |
| 0.999490048 | 0.10453669 | 0.62336379 | 0.92488423 | 0.01616816 | 0.93931145 |
| 0.999490048 | 0.02168033 | 0.62386957 | 0.92544162 | 0.01952124 | 0.65867652 |
| 0.999490048 | 0.04170418 | 0.62400649 | 0.92545941 | 0.08989748 | 0.29488012 |
| 0.999490048 | 0.05456429 | 0.62410323 | 0.92545941 | -0.064404 | 0.56335563 |
| 0.999490048 | -0.0660432 | 0.62460082 | 0.92571148 | -0.3219394 | 0.02333088 |
| 0.999490048 | -0.0612629 | 0.62477864 | 0.92571148 | 0.05517409 | 0.65944463 |
| 0.999490048 | 0.03718581 | 0.62479541 | 0.92571148 | -0.0271934 | 0.72030721 |
| 0.999490048 | 0.07544947 | 0.62482753 | 0.92571148 | 0.11527858 | 0.45824246 |
| 0.999490048 | 0.08943894 | 0.62442105 | 0.92571148 | 0.37651194 | 0.04721338 |
| 0.999490048 | -0.0518066 | 0.62507602 | 0.92591534 | 0.02078334 | 0.84426763 |
| 0.999490048 | -0.0306592 | 0.62535942 | 0.92617084 | -0.019411 | 0.75687979 |
| 0.999490048 | 0.04200342 | 0.62551165 | 0.926232 | 0.0110355 | 0.89774243 |
| 0.999490048 | 0.02443472 | 0.62579027 | 0.92648027 | 0.06231777 | 0.21953537 |
| 0.999490048 | 0.03190185 | 0.62643187 | 0.92664974 | 0.00528651 | 0.93556185 |
| 0.999490048 | 0.03659322 | 0.62645931 | 0.92664974 | 0.0883332 | 0.24536366 |
| 0.999490048 | 0.0411461 | 0.62645962 | 0.92664974 | 0.02879552 | 0.7330754 |
| 0.999490048 | 0.06838372 | 0.62645197 | 0.92664974 | 0.01425319 | 0.91900978 |
| 0.999490048 | 0.10977613 | 0.62644135 | 0.92664974 | 0.17048146 | 0.4512237 |
| 0.972489992 | 0.03122571 | 0.62670572 | 0.92684959 | -0.1152958 | 0.08098154 |
| 0.987721587 | -0.0443732 | 0.62851866 | 0.92692237 | 0.15708405 | 0.09517766 |
| 0.999490048 | -0.0405581 | 0.62787397 | 0.92692237 | -0.1124427 | 0.18574858 |
| 0.999490048 | -0.0365438 | 0.62877276 | 0.92692237 | -0.024871 | 0.7418176 |
| 0.999490048 | -0.0303052 | 0.62690699 | 0.92692237 | -0.0329428 | 0.59738629 |
| 0.999490048 | -0.0300484 | 0.62808173 | 0.92692237 | -0.0558096 | 0.37100463 |
| 0.999490048 | -0.0277996 | 0.62843261 | 0.92692237 | -0.0382783 | 0.50608653 |
| 0.999490048 | -0.0254503 | 0.62793079 | 0.92692237 | 0.0150309 | 0.77436432 |
| 0.999490048 | -0.0222819 | 0.62757344 | 0.92692237 | -0.0025639 | 0.95537798 |
| 0.999490048 | -0.0222636 | 0.62724494 | 0.92692237 | 0.0215625 | 0.63808012 |
| 0.999490048 | 0.0211604 | 0.6288641 | 0.92692237 | -0.0288503 | 0.51076584 |
| 0.999490048 | 0.02785465 | 0.62881185 | 0.92692237 | 0.04440899 | 0.44253872 |
| 0.999490048 | 0.03684437 | 0.62837083 | 0.92692237 | 0.08243218 | 0.28333058 |
| 0.999490048 | 0.03752438 | 0.62879369 | 0.92692237 | 0.03991787 | 0.60718114 |
| 0.999490048 | 0.03893685 | 0.62848957 | 0.92692237 | -0.0123533 | 0.87773773 |
| 0.999490048 | 0.03989404 | 0.62803325 | 0.92692237 | -0.0593982 | 0.47195437 |
| 0.999490048 | 0.0430472 | 0.62749663 | 0.92692237 | 0.04356909 | 0.62335007 |
| 0.999490048 | 0.05313606 | 0.62789286 | 0.92692237 | -0.0289422 | 0.7914371 |
| 0.999490048 | 0.06925441 | 0.62841306 | 0.92692237 | 0.06406934 | 0.65423984 |
| 0.999490048 | 0.27403621 | 0.62711991 | 0.92692237 | -0.3149616 | 0.5775357 |
| 0.999490048 | -0.0208986 | 0.62917919 | 0.92702081 | -0.0065248 | 0.87991067 |

| | | | | | |
|---|---|---|---|---|---|
| 0.999490048 | 0.02645609 | 0.6291775 | 0.92702081 | 0.01485725 | 0.78591659 |
| 0.999490048 | 0.02955641 | 0.62926395 | 0.92702081 | -5.52E-04 | 0.99279464 |
| 0.999490048 | -0.08226 | 0.63045589 | 0.92714142 | -0.3618378 | 0.04203965 |
| 0.999490048 | -0.059232 | 0.63051794 | 0.92714142 | 0.02928687 | 0.81168955 |
| 0.999490048 | -0.0381042 | 0.63063318 | 0.92714142 | -0.0031071 | 0.96865115 |
| 0.999490048 | -0.0161321 | 0.62993035 | 0.92714142 | -0.0396828 | 0.24137077 |
| 0.999490048 | 0.02269749 | 0.62959735 | 0.92714142 | -0.0120281 | 0.79793977 |
| 0.999490048 | 0.02309218 | 0.63001164 | 0.92714142 | 0.02004334 | 0.6756897 |
| 0.999490048 | 0.02425036 | 0.62972337 | 0.92714142 | 0.02092376 | 0.67724157 |
| 0.999490048 | 0.02525483 | 0.6298632 | 0.92714142 | 0.09365006 | 0.08240192 |
| 0.999490048 | 0.02655899 | 0.63067823 | 0.92714142 | 0.04914179 | 0.37636938 |
| 0.999490048 | 0.03026093 | 0.63021268 | 0.92714142 | 0.02942412 | 0.63966009 |
| 0.999490048 | 0.03068871 | 0.63064604 | 0.92714142 | 0.0295379 | 0.64345987 |
| 0.999490048 | 0.05423115 | 0.63025144 | 0.92714142 | 0.01687615 | 0.88067349 |
| 0.999490048 | -0.1303518 | 0.63127691 | 0.92761425 | -0.6590262 | 0.02128339 |
| 0.999490048 | -0.0512862 | 0.63166643 | 0.92761425 | 0.0254945 | 0.81132296 |
| 0.999490048 | -0.0400387 | 0.63156531 | 0.92761425 | -0.0747871 | 0.37315081 |
| 0.999490048 | 0.02836168 | 0.63134096 | 0.92761425 | 0.07806917 | 0.19315458 |
| 0.999490048 | 0.03915007 | 0.63163289 | 0.92761425 | 0.13170332 | 0.11498852 |
| 0.999490048 | 0.04959972 | 0.63119767 | 0.92761425 | -0.0735535 | 0.4777641 |
| 0.999490048 | 0.0267903 | 0.6318932 | 0.92778411 | 0.03775526 | 0.50056144 |
| 0.999490048 | -0.2162548 | 0.63239483 | 0.92811268 | -1.1016169 | 0.03471827 |
| 0.999490048 | -0.0575349 | 0.63288827 | 0.92811268 | -0.0676918 | 0.57489623 |
| 0.999490048 | -0.0472904 | 0.6323747 | 0.92811268 | 0.00858115 | 0.93067069 |
| 0.999490048 | -0.0436778 | 0.63397539 | 0.92811268 | -0.0105551 | 0.90819829 |
| 0.999490048 | -0.0333778 | 0.63277019 | 0.92811268 | 0.06151892 | 0.38108744 |
| 0.999490048 | -0.0294072 | 0.6341177 | 0.92811268 | -0.054278 | 0.38228742 |
| 0.999490048 | -0.0284694 | 0.63400846 | 0.92811268 | 0.00191124 | 0.97444442 |
| 0.999490048 | -0.0284337 | 0.63256297 | 0.92811268 | -0.0588853 | 0.32570888 |
| 0.999490048 | -0.0245219 | 0.63377334 | 0.92811268 | -0.0463692 | 0.37044547 |
| 0.999490048 | -0.0244149 | 0.63272257 | 0.92811268 | -0.0194975 | 0.70248794 |
| 0.999490048 | 0.02311567 | 0.63384916 | 0.92811268 | 0.03781357 | 0.43762144 |
| 0.999490048 | 0.0233874 | 0.63357286 | 0.92811268 | 0.10001729 | 0.04950829 |
| 0.999490048 | 0.02665606 | 0.63367705 | 0.92811268 | 0.02365831 | 0.67217145 |
| 0.999490048 | 0.02852482 | 0.63346974 | 0.92811268 | 0.02172162 | 0.71625369 |
| 0.999490048 | 0.02952437 | 0.63347331 | 0.92811268 | 0.02297772 | 0.71030549 |
| 0.999490048 | 0.04189143 | 0.63371982 | 0.92811268 | 0.06692378 | 0.44812359 |
| 0.999490048 | 0.04267959 | 0.63379166 | 0.92811268 | 0.06082834 | 0.49816317 |
| 0.999490048 | 0.04624427 | 0.63330805 | 0.92811268 | -0.0842609 | 0.38720438 |
| 0.999490048 | 0.04357225 | 0.63423728 | 0.928125 | 0.0556087 | 0.54429526 |
| 0.999490048 | -0.0852193 | 0.63465026 | 0.92856661 | -0.0253805 | 0.8872226 |
| 0.999490048 | -0.0370175 | 0.6351731 | 0.92868068 | 0.05341145 | 0.49466384 |

| | | | | | |
|---|---|---|---|---|---|
| 0.999490048 | -0.0336497 | 0.63511831 | 0.92868068 | -0.0124008 | 0.8609041 |
| 0.999490048 | -0.0214168 | 0.63483956 | 0.92868068 | 0.00263232 | 0.95334703 |
| 0.999490048 | 0.03370574 | 0.63502119 | 0.92868068 | 0.04084633 | 0.56554819 |
| 0.999490048 | -0.0560984 | 0.63549646 | 0.92899079 | -0.0331604 | 0.77901417 |
| 0.999490048 | 0.02505758 | 0.63569787 | 0.92912257 | 0.03067856 | 0.56236246 |
| 0.999490048 | -0.0283674 | 0.63602633 | 0.92943995 | -0.0374342 | 0.53296691 |
| 0.999490048 | -0.0256456 | 0.63621419 | 0.92955179 | 0.02110408 | 0.69689156 |
| 0.999490048 | -0.0273784 | 0.63663714 | 0.92995365 | -0.0314614 | 0.58749711 |
| 0.999490048 | 0.04206889 | 0.63671198 | 0.92995365 | 0.04279054 | 0.6309675 |
| 0.999490048 | -0.15134 | 0.63909987 | 0.92999071 | 0.13361303 | 0.67838221 |
| 0.999490048 | -0.0660201 | 0.63758776 | 0.92999071 | 0.04661031 | 0.73916969 |
| 0.999490048 | -0.0496563 | 0.639118 | 0.92999071 | 0.05043734 | 0.63387861 |
| 0.999490048 | -0.0383838 | 0.63906382 | 0.92999071 | -0.1197592 | 0.15096937 |
| 0.999490048 | -0.0365079 | 0.63798862 | 0.92999071 | 0.03371565 | 0.66380249 |
| 0.999490048 | -0.0272122 | 0.63839309 | 0.92999071 | -0.0506271 | 0.38452946 |
| 0.999490048 | -0.0240468 | 0.63999662 | 0.92999071 | -0.0171489 | 0.73845298 |
| 0.999490048 | -0.0228247 | 0.63817873 | 0.92999071 | -0.0456535 | 0.3501093 |
| 0.999490048 | -0.0220606 | 0.64007864 | 0.92999071 | -0.0231322 | 0.6239971 |
| 0.999490048 | -0.0209415 | 0.63905687 | 0.92999071 | -0.0345786 | 0.44040971 |
| 0.999490048 | -0.0209391 | 0.63962384 | 0.92999071 | -0.0441562 | 0.32709127 |
| 0.999490048 | -0.0209285 | 0.6380441 | 0.92999071 | 0.04164947 | 0.35231369 |
| 0.999490048 | 0.01604846 | 0.6369884 | 0.92999071 | 0.02693548 | 0.43019963 |
| 0.999490048 | 0.02227105 | 0.63772939 | 0.92999071 | -0.0024857 | 0.957994 |
| 0.999490048 | 0.02414523 | 0.63760073 | 0.92999071 | 0.03540394 | 0.49086195 |
| 0.999490048 | 0.02668737 | 0.63827596 | 0.92999071 | -0.0077892 | 0.89064211 |
| 0.999490048 | 0.02696212 | 0.63726582 | 0.92999071 | -0.023303 | 0.68344646 |
| 0.999490048 | 0.02762641 | 0.63989504 | 0.92999071 | 0.02955403 | 0.61684278 |
| 0.999490048 | 0.02879944 | 0.63834078 | 0.92999071 | 0.05706105 | 0.35482071 |
| 0.999490048 | 0.03082023 | 0.63854105 | 0.92999071 | -0.0010677 | 0.98698768 |
| 0.999490048 | 0.03502409 | 0.63884884 | 0.92999071 | -0.0235138 | 0.75241228 |
| 0.999490048 | 0.03596318 | 0.63828499 | 0.92999071 | 0.10552911 | 0.1748208 |
| 0.999490048 | 0.03725401 | 0.63968771 | 0.92999071 | 0.02945645 | 0.71104122 |
| 0.999490048 | 0.04039768 | 0.63834514 | 0.92999071 | 0.0368418 | 0.66806649 |
| 0.999490048 | 0.0470963 | 0.63984195 | 0.92999071 | -0.0850153 | 0.40063176 |
| 0.999490048 | 0.0587849 | 0.63968649 | 0.92999071 | -0.017382 | 0.88968519 |
| 0.999490048 | 0.06118492 | 0.63814213 | 0.92999071 | 0.06665533 | 0.60855786 |
| 0.999490048 | 0.08968989 | 0.63949178 | 0.92999071 | -0.0403462 | 0.83244227 |
| 0.999490048 | 0.08973701 | 0.63980218 | 0.92999071 | -0.1306449 | 0.49673647 |
| 0.999490048 | 0.12321001 | 0.63892451 | 0.92999071 | 0.1965564 | 0.45896112 |
| 0.999490048 | -0.021279 | 0.64034815 | 0.93005862 | 0.01065538 | 0.81471988 |
| 0.999490048 | 0.03175867 | 0.64032503 | 0.93005862 | -0.0013052 | 0.98464556 |
| 0.999490048 | -0.0271716 | 0.64064023 | 0.93032102 | 0.05198618 | 0.37456152 |

| | | | | | |
|---|---|---|---|---|---|
| 0.999490048 | 0.02210825 | 0.64077867 | 0.93036027 | 0.04204286 | 0.37760848 |
| 0.999490048 | -0.1796083 | 0.64133324 | 0.93042693 | -0.0432512 | 0.91009281 |
| 0.999490048 | -0.0536858 | 0.64107813 | 0.93042693 | -0.1730061 | 0.15046169 |
| 0.999490048 | 0.02874614 | 0.64130204 | 0.93042693 | 6.26E-04 | 0.99188916 |
| 0.999490048 | 0.03624497 | 0.64134757 | 0.93042693 | 0.07611579 | 0.33152401 |
| 0.999490048 | 0.03700092 | 0.64138173 | 0.93042693 | 0.00430991 | 0.95663972 |
| 0.999490048 | -0.0198012 | 0.64182658 | 0.93077462 | -0.0033505 | 0.93713316 |
| 0.999490048 | 0.04361048 | 0.64184434 | 0.93077462 | 0.06590373 | 0.48334583 |
| 0.999490048 | -0.0503864 | 0.64226427 | 0.93106019 | 0.00555387 | 0.9588807 |
| 0.999490048 | -0.019004 | 0.64223498 | 0.93106019 | 0.02166086 | 0.59667877 |
| 0.999490048 | -0.034184 | 0.64242807 | 0.93113598 | -0.0105277 | 0.88607241 |
| 0.999490048 | 0.0309019 | 0.64286164 | 0.93160269 | 0.11312109 | 0.09806558 |
| 0.999490048 | 0.04910814 | 0.64297418 | 0.9316041 | -0.1177469 | 0.27121958 |
| 0.999490048 | 0.02414074 | 0.64320839 | 0.93167374 | 0.01692576 | 0.74508036 |
| 0.999490048 | 0.02625457 | 0.6432454 | 0.93167374 | 0.03423587 | 0.54647213 |
| 0.999490048 | -0.0224909 | 0.64350246 | 0.93172751 | 0.03706996 | 0.44725178 |
| 0.999490048 | 0.02678134 | 0.6435057 | 0.93172751 | 0.02793644 | 0.62932859 |
| 0.827257028 | 0.03823992 | 0.6437289 | 0.93172756 | 0.25332874 | 0.00470987 |
| 0.999490048 | 0.09154027 | 0.64363062 | 0.93172756 | 0.37037109 | 0.06970559 |
| 0.999490048 | -0.045988 | 0.64398294 | 0.9318017 | 0.00970556 | 0.92214386 |
| 0.999490048 | -0.0226565 | 0.64422649 | 0.9318017 | 0.01081179 | 0.82529577 |
| 0.999490048 | 0.02494578 | 0.64419452 | 0.9318017 | -5.40E-05 | 0.99920078 |
| 0.999490048 | 0.09400524 | 0.64410544 | 0.9318017 | 0.18483853 | 0.37032161 |
| 0.999490048 | -0.0572458 | 0.64467836 | 0.93213235 | -0.0172234 | 0.88943256 |
| 0.999490048 | -0.0213399 | 0.6445763 | 0.93213235 | -0.021662 | 0.63961132 |
| 0.999490048 | -0.0215633 | 0.64497442 | 0.93239896 | -0.0072364 | 0.8768855 |
| 0.999490048 | -0.0240835 | 0.64514573 | 0.93248518 | 0.03490304 | 0.50551231 |
| 0.999490048 | -0.0460745 | 0.64541226 | 0.93254157 | -0.1184886 | 0.24219352 |
| 0.999490048 | -0.0411988 | 0.64575007 | 0.93254157 | 0.06401593 | 0.47639183 |
| 0.999490048 | -0.0225291 | 0.64579909 | 0.93254157 | -0.003159 | 0.94850852 |
| 0.999490048 | 0.02752078 | 0.64585484 | 0.93254157 | 0.00563134 | 0.92493277 |
| 0.999490048 | 0.0503048 | 0.6455307 | 0.93254157 | 0.0419396 | 0.70117159 |
| 0.999490048 | 0.08069721 | 0.64541626 | 0.93254157 | 0.17545636 | 0.32088788 |
| 0.999490048 | -0.0304857 | 0.6459709 | 0.93254789 | -0.0728345 | 0.27715559 |
| 0.999490048 | -0.023824 | 0.64630655 | 0.93270993 | -0.0143446 | 0.78202476 |
| 0.999490048 | 0.02577003 | 0.64627281 | 0.93270993 | -0.0316729 | 0.57312024 |
| 0.999490048 | -0.0863774 | 0.64731725 | 0.93275121 | -0.2256979 | 0.23757334 |
| 0.999490048 | -0.0306882 | 0.64720561 | 0.93275121 | 0.05345487 | 0.42732849 |
| 0.999490048 | -0.0301918 | 0.64731021 | 0.93275121 | 0.02212184 | 0.73721455 |
| 0.999490048 | -0.0223655 | 0.64753567 | 0.93275121 | 0.0552185 | 0.26403184 |
| 0.999490048 | -0.0223301 | 0.64688669 | 0.93275121 | -0.0106545 | 0.82670421 |
| 0.999490048 | -0.0214213 | 0.64755923 | 0.93275121 | 0.03923883 | 0.40468179 |

| | | | | | |
|---|---|---|---|---|---|
| 0.999490048 | 0.0243198 | 0.64701877 | 0.93275121 | -9.84E-04 | 0.98518656 |
| 0.999490048 | 0.03240026 | 0.64752088 | 0.93275121 | 0.05179258 | 0.46631949 |
| 0.999490048 | 0.04998715 | 0.64756393 | 0.93275121 | 0.13465411 | 0.22411007 |
| 0.999490048 | 0.06169025 | 0.64698259 | 0.93275121 | 0.21698841 | 0.11549186 |
| 0.999490048 | 0.11303781 | 0.64702003 | 0.93275121 | -0.220517 | 0.37805728 |
| 0.999490048 | -0.0743618 | 0.64859765 | 0.93290419 | 0.07750244 | 0.63487784 |
| 0.999490048 | -0.0446149 | 0.64852575 | 0.93290419 | -0.0978111 | 0.32146026 |
| 0.999490048 | -0.0386035 | 0.6488991 | 0.93290419 | 0.06544125 | 0.44199967 |
| 0.999490048 | -0.0265067 | 0.64827134 | 0.93290419 | -0.0159254 | 0.7837494 |
| 0.999490048 | -0.0206074 | 0.64817846 | 0.93290419 | -0.0073574 | 0.87035396 |
| 0.999490048 | -0.0201544 | 0.64881421 | 0.93290419 | -0.031138 | 0.48294706 |
| 0.999490048 | 0.01846113 | 0.6488511 | 0.93290419 | -0.0192396 | 0.63516192 |
| 0.999490048 | 0.02063122 | 0.64804263 | 0.93290419 | 0.01081405 | 0.81061271 |
| 0.999490048 | 0.02492976 | 0.64879747 | 0.93290419 | -0.0860001 | 0.12430245 |
| 0.999490048 | 0.04937204 | 0.64858806 | 0.93290419 | 0.07270524 | 0.50319803 |
| 0.999490048 | 0.06142717 | 0.64831125 | 0.93290419 | 0.10244355 | 0.44854349 |
| 0.999490048 | 0.02248745 | 0.64924646 | 0.93296053 | 0.00575318 | 0.90718813 |
| 0.999490048 | 0.02609723 | 0.6491462 | 0.93296053 | 0.05977571 | 0.30160426 |
| 0.999490048 | 0.08997105 | 0.64927349 | 0.93296053 | -0.0482003 | 0.80686446 |
| 0.999490048 | -0.0283678 | 0.64951359 | 0.93298443 | -0.0900039 | 0.15687465 |
| 0.999490048 | 0.0221708 | 0.64944102 | 0.93298443 | -0.0254867 | 0.60155184 |
| 0.999490048 | -0.0556475 | 0.65135132 | 0.93312476 | 0.16809597 | 0.17927736 |
| 0.999490048 | -0.0258911 | 0.65086281 | 0.93312476 | 0.0276757 | 0.62865909 |
| 0.999490048 | -0.0243691 | 0.65196723 | 0.93312476 | -0.0283201 | 0.60043077 |
| 0.999490048 | -0.0209718 | 0.65181148 | 0.93312476 | -0.0067346 | 0.88457319 |
| 0.999490048 | -0.0207886 | 0.65146574 | 0.93312476 | -0.0189563 | 0.68029995 |
| 0.999490048 | -0.0186253 | 0.65118072 | 0.93312476 | 0.02855847 | 0.48935926 |
| 0.999490048 | 0.01629949 | 0.65206982 | 0.93312476 | -0.0025238 | 0.9442294 |
| 0.999490048 | 0.02303696 | 0.65189807 | 0.93312476 | 0.03637712 | 0.47758305 |
| 0.999490048 | 0.02313837 | 0.65185544 | 0.93312476 | 0.06662169 | 0.200455 |
| 0.999490048 | 0.02329102 | 0.65187778 | 0.93312476 | 0.08263125 | 0.11772804 |
| 0.999490048 | 0.02476231 | 0.65096169 | 0.93312476 | -0.0113439 | 0.83554046 |
| 0.999490048 | 0.02518015 | 0.65187398 | 0.93312476 | 0.03113211 | 0.577391 |
| 0.999490048 | 0.02650146 | 0.65008103 | 0.93312476 | 0.02092915 | 0.71993664 |
| 0.999490048 | 0.02852865 | 0.6517989 | 0.93312476 | -0.0275994 | 0.66238444 |
| 0.999490048 | 0.0288195 | 0.6516562 | 0.93312476 | 0.01156115 | 0.85603714 |
| 0.999490048 | 0.02942682 | 0.6499593 | 0.93312476 | 0.00487328 | 0.93996981 |
| 0.999490048 | 0.03399131 | 0.65087256 | 0.93312476 | 0.02455074 | 0.74352809 |
| 0.999490048 | 0.03447224 | 0.65007973 | 0.93312476 | 0.12352803 | 0.11235986 |
| 0.999490048 | 0.04305808 | 0.6507933 | 0.93312476 | 0.02914549 | 0.75903815 |
| 0.999490048 | 0.05416296 | 0.65004844 | 0.93312476 | 0.01744124 | 0.88363006 |
| 0.999490048 | 0.08128349 | 0.65053929 | 0.93312476 | -0.2159741 | 0.24157972 |

| | | | | | |
|---|---|---|---|---|---|
| 0.999490048 | 0.13496103 | 0.65092985 | 0.93312476 | -0.21576 | 0.47390517 |
| 0.999490048 | -0.0467199 | 0.65258281 | 0.93332011 | -0.1218349 | 0.24588946 |
| 0.999490048 | -0.0385965 | 0.65277922 | 0.93332011 | -0.0511465 | 0.5516573 |
| 0.999490048 | -0.0238233 | 0.65301221 | 0.93332011 | -0.0473042 | 0.3748547 |
| 0.999490048 | -0.0214093 | 0.65240275 | 0.93332011 | -0.0350647 | 0.46221876 |
| 0.999490048 | 0.02212462 | 0.65304896 | 0.93332011 | -0.0473616 | 0.33938364 |
| 0.999490048 | 0.02232827 | 0.65243002 | 0.93332011 | -0.026012 | 0.60006137 |
| 0.999490048 | 0.02396829 | 0.65302579 | 0.93332011 | -0.0334249 | 0.53148914 |
| 0.999490048 | 0.06368836 | 0.65310053 | 0.93332011 | -7.01E-04 | 0.99603512 |
| 0.999490048 | 0.02287091 | 0.65331173 | 0.9333907 | -0.020629 | 0.68526419 |
| 0.999490048 | 0.03369211 | 0.65359706 | 0.9333907 | 0.05412327 | 0.47238061 |
| 0.999490048 | 0.03921099 | 0.653375 | 0.9333907 | 0.06845953 | 0.43490722 |
| 0.999490048 | 0.07580263 | 0.65356259 | 0.9333907 | 0.05312035 | 0.7528792 |
| 0.999490048 | 0.03063108 | 0.65384854 | 0.93359017 | -0.0280601 | 0.68113212 |
| 0.999490048 | -0.0262002 | 0.65398159 | 0.9336205 | -0.0734476 | 0.21512195 |
| 0.999490048 | -0.0267217 | 0.65430664 | 0.93365589 | -0.0243455 | 0.68319291 |
| 0.999490048 | 0.03146288 | 0.65434183 | 0.93365589 | 0.0771482 | 0.27704259 |
| 0.999490048 | 0.08207871 | 0.65421526 | 0.93365589 | 0.08145695 | 0.65665612 |
| 0.999490048 | -0.2624818 | 0.65471354 | 0.93371752 | 0.03597067 | 0.95103301 |
| 0.999490048 | -0.0284642 | 0.65472072 | 0.93371752 | 0.00284633 | 0.96425909 |
| 0.999490048 | -0.0281182 | 0.65482187 | 0.93371752 | 0.05521588 | 0.38270082 |
| 0.999490048 | -0.0181686 | 0.65494413 | 0.93371752 | 0.03350644 | 0.41208282 |
| 0.999490048 | 0.02227166 | 0.65490141 | 0.93371752 | 0.06571153 | 0.19396489 |
| 0.999490048 | 0.04102762 | 0.65507006 | 0.93373763 | -0.1175475 | 0.20697982 |
| 0.999490048 | -0.0684663 | 0.6561121 | 0.93374095 | 0.08324519 | 0.58904354 |
| 0.999490048 | -0.0317251 | 0.65614938 | 0.93374095 | 0.01062078 | 0.88130145 |
| 0.999490048 | -0.0215797 | 0.65596418 | 0.93374095 | 0.01412567 | 0.77032819 |
| 0.999490048 | 0.02116052 | 0.65523587 | 0.93374095 | 0.00929914 | 0.84418988 |
| 0.999490048 | 0.02350616 | 0.65619064 | 0.93374095 | 0.0893118 | 0.09924791 |
| 0.999490048 | 0.02725611 | 0.6557211 | 0.93374095 | 0.02714905 | 0.65698159 |
| 0.999490048 | 0.0298146 | 0.65535597 | 0.93374095 | 0.00434866 | 0.94799144 |
| 0.999490048 | 0.0411195 | 0.65609501 | 0.93374095 | 0.1258631 | 0.17990769 |
| 0.999490048 | 0.07530984 | 0.65603535 | 0.93374095 | -0.1857194 | 0.27683918 |
| 0.999490048 | 0.08083213 | 0.65611447 | 0.93374095 | -0.1418661 | 0.43640189 |
| 0.999490048 | -0.0680716 | 0.65659815 | 0.9339028 | 0.0316317 | 0.83606535 |
| 0.999490048 | -0.020228 | 0.65676336 | 0.9339028 | -0.002841 | 0.95013626 |
| 0.999490048 | 0.02450243 | 0.65686361 | 0.9339028 | 1.05E-06 | 0.99998478 |
| 0.999490048 | 0.03146305 | 0.65649323 | 0.9339028 | 0.04630318 | 0.51374019 |
| 0.999490048 | 0.07622544 | 0.65678163 | 0.9339028 | -0.0070111 | 0.96733345 |
| 0.999490048 | -0.0781303 | 0.65721825 | 0.93392995 | 0.04042057 | 0.81816032 |
| 0.999490048 | -0.0423305 | 0.65714741 | 0.93392995 | 0.00596607 | 0.95002188 |
| 0.999490048 | 0.0292381 | 0.65699815 | 0.93392995 | 0.08068539 | 0.22637255 |

| | | | | | |
|---|---|---|---|---|---|
| 0.999490048 | -0.0203126 | 0.65749806 | 0.93400966 | -0.0430584 | 0.35059451 |
| 0.999490048 | 0.02302814 | 0.65739631 | 0.93400966 | 0.04082895 | 0.43360764 |
| 0.999490048 | 0.04084619 | 0.65792161 | 0.93445236 | -0.079607 | 0.39075706 |
| 0.999490048 | -0.0214441 | 0.65834044 | 0.9345472 | 0.02301413 | 0.63518566 |
| 0.999490048 | -0.0200223 | 0.65843226 | 0.9345472 | -0.0221545 | 0.62488432 |
| 0.999490048 | 0.02068134 | 0.65843607 | 0.9345472 | -0.0076219 | 0.8703607 |
| 0.999490048 | 0.02548478 | 0.65838684 | 0.9345472 | -0.0122556 | 0.83136772 |
| 0.999490048 | 0.02279251 | 0.65858996 | 0.93460676 | 0.02385052 | 0.64387677 |
| 0.999490048 | -0.0290348 | 0.65905004 | 0.93462429 | -0.0242771 | 0.71202058 |
| 0.999490048 | -0.0223615 | 0.65902976 | 0.93462429 | -0.0549665 | 0.28276625 |
| 0.999490048 | -0.0199711 | 0.65904667 | 0.93462429 | -0.0162026 | 0.72018459 |
| 0.999490048 | 0.02900851 | 0.65894535 | 0.93462429 | 0.07049024 | 0.28805476 |
| 0.999490048 | -0.0311827 | 0.65982579 | 0.93556405 | -0.0194396 | 0.78352794 |
| 0.999490048 | -0.017184 | 0.65993679 | 0.93556405 | 0.00997338 | 0.79818051 |
| 0.999490048 | -0.0268362 | 0.66023811 | 0.93567349 | -0.0286373 | 0.63909778 |
| 0.999490048 | 0.05823196 | 0.66015941 | 0.93567349 | 0.17039695 | 0.20471453 |
| 0.999490048 | -0.2717043 | 0.66046928 | 0.93584227 | -1.0063855 | 0.12921463 |
| 0.999490048 | 0.05327662 | 0.66074165 | 0.93606935 | 0.0127164 | 0.91641585 |
| 0.999490048 | -0.0389903 | 0.66086949 | 0.93609165 | 0.06603839 | 0.45906947 |
| 0.999490048 | -0.122949 | 0.66188466 | 0.93692955 | 0.39452525 | 0.18288057 |
| 0.999490048 | 0.0437615 | 0.66190987 | 0.93692955 | 0.10554557 | 0.2960048 |
| 0.999490048 | 0.05945436 | 0.66171036 | 0.93692955 | 0.14469175 | 0.29145344 |
| 0.999490048 | 0.08261808 | 0.66175968 | 0.93692955 | 0.09621295 | 0.6106655 |
| 0.999490048 | 0.06089629 | 0.66210271 | 0.93704367 | -0.1301791 | 0.35346998 |
| 0.999490048 | -0.1226108 | 0.66232014 | 0.93714612 | -0.160043 | 0.56984642 |
| 0.999490048 | -0.114829 | 0.6628485 | 0.93714612 | 0.05909573 | 0.82167609 |
| 0.999490048 | -0.0306659 | 0.66263947 | 0.93714612 | -0.074698 | 0.29241588 |
| 0.999490048 | -0.022187 | 0.66277829 | 0.93714612 | -0.0667351 | 0.19633142 |
| 0.999490048 | 0.0256191 | 0.66257121 | 0.93714612 | -0.005225 | 0.92895327 |
| 0.999490048 | 0.02718865 | 0.66247017 | 0.93714612 | -4.26E-04 | 0.99453267 |
| 0.999490048 | 0.02727323 | 0.66321666 | 0.9375079 | 2.99E-04 | 0.99618856 |
| 0.999490048 | -0.0482161 | 0.66333649 | 0.93751857 | 0.00835223 | 0.93977935 |
| 0.999490048 | -0.0389808 | 0.66379291 | 0.93800488 | -0.1079631 | 0.23443127 |
| 0.999490048 | -0.0371913 | 0.66421571 | 0.93835142 | 0.06618004 | 0.44171148 |
| 0.999490048 | -0.037142 | 0.66426291 | 0.93835142 | -0.0599027 | 0.48524139 |
| 0.999490048 | -0.0215272 | 0.66462202 | 0.93869991 | -0.0271957 | 0.58432185 |
| 0.999490048 | 0.06186127 | 0.6648291 | 0.93883358 | 0.04146358 | 0.77077146 |
| 0.999490048 | -0.0150929 | 0.66523944 | 0.9392542 | 0.02978992 | 0.39564784 |
| 0.999490048 | -0.0226253 | 0.66542396 | 0.93935588 | -0.0789932 | 0.1390562 |
| 0.999490048 | -0.0696225 | 0.66584118 | 0.93939681 | -0.1449461 | 0.37157554 |
| 0.999490048 | -0.0679042 | 0.66606773 | 0.93939681 | -0.0569182 | 0.71740089 |
| 0.999490048 | -0.0508059 | 0.66572673 | 0.93939681 | 0.03131651 | 0.78977185 |

| | | | | | |
|---|---|---|---|---|---|
| 0.999490048 | -0.0422628 | 0.66733654 | 0.93939681 | -0.1262653 | 0.20561295 |
| 0.999490048 | -0.0291532 | 0.66614874 | 0.93939681 | -0.0552154 | 0.41609916 |
| 0.999490048 | -0.0277507 | 0.66726572 | 0.93939681 | 0.04625876 | 0.47506387 |
| 0.999490048 | -0.0259895 | 0.66724538 | 0.93939681 | -0.0513042 | 0.39857406 |
| 0.999490048 | -0.0215989 | 0.66735373 | 0.93939681 | 0.02639348 | 0.59978763 |
| 0.999490048 | 0.01817353 | 0.66650111 | 0.93939681 | 0.00406552 | 0.92306062 |
| 0.999490048 | 0.01845911 | 0.66719015 | 0.93939681 | 0.01771605 | 0.67978352 |
| 0.999490048 | 0.01921454 | 0.66697374 | 0.93939681 | 0.0054926 | 0.90193517 |
| 0.999490048 | 0.02152623 | 0.66563563 | 0.93939681 | -0.001943 | 0.9688267 |
| 0.999490048 | 0.02237984 | 0.66720004 | 0.93939681 | 0.05374522 | 0.30601255 |
| 0.999490048 | 0.03189308 | 0.66697132 | 0.93939681 | 0.012296 | 0.86802711 |
| 0.999490048 | 0.0334357 | 0.66736549 | 0.93939681 | -0.0660366 | 0.39851859 |
| 0.999490048 | 0.0394822 | 0.66707652 | 0.93939681 | 0.13066119 | 0.16206889 |
| 0.999490048 | 0.08781412 | 0.66593781 | 0.93939681 | 0.00946157 | 0.96282908 |
| 0.999490048 | -0.0362971 | 0.66808466 | 0.94009216 | -0.0857422 | 0.31517212 |
| 0.999490048 | -0.0205197 | 0.66804563 | 0.94009216 | -0.0347095 | 0.46974511 |
| 0.999490048 | -0.0528213 | 0.66847764 | 0.9401344 | -0.1998425 | 0.12396101 |
| 0.999490048 | -0.0293965 | 0.66864439 | 0.9401344 | -0.029124 | 0.67152507 |
| 0.999490048 | 0.02623476 | 0.66830746 | 0.9401344 | 0.01643238 | 0.78817585 |
| 0.999490048 | 0.06696469 | 0.66853732 | 0.9401344 | 0.02777589 | 0.85882884 |
| 0.999490048 | 0.0730706 | 0.66867763 | 0.9401344 | 0.17511455 | 0.31449152 |
| 0.999490048 | 0.0232425 | 0.6688005 | 0.94014884 | 0.11293347 | 0.04561412 |
| 0.999490048 | -0.0255032 | 0.66917565 | 0.9402332 | 0.0430556 | 0.47219447 |
| 0.999490048 | -0.0248179 | 0.66899504 | 0.9402332 | 0.02746759 | 0.63622919 |
| 0.999490048 | -0.0192456 | 0.66922687 | 0.9402332 | -0.0500046 | 0.27198587 |
| 0.999490048 | 0.0418656 | 0.66931092 | 0.9402332 | 0.03963135 | 0.68593449 |
| 0.999490048 | 0.03701621 | 0.66961266 | 0.94034068 | 0.00621939 | 0.9427465 |
| 0.999490048 | 0.05503841 | 0.66958039 | 0.94034068 | 0.02370144 | 0.85398373 |
| 0.999490048 | 0.08488074 | 0.66975324 | 0.94037995 | 0.05562948 | 0.77962654 |
| 0.999490048 | -0.0667845 | 0.66990356 | 0.94043287 | 0.00382558 | 0.98048317 |
| 0.999490048 | -0.0386437 | 0.67062739 | 0.94050028 | -0.1321041 | 0.15368072 |
| 0.999490048 | -0.0262469 | 0.67055776 | 0.94050028 | -0.0780887 | 0.2120348 |
| 0.999490048 | 0.0201263 | 0.67047251 | 0.94050028 | 0.02137571 | 0.6514071 |
| 0.999490048 | 0.02390673 | 0.67029975 | 0.94050028 | 0.02300891 | 0.68195036 |
| 0.999490048 | 0.0365582 | 0.67018773 | 0.94050028 | 0.04266267 | 0.619417 |
| 0.999490048 | 0.05199235 | 0.67044293 | 0.94050028 | -0.1226121 | 0.31957714 |
| 0.999490048 | -0.0245106 | 0.6712657 | 0.94123738 | 3.58E-05 | 0.999504 |
| 0.999490048 | -0.0214114 | 0.67176277 | 0.94177621 | -0.0413777 | 0.41515963 |
| 0.999490048 | 0.08466372 | 0.67190052 | 0.94181121 | -0.0236512 | 0.90531264 |
| 0.999490048 | 0.08262521 | 0.67202896 | 0.94183314 | -0.0311543 | 0.87297513 |
| 0.999490048 | 0.02283032 | 0.67220414 | 0.94192056 | -0.0035396 | 0.94760293 |
| 0.999490048 | -0.0315983 | 0.67251998 | 0.94220501 | 0.04030011 | 0.590239 |

| | | | | | |
|---|---|---|---|---|---|
| 0.999490048 | -0.024778 | 0.67296585 | 0.94261148 | -0.0235204 | 0.68861917 |
| 0.999490048 | 0.02571838 | 0.67310316 | 0.94261148 | -0.0096338 | 0.87423594 |
| 0.999490048 | 0.04971223 | 0.67314878 | 0.94261148 | -0.1287305 | 0.27952261 |
| 0.999490048 | 0.02199786 | 0.67344741 | 0.94274123 | 0.00777847 | 0.8813568 |
| 0.999490048 | 0.04102529 | 0.67346724 | 0.94274123 | -0.0459568 | 0.63704589 |
| 0.999490048 | -0.0522629 | 0.67430578 | 0.94328594 | -0.0578109 | 0.6421663 |
| 0.999490048 | -0.0424854 | 0.67399215 | 0.94328594 | -0.110861 | 0.27722513 |
| 0.999490048 | -0.0215121 | 0.67430824 | 0.94328594 | -0.0203747 | 0.69055949 |
| 0.999490048 | 0.03192819 | 0.674173 | 0.94328594 | -0.0080836 | 0.91508815 |
| 0.876151891 | 0.02816006 | 0.67462135 | 0.94356587 | -0.1432284 | 0.04046518 |
| 0.999490048 | 0.02933574 | 0.67505192 | 0.94400997 | -0.0025243 | 0.97117321 |
| 0.999490048 | -0.0621879 | 0.67520902 | 0.94407155 | -0.3601239 | 0.02134641 |
| 0.999490048 | -0.0196829 | 0.67544713 | 0.94424636 | -0.0287773 | 0.54125936 |
| 0.999490048 | -0.1823546 | 0.68049614 | 0.94447338 | 0.7421817 | 0.11983818 |
| 0.999490048 | -0.075911 | 0.67682102 | 0.94447338 | 0.11313307 | 0.53722366 |
| 0.999490048 | -0.0640368 | 0.67921352 | 0.94447338 | -0.0259119 | 0.86691681 |
| 0.999490048 | -0.058956 | 0.67774985 | 0.94447338 | 0.05290967 | 0.70911408 |
| 0.999490048 | -0.058011 | 0.67642608 | 0.94447338 | -0.2186184 | 0.12403877 |
| 0.999490048 | -0.0580081 | 0.67879996 | 0.94447338 | -0.0206914 | 0.88239548 |
| 0.999490048 | -0.0407761 | 0.67803477 | 0.94447338 | 0.02103104 | 0.8299616 |
| 0.999490048 | -0.0379928 | 0.68035486 | 0.94447338 | -0.1421843 | 0.13129248 |
| 0.999490048 | -0.0368518 | 0.68051572 | 0.94447338 | -0.0470953 | 0.59913469 |
| 0.999490048 | -0.0288985 | 0.67954009 | 0.94447338 | 0.01547208 | 0.82475128 |
| 0.999490048 | -0.0264659 | 0.68019527 | 0.94447338 | -0.0278391 | 0.66464881 |
| 0.999490048 | -0.0259897 | 0.68043618 | 0.94447338 | 0.04215509 | 0.5052871 |
| 0.999490048 | -0.0259688 | 0.677777 | 0.94447338 | -0.0385873 | 0.53782768 |
| 0.999490048 | -0.0245332 | 0.67817783 | 0.94447338 | -0.0071563 | 0.90350682 |
| 0.999490048 | -0.0245114 | 0.68061666 | 0.94447338 | -0.0453478 | 0.44820978 |
| 0.999490048 | -0.0240628 | 0.68092572 | 0.94447338 | 0.02667518 | 0.64863176 |
| 0.999490048 | -0.0236236 | 0.67799475 | 0.94447338 | -0.0026007 | 0.96347854 |
| 0.999490048 | -0.023329 | 0.67669605 | 0.94447338 | -0.1168624 | 0.04466654 |
| 0.999490048 | -0.0226647 | 0.67998815 | 0.94447338 | 0.00956646 | 0.86159162 |
| 0.999490048 | -0.0225693 | 0.67643559 | 0.94447338 | -0.0074232 | 0.89065151 |
| 0.999490048 | -0.0202578 | 0.68024762 | 0.94447338 | 0.01573346 | 0.74874047 |
| 0.881091405 | -0.0195195 | 0.6767623 | 0.94447338 | -0.1380847 | 0.00625458 |
| 0.999490048 | -0.0164548 | 0.67899444 | 0.94447338 | -0.083656 | 0.04326217 |
| 0.999490048 | -0.0161673 | 0.67792717 | 0.94447338 | -0.0180256 | 0.64348517 |
| 0.999490048 | 0.01676902 | 0.67946376 | 0.94447338 | 0.01175201 | 0.77194781 |
| 0.999490048 | 0.01835619 | 0.67753192 | 0.94447338 | 0.0104431 | 0.81277057 |
| 0.999490048 | 0.01994889 | 0.67626675 | 0.94447338 | -0.0341908 | 0.47569206 |
| 0.999490048 | 0.0202238 | 0.6801688 | 0.94447338 | -0.0190607 | 0.697573 |
| 0.999490048 | 0.02238435 | 0.67784305 | 0.94447338 | 0.0290367 | 0.59042251 |

| | | | | | |
|---|---|---|---|---|---|
| 0.999490048 | 0.0227021 | 0.67729586 | 0.94447338 | -0.0158238 | 0.77156431 |
| 0.999490048 | 0.02281271 | 0.67810814 | 0.94447338 | -0.0028372 | 0.95876236 |
| 0.999490048 | 0.0233104 | 0.67805586 | 0.94447338 | 0.03223975 | 0.56649184 |
| 0.999490048 | 0.02354633 | 0.68081995 | 0.94447338 | 0.06010148 | 0.29821874 |
| 0.999490048 | 0.02431966 | 0.67831903 | 0.94447338 | 0.01739276 | 0.76656151 |
| 0.999490048 | 0.02518318 | 0.67905907 | 0.94447338 | -7.56E-04 | 0.99006711 |
| 0.999490048 | 0.02586834 | 0.6764321 | 0.94447338 | 0.05369977 | 0.38903413 |
| 0.999490048 | 0.02715602 | 0.6801491 | 0.94447338 | 0.0752814 | 0.25844407 |
| 0.999490048 | 0.02840721 | 0.67852801 | 0.94447338 | -0.0318224 | 0.64257863 |
| 0.999490048 | 0.02864164 | 0.67781036 | 0.94447338 | 0.05101435 | 0.46100205 |
| 0.999490048 | 0.03058729 | 0.67957133 | 0.94447338 | -0.0112352 | 0.8792298 |
| 0.999490048 | 0.03259525 | 0.6798368 | 0.94447338 | 0.02705458 | 0.73180977 |
| 0.999490048 | 0.03307026 | 0.67904651 | 0.94447338 | 0.03018667 | 0.70558299 |
| 0.999490048 | 0.03419222 | 0.68039758 | 0.94447338 | -0.0871668 | 0.29814068 |
| 0.999490048 | 0.03837647 | 0.67928091 | 0.94447338 | 0.04641975 | 0.6172868 |
| 0.999490048 | 0.06922462 | 0.68076225 | 0.94447338 | 0.02831339 | 0.86617798 |
| 0.999490048 | 0.09532923 | 0.68002802 | 0.94447338 | -0.2213505 | 0.34184126 |
| 0.999490048 | 0.17908515 | 0.67767889 | 0.94447338 | -0.4235608 | 0.33740263 |
| 0.999490048 | -0.0611261 | 0.68181991 | 0.94458396 | -0.1473013 | 0.32705875 |
| 0.999490048 | -0.0404719 | 0.68199417 | 0.94458396 | 0.04374286 | 0.65795862 |
| 0.999490048 | -0.0335359 | 0.68224255 | 0.94458396 | -0.0179401 | 0.82643464 |
| 0.999490048 | -0.0286145 | 0.68195045 | 0.94458396 | -0.0368076 | 0.59850063 |
| 0.999490048 | -0.0174116 | 0.68205865 | 0.94458396 | 0.00852316 | 0.84086311 |
| 0.999490048 | 0.02721335 | 0.68156268 | 0.94458396 | -5.46E-04 | 0.9934252 |
| 0.999490048 | 0.02762192 | 0.68161074 | 0.94458396 | 0.04596487 | 0.49607716 |
| 0.999490048 | 0.04080402 | 0.68193766 | 0.94458396 | 0.10367312 | 0.30215708 |
| 0.999490048 | 0.04442587 | 0.68155942 | 0.94458396 | 0.10117577 | 0.3576988 |
| 0.999490048 | 0.06001862 | 0.68131193 | 0.94458396 | 0.04991942 | 0.73253928 |
| 0.999490048 | 0.07264287 | 0.6822498 | 0.94458396 | 0.0542129 | 0.75978885 |
| 0.999490048 | 0.0307622 | 0.68237234 | 0.94459699 | 0.07579207 | 0.31741757 |
| 0.999490048 | 0.03068222 | 0.68250512 | 0.94462419 | 0.05867278 | 0.43613543 |
| 0.999490048 | 0.03547703 | 0.68277371 | 0.94483932 | 0.14090827 | 0.11300171 |
| 0.999490048 | -0.0266764 | 0.68342282 | 0.94503229 | 3.59E-04 | 0.99561258 |
| 0.999490048 | 0.02349295 | 0.6832036 | 0.94503229 | -0.0624917 | 0.28250924 |
| 0.999490048 | 0.02623159 | 0.68347904 | 0.94503229 | -0.0643818 | 0.3209961 |
| 0.999490048 | 0.02905821 | 0.68339517 | 0.94503229 | 0.065992 | 0.35765719 |
| 0.999490048 | 0.03551431 | 0.68307671 | 0.94503229 | 0.07638457 | 0.38275915 |
| 0.999490048 | -0.034956 | 0.68387296 | 0.94508469 | 0.03217761 | 0.70772332 |
| 0.999490048 | -0.0347591 | 0.68381856 | 0.94508469 | 0.00892185 | 0.91661586 |
| 0.999490048 | 0.01504703 | 0.68396968 | 0.94508469 | 0.03903346 | 0.29554923 |
| 0.999490048 | 0.05322059 | 0.68391668 | 0.94508469 | 0.02429163 | 0.85238857 |
| 0.999490048 | -0.0545798 | 0.68423193 | 0.94529063 | -0.0179565 | 0.89340183 |

| | | | | | |
|---|---|---|---|---|---|
| 0.999490048 | -0.0550494 | 0.68470609 | 0.94542756 | -0.1301181 | 0.34081335 |
| 0.999490048 | -0.0227819 | 0.68461228 | 0.94542756 | -0.0150396 | 0.7883977 |
| 0.999490048 | -0.0182538 | 0.6846109 | 0.94542756 | 0.02715566 | 0.54647658 |
| 0.999490048 | -0.0173412 | 0.68489716 | 0.94542756 | 0.01046119 | 0.80641434 |
| 0.999490048 | 0.02758104 | 0.68486847 | 0.94542756 | -0.0615788 | 0.36803552 |
| 0.999490048 | -0.0187822 | 0.68517454 | 0.94565412 | 0.0483929 | 0.30067588 |
| 0.999490048 | -0.0353439 | 0.6853491 | 0.94570188 | 0.01171365 | 0.8930221 |
| 0.999490048 | -0.0220374 | 0.68543566 | 0.94570188 | -0.0078848 | 0.88460412 |
| 0.999490048 | -0.0184376 | 0.68572605 | 0.94594623 | -0.0286215 | 0.53084359 |
| 0.999490048 | -0.0486206 | 0.6865068 | 0.94655413 | -0.0195642 | 0.87081256 |
| 0.999490048 | -0.0267018 | 0.6863422 | 0.94655413 | -0.0875492 | 0.19239738 |
| 0.999490048 | 0.01759797 | 0.68648686 | 0.94655413 | 0.01473182 | 0.73532134 |
| 0.999490048 | 0.02371205 | 0.68673677 | 0.9465586 | 0.02099732 | 0.72091003 |
| 0.999490048 | 0.02814975 | 0.68666016 | 0.9465586 | 0.12491303 | 0.08212551 |
| 0.999490048 | -0.0600628 | 0.68694913 | 0.94669504 | -0.0277878 | 0.85190652 |
| 0.999490048 | -0.0355852 | 0.68724423 | 0.94678919 | -0.0647507 | 0.46550013 |
| 0.999490048 | 0.05335574 | 0.6872064 | 0.94678919 | 0.11354528 | 0.39422244 |
| 0.999490048 | 0.03452885 | 0.68750761 | 0.9469958 | -0.0689108 | 0.42434858 |
| 0.999490048 | -0.0497424 | 0.68817144 | 0.94711529 | -0.1851545 | 0.14318856 |
| 0.999490048 | -0.0496209 | 0.68781529 | 0.94711529 | -0.1803447 | 0.15198698 |
| 0.999490048 | -0.0412095 | 0.68810216 | 0.94711529 | -0.0278984 | 0.78561781 |
| 0.999490048 | -0.0328026 | 0.68827492 | 0.94711529 | -0.0950822 | 0.25043477 |
| 0.999490048 | -0.0213923 | 0.68796183 | 0.94711529 | -0.0881256 | 0.10658118 |
| 0.999490048 | 0.03238404 | 0.68819229 | 0.94711529 | 0.12755559 | 0.12229505 |
| 0.999490048 | -0.0371148 | 0.68945612 | 0.94742075 | 0.00903489 | 0.9223918 |
| 0.999490048 | -0.0246858 | 0.68906548 | 0.94742075 | 0.00729974 | 0.90567638 |
| 0.999490048 | -0.0201709 | 0.68973657 | 0.94742075 | 0.04073566 | 0.42239658 |
| 0.999490048 | 0.02124765 | 0.689745 | 0.94742075 | 0.00650202 | 0.90263079 |
| 0.999490048 | 0.02205824 | 0.6893768 | 0.94742075 | -0.0226907 | 0.68098203 |
| 0.999490048 | 0.02563537 | 0.68960342 | 0.94742075 | 0.08516575 | 0.19150114 |
| 0.999490048 | 0.02862615 | 0.68913767 | 0.94742075 | -0.0596067 | 0.40737713 |
| 0.950724435 | 0.0329404 | 0.68915243 | 0.94742075 | -0.1673942 | 0.05026942 |
| 0.999490048 | 0.03600471 | 0.68922061 | 0.94742075 | 0.04187674 | 0.64202417 |
| 0.999490048 | 0.04662189 | 0.68945717 | 0.94742075 | -0.1202611 | 0.30699851 |
| 0.999490048 | 0.08625358 | 0.68890158 | 0.94742075 | 0.02049359 | 0.92410305 |
| 0.999490048 | -0.0177392 | 0.69028475 | 0.9478503 | -0.0446396 | 0.32006976 |
| 0.999490048 | -0.0173164 | 0.69020447 | 0.9478503 | -0.027003 | 0.53517909 |
| 0.999490048 | 0.01946165 | 0.69041561 | 0.94787411 | 0.04123989 | 0.4012819 |
| 0.999490048 | -0.0242203 | 0.69150228 | 0.94897061 | 0.08860542 | 0.15436482 |
| 0.999490048 | -0.0195731 | 0.69215112 | 0.94897061 | 0.03993012 | 0.42154965 |
| 0.999490048 | -0.0175725 | 0.69162832 | 0.94897061 | -0.0223259 | 0.61464733 |
| 0.999490048 | -0.0148289 | 0.69233041 | 0.94897061 | 0.02447268 | 0.51487741 |

| | | | | | |
|---|---|---|---|---|---|
| 0.999490048 | 0.03563383 | 0.69174484 | 0.94897061 | 0.09571341 | 0.29156266 |
| 0.999490048 | 0.03786726 | 0.69191669 | 0.94897061 | 0.00337396 | 0.97179121 |
| 0.999490048 | 0.04283366 | 0.6917121 | 0.94897061 | 0.05051174 | 0.64026871 |
| 0.999490048 | 0.04776655 | 0.69235077 | 0.94897061 | -0.1018236 | 0.40159529 |
| 0.999490048 | 0.05662246 | 0.69215229 | 0.94897061 | 0.12172438 | 0.39735779 |
| 0.999490048 | 0.07739935 | 0.69187184 | 0.94897061 | 0.07210181 | 0.71180212 |
| 0.999490048 | 0.02234276 | 0.69257036 | 0.94897351 | 0.01171907 | 0.83553028 |
| 0.999490048 | 0.03356464 | 0.69258019 | 0.94897351 | -0.0641139 | 0.45201033 |
| 0.999490048 | -0.0276103 | 0.6930365 | 0.94897838 | -0.0247818 | 0.72303383 |
| 0.999490048 | -0.0266856 | 0.69360659 | 0.94897838 | 0.01601402 | 0.81292363 |
| 0.999490048 | -0.0199623 | 0.69348645 | 0.94897838 | 0.0396313 | 0.43604234 |
| 0.999490048 | 0.01303107 | 0.69289049 | 0.94897838 | 0.03331067 | 0.31686364 |
| 0.999490048 | 0.03872135 | 0.69333276 | 0.94897838 | 0.02949986 | 0.7637019 |
| 0.999490048 | 0.04095707 | 0.6935027 | 0.94897838 | 0.03304535 | 0.75037092 |
| 0.999490048 | 0.04179581 | 0.69353504 | 0.94897838 | 0.08752653 | 0.41170507 |
| 0.999490048 | 0.04938221 | 0.69342815 | 0.94897838 | 0.20378932 | 0.11228486 |
| 0.999490048 | 0.05333062 | 0.69321285 | 0.94897838 | -0.0173879 | 0.89751318 |
| 0.999490048 | -0.0196319 | 0.69382836 | 0.94912628 | 0.00385594 | 0.93827584 |
| 0.999490048 | -0.0739537 | 0.6968036 | 0.94917744 | 0.03231847 | 0.86462683 |
| 0.999490048 | -0.0703564 | 0.69671409 | 0.94917744 | -0.2530901 | 0.17767151 |
| 0.999490048 | -0.0645565 | 0.69494665 | 0.94917744 | -0.0198019 | 0.90412097 |
| 0.999490048 | -0.0529621 | 0.69407179 | 0.94917744 | -0.1316822 | 0.33194938 |
| 0.999490048 | -0.0511201 | 0.69495585 | 0.94917744 | -0.0668736 | 0.6083588 |
| 0.999490048 | -0.0407698 | 0.69643021 | 0.94917744 | 0.01500774 | 0.8856531 |
| 0.999490048 | -0.029016 | 0.69496529 | 0.94917744 | 0.00327954 | 0.96459498 |
| 0.999490048 | -0.0254347 | 0.69663215 | 0.94917744 | 0.02307825 | 0.72345463 |
| 0.999490048 | -0.0234094 | 0.69432944 | 0.94917744 | 0.03876597 | 0.51631956 |
| 0.999490048 | -0.020735 | 0.69565727 | 0.94917744 | -0.0343136 | 0.51851474 |
| 0.999490048 | -0.019783 | 0.69505204 | 0.94917744 | -0.0653604 | 0.202001 |
| 0.999490048 | -0.0192867 | 0.69642066 | 0.94917744 | 0.01300582 | 0.79230376 |
| 0.999490048 | -0.0181323 | 0.69592964 | 0.94917744 | 0.01834661 | 0.69252832 |
| 0.999490048 | -0.0175603 | 0.69584458 | 0.94917744 | -0.0550474 | 0.22655327 |
| 0.999490048 | 0.01514257 | 0.69550647 | 0.94917744 | 0.00541465 | 0.88854715 |
| 0.999490048 | 0.01830066 | 0.69584685 | 0.94917744 | 0.039814 | 0.39774306 |
| 0.999490048 | 0.01894439 | 0.69624799 | 0.94917744 | -0.0222042 | 0.64745328 |
| 0.999490048 | 0.01914043 | 0.69600329 | 0.94917744 | -0.0017295 | 0.97179345 |
| 0.999490048 | 0.02092409 | 0.69600941 | 0.94917744 | -0.0042813 | 0.93618799 |
| 0.999490048 | 0.02119545 | 0.69593344 | 0.94917744 | -7.42E-04 | 0.98907437 |
| 0.999490048 | 0.02286167 | 0.69621659 | 0.94917744 | 0.02515491 | 0.66759028 |
| 0.999490048 | 0.02410251 | 0.69675586 | 0.94917744 | -0.0100071 | 0.8713014 |
| 0.999490048 | 0.03340104 | 0.69682128 | 0.94917744 | 0.15227197 | 0.08440722 |
| 0.999490048 | 0.0348168 | 0.6955095 | 0.94917744 | -0.0664944 | 0.45659919 |

| | | | | | |
|---|---|---|---|---|---|
| 0.999490048 | 0.0404902 | 0.6953521 | 0.94917744 | -0.0274202 | 0.79068767 |
| 0.999490048 | 0.04927757 | 0.69491181 | 0.94917744 | 0.04178636 | 0.73934487 |
| 0.876151891 | -0.0558051 | 0.70085652 | 0.9492598 | -0.4269436 | 0.00642101 |
| 0.999490048 | -0.0479849 | 0.69856893 | 0.9492598 | 0.01337 | 0.9139551 |
| 0.999490048 | -0.0466775 | 0.70192873 | 0.9492598 | -0.0527937 | 0.66524163 |
| 0.999490048 | -0.0445714 | 0.70006658 | 0.9492598 | -0.0171583 | 0.88195704 |
| 0.999490048 | -0.0353588 | 0.70038144 | 0.9492598 | 0.04613565 | 0.61645293 |
| 0.999490048 | -0.0350094 | 0.69954536 | 0.9492598 | -0.0379656 | 0.67564574 |
| 0.999490048 | -0.0294706 | 0.69761887 | 0.9492598 | -0.0505217 | 0.50661441 |
| 0.999490048 | -0.0278504 | 0.6991348 | 0.9492598 | -0.0750266 | 0.30228537 |
| 0.999490048 | -0.0255996 | 0.69825158 | 0.9492598 | -0.0132973 | 0.84023124 |
| 0.999490048 | -0.0240032 | 0.70094306 | 0.9492598 | 0.04754646 | 0.44877566 |
| 0.999490048 | -0.0228895 | 0.6987854 | 0.9492598 | -0.053214 | 0.37147727 |
| 0.999490048 | -0.0216399 | 0.70199626 | 0.9492598 | 0.02282261 | 0.68660008 |
| 0.999490048 | -0.0206746 | 0.69762446 | 0.9492598 | -0.0052522 | 0.92126224 |
| 0.999490048 | -0.0201856 | 0.70208257 | 0.9492598 | -0.0363748 | 0.49206984 |
| 0.999490048 | -0.019846 | 0.70167616 | 0.9492598 | -0.0591748 | 0.25883456 |
| 0.999490048 | -0.0187978 | 0.69788386 | 0.9492598 | 8.01E-04 | 0.98678722 |
| 0.999490048 | -0.0187735 | 0.7001846 | 0.9492598 | -0.0304217 | 0.53362443 |
| 0.999490048 | -0.0177865 | 0.70167058 | 0.9492598 | -0.0099025 | 0.83094656 |
| 0.999490048 | -0.0174008 | 0.69985371 | 0.9492598 | -0.074732 | 0.10637989 |
| 0.999490048 | -0.0170349 | 0.69963988 | 0.9492598 | -0.0476238 | 0.28561906 |
| 0.999490048 | -0.015518 | 0.70022971 | 0.9492598 | -0.0242389 | 0.54844483 |
| 0.999490048 | 0.0129612 | 0.70016707 | 0.9492598 | 0.01493018 | 0.65749154 |
| 0.999490048 | 0.01843502 | 0.69913566 | 0.9492598 | -0.0072958 | 0.87827973 |
| 0.999490048 | 0.01887572 | 0.6988584 | 0.9492598 | -0.0113601 | 0.81572017 |
| 0.999490048 | 0.01993476 | 0.70033191 | 0.9492598 | 0.01590145 | 0.758714 |
| 0.999490048 | 0.0202276 | 0.70010876 | 0.9492598 | 0.05210957 | 0.32509244 |
| 0.999490048 | 0.02086613 | 0.70015696 | 0.9492598 | 0.05720626 | 0.29568831 |
| 0.999490048 | 0.0210496 | 0.69807766 | 0.9492598 | 0.01495651 | 0.78267357 |
| 0.999490048 | 0.02359074 | 0.70008332 | 0.9492598 | 0.0056926 | 0.92583627 |
| 0.999490048 | 0.02363746 | 0.69773464 | 0.9492598 | 0.08317927 | 0.17899106 |
| 0.999490048 | 0.02409726 | 0.6982617 | 0.9492598 | 0.07354914 | 0.24252184 |
| 0.999490048 | 0.02561273 | 0.69853871 | 0.9492598 | 0.02725872 | 0.6802659 |
| 0.999490048 | 0.02579512 | 0.70182496 | 0.9492598 | 0.02998569 | 0.65644638 |
| 0.999490048 | 0.02625773 | 0.69917544 | 0.9492598 | 0.0752026 | 0.27350921 |
| 0.999490048 | 0.0274766 | 0.69949717 | 0.9492598 | 0.0451587 | 0.52687136 |
| 0.999490048 | 0.03010788 | 0.70158508 | 0.9492598 | -0.014549 | 0.85293503 |
| 0.999490048 | 0.03158699 | 0.70183361 | 0.9492598 | -0.0152334 | 0.85335051 |
| 0.999490048 | 0.03253046 | 0.70068613 | 0.9492598 | -0.0209213 | 0.80457358 |
| 0.999490048 | 0.03283944 | 0.70183765 | 0.9492598 | 0.09423159 | 0.27700982 |
| 0.999490048 | 0.03726205 | 0.70111887 | 0.9492598 | -0.1562459 | 0.11601966 |

| | | | | | |
|---|---|---|---|---|---|
| 0.999490048 | 0.04164927 | 0.7021112 | 0.9492598 | -0.0115807 | 0.91519151 |
| 0.999490048 | 0.04333657 | 0.70126203 | 0.9492598 | 0.00538063 | 0.96195579 |
| 0.999490048 | 0.04482944 | 0.70197904 | 0.9492598 | -0.1112056 | 0.3461381 |
| 0.999490048 | 0.04801111 | 0.70078773 | 0.9492598 | 0.06188608 | 0.62117222 |
| 0.999490048 | 0.08281283 | 0.70127091 | 0.9492598 | -0.0060227 | 0.97771033 |
| 0.999490048 | 0.10127002 | 0.69730834 | 0.9492598 | 0.07320456 | 0.77842794 |
| 0.999490048 | -0.0493672 | 0.7023055 | 0.94936878 | 0.01774598 | 0.89058057 |
| 0.999490048 | -0.0222377 | 0.70246204 | 0.94942669 | 0.07496341 | 0.20455029 |
| 0.999490048 | -0.0387246 | 0.70305653 | 0.94979602 | -0.0258899 | 0.79867494 |
| 0.999490048 | -0.0221769 | 0.70317813 | 0.94979602 | -0.0143797 | 0.80469525 |
| 0.999490048 | -0.0204761 | 0.70351615 | 0.94979602 | 0.01043126 | 0.84610478 |
| 0.999490048 | -0.0169053 | 0.70299426 | 0.94979602 | 2.06E-04 | 0.99629435 |
| 0.999490048 | 0.02078897 | 0.70335155 | 0.94979602 | 0.00782067 | 0.88594675 |
| 0.999490048 | 0.02293745 | 0.70353154 | 0.94979602 | 0.00817661 | 0.89195374 |
| 0.999490048 | 0.05169147 | 0.70328806 | 0.94979602 | 0.02220606 | 0.86967166 |
| 0.999490048 | -0.0297989 | 0.70386291 | 0.95008432 | 0.06066252 | 0.44112754 |
| 0.999490048 | -0.0152638 | 0.70397266 | 0.95008432 | -0.0165096 | 0.6811641 |
| 0.999490048 | -0.015975 | 0.70419461 | 0.95009975 | 0.00401202 | 0.92393296 |
| 0.999490048 | 0.02704511 | 0.70421166 | 0.95009975 | 0.01643915 | 0.81733506 |
| 0.999490048 | -0.0759064 | 0.70444852 | 0.95026578 | -0.2572525 | 0.21360755 |
| 0.999490048 | 0.02083115 | 0.70460915 | 0.95032894 | 0.02253815 | 0.68175446 |
| 0.999490048 | -0.0738747 | 0.70893847 | 0.95035455 | 0.07217987 | 0.71530432 |
| 0.999490048 | -0.0597567 | 0.708105 | 0.95035455 | 0.08375727 | 0.60087494 |
| 0.999490048 | -0.0462828 | 0.70704047 | 0.95035455 | -0.1585841 | 0.20455635 |
| 0.999490048 | -0.0448687 | 0.7077579 | 0.95035455 | -0.0163811 | 0.89071386 |
| 0.999490048 | -0.044594 | 0.70625543 | 0.95035455 | 0.01379275 | 0.90708021 |
| 0.999490048 | -0.0419523 | 0.70931697 | 0.95035455 | 0.08736772 | 0.43968594 |
| 0.999490048 | -0.0404903 | 0.70514681 | 0.95035455 | -0.2085629 | 0.05978629 |
| 0.999490048 | -0.0392897 | 0.70656218 | 0.95035455 | 0.17154748 | 0.10881568 |
| 0.999490048 | -0.0361637 | 0.70611439 | 0.95035455 | -0.0235216 | 0.80609734 |
| 0.999490048 | -0.0314591 | 0.7084028 | 0.95035455 | -0.1210208 | 0.15799177 |
| 0.999490048 | -0.0309141 | 0.70619916 | 0.95035455 | 0.11147251 | 0.18130986 |
| 0.999490048 | -0.0303385 | 0.70716861 | 0.95035455 | 0.02928578 | 0.7168574 |
| 0.999490048 | -0.0283635 | 0.70896396 | 0.95035455 | 0.04481929 | 0.55615544 |
| 0.999490048 | -0.0262628 | 0.70574519 | 0.95035455 | -0.0781756 | 0.26634845 |
| 0.999490048 | -0.0254319 | 0.70939873 | 0.95035455 | -0.0404291 | 0.55441251 |
| 0.999490048 | -0.0241363 | 0.70606748 | 0.95035455 | -0.0736544 | 0.25532839 |
| 0.999490048 | -0.0229792 | 0.70778178 | 0.95035455 | 9.20E-04 | 0.98800972 |
| 0.999490048 | -0.0211769 | 0.70690094 | 0.95035455 | -0.0077173 | 0.89087247 |
| 0.999490048 | -0.0209928 | 0.70497038 | 0.95035455 | -0.0050343 | 0.92755528 |
| 0.999490048 | -0.0197982 | 0.70776234 | 0.95035455 | -0.0076842 | 0.88418229 |
| 0.999490048 | -0.0169412 | 0.71008151 | 0.95035455 | 0.00538873 | 0.90575422 |

| | | | | | |
|---|---|---|---|---|---|
| 0.999490048 | -0.0159023 | 0.70876281 | 0.95035455 | -0.019957 | 0.63951423 |
| 0.999490048 | -0.0143639 | 0.70644006 | 0.95035455 | -0.0332852 | 0.38572332 |
| 0.999490048 | 0.01410621 | 0.70986889 | 0.95035455 | -0.0409509 | 0.28504023 |
| 0.999490048 | 0.01915765 | 0.70961206 | 0.95035455 | 0.02044035 | 0.69119585 |
| 0.999490048 | 0.01917805 | 0.70865537 | 0.95035455 | 0.04948287 | 0.33878811 |
| 0.999490048 | 0.01939701 | 0.70951877 | 0.95035455 | -0.0192726 | 0.71129294 |
| 0.999490048 | 0.02350687 | 0.70509647 | 0.95035455 | 0.00454302 | 0.94161301 |
| 0.999490048 | 0.02395536 | 0.70915622 | 0.95035455 | 0.02465688 | 0.70106714 |
| 0.999490048 | 0.02417855 | 0.70651246 | 0.95035455 | -0.0186725 | 0.7710791 |
| 0.999490048 | 0.02448163 | 0.70907738 | 0.95035455 | 0.01866942 | 0.77588476 |
| 0.999490048 | 0.0245642 | 0.70888341 | 0.95035455 | 0.01485755 | 0.82118221 |
| 0.999490048 | 0.02491228 | 0.70714088 | 0.95035455 | -8.11E-05 | 0.9990223 |
| 0.999490048 | 0.02562875 | 0.70869939 | 0.95035455 | 0.05916422 | 0.3912921 |
| 0.999490048 | 0.02634483 | 0.7093448 | 0.95035455 | 0.01303998 | 0.85347214 |
| 0.999490048 | 0.02659037 | 0.70884168 | 0.95035455 | 0.03710329 | 0.60281674 |
| 0.999490048 | 0.03000835 | 0.70623153 | 0.95035455 | 0.03579431 | 0.65319728 |
| 0.999490048 | 0.03039826 | 0.70979266 | 0.95035455 | 0.01322614 | 0.87122879 |
| 0.999490048 | 0.03079788 | 0.7061984 | 0.95035455 | -0.0467779 | 0.56766465 |
| 0.999490048 | 0.03347656 | 0.70921171 | 0.95035455 | 0.16245931 | 0.07907437 |
| 0.999490048 | 0.03448874 | 0.70804484 | 0.95035455 | -0.0205882 | 0.82294954 |
| 0.999490048 | 0.03491046 | 0.70695115 | 0.95035455 | 0.05262491 | 0.57159903 |
| 0.999490048 | 0.03691646 | 0.70997353 | 0.95035455 | 0.14614433 | 0.14891885 |
| 0.999490048 | 0.03815958 | 0.70860088 | 0.95035455 | 0.09964472 | 0.33301546 |
| 0.999490048 | 0.05200776 | 0.70507405 | 0.95035455 | -0.0621475 | 0.65127503 |
| 0.999490048 | 0.06014923 | 0.71009126 | 0.95035455 | 0.08143479 | 0.61517582 |
| 0.999490048 | 0.06561526 | 0.70951724 | 0.95035455 | 0.04780853 | 0.78563444 |
| 0.999490048 | 0.11150644 | 0.70710097 | 0.95035455 | -0.0939604 | 0.75129881 |
| 0.999490048 | -0.1691094 | 0.71105083 | 0.95036473 | 0.05633595 | 0.9015062 |
| 0.999490048 | -0.0394857 | 0.71062887 | 0.95036473 | -0.1184661 | 0.27088084 |
| 0.999490048 | -0.0266509 | 0.71151683 | 0.95036473 | -0.0295881 | 0.68147924 |
| 0.999490048 | -0.0246046 | 0.71029586 | 0.95036473 | 0.01033382 | 0.87588294 |
| 0.999490048 | -0.0187692 | 0.71156086 | 0.95036473 | -0.0074989 | 0.88241995 |
| 0.999490048 | -0.0186831 | 0.71080992 | 0.95036473 | -0.0117317 | 0.81575181 |
| 0.999490048 | -0.0172306 | 0.71117145 | 0.95036473 | 0.01170562 | 0.80124263 |
| 0.999490048 | 0.01468412 | 0.71157848 | 0.95036473 | -0.0181365 | 0.64814824 |
| 0.999490048 | 0.01552583 | 0.71048801 | 0.95036473 | -0.0094124 | 0.82179599 |
| 0.999490048 | 0.02258808 | 0.71147603 | 0.95036473 | 0.02820036 | 0.64449827 |
| 0.999490048 | 0.02706029 | 0.71086418 | 0.95036473 | 0.05553711 | 0.44878646 |
| 0.999490048 | 0.05043658 | 0.71146334 | 0.95036473 | 8.09E-04 | 0.99526163 |
| 0.999490048 | 0.10257639 | 0.71088909 | 0.95036473 | 0.02695154 | 0.92209611 |
| 0.999490048 | -0.0973257 | 0.71341503 | 0.95062571 | 0.02372679 | 0.92856405 |
| 0.999490048 | -0.0579691 | 0.71214814 | 0.95062571 | 0.13215788 | 0.40861521 |

| | | | | | |
|---|---|---|---|---|---|
| 0.999490048 | -0.0413744 | 0.71411743 | 0.95062571 | -0.0640076 | 0.57162269 |
| 0.999490048 | -0.0370022 | 0.71416468 | 0.95062571 | 0.1594756 | 0.12286998 |
| 0.999490048 | -0.0321765 | 0.71282213 | 0.95062571 | -0.019748 | 0.82113022 |
| 0.999490048 | -0.0312588 | 0.71260376 | 0.95062571 | -0.0780956 | 0.36082518 |
| 0.999490048 | -0.0296777 | 0.7141177 | 0.95062571 | 0.09270286 | 0.25802943 |
| 0.999490048 | -0.0183936 | 0.71408508 | 0.95062571 | 0.04577065 | 0.36526712 |
| 0.999490048 | -0.0168845 | 0.71340727 | 0.95062571 | -0.0385379 | 0.40452903 |
| 0.999490048 | 0.01377946 | 0.71331899 | 0.95062571 | 0.00647907 | 0.8627022 |
| 0.999490048 | 0.01673327 | 0.71359198 | 0.95062571 | 0.02109929 | 0.6437713 |
| 0.999490048 | 0.01959587 | 0.71373021 | 0.95062571 | 0.04594817 | 0.39255047 |
| 0.999490048 | 0.02134609 | 0.71329167 | 0.95062571 | -0.0233577 | 0.68771481 |
| 0.999490048 | 0.02169034 | 0.71410336 | 0.95062571 | 0.02061422 | 0.72767438 |
| 0.999490048 | 0.02223866 | 0.71290694 | 0.95062571 | 0.06167535 | 0.31186616 |
| 0.999490048 | 0.02515676 | 0.71289007 | 0.95062571 | 0.00335811 | 0.96077073 |
| 0.999490048 | 0.03202996 | 0.71355037 | 0.95062571 | 0.07127951 | 0.41645526 |
| 0.999490048 | 0.03664049 | 0.71214562 | 0.95062571 | -0.015156 | 0.87855746 |
| 0.999490048 | 0.04384611 | 0.71378211 | 0.95062571 | 0.07326298 | 0.54095064 |
| 0.999490048 | 0.06872957 | 0.7138236 | 0.95062571 | -0.1374105 | 0.46909124 |
| 0.999490048 | 0.14726387 | 0.71298611 | 0.95062571 | 0.38777534 | 0.34468298 |
| 0.999490048 | -0.0243978 | 0.71431103 | 0.95066897 | 0.01226826 | 0.85380693 |
| 0.999490048 | 0.01777897 | 0.71454293 | 0.95067757 | 0.05048798 | 0.3034948 |
| 0.999490048 | 0.06507797 | 0.7145452 | 0.95067757 | 0.08298646 | 0.64156797 |
| 0.999490048 | 0.01726648 | 0.71488822 | 0.95078069 | -0.0414326 | 0.38373803 |
| 0.999490048 | 0.02134828 | 0.7149643 | 0.95078069 | 0.03390885 | 0.56266869 |
| 0.999490048 | 0.03557675 | 0.71487722 | 0.95078069 | 0.03028684 | 0.75571655 |
| 0.781057588 | 0.05249831 | 0.71516051 | 0.95089016 | -0.3671027 | 0.02370429 |
| 0.999490048 | -0.0299713 | 0.71547858 | 0.95096688 | -0.0979174 | 0.23966089 |
| 0.999490048 | 0.01926528 | 0.71547245 | 0.95096688 | -0.0137572 | 0.79450089 |
| 0.999490048 | 0.03190585 | 0.71555987 | 0.95096688 | -0.1163833 | 0.19085526 |
| 0.999490048 | -0.0209434 | 0.71590887 | 0.95127929 | -0.0015529 | 0.97844395 |
| 0.999490048 | -0.0911813 | 0.71651609 | 0.95158639 | 0.19260055 | 0.44958388 |
| 0.999490048 | -0.0387617 | 0.71693772 | 0.95158639 | 0.05865914 | 0.58387833 |
| 0.999490048 | -0.0227458 | 0.71676797 | 0.95158639 | 0.03196426 | 0.61064472 |
| 0.999490048 | -0.0192748 | 0.71667063 | 0.95158639 | -0.0123216 | 0.81639996 |
| 0.999490048 | -0.0174612 | 0.71651123 | 0.95158639 | -0.038357 | 0.42739163 |
| 0.999490048 | 0.02100731 | 0.71687161 | 0.95158639 | 0.05208633 | 0.37180521 |
| 0.999490048 | 0.03103856 | 0.71689082 | 0.95158639 | -0.0119013 | 0.88929328 |
| 0.999490048 | -0.0337369 | 0.71723095 | 0.95182429 | 0.07762598 | 0.4073156 |
| 0.999490048 | -0.0284671 | 0.71744756 | 0.95196045 | -0.02513 | 0.74931321 |
| 0.999490048 | 0.02196642 | 0.71782208 | 0.95215478 | 0.07041745 | 0.25235721 |
| 0.999490048 | 0.02851648 | 0.71777316 | 0.95215478 | 0.10415708 | 0.1938062 |
| 0.999490048 | -0.0296246 | 0.71794559 | 0.95216736 | -0.0182149 | 0.82410203 |

| | | | | | |
|---|---|---|---|---|---|
| 0.999490048 | -0.0213229 | 0.71820475 | 0.95220859 | -0.0453526 | 0.44486257 |
| 0.999490048 | 0.0150398 | 0.71819679 | 0.95220859 | -0.0152264 | 0.71486058 |
| 0.999490048 | 0.019211 | 0.71847941 | 0.95227032 | -0.0178093 | 0.73819097 |
| 0.999490048 | 0.02218125 | 0.71844811 | 0.95227032 | 0.01485748 | 0.80903731 |
| 0.999490048 | -0.0296534 | 0.71876249 | 0.95238628 | -0.0138167 | 0.86661184 |
| 0.999490048 | -0.0209589 | 0.71932954 | 0.95238628 | -0.0297482 | 0.61048507 |
| 0.999490048 | -0.0171786 | 0.71909945 | 0.95238628 | -0.0189914 | 0.69099832 |
| 0.999490048 | 0.02475957 | 0.71933844 | 0.95238628 | -0.0245226 | 0.72190561 |
| 0.999490048 | 0.02851083 | 0.7189613 | 0.95238628 | 0.04294402 | 0.58842535 |
| 0.999490048 | 0.04080084 | 0.71889871 | 0.95238628 | 0.00790802 | 0.94431318 |
| 0.987721587 | 0.0429763 | 0.7193653 | 0.95238628 | -0.2145818 | 0.08158127 |
| 0.999490048 | 0.01535991 | 0.71967849 | 0.95238884 | -0.04589 | 0.28849662 |
| 0.999490048 | 0.0175515 | 0.71966554 | 0.95238884 | -0.0037125 | 0.93941001 |
| 0.999490048 | 0.02783711 | 0.71970941 | 0.95238884 | -0.0187762 | 0.80861388 |
| 0.999490048 | -0.0391305 | 0.72050224 | 0.95254998 | 0.01516874 | 0.88957092 |
| 0.999490048 | -0.0247593 | 0.72077092 | 0.95254998 | -0.026466 | 0.70246446 |
| 0.999490048 | -0.0237353 | 0.72047604 | 0.95254998 | -0.0396021 | 0.55140656 |
| 0.999490048 | -0.0211376 | 0.72142827 | 0.95254998 | 7.25E-04 | 0.99022672 |
| 0.999490048 | -0.0210146 | 0.72101653 | 0.95254998 | 0.04419639 | 0.45461076 |
| 0.999490048 | -0.0190339 | 0.72109324 | 0.95254998 | -0.0820703 | 0.13204593 |
| 0.999490048 | -0.0189789 | 0.72031312 | 0.95254998 | -0.0295208 | 0.57828127 |
| 0.999490048 | 0.0149215 | 0.72136829 | 0.95254998 | 0.05735924 | 0.17780137 |
| 0.999490048 | 0.02115855 | 0.72015859 | 0.95254998 | 2.94E-05 | 0.99960254 |
| 0.999490048 | 0.02298884 | 0.72053285 | 0.95254998 | -0.0388271 | 0.5466556 |
| 0.999490048 | 0.02961023 | 0.72051142 | 0.95254998 | 0.03237904 | 0.69569999 |
| 0.999490048 | 0.03057629 | 0.72105544 | 0.95254998 | -0.0217628 | 0.79926951 |
| 0.999490048 | 0.05547991 | 0.72092276 | 0.95254998 | 0.08485155 | 0.58547423 |
| 0.999490048 | 0.0929266 | 0.72133879 | 0.95254998 | 0.00620646 | 0.98093874 |
| 0.999490048 | -0.0253868 | 0.72174913 | 0.95267234 | 0.02609769 | 0.71431435 |
| 0.999490048 | 0.05286876 | 0.72170552 | 0.95267234 | 0.27997456 | 0.06794328 |
| 0.999490048 | -0.1965333 | 0.72974927 | 0.95310259 | -1.2191388 | 0.05678857 |
| 0.999490048 | -0.0663556 | 0.7281687 | 0.95310259 | 0.00622756 | 0.97394676 |
| 0.999490048 | -0.0632258 | 0.72432428 | 0.95310259 | 0.12418644 | 0.49333543 |
| 0.999490048 | -0.0625528 | 0.72454511 | 0.95310259 | -0.3267987 | 0.07436985 |
| 0.999490048 | -0.0514398 | 0.72701583 | 0.95310259 | -0.2126164 | 0.16663773 |
| 0.999490048 | -0.0511812 | 0.722892 | 0.95310259 | -0.1168537 | 0.42063871 |
| 0.999490048 | -0.0395197 | 0.72878525 | 0.95310259 | -0.0867957 | 0.44842839 |
| 0.999490048 | -0.0372764 | 0.72557868 | 0.95310259 | 0.11031136 | 0.30351261 |
| 0.999490048 | -0.0363242 | 0.72435444 | 0.95310259 | 0.031929 | 0.75651259 |
| 0.999490048 | -0.0339066 | 0.73045192 | 0.95310259 | 0.00734386 | 0.94044836 |
| 0.999490048 | -0.0338305 | 0.72551258 | 0.95310259 | 0.04598303 | 0.63356112 |
| 0.999490048 | -0.0322674 | 0.72905522 | 0.95310259 | 0.0725209 | 0.43850793 |

| | | | | | |
|---|---|---|---|---|---|
| 0.999490048 | -0.0280918 | 0.72714863 | 0.95310259 | -0.0478366 | 0.55332539 |
| 0.999490048 | -0.02733 | 0.73007922 | 0.95310259 | 0.08276795 | 0.30074607 |
| 0.999490048 | -0.0272709 | 0.72656079 | 0.95310259 | -0.0026943 | 0.97240444 |
| 0.999490048 | -0.0270441 | 0.72963549 | 0.95310259 | -0.1488917 | 0.06573381 |
| 0.999490048 | -0.0269608 | 0.72872244 | 0.95310259 | -0.0163364 | 0.83342256 |
| 0.999490048 | -0.0224437 | 0.72830475 | 0.95310259 | 0.00204514 | 0.97471705 |
| 0.999490048 | -0.0218017 | 0.72751185 | 0.95310259 | -0.0707083 | 0.26390412 |
| 0.999490048 | -0.0212146 | 0.72697486 | 0.95310259 | -0.0266872 | 0.66072814 |
| 0.999490048 | -0.0204803 | 0.73061851 | 0.95310259 | 0.00371428 | 0.95015294 |
| 0.999490048 | -0.0204004 | 0.72908575 | 0.95310259 | -0.0092763 | 0.87474271 |
| 0.999490048 | -0.0198044 | 0.72487671 | 0.95310259 | -0.0360275 | 0.52320308 |
| 0.999490048 | -0.0184247 | 0.72842765 | 0.95310259 | -0.046664 | 0.38229048 |
| 0.999490048 | -0.017888 | 0.72592705 | 0.95310259 | -0.003455 | 0.94595366 |
| 0.999490048 | -0.017548 | 0.72767027 | 0.95310259 | -0.0319403 | 0.52735606 |
| 0.999490048 | -0.0174903 | 0.72938568 | 0.95310259 | -0.0217957 | 0.66660515 |
| 0.999490048 | -0.0173714 | 0.7266026 | 0.95310259 | -0.0042319 | 0.93204027 |
| 0.999490048 | -0.0164365 | 0.72699004 | 0.95310259 | 0.03303086 | 0.48454258 |
| 0.999490048 | -0.0159194 | 0.72663353 | 0.95310259 | -0.0293595 | 0.52031053 |
| 0.999490048 | -0.0152251 | 0.72523653 | 0.95310259 | 0.03336264 | 0.44334475 |
| 0.999490048 | -0.0151174 | 0.72700913 | 0.95310259 | -0.0236799 | 0.58515722 |
| 0.999490048 | -0.0150853 | 0.72291315 | 0.95310259 | -0.0183468 | 0.66649371 |
| 0.999490048 | -0.0141708 | 0.73051803 | 0.95310259 | -0.0277824 | 0.50094775 |
| 0.999490048 | -0.0125515 | 0.72339894 | 0.95310259 | 0.00792469 | 0.82304895 |
| 0.999490048 | 0.01671332 | 0.73086418 | 0.95310259 | 0.0228262 | 0.63885475 |
| 0.999490048 | 0.01721436 | 0.72769206 | 0.95310259 | -0.0443742 | 0.37268543 |
| 0.999490048 | 0.01731806 | 0.72671693 | 0.95310259 | 0.02785414 | 0.57477729 |
| 0.999490048 | 0.01786868 | 0.72973098 | 0.95310259 | -0.0083906 | 0.87100575 |
| 0.999490048 | 0.01807181 | 0.7268591 | 0.95310259 | -0.0544287 | 0.29750261 |
| 0.999490048 | 0.01899716 | 0.72562634 | 0.95310259 | 0.02769883 | 0.60935934 |
| 0.999490048 | 0.01901448 | 0.72513137 | 0.95310259 | 0.027824 | 0.6074049 |
| 0.999490048 | 0.0192222 | 0.72967331 | 0.95310259 | 0.01833253 | 0.74170047 |
| 0.999490048 | 0.02009819 | 0.72265435 | 0.95310259 | -0.0345906 | 0.54231349 |
| 0.999490048 | 0.02053152 | 0.73081216 | 0.95310259 | -0.024133 | 0.68605764 |
| 0.999490048 | 0.02068503 | 0.72640255 | 0.95310259 | 0.02410934 | 0.68352238 |
| 0.999490048 | 0.02198864 | 0.72818203 | 0.95310259 | 0.01619224 | 0.79789528 |
| 0.999490048 | 0.02235584 | 0.72923463 | 0.95310259 | -0.0181263 | 0.77888519 |
| 0.999490048 | 0.02275528 | 0.72752588 | 0.95310259 | 0.00697715 | 0.91482873 |
| 0.999490048 | 0.02327975 | 0.72638965 | 0.95310259 | 0.04445653 | 0.50535839 |
| 0.999490048 | 0.02358827 | 0.72828248 | 0.95310259 | -0.0030124 | 0.96456937 |
| 0.999490048 | 0.024092 | 0.72725446 | 0.95310259 | 0.11235253 | 0.11245952 |
| 0.999490048 | 0.0242173 | 0.72298191 | 0.95310259 | -0.0160146 | 0.81453633 |
| 0.999490048 | 0.02496606 | 0.72564831 | 0.95310259 | -0.0195202 | 0.78369518 |

| | | | | | |
|---|---|---|---|---|---|
| 0.999490048 | 0.02553323 | 0.72654366 | 0.95310259 | 0.01018011 | 0.88899073 |
| 0.999490048 | 0.02692119 | 0.72508928 | 0.95310259 | -0.0424988 | 0.57949537 |
| 0.999490048 | 0.02729779 | 0.72931866 | 0.95310259 | 0.06526911 | 0.41067615 |
| 0.999490048 | 0.02746638 | 0.72987794 | 0.95310259 | 0.061199 | 0.44384455 |
| 0.999490048 | 0.03006488 | 0.7289861 | 0.95310259 | 0.01839025 | 0.83203335 |
| 0.999490048 | 0.0339255 | 0.72651886 | 0.95310259 | 0.08115215 | 0.40550343 |
| 0.999490048 | 0.03425532 | 0.72620363 | 0.95310259 | 0.02228324 | 0.81968004 |
| 0.999490048 | 0.03560664 | 0.72901709 | 0.95310259 | -0.0083633 | 0.93507305 |
| 0.999490048 | 0.03850177 | 0.72531762 | 0.95310259 | -0.0664627 | 0.54515286 |
| 0.999490048 | 0.03893963 | 0.72565768 | 0.95310259 | -0.075647 | 0.4969058 |
| 0.999490048 | 0.03985421 | 0.72686499 | 0.95310259 | 0.06801185 | 0.55205242 |
| 0.999490048 | 0.04077268 | 0.723857 | 0.95310259 | -0.034179 | 0.76701348 |
| 0.448222939 | 0.04237506 | 0.72772064 | 0.95310259 | -0.3635271 | 0.00572868 |
| 0.999490048 | 0.04253578 | 0.73026104 | 0.95310259 | 0.07185145 | 0.56116591 |
| 0.999490048 | 0.05154199 | 0.72355581 | 0.95310259 | 0.07039929 | 0.62938452 |
| 0.999490048 | 0.05234189 | 0.73079619 | 0.95310259 | -0.1123632 | 0.46206036 |
| 0.999490048 | 0.05748707 | 0.72956322 | 0.95310259 | 0.17711187 | 0.29169432 |
| 0.999490048 | 0.06112499 | 0.73007235 | 0.95310259 | -0.0988646 | 0.57754313 |
| 0.999490048 | 0.06307759 | 0.72747947 | 0.95310259 | 0.22239833 | 0.22552893 |
| 0.999490048 | 0.07581023 | 0.72302013 | 0.95310259 | 0.07855108 | 0.71346611 |
| 0.999490048 | 0.07773402 | 0.72228177 | 0.95310259 | 0.00558423 | 0.97960482 |
| 0.999490048 | 0.08965759 | 0.72608238 | 0.95310259 | 0.35130373 | 0.17726424 |
| 0.999490048 | 0.09312931 | 0.73033758 | 0.95310259 | 0.10183619 | 0.70631277 |
| 0.999490048 | 0.02405233 | 0.73111526 | 0.95328113 | -0.0470887 | 0.50253027 |
| 0.999490048 | 0.02090327 | 0.73159957 | 0.95376369 | 0.03120172 | 0.60918317 |
| 0.999490048 | -0.0211458 | 0.73172621 | 0.95377987 | -0.0203803 | 0.74106208 |
| 0.999490048 | -0.0194654 | 0.73218142 | 0.95392648 | -0.0797918 | 0.16831537 |
| 0.999490048 | 0.01636159 | 0.73204607 | 0.95392648 | -0.0061821 | 0.89695693 |
| 0.999490048 | 0.04345281 | 0.73208438 | 0.95392648 | 0.05074679 | 0.68941457 |
| 0.999490048 | 0.01913371 | 0.73234059 | 0.95398501 | 0.02019925 | 0.71809089 |
| 0.987721587 | -0.0305865 | 0.73324217 | 0.95426625 | 0.16363553 | 0.07704276 |
| 0.999490048 | -0.0274997 | 0.73343791 | 0.95426625 | 0.06210752 | 0.44398277 |
| 0.999490048 | -0.0210996 | 0.73309703 | 0.95426625 | 0.01915711 | 0.75680538 |
| 0.999490048 | -0.0192276 | 0.73341127 | 0.95426625 | 0.01356528 | 0.80999506 |
| 0.999490048 | -0.0188781 | 0.73278705 | 0.95426625 | -0.0048211 | 0.93044227 |
| 0.999490048 | 0.0197064 | 0.73336589 | 0.95426625 | -0.0097023 | 0.86668856 |
| 0.999490048 | 0.02134179 | 0.73347076 | 0.95426625 | -0.0764708 | 0.22870979 |
| 0.999490048 | 0.03542736 | 0.73328365 | 0.95426625 | -0.0926729 | 0.37595945 |
| 0.999490048 | -0.0550951 | 0.73836793 | 0.95438206 | 0.02470582 | 0.88083193 |
| 0.999490048 | -0.0322081 | 0.73832579 | 0.95438206 | 0.11693466 | 0.23141948 |
| 0.999490048 | -0.0289499 | 0.73844526 | 0.95438206 | -0.0970361 | 0.26844515 |
| 0.999490048 | -0.0287378 | 0.73817033 | 0.95438206 | 0.03241371 | 0.70623322 |

| | | | | | |
|---|---|---|---|---|---|
| 0.999490048 | -0.0277172 | 0.7378069 | 0.95438206 | 0.01265174 | 0.87844507 |
| 0.999490048 | -0.0255101 | 0.73620599 | 0.95438206 | 0.02952505 | 0.69672022 |
| 0.999490048 | -0.025452 | 0.73726799 | 0.95438206 | -0.0194434 | 0.79764381 |
| 0.999490048 | -0.0236676 | 0.7361939 | 0.95438206 | -0.0373483 | 0.59560511 |
| 0.999490048 | -0.0231533 | 0.73802888 | 0.95438206 | -0.0359219 | 0.60439293 |
| 0.999490048 | -0.0229068 | 0.73721766 | 0.95438206 | -0.0264325 | 0.69872785 |
| 0.999490048 | -0.0228498 | 0.73825762 | 0.95438206 | 0.00799975 | 0.90677581 |
| 0.999490048 | -0.022407 | 0.73751723 | 0.95438206 | -0.001963 | 0.97655053 |
| 0.999490048 | -0.0211571 | 0.73631834 | 0.95438206 | -0.0609628 | 0.33589608 |
| 0.999490048 | -0.021049 | 0.73777737 | 0.95438206 | -0.0025869 | 0.96714232 |
| 0.999490048 | -0.0202162 | 0.73542734 | 0.95438206 | -0.0255869 | 0.66909431 |
| 0.999490048 | -0.0183436 | 0.7361877 | 0.95438206 | -0.0745428 | 0.17841706 |
| 0.999490048 | -0.0181027 | 0.73807294 | 0.95438206 | -0.0187011 | 0.72976877 |
| 0.999490048 | -0.0180895 | 0.73671529 | 0.95438206 | 0.0358554 | 0.50657137 |
| 0.999490048 | -0.0178573 | 0.73578141 | 0.95438206 | 0.04979182 | 0.35045979 |
| 0.999490048 | -0.0173372 | 0.73812315 | 0.95438206 | 0.00643712 | 0.90111114 |
| 0.999490048 | -0.0170351 | 0.73438719 | 0.95438206 | -0.0553601 | 0.27542136 |
| 0.999490048 | -0.0145087 | 0.73396314 | 0.95438206 | -0.0262765 | 0.53929253 |
| 0.999490048 | 0.01262373 | 0.73402012 | 0.95438206 | 0.00871285 | 0.81447255 |
| 0.999490048 | 0.01404904 | 0.73815792 | 0.95438206 | 0.0106449 | 0.79994783 |
| 0.999490048 | 0.01452742 | 0.73593181 | 0.95438206 | 0.01081968 | 0.80158132 |
| 0.999490048 | 0.01758741 | 0.73552313 | 0.95438206 | -0.0014034 | 0.97847106 |
| 0.999490048 | 0.01792403 | 0.73583961 | 0.95438206 | 0.05227215 | 0.32929111 |
| 0.999490048 | 0.02005521 | 0.73847455 | 0.95438206 | 0.03962264 | 0.5108673 |
| 0.999490048 | 0.02027381 | 0.73575703 | 0.95438206 | -0.0619601 | 0.30692457 |
| 0.999490048 | 0.02247904 | 0.73776697 | 0.95438206 | 0.08585478 | 0.20776906 |
| 0.999490048 | 0.02253898 | 0.73688898 | 0.95438206 | 0.05214381 | 0.43924748 |
| 0.999490048 | 0.02368127 | 0.73707249 | 0.95438206 | 0.01627674 | 0.81739538 |
| 0.999490048 | 0.02405169 | 0.73546778 | 0.95438206 | 0.01653914 | 0.81617183 |
| 0.999490048 | 0.02441655 | 0.73847096 | 0.95438206 | 0.09939818 | 0.1814694 |
| 0.999490048 | 0.02701092 | 0.73707726 | 0.95438206 | 0.02811339 | 0.72679189 |
| 0.999490048 | 0.0275258 | 0.73724187 | 0.95438206 | -0.0786694 | 0.34149546 |
| 0.999490048 | 0.02929418 | 0.7357458 | 0.95438206 | 0.01469355 | 0.86546055 |
| 0.999490048 | 0.03058543 | 0.73710094 | 0.95438206 | 0.1299817 | 0.16149754 |
| 0.999490048 | 0.03368948 | 0.73646141 | 0.95438206 | -0.0691401 | 0.49135014 |
| 0.999490048 | 0.03408859 | 0.73604393 | 0.95438206 | 0.11316524 | 0.26850101 |
| 0.999490048 | 0.0366287 | 0.7366341 | 0.95438206 | 0.16099444 | 0.14744556 |
| 0.999490048 | 0.04508481 | 0.73553627 | 0.95438206 | -0.0295756 | 0.82452963 |
| 0.999490048 | 0.0829542 | 0.73531931 | 0.95438206 | 0.01057379 | 0.96559014 |
| 0.999490048 | -0.1915111 | 0.7437033 | 0.95450483 | -0.0824334 | 0.88798325 |
| 0.984638128 | -0.0917139 | 0.74553405 | 0.95450483 | -0.7204184 | 0.01620046 |
| 0.999490048 | -0.0650701 | 0.743488 | 0.95450483 | -0.2317239 | 0.24972676 |

| | | | | | |
|---|---|---|---|---|---|
| 0.999490048 | -0.057636 | 0.74262543 | 0.95450483 | 0.25826672 | 0.16573448 |
| 0.999490048 | -0.0538686 | 0.74387811 | 0.95450483 | 0.08152341 | 0.62147049 |
| 0.999490048 | -0.0406728 | 0.7455828 | 0.95450483 | -0.1045215 | 0.40714838 |
| 0.436210265 | -0.0363584 | 0.74217077 | 0.95450483 | -0.4659139 | 0.00117615 |
| 0.999490048 | -0.0309379 | 0.74469237 | 0.95450483 | 0.0720506 | 0.45034873 |
| 0.999490048 | -0.0289792 | 0.74271421 | 0.95450483 | -0.0420448 | 0.6342948 |
| 0.999490048 | -0.0226014 | 0.74267792 | 0.95450483 | -0.0272926 | 0.69191511 |
| 0.999490048 | -0.022015 | 0.7439769 | 0.95450483 | 0.01349358 | 0.84123714 |
| 0.999490048 | -0.0214701 | 0.74255121 | 0.95450483 | -0.0548432 | 0.40423857 |
| 0.999490048 | -0.0202469 | 0.74341344 | 0.95450483 | 0.00212508 | 0.97256337 |
| 0.999490048 | -0.0198484 | 0.74352362 | 0.95450483 | 0.09127162 | 0.14067939 |
| 0.999490048 | -0.018683 | 0.74504781 | 0.95450483 | -0.0671898 | 0.24811722 |
| 0.999490048 | -0.0169152 | 0.74427817 | 0.95450483 | -0.0541537 | 0.30106919 |
| 0.999490048 | -0.0163992 | 0.74001008 | 0.95450483 | 0.00248318 | 0.95988158 |
| 0.999490048 | -0.0162003 | 0.74565689 | 0.95450483 | -0.0218882 | 0.66147773 |
| 0.999490048 | -0.0161163 | 0.74133802 | 0.95450483 | 0.02294908 | 0.63871513 |
| 0.999490048 | -0.0159894 | 0.74483574 | 0.95450483 | -0.0456913 | 0.35599971 |
| 0.999490048 | -0.0154573 | 0.74380541 | 0.95450483 | -0.0120319 | 0.79910796 |
| 0.999490048 | -0.0148925 | 0.7435974 | 0.95450483 | -0.0252254 | 0.58031471 |
| 0.999490048 | -0.014526 | 0.74061269 | 0.95450483 | -0.0297806 | 0.49884797 |
| 0.999490048 | -0.014151 | 0.74210936 | 0.95450483 | 0.00346006 | 0.93580634 |
| 0.999490048 | -0.0138073 | 0.74090269 | 0.95450483 | -0.0264792 | 0.52721537 |
| 0.999490048 | 0.01351948 | 0.74507111 | 0.95450483 | -5.49E-05 | 0.99894561 |
| 0.999490048 | 0.01386292 | 0.74501495 | 0.95450483 | 0.0063119 | 0.88218444 |
| 0.999490048 | 0.01541504 | 0.74059931 | 0.95450483 | -0.0019095 | 0.96725351 |
| 0.999490048 | 0.01640356 | 0.74039447 | 0.95450483 | 0.04577152 | 0.35878393 |
| 0.999490048 | 0.01695962 | 0.74490026 | 0.95450483 | -0.0111059 | 0.83117271 |
| 0.999490048 | 0.01709963 | 0.7414516 | 0.95450483 | 0.016915 | 0.74413909 |
| 0.999490048 | 0.01713577 | 0.74555613 | 0.95450483 | -0.0153364 | 0.77144053 |
| 0.999490048 | 0.017782 | 0.74151303 | 0.95450483 | 0.06222466 | 0.25413626 |
| 0.999490048 | 0.01816972 | 0.74212441 | 0.95450483 | 0.00991738 | 0.8573605 |
| 0.999490048 | 0.01854724 | 0.74426466 | 0.95450483 | -0.0568721 | 0.32148267 |
| 0.999490048 | 0.01879051 | 0.74457301 | 0.95450483 | 0.00129316 | 0.98209247 |
| 0.999490048 | 0.01915137 | 0.73956885 | 0.95450483 | -0.054308 | 0.34959117 |
| 0.999490048 | 0.01953029 | 0.7432254 | 0.95450483 | -0.0119623 | 0.84086898 |
| 0.999490048 | 0.02068148 | 0.74437738 | 0.95450483 | -0.0469869 | 0.46080997 |
| 0.999490048 | 0.02118871 | 0.73928795 | 0.95450483 | 0.0295381 | 0.64304441 |
| 0.999490048 | 0.02238442 | 0.739355 | 0.95450483 | 0.02514641 | 0.70866413 |
| 0.999490048 | 0.02248364 | 0.74028633 | 0.95450483 | 0.08402389 | 0.22190601 |
| 0.999490048 | 0.0234803 | 0.73932891 | 0.95450483 | 0.09863157 | 0.16984069 |
| 0.999490048 | 0.024848 | 0.7409224 | 0.95450483 | 0.00664047 | 0.9295189 |
| 0.999490048 | 0.02505249 | 0.74475008 | 0.95450483 | -0.0440519 | 0.56786319 |

| | | | | | |
|---|---|---|---|---|---|
| 0.999490048 | 0.02589979 | 0.73994319 | 0.95450483 | 0.01448442 | 0.85262296 |
| 0.999490048 | 0.02721831 | 0.74263579 | 0.95450483 | -0.0264353 | 0.74977287 |
| 0.999490048 | 0.02786158 | 0.74389092 | 0.95450483 | 0.06636826 | 0.43871964 |
| 0.999490048 | 0.03157646 | 0.74131535 | 0.95450483 | 3.07E-04 | 0.99743113 |
| 0.999490048 | 0.03216424 | 0.7436038 | 0.95450483 | -7.55E-04 | 0.99386457 |
| 0.999490048 | 0.03279254 | 0.7408808 | 0.95450483 | -0.0189261 | 0.84852888 |
| 0.999490048 | 0.03286267 | 0.73988833 | 0.95450483 | -0.035183 | 0.72230097 |
| 0.999490048 | 0.03612387 | 0.74128859 | 0.95450483 | -0.0211117 | 0.84689147 |
| 0.999490048 | 0.0376337 | 0.74135026 | 0.95450483 | 0.174345 | 0.1345851 |
| 0.999490048 | 0.03973462 | 0.7440343 | 0.95450483 | -0.0132615 | 0.91313964 |
| 0.999490048 | 0.04104848 | 0.7399824 | 0.95450483 | 0.07522138 | 0.54413055 |
| 0.999490048 | 0.04130286 | 0.73915299 | 0.95450483 | 0.06240221 | 0.61541864 |
| 0.999490048 | 0.04392198 | 0.74558851 | 0.95450483 | 0.15750648 | 0.25044977 |
| 0.999490048 | 0.05561954 | 0.74169834 | 0.95450483 | -0.1969059 | 0.25640415 |
| 0.999490048 | 0.07166906 | 0.740981 | 0.95450483 | 0.06034366 | 0.78069875 |
| 0.999490048 | 0.07235558 | 0.74305138 | 0.95450483 | 0.19234642 | 0.3866042 |
| 0.999490048 | 0.08724142 | 0.74510809 | 0.95450483 | -0.1878779 | 0.48913973 |
| 0.999490048 | -0.0159395 | 0.74578818 | 0.95452657 | -0.0296841 | 0.54705391 |
| 0.999490048 | -0.0343541 | 0.74607869 | 0.95475204 | 0.01648301 | 0.87643342 |
| 0.999490048 | 0.01318811 | 0.7463532 | 0.95495698 | -0.0099443 | 0.80722659 |
| 0.999490048 | 0.01493114 | 0.74647384 | 0.954965 | -0.0406391 | 0.38208372 |
| 0.999490048 | -0.0394218 | 0.74786988 | 0.95520837 | -0.0620033 | 0.61369465 |
| 0.999490048 | -0.0251094 | 0.74734678 | 0.95520837 | -0.1182247 | 0.13763794 |
| 0.999490048 | -0.0204497 | 0.74749167 | 0.95520837 | 0.04112607 | 0.51865604 |
| 0.999490048 | -0.018155 | 0.74728927 | 0.95520837 | -0.0801056 | 0.16292657 |
| 0.999490048 | -0.0162216 | 0.74689916 | 0.95520837 | 0.02726869 | 0.58818014 |
| 0.999490048 | -0.0157374 | 0.74748613 | 0.95520837 | -0.0634364 | 0.20133352 |
| 0.999490048 | -0.0121462 | 0.74792244 | 0.95520837 | -0.0640065 | 0.09915768 |
| 0.999490048 | 0.01625436 | 0.7469712 | 0.95520837 | -0.0418674 | 0.40870533 |
| 0.999490048 | 0.01687115 | 0.74775589 | 0.95520837 | 0.00957158 | 0.85512451 |
| 0.999490048 | 0.02389739 | 0.7476121 | 0.95520837 | -0.0612656 | 0.41208814 |
| 0.999490048 | 0.04179808 | 0.74729989 | 0.95520837 | 0.00719607 | 0.95571809 |
| 0.999490048 | -0.0389397 | 0.74842247 | 0.95570082 | 0.10024096 | 0.41172982 |
| 0.999490048 | -0.0238633 | 0.74885845 | 0.95596516 | 0.08276633 | 0.27220225 |
| 0.999490048 | 0.03392302 | 0.74879127 | 0.95596516 | -0.1066438 | 0.31845413 |
| 0.999490048 | 0.01956234 | 0.74898888 | 0.9559855 | 0.00881636 | 0.88524421 |
| 0.999490048 | 0.02069744 | 0.74939375 | 0.95635607 | 0.00599137 | 0.92625569 |
| 0.999490048 | -0.0415073 | 0.74992384 | 0.95644785 | -0.0619812 | 0.63507985 |
| 0.999490048 | 0.01638245 | 0.74961546 | 0.95644785 | -0.0223264 | 0.66388363 |
| 0.999490048 | 0.0360963 | 0.7498518 | 0.95644785 | -0.0013086 | 0.99075561 |
| 0.999490048 | 0.05939338 | 0.74979889 | 0.95644785 | 0.1092899 | 0.55828761 |
| 0.999490048 | 0.03171027 | 0.75005061 | 0.95646344 | 0.0204312 | 0.83727344 |

| | | | | | |
|---|---|---|---|---|---|
| 0.999490048 | -0.0155696 | 0.75019214 | 0.95649784 | 0.00185542 | 0.96970345 |
| 0.999490048 | -0.0279027 | 0.75092588 | 0.95678279 | -0.0832795 | 0.34727379 |
| 0.999490048 | -0.0216 | 0.7509288 | 0.95678279 | 0.00334251 | 0.96078582 |
| 0.999490048 | -0.0153108 | 0.75098855 | 0.95678279 | -0.0433148 | 0.37268288 |
| 0.999490048 | 0.02728948 | 0.75097152 | 0.95678279 | 0.09191638 | 0.2900866 |
| 0.999490048 | 0.02921433 | 0.75067597 | 0.95678279 | -0.0136503 | 0.88188078 |
| 0.999490048 | 0.0675938 | 0.75110905 | 0.95679033 | 0.00985018 | 0.96301661 |
| 0.999490048 | 0.02060018 | 0.7512582 | 0.95683435 | -0.0051017 | 0.93737011 |
| 0.999490048 | -0.0444363 | 0.7514247 | 0.95690045 | -0.1161465 | 0.41044391 |
| 0.999490048 | -0.2293445 | 0.75180427 | 0.95702028 | 0.06948451 | 0.92350416 |
| 0.999490048 | 0.01317694 | 0.75176607 | 0.95702028 | 0.03531783 | 0.39946551 |
| 0.999490048 | 0.01871922 | 0.75186264 | 0.95702028 | 0.03679353 | 0.53546164 |
| 0.999490048 | 0.0183146 | 0.75214957 | 0.95709371 | 0.00138922 | 0.98086944 |
| 0.999490048 | 0.0515813 | 0.75211082 | 0.95709371 | -0.0891544 | 0.5874734 |
| 0.999490048 | -0.041153 | 0.75251956 | 0.95722004 | 0.05581352 | 0.66916862 |
| 0.999490048 | -0.0186032 | 0.75258878 | 0.95722004 | -0.0436354 | 0.46167716 |
| 0.999490048 | -0.0160468 | 0.75259276 | 0.95722004 | -0.0354166 | 0.48828245 |
| 0.999490048 | -0.0134652 | 0.75286898 | 0.95742552 | -0.0406174 | 0.34609136 |
| 0.999490048 | -0.01974 | 0.75300189 | 0.95744873 | -0.0564735 | 0.37137466 |
| 0.999490048 | -0.0485395 | 0.75366124 | 0.95803597 | -0.129371 | 0.40933078 |
| 0.999490048 | 0.03444453 | 0.75369321 | 0.95803597 | 0.0677239 | 0.53843207 |
| 0.999490048 | 0.01806141 | 0.75398276 | 0.95811231 | -0.0049792 | 0.93108888 |
| 0.999490048 | 0.04205182 | 0.75389747 | 0.95811231 | 0.01630183 | 0.90318731 |
| 0.999490048 | -0.0253234 | 0.75456449 | 0.95846544 | 0.0465239 | 0.56663458 |
| 0.999490048 | -0.0159912 | 0.75462788 | 0.95846544 | 0.0145382 | 0.77626288 |
| 0.999490048 | 0.07307429 | 0.7547198 | 0.95846544 | -0.2042048 | 0.38578723 |
| 0.999490048 | 0.11995374 | 0.75451004 | 0.95846544 | 0.42986283 | 0.27929087 |
| 0.999490048 | -0.019252 | 0.75524824 | 0.95855843 | 0.10740522 | 0.09088434 |
| 0.999490048 | -0.0151868 | 0.75532219 | 0.95855843 | 0.01875695 | 0.70046784 |
| 0.999490048 | 0.01644441 | 0.75536701 | 0.95855843 | -0.0115999 | 0.82595674 |
| 0.999490048 | 0.01822472 | 0.75520127 | 0.95855843 | 0.00536817 | 0.9267607 |
| 0.999490048 | 0.06639402 | 0.75509717 | 0.95855843 | 0.03375126 | 0.87391915 |
| 0.999490048 | 0.03898938 | 0.75564097 | 0.95876038 | 0.1258783 | 0.31940962 |
| 0.999490048 | -0.0220858 | 0.75633185 | 0.95891063 | -0.0070984 | 0.92048881 |
| 0.999490048 | -0.021773 | 0.75608349 | 0.95891063 | -0.0275748 | 0.69419457 |
| 0.999490048 | -0.0152438 | 0.75636335 | 0.95891063 | -0.00977 | 0.84229155 |
| 0.999490048 | -0.0139709 | 0.75629193 | 0.95891063 | -0.0814447 | 0.07927374 |
| 0.999490048 | 0.01221411 | 0.75644842 | 0.95891063 | -0.0080848 | 0.83725517 |
| 0.999490048 | 0.01731527 | 0.75622595 | 0.95891063 | 0.0071909 | 0.89733586 |
| 0.999490048 | -0.0170322 | 0.75690497 | 0.95906056 | -0.034371 | 0.5334024 |
| 0.999490048 | -0.0126098 | 0.756997 | 0.95906056 | 0.00335811 | 0.93426728 |
| 0.956358655 | -0.0111424 | 0.75725584 | 0.95906056 | -0.0985377 | 0.01054845 |

| | | | | | |
|---|---|---|---|---|---|
| 0.999490048 | 0.01708167 | 0.7571665 | 0.95906056 | 0.00744364 | 0.89273316 |
| 0.999490048 | 0.01718708 | 0.75684843 | 0.95906056 | 0.02064212 | 0.71011279 |
| 0.999490048 | 0.02426146 | 0.75708952 | 0.95906056 | 0.0471605 | 0.54874865 |
| 0.999490048 | -0.0171427 | 0.75753414 | 0.95926752 | -0.0638673 | 0.25583451 |
| 0.999490048 | -0.0479518 | 0.75938352 | 0.95946707 | -0.018971 | 0.90347424 |
| 0.999490048 | -0.0413038 | 0.76101609 | 0.95946707 | 0.01456979 | 0.91449164 |
| 0.999490048 | -0.0353474 | 0.76136597 | 0.95946707 | -0.1694021 | 0.15360976 |
| 0.999490048 | -0.033506 | 0.76081358 | 0.95946707 | 0.13914401 | 0.21292651 |
| 0.999490048 | -0.0316966 | 0.75953453 | 0.95946707 | 0.00213746 | 0.98351621 |
| 0.972489992 | -0.0281093 | 0.75803454 | 0.95946707 | -0.2385069 | 0.01395897 |
| 0.999490048 | -0.0255409 | 0.75889281 | 0.95946707 | 9.09E-04 | 0.99127232 |
| 0.999490048 | -0.0243597 | 0.76063709 | 0.95946707 | 0.01900322 | 0.81208038 |
| 0.999490048 | -0.0238582 | 0.75885996 | 0.95946707 | -0.0035081 | 0.9639634 |
| 0.999490048 | -0.0222621 | 0.76012802 | 0.95946707 | 0.07444279 | 0.31184639 |
| 0.999490048 | -0.0213606 | 0.75954817 | 0.95946707 | 0.01685569 | 0.80908238 |
| 0.999490048 | -0.020195 | 0.75988979 | 0.95946707 | -0.0761369 | 0.25501549 |
| 0.999490048 | -0.0186996 | 0.76109079 | 0.95946707 | 0.02871471 | 0.64100073 |
| 0.999490048 | -0.0162357 | 0.76053227 | 0.95946707 | 0.04219544 | 0.4307673 |
| 0.999490048 | -0.0151302 | 0.76136872 | 0.95946707 | 0.03279915 | 0.51177672 |
| 0.999490048 | -0.0148961 | 0.76011652 | 0.95946707 | -0.0349258 | 0.47592179 |
| 0.999490048 | 0.01239369 | 0.76006482 | 0.95946707 | 0.04981546 | 0.22608098 |
| 0.999490048 | 0.01316666 | 0.75934188 | 0.95946707 | 0.01400591 | 0.74455221 |
| 0.999490048 | 0.01531662 | 0.75920724 | 0.95946707 | 0.03645547 | 0.46761681 |
| 0.999490048 | 0.01547053 | 0.76127556 | 0.95946707 | 0.07605651 | 0.14353476 |
| 0.999490048 | 0.01658862 | 0.76061214 | 0.95946707 | 0.0203882 | 0.70819346 |
| 0.999490048 | 0.01822885 | 0.75882545 | 0.95946707 | 0.05991991 | 0.31731903 |
| 0.999490048 | 0.01931171 | 0.75921154 | 0.95946707 | 0.04351524 | 0.4914362 |
| 0.999490048 | 0.02148486 | 0.75849172 | 0.95946707 | 0.10281033 | 0.14934522 |
| 0.999490048 | 0.02178268 | 0.75811368 | 0.95946707 | 0.02638544 | 0.70924901 |
| 0.999490048 | 0.02454208 | 0.76135325 | 0.95946707 | 0.01298249 | 0.87227922 |
| 0.999490048 | 0.0249332 | 0.75864798 | 0.95946707 | -0.0424886 | 0.60121875 |
| 0.999490048 | 0.02735174 | 0.75901296 | 0.95946707 | 0.00993268 | 0.91122227 |
| 0.999490048 | 0.0296125 | 0.75936482 | 0.95946707 | 0.14671799 | 0.1374501 |
| 0.999490048 | 0.03193245 | 0.75967569 | 0.95946707 | 0.11976085 | 0.25700914 |
| 0.999490048 | 0.04971258 | 0.75844319 | 0.95946707 | 0.12360868 | 0.44671172 |
| 0.999490048 | 0.06283377 | 0.76073956 | 0.95946707 | 0.33288338 | 0.13314051 |
| 0.999490048 | -0.0329192 | 0.76323856 | 0.95965095 | 0.06492369 | 0.55349758 |
| 0.999490048 | -0.0279562 | 0.76315927 | 0.95965095 | 0.03285428 | 0.72333409 |
| 0.999490048 | -0.0253467 | 0.7624551 | 0.95965095 | -0.0713337 | 0.3979469 |
| 0.999490048 | -0.0212619 | 0.76307935 | 0.95965095 | -0.0017954 | 0.97967381 |
| 0.999490048 | -0.0193367 | 0.76264706 | 0.95965095 | 0.03595768 | 0.57525263 |
| 0.999490048 | -0.0168689 | 0.76206812 | 0.95965095 | 0.00317972 | 0.95444865 |

| | | | | | |
|---|---|---|---|---|---|
| 0.999490048 | -0.0150667 | 0.76170755 | 0.95965095 | 0.02443718 | 0.62336196 |
| 0.999490048 | -0.0142926 | 0.76288811 | 0.95965095 | 0.02857909 | 0.5474398 |
| 0.999490048 | -0.0123423 | 0.76248396 | 0.95965095 | -4.82E-04 | 0.99056711 |
| 0.999490048 | 0.01290118 | 0.76318004 | 0.95965095 | 0.02461642 | 0.5662841 |
| 0.999490048 | 0.01719734 | 0.76257078 | 0.95965095 | 0.02122088 | 0.70943949 |
| 0.999490048 | 0.02123493 | 0.76319583 | 0.95965095 | 0.08257253 | 0.24735071 |
| 0.999490048 | 0.02945549 | 0.76219669 | 0.95965095 | -0.0465988 | 0.63261602 |
| 0.999490048 | 0.03015058 | 0.76263346 | 0.95965095 | -0.0306361 | 0.75893547 |
| 0.999490048 | 0.03847163 | 0.76276039 | 0.95965095 | -0.1948679 | 0.15184598 |
| 0.999490048 | -0.0124375 | 0.76353649 | 0.95988102 | -0.0375693 | 0.3670083 |
| 0.999490048 | 0.01307567 | 0.76385914 | 0.96014208 | 0.0207603 | 0.63385046 |
| 0.999490048 | -0.0391802 | 0.76452845 | 0.96034176 | 0.02661122 | 0.8387077 |
| 0.999490048 | -0.0340286 | 0.76484196 | 0.96034176 | -0.1308689 | 0.25577026 |
| 0.999490048 | -0.0303807 | 0.76471605 | 0.96034176 | 0.00542009 | 0.95737935 |
| 0.999490048 | -0.0275211 | 0.76489975 | 0.96034176 | 0.0756345 | 0.41389039 |
| 0.999490048 | -0.0242968 | 0.76544555 | 0.96034176 | 0.01117768 | 0.89075656 |
| 0.999490048 | -0.0227128 | 0.76620195 | 0.96034176 | -0.1294829 | 0.09890906 |
| 0.999490048 | -0.0177852 | 0.76620321 | 0.96034176 | 0.01966124 | 0.74242416 |
| 0.999490048 | -0.0166059 | 0.76538405 | 0.96034176 | -0.060174 | 0.284692 |
| 0.999490048 | 0.0145973 | 0.76564684 | 0.96034176 | 0.01695251 | 0.72929874 |
| 0.999490048 | 0.01617258 | 0.76586518 | 0.96034176 | -0.0324224 | 0.55158062 |
| 0.999490048 | 0.01805335 | 0.76443509 | 0.96034176 | -0.0257487 | 0.66938391 |
| 0.999490048 | 0.01811415 | 0.76442383 | 0.96034176 | -0.0456337 | 0.45245324 |
| 0.999490048 | 0.01934044 | 0.76612322 | 0.96034176 | 0.09705552 | 0.14377964 |
| 0.999490048 | 0.01942224 | 0.76576447 | 0.96034176 | 0.04038505 | 0.53681481 |
| 0.999490048 | 0.02066321 | 0.76546743 | 0.96034176 | 0.07298517 | 0.29693132 |
| 0.999490048 | 0.02125512 | 0.7651636 | 0.96034176 | 0.02172392 | 0.76015066 |
| 0.999490048 | 0.02366641 | 0.76563268 | 0.96034176 | 0.04547627 | 0.56771237 |
| 0.999490048 | 0.03117589 | 0.76579422 | 0.96034176 | -0.0039834 | 0.96961251 |
| 0.999490048 | 0.03520041 | 0.76556722 | 0.96034176 | 0.1076782 | 0.36974536 |
| 0.999490048 | 0.02065865 | 0.76644343 | 0.96049867 | 0.03437061 | 0.62172642 |
| 0.999490048 | 0.01950683 | 0.76658233 | 0.96052859 | -0.0343494 | 0.60184809 |
| 0.999490048 | -0.0174831 | 0.76697213 | 0.9606681 | -0.0798946 | 0.18312081 |
| 0.999490048 | 0.01542931 | 0.76698969 | 0.9606681 | 0.02194252 | 0.67374241 |
| 0.999490048 | 0.13703786 | 0.76703883 | 0.9606681 | -0.6798488 | 0.1664263 |
| 0.999490048 | 0.0334724 | 0.76765252 | 0.96129252 | 0.07002989 | 0.53769896 |
| 0.999490048 | -0.0230151 | 0.76787927 | 0.96143229 | -0.079783 | 0.31084996 |
| 0.999490048 | -0.0193964 | 0.76845871 | 0.96158103 | 0.00104349 | 0.98735283 |
| 0.999490048 | 0.01824306 | 0.76844457 | 0.96158103 | -0.0019421 | 0.9749755 |
| 0.999490048 | 0.0187914 | 0.76827542 | 0.96158103 | -0.0404058 | 0.52769618 |
| 0.999490048 | 0.02261348 | 0.76817036 | 0.96158103 | -2.94E-05 | 0.99969396 |
| 0.999490048 | -0.0244047 | 0.76873823 | 0.96174688 | -0.0188442 | 0.82033577 |

| | | | | | |
|---|---|---|---|---|---|
| 0.999490048 | -0.0124254 | 0.76905197 | 0.96174688 | 0.02194399 | 0.60474565 |
| 0.999490048 | 0.01957127 | 0.769049 | 0.96174688 | -0.0620856 | 0.3553739 |
| 0.999490048 | 0.03343255 | 0.76895412 | 0.96174688 | -0.143 | 0.21568671 |
| 0.999490048 | -0.0143038 | 0.76922007 | 0.96181306 | 7.44E-04 | 0.98782025 |
| 0.999490048 | -0.0409497 | 0.76983362 | 0.96186005 | -0.0729652 | 0.6028038 |
| 0.999490048 | -0.0216498 | 0.76978296 | 0.96186005 | -0.0945022 | 0.20833088 |
| 0.999490048 | -0.0169049 | 0.76962916 | 0.96186005 | -0.0270004 | 0.6404085 |
| 0.999490048 | 0.01447776 | 0.7696747 | 0.96186005 | -0.012165 | 0.80561233 |
| 0.835632585 | 0.07610991 | 0.76950807 | 0.96186005 | 0.85050775 | 0.00932303 |
| 0.999490048 | -0.032139 | 0.77117168 | 0.96200265 | 0.12674975 | 0.25715685 |
| 0.999490048 | -0.0197148 | 0.7704499 | 0.96200265 | 0.03722673 | 0.58248325 |
| 0.999490048 | -0.0164003 | 0.77110254 | 0.96200265 | -0.0899884 | 0.11908185 |
| 0.999490048 | -0.0125358 | 0.77150548 | 0.96200265 | 0.03258018 | 0.45259284 |
| 0.999490048 | 0.01144872 | 0.77025641 | 0.96200265 | -0.0238713 | 0.54373493 |
| 0.999490048 | 0.01297486 | 0.77142854 | 0.96200265 | 0.05263333 | 0.24468181 |
| 0.999490048 | 0.01629023 | 0.77099372 | 0.96200265 | 0.03436177 | 0.54041952 |
| 0.999490048 | 0.01836667 | 0.77094892 | 0.96200265 | 0.03566904 | 0.57271146 |
| 0.999490048 | 0.01953184 | 0.77104933 | 0.96200265 | 0.02536541 | 0.70565738 |
| 0.999490048 | 0.02138731 | 0.77032479 | 0.96200265 | -0.1088364 | 0.14560083 |
| 0.999490048 | 0.02235947 | 0.77132782 | 0.96200265 | 0.10272651 | 0.1891169 |
| 0.999490048 | 0.02936647 | 0.77018198 | 0.96200265 | 0.02602772 | 0.79565765 |
| 0.999490048 | 0.03460293 | 0.77156068 | 0.96200265 | -0.013717 | 0.90830651 |
| 0.999490048 | 0.03890842 | 0.7713654 | 0.96200265 | 0.21172895 | 0.12243296 |
| 0.999490048 | -0.0249848 | 0.77203199 | 0.96232248 | -0.0582708 | 0.50083798 |
| 0.999490048 | -0.0207551 | 0.77232202 | 0.96232248 | 0.0254933 | 0.72240954 |
| 0.999490048 | -0.0168449 | 0.77239344 | 0.96232248 | 0.03752132 | 0.5207819 |
| 0.999490048 | -0.0157618 | 0.77226974 | 0.96232248 | -0.0394962 | 0.47029934 |
| 0.999490048 | 0.01211047 | 0.77212263 | 0.96232248 | -0.0275879 | 0.51092942 |
| 0.999490048 | -0.0133828 | 0.77256795 | 0.9623963 | -0.0628238 | 0.18234766 |
| 0.999490048 | -0.026683 | 0.77272152 | 0.96244401 | -0.0035072 | 0.9696931 |
| 0.999490048 | -0.0677111 | 0.77723171 | 0.96255866 | 0.03706858 | 0.87683896 |
| 0.999490048 | -0.0506823 | 0.77594117 | 0.96255866 | 0.12303838 | 0.49133023 |
| 0.999490048 | -0.0338807 | 0.77513438 | 0.96255866 | -0.2402115 | 0.05121779 |
| 0.999490048 | -0.0316697 | 0.77612169 | 0.96255866 | 0.11960468 | 0.28795574 |
| 0.999490048 | -0.0315151 | 0.77603812 | 0.96255866 | -0.041728 | 0.70658531 |
| 0.999490048 | -0.0299855 | 0.77435391 | 0.96255866 | 0.01057092 | 0.91936346 |
| 0.999490048 | -0.024756 | 0.77749178 | 0.96255866 | 0.00405867 | 0.96301851 |
| 0.999490048 | -0.0245365 | 0.77772311 | 0.96255866 | 0.07024867 | 0.4216443 |
| 0.999490048 | -0.0213756 | 0.77522937 | 0.96255866 | 0.03882887 | 0.60466058 |
| 0.999490048 | -0.0205978 | 0.77461597 | 0.96255866 | -0.0271961 | 0.70555131 |
| 0.999490048 | -0.0203862 | 0.77568406 | 0.96255866 | 0.0215666 | 0.76309668 |
| 0.999490048 | -0.0182566 | 0.77688086 | 0.96255866 | -0.0024114 | 0.9701175 |

| | | | | | |
|---|---|---|---|---|---|
| 0.999490048 | -0.0182527 | 0.77567044 | 0.96255866 | -0.0461419 | 0.47323547 |
| 0.999490048 | -0.0179981 | 0.77352942 | 0.96255866 | -0.0238983 | 0.70257052 |
| 0.999490048 | -0.0172366 | 0.77442966 | 0.96255866 | -0.0010778 | 0.98569033 |
| 0.999490048 | -0.0153964 | 0.77482422 | 0.96255866 | -0.0285973 | 0.59592002 |
| 0.999490048 | -0.0137468 | 0.77715269 | 0.96255866 | -0.0313646 | 0.51984673 |
| 0.999490048 | -0.0132566 | 0.77681814 | 0.96255866 | 0.0255528 | 0.58561473 |
| 0.999490048 | -0.0131494 | 0.77434372 | 0.96255866 | 0.04668886 | 0.31328464 |
| 0.999490048 | -0.0128554 | 0.77675075 | 0.96255866 | 0.04219063 | 0.35582196 |
| 0.999490048 | -0.0116051 | 0.77363942 | 0.96255866 | -0.0167869 | 0.6776573 |
| 0.999490048 | -0.0110261 | 0.7751982 | 0.96255866 | 0.0330513 | 0.39505044 |
| 0.999490048 | 0.00972208 | 0.77761481 | 0.96255866 | -0.0155697 | 0.65146055 |
| 0.999490048 | 0.01436895 | 0.77768511 | 0.96255866 | 0.00648947 | 0.89847016 |
| 0.999490048 | 0.01551153 | 0.77403621 | 0.96255866 | -0.0107785 | 0.84180688 |
| 0.999490048 | 0.0156876 | 0.77738836 | 0.96255866 | -0.0494257 | 0.37646079 |
| 0.999490048 | 0.01703508 | 0.77374451 | 0.96255866 | 0.08628911 | 0.15343704 |
| 0.999490048 | 0.01726743 | 0.77443414 | 0.96255866 | 0.00392312 | 0.94804542 |
| 0.999490048 | 0.01738286 | 0.77619783 | 0.96255866 | 0.05260826 | 0.39271647 |
| 0.999490048 | 0.01855578 | 0.77777045 | 0.96255866 | -0.0128989 | 0.84439975 |
| 0.999490048 | 0.0188976 | 0.7761611 | 0.96255866 | 0.03336613 | 0.61626698 |
| 0.999490048 | 0.01943245 | 0.77666192 | 0.96255866 | -0.0058503 | 0.93189221 |
| 0.999490048 | 0.02010154 | 0.77452499 | 0.96255866 | 0.00810243 | 0.90801266 |
| 0.999490048 | 0.020483 | 0.77338825 | 0.96255866 | -0.0102303 | 0.88557513 |
| 0.999490048 | 0.02198273 | 0.77398402 | 0.96255866 | -0.0052935 | 0.94482663 |
| 0.999490048 | 0.02270637 | 0.77639108 | 0.96255866 | 0.16890956 | 0.04262878 |
| 0.999490048 | 0.02374555 | 0.77557212 | 0.96255866 | 0.07354417 | 0.38056691 |
| 0.999490048 | 0.02541842 | 0.77470773 | 0.96255866 | 0.13114378 | 0.1479539 |
| 0.999490048 | 0.02590009 | 0.77711758 | 0.96255866 | 0.0211759 | 0.81692174 |
| 0.999490048 | 0.02866893 | 0.77649712 | 0.96255866 | 0.07471136 | 0.46151267 |
| 0.999490048 | 0.03022927 | 0.77586789 | 0.96255866 | 0.12813352 | 0.23384083 |
| 0.999490048 | 0.03167248 | 0.77741814 | 0.96255866 | 0.10612739 | 0.34745704 |
| 0.999490048 | 0.06029585 | 0.77544069 | 0.96255866 | -0.1276345 | 0.5470847 |
| 0.999490048 | -0.0417655 | 0.77918315 | 0.96257231 | 0.19905474 | 0.18881403 |
| 0.999490048 | -0.0386551 | 0.77927948 | 0.96257231 | -0.0137756 | 0.92038795 |
| 0.999490048 | -0.0342464 | 0.77902441 | 0.96257231 | 0.0187296 | 0.87797037 |
| 0.999490048 | -0.0282884 | 0.77918305 | 0.96257231 | -0.1236259 | 0.22694961 |
| 0.999490048 | -0.0254209 | 0.77843463 | 0.96257231 | 0.02616831 | 0.77210842 |
| 0.999490048 | -0.0197886 | 0.77838004 | 0.96257231 | -0.0108811 | 0.87695118 |
| 0.999490048 | -0.0163179 | 0.77794902 | 0.96257231 | 0.03925106 | 0.49921764 |
| 0.999490048 | -0.0119938 | 0.77889373 | 0.96257231 | -0.0247793 | 0.56289023 |
| 0.987721587 | 0.01365805 | 0.7792801 | 0.96257231 | -0.0915995 | 0.06898003 |
| 0.999490048 | 0.01810671 | 0.77880946 | 0.96257231 | 0.01958083 | 0.76135925 |
| 0.999490048 | 0.02116347 | 0.77848676 | 0.96257231 | 0.09766749 | 0.20132377 |

| | | | | | |
|---|---|---|---|---|---|
| 0.999490048 | 0.03094116 | 0.77835163 | 0.96257231 | 0.09916289 | 0.37052737 |
| 0.999490048 | 0.03621885 | 0.77804849 | 0.96257231 | -0.0519519 | 0.68624288 |
| 0.999490048 | -0.018504 | 0.77971089 | 0.96288105 | 0.02608039 | 0.69364751 |
| 0.999490048 | -0.0131009 | 0.77976068 | 0.96288105 | -0.0575883 | 0.22553422 |
| 0.999490048 | 0.01695237 | 0.77988037 | 0.96288646 | 0.04412398 | 0.46897857 |
| 0.999490048 | -0.0724938 | 0.78156975 | 0.96297376 | 0.29556772 | 0.27101943 |
| 0.999490048 | -0.0631791 | 0.78996062 | 0.96297376 | -0.2200794 | 0.35727391 |
| 0.999490048 | -0.057963 | 0.79008381 | 0.96297376 | -0.1948859 | 0.37425164 |
| 0.999490048 | -0.046743 | 0.79321631 | 0.96297376 | 0.16246282 | 0.36588673 |
| 0.999490048 | -0.0460434 | 0.78356811 | 0.96297376 | 0.03374547 | 0.84040161 |
| 0.999490048 | -0.0390531 | 0.79131887 | 0.96297376 | -0.0399145 | 0.78683246 |
| 0.999490048 | -0.0368899 | 0.78380771 | 0.96297376 | 0.13929774 | 0.30499699 |
| 0.999490048 | -0.0367993 | 0.78711916 | 0.96297376 | -0.2192159 | 0.1272469 |
| 0.999490048 | -0.0326471 | 0.78240485 | 0.96297376 | -0.0351262 | 0.76637241 |
| 0.999490048 | -0.0323315 | 0.78720035 | 0.96297376 | -5.32E-04 | 0.99645363 |
| 0.999490048 | -0.0322779 | 0.78228781 | 0.96297376 | -0.0509928 | 0.66279323 |
| 0.999490048 | -0.0300285 | 0.78621075 | 0.96297376 | 0.01127651 | 0.9188194 |
| 0.999490048 | -0.0288394 | 0.79725002 | 0.96297376 | 0.04278286 | 0.7033413 |
| 0.999490048 | -0.0275054 | 0.79371287 | 0.96297376 | 0.02711568 | 0.79656703 |
| 0.999490048 | -0.0269461 | 0.79162063 | 0.96297376 | 0.04051345 | 0.6914547 |
| 0.999490048 | -0.0267721 | 0.78524294 | 0.96297376 | -0.0460212 | 0.63997782 |
| 0.999490048 | -0.0259942 | 0.78962068 | 0.96297376 | -0.0423055 | 0.66450032 |
| 0.999490048 | -0.0257579 | 0.787485 | 0.96297376 | -0.0739248 | 0.44152423 |
| 0.999490048 | -0.024078 | 0.79354477 | 0.96297376 | -0.1439302 | 0.12634734 |
| 0.999490048 | -0.0236457 | 0.79433859 | 0.96297376 | 0.03553729 | 0.69547067 |
| 0.999490048 | -0.0229624 | 0.79534875 | 0.96297376 | 0.06726426 | 0.44970224 |
| 0.999490048 | -0.0211909 | 0.78904365 | 0.96297376 | -0.0048386 | 0.95125282 |
| 0.999490048 | -0.0211429 | 0.78152559 | 0.96297376 | 0.04741778 | 0.5352279 |
| 0.999490048 | -0.0209747 | 0.78914355 | 0.96297376 | -0.0366306 | 0.64097007 |
| 0.999490048 | -0.0208624 | 0.78711061 | 0.96297376 | -0.1313326 | 0.09802596 |
| 0.999490048 | -0.0201821 | 0.7874258 | 0.96297376 | -0.0691146 | 0.35951456 |
| 0.999490048 | -0.0197656 | 0.79334319 | 0.96297376 | 0.00489019 | 0.94828839 |
| 0.999490048 | -0.0197252 | 0.79451781 | 0.96297376 | 0.03113856 | 0.68127154 |
| 0.999490048 | -0.0183954 | 0.78407411 | 0.96297376 | -0.046531 | 0.49000415 |
| 0.999490048 | -0.0183056 | 0.78488945 | 0.96297376 | -0.0748593 | 0.26988045 |
| 0.999490048 | -0.018091 | 0.78134909 | 0.96297376 | 0.04145898 | 0.52608312 |
| 0.999490048 | -0.0176028 | 0.79696555 | 0.96297376 | 0.01487217 | 0.82789044 |
| 0.999490048 | -0.0174098 | 0.79183004 | 0.96297376 | -0.0318364 | 0.62991159 |
| 0.999490048 | -0.0171409 | 0.78383103 | 0.96297376 | 0.00697555 | 0.91105316 |
| 0.999490048 | -0.0165172 | 0.78273454 | 0.96297376 | 0.01186674 | 0.84289634 |
| 0.999490048 | -0.0165166 | 0.79222717 | 0.96297376 | -0.0580322 | 0.35845735 |
| 0.999490048 | -0.0164216 | 0.79236332 | 0.96297376 | 0.01288433 | 0.8363257 |

| | | | | | |
|---|---|---|---|---|---|
| 0.999490048 | -0.0163405 | 0.79416968 | 0.96297376 | -0.0691305 | 0.2751584 |
| 0.999490048 | -0.015995 | 0.7963738 | 0.96297376 | 0.03269149 | 0.5986935 |
| 0.999490048 | -0.0159405 | 0.79659549 | 0.96297376 | -0.0127526 | 0.83659043 |
| 0.999490048 | -0.0156422 | 0.79349233 | 0.96297376 | 0.02366547 | 0.6923285 |
| 0.999490048 | -0.0155125 | 0.79523142 | 0.96297376 | -0.0108027 | 0.85653046 |
| 0.999490048 | -0.015404 | 0.78097034 | 0.96297376 | -0.0358333 | 0.51918027 |
| 0.999490048 | -0.0149008 | 0.78733719 | 0.96297376 | -0.0329305 | 0.55218082 |
| 0.999490048 | -0.0147946 | 0.78181164 | 0.96297376 | -0.0101856 | 0.84871795 |
| 0.999490048 | -0.0146913 | 0.78912375 | 0.96297376 | -0.0023414 | 0.96597758 |
| 0.999490048 | -0.0146347 | 0.7867124 | 0.96297376 | -0.0337208 | 0.53427635 |
| 0.999490048 | -0.0146262 | 0.79110514 | 0.96297376 | -0.0375201 | 0.4985241 |
| 0.999490048 | -0.0145137 | 0.79007552 | 0.96297376 | 0.0486504 | 0.37566139 |
| 0.999490048 | -0.0144907 | 0.78447122 | 0.96297376 | 0.00483233 | 0.92728099 |
| 0.999490048 | -0.0144843 | 0.78220789 | 0.96297376 | 0.0198761 | 0.70462104 |
| 0.999490048 | -0.0143944 | 0.78525713 | 0.96297376 | 0.07002514 | 0.19230556 |
| 0.999490048 | -0.0143483 | 0.78893786 | 0.96297376 | 7.87E-04 | 0.98827771 |
| 0.999490048 | -0.0141242 | 0.79433564 | 0.96297376 | -0.0084303 | 0.87629805 |
| 0.999490048 | -0.0136538 | 0.78090425 | 0.96297376 | 0.00896561 | 0.85501821 |
| 0.999490048 | -0.0135265 | 0.78903103 | 0.96297376 | -0.0215018 | 0.67093841 |
| 0.999490048 | -0.013511 | 0.79558349 | 0.96297376 | 0.03625285 | 0.48878769 |
| 0.999490048 | -0.0133749 | 0.78429142 | 0.96297376 | -0.0389604 | 0.42783442 |
| 0.999490048 | -0.0132379 | 0.78539224 | 0.96297376 | -0.0307439 | 0.52848283 |
| 0.999490048 | -0.0132023 | 0.78341445 | 0.96297376 | 0.02662272 | 0.58026633 |
| 0.999490048 | -0.0131126 | 0.79643601 | 0.96297376 | -0.0470751 | 0.35818521 |
| 0.999490048 | -0.0130583 | 0.78304667 | 0.96297376 | -0.0149369 | 0.75284524 |
| 0.999490048 | -0.0129244 | 0.7969301 | 0.96297376 | 0.01745127 | 0.72839659 |
| 0.999490048 | -0.0125409 | 0.79522334 | 0.96297376 | -0.0183096 | 0.70496653 |
| 0.999490048 | -0.0120687 | 0.794308 | 0.96297376 | 0.02481416 | 0.59273307 |
| 0.999490048 | -0.0119952 | 0.79333184 | 0.96297376 | 0.05168464 | 0.26465936 |
| 0.999490048 | -0.0118475 | 0.7897744 | 0.96297376 | 0.00974093 | 0.82645443 |
| 0.999490048 | -0.0113184 | 0.79461639 | 0.96297376 | -0.0675774 | 0.12871708 |
| 0.999490048 | -0.0111721 | 0.78902031 | 0.96297376 | -0.0605388 | 0.15517734 |
| 0.999490048 | -0.0110036 | 0.79294189 | 0.96297376 | 0.00105038 | 0.97999567 |
| 0.999490048 | -0.0107209 | 0.79323603 | 0.96297376 | 0.01938556 | 0.63606597 |
| 0.999490048 | -0.0106145 | 0.79309116 | 0.96297376 | 0.00278768 | 0.94504381 |
| 0.999490048 | 0.01034923 | 0.79008114 | 0.96297376 | 0.00852871 | 0.82631685 |
| 0.999490048 | 0.01036073 | 0.79536037 | 0.96297376 | -0.0122127 | 0.75988236 |
| 0.999490048 | 0.01145932 | 0.78576919 | 0.96297376 | -0.0317731 | 0.45330336 |
| 0.999490048 | 0.01161357 | 0.78294805 | 0.96297376 | -0.0073512 | 0.8615078 |
| 0.999490048 | 0.01172088 | 0.78762468 | 0.96297376 | 0.00840742 | 0.84671811 |
| 0.999490048 | 0.01205082 | 0.78365314 | 0.96297376 | 0.0166802 | 0.70411539 |
| 0.999490048 | 0.01208865 | 0.78909086 | 0.96297376 | -0.0131722 | 0.77072366 |

| | | | | | |
|---|---|---|---|---|---|
| 0.999490048 | 0.01227926 | 0.78469216 | 0.96297376 | 0.02505248 | 0.57815127 |
| 0.999490048 | 0.01244793 | 0.78718425 | 0.96297376 | -0.0605901 | 0.19605309 |
| 0.999490048 | 0.01262603 | 0.79287043 | 0.96297376 | 0.03506997 | 0.46785244 |
| 0.999490048 | 0.01291222 | 0.79668147 | 0.96297376 | 0.01609796 | 0.74814102 |
| 0.999490048 | 0.01314435 | 0.79607446 | 0.96297376 | 0.02256647 | 0.65769512 |
| 0.999490048 | 0.01346743 | 0.7946885 | 0.96297376 | 0.01211633 | 0.81486782 |
| 0.999490048 | 0.01364744 | 0.7828307 | 0.96297376 | 0.0048307 | 0.92222666 |
| 0.999490048 | 0.01371082 | 0.78898246 | 0.96297376 | 0.0244282 | 0.63401347 |
| 0.999490048 | 0.01378542 | 0.78979031 | 0.96297376 | 0.02003674 | 0.69863618 |
| 0.999490048 | 0.01384372 | 0.79007391 | 0.96297376 | -0.0436091 | 0.40466511 |
| 0.999490048 | 0.01392875 | 0.79667052 | 0.96297376 | -0.0101817 | 0.85056021 |
| 0.999490048 | 0.01400517 | 0.79440485 | 0.96297376 | 0.00399655 | 0.94068345 |
| 0.999490048 | 0.01402702 | 0.7858594 | 0.96297376 | 0.00597038 | 0.90788103 |
| 0.999490048 | 0.01411508 | 0.79087121 | 0.96297376 | 0.00447029 | 0.93302704 |
| 0.999490048 | 0.01497047 | 0.78261767 | 0.96297376 | -0.0106335 | 0.84456824 |
| 0.999490048 | 0.01500137 | 0.79132036 | 0.96297376 | -0.0251704 | 0.65748344 |
| 0.999490048 | 0.0152016 | 0.79666942 | 0.96297376 | 0.06620421 | 0.26745083 |
| 0.999490048 | 0.01521241 | 0.78292614 | 0.96297376 | 0.02356909 | 0.66981926 |
| 0.999490048 | 0.01541781 | 0.7855675 | 0.96297376 | 0.01253028 | 0.82495838 |
| 0.999490048 | 0.01546399 | 0.78384935 | 0.96297376 | -0.0011535 | 0.98366259 |
| 0.999490048 | 0.01556433 | 0.78575825 | 0.96297376 | -0.0015719 | 0.9780827 |
| 0.999490048 | 0.01561891 | 0.78627835 | 0.96297376 | 0.06071564 | 0.29684104 |
| 0.999490048 | 0.01573834 | 0.79449408 | 0.96297376 | 0.002411 | 0.96814868 |
| 0.999490048 | 0.01588262 | 0.79211111 | 0.96297376 | -0.1069541 | 0.08487004 |
| 0.999490048 | 0.01593002 | 0.79187008 | 0.96297376 | 0.04635186 | 0.44496391 |
| 0.999490048 | 0.0162553 | 0.78193968 | 0.96297376 | 0.01730882 | 0.76822242 |
| 0.999490048 | 0.01686299 | 0.78292354 | 0.96297376 | 0.08397288 | 0.17769684 |
| 0.999490048 | 0.01716224 | 0.78432736 | 0.96297376 | -0.0164846 | 0.7926325 |
| 0.999490048 | 0.01720461 | 0.79361802 | 0.96297376 | 0.03742447 | 0.57029246 |
| 0.999490048 | 0.01722742 | 0.79534253 | 0.96297376 | 0.07869678 | 0.24227127 |
| 0.999490048 | 0.01774213 | 0.78697853 | 0.96297376 | -0.0355799 | 0.58869549 |
| 0.999490048 | 0.01788598 | 0.79045686 | 0.96297376 | -0.0412988 | 0.54076321 |
| 0.999490048 | 0.01791947 | 0.78562599 | 0.96297376 | 0.05011006 | 0.44918574 |
| 0.999490048 | 0.01793013 | 0.78240518 | 0.96297376 | 0.04548981 | 0.48530737 |
| 0.999490048 | 0.01876476 | 0.79167903 | 0.96297376 | -0.0028242 | 0.96826811 |
| 0.999490048 | 0.02027727 | 0.78811734 | 0.96297376 | -0.1179831 | 0.12648647 |
| 0.999490048 | 0.02041208 | 0.79248209 | 0.96297376 | -0.0188727 | 0.80779419 |
| 0.999490048 | 0.02185989 | 0.79241365 | 0.96297376 | -0.0863027 | 0.3036449 |
| 0.999490048 | 0.02319675 | 0.78463303 | 0.96297376 | -0.0254877 | 0.7640022 |
| 0.999490048 | 0.0239812 | 0.78361502 | 0.96297376 | 0.06190823 | 0.48025832 |
| 0.999490048 | 0.02450649 | 0.79249283 | 0.96297376 | -0.0170691 | 0.85456137 |
| 0.999490048 | 0.02465156 | 0.79723835 | 0.96297376 | 0.03392681 | 0.7238031 |

| | | | | | |
|---|---|---|---|---|---|
| 0.999490048 | 0.0246595 | 0.78687425 | 0.96297376 | 0.02946803 | 0.74670175 |
| 0.999490048 | 0.02507897 | 0.78444491 | 0.96297376 | 0.05988921 | 0.5151137 |
| 0.999490048 | 0.02510205 | 0.79298746 | 0.96297376 | 0.02811793 | 0.76882578 |
| 0.999490048 | 0.02517477 | 0.79713825 | 0.96297376 | 0.06561306 | 0.50452452 |
| 0.999490048 | 0.02596167 | 0.79410068 | 0.96297376 | 0.09669423 | 0.33524273 |
| 0.999490048 | 0.0264096 | 0.79433426 | 0.96297376 | 0.01818438 | 0.85749715 |
| 0.999490048 | 0.0268129 | 0.78219495 | 0.96297376 | 0.02389059 | 0.80539747 |
| 0.999490048 | 0.02784205 | 0.78044704 | 0.96297376 | -0.1478065 | 0.14718095 |
| 0.999490048 | 0.0283109 | 0.79362649 | 0.96297376 | 0.02296068 | 0.83193825 |
| 0.999490048 | 0.0315668 | 0.79487761 | 0.96297376 | 0.04082741 | 0.73679731 |
| 0.999490048 | 0.03162444 | 0.78250152 | 0.96297376 | 0.05951165 | 0.60411933 |
| 0.999490048 | 0.03188712 | 0.79153781 | 0.96297376 | 0.24787836 | 0.0482041 |
| 0.999490048 | 0.0324724 | 0.78388277 | 0.96297376 | -0.0500432 | 0.67286036 |
| 0.999490048 | 0.0336347 | 0.79346763 | 0.96297376 | 0.03499111 | 0.78535054 |
| 0.999490048 | 0.03407319 | 0.79659305 | 0.96297376 | -0.0436038 | 0.74161484 |
| 0.999490048 | 0.03551904 | 0.78122569 | 0.96297376 | -0.2033751 | 0.12047248 |
| 0.999490048 | 0.03700417 | 0.79028676 | 0.96297376 | 0.10486735 | 0.45331093 |
| 0.999490048 | 0.03730855 | 0.78161524 | 0.96297376 | -0.0820401 | 0.54333223 |
| 0.999490048 | 0.03861753 | 0.79724706 | 0.96297376 | 0.04495142 | 0.76495652 |
| 0.999490048 | 0.03900709 | 0.79596768 | 0.96297376 | -0.1409023 | 0.35416493 |
| 0.999490048 | 0.04255783 | 0.79105399 | 0.96297376 | 0.08163484 | 0.61197721 |
| 0.999490048 | 0.04483087 | 0.79236951 | 0.96297376 | 0.00879477 | 0.9587874 |
| 0.999490048 | 0.0462472 | 0.78599505 | 0.96297376 | 0.35925804 | 0.04297222 |
| 0.999490048 | 0.05073919 | 0.78597831 | 0.96297376 | 0.28791515 | 0.14242239 |
| 0.999490048 | 0.05860383 | 0.79681517 | 0.96297376 | -0.0418528 | 0.85406264 |
| 0.781057588 | 0.06042519 | 0.79120001 | 0.96297376 | -0.6028421 | 0.02024806 |
| 0.999490048 | 0.06962368 | 0.78043323 | 0.96297376 | 0.27320962 | 0.27934511 |
| 0.999490048 | 0.06983134 | 0.7826329 | 0.96297376 | -0.2336647 | 0.35964273 |
| 0.999490048 | 0.07185819 | 0.78996212 | 0.96297376 | 0.11860311 | 0.66059676 |
| 0.999490048 | -0.0198161 | 0.79971546 | 0.96329548 | -0.0266653 | 0.73293133 |
| 0.999490048 | -0.0161342 | 0.79918155 | 0.96329548 | -0.0316038 | 0.61889745 |
| 0.999490048 | -0.0120996 | 0.79873758 | 0.96329548 | -0.0381548 | 0.42406364 |
| 0.999490048 | -0.0110995 | 0.79918034 | 0.96329548 | 0.00378885 | 0.93075637 |
| 0.999490048 | -0.0099058 | 0.79944522 | 0.96329548 | 0.00804055 | 0.83657916 |
| 0.999490048 | 0.01046033 | 0.79980076 | 0.96329548 | -0.0169444 | 0.68153053 |
| 0.999490048 | 0.01158012 | 0.79917213 | 0.96329548 | 0.00420369 | 0.92637325 |
| 0.999490048 | 0.01166278 | 0.79856334 | 0.96329548 | 0.04764897 | 0.30200969 |
| 0.999490048 | 0.01218766 | 0.79866623 | 0.96329548 | 0.00456069 | 0.92391456 |
| 0.999490048 | 0.01242 | 0.7986264 | 0.96329548 | 0.02545701 | 0.60178923 |
| 0.999490048 | 0.01278827 | 0.7977634 | 0.96329548 | 6.47E-04 | 0.98964722 |
| 0.999490048 | 0.01445593 | 0.7997664 | 0.96329548 | 0.01847529 | 0.74590672 |
| 0.999490048 | 0.01459064 | 0.79831504 | 0.96329548 | 0.02240468 | 0.69503874 |

| | | | | | |
|---|---|---|---|---|---|
| 0.999490048 | 0.0163005 | 0.79930368 | 0.96329548 | -0.0024545 | 0.96944173 |
| 0.999490048 | 0.01651794 | 0.79814022 | 0.96329548 | 0.03577633 | 0.5805392 |
| 0.999490048 | 0.01694716 | 0.79966519 | 0.96329548 | 0.01126908 | 0.86594331 |
| 0.999490048 | 0.02421592 | 0.79982366 | 0.96329548 | 0.04872156 | 0.61062706 |
| 0.999490048 | 0.02466903 | 0.79890951 | 0.96329548 | 0.05904111 | 0.54329106 |
| 0.999490048 | 0.0446799 | 0.7981472 | 0.96329548 | 0.08324713 | 0.63426856 |
| 0.999490048 | 0.06800852 | 0.79863934 | 0.96329548 | -0.2338484 | 0.3837282 |
| 0.999490048 | -0.0148946 | 0.80012508 | 0.96351953 | -0.0727104 | 0.22312676 |
| 0.999490048 | -0.0493786 | 0.80037963 | 0.96354814 | 0.35341668 | 0.09778899 |
| 0.999490048 | 0.02069056 | 0.80033511 | 0.96354814 | 0.06278644 | 0.44519342 |
| 0.999490048 | -0.0176334 | 0.80069492 | 0.96378875 | 0.1002061 | 0.15945924 |
| 0.999490048 | -0.0368127 | 0.80187817 | 0.9638526 | 0.09394386 | 0.52340967 |
| 0.999490048 | -0.0238655 | 0.8018414 | 0.9638526 | -0.0177277 | 0.852073 |
| 0.999490048 | -0.0218223 | 0.80175069 | 0.9638526 | -0.0531973 | 0.54175488 |
| 0.999490048 | -0.0193507 | 0.80393123 | 0.9638526 | 0.00575962 | 0.94106137 |
| 0.999490048 | -0.0191811 | 0.80321486 | 0.9638526 | 0.01686415 | 0.82655687 |
| 0.999490048 | -0.0178748 | 0.80125968 | 0.9638526 | -0.0404058 | 0.57036319 |
| 0.999490048 | -0.0170672 | 0.80247202 | 0.9638526 | 0.02990733 | 0.66155134 |
| 0.999490048 | -0.016853 | 0.80229296 | 0.9638526 | -0.0364135 | 0.5893792 |
| 0.999490048 | -0.0152158 | 0.80101188 | 0.9638526 | 6.87E-04 | 0.9909134 |
| 0.999490048 | -0.0150484 | 0.80249247 | 0.9638526 | -0.058347 | 0.3363725 |
| 0.999490048 | -0.0144207 | 0.80233725 | 0.9638526 | 0.00638105 | 0.91175726 |
| 0.999490048 | -0.0140749 | 0.80328735 | 0.9638526 | -0.0209644 | 0.71082963 |
| 0.999490048 | -0.0138451 | 0.80393025 | 0.9638526 | 0.04560257 | 0.41634654 |
| 0.999490048 | -0.0137734 | 0.80415416 | 0.9638526 | -0.1211194 | 0.03691718 |
| 0.999490048 | -0.0114621 | 0.80305672 | 0.9638526 | 0.04288338 | 0.35467006 |
| 0.999490048 | -0.0107281 | 0.8029298 | 0.9638526 | 0.0129559 | 0.76319101 |
| 0.999490048 | -0.0104162 | 0.80237482 | 0.9638526 | -0.0018737 | 0.96406848 |
| 0.999490048 | 0.00963078 | 0.80382106 | 0.9638526 | 0.02679851 | 0.49124472 |
| 0.999490048 | 0.01105558 | 0.80265476 | 0.9638526 | -0.0135112 | 0.76011727 |
| 0.999490048 | 0.01174502 | 0.80416915 | 0.9638526 | 0.01053915 | 0.82390567 |
| 0.999490048 | 0.01211398 | 0.80194606 | 0.9638526 | 0.01896722 | 0.69478527 |
| 0.999490048 | 0.01299893 | 0.80146781 | 0.9638526 | 0.07112445 | 0.17658122 |
| 0.999490048 | 0.016315 | 0.80423544 | 0.9638526 | 0.02638927 | 0.68877072 |
| 0.999490048 | 0.01651562 | 0.80432634 | 0.9638526 | -0.035681 | 0.59331302 |
| 0.999490048 | 0.0190552 | 0.8039018 | 0.9638526 | 0.090062 | 0.24670111 |
| 0.999490048 | 0.01949407 | 0.80173805 | 0.9638526 | -0.0191058 | 0.80560256 |
| 0.999490048 | 0.02499444 | 0.8042786 | 0.9638526 | 0.0298907 | 0.76701641 |
| 0.999490048 | 0.03367615 | 0.80388517 | 0.9638526 | -0.1082142 | 0.42755937 |
| 0.999490048 | 0.04535623 | 0.80380955 | 0.9638526 | -0.1312337 | 0.47859404 |
| 0.999490048 | 0.04537796 | 0.80371582 | 0.9638526 | -0.1168856 | 0.52535875 |
| 0.999490048 | 0.09392993 | 0.80307604 | 0.9638526 | -0.0149383 | 0.96830318 |

| | | | | | |
|---|---|---|---|---|---|
| 0.999490048 | -0.0293999 | 0.80455236 | 0.9639851 | -0.1109773 | 0.35414583 |
| 0.999490048 | 0.0111226 | 0.80514063 | 0.96441318 | 0.022346 | 0.6208024 |
| 0.999490048 | 0.02428126 | 0.80508007 | 0.96441318 | 0.04527804 | 0.64589254 |
| 0.999490048 | -0.0108895 | 0.8053592 | 0.96453663 | -0.0130457 | 0.76789557 |
| 0.999490048 | -0.0380481 | 0.80983247 | 0.96457186 | -0.0953688 | 0.54762255 |
| 0.999490048 | -0.0343311 | 0.8072826 | 0.96457186 | -0.0274261 | 0.8454611 |
| 0.999490048 | -0.0339948 | 0.80567587 | 0.96457186 | 0.03132644 | 0.82064238 |
| 0.999490048 | -0.0311626 | 0.81015998 | 0.96457186 | -0.1013242 | 0.43730014 |
| 0.999490048 | -0.0288463 | 0.80583913 | 0.96457186 | 0.03818795 | 0.74499336 |
| 0.999490048 | -0.0220774 | 0.80841936 | 0.96457186 | -0.1121978 | 0.2245166 |
| 0.999490048 | -0.0186467 | 0.80934506 | 0.96457186 | -0.1252804 | 0.11384377 |
| 0.999490048 | -0.0170893 | 0.81035588 | 0.96457186 | -0.0335638 | 0.63798396 |
| 0.999490048 | -0.0170201 | 0.80995816 | 0.96457186 | 0.05035051 | 0.47880834 |
| 0.999490048 | -0.0158883 | 0.80775343 | 0.96457186 | -0.0354936 | 0.5876217 |
| 0.999490048 | -0.0148378 | 0.80664094 | 0.96457186 | 0.07405672 | 0.22840276 |
| 0.987721587 | -0.0145986 | 0.80709341 | 0.96457186 | -0.1402533 | 0.02572653 |
| 0.999490048 | -0.0145662 | 0.80855663 | 0.96457186 | -0.0848883 | 0.16589567 |
| 0.999490048 | -0.0141267 | 0.8069036 | 0.96457186 | -0.0558148 | 0.3383861 |
| 0.999490048 | -0.0137069 | 0.80826053 | 0.96457186 | -0.0023991 | 0.96610117 |
| 0.999490048 | -0.0133386 | 0.80612087 | 0.96457186 | 0.00485208 | 0.92882227 |
| 0.999490048 | -0.0133079 | 0.8092123 | 0.96457186 | 0.00577505 | 0.91651402 |
| 0.999490048 | -0.0127774 | 0.80748048 | 0.96457186 | 0.00752409 | 0.88585745 |
| 0.999490048 | -0.0127499 | 0.80964078 | 0.96457186 | -0.036988 | 0.48654081 |
| 0.999490048 | -0.0126188 | 0.81029165 | 0.96457186 | 0.02990479 | 0.57046726 |
| 0.999490048 | -0.01216 | 0.80827132 | 0.96457186 | -0.0288201 | 0.5662755 |
| 0.999490048 | -0.0121031 | 0.80704298 | 0.96457186 | 0.01789772 | 0.7181575 |
| 0.999490048 | -0.0112308 | 0.80913899 | 0.96457186 | -0.0138622 | 0.76567457 |
| 0.999490048 | -0.0102069 | 0.81020727 | 0.96457186 | 0.01387196 | 0.74424907 |
| 0.999490048 | 0.01003218 | 0.8080112 | 0.96457186 | -0.0239804 | 0.56244488 |
| 0.999490048 | 0.01011297 | 0.80704237 | 0.96457186 | 0.0190094 | 0.64666023 |
| 0.999490048 | 0.0103281 | 0.80653488 | 0.96457186 | -0.0324965 | 0.44341128 |
| 0.999490048 | 0.0109718 | 0.80816935 | 0.96457186 | -0.0335855 | 0.45958598 |
| 0.999490048 | 0.01187745 | 0.80782942 | 0.96457186 | -0.0500424 | 0.31023503 |
| 0.999490048 | 0.01264802 | 0.80569198 | 0.96457186 | -5.63E-04 | 0.99125489 |
| 0.999490048 | 0.01323817 | 0.81007672 | 0.96457186 | 0.01498822 | 0.7855814 |
| 0.999490048 | 0.01331683 | 0.80641606 | 0.96457186 | 0.00932166 | 0.8637619 |
| 0.999490048 | 0.01402075 | 0.80668594 | 0.96457186 | -0.0066256 | 0.9078971 |
| 0.999490048 | 0.01439524 | 0.80725656 | 0.96457186 | 0.06378622 | 0.28497689 |
| 0.999490048 | 0.01455682 | 0.80649035 | 0.96457186 | 0.00682989 | 0.90845706 |
| 0.999490048 | 0.01479938 | 0.80877 | 0.96457186 | 0.04771043 | 0.43779604 |
| 0.999490048 | 0.01527105 | 0.80787454 | 0.96457186 | -3.98E-04 | 0.99494341 |
| 0.999490048 | 0.01689177 | 0.8096361 | 0.96457186 | -0.1268968 | 0.07931161 |

| | | | | | |
|---|---|---|---|---|---|
| 0.999490048 | 0.01972991 | 0.80718606 | 0.96457186 | 0.06822137 | 0.40180102 |
| 0.999490048 | 0.02481551 | 0.80866721 | 0.96457186 | -0.0877991 | 0.39478648 |
| 0.999490048 | 0.0251126 | 0.80895373 | 0.96457186 | 0.0017893 | 0.98624756 |
| 0.999490048 | 0.02646203 | 0.80907179 | 0.96457186 | 0.09032329 | 0.41242889 |
| 0.999490048 | 0.0455509 | 0.81003429 | 0.96457186 | -0.1228008 | 0.52173799 |
| 0.999490048 | 0.01726364 | 0.81049933 | 0.9646051 | -0.0142285 | 0.84330964 |
| 0.999490048 | -0.0276454 | 0.81069813 | 0.96462715 | 0.09721565 | 0.40269056 |
| 0.999490048 | -0.0227879 | 0.8107489 | 0.96462715 | 0.0317783 | 0.73856802 |
| 0.999490048 | 0.01185747 | 0.81089602 | 0.96466473 | 0.05329456 | 0.28743763 |
| 0.987721587 | -0.0522741 | 0.81169637 | 0.96477701 | -0.5337447 | 0.02119436 |
| 0.999490048 | -0.0412103 | 0.81175684 | 0.96477701 | 0.10642336 | 0.54152363 |
| 0.999490048 | -0.0107232 | 0.81134412 | 0.96477701 | -0.0528223 | 0.24584916 |
| 0.999490048 | 0.01229725 | 0.81112864 | 0.96477701 | 0.05468352 | 0.29307112 |
| 0.999490048 | 0.01251879 | 0.81128701 | 0.96477701 | -0.0068242 | 0.89640173 |
| 0.999490048 | 0.0159593 | 0.81185325 | 0.96477701 | 0.07027857 | 0.29953665 |
| 0.999490048 | 0.01713083 | 0.81191474 | 0.96477701 | -0.0819503 | 0.2608186 |
| 0.999490048 | 0.06851812 | 0.81176477 | 0.96477701 | 0.54507432 | 0.06699661 |
| 0.999490048 | 0.02545167 | 0.8120495 | 0.96479985 | 0.04574476 | 0.66950411 |
| 0.999490048 | -0.0080669 | 0.81264787 | 0.96524103 | -0.0183905 | 0.58985516 |
| 0.999490048 | 0.01022684 | 0.81265203 | 0.96524103 | 0.03271486 | 0.45070765 |
| 0.999490048 | 0.01837062 | 0.81277552 | 0.96525041 | 0.01550648 | 0.84151727 |
| 0.999490048 | -0.0281004 | 0.81297249 | 0.96534703 | -0.1484759 | 0.21806976 |
| 0.999490048 | 0.01683234 | 0.81332695 | 0.9656306 | -0.0408515 | 0.56764433 |
| 0.999490048 | -0.0263605 | 0.81347383 | 0.96566768 | 0.00245792 | 0.98243791 |
| 0.999490048 | -0.042675 | 0.81528285 | 0.96577644 | -0.2221951 | 0.23850439 |
| 0.999490048 | -0.0296482 | 0.81510556 | 0.96577644 | 0.11008068 | 0.38857737 |
| 0.999490048 | -0.0233994 | 0.81517439 | 0.96577644 | 0.03784195 | 0.70565491 |
| 0.999490048 | -0.0208422 | 0.8157371 | 0.96577644 | 0.06403531 | 0.4760432 |
| 0.999490048 | -0.0203011 | 0.81559514 | 0.96577644 | 0.0415646 | 0.63361122 |
| 0.999490048 | -0.0202278 | 0.81506626 | 0.96577644 | 0.10080101 | 0.24988033 |
| 0.999490048 | -0.0197254 | 0.81658695 | 0.96577644 | 0.00137398 | 0.98710324 |
| 0.987721587 | -0.0192002 | 0.81658577 | 0.96577644 | 0.15942555 | 0.06289533 |
| 0.999490048 | -0.0157282 | 0.81585574 | 0.96577644 | -0.065359 | 0.33742251 |
| 0.999490048 | -0.0146562 | 0.81456842 | 0.96577644 | -0.0066608 | 0.9150767 |
| 0.999490048 | -0.0143015 | 0.81499396 | 0.96577644 | 0.01581996 | 0.79579005 |
| 0.999490048 | -0.0141055 | 0.81696335 | 0.96577644 | 0.05004349 | 0.41446121 |
| 0.999490048 | -0.0135852 | 0.81424146 | 0.96577644 | 0.01833107 | 0.75132457 |
| 0.999490048 | -0.0124307 | 0.81518591 | 0.96577644 | -0.0047229 | 0.92920195 |
| 0.999490048 | -0.0103 | 0.81583468 | 0.96577644 | 0.02929365 | 0.50935371 |
| 0.999490048 | -0.0087872 | 0.81676038 | 0.96577644 | 0.01564003 | 0.68038565 |
| 0.999490048 | 0.00983761 | 0.81699146 | 0.96577644 | -0.0234691 | 0.58185052 |
| 0.999490048 | 0.01039955 | 0.81617206 | 0.96577644 | 0.04093463 | 0.36391412 |

| | | | | | |
|---|---|---|---|---|---|
| 0.999490048 | 0.0110819 | 0.81647477 | 0.96577644 | 0.01099586 | 0.81787317 |
| 0.999490048 | 0.01324647 | 0.81545357 | 0.96577644 | -0.0818949 | 0.15717519 |
| 0.999490048 | 0.01325098 | 0.8164399 | 0.96577644 | 0.00905693 | 0.87390334 |
| 0.999490048 | 0.01399816 | 0.81449597 | 0.96577644 | 0.02059025 | 0.73016945 |
| 0.999490048 | 0.01517689 | 0.8170353 | 0.96577644 | -0.020642 | 0.75312108 |
| 0.999490048 | 0.0171039 | 0.81423186 | 0.96577644 | -0.0142082 | 0.84521825 |
| 0.999490048 | 0.01837015 | 0.81476811 | 0.96577644 | 0.05911584 | 0.45322342 |
| 0.999490048 | 0.02018131 | 0.81390504 | 0.96577644 | -0.0036193 | 0.96630926 |
| 0.999490048 | 0.0211109 | 0.81666213 | 0.96577644 | -0.0617753 | 0.4992592 |
| 0.999490048 | 0.04423246 | 0.81570185 | 0.96577644 | -0.013293 | 0.94412948 |
| 0.999490048 | 0.04470019 | 0.81483514 | 0.96577644 | -0.070321 | 0.71278034 |
| 0.999490048 | 0.04661481 | 0.81612479 | 0.96577644 | -0.0625343 | 0.75518628 |
| 0.999490048 | -0.0169921 | 0.81716075 | 0.965788 | -0.0929573 | 0.21288016 |
| 0.999490048 | -0.0504775 | 0.81911492 | 0.96583976 | -0.389969 | 0.08628239 |
| 0.999490048 | -0.0460262 | 0.8214602 | 0.96583976 | 0.0963311 | 0.63855285 |
| 0.999490048 | -0.04405 | 0.81971102 | 0.96583976 | -0.2707044 | 0.17901091 |
| 0.999490048 | -0.0350063 | 0.82112364 | 0.96583976 | -0.1161361 | 0.4604165 |
| 0.999490048 | -0.0273537 | 0.82031233 | 0.96583976 | -0.113815 | 0.34864557 |
| 0.999490048 | -0.0259143 | 0.82027696 | 0.96583976 | -0.026299 | 0.81765869 |
| 0.999490048 | -0.0178483 | 0.82121713 | 0.96583976 | 0.10727548 | 0.1820268 |
| 0.999490048 | -0.0178254 | 0.82089758 | 0.96583976 | 0.03614202 | 0.64674811 |
| 0.999490048 | -0.0171811 | 0.8187107 | 0.96583976 | 0.11807617 | 0.12393391 |
| 0.999490048 | -0.0165241 | 0.81979245 | 0.96583976 | 0.02485238 | 0.73204695 |
| 0.999490048 | -0.0157049 | 0.81779444 | 0.96583976 | -0.0278532 | 0.68318355 |
| 0.999490048 | -0.015021 | 0.81973979 | 0.96583976 | -0.0011929 | 0.98555329 |
| 0.999490048 | -0.0135064 | 0.82159108 | 0.96583976 | -0.0769697 | 0.20587695 |
| 0.999490048 | -0.0133522 | 0.8206836 | 0.96583976 | -0.0924209 | 0.12536857 |
| 0.999490048 | -0.0130616 | 0.81853761 | 0.96583976 | 0.04253157 | 0.4573056 |
| 0.999490048 | -0.012964 | 0.81763396 | 0.96583976 | -0.0216066 | 0.70101808 |
| 0.999490048 | -0.0129498 | 0.82107945 | 0.96583976 | -0.0584493 | 0.31212833 |
| 0.999490048 | -0.0128134 | 0.8208915 | 0.96583976 | 0.07110675 | 0.21585803 |
| 0.999490048 | -0.012716 | 0.82159998 | 0.96583976 | 0.01916657 | 0.7341191 |
| 0.999490048 | -0.011804 | 0.82006815 | 0.96583976 | -0.052432 | 0.31699575 |
| 0.999490048 | 0.01048466 | 0.82093892 | 0.96583976 | -0.0049319 | 0.91517698 |
| 0.999490048 | 0.01048819 | 0.82151135 | 0.96583976 | 0.03184966 | 0.49510129 |
| 0.999490048 | 0.01073222 | 0.82071209 | 0.96583976 | 0.0050887 | 0.91439246 |
| 0.999490048 | 0.01089246 | 0.82025113 | 0.96583976 | -0.0099061 | 0.8362714 |
| 0.999490048 | 0.01162211 | 0.81879324 | 0.96583976 | 0.01050695 | 0.83590431 |
| 0.999490048 | 0.01216126 | 0.82098265 | 0.96583976 | -0.0072276 | 0.89298575 |
| 0.999490048 | 0.01586609 | 0.81827787 | 0.96583976 | 0.04690128 | 0.49878038 |
| 0.999490048 | 0.0160523 | 0.81742364 | 0.96583976 | -0.0238492 | 0.73177175 |
| 0.999490048 | 0.01753088 | 0.81986639 | 0.96583976 | 0.12361079 | 0.11716563 |

| | | | | | |
|---|---|---|---|---|---|
| 0.999490048 | 0.0233496 | 0.81796802 | 0.96583976 | 0.02423214 | 0.81122291 |
| 0.999490048 | 0.02336935 | 0.81929284 | 0.96583976 | -0.1837247 | 0.08148015 |
| 0.999490048 | 0.02483093 | 0.82137184 | 0.96583976 | -0.0301903 | 0.78374306 |
| 0.999490048 | 0.02788377 | 0.81798985 | 0.96583976 | 0.02926721 | 0.80913984 |
| 0.999490048 | 0.02900178 | 0.81979469 | 0.96583976 | 0.03112858 | 0.806847 |
| 0.999490048 | 0.04042227 | 0.81973366 | 0.96583976 | 0.09208811 | 0.60443959 |
| 0.999490048 | 0.04705839 | 0.81760283 | 0.96583976 | 0.04473607 | 0.82644821 |
| 0.999490048 | 0.04711911 | 0.81868646 | 0.96583976 | 0.12511468 | 0.54678735 |
| 0.999490048 | 0.0713655 | 0.81992226 | 0.96583976 | 0.21152602 | 0.50528275 |
| 0.999490048 | -0.0145442 | 0.82210078 | 0.96595113 | 0.06194329 | 0.34242833 |
| 0.999490048 | -0.0128899 | 0.82222201 | 0.96595113 | -0.0131457 | 0.81875871 |
| 0.999490048 | 0.0144934 | 0.82205367 | 0.96595113 | 0.00451215 | 0.94415337 |
| 0.999490048 | 0.01503367 | 0.82227313 | 0.96595113 | 0.00493088 | 0.9412433 |
| 0.999490048 | 0.01568562 | 0.82198645 | 0.96595113 | -0.0507595 | 0.46870297 |
| 0.999490048 | 0.01039846 | 0.8224568 | 0.96603099 | -0.0065828 | 0.88700674 |
| 0.999490048 | -0.0133662 | 0.82294922 | 0.96620163 | 0.01726547 | 0.77263542 |
| 0.999490048 | -0.0109907 | 0.82292786 | 0.96620163 | -0.0471534 | 0.34120579 |
| 0.999490048 | 0.01751342 | 0.82278657 | 0.96620163 | 0.08761618 | 0.26823137 |
| 0.999490048 | -0.0127306 | 0.82321976 | 0.96638338 | 0.03711026 | 0.51647123 |
| 0.999490048 | -0.0642698 | 0.82422673 | 0.96649203 | 0.03800541 | 0.89539536 |
| 0.999490048 | -0.046409 | 0.82628424 | 0.96649203 | -0.1889393 | 0.37516723 |
| 0.999490048 | -0.0330502 | 0.82703471 | 0.96649203 | 0.02058476 | 0.89171602 |
| 0.999490048 | -0.0303405 | 0.82504279 | 0.96649203 | 0.06963498 | 0.61264407 |
| 0.999490048 | -0.0279892 | 0.8274457 | 0.96649203 | 0.03154133 | 0.80598358 |
| 0.999490048 | -0.0233395 | 0.82673209 | 0.96649203 | -0.0492313 | 0.64482772 |
| 0.999490048 | -0.0208953 | 0.82722212 | 0.96649203 | 0.09977972 | 0.30227278 |
| 0.999490048 | -0.018881 | 0.82714887 | 0.96649203 | 0.03071076 | 0.72267116 |
| 0.999490048 | -0.0175824 | 0.82737578 | 0.96649203 | -0.1384895 | 0.09487327 |
| 0.999490048 | -0.0156273 | 0.82732548 | 0.96649203 | 0.04387626 | 0.54157506 |
| 0.999490048 | -0.0154031 | 0.82489405 | 0.96649203 | -0.0518666 | 0.45853558 |
| 0.999490048 | -0.0136096 | 0.82382128 | 0.96649203 | 0.01557349 | 0.79894012 |
| 0.999490048 | -0.013479 | 0.82541632 | 0.96649203 | -0.0234856 | 0.70100784 |
| 0.999490048 | -0.0133391 | 0.82747922 | 0.96649203 | -0.0662319 | 0.28456943 |
| 0.999490048 | -0.0119196 | 0.82455704 | 0.96649203 | 0.03860367 | 0.4748426 |
| 0.999490048 | -0.0116084 | 0.82657858 | 0.96649203 | -0.030093 | 0.5711246 |
| 0.999490048 | -0.0113985 | 0.82685908 | 0.96649203 | -0.0580586 | 0.27088605 |
| 0.999490048 | -0.0103623 | 0.82717262 | 0.96649203 | 0.00614728 | 0.89691336 |
| 0.999490048 | -0.0102935 | 0.82476411 | 0.96649203 | -0.0114251 | 0.80588658 |
| 0.999490048 | 0.00720461 | 0.82705763 | 0.96649203 | -0.0268084 | 0.41910011 |
| 0.999490048 | 0.01009417 | 0.82528264 | 0.96649203 | 0.02744373 | 0.54963456 |
| 0.999490048 | 0.01154799 | 0.82411143 | 0.96649203 | 0.0167429 | 0.74740147 |
| 0.999490048 | 0.01177863 | 0.82416073 | 0.96649203 | -0.0332869 | 0.53146961 |

| | | | | | |
|---|---|---|---|---|---|
| 0.999490048 | 0.01209289 | 0.82558835 | 0.96649203 | -0.0373728 | 0.49764823 |
| 0.999490048 | 0.0121766 | 0.8272681 | 0.96649203 | -0.0051857 | 0.92593114 |
| 0.999490048 | 0.01331149 | 0.82558229 | 0.96649203 | -0.0066411 | 0.91242253 |
| 0.999490048 | 0.01363658 | 0.82641274 | 0.96649203 | -0.0250697 | 0.68716331 |
| 0.999490048 | 0.01395308 | 0.82640497 | 0.96649203 | 0.0172869 | 0.78589505 |
| 0.999490048 | 0.01458098 | 0.82346782 | 0.96649203 | 0.02466316 | 0.70618061 |
| 0.999490048 | 0.01476496 | 0.82615853 | 0.96649203 | -0.0056357 | 0.93316274 |
| 0.999490048 | 0.01718275 | 0.82619079 | 0.96649203 | 0.10532432 | 0.18586528 |
| 0.999490048 | 0.01991524 | 0.82457954 | 0.96649203 | 0.11757897 | 0.19794135 |
| 0.999490048 | 0.02799231 | 0.82457622 | 0.96649203 | 0.12422084 | 0.32969923 |
| 0.999490048 | 0.03564518 | 0.8238524 | 0.96649203 | 0.11763427 | 0.46478523 |
| 0.999490048 | 0.03672754 | 0.82616071 | 0.96649203 | -0.0119586 | 0.94296586 |
| 0.999490048 | 0.05128167 | 0.82634867 | 0.96649203 | -0.033392 | 0.88633315 |
| 0.999490048 | -0.014667 | 0.82811041 | 0.96649403 | 0.0449701 | 0.50727857 |
| 0.999490048 | -0.0146226 | 0.82773094 | 0.96649403 | -0.0542357 | 0.4223965 |
| 0.999490048 | -0.0109501 | 0.82820847 | 0.96649403 | -0.0485162 | 0.34054473 |
| 0.999490048 | -0.0097664 | 0.82802761 | 0.96649403 | -0.0062713 | 0.88903089 |
| 0.999490048 | 0.00957935 | 0.82771745 | 0.96649403 | 0.01901169 | 0.66623976 |
| 0.999490048 | 0.01267277 | 0.82829117 | 0.96649403 | 0.00822694 | 0.88799448 |
| 0.999490048 | 0.04336972 | 0.82818247 | 0.96649403 | -0.1627979 | 0.41812738 |
| 0.999490048 | -0.0730861 | 0.82999229 | 0.9666693 | -0.6495609 | 0.08365575 |
| 0.999490048 | -0.0175041 | 0.83064099 | 0.9666693 | 0.01556127 | 0.8491848 |
| 0.999490048 | -0.0158063 | 0.8303425 | 0.9666693 | -0.0750463 | 0.31377093 |
| 0.999490048 | -0.0127146 | 0.82945816 | 0.9666693 | 0.02253734 | 0.70289076 |
| 0.999490048 | -0.0125801 | 0.82991024 | 0.9666693 | 0.09408992 | 0.11696174 |
| 0.999490048 | -0.0122476 | 0.8297791 | 0.9666693 | -0.0556299 | 0.33318913 |
| 0.999490048 | -0.0100558 | 0.8300438 | 0.9666693 | 0.01248126 | 0.78996933 |
| 0.999490048 | -0.0097845 | 0.82983237 | 0.9666693 | -0.0149751 | 0.74236386 |
| 0.999490048 | 0.01255445 | 0.82926101 | 0.9666693 | 0.02832833 | 0.62720276 |
| 0.999490048 | 0.01310981 | 0.82878002 | 0.9666693 | 0.05320814 | 0.38351828 |
| 0.999490048 | 0.01348975 | 0.82941906 | 0.9666693 | 0.00952984 | 0.87899913 |
| 0.999490048 | 0.01413848 | 0.82889782 | 0.9666693 | -3.14E-04 | 0.9961707 |
| 0.999490048 | 0.01429198 | 0.82955833 | 0.9666693 | -0.0349825 | 0.59910618 |
| 0.999490048 | 0.01486211 | 0.83014259 | 0.9666693 | 0.00786742 | 0.90955814 |
| 0.999490048 | 0.01635451 | 0.82886754 | 0.9666693 | -0.0461346 | 0.54334458 |
| 0.999490048 | 0.01731678 | 0.8302689 | 0.9666693 | 0.04486293 | 0.57966506 |
| 0.999490048 | 0.01769999 | 0.83055822 | 0.9666693 | -0.0331911 | 0.68856657 |
| 0.999490048 | 0.02705226 | 0.83057544 | 0.9666693 | 0.09774246 | 0.44200482 |
| 0.999490048 | 0.02738036 | 0.82969932 | 0.9666693 | 0.00506563 | 0.96824364 |
| 0.999490048 | 0.0084121 | 0.83093591 | 0.96673858 | -0.009173 | 0.81592151 |
| 0.999490048 | 0.0091566 | 0.83104785 | 0.96673858 | -0.0327319 | 0.44809118 |
| 0.999490048 | 0.04098743 | 0.83102477 | 0.96673858 | 0.05904161 | 0.75866934 |

| | | | | | |
|---|---|---|---|---|---|
| 0.999490048 | -0.0149795 | 0.83144189 | 0.96692755 | -0.1177708 | 0.10320529 |
| 0.999490048 | 0.08439949 | 0.83132961 | 0.96692755 | 0.3243859 | 0.42206851 |
| 0.999490048 | -0.0152314 | 0.83176374 | 0.96708077 | -0.0642435 | 0.37384764 |
| 0.999490048 | 0.03008255 | 0.83180528 | 0.96708077 | 0.1235398 | 0.38649151 |
| 0.999490048 | -0.018137 | 0.8322086 | 0.96724443 | -0.0633217 | 0.46173878 |
| 0.999490048 | -0.0105318 | 0.83209106 | 0.96724443 | -0.005943 | 0.90473855 |
| 0.999490048 | 0.01256268 | 0.83229356 | 0.96724443 | 0.06333839 | 0.29091526 |
| 0.999490048 | -0.0145356 | 0.83255256 | 0.96741078 | -0.0051899 | 0.93980078 |
| 0.999490048 | -0.0334077 | 0.83402165 | 0.96808241 | 0.0994057 | 0.53618275 |
| 0.999490048 | -0.019841 | 0.83631218 | 0.96808241 | -0.119411 | 0.22050586 |
| 0.999490048 | -0.0161202 | 0.83647553 | 0.96808241 | 0.00672525 | 0.93135405 |
| 0.999490048 | -0.0139712 | 0.83593266 | 0.96808241 | -0.0185189 | 0.78376199 |
| 0.999490048 | -0.0127771 | 0.83608115 | 0.96808241 | 0.03308811 | 0.59299335 |
| 0.999490048 | -0.0120519 | 0.83649276 | 0.96808241 | 0.01657685 | 0.77659483 |
| 0.999490048 | -0.0118657 | 0.8351279 | 0.96808241 | -0.0276946 | 0.62779481 |
| 0.999490048 | -0.0115287 | 0.83441459 | 0.96808241 | 0.03627881 | 0.512303 |
| 0.999490048 | -0.0111569 | 0.83493297 | 0.96808241 | -0.0050789 | 0.92440053 |
| 0.999490048 | -0.011128 | 0.83570175 | 0.96808241 | 0.02137951 | 0.69063717 |
| 0.999490048 | -0.0111121 | 0.83577196 | 0.96808241 | 0.05309936 | 0.32639437 |
| 0.999490048 | -0.0097264 | 0.83603933 | 0.96808241 | -0.0620338 | 0.19424769 |
| 0.999490048 | -0.0095868 | 0.83524261 | 0.96808241 | 0.02686142 | 0.56122777 |
| 0.999490048 | -0.0091039 | 0.83579356 | 0.96808241 | -0.0158701 | 0.71809099 |
| 0.999490048 | -0.0088939 | 0.83486145 | 0.96808241 | -0.0049712 | 0.90720975 |
| 0.999490048 | -0.0085813 | 0.8351467 | 0.96808241 | -0.0046992 | 0.90924163 |
| 0.999490048 | 0.00720417 | 0.83541357 | 0.96808241 | 0.03194198 | 0.36081193 |
| 0.999490048 | 0.00898977 | 0.83481033 | 0.96808241 | 0.02138275 | 0.62059573 |
| 0.999490048 | 0.01110334 | 0.83628019 | 0.96808241 | -0.0037402 | 0.94448068 |
| 0.999490048 | 0.01133746 | 0.83453079 | 0.96808241 | -0.0490825 | 0.36950873 |
| 0.999490048 | 0.01200046 | 0.83623732 | 0.96808241 | -0.009017 | 0.87654736 |
| 0.999490048 | 0.01405161 | 0.83536032 | 0.96808241 | 0.0165835 | 0.80627211 |
| 0.999490048 | 0.01408758 | 0.83590065 | 0.96808241 | 0.06126126 | 0.3713813 |
| 0.999490048 | 0.01410488 | 0.83346314 | 0.96808241 | 0.01074935 | 0.87266758 |
| 0.999490048 | 0.01494025 | 0.83644059 | 0.96808241 | -0.1178665 | 0.11227412 |
| 0.999490048 | 0.02630156 | 0.83481073 | 0.96808241 | 0.0547746 | 0.66454014 |
| 0.999490048 | 0.02698486 | 0.83568874 | 0.96808241 | 0.10794846 | 0.40972456 |
| 0.999490048 | 0.02815431 | 0.83625738 | 0.96808241 | 0.02428514 | 0.85848825 |
| 0.999490048 | 0.03712061 | 0.83514119 | 0.96808241 | 0.15654948 | 0.38357402 |
| 0.999490048 | -0.0142097 | 0.83705839 | 0.96820025 | 0.03495854 | 0.61365012 |
| 0.999490048 | -0.0109672 | 0.83703127 | 0.96820025 | -0.0278098 | 0.60280085 |
| 0.999490048 | 0.01486293 | 0.83690376 | 0.96820025 | 0.06613366 | 0.36349702 |
| 0.999490048 | 0.03813992 | 0.83682459 | 0.96820025 | -0.028505 | 0.87764357 |
| 0.999490048 | -0.0145733 | 0.83726332 | 0.96830315 | -0.0200459 | 0.77762273 |

| | | | | | |
|---|---|---|---|---|---|
| 0.999490048 | -0.0299595 | 0.8378496 | 0.96860451 | -0.1258904 | 0.39313171 |
| 0.999490048 | -0.0141139 | 0.8378719 | 0.96860451 | -0.056133 | 0.41865589 |
| 0.999490048 | -0.0132043 | 0.83774929 | 0.96860451 | 0.0298251 | 0.64427558 |
| 0.999490048 | -0.0153914 | 0.8384202 | 0.96896917 | 0.01847225 | 0.80669979 |
| 0.999490048 | 0.00999186 | 0.83861109 | 0.96896917 | 0.04513268 | 0.36144525 |
| 0.999490048 | 0.01019381 | 0.83865152 | 0.96896917 | -0.033993 | 0.49894602 |
| 0.987721587 | 0.01662218 | 0.83855444 | 0.96896917 | -0.1546984 | 0.06681296 |
| 0.999490048 | -0.0106269 | 0.83888302 | 0.96910255 | -0.0052882 | 0.91938447 |
| 0.999490048 | -0.0176024 | 0.83905159 | 0.9691632 | 0.02299496 | 0.79081633 |
| 0.999490048 | -0.0157074 | 0.83974266 | 0.96921498 | -0.0245251 | 0.75233867 |
| 0.999490048 | -0.0113254 | 0.83947202 | 0.96921498 | -0.0146089 | 0.79391907 |
| 0.999490048 | -0.0104312 | 0.83979286 | 0.96921498 | -0.0308656 | 0.55098987 |
| 0.999490048 | 0.00975366 | 0.83973025 | 0.96921498 | 0.03013284 | 0.53354494 |
| 0.999490048 | 0.01051059 | 0.83922581 | 0.96921498 | 0.02590872 | 0.61773703 |
| 0.999490048 | 0.01273389 | 0.83934901 | 0.96921498 | 0.03119801 | 0.62014025 |
| 0.999490048 | -0.0081212 | 0.84010478 | 0.96944098 | -0.0376366 | 0.35377757 |
| 0.999490048 | -0.0220425 | 0.84038219 | 0.96949314 | 0.10554278 | 0.34809087 |
| 0.999490048 | 0.02740362 | 0.84037161 | 0.96949314 | 0.23065312 | 0.09902993 |
| 0.999490048 | -0.0141162 | 0.84069823 | 0.96972375 | 0.04305949 | 0.54116089 |
| 0.999490048 | -0.0313886 | 0.84327109 | 0.9702627 | -0.3403199 | 0.04000359 |
| 0.999490048 | -0.0202883 | 0.84418665 | 0.9702627 | 0.03151269 | 0.76028416 |
| 0.999490048 | -0.0201648 | 0.84402448 | 0.9702627 | 0.06404317 | 0.53350405 |
| 0.999490048 | -0.0195434 | 0.84159244 | 0.9702627 | 0.09794958 | 0.32114328 |
| 0.999490048 | -0.0138824 | 0.84374473 | 0.9702627 | 0.04493837 | 0.52497844 |
| 0.999490048 | -0.0124948 | 0.84278578 | 0.9702627 | -0.0235967 | 0.70827604 |
| 0.999490048 | -0.0121974 | 0.84208351 | 0.9702627 | -0.0272856 | 0.65634771 |
| 0.999490048 | -0.012079 | 0.8430582 | 0.9702627 | -0.0468397 | 0.44515293 |
| 0.999490048 | -0.0112134 | 0.8434052 | 0.9702627 | -0.0297999 | 0.60047515 |
| 0.999490048 | -0.0107904 | 0.84284712 | 0.9702627 | 0.05032469 | 0.35907279 |
| 0.999490048 | -0.0102792 | 0.84353485 | 0.9702627 | -0.0209013 | 0.68853864 |
| 0.999490048 | -0.0095519 | 0.84393846 | 0.9702627 | 7.69E-04 | 0.98735801 |
| 0.999490048 | -0.0083438 | 0.84274215 | 0.9702627 | -0.0093873 | 0.82340001 |
| 0.999490048 | -0.0077382 | 0.84161007 | 0.9702627 | 0.01196686 | 0.75742528 |
| 0.999490048 | 0.01011234 | 0.84318811 | 0.9702627 | 0.0484387 | 0.34748211 |
| 0.999490048 | 0.01059045 | 0.84205057 | 0.9702627 | -0.004976 | 0.92538064 |
| 0.999490048 | 0.01113469 | 0.84368656 | 0.9702627 | -0.0646243 | 0.25843098 |
| 0.999490048 | 0.01138927 | 0.84416558 | 0.9702627 | 0.00622625 | 0.91440142 |
| 0.999490048 | 0.0117884 | 0.84270509 | 0.9702627 | -0.0146065 | 0.80582233 |
| 0.999490048 | 0.01388263 | 0.84199155 | 0.9702627 | 0.07322735 | 0.29813781 |
| 0.999490048 | 0.01538783 | 0.84385172 | 0.9702627 | 0.05071013 | 0.51787954 |
| 0.999490048 | 0.01551045 | 0.84279643 | 0.9702627 | 0.06215622 | 0.42950538 |
| 0.999490048 | 0.01653719 | 0.84164325 | 0.9702627 | 0.0713602 | 0.39194543 |

| | | | | | |
|---|---|---|---|---|---|
| 0.999490048 | 0.01884188 | 0.84378431 | 0.9702627 | 0.05835552 | 0.54301458 |
| 0.999490048 | 0.02224861 | 0.84318979 | 0.9702627 | 0.05975658 | 0.59610888 |
| 0.999490048 | 0.02912687 | 0.84141903 | 0.9702627 | 0.01693718 | 0.90735433 |
| 0.999490048 | -0.0309649 | 0.84821249 | 0.97082415 | -0.1086564 | 0.5035996 |
| 0.999490048 | -0.0271466 | 0.84829567 | 0.97082415 | -0.0339618 | 0.81089784 |
| 0.999490048 | -0.026873 | 0.84793552 | 0.97082415 | 0.00441173 | 0.97487765 |
| 0.999490048 | -0.0217437 | 0.85014869 | 0.97082415 | -0.0424007 | 0.71284041 |
| 0.999490048 | -0.0198741 | 0.84661358 | 0.97082415 | -0.192206 | 0.07031297 |
| 0.999490048 | -0.0194926 | 0.85071869 | 0.97082415 | 0.13098856 | 0.21299685 |
| 0.999490048 | -0.0189887 | 0.8461024 | 0.97082415 | -0.0170873 | 0.86133743 |
| 0.999490048 | -0.0189207 | 0.84578927 | 0.97082415 | -0.031603 | 0.74545493 |
| 0.999490048 | -0.0179166 | 0.84603444 | 0.97082415 | -0.0763442 | 0.41742328 |
| 0.999490048 | -0.0153116 | 0.85003335 | 0.97082415 | -0.0460069 | 0.57106773 |
| 0.999490048 | -0.0149006 | 0.84717647 | 0.97082415 | 0.05383208 | 0.48822436 |
| 0.999490048 | -0.0148842 | 0.84899361 | 0.97082415 | -0.0607 | 0.44005476 |
| 0.999490048 | -0.0142349 | 0.84564208 | 0.97082415 | -0.0089973 | 0.90204518 |
| 0.999490048 | -0.0141972 | 0.85083726 | 0.97082415 | 0.1220479 | 0.11486274 |
| 0.999490048 | -0.0140404 | 0.85079562 | 0.97082415 | -0.0696808 | 0.35455772 |
| 0.999490048 | -0.0130629 | 0.85078821 | 0.97082415 | 0.01596214 | 0.81822982 |
| 0.999490048 | -0.0122145 | 0.84788766 | 0.97082415 | 0.03760814 | 0.55603468 |
| 0.999490048 | -0.0120843 | 0.84709372 | 0.97082415 | -0.0116475 | 0.85255222 |
| 0.999490048 | -0.0119923 | 0.85011251 | 0.97082415 | -0.0085561 | 0.89273023 |
| 0.999490048 | -0.0117199 | 0.84503725 | 0.97082415 | -0.0845985 | 0.16630875 |
| 0.999490048 | -0.0114303 | 0.84903181 | 0.97082415 | 0.00395013 | 0.94753337 |
| 0.999490048 | -0.0114174 | 0.84812168 | 0.97082415 | 0.0305751 | 0.60886166 |
| 0.999490048 | -0.0106967 | 0.84977784 | 0.97082415 | 0.05360723 | 0.34667564 |
| 0.999490048 | -0.010584 | 0.84896209 | 0.97082415 | -0.0198969 | 0.72058134 |
| 0.999490048 | -0.0103834 | 0.84604779 | 0.97082415 | 0.04945898 | 0.35896286 |
| 0.999490048 | -0.0099448 | 0.84966085 | 0.97082415 | -0.0130654 | 0.80338887 |
| 0.999490048 | -0.0099244 | 0.84902772 | 0.97082415 | -0.0751951 | 0.15742591 |
| 0.999490048 | -0.0093957 | 0.84915699 | 0.97082415 | 0.03650526 | 0.46221969 |
| 0.999490048 | -0.0093525 | 0.84627621 | 0.97082415 | -0.0211821 | 0.66109178 |
| 0.999490048 | -0.0093419 | 0.84994402 | 0.97082415 | -4.14E-04 | 0.99331009 |
| 0.999490048 | -0.0090151 | 0.85040068 | 0.97082415 | -0.0239555 | 0.61701026 |
| 0.999490048 | -0.0083616 | 0.84863863 | 0.97082415 | -0.0094658 | 0.82895997 |
| 0.99974305 | 0.00867153 | 0.84519276 | 0.97082415 | 0.00869249 | 0.84482364 |
| 0.999490048 | 0.0090751 | 0.8478519 | 0.97082415 | 0.01247674 | 0.79202043 |
| 0.999490048 | 0.00941562 | 0.84771905 | 0.97082415 | -0.0037149 | 0.93958785 |
| 0.999490048 | 0.00949372 | 0.84702974 | 0.97082415 | 0.03374452 | 0.4947918 |
| 0.999490048 | 0.0100598 | 0.84706877 | 0.97082415 | -0.0264631 | 0.61271005 |
| 0.999490048 | 0.01015118 | 0.84923517 | 0.97082415 | -8.26E-04 | 0.98764922 |
| 0.999490048 | 0.01015946 | 0.84715015 | 0.97082415 | 0.03482884 | 0.51043244 |

| | | | | | |
|---|---|---|---|---|---|
| 0.999490048 | 0.01019505 | 0.84692681 | 0.97082415 | -0.0467466 | 0.37963507 |
| 0.999490048 | 0.01189563 | 0.84671099 | 0.97082415 | 0.09031102 | 0.15053727 |
| 0.999490048 | 0.01304725 | 0.84805399 | 0.97082415 | 0.0402942 | 0.55527901 |
| 0.999490048 | 0.01322701 | 0.84654005 | 0.97082415 | 0.07549282 | 0.27495352 |
| 0.999490048 | 0.01384796 | 0.84740758 | 0.97082415 | 0.00236426 | 0.97378222 |
| 0.999490048 | 0.01601102 | 0.84894412 | 0.97082415 | -0.1487338 | 0.09717504 |
| 0.999490048 | 0.01601162 | 0.84775787 | 0.97082415 | 0.08469249 | 0.31466584 |
| 0.999490048 | 0.01810902 | 0.85067413 | 0.97082415 | 0.03278851 | 0.73341513 |
| 0.999490048 | 0.01876275 | 0.85081036 | 0.97082415 | 0.09591781 | 0.34056982 |
| 0.999490048 | 0.0198876 | 0.84965266 | 0.97082415 | 0.08966748 | 0.39600935 |
| 0.999490048 | 0.02425643 | 0.84674101 | 0.97082415 | 0.07833727 | 0.53580103 |
| 0.999490048 | 0.02567064 | 0.84534833 | 0.97082415 | 0.06176843 | 0.63943908 |
| 0.999490048 | 0.03208165 | 0.84694108 | 0.97082415 | -0.0418482 | 0.80125929 |
| 0.999490048 | 0.04264482 | 0.84584179 | 0.97082415 | 0.05730433 | 0.79395204 |
| 0.999490048 | 0.01626719 | 0.85095968 | 0.97083117 | -0.0517863 | 0.55102732 |
| 0.999490048 | -0.0124467 | 0.85122085 | 0.97099646 | -0.0059258 | 0.92882655 |
| 0.999490048 | -0.0093342 | 0.85168101 | 0.97112751 | -0.0258733 | 0.60511893 |
| 0.999490048 | 0.01228199 | 0.85159828 | 0.97112751 | 0.07982411 | 0.23064929 |
| 0.999490048 | 0.01414963 | 0.85168464 | 0.97112751 | 0.01191884 | 0.87484412 |
| 0.999490048 | -0.0146213 | 0.85207867 | 0.97144414 | -0.0379745 | 0.62886877 |
| 0.999490048 | -0.0217148 | 0.85310658 | 0.97192495 | -0.0471048 | 0.6883204 |
| 0.999490048 | -0.0207998 | 0.85391475 | 0.97192495 | -0.0106033 | 0.92519903 |
| 0.999490048 | -0.0134056 | 0.85310157 | 0.97192495 | -0.0157499 | 0.82780705 |
| 0.999490048 | -0.0109538 | 0.85395775 | 0.97192495 | -0.0121145 | 0.83869716 |
| 0.999490048 | -0.0092536 | 0.85367691 | 0.97192495 | -0.0273161 | 0.58713535 |
| 0.999490048 | -0.0075582 | 0.85340917 | 0.97192495 | -0.0025451 | 0.95037323 |
| 0.999490048 | 0.00821322 | 0.85302618 | 0.97192495 | 0.02696348 | 0.54441835 |
| 0.999490048 | 0.00932335 | 0.85292951 | 0.97192495 | 0.00312428 | 0.95045037 |
| 0.999490048 | 0.01243331 | 0.85374265 | 0.97192495 | 0.06573307 | 0.33416306 |
| 0.999490048 | 0.01327735 | 0.85401358 | 0.97192495 | 0.01740628 | 0.80943112 |
| 0.999490048 | 0.01687175 | 0.85348883 | 0.97192495 | 0.15146182 | 0.10633975 |
| 0.999490048 | 0.01892453 | 0.85282389 | 0.97192495 | 0.172938 | 0.0990514 |
| 0.999490048 | 0.02104369 | 0.85388137 | 0.97192495 | -0.0536011 | 0.63962529 |
| 0.999490048 | -0.007288 | 0.85420699 | 0.97201258 | 0.02067485 | 0.60304019 |
| 0.999490048 | 0.01491322 | 0.85434341 | 0.97203536 | 0.17543288 | 0.03867458 |
| 0.999490048 | -0.02473 | 0.8545496 | 0.97211317 | -0.2050441 | 0.13711785 |
| 0.999490048 | -0.0107677 | 0.85511033 | 0.97211317 | 0.02810702 | 0.6342768 |
| 0.999490048 | -0.0097733 | 0.85506519 | 0.97211317 | 0.05816274 | 0.2824941 |
| 0.999490048 | -0.009044 | 0.85493368 | 0.97211317 | -0.0237644 | 0.6316206 |
| 0.999490048 | 0.01000746 | 0.85482747 | 0.97211317 | 0.02376915 | 0.66437399 |
| 0.999490048 | 0.01012565 | 0.85504464 | 0.97211317 | 0.00904503 | 0.87036069 |
| 0.999490048 | 0.02376035 | 0.85525275 | 0.97214272 | -0.1545693 | 0.25486299 |

| | | | | | |
|---|---|---|---|---|---|
| 0.999490048 | 0.00796563 | 0.85560336 | 0.97223909 | 0.02691885 | 0.53998961 |
| 0.999490048 | 0.01295757 | 0.85568684 | 0.97223909 | 0.00520553 | 0.94173986 |
| 0.999490048 | 0.02027869 | 0.85561945 | 0.97223909 | 0.07259704 | 0.51644633 |
| 0.999490048 | -0.0215502 | 0.85600997 | 0.97230807 | 0.0398393 | 0.73748761 |
| 0.999490048 | -0.0190361 | 0.8559261 | 0.97230807 | 0.06267266 | 0.55129892 |
| 0.999490048 | -0.011486 | 0.85672404 | 0.97230807 | 0.01283588 | 0.84011419 |
| 0.999490048 | -0.0104893 | 0.85679554 | 0.97230807 | -0.0343577 | 0.55572624 |
| 0.999490048 | -0.00965 | 0.85652987 | 0.97230807 | -0.0904705 | 0.09911968 |
| 0.999490048 | -0.0089556 | 0.85623671 | 0.97230807 | -0.020301 | 0.68172225 |
| 0.999490048 | -0.0086362 | 0.85668516 | 0.97230807 | -0.0579278 | 0.23235172 |
| 0.999490048 | 0.00610288 | 0.85673913 | 0.97230807 | 0.01562945 | 0.64445063 |
| 0.999490048 | 0.04877843 | 0.85621549 | 0.97230807 | -0.4447008 | 0.12592878 |
| 0.999490048 | 0.02238469 | 0.85696326 | 0.97236625 | 0.05844754 | 0.63855258 |
| 0.999490048 | -0.0354068 | 0.85784127 | 0.9725638 | 0.11746007 | 0.5536393 |
| 0.999490048 | -0.0149842 | 0.85791935 | 0.9725638 | -0.094337 | 0.26555614 |
| 0.999490048 | -0.0079867 | 0.85746257 | 0.9725638 | -8.00E-04 | 0.98564669 |
| 0.999490048 | 0.00746091 | 0.85812748 | 0.9725638 | -0.0046508 | 0.91125794 |
| 0.999490048 | 0.01034445 | 0.85780016 | 0.9725638 | 0.03431366 | 0.55356408 |
| 0.999490048 | 0.0105322 | 0.85807707 | 0.9725638 | -0.0744383 | 0.2133061 |
| 0.999490048 | 0.01254737 | 0.85739046 | 0.9725638 | -0.0204338 | 0.7699161 |
| 0.999490048 | 0.01663837 | 0.85788383 | 0.9725638 | 0.01751376 | 0.85049778 |
| 0.999490048 | 0.01701417 | 0.85818564 | 0.9725638 | 0.04282938 | 0.65341276 |
| 0.999490048 | -0.0156552 | 0.85865305 | 0.97296146 | 0.10075631 | 0.25774891 |
| 0.999490048 | 0.01896758 | 0.858822 | 0.97302085 | 0.00967193 | 0.92771287 |
| 0.999490048 | 0.01059209 | 0.85922417 | 0.97334443 | 0.01705039 | 0.77536813 |
| 0.999490048 | -0.0244975 | 0.85964982 | 0.97340742 | -0.1913754 | 0.17505174 |
| 0.999490048 | -0.0188762 | 0.85986265 | 0.97340742 | -0.031045 | 0.77165748 |
| 0.999490048 | -0.0128589 | 0.85960906 | 0.97340742 | 0.03969547 | 0.58606905 |
| 0.999490048 | -0.0083701 | 0.85980898 | 0.97340742 | -0.0349389 | 0.46326968 |
| 0.999490048 | -0.0068384 | 0.8598318 | 0.97340742 | -0.006907 | 0.85844087 |
| 0.999490048 | -0.0082126 | 0.86004943 | 0.97348688 | -0.0257203 | 0.58186905 |
| 0.999490048 | -0.0168737 | 0.86166818 | 0.97354239 | -0.0310625 | 0.7485579 |
| 0.999490048 | -0.0109187 | 0.86071328 | 0.97354239 | 0.00439954 | 0.94362042 |
| 0.999490048 | -0.010732 | 0.86127167 | 0.97354239 | 0.03494946 | 0.57042337 |
| 0.999490048 | -0.0100699 | 0.86096264 | 0.97354239 | 0.04801723 | 0.40674774 |
| 0.999490048 | -0.0098078 | 0.86068782 | 0.97354239 | 0.00681808 | 0.90288212 |
| 0.999490048 | -0.0091597 | 0.86155644 | 0.97354239 | -0.0246983 | 0.63883101 |
| 0.999490048 | -0.0085242 | 0.86046534 | 0.97354239 | -0.0305454 | 0.53027923 |
| 0.999490048 | -0.0081522 | 0.86056321 | 0.97354239 | -0.0326992 | 0.4831199 |
| 0.999490048 | -0.0055569 | 0.86184735 | 0.97354239 | -0.0049449 | 0.87692567 |
| 0.999490048 | 0.00804153 | 0.86153651 | 0.97354239 | 0.02115853 | 0.64688758 |
| 0.999490048 | 0.01372663 | 0.86146214 | 0.97354239 | 0.02661638 | 0.73528796 |

| | | | | | |
|---|---|---|---|---|---|
| 0.999490048 | 0.01874762 | 0.86088844 | 0.97354239 | 0.04161175 | 0.69765124 |
| 0.999490048 | 0.02072671 | 0.86131942 | 0.97354239 | -2.94E-04 | 0.99802443 |
| 0.999490048 | 0.0210264 | 0.86157584 | 0.97354239 | 0.08356308 | 0.49028495 |
| 0.999490048 | 0.02703527 | 0.86181328 | 0.97354239 | -0.0146957 | 0.92460171 |
| 0.999490048 | 0.01004303 | 0.86201627 | 0.9736015 | -0.0076859 | 0.89416838 |
| 0.999490048 | -0.0066976 | 0.86252466 | 0.97404394 | -0.0490347 | 0.21195256 |
| 0.999490048 | -0.0268306 | 0.86473623 | 0.97430224 | -0.1246312 | 0.43153185 |
| 0.999490048 | -0.0230086 | 0.86448952 | 0.97430224 | -0.0924863 | 0.49462619 |
| 0.999490048 | -0.0141795 | 0.86734199 | 0.97430224 | -0.0821838 | 0.33735472 |
| 0.999490048 | -0.0141272 | 0.86663612 | 0.97430224 | -0.0071364 | 0.9323811 |
| 0.999490048 | -0.0139193 | 0.86437397 | 0.97430224 | 0.07474825 | 0.36289165 |
| 0.999490048 | -0.0127393 | 0.86720205 | 0.97430224 | -0.0691146 | 0.36810949 |
| 0.999490048 | -0.0122219 | 0.86602771 | 0.97430224 | 0.00150597 | 0.98341019 |
| 0.999490048 | -0.0118628 | 0.86766552 | 0.97430224 | 0.0329066 | 0.64455571 |
| 0.999490048 | -0.0116805 | 0.86345626 | 0.97430224 | 0.0189804 | 0.77999977 |
| 0.999490048 | -0.0113512 | 0.86583659 | 0.97430224 | -0.059255 | 0.38137288 |
| 0.999490048 | -0.0112339 | 0.86835416 | 0.97430224 | -0.092362 | 0.18073145 |
| 0.999490048 | -0.011099 | 0.86389063 | 0.97430224 | 0.01147877 | 0.85928391 |
| 0.999490048 | -0.0108905 | 0.86635912 | 0.97430224 | -0.0270105 | 0.67685094 |
| 0.999490048 | -0.0108184 | 0.86785374 | 0.97430224 | 0.01491445 | 0.81861725 |
| 0.999490048 | -0.0107635 | 0.86569287 | 0.97430224 | -0.0269245 | 0.67270596 |
| 0.999490048 | -0.0104603 | 0.8675373 | 0.97430224 | 0.0427222 | 0.49763098 |
| 0.999490048 | -0.0104032 | 0.86725564 | 0.97430224 | 0.045133 | 0.47058211 |
| 0.999490048 | -0.0102337 | 0.86768411 | 0.97430224 | -0.0170594 | 0.78133371 |
| 0.999490048 | -0.0101474 | 0.86610309 | 0.97430224 | -0.0322508 | 0.5930023 |
| 0.999490048 | -0.0095883 | 0.86422857 | 0.97430224 | 0.04856187 | 0.38988653 |
| 0.999490048 | -0.0092412 | 0.8647604 | 0.97430224 | 0.0076529 | 0.88782458 |
| 0.999490048 | -0.0089192 | 0.86719525 | 0.97430224 | 0.00155029 | 0.97680592 |
| 0.999490048 | -0.0082901 | 0.8655639 | 0.97430224 | -0.0458238 | 0.35344586 |
| 0.999490048 | -0.0082157 | 0.86807454 | 0.97430224 | 9.98E-06 | 0.99983893 |
| 0.999490048 | 0.00700441 | 0.86391579 | 0.97430224 | -0.0080085 | 0.84465435 |
| 0.999490048 | 0.00730867 | 0.86782608 | 0.97430224 | 0.01655687 | 0.70649351 |
| 0.999490048 | 0.00780825 | 0.86599529 | 0.97430224 | 0.0361766 | 0.43699279 |
| 0.999490048 | 0.0080202 | 0.86648007 | 0.97430224 | -0.0053844 | 0.91011484 |
| 0.999490048 | 0.00877809 | 0.86755307 | 0.97430224 | 0.03807548 | 0.4716604 |
| 0.999490048 | 0.0089502 | 0.86811837 | 0.97430224 | -0.0031193 | 0.9538404 |
| 0.999490048 | 0.00910434 | 0.86663347 | 0.97430224 | 0.03549983 | 0.51427718 |
| 0.999490048 | 0.0092007 | 0.8639983 | 0.97430224 | -0.0310313 | 0.56465912 |
| 0.999490048 | 0.00947426 | 0.86436526 | 0.97430224 | -0.0594601 | 0.28904743 |
| 0.999490048 | 0.0102117 | 0.86767782 | 0.97430224 | 0.04956442 | 0.42161712 |
| 0.999490048 | 0.0105703 | 0.86707033 | 0.97430224 | 0.0287456 | 0.64956524 |
| 0.999490048 | 0.01100764 | 0.86406896 | 0.97430224 | 0.07678994 | 0.23883034 |

| | | | | | |
|---|---|---|---|---|---|
| 0.999490048 | 0.01137821 | 0.8666493 | 0.97430224 | -0.06308 | 0.35591918 |
| 0.999490048 | 0.01143249 | 0.86760586 | 0.97430224 | 0.00990046 | 0.8852071 |
| 0.999490048 | 0.01154789 | 0.86792705 | 0.97430224 | 0.07719562 | 0.27202587 |
| 0.999490048 | 0.01157435 | 0.8630932 | 0.97430224 | 0.03900857 | 0.5623437 |
| 0.999490048 | 0.0119349 | 0.86430765 | 0.97430224 | 0.03005404 | 0.6674419 |
| 0.999490048 | 0.01220332 | 0.86349865 | 0.97430224 | 0.04487013 | 0.52883042 |
| 0.999490048 | 0.01356262 | 0.86831031 | 0.97430224 | 0.02594523 | 0.75128125 |
| 0.999490048 | 0.01764252 | 0.86588924 | 0.97430224 | -0.0090793 | 0.93072714 |
| 0.999490048 | 0.017941 | 0.86546006 | 0.97430224 | 0.10079916 | 0.34536092 |
| 0.999490048 | 0.02306869 | 0.86714196 | 0.97430224 | 0.04183722 | 0.76173774 |
| 0.999490048 | 0.03794618 | 0.86822645 | 0.97430224 | 0.27221187 | 0.24040644 |
| 0.999490048 | 0.06534942 | 0.86535342 | 0.97430224 | 0.04305514 | 0.91103397 |
| 0.999490048 | -0.0161743 | 0.8690023 | 0.97459071 | 0.0283467 | 0.77267212 |
| 0.999490048 | -0.0146313 | 0.8700071 | 0.97459071 | 2.40E-04 | 0.99785452 |
| 0.999490048 | -0.0117893 | 0.87059547 | 0.97459071 | -0.0105247 | 0.88436611 |
| 0.999490048 | -0.0111077 | 0.8699981 | 0.97459071 | -0.0991302 | 0.15247414 |
| 0.999490048 | -0.0074738 | 0.87055124 | 0.97459071 | -0.012228 | 0.78985519 |
| 0.999490048 | -0.0074579 | 0.87044478 | 0.97459071 | -0.0540486 | 0.24358535 |
| 0.999490048 | -0.0063244 | 0.86972673 | 0.97459071 | 0.03808784 | 0.32786258 |
| 0.999490048 | 0.00696584 | 0.86969763 | 0.97459071 | -0.0432395 | 0.31340608 |
| 0.999490048 | 0.00726153 | 0.87032057 | 0.97459071 | -0.0264715 | 0.55306975 |
| 0.999490048 | 0.00820998 | 0.86889799 | 0.97459071 | 0.0149865 | 0.76333487 |
| 0.999490048 | 0.0088122 | 0.86942444 | 0.97459071 | 0.00556617 | 0.91728626 |
| 0.999490048 | 0.00883918 | 0.87033176 | 0.97459071 | 0.01502717 | 0.78149424 |
| 0.999490048 | 0.00949397 | 0.86974132 | 0.97459071 | 0.11710595 | 0.05161553 |
| 0.999490048 | 0.00984762 | 0.86878307 | 0.97459071 | -0.0263823 | 0.65860445 |
| 0.999490048 | 0.02034829 | 0.86950074 | 0.97459071 | 0.0525356 | 0.67191553 |
| 0.999490048 | 0.02166536 | 0.87011424 | 0.97459071 | 0.10439994 | 0.43347463 |
| 0.999490048 | 0.09377634 | 0.87040126 | 0.97459071 | 0.07545737 | 0.89553262 |
| 0.999490048 | -0.0096886 | 0.87089503 | 0.97466473 | -0.0407619 | 0.49603573 |
| 0.999490048 | -0.0070962 | 0.87087484 | 0.97466473 | -0.0090339 | 0.83608852 |
| 0.999490048 | -0.0148895 | 0.87142973 | 0.97490986 | -0.0800218 | 0.38787087 |
| 0.999490048 | -0.0061188 | 0.87155119 | 0.97490986 | -0.0101603 | 0.78842169 |
| 0.999490048 | 0.00867299 | 0.87158109 | 0.97490986 | -0.0724761 | 0.18444095 |
| 0.999490048 | 0.01076221 | 0.87130874 | 0.97490986 | -0.0544639 | 0.41534029 |
| 0.999490048 | -0.0134245 | 0.87193908 | 0.97491849 | -0.0545941 | 0.51382522 |
| 0.999490048 | -0.0047145 | 0.87177795 | 0.97491849 | -0.043267 | 0.14699261 |
| 0.999490048 | 0.0101497 | 0.8718703 | 0.97491849 | -0.0449026 | 0.47765769 |
| 0.999490048 | -0.0179095 | 0.87219669 | 0.97494544 | -0.0506462 | 0.6497634 |
| 0.999490048 | 0.01412071 | 0.87214983 | 0.97494544 | 0.10235173 | 0.24966125 |
| 0.999490048 | -0.014698 | 0.87248571 | 0.975114 | -0.0459874 | 0.61634311 |
| 0.999490048 | -0.0107596 | 0.87328174 | 0.975114 | 0.04189109 | 0.53610449 |

| | | | | | |
|---|---|---|---|---|---|
| 0.999490048 | -0.0093854 | 0.87315933 | 0.975114 | 0.03648181 | 0.53636312 |
| 0.999490048 | 0.00719777 | 0.87275209 | 0.975114 | 0.04775014 | 0.29327733 |
| 0.999490048 | 0.00858093 | 0.87327702 | 0.975114 | 0.03118087 | 0.56342215 |
| 0.999490048 | 0.00865009 | 0.87284328 | 0.975114 | -0.0714799 | 0.19348148 |
| 0.999490048 | 0.0165702 | 0.87278023 | 0.975114 | 0.05946048 | 0.56674886 |
| 0.999490048 | 0.01672288 | 0.87306895 | 0.975114 | 0.08028344 | 0.45110028 |
| 0.999490048 | -0.0117833 | 0.87356246 | 0.97516666 | 0.01516459 | 0.83775563 |
| 0.999490048 | -0.0074504 | 0.87355817 | 0.97516666 | -0.0031127 | 0.94697869 |
| 0.999490048 | -0.1086312 | 0.87429855 | 0.97556025 | -1.0900902 | 0.13936476 |
| 0.999490048 | -0.0083558 | 0.87461605 | 0.97556025 | -0.0566262 | 0.29025203 |
| 0.999490048 | -0.0081273 | 0.87446038 | 0.97556025 | -0.0134347 | 0.79404703 |
| 0.999490048 | 0.00816209 | 0.87461351 | 0.97556025 | -0.0605714 | 0.24785733 |
| 0.999490048 | 0.00840966 | 0.87450375 | 0.97556025 | 0.01721973 | 0.74658843 |
| 0.999490048 | 0.01234213 | 0.87450227 | 0.97556025 | -0.0343102 | 0.66114981 |
| 0.999490048 | 0.01753667 | 0.87479406 | 0.97562847 | 0.05150675 | 0.64413803 |
| 0.999490048 | 0.00984792 | 0.87504843 | 0.97565154 | 0.00696109 | 0.91148401 |
| 0.999490048 | 0.01855136 | 0.87503375 | 0.97565154 | 0.04582837 | 0.69800671 |
| 0.999490048 | 0.00737816 | 0.87518585 | 0.97567448 | -0.0195976 | 0.67697909 |
| 0.999490048 | -0.0276212 | 0.87633147 | 0.97582805 | 0.18326369 | 0.31724045 |
| 0.999490048 | -0.0152735 | 0.8759706 | 0.97582805 | 0.00620107 | 0.94946276 |
| 0.999490048 | -0.0072461 | 0.87620968 | 0.97582805 | 0.010323 | 0.82441779 |
| 0.999490048 | 0.00685643 | 0.87660839 | 0.97582805 | -4.03E-06 | 0.9999272 |
| 0.999490048 | 0.00812495 | 0.87595923 | 0.97582805 | -0.0334215 | 0.52244196 |
| 0.999490048 | 0.00819646 | 0.87544291 | 0.97582805 | -0.0014318 | 0.97814972 |
| 0.999490048 | 0.00859742 | 0.87635148 | 0.97582805 | -0.0406417 | 0.46433041 |
| 0.999490048 | 0.00981317 | 0.87623658 | 0.97582805 | 0.04926826 | 0.43695834 |
| 0.999490048 | 0.01427674 | 0.87642807 | 0.97582805 | 0.15562093 | 0.09914478 |
| 0.999490048 | 0.01474111 | 0.87660913 | 0.97582805 | -0.1788055 | 0.06860043 |
| 0.999490048 | 0.03556198 | 0.87634154 | 0.97582805 | 0.31677738 | 0.18345854 |
| 0.999490048 | -0.0082293 | 0.8767481 | 0.97585265 | 0.02011308 | 0.70498147 |
| 0.999490048 | -0.0128331 | 0.87695003 | 0.97594732 | -0.0974679 | 0.24592417 |
| 0.999490048 | 0.00955602 | 0.87712185 | 0.97600845 | -0.0880808 | 0.16227423 |
| 0.999490048 | -0.0070576 | 0.87728853 | 0.97606385 | 0.05650104 | 0.22324421 |
| 0.999490048 | -0.029612 | 0.87789331 | 0.97625432 | 0.06413532 | 0.73953431 |
| 0.999490048 | -0.0070725 | 0.87792738 | 0.97625432 | 0.0375407 | 0.41789395 |
| 0.999490048 | 0.00880856 | 0.8775987 | 0.97625432 | 0.03812142 | 0.50686307 |
| 0.999490048 | 0.01028656 | 0.87771307 | 0.97625432 | 0.01413303 | 0.83260602 |
| 0.999490048 | -0.0079229 | 0.87843924 | 0.97669342 | 0.01411449 | 0.78536605 |
| 0.999490048 | 0.00889069 | 0.87863265 | 0.97677841 | 0.05955034 | 0.31131374 |
| 0.999490048 | 0.00764703 | 0.87885544 | 0.97689602 | 0.01638334 | 0.7442145 |
| 0.999490048 | 0.01314536 | 0.87918574 | 0.9771331 | -0.0051241 | 0.9527435 |
| 0.999490048 | -0.0238791 | 0.87936895 | 0.97714893 | 0.08460643 | 0.59175204 |

| | | | | | |
|---|---|---|---|---|---|
| 0.999490048 | -0.0127673 | 0.87943404 | 0.97714893 | 0.07452774 | 0.37953352 |
| 0.999490048 | 0.00687356 | 0.87960391 | 0.97720765 | 0.00400834 | 0.92960285 |
| 0.999490048 | -0.1654401 | 0.8799436 | 0.97732496 | -0.0949449 | 0.93089087 |
| 0.999490048 | 0.00844053 | 0.87988744 | 0.97732496 | 0.03025857 | 0.58902477 |
| 0.999490048 | 0.00920649 | 0.88016794 | 0.97744412 | -0.0819705 | 0.18715862 |
| 0.999490048 | -0.006718 | 0.88044836 | 0.97752431 | 0.00108284 | 0.98065518 |
| 0.999490048 | 0.01054324 | 0.88059136 | 0.97752431 | -0.0043858 | 0.95016431 |
| 0.999490048 | 0.01122122 | 0.88056016 | 0.97752431 | 0.09710152 | 0.20086354 |
| 0.999490048 | 0.01091726 | 0.88085934 | 0.97756366 | 0.00472802 | 0.94823641 |
| 0.999490048 | 0.01300213 | 0.88086096 | 0.97756366 | 0.00154323 | 0.98580392 |
| 0.999490048 | -0.0059883 | 0.88123366 | 0.97784732 | -0.0348594 | 0.38792869 |
| 0.999490048 | 0.00847774 | 0.88136014 | 0.97785771 | 0.08219677 | 0.15618286 |
| 0.999490048 | -0.0312165 | 0.8816796 | 0.97803553 | -0.042642 | 0.83891799 |
| 0.999490048 | -0.0192637 | 0.88292597 | 0.97803553 | 0.02889353 | 0.82523564 |
| 0.999490048 | -0.008632 | 0.88288093 | 0.97803553 | 0.00760048 | 0.89678819 |
| 0.999490048 | -0.0081307 | 0.88232505 | 0.97803553 | 0.01536498 | 0.77981151 |
| 0.999490048 | -0.0063287 | 0.88240219 | 0.97803553 | -0.0183907 | 0.66781493 |
| 0.999490048 | 0.00787421 | 0.88245381 | 0.97803553 | 0.03572305 | 0.50418474 |
| 0.999490048 | 0.00794365 | 0.88199443 | 0.97803553 | 0.05644034 | 0.29679236 |
| 0.999490048 | 0.00894581 | 0.88196623 | 0.97803553 | 0.07795435 | 0.20302322 |
| 0.999490048 | 0.00935996 | 0.88246058 | 0.97803553 | -0.0010138 | 0.98721987 |
| 0.999490048 | 0.01797609 | 0.88270283 | 0.97803553 | 0.19061065 | 0.14448322 |
| 0.999490048 | 0.01891425 | 0.8825408 | 0.97803553 | 0.19285431 | 0.14052151 |
| 0.999490048 | 0.02006403 | 0.88278636 | 0.97803553 | -0.0159198 | 0.90686467 |
| 0.999490048 | -0.0129175 | 0.88361629 | 0.97835024 | -0.0083337 | 0.92474963 |
| 0.999490048 | -0.0058936 | 0.88358291 | 0.97835024 | -5.36E-04 | 0.98937617 |
| 0.999490048 | 0.00724234 | 0.88367874 | 0.97835024 | 7.16E-04 | 0.98845103 |
| 0.999490048 | 0.02801814 | 0.88348353 | 0.97835024 | -0.1002377 | 0.60299374 |
| 0.889155227 | 0.02783566 | 0.8839748 | 0.97854827 | -0.5231466 | 0.02177441 |
| 0.999490048 | -0.0404675 | 0.88423901 | 0.97858127 | 0.04648174 | 0.86721285 |
| 0.999490048 | -0.0080794 | 0.88420934 | 0.97858127 | 0.01094807 | 0.84358649 |
| 0.999490048 | -0.0066017 | 0.88464908 | 0.97890535 | -0.0297268 | 0.5152917 |
| 0.999490048 | -0.0390728 | 0.88557635 | 0.97910152 | 0.0528814 | 0.84566495 |
| 0.999490048 | -0.0257463 | 0.88519623 | 0.97910152 | 0.20857259 | 0.24841722 |
| 0.999490048 | -0.0158348 | 0.885598 | 0.97910152 | -0.1676055 | 0.13643907 |
| 0.999490048 | -0.010061 | 0.88558018 | 0.97910152 | 0.05959281 | 0.39735956 |
| 0.999490048 | -0.009878 | 0.88584268 | 0.97910152 | 0.07034783 | 0.31150199 |
| 0.999490048 | -0.005396 | 0.88497032 | 0.97910152 | -0.0294845 | 0.4320216 |
| 0.999490048 | 0.00703829 | 0.88507082 | 0.97910152 | -0.0396532 | 0.41843609 |
| 0.999490048 | 0.00916442 | 0.88599893 | 0.97910152 | 0.06848814 | 0.28936485 |
| 0.999490048 | 0.009439 | 0.88596472 | 0.97910152 | 0.06965364 | 0.29519052 |
| 0.999490048 | 0.01277707 | 0.88589783 | 0.97910152 | 0.03066665 | 0.73079301 |

| | | | | | |
|---|---|---|---|---|---|
| 0.999490048 | -0.0311617 | 0.88724548 | 0.97912875 | -0.2738202 | 0.22985759 |
| 0.999490048 | -0.0244568 | 0.88821354 | 0.97912875 | 0.04948001 | 0.77642393 |
| 0.999490048 | -0.0133407 | 0.8881061 | 0.97912875 | 0.03851672 | 0.68501373 |
| 0.999490048 | -0.0126218 | 0.88634022 | 0.97912875 | 0.11175971 | 0.21273513 |
| 0.999490048 | -0.0074221 | 0.88793921 | 0.97912875 | -0.0264519 | 0.61635277 |
| 0.999490048 | -0.0069131 | 0.88733741 | 0.97912875 | 0.04557257 | 0.35441867 |
| 0.999490048 | -0.0064728 | 0.88682221 | 0.97912875 | -0.0490761 | 0.28600508 |
| 0.999490048 | 0.00699545 | 0.88818925 | 0.97912875 | 0.02041228 | 0.68207304 |
| 0.999490048 | 0.00716503 | 0.88770821 | 0.97912875 | -0.0346682 | 0.49639889 |
| 0.999490048 | 0.0072225 | 0.88669413 | 0.97912875 | -0.0180336 | 0.72229525 |
| 0.999490048 | 0.00767644 | 0.88713563 | 0.97912875 | -0.0020097 | 0.97035429 |
| 0.999490048 | 0.0089246 | 0.8864384 | 0.97912875 | 0.05383794 | 0.39236735 |
| 0.999490048 | 0.01036089 | 0.88826339 | 0.97912875 | 0.04107963 | 0.57858903 |
| 0.999490048 | 0.01097144 | 0.88842381 | 0.97912875 | 0.06932905 | 0.37896666 |
| 0.999490048 | 0.01182242 | 0.8883286 | 0.97912875 | 0.10046249 | 0.23927554 |
| 0.999490048 | 0.0142217 | 0.88848605 | 0.97912875 | -0.0218004 | 0.82985635 |
| 0.999490048 | 0.01436644 | 0.88725776 | 0.97912875 | -0.0132134 | 0.89624952 |
| 0.999490048 | 0.01626323 | 0.88776964 | 0.97912875 | 0.05132332 | 0.6566328 |
| 0.999490048 | 0.01991875 | 0.88762227 | 0.97912875 | 0.10307376 | 0.47181626 |
| 0.999490048 | 0.02737915 | 0.8868403 | 0.97912875 | -0.0419686 | 0.8274758 |
| 0.999490048 | 0.0680403 | 0.88751812 | 0.97912875 | 0.15127104 | 0.75335468 |
| 0.999490048 | -0.0172246 | 0.88964743 | 0.97916927 | -0.0549745 | 0.65845191 |
| 0.999490048 | -0.0167824 | 0.88930179 | 0.97916927 | -0.0491094 | 0.68423141 |
| 0.999490048 | -0.0145373 | 0.88872998 | 0.97916927 | -0.0441988 | 0.67106111 |
| 0.999490048 | -0.0101042 | 0.8898005 | 0.97916927 | -0.1240318 | 0.09806605 |
| 0.999490048 | -0.0079187 | 0.88945487 | 0.97916927 | -0.0074978 | 0.89529375 |
| 0.999490048 | -0.0071094 | 0.88936958 | 0.97916927 | -0.0456308 | 0.3756691 |
| 0.999490048 | 0.00645713 | 0.88954354 | 0.97916927 | 0.00102971 | 0.98232715 |
| 0.999490048 | 0.00906991 | 0.88927554 | 0.97916927 | -0.0181253 | 0.78096992 |
| 0.999490048 | 0.00922593 | 0.88993001 | 0.97916927 | -0.0198792 | 0.76571395 |
| 0.999490048 | 0.01214395 | 0.88992529 | 0.97916927 | -0.0341456 | 0.69805243 |
| 0.999490048 | 0.01564075 | 0.88984016 | 0.97916927 | -0.1702426 | 0.14025543 |
| 0.999490048 | 0.02914483 | 0.88898692 | 0.97916927 | 0.05563217 | 0.79000342 |
| 0.999490048 | -0.0124073 | 0.8902314 | 0.97934774 | 0.13240373 | 0.14924448 |
| 0.999490048 | -0.0082593 | 0.89032679 | 0.97934774 | -0.1124171 | 0.0695196 |
| 0.999490048 | 0.00662226 | 0.89053884 | 0.97945196 | 0.02783606 | 0.56417609 |
| 0.999490048 | -0.0470154 | 0.89333389 | 0.97964694 | 0.03002885 | 0.93172797 |
| 0.999490048 | -0.0331492 | 0.89652371 | 0.97964694 | 0.29573307 | 0.25216947 |
| 0.999490048 | -0.0231382 | 0.89663828 | 0.97964694 | -0.0501726 | 0.77861206 |
| 0.999490048 | -0.0217297 | 0.89630634 | 0.97964694 | -0.1682046 | 0.31786394 |
| 0.999490048 | -0.0175744 | 0.89208333 | 0.97964694 | -0.0218441 | 0.86610158 |
| 0.999490048 | -0.0164933 | 0.89183 | 0.97964694 | -0.0274157 | 0.82122433 |

| | | | | | |
|---|---|---|---|---|---|
| 0.999490048 | -0.0142082 | 0.89365203 | 0.97964694 | 0.01369935 | 0.89743821 |
| 0.999490048 | -0.0140483 | 0.89669959 | 0.97964694 | -0.1406699 | 0.20095812 |
| 0.999490048 | -0.0138399 | 0.89542574 | 0.97964694 | 0.02719775 | 0.79627111 |
| 0.999490048 | -0.0138284 | 0.89625361 | 0.97964694 | -0.0072382 | 0.94557795 |
| 0.999490048 | -0.011989 | 0.89407638 | 0.97964694 | 0.0761317 | 0.40111142 |
| 0.999490048 | -0.0107021 | 0.89555003 | 0.97964694 | 3.08E-05 | 0.99969833 |
| 0.999490048 | -0.0089673 | 0.89284817 | 0.97964694 | -0.020352 | 0.75999715 |
| 0.999490048 | -0.0088255 | 0.89528622 | 0.97964694 | -0.032493 | 0.6287256 |
| 0.999490048 | -0.0084913 | 0.89329292 | 0.97964694 | 0.01278216 | 0.8400129 |
| 0.999490048 | -0.0082862 | 0.89265771 | 0.97964694 | -0.0161598 | 0.79252749 |
| 0.999490048 | -0.0080181 | 0.8965378 | 0.97964694 | 0.00419677 | 0.94572893 |
| 0.999490048 | -0.0077525 | 0.89506388 | 0.97964694 | -0.0481069 | 0.4161288 |
| 0.999490048 | -0.0075626 | 0.89449866 | 0.97964694 | -0.0238597 | 0.67614507 |
| 0.999490048 | -0.0075617 | 0.89412982 | 0.97964694 | 0.00489281 | 0.93137196 |
| 0.999490048 | -0.0074271 | 0.89302475 | 0.97964694 | 0.03260113 | 0.55631101 |
| 0.999490048 | -0.0072493 | 0.893283 | 0.97964694 | -0.0315496 | 0.56060162 |
| 0.999490048 | -0.0072002 | 0.89091435 | 0.97964694 | -0.0214571 | 0.6832009 |
| 0.999490048 | -0.0071165 | 0.89432156 | 0.97964694 | 0.00236456 | 0.96478943 |
| 0.999490048 | -0.0069145 | 0.89346722 | 0.97964694 | -0.0033304 | 0.94856274 |
| 0.999490048 | -0.006773 | 0.89572524 | 0.97964694 | -0.0114848 | 0.82418303 |
| 0.913389923 | -0.0065785 | 0.89096661 | 0.97964694 | 0.11153154 | 0.02705808 |
| 0.999490048 | -0.0063973 | 0.8961003 | 0.97964694 | 0.05050841 | 0.30755083 |
| 0.999490048 | -0.0061895 | 0.89141034 | 0.97964694 | 0.02108934 | 0.64248094 |
| 0.999490048 | -0.0061558 | 0.89420992 | 0.97964694 | -0.0633046 | 0.17930555 |
| 0.999490048 | -0.006149 | 0.89244472 | 0.97964694 | 0.01476025 | 0.7457253 |
| 0.999490048 | -0.006054 | 0.89385495 | 0.97964694 | -0.0774372 | 0.09697794 |
| 0.999490048 | -0.006046 | 0.89339911 | 0.97964694 | -0.0474194 | 0.29846077 |
| 0.999490048 | -0.0052496 | 0.89360634 | 0.97964694 | -0.0578867 | 0.14872845 |
| 0.999490048 | 0.00513771 | 0.89287061 | 0.97964694 | -7.63E-04 | 0.98403312 |
| 0.999490048 | 0.00562001 | 0.89319619 | 0.97964694 | 0.01033513 | 0.80506509 |
| 0.999490048 | 0.00603816 | 0.89225076 | 0.97964694 | -0.0122477 | 0.78362368 |
| 0.999490048 | 0.00618968 | 0.89519324 | 0.97964694 | -0.0549128 | 0.24881172 |
| 0.999490048 | 0.00644799 | 0.89307731 | 0.97964694 | -0.0123268 | 0.79730788 |
| 0.999490048 | 0.006836 | 0.89440458 | 0.97964694 | -0.0112398 | 0.8272929 |
| 0.999490048 | 0.00932047 | 0.89382987 | 0.97964694 | -0.0157941 | 0.8211395 |
| 0.999490048 | 0.00945892 | 0.89618656 | 0.97964694 | -0.0437143 | 0.54789569 |
| 0.999490048 | 0.00977757 | 0.89500193 | 0.97964694 | -0.0248667 | 0.73737679 |
| 0.999490048 | 0.01008839 | 0.89163503 | 0.97964694 | 0.06930688 | 0.35860651 |
| 0.999490048 | 0.01059533 | 0.89509207 | 0.97964694 | 0.0367582 | 0.64796637 |
| 0.999490048 | 0.01072403 | 0.8923995 | 0.97964694 | -0.0496791 | 0.53244353 |
| 0.999490048 | 0.0126605 | 0.89627155 | 0.97964694 | 0.07385328 | 0.4495004 |
| 0.999490048 | 0.01310614 | 0.89652208 | 0.97964694 | 0.05713547 | 0.57191194 |

| | | | | | |
|---|---|---|---|---|---|
| 0.999490048 | 0.01706381 | 0.89433829 | 0.97964694 | -0.035383 | 0.78312969 |
| 0.999490048 | 0.02302458 | 0.89498753 | 0.97964694 | 0.00843523 | 0.9614253 |
| 0.999490048 | 0.05138638 | 0.89459311 | 0.97964694 | -0.6253885 | 0.1341161 |
| 0.999490048 | -0.0119064 | 0.89690803 | 0.97974648 | -0.1127993 | 0.22643017 |
| 0.999490048 | -0.0222909 | 0.89743349 | 0.9799337 | 0.10778246 | 0.53462398 |
| 0.999490048 | -0.0112444 | 0.89749276 | 0.9799337 | -0.0594186 | 0.49794742 |
| 0.999490048 | -0.0102476 | 0.89739752 | 0.9799337 | 0.12076038 | 0.13727143 |
| 0.999490048 | 0.04572063 | 0.89754885 | 0.9799337 | -0.0177435 | 0.9601289 |
| 0.999490048 | -0.0063038 | 0.89805531 | 0.9802649 | -0.0250607 | 0.61138899 |
| 0.999490048 | 0.00520938 | 0.898087 | 0.9802649 | 0.01886451 | 0.64345277 |
| 0.999490048 | -0.0086093 | 0.89998456 | 0.98032834 | -0.0195169 | 0.77585476 |
| 0.999490048 | -0.0083462 | 0.89879811 | 0.98032834 | -0.0312531 | 0.63462531 |
| 0.999490048 | -0.0076208 | 0.89992651 | 0.98032834 | 0.03178838 | 0.60084406 |
| 0.999490048 | -0.0073324 | 0.89881293 | 0.98032834 | 0.03892834 | 0.50149766 |
| 0.999490048 | -0.0071955 | 0.89920674 | 0.98032834 | -0.0711645 | 0.21731769 |
| 0.999490048 | -0.0070128 | 0.89987002 | 0.98032834 | -0.0026194 | 0.96251019 |
| 0.999490048 | -0.0067532 | 0.89955381 | 0.98032834 | -0.0579944 | 0.28390165 |
| 0.999490048 | -0.0065323 | 0.9000236 | 0.98032834 | 0.01511019 | 0.7715066 |
| 0.999490048 | -0.0064223 | 0.89991441 | 0.98032834 | 0.00183934 | 0.971262 |
| 0.999490048 | -0.0061741 | 0.89972257 | 0.98032834 | -0.0152678 | 0.75553027 |
| 0.999490048 | -0.0052218 | 0.89855922 | 0.98032834 | 0.00679236 | 0.86831061 |
| 0.999490048 | -0.0051663 | 0.89882797 | 0.98032834 | -0.0250019 | 0.53984192 |
| 0.999490048 | 0.00548139 | 0.89916586 | 0.98032834 | -0.0555826 | 0.20608081 |
| 0.999490048 | 0.00793276 | 0.89892444 | 0.98032834 | 0.02252104 | 0.71870133 |
| 0.999490048 | 0.00860315 | 0.8993704 | 0.98032834 | 0.0741511 | 0.28132475 |
| 0.999490048 | 0.00957172 | 0.89909091 | 0.98032834 | 0.01673041 | 0.82464311 |
| 0.999490048 | 0.00517654 | 0.90023044 | 0.98042574 | -0.0475917 | 0.25522528 |
| 0.999490048 | 0.06956583 | 0.90036867 | 0.98044841 | -0.2948689 | 0.59789832 |
| 0.999490048 | 0.01870384 | 0.90050167 | 0.98046538 | -0.158879 | 0.29354158 |
| 0.999490048 | -0.0326938 | 0.90365716 | 0.98085702 | -0.0536906 | 0.84256008 |
| 0.999490048 | -0.0273174 | 0.90398134 | 0.98085702 | 0.16122506 | 0.4786576 |
| 0.999490048 | -0.0140245 | 0.90100425 | 0.98085702 | 0.01695343 | 0.88048 |
| 0.999490048 | -0.011903 | 0.90267024 | 0.98085702 | -0.0126523 | 0.89657825 |
| 0.999490048 | -0.0111847 | 0.90441537 | 0.98085702 | 0.15616315 | 0.10268702 |
| 0.999490048 | -0.0105504 | 0.90476501 | 0.98085702 | 0.01114861 | 0.89939447 |
| 0.999490048 | -0.009057 | 0.90325455 | 0.98085702 | -0.0267938 | 0.71946491 |
| 0.999490048 | -0.0086599 | 0.9023042 | 0.98085702 | 0.01612274 | 0.81929839 |
| 0.999490048 | -0.0079677 | 0.90259618 | 0.98085702 | -0.0016042 | 0.98033895 |
| 0.999490048 | -0.0078944 | 0.90169285 | 0.98085702 | -0.0401047 | 0.53188189 |
| 0.999490048 | -0.007524 | 0.90336437 | 0.98085702 | 0.07428618 | 0.23719691 |
| 0.999490048 | -0.0072628 | 0.90401625 | 0.98085702 | -0.0205979 | 0.73261706 |
| 0.999490048 | -0.0071197 | 0.90400505 | 0.98085702 | 0.00872244 | 0.88254642 |

| | | | | | |
|---|---|---|---|---|---|
| 0.999490048 | -0.0069885 | 0.90257693 | 0.98085702 | -0.0172802 | 0.76232157 |
| 0.999490048 | -0.0068407 | 0.90438755 | 0.98085702 | -0.0440839 | 0.44154915 |
| 0.999490048 | -0.0063596 | 0.90493535 | 0.98085702 | -0.0396339 | 0.45912791 |
| 0.999490048 | -0.0062953 | 0.90284502 | 0.98085702 | 0.00942706 | 0.85498533 |
| 0.999490048 | -0.0062497 | 0.90423515 | 0.98085702 | -0.0230946 | 0.65720437 |
| 0.999490048 | -0.0056413 | 0.90303164 | 0.98085702 | -0.0219828 | 0.6356781 |
| 0.999490048 | -0.0055141 | 0.90384781 | 0.98085702 | -0.0325089 | 0.47852008 |
| 0.999490048 | -0.0053569 | 0.90468718 | 0.98085702 | -0.0529137 | 0.24332594 |
| 0.999490048 | -0.0051195 | 0.90349897 | 0.98085702 | -0.0792364 | 0.06957176 |
| 0.999490048 | 0.00591416 | 0.90265871 | 0.98085702 | -0.0104736 | 0.82858055 |
| 0.999490048 | 0.00622201 | 0.90429055 | 0.98085702 | -0.0108787 | 0.83353365 |
| 0.999490048 | 0.00670014 | 0.90497276 | 0.98085702 | 0.07877732 | 0.16850644 |
| 0.999490048 | 0.0069913 | 0.90457691 | 0.98085702 | 0.03287612 | 0.57410391 |
| 0.999490048 | 0.00765837 | 0.9047077 | 0.98085702 | 0.01759572 | 0.78340226 |
| 0.999490048 | 0.00766573 | 0.90142738 | 0.98085702 | 0.06330769 | 0.31131623 |
| 0.999490048 | 0.00801969 | 0.90161097 | 0.98085702 | -0.0442855 | 0.49675909 |
| 0.999490048 | 0.00853158 | 0.90464972 | 0.98085702 | 0.04452977 | 0.53336973 |
| 0.999490048 | 0.00895218 | 0.90366063 | 0.98085702 | 0.09411673 | 0.21036362 |
| 0.999490048 | 0.00896706 | 0.90410531 | 0.98085702 | 0.04499278 | 0.5469262 |
| 0.999490048 | 0.01477378 | 0.9035866 | 0.98085702 | 0.07396987 | 0.54561857 |
| 0.999490048 | 0.02443646 | 0.90270494 | 0.98085702 | -0.080574 | 0.6873325 |
| 0.999490048 | 0.02482583 | 0.9045174 | 0.98085702 | 0.02003318 | 0.92288211 |
| 0.999490048 | 0.00780692 | 0.90509213 | 0.98085909 | 0.00443567 | 0.94598428 |
| 0.999490048 | 0.01640939 | 0.90544377 | 0.98111283 | 0.19229537 | 0.17192556 |
| 0.999490048 | 0.01047713 | 0.9056215 | 0.98117808 | 0.04010334 | 0.65140145 |
| 0.999490048 | -0.0082143 | 0.90639519 | 0.98188893 | -0.0423252 | 0.54601945 |
| 0.999490048 | -0.0078417 | 0.90670884 | 0.9819872 | 0.06627875 | 0.32658262 |
| 0.999490048 | 0.00848554 | 0.90672112 | 0.9819872 | 0.00324226 | 0.96428502 |
| 0.999490048 | -0.0184145 | 0.91101374 | 0.98207723 | -0.1891347 | 0.25714824 |
| 0.999490048 | -0.018214 | 0.90863258 | 0.98207723 | -0.176757 | 0.2712178 |
| 0.999490048 | -0.0170537 | 0.90718801 | 0.98207723 | 0.07129254 | 0.62676347 |
| 0.999490048 | -0.0165861 | 0.90946522 | 0.98207723 | 0.07383427 | 0.6154615 |
| 0.999490048 | -0.0130391 | 0.91160007 | 0.98207723 | 0.20133232 | 0.0956064 |
| 0.999490048 | -0.0105665 | 0.90872131 | 0.98207723 | -0.0903063 | 0.33171157 |
| 0.999490048 | -0.0104192 | 0.9074128 | 0.98207723 | 0.02054138 | 0.81871465 |
| 0.999490048 | -0.0088922 | 0.90798914 | 0.98207723 | -0.0968121 | 0.21534556 |
| 0.999490048 | -0.0082086 | 0.91163077 | 0.98207723 | 0.05810031 | 0.43492725 |
| 0.999490048 | -0.0079304 | 0.91002923 | 0.98207723 | -0.0486615 | 0.49010829 |
| 0.771925038 | -0.0072183 | 0.90973767 | 0.98207723 | 0.16856351 | 0.01297709 |
| 0.999490048 | -0.0068901 | 0.91012326 | 0.98207723 | 0.04028539 | 0.5110443 |
| 0.999490048 | -0.0068112 | 0.91099549 | 0.98207723 | -0.0627474 | 0.30814465 |
| 0.999490048 | -0.005957 | 0.90986124 | 0.98207723 | -0.0195899 | 0.71001509 |

| | | | | | |
|---|---|---|---|---|---|
| 0.999490048 | -0.0058378 | 0.91167312 | 0.98207723 | -0.0448871 | 0.39707478 |
| 0.999490048 | -0.0054122 | 0.90733527 | 0.98207723 | 0.02057198 | 0.65875404 |
| 0.999490048 | -0.0051913 | 0.90845541 | 0.98207723 | 0.01995103 | 0.65913697 |
| 0.999490048 | -0.0047575 | 0.90936695 | 0.98207723 | -0.0351866 | 0.40309158 |
| 0.999490048 | -0.0044852 | 0.9107189 | 0.98207723 | -0.0099574 | 0.80350276 |
| 0.999490048 | 0.00546984 | 0.90859183 | 0.98207723 | -0.0113325 | 0.812059 |
| 0.999490048 | 0.00547625 | 0.91084308 | 0.98207723 | -0.0266114 | 0.58741524 |
| 0.999490048 | 0.00548787 | 0.90728237 | 0.98207723 | 0.01261963 | 0.78895752 |
| 0.999490048 | 0.00594712 | 0.91080466 | 0.98207723 | -0.0107054 | 0.84023215 |
| 0.999490048 | 0.00635507 | 0.91174403 | 0.98207723 | 0.03556281 | 0.53661741 |
| 0.999490048 | 0.00662112 | 0.90961556 | 0.98207723 | -0.0055475 | 0.92422027 |
| 0.999490048 | 0.00692573 | 0.91108796 | 0.98207723 | 0.02174411 | 0.72618967 |
| 0.999490048 | 0.00696199 | 0.91110353 | 0.98207723 | 0.02774044 | 0.65701948 |
| 0.999490048 | 0.00696324 | 0.90819233 | 0.98207723 | -0.0133918 | 0.82454521 |
| 0.999490048 | 0.00709597 | 0.90929969 | 0.98207723 | -0.0111295 | 0.85821779 |
| 0.999490048 | 0.00724055 | 0.90708977 | 0.98207723 | -0.0292165 | 0.63838929 |
| 0.999490048 | 0.00748598 | 0.90818443 | 0.98207723 | -0.0181462 | 0.77995588 |
| 0.999490048 | 0.00765675 | 0.90837056 | 0.98207723 | 0.03777059 | 0.57139744 |
| 0.999490048 | 0.00803391 | 0.90937429 | 0.98207723 | 0.05210685 | 0.46273753 |
| 0.999490048 | 0.009398 | 0.91063421 | 0.98207723 | -0.0788344 | 0.35064214 |
| 0.999490048 | 0.00993755 | 0.90905399 | 0.98207723 | 0.03388038 | 0.69734291 |
| 0.999490048 | 0.01018347 | 0.90952312 | 0.98207723 | 0.09458693 | 0.29646366 |
| 0.999490048 | 0.01063854 | 0.90876423 | 0.98207723 | 0.07514026 | 0.42126739 |
| 0.999490048 | 0.0109926 | 0.91155771 | 0.98207723 | -0.0264399 | 0.78946843 |
| 0.999490048 | 0.01118644 | 0.90958483 | 0.98207723 | -0.0347373 | 0.7246519 |
| 0.999490048 | 0.01384898 | 0.91152222 | 0.98207723 | -0.0648105 | 0.60399451 |
| 0.999490048 | 0.01644205 | 0.90863579 | 0.98207723 | 0.03240023 | 0.8213063 |
| 0.999490048 | 0.02095628 | 0.91139761 | 0.98207723 | 0.08349927 | 0.6580906 |
| 0.999490048 | -0.0106067 | 0.91318204 | 0.98227203 | -0.063167 | 0.51787316 |
| 0.999490048 | -0.0071248 | 0.91345417 | 0.98227203 | 0.04212061 | 0.52221542 |
| 0.999490048 | -0.0060106 | 0.91264254 | 0.98227203 | -0.0221142 | 0.68693805 |
| 0.999490048 | -0.0057345 | 0.91333803 | 0.98227203 | 0.03730202 | 0.48115121 |
| 0.999490048 | -0.0055102 | 0.9131774 | 0.98227203 | 0.00984727 | 0.84555076 |
| 0.999490048 | -0.005129 | 0.91299165 | 0.98227203 | 0.03497575 | 0.45865786 |
| 0.999490048 | -0.0044441 | 0.91284988 | 0.98227203 | -0.0395669 | 0.33435857 |
| 0.999490048 | 0.00680777 | 0.91335151 | 0.98227203 | 0.07811786 | 0.21880723 |
| 0.999490048 | 0.01023409 | 0.91330282 | 0.98227203 | -0.0345307 | 0.71369914 |
| 0.999490048 | 0.01099869 | 0.91297728 | 0.98227203 | 0.07510627 | 0.45796624 |
| 0.999490048 | 0.01448057 | 0.91295406 | 0.98227203 | -0.0382141 | 0.77314084 |
| 0.999490048 | 0.01494336 | 0.91271157 | 0.98227203 | 0.01442209 | 0.91574423 |
| 0.999490048 | 0.01823399 | 0.91207467 | 0.98227203 | -0.2320011 | 0.16812987 |
| 0.992011001 | -0.0339534 | 0.91387685 | 0.98236952 | -0.7264737 | 0.0370809 |

| | | | | | |
|---|---|---|---|---|---|
| 0.999490048 | 0.00508324 | 0.91389778 | 0.98236952 | -4.01E-04 | 0.99319946 |
| 0.999490048 | 0.00936156 | 0.91387808 | 0.98236952 | -0.0034849 | 0.96787959 |
| 0.999490048 | 0.00637053 | 0.91402321 | 0.98237788 | -0.0377938 | 0.52350239 |
| 0.999490048 | 0.02244179 | 0.91417022 | 0.98240944 | 0.20320504 | 0.34308419 |
| 0.999490048 | 0.00821339 | 0.91436812 | 0.98249566 | -0.0109728 | 0.88578218 |
| 0.999490048 | -0.0105058 | 0.91474948 | 0.98258962 | -0.0650907 | 0.50899586 |
| 0.999490048 | 0.00705091 | 0.91480859 | 0.98258962 | 0.04817321 | 0.46719088 |
| 0.999490048 | 0.02718696 | 0.91461559 | 0.98258962 | 0.07901734 | 0.75597684 |
| 0.999490048 | -0.0057429 | 0.91594818 | 0.98368711 | -0.0162031 | 0.76605924 |
| 0.999490048 | 0.00990807 | 0.91609899 | 0.98372255 | -0.0538347 | 0.56828087 |
| 0.999490048 | -0.0098288 | 0.91673917 | 0.98393261 | 0.00798713 | 0.9322957 |
| 0.999490048 | -0.0077604 | 0.91676595 | 0.98393261 | 0.0838578 | 0.26474069 |
| 0.999490048 | -0.0074745 | 0.91645284 | 0.98393261 | 0.06473772 | 0.36746972 |
| 0.999490048 | 0.00500373 | 0.91666878 | 0.98393261 | -0.0328941 | 0.49356324 |
| 0.999490048 | -0.0069777 | 0.91746585 | 0.98399071 | 0.04028718 | 0.55102853 |
| 0.999490048 | 0.00590326 | 0.91744365 | 0.98399071 | 0.03495428 | 0.54086616 |
| 0.999490048 | 0.00685385 | 0.91738266 | 0.98399071 | 0.11741322 | 0.09607436 |
| 0.999490048 | 0.00696153 | 0.9173547 | 0.98399071 | 9.93E-04 | 0.98819132 |
| 0.999490048 | 0.00878848 | 0.91752715 | 0.98399071 | -0.0194586 | 0.8187352 |
| 0.999490048 | 0.01200209 | 0.91725894 | 0.98399071 | -0.0875659 | 0.45103988 |
| 0.999490048 | -0.0037351 | 0.91767368 | 0.98402148 | -0.015903 | 0.66045347 |
| 0.999490048 | -0.005724 | 0.91804628 | 0.98416824 | -0.0032905 | 0.95282943 |
| 0.999490048 | 0.02167416 | 0.9179948 | 0.98416824 | -0.1820546 | 0.39797904 |
| 0.999490048 | -0.0077609 | 0.91848251 | 0.98445728 | 0.02539921 | 0.7379242 |
| 0.999490048 | -0.0061583 | 0.91896625 | 0.98445728 | 0.02801566 | 0.64416989 |
| 0.999490048 | -0.0056665 | 0.91869857 | 0.98445728 | -0.0154074 | 0.78150706 |
| 0.999490048 | 0.00444086 | 0.91888652 | 0.98445728 | -0.0046137 | 0.91574108 |
| 0.999490048 | 0.00603985 | 0.91902329 | 0.98445728 | -0.0944908 | 0.12064331 |
| 0.999490048 | 0.00696899 | 0.91888664 | 0.98445728 | -0.0058753 | 0.93157973 |
| 0.999490048 | -0.0172873 | 0.91971194 | 0.98481594 | 0.02022378 | 0.90613611 |
| 0.999490048 | -0.0049729 | 0.91964803 | 0.98481594 | -0.0104987 | 0.8314092 |
| 0.999490048 | 0.00795396 | 0.91948258 | 0.98481594 | 0.0198219 | 0.80121298 |
| 0.999490048 | -0.0263974 | 0.92085663 | 0.98482354 | -0.0341024 | 0.89787496 |
| 0.578467169 | -0.0183539 | 0.92053596 | 0.98482354 | -0.5798829 | 0.0038511 |
| 0.999490048 | -0.0171559 | 0.92042181 | 0.98482354 | 0.22741087 | 0.2023875 |
| 0.999490048 | -0.0085051 | 0.91985723 | 0.98482354 | 0.0036759 | 0.96531198 |
| 0.999490048 | -0.0080074 | 0.92189072 | 0.98482354 | -0.0369166 | 0.65187199 |
| 0.999490048 | -0.0079664 | 0.9216906 | 0.98482354 | -0.0256078 | 0.75222202 |
| 0.999490048 | -0.007328 | 0.92188618 | 0.98482354 | -0.0751881 | 0.31918494 |
| 0.999490048 | -0.0067973 | 0.92154252 | 0.98482354 | 0.04782562 | 0.49037091 |
| 0.999490048 | -0.0051994 | 0.92147809 | 0.98482354 | 0.00773283 | 0.88346008 |
| 0.999490048 | -0.0047849 | 0.92187026 | 0.98482354 | -0.0473154 | 0.33660644 |

| | | | | | |
|---|---|---|---|---|---|
| 0.999490048 | -0.0044861 | 0.92111844 | 0.98482354 | 0.00410197 | 0.9278519 |
| 0.999490048 | 0.00384178 | 0.92169378 | 0.98482354 | -0.0400187 | 0.31084562 |
| 0.999490048 | 0.00438725 | 0.92175199 | 0.98482354 | -0.0606279 | 0.18245892 |
| 0.999490048 | 0.00529399 | 0.92041814 | 0.98482354 | 0.01760337 | 0.73998706 |
| 0.999490048 | 0.00565636 | 0.92199734 | 0.98482354 | -0.081636 | 0.16574917 |
| 0.999490048 | 0.00582807 | 0.92070007 | 0.98482354 | -0.0056014 | 0.92377412 |
| 0.967348048 | 0.00679426 | 0.92070065 | 0.98482354 | -0.1559906 | 0.02946359 |
| 0.999490048 | 0.0084811 | 0.92094673 | 0.98482354 | 0.02748228 | 0.74799879 |
| 0.999490048 | 0.01288078 | 0.92055873 | 0.98482354 | 0.06408517 | 0.62059786 |
| 0.999490048 | 0.01481876 | 0.9220779 | 0.98482354 | 0.09704936 | 0.52344977 |
| 0.999490048 | -0.0086883 | 0.92293889 | 0.9851131 | -0.0966842 | 0.28721952 |
| 0.999490048 | -0.0067764 | 0.92273999 | 0.9851131 | 0.00778184 | 0.91132342 |
| 0.999490048 | 0.00566912 | 0.92270717 | 0.9851131 | -0.0579796 | 0.32574857 |
| 0.999490048 | 0.00584822 | 0.92277526 | 0.9851131 | -0.043389 | 0.47426808 |
| 0.999490048 | 0.0062841 | 0.92286164 | 0.9851131 | 0.05317768 | 0.41564918 |
| 0.999490048 | -0.0064506 | 0.92397321 | 0.98524586 | 0.03655568 | 0.58970399 |
| 0.999490048 | -0.0064424 | 0.92337302 | 0.98524586 | 0.05782 | 0.39148227 |
| 0.999490048 | -0.0056678 | 0.92384886 | 0.98524586 | -0.0312666 | 0.59897065 |
| 0.999490048 | -0.0050556 | 0.92350572 | 0.98524586 | 0.03015512 | 0.5680726 |
| 0.999490048 | 0.00474665 | 0.92400723 | 0.98524586 | 0.02128503 | 0.66938757 |
| 0.999490048 | 0.00564161 | 0.92368557 | 0.98524586 | -0.009485 | 0.87207477 |
| 0.999490048 | 0.00575895 | 0.92330333 | 0.98524586 | 0.00980153 | 0.86986878 |
| 0.999490048 | 0.00788817 | 0.92362163 | 0.98524586 | -0.0402793 | 0.62525311 |
| 0.999490048 | 0.00370311 | 0.92437501 | 0.98551217 | -0.0642238 | 0.10875385 |
| 0.999490048 | 0.00402066 | 0.9247501 | 0.98566037 | -0.0221899 | 0.60313276 |
| 0.999490048 | 0.00804086 | 0.92474177 | 0.98566037 | 0.01332094 | 0.87566478 |
| 0.999490048 | 0.00767756 | 0.92493392 | 0.98573047 | -0.0418169 | 0.60873973 |
| 0.999490048 | -0.0056821 | 0.92529707 | 0.98599165 | -0.1075058 | 0.08519429 |
| 0.999490048 | 0.00578561 | 0.92554398 | 0.98601551 | 0.02620968 | 0.67256727 |
| 0.999490048 | 0.00604703 | 0.92555563 | 0.98601551 | 0.06371019 | 0.32952057 |
| 0.999490048 | -0.0314379 | 0.92584705 | 0.98612234 | -1.61E-04 | 0.9996193 |
| 0.999490048 | -0.0077864 | 0.92594942 | 0.98612234 | -0.1199142 | 0.16059265 |
| 0.999490048 | -0.0041251 | 0.92612385 | 0.98612234 | 0.00723573 | 0.87082373 |
| 0.999490048 | 0.00442176 | 0.92612831 | 0.98612234 | 0.08566493 | 0.08157534 |
| 0.999490048 | -0.0105076 | 0.92688321 | 0.9868003 | -0.0269952 | 0.81370589 |
| 0.999490048 | -0.0043493 | 0.9271465 | 0.98695477 | 0.02326481 | 0.62557409 |
| 0.999490048 | 0.0053295 | 0.92729203 | 0.98698387 | -0.009617 | 0.86923336 |
| 0.999490048 | 0.00433441 | 0.92773412 | 0.98723644 | -0.0294581 | 0.53914764 |
| 0.999490048 | 0.00577341 | 0.92776579 | 0.98723644 | 0.01445638 | 0.82050154 |
| 0.999490048 | -0.0038676 | 0.92804785 | 0.98741075 | -0.0099853 | 0.81573698 |
| 0.999490048 | -0.0091377 | 0.92854581 | 0.98768885 | -0.0606089 | 0.553357 |
| 0.999490048 | -0.0047515 | 0.92851791 | 0.98768885 | -0.0374259 | 0.48196546 |

| | | | | | |
|---|---|---|---|---|---|
| 0.999490048 | -0.0105305 | 0.92868292 | 0.98770887 | 0.14076245 | 0.23812396 |
| 0.999490048 | -0.0188632 | 0.92982318 | 0.98772918 | -0.0406715 | 0.84944207 |
| 0.999490048 | -0.0087222 | 0.92990628 | 0.98772918 | 0.07535388 | 0.44988127 |
| 0.999490048 | -0.003842 | 0.92937285 | 0.98772918 | -0.0170325 | 0.69479637 |
| 0.999490048 | 0.00453116 | 0.92891741 | 0.98772918 | -0.0231007 | 0.64991335 |
| 0.999490048 | 0.00466308 | 0.92984587 | 0.98772918 | -0.0055554 | 0.9164688 |
| 0.999490048 | 0.00537712 | 0.92912157 | 0.98772918 | -0.0239028 | 0.69297777 |
| 0.999490048 | 0.00538901 | 0.92980676 | 0.98772918 | -0.0084197 | 0.89054713 |
| 0.999490048 | 0.0059285 | 0.929396 | 0.98772918 | 0.0415068 | 0.53657665 |
| 0.999490048 | 0.0084805 | 0.92997525 | 0.98772918 | -0.112805 | 0.24878462 |
| 0.999490048 | 0.01321573 | 0.93000322 | 0.98772918 | 0.1802283 | 0.23754034 |
| 0.999490048 | 0.01462373 | 0.92922476 | 0.98772918 | -0.0234805 | 0.88660928 |
| 0.999490048 | -0.0054367 | 0.93030487 | 0.9879239 | -0.0127189 | 0.83793186 |
| 0.999490048 | -0.0158474 | 0.93087344 | 0.98802507 | 0.01095211 | 0.95219361 |
| 0.999490048 | -0.0139555 | 0.93059266 | 0.98802507 | 0.04412702 | 0.78315038 |
| 0.999490048 | -0.0038072 | 0.93079945 | 0.98802507 | 0.05463256 | 0.21971382 |
| 0.999490048 | 0.00284061 | 0.93080693 | 0.98802507 | -0.0022648 | 0.94480597 |
| 0.999490048 | -0.0059393 | 0.93126929 | 0.9883196 | 0.01858937 | 0.78733919 |
| 0.999490048 | -0.0055131 | 0.9316866 | 0.98863682 | 0.01005412 | 0.8757941 |
| 0.999490048 | 0.00458316 | 0.93190989 | 0.98873758 | -0.0354484 | 0.51053232 |
| 0.999490048 | 0.00623834 | 0.93201838 | 0.98873758 | -0.0572996 | 0.43610008 |
| 0.999490048 | -0.0041962 | 0.93233571 | 0.98894857 | -0.0148637 | 0.76379141 |
| 0.999490048 | -0.0047566 | 0.93258464 | 0.98896136 | 0.03388949 | 0.54813528 |
| 0.971241857 | 0.01183186 | 0.9324971 | 0.98896136 | -0.3570118 | 0.02946592 |
| 0.999490048 | -0.00482 | 0.9330208 | 0.98898739 | 0.00800257 | 0.88903757 |
| 0.999490048 | 0.00401847 | 0.93308295 | 0.98898739 | -0.0091103 | 0.84907024 |
| 0.999490048 | 0.00416188 | 0.93289621 | 0.98898739 | -0.0272931 | 0.58195538 |
| 0.999490048 | 0.00770039 | 0.93277796 | 0.98898739 | 0.02010124 | 0.82579521 |
| 0.999490048 | 0.00333501 | 0.93342241 | 0.98922162 | -0.0480373 | 0.23550178 |
| 0.999490048 | -0.0254171 | 0.93393587 | 0.98923113 | 0.0012539 | 0.9967364 |
| 0.999490048 | -0.005157 | 0.93396157 | 0.98923113 | -0.0637241 | 0.31088468 |
| 0.999490048 | -0.0041211 | 0.93357666 | 0.98923113 | -0.0149745 | 0.76220394 |
| 0.999490048 | 0.00385306 | 0.93369751 | 0.98923113 | -0.0021093 | 0.96367239 |
| 0.999490048 | 0.00959558 | 0.93402374 | 0.98923113 | 0.11935142 | 0.30822405 |
| 0.999490048 | -0.004622 | 0.93416993 | 0.98926048 | -0.0452337 | 0.42188186 |
| 0.999490048 | 0.00347006 | 0.93445747 | 0.98932867 | -0.0017176 | 0.96752816 |
| 0.999490048 | 0.00467309 | 0.93447128 | 0.98932867 | -0.0164925 | 0.77185412 |
| 0.999490048 | 0.00452865 | 0.93481603 | 0.98956819 | 0.02563185 | 0.64412261 |
| 0.999490048 | 0.00527542 | 0.93507287 | 0.98956828 | 0.02673696 | 0.68019535 |
| 0.972489992 | 0.02197283 | 0.93496699 | 0.98956828 | -0.5959953 | 0.01959081 |
| 0.999490048 | 0.02434011 | 0.93517165 | 0.98956828 | -0.0113686 | 0.96968917 |
| 0.999490048 | -0.0086701 | 0.93569331 | 0.98972644 | 0.11262233 | 0.29984665 |

| | | | | | |
|---|---|---|---|---|---|
| 0.999490048 | -0.0055029 | 0.93562176 | 0.98972644 | -0.0511318 | 0.45545263 |
| 0.999490048 | -0.003909 | 0.93579524 | 0.98972644 | -0.0113341 | 0.81540662 |
| 0.999490048 | 0.00392636 | 0.93570125 | 0.98972644 | 0.0664097 | 0.18027737 |
| 0.999490048 | 0.00446538 | 0.9361706 | 0.98974734 | -0.032682 | 0.55911994 |
| 0.999490048 | 0.00882039 | 0.9361308 | 0.98974734 | 0.11631378 | 0.29595893 |
| 0.999490048 | 0.01542914 | 0.93604524 | 0.98974734 | 0.16952977 | 0.3816249 |
| 0.999490048 | -0.0080599 | 0.93629193 | 0.9897503 | -0.1250247 | 0.22196412 |
| 0.999490048 | -0.0066656 | 0.93715557 | 0.98997268 | 0.01656128 | 0.84473985 |
| 0.999490048 | -0.0051566 | 0.93758955 | 0.98997268 | -0.031892 | 0.62898929 |
| 0.999490048 | -0.0037342 | 0.93718276 | 0.98997268 | -0.0214934 | 0.65075888 |
| 0.999490048 | 0.00336392 | 0.93667292 | 0.98997268 | 0.04781163 | 0.26478061 |
| 0.999490048 | 0.00360738 | 0.93761819 | 0.98997268 | 0.05945054 | 0.2044603 |
| 0.999490048 | 0.00394982 | 0.93681737 | 0.98997268 | -3.72E-04 | 0.99404011 |
| 0.999490048 | 0.00441176 | 0.9376879 | 0.98997268 | -0.0545221 | 0.33844038 |
| 0.999490048 | 0.0058027 | 0.93763669 | 0.98997268 | 0.00720323 | 0.92263006 |
| 0.999490048 | 0.01135134 | 0.9376729 | 0.98997268 | -0.0169135 | 0.90725494 |
| 0.999490048 | 0.02129386 | 0.93716987 | 0.98997268 | -0.143878 | 0.59532543 |
| 0.999490048 | -0.0095959 | 0.93849211 | 0.99026017 | -0.0286345 | 0.81797244 |
| 0.999490048 | -0.0068733 | 0.93914614 | 0.99026017 | 0.02737141 | 0.7613154 |
| 0.999490048 | -0.0062056 | 0.93904756 | 0.99026017 | 0.04686015 | 0.56493837 |
| 0.999490048 | -0.0061862 | 0.93897156 | 0.99026017 | 0.01183783 | 0.88353692 |
| 0.999490048 | -0.0058155 | 0.93868802 | 0.99026017 | -0.0068063 | 0.92827033 |
| 0.999490048 | -0.0047725 | 0.93862122 | 0.99026017 | -0.0342617 | 0.58153586 |
| 0.999490048 | -0.0047025 | 0.93876947 | 0.99026017 | -0.0228869 | 0.70888161 |
| 0.999490048 | -0.0041087 | 0.93906703 | 0.99026017 | -0.0143322 | 0.78987384 |
| 0.999490048 | 0.00493054 | 0.93905247 | 0.99026017 | -0.0480776 | 0.45840726 |
| 0.999490048 | 0.00517274 | 0.9385891 | 0.99026017 | 0.07363704 | 0.27844331 |
| 0.999490048 | -0.0071109 | 0.9394562 | 0.99046203 | -0.1744124 | 0.07150912 |
| 0.999490048 | -0.003875 | 0.93961295 | 0.99050223 | 0.00958845 | 0.85134677 |
| 0.999490048 | -0.0040757 | 0.93993141 | 0.99071286 | 0.0347547 | 0.52219395 |
| 0.31159876 | -0.0234768 | 0.94090712 | 0.99079036 | -1.4100855 | 7.99E-04 |
| 0.999490048 | -0.0060381 | 0.9409542 | 0.99079036 | 0.0606128 | 0.45961383 |
| 0.999490048 | 0.00343143 | 0.94072104 | 0.99079036 | 0.0363012 | 0.43428418 |
| 0.999490048 | 0.00447459 | 0.94064418 | 0.99079036 | 0.06760105 | 0.26657581 |
| 0.999490048 | 0.00463042 | 0.94082435 | 0.99079036 | -0.089336 | 0.16036303 |
| 0.999490048 | 0.00614547 | 0.94066164 | 0.99079036 | 0.14545551 | 0.08724162 |
| 0.999490048 | 0.00719809 | 0.94034166 | 0.99079036 | 0.01254767 | 0.89621328 |
| 0.999490048 | 0.00973138 | 0.94066043 | 0.99079036 | -0.0296907 | 0.82041398 |
| 0.999490048 | -0.0102196 | 0.94161105 | 0.99081919 | 0.19565088 | 0.16894925 |
| 0.999490048 | -0.0039635 | 0.94159095 | 0.99081919 | -0.0093358 | 0.86301218 |
| 0.999490048 | 0.0031703 | 0.94154461 | 0.99081919 | 0.07147944 | 0.10734284 |
| 0.999490048 | 0.00432207 | 0.94152556 | 0.99081919 | -0.0411798 | 0.48673498 |

| | | | | | |
|---|---|---|---|---|---|
| 0.999490048 | 0.00589395 | 0.94162005 | 0.99081919 | -0.0364784 | 0.65102356 |
| 0.999490048 | 0.00842417 | 0.94169355 | 0.99081919 | 0.20964672 | 0.07783155 |
| 0.999490048 | -0.0345481 | 0.94193795 | 0.99082188 | -0.1418012 | 0.76542659 |
| 0.999490048 | -0.0048503 | 0.94228941 | 0.99082188 | -0.0780209 | 0.25053914 |
| 0.999490048 | -0.0036088 | 0.94184385 | 0.99082188 | -0.0317437 | 0.52276773 |
| 0.999490048 | 0.00623733 | 0.94217079 | 0.99082188 | 6.29E-04 | 0.99416749 |
| 0.999490048 | 0.00628956 | 0.94208877 | 0.99082188 | 0.03295503 | 0.70387332 |
| 0.999490048 | -0.0164712 | 0.94605878 | 0.99091873 | 0.1570285 | 0.52285236 |
| 0.999490048 | -0.0138854 | 0.9441333 | 0.99091873 | 0.14539368 | 0.46548075 |
| 0.999490048 | -0.0079736 | 0.94368739 | 0.99091873 | -0.1153497 | 0.31184892 |
| 0.999490048 | -0.0059647 | 0.94569978 | 0.99091873 | -0.0640299 | 0.46707309 |
| 0.999490048 | -0.0054249 | 0.94700976 | 0.99091873 | 0.0080368 | 0.92156933 |
| 0.999490048 | -0.0043814 | 0.94352871 | 0.99091873 | -0.061772 | 0.32271763 |
| 0.999490048 | -0.004318 | 0.94675274 | 0.99091873 | -0.1111031 | 0.09487298 |
| 0.999490048 | -0.0042515 | 0.9439425 | 0.99091873 | 0.05300442 | 0.38431334 |
| 0.999490048 | -0.0040036 | 0.94463373 | 0.99091873 | -0.0035901 | 0.9503441 |
| 0.999490048 | -0.0037493 | 0.94682812 | 0.99091873 | -0.1104733 | 0.05818626 |
| 0.999490048 | -0.0036892 | 0.94649117 | 0.99091873 | -0.0260059 | 0.63689066 |
| 0.999490048 | -0.0036211 | 0.94334996 | 0.99091873 | 0.00368503 | 0.94235197 |
| 0.999490048 | -0.0036146 | 0.94481642 | 0.99091873 | -0.0101857 | 0.84540234 |
| 0.999490048 | -0.0033885 | 0.94383515 | 0.99091873 | -0.0382123 | 0.42981763 |
| 0.999490048 | -0.0033726 | 0.94517544 | 0.99091873 | -0.0286582 | 0.56031925 |
| 0.999490048 | -0.0031513 | 0.94543552 | 0.99091873 | -0.0675051 | 0.1510851 |
| 0.999490048 | 0.00256837 | 0.94571953 | 0.99091873 | -0.0031178 | 0.93413276 |
| 0.999490048 | 0.00296864 | 0.94402191 | 0.99091873 | -0.0060553 | 0.88613237 |
| 0.999490048 | 0.00350076 | 0.94700269 | 0.99091873 | -0.0487602 | 0.35861637 |
| 0.999490048 | 0.00357627 | 0.94566761 | 0.99091873 | 0.04557294 | 0.38871581 |
| 0.999490048 | 0.00395134 | 0.9455571 | 0.99091873 | -0.0209266 | 0.71795402 |
| 0.999490048 | 0.00396368 | 0.94397839 | 0.99091873 | -0.1041666 | 0.07385649 |
| 0.999490048 | 0.00398013 | 0.94641162 | 0.99091873 | 0.05369958 | 0.36839992 |
| 0.999490048 | 0.0041994 | 0.94590116 | 0.99091873 | 0.02060256 | 0.73947852 |
| 0.999490048 | 0.00441654 | 0.94479126 | 0.99091873 | -0.0332415 | 0.60319393 |
| 0.999490048 | 0.00445245 | 0.94406089 | 0.99091873 | -0.0553935 | 0.38628099 |
| 0.999490048 | 0.0044726 | 0.94513529 | 0.99091873 | -0.0723707 | 0.27134073 |
| 0.999490048 | 0.00476408 | 0.94638705 | 0.99091873 | -0.0299107 | 0.67342764 |
| 0.999490048 | 0.00477664 | 0.94380241 | 0.99091873 | -0.0095332 | 0.88813413 |
| 0.999490048 | 0.00488367 | 0.94392236 | 0.99091873 | 0.04847982 | 0.48711976 |
| 0.999490048 | 0.00498624 | 0.94615198 | 0.99091873 | -0.0194362 | 0.7924761 |
| 0.999490048 | 0.00541542 | 0.94289529 | 0.99091873 | -0.1013693 | 0.18768728 |
| 0.999490048 | 0.00564336 | 0.94491133 | 0.99091873 | 0.12080642 | 0.14761551 |
| 0.999490048 | 0.0057785 | 0.94646109 | 0.99091873 | 0.00493114 | 0.95430212 |
| 0.999490048 | 0.0061747 | 0.94257993 | 0.99091873 | 0.08313082 | 0.33668659 |

| | | | | | |
|---|---|---|---|---|---|
| 0.999490048 | 0.00834637 | 0.94552697 | 0.99091873 | 0.03765086 | 0.75813098 |
| 0.999490048 | 0.01009321 | 0.94410245 | 0.99091873 | 0.05156042 | 0.7205456 |
| 0.999490048 | 0.01129977 | 0.94602341 | 0.99091873 | 0.10411106 | 0.53435939 |
| 0.999490048 | 0.01317421 | 0.94681643 | 0.99091873 | 0.05121615 | 0.79551253 |
| 0.999490048 | -0.0076955 | 0.94745365 | 0.99100963 | -0.144807 | 0.22188298 |
| 0.999490048 | -0.007379 | 0.94816478 | 0.99100963 | -0.088451 | 0.43857721 |
| 0.999490048 | -0.0046531 | 0.94786386 | 0.99100963 | 0.07052057 | 0.32638525 |
| 0.999490048 | -0.0039512 | 0.94742227 | 0.99100963 | -0.0113273 | 0.85010193 |
| 0.999490048 | -0.003861 | 0.94760634 | 0.99100963 | 0.02237037 | 0.70380238 |
| 0.999490048 | -0.0030362 | 0.94761352 | 0.99100963 | -0.0587613 | 0.21072958 |
| 0.999490048 | 0.00299576 | 0.94778016 | 0.99100963 | -0.006248 | 0.89136605 |
| 0.999490048 | 0.00497856 | 0.94797881 | 0.99100963 | 0.07455417 | 0.33311848 |
| 0.999490048 | 0.00634951 | 0.94806178 | 0.99100963 | -0.0216295 | 0.82446924 |
| 0.999490048 | -0.0098169 | 0.9492688 | 0.99119322 | 0.05324167 | 0.73034369 |
| 0.999490048 | -0.0048371 | 0.94914169 | 0.99119322 | -0.0893664 | 0.24506782 |
| 0.999490048 | -0.0044017 | 0.94936232 | 0.99119322 | -0.0339611 | 0.62496812 |
| 0.999490048 | -0.0032214 | 0.94849506 | 0.99119322 | 0.01278258 | 0.79782671 |
| 0.999490048 | -0.0031824 | 0.94913764 | 0.99119322 | 0.05085562 | 0.3130018 |
| 0.999490048 | 0.00268942 | 0.94940879 | 0.99119322 | -0.0199969 | 0.63783114 |
| 0.999490048 | 0.00300166 | 0.94936452 | 0.99119322 | -0.0240488 | 0.61181387 |
| 0.999490048 | 0.00417384 | 0.94862106 | 0.99119322 | -0.0330182 | 0.61113968 |
| 0.999490048 | 0.01173671 | 0.94912878 | 0.99119322 | -0.0298451 | 0.87114686 |
| 0.999490048 | -0.0039934 | 0.94980289 | 0.99146453 | -0.011509 | 0.85607011 |
| 0.999490048 | 0.00267287 | 0.94990614 | 0.99146453 | 0.08277037 | 0.0605569 |
| 0.999490048 | 0.00278289 | 0.95030019 | 0.99175185 | 0.05879809 | 0.1954141 |
| 0.999490048 | -0.0042851 | 0.95042044 | 0.9917534 | -0.0100176 | 0.88444613 |
| 0.999490048 | -0.0060679 | 0.95065689 | 0.99178715 | 0.05581636 | 0.57044237 |
| 0.999490048 | -0.0025633 | 0.95069034 | 0.99178715 | -0.0130307 | 0.75347494 |
| 0.999490048 | 0.00323218 | 0.95092444 | 0.99190744 | -0.0084098 | 0.87280143 |
| 0.999490048 | -0.0035683 | 0.95107351 | 0.99193902 | 0.00283762 | 0.96108219 |
| 0.999490048 | 0.00308072 | 0.95121728 | 0.99196507 | 0.01092971 | 0.82824589 |
| 0.999490048 | -0.0056769 | 0.95141144 | 0.99204365 | 0.09083617 | 0.34362439 |
| 0.999490048 | -0.003381 | 0.95165314 | 0.9920479 | 0.02401482 | 0.66729501 |
| 0.999490048 | -0.0028931 | 0.95161648 | 0.9920479 | 0.02047496 | 0.66818769 |
| 0.999490048 | -0.0054502 | 0.95214102 | 0.99208686 | -0.1159851 | 0.20877332 |
| 0.999490048 | -0.0054166 | 0.95261358 | 0.99208686 | -0.0941182 | 0.30691122 |
| 0.999490048 | -0.003744 | 0.95275982 | 0.99208686 | -0.0816603 | 0.20369437 |
| 0.999490048 | -0.002492 | 0.95253764 | 0.99208686 | 0.02619809 | 0.53309723 |
| 0.999490048 | 0.0026822 | 0.95238036 | 0.99208686 | 0.05083511 | 0.26373985 |
| 0.999490048 | 0.00281674 | 0.95265045 | 0.99208686 | -0.0186807 | 0.69417226 |
| 0.999490048 | 0.0062628 | 0.95252814 | 0.99208686 | -0.0515857 | 0.62470712 |
| 0.999490048 | 0.00722878 | 0.95271656 | 0.99208686 | 0.08205355 | 0.50283613 |

| | | | | | |
|---|---|---|---|---|---|
| 0.999490048 | 0.00851782 | 0.95209085 | 0.99208686 | 0.03546626 | 0.80257505 |
| 0.999490048 | -0.0045182 | 0.95320734 | 0.99218166 | -0.1258667 | 0.11116522 |
| 0.999490048 | -0.0035759 | 0.95308983 | 0.99218166 | -0.0370429 | 0.54375412 |
| 0.999490048 | -0.0029592 | 0.95301255 | 0.99218166 | 0.04081313 | 0.41946469 |
| 0.999490048 | -0.0123999 | 0.95493513 | 0.99260066 | 0.33074908 | 0.15805811 |
| 0.999490048 | -0.0061801 | 0.95392825 | 0.99260066 | 0.01006631 | 0.92503951 |
| 0.999490048 | -0.0051784 | 0.95446233 | 0.99260066 | -0.0281852 | 0.75616888 |
| 0.999490048 | -0.0049461 | 0.95476728 | 0.99260066 | 0.06998727 | 0.4251725 |
| 0.999490048 | -0.0042789 | 0.95511906 | 0.99260066 | -0.0253394 | 0.73919637 |
| 0.999490048 | -0.0030854 | 0.95504529 | 0.99260066 | 0.00863497 | 0.87466849 |
| 0.999490048 | -0.0028873 | 0.95514018 | 0.99260066 | 0.02589225 | 0.61484128 |
| 0.999490048 | -0.0027154 | 0.95433748 | 0.99260066 | -9.71E-04 | 0.98366342 |
| 0.999490048 | -0.00263 | 0.9542678 | 0.99260066 | -0.0282446 | 0.53950322 |
| 0.999490048 | 0.00253289 | 0.95432459 | 0.99260066 | -0.0069763 | 0.87467516 |
| 0.999490048 | 0.00281245 | 0.95475206 | 0.99260066 | 0.01132539 | 0.81935571 |
| 0.999490048 | 0.00372467 | 0.95527412 | 0.99260066 | 0.03313915 | 0.61864968 |
| 0.999490048 | 0.00822945 | 0.95498916 | 0.99260066 | -0.010911 | 0.94034807 |
| 0.999490048 | 0.00961056 | 0.95526799 | 0.99260066 | -0.1078452 | 0.53068182 |
| 0.999490048 | -0.0048922 | 0.9558902 | 0.99275554 | 0.03727537 | 0.67397836 |
| 0.999490048 | -0.0024041 | 0.95589875 | 0.99275554 | -0.0435755 | 0.32098424 |
| 0.999490048 | 0.00234245 | 0.95581318 | 0.99275554 | -0.0381177 | 0.37111689 |
| 0.999490048 | 0.00412987 | 0.95559379 | 0.99275554 | 0.0531815 | 0.47560416 |
| 0.999490048 | -0.0028646 | 0.95606172 | 0.99280131 | -0.034022 | 0.51467801 |
| 0.999490048 | -0.0064075 | 0.95627184 | 0.99286095 | -0.1361509 | 0.25029596 |
| 0.999490048 | -0.0039011 | 0.95662652 | 0.99286095 | 0.02519995 | 0.72566576 |
| 0.999490048 | 0.00270476 | 0.95671368 | 0.99286095 | 0.02753287 | 0.58174298 |
| 0.999490048 | 0.00397763 | 0.9565689 | 0.99286095 | 0.00886481 | 0.90340728 |
| 0.999490048 | 0.00453336 | 0.95665785 | 0.99286095 | 0.04435676 | 0.59592129 |
| 0.999490048 | 0.00458854 | 0.95703247 | 0.99305936 | 0.08668243 | 0.31373511 |
| 0.999490048 | 0.00713367 | 0.95714272 | 0.99305936 | 0.07080665 | 0.59480038 |
| 0.99974305 | -0.0078909 | 0.95743344 | 0.99306682 | -0.0080076 | 0.95680434 |
| 0.999490048 | -0.0038765 | 0.95748101 | 0.99306682 | 0.01391869 | 0.84824104 |
| 0.999490048 | 0.00275711 | 0.9575067 | 0.99306682 | 0.01287853 | 0.80356456 |
| 0.999490048 | 0.00274037 | 0.95780268 | 0.99325042 | -0.0059287 | 0.90887385 |
| 0.999490048 | -0.0029883 | 0.95838635 | 0.99373228 | 0.03010543 | 0.60011822 |
| 0.999490048 | -0.0029739 | 0.95898356 | 0.99388249 | -0.0566002 | 0.33226843 |
| 0.999490048 | -0.001943 | 0.95904113 | 0.99388249 | 0.02642374 | 0.4870433 |
| 0.999490048 | 0.0021766 | 0.95871157 | 0.99388249 | -0.0516453 | 0.2261747 |
| 0.999490048 | 0.00318815 | 0.95912636 | 0.99388249 | 0.10313135 | 0.10643915 |
| 0.999490048 | 0.00908189 | 0.95901664 | 0.99388249 | 0.12583294 | 0.4815327 |
| 0.999490048 | 0.00264957 | 0.95940779 | 0.99405076 | -0.0372845 | 0.47611879 |
| 0.999490048 | 0.00408849 | 0.96027046 | 0.99474978 | -0.0057369 | 0.94427594 |

| | | | | | |
|---|---|---|---|---|---|
| 0.999490048 | 0.00843061 | 0.96035595 | 0.99474978 | -0.1832354 | 0.29249489 |
| 0.999490048 | 0.00847405 | 0.96043985 | 0.99474978 | 0.07720336 | 0.65200327 |
| 0.999490048 | 0.00213494 | 0.96065813 | 0.99485246 | -0.0297347 | 0.49410788 |
| 0.999490048 | -0.0042117 | 0.96080111 | 0.99487715 | -0.0517621 | 0.54727631 |
| 0.999490048 | -0.0046058 | 0.96107493 | 0.99489704 | 0.01306752 | 0.889889 |
| 0.999490048 | -0.0029424 | 0.96097012 | 0.99489704 | -0.0469142 | 0.43803096 |
| 0.999490048 | 0.00292919 | 0.96117777 | 0.99489704 | 0.06782259 | 0.2657165 |
| 0.999490048 | -0.0030582 | 0.96148017 | 0.99496337 | -0.0306713 | 0.62892016 |
| 0.999490048 | 0.00311084 | 0.96147366 | 0.99496337 | -0.0530248 | 0.41346347 |
| 0.999490048 | -0.0059268 | 0.961647 | 0.9950127 | -0.1486411 | 0.23450738 |
| 0.999490048 | -0.0025387 | 0.96197832 | 0.9952116 | -0.03105 | 0.56119337 |
| 0.999490048 | -0.0020656 | 0.96207761 | 0.9952116 | -0.001659 | 0.96953829 |
| 0.999490048 | -0.002572 | 0.96280778 | 0.99535038 | 0.04274701 | 0.44107344 |
| 0.999490048 | 0.00212852 | 0.96270285 | 0.99535038 | 0.01109099 | 0.80760253 |
| 0.999490048 | 0.00330209 | 0.96252433 | 0.99535038 | 0.00178865 | 0.97969485 |
| 0.999490048 | 0.00514386 | 0.9625398 | 0.99535038 | 0.04788539 | 0.66255281 |
| 0.999490048 | 0.00865224 | 0.96264894 | 0.99535038 | -0.0207341 | 0.91065669 |
| 0.999490048 | 0.00335461 | 0.96328944 | 0.9956699 | 0.06774396 | 0.35678054 |
| 0.999490048 | 0.00898953 | 0.96335534 | 0.9956699 | 0.01456551 | 0.94066606 |
| 0.999490048 | -0.0032918 | 0.96365763 | 0.99569108 | 0.01709723 | 0.81302518 |
| 0.999490048 | -0.0019482 | 0.96350128 | 0.99569108 | 0.08508094 | 0.05433007 |
| 0.999490048 | 0.00169108 | 0.96385281 | 0.99569108 | 0.0165086 | 0.65881442 |
| 0.999490048 | 0.00223101 | 0.96378703 | 0.99569108 | -0.0332433 | 0.50067649 |
| 0.999490048 | 0.00320168 | 0.96406522 | 0.99578731 | -0.0030392 | 0.9658878 |
| 0.999490048 | -0.0089796 | 0.9670209 | 0.9958413 | 0.00950211 | 0.96510301 |
| 0.999490048 | -0.0069146 | 0.96623114 | 0.9958413 | 0.12762719 | 0.43735646 |
| 0.999490048 | -0.0041919 | 0.96606954 | 0.9958413 | 0.05540412 | 0.57517305 |
| 0.999490048 | -0.0034297 | 0.96682211 | 0.9958413 | 0.07792281 | 0.3489225 |
| 0.999490048 | -0.0030099 | 0.9665449 | 0.9958413 | 0.07982482 | 0.27185654 |
| 0.999490048 | -0.0028498 | 0.96575112 | 0.9958413 | 0.02377876 | 0.72048111 |
| 0.999490048 | -0.0024736 | 0.96585068 | 0.9958413 | -0.0490211 | 0.3995749 |
| 0.999490048 | -0.0024628 | 0.96514606 | 0.9958413 | 0.03585606 | 0.52632532 |
| 0.999490048 | -0.0022037 | 0.96448173 | 0.9958413 | -0.0642436 | 0.20164523 |
| 0.999490048 | -0.0020211 | 0.96521368 | 0.9958413 | -0.0176574 | 0.70359542 |
| 0.999490048 | -0.0019565 | 0.96475103 | 0.9958413 | -0.022186 | 0.61716186 |
| 0.999490048 | -0.0017219 | 0.96709906 | 0.9958413 | 0.02907314 | 0.48827955 |
| 0.999490048 | 0.00184388 | 0.9649785 | 0.9958413 | 0.03982217 | 0.34731046 |
| 0.999490048 | 0.00194094 | 0.96611831 | 0.9958413 | -0.0122867 | 0.78816079 |
| 0.999490048 | 0.00201375 | 0.9647539 | 0.9958413 | -0.0375115 | 0.41358827 |
| 0.999490048 | 0.00201467 | 0.96590083 | 0.9958413 | 0.0340953 | 0.47170537 |
| 0.999490048 | 0.00211178 | 0.9664231 | 0.9958413 | 9.57E-04 | 0.98477439 |
| 0.999490048 | 0.00212899 | 0.96595552 | 0.9958413 | -0.0491106 | 0.32949959 |

| | | | | | |
|---|---|---|---|---|---|
| 0.999490048 | 0.00223105 | 0.96544835 | 0.9958413 | -0.0501413 | 0.33484027 |
| 0.999490048 | 0.00232228 | 0.96691917 | 0.9958413 | -0.0290113 | 0.60535877 |
| 0.999490048 | 0.002665 | 0.96449583 | 0.9958413 | -0.0311347 | 0.60401988 |
| 0.999490048 | 0.00334454 | 0.96648257 | 0.9958413 | 0.00486611 | 0.9512511 |
| 0.999490048 | 0.00347463 | 0.96672896 | 0.9958413 | 0.02312655 | 0.78146449 |
| 0.999490048 | 0.00353822 | 0.96537648 | 0.9958413 | 0.13659363 | 0.10290277 |
| 0.999490048 | 0.00462436 | 0.96681135 | 0.9958413 | -0.0790577 | 0.47910813 |
| 0.999490048 | -0.003965 | 0.96797343 | 0.99621345 | -0.091734 | 0.35706909 |
| 0.999490048 | -0.0021511 | 0.96803275 | 0.99621345 | 0.04009948 | 0.45754348 |
| 0.999490048 | -0.0020157 | 0.968057 | 0.99621345 | 0.03055832 | 0.54527063 |
| 0.999490048 | 0.00175021 | 0.96783417 | 0.99621345 | -0.0177818 | 0.68253407 |
| 0.999490048 | 0.00282213 | 0.96781799 | 0.99621345 | -0.0174956 | 0.80261653 |
| 0.999490048 | 0.00334288 | 0.96841003 | 0.99645395 | 0.01493603 | 0.85959387 |
| 0.999490048 | -0.0037153 | 0.9696487 | 0.99686859 | 0.02259637 | 0.817089 |
| 0.999490048 | -0.0025132 | 0.9693048 | 0.99686859 | 0.04241089 | 0.51787752 |
| 0.999490048 | -0.0018301 | 0.96957123 | 0.99686859 | 0.00377801 | 0.93723779 |
| 0.999490048 | 0.00182415 | 0.96922372 | 0.99686859 | -0.0651715 | 0.17605217 |
| 0.999490048 | 0.00245553 | 0.96897219 | 0.99686859 | 0.0028778 | 0.96364002 |
| 0.999490048 | 0.00312392 | 0.96941655 | 0.99686859 | -0.0179998 | 0.82524276 |
| 0.999490048 | 0.00323995 | 0.96931197 | 0.99686859 | 0.02421013 | 0.77393149 |
| 0.999490048 | 0.00225364 | 0.96996105 | 0.99694867 | -0.0645044 | 0.2866482 |
| 0.999490048 | 0.00736763 | 0.96996539 | 0.99694867 | -0.0021134 | 0.99138255 |
| 0.999490048 | 0.0022506 | 0.97032965 | 0.99709187 | -0.0333555 | 0.58260667 |
| 0.999490048 | 0.00333433 | 0.97034353 | 0.99709187 | 0.05671095 | 0.52882811 |
| 0.999490048 | -0.0042128 | 0.97068094 | 0.99719314 | -0.0627987 | 0.58490195 |
| 0.999490048 | 0.00213759 | 0.97065124 | 0.99719314 | 0.02717189 | 0.6407496 |
| 0.999490048 | -0.0049701 | 0.97099309 | 0.99734882 | 0.01625643 | 0.90533703 |
| 0.999490048 | -0.0032744 | 0.97107136 | 0.99734882 | 0.05543265 | 0.54078887 |
| 0.999490048 | -0.001566 | 0.9716328 | 0.99736801 | 0.03872016 | 0.38294393 |
| 0.999490048 | -0.0014983 | 0.97143862 | 0.99736801 | -0.0249402 | 0.55260996 |
| 0.999490048 | 0.00171313 | 0.97121323 | 0.99736801 | -0.0623679 | 0.19646536 |
| 0.999490048 | 0.00240303 | 0.97174264 | 0.99736801 | 0.04346698 | 0.52340027 |
| 0.999490048 | 0.003116 | 0.97157355 | 0.99736801 | -0.0622798 | 0.47855163 |
| 0.999490048 | 0.00369288 | 0.97180672 | 0.99736801 | 0.04358584 | 0.67711124 |
| 0.999490048 | -0.0091469 | 0.97564517 | 0.99745199 | -0.0085959 | 0.97711181 |
| 0.999490048 | -0.0067479 | 0.97703702 | 0.99745199 | 0.37752547 | 0.11631881 |
| 0.999490048 | -0.0053299 | 0.97433504 | 0.99745199 | 0.01419403 | 0.93172806 |
| 0.999490048 | -0.0050368 | 0.97469902 | 0.99745199 | 0.09329765 | 0.55824831 |
| 0.999490048 | -0.0048979 | 0.97937817 | 0.99745199 | 0.10840009 | 0.56853798 |
| 0.999490048 | -0.0048915 | 0.97332336 | 0.99745199 | 0.0969502 | 0.50932314 |
| 0.999490048 | -0.0034788 | 0.97354485 | 0.99745199 | 0.03656494 | 0.72773398 |
| 0.999490048 | -0.0031654 | 0.97621724 | 0.99745199 | 0.07169798 | 0.50147014 |

| | | | | | |
|---|---|---|---|---|---|
| 0.999490048 | -0.0029079 | 0.9743256 | 0.99745199 | -0.0608128 | 0.50284561 |
| 0.999490048 | -0.002895 | 0.97998487 | 0.99745199 | -0.0590881 | 0.60956055 |
| 0.999490048 | -0.0028803 | 0.98365408 | 0.99745199 | -0.0885967 | 0.5301988 |
| 0.999490048 | -0.0026935 | 0.97395105 | 0.99745199 | 0.04559892 | 0.58156074 |
| 0.999490048 | -0.0025955 | 0.97657502 | 0.99745199 | 0.0752706 | 0.3978931 |
| 0.999490048 | -0.0025148 | 0.97801163 | 0.99745199 | -0.0177574 | 0.8457502 |
| 0.999490048 | -0.00239 | 0.97923351 | 0.99745199 | 0.07206426 | 0.43536798 |
| 0.999490048 | -0.002337 | 0.98382789 | 0.99745199 | 0.07459395 | 0.52164176 |
| 0.999490048 | -0.0023244 | 0.97809523 | 0.99745199 | -0.005777 | 0.94559541 |
| 0.999490048 | -0.0022199 | 0.97943852 | 0.99745199 | -0.016775 | 0.84563882 |
| 0.999490048 | -0.0021429 | 0.97970638 | 0.99745199 | 0.044455 | 0.59873113 |
| 0.999490048 | -0.0021265 | 0.97228729 | 0.99745199 | 0.00278351 | 0.96373141 |
| 0.999490048 | -0.0020779 | 0.97768362 | 0.99745199 | 0.03284471 | 0.6590009 |
| 0.999490048 | -0.0019956 | 0.97641748 | 0.99745199 | 0.0388622 | 0.56613537 |
| 0.999490048 | -0.0019944 | 0.97237296 | 0.99745199 | -0.0107257 | 0.85229541 |
| 0.999490048 | -0.001965 | 0.98154279 | 0.99745199 | 0.05635366 | 0.50889608 |
| 0.999490048 | -0.0019365 | 0.97246514 | 0.99745199 | -0.0594488 | 0.29469595 |
| 0.999490048 | -0.0018762 | 0.9793929 | 0.99745199 | -0.0588662 | 0.42074967 |
| 0.999490048 | -0.0018169 | 0.97368463 | 0.99745199 | 0.10233687 | 0.0722346 |
| 0.999490048 | -0.0016447 | 0.97615271 | 0.99745199 | -0.0026567 | 0.96148955 |
| 0.999490048 | -0.0014418 | 0.97488335 | 0.99745199 | 0.00400593 | 0.9302994 |
| 0.999490048 | -0.001439 | 0.97641438 | 0.99745199 | 0.0031631 | 0.94818644 |
| 0.999490048 | -0.0013923 | 0.97567623 | 0.99745199 | -0.0446658 | 0.33259679 |
| 0.999490048 | -0.0013564 | 0.98053031 | 0.99745199 | 0.01191984 | 0.83026566 |
| 0.999490048 | -0.0013431 | 0.97558701 | 0.99745199 | -0.012177 | 0.78160011 |
| 0.999490048 | -0.0012922 | 0.98277139 | 0.99745199 | 0.044122 | 0.46335524 |
| 0.999490048 | -0.0012901 | 0.98322821 | 0.99745199 | -0.0595849 | 0.33610598 |
| 0.999490048 | -0.0012566 | 0.98269984 | 0.99745199 | -0.0122551 | 0.83258856 |
| 0.999490048 | -0.0012388 | 0.98293734 | 0.99745199 | -0.0240935 | 0.67796867 |
| 0.999490048 | -0.0011713 | 0.98119975 | 0.99745199 | 0.07296241 | 0.15062637 |
| 0.999490048 | -0.0011625 | 0.97872025 | 0.99745199 | 1.45E-04 | 0.99734738 |
| 0.999490048 | -0.0011567 | 0.97989609 | 0.99745199 | -0.0281456 | 0.54130203 |
| 0.999490048 | -0.0011355 | 0.98264272 | 0.99745199 | 0.0022597 | 0.96546753 |
| 0.999490048 | -0.0011338 | 0.98285052 | 0.99745199 | -0.0172631 | 0.74371258 |
| 0.999490048 | -0.0011127 | 0.98407299 | 0.99745199 | 0.00478844 | 0.93154378 |
| 0.999490048 | -0.0010864 | 0.9830529 | 0.99745199 | 0.00269937 | 0.95791095 |
| 0.999490048 | -0.0010853 | 0.98350253 | 0.99745199 | 0.02325528 | 0.65835351 |
| 0.999490048 | -0.0010046 | 0.98369953 | 0.99745199 | -0.0084301 | 0.86390652 |
| 0.999490048 | 8.27E-04 | 0.98364596 | 0.99745199 | -0.0420967 | 0.30201552 |
| 0.999490048 | 9.24E-04 | 0.98398316 | 0.99745199 | -0.0213436 | 0.64367589 |
| 0.999490048 | 9.35E-04 | 0.9814021 | 0.99745199 | -0.0118076 | 0.76871546 |
| 0.999490048 | 0.0010584 | 0.98258161 | 0.99745199 | 0.01913142 | 0.69356507 |

| | | | | | |
|---|---|---|---|---|---|
| 0.999490048 | 0.00107453 | 0.97893445 | 0.99745199 | 0.04529771 | 0.27153028 |
| 0.999490048 | 0.00107828 | 0.97894445 | 0.99745199 | 0.01961282 | 0.63198173 |
| 0.999490048 | 0.00121737 | 0.98117794 | 0.99745199 | -0.0390064 | 0.45228229 |
| 0.999490048 | 0.00121903 | 0.98048477 | 0.99745199 | 0.00671724 | 0.8927983 |
| 0.999490048 | 0.00126684 | 0.98011167 | 0.99745199 | -0.0060851 | 0.9047005 |
| 0.999490048 | 0.00136392 | 0.97843947 | 0.99745199 | -0.0173764 | 0.73092445 |
| 0.999490048 | 0.00137974 | 0.97498193 | 0.99745199 | -0.0217291 | 0.62223808 |
| 0.999490048 | 0.00142629 | 0.9782928 | 0.99745199 | -9.58E-04 | 0.9854207 |
| 0.999490048 | 0.00143002 | 0.97718948 | 0.99745199 | 0.00202541 | 0.96769708 |
| 0.999490048 | 0.00144149 | 0.97993032 | 0.99745199 | -0.038488 | 0.50372201 |
| 0.999490048 | 0.00147459 | 0.97977655 | 0.99745199 | 0.02404366 | 0.67988741 |
| 0.999490048 | 0.00148979 | 0.97912948 | 0.99745199 | -0.0078026 | 0.89104193 |
| 0.999490048 | 0.00151674 | 0.98319178 | 0.99745199 | -0.0707658 | 0.33028602 |
| 0.999490048 | 0.00159411 | 0.98134461 | 0.99745199 | -0.1319234 | 0.06185502 |
| 0.999490048 | 0.00159726 | 0.97339626 | 0.99745199 | 3.73E-04 | 0.99379084 |
| 0.999490048 | 0.00160292 | 0.9839177 | 0.99745199 | -0.0811322 | 0.31259925 |
| 0.999490048 | 0.00161232 | 0.97538565 | 0.99745199 | 0.07468547 | 0.16121964 |
| 0.999490048 | 0.00161447 | 0.97765137 | 0.99745199 | -0.0558636 | 0.33690401 |
| 0.999490048 | 0.00163185 | 0.98084155 | 0.99745199 | -0.0711167 | 0.30056533 |
| 0.999490048 | 0.0016575 | 0.9815313 | 0.99745199 | -0.0877737 | 0.22710754 |
| 0.999490048 | 0.00166916 | 0.98128692 | 0.99745199 | -0.0377438 | 0.59688289 |
| 0.999490048 | 0.00169747 | 0.97318556 | 0.99745199 | -0.0323954 | 0.52291708 |
| 0.999490048 | 0.00171415 | 0.97536525 | 0.99745199 | -0.0082869 | 0.88135288 |
| 0.999490048 | 0.0017533 | 0.97826426 | 0.99745199 | -0.0103313 | 0.87248646 |
| 0.999490048 | 0.00176497 | 0.97899935 | 0.99745199 | 0.02214342 | 0.74147907 |
| 0.999490048 | 0.00177567 | 0.97713268 | 0.99745199 | -0.0416438 | 0.50336534 |
| 0.999490048 | 0.0018219 | 0.98031168 | 0.99745199 | -0.0575982 | 0.4380734 |
| 0.999490048 | 0.00183824 | 0.98383474 | 0.99745199 | -0.0167384 | 0.85367471 |
| 0.999490048 | 0.00185744 | 0.97384336 | 0.99745199 | 0.09206436 | 0.11316531 |
| 0.999490048 | 0.00191189 | 0.97831042 | 0.99745199 | 0.01087069 | 0.87717952 |
| 0.999490048 | 0.00196068 | 0.97368255 | 0.99745199 | 0.01650363 | 0.78141655 |
| 0.999490048 | 0.0019653 | 0.9817694 | 0.99745199 | 0.01200121 | 0.88904567 |
| 0.999490048 | 0.00201199 | 0.97507746 | 0.99745199 | 0.11847569 | 0.07491761 |
| 0.999490048 | 0.00201374 | 0.98375614 | 0.99745199 | -0.0218953 | 0.82488545 |
| 0.999490048 | 0.00202202 | 0.98022125 | 0.99745199 | 0.0480411 | 0.55722157 |
| 0.999490048 | 0.00205867 | 0.97695548 | 0.99745199 | -0.0145654 | 0.8381287 |
| 0.999490048 | 0.00214235 | 0.97411729 | 0.99745199 | -0.0306138 | 0.64362684 |
| 0.999490048 | 0.00219619 | 0.98397181 | 0.99745199 | -0.0471362 | 0.6669229 |
| 0.999490048 | 0.00228566 | 0.97491628 | 0.99745199 | 0.12746993 | 0.08867043 |
| 0.999490048 | 0.00242292 | 0.97236377 | 0.99745199 | -0.0288858 | 0.68010811 |
| 0.999490048 | 0.00248343 | 0.97753824 | 0.99745199 | 0.03470459 | 0.69443336 |
| 0.999490048 | 0.00249809 | 0.98004402 | 0.99745199 | 0.10763714 | 0.28667286 |

| | | | | | |
|---|---|---|---|---|---|
| 0.999490048 | 0.0025483 | 0.97955625 | 0.99745199 | -0.0363502 | 0.71508062 |
| 0.999490048 | 0.00290095 | 0.97477196 | 0.99745199 | 0.01788362 | 0.8454862 |
| 0.999490048 | 0.003291 | 0.97677885 | 0.99745199 | -8.01E-04 | 0.99434836 |
| 0.999490048 | 0.00356806 | 0.97833354 | 0.99745199 | 0.05467564 | 0.67782029 |
| 0.999490048 | 0.00367783 | 0.97795496 | 0.99745199 | 0.11491876 | 0.39142852 |
| 0.999490048 | 0.00418572 | 0.9733917 | 0.99745199 | -0.0876504 | 0.48701826 |
| 0.971241857 | 0.00467498 | 0.98348815 | 0.99745199 | -0.5280055 | 0.02629317 |
| 0.999490048 | 0.00607138 | 0.974052 | 0.99745199 | 0.00846101 | 0.96384552 |
| 0.999490048 | 0.00975326 | 0.9742667 | 0.99745199 | -0.3803171 | 0.23018289 |
| 0.987721587 | 0.01100064 | 0.97287602 | 0.99745199 | 0.73704908 | 0.03961295 |
| 0.999490048 | -0.0030477 | 0.98504565 | 0.99747253 | -0.1640537 | 0.31790258 |
| 0.999490048 | -0.0022222 | 0.98442313 | 0.99747253 | -0.0553572 | 0.62752349 |
| 0.999490048 | -0.0015061 | 0.98537521 | 0.99747253 | -0.0016471 | 0.98400687 |
| 0.999490048 | -0.0014756 | 0.9845793 | 0.99747253 | 0.01124419 | 0.88293624 |
| 0.999490048 | -0.0010152 | 0.98449005 | 0.99747253 | 0.00402028 | 0.93863862 |
| 0.999490048 | -9.93E-04 | 0.9852872 | 0.99747253 | -0.0321786 | 0.55145139 |
| 0.999490048 | -9.74E-04 | 0.9845725 | 0.99747253 | -0.0170247 | 0.73569504 |
| 0.999490048 | -8.95E-04 | 0.98508023 | 0.99747253 | -0.0397781 | 0.40930138 |
| 0.999490048 | -8.27E-04 | 0.984941 | 0.99747253 | -0.0350555 | 0.42660917 |
| 0.999490048 | 0.00110718 | 0.9854073 | 0.99747253 | 0.01373825 | 0.82055231 |
| 0.999490048 | 0.00232471 | 0.98503293 | 0.99747253 | 0.00862348 | 0.94452388 |
| 0.999490048 | -0.0010182 | 0.98559246 | 0.99753904 | 0.04426377 | 0.4352557 |
| 0.999490048 | -0.0020856 | 0.98603265 | 0.99757436 | 0.11103329 | 0.35549362 |
| 0.999490048 | -0.0010089 | 0.9861636 | 0.99757436 | 5.21E-04 | 0.99284824 |
| 0.999490048 | 8.87E-04 | 0.98634418 | 0.99757436 | 0.00247051 | 0.96199894 |
| 0.999490048 | 0.00109526 | 0.98629768 | 0.99757436 | 0.00738825 | 0.90778318 |
| 0.999490048 | 0.00215385 | 0.98632506 | 0.99757436 | 0.10628538 | 0.4010344 |
| 0.999490048 | 0.00242015 | 0.9858534 | 0.99757436 | -0.0509645 | 0.70924932 |
| 0.999490048 | -6.67E-04 | 0.98679061 | 0.99766335 | -0.0669897 | 0.10524135 |
| 0.999490048 | 7.63E-04 | 0.98674201 | 0.99766335 | 5.68E-04 | 0.99012568 |
| 0.999490048 | 0.00096815 | 0.98670691 | 0.99766335 | 0.07041125 | 0.23235804 |
| 0.999490048 | -0.0011478 | 0.98696386 | 0.9977177 | -0.0722228 | 0.30898433 |
| 0.999490048 | -0.0033926 | 0.98793945 | 0.99773597 | -0.2617004 | 0.24994223 |
| 0.999490048 | -0.0025258 | 0.98798425 | 0.99773597 | -0.0429403 | 0.79805993 |
| 0.999490048 | -0.0021689 | 0.98712652 | 0.99773597 | -0.0403484 | 0.76425982 |
| 0.999490048 | -0.0020197 | 0.9884158 | 0.99773597 | -0.071693 | 0.60725196 |
| 0.999490048 | -0.001836 | 0.9883681 | 0.99773597 | 0.12224829 | 0.33621818 |
| 0.999490048 | -0.0010262 | 0.98774729 | 0.99773597 | 0.02829061 | 0.67258133 |
| 0.999490048 | -8.54E-04 | 0.98782088 | 0.99773597 | -0.0212467 | 0.7044623 |
| 0.999490048 | 8.51E-04 | 0.98777024 | 0.99773597 | 0.00273723 | 0.96066773 |
| 0.819178176 | 9.23E-04 | 0.98828724 | 0.99773597 | 0.16593926 | 0.01325264 |
| 0.999490048 | 9.83E-04 | 0.98832487 | 0.99773597 | 0.06982244 | 0.30382371 |

| | | | | | |
|---|---|---|---|---|---|
| 0.999490048 | 0.00104697 | 0.98773558 | 0.99773597 | -0.0488971 | 0.47509421 |
| 0.999490048 | 0.00195648 | 0.98776312 | 0.99773597 | -0.0191623 | 0.8806197 |
| 0.999490048 | -0.0010327 | 0.98870243 | 0.99790467 | -0.0737181 | 0.31704213 |
| 0.999490048 | -0.0023747 | 0.98896499 | 0.99800282 | 0.03159123 | 0.854066 |
| 0.999490048 | 5.06E-04 | 0.98903872 | 0.99800282 | 0.00544813 | 0.88238103 |
| 0.999490048 | -9.21E-04 | 0.98933403 | 0.99812394 | 0.03264937 | 0.63633555 |
| 0.999490048 | 8.71E-04 | 0.98939782 | 0.99812394 | -0.0110344 | 0.86635836 |
| 0.999490048 | -0.0020738 | 0.9904053 | 0.99818605 | -0.0277672 | 0.87211536 |
| 0.999490048 | -0.0020386 | 0.99053528 | 0.99818605 | -0.0463748 | 0.78742705 |
| 0.999490048 | -0.0010063 | 0.98963198 | 0.99818605 | 0.02983931 | 0.70041757 |
| 0.999490048 | -7.30E-04 | 0.99045315 | 0.99818605 | -0.0073001 | 0.90479868 |
| 0.999490048 | -5.43E-04 | 0.98990419 | 0.99818605 | 0.01804741 | 0.67476173 |
| 0.999490048 | 5.18E-04 | 0.99018205 | 0.99818605 | -0.0056032 | 0.89417533 |
| 0.999490048 | 7.25E-04 | 0.98976967 | 0.99818605 | -0.0015895 | 0.9775875 |
| 0.999490048 | 0.0012397 | 0.99043894 | 0.99818605 | 0.14495109 | 0.16929617 |
| 0.999490048 | 0.00203571 | 0.99042654 | 0.99818605 | 0.08528079 | 0.61610065 |
| 0.999490048 | -0.0012853 | 0.99066997 | 0.99820131 | 0.069513 | 0.528763 |
| 0.999490048 | 5.10E-04 | 0.99083439 | 0.99824652 | -0.0268012 | 0.54733196 |
| 0.999490048 | -0.001352 | 0.99159767 | 0.99841319 | -0.0249054 | 0.84624338 |
| 0.999490048 | -7.05E-04 | 0.99126068 | 0.99841319 | 0.01739828 | 0.78715262 |
| 0.999490048 | -4.95E-04 | 0.99123041 | 0.99841319 | 0.04299671 | 0.3444081 |
| 0.999490048 | 5.66E-04 | 0.99158556 | 0.99841319 | 0.06453785 | 0.23571902 |
| 0.999490048 | 9.56E-04 | 0.9914833 | 0.99841319 | -0.0205426 | 0.81871855 |
| 0.999490048 | -8.26E-04 | 0.9917544 | 0.9984506 | -0.0645388 | 0.42240626 |
| 0.999490048 | -0.0013898 | 0.9920492 | 0.99862698 | 0.00138974 | 0.99204966 |
| 0.999490048 | -0.002483 | 0.99267296 | 0.99870298 | 0.34960571 | 0.2125788 |
| 0.999490048 | -0.0010102 | 0.99334325 | 0.99870298 | 0.04488349 | 0.7112601 |
| 0.999490048 | -5.94E-04 | 0.99343363 | 0.99870298 | -0.0445692 | 0.53861257 |
| 0.999490048 | -5.80E-04 | 0.99321965 | 0.99870298 | 0.04116063 | 0.54788264 |
| 0.999490048 | -3.92E-04 | 0.9932272 | 0.99870298 | -0.0147577 | 0.74923679 |
| 0.999490048 | -3.49E-04 | 0.99324063 | 0.99870298 | -0.0471124 | 0.25915864 |
| 0.999490048 | 4.16E-04 | 0.99326231 | 0.99870298 | -0.0396388 | 0.42393736 |
| 0.999490048 | 4.56E-04 | 0.99348877 | 0.99870298 | -0.0486267 | 0.38792236 |
| 0.999490048 | 4.59E-04 | 0.99293973 | 0.99870298 | 0.07918775 | 0.13525472 |
| 0.999490048 | 5.18E-04 | 0.99355996 | 0.99870298 | -0.0155331 | 0.80872013 |
| 0.999490048 | 5.47E-04 | 0.99250433 | 0.99870298 | 0.08016372 | 0.17619793 |
| 0.999490048 | 7.70E-04 | 0.99231862 | 0.99870298 | 0.11828864 | 0.14779307 |
| 0.999490048 | 0.00157316 | 0.9940336 | 0.9990588 | 0.05322297 | 0.80040242 |
| 0.999490048 | -5.98E-04 | 0.99416178 | 0.99906738 | -0.0702385 | 0.39373065 |
| 0.999490048 | -4.25E-04 | 0.9946386 | 0.99924514 | 0.07646156 | 0.23361319 |
| 0.999490048 | -4.17E-04 | 0.99488451 | 0.99924514 | -0.0181377 | 0.78066217 |
| 0.999490048 | -3.84E-04 | 0.99493701 | 0.99924514 | -0.0291682 | 0.6304771 |

| | | | | | |
|---|---|---|---|---|---|
| 0.999490048 | -3.39E-04 | 0.99453618 | 0.99924514 | 0.06257308 | 0.21401972 |
| 0.999490048 | 3.27E-04 | 0.99489294 | 0.99924514 | -0.0568432 | 0.27221749 |
| 0.999490048 | 5.23E-04 | 0.99515013 | 0.99933897 | -0.0416688 | 0.62895951 |
| 0.999490048 | -3.33E-04 | 0.99538508 | 0.9994188 | 0.1007051 | 0.0891356 |
| 0.999490048 | -2.70E-04 | 0.99554524 | 0.9994188 | -0.0358808 | 0.46058696 |
| 0.999490048 | 3.24E-04 | 0.99558869 | 0.9994188 | -0.0447255 | 0.44803316 |
| 0.999490048 | 2.92E-04 | 0.99592512 | 0.9995162 | 0.03114154 | 0.58682479 |
| 0.999490048 | 0.00231651 | 0.99590824 | 0.9995162 | 0.50059629 | 0.2856622 |
| 0.999490048 | -4.81E-04 | 0.99626926 | 0.99968232 | 0.03549014 | 0.73058295 |
| 0.835632585 | -4.04E-04 | 0.99678722 | 0.99968232 | -0.2608296 | 0.01453157 |
| 0.999490048 | -3.76E-04 | 0.99812593 | 0.99968232 | -0.0437058 | 0.78549699 |
| 0.999490048 | -3.00E-04 | 0.99793054 | 0.99968232 | 0.07544605 | 0.52002926 |
| 0.999490048 | -2.93E-04 | 0.99770016 | 0.99968232 | 0.06488882 | 0.52516041 |
| 0.999490048 | -1.86E-04 | 0.99811695 | 0.99968232 | -0.0286272 | 0.71621844 |
| 0.999490048 | -1.81E-04 | 0.99734169 | 0.99968232 | -0.0155791 | 0.77419623 |
| 0.999490048 | -1.60E-04 | 0.99732302 | 0.99968232 | -0.0068956 | 0.88498494 |
| 0.999490048 | -1.53E-04 | 0.99744945 | 0.99968232 | 0.02734355 | 0.56854609 |
| 0.999490048 | -1.38E-04 | 0.99705735 | 0.99968232 | 0.00441266 | 0.9063822 |
| 0.999490048 | -1.23E-04 | 0.99749275 | 0.99968232 | 0.02585816 | 0.51114019 |
| 0.999490048 | 1.52E-04 | 0.99807883 | 0.99968232 | -7.17E-04 | 0.99095934 |
| 0.999490048 | 1.70E-04 | 0.99688152 | 0.99968232 | -0.0252313 | 0.56209452 |
| 0.999490048 | 3.36E-04 | 0.99725744 | 0.99968232 | 0.01330332 | 0.89167982 |
| 0.999490048 | 4.05E-04 | 0.99754502 | 0.99968232 | -0.050744 | 0.70033311 |
| 0.781057588 | 4.22E-04 | 0.9976081 | 0.99968232 | -0.3738131 | 0.01281076 |
| 0.999490048 | 8.59E-04 | 0.99686415 | 0.99968232 | -0.2973367 | 0.19149941 |
| 0.999490048 | -1.21E-04 | 0.99837867 | 0.99981553 | -0.0379025 | 0.52757012 |
| 0.999490048 | 8.56E-05 | 0.99851162 | 0.99982876 | -0.0093283 | 0.83895006 |
| 0.999490048 | -1.80E-05 | 0.99974548 | 0.999985 | -0.0357817 | 0.52769264 |
| 0.999490048 | 2.44E-05 | 0.99972243 | 0.999985 | 0.02152118 | 0.75930517 |
| 0.999490048 | 3.12E-05 | 0.99958731 | 0.999985 | 0.04221437 | 0.48624815 |
| 0.999490048 | 3.77E-05 | 0.99948932 | 0.999985 | -0.0245614 | 0.67694518 |
| 0.999490048 | 4.24E-05 | 0.99923468 | 0.999985 | -0.0161809 | 0.71440227 |
| 0.999490048 | 4.36E-05 | 0.99915849 | 0.999985 | -0.0111622 | 0.78727223 |
| 0.999490048 | 5.05E-05 | 0.99908583 | 0.999985 | -0.0158055 | 0.72010976 |
| 0.999490048 | 6.34E-05 | 0.99943427 | 0.999985 | -0.0619681 | 0.49061997 |
| 0.999490048 | 7.11E-05 | 0.99892834 | 0.999985 | 0.02054357 | 0.69823053 |
| 0.999490048 | 7.74E-06 | 0.99986559 | 0.99998535 | -0.0377082 | 0.41511337 |
| 0.999490048 | 2.59E-08 | 0.9999996 | 0.9999996 | 0.02058413 | 0.68806415 |

| AMPH-04647.6h - DMSO.6h__adj.P.Val |
| --- |
| 0.99252883 |
| 0.72975604 |
| 0.66733542 |
| 0.99122857 |
| 0.92548507 |
| 0.90265407 |
| 0.91267201 |
| 0.72975604 |
| 0.72204897 |
| 0.79151544 |
| 0.86545047 |
| 0.74240405 |
| 0.69921142 |
| 0.67021692 |
| 0.80984148 |
| 0.55521624 |
| 0.92861642 |
| 0.94171831 |
| 0.72975604 |
| 0.90265407 |
| 0.99998478 |
| 0.98924373 |
| 0.92672232 |
| 0.9829938 |
| 0.85141726 |
| 0.95585979 |
| 0.96709983 |
| 0.96484568 |
| 0.99252883 |
| 0.55521624 |
| 0.93752249 |
| 0.78307327 |
| 0.85669195 |
| 0.94171831 |
| 0.96181283 |
| 0.90265407 |
| 0.99183457 |
| 0.98956725 |

| |
|---|
| 0.74240405 |
| 0.98924373 |
| 0.96484568 |
| 0.94171831 |
| 0.98275443 |
| 0.99998478 |
| 0.80984148 |
| 0.96484568 |
| 0.90265407 |
| 0.98924373 |
| 0.55521624 |
| 0.69921142 |
| 0.96484568 |
| 0.93752249 |
| 0.91325047 |
| 0.99998478 |
| 0.98924373 |
| 0.926218 |
| 0.61773657 |
| 0.65445413 |
| 0.91325047 |
| 0.61079703 |
| 0.9520243 |
| 0.97078973 |
| 0.69921142 |
| 0.95951105 |
| 0.81286841 |
| 0.99998478 |
| 0.93752249 |
| 0.79151544 |
| 0.91325047 |
| 0.98732643 |
| 0.90297323 |
| 0.69921142 |
| 0.99998478 |
| 0.94171831 |
| 0.98924373 |
| 0.72975604 |
| 0.79151544 |
| 0.99998478 |
| 0.87052203 |
| 0.69921142 |

| |
|---|
| 0.90265407 |
| 0.80133949 |
| 0.926218 |
| 0.86079694 |
| 0.90265407 |
| 0.9467702 |
| 0.74240405 |
| 0.8773376 |
| 0.87052203 |
| 0.85669195 |
| 0.91325047 |
| 0.85141726 |
| 0.69921142 |
| 0.90265407 |
| 0.98924373 |
| 0.99969417 |
| 0.9403055 |
| 0.89958215 |
| 0.79151544 |
| 0.90297323 |
| 0.99122857 |
| 0.55521624 |
| 0.99122857 |
| 0.8773376 |
| 0.99122857 |
| 0.93752249 |
| 0.96484568 |
| 0.91325047 |
| 0.87669383 |
| 0.96181283 |
| 0.90297323 |
| 0.90265407 |
| 0.8773376 |
| 0.90297323 |
| 0.79151544 |
| 0.85141726 |
| 0.81286841 |
| 0.91325047 |
| 0.9403055 |
| 0.82029292 |
| 0.79151544 |
| 0.90265407 |

| |
|---|
| 0.94653188 |
| 0.72975604 |
| 0.93752249 |
| 0.65854402 |
| 0.80984148 |
| 0.95240191 |
| 0.90265407 |
| 0.99252883 |
| 0.71816793 |
| 0.58876713 |
| 0.99122857 |
| 0.90265407 |
| 0.85141726 |
| 0.8773376 |
| 0.96167371 |
| 0.93752249 |
| 0.87052203 |
| 0.92861642 |
| 0.96709089 |
| 0.7628002 |
| 0.91325047 |
| 0.99998478 |
| 0.99122857 |
| 0.97209533 |
| 0.93539424 |
| 0.80984148 |
| 0.79151544 |
| 0.72975604 |
| 0.93752249 |
| 0.74240405 |
| 0.9467702 |
| 0.61977276 |
| 0.81286841 |
| 0.80984148 |
| 0.90297323 |
| 0.99122857 |
| 0.91325047 |
| 0.94171831 |
| 0.93752249 |
| 0.98924373 |
| 0.61079703 |
| 0.99998478 |

| |
|---|
| 0.91325047 |
| 0.99998478 |
| 0.80984148 |
| 0.86327908 |
| 0.90265407 |
| 0.90265407 |
| 0.7958357 |
| 0.95034906 |
| 0.55521624 |
| 0.85141726 |
| 0.98924373 |
| 0.98924373 |
| 0.98924373 |
| 0.9403055 |
| 0.95240191 |
| 0.90265407 |
| 0.84320674 |
| 0.55521624 |
| 0.92861642 |
| 0.93752249 |
| 0.7958357 |
| 0.79151544 |
| 0.72975604 |
| 0.79151544 |
| 0.93427164 |
| 0.90265407 |
| 0.69921142 |
| 0.72975604 |
| 0.99998478 |
| 0.99969417 |
| 0.61773657 |
| 0.55521624 |
| 0.90265407 |
| 0.96484568 |
| 0.9467702 |
| 0.96181283 |
| 0.96484568 |
| 0.95859413 |
| 0.90297323 |
| 0.61773657 |
| 0.99969417 |
| 0.9467702 |

0.78927011
0.98924373
0.79151544
0.94412092
0.79151544
0.8773376
0.84586312
0.94747425
0.61977276
0.61773657
0.95951105
0.87669383
0.99101915
0.61079703
0.70718912
0.98924373
0.93752249
0.99830763
0.72975604
0.96865889
0.66733542
0.92861642
0.99969417
0.65854402
0.93752249
0.90265407
0.98924373
0.98924373
0.91325047
0.65854402
0.95034906
0.55521624
0.90265407
0.94171831
0.94653188
0.98924373
0.9403055
0.96484568
0.96484568
0.55521624
0.87669383
0.98924373

0.98924373
0.90265407
0.96167371
0.87669383
0.77062555
0.80618748
0.99122857
0.55521624
0.69921142
0.99122857
0.85971745
0.96484568
0.91325047
0.99998478
0.99998478
0.90265407
0.96181283
0.96484568
0.91267201
0.99252883
0.98924373
0.90265407
0.96181283
0.99969417
0.86327908
0.79777407
0.85141726
0.99122857
0.98924373
0.90265407
0.99122857
0.86537971
0.94171831
0.61079703
0.90265407
0.99998478
0.86537971
0.90265407
0.80984148
0.98924373
0.9403055
0.98924373

| |
|---|
| 0.74240405 |
| 0.95951105 |
| 0.79151544 |
| 0.98924373 |
| 0.95951105 |
| 0.96484568 |
| 0.926218 |
| 0.90265407 |
| 0.90297323 |
| 0.9403055 |
| 0.90297323 |
| 0.97815363 |
| 0.96484568 |
| 0.91105653 |
| 0.94171831 |
| 0.94171831 |
| 0.9865573 |
| 0.93752249 |
| 0.98325199 |
| 0.92861642 |
| 0.90265407 |
| 0.97368509 |
| 0.82029292 |
| 0.69921142 |
| 0.93752249 |
| 0.99969417 |
| 0.90265407 |
| 0.99998478 |
| 0.93928067 |
| 0.85141726 |
| 0.98924373 |
| 0.79151544 |
| 0.90265407 |
| 0.98362105 |
| 0.814903 |
| 0.90297323 |
| 0.926218 |
| 0.94653188 |
| 0.94171831 |
| 0.915492 |
| 0.85669195 |
| 0.94171831 |

| |
|---|
| 0.90265407 |
| 0.90265407 |
| 0.99969417 |
| 0.87944932 |
| 0.79151544 |
| 0.96181283 |
| 0.98732643 |
| 0.95951105 |
| 0.61773657 |
| 0.98924373 |
| 0.97954048 |
| 0.92075126 |
| 0.90265407 |
| 0.98924373 |
| 0.74240405 |
| 0.9467702 |
| 0.93752249 |
| 0.95859413 |
| 0.99122857 |
| 0.98924373 |
| 0.84320674 |
| 0.9403055 |
| 0.92861642 |
| 0.926218 |
| 0.98924373 |
| 0.98924373 |
| 0.8773376 |
| 0.85141726 |
| 0.89474679 |
| 0.79151544 |
| 0.90297323 |
| 0.80984148 |
| 0.85141726 |
| 0.91325047 |
| 0.74240405 |
| 0.93811476 |
| 0.92861642 |
| 0.90265407 |
| 0.94747425 |
| 0.99969417 |
| 0.98259014 |
| 0.90265407 |

| |
|---|
| 0.93054695 |
| 0.93752249 |
| 0.93752249 |
| 0.99998478 |
| 0.26622013 |
| 0.81293822 |
| 0.74240405 |
| 0.90265407 |
| 0.98924373 |
| 0.8773376 |
| 0.93594857 |
| 0.99998478 |
| 0.90265407 |
| 0.99122857 |
| 0.87669383 |
| 0.98924373 |
| 0.74240405 |
| 0.93752249 |
| 0.66733542 |
| 0.86327908 |
| 0.99998478 |
| 0.94171831 |
| 0.87669383 |
| 0.95240191 |
| 0.99998478 |
| 0.99122857 |
| 0.79151544 |
| 0.78307327 |
| 0.98259014 |
| 0.91325047 |
| 0.94402118 |
| 0.99998478 |
| 0.94171831 |
| 0.98259014 |
| 0.92861642 |
| 0.92861642 |
| 0.93752249 |
| 0.94171831 |
| 0.99969417 |
| 0.95951105 |
| 0.8773376 |
| 0.8773376 |

| |
|---|
| 0.84076524 |
| 0.97954048 |
| 0.98924373 |
| 0.69921142 |
| 0.99122857 |
| 0.79151544 |
| 0.94171831 |
| 0.90297323 |
| 0.93752249 |
| 0.91325047 |
| 0.99830763 |
| 0.9403055 |
| 0.9403055 |
| 0.61773657 |
| 0.95859413 |
| 0.96181283 |
| 0.93752249 |
| 0.84430401 |
| 0.93752249 |
| 0.98924373 |
| 0.74240405 |
| 0.61773657 |
| 0.94596453 |
| 0.94171831 |
| 0.87669383 |
| 0.9403055 |
| 0.96484568 |
| 0.87845 |
| 0.90297323 |
| 0.94641693 |
| 0.87052203 |
| 0.90265407 |
| 0.26622013 |
| 0.86537971 |
| 0.99186511 |
| 0.72975604 |
| 0.92548507 |
| 0.66733542 |
| 0.90265407 |
| 0.8773376 |
| 0.90265407 |
| 0.8773376 |

0.92861642
0.99998478
0.90297323
0.99998478
0.72975604
0.99969417
0.78307327
0.94002917
0.92548507
0.97368509
0.98924373
0.97126196
0.85141726
0.93752249
0.84625624
0.90265407
0.99969417
0.96181283
0.99846792
0.99998478
0.79151544
0.79151544
0.90265407
0.98735899
0.91325047
0.8773376
0.99969417
0.93752249
0.90265407
0.99998478
0.80984148
0.98924373
0.85141726
0.95951105
0.90265407
0.96484568
0.91325047
0.98924373
0.8773376
0.99969417
0.98924373
0.61773657

0.94171831
0.87669383
0.90297323
0.99969417
0.915492
0.99998478
0.85141726
0.85141726
0.8773376
0.90265407
0.97329503
0.98924373
0.91325047
0.87669383
0.90265407
0.96484568
0.85141726
0.90265407
0.86974476
0.9715425
0.55521624
0.90265407
0.90265407
0.99998478
0.9834625
0.97110689
0.91325047
0.79151544
0.94171831
0.96484568
0.91325047
0.99998478
0.91325047
0.95240191
0.99830763
0.99998478
0.9829938
0.96484568
0.72975604
0.90297323
0.99122857
0.98924373

0.99998478
0.94171831
0.92244927
0.61773657
0.92861642
0.85141726
0.92861642
0.98924373
0.99122857
0.61773657
0.99969417
0.98924373
0.95240191
0.99122857
0.96181283
0.95034906
0.93752249
0.99998478
0.92861642
0.9403055
0.99998478
0.9303776
0.9715425
0.90297323
0.85141726
0.91485898
0.8773376
0.93752249
0.96484568
0.98924373
0.94653188
0.90265407
0.99998478
0.91325047
0.96181283
0.94412092
0.98259014
0.98924373
0.99998478
0.93908842
0.99830763
0.93752249

0.9834625
0.8773376
0.98924373
0.85141726
0.90297323
0.90265407
0.90265407
0.93752249
0.99122857
0.99998478
0.93752249
0.84320674
0.91325047
0.98924373
0.95585979
0.9829938
0.99969417
0.85141726
0.87052203
0.99122857
0.72975604
0.90297323
0.86327908
0.96484568
0.98924373
0.99998478
0.90297323
0.90297323
0.99998478
0.90265407
0.99122857
0.95134169
0.9467702
0.98259014
0.90265407
0.97024001
0.8773376
0.98924373
0.787647
0.99122857
0.96484568
0.85141726

| |
|---|
| 0.95240191 |
| 0.93752249 |
| 0.95622281 |
| 0.90473298 |
| 0.93594857 |
| 0.90265407 |
| 0.79151544 |
| 0.99122857 |
| 0.92861642 |
| 0.91325047 |
| 0.98924373 |
| 0.99650199 |
| 0.98924373 |
| 0.99122857 |
| 0.99830763 |
| 0.99122857 |
| 0.90297323 |
| 0.72975604 |
| 0.80984148 |
| 0.99998478 |
| 0.61079703 |
| 0.80984148 |
| 0.926218 |
| 0.93752249 |
| 0.98259014 |
| 0.90297323 |
| 0.94171831 |
| 0.93752249 |
| 0.86327908 |
| 0.8773376 |
| 0.99969417 |
| 0.85141726 |
| 0.94897478 |
| 0.99122857 |
| 0.82029292 |
| 0.80984148 |
| 0.90265407 |
| 0.99122857 |
| 0.8971924 |
| 0.99998478 |
| 0.9403055 |
| 0.99122857 |

| |
|---|
| 0.89618249 |
| 0.92367831 |
| 0.90297323 |
| 0.99998478 |
| 0.90265407 |
| 0.93811476 |
| 0.91325047 |
| 0.85141726 |
| 0.915492 |
| 0.89060223 |
| 0.92367831 |
| 0.99122857 |
| 0.90265407 |
| 0.99998478 |
| 0.69921142 |
| 0.90265407 |
| 0.93151025 |
| 0.90297323 |
| 0.99122857 |
| 0.94683707 |
| 0.61773657 |
| 0.91325047 |
| 0.94171831 |
| 0.99998478 |
| 0.78307327 |
| 0.98924373 |
| 0.9403055 |
| 0.90265407 |
| 0.95654768 |
| 0.96353697 |
| 0.9403055 |
| 0.99122857 |
| 0.90265407 |
| 0.61773657 |
| 0.98924373 |
| 0.99998478 |
| 0.80984148 |
| 0.7958357 |
| 0.90265407 |
| 0.99998478 |
| 0.94653188 |
| 0.89077022 |

| |
|---|
| 0.99122857 |
| 0.99998478 |
| 0.99122857 |
| 0.9865573 |
| 0.94653188 |
| 0.69921142 |
| 0.98924373 |
| 0.99122857 |
| 0.96484568 |
| 0.79151544 |
| 0.95034906 |
| 0.95352171 |
| 0.94596453 |
| 0.98924373 |
| 0.99998478 |
| 0.95134169 |
| 0.99122857 |
| 0.97327383 |
| 0.9467702 |
| 0.91267201 |
| 0.91325047 |
| 0.92861642 |
| 0.99252883 |
| 0.85669195 |
| 0.90297323 |
| 0.99998478 |
| 0.90529035 |
| 0.94653188 |
| 0.95828764 |
| 0.93752249 |
| 0.99998478 |
| 0.915492 |
| 0.84320674 |
| 0.99969417 |
| 0.99998478 |
| 0.98924373 |
| 0.97110689 |
| 0.97329503 |
| 0.98924373 |
| 0.95859413 |
| 0.93594857 |
| 0.99122857 |

| |
|---|
| 0.91325047 |
| 0.96484568 |
| 0.99969417 |
| 0.98924373 |
| 0.92861642 |
| 0.98924373 |
| 0.96181283 |
| 0.90297323 |
| 0.97078973 |
| 0.98924373 |
| 0.99682234 |
| 0.94002917 |
| 0.69921142 |
| 0.80984148 |
| 0.99998478 |
| 0.99998478 |
| 0.90265407 |
| 0.74240405 |
| 0.98924373 |
| 0.87052203 |
| 0.98924373 |
| 0.90297323 |
| 0.90265407 |
| 0.99998478 |
| 0.91485898 |
| 0.9647026 |
| 0.99998478 |
| 0.99122857 |
| 0.98924373 |
| 0.90265407 |
| 0.99830763 |
| 0.98924373 |
| 0.90297323 |
| 0.98924373 |
| 0.61773657 |
| 0.93752249 |
| 0.96484568 |
| 0.79151544 |
| 0.7628002 |
| 0.8773376 |
| 0.96709089 |
| 0.93752249 |

0.97430523
0.96484568
0.94171831
0.98924373
0.94002917
0.94653188
0.98924373
0.99122857
0.93752249
0.91105653
0.9403055
0.93151025
0.9403055
0.94171831
0.99252883
0.99122857
0.92910003
0.94171831
0.915492
0.91325047
0.91325047
0.98924373
0.90265407
0.95951105
0.99122857
0.94171831
0.96484568
0.99122857
0.79151544
0.99998478
0.95951105
0.99998478
0.85801119
0.93752249
0.95240191
0.90265407
0.90265407
0.94653188
0.85801119
0.98924373
0.96181283
0.96484568

| |
|---|
| 0.92244927 |
| 0.96181283 |
| 0.99998478 |
| 0.99998478 |
| 0.97368509 |
| 0.69921142 |
| 0.99998478 |
| 0.98924373 |
| 0.98924373 |
| 0.92931319 |
| 0.96484568 |
| 0.98924373 |
| 0.69921142 |
| 0.96484568 |
| 0.8773376 |
| 0.90297323 |
| 0.99998478 |
| 0.72975604 |
| 0.90265407 |
| 0.98924373 |
| 0.99122857 |
| 0.90297323 |
| 0.9834625 |
| 0.96484568 |
| 0.98732643 |
| 0.8773376 |
| 0.61773657 |
| 0.99122857 |
| 0.90265407 |
| 0.99969417 |
| 0.81286841 |
| 0.99830763 |
| 0.9403055 |
| 0.91325047 |
| 0.99846792 |
| 0.98732643 |
| 0.96181283 |
| 0.95240191 |
| 0.99830763 |
| 0.91325047 |
| 0.93752249 |
| 0.99998478 |

0.93752249
0.90265407
0.99998478
0.98924373
0.93752249
0.9403055
0.99122857
0.90297323
0.99252883
0.90265407
0.96484568
0.98924373
0.90265407
0.79151544
0.99846792
0.9865573
0.98924373
0.99122857
0.99998478
0.99830763
0.99998478
0.99122857
0.98259014
0.92367831
0.99998478
0.98924373
0.98924373
0.99998478
0.93752249
0.90265407
0.96181283
0.85971745
0.94002917
0.90265407
0.90297323
0.915492
0.99998478
0.98924373
0.98732643
0.99122857
0.99969417
0.90297323

| |
|---|
| 0.99998478 |
| 0.90297323 |
| 0.9403055 |
| 0.93752249 |
| 0.94171831 |
| 0.94412092 |
| 0.8773376 |
| 0.98924373 |
| 0.90265407 |
| 0.90265407 |
| 0.915492 |
| 0.87052203 |
| 0.96181283 |
| 0.74281644 |
| 0.81249664 |
| 0.94171831 |
| 0.99969417 |
| 0.99122857 |
| 0.99969417 |
| 0.91267201 |
| 0.81249664 |
| 0.95951105 |
| 0.98924373 |
| 0.99998478 |
| 0.926218 |
| 0.90265407 |
| 0.93752249 |
| 0.99122857 |
| 0.98924373 |
| 0.90297323 |
| 0.90265407 |
| 0.9521979 |
| 0.85669195 |
| 0.95951105 |
| 0.94171831 |
| 0.90297323 |
| 0.99969417 |
| 0.66733542 |
| 0.98924373 |
| 0.93752249 |
| 0.8773376 |
| 0.78307327 |

| |
|---|
| 0.90265407 |
| 0.99969417 |
| 0.99122857 |
| 0.93054695 |
| 0.99252883 |
| 0.99998478 |
| 0.96484568 |
| 0.96167371 |
| 0.90297323 |
| 0.98924373 |
| 0.90297323 |
| 0.93811476 |
| 0.98956576 |
| 0.93752249 |
| 0.99252883 |
| 0.85801119 |
| 0.98924373 |
| 0.98924373 |
| 0.72225312 |
| 0.915492 |
| 0.96484568 |
| 0.915492 |
| 0.96181283 |
| 0.90265407 |
| 0.99122857 |
| 0.98924373 |
| 0.85904486 |
| 0.99998478 |
| 0.99969417 |
| 0.99969417 |
| 0.95951105 |
| 0.95585979 |
| 0.98924373 |
| 0.98924373 |
| 0.99122857 |
| 0.99830763 |
| 0.99998478 |
| 0.99969417 |
| 0.70655321 |
| 0.9557789 |
| 0.98924373 |
| 0.99948729 |

0.99969417
0.98924373
0.90265407
0.95240191
0.99122857
0.90265407
0.93752249
0.93752249
0.90297323
0.90265407
0.96484568
0.98924373
0.8773376
0.99252883
0.96484568
0.93427164
0.99865984
0.94171831
0.9403055
0.93752249
0.94653188
0.94171831
0.9403055
0.97126196
0.94897478
0.95240191
0.95034906
0.94171831
0.99122857
0.91325047
0.98924373
0.99252883
0.98785699
0.98888917
0.93752249
0.98259014
0.93752249
0.92672232
0.82029292
0.98924373
0.99998478
0.93752249

0.92367831
0.93752249
0.74240405
0.99122857
0.69921142
0.99998478
0.98924373
0.86004523
0.99998478
0.9384306
0.61773657
0.93752249
0.99252883
0.99122857
0.90297323
0.92861642
0.90297323
0.93752249
0.95034906
0.84430401
0.90265407
0.90297323
0.90265407
0.99252883
0.90265407
0.99617169
0.7958357
0.96920668
0.8773376
0.7958357
0.99998478
0.99122857
0.91267201
0.96181283
0.9829938
0.98924373
0.99998478
0.99122857
0.9403055
0.99998478
0.92861642
0.99969417

| |
|---|
| 0.9467702 |
| 0.99998478 |
| 0.85141726 |
| 0.99969417 |
| 0.89152698 |
| 0.96181283 |
| 0.96181283 |
| 0.93752249 |
| 0.95585979 |
| 0.99998478 |
| 0.94653188 |
| 0.94171831 |
| 0.93752249 |
| 0.99969417 |
| 0.91325047 |
| 0.82029292 |
| 0.99998478 |
| 0.93365848 |
| 0.99830763 |
| 0.99969417 |
| 0.86327908 |
| 0.99998478 |
| 0.99998478 |
| 0.99998478 |
| 0.96709089 |
| 0.9403055 |
| 0.95240191 |
| 0.99183457 |
| 0.99998478 |
| 0.90265407 |
| 0.99969417 |
| 0.915492 |
| 0.99969417 |
| 0.98924373 |
| 0.92548507 |
| 0.94846058 |
| 0.99998478 |
| 0.96484568 |
| 0.9985128 |
| 0.91325047 |
| 0.90265407 |
| 0.95835107 |

0.90265407
0.99998478
0.91267201
0.93752249
0.99998478
0.94171831
0.79151544
0.96484568
0.99969417
0.99122857
0.99969417
0.99808206
0.95951105
0.97327383
0.95951105
0.91325047
0.94171831
0.96484568
0.99122857
0.94171831
0.99122857
0.92861642
0.95034906
0.95034906
0.99998478
0.95951105
0.99122857
0.96181283
0.93752249
0.99998478
0.99941754
0.98924373
0.99122857
0.96181283
0.89618249
0.99969417
0.99252883
0.69921142
0.94171831
0.99122857
0.99122857
0.99122857

| |
|---|
| 0.93594857 |
| 0.95951105 |
| 0.90265407 |
| 0.9403055 |
| 0.85971745 |
| 0.98924373 |
| 0.99998478 |
| 0.96484568 |
| 0.78307327 |
| 0.95034906 |
| 0.9467702 |
| 0.99122857 |
| 0.58876713 |
| 0.90186583 |
| 0.90297323 |
| 0.99998478 |
| 0.87669383 |
| 0.99998478 |
| 0.98103056 |
| 0.94171831 |
| 0.90265407 |
| 0.787647 |
| 0.99122857 |
| 0.96484568 |
| 0.98924373 |
| 0.90265407 |
| 0.94653188 |
| 0.8773376 |
| 0.92861642 |
| 0.99122857 |
| 0.98924373 |
| 0.80984148 |
| 0.98924373 |
| 0.90297323 |
| 0.99830763 |
| 0.99969417 |
| 0.95170115 |
| 0.98924373 |
| 0.97815363 |
| 0.9338612 |
| 0.80984148 |
| 0.61079703 |

| |
|---|
| 0.99998478 |
| 0.84586312 |
| 0.96167371 |
| 0.99122857 |
| 0.98924373 |
| 0.98924373 |
| 0.9403055 |
| 0.926218 |
| 0.97126196 |
| 0.90297323 |
| 0.79151544 |
| 0.98924373 |
| 0.92861642 |
| 0.99122857 |
| 0.96874634 |
| 0.91325047 |
| 0.98924373 |
| 0.99122857 |
| 0.98924373 |
| 0.96101531 |
| 0.99998478 |
| 0.93752249 |
| 0.99998478 |
| 0.91325047 |
| 0.8773376 |
| 0.90265407 |
| 0.99122857 |
| 0.99122857 |
| 0.99998478 |
| 0.92548507 |
| 0.99998478 |
| 0.91325047 |
| 0.94653188 |
| 0.9403055 |
| 0.99846792 |
| 0.90297323 |
| 0.99309902 |
| 0.99122857 |
| 0.9944798 |
| 0.87052203 |
| 0.94897478 |
| 0.92548507 |

| |
|---|
| 0.99830763 |
| 0.90265407 |
| 0.99998478 |
| 0.93752249 |
| 0.96484568 |
| 0.92672232 |
| 0.95240191 |
| 0.91325047 |
| 0.91485898 |
| 0.90265407 |
| 0.94653188 |
| 0.99122857 |
| 0.99998478 |
| 0.99122857 |
| 0.95951105 |
| 0.86327908 |
| 0.787647 |
| 0.85141726 |
| 0.94171831 |
| 0.94171831 |
| 0.79151544 |
| 0.99969417 |
| 0.9985128 |
| 0.99846792 |
| 0.98732643 |
| 0.99680366 |
| 0.93752249 |
| 0.99122857 |
| 0.90265407 |
| 0.99122857 |
| 0.90297323 |
| 0.99998478 |
| 0.98924373 |
| 0.99998478 |
| 0.85141726 |
| 0.97078973 |
| 0.91325047 |
| 0.91333392 |
| 0.82029292 |
| 0.98924373 |
| 0.99998478 |
| 0.99998478 |

| |
|---|
| 0.98924373 |
| 0.99122857 |
| 0.95240191 |
| 0.99998478 |
| 0.97993742 |
| 0.99998478 |
| 0.90297323 |
| 0.99122857 |
| 0.99122857 |
| 0.94002917 |
| 0.99122857 |
| 0.98924373 |
| 0.90265407 |
| 0.96484568 |
| 0.99969417 |
| 0.90265407 |
| 0.98924373 |
| 0.99998478 |
| 0.99122857 |
| 0.99830763 |
| 0.99122857 |
| 0.94171831 |
| 0.98924373 |
| 0.99122857 |
| 0.90297323 |
| 0.9403055 |
| 0.94171831 |
| 0.93752249 |
| 0.99201427 |
| 0.99998478 |
| 0.99122857 |
| 0.94171831 |
| 0.99650199 |
| 0.95859413 |
| 0.9403055 |
| 0.86974476 |
| 0.96181283 |
| 0.94653188 |
| 0.9715425 |
| 0.99101915 |
| 0.87052203 |
| 0.96181283 |

| |
|---|
| 0.95859413 |
| 0.85141726 |
| 0.99998478 |
| 0.94171831 |
| 0.99122857 |
| 0.99002751 |
| 0.98924373 |
| 0.99122857 |
| 0.9467702 |
| 0.9998342 |
| 0.82029292 |
| 0.99969417 |
| 0.9403055 |
| 0.97479234 |
| 0.9865573 |
| 0.9520243 |
| 0.99969417 |
| 0.91325047 |
| 0.87052203 |
| 0.96167371 |
| 0.98924373 |
| 0.94171831 |
| 0.94762541 |
| 0.98924373 |
| 0.99122857 |
| 0.9829938 |
| 0.94171831 |
| 0.90265407 |
| 0.99998478 |
| 0.95951105 |
| 0.98924373 |
| 0.99122857 |
| 0.98924373 |
| 0.98924373 |
| 0.87052203 |
| 0.99998478 |
| 0.98924373 |
| 0.91267201 |
| 0.90265407 |
| 0.98924373 |
| 0.95951105 |
| 0.95951105 |

0.90265407
0.96181283
0.90265407
0.9834625
0.98924373
0.95134169
0.94171831
0.99122857
0.90297323
0.99969417
0.91325047
0.97024001
0.99122857
0.84076524
0.98924373
0.94412092
0.90265407
0.94171831
0.98924373
0.926218
0.9467702
0.98924373
0.99998478
0.82029292
0.99998478
0.96484568
0.99998478
0.98924373
0.98924373
0.96484568
0.99969417
0.96181283
0.96181283
0.99122857
0.93752249
0.96181283
0.80984148
0.90265407
0.79151544
0.91325047
0.99969417
0.99252883

| |
|---|
| 0.98924373 |
| 0.85141726 |
| 0.96484568 |
| 0.98924373 |
| 0.91267201 |
| 0.99969417 |
| 0.95859413 |
| 0.98924373 |
| 0.96484568 |
| 0.98924373 |
| 0.98924373 |
| 0.92672232 |
| 0.97085356 |
| 0.87944932 |
| 0.95240191 |
| 0.96484568 |
| 0.96484568 |
| 0.98732643 |
| 0.96484568 |
| 0.99122857 |
| 0.99998478 |
| 0.78307327 |
| 0.99650199 |
| 0.99998478 |
| 0.99998478 |
| 0.55521624 |
| 0.94653188 |
| 0.82029292 |
| 0.90265407 |
| 0.95951105 |
| 0.95859413 |
| 0.99998478 |
| 0.94171831 |
| 0.93752249 |
| 0.99658202 |
| 0.99122857 |
| 0.85141726 |
| 0.98924373 |
| 0.9403055 |
| 0.92548507 |
| 0.96181283 |
| 0.98924373 |

0.94171831
0.90265407
0.97329503
0.90265407
0.98924373
0.99969417
0.96484568
0.99122857
0.99830763
0.93752249
0.99969417
0.93151025
0.97110689
0.99998478
0.99122857
0.99122857
0.93752249
0.92861642
0.87052203
0.99122857
0.93752249
0.99122857
0.98924373
0.93752249
0.99969417
0.99969417
0.98924373
0.93752249
0.99122857
0.98924373
0.97993742
0.85141726
0.98924373
0.99122857
0.90265407
0.99122857
0.99830763
0.99998478
0.99122857
0.90297323
0.92861642
0.85669195

0.95240191
0.97209533
0.99122857
0.92861642
0.82029292
0.99122857
0.93365848
0.90265407
0.9647026
0.96920668
0.9403055
0.98924373
0.99998478
0.99998478
0.90297323
0.98924373
0.99122857
0.99998478
0.9467702
0.98924373
0.85141726
0.98924373
0.92861642
0.99122857
0.93752249
0.69921142
0.99680366
0.99252883
0.86974476
0.99998478
0.98924373
0.99998478
0.96484568
0.97993742
0.91267201
0.99122857
0.93811476
0.99122857
0.98924373
0.8773376
0.93752249
0.90297323

| |
|---|
| 0.926218 |
| 0.93594857 |
| 0.93752249 |
| 0.99122857 |
| 0.90297323 |
| 0.99998478 |
| 0.91325047 |
| 0.90265407 |
| 0.93752249 |
| 0.99252883 |
| 0.99332973 |
| 0.99998478 |
| 0.9829938 |
| 0.99998478 |
| 0.98924373 |
| 0.98924373 |
| 0.99122857 |
| 0.9467702 |
| 0.74240405 |
| 0.99002751 |
| 0.92861642 |
| 0.96484568 |
| 0.94596453 |
| 0.97327383 |
| 0.91325047 |
| 0.94412092 |
| 0.92548507 |
| 0.74240405 |
| 0.99122857 |
| 0.99122857 |
| 0.90297323 |
| 0.95240191 |
| 0.99998478 |
| 0.99969417 |
| 0.99998478 |
| 0.99998478 |
| 0.99969417 |
| 0.9338612 |
| 0.93427164 |
| 0.90265407 |
| 0.9829938 |
| 0.95951105 |

| |
|---|
| 0.90297323 |
| 0.93752249 |
| 0.98924373 |
| 0.96181283 |
| 0.90265407 |
| 0.80984148 |
| 0.80984148 |
| 0.79151544 |
| 0.93752249 |
| 0.90297323 |
| 0.9403055 |
| 0.9985128 |
| 0.96181283 |
| 0.98259014 |
| 0.99998478 |
| 0.99122857 |
| 0.91267201 |
| 0.89618249 |
| 0.91485898 |
| 0.91325047 |
| 0.98924373 |
| 0.93752249 |
| 0.98259014 |
| 0.99183457 |
| 0.99122857 |
| 0.99830763 |
| 0.9715425 |
| 0.98924373 |
| 0.95240191 |
| 0.95240191 |
| 0.90265407 |
| 0.90265407 |
| 0.90265407 |
| 0.99122857 |
| 0.93752249 |
| 0.9985128 |
| 0.97209533 |
| 0.98924373 |
| 0.94653188 |
| 0.99998478 |
| 0.90265407 |
| 0.96181283 |

| |
|---|
| 0.90297323 |
| 0.90297323 |
| 0.98924373 |
| 0.92861642 |
| 0.86545047 |
| 0.98924373 |
| 0.99998478 |
| 0.96484568 |
| 0.9467702 |
| 0.95951105 |
| 0.94653188 |
| 0.95859413 |
| 0.95951105 |
| 0.99998478 |
| 0.99998478 |
| 0.99122857 |
| 0.98924373 |
| 0.98924373 |
| 0.58876713 |
| 0.96484568 |
| 0.99998478 |
| 0.98924373 |
| 0.98924373 |
| 0.99998478 |
| 0.98924373 |
| 0.92861642 |
| 0.98424698 |
| 0.94171831 |
| 0.98924373 |
| 0.94171831 |
| 0.96484568 |
| 0.98924373 |
| 0.99998478 |
| 0.80984148 |
| 0.96484568 |
| 0.98924373 |
| 0.98924373 |
| 0.99998478 |
| 0.90297323 |
| 0.79151544 |
| 0.69921142 |
| 0.9403055 |

| |
|---|
| 0.93752249 |
| 0.98924373 |
| 0.98924373 |
| 0.96181283 |
| 0.97085356 |
| 0.98924373 |
| 0.94171831 |
| 0.98259014 |
| 0.96484568 |
| 0.99998478 |
| 0.99830763 |
| 0.99122857 |
| 0.98924373 |
| 0.98924373 |
| 0.99122857 |
| 0.99122857 |
| 0.91267201 |
| 0.99122857 |
| 0.94171831 |
| 0.98924373 |
| 0.65854402 |
| 0.94653188 |
| 0.90265407 |
| 0.98924373 |
| 0.99998478 |
| 0.72975604 |
| 0.98924373 |
| 0.80984148 |
| 0.99998478 |
| 0.61773657 |
| 0.95240191 |
| 0.99122857 |
| 0.90529035 |
| 0.98924373 |
| 0.93752249 |
| 0.99998478 |
| 0.98924373 |
| 0.9403055 |
| 0.98924373 |
| 0.98924373 |
| 0.95951105 |
| 0.93752249 |

0.99122857
0.92861642
0.98924373
0.98924373
0.98924373
0.99998478
0.99122857
0.93752249
0.98924373
0.99252883
0.96484568
0.98924373
0.98924373
0.98924373
0.92861642
0.98924373
0.72975604
0.99998478
0.93908842
0.90297323
0.98259014
0.96484568
0.98924373
0.94171831
0.90265407
0.96484568
0.72975604
0.96484568
0.90265407
0.99969417
0.94002917
0.96484568
0.91267201
0.96484568
0.90265407
0.92861642
0.95951105
0.99830763
0.77062555
0.99101915
0.99998478
0.91105653

| |
|---|
| 0.99969417 |
| 0.99998478 |
| 0.93752249 |
| 0.90297323 |
| 0.99969417 |
| 0.91325047 |
| 0.98924373 |
| 0.92861642 |
| 0.9865496 |
| 0.98924373 |
| 0.96484568 |
| 0.91325047 |
| 0.91105653 |
| 0.99998478 |
| 0.99998478 |
| 0.9403055 |
| 0.98924373 |
| 0.96167371 |
| 0.93752249 |
| 0.99998478 |
| 0.99998478 |
| 0.915492 |
| 0.94171831 |
| 0.93752249 |
| 0.98924373 |
| 0.90265407 |
| 0.93811476 |
| 0.7958357 |
| 0.99998478 |
| 0.90265407 |
| 0.91267201 |
| 0.99969417 |
| 0.98924373 |
| 0.93054695 |
| 0.91105653 |
| 0.90265407 |
| 0.99969417 |
| 0.99969417 |
| 0.98924373 |
| 0.97815363 |
| 0.99969417 |
| 0.96181283 |

| |
|---|
| 0.98924373 |
| 0.99122857 |
| 0.85141726 |
| 0.915492 |
| 0.98732643 |
| 0.90297323 |
| 0.74240405 |
| 0.92672232 |
| 0.99122857 |
| 0.96181283 |
| 0.92861642 |
| 0.90529035 |
| 0.95951105 |
| 0.98924373 |
| 0.98924373 |
| 0.91325047 |
| 0.9403055 |
| 0.92861642 |
| 0.98924373 |
| 0.99969417 |
| 0.99122857 |
| 0.92861642 |
| 0.93752249 |
| 0.98924373 |
| 0.9403055 |
| 0.98924373 |
| 0.84320674 |
| 0.87669383 |
| 0.99122857 |
| 0.93752249 |
| 0.98924373 |
| 0.99122857 |
| 0.86974476 |
| 0.69921142 |
| 0.96181283 |
| 0.99998478 |
| 0.94897478 |
| 0.99122857 |
| 0.80133949 |
| 0.98924373 |
| 0.94171831 |
| 0.9467702 |

0.99998478
0.99122857
0.95951105
0.96484568
0.99122857
0.7958357
0.98924373
0.99969417
0.94002917
0.96484568
0.99998478
0.98924373
0.92861642
0.93151025
0.99998478
0.98924373
0.99998478
0.96484568
0.98924373
0.95859413
0.95034906
0.99969417
0.97815363
0.74240405
0.98924373
0.98924373
0.90265407
0.98924373
0.90265407
0.96484568
0.98924373
0.94412092
0.99122857
0.99122857
0.99998478
0.95034906
0.9521979
0.99998478
0.90265407
0.98924373
0.92367831
0.96484568

| |
|---|
| 0.926218 |
| 0.84593444 |
| 0.99830763 |
| 0.99969417 |
| 0.99122857 |
| 0.94171831 |
| 0.90297323 |
| 0.91485898 |
| 0.94171831 |
| 0.72975604 |
| 0.96181283 |
| 0.93427164 |
| 0.95859413 |
| 0.90297323 |
| 0.99998478 |
| 0.98924373 |
| 0.87052203 |
| 0.98924373 |
| 0.96709089 |
| 0.99998478 |
| 0.9403055 |
| 0.91325047 |
| 0.93752249 |
| 0.98732643 |
| 0.96709089 |
| 0.97110689 |
| 0.98259014 |
| 0.96484568 |
| 0.98924373 |
| 0.95951105 |
| 0.99998478 |
| 0.95240191 |
| 0.99969417 |
| 0.90297323 |
| 0.69921142 |
| 0.98924373 |
| 0.96484568 |
| 0.91325047 |
| 0.99969417 |
| 0.90297323 |
| 0.98550175 |
| 0.99969417 |

| |
|---|
| 0.94171831 |
| 0.90265407 |
| 0.82029292 |
| 0.91325047 |
| 0.98924373 |
| 0.99252883 |
| 0.97815363 |
| 0.90297323 |
| 0.98924373 |
| 0.99122857 |
| 0.98924373 |
| 0.98924373 |
| 0.95951105 |
| 0.9403055 |
| 0.98924373 |
| 0.99998478 |
| 0.99998478 |
| 0.96181283 |
| 0.99437562 |
| 0.80984148 |
| 0.90265407 |
| 0.98924373 |
| 0.90297323 |
| 0.98924373 |
| 0.99998478 |
| 0.93752249 |
| 0.91267201 |
| 0.90265407 |
| 0.99122857 |
| 0.99122857 |
| 0.97993742 |
| 0.93752249 |
| 0.94171831 |
| 0.93752249 |
| 0.98924373 |
| 0.9829938 |
| 0.99998478 |
| 0.95034906 |
| 0.99998478 |
| 0.99998478 |
| 0.99998478 |
| 0.96181283 |

0.95240191
0.81286841
0.89618249
0.90265407
0.93427164
0.93752249
0.91325047
0.97954048
0.90297323
0.98924373
0.99998478
0.99998478
0.91267201
0.98924373
0.82029292
0.99969417
0.99252883
0.98924373
0.96484568
0.98924373
0.98924373
0.99998478
0.94897478
0.9865573
0.94171831
0.99122857
0.95240191
0.99122857
0.91325047
0.99969417
0.94897478
0.9403055
0.91485898
0.99122857
0.91325047
0.99122857
0.98924373
0.99969417
0.99998478
0.98924373
0.99948729
0.94897478

| |
|---|
| 0.99969417 |
| 0.9467702 |
| 0.98924373 |
| 0.98924373 |
| 0.90297323 |
| 0.93752249 |
| 0.99969417 |
| 0.99628097 |
| 0.9467702 |
| 0.90265407 |
| 0.94171831 |
| 0.99998478 |
| 0.98924373 |
| 0.90265407 |
| 0.98924373 |
| 0.96484568 |
| 0.99998478 |
| 0.95828764 |
| 0.92075126 |
| 0.99998478 |
| 0.99122857 |
| 0.95951105 |
| 0.90297323 |
| 0.91325047 |
| 0.95034906 |
| 0.92672232 |
| 0.99969417 |
| 0.97110689 |
| 0.96484568 |
| 0.91325047 |
| 0.99332973 |
| 0.98924373 |
| 0.99846792 |
| 0.99830763 |
| 0.95988499 |
| 0.99122857 |
| 0.99846792 |
| 0.94596453 |
| 0.95240191 |
| 0.99998478 |
| 0.96181283 |
| 0.95034906 |

| |
|---|
| 0.99252883 |
| 0.96816591 |
| 0.97815363 |
| 0.94171831 |
| 0.96484568 |
| 0.99998478 |
| 0.87669383 |
| 0.69921142 |
| 0.81286841 |
| 0.98924373 |
| 0.97992318 |
| 0.99998478 |
| 0.99998478 |
| 0.9403055 |
| 0.99998478 |
| 0.99969417 |
| 0.79151544 |
| 0.99122857 |
| 0.95034906 |
| 0.93752249 |
| 0.91325047 |
| 0.96484568 |
| 0.8773376 |
| 0.96181283 |
| 0.90265407 |
| 0.98924373 |
| 0.94747425 |
| 0.96484568 |
| 0.96484568 |
| 0.87052203 |
| 0.96709089 |
| 0.99122857 |
| 0.99122857 |
| 0.94653188 |
| 0.98924373 |
| 0.98924373 |
| 0.98924373 |
| 0.96816591 |
| 0.92861642 |
| 0.95240191 |
| 0.94171831 |
| 0.99998478 |

| |
|---|
| 0.93752249 |
| 0.95034906 |
| 0.99122857 |
| 0.96167371 |
| 0.99830763 |
| 0.8773376 |
| 0.99969417 |
| 0.95240191 |
| 0.96484568 |
| 0.96181283 |
| 0.99122857 |
| 0.98924373 |
| 0.93752249 |
| 0.99998478 |
| 0.9467702 |
| 0.99513459 |
| 0.94171831 |
| 0.98924373 |
| 0.99998478 |
| 0.94653188 |
| 0.80133949 |
| 0.99846792 |
| 0.97110689 |
| 0.94171831 |
| 0.81286841 |
| 0.99122857 |
| 0.98295353 |
| 0.95034906 |
| 0.96484568 |
| 0.90297323 |
| 0.98956576 |
| 0.99830763 |
| 0.96484568 |
| 0.98924373 |
| 0.99830763 |
| 0.94653188 |
| 0.98924373 |
| 0.97327383 |
| 0.99122857 |
| 0.87052203 |
| 0.98924373 |
| 0.98924373 |

| |
|---|
| 0.9985128 |
| 0.94171831 |
| 0.99122857 |
| 0.99122857 |
| 0.90265407 |
| 0.99998478 |
| 0.98924373 |
| 0.93752249 |
| 0.99101915 |
| 0.99998478 |
| 0.98924373 |
| 0.99998478 |
| 0.99998478 |
| 0.98924373 |
| 0.99998478 |
| 0.99830763 |
| 0.99969417 |
| 0.94171831 |
| 0.99998478 |
| 0.92672232 |
| 0.98924373 |
| 0.99830763 |
| 0.98924373 |
| 0.98924373 |
| 0.94653188 |
| 0.98924373 |
| 0.99122857 |
| 0.98924373 |
| 0.99998478 |
| 0.91485898 |
| 0.99998478 |
| 0.99830763 |
| 0.99846792 |
| 0.78307327 |
| 0.98924373 |
| 0.96484568 |
| 0.97993742 |
| 0.94171831 |
| 0.93752249 |
| 0.99969417 |
| 0.99122857 |
| 0.99252883 |

0.99969417
0.98259014
0.92672232
0.99998478
0.92861642
0.66733542
0.61773657
0.99969417
0.99998478
0.99969417
0.92861642
0.99998478
0.99122857
0.99252883
0.96181283
0.95951105
0.99252883
0.99998478
0.96181283
0.95240191
0.97815363
0.99122857
0.99122857
0.96484568
0.99998478
0.98924373
0.99122857
0.94171831
0.99643667
0.95951105
0.98924373
0.99122857
0.99830763
0.80984148
0.95134169
0.90265407
0.99969417
0.98924373
0.98924373
0.93752249
0.99122857
0.96484568

| |
|---|
| 0.9403055 |
| 0.79151544 |
| 0.93752249 |
| 0.90297323 |
| 0.90297323 |
| 0.90265407 |
| 0.99998478 |
| 0.98924373 |
| 0.98924373 |
| 0.98924373 |
| 0.92861642 |
| 0.99998478 |
| 0.85141726 |
| 0.93752249 |
| 0.99122857 |
| 0.99252883 |
| 0.87944932 |
| 0.99830763 |
| 0.99122857 |
| 0.96977284 |
| 0.95609471 |
| 0.97327383 |
| 0.99998478 |
| 0.9403055 |
| 0.94653188 |
| 0.96484568 |
| 0.99122857 |
| 0.80984148 |
| 0.99998478 |
| 0.7958357 |
| 0.99998478 |
| 0.99998478 |
| 0.92861642 |
| 0.95240191 |
| 0.98924373 |
| 0.99969417 |
| 0.98924373 |
| 0.99830763 |
| 0.98103056 |
| 0.96484568 |
| 0.94171831 |
| 0.96484568 |

0.94653188
0.99830763
0.72204897
0.91485898
0.92672232
0.99969417
0.96484568
0.99969417
0.99122857
0.91325047
0.99252883
0.99998478
0.99122857
0.94897478
0.98924373
0.96484568
0.98924373
0.92861642
0.87669383
0.99998478
0.94171831
0.98924373
0.98924373
0.98337226
0.9715425
0.93752249
0.99183457
0.99122857
0.91325047
0.98924373
0.99122857
0.94171831
0.93811476
0.99830763
0.94747425
0.93151025
0.85141726
0.8773376
0.99998478
0.94171831
0.95240191
0.94171831

| |
|---|
| 0.96484568 |
| 0.94653188 |
| 0.99998478 |
| 0.99969417 |
| 0.96484568 |
| 0.99998478 |
| 0.94653188 |
| 0.93427164 |
| 0.99122857 |
| 0.90297323 |
| 0.99122857 |
| 0.93752249 |
| 0.99650199 |
| 0.98325199 |
| 0.90297323 |
| 0.91325047 |
| 0.97327383 |
| 0.66733542 |
| 0.93752249 |
| 0.93030041 |
| 0.93752249 |
| 0.8773376 |
| 0.86079694 |
| 0.92861642 |
| 0.97110689 |
| 0.99252883 |
| 0.94171831 |
| 0.99998478 |
| 0.94596453 |
| 0.99122857 |
| 0.91325047 |
| 0.95951105 |
| 0.99998478 |
| 0.98568989 |
| 0.95034906 |
| 0.9829938 |
| 0.90297323 |
| 0.91325047 |
| 0.9467702 |
| 0.99122857 |
| 0.99122857 |
| 0.99998478 |

| |
|---|
| 0.98924373 |
| 0.93427164 |
| 0.98924373 |
| 0.95060469 |
| 0.90297323 |
| 0.94171831 |
| 0.99122857 |
| 0.98924373 |
| 0.99998478 |
| 0.94171831 |
| 0.98924373 |
| 0.98732643 |
| 0.90297323 |
| 0.90265407 |
| 0.99252883 |
| 0.99969417 |
| 0.99998478 |
| 0.99122857 |
| 0.94171831 |
| 0.99122857 |
| 0.90297323 |
| 0.96167371 |
| 0.99998478 |
| 0.96880292 |
| 0.90297323 |
| 0.9403055 |
| 0.99969417 |
| 0.98924373 |
| 0.96181283 |
| 0.95951105 |
| 0.90265407 |
| 0.90265407 |
| 0.9467702 |
| 0.99122857 |
| 0.97327383 |
| 0.99122857 |
| 0.99998478 |
| 0.91325047 |
| 0.926218 |
| 0.99122857 |
| 0.98924373 |
| 0.96484568 |

| |
|---|
| 0.99998478 |
| 0.93752249 |
| 0.99998478 |
| 0.91325047 |
| 0.99948729 |
| 0.96816591 |
| 0.98237613 |
| 0.97992318 |
| 0.93752249 |
| 0.96898185 |
| 0.94653188 |
| 0.98924373 |
| 0.98924373 |
| 0.98275443 |
| 0.99998478 |
| 0.99122857 |
| 0.98924373 |
| 0.98924373 |
| 0.87669383 |
| 0.99998478 |
| 0.99122857 |
| 0.96484568 |
| 0.99998478 |
| 0.99122857 |
| 0.99122857 |
| 0.96484568 |
| 0.99998478 |
| 0.99998478 |
| 0.96484568 |
| 0.99998478 |
| 0.96484568 |
| 0.96484568 |
| 0.94897478 |
| 0.99998478 |
| 0.99252883 |
| 0.99998478 |
| 0.99650199 |
| 0.98924373 |
| 0.99122857 |
| 0.926218 |
| 0.99122857 |
| 0.98924373 |

| |
|---|
| 0.98924373 |
| 0.98924373 |
| 0.92570173 |
| 0.99998478 |
| 0.98924373 |
| 0.98924373 |
| 0.9403055 |
| 0.96181283 |
| 0.99122857 |
| 0.90265407 |
| 0.90265407 |
| 0.98924373 |
| 0.99122857 |
| 0.8773376 |
| 0.79151544 |
| 0.99122857 |
| 0.99998478 |
| 0.85141726 |
| 0.99122857 |
| 0.98924373 |
| 0.98924373 |
| 0.99572919 |
| 0.99998478 |
| 0.99969417 |
| 0.99252883 |
| 0.92861642 |
| 0.95650711 |
| 0.93752249 |
| 0.86327908 |
| 0.96484568 |
| 0.96484568 |
| 0.99122857 |
| 0.99998478 |
| 0.99969417 |
| 0.93752249 |
| 0.98924373 |
| 0.99998478 |
| 0.98924373 |
| 0.98924373 |
| 0.98924373 |
| 0.99998478 |
| 0.93594857 |

| |
|---|
| 0.93752249 |
| 0.92861642 |
| 0.96484568 |
| 0.99969417 |
| 0.99998478 |
| 0.90297323 |
| 0.99846792 |
| 0.99998478 |
| 0.96709089 |
| 0.90265407 |
| 0.94653188 |
| 0.98924373 |
| 0.98237613 |
| 0.90265407 |
| 0.93594857 |
| 0.98924373 |
| 0.98237613 |
| 0.94171831 |
| 0.99998478 |
| 0.98924373 |
| 0.95846565 |
| 0.96484568 |
| 0.99122857 |
| 0.99969417 |
| 0.89084498 |
| 0.98924373 |
| 0.99969417 |
| 0.98924373 |
| 0.99122857 |
| 0.99122857 |
| 0.98924373 |
| 0.94171831 |
| 0.94171831 |
| 0.99998478 |
| 0.99998478 |
| 0.90297323 |
| 0.92861642 |
| 0.99998478 |
| 0.98924373 |
| 0.91267201 |
| 0.95240191 |
| 0.98924373 |

| |
|---|
| 0.98735899 |
| 0.9403055 |
| 0.96484568 |
| 0.99969417 |
| 0.98924373 |
| 0.99122857 |
| 0.93594857 |
| 0.96484568 |
| 0.90265407 |
| 0.97024001 |
| 0.98337226 |
| 0.926218 |
| 0.99998478 |
| 0.99998478 |
| 0.99998478 |
| 0.95240191 |
| 0.99122857 |
| 0.61079703 |
| 0.96484568 |
| 0.99122857 |
| 0.98918129 |
| 0.99122857 |
| 0.98924373 |
| 0.99969417 |
| 0.94653188 |
| 0.99969417 |
| 0.90297323 |
| 0.99830763 |
| 0.99122857 |
| 0.98924373 |
| 0.99969417 |
| 0.98924373 |
| 0.98732643 |
| 0.99969417 |
| 0.9403055 |
| 0.96484568 |
| 0.93752249 |
| 0.926218 |
| 0.93752249 |
| 0.99998478 |
| 0.98924373 |
| 0.93752249 |

| |
|---|
| 0.91325047 |
| 0.93752249 |
| 0.69921142 |
| 0.55555047 |
| 0.96484568 |
| 0.92672232 |
| 0.8773376 |
| 0.98924373 |
| 0.99998478 |
| 0.98259014 |
| 0.99122857 |
| 0.99998478 |
| 0.98924373 |
| 0.99948729 |
| 0.90265407 |
| 0.99846792 |
| 0.99998478 |
| 0.9403055 |
| 0.98924373 |
| 0.926218 |
| 0.95988499 |
| 0.9811568 |
| 0.99998478 |
| 0.90297323 |
| 0.98924373 |
| 0.99969417 |
| 0.85141726 |
| 0.99122857 |
| 0.99969417 |
| 0.94171831 |
| 0.99122857 |
| 0.79105523 |
| 0.94171831 |
| 0.99969417 |
| 0.99252883 |
| 0.99998478 |
| 0.93054695 |
| 0.98924373 |
| 0.96181283 |
| 0.99998478 |
| 0.99969417 |
| 0.93752249 |

| |
|---|
| 0.9403055 |
| 0.95240191 |
| 0.95650711 |
| 0.97327383 |
| 0.99122857 |
| 0.99998478 |
| 0.80984148 |
| 0.99969417 |
| 0.94171831 |
| 0.99830763 |
| 0.99122857 |
| 0.92548507 |
| 0.99998478 |
| 0.90265407 |
| 0.90265407 |
| 0.76761855 |
| 0.9467702 |
| 0.98924373 |
| 0.96484568 |
| 0.96484568 |
| 0.99122857 |
| 0.99998478 |
| 0.66733542 |
| 0.99122857 |
| 0.98924373 |
| 0.98924373 |
| 0.98924373 |
| 0.99998478 |
| 0.9467702 |
| 0.90265407 |
| 0.94002917 |
| 0.99122857 |
| 0.94653188 |
| 0.926218 |
| 0.96181283 |
| 0.91325047 |
| 0.99969417 |
| 0.95240191 |
| 0.99998478 |
| 0.92861642 |
| 0.98732643 |
| 0.98924373 |

| |
|---|
| 0.99122857 |
| 0.9338612 |
| 0.99998478 |
| 0.98466138 |
| 0.95240191 |
| 0.99969417 |
| 0.95034906 |
| 0.91485898 |
| 0.98924373 |
| 0.9865573 |
| 0.98924373 |
| 0.99998478 |
| 0.91564298 |
| 0.95951105 |
| 0.99998478 |
| 0.91325047 |
| 0.99808206 |
| 0.85141726 |
| 0.94653188 |
| 0.95951105 |
| 0.99998478 |
| 0.97327383 |
| 0.99998478 |
| 0.98924373 |
| 0.91325047 |
| 0.98924373 |
| 0.99969417 |
| 0.99998478 |
| 0.99122857 |
| 0.99969417 |
| 0.9936423 |
| 0.94897478 |
| 0.99998478 |
| 0.99969417 |
| 0.95240191 |
| 0.99969417 |
| 0.98924373 |
| 0.90265407 |
| 0.99830763 |
| 0.98924373 |
| 0.98924373 |
| 0.99122857 |

| |
|---|
| 0.99998478 |
| 0.98924373 |
| 0.99969417 |
| 0.98924373 |
| 0.98732643 |
| 0.99830763 |
| 0.98924373 |
| 0.99969417 |
| 0.99998478 |
| 0.99998478 |
| 0.99830763 |
| 0.9865573 |
| 0.99941754 |
| 0.96484568 |
| 0.99122857 |
| 0.98924373 |
| 0.99969417 |
| 0.95951105 |
| 0.98924373 |
| 0.99969417 |
| 0.99969417 |
| 0.91267201 |
| 0.95951105 |
| 0.98732643 |
| 0.98924373 |
| 0.93752249 |
| 0.99998478 |
| 0.98924373 |
| 0.94653188 |
| 0.787647 |
| 0.98924373 |
| 0.99846792 |
| 0.99969417 |
| 0.99122857 |
| 0.99998478 |
| 0.99572919 |
| 0.95034906 |
| 0.99808206 |
| 0.99969417 |
| 0.98924373 |
| 0.98924373 |
| 0.97329503 |

| |
|---|
| 0.99969417 |
| 0.99998478 |
| 0.99969417 |
| 0.99998478 |
| 0.96484568 |
| 0.99969417 |
| 0.99122857 |
| 0.926218 |
| 0.96484568 |
| 0.97329503 |
| 0.94171831 |
| 0.98924373 |
| 0.99998478 |
| 0.98924373 |
| 0.98924373 |
| 0.99572919 |
| 0.92861642 |
| 0.99122857 |
| 0.99122857 |
| 0.96484568 |
| 0.98924373 |
| 0.98259014 |
| 0.99808206 |
| 0.98924373 |
| 0.97992318 |
| 0.926218 |
| 0.95034906 |
| 0.99998478 |
| 0.99998478 |
| 0.99650199 |
| 0.94171831 |
| 0.90265407 |
| 0.99998478 |
| 0.85971745 |
| 0.96484568 |
| 0.99998478 |
| 0.99122857 |
| 0.79151544 |
| 0.99998478 |
| 0.99846792 |
| 0.99998478 |
| 0.99998478 |

| |
|---|
| 0.96181283 |
| 0.99122857 |
| 0.95951105 |
| 0.99122857 |
| 0.93752249 |
| 0.96181283 |
| 0.98732643 |
| 0.98924373 |
| 0.98924373 |
| 0.96484568 |
| 0.90265407 |
| 0.99846792 |
| 0.99998478 |
| 0.99969417 |
| 0.98924373 |
| 0.95352171 |
| 0.99969417 |
| 0.93752249 |
| 0.98924373 |
| 0.99998478 |
| 0.99969417 |
| 0.99998478 |
| 0.93811476 |
| 0.99122857 |
| 0.90297323 |
| 0.99998478 |
| 0.96484568 |
| 0.94002917 |
| 0.98924373 |
| 0.99122857 |
| 0.94653188 |
| 0.95034906 |
| 0.91485898 |
| 0.98924373 |
| 0.99969417 |
| 0.96484568 |
| 0.72204897 |
| 0.9715425 |
| 0.99122857 |
| 0.96484568 |
| 0.90265407 |
| 0.94171831 |

| |
|---|
| 0.99122857 |
| 0.93752249 |
| 0.94653188 |
| 0.98924373 |
| 0.99122857 |
| 0.80984148 |
| 0.99998478 |
| 0.93752249 |
| 0.98924373 |
| 0.99122857 |
| 0.8773376 |
| 0.99122857 |
| 0.95951105 |
| 0.98924373 |
| 0.94653188 |
| 0.93427164 |
| 0.98924373 |
| 0.89474679 |
| 0.9403055 |
| 0.99998478 |
| 0.99122857 |
| 0.9829938 |
| 0.98732643 |
| 0.99122857 |
| 0.96181283 |
| 0.96484568 |
| 0.85669195 |
| 0.99122857 |
| 0.99969417 |
| 0.95859413 |
| 0.98924373 |
| 0.90265407 |
| 0.9403055 |
| 0.99998478 |
| 0.96484568 |
| 0.99998478 |
| 0.99998478 |
| 0.84593444 |
| 0.94171831 |
| 0.95240191 |
| 0.99252883 |
| 0.99122857 |

| |
|---|
| 0.98732643 |
| 0.9865573 |
| 0.96484568 |
| 0.98924373 |
| 0.99122857 |
| 0.95988499 |
| 0.99998478 |
| 0.98924373 |
| 0.99998478 |
| 0.90265407 |
| 0.94846058 |
| 0.99998478 |
| 0.97329503 |
| 0.99969417 |
| 0.99998478 |
| 0.98924373 |
| 0.91325047 |
| 0.98924373 |
| 0.92861642 |
| 0.99969417 |
| 0.9403055 |
| 0.99998478 |
| 0.99998478 |
| 0.94653188 |
| 0.9520243 |
| 0.98924373 |
| 0.99781055 |
| 0.98924373 |
| 0.99998478 |
| 0.98924373 |
| 0.98732643 |
| 0.99998478 |
| 0.92690952 |
| 0.98924373 |
| 0.99998478 |
| 0.99998478 |
| 0.96484568 |
| 0.99998478 |
| 0.85141726 |
| 0.95951105 |
| 0.96181283 |
| 0.93752249 |

0.95951105
0.99998478
0.96209415
0.96181283
0.91325047
0.98924373
0.74240405
0.98732643
0.99969417
0.90265407
0.99680366
0.99998478
0.91325047
0.98924373
0.99122857
0.98924373
0.99998478
0.99122857
0.99998478
0.95240191
0.98924373
0.99998478
0.89618249
0.96181283
0.99969417
0.99998478
0.9936423
0.98924373
0.99122857
0.99122857
0.99998478
0.99998478
0.99969417
0.98924373
0.99998478
0.98924373
0.96484568
0.98924373
0.95240191
0.90297323
0.98924373
0.96484568

| |
|---|
| 0.98924373 |
| 0.98924373 |
| 0.96484568 |
| 0.99998478 |
| 0.96484568 |
| 0.99998478 |
| 0.97329503 |
| 0.98924373 |
| 0.99680366 |
| 0.99998478 |
| 0.99122857 |
| 0.96181283 |
| 0.90297323 |
| 0.9467702 |
| 0.99998478 |
| 0.96484568 |
| 0.95585979 |
| 0.99176357 |
| 0.94747425 |
| 0.98924373 |
| 0.98924373 |
| 0.99969417 |
| 0.93752249 |
| 0.98924373 |
| 0.98924373 |
| 0.91325047 |
| 0.98924373 |
| 0.99998478 |
| 0.98924373 |
| 0.94412092 |
| 0.98924373 |
| 0.94846058 |
| 0.99122857 |
| 0.96816591 |
| 0.94171831 |
| 0.91325047 |
| 0.99998478 |
| 0.99122857 |
| 0.96484568 |
| 0.99998478 |
| 0.98924373 |
| 0.91325047 |

| |
|---|
| 0.91564298 |
| 0.91325047 |
| 0.98924373 |
| 0.99998478 |
| 0.98924373 |
| 0.98924373 |
| 0.93752249 |
| 0.94171831 |
| 0.95951105 |
| 0.99998478 |
| 0.99969417 |
| 0.98924373 |
| 0.99998478 |
| 0.98924373 |
| 0.99830763 |
| 0.99122857 |
| 0.98924373 |
| 0.99969417 |
| 0.99998478 |
| 0.97110689 |
| 0.90265407 |
| 0.99122857 |
| 0.99998478 |
| 0.99998478 |
| 0.61773657 |
| 0.99998478 |
| 0.99122857 |
| 0.83011818 |
| 0.90297323 |
| 0.97815363 |
| 0.92861642 |
| 0.98924373 |
| 0.9647026 |
| 0.9865573 |
| 0.98924373 |
| 0.99998478 |
| 0.99122857 |
| 0.98924373 |
| 0.96484568 |
| 0.99998478 |
| 0.97711186 |
| 0.61773657 |

| |
|---|
| 0.98924373 |
| 0.98732643 |
| 0.9403055 |
| 0.99998478 |
| 0.99998478 |
| 0.99252883 |
| 0.99846792 |
| 0.98924373 |
| 0.9403055 |
| 0.94944907 |
| 0.96484568 |
| 0.99252883 |
| 0.96484568 |
| 0.99969417 |
| 0.91325047 |
| 0.93054695 |
| 0.99122857 |
| 0.97329503 |
| 0.94171831 |
| 0.9834625 |
| 0.85141726 |
| 0.99680366 |
| 0.96484568 |
| 0.99309902 |
| 0.99969417 |
| 0.99998478 |
| 0.90265407 |
| 0.96484568 |
| 0.90297323 |
| 0.98924373 |
| 0.92861642 |
| 0.93427164 |
| 0.98924373 |
| 0.99998478 |
| 0.99781055 |
| 0.99998478 |
| 0.9403055 |
| 0.93752249 |
| 0.96484568 |
| 0.99998478 |
| 0.98924373 |
| 0.98924373 |

| |
|---|
| 0.98924373 |
| 0.93594857 |
| 0.96181283 |
| 0.98924373 |
| 0.98732643 |
| 0.99998478 |
| 0.96181283 |
| 0.99122857 |
| 0.91267201 |
| 0.96484568 |
| 0.96484568 |
| 0.97815363 |
| 0.98924373 |
| 0.98924373 |
| 0.99998478 |
| 0.94171831 |
| 0.98924373 |
| 0.99969417 |
| 0.94171831 |
| 0.98924373 |
| 0.98924373 |
| 0.90265407 |
| 0.98924373 |
| 0.98259014 |
| 0.99969417 |
| 0.99998478 |
| 0.98924373 |
| 0.97110689 |
| 0.98924373 |
| 0.99252883 |
| 0.99969417 |
| 0.98924373 |
| 0.97327383 |
| 0.99122857 |
| 0.92075126 |
| 0.99998478 |
| 0.96484568 |
| 0.9985128 |
| 0.98924373 |
| 0.99122857 |
| 0.8773376 |
| 0.98103056 |

0.98732643
0.95585979
0.99998478
0.99998478
0.99122857
0.98732643
0.99969417
0.99122857
0.99998478
0.96709089
0.98924373
0.95240191
0.99122857
0.95951105
0.97327383
0.80984148
0.98924373
0.99122857
0.98924373
0.99998478
0.98924373
0.99998478
0.90265407
0.90297323
0.99122857
0.87669383
0.99122857
0.99122857
0.94596453
0.96484568
0.98924373
0.95859413
0.99998478
0.98924373
0.9403055
0.98924373
0.99998478
0.99998478
0.98924373
0.94171831
0.93752249
0.92861642

| |
|---|
| 0.99998478 |
| 0.96484568 |
| 0.91564298 |
| 0.90265407 |
| 0.90297323 |
| 0.93752249 |
| 0.98924373 |
| 0.99998478 |
| 0.9647026 |
| 0.90265407 |
| 0.99122857 |
| 0.98732643 |
| 0.99969417 |
| 0.98924373 |
| 0.95951105 |
| 0.99998478 |
| 0.96181283 |
| 0.99998478 |
| 0.99998478 |
| 0.90265407 |
| 0.95585979 |
| 0.92861642 |
| 0.99119687 |
| 0.99122857 |
| 0.99830763 |
| 0.96484568 |
| 0.99998478 |
| 0.9403055 |
| 0.91105653 |
| 0.94171831 |
| 0.88633644 |
| 0.98924373 |
| 0.90265407 |
| 0.98924373 |
| 0.90265407 |
| 0.99122857 |
| 0.96484568 |
| 0.99969417 |
| 0.99658202 |
| 0.90265407 |
| 0.926218 |
| 0.96484568 |

| |
|---|
| 0.95134169 |
| 0.99998478 |
| 0.95569044 |
| 0.90265407 |
| 0.99998478 |
| 0.94653188 |
| 0.99998478 |
| 0.80984148 |
| 0.99998478 |
| 0.99998478 |
| 0.93432929 |
| 0.90297323 |
| 0.93594857 |
| 0.98924373 |
| 0.99969417 |
| 0.99998478 |
| 0.95951105 |
| 0.99998478 |
| 0.90265407 |
| 0.98882544 |
| 0.99998478 |
| 0.99122857 |
| 0.98924373 |
| 0.95859413 |
| 0.99122857 |
| 0.99252883 |
| 0.98924373 |
| 0.96484568 |
| 0.8773376 |
| 0.98924373 |
| 0.72975604 |
| 0.94653188 |
| 0.98924373 |
| 0.99252883 |
| 0.99122857 |
| 0.61079703 |
| 0.91325047 |
| 0.99122857 |
| 0.99998478 |
| 0.99122857 |
| 0.93752249 |
| 0.99998478 |

| |
|---|
| 0.98732643 |
| 0.98924373 |
| 0.99998478 |
| 0.99969417 |
| 0.95240191 |
| 0.99998478 |
| 0.97126196 |
| 0.96484568 |
| 0.90265407 |
| 0.93752249 |
| 0.99969417 |
| 0.99122857 |
| 0.99998478 |
| 0.90265407 |
| 0.95034906 |
| 0.98924373 |
| 0.98924373 |
| 0.99998478 |
| 0.94171831 |
| 0.90265407 |
| 0.99969417 |
| 0.99122857 |
| 0.90297323 |
| 0.98924373 |
| 0.99969417 |
| 0.95951105 |
| 0.99998478 |
| 0.98924373 |
| 0.98924373 |
| 0.94653188 |
| 0.98924373 |
| 0.915492 |
| 0.67021692 |
| 0.94639331 |
| 0.99998478 |
| 0.93752249 |
| 0.9403055 |
| 0.89077022 |
| 0.92861642 |
| 0.93752249 |
| 0.99998478 |
| 0.99998478 |

| |
|---|
| 0.9647026 |
| 0.98924373 |
| 0.96181283 |
| 0.94171831 |
| 0.99830763 |
| 0.95240191 |
| 0.99122857 |
| 0.93752249 |
| 0.98924373 |
| 0.97815363 |
| 0.99122857 |
| 0.99998478 |
| 0.99998478 |
| 0.96484568 |
| 0.99998478 |
| 0.98924373 |
| 0.95134169 |
| 0.98924373 |
| 0.99513459 |
| 0.67021692 |
| 0.99998478 |
| 0.99969417 |
| 0.99998478 |
| 0.98924373 |
| 0.90297323 |
| 0.98924373 |
| 0.93594857 |
| 0.95240191 |
| 0.92861642 |
| 0.99122857 |
| 0.78307327 |
| 0.9865573 |
| 0.99252883 |
| 0.99122857 |
| 0.99998478 |
| 0.99998478 |
| 0.98924373 |
| 0.90297323 |
| 0.90265407 |
| 0.91325047 |
| 0.99969417 |
| 0.98924373 |

| |
|---|
| 0.90265407 |
| 0.99998478 |
| 0.97085356 |
| 0.99122857 |
| 0.93594857 |
| 0.99998478 |
| 0.99969417 |
| 0.98924373 |
| 0.80133949 |
| 0.99969417 |
| 0.99998478 |
| 0.98924373 |
| 0.99388898 |
| 0.99122857 |
| 0.99969417 |
| 0.96167371 |
| 0.98735899 |
| 0.99781055 |
| 0.90297323 |
| 0.99998478 |
| 0.98924373 |
| 0.96181283 |
| 0.97329503 |
| 0.98924373 |
| 0.99998478 |
| 0.99122857 |
| 0.99122857 |
| 0.99122857 |
| 0.9647026 |
| 0.99969417 |
| 0.9467702 |
| 0.9985128 |
| 0.98198365 |
| 0.91325047 |
| 0.99998478 |
| 0.99122857 |
| 0.97992318 |
| 0.99998478 |
| 0.93427164 |
| 0.93811476 |
| 0.91325047 |
| 0.95240191 |

| |
|---|
| 0.96484568 |
| 0.93752249 |
| 0.99122857 |
| 0.96181283 |
| 0.96484568 |
| 0.94171831 |
| 0.9403055 |
| 0.99998478 |
| 0.99122857 |
| 0.99122857 |
| 0.98924373 |
| 0.92075126 |
| 0.99122857 |
| 0.84430401 |
| 0.99998478 |
| 0.98924373 |
| 0.96484568 |
| 0.98924373 |
| 0.99948729 |
| 0.99122857 |
| 0.92548507 |
| 0.99969417 |
| 0.99998478 |
| 0.926218 |
| 0.99998478 |
| 0.99122857 |
| 0.99998478 |
| 0.99998478 |
| 0.99252883 |
| 0.99998478 |
| 0.98386625 |
| 0.90297323 |
| 0.94653188 |
| 0.99252883 |
| 0.97815363 |
| 0.98924373 |
| 0.99998478 |
| 0.95240191 |
| 0.98924373 |
| 0.98924373 |
| 0.8773376 |
| 0.90297323 |

| |
|---|
| 0.99998478 |
| 0.98924373 |
| 0.93594857 |
| 0.93752249 |
| 0.99998478 |
| 0.92367831 |
| 0.91267201 |
| 0.99122857 |
| 0.99969417 |
| 0.93752249 |
| 0.99969417 |
| 0.99998478 |
| 0.94747425 |
| 0.99122857 |
| 0.96181283 |
| 0.99122857 |
| 0.92672232 |
| 0.96181283 |
| 0.8773376 |
| 0.99998478 |
| 0.99969417 |
| 0.99969417 |
| 0.99846792 |
| 0.99846792 |
| 0.98924373 |
| 0.98732643 |
| 0.99998478 |
| 0.99367154 |
| 0.98924373 |
| 0.99998478 |
| 0.98924373 |
| 0.96484568 |
| 0.99969417 |
| 0.9865573 |
| 0.99513459 |
| 0.99513459 |
| 0.99122857 |
| 0.99998478 |
| 0.98924373 |
| 0.96484568 |
| 0.89618249 |
| 0.91325047 |

| |
|---|
| 0.9467702 |
| 0.99969417 |
| 0.90297323 |
| 0.99998478 |
| 0.99948729 |
| 0.99122857 |
| 0.98924373 |
| 0.99122857 |
| 0.99122857 |
| 0.98924373 |
| 0.99998478 |
| 0.99969417 |
| 0.96484568 |
| 0.98924373 |
| 0.96484568 |
| 0.99998478 |
| 0.99998478 |
| 0.99998478 |
| 0.94171831 |
| 0.94002917 |
| 0.99969417 |
| 0.69921142 |
| 0.99998478 |
| 0.93752249 |
| 0.99969417 |
| 0.99998478 |
| 0.99998478 |
| 0.99122857 |
| 0.99122857 |
| 0.99969417 |
| 0.99998478 |
| 0.9403055 |
| 0.99998478 |
| 0.99122857 |
| 0.96012745 |
| 0.99998478 |
| 0.91325047 |
| 0.99252883 |
| 0.99998478 |
| 0.99998478 |
| 0.90265407 |
| 0.98924373 |

| |
|---|
| 0.95951105 |
| 0.98924373 |
| 0.99998478 |
| 0.99998478 |
| 0.96484568 |
| 0.98924373 |
| 0.96167371 |
| 0.96484568 |
| 0.9403055 |
| 0.99122857 |
| 0.90297323 |
| 0.94171831 |
| 0.98924373 |
| 0.99122857 |
| 0.79151544 |
| 0.98924373 |
| 0.99969417 |
| 0.93752249 |
| 0.99122857 |
| 0.99122857 |
| 0.99998478 |
| 0.74240405 |
| 0.98924373 |
| 0.99122857 |
| 0.99969417 |
| 0.99122857 |
| 0.99998478 |
| 0.99998478 |
| 0.99998478 |
| 0.9715425 |
| 0.99998478 |
| 0.99998478 |
| 0.98924373 |
| 0.99998478 |
| 0.99998478 |
| 0.99969417 |
| 0.99998478 |
| 0.98924373 |
| 0.99998478 |
| 0.96816591 |
| 0.99998478 |
| 0.98259014 |

| |
|---|
| 0.99830763 |
| 0.98924373 |
| 0.96181283 |
| 0.99998478 |
| 0.90297323 |
| 0.99122857 |
| 0.96484568 |
| 0.99998478 |
| 0.99969417 |
| 0.98198365 |
| 0.99998478 |
| 0.94171831 |
| 0.99830763 |
| 0.9811568 |
| 0.99122857 |
| 0.98924373 |
| 0.98924373 |
| 0.99252883 |
| 0.99969417 |
| 0.93752249 |
| 0.99830763 |
| 0.94171831 |
| 0.92861642 |
| 0.98924373 |
| 0.99781055 |
| 0.99998478 |
| 0.98259014 |
| 0.94171831 |
| 0.99998478 |
| 0.99998478 |
| 0.93752249 |
| 0.98924373 |
| 0.94171831 |
| 0.9865573 |
| 0.95951105 |
| 0.98924373 |
| 0.95951105 |
| 0.92861642 |
| 0.98924373 |
| 0.9865573 |
| 0.90265407 |
| 0.99122857 |

| |
|---|
| 0.99830763 |
| 0.99998478 |
| 0.99122857 |
| 0.99122857 |
| 0.9467702 |
| 0.98924373 |
| 0.95951105 |
| 0.96484568 |
| 0.9985128 |
| 0.94747425 |
| 0.99122857 |
| 0.99998478 |
| 0.99122857 |
| 0.93752249 |
| 0.96181283 |
| 0.98924373 |
| 0.96816591 |
| 0.9403055 |
| 0.90265407 |
| 0.94171831 |
| 0.96181283 |
| 0.9985128 |
| 0.99252883 |
| 0.95654768 |
| 0.99122857 |
| 0.95240191 |
| 0.92672232 |
| 0.9557789 |
| 0.98924373 |
| 0.98924373 |
| 0.96181283 |
| 0.99122857 |
| 0.99122857 |
| 0.99122857 |
| 0.99658202 |
| 0.93594857 |
| 0.98732643 |
| 0.99998478 |
| 0.80984148 |
| 0.94653188 |
| 0.99998478 |
| 0.99122857 |

| |
|---|
| 0.99252883 |
| 0.94653188 |
| 0.99998478 |
| 0.99122857 |
| 0.98924373 |
| 0.98924373 |
| 0.99998478 |
| 0.98924373 |
| 0.99513459 |
| 0.99969417 |
| 0.99969417 |
| 0.98924373 |
| 0.99998478 |
| 0.91267201 |
| 0.91325047 |
| 0.65854402 |
| 0.90265407 |
| 0.9865573 |
| 0.8773376 |
| 0.96484568 |
| 0.99998478 |
| 0.96484568 |
| 0.96484568 |
| 0.98924373 |
| 0.94171831 |
| 0.98259014 |
| 0.99969417 |
| 0.97078973 |
| 0.95951105 |
| 0.98924373 |
| 0.915492 |
| 0.98924373 |
| 0.99122857 |
| 0.99002751 |
| 0.99998478 |
| 0.99998478 |
| 0.99998478 |
| 0.90297323 |
| 0.93752249 |
| 0.99830763 |
| 0.98924373 |
| 0.98924373 |

0.99998478
0.98924373
0.99998478
0.99998478
0.99122857
0.96484568
0.94641693
0.98924373
0.98924373
0.99998478
0.96181283
0.96181283
0.94171831
0.96181283
0.97085356
0.96484568
0.99252883
0.99998478
0.90265407
0.94653188
0.98924373
0.99998478
0.99122857
0.94897478
0.98924373
0.95951105
0.9403055
0.90265407
0.96484568
0.95585979
0.98732643
0.94653188
0.99122857
0.94641693
0.99201427
0.99998478
0.98924373
0.99998478
0.96181283
0.94171831
0.95859413
0.99998478

| |
|---|
| 0.99122857 |
| 0.99846792 |
| 0.99998478 |
| 0.96484568 |
| 0.99998478 |
| 0.99122857 |
| 0.99998478 |
| 0.91325047 |
| 0.91325047 |
| 0.99969417 |
| 0.98924373 |
| 0.99998478 |
| 0.99274826 |
| 0.92367831 |
| 0.99998478 |
| 0.99969417 |
| 0.99998478 |
| 0.90265407 |
| 0.92861642 |
| 0.9985128 |
| 0.94171831 |
| 0.91401174 |
| 0.99969417 |
| 0.99969417 |
| 0.98924373 |
| 0.82029292 |
| 0.92861642 |
| 0.96484568 |
| 0.98924373 |
| 0.99846792 |
| 0.98924373 |
| 0.93752249 |
| 0.98924373 |
| 0.90297323 |
| 0.99969417 |
| 0.99122857 |
| 0.98924373 |
| 0.99122857 |
| 0.99122857 |
| 0.99122857 |
| 0.94596453 |
| 0.99309902 |

0.98924373
0.78307327
0.92548507
0.99998478
0.99998478
0.97085356
0.98924373
0.98924373
0.99122857
0.92548507
0.99969417
0.79151544
0.90265407
0.99998478
0.99941754
0.90297323
0.99998478
0.91325047
0.98466138
0.91325047
0.99998478
0.92861642
0.94171831
0.92861642
0.98259014
0.98732643
0.96181283
0.98924373
0.99122857
0.98924373
0.99122857
0.99122857
0.99969417
0.99969417
0.99994907
0.99998478
0.96484568
0.99122857
0.99122857
0.98924373
0.98924373
0.97085356

| |
|---|
| 0.94171831 |
| 0.99998478 |
| 0.99122857 |
| 0.98924373 |
| 0.99969417 |
| 0.93811476 |
| 0.9467702 |
| 0.93365848 |
| 0.99998478 |
| 0.98924373 |
| 0.80984148 |
| 0.99122857 |
| 0.98924373 |
| 0.98924373 |
| 0.99969417 |
| 0.99998478 |
| 0.99830763 |
| 0.96484568 |
| 0.91325047 |
| 0.99122857 |
| 0.99998478 |
| 0.93594857 |
| 0.80984148 |
| 0.99998478 |
| 0.96484568 |
| 0.98924373 |
| 0.94928862 |
| 0.99969417 |
| 0.9865573 |
| 0.99122857 |
| 0.97209533 |
| 0.99658202 |
| 0.99998478 |
| 0.98924373 |
| 0.99969417 |
| 0.99998478 |
| 0.99998478 |
| 0.90265407 |
| 0.94171831 |
| 0.93365848 |
| 0.92548507 |
| 0.93752249 |

| |
|---|
| 0.93054695 |
| 0.98924373 |
| 0.96181283 |
| 0.99830763 |
| 0.99998478 |
| 0.99998478 |
| 0.99969417 |
| 0.98924373 |
| 0.96181283 |
| 0.94653188 |
| 0.96484568 |
| 0.93752249 |
| 0.85141726 |
| 0.98924373 |
| 0.95859413 |
| 0.99969417 |
| 0.99969417 |
| 0.99252883 |
| 0.98924373 |
| 0.99002751 |
| 0.99969417 |
| 0.926218 |
| 0.99122857 |
| 0.99122857 |
| 0.90297323 |
| 0.98924373 |
| 0.90297323 |
| 0.99998478 |
| 0.99998478 |
| 0.99998478 |
| 0.99998478 |
| 0.96943415 |
| 0.99969417 |
| 0.69921142 |
| 0.99252883 |
| 0.99122857 |
| 0.9829938 |
| 0.90297323 |
| 0.99998478 |
| 0.99998478 |
| 0.95951105 |
| 0.81249664 |

0.90297323
0.99998478
0.90265407
0.99998478
0.99969417
0.90265407
0.94171831
0.99969417
0.99122857
0.94897478
0.91325047
0.87669383
0.99998478
0.98924373
0.99604194
0.99252883
0.99122857
0.99998478
0.98924373
0.99969417
0.99998478
0.99617169
0.95951105
0.96484568
0.98924373
0.99122857
0.98924373
0.98924373
0.96484568
0.98924373
0.98924373
0.95951105
0.94653188
0.89618249
0.93752249
0.92861642
0.92672232
0.99122857
0.94171831
0.99969417
0.98924373
0.99122857

| |
|---|
| 0.92244927 |
| 0.98924373 |
| 0.94653188 |
| 0.98924373 |
| 0.97153778 |
| 0.98924373 |
| 0.93908842 |
| 0.99998478 |
| 0.99969417 |
| 0.90265407 |
| 0.99998478 |
| 0.99252883 |
| 0.99572919 |
| 0.99969417 |
| 0.91325047 |
| 0.98924373 |
| 0.99969417 |
| 0.99122857 |
| 0.80984148 |
| 0.98259014 |
| 0.98924373 |
| 0.99998478 |
| 0.9936085 |
| 0.98924373 |
| 0.96484568 |
| 0.98924373 |
| 0.99122857 |
| 0.98924373 |
| 0.99122857 |
| 0.99122857 |
| 0.98924373 |
| 0.99122857 |
| 0.93752249 |
| 0.915492 |
| 0.99998478 |
| 0.98924373 |
| 0.9467702 |
| 0.99122857 |
| 0.99969417 |
| 0.99122857 |
| 0.99969417 |
| 0.98924373 |

| |
|---|
| 0.96484568 |
| 0.96484568 |
| 0.99969417 |
| 0.99998478 |
| 0.99830763 |
| 0.98924373 |
| 0.99969417 |
| 0.98924373 |
| 0.94653188 |
| 0.99122857 |
| 0.99998478 |
| 0.93752249 |
| 0.99830763 |
| 0.96181283 |
| 0.94171831 |
| 0.99572919 |
| 0.98924373 |
| 0.98732643 |
| 0.99122857 |
| 0.99617169 |
| 0.93752249 |
| 0.93752249 |
| 0.90265407 |
| 0.96709089 |
| 0.98924373 |
| 0.98924373 |
| 0.99998478 |
| 0.99969417 |
| 0.99998478 |
| 0.99122857 |
| 0.96181283 |
| 0.93752249 |
| 0.98924373 |
| 0.99998478 |
| 0.95646366 |
| 0.91325047 |
| 0.98924373 |
| 0.63292047 |
| 0.99998478 |
| 0.91325047 |
| 0.98924373 |
| 0.99122857 |

| |
|---|
| 0.93752249 |
| 0.99969417 |
| 0.99969417 |
| 0.98924373 |
| 0.72204897 |
| 0.915492 |
| 0.99122857 |
| 0.99122857 |
| 0.98924373 |
| 0.99969417 |
| 0.93752249 |
| 0.99998478 |
| 0.99969417 |
| 0.96484568 |
| 0.99122857 |
| 0.99998478 |
| 0.99830763 |
| 0.92861642 |
| 0.98924373 |
| 0.98924373 |
| 0.99998478 |
| 0.99998478 |
| 0.98924373 |
| 0.99998478 |
| 0.95988499 |
| 0.99122857 |
| 0.99122857 |
| 0.99998478 |
| 0.98924373 |
| 0.99122857 |
| 0.99998478 |
| 0.99969417 |
| 0.91325047 |
| 0.96874634 |
| 0.95240191 |
| 0.95240191 |
| 0.99252883 |
| 0.99122857 |
| 0.99998478 |
| 0.99122857 |
| 0.99122857 |
| 0.9829938 |

| |
|---|
| 0.98924373 |
| 0.61773657 |
| 0.99122857 |
| 0.99998478 |
| 0.99969417 |
| 0.94653188 |
| 0.90265407 |
| 0.82029292 |
| 0.99998478 |
| 0.94171831 |
| 0.98924373 |
| 0.99998478 |
| 0.98259014 |
| 0.99122857 |
| 0.98924373 |
| 0.99969417 |
| 0.98924373 |
| 0.95951105 |
| 0.94171831 |
| 0.99998478 |
| 0.98924373 |
| 0.96181283 |
| 0.99122857 |
| 0.99658202 |
| 0.99998478 |
| 0.95951105 |
| 0.90265407 |
| 0.96484568 |
| 0.99122857 |
| 0.98259014 |
| 0.9829938 |
| 0.91267201 |
| 0.98924373 |
| 0.98924373 |
| 0.99969417 |
| 0.99323154 |
| 0.99969417 |
| 0.98924373 |
| 0.98259014 |
| 0.99998478 |
| 0.99122857 |
| 0.9467702 |

0.99969417
0.99998478
0.99122857
0.96484568
0.99846792
0.9403055
0.95951105
0.98924373
0.99122857
0.99122857
0.99781055
0.93752249
0.99998478
0.99658202
0.99122857
0.99998478
0.96484568
0.96484568
0.96484568
0.93752249
0.99998478
0.99122857
0.93594857
0.94171831
0.94380763
0.96181283
0.9403055
0.93752249
0.92548507
0.94653188
0.96484568
0.99122857
0.99122857
0.96816591
0.99998478
0.96484568
0.99122857
0.97327383
0.98924373
0.90297323
0.98924373
0.98924373

| |
|---|
| 0.98924373 |
| 0.98924373 |
| 0.93752249 |
| 0.97628191 |
| 0.9403055 |
| 0.86079694 |
| 0.99998478 |
| 0.98924373 |
| 0.98924373 |
| 0.90297323 |
| 0.99830763 |
| 0.99122857 |
| 0.61079703 |
| 0.98924373 |
| 0.90265407 |
| 0.9647026 |
| 0.99998478 |
| 0.96484568 |
| 0.99122857 |
| 0.99122857 |
| 0.99969417 |
| 0.99969417 |
| 0.99998478 |
| 0.99998478 |
| 0.98924373 |
| 0.99122857 |
| 0.98924373 |
| 0.99969417 |
| 0.99969417 |
| 0.99122857 |
| 0.99969417 |
| 0.99122857 |
| 0.98924373 |
| 0.98924373 |
| 0.99969417 |
| 0.98337226 |
| 0.98924373 |
| 0.99998478 |
| 0.96181283 |
| 0.95951105 |
| 0.99998478 |
| 0.98882544 |

| |
|---|
| 0.96484568 |
| 0.99650199 |
| 0.90297323 |
| 0.93594857 |
| 0.99969417 |
| 0.96484568 |
| 0.92548507 |
| 0.74240405 |
| 0.99969417 |
| 0.99122857 |
| 0.99998478 |
| 0.99998478 |
| 0.99846792 |
| 0.99830763 |
| 0.99998478 |
| 0.99122857 |
| 0.98924373 |
| 0.99252883 |
| 0.98924373 |
| 0.99969417 |
| 0.99122857 |
| 0.99969417 |
| 0.90265407 |
| 0.94641693 |
| 0.96484568 |
| 0.96484568 |
| 0.99252883 |
| 0.80984148 |
| 0.99388898 |
| 0.99998478 |
| 0.99998478 |
| 0.99122857 |
| 0.87052203 |
| 0.99969417 |
| 0.99252883 |
| 0.98924373 |
| 0.99252883 |
| 0.99157426 |
| 0.86327908 |
| 0.99969417 |
| 0.98924373 |
| 0.9403055 |

| |
|---|
| 0.98924373 |
| 0.99998478 |
| 0.99998478 |
| 0.90265407 |
| 0.95240191 |
| 0.99998478 |
| 0.99998478 |
| 0.99998478 |
| 0.90265407 |
| 0.99998478 |
| 0.99122857 |
| 0.99998478 |
| 0.98924373 |
| 0.99122857 |
| 0.98924373 |
| 0.90265407 |
| 0.99998478 |
| 0.99969417 |
| 0.95240191 |
| 0.99122857 |
| 0.98924373 |
| 0.99122857 |
| 0.99998478 |
| 0.99252883 |
| 0.96484568 |
| 0.97329503 |
| 0.98924373 |
| 0.99998478 |
| 0.99998478 |
| 0.93752249 |
| 0.98732643 |
| 0.98924373 |
| 0.91325047 |
| 0.99998478 |
| 0.96484568 |
| 0.98924373 |
| 0.99998478 |
| 0.99948729 |
| 0.99998478 |
| 0.99998478 |
| 0.99998478 |
| 0.96484568 |

| |
|---|
| 0.94653188 |
| 0.93752249 |
| 0.99122857 |
| 0.98924373 |
| 0.98924373 |
| 0.94171831 |
| 0.98924373 |
| 0.9403055 |
| 0.99969417 |
| 0.98924373 |
| 0.99969417 |
| 0.95240191 |
| 0.98924373 |
| 0.99122857 |
| 0.9467702 |
| 0.99122857 |
| 0.85801119 |
| 0.99998478 |
| 0.93151025 |
| 0.97368509 |
| 0.98924373 |
| 0.9985128 |
| 0.98924373 |
| 0.90297323 |
| 0.99122857 |
| 0.98924373 |
| 0.80984148 |
| 0.99252883 |
| 0.97209533 |
| 0.99998478 |
| 0.91325047 |
| 0.98705617 |
| 0.99122857 |
| 0.90265407 |
| 0.99998478 |
| 0.99969417 |
| 0.90265407 |
| 0.99122857 |
| 0.90297323 |
| 0.99969417 |
| 0.99122857 |
| 0.99183457 |

| |
|---|
| 0.99998478 |
| 0.99998478 |
| 0.99998478 |
| 0.93752249 |
| 0.92367831 |
| 0.85141726 |
| 0.98924373 |
| 0.96484568 |
| 0.99998478 |
| 0.99969417 |
| 0.98924373 |
| 0.95951105 |
| 0.99122857 |
| 0.98924373 |
| 0.98924373 |
| 0.99122857 |
| 0.99830763 |
| 0.98924373 |
| 0.98924373 |
| 0.99998478 |
| 0.99998478 |
| 0.9467702 |
| 0.96484568 |
| 0.94171831 |
| 0.99122857 |
| 0.99183457 |
| 0.98924373 |
| 0.93752249 |
| 0.99252883 |
| 0.96484568 |
| 0.96181283 |
| 0.99998478 |
| 0.99998478 |
| 0.99969417 |
| 0.99122857 |
| 0.99969417 |
| 0.99122857 |
| 0.99122857 |
| 0.93427164 |
| 0.99969417 |
| 0.99998478 |
| 0.98924373 |

| |
|---|
| 0.97024001 |
| 0.99998478 |
| 0.94171831 |
| 0.99122857 |
| 0.9829938 |
| 0.98924373 |
| 0.99998478 |
| 0.93752249 |
| 0.91267201 |
| 0.99998478 |
| 0.99998478 |
| 0.99998478 |
| 0.95988499 |
| 0.95643072 |
| 0.99998478 |
| 0.91325047 |
| 0.94653188 |
| 0.99846792 |
| 0.98924373 |
| 0.99969417 |
| 0.90297323 |
| 0.93054695 |
| 0.99998478 |
| 0.99122857 |
| 0.99122857 |
| 0.99969417 |
| 0.98924373 |
| 0.94171831 |
| 0.98924373 |
| 0.915492 |
| 0.93908842 |
| 0.96484568 |
| 0.98924373 |
| 0.99122857 |
| 0.99998478 |
| 0.94402118 |
| 0.99998478 |
| 0.99969417 |
| 0.99122857 |
| 0.99998478 |
| 0.8773376 |
| 0.93550274 |

| |
|---|
| 0.98924373 |
| 0.8773376 |
| 0.91325047 |
| 0.99122857 |
| 0.99969417 |
| 0.96484568 |
| 0.99998478 |
| 0.99998478 |
| 0.93594857 |
| 0.98924373 |
| 0.99122857 |
| 0.99122857 |
| 0.99252883 |
| 0.98924373 |
| 0.98924373 |
| 0.94653188 |
| 0.99998478 |
| 0.93752249 |
| 0.99998478 |
| 0.99969417 |
| 0.97327383 |
| 0.99122857 |
| 0.98924373 |
| 0.99969417 |
| 0.99680366 |
| 0.99830763 |
| 0.99252883 |
| 0.99998478 |
| 0.98924373 |
| 0.99969417 |
| 0.97085356 |
| 0.99122857 |
| 0.99969417 |
| 0.96484568 |
| 0.99969417 |
| 0.98924373 |
| 0.9715425 |
| 0.9403055 |
| 0.99969417 |
| 0.99122857 |
| 0.98466138 |
| 0.9467702 |

| |
|---|
| 0.99969417 |
| 0.97085356 |
| 0.99122857 |
| 0.99998478 |
| 0.9467702 |
| 0.95240191 |
| 0.99969417 |
| 0.99122857 |
| 0.94653188 |
| 0.99183457 |
| 0.98924373 |
| 0.98924373 |
| 0.98924373 |
| 0.96484568 |
| 0.99948729 |
| 0.87052203 |
| 0.99998478 |
| 0.99122857 |
| 0.93752249 |
| 0.95859413 |
| 0.99998478 |
| 0.99183457 |
| 0.92367831 |
| 0.99969417 |
| 0.99122857 |
| 0.99998478 |
| 0.99969417 |
| 0.94171831 |
| 0.99830763 |
| 0.97327383 |
| 0.99998478 |
| 0.91325047 |
| 0.99969417 |
| 0.98924373 |
| 0.9403055 |
| 0.99122857 |
| 0.98924373 |
| 0.99998478 |
| 0.90265407 |
| 0.99122857 |
| 0.99998478 |
| 0.99002751 |

| |
|---|
| 0.99122857 |
| 0.99998478 |
| 0.99998478 |
| 0.99969417 |
| 0.94683707 |
| 0.98924373 |
| 0.99998478 |
| 0.93488349 |
| 0.926218 |
| 0.87669383 |
| 0.99998478 |
| 0.93752249 |
| 0.98924373 |
| 0.96181283 |
| 0.9985128 |
| 0.99969417 |
| 0.96018853 |
| 0.98732643 |
| 0.98924373 |
| 0.98259014 |
| 0.93752249 |
| 0.80984148 |
| 0.9715425 |
| 0.98924373 |
| 0.96484568 |
| 0.99998478 |
| 0.99969417 |
| 0.98924373 |
| 0.91267201 |
| 0.98732643 |
| 0.92861642 |
| 0.94171831 |
| 0.99998478 |
| 0.98924373 |
| 0.98924373 |
| 0.99998478 |
| 0.99998478 |
| 0.98924373 |
| 0.98924373 |
| 0.98924373 |
| 0.99122857 |
| 0.80133949 |

| |
|---|
| 0.99998478 |
| 0.96181283 |
| 0.99998478 |
| 0.98705617 |
| 0.91267201 |
| 0.99122857 |
| 0.99252883 |
| 0.99808206 |
| 0.9403055 |
| 0.99122857 |
| 0.95951105 |
| 0.99998478 |
| 0.99998478 |
| 0.99998478 |
| 0.99122857 |
| 0.99998478 |
| 0.99998478 |
| 0.96181283 |
| 0.91812391 |
| 0.98924373 |
| 0.99122857 |
| 0.93427164 |
| 0.72975604 |
| 0.99998478 |
| 0.9403055 |
| 0.91325047 |
| 0.99122857 |
| 0.98924373 |
| 0.99998478 |
| 0.98924373 |
| 0.99969417 |
| 0.99002751 |
| 0.99969417 |
| 0.99122857 |
| 0.98924373 |
| 0.96484568 |
| 0.91267201 |
| 0.98924373 |
| 0.99122857 |
| 0.93811476 |
| 0.94653188 |
| 0.96484568 |

0.99998478
0.99122857
0.99122857
0.99122857
0.85141726
0.93594857
0.93752249
0.98237613
0.98924373
0.96167371
0.95988499
0.98924373
0.99998478
0.99122857
0.98924373
0.99998478
0.99998478
0.99252883
0.90529035
0.99998478
0.90297323
0.94171831
0.93752249
0.99969417
0.99969417
0.99969417
0.96484568
0.98924373
0.95240191
0.99969417
0.92861642
0.69921142
0.99252883
0.78307327
0.98924373
0.99122857
0.99969417
0.98924373
0.98924373
0.99122857
0.99122857
0.99122857

| |
|---|
| 0.98924373 |
| 0.96484568 |
| 0.95034906 |
| 0.99998478 |
| 0.99122857 |
| 0.9403055 |
| 0.99998478 |
| 0.98924373 |
| 0.99998478 |
| 0.93752249 |
| 0.98924373 |
| 0.99332973 |
| 0.99998478 |
| 0.98732643 |
| 0.93752249 |
| 0.98924373 |
| 0.93752249 |
| 0.99830763 |
| 0.95988499 |
| 0.99998478 |
| 0.99998478 |
| 0.99122857 |
| 0.99830763 |
| 0.99846792 |
| 0.98259014 |
| 0.98466138 |
| 0.95134169 |
| 0.79151544 |
| 0.98337226 |
| 0.99998478 |
| 0.93594857 |
| 0.94171831 |
| 0.94171831 |
| 0.97993742 |
| 0.90265407 |
| 0.99998478 |
| 0.99998478 |
| 0.99122857 |
| 0.95859413 |
| 0.91267201 |
| 0.94171831 |
| 0.99998478 |

| |
|---|
| 0.99122857 |
| 0.9715425 |
| 0.99122857 |
| 0.97329503 |
| 0.99969417 |
| 0.98237613 |
| 0.99998478 |
| 0.94171831 |
| 0.93752249 |
| 0.93752249 |
| 0.96484568 |
| 0.90265407 |
| 0.99998478 |
| 0.95034906 |
| 0.98924373 |
| 0.98924373 |
| 0.93752249 |
| 0.90265407 |
| 0.99122857 |
| 0.99122857 |
| 0.98924373 |
| 0.99122857 |
| 0.99122857 |
| 0.99650199 |
| 0.90297323 |
| 0.99998478 |
| 0.99998478 |
| 0.99998478 |
| 0.99998478 |
| 0.92367831 |
| 0.97078973 |
| 0.98924373 |
| 0.99122857 |
| 0.99808206 |
| 0.99122857 |
| 0.96181283 |
| 0.99183457 |
| 0.99998478 |
| 0.98924373 |
| 0.99998478 |
| 0.98924373 |
| 0.98924373 |

| |
|---|
| 0.99998478 |
| 0.99122857 |
| 0.99998478 |
| 0.98924373 |
| 0.94171831 |
| 0.98924373 |
| 0.99998478 |
| 0.99998478 |
| 0.99183457 |
| 0.9403055 |
| 0.99122857 |
| 0.99998478 |
| 0.90297323 |
| 0.92861642 |
| 0.98924373 |
| 0.99969417 |
| 0.99998478 |
| 0.98924373 |
| 0.96181283 |
| 0.98259014 |
| 0.99969417 |
| 0.9467702 |
| 0.98924373 |
| 0.99998478 |
| 0.99122857 |
| 0.99998478 |
| 0.99998478 |
| 0.94171831 |
| 0.98924373 |
| 0.98924373 |
| 0.90265407 |
| 0.99002751 |
| 0.98924373 |
| 0.99122857 |
| 0.92075126 |
| 0.99122857 |
| 0.98924373 |
| 0.99122857 |
| 0.99969417 |
| 0.96181283 |
| 0.94171831 |
| 0.99122857 |

| |
|---|
| 0.98924373 |
| 0.99680366 |
| 0.98732643 |
| 0.90297323 |
| 0.99781055 |
| 0.99998478 |
| 0.98924373 |
| 0.98924373 |
| 0.93752249 |
| 0.98924373 |
| 0.9403055 |
| 0.99998478 |
| 0.98924373 |
| 0.98924373 |
| 0.99122857 |
| 0.99998478 |
| 0.98924373 |
| 0.99969417 |
| 0.95240191 |
| 0.94653188 |
| 0.95951105 |
| 0.99252883 |
| 0.87822432 |
| 0.98924373 |
| 0.99998478 |
| 0.99122857 |
| 0.98924373 |
| 0.99122857 |
| 0.99122857 |
| 0.99122857 |
| 0.98924373 |
| 0.98924373 |
| 0.9985128 |
| 0.9467702 |
| 0.90297323 |
| 0.9403055 |
| 0.99998478 |
| 0.99122857 |
| 0.99998478 |
| 0.9811461 |
| 0.98924373 |
| 0.96484568 |

0.91325047
0.98924373
0.9985128
0.99998478
0.98924373
0.99122857
0.85669195
0.98924373
0.97954048
0.99998478
0.99969417
0.99998478
0.99658202
0.98259014
0.99846792
0.97327383
0.99998478
0.98386625
0.96181283
0.99122857
0.95240191
0.94171831
0.99998478
0.99998478
0.94171831
0.98924373
0.99947064
0.90297323
0.97209533
0.98924373
0.98924373
0.99572919
0.98924373
0.94171831
0.99122857
0.99122857
0.9403055
0.99998478
0.99998478
0.89474679
0.86327908
0.95134169

0.99122857
0.98924373
0.99998478
0.99122857
0.98924373
0.98924373
0.99998478
0.99969417
0.99998478
0.99252883
0.99998478
0.99122857
0.99085373
0.94171831
0.99998478
0.65854402
0.90297323
0.95951105
0.98924373
0.98924373
0.99998478
0.99998478
0.98924373
0.98924373
0.99998478
0.99998478
0.99969417
0.74240405
0.99122857
0.98924373
0.99122857
0.94171831
0.99998478
0.98924373
0.99830763
0.99969417
0.80133949
0.99122857
0.99969417
0.98924373
0.99846792
0.99998478

0.98924373
0.99122857
0.99998478
0.98924373
0.95951105
0.95951105
0.98924373
0.99998478
0.98924373
0.98924373
0.99969417
0.99998478
0.94639331
0.99969417
0.96484568
0.99998478
0.99252883
0.99122857
0.98924373
0.98924373
0.99969417
0.95951105
0.99122857
0.98924373
0.99998478
0.99998478
0.99998478
0.99998478
0.98924373
0.98924373
0.99998478
0.99830763
0.99998478
0.99998478
0.97329503
0.93594857
0.99998478
0.99808206
0.95951105
0.99998478
0.98924373
0.98924373

| |
|---|
| 0.95240191 |
| 0.98924373 |
| 0.99998478 |
| 0.98924373 |
| 0.90265407 |
| 0.96181283 |
| 0.93594857 |
| 0.99969417 |
| 0.98924373 |
| 0.99998478 |
| 0.99650199 |
| 0.94653188 |
| 0.98924373 |
| 0.98924373 |
| 0.99122857 |
| 0.99969417 |
| 0.99998478 |
| 0.92548507 |
| 0.99122857 |
| 0.99122857 |
| 0.96484568 |
| 0.99967514 |
| 0.97597835 |
| 0.99534532 |
| 0.96484568 |
| 0.99122857 |
| 0.98924373 |
| 0.99122857 |
| 0.99998478 |
| 0.98732643 |
| 0.99830763 |
| 0.9403055 |
| 0.99998478 |
| 0.99122857 |
| 0.99998478 |
| 0.99998478 |
| 0.9985128 |
| 0.98924373 |
| 0.98924373 |
| 0.91325047 |
| 0.99998478 |
| 0.94171831 |

| |
|---|
| 0.99969417 |
| 0.92672232 |
| 0.99998478 |
| 0.99122857 |
| 0.98924373 |
| 0.99658202 |
| 0.93752249 |
| 0.99830763 |
| 0.99122857 |
| 0.99998478 |
| 0.99998478 |
| 0.89618249 |
| 0.99998478 |
| 0.99998478 |
| 0.96816591 |
| 0.99998478 |
| 0.98924373 |
| 0.99998478 |
| 0.98924373 |
| 0.99122857 |
| 0.98924373 |
| 0.99998478 |
| 0.81286841 |
| 0.99101915 |
| 0.91485898 |
| 0.99122857 |
| 0.96484568 |
| 0.99301012 |
| 0.9403055 |
| 0.99969417 |
| 0.90265407 |
| 0.98924373 |
| 0.9834625 |
| 0.9467702 |
| 0.99998478 |
| 0.99969417 |
| 0.98924373 |
| 0.8773376 |
| 0.99998478 |
| 0.99998478 |
| 0.9467702 |
| 0.99969417 |

|
|---|
| 0.93594857 |
| 0.97954048 |
| 0.94171831 |
| 0.99998478 |
| 0.99998478 |
| 0.99122857 |
| 0.99830763 |
| 0.97085356 |
| 0.99830763 |
| 0.98732643 |
| 0.90265407 |
| 0.98880599 |
| 0.99998478 |
| 0.95240191 |
| 0.98924373 |
| 0.98924373 |
| 0.99830763 |
| 0.99969417 |
| 0.98924373 |
| 0.98924373 |
| 0.98924373 |
| 0.98924373 |
| 0.98924373 |
| 0.99969417 |
| 0.96484568 |
| 0.99122857 |
| 0.99998478 |
| 0.99969417 |
| 0.9985128 |
| 0.9403055 |
| 0.99998478 |
| 0.99969417 |
| 0.99122857 |
| 0.98924373 |
| 0.93752249 |
| 0.99122857 |
| 0.99969417 |
| 0.93752249 |
| 0.94171831 |
| 0.91325047 |
| 0.99122857 |
| 0.91325047 |

0.90265407
0.99122857
0.98924373
0.93752249
0.96484568
0.98732643
0.97110689
0.99122857
0.9467702
0.99998478
0.99998478
0.9985128
0.99969417
0.99122857
0.98924373
0.99998478
0.85141726
0.91325047
0.96484568
0.96816591
0.99998478
0.98924373
0.9403055
0.99969417
0.99830763
0.99948729
0.99846792
0.95240191
0.99969417
0.99830763
0.99998478
0.94171831
0.99969417
0.99998478
0.98924373
0.96484568
0.98924373
0.99998478
0.96484568
0.99830763
0.99122857
0.94653188

| |
|---|
| 0.91325047 |
| 0.98259014 |
| 0.99998478 |
| 0.98924373 |
| 0.99998478 |
| 0.98924373 |
| 0.96709089 |
| 0.94596453 |
| 0.98924373 |
| 0.90297323 |
| 0.99998478 |
| 0.97327383 |
| 0.9403055 |
| 0.80984148 |
| 0.86441249 |
| 0.9226577 |
| 0.96484568 |
| 0.96181283 |
| 0.98924373 |
| 0.99122857 |
| 0.78307327 |
| 0.9403055 |
| 0.94653188 |
| 0.99122857 |
| 0.99998478 |
| 0.99998478 |
| 0.9985128 |
| 0.91325047 |
| 0.99998478 |
| 0.99969417 |
| 0.99998478 |
| 0.99969417 |
| 0.97209533 |
| 0.99998478 |
| 0.99122857 |
| 0.99122857 |
| 0.99998478 |
| 0.99969417 |
| 0.99792604 |
| 0.99998478 |
| 0.99122857 |
| 0.98924373 |

| |
|---|
| 0.98924373 |
| 0.98924373 |
| 0.98924373 |
| 0.99830763 |
| 0.95240191 |
| 0.98924373 |
| 0.99998478 |
| 0.96484568 |
| 0.80984148 |
| 0.95414145 |
| 0.96484568 |
| 0.99122857 |
| 0.98924373 |
| 0.99122857 |
| 0.98924373 |
| 0.99122857 |
| 0.98924373 |
| 0.99122857 |
| 0.90297323 |
| 0.97110689 |
| 0.99969417 |
| 0.99846792 |
| 0.99252883 |
| 0.99998478 |
| 0.99969417 |
| 0.99998478 |
| 0.95240191 |
| 0.96484568 |
| 0.99998478 |
| 0.99969417 |
| 0.8773376 |
| 0.99998478 |
| 0.99122857 |
| 0.86327908 |
| 0.80984148 |
| 0.99122857 |
| 0.99122857 |
| 0.98924373 |
| 0.99122857 |
| 0.98924373 |
| 0.93752249 |
| 0.9715425 |

0.98924373
0.99969417
0.99252883
0.99830763
0.99969417
0.99122857
0.99998478
0.99998478
0.98924373
0.90265407
0.95240191
0.99998478
0.99002751
0.99477462
0.99969417
0.90904748
0.99998478
0.99998478
0.99830763
0.99998478
0.99998478
0.99122857
0.99998478
0.96484568
0.98924373
0.99998478
0.99998478
0.98924373
0.99998478
0.99122857
0.98924373
0.99998478
0.99969417
0.90265407
0.98924373
0.98924373
0.99998478
0.99969417
0.95240191
0.98924373
0.99998478
0.98924373

| |
|---|
| 0.99998478 |
| 0.93539424 |
| 0.99998478 |
| 0.99122857 |
| 0.93365848 |
| 0.92861642 |
| 0.99122857 |
| 0.90265407 |
| 0.99998478 |
| 0.99998478 |
| 0.90297323 |
| 0.99969417 |
| 0.98732643 |
| 0.98924373 |
| 0.98924373 |
| 0.91325047 |
| 0.90265407 |
| 0.99998478 |
| 0.99998478 |
| 0.99643667 |
| 0.99122857 |
| 0.95951105 |
| 0.98924373 |
| 0.99252883 |
| 0.94641693 |
| 0.99122857 |
| 0.99998478 |
| 0.99846792 |
| 0.99998478 |
| 0.98924373 |
| 0.97329503 |
| 0.99122857 |
| 0.99998478 |
| 0.9403055 |
| 0.99122857 |
| 0.99969417 |
| 0.98924373 |
| 0.99969417 |
| 0.89958215 |
| 0.99763077 |
| 0.99122857 |
| 0.92861642 |

| |
|---|
| 0.95585979 |
| 0.99998478 |
| 0.99122857 |
| 0.92861642 |
| 0.99998478 |
| 0.98924373 |
| 0.99998478 |
| 0.98924373 |
| 0.99122857 |
| 0.787647 |
| 0.98924373 |
| 0.90265407 |
| 0.99969417 |
| 0.98924373 |
| 0.99830763 |
| 0.98956576 |
| 0.98924373 |
| 0.98924373 |
| 0.99969417 |
| 0.99998478 |
| 0.9467702 |
| 0.91621901 |
| 0.9403055 |
| 0.93752249 |
| 0.98924373 |
| 0.99122857 |
| 0.98924373 |
| 0.99122857 |
| 0.98924373 |
| 0.99969417 |
| 0.98924373 |
| 0.95034906 |
| 0.94171831 |
| 0.96181283 |
| 0.97085356 |
| 0.99969417 |
| 0.98924373 |
| 0.98924373 |
| 0.99122857 |
| 0.95859413 |
| 0.94171831 |
| 0.92548507 |

| |
|---|
| 0.98924373 |
| 0.93752249 |
| 0.99998478 |
| 0.99846792 |
| 0.99830763 |
| 0.94641693 |
| 0.99998478 |
| 0.99998478 |
| 0.99998478 |
| 0.98924373 |
| 0.99122857 |
| 0.90297323 |
| 0.96484568 |
| 0.99846792 |
| 0.99998478 |
| 0.99122857 |
| 0.99998478 |
| 0.99998478 |
| 0.98924373 |
| 0.96484568 |
| 0.99969417 |
| 0.99998478 |
| 0.94002917 |
| 0.99969417 |
| 0.99122857 |
| 0.93752249 |
| 0.99122857 |
| 0.99998478 |
| 0.99122857 |
| 0.98924373 |
| 0.97110689 |
| 0.96484568 |
| 0.95240191 |
| 0.93752249 |
| 0.99998478 |
| 0.99122857 |
| 0.95212302 |
| 0.79151544 |
| 0.926218 |
| 0.99846792 |
| 0.99122857 |
| 0.98732643 |

| |
|---|
| 0.93752249 |
| 0.99122857 |
| 0.99122857 |
| 0.98924373 |
| 0.98924373 |
| 0.99830763 |
| 0.99122857 |
| 0.99830763 |
| 0.96167371 |
| 0.93432929 |
| 0.99998478 |
| 0.98924373 |
| 0.96181283 |
| 0.98259014 |
| 0.9985128 |
| 0.99122857 |
| 0.99969417 |
| 0.9403055 |
| 0.99998478 |
| 0.85141726 |
| 0.98924373 |
| 0.99969417 |
| 0.98924373 |
| 0.99969417 |
| 0.98924373 |
| 0.99998478 |
| 0.99830763 |
| 0.96484568 |
| 0.99122857 |
| 0.99122857 |
| 0.99969417 |
| 0.94171831 |
| 0.99998478 |
| 0.98924373 |
| 0.91485898 |
| 0.97430523 |
| 0.95240191 |
| 0.98924373 |
| 0.9985128 |
| 0.99969417 |
| 0.99122857 |
| 0.99122857 |

0.95240191
0.99969417
0.96484568
0.95240191
0.99998478
0.98924373
0.85141726
0.94897478
0.99998478
0.99998478
0.8773376
0.99969417
0.99122857
0.98924373
0.99998478
0.95859413
0.99998478
0.99998478
0.99998478
0.96484568
0.99846792
0.99969417
0.99998478
0.99969417
0.99122857
0.9403055
0.95240191
0.99830763
0.99998478
0.99998478
0.99969417
0.98924373
0.98924373
0.99998478
0.99998478
0.90297323
0.99998478
0.99122857
0.99998478
0.99122857
0.80984148
0.94653188

| |
|---|
| 0.95988499 |
| 0.99969417 |
| 0.98924373 |
| 0.98259014 |
| 0.99998478 |
| 0.99969417 |
| 0.96181283 |
| 0.99122857 |
| 0.9403055 |
| 0.98924373 |
| 0.97127035 |
| 0.99998478 |
| 0.98924373 |
| 0.99969417 |
| 0.99998478 |
| 0.98924373 |
| 0.90297323 |
| 0.92861642 |
| 0.93752249 |
| 0.98924373 |
| 0.96484568 |
| 0.91325047 |
| 0.95951105 |
| 0.96484568 |
| 0.99969417 |
| 0.91325047 |
| 0.99998478 |
| 0.9403055 |
| 0.98924373 |
| 0.95951105 |
| 0.99658202 |
| 0.98924373 |
| 0.94171831 |
| 0.99969417 |
| 0.96484568 |
| 0.99122857 |
| 0.98924373 |
| 0.98924373 |
| 0.98924373 |
| 0.96484568 |
| 0.99998478 |
| 0.99998478 |

| |
|---|
| 0.99969417 |
| 0.99998478 |
| 0.99998478 |
| 0.99122857 |
| 0.9865573 |
| 0.99998478 |
| 0.97711186 |
| 0.99122857 |
| 0.99969417 |
| 0.98924373 |
| 0.99998478 |
| 0.99122857 |
| 0.98924373 |
| 0.99998478 |
| 0.99998478 |
| 0.98924373 |
| 0.9403055 |
| 0.99969417 |
| 0.98924373 |
| 0.97110689 |
| 0.99998478 |
| 0.98924373 |
| 0.95240191 |
| 0.99969417 |
| 0.94171831 |
| 0.79151544 |
| 0.99969417 |
| 0.98924373 |
| 0.99122857 |
| 0.99122857 |
| 0.99969417 |
| 0.91325047 |
| 0.99998478 |
| 0.99998478 |
| 0.99969417 |
| 0.98924373 |
| 0.93908842 |
| 0.95951105 |
| 0.95951105 |
| 0.99122857 |
| 0.99998478 |
| 0.98924373 |

| |
|---|
| 0.98924373 |
| 0.96167371 |
| 0.99998478 |
| 0.94641693 |
| 0.926218 |
| 0.98924373 |
| 0.96181283 |
| 0.98924373 |
| 0.99252883 |
| 0.99122857 |
| 0.91325047 |
| 0.98924373 |
| 0.95951105 |
| 0.99122857 |
| 0.9985128 |
| 0.99998478 |
| 0.94171831 |
| 0.99998478 |
| 0.99998478 |
| 0.94171831 |
| 0.99122857 |
| 0.98924373 |
| 0.99998478 |
| 0.99969417 |
| 0.99998478 |
| 0.99969417 |
| 0.98924373 |
| 0.98732643 |
| 0.99998478 |
| 0.99122857 |
| 0.98924373 |
| 0.97711186 |
| 0.90265407 |
| 0.99122857 |
| 0.95034906 |
| 0.99998478 |
| 0.99122857 |
| 0.99998478 |
| 0.99998478 |
| 0.99572919 |
| 0.96484568 |
| 0.90265407 |

0.99998478
0.99969417
0.99998478
0.99865984
0.99122857
0.9467702
0.94897478
0.99513459
0.98924373
0.93752249
0.99969417
0.98924373
0.97327383
0.95240191
0.93752249
0.99998478
0.99122857
0.98924373
0.99513459
0.90265407
0.94171831
0.9467702
0.99998478
0.99122857
0.99122857
0.99998478
0.99122857
0.98924373
0.99122857
0.95859413
0.96181283
0.99650199
0.96181283
0.99998478
0.94002917
0.99998478
0.926218
0.9865573
0.98735899
0.98924373
0.99002751
0.93111203

| |
|---|
| 0.9985128 |
| 0.95951105 |
| 0.93811476 |
| 0.97992318 |
| 0.9834625 |
| 0.99122857 |
| 0.99969417 |
| 0.95240191 |
| 0.99969417 |
| 0.96181283 |
| 0.99998478 |
| 0.98924373 |
| 0.99781055 |
| 0.99122857 |
| 0.98924373 |
| 0.72975604 |
| 0.96181283 |
| 0.99998478 |
| 0.99830763 |
| 0.9467702 |
| 0.99122857 |
| 0.99830763 |
| 0.99998478 |
| 0.85141726 |
| 0.98924373 |
| 0.98924373 |
| 0.99122857 |
| 0.9865573 |
| 0.99998478 |
| 0.99948729 |
| 0.99998478 |
| 0.99122857 |
| 0.99998478 |
| 0.99998478 |
| 0.8773376 |
| 0.96484568 |
| 0.95240191 |
| 0.98924373 |
| 0.99969417 |
| 0.96484568 |
| 0.98924373 |
| 0.99998478 |

| |
|---|
| 0.96709089 |
| 0.99122857 |
| 0.90297323 |
| 0.98924373 |
| 0.99513459 |
| 0.98924373 |
| 0.98924373 |
| 0.9403055 |
| 0.94747425 |
| 0.99122857 |
| 0.98732643 |
| 0.93752249 |
| 0.99998478 |
| 0.95034906 |
| 0.94653188 |
| 0.99122857 |
| 0.91325047 |
| 0.98924373 |
| 0.98732643 |
| 0.99998478 |
| 0.99122857 |
| 0.9829938 |
| 0.90265407 |
| 0.99998478 |
| 0.9985128 |
| 0.98924373 |
| 0.99998478 |
| 0.98924373 |
| 0.98924373 |
| 0.98924373 |
| 0.99122857 |
| 0.95951105 |
| 0.91325047 |
| 0.99122857 |
| 0.97126196 |
| 0.9865573 |
| 0.99969417 |
| 0.99122857 |
| 0.99969417 |
| 0.94435487 |
| 0.98924373 |
| 0.99122857 |

0.97024001
0.99830763
0.99969417
0.99998478
0.99122857
0.95951105
0.98924373
0.80133949
0.99122857
0.99650199
0.98924373
0.87219413
0.99998478
0.99967514
0.99998478
0.94171831
0.99998478
0.9467702
0.99830763
0.99998478
0.98924373
0.90265407
0.90297323
0.93752249
0.99830763
0.99122857
0.97110689
0.98924373
0.99969417
0.99998478
0.99122857
0.98924373
0.98924373
0.95859413
0.99122857
0.99998478
0.98924373
0.99122857
0.99969417
0.99122857
0.99085373
0.99998478

0.99969417
0.99998478
0.86079694
0.99969417
0.99998478
0.94653188
0.99969417
0.99122857
0.99122857
0.90265407
0.97209533
0.99122857
0.99122857
0.99998478
0.79151544
0.99969417
0.9715425
0.94002917
0.91325047
0.98924373
0.98924373
0.85141726
0.98956576
0.99998478
0.99998478
0.97329503
0.97368509
0.99998478
0.96484568
0.97110689
0.99252883
0.98924373
0.8773376
0.99122857
0.99572919
0.9936423
0.98924373
0.98924373
0.97815363
0.98924373
0.99998478
0.98924373

| |
|---|
| 0.99998478 |
| 0.99998478 |
| 0.98924373 |
| 0.99969417 |
| 0.98924373 |
| 0.98924373 |
| 0.99252883 |
| 0.99122857 |
| 0.99122857 |
| 0.99122857 |
| 0.99830763 |
| 0.99122857 |
| 0.92861642 |
| 0.99122857 |
| 0.97504433 |
| 0.99830763 |
| 0.96484568 |
| 0.99122857 |
| 0.98924373 |
| 0.96484568 |
| 0.96484568 |
| 0.98924373 |
| 0.99998478 |
| 0.98924373 |
| 0.99998478 |
| 0.99122857 |
| 0.99122857 |
| 0.96484568 |
| 0.99998478 |
| 0.9985128 |
| 0.93752249 |
| 0.99388898 |
| 0.99122857 |
| 0.98259014 |
| 0.99998478 |
| 0.99122857 |
| 0.99998478 |
| 0.98924373 |
| 0.98924373 |
| 0.99969417 |
| 0.99998478 |
| 0.9715425 |

| |
|---|
| 0.97327383 |
| 0.99998478 |
| 0.92861642 |
| 0.99998478 |
| 0.96484568 |
| 0.99998478 |
| 0.99998478 |
| 0.98924373 |
| 0.99998478 |
| 0.99122857 |
| 0.99998478 |
| 0.90297323 |
| 0.95240191 |
| 0.99846792 |
| 0.98924373 |
| 0.98924373 |
| 0.99122857 |
| 0.65854402 |
| 0.90265407 |
| 0.99998478 |
| 0.99969417 |
| 0.99998478 |
| 0.97110689 |
| 0.99998478 |
| 0.99122857 |
| 0.99998478 |
| 0.98924373 |
| 0.94653188 |
| 0.98924373 |
| 0.99998478 |
| 0.99998478 |
| 0.99252883 |
| 0.9647026 |
| 0.95585979 |
| 0.99969417 |
| 0.98924373 |
| 0.94653188 |
| 0.98924373 |
| 0.99830763 |
| 0.95240191 |
| 0.99969417 |
| 0.9829938 |

| |
|---|
| 0.99998478 |
| 0.98924373 |
| 0.94171831 |
| 0.91325047 |
| 0.97327383 |
| 0.99122857 |
| 0.9647026 |
| 0.98924373 |
| 0.99969417 |
| 0.99998478 |
| 0.98924373 |
| 0.99122857 |
| 0.99969417 |
| 0.91325047 |
| 0.98924373 |
| 0.98924373 |
| 0.99998478 |
| 0.96167371 |
| 0.99969417 |
| 0.93427164 |
| 0.99122857 |
| 0.93752249 |
| 0.99122857 |
| 0.99122857 |
| 0.99998478 |
| 0.99122857 |
| 0.98924373 |
| 0.99998478 |
| 0.98924373 |
| 0.9403055 |
| 0.91325047 |
| 0.99998478 |
| 0.99085373 |
| 0.99650199 |
| 0.99122857 |
| 0.99998478 |
| 0.99998478 |
| 0.99846792 |
| 0.91325047 |
| 0.99969417 |
| 0.99998478 |
| 0.94653188 |

| |
|---|
| 0.98924373 |
| 0.9467702 |
| 0.98924373 |
| 0.9715425 |
| 0.98924373 |
| 0.96484568 |
| 0.99122857 |
| 0.98924373 |
| 0.99998478 |
| 0.99122857 |
| 0.98924373 |
| 0.98924373 |
| 0.99865984 |
| 0.99122857 |
| 0.94171831 |
| 0.99122857 |
| 0.95585979 |
| 0.99122857 |
| 0.99998478 |
| 0.99998478 |
| 0.97368509 |
| 0.9865573 |
| 0.94002917 |
| 0.9403055 |
| 0.99122857 |
| 0.99998478 |
| 0.99969417 |
| 0.99998478 |
| 0.90297323 |
| 0.99122857 |
| 0.99998478 |
| 0.93752249 |
| 0.95585979 |
| 0.98924373 |
| 0.99998478 |
| 0.99998478 |
| 0.99998478 |
| 0.98924373 |
| 0.99998478 |
| 0.99969417 |
| 0.99998478 |
| 0.94171831 |

| |
|---|
| 0.96484568 |
| 0.98924373 |
| 0.97992318 |
| 0.99122857 |
| 0.99122857 |
| 0.99998478 |
| 0.99998478 |
| 0.99122857 |
| 0.99437562 |
| 0.95859413 |
| 0.99650199 |
| 0.95951105 |
| 0.99969417 |
| 0.99969417 |
| 0.99122857 |
| 0.9403055 |
| 0.91591015 |
| 0.99998478 |
| 0.98924373 |
| 0.93752249 |
| 0.95951105 |
| 0.95951105 |
| 0.99122857 |
| 0.96484568 |
| 0.98924373 |
| 0.99969417 |
| 0.95951105 |
| 0.9403055 |
| 0.99998478 |
| 0.99998478 |
| 0.99998478 |
| 0.99998478 |
| 0.94641693 |
| 0.98924373 |
| 0.98924373 |
| 0.99122857 |
| 0.99969417 |
| 0.98237613 |
| 0.926218 |
| 0.97110689 |
| 0.99617169 |
| 0.99969417 |

| |
|---|
| 0.9403055 |
| 0.98732643 |
| 0.98924373 |
| 0.98259014 |
| 0.99122857 |
| 0.99998478 |
| 0.99122857 |
| 0.99998478 |
| 0.99998478 |
| 0.96181283 |
| 0.99998478 |
| 0.98259014 |
| 0.93594857 |
| 0.99998478 |
| 0.96181283 |
| 0.98924373 |
| 0.91325047 |
| 0.99122857 |
| 0.99969417 |
| 0.99998478 |
| 0.96181283 |
| 0.87052203 |
| 0.98924373 |
| 0.99122857 |
| 0.95240191 |
| 0.99122857 |
| 0.99998478 |
| 0.99998478 |
| 0.99969417 |
| 0.99998478 |
| 0.93054695 |
| 0.94171831 |
| 0.99122857 |
| 0.99122857 |
| 0.99122857 |
| 0.96369417 |
| 0.99998478 |
| 0.98732643 |
| 0.99998478 |
| 0.99998478 |
| 0.99998478 |
| 0.99122857 |

|  |
|---|
| 0.99122857 |
| 0.99998478 |
| 0.95734773 |
| 0.99998478 |
| 0.99122857 |
| 0.99122857 |
| 0.95585979 |
| 0.99183457 |
| 0.99998478 |
| 0.85757422 |
| 0.99998478 |
| 0.79151544 |
| 0.98924373 |
| 0.91325047 |
| 0.98924373 |
| 0.99998478 |
| 0.99332973 |
| 0.91325047 |
| 0.99998478 |
| 0.99998478 |
| 0.92244927 |
| 0.99122857 |
| 0.99969417 |
| 0.99122857 |
| 0.98924373 |
| 0.98924373 |
| 0.99998478 |
| 0.98924373 |
| 0.99122857 |
| 0.99998478 |
| 0.86938366 |
| 0.99998478 |
| 0.99998478 |
| 0.9985128 |
| 0.69921142 |
| 0.86327908 |
| 0.99122857 |
| 0.99969417 |
| 0.99969417 |
| 0.98924373 |
| 0.99252883 |
| 0.99122857 |

0.99969417
0.99998478
0.98924373
0.95988499
0.99969417
0.99998478
0.97815363
0.95034906
0.99122857
0.98924373
0.99998478
0.99830763
0.99252883
0.95988499
0.99122857
0.99998478
0.96484568
0.96484568
0.96484568
0.99122857
0.99969417
0.99122857
0.99998478
0.99998478
0.98924373
0.96167371
0.96484568
0.99830763
0.99969417
0.96181283
0.98924373
0.91325047
0.99998478
0.95859413
0.9647026
0.96484568
0.97368509
0.99309902
0.99998478
0.95951105
0.99998478
0.99998478

| |
|---|
| 0.96484568 |
| 0.99969417 |
| 0.98924373 |
| 0.99969417 |
| 0.97085356 |
| 0.96167371 |
| 0.99998478 |
| 0.99998478 |
| 0.98924373 |
| 0.99998478 |
| 0.94002917 |
| 0.99830763 |
| 0.99650199 |
| 0.90265407 |
| 0.99998478 |
| 0.98924373 |
| 0.9811568 |
| 0.98882544 |
| 0.92861642 |
| 0.92861642 |
| 0.99969417 |
| 0.9467702 |
| 0.91267201 |
| 0.91325047 |
| 0.99998478 |
| 0.99998478 |
| 0.98732643 |
| 0.99998478 |
| 0.99122857 |
| 0.93908842 |
| 0.98368972 |
| 0.8773376 |
| 0.99122857 |
| 0.96181283 |
| 0.99998478 |
| 0.9647026 |
| 0.98924373 |
| 0.98259014 |
| 0.93131953 |
| 0.98732643 |
| 0.99122857 |
| 0.98924373 |

0.95951105
0.99998478
0.99122857
0.98259014
0.98259014
0.99437562
0.99998478
0.98924373
0.99680366
0.99969417
0.98924373
0.96181283
0.99969417
0.9985128
0.9865573
0.91325047
0.99998478
0.99998478
0.99998478
0.93752249
0.99998478
0.96484568
0.99122857
0.99998478
0.99998478
0.99680366
0.98924373
0.98924373
0.9403055
0.99969417
0.99252883
0.94171831
0.99998478
0.98259014
0.99122857
0.99998478
0.99998478
0.99998478
0.99122857
0.99998478
0.90265407
0.98924373

0.99969417
0.99830763
0.69921142
0.99998478
0.99122857
0.99998478
0.99122857
0.99122857
0.98924373
0.96167371
0.99998478
0.98924373
0.97110689
0.99122857
0.99998478
0.98924373
0.95034906
0.99998478
0.98924373
0.99998478
0.91267201
0.95951105
0.98924373
0.99122857
0.99998478
0.99969417
0.99969417
0.96484568
0.95951105
0.99969417
0.99998478
0.93752249
0.94653188
0.99122857
0.99122857
0.95240191
0.98924373
0.99998478
0.99998478
0.99969417
0.95585979
0.91325047

| |
|---|
| 0.99998478 |
| 0.99998478 |
| 0.96484568 |
| 0.99122857 |
| 0.99998478 |
| 0.99969417 |
| 0.99998478 |
| 0.9403055 |
| 0.99969417 |
| 0.99969417 |
| 0.99998478 |
| 0.99998478 |
| 0.99998478 |
| 0.99998478 |
| 0.99998478 |
| 0.98924373 |
| 0.99122857 |
| 0.99998478 |
| 0.99969417 |
| 0.94171831 |
| 0.99122857 |
| 0.99563112 |
| 0.99122857 |
| 0.9403055 |
| 0.99998478 |
| 0.99998478 |
| 0.98924373 |
| 0.90265407 |
| 0.91267201 |
| 0.99969417 |
| 0.93427164 |
| 0.93752249 |
| 0.99604194 |
| 0.98924373 |
| 0.95240191 |
| 0.98924373 |
| 0.94897478 |
| 0.99998478 |
| 0.99998478 |
| 0.99998478 |
| 0.99998478 |
| 0.99998478 |

| |
|---|
| 0.99122857 |
| 0.97704683 |
| 0.95951105 |
| 0.99183457 |
| 0.96484568 |
| 0.99388898 |
| 0.99998478 |
| 0.99252883 |
| 0.99969417 |
| 0.99969417 |
| 0.99969417 |
| 0.99998478 |
| 0.97992318 |
| 0.99998478 |
| 0.99122857 |
| 0.99122857 |
| 0.99998478 |
| 0.98924373 |
| 0.90265407 |
| 0.99969417 |
| 0.98924373 |
| 0.92861642 |
| 0.96484568 |
| 0.99122857 |
| 0.99122857 |
| 0.99969417 |
| 0.9985128 |
| 0.99998478 |
| 0.95240191 |
| 0.99122857 |
| 0.99998478 |
| 0.99998478 |
| 0.99969417 |
| 0.99969417 |
| 0.99122857 |
| 0.99969417 |
| 0.99122857 |
| 0.98924373 |
| 0.99998478 |
| 0.99998478 |
| 0.99998478 |
| 0.98466138 |

| |
|---|
| 0.98924373 |
| 0.91325047 |
| 0.99969417 |
| 0.96709089 |
| 0.95034906 |
| 0.96977284 |
| 0.9829938 |
| 0.99998478 |
| 0.99122857 |
| 0.97993742 |
| 0.99122857 |
| 0.99808206 |
| 0.96181283 |
| 0.99998478 |
| 0.98732643 |
| 0.99998478 |
| 0.98924373 |
| 0.98924373 |
| 0.96484568 |
| 0.99998478 |
| 0.96181283 |
| 0.99122857 |
| 0.97430523 |
| 0.98924373 |
| 0.99948729 |
| 0.80133949 |
| 0.94653188 |
| 0.99969417 |
| 0.93908842 |
| 0.99998478 |
| 0.98924373 |
| 0.99122857 |
| 0.99122857 |
| 0.99969417 |
| 0.98924373 |
| 0.97110689 |
| 0.99998478 |
| 0.98368972 |
| 0.9985128 |
| 0.94747425 |
| 0.94002917 |
| 0.99969417 |

0.98924373
0.99534532
0.99830763
0.99969417
0.99998478
0.99122857
0.99183457
0.99658202
0.99122857
0.99998478
0.90265407
0.95951105
0.99998478
0.99969417
0.99998478
0.99252883
0.98924373
0.99998478
0.98924373
0.92367831
0.99101915
0.93752249
0.99998478
0.98924373
0.99252883
0.99969417
0.99122857
0.99998478
0.99513459
0.90265407
0.89132432
0.99998478
0.98924373
0.90265407
0.93752249
0.98732643
0.98924373
0.96181283
0.99966452
0.99998478
0.99122857
0.98924373

0.98924373
0.96167371
0.99998478
0.90265407
0.99998478
0.99998478
0.95240191
0.99122857
0.99998478
0.99998478
0.98924373
0.97368509
0.99998478
0.98924373
0.99122857
0.99998478
0.98924373
0.98924373
0.98924373
0.99122857
0.99122857
0.98924373
0.99969417
0.99122857
0.97126196
0.98956576
0.99998478
0.95988499
0.99122857
0.99122857
0.99830763
0.98924373
0.99122857
0.99122857
0.99969417
0.99969417
0.99998478
0.98924373
0.99998478
0.91325047
0.99969417
0.99969417

| |
|---|
| 0.99998478 |
| 0.99122857 |
| 0.9865573 |
| 0.98924373 |
| 0.99998478 |
| 0.98337226 |
| 0.99969417 |
| 0.99998478 |
| 0.98924373 |
| 0.98924373 |
| 0.98924373 |
| 0.99969417 |
| 0.67021692 |
| 0.98924373 |
| 0.99122857 |
| 0.98924373 |
| 0.95951105 |
| 0.99085373 |
| 0.94171831 |
| 0.99513459 |
| 0.99998478 |
| 0.93752249 |
| 0.99252883 |
| 0.98924373 |
| 0.99122857 |
| 0.99830763 |
| 0.93752249 |
| 0.99998478 |
| 0.99176357 |
| 0.99628097 |
| 0.90265407 |
| 0.98924373 |
| 0.99967514 |
| 0.99969417 |
| 0.99998478 |
| 0.99998478 |
| 0.94171831 |
| 0.97209533 |
| 0.99998478 |
| 0.94445388 |
| 0.95240191 |
| 0.99252883 |

| |
|---|
| 0.99998478 |
| 0.99252883 |
| 0.99969417 |
| 0.99122857 |
| 0.99122857 |
| 0.99252883 |
| 0.99998478 |
| 0.99998478 |
| 0.96484568 |
| 0.99998478 |
| 0.99122857 |
| 0.93752249 |
| 0.99830763 |
| 0.98924373 |
| 0.96484568 |
| 0.99998478 |
| 0.95465684 |
| 0.98924373 |
| 0.99969417 |
| 0.99969417 |
| 0.99969417 |
| 0.99998478 |
| 0.96484568 |
| 0.98924373 |
| 0.96181283 |
| 0.94171831 |
| 0.98924373 |
| 0.99969417 |
| 0.99969417 |
| 0.93752249 |
| 0.99808206 |
| 0.96484568 |
| 0.99998478 |
| 0.93594857 |
| 0.98924373 |
| 0.95240191 |
| 0.92861642 |
| 0.99969417 |
| 0.99998478 |
| 0.99998478 |
| 0.78307327 |
| 0.9467702 |

| |
|---|
| 0.93752249 |
| 0.99122857 |
| 0.98368972 |
| 0.5776957 |
| 0.98924373 |
| 0.99122857 |
| 0.99201427 |
| 0.99998478 |
| 0.9829938 |
| 0.99998478 |
| 0.92672232 |
| 0.9467702 |
| 0.96167371 |
| 0.99998478 |
| 0.99122857 |
| 0.99122857 |
| 0.96484568 |
| 0.99969417 |
| 0.99122857 |
| 0.98924373 |
| 0.99998478 |
| 0.98924373 |
| 0.99998478 |
| 0.99998478 |
| 0.99998478 |
| 0.96484568 |
| 0.99998478 |
| 0.99846792 |
| 0.99969417 |
| 0.94897478 |
| 0.99998478 |
| 0.9647026 |
| 0.99998478 |
| 0.96484568 |
| 0.99998478 |
| 0.98924373 |
| 0.99122857 |
| 0.99332973 |
| 0.94171831 |
| 0.93752249 |
| 0.99998478 |
| 0.98924373 |

| |
|---|
| 0.99998478 |
| 0.9985128 |
| 0.98924373 |
| 0.99998478 |
| 0.99998478 |
| 0.99998478 |
| 0.99658202 |
| 0.99998478 |
| 0.92548507 |
| 0.99998478 |
| 0.98924373 |
| 0.99122857 |
| 0.9467702 |
| 0.95034906 |
| 0.99969417 |
| 0.97733729 |
| 0.98924373 |
| 0.98924373 |
| 0.99998478 |
| 0.99969417 |
| 0.97368509 |
| 0.99122857 |
| 0.926218 |
| 0.98924373 |
| 0.93594857 |
| 0.99122857 |
| 0.9403055 |
| 0.90297323 |
| 0.98466138 |
| 0.99998478 |
| 0.9865573 |
| 0.99998478 |
| 0.9865573 |
| 0.95240191 |
| 0.96204485 |
| 0.99998478 |
| 0.99998478 |
| 0.99122857 |
| 0.99122857 |
| 0.99998478 |
| 0.98924373 |
| 0.99998478 |

| |
|---|
| 0.99998478 |
| 0.96484568 |
| 0.99998478 |
| 0.97126196 |
| 0.95951105 |
| 0.99998478 |
| 0.99998478 |
| 0.99998478 |
| 0.9865496 |
| 0.99998478 |
| 0.98259014 |
| 0.98924373 |
| 0.99998478 |
| 0.99122857 |
| 0.99122857 |
| 0.98924373 |
| 0.98924373 |
| 0.96484568 |
| 0.97110689 |
| 0.985273 |
| 0.98924373 |
| 0.99998478 |
| 0.99998478 |
| 0.98924373 |
| 0.99969417 |
| 0.97704683 |
| 0.95713239 |
| 0.90297323 |
| 0.99252883 |
| 0.99969417 |
| 0.99998478 |
| 0.99998478 |
| 0.96353697 |
| 0.99998478 |
| 0.99252883 |
| 0.99998478 |
| 0.90265407 |
| 0.99998478 |
| 0.99998478 |
| 0.98924373 |
| 0.99998478 |
| 0.72975604 |

| |
|---|
| 0.99998478 |
| 0.9936423 |
| 0.98924373 |
| 0.9494303 |
| 0.99998478 |
| 0.99998478 |
| 0.93054695 |
| 0.94171831 |
| 0.99998478 |
| 0.74240405 |
| 0.99998478 |
| 0.99969417 |
| 0.99998478 |
| 0.96181283 |
| 0.99969417 |
| 0.94897478 |
| 0.99122857 |
| 0.98924373 |
| 0.98924373 |
| 0.98924373 |
| 0.94171831 |
| 0.99846792 |
| 0.98924373 |
| 0.92861642 |
| 0.99329945 |
| 0.96181283 |
| 0.98924373 |
| 0.92861642 |
| 0.99332973 |
| 0.99998478 |
| 0.99122857 |
| 0.99998478 |
| 0.926218 |
| 0.95034906 |
| 0.98924373 |
| 0.92489453 |
| 0.98924373 |
| 0.99680366 |
| 0.98259014 |
| 0.99998478 |
| 0.98956725 |
| 0.99998478 |

| |
|---|
| 0.99122857 |
| 0.98924373 |
| 0.99998478 |
| 0.98924373 |
| 0.99342943 |
| 0.9467702 |
| 0.99122857 |
| 0.99969417 |
| 0.92861642 |
| 0.97078973 |
| 0.99122857 |
| 0.99998478 |
| 0.9494303 |
| 0.99998478 |
| 0.98686029 |
| 0.99998478 |
| 0.90297323 |
| 0.99842837 |
| 0.95951105 |
| 0.99830763 |
| 0.98924373 |
| 0.99122857 |
| 0.98924373 |
| 0.92861642 |
| 0.98924373 |
| 0.95951105 |
| 0.99969417 |
| 0.98924373 |
| 0.99998478 |
| 0.97110689 |
| 0.99122857 |
| 0.99122857 |
| 0.93752249 |
| 0.99122857 |
| 0.93752249 |
| 0.98924373 |
| 0.96181283 |
| 0.99998478 |
| 0.99998478 |
| 0.98924373 |
| 0.99998478 |
| 0.99969417 |

| |
|---|
| 0.99122857 |
| 0.96484568 |
| 0.94171831 |
| 0.99998478 |
| 0.99122857 |
| 0.94171831 |
| 0.99122857 |
| 0.99969417 |
| 0.72204897 |
| 0.95034906 |
| 0.99122857 |
| 0.91325047 |
| 0.98924373 |
| 0.98924373 |
| 0.9467702 |
| 0.98924373 |
| 0.98924373 |
| 0.99252883 |
| 0.92861642 |
| 0.93772337 |
| 0.99969417 |
| 0.99998478 |
| 0.91325047 |
| 0.98924373 |
| 0.99658202 |
| 0.98924373 |
| 0.98924373 |
| 0.98924373 |
| 0.93752249 |
| 0.99998478 |
| 0.99998478 |
| 0.98924373 |
| 0.8773376 |
| 0.95951105 |
| 0.99252883 |
| 0.99998478 |
| 0.99998478 |
| 0.98732643 |
| 0.99122857 |
| 0.99252883 |
| 0.99969417 |
| 0.99998478 |

| |
|---|
| 0.98924373 |
| 0.99252883 |
| 0.99998478 |
| 0.99122857 |
| 0.98924373 |
| 0.99122857 |
| 0.96181283 |
| 0.96484568 |
| 0.99122857 |
| 0.98203965 |
| 0.99122857 |
| 0.99998478 |
| 0.99998478 |
| 0.97209533 |
| 0.93054695 |
| 0.99998478 |
| 0.98003064 |
| 0.99998478 |
| 0.99122857 |
| 0.99998478 |
| 0.99998478 |
| 0.99998478 |
| 0.99998478 |
| 0.86327908 |
| 0.97329503 |
| 0.92861642 |
| 0.99969417 |
| 0.98924373 |
| 0.94641693 |
| 0.96484568 |
| 0.98924373 |
| 0.93752249 |
| 0.99998478 |
| 0.99998478 |
| 0.94171831 |
| 0.99969417 |
| 0.99998478 |
| 0.98924373 |
| 0.98924373 |
| 0.90265407 |
| 0.99969417 |
| 0.9403055 |

| |
|---|
| 0.97110689 |
| 0.99122857 |
| 0.99252883 |
| 0.94171831 |
| 0.98924373 |
| 0.95240191 |
| 0.96484568 |
| 0.9715425 |
| 0.97024001 |
| 0.99998478 |
| 0.99969417 |
| 0.96181283 |
| 0.915492 |
| 0.99969417 |
| 0.99998478 |
| 0.99122857 |
| 0.99998478 |
| 0.99252883 |
| 0.99969417 |
| 0.99186511 |
| 0.99122857 |
| 0.99122857 |
| 0.98924373 |
| 0.91485898 |
| 0.99252883 |
| 0.98924373 |
| 0.99998478 |
| 0.98924373 |
| 0.99122857 |
| 0.90297323 |
| 0.96587727 |
| 0.99998478 |
| 0.99122857 |
| 0.98924373 |
| 0.95240191 |
| 0.98924373 |
| 0.99969417 |
| 0.99122857 |
| 0.99998478 |
| 0.99998478 |
| 0.96484568 |
| 0.99998478 |

0.95414145
0.99122857
0.99998478
0.99243657
0.99998478
0.98924373
0.98924373
0.99998478
0.99998478
0.98924373
0.98924373
0.97209533
0.99998478
0.99252883
0.94002917
0.99998478
0.99998478
0.99998478
0.99122857
0.98924373
0.98924373
0.98924373
0.99122857
0.96484568
0.99865984
0.99830763
0.99252883
0.99122857
0.95240191
0.99969417
0.915492
0.93151025
0.99998478
0.99122857
0.99998478
0.99969417
0.99969417
0.98924373
0.99998478
0.99998478
0.99252883
0.99969417

| |
|---|
| 0.99101915 |
| 0.9403055 |
| 0.98924373 |
| 0.99846792 |
| 0.99122857 |
| 0.99969417 |
| 0.99998478 |
| 0.99122857 |
| 0.99252883 |
| 0.9829938 |
| 0.99998478 |
| 0.99998478 |
| 0.99998478 |
| 0.99998478 |
| 0.99998478 |
| 0.99122857 |
| 0.95240191 |
| 0.99122857 |
| 0.99969417 |
| 0.99998478 |
| 0.99998478 |
| 0.95951105 |
| 0.99998478 |
| 0.90265407 |
| 0.98924373 |
| 0.99969417 |
| 0.93752249 |
| 0.99969417 |
| 0.98924373 |
| 0.94653188 |
| 0.99122857 |
| 0.98924373 |
| 0.98924373 |
| 0.98924373 |
| 0.99998478 |
| 0.91485898 |
| 0.99969417 |
| 0.96181283 |
| 0.99969417 |
| 0.98924373 |
| 0.99998478 |
| 0.99680366 |

0.99846792
0.98924373
0.99969417
0.98924373
0.96484568
0.99998478
0.99969417
0.92861642
0.99998478
0.99830763
0.99122857
0.87669383
0.99122857
0.99969417
0.99830763
0.91325047
0.98924373
0.98924373
0.99969417
0.96484568
0.99122857
0.99998478
0.86327908
0.92861642
0.99998478
0.79151544
0.95713239
0.96591084
0.99122857
0.99830763
0.99122857
0.98880599
0.99998478
0.99998478
0.99122857
0.99998478
0.96167371
0.99998478
0.99122857
0.99998478
0.99846792
0.99252883

| |
|---|
| 0.99998478 |
| 0.99122857 |
| 0.99998478 |
| 0.99122857 |
| 0.98924373 |
| 0.99122857 |
| 0.97430523 |
| 0.94171831 |
| 0.90297323 |
| 0.98924373 |
| 0.93550274 |
| 0.98924373 |
| 0.99998478 |
| 0.98924373 |
| 0.99998478 |
| 0.99969417 |
| 0.98924373 |
| 0.99122857 |
| 0.99122857 |
| 0.99998478 |
| 0.96484568 |
| 0.99998478 |
| 0.99388898 |
| 0.98732643 |
| 0.85141726 |
| 0.96484568 |
| 0.99969417 |
| 0.99998478 |
| 0.98924373 |
| 0.99969417 |
| 0.99969417 |
| 0.99252883 |
| 0.93752249 |
| 0.99125052 |
| 0.99122857 |
| 0.9467702 |
| 0.99969417 |
| 0.99969417 |
| 0.98924373 |
| 0.98924373 |
| 0.98924373 |
| 0.99998478 |

0.96484568
0.99122857
0.99122857
0.99969417
0.98924373
0.99998478
0.99969417
0.98924373
0.99846792
0.94171831
0.91325047
0.99122857
0.98924373
0.99122857
0.94171831
0.80984148
0.93752249
0.96484568
0.99998478
0.99998478
0.99998478
0.99998478
0.98924373
0.98924373
0.98924373
0.99628097
0.99969417
0.99846792
0.98924373
0.99122857
0.98924373
0.98924373
0.96181283
0.99998478
0.99969417
0.99998478
0.99998478
0.95951105
0.99998478
0.98924373
0.99998478
0.90265407

0.98259014
0.98198365
0.99998478
0.9865573
0.98924373
0.99998478
0.9829938
0.99830763
0.95951105
0.79151544
0.98924373
0.9467702
0.95951105
0.99998478
0.96160364
0.95060469
0.90265407
0.99122857
0.99122857
0.98924373
0.99998478
0.94171831
0.98924373
0.99998478
0.94653188
0.97815363
0.99252883
0.98924373
0.99122857
0.9467702
0.99998478
0.90265407
0.96484568
0.99998478
0.99969417
0.98732643
0.9985128
0.99998478
0.98924373
0.99122857
0.99122857
0.96865889

| |
|---|
| 0.99969417 |
| 0.93427164 |
| 0.99998478 |
| 0.99830763 |
| 0.99865984 |
| 0.99998478 |
| 0.98924373 |
| 0.99998478 |
| 0.98924373 |
| 0.99998478 |
| 0.994855 |
| 0.99948729 |
| 0.94171831 |
| 0.90297323 |
| 0.99122857 |
| 0.93752249 |
| 0.98924373 |
| 0.96484568 |
| 0.99969417 |
| 0.93752249 |
| 0.99122857 |
| 0.91325047 |
| 0.99830763 |
| 0.99122857 |
| 0.99998478 |
| 0.9403055 |
| 0.91485898 |
| 0.98924373 |
| 0.99252883 |
| 0.96181283 |
| 0.94171831 |
| 0.99830763 |
| 0.96181283 |
| 0.99998478 |
| 0.98924373 |
| 0.99998478 |
| 0.99998478 |
| 0.99998478 |
| 0.99998478 |
| 0.98924373 |
| 0.99830763 |
| 0.91325047 |

0.99969417
0.90265407
0.99969417
0.99969417
0.99969417
0.99122857
0.99969417
0.98924373
0.98924373
0.96484568
0.99969417
0.99998478
0.99998478
0.98924373
0.99998478
0.99969417
0.96484568
0.95240191
0.98924373
0.99998478
0.97196599
0.99998478
0.99122857
0.99969417
0.99122857
0.96167371
0.99658202
0.90297323
0.98924373
0.98924373
0.99969417
0.99252883
0.95951105
0.98924373
0.98924373
0.95240191
0.99998478
0.99969417
0.98732643
0.98924373
0.99846792
0.98924373

| |
|---|
| 0.98924373 |
| 0.99998478 |
| 0.99998478 |
| 0.99122857 |
| 0.99969417 |
| 0.99252883 |
| 0.99998478 |
| 0.93752249 |
| 0.9403055 |
| 0.96484568 |
| 0.98924373 |
| 0.99998478 |
| 0.99998478 |
| 0.98924373 |
| 0.98732643 |
| 0.96484568 |
| 0.99998478 |
| 0.99122857 |
| 0.99998478 |
| 0.98732643 |
| 0.90265407 |
| 0.99998478 |
| 0.96181283 |
| 0.99252883 |
| 0.91325047 |
| 0.96484568 |
| 0.99969417 |
| 0.99842837 |
| 0.99122857 |
| 0.97430523 |
| 0.99998478 |
| 0.99998478 |
| 0.99122857 |
| 0.99998478 |
| 0.98924373 |
| 0.99122857 |
| 0.99122857 |
| 0.98924373 |
| 0.99998478 |
| 0.99969417 |
| 0.98924373 |
| 0.99969417 |

| |
|---|
| 0.90529035 |
| 0.98732643 |
| 0.9715425 |
| 0.97733729 |
| 0.98924373 |
| 0.99998478 |
| 0.95951105 |
| 0.99998478 |
| 0.98924373 |
| 0.94171831 |
| 0.99998478 |
| 0.99969417 |
| 0.99122857 |
| 0.99969417 |
| 0.99122857 |
| 0.98924373 |
| 0.99998478 |
| 0.99183457 |
| 0.96484568 |
| 0.9403055 |
| 0.98924373 |
| 0.99628097 |
| 0.99998478 |
| 0.99998478 |
| 0.96709089 |
| 0.99122857 |
| 0.99998478 |
| 0.97110689 |
| 0.99998478 |
| 0.99969417 |
| 0.97110689 |
| 0.99998478 |
| 0.91325047 |
| 0.99122857 |
| 0.98565258 |
| 0.99998478 |
| 0.97430523 |
| 0.99122857 |
| 0.99122857 |
| 0.96816591 |
| 0.99998478 |
| 0.99969417 |

0.98077375
0.98732643
0.99122857
0.99969417
0.96709089
0.98924373
0.90265407
0.99998478
0.99969417
0.9985128
0.99969417
0.98924373
0.98924373
0.99122857
0.99122857
0.96484568
0.96484568
0.90297323
0.98924373
0.85669195
0.99969417
0.98924373
0.9647026
0.98924373
0.99329945
0.99122857
0.98924373
0.99122857
0.9650009
0.99122857
0.99998478
0.99969417
0.99969417
0.96484568
0.99998478
0.95034906
0.99998478
0.99969417
0.95988499
0.98924373
0.98924373
0.97993742

| |
|---|
| 0.98924373 |
| 0.99122857 |
| 0.99998478 |
| 0.98924373 |
| 0.99969417 |
| 0.99998478 |
| 0.994855 |
| 0.90265407 |
| 0.94171831 |
| 0.99998478 |
| 0.99846792 |
| 0.98732643 |
| 0.98924373 |
| 0.98924373 |
| 0.98924373 |
| 0.99998478 |
| 0.91325047 |
| 0.96484568 |
| 0.99969417 |
| 0.98924373 |
| 0.99998478 |
| 0.99998478 |
| 0.93752249 |
| 0.99998478 |
| 0.99122857 |
| 0.96484568 |
| 0.99650199 |
| 0.9650009 |
| 0.99969417 |
| 0.93427164 |
| 0.98924373 |
| 0.99122857 |
| 0.99998478 |
| 0.99122857 |
| 0.99998478 |
| 0.99998478 |
| 0.99969417 |
| 0.99998478 |
| 0.98924373 |
| 0.99122857 |
| 0.99998478 |
| 0.98924373 |

| |
|---|
| 0.97327383 |
| 0.92861642 |
| 0.98924373 |
| 0.95414145 |
| 0.99998478 |
| 0.90297323 |
| 0.96181283 |
| 0.99830763 |
| 0.96484568 |
| 0.98237613 |
| 0.98924373 |
| 0.99122857 |
| 0.99969417 |
| 0.99969417 |
| 0.98924373 |
| 0.99998478 |
| 0.99122857 |
| 0.94412092 |
| 0.99998478 |
| 0.99122857 |
| 0.99122857 |
| 0.99998478 |
| 0.99969417 |
| 0.99998478 |
| 0.99122857 |
| 0.99998478 |
| 0.98924373 |
| 0.99998478 |
| 0.96484568 |
| 0.99969417 |
| 0.91267201 |
| 0.90297323 |
| 0.99122857 |
| 0.99122857 |
| 0.85141726 |
| 0.926218 |
| 0.99122857 |
| 0.95859413 |
| 0.99122857 |
| 0.99122857 |
| 0.99998478 |
| 0.94897478 |

| |
|---|
| 0.98924373 |
| 0.99998478 |
| 0.98924373 |
| 0.99830763 |
| 0.98924373 |
| 0.99998478 |
| 0.98924373 |
| 0.90297323 |
| 0.99122857 |
| 0.94596453 |
| 0.99122857 |
| 0.91485898 |
| 0.99122857 |
| 0.98924373 |
| 0.95240191 |
| 0.99998478 |
| 0.99998478 |
| 0.98924373 |
| 0.94171831 |
| 0.99969417 |
| 0.99998478 |
| 0.99122857 |
| 0.95034906 |
| 0.99998478 |
| 0.99969417 |
| 0.93752249 |
| 0.99969417 |
| 0.99122857 |
| 0.98924373 |
| 0.99998478 |
| 0.99122857 |
| 0.9985128 |
| 0.99998478 |
| 0.98924373 |
| 0.9834625 |
| 0.99998478 |
| 0.99122857 |
| 0.98924373 |
| 0.98924373 |
| 0.99998478 |
| 0.99122857 |
| 0.99830763 |

0.99252883
0.99998478
0.98924373
0.99998478
0.99998478
0.94171831
0.98924373
0.98924373
0.96484568
0.99998478
0.96816591
0.97085356
0.99998478
0.99122857
0.99969417
0.97329503
0.93752249
0.99998478
0.99122857
0.99969417
0.99122857
0.98924373
0.98924373
0.99969417
0.99122857
0.97954048
0.99998478
0.99998478
0.96484568
0.99998478
0.99998478
0.99252883
0.98924373
0.99998478
0.98924373
0.99998478
0.98924373
0.98924373
0.95951105
0.98732643
0.99122857
0.94653188

| |
|---|
| 0.96484568 |
| 0.99998478 |
| 0.95240191 |
| 0.98924373 |
| 0.99122857 |
| 0.98924373 |
| 0.9985128 |
| 0.99998478 |
| 0.96484568 |
| 0.99969417 |
| 0.94653188 |
| 0.99998478 |
| 0.99969417 |
| 0.99998478 |
| 0.99998478 |
| 0.92931319 |
| 0.99969417 |
| 0.94653188 |
| 0.96484568 |
| 0.96181283 |
| 0.98924373 |
| 0.99969417 |
| 0.99998478 |
| 0.99969417 |
| 0.8773376 |
| 0.99122857 |
| 0.99122857 |
| 0.98924373 |
| 0.99998478 |
| 0.98924373 |
| 0.99998478 |
| 0.97815363 |
| 0.99969417 |
| 0.93752249 |
| 0.98732643 |
| 0.98924373 |
| 0.92861642 |
| 0.98924373 |
| 0.99122857 |
| 0.9467702 |
| 0.99122857 |
| 0.98924373 |

0.98924373
0.95951105
0.98924373
0.94002917
0.98924373
0.98924373
0.99998478
0.99998478
0.92650934
0.95951105
0.99969417
0.9467702
0.99846792
0.99122857
0.99122857
0.99998478
0.99252883
0.99122857
0.96181283
0.99998478
0.99969417
0.99998478
0.98924373
0.99998478
0.98924373
0.98924373
0.90297323
0.90265407
0.93752249
0.99252883
0.9467702
0.93594857
0.94171831
0.99830763
0.98732643
0.98924373
0.99998478
0.99969417
0.96181283
0.99846792
0.99998478
0.99122857

| |
|---|
| 0.97327383 |
| 0.99998478 |
| 0.99998478 |
| 0.99122857 |
| 0.93752249 |
| 0.99998478 |
| 0.99998478 |
| 0.9403055 |
| 0.99998478 |
| 0.99998478 |
| 0.97815363 |
| 0.93365848 |
| 0.99998478 |
| 0.99969417 |
| 0.99998478 |
| 0.99969417 |
| 0.99122857 |
| 0.98924373 |
| 0.95951105 |
| 0.9403055 |
| 0.99998478 |
| 0.92861642 |
| 0.92672232 |
| 0.99998478 |
| 0.99998478 |
| 0.99998478 |
| 0.99998478 |
| 0.99122857 |
| 0.79151544 |
| 0.99998478 |
| 0.99998478 |
| 0.98924373 |
| 0.99998478 |
| 0.9467702 |
| 0.926218 |
| 0.98259014 |
| 0.96181283 |
| 0.98924373 |
| 0.98732643 |
| 0.95951105 |
| 0.95951105 |
| 0.99830763 |

0.94402118
0.99969417
0.99122857
0.94171831
0.99122857
0.96484568
0.95951105
0.99122857
0.98924373
0.99658202
0.99998478
0.97993742
0.99101915
0.97327383
0.94653188
0.99998478
0.99998478
0.99122857
0.98924373
0.99969417
0.99881099
0.99122857
0.99122857
0.99122857
0.90297323
0.99998478
0.97209533
0.99998478
0.99969417
0.99969417
0.99252883
0.92672232
0.99969417
0.92861642
0.90265407
0.98924373
0.99998478
0.94747425
0.99969417
0.96181283
0.99998478
0.99969417

0.99998478
0.9403055
0.99969417
0.99998478
0.98259014
0.99998478
0.99969417
0.99122857
0.99998478
0.99969417
0.99998478
0.98732643
0.99122857
0.99998478
0.98924373
0.98924373
0.99122857
0.99998478
0.99998478
0.99969417
0.80984148
0.96181283
0.99122857
0.93752249
0.99846792
0.90297323
0.95988499
0.92861642
0.99998478
0.99969417
0.99969417
0.9467702
0.99969417
0.99969417
0.99969417
0.98924373
0.99830763
0.96484568
0.99122857
0.98924373
0.98924373
0.98924373

| |
|---|
| 0.99969417 |
| 0.99998478 |
| 0.92548507 |
| 0.94171831 |
| 0.98924373 |
| 0.98924373 |
| 0.926218 |
| 0.99998478 |
| 0.99122857 |
| 0.99122857 |
| 0.99969417 |
| 0.99122857 |
| 0.99122857 |
| 0.98924373 |
| 0.94171831 |
| 0.99998478 |
| 0.95951105 |
| 0.99969417 |
| 0.99998478 |
| 0.99948729 |
| 0.99998478 |
| 0.98924373 |
| 0.9403055 |
| 0.99650199 |
| 0.95859413 |
| 0.99969417 |
| 0.94897478 |
| 0.99122857 |
| 0.95951105 |
| 0.99998478 |
| 0.98924373 |
| 0.99998478 |
| 0.99998478 |
| 0.90446904 |
| 0.99998478 |
| 0.99650199 |
| 0.99969417 |
| 0.99998478 |
| 0.98924373 |
| 0.94653188 |
| 0.99830763 |
| 0.99998478 |

| |
|---|
| 0.99969417 |
| 0.98924373 |
| 0.98924373 |
| 0.99998478 |
| 0.99122857 |
| 0.99122857 |
| 0.98924373 |
| 0.94653188 |
| 0.90265407 |
| 0.99994907 |
| 0.99998478 |
| 0.93752249 |
| 0.98924373 |
| 0.99969417 |
| 0.96181283 |
| 0.98924373 |
| 0.98924373 |
| 0.94171831 |
| 0.98924373 |
| 0.98924373 |
| 0.99122857 |
| 0.99998478 |
| 0.99998478 |
| 0.93752249 |
| 0.99122857 |
| 0.98924373 |
| 0.96484568 |
| 0.99998478 |
| 0.95034906 |
| 0.95240191 |
| 0.99122857 |
| 0.99122857 |
| 0.90297323 |
| 0.96484568 |
| 0.99969417 |
| 0.94171831 |
| 0.98924373 |
| 0.98924373 |
| 0.74240405 |
| 0.98924373 |
| 0.96181283 |
| 0.9936423 |

| |
|---|
| 0.98259014 |
| 0.99122857 |
| 0.99122857 |
| 0.9829938 |
| 0.99969417 |
| 0.99969417 |
| 0.99122857 |
| 0.99969417 |
| 0.99998478 |
| 0.98924373 |
| 0.99998478 |
| 0.99781055 |
| 0.99122857 |
| 0.99969417 |
| 0.99998478 |
| 0.99122857 |
| 0.99969417 |
| 0.98924373 |
| 0.98924373 |
| 0.96484568 |
| 0.99252883 |
| 0.95951105 |
| 0.98732643 |
| 0.99969417 |
| 0.9971726 |
| 0.99122857 |
| 0.99969417 |
| 0.99122857 |
| 0.98924373 |
| 0.98924373 |
| 0.99122857 |
| 0.98924373 |
| 0.99998478 |
| 0.98924373 |
| 0.96484568 |
| 0.94171831 |
| 0.99513459 |
| 0.98924373 |
| 0.99969417 |
| 0.99998478 |
| 0.93752249 |
| 0.85141726 |

| |
|---|
| 0.99998478 |
| 0.99998478 |
| 0.98924373 |
| 0.96484568 |
| 0.99998478 |
| 0.98924373 |
| 0.98924373 |
| 0.99948729 |
| 0.99969417 |
| 0.98924373 |
| 0.99998478 |
| 0.95240191 |
| 0.97085356 |
| 0.98924373 |
| 0.98924373 |
| 0.98924373 |
| 0.90297323 |
| 0.99998478 |
| 0.99969417 |
| 0.98924373 |
| 0.99122857 |
| 0.99998478 |
| 0.98259014 |
| 0.99830763 |
| 0.99122857 |
| 0.99969417 |
| 0.99998478 |
| 0.91325047 |
| 0.99998478 |
| 0.99998478 |
| 0.99998478 |
| 0.99969417 |
| 0.99998478 |
| 0.61977276 |
| 0.9403055 |
| 0.99998478 |
| 0.99122857 |
| 0.9985128 |
| 0.96353697 |
| 0.98924373 |
| 0.99998478 |
| 0.96484568 |

0.99998478
0.96181283
0.93752249
0.99830763
0.93752249
0.99998478
0.81286841
0.99846792
0.99122857
0.98924373
0.95951105
0.99998478
0.96484568
0.98924373
0.98732643
0.99122857
0.97992318
0.99122857
0.98924373
0.99122857
0.99998478
0.99998478
0.99122857
0.91267201
0.99122857
0.99998478
0.99122857
0.90265407
0.99122857
0.96484568
0.99998478
0.93594857
0.99998478
0.90265407
0.99969417
0.99122857
0.99998478
0.98924373
0.99969417
0.99969417
0.98924373
0.98924373

| |
|---|
| 0.94653188 |
| 0.99998478 |
| 0.98924373 |
| 0.99252883 |
| 0.99122857 |
| 0.99998478 |
| 0.99252883 |
| 0.99998478 |
| 0.98924373 |
| 0.9467702 |
| 0.94653188 |
| 0.99998478 |
| 0.99998478 |
| 0.99998478 |
| 0.99969417 |
| 0.94171831 |
| 0.99998478 |
| 0.99969417 |
| 0.99998478 |
| 0.98924373 |
| 0.98924373 |
| 0.99969417 |
| 0.98924373 |
| 0.81286841 |
| 0.99998478 |
| 0.99998478 |
| 0.99122857 |
| 0.99969417 |
| 0.94653188 |
| 0.99998478 |
| 0.96181283 |
| 0.99969417 |
| 0.99998478 |
| 0.96181283 |
| 0.98732643 |
| 0.99998478 |
| 0.99969417 |
| 0.99122857 |
| 0.99122857 |
| 0.79151544 |
| 0.99998478 |
| 0.96167371 |

| |
|---|
| 0.98924373 |
| 0.99969417 |
| 0.93752249 |
| 0.98924373 |
| 0.95951105 |
| 0.97329503 |
| 0.94171831 |
| 0.99998478 |
| 0.99122857 |
| 0.99122857 |
| 0.95240191 |
| 0.9403055 |
| 0.99998478 |
| 0.96484568 |
| 0.99998478 |
| 0.99998478 |
| 0.99122857 |
| 0.99969417 |
| 0.99969417 |
| 0.98924373 |
| 0.99998478 |
| 0.99998478 |
| 0.99122857 |
| 0.99332973 |
| 0.99969417 |
| 0.98924373 |
| 0.95646366 |
| 0.90265407 |
| 0.99998478 |
| 0.98924373 |
| 0.55521624 |
| 0.98924373 |
| 0.98924373 |
| 0.95240191 |
| 0.93594857 |
| 0.90297323 |
| 0.99998478 |
| 0.99969417 |
| 0.93752249 |
| 0.99998478 |
| 0.91267201 |
| 0.98924373 |

| |
|---|
| 0.99122857 |
| 0.90265407 |
| 0.99969417 |
| 0.9467702 |
| 0.98924373 |
| 0.99998478 |
| 0.99252883 |
| 0.98924373 |
| 0.98924373 |
| 0.96181283 |
| 0.98924373 |
| 0.99998478 |
| 0.96484568 |
| 0.90297323 |
| 0.97479234 |
| 0.99998478 |
| 0.89618249 |
| 0.99122857 |
| 0.99998478 |
| 0.99998478 |
| 0.98924373 |
| 0.98924373 |
| 0.92861642 |
| 0.99998478 |
| 0.99998478 |
| 0.96484568 |
| 0.97815363 |
| 0.99628097 |
| 0.90265407 |
| 0.97085356 |
| 0.99830763 |
| 0.99122857 |
| 0.97711186 |
| 0.95240191 |
| 0.99122857 |
| 0.99998478 |
| 0.98924373 |
| 0.99969417 |
| 0.93752249 |
| 0.92861642 |
| 0.99998478 |
| 0.96484568 |

| |
|---|
| 0.99969417 |
| 0.99650199 |
| 0.98924373 |
| 0.99969417 |
| 0.94171831 |
| 0.98924373 |
| 0.96484568 |
| 0.99998478 |
| 0.99252883 |
| 0.94171831 |
| 0.99998478 |
| 0.96484568 |
| 0.99969417 |
| 0.99830763 |
| 0.9467702 |
| 0.99122857 |
| 0.99969417 |
| 0.96181283 |
| 0.99122857 |
| 0.99122857 |
| 0.99122857 |
| 0.99998478 |
| 0.99998478 |
| 0.90265407 |
| 0.9403055 |
| 0.99998478 |
| 0.98924373 |
| 0.99881183 |
| 0.99998478 |
| 0.99998478 |
| 0.9998342 |
| 0.96484568 |
| 0.99122857 |
| 0.99122857 |
| 0.94171831 |
| 0.96181283 |
| 0.9403055 |
| 0.98924373 |
| 0.95240191 |
| 0.99252883 |
| 0.99122857 |
| 0.98924373 |

| |
|---|
| 0.99969417 |
| 0.91325047 |
| 0.98924373 |
| 0.98732643 |
| 0.93427164 |
| 0.99998478 |
| 0.99948729 |
| 0.98924373 |
| 0.99830763 |
| 0.99998478 |
| 0.99122857 |
| 0.99998478 |
| 0.98924373 |
| 0.99998478 |
| 0.99969417 |
| 0.99122857 |
| 0.99998478 |
| 0.98924373 |
| 0.99122857 |
| 0.9647026 |
| 0.97110689 |
| 0.98924373 |
| 0.98924373 |
| 0.9467702 |
| 0.99781055 |
| 0.99122857 |
| 0.99998478 |
| 0.99122857 |
| 0.96181283 |
| 0.99122857 |
| 0.99998478 |
| 0.99998478 |
| 0.99969417 |
| 0.99998478 |
| 0.99122857 |
| 0.96484568 |
| 0.98924373 |
| 0.94171831 |
| 0.91267201 |
| 0.98924373 |
| 0.98924373 |
| 0.99998478 |

| |
|---|
| 0.95951105 |
| 0.99122857 |
| 0.98924373 |
| 0.98924373 |
| 0.99998478 |
| 0.98924373 |
| 0.95240191 |
| 0.99122857 |
| 0.9865573 |
| 0.94641693 |
| 0.98924373 |
| 0.99998478 |
| 0.98924373 |
| 0.99969417 |
| 0.99998478 |
| 0.99122857 |
| 0.99998478 |
| 0.96484568 |
| 0.99998478 |
| 0.99969417 |
| 0.8773376 |
| 0.99122857 |
| 0.98924373 |
| 0.99998478 |
| 0.99998478 |
| 0.98924373 |
| 0.98956725 |
| 0.96484568 |
| 0.95240191 |
| 0.99650199 |
| 0.98259014 |
| 0.98924373 |
| 0.9403055 |
| 0.99252883 |
| 0.99122857 |
| 0.98924373 |
| 0.96484568 |
| 0.99969417 |
| 0.9865573 |
| 0.98924373 |
| 0.99998478 |
| 0.96484568 |

| |
|---|
| 0.96484568 |
| 0.99122857 |
| 0.99122857 |
| 0.99998478 |
| 0.99969417 |
| 0.90473298 |
| 0.98924373 |
| 0.96484568 |
| 0.98924373 |
| 0.98924373 |
| 0.99122857 |
| 0.99969417 |
| 0.99998478 |
| 0.99969417 |
| 0.98924373 |
| 0.99998478 |
| 0.93752249 |
| 0.99998478 |
| 0.99969417 |
| 0.99969417 |
| 0.95951105 |
| 0.99998478 |
| 0.99122857 |
| 0.98924373 |
| 0.99122857 |
| 0.99122857 |
| 0.99998478 |
| 0.98924373 |
| 0.97368509 |
| 0.98924373 |
| 0.9403055 |
| 0.98924373 |
| 0.98924373 |
| 0.99122857 |
| 0.99998478 |
| 0.91325047 |
| 0.99998478 |
| 0.98924373 |
| 0.98924373 |
| 0.98924373 |
| 0.99808206 |
| 0.98924373 |

0.98924373
0.99122857
0.98924373
0.99122857
0.98259014
0.99998478
0.98924373
0.98924373
0.99998478
0.99998478
0.99122857
0.99998478
0.99122857
0.98924373
0.99998478
0.98924373
0.95951105
0.98732643
0.90265407
0.99998478
0.99998478
0.99998478
0.96484568
0.99998478
0.99969417
0.98924373
0.96484568
0.99998478
0.99122857
0.92861642
0.99998478
0.98924373
0.99998478
0.99846792
0.99998478
0.99998478
0.99122857
0.99998478
0.96167371
0.99122857
0.99969417
0.99252883

0.95240191
0.99122857
0.98924373
0.99998478
0.99998478
0.99830763
0.99122857
0.99998478
0.99998478
0.98924373
0.99122857
0.99998478
0.96484568
0.90265407
0.99998478
0.96181283
0.93594857
0.96484568
0.96167371
0.94171831
0.99122857
0.98924373
0.99998478
0.99998478
0.99830763
0.98924373
0.98924373
0.99998478
0.91325047
0.99998478
0.99969417
0.99998478
0.90265407
0.99969417
0.98924373
0.99998478
0.99122857
0.99122857
0.90297323
0.99122857
0.99252883
0.95951105

| |
|---|
| 0.99548569 |
| 0.99998478 |
| 0.99998478 |
| 0.99122857 |
| 0.97992318 |
| 0.98924373 |
| 0.80984148 |
| 0.99998478 |
| 0.94402118 |
| 0.85669195 |
| 0.96181283 |
| 0.99122857 |
| 0.99998478 |
| 0.99998478 |
| 0.99998478 |
| 0.98924373 |
| 0.99830763 |
| 0.985273 |
| 0.98924373 |
| 0.99969417 |
| 0.99998478 |
| 0.98924373 |
| 0.96484568 |
| 0.99998478 |
| 0.99998478 |
| 0.99998478 |
| 0.98259014 |
| 0.99332973 |
| 0.91252886 |
| 0.99998478 |
| 0.94596453 |
| 0.96181283 |
| 0.9467702 |
| 0.99969417 |
| 0.99969417 |
| 0.99122857 |
| 0.96484568 |
| 0.99122857 |
| 0.99252883 |
| 0.99998478 |
| 0.74240405 |
| 0.96181283 |

| |
|---|
| 0.98924373 |
| 0.99998478 |
| 0.96181283 |
| 0.99998478 |
| 0.99998478 |
| 0.99122857 |
| 0.99998478 |
| 0.99998478 |
| 0.99969417 |
| 0.99252883 |
| 0.99998478 |
| 0.99122857 |
| 0.99998478 |
| 0.99998478 |
| 0.93752249 |
| 0.99122857 |
| 0.98924373 |
| 0.98924373 |
| 0.99998478 |
| 0.99969417 |
| 0.96484568 |
| 0.94653188 |
| 0.99969417 |
| 0.98732643 |
| 0.99998478 |
| 0.94171831 |
| 0.99388898 |
| 0.98924373 |
| 0.98924373 |
| 0.9985128 |
| 0.95034906 |
| 0.98879245 |
| 0.97815363 |
| 0.92548507 |
| 0.99969417 |
| 0.93752249 |
| 0.92861642 |
| 0.99969417 |
| 0.98109548 |
| 0.94641693 |
| 0.99969417 |
| 0.99122857 |

| |
|---|
| 0.94171831 |
| 0.95240191 |
| 0.99122857 |
| 0.90297323 |
| 0.98924373 |
| 0.98924373 |
| 0.99122857 |
| 0.95951105 |
| 0.99830763 |
| 0.7628002 |
| 0.99969417 |
| 0.98924373 |
| 0.98924373 |
| 0.99572919 |
| 0.99969417 |
| 0.99998478 |
| 0.98924373 |
| 0.99998478 |
| 0.98924373 |
| 0.99998478 |
| 0.98924373 |
| 0.99998478 |
| 0.99252883 |
| 0.74240405 |
| 0.93908842 |
| 0.98924373 |
| 0.99998478 |
| 0.98924373 |
| 0.99969417 |
| 0.98924373 |
| 0.99122857 |
| 0.99513459 |
| 0.99969417 |
| 0.99650199 |
| 0.98924373 |
| 0.99252883 |
| 0.98732643 |
| 0.99122857 |
