## Supplementary table 3 for "Selective degradation of BRD9 by a DCAF16-recruiting targeted glue: mode of action elucidation and in vivo proof of concept"

| Uniprot Accession | Protein name | DMSO_1_abundance_norm | DMSO_2_abundance_norm | DMSO_3_abundance_norm | AMPH-04647.10nM_1_abundance_norm | AMPH-04647.10nM_2_abundance_norm |
|---|---|---|---|---|---|---|
| Q9NXF7 | DCAF16 | 13.3022116 | 13.6229137 | 13.8432622 | 13.7719713 | 13.3621811 |
| Q16531 | DDB1 | 15.4747528 | 15.1329781 | 15.1419845 | 15.2305781 | 15.5732272 |
| Q96QD9 | FYTTD1 | 11.5652422 | 12.157579 | 12.88028 | 11.62933 | NA |
| P06454 | PTMA | 18.2969316 | 17.4315613 | 16.5026251 | 16.412588 | 18.2243155 |
| P06733 | ENO1 | 19.060339 | 18.2187 | 17.6754283 | 17.41902 | 19.0493183 |
| Q5T750 | KPLCE | 15.429414 | 16.3009957 | 15.0258834 | 14.026012 | 14.568147 |
| P62937 | PPIA | 20.2616775 | 19.4880677 | 18.8733667 | 19.0680398 | 20.4222128 |
| P04406 | GAPDH | 21.1488461 | 20.6750639 | 20.3708597 | 20.1599484 | 21.0329301 |
| Q96HC4 | PDLIM5 | 14.82412 | 15.1469653 | 15.2364324 | 15.4187079 | 14.3119319 |
| P05204 | HMGN2 | 18.5721788 | 16.2239234 | 15.7944126 | 15.4836522 | 16.9924612 |
| P61981 | YWHAG | 11.8103148 | 11.5818961 | 11.3123601 | 11.2760238 | 11.7313778 |
| P15259 | PGAM2 | 14.8990895 | 14.0915522 | 12.9516803 | 13.130221 | 14.4763927 |
| P60174 | TPI1 | 18.1211516 | 17.3895998 | 17.0284662 | 16.7299666 | 17.9644668 |
| Q8TA86 | RP9 | 14.2532777 | 14.3061473 | 14.6326548 | 14.5377218 | 14.3470311 |
| P50213 | IDH3A | 12.5014315 | 11.9500331 | 11.8825683 | 12.4272599 | 12.9556836 |
| P63241 | EIF5A | 12.6763179 | 11.8409189 | 11.2404398 | 11.1372025 | 12.4670309 |
| Q14697 | GANAB | 16.6165343 | 16.1455858 | 16.0504785 | 16.1166015 | 16.6074138 |
| P09429 | HMGB1 | 17.2495511 | 15.5554806 | 15.161357 | 15.1804158 | 16.3003055 |
| P31942 | HNRNPH3 | 14.3999687 | 14.6460296 | 14.1049744 | 14.0309788 | 14.5086779 |
| Q9BQ61 | TRIR | 13.3433935 | 13.6535297 | 12.9316217 | 12.3871095 | 13.2259121 |
| Q13427 | PPIG | 14.2445474 | 14.2972285 | 14.2413372 | 14.0987194 | 14.4385742 |
| Q9UJV9 | DDX41 | 13.8743371 | 13.5881594 | 14.5066795 | 14.6277402 | 14.3529904 |
| P61604 | HSPE1 | 15.9816493 | 15.0579278 | NA | NA | 15.1244288 |
| Q8WXF1 | PSPC1 | 16.1938257 | 16.2945069 | 15.7747694 | 15.9004428 | 16.1425577 |
| Q9UHB9 | SRP68 | 13.2939431 | 13.9842705 | 12.3925856 | 12.4509235 | 13.4752566 |
| Q58FF6 | HSP90AB4P | 14.9002089 | 14.5691927 | 14.1784484 | 13.8960696 | 14.4982763 |
| P23490 | LORICRIN | 16.6815096 | 17.5163123 | 16.3509128 | 14.4272814 | 15.5121538 |
| P27797 | CALR | 17.7572868 | 17.09923 | 16.5443015 | 16.8479677 | 17.6822935 |
| Q9UNZ5 | C19orf53 | 19.3785676 | 18.4827715 | 18.3382982 | 18.3030554 | 18.3533793 |
| O43809 | NUDT21 | 17.9074135 | 17.4435085 | 17.7477787 | 18.0035171 | 17.6371636 |
| P23284 | PPIB | 16.5939085 | 15.7188432 | 15.3007432 | 15.2400988 | 16.4365066 |
| P09234 | SNRPC | 16.0685663 | 14.9771444 | 14.919358 | 15.3787039 | 14.7583538 |
| P26368 | U2AF2 | 19.92232 | 19.5507449 | 19.8905432 | 20.0678045 | 19.8013 |
| Q9Y383 | LUC7L2 | 20.1160715 | 19.7297115 | 20.2203832 | 20.449992 | 20.2698468 |
| Q9BU76 | C1orf35 | 16.8943288 | 16.6509559 | 16.6241731 | 16.7379862 | 16.9511096 |
| Q9ULI4 | KIF26A | 17.2640712 | 17.3050778 | 17.8533077 | 17.9013705 | 17.6027765 |

| | | | | | | |
|---|---|---|---|---|---|---|
| O75400 | PRPF40A | 16.7318564 | 16.8704496 | 17.0790026 | 17.2953661 | 16.9049936 |
| Q12834 | CDC20 | 13.9871676 | 14.1743848 | 14.4323336 | 14.1662251 | 14.2981971 |
| Q8WXA9 | SREK1 | 14.1514156 | 14.4715764 | 15.284663 | 15.1873422 | 15.0553105 |
| Q9Y6A4 | CFAP20 | 15.5142325 | 15.7327675 | 14.7380307 | 15.4451337 | 15.0753477 |
| Q05519 | SRSF11 | 17.5851536 | 17.2854586 | 17.8644786 | 18.0378833 | 17.8209643 |
| P01860 | IGHG3 | 14.8899869 | 14.7770797 | 15.217291 | 14.3961299 | 15.0779687 |
| P00558 | PGK1 | 17.0773459 | 16.3252386 | 16.0707605 | 15.7401078 | 16.7991871 |
| Q96QA5 | GSDMA | 11.9861684 | 12.4637217 | 11.9058562 | 10.7159359 | 11.44265 |
| Q9NQ29 | LUC7L | 15.1226033 | 14.9633592 | 14.9060197 | 14.7147699 | 14.9959239 |
| Q01081 | U2AF1 | 18.9003343 | 18.4868212 | 18.9380547 | 19.0970196 | 19.071795 |
| O75152 | ZC3H11A | 13.0057701 | 12.7543142 | NA | 12.5150218 | 13.061808 |
| Q5XKP0 | MICOS13 | 15.4814634 | 15.9573331 | 16.2024771 | 15.796397 | 16.1056725 |
| P14618 | PKM | 18.9662572 | 18.2284596 | 17.9806749 | 18.0569397 | 18.6997028 |
| P0DI83 | RAB34 | 19.2628111 | 19.1235947 | 19.2365536 | 19.2444193 | 19.5122045 |
| P15954 | COX7C | 17.717122 | 17.3427505 | 17.0130507 | 17.2561511 | 17.7060119 |
| Q16629 | SRSF7 | 16.5288631 | 16.7638711 | 16.6688636 | 16.8301717 | 16.6933595 |
| O00425 | IGF2BP3 | 14.5723923 | 14.6218134 | 14.0708459 | 14.188603 | 14.7354873 |
| P12277 | CKB | 17.9377206 | 17.6089649 | 17.3235926 | 17.3624747 | 18.3502677 |
| Q53GQ0 | HSD17B12 | 12.197068 | 12.3355194 | 12.4690845 | 12.0849738 | 11.5599945 |
| Q9NYF8 | BCLAF1 | 15.0872741 | 15.6464747 | 15.3235719 | 15.5006946 | 15.6540784 |
| Q14739 | LBR | 16.5859168 | 16.7402219 | 16.5266431 | 16.7745257 | 16.6133911 |
| Q01518 | CAP1 | 15.8648889 | 15.1897384 | 14.5037425 | 14.718518 | 15.5577819 |
| P11387 | TOP1 | 17.7178136 | 17.5860627 | 17.9431628 | 17.831136 | 17.7296622 |
| Q16630 | CPSF6 | 17.8936522 | 17.4234033 | 17.8164942 | 17.9152867 | 17.8809574 |
| Q06830 | PRDX1 | 18.8337232 | 18.8150019 | 18.0754867 | 18.2673758 | 18.5878207 |
| Q7L014 | DDX46 | 19.9157811 | 19.4532998 | 20.0155562 | 20.0652638 | 19.9866029 |
| Q7Z589 | EMSY | 15.0503758 | 15.16326 | 15.4940105 | 15.4687137 | 15.1673847 |
| Q8IWX8 | CHERP | 17.4839309 | 17.4370452 | 17.4488791 | 17.6617731 | 17.5099503 |
| Q66PJ3 | ARL6IP4 | 17.3570186 | 17.1152548 | 17.5075087 | 17.5945779 | 17.5049562 |
| P84085 | ARF5 | 13.8139427 | 13.913856 | 13.7765002 | 14.1178822 | 13.9300251 |
| Q9NX58 | LYAR | 16.0308191 | 16.3753893 | 15.9391272 | 15.7521223 | 15.6867155 |
| P30101 | PDIA3 | 15.8274808 | 15.1916507 | 15.2695994 | 15.007474 | 15.7726089 |
| Q14498 | RBM39 | 20.2326649 | 20.0224702 | 20.6506353 | 20.7339293 | 20.4733744 |
| P61326 | MAGOH | 13.6404008 | 14.6501475 | 14.4730729 | 14.4786047 | 13.7660892 |
| P35268 | RPL22 | 18.0069915 | 17.4960662 | 17.2754391 | 17.1455405 | 17.1901645 |
| P01857 | IGHG1 | 19.4711444 | 19.4372087 | 19.697355 | 19.7503702 | 19.469066 |
| P00338 | LDHA | 18.9223116 | 18.1422686 | 18.4418981 | 18.0567039 | 18.8129738 |
| Q9Y3F4 | STRAP | 11.8729086 | 12.5270516 | 11.3947978 | 11.7508717 | 11.7658651 |
| P11279 | LAMP1 | 14.4845444 | 14.5755798 | 13.6314715 | 13.9454476 | 13.6665592 |
| O14925 | TIMM23 | 13.7979843 | 14.0068443 | 14.3546718 | 14.4497169 | 13.9589035 |
| P30048 | PRDX3 | 15.7847258 | 16.5533942 | 15.2248488 | 16.1257964 | 15.9221035 |
| Q9HAV4 | XPO5 | 13.6441681 | 13.2901835 | 13.1108693 | 13.1808283 | 12.9567323 |

| | | | | | | |
|---|---|---|---|---|---|---|
| Q9Y2W1 | THRAP3 | 14.8143387 | 15.2818774 | 14.8917271 | 14.8093678 | 14.8418481 |
| O15042 | U2SURP | 17.2929251 | 17.401995 | 17.6239105 | 17.5466999 | 17.3279634 |
| Q14318 | FKBP8 | 13.6441111 | 12.8942284 | 12.9971953 | 13.2045057 | 12.9797852 |
| P23246 | SFPQ | 19.2770106 | 19.0517209 | 19.3657501 | 19.3686614 | 19.4393305 |
| Q15233 | NONO | 17.2955149 | 17.1399926 | 17.2113499 | 16.9540694 | 17.3378411 |
| P62987 | UBA52 | 18.0479016 | 16.9760003 | 17.9911729 | 18.3183054 | 17.7068496 |
| P06576 | ATP5F1B | 18.0342781 | 17.9688921 | 17.4490736 | 17.7655265 | 18.0345164 |
| O95232 | LUC7L3 | 18.3737485 | 18.1200065 | 18.5131816 | 18.6986181 | 18.5272694 |
| P15531 | NME1 | 15.4715098 | 15.2125623 | 15.5540822 | 15.5472715 | 15.9359747 |
| Q07955 | SRSF1 | 15.7800931 | 16.2644168 | 16.2126773 | 16.3167433 | 15.9419353 |
| P52565 | ARHGDIA | 14.9598051 | 14.535089 | 14.4934891 | 14.5852772 | 14.9050532 |
| Q15084 | PDIA6 | 15.808398 | 15.5896437 | 15.5852106 | 15.4688656 | 15.9212543 |
| Q5T4T6 | SYCP2L | 13.5708411 | 13.2823773 | 14.5036862 | 14.4187233 | 14.2376127 |
| Q2TBE0 | CWF19L2 | 14.3157238 | 14.4020826 | 14.8440507 | 14.9567801 | 14.7592503 |
| P07900 | HSP90AA1 | 19.9380678 | 19.6364874 | 19.2062577 | 19.2650376 | 19.5541882 |
| Q96IZ7 | RSRC1 | 12.9837961 | 13.3437148 | 13.3340228 | 13.9698859 | 13.2404953 |
| P14866 | HNRNPL | 14.7998671 | 15.0933659 | 14.748302 | 15.4390793 | 14.7161823 |
| A0A8I5KQE6 | RPSA2 | 15.2674388 | 15.1942527 | 15.0074425 | 14.7528014 | 15.0717615 |
| Q13642 | FHL1 | 18.5637048 | 18.0270979 | 18.5051696 | 18.2901457 | 18.7197939 |
| Q53H12 | AGK | 17.2968124 | 16.785682 | 16.8256592 | 16.9633482 | 16.7832309 |
| Q15004 | PCLAF | 12.5073455 | 13.1001072 | 12.4163235 | 13.0736817 | 13.2909231 |
| P63244 | RACK1 | 18.5441113 | 18.1296427 | 18.1691082 | 18.3902195 | 18.3736516 |
| Q9H0G5 | NSRP1 | 14.3945087 | 14.4984964 | 14.8599614 | 14.9066262 | 14.5759109 |
| Q9UHX1 | PUF60 | 17.7202195 | 17.5468857 | 17.7820968 | 18.0757181 | 17.8255623 |
| P38159 | RBMX | 16.1563452 | 16.4509756 | 16.3460602 | 16.3585746 | 16.0604267 |
| Q16795 | NDUFA9 | 14.0509973 | 13.8053361 | 14.1188456 | 14.1424782 | 14.0172686 |
| Q14008 | CKAP5 | 11.4521147 | 12.0524268 | 11.0446154 | 11.15074 | 12.0853297 |
| Q13435 | SF3B2 | 17.1412825 | 17.0819259 | 16.7818594 | 17.0302578 | 16.750445 |
| Q5JTV8 | TOR1AIP1 | 14.2849463 | 14.8152955 | 14.2817618 | 14.293659 | 14.5830113 |
| Q5BKY9 | FAM133B | 19.4152279 | 18.871641 | 19.4753775 | 19.4462417 | 19.4376885 |
| O95373 | IPO7 | 14.1810395 | 14.3510942 | 14.4210788 | 14.3668727 | 13.9688981 |
| Q9Y277 | VDAC3 | 15.9740773 | 15.5489293 | 15.4531671 | 15.5508347 | 15.5204197 |
| Q8N5F7 | NKAP | 15.4640545 | 15.0108983 | 15.7053811 | 15.6185296 | 15.6959097 |
| Q15393 | SF3B3 | 14.0098303 | 14.4172302 | 13.9988643 | 14.0909787 | 14.1031609 |
| P62826 | RAN | 18.4203946 | 17.7388428 | 17.7069991 | 17.6987921 | 18.2225938 |
| O15235 | MRPS12 | 20.0894045 | 19.8614768 | 20.4802535 | 20.5902557 | 20.2254804 |
| P28799 | GRN | 22.0507783 | 21.4023759 | 22.156563 | 22.4789557 | 22.0368512 |
| Q9H9B4 | SFXN1 | 18.5596078 | 18.4576658 | 18.5569066 | 18.4523875 | 18.3947752 |
| Q15532 | SS18 | 14.3864288 | 14.5294606 | 14.990842 | 14.7332591 | 14.7865082 |
| P22626 | HNRNPA2B1 | 17.9715125 | 18.3752036 | 17.4686134 | 17.6142063 | 17.6783863 |
| Q8IYB3 | SRRM1 | 14.8384528 | 16.0294041 | 15.3051298 | 15.1508868 | 15.2330807 |
| P62258 | YWHAE | 18.8518059 | 18.4179948 | 18.2785291 | 18.4702896 | 18.6633605 |

| | | | | | | |
|---|---|---|---|---|---|---|
| P26639 | TARS1 | 15.7614207 | 15.538506 | 15.4012841 | 15.3468766 | 15.6222565 |
| P27348 | YWHAQ | 19.5069984 | 19.3110706 | 19.2181074 | 19.3110918 | 19.4103947 |
| P00505 | GOT2 | 12.1180704 | 11.7484396 | 12.9215609 | 12.4423225 | 12.4720266 |
| P14625 | HSP90B1 | 15.802331 | 15.6367329 | 15.4584658 | 15.4747094 | 15.5933896 |
| O75964 | ATP5MG | 16.9920832 | 17.0573475 | 17.1285081 | 17.1281189 | 16.8494281 |
| P60228 | EIF3E | 11.6192181 | 12.006347 | 11.5422965 | 11.4267304 | 11.6318342 |
| Q9Y295 | DRG1 | 13.570265 | 13.5445473 | 13.8056687 | 13.8762769 | 13.400237 |
| P53618 | COPB1 | 15.1202365 | 15.4748627 | 14.7206594 | 15.0078397 | 14.8532794 |
| Q96K21 | ZFYVE19 | 14.2891665 | 14.2793519 | 14.3131779 | 14.4957789 | 13.9561802 |
| P22314 | UBA1 | 17.7599512 | 17.5336071 | 17.4191908 | 17.4625829 | 17.5837008 |
| Q92945 | KHSRP | 17.262499 | 17.0380621 | 16.6159141 | 16.8859253 | 16.894091 |
| P84103 | SRSF3 | 18.749887 | 18.9242334 | 18.6867423 | 18.7109186 | 18.7348297 |
| P43243 | MATR3 | 15.5308349 | 16.2676313 | 15.615023 | 16.5014578 | 15.9873189 |
| P25490 | YY1 | 15.1501707 | 15.1536609 | 15.3609258 | 15.1947997 | 15.1910778 |
| P27816 | MAP4 | 16.3115168 | 16.4398502 | 16.2478937 | 16.5029574 | 16.2171026 |
| O75533 | SF3B1 | 15.9139312 | 16.602398 | 16.6845939 | 16.780084 | 16.2327392 |
| Q6UW68 | TMEM205 | 14.9106091 | 14.4503605 | 14.4587529 | 14.6488128 | 14.7088629 |
| Q9UN86 | G3BP2 | 16.1637099 | 16.6535794 | 15.9978038 | 16.0996501 | 16.0724437 |
| Q9UBQ5 | EIF3K | 14.2097714 | 14.7645288 | 14.9097135 | 14.7408833 | 14.4097914 |
| Q12874 | SF3A3 | 11.4395307 | 11.5356261 | 11.4253977 | 11.7303386 | 11.2229166 |
| Q13283 | G3BP1 | 15.5859609 | 16.0749158 | 15.5181755 | 15.3248496 | 15.5605006 |
| Q9Y266 | NUDC | 16.7722495 | 16.1265954 | 15.9471731 | 16.2348316 | 16.293145 |
| P08708 | RPS17 | 18.7000028 | 18.7041626 | 18.2126972 | 18.180956 | 18.0297296 |
| Q15365 | PCBP1 | 17.4602579 | 17.5635714 | 17.3247713 | 17.5240102 | 17.311732 |
| P55060 | CSE1L | 16.5085028 | 16.4566515 | 16.4505069 | 16.3528242 | 16.2145125 |
| P04040 | CAT | 17.2149997 | 17.5868259 | 17.0492482 | 16.7070727 | 16.8726229 |
| P45880 | VDAC2 | 18.5814632 | 18.0780806 | 17.9652565 | 18.2753575 | 18.2057007 |
| P08574 | CYC1 | 16.8520135 | 16.7558432 | 16.5730915 | 16.5328413 | 16.6914304 |
| P04083 | ANXA1 | 17.218469 | 14.180868 | 14.704906 | 14.178153 | 14.3855977 |
| P48643 | CCT5 | 17.9172237 | 17.7083277 | 17.3544891 | 17.4554748 | 17.469807 |
| Q8TCC3 | MRPL30 | 17.7004849 | 17.6736672 | 18.0610534 | 18.2216956 | 17.9302223 |
| Q8WWM7 | ATXN2L | 13.5630269 | 14.1909472 | 13.2280601 | 14.003489 | 13.248093 |
| Q9NTG7 | SIRT3 | 17.1810139 | 16.8196025 | 17.7483006 | 17.4479889 | 17.3451886 |
| Q7L4I2 | RSRC2 | 16.9891573 | 16.7435549 | 17.5158479 | 17.1225347 | 16.9644051 |
| P14406 | COX7A2 | 15.6785982 | 15.7971087 | 15.3456307 | 15.6672378 | 15.928494 |
| O14949 | UQCRQ | 17.0817741 | 16.9455802 | 17.1154501 | 16.9981518 | 16.9844714 |
| P56134 | ATP5MF | 15.61034 | 14.7910344 | 15.4957836 | 14.5037311 | 15.4193903 |
| P07737 | PFN1 | 15.8681442 | 15.144001 | 15.070356 | 14.8069839 | 15.8207707 |
| P35637 | FUS | 15.8280102 | 14.4765493 | 13.9231889 | 13.8494716 | 14.5498302 |
| Q13347 | EIF3I | 13.243214 | 13.963346 | 14.01619 | 13.3751091 | 14.0159611 |
| Q9NRP0 | OSTC | 14.9310798 | 14.8915806 | 14.5259934 | 14.1742281 | 14.4288887 |
| Q13867 | BLMH | 16.4354695 | 16.7083031 | 16.3725842 | 15.8594272 | 15.5892413 |

| | | | | | | |
|---|---|---|---|---|---|---|
| P11940 | PABPC1 | 17.9729794 | 18.6735751 | 17.8463121 | 18.1185047 | 17.9006469 |
| Q9Y597 | KCTD3 | 19.1463214 | 18.9873944 | 19.6398283 | 19.2842317 | 19.4766834 |
| Q99623 | PHB2 | 18.8824459 | 18.4273434 | 18.353141 | 18.251286 | 18.3061021 |
| Q99873 | PRMT1 | 11.379874 | 11.7878431 | 11.5833965 | 11.194363 | 11.2524694 |
| P13639 | EEF2 | 19.6331419 | 19.2551861 | 18.8820851 | 18.8174329 | 19.2521268 |
| P31943 | HNRNPH1 | 18.0067183 | 18.4346054 | 17.9415053 | 18.3646099 | 17.9044401 |
| P49257 | LMAN1 | 15.1403002 | 15.0839995 | 15.2647746 | 15.1170839 | 15.0403672 |
| O75534 | CSDE1 | 14.1050102 | 14.5544452 | 13.9529287 | 14.314829 | 14.5237706 |
| O43143 | DHX15 | 16.4020644 | 16.4687016 | 16.3927026 | 16.1779224 | 16.2705107 |
| A0A075B6K5 | IGLV3-9 | 13.8782894 | 14.9417241 | 15.5081145 | 15.1665907 | 14.9997335 |
| Q5VTL8 | PRPF38B | 14.3796698 | 14.2619321 | 14.5536968 | 14.5811299 | 14.51313 |
| P30153 | PPP2R1A | 12.9994844 | 12.4869854 | 12.173604 | 12.3068892 | 12.2812282 |
| Q96P63 | SERPINB12 | 15.3012727 | 16.0441708 | 15.6638421 | 15.0611382 | 15.1000241 |
| Q8N684 | CPSF7 | 14.6067466 | 14.1684261 | 14.3869081 | 14.2631081 | 13.8513903 |
| Q96TA2 | YME1L1 | 12.1505722 | 12.6503964 | 12.221274 | 12.0709504 | 12.0090383 |
| P62851 | RPS25 | 20.9078251 | 20.2705045 | 20.2677949 | 20.2491574 | 20.2846707 |
| P61978 | HNRNPK | 18.5988503 | 18.8687432 | 18.2464472 | 18.4071334 | 18.4580787 |
| P13804 | ETFA | 14.7570604 | 14.5114135 | 14.063838 | 14.1444198 | 14.526085 |
| Q8NC51 | SERBP1 | 18.5126911 | 17.5111526 | 17.1703123 | 17.0231979 | 17.6382472 |
| P46782 | RPS5 | 20.957368 | 20.3760194 | 20.6634226 | 20.4705524 | 20.4939168 |
| P07195 | LDHB | 17.9129163 | 17.3666924 | 17.7344941 | 17.5181764 | 18.2379792 |
| P49756 | RBM25 | 16.331404 | 16.5440718 | 16.6788705 | 16.6909531 | 16.517452 |
| Q14103 | HNRNPD | 15.1804674 | 15.5741693 | 14.702602 | 14.3304783 | 14.9587284 |
| Q12872 | SFSWAP | 17.082449 | 16.8585201 | 17.288725 | 17.4020635 | 17.2375178 |
| Q9BY77 | POLDIP3 | 19.2356753 | 19.0112555 | 19.4758883 | 19.4938543 | 19.3623244 |
| P63104 | YWHAZ | 16.0388131 | 15.3032062 | 14.9858229 | 14.9906468 | 15.5801128 |
| Q15637 | SF1 | 15.3075867 | 15.0256877 | 14.7153159 | 15.2000854 | 14.8717308 |
| Q86V81 | ALYREF | 19.1993589 | 17.7429416 | 17.1745848 | 17.2110848 | 17.5188513 |
| Q9NUQ3 | TXLNG | 13.9517209 | 14.7656117 | 13.5189388 | 13.6353728 | 13.7781263 |
| Q13428 | TCOF1 | 16.6508133 | 16.836116 | 16.8428109 | 16.8960674 | 16.6961429 |
| P35659 | DEK | 17.0635815 | 16.4933631 | 16.3410015 | 15.7535089 | 16.3804571 |
| P48556 | PSMD8 | 13.2403148 | 14.12525 | 13.774259 | 14.1859221 | 13.5866972 |
| P35606 | COPB2 | 14.3190194 | 14.5180757 | 13.9653789 | 14.2109096 | 14.3837208 |
| P17844 | DDX5 | 18.2384568 | 18.4149685 | 18.2318428 | 18.1278091 | 18.1381681 |
| Q16610 | ECM1 | 15.0925515 | 15.7107248 | 14.7920052 | 14.1760828 | 14.4517809 |
| Q9Y2R9 | MRPS7 | 14.9547511 | 15.3302194 | 14.9641591 | 15.2214126 | 14.6029888 |
| Q96PK6 | RBM14 | 17.0858913 | 16.8732207 | 17.2121515 | 17.2066762 | 17.2945151 |
| P42766 | RPL35 | 20.3096422 | 20.4335605 | 20.1661673 | 20.1392937 | 19.9796958 |
| Q9Y3A2 | UTP11 | 15.1953732 | 15.3492847 | 15.5633738 | 15.4481679 | 15.4199994 |
| Q8IWS0 | PHF6 | 14.4934538 | 14.6972535 | 14.2012252 | 14.2181867 | 14.2110465 |
| P68104 | EEF1A1 | 22.2962863 | 22.0167119 | 21.9942141 | 22.1293247 | 22.0608096 |
| Q15181 | PPA1 | 14.9831289 | 15.0038033 | 14.6637248 | 14.7679462 | 14.9915248 |

| | | | | | | |
|---|---|---|---|---|---|---|
| Q14703 | MBTPS1 | 15.2652656 | 14.8770926 | 15.6868553 | 15.1914307 | 15.5115925 |
| Q96FQ6 | S100A16 | 15.1143537 | 14.6689956 | 13.4989687 | 13.2847734 | 13.596523 |
| Q96RL1 | UIMC1 | 21.3361229 | 20.9710295 | 21.548467 | 21.8169161 | 21.283979 |
| P0CF74 | IGLC6 | 14.6412886 | 14.5069225 | 14.9686032 | 14.9477932 | 14.7439929 |
| P58107 | EPPK1 | 14.2365998 | 14.8844133 | 14.3488962 | 14.3541918 | 13.964099 |
| Q86VP6 | CAND1 | 12.4921557 | 12.7808977 | 12.1729328 | 11.8257459 | 12.1544527 |
| Q96K30 | RITA1 | 14.7825904 | 14.7509811 | 15.2920566 | 15.0592459 | 15.0575475 |
| O60841 | EIF5B | 15.3239124 | 15.1265644 | 14.6743159 | 14.8231351 | 14.8242929 |
| P25705 | ATP5F1A | 19.4484577 | 19.2376122 | 19.1383904 | 19.3080668 | 19.2465085 |
| P29508 | SERPINB3 | 11.7029174 | 12.0301844 | 11.7861705 | 11.8189733 | 11.2841466 |
| Q92878 | RAD50 | 13.8256901 | 14.3985995 | 13.3211099 | 14.253375 | 14.1018481 |
| Q00325 | SLC25A3 | 20.1072821 | 20.1947204 | 20.2059811 | 20.5068978 | 20.0468232 |
| Q96KM6 | ZNF512B | 19.0084111 | 18.9702298 | 19.8189064 | 19.7610471 | 19.6470425 |
| P06744 | GPI | 17.3999814 | 16.461867 | 16.3832081 | 15.9842035 | 17.415194 |
| P29966 | MARCKS | 15.2837483 | 14.7291927 | 14.1245287 | 14.3688585 | 14.7391538 |
| O43175 | PHGDH | 16.8066649 | 16.6807553 | 16.932456 | 17.1704195 | 16.8387667 |
| Q92841 | DDX17 | 15.719302 | 15.8557103 | 15.6046171 | 15.5830209 | 15.5063246 |
| Q08188 | TGM3 | 15.8347206 | 15.3882271 | 14.8606803 | 14.4949412 | 14.5253923 |
| Q92922 | SMARCC1 | 20.5212765 | 20.160869 | 20.8131045 | 20.9188581 | 20.644382 |
| Q9NZM4 | BICRA | 20.0462636 | 19.6633022 | 20.442598 | 20.4803754 | 20.2979861 |
| Q9BSD7 | NTPCR | 17.3398635 | 17.2180559 | 17.77854 | 17.6659881 | 17.4700409 |
| P01834 | IGKC | 20.5589707 | 20.4152357 | 21.0319946 | 21.004702 | 20.772211 |
| Q04637 | EIF4G1 | 10.6875718 | 11.3240012 | 11.491033 | 11.8629554 | 10.9063953 |
| P08238 | HSP90AB1 | 19.8616826 | 19.2770803 | 19.0222331 | 19.0866503 | 19.3591597 |
| P09651 | HNRNPA1 | 18.5114969 | 18.92366 | 18.2165363 | 18.294218 | 18.2381492 |
| P60842 | EIF4A1 | 16.8233441 | 16.3300888 | 16.3504683 | 16.1107273 | 16.678506 |
| Q9BTV4 | TMEM43 | 13.2912032 | 13.1814789 | 12.735083 | 12.8626753 | 12.9355815 |
| B9A064 | IGLL5 | 21.4362339 | 21.0545197 | 21.7893661 | 21.7471402 | 21.5077054 |
| P48444 | ARCN1 | 15.1654408 | 15.0579398 | 15.3041405 | 14.8760559 | 15.0414418 |
| Q96EM0 | L3HYPDH | 24.3119998 | 23.0201518 | 24.3915067 | 24.8258114 | 23.5483148 |
| Q14204 | DYNC1H1 | 14.66039 | 14.5052866 | 14.8265797 | 14.579884 | 14.2915009 |
| Q15029 | EFTUD2 | 14.9235736 | 15.7068459 | 15.2986066 | 15.3707503 | 15.0146466 |
| Q07666 | KHDRBS1 | 14.8604811 | 14.9181666 | 14.5303382 | 14.4947629 | 14.6712061 |
| Q9NWB6 | ARGLU1 | 21.6812205 | 21.2848456 | 21.8776209 | 22.1732294 | 21.9401912 |
| O60506 | SYNCRIP | 16.9491179 | 17.4068704 | 16.6762388 | 16.7067647 | 16.6566752 |
| P62269 | RPS18 | 20.2176429 | 19.9943014 | 20.1661811 | 20.2139459 | 20.1514546 |
| Q96GM5 | SMARCD1 | 19.6768809 | 19.389994 | 20.1266008 | 20.174865 | 19.984656 |
| Q9P021 | CRIPT | 15.2418248 | 14.6085596 | 14.9621494 | 14.769172 | 15.1138912 |
| P50990 | CCT8 | 18.3468047 | 17.8403338 | 17.6601101 | 17.9090176 | 17.9255103 |
| Q9H9J2 | MRPL44 | 13.735635 | 14.6071894 | 13.6382956 | 14.1072554 | 13.5831506 |
| Q01844 | EWSR1 | 15.9715012 | 15.0837175 | 15.1057254 | 15.0292207 | 15.0639125 |
| P55196 | AFDN | NA | 12.2134925 | 13.1783594 | NA | 12.0655431 |

| | | | | | | |
|---|---|---|---|---|---|---|
| P05114 | HMGN1 | 15.3075262 | 14.3278708 | 13.9561276 | 13.7363506 | 14.8026312 |
| Q6SPF0 | SAMD1 | 10.8131106 | 11.1574605 | 11.4955167 | 11.9507711 | 11.4400213 |
| P37802 | TAGLN2 | 16.1015714 | 15.5837119 | 15.6239641 | 15.9268342 | 16.39348 |
| P49755 | TMED10 | 16.0072683 | 15.8004592 | 15.403214 | 15.4549548 | 15.8381635 |
| Q92598 | HSPH1 | 13.4120611 | 13.5932151 | 12.960587 | 13.2492717 | 13.4752348 |
| P62249 | RPS16 | 21.4431176 | 21.0230389 | 21.2044776 | 21.3300585 | 21.0315046 |
| P54136 | RARS1 | 14.9278686 | 15.1745339 | 15.2125992 | 15.1472982 | 15.2448477 |
| O43674 | NDUFB5 | 15.7346494 | 14.9260141 | 15.2881098 | 15.2336338 | 15.4870587 |
| Q6AI39 | BICRAL | 18.6492124 | 18.47123 | 19.099487 | 19.0777178 | 18.9830682 |
| Q9H8M2 | BRD9 | 20.8466128 | 20.4143878 | 21.1848033 | 21.2438357 | 21.031976 |
| Q9BRR8 | GPATCH1 | 14.3956161 | 14.2154751 | 14.6185023 | 14.6801007 | 14.5439152 |
| P08579 | SNRPB2 | 15.9210171 | 16.023868 | 15.47051 | 15.9035608 | 15.6488235 |
| P24468 | NR2F2 | 13.0289504 | 13.3554533 | 13.5123122 | 13.0479483 | 13.5666291 |
| O95831 | AIFM1 | 15.2751528 | 16.3616442 | 15.9203332 | 15.8338494 | 15.7962617 |
| P46783 | RPS10 | 22.0494554 | 21.1958275 | 21.5738791 | 21.464052 | 21.4920994 |
| P27824 | CANX | 17.318291 | 16.8952452 | 16.4894435 | 16.7182225 | 17.0035816 |
| Q00839 | HNRNPU | 19.4651695 | 19.5453541 | 19.1169456 | 19.2038269 | 19.1089936 |
| P23396 | RPS3 | 19.9618723 | 19.9285405 | 19.9947079 | 20.0431106 | 19.8171301 |
| Q9UBB9 | TFIP11 | 15.1622181 | 14.0230659 | 15.2320449 | NA | 14.8944027 |
| Q86WX3 | RPS19BP1 | 13.3115375 | 14.1874885 | 14.0073868 | 14.2404735 | 14.1780268 |
| P62280 | RPS11 | 21.4557891 | 21.2050338 | 21.6024905 | 21.55314 | 21.2005418 |
| Q13885 | TUBB2A | 13.9904922 | 14.5234645 | 14.69473 | 14.6764075 | 13.9605271 |
| P31948 | STIP1 | 16.3770157 | 15.7668968 | 15.7960451 | 15.5823371 | 16.1118123 |
| Q9BXS6 | NUSAP1 | 15.5241579 | 15.3601184 | 15.9114207 | 15.9779112 | 15.6184217 |
| Q9UM00 | TMCO1 | 15.4559608 | 15.7092462 | 15.1263534 | 15.7766307 | 15.4864806 |
| O96019 | ACTL6A | 18.4149786 | 18.4525237 | 18.9490388 | 19.0540237 | 18.9081363 |
| P62273 | RPS29 | 21.8779973 | 21.4256611 | 21.9064622 | 21.862648 | 21.789546 |
| Q6P1L8 | MRPL14 | 19.0435878 | 18.6136235 | 19.0701868 | 19.2936625 | 18.9698936 |
| P51532 | SMARCA4 | 20.5598446 | 20.2161924 | 20.8156897 | 20.9562057 | 20.6916006 |
| P51116 | FXR2 | 14.46849 | 15.3137972 | 14.8094733 | 15.0556021 | 14.8077798 |
| P61081 | UBE2M | 14.0645469 | 13.150739 | NA | NA | NA |
| P62318 | SNRPD3 | 12.5159423 | 12.4492834 | 11.7860194 | 12.1686825 | 12.2957806 |
| Q4VC05 | BCL7A | 19.0293057 | 19.0572851 | 19.3917522 | 19.4907567 | 19.2498437 |
| Q96I25 | RBM17 | 16.5913855 | 16.5148812 | 16.8791677 | 17.0101856 | 16.8551282 |
| P07355 | ANXA2 | 17.9897102 | 17.7655511 | 17.6898596 | 17.201254 | 17.3157143 |
| Q9NZ01 | TECR | 18.542877 | 18.3018509 | 18.648615 | 18.7922935 | 18.4464911 |
| Q96EV2 | RBM33 | 13.1218036 | 12.6014993 | 13.4379609 | 13.282217 | 13.5539954 |
| Q8ND56 | LSM14A | 17.4818134 | 16.9829476 | 16.8664098 | 16.7694888 | 17.2102211 |
| P78371 | CCT2 | 15.0733371 | 14.8760604 | 14.4406328 | 14.3943941 | 14.6251814 |
| P10809 | HSPD1 | 19.5894874 | 19.3602362 | 18.7975575 | 19.0197856 | 19.3249062 |
| P49368 | CCT3 | 17.3181614 | 17.1708093 | 17.1529628 | 17.1619898 | 17.3198032 |
| P48047 | ATP5PO | 15.0799627 | 14.8079897 | 14.5720991 | 14.4086396 | 14.5414058 |

| | | | | | | |
|---|---|---|---|---|---|---|
| Q8TBC3 | SHKBP1 | 20.0581207 | 19.6611947 | 20.5605194 | 20.1987237 | 20.3213474 |
| Q14444 | CAPRIN1 | 16.6377101 | 17.4022769 | 16.6061913 | 16.6463312 | 16.4654525 |
| Q9P258 | RCC2 | 15.1114198 | 15.2335501 | 14.5750893 | 14.0766577 | 14.8380043 |
| O95292 | VAPB | 14.0351836 | 13.6796887 | 13.7285224 | 14.4398759 | 14.1617655 |
| O75376 | NCOR1 | 18.6323413 | 18.4435766 | 18.953197 | 19.0606507 | 18.5381244 |
| P09874 | PARP1 | 18.3711766 | 18.2775722 | 18.1216738 | 18.1432733 | 18.3029755 |
| P04844 | RPN2 | 15.3930039 | 15.369138 | 15.0979045 | 15.1904219 | 15.1801179 |
| P36542 | ATP5F1C | 16.7338302 | 16.6339759 | 16.4289516 | 16.4369332 | 16.6133953 |
| P60866 | RPS20 | 21.4964578 | 20.9345904 | 21.0971946 | 21.225998 | 21.1314964 |
| Q9UBM7 | DHCR7 | 16.1126673 | 16.4958001 | 15.9324734 | 16.1602224 | 16.1787313 |
| P49790 | NUP153 | 15.1472234 | 15.0435517 | 15.4597445 | 15.5116014 | 15.3977673 |
| P56192 | MARS1 | 14.3143858 | 14.8184317 | 13.9671296 | 14.1940714 | 14.1516499 |
| Q9UI09 | NDUFA12 | 14.9550835 | 14.5556324 | 14.7539333 | 14.3999056 | 14.2746364 |
| Q13823 | GNL2 | 13.1045653 | 13.5489193 | 13.209729 | 12.8918238 | 12.7137556 |
| Q13310 | PABPC4 | 15.448056 | 16.2434343 | 15.3945136 | 15.7658496 | 15.6802269 |
| Q9NR31 | SAR1A | 16.9882825 | 16.885128 | 15.5102396 | 15.3701042 | 15.8930389 |
| A6NHL2 | TUBAL3 | 14.6613185 | 14.6590072 | 14.4531896 | 14.6608101 | 14.4194957 |
| P21796 | VDAC1 | 17.8210614 | 17.6090484 | 17.4777152 | 17.6934759 | 17.7986614 |
| P49458 | SRP9 | 15.8245066 | 15.7034612 | 14.0859478 | 15.4687109 | 15.1336681 |
| Q9P2M4 | TBC1D14 | 13.30996 | 13.363818 | 13.9655847 | 13.9897071 | 13.6977286 |
| P05141 | SLC25A5 | 22.7128299 | 22.4930352 | 22.9321348 | 23.0179973 | 22.7077031 |
| P62861 | FAU | 18.7766681 | 17.8564219 | 17.8600778 | 17.7565441 | 17.7196276 |
| P12236 | SLC25A6 | 18.9487704 | 18.8528891 | 19.1184567 | 19.0524667 | 18.7704953 |
| Q3YEC7 | RABL6 | 11.3368228 | 12.0199726 | 11.7374685 | 11.5356968 | 11.4187889 |
| P22695 | UQCRC2 | 18.5262438 | 18.6096746 | 18.4624148 | 18.8149227 | 18.2839214 |
| P62829 | RPL23 | 20.6063535 | 20.4937284 | 20.6555739 | 20.6963567 | 20.4883582 |
| Q4L180 | FILIP1L | 16.9732854 | 14.3542009 | 15.4196206 | 16.2395704 | 16.5572465 |
| P63173 | RPL38 | 19.6698837 | 19.2224384 | 19.7785712 | 19.6014022 | 19.4932597 |
| P01861 | IGHG4 | 16.4065152 | 16.2105408 | 16.564374 | 16.6865749 | 16.4170934 |
| P62266 | RPS23 | 21.2507166 | 21.2074298 | 21.6962451 | 21.6739104 | 21.4241998 |
| P57088 | TMEM33 | 14.9821566 | 14.7124994 | 14.9540468 | 15.2589868 | 14.9605927 |
| P62081 | RPS7 | 19.5870577 | 19.3271617 | 18.7700594 | 18.6811224 | 18.9866686 |
| P02765 | AHSG | 15.4351118 | 15.3189097 | 15.4423711 | 15.3002727 | 15.4482209 |
| Q14152 | EIF3A | 18.5529462 | 15.8177044 | 14.6507365 | 14.9670163 | 15.2237217 |
| Q12906 | ILF3 | 17.5083694 | 17.9543803 | 17.4162508 | 17.6915877 | 17.6241299 |
| P53999 | SUB1 | 15.8358997 | 15.1530279 | 15.4618958 | 15.1971607 | 15.3165308 |
| Q14157 | UBAP2L | 16.2000926 | 16.0072329 | 14.7505138 | 15.3527752 | 15.679393 |
| P05089 | ARG1 | 16.6552452 | 17.9294783 | 16.9846931 | 16.3316944 | 16.0265413 |
| O75643 | SNRNP200 | 14.4838404 | 14.8336106 | 14.6755425 | 14.6842151 | 14.4604554 |
| Q9NZI8 | IGF2BP1 | 16.8436504 | 17.3573724 | 16.904683 | 17.1234953 | 17.1324165 |
| P58557 | YBEY | 11.3581871 | 11.2966064 | 11.5792783 | 11.5292248 | 11.5970045 |
| P17987 | TCP1 | 17.6739344 | 17.2322435 | 17.0814625 | 17.1919852 | 17.1885075 |

| | | | | | | |
|---|---|---|---|---|---|---|
| Q6P2Q9 | PRPF8 | 16.6018338 | 16.9210271 | 16.877628 | 17.0222632 | 16.6693522 |
| Q9BRP1 | PDCD2L | 12.5709006 | 13.3566198 | 12.8790468 | NA | 12.8560438 |
| O75934 | BCAS2 | 14.3881693 | 15.1167375 | 14.0694982 | 14.2202626 | 14.2091218 |
| P39019 | RPS19 | 20.599888 | 20.0392547 | 19.8562307 | 19.7686104 | 20.0069589 |
| P62701 | RPS4X | 21.1976874 | 21.0605252 | 20.8452056 | 20.9953905 | 20.6951427 |
| Q9UQ35 | SRRM2 | 16.580482 | 17.3857334 | 17.1394184 | 17.2539195 | 16.9488157 |
| P61626 | LYZ | 17.0913079 | 16.5536796 | 16.8265179 | 16.3221492 | 16.1526575 |
| Q96AG4 | LRRC59 | 19.2468479 | 18.544375 | 18.6019936 | 18.4883675 | 18.6677831 |
| P31944 | CASP14 | 16.2056395 | 16.6219314 | 16.7085348 | 16.2078323 | 15.3968216 |
| Q96IX5 | ATP5MK | 17.2703293 | 17.1965399 | 17.1775748 | 17.2887858 | 17.1892389 |
| Q9NP64 | ZCCHC17 | 14.4816115 | 14.6500084 | 15.3042093 | 14.9136337 | 15.175366 |
| P51531 | SMARCA2 | 18.8621347 | 18.6447989 | 19.1476008 | 19.1444366 | 19.0445313 |
| O75340 | PDCD6 | 13.3481465 | 13.3447247 | 13.6012215 | 13.4695917 | 13.3275331 |
| Q8NAV1 | PRPF38A | 15.9165426 | 15.8693009 | 15.9320589 | 16.1891823 | 16.0306818 |
| Q86TG7 | PEG10 | 11.6018018 | 12.3519559 | 11.587529 | 10.8783473 | 11.3184653 |
| Q86YT6 | MIB1 | 11.3882434 | 12.3502887 | 12.16254 | 11.3594185 | 12.003929 |
| P38646 | HSPA9 | 21.2990162 | 21.1267966 | 21.4623633 | 21.6945801 | 21.458266 |
| Q8WUA2 | PPIL4 | 11.4109709 | 11.3284762 | 11.3695753 | 11.8393437 | 11.2470079 |
| P04843 | RPN1 | 17.2287857 | 17.0672652 | 17.0093642 | 16.7859224 | 16.9173211 |
| Q3ZCQ8 | TIMM50 | 17.6023056 | 17.4708634 | 17.793365 | 17.4577747 | 17.2985932 |
| Q9P0L0 | VAPA | 16.5435013 | 16.3546541 | 16.451856 | 16.4627556 | 16.7257061 |
| P83881 | RPL36A | 21.873812 | 21.7636189 | 22.1446477 | 22.1921634 | 21.8308648 |
| Q13247 | SRSF6 | 18.178704 | 18.4413391 | 18.4106032 | 18.3941842 | 18.2665771 |
| P49411 | TUFM | 18.7122691 | 18.8010944 | 18.7410618 | 18.8810744 | 18.5659731 |
| Q9P035 | HACD3 | 16.0132916 | 16.0048871 | 15.9846917 | 16.0589693 | 16.1200779 |
| Q15366 | PCBP2 | 19.5003204 | 19.4129938 | 19.2259282 | 19.6198506 | 19.1361444 |
| Q14974 | KPNB1 | 17.2337202 | 17.1400922 | 16.8424574 | 16.8920878 | 16.9311876 |
| Q9Y2X3 | NOP58 | 12.4892295 | 13.0761275 | 12.8089671 | 12.8083988 | 12.5084153 |
| P32119 | PRDX2 | 21.6652309 | 21.8700768 | 21.1309082 | 21.1251704 | 21.4194111 |
| P35232 | PHB1 | 18.0333584 | 17.4193182 | 17.0473407 | 17.3302285 | 17.6291293 |
| P62195 | PSMC5 | 15.898835 | 16.0125769 | 15.8651498 | 15.892037 | 15.9333172 |
| P18085 | ARF4 | 16.3191576 | 16.1757118 | 16.4436016 | 16.4562543 | 16.2219583 |
| O60237 | PPP1R12B | 19.5022201 | 19.0991496 | 19.6558579 | 19.8357223 | 19.86392 |
| P25398 | RPS12 | 17.5674948 | 17.505967 | 17.1945362 | 17.5862755 | 17.4707321 |
| Q9BW19 | KIFC1 | 16.4723434 | 16.6784814 | 16.9574562 | 16.9340198 | 16.7997625 |
| Q15072 | ZNF146 | 13.7819443 | 13.8645874 | 13.5227131 | 14.1061903 | 14.4480077 |
| Q9NXE8 | CWC25 | 14.3403711 | 14.7369176 | 14.9106677 | 14.5695434 | 13.9495125 |
| Q8N6S5 | ARL6IP6 | 16.5772484 | 16.5306851 | 16.3393013 | 16.8818915 | 16.7621508 |
| P11388 | TOP2A | 14.9465294 | 15.5898801 | 15.4480036 | 15.2245223 | 14.909044 |
| Q8WWK9 | CKAP2 | 17.2349023 | 17.1536598 | 17.689316 | 17.5531791 | 17.4530564 |
| P12235 | SLC25A4 | 16.8103687 | 16.7344126 | 17.0688718 | 16.9741016 | 16.5704142 |
| P62750 | RPL23A | 19.4106052 | 19.8289558 | 18.7547415 | 18.9670682 | 18.8830348 |

| | | | | | | |
|---|---|---|---|---|---|---|
| Q8WUZ0 | BCL7C | 16.2845388 | 15.837708 | 16.5379719 | 16.2884637 | 16.2921248 |
| P24534 | EEF1B2 | 15.5738986 | 15.4076063 | 14.9721279 | 14.8308165 | 15.0026664 |
| P40227 | CCT6A | 16.3529015 | 16.1237924 | 15.7777685 | 16.0877546 | 16.1847451 |
| P0DOY2 | IGLC2 | 21.2930149 | 21.0806648 | 21.7043963 | 21.9218027 | 21.290627 |
| P61513 | RPL37A | 21.1089096 | 21.0009492 | 21.5881733 | 21.6044542 | 21.1195426 |
| P62857 | RPS28 | 14.7932617 | 14.576949 | 14.7888781 | 14.9631552 | 15.0166475 |
| Q9BY32 | ITPA | 13.7908183 | 13.447444 | 13.4143615 | 13.3369771 | 13.5092348 |
| P51148 | RAB5C | 16.5628319 | 16.2491201 | 16.4947449 | 16.541446 | 16.6908829 |
| P62244 | RPS15A | 19.9509678 | 20.2953684 | 19.6441927 | 19.8107926 | 19.6555645 |
| P55058 | PLTP | 17.0698127 | 17.443826 | 17.1302463 | 17.7041041 | 17.2155973 |
| P11021 | HSPA5 | 18.6786771 | 18.5306732 | 18.7072443 | 18.862014 | 18.7770758 |
| Q9H1K1 | ISCU | 14.5767733 | 14.3557614 | 14.6367743 | 14.4638115 | 14.8501699 |
| Q9ULV0 | MYO5B | 14.7504019 | 14.3310312 | 15.0325794 | 14.6633367 | 14.44449 |
| P61254 | RPL26 | 17.8751195 | 18.6164127 | 18.2815327 | 18.3570301 | 18.0313329 |
| O95573 | ACSL3 | 14.8648011 | 14.3589459 | 14.4028557 | 14.5679099 | 14.189091 |
| P49327 | FASN | 17.1034361 | 17.3644295 | 16.8748292 | 16.6060936 | 16.7515961 |
| P57721 | PCBP3 | 16.2643756 | 16.1525829 | 15.9295779 | 16.1316582 | 16.1375056 |
| O94915 | FRYL | 16.291136 | 15.9126039 | 16.4951174 | 16.5960905 | 16.60707 |
| P09211 | GSTP1 | 15.5786302 | 14.9411466 | 15.502452 | 15.0100497 | 15.1387373 |
| P61204 | ARF3 | 17.9850732 | 17.6624249 | 17.8168963 | 17.8539006 | 17.7484143 |
| Q13151 | HNRNPA0 | 14.3307703 | 14.6631546 | 13.8207268 | 14.3064489 | 14.4590824 |
| P07910 | HNRNPC | 18.3346112 | 19.01114 | 18.2368934 | 18.5184468 | 18.3095877 |
| Q9Y6Q3 | ZFP37 | 18.9603652 | 18.7546376 | 18.9415368 | 18.9734005 | 18.5462553 |
| Q8IWC1 | MAP7D3 | 13.5867836 | 14.2400222 | 14.2483839 | 14.2999811 | 13.9823115 |
| P26641 | EEF1G | 18.978896 | 18.7464706 | 18.5739609 | 18.7142434 | 18.6644844 |
| O75190 | DNAJB6 | 17.0093976 | 17.488635 | 17.4469487 | 17.748008 | 17.5052678 |
| P31153 | MAT2A | 15.2107902 | 15.4166454 | 14.9255657 | 15.0173954 | 15.2553205 |
| Q9Y3C6 | PPIL1 | 12.7279445 | 13.727542 | 13.838149 | 13.6702827 | 13.6839807 |
| O00483 | NDUFA4 | 17.6207869 | 17.4418152 | 17.6908442 | 18.0511198 | 17.5594255 |
| Q99729 | HNRNPAB | 15.8525378 | 16.4295989 | 15.3204242 | 15.8411423 | 15.4446429 |
| P84090 | ERH | 15.6239089 | 16.4636747 | 15.4444736 | 15.8960486 | 15.6858052 |
| P51153 | RAB13 | 16.8799955 | 16.5759674 | 16.547566 | 16.6618036 | 16.5956297 |
| P55209 | NAP1L1 | 14.2931397 | 13.939831 | 14.0866567 | 14.4448354 | 14.38263 |
| P15880 | RPS2 | 19.8899914 | 20.0586659 | 19.815553 | 19.8095671 | 19.6675313 |
| Q86U86 | PBRM1 | 15.9656628 | 15.9850807 | 16.4203116 | 16.3997032 | 16.3665125 |
| P07437 | TUBB | 19.7074152 | 19.9026358 | 19.7191612 | 19.897357 | 19.7093108 |
| P05023 | ATP1A1 | 17.0888243 | 16.8405986 | 16.5064869 | 16.8636007 | 16.7135271 |
| Q9UNL2 | SSR3 | 15.5226564 | 15.7194858 | 15.5626373 | 15.407988 | 15.6341299 |
| P27635 | RPL10 | 20.1779021 | 20.2994144 | 20.4163866 | 20.419319 | 20.1909722 |
| Q5M775 | SPECC1 | 15.6876005 | 15.3599611 | 15.3748653 | 15.344496 | 15.1954745 |
| Q9NTJ3 | SMC4 | 13.7413157 | 14.5264045 | 14.0662252 | 14.0553853 | 13.6207032 |
| P62263 | RPS14 | 21.4676372 | 21.3881374 | 21.4917312 | 21.3808469 | 21.2900073 |

| | | | | | | |
|---|---|---|---|---|---|---|
| P35613 | BSG | 11.7753358 | 11.786753 | 11.802602 | 11.9996943 | 11.9523053 |
| P37108 | SRP14 | 15.9124157 | 16.1450337 | 15.6643876 | 16.1449715 | 15.7717992 |
| Q9P0J0 | NDUFA13 | 13.119201 | 13.6613607 | 13.2885845 | 13.477399 | 13.147913 |
| Q15645 | TRIP13 | 12.8626981 | 13.4061077 | 13.0421781 | 13.3897224 | 12.5680732 |
| Q5T749 | KPRP | 17.120915 | 18.9396993 | 17.2009847 | 16.2790495 | 17.2021867 |
| Q15758 | SLC1A5 | 15.1507625 | 15.3502798 | 15.0519085 | 14.6150595 | 14.8813207 |
| Q9NX20 | MRPL16 | 16.2526726 | 16.2235073 | 16.2898711 | 16.4053917 | 16.2978874 |
| P16615 | ATP2A2 | 17.2022823 | 17.1026463 | 16.9780202 | 17.0208047 | 16.7474191 |
| P04080 | CSTB | 17.9173822 | 14.6918975 | 13.7584599 | 13.6433479 | 14.0550979 |
| P68363 | TUBA1B | 21.6228139 | 21.8637607 | 21.6302684 | 21.8583436 | 21.6482288 |
| O60716 | CTNND1 | 16.0093417 | 16.2770114 | 16.5856507 | 16.5553538 | 16.4731689 |
| P15311 | EZR | 13.7041461 | 13.3922604 | 12.9379988 | 12.0278182 | 13.2290067 |
| Q9H000 | MKRN2 | 11.4187926 | 11.711743 | 11.8128575 | 11.5460947 | 11.7931367 |
| O75475 | PSIP1 | 14.7546543 | 14.7560253 | 14.2677775 | 13.9961071 | 13.9045047 |
| P62491 | RAB11A | 17.6767656 | 17.2712084 | 17.2506884 | 17.2869193 | 17.6376368 |
| P62847 | RPS24 | 20.9505003 | 20.9729165 | 20.7266049 | 20.7260721 | 20.4724837 |
| P40616 | ARL1 | 13.5140782 | 13.9394069 | 13.7476634 | 13.7567109 | 13.7276104 |
| P46781 | RPS9 | 19.4234463 | 19.8378744 | 19.4235835 | 19.5403961 | 19.2019011 |
| P35998 | PSMC2 | 14.0755706 | 14.2642074 | 13.5591548 | 13.9105278 | 13.7072184 |
| Q8WUU5 | GATAD1 | 14.8611848 | 14.5550966 | 13.8040024 | 14.8431489 | 14.6034748 |
| O43242 | PSMD3 | 17.1104933 | 16.5618289 | 16.374512 | 16.7238067 | 16.642844 |
| Q9Y679 | AUP1 | 14.6274042 | 14.5942828 | 14.524923 | 14.0892626 | 15.0444687 |
| Q92616 | GCN1 | 13.9179623 | 13.6411719 | 13.4126596 | 13.3507115 | 13.5300526 |
| Q99615 | DNAJC7 | 15.530898 | 15.3755615 | 15.089676 | 15.3156715 | 15.2149247 |
| P61927 | RPL37 | 19.2247667 | 19.2737402 | 19.6967468 | 19.346869 | 19.1834674 |
| Q96GY0 | ZC2HC1A | 15.1989823 | 15.107817 | 15.2930788 | 15.5675381 | 15.0109219 |
| Q8IXM6 | NRM | 15.5291049 | 15.5520781 | 15.8090565 | 15.9694614 | 15.6461922 |
| P83731 | RPL24 | 19.5441947 | 19.7369486 | 19.7684954 | 19.7466734 | 19.4901099 |
| P25311 | AZGP1 | 14.875814 | 15.7547742 | 15.8190256 | 15.0161713 | 15.1058995 |
| P50395 | GDI2 | 12.0269091 | 11.5637938 | 11.4768913 | 11.250457 | 12.0629574 |
| P26599 | PTBP1 | 14.1668212 | 14.7724814 | 13.8316973 | 14.4350238 | 14.4315605 |
| Q14571 | ITPR2 | 18.4093304 | 18.1426971 | 19.0720185 | 19.2404092 | 19.1555051 |
| P0DMV8 | HSPA1A | 22.8628155 | 22.7373905 | 23.0184337 | 23.2187882 | 23.1745462 |
| P10599 | TXN | 16.9768194 | 16.5791381 | 17.0479495 | 17.0612877 | 16.7098197 |
| P46777 | RPL5 | 18.8564785 | 18.8018639 | 18.7686138 | 18.9620192 | 18.7860445 |
| O14980 | XPO1 | 11.6466294 | 11.8519112 | 11.6515653 | 11.4645508 | 11.5506141 |
| O00571 | DDX3X | 18.291828 | 17.9914569 | 18.082869 | 18.302854 | 18.3272774 |
| P51572 | BCAP31 | 18.2620558 | 17.5950981 | 17.4230302 | 17.95177 | 17.9992369 |
| P52292 | KPNA2 | 17.571569 | 17.6661484 | 17.5206543 | 17.8265788 | 17.3565273 |
| P18621 | RPL17 | 18.6576612 | 19.0662337 | 18.7644237 | 18.8290371 | 18.610823 |
| P33993 | MCM7 | 11.9262145 | 11.9473332 | 11.2508387 | 11.508827 | 11.8438215 |
| Q15369 | ELOC | 13.6732653 | 13.9569703 | 14.3789013 | 14.3264007 | 13.8734036 |

| | | | | | | |
|---|---|---|---|---|---|---|
| A0M8Q6 | IGLC7 | 18.5028305 | 18.4831727 | 19.0665494 | 19.0666628 | 18.8055738 |
| Q9BQE3 | TUBA1C | 17.0757888 | 17.5187909 | 17.4271108 | 17.3666439 | 17.2184557 |
| Q9Y5A9 | YTHDF2 | 14.4036286 | 15.3852501 | 14.4982391 | 14.426177 | 14.3324643 |
| P25205 | MCM3 | 12.4620173 | 12.9799073 | 13.2961875 | 13.5595434 | 13.4825726 |
| Q9NP72 | RAB18 | 14.1472431 | 14.0977391 | 14.1895261 | 13.9921405 | 13.8346307 |
| P39059 | COL15A1 | 17.8665937 | 17.7028972 | 17.7288949 | 17.6683591 | 17.9463365 |
| P62854 | RPS26 | 17.732971 | 18.471238 | 18.5451322 | 18.4924554 | 18.2292388 |
| Q8TF01 | PNISR | 16.2294002 | 15.9182895 | 16.5022674 | 16.4558619 | 16.461523 |
| Q96CX3 | ZNF501 | 11.6469701 | 11.3974562 | 11.3150825 | 11.9504173 | 11.3488981 |
| P34932 | HSPA4 | 14.2307888 | 13.8037152 | 13.2239189 | 13.2000766 | 13.6316332 |
| Q13263 | TRIM28 | 17.3778392 | 18.1980527 | 17.5098867 | 17.3834086 | 17.1753388 |
| Q15370 | ELOB | 15.5341708 | 16.0010663 | 16.1959667 | 15.8229355 | 16.1641509 |
| Q16186 | ADRM1 | 14.7616639 | 14.5016225 | 14.9219347 | 14.6360938 | 14.5810659 |
| Q9Y4W6 | AFG3L2 | 14.7106831 | 14.5698654 | 14.1898831 | 14.9017636 | 14.4156772 |
| Q6ZNB6 | NFXL1 | 11.2082838 | NA | 11.5430677 | 11.2441042 | NA |
| Q9P2N5 | RBM27 | 17.9506801 | 17.8423821 | 18.1981179 | 17.9949321 | 18.0378062 |
| P62753 | RPS6 | 19.8097613 | 20.3403425 | 19.9590901 | 20.0028598 | 19.8519244 |
| O60762 | DPM1 | 18.5363472 | 18.2557102 | 18.7286283 | 18.7905206 | 18.375981 |
| P84098 | RPL19 | 19.6959865 | 20.3507082 | 19.6856 | 19.7418388 | 19.4994796 |
| Q07021 | C1QBP | 16.4046178 | 16.1373984 | 15.512453 | 16.0746238 | 15.9818903 |
| Q9Y4W2 | LAS1L | 13.2961326 | 13.8525604 | 13.5603557 | 13.9946053 | 13.4473727 |
| P10412 | H1-4 | 19.9671969 | 20.5909684 | 19.5487123 | 19.6556656 | 19.5218794 |
| P24390 | KDELR1 | 16.7311712 | 16.7167697 | 16.303893 | 16.6538118 | 16.2308497 |
| Q09019 | DMWD | 15.3210392 | 15.2145109 | 15.2180441 | 15.2065725 | 14.4260961 |
| P62906 | RPL10A | 17.1536072 | 17.4781828 | 18.1540854 | 17.7833672 | 18.359217 |
| P42677 | RPS27 | 19.6984795 | 19.7188117 | 20.2238448 | 20.2544945 | 20.1521648 |
| P61353 | RPL27 | 18.6662793 | 18.777097 | 18.8625503 | 18.8934748 | 18.7435577 |
| Q13526 | PIN1 | 12.2058358 | 11.5869154 | 11.8266983 | 11.6014035 | 11.8167858 |
| P47914 | RPL29 | 16.7820997 | 17.0957088 | 16.8764901 | 17.0379565 | 16.5726324 |
| P22528 | SPRR1B | 17.7158028 | 19.2457732 | 17.2884857 | 16.2757704 | 16.9552361 |
| P68400 | CSNK2A1 | 14.4268985 | 14.5902679 | 14.4281963 | 14.5327868 | 14.3861707 |
| Q99832 | CCT7 | 13.8581623 | 13.4349559 | 12.9025412 | 13.499588 | 13.4693952 |
| P11142 | HSPA8 | 20.7951086 | 20.6134519 | 20.8822814 | 20.9912959 | 20.9340851 |
| Q9Y446 | PKP3 | 16.7735832 | 16.8483246 | 17.2486903 | 17.3723753 | 17.1111387 |
| Q12905 | ILF2 | 16.270655 | 16.838145 | 16.3699429 | 16.3256088 | 16.355324 |
| Q08211 | DHX9 | 14.8064129 | 15.7203138 | 15.3582681 | 15.0300625 | 15.3790955 |
| Q15382 | RHEB | 15.6661958 | 15.6897967 | 15.7886386 | 15.9436278 | 15.7275354 |
| P13010 | XRCC5 | 16.4504466 | 16.6962758 | 16.4578008 | 16.5658743 | 16.3556141 |
| Q9Y6X1 | SERP1 | 14.6142611 | 14.0737377 | 14.0860446 | 14.601636 | 14.1190537 |
| P78527 | PRKDC | 17.7139393 | 17.7629739 | 17.7379781 | 17.8461311 | 17.5482343 |
| Q9H0B6 | KLC2 | 15.457089 | 15.2332877 | 15.6692427 | 15.7181364 | 15.4401987 |
| O95197 | RTN3 | 13.7460257 | 13.8107064 | 13.6672611 | 14.0817486 | 13.5522658 |

| | | | | | | |
|---|---|---|---|---|---|---|
| O95071 | UBR5 | 13.5747711 | 13.6002103 | 13.30113 | 13.4470391 | 13.7481685 |
| Q69YL0 | NCBP2AS2 | 13.6149377 | 13.0113481 | 12.6858754 | 13.1898172 | 12.8684301 |
| P62277 | RPS13 | 20.473587 | 20.925387 | 20.7759726 | 20.7948294 | 20.6257023 |
| P68371 | TUBB4B | 19.84676 | 20.059153 | 20.0430381 | 20.1335903 | 20.0270496 |
| P61247 | RPS3A | 19.1207118 | 19.4941668 | 19.151215 | 19.1353116 | 19.0251249 |
| P62241 | RPS8 | 19.191676 | 19.7091402 | 19.4038434 | 19.5510736 | 19.2513627 |
| Q9BRJ6 | C7orf50 | 11.3552142 | 11.3862959 | 11.6601509 | 11.4812162 | 11.4656364 |
| Q06787 | FMR1 | 14.0818627 | 14.5069382 | 13.8978186 | 14.0560713 | 14.3467475 |
| Q9P107 | GMIP | 17.7771384 | 17.5043531 | 18.0400333 | 18.2272084 | 18.0549855 |
| P62899 | RPL31 | 19.4653426 | 19.7457097 | 19.4690528 | 19.4058379 | 19.0988743 |
| P22061 | PCMT1 | 19.6059712 | 19.4694526 | 20.023774 | 20.2344307 | 19.9217261 |
| Q8IXM3 | MRPL41 | 19.9747543 | 19.5149874 | 20.0664258 | 20.0599264 | 20.087278 |
| Q9NX63 | CHCHD3 | 16.0699715 | 16.573152 | 16.3136855 | 16.1453825 | 15.9518779 |
| Q6NXE6 | ARMC6 | 14.512832 | 14.1384858 | 14.5079318 | 13.636341 | 14.6835928 |
| Q9NYZ3 | GTSE1 | 17.1134235 | 17.1318174 | 17.6712243 | 17.6732411 | 17.6656964 |
| Q08554 | DSC1 | 18.5148762 | 19.0029079 | 18.7706449 | 18.2414586 | 18.2709431 |
| P53007 | SLC25A1 | 17.1987318 | 17.2779872 | 17.503255 | 17.5916876 | 17.3106217 |
| Q9P2E9 | RRBP1 | 18.4345656 | 18.561469 | 18.3397061 | 18.4241338 | 18.2842616 |
| P46776 | RPL27A | 19.3636992 | 19.7547144 | 19.5636146 | 19.4692856 | 19.1760113 |
| P32969 | RPL9 | 18.1583753 | 18.2824031 | 18.3051535 | 18.3371268 | 18.2471027 |
| P31689 | DNAJA1 | 17.1380045 | 17.6706834 | 17.1396134 | 17.489157 | 17.6161311 |
| P24539 | ATP5PB | 13.5730677 | 13.9106365 | 13.61472 | 13.8405794 | 13.3483842 |
| O60347 | TBC1D12 | 23.2023825 | 22.7620068 | 23.4868348 | 23.6827196 | 23.4337447 |
| Q71UM5 | RPS27L | 17.1474964 | 16.9673563 | 17.5221378 | 17.7159569 | 17.3731955 |
| Q7Z2W4 | ZC3HAV1 | 15.8245908 | 15.6968267 | 15.8897966 | 15.7431893 | 15.8621251 |
| P18077 | RPL35A | 19.3633545 | 19.4824344 | 19.8317548 | 19.9516704 | 19.4494631 |
| P29692 | EEF1D | 15.4349665 | 14.9825342 | 15.0362469 | 15.4534564 | 15.5394523 |
| P16402 | H1-3 | 14.7736743 | 15.5902969 | 14.6962241 | 15.0649063 | 14.6212069 |
| O00767 | SCD | 14.4623078 | 14.8477134 | 14.7282009 | 14.142565 | 14.6981247 |
| P48729 | CSNK1A1 | 18.7101363 | 18.34413 | 18.9766528 | 19.177399 | 19.0361057 |
| Q9Y5B6 | PAXBP1 | 12.0468575 | 12.1191012 | 12.5185784 | 12.6944873 | 12.4548801 |
| P14678 | SNRPB | 16.706108 | 16.904116 | 17.1244402 | 16.743344 | 16.9309373 |
| Q9Y6D6 | ARFGEF1 | 14.7359739 | 14.7220018 | 14.8295144 | 14.7128673 | 14.7296642 |
| P50991 | CCT4 | 14.9410838 | 14.9627298 | 14.2892728 | 14.8294727 | 14.7528236 |
| P11586 | MTHFD1 | 15.0861028 | 15.1963371 | 14.9523878 | 14.9615733 | 15.0588619 |
| P62979 | RPS27A | 17.9922086 | 18.0217 | 17.8510107 | 18.1345734 | 17.9153069 |
| Q9Y4E5 | ZNF451 | 16.0815675 | 16.1956965 | 16.6877648 | 16.2297066 | 15.9963296 |
| O60637 | TSPAN3 | 15.635733 | 15.5334286 | 15.7231425 | 15.3063647 | 15.2738019 |
| Q9Y230 | RUVBL2 | 15.1941442 | 15.7146677 | 15.0796586 | 15.310818 | 15.0286378 |
| Q96B49 | TOMM6 | 11.2573426 | 11.4631703 | 11.8447342 | 11.8064651 | 11.5662329 |
| P61313 | RPL15 | 19.2453654 | 19.5225739 | 19.3809515 | 19.6086081 | 19.1313152 |
| P23528 | CFL1 | 17.6446422 | 17.3568863 | 17.6578396 | 17.8146505 | 17.6570005 |

| | | | | | | |
|---|---|---|---|---|---|---|
| P21333 | FLNA | 16.8301399 | 17.4365383 | 17.1215465 | 17.3652639 | 17.2143467 |
| P52272 | HNRNPM | 19.2972739 | 19.8423347 | 19.3379173 | 19.5427497 | 19.5236265 |
| Q13148 | TARDBP | 12.6359409 | 12.7769884 | 12.6119757 | 12.9142736 | 13.0034338 |
| Q14980 | NUMA1 | 13.4556338 | 13.9975269 | 13.0881263 | 13.6750701 | 13.2180029 |
| P67870 | CSNK2B | 13.7491312 | 13.8298369 | 14.1049677 | 14.1532176 | 14.3930772 |
| Q14677 | CLINT1 | 15.3291326 | 15.6150259 | 15.0898433 | 15.5617234 | 15.5726396 |
| Q7Z6B7 | SRGAP1 | 13.7801819 | 13.9154329 | 14.2215186 | 13.9551256 | 14.2118484 |
| P22234 | PAICS | 19.2838063 | 19.0120774 | 19.5564171 | 19.6848039 | 19.4742736 |
| P51659 | HSD17B4 | 13.5785742 | 14.0963042 | 13.9580387 | 13.804083 | 13.6806923 |
| Q02790 | FKBP4 | 16.9561257 | 16.8422027 | 17.0020464 | 16.6778825 | 17.1439778 |
| P18583 | SON | 13.6067144 | 14.3181537 | 13.7055256 | 13.6781153 | 14.1623643 |
| P39023 | RPL3 | 20.162955 | 20.7790718 | 20.3858105 | 20.6563286 | 20.1496604 |
| P07948 | LYN | 14.892987 | 14.9506025 | 15.4030694 | 15.4867396 | 15.1901887 |
| Q9Y2R0 | COA3 | 15.3090301 | 15.1541066 | 15.7011979 | 15.6191587 | 15.7849614 |
| P56270 | MAZ | 16.8888169 | 16.4886964 | 17.1017793 | 16.797213 | 16.9679454 |
| P06748 | NPM1 | 19.6974914 | 20.3413136 | 19.7216338 | 19.6727799 | 19.5074406 |
| Q8N1F7 | NUP93 | 13.6455954 | 14.0128034 | 14.2279639 | 14.2220108 | 14.2464592 |
| Q5VUJ6 | LRCH2 | 11.5564567 | 11.9131166 | 12.0765418 | 11.9045907 | 11.676489 |
| Q92900 | UPF1 | 12.3033149 | 12.9875979 | 12.3565853 | 12.804896 | 12.785012 |
| Q9BXK1 | KLF16 | 13.1609023 | 11.9958753 | 12.4061349 | 12.6183908 | 12.103505 |
| P16989 | YBX3 | 14.6697671 | 15.6548037 | 15.1627558 | 15.3659047 | 15.110215 |
| O95816 | BAG2 | 16.9017421 | 17.8042287 | 17.2206418 | 17.5644584 | 17.0211187 |
| P46779 | RPL28 | 21.4285799 | 21.2433579 | 21.6449825 | 21.6595578 | 21.3978194 |
| Q15517 | CDSN | 15.2177379 | 15.9249288 | 15.2276958 | 15.2618556 | 14.7784713 |
| Q9NR30 | DDX21 | 15.0037999 | 16.0545247 | 15.4656944 | 15.0659678 | 15.0010712 |
| Q5T0W9 | FAM83B | 20.201604 | 19.7854997 | 20.5830985 | 20.8169463 | 20.7316 |
| P61163 | ACTR1A | 11.6467645 | 11.4837173 | 11.5021589 | 12.1096865 | 11.7255841 |
| O43663 | PRC1 | 16.2658185 | 16.3110958 | 16.5539919 | 16.609108 | 16.4564195 |
| Q9BVP2 | GNL3 | 15.814599 | 16.1409175 | 16.3693774 | 16.2907608 | 16.1891133 |
| P62917 | RPL8 | 19.0270255 | 19.642465 | 19.3395258 | 19.4304948 | 19.1977432 |
| P53985 | SLC16A1 | 17.0066397 | 17.0665087 | 17.7170528 | 17.7717488 | 17.7488794 |
| P40429 | RPL13A | 19.1023448 | 19.5934534 | 19.3055937 | 19.5246652 | 19.1762916 |
| P39656 | DDOST | 15.2688038 | 15.2760847 | 15.0381795 | 14.7986409 | 15.2219235 |
| Q6PGN9 | PSRC1 | 14.5237318 | 14.8249421 | 15.2362761 | 14.8328407 | 15.0571359 |
| Q5JSZ5 | PRRC2B | 12.9037911 | 12.9439407 | 13.0234015 | 12.4762634 | 13.1497465 |
| Q9P032 | NDUFAF4 | 11.5984102 | 11.8210114 | 11.6979151 | 12.0603021 | 11.4805861 |
| P40938 | RFC3 | 13.814404 | 14.2024086 | 13.3900963 | 13.4527279 | 13.1607303 |
| Q14257 | RCN2 | 17.8183015 | 18.1349832 | 18.0473726 | 18.1492743 | 17.8829581 |
| Q9UF83 | NA | 16.7885726 | 16.190766 | 16.8871937 | 17.2062609 | 16.9049679 |
| P46778 | RPL21 | 18.7545291 | 19.1022569 | 19.2122314 | 19.0598078 | 18.8704804 |
| Q02878 | RPL6 | 20.2234257 | 20.9166565 | 20.5375407 | 20.6614736 | 20.3109585 |
| P08047 | SP1 | 16.1254819 | 16.2507912 | 16.9318883 | 16.2623415 | 16.7339371 |

| | | | | | | |
|---|---|---|---|---|---|---|
| Q9H0U4 | RAB1B | 17.4805871 | 17.0438615 | 17.0163431 | 17.0618643 | 17.4994359 |
| P36578 | RPL4 | 19.6584349 | 20.2694087 | 20.025111 | 20.1268046 | 19.6022191 |
| P62913 | RPL11 | 19.8326577 | 19.5211244 | 19.9267909 | 19.7922646 | 19.5422107 |
| P42167 | TMPO | 16.7484416 | 16.8801146 | 16.7955881 | 16.7633576 | 17.1033636 |
| Q15459 | SF3A1 | 13.3218073 | 12.6182014 | 13.1650291 | 12.3030739 | 13.6078536 |
| P51149 | RAB7A | 16.0725813 | 15.6292425 | 16.0108479 | 16.0516208 | 16.0002371 |
| O15511 | ARPC5 | 16.957815 | 16.8976932 | 18.4824367 | 17.7678492 | 17.5594185 |
| Q01469 | FABP5 | 17.1601246 | 16.95039 | 16.9116892 | 17.2613441 | 16.6996205 |
| Q9BUF5 | TUBB6 | 13.8439104 | 14.2921623 | 14.5649007 | 14.6971656 | 14.5395511 |
| P41252 | IARS1 | 15.2102125 | 15.6648636 | 15.1989599 | 15.4429826 | 15.0326463 |
| Q9BQG0 | MYBBP1A | 12.5868426 | 13.1887353 | 12.869925 | 12.8068938 | 12.4448445 |
| Q02543 | RPL18A | 16.778669 | 17.3779731 | 17.2163009 | 17.1553065 | 17.0750031 |
| P62140 | PPP1CB | 19.680268 | 19.3330495 | 20.2352576 | 20.2621722 | 20.1708117 |
| Q15046 | KARS1 | 14.9588147 | 15.6849027 | 14.8866503 | 15.2384039 | 14.7969578 |
| P49207 | RPL34 | 19.3375631 | 19.5974643 | 19.6999871 | 19.7539471 | 19.1480567 |
| P52597 | HNRNPF | 14.8605196 | 15.3298137 | 15.1788581 | 15.3465613 | 14.8793231 |
| Q92522 | H1-10 | 17.783047 | 18.0914777 | 17.6208483 | 18.4777743 | 18.1432124 |
| Q00059 | TFAM | 17.1459758 | 16.6295029 | 16.7823685 | 16.6440436 | 16.7086922 |
| P31930 | UQCRC1 | 12.86813 | 13.1790681 | NA | 13.693394 | 13.5680455 |
| P16403 | H1-2 | 18.9503669 | 19.9803633 | 18.7317576 | 18.7997253 | 18.5559677 |
| P62910 | RPL32 | 19.707162 | 20.147644 | 20.2236645 | 20.1653473 | 19.8803539 |
| P49792 | RANBP2 | 15.7816138 | 16.2396501 | 15.5290263 | 15.5370873 | 16.002777 |
| Q01130 | SRSF2 | 17.7325863 | 17.5916017 | 17.8336726 | 18.3635786 | 17.8060461 |
| P02042 | HBD | 10.844541 | 10.7119641 | 11.6192401 | 11.7252183 | 11.0722662 |
| Q99959 | PKP2 | 16.6478156 | 16.0722747 | 16.3478888 | 16.4855985 | 16.7077996 |
| P82912 | MRPS11 | 14.232915 | 14.62885 | 14.2911079 | 14.5285328 | 13.7806389 |
| P62424 | RPL7A | 19.27689 | 19.8805379 | 19.5184371 | 19.7191894 | 19.2520435 |
| P06493 | CDK1 | 15.6784693 | 16.046001 | 16.4678167 | 16.3977395 | 16.3169773 |
| Q92974 | ARHGEF2 | 14.8153709 | 14.9256005 | 15.1164987 | 14.9467086 | 14.9517779 |
| P42704 | LRPPRC | 14.2468896 | 14.3611345 | 14.2095567 | 14.059462 | 14.1668749 |
| P05387 | RPLP2 | 18.122261 | 18.7062764 | 18.3600751 | 18.5719044 | 18.1960915 |
| Q92485 | SMPDL3B | 14.6482941 | 15.15825 | 15.5639113 | 14.8101017 | 14.8933565 |
| O60884 | DNAJA2 | 14.0090478 | 14.2530299 | 13.7929417 | 14.1866136 | 14.2759453 |
| Q02413 | DSG1 | 18.433423 | 19.2119519 | 18.598088 | 17.8683724 | 17.9333155 |
| P41212 | ETV6 | 23.1424075 | 22.8428322 | 23.4591621 | 23.6984301 | 23.5031409 |
| Q15070 | OXA1L | 14.6607073 | 14.6169103 | 15.194369 | 14.8448518 | 14.7344088 |
| P62888 | RPL30 | 17.7015918 | 18.3200079 | 17.83538 | 18.0138278 | 17.7765992 |
| P0C0S5 | H2AZ1 | 14.5608757 | 15.6448643 | 14.5054767 | 14.3677474 | 14.3537483 |
| P08134 | RHOC | 11.7421689 | 11.9922915 | 11.7218367 | 11.6036826 | 11.5532897 |
| P50914 | RPL14 | 18.2529929 | 19.1175444 | 18.61884 | 18.7832314 | 18.3835975 |
| P62633 | CNBP | 15.1142649 | 14.4419437 | 15.0138945 | 15.0062232 | 15.1823396 |
| P06241 | FYN | 16.7844643 | 16.6232862 | 16.9492255 | 17.2837978 | 16.9006637 |

| | | | | | | |
|---|---|---|---|---|---|---|
| P30050 | RPL12 | 19.2232674 | 19.6563185 | 19.624357 | 19.7809915 | 19.3111654 |
| Q14683 | SMC1A | 15.7109895 | 15.9426006 | 15.4161822 | 15.9061159 | 15.4019685 |
| P05388 | RPLP0 | 18.0926623 | 18.932357 | 18.7604422 | 18.7359617 | 18.2879332 |
| O15027 | SEC16A | 17.8994639 | 17.9901544 | 18.1988726 | 18.2807101 | 18.3024983 |
| Q6PID8 | KLHDC10 | 13.4904463 | 13.7796718 | 14.1947411 | 13.2354814 | 14.1493943 |
| Q9Y3U8 | RPL36 | 14.8558192 | 15.7790213 | 15.2526945 | 15.4931092 | 15.2774664 |
| O14974 | PPP1R12A | 20.65907 | 20.3668793 | 21.2510692 | 21.3235949 | 21.2043146 |
| P35611 | ADD1 | 14.3437531 | 14.6897112 | 14.7756027 | 14.7748908 | 14.6340838 |
| O14908 | GIPC1 | 16.5605568 | 16.4480515 | 17.2997724 | 17.2351988 | 17.158062 |
| E7ERA6 | RNF223 | 17.8429251 | 16.6894124 | 17.8466911 | 18.266741 | 17.7263931 |
| P26373 | RPL13 | 21.447963 | 21.3124768 | 21.5331996 | 21.6733052 | 21.3488841 |
| Q9BWS9 | CHID1 | 11.2407318 | 11.5581723 | 11.6182635 | 11.350957 | 11.6495765 |
| Q13835 | PKP1 | 18.4413341 | 18.7611002 | 17.7931991 | 16.6786788 | 17.1359045 |
| P63208 | SKP1 | 14.1624096 | 14.3374539 | 13.9496449 | 14.271497 | 14.1517325 |
| P02545 | LMNA | 16.0873766 | 16.2789347 | 16.3917776 | 16.5988683 | 16.3235878 |
| Q9NV79 | PCMTD2 | 12.4759687 | 12.5716361 | 12.8678301 | 12.7242738 | 12.2020207 |
| Q16891 | IMMT | 15.4444206 | 15.6003078 | 15.0913789 | 15.5103537 | 15.5682751 |
| O94901 | SUN1 | 19.7649751 | 19.4497206 | 19.9913972 | 20.4463716 | 20.2362617 |
| Q9NVI7 | ATAD3A | 19.7677785 | 19.6281952 | 20.1364002 | 20.0885773 | 19.9719902 |
| P55735 | SEC13 | 16.2685968 | 16.6547844 | 16.8139432 | 16.7278621 | 17.1086582 |
| Q07020 | RPL18 | 19.2267566 | 19.6883328 | 19.4131048 | 19.6059157 | 19.2957637 |
| P18124 | RPL7 | 19.3475974 | 20.0333055 | 19.6464181 | 19.8376007 | 19.5015314 |
| O43854 | EDIL3 | 13.5626398 | 13.5744438 | 14.2663322 | 14.4718134 | 13.59691 |
| P19338 | NCL | 19.5785645 | 19.8054306 | 19.6985276 | 19.7405014 | 19.4006429 |
| O95168 | NDUFB4 | 13.7174634 | 12.7591125 | 13.5045808 | 13.6280103 | 13.5173789 |
| P12268 | IMPDH2 | 14.0808545 | 15.3758612 | 13.684498 | 13.9920706 | 13.6025338 |
| P14868 | DARS1 | 16.2645695 | 16.8314965 | 15.6192488 | 16.063312 | 15.2873781 |
| Q9BRK4 | LZTS2 | 10.1484359 | 10.9710083 | 11.5195515 | 11.3624226 | 11.6888502 |
| P04908 | NA | 18.2858844 | 20.4569416 | 19.0175228 | 18.5876008 | 18.7415043 |
| Q12800 | TFCP2 | 22.4834193 | 22.1512401 | 22.8364838 | 22.9917825 | 23.030657 |
| Q02978 | SLC25A11 | 14.7413785 | 15.1758875 | 14.9954414 | 14.9959333 | 15.0545881 |
| Q8IY67 | RAVER1 | 12.2084429 | 12.4403399 | 12.4563228 | 12.5848442 | 12.9444358 |
| Q14160 | SCRIB | 14.5788511 | 15.2148859 | 15.2879723 | 15.2763737 | 14.9922411 |
| Q9BVC6 | TMEM109 | 13.5014961 | 14.5890273 | 13.9410027 | 14.2256562 | 13.9650667 |
| Q9NNW5 | WDR6 | 13.6891456 | 13.6457409 | 14.278709 | 14.2454022 | 13.5486376 |
| Q15154 | PCM1 | 14.7595417 | 14.7924883 | 15.2115393 | 15.3334063 | 15.0459277 |
| P15924 | DSP | 21.0258705 | 21.7113328 | 21.2377146 | 20.729907 | 20.6999009 |
| P62805 | NA | 19.5060043 | 21.3439957 | 20.1616298 | 19.592108 | 19.7974591 |
| Q8NCP5 | ZBTB44 | 13.4255851 | 13.53984 | 13.6722281 | 13.8554225 | 14.4388361 |
| Q5T9A4 | ATAD3B | 17.9787935 | 18.1948482 | 18.3048815 | 18.2950468 | 18.2194378 |
| Q8WVK2 | SNRNP27 | 14.1652694 | 13.879064 | 14.6559065 | 14.7665725 | 14.4838209 |
| Q14643 | ITPR1 | 15.457716 | 14.229143 | 15.1777142 | NA | 14.9448399 |

| | | | | | | |
|---|---|---|---|---|---|---|
| P67809 | YBX1 | 16.7389162 | 17.2318649 | 16.4641481 | 17.2906938 | 16.8700922 |
| P60953 | CDC42 | 15.9132953 | 15.5147713 | 15.8375778 | 16.0347382 | 15.9419626 |
| Q92859 | NEO1 | 14.2038132 | 13.4477758 | 13.7339248 | 14.1652178 | 13.3719861 |
| Q9P0V3 | SH3BP4 | 13.661761 | 14.1337586 | 14.2643368 | 14.4985902 | 14.1898292 |
| O60603 | TLR2 | 14.7600409 | 14.9310835 | 14.625029 | 14.6855118 | 15.3408878 |
| Q8WXH0 | SYNE2 | 17.8806768 | 17.5925346 | 18.1669282 | 18.446936 | 18.2552499 |
| Q9NZI7 | UBP1 | 20.9499025 | 20.5930796 | 21.2314486 | 21.4141093 | 21.4286053 |
| P05386 | RPLP1 | 15.0137409 | 15.804438 | 15.4908992 | 15.6989392 | 15.1416653 |
| Q9Y448 | KNSTRN | 15.1163119 | 15.3637931 | 15.3017353 | 15.7655898 | 15.36316 |
| P22087 | FBL | 15.3763321 | 15.7059609 | 15.7792045 | 15.5181166 | 15.6475515 |
| Q96N67 | DOCK7 | 14.9437192 | 15.1075693 | 15.5487674 | 15.6422861 | 15.5676954 |
| Q03252 | LMNB2 | 20.0862074 | 19.7744096 | 20.2747922 | 20.6464342 | 20.5271362 |
| O43491 | EPB41L2 | 12.3646935 | 11.7871524 | 12.8992043 | 12.3861401 | 13.3121518 |
| Q12792 | TWF1 | 11.4228883 | 11.3369151 | 11.9309204 | 11.9441884 | 12.0586561 |
| P61962 | DCAF7 | 13.7463305 | 14.0335391 | 14.0178426 | 14.2144861 | 13.8940515 |
| Q86V48 | LUZP1 | 15.7718419 | 15.8402084 | 16.3728827 | 16.3400113 | 16.1380344 |
| Q969Z0 | TBRG4 | 12.8832015 | 12.5097699 | 12.6149697 | 12.2872888 | 13.0689019 |
| P67936 | TPM4 | 15.3356821 | 15.6780942 | 16.4328366 | 16.0667973 | 15.8478713 |
| P33176 | KIF5B | 16.7330895 | 16.6246399 | 16.7779362 | 16.9218464 | 16.7330199 |
| Q9UKD2 | MRTO4 | 13.9715253 | 14.212528 | 14.4514824 | 14.4860302 | 14.3553111 |
| Q9Y265 | RUVBL1 | 14.7805723 | 15.3671241 | 14.6189539 | 14.8347331 | 14.6820145 |
| Q96HS1 | PGAM5 | 15.7373875 | 16.1407835 | 16.2080083 | 16.2987907 | 16.1378916 |
| P13682 | ZNF35 | 13.0793392 | 12.6872936 | 13.4719139 | 13.7786404 | 13.573776 |
| Q9H2D6 | TRIOBP | 14.8263074 | 14.9110169 | 14.9954889 | 15.4693125 | 15.2931835 |
| O60814 | H2BC12 | 19.0261844 | 21.2119781 | 19.9024952 | 19.3999374 | 19.6097746 |
| P14923 | JUP | 19.2199108 | 19.9300589 | 19.2377528 | 18.6869928 | 18.5258642 |
| O95235 | KIF20A | 13.5641097 | 13.5355735 | 13.9673946 | 14.0695436 | 13.8002303 |
| Q9H3U1 | UNC45A | 12.317348 | 12.2860859 | 12.3069552 | 12.5537684 | 11.8868643 |
| O43707 | ACTN4 | 16.6554899 | 16.2952151 | 16.8829004 | 17.0962119 | 16.9500859 |
| Q6Y7W6 | GIGYF2 | 13.6534864 | 14.2573694 | 13.7528637 | 14.2797235 | 13.3702322 |
| Q05682 | CALD1 | 13.0471203 | 13.0451893 | 12.2335296 | 12.4489751 | 12.5070063 |
| P50402 | EMD | 15.4600868 | 15.7286718 | 15.5600414 | 15.6427041 | 15.8927655 |
| P81605 | DCD | 20.0783248 | 19.6043836 | 19.5073173 | 19.9012013 | 18.8011742 |
| P07814 | EPRS1 | 14.0497122 | 14.8414607 | 14.2933836 | 13.9362472 | 14.1810005 |
| A0A075B6H9 | IGLV4-69 | 15.3339876 | 16.5108345 | 15.8849809 | 15.4272284 | 14.8289504 |
| Q8NF91 | SYNE1 | 14.8804567 | 14.6500121 | 14.9501364 | 15.3732986 | 15.3513883 |
| P13591 | NCAM1 | 13.3223535 | 13.177866 | 14.1750113 | 13.8999835 | 13.6812583 |
| P42166 | TMPO | 12.4375282 | 12.7119766 | 12.9594803 | 12.5277024 | 13.0191305 |
| Q92954 | PRG4 | 19.4998762 | 17.720987 | 16.8672929 | 16.7160118 | 16.5660845 |
| O15061 | SYNM | 14.9452301 | 14.9308732 | 15.540535 | 15.550401 | 15.5446695 |
| P35222 | CTNNB1 | 16.2223 | 16.5309289 | 16.8430883 | 16.6214786 | 16.3230297 |
| Q96KE9 | BTBD6 | 15.6930817 | 15.8150002 | 16.1735887 | 16.1284728 | 16.4302844 |

| | | | | | | |
|---|---|---|---|---|---|---|
| Q9Y2F9 | BTBD3 | 18.6520994 | 18.2894868 | 19.1379575 | 19.1710031 | 19.1842885 |
| Q9UQE7 | SMC3 | 14.2226388 | 14.2332967 | 14.0925491 | 14.4040578 | 13.6455245 |
| O75531 | BANF1 | 16.5201391 | 17.1890805 | 17.0825752 | 17.3872401 | 17.7269127 |
| Q92747 | ARPC1A | 18.4964274 | 18.4636923 | 19.2754697 | 19.333742 | 18.8894841 |
| Q69YQ0 | SPECC1L | 16.028026 | 16.0336296 | 16.881048 | 16.8267865 | 16.7072852 |
| P0DP23 | CALM1 | 18.1134609 | 17.5301452 | 19.1152367 | 19.0350536 | 18.2205063 |
| Q9NZN5 | ARHGEF12 | 10.7474828 | 11.4822205 | 11.5233418 | 11.5536946 | 11.588765 |
| Q8N137 | CNTROB | 15.0880782 | 14.5388598 | 15.4083903 | 15.3331437 | 15.7089879 |
| Q9H0E2 | TOLLIP | 13.425511 | 13.7299067 | 14.0990456 | 14.6282911 | 14.9195039 |
| Q9ULV4 | CORO1C | 18.6357123 | 18.4243084 | 19.3826666 | 19.6785498 | 18.9927369 |
| Q09666 | AHNAK | 17.7282245 | 17.4454959 | 17.4440753 | 17.675324 | 17.2657463 |
| Q9NQX4 | MYO5C | 15.191763 | 15.5937076 | 16.1698202 | 15.7394311 | 16.0054426 |
| O15143 | ARPC1B | 17.7350063 | 17.561176 | 18.5322251 | 18.4621312 | 18.4134847 |
| Q7Z406 | MYH14 | 18.3489844 | 18.5427181 | 19.5103827 | 19.220418 | 18.8844803 |
| Q9UM54 | MYO6 | 17.514096 | 17.3897255 | 18.3316843 | 18.3779683 | 18.0873323 |
| P35221 | CTNNA1 | 14.5403391 | 14.3532597 | 14.7329878 | 14.5606112 | 14.5858906 |
| Q01082 | SPTBN1 | 16.6708961 | 16.5419707 | 17.1989201 | 17.3985652 | 17.0772642 |
| P04792 | HSPB1 | 16.0998852 | 15.5435483 | 14.3924036 | 14.4854218 | 14.2566375 |
| O75955 | FLOT1 | 11.5287084 | 11.6786104 | 12.776826 | 12.5861507 | 12.6822734 |
| O75369 | FLNB | 15.6404106 | 15.9078935 | 16.2431345 | 16.4621813 | 16.3044929 |
| P54886 | ALDH18A1 | 15.8990576 | 16.1423305 | 16.2285606 | 16.5243229 | 16.7148994 |
| Q00610 | CLTC | 17.9647395 | 18.0627542 | 18.1941865 | 18.3329712 | 18.1979752 |
| P68431 | NA | 18.2972364 | 19.3897566 | 18.4663911 | 18.6350344 | 18.4000434 |
| P59998 | ARPC4 | 18.8984522 | 18.9535306 | 19.8139345 | 19.9826701 | 19.4979191 |
| Q9HD67 | MYO10 | 15.15924 | 15.2355708 | 16.1436182 | 15.7122459 | 15.7611428 |
| P78347 | GTF2I | 15.1119493 | 15.1790675 | 15.3199573 | 15.6281192 | 15.263349 |
| P38432 | COIL | 14.0715689 | 14.3988818 | 15.1110201 | 14.5497325 | 14.4765936 |
| P01706 | IGLV2-11 | 19.0277624 | 18.9898609 | 19.7257931 | 19.8664798 | 19.7263946 |
| Q9BZF9 | UACA | 17.4951909 | 17.375792 | 18.1708906 | 17.9957125 | 17.9954217 |
| P04746 | AMY2A | 15.8546702 | 14.029647 | 14.3709309 | 14.5055666 | 13.7516406 |
| O14654 | IRS4 | 18.3517828 | 18.2255922 | 18.6982846 | 18.8628484 | 18.4832423 |
| P49006 | MARCKSL1 | 15.785783 | 15.2103741 | 15.260512 | 15.3226943 | 15.8850839 |
| Q9Y2J2 | EPB41L3 | 15.5249625 | 15.8083706 | 15.9658725 | 15.9969452 | 15.9061103 |
| P06753 | TPM3 | 17.8689382 | 17.98122 | 18.8024473 | 18.9313606 | 18.1236701 |
| O43169 | CYB5B | 13.3917543 | 13.3430865 | 13.9051049 | 13.8778399 | 13.9852304 |
| Q13813 | SPTAN1 | 16.8503055 | 16.55121 | 17.1316359 | 17.6034398 | 17.2325158 |
| P20700 | LMNB1 | 18.3890271 | 18.6037263 | 19.1801662 | 19.5041805 | 19.1614563 |
| P04637 | TP53 | 16.1411955 | 15.7287941 | 16.5865105 | 16.3454424 | 16.3269338 |
| P55084 | HADHB | 15.0290184 | 14.7696112 | 15.5888828 | 15.5987229 | 15.5670489 |
| P29992 | GNA11 | 14.9838829 | 14.7187447 | 15.3436644 | 15.3390403 | 14.8876487 |
| Q9P2J3 | KLHL9 | 13.1085781 | 13.7829047 | 13.3889142 | 13.8561067 | 13.7829779 |
| Q0ZGT2 | NEXN | 12.6553508 | 13.4065461 | 13.6125917 | 14.287764 | 13.9476305 |

| | | | | | | |
|---|---|---|---|---|---|---|
| Q6UWP8 | SBSN | 16.35824 | 16.7674193 | 16.2776122 | 15.8446213 | 15.5726921 |
| Q9H2C0 | GAN | 14.5393113 | 14.273098 | 15.136391 | 14.9449864 | 15.3425656 |
| Q8WWI1 | LMO7 | 12.8204924 | 13.2392312 | 13.8573643 | 13.6498096 | 13.4517616 |
| P01040 | CSTA | 16.6231768 | 17.1751671 | 17.0600748 | 15.8513908 | 15.654751 |
| P08754 | GNAI3 | 17.3636594 | 16.7495914 | 17.6880322 | 17.8656057 | 17.3435292 |
| P34741 | SDC2 | 11.3533006 | 11.4131973 | 11.5366202 | 11.7412295 | 11.5840166 |
| Q9BPX5 | ARPC5L | 18.0731191 | 18.0339231 | 19.088147 | 19.4913292 | 19.152086 |
| O95425 | SVIL | 18.4454743 | 18.2260322 | 19.1462067 | 19.198249 | 19.0927689 |
| P63218 | GNG5 | 17.1286619 | 16.7643568 | 17.3857856 | 17.552786 | 17.3696796 |
| Q562R1 | ACTBL2 | 15.4907126 | 15.9155201 | 16.7594858 | 16.3288763 | 16.9005117 |
| P28290 | ITPRID2 | 16.924811 | 16.8174904 | 17.8006014 | 17.8339806 | 17.5132279 |
| Q9UHG3 | PCYOX1 | 18.1842781 | 18.2024009 | 18.7462101 | 18.7002335 | 19.259952 |
| P28289 | TMOD1 | 15.6083032 | 15.6756874 | 16.4826947 | 16.6798646 | 16.5259898 |
| P40939 | HADHA | 19.0506022 | 18.7886011 | 19.5156468 | 19.7050182 | 19.2432889 |
| Q12965 | MYO1E | 14.1927423 | 14.2067286 | 15.3356097 | 15.2831197 | 14.7147029 |
| Q9BUP0 | EFHD1 | 17.505778 | 17.3585731 | 18.4292841 | 18.789568 | 18.3340937 |
| P48681 | NES | 16.3335988 | 16.5990728 | 17.6279653 | 17.6114552 | 17.6762938 |
| O15145 | ARPC3 | 19.5547439 | 19.6484225 | 20.6927501 | 20.7598709 | 20.573533 |
| Q16352 | INA | 18.8357476 | 19.0119366 | 19.8964431 | 20.0912439 | 19.9356916 |
| P07196 | NEFL | 19.7103893 | 19.6968061 | 20.7232555 | 20.9079546 | 20.9462479 |
| P47756 | CAPZB | 19.6457687 | 19.7036137 | 20.4674054 | 20.7290334 | 20.4699889 |
| Q9Y4I1 | MYO5A | 16.7120295 | 16.9028996 | 17.876692 | 17.8121071 | 17.6130261 |
| Q96C19 | EFHD2 | 17.2618226 | 16.8958966 | 17.5959729 | 18.0492654 | 17.7254713 |
| Q08431 | MFGE8 | 17.8543029 | 17.7564618 | 18.4638382 | 18.6344737 | 18.2609824 |
| P61160 | ACTR2 | 17.8848554 | 18.0219465 | 19.1613806 | 19.2511835 | 18.9659252 |
| P08670 | VIM | 25.1009324 | 24.8577067 | 26.1111119 | 26.3212441 | 26.2371247 |
| Q15149 | PLEC | 24.1743396 | 23.9486613 | 25.0877685 | 25.3200185 | 25.2950577 |
| Q14126 | DSG2 | 17.3221334 | 17.0059468 | 17.6818281 | 17.8927331 | 17.669691 |
| Q6UVY6 | MOXD1 | 13.1574636 | 13.4707032 | 13.7356236 | 14.1910207 | 13.7283535 |
| O15144 | ARPC2 | 18.1559966 | 18.12239 | 19.1847698 | 19.2230776 | 18.9637872 |
| Q9UPN3 | MACF1 | 14.0003764 | 14.1270805 | 14.0853451 | 14.4159976 | 14.1795157 |
| P60660 | MYL6 | 21.9163997 | 21.8231199 | 22.8417882 | 22.9714549 | 22.3583132 |
| P08174 | CD55 | 13.1876262 | 13.821811 | 13.7483082 | 13.6557561 | 13.5922565 |
| Q6WCQ1 | MPRIP | 19.7554864 | 19.6439373 | 20.8528839 | 20.6534217 | 20.6363661 |
| Q14573 | ITPR3 | 16.9163288 | 17.0869626 | 17.6422274 | 17.7991777 | 17.6941594 |
| Q96SB3 | PPP1R9B | 14.9384775 | 14.6115836 | 15.5197055 | 15.8734177 | 15.5809337 |
| P35580 | MYH10 | 21.9234514 | 22.0589311 | 23.3280511 | 23.035261 | 22.8232557 |
| Q9P0K7 | RAI14 | 19.8006587 | 19.8991515 | 20.9739306 | 20.8887863 | 20.8797565 |
| Q9UBI6 | GNG12 | 17.4100897 | 17.3082794 | 17.8117867 | 18.1657095 | 17.6731376 |
| P47755 | CAPZA2 | 18.6694785 | 18.7206041 | 19.5304538 | 19.7616625 | 19.5534437 |
| P63096 | GNAI1 | 16.7591902 | 16.7428595 | 17.4095644 | 17.4674074 | 17.1385139 |
| Q9UHB6 | LIMA1 | 20.5456574 | 20.4177283 | 21.488035 | 21.7364653 | 21.3996395 |

| | | | | | | |
|---|---|---|---|---|---|---|
| Q14247 | CTTN | 18.0708663 | 17.9093824 | 19.0537004 | 19.2649063 | 19.0666965 |
| P60709 | ACTB | 25.6428593 | 25.4127215 | 26.7267882 | 27.0334962 | 26.5709311 |
| O00159 | MYO1C | 18.5158245 | 18.4982834 | 19.6196394 | 19.5847809 | 19.3514137 |
| P35579 | MYH9 | 23.2451705 | 23.2877532 | 24.6915874 | 24.4136137 | 24.1953457 |
| O00161 | SNAP23 | 16.7174857 | 16.4530613 | 17.1970698 | 17.2737171 | 17.2299615 |
| O94832 | MYO1D | 17.0792589 | 16.8266065 | 18.124376 | 18.1169659 | 17.8822233 |
| Q9NYL9 | TMOD3 | 18.1806434 | 17.9784192 | 19.0864496 | 19.3324278 | 18.9672102 |
| P13797 | PLS3 | 13.191744 | 11.2039609 | 11.0528286 | 11.2184703 | 11.2626894 |
| P52907 | CAPZA1 | 19.9939474 | 19.9977826 | 21.0570484 | 21.2564289 | 20.9762426 |
| P61158 | ACTR3 | 18.6437065 | 18.5995524 | 19.8270939 | 20.0003073 | 19.72418 |
| Q16643 | DBN1 | 18.4885902 | 18.3003006 | 19.3446286 | 19.8418174 | 19.3204239 |
| P02810 | PRH2 | 18.9088584 | 16.6736317 | 15.9275557 | 17.0090579 | 15.8570904 |
| P63261 | ACTG1 | 19.6158194 | 19.5195735 | 20.6376205 | 20.859045 | 20.549383 |
| P13987 | CD59 | 16.4327445 | 15.6725937 | 16.777959 | 16.7652569 | 16.8729972 |
| O14950 | MYL12B | 20.883064 | 20.9668705 | 22.1534842 | 22.1715176 | 21.6631903 |
| Q9HAV0 | GNB4 | 16.5770255 | 15.9643626 | 16.8187687 | 16.9350122 | 16.6200343 |
| P63092 | GNAS | 16.9427175 | 16.762776 | 17.2599073 | 17.4099792 | 16.9819029 |
| A8MTJ3 | GNAT3 | 17.4147603 | 16.8983631 | 17.6726058 | 17.8722145 | 17.2344564 |
| P14649 | MYL6B | 14.3474451 | 14.5233332 | 15.7647628 | 15.8788251 | 15.4727059 |
| Q9ULJ8 | PPP1R9A | 13.1732827 | 13.8896011 | 13.7405354 | 14.0074499 | 13.924532 |
| P68032 | ACTC1 | 23.4964056 | 23.3258571 | 24.7008433 | 24.8834093 | 24.4776753 |
| Q9BYX7 | POTEKP | 14.7584326 | 14.6587857 | 15.9063389 | 15.9095636 | 15.6650022 |
| O43795 | MYO1B | 18.9076634 | 18.9367572 | 20.1413266 | 20.1131672 | 19.8998185 |
| Q9Y6X9 | MORC2 | 17.5583144 | 17.5135612 | 18.8095105 | 18.8546834 | 18.6132561 |
| P62879 | GNB2 | 16.6713045 | 16.386393 | 17.2954388 | 17.1676056 | 16.9365689 |
| P62873 | GNB1 | 19.8916076 | 19.4709442 | 20.2489005 | 20.4493791 | 20.1039466 |
| P80723 | BASP1 | 18.2102964 | 17.6244646 | 18.4753957 | 18.7813571 | 18.3040772 |
| P50151 | GNG10 | 14.075985 | 13.695577 | 14.5114432 | 14.7320299 | 14.2168604 |
| A0A075B6P5 | IGKV2-28 | 15.6379555 | 14.2765637 | 16.1819512 | 15.4159882 | 14.9661009 |

| AMPH-04647.10nM_3_abundance_norm | AMPH-04647.300nM_1_abundance_norm | AMPH-04647.300nM_2_abundance_norm | AMPH-04647.300nM_3_abundance_norm | AMPH-04647.1000nM_1_abundance_norm | AMPH-03340.1nM_1_abundance_norm | AMPH-03340.10nM_1_abundance_norm |
|---|---|---|---|---|---|---|
| 12.8741228 | 13.8009504 | 13.2638229 | 14.4533342 | 14.5591761 | 14.9455216 | 16.5760719 |
| 15.3907256 | 15.0151384 | 15.0245599 | 15.8804746 | 14.9149881 | 15.5570256 | 15.8998813 |
| 11.7667683 | 12.3289295 | 11.5748201 | 12.314791 | 12.8242659 | 12.7790425 | 12.0441861 |
| 17.1957535 | 16.6647558 | 16.5826707 | 17.0059469 | 16.8683123 | 15.0761455 | 16.6263631 |
| 18.2344718 | 17.4127859 | 17.3411988 | 17.4140408 | 17.760227 | 16.989058 | 17.2949106 |
| 15.6255035 | 15.2099629 | 13.936533 | 14.5907661 | 14.7422324 | 14.1339244 | 14.8357984 |
| 19.6367039 | 19.0977672 | 18.9517256 | 19.0514159 | 19.3479412 | 18.7252528 | 18.8515998 |
| 20.6675716 | 20.3718584 | 19.9734043 | 19.9769299 | 20.5664459 | 19.3433889 | 20.2460908 |
| 14.8243483 | 14.0970298 | 13.8345218 | 14.4805811 | 15.1766916 | 14.868042 | 15.2051183 |
| 15.7341061 | 16.5080848 | 16.658306 | 16.2275797 | 16.120807 | 16.363787 | 16.5980102 |
| 11.3137615 | 11.4704626 | 11.3967175 | 10.5692187 | 13.0487017 | 10.9788937 | 11.369502 |
| 14.4375656 | 13.3981347 | 13.1038331 | 13.017241 | 12.4834013 | 11.6777773 | 12.635709 |
| 17.2626625 | 16.8278421 | 16.9204447 | 16.9687238 | 17.2956939 | 16.4376784 | 16.9774968 |
| 14.7495373 | 15.1556328 | 15.2844858 | 14.0754446 | 15.6209228 | 15.2722713 | 14.9777709 |
| 12.605204 | 11.5797946 | 12.7301361 | 12.0856302 | 12.5330492 | 12.7478171 | 12.7716105 |
| 11.6995222 | 11.2621403 | 11.207191 | 10.9572939 | 11.4017169 | 11.9507821 | 11.394183 |
| 16.2914748 | 16.0369671 | 15.8592149 | 15.8405978 | 16.1447527 | 15.8786615 | 15.7957164 |
| 15.6867628 | 15.5864241 | 14.8568377 | 14.9748095 | 15.2347851 | 15.1423632 | 15.3722037 |
| 13.8607557 | 13.1868891 | 14.4449956 | 13.0232501 | 13.6791464 | 15.2456076 | 14.3357045 |
| 12.3376582 | 13.8328283 | 11.9440182 | 12.3272681 | 14.3313247 | 13.0827746 | 11.7806812 |
| 14.266474 | 13.7650891 | 14.2241256 | 14.367969 | 14.0368692 | 14.2554228 | 13.6484853 |
| 14.2807853 | 14.7759858 | 14.5544105 | 14.5921458 | NA | 14.0430076 | 14.977107 |
| 15.2397903 | 14.2710442 | 14.5764895 | 14.9069508 | NA | NA | NA |
| 15.8167084 | 15.9473946 | 16.0582481 | 15.5514744 | 16.2590913 | 17.3583159 | 16.1586831 |
| 13.4689173 | 12.2176691 | 12.3378485 | 12.323752 | NA | 13.4632679 | 12.5508518 |
| 14.2694704 | 13.8316884 | 14.1602606 | 13.0788766 | 13.551602 | 14.4595806 | 13.5613628 |
| 16.0753654 | 17.6481795 | 15.1513257 | 14.6174097 | 15.448245 | 15.0850489 | 15.4896543 |
| 17.1006404 | 16.7205918 | 16.6679415 | 16.9310057 | 16.3865594 | 15.9429939 | 16.8365307 |
| 18.3935358 | 18.6080649 | 18.9035547 | 18.8440514 | 19.1574019 | 19.2014833 | 18.9629065 |
| 17.326351 | 17.6125857 | 17.5909482 | 17.6934826 | 18.1184302 | 18.083057 | 17.5288327 |
| 15.5189323 | 15.2838674 | 15.4157725 | 15.2398817 | 15.4612475 | 15.5122078 | 15.5389079 |
| 15.3017081 | 14.7560838 | 15.4223531 | 15.0351307 | 14.6206418 | 15.2276559 | 14.4247822 |
| 19.5337247 | 19.9949093 | 19.7159633 | 19.7779103 | 20.4760282 | 20.0304687 | 19.4609096 |
| 20.0582454 | 20.375327 | 20.1213364 | 20.1527318 | 20.8946141 | 20.5277137 | 20.1012976 |
| 17.0666971 | 16.5078105 | 16.9728036 | 16.9205231 | 16.8183325 | 16.4808424 | 16.9255875 |
| 17.8355161 | 17.6057886 | 18.2578323 | 18.4362805 | 19.0518596 | 18.1952247 | 18.6173012 |

| | | | | | | |
|---|---|---|---|---|---|---|
| 17.0629776 | 17.2409716 | 17.1021834 | 17.2493101 | 17.6296518 | 17.268804 | 16.9752506 |
| 13.898812 | 14.0982785 | 14.1273257 | 14.0390897 | 14.6418936 | 14.3118451 | 14.3744071 |
| 14.6831228 | 15.0051611 | 14.7967098 | 14.8175408 | 15.0078882 | 14.8362145 | 14.90324 |
| 15.0608454 | 15.9688716 | 15.898312 | 16.7981187 | 16.9982568 | 15.4939341 | 15.0648378 |
| 17.6038854 | 17.832006 | 17.8077227 | 17.7752635 | 18.1817765 | 18.2970393 | 17.5270866 |
| 15.3487514 | 14.6824565 | 14.4787891 | 15.8297958 | 15.5938177 | 15.6670881 | 15.6472319 |
| 16.4982817 | 16.2141054 | 16.1077022 | 15.9224227 | 16.6526378 | 15.4393568 | 16.0676066 |
| 11.6713298 | 11.1816345 | 10.8790243 | 10.2969837 | 11.5079676 | 10.7848389 | 11.1464733 |
| 14.8131912 | 15.0925915 | 14.4881319 | 14.880068 | 15.1517948 | 15.0461716 | 14.3690854 |
| 18.6579969 | 18.8690458 | 18.9901187 | 18.9849262 | 19.5912378 | 19.0831773 | 18.8928614 |
| 12.4770627 | 12.6925342 | NA | NA | 13.1135959 | 12.8749051 | 12.1409992 |
| 15.8229753 | 16.0110614 | 15.999686 | 15.5860409 | 16.236767 | 15.9331575 | 15.7557719 |
| 18.3446593 | 17.8950562 | 17.9096528 | 17.8036946 | 18.2537409 | 17.6158022 | 17.79889 |
| 19.0102552 | 19.2170353 | 19.2749483 | 19.308289 | 20.0144674 | 19.514241 | 19.8411722 |
| 17.2836759 | 17.0034025 | 17.0928573 | 16.5037864 | 15.8150773 | 17.0829844 | 16.8297712 |
| 16.5255879 | 16.3563328 | 16.4066608 | 16.5961126 | 16.7224811 | 16.7930748 | 16.3460703 |
| 14.0573998 | 13.9332797 | 14.0357963 | 13.3424024 | 12.6580011 | 15.5324899 | 13.8362574 |
| 17.7496663 | 17.1863404 | 17.343171 | 17.4749922 | 17.8318274 | 17.002418 | 17.1813577 |
| 12.2726295 | 12.1406555 | 12.1632803 | 12.0048687 | 12.3225125 | 12.3603908 | 12.7769345 |
| 15.3612651 | 14.7682252 | 15.0065118 | 15.3554102 | 14.9885339 | 16.1858743 | 15.1070342 |
| 16.4613537 | 16.5968534 | 16.4852019 | 16.8386147 | 17.3652971 | 16.9272471 | 16.5062281 |
| 15.4110526 | 14.8388725 | 14.4168737 | 14.3016719 | 14.4828125 | 14.4217198 | 14.414081 |
| 17.672364 | 17.8898617 | 17.9548274 | 17.7722467 | 18.2709388 | 18.2090723 | 17.9284682 |
| 17.6289594 | 17.8762402 | 17.7495488 | 17.8242179 | 18.3664344 | 18.1710508 | 17.6745416 |
| 18.4232856 | 18.2284493 | 17.850034 | 17.7716677 | 18.27919 | 18.6704631 | 17.8018187 |
| 19.7447113 | 20.0879715 | 20.050636 | 20.125069 | 20.7152247 | 20.2091647 | 19.8664435 |
| 15.2509169 | 15.2467531 | 15.4425867 | 15.5668015 | 15.0039493 | 15.5742289 | 15.6232377 |
| 17.5271023 | 17.5546771 | 17.3613112 | 17.3499395 | 17.9746454 | 17.8523724 | 17.3469605 |
| 17.5149374 | 17.5649248 | 17.5476389 | 17.6427172 | 18.2624879 | 17.7566944 | 17.55868 |
| 13.8501455 | 13.4931617 | 13.6732111 | 13.3993987 | 14.3693204 | 14.1668605 | 13.6875111 |
| 15.9760565 | 15.9713358 | 15.692005 | 15.3380131 | 15.6725782 | 16.0186844 | 15.8403503 |
| 15.5968388 | 15.2214755 | 14.8135087 | 14.8730356 | 14.9122059 | 14.953617 | 14.8619146 |
| 20.4102811 | 20.8019129 | 20.7735788 | 20.5709916 | 21.3215886 | 20.7035852 | 20.4931685 |
| 14.2684752 | 14.4045525 | 14.011163 | 13.6307439 | 13.7194146 | 14.7570712 | 13.8158229 |
| 17.076595 | 16.7814351 | 16.9243387 | 16.8902242 | 17.3447232 | 17.5528518 | 16.9442776 |
| 20.0692543 | 19.77547 | 19.825476 | 19.8366921 | 20.6575587 | 19.9593453 | 20.1053334 |
| 18.7647999 | 18.2022074 | 18.4490278 | 18.412993 | 18.705877 | 18.0098641 | 18.0555006 |
| 11.8237351 | 11.0405693 | 11.4588453 | 10.665061 | 11.5401876 | 12.4286957 | 11.3007136 |
| 14.0975861 | 14.1421538 | 13.6491 | 12.951387 | 13.5317679 | 13.5888886 | 13.5551762 |
| 14.0662149 | 14.0129228 | 13.7228562 | 13.9742452 | 14.4716189 | 14.3540743 | 14.0862591 |
| 15.7619435 | 15.1888304 | 15.3111807 | 15.2998635 | 15.6636871 | 15.8153178 | 15.0538774 |
| 12.9727568 | 13.0247698 | 12.5739464 | 12.5524784 | 13.1258914 | 13.2874189 | 12.5928794 |

| | | | | | | |
|---|---|---|---|---|---|---|
| 14.7371287 | 14.5441532 | 14.6896956 | 14.2070224 | 14.6797504 | 15.8465709 | 14.6374182 |
| 17.3819385 | 17.577683 | 17.808388 | 17.8229076 | 18.5372729 | 18.2617084 | 17.9702822 |
| 12.9805489 | NA | 13.2304353 | 12.4733894 | 13.1741376 | 14.0575394 | 13.1548649 |
| 18.8257239 | 18.7886296 | 19.3207201 | 19.0139178 | 19.3979541 | 20.9090766 | 19.0644921 |
| 16.9110258 | 16.6190393 | 17.2927845 | 16.7015841 | 16.8473997 | 18.8458011 | 17.1081807 |
| 18.0572818 | 17.8414737 | 18.1343147 | 18.1451625 | 17.9052529 | 18.4215147 | 18.4227243 |
| 17.6977042 | 17.5237073 | 17.5199518 | 17.2405111 | 17.6375689 | 17.7289505 | 17.1610183 |
| 18.4172852 | 18.6839874 | 18.6466773 | 18.5333113 | 19.3092662 | 18.7438493 | 18.3890236 |
| 15.6521552 | 15.3913876 | 15.3530144 | 15.5968558 | 15.932088 | 15.6009474 | 15.4632952 |
| 15.5142302 | 15.9954165 | 15.9045209 | 16.0706866 | 16.4639832 | 16.6897574 | 16.2177448 |
| 14.6103369 | 14.3511841 | 14.8922317 | 14.6223 | 14.9472147 | 14.165766 | 14.6511203 |
| 15.5757919 | 15.3344105 | 15.4379337 | 15.5919326 | 15.9250227 | 15.1073136 | 15.6671055 |
| 14.3066626 | 14.7456343 | 14.3654071 | 14.1828925 | 14.9561622 | 14.6417251 | 14.5933892 |
| 14.9187505 | 15.0143558 | 14.6909791 | 14.7317979 | 15.000506 | 14.8308708 | 14.7070108 |
| 19.4753396 | 19.0959445 | 19.1154097 | 18.8928203 | 19.4763595 | 19.7978368 | 18.9604199 |
| 12.787496 | 13.5436122 | 13.3687909 | 13.5998045 | 13.6827442 | 13.1519204 | 13.1957318 |
| 14.9507671 | 15.2070371 | 15.4606645 | 14.618749 | 18.3950638 | 15.7945698 | 15.0777779 |
| 14.9796248 | 14.8540196 | 14.4728343 | 14.1515063 | 14.5883047 | 15.2369961 | 14.28461 |
| 18.6653185 | 18.5098969 | 18.9435509 | 18.8892935 | 19.2286361 | 19.4541237 | 18.8823544 |
| 16.7532258 | 16.9205128 | 16.5953464 | 17.0849188 | 17.4272678 | 17.4097929 | 16.8897507 |
| 13.0804884 | 14.1065808 | 13.9206979 | 13.6393022 | 14.2684539 | 13.1032911 | 14.1722763 |
| 18.0572635 | 18.2387805 | 18.1739384 | 18.0819553 | 19.2043482 | 18.5505803 | 18.2869266 |
| 14.4519521 | 14.748865 | 14.7437578 | 14.7373472 | 14.9896141 | 15.6889464 | 14.8199729 |
| 17.7885353 | 18.0630364 | 17.9326098 | 17.9295454 | 18.5251145 | 18.1471176 | 17.9285973 |
| 15.6992845 | 15.7800811 | 16.1134758 | 15.8406729 | 15.9326521 | 16.6635034 | 15.9423605 |
| 14.0998702 | 13.4926591 | 13.9351585 | 13.9036104 | 13.6956812 | 14.3901426 | 13.7745428 |
| 11.9181955 | 11.7575697 | 11.1406064 | 10.8057636 | 11.6427836 | 11.8044887 | 12.0064677 |
| 16.7536303 | 16.8493644 | 16.8149651 | 16.8546034 | 17.1257055 | 17.459354 | 16.8738236 |
| 14.1845356 | 13.9492943 | 14.0712049 | 13.5194362 | 14.1622461 | 14.0085296 | 14.1405648 |
| 19.4091531 | 19.4139105 | 19.7902638 | 19.6686433 | 20.3822759 | 19.9966152 | 19.4987281 |
| 14.0699087 | 13.9085615 | 14.087157 | 13.9331367 | 15.3900217 | 14.3710139 | 14.0752422 |
| 15.460451 | 15.4797211 | 15.8232898 | 15.6310438 | 15.9535897 | 15.5798689 | 15.4055097 |
| 15.5483488 | 16.0568106 | 16.1166802 | 16.0005423 | 16.2399042 | 15.9149682 | 15.8173338 |
| 14.0867226 | 13.7235725 | 14.00484 | 13.872848 | 14.3651334 | 14.3982768 | 13.6539106 |
| 17.9357969 | 17.7036662 | 17.6789626 | 17.6147251 | 18.0644117 | 17.9998765 | 17.4219363 |
| 20.3417775 | 20.497073 | 20.6461203 | 20.6074138 | 21.1651497 | 20.6335268 | 20.7744706 |
| 21.9833094 | 22.138365 | 22.5860987 | 22.4725544 | 22.8022853 | 22.788707 | 22.4438924 |
| 18.4452813 | 18.2680301 | 18.5749165 | 18.5551488 | 19.0571772 | 19.0950341 | 18.3512828 |
| 14.8107678 | 15.1925159 | 15.448367 | 15.5487984 | 15.9088354 | 15.5186178 | 15.1218856 |
| 17.4778531 | 17.6299983 | 17.4238072 | 16.8219339 | 17.0450996 | 18.4306646 | 17.0672844 |
| 14.9823007 | 15.3815445 | 14.5856151 | 15.3814064 | 15.7663639 | 15.1864882 | 14.9306831 |
| 18.6438131 | 18.0904665 | 18.0115763 | 18.3170667 | 18.3062234 | 18.1530693 | 18.1732066 |

| | | | | | | |
|---|---|---|---|---|---|---|
| 15.7944652 | 15.5582415 | 15.5784821 | 15.8023941 | 16.0396926 | 15.5946209 | 15.7286572 |
| 19.4399586 | 19.120042 | 19.1940839 | 19.0429667 | 19.6512998 | 19.3735057 | 19.2109461 |
| 12.9055388 | 12.9433062 | 12.99618 | 13.1598458 | 13.3406864 | 11.9943089 | 13.1968607 |
| 15.3570079 | 15.4229817 | 15.4889698 | 15.5437011 | 15.4997572 | 15.4348976 | 15.433573 |
| 16.7916547 | 17.0010703 | 17.2604202 | 16.9248259 | 17.7791122 | 17.848315 | 17.1166924 |
| 11.4090029 | 11.464926 | 11.6570632 | 10.8310157 | 12.075454 | 12.0190717 | 11.7091045 |
| 13.5911125 | 13.2677746 | 13.4099312 | 13.5940846 | 13.7342102 | 14.3310024 | 13.6809789 |
| 15.1311758 | 14.9689829 | 14.3080316 | 14.6839845 | 15.0316309 | 15.4389211 | 15.0474434 |
| 13.8750956 | 13.6257262 | 14.8175803 | 14.5450111 | 15.536189 | 14.2070468 | 14.9667568 |
| 17.2719169 | 17.0054296 | 17.1475991 | 17.0266787 | 17.0737281 | 17.3444683 | 17.2647849 |
| 16.5971159 | 17.0014389 | 16.8047535 | 16.4748126 | 17.2109502 | 17.1802485 | 16.3849788 |
| 18.2430299 | 18.5066308 | 18.5168038 | 18.3708233 | 18.4596794 | 19.065665 | 18.3480281 |
| 15.4141636 | 15.7586308 | 15.7757251 | 15.8446306 | 15.8968463 | 16.3242735 | 15.6982058 |
| 15.2135882 | 15.2241722 | 15.2439474 | 15.3009847 | 15.6481945 | 15.4642818 | 15.3380923 |
| 16.3946173 | 16.5645109 | 16.6873781 | 16.7406138 | 17.0598139 | 17.2323029 | 16.680173 |
| 16.1890634 | 16.56487 | 16.5192421 | 16.4523327 | 17.0995071 | 16.9683064 | 16.7990473 |
| 14.8339035 | 14.3792697 | 14.3335709 | 14.0607717 | 14.853967 | 14.2356433 | 13.9888711 |
| 16.0400382 | 15.8024038 | 16.3059381 | 15.9139224 | 16.5468755 | 17.0136599 | 16.2955095 |
| 14.3672132 | 14.1086738 | 14.4675711 | 14.311559 | 14.094754 | 14.7363825 | 14.2557757 |
| 11.6821132 | 11.3029509 | 11.5471588 | 11.5219256 | 11.959128 | 12.1495771 | 11.6677562 |
| 15.2642279 | 14.9590843 | 15.3026043 | 14.823151 | 14.9049265 | 16.2293988 | 14.8651201 |
| 16.0918166 | 15.9585755 | 16.0326827 | 15.9357688 | 16.6006771 | 16.2922091 | 15.749147 |
| 18.1212288 | 18.0094726 | 18.1245122 | 17.9142441 | 18.204296 | 18.6228876 | 17.9282181 |
| 17.35969 | 17.396035 | 17.2182156 | 17.1179059 | 17.5894716 | 17.6664302 | 17.1493167 |
| 16.2751804 | 16.2552656 | 16.4098763 | 16.2384824 | 16.7755505 | 16.9342019 | 16.1971776 |
| 16.7563686 | 17.359173 | 16.6883851 | 16.8102875 | 17.1947158 | 16.6722769 | 17.0409398 |
| 18.0332985 | 18.071581 | 18.0284561 | 17.7529424 | 18.203256 | 17.951854 | 17.8655004 |
| 16.6660926 | 16.9238777 | 17.1197667 | 17.2432436 | 17.1708435 | 17.1608527 | 16.7461388 |
| 13.9392678 | 13.8625475 | 14.8153164 | 14.9690789 | 16.6431944 | 14.9710205 | 14.8877651 |
| 17.5778147 | 17.2442444 | 17.4831172 | 17.261097 | 17.7845757 | 17.725854 | 17.1513768 |
| 17.7731795 | 18.1406772 | 18.2816903 | 18.2512501 | 18.9400595 | 18.6401375 | 18.4470211 |
| 13.0681722 | 13.1202756 | 13.38904 | 12.5628681 | 13.4654881 | 14.8608699 | 13.0303892 |
| 17.6014172 | 17.3432769 | 18.0138866 | 17.3804194 | 18.4842393 | 18.0414847 | 17.5908259 |
| 16.7516245 | 17.0276057 | 17.1646642 | 17.2629296 | 17.7270836 | 17.3424259 | 17.0407389 |
| 15.7639078 | 15.4216249 | 15.5424604 | 14.7607723 | 15.2286236 | 15.5856164 | 15.0729226 |
| 16.8965852 | 17.2669331 | 17.1430656 | 16.9618166 | 17.5323542 | 17.604393 | 17.1127979 |
| 15.104907 | 15.1568037 | 15.287103 | 14.710768 | 15.6472495 | 15.762638 | 15.4545926 |
| 15.2702306 | 15.1965565 | 15.1567251 | 14.8463008 | 15.0311966 | 14.8657564 | 14.6360088 |
| 13.7503817 | 13.6624153 | 13.5634769 | 13.9156204 | 13.5285114 | 15.4226484 | 13.9445486 |
| 13.9138485 | 13.4673576 | 13.3044177 | 13.6736904 | 13.6804917 | 13.8761575 | 13.9268904 |
| 14.5834955 | 14.2928217 | 14.2370132 | 13.4120139 | 13.8276078 | 14.7036626 | 14.3322446 |
| 16.4998771 | 16.9600892 | 16.1001428 | 15.7966585 | 16.3693308 | 15.6927963 | 16.3379791 |

| | | | | | | |
|---|---|---|---|---|---|---|
| 17.6747506 | 17.9902085 | 18.0759865 | 17.8207531 | 18.1949928 | 18.7894527 | 17.9040386 |
| 19.1475608 | 19.3866624 | 19.3555986 | 19.3787194 | 19.503573 | 19.4420282 | 19.6592657 |
| 18.2958759 | 18.2399623 | 18.256638 | 17.9453947 | 18.5806374 | 18.0559944 | 18.2519087 |
| NA | 11.4142764 | 11.9281816 | NA | 11.328044 | 11.4998662 | 11.3988562 |
| 18.9554455 | 18.64113 | 18.9224174 | 18.6705481 | 19.055936 | 18.9491761 | 18.6640154 |
| 17.7577086 | 17.5877231 | 17.9541587 | 17.6218284 | 17.8073072 | 18.9131552 | 17.6047056 |
| 15.0552186 | 14.7607602 | 14.7845653 | 14.5515343 | 13.8903221 | 15.2625637 | 14.584551 |
| 14.439028 | 14.3372839 | 14.9011413 | 14.3950246 | 14.358085 | 15.6994494 | 14.5300332 |
| 16.4797887 | 16.3537727 | 16.6283709 | 16.0776637 | 16.6579623 | 16.8239356 | 16.2384794 |
| 14.7082127 | 15.5393095 | 16.0080004 | 14.9486527 | 16.5008104 | 15.6733229 | 14.915754 |
| 14.418042 | 14.7827118 | 14.371101 | 14.4843191 | 14.8244302 | 14.8902085 | 14.4095863 |
| 12.8894694 | 12.5167655 | 12.9590955 | 12.8005708 | 13.041042 | 12.232739 | 12.2516717 |
| 15.3223959 | 15.3905086 | 14.984878 | 14.8727753 | 15.779922 | 14.9318042 | 15.4366531 |
| 13.501136 | 14.2294606 | 14.5553299 | 14.4138406 | 14.937059 | 14.4301045 | 14.3258028 |
| 11.8990731 | 11.588144 | 12.4128814 | 11.8119295 | 11.6864175 | 12.5540041 | 11.1022959 |
| 20.0390862 | 20.1422704 | 20.2035352 | 20.1041325 | 20.6050453 | 20.5849182 | 20.0959494 |
| 18.1357558 | 18.1513093 | 18.3435585 | 17.7877905 | 18.162794 | 18.9614202 | 17.8967659 |
| 14.4567262 | 14.5163864 | 14.6228536 | 14.7241617 | 15.1054409 | 14.3886045 | 14.9478664 |
| 17.0041465 | 17.0523413 | 17.0401514 | 17.1708331 | 17.0043299 | 17.8740412 | 17.1328724 |
| 20.4344821 | 20.5829736 | 20.7152936 | 20.9497359 | 21.1341853 | 21.2258655 | 20.4110292 |
| 18.061514 | 17.5491638 | 17.8832443 | 17.9102901 | 17.9477479 | 17.5889 | 17.7686502 |
| 16.5856685 | 16.7000347 | 16.8046033 | 16.7317528 | 18.1472066 | 16.8258069 | 16.0565865 |
| 14.3507577 | 14.6510095 | 14.756447 | 14.3751244 | 14.1593484 | 15.5354467 | 14.0060336 |
| 17.0321339 | 17.3756069 | 17.5799861 | 17.4839008 | 18.1300552 | 17.5305348 | 17.3920901 |
| 19.1912295 | 19.3803519 | 19.5563669 | 19.5967998 | 20.1849151 | 19.9698223 | 19.7333659 |
| 15.5573888 | 14.9085776 | 14.8243094 | 14.897851 | 14.8829915 | 14.8144747 | 14.8433094 |
| 14.9008073 | 14.8394339 | 14.6753556 | 14.9769129 | 14.7089136 | 15.3644939 | 15.2020266 |
| 16.9262736 | 16.5857582 | 17.0102707 | 17.1523559 | 16.6741375 | 17.7268123 | 16.4708686 |
| 13.4145833 | 13.3510245 | 13.1639589 | 12.872026 | 13.162812 | 14.8176267 | 13.6841809 |
| 16.5498903 | 17.0139438 | 16.7490532 | 16.5919415 | 17.8901992 | 16.8717121 | 16.8424706 |
| 15.9731805 | 16.655644 | 16.3131497 | 16.2501095 | 16.6247658 | 16.751644 | 16.4365909 |
| 13.3025989 | 13.7929957 | 13.7388802 | 13.8245681 | 14.325546 | 13.2249233 | 14.1789989 |
| 14.092049 | 13.7796375 | 13.9223778 | 13.8041854 | 13.8409225 | 14.5669549 | 13.531009 |
| 17.953223 | 17.9730636 | 18.2077472 | 17.9520885 | 18.5234497 | 18.6767917 | 18.1505946 |
| 14.5769284 | 15.9537383 | 14.0226642 | 13.967419 | 14.9755267 | 14.4828017 | 14.3436332 |
| 14.8603085 | 14.8632792 | 14.7932772 | 14.9806371 | 14.6896162 | 15.6917666 | 14.7844199 |
| 16.8475081 | 16.9000412 | 17.2469114 | 16.961673 | 17.7686286 | 17.8465345 | 16.7572925 |
| 20.0113154 | 20.0392914 | 20.1576055 | 19.9482322 | 20.4495654 | 20.4318184 | 19.9287187 |
| 15.4764489 | 15.4208401 | 15.7866681 | 15.5892165 | 16.0266154 | 15.7886451 | 15.8248073 |
| 14.1807467 | 13.8785128 | 14.0386967 | 13.8095739 | 14.0604905 | 14.4258201 | 13.8689457 |
| 21.89368 | 21.8785203 | 22.1250774 | 21.9469767 | 22.5565653 | 22.463681 | 21.909026 |
| 14.7938248 | 14.7592769 | 14.9110724 | 14.4057102 | 14.7168418 | 15.3409859 | 14.4001839 |

| | | | | | | |
|---|---|---|---|---|---|---|
| 15.2452554 | 16.1308208 | 17.0501133 | 17.3354156 | 17.4132252 | 15.9685311 | 16.6353847 |
| 14.5477381 | 13.9151666 | 12.969894 | 13.8015814 | 13.5028179 | 13.3340079 | 13.9766494 |
| 21.3716197 | 21.3248952 | 21.8774506 | 21.7663436 | 21.0638486 | 22.2281966 | 21.6836322 |
| 14.9839207 | 14.9137486 | 15.2362465 | 15.0579097 | 15.7683726 | 15.1591663 | 15.1054279 |
| 14.5427994 | 14.0260421 | 13.8156422 | 13.9096314 | 14.8921341 | 15.0677962 | 14.0025026 |
| 12.0529764 | 11.9197595 | 11.8017636 | 11.6932457 | 12.3530423 | 12.1627028 | 11.4939958 |
| 14.8288711 | 15.2853286 | 15.3815931 | 15.3157938 | 15.7266917 | 15.4090576 | 15.2941973 |
| 14.8484981 | 15.0049085 | 14.6276131 | 14.6502887 | 14.8852915 | 15.2995406 | 15.6891197 |
| 19.115337 | 19.014657 | 19.1654879 | 18.9141265 | 19.4034215 | 19.4049251 | 19.0196413 |
| 11.7529524 | 11.3251441 | 11.3916396 | 11.5595995 | 11.5551971 | 11.4435562 | 11.7486845 |
| 13.668919 | 13.2814547 | 13.6503908 | 12.9664378 | 14.0751498 | 13.4979169 | 13.6652611 |
| 20.0876122 | 20.3211359 | 20.265323 | 20.329571 | 20.8235652 | 20.7879344 | 20.1721002 |
| 19.9012849 | 19.9029281 | 20.1161627 | 20.2610858 | 20.6285294 | 20.261121 | 20.1940716 |
| 17.1303966 | 16.6074825 | 16.8717107 | 16.3195692 | 16.7397524 | 15.8306883 | 16.9576905 |
| 14.5277361 | 14.2568042 | 13.9361392 | 13.9633955 | 13.2349656 | 14.6143855 | 14.0325312 |
| 16.8220226 | 16.8358741 | 16.3449897 | 16.7970179 | 17.6227037 | 17.3580505 | 16.4856264 |
| 15.2341544 | 15.3095122 | 15.3718031 | 15.1407836 | 15.4942103 | 16.2181468 | 15.2144226 |
| 14.1959997 | 15.1974955 | 13.8835054 | 13.7292924 | 14.9682857 | 13.531742 | 14.1422121 |
| 20.5757873 | 20.6549329 | 20.6910719 | 20.7288006 | 21.2874627 | 21.2293745 | 20.9283727 |
| 20.2593878 | 20.4023334 | 20.352437 | 20.2885266 | 20.8802767 | 20.8296813 | 20.6354301 |
| 17.4243431 | 17.552571 | 17.7278318 | 17.5787508 | 17.9777221 | 17.8854781 | 17.7566841 |
| 20.8675341 | 21.094071 | 21.1815341 | 21.1035008 | 21.9407434 | 21.4365767 | 21.1119089 |
| 11.0498252 | 11.6161117 | 12.0416933 | 11.0550169 | 11.3577795 | 12.302619 | 11.5875465 |
| 19.2585855 | 18.8791231 | 19.0390562 | 18.9414981 | 19.1263839 | 19.6817996 | 18.8292894 |
| 17.9635302 | 17.8713112 | 18.4322574 | 17.8696821 | 17.9205749 | 19.0124803 | 17.9675732 |
| 16.4899982 | 16.1469546 | 16.5788499 | 16.2266701 | 16.7356097 | 16.6357727 | 16.3246874 |
| 12.6236643 | 12.7455188 | 12.7445025 | 12.5800446 | 12.4267215 | 12.5127414 | 12.5069488 |
| 21.5573876 | 21.821116 | 22.0278895 | 21.9064803 | 22.8736164 | 22.2877534 | 22.0024027 |
| 14.9038748 | 15.3319982 | 14.7000824 | 14.9649967 | 15.4844448 | 15.522359 | 15.1124212 |
| 24.0824491 | 23.7009663 | 25.603966 | 24.9610303 | 26.3585665 | 24.0676426 | 25.1389447 |
| 13.9764756 | 14.2095763 | 14.5551488 | 14.3570652 | 14.9921988 | 15.055342 | 14.2278038 |
| 15.0724671 | 14.9727104 | 14.7206535 | 15.0151265 | 14.5940924 | 15.5682059 | 14.9584166 |
| 14.4211661 | 14.1892582 | 14.2133272 | 13.7844693 | 14.5387163 | 15.1894159 | 13.4506655 |
| 21.7624255 | 22.1876897 | 22.3130014 | 22.3077917 | 23.2315554 | 22.4351869 | 22.1879713 |
| 16.4250713 | 16.4583386 | 16.6464742 | 16.2088469 | 16.8139578 | 17.2445915 | 16.3585969 |
| 19.8510627 | 19.9944414 | 20.1656257 | 20.2508868 | 20.4306658 | 20.7352428 | 19.9245207 |
| 19.9144369 | 20.0138631 | 19.9668405 | 20.0431525 | 20.5510838 | 20.5524716 | 20.2408164 |
| 14.982575 | 14.940617 | 15.1546235 | 15.3027121 | 15.6285093 | 15.1316879 | 15.2617638 |
| 17.8541843 | 17.5488832 | 17.5388215 | 17.5559786 | 17.4145386 | 17.7455851 | 17.3901153 |
| 13.6817054 | 13.9995608 | 13.7559344 | 13.7753883 | 14.1064613 | 14.5803465 | 13.580743 |
| 14.6631415 | 14.7978018 | 15.0386247 | 14.7504254 | 15.5477008 | 15.57889 | 14.9185209 |
| 12.9142063 | 12.552831 | 13.4215865 | 13.3565664 | 13.8543762 | 12.6150948 | 13.4823622 |

| | | | | | | |
|---|---|---|---|---|---|---|
| 13.435817 | 13.8048948 | 14.027979 | 13.5601152 | 14.5401513 | 13.2939495 | 13.4024096 |
| 11.1368733 | 11.1194955 | 11.9544461 | 11.4725814 | 11.8840786 | 11.1803877 | 12.1694208 |
| 15.9724282 | 15.8972116 | 16.0206615 | 15.8495103 | 15.4810915 | 15.768243 | 15.3947028 |
| 15.7262079 | 15.4351362 | 15.3564546 | 14.6205715 | 15.1841541 | 14.7474442 | 14.8113692 |
| 13.4966139 | 12.7339779 | 13.1231478 | 13.430708 | 12.7170588 | 13.4243568 | 12.7500272 |
| 20.9045805 | 21.1326023 | 21.2316424 | 21.1858267 | 21.752897 | 21.6230685 | 21.1454545 |
| 15.1216849 | 14.8599096 | 15.2971331 | 14.8443752 | 15.2407677 | 15.4907994 | 15.4687551 |
| 14.6200751 | 14.8510723 | 15.3548543 | 15.470247 | 15.4582986 | 15.5399617 | 14.863756 |
| 18.8712992 | 18.9698169 | 19.0591762 | 19.156519 | 19.6006769 | 19.4587438 | 19.2433408 |
| 20.9721422 | 21.1162702 | 21.0721365 | 21.0216016 | 21.5403594 | 21.6189542 | 21.2791575 |
| 14.825224 | 14.8360909 | 14.7184411 | 14.7967038 | 15.6069312 | 14.8560414 | 14.7487002 |
| 15.6890151 | 15.888533 | 15.5536256 | 15.870318 | 15.8834295 | 16.0474749 | 15.4701245 |
| 13.0038882 | 13.4511664 | 12.4605709 | 12.9512429 | 13.9523626 | 13.2332751 | 12.773004 |
| 16.3619808 | 14.5146526 | 15.5032001 | 15.890347 | 16.2740385 | 16.2812859 | 15.8738648 |
| 21.3464962 | 21.4253213 | 21.6846298 | 21.4541092 | 21.932046 | 22.0746792 | 21.5157476 |
| 16.5684976 | 16.682694 | 16.6452532 | 16.3407079 | 16.5723831 | 16.4134604 | 16.2612267 |
| 18.9312717 | 18.7971835 | 18.9713023 | 18.777815 | 18.7842714 | 19.6277375 | 18.9468372 |
| 19.8160373 | 19.7352174 | 20.0304481 | 20.1074473 | 20.4270789 | 20.4840041 | 19.8663069 |
| 14.8413962 | 14.9728627 | 14.2409508 | 15.2999705 | 15.3716522 | 14.5484243 | 14.327952 |
| 13.9023523 | 14.1782632 | 14.2914149 | 13.7644355 | 14.6087981 | 14.2431103 | 14.1101405 |
| 21.2808243 | 21.4260141 | 21.6958238 | 21.4730292 | 22.090731 | 21.8769788 | 21.5244386 |
| 13.9706253 | 13.8875812 | 13.6777008 | 14.1047825 | 13.7977348 | 14.3467409 | 13.7257753 |
| 15.2933069 | 15.5932827 | 15.8026922 | 15.577549 | 15.6994454 | 15.6119858 | 15.6251071 |
| 16.0214992 | 15.7491197 | 16.1116838 | 15.8148802 | 16.5112971 | 16.3602696 | 15.7338799 |
| 15.8463179 | 15.9623821 | 15.6008869 | 15.1474826 | 15.848147 | 16.0784715 | 15.6662299 |
| 18.6693288 | 18.9188214 | 19.0043946 | 18.9085919 | 19.2332357 | 19.3982212 | 19.0833227 |
| 21.8049153 | 21.8002441 | 21.9673508 | 21.6892197 | 22.4209628 | 22.1858604 | 21.9284671 |
| 18.8063404 | 19.1387795 | 19.506753 | 19.482947 | 19.8349807 | 19.3744718 | 19.3931636 |
| 20.6208636 | 20.7780732 | 20.690364 | 20.6936724 | 21.2910425 | 21.2212336 | 20.9934311 |
| 14.6887115 | 14.7276046 | 14.9072412 | 14.769306 | 15.12389 | 15.966098 | 14.9201012 |
| 14.1385852 | NA | 15.0596482 | 13.8122362 | 14.4881595 | NA | 13.4914877 |
| 11.8344384 | 11.4436349 | 11.6726112 | 12.031719 | 11.2532898 | 13.0192049 | 11.5730177 |
| 19.2673234 | 19.5292879 | 19.7801547 | 19.3907954 | 19.7856983 | 20.1696281 | 19.7097081 |
| 16.8676521 | 16.9453517 | 16.7769827 | 17.0961433 | 17.318784 | 17.1804873 | 16.7647674 |
| 17.4704176 | 17.5043363 | 17.3515757 | 17.1620107 | 17.6551646 | 16.8506112 | 17.5276653 |
| 18.4756943 | 18.7076955 | 18.5977875 | 18.5640816 | 19.0460907 | 18.8367782 | 18.4870476 |
| 13.4924475 | 13.5516355 | 13.2115739 | 13.4823147 | 13.2660714 | 13.5534188 | 13.7187552 |
| 16.5003731 | 16.9313756 | 17.2175471 | 17.3103528 | 18.0170154 | 17.1339992 | 17.2056037 |
| 14.6229535 | 14.2973446 | 14.3911619 | 14.3260425 | 14.5962599 | 14.6930359 | 14.4792679 |
| 19.0310585 | 18.7502727 | 18.7737484 | 18.6745064 | 18.6353902 | 19.0104462 | 18.6221175 |
| 17.2223113 | 17.1936965 | 17.1258198 | 16.9538437 | 17.4323758 | 17.6426253 | 17.199578 |
| 14.7032913 | 14.7238926 | 14.3745145 | 14.5843213 | 14.6813837 | 14.8965037 | 14.4511783 |

| | | | | | | |
|---|---|---|---|---|---|---|
| 19.9228809 | 20.2089807 | 20.1035967 | 20.1862431 | 20.5271667 | 20.1903321 | 20.3425549 |
| 16.1277486 | 16.1481026 | 16.680184 | 16.0186024 | 16.2597802 | 17.7485337 | 16.1636132 |
| 14.8210756 | 14.7965113 | 13.7746306 | 13.9031279 | 14.5792998 | 14.8658258 | 13.9235251 |
| 13.9884421 | 13.7001208 | 13.6426065 | 13.2498392 | 13.1893033 | 14.0105772 | 13.6860493 |
| 18.7153731 | 18.732615 | 19.1595157 | 19.0833026 | 19.0366527 | 19.3974815 | 19.0458487 |
| 18.1131092 | 18.0202032 | 18.0923961 | 18.0271844 | 18.1146748 | 18.746162 | 18.0289216 |
| 15.163094 | 15.2475975 | 15.198874 | 14.9368357 | 15.197321 | 15.3319019 | 15.0806366 |
| 16.3010011 | 16.3830918 | 17.0262062 | 16.6692263 | 17.0190619 | 16.5257003 | 16.7337034 |
| 20.9532193 | 21.0590718 | 21.1368838 | 21.1989661 | 21.6769209 | 21.5894787 | 21.1132379 |
| 16.2044689 | 16.3654547 | 16.2969721 | 16.222451 | 16.5806602 | 15.9613017 | 15.7360462 |
| 15.4147981 | 15.5961381 | 14.9761449 | 15.4762298 | 15.4950764 | 15.5194579 | 15.4708865 |
| 14.3936463 | 13.7053722 | 13.8216838 | 13.8621759 | 14.2265234 | 14.7338766 | 13.6687323 |
| 14.656849 | 14.2031846 | 14.914885 | 14.9005841 | 15.2496842 | 14.8153747 | 14.3898069 |
| 12.2804853 | 12.6371376 | 12.6282095 | 12.504803 | 12.5768354 | 13.1181344 | 12.368155 |
| 15.2329122 | 15.1657505 | 15.5576825 | 15.1948915 | 15.4348677 | 16.3966448 | 15.217893 |
| 16.151771 | 15.2200448 | 16.1586679 | 14.6587084 | 15.9317434 | 15.1668552 | 15.4172061 |
| 14.4798617 | 14.2155856 | 14.9789723 | 14.643443 | 15.5853939 | 15.0934356 | 14.8368999 |
| 17.4547248 | 17.5541543 | 17.7326283 | 17.2923949 | 17.5008662 | 17.4349867 | 17.3365802 |
| 14.6076213 | 14.6731101 | 14.7532487 | 13.9834501 | 13.7706344 | 16.0065235 | 14.8151004 |
| 13.6654723 | 13.8546894 | 14.325432 | 13.7661377 | 13.8739118 | 14.0749341 | 14.8211029 |
| 22.5508736 | 22.8730965 | 22.7541881 | 22.8128133 | 23.3777207 | 23.3558471 | 22.804455 |
| 17.7697173 | 17.6499367 | 17.7550614 | 17.7617901 | 17.9353653 | 18.151865 | 17.6760402 |
| 18.8340333 | 19.0110865 | 18.6422006 | 18.8647319 | 19.4176219 | 19.4947455 | 18.9014644 |
| 11.4097083 | 12.0352032 | 11.3334944 | 10.9644137 | 11.6941498 | 11.9028893 | 11.6858521 |
| 18.2431642 | 18.45783 | 18.8359884 | 18.4119429 | 18.6563329 | 19.0338036 | 18.4724565 |
| 20.3051629 | 20.6828881 | 20.6558085 | 20.7014831 | 21.0147742 | 20.9722187 | 20.5889766 |
| 15.6559382 | 14.7531542 | 17.5603715 | 14.8144167 | 16.0693576 | 13.1641448 | 15.0950022 |
| 19.5467813 | 19.8207005 | 19.687807 | 19.4697955 | 20.0221258 | 19.9011499 | 19.4686719 |
| 16.3957603 | 16.4832964 | 16.7181583 | 16.6785341 | 17.1013399 | 16.7283271 | 16.9030635 |
| 21.5029508 | 21.6233675 | 21.6095727 | 21.6939356 | 21.8069714 | 22.072089 | 21.5057579 |
| 15.0194421 | 15.3194272 | 15.0982322 | 15.2667284 | 15.7949294 | 15.4117322 | 15.3595335 |
| 18.7207904 | 18.4675402 | 18.3781651 | 18.4611647 | 18.63218 | 19.0725184 | 18.4241778 |
| 15.4527727 | 15.4421658 | 15.317204 | 15.3094732 | 15.7214161 | 16.3062685 | 15.815037 |
| 15.1912801 | 14.8635571 | 14.7416026 | 14.6525127 | 15.0453547 | 15.9829207 | 15.134379 |
| 17.4508791 | 17.1223359 | 17.3771477 | 17.344015 | 17.4886091 | 17.8539116 | 17.2072132 |
| 14.9568701 | 15.3996885 | 15.1539548 | 15.1405445 | 15.5825124 | 15.3113072 | 15.7619036 |
| 15.2756448 | 15.440742 | 15.4031946 | 15.4868652 | 15.8601216 | 16.266252 | 15.4729619 |
| 16.4960855 | 18.0744962 | 16.0901187 | 15.883667 | 16.8048204 | 16.0759363 | 16.5249783 |
| 14.446438 | 14.5881747 | 14.7543904 | 14.6013338 | 14.8879471 | 14.9476907 | 14.7674094 |
| 16.7901797 | 16.8980948 | 17.4348669 | 16.7166632 | 17.1612926 | 17.8696842 | 16.9443027 |
| 11.6583911 | 11.8844537 | 11.3271598 | 11.4701213 | 12.0078982 | 12.0491363 | 11.8610823 |
| 17.029688 | 16.8368185 | 16.9536396 | 17.0211693 | 17.148014 | 17.4059628 | 17.0211044 |

| | | | | | | |
|---|---|---|---|---|---|---|
| 16.5480927 | 16.7802677 | 16.6050864 | 16.5972111 | 16.7729914 | 17.170311 | 16.5304998 |
| 13.4140331 | 13.9927006 | 13.1083151 | 13.2084666 | 13.663304 | 12.4191374 | 13.9041615 |
| 14.128439 | 13.7974279 | 13.9094493 | 13.7536201 | 13.6418601 | 14.8680127 | 14.083823 |
| 19.5306529 | 19.5982869 | 19.6001358 | 19.4612594 | 19.5251533 | 20.0597799 | 19.6155496 |
| 20.5213187 | 20.6546898 | 20.700451 | 20.5570595 | 20.4540783 | 21.1500254 | 20.4258342 |
| 16.8475424 | 17.1085518 | 17.353537 | 17.0257354 | 17.4528863 | 17.4852395 | 16.9046873 |
| 16.3962383 | 16.3858896 | 16.8052361 | 16.1217269 | 17.2387025 | 17.1317452 | 16.469272 |
| 18.4444574 | 18.4167711 | 18.3158811 | 18.3427123 | 18.565669 | 18.7404902 | 18.2789894 |
| 15.9248587 | 16.9680851 | 16.1137093 | 15.7781584 | 16.609266 | 15.3412935 | 15.9743389 |
| 17.2620027 | 17.1701981 | 17.3753497 | 17.3005505 | 17.8566325 | 17.8664315 | 17.2784612 |
| 15.1419451 | 15.1766298 | 15.8391248 | 15.0015063 | 15.5217624 | 14.9887693 | 14.8756447 |
| 18.9939181 | 18.9979753 | 19.104684 | 19.057205 | 19.5547961 | 19.5389728 | 19.2078151 |
| 13.6567553 | 13.7517314 | 13.3442462 | 13.150664 | 13.9701197 | 13.7665676 | 13.5093768 |
| 15.9631149 | 16.1680461 | 16.0086741 | 16.4661422 | 16.6388291 | 16.544586 | 15.9158186 |
| 11.5208204 | 11.5801761 | 11.501626 | 11.3587313 | 11.5817296 | 12.4406251 | 11.4045948 |
| 12.7482071 | 11.7365429 | 12.4192689 | 11.6312289 | 12.4465315 | 11.1053018 | 12.0909306 |
| 21.364491 | 21.5427107 | 21.7173195 | 21.7207662 | 22.1726893 | 21.9596755 | 21.7125599 |
| 11.4304792 | 11.5643104 | 11.5246147 | 11.9099311 | 11.5372674 | 11.6656201 | 10.9616371 |
| 16.956498 | 17.0736495 | 16.7006588 | 17.0421524 | 17.3111309 | 16.8037487 | 16.7874202 |
| 17.5216577 | 17.7437752 | 17.4714041 | 17.9218798 | 19.0718115 | 18.1139731 | 17.7003618 |
| 16.4384219 | 16.3335566 | 16.2983063 | 16.0469901 | 16.0678307 | 16.5176427 | 16.0790123 |
| 21.8043742 | 21.9806622 | 22.1049196 | 22.0371011 | 22.8301528 | 22.3940822 | 22.0365634 |
| 18.0879858 | 17.8667745 | 18.108725 | 18.2921391 | 18.1561674 | 18.6614369 | 18.4039718 |
| 18.5720448 | 18.5222611 | 18.6471838 | 18.4620834 | 18.9634064 | 19.005093 | 18.6218614 |
| 15.9970718 | 16.1021219 | 15.8736116 | 15.9245764 | 16.1547272 | 16.3394727 | 16.0388221 |
| 19.1330946 | 19.2006109 | 19.7418484 | 19.3525728 | 19.4669839 | 19.9665711 | 19.0723999 |
| 16.8540688 | 16.6811201 | 16.7251667 | 16.4503793 | 16.5325432 | 17.0933385 | 16.5090976 |
| 11.9856639 | 12.9751087 | 12.4642907 | 12.348436 | 13.7393034 | 13.065883 | 13.019929 |
| 21.2024765 | 21.9189412 | 21.1075873 | 20.9700918 | 21.6115803 | 21.3425152 | 21.1670358 |
| 17.3471672 | 17.056991 | 17.4586962 | 17.0957722 | 17.1748845 | 17.2854417 | 16.9896838 |
| 15.8890953 | 15.7880793 | 15.6936439 | 15.9387744 | 15.6664764 | 16.0994842 | 15.7651364 |
| 16.0533922 | 16.3595625 | 16.1827471 | 16.1993474 | 16.6136756 | 16.799896 | 16.1269702 |
| 19.5847163 | 19.7894794 | 19.4800648 | 19.7308307 | 19.6292631 | 19.6303167 | 19.6913359 |
| 17.3418237 | 17.1918223 | 17.4261306 | 17.3226439 | 17.759006 | 18.2351962 | 17.3066995 |
| 16.7610242 | 16.8226907 | 17.046921 | 17.2549262 | 17.6168354 | 17.2700116 | 17.186239 |
| 14.2735574 | 13.9747335 | 14.2060856 | 13.8370655 | 13.9920189 | 13.9565347 | 13.3392848 |
| 14.236578 | 14.3723717 | 15.2279762 | 14.6513441 | 15.0823702 | 15.9558606 | 14.6618019 |
| 16.7236005 | 17.1526879 | 17.1823851 | 17.3752806 | 17.8479422 | 17.2888904 | 17.2460788 |
| 14.813362 | 14.8498197 | 15.2521931 | 14.5014354 | 15.2855571 | 15.4479334 | 15.2546925 |
| 17.5752633 | 17.7037629 | 17.7864101 | 17.7280824 | 18.3403435 | 18.0312339 | 17.6578292 |
| 16.5928445 | 16.8821509 | 16.9974399 | 16.8475623 | 17.4581397 | 17.1366728 | 17.050184 |
| 18.9119795 | 18.4756646 | 18.5703707 | 18.7695658 | 18.5480996 | 18.9412476 | 18.3946394 |

| | | | | | | |
|---|---|---|---|---|---|---|
| 16.2156844 | 16.437789 | 16.6802216 | 16.5640128 | 16.6741483 | 16.834999 | 16.5846647 |
| 15.0658025 | 14.9357726 | 14.4605853 | 14.7354701 | 14.8984346 | 15.5682006 | 14.6954702 |
| 15.9103705 | 15.7360757 | 15.6946503 | 15.8789034 | 15.9250017 | 16.1085992 | 15.7402379 |
| 21.5481643 | 21.7478708 | 22.0280117 | 21.8546332 | 22.740632 | 22.0705355 | 21.6750425 |
| 21.1961492 | 21.4909982 | 21.5815003 | 21.4957908 | 22.0518148 | 21.9505458 | 21.475449 |
| 14.992363 | 14.8307448 | 14.5422931 | 14.9378618 | 15.2893598 | 15.4584507 | 14.5005061 |
| 13.6170897 | 13.395804 | 13.9698624 | 13.6823856 | 13.9780723 | 13.6328232 | 13.4182994 |
| 16.6070692 | 16.2606947 | 16.3917597 | 16.3975255 | 16.5800708 | 16.5864485 | 16.2517012 |
| 19.6112083 | 19.5504561 | 19.6089564 | 19.4838924 | 19.5564059 | 20.1124942 | 19.2092934 |
| 17.2172679 | 17.2832713 | 17.886445 | 17.7803014 | 18.02385 | 17.6321342 | 17.9435899 |
| 18.7363381 | 18.664941 | 18.7020955 | 18.7684194 | 19.1309771 | 18.8591765 | 18.7710545 |
| 14.4249091 | 14.6702151 | 15.1096725 | 15.0022851 | 15.356508 | 15.1100439 | 15.146221 |
| 14.6622184 | 14.7115733 | 14.4187898 | 15.0197637 | 15.7977543 | 15.0166445 | 15.0177713 |
| 18.156022 | 17.9338064 | 18.2808519 | 18.3431432 | 18.7042156 | 18.4698258 | 18.1660316 |
| 13.489375 | 14.0091608 | 14.8616225 | 14.4330248 | 14.0699362 | 14.8590209 | 14.545759 |
| 16.8935882 | 16.8441034 | 16.7339742 | 16.8163157 | 17.0736786 | 17.4379468 | 16.5594552 |
| 15.7126406 | 15.6978494 | 16.5908167 | 15.9569258 | 15.9600449 | 16.502377 | 15.8561065 |
| 15.9971737 | 16.250978 | 15.9857157 | 16.0249032 | 16.3529012 | 16.4182086 | 16.4241547 |
| 15.1803922 | 15.4052546 | 14.9301176 | 15.0970877 | 14.7702242 | 15.2967368 | 15.1014985 |
| 17.683422 | 17.7461915 | 17.7332929 | 17.5622086 | 17.8587419 | 18.1733355 | 17.4200166 |
| 13.9175511 | 14.21664 | 14.3471633 | 13.9808006 | 14.2722378 | 14.5383377 | 14.161132 |
| 17.9990919 | 18.3237887 | 18.4631048 | 17.9401609 | 18.0919118 | 18.9689634 | 18.0993606 |
| 18.8921727 | 18.7916133 | 19.0910331 | 18.9132201 | 17.9101118 | 19.1779664 | 19.1172808 |
| 14.5015378 | 13.9413114 | 13.6230937 | 14.5105885 | 13.8645532 | 14.5382008 | 14.7017586 |
| 18.6164555 | 18.4929276 | 18.595151 | 18.4287529 | 18.9736468 | 19.0709761 | 18.3915195 |
| 17.4490426 | 17.3429838 | 17.7228384 | 17.5724423 | 18.2510397 | 17.9858837 | 17.6195361 |
| 14.8818704 | 14.7929656 | 14.9057957 | 14.7788844 | 15.4182927 | 15.6105604 | 15.9136069 |
| 13.5989565 | 14.2152552 | 13.5564931 | 13.7799424 | 13.9149787 | 13.3022536 | 14.720854 |
| 17.3498834 | 17.5429998 | 17.7401768 | 17.8867176 | 18.2878171 | 17.9580212 | 17.622084 |
| 15.2938928 | 14.8859451 | 15.6201642 | 15.4000352 | 15.1478537 | 16.0737587 | 15.2482113 |
| 15.2513493 | 15.3865937 | 15.7232196 | 15.3979382 | 14.9206112 | 16.9442456 | 15.4201265 |
| 16.4059787 | 16.8254607 | 16.7241996 | 16.601598 | 17.2255705 | 16.9074895 | 16.2834066 |
| 14.2509263 | 14.0825648 | 14.4421695 | 14.3588236 | 14.7288648 | 14.5610057 | 14.4248369 |
| 19.5757659 | 19.5628927 | 19.7028278 | 19.5531282 | 19.5833867 | 20.2933298 | 19.4372118 |
| 16.1628769 | 16.2789729 | 16.1576229 | 16.4596442 | 16.6009075 | 16.5055053 | 16.3243525 |
| 19.6958202 | 19.7376717 | 19.7206319 | 19.6053166 | 20.2500443 | 20.2616559 | 19.5324044 |
| 16.5482985 | 16.8771329 | 16.87315 | 16.2767013 | 17.0680822 | 16.9405822 | 16.6756869 |
| 15.7121726 | 15.6695447 | 15.5184543 | 15.6171015 | 15.6894878 | 16.0711529 | 15.764403 |
| 20.3315266 | 20.2822442 | 20.4660131 | 20.4002628 | 20.6381204 | 20.8769663 | 20.343928 |
| 15.0553828 | 15.1034694 | 15.2385992 | 15.7053264 | 15.7226915 | 15.3434224 | 15.7131616 |
| 13.51177 | 13.9291844 | 13.6276335 | 13.7436306 | 14.200743 | 14.2099042 | 14.6711578 |
| 21.072412 | 21.2500791 | 21.2916888 | 21.19477 | 21.6349581 | 21.832644 | 21.2043929 |

| | | | | | | |
|---|---|---|---|---|---|---|
| 11.6012992 | 11.8675786 | 11.2233185 | 11.4258284 | 11.8073876 | 11.592671 | 11.5378825 |
| 15.5115279 | 15.5317962 | 15.7837094 | 15.614508 | 15.7417159 | 15.9760532 | 15.9828025 |
| 13.041547 | 12.7986957 | 13.2080845 | 12.6580759 | 13.0313931 | 12.777327 | 12.3790798 |
| 13.1155875 | 12.9913066 | 12.4047483 | 13.0271947 | 12.9486942 | 13.4870398 | 12.4461127 |
| 17.8761751 | 17.9297164 | 17.2467804 | 16.7298933 | 16.9315143 | 16.6344353 | 17.0427798 |
| 15.0229691 | 14.9356801 | 15.0569254 | 14.6638273 | 15.3650664 | 15.3685013 | 14.9386098 |
| 16.1225651 | 16.200787 | 16.5757209 | 16.411763 | 16.8359048 | 16.8359052 | 16.3842114 |
| 16.9165433 | 16.789957 | 16.7710196 | 16.6559609 | 16.7779056 | 17.2668707 | 16.6128484 |
| 14.4669027 | 14.3471777 | 13.3872921 | 13.1894323 | 13.226048 | 13.1711038 | 13.7401368 |
| 21.6582771 | 21.5047883 | 21.6720929 | 21.5874618 | 22.069445 | 22.2244934 | 21.4420706 |
| 16.2007094 | 16.6703622 | 16.6887176 | 16.6298733 | 17.0935472 | 16.584932 | 16.8508461 |
| 13.7779921 | 13.4953406 | 13.2215072 | 12.9997536 | 13.4750733 | 12.4599091 | 13.0580618 |
| 11.4357575 | 11.4665062 | 11.7339482 | 11.7005277 | 11.5537294 | 12.0484296 | 11.374132 |
| 14.0518357 | 13.6048094 | 13.6977695 | 14.0557634 | 14.1337869 | 14.5010412 | 13.8496029 |
| 17.2020863 | 17.3280673 | 17.203615 | 16.9696364 | 17.0010813 | 17.1652392 | 16.8876463 |
| 20.4431457 | 20.6175884 | 20.4546033 | 20.2350598 | 20.3149105 | 21.1922327 | 20.3590474 |
| 13.7089385 | 13.5872167 | 13.6207779 | 13.9061009 | 13.9931272 | 14.3153283 | 13.6778781 |
| 19.2957777 | 19.2047159 | 19.4451404 | 19.2044442 | 19.2326334 | 19.9698821 | 19.2218617 |
| 14.027353 | 13.6899972 | 13.5010241 | 13.4791065 | 13.6711115 | 14.1061485 | 13.6621706 |
| 14.1895517 | 14.2597196 | 14.8144324 | 14.3690864 | 14.4321004 | 14.8395629 | 14.6705069 |
| 16.4903117 | 16.5210176 | 16.4851852 | 16.4357703 | 16.4895816 | 16.7162612 | 15.8875873 |
| 15.3104946 | 15.1130083 | 14.0105304 | 14.630608 | 14.011417 | 14.3470188 | 14.3587249 |
| 13.4368379 | 13.4106581 | 13.2255476 | 13.3633186 | 13.8004663 | 13.7256645 | 12.8098737 |
| 15.194777 | 15.2955196 | 15.2248252 | 15.0908059 | 15.5300639 | 15.42831 | 15.0317404 |
| 19.3656926 | 19.5104866 | 19.144762 | 19.0932558 | 19.5635263 | 19.9526527 | 19.139792 |
| 15.2281822 | 15.3146361 | 15.3256267 | 15.498844 | 15.888039 | 15.4650264 | 15.5127392 |
| 15.6528698 | 15.8362327 | 16.0075236 | 15.9635045 | 15.7409683 | 15.8156187 | 16.012874 |
| 19.6265545 | 19.650941 | 19.7068782 | 19.6056147 | 19.9024631 | 20.1357213 | 19.649522 |
| 15.2181997 | 15.6120932 | 15.7031184 | 16.1987734 | 15.6388446 | 14.9537008 | 14.4433337 |
| 11.7512238 | 11.8031259 | 11.6875405 | 11.3777346 | 11.9481082 | 11.6426447 | 11.600919 |
| 14.2241129 | 14.2430383 | 14.3672448 | 13.5053386 | 13.2483461 | 14.9452694 | 13.7711181 |
| 19.2363423 | 19.0125083 | 18.92876 | 19.1776008 | 19.2166248 | 18.5280602 | 19.1388196 |
| 23.0571819 | 22.9447234 | 23.0212567 | 23.2002865 | 23.361171 | 23.4047135 | 23.0411748 |
| 16.5535297 | 16.9635996 | 16.5105861 | 16.4257636 | 17.1089873 | 16.9573524 | 16.6231417 |
| 18.8103482 | 18.6230116 | 18.6569024 | 18.6961058 | 18.9280281 | 19.0718168 | 18.7701687 |
| 11.5402492 | 11.459165 | 11.6866234 | 11.6108522 | 11.8014685 | 11.7259646 | 11.5336376 |
| 17.8638227 | 18.0899241 | 18.1667332 | 17.7841446 | 18.3087773 | 18.6913272 | 17.9747023 |
| 17.6995468 | 17.4953837 | 17.5609591 | 17.2592594 | 16.9883831 | 17.1272347 | 17.2145406 |
| 17.3063826 | 17.4636246 | 17.723787 | 17.5685466 | 17.6436162 | 18.1580252 | 17.5237307 |
| 18.7450375 | 18.6886615 | 18.590614 | 18.5068273 | 18.5117697 | 19.2265657 | 18.5305864 |
| 11.2922112 | 11.8484283 | 11.3103469 | 11.3510139 | 11.7608978 | 12.1842341 | 11.4676077 |
| 14.0756241 | 14.0729803 | 14.6292271 | 14.1434895 | 15.2164649 | 14.1231549 | 14.3537538 |

| | | | | | | |
|---|---|---|---|---|---|---|
| 18.8392397 | 18.99719 | 18.7561918 | 18.5597108 | 19.1685243 | 19.4715038 | 18.6722445 |
| 17.279771 | 17.1265602 | 17.2060738 | 17.2442717 | 17.4325193 | 17.8093841 | 17.305504 |
| 14.5670791 | 14.2243014 | 13.9898556 | 13.6742555 | 14.4993125 | 14.1172625 | 14.0526333 |
| 12.8661534 | 13.0836148 | 13.5057731 | 13.3369363 | 13.6389909 | 13.5314691 | 13.2828789 |
| 13.8916019 | 13.9804596 | 14.2819106 | 14.1967507 | 14.411536 | 14.2417427 | 14.0779479 |
| 17.6344855 | 17.2910091 | 18.0073234 | 17.7584253 | 18.2443054 | 17.6760443 | 18.098992 |
| 18.2754959 | 18.2176463 | 18.3182008 | 18.3418438 | 18.4004277 | 18.8044977 | 18.3280939 |
| 16.4490502 | 16.4573779 | 16.642176 | 16.4996308 | 16.7167962 | 16.4302852 | 16.5918662 |
| 11.2677538 | 11.5923534 | 11.8153162 | 11.6425918 | 12.1039696 | 11.8331688 | 11.2106359 |
| 13.8220658 | 13.3271737 | 13.54408 | 13.7513605 | 11.8389785 | 12.6436952 | 13.5206047 |
| 17.0331334 | 16.9286571 | 17.2961916 | 16.9558622 | 17.7429824 | 17.8670481 | 16.9958821 |
| 15.7628565 | 16.1278036 | 16.1982972 | 16.3539715 | 16.2256477 | 16.1907031 | 15.9030911 |
| 14.554974 | 14.6985899 | 14.712282 | 14.9183558 | 15.1609891 | 14.7769386 | 14.8864711 |
| 14.0984619 | 14.4392423 | 14.9525243 | 14.7413897 | 15.2694887 | 14.6753485 | 14.4829969 |
| 11.2796205 | 12.0588472 | 11.9594934 | 11.8361364 | 11.4764831 | NA | 11.8467227 |
| 17.7528443 | 17.990193 | 18.111278 | 17.8690315 | 18.4469397 | 18.0968784 | 17.9805518 |
| 19.7290027 | 19.7840279 | 19.8600236 | 19.5914356 | 19.7384266 | 20.333808 | 19.5373062 |
| 18.4045557 | 18.8161653 | 18.8811476 | 18.7370458 | 19.3264074 | 18.9020783 | 18.7401652 |
| 19.8449387 | 19.600162 | 19.5097678 | 19.5572795 | 19.3595776 | 20.390523 | 19.4637466 |
| 15.9860382 | 15.6691921 | 15.5088189 | 15.8871242 | 15.4770686 | 16.2589431 | 15.5832555 |
| 13.4226375 | 13.5027225 | 13.7825629 | 14.1031711 | 14.3462851 | 14.0238892 | 13.7021003 |
| 19.6493078 | 19.6461943 | 19.7478831 | 19.3684758 | 19.1567507 | 19.8937693 | 19.290893 |
| 16.6341239 | 16.584222 | 16.2773155 | 16.2588419 | 16.7275066 | 16.7977259 | 16.366007 |
| 14.9660233 | 15.5304557 | 15.5710469 | 15.8224003 | 16.7995985 | 15.9050269 | 15.5481725 |
| 18.3027572 | 17.8758859 | 17.9989933 | 18.0931141 | 18.3475122 | 19.2416053 | 18.2851537 |
| 20.1354731 | 20.4307918 | 20.3718188 | 20.2294117 | 21.3451285 | 20.4835376 | 20.1757079 |
| 18.9496369 | 18.8628835 | 18.8863797 | 18.9063628 | 19.4113302 | 19.431507 | 18.6175679 |
| 11.4171878 | 11.6317862 | 11.3006567 | 11.4580816 | 12.2922079 | 11.6472242 | 11.5043454 |
| 16.6020332 | 16.6298442 | 16.469005 | 16.4588566 | 18.3422198 | 17.0524467 | 16.6560363 |
| 18.631823 | 18.1926138 | 16.8566418 | 16.6172167 | 17.4790239 | 16.5393158 | 17.3482574 |
| 14.9541588 | 14.2705175 | 14.0562292 | 14.320947 | 13.964725 | 14.4281176 | 14.4194626 |
| 13.0189545 | 13.719438 | 12.9334394 | 13.0072821 | 14.1180362 | 13.3385388 | 12.5613926 |
| 20.8799516 | 20.7439077 | 20.7777268 | 20.9673344 | 21.0399817 | 21.1218327 | 20.8186358 |
| 17.1544367 | 17.3509704 | 17.5382251 | 17.3844928 | 17.6969293 | 17.1241979 | 17.4275967 |
| 16.0652831 | 16.0878656 | 16.2873711 | 16.2629447 | 16.1544796 | 16.8223552 | 16.0542817 |
| 14.8898738 | 14.827384 | 14.9544741 | 14.4030973 | 14.5836574 | 15.6033457 | 14.63412 |
| 15.8667115 | 15.8940236 | 16.0307324 | 15.839178 | 16.3440473 | 15.781477 | 15.9808448 |
| 16.3215041 | 16.3323343 | 16.4945289 | 16.2109811 | 16.5236174 | 16.8154362 | 16.1058932 |
| 13.7980682 | 14.0100791 | 14.1905259 | 13.4865706 | 14.3461699 | 13.9702874 | 13.7746189 |
| 17.6172212 | 17.6681818 | 17.7069726 | 17.6846157 | 17.8761756 | 18.0509612 | 17.6646208 |
| 15.115953 | 15.7965069 | 15.7040567 | 15.793469 | 15.9906062 | 15.3665257 | 15.4238693 |
| 13.7690967 | 13.8764145 | 14.0857903 | 13.9054277 | 14.0693428 | 13.7154577 | 13.5774827 |

| | | | | | | |
|---|---|---|---|---|---|---|
| 13.6372233 | 13.8578818 | 13.7509204 | 14.1929682 | 13.6547149 | 13.5941489 | 13.3150796 |
| 13.1674044 | 12.776889 | 12.7555533 | 13.6666653 | 12.9147385 | 13.7121181 | 12.4112037 |
| 20.475636 | 20.5720558 | 20.9351961 | 20.7703324 | 20.7454412 | 21.043667 | 20.4990434 |
| 19.9180103 | 20.0484035 | 19.9850915 | 19.921869 | 20.1682757 | 20.3906398 | 19.8243771 |
| 19.0007195 | 18.9940873 | 18.8305055 | 18.7629108 | 18.7189106 | 19.4621548 | 18.9214288 |
| 19.2446302 | 19.3241578 | 19.2512897 | 19.2565154 | 19.1873489 | 19.855842 | 19.155361 |
| 11.6856526 | 11.5980056 | 11.3908841 | 11.9465577 | 12.1723065 | 11.6834735 | 11.4154586 |
| 13.9840088 | 13.8516784 | 13.1934533 | 13.9479354 | 13.5191363 | 14.5706998 | 14.038257 |
| 17.8338758 | 18.1013451 | 18.0183356 | 18.0072444 | 18.402144 | 18.3837635 | 18.0787522 |
| 18.9508486 | 18.993904 | 19.7294284 | 18.9657877 | 18.8717816 | 19.5406467 | 19.0119809 |
| 19.9822284 | 20.0408458 | 20.4006098 | 20.3802543 | 20.8209823 | 20.5164646 | 20.1725747 |
| 19.7463791 | 20.2151474 | 20.1642675 | 20.0794265 | 20.6844412 | 20.142891 | 20.233771 |
| 15.9394011 | 15.7186693 | 16.0238969 | 15.7329695 | 15.9301793 | 16.3578183 | 16.186971 |
| 14.0419487 | 14.5485997 | 14.5851589 | 14.4387522 | 14.9197605 | 14.4412756 | 13.9552683 |
| 17.6732094 | 17.6647727 | 17.9817517 | 17.8783219 | 18.4572081 | 17.691397 | 18.0247966 |
| 18.3209494 | 19.6212705 | 18.0356233 | 17.6367334 | 19.3011574 | 17.9735918 | 18.2277673 |
| 17.2291346 | 17.4800655 | 17.3576553 | 17.5769548 | 18.1688575 | 17.6994681 | 17.370754 |
| 18.2278447 | 18.2666513 | 18.4594465 | 18.2449897 | 18.6445228 | 18.575028 | 18.4723014 |
| 19.3115796 | 19.3581166 | 19.4906418 | 19.297718 | 19.4669327 | 19.9006887 | 19.1959444 |
| 18.1531542 | 18.1622112 | 18.286427 | 18.1748396 | 18.3411928 | 18.3610499 | 18.1517986 |
| 17.665686 | 17.2179371 | 17.657946 | 17.9321301 | 17.7299395 | 17.7787692 | 17.3826119 |
| 13.3121468 | 13.6237348 | 13.9630796 | 14.0335355 | 14.1442909 | 13.4172837 | 13.6432399 |
| 23.4117619 | 23.4964948 | 23.6818904 | 23.6787281 | 24.054751 | 23.9707527 | 23.5328559 |
| 17.4245701 | 17.4046238 | 17.7099185 | 17.392245 | 18.3411103 | 17.3236736 | 17.3958012 |
| 15.6558439 | 15.8503547 | 15.4833609 | 15.7516139 | 16.1138057 | 16.4950277 | 16.1789639 |
| 19.7151204 | 19.5647472 | 19.6302441 | 19.5868995 | 19.5872509 | 19.7056999 | 19.7063727 |
| 15.0803703 | 14.6142356 | 15.1370948 | 14.9865013 | 15.0769657 | 15.0025761 | 14.8991366 |
| 14.8182716 | 15.1853773 | 14.8135707 | 14.3140156 | 14.8795456 | 14.6383668 | 15.0082686 |
| 14.4897172 | 14.6414952 | 14.4325013 | 14.532167 | 14.5253659 | 15.0245778 | 14.7025008 |
| 18.8549353 | 19.0880312 | 18.8563247 | 19.092004 | 19.3940722 | 18.9744661 | 19.2210061 |
| 12.800107 | 12.6431357 | 12.3784917 | 12.7258898 | 12.5412637 | 12.9619823 | 12.6583979 |
| 16.7384738 | 16.3720077 | 16.8743585 | 16.8543974 | 16.8155975 | 17.562953 | 17.0893706 |
| 14.5654188 | 14.8429767 | 14.6283793 | 14.6100148 | 15.2684314 | 14.9046383 | 14.7258817 |
| 14.7952937 | 14.5022845 | 14.2512833 | 13.7919675 | 13.9321438 | 15.1782076 | 14.0952021 |
| 14.9387027 | 15.2299354 | 14.8469306 | 14.6299142 | 15.37427 | 15.0351592 | 14.6740207 |
| 17.9689758 | 18.0892209 | 17.8857539 | 18.0755642 | 18.0685589 | 17.9526253 | 18.0844679 |
| 15.8724042 | 16.2242531 | 16.5176342 | 17.1744425 | 17.5158024 | 17.1590044 | 16.5003983 |
| 15.2134705 | 15.6271527 | 15.6662789 | 15.4501277 | 14.191697 | 15.214056 | 14.9989383 |
| 14.9579932 | 15.0955212 | 14.7234127 | 14.6869584 | 14.8509125 | 15.5429724 | 14.6010491 |
| 11.4988264 | 11.5691136 | 12.3393699 | 11.5685612 | 10.791686 | 11.2111966 | 12.3432822 |
| 19.31615 | 19.3088004 | 19.4369759 | 19.4101852 | 19.4975288 | 19.7676691 | 19.4546109 |
| 17.6326037 | 17.2652606 | 17.3094444 | 17.9308751 | 17.1337748 | 16.7808408 | 17.7028082 |

| | | | | | | |
|---|---|---|---|---|---|---|
| 17.2393965 | 16.6971111 | 16.8722825 | 17.3736034 | 16.9318905 | 17.5711155 | 17.0331611 |
| 19.2018299 | 19.1138753 | 19.3100271 | 19.0420481 | 19.238632 | 20.1495093 | 19.1572071 |
| 12.6591723 | 12.4810878 | 12.1918088 | 11.9610276 | 11.3048671 | 11.9620855 | 12.8011349 |
| 13.2029014 | 13.361991 | 13.343927 | 12.7868431 | 14.0252118 | 13.6815971 | 12.8970101 |
| 14.5256469 | 13.7960316 | 13.9685729 | 14.3289895 | 14.0108486 | 13.8816987 | 14.3797066 |
| 15.286031 | 15.0411312 | 15.1950984 | 15.0526394 | 15.2297052 | 15.4127725 | 14.9954067 |
| 13.8343897 | 14.2093623 | 14.0763901 | 14.3338105 | 14.7057829 | 13.9222455 | 14.2546175 |
| 19.5363123 | 19.5170525 | 19.8027989 | 19.6300549 | 20.1801011 | 19.6909277 | 19.628005 |
| 13.9693029 | 13.9871404 | 13.7932348 | 13.7027762 | 14.045066 | 14.3203182 | 13.7420347 |
| 16.5665647 | 16.9052574 | 16.6272679 | 16.6627325 | 16.7028463 | 16.5583208 | 16.6236281 |
| 13.9262487 | 14.2578086 | 13.4912698 | 14.0037018 | 14.4996433 | 14.3436986 | 14.1488827 |
| 20.3171408 | 20.3014064 | 20.325075 | 20.6395067 | 20.4549935 | 20.8797599 | 20.2631333 |
| 15.3763742 | 15.4242681 | 15.2532894 | 15.7040359 | 15.7696684 | 15.4268197 | 15.5586963 |
| 15.3011194 | 15.5297861 | 15.298404 | 15.5972645 | 15.7048763 | 15.3373326 | 15.5154166 |
| 16.98399 | 17.4395166 | 17.2947178 | 17.0046348 | 17.2705324 | 16.9571174 | 17.4130197 |
| 19.4271573 | 19.4492126 | 19.4878645 | 19.6669543 | 19.5034546 | 19.605971 | 19.2864816 |
| 14.5171192 | 14.501698 | 14.2977166 | 14.4263645 | 14.3396965 | 14.3351909 | 14.3885599 |
| 11.9674375 | 11.9112707 | 11.4830402 | 11.7697913 | 10.4851207 | 11.5360767 | 11.7284335 |
| 12.3616997 | 12.1860077 | 12.4990031 | 11.7891738 | 11.3560762 | 13.5142592 | 12.2497995 |
| 12.5763096 | 12.3799424 | 12.9191594 | 12.7157932 | 12.8679386 | 12.8229046 | 12.9236476 |
| 14.81662 | 15.076589 | 14.864036 | 14.6902703 | 15.0188344 | 16.2476686 | 15.1357232 |
| 17.2467893 | 16.9038391 | 17.2031939 | 16.9064695 | 17.3223201 | 17.6003196 | 17.2395441 |
| 21.4291125 | 21.5399364 | 21.6195881 | 21.6690397 | 22.1365654 | 22.249414 | 21.5443533 |
| 14.8879462 | 16.5441826 | 14.8130639 | 14.4990621 | 15.6625373 | 14.8486195 | 14.8966518 |
| 14.8853676 | 14.8470529 | 15.1902756 | 14.9687422 | 14.7560953 | 15.2574118 | 15.1339695 |
| 20.4234727 | 20.6615416 | 20.5837733 | 20.717712 | 20.9402634 | 20.6395318 | 20.8465071 |
| 11.3241145 | 12.1559849 | 10.897768 | 11.6970757 | 11.7130673 | 11.7100983 | 12.1081327 |
| 16.0869956 | 16.5005224 | 16.4499652 | 16.634222 | 16.7910868 | 16.2823012 | 16.4127975 |
| 16.1546828 | 16.1848478 | 16.5446763 | 16.4842643 | 16.5185441 | 16.3877542 | 16.1136333 |
| 19.3667864 | 19.1598736 | 19.0819912 | 19.0611666 | 19.278276 | 19.635081 | 19.3097167 |
| 17.5928475 | 18.037754 | 18.2417107 | 17.2938244 | 16.9516487 | 18.2724926 | 18.2871889 |
| 19.288985 | 19.1086333 | 19.1734437 | 19.3262521 | 19.334803 | 19.5139127 | 19.151195 |
| 15.0417252 | 15.258337 | 14.9410884 | 14.9919311 | 15.3646375 | 15.2126182 | 14.9651266 |
| 14.9874529 | 14.9434774 | 15.2834561 | 15.7772686 | 16.3905687 | 15.2486273 | 15.2593624 |
| 12.3289901 | 13.3501194 | 12.5411897 | 12.90888 | 12.420492 | 13.1307649 | 12.7649261 |
| 11.9080694 | 11.5303963 | 11.7058732 | 11.4163431 | 11.4308649 | 12.0462905 | 11.8987426 |
| 13.4256961 | 13.4226954 | 13.1396173 | 13.6844086 | 13.3958285 | 13.0685534 | 13.3605125 |
| 17.9266326 | 17.7685923 | 17.8933504 | 18.0172667 | 17.9653204 | 18.2899875 | 17.9923708 |
| 16.8356636 | 17.1614415 | 17.1579688 | 16.9497414 | 17.198674 | 16.9920123 | 17.2451593 |
| 18.8525977 | 19.1429059 | 19.0478174 | 18.9326553 | 19.0764077 | 19.516323 | 19.0615233 |
| 20.2841166 | 20.1531211 | 20.1908017 | 20.1692343 | 20.1432112 | 20.8552379 | 20.1427658 |
| 16.3183369 | 16.8036181 | 16.751984 | 16.2971172 | 17.9406425 | 17.2920138 | 16.7182156 |

| | | | | | | |
|---|---|---|---|---|---|---|
| 17.1495915 | 17.2515503 | 17.3466362 | 17.1497274 | 17.3480569 | 17.1021466 | 16.8262628 |
| 19.6799231 | 19.7041871 | 19.5832797 | 19.7872208 | 19.6347499 | 20.3209351 | 19.6623638 |
| 19.5103718 | 19.6417026 | 19.663106 | 19.5751028 | 19.7622559 | 19.7311296 | 19.5775226 |
| 17.0682039 | 16.9962284 | 17.1191341 | 17.2150063 | 17.0282047 | 16.9923003 | 16.9636428 |
| 11.983742 | 12.8895009 | 13.7610445 | 13.0441802 | 13.2618666 | 13.5423195 | 13.0660835 |
| 15.6977359 | 16.289076 | 15.3517183 | 15.8651162 | 15.403354 | 15.7028274 | 15.6745408 |
| 17.5554838 | 17.5173151 | 17.7216636 | 18.1257053 | 17.961739 | 16.8275859 | 17.7760934 |
| 17.3295869 | 16.8761107 | 17.1457839 | 16.3349496 | 18.3504045 | 16.4695077 | 16.3679926 |
| 14.3331429 | 14.4104244 | 14.8541938 | 14.6545173 | 14.7865932 | 14.6394989 | 14.7400491 |
| 15.095246 | 15.5212304 | 15.4591124 | 15.4212137 | 15.6716803 | 15.5075378 | 15.4861361 |
| 12.6506521 | 12.9055497 | 13.1741099 | 12.518072 | 12.4942889 | 12.9470489 | 12.5969085 |
| 17.0600683 | 16.9678271 | 17.105647 | 17.0822409 | 16.9935764 | 17.5904551 | 17.0437166 |
| 20.2205757 | 20.0001228 | 19.8114232 | 20.2541783 | 19.6585249 | 19.7299621 | 20.0454621 |
| 14.5893274 | 14.6536854 | 15.4446268 | 15.097386 | 15.8111481 | 15.0904118 | 15.0894699 |
| 19.372877 | 19.3911164 | 19.3394353 | 19.4705562 | 19.6433265 | 20.1689605 | 19.3474608 |
| 14.8569053 | 15.1231737 | 15.2284494 | 14.8918916 | 15.0086821 | 15.5345371 | 14.9882939 |
| 18.0195591 | 17.7469073 | 18.3014351 | 18.2678964 | 18.0718336 | 17.6957674 | 18.3589099 |
| 16.67954 | 16.7608125 | 16.7511553 | 16.7406975 | 16.4654496 | 16.801139 | 16.7833906 |
| 12.9984521 | 13.5240466 | 13.7950694 | 13.3286963 | 14.298676 | 13.2044377 | 13.7355406 |
| 18.7965154 | 18.9337825 | 19.011655 | 18.4606192 | 18.0019554 | 19.030679 | 18.4786783 |
| 19.9944808 | 19.88371 | 19.9658547 | 19.9942231 | 20.6551083 | 20.4550095 | 19.8038833 |
| 15.7700377 | 16.2505089 | 15.5954162 | 15.1062689 | 15.9286414 | 15.3885235 | 15.6789395 |
| 17.6773285 | 17.8479016 | 17.9405391 | 18.0361896 | 18.5437129 | 17.8597436 | 17.9681365 |
| 12.0288416 | 12.3525891 | 11.8861253 | 11.655523 | 11.3652027 | 12.4250667 | 11.5267154 |
| 16.0452284 | 16.5039145 | 16.4475194 | 16.8564069 | 16.9711983 | 16.0978079 | 16.7572748 |
| 14.0967948 | 14.5360605 | 14.958665 | 14.0144452 | 14.3678401 | 14.7539279 | 14.8759361 |
| 19.2560916 | 19.2794691 | 19.0896422 | 19.0689131 | 19.0047701 | 19.9526664 | 19.1517243 |
| 16.2846013 | 16.6276675 | 16.3029526 | 16.2061744 | 16.6024319 | 16.9202027 | 16.3326497 |
| 14.799023 | 14.7633881 | 14.6153789 | 14.5777397 | 14.5221719 | 15.0346415 | 14.5662771 |
| 14.2794522 | 13.9865675 | 14.0738022 | 13.8388723 | 13.7516441 | 14.199571 | 13.9546958 |
| 18.0144535 | 18.0874874 | 18.0681469 | 18.0181666 | 17.9062623 | 18.7510474 | 17.9368594 |
| 15.1085312 | 15.5102646 | 13.5423383 | 14.9618179 | NA | 14.4408938 | 14.9991414 |
| 14.2110815 | 14.1756801 | 14.1060466 | 14.1416229 | 13.8764026 | 14.346425 | 14.020463 |
| 18.2500171 | 19.1178757 | 18.2391593 | 17.7265746 | 18.5683917 | 17.9738536 | 18.1343706 |
| 23.4544544 | 23.6291107 | 23.7419799 | 23.6796021 | 24.1103682 | 23.7570159 | 23.4644315 |
| 14.3117274 | 15.1790082 | 14.926968 | 15.6791697 | 14.2327486 | 14.8737641 | 14.8365849 |
| 17.7040745 | 17.604571 | 17.5833371 | 17.6311963 | 17.6165268 | 18.0962395 | 17.6480215 |
| 14.3984957 | 14.9919248 | 15.0447014 | 14.3253751 | 14.394702 | 15.0142663 | 14.6185094 |
| 11.6490961 | 11.480033 | 12.0000797 | 11.912119 | 11.2346906 | 11.5630085 | 11.7421724 |
| 18.3380683 | 18.4643571 | 18.2812265 | 18.2828254 | 17.8977756 | 19.0981584 | 18.4423461 |
| 15.073325 | 15.193092 | 15.0425663 | 15.2536579 | 15.1192292 | 14.5984692 | 15.0355123 |
| 17.4438424 | 16.9191268 | 16.9407908 | 17.276677 | 17.2797224 | 16.2808619 | 17.1859477 |

| | | | | | | |
|---|---|---|---|---|---|---|
| 19.2256355 | 19.2188884 | 19.2237909 | 19.3059327 | 19.5030018 | 19.8235811 | 19.256377 |
| 15.781703 | 15.6341936 | 15.58602 | 15.7568036 | 15.5887005 | 15.8124167 | 15.5681702 |
| 18.4030422 | 18.2729611 | 18.2437063 | 18.281647 | 18.4536779 | 18.8622676 | 18.2605175 |
| 18.1447601 | 18.0792686 | 18.0774806 | 18.3125414 | 18.3143968 | 18.1177287 | 18.2012584 |
| 13.9974638 | 13.5254473 | 13.4758786 | 13.9377231 | 14.3532619 | 14.6250478 | 14.2750104 |
| 15.333179 | 15.3596348 | 15.1467867 | 15.2209764 | 15.5852755 | 15.5102533 | 15.2580954 |
| 21.1810848 | 21.1461751 | 20.9970203 | 21.4195409 | 21.1083856 | 20.8143475 | 21.2530083 |
| 14.6218965 | 14.4489914 | 14.3278311 | 14.9642141 | 15.0215629 | 14.2911308 | 15.0101732 |
| 16.8794579 | 17.2690795 | 17.0847258 | 17.4907801 | 17.2645219 | 16.978357 | 17.1990335 |
| 17.857114 | 17.2784491 | 18.1786231 | 17.0768733 | 18.1814216 | 16.4369096 | 17.6248186 |
| 21.4747139 | 21.586105 | 21.8183343 | 21.7728018 | 22.320616 | 22.1009004 | 21.5528543 |
| 11.5173939 | 11.5166842 | 11.4660873 | 11.9907036 | 12.2994514 | 11.3521381 | 11.9717029 |
| 17.8625689 | 18.9882963 | 17.0509213 | 16.8401125 | 18.5366185 | 17.0632039 | 17.6008642 |
| 14.0405186 | 13.762743 | 13.7017954 | 14.1493777 | 13.5142559 | 13.9511116 | 13.8866558 |
| 16.8060698 | 16.3729334 | 16.431929 | 16.5936764 | 16.6075064 | 16.5749335 | 16.5394017 |
| 12.4142893 | 12.5098954 | 14.6615057 | 13.9708566 | 12.7101902 | 13.1810456 | 13.4834292 |
| 15.1536513 | 15.355064 | 15.0088069 | 15.3970833 | 15.010216 | 15.4083148 | 15.2920617 |
| 19.8862536 | 20.1205352 | 19.994484 | 20.0231896 | 20.1309572 | 19.6159698 | 20.1729422 |
| 19.8629646 | 20.0532786 | 19.8568222 | 20.0213543 | 19.9899431 | 19.786962 | 19.9699063 |
| 16.8870558 | 17.0921514 | 16.9077239 | 16.8235266 | 17.0683721 | 16.9704575 | 16.8747337 |
| 19.1642105 | 19.2381986 | 19.3695135 | 19.1149817 | 19.2789293 | 19.6975176 | 19.2206364 |
| 19.4444308 | 19.378976 | 19.2947031 | 19.3485407 | 19.0399893 | 20.0121115 | 19.3215297 |
| 14.0171454 | 14.3706813 | 14.0369059 | 13.7099671 | 14.3577256 | 13.6451844 | 13.6081409 |
| 19.4475744 | 19.3951122 | 19.3238387 | 19.6118631 | 19.3123282 | 19.3444654 | 19.421864 |
| 13.3790066 | 13.5582313 | 14.1347502 | 14.0587469 | 13.951659 | 13.7913751 | 13.886622 |
| 13.3350895 | 15.8168311 | 13.9059621 | 13.0956779 | 16.2336021 | 14.3819193 | 13.4422946 |
| 15.4055626 | 15.5023497 | 16.0054679 | 15.3377964 | 15.2591055 | 16.0478867 | 15.3845071 |
| 11.1327713 | 11.0043191 | 12.3494019 | 12.4508358 | 12.1441039 | 10.6742577 | 11.9509762 |
| 19.127053 | 19.547654 | 19.5193053 | 18.3852986 | 18.0286327 | 19.9809405 | 19.178374 |
| 22.8221794 | 22.9725915 | 22.9678927 | 22.9533731 | 23.6446248 | 23.3506196 | 22.9875695 |
| 14.9079939 | 14.9657384 | 15.0855414 | 15.3189101 | 15.6250964 | 15.2178305 | 15.0505061 |
| 13.1929693 | NA | 13.2468727 | 13.8536906 | 12.889959 | NA | 12.9943498 |
| 15.0506388 | 15.4807452 | 15.3776079 | 15.6916518 | 16.1201314 | 15.0470833 | 15.8830062 |
| 13.9655753 | 14.1151738 | 14.3891582 | 14.0938241 | 14.9736129 | 13.3549736 | 13.9108575 |
| 14.0378145 | 13.5942786 | 13.4265011 | 13.8762888 | 12.9451922 | 13.0521509 | 13.7037733 |
| 14.9657107 | 14.8400661 | 15.020051 | 14.9392484 | 14.327206 | 14.9144305 | 15.1335743 |
| 21.0694724 | 21.6272907 | 20.8432169 | 20.8100979 | 21.5490964 | 20.5976006 | 20.8245443 |
| 19.9034163 | 20.7542283 | 20.4586463 | 19.5467898 | 19.7429407 | 21.0506405 | 20.2880624 |
| 13.80592 | 14.1329964 | 13.9955902 | 14.170926 | 13.8864648 | 13.869036 | 14.3421733 |
| 18.0649572 | 18.2439271 | 17.9116155 | 18.1304236 | 17.9804851 | 18.011208 | 18.0735189 |
| 14.853693 | 14.4024182 | 14.449629 | 15.3229646 | 14.7055085 | 14.0750227 | 14.887627 |
| 15.5944325 | 16.0727632 | 14.8696721 | 14.6823365 | NA | NA | 15.4785456 |

| | | | | | | |
|---|---|---|---|---|---|---|
| 16.0382761 | 16.5263803 | 16.7365611 | 16.4787855 | 16.3612327 | 17.628198 | 16.5493853 |
| 16.2177753 | 15.7939853 | 15.5397498 | 15.8998501 | 14.7713564 | 15.4363456 | 15.5467939 |
| 13.6687718 | 13.9812985 | 13.7128493 | 13.8103431 | 13.2024924 | 13.8051354 | 13.7783776 |
| 14.2572578 | 14.2282422 | 14.2491753 | 14.2015111 | 13.5867682 | 13.7117266 | 14.1665215 |
| 14.7990395 | 14.5136429 | 16.0922672 | 14.0949606 | 15.1030818 | 14.6299486 | 13.96304 |
| 18.079213 | 18.2910927 | 17.9652503 | 18.0872261 | 17.9942614 | 17.7045319 | 18.1452436 |
| 21.1980299 | 21.4570031 | 21.4205693 | 21.3439662 | 22.0965126 | 21.7044717 | 21.4545693 |
| 15.2444141 | 15.2023629 | 14.9456276 | 15.2723727 | 14.7384673 | 15.9059569 | 15.2406573 |
| 15.2081152 | 15.5794181 | 15.523193 | 15.4941151 | 15.8319194 | 15.5029669 | 15.2454478 |
| 15.4762429 | 15.8089043 | 15.9215405 | 15.7598497 | 15.4128569 | 16.075799 | 15.6827778 |
| 15.5406077 | 15.0831432 | 15.2747578 | 15.6978999 | 15.4385336 | 15.3694666 | 15.5766548 |
| 20.3709636 | 20.2274546 | 20.1092001 | 20.2706699 | 19.8262955 | 19.7422799 | 20.4328266 |
| 12.3495991 | 12.8083669 | 12.0622589 | 13.5196524 | 12.3410861 | 12.7754588 | 11.8845454 |
| 12.0478224 | 11.6955235 | 11.827908 | 12.1290142 | 11.3340052 | 11.1127637 | 11.7557891 |
| 13.9424471 | 13.9713891 | 14.1180178 | 14.5250952 | 14.3832058 | 14.5188328 | 13.7300791 |
| 16.1032941 | 16.1134318 | 15.970409 | 16.3137301 | 16.0106644 | 15.6783477 | 16.1919185 |
| 11.9144562 | 13.0505392 | 12.8639298 | 12.4287494 | 12.6562149 | 13.1107766 | 12.7819399 |
| 16.2490486 | 16.0757794 | 15.7416808 | 16.3411758 | 16.1842112 | 15.5886789 | 16.4187017 |
| 16.5921552 | 16.8479196 | 16.8474156 | 16.7808645 | 16.3641958 | 16.5179133 | 16.72496 |
| 14.8917251 | 14.310716 | 14.5641398 | 14.9672827 | 14.3855416 | 13.6976616 | 14.9640003 |
| 14.8184587 | 15.020536 | 14.5363026 | 14.5575879 | 14.0842952 | 14.7058278 | 14.1016512 |
| 16.2705216 | 16.0486346 | 15.8049464 | 16.1126886 | 15.5539234 | 15.9424052 | 16.0313754 |
| 13.671753 | 13.4867344 | 13.594716 | 13.8563288 | 13.5954001 | 13.4990702 | 13.9345846 |
| 15.2899651 | 15.5562395 | 15.1955907 | 15.3535494 | 15.5924156 | 14.6963347 | 15.1534604 |
| 19.8616049 | 20.758328 | 20.4977448 | 19.3438486 | 19.7632115 | 21.0055189 | 20.2001123 |
| 19.1154499 | 20.011831 | 18.7405408 | 18.7277495 | 19.4694416 | 17.8933042 | 18.8552042 |
| 14.0552755 | 13.9018534 | 13.5052245 | 14.3118898 | 14.0400249 | 13.6042372 | 13.9192008 |
| 11.6314391 | 12.48587 | 12.2870377 | 12.973681 | 12.900489 | 11.6870052 | 10.9780744 |
| 17.1043816 | 16.5193913 | 16.6930801 | 17.4047155 | 16.7473158 | 16.0396151 | 17.0698959 |
| 14.0903372 | 14.1588435 | 14.0047947 | 14.0761639 | 14.7747779 | 14.5017917 | 13.6930084 |
| 12.5994491 | 12.1895175 | 13.0727243 | 13.2864089 | 12.9355623 | 11.9400686 | 12.5576478 |
| 15.7155531 | 16.9663416 | 15.4650716 | 15.3693318 | 15.5744485 | 15.7270717 | 15.4629764 |
| 18.8878003 | 19.2667584 | 20.1541852 | 18.9664274 | 20.1793656 | 19.3298043 | 19.0160523 |
| 14.0771542 | 14.0482864 | 14.0148859 | 14.0117151 | 13.8580599 | 14.1232789 | 13.9521053 |
| 15.2331993 | 16.2450009 | 15.4176463 | 14.6963027 | 16.3708574 | 15.0007033 | 15.1633474 |
| 15.0448725 | 15.0080421 | 15.071624 | 15.2408643 | 14.7430386 | 14.9172286 | 15.0360711 |
| 13.7116496 | 14.1837143 | 13.9861826 | 13.9473323 | 13.3746566 | 12.9235963 | 14.0705476 |
| 12.9213936 | 12.9064885 | 13.0045736 | 13.0044643 | 13.0587562 | 12.7787292 | 12.7989729 |
| 16.5698551 | 19.5794995 | 17.0189685 | 17.3936468 | 17.2743338 | 17.0814657 | 17.7618195 |
| 15.810366 | 15.2526253 | 15.3694399 | 16.1121424 | 15.5989929 | 14.9959372 | 15.968284 |
| 16.4676256 | 16.3568075 | 16.3244309 | 16.5176692 | 15.6463158 | 16.0916817 | 16.4245832 |
| 16.247537 | 16.3213829 | 15.9422149 | 15.9729952 | 15.9170285 | 15.907382 | 16.0234876 |

| | | | | | | |
|---|---|---|---|---|---|---|
| 19.1662994 | 19.1839025 | 18.9445721 | 19.0789825 | 18.9357363 | 18.8685078 | 19.0830858 |
| 13.9899754 | 14.5388263 | 13.8836184 | 13.1687562 | 13.6128919 | 14.1253846 | 13.7489029 |
| 17.645233 | 17.2848552 | 17.2754149 | 16.8878659 | 16.4361612 | 17.2706703 | 16.9741378 |
| 19.4192134 | 19.1755247 | 19.10962 | 19.5514161 | 18.6410606 | 17.8974481 | 19.1100439 |
| 16.6569798 | 16.8471134 | 16.3271801 | 17.0977064 | 16.3648294 | 15.7664911 | 16.5269611 |
| 18.6524464 | 18.5567678 | 18.4993102 | 19.2651559 | 18.5522081 | 17.7696308 | 18.9225826 |
| 12.2277243 | 11.979031 | 11.8066662 | 11.6233224 | 11.7934613 | 11.3701113 | 11.5336665 |
| 15.8590027 | 16.1601613 | 16.0608686 | 15.6706207 | 15.9971717 | 16.1316867 | 15.9020237 |
| 13.7112865 | 13.9873986 | 14.5654329 | 14.5895774 | 14.7761506 | 13.2060053 | 14.9572759 |
| 19.0792199 | 19.2634633 | 19.213054 | 19.8626318 | 19.1566419 | 18.7793756 | 19.3926688 |
| 18.0621724 | 17.3806646 | 17.1981553 | 17.5515291 | 17.0653426 | 17.2239473 | 17.4099409 |
| 15.7074747 | 15.5596882 | 15.7940827 | 15.9150747 | 15.622561 | 14.7594844 | 15.9360359 |
| 18.4666626 | 18.2113352 | 18.3170015 | 18.6782906 | 18.3470533 | 17.7566462 | 18.390734 |
| 18.9767922 | 19.1379955 | 18.5970189 | 18.9584962 | 18.8694571 | 18.1131235 | 19.1434154 |
| 18.2480031 | 18.0272183 | 17.9224493 | 18.4405123 | 17.7371244 | 17.2992125 | 18.3765268 |
| 14.9417428 | 14.6647989 | 14.063781 | 14.8543063 | 14.4263922 | 14.0462366 | 14.7361557 |
| 17.2946399 | 16.8748404 | 17.0058359 | 17.6970068 | 16.7237777 | 16.7246595 | 17.2133012 |
| 15.0735601 | 14.5548321 | 15.3441839 | 13.7793727 | 15.7659964 | 14.5664756 | 14.5779945 |
| 12.762056 | 12.5192488 | 12.7209206 | 12.932566 | 11.658303 | 12.111214 | 13.0158294 |
| 16.5196022 | 16.068116 | 15.8667606 | 16.5454473 | 15.7781812 | 15.877863 | 16.303154 |
| 16.5811459 | 16.1970058 | 16.1686107 | 16.1090063 | 15.7751142 | 15.8743949 | 16.2412006 |
| 18.1996289 | 18.0209463 | 17.9085972 | 18.1950153 | 17.3763088 | 18.2766486 | 17.8647372 |
| 18.8445044 | 19.0340795 | 18.8412865 | 18.293446 | 17.511608 | 19.3480741 | 18.8017289 |
| 19.6095477 | 19.5140886 | 19.4992067 | 20.0819663 | 19.1715475 | 19.010555 | 19.6997511 |
| 15.7664769 | 15.7976021 | 15.3479427 | 15.9676473 | 15.5714262 | 14.9589072 | 15.830329 |
| 15.4663747 | 14.9272967 | 14.9299996 | 15.4754441 | 14.2188235 | 14.9839573 | 15.3130575 |
| 14.3834082 | 14.7303259 | 14.3892441 | 14.8907141 | 13.7959071 | 13.5452647 | 14.810229 |
| 19.8068818 | 19.7099886 | 19.6971813 | 20.281752 | 19.4203722 | 19.0625228 | 19.888918 |
| 18.0830509 | 18.0054873 | 17.9361868 | 18.5517753 | 18.0134389 | 17.1060831 | 18.0834885 |
| 13.6161973 | 15.3345736 | 13.9189063 | 13.8049951 | 13.2176544 | 13.5132921 | 13.8656176 |
| 18.937863 | 18.5683608 | 18.2874225 | 18.9607991 | 18.2696464 | 17.9280873 | 18.5465404 |
| 15.6623118 | 15.2247387 | 14.695885 | 15.6995105 | 14.371307 | 14.5628644 | 14.8563597 |
| 16.2340549 | 15.6628644 | 15.4552062 | 15.8277257 | 15.4402742 | 15.0927621 | 15.9346179 |
| 18.5139937 | 18.5269992 | 18.4323241 | 18.9365992 | 18.7933835 | 17.9030601 | 19.2105836 |
| 14.14477 | 13.7020864 | 13.4488345 | 14.3358006 | 13.5323765 | 12.3250882 | 13.6964132 |
| 17.501383 | 16.8427128 | 16.8865232 | 17.9594549 | 16.7164558 | 16.4058079 | 17.2078746 |
| 19.6583698 | 19.013716 | 18.756165 | 19.4911448 | 18.030027 | 18.2107727 | 19.0814223 |
| 16.2444923 | 16.2212881 | 16.3700921 | 16.6118929 | 16.0963008 | 15.6147806 | 16.6991705 |
| 15.6443604 | 15.7444355 | 14.9780595 | 15.8467648 | 15.3338613 | 14.7826614 | 15.7926109 |
| 15.5866944 | 15.3409844 | 14.5668154 | 15.270903 | 15.0975442 | 14.6759358 | 14.8660098 |
| 13.6935816 | 13.5245378 | 13.9072959 | 14.3330365 | 13.1104236 | 13.4898823 | 13.1179173 |
| 13.9415641 | 14.296565 | 13.7156111 | 14.8083175 | 12.7983548 | 12.5590318 | 14.1229587 |

| | | | | | | |
|---|---|---|---|---|---|---|
| 15.5318947 | 18.2623891 | 15.5643179 | 15.551685 | 16.5875022 | 14.9614577 | 15.962493 |
| 15.5220986 | 14.7122816 | 14.549566 | 15.4196362 | NA | 14.1625207 | 15.0532273 |
| 13.6991755 | 13.4911314 | 13.6777744 | 13.9943054 | 12.788634 | 12.8476579 | 13.8719242 |
| 15.6602387 | 16.9470094 | 17.0652134 | 15.8167732 | 15.9184923 | 15.8208099 | 15.3724702 |
| 17.8763399 | 17.4899166 | 17.0034217 | 17.7196077 | 17.1080512 | 16.2472929 | 17.4333023 |
| 11.7143543 | 11.6345755 | 11.6877402 | 12.4540463 | 10.7256744 | 10.6128647 | 12.4659119 |
| 19.1567225 | 19.1783309 | 19.0000446 | 19.6802551 | 18.635526 | 17.7764034 | 19.1975723 |
| 19.298883 | 19.1650748 | 18.9078436 | 19.6039875 | 18.5224859 | 18.1217895 | 19.3860593 |
| 17.5920783 | 17.1426986 | 17.2788128 | 17.5456902 | 16.7706543 | 16.1993952 | 17.2057246 |
| 16.8510922 | 16.775039 | 15.9946891 | 16.9276738 | 15.3395489 | 15.37754 | 16.4245958 |
| 18.0954694 | 17.5271255 | 17.6309414 | 18.1909032 | 17.6186221 | 16.8739491 | 17.6593355 |
| 18.9376868 | 18.5896663 | 18.7768509 | 19.6917372 | 18.9017294 | 17.0245769 | 19.1679908 |
| 16.7098821 | 16.3941289 | 16.3432522 | 16.8518567 | 15.9966343 | 15.5275734 | 16.8447921 |
| 19.8305884 | 19.3610829 | 19.2136352 | 19.6989828 | 18.5397697 | 18.0073624 | 19.3981253 |
| 15.0248174 | 14.9637935 | 14.636076 | 14.8287717 | 14.285123 | 13.8569154 | 15.1903905 |
| 18.7559289 | 18.3017978 | 18.1737439 | 19.3719593 | 17.4854067 | 16.8392073 | 18.4420156 |
| 17.8872634 | 17.370064 | 17.5762973 | 18.2252878 | 17.2413522 | 16.2306224 | 17.7020753 |
| 20.7088554 | 20.4245845 | 20.2586308 | 20.9700598 | 20.1340081 | 19.3799455 | 20.5908063 |
| 20.262591 | 19.7134361 | 19.7521897 | 20.5349113 | 19.3925922 | 18.6566342 | 19.9798136 |
| 21.0764485 | 20.4738132 | 20.7008457 | 21.3988699 | 20.2565765 | 19.5459709 | 20.8690612 |
| 20.614673 | 20.482128 | 20.480384 | 21.2786386 | 20.4943623 | 19.4439764 | 20.5945003 |
| 17.8281834 | 17.5551343 | 17.4314564 | 17.8795184 | 16.7655823 | 16.2235501 | 17.8493662 |
| 18.0922421 | 17.6117558 | 17.4924539 | 18.6757289 | 17.2621382 | 16.5424163 | 17.968585 |
| 18.7987523 | 18.6984519 | 18.2866267 | 18.8825688 | 17.9056258 | 17.5948895 | 18.4767812 |
| 19.0355735 | 18.9181158 | 18.6773388 | 19.4859326 | 18.4265593 | 17.633776 | 19.130308 |
| 26.3891 | 25.9400223 | 25.9417506 | 26.6896698 | 25.534674 | 24.8610655 | 26.1403028 |
| 25.5202446 | 25.0065858 | 24.8730919 | 25.7054789 | 24.5941597 | 23.8787052 | 25.3386163 |
| 18.0774066 | 17.6856397 | 17.3998404 | 17.9361515 | 16.6924649 | 16.5723729 | 17.7198206 |
| 14.3438623 | 14.0643445 | 13.8139748 | 14.1334995 | 12.312483 | 12.3854362 | 13.9171037 |
| 19.1846248 | 18.91382 | 18.7787147 | 19.4163929 | 18.5091831 | 17.3861049 | 18.9678559 |
| 14.2111017 | 14.3691671 | 15.0445656 | 14.3169643 | 13.911846 | 13.8007009 | 13.9482757 |
| 22.3985673 | 22.7271078 | 22.4546096 | 22.8549333 | 22.6711435 | 21.465048 | 22.7589383 |
| 13.8073521 | 14.1609077 | 12.9599233 | 13.3452182 | 12.6484398 | 12.1711638 | 13.0777598 |
| 20.7886229 | 20.6259216 | 20.2511604 | 20.9499398 | 20.0263704 | 19.5684994 | 20.7784933 |
| 17.9192319 | 17.7781504 | 17.3879574 | 18.1707914 | 16.6196533 | 16.8860647 | 17.8337927 |
| 16.1061533 | 15.2918546 | 15.0177785 | 16.4011273 | 15.0569514 | 14.1319645 | 15.6939621 |
| 22.779545 | 22.9410959 | 22.5894447 | 22.9162194 | 22.4160168 | 21.8203502 | 22.9961793 |
| 21.0286463 | 20.8203756 | 20.5029594 | 21.1587239 | 20.1180292 | 19.6090216 | 20.9271234 |
| 18.1366108 | 17.8769473 | 17.6786914 | 18.1036839 | 16.7825134 | 15.8124429 | 17.6625175 |
| 19.7281371 | 19.4457486 | 19.0004039 | 20.1041992 | 18.7877221 | 18.0335398 | 19.4390105 |
| 17.6628725 | 17.3099171 | 16.8079303 | 17.5232045 | 16.1263518 | 16.0109053 | 17.1309387 |
| 21.7689472 | 21.4343318 | 21.1991066 | 22.0657861 | 20.8418114 | 20.183904 | 21.5811622 |

| | | | | | | |
|---|---|---|---|---|---|---|
| 19.287521 | 18.8193994 | 18.9170673 | 19.7055267 | 18.5608616 | 18.1110383 | 19.2201087 |
| 26.7331002 | 26.5657528 | 26.3462269 | 27.2682497 | 26.3049039 | 25.1599471 | 26.6496539 |
| 19.5989969 | 19.4421275 | 19.0773623 | 19.6204025 | 18.6591238 | 18.0769753 | 19.5290518 |
| 24.2028059 | 24.3526792 | 23.9461691 | 24.2582448 | 23.7397067 | 23.0012068 | 24.3853517 |
| 17.6941571 | 17.3651093 | 16.7913798 | 17.4866261 | 16.0640559 | 15.8695375 | 16.9677912 |
| 18.0485537 | 17.9004879 | 17.725829 | 18.2431603 | 17.1911624 | 16.4409938 | 18.0280637 |
| 19.2595244 | 18.9443348 | 18.6112471 | 19.6826258 | 18.2984691 | 17.5474649 | 18.8804015 |
| 11.5272703 | 13.7152276 | 11.328905 | 11.5906194 | 10.5773116 | 10.7728571 | 11.4781668 |
| 21.1320325 | 20.7625129 | 20.7336246 | 21.6421829 | 20.2882915 | 19.6267046 | 21.0617189 |
| 19.8810365 | 19.6047019 | 19.3515797 | 20.234417 | 19.103083 | 17.8477973 | 19.8005188 |
| 19.748823 | 19.1863579 | 19.1066817 | 20.3721681 | 18.8002239 | 17.9109478 | 19.522948 |
| 14.923039 | 17.9339482 | 15.9410763 | 16.6624501 | NA | 15.2833634 | 14.9537886 |
| 20.8710525 | 20.540377 | 20.3522865 | 21.2525915 | 20.1452451 | 19.1760232 | 20.8041168 |
| 17.089844 | 17.0412522 | 16.179127 | 17.1379877 | 16.1999713 | 15.352009 | 16.8445005 |
| 21.6853536 | 21.9742534 | 21.550962 | 21.9968346 | 21.5628133 | 20.4792894 | 21.8142301 |
| 17.0176543 | 16.9234351 | 16.2080269 | 17.0099454 | 15.5867379 | 14.7841817 | 16.7813967 |
| 17.7520713 | 17.3770051 | 16.8327897 | 17.4333094 | 16.0330236 | 15.439763 | 17.0178516 |
| 18.0819511 | 17.5904819 | 17.1558533 | 17.679214 | 16.1735189 | 16.0456746 | 17.5266286 |
| 15.8439893 | 16.1895078 | 16.0498013 | 16.3098883 | 16.003134 | 14.5979618 | 16.3889905 |
| 13.6669453 | 15.9362896 | 14.913568 | 14.2786811 | NA | 13.8003194 | 14.8685409 |
| 24.6064254 | 24.4993529 | 24.2756517 | 25.2121248 | 24.1480896 | 22.9027263 | 24.5335239 |
| 15.9888005 | 15.7202318 | 15.3330176 | 16.4035146 | 14.8545324 | 14.0903469 | 15.6700387 |
| 20.0948281 | 20.0172368 | 19.6267309 | 20.1221822 | 18.9714335 | 18.2883961 | 19.9615872 |
| 18.7547383 | 18.3839804 | 18.2163013 | 19.0132234 | 17.9000853 | 16.4188634 | 18.6653369 |
| 17.4009349 | 17.2759639 | 16.4824053 | 17.2390173 | 15.4279022 | 15.1717446 | 16.9689445 |
| 20.6505023 | 20.2073599 | 19.623955 | 20.4174159 | 18.9431871 | 18.3106067 | 19.9627331 |
| 19.1556309 | 18.3446826 | 18.0338914 | 19.1719748 | 16.9880468 | 16.6123891 | 18.2490697 |
| 15.1121922 | 14.3145049 | 14.1061092 | 14.9108121 | 13.056612 | 12.6555821 | 14.2347798 |
| 14.5979224 | 18.4166559 | 17.0614797 | 17.5076013 | 15.9231014 | 17.6616733 | 20.5801431 |

| AMPH-03340.10nM_2_abundance_norm | AMPH-03340.10nM_3_abundance_norm | AMPH-03340.300nM_1_abundance_norm | AMPH-03340.300nM_2_abundance_norm | AMPH-03340.300nM_3_abundance_norm | AMPH-03340.1000nM_1_abundance_norm | AMPH-03340.1nM - AMPH-04647.10nM__logFC |
|---|---|---|---|---|---|---|
| 16.269552 | 16.5640638 | 18.1006316 | 18.443451 | 17.3692854 | 19.4094112 | 1.609429863 |
| 15.8507383 | 16.0390059 | 16.9702539 | 17.5505126 | 16.4876615 | 18.2369899 | 0.15884868 |
| 11.8557663 | 12.4047141 | 13.2707377 | 15.0324024 | 12.831805 | 12.7465245 | 1.080993298 |
| 16.6810571 | 16.7150936 | 17.988018 | 17.548235 | 18.7731456 | 16.0262989 | -2.2014068 |
| 17.1578104 | 17.722775 | 18.50694 | 18.101803 | 19.4000916 | 17.1542055 | -1.24521202 |
| 14.2316034 | 14.8019348 | 17.6065781 | 14.8606911 | 14.8532881 | 15.3318859 | -0.605963063 |
| 18.8091725 | 19.0602191 | 19.9683426 | 19.7563418 | 20.6538592 | 19.8478083 | -0.983732682 |
| 19.6507686 | 20.3467444 | 21.0687562 | 20.8367675 | 21.4959895 | 19.9155426 | -1.276761091 |
| 14.8551095 | 14.561272 | 15.0565646 | 15.0791676 | 15.3313846 | 15.5410231 | 0.016379311 |
| 16.9576187 | 17.3527288 | 16.9407459 | 17.6350303 | 17.7369222 | 16.1951257 | 0.293713841 |
| 11.402799 | 11.4929567 | 11.5617922 | 12.4624606 | 12.3219954 | 11.9368321 | -0.461493966 |
| 12.6945674 | 12.4147548 | 14.0277073 | 13.7173838 | 14.6188641 | 12.372925 | -2.336949136 |
| 16.978751 | 17.361107 | 17.6010382 | 17.6521403 | 18.2486554 | 17.2462979 | -0.881353557 |
| 15.5460232 | 15.4828556 | 15.5480533 | 16.0167333 | 15.6963944 | 15.6253418 | 0.727507857 |
| 12.7214028 | 12.0288878 | 13.0298464 | 12.8661364 | 13.0928118 | 12.1243797 | 0.08510126 |
| 10.697244 | 11.1963435 | 11.6608729 | 11.7140224 | 12.5855875 | 10.7221084 | 0.18286361 |
| 15.9170243 | 16.4529791 | 16.6290129 | 16.6340421 | 16.9003452 | 16.2979288 | -0.459835158 |
| 15.2058492 | 15.2445282 | 16.0990614 | 15.3375281 | 16.396834 | 15.2313778 | -0.580131502 |
| 14.8922543 | 13.7153837 | 13.7629159 | 14.1349573 | 15.0446941 | 14.8403486 | 1.112136789 |
| 12.3952042 | 13.3094544 | 13.7074508 | 13.2462217 | 13.4064681 | 14.1550407 | 0.432547966 |
| 15.0570955 | 14.5909936 | 14.5761755 | 14.4229689 | 15.5902073 | 14.7060085 | -0.012499684 |
| 14.8095387 | 14.7999236 | 15.4113585 | 15.4499261 | 15.2461625 | 14.882623 | -0.377497737 |
| 14.9729526 | 14.8985859 | 14.987887 | 15.0080008 | 15.9365976 | NA | NA |
| 16.9166375 | 16.1924368 | 16.3153663 | 16.4155548 | 16.9771127 | 17.4050671 | 1.405079614 |
| 11.957762 | NA | NA | 12.6140042 | 13.401103 | 13.4965202 | 0.331568799 |
| 13.703253 | 13.5886405 | 14.4915377 | 13.6580364 | 15.0507142 | 13.817654 | 0.238308522 |
| 15.7650302 | 17.1302498 | 17.995407 | 16.3341197 | 15.1832205 | 15.8148131 | -0.253217888 |
| 16.7150264 | 16.6646423 | 17.4322738 | 17.0515978 | 17.9129229 | 16.1896759 | -1.267306693 |
| 18.9076693 | 19.0360826 | 19.0792338 | 19.309545 | 20.0343185 | 19.8035325 | 0.851493109 |
| 18.1939507 | 18.0122446 | 17.7255171 | 17.8944957 | 19.2875537 | 18.5885086 | 0.427379791 |
| 15.050228 | 15.5053112 | 15.7865499 | 15.7197581 | 16.4436687 | 15.5386912 | -0.219638036 |
| 15.2194716 | 15.5900533 | 15.4230233 | 15.4153117 | 16.3448416 | 15.994598 | 0.08140066 |
| 20.3852588 | 20.4343882 | 20.0018641 | 20.119341 | 21.3088041 | 20.5490053 | 0.229525587 |
| 20.5408243 | 20.7550954 | 20.3407222 | 20.6955412 | 21.5263553 | 20.8704719 | 0.26835236 |
| 17.048114 | 17.6487442 | 17.102503 | 17.3781076 | 17.8228379 | 17.0850443 | -0.437755186 |
| 18.4453558 | 18.5894222 | 18.6581027 | 19.1481901 | 18.3942178 | 19.5341648 | 0.415337009 |

| | | | | | | |
|---|---|---|---|---|---|---|
| 17.9393184 | 17.8362495 | 17.462345 | 17.6343345 | 18.3806951 | 17.7889923 | 0.181024872 |
| 14.4286804 | 14.492321 | 14.5954566 | 14.9372878 | 14.6111791 | 14.5293608 | 0.190767055 |
| 15.2858545 | 14.8320642 | 15.2168274 | 15.1952832 | 16.084053 | 15.0697419 | -0.139043957 |
| 15.8940418 | 17.1274817 | 16.8280029 | 16.4950598 | 17.2143984 | 16.2953166 | 0.300158559 |
| 18.1850553 | 18.242626 | 17.8970328 | 18.1418988 | 19.2476641 | 18.3065719 | 0.476128275 |
| 15.7721044 | 15.8973961 | 15.9653581 | 15.1627197 | 15.725442 | 16.2648756 | 0.726138064 |
| 15.8396143 | 16.3074134 | 16.6862598 | 16.3657894 | 17.0545098 | 16.0454427 | -0.906502066 |
| 10.8997106 | 13.2341244 | 11.0442046 | 12.6466958 | 10.5182057 | 10.9369701 | -0.491799677 |
| 15.5943896 | 15.4656915 | 15.1334159 | 15.3291828 | 15.8494749 | 15.3994318 | 0.204876636 |
| 19.3712451 | 19.5815284 | 19.2463846 | 19.550283 | 19.8908382 | 19.4503304 | 0.14090675 |
| 12.6054031 | NA | 13.3491509 | NA | 13.2495192 | 14.4471858 | 0.190274269 |
| 16.2571284 | 16.2966179 | 16.4185473 | 16.4202496 | 16.5701856 | 16.0034896 | 0.024809202 |
| 17.3977545 | 17.7780677 | 18.1852948 | 18.1336341 | 19.0907366 | 17.7449884 | -0.751298337 |
| 19.3529481 | 19.8676234 | 19.7115092 | 19.8913942 | 19.9954852 | 20.143384 | 0.25861463 |
| 16.87159 | 16.992485 | 17.3775596 | 17.2537399 | 17.7657338 | 16.0047604 | -0.332295265 |
| 17.2491783 | 17.0074976 | 16.617301 | 16.6577575 | 17.8727622 | 17.2716714 | 0.110035073 |
| 14.9083581 | 13.6789691 | 14.5887283 | 13.7354804 | 14.7741452 | 14.3170334 | 1.205326528 |
| 17.2266337 | 17.5320799 | 17.8943057 | 17.4853394 | 18.408805 | 17.1889811 | -0.8183849 |
| 12.5175617 | 12.2827461 | 12.9027002 | 12.3444437 | 12.8429361 | 13.1405289 | 0.387858205 |
| 15.6789791 | 15.1708997 | 15.3518795 | 15.3953015 | 16.158714 | 16.5483187 | 0.68052819 |
| 17.0444974 | 17.1855276 | 16.7947598 | 17.0998607 | 17.7991739 | 17.9792953 | 0.310823583 |
| 14.076364 | 14.3895449 | 15.2230587 | 14.5988097 | 15.5072537 | 14.6515288 | -0.807397714 |
| 18.0334322 | 18.4527884 | 18.0979064 | 18.1411178 | 19.138498 | 18.3660225 | 0.46468495 |
| 18.1459163 | 18.0756767 | 17.916764 | 18.2325454 | 19.0498095 | 18.7645965 | 0.362649635 |
| 17.68987 | 17.7458875 | 18.4190659 | 18.1875136 | 18.9857199 | 18.6123431 | 0.244302368 |
| 20.4679628 | 20.4381491 | 20.179382 | 20.3982147 | 21.427828 | 20.9844438 | 0.276971996 |
| 15.1733698 | 15.5007557 | 15.398463 | 15.6976522 | 16.8755319 | 15.3513847 | 0.278557199 |
| 17.7046094 | 17.7383749 | 17.6624723 | 17.7741905 | 18.520815 | 18.2562929 | 0.286097164 |
| 18.0938098 | 18.1359519 | 17.6317445 | 18.0127486 | 18.7974809 | 18.2426223 | 0.218537271 |
| 13.5384097 | 14.2364604 | 13.9119493 | 14.1932586 | 14.137391 | 14.6429748 | 0.200842948 |
| 16.2375358 | 16.1685049 | 16.1859011 | 15.7434232 | 16.7487634 | 16.6005815 | 0.213719552 |
| 15.2762589 | 14.8798829 | 15.5923818 | 15.0558688 | 15.9348848 | 15.4826261 | -0.505356917 |
| 21.2377236 | 21.220236 | 20.834512 | 21.1847316 | 21.7969214 | 21.1855974 | 0.164390226 |
| 14.2828474 | 14.472727 | 14.6026748 | 14.4977758 | 14.6018003 | 14.434479 | 0.586014857 |
| 17.0819966 | 17.4906682 | 17.2236458 | 17.3588197 | 17.6313065 | 18.1758907 | 0.415418484 |
| 19.5790678 | 19.9580748 | 20.0201519 | 20.1852495 | 20.8337183 | 20.5262892 | 0.19644849 |
| 18.0280737 | 18.5181686 | 18.5710296 | 18.6295249 | 19.4642144 | 18.4882583 | -0.534961769 |
| 11.202009 | 10.8567958 | 11.3602053 | 11.5809235 | 11.8235139 | 12.0643611 | 0.648538434 |
| 13.8870473 | 13.2029193 | 14.9749639 | 13.7120771 | 13.6466741 | 13.7808426 | -0.31430906 |
| 14.3919802 | 14.3001513 | 14.2773016 | 14.3840888 | 14.6336661 | 14.9052054 | 0.195795839 |
| 15.1582618 | 15.3983216 | 15.8224965 | 15.662155 | 15.8979176 | 15.7803022 | -0.121296673 |
| 13.4061453 | 12.8811806 | 12.9310194 | 13.1447617 | 13.6452968 | 13.3343152 | 0.250646441 |

| | | | | | | |
|---|---|---|---|---|---|---|
| 15.3046313 | 14.9213665 | 14.9401292 | 14.838107 | 15.2316786 | 15.8301699 | 1.050456065 |
| 17.9505731 | 18.2050759 | 17.9350283 | 18.2118599 | 18.6276393 | 18.5159506 | 0.842841128 |
| 13.6275851 | 12.9030037 | 13.2157972 | 13.9331235 | 12.9713654 | 13.5637643 | 1.002592828 |
| 20.4985784 | 19.0583001 | 19.5977102 | 19.1660571 | 19.9224357 | 20.9552456 | 1.697837926 |
| 18.3468915 | 16.9209603 | 17.4364187 | 16.7389139 | 17.9957613 | 18.8626154 | 1.778155613 |
| 17.9288473 | 17.8064767 | 18.251404 | 18.6169208 | 18.7926987 | 18.1248982 | 0.394035759 |
| 17.2670951 | 17.1994753 | 17.7983143 | 17.8332556 | 18.1839237 | 17.926721 | -0.103631874 |
| 19.0203968 | 19.0313742 | 18.6664037 | 19.0189959 | 19.6849999 | 19.0241604 | 0.196125058 |
| 15.4536259 | 15.4910442 | 15.8374655 | 15.6237879 | 16.377621 | 15.8561344 | -0.110853086 |
| 16.4438052 | 15.6156197 | 16.1868472 | 16.4903029 | 16.7812654 | 17.0702659 | 0.765454441 |
| 14.5541777 | 15.1695605 | 14.9770507 | 14.9491459 | 15.4235624 | 14.7458495 | -0.534456417 |
| 15.5920787 | 15.7209531 | 15.8520681 | 15.926579 | 16.0663909 | 15.9803615 | -0.547990338 |
| 13.8468204 | 13.8125807 | 14.6635298 | 14.8179336 | 15.2859477 | 14.6873436 | 0.320725529 |
| 15.4311193 | 15.2059048 | 15.0405471 | 15.1001206 | 15.7612052 | 15.427512 | -0.047389457 |
| 18.9515896 | 19.0260156 | 19.280505 | 19.2523547 | 20.0325406 | 19.5379288 | 0.366314943 |
| 13.8822415 | 13.6162617 | 13.5232952 | 14.0628229 | 14.3757806 | 13.9529772 | -0.180705356 |
| 15.4408077 | 15.5882855 | 15.270182 | 15.7537826 | 15.7067564 | 18.7504499 | 0.759226902 |
| 14.4424616 | 14.8134511 | 14.9180099 | 14.5566225 | 15.4429379 | 14.7595229 | 0.302266863 |
| 18.9434226 | 18.7397038 | 18.7428391 | 19.1999176 | 19.8391397 | 19.6396793 | 0.895704311 |
| 17.5973404 | 17.5091689 | 17.259304 | 17.1735565 | 17.6064271 | 17.5671026 | 0.576524588 |
| 13.8271985 | 14.2636443 | 14.2080141 | 14.4537787 | 14.4386516 | 13.9006146 | -0.045073265 |
| 17.9645122 | 18.8017288 | 18.493556 | 18.4459352 | 18.9834868 | 19.1497141 | 0.276868778 |
| 15.1640732 | 15.4014027 | 14.9100469 | 15.0869409 | 15.6419079 | 16.0295792 | 1.04411668 |
| 18.2034849 | 18.2389266 | 17.9731836 | 18.181886 | 19.1649966 | 18.584978 | 0.250512348 |
| 16.4491362 | 16.0587871 | 16.0969002 | 16.0392216 | 16.9816942 | 16.828545 | 0.624074749 |
| 14.1875401 | 13.7303552 | 14.1795367 | 13.841487 | 14.692366 | 14.063233 | 0.303603589 |
| 11.8338834 | 11.4858116 | 11.7946495 | 11.5639149 | 11.7236249 | 11.5549152 | 0.08640025 |
| 17.1163761 | 17.3185293 | 16.956221 | 17.1291842 | 17.8073245 | 17.4513004 | 0.614576334 |
| 14.283445 | 15.3319869 | 14.1883339 | 14.9029751 | 13.8188622 | 14.2008106 | -0.345205737 |
| 20.0978826 | 20.1655213 | 19.7817693 | 20.2298127 | 20.2230692 | 19.9589349 | 0.565587483 |
| 14.1165847 | 14.3503189 | 14.2387192 | 14.4596356 | 14.5885955 | 15.0819972 | 0.235787355 |
| 16.055612 | 15.936033 | 15.9297166 | 15.9418812 | 16.416123 | 16.0244752 | 0.069300444 |
| 16.6649755 | 16.3338446 | 16.1282254 | 16.4825101 | 16.9073235 | 15.7848641 | 0.294038812 |
| 14.3971101 | 13.9943994 | 13.9547721 | 14.2677719 | 14.721787 | 14.6038387 | 0.30465608 |
| 17.5800072 | 17.6565029 | 17.905997 | 17.8864408 | 18.5460751 | 17.9698711 | 0.047482255 |
| 20.4526162 | 20.6332616 | 20.6701933 | 21.1218281 | 21.2930749 | 20.4885826 | 0.2476889 |
| 22.2293144 | 22.2248627 | 22.5444376 | 22.705973 | 23.2738372 | 23.278249 | 0.622334936 |
| 18.7963717 | 18.5914182 | 18.6446425 | 18.6832512 | 19.3929863 | 19.4162622 | 0.664219423 |
| 15.4508549 | 15.5153163 | 15.6090046 | 15.9480472 | 15.9520086 | 17.1898019 | 0.741772732 |
| 18.1931017 | 17.0406552 | 17.8208565 | 17.3588641 | 18.0078395 | 18.8403574 | 0.840516097 |
| 15.8073291 | 15.5540814 | 15.1995412 | 15.3967194 | 16.0622486 | 18.6533248 | 0.064398751 |
| 17.9167682 | 18.0997096 | 18.2804611 | 18.5579317 | 18.8885199 | 18.2991082 | -0.439418484 |

| | | | | | | |
|---|---|---|---|---|---|---|
| 15.7851365 | 16.0351039 | 16.0729878 | 15.827821 | 16.3395265 | 16.2235244 | 0.006754773 |
| 19.0139569 | 18.9849896 | 19.4714629 | 19.424282 | 19.760006 | 19.5337684 | -0.01364269 |
| 13.2144088 | 13.897522 | 13.1264508 | 13.2892683 | 13.9820917 | 13.4332402 | -0.612320394 |
| 15.4704633 | 15.5130285 | 15.6697746 | 15.7669778 | 16.314453 | 15.8433393 | -0.040138029 |
| 17.0853859 | 17.1642337 | 17.1932425 | 17.6248043 | 17.657966 | 18.2279448 | 0.92524777 |
| 11.8592004 | 11.6008831 | 11.6141832 | 11.5183078 | 12.1100813 | 11.308136 | 0.529882516 |
| 13.577823 | 13.4468063 | 13.7491741 | 14.0063918 | 13.803695 | 13.8786861 | 0.70846027 |
| 14.4915754 | 14.6726896 | 14.7358851 | 15.089316 | 15.4093118 | 15.088389 | 0.441489511 |
| 14.8195324 | 15.1432191 | 14.4244669 | 15.1209312 | 14.7100402 | 14.8476081 | 0.098028536 |
| 16.964525 | 16.9841784 | 17.1266812 | 17.4202427 | 17.8993219 | 17.5079059 | -0.094931856 |
| 16.8878831 | 16.8651255 | 16.8784544 | 16.9061418 | 17.7623178 | 17.3358445 | 0.387871137 |
| 19.1312344 | 18.820172 | 18.6318694 | 18.6153486 | 19.4102305 | 19.2541784 | 0.502738899 |
| 16.308613 | 15.6435378 | 16.134593 | 16.4039152 | 16.1026936 | 16.5924541 | 0.356626702 |
| 15.5374666 | 15.3798835 | 15.7192226 | 15.8691407 | 15.4408707 | 15.921741 | 0.264459941 |
| 16.4842311 | 16.6011577 | 16.6686631 | 17.076734 | 17.5043233 | 16.8142322 | 0.860743826 |
| 16.8714802 | 17.0922031 | 16.6162878 | 16.9033227 | 17.273824 | 16.912048 | 0.567677599 |
| 14.6523616 | 14.4515543 | 14.487791 | 14.7510914 | 14.791718 | 15.5096599 | -0.494883146 |
| 16.7157657 | 16.3278062 | 16.3220555 | 16.3801433 | 16.5663696 | 17.2051122 | 0.942949278 |
| 14.6241625 | 14.3801914 | 14.1342602 | 14.8860326 | 15.1095878 | 14.7950956 | 0.230419849 |
| 11.5907335 | 11.600661 | 11.7303985 | 11.9514898 | 11.9232054 | 12.0738913 | 0.604454262 |
| 15.2652219 | 14.9732884 | 15.3551325 | 15.1502067 | 15.8119224 | 16.4465033 | 0.846206108 |
| 16.191211 | 16.0587106 | 16.3804344 | 16.2225361 | 16.5557764 | 16.8410074 | 0.085611359 |
| 18.1240863 | 18.3501187 | 18.1855028 | 18.2579636 | 18.8359068 | 18.680222 | 0.512249491 |
| 17.5850913 | 17.3822915 | 17.4148284 | 17.491713 | 18.0518851 | 17.9834988 | 0.267952864 |
| 16.3516935 | 16.4415221 | 16.4152307 | 16.8086421 | 16.9047445 | 16.7290007 | 0.653362847 |
| 16.8872432 | 17.3784322 | 17.6573349 | 17.6387316 | 16.7864732 | 17.105914 | -0.106411132 |
| 18.1896286 | 17.9782428 | 18.1771281 | 18.1953619 | 18.6994051 | 18.6618833 | -0.21959822 |
| 17.1334515 | 17.3356085 | 17.2522473 | 17.383587 | 17.8689504 | 17.8524474 | 0.530731294 |
| 14.6248213 | 13.8752575 | 14.9571811 | 14.9142502 | 14.9906094 | 15.1910383 | 0.803347701 |
| 17.2913546 | 17.4114354 | 17.5970931 | 17.6020793 | 18.0031512 | 17.776817 | 0.22482186 |
| 18.2136941 | 18.3819194 | 18.4731548 | 18.6626483 | 18.7509565 | 18.7260458 | 0.665105005 |
| 14.1924763 | 12.9566884 | 13.7545464 | 12.7404454 | 13.7893899 | 14.4849461 | 1.420951801 |
| 17.319878 | 17.8331111 | 17.6571002 | 18.1839952 | 18.1077358 | 18.1773985 | 0.576619816 |
| 17.4854777 | 17.5281231 | 17.090141 | 17.3178521 | 18.2546222 | 17.3981852 | 0.396237798 |
| 15.2082251 | 15.0873966 | 15.418869 | 15.7213763 | 15.7910844 | 15.9566972 | -0.200930102 |
| 17.2849711 | 17.2124521 | 17.2429125 | 17.5720109 | 17.7589187 | 18.214969 | 0.644656874 |
| 15.3291301 | 15.6820052 | 15.5246437 | 15.0444719 | 15.7866972 | 15.7006057 | 0.75329517 |
| 14.9033823 | 14.9909694 | 15.3964239 | 15.0830086 | 15.9200098 | 15.2337768 | -0.433572016 |
| 14.3356678 | 14.0579842 | 13.4861018 | 14.0842003 | 14.7708992 | 14.9511138 | 1.372753833 |
| 13.8781096 | 14.618886 | 13.7666952 | 14.281118 | 13.5922106 | 14.0754673 | 0.107851274 |
| 14.144581 | 13.881829 | 14.2234844 | 14.4739185 | 14.4347035 | 14.8578447 | 0.30812519 |
| 15.4851147 | 17.2627598 | 17.6205586 | 16.8098722 | 15.6150548 | 16.3483652 | -0.290052226 |

| | | | | | | |
|---|---|---|---|---|---|---|
| 18.2591495 | 18.19598 | 18.1646907 | 18.4282845 | 18.4801445 | 19.1995114 | 0.891485315 |
| 18.9385093 | 19.2274256 | 19.1820418 | 19.8627508 | 20.259989 | 19.4204749 | 0.139202907 |
| 18.1596229 | 18.1330154 | 18.369783 | 18.4038743 | 18.8502306 | 19.0878907 | -0.228426924 |
| 10.9073977 | 11.7670327 | 12.0464301 | 11.8622065 | 12.2859238 | 12.7134165 | 0.276449967 |
| 18.5528703 | 18.7596158 | 18.9113068 | 18.8442954 | 19.6444944 | 19.1499864 | -0.059158972 |
| 18.3558276 | 17.8652862 | 17.9485455 | 17.9104333 | 18.4693462 | 18.5715123 | 0.904235656 |
| 14.5044616 | 14.9728887 | 15.1804289 | 14.5830299 | 15.4973788 | 14.3722276 | 0.191673861 |
| 14.6266049 | 14.4598791 | 14.9745864 | 14.54494 | 15.2759289 | 15.1451452 | 1.273573579 |
| 16.5238561 | 16.3570878 | 16.8056388 | 16.5286362 | 16.8873342 | 16.6060984 | 0.514528352 |
| 15.3010259 | 15.487248 | 15.9941478 | 16.0999409 | 15.5613994 | 15.1904463 | 0.715143898 |
| 14.9234793 | 14.8550825 | 14.5411201 | 14.6589446 | 15.5917238 | 15.2197739 | 0.386107814 |
| 12.8722679 | NA | 13.2595325 | 13.0237364 | NA | 13.1794671 | -0.25978993 |
| 15.0244613 | 15.7248757 | 15.6208104 | 15.9494895 | 14.8232092 | 14.990167 | -0.229381864 |
| 14.4203091 | 14.9041209 | 14.2709469 | 15.2267783 | 14.8461875 | 14.7411582 | 0.558226351 |
| 11.529836 | 12.2054716 | 12.3669513 | 12.1708581 | 12.4201198 | 12.2813269 | 0.560983489 |
| 20.2151057 | 20.4797861 | 20.2756584 | 20.2527344 | 21.0621906 | 20.9507735 | 0.39394671 |
| 18.2070826 | 18.2555431 | 18.2222993 | 18.3040306 | 18.8964841 | 19.2059711 | 0.627764251 |
| 14.5305488 | 14.8254029 | 14.707364 | 15.0109493 | 15.2838775 | 14.6780335 | 0.012860836 |
| 17.1375184 | 17.3054696 | 16.9461954 | 17.5132278 | 17.9407045 | 18.0563562 | 0.652177352 |
| 20.759907 | 20.9199438 | 20.772306 | 20.9415992 | 21.6620642 | 21.8213948 | 0.759548369 |
| 17.4735173 | 17.7920256 | 17.8639163 | 17.9545238 | 18.6491407 | 17.9881517 | -0.350323158 |
| 17.2489087 | 17.2326817 | 16.5939851 | 17.0209055 | 17.7461291 | 17.3303538 | 0.22778233 |
| 15.0781501 | 14.5896539 | 14.7213728 | 14.5548032 | 15.6309444 | 15.8626152 | 0.98879191 |
| 17.6388862 | 17.7017931 | 17.5486262 | 17.8687669 | 18.1422735 | 18.1069416 | 0.306629743 |
| 19.4816035 | 19.7078307 | 19.7151522 | 19.9919544 | 19.9416923 | 20.1802197 | 0.6206862 |
| 14.6944366 | 14.570598 | 15.2497901 | 14.8739979 | 15.6190181 | 14.3713136 | -0.561574731 |
| 15.2216793 | 15.3270517 | 14.7446716 | 15.2557024 | 15.5947518 | 15.1096985 | 0.373619458 |
| 17.2769599 | 17.0485564 | 17.1437432 | 16.7311229 | 17.9748806 | 18.1345415 | 0.508075677 |
| 13.3615671 | 13.6307176 | 13.2815189 | 13.3256863 | 13.8810015 | 14.3623897 | 1.208265965 |
| 16.8527747 | 17.1960343 | 17.0659597 | 16.9516289 | 17.437817 | 17.3212704 | 0.157678604 |
| 16.7588585 | 17.0737663 | 16.7576668 | 16.6605995 | 16.8960339 | 17.347705 | 0.715928447 |
| 13.3623165 | 14.1175987 | 14.1403019 | 14.4374968 | 13.8663363 | 14.4967712 | -0.466816036 |
| 13.8969142 | 14.891373 | 13.9611504 | 14.2519462 | 14.3800037 | 14.6566843 | 0.338061758 |
| 18.4686584 | 18.3075407 | 18.2503495 | 18.2477602 | 18.7181913 | 18.9349702 | 0.603724965 |
| 14.8068753 | 15.3814469 | 15.3195004 | 15.2726078 | 14.430995 | 14.8548501 | 0.081204267 |
| 14.6439793 | 14.975448 | 14.8086608 | 15.474455 | 15.4321924 | 15.0431073 | 0.796863275 |
| 17.9779258 | 17.4433601 | 17.394275 | 17.2854184 | 17.5039694 | 18.4390837 | 0.730301315 |
| 19.9585061 | 20.371168 | 20.1196354 | 20.2385559 | 20.8574043 | 20.7572628 | 0.388383463 |
| 15.5734109 | 15.7734292 | 15.8062624 | 15.8973578 | 16.1613124 | 15.8461413 | 0.340439702 |
| 14.1890172 | 14.1421212 | 14.5357278 | 14.224828 | 14.0303898 | 14.419349 | 0.222493487 |
| 21.8343274 | 21.9946934 | 21.9906837 | 22.2622512 | 22.759504 | 22.7315365 | 0.435742903 |
| 14.6985885 | 14.6542723 | 14.7815273 | 14.9414806 | 15.4147986 | 15.0769115 | 0.489887308 |

| | | | | | | |
|---|---|---|---|---|---|---|
| 16.842801 | 17.1856826 | 16.9357442 | 17.4210619 | 17.2113707 | 17.3255063 | 0.652438214 |
| 12.9250879 | 14.0440057 | 15.0531487 | 13.5637565 | 13.1183162 | 13.931284 | -0.47567024 |
| 21.2982003 | 21.88165 | 21.687209 | 22.2686397 | 22.0602793 | 20.8326984 | 0.737358289 |
| 15.1795251 | 14.8954528 | 15.1138036 | 15.6087393 | 15.5315386 | 15.6199183 | 0.267263992 |
| 14.0704246 | 14.3091284 | 14.4066119 | 14.0189974 | 14.3681571 | 14.1881513 | 0.78076609 |
| 11.9408463 | 11.7962826 | 12.1150355 | 11.9044302 | 12.4366887 | 12.0658925 | 0.15164444 |
| 15.1182488 | 15.3232109 | 15.5352424 | 15.5864842 | 15.8981016 | 15.2812045 | 0.42716948 |
| 14.9343664 | 15.9086565 | 14.7305951 | 14.8157345 | 15.7728982 | 15.6518467 | 0.467565289 |
| 18.9379287 | 18.8528082 | 19.157053 | 19.3207644 | 19.6376721 | 19.5502274 | 0.181621086 |
| 11.4385538 | 12.0841215 | 11.4424561 | 12.0579131 | 11.7962843 | 11.072725 | -0.175134531 |
| 14.0534903 | 12.8260148 | 13.4468342 | 13.7591113 | 13.7112001 | 13.3948544 | -0.510130506 |
| 20.5875802 | 20.7234528 | 20.4762313 | 20.6483906 | 20.8064829 | 21.0747467 | 0.574156703 |
| 19.901993 | 20.0348447 | 20.1354888 | 20.2955638 | 20.8621099 | 20.2591805 | 0.491329544 |
| 16.9674954 | 16.7926399 | 16.6690197 | 16.7118959 | 17.4270524 | NA | -1.012576419 |
| 13.9583691 | 13.781776 | 14.3622708 | 13.9636752 | 14.8340537 | 15.0370115 | 0.069136037 |
| 16.7147749 | 16.8907762 | 16.6422554 | 16.9534618 | 17.3852938 | 17.5994665 | 0.414314198 |
| 15.5185137 | 15.209893 | 15.4867937 | 15.4587888 | 15.8791061 | 16.323229 | 0.776980119 |
| 14.2453926 | 15.4155521 | 15.5454284 | 14.8789032 | 13.3875893 | 14.0572979 | -0.873702397 |
| 20.6428463 | 20.912664 | 21.0354709 | 21.4797975 | 20.5566264 | 21.7683364 | 0.516365378 |
| 20.2185498 | 20.4844993 | 20.7334049 | 21.0955044 | 20.2103799 | 21.546754 | 0.483764878 |
| 17.465752 | 17.7307078 | 17.6334098 | 17.968891 | 18.2505901 | 17.8139004 | 0.365354079 |
| 20.7868957 | 21.0133335 | 21.0815808 | 21.536505 | 21.7541224 | 21.6148052 | 0.555094277 |
| 12.2292699 | 12.2039523 | 11.8373148 | 11.9298366 | 11.9358009 | 11.2729569 | 1.029560339 |
| 18.7479174 | 18.9042379 | 19.1449042 | 19.0721282 | 19.6283591 | 19.5389277 | 0.44700115 |
| 18.6278844 | 18.1502521 | 18.2769668 | 18.0262691 | 18.8492843 | 19.5521005 | 0.847181178 |
| 16.1746592 | 16.2430809 | 16.5334867 | 16.4563928 | 16.9410496 | 16.724899 | 0.209362253 |
| 12.8554704 | 13.012558 | 12.9506375 | 13.1095198 | 12.9839268 | 12.7494734 | -0.294565691 |
| 21.5966166 | 21.9873762 | 21.7884527 | 22.4253551 | 22.511906 | 22.4287569 | 0.683675673 |
| 15.1551121 | 15.2998051 | 15.0168766 | 15.5657795 | 15.3722769 | 15.2711905 | 0.581901501 |
| 24.5812533 | 25.2572268 | 24.7808613 | 25.6968251 | 24.7460823 | 25.599036 | -0.084549191 |
| 14.3433547 | 13.9318282 | 14.4378662 | 14.9918338 | 14.6436713 | 15.0507718 | 0.772721839 |
| 15.0914747 | 15.0315096 | 15.0527226 | 15.0096514 | 15.5949495 | 15.8807067 | 0.415584547 |
| 14.9558341 | 13.683844 | 14.5376853 | 14.2752952 | 14.3228931 | 15.0565539 | 0.660370906 |
| 22.38211 | 22.3865479 | 22.338396 | 22.7034486 | 22.7104389 | 23.0367499 | 0.476571577 |
| 16.9631354 | 16.6150801 | 16.7228741 | 16.4475217 | 17.0870136 | 17.6760492 | 0.648421136 |
| 20.1919019 | 20.3134153 | 20.1038081 | 20.362122 | 20.8877641 | 20.9418428 | 0.663088349 |
| 19.9212267 | 20.1141225 | 20.4071282 | 20.722622 | 19.8323769 | 21.1656523 | 0.527818981 |
| 15.3422739 | 15.4016946 | 15.2477542 | 15.4784161 | 15.6030248 | 14.9041493 | 0.176475103 |
| 17.3194744 | 17.400061 | 17.7842393 | 17.5446545 | 18.244688 | 18.0060044 | -0.15065235 |
| 13.7512099 | 14.1635939 | 14.0890076 | 14.5789466 | 13.7922187 | 13.8114106 | 0.789642742 |
| 15.0146651 | 15.3106552 | 14.8667847 | 15.1265517 | 15.5223524 | 15.5941689 | 0.66013182 |
| 13.3160251 | 14.0553304 | 13.8004087 | 12.5485415 | 13.9085639 | 13.1936688 | 0.125220115 |

| | | | | | | |
|---|---|---|---|---|---|---|
| 13.4568559 | 14.2918401 | 14.158964 | 14.5012126 | 13.6584987 | 13.4873057 | -0.697650113 |
| 11.5819403 | 12.5658513 | 11.5812066 | 12.3230014 | 11.5649827 | 11.6552975 | -0.328834191 |
| 15.4762416 | 15.7543783 | 15.913099 | 16.0987138 | 16.675681 | 16.3292516 | -0.329337768 |
| 14.979451 | 15.0286043 | 15.446747 | 15.300164 | 15.5850933 | 15.6741168 | -0.925664576 |
| 12.6938792 | 13.0005062 | 13.557141 | 13.0003006 | 13.6453007 | 13.1054942 | 0.017316663 |
| 21.070256 | 21.2818725 | 21.1747111 | 21.5334494 | 21.7526443 | 21.9205999 | 0.534353932 |
| 14.9792856 | 15.1866422 | 15.3362769 | 15.2895244 | 15.2830925 | 15.1472858 | 0.319522511 |
| 14.9111367 | 15.4128283 | 15.5839162 | 15.598579 | 15.3999142 | 14.84977 | 0.426372467 |
| 19.0206299 | 19.1861362 | 19.3661308 | 19.8057067 | 18.919799 | 20.3161606 | 0.481382064 |
| 20.9108266 | 21.2007245 | 21.3881929 | 21.7341298 | 20.9915643 | 22.0623687 | 0.536302924 |
| 15.0562775 | 15.0333973 | 14.7923249 | 15.0524193 | 15.4021724 | 15.3515508 | 0.172961401 |
| 16.0171044 | 15.9747402 | 15.8369831 | 15.9530132 | 16.4160895 | 16.7634172 | 0.300341746 |
| 13.7403706 | 12.8639089 | 13.3646495 | 13.1778341 | 13.2118274 | 13.9316325 | 0.027119941 |
| 15.2685192 | 15.4744267 | 15.5940029 | 15.7966552 | 15.40622 | 15.9125394 | 0.283921894 |
| 21.3463019 | 21.632086 | 21.560264 | 21.6787848 | 22.2111168 | 22.2816831 | 0.640463293 |
| 16.5281699 | 16.4977032 | 16.7073487 | 16.7201817 | 17.126562 | 16.9674749 | -0.349973547 |
| 19.0123896 | 18.8855542 | 19.074749 | 18.7470864 | 19.6058176 | 19.8866503 | 0.546373416 |
| 19.943319 | 20.1154031 | 19.8881451 | 20.3108981 | 20.5530135 | 20.6535073 | 0.591911454 |
| 15.3045701 | 15.0426279 | 15.0041168 | 15.283081 | 15.0979698 | 15.287005 | -0.319475133 |
| 14.483565 | 13.9943393 | 14.39861 | 14.3342871 | 14.3718318 | 13.919282 | 0.136159488 |
| 21.3836327 | 21.6640712 | 21.5597095 | 21.7889529 | 22.1131535 | 21.9630928 | 0.532143422 |
| 13.9581516 | 13.6466282 | 14.1663295 | 13.8544022 | 14.5131141 | 13.8836872 | 0.144220881 |
| 15.7032072 | 15.8766659 | 15.7857117 | 15.8889145 | 16.1614119 | 16.3044962 | -0.050499626 |
| 15.7693644 | 15.9892029 | 16.1346491 | 16.1350704 | 16.2683784 | 16.0189328 | 0.487658846 |
| 15.8450387 | 15.6211348 | 15.5805389 | 15.9052932 | 16.0855421 | 15.8811209 | 0.375328413 |
| 18.7830275 | 19.0008694 | 19.2187747 | 19.6726606 | 18.7959785 | 20.0119622 | 0.521058258 |
| 21.5757386 | 21.9831845 | 21.7897638 | 22.1548736 | 22.3661319 | 22.3884229 | 0.366824034 |
| 19.2518923 | 19.4388275 | 19.4153834 | 19.6102491 | 19.9564746 | 19.4996585 | 0.351172975 |
| 20.6135232 | 20.8978612 | 21.0170178 | 21.3858794 | 20.6083548 | 21.8931353 | 0.465010306 |
| 15.6767852 | 14.8339861 | 15.3463689 | 14.7146921 | 15.1909763 | 16.0348258 | 1.115400228 |
| 14.0478448 | 14.8422836 | 14.5121492 | 14.587486 | 15.051112 | NA | NA |
| 11.9604301 | 11.6218175 | 11.6723128 | 11.5029317 | 12.8120909 | 12.8923523 | 0.919571028 |
| 19.2159606 | 19.8267665 | 19.8020171 | 20.1870884 | 19.5484691 | 21.281501 | 0.833653509 |
| 17.0375498 | 17.1069627 | 16.8886168 | 17.0310693 | 17.7327574 | 17.3649866 | 0.269498687 |
| 17.1014927 | 17.8676584 | 17.9698931 | 17.584985 | 17.2955814 | 16.9907375 | -0.478517427 |
| 19.0650504 | 19.0707918 | 18.7330727 | 18.831106 | 19.1348892 | 19.2183568 | 0.265285219 |
| 13.4440349 | 13.4839544 | 13.150303 | 13.2621283 | 14.6547038 | 13.3244795 | 0.110532213 |
| 16.9946301 | 17.6979164 | 16.9585186 | 17.7525348 | 17.5689548 | 18.2430293 | 0.30730484 |
| 14.2596966 | 14.0318465 | 14.6142218 | 14.3769681 | 14.8380039 | 14.6893302 | 0.145526229 |
| 18.4572895 | 18.3891617 | 18.8628916 | 18.7913062 | 19.3581727 | 18.9855905 | -0.114803903 |
| 16.9887727 | 16.9885445 | 17.2096444 | 17.1765822 | 17.6992869 | 17.7696196 | 0.407923922 |
| 14.6202141 | 13.8603465 | 14.8563323 | 14.5301464 | 15.1071834 | 14.7661551 | 0.345391423 |

| | | | | | | |
|---|---|---|---|---|---|---|
| 19.7882783 | 19.9166377 | 19.8936211 | 20.4631438 | 20.9513347 | 20.0054247 | 0.042681446 |
| 16.6619778 | 16.3996483 | 16.4392216 | 16.4339666 | 16.7821316 | 17.8466815 | 1.335356221 |
| 14.3918413 | 14.3338017 | 14.2148367 | 14.3495665 | 14.7180043 | 15.374732 | 0.287246563 |
| 13.7669381 | 13.4585924 | 13.9042759 | 13.4193527 | 14.0762284 | 13.6615536 | -0.186117316 |
| 18.7148691 | 19.2592342 | 19.0454682 | 19.3718485 | 19.3631161 | 18.365025 | 0.626098746 |
| 18.2254378 | 18.5099414 | 18.177813 | 18.0967313 | 18.6635691 | 18.8632381 | 0.559709344 |
| 15.4507457 | 15.3532925 | 15.2218022 | 15.3148303 | 15.6394563 | 15.4177472 | 0.154023997 |
| 16.4966316 | 16.633666 | 16.8277829 | 17.0431762 | 17.0002172 | 17.278665 | 0.075257078 |
| 20.9714403 | 21.2077289 | 21.1697133 | 21.3731213 | 21.6443942 | 22.0501171 | 0.485907425 |
| 16.661663 | 16.4887508 | 16.5088161 | 16.3818135 | 16.7817975 | 16.5966542 | -0.219839207 |
| 15.4610101 | 15.3466848 | 15.3758961 | 15.6914368 | 15.7660862 | 15.3158669 | 0.078068973 |
| 13.9871364 | 14.0752615 | 13.9472435 | 13.9972641 | 14.2274553 | 14.5316561 | 0.487420757 |
| 14.8978308 | 14.7392386 | 14.7704949 | 14.9094597 | 15.1159245 | 15.1520739 | 0.371577665 |
| 12.5341108 | 12.9705945 | 13.2308486 | 12.6460806 | 12.6703119 | 12.8925171 | 0.489446127 |
| 15.7604719 | 15.4434216 | 15.412174 | 15.4443728 | 15.8378628 | 16.1456723 | 0.836981939 |
| 15.1831044 | 15.3181054 | 15.2475767 | 15.519744 | 16.0436007 | 16.0086449 | -0.638116141 |
| 15.0917669 | 15.1336347 | 14.7409839 | 15.1021852 | 14.7681426 | 15.3815606 | 0.573379741 |
| 17.6135927 | 17.369669 | 17.5647568 | 17.7172332 | 18.0687846 | 18.1616055 | -0.213967292 |
| 14.5122734 | 15.6682578 | 14.6773361 | 14.5342759 | 14.9676536 | 15.5120255 | 0.93652343 |
| 13.9686067 | 14.1418813 | 14.1106064 | 14.1223816 | 14.4765466 | 13.9273406 | 0.290631376 |
| 22.8046003 | 22.8354726 | 22.8576732 | 23.1413019 | 23.2025653 | 23.5168156 | 0.596989163 |
| 17.5993502 | 17.7462867 | 17.5973905 | 17.831337 | 18.4994808 | 18.0487395 | 0.40323527 |
| 18.9141924 | 18.9322419 | 18.9302614 | 19.1586937 | 19.1884033 | 19.3376077 | 0.609080351 |
| 11.8753866 | 11.5118092 | 11.3604411 | 11.4365201 | 12.2851929 | 11.8076841 | 0.448157981 |
| 18.3882983 | 18.4735993 | 18.4965932 | 19.0491066 | 18.908813 | 19.1270053 | 0.586467556 |
| 20.6477914 | 20.760768 | 20.5995048 | 20.9135722 | 21.2732016 | 20.8663253 | 0.475592823 |
| 15.5466707 | 17.0687951 | 15.9721614 | 16.4700601 | 15.4264841 | 13.3802397 | -2.986773629 |
| 19.4692659 | 19.4942491 | 19.6669012 | 19.8738873 | 20.1751719 | 19.9573911 | 0.354002179 |
| 16.5080495 | 16.6066289 | 16.6055815 | 16.9979994 | 17.01377 | 17.0991476 | 0.228517551 |
| 21.4400932 | 21.8211616 | 21.4831314 | 21.8997498 | 22.2793393 | 21.9786433 | 0.538402047 |
| 15.9664488 | 15.5972139 | 14.8815166 | 15.4626662 | 16.068212 | 16.1242099 | 0.33205837 |
| 18.4648458 | 18.6300592 | 18.5733608 | 18.5335078 | 18.926088 | 19.6206209 | 0.276324561 |
| 15.7418448 | 15.6567178 | 15.3833237 | 15.597013 | 15.812106 | 15.4243029 | 0.9058464 |
| 14.4738398 | 14.8025601 | 14.950797 | 14.7934029 | 15.2355794 | 15.3679013 | 0.855581323 |
| 17.4874802 | 17.2867228 | 17.4450065 | 17.4422296 | 17.6761013 | 18.3739905 | 0.265045976 |
| 15.6019969 | 15.522836 | 15.1932319 | 15.4772009 | 15.7381451 | 15.681237 | 0.154453332 |
| 15.256686 | 15.2604948 | 15.5528679 | 15.8384578 | 15.6534992 | 15.7316336 | 0.830314364 |
| 16.6189194 | 17.1543337 | 18.1663769 | 17.1941591 | 15.4008971 | 16.2239268 | -0.208837412 |
| 14.5807334 | 14.7909889 | 14.71753 | 14.8826483 | 15.051773 | 15.0940959 | 0.417321156 |
| 17.528389 | 17.0759598 | 17.1866385 | 17.1004341 | 17.4693413 | 17.9899317 | 0.854320411 |
| 11.6694675 | 11.6722863 | 11.6430964 | 11.99327 | 11.746553 | 11.5244592 | 0.454262799 |
| 16.8803627 | 16.99517 | 16.7966373 | 17.2272911 | 17.4879341 | 17.4757559 | 0.269235913 |

| | | | | | | |
|---|---|---|---|---|---|---|
| 16.7804408 | 16.7580727 | 16.7913343 | 16.6896196 | 17.1967442 | 17.4123003 | 0.423741615 |
| 12.5672772 | 13.5195856 | 14.1522293 | 13.92163 | 12.9278569 | NA | -0.715901087 |
| 14.2305099 | 14.4263488 | 14.4021971 | 13.2378977 | 14.5115452 | 14.551205 | 0.682071555 |
| 19.5207559 | 19.7124088 | 19.6072496 | 19.6704446 | 20.0689094 | 20.4657626 | 0.29103912 |
| 20.6957072 | 20.6480153 | 20.6857956 | 20.5371133 | 21.3659982 | 21.2993403 | 0.412741435 |
| 17.3048724 | 17.3256448 | 17.040051 | 17.4534316 | 17.6704459 | 17.562793 | 0.468480334 |
| 16.8623985 | 16.3996811 | 16.8913233 | 16.2637682 | 16.8313572 | 17.4880762 | 0.841396855 |
| 18.3143563 | 18.6124826 | 18.37964 | 18.4276605 | 18.9402142 | 19.076489 | 0.206954213 |
| 16.3021048 | 17.313223 | 16.982685 | 16.6909645 | 15.8574527 | 16.3995162 | -0.501877405 |
| 17.1626058 | 17.3665019 | 17.2008879 | 17.6182836 | 17.6864352 | 17.9155184 | 0.619755693 |
| 15.5575677 | 15.8709489 | 15.3569977 | 15.5905214 | 15.727255 | 14.9566023 | -0.088212297 |
| 18.9069993 | 19.2138384 | 19.1993459 | 19.6057556 | 18.9991954 | 20.2235964 | 0.47801079 |
| 13.4887566 | 13.447812 | 13.4503213 | 13.6242518 | 13.8161887 | 14.0473884 | 0.281940947 |
| 16.4310409 | 16.2366621 | 15.9019776 | 16.5180903 | 16.8634128 | 16.3156751 | 0.483592971 |
| 11.5748548 | 11.2437707 | 11.784202 | 11.4095093 | 11.8870929 | 11.9581724 | 1.201414117 |
| 12.183316 | 12.4584455 | 12.0781198 | 11.6910447 | 12.6521607 | 13.2461985 | -0.931883078 |
| 21.2730214 | 21.6487623 | 21.703396 | 21.9985149 | 21.9099386 | 22.4313661 | 0.453896481 |
| 12.0644769 | 12.0853225 | 11.5373645 | 11.5302936 | 12.5594111 | 11.9725338 | 0.160009883 |
| 16.6613833 | 16.8888604 | 16.9236951 | 17.12788 | 17.3899633 | 17.4291742 | -0.082831747 |
| 17.498695 | 17.9776234 | 17.7616885 | 18.1167723 | 17.8813343 | 18.1520826 | 0.687964641 |
| 16.1380267 | 16.1212616 | 16.4450271 | 16.3013679 | 16.5493145 | 16.7113746 | -0.024651843 |
| 22.0421842 | 22.3395592 | 21.974023 | 22.4980122 | 22.262735 | 22.2727879 | 0.451614723 |
| 18.2532615 | 18.4559466 | 18.1064976 | 18.1324885 | 18.6369789 | 18.8625106 | 0.411854501 |
| 18.5739734 | 18.6090333 | 18.5842549 | 18.7760844 | 18.8748469 | 19.0951169 | 0.332062269 |
| 16.2164385 | 16.280044 | 16.0852271 | 15.8515774 | 16.5665254 | 16.3170404 | 0.280766392 |
| 19.3169796 | 19.2630603 | 19.2996929 | 19.5393397 | 20.0527441 | 19.509437 | 0.670207949 |
| 16.4932139 | 16.2588551 | 16.6946875 | 16.5072013 | 17.2436613 | 17.0265477 | 0.200890443 |
| 13.1952507 | 12.353106 | 13.0016093 | 12.6477045 | 12.7236607 | 12.8620453 | 0.631723643 |
| 20.9610151 | 21.1432362 | 21.501529 | 21.5258836 | 21.5463674 | 21.6105429 | 0.093495932 |
| 17.2843226 | 17.3944734 | 17.3316777 | 17.3029292 | 17.552412 | 17.6510046 | -0.150066652 |
| 15.6905975 | 15.7184224 | 15.9132795 | 15.8689912 | 16.212215 | 16.1402237 | 0.194667716 |
| 16.081065 | 16.1462537 | 16.1340749 | 16.4896568 | 16.6915678 | 16.9496761 | 0.556027731 |
| 19.2526597 | 19.4208449 | 19.3784733 | 19.9675912 | 20.225724 | 19.3228079 | -0.131136145 |
| 17.1616302 | 17.1671876 | 17.1958209 | 17.3867832 | 17.9294027 | 18.4862455 | 0.768919139 |
| 16.9410642 | 17.028854 | 17.0859079 | 17.4943393 | 17.1119513 | 17.3331399 | 0.438409427 |
| 14.2741059 | 14.8842961 | 14.2738118 | 14.3243326 | 13.9840139 | NA | -0.319383733 |
| 14.8159498 | 15.1777418 | 14.7642837 | 14.8894121 | 15.1584572 | 15.0948044 | 1.703982637 |
| 17.03245 | 17.2018202 | 17.2680828 | 17.5137758 | 17.481982 | 17.5735607 | 0.499676161 |
| 15.1863231 | 14.9119276 | 14.9989601 | 15.0383263 | 15.1186951 | 15.6210353 | 0.46562398 |
| 17.6280734 | 17.759234 | 17.7376312 | 17.9568779 | 18.0754004 | 18.3008936 | 0.504067613 |
| 17.0052213 | 17.0037911 | 16.9320189 | 17.0881941 | 17.2553073 | 17.6506266 | 0.424219414 |
| 18.6150702 | 18.9164408 | 18.5840653 | 18.5413429 | 19.2335079 | 19.650281 | 0.02055349 |

| | | | | | | |
|---|---|---|---|---|---|---|
| 16.4364966 | 16.5411841 | 16.7542447 | 17.0568447 | 16.4139898 | 17.7013154 | 0.569574651 |
| 14.673293 | 14.1865041 | 14.7254539 | 14.8605688 | 15.0864842 | 15.4717388 | 0.601772179 |
| 15.6814644 | 15.6304662 | 15.7110079 | 15.8128006 | 16.3254056 | 16.2080122 | 0.047642498 |
| 21.4182923 | 21.8160898 | 21.7058615 | 22.1424242 | 22.3184762 | 22.47394 | 0.483670813 |
| 21.1065623 | 21.5491703 | 21.3974862 | 21.9116524 | 21.7935334 | 22.0005094 | 0.643830454 |
| 14.5343195 | 14.769359 | 14.6757354 | 14.7070014 | 15.4616948 | 15.5418044 | 0.467728762 |
| 13.5534585 | 13.7058129 | 13.6052798 | 13.7084575 | 14.2613024 | 13.5316477 | 0.145056024 |
| 16.0561304 | 15.9451849 | 16.3747818 | 16.563956 | 16.6343475 | 17.078366 | -0.026684224 |
| 19.6189741 | 19.5892105 | 19.5557619 | 19.7202463 | 19.8889595 | 20.1192488 | 0.419972402 |
| 17.5915712 | 17.7000071 | 17.8412404 | 18.2546609 | 17.3704475 | 17.9939149 | 0.2531444 |
| 18.4782487 | 18.7386478 | 18.7565258 | 18.9664905 | 18.9275023 | 19.0767683 | 0.06736723 |
| 14.9321441 | 14.8908986 | 15.055208 | 15.2731767 | 14.9678786 | 15.4833873 | 0.530413702 |
| 14.8610935 | 15.3131534 | 14.8602634 | 14.5677373 | 15.2336149 | 15.1472241 | 0.426629451 |
| 18.2070869 | 18.4827181 | 18.0424051 | 18.4473753 | 18.5775192 | 18.7168843 | 0.288364108 |
| 14.4444989 | 14.9902779 | 14.945443 | 14.248786 | 14.6178094 | 14.3407081 | 0.776895655 |
| 16.6375553 | 16.7269182 | 16.7762685 | 16.8345887 | 17.2855663 | 16.8733881 | 0.687520855 |
| 16.1655357 | 15.956649 | 15.9571304 | 16.1924952 | 16.5907331 | 16.6639576 | 0.508442161 |
| 15.9315264 | 15.8434742 | 15.8506405 | 16.2431552 | 16.658913 | 15.8107419 | 0.018097226 |
| 14.8934071 | 15.4499911 | 15.2574273 | 15.5215251 | 15.1362759 | 14.3684726 | 0.187010376 |
| 17.4263481 | 17.4602466 | 17.5225118 | 17.8125299 | 18.1862147 | 18.255719 | 0.411423211 |
| 14.4592731 | 14.2706292 | 14.3014891 | 14.0782393 | 14.637889 | 14.677066 | 0.310643544 |
| 18.669737 | 18.1484632 | 18.2934234 | 18.1465971 | 18.7572609 | 19.0430588 | 0.693254545 |
| 18.8509858 | 19.1506797 | 19.1348408 | 19.3553625 | 18.7738233 | 18.2209214 | 0.374023599 |
| 13.8253251 | 14.3612185 | 14.4095558 | 13.8194503 | 14.313255 | 14.0476689 | 0.276923952 |
| 18.3847286 | 18.5286472 | 18.3956065 | 18.52151 | 19.0649905 | 19.172114 | 0.405914948 |
| 17.3459279 | 17.3854788 | 17.5115368 | 17.8368594 | 17.7464632 | 18.147133 | 0.418444254 |
| 14.9581095 | 15.2346295 | 14.8590872 | 14.8601645 | 15.212828 | 15.609518 | 0.559031625 |
| 13.6098544 | 14.1363245 | 13.4416724 | 14.0123537 | 14.5508449 | 14.6503885 | -0.348819712 |
| 17.4405326 | 17.6624197 | 17.5688801 | 18.0330881 | 18.0186987 | 18.3690875 | 0.304544913 |
| 15.7200877 | 15.0427129 | 15.1542393 | 15.1680412 | 16.0287081 | 16.7255898 | 0.547199342 |
| 15.7036659 | 15.6140316 | 15.4982578 | 15.322177 | 16.1300245 | 17.1880647 | 1.333177833 |
| 16.727694 | 16.9397427 | 16.6402613 | 16.9682125 | 16.9820295 | 17.3206622 | 0.353018783 |
| 14.2545151 | 14.3416569 | 14.3093033 | 14.4270541 | 14.5807501 | 14.5410422 | 0.201541838 |
| 19.6842762 | 19.6730765 | 19.5244958 | 19.5214689 | 20.2004743 | 20.2594806 | 0.609041686 |
| 15.9982989 | 16.2750384 | 16.3333942 | 16.6355646 | 16.353595 | 16.7265253 | 0.195807775 |
| 19.6548026 | 19.6158491 | 19.6236392 | 19.8334964 | 20.0299061 | 20.4606535 | 0.494159838 |
| 16.7505094 | 16.7328285 | 16.9265945 | 16.8642183 | 16.6580716 | 16.6012571 | 0.232106766 |
| 15.7973631 | 15.708859 | 15.6293428 | 15.7375692 | 15.8589127 | 16.322717 | 0.486389442 |
| 20.3922332 | 20.6128366 | 20.2681593 | 20.4898833 | 20.8018971 | 20.8318559 | 0.563027076 |
| 14.9271097 | 15.7352839 | 15.2094943 | 15.7831054 | 15.4644685 | 15.4561241 | 0.144971241 |
| 13.3846044 | 13.8876603 | 13.8314013 | 13.7430386 | 14.1252159 | 13.4750291 | 0.480618034 |
| 21.1301462 | 21.3165297 | 21.1847014 | 21.3533801 | 21.597086 | 21.9334949 | 0.584888574 |

| | | | | | | |
|---|---|---|---|---|---|---|
| 11.447678 | 11.2130396 | 11.5679035 | 11.4593096 | 11.8820432 | 11.7818193 | -0.258428543 |
| 15.7710061 | 15.9934047 | 15.5288988 | 15.8437256 | 15.9488781 | 16.0068093 | 0.166620364 |
| 13.5388452 | 13.1518413 | 12.4800866 | 13.556822 | NA | 13.9159959 | -0.444959386 |
| 12.9819446 | 12.780055 | 12.5562711 | 12.7716221 | 13.4834417 | 13.2456584 | 0.462578753 |
| 16.7936518 | 17.2119101 | 18.7116158 | 16.9517987 | 16.6297302 | 17.8362443 | -0.484701777 |
| 15.1084347 | 15.0697866 | 14.7726903 | 14.9362772 | 15.3265537 | 15.5259061 | 0.528718221 |
| 16.4649435 | 16.3974299 | 16.3636963 | 16.7038472 | 16.4991457 | 16.9564082 | 0.560623764 |
| 16.7720983 | 17.0033787 | 16.7964264 | 16.8428812 | 16.9516492 | 17.1063257 | 0.371948371 |
| 13.2794285 | 13.7266343 | 13.354118 | 13.4707617 | 14.4599785 | 13.1521373 | -0.884012375 |
| 21.4681851 | 21.5874959 | 21.4565046 | 21.7565109 | 21.9084741 | 22.4311472 | 0.502876899 |
| 16.4840465 | 16.6730945 | 16.4629788 | 16.9634278 | 16.9196857 | 16.4796774 | 0.175187918 |
| 12.8418084 | 13.598033 | 13.4963115 | 12.9377862 | 13.6379224 | 13.3633512 | -0.551696582 |
| 11.9819709 | 11.5709375 | 11.8441319 | 11.715391 | 11.6926295 | 11.0045261 | 0.456766642 |
| 13.3784745 | 14.7679811 | 13.3661778 | 13.9419636 | 14.3992683 | 14.8286577 | 0.516892066 |
| 17.0487282 | 17.0457912 | 17.1141113 | 17.3476533 | 17.385053 | 17.543824 | -0.210308284 |
| 20.4132491 | 20.4775592 | 20.3168249 | 20.2984882 | 21.0350945 | 20.9506441 | 0.644998896 |
| 13.9478506 | 13.6528177 | 13.6982522 | 13.8683642 | 13.8824228 | 14.197404 | 0.584241649 |
| 19.2155482 | 19.3352341 | 19.1848328 | 19.2400033 | 19.7574341 | 19.9726519 | 0.623857144 |
| 13.4586447 | 13.4513946 | 13.7065634 | 13.6020306 | 13.6833861 | 13.969971 | 0.2244487 |
| 14.3685681 | 14.8719019 | 14.0774342 | 14.7527959 | 14.9265623 | 14.2711975 | 0.294171073 |
| 16.302582 | 16.5143124 | 16.8106434 | 16.2587571 | 16.6857741 | 16.5517978 | 0.097273662 |
| 14.5599331 | 15.2347296 | 14.6321165 | 14.6896558 | 14.7448409 | 15.1419378 | -0.467723143 |
| 13.732779 | 13.3925173 | 13.4247303 | 13.4981823 | 13.3868073 | 14.2655601 | 0.286463861 |
| 15.0667828 | 14.9801701 | 15.2733931 | 15.3705805 | 15.2709953 | 15.7516848 | 0.186518918 |
| 19.2066744 | 19.3868676 | 19.0419838 | 19.4770887 | 19.5274246 | 20.0240897 | 0.653976326 |
| 15.5522604 | 15.5171755 | 15.3880574 | 15.8289374 | 15.2122333 | 15.9393664 | 0.196145677 |
| 15.7915114 | 15.8758557 | 15.7203889 | 16.338585 | 16.0377658 | 15.1599001 | 0.059444251 |
| 19.5195831 | 19.7081054 | 19.6485269 | 19.7489372 | 19.8543693 | 20.0171768 | 0.514608661 |
| 15.162065 | 16.9044578 | 16.5494141 | 16.0853992 | 15.1665221 | 15.9658 | -0.159722712 |
| 11.2126554 | 11.1995864 | 11.9571761 | 11.4555146 | 11.7381581 | 11.6372318 | -0.045568056 |
| 14.3364676 | 13.2592777 | 13.8552776 | 13.8592624 | 14.6734988 | 14.9671477 | 0.581703654 |
| 19.1893389 | 19.349899 | 19.1419909 | 19.5060244 | 18.7397843 | 19.2790638 | -0.682691994 |
| 22.779117 | 22.9215459 | 22.9939813 | 23.2025942 | 23.2378442 | 23.6510822 | 0.254541433 |
| 16.438836 | 16.98973 | 16.5701266 | 16.9353455 | 16.659883 | 16.5399456 | 0.182473318 |
| 18.4450741 | 18.5048724 | 18.5857803 | 18.730367 | 18.9232634 | 18.7895741 | 0.21901285 |
| 11.7903697 | 11.5796487 | 11.3233121 | 11.836963 | 11.853063 | 11.9787047 | 0.207493228 |
| 18.1099354 | 18.0402133 | 17.8816373 | 18.0468596 | 18.3686025 | 18.7249079 | 0.526675845 |
| 17.0301511 | 17.2431962 | 17.3924444 | 17.535192 | 17.6427096 | 17.8682403 | -0.756283166 |
| 17.5516803 | 17.6195491 | 17.562724 | 17.5720465 | 17.8735928 | 17.9163498 | 0.661528969 |
| 18.5899547 | 18.7244706 | 18.5359674 | 18.3841101 | 19.1172134 | 19.0536496 | 0.498266533 |
| 11.5645755 | 11.7000567 | 11.8674039 | 11.3557673 | 11.5352852 | 11.7792291 | 0.635947483 |
| 14.2471628 | 13.9793543 | 14.054479 | 14.59268 | 14.4443957 | 14.8201231 | 0.031345438 |

| | | | | | | |
|---|---|---|---|---|---|---|
| 18.0945892 | 18.2530732 | 18.5292412 | 18.7907651 | 19.2387141 | 19.0477002 | 0.567678347 |
| 17.0447446 | 17.2405464 | 17.1291878 | 17.2633323 | 17.4212755 | 17.9180314 | 0.521093903 |
| 14.1097611 | 13.9483295 | 14.1578693 | 13.4638328 | 14.5000021 | 15.0005088 | -0.32464428 |
| 13.3319838 | 13.1166087 | 13.2473245 | 13.5584349 | 13.3517222 | 13.3573354 | 0.228712668 |
| 14.0927674 | 14.179121 | 14.0707468 | 13.9672978 | 14.6521989 | 14.3438242 | 0.335618326 |
| 17.8300677 | 18.0755894 | 17.6918539 | 18.0334528 | 17.5615654 | 18.0517707 | -0.073682696 |
| 18.212524 | 18.2366259 | 18.3939881 | 18.353522 | 18.3567558 | 18.6060654 | 0.472100994 |
| 16.6904354 | 16.8994162 | 16.4771561 | 16.4997055 | 16.845395 | 16.5211831 | -0.025193166 |
| 11.8229324 | 11.8090872 | 11.9200547 | 12.0235452 | 11.3295254 | 11.873502 | 0.310812405 |
| 12.9011287 | 13.7213127 | 13.4156864 | 13.7893674 | 13.6393679 | 13.4759135 | -0.907563314 |
| 16.7073084 | 17.0893484 | 16.9159966 | 17.3999066 | 17.0827188 | 17.6175029 | 0.669754497 |
| 15.8753061 | 15.7644783 | 15.933564 | 16.3555525 | 16.6072943 | 16.3877803 | 0.274055416 |
| 14.6338088 | 14.7668401 | 14.8322208 | 14.9607912 | 14.7502439 | 14.9904041 | 0.186227393 |
| 14.4436367 | 14.3473751 | 14.8086586 | 14.5535606 | 14.9836041 | 14.950737 | 0.203380895 |
| 11.6653835 | 10.8772047 | 11.8093898 | NA | 12.2332443 | 12.1431294 | NA |
| 17.8828586 | 18.064411 | 17.7851753 | 18.2330093 | 18.154575 | 17.814675 | 0.168350864 |
| 19.678013 | 19.812469 | 19.6954351 | 19.6569685 | 20.0840357 | 20.2568688 | 0.472545684 |
| 18.6464816 | 18.871418 | 18.8499286 | 19.0046763 | 18.7793084 | 19.2744087 | 0.378392534 |
| 19.4003351 | 19.4774062 | 19.4776116 | 19.3506716 | 20.0357873 | 20.4756587 | 0.695103954 |
| 15.6638212 | 15.5200062 | 15.8271043 | 15.365803 | 16.0668183 | 16.2790553 | 0.244759011 |
| 13.7728361 | 13.9855726 | 13.7361398 | 13.8575028 | 13.9848328 | 14.7694751 | 0.402350716 |
| 19.7832986 | 19.8421914 | 19.4100882 | 19.4522041 | 20.0899367 | 20.2882164 | 0.284818304 |
| 16.6684505 | 16.5218929 | 16.2184114 | 16.4994648 | 16.5913698 | 17.1448415 | 0.29146406 |
| 15.4932033 | 16.0235613 | 15.360077 | 15.8534005 | 15.8972524 | 16.1573033 | 1.038796282 |
| 18.1754306 | 17.6271539 | 17.6709415 | 17.9849605 | 18.4986403 | 18.2642334 | 1.093158099 |
| 20.0513931 | 20.2502169 | 20.2991186 | 20.4886939 | 20.4255054 | 20.5582648 | 0.302826817 |
| 18.7699054 | 18.6580792 | 18.6994517 | 18.9178726 | 19.2180836 | 19.031756 | 0.569283802 |
| 11.2950305 | 11.7975263 | 11.4054934 | 11.2192435 | 11.9430865 | 11.8707995 | 0.035431896 |
| 16.3033368 | 16.3587142 | 16.2779311 | 16.7420283 | 16.7146035 | 17.5556304 | 0.314906039 |
| 16.7914211 | 17.6254787 | 18.3492324 | 16.8452768 | 16.6475813 | 17.7034379 | -0.748293991 |
| 14.7342818 | 15.1772775 | 14.5463728 | 13.8792444 | 14.3945684 | 14.8467673 | -0.196254503 |
| 12.8379839 | 12.8986958 | 13.1421031 | 13.2967276 | 13.3924158 | 13.828114 | 0.009226271 |
| 20.5674327 | 20.6638275 | 20.7561803 | 21.0056698 | 20.8972038 | 21.2645602 | 0.186721829 |
| 17.0919173 | 17.3680146 | 17.1621678 | 17.6996409 | 17.5817395 | 16.7427031 | -0.088452361 |
| 16.3714157 | 15.9745735 | 16.1403349 | 16.0172924 | 16.6499504 | 17.068322 | 0.573616567 |
| 15.3633936 | 14.5432278 | 14.8356185 | 14.3978852 | 15.119893 | 15.5992407 | 0.503668501 |
| 15.9401821 | 16.1575588 | 15.8699333 | 16.277678 | 15.7823828 | 16.0614493 | -0.064481218 |
| 16.4080737 | 15.992124 | 16.450815 | 16.0352359 | 16.7100222 | 17.1071674 | 0.401105349 |
| 13.5216206 | 13.9017308 | 14.321306 | 13.9921764 | 13.5287433 | 14.0332665 | -0.202631887 |
| 17.6463789 | 17.7411489 | 17.5957491 | 17.7260055 | 17.8911687 | 18.2229096 | 0.380432301 |
| 15.7518525 | 15.9007619 | 15.602261 | 16.2706651 | 15.5712208 | 15.6596643 | -0.058236996 |
| 13.8431587 | 14.180395 | 13.902846 | 14.0195926 | 14.0942064 | 14.2017243 | -0.085579324 |

| | | | | | | |
|---|---|---|---|---|---|---|
| 13.5906512 | 13.4151693 | 13.6147179 | 14.2351311 | 14.1008388 | 14.4106246 | -0.016661448 |
| 13.4975612 | 13.3565545 | 13.0343789 | 12.9201587 | 13.3919065 | 13.6605154 | 0.636900872 |
| 20.5095468 | 20.706657 | 20.5385783 | 20.5949955 | 21.2888096 | 21.0667275 | 0.411611144 |
| 19.7678162 | 19.8619607 | 19.8615565 | 20.0436157 | 20.1942213 | 20.5052344 | 0.364423038 |
| 18.8513506 | 18.9292155 | 18.8245558 | 18.7209158 | 19.1837326 | 19.6443714 | 0.408436172 |
| 19.0488873 | 19.1333587 | 19.0401946 | 19.3067546 | 19.6174619 | 19.8428819 | 0.506819784 |
| 11.4999251 | 11.7207289 | 11.8216983 | 11.634479 | 11.6094313 | 12.5141709 | 0.139305141 |
| 14.3067156 | 13.6729709 | 14.2540222 | 12.7822738 | 14.0865076 | 15.1360747 | 0.44175731 |
| 17.5817357 | 17.8293881 | 17.6881292 | 18.253487 | 18.3128546 | 18.4738861 | 0.345073598 |
| 18.9029846 | 19.292626 | 18.8949269 | 19.4992671 | 19.4176821 | 19.8258587 | 0.388793115 |
| 19.9284467 | 20.0049574 | 20.0865528 | 20.40873 | 20.443643 | 20.9982953 | 0.470336259 |
| 19.7514806 | 20.0212761 | 19.9827701 | 20.3711338 | 20.2196606 | 20.1860327 | 0.178363163 |
| 16.0910781 | 15.8174296 | 15.9432907 | 15.8975146 | 15.7481395 | 16.3338908 | 0.345597847 |
| 14.3698775 | 14.3544469 | 14.7911157 | 14.3481978 | 14.5405292 | 14.3648591 | 0.320648074 |
| 17.7759465 | 17.9097514 | 17.6289315 | 18.1182913 | 17.88228 | 17.8001054 | 0.020681422 |
| 18.6892922 | 18.821601 | 18.7108865 | 18.9704143 | 17.7124241 | 18.8431147 | -0.304191894 |
| 17.5050759 | 17.5945778 | 17.3227692 | 17.6470284 | 17.5420335 | 18.1913285 | 0.322320119 |
| 18.1459903 | 18.1911569 | 18.3361493 | 18.1624323 | 18.559219 | 19.1078653 | 0.262947994 |
| 19.2034241 | 19.5259803 | 19.1619843 | 19.3081357 | 19.7535156 | 19.432286 | 0.581729825 |
| 18.0589519 | 17.967796 | 18.1612818 | 18.2154228 | 18.3227532 | 18.6241568 | 0.115255259 |
| 17.1933384 | 17.2376543 | 17.2876931 | 17.8906126 | 17.7052319 | 17.8286249 | 0.188444546 |
| 13.7524711 | 13.6864349 | 13.7701844 | 14.0565893 | 13.8596842 | 14.3139348 | -0.08308644 |
| 23.1950986 | 23.5455903 | 23.4188117 | 23.8852763 | 23.6188565 | 24.256612 | 0.461343971 |
| 17.4389708 | 17.7553407 | 17.5768889 | 17.6621464 | 17.3294618 | 18.1779627 | -0.180900605 |
| 15.8912881 | 16.2441375 | 15.5495529 | 15.8004453 | 15.7958494 | 16.1817482 | 0.74130832 |
| 19.3889244 | 19.7619647 | 19.6423692 | 19.7550108 | 19.4435837 | 19.691553 | 2.82E-04 |
| 14.6782137 | 15.0767519 | 15.1074527 | 14.361301 | 15.3216621 | 15.0487896 | -0.355183637 |
| 14.7780811 | 14.7000357 | 14.572846 | 14.9238042 | 14.8681184 | 15.3410302 | -0.196428104 |
| 14.9406031 | 14.7302262 | 14.5084223 | 14.2020684 | 14.9401855 | 15.1119826 | 0.581108863 |
| 18.8904027 | 19.1250121 | 18.9528523 | 19.4865669 | 18.6298913 | 19.0297967 | -0.0483472 |
| 12.9916353 | 12.7015026 | 12.1902171 | 12.0611109 | 13.5237872 | 12.6458807 | 0.312157523 |
| 16.7054024 | 16.9155236 | 16.4784124 | 16.4381155 | 17.208215 | 17.5118249 | 0.758701312 |
| 14.6377014 | 14.8986198 | 14.2899986 | 15.0289314 | 14.7765409 | 15.3518428 | 0.235321527 |
| 14.3125408 | 14.0695422 | 13.9907475 | 13.8967713 | 14.6683124 | 14.5480691 | 0.385677545 |
| 14.7509281 | 15.0156134 | 15.0545311 | 14.7090547 | 14.9493383 | 14.8393443 | 0.048779884 |
| 17.822175 | 18.052114 | 17.8236614 | 18.1788954 | 18.0481059 | 17.9952011 | -0.053660068 |
| 16.3957787 | 16.8433028 | 16.659665 | 16.4561835 | 16.7995563 | 16.6705791 | 1.126190955 |
| 15.0747584 | 15.7061553 | 15.626991 | 15.4157529 | 15.6988599 | 14.3840644 | -0.050489693 |
| 14.8625257 | 14.6613983 | 14.775755 | 14.8365407 | 14.8892629 | 15.2520573 | 0.443822732 |
| 11.7853106 | 11.6647147 | 11.6012156 | 11.9285009 | 11.9424847 | 9.62893531 | -0.41264489 |
| 19.2022664 | 19.30257 | 19.2025446 | 19.3777977 | 19.5649726 | 19.946041 | 0.415644641 |
| 16.9454822 | 17.4052165 | 17.4426297 | 17.2951638 | 17.7433514 | 17.3423501 | -0.920577484 |

| | | | | | | |
|---|---|---|---|---|---|---|
| 16.8595598 | 16.9826011 | 16.9007081 | 16.8831055 | 17.1271502 | 17.0036821 | 0.298113103 |
| 19.4393917 | 19.0230098 | 19.1690191 | 18.8133846 | 19.449014 | 20.0097863 | 0.726773898 |
| 12.7878915 | 12.1032026 | 12.6656467 | 11.9437081 | 11.9777446 | 12.1862914 | -0.896874388 |
| 13.1396876 | 13.4217759 | 12.9386134 | 13.2276208 | 13.2753022 | 13.7750142 | 0.316272253 |
| 14.1069062 | 14.0458332 | 14.070205 | 13.8220118 | 14.1467944 | 14.0870995 | -0.475615236 |
| 14.6187856 | 14.7280669 | 14.8267347 | 14.9715296 | 15.4343726 | 15.487683 | -0.060692144 |
| 14.177202 | 14.0298228 | 13.9246362 | 14.037687 | 14.5971217 | 14.5034791 | -0.078209052 |
| 19.2616221 | 19.5440704 | 19.4851374 | 19.7570819 | 19.6468677 | 20.0219061 | 0.125797794 |
| 13.9063694 | 13.7433407 | 13.3803563 | 14.0378489 | 13.9805348 | 13.9443406 | 0.502292171 |
| 16.4448096 | 16.6866053 | 16.6041659 | 16.7791793 | 16.7220234 | 16.6660407 | -0.237820875 |
| 13.7627199 | 14.3416984 | 13.5900808 | 13.759353 | 14.312891 | 13.1475963 | 0.421455796 |
| 20.2214957 | 20.2190121 | 20.0755548 | 20.1772424 | 20.9162618 | 20.8567099 | 0.50538333 |
| 15.5265852 | 15.4551413 | 15.404511 | 15.3326813 | 15.54586 | 15.2247014 | 0.075718813 |
| 15.1636304 | 15.4813397 | 14.9458246 | 15.5803442 | 15.796419 | 15.1174145 | -0.231080528 |
| 17.3398206 | 17.1937195 | 17.2517873 | 17.5436859 | 16.8396632 | 17.0996963 | 0.040734594 |
| 19.1935512 | 19.4401534 | 19.5224047 | 19.3688944 | 19.6025136 | 20.2294608 | 0.070178428 |
| 14.6108198 | 14.3833245 | 14.4906113 | 14.44296 | 14.1786267 | 14.5928287 | 0.006661206 |
| 11.5224947 | 11.4810468 | 11.5923622 | 11.7046124 | 11.736767 | 9.94009735 | -0.313428994 |
| 12.9011628 | 12.053177 | 12.1689068 | 11.8532628 | 12.3211269 | 11.5925462 | 0.863723317 |
| 12.7710124 | 12.9534745 | 12.3875668 | 13.0818922 | 12.4136545 | 12.5749404 | 0.390169482 |
| 15.2461235 | 14.9366622 | 14.1338892 | 14.9101637 | 15.4449267 | 16.0614867 | 1.150088717 |
| 16.9449021 | 16.6966998 | 17.0395653 | 16.8398825 | 16.988608 | 17.2043236 | 0.322864149 |
| 21.3298439 | 21.8203118 | 21.4396192 | 21.8579792 | 21.384193 | 22.5891634 | 0.753917378 |
| 15.0381545 | 15.289992 | 15.1801436 | 15.5627028 | 14.9606298 | 15.2620719 | -0.127471595 |
| 14.8788792 | 14.9559255 | 14.9653069 | 14.6423109 | 15.2408006 | 15.7892366 | 0.27327628 |
| 20.4899549 | 20.7203322 | 20.513579 | 21.0872558 | 20.2013704 | 20.6247765 | -0.017807908 |
| 11.550939 | 11.9816321 | 11.4072181 | 11.4574861 | 11.7213155 | 10.9341786 | -0.009696757 |
| 16.1338993 | 16.4039172 | 16.3508712 | 16.8653649 | 16.2029439 | 16.6448016 | -0.101873152 |
| 16.3641388 | 16.8123219 | 16.1101722 | 16.3028801 | 16.6198565 | 16.270952 | 0.176235217 |
| 19.0795688 | 19.4151421 | 18.9853124 | 19.1384512 | 18.9962153 | 19.5403077 | 0.303406219 |
| 17.9066467 | 17.3578173 | 17.5565654 | 17.8987256 | 17.9235196 | 17.8358934 | 0.568000648 |
| 18.982678 | 18.8684458 | 18.9994292 | 19.031355 | 19.3809862 | 19.8309845 | 0.183932179 |
| 14.905184 | 15.1415758 | 14.69465 | 14.85648 | 15.4399291 | 15.328247 | 0.191854973 |
| 15.4329345 | 15.1346864 | 15.1292586 | 15.2909484 | 15.3755656 | 15.4641415 | 0.289484088 |
| 12.6182555 | 12.68208 | 12.8522835 | 13.1597299 | 12.5785086 | 12.5483884 | 0.479098252 |
| 11.5457184 | 11.6317325 | 11.1337307 | 11.6778313 | 11.6297102 | 11.1822479 | 0.22997132 |
| 13.5606057 | 13.3973024 | 13.4616519 | 13.3099765 | 13.2602637 | 13.7479205 | -0.277831299 |
| 17.9226638 | 18.0016763 | 17.7892085 | 17.863017 | 17.8108552 | 18.191168 | 0.303699163 |
| 16.9496874 | 16.9106372 | 17.075522 | 17.1933445 | 16.7817699 | 17.3050245 | 0.009714812 |
| 18.6991645 | 18.924238 | 18.8999563 | 18.9617383 | 19.03644 | 19.5002129 | 0.588694317 |
| 20.0497584 | 19.9910179 | 20.031595 | 19.9335497 | 20.3161337 | 20.7515141 | 0.436388326 |
| 16.7215517 | 16.6657176 | 16.5296293 | 16.7822768 | 16.3059756 | 16.9962616 | 0.85380865 |

| | | | | | | |
|---|---|---|---|---|---|---|
| 17.0599411 | 17.1505376 | 17.0535911 | 17.2102322 | 17.2466367 | 17.4127181 | -0.134817244 |
| 19.4658978 | 19.5152847 | 19.4545679 | 19.4707632 | 19.9115505 | 20.173154 | 0.517952898 |
| 19.3893341 | 19.5125433 | 19.3136242 | 19.5750778 | 19.7529186 | 20.0525706 | 0.116180596 |
| 17.2820818 | 16.8597816 | 16.9473265 | 17.054358 | 17.0749156 | 17.2943618 | 0.013991974 |
| 13.1346311 | 12.2561705 | 12.6861618 | 12.9498331 | 13.8038338 | 13.816717 | 0.910763022 |
| 15.4057971 | 15.6508229 | 15.8733167 | 15.6436335 | 15.7324621 | 15.9487323 | -0.213703858 |
| 17.5255915 | 17.6870578 | 17.4929516 | 17.5150478 | 18.094101 | 17.0584779 | -0.799997942 |
| 16.8065635 | 17.8858565 | 16.956155 | 17.0971866 | 16.0231624 | 16.4186861 | -0.62734285 |
| 14.6291565 | 14.6164711 | 14.3104203 | 14.5831808 | 14.7445639 | 14.6391637 | 0.116212418 |
| 15.5597852 | 15.295056 | 15.3421787 | 15.4168123 | 15.3529385 | 15.5446732 | 0.317246185 |
| 12.8887931 | 12.8496951 | 12.9444017 | 12.5016085 | 12.8556447 | 12.9855344 | 0.312918765 |
| 16.8408977 | 16.7079435 | 16.8071084 | 16.8610955 | 17.1844263 | 17.3383714 | 0.49366251 |
| 19.710393 | 19.7270705 | 19.6770025 | 20.0993463 | 19.9857341 | 19.4170631 | -0.48789112 |
| 14.7054655 | 15.013234 | 14.7908638 | 14.7967526 | 15.2943739 | 15.8313911 | 0.215515416 |
| 19.1098446 | 19.340407 | 19.1681169 | 19.4300982 | 19.2830451 | 20.1795335 | 0.744000251 |
| 15.0670276 | 14.8917048 | 14.8309785 | 14.8241504 | 15.2603023 | 15.2451836 | 0.506940491 |
| 18.359352 | 18.327552 | 18.1331802 | 18.1243915 | 17.7264032 | 18.0948517 | -0.51774783 |
| 16.9991504 | 16.6689752 | 16.4155942 | 16.689807 | 16.8114336 | 17.1527647 | 0.123713701 |
| 13.30765 | 13.8823638 | 13.8058115 | 13.2131708 | 13.2841794 | NA | -0.215526148 |
| 18.9409388 | 18.7979321 | 18.7012272 | 18.368948 | 18.991092 | 19.5543974 | 0.313276209 |
| 19.757783 | 19.8646316 | 19.8360728 | 19.8387636 | 19.8130996 | 20.6556589 | 0.441615498 |
| 16.0972682 | 15.7197306 | 16.0254251 | 15.3575404 | 15.1883154 | 16.0982717 | -0.381443876 |
| 18.1568692 | 18.2742738 | 17.6140752 | 17.8281809 | 17.9981628 | 18.0600652 | -0.089240766 |
| 11.5923967 | 11.3019436 | 11.9374029 | 11.6789629 | 11.8869029 | 11.193397 | 0.816291322 |
| 16.0391446 | 16.3981953 | 16.3281654 | 16.5741107 | 16.509195 | 16.8388509 | -0.315067627 |
| 14.3923239 | 14.622573 | 14.0263682 | 14.7197232 | 14.362433 | 14.8462983 | 0.618605702 |
| 18.9740221 | 19.0357144 | 19.0137292 | 18.7428817 | 19.2652439 | 19.6847749 | 0.543558184 |
| 16.3625205 | 16.2269239 | 16.2799418 | 16.0498619 | 16.3896663 | 16.9567732 | 0.587096676 |
| 14.4070503 | 14.7818693 | 14.6399926 | 14.6182626 | 14.2740096 | 15.0997643 | 0.135471677 |
| 13.7433171 | 14.3887708 | 14.1348494 | 13.5919336 | 13.7468006 | 13.1028201 | 0.030974643 |
| 17.6063539 | 17.9065371 | 17.8254854 | 17.9960239 | 17.9262944 | 18.8506691 | 0.490230875 |
| 14.6742229 | 14.7756631 | 14.5669277 | 14.5120981 | 14.4760895 | NA | -0.496436079 |
| 13.8165973 | 13.9870571 | 13.7768258 | 13.9397526 | 14.2444437 | 14.3080918 | 0.121878185 |
| 18.1331659 | 18.4484199 | 18.2293647 | 18.3749856 | 18.0004988 | 18.5638993 | -0.043381472 |
| 23.2043448 | 23.4587593 | 23.3289216 | 23.8133541 | 23.4250367 | 24.4187091 | 0.205007389 |
| 15.2306688 | 15.246221 | 14.9024608 | 14.7865228 | 15.6109177 | 15.4535873 | 0.24343472 |
| 17.4478771 | 17.3104137 | 17.2309723 | 17.3373218 | 17.7596534 | 18.0351305 | 0.264739031 |
| 15.1643408 | 14.3453488 | 14.4968856 | 14.3624834 | 14.9862964 | 14.3887942 | 0.640935844 |
| 11.6003549 | 11.5372239 | 11.4663204 | 11.758463 | 11.6472872 | 11.1860726 | -0.039014289 |
| 18.1880214 | 18.0341145 | 18.1235994 | 18.0057736 | 18.3732187 | 18.7256479 | 0.59652599 |
| 15.0497185 | 14.7157299 | 14.7477773 | 14.9536946 | 15.2516136 | 14.5377085 | -0.488826767 |
| 16.463748 | 16.4043715 | 16.2802095 | 15.9110884 | 18.4050751 | 15.9335738 | -0.928572674 |

| | | | | | | |
|---|---|---|---|---|---|---|
| 18.9697501 | 18.9849859 | 19.0226779 | 19.0444841 | 19.1317221 | 19.5886797 | 0.384316947 |
| 15.6199776 | 15.5626313 | 15.4566184 | 15.454026 | 15.5130641 | 16.0629176 | 0.115820898 |
| 17.8333709 | 18.1157038 | 18.0559146 | 18.0885164 | 18.0989198 | 18.7403821 | 0.3866219 |
| 17.9218326 | 18.0612415 | 17.8142606 | 18.3254208 | 17.773838 | 17.8583346 | -0.124927518 |
| 13.7326548 | 13.9102332 | 13.1987505 | 13.7224314 | 13.4440915 | 14.9226428 | 0.830934641 |
| 15.1940065 | 15.4372959 | 14.5186489 | 15.0507018 | 15.5737943 | 15.2877581 | 0.142335079 |
| 20.7918411 | 20.9309422 | 20.739594 | 21.1523729 | 21.0729696 | 20.8319903 | -0.421983957 |
| 14.6051015 | 14.1719307 | 14.618259 | 14.4118162 | 14.1128929 | 14.7226326 | -0.385826201 |
| 17.1313596 | 17.1028893 | 16.9910267 | 17.2058818 | 17.0298687 | 17.3203287 | -0.112549272 |
| 17.3533039 | 18.3320256 | 17.922104 | 17.4852633 | 16.5076058 | 16.4432096 | -1.513173131 |
| 21.3731115 | 21.8169983 | 21.3556685 | 21.8945012 | 21.305966 | 22.4601763 | 0.601932633 |
| 11.9151284 | 11.857005 | 11.5874389 | 11.4862397 | 11.2770231 | 11.9766604 | -0.153837703 |
| 17.210673 | 18.2293745 | 18.1632841 | 17.9077724 | 16.1808118 | 17.6440426 | -0.162513475 |
| 13.6540722 | 13.5501087 | 13.7866975 | 13.5712437 | 13.6172517 | 14.1189295 | -0.203471114 |
| 16.3698697 | 16.5177732 | 16.2293575 | 16.5629343 | 15.9381285 | 16.8636716 | -0.001241854 |
| 13.4103022 | 14.1109571 | 13.419926 | 13.6206505 | 13.4307497 | 13.39832 | 0.734184397 |
| 15.18859 | 14.9592122 | 15.2053341 | 14.7537252 | 15.1175871 | 15.8732464 | -0.002445274 |
| 19.7360874 | 19.8811439 | 19.7520349 | 20.1269121 | 19.5621367 | 19.661546 | -0.57365914 |
| 19.7590486 | 19.8047523 | 19.6709679 | 19.747166 | 19.7967993 | 19.8562295 | -0.187548672 |
| 16.6306478 | 16.5775173 | 16.5740034 | 16.9279688 | 16.593393 | 16.5954518 | 0.062598779 |
| 19.025851 | 19.0226775 | 18.808456 | 18.7529051 | 19.4317922 | 19.6506609 | 0.342221012 |
| 19.184183 | 19.1275789 | 19.0358978 | 18.9113182 | 19.3436746 | 19.9679469 | 0.417590536 |
| 13.9748867 | 13.2901862 | 13.6255284 | 13.8388151 | 13.9212143 | 13.6082269 | -0.383438536 |
| 19.1719544 | 19.1176096 | 19.1644967 | 19.2942747 | 19.1219307 | 19.8847809 | -0.185107467 |
| 13.7712304 | 13.4470234 | 13.3943834 | 13.8078622 | 13.779746 | 13.5517362 | 0.283243181 |
| 13.8348705 | 13.6489289 | 13.9179969 | 14.2611169 | 13.867461 | 14.0976053 | 0.73868795 |
| 15.329583 | 15.3063848 | 15.1065896 | 15.4085583 | 15.5540676 | 16.7473451 | 0.46246915 |
| 12.1583404 | 12.0062866 | 11.9838744 | 11.5085496 | 11.5271953 | 11.5750949 | -0.720423644 |
| 19.8673099 | 18.4375833 | 18.7659132 | 18.3133139 | 19.5649559 | 18.9051747 | 1.16222112 |
| 22.5674964 | 22.8325011 | 22.6852504 | 23.171189 | 22.2253095 | 23.5038667 | 0.402413243 |
| 15.1482024 | 15.1459297 | 14.9853482 | 14.6883941 | 14.8558341 | 15.36877 | 0.231658717 |
| 12.3963655 | 12.900247 | NA | 13.4848792 | 13.0537157 | NA | NA |
| 15.4814255 | 15.5133803 | 15.2801653 | 15.5159393 | 14.9058473 | 15.2384246 | -0.059334606 |
| 13.136552 | 13.1362118 | 13.8563407 | 14.0028055 | 13.8896068 | 14.4691318 | -0.697125792 |
| 14.0922883 | 13.3134651 | 13.4083098 | 13.3081378 | 13.3278282 | 12.0111705 | -0.891800534 |
| 15.0898521 | 14.7036473 | 14.9415027 | 14.7827226 | 14.2167995 | 14.7926523 | -0.200584431 |
| 20.8006611 | 20.9358795 | 20.9889279 | 21.0778497 | 20.3517426 | 21.1244746 | -0.235492874 |
| 20.8765096 | 19.6686906 | 19.8494598 | 19.662766 | 20.3742755 | 20.1849189 | 1.286312724 |
| 13.8579743 | 13.4202346 | 13.9213387 | 13.9257806 | 13.5774581 | 13.5501864 | -0.164356888 |
| 17.9785129 | 17.6742229 | 17.6973595 | 17.8203551 | 17.8824302 | 17.9636909 | -0.181939218 |
| 14.7168365 | 14.8470604 | 14.3640357 | 14.4070588 | 14.5026518 | 14.5904169 | -0.626339419 |
| 15.2694055 | 14.8878785 | 14.8479858 | 14.9539958 | NA | NA | NA |

| | | | | | | |
|---|---|---|---|---|---|---|
| 16.7163102 | 16.5247405 | 15.4790734 | 16.3642919 | 16.9643779 | 18.0466168 | 0.895177333 |
| 15.4487335 | 15.3239639 | 15.6219341 | 15.3998051 | 15.2678576 | 15.3380414 | -0.628479709 |
| 13.9799363 | 13.8510136 | 13.6445584 | 13.251156 | 13.652937 | 13.3717335 | 0.069810161 |
| 14.2107418 | 14.311372 | 14.0610243 | 14.2028657 | 13.4534734 | 14.3044843 | -0.603499121 |
| 14.6118716 | 15.610787 | 14.8968046 | 14.9251753 | 13.9143012 | NA | -0.311864412 |
| 17.7961254 | 17.8182993 | 17.7837842 | 18.1578865 | 17.4309256 | 17.5173132 | -0.555934351 |
| 20.9687365 | 21.1860137 | 21.0597419 | 21.5349897 | 20.6446811 | 21.8092934 | 0.357556873 |
| 14.8648022 | 14.7042848 | 14.68842 | 14.9183437 | 14.82743 | 15.3150288 | 0.544284033 |
| 15.3461825 | 15.3526852 | 15.1819712 | 15.5419629 | 14.8808299 | 15.0329042 | 0.057345258 |
| 15.5067989 | 15.9931944 | 15.2924088 | 15.5722657 | 15.6228053 | 15.5064149 | 0.528495356 |
| 15.4393343 | 15.2813001 | 14.9204297 | 15.0861291 | 15.0448874 | 15.1804102 | -0.214063125 |
| 19.9993486 | 20.008568 | 19.9251558 | 20.1877166 | 19.4567482 | 19.3879615 | -0.772564831 |
| 13.1755434 | 12.9184863 | 13.173144 | 12.0174318 | 12.1613184 | NA | 0.092828507 |
| 11.4558289 | 11.4872081 | 11.64319 | 11.4066027 | 11.5136906 | 10.8711274 | -0.904125199 |
| 14.1354975 | 13.3477994 | 14.1772006 | 13.3077258 | 14.0394742 | 14.5548284 | 0.501837867 |
| 16.0036011 | 16.081601 | 15.7027413 | 15.9594707 | 15.6378397 | 15.7572324 | -0.515432186 |
| 12.4528838 | 12.1967126 | 12.4365685 | 12.3383573 | 12.4694623 | 12.8803368 | 0.687227579 |
| 15.9013949 | 15.8219968 | 15.7490374 | 15.8354729 | 15.465261 | 15.5575392 | -0.465893486 |
| 16.4942598 | 16.6277393 | 16.171335 | 16.7254814 | 16.4451633 | 16.5888731 | -0.231093877 |
| 14.3041303 | 14.4265782 | 14.6785163 | 14.6282698 | 13.3972278 | 13.6865298 | -0.880027237 |
| 14.2633835 | 14.1106901 | 14.3789108 | 14.6339827 | 13.9483734 | 14.2489665 | -0.072574319 |
| 16.0324286 | 15.6520143 | 15.795944 | 15.6201088 | 15.3970422 | 15.7569325 | -0.293329399 |
| 13.6706843 | 13.5172905 | 13.4492251 | 13.4491449 | 12.8791503 | 13.1580841 | -0.175652983 |
| 15.3971048 | 15.0253142 | 14.9184491 | 14.984111 | 15.0363597 | 14.7858834 | -0.654485718 |
| 20.9681349 | 19.6934433 | 19.6522174 | 19.2354887 | 20.5182943 | 20.1015898 | 1.38174659 |
| 18.7902039 | 18.7525786 | 19.4651058 | 18.9088411 | 17.9074318 | 18.9413083 | -0.882798152 |
| 13.4967611 | 13.6320142 | 13.9991929 | 13.2064089 | 13.3005747 | 14.360154 | -0.370779324 |
| 12.0581198 | 12.7551421 | 12.1429867 | 11.9400831 | 12.4484653 | 12.7999278 | -0.337018712 |
| 16.4138528 | 16.7361458 | 16.5456115 | 16.5587875 | 16.2845058 | 16.1184908 | -1.010611382 |
| 13.111251 | 14.9788089 | 13.3375582 | 13.8389748 | 13.8275542 | 14.8634024 | 0.588360691 |
| 12.200604 | 12.9498171 | 12.5229096 | 12.4586154 | 12.3257173 | 12.3093918 | -0.578408275 |
| 15.5864926 | 15.4775239 | 15.8167464 | 15.6670487 | 15.04644 | 15.8316502 | -0.023269217 |
| 18.7353776 | 19.5021136 | 18.8211783 | 19.7643866 | 18.4918928 | 20.0051839 | 0.13307907 |
| 13.3097054 | 13.463173 | 13.9940904 | 13.3813393 | 13.3813726 | 13.6512449 | 0.058478221 |
| 15.2806277 | 15.3902498 | 14.739743 | 15.3550468 | 14.9441142 | 15.5102398 | -0.162422766 |
| 14.8350413 | 14.9186373 | 14.7503607 | 14.9871192 | 14.2620796 | 14.0327409 | -0.339291165 |
| 13.4556168 | 14.4535619 | 13.6368697 | 13.6946046 | 13.4646885 | 13.7134174 | -0.840700821 |
| 13.5383907 | 12.6302974 | 12.4317048 | 12.7620454 | 12.3774745 | 13.214937 | -0.044012984 |
| 17.1322777 | 17.0573567 | 16.8972409 | 17.0855396 | 18.6648444 | 16.8921595 | 0.464148593 |
| 15.2355063 | 15.865695 | 15.1551127 | 15.2600366 | 14.9497847 | 14.9906721 | -0.639208263 |
| 16.2412602 | 16.2755629 | 16.0941217 | 15.9994248 | 15.7304321 | 15.9787258 | -0.379029641 |
| 15.8812336 | 15.6580293 | 15.6919887 | 15.9614917 | 15.204896 | 15.2883975 | -0.361382749 |

| | | | | | | |
|---|---|---|---|---|---|---|
| 18.6848117 | 18.7210652 | 18.4043774 | 19.0848506 | 18.3210927 | 18.2679311 | -0.305355908 |
| 13.7208238 | 13.719623 | 13.2703634 | 13.7665506 | 13.1282909 | 13.4104099 | 0.112198714 |
| 17.7332584 | 16.9922931 | 17.1169853 | 16.9713814 | 15.9285645 | 16.9425508 | -0.315791644 |
| 18.7383008 | 18.8714937 | 19.003569 | 18.9002958 | 18.4859624 | 18.55938 | -1.316698386 |
| 16.6182549 | 16.1075132 | 16.4466456 | 16.2369197 | 16.0856881 | 15.1327412 | -0.963859452 |
| 18.2824935 | 18.2540441 | 18.3290408 | 18.2770085 | 18.2057959 | 17.6994462 | -0.8663713 |
| 12.2282362 | 11.8684144 | 11.4077754 | 11.5988957 | 10.8655189 | 11.6289126 | -0.419949986 |
| 15.9729045 | 15.9052776 | 15.3036568 | 15.9284994 | 15.0978128 | 15.4967116 | 0.497975254 |
| 13.9464811 | 13.8198341 | 14.7920182 | 13.8328044 | 12.952167 | 14.0102984 | -1.213688478 |
| 18.9936961 | 18.9671265 | 19.0425965 | 19.1155125 | 18.5837243 | 19.0103176 | -0.47079324 |
| 17.1115117 | 17.1590626 | 16.9487483 | 17.1064533 | 16.4690833 | 16.8439427 | -0.44380026 |
| 15.6110549 | 15.8376369 | 15.6100667 | 15.4375991 | 14.5364854 | 14.7156054 | -1.05796504 |
| 18.1877985 | 18.3495193 | 18.2281946 | 17.934571 | 17.340555 | 17.7589464 | -0.690779998 |
| 18.8195997 | 18.7564145 | 18.5916408 | 18.3135121 | 18.0688059 | 17.7709824 | -0.91410664 |
| 18.0877945 | 17.8876466 | 17.773721 | 17.6415896 | 17.2514387 | 17.1050812 | -0.938555352 |
| 14.4053513 | 14.1520494 | 14.2423206 | 13.7577789 | 13.84444 | 14.0376057 | -0.649844956 |
| 16.9626306 | 16.7091115 | 16.8676994 | 16.7410482 | 16.226845 | 16.1465851 | -0.532163604 |
| 13.7846892 | 14.4416736 | 14.1438663 | 14.4379542 | 13.3099022 | 14.0474924 | -0.038730847 |
| 11.8255071 | 12.5543974 | 12.1344341 | 12.5820465 | 11.6650886 | NA | -0.565612744 |
| 16.0616091 | 15.8554981 | 15.8737743 | 15.6364643 | 15.1733901 | 15.420577 | -0.550895774 |
| 15.8597636 | 15.2991231 | 15.4357312 | 15.5770794 | 15.6465982 | 15.4359534 | -0.732394469 |
| 17.5834368 | 17.6715316 | 17.5632162 | 17.382488 | 17.3611319 | 17.5289479 | 0.033123531 |
| 19.2712254 | 18.0762785 | 18.2328211 | 17.6694472 | 18.4468881 | 18.6092627 | 0.721546727 |
| 19.3837863 | 19.5590089 | 19.3334163 | 19.3085207 | 18.6049127 | 18.8383151 | -0.686157351 |
| 15.7006729 | 15.564156 | 15.4829066 | 15.1913828 | 14.5769961 | 15.0604211 | -0.787714676 |
| 14.9232832 | 14.806053 | 14.970037 | 14.3341925 | 14.0891757 | 14.718804 | -0.468656951 |
| 14.5066783 | 13.3686772 | 14.2469537 | 13.7219379 | 14.098025 | 13.295676 | -0.924646714 |
| 19.4377986 | 19.5338791 | 19.683479 | 19.598138 | 18.4429848 | 19.2294926 | -0.737395937 |
| 17.9432085 | 18.0631064 | 17.8262227 | 17.752665 | 16.9368192 | 17.3690496 | -0.918645302 |
| 13.7516499 | 13.6325192 | 13.4966904 | 13.8463296 | 13.731374 | 13.9461524 | -0.444509371 |
| 18.3447837 | 18.1813817 | 18.1727531 | 17.9395294 | 17.7063414 | 18.2874463 | -0.83323065 |
| 14.5735188 | 13.8625765 | 14.0134121 | 14.4091794 | 15.1654961 | 15.7537971 | -1.060498946 |
| 15.5285989 | 15.6396538 | 15.4240882 | 15.1194543 | 14.3428003 | 14.6777529 | -0.952941396 |
| 18.2671537 | 18.3127869 | 18.3646403 | 18.3698224 | 17.0820364 | 18.3154124 | -0.619947955 |
| 13.7119973 | 13.2841039 | 13.3576587 | 13.3471198 | 12.6968283 | 13.2965449 | -1.677525213 |
| 16.9564479 | 16.9851058 | 16.8922406 | 16.6581473 | 16.0494865 | 16.6151471 | -1.039971659 |
| 19.0811917 | 18.7091951 | 18.8878218 | 18.4544626 | 17.8167059 | 18.1440938 | -1.230562789 |
| 16.1613682 | 16.2277274 | 16.0528091 | 15.6408506 | 15.3820781 | 16.1411714 | -0.69084228 |
| 15.3990708 | 15.4816607 | 14.9801792 | 14.679652 | 14.7636493 | 15.2586296 | -0.820715962 |
| 14.8340066 | 14.7199305 | 14.6903336 | 14.2433548 | 14.0606814 | 14.6093676 | -0.595191959 |
| 13.4776215 | 13.3569997 | 13.0762588 | NA | 13.2704214 | 13.432619 | -0.287673064 |
| 13.9309371 | 13.9151626 | 13.8136822 | 13.199779 | NA | 12.9480689 | -1.499954402 |

| | | | | | | |
|---|---|---|---|---|---|---|
| 16.1471916 | 15.9753331 | 15.5102738 | 16.1423062 | 15.4226763 | 16.4586894 | -0.688278338 |
| 14.3463214 | 15.2372537 | 14.3577576 | 14.4881834 | 13.5222763 | 13.7517542 | -1.107362821 |
| 13.5754226 | 13.3541419 | 13.4948815 | 13.1879849 | 12.1643583 | 12.3718292 | -0.752590945 |
| 15.205162 | 16.8409786 | 15.7773042 | 16.7945077 | 14.9126543 | 16.0708362 | 0.098683061 |
| 17.1744181 | 16.8233481 | 16.6676178 | 16.7836997 | 16.3948275 | 16.7010858 | -1.447865381 |
| 12.2403653 | 11.9043372 | 11.5100047 | 11.0115557 | 10.7986736 | 11.1345601 | -1.067002124 |
| 18.8318368 | 18.831823 | 18.8434529 | 18.5979991 | 17.9191462 | 17.5555316 | -1.49030914 |
| 19.0909135 | 19.0026085 | 18.8669294 | 18.3528065 | 17.952804 | 17.6475227 | -1.074844167 |
| 17.0316083 | 16.5968516 | 16.8804462 | 16.329543 | 16.248783 | 16.4123666 | -1.305452737 |
| 16.3767262 | 15.6433561 | 15.9187409 | 15.5348853 | NA | 15.1732922 | -1.315953389 |
| 17.5478559 | 17.5357643 | 17.2123662 | 16.9055648 | 16.7028622 | 17.2924125 | -0.94027687 |
| 18.9482254 | 18.4511664 | 18.8803127 | 18.6146206 | 16.9795238 | 16.9410247 | -1.941380545 |
| 16.4506246 | 16.0423232 | 16.2343418 | 15.8725509 | 14.8838847 | 15.2630118 | -1.11100543 |
| 19.3306273 | 18.7577389 | 18.9570875 | 18.3768933 | 18.3064725 | 18.7559538 | -1.585602786 |
| 14.9940334 | 14.6820438 | 14.746416 | 14.2595745 | 12.7872248 | 13.3737337 | -1.150631301 |
| 18.0503167 | 18.0269716 | 18.1111562 | 17.6722531 | 17.4059774 | 16.5673296 | -1.787322872 |
| 17.4727033 | 17.553385 | 17.3505685 | 17.2920479 | 15.8706201 | 16.3181385 | -1.494381713 |
| 20.2031493 | 20.1791613 | 20.1180341 | 20.1081517 | 18.7207378 | 19.1883556 | -1.300807578 |
| 19.6585441 | 19.6901367 | 19.5889031 | 19.3703469 | 18.3151771 | 18.7115428 | -1.439874609 |
| 20.6378077 | 20.5281552 | 20.4831252 | 20.263256 | 19.0961251 | 19.4031521 | -1.430912801 |
| 20.3111841 | 20.2528795 | 20.2662718 | 20.2016806 | 18.9810616 | 20.0523072 | -1.16058877 |
| 17.5403824 | 17.3672547 | 17.2981026 | 16.8973529 | 15.869863 | 16.4160279 | -1.527555425 |
| 17.5335954 | 17.7672036 | 17.5366973 | 17.534875 | 15.8090752 | 16.9102861 | -1.413243355 |
| 18.3347015 | 17.7946429 | 17.8842876 | 17.7773647 | 17.278102 | 17.4542009 | -0.969846589 |
| 18.6345963 | 18.7269298 | 18.6139283 | 18.5547992 | 16.9488514 | 17.5570465 | -1.450451439 |
| 25.7283392 | 25.7779477 | 25.7427644 | 25.5596561 | 24.2963938 | 24.9591561 | -1.454757425 |
| 24.8890423 | 24.9295476 | 24.7489432 | 24.482114 | 23.3602623 | 23.8244901 | -1.499735055 |
| 17.4725614 | 16.9533565 | 17.013721 | 16.9424258 | 16.042682 | 16.2809598 | -1.307570679 |
| 13.7040058 | 13.0365146 | 12.7954154 | 13.6409368 | 12.5139212 | 12.6787805 | -1.702309258 |
| 18.5799158 | 18.591966 | 18.6779067 | 18.3763153 | 16.9796035 | 17.1775078 | -1.737725016 |
| 14.078827 | 13.9650348 | 13.7216191 | 13.7796562 | 13.1192165 | 13.1771547 | -0.468170779 |
| 22.2957008 | 22.3400061 | 22.3697761 | 22.1940169 | 20.331149 | 21.8572145 | -1.111063778 |
| 13.4143418 | 13.6484109 | 13.1035424 | 12.8213382 | 11.3669697 | 11.2246355 | -1.513957755 |
| 20.4359353 | 20.1616595 | 20.1433056 | 19.7636546 | 18.6902729 | 19.4529286 | -1.124304162 |
| 17.5043256 | 17.1136052 | 17.4039869 | 16.5617117 | 16.1176561 | 16.5771307 | -0.918124937 |
| 15.4164554 | 15.2617122 | 15.2236212 | 14.4321147 | 13.7187203 | 14.110872 | -1.721537043 |
| 22.7452954 | 22.4484411 | 22.4887941 | 22.0840269 | 20.5205595 | 21.6083247 | -1.059003638 |
| 20.6031108 | 20.3364113 | 20.3551486 | 19.856003 | 18.7342711 | 19.4135081 | -1.323374781 |
| 17.4965415 | 16.5074524 | 17.211498 | 16.4810741 | 16.4186379 | 16.3102571 | -2.1793764 |
| 19.1157662 | 18.9966632 | 19.0574056 | 18.71727 | 17.2016463 | 17.7477811 | -1.647541354 |
| 17.2559172 | 16.310884 | 16.6470729 | 16.2068732 | 15.1408604 | 16.5909416 | -1.412026018 |
| 21.2210118 | 21.1652858 | 20.9671363 | 20.5297391 | 19.5424002 | 20.2505379 | -1.451113344 |

| | | | | | | |
|---|---|---|---|---|---|---|
| 18.7301245 | 18.7613894 | 18.656923 | 18.3179055 | 16.7646525 | 18.038315 | -1.0953363 |
| 26.1545513 | 26.1536097 | 26.1780108 | 26.0060497 | 24.2728556 | 25.3123216 | -1.619228726 |
| 19.1731022 | 18.9146019 | 18.8500516 | 18.2618308 | 17.2979876 | 17.9204829 | -1.434755184 |
| 24.0755043 | 23.8512041 | 23.8143574 | 23.3708272 | 21.6230924 | 22.7829567 | -1.269381675 |
| 16.8563941 | 16.337863 | 16.5229462 | 15.8710122 | 15.4744868 | 15.5504593 | -1.529741041 |
| 17.8084829 | 17.4674448 | 17.4861838 | 16.77197 | 15.7659822 | 16.5707892 | -1.574920526 |
| 18.7602838 | 18.3629475 | 18.5762728 | 18.3164034 | 16.4538677 | 17.1421751 | -1.638922533 |
| 10.7535526 | 10.481355 | 10.9407235 | 10.3180954 | 11.3328176 | 10.0977823 | -0.563286229 |
| 20.6829465 | 20.5372316 | 20.5145017 | 20.3117403 | 18.2380085 | 18.9312151 | -1.494863392 |
| 19.3032259 | 19.3191402 | 19.3269212 | 19.0296133 | 16.7088809 | 17.7209668 | -2.020710671 |
| 18.9443074 | 19.095241 | 18.946585 | 18.5381478 | 17.0169077 | 18.2780822 | -1.72607362 |
| 15.6303517 | 15.2411311 | 15.4566184 | NA | NA | 15.8908856 | -0.646365736 |
| 20.2855967 | 20.1381761 | 20.2396918 | 19.8625843 | 17.8299815 | 19.2130817 | -1.583803602 |
| 16.1509107 | 15.9759792 | 15.8949296 | 14.9217241 | 15.3270296 | 16.1316955 | -1.557357066 |
| 21.5377345 | 21.4955857 | 21.40891 | 21.0597611 | 18.7462292 | 20.8214811 | -1.360731054 |
| 16.2759467 | 16.0299205 | 16.1858244 | 15.2863429 | 14.3500183 | 15.0858302 | -2.073385214 |
| 16.9937991 | 16.0342855 | 16.25832 | 15.5251944 | 15.4686095 | 15.9295115 | -1.941554801 |
| 17.2364273 | 16.4934457 | 16.6764064 | 16.024185 | 15.2828706 | 15.8496465 | -1.68386607 |
| 15.6620538 | 15.8801548 | 15.7906121 | 15.5607168 | 12.7019442 | 15.1147017 | -1.133878259 |
| 12.9359875 | 14.4389345 | 13.3311399 | 13.7433498 | 13.2505381 | NA | -0.065989647 |
| 24.1870942 | 24.1199873 | 24.1062605 | 23.9036905 | 21.148317 | 22.7146468 | -1.753110303 |
| 15.3795945 | 15.2265325 | 15.0796287 | 14.87374 | 12.6260645 | 14.2785099 | -1.764108506 |
| 19.6946614 | 19.3280728 | 19.2309731 | 18.558766 | 16.9239474 | 18.0044444 | -1.747541819 |
| 18.1684807 | 18.1398406 | 18.0989024 | 17.8198799 | 14.6156993 | 16.5406684 | -2.322029203 |
| 16.8559668 | 15.8519716 | 16.2332214 | 15.3136331 | 14.2967731 | 14.6396407 | -1.996625166 |
| 19.8728962 | 19.021909 | 19.225451 | 18.3571687 | 17.5061977 | 18.3644 | -2.090669278 |
| 17.7338882 | 17.0450557 | 17.4064615 | 16.6981818 | 15.8882614 | 16.6895053 | -2.134632658 |
| 13.9441333 | 13.0304268 | 13.0602221 | 12.964661 | 11.1042906 | 12.5058684 | -2.031445395 |
| 16.6680291 | 18.8491962 | 16.0836142 | 15.1563887 | 14.3937831 | 16.9642588 | 2.668336128 |

| AMPH-03340.1nM - AMPH-04647.10nM__P.Value | AMPH-03340.1nM - AMPH-04647.10nM__adj.P.Val | AMPH-03340.10nM - AMPH-04647.10nM__logFC | AMPH-03340.10nM - AMPH-04647.10nM__P.Value | AMPH-03340.10nM - AMPH-04647.10nM__adj.P.Val | AMPH-03340.300nM - AMPH-04647.300nM__logFC |
|---|---|---|---|---|---|
| 0.003260271 | 0.098532628 | 3.13380421 | 1.09E-07 | 8.98E-05 | 4.13175351 |
| 0.68026888 | 0.79164255 | 0.53169822 | 0.06586937 | 0.49336544 | 1.69608503 |
| 0.14918866 | 0.33261734 | 0.40350631 | 0.45667582 | 0.82221677 | 1.63880151 |
| 0.004011056 | 0.10782501 | -0.603381 | 0.20558904 | 0.67246456 | 1.35200842 |
| 0.051076945 | 0.20944114 | -0.842438 | 0.0605914 | 0.49336544 | 1.28026969 |
| 0.508038793 | 0.660585639 | -0.1167753 | 0.85581266 | 0.95510089 | 1.1944318 |
| 0.069247854 | 0.236001344 | -0.8019884 | 0.03956242 | 0.43746034 | 1.09254499 |
| 0.005254928 | 0.10782501 | -0.5389488 | 0.06947358 | 0.49336544 | 1.02644021 |
| 0.965542971 | 0.976490096 | 0.02217063 | 0.93410128 | 0.98353605 | 1.01832802 |
| 0.718251014 | 0.822009576 | 0.89937938 | 0.13281492 | 0.57964125 | 0.97290929 |
| 0.243820573 | 0.452176335 | -0.0186351 | 0.94562249 | 0.98883075 | 0.9699498 |
| 0.001674297 | 0.08174711 | -1.4330494 | 0.00467822 | 0.25605458 | 0.94824879 |
| 0.063941064 | 0.231625316 | -0.213247 | 0.50314009 | 0.83389062 | 0.92827447 |
| 0.086085044 | 0.251982331 | 0.79078651 | 0.01303003 | 0.38205905 | 0.9152059 |
| 0.835027987 | 0.90130005 | -0.1554155 | 0.59229126 | 0.86921425 | 0.86441126 |
| 0.738342494 | 0.837509499 | -0.671995 | 0.09794843 | 0.51403768 | 0.84461919 |
| 0.142456166 | 0.324226173 | -0.2832567 | 0.19731911 | 0.65322174 | 0.80887348 |
| 0.375548442 | 0.560049355 | -0.448301 | 0.33413654 | 0.7414219 | 0.80511741 |
| 0.069765684 | 0.236001344 | 0.18097669 | 0.65875711 | 0.8954615 | 0.7624775 |
| 0.507992012 | 0.660585639 | -0.1551134 | 0.73560791 | 0.91801006 | 0.75200866 |
| 0.979306235 | 0.984399776 | 0.16426896 | 0.63115954 | 0.89226287 | 0.74405604 |
| 0.212599946 | 0.412070204 | 0.44168442 | 0.04828477 | 0.4663741 | 0.72830172 |
| NA | NA | -0.2463403 | 0.51794529 | 0.83389062 | 0.72600028 |
| 0.001131292 | 0.08174711 | 0.46934955 | 0.07567728 | 0.49336544 | 0.71697225 |
| 0.556316481 | 0.688853184 | -0.8773923 | 0.06514713 | 0.49336544 | 0.7144637 |
| 0.619671208 | 0.73819734 | -0.6035201 | 0.0901894 | 0.51399895 | 0.70982088 |
| 0.822517252 | 0.895427669 | 0.79004459 | 0.33013991 | 0.74055975 | 0.69861078 |
| 0.008356766 | 0.128662666 | -0.4715675 | 0.12963956 | 0.57222625 | 0.69241851 |
| 0.039109346 | 0.185542013 | 0.61889597 | 0.03465379 | 0.42996019 | 0.68914209 |
| 0.377850708 | 0.560049355 | 0.25599876 | 0.45328765 | 0.81848384 | 0.67018338 |
| 0.652927075 | 0.766602148 | -0.3670302 | 0.29566558 | 0.70977031 | 0.67015168 |
| 0.877412951 | 0.926221175 | -0.0681529 | 0.85510522 | 0.95510089 | 0.65653632 |
| 0.622404393 | 0.73970828 | 0.29257579 | 0.37925298 | 0.77262208 | 0.64707546 |
| 0.48842312 | 0.650259264 | 0.20637771 | 0.45184242 | 0.81848384 | 0.63774119 |
| 0.194711052 | 0.390373709 | 0.28888431 | 0.22444464 | 0.67389428 | 0.63410375 |
| 0.247729493 | 0.45426352 | 0.7708054 | 0.00674209 | 0.27273696 | 0.63353635 |

| | | | | | |
|---|---|---|---|---|---|
| 0.636183787 | 0.750688924 | 0.49582709 | 0.0817268 | 0.49336544 | 0.62830314 |
| 0.428239979 | 0.602489349 | 0.31072477 | 0.08085738 | 0.49336544 | 0.62640985 |
| 0.738788685 | 0.837509499 | 0.03179443 | 0.91397465 | 0.97577788 | 0.62558398 |
| 0.624582114 | 0.73970828 | 0.83501149 | 0.0687617 | 0.49336544 | 0.62405294 |
| 0.280066568 | 0.481124883 | 0.16401166 | 0.5928867 | 0.86921425 | 0.62386782 |
| 0.141124348 | 0.323476034 | 0.83129413 | 0.02407695 | 0.40341172 | 0.62082611 |
| 0.046077899 | 0.201067195 | -0.2743141 | 0.3655604 | 0.76393645 | 0.62077627 |
| 0.555753284 | 0.688853184 | 0.48346425 | 0.41540428 | 0.81100992 | 0.61715456 |
| 0.619687718 | 0.73819734 | 0.30176048 | 0.30816562 | 0.72563321 | 0.61709408 |
| 0.656243605 | 0.769389055 | 0.33960777 | 0.14308434 | 0.58443903 | 0.6144717 |
| 0.555620239 | 0.688853184 | -0.3114296 | 0.23634105 | 0.67596142 | 0.60680081 |
| 0.93504182 | 0.96094978 | 0.19482442 | 0.37219802 | 0.76393645 | 0.60406477 |
| 0.082416249 | 0.251982331 | -0.7088632 | 0.02552529 | 0.40459153 | 0.60042067 |
| 0.317702017 | 0.521619408 | 0.43162153 | 0.02803249 | 0.41491024 | 0.59937196 |
| 0.306058895 | 0.509681752 | -0.5173309 | 0.03442459 | 0.42996019 | 0.59899571 |
| 0.796224505 | 0.879187004 | 0.18454235 | 0.54242802 | 0.83875173 | 0.59623818 |
| 0.026418201 | 0.158357662 | -0.1859685 | 0.59760372 | 0.87351067 | 0.59562518 |
| 0.048353899 | 0.206580009 | -0.5074458 | 0.07882809 | 0.49336544 | 0.59464881 |
| 0.213578146 | 0.412985231 | 0.55321488 | 0.01964908 | 0.39548458 | 0.59375846 |
| 0.070368596 | 0.236299482 | -0.1863751 | 0.46122685 | 0.82452895 | 0.59191595 |
| 0.376261632 | 0.560049355 | 0.2956609 | 0.23907656 | 0.67673222 | 0.59104145 |
| 0.10289442 | 0.271199549 | -0.9357875 | 0.01246633 | 0.3790688 | 0.590568 |
| 0.195973585 | 0.390373709 | 0.39384222 | 0.12578198 | 0.57222625 | 0.58686214 |
| 0.317281362 | 0.521619408 | 0.15697702 | 0.53587703 | 0.83875173 | 0.58303732 |
| 0.477847131 | 0.646064477 | -0.6803019 | 0.01210323 | 0.3790688 | 0.58071611 |
| 0.494286255 | 0.654768806 | 0.32532575 | 0.26299646 | 0.68489683 | 0.58058273 |
| 0.512963791 | 0.66546654 | 0.13678264 | 0.64830213 | 0.8954615 | 0.57183523 |
| 0.336844292 | 0.53494449 | 0.03037304 | 0.88348214 | 0.96927913 | 0.56385002 |
| 0.542876027 | 0.685694079 | 0.39132339 | 0.13626159 | 0.57964125 | 0.56223105 |
| 0.450761336 | 0.622370982 | -0.1452238 | 0.44081401 | 0.81757951 | 0.55894244 |
| 0.543947977 | 0.685694079 | 0.27716553 | 0.27293298 | 0.69098729 | 0.55891129 |
| 0.194276383 | 0.390373709 | -0.4529551 | 0.10578642 | 0.52006376 | 0.55837183 |
| 0.661130805 | 0.773236608 | 0.44451441 | 0.10848627 | 0.52086098 | 0.55656056 |
| 0.162618103 | 0.350122354 | 0.01940937 | 0.94598303 | 0.98883075 | 0.55193051 |
| 0.165104506 | 0.353609651 | 0.03488078 | 0.86453651 | 0.95658285 | 0.53925798 |
| 0.541321779 | 0.684834995 | 0.1179285 | 0.60339866 | 0.8783516 | 0.53382722 |
| 0.196143653 | 0.390373709 | -0.3442449 | 0.23712167 | 0.67596142 | 0.5335136 |
| 0.103927404 | 0.271810133 | -0.6603178 | 0.02511132 | 0.40459153 | 0.53338904 |
| 0.562839469 | 0.6927255 | -0.3548167 | 0.35997459 | 0.76002696 | 0.53035809 |
| 0.478214392 | 0.646064477 | 0.1011851 | 0.60273403 | 0.8783516 | 0.52834407 |
| 0.739517895 | 0.837509499 | -0.7331275 | 0.01145669 | 0.37623771 | 0.52756482 |
| 0.474024273 | 0.643600344 | -0.076704 | 0.75490846 | 0.92263129 | 0.52329445 |

| | | | | | |
|---|---|---|---|---|---|
| 0.00243982 | 0.090495129 | 0.15835714 | 0.44692107 | 0.81757951 | 0.52301452 |
| 0.005681955 | 0.10782501 | 0.62310978 | 0.00422033 | 0.25605458 | 0.52184964 |
| 0.03142123 | 0.167945251 | 0.17353796 | 0.56610392 | 0.85122952 | 0.52151634 |
| 0.002850725 | 0.098532628 | 0.32921823 | 0.34113184 | 0.74732161 | 0.52097851 |
| 0.003122839 | 0.098532628 | 0.39103206 | 0.28837947 | 0.7065936 | 0.51922863 |
| 0.332560425 | 0.5348071 | 0.02520382 | 0.92897508 | 0.98284606 | 0.51335755 |
| 0.698430238 | 0.804970444 | -0.6233862 | 0.00450516 | 0.25605458 | 0.51044112 |
| 0.575178383 | 0.703666508 | 0.26587394 | 0.28965095 | 0.7065936 | 0.50214114 |
| 0.690754944 | 0.798379652 | -0.2424787 | 0.22923868 | 0.67475571 | 0.49920553 |
| 0.047461709 | 0.203835552 | 0.16808696 | 0.51153732 | 0.83389062 | 0.49593046 |
| 0.104664189 | 0.272862551 | 0.09139706 | 0.6814727 | 0.90170221 | 0.49468103 |
| 0.026781075 | 0.158357662 | 0.00474185 | 0.97633217 | 0.99336075 | 0.49358705 |
| 0.459606699 | 0.6313789 | -0.2367361 | 0.44054871 | 0.81757951 | 0.49115906 |
| 0.887248547 | 0.930584594 | 0.236418 | 0.32540827 | 0.73395656 | 0.48824671 |
| 0.262498914 | 0.465650248 | -0.4521801 | 0.0606378 | 0.49336544 | 0.48707529 |
| 0.66142053 | 0.773236608 | 0.23211925 | 0.42986754 | 0.81724722 | 0.48323042 |
| 0.047126669 | 0.203467522 | 0.33361415 | 0.19811941 | 0.6532371 | 0.48142344 |
| 0.380851164 | 0.561979295 | -0.4212217 | 0.0959415 | 0.51403768 | 0.47973675 |
| 0.02346221 | 0.149403738 | 0.29674087 | 0.25426021 | 0.68489683 | 0.47971843 |
| 0.0836517 | 0.251982331 | 0.49881835 | 0.03876263 | 0.43746034 | 0.47950319 |
| 0.877077419 | 0.926221175 | 0.939342 | 3.58E-04 | 0.073548 | 0.47795453 |
| 0.38829927 | 0.567835491 | 0.07734435 | 0.73032826 | 0.91801006 | 0.47610128 |
| 0.004160963 | 0.10782501 | 0.48365316 | 0.04228672 | 0.43868947 | 0.46964188 |
| 0.479113412 | 0.6462049 | 0.22706437 | 0.36701234 | 0.76393645 | 0.46495821 |
| 0.098855227 | 0.268369116 | 0.11066599 | 0.66466334 | 0.8954615 | 0.46119544 |
| 0.335680941 | 0.53494449 | -0.1890596 | 0.39479367 | 0.78538443 | 0.46065392 |
| 0.842663377 | 0.907141578 | 0.05729914 | 0.85230834 | 0.95510089 | 0.45941653 |
| 0.057299462 | 0.217471448 | 0.25813196 | 0.23934969 | 0.67673222 | 0.45793227 |
| 0.459165382 | 0.6313789 | 0.23159693 | 0.48205848 | 0.82809008 | 0.45674526 |
| 0.086698298 | 0.252663612 | 0.48968291 | 0.04049572 | 0.43746034 | 0.45394451 |
| 0.319425439 | 0.522347011 | 0.04548874 | 0.78242052 | 0.93096702 | 0.4526984 |
| 0.817119619 | 0.892596532 | 0.2884831 | 0.18672071 | 0.65322174 | 0.45122206 |
| 0.403653198 | 0.584008883 | 0.65112193 | 0.0171372 | 0.39548458 | 0.4480086 |
| 0.34526206 | 0.540404471 | -0.0784807 | 0.72735651 | 0.91801006 | 0.44769015 |
| 0.883894422 | 0.930057686 | -0.3995788 | 0.09810782 | 0.51403768 | 0.44705304 |
| 0.394660846 | 0.57405214 | 0.23427828 | 0.25958755 | 0.68489683 | 0.44482974 |
| 0.087066592 | 0.252833946 | 0.1329844 | 0.587428 | 0.86897007 | 0.44240988 |
| 0.032104027 | 0.167945251 | 0.14887622 | 0.46406849 | 0.82452895 | 0.44092821 |
| 0.014042884 | 0.136070839 | 0.58584059 | 0.00730842 | 0.27273696 | 0.43979304 |
| 0.08338351 | 0.251982331 | -0.1564681 | 0.63161587 | 0.89226287 | 0.43727359 |
| 0.891425949 | 0.931452308 | 0.30860842 | 0.36196168 | 0.76002696 | 0.43664775 |
| 0.127228872 | 0.307171997 | -0.5292596 | 0.01513669 | 0.39548458 | 0.43593443 |

| | | | | | |
|---|---|---|---|---|---|
| 0.979380703 | 0.984399776 | 0.26176643 | 0.17095482 | 0.62657995 | 0.4337392 |
| 0.949455434 | 0.971774232 | -0.3171842 | 0.05179609 | 0.48878836 | 0.43288612 |
| 0.169948144 | 0.361378298 | 0.82963452 | 0.0148056 | 0.39548458 | 0.43282632 |
| 0.873518351 | 0.925702564 | -0.0026807 | 0.98799836 | 0.99435417 | 0.43185094 |
| 0.00142665 | 0.08174711 | 0.19903675 | 0.25049508 | 0.68324405 | 0.4298988 |
| 0.116739842 | 0.294011453 | 0.23387347 | 0.31469496 | 0.72670454 | 0.42985584 |
| 0.009099615 | 0.128995227 | -0.0540061 | 0.75001191 | 0.92263129 | 0.42915682 |
| 0.21597398 | 0.416630656 | -0.2601955 | 0.2984577 | 0.71438418 | 0.42450463 |
| 0.804517517 | 0.887143641 | 0.86748453 | 0.00683858 | 0.27273696 | 0.42237357 |
| 0.733229262 | 0.835635584 | -0.3682375 | 0.07681053 | 0.49336544 | 0.42217948 |
| 0.300229541 | 0.502023168 | -0.0797149 | 0.75925125 | 0.92263129 | 0.42196966 |
| 0.160522082 | 0.34744302 | 0.20355211 | 0.41069695 | 0.80473078 | 0.42106356 |
| 0.365825098 | 0.555890652 | -0.0841946 | 0.7596818 | 0.92263129 | 0.42073847 |
| 0.225948595 | 0.429171026 | 0.2186589 | 0.16064401 | 0.61919592 | 0.42004324 |
| 0.005948703 | 0.107869808 | 0.21696153 | 0.26914315 | 0.69052039 | 0.41907252 |
| 0.106127892 | 0.275338679 | 0.52028139 | 0.04180922 | 0.43868947 | 0.41899662 |
| 0.094612511 | 0.26622003 | -0.3662641 | 0.08154474 | 0.49336544 | 0.41899601 |
| 0.004345 | 0.10782501 | 0.37564981 | 0.07705231 | 0.49336544 | 0.41543471 |
| 0.523274124 | 0.670737759 | -0.0859194 | 0.73507739 | 0.91801006 | 0.41402553 |
| 0.018176129 | 0.136070839 | 0.07459412 | 0.64883428 | 0.8954615 | 0.41101947 |
| 0.014266735 | 0.136070839 | -0.3486492 | 0.12609551 | 0.57222625 | 0.41080729 |
| 0.769279905 | 0.858732332 | -0.2069082 | 0.32383378 | 0.73241746 | 0.41057327 |
| 0.090556277 | 0.258370357 | 0.02350292 | 0.90790779 | 0.97302755 | 0.41038145 |
| 0.327252004 | 0.529836578 | -0.0262442 | 0.89022662 | 0.97256731 | 0.40875663 |
| 0.009623039 | 0.13087333 | 0.04929203 | 0.75471153 | 0.92263129 | 0.40833096 |
| 0.769273465 | 0.858732332 | 0.32351701 | 0.21904419 | 0.67389428 | 0.40823136 |
| 0.449093509 | 0.621119158 | -0.1603283 | 0.434793 | 0.81757951 | 0.40630518 |
| 0.074283609 | 0.243630549 | 0.44161148 | 0.03946721 | 0.43746034 | 0.40596559 |
| 0.334610216 | 0.53494449 | 0.29494182 | 0.61207956 | 0.88540816 | 0.40503263 |
| 0.382371674 | 0.562517969 | -0.2163099 | 0.23986489 | 0.67673222 | 0.40462168 |
| 0.013363894 | 0.136070839 | 0.37251242 | 0.041885 | 0.43868947 | 0.40438068 |
| 0.022369816 | 0.148887061 | -0.0467335 | 0.90689634 | 0.97302755 | 0.40406603 |
| 0.127918805 | 0.307910751 | 0.11640677 | 0.65150629 | 0.8954615 | 0.40374946 |
| 0.326485779 | 0.52964691 | 0.40525845 | 0.16329122 | 0.61988865 | 0.40247196 |
| 0.50035405 | 0.658530492 | -0.6636984 | 0.00588915 | 0.27273696 | 0.40215735 |
| 0.012031953 | 0.136070839 | 0.24367097 | 0.14633499 | 0.59222785 | 0.4006756 |
| 0.076612963 | 0.245345666 | 0.47923313 | 0.10755779 | 0.52086098 | 0.40037938 |
| 0.296613587 | 0.499044716 | -0.4558749 | 0.12904313 | 0.57222625 | 0.39995329 |
| 0.031974518 | 0.167945251 | 0.06283899 | 0.87993506 | 0.96927913 | 0.39989621 |
| 0.783652066 | 0.870890035 | 0.37298907 | 0.19226764 | 0.65322174 | 0.39818605 |
| 0.357679693 | 0.548671761 | -0.2759859 | 0.24815751 | 0.68324405 | 0.39675254 |
| 0.681699537 | 0.79164255 | 0.37910265 | 0.45163434 | 0.81848384 | 0.39619837 |

| | | | | | |
|---|---|---|---|---|---|
| 0.010210833 | 0.134288599 | 0.22175526 | 0.31571182 | 0.72670454 | 0.39539055 |
| 0.711497405 | 0.816570861 | -0.0277585 | 0.91674442 | 0.97578123 | 0.39460042 |
| 0.385147851 | 0.564238144 | -0.1029057 | 0.57702852 | 0.85978296 | 0.39396428 |
| 0.442678432 | 0.616748597 | 0.13434599 | 0.61421414 | 0.88540816 | 0.3936245 |
| 0.859822726 | 0.916041167 | -0.3495012 | 0.15472134 | 0.60488676 | 0.38866704 |
| 0.017943116 | 0.136070839 | -0.0669797 | 0.7835795 | 0.93099677 | 0.38820492 |
| 0.524904216 | 0.670737759 | -0.3835894 | 0.08586501 | 0.50715949 | 0.3879926 |
| 9.40E-04 | 0.08174711 | 0.11296321 | 0.61088446 | 0.88540816 | 0.38733514 |
| 0.052183727 | 0.212909608 | 0.06373379 | 0.71616574 | 0.91801006 | 0.38726733 |
| 0.174326816 | 0.364968222 | 0.27649701 | 0.44720574 | 0.81757951 | 0.38650849 |
| 0.28241184 | 0.482990679 | 0.22528207 | 0.37198979 | 0.76393645 | 0.38455218 |
| 0.488491335 | 0.650259264 | 0.06944089 | 0.813196 | 0.94565711 | 0.38282386 |
| 0.573217227 | 0.702320206 | 0.23414391 | 0.41892872 | 0.81171413 | 0.38178238 |
| 0.145701554 | 0.329342017 | 0.67819949 | 0.01897191 | 0.39548458 | 0.38176056 |
| 0.154554383 | 0.339936324 | -0.3804861 | 0.17089643 | 0.62657995 | 0.38165807 |
| 0.244730961 | 0.452835519 | 0.07264229 | 0.7563189 | 0.92263129 | 0.3802151 |
| 0.067193379 | 0.232947913 | -0.2138588 | 0.35588259 | 0.76002696 | 0.38005189 |
| 0.965719985 | 0.976490096 | 0.3921957 | 0.07969083 | 0.49336544 | 0.37959634 |
| 0.159808369 | 0.34744302 | -0.0299104 | 0.92472359 | 0.97961041 | 0.37893397 |
| 0.034811859 | 0.175348622 | 0.2306429 | 0.3334022 | 0.7414219 | 0.3759888 |
| 0.32483585 | 0.52907396 | -0.2611588 | 0.30018251 | 0.71544499 | 0.37496087 |
| 0.606812197 | 0.730322644 | 0.24803445 | 0.43102076 | 0.81724722 | 0.37487631 |
| 0.049296112 | 0.209024447 | 0.01129104 | 0.97281332 | 0.99336075 | 0.37484652 |
| 0.259271565 | 0.462944414 | 0.35368472 | 0.07549328 | 0.49336544 | 0.37339093 |
| 0.018148486 | 0.136070839 | 0.29179727 | 0.09756062 | 0.51403768 | 0.37176008 |
| 0.147338867 | 0.330789206 | -0.6732681 | 0.02077638 | 0.39548458 | 0.37068937 |
| 0.236334745 | 0.444352884 | 0.25937805 | 0.24456344 | 0.68324405 | 0.3678078 |
| 0.418257199 | 0.596225211 | -0.2866083 | 0.51666793 | 0.83389062 | 0.3671206 |
| 0.006988816 | 0.116385189 | -0.0505389 | 0.8550423 | 0.95510089 | 0.36706572 |
| 0.547883014 | 0.685694079 | 0.24972636 | 0.18915618 | 0.65322174 | 0.36682239 |
| 0.044667885 | 0.199157639 | 0.72068973 | 0.00716938 | 0.27273696 | 0.36513238 |
| 0.260033764 | 0.463291597 | 0.19456533 | 0.50046849 | 0.83389062 | 0.36256365 |
| 0.397294534 | 0.576854697 | -0.122461 | 0.66156203 | 0.8954615 | 0.36229985 |
| 0.022644088 | 0.149012709 | 0.23586446 | 0.17967677 | 0.63969603 | 0.36113391 |
| 0.901189104 | 0.938297297 | 0.44238775 | 0.34636262 | 0.75030004 | 0.35976054 |
| 0.017916177 | 0.136070839 | -0.0936209 | 0.6636595 | 0.8954615 | 0.35937157 |
| 0.056419973 | 0.216144123 | 0.27662629 | 0.28459977 | 0.70378436 | 0.35834579 |
| 0.183014029 | 0.375101655 | 0.04269595 | 0.83086468 | 0.95252044 | 0.35682216 |
| 0.157684381 | 0.344961006 | 0.27567708 | 0.10932883 | 0.52134082 | 0.35606927 |
| 0.384987396 | 0.564238144 | -0.1366319 | 0.44907929 | 0.81757951 | 0.3547207 |
| 0.121964943 | 0.299769257 | -0.1152558 | 0.54828357 | 0.84024713 | 0.35395479 |
| 0.098413541 | 0.268369116 | -0.2667504 | 0.19450763 | 0.65322174 | 0.35391566 |

| | | | | | |
|---|---|---|---|---|---|
| 0.129761052 | 0.309605316 | 1.57186325 | 7.47E-05 | 0.03067894 | 0.35060908 |
| 0.532510107 | 0.677891182 | -0.1610972 | 0.76374387 | 0.92561984 | 0.34952649 |
| 0.042102573 | 0.194099999 | 0.13032256 | 0.58565338 | 0.86790871 | 0.34914622 |
| 0.313553135 | 0.517933924 | 0.168233 | 0.36786986 | 0.76393645 | 0.34872558 |
| 0.016976928 | 0.136070839 | -0.1596782 | 0.44761232 | 0.81757951 | 0.3474836 |
| 0.594731435 | 0.720385949 | -0.2673501 | 0.19587796 | 0.65322174 | 0.3471285 |
| 0.102895446 | 0.271199549 | 0.26333081 | 0.15057035 | 0.59596336 | 0.34570424 |
| 0.274811083 | 0.477118817 | 0.67873885 | 0.03481565 | 0.42996019 | 0.3454725 |
| 0.437240436 | 0.61409328 | -0.2865113 | 0.09577435 | 0.51403768 | 0.34040604 |
| 0.570163517 | 0.700682877 | 0.13842917 | 0.52620941 | 0.83603958 | 0.34009009 |
| 0.273959329 | 0.476654184 | -0.4931253 | 0.14170391 | 0.58272509 | 0.33962079 |
| 0.037900508 | 0.181922439 | 0.28060002 | 0.13581836 | 0.57964125 | 0.33835829 |
| 0.164758489 | 0.353609651 | 0.27384492 | 0.26738143 | 0.68815095 | 0.33766198 |
| 0.059836035 | 0.221937292 | 0.06267721 | 0.86054398 | 0.95510089 | 0.33640188 |
| 0.85746639 | 0.915827977 | -0.621024 | 0.03535283 | 0.42996019 | 0.3345536 |
| 0.182176548 | 0.374448523 | -0.2466771 | 0.25700578 | 0.68489683 | 0.33437643 |
| 0.005846039 | 0.107869808 | -0.1268902 | 0.46726105 | 0.82452895 | 0.33419657 |
| 0.245648264 | 0.453504487 | 0.19560783 | 0.70713896 | 0.91602089 | 0.33387589 |
| 0.125001451 | 0.305392768 | 0.11495184 | 0.61579545 | 0.88540816 | 0.33236315 |
| 0.159530348 | 0.34744302 | 0.10024329 | 0.67018976 | 0.89845836 | 0.3319974 |
| 0.197978631 | 0.392113016 | 0.13092395 | 0.5048289 | 0.83389062 | 0.33124578 |
| 0.068137052 | 0.234598458 | 0.08923032 | 0.66054876 | 0.8954615 | 0.3310341 |
| 0.030210759 | 0.165449526 | 0.73386427 | 0.02910268 | 0.41918076 | 0.33004349 |
| 0.151605041 | 0.336167699 | -0.4076503 | 0.07056457 | 0.49336544 | 0.32857137 |
| 0.035533681 | 0.176173628 | 0.08327077 | 0.75170268 | 0.92263129 | 0.32642319 |
| 0.479902168 | 0.6462049 | -0.1789347 | 0.39508376 | 0.78538443 | 0.32615153 |
| 0.271469675 | 0.475363208 | -0.015648 | 0.93271598 | 0.98353605 | 0.32467273 |
| 0.046753583 | 0.202930445 | 0.25805408 | 0.26444859 | 0.68489683 | 0.32340931 |
| 0.043425239 | 0.197960866 | 0.2486553 | 0.2015691 | 0.66195291 | 0.31928522 |
| 0.903439071 | 0.939116282 | 0.84028315 | 0.10406187 | 0.52006376 | 0.31926872 |
| 0.015969051 | 0.136070839 | -0.114958 | 0.57453404 | 0.85850096 | 0.31719366 |
| 0.188144416 | 0.380957429 | -0.1254877 | 0.56430063 | 0.8500749 | 0.31627767 |
| 0.138712012 | 0.322965105 | -0.4989305 | 0.11554934 | 0.53876861 | 0.31627296 |
| 0.085095141 | 0.251982331 | 0.36026104 | 0.06755874 | 0.49336544 | 0.31460026 |
| 0.064151129 | 0.231625316 | 0.04943378 | 0.83186012 | 0.95252044 | 0.31458323 |
| 0.033501739 | 0.171933454 | 0.07112489 | 0.7265595 | 0.91801006 | 0.31424679 |
| 0.119719997 | 0.296061492 | 0.06740257 | 0.769376 | 0.92754434 | 0.31275701 |
| 0.507113766 | 0.660585639 | 0.38003135 | 0.05674525 | 0.49336544 | 0.31041417 |
| 0.59679032 | 0.720385949 | -0.5263538 | 0.01788415 | 0.39548458 | 0.30996619 |
| 0.057740758 | 0.218131754 | 0.04114512 | 0.88131187 | 0.96927913 | 0.30976308 |
| 0.075924553 | 0.245345666 | 0.16252219 | 0.51595456 | 0.83389062 | 0.30961229 |
| 0.840571566 | 0.90608507 | 1.12803122 | 0.02830082 | 0.41491024 | 0.30884339 |

| | | | | | |
|---|---|---|---|---|---|
| 0.227371933 | 0.430476791 | -0.2745645 | 0.49395564 | 0.83389062 | 0.30856207 |
| 0.469543614 | 0.640234876 | 0.59651558 | 0.07690858 | 0.49336544 | 0.30755588 |
| 0.302928183 | 0.50549979 | -0.5558066 | 0.02276842 | 0.40105221 | 0.30670344 |
| 0.010256948 | 0.134288599 | -0.7333006 | 0.00502727 | 0.25796202 | 0.30661397 |
| 0.957791807 | 0.973553946 | -0.5922359 | 0.02045793 | 0.39548458 | 0.30496953 |
| 0.054861677 | 0.214196787 | 0.07714646 | 0.67589103 | 0.89845836 | 0.30357782 |
| 0.207912581 | 0.403944444 | 0.04028403 | 0.81742988 | 0.94636732 | 0.30249197 |
| 0.267967267 | 0.47125278 | -0.0510155 | 0.84811931 | 0.95510089 | 0.30207861 |
| 0.134022802 | 0.315166013 | 0.17267389 | 0.43357845 | 0.81757951 | 0.30204144 |
| 0.102256877 | 0.271199549 | 0.04758491 | 0.82971134 | 0.95252044 | 0.3012929 |
| 0.507040218 | 0.660585639 | 0.26304507 | 0.16488028 | 0.61988865 | 0.29856023 |
| 0.335043401 | 0.53494449 | 0.07352326 | 0.73485321 | 0.91801006 | 0.29786974 |
| 0.946845957 | 0.970636057 | -0.080394 | 0.7801505 | 0.93096702 | 0.29711028 |
| 0.556292724 | 0.688853184 | -0.458427 | 0.19004249 | 0.65322174 | 0.29622612 |
| 0.055963401 | 0.215907026 | 0.06382929 | 0.773656 | 0.93014087 | 0.29536845 |
| 0.267173496 | 0.470871648 | -0.3344006 | 0.14053481 | 0.58272509 | 0.29514574 |
| 0.074645263 | 0.243642138 | -0.1331037 | 0.51828615 | 0.83389062 | 0.29378407 |
| 0.034775267 | 0.175348622 | 0.082917 | 0.65108077 | 0.8954615 | 0.2929813 |
| 0.547377098 | 0.685694079 | 0.02381725 | 0.95167691 | 0.98883675 | 0.29046123 |
| 0.676568899 | 0.788686031 | 0.08906406 | 0.69945121 | 0.91602089 | 0.29020504 |
| 0.056093492 | 0.215907026 | 0.17921211 | 0.33867835 | 0.74732161 | 0.28898294 |
| 0.682985729 | 0.79164255 | -0.4256683 | 0.10329086 | 0.52006376 | 0.28792708 |
| 0.873373273 | 0.925702564 | 0.07250795 | 0.74657421 | 0.92078002 | 0.28750475 |
| 0.076771976 | 0.245345666 | -0.041795 | 0.82048987 | 0.94636732 | 0.28747139 |
| 0.250508458 | 0.456283262 | 0.00765803 | 0.97291094 | 0.99336075 | 0.28687421 |
| 0.110894037 | 0.283826027 | 0.07857689 | 0.72238456 | 0.91801006 | 0.28520195 |
| 0.193096666 | 0.390017028 | 0.01009366 | 0.95835309 | 0.99051129 | 0.28465158 |
| 0.226028217 | 0.429171026 | 0.33799561 | 0.10657239 | 0.52080911 | 0.28454253 |
| 0.139347103 | 0.32303192 | 0.0787152 | 0.71338728 | 0.91801006 | 0.28304747 |
| 0.008056255 | 0.128662666 | 0.29292637 | 0.27232527 | 0.69098729 | 0.28262849 |
| NA | NA | -0.0113798 | 0.98471176 | 0.99336075 | 0.28097356 |
| 0.052608514 | 0.213574865 | -0.3812121 | 0.23537598 | 0.67596142 | 0.2797901 |
| 0.013087727 | 0.136070839 | 0.24817047 | 0.25268657 | 0.68467219 | 0.27911218 |
| 0.375525242 | 0.560049355 | 0.05877137 | 0.7819229 | 0.93096702 | 0.27798861 |
| 0.136152491 | 0.318339349 | 0.16981019 | 0.44105108 | 0.81757951 | 0.27751228 |
| 0.341278925 | 0.536580373 | 0.30280362 | 0.13387598 | 0.57964125 | 0.27650108 |
| 0.811664039 | 0.890602236 | 0.10602819 | 0.74671676 | 0.92078002 | 0.2738703 |
| 0.420195723 | 0.596225211 | 0.47268908 | 0.09205357 | 0.51399895 | 0.27357756 |
| 0.596362807 | 0.720385949 | -0.2905727 | 0.14787663 | 0.59222785 | 0.27154828 |
| 0.703026973 | 0.809125543 | -0.6357273 | 0.00851753 | 0.30403894 | 0.27128101 |
| 0.100654979 | 0.270279569 | -0.1757363 | 0.30273666 | 0.71741699 | 0.27071783 |
| 0.285111648 | 0.483682131 | -0.2405326 | 0.29217531 | 0.7065936 | 0.27031121 |

| | | | | | |
|---|---|---|---|---|---|
| 0.911885568 | 0.943090778 | -0.1318271 | 0.63018895 | 0.89226287 | 0.26975971 |
| 0.001803245 | 0.08174711 | -0.0047644 | 0.98489117 | 0.99336075 | 0.26947692 |
| 0.497193907 | 0.656489042 | -0.3621898 | 0.23410466 | 0.67596142 | 0.26937924 |
| 0.530506517 | 0.67639581 | -0.5595012 | 0.01590181 | 0.39548458 | 0.2690969 |
| 0.049675885 | 0.209024447 | 0.23526793 | 0.27335343 | 0.69098729 | 0.26833313 |
| 0.042688611 | 0.195696105 | 0.06831426 | 0.70670853 | 0.91602089 | 0.26610988 |
| 0.524770839 | 0.670737759 | 0.11701367 | 0.49488009 | 0.83389062 | 0.26426053 |
| 0.76899241 | 0.858732332 | 0.17089049 | 0.35237184 | 0.75931045 | 0.26421733 |
| 0.072851923 | 0.241655161 | -0.0061022 | 0.97304838 | 0.99336075 | 0.26410235 |
| 0.495963683 | 0.655926038 | 0.11434578 | 0.61515575 | 0.88540816 | 0.26251641 |
| 0.769281047 | 0.858732332 | -0.0151951 | 0.93555902 | 0.98353605 | 0.26163545 |
| 0.108024527 | 0.278949411 | -0.3360791 | 0.11629851 | 0.53876861 | 0.26091034 |
| 0.243246197 | 0.452138718 | 0.23182842 | 0.30064375 | 0.71544499 | 0.25907511 |
| 0.13892249 | 0.322965105 | -0.0044014 | 0.984376 | 0.99336075 | 0.25903036 |
| 0.029641581 | 0.163429259 | -0.0857341 | 0.7315884 | 0.91801006 | 0.25869504 |
| 0.281774774 | 0.482990679 | -0.4988327 | 0.23616144 | 0.67596142 | 0.25783348 |
| 0.054406332 | 0.214196787 | 0.50071137 | 0.02119528 | 0.39548458 | 0.25777024 |
| 0.421844424 | 0.596225211 | -0.2090068 | 0.27204398 | 0.69098729 | 0.25719904 |
| 0.134718718 | 0.315892165 | -0.0714562 | 0.86657856 | 0.95755181 | 0.25648557 |
| 0.42073305 | 0.596225211 | 0.52622764 | 0.05165109 | 0.48878836 | 0.25442518 |
| 0.024339854 | 0.151613136 | 0.05598467 | 0.74344141 | 0.92078002 | 0.25381417 |
| 0.27213046 | 0.475499904 | -0.0747373 | 0.76896417 | 0.92754434 | 0.25380673 |
| 0.013997461 | 0.136070839 | 0.03030113 | 0.8467544 | 0.95510089 | 0.25311309 |
| 0.284874784 | 0.483682131 | 0.23628462 | 0.42105275 | 0.81171413 | 0.24968092 |
| 0.050636272 | 0.20944114 | -0.0025513 | 0.98970979 | 0.99455537 | 0.24958386 |
| 0.073905706 | 0.243630549 | 0.1692194 | 0.34808485 | 0.75204649 | 0.24869969 |
| 0.013251298 | 0.136070839 | -0.2474291 | 0.74548951 | 0.92078002 | 0.24692108 |
| 0.184662399 | 0.375771863 | -0.0697521 | 0.70347987 | 0.91602089 | 0.24588574 |
| 0.371211362 | 0.558871718 | 0.17277109 | 0.33999014 | 0.74732161 | 0.24578734 |
| 0.083905887 | 0.251982331 | 0.05531724 | 0.79110099 | 0.93650527 | 0.24511491 |
| 0.361013934 | 0.551661742 | 0.56139158 | 0.04016874 | 0.43746034 | 0.242669 |
| 0.340900314 | 0.536580373 | -0.2898329 | 0.16550318 | 0.61988865 | 0.24202889 |
| 6.00E-04 | 0.081545586 | 0.33744439 | 0.03657812 | 0.42996019 | 0.2411999 |
| 0.357712472 | 0.548671761 | -0.3237463 | 0.61882314 | 0.88820594 | 0.24070236 |
| 0.292705469 | 0.494508618 | -0.2617269 | 0.14889735 | 0.59342099 | 0.23994629 |
| 0.589485771 | 0.71794088 | 0.47205828 | 0.03134296 | 0.4288761 | 0.23813004 |
| 0.056742562 | 0.216364162 | -0.1058901 | 0.71409606 | 0.91801006 | 0.23800768 |
| 0.816505378 | 0.892596532 | 0.48130344 | 0.45360554 | 0.81848384 | 0.23771709 |
| 0.077271858 | 0.245345666 | 0.18267439 | 0.25790272 | 0.68489683 | 0.23601742 |
| 0.017738621 | 0.136070839 | 0.16752002 | 0.46979483 | 0.82452895 | 0.23559632 |
| 0.080919027 | 0.251982331 | 0.13940524 | 0.42847265 | 0.81724722 | 0.23372824 |
| 0.339816796 | 0.536580373 | -0.1711812 | 0.3892825 | 0.77951446 | 0.2334117 |

| | | | | | |
|---|---|---|---|---|---|
| 0.11973075 | 0.296061492 | -0.0568982 | 0.75790837 | 0.92263129 | 0.23171095 |
| 0.242745508 | 0.452138718 | 0.19530298 | 0.66178474 | 0.8954615 | 0.23074467 |
| 0.132754174 | 0.313084988 | 0.06095279 | 0.84318995 | 0.95510089 | 0.23038091 |
| 0.328554995 | 0.530892824 | -0.1525026 | 0.46565475 | 0.82452895 | 0.22897383 |
| 0.184211357 | 0.375771863 | -0.1474318 | 0.49225583 | 0.83389062 | 0.22556895 |
| 0.169165917 | 0.361359655 | 0.16164229 | 0.49121337 | 0.83389062 | 0.22536807 |
| 0.01934374 | 0.140932966 | 0.28676883 | 0.2245403 | 0.67389428 | 0.22453203 |
| 0.490128952 | 0.651376587 | -0.1315932 | 0.5342226 | 0.83875173 | 0.22405011 |
| 0.37354983 | 0.559296626 | 0.686718 | 0.0967101 | 0.51403768 | 0.22371649 |
| 0.012802279 | 0.136070839 | 0.02251383 | 0.8859516 | 0.96982168 | 0.21983613 |
| 0.82835615 | 0.897660848 | 0.35773887 | 0.22555435 | 0.67389428 | 0.2191711 |
| 0.084963184 | 0.251982331 | 0.04858896 | 0.79403175 | 0.93650527 | 0.21481089 |
| 0.279091772 | 0.480901287 | -0.0026449 | 0.98829842 | 0.99435417 | 0.21470673 |
| 0.146453964 | 0.330128272 | 0.13351419 | 0.55820974 | 0.84787715 | 0.21353942 |
| 0.0022213 | 0.090495129 | 0.16852908 | 0.4730273 | 0.82586251 | 0.21342361 |
| 0.084149802 | 0.251982331 | 0.20704577 | 0.56888934 | 0.85229588 | 0.21142815 |
| 0.077809803 | 0.246096122 | 0.03900216 | 0.82072294 | 0.94636732 | 0.21035102 |
| 0.724070758 | 0.827509437 | 0.19820197 | 0.53816329 | 0.83875173 | 0.20940434 |
| 0.727251635 | 0.829982285 | -0.1073592 | 0.52473194 | 0.83603958 | 0.20835923 |
| 0.016013075 | 0.136070839 | 0.29955155 | 0.11448171 | 0.53708275 | 0.2075787 |
| 0.907858931 | 0.942246165 | -0.4295277 | 0.01134205 | 0.37623771 | 0.20561883 |
| 0.097254706 | 0.268369116 | 0.19696813 | 0.29186245 | 0.7065936 | 0.20402909 |
| 0.128510932 | 0.308426236 | 0.12147754 | 0.51174636 | 0.83389062 | 0.20277546 |
| 0.12723546 | 0.307171997 | -0.071408 | 0.62993208 | 0.89226287 | 0.20121928 |
| 0.273680634 | 0.476654184 | 0.11972851 | 0.50339054 | 0.83389062 | 0.20100664 |
| 0.044975995 | 0.199157639 | -0.0788832 | 0.72001788 | 0.91801006 | 0.19891485 |
| 0.468502146 | 0.640234876 | -0.4720592 | 0.02638539 | 0.40459153 | 0.19629471 |
| 0.115169552 | 0.291511526 | 0.4219359 | 0.13495421 | 0.57964125 | 0.19504633 |
| 0.784441391 | 0.870890035 | -0.1585903 | 0.51437284 | 0.83389062 | 0.1923866 |
| 0.64529275 | 0.75982523 | -0.2126817 | 0.36159538 | 0.76002696 | 0.19185316 |
| 0.350179056 | 0.543464079 | -0.1800978 | 0.22654668 | 0.67389428 | 0.19132941 |
| 0.035718241 | 0.176173628 | -0.125772 | 0.47042997 | 0.82452895 | 0.19121415 |
| 0.682629795 | 0.79164255 | -0.306506 | 0.18889261 | 0.65322174 | 0.19047118 |
| 0.011624934 | 0.136070839 | -0.254438 | 0.19711552 | 0.65322174 | 0.19046998 |
| 0.103649518 | 0.271810133 | 0.22045027 | 0.23638392 | 0.67596142 | 0.18922017 |
| 0.446424403 | 0.620370874 | -0.1100228 | 0.70849362 | 0.91602089 | 0.18809123 |
| 2.07E-04 | 0.081545586 | 0.63328654 | 0.02105656 | 0.39548458 | 0.18682033 |
| 0.029030245 | 0.163429259 | 0.37090206 | 0.02295914 | 0.40105221 | 0.18449565 |
| 0.147931242 | 0.330789206 | 0.13533831 | 0.53947091 | 0.83875173 | 0.18417777 |
| 0.045152161 | 0.199157639 | 0.15454595 | 0.35716053 | 0.76002696 | 0.18388469 |
| 0.081423574 | 0.251982331 | 0.3072787 | 0.07499209 | 0.49336544 | 0.18278903 |
| 0.955248495 | 0.973137043 | -0.278644 | 0.29161419 | 0.7065936 | 0.181105 |

| | | | | | |
|---|---|---|---|---|---|
| 0.058188484 | 0.218810153 | 0.25535746 | 0.21240036 | 0.67389428 | 0.18101861 |
| 0.054565171 | 0.214196787 | -0.448006 | 0.04434985 | 0.43868947 | 0.18022629 |
| 0.861033793 | 0.916041167 | -0.3769005 | 0.0655108 | 0.49336544 | 0.17986158 |
| 0.142189957 | 0.324226173 | 0.04961021 | 0.82494163 | 0.94989772 | 0.17874869 |
| 0.043987836 | 0.199157639 | 0.07034521 | 0.73791366 | 0.91931277 | 0.17812755 |
| 0.122801639 | 0.300919332 | -0.3893271 | 0.07359256 | 0.49336544 | 0.17784396 |
| 0.624100908 | 0.73970828 | 0.0714231 | 0.73235652 | 0.91801006 | 0.17566255 |
| 0.901501325 | 0.938297297 | -0.5287939 | 0.00321668 | 0.22007459 | 0.17436846 |
| 0.126908379 | 0.307171997 | -0.2200292 | 0.24935424 | 0.68324405 | 0.17388758 |
| 0.465009423 | 0.636657197 | 0.36606628 | 0.14651587 | 0.59222785 | 0.17211031 |
| 0.740741413 | 0.837509499 | -0.129159 | 0.37547965 | 0.7687501 | 0.17168757 |
| 0.050932644 | 0.20944114 | 0.41012439 | 0.03493463 | 0.42996019 | 0.17136351 |
| 0.204929235 | 0.40119559 | 0.47399104 | 0.05523474 | 0.49336544 | 0.17049623 |
| 0.351011581 | 0.543464079 | 0.10381721 | 0.63097647 | 0.89226287 | 0.16983269 |
| 0.081591316 | 0.251982331 | 0.5780533 | 0.06834952 | 0.49336544 | 0.1694101 |
| 0.014486869 | 0.136070839 | -0.1091164 | 0.54248132 | 0.83875173 | 0.16734344 |
| 0.145602757 | 0.329342017 | -0.0011711 | 0.99606965 | 0.99850205 | 0.16492228 |
| 0.959236976 | 0.973553946 | -0.3337263 | 0.1959289 | 0.65322174 | 0.16370392 |
| 0.534939696 | 0.678865929 | 0.03857247 | 0.85547356 | 0.95510089 | 0.16092283 |
| 0.110956498 | 0.283826027 | -0.3263752 | 0.07692898 | 0.49336544 | 0.15985443 |
| 0.349474714 | 0.543464079 | 0.06931727 | 0.7645879 | 0.92561984 | 0.15767119 |
| 0.07123346 | 0.238223373 | 0.03014476 | 0.90630414 | 0.97302755 | 0.15674235 |
| 0.170060376 | 0.361378298 | 0.23570596 | 0.21832225 | 0.67389428 | 0.15605337 |
| 0.502160503 | 0.659843752 | 0.03482392 | 0.90425567 | 0.97302755 | 0.15575581 |
| 0.130371454 | 0.31015483 | -0.230096 | 0.21870788 | 0.67389428 | 0.15509184 |
| 0.119080137 | 0.296061492 | -0.1171252 | 0.52209333 | 0.83555287 | 0.15219832 |
| 0.100358851 | 0.270279569 | 0.31725319 | 0.17998756 | 0.63969603 | 0.15147801 |
| 0.483479708 | 0.649949656 | 0.50460435 | 0.16270688 | 0.61988865 | 0.15106009 |
| 0.293619683 | 0.495028225 | -0.0784641 | 0.6970601 | 0.91565815 | 0.15025756 |
| 0.237779618 | 0.446041766 | -0.1895554 | 0.55558207 | 0.84625767 | 0.14828137 |
| 0.003766251 | 0.105974512 | -0.0317931 | 0.90893406 | 0.97302755 | 0.14756924 |
| 0.201540218 | 0.398082696 | 0.09581042 | 0.61523651 | 0.88540816 | 0.14641497 |
| 0.377048512 | 0.560049355 | -0.0191276 | 0.90431833 | 0.97302755 | 0.1445165 |
| 0.035242471 | 0.176173628 | -0.0860999 | 0.6490885 | 0.8954615 | 0.14253012 |
| 0.438966135 | 0.614685052 | -0.1104676 | 0.53533534 | 0.83875173 | 0.14210459 |
| 0.032812225 | 0.170539973 | -0.1664773 | 0.27895269 | 0.69610989 | 0.14114051 |
| 0.415007489 | 0.596207942 | 0.01119949 | 0.95511272 | 0.98883675 | 0.1406334 |
| 0.029330928 | 0.163429259 | 0.17211152 | 0.24545067 | 0.68324405 | 0.14024139 |
| 0.028825843 | 0.163429259 | 0.13572667 | 0.42118175 | 0.81171413 | 0.13713984 |
| 0.673139803 | 0.785811272 | 0.26006728 | 0.29262098 | 0.7065936 | 0.1365577 |
| 0.254547583 | 0.459537229 | 0.25185468 | 0.39357713 | 0.78538443 | 0.13306911 |
| 0.015247467 | 0.136070839 | -0.0307325 | 0.84072856 | 0.95510089 | 0.13287654 |

| | | | | | |
|---|---|---|---|---|---|
| 0.354719205 | 0.547166108 | -0.4515662 | 0.03271712 | 0.42996019 | 0.13084361 |
| 0.550498629 | 0.687912529 | 0.10630494 | 0.58967443 | 0.86921425 | 0.1304963 |
| 0.339303905 | 0.536580373 | -0.1990309 | 0.54130275 | 0.83875173 | 0.13016892 |
| 0.261562614 | 0.465000202 | -0.2884236 | 0.31980462 | 0.72670454 | 0.12936174 |
| 0.558423247 | 0.690414197 | -0.1030232 | 0.85959677 | 0.95510089 | 0.12891818 |
| 0.058468596 | 0.21885493 | 0.19916056 | 0.29148991 | 0.7065936 | 0.12636281 |
| 0.02030659 | 0.142846354 | 0.14024685 | 0.37131851 | 0.76393645 | 0.12613943 |
| 0.100986095 | 0.270279569 | -0.0988139 | 0.52064492 | 0.83486227 | 0.12467307 |
| 0.412899955 | 0.59428954 | -0.4730496 | 0.53340943 | 0.83875173 | 0.12031869 |
| 0.037568336 | 0.181395044 | -0.222366 | 0.1733538 | 0.62990676 | 0.11904886 |
| 0.525246848 | 0.670737759 | 0.259585 | 0.19343547 | 0.65322174 | 0.11904637 |
| 0.307113289 | 0.510396016 | 0.15436204 | 0.68140279 | 0.90170221 | 0.1184729 |
| 0.098657428 | 0.268369116 | 0.05068384 | 0.78563619 | 0.93209149 | 0.11705677 |
| 0.265167167 | 0.469363141 | 0.01453699 | 0.96385148 | 0.99287587 | 0.1163558 |
| 0.412943835 | 0.59428954 | -0.3814922 | 0.04786847 | 0.4663741 | 0.11516627 |
| 0.036347698 | 0.177603121 | -0.1306153 | 0.51902327 | 0.83389062 | 0.11438537 |
| 0.021185707 | 0.147522333 | 0.02842883 | 0.86119015 | 0.95510089 | 0.1116479 |
| 0.03210718 | 0.167945251 | -0.088477 | 0.64126218 | 0.8954615 | 0.10932326 |
| 0.393820363 | 0.573852529 | -0.3576298 | 0.06709284 | 0.49336544 | 0.10728405 |
| 0.487624072 | 0.650259264 | 0.09160049 | 0.75824222 | 0.92263129 | 0.10451799 |
| 0.756132552 | 0.849867993 | -0.3841603 | 0.09829953 | 0.51403768 | 0.1044005 |
| 0.316922676 | 0.521619408 | -0.0969461 | 0.76552481 | 0.92561984 | 0.1041555 |
| 0.344880868 | 0.540404471 | -0.1274773 | 0.54838497 | 0.84024713 | 0.10339859 |
| 0.382594942 | 0.562517969 | -0.21556 | 0.16256459 | 0.61988865 | 0.10127273 |
| 0.026624077 | 0.158357662 | -0.0542317 | 0.77621499 | 0.93032483 | 0.09933089 |
| 0.460444148 | 0.63146626 | 0.25851098 | 0.17877752 | 0.63969603 | 0.09670713 |
| 0.814994246 | 0.892596532 | 0.13723925 | 0.44912102 | 0.81757951 | 0.09649299 |
| 0.021513674 | 0.147522333 | 0.00462423 | 0.97430418 | 0.99336075 | 0.09613318 |
| 0.823003372 | 0.895427669 | 0.38986201 | 0.44442642 | 0.81757951 | 0.09578342 |
| 0.891500309 | 0.931452308 | -0.3504925 | 0.15266265 | 0.59969395 | 0.09414924 |
| 0.224877276 | 0.429171026 | -0.5746113 | 0.09765394 | 0.51403768 | 0.09080566 |
| 0.053525601 | 0.214196787 | 0.01526695 | 0.94788039 | 0.98883075 | 0.08964348 |
| 0.246747719 | 0.45426352 | -0.2362262 | 0.13478431 | 0.57964125 | 0.08938436 |
| 0.555000042 | 0.688853184 | -0.0909765 | 0.67630605 | 0.89845836 | 0.08846861 |
| 0.308484275 | 0.510831547 | -0.2794322 | 0.0769987 | 0.49336544 | 0.08779697 |
| 0.393522662 | 0.573852529 | 0.11608067 | 0.49755756 | 0.83389062 | 0.08556583 |
| 0.060336681 | 0.22278159 | -0.1230344 | 0.51130326 | 0.83389062 | 0.08543251 |
| 0.018597696 | 0.13796109 | -0.7208886 | 0.00293034 | 0.22007459 | 0.08491461 |
| 0.01375801 | 0.136070839 | 0.06849045 | 0.68728172 | 0.90571154 | 0.08413502 |
| 0.083233804 | 0.251982331 | -0.1132953 | 0.55871077 | 0.84787715 | 0.08372933 |
| 0.069484403 | 0.236001344 | 0.02912668 | 0.90065016 | 0.97302755 | 0.08288912 |
| 0.922852169 | 0.949618373 | 0.10161415 | 0.65813465 | 0.8954615 | 0.08195262 |

| | | | | | |
|---|---|---|---|---|---|
| 0.098717792 | 0.268369116 | -0.5638565 | 0.02610168 | 0.40459153 | 0.08187589 |
| 0.033439854 | 0.171933454 | -0.0913586 | 0.56884745 | 0.85229588 | 0.07896333 |
| 0.421791253 | 0.596225211 | -0.4049988 | 0.16610902 | 0.61988865 | 0.07776386 |
| 0.491691665 | 0.652390892 | -0.0589327 | 0.80071561 | 0.93651857 | 0.07705245 |
| 0.197834012 | 0.392113016 | 0.21048775 | 0.2504673 | 0.68324405 | 0.07704086 |
| 0.793650155 | 0.877531879 | 0.25182267 | 0.21811333 | 0.67389428 | 0.07670477 |
| 0.101023613 | 0.270279569 | -0.0733154 | 0.70596999 | 0.91602089 | 0.07552495 |
| 0.919468398 | 0.947331077 | 0.27176087 | 0.13826126 | 0.58211534 | 0.07435727 |
| 0.369478287 | 0.55776197 | 0.09186206 | 0.70423219 | 0.91602089 | 0.07428792 |
| 0.03150991 | 0.167945251 | -0.1702432 | 0.53724343 | 0.83875173 | 0.07393581 |
| 0.050076161 | 0.209411274 | -0.2664473 | 0.24827131 | 0.68324405 | 0.07263706 |
| 0.362916118 | 0.553531874 | -0.0690225 | 0.74269041 | 0.92078002 | 0.07211286 |
| 0.401456429 | 0.581862249 | 0.17166212 | 0.2781719 | 0.69610989 | 0.07134272 |
| 0.521308071 | 0.670737759 | -0.047298 | 0.8318581 | 0.95252044 | 0.07088901 |
| NA | NA | 0.20124124 | 0.47783555 | 0.82764344 | 0.0698247 |
| 0.48548105 | 0.650259264 | 0.04741294 | 0.77958592 | 0.93096702 | 0.06741899 |
| 0.083817538 | 0.251982331 | -0.1853329 | 0.31982952 | 0.72670454 | 0.06698408 |
| 0.13195641 | 0.312522442 | 0.22900249 | 0.19260263 | 0.65322174 | 0.06651818 |
| 0.03403767 | 0.173592119 | -0.2482564 | 0.25593286 | 0.68489683 | 0.06562038 |
| 0.452726148 | 0.624027933 | -0.4251565 | 0.07826623 | 0.49336544 | 0.06486345 |
| 0.194386362 | 0.390373709 | 0.19863111 | 0.35760804 | 0.76002696 | 0.06333964 |
| 0.447792211 | 0.620370874 | 0.02984336 | 0.90953048 | 0.97302755 | 0.06322529 |
| 0.284870263 | 0.483682131 | 0.01252167 | 0.9471186 | 0.98883075 | 0.06295549 |
| 0.004792047 | 0.10782501 | 0.82208176 | 0.00215784 | 0.22007459 | 0.06227571 |
| 0.014500975 | 0.136070839 | -0.1192011 | 0.67459019 | 0.89845836 | 0.06218299 |
| 0.20762665 | 0.403944444 | -0.0216049 | 0.8959282 | 0.97256731 | 0.06043185 |
| 0.020262298 | 0.142846354 | -0.1803723 | 0.26130585 | 0.68489683 | 0.05992729 |
| 0.9117003 | 0.943090778 | -0.0794916 | 0.72548021 | 0.91801006 | 0.05909964 |
| 0.247566525 | 0.45426352 | -0.2981783 | 0.12827741 | 0.57222625 | 0.05895237 |
| 0.424418862 | 0.598144717 | -0.0325574 | 0.96034955 | 0.99051129 | 0.05853936 |
| 0.546472137 | 0.685694079 | 0.15263521 | 0.50771577 | 0.83389062 | 0.05749736 |
| 0.97992908 | 0.984399776 | -0.5632885 | 0.04379616 | 0.43868947 | 0.05702897 |
| 0.379974679 | 0.5617017 | -0.2518122 | 0.10550425 | 0.52006376 | 0.05669501 |
| 0.739821695 | 0.837509499 | 0.08319261 | 0.65917647 | 0.8954615 | 0.05661995 |
| 0.065765694 | 0.232947913 | -0.115315 | 0.57993991 | 0.86255556 | 0.05646545 |
| 0.2261563 | 0.429171026 | -0.2527634 | 0.38424742 | 0.77893119 | 0.05614707 |
| 0.77744336 | 0.866658172 | 0.18023698 | 0.27353334 | 0.69098729 | 0.05535336 |
| 0.152853581 | 0.337104114 | -0.2456338 | 0.21164681 | 0.67389428 | 0.05274295 |
| 0.595380361 | 0.720385949 | -0.4402625 | 0.11680976 | 0.53876861 | 0.05168341 |
| 0.074343145 | 0.243630549 | 0.01352069 | 0.92427732 | 0.97961041 | 0.05105104 |
| 0.853642594 | 0.912938869 | 0.26739852 | 0.24226794 | 0.68117115 | 0.05003814 |
| 0.742058049 | 0.837509499 | 0.06597509 | 0.71981797 | 0.91801006 | 0.04967086 |

| | | | | | |
|---|---|---|---|---|---|
| 0.951528936 | 0.971774232 | -0.1705103 | 0.38541684 | 0.77937741 | 0.04963913 |
| 0.179267661 | 0.37033522 | 0.01322259 | 0.96750322 | 0.99336075 | 0.04911213 |
| 0.180613337 | 0.372172936 | -0.0603068 | 0.7747603 | 0.93014087 | 0.04826636 |
| 0.092611998 | 0.2614927 | -0.2081654 | 0.16703764 | 0.62053348 | 0.04800986 |
| 0.096411253 | 0.268369116 | -0.1530536 | 0.36137757 | 0.76002696 | 0.04723355 |
| 0.065324255 | 0.232947913 | -0.2364865 | 0.20853306 | 0.67389428 | 0.04414941 |
| 0.57720068 | 0.705083465 | 0.00120247 | 0.99453979 | 0.99818725 | 0.04338708 |
| 0.367718386 | 0.556841873 | -0.1229614 | 0.71962213 | 0.91801006 | 0.04324547 |
| 0.25434156 | 0.459537229 | -0.2087312 | 0.32655575 | 0.73452676 | 0.04251525 |
| 0.253855331 | 0.459537229 | -0.0826565 | 0.72599821 | 0.91801006 | 0.04091864 |
| 0.092598334 | 0.2614927 | -0.0108021 | 0.95413653 | 0.98883675 | 0.03907195 |
| 0.515541504 | 0.666509036 | 0.03764804 | 0.84504462 | 0.95510089 | 0.03824103 |
| 0.173954109 | 0.364968222 | 0.01960572 | 0.91021335 | 0.97302755 | 0.03780304 |
| 0.351194814 | 0.543464079 | 0.10590341 | 0.65936799 | 0.8954615 | 0.03577734 |
| 0.937588036 | 0.962354512 | 0.23278254 | 0.22483878 | 0.67389428 | 0.0348855 |
| 0.612946131 | 0.735535357 | 0.30176982 | 0.47980807 | 0.82809008 | 0.03336586 |
| 0.174433341 | 0.364968222 | 0.11298795 | 0.49005616 | 0.83389062 | 0.0323852 |
| 0.257755454 | 0.46226033 | -0.0422638 | 0.79259558 | 0.93650527 | 0.02890436 |
| 0.040809799 | 0.189619128 | -0.0105092 | 0.95499989 | 0.98883675 | 0.0257197 |
| 0.554446932 | 0.688853184 | -0.1862791 | 0.18766304 | 0.65322174 | 0.02532666 |
| 0.545361788 | 0.685694079 | -0.3191232 | 0.15959444 | 0.61911421 | 0.02517478 |
| 0.750666113 | 0.844887653 | 0.19367849 | 0.30322009 | 0.71741699 | 0.02203598 |
| 0.121216795 | 0.298830528 | -0.0848938 | 0.67452156 | 0.89845836 | 0.02194372 |
| 0.504407944 | 0.660585639 | 0.02546341 | 0.89347002 | 0.97256731 | 0.02056992 |
| 0.005191183 | 0.10782501 | 0.35107706 | 0.04422762 | 0.43868947 | 0.02017269 |
| 0.999132447 | 0.999132447 | -0.0863307 | 0.63905623 | 0.8954615 | 0.01969096 |
| 0.322863295 | 0.526912898 | -0.4730589 | 0.07382261 | 0.49336544 | 0.01752802 |
| 0.596381997 | 0.720385949 | -0.0059998 | 0.98165624 | 0.99336075 | 0.01726831 |
| 0.062006609 | 0.227916186 | 0.34764108 | 0.10834984 | 0.52086098 | 0.01483757 |
| 0.878779408 | 0.926465112 | 0.05599363 | 0.80287409 | 0.93763816 | 0.01098355 |
| 0.4757964 | 0.644933327 | 0.13402045 | 0.66323203 | 0.8954615 | 0.00919939 |
| 0.028451687 | 0.163429259 | 0.09918047 | 0.65922415 | 0.8954615 | 0.00799311 |
| 0.369790718 | 0.55776197 | 0.08475089 | 0.64429939 | 0.8954615 | 0.00470004 |
| 0.279347071 | 0.480901287 | -0.633435 | 0.02019834 | 0.39548458 | 0.00343198 |
| 0.847802796 | 0.909076322 | -0.1728586 | 0.34363002 | 0.74832957 | 0.00204795 |
| 0.806000467 | 0.887579461 | -0.0200331 | 0.89675207 | 0.97256731 | 4.12E-05 |
| 0.004893893 | 0.10782501 | 0.54701318 | 0.03824567 | 0.43746034 | -3.08E-04 |
| 0.847433057 | 0.909076322 | -0.004595 | 0.98023147 | 0.99336075 | -6.52E-04 |
| 0.117872439 | 0.295911501 | -0.3908253 | 0.05676915 | 0.49336544 | -0.0014446 |
| 0.251501884 | 0.457072466 | 0.307261 | 0.22837884 | 0.67475571 | -0.0016145 |
| 0.095120781 | 0.266564585 | -0.0322087 | 0.84750948 | 0.95510089 | -0.0035489 |
| 0.010995256 | 0.135941346 | -0.3502493 | 0.13814996 | 0.58211534 | -0.0081451 |

| | | | | | |
|---|---|---|---|---|---|
| 0.297277517 | 0.499132621 | -0.3145617 | 0.12829702 | 0.57222625 | -0.0106777 |
| 0.025184087 | 0.153359812 | -0.2161992 | 0.31057433 | 0.7264431 | -0.011511 |
| 0.016947988 | 0.136070839 | -0.2948836 | 0.22930188 | 0.67475571 | -0.0156083 |
| 0.378170582 | 0.560049355 | -0.2125003 | 0.4017519 | 0.79479112 | -0.0170749 |
| 0.088829336 | 0.255487457 | -0.1798319 | 0.34520698 | 0.74977495 | -0.0181943 |
| 0.827330309 | 0.897660848 | -0.6927116 | 0.00287839 | 0.22007459 | -0.018744 |
| 0.780464959 | 0.868839573 | 0.15342625 | 0.4435666 | 0.81757951 | -0.0200393 |
| 0.61520238 | 0.737158799 | -0.0872308 | 0.62200037 | 0.89120821 | -0.0202731 |
| 0.091850181 | 0.261148945 | -0.0207778 | 0.91729846 | 0.97578123 | -0.0281371 |
| 0.336449632 | 0.53494449 | -0.2111273 | 0.23180335 | 0.67596142 | -0.0299631 |
| 0.277448804 | 0.480217643 | 0.1621909 | 0.54856573 | 0.84024713 | -0.0301518 |
| 0.142643623 | 0.324226173 | -0.1398296 | 0.55338014 | 0.84447044 | -0.0323097 |
| 0.76326271 | 0.856702024 | 0.16237343 | 0.36725586 | 0.76393645 | -0.032847 |
| 0.484585452 | 0.650259264 | -0.1816176 | 0.43816072 | 0.81757951 | -0.0342889 |
| 0.890507899 | 0.931452308 | 0.39913716 | 0.07206238 | 0.49336544 | -0.0345776 |
| 0.787836654 | 0.872285902 | -0.2290639 | 0.225776 | 0.67389428 | -0.0367395 |
| 0.978904642 | 0.984399776 | 0.13237172 | 0.46178048 | 0.82452895 | -0.0378604 |
| 0.222841688 | 0.427856041 | -0.2721807 | 0.13940016 | 0.58272509 | -0.0434535 |
| 0.035839243 | 0.176173628 | -0.2491561 | 0.36042856 | 0.76002696 | -0.0436294 |
| 0.336962583 | 0.53494449 | 0.4499764 | 0.12693776 | 0.57222625 | -0.0439272 |
| 0.016910653 | 0.136070839 | 0.00858976 | 0.97762836 | 0.99336075 | -0.0473052 |
| 0.332943636 | 0.5348071 | -0.3170735 | 0.18523443 | 0.65322174 | -0.0484822 |
| 0.010466276 | 0.134288599 | 0.06933975 | 0.70748778 | 0.91602089 | -0.0489243 |
| 0.825521662 | 0.896971606 | 0.09884174 | 0.80901415 | 0.94345449 | -0.0509442 |
| 0.420141791 | 0.596225211 | 0.00545586 | 0.98162687 | 0.99336075 | -0.0525508 |
| 0.958259558 | 0.973553946 | 0.02825839 | 0.90651939 | 0.97302755 | -0.0536072 |
| 0.980780659 | 0.984399776 | 0.16043956 | 0.57512244 | 0.85850096 | -0.0549364 |
| 0.718017934 | 0.822009576 | -0.067303 | 0.73574984 | 0.91801006 | -0.0551765 |
| 0.561039354 | 0.69155304 | 0.21851235 | 0.31397647 | 0.72670454 | -0.0602932 |
| 0.225729984 | 0.429171026 | -0.0635323 | 0.71324609 | 0.91801006 | -0.0610175 |
| 0.170672051 | 0.361736087 | 0.14605906 | 0.60792088 | 0.88336822 | -0.0648262 |
| 0.469976337 | 0.640234876 | -0.3292076 | 0.08078089 | 0.49336544 | -0.0655196 |
| 0.504073882 | 0.660585639 | -0.0168011 | 0.93353641 | 0.98353605 | -0.0667658 |
| 0.365112572 | 0.55584302 | 0.31651794 | 0.16960261 | 0.62657995 | -0.0694765 |
| 0.176020011 | 0.366409002 | 0.03675388 | 0.8794312 | 0.96927913 | -0.069889 |
| 0.417951943 | 0.596225211 | -0.1242547 | 0.5339915 | 0.83875173 | -0.0704468 |
| 0.351653227 | 0.543464079 | 0.09308878 | 0.65510694 | 0.8954615 | -0.0716097 |
| 0.160346894 | 0.34744302 | -0.0140514 | 0.92416548 | 0.97961041 | -0.0720429 |
| 0.973196011 | 0.982831615 | 0.05286384 | 0.79620058 | 0.93650527 | -0.0728384 |
| 0.021449983 | 0.147522333 | -0.0326534 | 0.84241516 | 0.95510089 | -0.0750813 |
| 0.118219546 | 0.295911501 | -0.3576688 | 0.07411166 | 0.49336544 | -0.0772929 |
| 0.018031221 | 0.136070839 | 0.26362312 | 0.26284178 | 0.68489683 | -0.0782792 |

| | | | | | |
|---|---|---|---|---|---|
| 0.594605163 | 0.720385949 | -0.2247167 | 0.2198259 | 0.67389428 | -0.0791513 |
| 0.085554539 | 0.251982331 | -0.2551335 | 0.21872032 | 0.67389428 | -0.0792687 |
| 0.624040508 | 0.73970828 | -0.1218157 | 0.46963003 | 0.82452895 | -0.0794303 |
| 0.950602158 | 0.971774232 | 0.05686039 | 0.72240001 | 0.91801006 | -0.0845895 |
| 0.140481063 | 0.323047305 | 0.18740519 | 0.65659995 | 0.8954615 | -0.0849656 |
| 0.498917185 | 0.657700198 | -0.3394777 | 0.14088399 | 0.58272509 | -0.0854994 |
| 0.113201024 | 0.287763351 | 0.03533038 | 0.91754338 | 0.97578123 | -0.0875279 |
| 0.24155973 | 0.451417847 | -0.076713 | 0.83566276 | 0.95288767 | -0.0934467 |
| 0.686564743 | 0.794662171 | 0.13860572 | 0.49839428 | 0.83389062 | -0.0936568 |
| 0.201968427 | 0.398082696 | 0.25670077 | 0.14771489 | 0.59222785 | -0.0965424 |
| 0.308627393 | 0.510831547 | 0.14433541 | 0.50209255 | 0.83389062 | -0.0986922 |
| 0.06737219 | 0.232947913 | -0.2326067 | 0.20798663 | 0.67389428 | -0.1010283 |
| 0.13213265 | 0.312522442 | -0.3902113 | 0.09194989 | 0.51399895 | -0.1012138 |
| 0.572028676 | 0.701917895 | 0.06116008 | 0.81974585 | 0.94636732 | -0.1045692 |
| 0.009181934 | 0.128995227 | -0.1590561 | 0.3780799 | 0.77214825 | -0.1066159 |
| 0.081557581 | 0.251982331 | -0.0452544 | 0.81640099 | 0.94636732 | -0.1093612 |
| 0.086143024 | 0.251982331 | 0.13508936 | 0.50738541 | 0.83389062 | -0.1107546 |
| 0.609060132 | 0.731948554 | 0.13974682 | 0.41727081 | 0.81171413 | -0.1119435 |
| 0.53401949 | 0.678753744 | 0.22188762 | 0.36915022 | 0.76393645 | -0.1148836 |
| 0.447677944 | 0.620370874 | 0.02178032 | 0.9398758 | 0.98548918 | -0.1149299 |
| 0.06609041 | 0.232947913 | -0.2046281 | 0.21292562 | 0.67389428 | -0.1186172 |
| 0.372320458 | 0.559296626 | 0.0620121 | 0.835297 | 0.95288767 | -0.126971 |
| 0.741248109 | 0.837509499 | 0.18410876 | 0.34225691 | 0.74732161 | -0.1280705 |
| 0.053165446 | 0.214196787 | -0.1350901 | 0.62938068 | 0.89226287 | -0.1303229 |
| 0.341280899 | 0.536580373 | -0.0146706 | 0.94921509 | 0.98883675 | -0.1321232 |
| 0.119125393 | 0.296061492 | 0.49495551 | 0.08122739 | 0.49336544 | -0.1335488 |
| 0.069990595 | 0.236001344 | -0.3552879 | 0.09132505 | 0.51399895 | -0.1387232 |
| 0.049694538 | 0.209024447 | -0.0257413 | 0.8961261 | 0.97256731 | -0.1391082 |
| 0.560743011 | 0.69155304 | -0.3141043 | 0.07062375 | 0.49336544 | -0.141414 |
| 0.908759476 | 0.942246165 | -0.1396684 | 0.4689915 | 0.82452895 | -0.1418861 |
| 0.08194115 | 0.251982331 | -0.444233 | 0.03065342 | 0.42655014 | -0.141999 |
| 0.355668952 | 0.547595972 | -0.1209874 | 0.74694214 | 0.92078002 | -0.1531018 |
| 0.604581648 | 0.728712887 | -0.2831743 | 0.1033053 | 0.52006376 | -0.1541092 |
| 0.918521291 | 0.947331077 | 0.22141713 | 0.46450383 | 0.82452895 | -0.1595868 |
| 0.442910144 | 0.616748597 | -0.1761633 | 0.3534808 | 0.75970611 | -0.1611268 |
| 0.52266356 | 0.670737759 | 0.47416221 | 0.09209615 | 0.51399895 | -0.1617482 |
| 0.347754794 | 0.542577269 | -0.3627298 | 0.08042265 | 0.49336544 | -0.163719 |
| 0.161833989 | 0.349355913 | 0.33606917 | 0.29210621 | 0.7065936 | -0.172112 |
| 0.870148466 | 0.924532745 | 0.02456093 | 0.88427536 | 0.96927913 | -0.1733871 |
| 0.069504431 | 0.236001344 | -0.2801385 | 0.21316641 | 0.67389428 | -0.1752724 |
| 0.098398315 | 0.268369116 | -0.1536424 | 0.44485652 | 0.81757951 | -0.1787436 |
| 0.177432601 | 0.367474625 | -0.5247455 | 0.2755685 | 0.69399306 | -0.1800739 |

| | | | | | |
|---|---|---|---|---|---|
| 0.147963309 | 0.330789206 | -0.3688931 | 0.05634451 | 0.49336544 | -0.1832426 |
| 0.636134953 | 0.750688924 | -0.1130028 | 0.51539432 | 0.83389062 | -0.1844362 |
| 0.206509568 | 0.403138295 | -0.4057816 | 0.06922644 | 0.49336544 | -0.1849879 |
| 0.618253837 | 0.73819734 | -0.181212 | 0.31337403 | 0.72670454 | -0.1852571 |
| 0.040762011 | 0.189619128 | 0.17851962 | 0.50542956 | 0.83389062 | -0.1912585 |
| 0.697565772 | 0.804970444 | -0.0714523 | 0.78235037 | 0.93096702 | -0.194751 |
| 0.195249751 | 0.390373709 | -0.2444009 | 0.2840951 | 0.70378436 | -0.1992666 |
| 0.256867423 | 0.46226033 | -0.0812219 | 0.73028328 | 0.91801006 | -0.1993562 |
| 0.707874714 | 0.813557418 | 0.05352122 | 0.80077471 | 0.93651857 | -0.2059361 |
| 0.020259761 | 0.142846354 | -0.1800333 | 0.66710832 | 0.89639268 | -0.2063241 |
| 0.037515036 | 0.181395044 | 0.08202033 | 0.66532462 | 0.8954615 | -0.2070352 |
| 0.547860934 | 0.685694079 | 0.40863633 | 0.03608864 | 0.42996019 | -0.2075911 |
| 0.845066745 | 0.908530256 | 0.45458649 | 0.44399724 | 0.81757951 | -0.209154 |
| 0.412752015 | 0.59428954 | -0.4576371 | 0.01753504 | 0.39548458 | -0.2129077 |
| 0.996246196 | 0.997468584 | -0.1004938 | 0.59220236 | 0.86921425 | -0.2227061 |
| 0.204186082 | 0.40119559 | 1.22136827 | 0.0072006 | 0.27273696 | -0.2236438 |
| 0.993059602 | 0.995499552 | -0.2641387 | 0.19724923 | 0.65322174 | -0.2281026 |
| 0.067037627 | 0.232947913 | -0.2595711 | 0.22501008 | 0.67389428 | -0.232375 |
| 0.415844625 | 0.596225211 | -0.1299417 | 0.42509672 | 0.81626642 | -0.2388406 |
| 0.812019686 | 0.890602236 | -0.2135591 | 0.26155719 | 0.68489683 | -0.2426789 |
| 0.249249228 | 0.455005301 | -0.265575 | 0.20818049 | 0.67389428 | -0.2431802 |
| 0.140322276 | 0.323047305 | -0.3834238 | 0.06167692 | 0.49336544 | -0.2437764 |
| 0.325750556 | 0.52950688 | -0.4042183 | 0.15098706 | 0.59596336 | -0.2439988 |
| 0.417192813 | 0.596225211 | -0.2924303 | 0.08255704 | 0.49473962 | -0.2500373 |
| 0.422326191 | 0.596225211 | 0.19349338 | 0.4378798 | 0.81757951 | -0.2565789 |
| 0.345699919 | 0.540404471 | -0.0012 | 0.99824359 | 0.99938403 | -0.2572987 |
| 0.270953549 | 0.475363208 | -0.2452592 | 0.40460501 | 0.79851133 | -0.2587995 |
| 0.186474752 | 0.378515914 | 0.64385301 | 0.10082302 | 0.52006376 | -0.2616458 |
| 0.129560875 | 0.309605316 | 0.34236969 | 0.51350379 | 0.83389062 | -0.2693583 |
| 0.231889394 | 0.437001722 | -0.1523507 | 0.51450118 | 0.83389062 | -0.2707028 |
| 0.319145493 | 0.522347011 | 0.12870769 | 0.43067798 | 0.81724722 | -0.2802045 |
| NA | NA | -0.1437623 | 0.5531883 | 0.84447044 | -0.2809842 |
| 0.849172444 | 0.909350019 | 0.51951942 | 0.03061897 | 0.42655014 | -0.2826843 |
| 0.076053617 | 0.245345666 | -0.657559 | 0.02275774 | 0.40105221 | -0.2831344 |
| 0.019546245 | 0.141148103 | -0.2407759 | 0.33226199 | 0.7414219 | -0.2842642 |
| 0.508391889 | 0.660585639 | -0.1393237 | 0.51581317 | 0.83389062 | -0.2861135 |
| 0.524658142 | 0.670737759 | 0.02060153 | 0.93681509 | 0.98353605 | -0.2873618 |
| 0.050299767 | 0.209411274 | 0.5134264 | 0.24728249 | 0.68324405 | -0.2910544 |
| 0.618394688 | 0.73819734 | -0.1599321 | 0.49472949 | 0.83389062 | -0.2916451 |
| 0.440254924 | 0.615150716 | -0.2843957 | 0.10062466 | 0.52006376 | -0.2952738 |
| 0.088919654 | 0.255487457 | 0.11581249 | 0.64042808 | 0.8954615 | -0.3004218 |
| NA | NA | -0.057693 | 0.8943643 | 0.97256731 | -0.3072664 |

| | | | | | |
|---|---|---|---|---|---|
| 0.090146692 | 0.258104213 | -0.1362087 | 0.70153815 | 0.91602089 | -0.3113279 |
| 0.018896469 | 0.138914584 | -0.6249949 | 0.0021976 | 0.22007459 | -0.3146628 |
| 0.8299867 | 0.898222566 | 0.1344506 | 0.56072535 | 0.84854103 | -0.3186132 |
| 0.055644424 | 0.215907026 | -0.0856806 | 0.68204272 | 0.90170221 | -0.3205218 |
| 0.649974651 | 0.764235324 | -0.2132468 | 0.66071072 | 0.8954615 | -0.3215299 |
| 0.08164588 | 0.251982331 | -0.3405769 | 0.12648573 | 0.57222625 | -0.3236576 |
| 0.27777295 | 0.480217643 | -0.1438083 | 0.531202 | 0.83875173 | -0.3273753 |
| 0.10302924 | 0.271199549 | -0.4250914 | 0.07452445 | 0.49336544 | -0.3287232 |
| 0.831076026 | 0.898222566 | -0.1308498 | 0.49449256 | 0.83389062 | -0.330654 |
| 0.04561814 | 0.200131196 | 0.18028669 | 0.3084604 | 0.72563321 | -0.3342715 |
| 0.439168364 | 0.614685052 | -0.1511 | 0.43995459 | 0.81757951 | -0.3347849 |
| 0.017825467 | 0.136070839 | -0.3679303 | 0.09265755 | 0.51399895 | -0.3459013 |
| 0.8856064 | 0.930057686 | -0.0231053 | 0.95959843 | 0.99051129 | -0.346128 |
| 0.003387603 | 0.098724422 | -0.4506136 | 0.02661138 | 0.40459153 | -0.3629875 |
| 0.177028374 | 0.367474625 | -0.2792029 | 0.28230369 | 0.70233735 | -0.3633672 |
| 0.063253022 | 0.231455004 | -0.1014064 | 0.58403591 | 0.86707682 | -0.3658398 |
| 0.084001976 | 0.251982331 | 0.05362977 | 0.84052181 | 0.95510089 | -0.3662767 |
| 0.231650653 | 0.437001722 | -0.0072079 | 0.97857462 | 0.99336075 | -0.3696216 |
| 0.33256486 | 0.5348071 | -0.1333541 | 0.42652655 | 0.81626642 | -0.3780733 |
| 0.066387136 | 0.232947913 | -0.0127859 | 0.96800038 | 0.99336075 | -0.3793748 |
| 0.819674261 | 0.894190103 | -0.6198271 | 0.01379506 | 0.39054282 | -0.3843866 |
| 0.266992566 | 0.470871648 | -0.3304618 | 0.08653277 | 0.50745289 | -0.3843915 |
| 0.578550827 | 0.705676345 | 0.03279662 | 0.88278542 | 0.96927913 | -0.386753 |
| 0.008685566 | 0.128995227 | -0.1588606 | 0.31455435 | 0.72670454 | -0.38882 |
| 0.077008499 | 0.245345666 | 0.66345788 | 0.21622622 | 0.67389428 | -0.3979737 |
| 0.125827885 | 0.306494192 | 0.02322656 | 0.95260857 | 0.98883675 | -0.3995809 |
| 0.299549635 | 0.501914789 | -0.2923578 | 0.24950144 | 0.68324405 | -0.4042638 |
| 0.516217783 | 0.666509036 | -0.0935785 | 0.79752087 | 0.93651857 | -0.4050179 |
| 0.009191861 | 0.128995227 | -0.3102616 | 0.21212513 | 0.67389428 | -0.4094274 |
| 0.282928364 | 0.482990679 | 0.01425846 | 0.97002781 | 0.99336075 | -0.411905 |
| 0.172911641 | 0.364588886 | 0.05087947 | 0.86087886 | 0.95510089 | -0.4138028 |
| 0.960872402 | 0.974002335 | -0.2413433 | 0.47580142 | 0.82586251 | -0.4235033 |
| 0.810414294 | 0.890602236 | -0.1122107 | 0.77492857 | 0.93014087 | -0.4366378 |
| 0.860897291 | 0.916041167 | -0.4898061 | 0.05254839 | 0.49025257 | -0.4393617 |
| 0.746953041 | 0.841869726 | 0.11494891 | 0.74674388 | 0.92078002 | -0.4400153 |
| 0.222111998 | 0.427460826 | -0.3266033 | 0.10370516 | 0.52006376 | -0.4403236 |
| 0.04033582 | 0.189619128 | 0.22894493 | 0.39968122 | 0.79260454 | -0.4403554 |
| 0.894872252 | 0.933779742 | 0.16647814 | 0.4833906 | 0.82852543 | -0.4481005 |
| 0.636613646 | 0.750688924 | 0.69983416 | 0.32042271 | 0.72670454 | -0.4481633 |
| 0.098499228 | 0.268369116 | 0.05468293 | 0.83373858 | 0.95288767 | -0.4564245 |
| 0.150426859 | 0.334464079 | -0.1569092 | 0.38843684 | 0.77951446 | -0.4583096 |
| 0.24175196 | 0.451417847 | -0.4145146 | 0.06712616 | 0.49336544 | -0.4594055 |

| | | | | | |
|---|---|---|---|---|---|
| 0.367815894 | 0.556841873 | -0.3442094 | 0.15986871 | 0.61911421 | -0.4657121 |
| 0.78692161 | 0.872285902 | -0.2834027 | 0.34149419 | 0.74732161 | -0.475332 |
| 0.471289698 | 0.640953989 | -0.3532322 | 0.26089328 | 0.68489683 | -0.4770683 |
| 0.00159832 | 0.08174711 | -0.3075337 | 0.22036312 | 0.67389428 | -0.4822445 |
| 0.015975259 | 0.136070839 | -0.3127741 | 0.23023958 | 0.67509534 | -0.5009155 |
| 0.098994 | 0.268369116 | -0.1496287 | 0.67297301 | 0.89845836 | -0.5031295 |
| 0.291734638 | 0.493891005 | 0.08671109 | 0.75385976 | 0.92263129 | -0.5122766 |
| 0.183413677 | 0.375101655 | 0.29302382 | 0.26373227 | 0.68489683 | -0.5205605 |
| 0.067303861 | 0.232947913 | -0.1784968 | 0.68667035 | 0.90571154 | -0.5218064 |
| 0.248422524 | 0.454512958 | -0.1323384 | 0.63967109 | 0.8954615 | -0.5324386 |
| 0.172080325 | 0.363776024 | -0.4409092 | 0.06252843 | 0.49336544 | -0.5353547 |
| 0.016719926 | 0.136070839 | -0.0225402 | 0.93611406 | 0.98353605 | -0.5615648 |
| 0.076625442 | 0.245345666 | -0.1380756 | 0.59794519 | 0.87351067 | -0.5677689 |
| 0.029101667 | 0.163429259 | -0.1207536 | 0.65742321 | 0.8954615 | -0.573184 |
| 0.0171389 | 0.136070839 | -0.1204453 | 0.63218457 | 0.89226287 | -0.5744769 |
| 0.064123085 | 0.231625316 | -0.2648961 | 0.26647305 | 0.68796973 | -0.5794489 |
| 0.156748474 | 0.343835362 | -0.295142 | 0.26026395 | 0.68489683 | -0.5806968 |
| 0.952864403 | 0.971921691 | -0.3370874 | 0.47101593 | 0.82452895 | -0.5955553 |
| 0.258917335 | 0.462944414 | -0.2115821 | 0.54377246 | 0.83916765 | -0.5970554 |
| 0.110006939 | 0.283172436 | -0.3553384 | 0.14195496 | 0.58272509 | -0.5988984 |
| 0.021954104 | 0.148054125 | -0.8067603 | 0.00125279 | 0.20570872 | -0.6050713 |
| 0.875909374 | 0.926221175 | -0.5369565 | 0.00255835 | 0.22007459 | -0.6059076 |
| 0.152493844 | 0.337104114 | 0.08988351 | 0.79387825 | 0.93650527 | -0.6065518 |
| 0.095388307 | 0.266564585 | -0.1491969 | 0.59149123 | 0.86921425 | -0.6161373 |
| 0.059222058 | 0.220663012 | -0.0482359 | 0.86164913 | 0.95510089 | -0.6206355 |
| 0.175532886 | 0.3663295 | -0.4384831 | 0.07976483 | 0.49336544 | -0.6464451 |
| 0.066457895 | 0.232947913 | -0.2413833 | 0.47520325 | 0.82586251 | -0.6477891 |
| 0.106288828 | 0.275338679 | -0.1797202 | 0.56175434 | 0.84854103 | -0.6547734 |
| 0.023732618 | 0.149403738 | 0.00520609 | 0.98410798 | 0.99336075 | -0.6592475 |
| 0.48845314 | 0.650259264 | -0.2078726 | 0.64524354 | 0.8954615 | -0.6613604 |
| 0.014366939 | 0.136070839 | -0.4037493 | 0.07664191 | 0.49336544 | -0.6659862 |
| 0.040898243 | 0.189619128 | -1.192545 | 0.0029552 | 0.22007459 | -0.6773489 |
| 0.014261375 | 0.136070839 | -0.3447466 | 0.17520153 | 0.63365841 | -0.6864845 |
| 0.257730074 | 0.46226033 | 0.07383331 | 0.84543167 | 0.95510089 | -0.6931412 |
| 3.22E-04 | 0.081545586 | -0.438442 | 0.10396157 | 0.52006376 | -0.6950383 |
| 0.022442535 | 0.148887061 | -0.3959701 | 0.18955847 | 0.65322174 | -0.6962722 |
| 0.008218533 | 0.128662666 | -0.4840658 | 0.10985623 | 0.52134082 | -0.7006785 |
| 0.054646569 | 0.214196787 | 0.05713253 | 0.81015276 | 0.94345449 | -0.7091785 |
| 0.03055209 | 0.166203369 | -0.0455966 | 0.85314092 | 0.95510089 | -0.7152598 |
| 0.115389979 | 0.291511526 | -0.4644788 | 0.08506732 | 0.50608889 | -0.728111 |
| 0.373393623 | 0.559296626 | -0.4600426 | 0.05701874 | 0.49336544 | -0.7482833 |
| 0.002996678 | 0.098532628 | -0.0693001 | 0.81667995 | 0.94636732 | -0.7667672 |

| | | | | | |
|---|---|---|---|---|---|
| 0.360832649 | 0.551661742 | 0.37860322 | 0.47459738 | 0.82586251 | -0.7677119 |
| 0.030945344 | 0.16722782 | -0.3909494 | 0.25168967 | 0.6842292 | -0.7710889 |
| 0.112654874 | 0.28726993 | 2.47E-04 | 0.99938403 | 0.99938403 | -0.7719954 |
| 0.88529692 | 0.930057686 | 0.08407673 | 0.86203381 | 0.95510089 | -0.7815099 |
| 0.001583554 | 0.08174711 | -0.5514688 | 0.05465341 | 0.49336544 | -0.7889337 |
| 0.006300574 | 0.109388686 | 0.52367137 | 0.04313705 | 0.43868947 | -0.8187093 |
| 0.003155911 | 0.098532628 | -0.3129686 | 0.31050052 | 0.7264431 | -0.8326774 |
| 0.013114375 | 0.136070839 | -0.0367732 | 0.89299836 | 0.97256731 | -0.8347887 |
| 9.21E-04 | 0.08174711 | -0.5601198 | 0.02403578 | 0.40341172 | -0.8361431 |
| 0.016607031 | 0.136070839 | -0.5452674 | 0.13208718 | 0.57964125 | -0.8389875 |
| 0.022987143 | 0.149291216 | -0.2332408 | 0.38662888 | 0.77951446 | -0.8427256 |
| 0.005307618 | 0.10782501 | -0.1101632 | 0.79603908 | 0.93650527 | -0.8612657 |
| 0.029388092 | 0.163429259 | -0.1926655 | 0.56224887 | 0.84854103 | -0.8661535 |
| 5.93E-04 | 0.081545586 | -0.4308013 | 0.11394488 | 0.53708275 | -0.8777492 |
| 0.066392337 | 0.232947913 | -0.0520574 | 0.90061338 | 0.97302755 | -0.8784753 |
| 0.001787037 | 0.08174711 | -0.4534289 | 0.19119255 | 0.65322174 | -0.8860381 |
| 0.014706977 | 0.136070839 | -0.1489496 | 0.70171908 | 0.91602089 | -0.8861375 |
| 0.02305232 | 0.149291216 | -0.3563808 | 0.3406378 | 0.74732161 | -0.9021172 |
| 0.008909067 | 0.128995227 | -0.320344 | 0.35720451 | 0.76002696 | -0.9087033 |
| 0.011995269 | 0.136070839 | -0.2985423 | 0.41026263 | 0.80473078 | -0.9103408 |
| 0.024986674 | 0.153359812 | -0.2183771 | 0.51601112 | 0.83389062 | -0.9307122 |
| 0.006165453 | 0.109369766 | -0.1654378 | 0.6307278 | 0.89226287 | -0.9335969 |
| 0.027618354 | 0.160975551 | -0.1991983 | 0.63251765 | 0.89226287 | -0.9664303 |
| 0.018044355 | 0.136070839 | -0.3626942 | 0.17943974 | 0.63969603 | -0.9759644 |
| 0.025003921 | 0.153359812 | -0.253616 | 0.54577437 | 0.84024713 | -0.9879361 |
| 0.014268101 | 0.136070839 | -0.4336263 | 0.25912586 | 0.68489683 | -0.9908761 |
| 0.009326861 | 0.128995227 | -0.3260382 | 0.37150846 | 0.76393645 | -0.9979457 |
| 0.004907614 | 0.10782501 | -0.4980307 | 0.09481954 | 0.51403768 | -1.0076009 |
| 0.001043371 | 0.08174711 | -0.5352042 | 0.08962771 | 0.51399895 | -1.0205151 |
| 0.005521643 | 0.10782501 | -0.410584 | 0.29372076 | 0.70716935 | -1.0250341 |
| 0.140541413 | 0.323047305 | -0.2714925 | 0.22107447 | 0.67389428 | -1.0367351 |
| 0.087736493 | 0.253875809 | -0.1112301 | 0.79899856 | 0.93651857 | -1.0472363 |
| 0.015080251 | 0.136070839 | -0.3049507 | 0.44461904 | 0.81757951 | -1.0580663 |
| 0.044274979 | 0.199157639 | -0.2341076 | 0.52647072 | 0.83603958 | -1.0765963 |
| 0.053735041 | 0.214196787 | -0.3202818 | 0.3170516 | 0.72670454 | -1.0845148 |
| 0.006713149 | 0.114123535 | -0.396125 | 0.32016053 | 0.72670454 | -1.1121014 |
| 0.101698118 | 0.271194981 | -0.149382 | 0.73227641 | 0.91801006 | -1.1177931 |
| 0.023802066 | 0.149403738 | -0.3101812 | 0.41587719 | 0.81100992 | -1.1788787 |
| 1.19E-04 | 0.081545586 | -0.7696488 | 0.02033469 | 0.39548458 | -1.1827042 |
| 0.011524335 | 0.136070839 | -0.4972678 | 0.23645716 | 0.67596142 | -1.1913433 |
| 0.014057676 | 0.136070839 | -0.5236846 | 0.1642451 | 0.61988865 | -1.2154152 |
| 0.010945763 | 0.135941346 | -0.3125307 | 0.38837999 | 0.77951446 | -1.2199829 |

| | | | | | |
|---|---|---|---|---|---|
| 0.085147897 | 0.251982331 | -0.3025004 | 0.48177738 | 0.82809008 | -1.2341707 |
| 0.021808904 | 0.148054125 | -0.4599042 | 0.3186052 | 0.72670454 | -1.2411045 |
| 0.012981523 | 0.136070839 | -0.3061452 | 0.40634505 | 0.80002227 | -1.2433408 |
| 0.072777437 | 0.241655161 | -0.1665684 | 0.72458233 | 0.91801006 | -1.2496053 |
| 0.002106448 | 0.090466412 | -0.6785958 | 0.03374994 | 0.42996019 | -1.2582233 |
| 0.010532439 | 0.134288599 | -0.2479172 | 0.52316897 | 0.83564537 | -1.2817804 |
| 0.023740945 | 0.149403738 | -0.5185098 | 0.27642088 | 0.69401084 | -1.2972212 |
| 0.516190211 | 0.666509036 | -0.4317852 | 0.48212796 | 0.82809008 | -1.3477051 |
| 0.044599263 | 0.199157639 | -0.3609357 | 0.46452273 | 0.82452895 | -1.3580233 |
| 0.016838857 | 0.136070839 | -0.394213 | 0.46774778 | 0.82452895 | -1.3750944 |
| 0.016451193 | 0.136070839 | -0.4495226 | 0.33381254 | 0.7414219 | -1.3878557 |
| 0.53676856 | 0.680129107 | -0.6546387 | 0.3801991 | 0.77263233 | -1.3892064 |
| 0.038661046 | 0.1844878 | -0.3505303 | 0.48763629 | 0.83389062 | -1.4043325 |
| 0.005663834 | 0.10782501 | -0.5855693 | 0.10549139 | 0.52006376 | -1.4048945 |
| 0.086155723 | 0.251982331 | -0.2241703 | 0.67410092 | 0.89845836 | -1.4357165 |
| 0.001784193 | 0.08174711 | -0.4951456 | 0.21691174 | 0.67389428 | -1.4397406 |
| 4.50E-04 | 0.081545586 | -0.699339 | 0.03665921 | 0.42996019 | -1.4636601 |
| 0.004620606 | 0.10782501 | -0.6440401 | 0.09092769 | 0.51399895 | -1.4806957 |
| 0.205022746 | 0.40119559 | 0.24522626 | 0.69067946 | 0.90873051 | -1.4986414 |
| 0.917991711 | 0.947331077 | 0.21484523 | 0.63673213 | 0.8954615 | -1.6011703 |
| 0.054815963 | 0.214196787 | -0.3756348 | 0.53637912 | 0.83875173 | -1.6096205 |
| 0.029499983 | 0.163429259 | -0.4290669 | 0.41938245 | 0.81171413 | -1.6257769 |
| 0.016822463 | 0.136070839 | -0.3744975 | 0.42587325 | 0.81626642 | -1.6841545 |
| 0.026229438 | 0.158357662 | -0.4163399 | 0.53970744 | 0.83875173 | -1.6930079 |
| 0.004921897 | 0.10782501 | -0.6094088 | 0.17339699 | 0.62990676 | -1.7179196 |
| 0.001611514 | 0.08174711 | -0.7820966 | 0.05907455 | 0.49336544 | -1.7199711 |
| 0.002429496 | 0.090495129 | -1.0710172 | 0.02048466 | 0.39548458 | -1.852548 |
| 0.007679406 | 0.125327911 | -0.9505809 | 0.05911664 | 0.49336544 | -2.0674175 |
| 0.027109173 | 0.1591445 | 3.70578563 | 2.48E-04 | 0.06786138 | -2.4506503 |

| AMPH-03340.300nM - AMPH-04647.300nM__P.Value | AMPH-03340.300nM - AMPH-04647.300nM__adj.P.Val | AMPH-03340.1000nM - AMPH-04647.1000nM__logFC | AMPH-03340.1000nM - AMPH-04647.1000nM__P.Value | AMPH-03340.1000nM - AMPH-04647.1000nM__adj.P.Val | AMPH-03340.10nM - DMSO__logFC | AMPH-03340.10nM - DMSO__P.Value |
|---|---|---|---|---|---|---|
| 3.06E-09 | 2.51E-06 | 4.85023512 | 4.41E-07 | 3.54E-04 | 2.88043342 | 3.12E-07 |
| 1.60E-05 | 0.00656621 | 3.32200179 | 4.08E-06 | 0.00163947 | 0.67997002 | 0.0229466 |
| 0.00391557 | 0.10106781 | -0.0777414 | 0.92546504 | 0.96884622 | -0.0994783 | 0.83584884 |
| 0.00989583 | 0.14669425 | -0.8420134 | 0.30304088 | 0.67301027 | -0.7362014 | 0.12747389 |
| 0.00772072 | 0.14078773 | -0.6060216 | 0.41154183 | 0.73080926 | -0.9263238 | 0.04146171 |
| 0.07879673 | 0.26820895 | 0.58965347 | 0.59791204 | 0.83776197 | -0.9623188 | 0.14909094 |
| 0.00788823 | 0.14078773 | 0.49986709 | 0.42855935 | 0.73624298 | -0.6340402 | 0.09459394 |
| 0.00212746 | 0.08733222 | -0.6509032 | 0.19236121 | 0.58361665 | -0.6503886 | 0.03245379 |
| 0.00164502 | 0.08733222 | 0.36433146 | 0.43767121 | 0.7387887 | -0.1953393 | 0.47040743 |
| 0.10620153 | 0.31057514 | 0.07431871 | 0.94044769 | 0.97689915 | 0.10594761 | 0.85372174 |
| 0.00272668 | 0.09647579 | -1.1118697 | 0.031022 | 0.33613674 | -0.1464377 | 0.59384079 |
| 0.04358984 | 0.22218971 | -0.1104763 | 0.88374737 | 0.95710625 | -1.399097 | 0.00548651 |
| 0.00953062 | 0.14616752 | -0.0493959 | 0.92808444 | 0.97032495 | -0.4072876 | 0.21018868 |
| 0.00533553 | 0.12167965 | 0.00441892 | 0.9928332 | 0.99805425 | 0.93818997 | 0.00452106 |
| 0.00850795 | 0.14515769 | -0.4086695 | 0.41928499 | 0.73281212 | 0.39595608 | 0.18409113 |
| 0.04261529 | 0.2217997 | -0.6796085 | 0.3184578 | 0.68160393 | -0.823302 | 0.04737566 |
| 0.00165957 | 0.08733222 | 0.15317612 | 0.67921264 | 0.87082935 | -0.2156263 | 0.32044369 |
| 0.09375123 | 0.29603753 | -0.0034074 | 0.99656107 | 0.99841214 | -0.7146025 | 0.13291285 |
| 0.07772335 | 0.26699109 | 1.16120225 | 0.11646642 | 0.51041692 | -0.0692101 | 0.86548549 |
| 0.11673039 | 0.32708413 | -0.176284 | 0.8244513 | 0.94229238 | -0.8144017 | 0.09164897 |
| 0.04284441 | 0.2217997 | 0.66913924 | 0.26748534 | 0.64783091 | 0.17115378 | 0.61704599 |
| 0.00303572 | 0.099693 | NA | NA | NA | 0.87246441 | 7.47E-04 |
| 0.03407631 | 0.20272937 | NA | NA | NA | -0.5840193 | 0.14111662 |
| 0.0108501 | 0.14982855 | 1.14597581 | 0.01699167 | 0.27645753 | 0.33488516 | 0.19291074 |
| 0.12449532 | 0.3318528 | NA | NA | NA | -0.9692929 | 0.04452055 |
| 0.05028998 | 0.2231788 | 0.26605203 | 0.65072553 | 0.85908592 | -0.9315313 | 0.0137988 |
| 0.3874374 | 0.59566686 | 0.36656811 | 0.79098161 | 0.92824212 | -0.7212668 | 0.37264477 |
| 0.03287154 | 0.20187361 | -0.1968835 | 0.70402032 | 0.88580961 | -0.394873 | 0.19906883 |
| 0.02072745 | 0.1660206 | 0.64613061 | 0.18087595 | 0.57346985 | 0.23567371 | 0.38890321 |
| 0.06268017 | 0.24250524 | 0.47007839 | 0.42714598 | 0.73587133 | 0.21210911 | 0.53307276 |
| 0.06700296 | 0.24885909 | 0.07744363 | 0.89664766 | 0.95992639 | -0.5063493 | 0.15597853 |
| 0.09439056 | 0.29691438 | 1.37395612 | 0.04778502 | 0.38419153 | -0.2435872 | 0.51689179 |
| 0.0640235 | 0.24263403 | 0.07297702 | 0.89786393 | 0.95995026 | 0.30564953 | 0.35892001 |
| 0.03108472 | 0.19846802 | -0.0241422 | 0.95906237 | 0.98352825 | 0.44368372 | 0.11805454 |
| 0.01428743 | 0.14982855 | 0.26671183 | 0.50934376 | 0.7800336 | 0.48432931 | 0.05107825 |
| 0.02068584 | 0.1660206 | 0.48230525 | 0.27197417 | 0.64783091 | 1.07654083 | 5.53E-04 |

| | | | | | | |
|---|---|---|---|---|---|---|
| 0.03216394 | 0.20187361 | 0.15934045 | 0.73338281 | 0.90574466 | 0.68983663 | 0.02047238 |
| 0.0019267 | 0.08733222 | -0.1125328 | 0.70033013 | 0.88393316 | 0.23384085 | 0.17889012 |
| 0.04785332 | 0.22218971 | 0.06185364 | 0.9034314 | 0.96079213 | 0.37116791 | 0.21965208 |
| 0.16302723 | 0.3824153 | -0.7029402 | 0.35476602 | 0.70018966 | 0.70044354 | 0.12056233 |
| 0.05589538 | 0.23446999 | 0.12479537 | 0.81354933 | 0.94081146 | 0.40655907 | 0.19619552 |
| 0.08010208 | 0.2695238 | 0.67105793 | 0.25903028 | 0.64476888 | 0.8107916 | 0.02717359 |
| 0.05242567 | 0.22894403 | -0.6071951 | 0.25179742 | 0.6390225 | -0.4195703 | 0.17436159 |
| 0.30203094 | 0.5091733 | -0.5709975 | 0.57625447 | 0.82660601 | -0.3584793 | 0.5437875 |
| 0.04808783 | 0.22218971 | 0.24763706 | 0.6242383 | 0.8480478 | 0.1457281 | 0.61763303 |
| 0.01382713 | 0.14982855 | -0.1409075 | 0.71596384 | 0.89384306 | 0.50680824 | 0.03609784 |
| 0.09802467 | 0.30097498 | 1.33358994 | 0.00636356 | 0.24466371 | -0.506841 | 0.09157686 |
| 0.01244634 | 0.14982855 | -0.2332774 | 0.534285 | 0.79644543 | 0.2227482 | 0.30967012 |
| 0.05282416 | 0.22946366 | -0.5087525 | 0.31894714 | 0.68160393 | -0.7335599 | 0.02154558 |
| 0.00424023 | 0.10135281 | 0.12891661 | 0.67974573 | 0.87082935 | 0.47959475 | 0.01646877 |
| 0.01677286 | 0.1555914 | 0.18968312 | 0.62831232 | 0.8480478 | -0.4596924 | 0.056273 |
| 0.06291549 | 0.24250524 | 0.54919033 | 0.30136672 | 0.67301027 | 0.21371611 | 0.48151734 |
| 0.10526674 | 0.31057514 | 1.65903235 | 0.01446872 | 0.27094123 | -0.280489 | 0.42884699 |
| 0.04322528 | 0.2217997 | -0.6428463 | 0.1875364 | 0.57992024 | -0.3100689 | 0.26650298 |
| 0.01343098 | 0.14982855 | 0.81801644 | 0.04130087 | 0.37310001 | 0.19185682 | 0.37759877 |
| 0.03021752 | 0.19534319 | 1.55978482 | 0.00248791 | 0.18548002 | -0.0334693 | 0.8937383 |
| 0.02737901 | 0.18935579 | 0.61399826 | 0.1624021 | 0.55487874 | 0.29449044 | 0.24085186 |
| 0.09259475 | 0.29577256 | 0.16871625 | 0.77056261 | 0.92170206 | -0.8927933 | 0.01617408 |
| 0.02919828 | 0.19358394 | 0.09508369 | 0.82393693 | 0.94229238 | 0.3892166 | 0.12993072 |
| 0.03319178 | 0.20187361 | 0.39816215 | 0.36832716 | 0.7067662 | 0.25419497 | 0.32137432 |
| 0.02767685 | 0.18935579 | 0.33315311 | 0.4303404 | 0.73772639 | -0.8288785 | 0.00344039 |
| 0.05598928 | 0.23446999 | 0.26921908 | 0.58634923 | 0.83438014 | 0.46263942 | 0.11925216 |
| 0.07129901 | 0.25579425 | 0.34743531 | 0.50536442 | 0.77910145 | 0.19657228 | 0.51380155 |
| 0.01478211 | 0.14982855 | 0.2816475 | 0.43747877 | 0.7387887 | 0.14002988 | 0.50254325 |
| 0.03928007 | 0.21532576 | -0.0198655 | 0.96373966 | 0.98461483 | 0.60288652 | 0.02867347 |
| 0.00842611 | 0.14515769 | 0.27365438 | 0.40259662 | 0.72251714 | -0.0139726 | 0.94025199 |
| 0.0370108 | 0.21101298 | 0.9280033 | 0.04433745 | 0.38419153 | -0.0329815 | 0.89397283 |
| 0.05114145 | 0.22573725 | 0.57042014 | 0.22953104 | 0.61067965 | -0.4235582 | 0.12827822 |
| 0.04970476 | 0.2231788 | -0.1359912 | 0.76674373 | 0.92170206 | 0.68178591 | 0.01965578 |
| 0.06963295 | 0.25350939 | 0.7150644 | 0.16407412 | 0.55487874 | -0.0640747 | 0.8231598 |
| 0.01746504 | 0.1555914 | 0.8311675 | 0.03110746 | 0.33613674 | -0.4205182 | 0.05447312 |
| 0.03021458 | 0.19534319 | -0.1312695 | 0.7377573 | 0.9097498 | 0.34558932 | 0.14131414 |
| 0.07604151 | 0.26453425 | -0.2176188 | 0.65922686 | 0.86181853 | -0.3015785 | 0.29759996 |
| 0.06251925 | 0.24250524 | 0.52417348 | 0.27062533 | 0.64783091 | -0.8117466 | 0.00806119 |
| 0.17882992 | 0.39172402 | 0.2490747 | 0.70714095 | 0.8878476 | -0.6821509 | 0.08998566 |
| 0.0145105 | 0.14982855 | 0.43358647 | 0.2085984 | 0.59814652 | 0.20629679 | 0.29579559 |
| 0.05536978 | 0.23446999 | 0.11661511 | 0.79393916 | 0.92824212 | -0.650836 | 0.02180244 |
| 0.04717281 | 0.22218971 | 0.20842381 | 0.6251966 | 0.8480478 | -0.3883385 | 0.12894037 |

| | | | | | | |
|---|---|---|---|---|---|---|
| 0.02136378 | 0.1660206 | 1.15041949 | 0.00532302 | 0.24466371 | -0.0415091 | 0.84046456 |
| 0.01275992 | 0.14982855 | -0.0213222 | 0.9474436 | 0.97721252 | 0.60236686 | 0.00529779 |
| 0.13715658 | 0.34623221 | 0.38962669 | 0.45891543 | 0.75146233 | 0.04997295 | 0.86799807 |
| 0.14098588 | 0.35182307 | 1.55729143 | 0.01733693 | 0.27645753 | 0.30896296 | 0.37075277 |
| 0.16475896 | 0.38393104 | 2.01521566 | 0.00531747 | 0.24466371 | 0.24305837 | 0.50406782 |
| 0.08536908 | 0.27971896 | 0.21964532 | 0.6549216 | 0.86143329 | 0.38099116 | 0.19136159 |
| 0.01525632 | 0.15038994 | 0.28915207 | 0.3826773 | 0.71114006 | -0.6082184 | 0.00531123 |
| 0.05626137 | 0.23446999 | -0.2851059 | 0.50688506 | 0.77922675 | 0.47795266 | 0.06763767 |
| 0.02125186 | 0.1660206 | -0.0759536 | 0.82353766 | 0.94229238 | 0.05660371 | 0.77359695 |
| 0.0664622 | 0.24802486 | 0.60628273 | 0.1822169 | 0.57346985 | 0.00666084 | 0.97908573 |
| 0.03930405 | 0.21532576 | -0.2013653 | 0.60227167 | 0.83776197 | 0.12882508 | 0.56405222 |
| 0.0070182 | 0.13718908 | 0.05533879 | 0.84167165 | 0.94578917 | -0.0010383 | 0.9948166 |
| 0.12152876 | 0.32823428 | -0.2688186 | 0.61095653 | 0.84111139 | 0.2986286 | 0.33357416 |
| 0.05357232 | 0.23027682 | 0.42700604 | 0.30585335 | 0.67301027 | 0.59405928 | 0.02239655 |
| 0.04523512 | 0.22218971 | 0.06156932 | 0.87506406 | 0.95332183 | -0.6142626 | 0.01488912 |
| 0.11240407 | 0.3193209 | 0.27023302 | 0.5934436 | 0.83776197 | 0.34423374 | 0.24781122 |
| 0.07134821 | 0.25579425 | 0.35538615 | 0.42018208 | 0.73281212 | 0.48844537 | 0.06775025 |
| 0.06139161 | 0.24250524 | 0.17121816 | 0.68201675 | 0.87082935 | -0.6428705 | 0.01632329 |
| 0.07496352 | 0.26414183 | 0.41104321 | 0.35793277 | 0.70018966 | 0.48983612 | 0.06965065 |
| 0.04579935 | 0.22218971 | 0.13983485 | 0.7181276 | 0.8951544 | 0.36270211 | 0.11983085 |
| 0.0328453 | 0.20187361 | -0.3678392 | 0.31143391 | 0.67733577 | 1.41311433 | 5.63E-06 |
| 0.0478229 | 0.22218971 | -0.0546341 | 0.88800983 | 0.95710625 | 0.07010178 | 0.75463732 |
| 0.04776262 | 0.22218971 | 1.03996509 | 0.01485655 | 0.27146966 | 0.54416073 | 0.02473908 |
| 0.07668837 | 0.26539136 | 0.0598636 | 0.88922201 | 0.95710625 | 0.44060231 | 0.09169328 |
| 0.08585771 | 0.27971896 | 0.89589292 | 0.05709853 | 0.39354925 | -0.1676991 | 0.513076 |
| 0.05022423 | 0.2231788 | 0.36755182 | 0.34109994 | 0.69611043 | -0.094247 | 0.66838957 |
| 0.15026664 | 0.36936801 | -0.0878684 | 0.86905836 | 0.95318114 | 0.25900193 | 0.4056112 |
| 0.046565 | 0.22218971 | 0.3255949 | 0.38607128 | 0.71114006 | 0.1012204 | 0.63750448 |
| 0.17601069 | 0.38876695 | 0.0385645 | 0.94564069 | 0.97721252 | 0.12466437 | 0.70335516 |
| 0.05514596 | 0.23446999 | -0.423341 | 0.27950847 | 0.65094491 | 0.66662854 | 0.00819215 |
| 0.01390228 | 0.14982855 | -0.3080245 | 0.28938746 | 0.66583113 | -0.1370222 | 0.41055376 |
| 0.04735646 | 0.22218971 | 0.07088544 | 0.84680992 | 0.94578917 | 0.14032698 | 0.51071561 |
| 0.08437723 | 0.27847077 | -0.4550401 | 0.29482075 | 0.66583113 | 0.87860666 | 0.0026002 |
| 0.06152768 | 0.24250524 | 0.23870529 | 0.54219734 | 0.80235959 | -0.1268349 | 0.5744601 |
| 0.06729198 | 0.24885909 | -0.0945406 | 0.81245559 | 0.94081146 | -0.4025967 | 0.095827 |
| 0.04229843 | 0.2217997 | -0.6765671 | 0.0700789 | 0.41761531 | 0.47640454 | 0.03126403 |
| 0.08555534 | 0.27971896 | 0.47596369 | 0.2701904 | 0.64783091 | 0.42945077 | 0.09417952 |
| 0.04236383 | 0.2217997 | 0.359085 | 0.31217854 | 0.67733577 | 0.05496417 | 0.78518697 |
| 0.03396195 | 0.20272937 | 1.28096659 | 0.00141239 | 0.15677717 | 0.72710849 | 0.00161224 |
| 0.19202523 | 0.4006729 | 1.79525782 | 0.00570873 | 0.24466371 | -0.5047628 | 0.13580564 |
| 0.20352904 | 0.41195658 | 2.88696086 | 1.54E-04 | 0.04121815 | 0.03970224 | 0.90525943 |
| 0.03901221 | 0.21532576 | -0.0071152 | 0.9832186 | 0.9931002 | -0.4528818 | 0.03295012 |

| | | | | | | |
|---|---|---|---|---|---|---|
| 0.03118438 | 0.19846802 | 0.18383176 | 0.56802462 | 0.81844408 | 0.28256225 | 0.14152003 |
| 0.01152438 | 0.14982855 | -0.1175314 | 0.65678931 | 0.86143329 | -0.275428 | 0.08626757 |
| 0.17008619 | 0.3846853 | 0.09255375 | 0.86095564 | 0.95286706 | 1.17357358 | 0.00148439 |
| 0.02683723 | 0.18935579 | 0.34358205 | 0.2759227 | 0.65094491 | -0.160155 | 0.37557392 |
| 0.0212041 | 0.1660206 | 0.44883262 | 0.1407372 | 0.54092474 | 0.06279105 | 0.71110813 |
| 0.07564191 | 0.26453425 | -0.767318 | 0.06800271 | 0.41108407 | 4.42E-04 | 0.99845565 |
| 0.02143506 | 0.1660206 | 0.14447592 | 0.62350604 | 0.8480478 | -0.0716242 | 0.67303514 |
| 0.09967807 | 0.30097498 | 0.05675812 | 0.89381093 | 0.95816532 | -0.3680167 | 0.14876944 |
| 0.14621353 | 0.36157019 | -0.6885808 | 0.16965488 | 0.56174159 | 0.68260397 | 0.02596548 |
| 0.04587187 | 0.22218971 | 0.4341778 | 0.21482108 | 0.59970885 | -0.4997536 | 0.0211708 |
| 0.11992427 | 0.32823428 | 0.1248943 | 0.7815473 | 0.92594338 | -0.2594959 | 0.32605746 |
| 0.10095444 | 0.30249486 | 0.79449903 | 0.07638847 | 0.42948485 | -0.020476 | 0.93324811 |
| 0.14098635 | 0.35182307 | 0.69560781 | 0.1585878 | 0.55397439 | 0.07895579 | 0.77417137 |
| 0.01280128 | 0.14982855 | 0.2735465 | 0.30291856 | 0.67301027 | 0.19689496 | 0.20357592 |
| 0.04297504 | 0.2217997 | -0.2455817 | 0.46466319 | 0.75509356 | 0.2554337 | 0.1968803 |
| 0.0929468 | 0.29577256 | -0.1874591 | 0.64897857 | 0.85908592 | 0.52060252 | 0.04170007 |
| 0.04971311 | 0.2231788 | 0.65569291 | 0.07279563 | 0.42411369 | -0.2423118 | 0.23511829 |
| 0.05321749 | 0.22995556 | 0.65823675 | 0.07401788 | 0.42507413 | 0.17466278 | 0.39041037 |
| 0.11806954 | 0.32807533 | 0.70034161 | 0.12618947 | 0.51763437 | -0.2079613 | 0.41733943 |
| 0.02221749 | 0.16889405 | 0.11476333 | 0.68561564 | 0.87082935 | 0.15286543 | 0.3562986 |
| 0.07581498 | 0.26453425 | 1.5415768 | 9.41E-04 | 0.12611263 | -0.6918073 | 0.00597206 |
| 0.06161375 | 0.24250524 | 0.24033039 | 0.50415409 | 0.77910145 | -0.2823165 | 0.18450593 |
| 0.05846903 | 0.24001535 | 0.47592599 | 0.18977052 | 0.58231375 | -0.4048131 | 0.06155773 |
| 0.04566914 | 0.22218971 | 0.39402716 | 0.24294736 | 0.62405647 | -0.0773004 | 0.68513847 |
| 0.01915982 | 0.16558122 | -0.0465498 | 0.86457155 | 0.95286706 | -0.141756 | 0.37486117 |
| 0.12659129 | 0.33526275 | -0.0888018 | 0.84145636 | 0.94578917 | -0.1814862 | 0.48247538 |
| 0.06061938 | 0.24250524 | 0.45862732 | 0.2052324 | 0.59482251 | -0.1971428 | 0.33951901 |
| 0.05565755 | 0.23446999 | 0.68160397 | 0.0626237 | 0.39959887 | 0.34475017 | 0.09824051 |
| 0.48793983 | 0.67807688 | -1.452156 | 0.16218361 | 0.55487874 | -0.9054663 | 0.13328482 |
| 0.03738516 | 0.21167737 | -0.0077588 | 0.98009106 | 0.9917316 | -0.3752912 | 0.05117959 |
| 0.02904715 | 0.19358394 | -0.2140137 | 0.47065762 | 0.75759091 | 0.53580973 | 0.006088 |
| 0.32041462 | 0.52627686 | 1.01945801 | 0.15549437 | 0.55397439 | -0.2674935 | 0.50650045 |
| 0.13138937 | 0.34375786 | -0.3068409 | 0.49389949 | 0.77557654 | 0.33163265 | 0.20940929 |
| 0.1660116 | 0.38393104 | -0.3288984 | 0.50209758 | 0.77910145 | 0.26859316 | 0.34623369 |
| 0.07009334 | 0.25350939 | 0.72807355 | 0.05958838 | 0.39354925 | -0.4842644 | 0.03314633 |
| 0.02390132 | 0.1752052 | 0.68261488 | 0.02590828 | 0.33063899 | 0.15580562 | 0.34235968 |
| 0.17297047 | 0.38800208 | 0.0533562 | 0.9136673 | 0.96361813 | 0.18952328 | 0.50803293 |
| 0.17897513 | 0.39172402 | 0.20258021 | 0.68556012 | 0.87082935 | -0.5173802 | 0.08843229 |
| 0.34401457 | 0.55379601 | 1.42260237 | 0.06363966 | 0.40136977 | -0.6298493 | 0.14501749 |
| 0.16564221 | 0.38393104 | 0.39497562 | 0.41656283 | 0.73125877 | 0.40037865 | 0.16347755 |
| 0.10497305 | 0.31057514 | 1.03023686 | 0.02088294 | 0.31298919 | -0.663333 | 0.01155718 |
| 0.43191993 | 0.62985126 | -0.0209656 | 0.98063013 | 0.9917316 | -0.1435011 | 0.77379984 |

| | | | | | | |
|---|---|---|---|---|---|---|
| 0.08445703 | 0.27847077 | 1.00451865 | 0.01633929 | 0.27645753 | -0.0445662 | 0.8374359 |
| 0.15213641 | 0.37173808 | -0.0830981 | 0.85667768 | 0.95286706 | 0.01721884 | 0.94829169 |
| 0.04594956 | 0.22218971 | 0.50725325 | 0.12609163 | 0.51763437 | -0.3727944 | 0.05721853 |
| 0.15491515 | 0.37483905 | 1.38537246 | 0.00468773 | 0.24466371 | -0.2259423 | 0.34954816 |
| 0.11646502 | 0.32708413 | 0.09405046 | 0.81875276 | 0.94229238 | -0.5979705 | 0.02189686 |
| 0.12655736 | 0.33526275 | 0.76420515 | 0.08600339 | 0.45193939 | -0.1856698 | 0.45047774 |
| 0.08267566 | 0.2757875 | 0.48190549 | 0.20177714 | 0.59207598 | -0.4757243 | 0.03783474 |
| 0.09565287 | 0.29788726 | 0.7870602 | 0.05463149 | 0.39354925 | 0.33471105 | 0.14501868 |
| 0.04036417 | 0.21704662 | -0.0518639 | 0.86411528 | 0.95286706 | -0.0480151 | 0.78393337 |
| 0.29263563 | 0.50025656 | -1.3103642 | 0.05014593 | 0.39354925 | 0.45863331 | 0.21532861 |
| 0.13748076 | 0.34623221 | 0.39534369 | 0.36589753 | 0.7058246 | 0.33094982 | 0.19673212 |
| 0.20727048 | 0.41606128 | 0.13842505 | 0.76125519 | 0.92154871 | 0.0086119 | 0.97659363 |
| 0.19576079 | 0.40516661 | -0.789755 | 0.1268618 | 0.5177507 | -0.2744318 | 0.34552096 |
| 0.1584235 | 0.37483905 | -0.1959008 | 0.6659165 | 0.86915076 | 0.16271732 | 0.53594233 |
| 0.16966701 | 0.3846853 | 0.5949094 | 0.21365901 | 0.59970885 | -0.728213 | 0.01509478 |
| 0.11949851 | 0.32823428 | 0.34572823 | 0.39906128 | 0.71938402 | -0.2184278 | 0.35728856 |
| 0.11158839 | 0.3193209 | 1.04317714 | 0.01741539 | 0.27645753 | -0.4515497 | 0.06311821 |
| 0.08880939 | 0.28819174 | -0.4274074 | 0.25462634 | 0.63999508 | 0.32383539 | 0.14116923 |
| 0.2427812 | 0.45300765 | 1.05202629 | 0.0706414 | 0.41761531 | -0.5394319 | 0.10429641 |
| 0.12445992 | 0.3318528 | 0.68720951 | 0.10654874 | 0.4883708 | 0.03135664 | 0.89364018 |
| 0.14453749 | 0.35938533 | 0.04040373 | 0.92484265 | 0.96884622 | 0.00669676 | 0.97839125 |
| 0.24044074 | 0.45172048 | -0.8168527 | 0.14516541 | 0.54686998 | 0.32794357 | 0.30171923 |
| 0.26843105 | 0.47496098 | 1.70326687 | 0.00896513 | 0.24466371 | -0.594467 | 0.08875224 |
| 0.06213432 | 0.24250524 | -0.0231136 | 0.94336554 | 0.97721252 | 0.50102507 | 0.01649069 |
| 0.04001227 | 0.21704662 | -0.0046954 | 0.98708969 | 0.99575923 | 0.39999366 | 0.02879631 |
| 0.17416679 | 0.38876695 | -0.5116779 | 0.27315258 | 0.64783091 | -0.7398327 | 0.01250412 |
| 0.10679003 | 0.31090289 | 0.4007849 | 0.29688341 | 0.66674376 | 0.23405576 | 0.29157213 |
| 0.40839119 | 0.60844391 | 1.46040403 | 0.07021925 | 0.41761531 | -1.1068335 | 0.02200554 |
| 0.19761913 | 0.40662984 | 1.19957774 | 0.02282327 | 0.31637777 | -0.5199352 | 0.07583031 |
| 0.06189903 | 0.24250524 | -0.5689287 | 0.09076728 | 0.46187909 | 0.18717982 | 0.31858539 |
| 0.13436308 | 0.34623221 | 0.72293919 | 0.09067134 | 0.46187909 | 0.12375658 | 0.59896192 |
| 0.21812102 | 0.42535638 | 0.17122523 | 0.73054602 | 0.9045453 | 0.17303012 | 0.54837842 |
| 0.20673029 | 0.41599403 | 0.81576179 | 0.10709046 | 0.4883708 | -0.1610592 | 0.56576193 |
| 0.04817268 | 0.22218971 | 0.41152043 | 0.17670778 | 0.56670496 | 0.01384185 | 0.93516946 |
| 0.44137664 | 0.64136322 | -0.1206766 | 0.88026488 | 0.95510521 | -0.354442 | 0.44801849 |
| 0.10986143 | 0.31759237 | 0.35349116 | 0.34912049 | 0.69650836 | -0.2817608 | 0.20226173 |
| 0.17134175 | 0.38540157 | 0.67045509 | 0.14161594 | 0.54092474 | 0.33577161 | 0.1981662 |
| 0.09001528 | 0.28981389 | 0.30769739 | 0.38037729 | 0.71114006 | -0.2169924 | 0.28713254 |
| 0.04423973 | 0.22218971 | -0.1804741 | 0.5289479 | 0.79423262 | 0.35453855 | 0.04503808 |
| 0.06241308 | 0.24250524 | 0.35885849 | 0.25715792 | 0.64209618 | -0.3972828 | 0.03964825 |
| 0.07938462 | 0.26820895 | 0.17497113 | 0.59821745 | 0.83776197 | -0.1897219 | 0.32825607 |
| 0.09197823 | 0.29497707 | 0.36006967 | 0.30637035 | 0.67301027 | -0.2992041 | 0.14860315 |

| | | | | | | |
|---|---|---|---|---|---|---|
| 0.24158081 | 0.45268389 | -0.0877189 | 0.862411 | 0.95286706 | 1.61155159 | 5.82E-05 |
| 0.51680802 | 0.69786084 | 0.42846607 | 0.64513347 | 0.85681478 | -0.7788583 | 0.16020788 |
| 0.15681197 | 0.37483905 | -0.2311502 | 0.5767736 | 0.82660601 | 0.33595435 | 0.17198323 |
| 0.07397808 | 0.26292643 | -0.1484543 | 0.64284679 | 0.85570997 | 0.35453054 | 0.06979055 |
| 0.1108617 | 0.3193209 | -0.7039828 | 0.06630468 | 0.40385579 | -0.3626179 | 0.09742455 |
| 0.09945858 | 0.30097498 | -0.2871498 | 0.41402389 | 0.73080926 | -0.7382872 | 0.00208636 |
| 0.06554721 | 0.2468544 | -0.4454872 | 0.15962037 | 0.55397439 | 0.30334292 | 0.10156643 |
| 0.25472662 | 0.46267823 | 0.76655529 | 0.15026483 | 0.55181975 | 0.46911663 | 0.12894088 |
| 0.05206913 | 0.22860296 | 0.14680587 | 0.6059653 | 0.83973408 | -0.3380274 | 0.05352594 |
| 0.1322841 | 0.34375786 | -0.4824721 | 0.21171207 | 0.59970885 | -0.0826375 | 0.70386123 |
| 0.30185621 | 0.5091733 | -0.6802954 | 0.23537522 | 0.61471277 | -0.3335444 | 0.31026818 |
| 0.0769344 | 0.26539136 | 0.25118148 | 0.42742775 | 0.73587133 | 0.32504986 | 0.08799477 |
| 0.17615262 | 0.38876695 | -0.3693489 | 0.38370806 | 0.71114006 | 0.7777873 | 0.00534977 |
| 0.3529476 | 0.56362207 | NA | NA | NA | 0.1575898 | 0.6596284 |
| 0.23080691 | 0.43863999 | 1.80204585 | 0.00155997 | 0.15677717 | -0.7882645 | 0.01035105 |
| 0.1313394 | 0.34375786 | -0.0232372 | 0.94969454 | 0.97766249 | -0.1095662 | 0.60805099 |
| 0.06885083 | 0.2523506 | 0.82901868 | 0.01345692 | 0.27094123 | -0.4122667 | 0.02897766 |
| 0.52333825 | 0.70272385 | -0.9109878 | 0.31981006 | 0.68160393 | -0.7601571 | 0.15802211 |
| 0.15967891 | 0.37563434 | 0.48087368 | 0.23527694 | 0.61471277 | 0.32954431 | 0.16300866 |
| 0.17133969 | 0.38540157 | 0.66647728 | 0.11679448 | 0.51041692 | 0.39543843 | 0.10782925 |
| 0.10493488 | 0.31057514 | -0.1638217 | 0.62865235 | 0.8480478 | 0.20556149 | 0.3005309 |
| 0.11787517 | 0.32807533 | -0.3259381 | 0.35996454 | 0.70022229 | 0.30197902 | 0.15084415 |
| 0.29360587 | 0.50025656 | -0.0848226 | 0.87374938 | 0.95318114 | 0.83938753 | 0.01469283 |
| 0.13718101 | 0.34623221 | 0.41254385 | 0.27244125 | 0.64783091 | -0.5598504 | 0.01755614 |
| 0.22626449 | 0.43200732 | 1.63152563 | 0.00253766 | 0.18548002 | -0.3019945 | 0.26118179 |
| 0.13181975 | 0.34375786 | -0.0107107 | 0.9762465 | 0.99053243 | -0.2538246 | 0.23356496 |
| 0.09578835 | 0.29788726 | 0.32275189 | 0.32316243 | 0.68204616 | -0.277596 | 0.14917738 |
| 0.16706 | 0.38411987 | -0.4448595 | 0.26660578 | 0.64783091 | 0.43542525 | 0.06975908 |
| 0.10716946 | 0.31090504 | -0.2132543 | 0.51800074 | 0.7902406 | 0.01327246 | 0.94403159 |
| 0.52006898 | 0.70024461 | -0.7595304 | 0.38004419 | 0.71114006 | 1.08458883 | 0.04139545 |
| 0.13472055 | 0.34623221 | 0.05857304 | 0.86815813 | 0.95318114 | -0.4964232 | 0.02606912 |
| 0.15858148 | 0.37483905 | 1.28661425 | 0.00346707 | 0.2144247 | -0.2825417 | 0.20470894 |
| 0.30583318 | 0.5132799 | 0.51783763 | 0.33210107 | 0.68994641 | -0.7395474 | 0.02595712 |
| 0.10569638 | 0.31057514 | -0.1948055 | 0.54673083 | 0.80655337 | 0.704314 | 0.00164378 |
| 0.18992896 | 0.39880225 | 0.86209149 | 0.04663418 | 0.38419153 | -0.3651382 | 0.13202753 |
| 0.13688656 | 0.34623221 | 0.51117696 | 0.16054233 | 0.55397439 | 0.0172375 | 0.93231839 |
| 0.18676716 | 0.39519545 | 0.61456851 | 0.13751514 | 0.53765116 | 0.36089663 | 0.13140496 |
| 0.11215988 | 0.3193209 | -0.7243601 | 0.03836554 | 0.367213 | 0.39773285 | 0.04739043 |
| 0.13721708 | 0.34623221 | 0.5914658 | 0.1042848 | 0.48235751 | -0.5791993 | 0.01049469 |
| 0.27124582 | 0.47890928 | -0.2950507 | 0.53906863 | 0.80112971 | -0.1618578 | 0.55922637 |
| 0.2246996 | 0.43200732 | 0.04646802 | 0.91397551 | 0.96361813 | -0.305701 | 0.23031146 |
| 0.46147828 | 0.65299351 | -0.6607074 | 0.36584356 | 0.7058246 | 0.92197996 | 0.06452993 |

| | | | | | | |
|---|---|---|---|---|---|---|
| 0.44304521 | 0.64265038 | -1.0528456 | 0.1419591 | 0.54092474 | -0.813473 | 0.05594268 |
| 0.34207935 | 0.55176257 | -0.2287812 | 0.67927749 | 0.87082935 | 0.95037486 | 0.00868256 |
| 0.18063673 | 0.39172402 | 0.84816015 | 0.04086913 | 0.37310001 | -0.2279749 | 0.31284526 |
| 0.18770724 | 0.39616361 | 0.48996271 | 0.22213544 | 0.60713403 | -0.7971723 | 0.00281635 |
| 0.20064481 | 0.40940549 | 0.38843541 | 0.34028066 | 0.69611043 | -0.5071502 | 0.04214606 |
| 0.11472364 | 0.32478659 | 0.16770289 | 0.60043682 | 0.83776197 | -0.0576837 | 0.75425458 |
| 0.09872521 | 0.30097498 | -0.0934819 | 0.75727828 | 0.92110701 | 0.10656042 | 0.54371584 |
| 0.26707536 | 0.47460795 | -0.6085286 | 0.20013051 | 0.59156226 | -0.2536841 | 0.34824663 |
| 0.18005142 | 0.39172402 | 0.71548371 | 0.07396246 | 0.42507413 | 0.41005919 | 0.07587568 |
| 0.18666019 | 0.39519545 | 0.52200922 | 0.18653752 | 0.57905856 | 0.31496823 | 0.1686235 |
| 0.11837044 | 0.32807533 | -0.2553804 | 0.4254338 | 0.73587133 | 0.53626053 | 0.00964841 |
| 0.18303291 | 0.39172402 | 0.8799877 | 0.03130014 | 0.33613674 | 0.01552466 | 0.94287778 |
| 0.31057881 | 0.51721136 | -0.0207301 | 0.966809 | 0.98461483 | -0.1731441 | 0.54975394 |
| 0.388727 | 0.59653246 | -0.3614992 | 0.54096967 | 0.80235959 | -0.3134398 | 0.36251014 |
| 0.19598496 | 0.40516661 | 0.34963712 | 0.36931554 | 0.70697546 | -0.1083422 | 0.62639504 |
| 0.18965007 | 0.39880225 | 0.39509182 | 0.30484533 | 0.67301027 | -0.47196 | 0.04451253 |
| 0.16535462 | 0.38393104 | 1.10237893 | 0.00668262 | 0.24466371 | -0.4275627 | 0.05118008 |
| 0.12440732 | 0.3318528 | 0.22642839 | 0.47818659 | 0.76433801 | 0.01330275 | 0.94195068 |
| 0.41464287 | 0.61347994 | -0.0846472 | 0.88946293 | 0.95710625 | 0.08594036 | 0.8070648 |
| 0.21967248 | 0.42535638 | -0.6895161 | 0.09959474 | 0.47663196 | 0.36054399 | 0.1325987 |
| 0.13226154 | 0.34375786 | -0.1276382 | 0.69001596 | 0.87471896 | 0.10294302 | 0.57845156 |
| 0.25842284 | 0.46579227 | 0.08595232 | 0.84211957 | 0.94578917 | -0.6260439 | 0.02239286 |
| 0.21199408 | 0.42142164 | 0.60505083 | 0.13424767 | 0.5343323 | -0.2449925 | 0.28393876 |
| 0.13370315 | 0.34623221 | -0.4923642 | 0.13775639 | 0.53765116 | 0.23225008 | 0.21939517 |
| 0.21595808 | 0.42315414 | 0.03297388 | 0.93273215 | 0.97391772 | 0.28028096 | 0.22613813 |
| 0.20909897 | 0.41821975 | 0.77872643 | 0.05661754 | 0.39354925 | 0.35022615 | 0.12808731 |
| 0.1556508 | 0.37483905 | -0.0325399 | 0.92258022 | 0.9683479 | 0.0924232 | 0.6339039 |
| 0.16868699 | 0.38411987 | -0.3353221 | 0.34034925 | 0.69611043 | 0.45216177 | 0.03669493 |
| 0.19879192 | 0.40802042 | 0.60209286 | 0.11970826 | 0.51041692 | 0.30436295 | 0.16891984 |
| 0.28883843 | 0.49610116 | 0.91093585 | 0.05914185 | 0.39354925 | 0.27970401 | 0.29366172 |
| 0.55306155 | 0.73118121 | NA | NA | NA | 0.5195624 | 0.28251241 |
| 0.37833235 | 0.58716608 | 1.63906255 | 0.00804064 | 0.24466371 | -0.5319933 | 0.10534676 |
| 0.20096274 | 0.40940549 | 1.49580271 | 9.39E-04 | 0.12611263 | 0.42469739 | 0.06025575 |
| 0.20294018 | 0.41195658 | 0.04620263 | 0.89990824 | 0.96079213 | 0.30794852 | 0.16103441 |
| 0.21579462 | 0.42315414 | -0.6644271 | 0.09457409 | 0.46546266 | -0.3161015 | 0.16181709 |
| 0.16832999 | 0.38411987 | 0.17226608 | 0.6097211 | 0.84085037 | 0.37651561 | 0.06775002 |
| 0.40902889 | 0.60844391 | 0.05840814 | 0.91804784 | 0.96485028 | 0.49516022 | 0.14588499 |
| 0.31338515 | 0.5192185 | 0.22601391 | 0.62578543 | 0.8480478 | 0.18899313 | 0.48202124 |
| 0.17430286 | 0.38876695 | 0.09307031 | 0.78133326 | 0.92594338 | -0.5397398 | 0.01282348 |
| 0.21417231 | 0.42311413 | 0.35020033 | 0.34872498 | 0.69650836 | -0.7595708 | 0.00258489 |
| 0.12133229 | 0.32823428 | 0.3372438 | 0.25551409 | 0.63999508 | -0.1550128 | 0.36133147 |
| 0.2388737 | 0.44980575 | 0.08477135 | 0.82705323 | 0.94229238 | -0.5094375 | 0.03582063 |

| | | | | | | |
|---|---|---|---|---|---|---|
| 0.33075239 | 0.53878515 | -0.521742 | 0.27940876 | 0.65094491 | -0.0774547 | 0.77667772 |
| 0.2937113 | 0.50025656 | 1.5869013 | 0.00226275 | 0.18548002 | -0.4736464 | 0.07554455 |
| 0.37084436 | 0.57992995 | 0.79543226 | 0.1370456 | 0.53765116 | -0.7569637 | 0.02081666 |
| 0.20936458 | 0.41821975 | 0.47225038 | 0.20373288 | 0.59482251 | -0.1772717 | 0.40083379 |
| 0.21439606 | 0.42311413 | -0.6716277 | 0.08095496 | 0.43431059 | 0.33027904 | 0.13160019 |
| 0.15650431 | 0.37483905 | 0.7485633 | 0.02885466 | 0.33613674 | -0.0020406 | 0.9910094 |
| 0.13549047 | 0.34623221 | 0.22042616 | 0.4585496 | 0.75146233 | 0.0082095 | 0.96148618 |
| 0.15888428 | 0.37483905 | 0.25960317 | 0.41304361 | 0.73080926 | 0.02241443 | 0.90142374 |
| 0.15841233 | 0.37483905 | 0.37319624 | 0.24438261 | 0.62430627 | -0.0786119 | 0.66444646 |
| 0.25727805 | 0.46525392 | 0.015994 | 0.9674698 | 0.98461483 | 0.11517306 | 0.61262377 |
| 0.17789391 | 0.39155736 | -0.1792095 | 0.58393276 | 0.83241479 | 0.20935394 | 0.27552201 |
| 0.21490694 | 0.42311413 | 0.3051327 | 0.39538975 | 0.71772396 | -0.4562723 | 0.03917431 |
| 0.24958111 | 0.456361 | -0.0976103 | 0.79778023 | 0.92824212 | -0.0792576 | 0.71889677 |
| 0.26000374 | 0.46607656 | 0.31568177 | 0.42278772 | 0.73576045 | -0.6634511 | 0.00926897 |
| 0.30868226 | 0.51509784 | 0.71080459 | 0.11572692 | 0.51041692 | -0.2214058 | 0.38128931 |
| 0.53284779 | 0.71133013 | 0.07690145 | 0.91389568 | 0.96361813 | -1.155078 | 0.01227662 |
| 0.20372031 | 0.41195658 | -0.2038333 | 0.5526979 | 0.80794383 | 0.42959544 | 0.0430504 |
| 0.18110872 | 0.39172402 | 0.66073933 | 0.05540276 | 0.39354925 | -0.1959944 | 0.3017861 |
| 0.54886326 | 0.72797534 | 1.74139114 | 0.03010116 | 0.33613674 | -0.2060947 | 0.6292483 |
| 0.32178068 | 0.52730926 | 0.0534288 | 0.90270858 | 0.96079213 | 0.7640761 | 0.00792524 |
| 0.15203479 | 0.37173808 | 0.13909483 | 0.63939628 | 0.85252837 | 0.10217605 | 0.55204195 |
| 0.32618259 | 0.53345798 | 0.11337422 | 0.79691013 | 0.92824212 | -0.4904969 | 0.06912548 |
| 0.12193843 | 0.32823428 | -0.0800142 | 0.76846761 | 0.92170206 | -0.0574058 | 0.7147355 |
| 0.39583483 | 0.60070313 | 0.11353425 | 0.82149904 | 0.94229238 | -0.007072 | 0.98056271 |
| 0.21954711 | 0.42535638 | 0.47067237 | 0.1834063 | 0.57346985 | -0.087993 | 0.65758766 |
| 0.17534946 | 0.38876695 | -0.1484489 | 0.63062971 | 0.84840812 | 0.08062668 | 0.65087047 |
| 0.74599209 | 0.85658672 | -2.6891179 | 0.05627655 | 0.39354925 | 0.32112037 | 0.673957 |
| 0.19208458 | 0.4006729 | -0.0647347 | 0.83809899 | 0.94578917 | -0.0795688 | 0.6643919 |
| 0.18155104 | 0.39172402 | -0.0021923 | 0.99433017 | 0.99805425 | 0.2787706 | 0.13304175 |
| 0.25115244 | 0.45821367 | 0.17167186 | 0.63600496 | 0.85082859 | 0.20420705 | 0.33563258 |
| 0.34562817 | 0.55530475 | 0.32928049 | 0.45726751 | 0.75146233 | 0.75816445 | 0.00850403 |
| 0.24205631 | 0.45268389 | 0.98844083 | 0.01193537 | 0.26655653 | -0.721732 | 0.00258789 |
| 0.12113695 | 0.32823428 | -0.2971133 | 0.26036708 | 0.6460961 | 0.339069 | 0.03581029 |
| 0.71091043 | 0.84321225 | 0.32254659 | 0.77409049 | 0.92202779 | -1.5368694 | 0.02967457 |
| 0.18321805 | 0.39172402 | 0.88538141 | 0.00973458 | 0.24466371 | -0.2991948 | 0.10257667 |
| 0.24828901 | 0.4550131 | 0.09872468 | 0.77742552 | 0.92463035 | 0.14530434 | 0.47458366 |
| 0.41471543 | 0.61347994 | -0.128488 | 0.79719997 | 0.92824212 | -0.3225655 | 0.27362162 |
| 0.70917387 | 0.84259297 | -0.5808936 | 0.59965009 | 0.83776197 | -0.4237284 | 0.50845109 |
| 0.14968903 | 0.36905314 | 0.20614879 | 0.45515933 | 0.75143348 | 0.04871275 | 0.75791095 |
| 0.31368134 | 0.5192185 | 0.82863908 | 0.05186134 | 0.39354925 | 0.14764855 | 0.52321793 |
| 0.19290697 | 0.40095347 | -0.483439 | 0.12453376 | 0.51763437 | 0.32292143 | 0.07947935 |
| 0.24562684 | 0.45316772 | 0.32774184 | 0.34266559 | 0.69611043 | -0.3636677 | 0.07968443 |

| | | | | | | |
|---|---|---|---|---|---|---|
| 0.22104191 | 0.42700097 | 0.63930882 | 0.06041242 | 0.39354925 | -0.1104919 | 0.55133215 |
| 0.56443599 | 0.7402587 | NA | NA | NA | 0.394819 | 0.329932 |
| 0.45878596 | 0.6522349 | 0.9093449 | 0.10415879 | 0.48235751 | -0.2779078 | 0.37359002 |
| 0.27890691 | 0.48313201 | 0.94060934 | 0.01803199 | 0.27880225 | -0.5488863 | 0.01704155 |
| 0.29872023 | 0.50566867 | 0.84526196 | 0.03469557 | 0.3531043 | -0.4446205 | 0.05138111 |
| 0.34095246 | 0.5510275 | 0.10990675 | 0.78548165 | 0.92599303 | 0.14319022 | 0.54126172 |
| 0.3367767 | 0.54751221 | 0.24937365 | 0.53399119 | 0.79644543 | -0.246718 | 0.29282288 |
| 0.2960819 | 0.5022381 | 0.51082002 | 0.17486698 | 0.56670496 | -0.3957961 | 0.07555866 |
| 0.57156205 | 0.74421521 | -0.2097498 | 0.75847107 | 0.92116426 | 0.01785363 | 0.9637818 |
| 0.17536515 | 0.38876695 | 0.0588859 | 0.82860785 | 0.94229238 | 0.05437493 | 0.72949963 |
| 0.45043836 | 0.64992951 | -0.5651602 | 0.26693956 | 0.64783091 | 0.6227774 | 0.04427495 |
| 0.25876403 | 0.46579227 | 0.66880025 | 0.05249927 | 0.39354925 | 0.22470618 | 0.23844556 |
| 0.24528049 | 0.45316772 | 0.07726863 | 0.80482707 | 0.9350881 | 0.05061755 | 0.77925708 |
| 0.35355099 | 0.56362207 | -0.3231539 | 0.41593741 | 0.73125877 | 0.28853978 | 0.21559617 |
| 0.36611741 | 0.57582834 | 0.37644274 | 0.35764455 | 0.70018966 | -0.4393555 | 0.07479186 |
| 0.56083445 | 0.73770118 | 0.79966693 | 0.21402741 | 0.59970885 | 0.27720661 | 0.44768514 |
| 0.23314603 | 0.44104352 | 0.25867682 | 0.39136762 | 0.71513537 | 0.2487225 | 0.1626405 |
| 0.51577104 | 0.69786084 | 0.43526642 | 0.43695347 | 0.7387887 | 0.33413805 | 0.30524129 |
| 0.22592246 | 0.43200732 | 0.11804322 | 0.68508932 | 0.87082935 | -0.3225837 | 0.06984283 |
| 0.26319477 | 0.46901585 | -0.9197289 | 0.00973786 | 0.24466371 | 0.10338202 | 0.57091483 |
| 0.18584902 | 0.39519545 | 0.64354384 | 0.02458629 | 0.3240553 | -0.337237 | 0.03844577 |
| 0.27558565 | 0.48197752 | -0.5573649 | 0.09494512 | 0.46546266 | 0.21207609 | 0.25788281 |
| 0.27982829 | 0.4836611 | 0.70634321 | 0.03995872 | 0.37310001 | 0.02751116 | 0.88101185 |
| 0.18651115 | 0.39519545 | 0.13171051 | 0.60808219 | 0.84003107 | -0.1498524 | 0.31862117 |
| 0.2679952 | 0.47496098 | 0.16231322 | 0.59932056 | 0.83776197 | 0.17747805 | 0.32576304 |
| 0.37185294 | 0.58040164 | 0.04245309 | 0.91113762 | 0.96361813 | -0.1622675 | 0.46420708 |
| 0.32050966 | 0.52627686 | 0.49400451 | 0.15654408 | 0.55397439 | -0.6517011 | 0.00404613 |
| 0.47579003 | 0.66924474 | -0.8772581 | 0.07750191 | 0.4297347 | 0.06465384 | 0.81164585 |
| 0.43057731 | 0.62901063 | -0.0010374 | 0.99802016 | 0.99841214 | -0.4649763 | 0.06967933 |
| 0.40923648 | 0.60844391 | 0.47612009 | 0.24286919 | 0.62405647 | -0.2771791 | 0.23915056 |
| 0.20034649 | 0.40940549 | 0.47374721 | 0.07510626 | 0.42524954 | -0.2008018 | 0.18020243 |
| 0.27818699 | 0.48313201 | 0.33600042 | 0.27155685 | 0.64783091 | -0.1947273 | 0.26981579 |
| 0.40542762 | 0.60844391 | -0.3064552 | 0.43885585 | 0.7387887 | 0.03587099 | 0.87400331 |
| 0.32804934 | 0.53544435 | 0.72723946 | 0.042034 | 0.37550369 | -0.2108269 | 0.28079786 |
| 0.30634245 | 0.5132799 | -0.2836956 | 0.37370077 | 0.71038272 | 0.34929208 | 0.0699582 |
| 0.52468906 | 0.70272385 | NA | NA | NA | 0.44281402 | 0.14652406 |
| 0.45718599 | 0.65164878 | 0.01243418 | 0.97698037 | 0.99053243 | 0.22251238 | 0.3778101 |
| 0.22552937 | 0.43200732 | -0.2743815 | 0.29478604 | 0.66583113 | 0.67770477 | 3.51E-04 |
| 0.40634326 | 0.60844391 | 0.33547821 | 0.38315277 | 0.71114006 | -0.21049 | 0.34442029 |
| 0.27620006 | 0.48197752 | -0.0394499 | 0.89044213 | 0.95710625 | 0.3224195 | 0.06662405 |
| 0.27215246 | 0.47947891 | 0.19248694 | 0.4985723 | 0.7789967 | 0.1485144 | 0.3688227 |
| 0.48811624 | 0.67807688 | 1.10218143 | 0.02509646 | 0.32544444 | -0.689384 | 0.0166721 |

| | | | | | | |
|---|---|---|---|---|---|---|
| 0.37017971 | 0.57992995 | 1.02716712 | 0.00877731 | 0.24466371 | 0.30070889 | 0.14607559 |
| 0.38995908 | 0.59730673 | 0.57330421 | 0.12524728 | 0.51763437 | -0.7994552 | 0.00143826 |
| 0.35706908 | 0.56593381 | 0.28301052 | 0.40155611 | 0.722262 | -0.4007646 | 0.05186775 |
| 0.43023312 | 0.62901063 | -0.266692 | 0.49561387 | 0.77675156 | 0.27711619 | 0.22835111 |
| 0.40183754 | 0.60729853 | -0.0513054 | 0.88776355 | 0.95710625 | 0.14438318 | 0.49495864 |
| 0.39246928 | 0.59891687 | 0.25244457 | 0.48154867 | 0.76524896 | -0.1183014 | 0.56664944 |
| 0.40516321 | 0.60844391 | -0.4464245 | 0.2283185 | 0.61067965 | 0.00831563 | 0.96817289 |
| 0.26335725 | 0.46901585 | 0.49829517 | 0.07486289 | 0.42524954 | -0.3512268 | 0.03389522 |
| 0.35841927 | 0.56697922 | 0.56284299 | 0.09742437 | 0.47136383 | -0.491017 | 0.01767285 |
| 0.48205391 | 0.67606246 | -0.0299351 | 0.94322417 | 0.97721252 | 0.53042774 | 0.04265349 |
| 0.243668 | 0.45316772 | -0.0542088 | 0.82771709 | 0.94229238 | 0.02378542 | 0.86858684 |
| 0.3465762 | 0.55574036 | 0.12687927 | 0.68355179 | 0.87082935 | 0.46665154 | 0.01871468 |
| 0.46507043 | 0.65605295 | -0.6505302 | 0.11997005 | 0.51041692 | 0.35933524 | 0.13548345 |
| 0.43511935 | 0.63339182 | 0.01266869 | 0.9728913 | 0.98888066 | 0.02759056 | 0.89802926 |
| 0.57249532 | 0.74421521 | 0.27077188 | 0.6021965 | 0.83776197 | 0.11797772 | 0.69344019 |
| 0.35449363 | 0.56402959 | -0.2002905 | 0.51896398 | 0.7902406 | -0.472922 | 0.01682729 |
| 0.4915533 | 0.6805485 | 0.70391265 | 0.10334488 | 0.48235751 | -0.1227484 | 0.60733793 |
| 0.51637161 | 0.69786084 | -0.5421593 | 0.22352198 | 0.60713403 | -0.1665673 | 0.5091733 |
| 0.45167778 | 0.65057448 | -0.4017516 | 0.28310561 | 0.65595652 | -0.192444 | 0.37014605 |
| 0.36607222 | 0.57582834 | 0.39697714 | 0.20164077 | 0.59207598 | -0.3859277 | 0.0403676 |
| 0.49861671 | 0.68685289 | 0.40482817 | 0.32045558 | 0.68160393 | 0.02546089 | 0.91229184 |
| 0.54311766 | 0.72268979 | 0.95114701 | 0.04616898 | 0.38419153 | -0.2216946 | 0.39281617 |
| 0.40801241 | 0.60844391 | 0.3108096 | 0.34325519 | 0.69611043 | 0.15413557 | 0.41366021 |
| 0.59238659 | 0.76032926 | 0.18311567 | 0.71555562 | 0.89384306 | 0.2710375 | 0.35642096 |
| 0.4002073 | 0.60621807 | 0.19846719 | 0.53190611 | 0.7963734 | -0.3314774 | 0.08455208 |
| 0.4078494 | 0.60844391 | -0.1039067 | 0.74166796 | 0.91068668 | 0.13532048 | 0.4606529 |
| 0.51164414 | 0.69546331 | 0.19122537 | 0.6311893 | 0.84840812 | 0.1844482 | 0.42591661 |
| 0.66609874 | 0.8162195 | 0.73540983 | 0.23548698 | 0.61471277 | 0.7244658 | 0.05259075 |
| 0.45918608 | 0.6522349 | 0.08127039 | 0.81560396 | 0.94081146 | -0.00947 | 0.96242513 |
| 0.64394709 | 0.80001162 | 1.57773614 | 0.01139849 | 0.26183949 | -0.5305163 | 0.11280189 |
| 0.59723258 | 0.76373519 | 2.26745353 | 2.67E-04 | 0.05371205 | -0.2647444 | 0.34838535 |
| 0.44515239 | 0.6445681 | 0.09509166 | 0.77267063 | 0.92170206 | -0.0175618 | 0.92627508 |
| 0.3705591 | 0.57992995 | -0.1878227 | 0.4989842 | 0.7789967 | 0.23379381 | 0.15645649 |
| 0.4541707 | 0.65101331 | 0.67609396 | 0.05317124 | 0.39354925 | -0.3232152 | 0.10242057 |
| 0.42726473 | 0.62752118 | 0.1256178 | 0.68285794 | 0.87082935 | 0.07554488 | 0.67049541 |
| 0.35588228 | 0.56514381 | 0.21060927 | 0.42458899 | 0.73587133 | -0.1753854 | 0.2550958 |
| 0.48352872 | 0.67606246 | -0.466825 | 0.18917521 | 0.58231375 | -0.092295 | 0.64399724 |
| 0.34009566 | 0.5507269 | 0.63322923 | 0.02180091 | 0.31637777 | 0.15528184 | 0.29256767 |
| 0.4164741 | 0.61497344 | 0.19373553 | 0.50583452 | 0.77910145 | 0.15176487 | 0.36979029 |
| 0.57508573 | 0.74421521 | -0.2665674 | 0.52815266 | 0.79423262 | -0.0156239 | 0.94857588 |
| 0.64909557 | 0.80016136 | -0.7257139 | 0.1649453 | 0.55487874 | -0.1301743 | 0.6561816 |
| 0.39085637 | 0.59756626 | 0.29853681 | 0.26986793 | 0.64783091 | -0.2321457 | 0.14391971 |

| | | | | | | |
|---|---|---|---|---|---|---|
| 0.50439042 | 0.69145008 | -0.0255683 | 0.93948528 | 0.97689915 | -0.3886969 | 0.06083768 |
| 0.50905635 | 0.69424462 | 0.26509341 | 0.43999493 | 0.7387887 | 0.00845878 | 0.96559269 |
| 0.71958093 | 0.84321225 | 0.88460277 | 0.1310774 | 0.52430959 | -0.3331266 | 0.31268449 |
| 0.65085394 | 0.80112607 | 0.29696419 | 0.5497179 | 0.80794383 | -0.3676239 | 0.20961258 |
| 0.82487591 | 0.89727729 | 0.90473002 | 0.37627017 | 0.71114006 | -0.7377525 | 0.21756621 |
| 0.49824619 | 0.68685289 | 0.16083974 | 0.61737601 | 0.8456053 | -0.1453732 | 0.43709913 |
| 0.42009836 | 0.61810172 | 0.12050335 | 0.65393428 | 0.86143329 | 0.16017795 | 0.30928296 |
| 0.41968318 | 0.61810172 | 0.32842008 | 0.22652178 | 0.61067965 | -0.2982078 | 0.06632615 |
| 0.87331816 | 0.92158639 | -0.0739107 | 0.95488653 | 0.98049651 | -1.8738466 | 0.02378172 |
| 0.45548332 | 0.65101331 | 0.36170227 | 0.1993312 | 0.59137373 | -0.2063638 | 0.20438569 |
| 0.5413038 | 0.72144549 | -0.6138698 | 0.08306647 | 0.44228769 | 0.37866112 | 0.06594857 |
| 0.75239136 | 0.85912838 | -0.1117221 | 0.86345042 | 0.95286706 | -0.1788341 | 0.63468155 |
| 0.53223743 | 0.71133013 | -0.5492033 | 0.1043908 | 0.48235751 | -0.0054509 | 0.97663397 |
| 0.7174148 | 0.84321225 | 0.69487073 | 0.22334105 | 0.60713403 | -0.5941329 | 0.07986115 |
| 0.52407302 | 0.70272385 | 0.54274279 | 0.09680364 | 0.47136383 | -0.4054989 | 0.03699043 |
| 0.57159813 | 0.74421521 | 0.6357336 | 0.08384329 | 0.44348685 | -0.466722 | 0.03278329 |
| 0.49562712 | 0.68503344 | 0.20427685 | 0.4720809 | 0.75759091 | 0.02579926 | 0.87390456 |
| 0.56559854 | 0.74060032 | 0.74001842 | 0.03704902 | 0.36326108 | -0.3040867 | 0.12360727 |
| 0.56158738 | 0.73770118 | 0.29885958 | 0.35501544 | 0.70018966 | -0.442241 | 0.02769464 |
| 0.72557719 | 0.84376611 | -0.1609029 | 0.75493995 | 0.92104965 | 0.23023102 | 0.44321899 |
| 0.6381219 | 0.79758002 | 0.06221621 | 0.87099882 | 0.95318114 | -0.4474508 | 0.05808792 |
| 0.74871569 | 0.85731602 | 1.13052077 | 0.05948999 | 0.39354925 | 0.13559248 | 0.67705267 |
| 0.62562973 | 0.78779449 | 0.46509386 | 0.21583746 | 0.60046131 | -0.3455412 | 0.11734918 |
| 0.50019756 | 0.68787638 | 0.22162094 | 0.39658369 | 0.71813803 | -0.3058141 | 0.05502258 |
| 0.6038427 | 0.76980568 | 0.46056338 | 0.17688828 | 0.56670496 | -0.1539733 | 0.42429745 |
| 0.6049915 | 0.77007445 | 0.05132736 | 0.87352152 | 0.95318114 | 0.32743236 | 0.09447085 |
| 0.5927049 | 0.76032926 | -0.5810682 | 0.07741201 | 0.4297347 | 0.2633339 | 0.15706674 |
| 0.50646995 | 0.69186661 | 0.11471376 | 0.64580816 | 0.85681478 | -0.0574761 | 0.68973237 |
| 0.84950481 | 0.90694857 | 0.32695539 | 0.70895294 | 0.8878476 | 0.0200809 | 0.96824817 |
| 0.69094437 | 0.83479344 | -0.3108764 | 0.45182243 | 0.7503289 | -0.3514778 | 0.15159278 |
| 0.78347608 | 0.87634041 | 1.71880159 | 0.00828239 | 0.24466371 | -0.4680454 | 0.17043719 |
| 0.7018316 | 0.83686678 | 0.06243907 | 0.87737438 | 0.95454533 | 0.68467052 | 0.00965578 |
| 0.55806748 | 0.73636337 | 0.28991123 | 0.27989033 | 0.65094491 | 0.04106598 | 0.78684918 |
| 0.68469203 | 0.83032815 | -0.5690417 | 0.14555992 | 0.54686998 | -0.1840664 | 0.40273309 |
| 0.55877513 | 0.73636337 | -0.138454 | 0.59412258 | 0.83776197 | -0.2356137 | 0.12996161 |
| 0.61573981 | 0.7813329 | 0.1772362 | 0.54913292 | 0.80794383 | -0.0821499 | 0.62978447 |
| 0.64702523 | 0.80001162 | 0.41613063 | 0.2090537 | 0.59814652 | -0.0804343 | 0.66625547 |
| 0.67976627 | 0.8280239 | 0.87985722 | 0.02413272 | 0.32337839 | -0.5974321 | 0.01003016 |
| 0.6214685 | 0.78495745 | 0.27273359 | 0.36056222 | 0.70022229 | -0.0211372 | 0.90087262 |
| 0.66475569 | 0.81579136 | 0.5418799 | 0.11998607 | 0.51041692 | -0.2144357 | 0.27572763 |
| 0.72291162 | 0.84321225 | 0.01833128 | 0.96380975 | 0.98461483 | -0.1307155 | 0.57732274 |
| 0.72086067 | 0.84321225 | -0.3963418 | 0.32575339 | 0.68204616 | 0.19037796 | 0.41114189 |

| | | | | | | |
|---|---|---|---|---|---|---|
| 0.72398957 | 0.84321225 | -0.1208241 | 0.7633759 | 0.92154871 | -0.3442152 | 0.15172262 |
| 0.62189278 | 0.78495745 | 0.48551212 | 0.09469894 | 0.46546266 | -0.1436318 | 0.37439479 |
| 0.78336738 | 0.87634041 | 0.50119633 | 0.31445809 | 0.67963522 | -0.7254646 | 0.02011618 |
| 0.74157731 | 0.85314094 | -0.2816555 | 0.4893524 | 0.77144966 | 0.33111974 | 0.16999217 |
| 0.6676608 | 0.81691434 | -0.0677118 | 0.82711049 | 0.94229238 | -0.028224 | 0.87465561 |
| 0.70057509 | 0.83686678 | -0.1925347 | 0.57843717 | 0.8275151 | 0.23542107 | 0.24801695 |
| 0.69758429 | 0.83686678 | 0.20563772 | 0.54289007 | 0.80235959 | 0.00930086 | 0.96173079 |
| 0.67393987 | 0.82214655 | -0.195613 | 0.5244939 | 0.79414895 | 0.51058686 | 0.01033612 |
| 0.75864256 | 0.86182872 | -0.2304675 | 0.58347348 | 0.83241479 | 0.16104888 | 0.50796436 |
| 0.78758906 | 0.87735498 | 1.63693501 | 0.00335674 | 0.2144247 | -0.3717923 | 0.18855762 |
| 0.74758687 | 0.85721902 | -0.1254794 | 0.74822735 | 0.91563895 | -0.7644132 | 0.0037609 |
| 0.73165036 | 0.84724377 | 0.16213258 | 0.65667526 | 0.86143329 | -0.0627761 | 0.76518233 |
| 0.64650222 | 0.80001162 | -0.170585 | 0.52804743 | 0.79423262 | 0.03396631 | 0.82665776 |
| 0.75056182 | 0.85823294 | -0.3187517 | 0.41398194 | 0.73080926 | -0.0654743 | 0.76899124 |
| 0.80352773 | 0.88668853 | 0.66664639 | 0.14396487 | 0.54597997 | 0.08742783 | 0.75560701 |
| 0.69108889 | 0.83479344 | -0.6322646 | 0.04503228 | 0.38419153 | -0.0211195 | 0.90068181 |
| 0.71500009 | 0.84321225 | 0.5184422 | 0.1177237 | 0.51041692 | -0.3604686 | 0.06428603 |
| 0.69711879 | 0.83686678 | -0.0519988 | 0.86026032 | 0.95286706 | 0.24579301 | 0.16383337 |
| 0.75895513 | 0.86182872 | 1.11608108 | 0.0080951 | 0.24466371 | -0.4636023 | 0.04384946 |
| 0.77650834 | 0.87564014 | 0.80198675 | 0.05748331 | 0.39354925 | -0.4291288 | 0.07580425 |
| 0.76617322 | 0.86762512 | 0.42319006 | 0.26140229 | 0.64611119 | 0.25048673 | 0.25010201 |
| 0.80991093 | 0.88895304 | 1.1314657 | 0.02361483 | 0.32180214 | -0.3968316 | 0.14583879 |
| 0.73926199 | 0.85314094 | 0.41733496 | 0.2145009 | 0.59970885 | -0.0651612 | 0.73051308 |
| 0.78158549 | 0.87634041 | -0.6422952 | 0.11414396 | 0.51041692 | 0.43711435 | 0.06685075 |
| 0.82623828 | 0.89727729 | -0.0832788 | 0.86516536 | 0.95286706 | 0.43395428 | 0.14047471 |
| 0.7150199 | 0.84321225 | -0.7868637 | 0.01392346 | 0.27094123 | 0.27872729 | 0.10734224 |
| 0.70333505 | 0.83686678 | -0.3795742 | 0.17674739 | 0.56670496 | -0.0867914 | 0.58233467 |
| 0.79384336 | 0.88192883 | -0.4214084 | 0.29111408 | 0.66583113 | -0.3408491 | 0.14641449 |
| 0.75423526 | 0.86003771 | -0.7865894 | 0.02723334 | 0.33604738 | -0.4787371 | 0.02100296 |
| 0.92877966 | 0.95316012 | 0.22441403 | 0.84326753 | 0.94578917 | -0.8283015 | 0.21844149 |
| 0.80164744 | 0.88580424 | 0.88204234 | 0.03936343 | 0.37233171 | 0.29521976 | 0.20946456 |
| 0.82608269 | 0.89727729 | -0.2899223 | 0.52169052 | 0.79289069 | -0.632529 | 0.02600737 |
| 0.70300637 | 0.83686678 | 0.22457855 | 0.38836362 | 0.71126277 | -0.0803153 | 0.5900912 |
| 0.76361612 | 0.8659238 | -0.9542263 | 0.00967201 | 0.24466371 | 0.33897682 | 0.08730826 |
| 0.78550795 | 0.87735498 | 0.91384247 | 0.02102166 | 0.31298919 | -0.3594907 | 0.09875475 |
| 0.84476997 | 0.90542578 | 1.01558323 | 0.05569362 | 0.39354925 | -0.4480845 | 0.13337274 |
| 0.73166362 | 0.84724377 | -0.282598 | 0.31974908 | 0.68160393 | 0.31131818 | 0.06887388 |
| 0.78293085 | 0.87634041 | 0.58354998 | 0.09403803 | 0.46546266 | -0.3661441 | 0.07114804 |
| 0.8474273 | 0.90590861 | -0.3129034 | 0.50438848 | 0.77910145 | -0.5253577 | 0.06582353 |
| 0.72026124 | 0.84321225 | 0.34673399 | 0.17350592 | 0.56670496 | -0.0542476 | 0.70363382 |
| 0.82268693 | 0.89727729 | -0.3309419 | 0.39783089 | 0.7187776 | 0.23895481 | 0.29371893 |
| 0.7869089 | 0.87735498 | 0.13238145 | 0.6779802 | 0.87082935 | 0.12568107 | 0.49698502 |

| | | | | | | |
|---|---|---|---|---|---|---|
| 0.79808042 | 0.88424295 | 0.75590976 | 0.03754519 | 0.36369072 | -0.0517371 | 0.78976346 |
| 0.87976972 | 0.92482835 | 0.7457769 | 0.19793504 | 0.58940656 | -0.0156139 | 0.96163069 |
| 0.81875785 | 0.89675319 | 0.32128625 | 0.3845529 | 0.71114006 | -0.1532332 | 0.47068088 |
| 0.74195205 | 0.85314094 | 0.33695872 | 0.19471472 | 0.58414417 | -0.1649324 | 0.26764719 |
| 0.77524432 | 0.87548223 | 0.92546082 | 0.00520199 | 0.24466371 | -0.3546996 | 0.04594253 |
| 0.80928692 | 0.88895304 | 0.65553297 | 0.05312022 | 0.39354925 | -0.3223508 | 0.09374382 |
| 0.8051766 | 0.88731542 | 0.34186442 | 0.27174237 | 0.64783091 | 0.07815053 | 0.65762023 |
| 0.89933345 | 0.93747091 | 1.61693836 | 0.01449064 | 0.27094123 | -0.1562254 | 0.64880066 |
| 0.83899641 | 0.90432235 | 0.07174215 | 0.84308233 | 0.94578917 | 0.05611706 | 0.78867434 |
| 0.86203923 | 0.91320543 | 0.95407713 | 0.03157829 | 0.33613674 | -0.4908379 | 0.05167052 |
| 0.83532303 | 0.90236869 | 0.17731303 | 0.58767178 | 0.83478465 | 0.33559367 | 0.08995503 |
| 0.84263897 | 0.90432235 | -0.4984085 | 0.1499552 | 0.55181975 | 0.15012005 | 0.44037838 |
| 0.82797194 | 0.89797221 | 0.40371152 | 0.19342251 | 0.58414417 | -0.2871101 | 0.11443038 |
| 0.88126076 | 0.9252111 | -0.5549015 | 0.19432189 | 0.58414417 | -0.1598856 | 0.50762248 |
| 0.85190162 | 0.90760167 | -0.6571027 | 0.05724613 | 0.39354925 | 0.59800976 | 0.00555755 |
| 0.93717265 | 0.96057272 | -0.4580427 | 0.53492603 | 0.79644543 | -0.1832561 | 0.66608298 |
| 0.84195451 | 0.90432235 | 0.02247107 | 0.93629603 | 0.97637096 | 0.16347793 | 0.3223642 |
| 0.85719554 | 0.91042373 | 0.46334252 | 0.11153551 | 0.50378959 | -0.1754307 | 0.28443193 |
| 0.890191 | 0.9300694 | -0.0346467 | 0.9144784 | 0.96361813 | -0.2522265 | 0.18925529 |
| 0.853433 | 0.90760167 | 0.28296408 | 0.24459761 | 0.62430627 | -0.1891284 | 0.18140278 |
| 0.90846442 | 0.9417289 | 0.09868538 | 0.79489358 | 0.92824212 | -0.0448989 | 0.83760981 |
| 0.90498255 | 0.94049452 | 0.16964388 | 0.59746459 | 0.83776197 | -0.0054261 | 0.97654526 |
| 0.91330428 | 0.9432101 | 0.20186098 | 0.56534335 | 0.81844408 | 0.27410688 | 0.18746822 |
| 0.91384386 | 0.9432101 | -0.1631475 | 0.62175094 | 0.8480478 | 0.31770743 | 0.110552 |
| 0.90093124 | 0.93747091 | 0.06794249 | 0.80887353 | 0.93843336 | 0.30105848 | 0.07899124 |
| 0.91448872 | 0.9432101 | 0.10430212 | 0.74304784 | 0.91068668 | 0.05990602 | 0.74433101 |
| 0.94402289 | 0.96518405 | -0.0281761 | 0.94804199 | 0.97721252 | -0.2665485 | 0.29513046 |
| 0.94724124 | 0.96556831 | 0.46148459 | 0.31599905 | 0.68113467 | -0.19127 | 0.46778964 |
| 0.94274829 | 0.96508273 | 0.58661672 | 0.1169378 | 0.51041692 | 0.11170266 | 0.59064267 |
| 0.96090399 | 0.9751572 | -0.3642755 | 0.35531646 | 0.70018966 | 0.40183394 | 0.08889432 |
| 0.97607359 | 0.98689214 | 0.10461701 | 0.84399623 | 0.94578917 | 0.55566622 | 0.08601645 |
| 0.97155097 | 0.98353064 | 0.69622738 | 0.08888443 | 0.46105216 | -0.0081226 | 0.97109049 |
| 0.97949489 | 0.98913321 | 0.08341146 | 0.79252996 | 0.92824212 | -0.0084291 | 0.96323605 |
| 0.98890628 | 0.99618657 | 0.61592521 | 0.16421568 | 0.55487874 | -0.5719337 | 0.03306411 |
| 0.9909022 | 0.9969739 | -0.5349256 | 0.10150315 | 0.48235751 | -0.2647552 | 0.15529155 |
| 0.99978678 | 0.99978678 | -0.0733577 | 0.78404001 | 0.92599303 | 0.0312792 | 0.83950636 |
| 0.99899154 | 0.99978678 | -0.8452233 | 0.06069665 | 0.39354925 | 0.25815038 | 0.29933075 |
| 0.99719526 | 0.99963042 | 0.19236733 | 0.55176677 | 0.80794383 | -0.3708174 | 0.06078482 |
| 0.99399572 | 0.99846423 | 0.40114484 | 0.23931867 | 0.62269324 | -0.6211658 | 0.00518795 |
| 0.99481576 | 0.99846423 | -1.1627507 | 0.01539213 | 0.27500607 | 0.40935346 | 0.11530896 |
| 0.98308456 | 0.9915386 | 0.4485122 | 0.13722401 | 0.53765116 | -0.0631479 | 0.7066882 |
| 0.97137443 | 0.98353064 | 0.20857526 | 0.59757161 | 0.83776197 | -0.2019537 | 0.38021363 |

| | | | | | | |
|---|---|---|---|---|---|---|
| 0.95706406 | 0.97246237 | 0.07179159 | 0.83457496 | 0.94578917 | -0.1709676 | 0.39481552 |
| 0.95614128 | 0.97246237 | 0.77115423 | 0.04773204 | 0.38419153 | -0.2859724 | 0.18560542 |
| 0.94792699 | 0.96556831 | 0.88142429 | 0.04752797 | 0.38419153 | -0.110892 | 0.64388881 |
| 0.94560422 | 0.96556831 | -0.2501976 | 0.56607458 | 0.81844408 | -0.3609378 | 0.16376579 |
| 0.92270222 | 0.9492964 | 0.07625085 | 0.81453037 | 0.94081146 | 0.28283674 | 0.14650531 |
| 0.92404853 | 0.94949167 | 0.2579778 | 0.4532262 | 0.7503289 | -0.5639142 | 0.01097902 |
| 0.91945774 | 0.9471453 | -0.2023038 | 0.55797677 | 0.81136791 | 0.18150297 | 0.36670092 |
| 0.90840164 | 0.9417289 | -0.1581951 | 0.60585934 | 0.83973408 | 0.19379891 | 0.28134218 |
| 0.88819139 | 0.9300694 | -0.1007254 | 0.77163109 | 0.92170206 | -0.0803907 | 0.68865899 |
| 0.86182928 | 0.91320543 | -0.0368056 | 0.90173788 | 0.96079213 | -0.3484439 | 0.05794528 |
| 0.91064631 | 0.94280028 | -1.352047 | 0.01017547 | 0.24782959 | 0.20763576 | 0.44433678 |
| 0.89041967 | 0.9300694 | 0.40171638 | 0.33075561 | 0.68893138 | -0.2080654 | 0.38137959 |
| 0.85325254 | 0.90760167 | -0.544967 | 0.09225568 | 0.46358477 | 0.43125463 | 0.02648502 |
| 0.88241468 | 0.92523941 | -0.5874618 | 0.15804909 | 0.55397439 | -0.001316 | 0.99546901 |
| 0.86876141 | 0.91914062 | -0.170836 | 0.63857709 | 0.85252837 | 0.48908911 | 0.03172328 |
| 0.84197785 | 0.90432235 | 0.72600618 | 0.03581629 | 0.35995375 | -0.6134175 | 0.00427016 |
| 0.83181637 | 0.9009515 | 0.25313227 | 0.4174813 | 0.73127444 | 0.49878055 | 0.01261633 |
| 0.80626243 | 0.88732099 | -0.5450233 | 0.09142541 | 0.46230206 | -0.27138 | 0.14048091 |
| 0.87091197 | 0.92023002 | 0.23646995 | 0.61257429 | 0.84189697 | -0.1477863 | 0.58387088 |
| 0.8765262 | 0.92343323 | -0.2929982 | 0.55197099 | 0.80794383 | 0.36174068 | 0.21324864 |
| 0.87731781 | 0.92343323 | 1.0426523 | 0.0648244 | 0.40136977 | -0.0562726 | 0.85432682 |
| 0.83448689 | 0.90236869 | -0.1179965 | 0.76922749 | 0.92170206 | -0.3484889 | 0.14788197 |
| 0.7909408 | 0.87989485 | 0.45259796 | 0.17062164 | 0.56221227 | 0.12586293 | 0.4982642 |
| 0.90078499 | 0.93747091 | -0.4004655 | 0.57359981 | 0.82499865 | -0.3818547 | 0.35727788 |
| 0.82460563 | 0.89727729 | 1.03314135 | 0.02225056 | 0.31637777 | -0.5184149 | 0.04246191 |
| 0.82382661 | 0.89727729 | -0.3154869 | 0.45355702 | 0.7503289 | 0.49553066 | 0.05427839 |
| 0.84704727 | 0.90590861 | -0.7788888 | 0.12967208 | 0.52390128 | 0.33602103 | 0.24907879 |
| 0.78188446 | 0.87634041 | -0.1462852 | 0.67228506 | 0.87082935 | -0.0600974 | 0.76304896 |
| 0.77751725 | 0.87564014 | -0.2475921 | 0.5057034 | 0.77910145 | 0.32173333 | 0.14618435 |
| 0.72407385 | 0.84321225 | 0.26203167 | 0.38678276 | 0.71114006 | -0.0681963 | 0.69333452 |
| 0.81920207 | 0.89675319 | 0.88424466 | 0.0875711 | 0.45718939 | 0.58715058 | 0.05301218 |
| 0.71397102 | 0.84321225 | 0.49618152 | 0.12386331 | 0.51763437 | -0.3330244 | 0.0776768 |
| 0.74079567 | 0.85314094 | -0.0363905 | 0.91694594 | 0.96485028 | -0.1903939 | 0.35208622 |
| 0.7554748 | 0.86025633 | -0.9264272 | 0.02800395 | 0.33604738 | 0.41401111 | 0.07889011 |
| 0.77324341 | 0.8744254 | 0.12789644 | 0.76084177 | 0.92154871 | -0.2686239 | 0.2775837 |
| 0.72314094 | 0.84321225 | -0.248617 | 0.47349451 | 0.75834579 | -0.0137144 | 0.94489677 |
| 0.73076021 | 0.84724377 | 0.35209199 | 0.33569591 | 0.69382909 | -0.3628294 | 0.09660415 |
| 0.62693033 | 0.78822328 | 0.22584762 | 0.38350175 | 0.71114006 | -0.0279821 | 0.84972646 |
| 0.72223561 | 0.84321225 | 0.10635051 | 0.76428902 | 0.92154871 | 0.41298384 | 0.05849828 |
| 0.64857477 | 0.80016136 | 0.42380516 | 0.15099545 | 0.55181975 | -0.1280305 | 0.44027245 |
| 0.68336795 | 0.8299521 | 0.60830295 | 0.07892002 | 0.43164421 | -0.4980269 | 0.01756937 |
| 0.73431466 | 0.84911596 | -0.9443809 | 0.02987681 | 0.33613674 | 0.26577448 | 0.25913668 |

| | | | | | | |
|---|---|---|---|---|---|---|
| 0.65809024 | 0.80882049 | 0.06466121 | 0.83421203 | 0.94578917 | -0.1680167 | 0.35330294 |
| 0.69531176 | 0.83686678 | 0.53840414 | 0.13886402 | 0.53935592 | -0.4364694 | 0.04462011 |
| 0.63548165 | 0.79758002 | 0.29031474 | 0.32371803 | 0.68204616 | -0.2670576 | 0.12521233 |
| 0.5981507 | 0.76373519 | 0.26615707 | 0.34372617 | 0.69611043 | 0.22712064 | 0.16933295 |
| 0.8398023 | 0.90432235 | 0.55485037 | 0.45029429 | 0.7495582 | -0.2160509 | 0.60866518 |
| 0.70043894 | 0.83686678 | 0.54537823 | 0.16978011 | 0.56174159 | -0.3271703 | 0.15477268 |
| 0.79777283 | 0.88424295 | -0.9032612 | 0.14165351 | 0.54092474 | 0.21693261 | 0.52786856 |
| 0.80057644 | 0.88580424 | -1.9317183 | 0.00809616 | 0.24466371 | 0.01273628 | 0.97250487 |
| 0.64587079 | 0.80001162 | -0.1474295 | 0.67602851 | 0.87082935 | 0.42823443 | 0.04942655 |
| 0.57387808 | 0.74421521 | -0.1270071 | 0.66854342 | 0.87082935 | 0.08898039 | 0.60394108 |
| 0.64485748 | 0.80001162 | 0.49124554 | 0.19699137 | 0.58877719 | -0.1033688 | 0.62939313 |
| 0.57572858 | 0.74421521 | 0.34479505 | 0.27766118 | 0.65094491 | -0.2601284 | 0.16193904 |
| 0.64652277 | 0.80001162 | -0.2414618 | 0.52899219 | 0.79423262 | 0.07811687 | 0.72297533 |
| 0.69734064 | 0.83686678 | 0.02024299 | 0.96523607 | 0.98461483 | -0.2407328 | 0.37605819 |
| 0.5516063 | 0.7304335 | 0.53620705 | 0.09790766 | 0.47136383 | -0.2791007 | 0.13226389 |
| 0.57651751 | 0.74421521 | 0.23650144 | 0.48696351 | 0.76919187 | -0.1407217 | 0.47397067 |
| 0.58582565 | 0.75386028 | 0.0230181 | 0.94759536 | 0.97721252 | 0.51681362 | 0.02060905 |
| 0.51407319 | 0.69761007 | 0.68731508 | 0.0321922 | 0.33613674 | -0.0354436 | 0.83521068 |
| 0.63814078 | 0.79758002 | NA | NA | NA | 0.61825246 | 0.03702425 |
| 0.69142453 | 0.83479344 | 1.55244203 | 0.00679176 | 0.24466371 | -0.4816462 | 0.11119034 |
| 0.46210625 | 0.65299351 | 5.51E-04 | 0.99841214 | 0.99841214 | -0.2173909 | 0.18726311 |
| 0.6710851 | 0.81988224 | 0.16963034 | 0.74292735 | 0.91068668 | -0.0181173 | 0.95152926 |
| 0.50532284 | 0.69145008 | -0.4836477 | 0.15794337 | 0.55397439 | 0.4138063 | 0.04401586 |
| 0.64142851 | 0.80001162 | -0.1718057 | 0.72256204 | 0.89789781 | 0.41510351 | 0.15141395 |
| 0.56846507 | 0.74316851 | -0.1323474 | 0.74056112 | 0.91068668 | 0.04221189 | 0.85466753 |
| 0.62052001 | 0.78495745 | 0.47845817 | 0.3125517 | 0.67733577 | 0.24598675 | 0.36671828 |
| 0.49111831 | 0.6805485 | 0.68000477 | 0.06481945 | 0.40136977 | -0.5048014 | 0.02184877 |
| 0.48419577 | 0.67606246 | 0.35434127 | 0.30836124 | 0.67529376 | 0.24326901 | 0.2292239 |
| 0.3937783 | 0.5997996 | 0.57759233 | 0.05660788 | 0.39354925 | -0.3674245 | 0.03805684 |
| 0.46209575 | 0.65299351 | -0.648824 | 0.06539731 | 0.40136977 | -0.243599 | 0.21499395 |
| 0.45594721 | 0.65101331 | 0.94440682 | 0.01048036 | 0.24782959 | -0.5796207 | 0.00722826 |
| 0.68337103 | 0.8299521 | NA | NA | NA | -0.3071427 | 0.41735192 |
| 0.35942109 | 0.56747061 | 0.43168923 | 0.14746395 | 0.55144658 | -0.0769673 | 0.6434281 |
| 0.59637706 | 0.76373519 | -0.0044924 | 0.99309746 | 0.99805425 | -0.5091688 | 0.10541442 |
| 0.3948975 | 0.60039046 | 0.30834084 | 0.34858295 | 0.69650836 | 0.22771127 | 0.23504706 |
| 0.54771642 | 0.72762974 | 1.22083869 | 0.01753649 | 0.27645753 | 0.28049605 | 0.30326033 |
| 0.40982885 | 0.60844391 | 0.41860376 | 0.2301442 | 0.61067965 | -0.4835558 | 0.02474764 |
| 0.58403658 | 0.75273789 | -0.0059078 | 0.99129623 | 0.99805425 | -0.1943393 | 0.53703671 |
| 0.31288041 | 0.5192185 | -0.048618 | 0.86790107 | 0.95318114 | -0.1921819 | 0.26524674 |
| 0.42833235 | 0.62796582 | 0.82787225 | 0.04277437 | 0.37791861 | -0.4416318 | 0.05868377 |
| 0.37582125 | 0.58437358 | -0.5815207 | 0.10751447 | 0.4883708 | 0.07695248 | 0.69974039 |
| 0.7029929 | 0.83686678 | -1.3461487 | 0.11480767 | 0.51041692 | -0.1009696 | 0.8304126 |

| | | | | | | |
|---|---|---|---|---|---|---|
| 0.31955268 | 0.52627686 | 0.08567788 | 0.7847153 | 0.92599303 | -0.4309433 | 0.02905186 |
| 0.29430441 | 0.50025656 | 0.47421713 | 0.12788395 | 0.51928633 | -0.1063311 | 0.54015048 |
| 0.38541499 | 0.59478516 | 0.28670421 | 0.43614142 | 0.7387887 | -0.5252898 | 0.02312981 |
| 0.3030823 | 0.5098987 | -0.4560622 | 0.1506827 | 0.55181975 | 0.03194723 | 0.85643059 |
| 0.47605229 | 0.66924474 | 0.5693809 | 0.22852438 | 0.61067965 | 0.15101311 | 0.5723293 |
| 0.45537697 | 0.65101331 | -0.2975174 | 0.5093503 | 0.7800336 | 6.21E-04 | 0.99808189 |
| 0.37924078 | 0.58726281 | -0.2763953 | 0.47917353 | 0.76439587 | 0.23292443 | 0.30642728 |
| 0.4024 | 0.60729853 | -0.2989302 | 0.46712319 | 0.75719162 | -0.0072872 | 0.97527245 |
| 0.33898207 | 0.55000846 | 0.05580683 | 0.87916836 | 0.95510521 | 0.37496721 | 0.09274749 |
| 0.62242058 | 0.78495745 | -1.738212 | 0.02779262 | 0.33604738 | 0.31037314 | 0.46127169 |
| 0.2832544 | 0.48855433 | 0.13956035 | 0.67083554 | 0.87082935 | 0.14977493 | 0.43312651 |
| 0.25927779 | 0.46579227 | -0.322791 | 0.30908968 | 0.67529376 | 0.44222289 | 0.02500609 |
| 0.72250401 | 0.84321225 | -0.8925759 | 0.38693111 | 0.71114006 | -0.6515739 | 0.27772167 |
| 0.23189981 | 0.43969919 | 0.60467361 | 0.05944046 | 0.39354925 | -0.4528906 | 0.01852399 |
| 0.244297 | 0.45316772 | 0.25616515 | 0.43322927 | 0.7387887 | 0.22298519 | 0.24373431 |
| 0.57546999 | 0.74421521 | 0.68812983 | 0.32552385 | 0.68204616 | 1.02975124 | 0.01914428 |
| 0.26196734 | 0.46857338 | 0.86303045 | 0.02274749 | 0.31637777 | -0.2320811 | 0.25410271 |
| 0.27494423 | 0.48197752 | -0.4694112 | 0.20626256 | 0.59482251 | 0.19469353 | 0.3573377 |
| 0.15300456 | 0.37274997 | -0.1337135 | 0.63310069 | 0.84840812 | 4.44E-04 | 0.99779852 |
| 0.20488115 | 0.41328606 | -0.4729204 | 0.15676805 | 0.55397439 | 0.11519143 | 0.53833904 |
| 0.24703014 | 0.45473486 | 0.37173163 | 0.30444316 | 0.67301027 | -0.3530098 | 0.10113766 |
| 0.21790867 | 0.42535638 | 0.92795768 | 0.01305456 | 0.27094123 | -0.4646765 | 0.02733605 |
| 0.37495467 | 0.58413242 | -0.7494987 | 0.12613883 | 0.51763437 | -0.176734 | 0.51767477 |
| 0.13231119 | 0.34375786 | 0.57245268 | 0.05293101 | 0.39354925 | -0.4570316 | 0.01102485 |
| 0.30746603 | 0.51411326 | -0.3999229 | 0.35673616 | 0.70018966 | 0.37457306 | 0.14415908 |
| 0.63825831 | 0.79758002 | -2.1359968 | 0.03679543 | 0.36326108 | -0.7383732 | 0.1892212 |
| 0.37982528 | 0.58726281 | 1.48823964 | 0.00918112 | 0.24466371 | -0.89828 | 0.00697963 |
| 0.48750665 | 0.67807688 | -0.569009 | 0.3856809 | 0.71114006 | 1.15886911 | 0.0068252 |
| 0.60617063 | 0.77038095 | 0.87654207 | 0.33841513 | 0.69611043 | -0.0923605 | 0.85909842 |
| 0.25427798 | 0.46267823 | -0.140758 | 0.72663551 | 0.90156628 | 0.30547459 | 0.20103269 |
| 0.09876869 | 0.30097498 | -0.2563264 | 0.36657116 | 0.7058246 | 0.14397698 | 0.37934462 |
| 0.34932112 | 0.55904998 | NA | NA | NA | 0.39528557 | 0.11883216 |
| 0.21253987 | 0.42148606 | -0.8817068 | 0.03360154 | 0.3463543 | 0.59870088 | 0.01496937 |
| 0.29020199 | 0.49740258 | -0.5044811 | 0.27712207 | 0.65094491 | -0.6159683 | 0.03112907 |
| 0.25538719 | 0.46285405 | -0.9340217 | 0.04082689 | 0.37310001 | -0.1680229 | 0.49496729 |
| 0.19228395 | 0.4006729 | 0.46544628 | 0.21912106 | 0.60147024 | 0.05450149 | 0.79807908 |
| 0.2789337 | 0.48313201 | -0.4246218 | 0.3529441 | 0.70018966 | -0.4712777 | 0.08569793 |
| 0.50503664 | 0.69145008 | 0.44197821 | 0.55822092 | 0.81136791 | -0.0594557 | 0.8908617 |
| 0.22165144 | 0.42717331 | -0.3362784 | 0.40898712 | 0.7302279 | 0.32757633 | 0.17270179 |
| 0.08938458 | 0.28891629 | -0.0167942 | 0.9531198 | 0.97993391 | -0.2507562 | 0.14358811 |
| 0.23566381 | 0.44478158 | -0.1150916 | 0.78815631 | 0.92778576 | 0.58376133 | 0.03018146 |
| 0.48379848 | 0.67606246 | NA | NA | NA | 0.25708549 | 0.51193105 |

| | | | | | | |
|---|---|---|---|---|---|---|
| 0.38622298 | 0.59491382 | 1.68538414 | 0.01410189 | 0.27094123 | -0.2148311 | 0.54709311 |
| 0.08158309 | 0.27338661 | 0.56668494 | 0.07131601 | 0.41852607 | -0.3153844 | 0.08095672 |
| 0.17944121 | 0.39172402 | 0.16924107 | 0.67156365 | 0.87082935 | 0.07460457 | 0.74580992 |
| 0.1395571 | 0.35038649 | 0.71771605 | 0.06220999 | 0.39959887 | 0.20959296 | 0.32332212 |
| 0.50993196 | 0.69428547 | NA | NA | NA | -0.043485 | 0.92844342 |
| 0.14489226 | 0.35938533 | -0.4769483 | 0.21021359 | 0.59933238 | 0.03984289 | 0.85212954 |
| 0.16563223 | 0.38393104 | -0.2872191 | 0.47128335 | 0.75759091 | 0.27829628 | 0.23423904 |
| 0.15859576 | 0.37483905 | 0.57656149 | 0.15375381 | 0.55397439 | -0.4997779 | 0.03977716 |
| 0.09770483 | 0.30097498 | -0.7990153 | 0.0265448 | 0.33346907 | 0.0541584 | 0.775847 |
| 0.06999206 | 0.25350939 | 0.09355801 | 0.75628197 | 0.92110701 | 0.10709119 | 0.54036763 |
| 0.099714 | 0.30097498 | -0.2581234 | 0.44612154 | 0.74570004 | 0.23241113 | 0.24149376 |
| 0.11190851 | 0.3193209 | -0.438334 | 0.23517038 | 0.61471277 | 0.10177801 | 0.6257856 |
| 0.45250687 | 0.65062722 | NA | NA | NA | 0.30917496 | 0.50132994 |
| 0.06601145 | 0.24746759 | -0.4628778 | 0.16448681 | 0.55487874 | 0.00270079 | 0.98839127 |
| 0.16750701 | 0.38411987 | 0.1716226 | 0.69760559 | 0.88187877 | -0.1947788 | 0.44845286 |
| 0.06237654 | 0.24250524 | -0.253432 | 0.43217197 | 0.7387887 | 0.09739588 | 0.59884392 |
| 0.18291729 | 0.39172402 | 0.22412192 | 0.62850716 | 0.8480478 | -0.1921349 | 0.47470782 |
| 0.18235857 | 0.39172402 | -0.626672 | 0.1912075 | 0.58231375 | 0.23182685 | 0.393913 |
| 0.03554037 | 0.20548339 | 0.22467736 | 0.43897178 | 0.7387887 | -0.0962355 | 0.56398742 |
| 0.24537148 | 0.45316772 | -0.6990118 | 0.21795641 | 0.60147024 | 0.35305772 | 0.27814507 |
| 0.10340981 | 0.3087253 | 0.16467131 | 0.67347601 | 0.87082935 | -0.7636418 | 0.00374296 |
| 0.04995644 | 0.2231788 | 0.20300906 | 0.52431303 | 0.79414895 | -0.1234537 | 0.50278386 |
| 0.09798141 | 0.30097498 | -0.437316 | 0.26678689 | 0.64783091 | 0.62800421 | 0.01201191 |
| 0.02274087 | 0.16972958 | -0.8065322 | 0.008365 | 0.24466371 | 0.28102208 | 0.08598004 |
| 0.45035812 | 0.64992951 | 0.33837823 | 0.70883784 | 0.8878476 | 0.24034429 | 0.64637128 |
| 0.3153428 | 0.52091838 | -0.5281332 | 0.44014899 | 0.7387887 | -0.6632453 | 0.10562128 |
| 0.11868255 | 0.32807533 | 0.32012914 | 0.46027923 | 0.75216362 | -0.0063673 | 0.97950641 |
| 0.27650598 | 0.48197752 | -0.1005612 | 0.87343921 | 0.95318114 | -0.3730176 | 0.31471186 |
| 0.10629917 | 0.31057514 | -0.628825 | 0.14821982 | 0.55170711 | 0.12876303 | 0.59614955 |
| 0.28754147 | 0.4949089 | 0.08862455 | 0.89277465 | 0.95816532 | 0.03978293 | 0.9165213 |
| 0.16823815 | 0.38411987 | -0.6261704 | 0.22504531 | 0.60921357 | -0.2059234 | 0.48179882 |
| 0.21916378 | 0.42535638 | 0.25720167 | 0.65904676 | 0.86181853 | -0.0739357 | 0.82566478 |
| 0.27545108 | 0.48197752 | -0.1741817 | 0.79769637 | 0.92824212 | -0.6454941 | 0.11534549 |
| 0.07833007 | 0.26795412 | -0.206815 | 0.61427479 | 0.8427934 | -0.8198576 | 0.00317238 |
| 0.22769309 | 0.43372628 | -0.8606176 | 0.1760852 | 0.56670496 | -0.6318593 | 0.09133354 |
| 0.0340537 | 0.20272937 | -0.7102978 | 0.04623134 | 0.38419153 | 0.10304814 | 0.59197785 |
| 0.11667819 | 0.32708413 | 0.33876073 | 0.47034549 | 0.75759091 | 0.43483181 | 0.12094917 |
| 0.07273636 | 0.25963717 | 0.15618082 | 0.70247674 | 0.88525282 | 0.28622528 | 0.23594044 |
| 0.52027919 | 0.70024461 | -0.3821744 | 0.75024703 | 0.91671522 | -0.7122341 | 0.31222213 |
| 0.09553788 | 0.29788726 | -0.6083209 | 0.19084307 | 0.58231375 | 0.55094901 | 0.04871875 |
| 0.02073929 | 0.1660206 | 0.33241005 | 0.29458463 | 0.66583113 | -0.2183037 | 0.23578769 |
| 0.04506431 | 0.22218971 | -0.628631 | 0.10429194 | 0.48235751 | -0.03964 | 0.85242088 |

| | | | | | | |
|---|---|---|---|---|---|---|
| 0.06413104 | 0.24263403 | -0.6678052 | 0.11851265 | 0.51041692 | 0.13647299 | 0.56584682 |
| 0.12062782 | 0.32823428 | -0.202482 | 0.6908539 | 0.87471896 | -0.453045 | 0.13756018 |
| 0.1357135 | 0.34623221 | 0.50638966 | 0.34871184 | 0.69650836 | 0.30263149 | 0.33257837 |
| 0.06368693 | 0.24263403 | -0.0816806 | 0.84697538 | 0.94578917 | 0.16141635 | 0.51189496 |
| 0.06400386 | 0.24263403 | -1.2320882 | 0.01260537 | 0.27094123 | 0.10334187 | 0.68499836 |
| 0.16890045 | 0.38411987 | -0.852762 | 0.17762394 | 0.56670496 | 0.2334258 | 0.5121638 |
| 0.0793734 | 0.26820895 | -0.1645487 | 0.73128662 | 0.9045453 | 0.62575732 | 0.03647567 |
| 0.05733152 | 0.2377231 | -0.5004601 | 0.27015585 | 0.64783091 | 0.91495919 | 0.00262283 |
| 0.24828973 | 0.4550131 | -0.7658522 | 0.32473184 | 0.68204616 | 0.48970926 | 0.27725518 |
| 0.07439538 | 0.26326986 | -0.1463242 | 0.76451865 | 0.92154871 | 0.30360138 | 0.29063112 |
| 0.02757034 | 0.18935579 | -0.2213999 | 0.56733798 | 0.81844408 | -0.3124268 | 0.17386265 |
| 0.06106453 | 0.24250524 | -0.9069556 | 0.07842154 | 0.43164421 | 0.14314565 | 0.61232735 |
| 0.04319622 | 0.2217997 | -0.5881069 | 0.20549152 | 0.59482251 | 0.3665481 | 0.17365258 |
| 0.04910007 | 0.2231788 | -1.0984748 | 0.03178007 | 0.33613674 | 0.10578145 | 0.6974202 |
| 0.03470056 | 0.20495797 | -0.6320432 | 0.16003961 | 0.55397439 | 0.37215403 | 0.15246081 |
| 0.02372373 | 0.1752052 | -0.3887865 | 0.34343377 | 0.69611043 | -0.11101 | 0.63531298 |
| 0.03651291 | 0.20963009 | -0.5771927 | 0.20641229 | 0.59482251 | 0.15775212 | 0.54085002 |
| 0.21141812 | 0.42129679 | -1.7185039 | 0.04644397 | 0.38419153 | -1.0771599 | 0.03255868 |
| 0.10059936 | 0.30249486 | NA | NA | NA | 0.47052968 | 0.18781122 |
| 0.01989124 | 0.1660206 | -0.3576042 | 0.3814704 | 0.71114006 | 0.14294091 | 0.54170147 |
| 0.0093125 | 0.14616752 | -0.3391608 | 0.34748343 | 0.69650836 | -0.2899538 | 0.17180829 |
| 0.00100611 | 0.07509234 | 0.15263907 | 0.55907689 | 0.81136791 | -0.3673248 | 0.02545346 |
| 0.09352181 | 0.29603753 | 1.09765468 | 0.08102809 | 0.43431059 | -0.0013838 | 0.99678662 |
| 0.03934088 | 0.21532576 | -0.3332324 | 0.4903719 | 0.77154404 | 0.32554302 | 0.25042292 |
| 0.03815781 | 0.21457232 | -0.5110051 | 0.29571057 | 0.66597003 | 0.18557629 | 0.50571302 |
| 0.01448355 | 0.14982855 | 0.49998043 | 0.23440197 | 0.61471277 | -0.1895268 | 0.42839298 |
| 0.06872096 | 0.2523506 | -0.5002311 | 0.39475463 | 0.71772396 | -0.2986287 | 0.37929924 |
| 0.04777679 | 0.22218971 | -0.1908796 | 0.72098549 | 0.8973256 | 0.37239305 | 0.23818058 |
| 0.02204939 | 0.16889405 | -0.6443893 | 0.16898513 | 0.56174159 | 0.34930996 | 0.19482272 |
| 0.15628647 | 0.37483905 | 0.72849796 | 0.35711307 | 0.70018966 | -1.0018204 | 0.0393859 |
| 0.00695684 | 0.13718908 | 0.01779988 | 0.961941 | 0.98461483 | -0.0676513 | 0.75383735 |
| 0.06141287 | 0.24250524 | 1.38249009 | 0.03100954 | 0.33613674 | -0.9880714 | 0.01011502 |
| 0.01285237 | 0.14982855 | -0.7625213 | 0.08947693 | 0.46115034 | -0.065445 | 0.79044357 |
| 0.08297139 | 0.2757875 | -0.4779711 | 0.47009362 | 0.75759091 | 0.37930624 | 0.32470609 |
| 0.01530157 | 0.15038994 | -0.2358316 | 0.5980583 | 0.83776197 | 0.01752287 | 0.94564114 |
| 0.02923801 | 0.19358394 | -0.1013087 | 0.84165498 | 0.94578917 | 0.2054256 | 0.48639648 |
| 0.02677421 | 0.18935579 | 0.11406679 | 0.81990591 | 0.94229238 | 0.23296316 | 0.42548495 |
| 0.00866349 | 0.14515769 | 0.04487061 | 0.91318782 | 0.96361813 | 0.21058867 | 0.38196889 |
| 0.01018462 | 0.14669425 | -0.0752317 | 0.86002144 | 0.95286706 | 0.42861001 | 0.09766885 |
| 0.01141553 | 0.14982855 | -0.4881766 | 0.28013301 | 0.65094491 | -0.2087817 | 0.41936411 |
| 0.00941265 | 0.14616752 | 0.32219545 | 0.41448883 | 0.73080926 | -0.1092862 | 0.62877528 |
| 0.03542734 | 0.20548339 | 0.14971413 | 0.77259849 | 0.92170206 | 0.76485659 | 0.02123392 |

| | | | | | | |
|---|---|---|---|---|---|---|
| 0.1581429 | 0.37483905 | -0.1288129 | 0.88731732 | 0.95710625 | -0.4394179 | 0.40807756 |
| 0.03319481 | 0.20187361 | NA | NA | NA | 0.22933401 | 0.49472813 |
| 0.02759904 | 0.18935579 | -0.4168048 | 0.45716058 | 0.75146233 | 0.29480032 | 0.364697 |
| 0.1217673 | 0.32823428 | 0.15234392 | 0.85574399 | 0.95286706 | -1.1466027 | 0.02973369 |
| 0.00946782 | 0.14616752 | -0.4069654 | 0.38741212 | 0.71114006 | -0.1234048 | 0.64668788 |
| 0.0036453 | 0.09975985 | 0.40888567 | 0.333861 | 0.69181507 | 0.76916547 | 0.00556007 |
| 0.01399585 | 0.14982855 | -1.0799944 | 0.05441483 | 0.39354925 | 0.55534761 | 0.08268881 |
| 0.00751782 | 0.14078773 | -0.8749631 | 0.0804401 | 0.43431059 | 0.55395603 | 0.05774388 |
| 0.00200677 | 0.08733222 | -0.3582877 | 0.36695856 | 0.7058246 | -0.1482066 | 0.51507416 |
| 0.04534491 | 0.22218971 | -0.1662566 | 0.78198452 | 0.92594338 | 0.09298652 | 0.78861927 |
| 0.00593054 | 0.13159379 | -0.3262096 | 0.4824819 | 0.76524896 | 0.40001763 | 0.14741597 |
| 0.05815142 | 0.23991116 | -1.9607047 | 0.0167656 | 0.27645753 | 0.47816452 | 0.27175419 |
| 0.01810345 | 0.15811634 | -0.7336225 | 0.2126968 | 0.59970885 | 0.52368487 | 0.1286199 |
| 0.00393931 | 0.10106781 | 0.21618407 | 0.63314039 | 0.84840812 | 0.04388049 | 0.86626137 |
| 0.04951625 | 0.2231788 | -0.9113892 | 0.21919251 | 0.60147024 | 0.37712902 | 0.37233449 |
| 0.01762485 | 0.1555914 | -0.9180771 | 0.13057539 | 0.52430959 | 0.40855622 | 0.23629331 |
| 0.03523438 | 0.20548339 | -0.9232136 | 0.18325604 | 0.57346985 | 0.72250887 | 0.07836019 |
| 0.02536049 | 0.18425631 | -0.9456526 | 0.15274757 | 0.55325116 | 0.35906681 | 0.33712743 |
| 0.01701407 | 0.1555914 | -0.6810494 | 0.26198041 | 0.64611119 | 0.52812239 | 0.13859084 |
| 0.02121518 | 0.1660206 | -0.8534244 | 0.18279997 | 0.57346985 | 0.63485772 | 0.0923115 |
| 0.0128972 | 0.14982855 | -0.4420551 | 0.44903762 | 0.74901711 | 0.44725872 | 0.19360593 |
| 0.01471575 | 0.14982855 | -0.3495544 | 0.55839402 | 0.81136791 | 0.42179404 | 0.23049094 |
| 0.03227385 | 0.20187361 | -0.3518521 | 0.62585191 | 0.8480478 | 0.50523062 | 0.23515446 |
| 0.00188116 | 0.08733222 | -0.4514249 | 0.32727132 | 0.68344451 | 0.17717424 | 0.5015301 |
| 0.02988547 | 0.19534319 | -0.8695128 | 0.24038347 | 0.62344616 | 0.47455055 | 0.265862 |
| 0.01747403 | 0.1555914 | -0.5755179 | 0.38278767 | 0.71114006 | 0.52561292 | 0.17575615 |
| 0.01327558 | 0.14982855 | -0.7696696 | 0.22871082 | 0.61067965 | 0.64881228 | 0.08719231 |
| 0.00269515 | 0.09647579 | -0.4115051 | 0.40756526 | 0.72980505 | 0.04527673 | 0.87310374 |
| 0.00362515 | 0.09975985 | 0.3662975 | 0.48351551 | 0.76524896 | 0.09794457 | 0.74385674 |
| 0.01627452 | 0.15536487 | -1.3316753 | 0.06009992 | 0.39354925 | 0.2255271 | 0.55872845 |
| 2.24E-04 | 0.05196388 | -0.7346913 | 0.06493596 | 0.40136977 | -0.0735548 | 0.73391384 |
| 0.02808558 | 0.19056413 | -0.813929 | 0.29114228 | 0.66583113 | 0.27111248 | 0.53713441 |
| 0.01607075 | 0.15522458 | -1.4238043 | 0.05199802 | 0.39354925 | -0.2057443 | 0.60398698 |
| 0.00961394 | 0.14616752 | -0.5734419 | 0.37374613 | 0.71038272 | 0.37459349 | 0.3161024 |
| 0.00334973 | 0.09975985 | -0.0425226 | 0.93776261 | 0.9766336 | 0.26873489 | 0.39871958 |
| 0.01160894 | 0.14982855 | -0.9460794 | 0.17697796 | 0.56670496 | 0.43412101 | 0.27760199 |
| 0.02024198 | 0.1660206 | -0.8076921 | 0.29412335 | 0.66583113 | 0.29316072 | 0.50450939 |
| 0.00645365 | 0.13585766 | -0.7045211 | 0.29001139 | 0.66583113 | 0.3976349 | 0.3005432 |
| 0.00123849 | 0.07821557 | -0.4722562 | 0.37081477 | 0.70815932 | -0.2878815 | 0.34547828 |
| 0.01011175 | 0.14669425 | -1.0399409 | 0.15738717 | 0.55397439 | 0.21030111 | 0.60926266 |
| 0.00416283 | 0.10135281 | 0.46458983 | 0.46488969 | 0.75509356 | -0.0712914 | 0.84462618 |
| 0.00361458 | 0.09975985 | -0.5912735 | 0.34732663 | 0.69650836 | 0.50534637 | 0.17178674 |

| | | | | | | |
|---|---|---|---|---|---|---|
| 0.01039265 | 0.14710979 | -0.5225466 | 0.48292833 | 0.76524896 | 0.55922449 | 0.20263626 |
| 0.01424381 | 0.14982855 | -0.9925823 | 0.21831833 | 0.60147024 | 0.39181529 | 0.39323091 |
| 0.00360054 | 0.09975985 | -0.7386409 | 0.25272995 | 0.6390225 | 0.32766956 | 0.37499537 |
| 0.01710793 | 0.1555914 | -0.95675 | 0.25274771 | 0.6390225 | 0.3625163 | 0.44690691 |
| 6.28E-04 | 0.06443541 | -0.5135966 | 0.32204606 | 0.68204616 | -0.0685228 | 0.81600031 |
| 0.00432077 | 0.10135281 | -0.6203731 | 0.36004746 | 0.70022229 | 0.42458331 | 0.28077022 |
| 0.01309169 | 0.14982855 | -1.1562941 | 0.16671703 | 0.55850205 | 0.25270691 | 0.589819 |
| 0.0404484 | 0.21704662 | -0.4795293 | 0.65052587 | 0.85908592 | -0.9118197 | 0.14927683 |
| 0.01317827 | 0.14982855 | -1.3570763 | 0.12458351 | 0.51763437 | 0.41103955 | 0.40618526 |
| 0.02057404 | 0.1660206 | -1.3821162 | 0.15276338 | 0.55325116 | 0.45084402 | 0.4076188 |
| 0.00783118 | 0.14078773 | -0.5221417 | 0.51305648 | 0.78421561 | 0.47632565 | 0.30676281 |
| 0.19641451 | 0.40516661 | NA | NA | NA | -1.8949248 | 0.0213621 |
| 0.01249536 | 0.14982855 | -0.9321634 | 0.29206929 | 0.66583113 | 0.48495873 | 0.34065072 |
| 9.43E-04 | 0.07509234 | -0.0682759 | 0.90902289 | 0.96361813 | 0.02936439 | 0.93215595 |
| 0.01538703 | 0.15038994 | -0.7413323 | 0.42584397 | 0.73587133 | 0.28137723 | 0.5982722 |
| 0.00204728 | 0.08733222 | -0.5009078 | 0.46281907 | 0.75477999 | -0.0909643 | 0.81579139 |
| 2.53E-04 | 0.05196388 | -0.103512 | 0.84667725 | 0.94578917 | -0.3064882 | 0.32935314 |
| 9.04E-04 | 0.07509234 | -0.3238723 | 0.60682276 | 0.83973408 | -0.2430758 | 0.50489151 |
| 0.02602821 | 0.18744874 | -0.8884323 | 0.40961789 | 0.7302279 | 1.09855267 | 0.0898205 |
| 0.00282024 | 0.09647579 | NA | NA | NA | 0.48001456 | 0.29881996 |
| 0.0164822 | 0.1555389 | -1.4334429 | 0.18402391 | 0.57346985 | 0.43916648 | 0.47086044 |
| 0.00691009 | 0.13718908 | -0.5760225 | 0.52948841 | 0.79423262 | 0.31753622 | 0.54801202 |
| 0.00235542 | 0.09208585 | -0.9669892 | 0.2414857 | 0.62405647 | 0.33285807 | 0.47806985 |
| 0.02256934 | 0.16972958 | -1.3594169 | 0.25552043 | 0.63999508 | 0.36409068 | 0.59117092 |
| 0.00116481 | 0.07821557 | -0.7882615 | 0.30235045 | 0.67301027 | -0.2254178 | 0.60427604 |
| 4.62E-04 | 0.05985127 | -0.5787871 | 0.39546233 | 0.71772396 | -0.2513047 | 0.52046686 |
| 4.66E-04 | 0.05985127 | -0.2985415 | 0.68134493 | 0.87082935 | -0.427381 | 0.31591058 |
| 5.10E-04 | 0.05985127 | -0.5507436 | 0.50384423 | 0.77910145 | -0.3578884 | 0.45278296 |
| 0.00630185 | 0.13585766 | 1.04115744 | 0.44582265 | 0.74570004 | 3.33363268 | 6.33E-04 |

| AMPH-03340.10nM - DMSO__adj.P.Val | AMPH-03340.300nM - DMSO__logFC | AMPH-03340.300nM - DMSO__P.Value | AMPH-03340.300nM - DMSO__adj.P.Val | AMPH-03340.1000nM - DMSO__logFC | AMPH-03340.1000nM - DMSO__P.Value | AMPH-03340.1000nM - DMSO__adj.P.Val |
|---|---|---|---|---|---|---|
| 2.56E-04 | 4.38166016 | 1.40E-09 | 1.15E-06 | 5.81994868 | 3.22E-09 | 2.60E-06 |
| 0.204189 | 1.75290419 | 1.12E-05 | 0.00172842 | 2.98708478 | 1.33E-06 | 2.68E-04 |
| 0.91935516 | 1.51061463 | 0.00665213 | 0.08504841 | 0.5454908 | 0.42701109 | 0.59717373 |
| 0.38020494 | 0.69276017 | 0.14965797 | 0.36677372 | -1.3840738 | 0.04900358 | 0.16993516 |
| 0.2426034 | 0.35145573 | 0.40949784 | 0.64653409 | -1.1639503 | 0.06604366 | 0.19546987 |
| 0.38661287 | 0.18808808 | 0.76998298 | 0.8601561 | -0.2535452 | 0.78043764 | 0.87098565 |
| 0.3235901 | 0.58514389 | 0.12027514 | 0.33670211 | 0.30677102 | 0.54988502 | 0.71089136 |
| 0.22305848 | 0.40224783 | 0.16468687 | 0.38576418 | -0.8160473 | 0.05381518 | 0.180426 |
| 0.67939616 | 0.08653306 | 0.74732675 | 0.84901121 | 0.47185055 | 0.22555633 | 0.41308969 |
| 0.92692476 | 0.57406117 | 0.32592402 | 0.57175988 | -0.6683793 | 0.41609355 | 0.58776851 |
| 0.75354449 | 0.54722576 | 0.06040124 | 0.23204241 | 0.36864175 | 0.34768579 | 0.52412336 |
| 0.14725224 | 0.1405444 | 0.7476676 | 0.84901121 | -1.607849 | 0.01860659 | 0.09826227 |
| 0.45292626 | 0.3208721 | 0.31848887 | 0.56232121 | -0.2667746 | 0.55296843 | 0.71339405 |
| 0.14725224 | 1.35636703 | 2.36E-04 | 0.01613401 | 1.22798181 | 0.00747804 | 0.06165567 |
| 0.42670244 | 0.88492057 | 0.00735972 | 0.08504841 | 0.01303543 | 0.97453252 | 0.98427785 |
| 0.25101641 | 0.06760206 | 0.86115839 | 0.92178753 | -1.1971171 | 0.04221966 | 0.15506128 |
| 0.55975377 | 0.45026717 | 0.04914872 | 0.21402119 | 0.02706261 | 0.92848992 | 0.9615595 |
| 0.38020494 | -0.0443217 | 0.92265555 | 0.96007631 | -0.7574185 | 0.25183818 | 0.43294733 |
| 0.93456055 | -0.0694685 | 0.8649886 | 0.92361698 | 0.45669104 | 0.43413388 | 0.60168125 |
| 0.32171019 | 0.14386524 | 0.75406959 | 0.85274261 | 0.84552579 | 0.20519858 | 0.39523805 |
| 0.76597692 | 0.60207956 | 0.09324788 | 0.29558497 | 0.44497076 | 0.36296763 | 0.5399595 |
| 0.08755387 | 1.37942372 | 8.32E-06 | 0.00172842 | 0.8928977 | 0.00786718 | 0.06412219 |
| 0.38661287 | -0.2089601 | 0.54748655 | 0.74050487 | NA | NA | NA |
| 0.43510911 | 0.48164394 | 0.06909183 | 0.24572073 | 1.31736644 | 0.00187262 | 0.0290977 |
| 0.2426034 | -0.2160461 | 0.62695574 | 0.78946421 | 0.27292049 | 0.62740363 | 0.76003319 |
| 0.19532438 | -0.1491872 | 0.6600648 | 0.80425909 | -0.7316293 | 0.14115905 | 0.32078714 |
| 0.59603045 | -0.3453292 | 0.66606232 | 0.80773584 | -1.0347651 | 0.36597127 | 0.54158385 |
| 0.44052698 | 0.33199207 | 0.27621183 | 0.51421748 | -0.9439301 | 0.03868371 | 0.14549291 |
| 0.6058625 | 0.74115334 | 0.01407346 | 0.11606357 | 1.07032005 | 0.01251307 | 0.0815368 |
| 0.71746351 | 0.60295529 | 0.09037424 | 0.28983302 | 0.88894174 | 0.07873203 | 0.21677029 |
| 0.39402079 | 0.11216057 | 0.74493247 | 0.84901121 | -0.3324738 | 0.49803446 | 0.66015551 |
| 0.70133826 | 0.40603595 | 0.28621481 | 0.5215385 | 0.67290837 | 0.21474075 | 0.40257664 |
| 0.58817032 | 0.68880042 | 0.05037205 | 0.21402119 | 0.76113593 | 0.11679286 | 0.28254082 |
| 0.36713172 | 0.83215086 | 0.00736416 | 0.08504841 | 0.8484165 | 0.04076689 | 0.15040936 |
| 0.26124802 | 0.7113302 | 0.00725227 | 0.08504841 | 0.36189172 | 0.27922404 | 0.45394975 |
| 0.08665997 | 1.25935126 | 1.32E-04 | 0.01205938 | 2.0600126 | 3.06E-05 | 0.00308814 |

| | | | | | | |
|---|---|---|---|---|---|---|
| 0.20206115 | 0.93202198 | 0.00329192 | 0.06285274 | 0.89522274 | 0.03107996 | 0.1255811 |
| 0.41842959 | 0.51667917 | 0.00730106 | 0.08504841 | 0.33139876 | 0.17803169 | 0.36325658 |
| 0.46358447 | 0.8628362 | 0.00964707 | 0.09770636 | 0.43385686 | 0.306279 | 0.48524202 |
| 0.36914225 | 1.51747684 | 0.00293745 | 0.05882069 | 0.96697308 | 0.12892623 | 0.30522724 |
| 0.43804532 | 0.85050163 | 0.01296087 | 0.11325581 | 0.7282083 | 0.10750469 | 0.26976332 |
| 0.21374185 | 0.65638739 | 0.06586436 | 0.24367421 | 1.3034231 | 0.01403296 | 0.08721575 |
| 0.41017441 | 0.21107133 | 0.48370497 | 0.69386761 | -0.4456723 | 0.30079045 | 0.48031361 |
| 0.71892035 | -0.7155467 | 0.23441333 | 0.46678674 | -1.181612 | 0.16872638 | 0.35362707 |
| 0.76597692 | 0.44003047 | 0.14514291 | 0.36277763 | 0.40210446 | 0.33595831 | 0.51705583 |
| 0.23382239 | 0.78743188 | 0.00284095 | 0.05882069 | 0.6752603 | 0.04652738 | 0.16561289 |
| 0.32171019 | 0.41929288 | 0.1530457 | 0.3706505 | 1.56714367 | 9.45E-04 | 0.0212038 |
| 0.55256004 | 0.58923634 | 0.01430779 | 0.11606357 | 0.12306509 | 0.6868279 | 0.8019609 |
| 0.20206115 | 0.07809126 | 0.78761986 | 0.87415549 | -0.6468089 | 0.12969035 | 0.30522724 |
| 0.20151958 | 0.65847639 | 0.00216533 | 0.05228638 | 0.93573088 | 0.00208853 | 0.03184025 |
| 0.27017623 | 0.10803669 | 0.63291468 | 0.78970053 | -1.3528806 | 6.67E-04 | 0.01759982 |
| 0.68650309 | 0.39540762 | 0.20203804 | 0.43612502 | 0.61780549 | 0.16125027 | 0.34537831 |
| 0.64015159 | -0.0555659 | 0.87411232 | 0.92959354 | -0.1046504 | 0.83294749 | 0.89976146 |
| 0.51971247 | 0.30605733 | 0.27247411 | 0.51086349 | -0.4344449 | 0.27076462 | 0.44923576 |
| 0.59619098 | 0.36280267 | 0.10658865 | 0.31821558 | 0.80663825 | 0.01673456 | 0.09452812 |
| 0.95071462 | 0.28285812 | 0.26931808 | 0.51064699 | 1.19587845 | 0.00389535 | 0.04526999 |
| 0.48704279 | 0.61367084 | 0.02279842 | 0.13329911 | 1.36170137 | 0.00125639 | 0.0231817 |
| 0.20151958 | -0.0764159 | 0.81892955 | 0.89543159 | -0.5345944 | 0.26759474 | 0.44765757 |
| 0.38020494 | 0.71016106 | 0.01071341 | 0.10471083 | 0.61700949 | 0.09277548 | 0.24259738 |
| 0.56018751 | 0.68852307 | 0.01441959 | 0.11606357 | 1.05341333 | 0.00915734 | 0.06915073 |
| 0.14703314 | -0.0439708 | 0.8554269 | 0.91865078 | 0.03760582 | 0.91223005 | 0.94862533 |
| 0.36806776 | 0.87359587 | 0.00721668 | 0.08504841 | 1.18956479 | 0.00911015 | 0.06915073 |
| 0.69955402 | 0.7546669 | 0.02189173 | 0.13153685 | 0.11550255 | 0.78485053 | 0.87470239 |
| 0.69955402 | 0.52920754 | 0.02067969 | 0.12960326 | 0.79967447 | 0.01455275 | 0.08841069 |
| 0.21683252 | 0.82073064 | 0.00500715 | 0.07612725 | 0.9160283 | 0.02015025 | 0.10304685 |
| 0.98150616 | 0.24609997 | 0.19997934 | 0.43549877 | 0.80820849 | 0.00734441 | 0.06117817 |
| 0.95071462 | 0.11091736 | 0.6550499 | 0.8014843 | 0.48546964 | 0.17926562 | 0.36485295 |
| 0.38020494 | 0.0981348 | 0.71389563 | 0.83552843 | 0.05304907 | 0.88832003 | 0.93337137 |
| 0.20206115 | 0.97013153 | 0.0021313 | 0.05228638 | 0.88367391 | 0.03013431 | 0.12550787 |
| 0.91202995 | 0.31287655 | 0.28466098 | 0.52050483 | 0.17993858 | 0.65804237 | 0.7802301 |
| 0.26620493 | -0.1882416 | 0.36408851 | 0.61128152 | 0.5830584 | 0.05877581 | 0.18628897 |
| 0.38661287 | 0.81113722 | 0.00251076 | 0.05571167 | 0.99105322 | 0.00682016 | 0.05800726 |
| 0.5453785 | 0.3860969 | 0.18758327 | 0.42150243 | -0.0139012 | 0.97238276 | 0.98427785 |
| 0.17245527 | -0.3433718 | 0.21407799 | 0.44709742 | 0.13277508 | 0.72741168 | 0.83250515 |
| 0.32120969 | -0.1192935 | 0.75510875 | 0.85274317 | -0.4496893 | 0.41064148 | 0.58415196 |
| 0.54328451 | 0.37851873 | 0.06588998 | 0.24367421 | 0.8520386 | 0.00665569 | 0.05721062 |
| 0.20206115 | -0.0601332 | 0.81549336 | 0.89543159 | -0.0740207 | 0.8390147 | 0.90351864 |
| 0.38020494 | -0.1080477 | 0.66067171 | 0.80425909 | -0.0140918 | 0.9675961 | 0.98218297 |

| | | | | | | |
|---|---|---|---|---|---|---|
| 0.92002854 | 0.00732382 | 0.97164752 | 0.98511131 | 0.83418881 | 0.01111395 | 0.07675272 |
| 0.14725224 | 0.81856562 | 5.08E-04 | 0.02937099 | 1.07634042 | 9.39E-04 | 0.0212038 |
| 0.9346131 | 0.1949171 | 0.51994556 | 0.71743749 | 0.3852527 | 0.3720823 | 0.54662272 |
| 0.59603045 | 0.33057377 | 0.33921097 | 0.59037865 | 1.72375167 | 0.00255375 | 0.03620056 |
| 0.69955402 | 0.17474548 | 0.62967682 | 0.78970053 | 1.64699625 | 0.0052839 | 0.0520657 |
| 0.43280403 | 0.88198292 | 0.00658165 | 0.08504841 | 0.45320657 | 0.26759411 | 0.44765757 |
| 0.14725224 | 0.12108327 | 0.52396107 | 0.71849817 | 0.10930638 | 0.68288318 | 0.7996661 |
| 0.28717843 | 0.78782095 | 0.00556359 | 0.08013522 | 0.68851487 | 0.06322717 | 0.19133914 |
| 0.87668234 | 0.53357337 | 0.01496131 | 0.11706618 | 0.4434163 | 0.12615591 | 0.29980582 |
| 0.99021155 | 0.40040944 | 0.13058155 | 0.34533555 | 0.9845369 | 0.01424812 | 0.08721575 |
| 0.73378421 | 0.45379189 | 0.05590568 | 0.22280857 | 0.08305503 | 0.79161638 | 0.87860719 |
| 0.99845565 | 0.28726188 | 0.08827799 | 0.28679553 | 0.3192774 | 0.17207314 | 0.35399507 |
| 0.56936462 | 1.13683552 | 0.00183875 | 0.05205559 | 0.90170873 | 0.05031962 | 0.17301383 |
| 0.20206115 | 0.78000529 | 0.0045355 | 0.07610054 | 0.90689301 | 0.01500423 | 0.08971656 |
| 0.19988415 | -0.0718042 | 0.75111663 | 0.85174965 | -0.0556755 | 0.86177532 | 0.91620323 |
| 0.49663882 | 0.76678835 | 0.01752476 | 0.12181362 | 0.73246599 | 0.09086567 | 0.24071956 |
| 0.28717843 | 0.6963953 | 0.01345976 | 0.11392227 | 3.86993826 | 2.04E-08 | 8.24E-06 |
| 0.20151958 | -0.1838546 | 0.44935693 | 0.67444614 | -0.3968551 | 0.25448069 | 0.43515202 |
| 0.28717843 | 0.89530807 | 0.0028924 | 0.05882069 | 1.2743552 | 0.00276072 | 0.03786635 |
| 0.36846864 | 0.37704464 | 0.10702942 | 0.31837375 | 0.59771809 | 0.07396759 | 0.20834169 |
| 0.00230941 | 1.69222273 | 6.85E-07 | 2.81E-04 | 1.22602259 | 7.19E-04 | 0.01759982 |
| 0.86400189 | 0.36003857 | 0.12353689 | 0.34012823 | 0.86875996 | 0.01414612 | 0.08721575 |
| 0.20983084 | 0.62864305 | 0.01147073 | 0.10824682 | 1.44525701 | 3.14E-04 | 0.01102751 |
| 0.32171019 | 0.75695475 | 0.00756216 | 0.08504841 | 0.90191076 | 0.02000737 | 0.10304685 |
| 0.69955402 | 0.05481165 | 0.82957534 | 0.89971117 | 0.51075128 | 0.17011443 | 0.35368215 |
| 0.7975986 | 0.24607025 | 0.2722324 | 0.51086349 | 0.07150671 | 0.81781924 | 0.89383192 |
| 0.62100204 | 0.17767745 | 0.56579927 | 0.75092333 | 0.03852955 | 0.92942818 | 0.9615595 |
| 0.77885591 | 0.2958873 | 0.18066682 | 0.41432251 | 0.44961115 | 0.15224957 | 0.33519797 |
| 0.82200579 | -0.1572775 | 0.63145483 | 0.78970053 | -0.2598572 | 0.57577105 | 0.72918967 |
| 0.17245527 | 0.82413494 | 0.00191264 | 0.05228638 | 0.70485277 | 0.03751925 | 0.14299789 |
| 0.62393252 | 0.11124592 | 0.50231078 | 0.70495239 | 0.76425966 | 0.00468843 | 0.0489601 |
| 0.69955402 | 0.43718234 | 0.05372811 | 0.21651549 | 0.36575061 | 0.23375535 | 0.41467388 |
| 0.13458393 | 1.11257497 | 3.84E-04 | 0.02426027 | 0.39141948 | 0.27074537 | 0.44923576 |
| 0.7403952 | 0.17280205 | 0.44644583 | 0.67130408 | 0.46186377 | 0.16061348 | 0.34537831 |
| 0.32644798 | 0.15742547 | 0.49693363 | 0.70161204 | 0.01445891 | 0.96451816 | 0.98152478 |
| 0.22305848 | 0.88465384 | 5.37E-04 | 0.02937099 | 0.34487099 | 0.24146986 | 0.42231525 |
| 0.3235901 | 0.97151015 | 0.00112799 | 0.04497455 | 1.40834329 | 9.23E-04 | 0.0212038 |
| 0.88293084 | 0.3822333 | 0.07343652 | 0.25547196 | 0.89153552 | 0.00643865 | 0.05623739 |
| 0.1226858 | 1.20077634 | 1.47E-05 | 0.00172842 | 2.5542248 | 1.21E-07 | 3.26E-05 |
| 0.38447045 | -0.2092565 | 0.52272474 | 0.71849817 | 0.90191423 | 0.06518361 | 0.19363367 |
| 0.95652252 | 0.1618408 | 0.62890981 | 0.78970053 | 3.26232921 | 5.13E-06 | 5.92E-04 |
| 0.22305848 | 0.05952765 | 0.76090208 | 0.85575426 | -0.2170017 | 0.43704642 | 0.60364702 |

| | | | | | | |
|---|---|---|---|---|---|---|
| 0.38661287 | 0.51304148 | 0.01324307 | 0.11325581 | 0.65645409 | 0.02251913 | 0.10895484 |
| 0.32047815 | 0.20652484 | 0.18839845 | 0.42150243 | 0.18837619 | 0.38776846 | 0.56555399 |
| 0.1226858 | 1.20324666 | 0.00121896 | 0.04497455 | 1.17054989 | 0.01499594 | 0.08971656 |
| 0.59603045 | 0.28455854 | 0.12595455 | 0.34201219 | 0.21082938 | 0.4086178 | 0.58248851 |
| 0.82811316 | 0.43269137 | 0.02051663 | 0.12960326 | 1.1686319 | 1.90E-04 | 0.00810113 |
| 0.99845565 | 0.02490356 | 0.91319713 | 0.95408514 | -0.4144846 | 0.21220577 | 0.40257664 |
| 0.80075065 | 0.21292664 | 0.22064348 | 0.45506071 | 0.23852578 | 0.32719399 | 0.50938871 |
| 0.38661287 | -0.0270819 | 0.91214971 | 0.95408514 | -0.0168638 | 0.96124825 | 0.97943076 |
| 0.20983084 | 0.45791397 | 0.11746811 | 0.33451359 | 0.55370935 | 0.17578981 | 0.35959029 |
| 0.20206115 | -0.0888344 | 0.65245829 | 0.80069994 | -0.0630104 | 0.8208209 | 0.89383192 |
| 0.56238061 | 0.21014627 | 0.42370947 | 0.65832537 | 0.36368614 | 0.33027573 | 0.51221265 |
| 0.97729171 | 0.09886197 | 0.68674577 | 0.81950331 | 0.46722418 | 0.19011746 | 0.37929606 |
| 0.87668234 | 0.40923755 | 0.1513588 | 0.36983803 | 0.78795766 | 0.05765898 | 0.18561139 |
| 0.44569554 | 0.45482553 | 0.00799815 | 0.08544939 | 0.70015517 | 0.00462653 | 0.0489601 |
| 0.43804532 | 0.75015323 | 0.00132272 | 0.04497455 | 0.48114533 | 0.09249248 | 0.24259738 |
| 0.2426034 | 0.53083715 | 0.03835803 | 0.18416343 | 0.51174027 | 0.14187428 | 0.32110482 |
| 0.48257912 | 0.07029263 | 0.72444686 | 0.83865842 | 0.90308578 | 0.0054745 | 0.05220092 |
| 0.60705855 | 0.15115845 | 0.45582645 | 0.67551175 | 0.93341452 | 0.00465435 | 0.0489601 |
| 0.62986383 | 0.08195562 | 0.74682536 | 0.84901121 | 0.16709103 | 0.64230864 | 0.77166992 |
| 0.5867485 | 0.40151309 | 0.02494318 | 0.14188246 | 0.60703983 | 0.01777216 | 0.09563282 |
| 0.15146213 | -0.2872635 | 0.20164888 | 0.43612502 | 0.72015261 | 0.03216322 | 0.12739159 |
| 0.42670244 | 0.10424295 | 0.61456336 | 0.78670568 | 0.55900143 | 0.07075066 | 0.20416618 |
| 0.27922042 | -0.1124964 | 0.58176123 | 0.76565751 | 0.14126783 | 0.62437698 | 0.76003319 |
| 0.81051683 | 0.20327526 | 0.29446217 | 0.53016106 | 0.53396524 | 0.06232316 | 0.19074665 |
| 0.59603045 | 0.23765204 | 0.14641875 | 0.36277763 | 0.25711363 | 0.2592572 | 0.43916104 |
| 0.68650309 | 0.07715532 | 0.76360334 | 0.85761743 | -0.1777772 | 0.6250507 | 0.76003319 |
| 0.57591964 | 0.1490316 | 0.46719349 | 0.68371811 | 0.45361657 | 0.12971898 | 0.30522724 |
| 0.32920596 | 0.77461218 | 0.00132569 | 0.04497455 | 1.12546471 | 0.00106818 | 0.02259002 |
| 0.38020494 | -0.4140674 | 0.4784559 | 0.68914438 | -0.1770427 | 0.82894331 | 0.89723855 |
| 0.26124802 | 0.07409436 | 0.68069171 | 0.81367954 | 0.11680345 | 0.64667793 | 0.77355455 |
| 0.15146213 | 0.81718472 | 2.18E-04 | 0.01613401 | 0.91431069 | 0.00160598 | 0.02648228 |
| 0.69955402 | -0.2325508 | 0.56283996 | 0.75092333 | 0.82426805 | 0.15928437 | 0.34504496 |
| 0.4528735 | 0.73330473 | 0.01127031 | 0.10759217 | 0.92775945 | 0.02066885 | 0.1043777 |
| 0.58011808 | 0.47135175 | 0.10896029 | 0.3217856 | 0.31533182 | 0.43200683 | 0.59976206 |
| 0.22305848 | 0.03666404 | 0.86089086 | 0.92178753 | 0.34958463 | 0.2484862 | 0.42809564 |
| 0.57834835 | 0.47701259 | 0.00921669 | 0.09458631 | 1.1673676 | 1.24E-04 | 0.00648765 |
| 0.69955402 | 0.15288494 | 0.59237072 | 0.77147967 | 0.40155303 | 0.32597341 | 0.50846817 |
| 0.32120969 | 0.10564701 | 0.7144226 | 0.83552843 | -0.1270569 | 0.75550461 | 0.85139152 |
| 0.38661287 | -0.6288491 | 0.14561079 | 0.36277763 | 0.20853102 | 0.7234912 | 0.83037059 |
| 0.39913114 | 0.13909128 | 0.61763783 | 0.78670568 | 0.33455064 | 0.39977738 | 0.57964261 |
| 0.18247005 | -0.4055158 | 0.09817755 | 0.30416515 | 0.0749601 | 0.82042878 | 0.89383192 |
| 0.87668234 | 0.17637629 | 0.72409939 | 0.83865842 | -0.157087 | 0.82384615 | 0.89471463 |

| | | | | | | |
|---|---|---|---|---|---|---|
| 0.91935516 | 0.19341772 | 0.37949826 | 0.62189235 | 1.03522254 | 0.0039219 | 0.04526999 |
| 0.9820691 | 0.51041252 | 0.07018197 | 0.24835948 | 0.16262692 | 0.66596237 | 0.78669239 |
| 0.27219985 | -0.0130141 | 0.943459 | 0.97291136 | 0.53358061 | 0.05462632 | 0.18097774 |
| 0.58210759 | 0.48114894 | 0.05997664 | 0.23204241 | 1.12971196 | 0.00472635 | 0.0489601 |
| 0.20206115 | -0.1234388 | 0.60383411 | 0.7807052 | -0.1068179 | 0.75012956 | 0.8466764 |
| 0.6592553 | -0.018168 | 0.9405384 | 0.97291136 | 0.44390268 | 0.21028591 | 0.40073353 |
| 0.23670201 | -0.0760789 | 0.71984357 | 0.83859556 | -0.7907972 | 0.01734625 | 0.09555675 |
| 0.38661287 | 0.72769039 | 0.00461637 | 0.07610054 | 0.94101719 | 0.00821513 | 0.06572105 |
| 0.88286598 | 0.31938022 | 0.08374615 | 0.27836272 | 0.18494216 | 0.45901481 | 0.62438379 |
| 0.45975183 | 1.10912002 | 0.00712516 | 0.08504841 | 0.4144036 | 0.42110066 | 0.59276888 |
| 0.43804532 | 0.53216326 | 0.04664262 | 0.20587954 | 0.82134099 | 0.03194192 | 0.12739159 |
| 0.99021155 | 0.58827653 | 0.06275431 | 0.23852449 | 0.62610914 | 0.11021656 | 0.27486106 |
| 0.58010779 | -0.2052588 | 0.47741244 | 0.68914438 | -0.6795948 | 0.10917415 | 0.27310437 |
| 0.71847643 | 0.39394395 | 0.14644334 | 0.36277763 | 0.35379792 | 0.34569472 | 0.52331084 |
| 0.19988415 | -0.0214378 | 0.93638786 | 0.97291136 | -0.0594206 | 0.87573911 | 0.92444603 |
| 0.5867485 | 0.04815296 | 0.83686731 | 0.90403692 | 0.468732 | 0.17042759 | 0.35368215 |
| 0.28472554 | -0.0970756 | 0.67142393 | 0.81234619 | 0.6346242 | 0.0645836 | 0.19327239 |
| 0.38661287 | 0.55662624 | 0.01775748 | 0.12181362 | 0.23392952 | 0.43929957 | 0.60572364 |
| 0.34114483 | -0.2646761 | 0.40879087 | 0.64653409 | 0.32497084 | 0.47196595 | 0.63712067 |
| 0.95071462 | 0.4597198 | 0.06536543 | 0.24367421 | 1.15579147 | 0.003111 | 0.03927641 |
| 0.99021155 | 0.48449266 | 0.06550961 | 0.24367421 | 0.31678406 | 0.37181247 | 0.54662272 |
| 0.54574095 | 0.60222449 | 0.06878743 | 0.24572073 | 0.81223839 | 0.08111364 | 0.2206728 |
| 0.32120969 | -0.1833727 | 0.58198633 | 0.76565751 | 0.71020234 | 0.1447325 | 0.32318159 |
| 0.20151958 | 0.77665748 | 8.34E-04 | 0.0380313 | 1.03037691 | 0.00139678 | 0.0245348 |
| 0.21683252 | 0.64199328 | 0.0015361 | 0.04670897 | 0.93927998 | 0.00117501 | 0.02315628 |
| 0.18470305 | -0.195012 | 0.46407861 | 0.68158952 | -1.0713004 | 0.01091356 | 0.07601856 |
| 0.54189492 | 0.18217848 | 0.40806464 | 0.64653409 | 0.09350168 | 0.76153681 | 0.85699407 |
| 0.20206115 | -0.7557129 | 0.10090675 | 0.30797189 | 0.09557976 | 0.87759771 | 0.92450971 |
| 0.30093686 | -0.5826882 | 0.04956289 | 0.21402119 | 0.2836326 | 0.4723209 | 0.63712067 |
| 0.55775689 | 0.37522183 | 0.05681821 | 0.22426804 | 0.54469038 | 0.05130658 | 0.17491862 |
| 0.75537286 | 0.13878479 | 0.5558554 | 0.74690227 | 0.71505629 | 0.04489427 | 0.16235084 |
| 0.72150431 | 0.43477042 | 0.14422057 | 0.36277763 | 0.78349667 | 0.06868051 | 0.20178568 |
| 0.73378421 | -0.0697912 | 0.80254125 | 0.88441123 | 0.38919295 | 0.33178726 | 0.51357108 |
| 0.97805622 | 0.11034429 | 0.51976267 | 0.71743749 | 0.6398808 | 0.01676494 | 0.09452812 |
| 0.65746392 | -0.1907261 | 0.68087435 | 0.81367954 | -0.3435771 | 0.60104918 | 0.74713098 |
| 0.44482451 | 0.1553929 | 0.4730202 | 0.68613001 | -0.0399359 | 0.89532118 | 0.9370719 |
| 0.43971474 | 0.33746647 | 0.19604021 | 0.43088465 | 1.38199591 | 0.00145451 | 0.02492591 |
| 0.53698363 | 0.10207517 | 0.61096636 | 0.78670568 | 0.45413946 | 0.12357208 | 0.29540308 |
| 0.24326489 | 0.5856336 | 0.00265151 | 0.05728654 | 0.47679738 | 0.05509966 | 0.18097774 |
| 0.23837259 | -0.200329 | 0.27254349 | 0.51086349 | -0.0446285 | 0.85986856 | 0.9153805 |
| 0.56498582 | 0.23507548 | 0.22987341 | 0.46491584 | 0.62913234 | 0.03210322 | 0.12739159 |
| 0.38661287 | 0.16238316 | 0.42095507 | 0.65832537 | 0.19335916 | 0.49658144 | 0.65993059 |

| | | | | | | |
|---|---|---|---|---|---|---|
| 0.01591854 | 1.91298778 | 9.45E-06 | 0.00172842 | 2.04910182 | 1.65E-04 | 0.00784582 |
| 0.39913114 | -0.5156989 | 0.34300928 | 0.59037865 | -0.4961554 | 0.51526357 | 0.67037445 |
| 0.40927024 | 0.72016949 | 0.00792607 | 0.08544939 | -0.4525081 | 0.19197131 | 0.38091616 |
| 0.28717843 | 0.71242243 | 0.00142429 | 0.04497455 | 0.91431353 | 0.00295233 | 0.0384748 |
| 0.32863165 | -0.2253809 | 0.28857266 | 0.52299813 | -0.3018184 | 0.31394568 | 0.49516293 |
| 0.13458393 | -0.329944 | 0.11574433 | 0.3334249 | -0.4161029 | 0.15710685 | 0.34216263 |
| 0.33623404 | 0.73140003 | 8.17E-04 | 0.0380313 | 0.33932847 | 0.18757673 | 0.37796009 |
| 0.38020494 | 0.06481174 | 0.8270016 | 0.89929578 | 0.6102492 | 0.15999116 | 0.34537831 |
| 0.26472769 | 0.09700972 | 0.55550037 | 0.74690227 | 0.27540728 | 0.24516761 | 0.42509749 |
| 0.82200579 | -0.0742063 | 0.73272529 | 0.8397918 | -0.7670324 | 0.02299468 | 0.10929234 |
| 0.55256004 | -0.2094179 | 0.51945936 | 0.71743749 | -0.4536121 | 0.32853413 | 0.51049149 |
| 0.32120969 | 0.47437373 | 0.01793959 | 0.12181362 | 0.90541884 | 0.00276499 | 0.03786635 |
| 0.14725224 | 1.16520506 | 2.14E-04 | 0.01613401 | 0.99333142 | 0.0101204 | 0.07236537 |
| 0.79290617 | 0.18763721 | 0.60052491 | 0.78011227 | NA | NA | NA |
| 0.1758396 | -0.3258234 | 0.24259991 | 0.47763675 | 0.32452155 | 0.40481627 | 0.58141958 |
| 0.76018942 | 0.18704494 | 0.38549572 | 0.62547824 | 0.79284112 | 0.01754602 | 0.09555675 |
| 0.21683252 | -0.1183136 | 0.49736688 | 0.70161204 | 0.59668586 | 0.02595444 | 0.11848128 |
| 0.39553705 | -0.7572357 | 0.15951719 | 0.38070819 | -1.3039115 | 0.0918434 | 0.24229341 |
| 0.39913114 | 0.52554825 | 0.03388063 | 0.1713177 | 1.26991971 | 0.00124003 | 0.0231817 |
| 0.34581179 | 0.6290418 | 0.01604004 | 0.11706618 | 1.49603273 | 3.97E-04 | 0.01237993 |
| 0.54574095 | 0.50547716 | 0.01905244 | 0.12414327 | 0.36841391 | 0.19446252 | 0.38230102 |
| 0.38805069 | 0.78866906 | 0.0013523 | 0.04497455 | 0.94607156 | 0.00451466 | 0.0489601 |
| 0.19988415 | 0.73344875 | 0.02918003 | 0.15865433 | 0.10542158 | 0.8089926 | 0.89055316 |
| 0.20151958 | -0.1052015 | 0.62185125 | 0.78793079 | 0.15192905 | 0.6145542 | 0.75465014 |
| 0.51299103 | -0.166391 | 0.5294348 | 0.72444329 | 1.00153611 | 0.01557969 | 0.09188608 |
| 0.48257912 | 0.14234261 | 0.49668247 | 0.70161204 | 0.22359861 | 0.45111187 | 0.61779388 |
| 0.38661287 | -0.0545603 | 0.76878848 | 0.8601561 | -0.3197817 | 0.23429479 | 0.41467388 |
| 0.28717843 | 0.81519804 | 0.00243638 | 0.05556308 | 1.00205038 | 0.00639345 | 0.05623739 |
| 0.98150616 | 0.14247064 | 0.4555806 | 0.67551175 | 0.09535015 | 0.72195342 | 0.83037059 |
| 0.2426034 | 1.1667035 | 0.02991932 | 0.16127571 | 1.69114992 | 0.02662956 | 0.12020496 |
| 0.20983084 | 0.02703832 | 0.89439362 | 0.94625923 | 0.38668639 | 0.1928361 | 0.38095738 |
| 0.44579852 | -0.0905675 | 0.67650678 | 0.81292206 | 0.57103134 | 0.07789103 | 0.21634061 |
| 0.20983084 | -0.3910374 | 0.20982386 | 0.44075808 | 0.28689193 | 0.50664377 | 0.66455871 |
| 0.1226858 | 0.96953214 | 9.93E-05 | 0.01019503 | 1.42218752 | 6.91E-05 | 0.00544275 |
| 0.38020494 | -0.2582726 | 0.27714787 | 0.51479277 | 0.66530687 | 0.05826585 | 0.18628897 |
| 0.97729171 | 0.32518962 | 0.12473416 | 0.34022176 | 0.81580098 | 0.01157682 | 0.07860565 |
| 0.38020494 | 0.58955049 | 0.0201137 | 0.12901053 | 1.43449369 | 4.74E-04 | 0.01384725 |
| 0.25101641 | 0.50555377 | 0.0151746 | 0.11706618 | -0.033362 | 0.89944578 | 0.93852637 |
| 0.1758396 | -0.0912223 | 0.65010243 | 0.79986567 | 0.05692152 | 0.84089854 | 0.90351864 |
| 0.72992821 | 0.15968424 | 0.5644516 | 0.75092333 | -0.1822961 | 0.64112635 | 0.77166992 |
| 0.48028696 | -0.2150851 | 0.39260347 | 0.63360184 | 0.20718751 | 0.55761135 | 0.71504275 |
| 0.28717843 | 0.72324542 | 0.13664286 | 0.35389208 | 0.49774284 | 0.42960902 | 0.59854313 |

| | | | | | | |
|---|---|---|---|---|---|---|
| 0.27017024 | -0.4242831 | 0.29595187 | 0.53167721 | -1.0432025 | 0.07972758 | 0.21791411 |
| 0.17386306 | 0.66770097 | 0.05062719 | 0.21402119 | 0.49993485 | 0.27724954 | 0.45347698 |
| 0.55354732 | 0.45941545 | 0.05307115 | 0.21651549 | 0.55950248 | 0.09043075 | 0.24035543 |
| 0.13601303 | -0.2929791 | 0.20691465 | 0.43922288 | -0.0628637 | 0.84386625 | 0.90550323 |
| 0.2426034 | 0.07895968 | 0.73341135 | 0.8397918 | -0.2164602 | 0.51150658 | 0.66879835 |
| 0.86400189 | 0.26339024 | 0.16664715 | 0.38868554 | 0.69705524 | 0.01609563 | 0.09318167 |
| 0.71892035 | 0.19796406 | 0.26677189 | 0.5081664 | 0.0422853 | 0.86389362 | 0.91716526 |
| 0.58135034 | 0.21121202 | 0.43263271 | 0.66552154 | -0.4664878 | 0.22746159 | 0.41308969 |
| 0.30093686 | 0.62390239 | 0.01115473 | 0.10759217 | 1.57618417 | 1.24E-04 | 0.00648765 |
| 0.40888927 | 0.55602769 | 0.0223666 | 0.13210774 | 1.24710067 | 0.00111832 | 0.02259002 |
| 0.1758396 | 0.67244099 | 0.00210661 | 0.05228638 | 0.94168632 | 0.00226663 | 0.03391544 |
| 0.98150616 | 0.26356353 | 0.23561013 | 0.46678674 | 0.95828549 | 0.00647287 | 0.05623739 |
| 0.72215678 | -0.0474683 | 0.8689779 | 0.92533185 | 0.63272724 | 0.13524023 | 0.31582112 |
| 0.5916915 | -0.2534174 | 0.45931535 | 0.67767053 | 0.06016264 | 0.90019547 | 0.93852637 |
| 0.77403151 | 0.21033451 | 0.35014999 | 0.59890238 | 0.67529577 | 0.04529215 | 0.16235084 |
| 0.2426034 | -0.0496291 | 0.82017607 | 0.89543159 | 0.0664817 | 0.82950148 | 0.89723855 |
| 0.26124802 | -0.233272 | 0.26471448 | 0.50659811 | 0.51082729 | 0.0933649 | 0.24335109 |
| 0.98150616 | 0.28897866 | 0.12921811 | 0.34452754 | 0.6918004 | 0.01614534 | 0.09318167 |
| 0.89905048 | 0.32261291 | 0.3663189 | 0.61358533 | 0.48122874 | 0.34149589 | 0.52252085 |
| 0.38020494 | 0.532772 | 0.03316652 | 0.16912863 | 0.0838111 | 0.79697159 | 0.88333751 |
| 0.74320615 | 0.39950081 | 0.04412063 | 0.19902766 | 0.5419883 | 0.05222074 | 0.1772872 |
| 0.20206115 | -0.224947 | 0.37308308 | 0.61506267 | -0.5192084 | 0.15515055 | 0.33881525 |
| 0.53314753 | -0.0346398 | 0.87712958 | 0.93159558 | 0.32451039 | 0.31437944 | 0.49516293 |
| 0.46358447 | 0.5808003 | 0.006107 | 0.08498049 | 0.42036718 | 0.12196796 | 0.29330391 |
| 0.47604976 | 0.4266046 | 0.07429464 | 0.25736667 | 0.45060075 | 0.17196979 | 0.35399507 |
| 0.38020494 | 0.62362423 | 0.01197091 | 0.10920126 | 1.40644848 | 3.98E-04 | 0.01237993 |
| 0.77733525 | 0.36688288 | 0.07341215 | 0.25547196 | 0.65171601 | 0.02898453 | 0.12391271 |
| 0.23382239 | 0.75156969 | 0.00176876 | 0.0518627 | 0.59052582 | 0.05120061 | 0.17491862 |
| 0.40888927 | 0.47317512 | 0.04045671 | 0.1906336 | 1.36255977 | 3.98E-04 | 0.01237993 |
| 0.54189492 | 0.22009225 | 0.40514188 | 0.64653409 | 1.17090564 | 0.00592559 | 0.05440764 |
| 0.53197865 | 1.10927276 | 0.03527046 | 0.17444007 | NA | NA | NA |
| 0.34139791 | -0.2546366 | 0.42159188 | 0.65832537 | 0.64193729 | 0.16186135 | 0.34537831 |
| 0.27748741 | 0.68641056 | 0.00515088 | 0.07688862 | 2.12205335 | 3.93E-06 | 5.29E-04 |
| 0.39913114 | 0.55566973 | 0.01811043 | 0.12181362 | 0.70317515 | 0.03130103 | 0.12582701 |
| 0.39913114 | -0.1982205 | 0.37029441 | 0.61506267 | -0.8243028 | 0.01631293 | 0.09348117 |
| 0.28717843 | 0.40190833 | 0.05298164 | 0.21651549 | 0.72057587 | 0.01787197 | 0.09563282 |
| 0.38661287 | 0.63529042 | 0.06809934 | 0.24572073 | 0.27072491 | 0.56145509 | 0.71780967 |
| 0.68650309 | 0.31627912 | 0.24661333 | 0.48322086 | 1.13263898 | 0.00830163 | 0.06576192 |
| 0.18470305 | -0.1869455 | 0.34139381 | 0.59037865 | -0.1073466 | 0.69536953 | 0.8095945 |
| 0.13458393 | -0.2449702 | 0.25962488 | 0.49918508 | -0.2635032 | 0.38673049 | 0.56506011 |
| 0.5909425 | 0.14785997 | 0.38328319 | 0.6231198 | 0.55564174 | 0.03108443 | 0.1255811 |
| 0.23382239 | 0.0112035 | 0.96007977 | 0.98090261 | -0.0538621 | 0.86495041 | 0.91716526 |

| | | | | | | |
|---|---|---|---|---|---|---|
| 0.87710098 | 0.34275495 | 0.22115618 | 0.45506071 | -0.0878535 | 0.81994405 | 0.89383192 |
| 0.30093686 | -0.3302862 | 0.20239176 | 0.43612502 | 0.96462203 | 0.01510081 | 0.08971656 |
| 0.20206115 | -0.5458839 | 0.08171302 | 0.27494421 | 0.40137893 | 0.34649914 | 0.52331084 |
| 0.61626318 | -0.0145126 | 0.94447059 | 0.97291136 | -0.1529113 | 0.6055041 | 0.74923018 |
| 0.38020494 | 0.58377262 | 0.01320756 | 0.11325581 | -0.3113466 | 0.30409588 | 0.48324381 |
| 0.99830517 | 0.05589693 | 0.75797381 | 0.85363032 | 0.60643063 | 0.02993105 | 0.12545085 |
| 0.98908107 | 0.10534747 | 0.53823712 | 0.7316104 | 0.13106506 | 0.58762441 | 0.7384145 |
| 0.95369702 | 0.3581395 | 0.06310617 | 0.23875651 | 0.6797458 | 0.0168466 | 0.09452812 |
| 0.79620923 | 0.21966197 | 0.23574802 | 0.46678674 | 0.87403616 | 0.00354563 | 0.04292091 |
| 0.76206684 | 0.37716208 | 0.11165546 | 0.32738975 | 0.41634061 | 0.20651348 | 0.39571993 |
| 0.5286044 | 0.39429986 | 0.05044542 | 0.21402119 | 0.09902708 | 0.71010897 | 0.82267315 |
| 0.23837259 | -0.3093281 | 0.14571488 | 0.36277763 | 0.16500707 | 0.57073514 | 0.72611375 |
| 0.83481506 | 0.17707667 | 0.42552467 | 0.65832537 | 0.39719088 | 0.21380367 | 0.40257664 |
| 0.1758396 | -0.4386575 | 0.06648226 | 0.24476204 | -0.3952207 | 0.22594514 | 0.41308969 |
| 0.59619098 | -0.1305315 | 0.60250346 | 0.78056529 | 0.45033763 | 0.21438055 | 0.40257664 |
| 0.18470305 | -0.8575762 | 0.05135466 | 0.21402119 | -0.4525718 | 0.44058257 | 0.60645778 |
| 0.2426034 | 0.27926548 | 0.17055749 | 0.39555847 | 0.7903888 | 0.01169387 | 0.07873872 |
| 0.54574095 | 0.14764989 | 0.43287272 | 0.66552154 | 0.52566385 | 0.06115965 | 0.18887265 |
| 0.77403151 | -0.4782167 | 0.27107027 | 0.51086349 | 0.30738697 | 0.61080366 | 0.75298711 |
| 0.17245527 | 0.69005731 | 0.01438373 | 0.11606357 | 0.38088642 | 0.29524984 | 0.47333705 |
| 0.72284918 | 0.35451354 | 0.05257417 | 0.21651549 | 0.80414895 | 0.0042733 | 0.04795587 |
| 0.28717843 | -0.1883198 | 0.46287353 | 0.68103793 | -0.1156497 | 0.7479844 | 0.84645854 |
| 0.83115842 | 0.11908072 | 0.45186516 | 0.67475057 | 0.36423558 | 0.11604991 | 0.28158656 |
| 0.99021155 | -0.0040366 | 0.9889047 | 0.99496417 | 0.10959616 | 0.7897445 | 0.87773529 |
| 0.79165133 | 0.28539319 | 0.1636746 | 0.38576418 | 0.5942275 | 0.04802793 | 0.169461 |
| 0.78698771 | 0.34354093 | 0.0685304 | 0.24572073 | 0.28110667 | 0.27317013 | 0.45045197 |
| 0.80075065 | 0.37386625 | 0.62464291 | 0.78793079 | -2.2021292 | 0.05564074 | 0.18177589 |
| 0.79620923 | 0.34835567 | 0.07237848 | 0.25503319 | 0.40042665 | 0.13674073 | 0.31748996 |
| 0.38020494 | 0.47864025 | 0.01584854 | 0.11706618 | 0.70533753 | 0.01261691 | 0.08155568 |
| 0.57168952 | 0.50260965 | 0.02756653 | 0.1518934 | 0.59384607 | 0.05925281 | 0.18628897 |
| 0.17386306 | 0.58789728 | 0.03277018 | 0.16815196 | 1.24130892 | 0.00323735 | 0.0402427 |
| 0.13458393 | -0.5504407 | 0.01461961 | 0.11653107 | 0.39252792 | 0.18295271 | 0.36956447 |
| 0.23382239 | 0.19868338 | 0.19567329 | 0.43088465 | 0.02550536 | 0.90365835 | 0.94092261 |
| 0.21795857 | -1.3472025 | 0.05227695 | 0.21651549 | -0.9725611 | 0.29788191 | 0.47661105 |
| 0.33686179 | -0.105221 | 0.54931168 | 0.74117427 | 0.74765695 | 0.00793591 | 0.06412219 |
| 0.6813551 | -0.0140818 | 0.94424892 | 0.97291136 | 0.19762923 | 0.49135736 | 0.6573125 |
| 0.52857259 | 0.02899521 | 0.91991418 | 0.9584385 | 0.07902055 | 0.84644481 | 0.90706552 |
| 0.69955402 | -0.2693278 | 0.67283241 | 0.81234619 | -0.9658787 | 0.29236062 | 0.47218612 |
| 0.86543101 | 0.21965258 | 0.17792786 | 0.41033363 | 0.42976471 | 0.06971863 | 0.20263543 |
| 0.70651631 | 0.21690266 | 0.35231421 | 0.5991018 | 0.95469637 | 0.00949011 | 0.0710001 |
| 0.30638319 | 0.38294919 | 0.04152876 | 0.19262773 | 0.11310192 | 0.64718178 | 0.77355455 |
| 0.30638319 | -0.1585927 | 0.42423871 | 0.65832537 | 0.1465424 | 0.59921499 | 0.74713098 |

| | | | | | | |
|---|---|---|---|---|---|---|
| 0.72284918 | 0.09240301 | 0.6176525 | 0.78670568 | 0.61213726 | 0.03107614 | 0.1255811 |
| 0.56549932 | 0.73171631 | 0.08299465 | 0.27792865 | NA | NA | NA |
| 0.59603045 | -0.474255 | 0.13883506 | 0.35722845 | 0.02640336 | 0.9516541 | 0.97580776 |
| 0.20151958 | -0.3829233 | 0.08027712 | 0.27395812 | 0.30063817 | 0.3133951 | 0.49516293 |
| 0.26124802 | -0.1715037 | 0.42573768 | 0.65832537 | 0.26486752 | 0.38542024 | 0.56416586 |
| 0.71847643 | 0.35276487 | 0.14488976 | 0.36277763 | 0.52758174 | 0.12475208 | 0.2973442 |
| 0.54189492 | -0.1616856 | 0.48569272 | 0.69386761 | 0.66424102 | 0.05601757 | 0.18177589 |
| 0.30093686 | -0.2152339 | 0.31478173 | 0.55697371 | 0.27875013 | 0.35589407 | 0.53153865 |
| 0.98908107 | -0.0016678 | 0.9966153 | 0.99783068 | -0.112519 | 0.83974692 | 0.90351864 |
| 0.84472005 | 0.2870542 | 0.08327704 | 0.27792865 | 0.70070369 | 0.00608892 | 0.05527922 |
| 0.2426034 | 0.74631499 | 0.01902253 | 0.12414327 | 0.14465921 | 0.72255749 | 0.83037059 |
| 0.48456387 | 0.38325418 | 0.05406294 | 0.21651549 | 1.33875161 | 1.28E-04 | 0.00648765 |
| 0.87880503 | 0.19888968 | 0.28018667 | 0.51633949 | 0.61602412 | 0.02725728 | 0.12101033 |
| 0.45975183 | 0.52185946 | 0.03401314 | 0.1713177 | 0.40970772 | 0.21384345 | 0.40257664 |
| 0.30093686 | -0.1534941 | 0.5127474 | 0.71350105 | 0.11107682 | 0.73623346 | 0.83710001 |
| 0.65746392 | 0.17341767 | 0.63266062 | 0.78970053 | 1.2791744 | 0.02290089 | 0.10929234 |
| 0.39913114 | 0.57455778 | 0.00418013 | 0.07301892 | 1.13530742 | 2.90E-04 | 0.0106547 |
| 0.5495682 | 0.50601559 | 0.12913624 | 0.34452754 | 0.60285965 | 0.19591413 | 0.38328963 |
| 0.28717843 | 0.04537443 | 0.78682517 | 0.87415549 | 0.32736913 | 0.18108253 | 0.36762483 |
| 0.73814343 | 0.2977537 | 0.11648875 | 0.33439604 | 0.52990459 | 0.05365177 | 0.180426 |
| 0.23732313 | -0.0181006 | 0.90434034 | 0.95065739 | 0.26137076 | 0.23166504 | 0.41451545 |
| 0.50894661 | 0.31756389 | 0.09895635 | 0.30428152 | 0.34542839 | 0.19574972 | 0.38328963 |
| 0.94181084 | -0.0515604 | 0.77928194 | 0.86928053 | 0.51896185 | 0.06103961 | 0.18887265 |
| 0.55775689 | -0.006413 | 0.96534208 | 0.9833075 | 0.34364177 | 0.11551412 | 0.28158656 |
| 0.56238061 | 0.16681982 | 0.35470677 | 0.60168236 | 0.31608362 | 0.22037938 | 0.40747491 |
| 0.67453807 | 0.25084474 | 0.2640026 | 0.5064162 | 0.12968953 | 0.67714899 | 0.7967137 |
| 0.14725224 | -0.2569066 | 0.19894886 | 0.43440695 | -0.0455423 | 0.86829918 | 0.91950948 |
| 0.90292851 | -4.50E-04 | 0.99867558 | 0.99867558 | 0.0706039 | 0.85394363 | 0.91138614 |
| 0.28717843 | -0.030812 | 0.89845134 | 0.9493289 | 0.0551376 | 0.87173572 | 0.9219404 |
| 0.48479656 | -0.1043328 | 0.65080422 | 0.79986567 | 0.15099888 | 0.64325754 | 0.77166992 |
| 0.42030169 | 0.07264135 | 0.61805745 | 0.78670568 | 0.21470308 | 0.30441968 | 0.48324381 |
| 0.52368501 | 0.12560952 | 0.47099294 | 0.68613001 | 0.63685241 | 0.01855798 | 0.09826227 |
| 0.93623502 | 0.43818698 | 0.06820102 | 0.24572073 | -0.096268 | 0.7637393 | 0.85827727 |
| 0.53099293 | 0.08133627 | 0.67188944 | 0.81234619 | 1.06357949 | 0.00126237 | 0.0231817 |
| 0.28717843 | 0.52797247 | 0.01012203 | 0.10012272 | 0.63037952 | 0.02517521 | 0.11623755 |
| 0.38661287 | 0.47097114 | 0.12428559 | 0.34012823 | NA | NA | NA |
| 0.59619098 | 0.27473222 | 0.27960083 | 0.51633949 | 0.43215229 | 0.23136223 | 0.41451545 |
| 0.0719728 | 0.9388686 | 1.37E-05 | 0.00172842 | 1.09114916 | 1.04E-04 | 0.00646322 |
| 0.58010779 | -0.2761439 | 0.21994687 | 0.45506071 | 0.29289762 | 0.35198271 | 0.52862831 |
| 0.28717843 | 0.56401051 | 0.00361713 | 0.06455795 | 0.94160093 | 0.0010337 | 0.02257386 |
| 0.59603045 | 0.22062236 | 0.18920145 | 0.42150243 | 0.77940886 | 0.00383291 | 0.04526999 |
| 0.20151958 | -0.5451288 | 0.04980787 | 0.21402119 | 0.31884685 | 0.39028456 | 0.56782173 |

| | | | | | | |
|---|---|---|---|---|---|---|
| 0.38661287 | 0.52162019 | 0.01814282 | 0.12181362 | 1.4812425 | 9.36E-05 | 0.00630528 |
| 0.1226858 | -0.427042 | 0.05380901 | 0.21651549 | 0.15386121 | 0.60073881 | 0.74713098 |
| 0.26124802 | -0.1350828 | 0.48628108 | 0.69386761 | 0.12319145 | 0.65180732 | 0.77723958 |
| 0.47947892 | 0.69622861 | 0.00675424 | 0.08504841 | 1.11458132 | 0.00291849 | 0.0384748 |
| 0.69942883 | 0.46821331 | 0.03904731 | 0.18638281 | 0.76783204 | 0.01934719 | 0.10085504 |
| 0.73378421 | 0.22844762 | 0.27596213 | 0.51421748 | 0.82210814 | 0.01182002 | 0.07893038 |
| 0.98995236 | 0.30747193 | 0.1549703 | 0.3731103 | -0.0192269 | 0.94799049 | 0.97580776 |
| 0.22624369 | 0.08879611 | 0.56252041 | 0.75092333 | 0.64280032 | 0.00871317 | 0.06835184 |
| 0.20151958 | -0.2418538 | 0.20761039 | 0.43929931 | 0.15573918 | 0.55728364 | 0.71504275 |
| 0.2426034 | 0.60748791 | 0.02289303 | 0.13329911 | 0.77928654 | 0.03618691 | 0.13989965 |
| 0.9346131 | 0.2446413 | 0.10461641 | 0.31461566 | 0.43790339 | 0.04526801 | 0.16235084 |
| 0.20206115 | 0.57565139 | 0.0054396 | 0.07974841 | 0.96028423 | 0.00167411 | 0.02705359 |
| 0.38447045 | 0.18253435 | 0.43478726 | 0.66597079 | 0.44255331 | 0.18945711 | 0.37929606 |
| 0.95369702 | 0.09807818 | 0.64982892 | 0.79986567 | 0.45919592 | 0.14651265 | 0.32612182 |
| 0.81563667 | 0.06181193 | 0.83605229 | 0.90403692 | -0.2014928 | 0.63447036 | 0.7674432 |
| 0.20151958 | -0.1487571 | 0.40898079 | 0.64653409 | -0.2408435 | 0.34630888 | 0.52331084 |
| 0.76018942 | 0.13127408 | 0.5828696 | 0.76565751 | 0.5484454 | 0.11862209 | 0.28610939 |
| 0.69955402 | 0.01795046 | 0.94284656 | 0.97291136 | -0.4222105 | 0.24456313 | 0.42496131 |
| 0.59603045 | -0.0356668 | 0.86624248 | 0.92361698 | -0.9722703 | 0.00505821 | 0.05108794 |
| 0.24015795 | 0.01895399 | 0.91341159 | 0.95408514 | 0.43425426 | 0.09408825 | 0.24444794 |
| 0.96271414 | 0.06765526 | 0.77005449 | 0.8601561 | 0.40551542 | 0.22688117 | 0.41308969 |
| 0.6090444 | -0.1284544 | 0.61747054 | 0.78670568 | 0.51551063 | 0.1689357 | 0.35362707 |
| 0.626596 | 0.20249567 | 0.28690997 | 0.5215385 | -0.6645918 | 0.02204302 | 0.10835435 |
| 0.5867485 | 0.15569046 | 0.59254031 | 0.77147967 | 0.02260564 | 0.95595955 | 0.97740909 |
| 0.31989521 | -0.1057402 | 0.56364052 | 0.75092333 | 0.40567147 | 0.13058428 | 0.30583218 |
| 0.6714611 | 0.38329269 | 0.04930695 | 0.21402119 | 0.83213919 | 0.00514906 | 0.05136351 |
| 0.6380977 | -0.2069738 | 0.37300535 | 0.61506267 | 0.42518428 | 0.20243207 | 0.39224248 |
| 0.26327441 | 0.57041181 | 0.11787725 | 0.33451359 | 1.21917671 | 0.02442784 | 0.11409073 |
| 0.98908107 | 0.28907351 | 0.16492476 | 0.38576418 | 0.78460536 | 0.01368072 | 0.08703956 |
| 0.35756893 | -0.4171908 | 0.20482305 | 0.43677851 | 0.85806952 | 0.0733742 | 0.20834169 |
| 0.58135034 | -0.193866 | 0.48914603 | 0.69560876 | 1.34404568 | 0.00355904 | 0.04292091 |
| 0.9749639 | 0.19565816 | 0.31144617 | 0.55345737 | 0.65281925 | 0.02634952 | 0.11960907 |
| 0.39402079 | 0.33249336 | 0.05123828 | 0.21402119 | 0.43449968 | 0.0690454 | 0.20178568 |
| 0.33686179 | -0.1725904 | 0.36695541 | 0.61358533 | 0.33807723 | 0.21733045 | 0.40554967 |
| 0.79895026 | 0.31716624 | 0.08911277 | 0.28690819 | 0.60284031 | 0.02762922 | 0.1213283 |
| 0.5046594 | 0.05260984 | 0.727314 | 0.83865842 | 0.6842495 | 0.00542714 | 0.05220092 |
| 0.78329146 | 0.00432489 | 0.98265732 | 0.99185709 | -0.2107128 | 0.45838171 | 0.62438379 |
| 0.54189492 | 0.1403484 | 0.33973911 | 0.59037865 | 0.72112386 | 0.00286572 | 0.0384748 |
| 0.59603045 | 0.22207882 | 0.19638743 | 0.43088465 | 0.53395488 | 0.03668734 | 0.14115889 |
| 0.9820691 | 0.01154708 | 0.96198097 | 0.98110109 | -0.0180182 | 0.95805445 | 0.97740909 |
| 0.79165133 | -0.2114299 | 0.47217482 | 0.68613001 | -0.636286 | 0.13789238 | 0.31812894 |
| 0.38661287 | -0.0707794 | 0.64451183 | 0.79703317 | 0.48432631 | 0.03834356 | 0.14545349 |

| | | | | | | |
|---|---|---|---|---|---|---|
| 0.27748741 | -0.1518115 | 0.43983389 | 0.66752555 | -0.006411 | 0.98139847 | 0.98845389 |
| 0.98970237 | -0.1334448 | 0.49968293 | 0.70324205 | 0.09953032 | 0.72027534 | 0.83021751 |
| 0.55354732 | -0.3379278 | 0.35801148 | 0.60456228 | 0.55961382 | 0.23405724 | 0.41467388 |
| 0.4528735 | -0.1665497 | 0.56104744 | 0.75092333 | 0.14199709 | 0.72499617 | 0.83091759 |
| 0.46275093 | -0.3228181 | 0.58125196 | 0.76565751 | 0.08237795 | 0.92030343 | 0.95579071 |
| 0.65010577 | -0.1724765 | 0.35861368 | 0.60456228 | 0.3415892 | 0.2048528 | 0.39523805 |
| 0.55256004 | 0.26687939 | 0.10036437 | 0.30745952 | 0.70105785 | 0.00551766 | 0.05220092 |
| 0.28717843 | -0.230664 | 0.1460108 | 0.36277763 | 0.01200943 | 0.95565189 | 0.97740909 |
| 0.20771056 | -1.6942938 | 0.03795688 | 0.18416343 | -2.3037759 | 0.04485604 | 0.16235084 |
| 0.44579852 | 0.00154887 | 0.99217492 | 0.99703257 | 0.72553289 | 0.00505603 | 0.05108794 |
| 0.28717843 | 0.49136281 | 0.02136398 | 0.13044095 | 0.18900951 | 0.49359912 | 0.65921998 |
| 0.77733525 | 0.01253825 | 0.97331751 | 0.98531896 | 0.0185494 | 0.97208769 | 0.98427785 |
| 0.99021155 | 0.1029198 | 0.58229986 | 0.76565751 | -0.6432716 | 0.02575447 | 0.11823642 |
| 0.30638319 | -0.6903492 | 0.04549803 | 0.20411958 | 0.23583862 | 0.60485545 | 0.74923018 |
| 0.23382239 | -0.1172816 | 0.51659177 | 0.71642203 | 0.14426991 | 0.57193531 | 0.72611375 |
| 0.22305848 | -0.3332047 | 0.11331247 | 0.33043208 | 0.06730349 | 0.81305834 | 0.89220458 |
| 0.93623502 | 0.08263021 | 0.61272723 | 0.78670568 | 0.46368782 | 0.05874682 | 0.18628897 |
| 0.37585765 | -0.1675447 | 0.38229757 | 0.62275061 | 0.41101713 | 0.13971949 | 0.31981118 |
| 0.21450286 | -0.3023176 | 0.11566194 | 0.3334249 | 0.00366013 | 0.98875938 | 0.99147625 |
| 0.65564467 | 0.1788362 | 0.549787 | 0.74117427 | -0.1355638 | 0.74741932 | 0.84645854 |
| 0.27219985 | -0.0972198 | 0.66123615 | 0.80425909 | -0.1304802 | 0.67740385 | 0.7967137 |
| 0.80326625 | 0.10666768 | 0.74288883 | 0.84827779 | 0.55973442 | 0.23453708 | 0.41467388 |
| 0.36632577 | -0.2206913 | 0.30483673 | 0.54406729 | 0.60829551 | 0.05652513 | 0.18268921 |
| 0.26729905 | -0.0270555 | 0.85599007 | 0.91865078 | 0.41963963 | 0.06171378 | 0.18959974 |
| 0.63800038 | -0.0495856 | 0.79489141 | 0.8771584 | 0.62567177 | 0.03273331 | 0.12901716 |
| 0.3235901 | 0.27645002 | 0.15229227 | 0.36991702 | 0.73940702 | 0.01236937 | 0.08125569 |
| 0.39434799 | 0.40216675 | 0.0383519 | 0.18416343 | -0.4701797 | 0.07982992 | 0.21791411 |
| 0.81360672 | 0.06739823 | 0.64004402 | 0.79469368 | 0.33396395 | 0.1158144 | 0.28158656 |
| 0.98995236 | 0.45057382 | 0.37843409 | 0.62189235 | 0.48259537 | 0.50221776 | 0.66089894 |
| 0.38805069 | 0.0277515 | 0.9064694 | 0.95167695 | -0.0519662 | 0.87639314 | 0.92444603 |
| 0.40914893 | -0.1276537 | 0.69965112 | 0.82822293 | 0.71014778 | 0.14330217 | 0.32178236 |
| 0.1758396 | 0.58791789 | 0.02226439 | 0.13210774 | 0.73771519 | 0.03889414 | 0.14549291 |
| 0.88293084 | 0.27192667 | 0.08898458 | 0.28690819 | 0.77820231 | 0.00232641 | 0.0341771 |
| 0.61802592 | -0.1461839 | 0.50438958 | 0.70666185 | -0.3280234 | 0.29515532 | 0.47333705 |
| 0.38020494 | -0.0625152 | 0.67626902 | 0.81292206 | -0.0194113 | 0.92673545 | 0.96123523 |
| 0.77403151 | -0.0455893 | 0.78848742 | 0.87415549 | 0.26200272 | 0.28492717 | 0.46136503 |
| 0.79620923 | -0.0230182 | 0.90147096 | 0.94938929 | 0.60285661 | 0.0346332 | 0.13518661 |
| 0.1758396 | -0.2366127 | 0.25951512 | 0.49918508 | 0.1081789 | 0.70984668 | 0.82267315 |
| 0.95369702 | 0.08333059 | 0.62477826 | 0.78793079 | 0.33022592 | 0.18259325 | 0.36956447 |
| 0.5286044 | -0.1503426 | 0.43986763 | 0.66752555 | 0.2242101 | 0.41588829 | 0.58776851 |
| 0.74291845 | -0.1219767 | 0.60277515 | 0.78056529 | 0.07110024 | 0.82948118 | 0.89723855 |
| 0.62393252 | 0.36080593 | 0.13040499 | 0.34533555 | 0.81707743 | 0.02194343 | 0.10835435 |

| | | | | | | |
|---|---|---|---|---|---|---|
| 0.38805069 | 0.16872259 | 0.46997636 | 0.68613001 | 0.36351604 | 0.27675342 | 0.45347698 |
| 0.59603045 | -0.0692983 | 0.6648416 | 0.80744815 | 0.57746792 | 0.02039692 | 0.10365227 |
| 0.20206115 | -0.7218045 | 0.02064369 | 0.12960326 | 0.23813623 | 0.55358672 | 0.71339405 |
| 0.40914893 | 0.47312315 | 0.05754262 | 0.22604064 | 0.44463135 | 0.19117386 | 0.38046423 |
| 0.93623502 | 0.08524504 | 0.63497685 | 0.79107131 | 0.1989881 | 0.43642368 | 0.60364702 |
| 0.49663882 | -0.0038379 | 0.98460844 | 0.99185709 | 0.2856421 | 0.31865355 | 0.49994577 |
| 0.98908107 | 0.1183082 | 0.54428872 | 0.73739445 | 0.35628503 | 0.20667551 | 0.39571993 |
| 0.1758396 | 0.39076647 | 0.04008087 | 0.19021037 | 0.30453076 | 0.2335056 | 0.41467388 |
| 0.69955402 | 0.30453882 | 0.21952051 | 0.45506071 | 0.42033243 | 0.23022318 | 0.41451545 |
| 0.42922264 | -0.1380004 | 0.61615998 | 0.78670568 | -0.2768942 | 0.47891761 | 0.64279972 |
| 0.14703314 | -0.5623855 | 0.02322229 | 0.1342641 | -0.0777566 | 0.80738961 | 0.8900011 |
| 0.87131025 | 0.38840234 | 0.08006806 | 0.27395812 | 0.47737902 | 0.12338974 | 0.29540308 |
| 0.91344014 | 0.11934492 | 0.44599669 | 0.67130408 | 0.26199711 | 0.24344736 | 0.42484983 |
| 0.87443464 | 0.29179725 | 0.20300682 | 0.43630523 | 0.46059313 | 0.1581651 | 0.34354141 |
| 0.86400189 | 0.64564126 | 0.05393068 | 0.21651549 | 0.76745366 | 0.06025294 | 0.18797057 |
| 0.95369702 | 0.06052649 | 0.7211309 | 0.83859556 | -0.182385 | 0.45052942 | 0.61779388 |
| 0.28717843 | -0.2242515 | 0.23240139 | 0.46536961 | 0.2204708 | 0.40029774 | 0.57964261 |
| 0.39913114 | 0.37107586 | 0.04337961 | 0.19785924 | 0.76751346 | 0.00576741 | 0.053564 |
| 0.2426034 | -0.2894081 | 0.18888439 | 0.42150243 | 0.5648938 | 0.07717655 | 0.21577388 |
| 0.30093686 | -0.2649145 | 0.25663319 | 0.49809893 | 0.26089896 | 0.42411176 | 0.59596921 |
| 0.49781409 | 0.28980891 | 0.18667964 | 0.42150243 | 1.19979221 | 0.00111705 | 0.02259002 |
| 0.38661287 | -0.3848829 | 0.15747151 | 0.37748779 | 0.25259051 | 0.49988004 | 0.66037314 |
| 0.84472005 | -0.1475294 | 0.43943699 | 0.66752555 | 0.56089687 | 0.05018941 | 0.17301383 |
| 0.28717843 | 0.45237861 | 0.05885963 | 0.23011311 | 0.90610527 | 0.01125494 | 0.07706774 |
| 0.38661287 | 0.4562223 | 0.12266588 | 0.3390865 | 0.66894158 | 0.11055994 | 0.27486902 |
| 0.34559993 | 0.52406062 | 0.00589006 | 0.08337475 | 0.67788615 | 0.01023742 | 0.07255997 |
| 0.74702619 | 0.17649377 | 0.27125193 | 0.51086349 | 0.26311382 | 0.24725391 | 0.42688283 |
| 0.38661287 | -0.350542 | 0.13608521 | 0.35389208 | -0.0023504 | 0.9941294 | 0.9941294 |
| 0.20206115 | -0.3399119 | 0.08654413 | 0.28421091 | 0.6375309 | 0.02820327 | 0.12317967 |
| 0.46341205 | -0.8026571 | 0.23228586 | 0.46536961 | -0.379916 | 0.68253861 | 0.7996661 |
| 0.4528735 | -0.2083923 | 0.36898582 | 0.61506267 | 0.36497975 | 0.26947028 | 0.44923576 |
| 0.20983084 | -0.121471 | 0.64078833 | 0.79469368 | 0.42956081 | 0.25264323 | 0.43340919 |
| 0.75234336 | 0.12273731 | 0.41355902 | 0.65169281 | 0.50094625 | 0.02875488 | 0.12391271 |
| 0.32120969 | 0.5243167 | 0.01289235 | 0.11325581 | -0.214163 | 0.42573615 | 0.59717373 |
| 0.32958395 | -0.2237217 | 0.2900033 | 0.52328068 | 0.57540771 | 0.06502059 | 0.19363367 |
| 0.38020494 | -0.5105327 | 0.09082564 | 0.29014726 | 0.30424239 | 0.4573008 | 0.62438379 |
| 0.28717843 | 0.26178765 | 0.119883 | 0.33670211 | 0.34657228 | 0.14336838 | 0.32178236 |
| 0.29060966 | -0.13615 | 0.4803158 | 0.69061169 | 0.57232634 | 0.04869809 | 0.16972287 |
| 0.28717843 | -0.3106059 | 0.25820323 | 0.49918508 | -0.224748 | 0.55630564 | 0.71504275 |
| 0.82200579 | -6.56E-04 | 0.99631991 | 0.99783068 | 0.48461245 | 0.02758995 | 0.1213283 |
| 0.54189492 | 0.36150923 | 0.1208817 | 0.33670211 | 0.20645792 | 0.51605558 | 0.67037445 |
| 0.69955402 | 0.26421728 | 0.16444906 | 0.38576418 | 0.46039322 | 0.0920595 | 0.24229341 |

| | | | | | | |
|---|---|---|---|---|---|---|
| 0.88293084 | 0.49152546 | 0.02144888 | 0.13044095 | 0.91858748 | 0.0040945 | 0.04659655 |
| 0.98908107 | 0.01142762 | 0.9719125 | 0.98511131 | 0.55646161 | 0.23715306 | 0.4165645 |
| 0.67939616 | 0.08247892 | 0.69589946 | 0.82682121 | 0.34174526 | 0.26166245 | 0.44230808 |
| 0.52070698 | 0.05014747 | 0.73094463 | 0.83930844 | 0.52225073 | 0.02138828 | 0.10733992 |
| 0.2465282 | -0.3456298 | 0.05101973 | 0.21402119 | 0.38900682 | 0.11176517 | 0.27616591 |
| 0.3235901 | -0.1134161 | 0.53761576 | 0.7316104 | 0.40799538 | 0.12994823 | 0.30522724 |
| 0.79165133 | 0.22131586 | 0.22028419 | 0.45506071 | 1.04695061 | 7.13E-04 | 0.01759982 |
| 0.78564209 | -0.4546053 | 0.1968109 | 0.43088465 | 0.97386814 | 0.05905923 | 0.18628897 |
| 0.88293084 | 0.31098201 | 0.15189389 | 0.36991702 | 0.70004452 | 0.02996537 | 0.12545085 |
| 0.26124802 | -0.2894097 | 0.23081751 | 0.46536961 | 0.2658237 | 0.4296473 | 0.59854313 |
| 0.32120969 | 0.61324267 | 0.00488579 | 0.07612725 | 1.29856269 | 1.89E-04 | 0.00810113 |
| 0.6526185 | 0.33913233 | 0.09412015 | 0.29720247 | 0.3339769 | 0.23188241 | 0.41451545 |
| 0.36133594 | -0.4559547 | 0.01802438 | 0.12181362 | 0.01495448 | 0.95149269 | 0.97580776 |
| 0.69955402 | 0.17353104 | 0.47269859 | 0.68613001 | -0.0215575 | 0.94923894 | 0.97580776 |
| 0.14725224 | 0.57101254 | 0.00746883 | 0.08504841 | 0.49461701 | 0.0769141 | 0.21577388 |
| 0.79620923 | -0.2982347 | 0.48487855 | 0.69386761 | 0.08030503 | 0.89330809 | 0.93617761 |
| 0.56072247 | 0.17728572 | 0.28462979 | 0.52050483 | 0.86467055 | 0.00175273 | 0.02776871 |
| 0.53314753 | -0.0926467 | 0.56616509 | 0.75092333 | 0.66261843 | 0.00991809 | 0.07155191 |
| 0.42922264 | -0.1527976 | 0.4174573 | 0.65657557 | -0.12839 | 0.62736302 | 0.76003319 |
| 0.42190279 | -0.0154914 | 0.90999751 | 0.95408514 | 0.37551288 | 0.06819504 | 0.20110072 |
| 0.91935516 | 0.31174542 | 0.1688052 | 0.3926036 | 0.51252448 | 0.1136779 | 0.28003581 |
| 0.99021155 | 0.19601124 | 0.29765024 | 0.53239835 | 0.61446009 | 0.03074682 | 0.1255811 |
| 0.42922264 | 0.49057346 | 0.02625277 | 0.14662261 | 1.10620397 | 0.00138926 | 0.0245348 |
| 0.35316416 | 0.31050223 | 0.11815949 | 0.33451359 | 0.9656326 | 0.00250373 | 0.0361253 |
| 0.30638319 | -0.0884555 | 0.58704542 | 0.76868308 | 0.37801019 | 0.11480781 | 0.28158656 |
| 0.8570768 | 0.05447331 | 0.76673688 | 0.85996035 | 0.13237178 | 0.61133607 | 0.75298711 |
| 0.54327827 | -0.2211106 | 0.38227206 | 0.62275061 | -0.1024596 | 0.77196036 | 0.86391131 |
| 0.67854292 | -0.2318089 | 0.38094542 | 0.62275061 | 0.32096508 | 0.39072882 | 0.56782173 |
| 0.75234336 | -0.129182 | 0.53459999 | 0.72908071 | 0.43257522 | 0.1536009 | 0.33634019 |
| 0.32120969 | 0.3461305 | 0.13774291 | 0.35561926 | 0.35282365 | 0.27577418 | 0.45289743 |
| 0.32047815 | 0.36352607 | 0.24732714 | 0.48346567 | 0.41770168 | 0.34338932 | 0.52252085 |
| 0.99021155 | -0.2033071 | 0.37124104 | 0.61506267 | 0.60027018 | 0.07400255 | 0.20834169 |
| 0.98908107 | -0.0640064 | 0.7268955 | 0.83865842 | 0.58934615 | 0.03562429 | 0.13838666 |
| 0.22305848 | -0.5457517 | 0.04063445 | 0.1906336 | -0.1829597 | 0.60195825 | 0.74713098 |
| 0.39350111 | -0.1739679 | 0.34063983 | 0.59037865 | -0.2389316 | 0.35431245 | 0.53015641 |
| 0.92002854 | 0.06191447 | 0.689138 | 0.82116444 | 0.04022802 | 0.85385585 | 0.91138614 |
| 0.54574095 | 0.31679205 | 0.20703929 | 0.43922288 | 0.34890286 | 0.32046418 | 0.50181213 |
| 0.27748741 | -0.0502334 | 0.78675208 | 0.87415549 | -1.2467037 | 2.46E-04 | 0.00946159 |
| 0.14725224 | -0.4956373 | 0.01957493 | 0.12654344 | -0.0774328 | 0.77578269 | 0.86698813 |
| 0.36144521 | 0.30231803 | 0.23557504 | 0.46678674 | -1.8928137 | 7.41E-05 | 0.00544275 |
| 0.82413495 | -0.001192 | 0.99431802 | 0.99783068 | 0.56307736 | 0.02930375 | 0.12396558 |
| 0.59619098 | -0.0594078 | 0.79377245 | 0.87710253 | -0.2107727 | 0.51463691 | 0.67037445 |

| | | | | | | |
|---|---|---|---|---|---|---|
| 0.60929237 | -0.159087 | 0.42769238 | 0.66002903 | -0.1257262 | 0.65511037 | 0.77957169 |
| 0.42803946 | -0.3487028 | 0.11158743 | 0.32738975 | 0.51727763 | 0.09653514 | 0.24920254 |
| 0.78329146 | -0.4792685 | 0.06011246 | 0.23204241 | -0.488677 | 0.16277469 | 0.34611039 |
| 0.39913114 | -0.3665835 | 0.15770805 | 0.37748779 | 0.26125193 | 0.46468621 | 0.62997728 |
| 0.38661287 | 0.11835846 | 0.53074377 | 0.72502601 | 0.19245419 | 0.47204912 | 0.63712067 |
| 0.17747854 | -0.2671216 | 0.18789399 | 0.42150243 | 0.14301567 | 0.60863763 | 0.75195598 |
| 0.59501127 | 0.21410382 | 0.28964373 | 0.52328068 | 0.53110131 | 0.07380172 | 0.20834169 |
| 0.53099293 | 0.34559538 | 0.06524275 | 0.24367421 | 0.73780578 | 0.00908846 | 0.06915073 |
| 0.8135094 | -0.078059 | 0.69718898 | 0.82715629 | 0.06670157 | 0.81380542 | 0.89220458 |
| 0.27219985 | -0.2316687 | 0.19172195 | 0.42426879 | -0.2674175 | 0.28182098 | 0.45725171 |
| 0.65611599 | 0.0106437 | 0.96839357 | 0.98397416 | -0.7292016 | 0.0705702 | 0.20416618 |
| 0.59619098 | -0.0529261 | 0.82154742 | 0.89573763 | 0.41409748 | 0.22396546 | 0.41308969 |
| 0.21110874 | 0.34546447 | 0.06711741 | 0.24572073 | 0.14248179 | 0.57244364 | 0.72611375 |
| 0.99845565 | 0.05275108 | 0.82009291 | 0.89543159 | -0.270697 | 0.41441225 | 0.58744755 |
| 0.22305848 | 0.38528127 | 0.08138666 | 0.27494421 | 0.27326549 | 0.36275751 | 0.5399595 |
| 0.14725224 | -0.4222087 | 0.03467952 | 0.17255689 | 0.30931454 | 0.24637052 | 0.42626849 |
| 0.18470305 | 0.40861177 | 0.03459901 | 0.17255689 | 0.63070784 | 0.02288496 | 0.10929234 |
| 0.38661287 | -0.1707911 | 0.34240225 | 0.59037865 | -1.9086077 | 1.67E-06 | 2.69E-04 |
| 0.74782838 | -0.4347339 | 0.1209831 | 0.33670211 | -0.9566199 | 0.02212682 | 0.10835435 |
| 0.45831711 | 0.10673368 | 0.70637465 | 0.83204245 | 0.05396961 | 0.89261364 | 0.93617761 |
| 0.92692476 | -0.3327823 | 0.28713204 | 0.5215385 | 0.89904449 | 0.05268759 | 0.17812373 |
| 0.38661287 | -0.3528522 | 0.14323339 | 0.36277763 | -0.1045473 | 0.75027265 | 0.8466764 |
| 0.69955402 | 0.12162373 | 0.51261798 | 0.71350105 | 1.15018999 | 4.80E-04 | 0.01384725 |
| 0.5867485 | -0.2222954 | 0.5883053 | 0.76910613 | -0.1947156 | 0.73660656 | 0.83710001 |
| 0.2426034 | -0.5585335 | 0.03054421 | 0.16192 | 0.28123031 | 0.4070599 | 0.58213169 |
| 0.26620493 | 0.4106677 | 0.10380235 | 0.31331518 | 0.43470911 | 0.21405626 | 0.40257664 |
| 0.49755154 | -0.0155404 | 0.95645925 | 0.98034088 | -0.610035 | 0.14479175 | 0.32318159 |
| 0.87008778 | 0.09609129 | 0.63063773 | 0.78970053 | 0.26783287 | 0.34892289 | 0.52500875 |
| 0.38661287 | 0.23600494 | 0.27821972 | 0.51561713 | 0.16265402 | 0.59128102 | 0.74185569 |
| 0.81563667 | -0.2963458 | 0.10172227 | 0.30931106 | 0.20396889 | 0.40875122 | 0.58248851 |
| 0.26377575 | 0.5295365 | 0.07748382 | 0.26728661 | 0.57249299 | 0.16753171 | 0.35251464 |
| 0.30638319 | -0.1965405 | 0.28049624 | 0.51633949 | 0.49718722 | 0.06407778 | 0.19318973 |
| 0.58514735 | -0.1973363 | 0.33534934 | 0.58704011 | 0.13389096 | 0.63967239 | 0.77142581 |
| 0.30638319 | 0.40360754 | 0.0859052 | 0.28324566 | 0.60249147 | 0.07143333 | 0.20467422 |
| 0.5286044 | -0.0935371 | 0.70010562 | 0.82822293 | -0.408656 | 0.2443313 | 0.42496131 |
| 0.98150616 | -0.2253548 | 0.26665732 | 0.5081664 | -0.523531 | 0.07791475 | 0.21634061 |
| 0.32773555 | -0.4583389 | 0.04093996 | 0.19097563 | -0.0543825 | 0.85317584 | 0.91138614 |
| 0.92692476 | -0.1791921 | 0.23652044 | 0.46678674 | 0.19094892 | 0.36731 | 0.54257126 |
| 0.27219985 | 0.39470138 | 0.06913702 | 0.24572073 | 0.6828471 | 0.03029532 | 0.12553136 |
| 0.6526185 | -0.0569609 | 0.72911866 | 0.83873656 | 0.47720704 | 0.05467391 | 0.18097774 |
| 0.20151958 | -0.4654482 | 0.02478562 | 0.14188246 | 0.19230649 | 0.47574643 | 0.64067185 |
| 0.51019476 | 0.10324009 | 0.65492263 | 0.8014843 | 0.5602078 | 0.10124094 | 0.25643473 |

| | | | | | | |
|---|---|---|---|---|---|---|
| 0.58598326 | -0.0101106 | 0.95476335 | 0.98034088 | 0.23245418 | 0.36353709 | 0.5399595 |
| 0.2426034 | -0.3720243 | 0.08120952 | 0.27494421 | 0.18883584 | 0.51153141 | 0.66879835 |
| 0.37933328 | -0.2129841 | 0.21456319 | 0.44709742 | 0.29237967 | 0.22758303 | 0.41308969 |
| 0.40888927 | 0.21748528 | 0.1869674 | 0.42150243 | 0.48631369 | 0.04541001 | 0.16235084 |
| 0.76018942 | 0.11159694 | 0.79072779 | 0.87491579 | 0.78170441 | 0.20134938 | 0.39108246 |
| 0.39340052 | -0.1544198 | 0.48972212 | 0.69560876 | 0.04450839 | 0.88712495 | 0.93332938 |
| 0.71162576 | 0.2547185 | 0.45975942 | 0.67767053 | -0.3875038 | 0.42718615 | 0.59717373 |
| 0.99021155 | -0.3152332 | 0.39963504 | 0.64207508 | -0.5887151 | 0.27040558 | 0.44923576 |
| 0.25846624 | 0.31239721 | 0.13923642 | 0.35722845 | 0.40550595 | 0.17217829 | 0.35399507 |
| 0.75857894 | 0.01263116 | 0.94102065 | 0.97291136 | 0.18666114 | 0.44420987 | 0.61041084 |
| 0.77403151 | -0.114616 | 0.59293992 | 0.77147967 | 0.10370015 | 0.73156853 | 0.83554615 |
| 0.39913114 | -0.1734376 | 0.34183267 | 0.59037865 | 0.21405709 | 0.40500369 | 0.58141958 |
| 0.83836546 | 0.17116932 | 0.44116122 | 0.66825344 | -0.3324619 | 0.29454171 | 0.47333705 |
| 0.59603045 | -0.2161258 | 0.4255442 | 0.65832537 | 0.65460185 | 0.10031176 | 0.25596503 |
| 0.38020494 | -0.2512514 | 0.1722264 | 0.39830387 | 0.63452869 | 0.02208812 | 0.10835435 |
| 0.6813551 | -0.1512534 | 0.44216635 | 0.66854249 | 0.12211978 | 0.65856055 | 0.7802301 |
| 0.20206115 | 0.1628673 | 0.42578657 | 0.65832537 | 0.26306072 | 0.36465988 | 0.54063336 |
| 0.91935516 | -0.2136708 | 0.22185376 | 0.45535484 | 0.30014898 | 0.22480977 | 0.41308969 |
| 0.23382239 | 0.4107882 | 0.14714367 | 0.36277763 | NA | NA | NA |
| 0.35382662 | -0.5337402 | 0.08041889 | 0.27395812 | 0.33356814 | 0.41945065 | 0.59147666 |
| 0.42922264 | -0.1968448 | 0.22990966 | 0.46491584 | 0.6295021 | 0.01296649 | 0.0831502 |
| 0.98388605 | -0.3263365 | 0.28350336 | 0.52050483 | 0.24817496 | 0.55840591 | 0.71504275 |
| 0.2426034 | 0.0941861 | 0.62294658 | 0.78793079 | 0.34077832 | 0.21895208 | 0.4057644 |
| 0.38805069 | 0.77584117 | 0.01305448 | 0.11325581 | 0.13481527 | 0.73290206 | 0.83554615 |
| 0.92692476 | 0.11449734 | 0.62063874 | 0.78793079 | 0.48285789 | 0.15313339 | 0.33622767 |
| 0.59501127 | -0.0147828 | 0.95610343 | 0.98034088 | 0.46200734 | 0.23564035 | 0.41495416 |
| 0.20206115 | -0.5513367 | 0.01368572 | 0.11465285 | 0.12615318 | 0.65635219 | 0.77990084 |
| 0.48008373 | 0.17572764 | 0.37928609 | 0.62189235 | 0.89267748 | 0.00555604 | 0.05220092 |
| 0.23670201 | -0.4417351 | 0.01547712 | 0.11706618 | 0.14727423 | 0.52756339 | 0.68312696 |
| 0.45975183 | -0.4479991 | 0.03132826 | 0.16382485 | -1.1697068 | 5.66E-04 | 0.01577633 |
| 0.16484455 | -0.4802696 | 0.02099695 | 0.12961275 | 0.45446497 | 0.10432338 | 0.26259592 |
| 0.62986383 | -0.6051134 | 0.12163806 | 0.33738124 | NA | NA | NA |
| 0.78329146 | -0.0313325 | 0.85005321 | 0.91587098 | 0.28975201 | 0.22822027 | 0.41308969 |
| 0.34139791 | -0.5462046 | 0.08443338 | 0.27951535 | -0.1839217 | 0.66550249 | 0.78669239 |
| 0.48257912 | 0.37430355 | 0.06048365 | 0.23204241 | 1.27057517 | 2.22E-04 | 0.00897964 |
| 0.54720161 | 0.27597161 | 0.31083924 | 0.55345737 | 0.62959181 | 0.11171435 | 0.27616591 |
| 0.20983084 | -0.5096774 | 0.01898091 | 0.12414327 | 0.08280397 | 0.76577118 | 0.85936544 |
| 0.71847643 | -0.2885171 | 0.36329421 | 0.61119783 | -0.5149448 | 0.25527482 | 0.43515202 |
| 0.5196969 | -0.1947422 | 0.25920846 | 0.49918508 | -0.6326931 | 0.01700247 | 0.09474482 |
| 0.27219985 | -0.4955952 | 0.0366022 | 0.17994254 | 0.06252212 | 0.83999268 | 0.90351864 |
| 0.82069552 | 0.12766079 | 0.52421486 | 0.71849817 | -0.3189925 | 0.26759728 | 0.44765757 |
| 0.91635584 | 0.079799 | 0.86554611 | 0.92361698 | -0.8520849 | 0.2135813 | 0.40257664 |

| | | | | | | |
|---|---|---|---|---|---|---|
| 0.21683252 | -0.4350196 | 0.02778989 | 0.15210335 | 0.08736542 | 0.73317106 | 0.83554615 |
| 0.71847643 | -0.2153546 | 0.22394253 | 0.4584958 | 0.37299352 | 0.14133691 | 0.32078714 |
| 0.204189 | -0.5140369 | 0.02572048 | 0.14463364 | 0.14522831 | 0.6267247 | 0.76003319 |
| 0.9276115 | -0.0583238 | 0.7415195 | 0.84789347 | -0.1711623 | 0.49642515 | 0.65993059 |
| 0.73880874 | -0.3665286 | 0.18203128 | 0.4162888 | 1.10102311 | 0.00973826 | 0.0710116 |
| 0.99845565 | -0.24813 | 0.34449802 | 0.59170058 | -0.0080869 | 0.98233722 | 0.98845389 |
| 0.54989578 | 0.22930603 | 0.31372923 | 0.55631036 | 0.07298417 | 0.81754787 | 0.89383192 |
| 0.99021155 | -0.222033 | 0.35245575 | 0.5991018 | 0.11961029 | 0.7196502 | 0.83021751 |
| 0.32265122 | 0.30613214 | 0.16304037 | 0.38576418 | 0.5508685 | 0.08188718 | 0.22202967 |
| 0.6714611 | -0.1546852 | 0.71147156 | 0.83552843 | -1.0164666 | 0.10090481 | 0.25638706 |
| 0.64536636 | 0.08749875 | 0.64461635 | 0.79703317 | 1.02896316 | 0.00148075 | 0.02492591 |
| 0.20983084 | -0.0221553 | 0.90197764 | 0.94938929 | 0.50427123 | 0.06273796 | 0.19128335 |
| 0.5286044 | -0.9145884 | 0.13504901 | 0.35389208 | -0.6878352 | 0.41349379 | 0.58717572 |
| 0.20206115 | -0.4914385 | 0.01183458 | 0.10917072 | -0.0309066 | 0.89982027 | 0.93852637 |
| 0.49045556 | -0.0092229 | 0.9605916 | 0.98090261 | 0.61097525 | 0.03322552 | 0.13032145 |
| 0.20206115 | 0.85196379 | 0.04610596 | 0.20572281 | 0.75984176 | 0.18968905 | 0.37929606 |
| 0.503909 | -0.3531536 | 0.0916325 | 0.29159024 | 0.49454398 | 0.09454693 | 0.2448523 |
| 0.5867485 | 0.07833025 | 0.7076087 | 0.83230193 | -0.0738183 | 0.80236482 | 0.88734156 |
| 0.99845565 | -0.1058136 | 0.51441076 | 0.71460446 | 0.01210486 | 0.95761879 | 0.97740909 |
| 0.71847643 | 0.11934691 | 0.52398834 | 0.71849817 | 0.01634361 | 0.95044311 | 0.97580776 |
| 0.33617013 | -0.4450137 | 0.04395637 | 0.19902766 | 0.20792952 | 0.47735555 | 0.6417692 |
| 0.21374185 | -0.5788102 | 0.00828531 | 0.08720817 | 0.29217326 | 0.29277877 | 0.47218612 |
| 0.70133826 | -0.0059526 | 0.98248468 | 0.99185709 | -0.1929117 | 0.61648759 | 0.75587553 |
| 0.17747854 | -0.5006069 | 0.00631432 | 0.08498447 | 0.19060665 | 0.40381966 | 0.58141958 |
| 0.38661287 | 0.33361166 | 0.18995847 | 0.42150243 | 0.22468395 | 0.52268747 | 0.67789964 |
| 0.42922264 | -0.3648796 | 0.50664405 | 0.70740606 | -0.2827992 | 0.71439836 | 0.82579953 |
| 0.1637222 | -0.8820331 | 0.00782384 | 0.08544939 | 0.50890688 | 0.22779097 | 0.41308969 |
| 0.1637222 | 0.79354116 | 0.04803325 | 0.21088396 | 0.69542962 | 0.20129592 | 0.39108246 |
| 0.92927511 | -0.3720552 | 0.47827837 | 0.68914438 | -0.3482748 | 0.63712754 | 0.76950531 |
| 0.44367699 | 0.20353522 | 0.3866253 | 0.62607372 | 1.01348568 | 0.00699632 | 0.05888567 |
| 0.59619098 | -0.1277103 | 0.43418049 | 0.66597079 | 0.39786753 | 0.09753996 | 0.25099456 |
| 0.36806776 | 0.90092891 | 0.00491468 | 0.07612725 | NA | NA | NA |
| 0.19988415 | 0.20674757 | 0.35577901 | 0.60225684 | 0.21118819 | 0.50168033 | 0.66089894 |
| 0.22305848 | -0.0942577 | 0.7199966 | 0.83859556 | 0.45862306 | 0.22852858 | 0.41308969 |
| 0.69942883 | -0.5231066 | 0.04636552 | 0.20576266 | -1.860028 | 7.41E-05 | 0.00544275 |
| 0.89024854 | -0.2741815 | 0.2103956 | 0.44075808 | -0.1285375 | 0.6702853 | 0.7906431 |
| 0.32047815 | -0.5187993 | 0.06114091 | 0.23347295 | -0.2004981 | 0.58732338 | 0.7384145 |
| 0.95071462 | -0.3750428 | 0.39281771 | 0.63360184 | -0.152291 | 0.80387874 | 0.88734156 |
| 0.40979239 | 0.2623081 | 0.26929114 | 0.51064699 | 0.00430205 | 0.98955067 | 0.99147625 |
| 0.38661287 | -0.3594594 | 0.04317637 | 0.19785924 | -0.1958168 | 0.40696298 | 0.58213169 |
| 0.21928298 | 0.19116879 | 0.44371802 | 0.66965532 | 0.35700364 | 0.31550313 | 0.49596601 |
| 0.69955402 | -0.0538669 | 0.9013282 | 0.94938929 | NA | NA | NA |

| | | | | | | |
|---|---|---|---|---|---|---|
| 0.72150431 | -0.5423953 | 0.14124282 | 0.36069402 | 1.23497375 | 0.02478362 | 0.11508714 |
| 0.30914171 | -0.3253492 | 0.07273795 | 0.25520451 | -0.4171735 | 0.10042192 | 0.25596503 |
| 0.85757696 | -0.2789541 | 0.23641253 | 0.46678674 | -0.4234378 | 0.20544552 | 0.39523805 |
| 0.5611997 | -0.1141644 | 0.58602286 | 0.76856991 | 0.28453214 | 0.34244025 | 0.52252085 |
| 0.97599494 | -0.1932908 | 0.69055619 | 0.82166178 | NA | NA | NA |
| 0.92692476 | -0.0891811 | 0.67727002 | 0.81292206 | -0.3627334 | 0.24147234 | 0.42231525 |
| 0.48257912 | 0.15499401 | 0.50023552 | 0.70324205 | 0.88448319 | 0.0141759 | 0.08721575 |
| 0.23837259 | -0.6249615 | 0.01317533 | 0.11325581 | -0.1213306 | 0.70360055 | 0.81799891 |
| 0.87710098 | -0.0590254 | 0.75637932 | 0.85300471 | -0.2277093 | 0.40251907 | 0.58141958 |
| 0.71847643 | -0.1246725 | 0.47702588 | 0.68914438 | -0.1140843 | 0.64369496 | 0.77166992 |
| 0.48714097 | -0.1828699 | 0.35231153 | 0.5991018 | -0.0196084 | 0.94290184 | 0.97300727 |
| 0.77403151 | -0.1885962 | 0.3710364 | 0.61506267 | -0.6571749 | 0.03871428 | 0.14549291 |
| 0.69955402 | 0.10028132 | 0.82611796 | 0.89929578 | NA | NA | NA |
| 0.99689095 | -0.0424135 | 0.81942827 | 0.89543159 | -0.6924472 | 0.01750337 | 0.09555675 |
| 0.65746392 | -0.0911039 | 0.72079118 | 0.83859556 | 0.62225765 | 0.09961531 | 0.25552116 |
| 0.75537286 | -0.2282938 | 0.22743015 | 0.4621786 | -0.2377452 | 0.36847036 | 0.54329207 |
| 0.6813551 | -0.2545177 | 0.34692976 | 0.59463326 | 0.21102314 | 0.57741197 | 0.73012342 |
| 0.6090444 | -0.1322805 | 0.62360811 | 0.78793079 | -0.2579984 | 0.50018361 | 0.66037314 |
| 0.73378421 | -0.264562 | 0.12622374 | 0.34201219 | -0.1230154 | 0.6016432 | 0.74713098 |
| 0.5286044 | 0.02282607 | 0.94290949 | 0.97291136 | -0.5253154 | 0.25490558 | 0.43515202 |
| 0.14703314 | -0.6017945 | 0.01621662 | 0.11706618 | -0.6732503 | 0.04870501 | 0.16972287 |
| 0.69955402 | -0.4243615 | 0.03274295 | 0.16815196 | -0.271794 | 0.30223062 | 0.48166142 |
| 0.18470305 | 0.17965786 | 0.42443369 | 0.65832537 | 0.07856853 | 0.80291589 | 0.88734156 |
| 0.32047815 | 0.0687022 | 0.65892378 | 0.80425909 | -0.1250543 | 0.570826 | 0.72611375 |
| 0.7842404 | -0.2448858 | 0.64020539 | 0.79469368 | 0.05470387 | 0.94091975 | 0.97220353 |
| 0.34139791 | -0.7021146 | 0.08837913 | 0.28679553 | -0.5212658 | 0.35306957 | 0.52927683 |
| 0.99021155 | -0.1869671 | 0.45520202 | 0.67551175 | 0.67112805 | 0.07122583 | 0.20467422 |
| 0.55565256 | -0.1262846 | 0.72942497 | 0.83873656 | 0.49646479 | 0.3430751 | 0.52252085 |
| 0.75530676 | -0.1482335 | 0.54246426 | 0.73613745 | -0.492711 | 0.1640951 | 0.3480022 |
| 0.9659358 | -0.2198774 | 0.56462528 | 0.75092333 | 0.97549593 | 0.0850655 | 0.22910975 |
| 0.68650309 | -0.3395323 | 0.25308848 | 0.49291016 | -0.4658879 | 0.2668427 | 0.44765757 |
| 0.91344014 | -0.072855 | 0.82816869 | 0.89937368 | 0.24871686 | 0.60178323 | 0.74713098 |
| 0.36144521 | -0.7041894 | 0.08831829 | 0.28679553 | 0.27517529 | 0.62104093 | 0.75915442 |
| 0.1446959 | -0.8092514 | 0.00347805 | 0.06345508 | -0.7436073 | 0.03921391 | 0.14601309 |
| 0.32171019 | -0.8969663 | 0.02194951 | 0.13153685 | -0.3996945 | 0.43146677 | 0.59976206 |
| 0.75234336 | -0.1603486 | 0.40765215 | 0.64653409 | -0.7941275 | 0.00957841 | 0.07100325 |
| 0.36914225 | 0.04031065 | 0.8806614 | 0.93413825 | 0.15500709 | 0.68388717 | 0.79968282 |
| 0.48257912 | -0.1792535 | 0.45108999 | 0.67475057 | 0.51194201 | 0.13948059 | 0.31981118 |
| 0.55354732 | -0.4801771 | 0.49139332 | 0.69677705 | -1.1372259 | 0.25619468 | 0.43580063 |
| 0.25639803 | -0.0172347 | 0.94721434 | 0.974515 | -0.1482074 | 0.68809566 | 0.8022818 |
| 0.48257912 | -0.5907795 | 0.00463462 | 0.07610054 | -0.5533799 | 0.04316078 | 0.15780048 |
| 0.92692476 | -0.2744314 | 0.21034035 | 0.44075808 | -0.6054927 | 0.05955413 | 0.18651062 |

| | | | | | | |
|---|---|---|---|---|---|---|
| 0.73378421 | -0.089741 | 0.70481375 | 0.83139668 | -0.4252502 | 0.21577624 | 0.40358148 |
| 0.38661287 | -0.7944266 | 0.01512825 | 0.11706618 | -0.7724183 | 0.07826812 | 0.21657753 |
| 0.56884759 | -0.2582879 | 0.4061481 | 0.64653409 | 0.01195255 | 0.97804388 | 0.98659108 |
| 0.69955402 | 0.05141259 | 0.8333854 | 0.90265095 | -0.1858164 | 0.59246049 | 0.74218307 |
| 0.81051683 | -0.0578167 | 0.82010449 | 0.89543159 | -1.1814934 | 0.00465299 | 0.0489601 |
| 0.69955402 | 0.01766747 | 0.96012749 | 0.98090261 | -0.5535015 | 0.27823648 | 0.45355218 |
| 0.23382239 | 0.03971492 | 0.88564859 | 0.93821612 | 0.37789756 | 0.34095586 | 0.52252085 |
| 0.13458393 | 0.43154692 | 0.10821333 | 0.32073338 | 0.48493552 | 0.19445025 | 0.38230102 |
| 0.5286044 | 0.10750878 | 0.80769048 | 0.8888926 | 0.25881067 | 0.67923429 | 0.79770539 |
| 0.54189492 | 0.09971535 | 0.723826 | 0.83865842 | 0.19608851 | 0.62390832 | 0.76003319 |
| 0.41017441 | -0.6978369 | 0.00630009 | 0.08498447 | -0.6953225 | 0.0404819 | 0.15004299 |
| 0.76206684 | -0.4570465 | 0.11982325 | 0.33670211 | -0.9361582 | 0.03074518 | 0.1255811 |
| 0.41017441 | -0.1083623 | 0.67838857 | 0.81307593 | -0.183856 | 0.61931879 | 0.75819634 |
| 0.81914447 | -0.4760422 | 0.09536185 | 0.29996966 | -1.0297127 | 0.01596665 | 0.09318167 |
| 0.38872772 | -0.1895855 | 0.45389889 | 0.67551175 | -0.6400874 | 0.08687985 | 0.23244674 |
| 0.77733525 | -0.5940157 | 0.02094864 | 0.12961275 | -0.5045898 | 0.14115702 | 0.32078714 |
| 0.71847643 | -0.1920648 | 0.45798341 | 0.67748537 | -0.6573439 | 0.08564394 | 0.22990134 |
| 0.22305848 | -1.3813714 | 0.00872308 | 0.09065383 | -1.2977866 | 0.06297199 | 0.19128335 |
| 0.42922264 | 0.13247481 | 0.70271866 | 0.83011801 | NA | NA | NA |
| 0.71847643 | -0.36927 | 0.12809107 | 0.34452754 | -0.5099025 | 0.13662009 | 0.31748996 |
| 0.40927024 | -0.5368467 | 0.01824979 | 0.12181362 | -0.6540295 | 0.03736261 | 0.14299789 |
| 0.20983084 | -0.6382814 | 6.53E-04 | 0.03349417 | -0.5449455 | 0.02004195 | 0.10304685 |
| 0.99845565 | -0.6014092 | 0.09606984 | 0.30050264 | -0.108532 | 0.82341376 | 0.89471463 |
| 0.49781409 | -0.1396892 | 0.61510873 | 0.78670568 | -0.3836574 | 0.3347499 | 0.51617924 |
| 0.69955402 | -0.4290478 | 0.13630005 | 0.35389208 | -0.4523885 | 0.25847374 | 0.43875374 |
| 0.64015159 | -0.7391897 | 0.00652572 | 0.08504841 | -0.4848541 | 0.16200294 | 0.34537831 |
| 0.59619098 | -0.5048514 | 0.14686931 | 0.36277763 | -1.2314809 | 0.01889643 | 0.09914488 |
| 0.48456387 | -0.0062716 | 0.98374749 | 0.99185709 | -0.0183129 | 0.96645117 | 0.98218297 |
| 0.43702037 | -0.1753889 | 0.50558077 | 0.70712405 | -0.3115749 | 0.4051228 | 0.58141958 |
| 0.23837259 | -1.0602847 | 0.03056955 | 0.16192 | -0.8055969 | 0.21790931 | 0.4056929 |
| 0.86400189 | -0.4856786 | 0.0373031 | 0.18229668 | -0.1377736 | 0.65218866 | 0.77723958 |
| 0.1758396 | -0.8895272 | 0.0181824 | 0.12181362 | 0.33490734 | 0.48941609 | 0.65580133 |
| 0.88293084 | -0.8042876 | 0.00483954 | 0.07612725 | -1.088649 | 0.0064481 | 0.05623739 |
| 0.56238061 | -0.2787022 | 0.46577026 | 0.68285247 | 0.09787724 | 0.85498848 | 0.91138614 |
| 0.98150616 | -0.4127797 | 0.12390099 | 0.34012823 | -0.2501037 | 0.49495914 | 0.65993059 |
| 0.69088496 | -0.3110924 | 0.2971084 | 0.53239835 | -0.2292367 | 0.58161684 | 0.73429127 |
| 0.6380977 | -0.3379764 | 0.25335943 | 0.49291016 | -0.5802128 | 0.17006143 | 0.35368215 |
| 0.59619098 | -0.4602541 | 0.0682909 | 0.24572073 | -0.0109953 | 0.97389427 | 0.98427785 |
| 0.32863165 | -0.321344 | 0.20481465 | 0.43677851 | 0.12945881 | 0.71067557 | 0.82267315 |
| 0.63173933 | -0.6839741 | 0.01618663 | 0.11706618 | -0.4060631 | 0.27158829 | 0.44967897 |
| 0.77403151 | -0.2534589 | 0.32292922 | 0.56771924 | 0.00581999 | 0.98540946 | 0.990312 |
| 0.20206115 | 0.2819011 | 0.40483023 | 0.64653409 | -0.2767607 | 0.51570606 | 0.67037445 |

| | | | | | | |
|---|---|---|---|---|---|---|
| 0.62158011 | -0.7760051 | 0.15400192 | 0.37186935 | -0.0090678 | 0.99024918 | 0.99147625 |
| 0.69942883 | -0.526861 | 0.12925028 | 0.34452754 | -0.8978459 | 0.07234156 | 0.2065441 |
| 0.59407983 | -0.356621 | 0.27602051 | 0.51421748 | -0.9338667 | 0.0541779 | 0.18089152 |
| 0.21795857 | -1.1246508 | 0.03250271 | 0.16815196 | -0.88197 | 0.20998372 | 0.40073353 |
| 0.7842404 | -0.6517127 | 0.02649345 | 0.14696706 | -0.5660085 | 0.15063256 | 0.33345508 |
| 0.14725224 | -0.327628 | 0.18634579 | 0.42150243 | -0.2998126 | 0.38397245 | 0.56306667 |
| 0.31429405 | 0.05513635 | 0.85565591 | 0.91865078 | -0.8428648 | 0.06455514 | 0.19327239 |
| 0.27219985 | -0.2150578 | 0.43634847 | 0.66711749 | -0.9583817 | 0.02400373 | 0.11292232 |
| 0.70012564 | -0.6066774 | 0.01591733 | 0.11706618 | -0.6805682 | 0.04749909 | 0.1683301 |
| 0.88293084 | -0.3284264 | 0.40275952 | 0.64583119 | -0.8819473 | 0.08894365 | 0.23718307 |
| 0.38661287 | -0.2407032 | 0.37202806 | 0.61506267 | 0.11144486 | 0.76722 | 0.85979717 |
| 0.52620327 | -0.2194773 | 0.60784908 | 0.78466053 | -1.436605 | 0.02886841 | 0.12391271 |
| 0.38020494 | -0.258636 | 0.4387868 | 0.66752555 | -0.6592166 | 0.17263548 | 0.35403418 |
| 0.93456055 | -0.5714656 | 0.04196727 | 0.19356814 | -0.3623296 | 0.33337594 | 0.51504352 |
| 0.59603045 | -0.6472885 | 0.13576275 | 0.35389208 | -1.2046265 | 0.0559111 | 0.18177589 |
| 0.48257912 | -0.0347495 | 0.91772729 | 0.95737497 | -1.1972155 | 0.02228712 | 0.1084819 |
| 0.30638319 | -0.0158001 | 0.96750844 | 0.98397416 | -0.5354071 | 0.33706994 | 0.51778044 |
| 0.57304684 | -0.316331 | 0.39604925 | 0.63756164 | -0.7769499 | 0.15037599 | 0.33345508 |
| 0.38661287 | -0.1565667 | 0.64895088 | 0.79986567 | -0.5364996 | 0.27841817 | 0.45355218 |
| 0.32250102 | -0.0959815 | 0.7889759 | 0.87415549 | -0.6403316 | 0.2186748 | 0.4057644 |
| 0.43548074 | -0.1225913 | 0.71398692 | 0.83552843 | 0.11337793 | 0.81032042 | 0.89080123 |
| 0.48028696 | -0.4754342 | 0.1793987 | 0.41256675 | -0.7478459 | 0.1381971 | 0.31812894 |
| 0.48257912 | -0.2910149 | 0.48680602 | 0.69386761 | -0.3409446 | 0.56350491 | 0.7192922 |
| 0.69955402 | -0.3782829 | 0.16259444 | 0.38576418 | -0.5706667 | 0.13813283 | 0.31812894 |
| 0.5196969 | -0.3168678 | 0.45202535 | 0.67475057 | -0.7990143 | 0.18918473 | 0.37929606 |
| 0.41227372 | -0.1569789 | 0.67688071 | 0.81292206 | -0.3974276 | 0.45869782 | 0.62438379 |
| 0.32120969 | -0.2064833 | 0.56812958 | 0.75231353 | -0.5790997 | 0.26552468 | 0.44765757 |
| 0.93623502 | -0.6703598 | 0.03005504 | 0.16127571 | -1.0556763 | 0.01762123 | 0.09555675 |
| 0.8570768 | -0.4711723 | 0.13081529 | 0.34533555 | -0.7758162 | 0.08263921 | 0.22331934 |
| 0.72992821 | -0.4764436 | 0.22612363 | 0.460755 | -1.310211 | 0.02718119 | 0.12101033 |
| 0.84745888 | -0.5307701 | 0.02505841 | 0.14188246 | -0.8937793 | 0.00975531 | 0.0710116 |
| 0.71847643 | -0.5621219 | 0.21044722 | 0.44075808 | -0.3365548 | 0.58742259 | 0.7384145 |
| 0.75857894 | -1.1552984 | 0.00975873 | 0.09770636 | -2.3612796 | 6.90E-04 | 0.01759982 |
| 0.5557175 | -0.5516915 | 0.14779307 | 0.36328775 | -0.631174 | 0.23572272 | 0.41495416 |
| 0.61416281 | -0.5207214 | 0.11352278 | 0.33043208 | -0.6380422 | 0.16575995 | 0.34969724 |
| 0.5286044 | -0.5651035 | 0.16339737 | 0.38576418 | -0.9123835 | 0.11515221 | 0.28158656 |
| 0.69955402 | -0.7390177 | 0.10584966 | 0.31716266 | -0.8284864 | 0.1923438 | 0.38091616 |
| 0.54574095 | -0.576106 | 0.14146586 | 0.36069402 | -0.8110722 | 0.14313265 | 0.32178236 |
| 0.58010779 | -0.8063153 | 0.01591706 | 0.11706618 | -1.1997948 | 0.01199238 | 0.07942496 |
| 0.76018942 | -0.6480715 | 0.12910369 | 0.34452754 | -1.225731 | 0.04873231 | 0.16972287 |
| 0.92335298 | -0.9722692 | 0.01625523 | 0.11706618 | -0.3795964 | 0.46458905 | 0.62997728 |
| 0.40927024 | -0.470715 | 0.20114924 | 0.43612502 | -0.5666023 | 0.27281625 | 0.45045197 |

| | | | | | | |
|---|---|---|---|---|---|---|
| 0.44482451 | -0.4314893 | 0.31999154 | 0.56376193 | -0.3063346 | 0.61293482 | 0.7538072 |
| 0.6090444 | -0.4418177 | 0.33735885 | 0.5893013 | -0.6151347 | 0.3446649 | 0.52331084 |
| 0.59603045 | -0.7412924 | 0.0568012 | 0.22426804 | -0.9574328 | 0.07887434 | 0.21677029 |
| 0.65746392 | -0.8054114 | 0.10371038 | 0.31331518 | -0.958547 | 0.16524937 | 0.34953271 |
| 0.90531927 | -0.8330572 | 0.01183434 | 0.10917072 | -1.2387463 | 0.00879924 | 0.06836334 |
| 0.53099293 | -0.6687018 | 0.09878338 | 0.30428152 | -0.7726246 | 0.17071292 | 0.35368215 |
| 0.75234336 | -0.6329894 | 0.18833094 | 0.42150243 | -1.2729956 | 0.06917653 | 0.20178568 |
| 0.38661287 | -0.952299 | 0.13327963 | 0.35071339 | -1.7183955 | 0.06124336 | 0.18887265 |
| 0.62100204 | -0.6615093 | 0.1895402 | 0.42150243 | -1.4183777 | 0.05506987 | 0.18097774 |
| 0.62158011 | -0.6683125 | 0.22616841 | 0.460755 | -1.3024842 | 0.1027602 | 0.2594695 |
| 0.54989578 | -0.5439596 | 0.24574307 | 0.48266761 | -0.4330909 | 0.5064969 | 0.66455871 |
| 0.20206115 | -1.7133969 | 0.11720727 | 0.33451359 | -1.2791296 | 0.23186247 | 0.41451545 |
| 0.57664793 | -0.6135853 | 0.23235709 | 0.46536961 | -0.7112561 | 0.3236358 | 0.5057983 |
| 0.97729171 | -0.9132046 | 0.01715235 | 0.12181362 | -0.1627369 | 0.73913759 | 0.83879659 |
| 0.75537286 | -0.9295061 | 0.09626333 | 0.30050264 | -0.5129918 | 0.49838473 | 0.66015551 |
| 0.90531927 | -1.1793238 | 0.00801413 | 0.08544939 | -1.3675554 | 0.02403792 | 0.11292232 |
| 0.56549932 | -1.237759 | 0.00107751 | 0.04497455 | -1.0589554 | 0.02678378 | 0.1202294 |
| 0.69955402 | -1.3340891 | 0.00205443 | 0.05228638 | -1.4789298 | 0.01047442 | 0.07359419 |
| 0.32120969 | -0.1940893 | 0.75254462 | 0.85219191 | 0.23618798 | 0.7860659 | 0.87485021 |
| 0.54574095 | -0.1594638 | 0.72546687 | 0.83865842 | NA | NA | NA |
| 0.67939616 | -0.7882793 | 0.20446468 | 0.43677851 | -1.1263886 | 0.20005803 | 0.39045141 |
| 0.72150431 | -0.914708 | 0.09755605 | 0.30338453 | -0.8293425 | 0.2744555 | 0.45164978 |
| 0.68498316 | -1.0906869 | 0.03126749 | 0.16382485 | -1.324138 | 0.05920577 | 0.18628897 |
| 0.75234336 | -1.1156349 | 0.11390046 | 0.33043208 | -1.4197936 | 0.15149837 | 0.33445542 |
| 0.75857894 | -1.5031695 | 0.00319576 | 0.06246942 | -2.144738 | 0.00299989 | 0.0384748 |
| 0.7039593 | -1.5075449 | 0.00138338 | 0.04497455 | -1.5060841 | 0.01414502 | 0.08721575 |
| 0.5557175 | -1.439084 | 0.00342507 | 0.06345508 | -1.4138803 | 0.02871583 | 0.12391271 |
| 0.66144984 | -1.7179439 | 0.00226756 | 0.05319046 | -1.5884667 | 0.02921134 | 0.12396558 |
| 0.08665997 | -0.1542282 | 0.84341488 | 0.90991277 | 1.59876867 | 0.1619884 | 0.34537831 |

| AMPH-04647.10nM - DMSO__logFC | AMPH-04647.10nM - DMSO__P.Value | AMPH-04647.10nM - DMSO__adj.P.Val | AMPH-04647.300nM - DMSO__logFC | AMPH-04647.300nM - DMSO__P.Value | AMPH-04647.300nM - DMSO__adj.P.Val | AMPH-04647.1000nM - DMSO__logFC |
|---|---|---|---|---|---|---|
| -0.2533708 | 0.44549873 | 0.83552847 | 0.24990665 | 0.45161812 | 0.65311727 | 0.96971356 |
| 0.14827181 | 0.58728944 | 0.89531116 | 0.05681916 | 0.83449996 | 0.91106976 | -0.334917 |
| -0.5029846 | 0.35627707 | 0.7948464 | -0.1281869 | 0.78957828 | 0.88800516 | 0.62323219 |
| -0.1328204 | 0.77456407 | 0.92229021 | -0.6592482 | 0.16892591 | 0.41773545 | -0.5420604 |
| -0.0838858 | 0.84215126 | 0.94068869 | -0.928814 | 0.04099103 | 0.29664784 | -0.5579288 |
| -0.8455435 | 0.20120703 | 0.70594431 | -1.0063437 | 0.1326394 | 0.38619401 | -0.8431986 |
| 0.16794818 | 0.64255557 | 0.9033187 | -0.5074011 | 0.17346193 | 0.4244102 | -0.1930961 |
| -0.1114399 | 0.69097594 | 0.91646405 | -0.6241924 | 0.03895016 | 0.29664784 | -0.165144 |
| -0.2175099 | 0.42256982 | 0.82524955 | -0.931795 | 0.00317679 | 0.2057742 | 0.10751909 |
| -0.7934318 | 0.18105171 | 0.69136491 | -0.3988481 | 0.491069 | 0.67021392 | -0.742698 |
| -0.1278026 | 0.64123082 | 0.9033187 | -0.422724 | 0.13716156 | 0.38965274 | 1.48051142 |
| 0.03395245 | 0.93794291 | 0.97474826 | -0.8077044 | 0.07986466 | 0.33574662 | -1.4973727 |
| -0.1940406 | 0.5418033 | 0.87657089 | -0.6074024 | 0.07017814 | 0.32210016 | -0.2173787 |
| 0.14740346 | 0.60551065 | 0.90211763 | 0.44116113 | 0.13577788 | 0.38840989 | 1.2235629 |
| 0.55137155 | 0.07187088 | 0.59481158 | 0.02050931 | 0.94337554 | 0.97178333 | 0.42170492 |
| -0.151307 | 0.69608945 | 0.91708348 | -0.7770171 | 0.05945851 | 0.30156336 | -0.5175086 |
| 0.06763045 | 0.75150954 | 0.92229021 | -0.3586063 | 0.10849411 | 0.36895227 | -0.1261135 |
| -0.2663015 | 0.56196028 | 0.8791092 | -0.8494391 | 0.07859866 | 0.33574662 | -0.7540111 |
| -0.2501868 | 0.54279136 | 0.87657089 | -0.831946 | 0.05662996 | 0.2980333 | -0.7045112 |
| -0.6592883 | 0.16490662 | 0.67483444 | -0.6081434 | 0.19795507 | 0.44702822 | 1.0218098 |
| 0.00688482 | 0.98387985 | 0.99680173 | -0.1419765 | 0.67786706 | 0.8071126 | -0.2241685 |
| 0.43077998 | 0.0533437 | 0.59481158 | 0.651122 | 0.00648137 | 0.25347982 | NA |
| -0.337679 | 0.37939088 | 0.80918 | -0.9349603 | 0.01761994 | 0.29664784 | NA |
| -0.1344644 | 0.59174424 | 0.89800743 | -0.2353283 | 0.35293612 | 0.57864578 | 0.17139063 |
| -0.0919006 | 0.81651703 | 0.93201484 | -0.9305098 | 0.03298823 | 0.29664784 | NA |
| -0.3280112 | 0.33979249 | 0.77971341 | -0.8590081 | 0.02123358 | 0.29664784 | -0.9976813 |
| -1.5113114 | 0.07384375 | 0.59481158 | -1.0439399 | 0.20364014 | 0.44702822 | -1.4013333 |
| 0.07669449 | 0.79742433 | 0.92439097 | -0.3604264 | 0.23887678 | 0.47506421 | -0.7470466 |
| -0.3832223 | 0.16998809 | 0.67483444 | 0.05201124 | 0.84728728 | 0.91537256 | 0.42418944 |
| -0.0438897 | 0.89669111 | 0.96815134 | -0.0672281 | 0.84241479 | 0.9149855 | 0.41886335 |
| -0.1393191 | 0.68644138 | 0.91488372 | -0.5579911 | 0.1206451 | 0.38619401 | -0.4099174 |
| -0.1754343 | 0.63943478 | 0.9033187 | -0.2505004 | 0.50524946 | 0.6780296 | -0.7010478 |
| 0.01307374 | 0.96822058 | 0.99292113 | 0.04172496 | 0.89884787 | 0.94776972 | 0.68815891 |
| 0.237306 | 0.38850897 | 0.81368843 | 0.19440967 | 0.47800265 | 0.65956332 | 0.8725587 |
| 0.19544499 | 0.40450325 | 0.81802578 | 0.07722645 | 0.73923393 | 0.84727753 | 0.09517988 |
| 0.30573543 | 0.22976233 | 0.72189217 | 0.62581491 | 0.02201086 | 0.29664784 | 1.57770735 |

| | | | | | | |
|---|---|---|---|---|---|---|
| 0.19400954 | 0.47566261 | 0.84158705 | 0.30371884 | 0.27026424 | 0.50659118 | 0.7358823 |
| -0.0768839 | 0.64913512 | 0.90945382 | -0.1097307 | 0.51771607 | 0.68579228 | 0.44393161 |
| 0.33937349 | 0.25967924 | 0.72763363 | 0.23725221 | 0.42539119 | 0.63499303 | 0.37200322 |
| -0.1345679 | 0.75574974 | 0.92229021 | 0.89342391 | 0.05335076 | 0.2969548 | 1.66991328 |
| 0.24254741 | 0.43189178 | 0.827419 | 0.22663381 | 0.46208875 | 0.65690101 | 0.60341292 |
| -0.0205025 | 0.9512678 | 0.98274336 | 0.03556128 | 0.91559167 | 0.95825396 | 0.63236517 |
| -0.1452561 | 0.62824261 | 0.90211763 | -0.4097049 | 0.18406543 | 0.43424633 | 0.16152277 |
| -0.8419435 | 0.16569019 | 0.67483444 | -1.3327013 | 0.03610968 | 0.29664784 | -0.6106145 |
| -0.1560324 | 0.59323144 | 0.89860335 | -0.1770636 | 0.54499715 | 0.70797889 | 0.1544674 |
| 0.16720046 | 0.45796242 | 0.84158705 | 0.17296018 | 0.44293288 | 0.64696892 | 0.81616777 |
| -0.1954113 | 0.44687052 | 0.83552847 | -0.1875079 | 0.58348989 | 0.73586053 | 0.23355373 |
| 0.02792378 | 0.89679634 | 0.96815134 | -0.0148284 | 0.94507333 | 0.97231229 | 0.35634251 |
| -0.0246967 | 0.93202325 | 0.97474826 | -0.5223294 | 0.08716424 | 0.34908216 | -0.1380564 |
| 0.04797322 | 0.78981702 | 0.92229021 | 0.05910442 | 0.7427761 | 0.84814907 | 0.80681426 |
| 0.05763855 | 0.79828648 | 0.92439097 | -0.490959 | 0.04319042 | 0.29664784 | -1.5425637 |
| 0.02917377 | 0.9227852 | 0.97342657 | -0.2008306 | 0.50790281 | 0.6780296 | 0.06861516 |
| -0.0945205 | 0.78771371 | 0.92229021 | -0.6511911 | 0.07909797 | 0.33574662 | -1.7636828 |
| 0.19737691 | 0.47361331 | 0.84158705 | -0.2885915 | 0.29965336 | 0.53950747 | 0.20840139 |
| -0.3613581 | 0.10786731 | 0.59594639 | -0.2309558 | 0.29110911 | 0.52876235 | -0.0113782 |
| 0.15290583 | 0.54423568 | 0.87657089 | -0.3090578 | 0.22935427 | 0.46608875 | -0.3639064 |
| -0.0011705 | 0.996187 | 0.9983007 | 0.02262938 | 0.9263959 | 0.96351206 | 0.74770311 |
| 0.04299423 | 0.89745089 | 0.96815134 | -0.6669839 | 0.06074614 | 0.30156336 | -0.7033107 |
| -0.0046256 | 0.98503001 | 0.99680173 | 0.12329892 | 0.61853584 | 0.76248938 | 0.5219258 |
| 0.09721795 | 0.70022569 | 0.91708348 | 0.10548575 | 0.67626659 | 0.8071126 | 0.65525118 |
| -0.1485766 | 0.54061592 | 0.87657089 | -0.6246869 | 0.01926341 | 0.29664784 | -0.2955473 |
| 0.13731368 | 0.63033304 | 0.90211763 | 0.29301313 | 0.31137255 | 0.55221047 | 0.92034571 |
| 0.05978964 | 0.84149617 | 0.94068869 | 0.18283168 | 0.54324677 | 0.70682346 | -0.2319328 |
| 0.10965684 | 0.59840821 | 0.90084238 | -0.0346425 | 0.86726146 | 0.9250153 | 0.51802697 |
| 0.21156313 | 0.40749428 | 0.81802578 | 0.25849959 | 0.31435949 | 0.55622661 | 0.93589383 |
| 0.13125127 | 0.48517623 | 0.8455316 | -0.3128425 | 0.10920268 | 0.36895227 | 0.53455411 |
| -0.3101471 | 0.22231787 | 0.71226066 | -0.4479939 | 0.08612319 | 0.34660363 | -0.4425337 |
| 0.02939693 | 0.91235584 | 0.97342657 | -0.460237 | 0.10077922 | 0.36186717 | -0.5173711 |
| 0.2372715 | 0.37601151 | 0.80918 | 0.41357097 | 0.13306277 | 0.38619401 | 1.01966515 |
| -0.083484 | 0.77105265 | 0.92229021 | -0.239054 | 0.40972153 | 0.62062985 | -0.5351258 |
| -0.455399 | 0.03928601 | 0.59481158 | -0.7274996 | 0.00267298 | 0.2057742 | -0.2481091 |
| 0.22766083 | 0.32210391 | 0.762096 | 0.27731 | 0.23160411 | 0.46834229 | 1.12232269 |
| 0.04266643 | 0.88059573 | 0.96395879 | -0.1474167 | 0.60534418 | 0.75490144 | 0.2037176 |
| -0.1514287 | 0.57529699 | 0.88948932 | -0.8767608 | 0.00491583 | 0.24305283 | -0.3913984 |
| -0.3273343 | 0.39734606 | 0.81759677 | -0.6496516 | 0.104804 | 0.36463493 | -0.698764 |
| 0.10511169 | 0.58887701 | 0.89531116 | -0.1498253 | 0.44350565 | 0.64696892 | 0.41845213 |
| 0.08229152 | 0.74968209 | 0.92229021 | -0.587698 | 0.03535802 | 0.29664784 | -0.1906358 |
| -0.3116345 | 0.21649634 | 0.71226066 | -0.6313421 | 0.01989185 | 0.29664784 | -0.2225156 |

| | | | | | | |
|---|---|---|---|---|---|---|
| -0.1998663 | 0.33999683 | 0.77971341 | -0.5156907 | 0.02293079 | 0.29664784 | -0.3162307 |
| -0.0207429 | 0.91157326 | 0.97342657 | 0.29671598 | 0.12768126 | 0.38619401 | 1.09766266 |
| -0.123565 | 0.68192097 | 0.91434239 | -0.3265992 | 0.33958271 | 0.57602769 | -0.004374 |
| -0.0202553 | 0.95253105 | 0.98274336 | -0.1904047 | 0.57785859 | 0.73213257 | 0.16646024 |
| -0.1479737 | 0.68269414 | 0.91434239 | -0.3444832 | 0.34749539 | 0.57788722 | -0.3682194 |
| 0.35578733 | 0.22066854 | 0.71226066 | 0.36862537 | 0.20532104 | 0.44832067 | 0.23356125 |
| 0.0151678 | 0.93591725 | 0.97474826 | -0.3893578 | 0.05382344 | 0.2969548 | -0.1798457 |
| 0.21207872 | 0.39491067 | 0.81759677 | 0.28567981 | 0.25664811 | 0.48903939 | 0.97362072 |
| 0.29908241 | 0.14315108 | 0.6275099 | 0.03436784 | 0.86121405 | 0.92255462 | 0.51936989 |
| -0.1614261 | 0.52816632 | 0.87657089 | -0.095521 | 0.70766001 | 0.82422923 | 0.37825417 |
| 0.03742802 | 0.86618333 | 0.95582865 | -0.0408891 | 0.85395627 | 0.92007624 | 0.28442028 |
| -0.0057802 | 0.97115185 | 0.99292113 | -0.2063252 | 0.20960594 | 0.45028669 | 0.26393861 |
| 0.53536471 | 0.09392164 | 0.59481158 | 0.64567647 | 0.04783204 | 0.29664784 | 1.17052737 |
| 0.35764127 | 0.14522154 | 0.6275099 | 0.29175858 | 0.22895766 | 0.46608875 | 0.47988696 |
| -0.1620825 | 0.47724281 | 0.84158705 | -0.5588795 | 0.02431832 | 0.29664784 | -0.1172448 |
| 0.11211449 | 0.70055591 | 0.91708348 | 0.28355792 | 0.33746234 | 0.57480619 | 0.46223297 |
| 0.15483121 | 0.54098343 | 0.87657089 | 0.21497186 | 0.39888079 | 0.60958466 | 3.51455211 |
| -0.2216488 | 0.36395646 | 0.79801576 | -0.6635913 | 0.01371989 | 0.28790926 | -0.5680733 |
| 0.19309525 | 0.45216846 | 0.83988758 | 0.41558964 | 0.11792806 | 0.38619401 | 0.86331199 |
| -0.1361162 | 0.54452028 | 0.87657089 | -0.1024585 | 0.64735101 | 0.78853884 | 0.45788323 |
| 0.47377232 | 0.03417442 | 0.59481158 | 1.21426821 | 2.94E-05 | 0.02412214 | 1.59386183 |
| -0.0072426 | 0.97419681 | 0.99479549 | -0.1160627 | 0.60594403 | 0.75490144 | 0.92339406 |
| 0.06050757 | 0.78434262 | 0.92229021 | 0.15900117 | 0.47551433 | 0.65793667 | 0.40529192 |
| 0.21353794 | 0.39546186 | 0.81759677 | 0.29199655 | 0.2503866 | 0.48449947 | 0.84204716 |
| -0.2783651 | 0.28389603 | 0.74734577 | -0.4063838 | 0.12586364 | 0.38619401 | -0.3851416 |
| 0.09481265 | 0.66653143 | 0.91189286 | -0.2145837 | 0.33589113 | 0.57462107 | -0.2960451 |
| 0.2017028 | 0.51517668 | 0.87270105 | -0.2817391 | 0.36671859 | 0.58689271 | 0.12639796 |
| -0.1569116 | 0.46757607 | 0.84158705 | -0.162045 | 0.45341824 | 0.65311727 | 0.12401625 |
| -0.1069326 | 0.74377657 | 0.92229021 | -0.6140228 | 0.07584917 | 0.33574662 | -0.2984217 |
| 0.17694563 | 0.42914273 | 0.82727435 | 0.37019043 | 0.11026567 | 0.37034058 | 1.12819373 |
| -0.182511 | 0.27740218 | 0.73791976 | -0.3414525 | 0.05264713 | 0.2969548 | 1.07228419 |
| -0.1481561 | 0.48780177 | 0.84669187 | -0.0140397 | 0.94712801 | 0.97320663 | 0.29486517 |
| 0.22748472 | 0.36202793 | 0.79801576 | 0.66456637 | 0.0153513 | 0.29310264 | 0.84645953 |
| -0.0483542 | 0.82968921 | 0.93720094 | -0.2748881 | 0.2329643 | 0.46926845 | 0.22315847 |
| -0.0030179 | 0.989523 | 0.99803241 | -0.2896276 | 0.22005133 | 0.45621753 | 0.10899949 |
| 0.24212626 | 0.24467418 | 0.72763363 | 0.43982409 | 0.04435232 | 0.29664784 | 1.02143814 |
| 0.29646637 | 0.23583115 | 0.72352382 | 0.52910027 | 0.04396932 | 0.29664784 | 0.9323796 |
| -0.093912 | 0.64227146 | 0.9033187 | -0.0586949 | 0.771042 | 0.87434458 | 0.53245053 |
| 0.1412679 | 0.46369021 | 0.84158705 | 0.76098331 | 0.00112578 | 0.1736455 | 1.27325822 |
| -0.3482946 | 0.29347208 | 0.75163014 | -0.6465301 | 0.06197443 | 0.30286315 | -0.8933436 |
| -0.2689062 | 0.42535386 | 0.82524955 | -0.2748069 | 0.41549895 | 0.62591677 | 0.37536834 |
| 0.07637781 | 0.69652553 | 0.91708348 | -0.3764068 | 0.0695609 | 0.32210016 | -0.2098865 |

| | | | | | | |
|---|---|---|---|---|---|---|
| 0.02079582 | 0.91042358 | 0.97342657 | 0.07930228 | 0.66886099 | 0.80582093 | 0.47262234 |
| 0.04175622 | 0.78405696 | 0.92229021 | -0.2263613 | 0.1518183 | 0.40854872 | 0.30590761 |
| 0.34393906 | 0.26981608 | 0.73106755 | 0.77042035 | 0.02185745 | 0.29664784 | 1.07799614 |
| -0.1574743 | 0.38339954 | 0.80918 | -0.1472924 | 0.41412244 | 0.62546618 | -0.1327527 |
| -0.1362457 | 0.42575863 | 0.82524955 | 0.00279257 | 0.98682571 | 0.99360656 | 0.71979928 |
| -0.2334314 | 0.31557829 | 0.76052831 | -0.4049523 | 0.09225082 | 0.35527307 | 0.35283341 |
| -0.0176181 | 0.91708023 | 0.97342657 | -0.2162302 | 0.21393627 | 0.45221078 | 0.09404986 |
| -0.1078212 | 0.66150692 | 0.91189286 | -0.4515865 | 0.08167027 | 0.33574662 | -0.073622 |
| -0.1848806 | 0.51196638 | 0.87105869 | 0.0355404 | 0.89893659 | 0.94776972 | 1.24229017 |
| -0.1315161 | 0.50676856 | 0.86656178 | -0.5110139 | 0.01887982 | 0.29664784 | -0.4971882 |
| -0.179781 | 0.4924026 | 0.85287454 | -0.2118234 | 0.42010394 | 0.63053992 | 0.23879184 |
| -0.2240281 | 0.36644573 | 0.79801576 | -0.3222016 | 0.20066523 | 0.44702822 | -0.3272748 |
| 0.16315036 | 0.55515252 | 0.8791092 | -0.0115009 | 0.96661473 | 0.97943329 | 0.09234986 |
| -0.0217639 | 0.88498183 | 0.96747015 | 0.03478229 | 0.81727558 | 0.90064866 | 0.42660867 |
| 0.03847217 | 0.8413452 | 0.94068869 | 0.33108071 | 0.10072793 | 0.36186717 | 0.72672699 |
| 3.21E-04 | 0.99891836 | 0.99891836 | 0.11184054 | 0.63817611 | 0.77967647 | 0.69919938 |
| 0.12395225 | 0.53608476 | 0.87657089 | -0.3487034 | 0.09567002 | 0.36029856 | 0.24739287 |
| -0.200987 | 0.32502061 | 0.76336295 | -0.2642763 | 0.20106801 | 0.44702822 | 0.27517777 |
| -0.1220419 | 0.63147985 | 0.90211763 | -0.3320699 | 0.2031484 | 0.44702822 | -0.5332506 |
| 0.07827131 | 0.63289771 | 0.90211763 | -0.0095064 | 0.9535515 | 0.97614187 | 0.4922765 |
| -0.343158 | 0.13169243 | 0.62137637 | -0.6980708 | 0.00563178 | 0.25039365 | -0.8214242 |
| -0.0754083 | 0.71506193 | 0.92229021 | -0.3063303 | 0.15207489 | 0.40854872 | 0.31867104 |
| -0.428316 | 0.04946429 | 0.59481158 | -0.5228779 | 0.01988458 | 0.29664784 | -0.3346582 |
| -0.0510562 | 0.78850818 | 0.92229021 | -0.2054814 | 0.28945525 | 0.52750821 | 0.13993808 |
| -0.191048 | 0.23690011 | 0.72352382 | -0.1706789 | 0.28828313 | 0.52750821 | 0.30366344 |
| -0.5050032 | 0.06404855 | 0.59481158 | -0.331076 | 0.20902827 | 0.45028669 | -0.0889755 |
| -0.0368145 | 0.85620337 | 0.94992292 | -0.2572736 | 0.21767095 | 0.45543404 | -0.0050107 |
| -0.0968613 | 0.62682114 | 0.90211763 | 0.36864659 | 0.07899202 | 0.33574662 | 0.44386075 |
| -1.2004082 | 0.05288281 | 0.59481158 | -0.8191001 | 0.17141565 | 0.42135405 | 1.27511337 |
| -0.1589813 | 0.38234757 | 0.80918 | -0.3305273 | 0.08148578 | 0.33574662 | 0.12456223 |
| 0.16329731 | 0.34366581 | 0.7836255 | 0.41280404 | 0.02633755 | 0.29664784 | 1.12832435 |
| -0.22076 | 0.58258394 | 0.89402134 | -0.6366169 | 0.12665727 | 0.38619401 | -0.19519 |
| 0.21522588 | 0.40773565 | 0.81802578 | 0.32955527 | 0.21212158 | 0.45117052 | 1.23460032 |
| -0.1366653 | 0.6277189 | 0.90211763 | 0.06887979 | 0.80631614 | 0.89336782 | 0.64423023 |
| 0.17943398 | 0.39694842 | 0.81759677 | -0.3654933 | 0.09649546 | 0.36085978 | -0.3784889 |
| -0.0878653 | 0.58824174 | 0.89531116 | 0.07633698 | 0.63767158 | 0.77967647 | 0.48475272 |
| -0.2897098 | 0.31657578 | 0.76052831 | -0.2474944 | 0.38996404 | 0.60505428 | 0.34819684 |
| -0.0615053 | 0.83103811 | 0.93720094 | -0.2943063 | 0.31565169 | 0.55731191 | -0.3296372 |
| -0.6926883 | 0.11165561 | 0.59594639 | -1.0287453 | 0.02427966 | 0.29664784 | -1.2140714 |
| 0.02738958 | 0.92135639 | 0.97342657 | -0.2590948 | 0.35760918 | 0.58094699 | -0.060425 |
| -0.3873472 | 0.11272271 | 0.59594639 | -0.8022683 | 0.00342431 | 0.2057742 | -0.9552768 |
| -0.5226037 | 0.30374953 | 0.75340896 | -0.2198221 | 0.66037638 | 0.7984816 | -0.1361214 |

| | | | | | | |
|---|---|---|---|---|---|---|
| -0.2663214 | 0.23181024 | 0.72352382 | -0.2019728 | 0.35936929 | 0.58094699 | 0.0307039 |
| 0.0449773 | 0.86554027 | 0.95582865 | 0.11581209 | 0.66374805 | 0.80137816 | 0.24572499 |
| -0.2698887 | 0.15612083 | 0.66069691 | -0.4069784 | 0.04008552 | 0.29664784 | 0.02632736 |
| -0.3602883 | 0.19002925 | 0.69198449 | 0.08752445 | 0.7418556 | 0.84814907 | -0.2556605 |
| -0.2484693 | 0.30317021 | 0.75340896 | -0.5121059 | 0.04456307 | 0.29664784 | -0.2008684 |
| -0.1186901 | 0.62746117 | 0.90211763 | -0.4063729 | 0.11109174 | 0.37075738 | -0.3203025 |
| -0.0921349 | 0.66436281 | 0.91189286 | -0.4640715 | 0.04208582 | 0.29664784 | -1.2727027 |
| 0.22174784 | 0.32412482 | 0.76336295 | 0.34035525 | 0.13884402 | 0.39198813 | 0.15395699 |
| -0.1117489 | 0.52581161 | 0.87657089 | -0.0678871 | 0.69863874 | 0.82128384 | 0.23680606 |
| 0.18213631 | 0.61452892 | 0.90211763 | 0.72261153 | 0.05993036 | 0.30156336 | 1.72476776 |
| 0.10566776 | 0.67197951 | 0.91189286 | 0.14761108 | 0.55537914 | 0.71580263 | 0.42599731 |
| -0.060829 | 0.81697069 | 0.93201484 | 0.20545268 | 0.43962602 | 0.64696892 | 0.48768409 |
| -0.5085757 | 0.09167263 | 0.59481158 | -0.5870412 | 0.05523621 | 0.2969548 | 0.11016013 |
| -0.5154822 | 0.0637282 | 0.59481158 | 0.01218339 | 0.962782 | 0.97795176 | 0.54969868 |
| -0.3477269 | 0.20834189 | 0.71226066 | -0.4030959 | 0.14844672 | 0.40628357 | -0.65433 |
| -0.2910701 | 0.22518172 | 0.71226066 | -0.3320621 | 0.16968264 | 0.4183467 | 0.12300377 |
| -0.237691 | 0.3065273 | 0.75800877 | -0.4771275 | 0.05108664 | 0.29664784 | -0.4085529 |
| -0.0683603 | 0.74709707 | 0.92229021 | 0.1770299 | 0.4085648 | 0.62002163 | 0.66133687 |
| -0.5095215 | 0.12313909 | 0.61712325 | -0.6436101 | 0.05704018 | 0.29828017 | -0.7270554 |
| -0.1992863 | 0.4013098 | 0.81802578 | 0.083731 | 0.72160327 | 0.83207343 | 0.46858195 |
| 0.26785557 | 0.28842835 | 0.74734577 | 0.10953179 | 0.65887041 | 0.7984816 | 0.27638033 |
| 0.07990912 | 0.79776878 | 0.92439097 | 0.22734818 | 0.46965468 | 0.65793667 | 1.62909113 |
| -0.6057581 | 0.08342824 | 0.59481158 | -0.5582193 | 0.10794217 | 0.36895227 | -0.9930645 |
| 0.14734035 | 0.43775724 | 0.83194141 | 0.40326655 | 0.04596635 | 0.29664784 | 1.05349053 |
| 0.10819639 | 0.52147265 | 0.87657089 | 0.27023319 | 0.12244999 | 0.38619401 | 0.94397539 |
| -0.0665646 | 0.80100859 | 0.92584273 | -0.5657014 | 0.0461924 | 0.29664784 | -0.5596226 |
| -0.0253223 | 0.90734323 | 0.97252842 | -0.1856293 | 0.39945986 | 0.60958466 | -0.3072832 |
| -0.8202252 | 0.07732718 | 0.59481158 | -1.1228335 | 0.02046127 | 0.29664784 | -1.3648243 |
| -0.4693963 | 0.10550912 | 0.59481158 | -0.949754 | 0.00345004 | 0.2057742 | -0.9159451 |
| -0.0625465 | 0.73490464 | 0.92229021 | 0.00839945 | 0.96365771 | 0.97795176 | 1.11361912 |
| -0.5969332 | 0.02089847 | 0.59481158 | -0.2263476 | 0.34160173 | 0.57641717 | -0.0078829 |
| -0.0215352 | 0.94007395 | 0.97572783 | 0.07220676 | 0.80119224 | 0.89336782 | 0.61227143 |
| -0.0385982 | 0.88992133 | 0.96815134 | -0.4320911 | 0.1365524 | 0.38926916 | -0.4265688 |
| -0.2220226 | 0.20497925 | 0.71007579 | -0.2507896 | 0.15533274 | 0.41284748 | 0.22836037 |
| -0.7968298 | 0.10080198 | 0.59481158 | -0.5504866 | 0.2452573 | 0.47975626 | -0.2229005 |
| -0.1881399 | 0.38696444 | 0.81312991 | -0.2039787 | 0.34937126 | 0.57829396 | -0.393427 |
| 0.05914532 | 0.81550122 | 0.93201484 | -0.0208793 | 0.93429543 | 0.96691378 | 0.71154082 |
| -0.2596884 | 0.20652314 | 0.71195432 | -0.254747 | 0.21481389 | 0.45221078 | 0.14644206 |
| 0.07886147 | 0.63267447 | 0.90211763 | 0.22956433 | 0.17651048 | 0.42561576 | 0.65727148 |
| -0.2606509 | 0.1592809 | 0.66695142 | -0.5550497 | 0.00675528 | 0.25347982 | -0.403487 |
| -0.074466 | 0.69703115 | 0.91708348 | -0.1188793 | 0.53594627 | 0.70065588 | 0.4541612 |
| -0.0324537 | 0.87079547 | 0.95959652 | -0.1915325 | 0.34464701 | 0.57788722 | -0.1667105 |

| | | | | | | |
|---|---|---|---|---|---|---|
| 0.03968834 | 0.89194623 | 0.96815134 | 1.5623787 | 7.94E-05 | 0.03258502 | 2.13682074 |
| -0.6177612 | 0.25922705 | 0.72763363 | -0.8652254 | 0.12162538 | 0.38619401 | -0.9246214 |
| 0.20563178 | 0.39338093 | 0.81759677 | 0.37102327 | 0.13413536 | 0.38619401 | -0.2213579 |
| 0.18629754 | 0.32023922 | 0.76207652 | 0.36369685 | 0.06361509 | 0.30542686 | 1.06276782 |
| -0.2029397 | 0.33747253 | 0.77951561 | -0.5728645 | 0.01388194 | 0.28790926 | 0.40216434 |
| -0.4709371 | 0.03110386 | 0.59481158 | -0.6770725 | 0.00389257 | 0.21305333 | -0.1289531 |
| 0.0400121 | 0.82076337 | 0.93201484 | 0.38569578 | 0.04265071 | 0.29664784 | 0.78481562 |
| -0.2096222 | 0.48315616 | 0.84398129 | -0.2806608 | 0.35108485 | 0.57864578 | -0.1563061 |
| -0.051516 | 0.75316337 | 0.92229021 | -0.2433963 | 0.15160005 | 0.40854872 | 0.12860142 |
| -0.2210667 | 0.3167161 | 0.76052831 | -0.4142964 | 0.07174734 | 0.32543958 | -0.2845603 |
| 0.15958092 | 0.62243332 | 0.90211763 | -0.5490387 | 0.10481589 | 0.36463493 | 0.22668328 |
| 0.04444984 | 0.80587409 | 0.92664234 | 0.13601543 | 0.45607501 | 0.65461116 | 0.65423736 |
| 0.50394238 | 0.05155231 | 0.59481158 | 0.82754308 | 0.00350894 | 0.2057742 | 1.36268027 |
| 0.09491259 | 0.79035761 | 0.92229021 | -0.1487647 | 0.67746932 | 0.8071126 | -0.0085998 |
| -0.1672405 | 0.54142428 | 0.87657089 | -0.660377 | 0.02661553 | 0.29664784 | -1.4775243 |
| 0.13711089 | 0.52211016 | 0.87657089 | -0.1473315 | 0.49205839 | 0.67021392 | 0.8160783 |
| -0.2853765 | 0.11468883 | 0.59594639 | -0.4525102 | 0.01824406 | 0.29664784 | -0.2323328 |
| -0.9557649 | 0.0816277 | 0.59481158 | -1.0911116 | 0.0501873 | 0.29664784 | -0.3929237 |
| 0.21459247 | 0.35413746 | 0.79222575 | 0.1931851 | 0.40284114 | 0.61360404 | 0.78904603 |
| 0.29519514 | 0.22073762 | 0.71226066 | 0.2970444 | 0.21800923 | 0.45543404 | 0.82955545 |
| 0.07463754 | 0.70228087 | 0.91708348 | 0.17423139 | 0.37767655 | 0.59948289 | 0.53223562 |
| 0.2127487 | 0.30265982 | 0.75340896 | 0.45763496 | 0.03695208 | 0.29664784 | 1.2720097 |
| 0.10552326 | 0.73248401 | 0.92229021 | 0.40340526 | 0.20347161 | 0.44702822 | 0.19024417 |
| -0.1522002 | 0.47756171 | 0.84158705 | -0.4337729 | 0.05606333 | 0.2969548 | -0.2606148 |
| -0.3852653 | 0.15729383 | 0.66224735 | -0.4928142 | 0.07649304 | 0.33574662 | -0.6299895 |
| -0.0748899 | 0.71902418 | 0.92229021 | -0.1838089 | 0.38267049 | 0.600166 | 0.23430929 |
| -0.261948 | 0.17179098 | 0.67483444 | -0.3792331 | 0.05566351 | 0.2969548 | -0.6425336 |
| 0.17737117 | 0.43773561 | 0.83194141 | 0.49178873 | 0.04355308 | 0.29664784 | 1.44690988 |
| -0.2353828 | 0.22532806 | 0.71226066 | -0.1768146 | 0.35697857 | 0.58094699 | 0.30860443 |
| 0.24430568 | 0.62150572 | 0.90211763 | 0.84743478 | 0.10140867 | 0.36198487 | 2.45068037 |
| -0.3814652 | 0.07683567 | 0.59481158 | -0.2901553 | 0.16855851 | 0.41773545 | 0.32811335 |
| -0.157054 | 0.47209368 | 0.84158705 | -0.4068452 | 0.07589128 | 0.33574662 | -0.7155829 |
| -0.240617 | 0.43235414 | 0.827419 | -0.7073104 | 0.03200348 | 0.29664784 | -0.2309457 |
| 0.34405296 | 0.07938115 | 0.59481158 | 0.65493188 | 0.00282856 | 0.2057742 | 1.61699301 |
| -0.414572 | 0.09075003 | 0.59481158 | -0.5728558 | 0.02484019 | 0.29664784 | -0.1967846 |
| -0.0538874 | 0.79085543 | 0.92229021 | 0.01094283 | 0.95699976 | 0.97723483 | 0.30462402 |
| 0.29349406 | 0.21373561 | 0.71226066 | 0.27679348 | 0.23955735 | 0.47506421 | 0.81992518 |
| 0.0177015 | 0.92445047 | 0.97342657 | 0.1951396 | 0.30494263 | 0.54426858 | 0.69099809 |
| -0.0528455 | 0.79222034 | 0.92257149 | -0.4011884 | 0.06047543 | 0.30156336 | -0.5345443 |
| -0.2030029 | 0.46544024 | 0.84158705 | -0.1500788 | 0.58785056 | 0.73795919 | 0.11275461 |
| -0.4682231 | 0.07511923 | 0.59481158 | -0.5246974 | 0.0490375 | 0.29664784 | 0.16071949 |
| -0.2060513 | 0.68594427 | 0.91488372 | 0.41440202 | 0.37913584 | 0.59948289 | 1.15845022 |

| | | | | | | |
|---|---|---|---|---|---|---|
| -0.5389086 | 0.18939809 | 0.69198449 | -0.7328452 | 0.08150808 | 0.33574662 | 0.00964313 |
| 0.35385928 | 0.27679118 | 0.73791976 | 0.36014508 | 0.26870128 | 0.50481408 | 0.72871601 |
| 0.32783166 | 0.1542141 | 0.65600922 | 0.152712 | 0.49469528 | 0.67042818 | -0.2886577 |
| -0.0638717 | 0.77733963 | 0.92229021 | -0.599593 | 0.0167915 | 0.29664784 | -0.5528264 |
| 0.08508571 | 0.71369874 | 0.92229021 | -0.2260099 | 0.33665376 | 0.57462107 | -0.6048956 |
| -0.1348302 | 0.46780276 | 0.84158705 | -0.0401876 | 0.82720735 | 0.90672528 | 0.52935235 |
| 0.06627639 | 0.7045487 | 0.91708348 | -0.1045279 | 0.5513293 | 0.71282103 | 0.13576721 |
| -0.2026686 | 0.45108679 | 0.83977835 | -0.0908666 | 0.7333755 | 0.84327911 | 0.14204076 |
| 0.2373853 | 0.28625065 | 0.74734577 | 0.32186095 | 0.15484037 | 0.41284748 | 0.86070047 |
| 0.26738332 | 0.2382101 | 0.72352382 | 0.25473479 | 0.26008343 | 0.49200115 | 0.72509145 |
| 0.27321546 | 0.15020509 | 0.64564595 | 0.37388077 | 0.05588418 | 0.2969548 | 1.19706673 |
| -0.0579986 | 0.78918313 | 0.92229021 | -0.0343062 | 0.87423445 | 0.92852068 | 0.07829779 |
| -0.0927501 | 0.74754813 | 0.92229021 | -0.3445786 | 0.24248006 | 0.47625868 | 0.65345732 |
| 0.14498723 | 0.66999167 | 0.91189286 | -0.5496435 | 0.12079803 | 0.38619401 | 0.42166182 |
| -0.1721715 | 0.4420773 | 0.8343574 | -0.0850339 | 0.70194248 | 0.82327825 | 0.32565865 |
| -0.1375593 | 0.53116772 | 0.87657089 | -0.3447749 | 0.12951895 | 0.38619401 | -0.3286101 |
| -0.2944589 | 0.16444933 | 0.67483444 | -0.5270561 | 0.01976318 | 0.29664784 | -0.5915516 |
| -0.0696143 | 0.70386164 | 0.91708348 | -0.0040026 | 0.9825181 | 0.99340808 | 0.46537201 |
| 0.06212311 | 0.87445504 | 0.95979624 | 0.03215168 | 0.92713097 | 0.96351206 | 0.56587592 |
| 0.27147993 | 0.24933525 | 0.72763363 | 0.24256695 | 0.3010837 | 0.54089654 | 0.77332721 |
| -0.0762691 | 0.67982795 | 0.91434239 | 0.11051786 | 0.55115194 | 0.71282103 | 0.66962648 |
| -0.2003756 | 0.42619465 | 0.82524955 | -0.5128741 | 0.05424626 | 0.2969548 | -0.6051607 |
| -0.3175004 | 0.17060079 | 0.67483444 | -0.3221446 | 0.16483183 | 0.41773545 | -0.2805404 |
| 0.27404506 | 0.15139024 | 0.64735096 | 0.29332891 | 0.12655849 | 0.38619401 | 0.91273143 |
| 0.27262292 | 0.23843382 | 0.72352382 | 0.13973039 | 0.53828434 | 0.70147848 | 0.41762687 |
| 0.27164926 | 0.23037241 | 0.72189217 | 0.33842227 | 0.14041389 | 0.39210818 | 0.62772205 |
| 0.08232954 | 0.67114029 | 0.91189286 | 0.08223131 | 0.67150732 | 0.80600513 | 0.68425593 |
| 0.11416616 | 0.56986212 | 0.88441739 | 0.46702716 | 0.03173428 | 0.29664784 | 0.92584797 |
| 0.22564775 | 0.30053702 | 0.75340896 | 0.19012765 | 0.38042702 | 0.59948289 | 0.76046691 |
| -0.0132224 | 0.95961431 | 0.98603674 | -0.0625362 | 0.81090413 | 0.89724028 | 0.25996979 |
| 0.53094223 | 0.40743336 | 0.81802578 | 0.82829921 | 0.12895604 | 0.38619401 | 0.88051652 |
| -0.1507812 | 0.63160857 | 0.90211763 | -0.5344267 | 0.10390361 | 0.36463493 | -0.9971253 |
| 0.17652692 | 0.41050746 | 0.81802578 | 0.40729837 | 0.07022647 | 0.32210016 | 0.62625064 |
| 0.24917716 | 0.25113102 | 0.72763363 | 0.27768112 | 0.20341229 | 0.44702822 | 0.65697252 |
| -0.4859117 | 0.0393088 | 0.59481158 | -0.4757327 | 0.04301484 | 0.29664784 | -0.1598757 |
| 0.07371199 | 0.7045949 | 0.91708348 | 0.12540724 | 0.52093466 | 0.68649656 | 0.54830978 |
| 0.38913203 | 0.2464282 | 0.72763363 | 0.36142012 | 0.28014264 | 0.52035545 | 0.21231676 |
| -0.283696 | 0.29646947 | 0.75163014 | 0.04270156 | 0.87272788 | 0.92812123 | 0.90662507 |
| -0.2491671 | 0.21026255 | 0.71226066 | -0.4584938 | 0.02970567 | 0.29664784 | -0.2004169 |
| -0.1238436 | 0.56217481 | 0.8791092 | -0.5162512 | 0.02644331 | 0.29664784 | -0.6137035 |
| 0.02072357 | 0.90143576 | 0.96868948 | -0.1228579 | 0.46694088 | 0.65789131 | 0.21839794 |
| -0.2689049 | 0.24120834 | 0.72440603 | -0.2591077 | 0.25796891 | 0.49026035 | -0.1386334 |

| | | | | | | |
|---|---|---|---|---|---|---|
| 0.05437245 | 0.84205457 | 0.94068869 | 0.07299524 | 0.78918957 | 0.88800516 | 0.43388842 |
| -0.468882 | 0.07821772 | 0.59481158 | -0.5997631 | 0.02900387 | 0.29664784 | -0.6222793 |
| -0.3947739 | 0.19671381 | 0.70594431 | -0.8152632 | 0.01402725 | 0.28790926 | -0.3940533 |
| 0.38222959 | 0.08248624 | 0.59481158 | -0.2836095 | 0.1871515 | 0.43587344 | -0.6251617 |
| 0.09501112 | 0.65242028 | 0.91189286 | 0.31543949 | 0.14845928 | 0.40628357 | 0.36028109 |
| -0.0703548 | 0.69841526 | 0.91708348 | -0.2102129 | 0.25673079 | 0.48903939 | -0.1421327 |
| -0.1088042 | 0.52517322 | 0.87657089 | -0.1589131 | 0.35725574 | 0.58094699 | -0.0893611 |
| -0.1484761 | 0.41733676 | 0.82524955 | 0.09392218 | 0.60538132 | 0.75490144 | 0.42014262 |
| -0.0725097 | 0.68892738 | 0.91646405 | -0.0444404 | 0.80584764 | 0.89336782 | 0.50083992 |
| 8.27E-04 | 0.99708474 | 0.9983007 | 0.11464566 | 0.61423734 | 0.75858715 | 0.40034661 |
| 0.22454906 | 0.2436316 | 0.72735106 | 0.1326644 | 0.48405439 | 0.66456296 | 0.27823656 |
| -0.1201932 | 0.55932262 | 0.8791092 | -0.5702384 | 0.01295576 | 0.28747783 | -0.1401256 |
| -0.311086 | 0.17107834 | 0.67483444 | -0.0819984 | 0.70965463 | 0.82422923 | 0.49480117 |
| -0.6590496 | 0.00964606 | 0.59481158 | -0.6976878 | 0.00679239 | 0.25347982 | -0.7109025 |
| -0.1356718 | 0.58843653 | 0.89531116 | -0.3892265 | 0.1340819 | 0.38619401 | -0.260467 |
| -0.6562453 | 0.12557983 | 0.61712325 | -1.1154097 | 0.01492573 | 0.29310264 | -0.5294733 |
| -0.0711159 | 0.71867807 | 0.92229021 | 0.02149524 | 0.91311434 | 0.95742896 | 0.9942221 |
| 0.0130124 | 0.94428732 | 0.9788635 | -0.1095491 | 0.55869899 | 0.71696984 | -0.1350755 |
| -0.1346384 | 0.75188766 | 0.92229021 | -0.7347022 | 0.09993746 | 0.36186717 | -1.4340042 |
| 0.23784846 | 0.35326879 | 0.79222575 | 0.43563214 | 0.10020709 | 0.36186717 | 0.32745762 |
| 0.04619138 | 0.78699793 | 0.92229021 | 0.10069937 | 0.5577317 | 0.71696984 | 0.66505412 |
| -0.4157596 | 0.11753163 | 0.60308416 | -0.4421265 | 0.09781087 | 0.36085978 | -0.2290239 |
| -0.087707 | 0.57775674 | 0.89117535 | -0.1340324 | 0.39837942 | 0.60958466 | 0.44424981 |
| -0.2433566 | 0.4076121 | 0.81802578 | -0.2537175 | 0.38843797 | 0.60399162 | -0.0039381 |
| -0.0854417 | 0.66684606 | 0.91189286 | 0.03580933 | 0.85643583 | 0.92118155 | 0.12355513 |
| -0.0885927 | 0.61926657 | 0.90211763 | 0.09484125 | 0.59500668 | 0.74566298 | 0.4295556 |
| 0.56854944 | 0.45933309 | 0.84158705 | 0.12694517 | 0.86755393 | 0.9250153 | 0.48698863 |
| -0.0098167 | 0.95717017 | 0.98475778 | 0.10246993 | 0.57718719 | 0.73213257 | 0.46516137 |
| 0.10599951 | 0.5542683 | 0.8791092 | 0.2328529 | 0.20419394 | 0.44704859 | 0.70752986 |
| 0.14888981 | 0.47928918 | 0.84260475 | 0.25749474 | 0.22910345 | 0.46608875 | 0.42217421 |
| 0.19677287 | 0.44197048 | 0.8343574 | 0.34522828 | 0.1865172 | 0.43587344 | 0.91202843 |
| -0.4318991 | 0.04659524 | 0.59481158 | -0.7924696 | 0.00126903 | 0.1736455 | -0.5959129 |
| 0.00162461 | 0.99129815 | 0.9983007 | -0.0425165 | 0.77563862 | 0.87834387 | 0.32261862 |
| -1.213123 | 0.07695545 | 0.59481158 | -1.5879049 | 0.02540235 | 0.29664784 | -1.2951077 |
| -0.0374679 | 0.83021669 | 0.93720094 | -0.3451673 | 0.06342117 | 0.30542686 | -0.1377245 |
| -0.3267539 | 0.12038399 | 0.61009415 | -0.2522119 | 0.22270692 | 0.45710595 | 0.09890454 |
| -0.2166754 | 0.45686469 | 0.84158705 | -0.2090125 | 0.47263669 | 0.65793667 | 0.20750852 |
| -0.9050318 | 0.16897637 | 0.67483444 | -0.5070449 | 0.43032295 | 0.64002744 | -0.3849851 |
| -0.1339616 | 0.40180732 | 0.81802578 | -0.0163648 | 0.91740458 | 0.95825592 | 0.22361592 |
| -0.0198715 | 0.93103797 | 0.97474826 | -0.0186937 | 0.93511515 | 0.96691378 | 0.12605729 |
| 0.18351619 | 0.30111782 | 0.75340896 | 0.14922095 | 0.39744293 | 0.60958466 | 0.59654088 |
| -0.1924866 | 0.33466831 | 0.77836454 | -0.3920044 | 0.06097383 | 0.30156336 | -0.1811994 |

| | | | | | | |
|---|---|---|---|---|---|---|
| -0.0535936 | 0.7715235 | 0.92229021 | -0.1393079 | 0.45423868 | 0.65311727 | -0.0271716 |
| 0.19951602 | 0.65500839 | 0.91189286 | 0.50097164 | 0.2213253 | 0.45710595 | 0.7277816 |
| -0.3388606 | 0.28118088 | 0.74467581 | -0.7046359 | 0.03497149 | 0.29664784 | -0.8829415 |
| -0.3963837 | 0.07123018 | 0.59481158 | -0.6118971 | 0.00920152 | 0.28367628 | -0.6399712 |
| -0.2971888 | 0.17680085 | 0.68793127 | -0.3970726 | 0.07799258 | 0.33574662 | -0.5803944 |
| -0.0184521 | 0.93683804 | 0.97474826 | 0.1273968 | 0.58624967 | 0.73707654 | 0.41767499 |
| -0.5334868 | 0.03284492 | 0.59481158 | -0.3862176 | 0.10889317 | 0.36895227 | 0.41486737 |
| -0.2642028 | 0.22129348 | 0.71226066 | -0.439284 | 0.05130815 | 0.29664784 | -0.2320699 |
| -0.6688644 | 0.10486782 | 0.59481158 | -0.2253843 | 0.5687308 | 0.72504346 | 0.09723075 |
| 0.0318611 | 0.83921133 | 0.94068869 | 0.06721807 | 0.66939083 | 0.80582093 | 0.64181779 |
| 0.26503853 | 0.36365934 | 0.79801576 | 0.52714389 | 0.08260757 | 0.33574662 | 0.70981936 |
| 0.17611722 | 0.35095752 | 0.78941403 | 0.16844329 | 0.37171966 | 0.59143768 | 0.66995135 |
| 0.05326244 | 0.76807901 | 0.92229021 | -0.0158171 | 0.93011824 | 0.9641756 | 0.53875549 |
| 0.15502558 | 0.4975177 | 0.8581135 | 0.30832004 | 0.18740965 | 0.43587344 | 0.73286164 |
| -0.6078845 | 0.01842127 | 0.59481158 | -0.3669177 | 0.1303841 | 0.38619401 | -0.2653659 |
| 0.07016084 | 0.84614001 | 0.94385998 | -0.0380105 | 0.91623551 | 0.95825396 | 0.47950747 |
| 0.20972034 | 0.23448117 | 0.72352382 | 0.36420677 | 0.04857651 | 0.29664784 | 0.87663059 |
| 0.13593609 | 0.67177049 | 0.91189286 | 0.29661125 | 0.36088059 | 0.58094699 | 0.16759322 |
| -0.2152245 | 0.21177127 | 0.71226066 | -0.1629848 | 0.3386882 | 0.57569982 | 0.20932591 |
| -0.1961695 | 0.28921288 | 0.74734577 | 0.090175 | 0.62061007 | 0.76365175 | 1.44963348 |
| 0.09229073 | 0.54261773 | 0.87657089 | -0.2237195 | 0.15227273 | 0.40854872 | -0.3821731 |
| 0.01510796 | 0.93426881 | 0.97474826 | 0.1135348 | 0.53811832 | 0.70147848 | 0.90279325 |
| -0.0939664 | 0.61068486 | 0.90211763 | -0.2543359 | 0.18022425 | 0.42885589 | -0.1873814 |
| -0.0784443 | 0.59691134 | 0.90084238 | -0.2076323 | 0.17369285 | 0.4244102 | 0.21193127 |
| 0.05774954 | 0.74536523 | 0.92229021 | -0.0341868 | 0.84736071 | 0.91537256 | 0.15377041 |
| -0.0833843 | 0.70484735 | 0.91708348 | 0.0519299 | 0.81322274 | 0.89859471 | 0.08723643 |
| -0.1796419 | 0.36192543 | 0.79801576 | -0.4532013 | 0.03194444 | 0.29664784 | -0.5395468 |
| -0.3572821 | 0.20064399 | 0.70594431 | -0.1954962 | 0.47479126 | 0.65793667 | 0.94786204 |
| -0.306386 | 0.21689219 | 0.71226066 | -0.2231985 | 0.36229721 | 0.58094923 | 0.05617497 |
| -0.0644974 | 0.77912805 | 0.92229021 | -0.296186 | 0.20999715 | 0.45028669 | -0.3251212 |
| -0.0207041 | 0.88653108 | 0.96749122 | -0.1186881 | 0.41866765 | 0.62953506 | -0.2590441 |
| -0.0689554 | 0.69040161 | 0.91646405 | -0.0656046 | 0.70464551 | 0.82422923 | 0.30085199 |
| 0.342377 | 0.14511806 | 0.6275099 | 0.2477158 | 0.28325327 | 0.52183303 | 0.21018727 |
| 0.04361111 | 0.8199382 | 0.93201484 | -0.1091337 | 0.57080792 | 0.72656326 | 0.33634003 |
| 0.12884181 | 0.4817414 | 0.84330424 | 0.3387523 | 0.07786184 | 0.33574662 | 0.91407511 |
| 0.55283687 | 0.0754441 | 0.59481158 | 0.28287992 | 0.34302456 | 0.57709664 | 0.2689373 |
| -0.4107742 | 0.11457131 | 0.59594639 | 0.0879119 | 0.72440952 | 0.83413775 | 0.41971812 |
| 0.30680271 | 0.05326788 | 0.59481158 | 0.75437294 | 1.29E-04 | 0.03541381 | 1.36553067 |
| -0.3458283 | 0.12997783 | 0.62041745 | -0.4603216 | 0.05016506 | 0.29664784 | -0.0425806 |
| 0.16787355 | 0.31850712 | 0.76052831 | 0.38012581 | 0.03421244 | 0.29664784 | 0.98105083 |
| -0.1587643 | 0.33767739 | 0.77951561 | 0.03783334 | 0.81648704 | 0.90064866 | 0.58692192 |
| -0.41074 | 0.12831725 | 0.61969684 | -0.7262338 | 0.01250938 | 0.2852833 | -0.7833346 |

| | | | | | | |
|---|---|---|---|---|---|---|
| 0.04535143 | 0.81997791 | 0.93201484 | 0.34060158 | 0.10311384 | 0.36463493 | 0.45407538 |
| -0.3514492 | 0.10530323 | 0.59481158 | -0.6072683 | 0.00958273 | 0.28367628 | -0.419443 |
| -0.023864 | 0.9012826 | 0.96868948 | -0.3149443 | 0.11733768 | 0.38619401 | -0.1598191 |
| 0.22750598 | 0.31866229 | 0.76052831 | 0.51747992 | 0.03357643 | 0.29664784 | 1.38127336 |
| 0.07403797 | 0.72475784 | 0.92229021 | 0.29008576 | 0.18073586 | 0.42885589 | 0.81913742 |
| 0.27102567 | 0.19991853 | 0.70594431 | 0.05060366 | 0.80546493 | 0.89336782 | 0.56966357 |
| -0.0631075 | 0.76241286 | 0.92229021 | 0.13180939 | 0.530006 | 0.69621588 | 0.42719762 |
| 0.17756704 | 0.255082 | 0.72763363 | -0.0855724 | 0.57662929 | 0.73213257 | 0.14450515 |
| -0.2709878 | 0.16084821 | 0.66695142 | -0.4157414 | 0.03922338 | 0.29664784 | -0.4071038 |
| 0.16436146 | 0.50166632 | 0.86164864 | 0.43537759 | 0.08877016 | 0.34914562 | 0.80922164 |
| 0.15294442 | 0.29662383 | 0.75163014 | 0.07295373 | 0.61327858 | 0.75858715 | 0.49211223 |
| 0.05652714 | 0.75281915 | 0.92229021 | 0.40428788 | 0.03722033 | 0.29664784 | 0.83340496 |
| -0.1146558 | 0.62142248 | 0.90211763 | 0.01203812 | 0.95846929 | 0.97751961 | 1.09308347 |
| -0.0762266 | 0.7238145 | 0.92229021 | -0.0717545 | 0.73934682 | 0.84727753 | 0.44652723 |
| -0.4600756 | 0.13852816 | 0.62582794 | -0.1075982 | 0.71908769 | 0.83150844 | -0.4722647 |
| -0.3638056 | 0.05588802 | 0.59481158 | -0.3161005 | 0.09170525 | 0.35514156 | -0.040553 |
| -0.1215773 | 0.61073828 | 0.90211763 | -0.0336482 | 0.88748724 | 0.9401639 | -0.1554673 |
| 0.16715894 | 0.50769332 | 0.86656178 | -0.1457535 | 0.56279949 | 0.7197171 | 0.11994877 |
| -0.2310165 | 0.28497248 | 0.74734577 | -0.1965897 | 0.36022142 | 0.58094699 | -0.5705188 |
| -0.0595525 | 0.73310025 | 0.92229021 | -0.1409004 | 0.42414239 | 0.63428215 | 0.03727711 |
| -0.0438564 | 0.84958404 | 0.94641587 | -0.0900159 | 0.69767877 | 0.82128384 | 6.87E-04 |
| -0.2518394 | 0.3334859 | 0.77836454 | -0.2851967 | 0.27562808 | 0.51429694 | -0.4356364 |
| -0.0815704 | 0.66257848 | 0.91189286 | 0.0464423 | 0.80335008 | 0.89336782 | -0.9754014 |
| 0.23621358 | 0.41988094 | 0.82524955 | -6.53E-05 | 0.99981985 | 0.99981985 | -0.16051 |
| -0.1013814 | 0.5796439 | 0.89117535 | -0.260832 | 0.16634692 | 0.41773545 | 0.20720428 |
| 0.25244569 | 0.17860581 | 0.68982571 | 0.23109438 | 0.21587833 | 0.45328928 | 0.93604588 |
| -0.132805 | 0.56434481 | 0.8791092 | -0.3584518 | 0.13304454 | 0.38619401 | 0.2339589 |
| 0.21986145 | 0.53149464 | 0.87657089 | 0.41935172 | 0.24103213 | 0.47562302 | 0.48376687 |
| 0.06899413 | 0.73196102 | 0.92229021 | 0.13881595 | 0.49347069 | 0.67042818 | 0.70333497 |
| -0.340961 | 0.29556595 | 0.75163014 | -0.5654721 | 0.09284914 | 0.35527307 | -0.7196666 |
| -0.2329513 | 0.40766369 | 0.81802578 | -0.3414352 | 0.23120943 | 0.46834229 | -0.9234079 |
| -0.1133722 | 0.55273075 | 0.8791092 | 0.04924318 | 0.79549074 | 0.89221024 | 0.55772759 |
| 0.25292142 | 0.12746438 | 0.61922046 | 0.18797686 | 0.24867355 | 0.48265008 | 0.62232235 |
| -0.2371153 | 0.22089175 | 0.71226066 | -0.3151205 | 0.11051577 | 0.37034058 | -0.3380167 |
| 0.18601249 | 0.30251098 | 0.75340896 | 0.17506165 | 0.33083504 | 0.57365826 | 0.47722251 |
| -0.008908 | 0.95281817 | 0.98274336 | -0.0885307 | 0.55890463 | 0.71696984 | 0.47364023 |
| -0.1034945 | 0.60467768 | 0.90211763 | -0.1363085 | 0.49685288 | 0.67136068 | 0.25611227 |
| -0.0168297 | 0.90737731 | 0.97252842 | 1.07E-04 | 0.99940964 | 0.99981985 | 0.08789463 |
| 0.0160382 | 0.92340427 | 0.97342657 | 0.08493898 | 0.61220031 | 0.75858715 | 0.34021935 |
| -0.2756912 | 0.26572698 | 0.72971906 | -0.1250106 | 0.60750261 | 0.75569643 | 0.24854923 |
| -0.382029 | 0.20292397 | 0.70893863 | -0.344499 | 0.24839858 | 0.48265008 | 0.08942784 |
| -0.2014132 | 0.20071363 | 0.70594431 | -0.203656 | 0.19602765 | 0.44702822 | 0.1857895 |

| | | | | | | |
|---|---|---|---|---|---|---|
| 0.06286933 | 0.74688297 | 0.92229021 | -0.2826551 | 0.16067354 | 0.41712905 | 0.01915729 |
| -0.0978462 | 0.61933843 | 0.90211763 | -0.2639411 | 0.19183966 | 0.43994515 | -0.1655631 |
| -0.1340957 | 0.67948221 | 0.91434239 | -0.4680967 | 0.16356899 | 0.41773545 | -0.324989 |
| -0.0792003 | 0.7812266 | 0.92229021 | -0.2959114 | 0.30784288 | 0.5482408 | -0.1549671 |
| -0.6347292 | 0.28543364 | 0.74734577 | -0.4517363 | 0.44256893 | 0.64696892 | -0.8223521 |
| -0.3445338 | 0.07848516 | 0.59481158 | -0.2988393 | 0.1220501 | 0.38619401 | 0.18074945 |
| 0.0199311 | 0.89747214 | 0.96815134 | 0.14073997 | 0.36968644 | 0.58934478 | 0.5805545 |
| -0.1993939 | 0.20466953 | 0.71007579 | -0.3553371 | 0.03242623 | 0.29664784 | -0.3164106 |
| -1.400797 | 0.07916058 | 0.59481158 | -1.8146125 | 0.02778077 | 0.29664784 | -2.2298652 |
| 0.01600221 | 0.91930623 | 0.97342657 | -0.1175 | 0.46125442 | 0.65690101 | 0.36383063 |
| 0.11907612 | 0.54120408 | 0.87657089 | 0.37231644 | 0.07008844 | 0.32210016 | 0.80287928 |
| -0.3331961 | 0.38067542 | 0.80918 | -0.1059346 | 0.77779037 | 0.87956735 | 0.13027149 |
| -0.0561347 | 0.76330621 | 0.92229021 | -0.014137 | 0.93945472 | 0.97003558 | -0.0940683 |
| -0.6086699 | 0.07348193 | 0.59481158 | -0.806705 | 0.02237065 | 0.29664784 | -0.4590321 |
| -0.0240067 | 0.89361595 | 0.96815134 | -0.2324479 | 0.2081502 | 0.45028669 | -0.3984729 |
| -0.3361067 | 0.11046915 | 0.59594639 | -0.4475901 | 0.03945208 | 0.29664784 | -0.5684301 |
| -0.0026296 | 0.98708868 | 0.99680173 | -0.0290177 | 0.85834821 | 0.92118155 | 0.25941097 |
| -0.2156098 | 0.26502965 | 0.72971906 | -0.2768679 | 0.15802181 | 0.41449172 | -0.3290013 |
| -0.0846112 | 0.64634145 | 0.90708774 | -0.4096016 | 0.03919649 | 0.29664784 | -0.2951995 |
| 0.13863053 | 0.64207245 | 0.9033187 | 0.07431821 | 0.80269259 | 0.89336782 | 0.02533909 |
| -0.0632906 | 0.77500025 | 0.92229021 | -0.2016203 | 0.36878882 | 0.58905763 | -0.1926964 |
| 0.23253856 | 0.47768522 | 0.84158705 | 0.00251218 | 0.99382402 | 0.99746885 | -0.5707863 |
| -0.2180639 | 0.31041935 | 0.76052831 | -0.3240899 | 0.13989345 | 0.39198813 | 0.14320165 |
| -0.0902541 | 0.54728899 | 0.8791092 | -0.1283283 | 0.39523504 | 0.60879544 | 0.19801868 |
| -0.0997416 | 0.60236178 | 0.90211763 | -0.1489165 | 0.43933828 | 0.64696892 | 0.16510839 |
| 0.06892139 | 0.71175995 | 0.92169545 | 0.17974289 | 0.34191859 | 0.57641717 | 0.68807965 |
| 0.12609464 | 0.48610343 | 0.8455316 | 0.30567376 | 0.10448922 | 0.36463493 | 0.11088851 |
| -0.0621003 | 0.66637235 | 0.91189286 | -0.0287349 | 0.84146818 | 0.9149855 | 0.21925019 |
| -0.3697811 | 0.46777482 | 0.84158705 | 0.3547904 | 0.48568648 | 0.66569049 | 0.15563998 |
| -9.85E-04 | 0.99667076 | 0.9983007 | -0.0663977 | 0.77887625 | 0.87958377 | 0.25891014 |
| 0.10656584 | 0.74720994 | 0.92229021 | -0.2184593 | 0.51126255 | 0.68140674 | -1.0086538 |
| 0.66940357 | 0.01102858 | 0.59481158 | 0.4982744 | 0.04715678 | 0.29664784 | 0.67527613 |
| 0.27729219 | 0.08345099 | 0.59481158 | 0.18254231 | 0.24041824 | 0.47562259 | 0.48829108 |
| -0.0930899 | 0.66927318 | 0.91189286 | -0.2346525 | 0.28977613 | 0.52750821 | 0.24101832 |
| 0.04381857 | 0.76941202 | 0.92229021 | -0.1503121 | 0.32241578 | 0.56560546 | 0.11904273 |
| -0.1982306 | 0.25392315 | 0.72763363 | -0.1311551 | 0.44444637 | 0.64696892 | 0.08476652 |
| 0.04260006 | 0.81889052 | 0.93201484 | -0.1084507 | 0.56194974 | 0.7197171 | 0.18672598 |
| 0.12345652 | 0.54975927 | 0.8791092 | -0.3215273 | 0.13242525 | 0.38619401 | -0.7716783 |
| -0.0896276 | 0.59909859 | 0.90084238 | -8.04E-04 | 0.99621666 | 0.99864943 | 0.05749233 |
| -0.1011403 | 0.60120562 | 0.90211763 | -0.2340719 | 0.23597809 | 0.47253174 | -0.3176698 |
| -0.1598422 | 0.49674184 | 0.8581135 | -0.2048658 | 0.38620229 | 0.60301822 | 0.05276896 |
| 0.08876381 | 0.69886549 | 0.91708348 | 0.27885331 | 0.23488027 | 0.47148338 | 1.21341921 |

| | | | | | | |
|---|---|---|---|---|---|---|
| 0.21964128 | 0.34998987 | 0.78940024 | 0.0868467 | 0.70805717 | 0.82422923 | 0.48434015 |
| -0.0522733 | 0.74346 | 0.92229021 | -0.1482616 | 0.35962937 | 0.58094699 | 0.09195581 |
| -0.3204658 | 0.26717426 | 0.72971906 | -0.7995684 | 0.01185771 | 0.2852833 | -0.2630601 |
| 0.39005242 | 0.1103195 | 0.59594639 | 0.3960707 | 0.10539721 | 0.36501668 | 0.72628685 |
| -0.2387118 | 0.19536361 | 0.70594431 | 0.00820418 | 0.9634115 | 0.97795176 | 0.26669992 |
| -0.0164016 | 0.9343011 | 0.97474826 | -0.0805427 | 0.68645131 | 0.81441694 | 0.4781768 |
| 0.0826163 | 0.67095039 | 0.91189286 | 0.04278325 | 0.82545918 | 0.90672528 | 0.15064731 |
| 0.23882599 | 0.18886243 | 0.69198449 | 0.31640919 | 0.08847145 | 0.34914562 | 0.5001438 |
| 0.06918682 | 0.7747026 | 0.92229021 | 0.23025089 | 0.34777042 | 0.57788722 | 0.65079996 |
| -0.2015491 | 0.46632488 | 0.84158705 | -0.2119362 | 0.44411943 | 0.64696892 | -1.9138292 |
| -0.4979659 | 0.04068945 | 0.59481158 | -0.6350226 | 0.01214887 | 0.2852833 | 0.04772285 |
| 0.00624637 | 0.9762531 | 0.99539868 | 0.31628948 | 0.14683907 | 0.40454655 | 0.31524645 |
| -0.1376958 | 0.38085525 | 0.80918 | 0.0480022 | 0.75715516 | 0.86216974 | 0.43258206 |
| -0.0181763 | 0.93492592 | 0.97474826 | 0.22090825 | 0.3292795 | 0.57365826 | 0.77934488 |
| -0.1138134 | 0.71171691 | 0.92169545 | 0.57581655 | 0.05881572 | 0.30156336 | 0.10080727 |
| -0.0685325 | 0.68628421 | 0.91488372 | -0.0068925 | 0.96750353 | 0.97943329 | 0.44987967 |
| -0.1751356 | 0.34628344 | 0.78535554 | -0.2912356 | 0.12713257 | 0.38619401 | -0.2979714 |
| 0.01679053 | 0.9215389 | 0.97342657 | 0.30455768 | 0.08995509 | 0.35001482 | 0.81951221 |
| -0.2153459 | 0.32168888 | 0.762096 | -0.3550285 | 0.11220141 | 0.37294477 | -0.5511873 |
| -0.0039723 | 0.98610992 | 0.99680173 | -0.329778 | 0.1629507 | 0.41773545 | -0.5410878 |
| 0.05185562 | 0.807546 | 0.92726611 | 0.22646927 | 0.29637988 | 0.53478656 | 0.77660215 |
| -0.4266749 | 0.11992987 | 0.61009415 | -0.4481082 | 0.103873 | 0.36463493 | -0.8788752 |
| -0.0776828 | 0.68161034 | 0.91434239 | -0.2104848 | 0.27515004 | 0.51429694 | 0.14356191 |
| -0.3849674 | 0.10210494 | 0.59481158 | 0.3901029 | 0.09801672 | 0.36085978 | 1.54840048 |
| 0.55315537 | 0.06613338 | 0.59481158 | 0.39403931 | 0.17789755 | 0.42579976 | 0.75222042 |
| 0.30033215 | 0.08488179 | 0.59481158 | 0.46362877 | 0.01241861 | 0.2852833 | 1.46474984 |
| 0.09358094 | 0.55352677 | 0.8791092 | 0.11656649 | 0.46204179 | 0.65690101 | 0.64268798 |
| -0.2613575 | 0.25819969 | 0.72763363 | -0.4096417 | 0.08571429 | 0.34660363 | 0.41905806 |
| -0.1805588 | 0.34456615 | 0.7836255 | -0.3988643 | 0.04823083 | 0.29664784 | 1.42412033 |
| -0.7957441 | 0.23613367 | 0.72352382 | -0.8611965 | 0.20165319 | 0.44702822 | -0.60433 |
| 0.14258456 | 0.53565542 | 0.87657089 | -0.2658897 | 0.25586408 | 0.48903939 | -0.5170626 |
| -0.0692406 | 0.78972361 | 0.92229021 | -0.1785 | 0.49485929 | 0.67042818 | 0.71948309 |
| 0.17149687 | 0.2585017 | 0.72763363 | 0.0660423 | 0.6572497 | 0.7984816 | 0.2763677 |
| 0.2557842 | 0.18729923 | 0.69198449 | 0.46769675 | 0.02360014 | 0.29664784 | 0.74006328 |
| -0.2441757 | 0.24990817 | 0.72763363 | -0.2801872 | 0.18992743 | 0.43800679 | -0.3384348 |
| -0.1953211 | 0.49903165 | 0.8589203 | -0.5666798 | 0.06336092 | 0.30542686 | -0.7113408 |
| 0.1310812 | 0.42120013 | 0.82524955 | 0.20643429 | 0.21274308 | 0.45132317 | 0.62917026 |
| -0.1205102 | 0.53136891 | 0.87657089 | -0.188893 | 0.33137038 | 0.57365826 | -0.0112236 |
| -0.0850952 | 0.75167947 | 0.92229021 | -0.3622893 | 0.19073979 | 0.43864808 | 0.08815546 |
| -0.0677682 | 0.63510839 | 0.90211763 | -0.051707 | 0.71683655 | 0.83007448 | 0.13787846 |
| -0.0284437 | 0.89857652 | 0.96815134 | 0.31147108 | 0.17677829 | 0.42561576 | 0.53739981 |
| 0.05970598 | 0.74536831 | 0.92229021 | 0.21454642 | 0.25347615 | 0.4873628 | 0.32801177 |

| | | | | | | |
|---|---|---|---|---|---|---|
| 0.11877317 | 0.54266421 | 0.87657089 | 0.44188633 | 0.03554347 | 0.29664784 | 0.16267772 |
| -0.0288365 | 0.92921071 | 0.97474826 | -0.0376845 | 0.90758546 | 0.95406871 | -0.1893153 |
| -0.0929263 | 0.65990241 | 0.91189286 | 0.03421256 | 0.87091287 | 0.9273923 | 0.020459 |
| 0.04323303 | 0.76675899 | 0.92229021 | 0.00213761 | 0.98828031 | 0.99360656 | 0.18529201 |
| -0.2016459 | 0.23409962 | 0.72352382 | -0.3928634 | 0.0293276 | 0.29664784 | -0.536454 |
| -0.0858643 | 0.63976306 | 0.9033187 | -0.1575656 | 0.39472962 | 0.60879544 | -0.2475376 |
| 0.07694807 | 0.66252582 | 0.91189286 | 0.17792878 | 0.31987671 | 0.56235284 | 0.70508619 |
| -0.033264 | 0.92247495 | 0.97342657 | -0.4978508 | 0.15987765 | 0.416697 | -0.6430702 |
| 0.26484828 | 0.21783424 | 0.71226066 | 0.26846676 | 0.2119421 | 0.45117052 | 0.62830236 |
| -0.4081814 | 0.09884177 | 0.59481158 | -0.3303283 | 0.17461074 | 0.42538699 | -0.6882534 |
| 0.34639578 | 0.08103039 | 0.59481158 | 0.57417072 | 0.0074769 | 0.26689273 | 1.12124966 |
| 0.11247201 | 0.56140599 | 0.8791092 | 0.30089131 | 0.13349291 | 0.38619401 | 0.83238542 |
| -0.3067158 | 0.09364391 | 0.59481158 | -0.4937578 | 0.01162024 | 0.2852833 | -0.388757 |
| -0.2657891 | 0.27716883 | 0.73791976 | 0.1377537 | 0.56733343 | 0.72438686 | 0.533344 |
| 0.36522721 | 0.06598118 | 0.59481158 | 0.53612704 | 0.01092811 | 0.2852833 | 1.15171972 |
| -0.485026 | 0.26259658 | 0.72971906 | -0.3316006 | 0.43831462 | 0.64696892 | 0.53834772 |
| 0.05048998 | 0.7561441 | 0.92229021 | 0.14490052 | 0.37864053 | 0.59948289 | 0.84219948 |
| -0.1331669 | 0.41247354 | 0.81995345 | -0.1215511 | 0.45352774 | 0.65311727 | 0.19927591 |
| -0.2417172 | 0.20725589 | 0.71195432 | -0.1785173 | 0.34550202 | 0.57788722 | -0.0937433 |
| -0.0028494 | 0.98340786 | 0.99680173 | -0.040818 | 0.76612438 | 0.87117467 | 0.09254881 |
| 0.27422426 | 0.22266884 | 0.71226066 | 0.28657064 | 0.20359988 | 0.44702822 | 0.4138391 |
| -0.1991046 | 0.29038161 | 0.74734577 | 0.17397527 | 0.3533301 | 0.57864578 | 0.44481621 |
| 0.35900069 | 0.09082991 | 0.59481158 | 0.46862974 | 0.03252474 | 0.29664784 | 0.90434299 |
| 0.29224402 | 0.13949636 | 0.62582794 | 0.28993231 | 0.14241958 | 0.39636092 | 1.12878013 |
| -0.0500186 | 0.75791259 | 0.92229021 | -0.1086282 | 0.50591324 | 0.6780296 | 0.3100677 |
| 0.14623671 | 0.43026328 | 0.82727435 | 0.03478235 | 0.84962916 | 0.91661701 | 0.02806966 |
| 0.20651049 | 0.41368219 | 0.82036975 | -0.2386386 | 0.34675576 | 0.57788722 | -0.0742835 |
| -0.1852702 | 0.48161322 | 0.84330424 | -0.2490772 | 0.34749116 | 0.57788722 | -0.1405195 |
| -0.2359384 | 0.26414391 | 0.72971906 | -0.1440196 | 0.48945359 | 0.66973566 | -0.1540415 |
| 0.34584031 | 0.13804846 | 0.62582794 | 0.33514695 | 0.14971718 | 0.40836481 | 0.71709919 |
| 0.42164577 | 0.18311095 | 0.69198449 | 0.35432668 | 0.25893874 | 0.49096699 | 0.31308467 |
| -0.107303 | 0.63349186 | 0.90211763 | -0.2113002 | 0.35327278 | 0.57864578 | -0.0959572 |
| -0.0931799 | 0.61200581 | 0.90211763 | -0.0687064 | 0.70782404 | 0.82422923 | 0.50593469 |
| 0.06150124 | 0.80344516 | 0.92644449 | -0.5491837 | 0.03955668 | 0.29664784 | -0.7988849 |
| -0.0918966 | 0.61054986 | 0.90211763 | -0.1760159 | 0.33516831 | 0.57462107 | 0.29599403 |
| 0.05131227 | 0.7400234 | 0.92229021 | 0.06187323 | 0.68933315 | 0.81547913 | 0.11358576 |
| -0.2888628 | 0.24772552 | 0.72763363 | 0.31710038 | 0.20662189 | 0.44996438 | 1.19412619 |
| -0.3662223 | 0.06367107 | 0.59481158 | -0.0495816 | 0.78944682 | 0.88800516 | -1.439071 |
| -0.2303405 | 0.24176279 | 0.72440603 | -0.4941927 | 0.01987177 | 0.29664784 | -0.4785776 |
| 0.10209245 | 0.68207469 | 0.91434239 | 0.30393253 | 0.23320527 | 0.46926845 | -0.7300631 |
| -0.0309392 | 0.85344288 | 0.94814155 | 0.00235688 | 0.9887656 | 0.99360656 | 0.11456516 |
| 0.14829554 | 0.51670455 | 0.8728692 | -0.0512626 | 0.82147818 | 0.90406647 | -0.4193479 |

| | | | | | | |
|---|---|---|---|---|---|---|
| 0.14359411 | 0.47313629 | 0.84158705 | -0.1484093 | 0.45870462 | 0.65690101 | -0.1975178 |
| -0.0697732 | 0.73934919 | 0.92229021 | -0.3371918 | 0.12286233 | 0.38619401 | -0.2538766 |
| 0.18399157 | 0.44609468 | 0.83552847 | -0.4636603 | 0.06791206 | 0.32210016 | -1.3701013 |
| -0.1484375 | 0.55553737 | 0.8791092 | -0.3495086 | 0.17662332 | 0.42561576 | 0.51144955 |
| 0.46266867 | 0.02457334 | 0.59481158 | 0.13655276 | 0.47055767 | 0.65793667 | 0.11620334 |
| 0.12879739 | 0.51554203 | 0.87270105 | -0.2483776 | 0.21891385 | 0.4561118 | -0.1149621 |
| 0.02807672 | 0.88735797 | 0.96749122 | 0.23414311 | 0.24866084 | 0.48265008 | 0.73340509 |
| 0.28102968 | 0.12628101 | 0.61712325 | 0.36586848 | 0.05254114 | 0.2969548 | 0.89600086 |
| -0.0596129 | 0.76608361 | 0.92229021 | -0.0499219 | 0.80316296 | 0.89336782 | 0.16742697 |
| -0.1373166 | 0.43001361 | 0.82727435 | -0.2017057 | 0.25227235 | 0.48618684 | -0.230612 |
| 0.04544486 | 0.86572194 | 0.95582865 | 0.04079547 | 0.87933663 | 0.93273304 | 0.6228454 |
| -0.0682358 | 0.77133795 | 0.92229021 | -0.0206164 | 0.92993328 | 0.9641756 | 0.0123811 |
| 0.2688812 | 0.14491289 | 0.6275099 | 0.3783115 | 0.0473337 | 0.29664784 | 0.68744878 |
| 0.18030164 | 0.44141661 | 0.8343574 | 0.08704001 | 0.70791444 | 0.82422923 | 0.31676477 |
| 0.08995195 | 0.66818703 | 0.91189286 | 0.41985887 | 0.05991659 | 0.30156336 | 0.4441015 |
| -0.3843537 | 0.05152685 | 0.59481158 | -0.3854691 | 0.05093683 | 0.29664784 | -0.4166916 |
| 0.36640883 | 0.05456065 | 0.59481158 | 0.44647213 | 0.02275898 | 0.29664784 | 0.37757557 |
| 8.01E-04 | 0.99639043 | 0.9983007 | -0.1273376 | 0.47589514 | 0.65793667 | -1.3635844 |
| 0.10136987 | 0.70634039 | 0.91757193 | -0.3911045 | 0.15970948 | 0.416697 | -1.1930898 |
| -0.0882357 | 0.75527676 | 0.92229021 | 0.15066084 | 0.59580379 | 0.74566298 | 0.34696777 |
| -0.0648623 | 0.83243129 | 0.93748434 | -0.2854771 | 0.35864703 | 0.58094699 | -0.1436078 |
| -0.0314154 | 0.89225077 | 0.96815134 | -0.30437 | 0.2023942 | 0.44702822 | 0.01344923 |
| 0.05652318 | 0.75950265 | 0.92229021 | 0.17054798 | 0.36202785 | 0.58094923 | 0.69759203 |
| -0.4806964 | 0.25057794 | 0.72763363 | -0.1713513 | 0.67585139 | 0.8071126 | 0.20574983 |
| -0.5238707 | 0.04061685 | 0.59481158 | -0.5059828 | 0.04696272 | 0.29664784 | -0.751911 |
| 0.46727227 | 0.06765946 | 0.59481158 | 0.46427492 | 0.06924266 | 0.32210016 | 0.75019599 |
| 0.17558147 | 0.54004294 | 0.87657089 | 0.03939599 | 0.88994941 | 0.9415573 | 0.16885376 |
| 0.0072056 | 0.9711188 | 0.99292113 | 0.15126777 | 0.45182634 | 0.65311727 | 0.41411808 |
| 0.10322098 | 0.62951866 | 0.90211763 | 0.29629816 | 0.17841062 | 0.42579976 | 0.41024616 |
| -0.004664 | 0.97842477 | 0.99539868 | -0.2353284 | 0.18615855 | 0.43587344 | -0.0580628 |
| 0.44109153 | 0.13496859 | 0.6256866 | 0.59436266 | 0.05050962 | 0.29664784 | -0.3117517 |
| -0.0038167 | 0.98292435 | 0.99680173 | -0.1310209 | 0.46676196 | 0.65789131 | 0.0010057 |
| -0.1735928 | 0.39497841 | 0.81759677 | -0.1305705 | 0.5199677 | 0.68632392 | 0.17028151 |
| 0.09749318 | 0.66263261 | 0.91189286 | 0.47308403 | 0.04800627 | 0.29664784 | 1.52891871 |
| -0.3053778 | 0.21982265 | 0.71226066 | -0.0236481 | 0.92223112 | 0.96183028 | -0.5365525 |
| 0.11054028 | 0.57952676 | 0.89117535 | -0.154908 | 0.43986371 | 0.64696892 | -0.274914 |
| -0.4559182 | 0.0418712 | 0.59481158 | -0.3867292 | 0.07841334 | 0.33574662 | -0.4064745 |
| -0.0139307 | 0.92481452 | 0.97342657 | -0.1071493 | 0.47198426 | 0.65793667 | -0.0348987 |
| 0.36012001 | 0.09416896 | 0.59481158 | 0.46753976 | 0.03507423 | 0.29664784 | 0.57649658 |
| -0.0953772 | 0.56351486 | 0.8791092 | 0.01812037 | 0.91211013 | 0.95742896 | 0.05340188 |
| -0.1403581 | 0.46192426 | 0.84158705 | -0.3881553 | 0.05482749 | 0.2969548 | -0.4159965 |
| 0.00215136 | 0.99254241 | 0.9983007 | 0.1815193 | 0.43536389 | 0.64635399 | 1.50458868 |

| | | | | | | |
|---|---|---|---|---|---|---|
| 0.05669998 | 0.75081652 | 0.92229021 | 0.06904071 | 0.69924166 | 0.82128384 | 0.16779297 |
| -0.1813359 | 0.37571481 | 0.80918 | -0.2927556 | 0.16156765 | 0.41712905 | -0.3495683 |
| -0.145242 | 0.39045109 | 0.81567518 | -0.1335538 | 0.42878027 | 0.63889038 | 0.00206493 |
| 0.17026026 | 0.29586191 | 0.75163014 | 0.30207483 | 0.07431164 | 0.33521897 | 0.22015662 |
| -0.4034561 | 0.344535 | 0.7836255 | 0.19656256 | 0.64109148 | 0.78207445 | 0.22685404 |
| 0.01230738 | 0.95572048 | 0.98450002 | -0.0689204 | 0.75624613 | 0.86216974 | -0.5008698 |
| 0.18160222 | 0.5964049 | 0.90084238 | 0.34224639 | 0.32430074 | 0.56769916 | 0.51575742 |
| 0.08944925 | 0.80892083 | 0.92754749 | -0.2217865 | 0.55091696 | 0.71282103 | 1.34300321 |
| 0.28962871 | 0.16816675 | 0.67483444 | 0.40605402 | 0.06076998 | 0.30156336 | 0.55293544 |
| -0.1677204 | 0.33398188 | 0.77836454 | 0.10917352 | 0.52547394 | 0.69136875 | 0.31366828 |
| -0.2477042 | 0.25655573 | 0.72763363 | -0.0159238 | 0.94049735 | 0.97003558 | -0.3875454 |
| -0.0275217 | 0.87820192 | 0.96262186 | -0.0724093 | 0.6875392 | 0.81453057 | -0.130738 |
| 0.46832822 | 0.0474842 | 0.59481158 | 0.27238308 | 0.22757018 | 0.46592298 | -0.0910001 |
| -0.3018929 | 0.2706998 | 0.73106755 | -0.1115565 | 0.6783285 | 0.8071126 | 0.63435886 |
| -0.1200446 | 0.5033582 | 0.86274965 | -0.1446355 | 0.42172495 | 0.63181786 | 0.09832164 |
| -0.0954672 | 0.62542809 | 0.90211763 | -0.0418922 | 0.82980127 | 0.90714626 | -0.1143817 |
| 0.38172426 | 0.0747951 | 0.59481158 | 0.27362193 | 0.18957262 | 0.43800679 | 0.24004262 |
| -0.1751905 | 0.31237813 | 0.76052831 | -0.1017273 | 0.55268493 | 0.7134502 | -0.3871661 |
| 0.39636484 | 0.16075815 | 0.66695142 | 0.52567175 | 0.07000432 | 0.32210016 | 1.27507697 |
| -0.5034265 | 0.09724549 | 0.59481158 | -0.4188104 | 0.16155559 | 0.41712905 | -1.2188739 |
| -0.0127628 | 0.93632559 | 0.97474826 | -0.0782276 | 0.62580764 | 0.76799413 | 0.6289515 |
| -0.0801294 | 0.7883013 | 0.92229021 | -0.1993654 | 0.50690099 | 0.6780296 | 0.07854463 |
| 0.22969754 | 0.24013859 | 0.72440603 | 0.22225658 | 0.25498348 | 0.48903939 | 0.82442605 |
| 0.55019365 | 0.06380328 | 0.59481158 | 0.90616411 | 0.00503276 | 0.24305283 | 0.30662098 |
| 0.05688248 | 0.80513716 | 0.92664234 | 0.24662056 | 0.29385538 | 0.53158737 | 0.61520532 |
| -0.2489688 | 0.36110533 | 0.79801576 | 0.11876598 | 0.6594219 | 0.7984816 | -0.0164508 |
| -0.1495135 | 0.45864345 | 0.84158705 | -0.4126136 | 0.05369048 | 0.2969548 | -0.5538516 |
| 0.26901034 | 0.18608259 | 0.69198449 | 0.31483583 | 0.12584379 | 0.38619401 | 0.53833621 |
| -0.0533202 | 0.74505697 | 0.92229021 | -0.3003211 | 0.08244277 | 0.33574662 | -0.4303181 |
| -0.1039306 | 0.5884468 | 0.89531116 | -0.306113 | 0.12479859 | 0.38619401 | -0.5208828 |
| -0.1353877 | 0.47679779 | 0.84158705 | -0.3382705 | 0.08882701 | 0.34914562 | -0.4899418 |
| -0.1861553 | 0.6204448 | 0.90211763 | -0.4520116 | 0.23885227 | 0.47506421 | NA |
| 0.20620697 | 0.225385 | 0.71226066 | 0.1227767 | 0.46287483 | 0.65690101 | -0.1419372 |
| -0.7305859 | 0.02612972 | 0.59481158 | -0.3866178 | 0.21006066 | 0.45028669 | -0.1794293 |
| 0.40387455 | 0.04479213 | 0.59481158 | 0.53543032 | 0.01107776 | 0.2852833 | 0.96223433 |
| -0.1936662 | 0.4728651 | 0.84158705 | 0.43771979 | 0.11733 | 0.38619401 | -0.5912469 |
| -0.1208261 | 0.54078802 | 0.87657089 | -0.3459585 | 0.09390287 | 0.35527307 | -0.3357998 |
| -0.5304085 | 0.10577648 | 0.59481158 | -0.1164051 | 0.71036175 | 0.82422923 | -0.5090369 |
| -0.2167429 | 0.21160426 | 0.71226066 | -0.0213551 | 0.89928455 | 0.94776972 | -0.5840751 |
| -0.1614933 | 0.46477469 | 0.84158705 | -0.3203227 | 0.15796041 | 0.41449172 | -0.7653501 |
| 0.23059487 | 0.25752604 | 0.72763363 | 0.30640438 | 0.13894733 | 0.39198813 | 0.26252816 |
| 0.42377596 | 0.37509157 | 0.80918 | 0.25987289 | 0.58320475 | 0.73586053 | 0.49406376 |

| | | | | | | |
|---|---|---|---|---|---|---|
| -0.0620501 | 0.73204007 | 0.92229021 | -0.2517769 | 0.17789497 | 0.42579976 | 0.00168754 |
| 0.00667173 | 0.96913306 | 0.99292113 | -0.0309184 | 0.85776002 | 0.92118155 | -0.1012236 |
| -0.1195081 | 0.5719789 | 0.8860277 | -0.329049 | 0.13306274 | 0.38619401 | -0.1414759 |
| 0.21315924 | 0.23882455 | 0.72352382 | 0.12693325 | 0.47602239 | 0.65793667 | 0.28489986 |
| -0.0275065 | 0.91763345 | 0.97342657 | -0.17527 | 0.51308981 | 0.68273376 | 0.53164221 |
| 0.07207321 | 0.78051231 | 0.92229021 | -0.053379 | 0.83639642 | 0.91192757 | 0.28943055 |
| 0.47732529 | 0.04696073 | 0.59481158 | 0.42857262 | 0.07081778 | 0.32300776 | 0.34937943 |
| 0.07393466 | 0.75357249 | 0.92229021 | -0.0226768 | 0.92316963 | 0.96183028 | 0.41854052 |
| 0.32144599 | 0.14436388 | 0.6275099 | 0.51206823 | 0.02716915 | 0.29664784 | 0.49506167 |
| 0.49040648 | 0.25101892 | 0.72763363 | 0.05163895 | 0.90151505 | 0.94890238 | 0.7217454 |
| 0.0677546 | 0.72055562 | 0.92229021 | 0.29453391 | 0.13453287 | 0.38619401 | 0.88940281 |
| 0.03358656 | 0.85193892 | 0.94775318 | 0.18543581 | 0.31141711 | 0.55221047 | 0.82706223 |
| -1.1061604 | 0.07558436 | 0.59481158 | -0.7054344 | 0.2415771 | 0.47562302 | 0.20474069 |
| 0.00474657 | 0.97818113 | 0.99539868 | -0.2785308 | 0.12419564 | 0.38619401 | -0.6355802 |
| 0.32347898 | 0.09908216 | 0.59481158 | 0.21348326 | 0.2634667 | 0.49725555 | 0.3548101 |
| -0.191617 | 0.63084936 | 0.90211763 | 1.07560762 | 0.01517976 | 0.29310264 | 0.07171193 |
| 0.0320576 | 0.87193545 | 0.95959652 | -0.125051 | 0.53216078 | 0.69792971 | -0.3684865 |
| 0.45426463 | 0.04308847 | 0.59481158 | 0.31070529 | 0.15080821 | 0.40854872 | 0.39559293 |
| 0.13038606 | 0.4235507 | 0.82524955 | 0.13302705 | 0.41443801 | 0.62546618 | 0.1458184 |
| 0.32875056 | 0.0930543 | 0.59481158 | 0.36202584 | 0.06705866 | 0.32008814 | 0.48926398 |
| -0.0874348 | 0.67081416 | 0.91189286 | -0.2018335 | 0.3331246 | 0.57456995 | -0.1638021 |
| -0.0812527 | 0.67392555 | 0.91231039 | -0.3350337 | 0.09778119 | 0.36085978 | -0.6357844 |
| 0.22748433 | 0.40728284 | 0.81802578 | 0.23804621 | 0.38640005 | 0.60301822 | 0.55658698 |
| -0.1646013 | 0.31084163 | 0.76052831 | -0.2505696 | 0.13155285 | 0.38619401 | -0.381846 |
| 0.18107968 | 0.46724455 | 0.84158705 | 0.59019058 | 0.02854517 | 0.29664784 | 0.62460683 |
| -0.7371733 | 0.18989862 | 0.69198449 | -0.1075809 | 0.84365898 | 0.9149855 | 1.85319754 |
| -0.6530207 | 0.03790585 | 0.59481158 | -0.6232336 | 0.04618747 | 0.29664784 | -0.9793328 |
| 0.5150161 | 0.1819691 | 0.69165106 | 1.055187 | 0.0120121 | 0.2852833 | 1.26443862 |
| -0.4347302 | 0.40889177 | 0.81802578 | -0.1026969 | 0.84354368 | 0.9149855 | -1.2248169 |
| 0.45782525 | 0.06379111 | 0.59481158 | 0.47423805 | 0.05585065 | 0.2969548 | 1.1542437 |
| 0.01526929 | 0.92469292 | 0.97342657 | 0.15249419 | 0.35259261 | 0.57864578 | 0.65419397 |
| 0.53904787 | 0.04067896 | 0.59481158 | 1.18191312 | 6.98E-04 | 0.14335853 | 0.52159041 |
| 0.07918146 | 0.72010838 | 0.92229021 | 0.4894319 | 0.03994716 | 0.29664784 | 1.09289496 |
| 0.04159072 | 0.87409128 | 0.95979624 | 0.18887668 | 0.47559863 | 0.65793667 | 0.96310415 |
| 0.07275294 | 0.76608492 | 0.92229021 | -0.2388424 | 0.33603182 | 0.57462107 | -0.9260062 |
| 0.19382515 | 0.36930949 | 0.80212458 | 0.01193208 | 0.95528392 | 0.97669751 | -0.5939837 |
| -0.4918792 | 0.07412706 | 0.59481158 | -0.2314375 | 0.37974464 | 0.59948289 | 0.22412374 |
| -0.5728821 | 0.19924974 | 0.70594431 | -0.0839884 | 0.84635892 | 0.91537256 | -0.5942692 |
| 0.48750845 | 0.05039173 | 0.59481158 | 0.55395315 | 0.02894731 | 0.29664784 | 0.34058046 |
| 0.03363956 | 0.83849171 | 0.94068869 | -0.0641856 | 0.6979111 | 0.82128384 | -0.1790226 |
| 0.46794884 | 0.07387814 | 0.59481158 | 0.49159063 | 0.06183759 | 0.30286315 | 0.47209526 |
| 0.31477849 | 0.47339509 | 0.84158705 | 0.25339957 | 0.51789429 | 0.68579228 | NA |

| | | | | | | |
|---|---|---|---|---|---|---|
| -0.0786224 | 0.82462114 | 0.93381235 | -0.2310674 | 0.51764796 | 0.68579228 | -0.4504104 |
| 0.30961054 | 0.08609207 | 0.59481158 | -0.0106864 | 0.95015849 | 0.97390509 | -0.9838584 |
| -0.059846 | 0.79469757 | 0.92414548 | 0.03965905 | 0.86299872 | 0.92255462 | -0.5926789 |
| 0.29527359 | 0.17105948 | 0.67483444 | 0.20635739 | 0.33059467 | 0.57365826 | -0.4331839 |
| 0.16976189 | 0.72643847 | 0.92229021 | 0.1282391 | 0.79136353 | 0.88879543 | 0.33103062 |
| 0.38041975 | 0.09093669 | 0.59481158 | 0.23447645 | 0.28237323 | 0.52183303 | 0.11421488 |
| 0.42210463 | 0.08006946 | 0.59481158 | 0.48236932 | 0.04884781 | 0.29664784 | 1.17170233 |
| -0.0746865 | 0.74033027 | 0.92229021 | -0.2962383 | 0.20099717 | 0.44702822 | -0.6978921 |
| 0.18500821 | 0.33801164 | 0.77951561 | 0.27162863 | 0.16712147 | 0.41773545 | 0.57130598 |
| -0.0731955 | 0.67450963 | 0.91231039 | 0.20959901 | 0.23935677 | 0.47506421 | -0.2076423 |
| 0.38351109 | 0.06291409 | 0.59481158 | 0.15191496 | 0.43754421 | 0.64696892 | 0.23851492 |
| 0.4697083 | 0.03694326 | 0.59481158 | 0.15730514 | 0.45365112 | 0.65311727 | -0.2188409 |
| 0.33228024 | 0.47044676 | 0.84158705 | 0.44640932 | 0.33576405 | 0.57462107 | -0.009264 |
| 0.45331434 | 0.02585664 | 0.59481158 | 0.32057399 | 0.10016281 | 0.36186717 | -0.2295694 |
| 0.08442416 | 0.74041497 | 0.92229021 | 0.2722633 | 0.2939594 | 0.53158737 | 0.45063505 |
| 0.19880228 | 0.29024923 | 0.74734577 | 0.13754601 | 0.45971082 | 0.65690101 | 0.01568678 |
| -0.2457647 | 0.36326011 | 0.79801576 | 0.11175908 | 0.67571053 | 0.8071126 | -0.0130988 |
| 0.23903477 | 0.37975921 | 0.80918 | 0.23734107 | 0.38305358 | 0.600166 | 0.36867355 |
| 0.03711863 | 0.8230227 | 0.93328955 | 0.11351135 | 0.49718306 | 0.67136068 | -0.3476928 |
| 0.36584359 | 0.26182972 | 0.72971906 | 0.40220092 | 0.21944478 | 0.4561118 | 0.1736964 |
| -0.1438147 | 0.52553632 | 0.87657089 | -0.2174079 | 0.34158501 | 0.57641717 | -0.8379216 |
| 0.20700819 | 0.26789207 | 0.72971906 | -0.0399699 | 0.82701525 | 0.90672528 | -0.474803 |
| 0.59520759 | 0.01618241 | 0.59481158 | 0.56641081 | 0.02097893 | 0.29664784 | 0.51588455 |
| 0.43988266 | 0.0117293 | 0.59481158 | 0.45752215 | 0.00930815 | 0.28367628 | 0.6814779 |
| -0.4231136 | 0.42287184 | 0.82524955 | 0.15308793 | 0.76960239 | 0.87391917 | -0.2836744 |
| -0.6864718 | 0.09499267 | 0.59481158 | -0.3025337 | 0.44365756 | 0.64696892 | 0.00686743 |
| 0.28599053 | 0.25948821 | 0.72763363 | 0.21729664 | 0.38707747 | 0.60301822 | 0.35099892 |
| -0.2794391 | 0.44778627 | 0.83552847 | 0.27873321 | 0.44891078 | 0.65231107 | 0.59702598 |
| 0.43902463 | 0.08533093 | 0.59481158 | 0.26119384 | 0.28969853 | 0.52750821 | 0.13611395 |
| 0.02552448 | 0.94637672 | 0.97979229 | 0.19202756 | 0.61444635 | 0.75858715 | 0.88687138 |
| -0.2568029 | 0.38267074 | 0.80918 | 0.07427051 | 0.79820757 | 0.89336782 | 0.16028254 |
| 0.16740758 | 0.61921966 | 0.90211763 | 0.35064834 | 0.30494951 | 0.54426858 | -0.0084848 |
| -0.5332833 | 0.1872679 | 0.69198449 | -0.2675516 | 0.49830008 | 0.67176414 | 0.44935702 |
| -0.3300515 | 0.17582243 | 0.68738197 | -0.3698897 | 0.13229257 | 0.38619401 | -0.5367923 |
| -0.7468082 | 0.05009153 | 0.59481158 | -0.456951 | 0.21119489 | 0.45117052 | 0.46092312 |
| 0.42965142 | 0.03795915 | 0.59481158 | 0.27997507 | 0.15802016 | 0.41449172 | -0.0838297 |
| 0.20588688 | 0.4477385 | 0.83552847 | 0.4806661 | 0.08930643 | 0.34914562 | -0.1837536 |
| 0.11974713 | 0.61267205 | 0.90211763 | 0.26884708 | 0.26423064 | 0.49755357 | 0.35576119 |
| -1.4120683 | 0.0562558 | 0.59481158 | -0.0320138 | 0.96309809 | 0.97795176 | -0.7550516 |
| 0.49626608 | 0.07229466 | 0.59481158 | 0.4391898 | 0.10750193 | 0.36895227 | 0.46011351 |
| -0.0613944 | 0.73288324 | 0.92229021 | -0.1324699 | 0.46484837 | 0.65789131 | -0.8857899 |
| 0.37487454 | 0.09435593 | 0.59481158 | 0.18497414 | 0.3913323 | 0.60505428 | 0.02313835 |

| | | | | | | |
|---|---|---|---|---|---|---|
| 0.48068243 | 0.0569415 | 0.59481158 | 0.37597111 | 0.12717703 | 0.38619401 | 0.24255503 |
| -0.1696423 | 0.56507395 | 0.8791092 | -0.3190946 | 0.28619627 | 0.52565356 | -0.5699363 |
| 0.65586368 | 0.04696897 | 0.59481158 | 0.21878039 | 0.48014669 | 0.66141012 | -0.4944371 |
| 0.46895003 | 0.07047371 | 0.59481158 | 0.53365714 | 0.04270807 | 0.29664784 | -0.1041358 |
| 0.41611596 | 0.1171841 | 0.60308416 | 0.44309873 | 0.09708943 | 0.36085978 | 0.05059483 |
| 0.38305451 | 0.28817695 | 0.74734577 | 0.52079701 | 0.15538352 | 0.41284748 | 0.29926054 |
| 0.53904623 | 0.06636212 | 0.59481158 | 0.55199152 | 0.06079399 | 0.30156336 | 0.54244627 |
| 0.62193538 | 0.02668302 | 0.59481158 | 0.95210747 | 0.00195177 | 0.2057742 | 0.98539557 |
| 0.66820605 | 0.14506045 | 0.6275099 | 0.62931516 | 0.16818666 | 0.41773545 | 1.02466286 |
| 0.43593978 | 0.13693872 | 0.62582794 | 0.63215392 | 0.038043 | 0.29664784 | 0.34241275 |
| 0.12848235 | 0.56537108 | 0.8791092 | -0.1624822 | 0.46877761 | 0.65789131 | -0.4739226 |
| 0.16568588 | 0.55811703 | 0.8791092 | 0.10451825 | 0.71078477 | 0.82422923 | -0.0292026 |
| 0.50462371 | 0.06832706 | 0.59481158 | 0.45940658 | 0.09384181 | 0.35527307 | 0.40425082 |
| 0.22653508 | 0.40951772 | 0.81802578 | 0.09714181 | 0.72094132 | 0.83207343 | 0.06876206 |
| 0.4925993 | 0.06477312 | 0.59481158 | 0.38489138 | 0.13986777 | 0.39198813 | -0.0080442 |
| 0.15388603 | 0.51244987 | 0.87105869 | -0.0145668 | 0.95018024 | 0.97390509 | -0.1158033 |
| 0.45289412 | 0.093157 | 0.59481158 | 0.38863208 | 0.14451283 | 0.40082782 | -0.0801512 |
| -0.7400725 | 0.12581441 | 0.61712325 | -0.7858161 | 0.10581482 | 0.36501668 | 0.42071737 |
| 0.68211179 | 0.06422011 | 0.59481158 | 0.7295302 | 0.04959406 | 0.29664784 | -0.336412 |
| 0.49827929 | 0.04644936 | 0.59481158 | 0.22962845 | 0.33189729 | 0.57365826 | -0.1522984 |
| 0.51680647 | 0.02218437 | 0.59481158 | 0.06822467 | 0.73994883 | 0.84727753 | -0.3148687 |
| 0.16963171 | 0.26842285 | 0.72971906 | -0.0323738 | 0.82917892 | 0.90714626 | -0.6975846 |
| -0.0912673 | 0.79078588 | 0.92229021 | 0.00514264 | 0.98805841 | 0.99360656 | -1.2061867 |
| 0.47473991 | 0.10205186 | 0.59481158 | 0.47644811 | 0.10093494 | 0.36186717 | -0.0504249 |
| 0.23381222 | 0.40389621 | 0.81802578 | 0.19158769 | 0.49225212 | 0.67021392 | 0.05861655 |
| 0.24895625 | 0.30204617 | 0.75340896 | -0.0927445 | 0.69589917 | 0.82128384 | -0.9848345 |
| -0.0572455 | 0.86437406 | 0.95582865 | 0.14293778 | 0.67059084 | 0.80600513 | -0.7312498 |
| 0.55211323 | 0.08891075 | 0.59481158 | 0.64850179 | 0.04967064 | 0.29664784 | 0.17256669 |
| 0.34410387 | 0.20117283 | 0.70594431 | 0.48385864 | 0.0800147 | 0.33574662 | 0.33281443 |
| -0.7939479 | 0.0935619 | 0.59481158 | -0.3989244 | 0.38171738 | 0.600166 | -1.5340949 |
| 0.33609803 | 0.13405501 | 0.6256866 | 0.18030756 | 0.40823211 | 0.62002163 | -0.1555734 |
| 0.20447363 | 0.54975413 | 0.8791092 | -0.2121783 | 0.53501842 | 0.70055841 | -1.0475828 |
| 0.27930161 | 0.26677905 | 0.72971906 | -0.1178031 | 0.63337299 | 0.77611825 | -0.3261276 |
| 0.30547293 | 0.42491154 | 0.82524955 | 0.41443899 | 0.28348055 | 0.52183303 | 0.57584829 |
| 0.45596485 | 0.09201996 | 0.59481158 | 0.28225864 | 0.28208057 | 0.52183303 | -0.0142721 |
| 0.60139575 | 0.0547076 | 0.59481158 | 0.38517984 | 0.2011875 | 0.44702822 | -0.127928 |
| 0.71702899 | 0.02393615 | 0.59481158 | 0.36270203 | 0.22185659 | 0.45710595 | -0.6942796 |
| 0.15345613 | 0.52141837 | 0.87657089 | 0.24892434 | 0.30396527 | 0.54426858 | -0.0558659 |
| 0.4742066 | 0.06968656 | 0.59481158 | 0.39391579 | 0.12521189 | 0.38619401 | 0.20469047 |
| 0.2556971 | 0.32542878 | 0.76336295 | 0.04413693 | 0.86293774 | 0.92255462 | 0.08211351 |
| 0.35075639 | 0.13565434 | 0.6256866 | 0.49482441 | 0.04266382 | 0.29664784 | -0.3163755 |
| 0.83415667 | 0.01346224 | 0.59481158 | 1.04866833 | 0.00324332 | 0.2057742 | -0.4264748 |

| | | | | | | |
|---|---|---|---|---|---|---|
| -0.8180212 | 0.13440161 | 0.6256866 | -0.0082932 | 0.98738851 | 0.99360656 | 0.11974506 |
| 0.62028343 | 0.07841318 | 0.59481158 | 0.24422787 | 0.46762537 | 0.65789131 | NA |
| 0.29455293 | 0.36508797 | 0.79801576 | 0.41537441 | 0.20783022 | 0.45028669 | -0.5170619 |
| -1.2306794 | 0.02106893 | 0.59481158 | -0.3431409 | 0.48182654 | 0.66261237 | -1.0343139 |
| 0.42806393 | 0.12612921 | 0.61712325 | 0.13722097 | 0.61058729 | 0.75838452 | -0.1590431 |
| 0.2454941 | 0.31556802 | 0.76052831 | 0.49108131 | 0.05587436 | 0.2969548 | -0.7086983 |
| 0.86831618 | 0.01102753 | 0.59481158 | 0.8878138 | 0.00967471 | 0.28367628 | 0.23712959 |
| 0.59072923 | 0.04470429 | 0.59481158 | 0.6197309 | 0.03641601 | 0.29664784 | -0.0834186 |
| 0.41191318 | 0.0842468 | 0.59481158 | 0.22946574 | 0.31851717 | 0.56116436 | -0.3222805 |
| 0.63825389 | 0.08245475 | 0.59481158 | 0.51056113 | 0.15638617 | 0.41417112 | -0.7156906 |
| 0.6332584 | 0.02908691 | 0.59481158 | 0.60202244 | 0.03660478 | 0.29664784 | 0.43765447 |
| 0.58832775 | 0.18091355 | 0.69136491 | 0.64178841 | 0.1467512 | 0.40454655 | 0.52409975 |
| 0.71635039 | 0.0441759 | 0.59481158 | 0.60751748 | 0.08192848 | 0.33574662 | 0.07440588 |
| 0.47468179 | 0.08429864 | 0.59481158 | 0.30628362 | 0.25080667 | 0.48449947 | -0.5785136 |
| 0.42918646 | 0.31197107 | 0.76052831 | 0.23118686 | 0.58111946 | 0.73512954 | -0.2932372 |
| 0.86198513 | 0.0203389 | 0.59481158 | 0.85128858 | 0.0216715 | 0.29664784 | -0.2791384 |
| 0.8714585 | 0.03792437 | 0.59481158 | 0.87033738 | 0.03813757 | 0.29664784 | 0.38780653 |
| 0.7154476 | 0.0673712 | 0.59481158 | 0.58578618 | 0.12697056 | 0.38619401 | 0.16870264 |
| 0.84846641 | 0.02417259 | 0.59481158 | 0.7521366 | 0.04191454 | 0.29664784 | 0.14454983 |
| 0.93340001 | 0.01865275 | 0.59481158 | 0.81435929 | 0.03598615 | 0.29664784 | 0.21309286 |
| 0.66563587 | 0.06133642 | 0.59481158 | 0.80812094 | 0.02691514 | 0.29664784 | 0.55543303 |
| 0.58723182 | 0.10262856 | 0.59481158 | 0.45816266 | 0.19473256 | 0.44533546 | -0.3982914 |
| 0.70442892 | 0.10547539 | 0.59481158 | 0.67541546 | 0.11928926 | 0.38619401 | 0.01090753 |
| 0.53986847 | 0.05377868 | 0.59481158 | 0.5976815 | 0.03515686 | 0.29664784 | -0.1192418 |
| 0.72816656 | 0.09684272 | 0.59481158 | 0.67106823 | 0.12331749 | 0.38619401 | 0.07049849 |
| 0.95923926 | 0.02068517 | 0.59481158 | 0.83389723 | 0.03989196 | 0.29664784 | 0.17809035 |
| 0.9748505 | 0.01511437 | 0.59481158 | 0.79146244 | 0.04144937 | 0.29664784 | 0.19056988 |
| 0.54330747 | 0.07084762 | 0.59481158 | 0.33724109 | 0.24542951 | 0.47975626 | -0.6441712 |
| 0.63314872 | 0.04874635 | 0.59481158 | 0.54934285 | 0.08225837 | 0.33574662 | -1.1421137 |
| 0.63611109 | 0.11290601 | 0.59594639 | 0.54859044 | 0.16685571 | 0.41773545 | 0.02146429 |
| 0.19793767 | 0.36638489 | 0.79801576 | 0.50596499 | 0.031426 | 0.29664784 | -0.159088 |
| 0.38234256 | 0.38725188 | 0.81312991 | 0.48511433 | 0.27643736 | 0.51463735 | 0.47737421 |
| 0.09920641 | 0.8017956 | 0.92584273 | -0.0972321 | 0.80565194 | 0.89336782 | -0.9374753 |
| 0.60870104 | 0.11302616 | 0.59594639 | 0.52490476 | 0.16703571 | 0.41773545 | -0.0577321 |
| 0.58901668 | 0.07681612 | 0.59481158 | 0.56379343 | 0.08891274 | 0.34914562 | -0.5955197 |
| 0.83024603 | 0.04830176 | 0.59481158 | 0.54699797 | 0.17640002 | 0.42561576 | 0.0336959 |
| 0.44254269 | 0.31850621 | 0.76052831 | 0.37877544 | 0.39099572 | 0.60505428 | -0.0207944 |
| 0.70781611 | 0.07607658 | 0.59481158 | 0.6027727 | 0.12525399 | 0.38619401 | -0.106551 |
| 0.48176737 | 0.12432441 | 0.61712325 | 0.37638894 | 0.22243158 | 0.45710595 | -0.7275386 |
| 0.70756896 | 0.10002492 | 0.59481158 | 0.54327177 | 0.19795825 | 0.44702822 | -0.1857901 |
| 0.45239324 | 0.22556367 | 0.71226066 | 0.24314597 | 0.50695067 | 0.6780296 | -0.8441863 |
| 0.81787711 | 0.03501885 | 0.59481158 | 0.74926792 | 0.05068535 | 0.29664784 | 0.02467113 |

| | | | | | | |
|---|---|---|---|---|---|---|
| 0.86172494 | 0.05830655 | 0.59481158 | 0.80268141 | 0.07545783 | 0.33574662 | 0.21621192 |
| 0.85171945 | 0.07585274 | 0.59481158 | 0.79928679 | 0.09363009 | 0.35527307 | 0.37744755 |
| 0.6338148 | 0.09777356 | 0.59481158 | 0.50204839 | 0.18191217 | 0.43040314 | -0.2187919 |
| 0.52908471 | 0.27240868 | 0.73327058 | 0.44419396 | 0.35381264 | 0.57864578 | -0.001797 |
| 0.61007294 | 0.05285774 | 0.59481158 | 0.42516612 | 0.16299141 | 0.41773545 | -0.7251497 |
| 0.67250051 | 0.09705677 | 0.59481158 | 0.61307859 | 0.12733553 | 0.38619401 | -0.1522514 |
| 0.77121674 | 0.11401954 | 0.59594639 | 0.66423182 | 0.16871862 | 0.41773545 | -0.1167016 |
| -0.4800345 | 0.43549678 | 0.83149501 | 0.39540614 | 0.51920168 | 0.68632392 | -1.2388662 |
| 0.7719752 | 0.12977714 | 0.62041745 | 0.69651399 | 0.16852774 | 0.41773545 | -0.0613013 |
| 0.84505702 | 0.13159176 | 0.62137637 | 0.70678194 | 0.20190777 | 0.44702822 | 0.07963202 |
| 0.92584829 | 0.05802681 | 0.59481158 | 0.84389609 | 0.08091725 | 0.33574662 | 0.08905075 |
| -1.2402861 | 0.10958072 | 0.59594639 | -0.3241904 | 0.66002306 | 0.7984816 | NA |
| 0.83548904 | 0.11107502 | 0.59594639 | 0.79074719 | 0.12980971 | 0.38619401 | 0.22090733 |
| 0.61493364 | 0.09058295 | 0.59481158 | 0.49168991 | 0.1683655 | 0.41773545 | -0.094461 |
| 0.50554758 | 0.34903369 | 0.78940024 | 0.5062104 | 0.34842165 | 0.57788722 | 0.22834045 |
| 0.40418136 | 0.30916593 | 0.76052831 | 0.2604169 | 0.50775426 | 0.6780296 | -0.8666477 |
| 0.39285086 | 0.21602395 | 0.71226066 | 0.22590113 | 0.46875173 | 0.65789131 | -0.9554433 |
| 0.4009643 | 0.27773107 | 0.73791976 | 0.14660668 | 0.68605352 | 0.81441694 | -1.1550575 |
| 0.85332642 | 0.17896818 | 0.68982571 | 1.30455212 | 0.04813007 | 0.29664784 | 1.12462032 |
| 0.26516933 | 0.56077741 | 0.8791092 | 1.44170649 | 0.00579474 | 0.25039365 | NA |
| 0.81480129 | 0.19048538 | 0.69198449 | 0.82134114 | 0.18716059 | 0.43587344 | 0.30705427 |
| 0.74660307 | 0.16948691 | 0.67483444 | 0.71106894 | 0.18937662 | 0.43800679 | -0.2533199 |
| 0.70735554 | 0.14339305 | 0.6275099 | 0.59346757 | 0.21451224 | 0.45221078 | -0.3571489 |
| 0.78043057 | 0.25809201 | 0.72763363 | 0.57737303 | 0.39795745 | 0.60958466 | -0.0603768 |
| 0.38399103 | 0.38159555 | 0.80918 | 0.21475009 | 0.62133906 | 0.76365175 | -1.3564765 |
| 0.53079192 | 0.1852868 | 0.69198449 | 0.21242615 | 0.58620103 | 0.73707654 | -0.927297 |
| 0.64363617 | 0.13935902 | 0.62582794 | 0.41346404 | 0.3313809 | 0.57365826 | -1.1153388 |
| 0.59269245 | 0.22150645 | 0.71226066 | 0.34947368 | 0.46327124 | 0.65690101 | -1.0377231 |
| -0.3721529 | 0.63475911 | 0.90211763 | 2.29642218 | 0.00943414 | 0.28367628 | 0.55761124 |

| AMPH-04647.1000 nM - DMSO__P.Value | AMPH-04647.1000 nM - DMSO__adj.P.Val |
|---|---|
| 0.05159427 | 0.20983078 |
| 0.38979616 | 0.61127455 |
| 0.36587672 | 0.59634507 |
| 0.4134506 | 0.63421761 |
| 0.3560292 | 0.59071784 |
| 0.36049186 | 0.59358189 |
| 0.70556198 | 0.83364717 |
| 0.67708404 | 0.8262441 |
| 0.77704385 | 0.86896376 |
| 0.36749278 | 0.59754326 |
| 0.00153665 | 0.03287626 |
| 0.02658018 | 0.15093977 |
| 0.62804477 | 0.7980417 |
| 0.00764793 | 0.07870589 |
| 0.31067056 | 0.55268089 |
| 0.35106144 | 0.58889956 |
| 0.67669734 | 0.8262441 |
| 0.25388042 | 0.49432195 |
| 0.23439086 | 0.46820583 |
| 0.13053228 | 0.34261012 |
| 0.64307877 | 0.80929737 |
| NA | NA |
| NA | NA |
| 0.62850169 | 0.7980417 |
| NA | NA |
| 0.0515373 | 0.20983078 |
| 0.22627808 | 0.45876328 |
| 0.0927341 | 0.28982858 |
| 0.27665988 | 0.51588183 |
| 0.38717183 | 0.61127455 |
| 0.40545915 | 0.62431494 |
| 0.19719176 | 0.43095941 |
| 0.15301859 | 0.37144594 |
| 0.03608936 | 0.17569251 |
| 0.77163981 | 0.86516094 |
| 4.05E-04 | 0.01568767 |

| | |
|---|---|
| 0.0691286 | 0.24351511 |
| 0.07810813 | 0.26025372 |
| 0.37804588 | 0.60502224 |
| 0.01441242 | 0.1157844 |
| 0.1762188 | 0.40356587 |
| 0.19593698 | 0.43053179 |
| 0.70295538 | 0.83364717 |
| 0.46595882 | 0.68503529 |
| 0.70773964 | 0.83364717 |
| 0.01936629 | 0.13012228 |
| 0.49660543 | 0.70834897 |
| 0.25284702 | 0.49432195 |
| 0.73647115 | 0.84373188 |
| 0.00589043 | 0.06658637 |
| 2.07E-04 | 0.01204292 |
| 0.87197722 | 0.93202857 |
| 0.00268682 | 0.04211205 |
| 0.5911316 | 0.78092466 |
| 0.9700685 | 0.99217835 |
| 0.31315948 | 0.55468117 |
| 0.04494783 | 0.1975275 |
| 0.15088089 | 0.37033984 |
| 0.14940745 | 0.36808561 |
| 0.08170816 | 0.26594675 |
| 0.39247886 | 0.61127455 |
| 0.03483823 | 0.17293675 |
| 0.5850649 | 0.77595067 |
| 0.0930446 | 0.28982858 |
| 0.01802438 | 0.12628845 |
| 0.0576469 | 0.22326879 |
| 0.21842601 | 0.45214166 |
| 0.1844951 | 0.41665142 |
| 0.01458584 | 0.1157844 |
| 0.19976595 | 0.43541479 |
| 0.39672001 | 0.6138345 |
| 0.00294219 | 0.04265434 |
| 0.61353184 | 0.79049349 |
| 0.31203572 | 0.55389748 |
| 0.20852235 | 0.44151844 |
| 0.14141867 | 0.35706019 |
| 0.602331 | 0.78689746 |
| 0.5242412 | 0.72969998 |

| | |
|---|---|
| 0.2876616 | 0.52911511 |
| 7.99E-04 | 0.02164517 |
| 0.99178842 | 0.99792926 |
| 0.72993706 | 0.84215114 |
| 0.47469835 | 0.68937707 |
| 0.56149274 | 0.75578411 |
| 0.50362985 | 0.71707718 |
| 0.01267198 | 0.11154155 |
| 0.07743091 | 0.25905897 |
| 0.30177961 | 0.54160447 |
| 0.37185592 | 0.60103154 |
| 0.25404918 | 0.49432195 |
| 0.01466887 | 0.1157844 |
| 0.16542007 | 0.38645551 |
| 0.71438002 | 0.83567044 |
| 0.2714053 | 0.50841592 |
| 7.01E-08 | 5.70E-05 |
| 0.1108453 | 0.31843544 |
| 0.02804208 | 0.15218014 |
| 0.16142605 | 0.38248725 |
| 6.33E-05 | 0.00672269 |
| 0.00996988 | 0.09649423 |
| 0.20727684 | 0.44151844 |
| 0.02807752 | 0.15218014 |
| 0.29403967 | 0.53510788 |
| 0.34746506 | 0.58851894 |
| 0.7717079 | 0.86516094 |
| 0.6828169 | 0.82731765 |
| 0.52121125 | 0.72808376 |
| 0.00243871 | 0.04202677 |
| 3.26E-04 | 0.01393391 |
| 0.33285479 | 0.57576797 |
| 0.02625286 | 0.15030687 |
| 0.48615949 | 0.69831743 |
| 0.73786007 | 0.84373188 |
| 0.00269351 | 0.04211205 |
| 0.01531686 | 0.11747744 |
| 0.07738576 | 0.25905897 |
| 2.65E-04 | 0.01381618 |
| 0.06748631 | 0.24170207 |
| 0.43123728 | 0.65287879 |
| 0.45191525 | 0.66922968 |

| | |
|---|---|
| 0.08652134 | 0.27693642 |
| 0.16953633 | 0.39268671 |
| 0.02305362 | 0.14092176 |
| 0.60017018 | 0.78689746 |
| 0.00824072 | 0.08374629 |
| 0.28464021 | 0.52593748 |
| 0.69506772 | 0.83364717 |
| 0.8321429 | 0.90687959 |
| 0.00631279 | 0.06843065 |
| 0.08964858 | 0.28140654 |
| 0.51867185 | 0.72703485 |
| 0.35131155 | 0.58889956 |
| 0.8123363 | 0.89489081 |
| 0.06009151 | 0.22866833 |
| 0.0162214 | 0.11989087 |
| 0.05153335 | 0.20983078 |
| 0.38570914 | 0.61127455 |
| 0.34016661 | 0.58099884 |
| 0.15168901 | 0.37033984 |
| 0.04721775 | 0.20513796 |
| 0.01677011 | 0.12170861 |
| 0.28412982 | 0.52593748 |
| 0.25510356 | 0.49432195 |
| 0.60441898 | 0.78689746 |
| 0.18651269 | 0.41888072 |
| 0.80616442 | 0.89171657 |
| 0.98607742 | 0.99743524 |
| 0.12918652 | 0.34137607 |
| 0.13480627 | 0.34604748 |
| 0.62513952 | 0.7980417 |
| 2.72E-04 | 0.01381618 |
| 0.73027866 | 0.84215114 |
| 0.00370998 | 0.04787642 |
| 0.12028739 | 0.33263146 |
| 0.21330913 | 0.44811452 |
| 0.04811123 | 0.20513796 |
| 0.39236058 | 0.61127455 |
| 0.42370591 | 0.64629063 |
| 0.05379561 | 0.21439131 |
| 0.8776335 | 0.93503426 |
| 0.01039703 | 0.09777925 |
| 0.84701053 | 0.91815941 |

| | |
|---|---|
| 0.92033588 | 0.96173916 |
| 0.51600054 | 0.7262389 |
| 0.91919847 | 0.96173916 |
| 0.45123413 | 0.66922968 |
| 0.5510792 | 0.75434241 |
| 0.3596461 | 0.59358189 |
| 6.59E-04 | 0.02061083 |
| 0.62377373 | 0.7980417 |
| 0.34603017 | 0.58731216 |
| 0.00380285 | 0.04830809 |
| 0.23767258 | 0.47194886 |
| 0.20426144 | 0.43863552 |
| 0.78583774 | 0.87518641 |
| 0.1515069 | 0.37033984 |
| 0.10093706 | 0.30620086 |
| 0.71043169 | 0.83364717 |
| 0.2178927 | 0.45214166 |
| 0.04071455 | 0.18914816 |
| 0.12007201 | 0.33263146 |
| 0.17187355 | 0.39472655 |
| 0.43432844 | 0.65511878 |
| 0.00201682 | 0.03999209 |
| 0.04844587 | 0.20513796 |
| 0.00117138 | 0.02645363 |
| 0.00112893 | 0.02622343 |
| 0.14864972 | 0.36808509 |
| 0.32622901 | 0.56671835 |
| 0.04149743 | 0.18953599 |
| 0.03145625 | 0.16186031 |
| 6.34E-04 | 0.02061083 |
| 0.98100881 | 0.99743524 |
| 0.14579294 | 0.36470664 |
| 0.28904753 | 0.5304642 |
| 0.35018895 | 0.58889956 |
| 0.7337086 | 0.843713 |
| 0.20771056 | 0.44151844 |
| 0.06225845 | 0.23218404 |
| 0.60587116 | 0.78689746 |
| 0.01191283 | 0.10761257 |
| 0.12590625 | 0.33671637 |
| 0.10819596 | 0.31528071 |
| 0.5569239 | 0.75463278 |

| | |
|---|---|
| 1.10E-04 | 0.00897043 |
| 0.23369584 | 0.46796728 |
| 0.51361386 | 0.72576789 |
| 9.28E-04 | 0.02415382 |
| 0.18555805 | 0.41789113 |
| 0.65053395 | 0.81241797 |
| 0.00624597 | 0.06843065 |
| 0.7098255 | 0.83364717 |
| 0.58023876 | 0.7755903 |
| 0.36067583 | 0.59358189 |
| 0.62090272 | 0.79620491 |
| 0.02043597 | 0.13291553 |
| 0.00111548 | 0.02622343 |
| 0.98639352 | 0.99743524 |
| 0.00151011 | 0.03287626 |
| 0.01501897 | 0.11628977 |
| 0.34963635 | 0.58889956 |
| 0.59453235 | 0.78212751 |
| 0.02562392 | 0.14774643 |
| 0.02304128 | 0.14092176 |
| 0.06890286 | 0.24351511 |
| 4.52E-04 | 0.01672164 |
| 0.66342111 | 0.82094575 |
| 0.3917819 | 0.61127455 |
| 0.10591191 | 0.31197966 |
| 0.43018038 | 0.65249375 |
| 0.02537237 | 0.14774643 |
| 3.84E-04 | 0.01562027 |
| 0.25927505 | 0.49832297 |
| 0.00291666 | 0.04265434 |
| 0.26520386 | 0.50265057 |
| 0.03171993 | 0.16208257 |
| 0.59192112 | 0.78092466 |
| 1.82E-05 | 0.00370229 |
| 0.55228851 | 0.75434241 |
| 0.29793926 | 0.5370834 |
| 0.02189785 | 0.13800737 |
| 0.01821415 | 0.12628845 |
| 0.0750064 | 0.25607259 |
| 0.77257914 | 0.86516094 |
| 0.64839683 | 0.81099481 |
| 0.08094016 | 0.26534013 |

| | |
|---|---|
| 0.98633202 | 0.99743524 |
| 0.12142813 | 0.33464769 |
| 0.36448883 | 0.59634507 |
| 0.09906472 | 0.30164652 |
| 0.08037724 | 0.26456152 |
| 0.05689051 | 0.2223653 |
| 0.58388699 | 0.77595067 |
| 0.70665908 | 0.83364717 |
| 0.01285941 | 0.11154155 |
| 0.03292675 | 0.16626989 |
| 3.14E-04 | 0.01393391 |
| 0.7985229 | 0.88688404 |
| 0.12384917 | 0.33563126 |
| 0.38571937 | 0.61127455 |
| 0.3078318 | 0.55003792 |
| 0.29624496 | 0.53640791 |
| 0.05577266 | 0.21990773 |
| 0.08761364 | 0.27921613 |
| 0.26642914 | 0.50265057 |
| 0.02942304 | 0.15533072 |
| 0.02005107 | 0.13146387 |
| 0.10164644 | 0.30720651 |
| 0.38228939 | 0.61061154 |
| 0.00297752 | 0.04265434 |
| 0.20347437 | 0.43863552 |
| 0.05945536 | 0.22866833 |
| 0.02290121 | 0.14092176 |
| 0.00476423 | 0.05613507 |
| 0.02236125 | 0.13984379 |
| 0.48524468 | 0.69823703 |
| 0.18163994 | 0.41249517 |
| 0.03758831 | 0.17976057 |
| 0.05126195 | 0.20983078 |
| 0.04223514 | 0.19061742 |
| 0.60590136 | 0.78689746 |
| 0.06082577 | 0.22894481 |
| 0.64806085 | 0.81099481 |
| 0.02776368 | 0.15218014 |
| 0.46773153 | 0.68640024 |
| 0.05599127 | 0.21990773 |
| 0.36308468 | 0.59633908 |
| 0.66248607 | 0.82094575 |

| | |
|---|---|
| 0.27096718 | 0.50841592 |
| 0.09630568 | 0.29657773 |
| 0.35510088 | 0.59071784 |
| 0.04823517 | 0.20513796 |
| 0.2372585 | 0.47194886 |
| 0.58097724 | 0.7755903 |
| 0.71080598 | 0.83364717 |
| 0.11638029 | 0.32661392 |
| 0.06536449 | 0.23634187 |
| 0.22352218 | 0.45686451 |
| 0.3043061 | 0.54493582 |
| 0.62963101 | 0.7980417 |
| 0.12687101 | 0.33727918 |
| 0.03867833 | 0.18282256 |
| 0.4645566 | 0.68421108 |
| 0.36876651 | 0.59841751 |
| 0.00262282 | 0.04211205 |
| 0.60963488 | 0.78922477 |
| 0.02892696 | 0.15446365 |
| 0.36578725 | 0.59634507 |
| 0.01383416 | 0.11360779 |
| 0.52681082 | 0.73088259 |
| 0.06019068 | 0.22866833 |
| 0.9923457 | 0.99792926 |
| 0.65985522 | 0.82094575 |
| 0.10303335 | 0.30910005 |
| 0.65202654 | 0.8130331 |
| 0.08792046 | 0.27921613 |
| 0.0123964 | 0.11075023 |
| 0.16681893 | 0.38749654 |
| 0.02099959 | 0.1333802 |
| 0.05146862 | 0.20983078 |
| 0.14092645 | 0.35692586 |
| 0.17176655 | 0.39472655 |
| 0.57909221 | 0.7755903 |
| 0.72890243 | 0.84215114 |
| 0.61247321 | 0.7903821 |
| 0.66954118 | 0.82600452 |
| 0.32471102 | 0.56671835 |
| 0.69861524 | 0.83364717 |
| 0.02684988 | 0.15093977 |
| 0.51689736 | 0.7262389 |

| | |
|---|---|
| 0.91697987 | 0.96070184 |
| 0.20969405 | 0.44280848 |
| 0.05767091 | 0.22326879 |
| 0.04267204 | 0.19061742 |
| 0.06948893 | 0.24351511 |
| 0.2171841 | 0.45214166 |
| 0.21461696 | 0.44969996 |
| 0.44000793 | 0.66112877 |
| 0.86125738 | 0.9286502 |
| 0.01046346 | 0.09777925 |
| 0.09680713 | 0.29699697 |
| 0.02092773 | 0.1333802 |
| 0.04910179 | 0.2068381 |
| 0.03509786 | 0.17293675 |
| 0.42533297 | 0.64755751 |
| 0.35545455 | 0.59071784 |
| 0.0024388 | 0.04202677 |
| 0.71162502 | 0.83364717 |
| 0.38356392 | 0.61127455 |
| 4.56E-05 | 0.00672269 |
| 0.0886899 | 0.28056378 |
| 0.00311406 | 0.04291076 |
| 0.47484767 | 0.68937707 |
| 0.31860358 | 0.56309719 |
| 0.54282019 | 0.74925776 |
| 0.77906228 | 0.87002423 |
| 0.06479985 | 0.23634187 |
| 0.02432724 | 0.14331915 |
| 0.86934632 | 0.93202857 |
| 0.32523694 | 0.56671835 |
| 0.21911908 | 0.45214166 |
| 0.22989588 | 0.46493867 |
| 0.51419717 | 0.72576789 |
| 0.2262738 | 0.45876328 |
| 0.00266922 | 0.04211205 |
| 0.5201896 | 0.72790731 |
| 0.24439905 | 0.48227288 |
| 1.00E-05 | 0.00370229 |
| 0.89072077 | 0.94168529 |
| 7.37E-04 | 0.02065337 |
| 0.02094544 | 0.1333802 |
| 0.04668787 | 0.20407118 |

| | |
|---|---|
| 0.12252695 | 0.33563126 |
| 0.16613615 | 0.38701629 |
| 0.55923679 | 0.75463278 |
| 5.34E-04 | 0.01808044 |
| 0.01366202 | 0.11333901 |
| 0.06515017 | 0.23634187 |
| 0.16183962 | 0.38248725 |
| 0.50598132 | 0.71916575 |
| 0.13802584 | 0.35067189 |
| 0.03051481 | 0.15902911 |
| 0.02692038 | 0.15093977 |
| 0.00465394 | 0.055642 |
| 0.00416094 | 0.05204377 |
| 0.15716359 | 0.37689552 |
| 0.27354664 | 0.51124923 |
| 0.87204987 | 0.93202857 |
| 0.64504883 | 0.80929737 |
| 0.73529741 | 0.84373188 |
| 0.07235232 | 0.24924762 |
| 0.87982792 | 0.93503426 |
| 0.99832219 | 0.99832219 |
| 0.24060108 | 0.47593352 |
| 0.00200024 | 0.03999209 |
| 0.69574883 | 0.83364717 |
| 0.42614785 | 0.64758543 |
| 0.0022516 | 0.04160725 |
| 0.47418196 | 0.68937707 |
| 0.33498205 | 0.57752373 |
| 0.02426557 | 0.14331915 |
| 0.12694641 | 0.33727918 |
| 0.03102272 | 0.16064634 |
| 0.05240531 | 0.21119191 |
| 0.01351212 | 0.11333901 |
| 0.21740844 | 0.45214166 |
| 0.07227022 | 0.24924762 |
| 0.03957959 | 0.18600118 |
| 0.36964718 | 0.59865171 |
| 0.66839342 | 0.82584171 |
| 0.16374577 | 0.38475524 |
| 0.47223942 | 0.68937707 |
| 0.82832582 | 0.90494986 |
| 0.39610174 | 0.6138345 |

| | |
|---|---|
| 0.94445921 | 0.98141178 |
| 0.55299886 | 0.75434241 |
| 0.48167267 | 0.69679694 |
| 0.70115647 | 0.83364717 |
| 0.32622325 | 0.56671835 |
| 0.49342251 | 0.7075 |
| 0.01691645 | 0.12170861 |
| 0.15761928 | 0.37689552 |
| 0.05120478 | 0.20983078 |
| 0.11908549 | 0.33156337 |
| 0.00963889 | 0.09441471 |
| 0.8060901 | 0.89171657 |
| 0.72138989 | 0.83911345 |
| 0.3201757 | 0.56464825 |
| 0.13190279 | 0.34261012 |
| 0.06082667 | 0.22894481 |
| 0.26947303 | 0.50713327 |
| 0.23077044 | 0.4655493 |
| 0.26647281 | 0.50265057 |
| 0.95191393 | 0.98344156 |
| 0.5404405 | 0.7485147 |
| 0.22587904 | 0.45876328 |
| 0.63273991 | 0.80002728 |
| 0.35476737 | 0.59071784 |
| 0.54268306 | 0.74925776 |
| 0.01832969 | 0.12628845 |
| 0.66325002 | 0.82094575 |
| 0.28991521 | 0.53085825 |
| 0.82744784 | 0.90494986 |
| 0.44300457 | 0.66206382 |
| 0.04472558 | 0.1975275 |
| 0.05584846 | 0.21990773 |
| 0.03579354 | 0.17530212 |
| 0.43768359 | 0.65895695 |
| 0.57491757 | 0.7713003 |
| 0.72454763 | 0.84030987 |
| 0.48140557 | 0.69679694 |
| 0.01671792 | 0.12170861 |
| 0.81080666 | 0.89489081 |
| 0.25450453 | 0.49432195 |
| 0.87295974 | 0.93202857 |
| 0.00181676 | 0.03787255 |

| | |
|---|---|
| 0.15365157 | 0.37178192 |
| 0.6842501 | 0.82782043 |
| 0.51343486 | 0.72576789 |
| 0.04133553 | 0.18953599 |
| 0.30097769 | 0.54136031 |
| 0.10516964 | 0.31141059 |
| 0.58463911 | 0.77595067 |
| 0.05989465 | 0.22866833 |
| 0.07228063 | 0.24924762 |
| 1.72E-04 | 0.01204292 |
| 0.88097939 | 0.93503426 |
| 0.29755149 | 0.5370834 |
| 0.06379844 | 0.23576424 |
| 0.02418911 | 0.14331915 |
| 0.78896624 | 0.8774686 |
| 0.07586749 | 0.25635962 |
| 0.26046682 | 0.49836463 |
| 0.00370866 | 0.04787642 |
| 0.08385406 | 0.27003759 |
| 0.10948603 | 0.3167692 |
| 0.01956103 | 0.13035345 |
| 0.029998 | 0.15734437 |
| 0.59265746 | 0.78092466 |
| 1.92E-04 | 0.01204292 |
| 0.07599344 | 0.25635962 |
| 1.52E-05 | 0.00370229 |
| 0.01039711 | 0.09777925 |
| 0.20270669 | 0.43863552 |
| 7.83E-05 | 0.0070694 |
| 0.51721073 | 0.7262389 |
| 0.12541598 | 0.33671637 |
| 0.06523372 | 0.23634187 |
| 0.20084121 | 0.43658797 |
| 0.01303376 | 0.11154155 |
| 0.25903795 | 0.49832297 |
| 0.09522955 | 0.29437881 |
| 0.01363016 | 0.11333901 |
| 0.96688902 | 0.99217835 |
| 0.81654887 | 0.89831425 |
| 0.49662843 | 0.70834897 |
| 0.10448023 | 0.31114442 |
| 0.21875663 | 0.45214166 |

| | |
|---|---|
| 0.55529341 | 0.75463278 |
| 0.68091458 | 0.8262441 |
| 0.94519733 | 0.98141178 |
| 0.37472711 | 0.60243265 |
| 0.03443336 | 0.17280443 |
| 0.3458715 | 0.58731216 |
| 0.01169662 | 0.10684664 |
| 0.19670511 | 0.43095941 |
| 0.04796536 | 0.20513796 |
| 0.05343719 | 0.21401199 |
| 7.01E-04 | 0.02065337 |
| 0.00746993 | 0.07805324 |
| 0.1293282 | 0.34137607 |
| 0.13075186 | 0.34261012 |
| 5.31E-04 | 0.01808044 |
| 0.37521397 | 0.60243265 |
| 0.00213924 | 0.04140956 |
| 0.38650555 | 0.61127455 |
| 0.72248391 | 0.83911345 |
| 0.63428272 | 0.80073269 |
| 0.19473625 | 0.42905304 |
| 0.10430411 | 0.31114442 |
| 0.00589695 | 0.06658637 |
| 7.33E-04 | 0.02065337 |
| 0.18957504 | 0.42341898 |
| 0.91380861 | 0.95985323 |
| 0.83346141 | 0.90710058 |
| 0.70409504 | 0.83364717 |
| 0.59982225 | 0.78689746 |
| 0.03673665 | 0.17777917 |
| 0.47451056 | 0.68937707 |
| 0.76246267 | 0.86079421 |
| 0.06590573 | 0.23708567 |
| 0.03495633 | 0.17293675 |
| 0.25496151 | 0.49432195 |
| 0.60451699 | 0.78689746 |
| 0.00327057 | 0.04431617 |
| 6.15E-05 | 0.00672269 |
| 0.09392824 | 0.29146434 |
| 0.05247327 | 0.21119191 |
| 0.62983181 | 0.7980417 |
| 0.20448077 | 0.43863552 |

| | |
|---|---|
| 0.4851114 | 0.69823703 |
| 0.39714262 | 0.6138345 |
| 9.81E-04 | 0.02416157 |
| 0.16299164 | 0.38409334 |
| 0.66227119 | 0.82094575 |
| 0.68014474 | 0.8262441 |
| 0.01823934 | 0.12628845 |
| 0.00248129 | 0.04202677 |
| 0.55646629 | 0.75463278 |
| 0.35083488 | 0.58889956 |
| 0.11690609 | 0.32661392 |
| 0.9702113 | 0.99217835 |
| 0.01427518 | 0.1157844 |
| 0.34163436 | 0.58228245 |
| 0.14832463 | 0.36808509 |
| 0.12525232 | 0.33671637 |
| 0.14895448 | 0.36808509 |
| 6.62E-05 | 0.00672269 |
| 0.00626563 | 0.06843065 |
| 0.39156838 | 0.61127455 |
| 0.74074697 | 0.84582484 |
| 0.96728187 | 0.99217835 |
| 0.01620648 | 0.11989087 |
| 0.72234452 | 0.83911345 |
| 0.03810589 | 0.18117011 |
| 0.04111779 | 0.18953599 |
| 0.6758136 | 0.8262441 |
| 0.15601203 | 0.37637322 |
| 0.18711428 | 0.41907413 |
| 0.81201674 | 0.89489081 |
| 0.44145574 | 0.66112877 |
| 0.99681817 | 0.99832219 |
| 0.55253301 | 0.75434241 |
| 2.02E-04 | 0.01204292 |
| 0.13271623 | 0.34362513 |
| 0.33529053 | 0.57752373 |
| 0.18030368 | 0.41060754 |
| 0.86728101 | 0.93202857 |
| 0.06143888 | 0.23018345 |
| 0.81821114 | 0.89892656 |
| 0.13492872 | 0.34604748 |
| 3.18E-04 | 0.01393391 |

| | |
|---|---|
| 0.50892081 | 0.72208136 |
| 0.23266356 | 0.46796728 |
| 0.9930194 | 0.99792926 |
| 0.33765613 | 0.58036878 |
| 0.70325751 | 0.83364717 |
| 0.12570778 | 0.33671637 |
| 0.29468394 | 0.53510788 |
| 0.02005076 | 0.13146387 |
| 0.06978969 | 0.24351511 |
| 0.20681616 | 0.44151844 |
| 0.21168129 | 0.44584686 |
| 0.60826072 | 0.7887017 |
| 0.77012657 | 0.86516094 |
| 0.11027579 | 0.31792275 |
| 0.69679808 | 0.83364717 |
| 0.67881735 | 0.8262441 |
| 0.40698546 | 0.62548048 |
| 0.12352128 | 0.33563126 |
| 0.00335272 | 0.04468455 |
| 0.00866376 | 0.0869585 |
| 0.01303062 | 0.11154155 |
| 0.85225655 | 0.92261595 |
| 0.0074885 | 0.07805324 |
| 0.44156571 | 0.66112877 |
| 0.07469915 | 0.25607259 |
| 0.96545071 | 0.99217835 |
| 0.06540827 | 0.23634187 |
| 0.06905724 | 0.24351511 |
| 0.07887715 | 0.26067935 |
| 0.06951709 | 0.24351511 |
| 0.08210332 | 0.26594675 |
| NA | NA |
| 0.54715722 | 0.75268836 |
| 0.67314422 | 0.8262441 |
| 0.00227182 | 0.04160725 |
| 0.13328313 | 0.34399742 |
| 0.23800619 | 0.47194886 |
| 0.26052271 | 0.49836463 |
| 0.02553565 | 0.14774643 |
| 0.02430574 | 0.14331915 |
| 0.35821392 | 0.59313221 |
| 0.46265198 | 0.68264258 |

| | |
|---|---|
| 0.99473502 | 0.99832219 |
| 0.67896732 | 0.8262441 |
| 0.63558702 | 0.80113527 |
| 0.26436785 | 0.50265057 |
| 0.17175671 | 0.39472655 |
| 0.43324517 | 0.65469948 |
| 0.27912009 | 0.51927833 |
| 0.22052052 | 0.45388147 |
| 0.11431402 | 0.32495559 |
| 0.23314305 | 0.46796728 |
| 0.00429695 | 0.05293059 |
| 0.00502044 | 0.0583088 |
| 0.80560898 | 0.89171657 |
| 0.01928032 | 0.13012228 |
| 0.19241833 | 0.4272102 |
| 0.89842196 | 0.94736323 |
| 0.20304024 | 0.43863552 |
| 0.19284888 | 0.4272102 |
| 0.52506087 | 0.72969998 |
| 0.07865502 | 0.26067935 |
| 0.57433532 | 0.7713003 |
| 0.03189817 | 0.16208257 |
| 0.1612796 | 0.38248725 |
| 0.10631093 | 0.3120245 |
| 0.08944146 | 0.28140654 |
| 0.02788555 | 0.15218014 |
| 0.02906881 | 0.15446365 |
| 0.02848303 | 0.15335563 |
| 0.11168894 | 0.31972925 |
| 0.00290789 | 0.04265434 |
| 0.01108119 | 0.10237506 |
| 0.14309809 | 0.36018187 |
| 0.00299053 | 0.04265434 |
| 0.01903768 | 0.13006417 |
| 0.0160218 | 0.11989087 |
| 0.06378976 | 0.23576424 |
| 0.54459544 | 0.75043406 |
| 0.33992639 | 0.58099884 |
| 0.30884958 | 0.5506463 |
| 0.44746662 | 0.66506465 |
| 0.19009787 | 0.42342347 |
| NA | NA |

| | |
|---|---|
| 0.37568678 | 0.60243265 |
| 9.51E-04 | 0.02415382 |
| 0.08403384 | 0.27003759 |
| 0.15662857 | 0.37674269 |
| 0.63018791 | 0.7980417 |
| 0.70609081 | 0.83364717 |
| 0.00230298 | 0.04160725 |
| 0.04198563 | 0.19061742 |
| 0.04776168 | 0.20513796 |
| 0.40396228 | 0.62319039 |
| 0.38991337 | 0.61127455 |
| 0.4608926 | 0.68128306 |
| 0.98854069 | 0.99792926 |
| 0.38820784 | 0.61127455 |
| 0.22259203 | 0.45686451 |
| 0.95199079 | 0.98344156 |
| 0.97225705 | 0.99261671 |
| 0.33934084 | 0.58099884 |
| 0.15305583 | 0.37144594 |
| 0.70071823 | 0.83364717 |
| 0.01762126 | 0.12566737 |
| 0.08210656 | 0.26594675 |
| 0.11670786 | 0.32661392 |
| 0.00675599 | 0.07227137 |
| 0.70145786 | 0.83364717 |
| 0.99008564 | 0.99792926 |
| 0.32512428 | 0.56671835 |
| 0.25753079 | 0.49732193 |
| 0.69136782 | 0.83364717 |
| 0.11427107 | 0.32495559 |
| 0.69687353 | 0.83364717 |
| 0.98572716 | 0.99743524 |
| 0.42282103 | 0.64615319 |
| 0.12322613 | 0.33563126 |
| 0.36602237 | 0.59634507 |
| 0.75703902 | 0.85781806 |
| 0.6297372 | 0.7980417 |
| 0.29486879 | 0.53510788 |
| 0.4451226 | 0.66279245 |
| 0.22365569 | 0.45686451 |
| 0.00308975 | 0.04291076 |
| 0.93876438 | 0.97848134 |

| | |
|---|---|
| 0.47203952 | 0.68937707 |
| 0.18304307 | 0.41452373 |
| 0.26558686 | 0.50265057 |
| 0.76338576 | 0.86079421 |
| 0.88802948 | 0.94128809 |
| 0.55183041 | 0.75434241 |
| 0.17873078 | 0.40816889 |
| 0.01488817 | 0.11628977 |
| 0.1163798 | 0.32661392 |
| 0.39598009 | 0.6138345 |
| 0.14657054 | 0.36552716 |
| 0.94146225 | 0.98003688 |
| 0.28256056 | 0.52447884 |
| 0.85786264 | 0.9274499 |
| 0.98189148 | 0.99743524 |
| 0.72598128 | 0.84077319 |
| 0.82509949 | 0.90405106 |
| 0.52382746 | 0.72969998 |
| 0.49561874 | 0.70834897 |
| 0.6447426 | 0.80929737 |
| 0.28759444 | 0.52911511 |
| 0.00464325 | 0.055642 |
| 0.02387814 | 0.14331915 |
| 0.89744918 | 0.94736323 |
| 0.88094406 | 0.93503426 |
| 0.00944 | 0.09359419 |
| 0.13802368 | 0.35067189 |
| 0.69266612 | 0.83364717 |
| 0.3746854 | 0.60243265 |
| 0.0274638 | 0.15218014 |
| 0.61113825 | 0.78991319 |
| 0.04307404 | 0.19136171 |
| 0.35587228 | 0.59071784 |
| 0.29165478 | 0.53284345 |
| 0.96867535 | 0.99217835 |
| 0.75758101 | 0.85781806 |
| 0.10533569 | 0.31141059 |
| 0.86800655 | 0.93202857 |
| 0.55897131 | 0.75463278 |
| 0.8204103 | 0.90012628 |
| 0.32911749 | 0.57051709 |
| 0.3218156 | 0.56508872 |

| | |
|---|---|
| 0.87184781 | 0.93202857 |
| NA | NA |
| 0.26459059 | 0.50265057 |
| 0.14553249 | 0.36470664 |
| 0.67595973 | 0.8262441 |
| 0.05161889 | 0.20983078 |
| 0.58196107 | 0.77563008 |
| 0.8292591 | 0.90494986 |
| 0.32174306 | 0.56508872 |
| 0.15979224 | 0.38097094 |
| 0.25536927 | 0.49432195 |
| 0.39029786 | 0.61127455 |
| 0.87356184 | 0.93202857 |
| 0.13174383 | 0.34261012 |
| 0.62032699 | 0.79620491 |
| 0.55970918 | 0.75463278 |
| 0.48345329 | 0.6981306 |
| 0.74622597 | 0.85088599 |
| 0.76584298 | 0.8623689 |
| 0.67494004 | 0.8262441 |
| 0.25044534 | 0.49300741 |
| 0.41677122 | 0.63810734 |
| 0.98516737 | 0.99743524 |
| 0.7475648 | 0.85121875 |
| 0.90488604 | 0.95294346 |
| 0.73787622 | 0.84373188 |
| 0.70857955 | 0.83364717 |
| 0.1236256 | 0.33563126 |
| 0.01546445 | 0.11750091 |
| 0.96841788 | 0.99217835 |
| 0.60415264 | 0.78689746 |
| 0.44393771 | 0.66224102 |
| 0.10903296 | 0.316585 |
| 0.91143798 | 0.95860165 |
| 0.19390165 | 0.42837512 |
| 0.95146493 | 0.98344156 |
| 0.97308182 | 0.99261671 |
| 0.84159986 | 0.91473354 |
| 0.10267562 | 0.30910005 |
| 0.74881037 | 0.85144452 |
| 0.11639498 | 0.32661392 |
| 0.96108311 | 0.99157432 |

| | |
|---|---|
| 0.72043384 | 0.83911345 |
| 0.55812097 | 0.75463278 |
| 0.67194095 | 0.8262441 |
| 0.99785065 | 0.99832219 |
| 0.09735446 | 0.29755328 |
| 0.78031842 | 0.87023166 |
| 0.85961631 | 0.92811164 |
| 0.16475236 | 0.38600482 |
| 0.92933884 | 0.96990049 |
| 0.91662268 | 0.96070184 |
| 0.89053062 | 0.94168529 |
| NA | NA |
| 0.75546256 | 0.85780874 |
| 0.84653373 | 0.91815941 |
| 0.76162285 | 0.86079421 |
| 0.13170318 | 0.34261012 |
| 0.04253747 | 0.19061742 |
| 0.03706662 | 0.17831457 |
| 0.20854008 | 0.44151844 |
| NA | NA |
| 0.71958479 | 0.83911345 |
| 0.73357139 | 0.843713 |
| 0.5889977 | 0.77989434 |
| 0.949522 | 0.98344156 |
| 0.04025613 | 0.18809331 |
| 0.10712537 | 0.3132839 |
| 0.07527841 | 0.25607259 |
| 0.13535896 | 0.34605923 |
| 0.61492967 | 0.79104086 |
