## Supplementary table 4 for "Selective degradation of BRD9 by a DCAF16-recruiting targeted glue: mode of action elucidation and in vivo proof of concept"

| Component name | Protein name | Peptide |
|---|---|---|
| 1:T3&:AMPTX-1-ent-1 [1]+$H^+$ | **DCAF16** | **CLLK** |
| 1:T7&:AMPTX-1-ent-1 [?]+$H^+$ | **DCAF16** | **LSHCSHCVPK** |
| 1:T28-30&:AMPTX-1-ent-1 [6]+$H^+$ | **His8-Avi-DDB1deltaB** | **QSTIVCHNRVDPNGSRYLLGDMEGR** |
| 1:T43-44&:AMPTX-1-ent-1 [?]+$H^+$ | **His8-Avi-DDB1deltaB** | **KICYQEVSQCFGVLSSR** |
| 1:T44&:AMPTX-1-ent-1 [2]+$H^+$ | **His8-Avi-DDB1deltaB** | **ICYQEVSQCFGVLSSR** |

| Modifiers | Response | Sequence start | Sequence end | Observed mass (Da) | Expected mass (Da) | Mass error (ppm) |
|---|---|---|---|---|---|---|
| **AMPTX-1-*ent-1* [1]** | **3135110** | **58** | **61** | 1167.7035 | 1167.6999 | 3.1 |
| **AMPTX-1-*ent-1* [?]** | **1569880** | **97** | **106** | 1801.9385 | 1801.9281 | 5.8 |
| **AMPTX-1-*ent-1* [6]** | **1416422** | **301** | **325** | 3508.7525 | 3508.7617 | -2.6 |
| **AMPTX-1-*ent-1* [?]** | **5599026** | **465** | **481** | 2638.3358 | 2638.356 | -7.7 |
| **AMPTX-1-*ent-1* [2]** | **2628654** | **466** | **481** | 2510.2432 | 2510.2611 | -7.1 |

| Observed m/z | Charge | Observed RT ( min) |
|---|---|---|
| 584.3554 | 2 | 34.28 |
| 601.3177 | 3 | 41.02 |
| 877.9436 | 4 | 28.51 |
| 880.1168 | 3 | 30.53 |
| 837.4192 | 3 | 35.18 |

| Component name | Protein name | Peptide |
|---|---|---|
| 1:T3&:AMPTX-1-ent-1 [1]+$H^+$ | **DCAF16** | **CLLK** |
| 1:T4-5&:AMPTX-1-ent-1 [7]+$H^+$ | **DDA1** | **FHADSVCKASNR** |
| 1:T43-44&:AMPTX-1-ent-1 [?]+$H^+$ | **His8-Avi-DDB1deltaB** | **KICYQEVSQCFGVLSSR** |
| 1:T44&:AMPTX-1-ent-1 [2]+$H^+$ | **His8-Avi-DDB1deltaB** | **ICYQEVSQCFGVLSSR** |
| 1:T57-58&:AMPTX-1-ent-1 [3]+$H^+$ | **His8-Avi-DDB1deltaB** | **TECNHYNNIMALYLKTK** |

| Modifiers | Response | Sequence start | Sequence end | Observed mass (Da) | Expected mass (Da) |
|---|---|---|---|---|---|
| **AMPTX-1-*ent-1* [1]** | **70251** | **58** | **61** | 1167.7031 | 1167.6999 |
| **AMPTX-1-*ent-1* [7]** | **114328** | **19** | **30** | 2026.0344 | 2026.0368 |
| **AMPTX-1-*ent-1* [?]** | **538628** | **465** | **481** | 2638.3387 | 2638.356 |
| **AMPTX-1-*ent-1* [2]** | **198198** | **466** | **481** | 2510.2436 | 2510.2611 |
| **AMPTX-1-*ent-1* [3]** | **295905** | **643** | **659** | 2747.4008 | 2747.4088 |

| Mass error ( ppm) | Observed m/z | Charge | Observed RT (min) |
|---|---|---|---|
| 2.7 | 584.3552 | 2 | 34.31 |
| -1.2 | 1013.521 | 2 | 26.95 |
| -6.6 | 660.3401 | 4 | 30.56 |
| -7 | 837.4194 | 3 | 35.18 |
| -2.9 | 458.7395 | 6 | 56.82 |
